# Network inference from temporal phosphoproteomics informed by protein-protein interactions

**DOI:** 10.1101/2023.04.26.538385

**Authors:** Michael Plank

## Abstract

Network inference from time-course data holds the promise to overcome challenges associated with other methods for deciphering cell signaling networks. Integration of protein-protein interactions in this process is frequently employed to limit wiring options.

In this study, a graph approach for the analysis of data of high temporal resolution is introduced and applied to a 5 s resolution phosphoproteomics dataset. Steiner trees informed by protein-protein interactions are constructed on individual time slices, which are then stitched together into a temporal signaling network.

Systematic benchmarking against existing knowledge indicates that the approach enriches signaling-relevant edges. The workflow is compatible with future extensions for reliably extracting extended signaling paths.

## Introduction

The first step in understanding any system is the cataloging of its components and their functionally relevant interactions. In the context of cell signaling, this entails defining which proteins participate in a signaling network and which regulatory interactions exist between them. This process of causal network inference is often based on manipulation of the network, for example through gene deletions, and the reconstruction of regulatory relationships from the observed consequences ^1–4^. However, networks from artificial manipulations may no longer represent the unmanipulated system. Therefore, inference based on the intact network is preferable. An alternative option for network inference is the physical mapping of interactions between components of the signaling pathway. This has been a successful strategy in the delineation of transcriptional networks based on transcription factor – promoter interactions ^5^.

Finally, signaling dynamics may be reconstructed from time course data, based on the reasoning that the ordering of observable changes (e.g. in transcription or protein phosphorylation) should reflect the ordering of the components in a pathway. This strategy is limited by the completeness and the temporal resolution of the available data. Even with high-quality data, however, the number of combinations in which signaling components can be arranged consistent with the data is generally too high to render this approach useful in practice. Therefore, in the past, inference from temporal data has mainly been employed to prioritize between a limited number of network models ^6,7^. An alternative that retains the option of *de novo* inference, in contrast to prioritization alone, is the combination of the time course and physical interaction-based approaches. In the case of phospho-signaling, this may be achieved by restricting signaling edges to previously observed protein-protein interactions (PPIs).

Unlike in transcriptional networks, where it is comparably straightforward to measure if the binding of a transcription factor to a promoter causes induction or repression of the corresponding gene, the signaling consequence of PPIs is much harder to determine. The challenge, therefore, is to distinguish the interactions that are signaling-relevant (in the given context) from all other PPIs.

Graph algorithms, such as Steiner trees, have been widely employed for network inference from static snapshots of signaling networks ^8–11^. These aim to determine the most likely trees connecting required nodes, such as analytes that change in abundance upon a treatment. Unaffected analytes are included in the network where required and are referred to as “Steiner nodes” in the context of Steiner trees. The reasoning behind these methods is that some connection should exist between analytes affected by the same treatment. The signaling relevance of the inferred connections here stems from the selection of required nodes, whereas only a subset of edges in the underlying network is signaling-relevant. For example, in the case of inferring phospho-signaling in a PPI network, an interaction between two proteins may or may not be relevant for transduction of the signal, but if the phosphorylation levels of both proteins change upon treatment, confidence in its signaling-relevance increases.

Time course data add a further layer of information, as they provide an order in which analytes need to be connected. Network inference on temporal phosphoproteomics data, informed by PPI has been performed previously ^12,13^. Using a graph-based method, Budak, et al. employed a two-step approach: First, graphs were built on PPI networks around proteins significantly differentially phosphorylated (in the following referred to as “significant proteins”) at each time point (TP) in the time course ^12^. Second, selected proteins were connected in the order in which they entered graphs, again through edges of the underlying PPI network (Fig. 1A). As graphs are built first for individual TPs, it is inherently assumed that the experimental sampling rate is slow compared to the speed of signal transduction, which implies that multiple signaling steps are represented within each TP. This in turn means that the order in which the signal is transduced within each TP cannot be resolved.

**Fig. 1.**
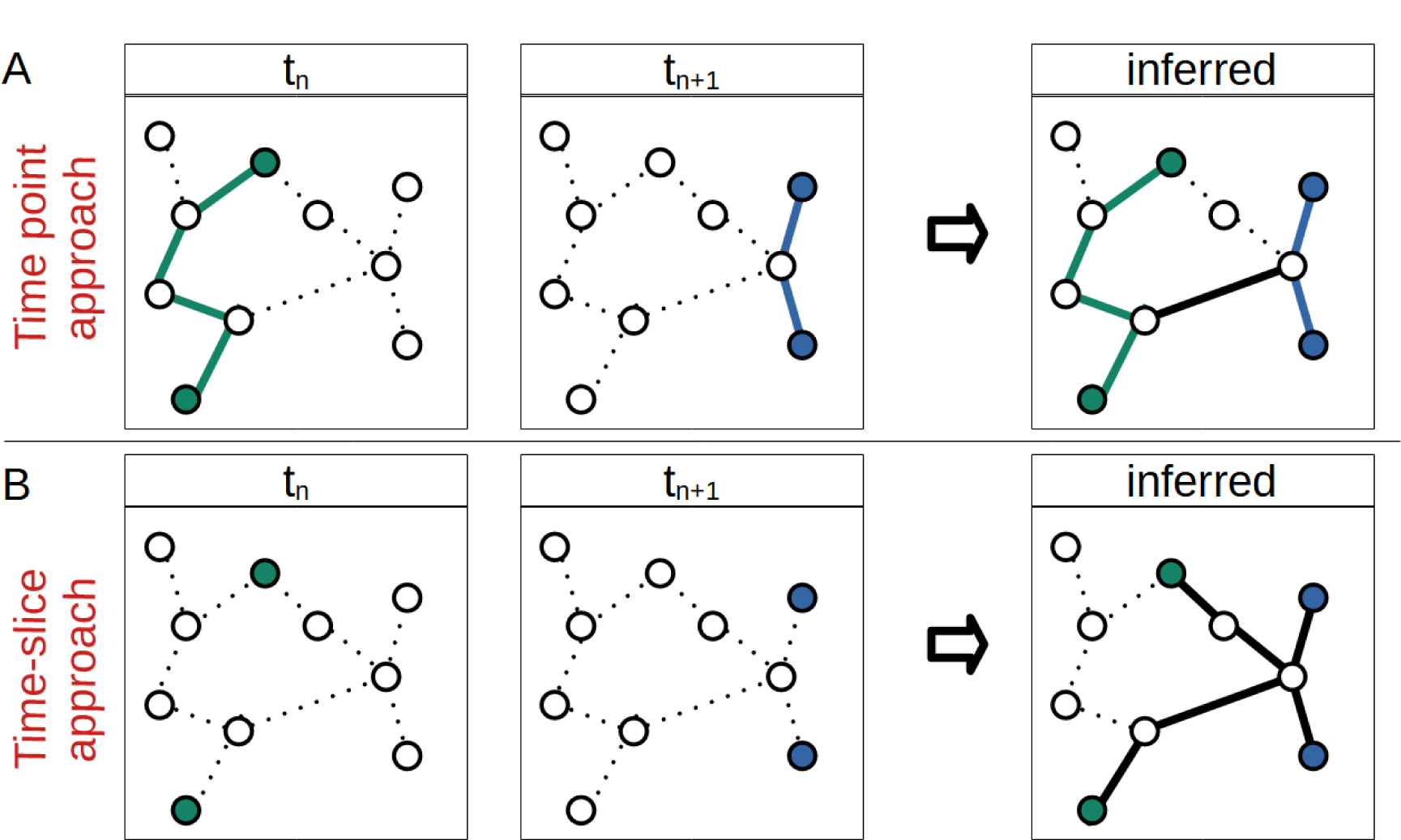
Comparison of time point and time-slice based approach to network inference from temporal data. A, Time point approach: Graphs are constructed at each TP, choosing significant proteins at the given TP as terminal nodes. The TP graphs are then connected into one graph via an edge linking selected nodes. B, Time slice approach: Graphs are constructed on time-slices of two adjacent TPs, treating significant proteins from the first and second TP as outgoing and incoming terminals respectively. Nodes that are not chosen in the at individual TPs can be chosen in a time-slice and nodes necessary for a TP-graph may be rejected for a time-slice graph. Colored nodes represent proteins significant at the TP. Dotted lines represent edges in the PPI network. Full lines represent edges of the inferred graph.

Here I explore an approach that immediately searches for the most likely connections between TPs, without first searching for connections within TPs (Fig. 1B). The approach is aimed at high-resolution data so that individual signaling steps are resolved between TPs. The underlying assumption of being able to trace signaling flow (rather than having to reconstruct multiple layers of signaling in each TP) justifies simplifying assumptions that ease computational burdens; in particular in the present analysis, a sub-network close to the significant proteins, instead of the complete proteome served as the underlying PPI network. The dataset used for this study is a phosphoproteomics time course of the yeast osmotic stress response with samples taken every 5 s for the first minute after stimulation ^14^. The goal of this work is the introduction of a PPI-informed network inference approach for high-resolution time course data and evaluation of its capacity to identify cellular signaling pathways.

## Methods

### Workflow

An overview of the computational analysis is outlined in this section before the detailed discussion below. The workflow is shown in fig. 2. Network inference was guided by an underlying unexpanded (fig. 2 left) or expanded (right) PPI network. In each case, significantly differentially phosphorylated proteins and their TPs of significance were extracted from the study by Kanshin et al. (fig. 2, steps A, B) ^14^. The PPI network for these proteins was obtained from STRING-db with or without expansion by one shell of interacting proteins (C) and filtered to yield high-confidence interactions (D) ^15^. Significant proteins were flagged in the network and the TP of significance retained (E). In the case of the expanded network, Steiner trees were built on slices of two neighboring TPs (F). This was done once, as Classical Steiner Trees on fully connected subgraphs, and once as Prize Collecting Steiner Trees via integer linear programming (ILP). (A Classical Steiner Tree here refers to a Steiner Tree in which a node is either required or not, without assigning node prizes or edge costs.) Time slices were combined into a graph representing the full time course (G). From this expanded network, or from the original unexpanded network, paths through the network over time were extracted (H) and plotted (I). The derived temporal network was evaluated via the betweenness centrality of its nodes (J), its edge confidence scores, and connections to kinases (K).

**Fig. 2.**
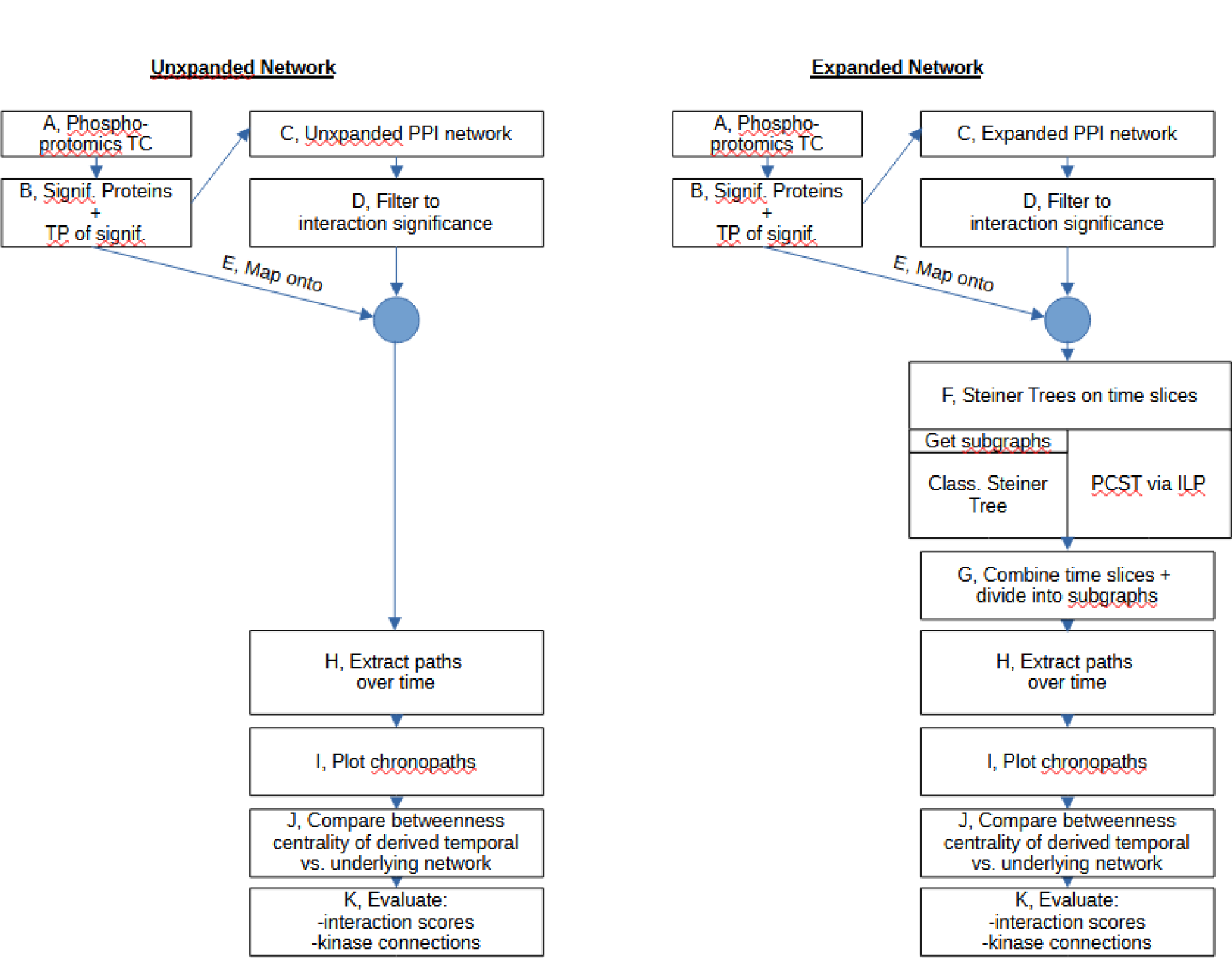
Outline of the workflow used in this study with unexpanded (left) and expanded (right) underlying PPI network. See text for details. TC: time course, TP: timepoint, PPI: protein-protein interaction, PCST: Prize-Collecting Steiner Tree, ILP: Integer Linear Programming

Analysis was performed in R (version 4.2.0) using the igraph package ^16,17^. Gurobi (version 9.5.2) was used for ILP ^18^.

### Classification of phosphosites as significant

As the phosphoproteomics dataset does not include replicate measurements, phosphosites were defined as significantly changed in abundance if their intensity changed by at least three-fold in either direction at any TP after hyperosmotic shock. In case a phosphosite exhibited a significant change at more than one TP, the first significant TP was selected. For simplicity, significant phosphosites on the same protein were collapsed, keeping the first TP of significance.

### Construction of the underlying network

To construct an underlying PPI network, significant proteins were imported into STRING-db ^15^. The network was expanded by nodes of the first neighboring shell and edges with a combined confidence score of at least 0.7 from experimentally determined interactions, curated databases, and text mining were retained. As in subsequent steps, a maximum of two Steiner nodes was allowed between terminals, the reduced underlying network compared to the full yeast interactome does not pose notable constraints on the further procedure. The network obtained from STRING-db was further filtered to a score threshold of 0.7 on experimentally determined interactions. The other types of evidence were used only for evaluation of inferences (see below). A network of limited size (574 nodes, 6092 edges) was obtained in this way to restrict computational requirements.

For the Classical Steiner Tree approach, as the corresponding algorithm requires a fully connected subgraph, the graph was split into fully connected subgraphs, and subgraphs of less than ten nodes were removed. A single subgraph of 548 nodes remained. Underlying networks for the Prize Collecting Steiner Tree approach were constructed in the same way, with the exception that no division into subgraphs was performed. Unexpanded networks were obtained from STRING-db in the same way but without network expansion. Paths from the unexpanded network were obtained by linking proteins significant at adjacent TPs.

Network communities were derived based on edge betweenness ^19^. For their annotations, a keyword was selected to represent process and function GO-terms found enriched by the Gene Ontology Finder tool of the Saccharomyces Genome Database ^20^. Within-cluster sums of squares were calculated as Σ(d ^2^)/(n-1), where d is the number of edges between node i and the center of the cluster and n the number of nodes in the cluster. Centers were defined as the node with the shortest total distance to all other nodes in the cluster.

### Construction of Steiner Trees

Steiner trees were constructed on time slices of the phosphoproteomic data; pairs of adjacent measurement TPs were selected and proteins significant at these TPs were defined as required terminal nodes. A modified version of Steiner trees was constructed at each time-slice, where terminals of the first TP (“out-terminals”) were only assigned outgoing edges and terminals from the second TP (“in-terminals”) only incoming edges from the underlying PPI network. All edges not linked to terminal nodes were considered in both directions. Steiner trees were obtained on the resulting directed graphs.

### Classical Steiner Trees (CSTs)

Classical Steiner Trees (CSTs) were built on the underlying network between the terminal nodes using the minimum spanning tree algorithm of the SteinerNet R-package ^21^. Steiner nodes were assigned a TP in the middle of the TPs of the terminals. Subsequently, all paths between terminals containing maximally two consecutive Steiner nodes were retained. Terminal nodes that could not be connected to any other terminal node in this way were discarded.

### Prize Collecting Steiner Trees (PCSTs)

For the construction of Prize Collecting Steiner Trees (PCSTs), prizes were assigned to nodes and costs to edges: Absolute values of the log_3_ of each protein’s maximum phosphorylation change (in either direction) was selected as node prizes and 1-e_S_, where e_S_ is the “experimentally determined interaction” confidence score obtained from STRING-db, as edge costs. PCSTs were constructed as trees for each time slice that contained the required nodes while minimizing the sum of node prizes minus edge costs.

Node prizes and edge costs were chosen in this way based on the following rationales: At these settings, node prizes and edge costs are always positive. It was reasoned, that even a protein without observable phosphorylation change should not contribute negatively to the objective function, as it may constitute a missing value or signal via a mechanism other than phosphorylation. Further, edges should not contribute positively, as even a highly confident PPI may not be signaling relevant. Proteins with no observed phosphorylation change should contribute no prize. Edges with the maximum confidence of 1 should contribute no cost, while edges of zero confidence incur a cost of 1. Proteins with a fold-change above 3 were included in trees as required nodes. At the given settings, nodes with a fold-change close to 3 compensate for the inclusion of one edge of very low confidence.

The algorithm was implemented as an Integer Linear Program with binary variables y_i_, indicating if a node i was selected as part of the tree, and x_ij_, indicating if an edge from node i to node j was selected, minimizing the objective function in eq. (1) ^10^.

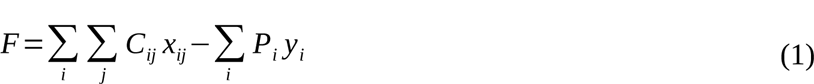

The following constraints were imposed: 1, out-terminals need to have at least one outgoing edge; 2, in-terminals need to have at least one incoming edge; 3, Steiner nodes need to have at least one incoming and one outgoing edge; 4, all terminal nodes need to be selected; 5, nodes with at least one incoming or outgoing edge need to be selected; 6, no circles are allowed. Therefore, the algorithm was allowed to return multiple disjoint trees, as long as each connected at least one out-terminal to at least one in-terminal.

Subsequently, all paths between terminals containing maximally two consecutive Steiner nodes were retained. Terminal nodes that could not be connected to any other terminal node in this way were discarded.

### Extraction of paths from temporal signaling networks

The temporal signaling network was constructed by concatenation of slices with significant proteins constituting an in-terminal in the earlier TP and an out-terminal in the subsequent TP. Paths were extracted by designating each out-terminal as the starting node of a path. For each path, the network was then searched for nodes that had an input from the starting node and the TP of the starting node. For each such node, a copy of the path was created, and the new node appended. The process was then repeated for each newly created path. Path extraction was terminated if more than two Steiner nodes were encountered in a row or if the total path length exceeded ten nodes. Once no further node could be connected, a path was stored. Steiner nodes were then trimmed off the ends of each path so that each path ended in an in-terminal, and only paths with a maximum of two Steiner nodes between significant proteins were retained. Paths that constituted subpaths or duplicates of other paths were removed if both their node sequence and TP-sequence were matched.

### Evaluation of inferred networks

Paths inferred via Steiner Tree approaches were evaluated via the ‘curated database’ and ‘text mining’ scores of their edges as obtained from STRING-db. Note that the underlying PPI network had been built based on edge selection only via ‘experimentally determined interaction’ scores. Scores of edges on Steiner trees and on inferred paths were compared to those of the underlying network via a one-sided Wilcoxon rank test.

Additionally, the enrichment of edges between any upstream kinase and terminal nodes was compared between inferred paths, underlying networks, and Steiner trees. Kinase identities were obtained from the Biogrid kinome project ^22^. Edges from a kinase to a terminal node were counted and compared to all connections to a terminal node. As the underlying network is undirected, any edge between a kinase and significant protein and any edge associated with a significant protein, respectively, were counted. Enrichment analysis was performed using Fisher’s Exact test.

GO analysis of the inferred networks was performed using the Gene Ontology Finder tool of the Saccharomyces Genome Database with all proteins in the underlying PPI network as background ^20^. The betweenness centrality of a node n was calculated as the number of shortest paths traversing the node ^23^. In case several shortest paths existed between a pair of nodes, the fraction of these paths traversing n was used for the count.

## Results

### Unexpanded underlying PPI network

In a first step, I explored if signaling information could be extracted by connecting the 138 proteins differentially phosphorylated after hyperosmotic shock (>3x change) based on their PPIs. Interaction data were obtained from STRING-db ^15^. PPIs are associated with confidence scores for different types of evidence. Interactions with an “experimental evidence” score above 0.7 (“high confidence”) were selected.

The 32 proteins that had at least one interaction with another protein formed a sparsely connected network with 23 edges (fig. 3A). The network consisted of nine disjoint subgraphs with two to nine nodes each. The largest subgraph contained the well-known effectors of the osmotic stress response Hog1, Pbs2, Ssk2, and Sko1.

**Fig. 3.**
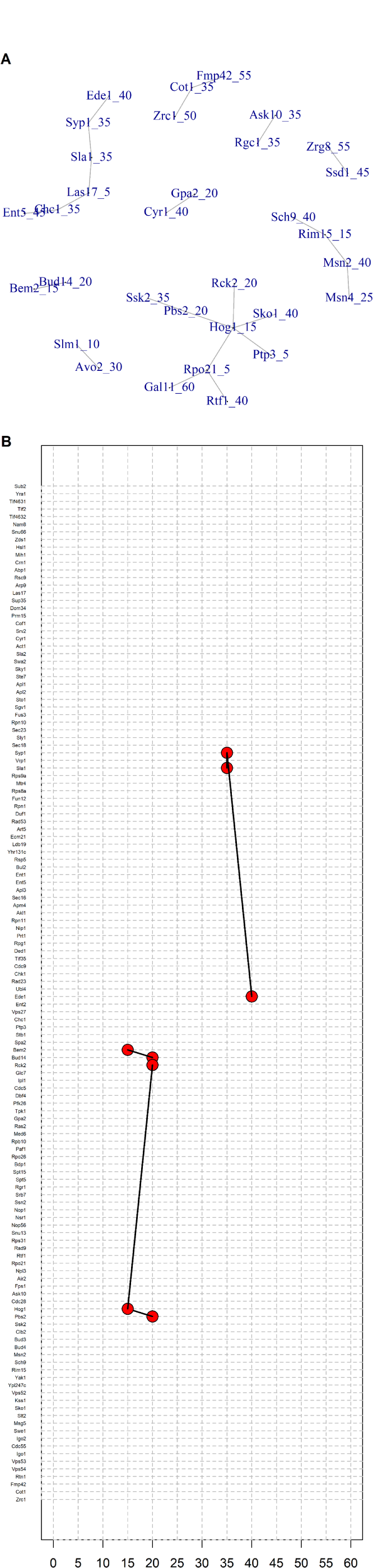
Network of significant proteins. A, Network of significantly differentially phosphorylated proteins with interaction score above 0.7. No network expansion was applied. Numbers after the protein name indicate the TP of significance in s. Only proteins for which interactions were detected are shown. B, Signaling paths among proteins significantly differentially phosphorylated at the indicated TPs (in s) after hyperosmotic stress. Interacting proteins with an experimental evidence interaction score > 0.7 and significant at the same or subsequent TPs are connected by edges.

Even more edges were removed when extracting signaling paths between proteins. Signaling paths were defined as sequences of neighbors in the underlying network that exhibited significant changes at the same or subsequent TPs. At the interaction confidence cutoff of 0.7, only one- and two-edge paths were found (fig. 3B).

Relaxing the interaction confidence threshold applied in STRING from 0.7 to 0.4 and 0.15 increased the number of edges in the PPI network to 53 and 90, respectively. Therefore, even at low interaction confidence, the proteins remained sparsely connected.

In summary, in the case of *S. cerevisiae* hyperosmotic stress-responsive signaling, differential phosphorylation alone appears to yield insufficient information to place proteins into a network that is sufficiently connected to extract signaling information. As signals can be passed on in ways other than protein phosphorylation and, as not all phosphorylation events are detected in mass spectrometry experiments (see section ‘Evaluation by comparison to published studies’ below), it may be overly restrictive to build a network based solely on differentially phosphorylated proteins.

### Underlying network with 1-shell expansion

To allow for signal transmission via means other than protein phosphorylation and for missing values in the phosphoproteomics experiment, one shell of interacting nodes was added to the network of significantly differentially phosphorylated proteins with an ‘experimental evidence’ interaction confidence cutoff of 0.7. This yielded a graph of 574 nodes and 6092 edges (fig. 4A). The median and mean node degree were 8 and 21 respectively and the median and mean node betweenness centrality were 73 and 864 (fig. 4B). Importantly, the vast majority (548) of nodes formed a fully connected subgraph, while only 26 nodes were part of smaller subgraphs of up to 4 nodes (fig. 4A). The obtained network retained 88 significant proteins with at least one confident PPI (fig. 4C). Unlike the network of only the significant proteins, the expanded network in theory offers the possibility for extracting signaling routes that connect most of the significant proteins.

**Fig. 4.**
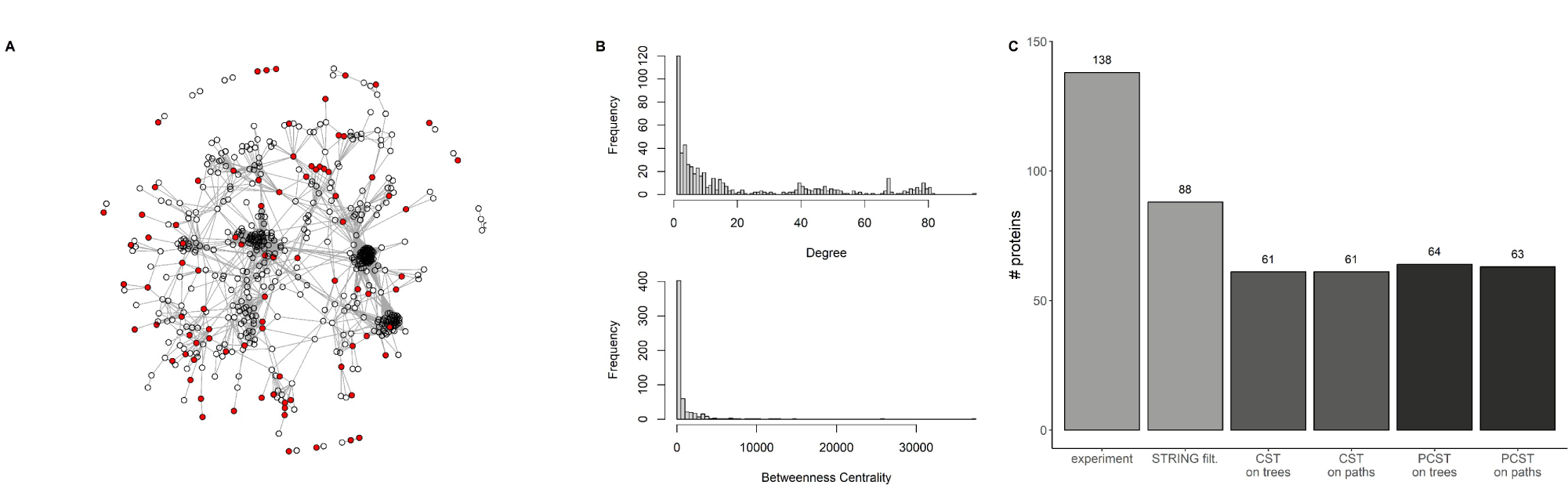
Protein-Protein interaction network via 1-shell expansion from significant proteins. A, PPI network obtained through expansion of the network of differentially phosphorylated proteins by one shell of interactors and trimming to an interaction confidence score above 0.7. Significant proteins are shown in red. (Three subgraphs do not contain significant nodes, because edges were removed when thresholding on edge confidence scores.) B, Distirbutions of node degrees (top) and betweenness centrality scores of the 1-shell expanded network. C, Number of significant proteins at different steps of network inference. 138 significant proteins were found experimentally, of which 88 were part of the PPI network obtained from STRING-db with at least one confident interaction. 61 proteins each were on CST trees and paths extracted over time. 64 proteins were on PCST trees and 63 on paths

Based on edge betweenness, the underlying network could be divided into 30 communities ^19^. Representative keywords based on significantly enriched GO-terms were associated with each of the eleven communities that encompassed at least ten members. These communities are indicated by color in the PPI network in fig. 5A. Two of them were associated with “signal transduction” gene ontologies and manual inspection revealed that one mainly contained proteins of the MAPK (“Signal transduction (I)”, dark blue in fig. 5*A*) and the other of the Protein Kinase A (“Signal transduction (II)” light blue fig. 5*A*) pathway. A third community was strongly enriched in members of the TOR signaling pathway (purple in fig. 5*A*).

**Fig. 5.**
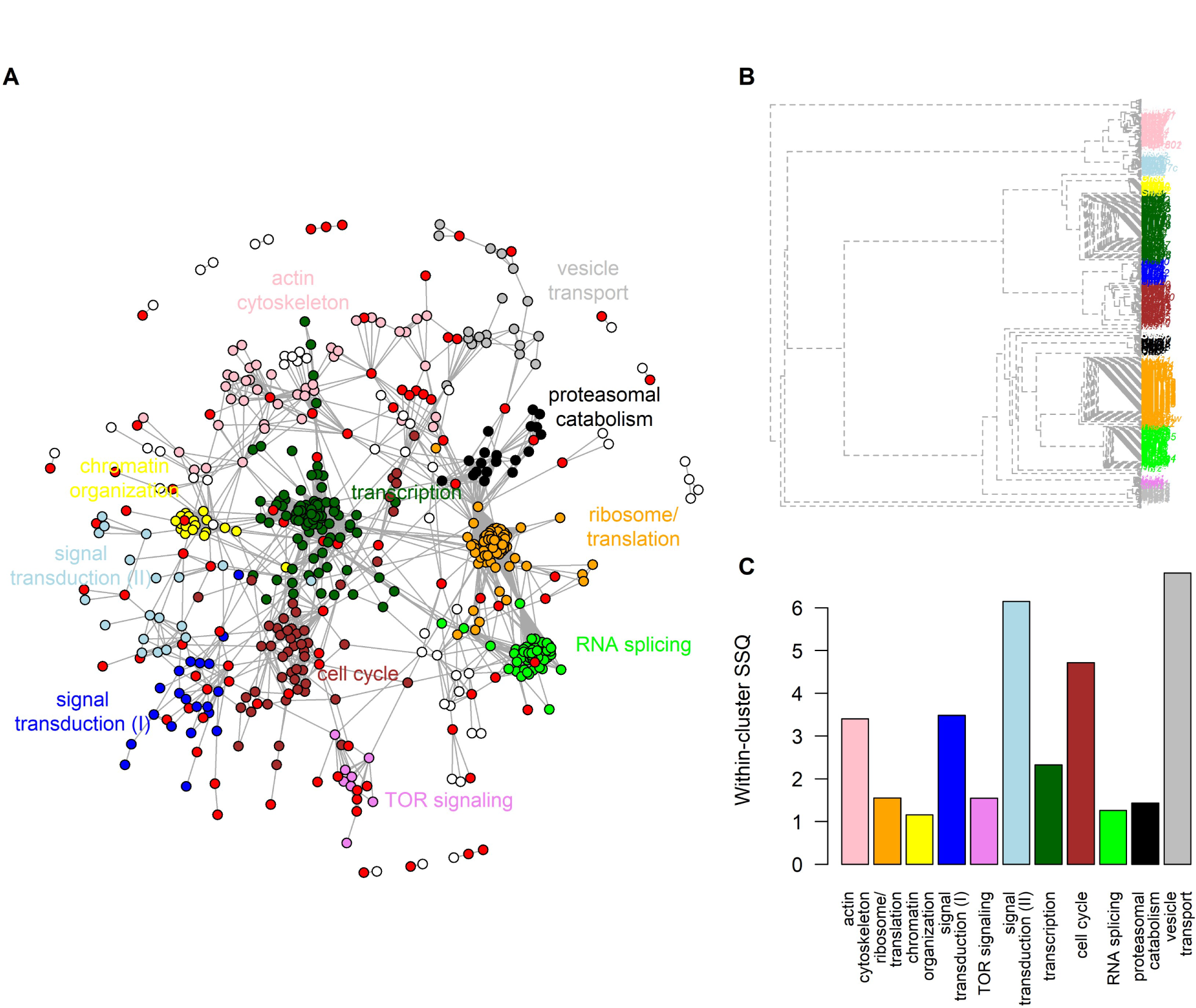
Communities in the PPI network. A: PPI network with communities defined by edge betweenness indicated by color coding. Communities are labelled in the corresponding color by a keyword summarizing the main functional category of its members. Nodes corresponding to significantly differentially phosphorylated proteins are shown in red. B: Dendrogram of communities in the underlying networks defined by edge betweenness. Color coding as in A. Branch length is proportional to the number of edge cuts necessary for separation of communities. C: Density of network communities shown as within-cluster sum-of-squares (SSQ). Color coding as in A.

Unsurprisingly, the clusters associated with chromatin organization and transcription, as well as ribosome/translation and RNA-splicing, respectively, were closely connected. Furthermore, one signal transduction cluster was closely associated with the cell cycle cluster, while the other signaling cluster was closer to the actin cytoskeleton cluster (fig. 5B). The clusters corresponding to the “ribosome/translation”, “chromatin organization”, “TOR signaling”, “RNA splicing” and “proteasomal catabolism” categories were highly compact, consistent with many of the cluster members existing in protein complexes (fig. 5C).

The expansion of the network affords the possibility of connecting significant proteins via interactors, which may have been missed in the phosphoproteomic analysis, or which may transmit information via means other than phosphorylation. As these proteins are not differentially phosphorylated, the straightforward extraction of signaling paths by connecting proteins reaching significance at subsequent TPs is no longer possible.

Therefore, the network according to which the signal was transmitted over time was extracted via graph-based approaches (fig. 2). In a time-slice-based approach, proteins significant at two subsequent TPs were first connected via interactors according to the underlying PPI network. The time-slices were then stitched together to obtain the signaling flow over the course of the experiment.

Two Steiner Tree approaches were evaluated for the extraction of high-confidence interactions between significant proteins. In the first, subsequently referred to as Classical Steiner Tree (CST), the tree with the minimum number of nodes containing all significant proteins of the time-slice was constructed. Second, a Prize-collecting Steiner Tree (PCST) was built based on the maximization of node prizes minus edge costs.

### Classical Steiner Tree approach

#### Description of the inferred network

The inference of signaling networks based on underlying PPI graphs has recently emerged as a potential way of network modeling in contexts in which little is known about the underlying network structure ^10,12,13^. Here, CSTs were constructed for each 5 s slice of adjacent TPs, where proteins significant at the first TP were treated as outgoing and of the second TP as incoming terminals (fig. 6A). A minimum-spanning tree algorithm was applied, then trees were trimmed to a maximum of two Steiner nodes between outgoing and incoming terminals, which at some time slices resulted in splitting into subtrees ^21^. A median of 51% of significant nodes in the input were retained in the Steiner trees, and only for the last TP, no tree was obtained (fig. 6B).

**Fig. 6.**
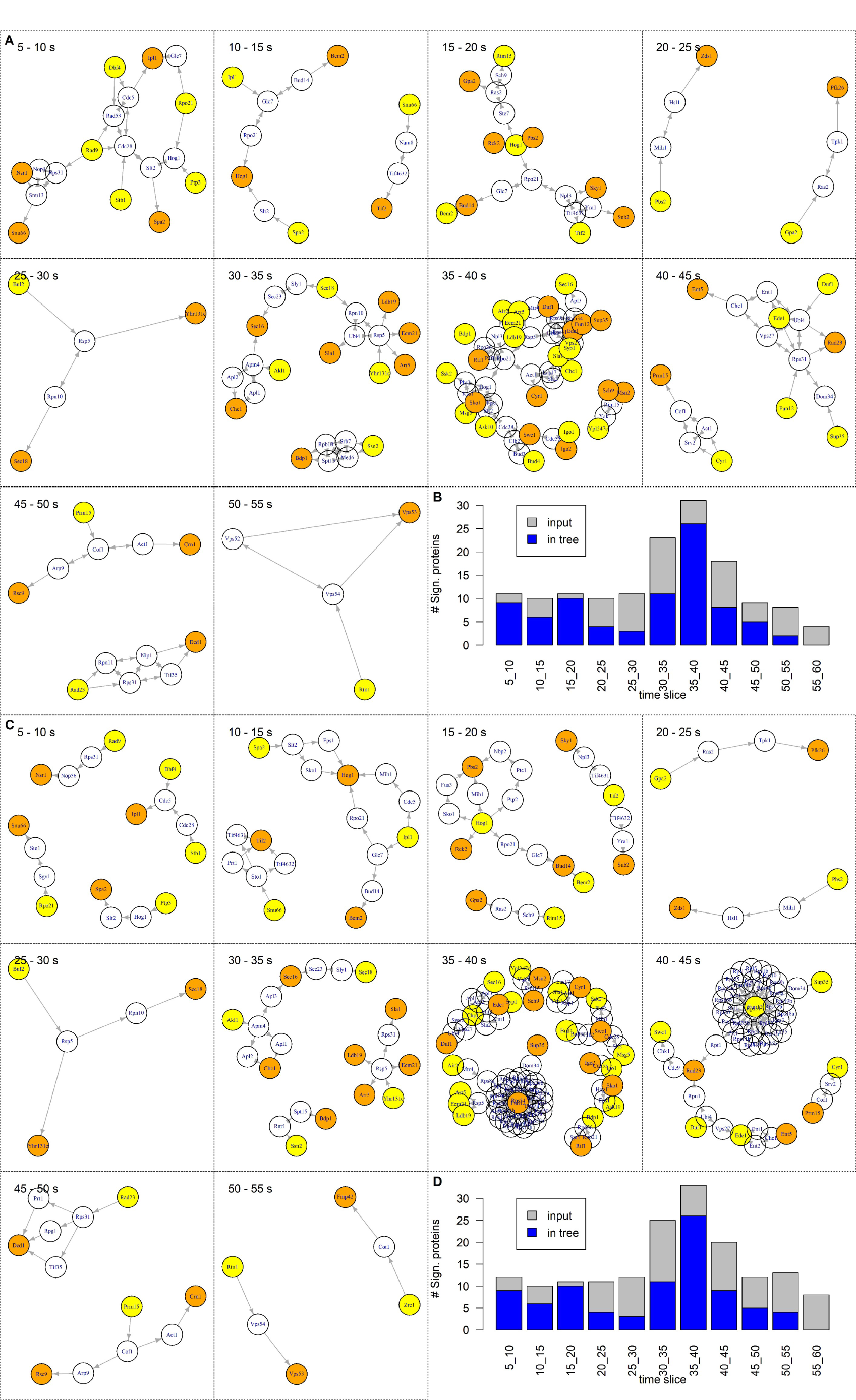
Steiner trees for time-slices of the phosphoproteomics experiment. Steiner trees created via CST (A) and PCST (C) approach at each time-slice. Incoming terminals are shown in yellow, outgoing terminals in orange and Steiner nodes in white. The 55 - 60 s time slice not containing inferred trees are omitted. Number of significant nodes in the input and output of the creation of CST (B) and PCST (D) at each time-slice.

Through linking up the Steiner trees of the time slices, the signaling flow through the network over time was obtained and is shown in fig. 7A and suppl. video 1. The network contained 258 edges and 115 nodes, of which 61 were significant. Only the tree obtained for the 50 to 55 min time slice was unconnected, while all other TPs form a connected graph.

**Fig. 7.**
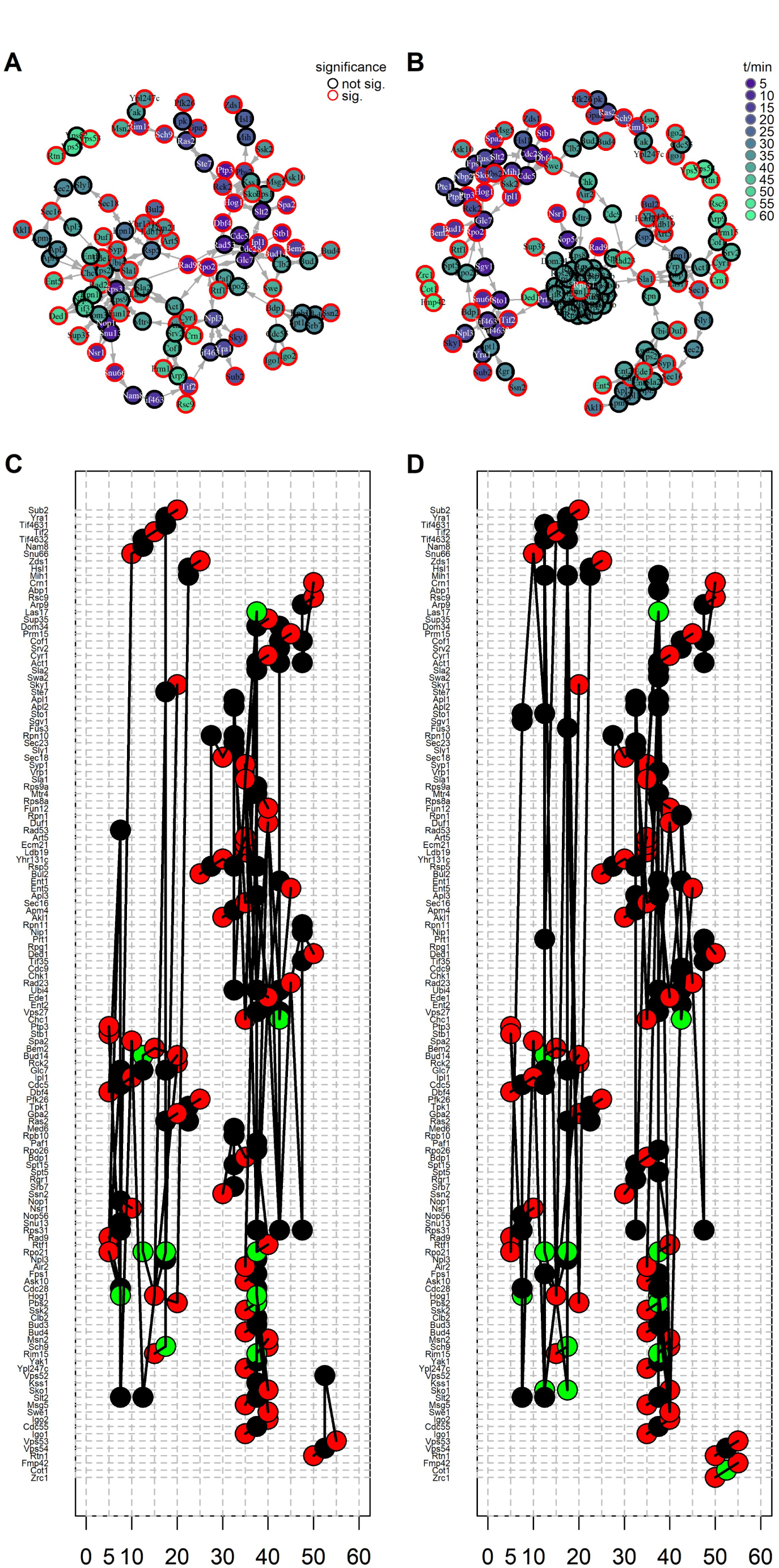
Time development of inferred signaling networks. Signaling networks representing the full phosphoproteomics time course, obtained via the CST (A) and PCST (B) approach. Node fill colors indicate the (first) TP at which a node was part of a Steiner tree. Node frame color indicates if a protein was significantly differentially phosphorylated. Chronopath plot formed by the extracted paths obtained via the CST (B) and PCST (D) approach. Red nodes indicate terminals. Significant proteins that became part of the network at a different TP than the one of their significance are shown in green. Chronopath plots are show in high-resolution in suppl. files 1 and 5.

Subsequently, paths through trees of adjacent time slices were extracted. After eliminating subpaths contained in longer paths, 1127 such paths were obtained.

Individual paths are shown in suppl. file 1. To simultaneously visualize all extracted signaling paths over time, they were presented in a graph referred to as ‘chronopath plot’ here (fig. 7B and suppl. file 2). Paths spanning the first significant proteins at 5 s to the last ones at 55 s were not obtained. Instead, individual sections are disconnected at 25 and 50 s. (The higher number of sections here, compared to the number of subgraphs above is due to the application of a threshold on the maximum number of subsequent Steiner nodes in a path.) Within each section, several nodes are connected by more than one alternative path. In other words, paths are entangled and threaded into networks with multiple inputs and outputs.

Many of the significant proteins are connecting nodes (e.g. Hog1, Yhr131c, and Prm15), whereas others either receive only input (e.g. Msn2, Pfk26, and Zds1) or only provide output (e.g. Ypl247c, Bul2, and Ssk2). It cannot be distinguished if these are truely nodes that connect to biological output or input, respectively, that is not captured by PPI or if connections were missed at the chosen analysis parameters. It is not unexpected, that the detectable signaling ends on certain proteins, such as transcription factors, while other proteins may directly sense the change in osmolarity without input from other proteins.

A further noteworthy feature is that many proteins significant at a given TP additionally enter the network as Steiner nodes at other TPs (green nodes in chronopath plots). This may be an artifact, or could reflect true multiple participation points of these proteins in the transduction path of the signal.

#### Hypothesis generation

I next attempted to derive novel hypotheses about the osmotic stress response from the inferred network.

First, GO analysis was used as a means of systematic hypothesis generation. The GO-term “positive regulation of cell cycle process” was found to be significantly enriched (corrected p = 0.008) compared to the underlying PPI network. Proteins belonging to this category are highlighted in suppl. file 3. Notably, five of the nodes were Steiner nodes in the 35 to 40 s time-slice, indicating that signal transduction started to impact cell cycle regulation around this time after stimulation.

As a second way to systematically generate hypotheses about novel signaling paths, it was reasoned that proteins with a central role in the signal transduction should preferentially be located on many paths through the network. Therefore, the betweenness centrality of nodes in the inferred network was calculated, and the results are shown in fig. 8A. Hog1 was identified as a hub node, which is unsurprising given its central role in the osmotic stress response. The identification of Hog1 as a hub lends confidence to the method’s ability to identify other signaling hubs in the network. Another hub node with betweenness centrality similar to Hog1 was the RNA-Pol II subunit Rpo21. The chronopath plot reveals an intimate connection between Rpo21 and Hog1 (fig. 7*B*). Although Rpo21 is differentially phosphorylated 5 s after salt addition and Hog1 is phosphorylated 10 s later, the network inference algorithm found a way of establishing direct connections between the nodes by either introducing Hog1 or Rpo21 as Steiner nodes. Recruitment of Rpo21 and Hog1 to stress-responsive genes after osmotic stress, which in turn leads to the accumulation of DNA remodelers, has previously been shown ^24,25^. The possibility that the recruitment of Hog1 is influenced by the observed early phosphorylation of Rpo21 arises as a hypothesis for future research.

**Fig. 8.**
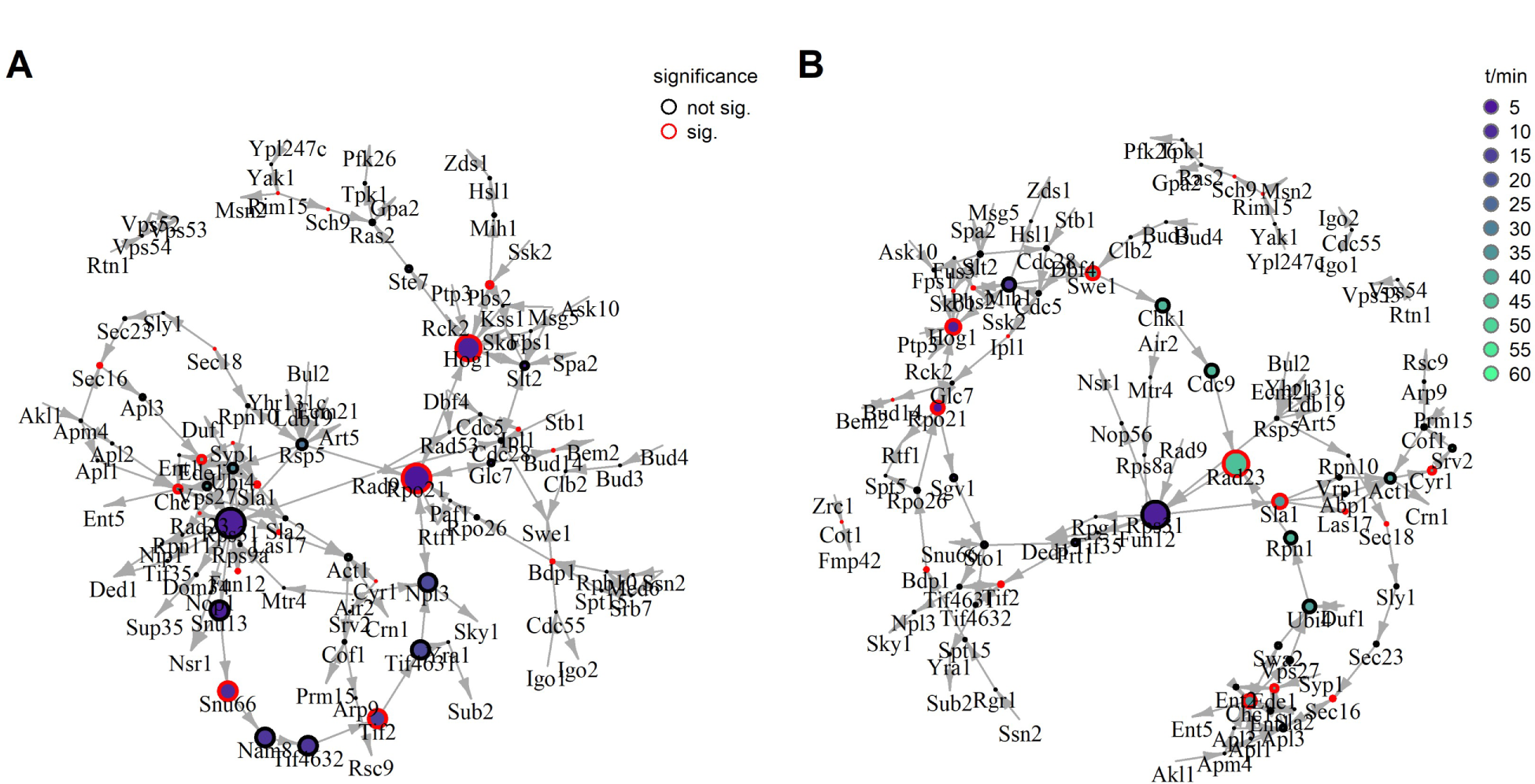
Inferred signaling network over time after path extraction and trimming, with node sizes indicating the betweenness centrality of nodes. Networks derived via CST (A) and PCST (B). Node fill colors indicate the (first) TP at which a node was part of a Steiner tree. Node frame color indicates if a protein is significantly differentially phosphorylated. (as fig. 7A and C)

Finally, it was explored how the different functional communities in the underlying PPI network are connected by signaling over time. Inspection of the chronopath plot with respect to communities revealed an extended path of nodes of the “Signal Transduction I” community (suppl. file 4; path #1126 in suppl. file 1). While the specific node ordering may not reflect the true signaling flow, the finding that a specific functional community is involved in a signaling process allows the generation of hypotheses for future experiments. In the case of the “Signal Transduction I” nodes, the involvement of the High Osmolarity Glycerol (HOG) pathway in osmosensing is already well-established (see below). An interesting question that arises is: does the observed differential phosphorylation of Spa2 indeed play a role in HOG-signaling? Spa2 is not a core member of the HOG-pathway but instead acts as an adapter for the MAPKK Mkk1 and the MAPK Slt2/Mpk1 ^26^. Therefore, it may be hypothesized that the observed phosphorylation of Spa2 directs it to the MAPKK Pbs1 and MAPK Hog1 instead. The path continues into nodes of the cell cycle regulation community. Connections between different communities are some of the most valuable results of network inference. In this case, the edge suggests a hypothesis for cell cycle regulation via the HOG-pathway. The known mechanism of regulation of the G2/M-transition in response to hyperosmotic stress is the phosphorylation of Hsl1 by Hog1, which in turn causes the accumulation of the Swe1 kinase ^27^. It is conceivable, that the HOG-pathway regulates Mih1 as well; Mih is the phosphatase counteracting Swe1 phosphorylation of Cdc28 ^28^ (path #1126 in suppl. file 1).

Another community that stands out in the inferred paths comprises members of the mediator complex (Ssn20, Rgr1, Med6) and other proteins involved in transcription and DNA remodeling (Bdp1, Spt15, Rpo21, Rpo26, and Paf1; paths #20 to 27 in suppl. file 1). Determining the exact regulation of this function, possibly via Hog1 as a kinase and Glc7 as phosphatase (path #32 in suppl. file 1) warrants further investigation.

Finally, a subpath of proteins of the ‘Signaling Transduction II’ community (paths #28, 191, and 1097 in suppl. file 1) accurately reflects knowledge gleaned from the literature of the Protein Kinase A (PKA) pathway, according to which the small GTPase Ras2 activates PKA subunit Tpk1, which in turn phosphorylates Pfk26 ^29,30^. The G-alpha subunit Gpa2, which on the inferred path is linked to Ras2 both as its upstream and downstream node, in reality, forms an independent input branch to PKA. An interesting question for future research is if the differential phosphorylation of Gpa2 that led to its integration in this path is physiologically relevant. A potential role of Sch9, which was linked upstream of Ras2 will also be interesting to explore. Finally, the inferred paths predict input to Ras2 via Ste7 also from the HOG-pathway, however, this is based on interaction evidence only of orthologues in other species and has not been observed in *S. cerevisiae* ^15^.

In summary, the network inferred via the CST approach supports the generation of novel hypotheses for experimental testing. To add credibility to these inferences, a systematic benchmarking of the network was addressed next.

#### Systematic evaluation

Systematic evaluation of the network inference approach was performed via edge confidence scores that had not been used in constructing the network and via the number of kinase-substrate edges in the network.

First, edge confidence scores from STRING-db were used to evaluate if the network inference approach extracted paths of biological relevance. In the construction of the underlying network, edges were filtered based on the ‘experimental evidence’ (“expScore”) score. The ‘curated database’ (“DBscore”) and ‘textmining’ (“textmScore”) scores, which had not been used in network inference, were used for evaluation. If biologically meaningful information were extracted, it is expected that the distribution of these two scores should increase over the process of network inference. The score distributions in the underlying network, Steiner Trees, and extracted paths are shown in fig. 9A. No increase in DBscore was observed in the process of network inference. In contrast, the text mining score of edges in inferred Steiner trees (p = 0.0002; median = 0.78) and extracted paths (p = 0.0015; median= 0.78) was significantly higher than in the underlying network (median = 0.66). This suggests that the time-slice-based CST approach was able to extract signaling-relevant connections. Based on the same argument as for interaction scores, previous knowledge on kinase substrate relationships was not used for inference but was reserved for evaluation of the inferred network.

**Fig. 9.**
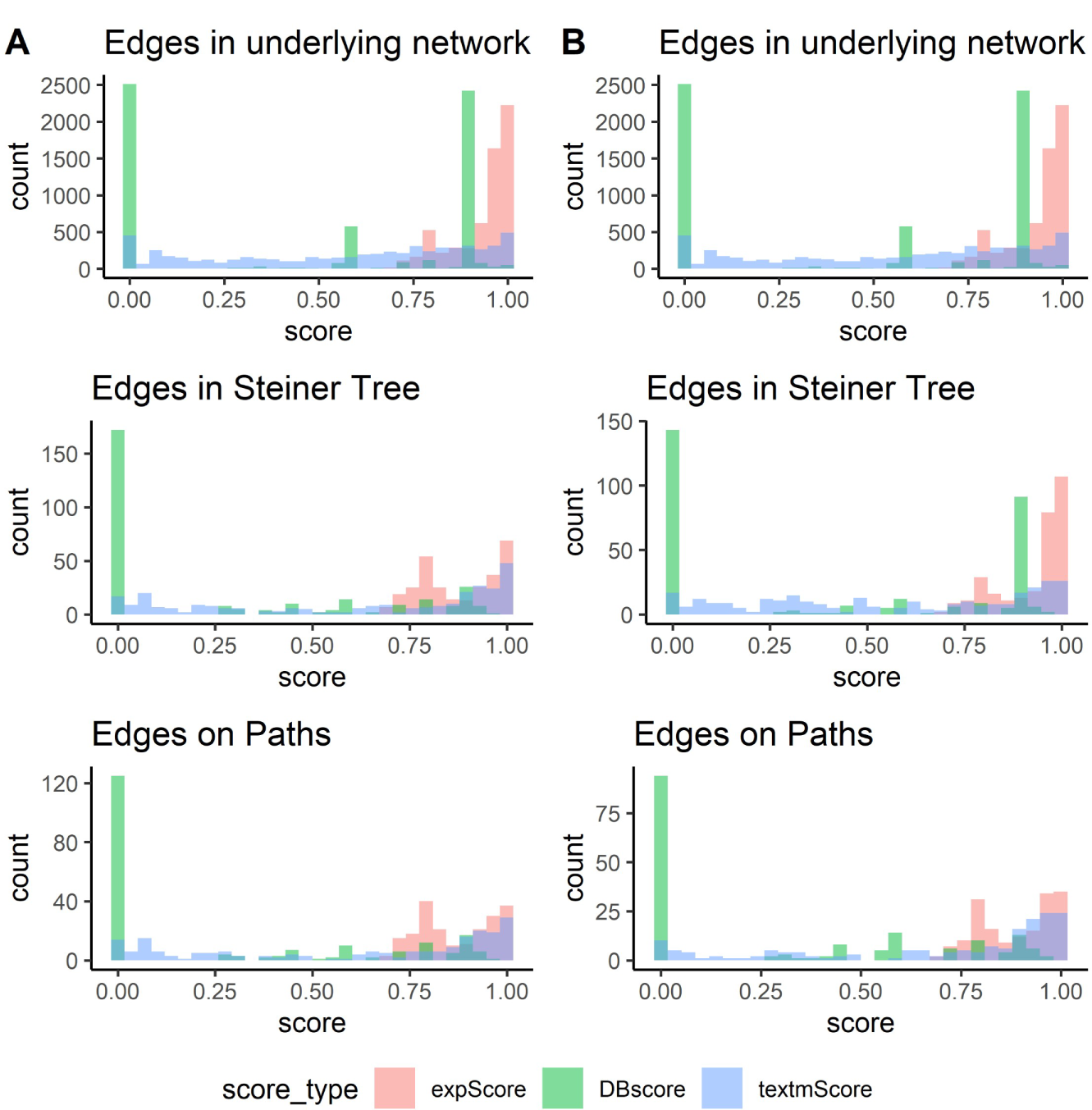
Evaluation of network inference via edge scores. Distribution of edge scores of type ‘experimental evidence’ (expScore’), ‘database’ (Dbscore) and ‘textmining’ (textmScore). Edge scores are shown as histograms for the underlying network, Steiner trees and exptracted paths for the CST (A) and PCST (B) approach.

It is expected that correct networks are enriched in edges from kinases to proteins with significant phosphorylation change.

In the underlying PPI network, 614 edges connected to significant proteins, of which 70 originated from kinases (11%). In the inferred network, 15 of the 66 edges connecting to significant proteins were from kinases (23%). This constitutes a significant enrichment of edges from kinases during network inference (p = 0.025). Therefore this test also confirms the potential of the approach for enriching signaling-relevant edges.

#### Evaluation by comparison to published studies

In addition, the results of network inference were also evaluated by comparison of the inferred network with published studies. The major pathway associated with the response to osmotic stress in *S. cerevisiae* is the HOG pathway (Fig. 10) ^31^. The central kinase, Hog1, is activated via phosphorylation by Pbs2, which is itself regulated via at least one of two branches: In the first branch, the membrane sensor Sln1 transmits information via a phosphorylation cascade of Ypd1, Ssk1, and the MAPKKKs Ssk2 / Ssk22. In the second branch, Ste50, Ste20, and/or Cla4 phosphorylate Ste11, which also serves as a kinase for Pbs2. Ste50 and Ste20 are anchored to the plasma membrane by Opy2 and Cdc42, respectively, and activated via the osmosensor Sho1 and its partners Hkr1 and Msb2. Further, Rps31 also exhibited an above-average betweenness centrality, even though it was not found differentially phosphorylated. This ribosomal protein entered the inferred network at five TPs and, therefore, appears central to the connection between the hyperosmotic stress response and translation regulation.

**Fig. 10.**
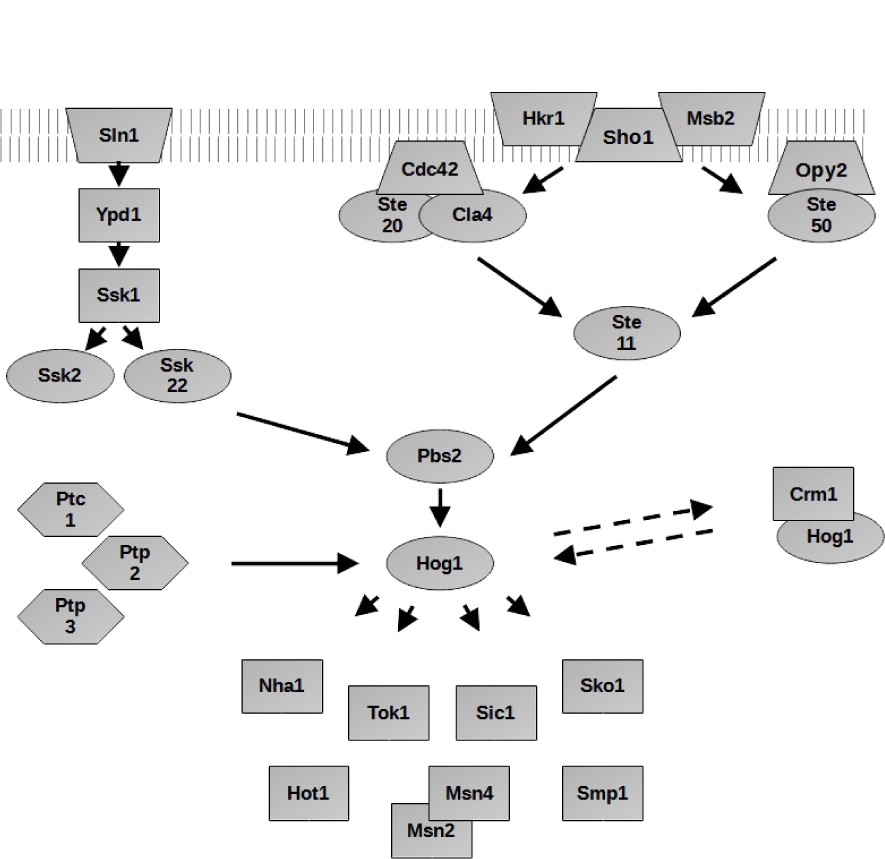
Model of the HOG-pathway assembled manually from literature. See text for details.

Notably, the Sln1 - Ypd1 - Ssk1 branch is a two-component system, which is less frequent in eukaryotic than bacterial signal transduction ^32^. Due to their lability, the histidine- and aspartate-phosphorylations on these proteins are not detectable by standard proteomics methods. The HOG-pathway therefore serves as an example of a signaling network that could, even in theory, only be partially reconstructed from the phosphoproteomics data (without PPI information) alone. Pbs2-dependent phosphorylation of Hog1 leads to its nuclear import. Targets of Hog1 include Nha1, Tok1, Sic1, Sho1, Ste50 and the transcription factors Msn2/4, Sko1, Hot1 and Smp1 ^27,31,33–37^. The phosphatases Ptc1, Ptp2, and Ptp3 counteract Hog1 phosphorylation and the karyopherin Crm1 is responsible for its re-export from the nucleus ^33^.

Of the 28 proteins mentioned that constitute the HOG-pathway, 17 were part of the underlying PPI network and Hog1, Ptp3, Pbs2, Ssk2, Msn2, and Sko1 were in the inferred network. A direct connection between Pbs2 and Hog1 was identified, however in the reverse direction than reported in the literature, as Hog1 reached significance after 15 and Pbs2 after 20 s. Despite the significance of Ptp3 already after 5 s, the reported direct Ptp3 – Hog1 connection was identified by recruiting Hog1 into the network between 5 and 10 s as a Steiner node (see e.g. path #36 in suppl. file 1). Similarly, the Ssk2-Pbs2 edge was found via the inclusion of Pbs2 as a Steiner node between 35 and 40 s, after it had already entered the network as a significant node at 20 s. Ssk2 did not have an input in the network, which is reasonable, as the Asp-phosphorylation of Ssk1 is undetectable by standard phosphoproteomics. Finally, the known Hog1 targets Msn2 and Sko1 both reached significance at 40 s. The inferred edge from upstream kinase Rim15 to Msn2 is expected as part of the general stress response to the increase in osmolarity, however, phosphorylation of Msn2 by Hog1 on a different site was not inferred ^39,40^.

For 13 of the listed proteins in the HOG-pathway, an observable Ser/Thr/Tyr phosphorylation difference is expected (Ssk2, Ssk22, Ste11, Pbs2, Hog1, Nha1, Tok1, Sic1, Msn2, Msn4, Sko1, Hot1, and Smp1) and for six of them, differential phosphorylation was observed by phosphoproteomics (Ssk2, Pbs2, Hog1, Msn2, Msn4, and Sko1). Out of these, in turn, all except Msn4 were mapped on the inferred pathway. The network inference approach therefore preferentially integrates signaling-relevant significant proteins in the network as the selection into the network among all significant proteins was only 51% (61 of 120).

In contrast, Ptp3 was the only one of the 13 proteins participating in HOG-signaling without an expected observable phosphorylation change that was part of the inferred network. Similarly, none of the proteins with an expected, but unobserved, phosphorylation change was part of the inferred network. Therefore, the ability of the CST approach to map proteins signaling through mechanisms other than phosphorylation, or with unobserved phosphorylation changes, appears limited.

In summary, comparison to published studies suggests that network inference based on CST efficiently mapped signaling relevant differentially phosphorylated proteins to the inferred network. However, the network is most valuable with respect to the identities of proteins involved in a signaling process, rather than their exact order on the pathway. It was next explored if an algorithm of constructing trees at time-slices that encodes varying degrees of confidence in differential phosphorylation and PPIs would efficiently prioritize between alternative paths and contribute additional signaling-relevant connections.

### Prize Collecting Steiner Tree approaches

#### Description of the inferred network

In the previous section, I introduced an approach for reconstructing temporal signaling networks based on trees built on individual time slices. The CSTs used for this purpose rely on the premise that signals are transmitted through shortest paths. As an alternative, trees were next built based on the degree of confidence in significant phosphorylation changes at nodes and in the interactions represented by edges, using PCSTs. PCSTs have been used for this purpose, however not in a time-slice manner ^12^.

From a high-level perspective, the Classical and Prize-collecting Steiner Trees were largely similar across time slices (fig. 6C, D). For example in the first slice, a similar number of nodes was chosen and there is a large overlap between the sets of nodes. However, unlike the CST, the PCST consisted of four disjoint subgraphs. With the exception of Rpo21 and Snu66, which have been completely rewired, nodes that are neighbors in the PCST are also neighbors in the CST. However, several edges present in the CST are lacking in the PCST. This is because edges between already selected nodes are obtained “for free” in a CST, while they incur a cost in a PCST. Also at the second time-slice, the CST and PCST resemble each other through a largely overlapping set of selected nodes and, by each being split into two subgraphs with the same terminal nodes. Unlike at the first time-slice, the PCST has more nodes (and therefore more edges) than the CST here. This is due to the inclusion of nodes that contribute higher prizes than the associated edge costs and, apparently, the longer paths selected in the PCST (e.g. Slt2 – Rps1 – Hog1) come with a higher cost-benefit than the shorter paths in the CST (Slt2 – Hog1). The trend of largely similar selected nodes with subtly distinct wiring is also observed at other TPs. At the 35 – 40 s and 40 – 45 s time-slices, distinct clusters of ribosomal and translation-associated proteins were observed only in the PCST results. Many of the proteins in these clusters were discarded during path extraction as they are on paths with a number of subsequent Steiner nodes above the specified limit (see below). The Steiner nodes in these clusters are associated with very low node prizes, which likely only stem from random variation in protein phosphorylation. They are included due to low edge costs originating from a high confidence in their interactions. In future work, it may be useful to represent stable protein complexes by single nodes where appropriate.

Upon linking the trees obtained at each time-slice, a network encompassing the full time course, consisting of 150 nodes and 285 edges was obtained (fig. 7C and suppl. video 2). Notably, the network was split into one major and four minor disjoint subgraphs, while in the case of CSTs, only one small subgraph was separate from the major network. This is again a consequence of the fact, that in a CST edges between selected nodes are obtained “for free” (e.g. Cdc55 is connected to the main CST network, even though it would also be connected to an outgoing and an incoming terminal without this connection; fig. 7A), while in a PCST additional edges are associated with a cost and, therefore, are only selected to introduce additional nodes. In other words, a CST cannot select between alternative, equal-length routes connecting two nodes, while a PCST does so via its prize-vs-cost calculation. A noteworthy subgraph inferred via the PCST approach is, for example, formed by growth-regulatory proteins of the PKA and TOR pathways (Gpa2, Ras2, Tpk1, Sch9, Rim15, Yak1, Msn2).

The chronopath plot for the PCST-derived paths confirmed the similarity to the solution obtained via CST (fig. 7D and suppl. file 5). Only a small number of nodes are present in one, but not the other solution, whereas there is a notable difference in the wiring of the edges. This difference becomes particularly obvious in the fact that 108 paths were extracted via the PCST, compared to 1127 via the CST approach (suppl. file 6). This again reflects the algorithmic difference causing CST to include all equal-length paths between required nodes, while PCST selects the path with the highest prize-to-cost-ratio.

As for CST, the chronopath is divided into sections lacking connections at the 25 and 50 s TPs. The re-use of significant proteins at TPs other than the TP of significance is also similar between both approaches and, as in the CST chronopath, Rpo21 re-appears three times as a Steiner node in the PCST results.

However, it is important to keep in mind that the PCST approach is dependent on multiple parameter choices. These parameters had been set to subjectively reasonable values largely arbitrarily (see methods). However, a higher prize-to-cost ratio would, for example, lead to a larger network and therefore likely integrate all significant proteins into one graph.

#### Hypothesis generation

As for the CST approach, the inferred network was analyzed with respect to GO enrichment, betweenness centrality scores, and network communities.

No significant ‘GO-function’ or ‘GO-process’ ontologies were obtained at an FDR of 0.05.

Fig. 8B reveals that the major subgraph consists largely of two parts that are linked by only a few connections. Connecting nodes between the parts are Rps31 and Rad23, which is reflected by their high betweenness centrality. In the network, Rps31 signals towards proteins associated with translation (e.g. Ded1, Tif2, Tif35) on one branch and a set of proteins associated with diverse functions through Sla1. Rad23 receives input from cell cycle-associated proteins and a branch rich in proteins involved in endocytosis (e.g. Swa2, Ent1, Ede1). It can be speculated that Rad23, which entered the network at a later TP, provides feedback to Rps31. Though, a high-confidence physical interaction has been observed between the two proteins, no information exists about their functional relationship.

Similar to the network inferred via CSTs, extended paths enriched in members of the HOG-pathway were observed (“Signal Transduction I”, e.g. path #96 in suppl. file 6: Ptp3 – Hog1 – Slt2 – Spa2 – Slt2 – Fps1 – Hog1 – Sko1 – Fus3 – Pbs2). The path further suggests a link between HOG-signaling and the cell cycle via a Pbs2 – Mih1 edge (suppl. file 7). A path collection of transcriptional proteins between Ssn2 and Rtf1 was also detected, similar to the one observed based on CSTs (paths #11 and 12 in suppl. file 6). The number of connections via the PCST approach was however smaller and only Rgr1, Spt15, Rpo26, Rpo21, and Spt5 were selected as Steiner nodes.

Finally, the subpath of proteins from the ‘Signaling Transduction II’ community consisting of members of the PKA pathway was identical to the one inferred via CST, however, without the upstream connection to the HOG-pathway (path #16 in suppl. file 6). Indeed, these nodes are part of a subgraph that is disjoint from the core temporal network (fig. 7B). This may be interpreted as PKA-signaling being regulated independently of the HOG-pathway in the response to hyperosmotic stress. The regulation of Ras2 occurs via Sch9, but not via Ste7.

#### Systematic evaluation

As for the CST approach, systematic evaluation of the inferred network was performed by asking if an enrichment of edges with high confidence scores was observed. As only the ‘experimental evidence’ (expScore) score had been used for the construction of the underlying network and for edge costs of PCSTs, the ‘database’ (DBscore) and ‘textmining scores’ (textmScore) remained unbiased for evaluation. No significant increase in DBscore was observed over the inference process. In contrast, there was a significant increase in textmScore from the edges in the underlying network to the ones on extracted paths (p < 0.0001), as was the case in the CST approach (fig. 9B). In contrast to CSTs, this increase is only visible at the step of path extraction and not upon inferring Steiner trees (p = 0.49). Further investigation of this phenomenon revealed that edges in the ribosomal protein/translation cluster, which was present in the Steiner trees, but was largely dropped upon path extraction, were associated with low textmScores (suppl. fig. 1). This is probably because the relationship between proteins stably associated in a complex is rarely functionally investigated and therefore is rarely found via text mining. Therefore, network inference based on PCSTs is able to extract signaling-relevant paths from an underlying PPI network but is vulnerable to the inclusion of high-scoring interactors without signaling-relevance.

A significantly increased frequency of edges between kinases and significant proteins compared to the underlying network was observed neither in inferred trees nor in extracted paths.

#### Evaluation by comparison to published studies

The paths extracted based on the PCST approach were also evaluated in the light of existing knowledge about the HOG-pathway. Of the 28 proteins defined as members of the HOG-pathway (see above) eight were recovered on the inferred trees (Ssk2, Pbs2, Hog1, Ptc1, Ptp2, Ptp3, Msn2, Sko1) and six on extracted paths (Ssk2, Pbs2, Hog1, Ptp3, Msn2, Sko1) (Fig. 10). Out of the eight differentially phosphorylated HOG-pathway proteins, all except two were mapped to the inferred tree and extracted paths. Therefore, like for CSTs, the approach based on PCSTs was highly efficient in recovering expected signaling-relevant proteins from significant proteins. Two proteins of the HOG-pathway which were not significant in the phosphoproteomics experiment, Ptc1, and Ptp2, were mapped to the inferred trees, but discarded during path extraction. Therefore PCSTs were similar to CSTs in recovering significant proteins and outperformed CSTs in recovering HOG-pathway proteins before path extraction. The phosphatases Ptc1 and Ptp2 had originally been placed on a connection from Hog1 to Pbs1 in the tree for the 15 – 20 s time-slice (fig. 6). The path was later eliminated as it contained a third Steiner node on the connection. The dropping of paths exceeding a limit on consecutive Steiner nodes is, therefore, a trade-off between the elimination of false positives and difficult-to-interpret paths on one hand and the loss of true positive connections on the other.

Ssk2 served as an input for Pbs2 in the inferred path, consistent with literature knowledge. Note that the Pbs2 node which received the input was a Steiner node, which is present in addition to the Pbs2 node at its TP of significance (fig. 7D). The connection from Hog1 to Sko1 reported in the literature was also recovered ^41^. Notably, this connection existed with Hog1 as significant and Sko1 as Steiner node and *vice versa* as the TPs of significance of the two proteins did not place them into a common time-slice. Therefore, as for CSTs, the inference algorithm introduced significant proteins as Steiner nodes at other TPs to permit the recovery of signaling-relevant connections that may otherwise be missed, possibly due to inaccurate determination of TPs of significance.

## Discussion

### Goal of the study

Mapping the structure of cellular signaling networks is a major prerequisite in enhancing our understanding of physiology and pathology. In this work, the systematic construction of signaling networks from temporal phosphoproteomics data has been explored. Other existing network inference approaches are based on mechanistic data, such as the binding of transcription factors to promoters or on observed changes as a consequence of manipulating the system, e.g. through gene deletions (see mini-review submitted with this article). The former does not readily translate to phosphorylation networks as, in this context, signaling-relevant connections are less easily identified than for transcription factor – target gene pairs. Inference based on system manipulation suffers from the difficulty of distinguishing its direct from indirect consequences. Therefore, inference from time course data is likely to gain importance with the increasing availability and quality of proteomics and phosphoproteomics data.

However, for guiding further experiments, too many alternative options for network wiring are expected when basing inference on these data alone. Therefore, inference was guided by underlying PPI networks in the current study.

When mapping only significant proteins to a PPI network, very few edges connecting proteins with phosphorylation changes at subsequent TPs were found. Therefore, network expansion allowing the inclusion of non-significant interacting nodes was performed. This expansion pursues the aim of adding missing connections that may have been missed due to incomplete data or signaling not involving phosphorylation but comes with the challenge of distinguishing signaling-relevant from other interactions.

The approach presented here sought to connect phosphorylation changes at a given TP to observations at the previous TP through the construction of trees that connect signaling nodes in slices of adjacent TPs. Both a CST and PCST algorithm were explored for this task. The goal of this work was the introduction and evaluation of the approach.

### Comparison to existing literature

Network inference from temporal phosphoproteomics data based on an underlying PPI network had been performed previously in a graph approach by Budak, et al. ^12^. In this study, in a first step, networks from significant proteins and their interactors were constructed at each TP via a Prize-Collecting Steiner Forest method. The TPs at which proteins were members of Steiner trees were mapped to a PPI network and nodes connected according to this temporal order. In contrast, in the current study, Steiner trees were built on slices of neighbouring TPs, which were subsequently stitched together. While this difference may appear subtle, it relies on a fundamentally different concept (Fig. 1): I argue that the signaling relevance of an edge changes during the signaling time course and that it should be possible to infer this relevance by selecting paths that connect proteins significant at a later TP to those significant at an earlier TP. Therefore, node and edge selection are performed for every time-slice. This is unlike in studies, where signaling-relevant nodes are selected via the plausibility of networks at every TP, but without explicit consideration of signal transfer from one TP to the next. In other words, TP-based approaches ask what are the most plausible proteins to take part in signaling at every TP and then connect the proteins over time, while the time-slice approach used here directly searches for the most plausible connections over time.

A second major difference to previous studies is that the inference here was applied to a tightly-resolved phosphoproteomics time course, while in previous network inference studies TPs were separated by minutes to hours ^6,7,12^. While further work is needed in comparing approaches, the algorithm introduced in the current study is designed for well-resolved signaling events and may not be optimal for the datasets in these previous studies. This is because too many signaling steps may occur over several minutes. An approach such as the one by Budak, et al. may be better suited for these cases ^12^. Based on the concept of directly tracing signaling flow (rather than reconstructing multiple signaling steps at a TP), it was assumed that few signaling steps should occur between observed phosphorylation changes. This justified algorithmic shortcuts, reducing computational requirements, namely the limitation of the underlying PPI network to a 1-shell expansion around significant proteins and the restriction to maximally two Steiner nodes between terminals. The distinction of significant proteins into outgoing and incoming terminals further rendered computations more efficient.

The phosphoproteomics dataset chosen for this study captured very early TPs after stimulation ^14^. This study and our own work showed that at least a wave of first responders reaches near-complete phosphorylation change on a timescale of seconds after stimulation ^42^. If, to which extent and why signaling may be faster in certain organisms or along certain pathways are exciting questions for future research. In any case, the high temporal resolution of early signaling is an important prerequisite for meaningful output from the time-slice approach applied here, but likely also for other inference efforts.

### Summary and interpretation of results

Compared to connecting only significant proteins via PPIs, which extracted only disconnected edges, the time-slice-based approaches on expanded underlying networks recovered paths spanning experimental results from 5 to 25 s and 25 to 50 s, providing new candidate models for signaling flow.

In comparison to CST, the PCST approach was more selective in choosing from connections between required nodes. This is reflected in the less densely connected network originating from this approach and in the more than ten times lower number of extracted paths.

On the other, the PCST network expanded beyond the selections of CST in the context of low-cost connections, such as protein complexes, thereby including proteins in the network that are likely not central to the signaling process. Therefore, the intersection of edges provided by the two approaches may be a viable way to limit the size of inferred networks. Alternatively, for future applications, PCSTs may be the preferred option if signaling-relevance could be encoded in edge scores (see below).

Overall, both approaches show promise in selecting signaling-relevant nodes and edges, as demonstrated by the enrichment of edges with high ‘textmining’ confidence scores in the inferred network.

### Study limitations and future work

On the other hand, only fragments of signaling paths could be extracted, and ambiguity about their temporal order remained. Tuning the multiple parameters of the inference algorithm is a potential starting point for improving the performance of path extraction. Even though no phospho-signaling network is defined with both sufficient completeness and edge confidence to serve as a ground truth for parameter selection, available literature is an important source for benchmarking inferred models ^43^. The networks derived in this study were therefore evaluated via comparison to published studies about HOG-signaling. In addition, a second, more systematic approach was applied: Networks were inferred on an experimentally determined PPI score but evaluated on independent scores from curated databases and text mining. Optimizing parameter choices based on this strategy will be the topic of a subsequent study.

Inspection of the inferred paths indicated that the correct determination of the TP of significance was a major challenge in inferring the correct order of signal propagation. In this study, time-slices consisted of two subsequent TPs. In future work, the approach may be extended to slices encompassing more TPs, in which paths from an earlier to any later significant protein are allowed. This alteration would shift the burden from accurate experimental results to the underlying network. This study introduced a network inference method for high-resolution temporal data and emphasized the evaluation of the approach. Edge confidence scores from curated databases and text mining were therefore employed only for the evaluation, but not the inference of networks. However, it is obvious that edge selection based on curated knowledge would be tremendously beneficial for network construction, especially as it may include signaling mechanisms not observable by the experimental method used in generating the data for network inference. Further, integration of known kinase-substrate relationships or other information on canonical signaling pathways would provide a scaffold *ab initio,* and the inference challenge would be shifted towards filling in the less studied peripheral parts ^3,4^. Different interaction databases have been compared recently in the context of network inference ^44^. PCST is an approach well suited for incorporating several types of interaction information via the capacity of encoding edge confidence, . Three requirements remain to be addressed for efficiently employing this approach: 1) systematic cataloging of curated knowledge. 2) means to account for the context-specificity of certain edges. 3) Means of evaluating inferred networks independent from the curated knowledge used as input. If these limitations can be overcome, network inference on knowledge-based networks offers promises beyond phospho-signaling. Even, and maybe especially, in cases where no agreement on accepted knowledge exists, this approach suggests a framework for comparing models systematically.

Even with these improvements, graph-based network inference methods remain limited by the discretization of phosphorylation differences and TPs. Therefore, rather than as a stand-alone tool, they should be used in conjunction with other inference methods that do not suffer from this limitation, such as Dynamic Bayesian Network and Ordinary Differential Equation models (see mini-review submitted with this article). These tools can build on the hypotheses generated by the graph approach through prioritization among significant proteins alongside with integration of proteins without observable change in the underlying data.

## Acknowledgements

I thank the members of the Stack Overflow community for advice and Carol Dieckmann (University of Arizona) for helpful comments on the manuscript.

## Supplement

Suppl. file 1: Individual chronopath plots for CST approach

Suppl. file 2: Combined charonopath plot for CST approach (Version of fig. 7B in .pdf format) Suppl. file 3: Combined charonopath plot for CST approach, with nodes of GO-term ‘positive regulation of cell cycle process’ highlighted

Suppl. file 4: Combined charonopath plot for CST approach, with nodes colored according to communities

Suppl. file 5: Combined charonopath plot for PCST approach (Version of fig. 7D in .pdf format) Suppl. file 6: Individual chronopath plots for PCST approach

Suppl. file 7: Combined charonopath plot for PCST approach, with nodes colored according to communities

Suppl. Fig. 1. **Distribution of experimental evidence (‘expScore’) vs. text mining scores (‘textmScore’) in Steiner trees inferred via the PCST approach.** Nodes in the cluster of ribosomal / translation-associated proteins are represented by edges involving the hub proteins Fun12 and Rps31 and shown in blue. Boxplots show the textmScore distribution of edges from/to Fun12 or Rps31 (blue) and all other edges (grey).

