## Supplementary figures and images for "Network inference from temporal phosphoproteomics informed by protein-protein interactions"

### Supplemental Figure 1

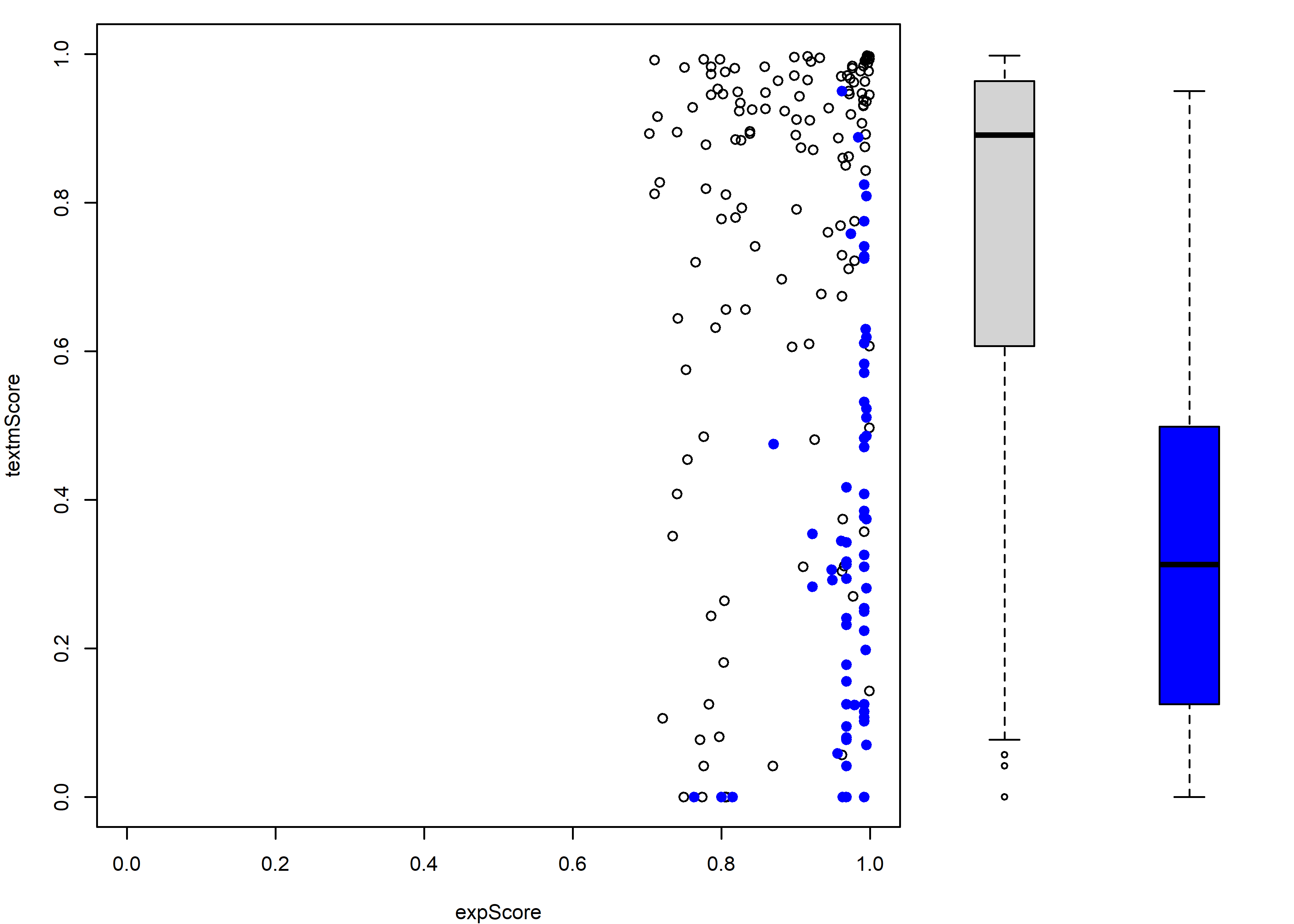

### Supplemental File 2

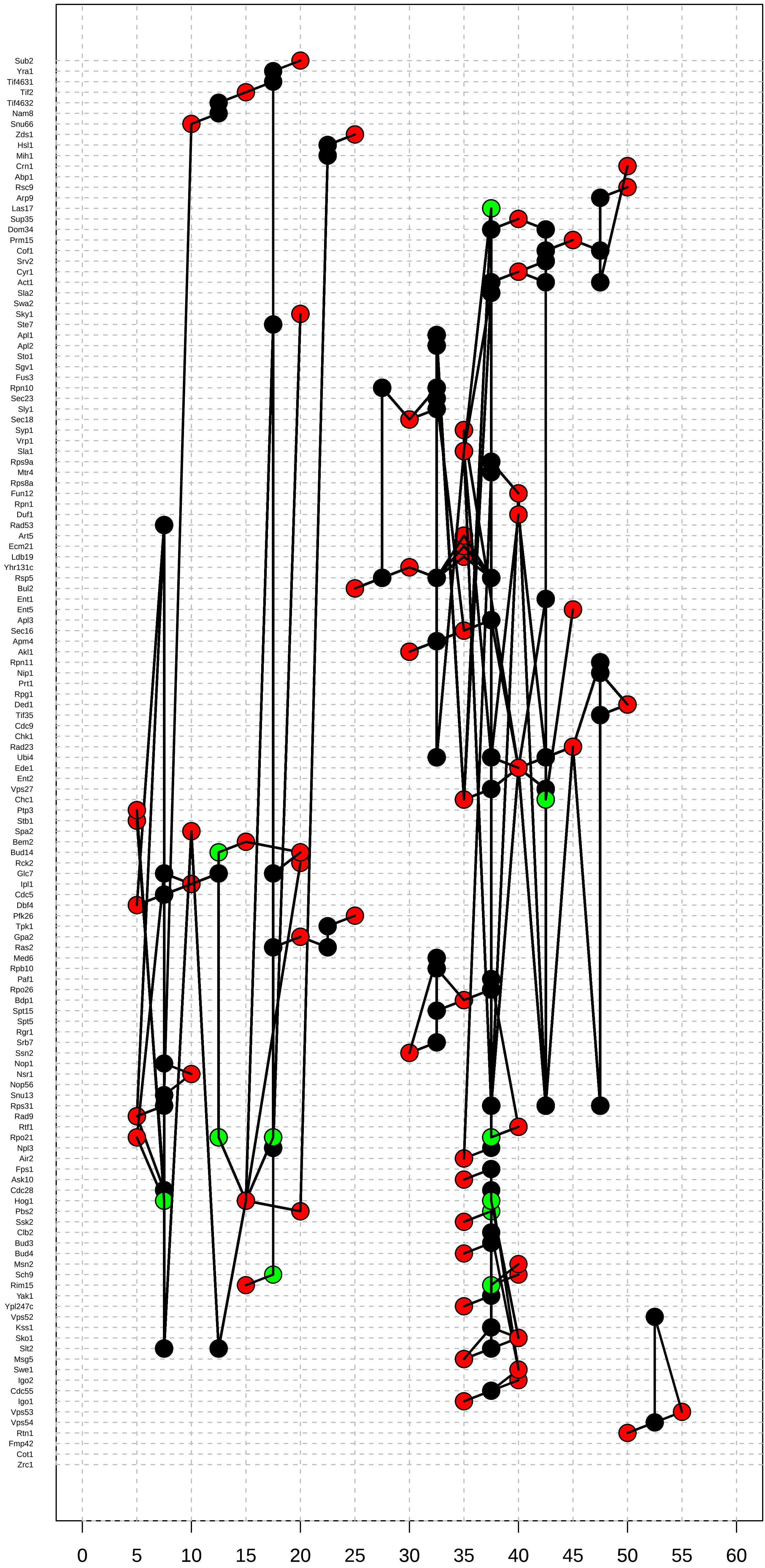

### Supplemental File 3

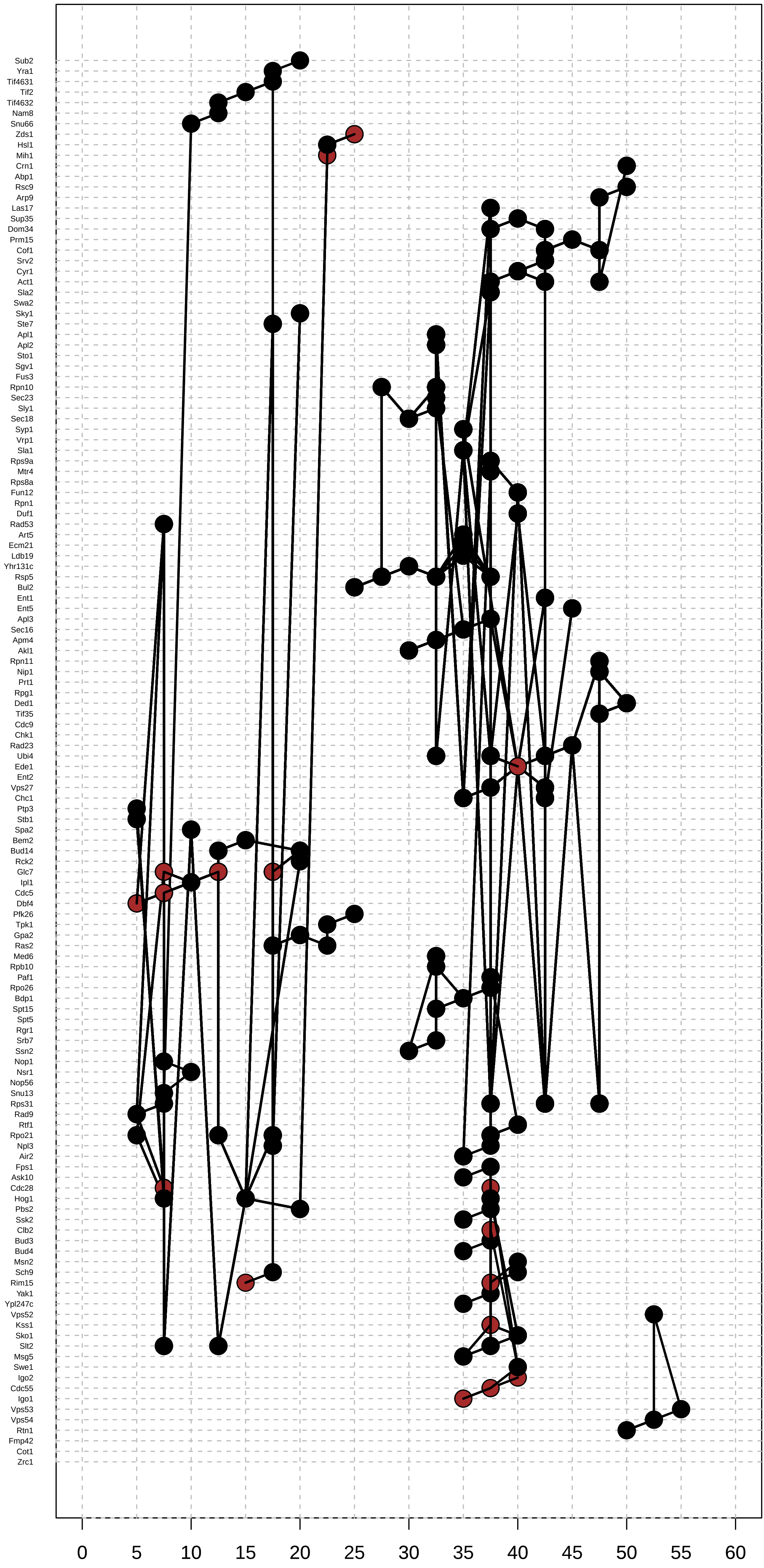

### Supplemental File 4

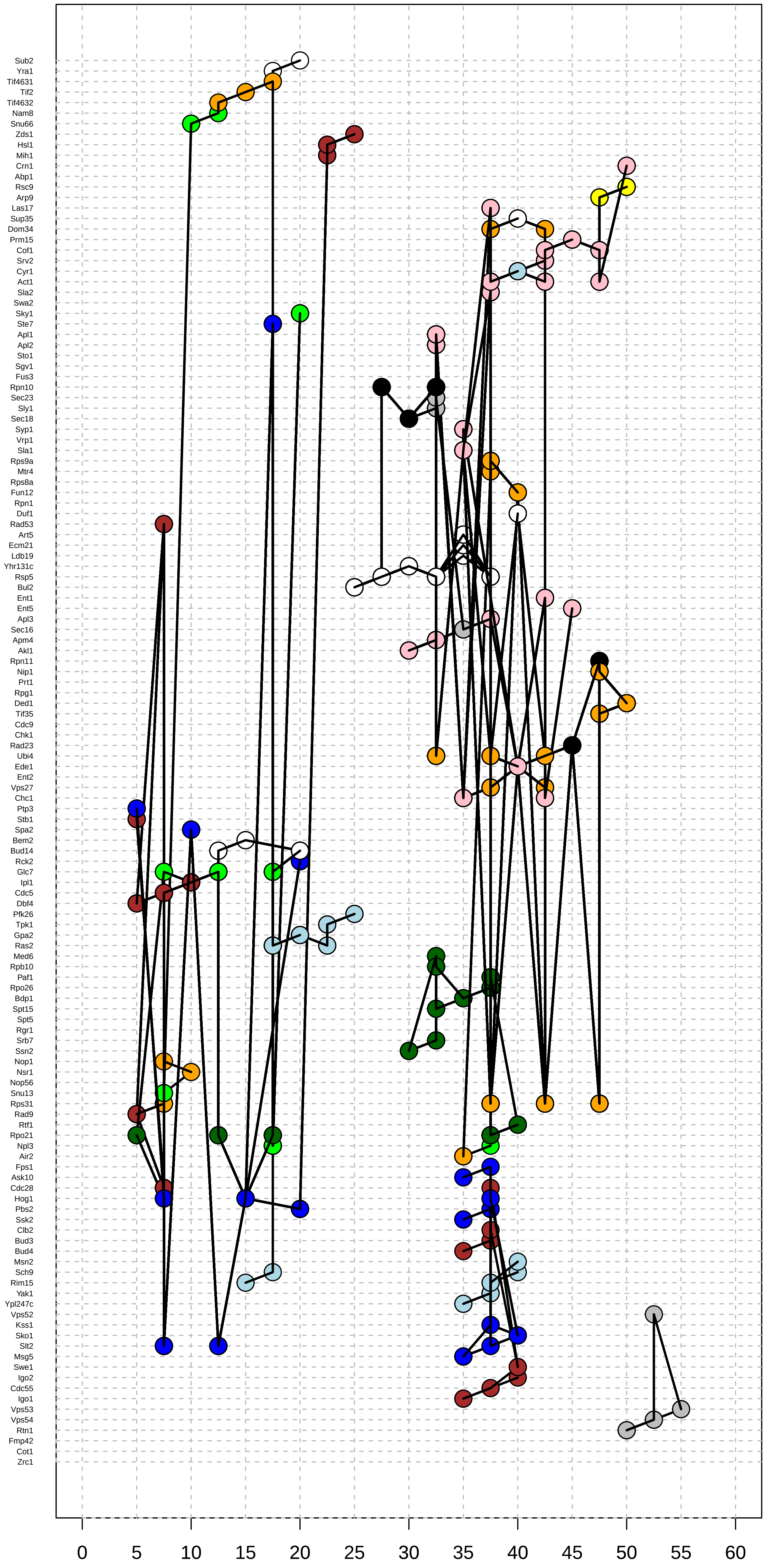

### Supplemental File 5

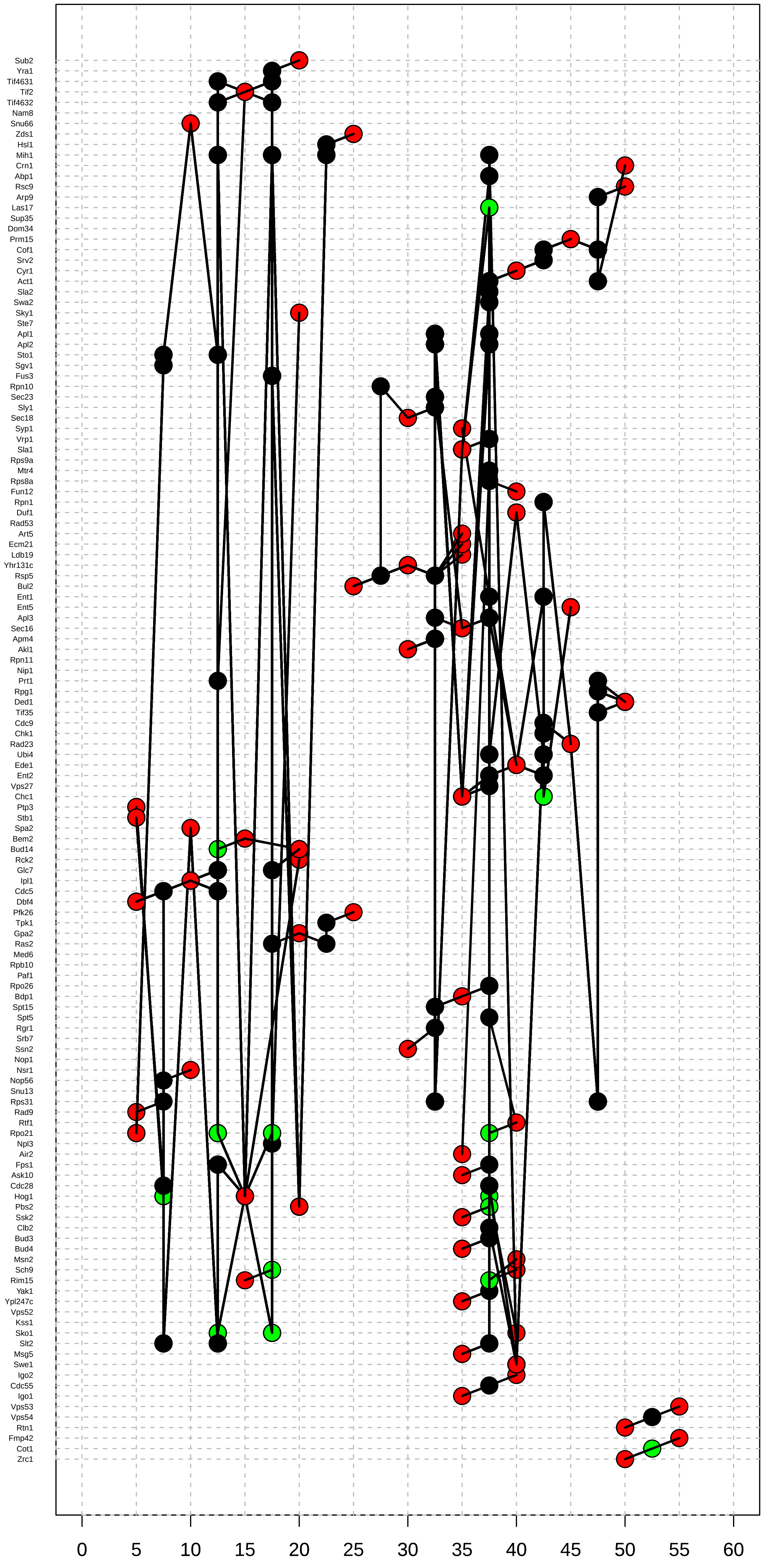

### Supplemental File 7

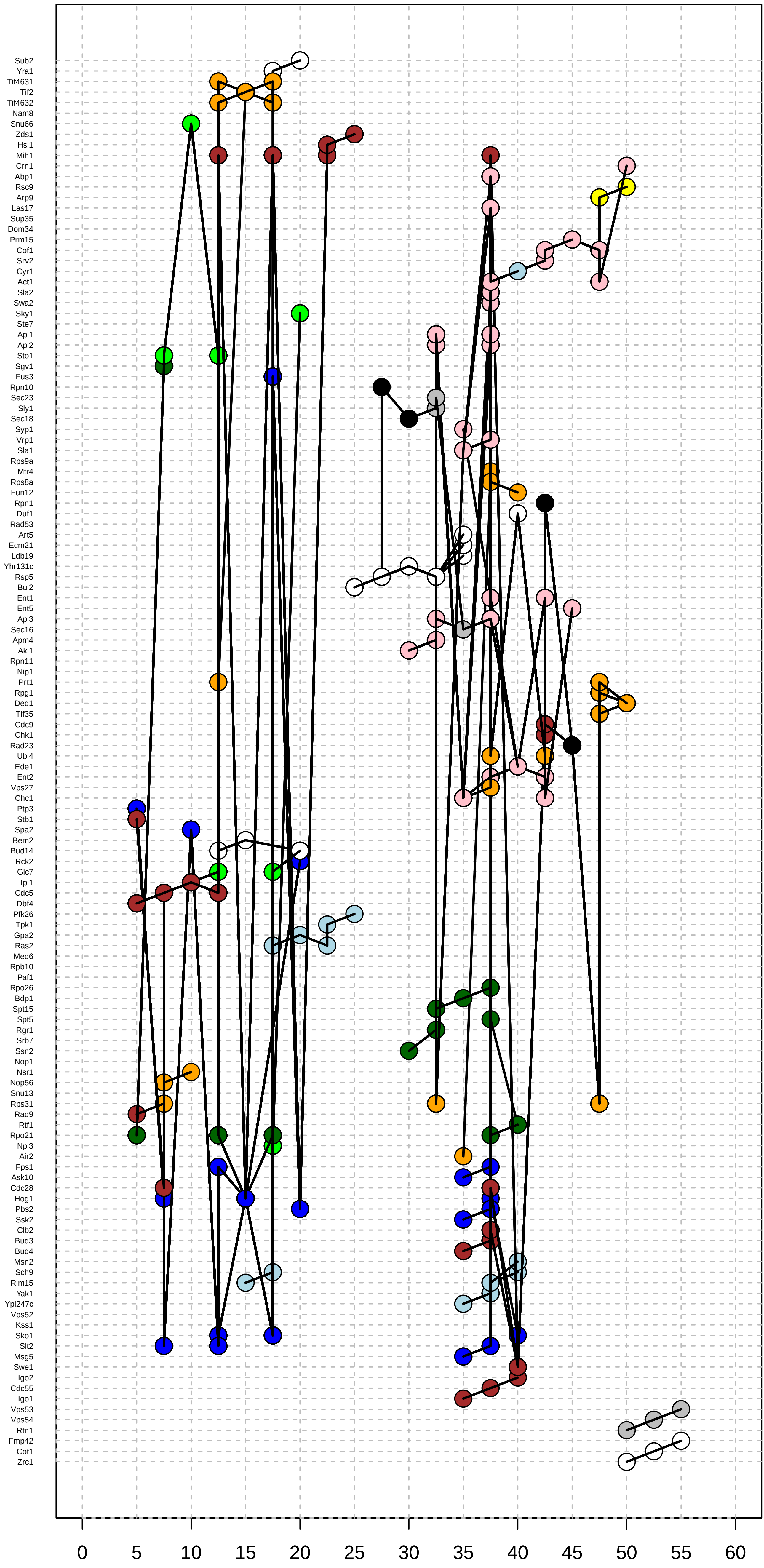

### Supplemental Video 1

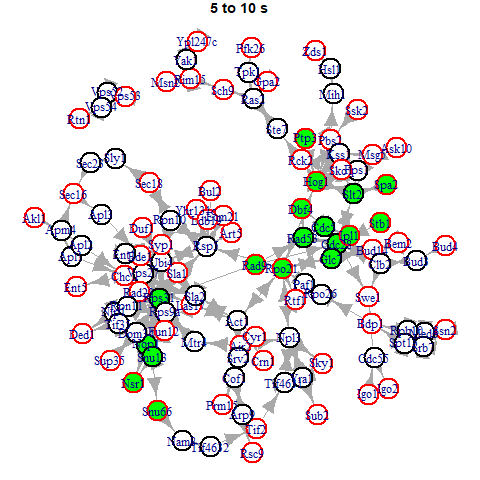

### Supplemental Video 2

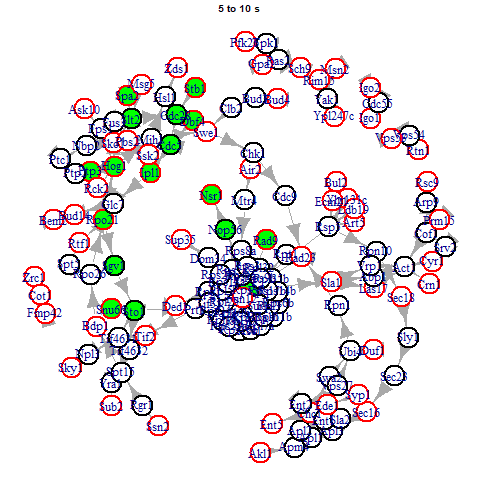
