## Supplemental File 1 for "Network inference from temporal phosphoproteomics informed by protein-protein interactions"

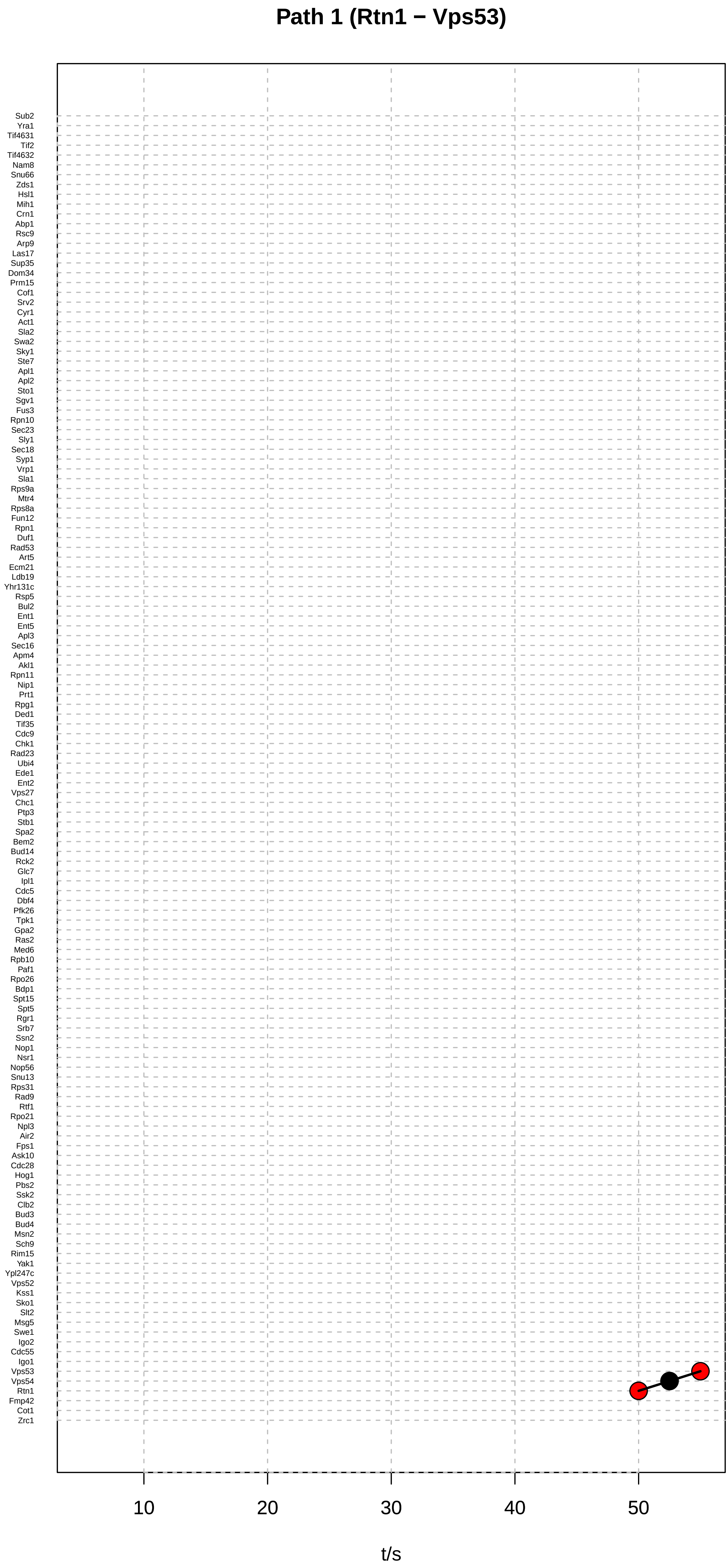
Path 1 (Rtn1 − Vps53)
Sub2
Yra1
Tif4631
Tif2
Tif4632
Nam8
Snu66
Zds1
Hsl1
Mih1
Crn1
Abp1
Rsc9
Arp9
Las17
Sup35
Dom34
Prm15
Cof1
Srv2
Cyr1
Act1
Sla2
Swa2
Sky1
Ste7
Apl1
Apl2
Sto1
Sgv1
Fus3
Rpn10
Sec23
Sly1
Sec18
Syp1
Vrp1
Sla1
Rps9a
Mtr4
Rps8a
Fun12
Rpn1
Duf1
Rad53
Art5
Ecm21
Ldb19
Yhr131c
Rsp5
Bul2
Ent1
Ent5
Apl3
Sec16
Apm4
Akl1
Rpn11
Nip1
Prt1
Rpg1
Ded1
Tif35
Cdc9
Chk1
Rad23
Ubi4
Ede1
Ent2
Vps27
Chc1
Ptp3
Stb1
Spa2
Bem2
Bud14
Rck2
Glc7
Ipl1
Cdc5
Dbf4
Pfk26
Tpk1
Gpa2
Ras2
Med6
Rpb10
Paf1
Rpo26
Bdp1
Spt15
Spt5
Rgr1
Srb7
Ssn2
Nop1
Nsr1
Nop56
Snu13
Rps31
Rad9
Rtf1
Rpo21
Npl3
Air2
Fps1
Ask10
Cdc28
Hog1
Pbs2
Ssk2
Clb2
Bud3
Bud4
Msn2
Sch9
Rim15
Yak1
Ypl247c
Vps52
Kss1
Sko1
Slt2
Msg5
Swe1
Igo2
Cdc55
Igo1
Vps53
Vps54
Rtn1
Fmp42
Cot1
Zrc1
10
20
30
40
50
t/s

### Path 2 (Igo1 − Igo2)

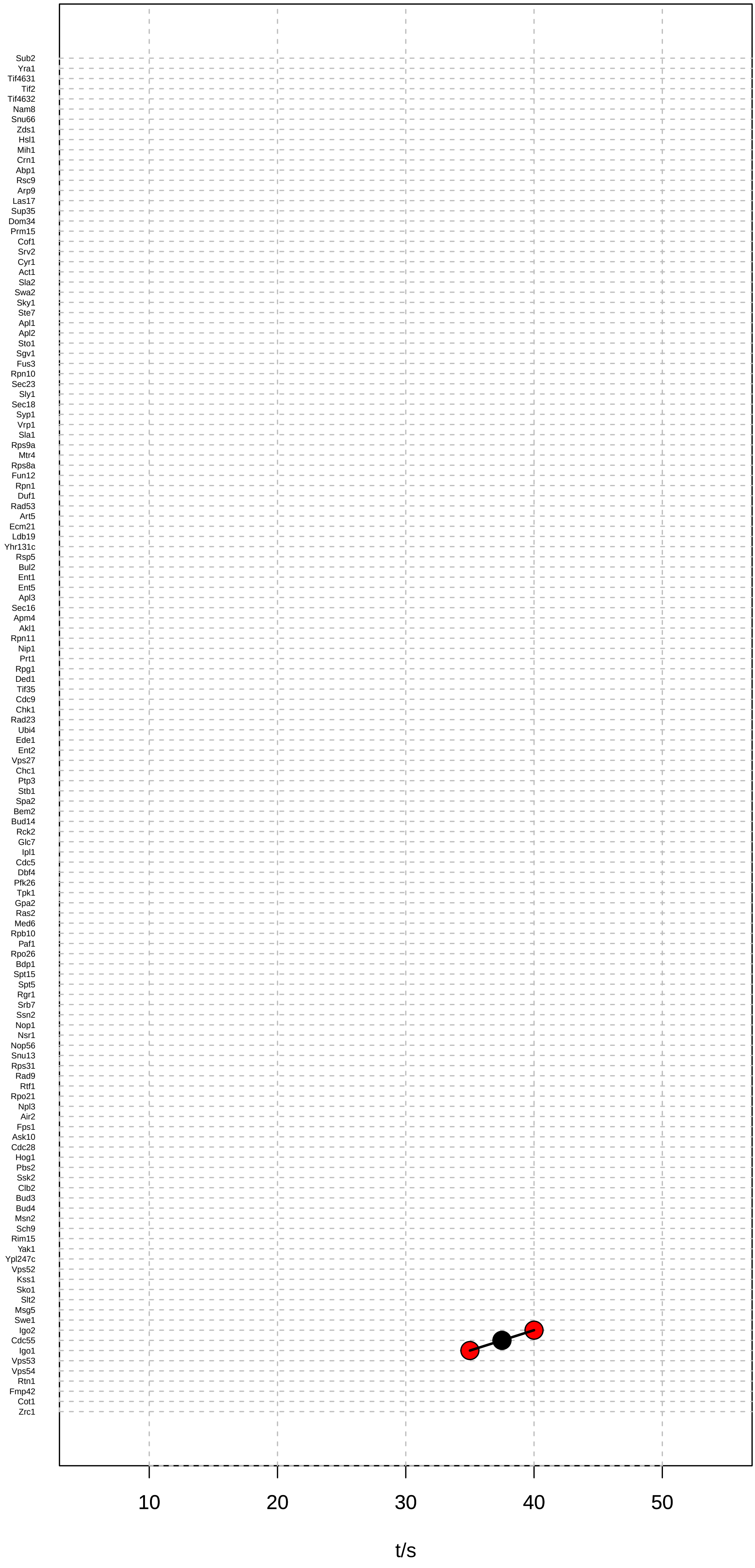

#### Path 3 (Igo1 − Swe1)

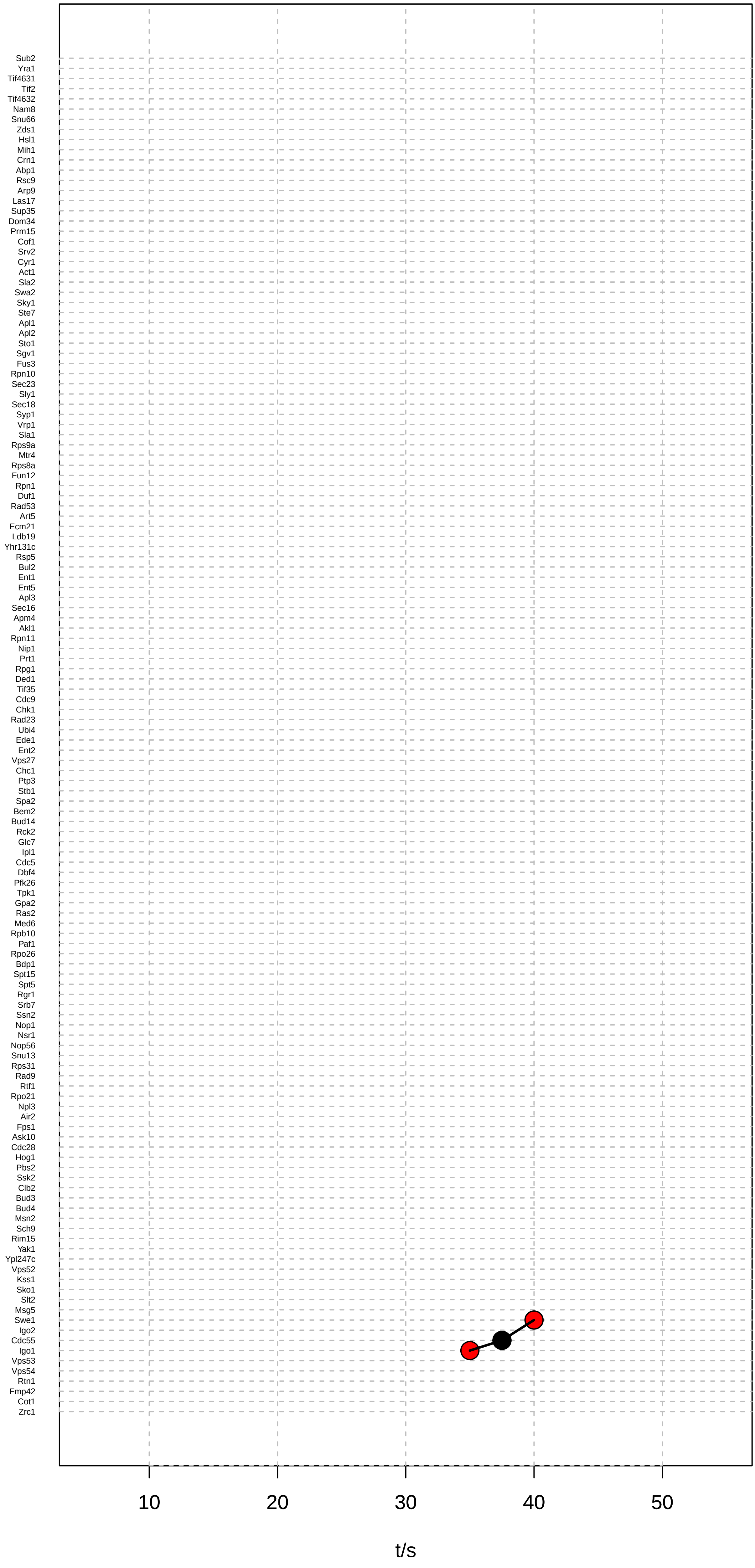

### Path 4 (Msg5 − Sko1)

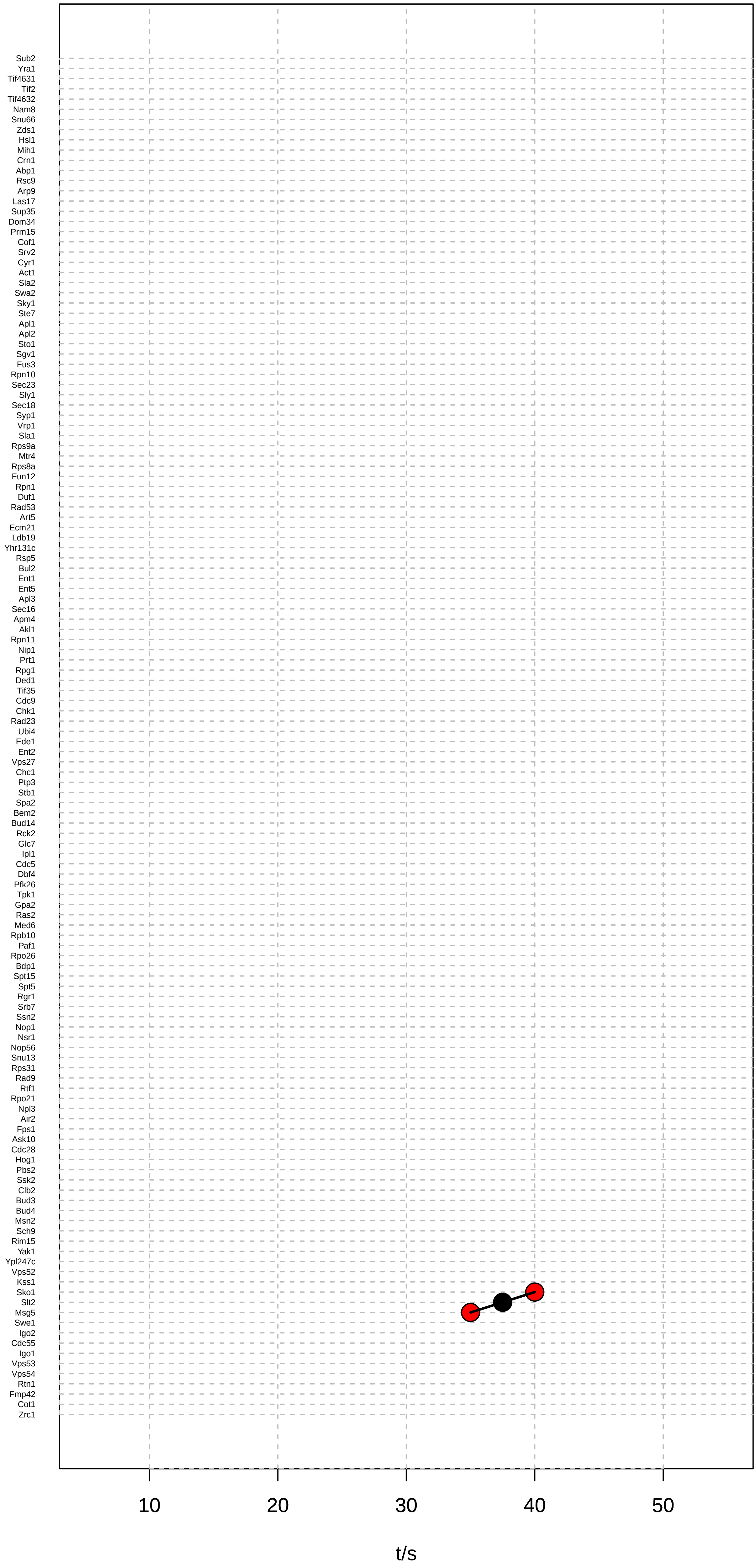

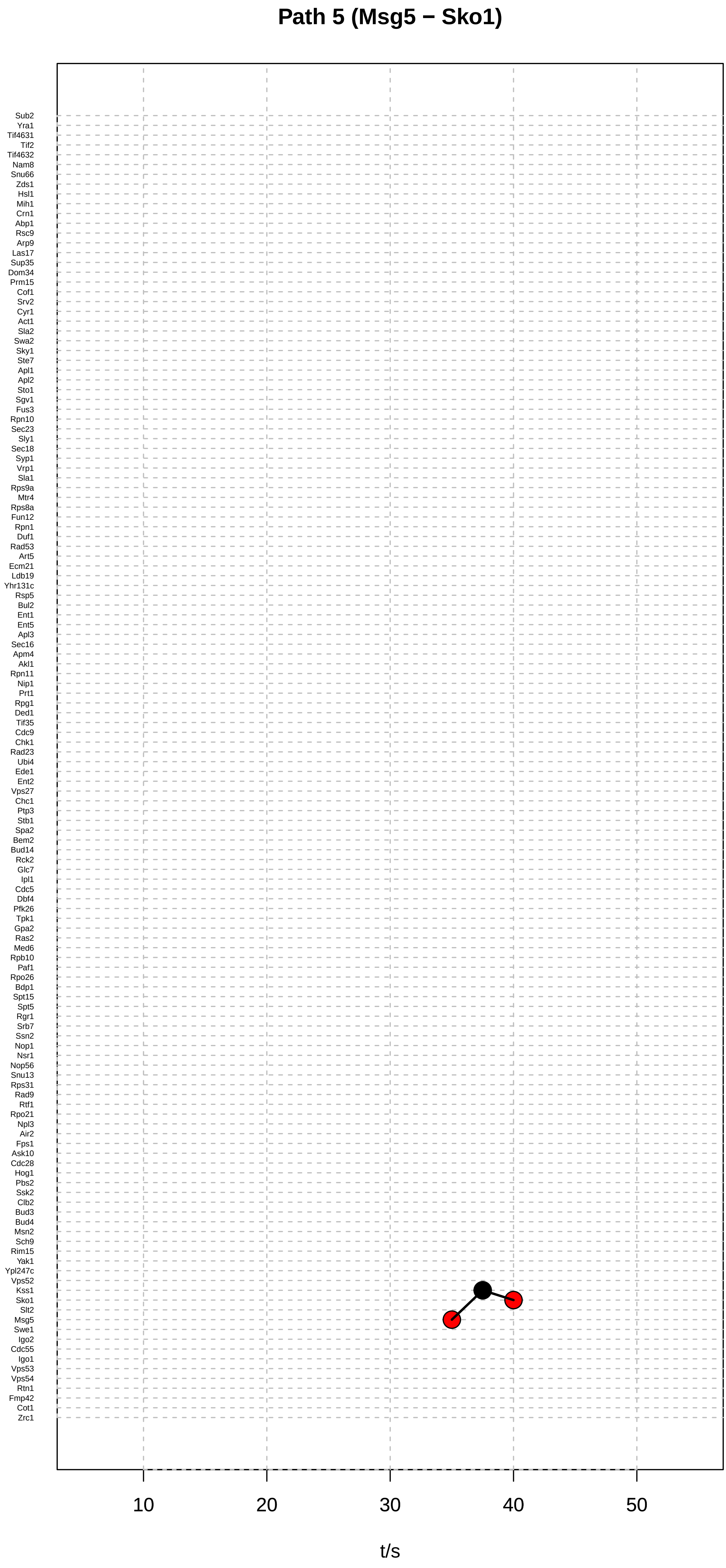

Path 5 (Msg5 − Sko1)
Sub2
Yra1
Tif4631
Tif2
Tif4632
Nam8
Snu66
Zds1
Hsl1
Mih1
Crn1
Abp1
Rsc9
Arp9
Las17
Sup35
Dom34
Prm15
Cof1
Srv2
Cyr1
Act1
Sla2
Swa2
Sky1
Ste7
Apl1
Apl2
Sto1
Sgv1
Fus3
Rpn10
Sec23
Sly1
Sec18
Syp1
Vrp1
Sla1
Rps9a
Mtr4
Rps8a
Fun12
Rpn1
Duf1
Rad53
Art5
Ecm21
Ldb19
Yhr131c
Rsp5
Bul2
Ent1
Ent5
Apl3
Sec16
Apm4
Akl1
Rpn11
Nip1
Prt1
Rpg1
Ded1
Tif35
Cdc9
Chk1
Rad23
Ubi4
Ede1
Ent2
Vps27
Chc1
Ptp3
Stb1
Spa2
Bem2
Bud14
Rck2
Glc7
Ipl1
Cdc5
Dbf4
Pfk26
Tpk1
Gpa2
Ras2
Med6
Rpb10
Paf1
Rpo26
Bdp1
Spt15
Spt5
Rgr1
Srb7
Ssn2
Nop1
Nsr1
Nop56
Snu13
Rps31
Rad9
Rtf1
Rpo21
Npl3
Air2
Fps1
Ask10
Cdc28
Hog1
Pbs2
Ssk2
Clb2
Bud3
Bud4
Msn2
Sch9
Rim15
Yak1
Ypl247c
Vps52
Kss1
Sko1
Slt2
Msg5
Swe1
Igo2
Cdc55
Igo1
Vps53
Vps54
Rtn1
Fmp42
Cot1
Zrc1
10
20
30
40
50
t/s

### Path 6 (Rtn1 − Vps53)

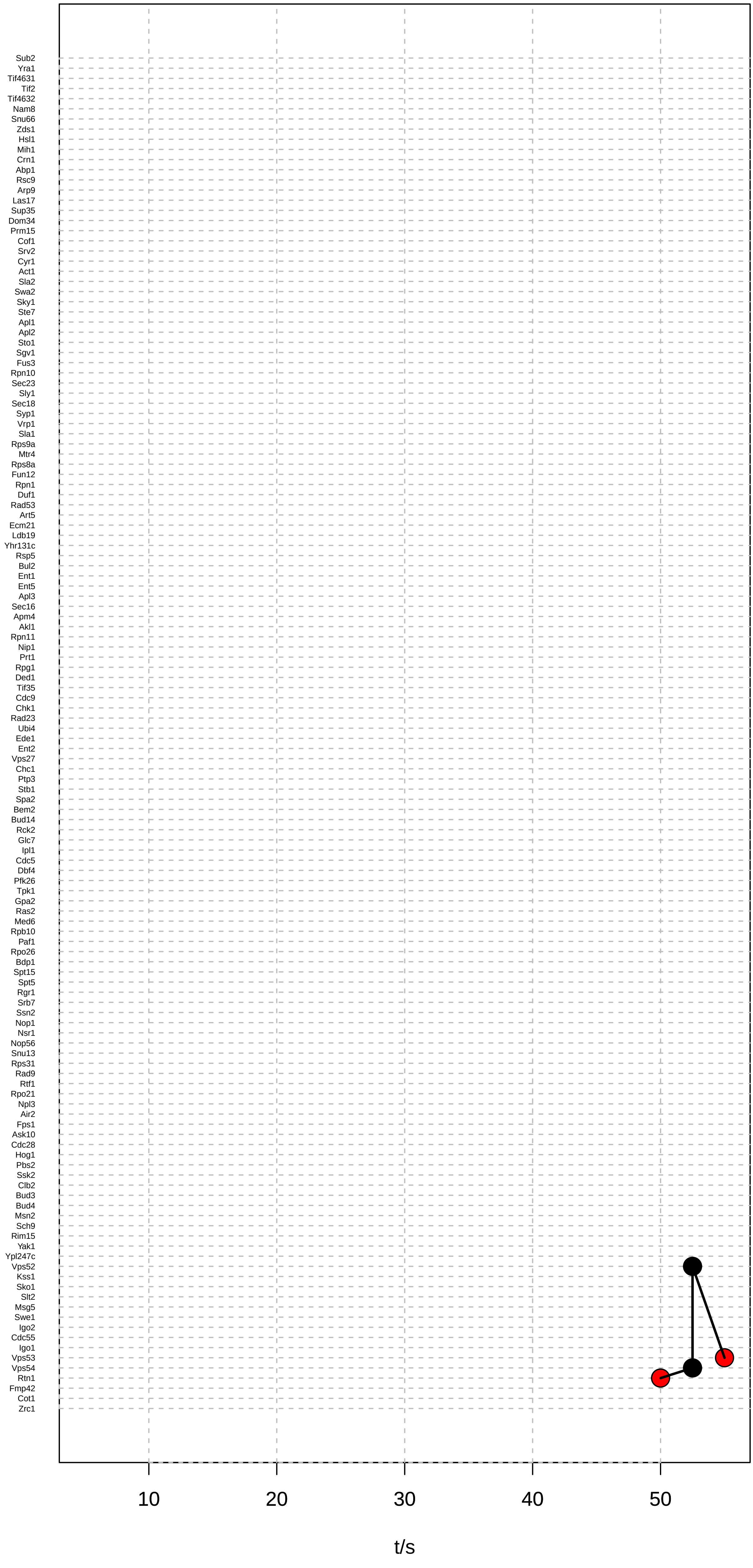

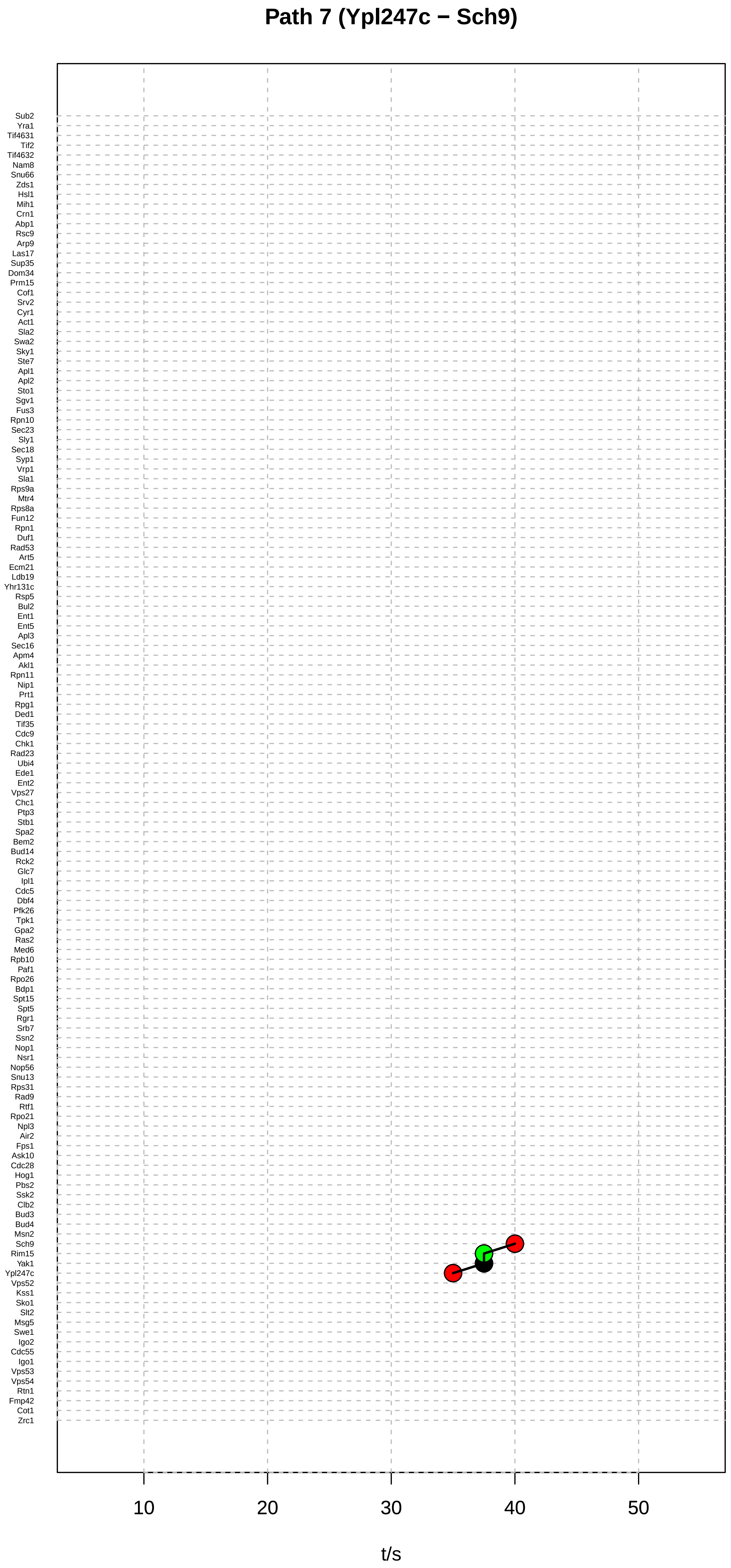
Path 7 (Ypl247c − Sch9)
Sub2
Yra1
Tif4631
Tif2
Tif4632
Nam8
Snu66
Zds1
Hsl1
Mih1
Crn1
Abp1
Rsc9
Arp9
Las17
Sup35
Dom34
Prm15
Cof1
Srv2
Cyr1
Act1
Sla2
Swa2
Sky1
Ste7
Apl1
Apl2
Sto1
Sgv1
Fus3
Rpn10
Sec23
Sly1
Sec18
Syp1
Vrp1
Sla1
Rps9a
Mtr4
Rps8a
Fun12
Rpn1
Duf1
Rad53
Art5
Ecm21
Ldb19
Yhr131c
Rsp5
Bul2
Ent1
Ent5
Apl3
Sec16
Apm4
Akl1
Rpn11
Nip1
Prt1
Rpg1
Ded1
Tif35
Cdc9
Chk1
Rad23
Ubi4
Ede1
Ent2
Vps27
Chc1
Ptp3
Stb1
Spa2
Bem2
Bud14
Rck2
Glc7
Ipl1
Cdc5
Dbf4
Pfk26
Tpk1
Gpa2
Ras2
Med6
Rpb10
Paf1
Rpo26
Bdp1
Spt15
Spt5
Rgr1
Srb7
Ssn2
Nop1
Nsr1
Nop56
Snu13
Rps31
Rad9
Rtf1
Rpo21
Npl3
Air2
Fps1
Ask10
Cdc28
Hog1
Pbs2
Ssk2
Clb2
Bud3
Bud4
Msn2
Sch9
Rim15
Yak1
Ypl247c
Vps52
Kss1
Sko1
Slt2
Msg5
Swe1
Igo2
Cdc55
Igo1
Vps53
Vps54
Rtn1
Fmp42
Cot1
Zrc1
10
20
30
40
50
t/s

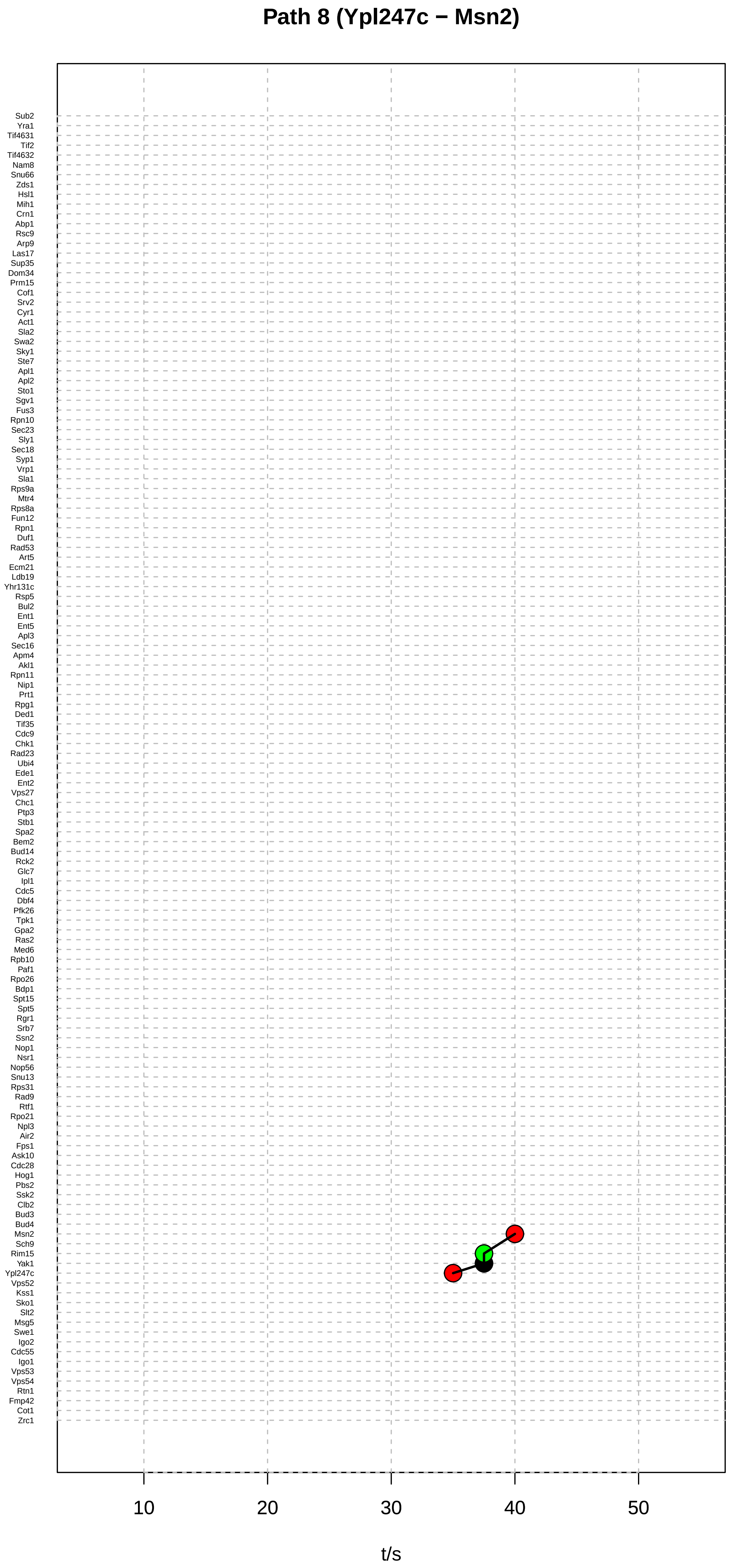

Path 8 (Ypl247c − Msn2)
Sub2
Yra1
Tif4631
Tif2
Tif4632
Nam8
Snu66
Zds1
Hsl1
Mih1
Crn1
Abp1
Rsc9
Arp9
Las17
Sup35
Dom34
Prm15
Cof1
Srv2
Cyr1
Act1
Sla2
Swa2
Sky1
Ste7
Apl1
Apl2
Sto1
Sgv1
Fus3
Rpn10
Sec23
Sly1
Sec18
Syp1
Vrp1
Sla1
Rps9a
Mtr4
Rps8a
Fun12
Rpn1
Duf1
Rad53
Art5
Ecm21
Ldb19
Yhr131c
Rsp5
Bul2
Ent1
Ent5
Apl3
Sec16
Apm4
Akl1
Rpn11
Nip1
Prt1
Rpg1
Ded1
Tif35
Cdc9
Chk1
Rad23
Ubi4
Ede1
Ent2
Vps27
Chc1
Ptp3
Stb1
Spa2
Bem2
Bud14
Rck2
Glc7
Ipl1
Cdc5
Dbf4
Pfk26
Tpk1
Gpa2
Ras2
Med6
Rpb10
Paf1
Rpo26
Bdp1
Spt15
Spt5
Rgr1
Srb7
Ssn2
Nop1
Nsr1
Nop56
Snu13
Rps31
Rad9
Rtf1
Rpo21
Npl3
Air2
Fps1
Ask10
Cdc28
Hog1
Pbs2
Ssk2
Clb2
Bud3
Bud4
Msn2
Sch9
Rim15
Yak1
Ypl247c
Vps52
Kss1
Sko1
Slt2
Msg5
Swe1
Igo2
Cdc55
Igo1
Vps53
Vps54
Rtn1
Fmp42
Cot1
Zrc1
10
20
30
40
50
t/s

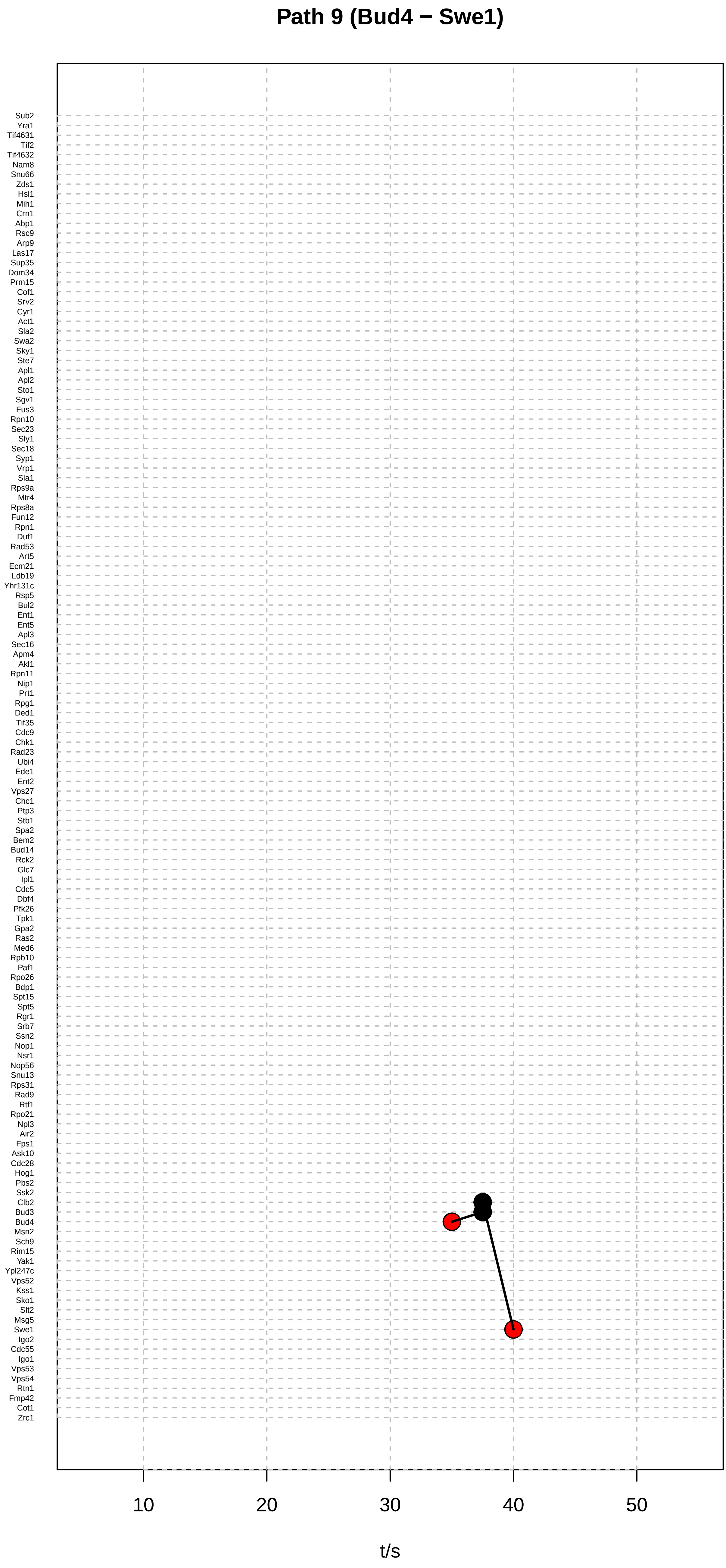
Path 9 (Bud4 − Swe1)
Sub2
Yra1
Tif4631
Tif2
Tif4632
Nam8
Snu66
Zds1
Hsl1
Mih1
Crn1
Abp1
Rsc9
Arp9
Las17
Sup35
Dom34
Prm15
Cof1
Srv2
Cyr1
Act1
Sla2
Swa2
Sky1
Ste7
Apl1
Apl2
Sto1
Sgv1
Fus3
Rpn10
Sec23
Sly1
Sec18
Syp1
Vrp1
Sla1
Rps9a
Mtr4
Rps8a
Fun12
Rpn1
Duf1
Rad53
Art5
Ecm21
Ldb19
Yhr131c
Rsp5
Bul2
Ent1
Ent5
Apl3
Sec16
Apm4
Akl1
Rpn11
Nip1
Prt1
Rpg1
Ded1
Tif35
Cdc9
Chk1
Rad23
Ubi4
Ede1
Ent2
Vps27
Chc1
Ptp3
Stb1
Spa2
Bem2
Bud14
Rck2
Glc7
Ipl1
Cdc5
Dbf4
Pfk26
Tpk1
Gpa2
Ras2
Med6
Rpb10
Paf1
Rpo26
Bdp1
Spt15
Spt5
Rgr1
Srb7
Ssn2
Nop1
Nsr1
Nop56
Snu13
Rps31
Rad9
Rtf1
Rpo21
Npl3
Air2
Fps1
Ask10
Cdc28
Hog1
Pbs2
Ssk2
Clb2
Bud3
Bud4
Msn2
Sch9
Rim15
Yak1
Ypl247c
Vps52
Kss1
Sko1
Slt2
Msg5
Swe1
Igo2
Cdc55
Igo1
Vps53
Vps54
Rtn1
Fmp42
Cot1
Zrc1
10
20
30
40
50
t/s

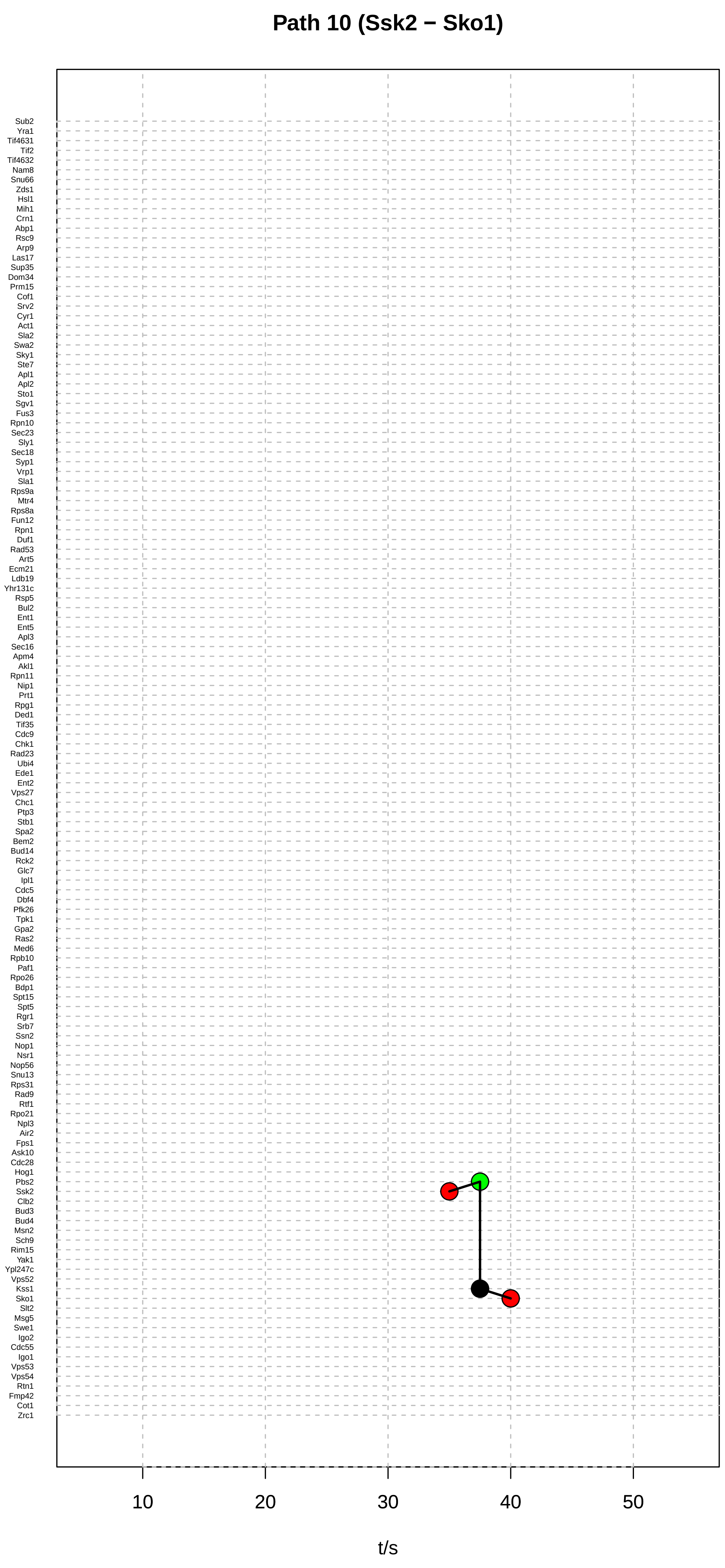
Path 10 (Ssk2 − Sko1)
Sub2
Yra1
Tif4631
Tif2
Tif4632
Nam8
Snu66
Zds1
Hsl1
Mih1
Crn1
Abp1
Rsc9
Arp9
Las17
Sup35
Dom34
Prm15
Cof1
Srv2
Cyr1
Act1
Sla2
Swa2
Sky1
Ste7
Apl1
Apl2
Sto1
Sgv1
Fus3
Rpn10
Sec23
Sly1
Sec18
Syp1
Vrp1
Sla1
Rps9a
Mtr4
Rps8a
Fun12
Rpn1
Duf1
Rad53
Art5
Ecm21
Ldb19
Yhr131c
Rsp5
Bul2
Ent1
Ent5
Apl3
Sec16
Apm4
Akl1
Rpn11
Nip1
Prt1
Rpg1
Ded1
Tif35
Cdc9
Chk1
Rad23
Ubi4
Ede1
Ent2
Vps27
Chc1
Ptp3
Stb1
Spa2
Bem2
Bud14
Rck2
Glc7
Ipl1
Cdc5
Dbf4
Pfk26
Tpk1
Gpa2
Ras2
Med6
Rpb10
Paf1
Rpo26
Bdp1
Spt15
Spt5
Rgr1
Srb7
Ssn2
Nop1
Nsr1
Nop56
Snu13
Rps31
Rad9
Rtf1
Rpo21
Npl3
Air2
Fps1
Ask10
Cdc28
Hog1
Pbs2
Ssk2
Clb2
Bud3
Bud4
Msn2
Sch9
Rim15
Yak1
Ypl247c
Vps52
Kss1
Sko1
Slt2
Msg5
Swe1
Igo2
Cdc55
Igo1
Vps53
Vps54
Rtn1
Fmp42
Cot1
Zrc1
10
20
30
40
50
t/s

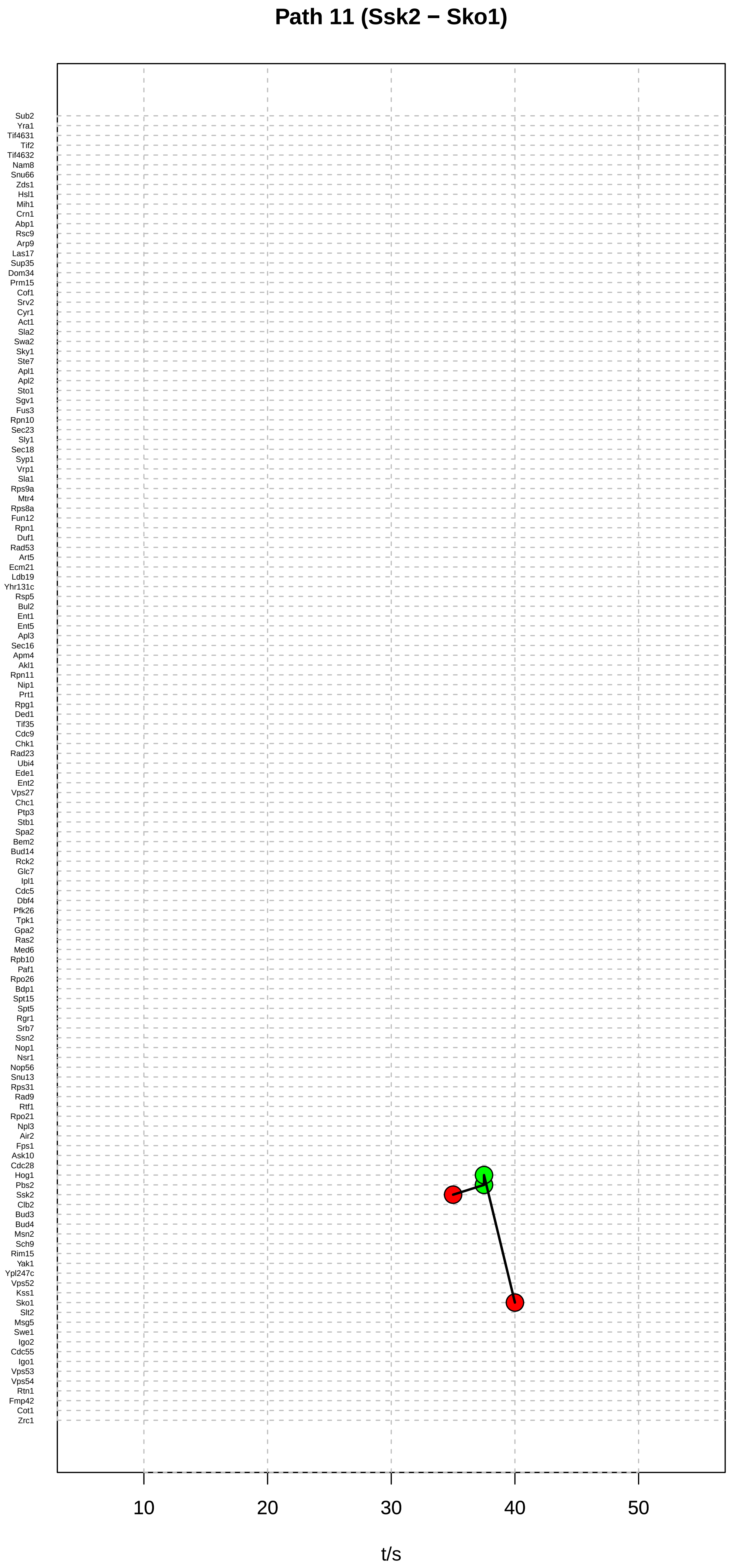

Path 11 (Ssk2 − Sko1)
Sub2
Yra1
Tif4631
Tif2
Tif4632
Nam8
Snu66
Zds1
Hsl1
Mih1
Crn1
Abp1
Rsc9
Arp9
Las17
Sup35
Dom34
Prm15
Cof1
Srv2
Cyr1
Act1
Sla2
Swa2
Sky1
Ste7
Apl1
Apl2
Sto1
Sgv1
Fus3
Rpn10
Sec23
Sly1
Sec18
Syp1
Vrp1
Sla1
Rps9a
Mtr4
Rps8a
Fun12
Rpn1
Duf1
Rad53
Art5
Ecm21
Ldb19
Yhr131c
Rsp5
Bul2
Ent1
Ent5
Apl3
Sec16
Apm4
Akl1
Rpn11
Nip1
Prt1
Rpg1
Ded1
Tif35
Cdc9
Chk1
Rad23
Ubi4
Ede1
Ent2
Vps27
Chc1
Ptp3
Stb1
Spa2
Bem2
Bud14
Rck2
Glc7
Ipl1
Cdc5
Dbf4
Pfk26
Tpk1
Gpa2
Ras2
Med6
Rpb10
Paf1
Rpo26
Bdp1
Spt15
Spt5
Rgr1
Srb7
Ssn2
Nop1
Nsr1
Nop56
Snu13
Rps31
Rad9
Rtf1
Rpo21
Npl3
Air2
Fps1
Ask10
Cdc28
Hog1
Pbs2
Ssk2
Clb2
Bud3
Bud4
Msn2
Sch9
Rim15
Yak1
Ypl247c
Vps52
Kss1
Sko1
Slt2
Msg5
Swe1
Igo2
Cdc55
Igo1
Vps53
Vps54
Rtn1
Fmp42
Cot1
Zrc1
10
20
30
40
50
t/s

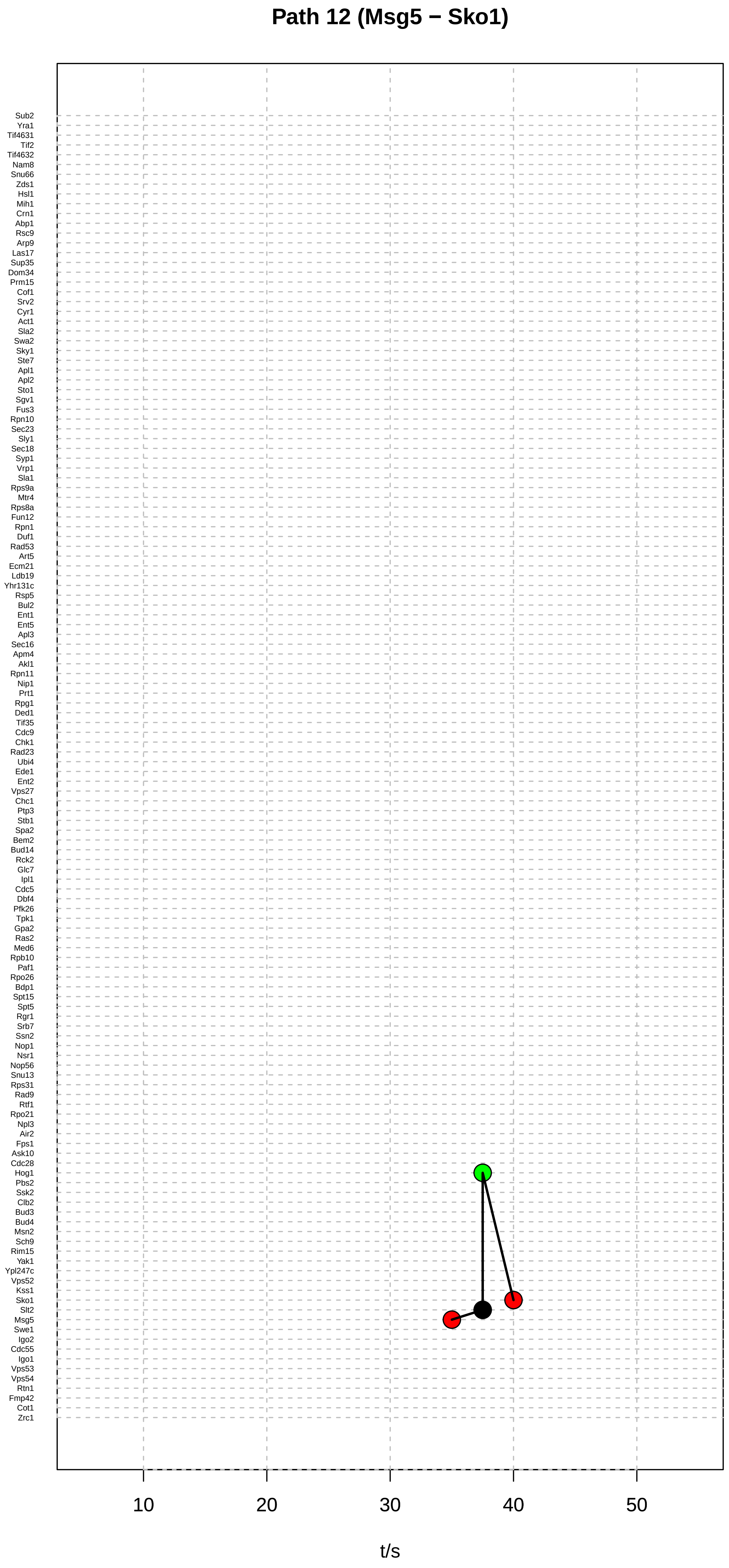

Path 12 (Msg5 − Sko1)
Sub2
Yra1
Tif4631
Tif2
Tif4632
Nam8
Snu66
Zds1
Hsl1
Mih1
Crn1
Abp1
Rsc9
Arp9
Las17
Sup35
Dom34
Prm15
Cof1
Srv2
Cyr1
Act1
Sla2
Swa2
Sky1
Ste7
Apl1
Apl2
Sto1
Sgv1
Fus3
Rpn10
Sec23
Sly1
Sec18
Syp1
Vrp1
Sla1
Rps9a
Mtr4
Rps8a
Fun12
Rpn1
Duf1
Rad53
Art5
Ecm21
Ldb19
Yhr131c
Rsp5
Bul2
Ent1
Ent5
Apl3
Sec16
Apm4
Akl1
Rpn11
Nip1
Prt1
Rpg1
Ded1
Tif35
Cdc9
Chk1
Rad23
Ubi4
Ede1
Ent2
Vps27
Chc1
Ptp3
Stb1
Spa2
Bem2
Bud14
Rck2
Glc7
Ipl1
Cdc5
Dbf4
Pfk26
Tpk1
Gpa2
Ras2
Med6
Rpb10
Paf1
Rpo26
Bdp1
Spt15
Spt5
Rgr1
Srb7
Ssn2
Nop1
Nsr1
Nop56
Snu13
Rps31
Rad9
Rtf1
Rpo21
Npl3
Air2
Fps1
Ask10
Cdc28
Hog1
Pbs2
Ssk2
Clb2
Bud3
Bud4
Msn2
Sch9
Rim15
Yak1
Ypl247c
Vps52
Kss1
Sko1
Slt2
Msg5
Swe1
Igo2
Cdc55
Igo1
Vps53
Vps54
Rtn1
Fmp42
Cot1
Zrc1
10
20
30
40
50
t/s

#### Path 13 (Msg5 − Swe1)

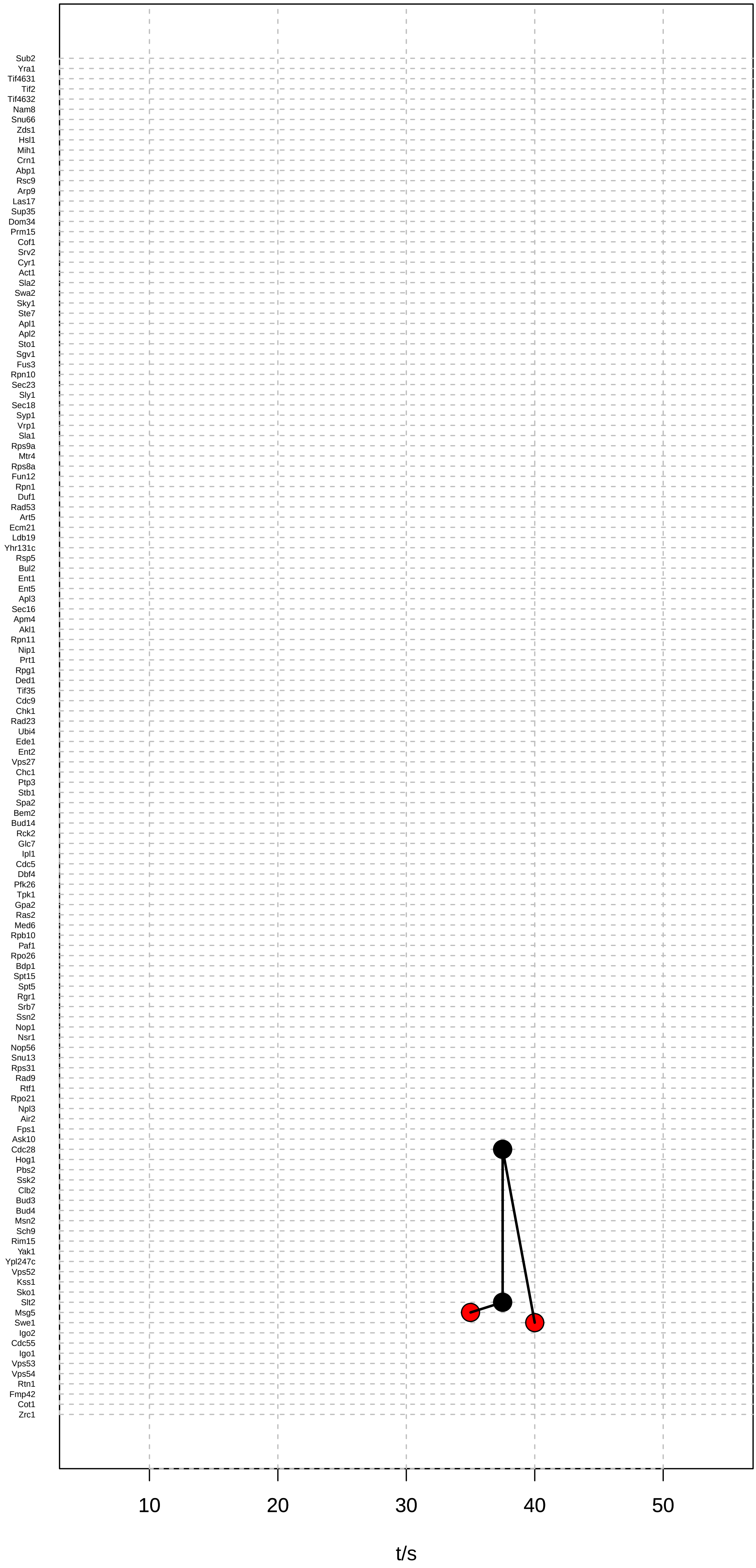

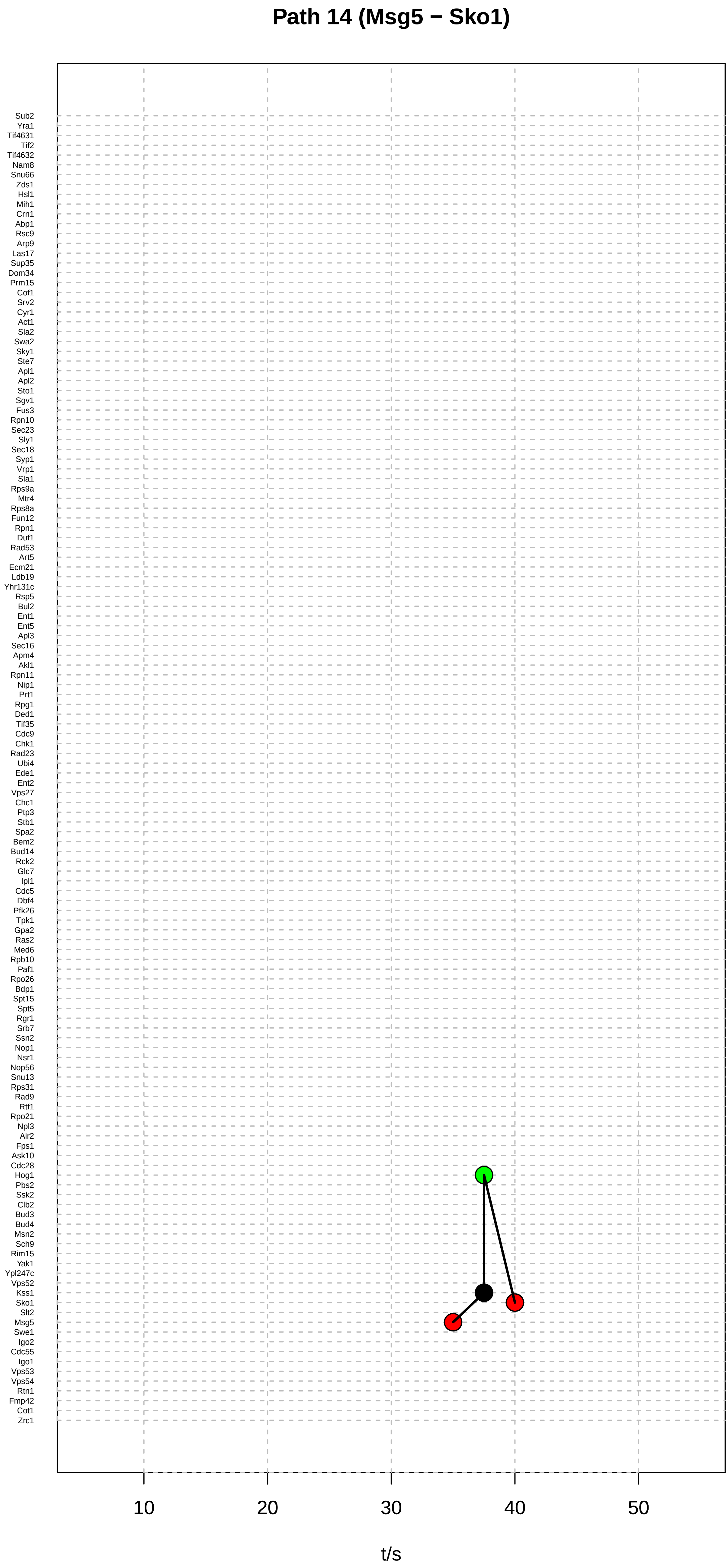
Path 14 (Msg5 − Sko1)
Sub2
Yra1
Tif4631
Tif2
Tif4632
Nam8
Snu66
Zds1
Hsl1
Mih1
Crn1
Abp1
Rsc9
Arp9
Las17
Sup35
Dom34
Prm15
Cof1
Srv2
Cyr1
Act1
Sla2
Swa2
Sky1
Ste7
Apl1
Apl2
Sto1
Sgv1
Fus3
Rpn10
Sec23
Sly1
Sec18
Syp1
Vrp1
Sla1
Rps9a
Mtr4
Rps8a
Fun12
Rpn1
Duf1
Rad53
Art5
Ecm21
Ldb19
Yhr131c
Rsp5
Bul2
Ent1
Ent5
Apl3
Sec16
Apm4
Akl1
Rpn11
Nip1
Prt1
Rpg1
Ded1
Tif35
Cdc9
Chk1
Rad23
Ubi4
Ede1
Ent2
Vps27
Chc1
Ptp3
Stb1
Spa2
Bem2
Bud14
Rck2
Glc7
Ipl1
Cdc5
Dbf4
Pfk26
Tpk1
Gpa2
Ras2
Med6
Rpb10
Paf1
Rpo26
Bdp1
Spt15
Spt5
Rgr1
Srb7
Ssn2
Nop1
Nsr1
Nop56
Snu13
Rps31
Rad9
Rtf1
Rpo21
Npl3
Air2
Fps1
Ask10
Cdc28
Hog1
Pbs2
Ssk2
Clb2
Bud3
Bud4
Msn2
Sch9
Rim15
Yak1
Ypl247c
Vps52
Kss1
Sko1
Slt2
Msg5
Swe1
Igo2
Cdc55
Igo1
Vps53
Vps54
Rtn1
Fmp42
Cot1
Zrc1
10
20
30
40
50
t/s

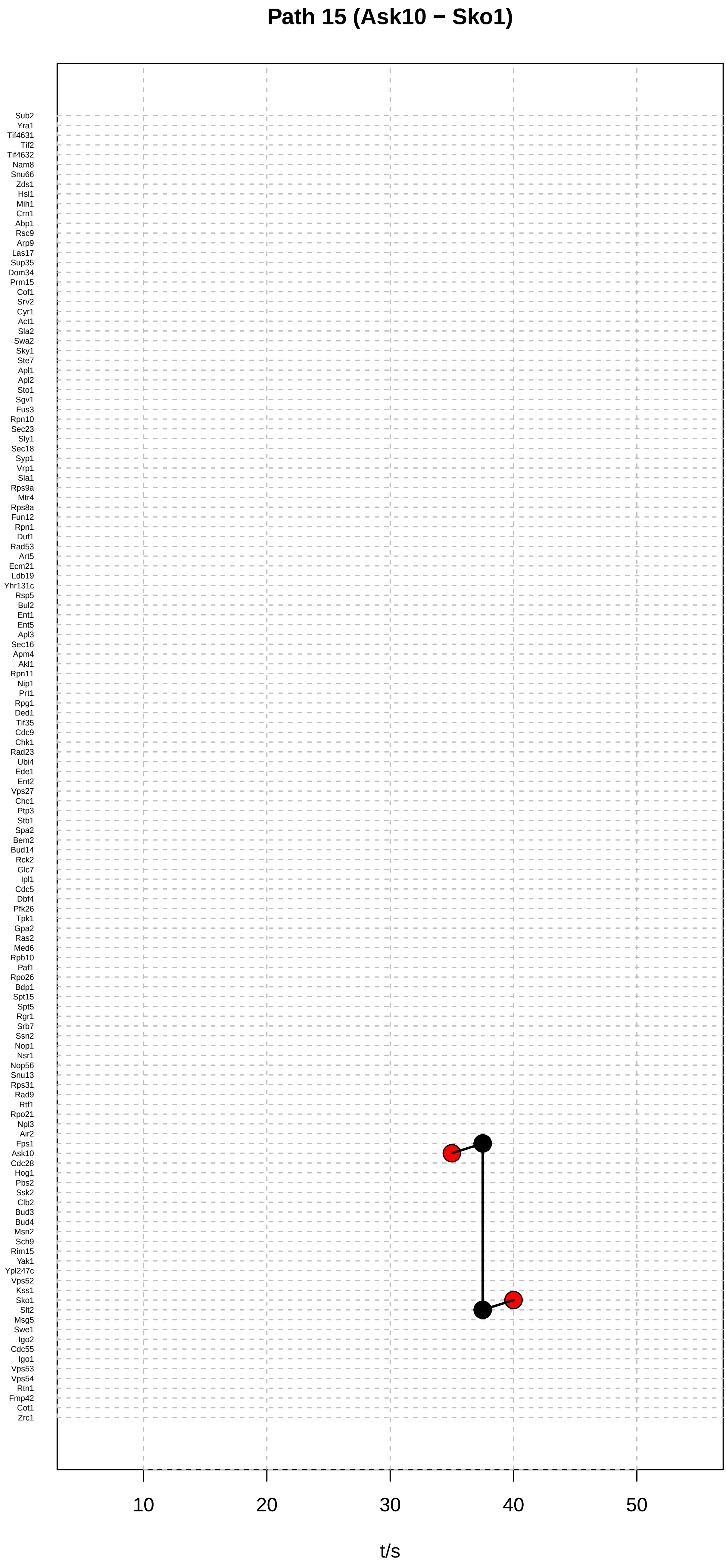

Path 15 (Ask10 − Sko1)
Sub2
Yra1
Tif4631
Tif2
Tif4632
Nam8
Snu66
Zds1
Hsl1
Mih1
Crn1
Abp1
Rsc9
Arp9
Las17
Sup35
Dom34
Prm15
Cof1
Srv2
Cyr1
Act1
Sla2
Swa2
Sky1
Ste7
Apl1
Apl2
Sto1
Sgv1
Fus3
Rpn10
Sec23
Sly1
Sec18
Syp1
Vrp1
Sla1
Rps9a
Mtr4
Rps8a
Fun12
Rpn1
Duf1
Rad53
Art5
Ecm21
Ldb19
Yhr131c
Rsp5
Bul2
Ent1
Ent5
Apl3
Sec16
Apm4
Akl1
Rpn11
Nip1
Prt1
Rpg1
Ded1
Tif35
Cdc9
Chk1
Rad23
Ubi4
Ede1
Ent2
Vps27
Chc1
Ptp3
Stb1
Spa2
Bem2
Bud14
Rck2
Glc7
Ipl1
Cdc5
Dbf4
Pfk26
Tpk1
Gpa2
Ras2
Med6
Rpb10
Paf1
Rpo26
Bdp1
Spt15
Spt5
Rgr1
Srb7
Ssn2
Nop1
Nsr1
Nop56
Snu13
Rps31
Rad9
Rtf1
Rpo21
Npl3
Air2
Fps1
Ask10
Cdc28
Hog1
Pbs2
Ssk2
Clb2
Bud3
Bud4
Msn2
Sch9
Rim15
Yak1
Ypl247c
Vps52
Kss1
Sko1
Slt2
Msg5
Swe1
Igo2
Cdc55
Igo1
Vps53
Vps54
Rtn1
Fmp42
Cot1
Zrc1
10
20
30
40
50
t/s

### Path 16 (Ask10 − Sko1)

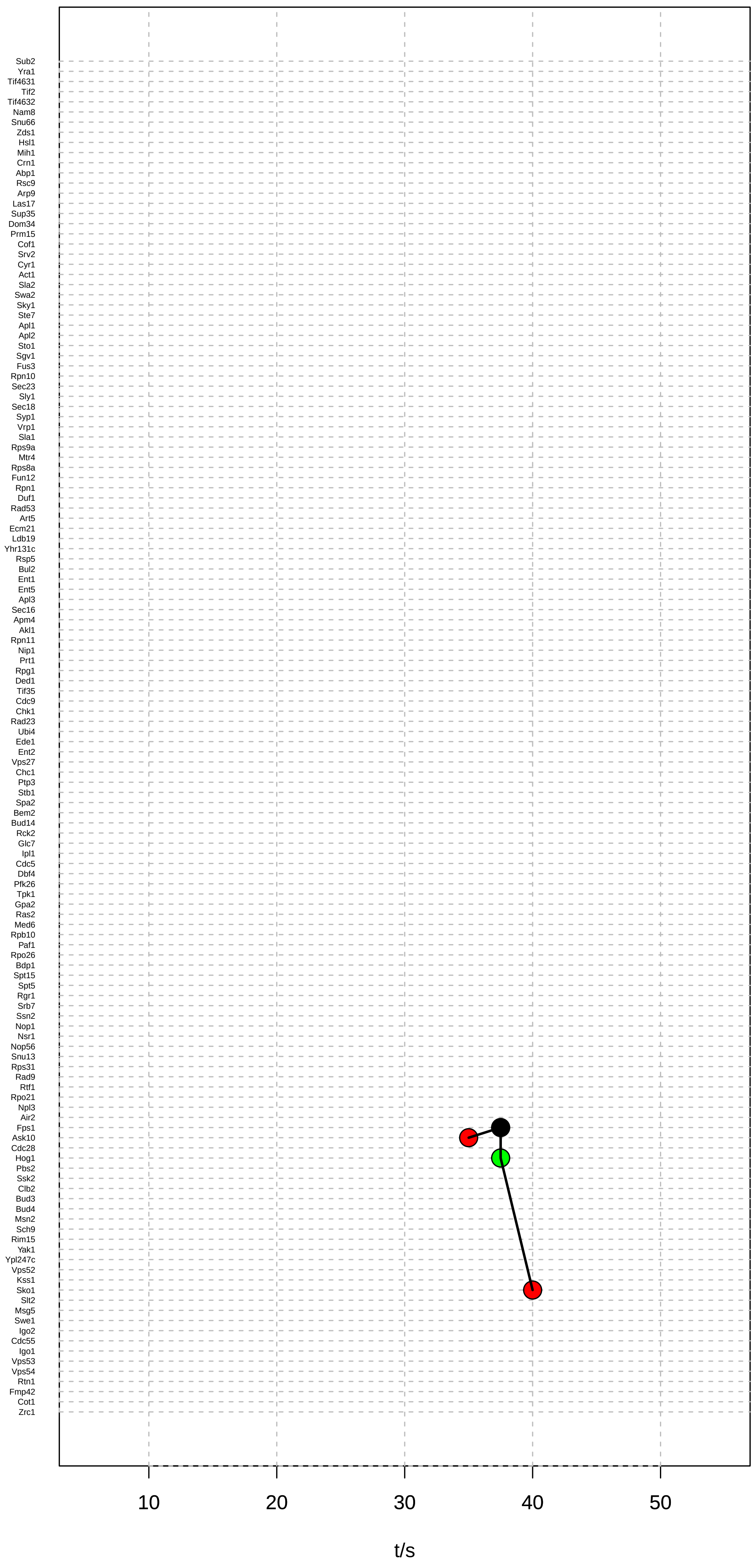

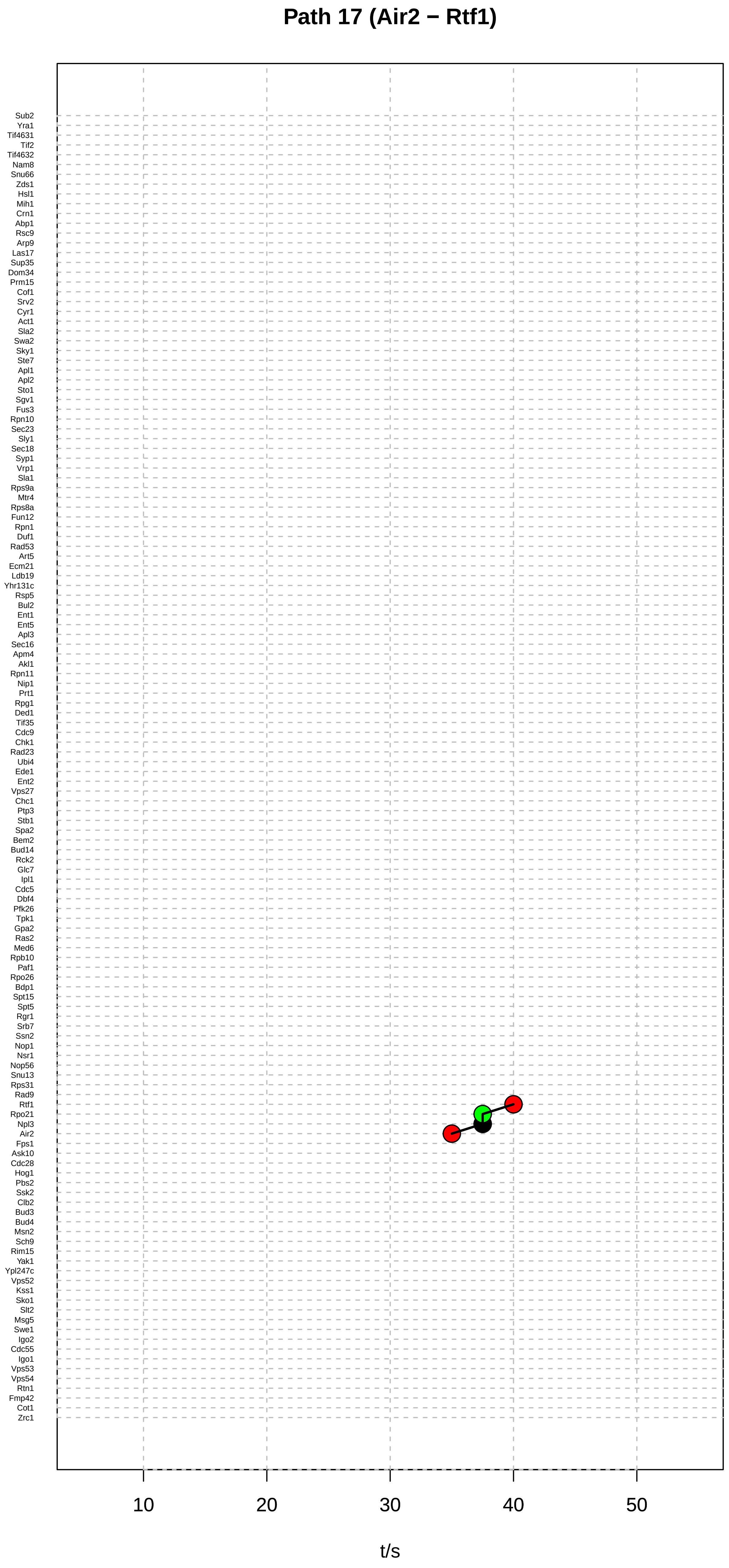
Path 17 (Air2 − Rtf1)
Sub2
Yra1
Tif4631
Tif2
Tif4632
Nam8
Snu66
Zds1
Hsl1
Mih1
Crn1
Abp1
Rsc9
Arp9
Las17
Sup35
Dom34
Prm15
Cof1
Srv2
Cyr1
Act1
Sla2
Swa2
Sky1
Ste7
Apl1
Apl2
Sto1
Sgv1
Fus3
Rpn10
Sec23
Sly1
Sec18
Syp1
Vrp1
Sla1
Rps9a
Mtr4
Rps8a
Fun12
Rpn1
Duf1
Rad53
Art5
Ecm21
Ldb19
Yhr131c
Rsp5
Bul2
Ent1
Ent5
Apl3
Sec16
Apm4
Akl1
Rpn11
Nip1
Prt1
Rpg1
Ded1
Tif35
Cdc9
Chk1
Rad23
Ubi4
Ede1
Ent2
Vps27
Chc1
Ptp3
Stb1
Spa2
Bem2
Bud14
Rck2
Glc7
Ipl1
Cdc5
Dbf4
Pfk26
Tpk1
Gpa2
Ras2
Med6
Rpb10
Paf1
Rpo26
Bdp1
Spt15
Spt5
Rgr1
Srb7
Ssn2
Nop1
Nsr1
Nop56
Snu13
Rps31
Rad9
Rtf1
Rpo21
Npl3
Air2
Fps1
Ask10
Cdc28
Hog1
Pbs2
Ssk2
Clb2
Bud3
Bud4
Msn2
Sch9
Rim15
Yak1
Ypl247c
Vps52
Kss1
Sko1
Slt2
Msg5
Swe1
Igo2
Cdc55
Igo1
Vps53
Vps54
Rtn1
Fmp42
Cot1
Zrc1
10
20
30
40
50
t/s

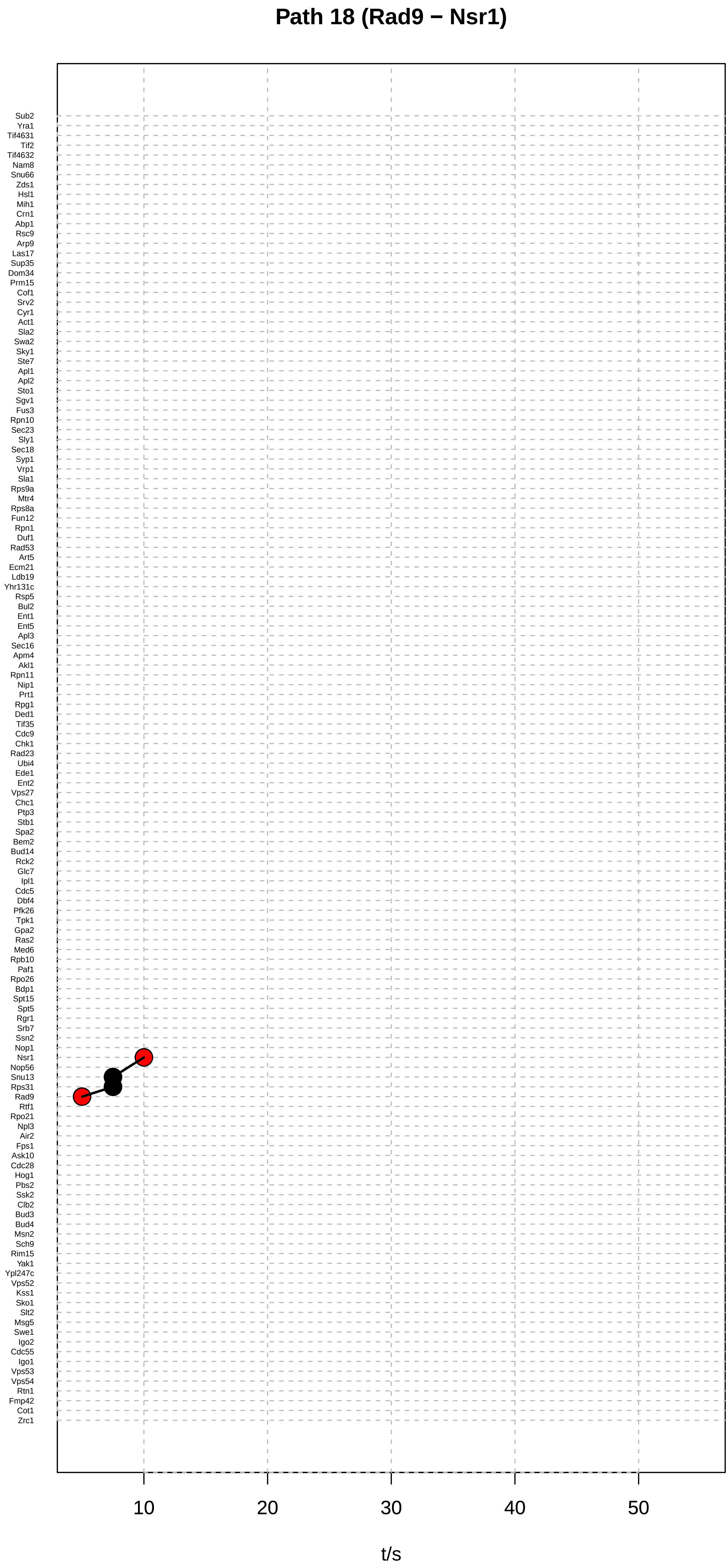
Path 18 (Rad9 − Nsr1)
Sub2
Yra1
Tif4631
Tif2
Tif4632
Nam8
Snu66
Zds1
Hsl1
Mih1
Crn1
Abp1
Rsc9
Arp9
Las17
Sup35
Dom34
Prm15
Cof1
Srv2
Cyr1
Act1
Sla2
Swa2
Sky1
Ste7
Apl1
Apl2
Sto1
Sgv1
Fus3
Rpn10
Sec23
Sly1
Sec18
Syp1
Vrp1
Sla1
Rps9a
Mtr4
Rps8a
Fun12
Rpn1
Duf1
Rad53
Art5
Ecm21
Ldb19
Yhr131c
Rsp5
Bul2
Ent1
Ent5
Apl3
Sec16
Apm4
Akl1
Rpn11
Nip1
Prt1
Rpg1
Ded1
Tif35
Cdc9
Chk1
Rad23
Ubi4
Ede1
Ent2
Vps27
Chc1
Ptp3
Stb1
Spa2
Bem2
Bud14
Rck2
Glc7
Ipl1
Cdc5
Dbf4
Pfk26
Tpk1
Gpa2
Ras2
Med6
Rpb10
Paf1
Rpo26
Bdp1
Spt15
Spt5
Rgr1
Srb7
Ssn2
Nop1
Nsr1
Nop56
Snu13
Rps31
Rad9
Rtf1
Rpo21
Npl3
Air2
Fps1
Ask10
Cdc28
Hog1
Pbs2
Ssk2
Clb2
Bud3
Bud4
Msn2
Sch9
Rim15
Yak1
Ypl247c
Vps52
Kss1
Sko1
Slt2
Msg5
Swe1
Igo2
Cdc55
Igo1
Vps53
Vps54
Rtn1
Fmp42
Cot1
Zrc1
10
20
30
40
50
t/s

### Path 19 (Rad9 − Nsr1)

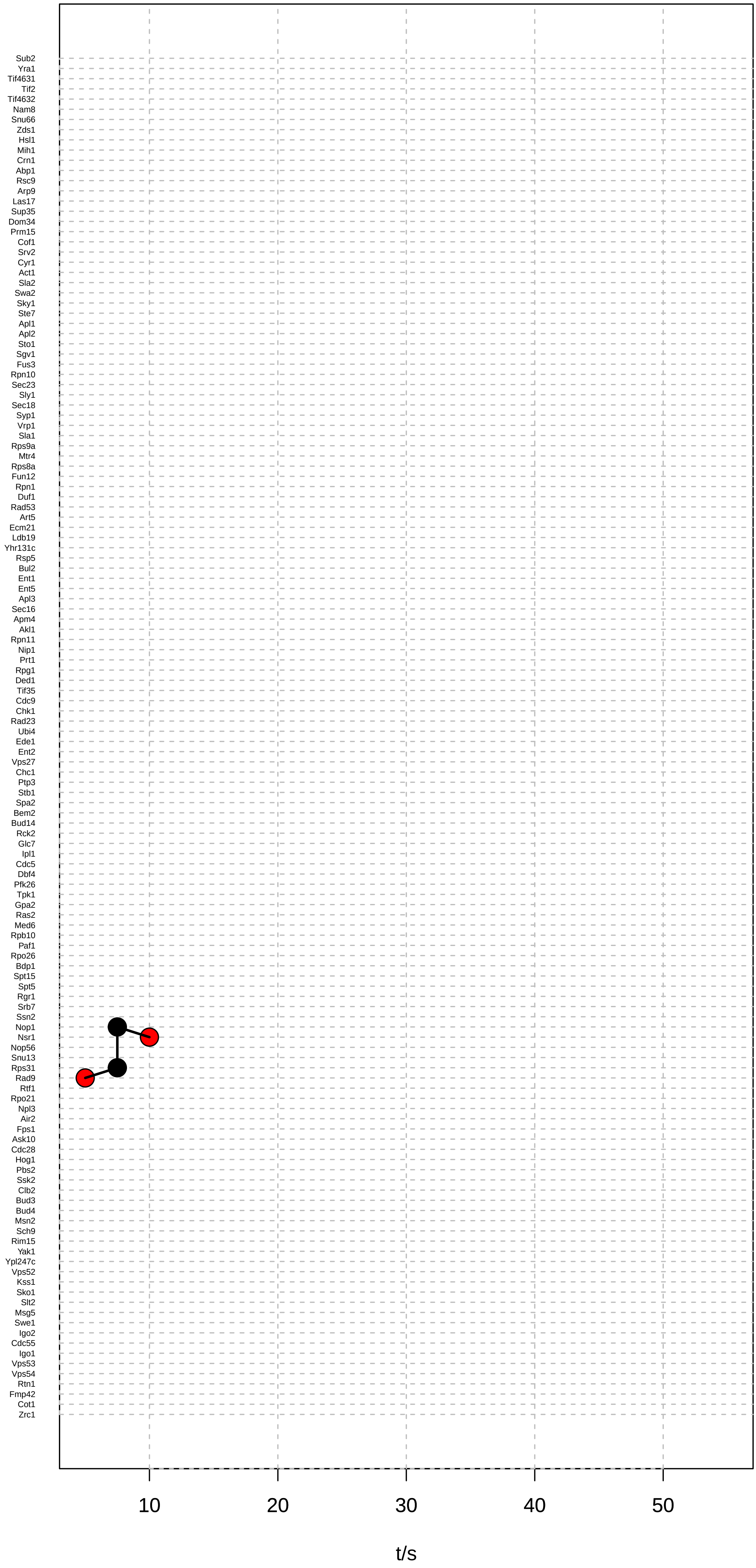

#### Path 20 (Ssn2 − Rtf1)

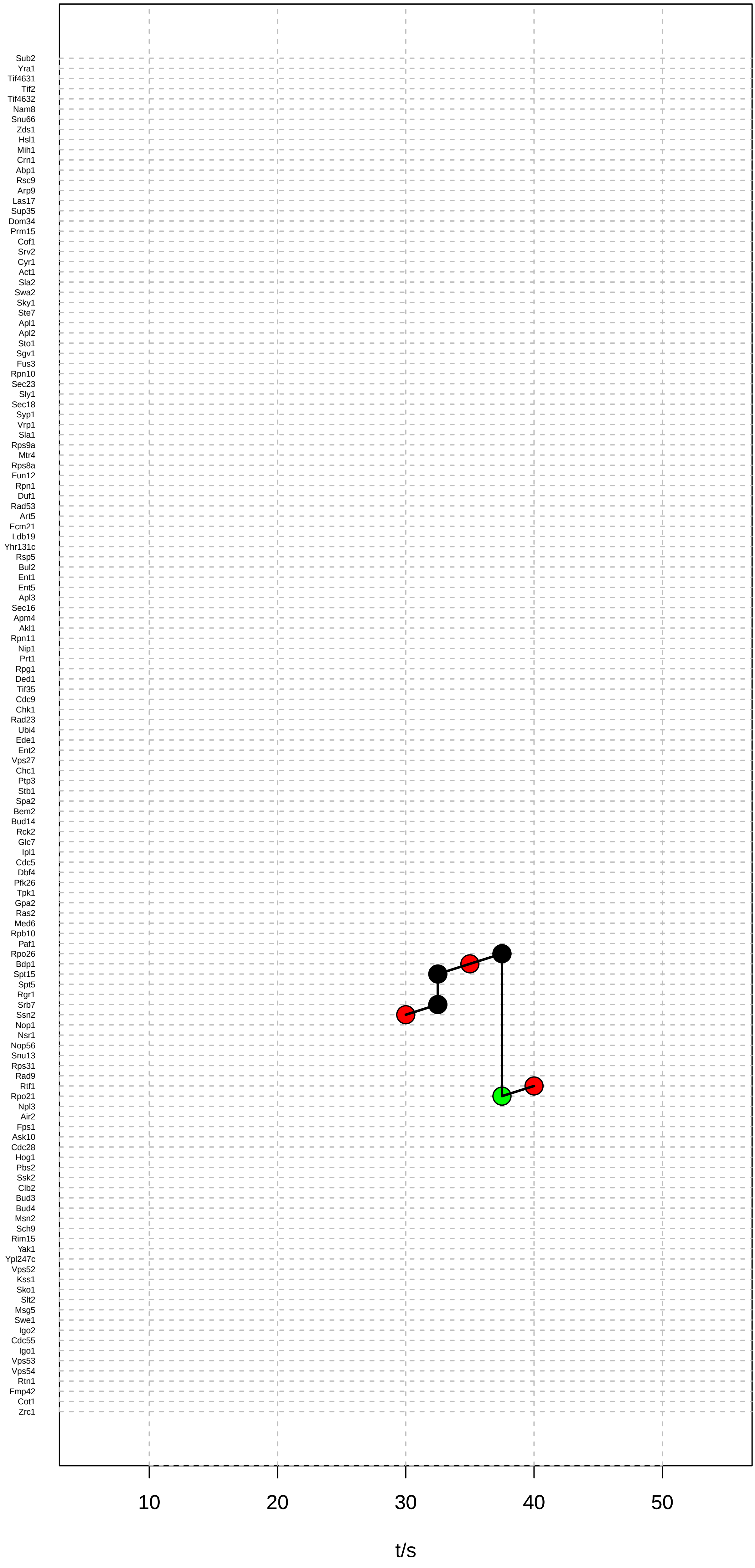

### Path 21 (Ssn2 − Rtf1)

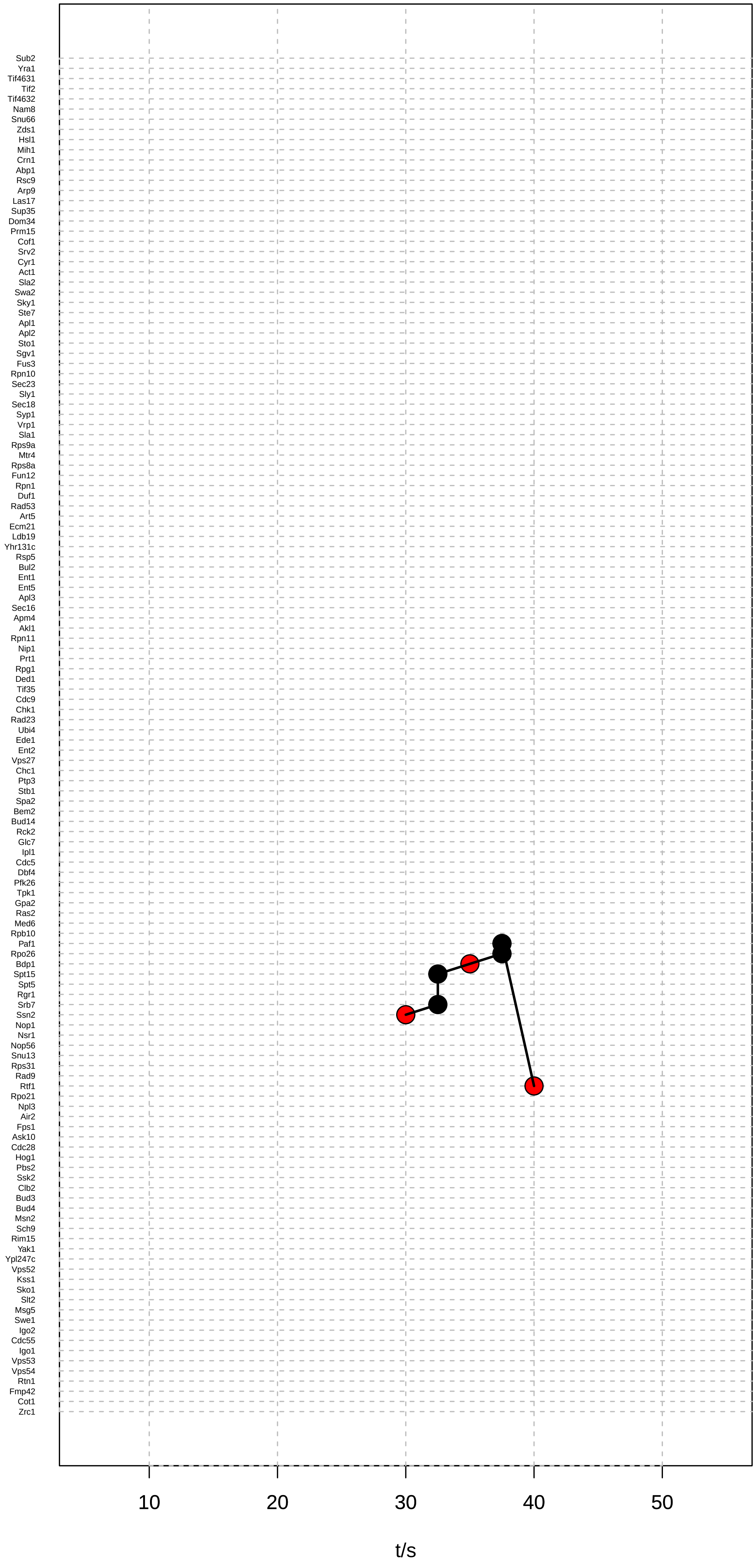

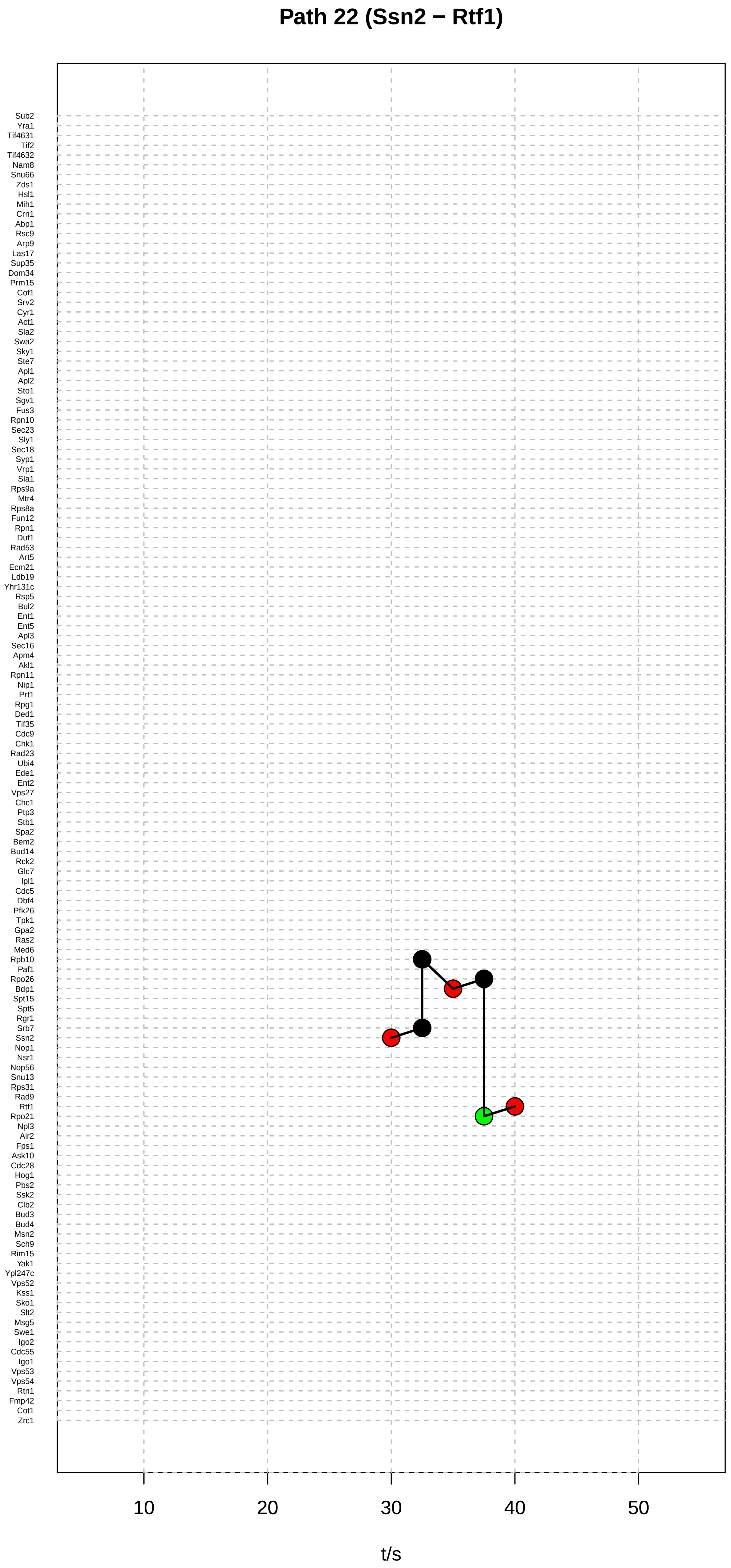

Path 22 (Ssn2 − Rtf1)
Sub2
Yra1
Tif4631
Tif2
Tif4632
Nam8
Snu66
Zds1
Hsl1
Mih1
Crn1
Abp1
Rsc9
Arp9
Las17
Sup35
Dom34
Prm15
Cof1
Srv2
Cyr1
Act1
Sla2
Swa2
Sky1
Ste7
Apl1
Apl2
Sto1
Sgv1
Fus3
Rpn10
Sec23
Sly1
Sec18
Syp1
Vrp1
Sla1
Rps9a
Mtr4
Rps8a
Fun12
Rpn1
Duf1
Rad53
Art5
Ecm21
Ldb19
Yhr131c
Rsp5
Bul2
Ent1
Ent5
Apl3
Sec16
Apm4
Akl1
Rpn11
Nip1
Prt1
Rpg1
Ded1
Tif35
Cdc9
Chk1
Rad23
Ubi4
Ede1
Ent2
Vps27
Chc1
Ptp3
Stb1
Spa2
Bem2
Bud14
Rck2
Glc7
Ipl1
Cdc5
Dbf4
Pfk26
Tpk1
Gpa2
Ras2
Med6
Rpb10
Paf1
Rpo26
Bdp1
Spt15
Spt5
Rgr1
Srb7
Ssn2
Nop1
Nsr1
Nop56
Snu13
Rps31
Rad9
Rtf1
Rpo21
Npl3
Air2
Fps1
Ask10
Cdc28
Hog1
Pbs2
Ssk2
Clb2
Bud3
Bud4
Msn2
Sch9
Rim15
Yak1
Ypl247c
Vps52
Kss1
Sko1
Slt2
Msg5
Swe1
Igo2
Cdc55
Igo1
Vps53
Vps54
Rtn1
Fmp42
Cot1
Zrc1
10
20
30
40
50
t/s

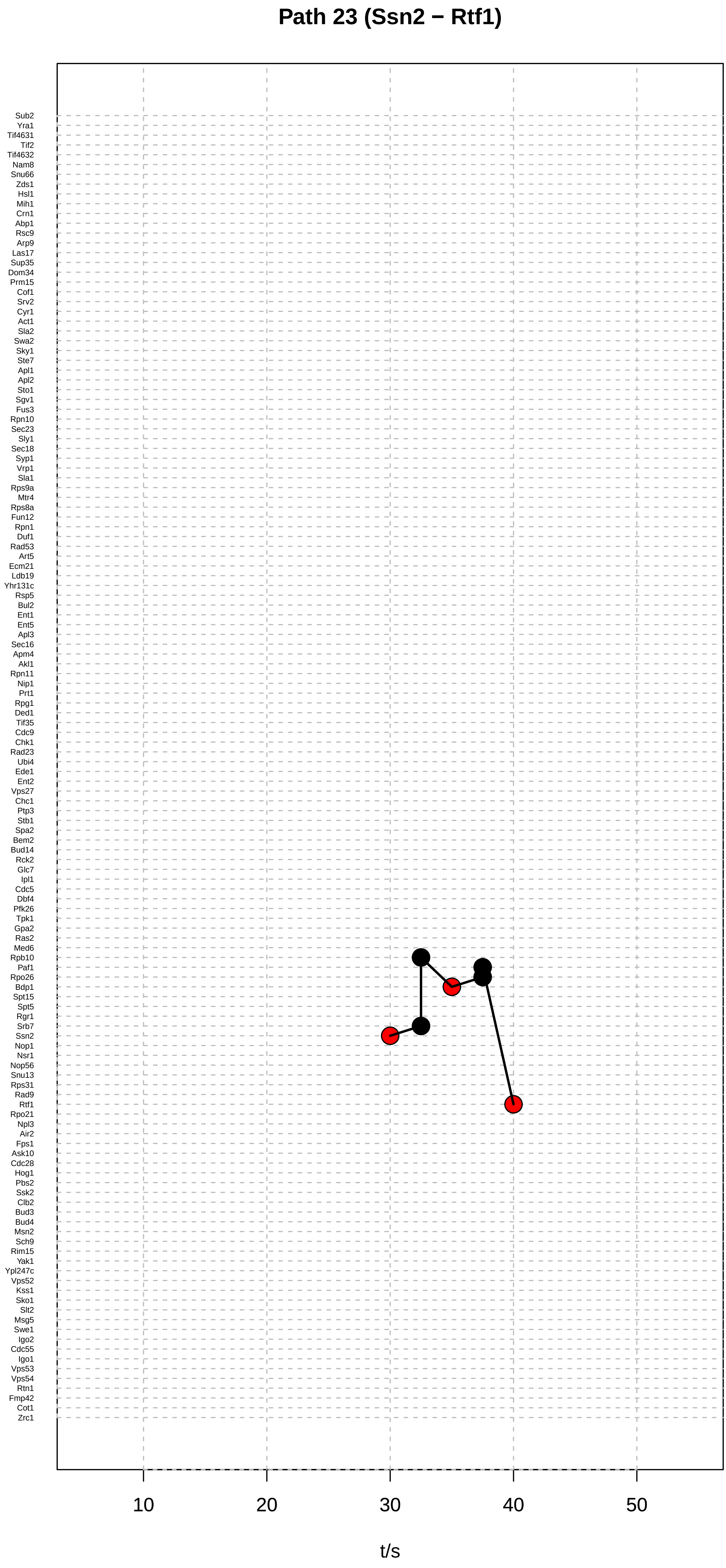
Path 23 (Ssn2 − Rtf1)
Sub2
Yra1
Tif4631
Tif2
Tif4632
Nam8
Snu66
Zds1
Hsl1
Mih1
Crn1
Abp1
Rsc9
Arp9
Las17
Sup35
Dom34
Prm15
Cof1
Srv2
Cyr1
Act1
Sla2
Swa2
Sky1
Ste7
Apl1
Apl2
Sto1
Sgv1
Fus3
Rpn10
Sec23
Sly1
Sec18
Syp1
Vrp1
Sla1
Rps9a
Mtr4
Rps8a
Fun12
Rpn1
Duf1
Rad53
Art5
Ecm21
Ldb19
Yhr131c
Rsp5
Bul2
Ent1
Ent5
Apl3
Sec16
Apm4
Akl1
Rpn11
Nip1
Prt1
Rpg1
Ded1
Tif35
Cdc9
Chk1
Rad23
Ubi4
Ede1
Ent2
Vps27
Chc1
Ptp3
Stb1
Spa2
Bem2
Bud14
Rck2
Glc7
Ipl1
Cdc5
Dbf4
Pfk26
Tpk1
Gpa2
Ras2
Med6
Rpb10
Paf1
Rpo26
Bdp1
Spt15
Spt5
Rgr1
Srb7
Ssn2
Nop1
Nsr1
Nop56
Snu13
Rps31
Rad9
Rtf1
Rpo21
Npl3
Air2
Fps1
Ask10
Cdc28
Hog1
Pbs2
Ssk2
Clb2
Bud3
Bud4
Msn2
Sch9
Rim15
Yak1
Ypl247c
Vps52
Kss1
Sko1
Slt2
Msg5
Swe1
Igo2
Cdc55
Igo1
Vps53
Vps54
Rtn1
Fmp42
Cot1
Zrc1
10
20
30
40
50
t/s

### Path 24 (Ssn2 − Rtf1)

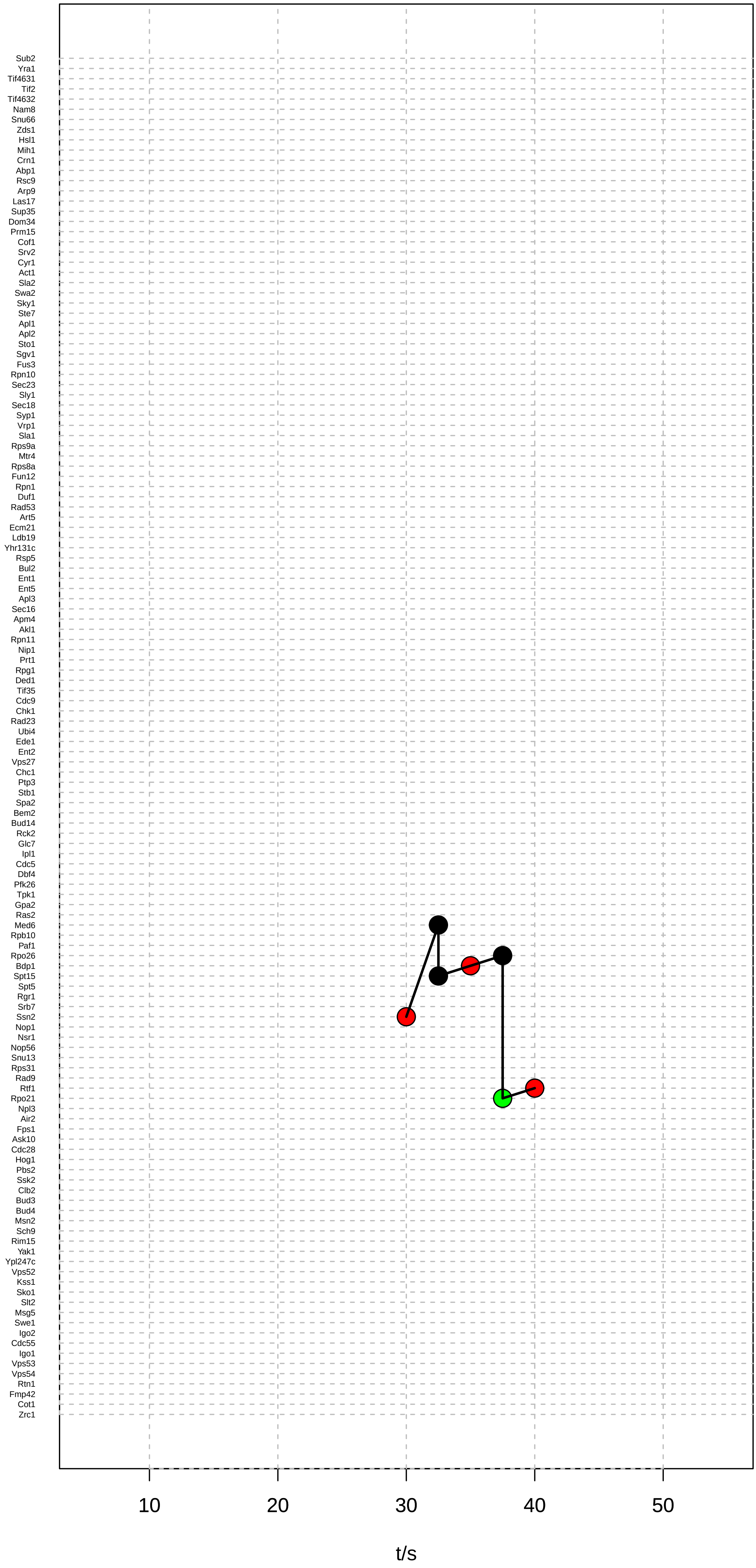

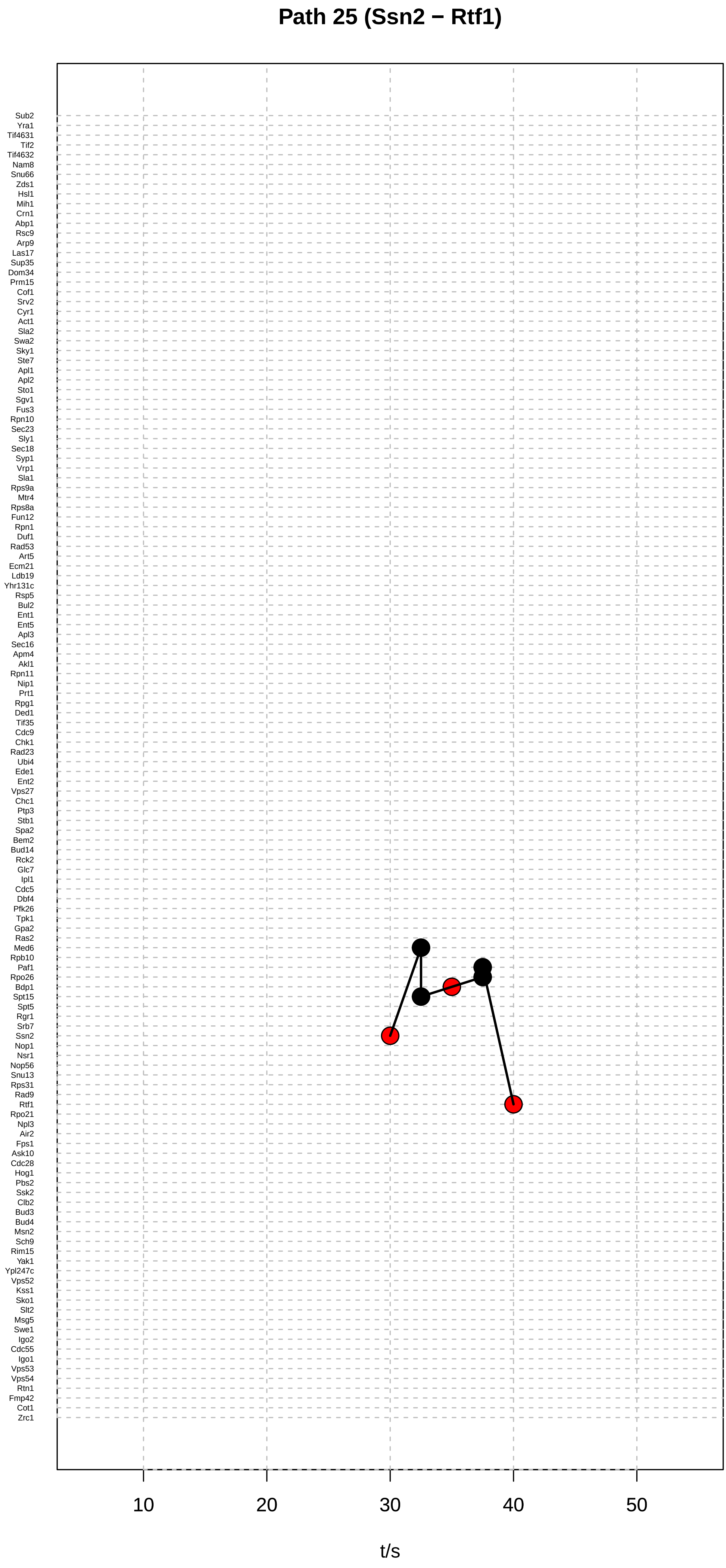

Path 25 (Ssn2 − Rtf1)
Sub2
Yra1
Tif4631
Tif2
Tif4632
Nam8
Snu66
Zds1
Hsl1
Mih1
Crn1
Abp1
Rsc9
Arp9
Las17
Sup35
Dom34
Prm15
Cof1
Srv2
Cyr1
Act1
Sla2
Swa2
Sky1
Ste7
Apl1
Apl2
Sto1
Sgv1
Fus3
Rpn10
Sec23
Sly1
Sec18
Syp1
Vrp1
Sla1
Rps9a
Mtr4
Rps8a
Fun12
Rpn1
Duf1
Rad53
Art5
Ecm21
Ldb19
Yhr131c
Rsp5
Bul2
Ent1
Ent5
Apl3
Sec16
Apm4
Akl1
Rpn11
Nip1
Prt1
Rpg1
Ded1
Tif35
Cdc9
Chk1
Rad23
Ubi4
Ede1
Ent2
Vps27
Chc1
Ptp3
Stb1
Spa2
Bem2
Bud14
Rck2
Glc7
Ipl1
Cdc5
Dbf4
Pfk26
Tpk1
Gpa2
Ras2
Med6
Rpb10
Paf1
Rpo26
Bdp1
Spt15
Spt5
Rgr1
Srb7
Ssn2
Nop1
Nsr1
Nop56
Snu13
Rps31
Rad9
Rtf1
Rpo21
Npl3
Air2
Fps1
Ask10
Cdc28
Hog1
Pbs2
Ssk2
Clb2
Bud3
Bud4
Msn2
Sch9
Rim15
Yak1
Ypl247c
Vps52
Kss1
Sko1
Slt2
Msg5
Swe1
Igo2
Cdc55
Igo1
Vps53
Vps54
Rtn1
Fmp42
Cot1
Zrc1
10
20
30
40
50
t/s

#### Path 26 (Ssn2 − Rtf1)

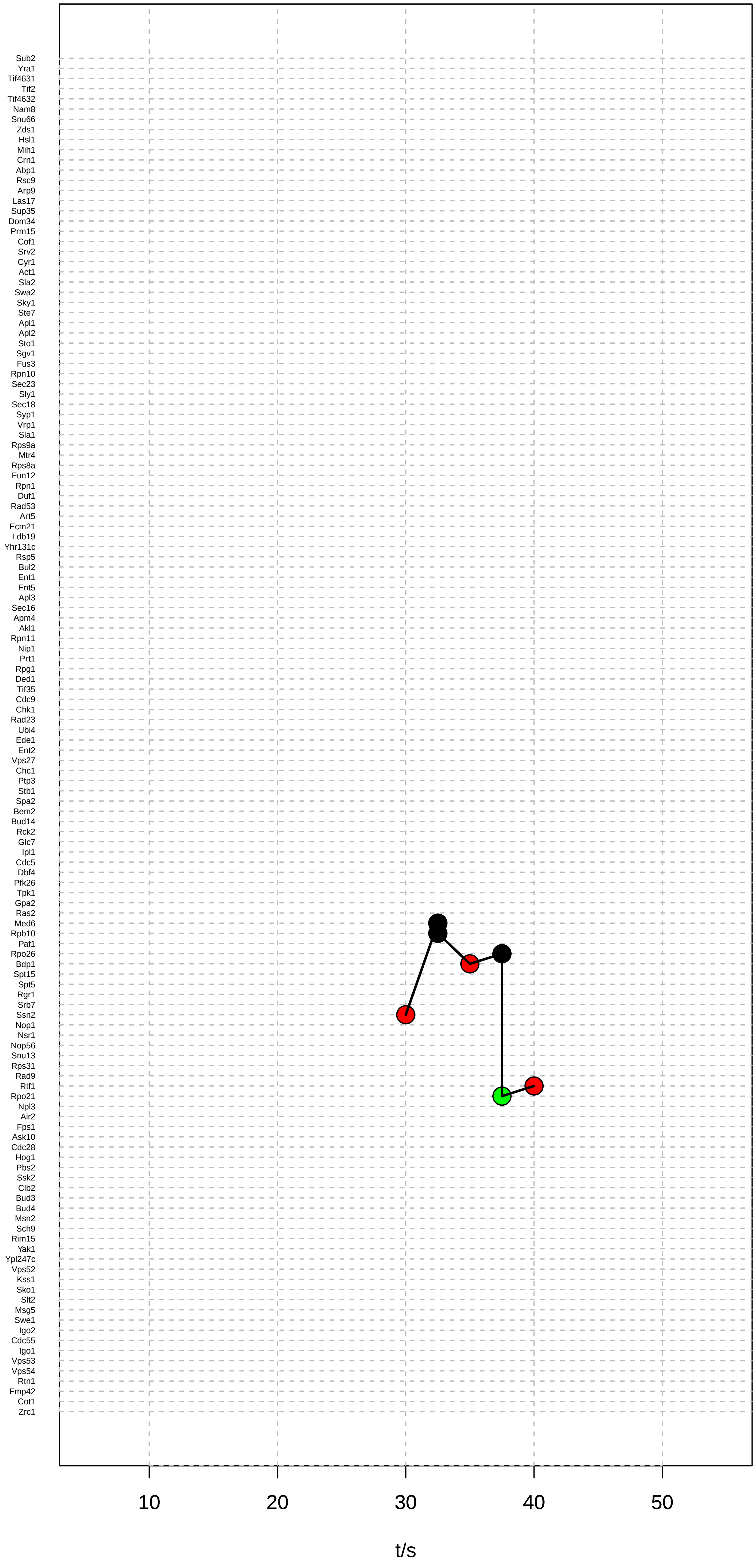

#### Path 27 (Ssn2 − Rtf1)

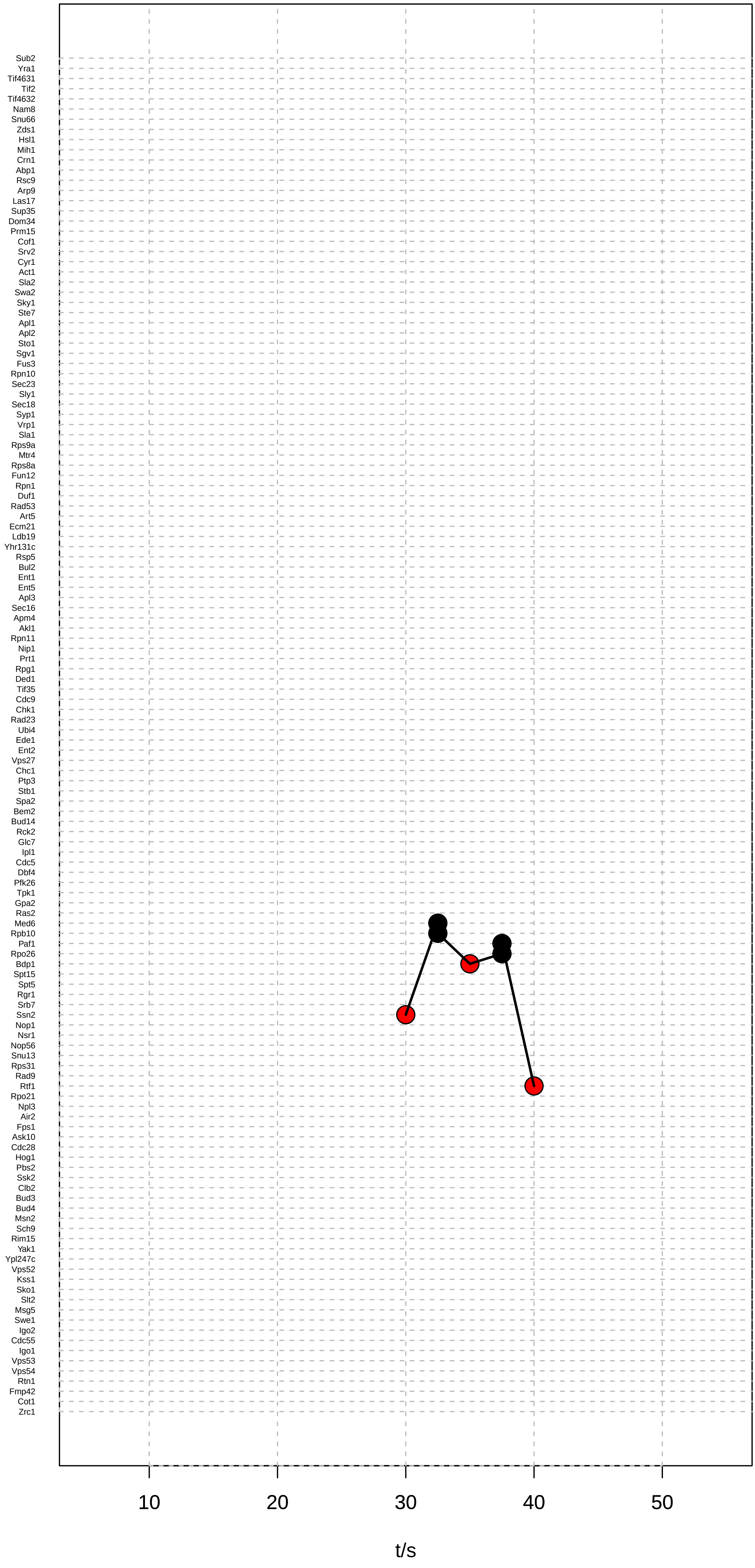

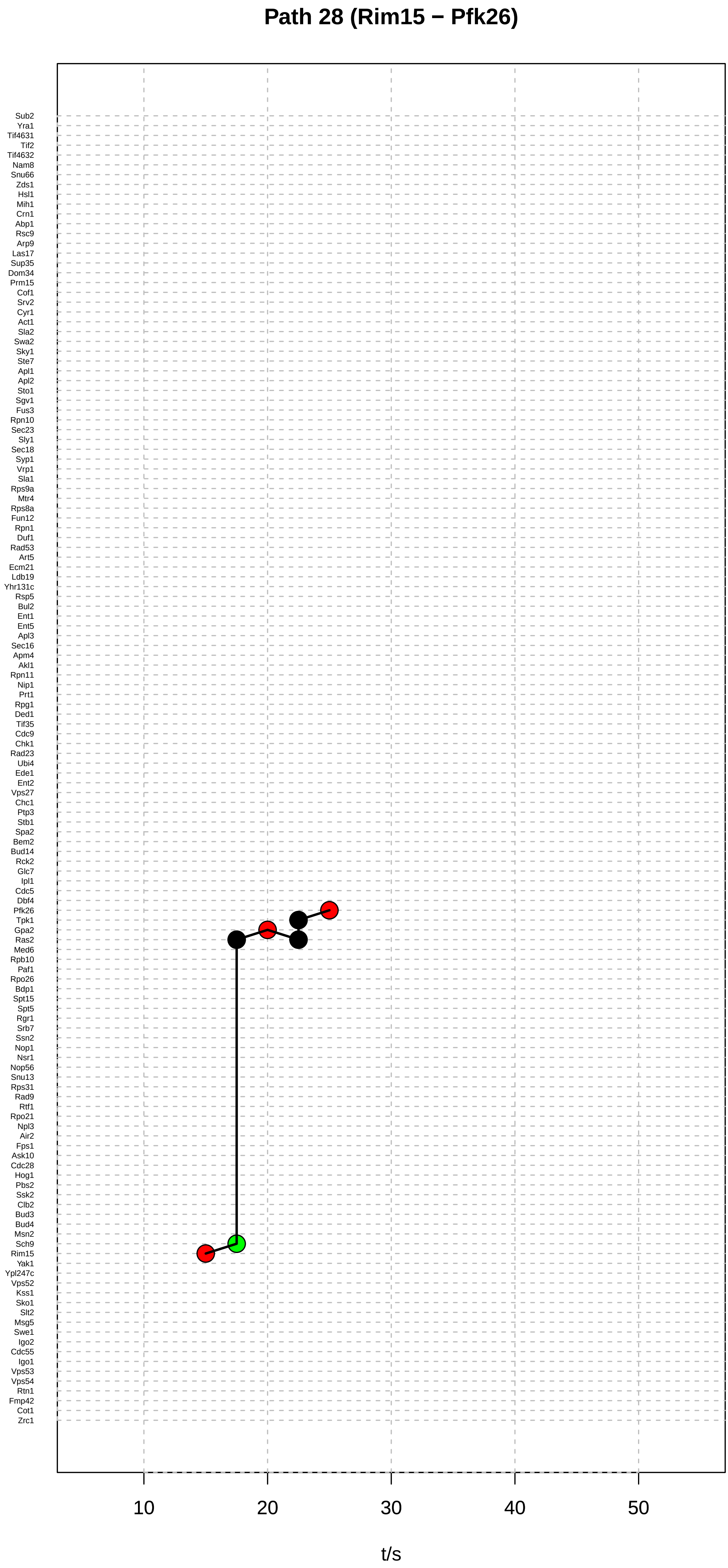

Path 28 (Rim15 − Pfk26)
Sub2
Yra1
Tif4631
Tif2
Tif4632
Nam8
Snu66
Zds1
Hsl1
Mih1
Crn1
Abp1
Rsc9
Arp9
Las17
Sup35
Dom34
Prm15
Cof1
Srv2
Cyr1
Act1
Sla2
Swa2
Sky1
Ste7
Apl1
Apl2
Sto1
Sgv1
Fus3
Rpn10
Sec23
Sly1
Sec18
Syp1
Vrp1
Sla1
Rps9a
Mtr4
Rps8a
Fun12
Rpn1
Duf1
Rad53
Art5
Ecm21
Ldb19
Yhr131c
Rsp5
Bul2
Ent1
Ent5
Apl3
Sec16
Apm4
Akl1
Rpn11
Nip1
Prt1
Rpg1
Ded1
Tif35
Cdc9
Chk1
Rad23
Ubi4
Ede1
Ent2
Vps27
Chc1
Ptp3
Stb1
Spa2
Bem2
Bud14
Rck2
Glc7
Ipl1
Cdc5
Dbf4
Pfk26
Tpk1
Gpa2
Ras2
Med6
Rpb10
Paf1
Rpo26
Bdp1
Spt15
Spt5
Rgr1
Srb7
Ssn2
Nop1
Nsr1
Nop56
Snu13
Rps31
Rad9
Rtf1
Rpo21
Npl3
Air2
Fps1
Ask10
Cdc28
Hog1
Pbs2
Ssk2
Clb2
Bud3
Bud4
Msn2
Sch9
Rim15
Yak1
Ypl247c
Vps52
Kss1
Sko1
Slt2
Msg5
Swe1
Igo2
Cdc55
Igo1
Vps53
Vps54
Rtn1
Fmp42
Cot1
Zrc1
10
20
30
40
50
t/s

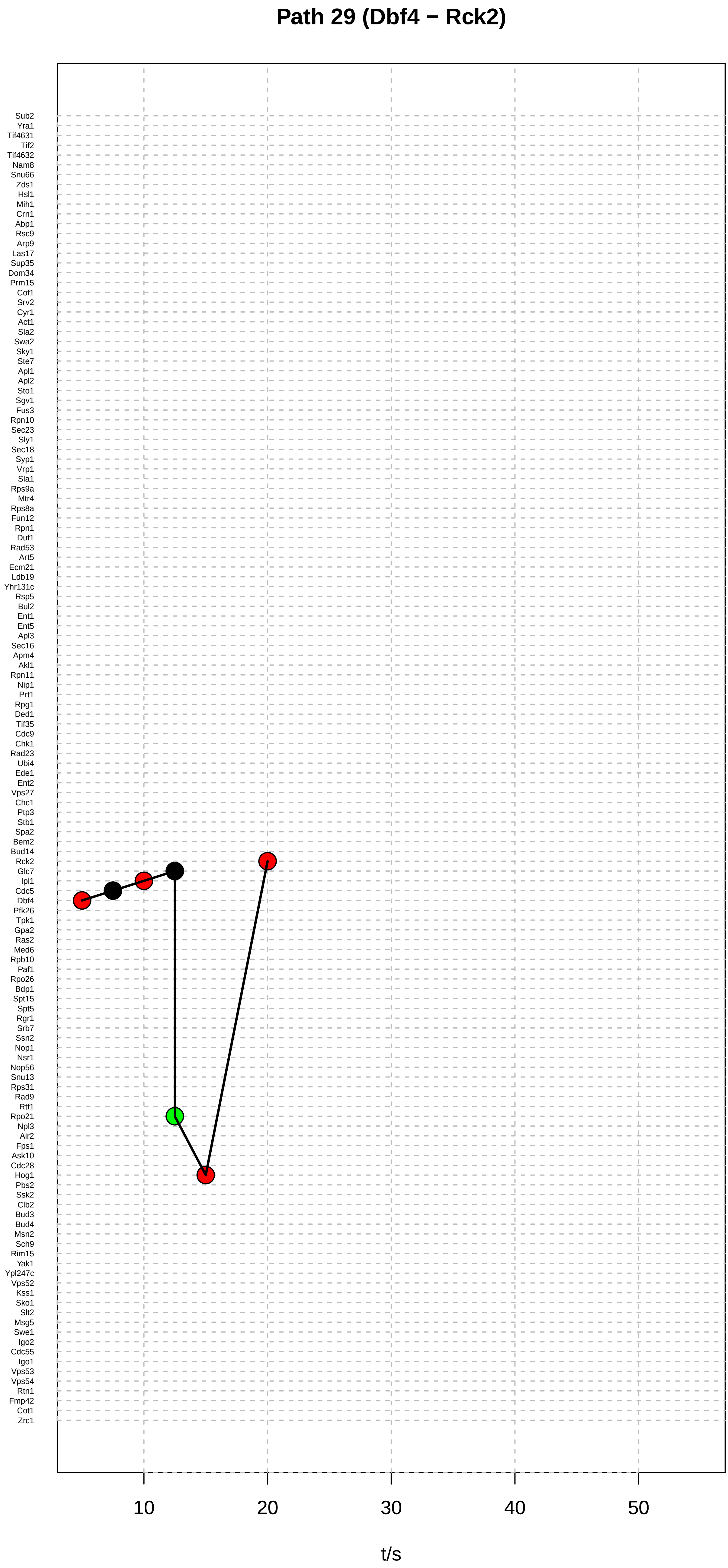

Path 29 (Dbf4 − Rck2)
10
20
30
40
50
t/s

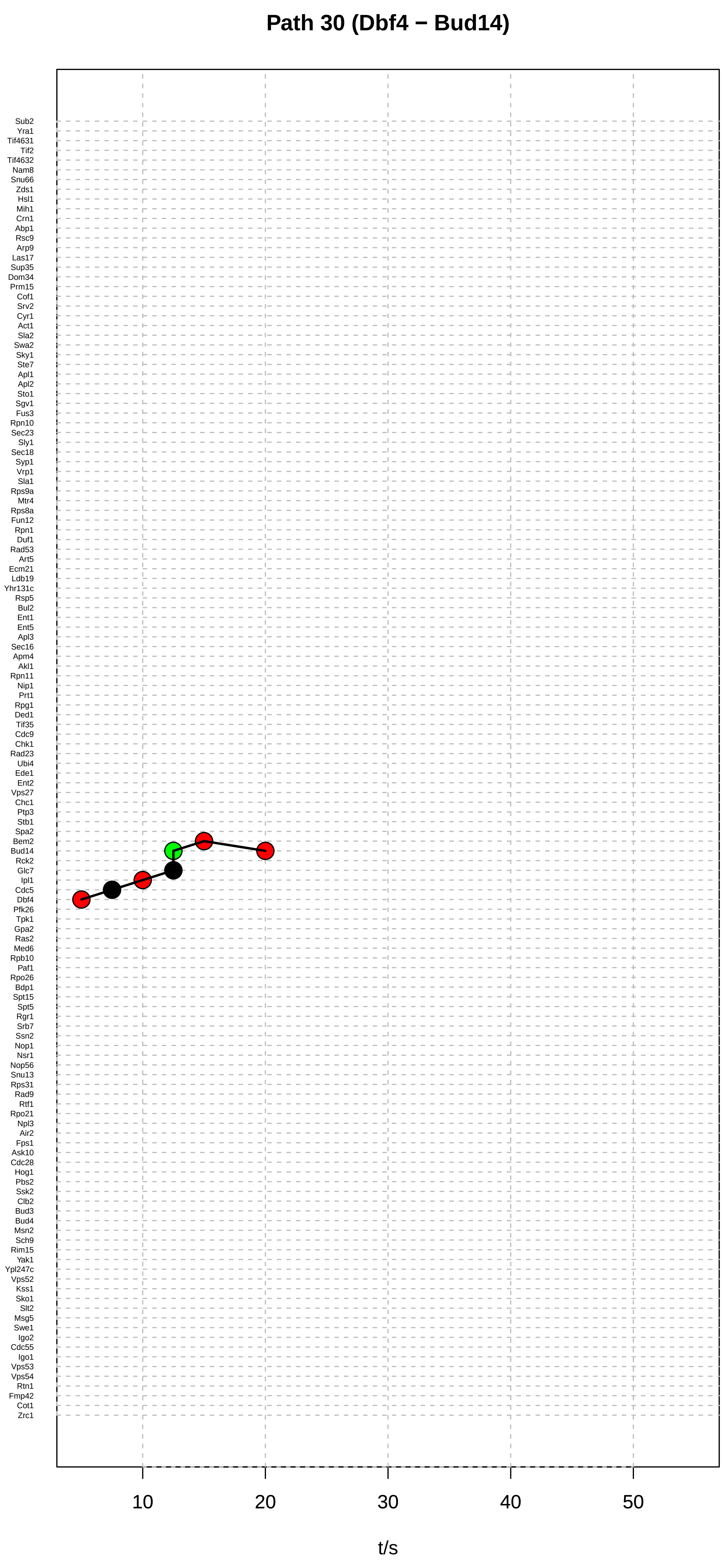
Path 30 (Dbf4 − Bud14)
Sub2
Yra1
Tif4631
Tif2
Tif4632
Nam8
Snu66
Zds1
Hsl1
Mih1
Crn1
Abp1
Rsc9
Arp9
Las17
Sup35
Dom34
Prm15
Cof1
Srv2
Cyr1
Act1
Sla2
Swa2
Sky1
Ste7
Apl1
Apl2
Sto1
Sgv1
Fus3
Rpn10
Sec23
Sly1
Sec18
Syp1
Vrp1
Sla1
Rps9a
Mtr4
Rps8a
Fun12
Rpn1
Duf1
Rad53
Art5
Ecm21
Ldb19
Yhr131c
Rsp5
Bul2
Ent1
Ent5
Apl3
Sec16
Apm4
Akl1
Rpn11
Nip1
Prt1
Rpg1
Ded1
Tif35
Cdc9
Chk1
Rad23
Ubi4
Ede1
Ent2
Vps27
Chc1
Ptp3
Stb1
Spa2
Bem2
Bud14
Rck2
Glc7
Ipl1
Cdc5
Dbf4
Pfk26
Tpk1
Gpa2
Ras2
Med6
Rpb10
Paf1
Rpo26
Bdp1
Spt15
Spt5
Rgr1
Srb7
Ssn2
Nop1
Nsr1
Nop56
Snu13
Rps31
Rad9
Rtf1
Rpo21
Npl3
Air2
Fps1
Ask10
Cdc28
Hog1
Pbs2
Ssk2
Clb2
Bud3
Bud4
Msn2
Sch9
Rim15
Yak1
Ypl247c
Vps52
Kss1
Sko1
Slt2
Msg5
Swe1
Igo2
Cdc55
Igo1
Vps53
Vps54
Rtn1
Fmp42
Cot1
Zrc1
10
20
30
40
50
t/s

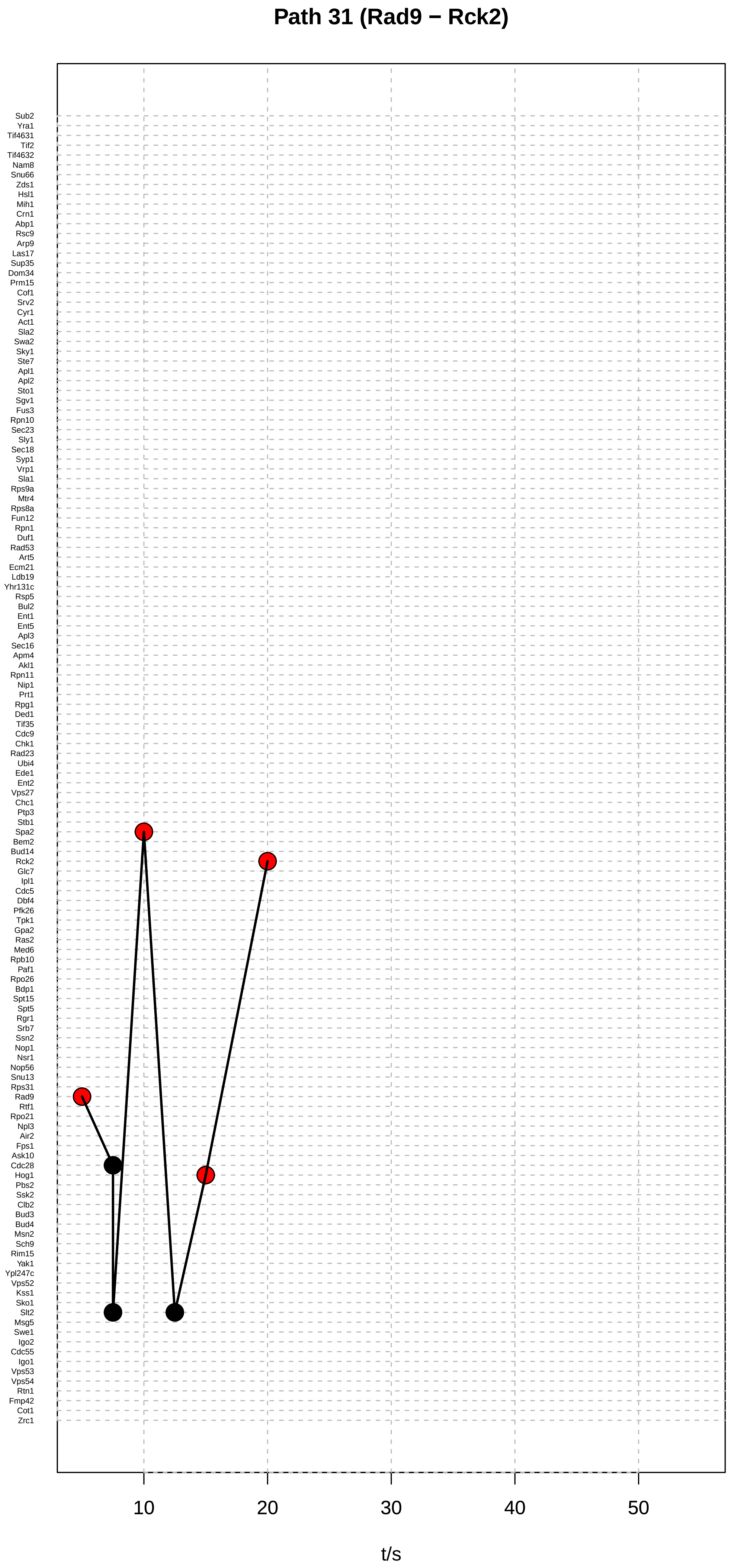
Path 31 (Rad9 − Rck2)
Sub2
Yra1
Tif4631
Tif2
Tif4632
Nam8
Snu66
Zds1
Hsl1
Mih1
Crn1
Abp1
Rsc9
Arp9
Las17
Sup35
Dom34
Prm15
Cof1
Srv2
Cyr1
Act1
Sla2
Swa2
Sky1
Ste7
Apl1
Apl2
Sto1
Sgv1
Fus3
Rpn10
Sec23
Sly1
Sec18
Syp1
Vrp1
Sla1
Rps9a
Mtr4
Rps8a
Fun12
Rpn1
Duf1
Rad53
Art5
Ecm21
Ldb19
Yhr131c
Rsp5
Bul2
Ent1
Ent5
Apl3
Sec16
Apm4
Akl1
Rpn11
Nip1
Prt1
Rpg1
Ded1
Tif35
Cdc9
Chk1
Rad23
Ubi4
Ede1
Ent2
Vps27
Chc1
Ptp3
Stb1
Spa2
Bem2
Bud14
Rck2
Glc7
Ipl1
Cdc5
Dbf4
Pfk26
Tpk1
Gpa2
Ras2
Med6
Rpb10
Paf1
Rpo26
Bdp1
Spt15
Spt5
Rgr1
Srb7
Ssn2
Nop1
Nsr1
Nop56
Snu13
Rps31
Rad9
Rtf1
Rpo21
Npl3
Air2
Fps1
Ask10
Cdc28
Hog1
Pbs2
Ssk2
Clb2
Bud3
Bud4
Msn2
Sch9
Rim15
Yak1
Ypl247c
Vps52
Kss1
Sko1
Slt2
Msg5
Swe1
Igo2
Cdc55
Igo1
Vps53
Vps54
Rtn1
Fmp42
Cot1
Zrc1
10
20
30
40
50
t/s

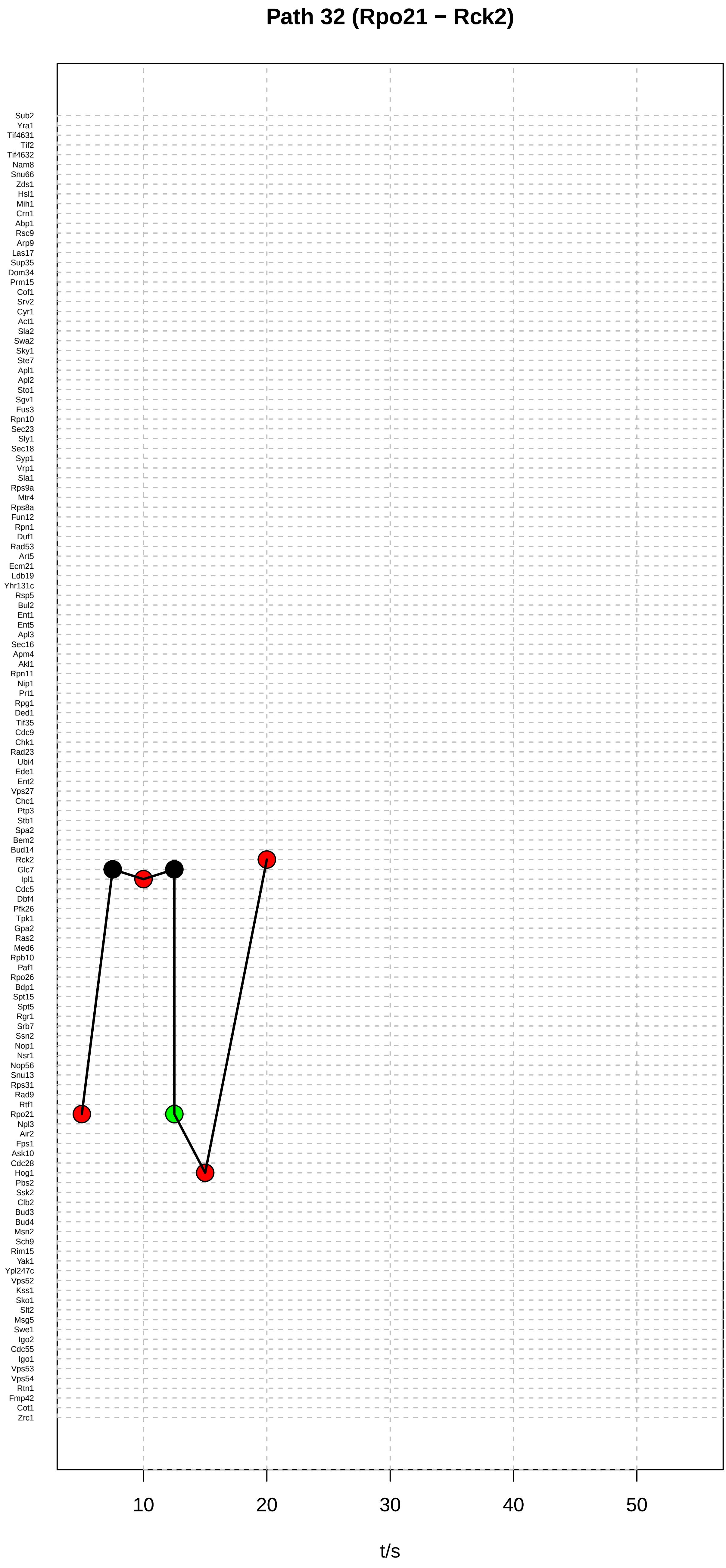

Path 32 (Rpo21 − Rck2)
Sub2
Yra1
Tif4631
Tif2
Tif4632
Nam8
Snu66
Zds1
Hsl1
Mih1
Crn1
Abp1
Rsc9
Arp9
Las17
Sup35
Dom34
Prm15
Cof1
Srv2
Cyr1
Act1
Sla2
Swa2
Sky1
Ste7
Apl1
Apl2
Sto1
Sgv1
Fus3
Rpn10
Sec23
Sly1
Sec18
Syp1
Vrp1
Sla1
Rps9a
Mtr4
Rps8a
Fun12
Rpn1
Duf1
Rad53
Art5
Ecm21
Ldb19
Yhr131c
Rsp5
Bul2
Ent1
Ent5
Apl3
Sec16
Apm4
Akl1
Rpn11
Nip1
Prt1
Rpg1
Ded1
Tif35
Cdc9
Chk1
Rad23
Ubi4
Ede1
Ent2
Vps27
Chc1
Ptp3
Stb1
Spa2
Bem2
Bud14
Rck2
Glc7
Ipl1
Cdc5
Dbf4
Pfk26
Tpk1
Gpa2
Ras2
Med6
Rpb10
Paf1
Rpo26
Bdp1
Spt15
Spt5
Rgr1
Srb7
Ssn2
Nop1
Nsr1
Nop56
Snu13
Rps31
Rad9
Rtf1
Rpo21
Npl3
Air2
Fps1
Ask10
Cdc28
Hog1
Pbs2
Ssk2
Clb2
Bud3
Bud4
Msn2
Sch9
Rim15
Yak1
Ypl247c
Vps52
Kss1
Sko1
Slt2
Msg5
Swe1
Igo2
Cdc55
Igo1
Vps53
Vps54
Rtn1
Fmp42
Cot1
Zrc1
10
20
30
40
50
t/s

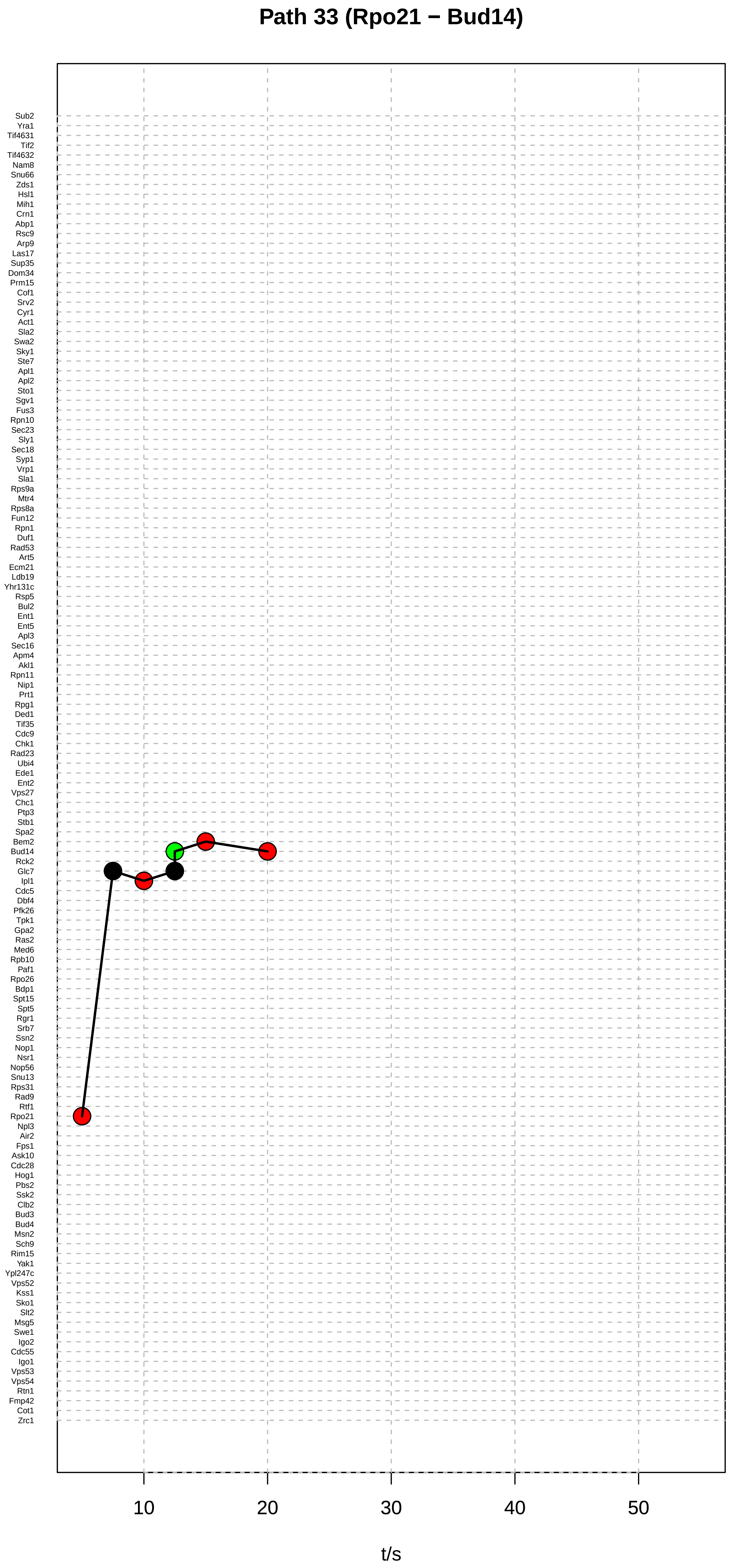

Path 33 (Rpo21 − Bud14)
Sub2
Yra1
Tif4631
Tif2
Tif4632
Nam8
Snu66
Zds1
Hsl1
Mih1
Crn1
Abp1
Rsc9
Arp9
Las17
Sup35
Dom34
Prm15
Cof1
Srv2
Cyr1
Act1
Sla2
Swa2
Sky1
Ste7
Apl1
Apl2
Sto1
Sgv1
Fus3
Rpn10
Sec23
Sly1
Sec18
Syp1
Vrp1
Sla1
Rps9a
Mtr4
Rps8a
Fun12
Rpn1
Duf1
Rad53
Art5
Ecm21
Ldb19
Yhr131c
Rsp5
Bul2
Ent1
Ent5
Apl3
Sec16
Apm4
Akl1
Rpn11
Nip1
Prt1
Rpg1
Ded1
Tif35
Cdc9
Chk1
Rad23
Ubi4
Ede1
Ent2
Vps27
Chc1
Ptp3
Stb1
Spa2
Bem2
Bud14
Rck2
Glc7
Ipl1
Cdc5
Dbf4
Pfk26
Tpk1
Gpa2
Ras2
Med6
Rpb10
Paf1
Rpo26
Bdp1
Spt15
Spt5
Rgr1
Srb7
Ssn2
Nop1
Nsr1
Nop56
Snu13
Rps31
Rad9
Rtf1
Rpo21
Npl3
Air2
Fps1
Ask10
Cdc28
Hog1
Pbs2
Ssk2
Clb2
Bud3
Bud4
Msn2
Sch9
Rim15
Yak1
Ypl247c
Vps52
Kss1
Sko1
Slt2
Msg5
Swe1
Igo2
Cdc55
Igo1
Vps53
Vps54
Rtn1
Fmp42
Cot1
Zrc1
10
20
30
40
50
t/s

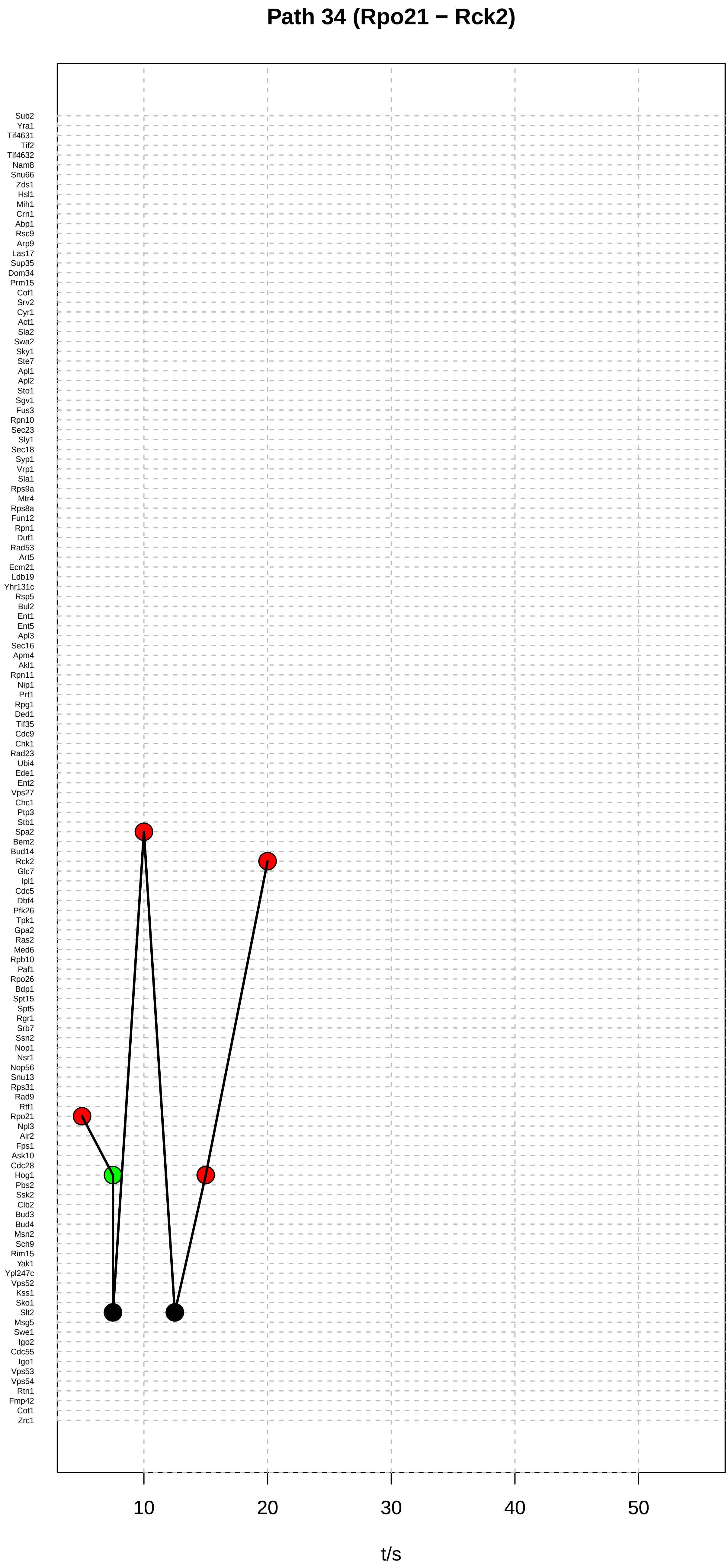

Path 34 (Rpo21 − Rck2)
Sub2
Yra1
Tif4631
Tif2
Tif4632
Nam8
Snu66
Zds1
Hsl1
Mih1
Crn1
Abp1
Rsc9
Arp9
Las17
Sup35
Dom34
Prm15
Cof1
Srv2
Cyr1
Act1
Sla2
Swa2
Sky1
Ste7
Apl1
Apl2
Sto1
Sgv1
Fus3
Rpn10
Sec23
Sly1
Sec18
Syp1
Vrp1
Sla1
Rps9a
Mtr4
Rps8a
Fun12
Rpn1
Duf1
Rad53
Art5
Ecm21
Ldb19
Yhr131c
Rsp5
Bul2
Ent1
Ent5
Apl3
Sec16
Apm4
Akl1
Rpn11
Nip1
Prt1
Rpg1
Ded1
Tif35
Cdc9
Chk1
Rad23
Ubi4
Ede1
Ent2
Vps27
Chc1
Ptp3
Stb1
Spa2
Bem2
Bud14
Rck2
Glc7
Ipl1
Cdc5
Dbf4
Pfk26
Tpk1
Gpa2
Ras2
Med6
Rpb10
Paf1
Rpo26
Bdp1
Spt15
Spt5
Rgr1
Srb7
Ssn2
Nop1
Nsr1
Nop56
Snu13
Rps31
Rad9
Rtf1
Rpo21
Npl3
Air2
Fps1
Ask10
Cdc28
Hog1
Pbs2
Ssk2
Clb2
Bud3
Bud4
Msn2
Sch9
Rim15
Yak1
Ypl247c
Vps52
Kss1
Sko1
Slt2
Msg5
Swe1
Igo2
Cdc55
Igo1
Vps53
Vps54
Rtn1
Fmp42
Cot1
Zrc1
10
20
30
40
50
t/s

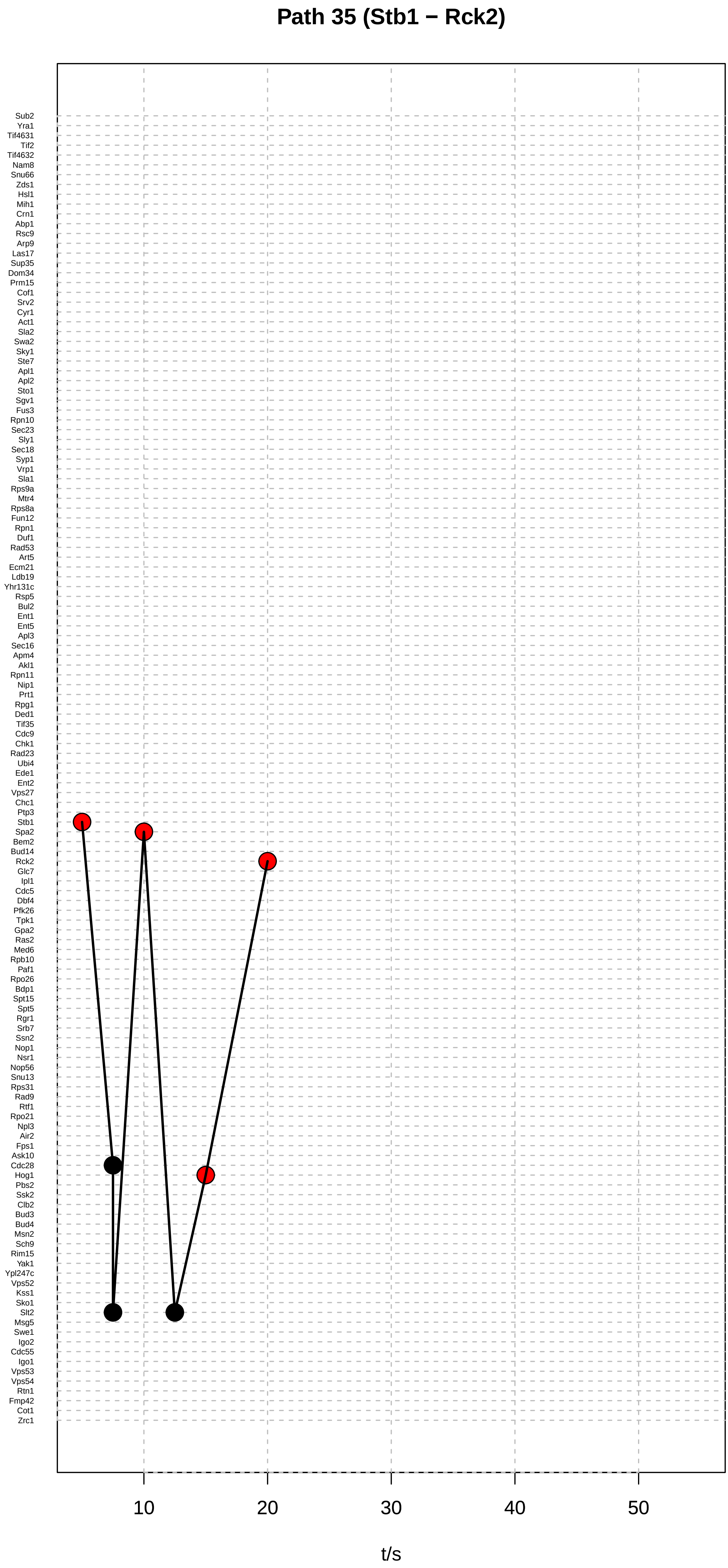

Path 35 (Stb1 − Rck2)
Sub2
Yra1
Tif4631
Tif2
Tif4632
Nam8
Snu66
Zds1
Hsl1
Mih1
Crn1
Abp1
Rsc9
Arp9
Las17
Sup35
Dom34
Prm15
Cof1
Srv2
Cyr1
Act1
Sla2
Swa2
Sky1
Ste7
Apl1
Apl2
Sto1
Sgv1
Fus3
Rpn10
Sec23
Sly1
Sec18
Syp1
Vrp1
Sla1
Rps9a
Mtr4
Rps8a
Fun12
Rpn1
Duf1
Rad53
Art5
Ecm21
Ldb19
Yhr131c
Rsp5
Bul2
Ent1
Ent5
Apl3
Sec16
Apm4
Akl1
Rpn11
Nip1
Prt1
Rpg1
Ded1
Tif35
Cdc9
Chk1
Rad23
Ubi4
Ede1
Ent2
Vps27
Chc1
Ptp3
Stb1
Spa2
Bem2
Bud14
Rck2
Glc7
Ipl1
Cdc5
Dbf4
Pfk26
Tpk1
Gpa2
Ras2
Med6
Rpb10
Paf1
Rpo26
Bdp1
Spt15
Spt5
Rgr1
Srb7
Ssn2
Nop1
Nsr1
Nop56
Snu13
Rps31
Rad9
Rtf1
Rpo21
Npl3
Air2
Fps1
Ask10
Cdc28
Hog1
Pbs2
Ssk2
Clb2
Bud3
Bud4
Msn2
Sch9
Rim15
Yak1
Ypl247c
Vps52
Kss1
Sko1
Slt2
Msg5
Swe1
Igo2
Cdc55
Igo1
Vps53
Vps54
Rtn1
Fmp42
Cot1
Zrc1
10
20
30
40
50
t/s

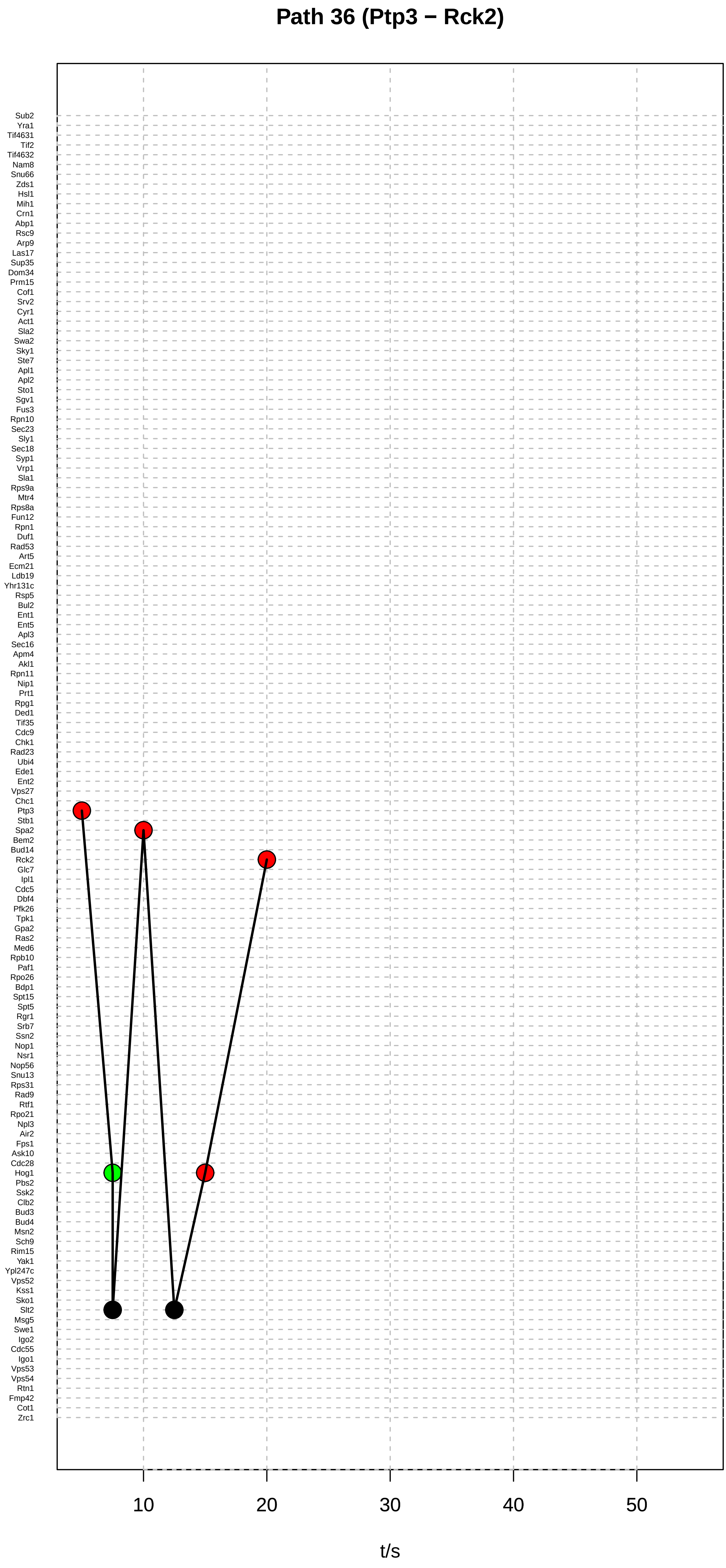

Path 36 (Ptp3 − Rck2)
Sub2
Yra1
Tif4631
Tif2
Tif4632
Nam8
Snu66
Zds1
Hsl1
Mih1
Crn1
Abp1
Rsc9
Arp9
Las17
Sup35
Dom34
Prm15
Cof1
Srv2
Cyr1
Act1
Sla2
Swa2
Sky1
Ste7
Apl1
Apl2
Sto1
Sgv1
Fus3
Rpn10
Sec23
Sly1
Sec18
Syp1
Vrp1
Sla1
Rps9a
Mtr4
Rps8a
Fun12
Rpn1
Duf1
Rad53
Art5
Ecm21
Ldb19
Yhr131c
Rsp5
Bul2
Ent1
Ent5
Apl3
Sec16
Apm4
Akl1
Rpn11
Nip1
Prt1
Rpg1
Ded1
Tif35
Cdc9
Chk1
Rad23
Ubi4
Ede1
Ent2
Vps27
Chc1
Ptp3
Stb1
Spa2
Bem2
Bud14
Rck2
Glc7
Ipl1
Cdc5
Dbf4
Pfk26
Tpk1
Gpa2
Ras2
Med6
Rpb10
Paf1
Rpo26
Bdp1
Spt15
Spt5
Rgr1
Srb7
Ssn2
Nop1
Nsr1
Nop56
Snu13
Rps31
Rad9
Rtf1
Rpo21
Npl3
Air2
Fps1
Ask10
Cdc28
Hog1
Pbs2
Ssk2
Clb2
Bud3
Bud4
Msn2
Sch9
Rim15
Yak1
Ypl247c
Vps52
Kss1
Sko1
Slt2
Msg5
Swe1
Igo2
Cdc55
Igo1
Vps53
Vps54
Rtn1
Fmp42
Cot1
Zrc1
10
20
30
40
50
t/s

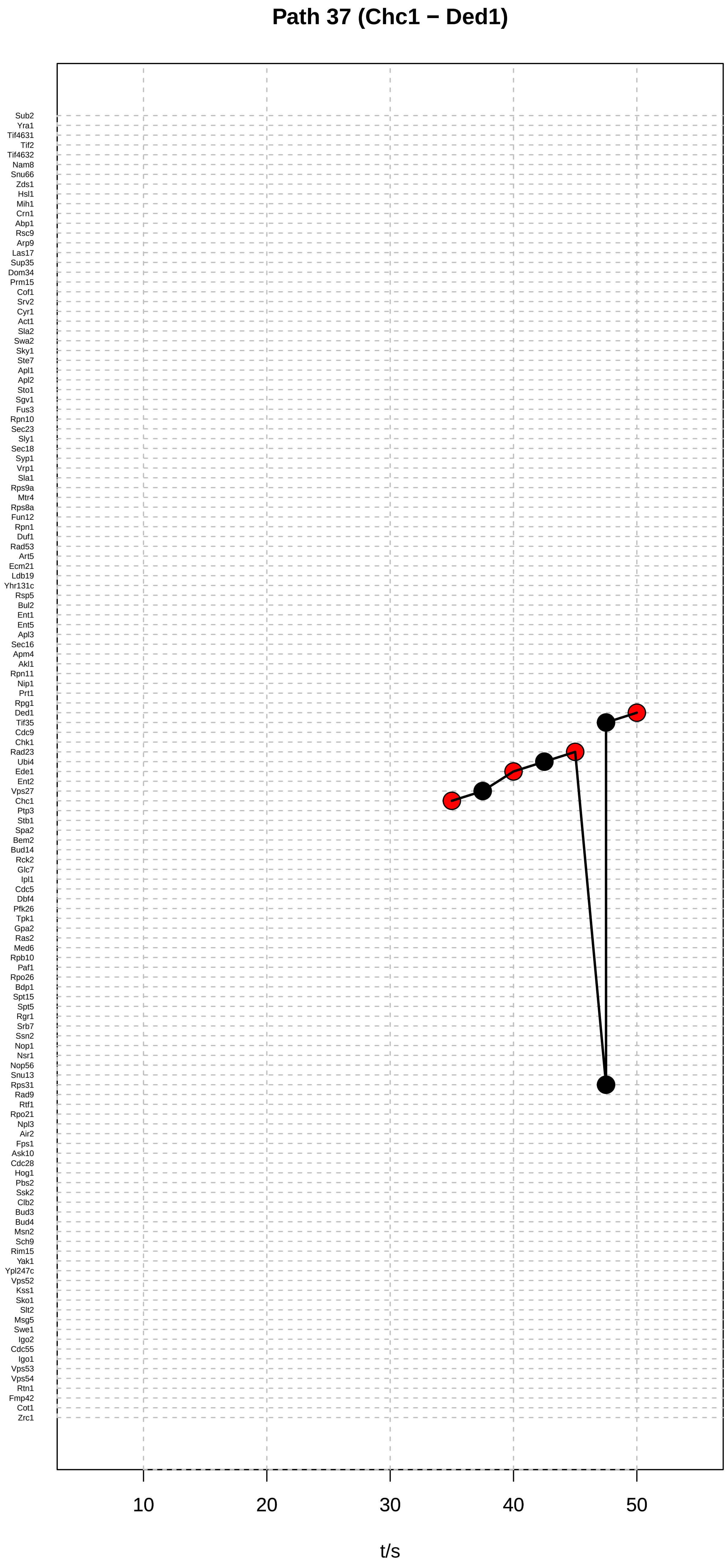
Path 37 (Chc1 − Ded1)
Sub2
Yra1
Tif4631
Tif2
Tif4632
Nam8
Snu66
Zds1
Hsl1
Mih1
Crn1
Abp1
Rsc9
Arp9
Las17
Sup35
Dom34
Prm15
Cof1
Srv2
Cyr1
Act1
Sla2
Swa2
Sky1
Ste7
Apl1
Apl2
Sto1
Sgv1
Fus3
Rpn10
Sec23
Sly1
Sec18
Syp1
Vrp1
Sla1
Rps9a
Mtr4
Rps8a
Fun12
Rpn1
Duf1
Rad53
Art5
Ecm21
Ldb19
Yhr131c
Rsp5
Bul2
Ent1
Ent5
Apl3
Sec16
Apm4
Akl1
Rpn11
Nip1
Prt1
Rpg1
Ded1
Tif35
Cdc9
Chk1
Rad23
Ubi4
Ede1
Ent2
Vps27
Chc1
Ptp3
Stb1
Spa2
Bem2
Bud14
Rck2
Glc7
Ipl1
Cdc5
Dbf4
Pfk26
Tpk1
Gpa2
Ras2
Med6
Rpb10
Paf1
Rpo26
Bdp1
Spt15
Spt5
Rgr1
Srb7
Ssn2
Nop1
Nsr1
Nop56
Snu13
Rps31
Rad9
Rtf1
Rpo21
Npl3
Air2
Fps1
Ask10
Cdc28
Hog1
Pbs2
Ssk2
Clb2
Bud3
Bud4
Msn2
Sch9
Rim15
Yak1
Ypl247c
Vps52
Kss1
Sko1
Slt2
Msg5
Swe1
Igo2
Cdc55
Igo1
Vps53
Vps54
Rtn1
Fmp42
Cot1
Zrc1
10
20
30
40
50
t/s

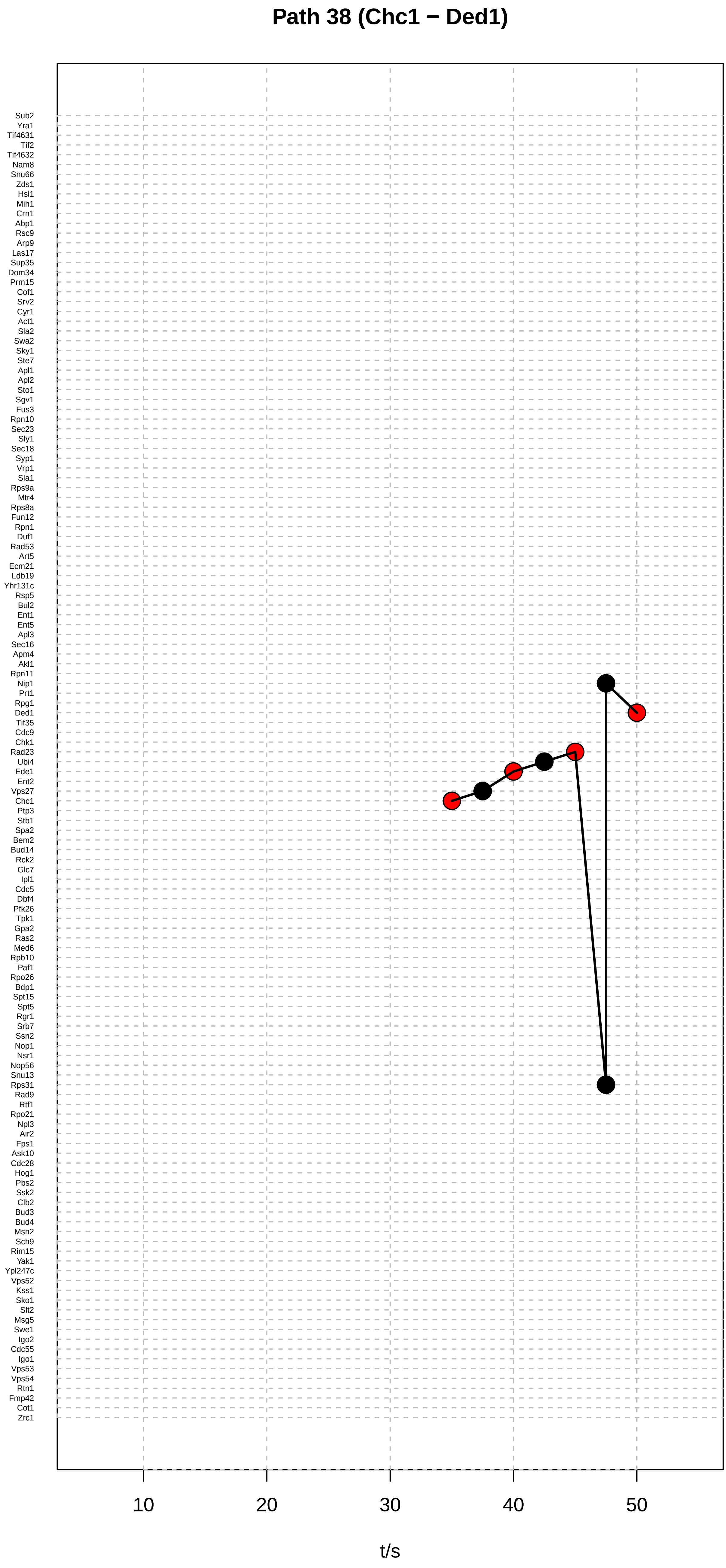

Path 38 (Chc1 − Ded1)
Sub2
Yra1
Tif4631
Tif2
Tif4632
Nam8
Snu66
Zds1
Hsl1
Mih1
Crn1
Abp1
Rsc9
Arp9
Las17
Sup35
Dom34
Prm15
Cof1
Srv2
Cyr1
Act1
Sla2
Swa2
Sky1
Ste7
Apl1
Apl2
Sto1
Sgv1
Fus3
Rpn10
Sec23
Sly1
Sec18
Syp1
Vrp1
Sla1
Rps9a
Mtr4
Rps8a
Fun12
Rpn1
Duf1
Rad53
Art5
Ecm21
Ldb19
Yhr131c
Rsp5
Bul2
Ent1
Ent5
Apl3
Sec16
Apm4
Akl1
Rpn11
Nip1
Prt1
Rpg1
Ded1
Tif35
Cdc9
Chk1
Rad23
Ubi4
Ede1
Ent2
Vps27
Chc1
Ptp3
Stb1
Spa2
Bem2
Bud14
Rck2
Glc7
Ipl1
Cdc5
Dbf4
Pfk26
Tpk1
Gpa2
Ras2
Med6
Rpb10
Paf1
Rpo26
Bdp1
Spt15
Spt5
Rgr1
Srb7
Ssn2
Nop1
Nsr1
Nop56
Snu13
Rps31
Rad9
Rtf1
Rpo21
Npl3
Air2
Fps1
Ask10
Cdc28
Hog1
Pbs2
Ssk2
Clb2
Bud3
Bud4
Msn2
Sch9
Rim15
Yak1
Ypl247c
Vps52
Kss1
Sko1
Slt2
Msg5
Swe1
Igo2
Cdc55
Igo1
Vps53
Vps54
Rtn1
Fmp42
Cot1
Zrc1
10
20
30
40
50
t/s

### Path 39 (Chc1 − Ded1)

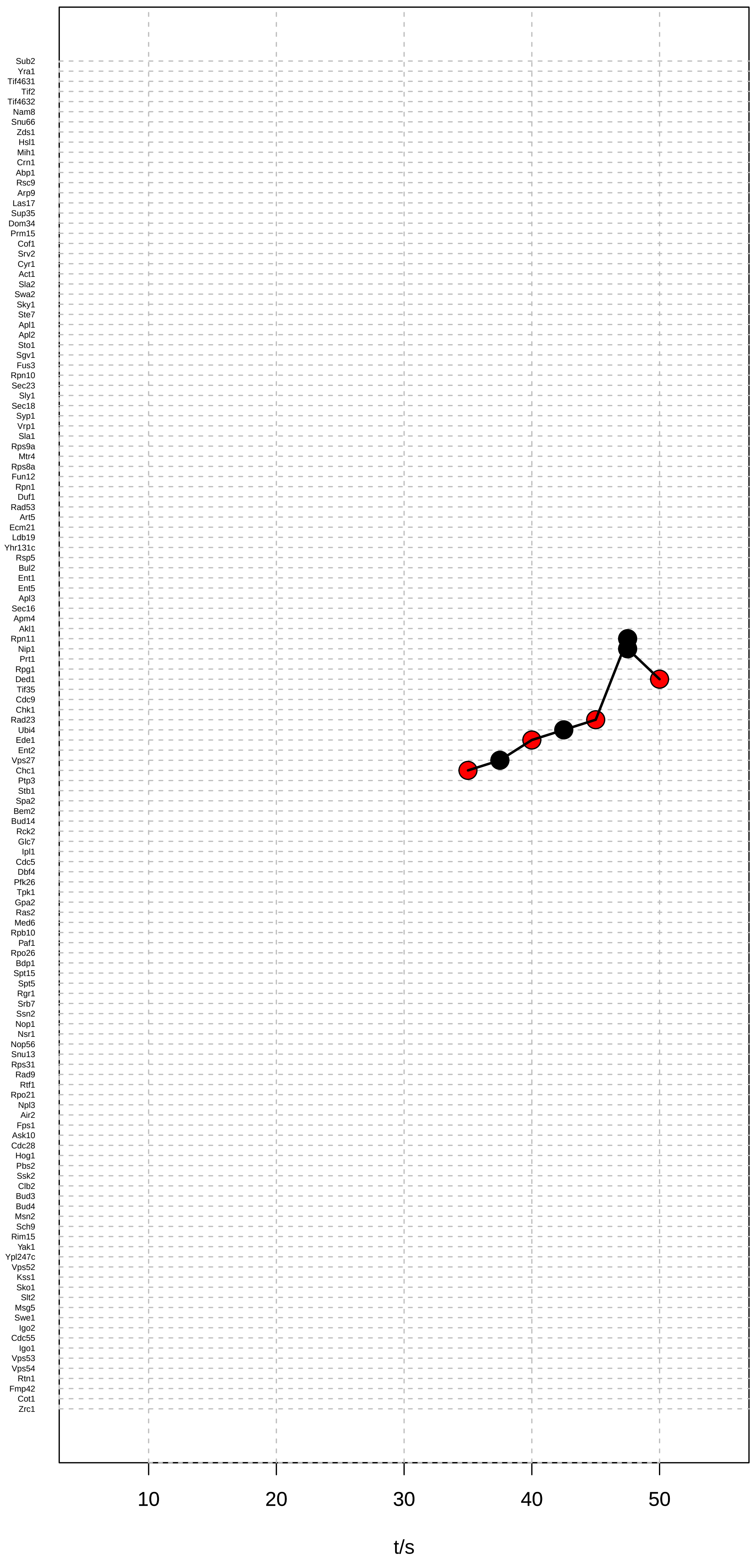

**Path 40 (Chc1 − Ded1)**

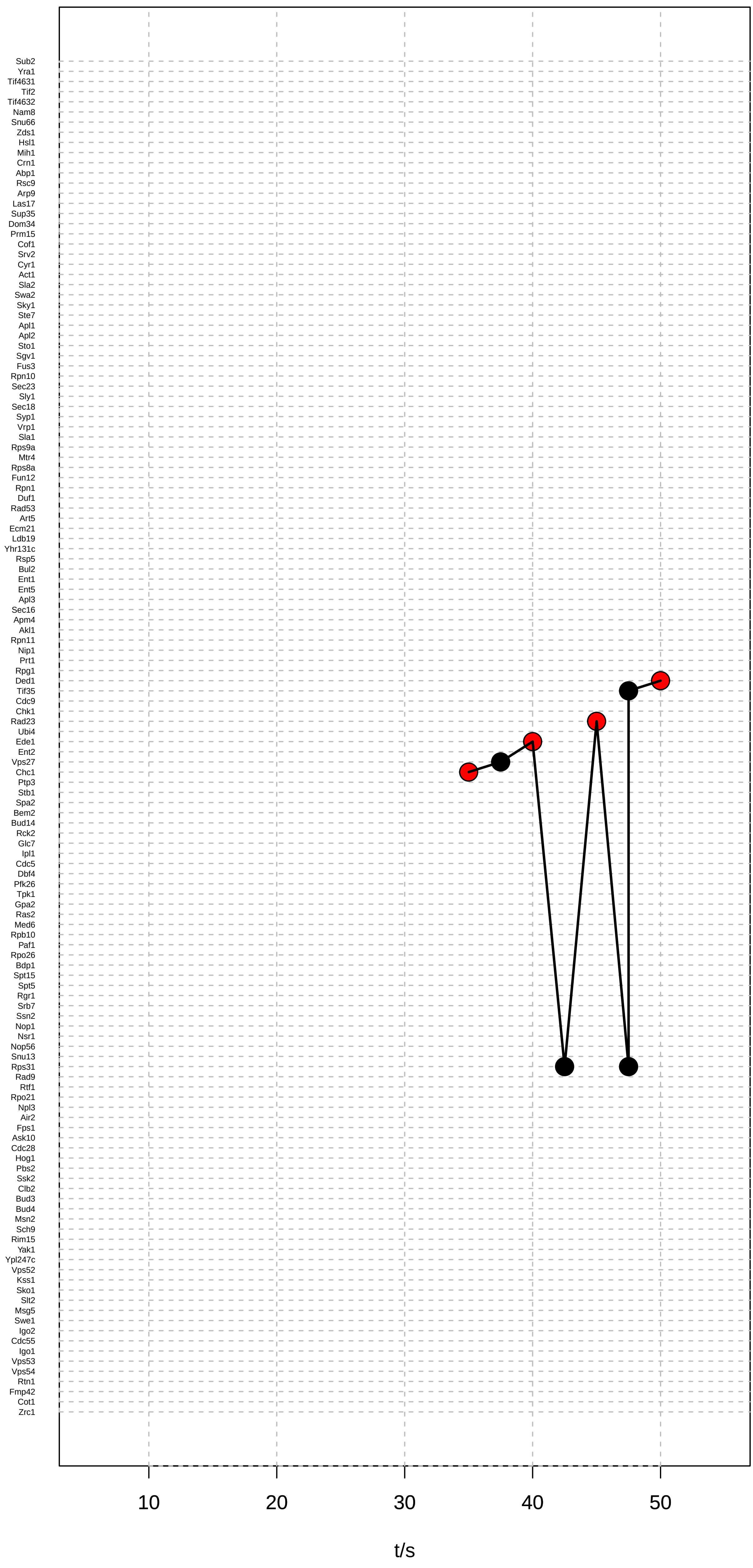

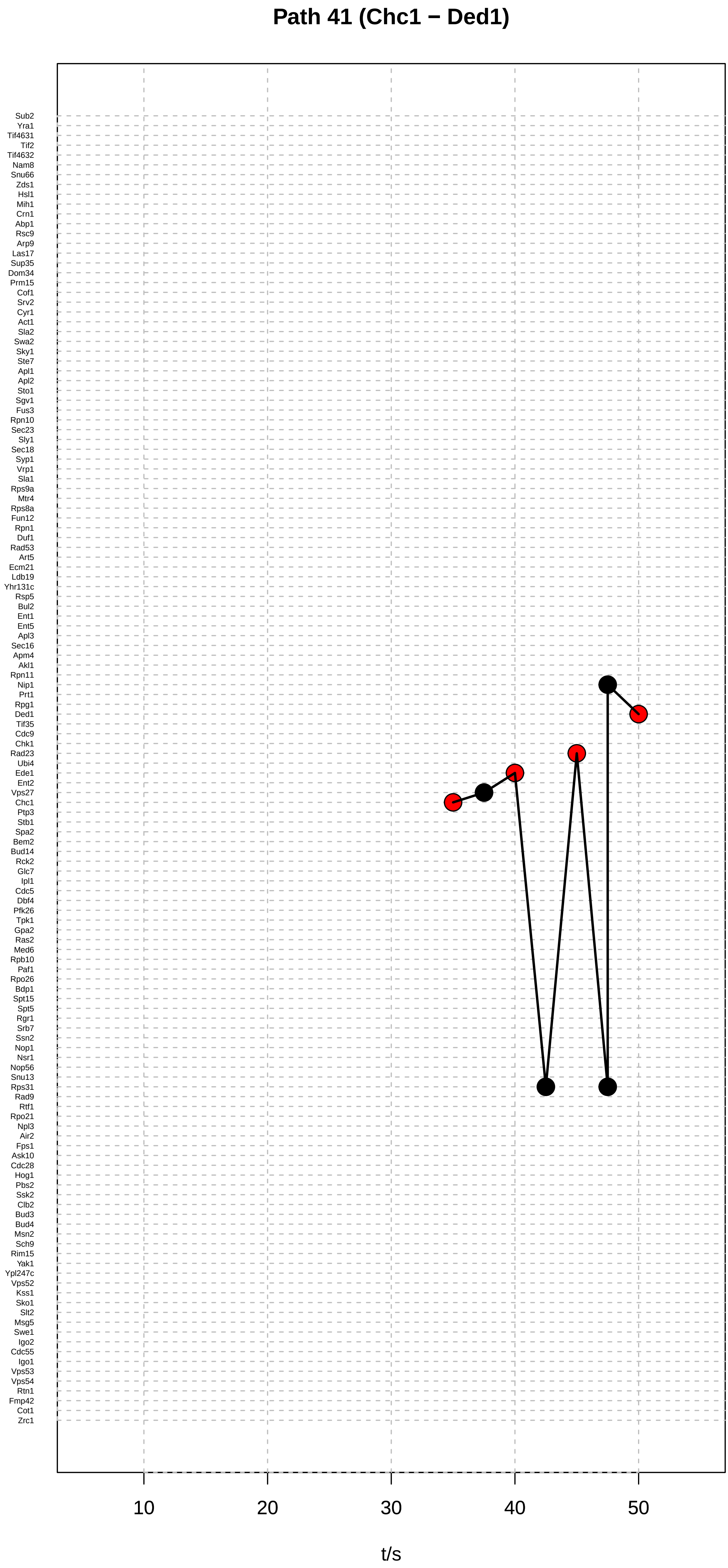

Path 41 (Chc1 − Ded1)
Sub2
Yra1
Tif4631
Tif2
Tif4632
Nam8
Snu66
Zds1
Hsl1
Mih1
Crn1
Abp1
Rsc9
Arp9
Las17
Sup35
Dom34
Prm15
Cof1
Srv2
Cyr1
Act1
Sla2
Swa2
Sky1
Ste7
Apl1
Apl2
Sto1
Sgv1
Fus3
Rpn10
Sec23
Sly1
Sec18
Syp1
Vrp1
Sla1
Rps9a
Mtr4
Rps8a
Fun12
Rpn1
Duf1
Rad53
Art5
Ecm21
Ldb19
Yhr131c
Rsp5
Bul2
Ent1
Ent5
Apl3
Sec16
Apm4
Akl1
Rpn11
Nip1
Prt1
Rpg1
Ded1
Tif35
Cdc9
Chk1
Rad23
Ubi4
Ede1
Ent2
Vps27
Chc1
Ptp3
Stb1
Spa2
Bem2
Bud14
Rck2
Glc7
Ipl1
Cdc5
Dbf4
Pfk26
Tpk1
Gpa2
Ras2
Med6
Rpb10
Paf1
Rpo26
Bdp1
Spt15
Spt5
Rgr1
Srb7
Ssn2
Nop1
Nsr1
Nop56
Snu13
Rps31
Rad9
Rtf1
Rpo21
Npl3
Air2
Fps1
Ask10
Cdc28
Hog1
Pbs2
Ssk2
Clb2
Bud3
Bud4
Msn2
Sch9
Rim15
Yak1
Ypl247c
Vps52
Kss1
Sko1
Slt2
Msg5
Swe1
Igo2
Cdc55
Igo1
Vps53
Vps54
Rtn1
Fmp42
Cot1
Zrc1
10
20
30
40
50
t/s

#### Path 42 (Chc1 − Ded1)

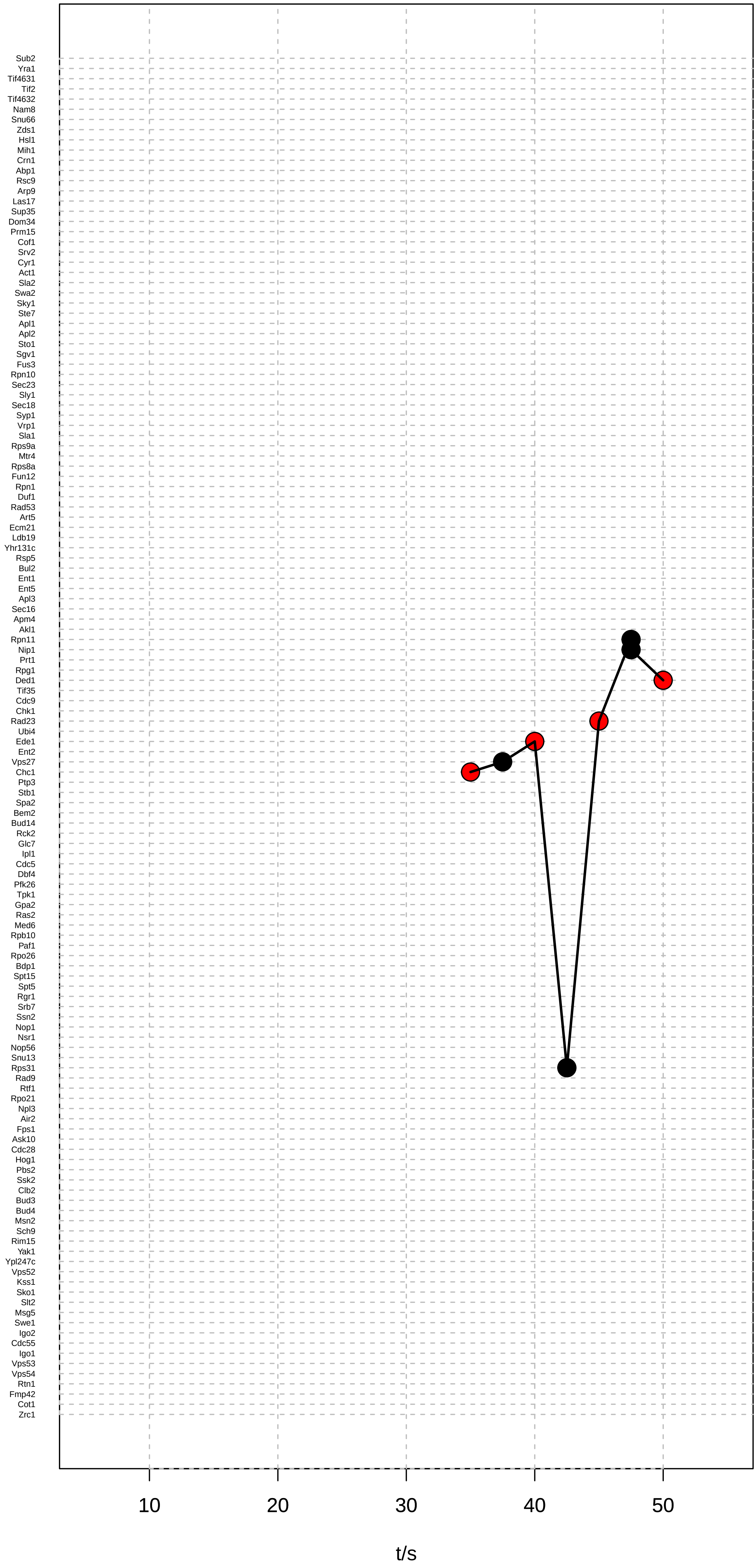

#### Path 43 (Akl1 − Ent5)

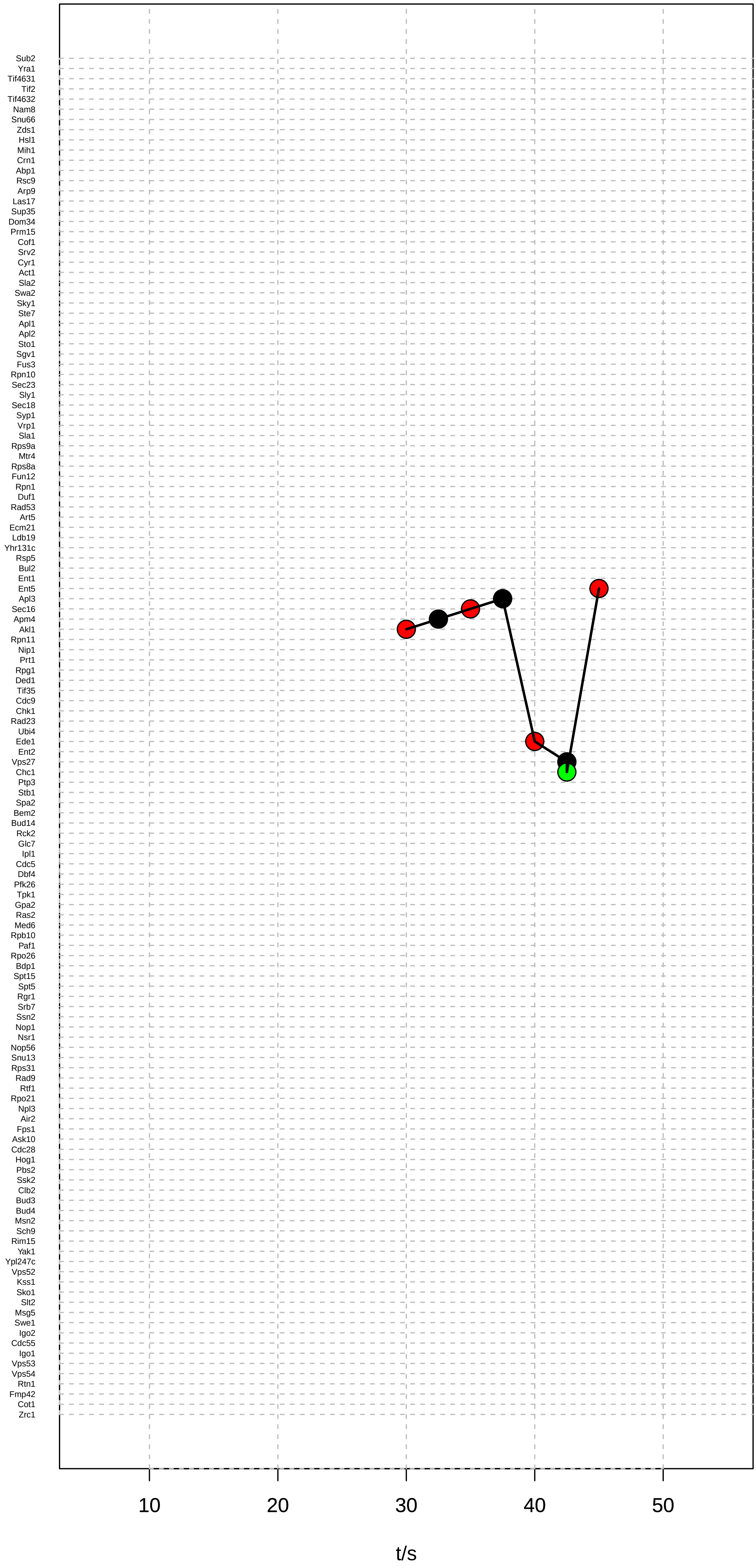

### Path 44 (Akl1 − Ent5)

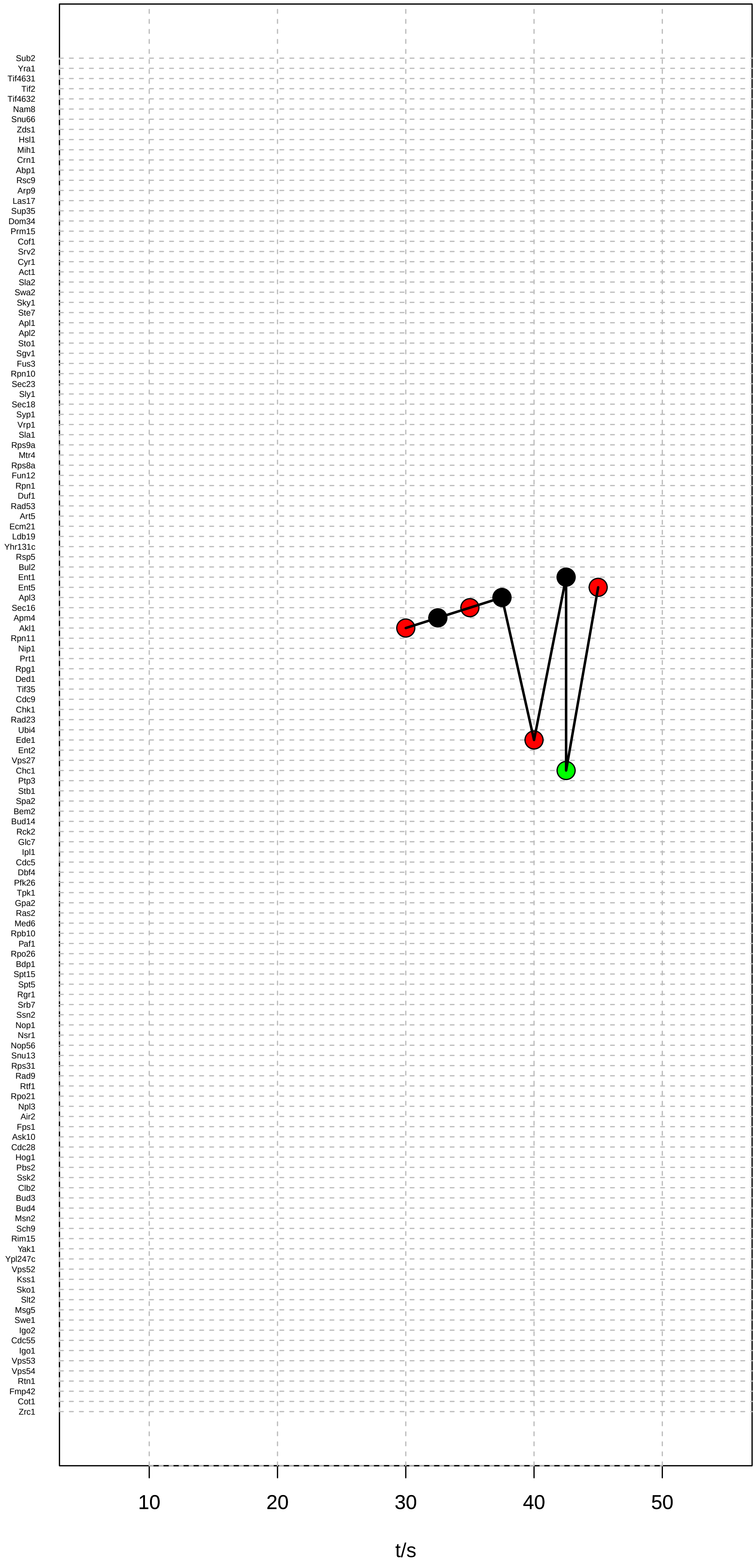

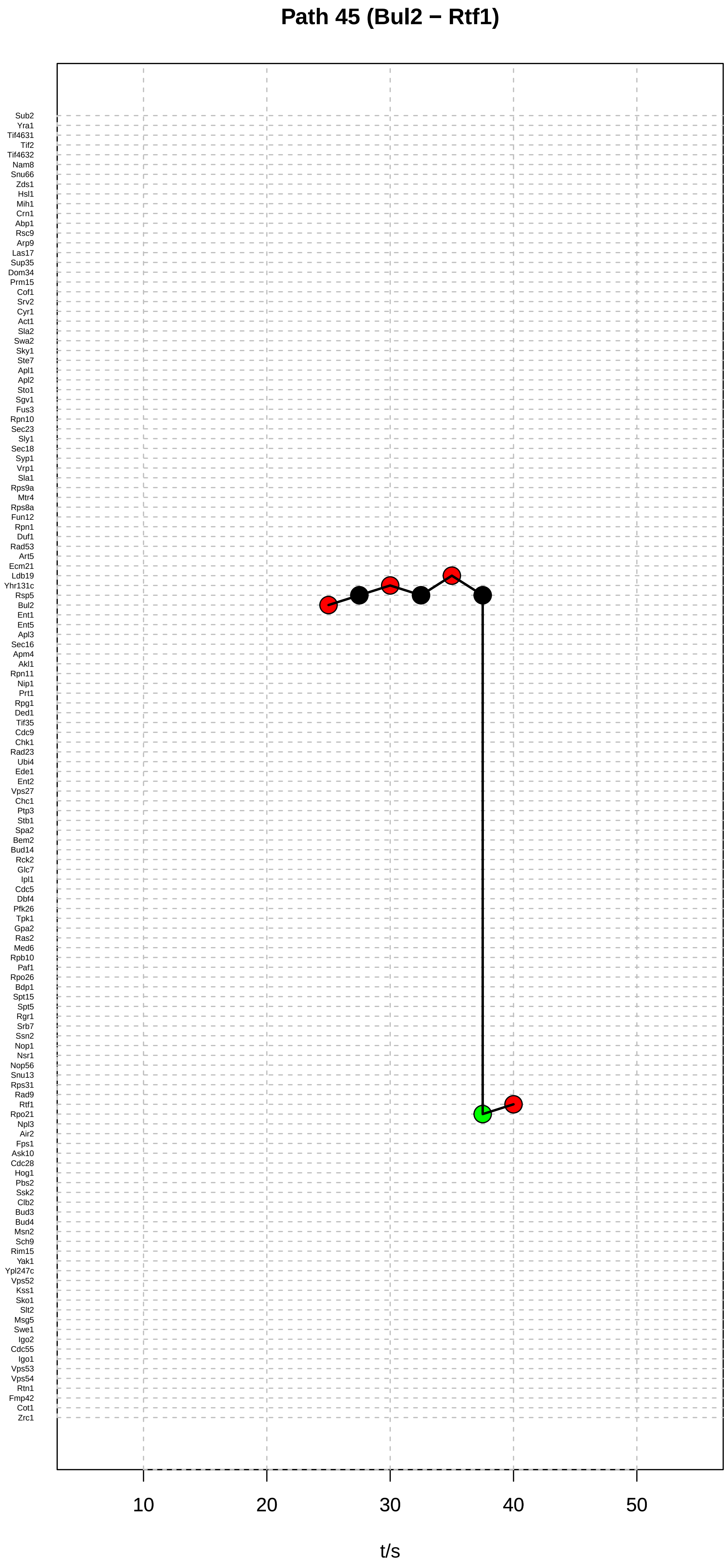

Path 45 (Bul2 − Rtf1)
Sub2
Yra1
Tif4631
Tif2
Tif4632
Nam8
Snu66
Zds1
Hsl1
Mih1
Crn1
Abp1
Rsc9
Arp9
Las17
Sup35
Dom34
Prm15
Cof1
Srv2
Cyr1
Act1
Sla2
Swa2
Sky1
Ste7
Apl1
Apl2
Sto1
Sgv1
Fus3
Rpn10
Sec23
Sly1
Sec18
Syp1
Vrp1
Sla1
Rps9a
Mtr4
Rps8a
Fun12
Rpn1
Duf1
Rad53
Art5
Ecm21
Ldb19
Yhr131c
Rsp5
Bul2
Ent1
Ent5
Apl3
Sec16
Apm4
Akl1
Rpn11
Nip1
Prt1
Rpg1
Ded1
Tif35
Cdc9
Chk1
Rad23
Ubi4
Ede1
Ent2
Vps27
Chc1
Ptp3
Stb1
Spa2
Bem2
Bud14
Rck2
Glc7
Ipl1
Cdc5
Dbf4
Pfk26
Tpk1
Gpa2
Ras2
Med6
Rpb10
Paf1
Rpo26
Bdp1
Spt15
Spt5
Rgr1
Srb7
Ssn2
Nop1
Nsr1
Nop56
Snu13
Rps31
Rad9
Rtf1
Rpo21
Npl3
Air2
Fps1
Ask10
Cdc28
Hog1
Pbs2
Ssk2
Clb2
Bud3
Bud4
Msn2
Sch9
Rim15
Yak1
Ypl247c
Vps52
Kss1
Sko1
Slt2
Msg5
Swe1
Igo2
Cdc55
Igo1
Vps53
Vps54
Rtn1
Fmp42
Cot1
Zrc1
10
20
30
40
50
t/s

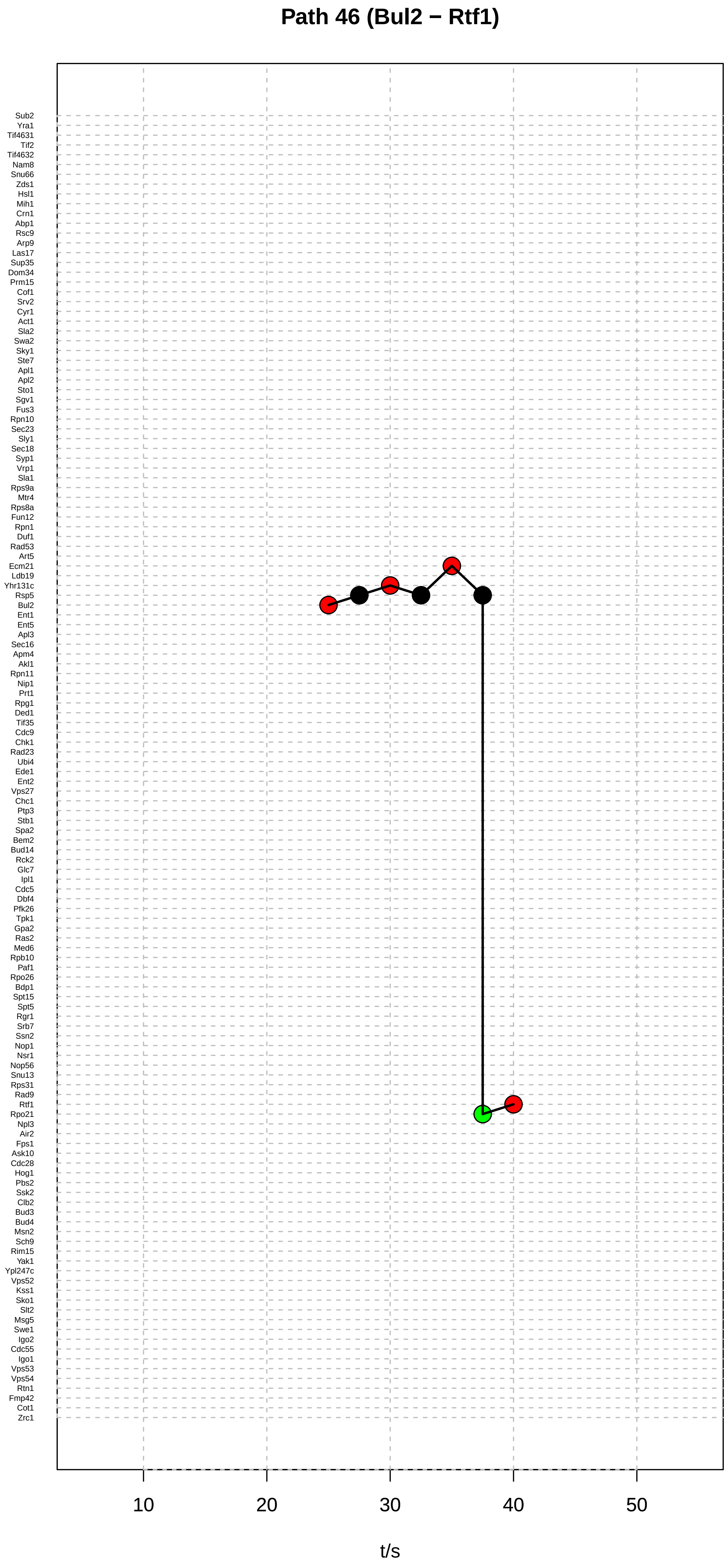

Path 46 (Bul2 − Rtf1)
Sub2
Yra1
Tif4631
Tif2
Tif4632
Nam8
Snu66
Zds1
Hsl1
Mih1
Crn1
Abp1
Rsc9
Arp9
Las17
Sup35
Dom34
Prm15
Cof1
Srv2
Cyr1
Act1
Sla2
Swa2
Sky1
Ste7
Apl1
Apl2
Sto1
Sgv1
Fus3
Rpn10
Sec23
Sly1
Sec18
Syp1
Vrp1
Sla1
Rps9a
Mtr4
Rps8a
Fun12
Rpn1
Duf1
Rad53
Art5
Ecm21
Ldb19
Yhr131c
Rsp5
Bul2
Ent1
Ent5
Apl3
Sec16
Apm4
Akl1
Rpn11
Nip1
Prt1
Rpg1
Ded1
Tif35
Cdc9
Chk1
Rad23
Ubi4
Ede1
Ent2
Vps27
Chc1
Ptp3
Stb1
Spa2
Bem2
Bud14
Rck2
Glc7
Ipl1
Cdc5
Dbf4
Pfk26
Tpk1
Gpa2
Ras2
Med6
Rpb10
Paf1
Rpo26
Bdp1
Spt15
Spt5
Rgr1
Srb7
Ssn2
Nop1
Nsr1
Nop56
Snu13
Rps31
Rad9
Rtf1
Rpo21
Npl3
Air2
Fps1
Ask10
Cdc28
Hog1
Pbs2
Ssk2
Clb2
Bud3
Bud4
Msn2
Sch9
Rim15
Yak1
Ypl247c
Vps52
Kss1
Sko1
Slt2
Msg5
Swe1
Igo2
Cdc55
Igo1
Vps53
Vps54
Rtn1
Fmp42
Cot1
Zrc1
10
20
30
40
50
t/s

### Path 47 (Bul2 − Rtf1)

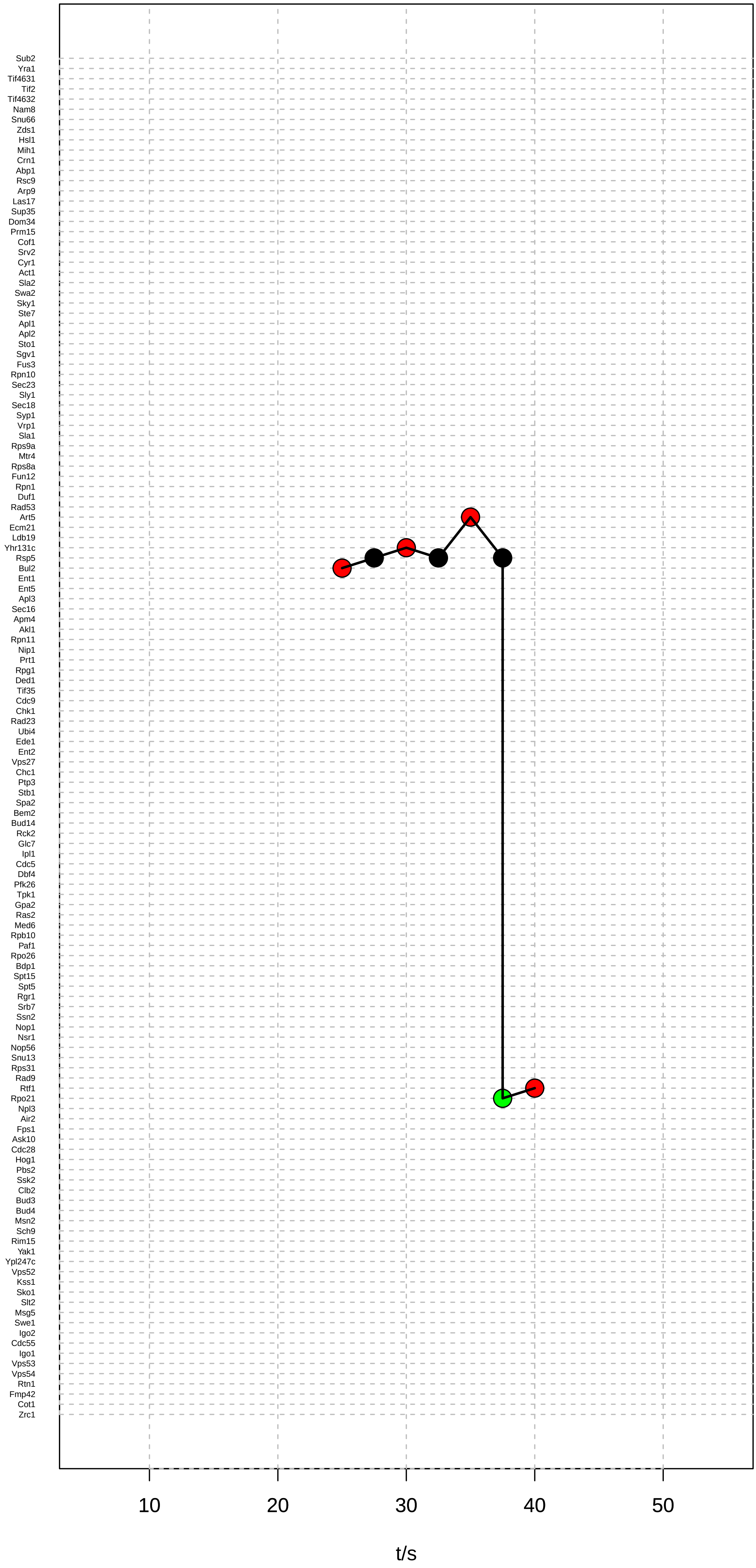

### Path 48 (Dbf4 − Rck2)

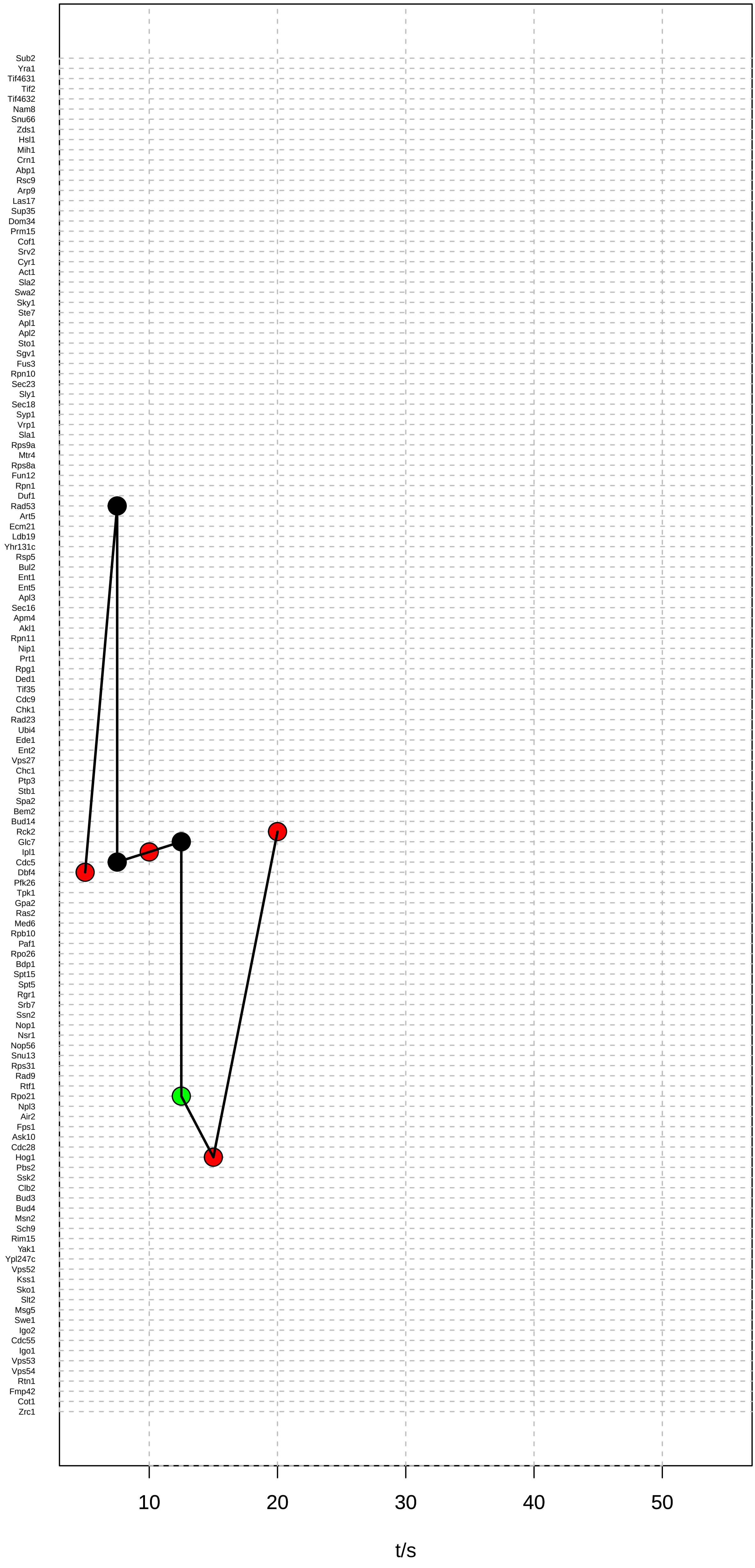

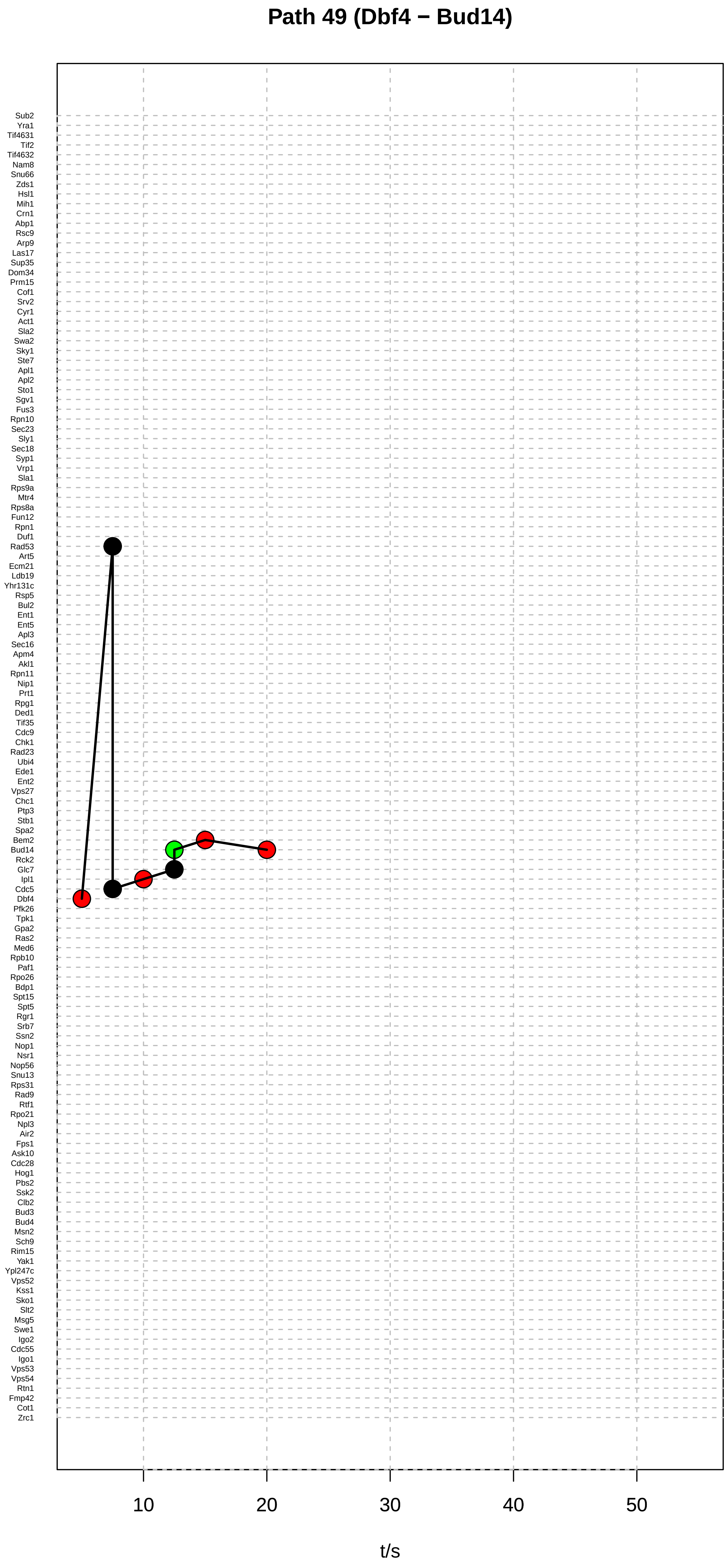

Path 49 (Dbf4 − Bud14)
Sub2
Yra1
Tif4631
Tif2
Tif4632
Nam8
Snu66
Zds1
Hsl1
Mih1
Crn1
Abp1
Rsc9
Arp9
Las17
Sup35
Dom34
Prm15
Cof1
Srv2
Cyr1
Act1
Sla2
Swa2
Sky1
Ste7
Apl1
Apl2
Sto1
Sgv1
Fus3
Rpn10
Sec23
Sly1
Sec18
Syp1
Vrp1
Sla1
Rps9a
Mtr4
Rps8a
Fun12
Rpn1
Duf1
Rad53
Art5
Ecm21
Ldb19
Yhr131c
Rsp5
Bul2
Ent1
Ent5
Apl3
Sec16
Apm4
Akl1
Rpn11
Nip1
Prt1
Rpg1
Ded1
Tif35
Cdc9
Chk1
Rad23
Ubi4
Ede1
Ent2
Vps27
Chc1
Ptp3
Stb1
Spa2
Bem2
Bud14
Rck2
Glc7
Ipl1
Cdc5
Dbf4
Pfk26
Tpk1
Gpa2
Ras2
Med6
Rpb10
Paf1
Rpo26
Bdp1
Spt15
Spt5
Rgr1
Srb7
Ssn2
Nop1
Nsr1
Nop56
Snu13
Rps31
Rad9
Rtf1
Rpo21
Npl3
Air2
Fps1
Ask10
Cdc28
Hog1
Pbs2
Ssk2
Clb2
Bud3
Bud4
Msn2
Sch9
Rim15
Yak1
Ypl247c
Vps52
Kss1
Sko1
Slt2
Msg5
Swe1
Igo2
Cdc55
Igo1
Vps53
Vps54
Rtn1
Fmp42
Cot1
Zrc1
10
20
30
40
50
t/s

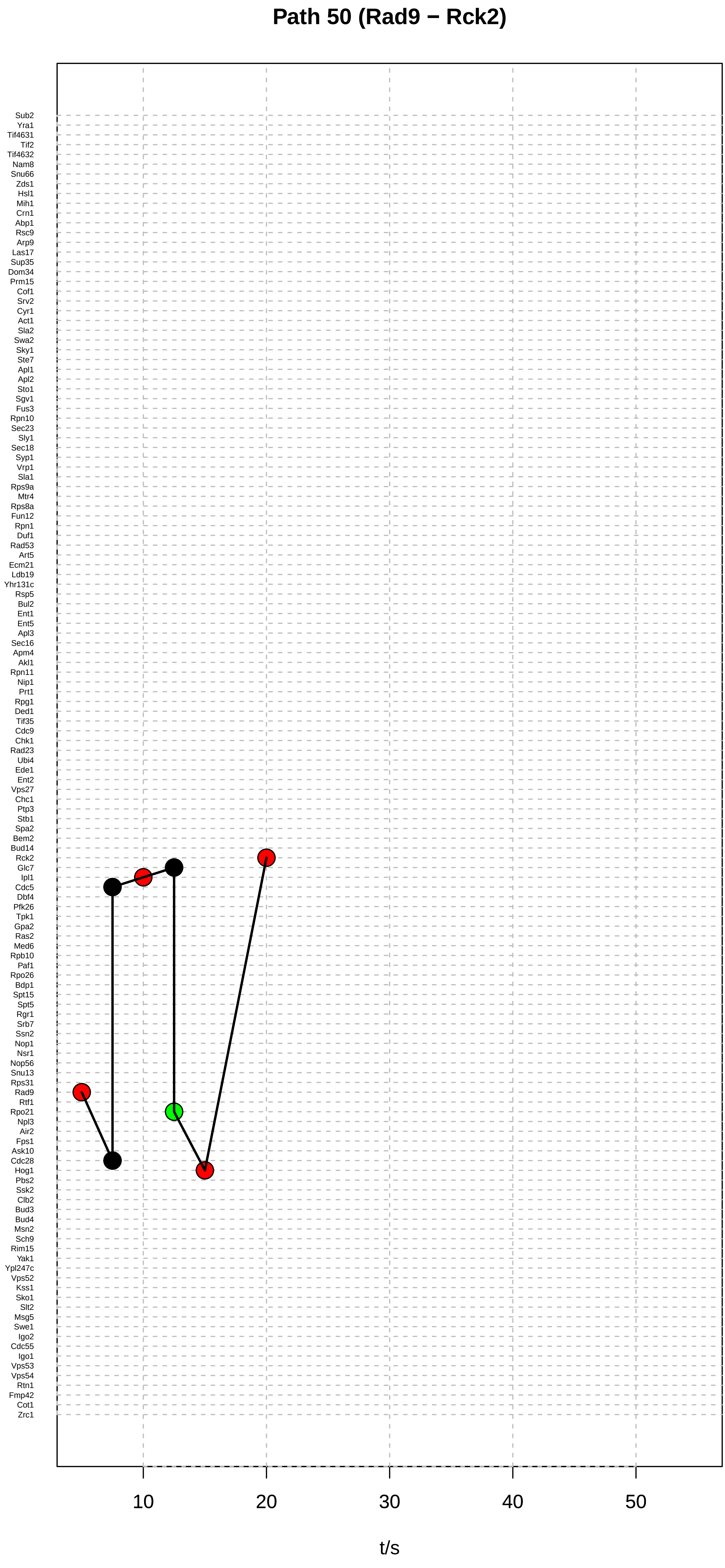

Path 50 (Rad9 − Rck2)
t/s
10
20
30
40
50
Sub2
Yra1
Tif4631
Tif2
Tif4632
Nam8
Snu66
Zds1
Hsl1
Mih1
Crn1
Abp1
Rsc9
Arp9
Las17
Sup35
Dom34
Prm15
Cof1
Srv2
Cyr1
Act1
Sla2
Swa2
Sky1
Ste7
Apl1
Apl2
Sto1
Sgv1
Fus3
Rpn10
Sec23
Sly1
Sec18
Syp1
Vrp1
Sla1
Rps9a
Mtr4
Rps8a
Fun12
Rpn1
Duf1
Rad53
Art5
Ecm21
Ldb19
Yhr131c
Rsp5
Bul2
Ent1
Ent5
Apl3
Sec16
Apm4
Akl1
Rpn11
Nip1
Prt1
Rpg1
Ded1
Tif35
Cdc9
Chk1
Rad23
Ubi4
Ede1
Ent2
Vps27
Chc1
Ptp3
Stb1
Spa2
Bem2
Bud14
Rck2
Glc7
Ipl1
Cdc5
Dbf4
Pfk26
Tpk1
Gpa2
Ras2
Med6
Rpb10
Paf1
Rpo26
Bdp1
Spt15
Spt5
Rgr1
Srb7
Ssn2
Nop1
Nsr1
Nop56
Snu13
Rps31
Rad9
Rtf1
Rpo21
Npl3
Air2
Fps1
Ask10
Cdc28
Hog1
Pbs2
Ssk2
Clb2
Bud3
Bud4
Msn2
Sch9
Rim15
Yak1
Ypl247c
Vps52
Kss1
Sko1
Slt2
Msg5
Swe1
Igo2
Cdc55
Igo1
Vps53
Vps54
Rtn1
Fmp42
Cot1
Zrc1

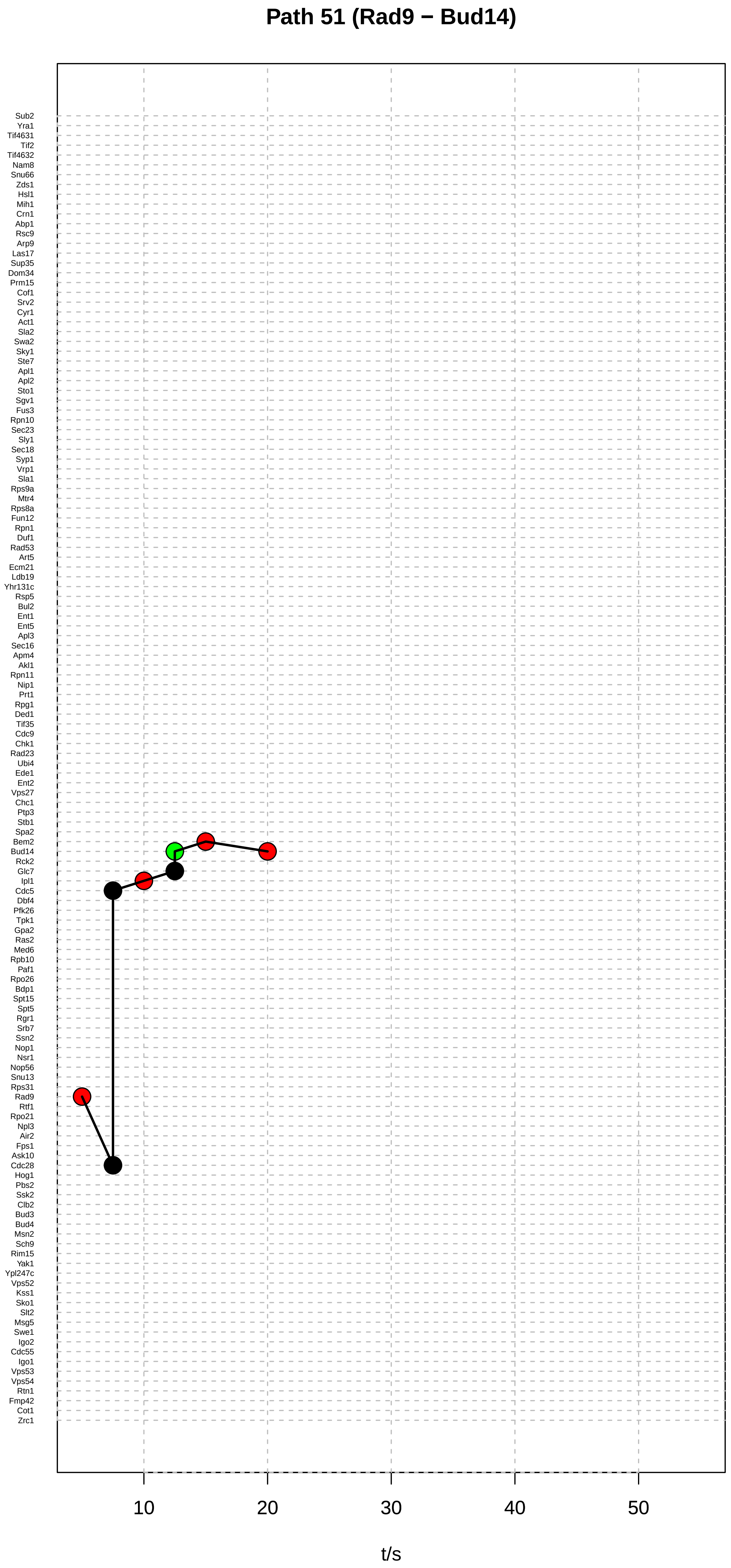
Path 51 (Rad9 − Bud14)
Sub2
Yra1
Tif4631
Tif2
Tif4632
Nam8
Snu66
Zds1
Hsl1
Mih1
Crn1
Abp1
Rsc9
Arp9
Las17
Sup35
Dom34
Prm15
Cof1
Srv2
Cyr1
Act1
Sla2
Swa2
Sky1
Ste7
Apl1
Apl2
Sto1
Sgv1
Fus3
Rpn10
Sec23
Sly1
Sec18
Syp1
Vrp1
Sla1
Rps9a
Mtr4
Rps8a
Fun12
Rpn1
Duf1
Rad53
Art5
Ecm21
Ldb19
Yhr131c
Rsp5
Bul2
Ent1
Ent5
Apl3
Sec16
Apm4
Akl1
Rpn11
Nip1
Prt1
Rpg1
Ded1
Tif35
Cdc9
Chk1
Rad23
Ubi4
Ede1
Ent2
Vps27
Chc1
Ptp3
Stb1
Spa2
Bem2
Bud14
Rck2
Glc7
Ipl1
Cdc5
Dbf4
Pfk26
Tpk1
Gpa2
Ras2
Med6
Rpb10
Paf1
Rpo26
Bdp1
Spt15
Spt5
Rgr1
Srb7
Ssn2
Nop1
Nsr1
Nop56
Snu13
Rps31
Rad9
Rtf1
Rpo21
Npl3
Air2
Fps1
Ask10
Cdc28
Hog1
Pbs2
Ssk2
Clb2
Bud3
Bud4
Msn2
Sch9
Rim15
Yak1
Ypl247c
Vps52
Kss1
Sko1
Slt2
Msg5
Swe1
Igo2
Cdc55
Igo1
Vps53
Vps54
Rtn1
Fmp42
Cot1
Zrc1
10
20
30
40
50
t/s

#### Path 52 (Rad9 − Rck2)

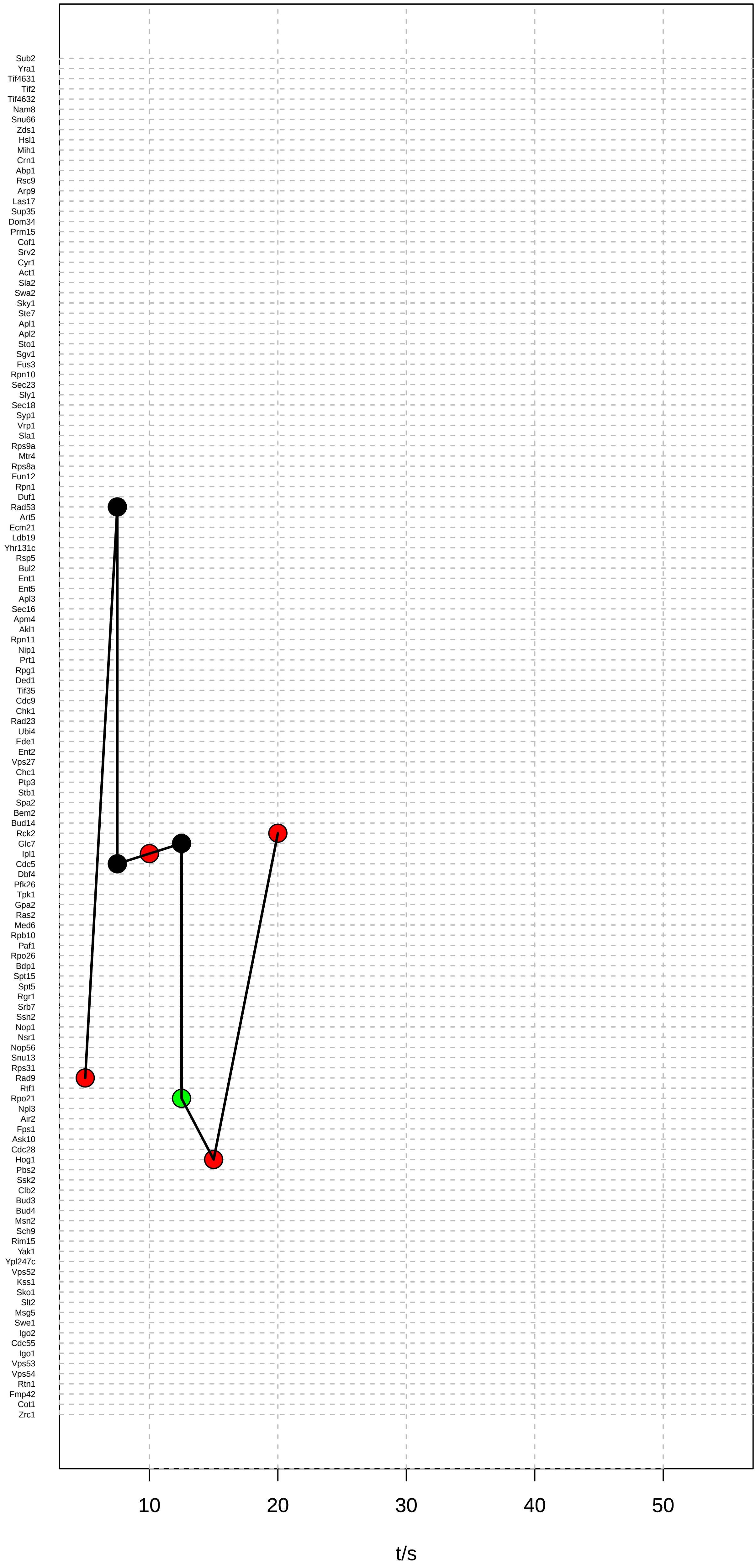

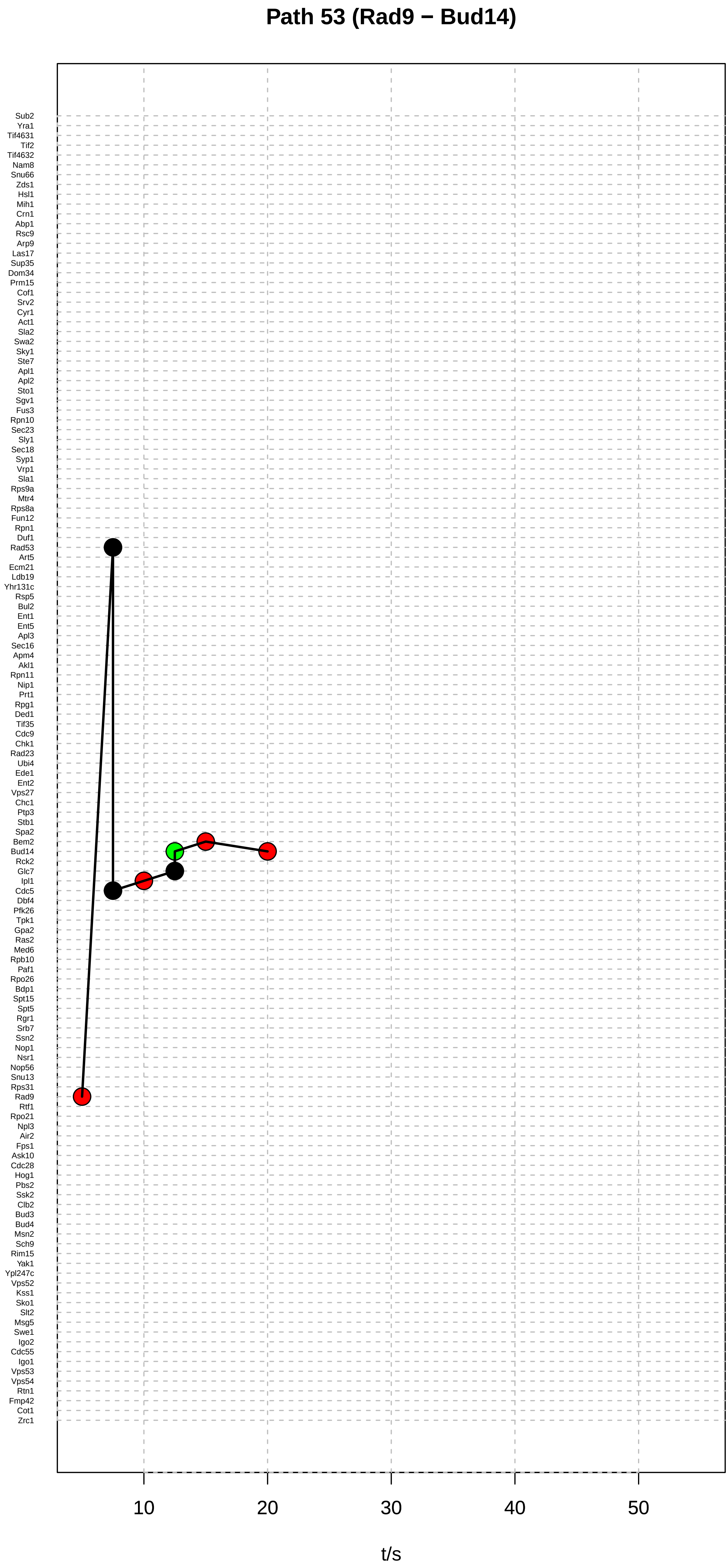

Path 53 (Rad9 − Bud14)
Sub2
Yra1
Tif4631
Tif2
Tif4632
Nam8
Snu66
Zds1
Hsl1
Mih1
Crn1
Abp1
Rsc9
Arp9
Las17
Sup35
Dom34
Prm15
Cof1
Srv2
Cyr1
Act1
Sla2
Swa2
Sky1
Ste7
Apl1
Apl2
Sto1
Sgv1
Fus3
Rpn10
Sec23
Sly1
Sec18
Syp1
Vrp1
Sla1
Rps9a
Mtr4
Rps8a
Fun12
Rpn1
Duf1
Rad53
Art5
Ecm21
Ldb19
Yhr131c
Rsp5
Bul2
Ent1
Ent5
Apl3
Sec16
Apm4
Akl1
Rpn11
Nip1
Prt1
Rpg1
Ded1
Tif35
Cdc9
Chk1
Rad23
Ubi4
Ede1
Ent2
Vps27
Chc1
Ptp3
Stb1
Spa2
Bem2
Bud14
Rck2
Glc7
Ipl1
Cdc5
Dbf4
Pfk26
Tpk1
Gpa2
Ras2
Med6
Rpb10
Paf1
Rpo26
Bdp1
Spt15
Spt5
Rgr1
Srb7
Ssn2
Nop1
Nsr1
Nop56
Snu13
Rps31
Rad9
Rtf1
Rpo21
Npl3
Air2
Fps1
Ask10
Cdc28
Hog1
Pbs2
Ssk2
Clb2
Bud3
Bud4
Msn2
Sch9
Rim15
Yak1
Ypl247c
Vps52
Kss1
Sko1
Slt2
Msg5
Swe1
Igo2
Cdc55
Igo1
Vps53
Vps54
Rtn1
Fmp42
Cot1
Zrc1
10
20
30
40
50
t/s

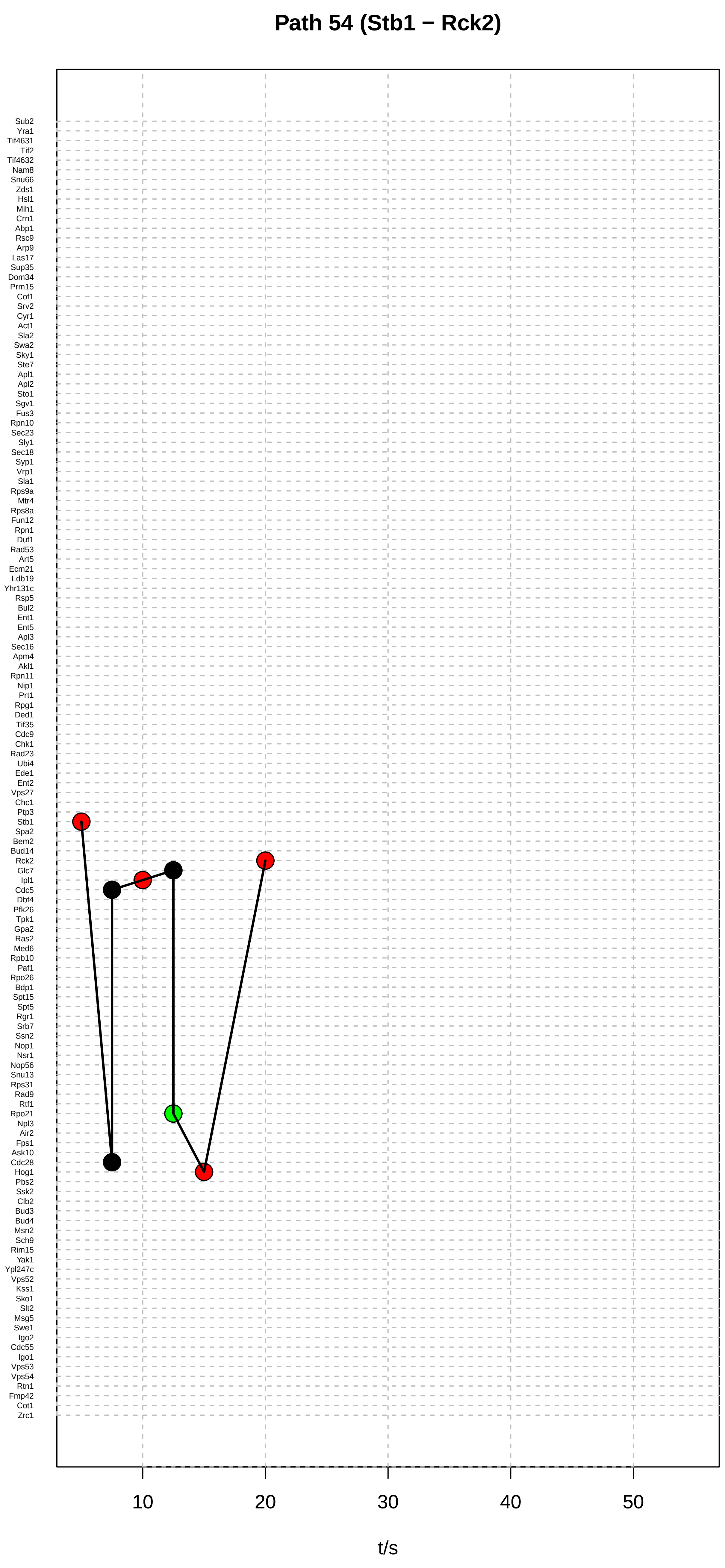

Path 54 (Stb1 − Rck2)
Sub2
Yra1
Tif4631
Tif2
Tif4632
Nam8
Snu66
Zds1
Hsl1
Mih1
Crn1
Abp1
Rsc9
Arp9
Las17
Sup35
Dom34
Prm15
Cof1
Srv2
Cyr1
Act1
Sla2
Swa2
Sky1
Ste7
Apl1
Apl2
Sto1
Sgv1
Fus3
Rpn10
Sec23
Sly1
Sec18
Syp1
Vrp1
Sla1
Rps9a
Mtr4
Rps8a
Fun12
Rpn1
Duf1
Rad53
Art5
Ecm21
Ldb19
Yhr131c
Rsp5
Bul2
Ent1
Ent5
Apl3
Sec16
Apm4
Akl1
Rpn11
Nip1
Prt1
Rpg1
Ded1
Tif35
Cdc9
Chk1
Rad23
Ubi4
Ede1
Ent2
Vps27
Chc1
Ptp3
Stb1
Spa2
Bem2
Bud14
Rck2
Glc7
Ipl1
Cdc5
Dbf4
Pfk26
Tpk1
Gpa2
Ras2
Med6
Rpb10
Paf1
Rpo26
Bdp1
Spt15
Spt5
Rgr1
Srb7
Ssn2
Nop1
Nsr1
Nop56
Snu13
Rps31
Rad9
Rtf1
Rpo21
Npl3
Air2
Fps1
Ask10
Cdc28
Hog1
Pbs2
Ssk2
Clb2
Bud3
Bud4
Msn2
Sch9
Rim15
Yak1
Ypl247c
Vps52
Kss1
Sko1
Slt2
Msg5
Swe1
Igo2
Cdc55
Igo1
Vps53
Vps54
Rtn1
Fmp42
Cot1
Zrc1
10
20
30
40
50
t/s

#### Path 55 (Stb1 − Bud14)

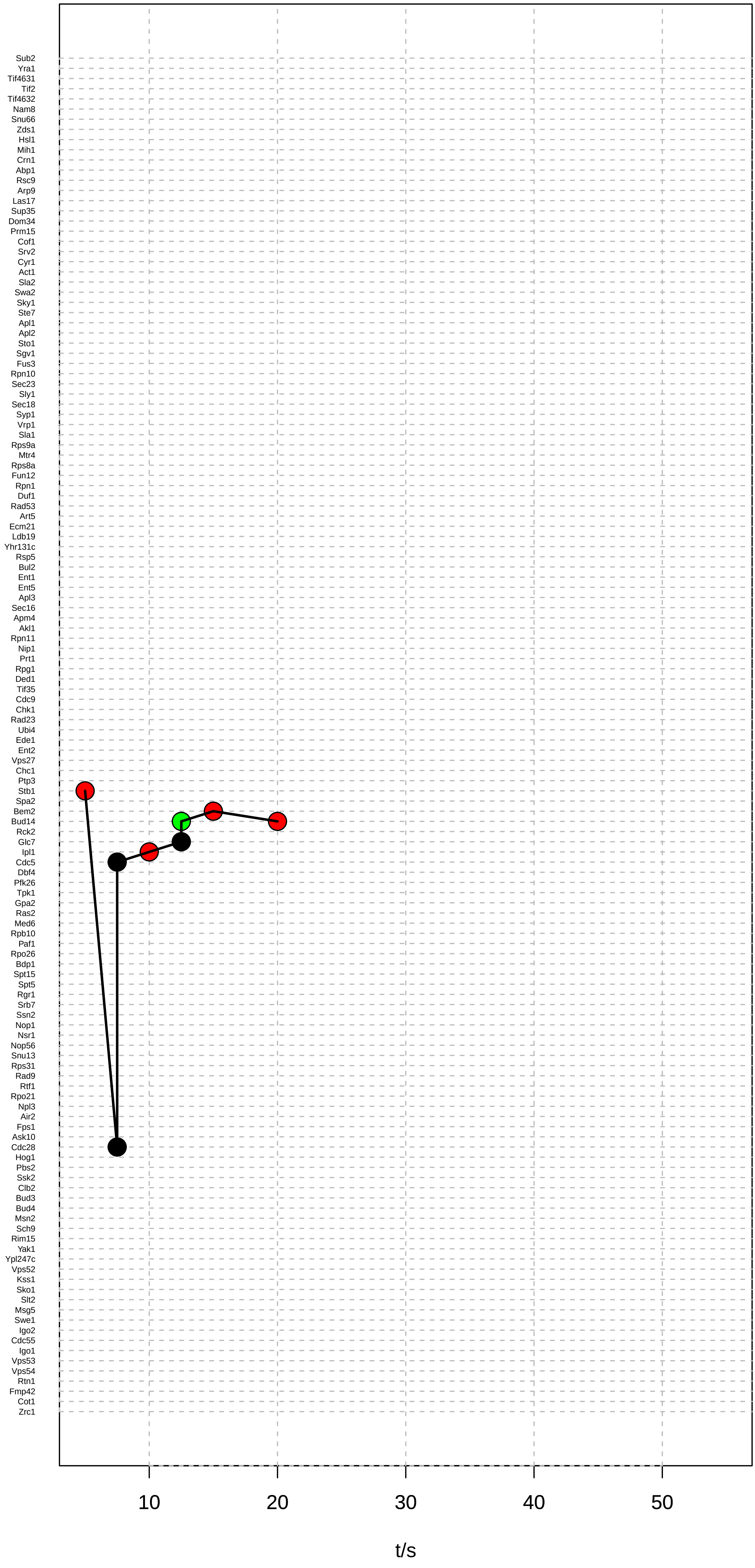

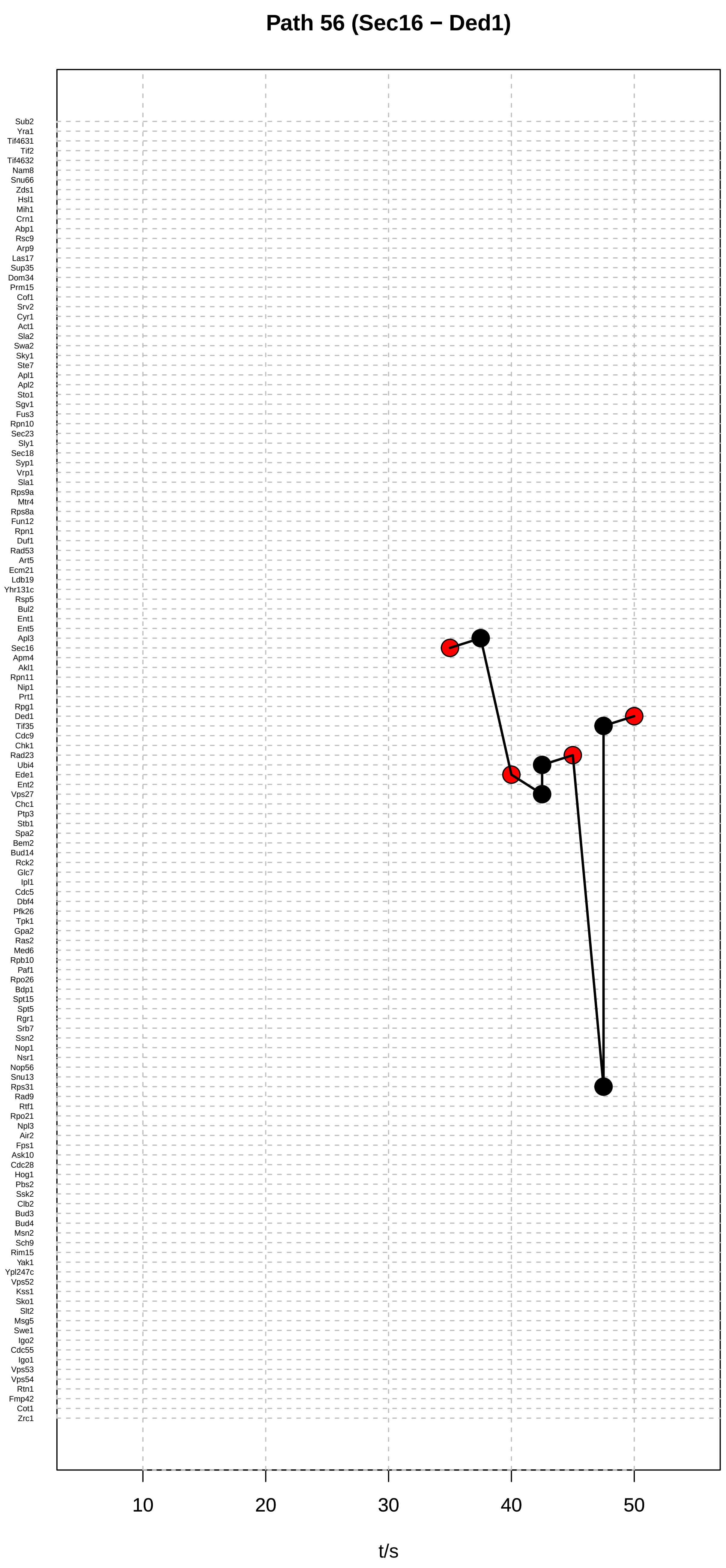

Path 56 (Sec16 − Ded1)
Sub2
Yra1
Tif4631
Tif2
Tif4632
Nam8
Snu66
Zds1
Hsl1
Mih1
Crn1
Abp1
Rsc9
Arp9
Las17
Sup35
Dom34
Prm15
Cof1
Srv2
Cyr1
Act1
Sla2
Swa2
Sky1
Ste7
Apl1
Apl2
Sto1
Sgv1
Fus3
Rpn10
Sec23
Sly1
Sec18
Syp1
Vrp1
Sla1
Rps9a
Mtr4
Rps8a
Fun12
Rpn1
Duf1
Rad53
Art5
Ecm21
Ldb19
Yhr131c
Rsp5
Bul2
Ent1
Ent5
Apl3
Sec16
Apm4
Akl1
Rpn11
Nip1
Prt1
Rpg1
Ded1
Tif35
Cdc9
Chk1
Rad23
Ubi4
Ede1
Ent2
Vps27
Chc1
Ptp3
Stb1
Spa2
Bem2
Bud14
Rck2
Glc7
Ipl1
Cdc5
Dbf4
Pfk26
Tpk1
Gpa2
Ras2
Med6
Rpb10
Paf1
Rpo26
Bdp1
Spt15
Spt5
Rgr1
Srb7
Ssn2
Nop1
Nsr1
Nop56
Snu13
Rps31
Rad9
Rtf1
Rpo21
Npl3
Air2
Fps1
Ask10
Cdc28
Hog1
Pbs2
Ssk2
Clb2
Bud3
Bud4
Msn2
Sch9
Rim15
Yak1
Ypl247c
Vps52
Kss1
Sko1
Slt2
Msg5
Swe1
Igo2
Cdc55
Igo1
Vps53
Vps54
Rtn1
Fmp42
Cot1
Zrc1
10
20
30
40
50
t/s

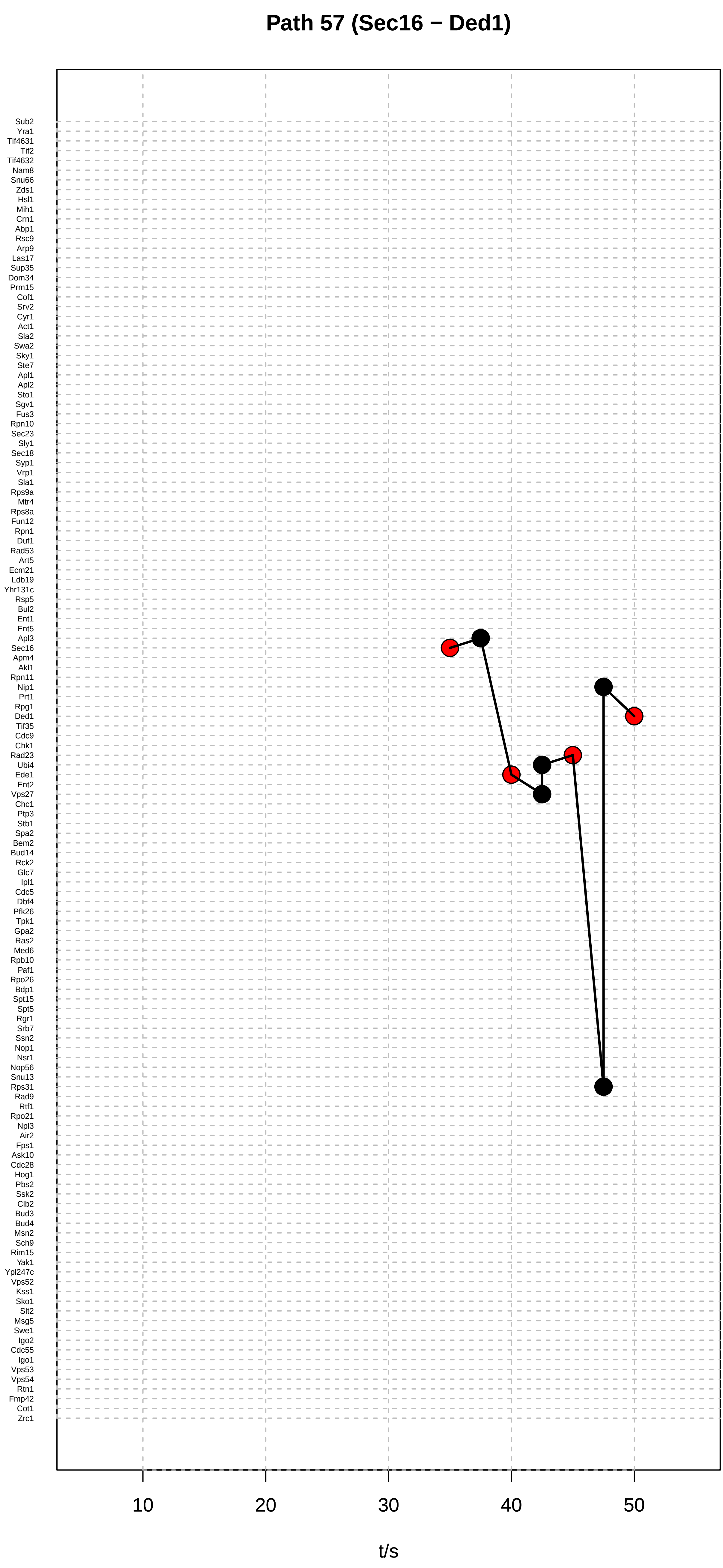

Path 57 (Sec16 − Ded1)
Sub2
Yra1
Tif4631
Tif2
Tif4632
Nam8
Snu66
Zds1
Hsl1
Mih1
Crn1
Abp1
Rsc9
Arp9
Las17
Sup35
Dom34
Prm15
Cof1
Srv2
Cyr1
Act1
Sla2
Swa2
Sky1
Ste7
Apl1
Apl2
Sto1
Sgv1
Fus3
Rpn10
Sec23
Sly1
Sec18
Syp1
Vrp1
Sla1
Rps9a
Mtr4
Rps8a
Fun12
Rpn1
Duf1
Rad53
Art5
Ecm21
Ldb19
Yhr131c
Rsp5
Bul2
Ent1
Ent5
Apl3
Sec16
Apm4
Akl1
Rpn11
Nip1
Prt1
Rpg1
Ded1
Tif35
Cdc9
Chk1
Rad23
Ubi4
Ede1
Ent2
Vps27
Chc1
Ptp3
Stb1
Spa2
Bem2
Bud14
Rck2
Glc7
Ipl1
Cdc5
Dbf4
Pfk26
Tpk1
Gpa2
Ras2
Med6
Rpb10
Paf1
Rpo26
Bdp1
Spt15
Spt5
Rgr1
Srb7
Ssn2
Nop1
Nsr1
Nop56
Snu13
Rps31
Rad9
Rtf1
Rpo21
Npl3
Air2
Fps1
Ask10
Cdc28
Hog1
Pbs2
Ssk2
Clb2
Bud3
Bud4
Msn2
Sch9
Rim15
Yak1
Ypl247c
Vps52
Kss1
Sko1
Slt2
Msg5
Swe1
Igo2
Cdc55
Igo1
Vps53
Vps54
Rtn1
Fmp42
Cot1
Zrc1
10
20
30
40
50
t/s

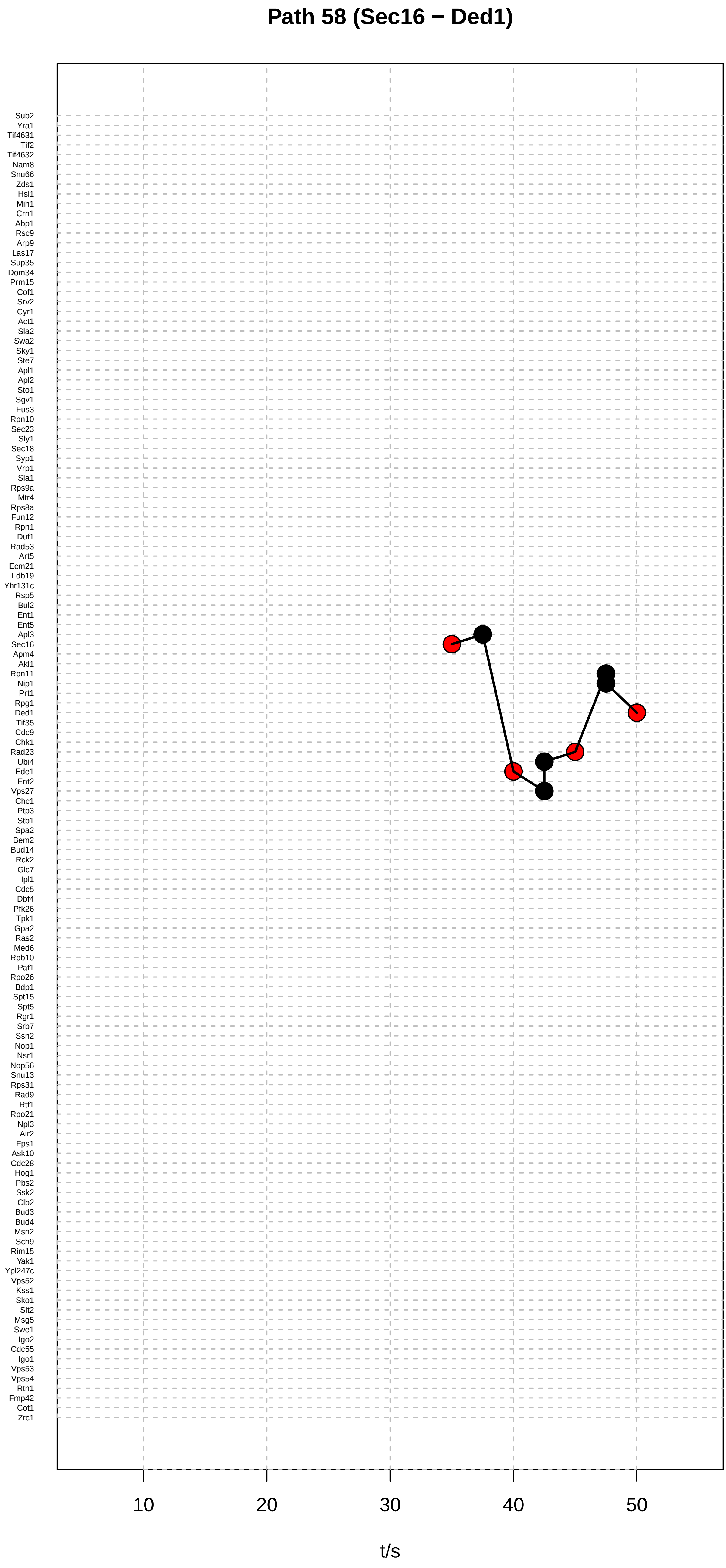

Path 58 (Sec16 − Ded1)
Sub2
Yra1
Tif4631
Tif2
Tif4632
Nam8
Snu66
Zds1
Hsl1
Mih1
Crn1
Abp1
Rsc9
Arp9
Las17
Sup35
Dom34
Prm15
Cof1
Srv2
Cyr1
Act1
Sla2
Swa2
Sky1
Ste7
Apl1
Apl2
Sto1
Sgv1
Fus3
Rpn10
Sec23
Sly1
Sec18
Syp1
Vrp1
Sla1
Rps9a
Mtr4
Rps8a
Fun12
Rpn1
Duf1
Rad53
Art5
Ecm21
Ldb19
Yhr131c
Rsp5
Bul2
Ent1
Ent5
Apl3
Sec16
Apm4
Akl1
Rpn11
Nip1
Prt1
Rpg1
Ded1
Tif35
Cdc9
Chk1
Rad23
Ubi4
Ede1
Ent2
Vps27
Chc1
Ptp3
Stb1
Spa2
Bem2
Bud14
Rck2
Glc7
Ipl1
Cdc5
Dbf4
Pfk26
Tpk1
Gpa2
Ras2
Med6
Rpb10
Paf1
Rpo26
Bdp1
Spt15
Spt5
Rgr1
Srb7
Ssn2
Nop1
Nsr1
Nop56
Snu13
Rps31
Rad9
Rtf1
Rpo21
Npl3
Air2
Fps1
Ask10
Cdc28
Hog1
Pbs2
Ssk2
Clb2
Bud3
Bud4
Msn2
Sch9
Rim15
Yak1
Ypl247c
Vps52
Kss1
Sko1
Slt2
Msg5
Swe1
Igo2
Cdc55
Igo1
Vps53
Vps54
Rtn1
Fmp42
Cot1
Zrc1
10
20
30
40
50
t/s

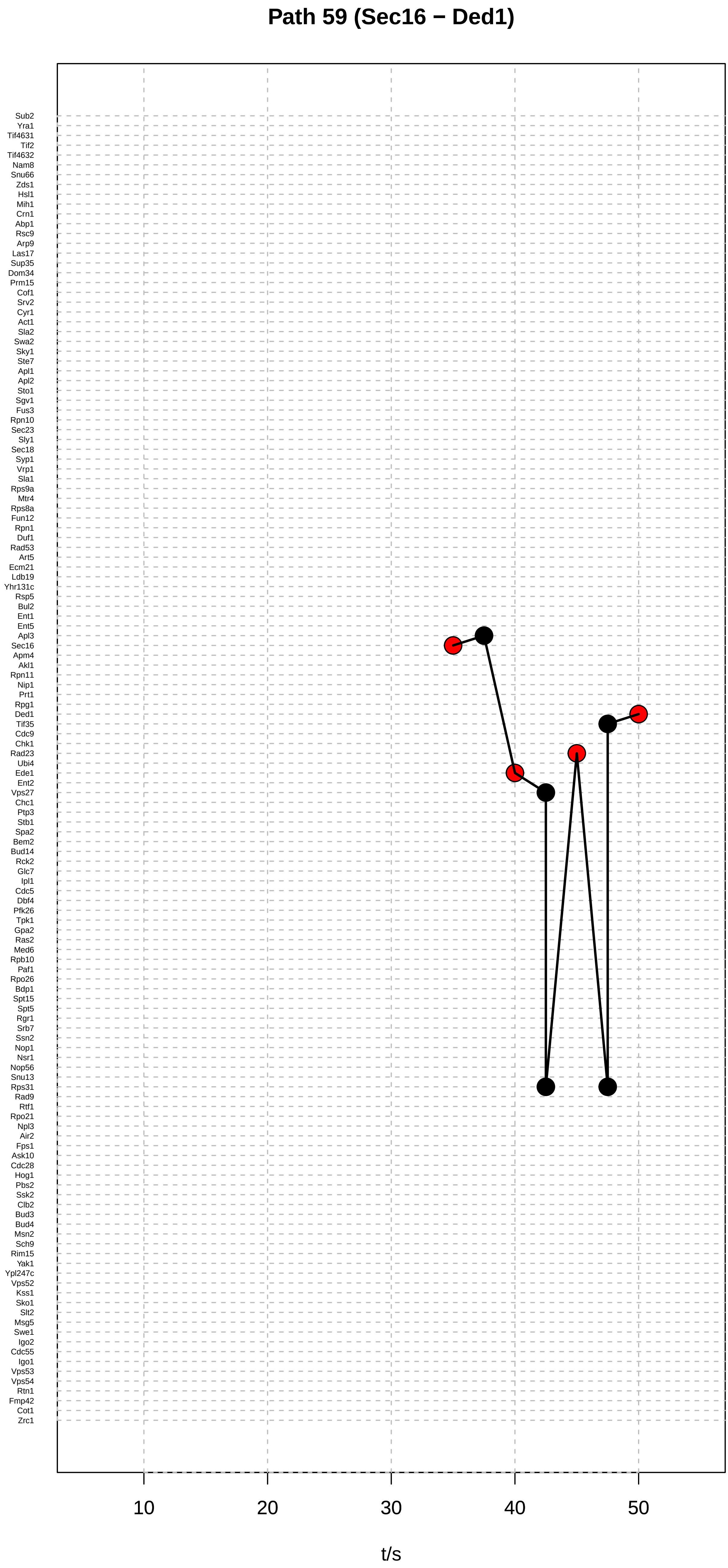
Path 59 (Sec16 − Ded1)
Sub2
Yra1
Tif4631
Tif2
Tif4632
Nam8
Snu66
Zds1
Hsl1
Mih1
Crn1
Abp1
Rsc9
Arp9
Las17
Sup35
Dom34
Prm15
Cof1
Srv2
Cyr1
Act1
Sla2
Swa2
Sky1
Ste7
Apl1
Apl2
Sto1
Sgv1
Fus3
Rpn10
Sec23
Sly1
Sec18
Syp1
Vrp1
Sla1
Rps9a
Mtr4
Rps8a
Fun12
Rpn1
Duf1
Rad53
Art5
Ecm21
Ldb19
Yhr131c
Rsp5
Bul2
Ent1
Ent5
Apl3
Sec16
Apm4
Akl1
Rpn11
Nip1
Prt1
Rpg1
Ded1
Tif35
Cdc9
Chk1
Rad23
Ubi4
Ede1
Ent2
Vps27
Chc1
Ptp3
Stb1
Spa2
Bem2
Bud14
Rck2
Glc7
Ipl1
Cdc5
Dbf4
Pfk26
Tpk1
Gpa2
Ras2
Med6
Rpb10
Paf1
Rpo26
Bdp1
Spt15
Spt5
Rgr1
Srb7
Ssn2
Nop1
Nsr1
Nop56
Snu13
Rps31
Rad9
Rtf1
Rpo21
Npl3
Air2
Fps1
Ask10
Cdc28
Hog1
Pbs2
Ssk2
Clb2
Bud3
Bud4
Msn2
Sch9
Rim15
Yak1
Ypl247c
Vps52
Kss1
Sko1
Slt2
Msg5
Swe1
Igo2
Cdc55
Igo1
Vps53
Vps54
Rtn1
Fmp42
Cot1
Zrc1
10
20
30
40
50
t/s

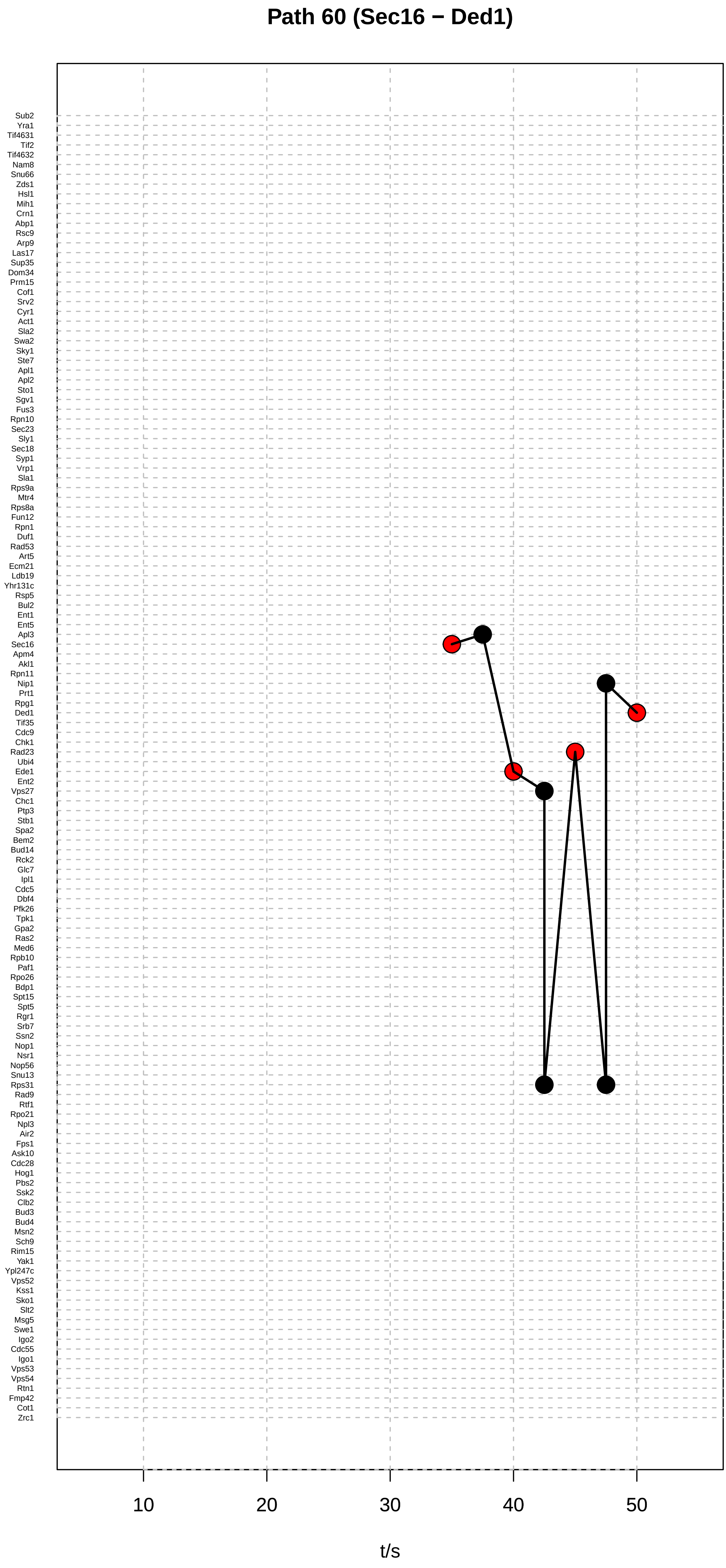
Path 60 (Sec16 − Ded1)
Sub2
Yra1
Tif4631
Tif2
Tif4632
Nam8
Snu66
Zds1
Hsl1
Mih1
Crn1
Abp1
Rsc9
Arp9
Las17
Sup35
Dom34
Prm15
Cof1
Srv2
Cyr1
Act1
Sla2
Swa2
Sky1
Ste7
Apl1
Apl2
Sto1
Sgv1
Fus3
Rpn10
Sec23
Sly1
Sec18
Syp1
Vrp1
Sla1
Rps9a
Mtr4
Rps8a
Fun12
Rpn1
Duf1
Rad53
Art5
Ecm21
Ldb19
Yhr131c
Rsp5
Bul2
Ent1
Ent5
Apl3
Sec16
Apm4
Akl1
Rpn11
Nip1
Prt1
Rpg1
Ded1
Tif35
Cdc9
Chk1
Rad23
Ubi4
Ede1
Ent2
Vps27
Chc1
Ptp3
Stb1
Spa2
Bem2
Bud14
Rck2
Glc7
Ipl1
Cdc5
Dbf4
Pfk26
Tpk1
Gpa2
Ras2
Med6
Rpb10
Paf1
Rpo26
Bdp1
Spt15
Spt5
Rgr1
Srb7
Ssn2
Nop1
Nsr1
Nop56
Snu13
Rps31
Rad9
Rtf1
Rpo21
Npl3
Air2
Fps1
Ask10
Cdc28
Hog1
Pbs2
Ssk2
Clb2
Bud3
Bud4
Msn2
Sch9
Rim15
Yak1
Ypl247c
Vps52
Kss1
Sko1
Slt2
Msg5
Swe1
Igo2
Cdc55
Igo1
Vps53
Vps54
Rtn1
Fmp42
Cot1
Zrc1
10
20
30
40
50
t/s

### Path 61 (Sec16 − Ded1)

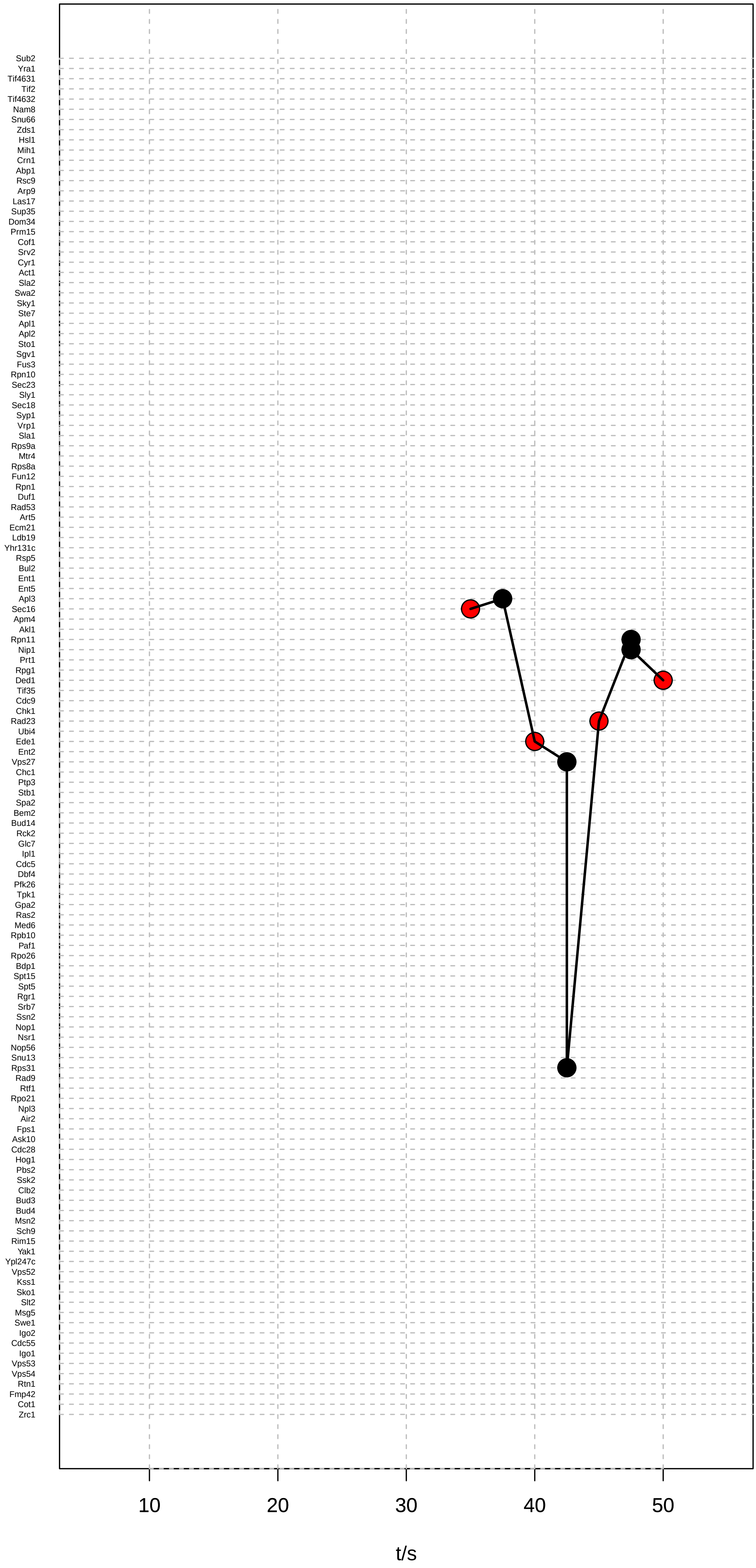

### Path 62 (Sec16 − Ded1)

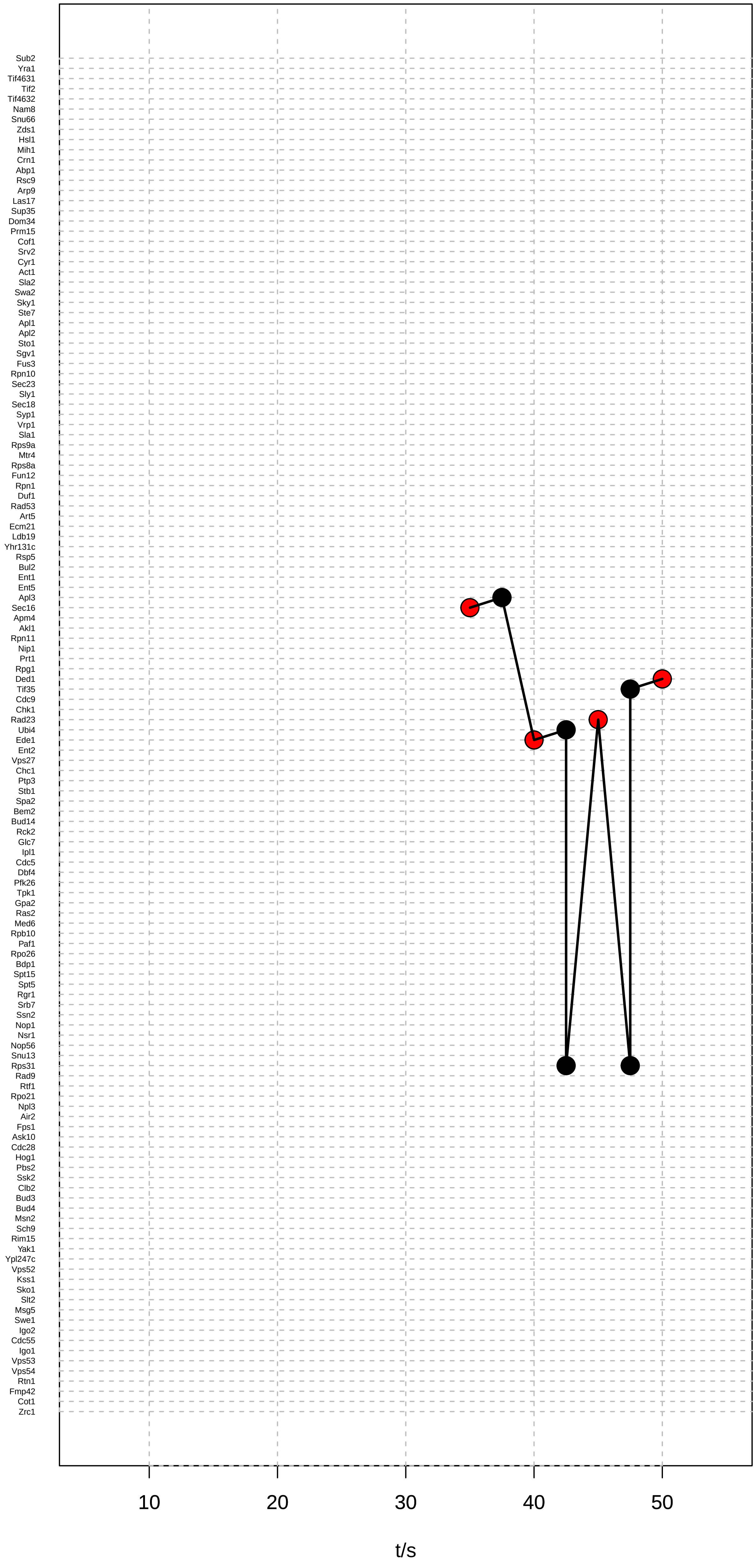

### Path 63 (Sec16 − Ded1)

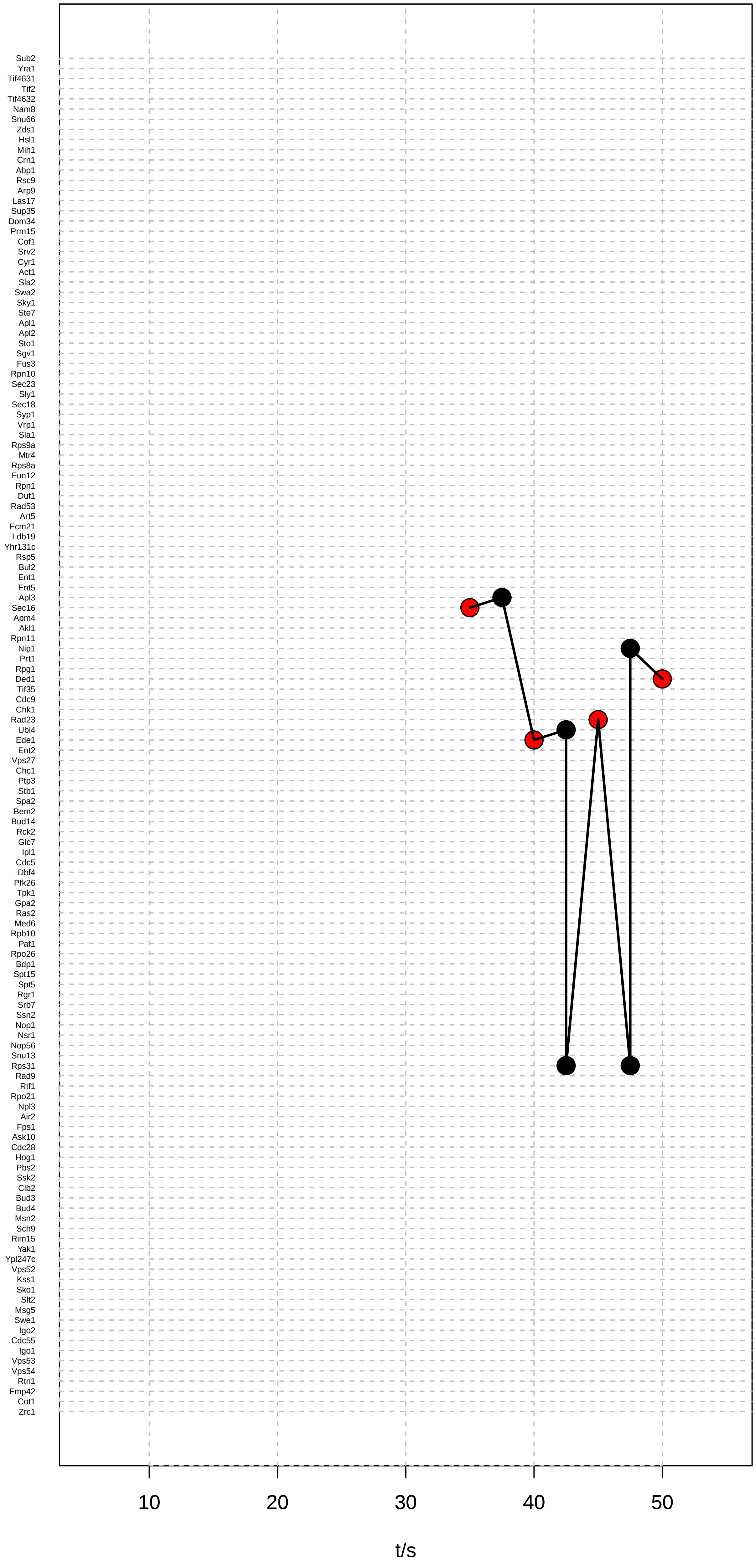

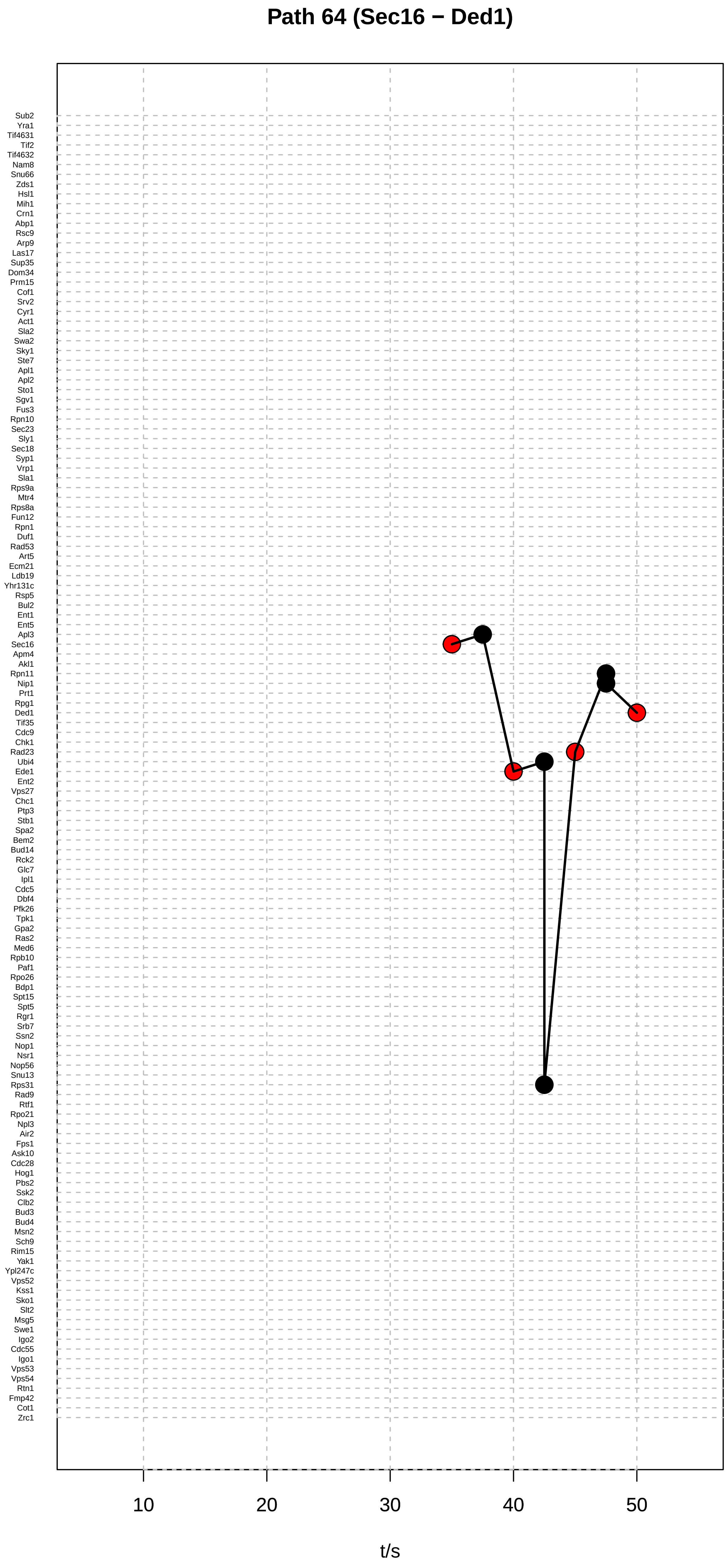
Path 64 (Sec16 − Ded1)
Sub2
Yra1
Tif4631
Tif2
Tif4632
Nam8
Snu66
Zds1
Hsl1
Mih1
Crn1
Abp1
Rsc9
Arp9
Las17
Sup35
Dom34
Prm15
Cof1
Srv2
Cyr1
Act1
Sla2
Swa2
Sky1
Ste7
Apl1
Apl2
Sto1
Sgv1
Fus3
Rpn10
Sec23
Sly1
Sec18
Syp1
Vrp1
Sla1
Rps9a
Mtr4
Rps8a
Fun12
Rpn1
Duf1
Rad53
Art5
Ecm21
Ldb19
Yhr131c
Rsp5
Bul2
Ent1
Ent5
Apl3
Sec16
Apm4
Akl1
Rpn11
Nip1
Prt1
Rpg1
Ded1
Tif35
Cdc9
Chk1
Rad23
Ubi4
Ede1
Ent2
Vps27
Chc1
Ptp3
Stb1
Spa2
Bem2
Bud14
Rck2
Glc7
Ipl1
Cdc5
Dbf4
Pfk26
Tpk1
Gpa2
Ras2
Med6
Rpb10
Paf1
Rpo26
Bdp1
Spt15
Spt5
Rgr1
Srb7
Ssn2
Nop1
Nsr1
Nop56
Snu13
Rps31
Rad9
Rtf1
Rpo21
Npl3
Air2
Fps1
Ask10
Cdc28
Hog1
Pbs2
Ssk2
Clb2
Bud3
Bud4
Msn2
Sch9
Rim15
Yak1
Ypl247c
Vps52
Kss1
Sko1
Slt2
Msg5
Swe1
Igo2
Cdc55
Igo1
Vps53
Vps54
Rtn1
Fmp42
Cot1
Zrc1
10
20
30
40
50
t/s

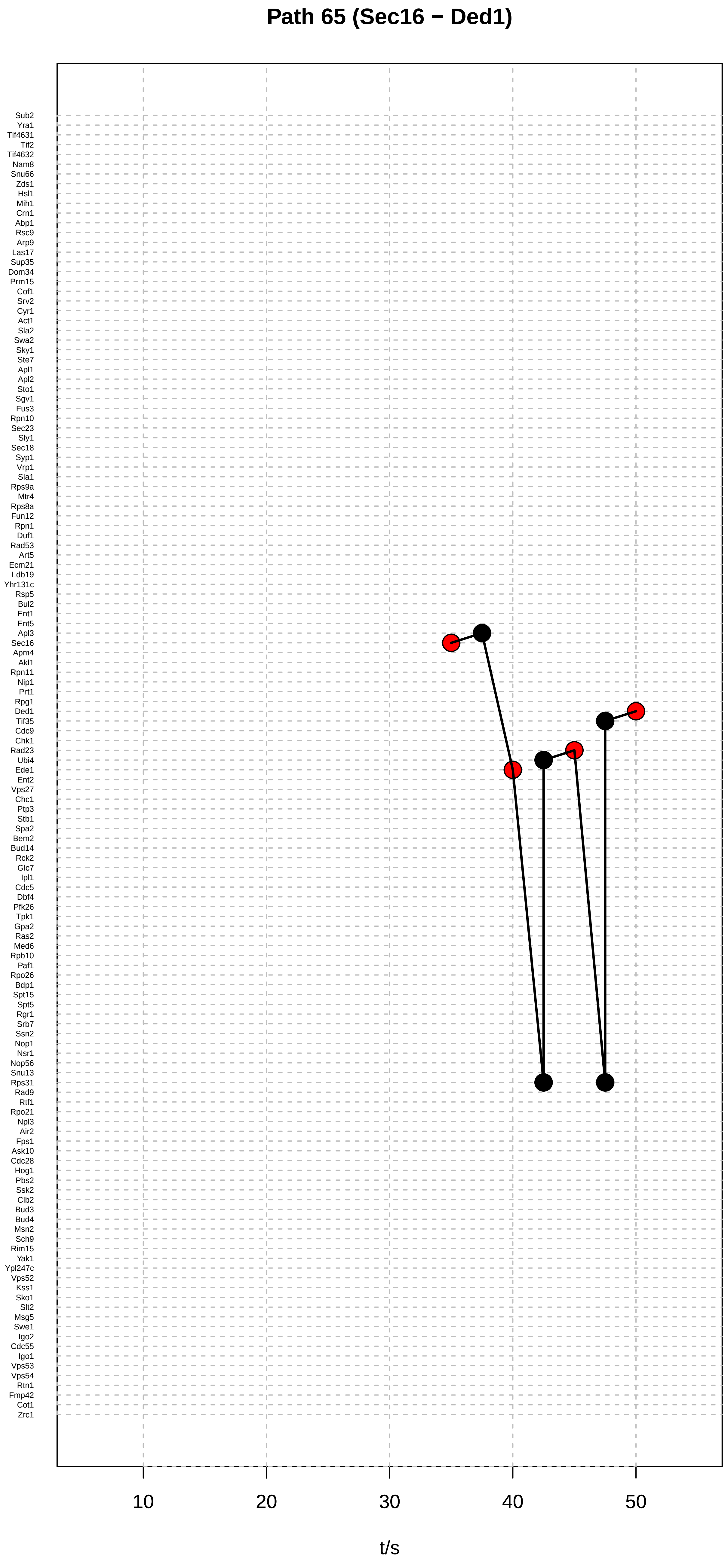
Path 65 (Sec16 − Ded1)
Sub2
Yra1
Tif4631
Tif2
Tif4632
Nam8
Snu66
Zds1
Hsl1
Mih1
Crn1
Abp1
Rsc9
Arp9
Las17
Sup35
Dom34
Prm15
Cof1
Srv2
Cyr1
Act1
Sla2
Swa2
Sky1
Ste7
Apl1
Apl2
Sto1
Sgv1
Fus3
Rpn10
Sec23
Sly1
Sec18
Syp1
Vrp1
Sla1
Rps9a
Mtr4
Rps8a
Fun12
Rpn1
Duf1
Rad53
Art5
Ecm21
Ldb19
Yhr131c
Rsp5
Bul2
Ent1
Ent5
Apl3
Sec16
Apm4
Akl1
Rpn11
Nip1
Prt1
Rpg1
Ded1
Tif35
Cdc9
Chk1
Rad23
Ubi4
Ede1
Ent2
Vps27
Chc1
Ptp3
Stb1
Spa2
Bem2
Bud14
Rck2
Glc7
Ipl1
Cdc5
Dbf4
Pfk26
Tpk1
Gpa2
Ras2
Med6
Rpb10
Paf1
Rpo26
Bdp1
Spt15
Spt5
Rgr1
Srb7
Ssn2
Nop1
Nsr1
Nop56
Snu13
Rps31
Rad9
Rtf1
Rpo21
Npl3
Air2
Fps1
Ask10
Cdc28
Hog1
Pbs2
Ssk2
Clb2
Bud3
Bud4
Msn2
Sch9
Rim15
Yak1
Ypl247c
Vps52
Kss1
Sko1
Slt2
Msg5
Swe1
Igo2
Cdc55
Igo1
Vps53
Vps54
Rtn1
Fmp42
Cot1
Zrc1
10
20
30
40
50
t/s

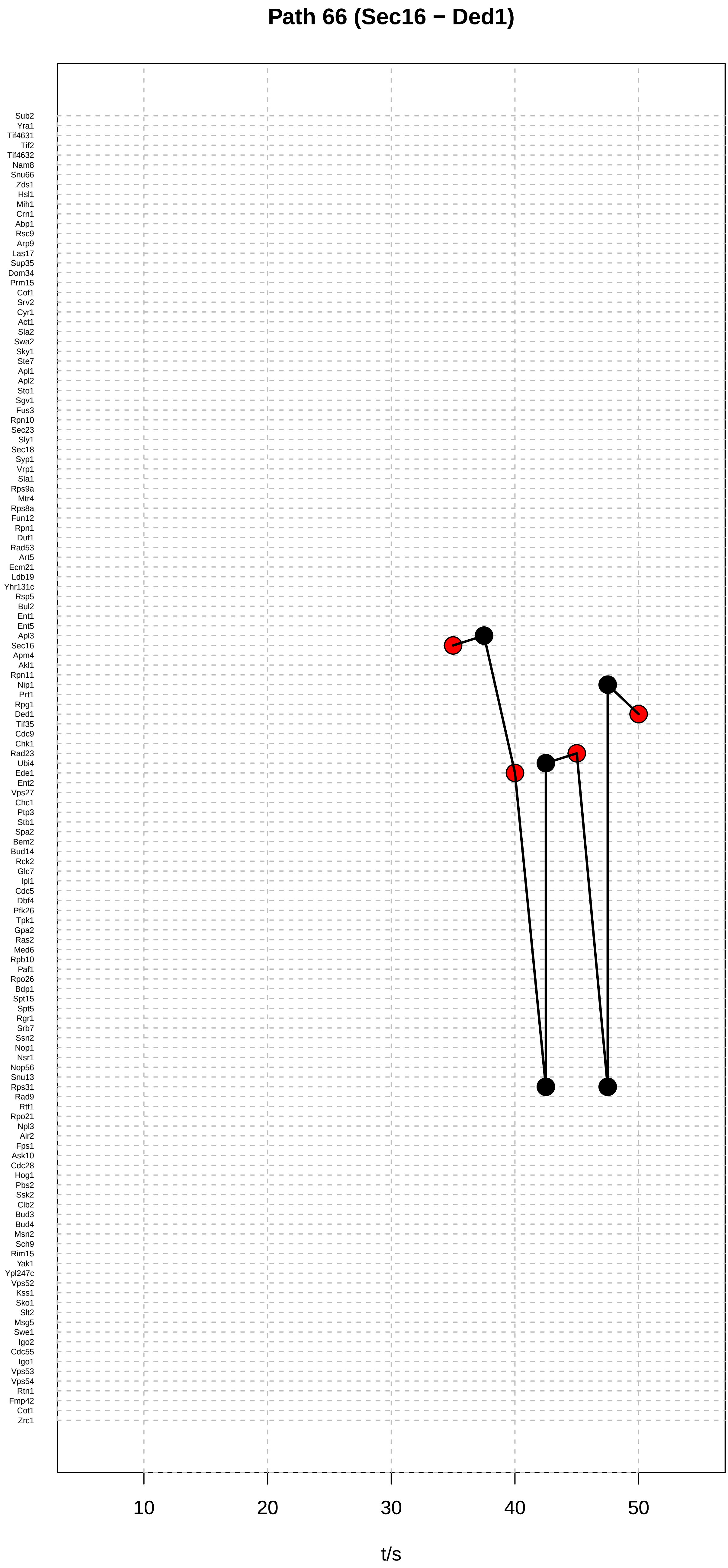

Path 66 (Sec16 − Ded1)
Sub2
Yra1
Tif4631
Tif2
Tif4632
Nam8
Snu66
Zds1
Hsl1
Mih1
Crn1
Abp1
Rsc9
Arp9
Las17
Sup35
Dom34
Prm15
Cof1
Srv2
Cyr1
Act1
Sla2
Swa2
Sky1
Ste7
Apl1
Apl2
Sto1
Sgv1
Fus3
Rpn10
Sec23
Sly1
Sec18
Syp1
Vrp1
Sla1
Rps9a
Mtr4
Rps8a
Fun12
Rpn1
Duf1
Rad53
Art5
Ecm21
Ldb19
Yhr131c
Rsp5
Bul2
Ent1
Ent5
Apl3
Sec16
Apm4
Akl1
Rpn11
Nip1
Prt1
Rpg1
Ded1
Tif35
Cdc9
Chk1
Rad23
Ubi4
Ede1
Ent2
Vps27
Chc1
Ptp3
Stb1
Spa2
Bem2
Bud14
Rck2
Glc7
Ipl1
Cdc5
Dbf4
Pfk26
Tpk1
Gpa2
Ras2
Med6
Rpb10
Paf1
Rpo26
Bdp1
Spt15
Spt5
Rgr1
Srb7
Ssn2
Nop1
Nsr1
Nop56
Snu13
Rps31
Rad9
Rtf1
Rpo21
Npl3
Air2
Fps1
Ask10
Cdc28
Hog1
Pbs2
Ssk2
Clb2
Bud3
Bud4
Msn2
Sch9
Rim15
Yak1
Ypl247c
Vps52
Kss1
Sko1
Slt2
Msg5
Swe1
Igo2
Cdc55
Igo1
Vps53
Vps54
Rtn1
Fmp42
Cot1
Zrc1
10
20
30
40
50
t/s

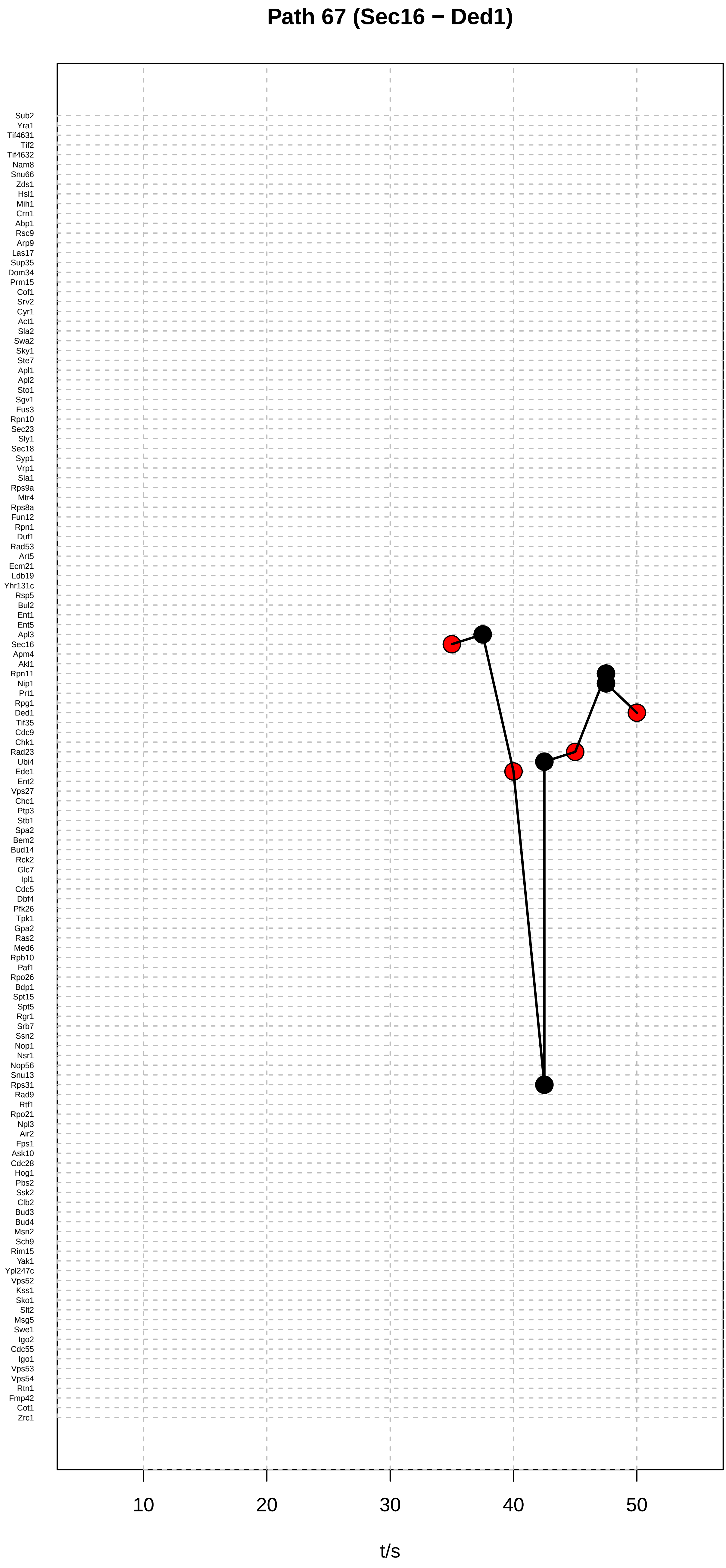
Path 67 (Sec16 − Ded1)
Sub2
Yra1
Tif4631
Tif2
Tif4632
Nam8
Snu66
Zds1
Hsl1
Mih1
Crn1
Abp1
Rsc9
Arp9
Las17
Sup35
Dom34
Prm15
Cof1
Srv2
Cyr1
Act1
Sla2
Swa2
Sky1
Ste7
Apl1
Apl2
Sto1
Sgv1
Fus3
Rpn10
Sec23
Sly1
Sec18
Syp1
Vrp1
Sla1
Rps9a
Mtr4
Rps8a
Fun12
Rpn1
Duf1
Rad53
Art5
Ecm21
Ldb19
Yhr131c
Rsp5
Bul2
Ent1
Ent5
Apl3
Sec16
Apm4
Akl1
Rpn11
Nip1
Prt1
Rpg1
Ded1
Tif35
Cdc9
Chk1
Rad23
Ubi4
Ede1
Ent2
Vps27
Chc1
Ptp3
Stb1
Spa2
Bem2
Bud14
Rck2
Glc7
Ipl1
Cdc5
Dbf4
Pfk26
Tpk1
Gpa2
Ras2
Med6
Rpb10
Paf1
Rpo26
Bdp1
Spt15
Spt5
Rgr1
Srb7
Ssn2
Nop1
Nsr1
Nop56
Snu13
Rps31
Rad9
Rtf1
Rpo21
Npl3
Air2
Fps1
Ask10
Cdc28
Hog1
Pbs2
Ssk2
Clb2
Bud3
Bud4
Msn2
Sch9
Rim15
Yak1
Ypl247c
Vps52
Kss1
Sko1
Slt2
Msg5
Swe1
Igo2
Cdc55
Igo1
Vps53
Vps54
Rtn1
Fmp42
Cot1
Zrc1
10
20
30
40
50
t/s

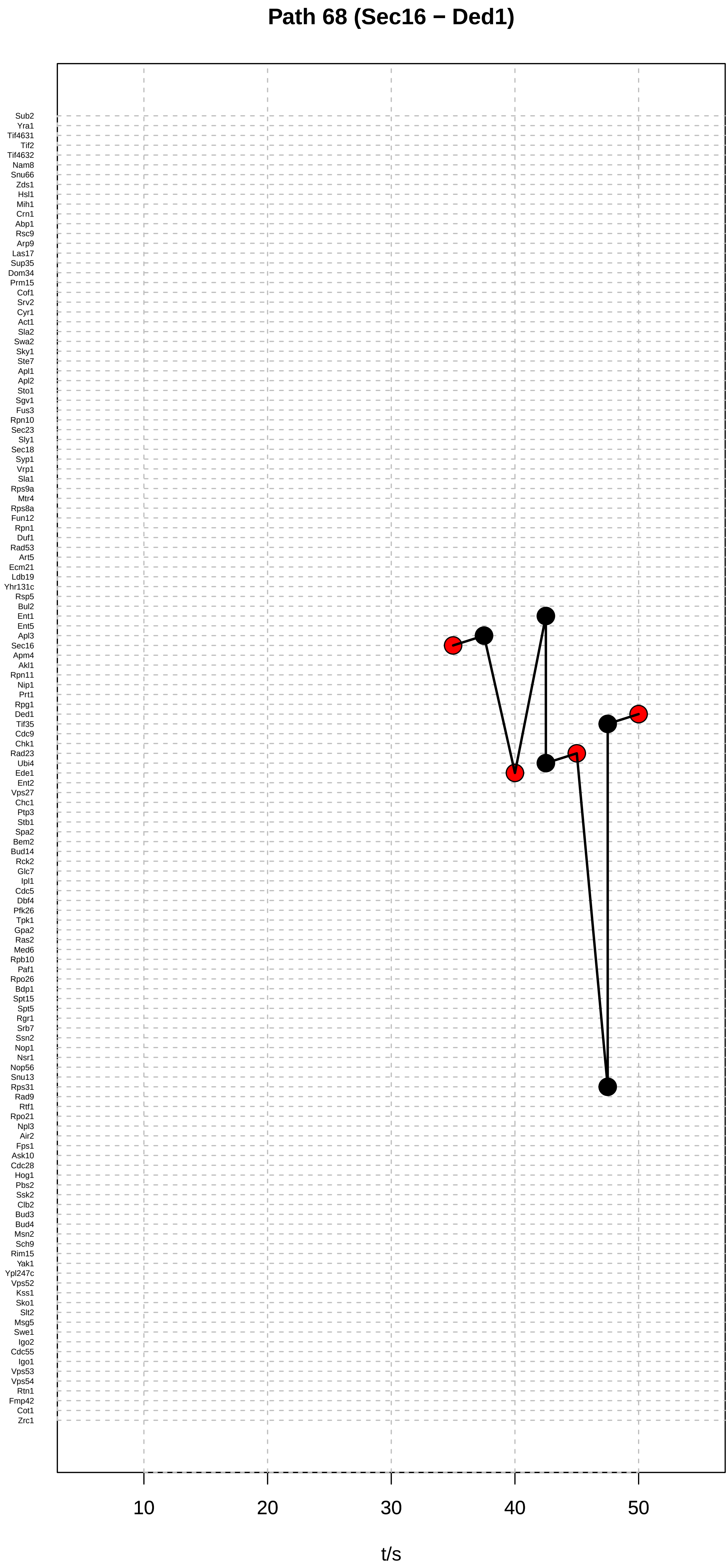

Path 68 (Sec16 − Ded1)
Sub2
Yra1
Tif4631
Tif2
Tif4632
Nam8
Snu66
Zds1
Hsl1
Mih1
Crn1
Abp1
Rsc9
Arp9
Las17
Sup35
Dom34
Prm15
Cof1
Srv2
Cyr1
Act1
Sla2
Swa2
Sky1
Ste7
Apl1
Apl2
Sto1
Sgv1
Fus3
Rpn10
Sec23
Sly1
Sec18
Syp1
Vrp1
Sla1
Rps9a
Mtr4
Rps8a
Fun12
Rpn1
Duf1
Rad53
Art5
Ecm21
Ldb19
Yhr131c
Rsp5
Bul2
Ent1
Ent5
Apl3
Sec16
Apm4
Akl1
Rpn11
Nip1
Prt1
Rpg1
Ded1
Tif35
Cdc9
Chk1
Rad23
Ubi4
Ede1
Ent2
Vps27
Chc1
Ptp3
Stb1
Spa2
Bem2
Bud14
Rck2
Glc7
Ipl1
Cdc5
Dbf4
Pfk26
Tpk1
Gpa2
Ras2
Med6
Rpb10
Paf1
Rpo26
Bdp1
Spt15
Spt5
Rgr1
Srb7
Ssn2
Nop1
Nsr1
Nop56
Snu13
Rps31
Rad9
Rtf1
Rpo21
Npl3
Air2
Fps1
Ask10
Cdc28
Hog1
Pbs2
Ssk2
Clb2
Bud3
Bud4
Msn2
Sch9
Rim15
Yak1
Ypl247c
Vps52
Kss1
Sko1
Slt2
Msg5
Swe1
Igo2
Cdc55
Igo1
Vps53
Vps54
Rtn1
Fmp42
Cot1
Zrc1
10
20
30
40
50
t/s

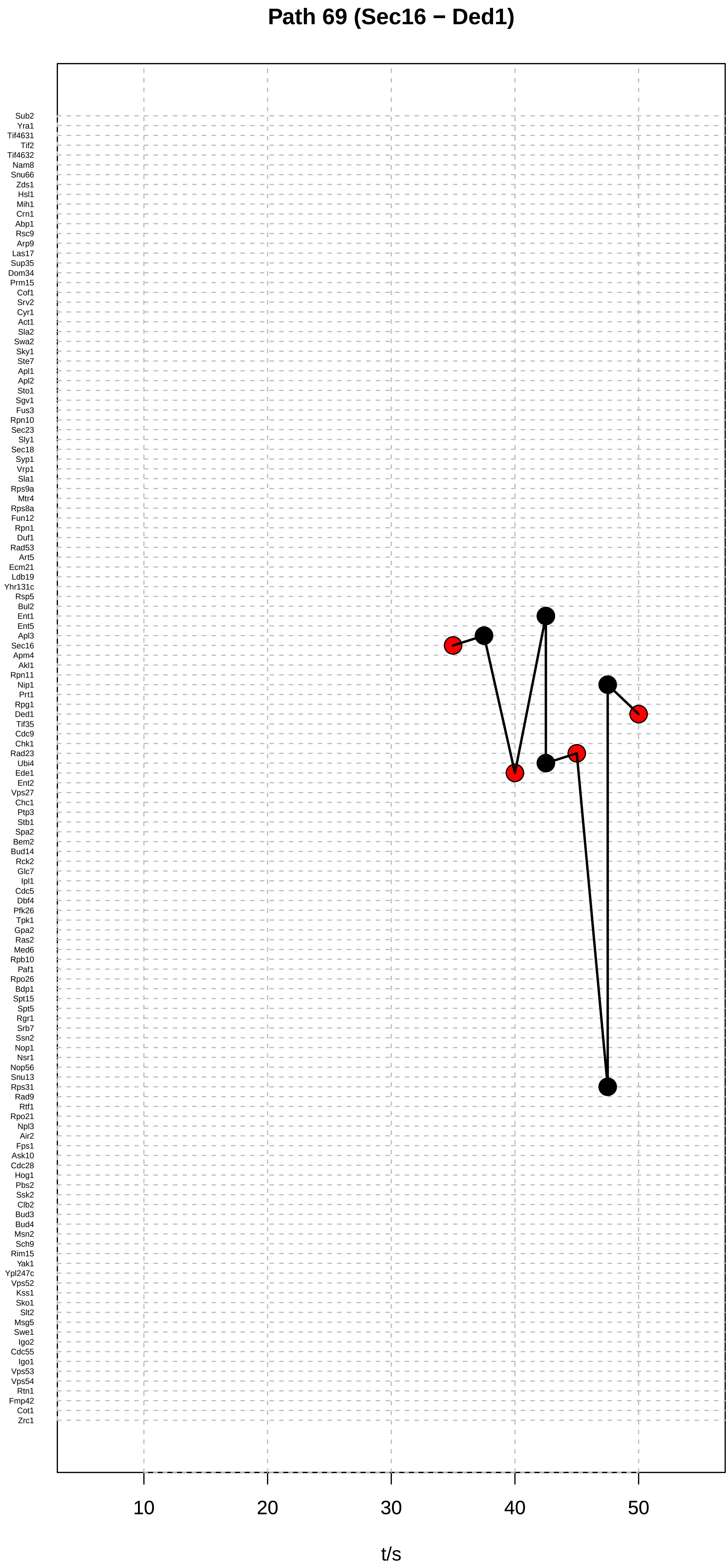
Path 69 (Sec16 − Ded1)
Sub2
Yra1
Tif4631
Tif2
Tif4632
Nam8
Snu66
Zds1
Hsl1
Mih1
Crn1
Abp1
Rsc9
Arp9
Las17
Sup35
Dom34
Prm15
Cof1
Srv2
Cyr1
Act1
Sla2
Swa2
Sky1
Ste7
Apl1
Apl2
Sto1
Sgv1
Fus3
Rpn10
Sec23
Sly1
Sec18
Syp1
Vrp1
Sla1
Rps9a
Mtr4
Rps8a
Fun12
Rpn1
Duf1
Rad53
Art5
Ecm21
Ldb19
Yhr131c
Rsp5
Bul2
Ent1
Ent5
Apl3
Sec16
Apm4
Akl1
Rpn11
Nip1
Prt1
Rpg1
Ded1
Tif35
Cdc9
Chk1
Rad23
Ubi4
Ede1
Ent2
Vps27
Chc1
Ptp3
Stb1
Spa2
Bem2
Bud14
Rck2
Glc7
Ipl1
Cdc5
Dbf4
Pfk26
Tpk1
Gpa2
Ras2
Med6
Rpb10
Paf1
Rpo26
Bdp1
Spt15
Spt5
Rgr1
Srb7
Ssn2
Nop1
Nsr1
Nop56
Snu13
Rps31
Rad9
Rtf1
Rpo21
Npl3
Air2
Fps1
Ask10
Cdc28
Hog1
Pbs2
Ssk2
Clb2
Bud3
Bud4
Msn2
Sch9
Rim15
Yak1
Ypl247c
Vps52
Kss1
Sko1
Slt2
Msg5
Swe1
Igo2
Cdc55
Igo1
Vps53
Vps54
Rtn1
Fmp42
Cot1
Zrc1
10
20
30
40
50
t/s

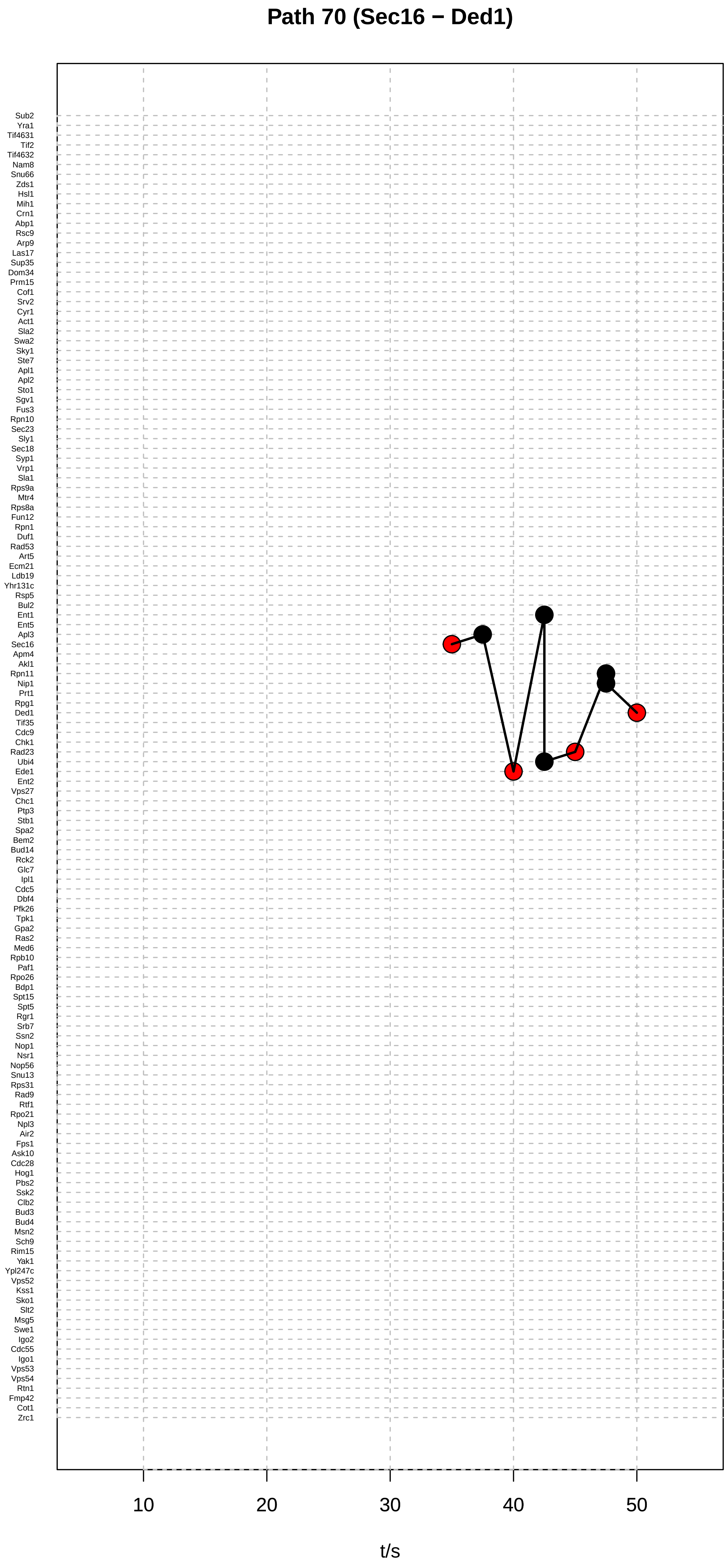
Path 70 (Sec16 − Ded1)
Sub2
Yra1
Tif4631
Tif2
Tif4632
Nam8
Snu66
Zds1
Hsl1
Mih1
Crn1
Abp1
Rsc9
Arp9
Las17
Sup35
Dom34
Prm15
Cof1
Srv2
Cyr1
Act1
Sla2
Swa2
Sky1
Ste7
Apl1
Apl2
Sto1
Sgv1
Fus3
Rpn10
Sec23
Sly1
Sec18
Syp1
Vrp1
Sla1
Rps9a
Mtr4
Rps8a
Fun12
Rpn1
Duf1
Rad53
Art5
Ecm21
Ldb19
Yhr131c
Rsp5
Bul2
Ent1
Ent5
Apl3
Sec16
Apm4
Akl1
Rpn11
Nip1
Prt1
Rpg1
Ded1
Tif35
Cdc9
Chk1
Rad23
Ubi4
Ede1
Ent2
Vps27
Chc1
Ptp3
Stb1
Spa2
Bem2
Bud14
Rck2
Glc7
Ipl1
Cdc5
Dbf4
Pfk26
Tpk1
Gpa2
Ras2
Med6
Rpb10
Paf1
Rpo26
Bdp1
Spt15
Spt5
Rgr1
Srb7
Ssn2
Nop1
Nsr1
Nop56
Snu13
Rps31
Rad9
Rtf1
Rpo21
Npl3
Air2
Fps1
Ask10
Cdc28
Hog1
Pbs2
Ssk2
Clb2
Bud3
Bud4
Msn2
Sch9
Rim15
Yak1
Ypl247c
Vps52
Kss1
Sko1
Slt2
Msg5
Swe1
Igo2
Cdc55
Igo1
Vps53
Vps54
Rtn1
Fmp42
Cot1
Zrc1
10
20
30
40
50
t/s

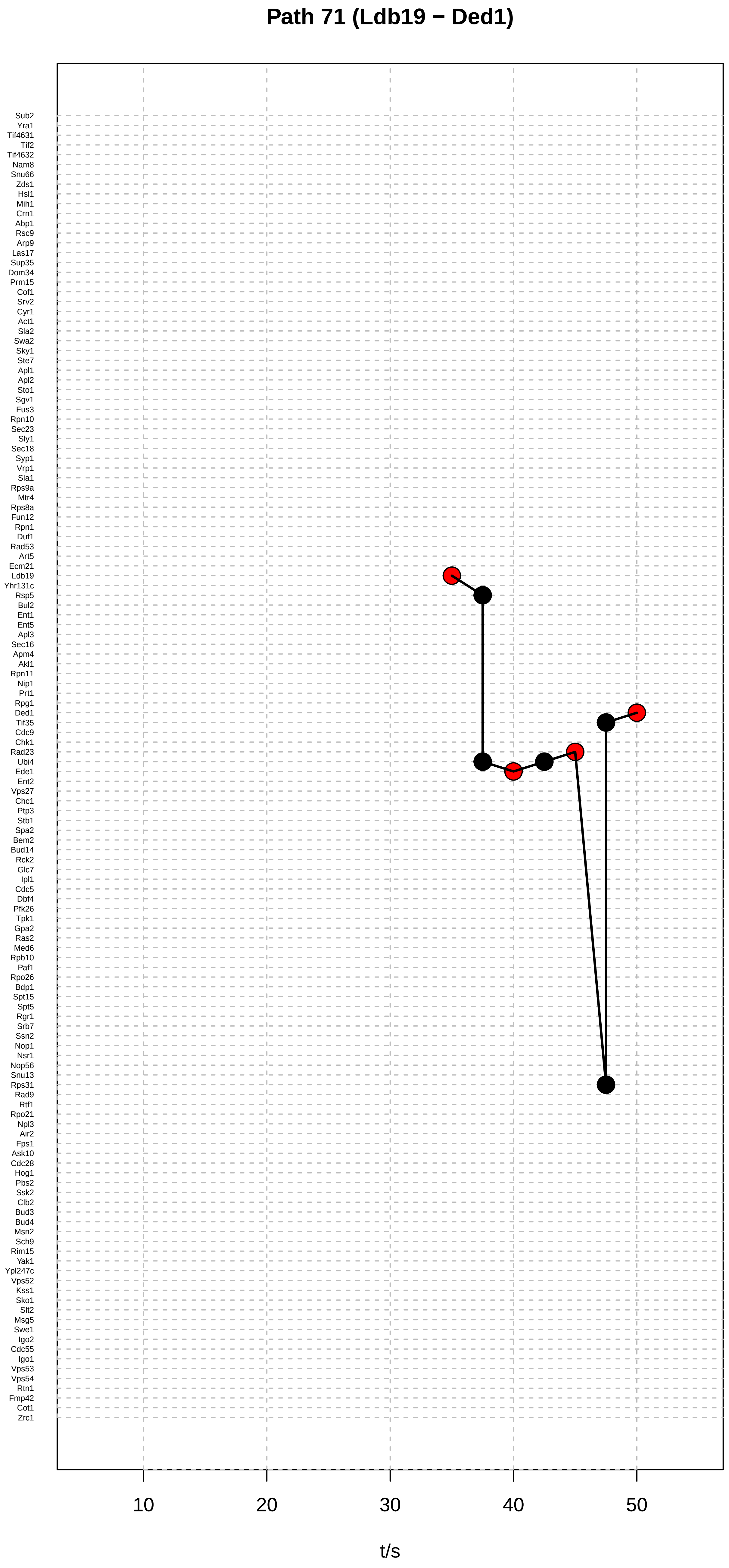
Path 71 (Ldb19 − Ded1)
Sub2
Yra1
Tif4631
Tif2
Tif4632
Nam8
Snu66
Zds1
Hsl1
Mih1
Crn1
Abp1
Rsc9
Arp9
Las17
Sup35
Dom34
Prm15
Cof1
Srv2
Cyr1
Act1
Sla2
Swa2
Sky1
Ste7
Apl1
Apl2
Sto1
Sgv1
Fus3
Rpn10
Sec23
Sly1
Sec18
Syp1
Vrp1
Sla1
Rps9a
Mtr4
Rps8a
Fun12
Rpn1
Duf1
Rad53
Art5
Ecm21
Ldb19
Yhr131c
Rsp5
Bul2
Ent1
Ent5
Apl3
Sec16
Apm4
Akl1
Rpn11
Nip1
Prt1
Rpg1
Ded1
Tif35
Cdc9
Chk1
Rad23
Ubi4
Ede1
Ent2
Vps27
Chc1
Ptp3
Stb1
Spa2
Bem2
Bud14
Rck2
Glc7
Ipl1
Cdc5
Dbf4
Pfk26
Tpk1
Gpa2
Ras2
Med6
Rpb10
Paf1
Rpo26
Bdp1
Spt15
Spt5
Rgr1
Srb7
Ssn2
Nop1
Nsr1
Nop56
Snu13
Rps31
Rad9
Rtf1
Rpo21
Npl3
Air2
Fps1
Ask10
Cdc28
Hog1
Pbs2
Ssk2
Clb2
Bud3
Bud4
Msn2
Sch9
Rim15
Yak1
Ypl247c
Vps52
Kss1
Sko1
Slt2
Msg5
Swe1
Igo2
Cdc55
Igo1
Vps53
Vps54
Rtn1
Fmp42
Cot1
Zrc1
10
20
30
40
50
t/s

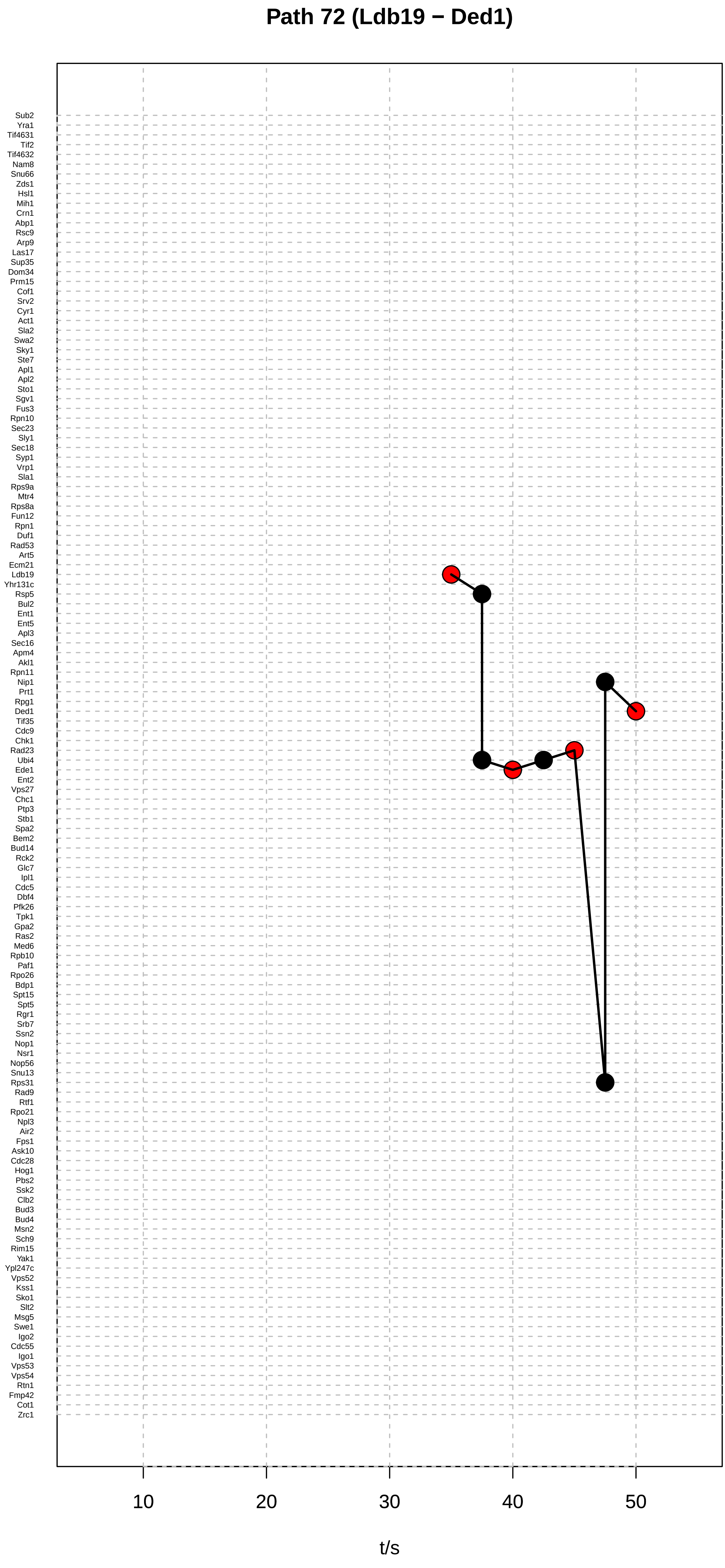

Path 72 (Ldb19 − Ded1)
Sub2
Yra1
Tif4631
Tif2
Tif4632
Nam8
Snu66
Zds1
Hsl1
Mih1
Crn1
Abp1
Rsc9
Arp9
Las17
Sup35
Dom34
Prm15
Cof1
Srv2
Cyr1
Act1
Sla2
Swa2
Sky1
Ste7
Apl1
Apl2
Sto1
Sgv1
Fus3
Rpn10
Sec23
Sly1
Sec18
Syp1
Vrp1
Sla1
Rps9a
Mtr4
Rps8a
Fun12
Rpn1
Duf1
Rad53
Art5
Ecm21
Ldb19
Yhr131c
Rsp5
Bul2
Ent1
Ent5
Apl3
Sec16
Apm4
Akl1
Rpn11
Nip1
Prt1
Rpg1
Ded1
Tif35
Cdc9
Chk1
Rad23
Ubi4
Ede1
Ent2
Vps27
Chc1
Ptp3
Stb1
Spa2
Bem2
Bud14
Rck2
Glc7
Ipl1
Cdc5
Dbf4
Pfk26
Tpk1
Gpa2
Ras2
Med6
Rpb10
Paf1
Rpo26
Bdp1
Spt15
Spt5
Rgr1
Srb7
Ssn2
Nop1
Nsr1
Nop56
Snu13
Rps31
Rad9
Rtf1
Rpo21
Npl3
Air2
Fps1
Ask10
Cdc28
Hog1
Pbs2
Ssk2
Clb2
Bud3
Bud4
Msn2
Sch9
Rim15
Yak1
Ypl247c
Vps52
Kss1
Sko1
Slt2
Msg5
Swe1
Igo2
Cdc55
Igo1
Vps53
Vps54
Rtn1
Fmp42
Cot1
Zrc1
10
20
30
40
50
t/s

### Path 73 (Ldb19 − Ded1)

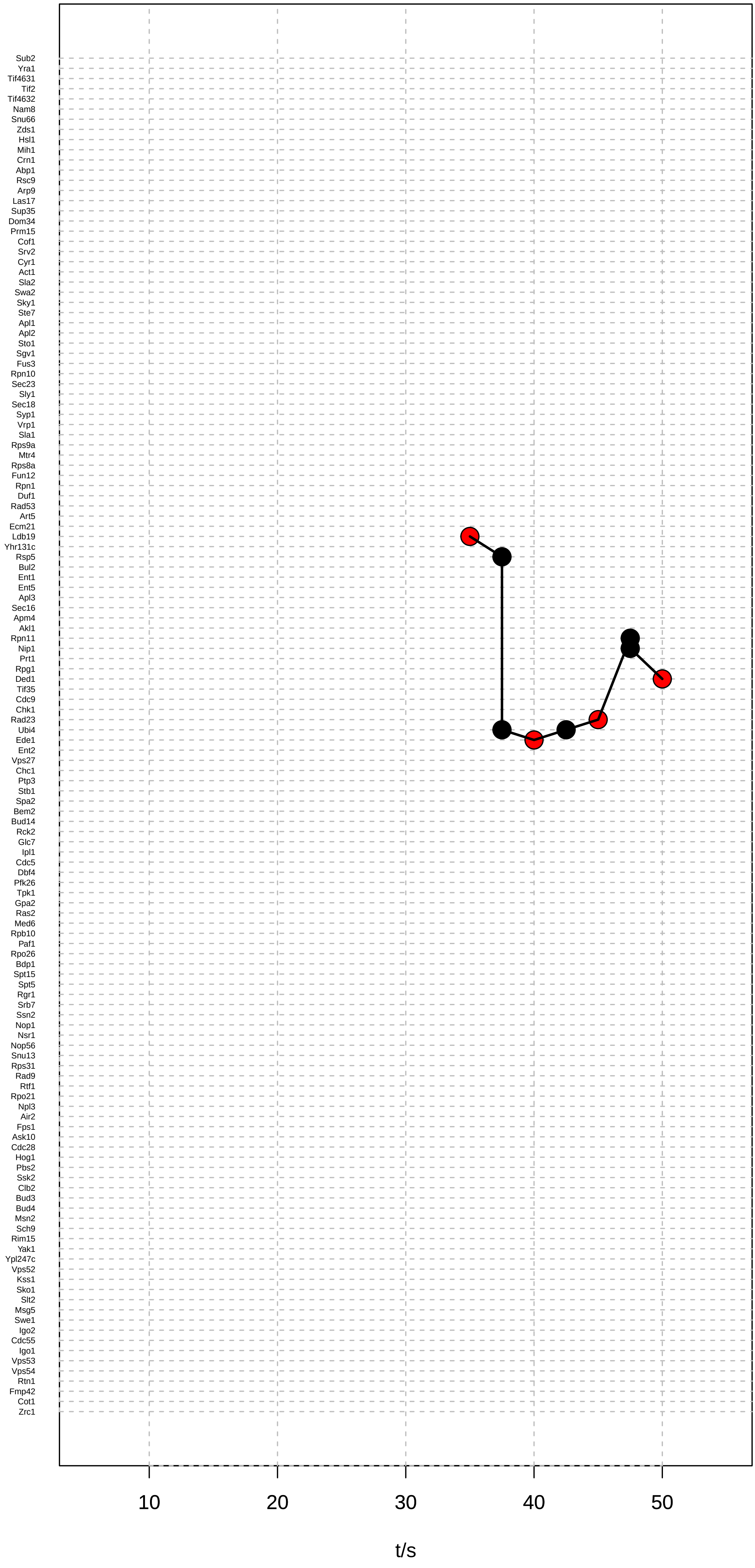

#### Path 74 (Ldb19 − Ded1)

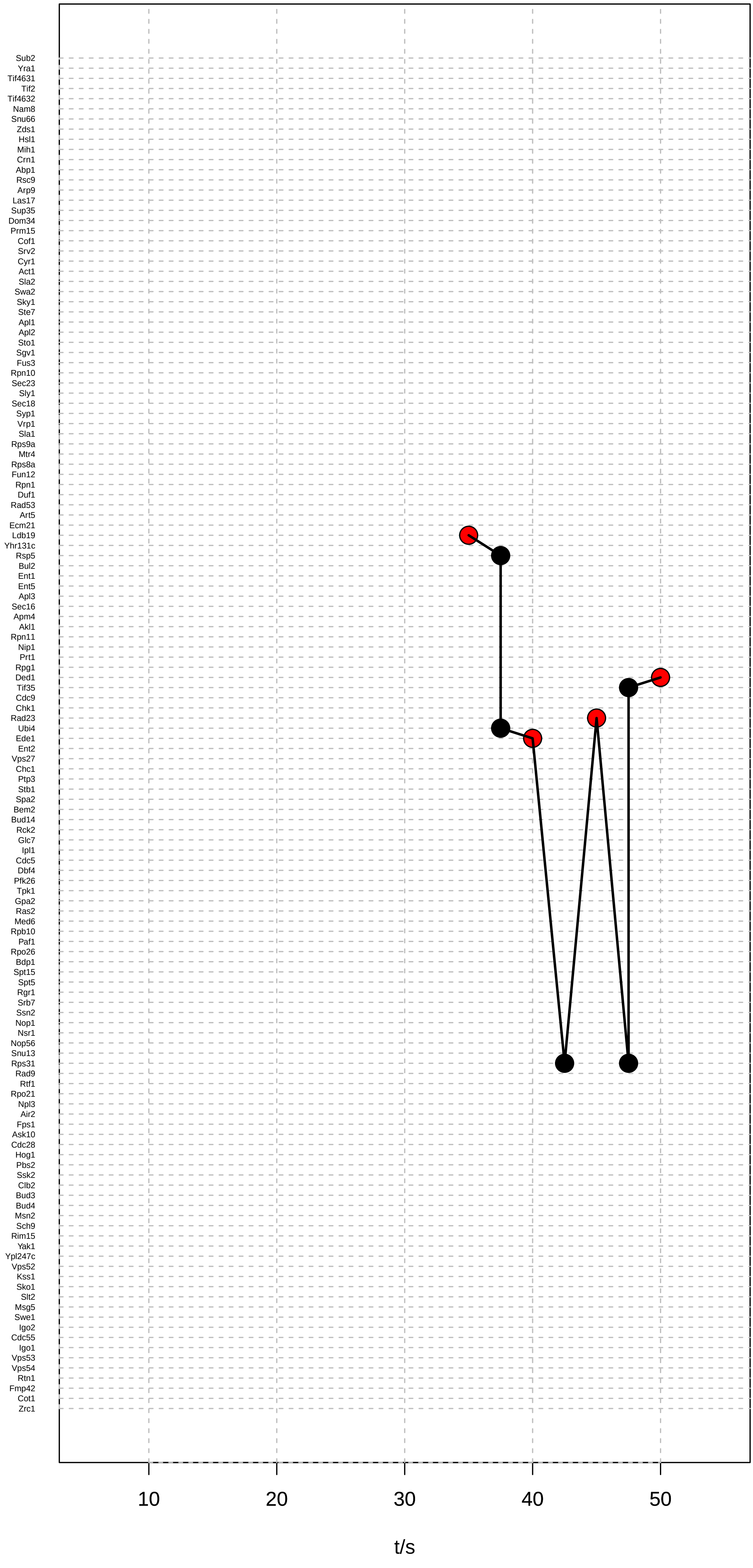

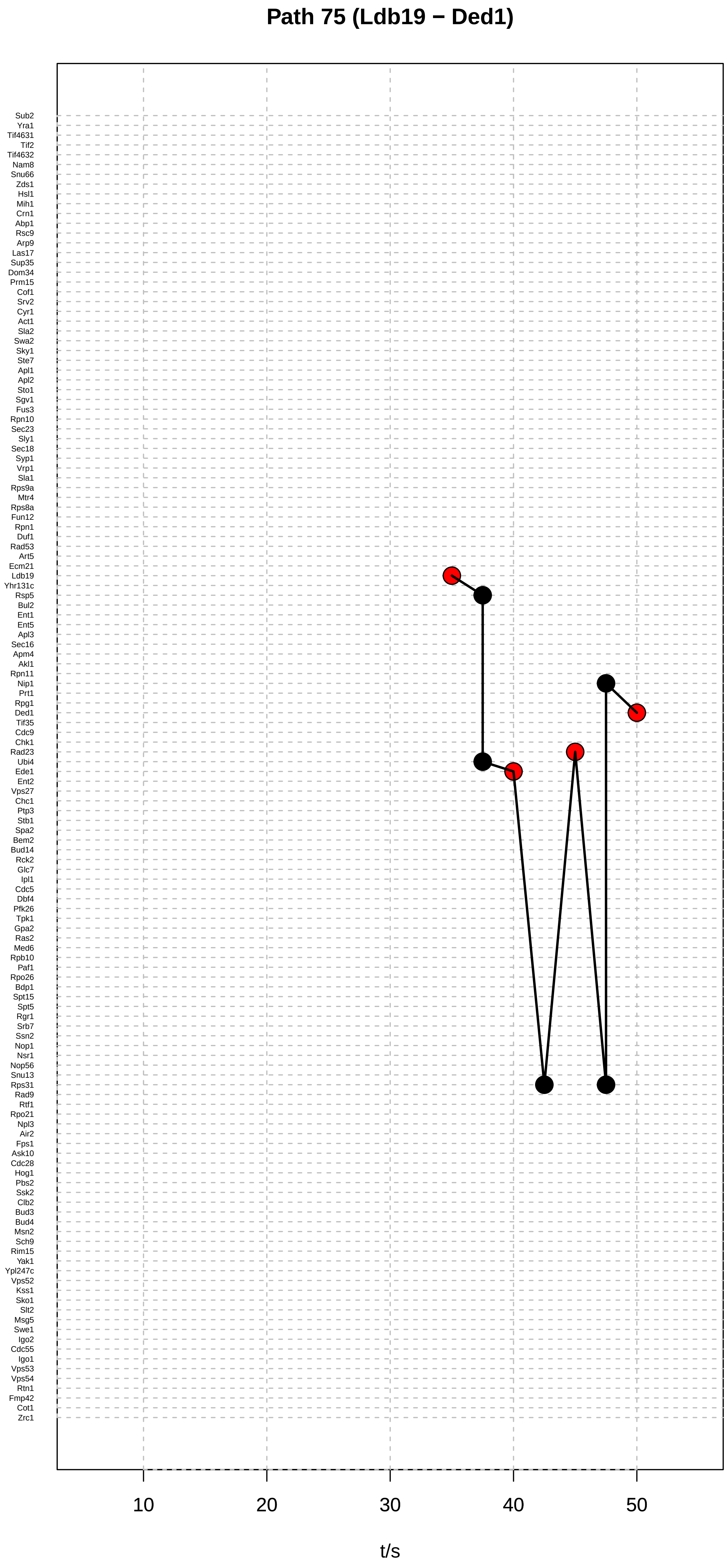

Path 75 (Ldb19 – Ded1)
Sub2
Yra1
Tif4631
Tif2
Tif4632
Nam8
Snu66
Zds1
Hsl1
Mih1
Crn1
Abp1
Rsc9
Arp9
Las17
Sup35
Dom34
Prm15
Cof1
Srv2
Cyr1
Act1
Sla2
Swa2
Sky1
Ste7
Apl1
Apl2
Sto1
Sgv1
Fus3
Rpn10
Sec23
Sly1
Sec18
Syp1
Vrp1
Sla1
Rps9a
Mtr4
Rps8a
Fun12
Rpn1
Duf1
Rad53
Art5
Ecm21
Ldb19
Yhr131c
Rsp5
Bul2
Ent1
Ent5
Apl3
Sec16
Apm4
Akl1
Rpn11
Nip1
Prt1
Rpg1
Ded1
Tif35
Cdc9
Chk1
Rad23
Ubi4
Ede1
Ent2
Vps27
Chc1
Ptp3
Stb1
Spa2
Bem2
Bud14
Rck2
Glc7
Ipl1
Cdc5
Dbf4
Pfk26
Tpk1
Gpa2
Ras2
Med6
Rpb10
Paf1
Rpo26
Bdp1
Spt15
Spt5
Rgr1
Srb7
Ssn2
Nop1
Nsr1
Nop56
Snu13
Rps31
Rad9
Rtf1
Rpo21
Npl3
Air2
Fps1
Ask10
Cdc28
Hog1
Pbs2
Ssk2
Clb2
Bud3
Bud4
Msn2
Sch9
Rim15
Yak1
Ypl247c
Vps52
Kss1
Sko1
Slt2
Msg5
Swe1
Igo2
Cdc55
Igo1
Vps53
Vps54
Rtn1
Fmp42
Cot1
Zrc1
10
20
30
40
50
t/s

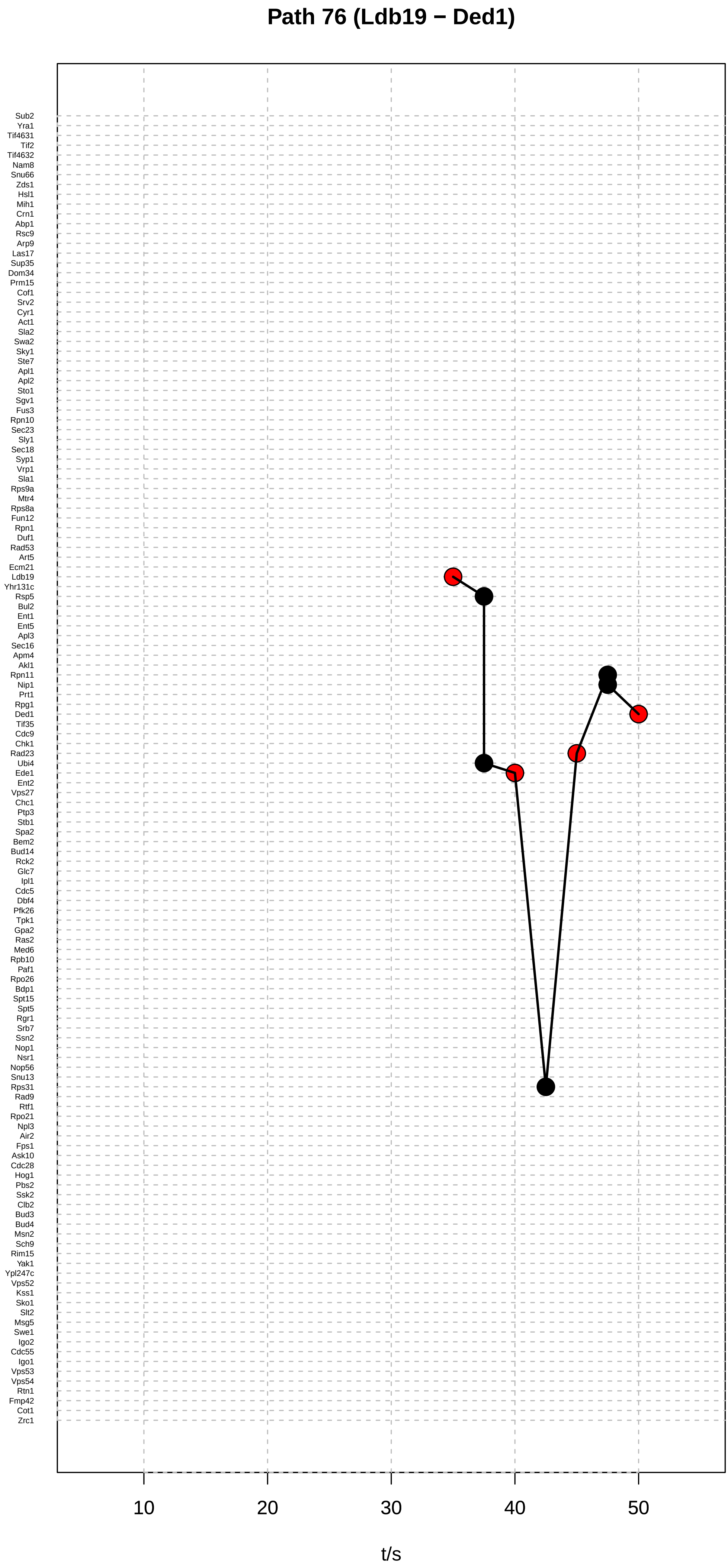

Path 76 (Ldb19 − Ded1)
Sub2
Yra1
Tif4631
Tif2
Tif4632
Nam8
Snu66
Zds1
Hsl1
Mih1
Crn1
Abp1
Rsc9
Arp9
Las17
Sup35
Dom34
Prm15
Cof1
Srv2
Cyr1
Act1
Sla2
Swa2
Sky1
Ste7
Apl1
Apl2
Sto1
Sgv1
Fus3
Rpn10
Sec23
Sly1
Sec18
Syp1
Vrp1
Sla1
Rps9a
Mtr4
Rps8a
Fun12
Rpn1
Duf1
Rad53
Art5
Ecm21
Ldb19
Yhr131c
Rsp5
Bul2
Ent1
Ent5
Apl3
Sec16
Apm4
Akl1
Rpn11
Nip1
Prt1
Rpg1
Ded1
Tif35
Cdc9
Chk1
Rad23
Ubi4
Ede1
Ent2
Vps27
Chc1
Ptp3
Stb1
Spa2
Bem2
Bud14
Rck2
Glc7
Ipl1
Cdc5
Dbf4
Pfk26
Tpk1
Gpa2
Ras2
Med6
Rpb10
Paf1
Rpo26
Bdp1
Spt15
Spt5
Rgr1
Srb7
Ssn2
Nop1
Nsr1
Nop56
Snu13
Rps31
Rad9
Rtf1
Rpo21
Npl3
Air2
Fps1
Ask10
Cdc28
Hog1
Pbs2
Ssk2
Clb2
Bud3
Bud4
Msn2
Sch9
Rim15
Yak1
Ypl247c
Vps52
Kss1
Sko1
Slt2
Msg5
Swe1
Igo2
Cdc55
Igo1
Vps53
Vps54
Rtn1
Fmp42
Cot1
Zrc1
10
20
30
40
50
t/s

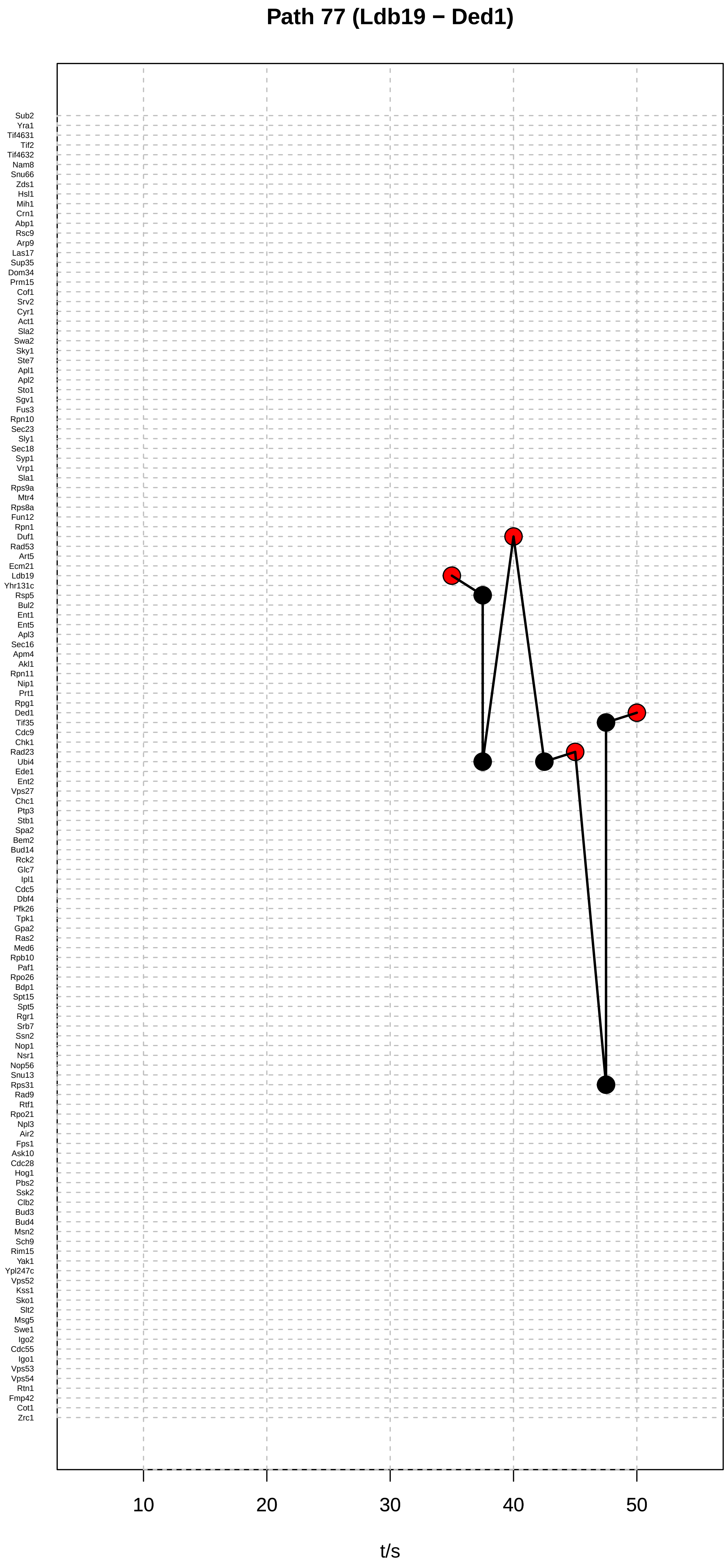
Path 77 (Ldb19 – Ded1)
Sub2
Yra1
Tif4631
Tif2
Tif4632
Nam8
Snu66
Zds1
Hsl1
Mih1
Crn1
Abp1
Rsc9
Arp9
Las17
Sup35
Dom34
Prm15
Cof1
Srv2
Cyr1
Act1
Sla2
Swa2
Sky1
Ste7
Apl1
Apl2
Sto1
Sgv1
Fus3
Rpn10
Sec23
Sly1
Sec18
Syp1
Vrp1
Sla1
Rps9a
Mtr4
Rps8a
Fun12
Rpn1
Duf1
Rad53
Art5
Ecm21
Ldb19
Yhr131c
Rsp5
Bul2
Ent1
Ent5
Apl3
Sec16
Apm4
Akl1
Rpn11
Nip1
Prt1
Rpg1
Ded1
Tif35
Cdc9
Chk1
Rad23
Ubi4
Ede1
Ent2
Vps27
Chc1
Ptp3
Stb1
Spa2
Bem2
Bud14
Rck2
Glc7
Ipl1
Cdc5
Dbf4
Pfk26
Tpk1
Gpa2
Ras2
Med6
Rpb10
Paf1
Rpo26
Bdp1
Spt15
Spt5
Rgr1
Srb7
Ssn2
Nop1
Nsr1
Nop56
Snu13
Rps31
Rad9
Rtf1
Rpo21
Npl3
Air2
Fps1
Ask10
Cdc28
Hog1
Pbs2
Ssk2
Clb2
Bud3
Bud4
Msn2
Sch9
Rim15
Yak1
Ypl247c
Vps52
Kss1
Sko1
Slt2
Msg5
Swe1
Igo2
Cdc55
Igo1
Vps53
Vps54
Rtn1
Fmp42
Cot1
Zrc1
10
20
30
40
50
t/s

#### Path 78 (Ldb19 − Ded1)

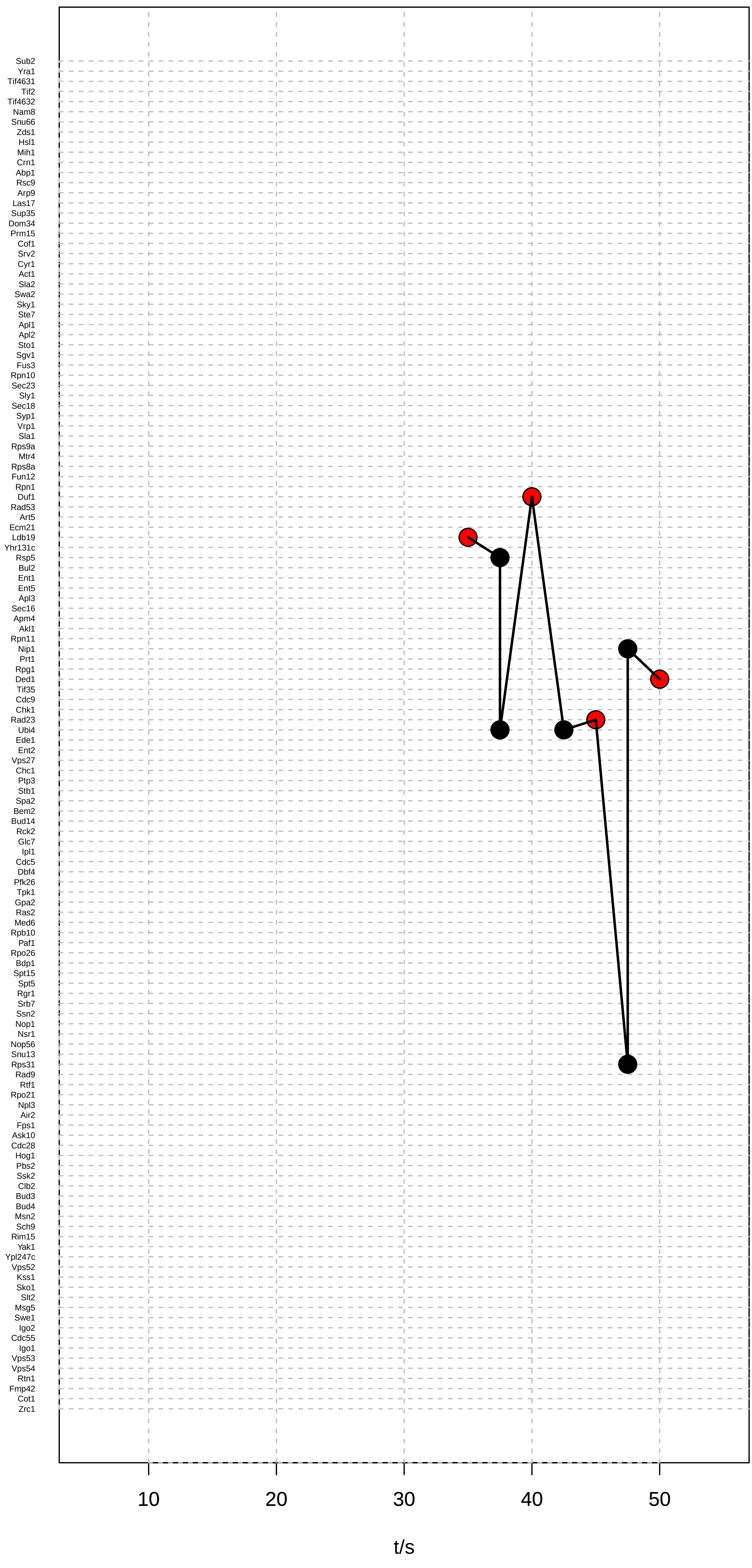

#### Path 79 (Ldb19 − Ded1)

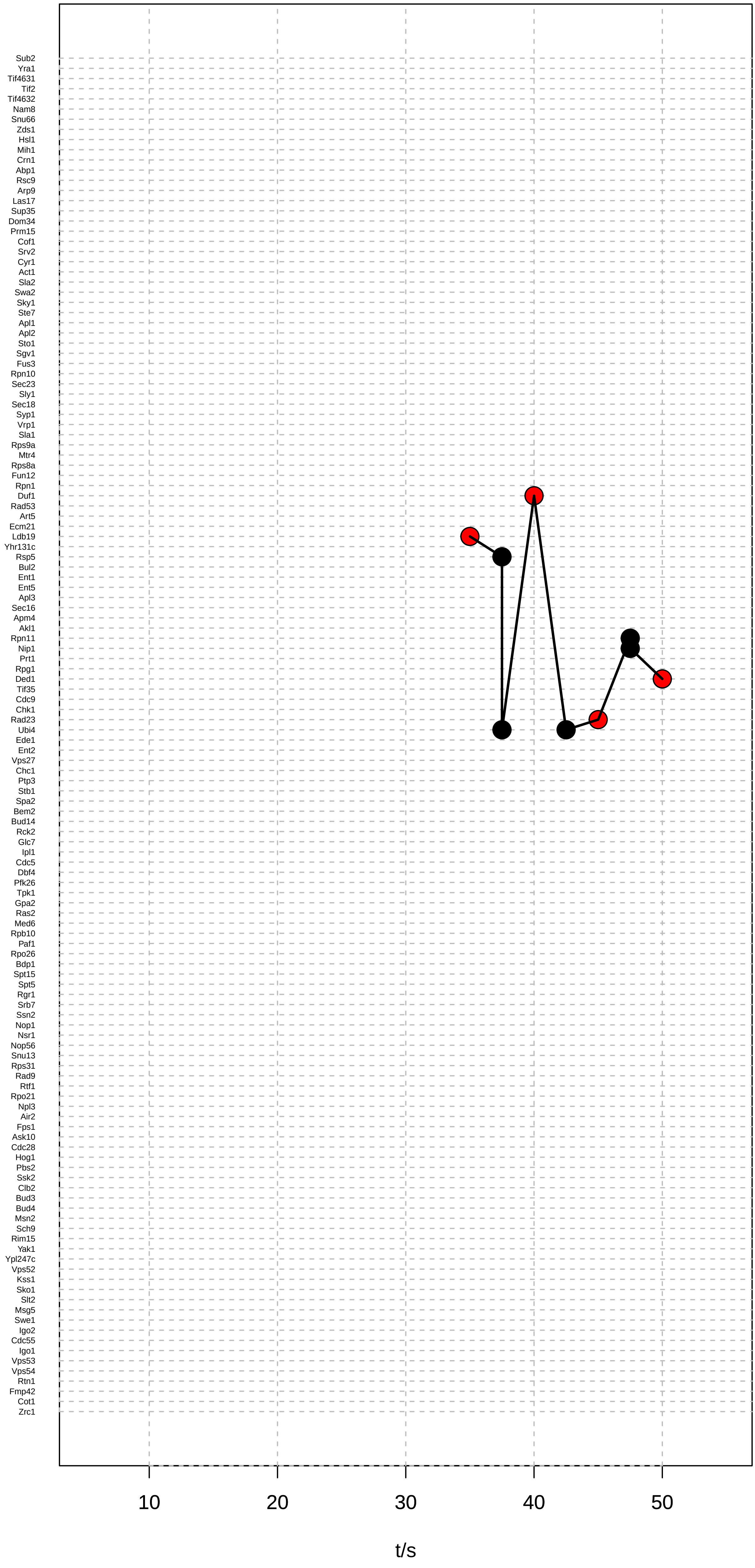

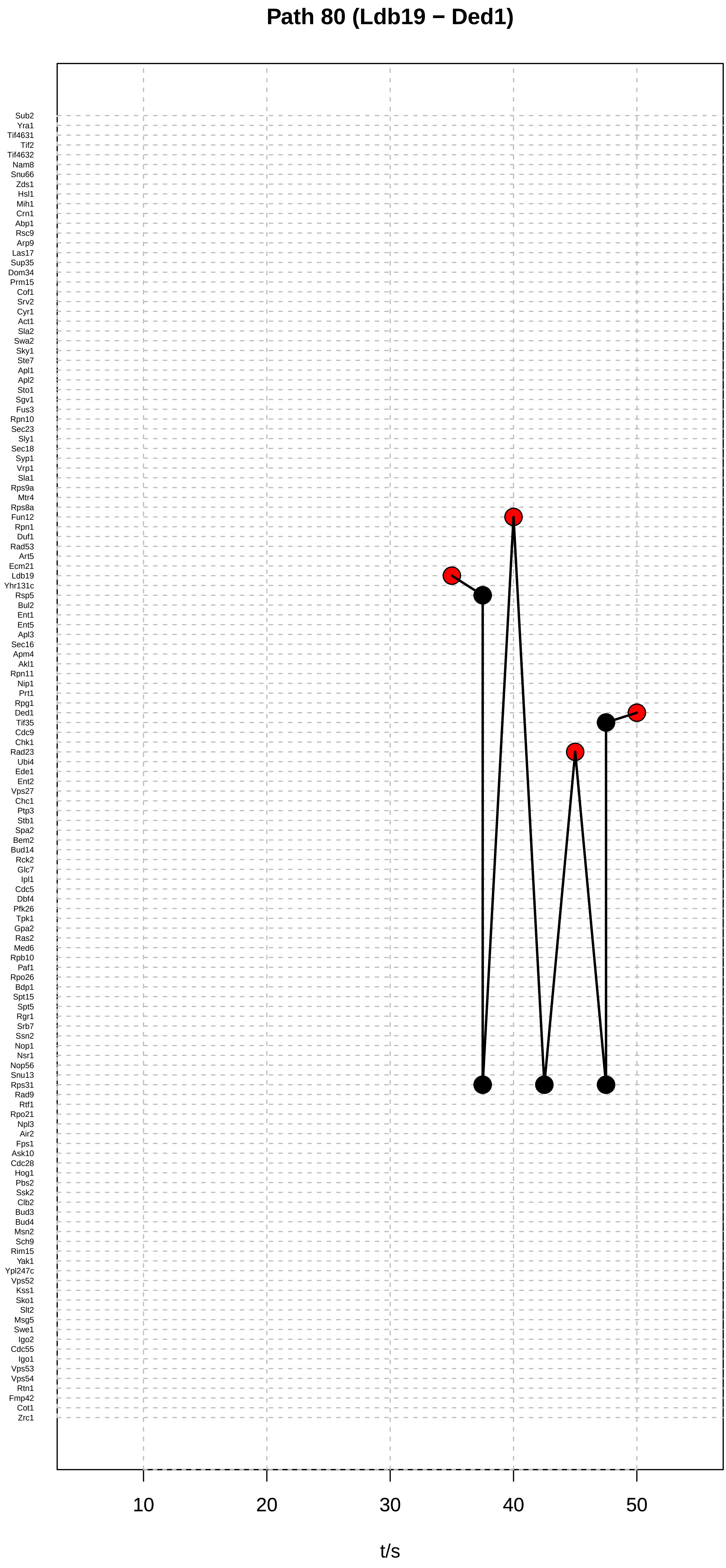

Path 80 (Ldb19 − Ded1)
Sub2
Yra1
Tif4631
Tif2
Tif4632
Nam8
Snu66
Zds1
Hsl1
Mih1
Crn1
Abp1
Rsc9
Arp9
Las17
Sup35
Dom34
Prm15
Cof1
Srv2
Cyr1
Act1
Sla2
Swa2
Sky1
Ste7
Apl1
Apl2
Sto1
Sgv1
Fus3
Rpn10
Sec23
Sly1
Sec18
Syp1
Vrp1
Sla1
Rps9a
Mtr4
Rps8a
Fun12
Rpn1
Duf1
Rad53
Art5
Ecm21
Ldb19
Yhr131c
Rsp5
Bul2
Ent1
Ent5
Apl3
Sec16
Apm4
Akl1
Rpn11
Nip1
Prt1
Rpg1
Ded1
Tif35
Cdc9
Chk1
Rad23
Ubi4
Ede1
Ent2
Vps27
Chc1
Ptp3
Stb1
Spa2
Bem2
Bud14
Rck2
Glc7
Ipl1
Cdc5
Dbf4
Pfk26
Tpk1
Gpa2
Ras2
Med6
Rpb10
Paf1
Rpo26
Bdp1
Spt15
Spt5
Rgr1
Srb7
Ssn2
Nop1
Nsr1
Nop56
Snu13
Rps31
Rad9
Rtf1
Rpo21
Npl3
Air2
Fps1
Ask10
Cdc28
Hog1
Pbs2
Ssk2
Clb2
Bud3
Bud4
Msn2
Sch9
Rim15
Yak1
Ypl247c
Vps52
Kss1
Sko1
Slt2
Msg5
Swe1
Igo2
Cdc55
Igo1
Vps53
Vps54
Rtn1
Fmp42
Cot1
Zrc1
10
20
30
40
50
t/s

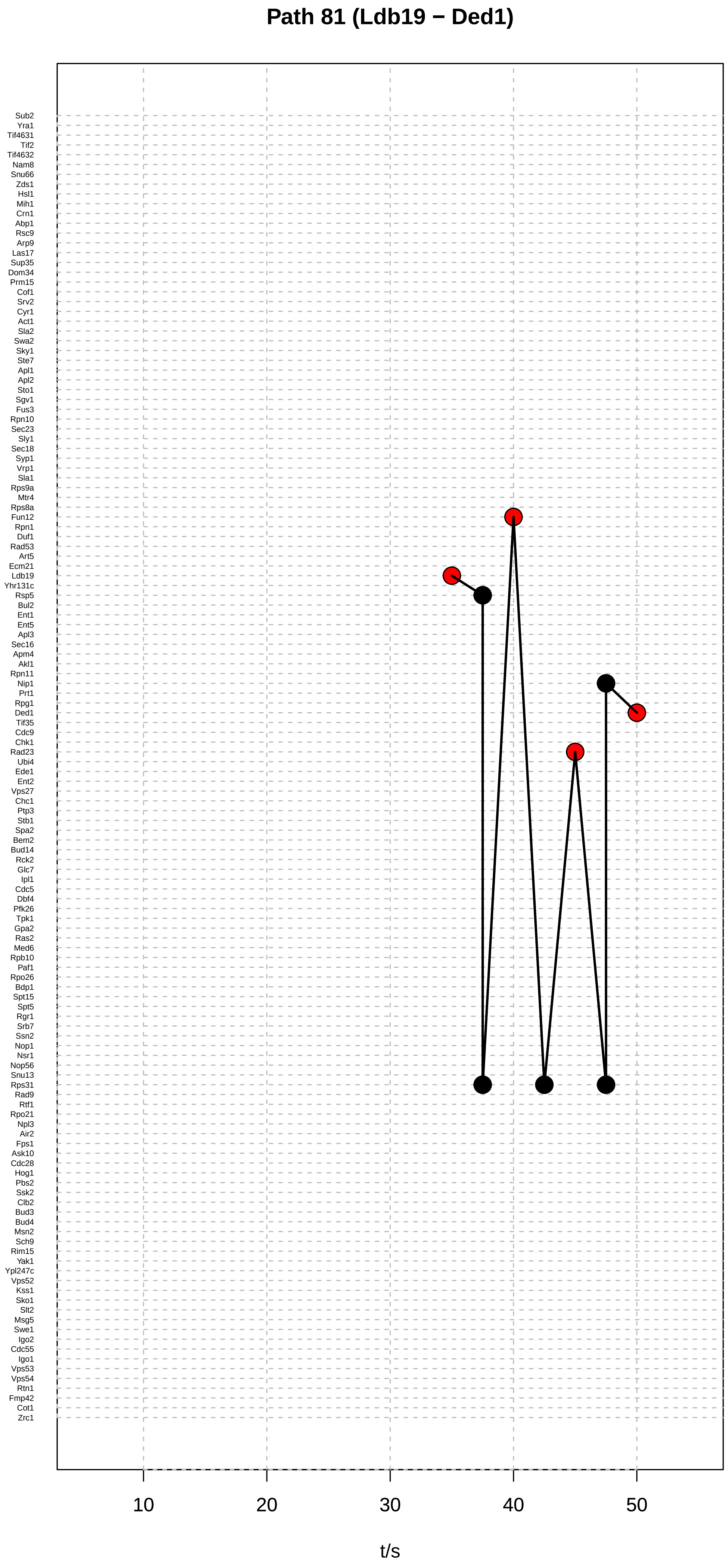

Path 81 (Ldb19 − Ded1)
Sub2
Yra1
Tif4631
Tif2
Tif4632
Nam8
Snu66
Zds1
Hsl1
Mih1
Crn1
Abp1
Rsc9
Arp9
Las17
Sup35
Dom34
Prm15
Cof1
Srv2
Cyr1
Act1
Sla2
Swa2
Sky1
Ste7
Apl1
Apl2
Sto1
Sgv1
Fus3
Rpn10
Sec23
Sly1
Sec18
Syp1
Vrp1
Sla1
Rps9a
Mtr4
Rps8a
Fun12
Rpn1
Duf1
Rad53
Art5
Ecm21
Ldb19
Yhr131c
Rsp5
Bul2
Ent1
Ent5
Apl3
Sec16
Apm4
Akl1
Rpn11
Nip1
Prt1
Rpg1
Ded1
Tif35
Cdc9
Chk1
Rad23
Ubi4
Ede1
Ent2
Vps27
Chc1
Ptp3
Stb1
Spa2
Bem2
Bud14
Rck2
Glc7
Ipl1
Cdc5
Dbf4
Pfk26
Tpk1
Gpa2
Ras2
Med6
Rpb10
Paf1
Rpo26
Bdp1
Spt15
Spt5
Rgr1
Srb7
Ssn2
Nop1
Nsr1
Nop56
Snu13
Rps31
Rad9
Rtf1
Rpo21
Npl3
Air2
Fps1
Ask10
Cdc28
Hog1
Pbs2
Ssk2
Clb2
Bud3
Bud4
Msn2
Sch9
Rim15
Yak1
Ypl247c
Vps52
Kss1
Sko1
Slt2
Msg5
Swe1
Igo2
Cdc55
Igo1
Vps53
Vps54
Rtn1
Fmp42
Cot1
Zrc1
10
20
30
40
50
t/s

### Path 82 (Ldb19 − Ded1)

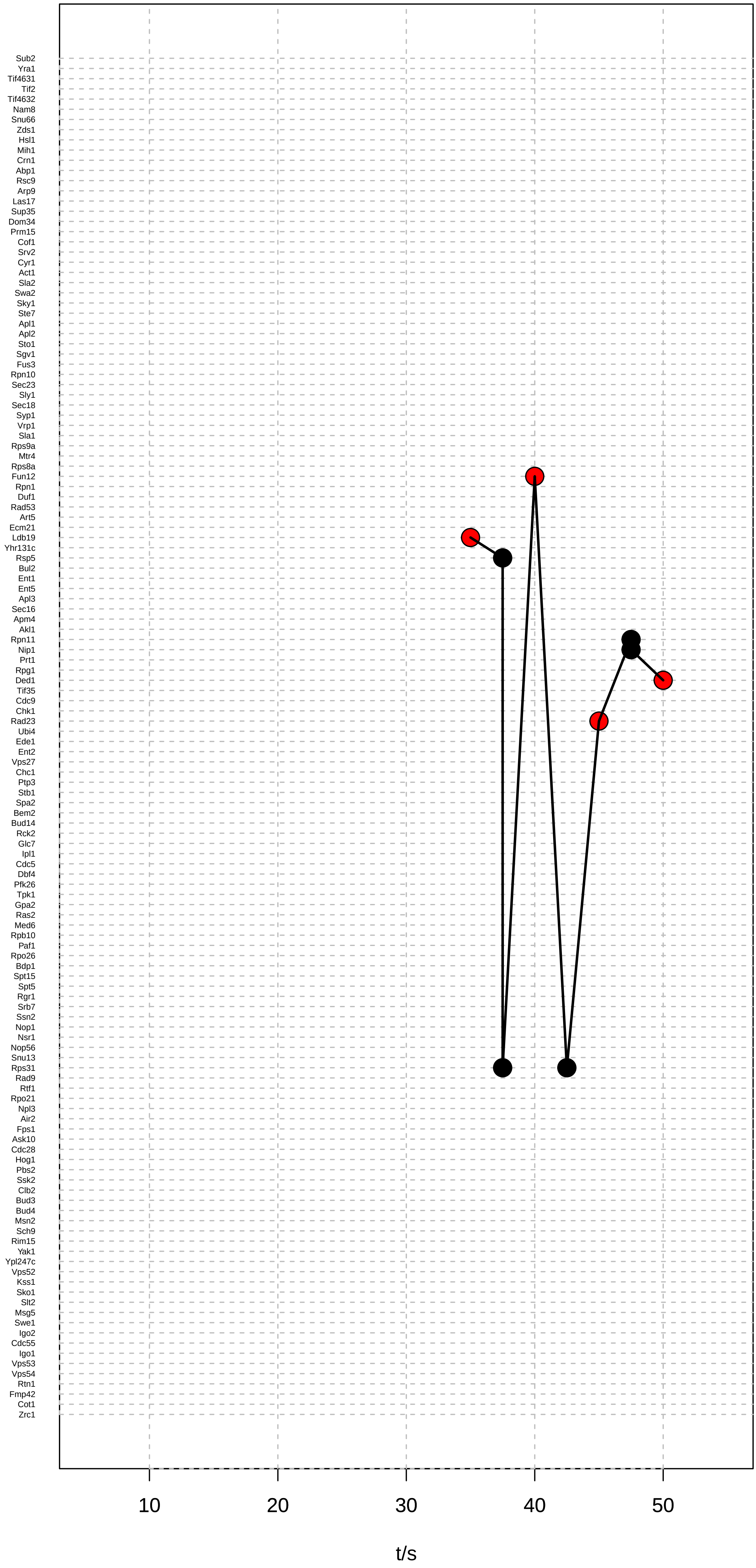

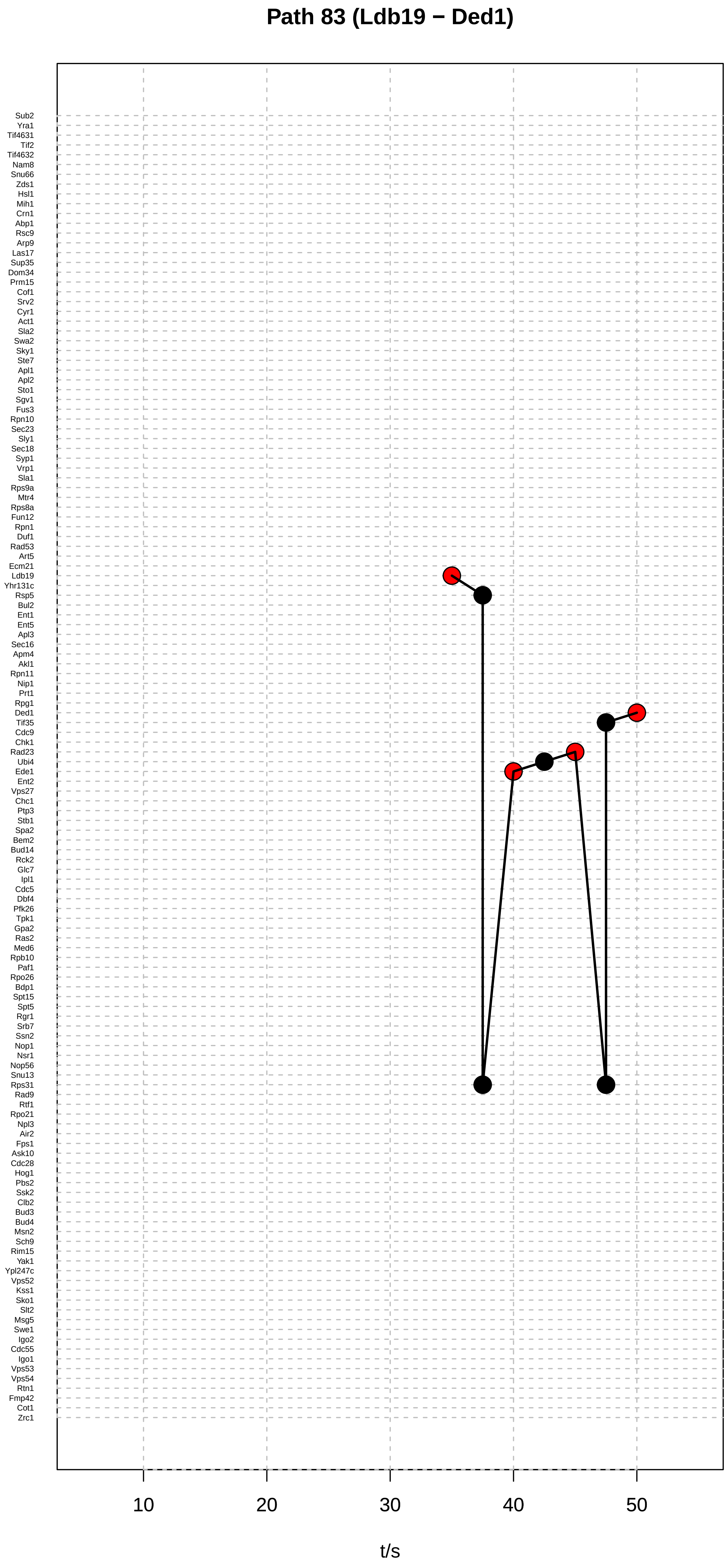
Path 83 (Ldb19 − Ded1)
Sub2
Yra1
Tif4631
Tif2
Tif4632
Nam8
Snu66
Zds1
Hsl1
Mih1
Crn1
Abp1
Rsc9
Arp9
Las17
Sup35
Dom34
Prm15
Cof1
Srv2
Cyr1
Act1
Sla2
Swa2
Sky1
Ste7
Apl1
Apl2
Sto1
Sgv1
Fus3
Rpn10
Sec23
Sly1
Sec18
Syp1
Vrp1
Sla1
Rps9a
Mtr4
Rps8a
Fun12
Rpn1
Duf1
Rad53
Art5
Ecm21
Ldb19
Yhr131c
Rsp5
Bul2
Ent1
Ent5
Apl3
Sec16
Apm4
Akl1
Rpn11
Nip1
Prt1
Rpg1
Ded1
Tif35
Cdc9
Chk1
Rad23
Ubi4
Ede1
Ent2
Vps27
Chc1
Ptp3
Stb1
Spa2
Bem2
Bud14
Rck2
Glc7
Ipl1
Cdc5
Dbf4
Pfk26
Tpk1
Gpa2
Ras2
Med6
Rpb10
Paf1
Rpo26
Bdp1
Spt15
Spt5
Rgr1
Srb7
Ssn2
Nop1
Nsr1
Nop56
Snu13
Rps31
Rad9
Rtf1
Rpo21
Npl3
Air2
Fps1
Ask10
Cdc28
Hog1
Pbs2
Ssk2
Clb2
Bud3
Bud4
Msn2
Sch9
Rim15
Yak1
Ypl247c
Vps52
Kss1
Sko1
Slt2
Msg5
Swe1
Igo2
Cdc55
Igo1
Vps53
Vps54
Rtn1
Fmp42
Cot1
Zrc1
10
20
30
40
50
t/s

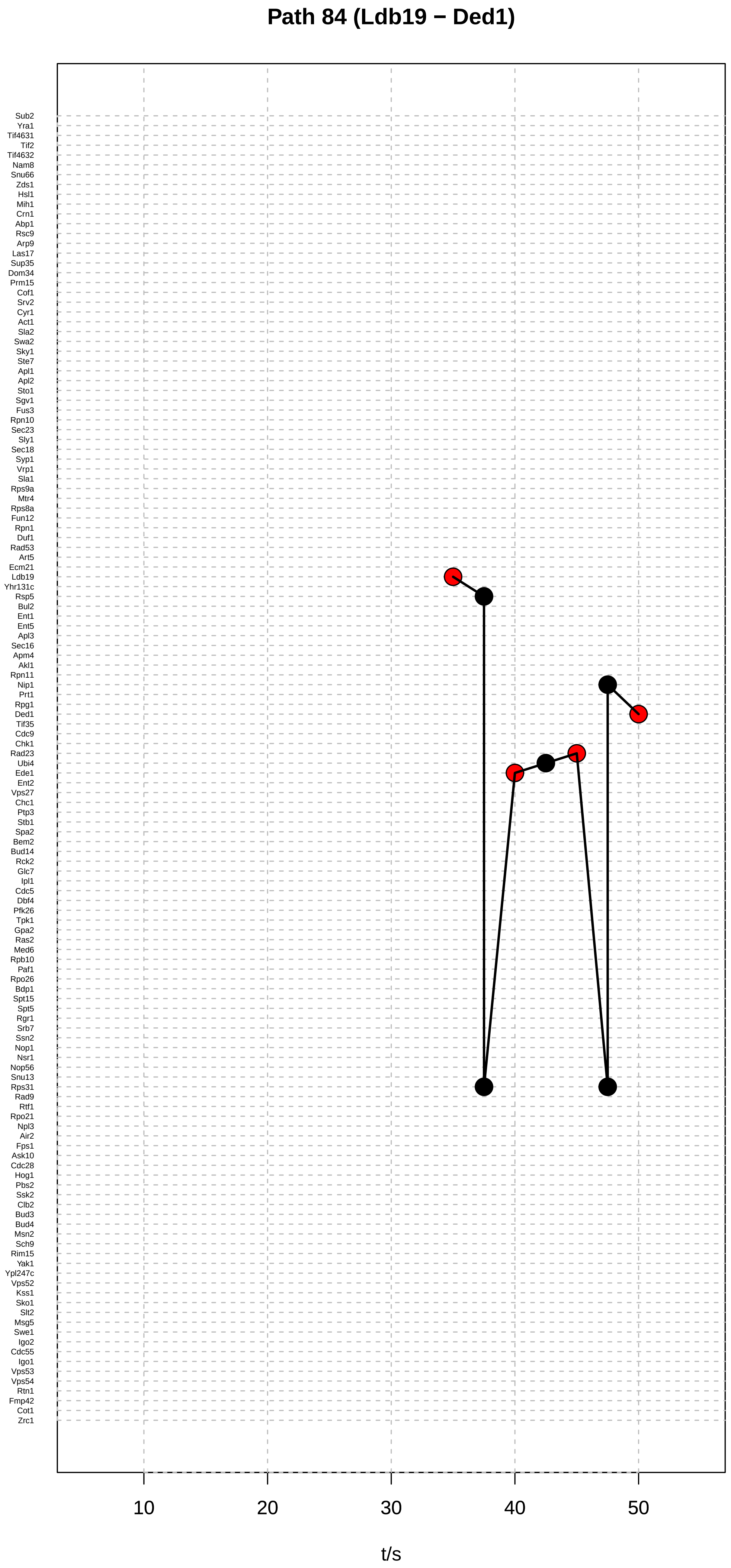

Path 84 (Ldb19 − Ded1)
Sub2
Yra1
Tif4631
Tif2
Tif4632
Nam8
Snu66
Zds1
Hsl1
Mih1
Crn1
Abp1
Rsc9
Arp9
Las17
Sup35
Dom34
Prm15
Cof1
Srv2
Cyr1
Act1
Sla2
Swa2
Sky1
Ste7
Apl1
Apl2
Sto1
Sgv1
Fus3
Rpn10
Sec23
Sly1
Sec18
Syp1
Vrp1
Sla1
Rps9a
Mtr4
Rps8a
Fun12
Rpn1
Duf1
Rad53
Art5
Ecm21
Ldb19
Yhr131c
Rsp5
Bul2
Ent1
Ent5
Apl3
Sec16
Apm4
Akl1
Rpn11
Nip1
Prt1
Rpg1
Ded1
Tif35
Cdc9
Chk1
Rad23
Ubi4
Ede1
Ent2
Vps27
Chc1
Ptp3
Stb1
Spa2
Bem2
Bud14
Rck2
Glc7
Ipl1
Cdc5
Dbf4
Pfk26
Tpk1
Gpa2
Ras2
Med6
Rpb10
Paf1
Rpo26
Bdp1
Spt15
Spt5
Rgr1
Srb7
Ssn2
Nop1
Nsr1
Nop56
Snu13
Rps31
Rad9
Rtf1
Rpo21
Npl3
Air2
Fps1
Ask10
Cdc28
Hog1
Pbs2
Ssk2
Clb2
Bud3
Bud4
Msn2
Sch9
Rim15
Yak1
Ypl247c
Vps52
Kss1
Sko1
Slt2
Msg5
Swe1
Igo2
Cdc55
Igo1
Vps53
Vps54
Rtn1
Fmp42
Cot1
Zrc1
10
20
30
40
50
t/s

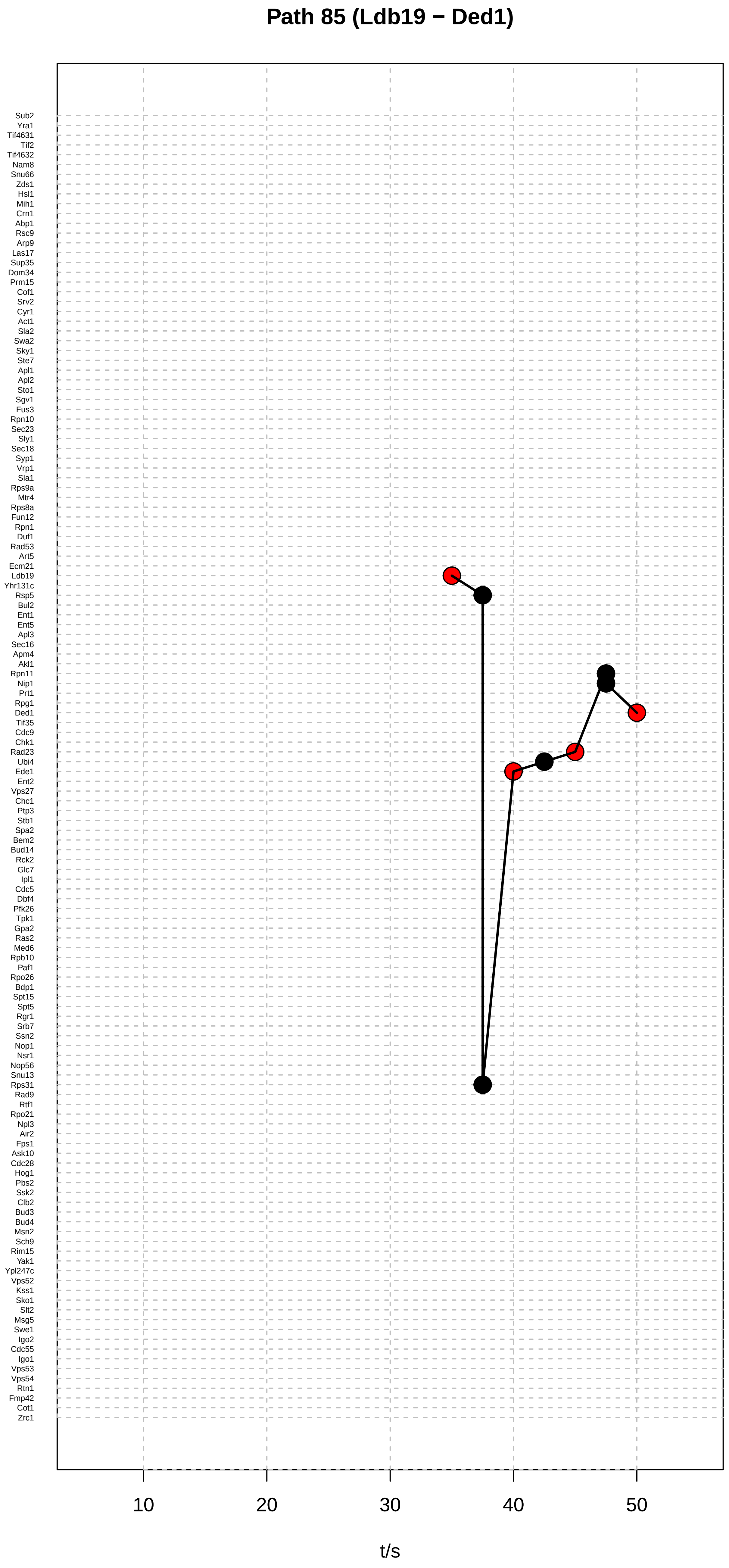
Path 85 (Ldb19 – Ded1)
Sub2
Yra1
Tif4631
Tif2
Tif4632
Nam8
Snu66
Zds1
Hsl1
Mih1
Crn1
Abp1
Rsc9
Arp9
Las17
Sup35
Dom34
Prm15
Cof1
Srv2
Cyr1
Act1
Sla2
Swa2
Sky1
Ste7
Apl1
Apl2
Sto1
Sgv1
Fus3
Rpn10
Sec23
Sly1
Sec18
Syp1
Vrp1
Sla1
Rps9a
Mtr4
Rps8a
Fun12
Rpn1
Duf1
Rad53
Art5
Ecm21
Ldb19
Yhr131c
Rsp5
Bul2
Ent1
Ent5
Apl3
Sec16
Apm4
Akl1
Rpn11
Nip1
Prt1
Rpg1
Ded1
Tif35
Cdc9
Chk1
Rad23
Ubi4
Ede1
Ent2
Vps27
Chc1
Ptp3
Stb1
Spa2
Bem2
Bud14
Rck2
Glc7
Ipl1
Cdc5
Dbf4
Pfk26
Tpk1
Gpa2
Ras2
Med6
Rpb10
Paf1
Rpo26
Bdp1
Spt15
Spt5
Rgr1
Srb7
Ssn2
Nop1
Nsr1
Nop56
Snu13
Rps31
Rad9
Rtf1
Rpo21
Npl3
Air2
Fps1
Ask10
Cdc28
Hog1
Pbs2
Ssk2
Clb2
Bud3
Bud4
Msn2
Sch9
Rim15
Yak1
Ypl247c
Vps52
Kss1
Sko1
Slt2
Msg5
Swe1
Igo2
Cdc55
Igo1
Vps53
Vps54
Rtn1
Fmp42
Cot1
Zrc1
10
20
30
40
50
t/s

### Path 86 (Ldb19 − Ded1)

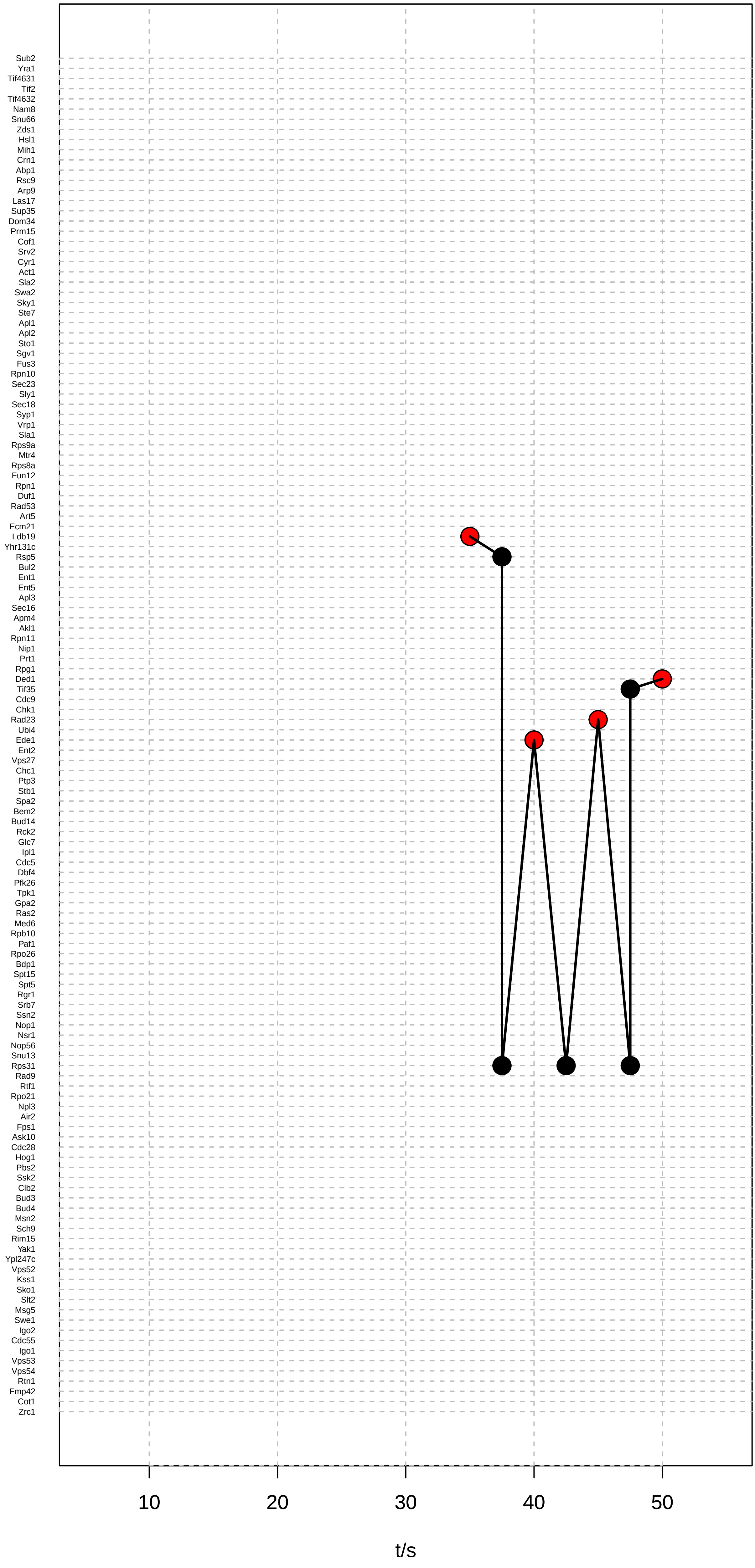

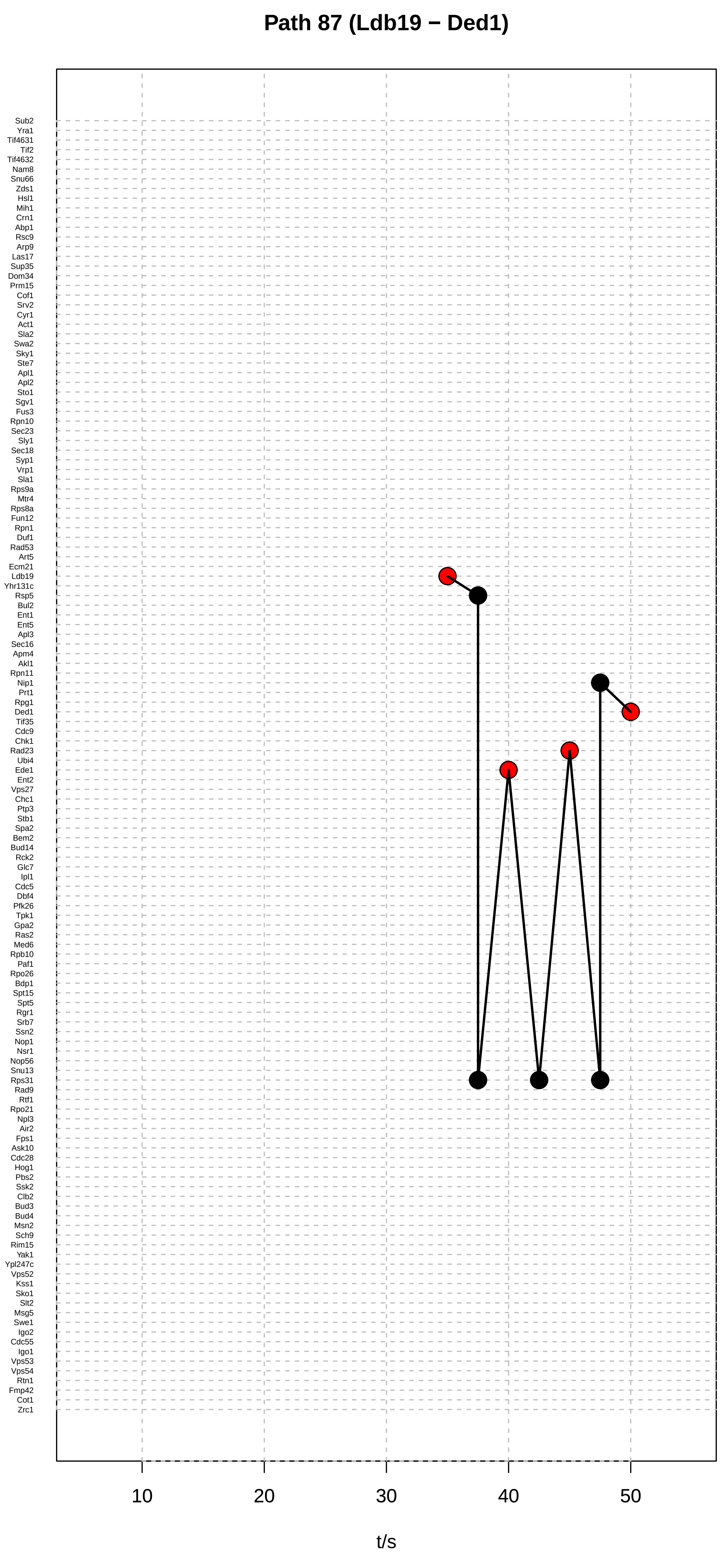

Path 87 (Ldb19 – Ded1)
Sub2
Yra1
Tif4631
Tif2
Tif4632
Nam8
Snu66
Zds1
Hsl1
Mih1
Crn1
Abp1
Rsc9
Arp9
Las17
Sup35
Dom34
Prm15
Cof1
Srv2
Cyr1
Act1
Sla2
Swa2
Sky1
Ste7
Apl1
Apl2
Sto1
Sgv1
Fus3
Rpn10
Sec23
Sly1
Sec18
Syp1
Vrp1
Sla1
Rps9a
Mtr4
Rps8a
Fun12
Rpn1
Duf1
Rad53
Art5
Ecm21
Ldb19
Yhr131c
Rsp5
Bul2
Ent1
Ent5
Apl3
Sec16
Apm4
Akl1
Rpn11
Nip1
Prt1
Rpg1
Ded1
Tif35
Cdc9
Chk1
Rad23
Ubi4
Ede1
Ent2
Vps27
Chc1
Ptp3
Stb1
Spa2
Bem2
Bud14
Rck2
Glc7
Ipl1
Cdc5
Dbf4
Pfk26
Tpk1
Gpa2
Ras2
Med6
Rpb10
Paf1
Rpo26
Bdp1
Spt15
Spt5
Rgr1
Srb7
Ssn2
Nop1
Nsr1
Nop56
Snu13
Rps31
Rad9
Rtf1
Rpo21
Npl3
Air2
Fps1
Ask10
Cdc28
Hog1
Pbs2
Ssk2
Clb2
Bud3
Bud4
Msn2
Sch9
Rim15
Yak1
Ypl247c
Vps52
Kss1
Sko1
Slt2
Msg5
Swe1
Igo2
Cdc55
Igo1
Vps53
Vps54
Rtn1
Fmp42
Cot1
Zrc1
10
20
30
40
50
t/s

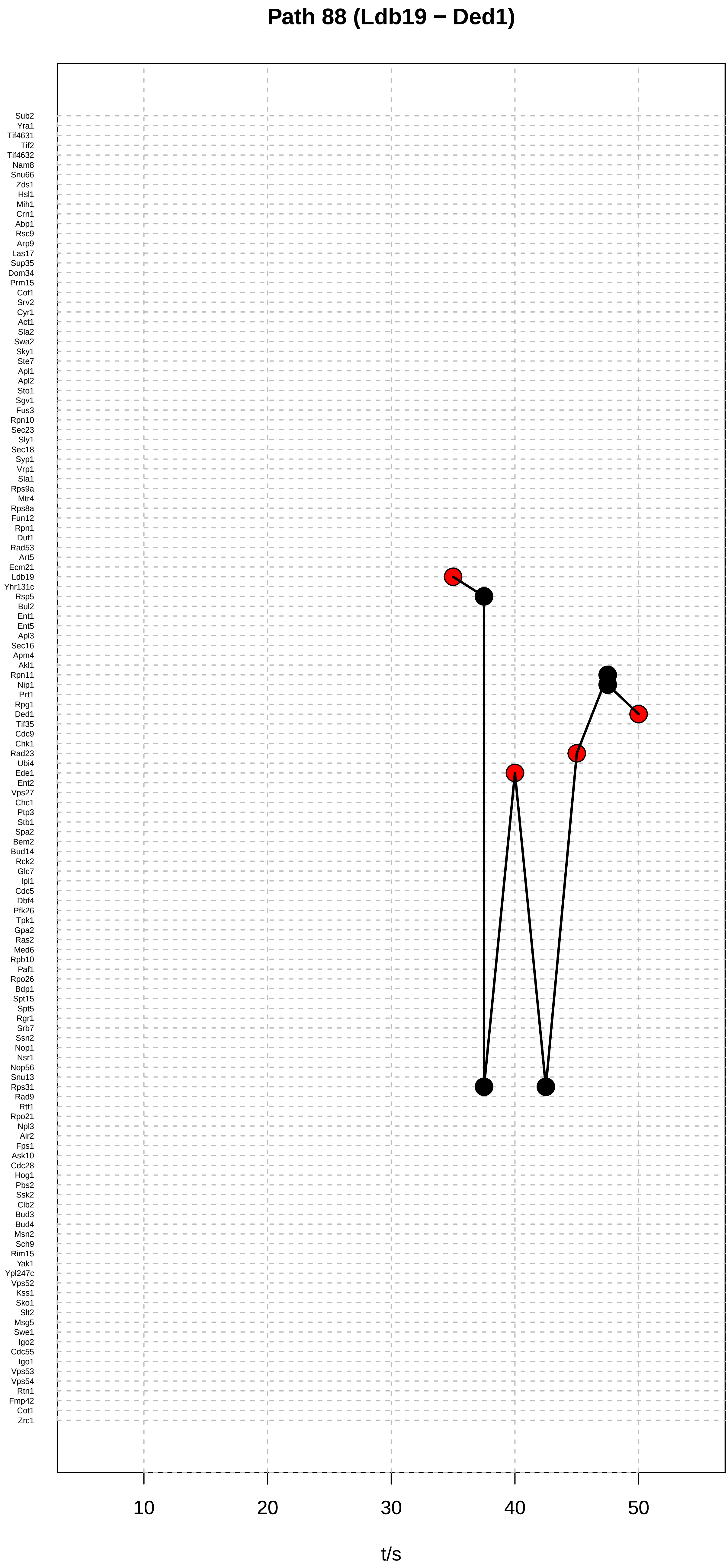

Path 88 (Ldb19 − Ded1)
Sub2
Yra1
Tif4631
Tif2
Tif4632
Nam8
Snu66
Zds1
Hsl1
Mih1
Crn1
Abp1
Rsc9
Arp9
Las17
Sup35
Dom34
Prm15
Cof1
Srv2
Cyr1
Act1
Sla2
Swa2
Sky1
Ste7
Apl1
Apl2
Sto1
Sgv1
Fus3
Rpn10
Sec23
Sly1
Sec18
Syp1
Vrp1
Sla1
Rps9a
Mtr4
Rps8a
Fun12
Rpn1
Duf1
Rad53
Art5
Ecm21
Ldb19
Yhr131c
Rsp5
Bul2
Ent1
Ent5
Apl3
Sec16
Apm4
Akl1
Rpn11
Nip1
Prt1
Rpg1
Ded1
Tif35
Cdc9
Chk1
Rad23
Ubi4
Ede1
Ent2
Vps27
Chc1
Ptp3
Stb1
Spa2
Bem2
Bud14
Rck2
Glc7
Ipl1
Cdc5
Dbf4
Pfk26
Tpk1
Gpa2
Ras2
Med6
Rpb10
Paf1
Rpo26
Bdp1
Spt15
Spt5
Rgr1
Srb7
Ssn2
Nop1
Nsr1
Nop56
Snu13
Rps31
Rad9
Rtf1
Rpo21
Npl3
Air2
Fps1
Ask10
Cdc28
Hog1
Pbs2
Ssk2
Clb2
Bud3
Bud4
Msn2
Sch9
Rim15
Yak1
Ypl247c
Vps52
Kss1
Sko1
Slt2
Msg5
Swe1
Igo2
Cdc55
Igo1
Vps53
Vps54
Rtn1
Fmp42
Cot1
Zrc1
10
20
30
40
50
t/s

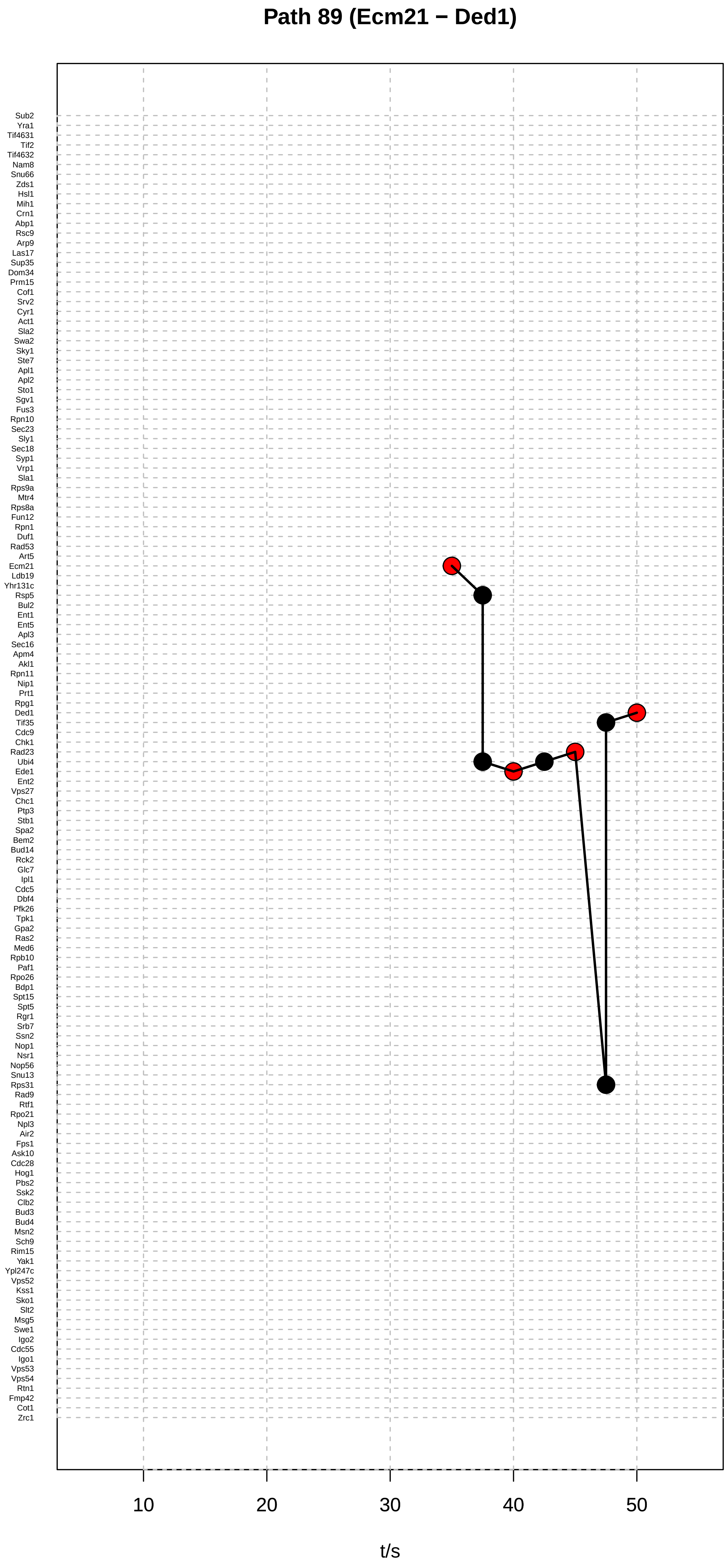

Path 89 (Ecm21 − Ded1)
Sub2
Yra1
Tif4631
Tif2
Tif4632
Nam8
Snu66
Zds1
Hsl1
Mih1
Crn1
Abp1
Rsc9
Arp9
Las17
Sup35
Dom34
Prm15
Cof1
Srv2
Cyr1
Act1
Sla2
Swa2
Sky1
Ste7
Apl1
Apl2
Sto1
Sgv1
Fus3
Rpn10
Sec23
Sly1
Sec18
Syp1
Vrp1
Sla1
Rps9a
Mtr4
Rps8a
Fun12
Rpn1
Duf1
Rad53
Art5
Ecm21
Ldb19
Yhr131c
Rsp5
Bul2
Ent1
Ent5
Apl3
Sec16
Apm4
Akl1
Rpn11
Nip1
Prt1
Rpg1
Ded1
Tif35
Cdc9
Chk1
Rad23
Ubi4
Ede1
Ent2
Vps27
Chc1
Ptp3
Stb1
Spa2
Bem2
Bud14
Rck2
Glc7
Ipl1
Cdc5
Dbf4
Pfk26
Tpk1
Gpa2
Ras2
Med6
Rpb10
Paf1
Rpo26
Bdp1
Spt15
Spt5
Rgr1
Srb7
Ssn2
Nop1
Nsr1
Nop56
Snu13
Rps31
Rad9
Rtf1
Rpo21
Npl3
Air2
Fps1
Ask10
Cdc28
Hog1
Pbs2
Ssk2
Clb2
Bud3
Bud4
Msn2
Sch9
Rim15
Yak1
Ypl247c
Vps52
Kss1
Sko1
Slt2
Msg5
Swe1
Igo2
Cdc55
Igo1
Vps53
Vps54
Rtn1
Fmp42
Cot1
Zrc1
10
20
30
40
50
t/s

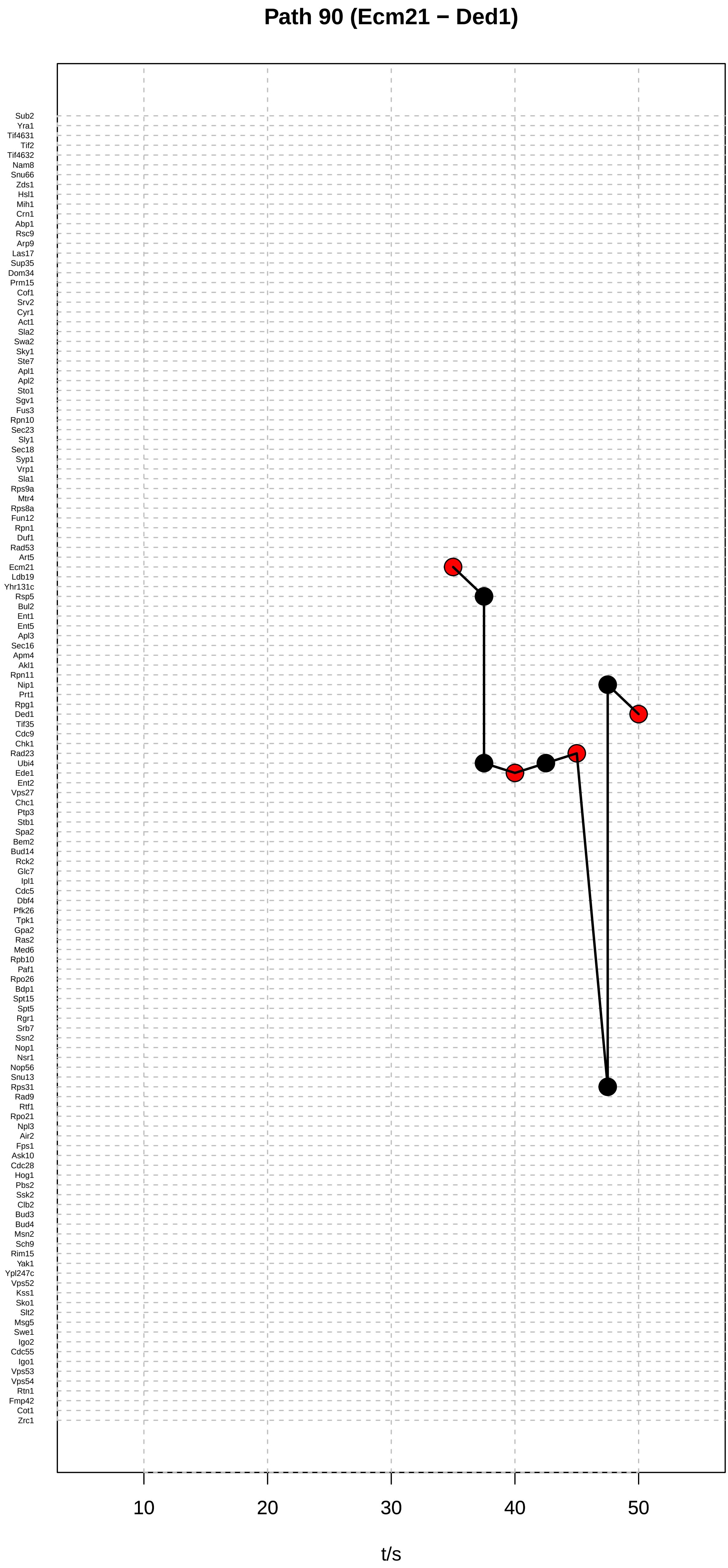

Path 90 (Ecm21 − Ded1)
Sub2
Yra1
Tif4631
Tif2
Tif4632
Nam8
Snu66
Zds1
Hsl1
Mih1
Crn1
Abp1
Rsc9
Arp9
Las17
Sup35
Dom34
Prm15
Cof1
Srv2
Cyr1
Act1
Sla2
Swa2
Sky1
Ste7
Apl1
Apl2
Sto1
Sgv1
Fus3
Rpn10
Sec23
Sly1
Sec18
Syp1
Vrp1
Sla1
Rps9a
Mtr4
Rps8a
Fun12
Rpn1
Duf1
Rad53
Art5
Ecm21
Ldb19
Yhr131c
Rsp5
Bul2
Ent1
Ent5
Apl3
Sec16
Apm4
Akl1
Rpn11
Nip1
Prt1
Rpg1
Ded1
Tif35
Cdc9
Chk1
Rad23
Ubi4
Ede1
Ent2
Vps27
Chc1
Ptp3
Stb1
Spa2
Bem2
Bud14
Rck2
Glc7
Ipl1
Cdc5
Dbf4
Pfk26
Tpk1
Gpa2
Ras2
Med6
Rpb10
Paf1
Rpo26
Bdp1
Spt15
Spt5
Rgr1
Srb7
Ssn2
Nop1
Nsr1
Nop56
Snu13
Rps31
Rad9
Rtf1
Rpo21
Npl3
Air2
Fps1
Ask10
Cdc28
Hog1
Pbs2
Ssk2
Clb2
Bud3
Bud4
Msn2
Sch9
Rim15
Yak1
Ypl247c
Vps52
Kss1
Sko1
Slt2
Msg5
Swe1
Igo2
Cdc55
Igo1
Vps53
Vps54
Rtn1
Fmp42
Cot1
Zrc1
10
20
30
40
50
t/s

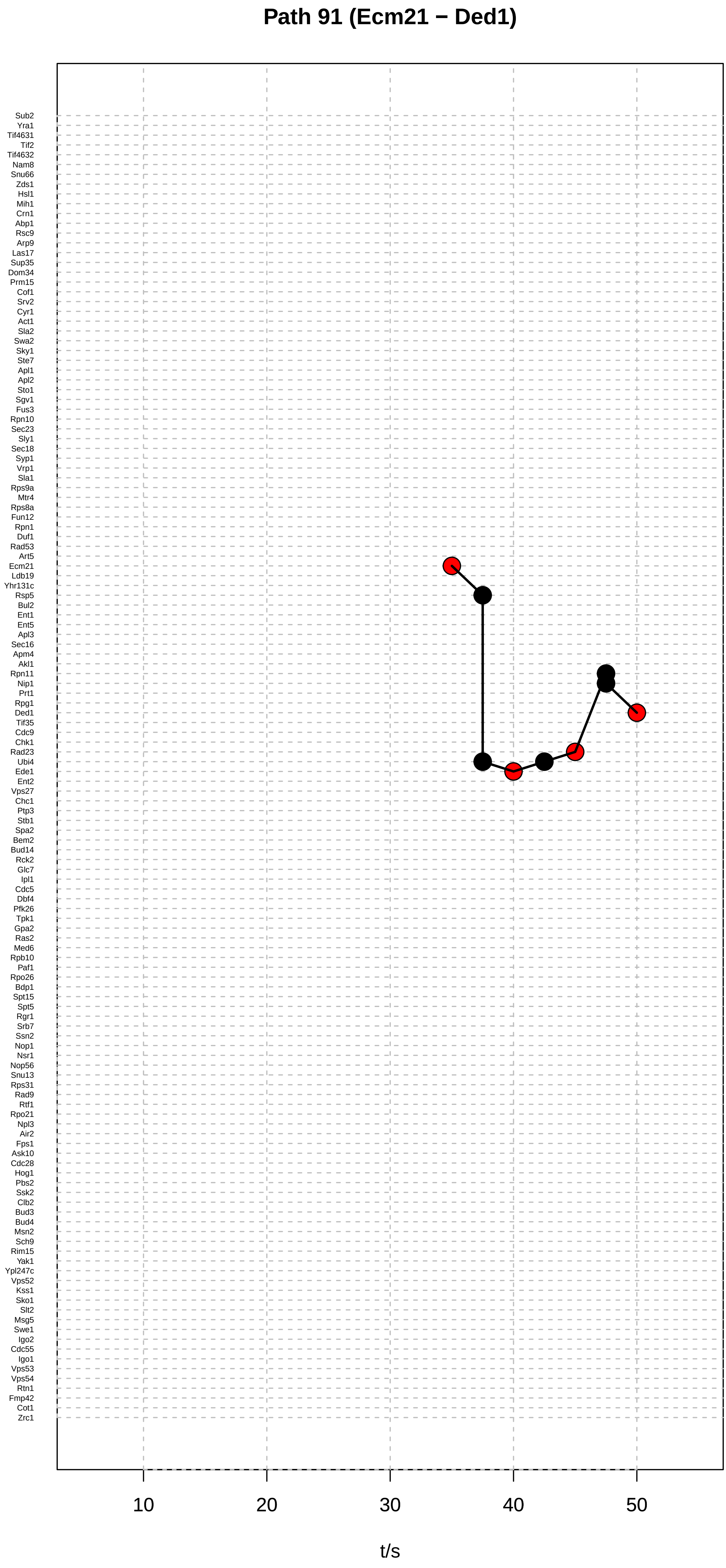
Path 91 (Ecm21 − Ded1)
Sub2
Yra1
Tif4631
Tif2
Tif4632
Nam8
Snu66
Zds1
Hsl1
Mih1
Crn1
Abp1
Rsc9
Arp9
Las17
Sup35
Dom34
Prm15
Cof1
Srv2
Cyr1
Act1
Sla2
Swa2
Sky1
Ste7
Apl1
Apl2
Sto1
Sgv1
Fus3
Rpn10
Sec23
Sly1
Sec18
Syp1
Vrp1
Sla1
Rps9a
Mtr4
Rps8a
Fun12
Rpn1
Duf1
Rad53
Art5
Ecm21
Ldb19
Yhr131c
Rsp5
Bul2
Ent1
Ent5
Apl3
Sec16
Apm4
Akl1
Rpn11
Nip1
Prt1
Rpg1
Ded1
Tif35
Cdc9
Chk1
Rad23
Ubi4
Ede1
Ent2
Vps27
Chc1
Ptp3
Stb1
Spa2
Bem2
Bud14
Rck2
Glc7
Ipl1
Cdc5
Dbf4
Pfk26
Tpk1
Gpa2
Ras2
Med6
Rpb10
Paf1
Rpo26
Bdp1
Spt15
Spt5
Rgr1
Srb7
Ssn2
Nop1
Nsr1
Nop56
Snu13
Rps31
Rad9
Rtf1
Rpo21
Npl3
Air2
Fps1
Ask10
Cdc28
Hog1
Pbs2
Ssk2
Clb2
Bud3
Bud4
Msn2
Sch9
Rim15
Yak1
Ypl247c
Vps52
Kss1
Sko1
Slt2
Msg5
Swe1
Igo2
Cdc55
Igo1
Vps53
Vps54
Rtn1
Fmp42
Cot1
Zrc1
10
20
30
40
50
t/s

### Path 92 (Ecm21 − Ded1)

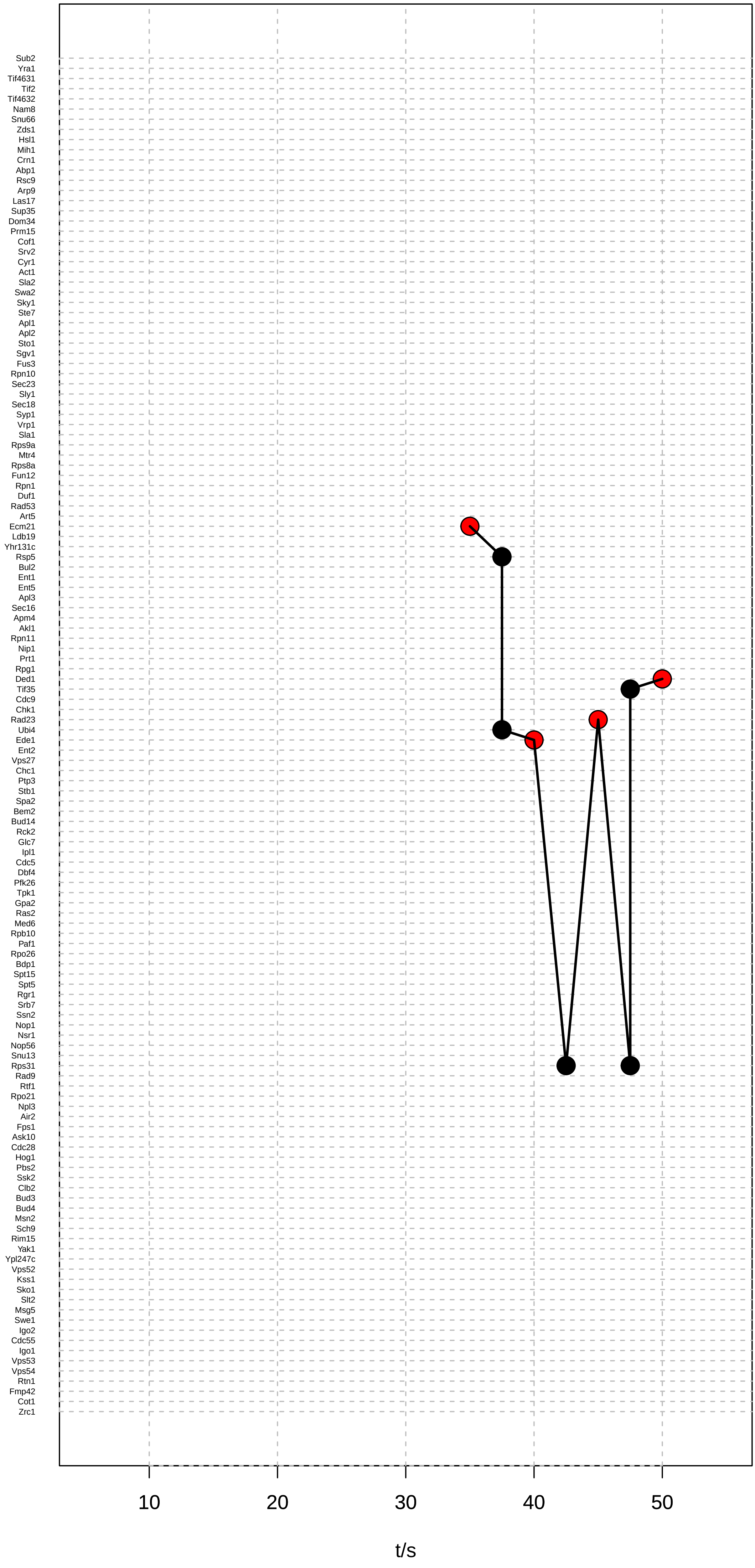

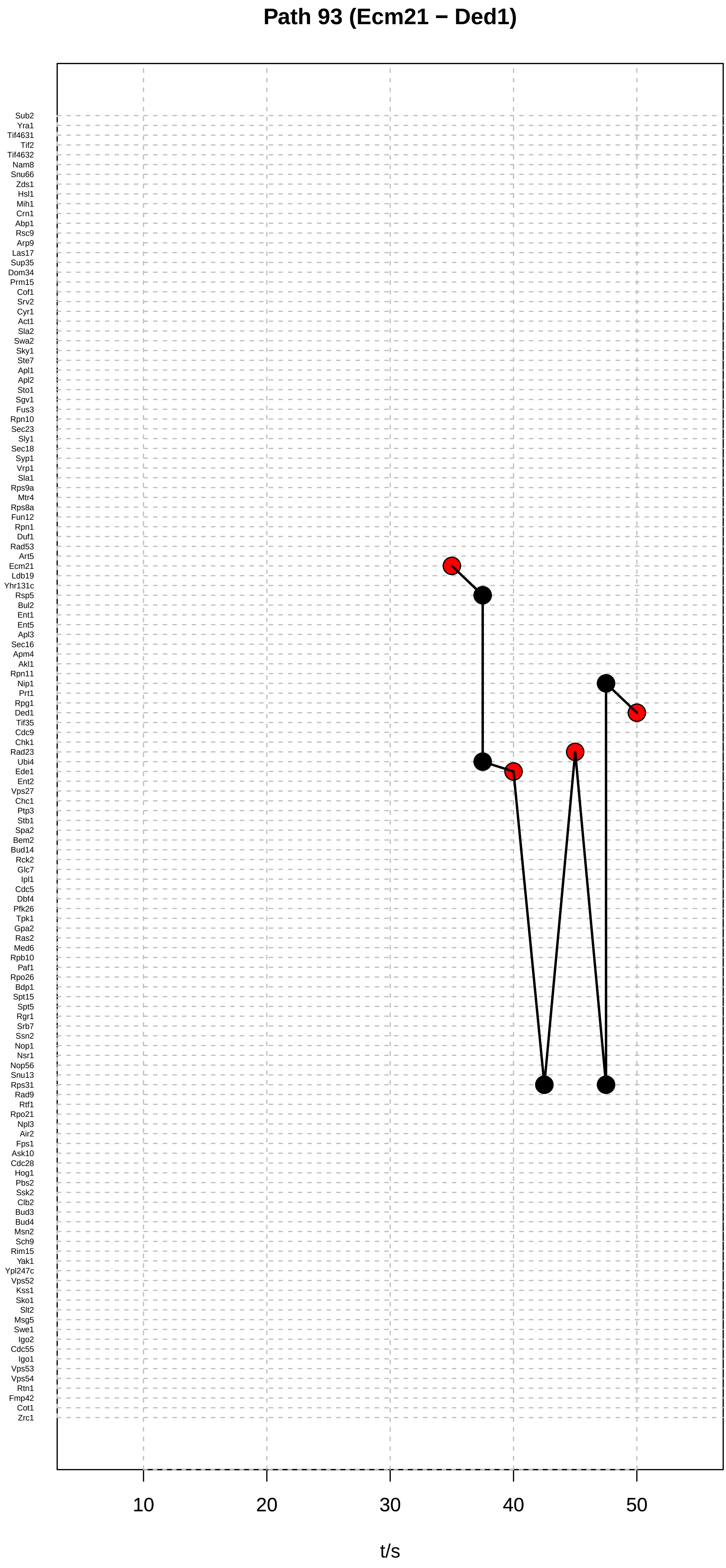
Path 93 (Ecm21 − Ded1)
Sub2
Yra1
Tif4631
Tif2
Tif4632
Nam8
Snu66
Zds1
Hsl1
Mih1
Crn1
Abp1
Rsc9
Arp9
Las17
Sup35
Dom34
Prm15
Cof1
Srv2
Cyr1
Act1
Sla2
Swa2
Sky1
Ste7
Apl1
Apl2
Sto1
Sgv1
Fus3
Rpn10
Sec23
Sly1
Sec18
Syp1
Vrp1
Sla1
Rps9a
Mtr4
Rps8a
Fun12
Rpn1
Duf1
Rad53
Art5
Ecm21
Ldb19
Yhr131c
Rsp5
Bul2
Ent1
Ent5
Apl3
Sec16
Apm4
Akl1
Rpn11
Nip1
Prt1
Rpg1
Ded1
Tif35
Cdc9
Chk1
Rad23
Ubi4
Ede1
Ent2
Vps27
Chc1
Ptp3
Stb1
Spa2
Bem2
Bud14
Rck2
Glc7
Ipl1
Cdc5
Dbf4
Pfk26
Tpk1
Gpa2
Ras2
Med6
Rpb10
Paf1
Rpo26
Bdp1
Spt15
Spt5
Rgr1
Srb7
Ssn2
Nop1
Nsr1
Nop56
Snu13
Rps31
Rad9
Rtf1
Rpo21
Npl3
Air2
Fps1
Ask10
Cdc28
Hog1
Pbs2
Ssk2
Clb2
Bud3
Bud4
Msn2
Sch9
Rim15
Yak1
Ypl247c
Vps52
Kss1
Sko1
Slt2
Msg5
Swe1
Igo2
Cdc55
Igo1
Vps53
Vps54
Rtn1
Fmp42
Cot1
Zrc1
10
20
30
40
50
t/s

### Path 94 (Ecm21 − Ded1)

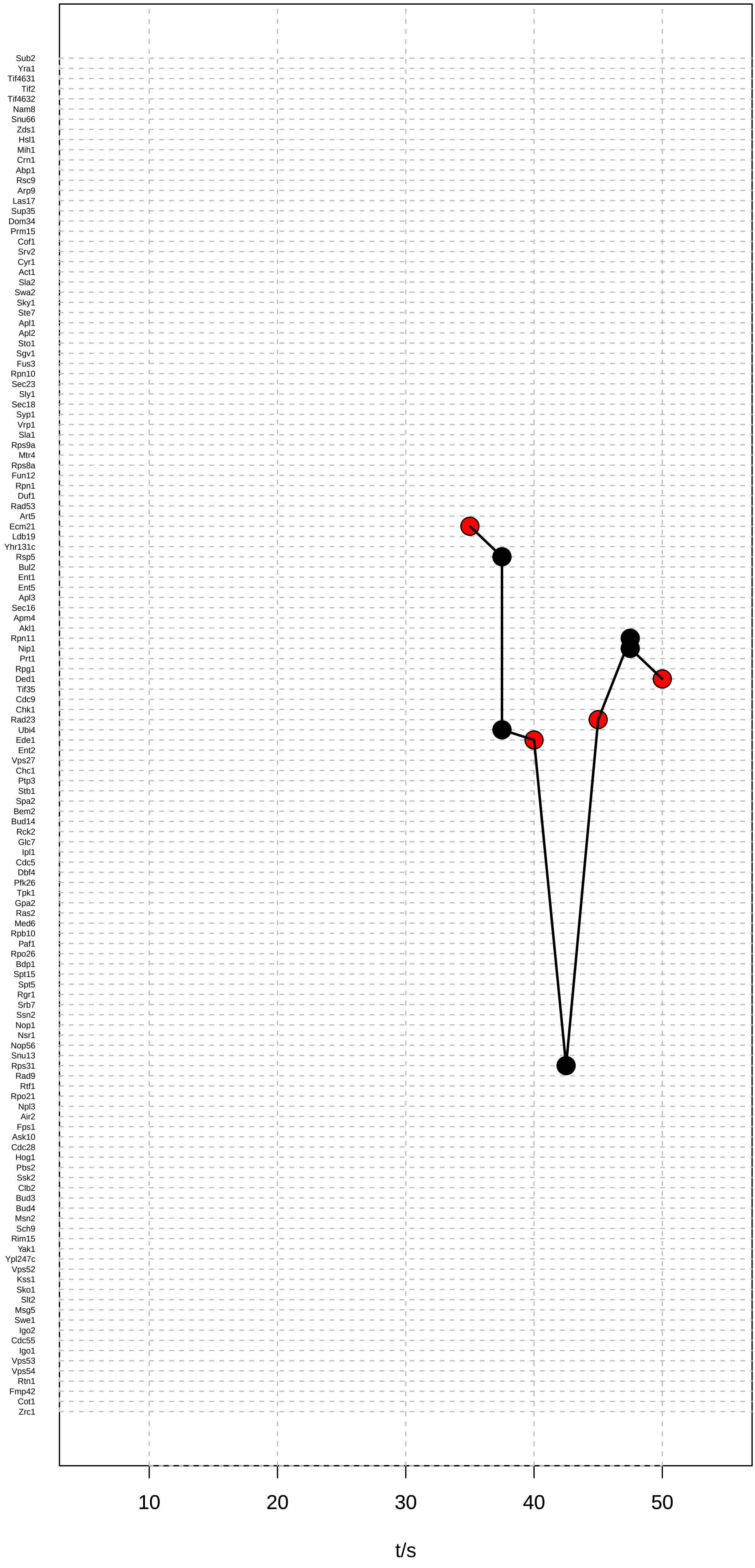

Path 95 (Ecm21 − Ded1)

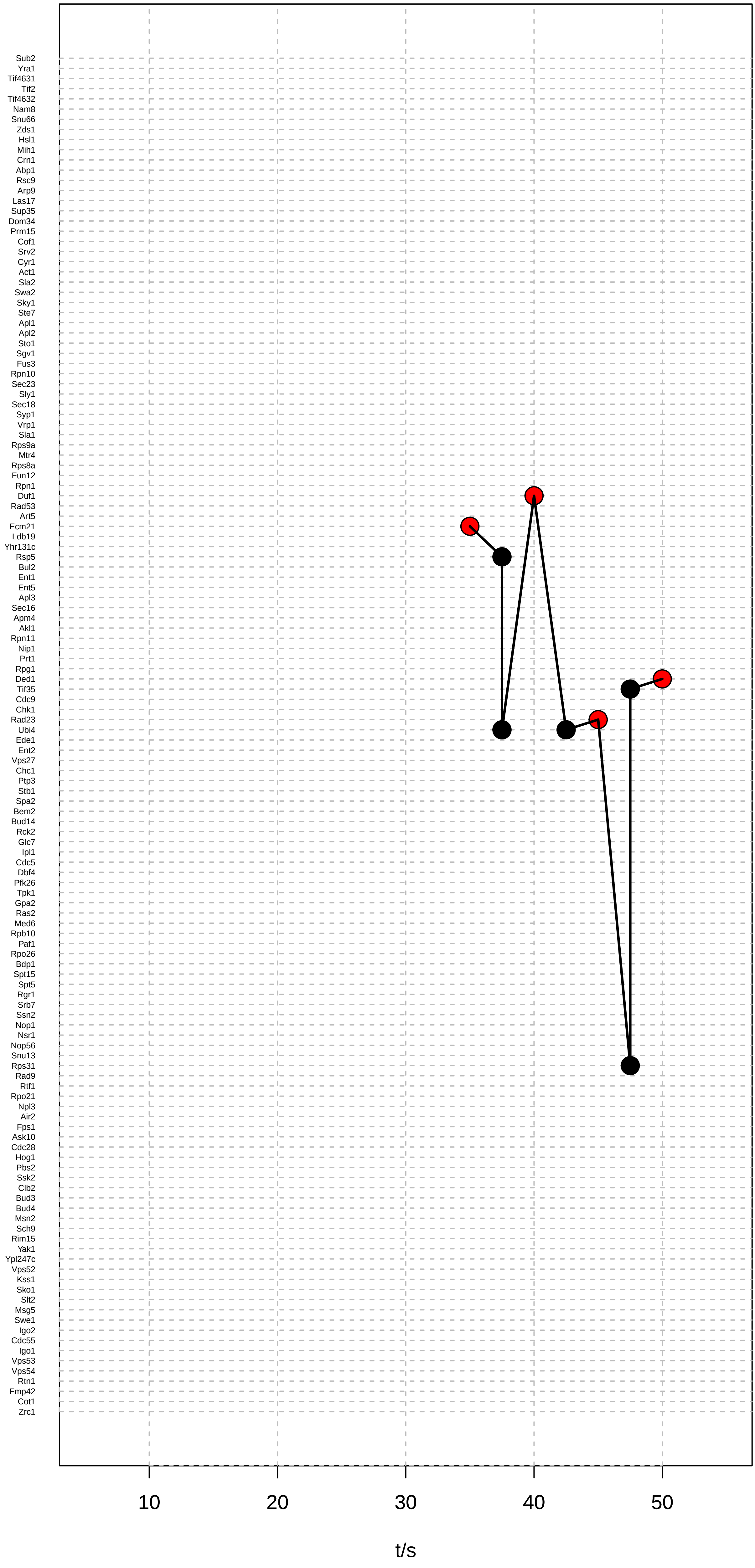

### Path 96 (Ecm21 − Ded1)

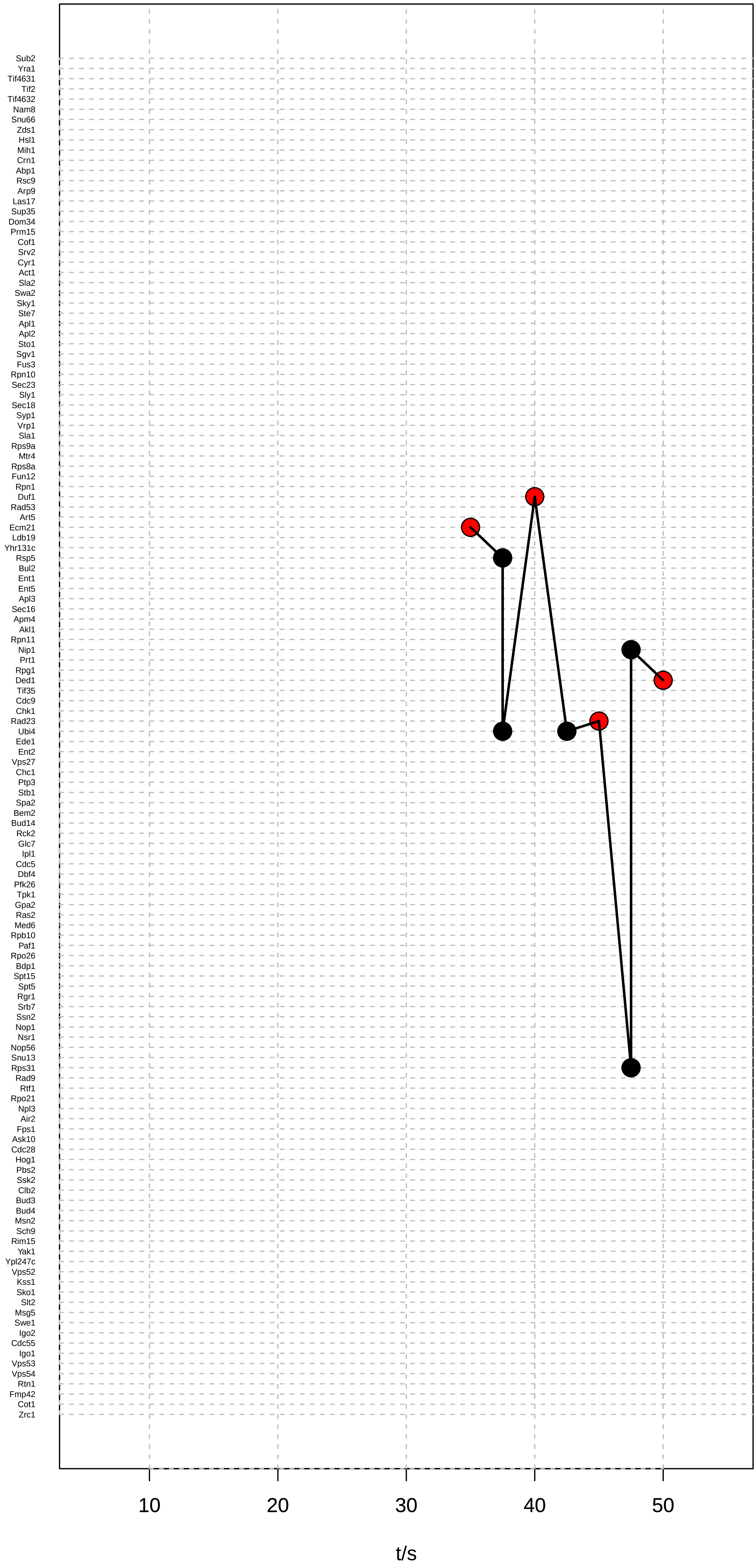

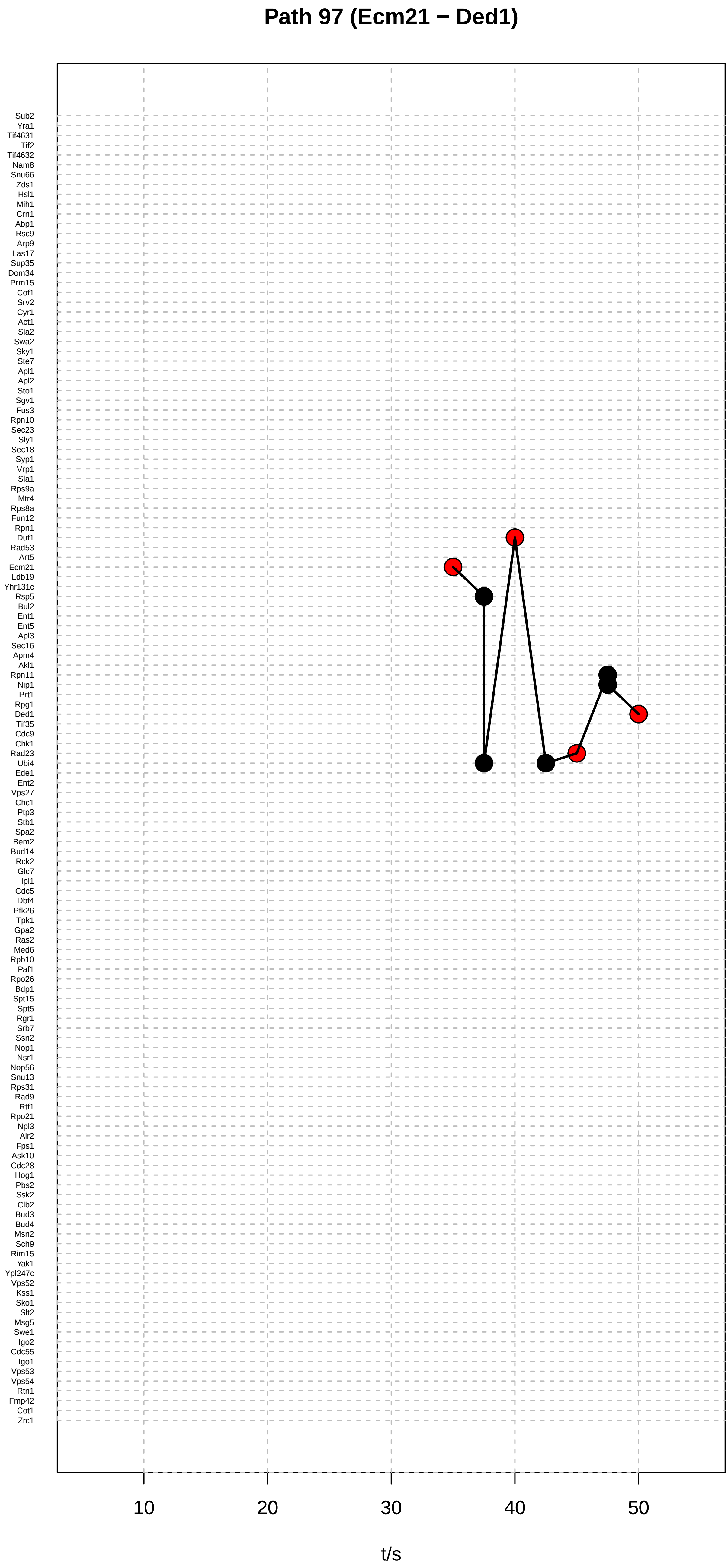

Path 97 (Ecm21 − Ded1)
Sub2
Yra1
Tif4631
Tif2
Tif4632
Nam8
Snu66
Zds1
Hsl1
Mih1
Crn1
Abp1
Rsc9
Arp9
Las17
Sup35
Dom34
Prm15
Cof1
Srv2
Cyr1
Act1
Sla2
Swa2
Sky1
Ste7
Apl1
Apl2
Sto1
Sgv1
Fus3
Rpn10
Sec23
Sly1
Sec18
Syp1
Vrp1
Sla1
Rps9a
Mtr4
Rps8a
Fun12
Rpn1
Duf1
Rad53
Art5
Ecm21
Ldb19
Yhr131c
Rsp5
Bul2
Ent1
Ent5
Apl3
Sec16
Apm4
Akl1
Rpn11
Nip1
Prt1
Rpg1
Ded1
Tif35
Cdc9
Chk1
Rad23
Ubi4
Ede1
Ent2
Vps27
Chc1
Ptp3
Stb1
Spa2
Bem2
Bud14
Rck2
Glc7
Ipl1
Cdc5
Dbf4
Pfk26
Tpk1
Gpa2
Ras2
Med6
Rpb10
Paf1
Rpo26
Bdp1
Spt15
Spt5
Rgr1
Srb7
Ssn2
Nop1
Nsr1
Nop56
Snu13
Rps31
Rad9
Rtf1
Rpo21
Npl3
Air2
Fps1
Ask10
Cdc28
Hog1
Pbs2
Ssk2
Clb2
Bud3
Bud4
Msn2
Sch9
Rim15
Yak1
Ypl247c
Vps52
Kss1
Sko1
Slt2
Msg5
Swe1
Igo2
Cdc55
Igo1
Vps53
Vps54
Rtn1
Fmp42
Cot1
Zrc1
10
20
30
40
50
t/s

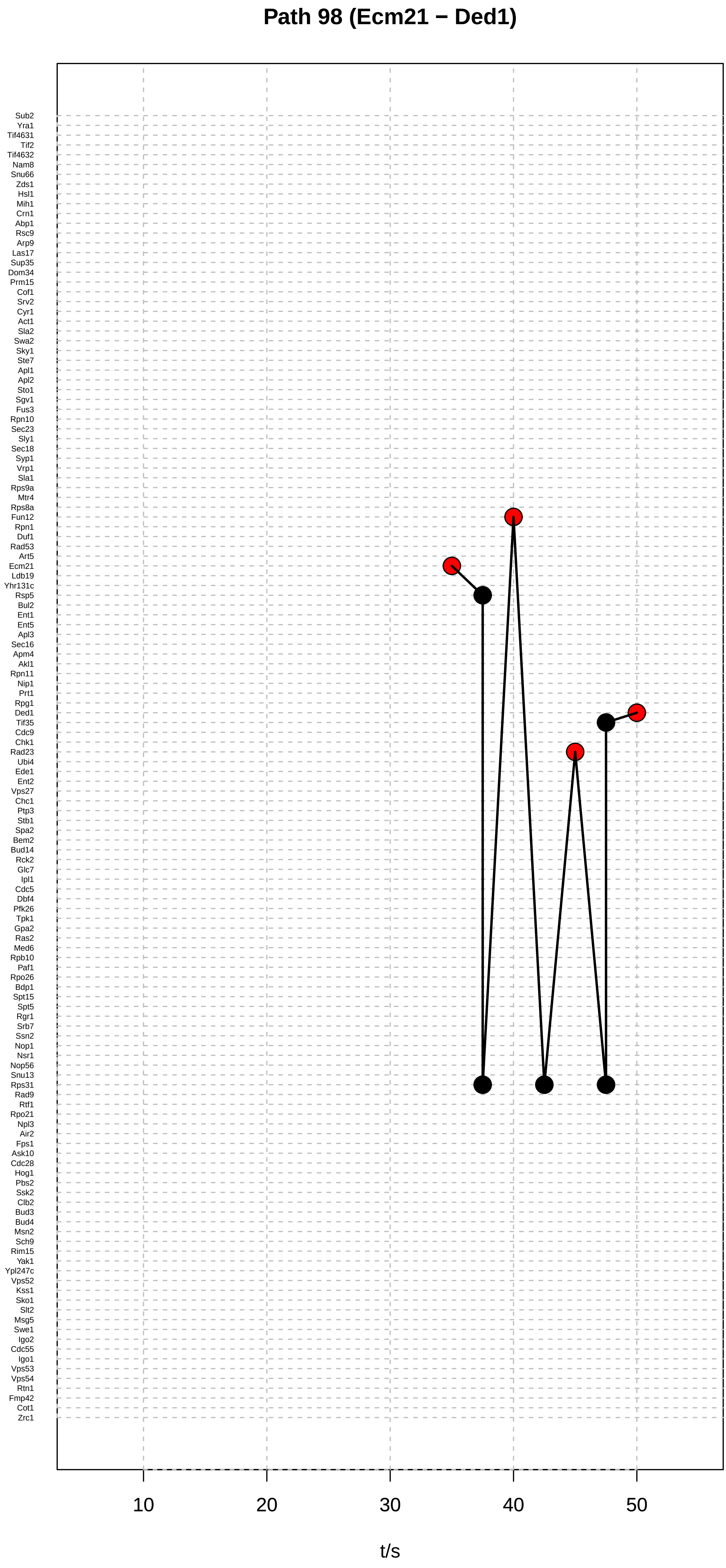
Path 98 (Ecm21 − Ded1)
Sub2
Yra1
Tif4631
Tif2
Tif4632
Nam8
Snu66
Zds1
Hsl1
Mih1
Crn1
Abp1
Rsc9
Arp9
Las17
Sup35
Dom34
Prm15
Cof1
Srv2
Cyr1
Act1
Sla2
Swa2
Sky1
Ste7
Apl1
Apl2
Sto1
Sgv1
Fus3
Rpn10
Sec23
Sly1
Sec18
Syp1
Vrp1
Sla1
Rps9a
Mtr4
Rps8a
Fun12
Rpn1
Duf1
Rad53
Art5
Ecm21
Ldb19
Yhr131c
Rsp5
Bul2
Ent1
Ent5
Apl3
Sec16
Apm4
Akl1
Rpn11
Nip1
Prt1
Rpg1
Ded1
Tif35
Cdc9
Chk1
Rad23
Ubi4
Ede1
Ent2
Vps27
Chc1
Ptp3
Stb1
Spa2
Bem2
Bud14
Rck2
Glc7
Ipl1
Cdc5
Dbf4
Pfk26
Tpk1
Gpa2
Ras2
Med6
Rpb10
Paf1
Rpo26
Bdp1
Spt15
Spt5
Rgr1
Srb7
Ssn2
Nop1
Nsr1
Nop56
Snu13
Rps31
Rad9
Rtf1
Rpo21
Npl3
Air2
Fps1
Ask10
Cdc28
Hog1
Pbs2
Ssk2
Clb2
Bud3
Bud4
Msn2
Sch9
Rim15
Yak1
Ypl247c
Vps52
Kss1
Sko1
Slt2
Msg5
Swe1
Igo2
Cdc55
Igo1
Vps53
Vps54
Rtn1
Fmp42
Cot1
Zrc1
10
20
30
40
50
t/s

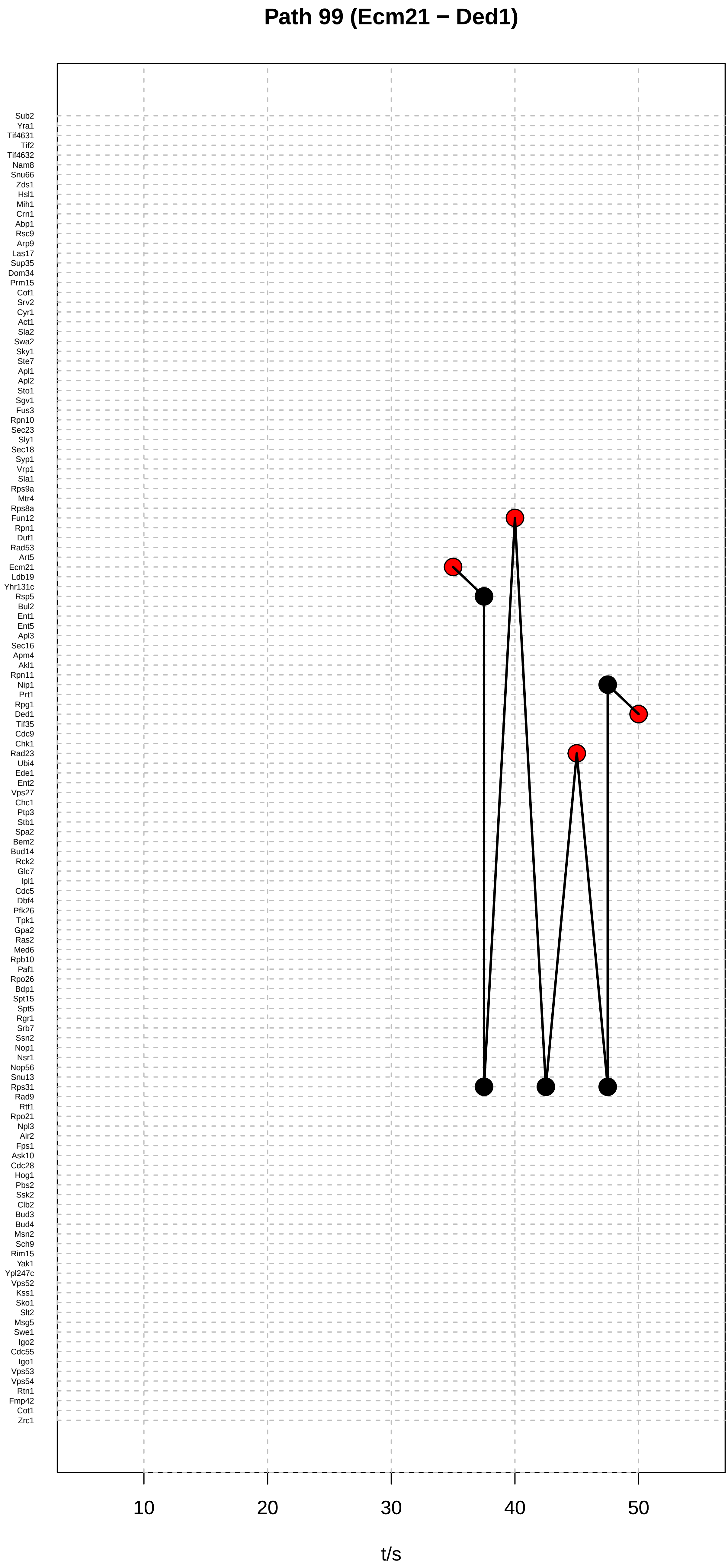
Path 99 (Ecm21 − Ded1)
Sub2
Yra1
Tif4631
Tif2
Tif4632
Nam8
Snu66
Zds1
Hsl1
Mih1
Crn1
Abp1
Rsc9
Arp9
Las17
Sup35
Dom34
Prm15
Cof1
Srv2
Cyr1
Act1
Sla2
Swa2
Sky1
Ste7
Apl1
Apl2
Sto1
Sgv1
Fus3
Rpn10
Sec23
Sly1
Sec18
Syp1
Vrp1
Sla1
Rps9a
Mtr4
Rps8a
Fun12
Rpn1
Duf1
Rad53
Art5
Ecm21
Ldb19
Yhr131c
Rsp5
Bul2
Ent1
Ent5
Apl3
Sec16
Apm4
Akl1
Rpn11
Nip1
Prt1
Rpg1
Ded1
Tif35
Cdc9
Chk1
Rad23
Ubi4
Ede1
Ent2
Vps27
Chc1
Ptp3
Stb1
Spa2
Bem2
Bud14
Rck2
Glc7
Ipl1
Cdc5
Dbf4
Pfk26
Tpk1
Gpa2
Ras2
Med6
Rpb10
Paf1
Rpo26
Bdp1
Spt15
Spt5
Rgr1
Srb7
Ssn2
Nop1
Nsr1
Nop56
Snu13
Rps31
Rad9
Rtf1
Rpo21
Npl3
Air2
Fps1
Ask10
Cdc28
Hog1
Pbs2
Ssk2
Clb2
Bud3
Bud4
Msn2
Sch9
Rim15
Yak1
Ypl247c
Vps52
Kss1
Sko1
Slt2
Msg5
Swe1
Igo2
Cdc55
Igo1
Vps53
Vps54
Rtn1
Fmp42
Cot1
Zrc1
10
20
30
40
50
t/s

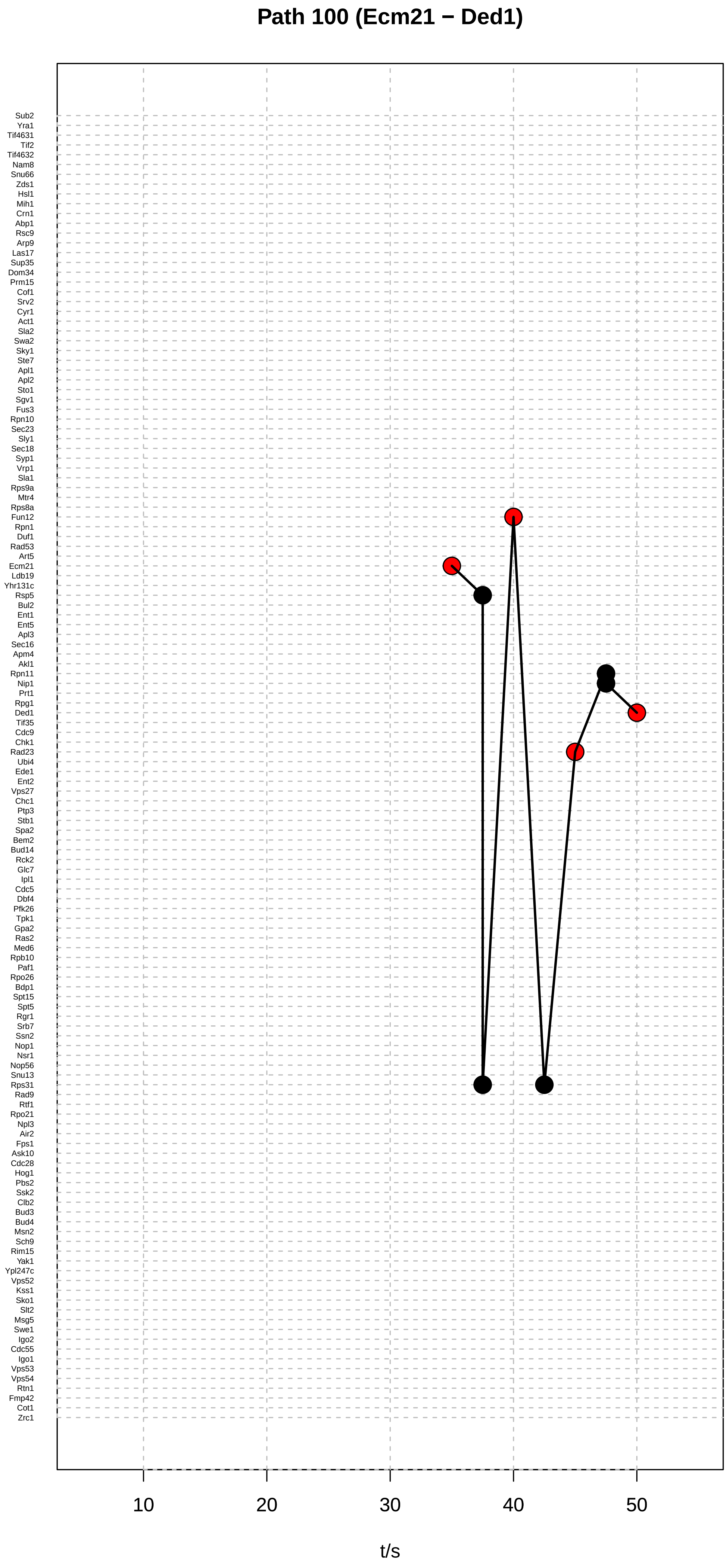

Path 100 (Ecm21 − Ded1)
Sub2
Yra1
Tif4631
Tif2
Tif4632
Nam8
Snu66
Zds1
Hsl1
Mih1
Crn1
Abp1
Rsc9
Arp9
Las17
Sup35
Dom34
Prm15
Cof1
Srv2
Cyr1
Act1
Sla2
Swa2
Sky1
Ste7
Apl1
Apl2
Sto1
Sgv1
Fus3
Rpn10
Sec23
Sly1
Sec18
Syp1
Vrp1
Sla1
Rps9a
Mtr4
Rps8a
Fun12
Rpn1
Duf1
Rad53
Art5
Ecm21
Ldb19
Yhr131c
Rsp5
Bul2
Ent1
Ent5
Apl3
Sec16
Apm4
Akl1
Rpn11
Nip1
Prt1
Rpg1
Ded1
Tif35
Cdc9
Chk1
Rad23
Ubi4
Ede1
Ent2
Vps27
Chc1
Ptp3
Stb1
Spa2
Bem2
Bud14
Rck2
Glc7
Ipl1
Cdc5
Dbf4
Pfk26
Tpk1
Gpa2
Ras2
Med6
Rpb10
Paf1
Rpo26
Bdp1
Spt15
Spt5
Rgr1
Srb7
Ssn2
Nop1
Nsr1
Nop56
Snu13
Rps31
Rad9
Rtf1
Rpo21
Npl3
Air2
Fps1
Ask10
Cdc28
Hog1
Pbs2
Ssk2
Clb2
Bud3
Bud4
Msn2
Sch9
Rim15
Yak1
Ypl247c
Vps52
Kss1
Sko1
Slt2
Msg5
Swe1
Igo2
Cdc55
Igo1
Vps53
Vps54
Rtn1
Fmp42
Cot1
Zrc1
10
20
30
40
50
t/s

#### Path 101 (Ecm21 − Ded1)

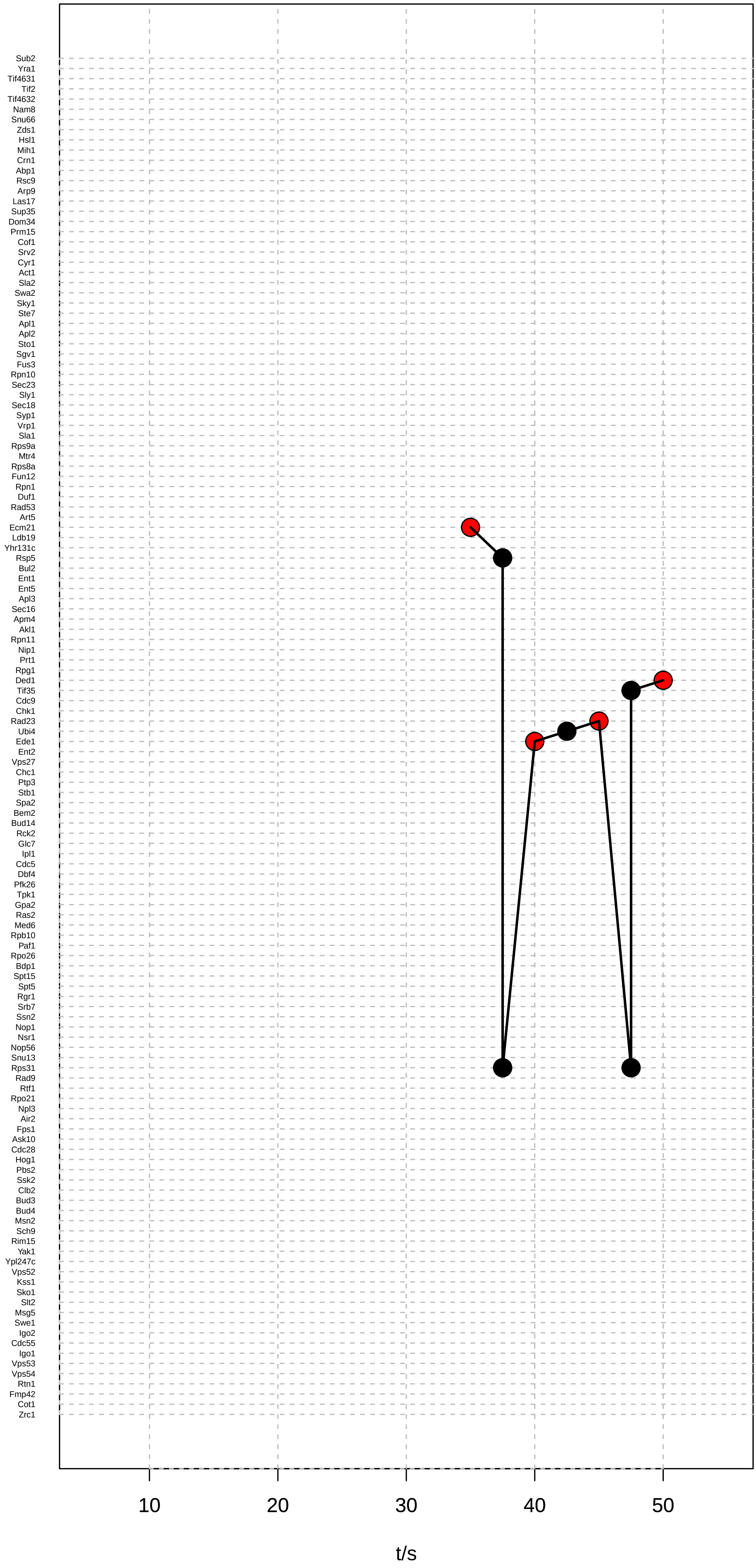

### Path 102 (Ecm21 − Ded1)

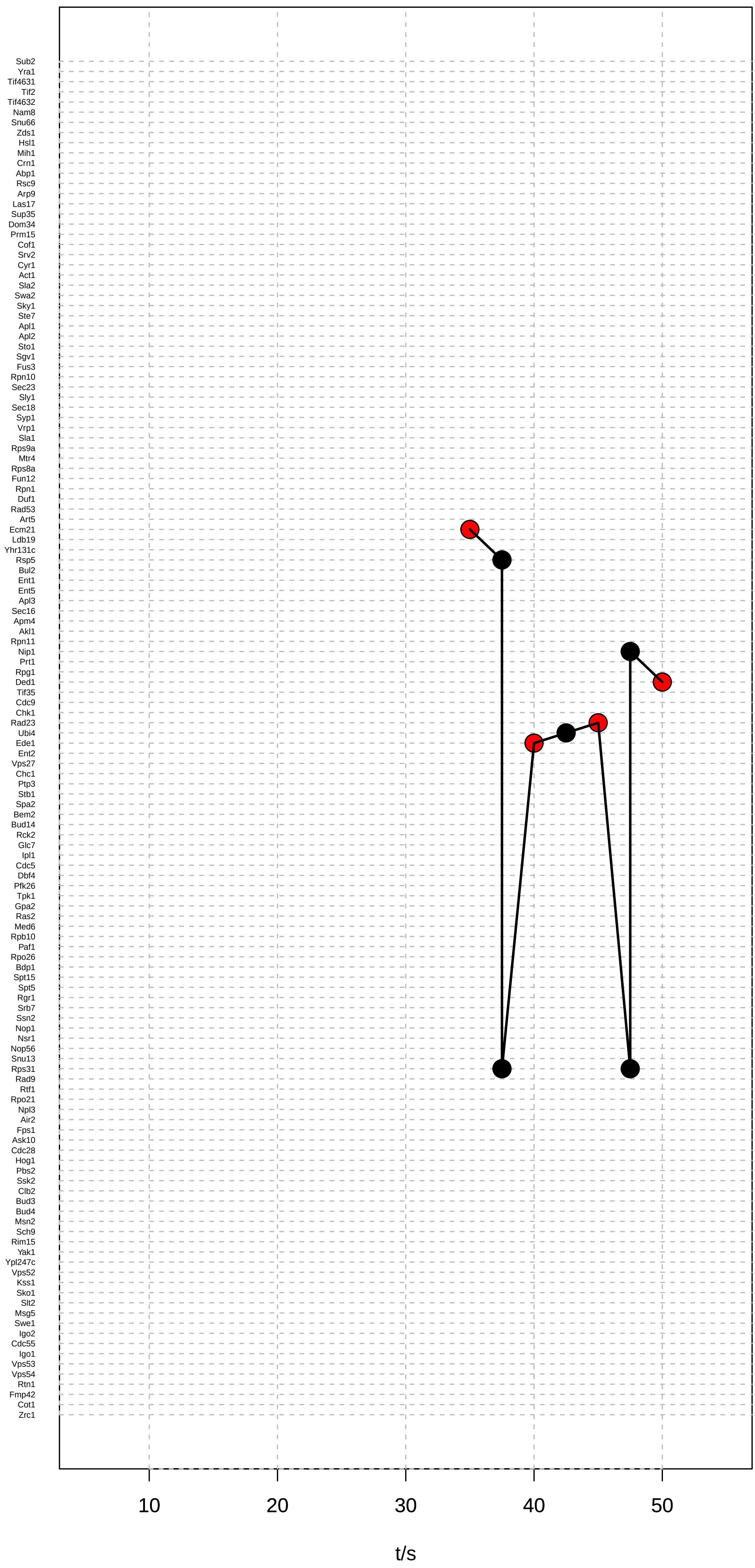

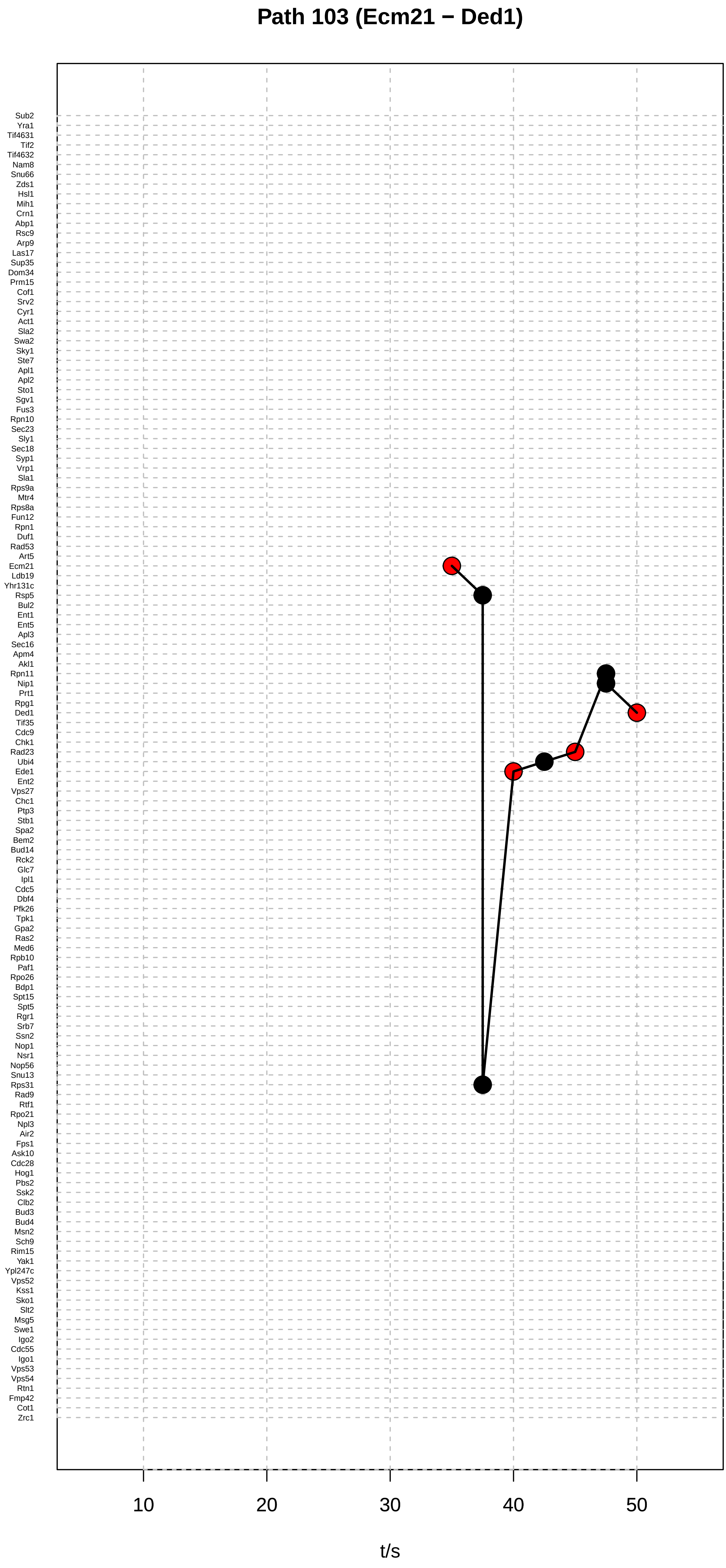

Path 103 (Ecm21 − Ded1)
Sub2
Yra1
Tif4631
Tif2
Tif4632
Nam8
Snu66
Zds1
Hsl1
Mih1
Crn1
Abp1
Rsc9
Arp9
Las17
Sup35
Dom34
Prm15
Cof1
Srv2
Cyr1
Act1
Sla2
Swa2
Sky1
Ste7
Apl1
Apl2
Sto1
Sgv1
Fus3
Rpn10
Sec23
Sly1
Sec18
Syp1
Vrp1
Sla1
Rps9a
Mtr4
Rps8a
Fun12
Rpn1
Duf1
Rad53
Art5
Ecm21
Ldb19
Yhr131c
Rsp5
Bul2
Ent1
Ent5
Apl3
Sec16
Apm4
Akl1
Rpn11
Nip1
Prt1
Rpg1
Ded1
Tif35
Cdc9
Chk1
Rad23
Ubi4
Ede1
Ent2
Vps27
Chc1
Ptp3
Stb1
Spa2
Bem2
Bud14
Rck2
Glc7
Ipl1
Cdc5
Dbf4
Pfk26
Tpk1
Gpa2
Ras2
Med6
Rpb10
Paf1
Rpo26
Bdp1
Spt15
Spt5
Rgr1
Srb7
Ssn2
Nop1
Nsr1
Nop56
Snu13
Rps31
Rad9
Rtf1
Rpo21
Npl3
Air2
Fps1
Ask10
Cdc28
Hog1
Pbs2
Ssk2
Clb2
Bud3
Bud4
Msn2
Sch9
Rim15
Yak1
Ypl247c
Vps52
Kss1
Sko1
Slt2
Msg5
Swe1
Igo2
Cdc55
Igo1
Vps53
Vps54
Rtn1
Fmp42
Cot1
Zrc1
10
20
30
40
50
t/s

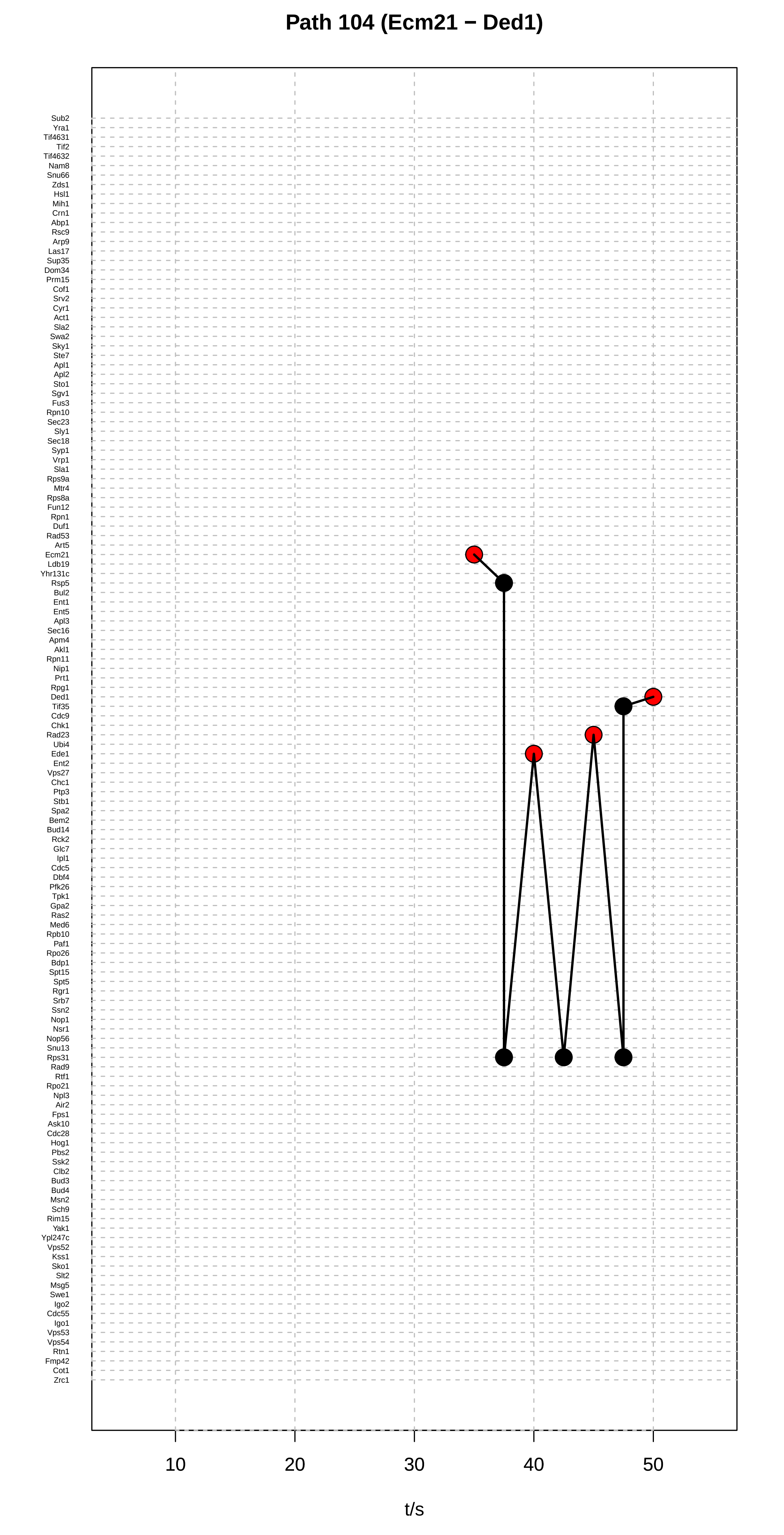
Path 104 (Ecm21 − Ded1)
Sub2
Yra1
Tif4631
Tif2
Tif4632
Nam8
Snu66
Zds1
Hsl1
Mih1
Crn1
Abp1
Rsc9
Arp9
Las17
Sup35
Dom34
Prm15
Cof1
Srv2
Cyr1
Act1
Sla2
Swa2
Sky1
Ste7
Apl1
Apl2
Sto1
Sgv1
Fus3
Rpn10
Sec23
Sly1
Sec18
Syp1
Vrp1
Sla1
Rps9a
Mtr4
Rps8a
Fun12
Rpn1
Duf1
Rad53
Art5
Ecm21
Ldb19
Yhr131c
Rsp5
Bul2
Ent1
Ent5
Apl3
Sec16
Apm4
Akl1
Rpn11
Nip1
Prt1
Rpg1
Ded1
Tif35
Cdc9
Chk1
Rad23
Ubi4
Ede1
Ent2
Vps27
Chc1
Ptp3
Stb1
Spa2
Bem2
Bud14
Rck2
Glc7
Ipl1
Cdc5
Dbf4
Pfk26
Tpk1
Gpa2
Ras2
Med6
Rpb10
Paf1
Rpo26
Bdp1
Spt15
Spt5
Rgr1
Srb7
Ssn2
Nop1
Nsr1
Nop56
Snu13
Rps31
Rad9
Rtf1
Rpo21
Npl3
Air2
Fps1
Ask10
Cdc28
Hog1
Pbs2
Ssk2
Clb2
Bud3
Bud4
Msn2
Sch9
Rim15
Yak1
Ypl247c
Vps52
Kss1
Sko1
Slt2
Msg5
Swe1
Igo2
Cdc55
Igo1
Vps53
Vps54
Rtn1
Fmp42
Cot1
Zrc1
10
20
30
40
50
t/s

### Path 105 (Ecm21 − Ded1)

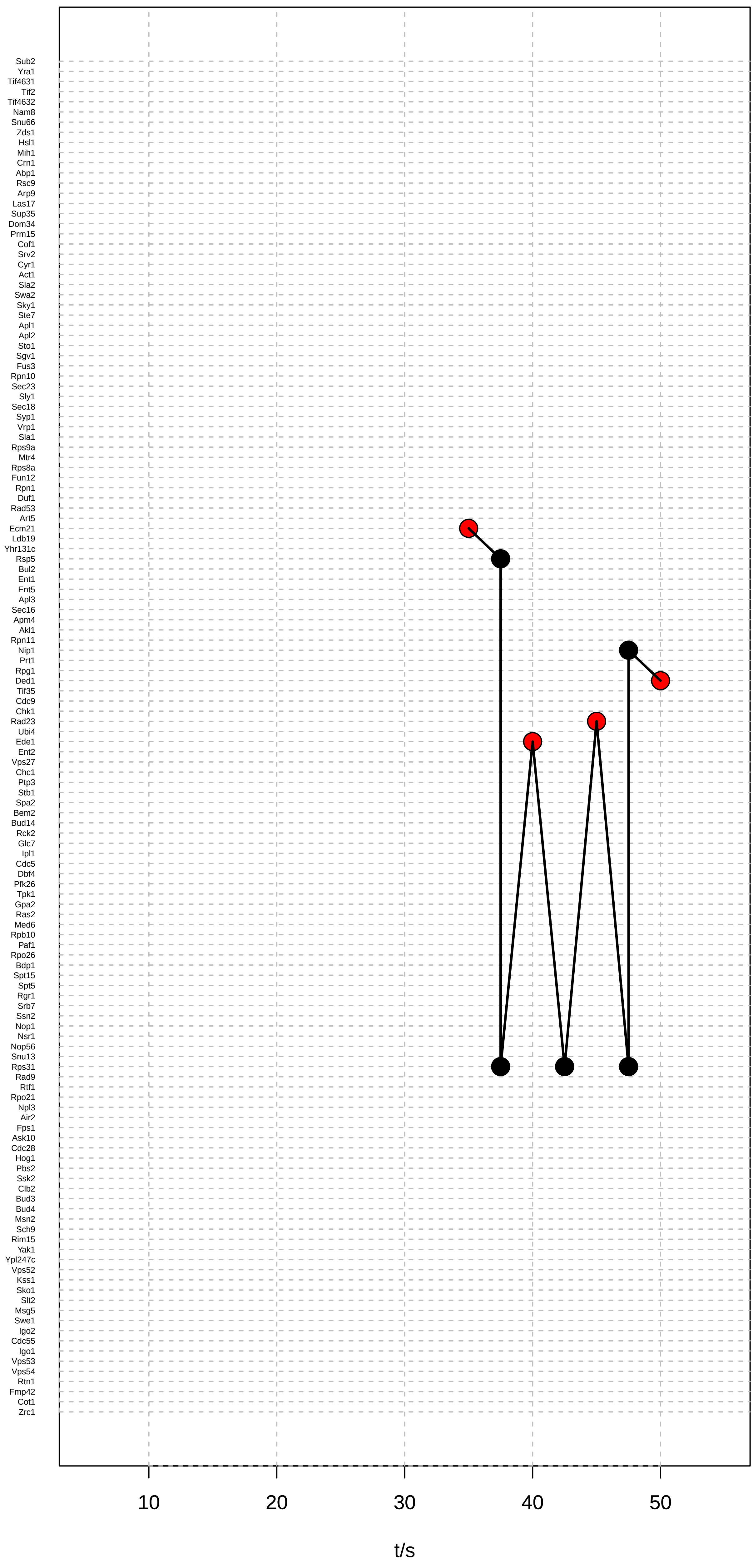

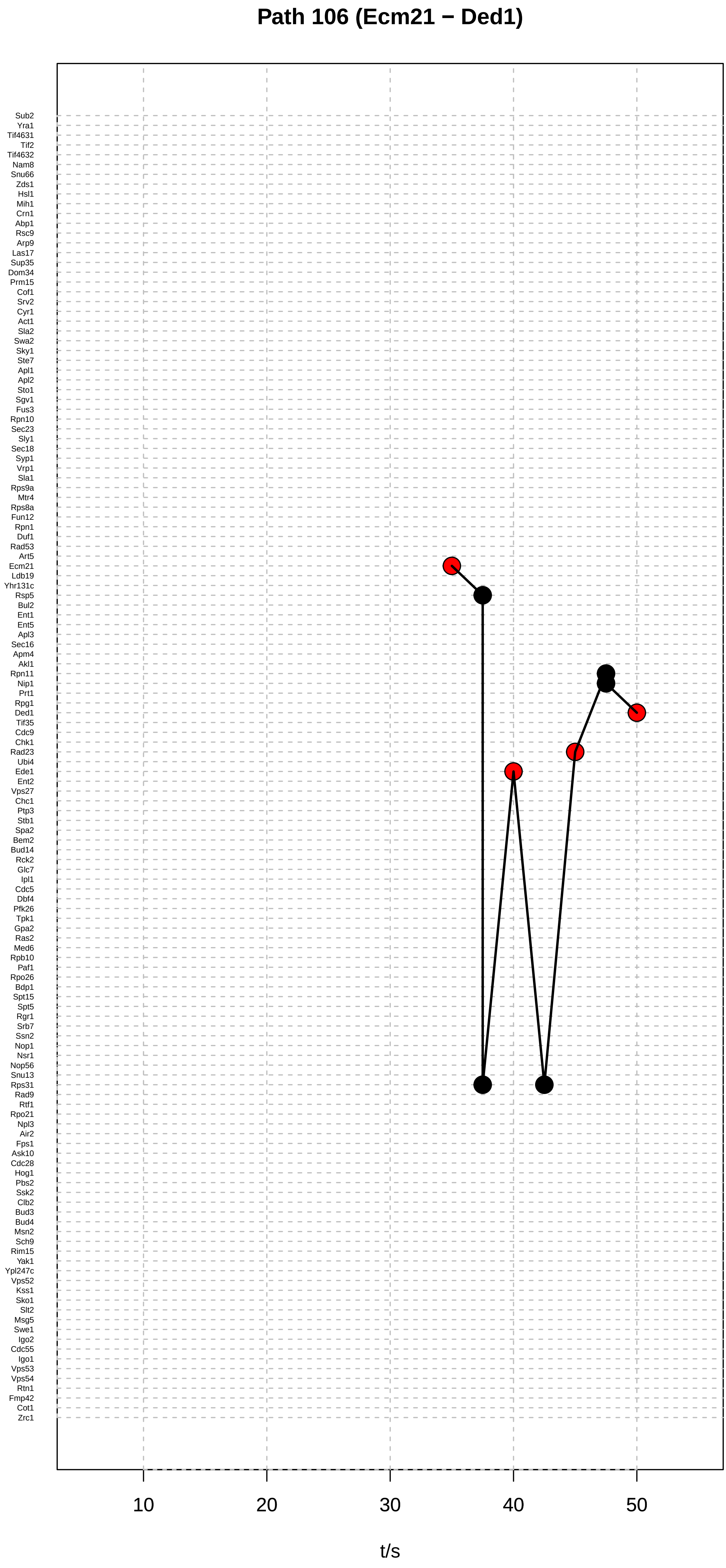

Path 106 (Ecm21 − Ded1)
Sub2
Yra1
Tif4631
Tif2
Tif4632
Nam8
Snu66
Zds1
Hsl1
Mih1
Crn1
Abp1
Rsc9
Arp9
Las17
Sup35
Dom34
Prm15
Cof1
Srv2
Cyr1
Act1
Sla2
Swa2
Sky1
Ste7
Apl1
Apl2
Sto1
Sgv1
Fus3
Rpn10
Sec23
Sly1
Sec18
Syp1
Vrp1
Sla1
Rps9a
Mtr4
Rps8a
Fun12
Rpn1
Duf1
Rad53
Art5
Ecm21
Ldb19
Yhr131c
Rsp5
Bul2
Ent1
Ent5
Apl3
Sec16
Apm4
Akl1
Rpn11
Nip1
Prt1
Rpg1
Ded1
Tif35
Cdc9
Chk1
Rad23
Ubi4
Ede1
Ent2
Vps27
Chc1
Ptp3
Stb1
Spa2
Bem2
Bud14
Rck2
Glc7
Ipl1
Cdc5
Dbf4
Pfk26
Tpk1
Gpa2
Ras2
Med6
Rpb10
Paf1
Rpo26
Bdp1
Spt15
Spt5
Rgr1
Srb7
Ssn2
Nop1
Nsr1
Nop56
Snu13
Rps31
Rad9
Rtf1
Rpo21
Npl3
Air2
Fps1
Ask10
Cdc28
Hog1
Pbs2
Ssk2
Clb2
Bud3
Bud4
Msn2
Sch9
Rim15
Yak1
Ypl247c
Vps52
Kss1
Sko1
Slt2
Msg5
Swe1
Igo2
Cdc55
Igo1
Vps53
Vps54
Rtn1
Fmp42
Cot1
Zrc1
10
20
30
40
50
t/s

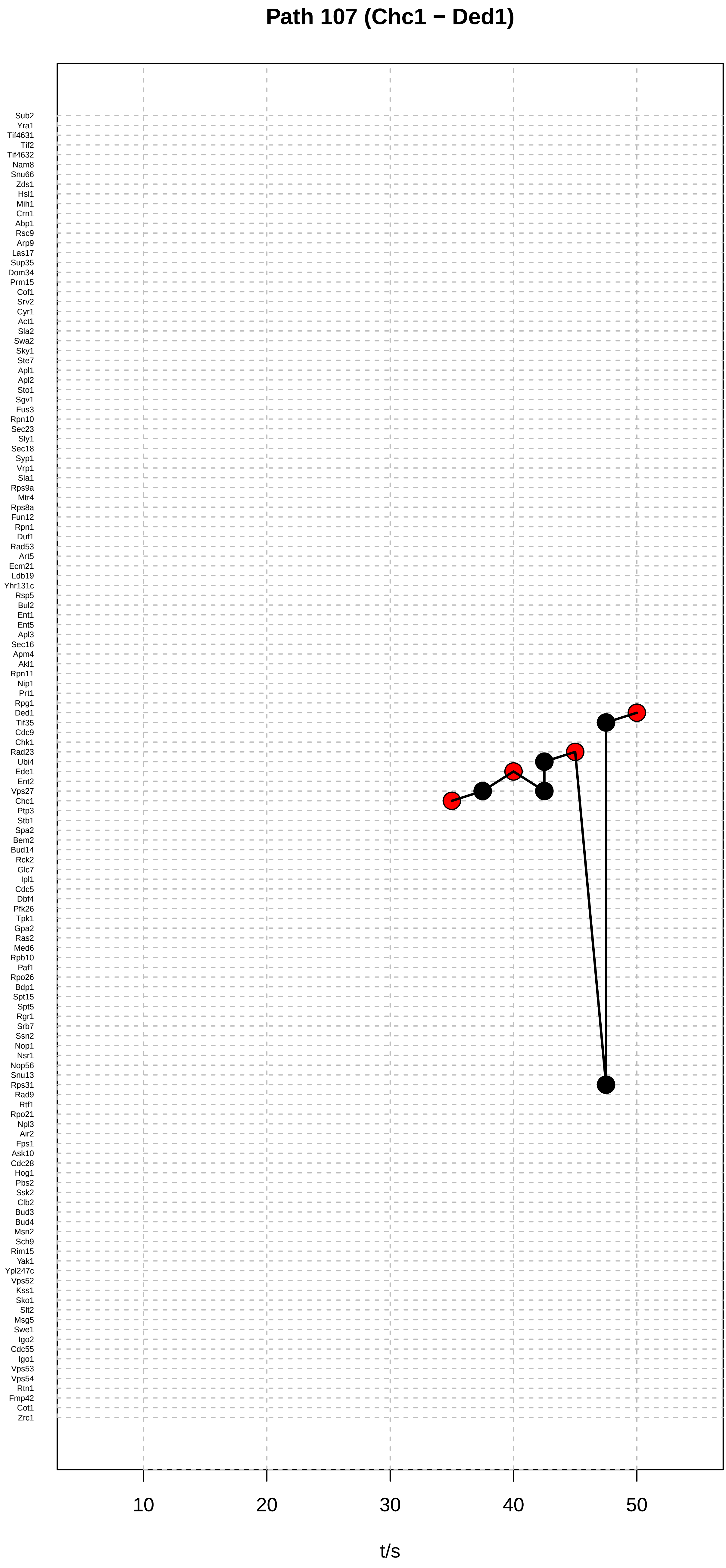
Path 107 (Chc1 − Ded1)
Sub2
Yra1
Tif4631
Tif2
Tif4632
Nam8
Snu66
Zds1
Hsl1
Mih1
Crn1
Abp1
Rsc9
Arp9
Las17
Sup35
Dom34
Prm15
Cof1
Srv2
Cyr1
Act1
Sla2
Swa2
Sky1
Ste7
Apl1
Apl2
Sto1
Sgv1
Fus3
Rpn10
Sec23
Sly1
Sec18
Syp1
Vrp1
Sla1
Rps9a
Mtr4
Rps8a
Fun12
Rpn1
Duf1
Rad53
Art5
Ecm21
Ldb19
Yhr131c
Rsp5
Bul2
Ent1
Ent5
Apl3
Sec16
Apm4
Akl1
Rpn11
Nip1
Prt1
Rpg1
Ded1
Tif35
Cdc9
Chk1
Rad23
Ubi4
Ede1
Ent2
Vps27
Chc1
Ptp3
Stb1
Spa2
Bem2
Bud14
Rck2
Glc7
Ipl1
Cdc5
Dbf4
Pfk26
Tpk1
Gpa2
Ras2
Med6
Rpb10
Paf1
Rpo26
Bdp1
Spt15
Spt5
Rgr1
Srb7
Ssn2
Nop1
Nsr1
Nop56
Snu13
Rps31
Rad9
Rtf1
Rpo21
Npl3
Air2
Fps1
Ask10
Cdc28
Hog1
Pbs2
Ssk2
Clb2
Bud3
Bud4
Msn2
Sch9
Rim15
Yak1
Ypl247c
Vps52
Kss1
Sko1
Slt2
Msg5
Swe1
Igo2
Cdc55
Igo1
Vps53
Vps54
Rtn1
Fmp42
Cot1
Zrc1
10
20
30
40
50
t/s

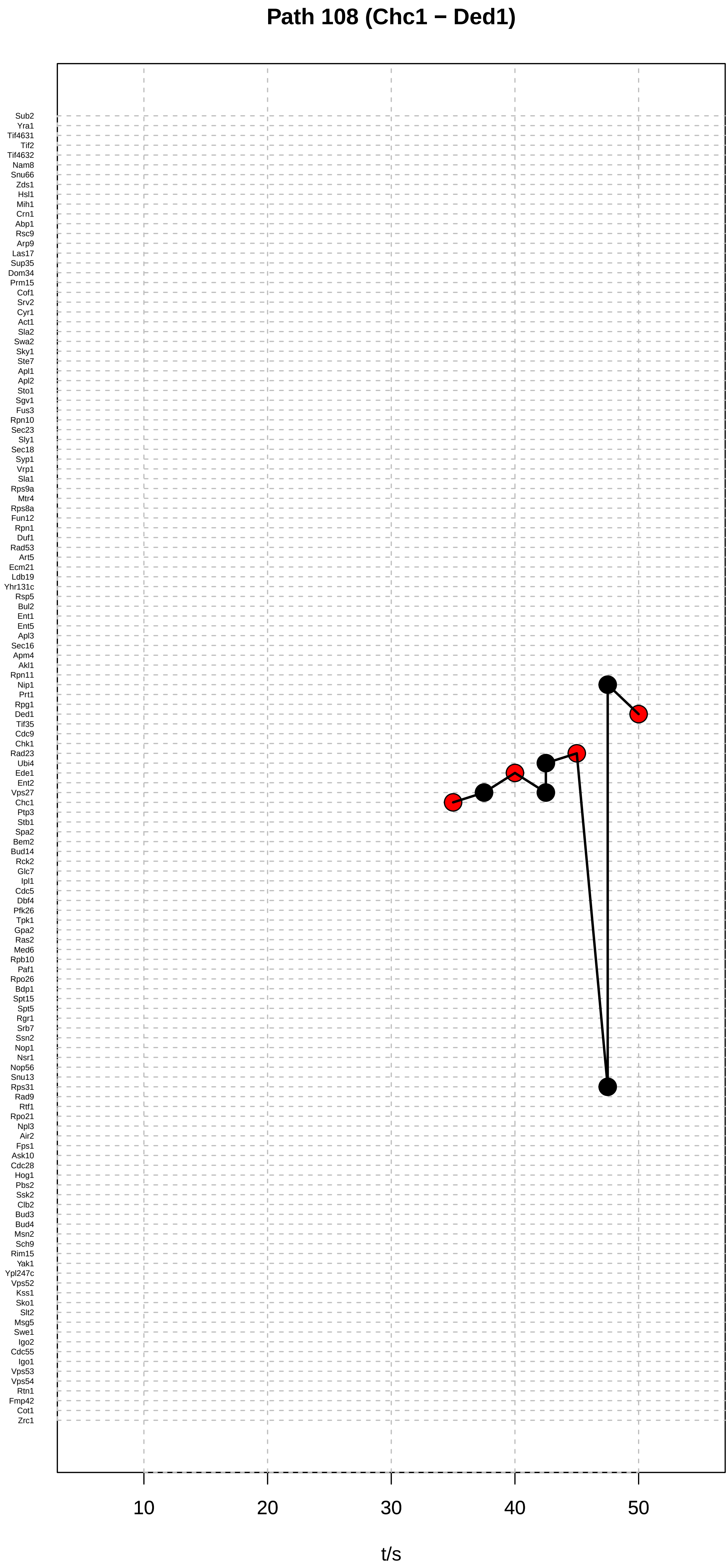
Path 108 (Chc1 − Ded1)
Sub2
Yra1
Tif4631
Tif2
Tif4632
Nam8
Snu66
Zds1
Hsl1
Mih1
Crn1
Abp1
Rsc9
Arp9
Las17
Sup35
Dom34
Prm15
Cof1
Srv2
Cyr1
Act1
Sla2
Swa2
Sky1
Ste7
Apl1
Apl2
Sto1
Sgv1
Fus3
Rpn10
Sec23
Sly1
Sec18
Syp1
Vrp1
Sla1
Rps9a
Mtr4
Rps8a
Fun12
Rpn1
Duf1
Rad53
Art5
Ecm21
Ldb19
Yhr131c
Rsp5
Bul2
Ent1
Ent5
Apl3
Sec16
Apm4
Akl1
Rpn11
Nip1
Prt1
Rpg1
Ded1
Tif35
Cdc9
Chk1
Rad23
Ubi4
Ede1
Ent2
Vps27
Chc1
Ptp3
Stb1
Spa2
Bem2
Bud14
Rck2
Glc7
Ipl1
Cdc5
Dbf4
Pfk26
Tpk1
Gpa2
Ras2
Med6
Rpb10
Paf1
Rpo26
Bdp1
Spt15
Spt5
Rgr1
Srb7
Ssn2
Nop1
Nsr1
Nop56
Snu13
Rps31
Rad9
Rtf1
Rpo21
Npl3
Air2
Fps1
Ask10
Cdc28
Hog1
Pbs2
Ssk2
Clb2
Bud3
Bud4
Msn2
Sch9
Rim15
Yak1
Ypl247c
Vps52
Kss1
Sko1
Slt2
Msg5
Swe1
Igo2
Cdc55
Igo1
Vps53
Vps54
Rtn1
Fmp42
Cot1
Zrc1
10
20
30
40
50
t/s

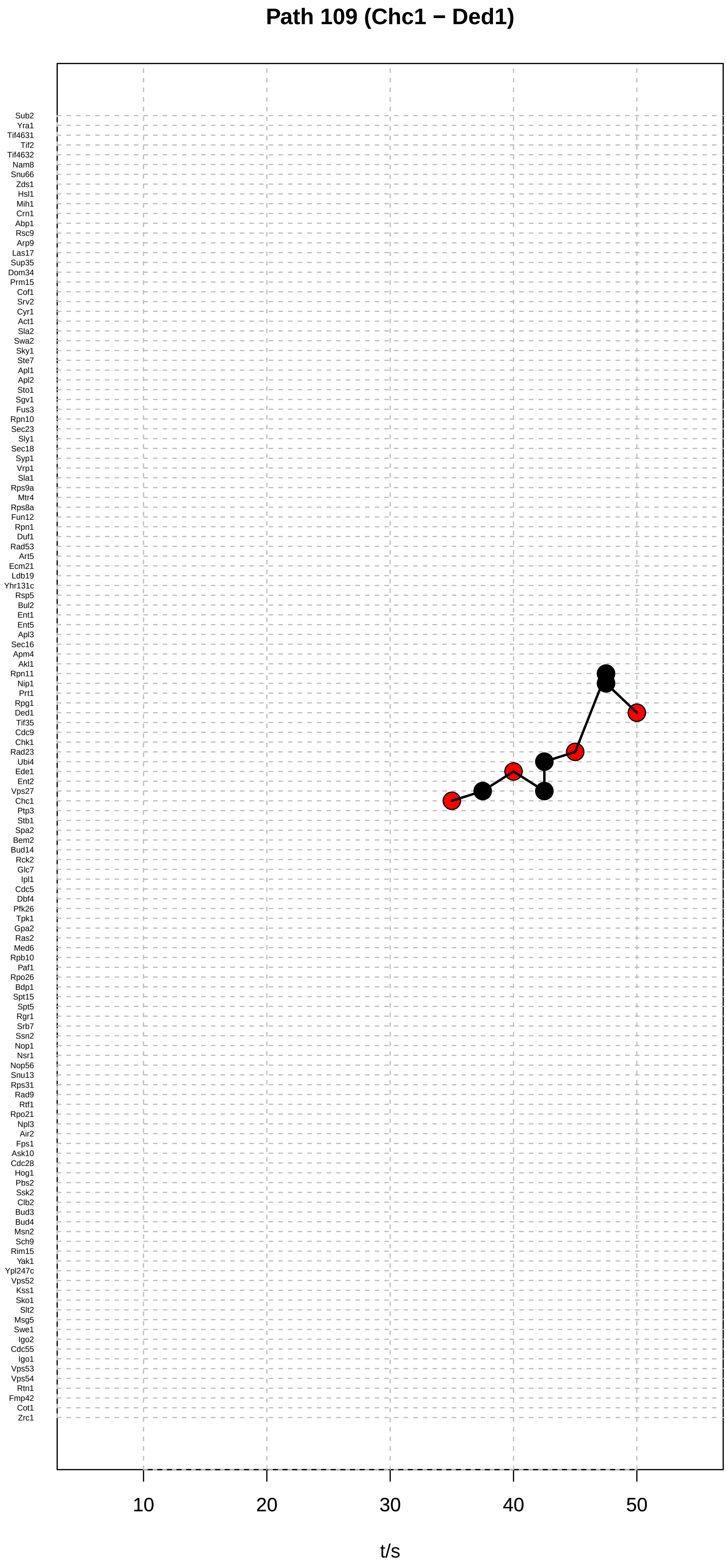

Path 109 (Chc1 − Ded1)
Sub2
Yra1
Tif4631
Tif2
Tif4632
Nam8
Snu66
Zds1
Hsl1
Mih1
Crn1
Abp1
Rsc9
Arp9
Las17
Sup35
Dom34
Prm15
Cof1
Srv2
Cyr1
Act1
Sla2
Swa2
Sky1
Ste7
Apl1
Apl2
Sto1
Sgv1
Fus3
Rpn10
Sec23
Sly1
Sec18
Syp1
Vrp1
Sla1
Rps9a
Mtr4
Rps8a
Fun12
Rpn1
Duf1
Rad53
Art5
Ecm21
Ldb19
Yhr131c
Rsp5
Bul2
Ent1
Ent5
Apl3
Sec16
Apm4
Akl1
Rpn11
Nip1
Prt1
Rpg1
Ded1
Tif35
Cdc9
Chk1
Rad23
Ubi4
Ede1
Ent2
Vps27
Chc1
Ptp3
Stb1
Spa2
Bem2
Bud14
Rck2
Glc7
Ipl1
Cdc5
Dbf4
Pfk26
Tpk1
Gpa2
Ras2
Med6
Rpb10
Paf1
Rpo26
Bdp1
Spt15
Spt5
Rgr1
Srb7
Ssn2
Nop1
Nsr1
Nop56
Snu13
Rps31
Rad9
Rtf1
Rpo21
Npl3
Air2
Fps1
Ask10
Cdc28
Hog1
Pbs2
Ssk2
Clb2
Bud3
Bud4
Msn2
Sch9
Rim15
Yak1
Ypl247c
Vps52
Kss1
Sko1
Slt2
Msg5
Swe1
Igo2
Cdc55
Igo1
Vps53
Vps54
Rtn1
Fmp42
Cot1
Zrc1
10
20
30
40
50
t/s

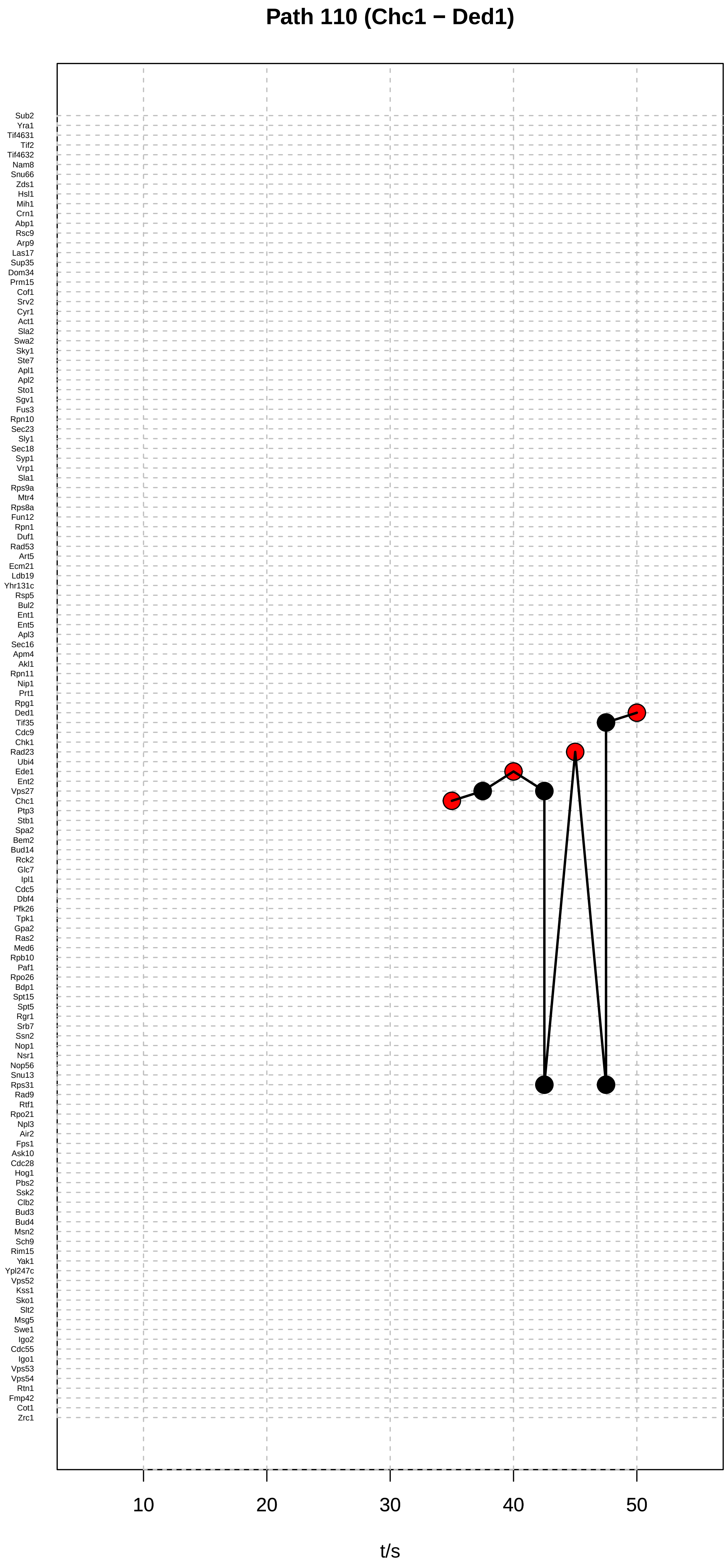
Path 110 (Chc1 − Ded1)
Sub2
Yra1
Tif4631
Tif2
Tif4632
Nam8
Snu66
Zds1
Hsl1
Mih1
Crn1
Abp1
Rsc9
Arp9
Las17
Sup35
Dom34
Prm15
Cof1
Srv2
Cyr1
Act1
Sla2
Swa2
Sky1
Ste7
Apl1
Apl2
Sto1
Sgv1
Fus3
Rpn10
Sec23
Sly1
Sec18
Syp1
Vrp1
Sla1
Rps9a
Mtr4
Rps8a
Fun12
Rpn1
Duf1
Rad53
Art5
Ecm21
Ldb19
Yhr131c
Rsp5
Bul2
Ent1
Ent5
Apl3
Sec16
Apm4
Akl1
Rpn11
Nip1
Prt1
Rpg1
Ded1
Tif35
Cdc9
Chk1
Rad23
Ubi4
Ede1
Ent2
Vps27
Chc1
Ptp3
Stb1
Spa2
Bem2
Bud14
Rck2
Glc7
Ipl1
Cdc5
Dbf4
Pfk26
Tpk1
Gpa2
Ras2
Med6
Rpb10
Paf1
Rpo26
Bdp1
Spt15
Spt5
Rgr1
Srb7
Ssn2
Nop1
Nsr1
Nop56
Snu13
Rps31
Rad9
Rtf1
Rpo21
Npl3
Air2
Fps1
Ask10
Cdc28
Hog1
Pbs2
Ssk2
Clb2
Bud3
Bud4
Msn2
Sch9
Rim15
Yak1
Ypl247c
Vps52
Kss1
Sko1
Slt2
Msg5
Swe1
Igo2
Cdc55
Igo1
Vps53
Vps54
Rtn1
Fmp42
Cot1
Zrc1
10
20
30
40
50
t/s

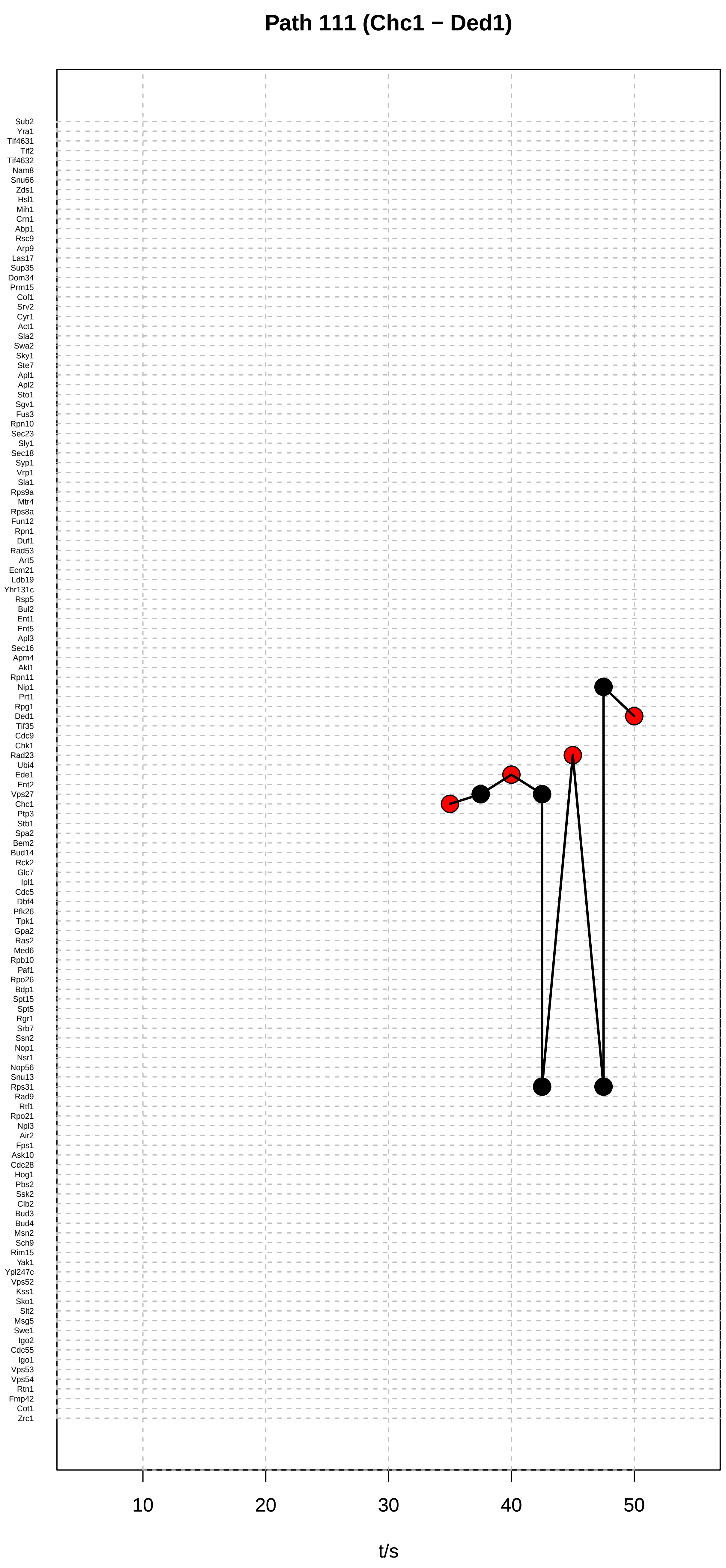

Path 111 (Chc1 − Ded1)
Sub2
Yra1
Tif4631
Tif2
Tif4632
Nam8
Snu66
Zds1
Hsl1
Mih1
Crn1
Abp1
Rsc9
Arp9
Las17
Sup35
Dom34
Prm15
Cof1
Srv2
Cyr1
Act1
Sla2
Swa2
Sky1
Ste7
Apl1
Apl2
Sto1
Sgv1
Fus3
Rpn10
Sec23
Sly1
Sec18
Syp1
Vrp1
Sla1
Rps9a
Mtr4
Rps8a
Fun12
Rpn1
Duf1
Rad53
Art5
Ecm21
Ldb19
Yhr131c
Rsp5
Bul2
Ent1
Ent5
Apl3
Sec16
Apm4
Akl1
Rpn11
Nip1
Prt1
Rpg1
Ded1
Tif35
Cdc9
Chk1
Rad23
Ubi4
Ede1
Ent2
Vps27
Chc1
Ptp3
Stb1
Spa2
Bem2
Bud14
Rck2
Glc7
Ipl1
Cdc5
Dbf4
Pfk26
Tpk1
Gpa2
Ras2
Med6
Rpb10
Paf1
Rpo26
Bdp1
Spt15
Spt5
Rgr1
Srb7
Ssn2
Nop1
Nsr1
Nop56
Snu13
Rps31
Rad9
Rtf1
Rpo21
Npl3
Air2
Fps1
Ask10
Cdc28
Hog1
Pbs2
Ssk2
Clb2
Bud3
Bud4
Msn2
Sch9
Rim15
Yak1
Ypl247c
Vps52
Kss1
Sko1
Slt2
Msg5
Swe1
Igo2
Cdc55
Igo1
Vps53
Vps54
Rtn1
Fmp42
Cot1
Zrc1
10
20
30
40
50
t/s

#### Path 112 (Chc1 − Ded1)

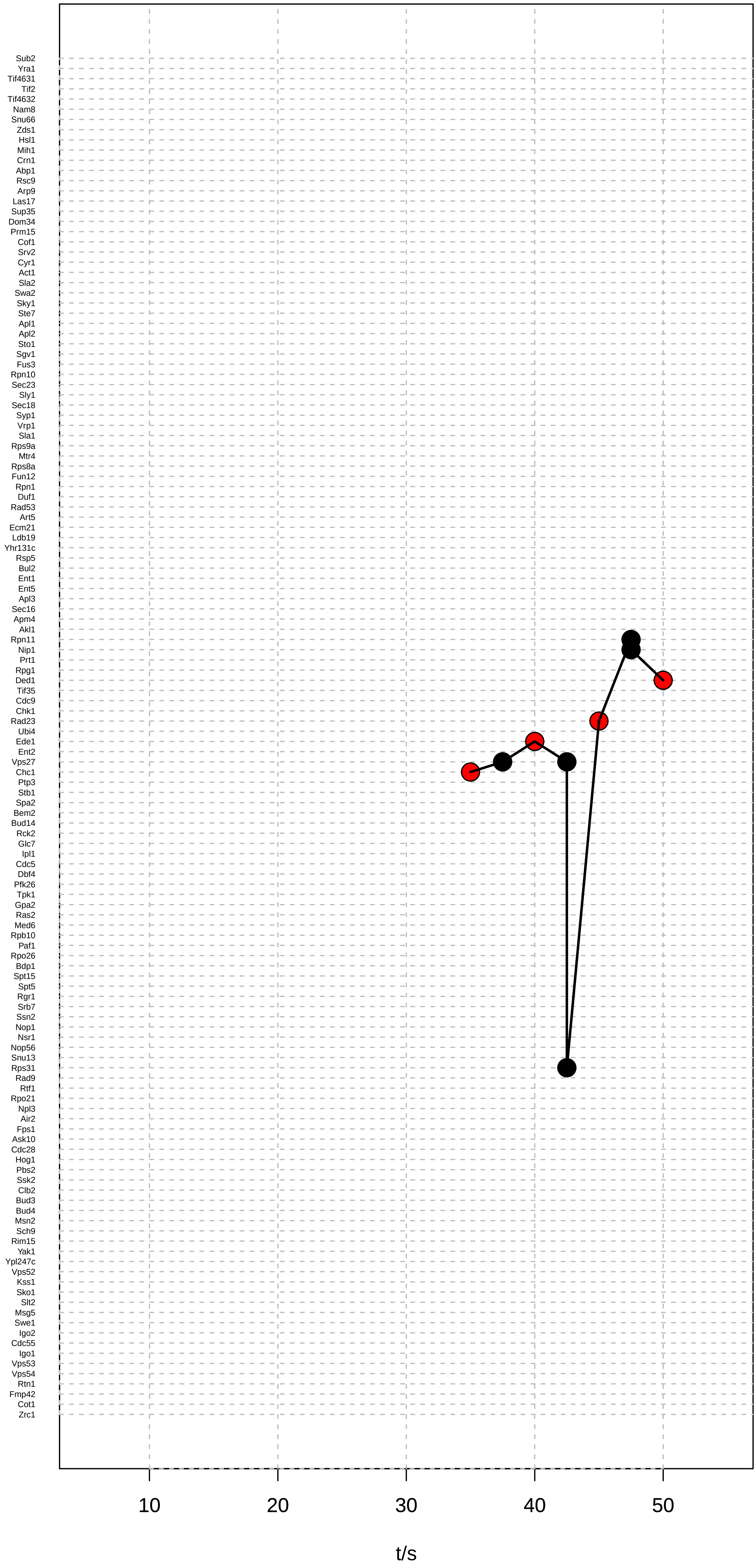

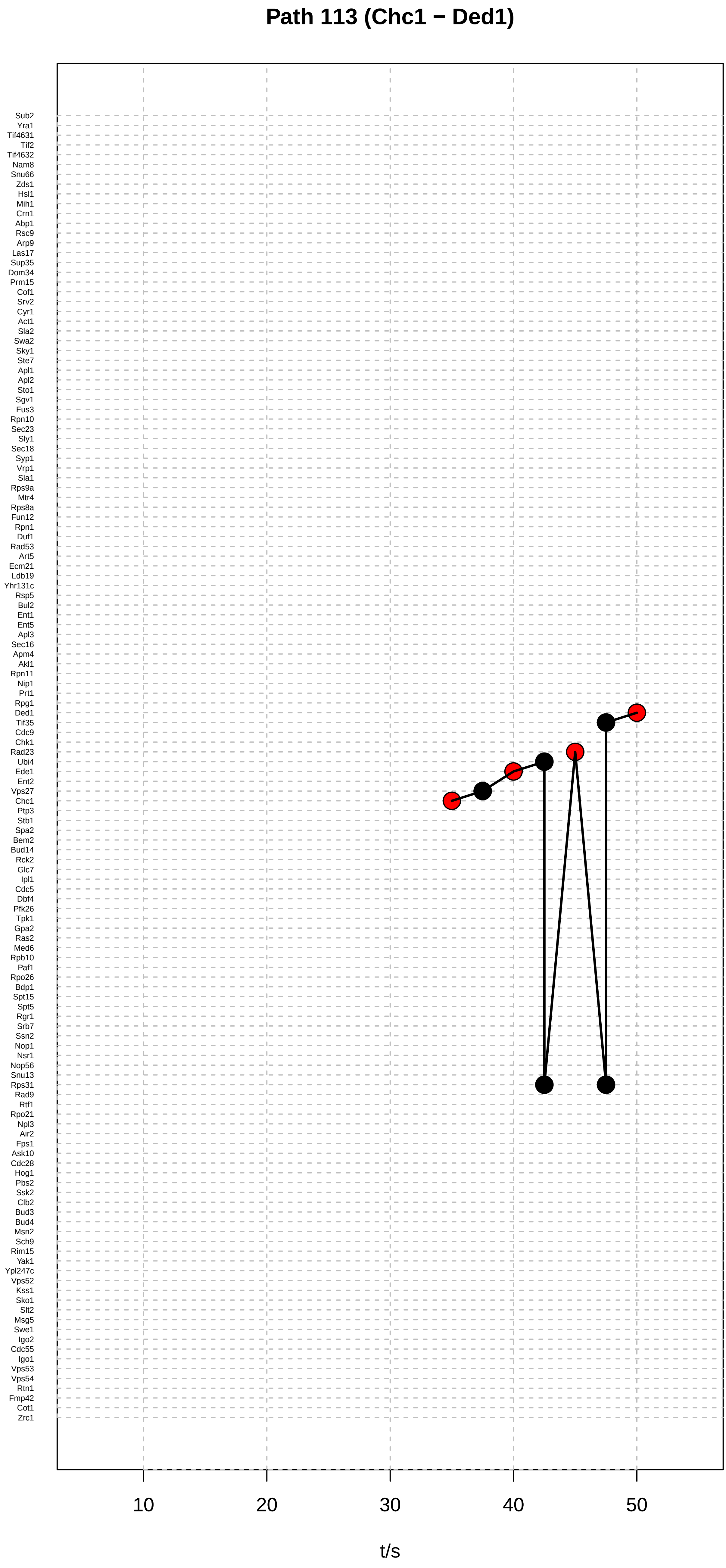

Path 113 (Chc1 − Ded1)
Sub2
Yra1
Tif4631
Tif2
Tif4632
Nam8
Snu66
Zds1
Hsl1
Mih1
Crn1
Abp1
Rsc9
Arp9
Las17
Sup35
Dom34
Prm15
Cof1
Srv2
Cyr1
Act1
Sla2
Swa2
Sky1
Ste7
Apl1
Apl2
Sto1
Sgv1
Fus3
Rpn10
Sec23
Sly1
Sec18
Syp1
Vrp1
Sla1
Rps9a
Mtr4
Rps8a
Fun12
Rpn1
Duf1
Rad53
Art5
Ecm21
Ldb19
Yhr131c
Rsp5
Bul2
Ent1
Ent5
Apl3
Sec16
Apm4
Akl1
Rpn11
Nip1
Prt1
Rpg1
Ded1
Tif35
Cdc9
Chk1
Rad23
Ubi4
Ede1
Ent2
Vps27
Chc1
Ptp3
Stb1
Spa2
Bem2
Bud14
Rck2
Glc7
Ipl1
Cdc5
Dbf4
Pfk26
Tpk1
Gpa2
Ras2
Med6
Rpb10
Paf1
Rpo26
Bdp1
Spt15
Spt5
Rgr1
Srb7
Ssn2
Nop1
Nsr1
Nop56
Snu13
Rps31
Rad9
Rtf1
Rpo21
Npl3
Air2
Fps1
Ask10
Cdc28
Hog1
Pbs2
Ssk2
Clb2
Bud3
Bud4
Msn2
Sch9
Rim15
Yak1
Ypl247c
Vps52
Kss1
Sko1
Slt2
Msg5
Swe1
Igo2
Cdc55
Igo1
Vps53
Vps54
Rtn1
Fmp42
Cot1
Zrc1
10
20
30
40
50
t/s

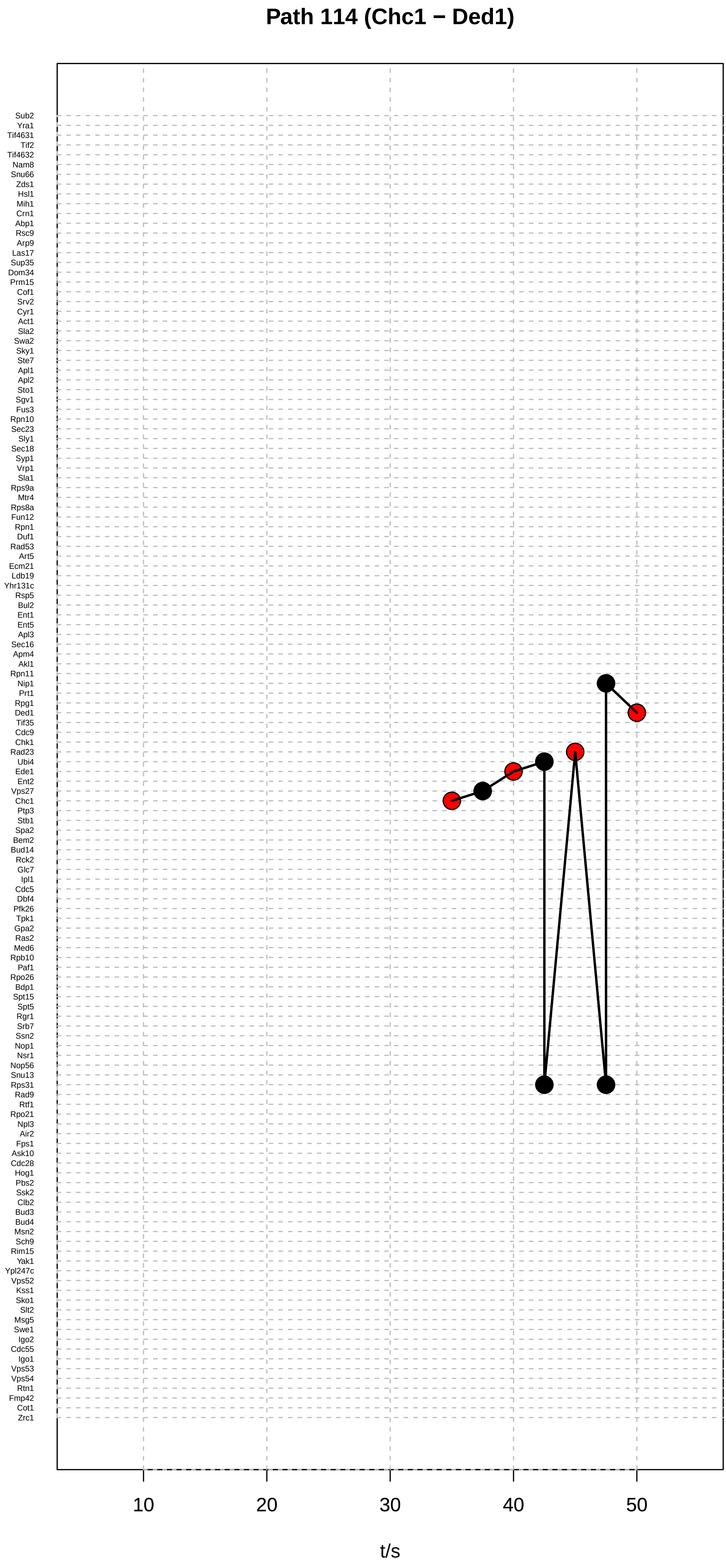
Path 114 (Chc1 − Ded1)
Sub2
Yra1
Tif4631
Tif2
Tif4632
Nam8
Snu66
Zds1
Hsl1
Mih1
Crn1
Abp1
Rsc9
Arp9
Las17
Sup35
Dom34
Prm15
Cof1
Srv2
Cyr1
Act1
Sla2
Swa2
Sky1
Ste7
Apl1
Apl2
Sto1
Sgv1
Fus3
Rpn10
Sec23
Sly1
Sec18
Syp1
Vrp1
Sla1
Rps9a
Mtr4
Rps8a
Fun12
Rpn1
Duf1
Rad53
Art5
Ecm21
Ldb19
Yhr131c
Rsp5
Bul2
Ent1
Ent5
Apl3
Sec16
Apm4
Akl1
Rpn11
Nip1
Prt1
Rpg1
Ded1
Tif35
Cdc9
Chk1
Rad23
Ubi4
Ede1
Ent2
Vps27
Chc1
Ptp3
Stb1
Spa2
Bem2
Bud14
Rck2
Glc7
Ipl1
Cdc5
Dbf4
Pfk26
Tpk1
Gpa2
Ras2
Med6
Rpb10
Paf1
Rpo26
Bdp1
Spt15
Spt5
Rgr1
Srb7
Ssn2
Nop1
Nsr1
Nop56
Snu13
Rps31
Rad9
Rtf1
Rpo21
Npl3
Air2
Fps1
Ask10
Cdc28
Hog1
Pbs2
Ssk2
Clb2
Bud3
Bud4
Msn2
Sch9
Rim15
Yak1
Ypl247c
Vps52
Kss1
Sko1
Slt2
Msg5
Swe1
Igo2
Cdc55
Igo1
Vps53
Vps54
Rtn1
Fmp42
Cot1
Zrc1
10
20
30
40
50
t/s

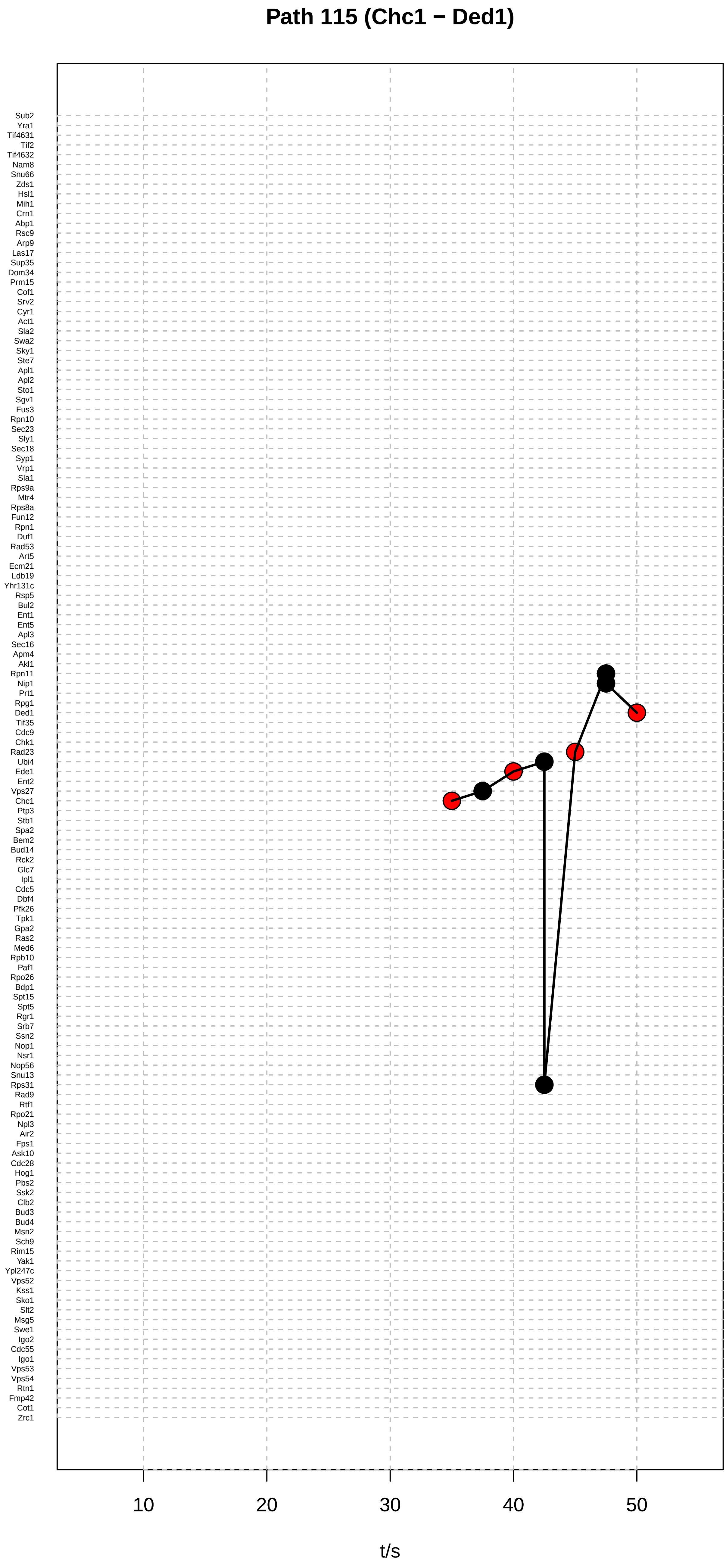

Path 115 (Chc1 − Ded1)
Sub2
Yra1
Tif4631
Tif2
Tif4632
Nam8
Snu66
Zds1
Hsl1
Mih1
Crn1
Abp1
Rsc9
Arp9
Las17
Sup35
Dom34
Prm15
Cof1
Srv2
Cyr1
Act1
Sla2
Swa2
Sky1
Ste7
Apl1
Apl2
Sto1
Sgv1
Fus3
Rpn10
Sec23
Sly1
Sec18
Syp1
Vrp1
Sla1
Rps9a
Mtr4
Rps8a
Fun12
Rpn1
Duf1
Rad53
Art5
Ecm21
Ldb19
Yhr131c
Rsp5
Bul2
Ent1
Ent5
Apl3
Sec16
Apm4
Akl1
Rpn11
Nip1
Prt1
Rpg1
Ded1
Tif35
Cdc9
Chk1
Rad23
Ubi4
Ede1
Ent2
Vps27
Chc1
Ptp3
Stb1
Spa2
Bem2
Bud14
Rck2
Glc7
Ipl1
Cdc5
Dbf4
Pfk26
Tpk1
Gpa2
Ras2
Med6
Rpb10
Paf1
Rpo26
Bdp1
Spt15
Spt5
Rgr1
Srb7
Ssn2
Nop1
Nsr1
Nop56
Snu13
Rps31
Rad9
Rtf1
Rpo21
Npl3
Air2
Fps1
Ask10
Cdc28
Hog1
Pbs2
Ssk2
Clb2
Bud3
Bud4
Msn2
Sch9
Rim15
Yak1
Ypl247c
Vps52
Kss1
Sko1
Slt2
Msg5
Swe1
Igo2
Cdc55
Igo1
Vps53
Vps54
Rtn1
Fmp42
Cot1
Zrc1
10
20
30
40
50
t/s

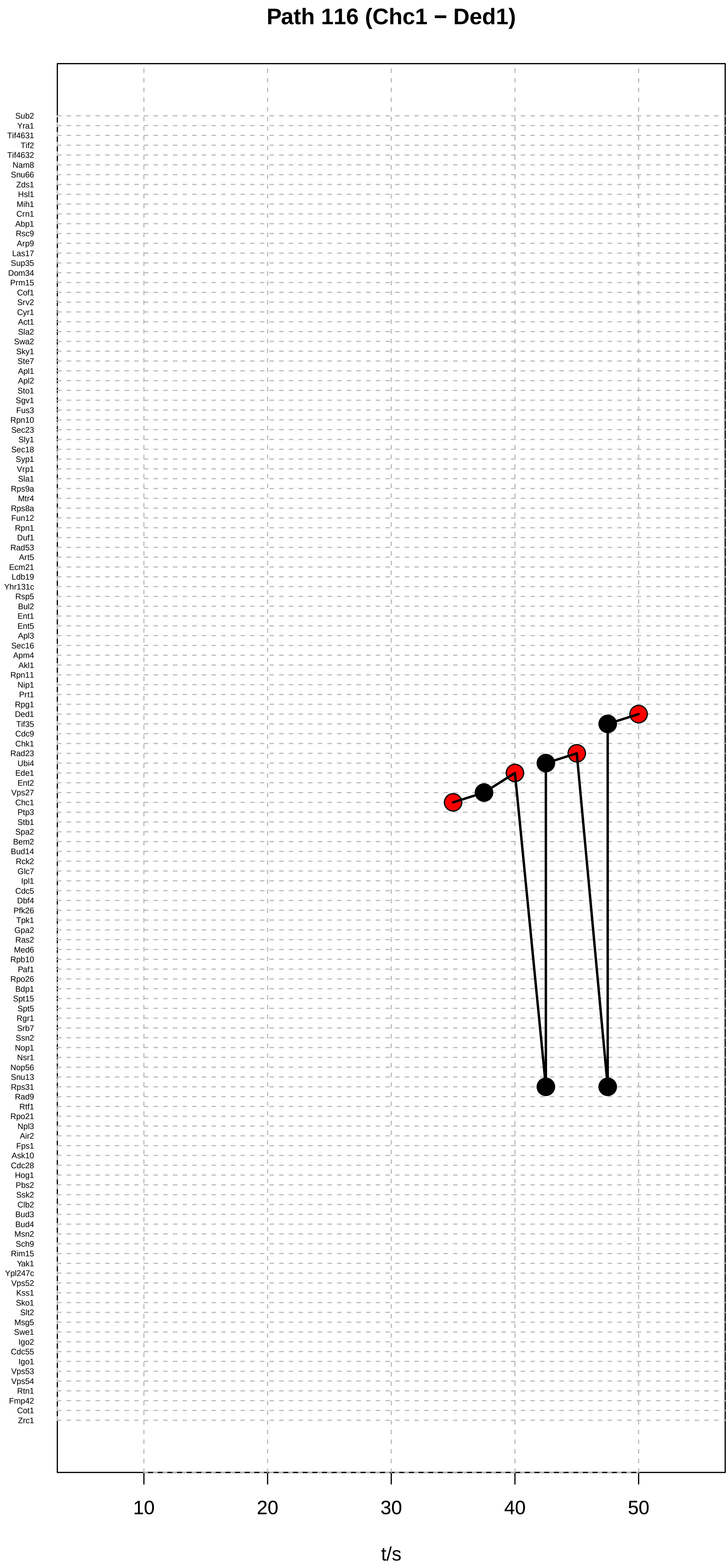

Path 116 (Chc1 − Ded1)
Sub2
Yra1
Tif4631
Tif2
Tif4632
Nam8
Snu66
Zds1
Hsl1
Mih1
Crn1
Abp1
Rsc9
Arp9
Las17
Sup35
Dom34
Prm15
Cof1
Srv2
Cyr1
Act1
Sla2
Swa2
Sky1
Ste7
Apl1
Apl2
Sto1
Sgv1
Fus3
Rpn10
Sec23
Sly1
Sec18
Syp1
Vrp1
Sla1
Rps9a
Mtr4
Rps8a
Fun12
Rpn1
Duf1
Rad53
Art5
Ecm21
Ldb19
Yhr131c
Rsp5
Bul2
Ent1
Ent5
Apl3
Sec16
Apm4
Akl1
Rpn11
Nip1
Prt1
Rpg1
Ded1
Tif35
Cdc9
Chk1
Rad23
Ubi4
Ede1
Ent2
Vps27
Chc1
Ptp3
Stb1
Spa2
Bem2
Bud14
Rck2
Glc7
Ipl1
Cdc5
Dbf4
Pfk26
Tpk1
Gpa2
Ras2
Med6
Rpb10
Paf1
Rpo26
Bdp1
Spt15
Spt5
Rgr1
Srb7
Ssn2
Nop1
Nsr1
Nop56
Snu13
Rps31
Rad9
Rtf1
Rpo21
Npl3
Air2
Fps1
Ask10
Cdc28
Hog1
Pbs2
Ssk2
Clb2
Bud3
Bud4
Msn2
Sch9
Rim15
Yak1
Ypl247c
Vps52
Kss1
Sko1
Slt2
Msg5
Swe1
Igo2
Cdc55
Igo1
Vps53
Vps54
Rtn1
Fmp42
Cot1
Zrc1
10
20
30
40
50
t/s

#### Path 117 (Chc1 − Ded1)

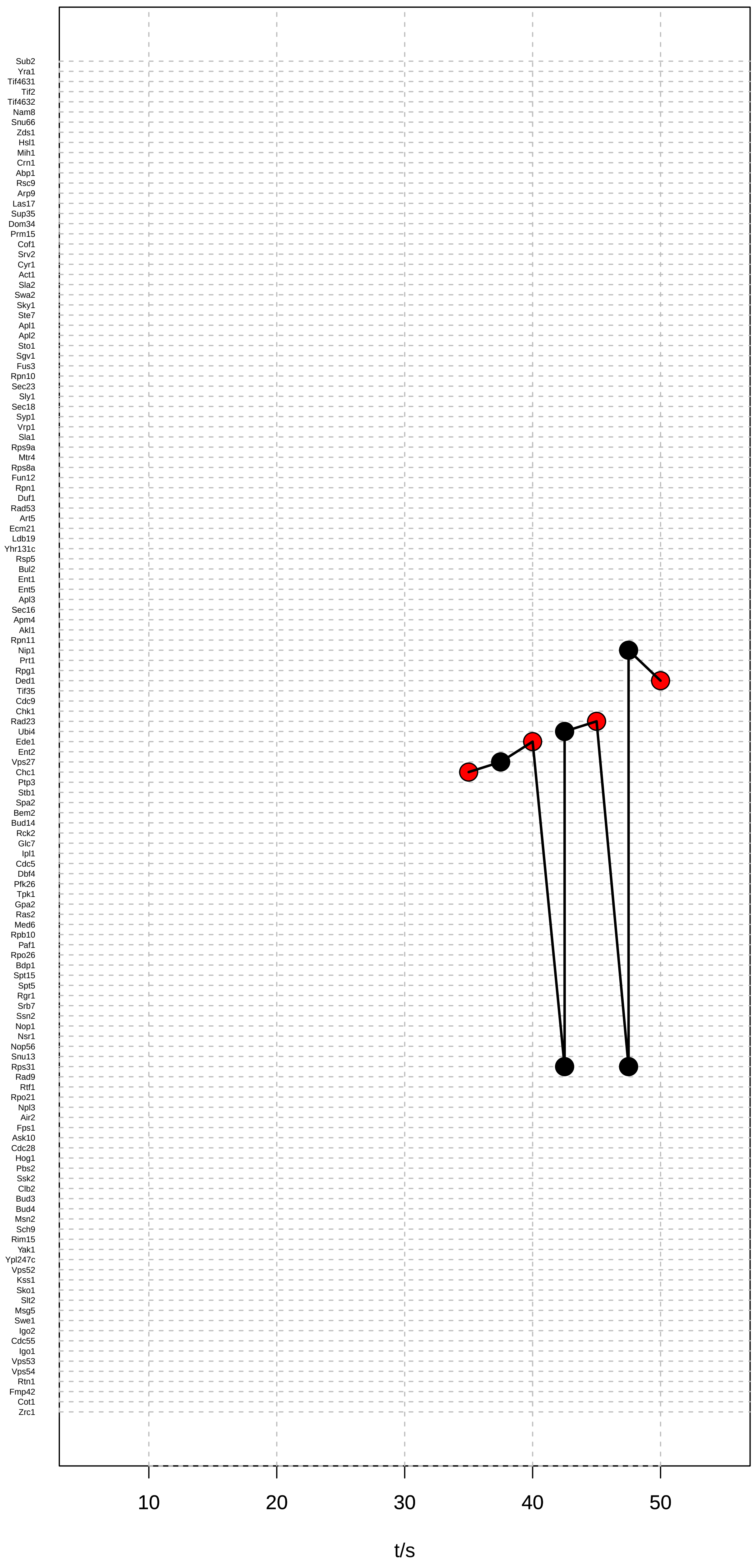

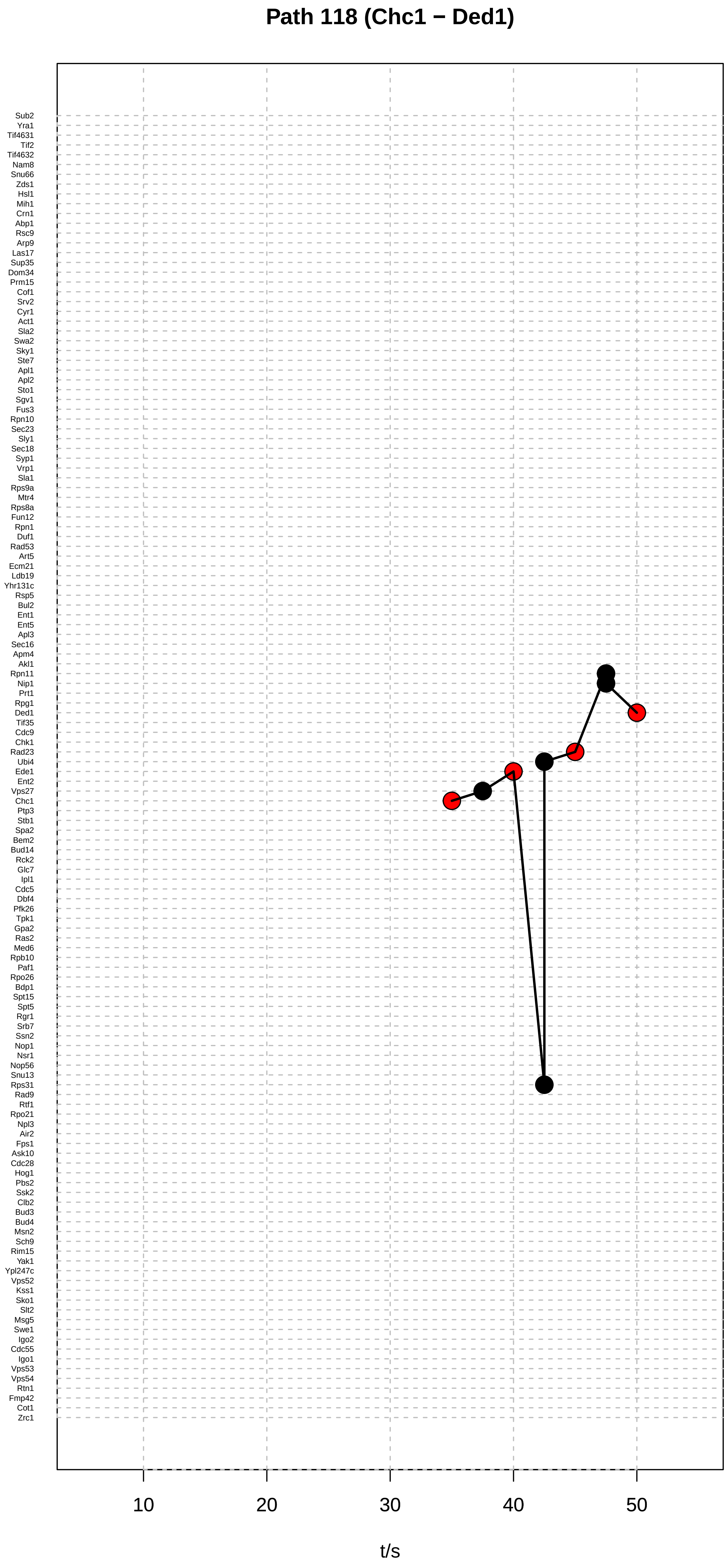
Path 118 (Chc1 − Ded1)
Sub2
Yra1
Tif4631
Tif2
Tif4632
Nam8
Snu66
Zds1
Hsl1
Mih1
Crn1
Abp1
Rsc9
Arp9
Las17
Sup35
Dom34
Prm15
Cof1
Srv2
Cyr1
Act1
Sla2
Swa2
Sky1
Ste7
Apl1
Apl2
Sto1
Sgv1
Fus3
Rpn10
Sec23
Sly1
Sec18
Syp1
Vrp1
Sla1
Rps9a
Mtr4
Rps8a
Fun12
Rpn1
Duf1
Rad53
Art5
Ecm21
Ldb19
Yhr131c
Rsp5
Bul2
Ent1
Ent5
Apl3
Sec16
Apm4
Akl1
Rpn11
Nip1
Prt1
Rpg1
Ded1
Tif35
Cdc9
Chk1
Rad23
Ubi4
Ede1
Ent2
Vps27
Chc1
Ptp3
Stb1
Spa2
Bem2
Bud14
Rck2
Glc7
Ipl1
Cdc5
Dbf4
Pfk26
Tpk1
Gpa2
Ras2
Med6
Rpb10
Paf1
Rpo26
Bdp1
Spt15
Spt5
Rgr1
Srb7
Ssn2
Nop1
Nsr1
Nop56
Snu13
Rps31
Rad9
Rtf1
Rpo21
Npl3
Air2
Fps1
Ask10
Cdc28
Hog1
Pbs2
Ssk2
Clb2
Bud3
Bud4
Msn2
Sch9
Rim15
Yak1
Ypl247c
Vps52
Kss1
Sko1
Slt2
Msg5
Swe1
Igo2
Cdc55
Igo1
Vps53
Vps54
Rtn1
Fmp42
Cot1
Zrc1
10
20
30
40
50
t/s

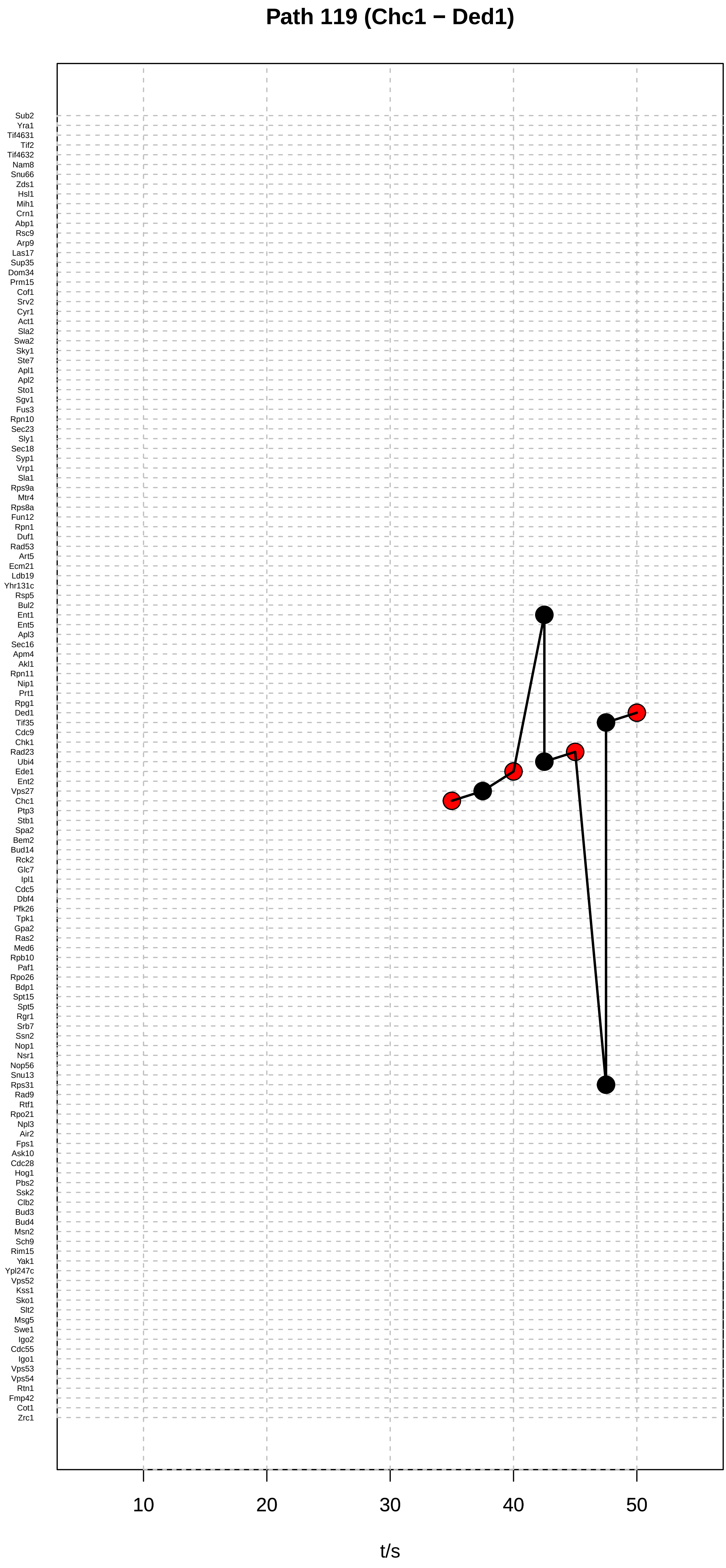
Path 119 (Chc1 − Ded1)
Sub2
Yra1
Tif4631
Tif2
Tif4632
Nam8
Snu66
Zds1
Hsl1
Mih1
Crn1
Abp1
Rsc9
Arp9
Las17
Sup35
Dom34
Prm15
Cof1
Srv2
Cyr1
Act1
Sla2
Swa2
Sky1
Ste7
Apl1
Apl2
Sto1
Sgv1
Fus3
Rpn10
Sec23
Sly1
Sec18
Syp1
Vrp1
Sla1
Rps9a
Mtr4
Rps8a
Fun12
Rpn1
Duf1
Rad53
Art5
Ecm21
Ldb19
Yhr131c
Rsp5
Bul2
Ent1
Ent5
Apl3
Sec16
Apm4
Akl1
Rpn11
Nip1
Prt1
Rpg1
Ded1
Tif35
Cdc9
Chk1
Rad23
Ubi4
Ede1
Ent2
Vps27
Chc1
Ptp3
Stb1
Spa2
Bem2
Bud14
Rck2
Glc7
Ipl1
Cdc5
Dbf4
Pfk26
Tpk1
Gpa2
Ras2
Med6
Rpb10
Paf1
Rpo26
Bdp1
Spt15
Spt5
Rgr1
Srb7
Ssn2
Nop1
Nsr1
Nop56
Snu13
Rps31
Rad9
Rtf1
Rpo21
Npl3
Air2
Fps1
Ask10
Cdc28
Hog1
Pbs2
Ssk2
Clb2
Bud3
Bud4
Msn2
Sch9
Rim15
Yak1
Ypl247c
Vps52
Kss1
Sko1
Slt2
Msg5
Swe1
Igo2
Cdc55
Igo1
Vps53
Vps54
Rtn1
Fmp42
Cot1
Zrc1
10
20
30
40
50
t/s

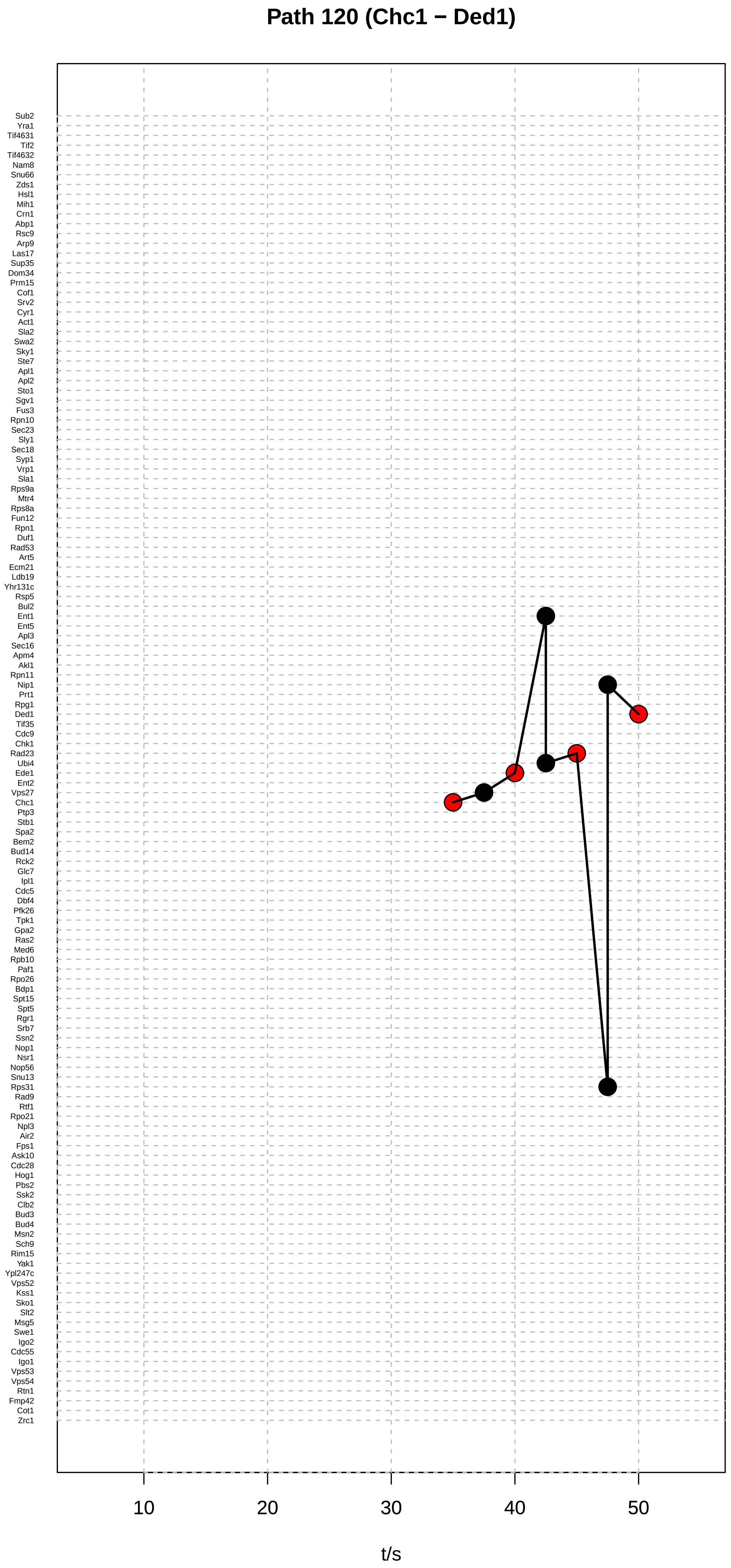

Path 120 (Chc1 − Ded1)
Sub2
Yra1
Tif4631
Tif2
Tif4632
Nam8
Snu66
Zds1
Hsl1
Mih1
Crn1
Abp1
Rsc9
Arp9
Las17
Sup35
Dom34
Prm15
Cof1
Srv2
Cyr1
Act1
Sla2
Swa2
Sky1
Ste7
Apl1
Apl2
Sto1
Sgv1
Fus3
Rpn10
Sec23
Sly1
Sec18
Syp1
Vrp1
Sla1
Rps9a
Mtr4
Rps8a
Fun12
Rpn1
Duf1
Rad53
Art5
Ecm21
Ldb19
Yhr131c
Rsp5
Bul2
Ent1
Ent5
Apl3
Sec16
Apm4
Akl1
Rpn11
Nip1
Prt1
Rpg1
Ded1
Tif35
Cdc9
Chk1
Rad23
Ubi4
Ede1
Ent2
Vps27
Chc1
Ptp3
Stb1
Spa2
Bem2
Bud14
Rck2
Glc7
Ipl1
Cdc5
Dbf4
Pfk26
Tpk1
Gpa2
Ras2
Med6
Rpb10
Paf1
Rpo26
Bdp1
Spt15
Spt5
Rgr1
Srb7
Ssn2
Nop1
Nsr1
Nop56
Snu13
Rps31
Rad9
Rtf1
Rpo21
Npl3
Air2
Fps1
Ask10
Cdc28
Hog1
Pbs2
Ssk2
Clb2
Bud3
Bud4
Msn2
Sch9
Rim15
Yak1
Ypl247c
Vps52
Kss1
Sko1
Slt2
Msg5
Swe1
Igo2
Cdc55
Igo1
Vps53
Vps54
Rtn1
Fmp42
Cot1
Zrc1
10
20
30
40
50
t/s

#### Path 121 (Chc1 − Ded1)

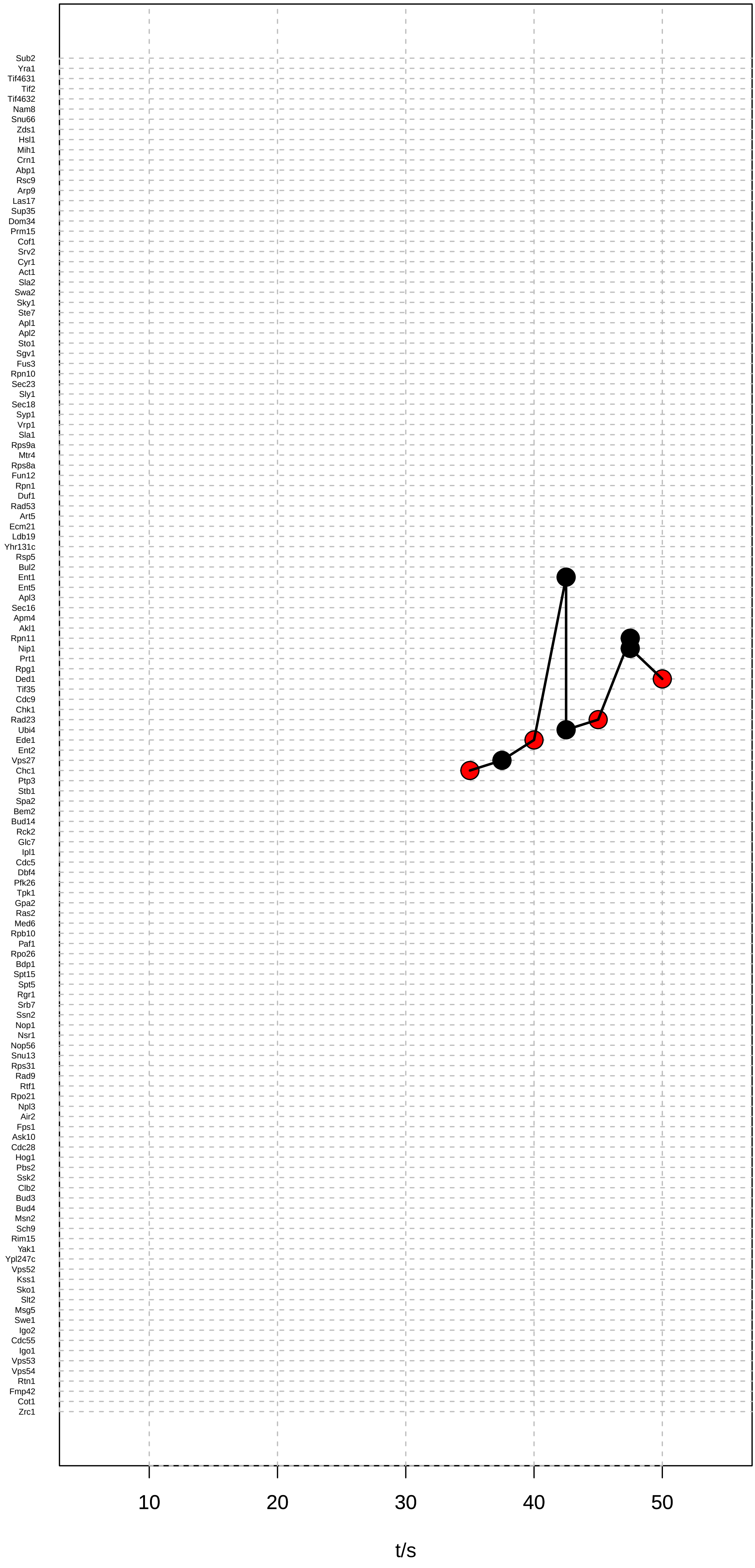

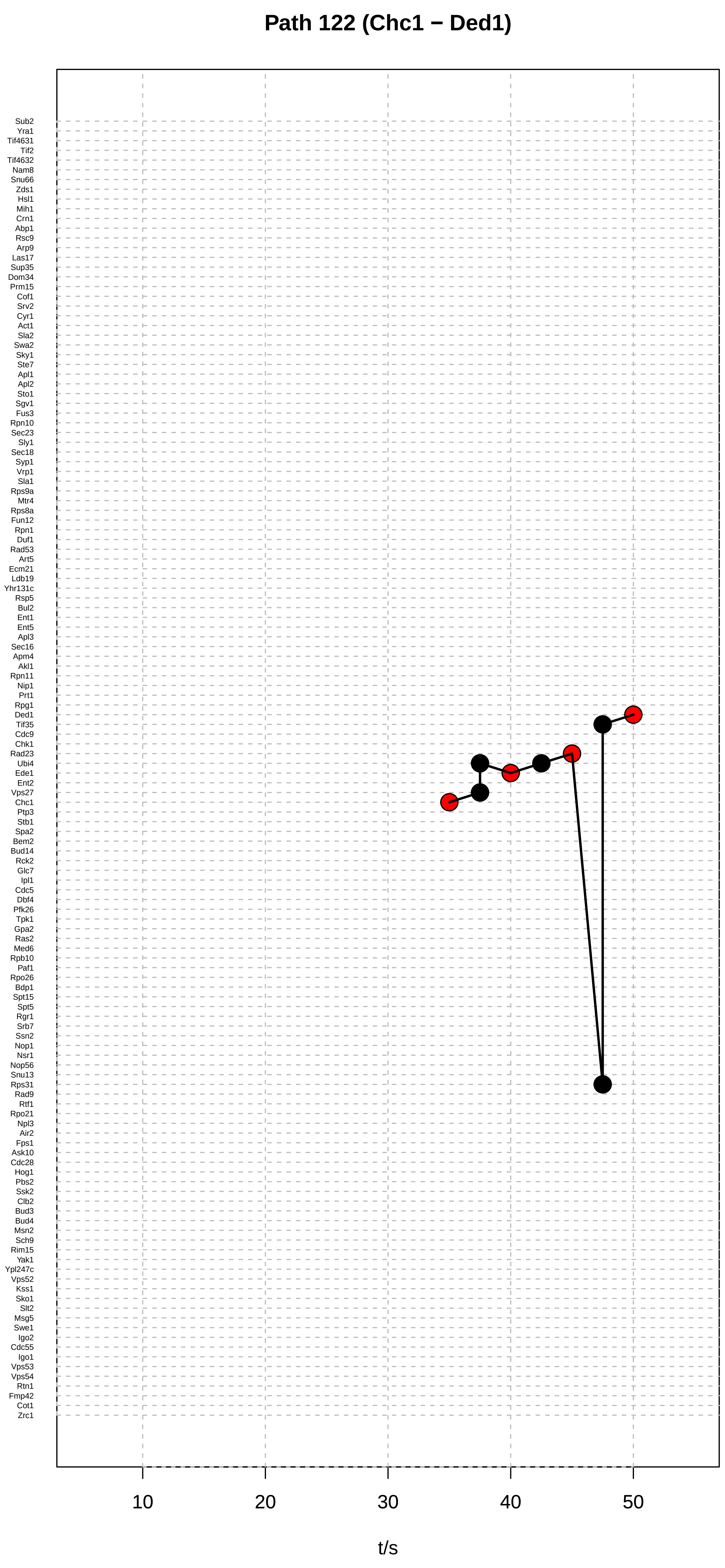
Path 122 (Chc1 − Ded1)
t/s
10
20
30
40
50

#### Path 123 (Chhc1 − Ded1)

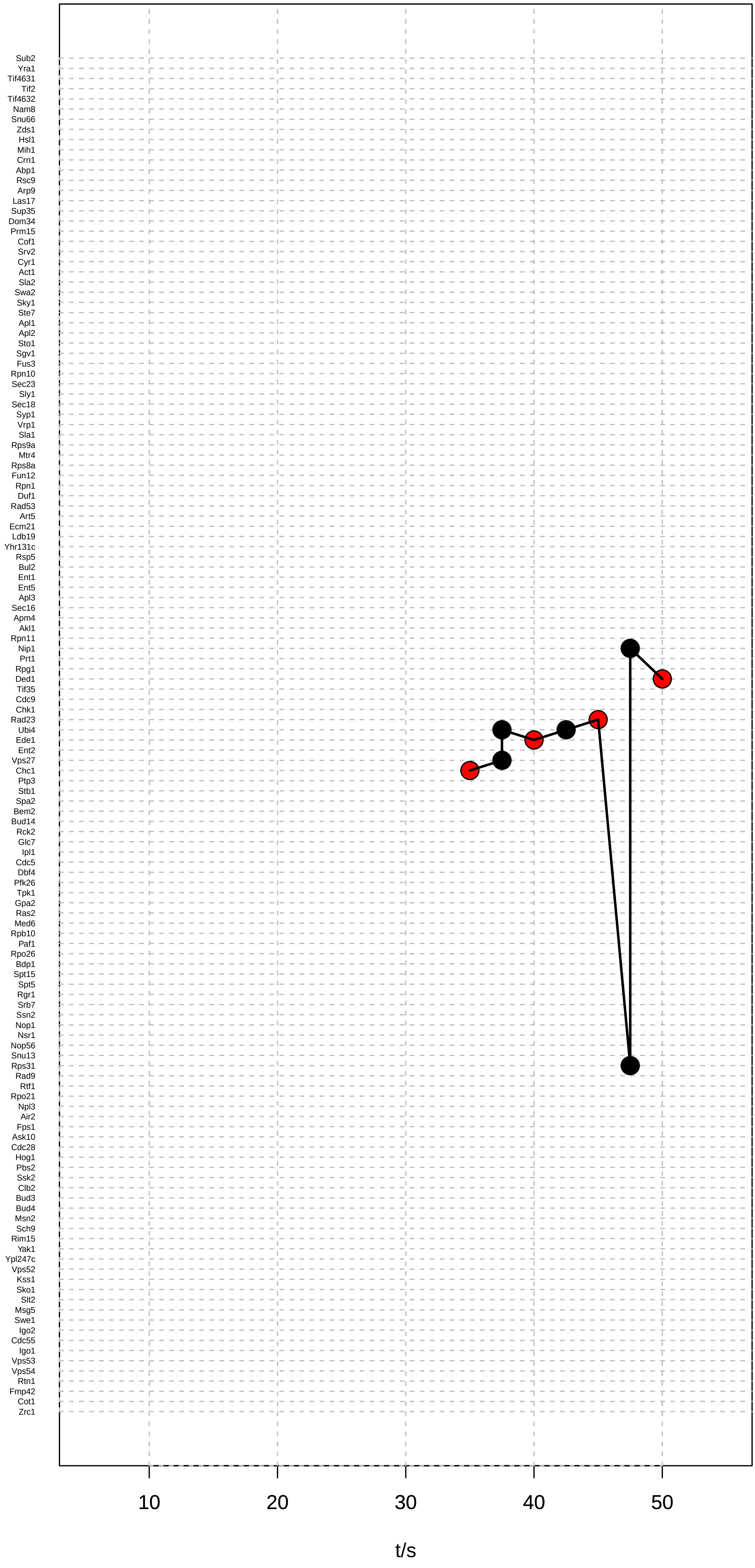

### Path 124 (Chc1 − Ded1)

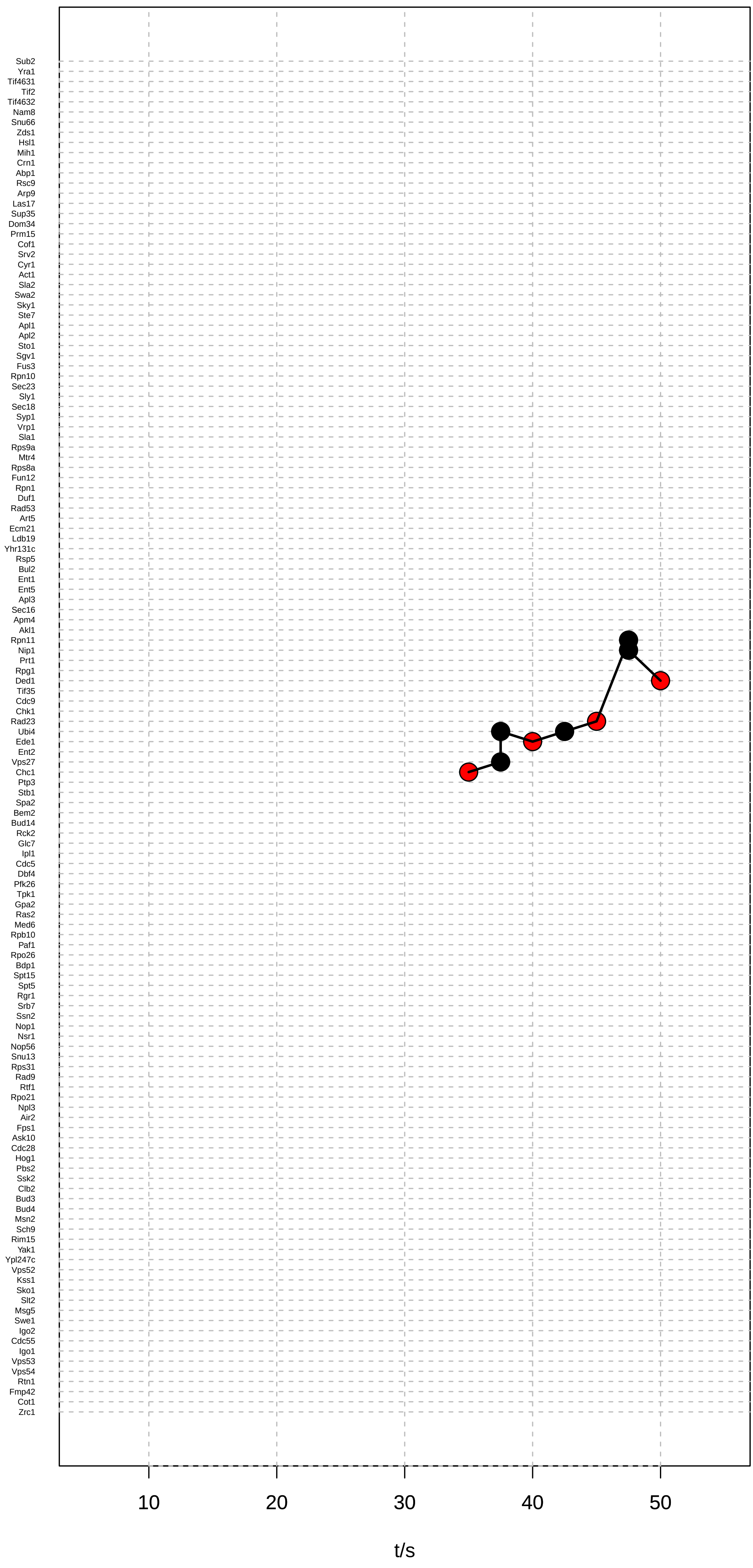

### Path 125 (Chc1 − Ded1)

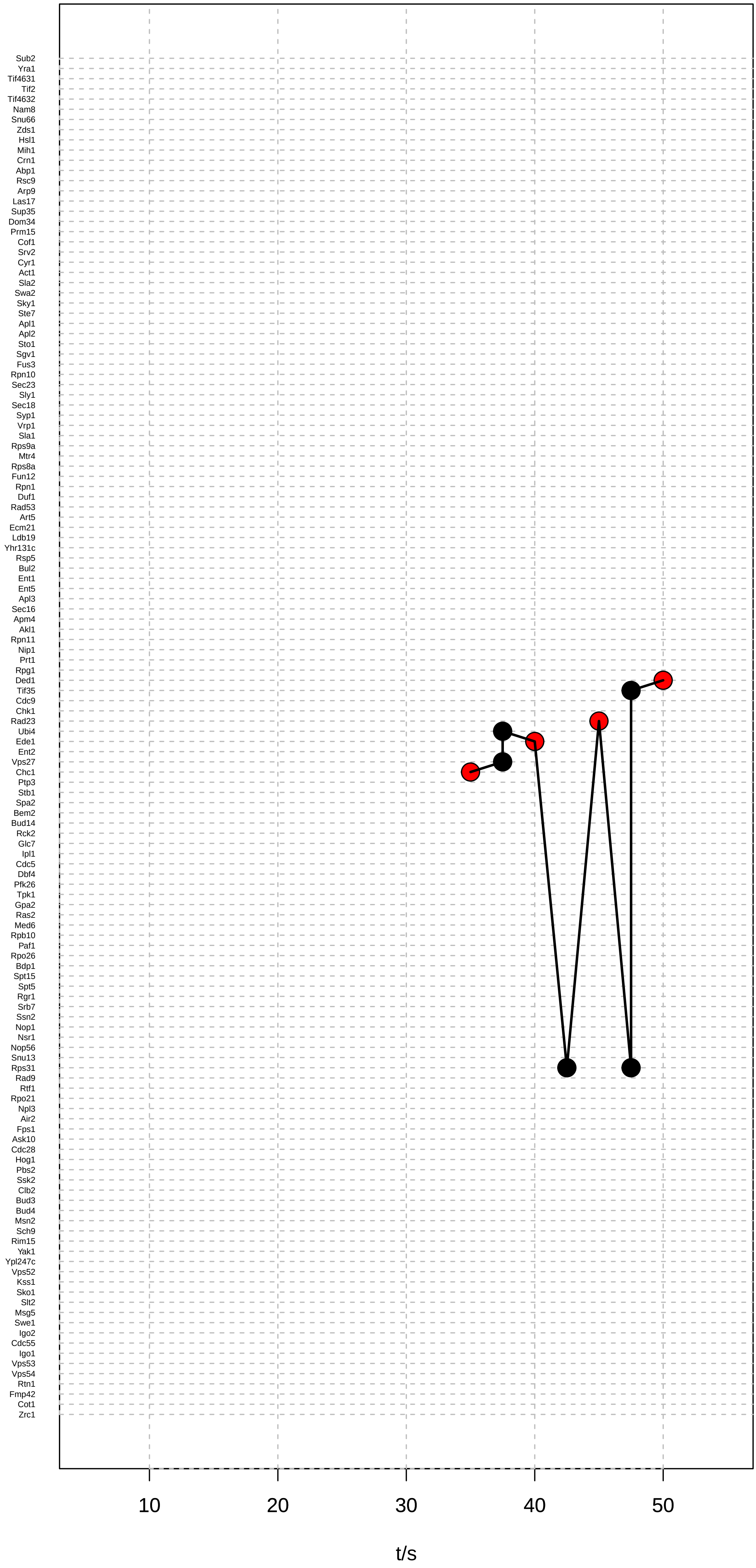

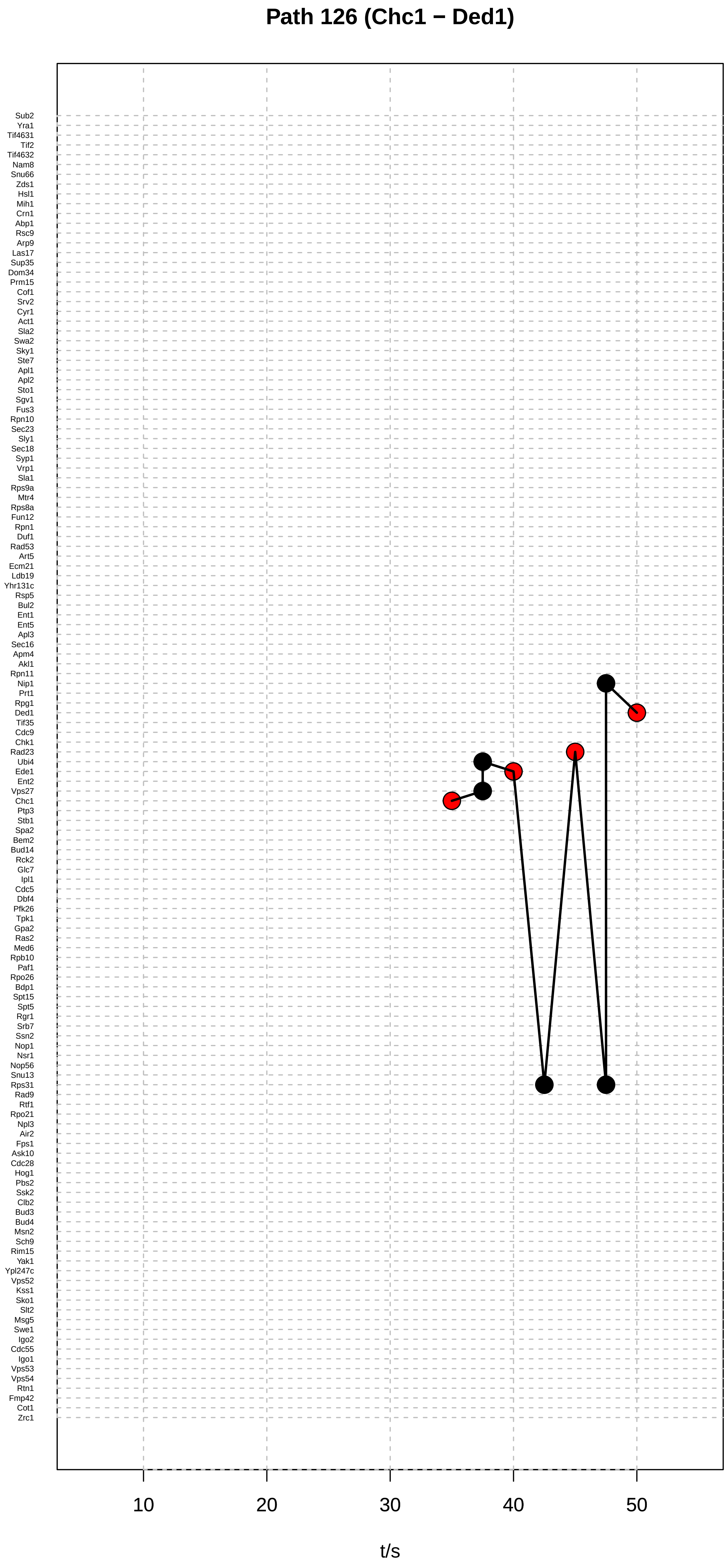

Path 126 (Chc1 − Ded1)
Sub2
Yra1
Tif4631
Tif2
Tif4632
Nam8
Snu66
Zds1
Hsl1
Mih1
Crn1
Abp1
Rsc9
Arp9
Las17
Sup35
Dom34
Prm15
Cof1
Srv2
Cyr1
Act1
Sla2
Swa2
Sky1
Ste7
Apl1
Apl2
Sto1
Sgv1
Fus3
Rpn10
Sec23
Sly1
Sec18
Syp1
Vrp1
Sla1
Rps9a
Mtr4
Rps8a
Fun12
Rpn1
Duf1
Rad53
Art5
Ecm21
Ldb19
Yhr131c
Rsp5
Bul2
Ent1
Ent5
Apl3
Sec16
Apm4
Akl1
Rpn11
Nip1
Prt1
Rpg1
Ded1
Tif35
Cdc9
Chk1
Rad23
Ubi4
Ede1
Ent2
Vps27
Chc1
Ptp3
Stb1
Spa2
Bem2
Bud14
Rck2
Glc7
Ipl1
Cdc5
Dbf4
Pfk26
Tpk1
Gpa2
Ras2
Med6
Rpb10
Paf1
Rpo26
Bdp1
Spt15
Spt5
Rgr1
Srb7
Ssn2
Nop1
Nsr1
Nop56
Snu13
Rps31
Rad9
Rtf1
Rpo21
Npl3
Air2
Fps1
Ask10
Cdc28
Hog1
Pbs2
Ssk2
Clb2
Bud3
Bud4
Msn2
Sch9
Rim15
Yak1
Ypl247c
Vps52
Kss1
Sko1
Slt2
Msg5
Swe1
Igo2
Cdc55
Igo1
Vps53
Vps54
Rtn1
Fmp42
Cot1
Zrc1
10
20
30
40
50
t/s

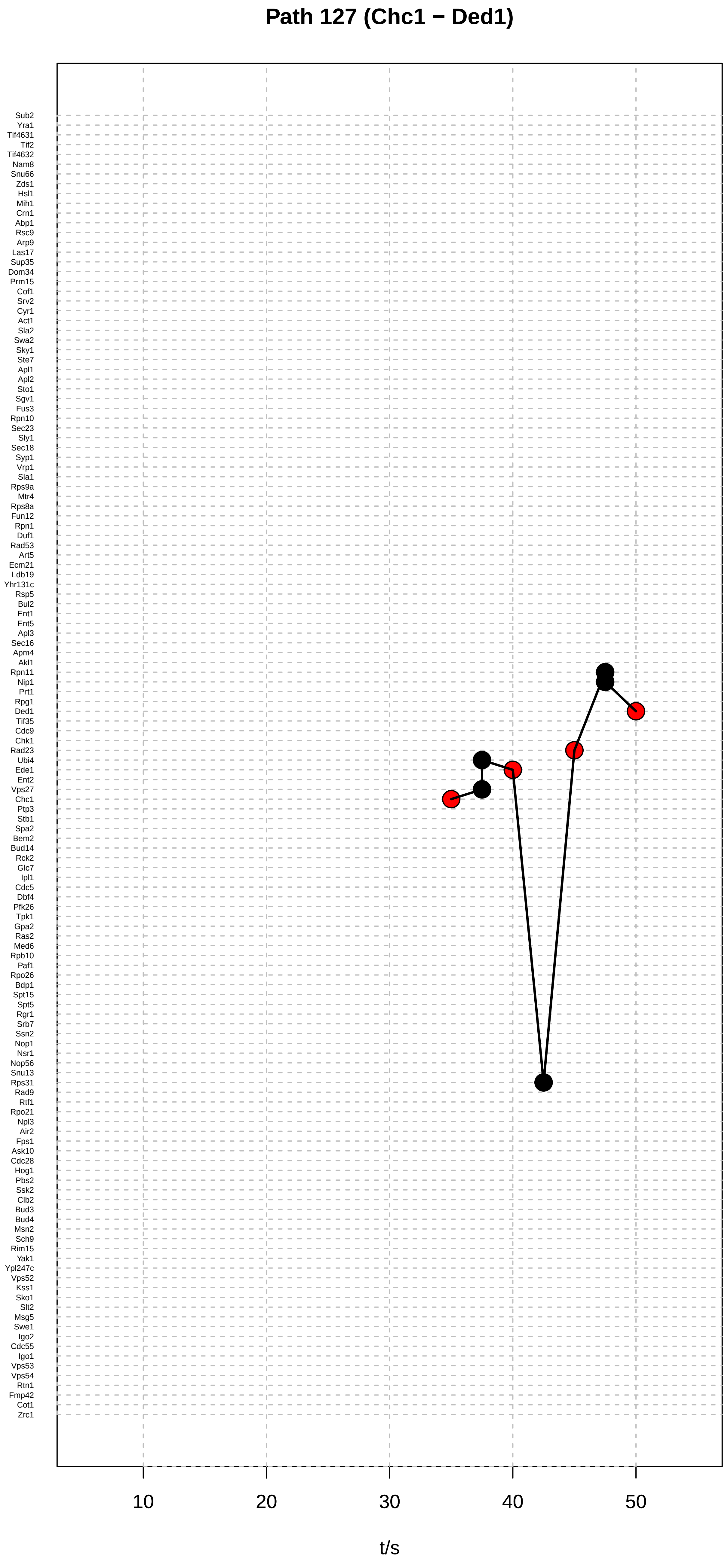

Path 127 (Chc1 − Ded1)
Sub2
Yra1
Tif4631
Tif2
Tif4632
Nam8
Snu66
Zds1
Hsl1
Mih1
Crn1
Abp1
Rsc9
Arp9
Las17
Sup35
Dom34
Prm15
Cof1
Srv2
Cyr1
Act1
Sla2
Swa2
Sky1
Ste7
Apl1
Apl2
Sto1
Sgv1
Fus3
Rpn10
Sec23
Sly1
Sec18
Syp1
Vrp1
Sla1
Rps9a
Mtr4
Rps8a
Fun12
Rpn1
Duf1
Rad53
Art5
Ecm21
Ldb19
Yhr131c
Rsp5
Bul2
Ent1
Ent5
Apl3
Sec16
Apm4
Akl1
Rpn11
Nip1
Prt1
Rpg1
Ded1
Tif35
Cdc9
Chk1
Rad23
Ubi4
Ede1
Ent2
Vps27
Chc1
Ptp3
Stb1
Spa2
Bem2
Bud14
Rck2
Glc7
Ipl1
Cdc5
Dbf4
Pfk26
Tpk1
Gpa2
Ras2
Med6
Rpb10
Paf1
Rpo26
Bdp1
Spt15
Spt5
Rgr1
Srb7
Ssn2
Nop1
Nsr1
Nop56
Snu13
Rps31
Rad9
Rtf1
Rpo21
Npl3
Air2
Fps1
Ask10
Cdc28
Hog1
Pbs2
Ssk2
Clb2
Bud3
Bud4
Msn2
Sch9
Rim15
Yak1
Ypl247c
Vps52
Kss1
Sko1
Slt2
Msg5
Swe1
Igo2
Cdc55
Igo1
Vps53
Vps54
Rtn1
Fmp42
Cot1
Zrc1
10
20
30
40
50
t/s

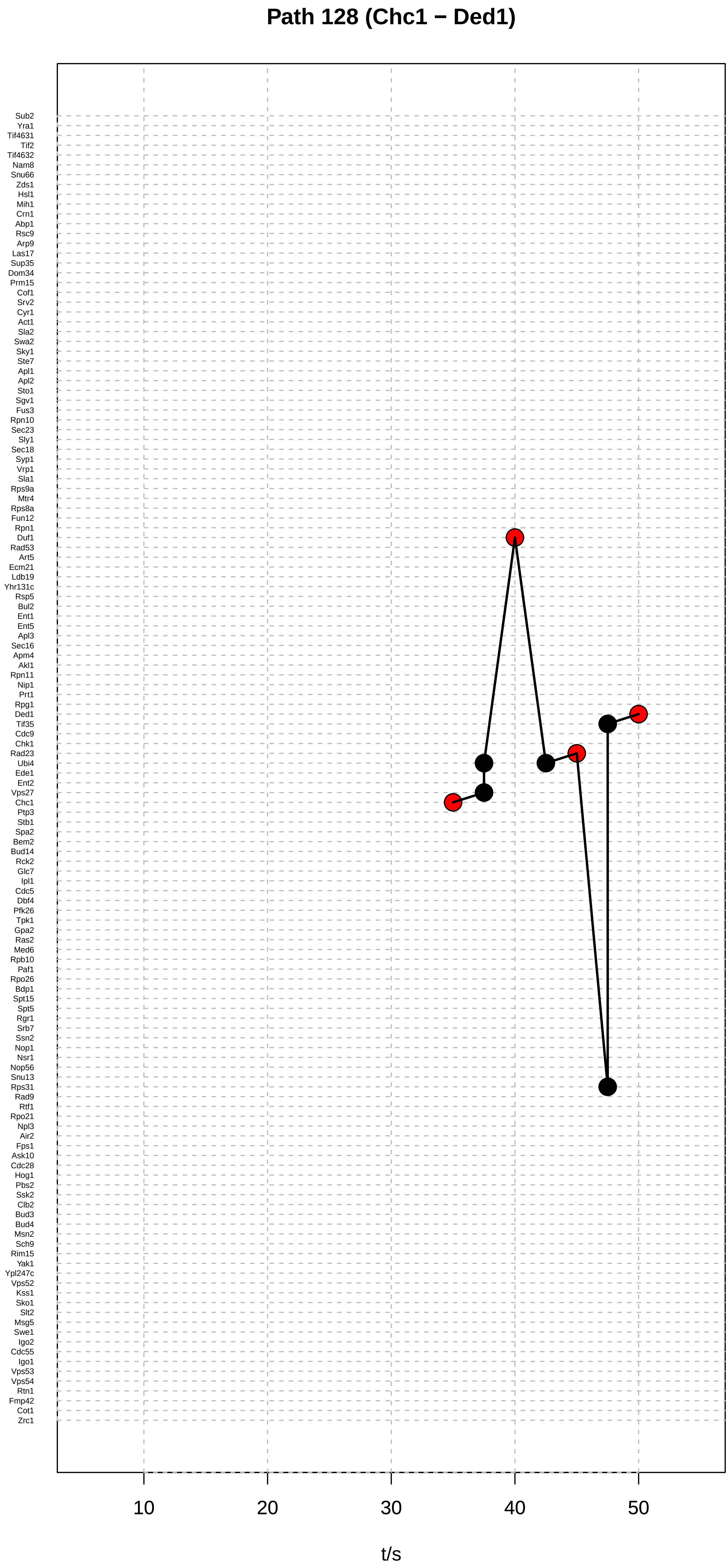
Path 128 (Chc1 − Ded1)
t/s
10
20
30
40
50
Sub2
Yra1
Tif4631
Tif2
Tif4632
Nam8
Snu66
Zds1
Hsl1
Mih1
Crn1
Abp1
Rsc9
Arp9
Las17
Sup35
Dom34
Prm15
Cof1
Srv2
Cyr1
Act1
Sla2
Swa2
Sky1
Ste7
Apl1
Apl2
Sto1
Sgv1
Fus3
Rpn10
Sec23
Sly1
Sec18
Syp1
Vrp1
Sla1
Rps9a
Mtr4
Rps8a
Fun12
Rpn1
Duf1
Rad53
Art5
Ecm21
Ldb19
Yhr131c
Rsp5
Bul2
Ent1
Ent5
Apl3
Sec16
Apm4
Akl1
Rpn11
Nip1
Prt1
Rpg1
Ded1
Tif35
Cdc9
Chk1
Rad23
Ubi4
Ede1
Ent2
Vps27
Chc1
Ptp3
Stb1
Spa2
Bem2
Bud14
Rck2
Glc7
Ipl1
Cdc5
Dbf4
Pfk26
Tpk1
Gpa2
Ras2
Med6
Rpb10
Paf1
Rpo26
Bdp1
Spt15
Spt5
Rgr1
Srb7
Ssn2
Nop1
Nsr1
Nop56
Snu13
Rps31
Rad9
Rtf1
Rpo21
Npl3
Air2
Fps1
Ask10
Cdc28
Hog1
Pbs2
Ssk2
Clb2
Bud3
Bud4
Msn2
Sch9
Rim15
Yak1
Ypl247c
Vps52
Kss1
Sko1
Slt2
Msg5
Swe1
Igo2
Cdc55
Igo1
Vps53
Vps54
Rtn1
Fmp42
Cot1
Zrc1

Path 129 (Chc1 − Ded1)

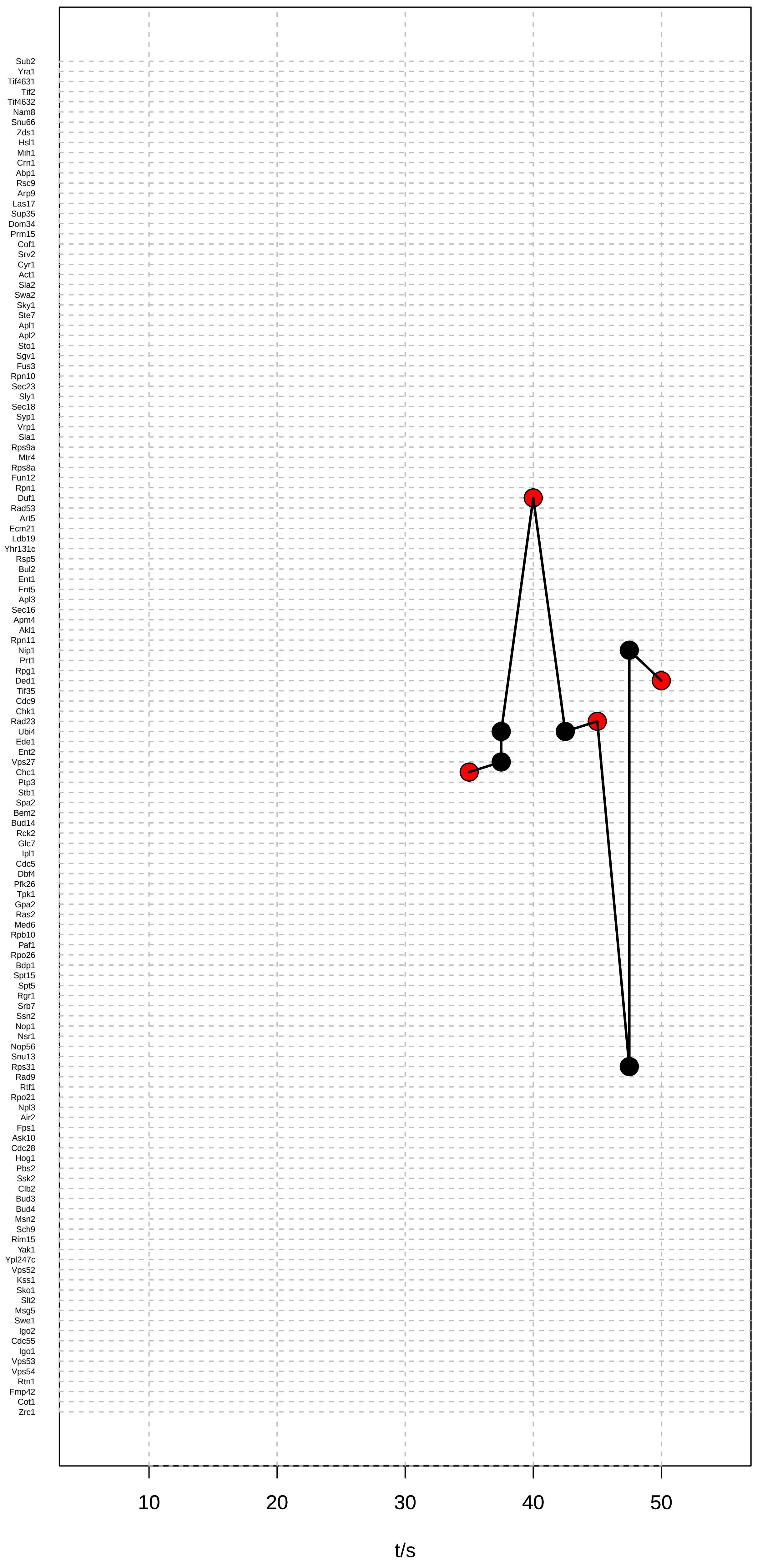

#### Path 130 (Chc1 − Ded1)

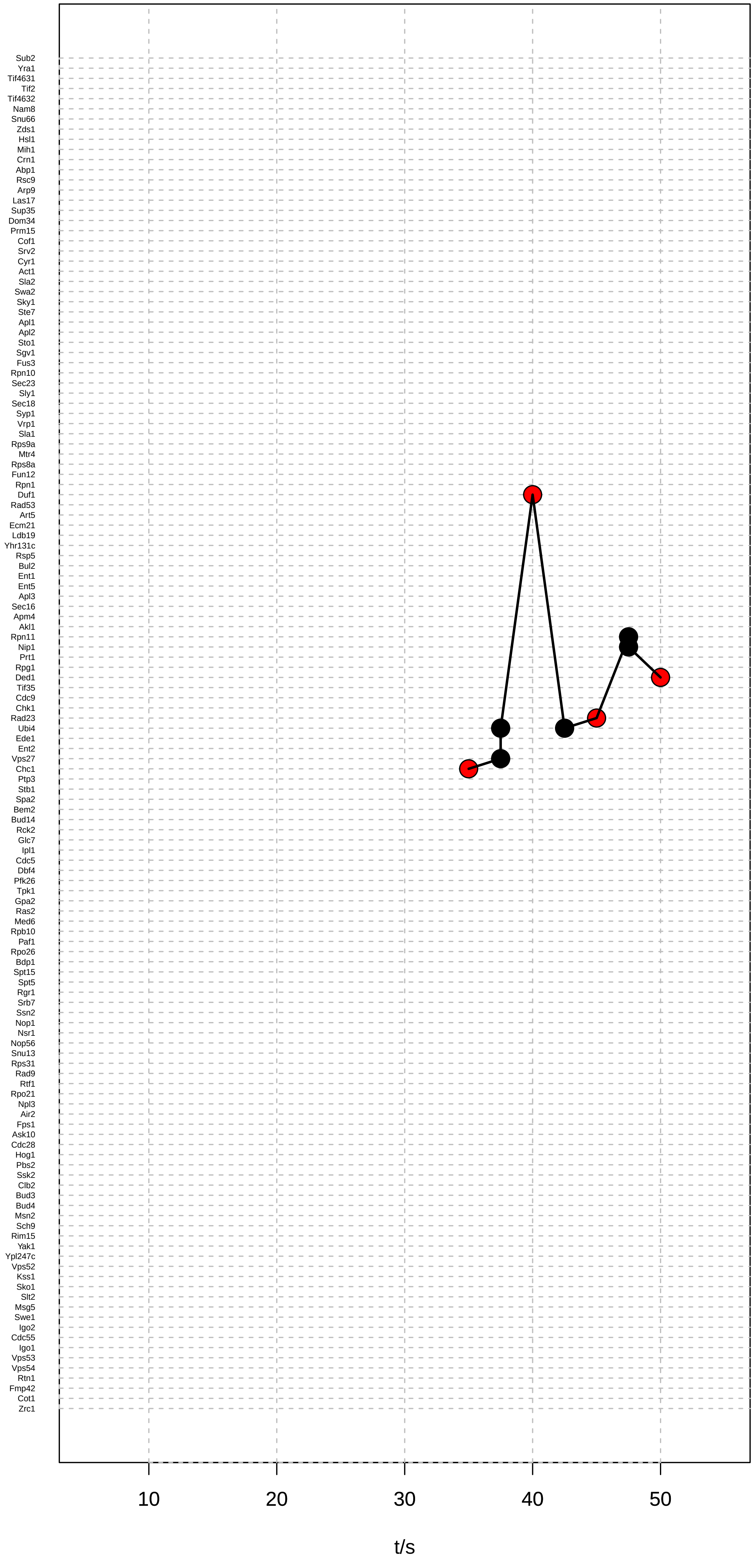

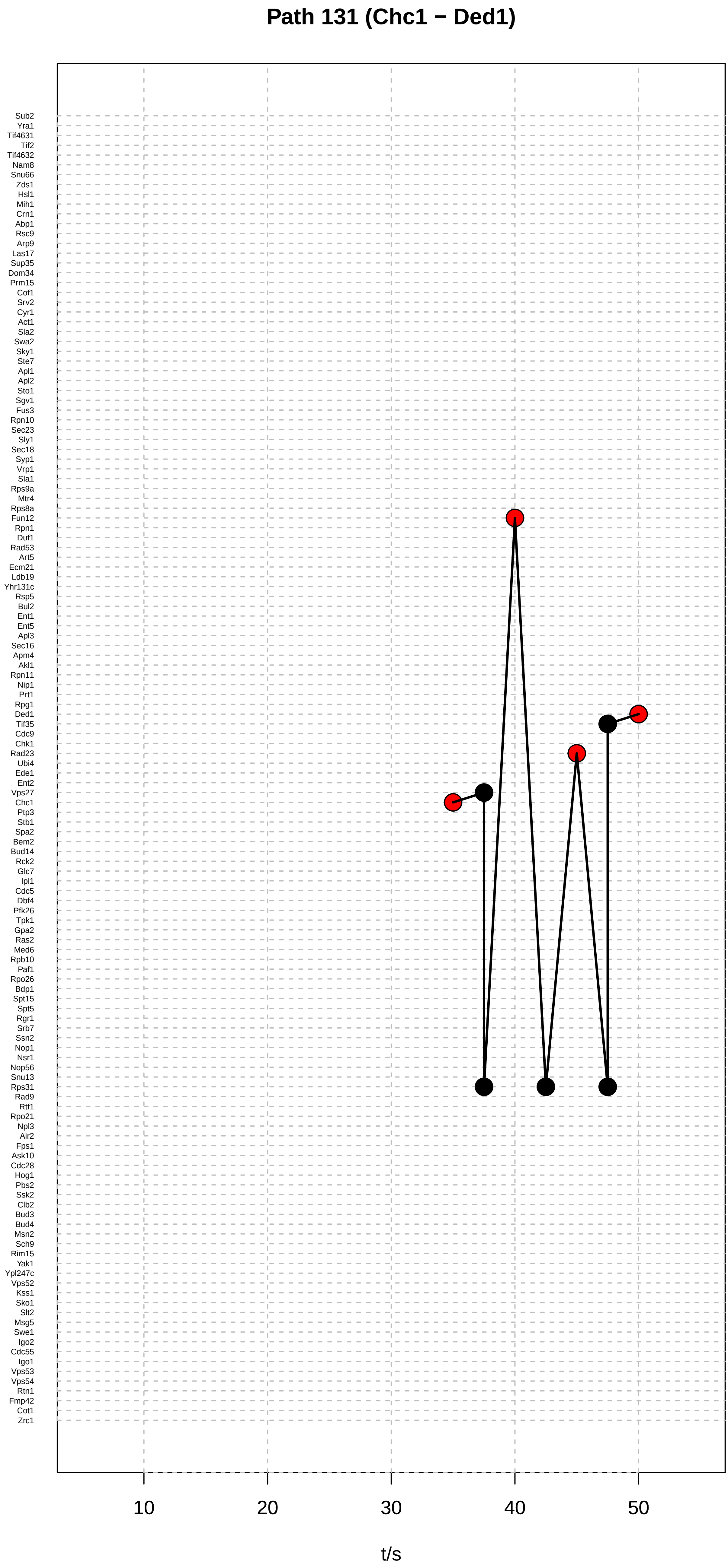
Path 131 (Chc1 − Ded1)
Sub2
Yra1
Tif4631
Tif2
Tif4632
Nam8
Snu66
Zds1
Hsl1
Mih1
Crn1
Abp1
Rsc9
Arp9
Las17
Sup35
Dom34
Prm15
Cof1
Srv2
Cyr1
Act1
Sla2
Swa2
Sky1
Ste7
Apl1
Apl2
Sto1
Sgv1
Fus3
Rpn10
Sec23
Sly1
Sec18
Syp1
Vrp1
Sla1
Rps9a
Mtr4
Rps8a
Fun12
Rpn1
Duf1
Rad53
Art5
Ecm21
Ldb19
Yhr131c
Rsp5
Bul2
Ent1
Ent5
Apl3
Sec16
Apm4
Akl1
Rpn11
Nip1
Prt1
Rpg1
Ded1
Tif35
Cdc9
Chk1
Rad23
Ubi4
Ede1
Ent2
Vps27
Chc1
Ptp3
Stb1
Spa2
Bem2
Bud14
Rck2
Glc7
Ipl1
Cdc5
Dbf4
Pfk26
Tpk1
Gpa2
Ras2
Med6
Rpb10
Paf1
Rpo26
Bdp1
Spt15
Spt5
Rgr1
Srb7
Ssn2
Nop1
Nsr1
Nop56
Snu13
Rps31
Rad9
Rtf1
Rpo21
Npl3
Air2
Fps1
Ask10
Cdc28
Hog1
Pbs2
Ssk2
Clb2
Bud3
Bud4
Msn2
Sch9
Rim15
Yak1
Ypl247c
Vps52
Kss1
Sko1
Slt2
Msg5
Swe1
Igo2
Cdc55
Igo1
Vps53
Vps54
Rtn1
Fmp42
Cot1
Zrc1
10
20
30
40
50
t/s

#### Path 132 (Chc1 − Ded1)

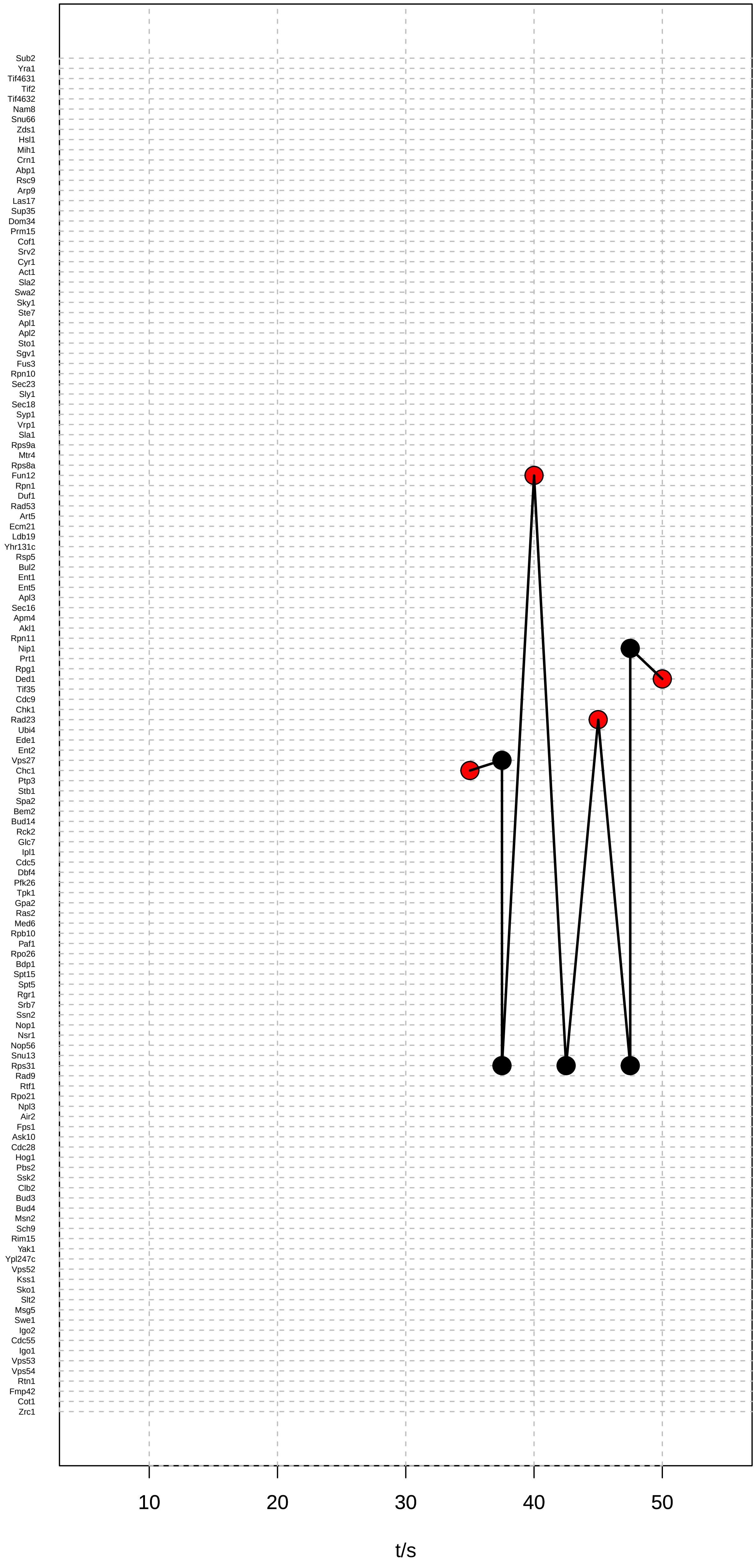

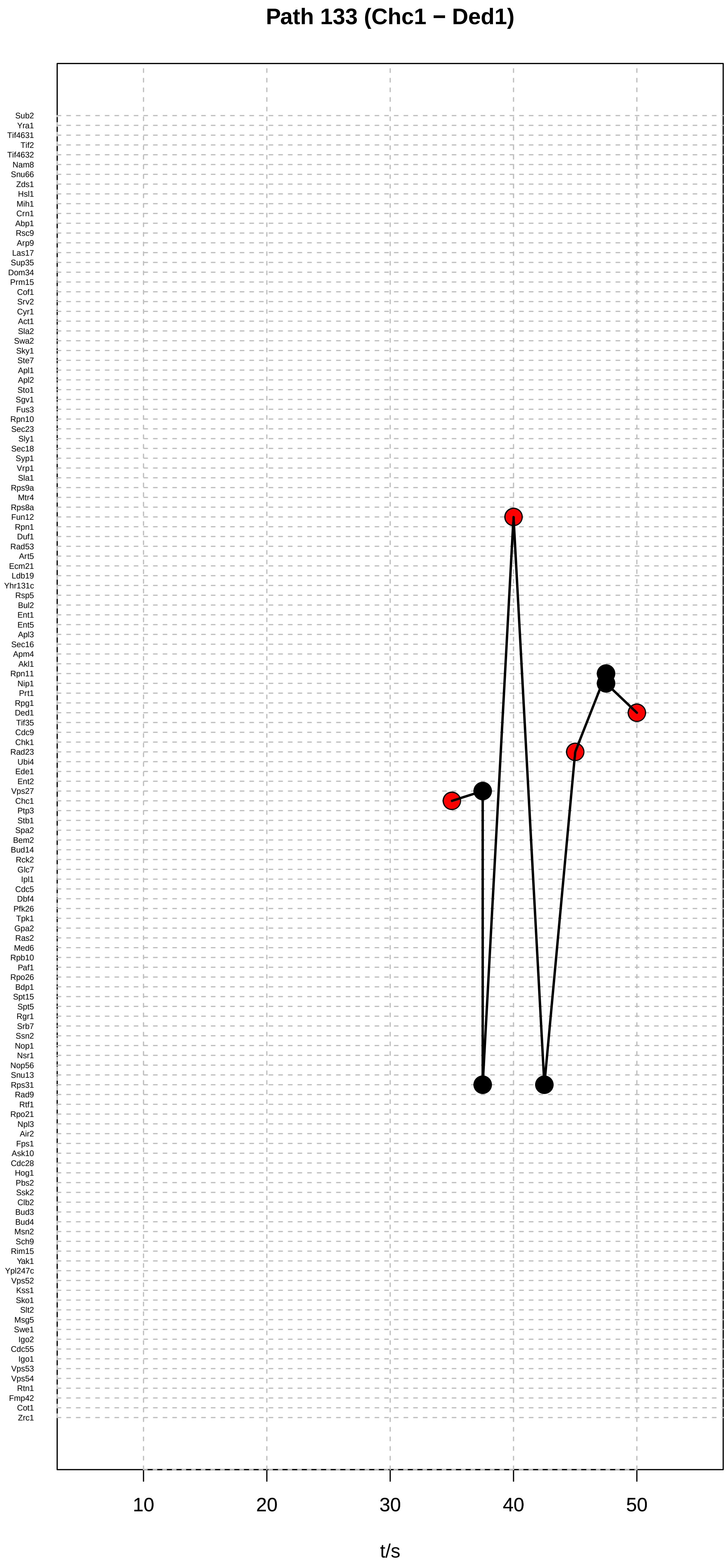

Path 133 (Chc1 − Ded1)
Sub2
Yra1
Tif4631
Tif2
Tif4632
Nam8
Snu66
Zds1
Hsl1
Mih1
Crn1
Abp1
Rsc9
Arp9
Las17
Sup35
Dom34
Prm15
Cof1
Srv2
Cyr1
Act1
Sla2
Swa2
Sky1
Ste7
Apl1
Apl2
Sto1
Sgv1
Fus3
Rpn10
Sec23
Sly1
Sec18
Syp1
Vrp1
Sla1
Rps9a
Mtr4
Rps8a
Fun12
Rpn1
Duf1
Rad53
Art5
Ecm21
Ldb19
Yhr131c
Rsp5
Bul2
Ent1
Ent5
Apl3
Sec16
Apm4
Akl1
Rpn11
Nip1
Prt1
Rpg1
Ded1
Tif35
Cdc9
Chk1
Rad23
Ubi4
Ede1
Ent2
Vps27
Chc1
Ptp3
Stb1
Spa2
Bem2
Bud14
Rck2
Glc7
Ipl1
Cdc5
Dbf4
Pfk26
Tpk1
Gpa2
Ras2
Med6
Rpb10
Paf1
Rpo26
Bdp1
Spt15
Spt5
Rgr1
Srb7
Ssn2
Nop1
Nsr1
Nop56
Snu13
Rps31
Rad9
Rtf1
Rpo21
Npl3
Air2
Fps1
Ask10
Cdc28
Hog1
Pbs2
Ssk2
Clb2
Bud3
Bud4
Msn2
Sch9
Rim15
Yak1
Ypl247c
Vps52
Kss1
Sko1
Slt2
Msg5
Swe1
Igo2
Cdc55
Igo1
Vps53
Vps54
Rtn1
Fmp42
Cot1
Zrc1
10
20
30
40
50
t/s

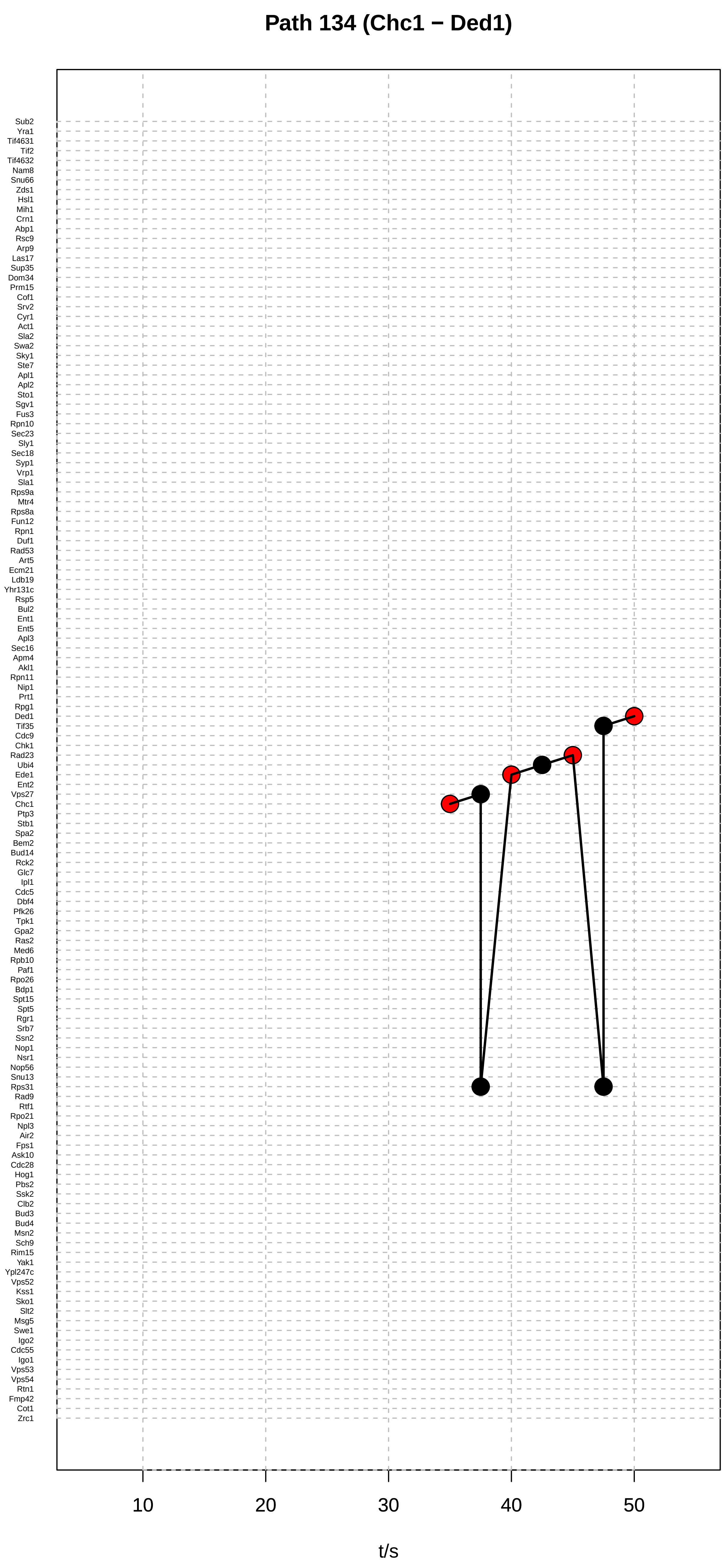

Path 134 (Chc1 − Ded1)
Sub2
Yra1
Tif4631
Tif2
Tif4632
Nam8
Snu66
Zds1
Hsl1
Mih1
Crn1
Abp1
Rsc9
Arp9
Las17
Sup35
Dom34
Prm15
Cof1
Srv2
Cyr1
Act1
Sla2
Swa2
Sky1
Ste7
Apl1
Apl2
Sto1
Sgv1
Fus3
Rpn10
Sec23
Sly1
Sec18
Syp1
Vrp1
Sla1
Rps9a
Mtr4
Rps8a
Fun12
Rpn1
Duf1
Rad53
Art5
Ecm21
Ldb19
Yhr131c
Rsp5
Bul2
Ent1
Ent5
Apl3
Sec16
Apm4
Akl1
Rpn11
Nip1
Prt1
Rpg1
Ded1
Tif35
Cdc9
Chk1
Rad23
Ubi4
Ede1
Ent2
Vps27
Chc1
Ptp3
Stb1
Spa2
Bem2
Bud14
Rck2
Glc7
Ipl1
Cdc5
Dbf4
Pfk26
Tpk1
Gpa2
Ras2
Med6
Rpb10
Paf1
Rpo26
Bdp1
Spt15
Spt5
Rgr1
Srb7
Ssn2
Nop1
Nsr1
Nop56
Snu13
Rps31
Rad9
Rtf1
Rpo21
Npl3
Air2
Fps1
Ask10
Cdc28
Hog1
Pbs2
Ssk2
Clb2
Bud3
Bud4
Msn2
Sch9
Rim15
Yak1
Ypl247c
Vps52
Kss1
Sko1
Slt2
Msg5
Swe1
Igo2
Cdc55
Igo1
Vps53
Vps54
Rtn1
Fmp42
Cot1
Zrc1
10
20
30
40
50
t/s

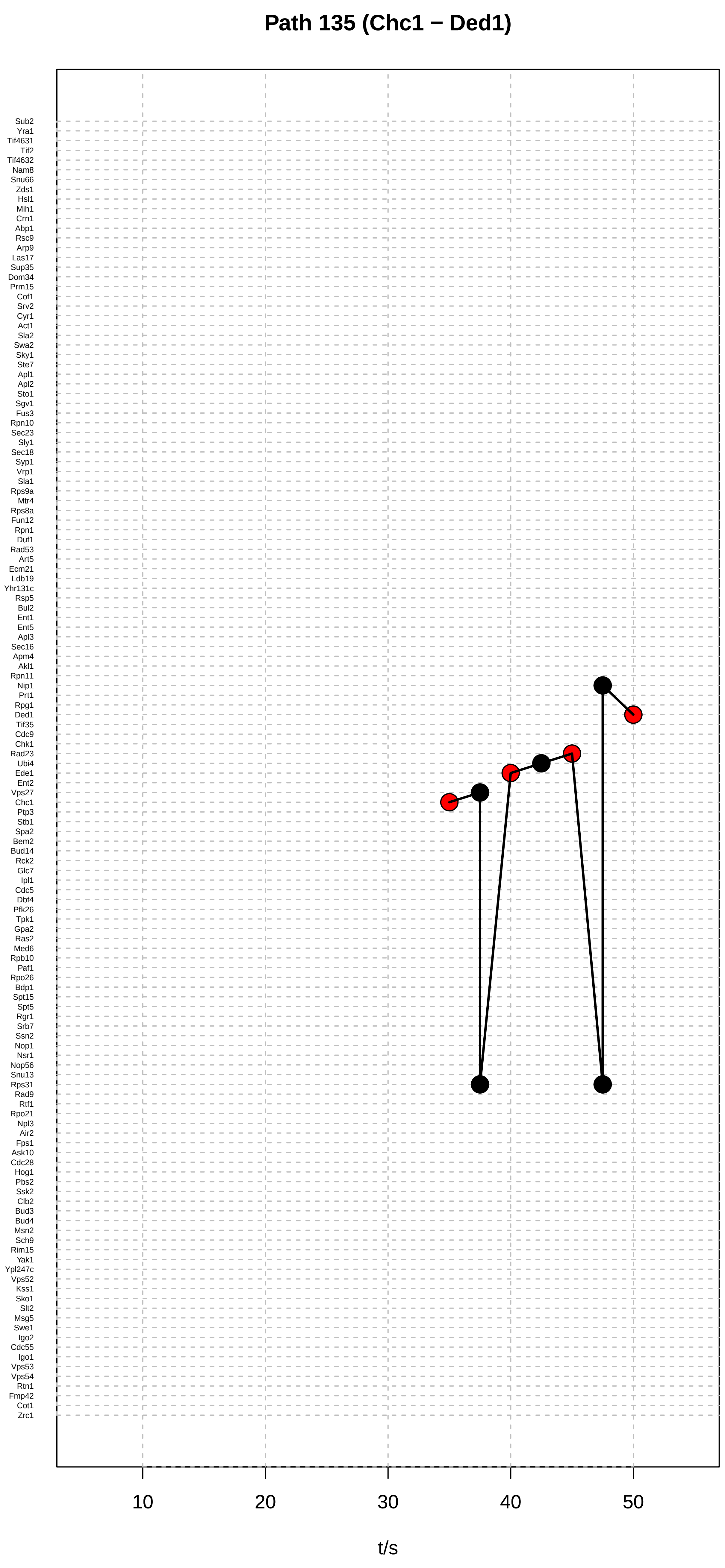

Path 135 (Chc1 − Ded1)
Sub2
Yra1
Tif4631
Tif2
Tif4632
Nam8
Snu66
Zds1
Hsl1
Mih1
Crn1
Abp1
Rsc9
Arp9
Las17
Sup35
Dom34
Prm15
Cof1
Srv2
Cyr1
Act1
Sla2
Swa2
Sky1
Ste7
Apl1
Apl2
Sto1
Sgv1
Fus3
Rpn10
Sec23
Sly1
Sec18
Syp1
Vrp1
Sla1
Rps9a
Mtr4
Rps8a
Fun12
Rpn1
Duf1
Rad53
Art5
Ecm21
Ldb19
Yhr131c
Rsp5
Bul2
Ent1
Ent5
Apl3
Sec16
Apm4
Akl1
Rpn11
Nip1
Prt1
Rpg1
Ded1
Tif35
Cdc9
Chk1
Rad23
Ubi4
Ede1
Ent2
Vps27
Chc1
Ptp3
Stb1
Spa2
Bem2
Bud14
Rck2
Glc7
Ipl1
Cdc5
Dbf4
Pfk26
Tpk1
Gpa2
Ras2
Med6
Rpb10
Paf1
Rpo26
Bdp1
Spt15
Spt5
Rgr1
Srb7
Ssn2
Nop1
Nsr1
Nop56
Snu13
Rps31
Rad9
Rtf1
Rpo21
Npl3
Air2
Fps1
Ask10
Cdc28
Hog1
Pbs2
Ssk2
Clb2
Bud3
Bud4
Msn2
Sch9
Rim15
Yak1
Ypl247c
Vps52
Kss1
Sko1
Slt2
Msg5
Swe1
Igo2
Cdc55
Igo1
Vps53
Vps54
Rtn1
Fmp42
Cot1
Zrc1
10
20
30
40
50
t/s

#### Path 136 (Chc1 − Ded1)

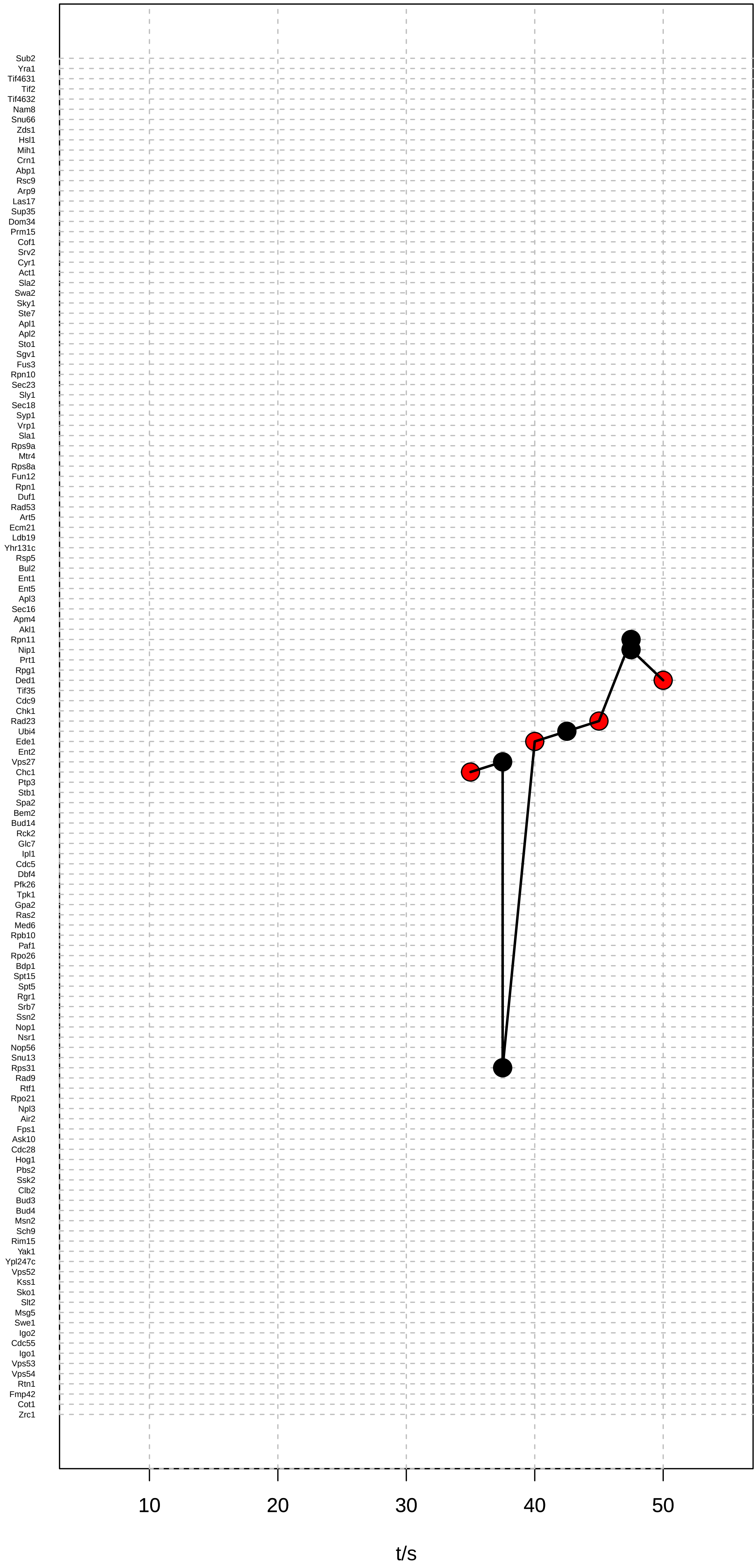

### Path 137 (Chc1 − Ded1)

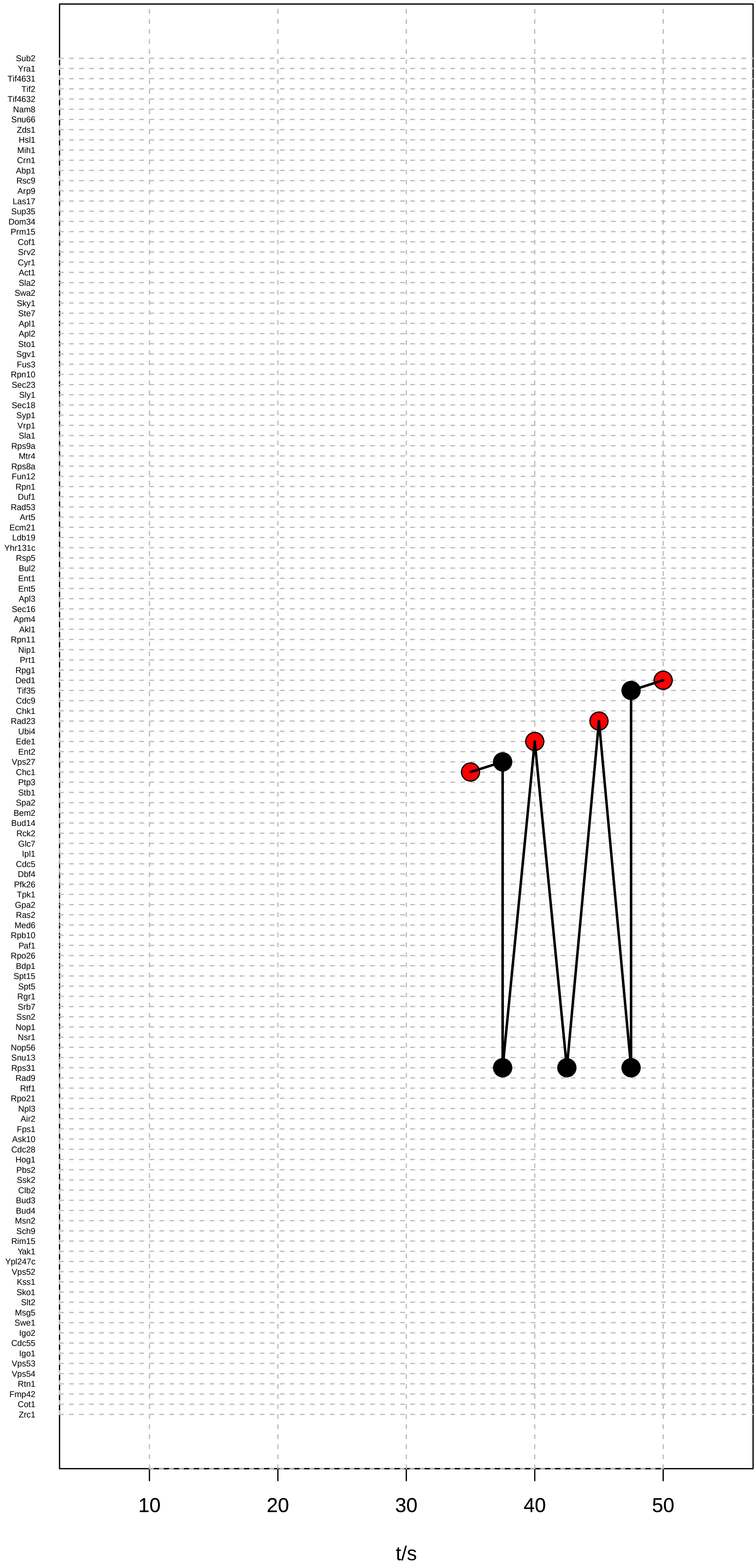

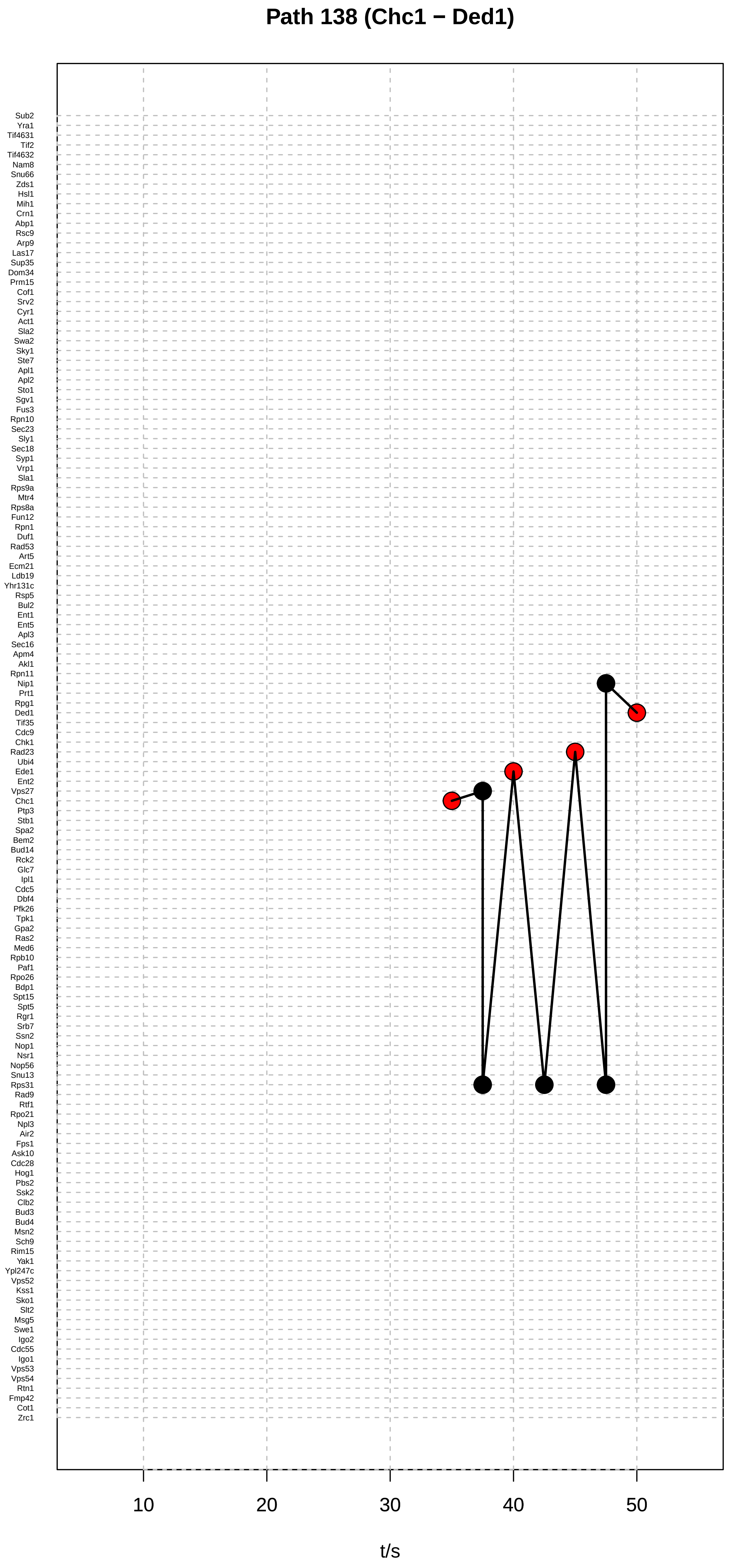
Path 138 (Chc1 − Ded1)
Sub2
Yra1
Tif4631
Tif2
Tif4632
Nam8
Snu66
Zds1
Hsl1
Mih1
Crn1
Abp1
Rsc9
Arp9
Las17
Sup35
Dom34
Prm15
Cof1
Srv2
Cyr1
Act1
Sla2
Swa2
Sky1
Ste7
Apl1
Apl2
Sto1
Sgv1
Fus3
Rpn10
Sec23
Sly1
Sec18
Syp1
Vrp1
Sla1
Rps9a
Mtr4
Rps8a
Fun12
Rpn1
Duf1
Rad53
Art5
Ecm21
Ldb19
Yhr131c
Rsp5
Bul2
Ent1
Ent5
Apl3
Sec16
Apm4
Akl1
Rpn11
Nip1
Prt1
Rpg1
Ded1
Tif35
Cdc9
Chk1
Rad23
Ubi4
Ede1
Ent2
Vps27
Chc1
Ptp3
Stb1
Spa2
Bem2
Bud14
Rck2
Glc7
Ipl1
Cdc5
Dbf4
Pfk26
Tpk1
Gpa2
Ras2
Med6
Rpb10
Paf1
Rpo26
Bdp1
Spt15
Spt5
Rgr1
Srb7
Ssn2
Nop1
Nsr1
Nop56
Snu13
Rps31
Rad9
Rtf1
Rpo21
Npl3
Air2
Fps1
Ask10
Cdc28
Hog1
Pbs2
Ssk2
Clb2
Bud3
Bud4
Msn2
Sch9
Rim15
Yak1
Ypl247c
Vps52
Kss1
Sko1
Slt2
Msg5
Swe1
Igo2
Cdc55
Igo1
Vps53
Vps54
Rtn1
Fmp42
Cot1
Zrc1
10
20
30
40
50
t/s

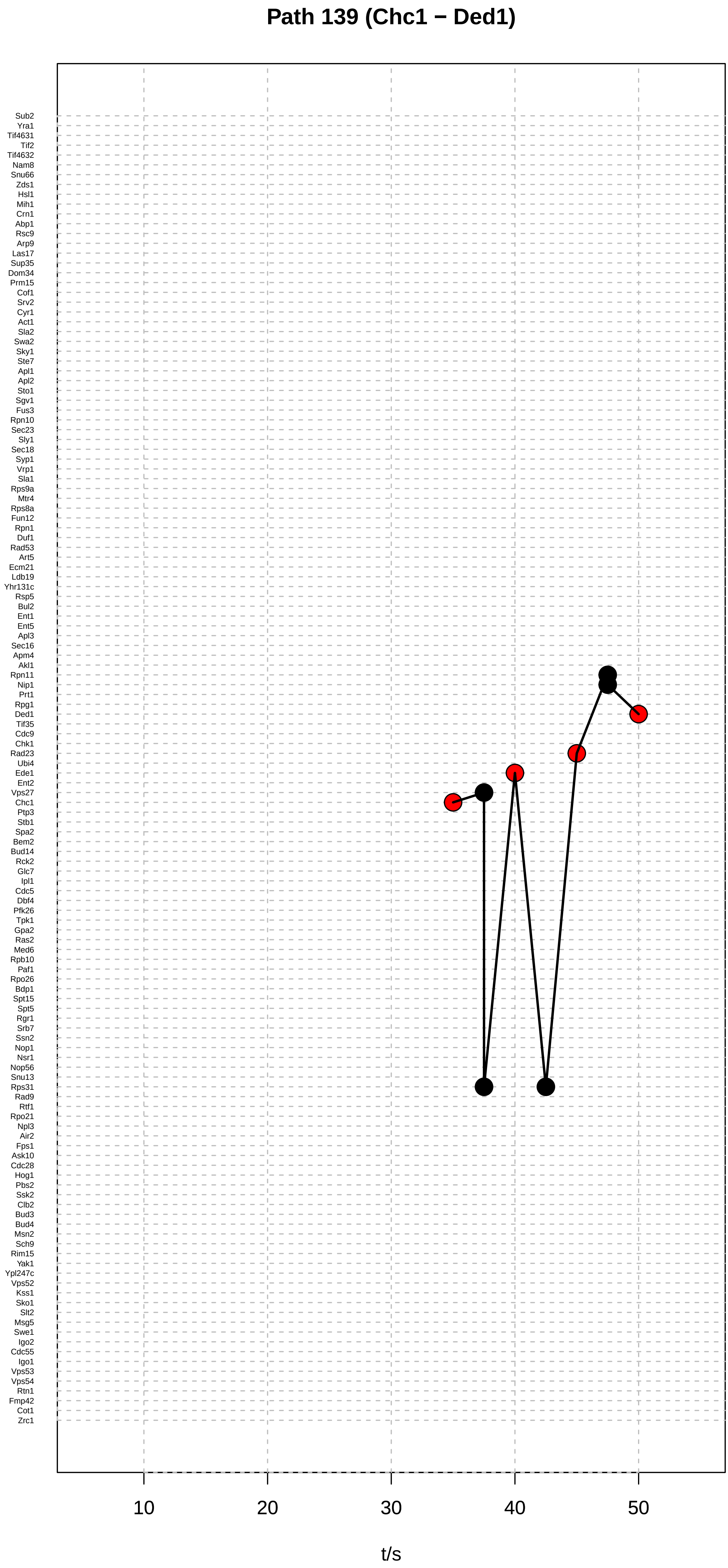

Path 139 (Chc1 − Ded1)
Sub2
Yra1
Tif4631
Tif2
Tif4632
Nam8
Snu66
Zds1
Hsl1
Mih1
Crn1
Abp1
Rsc9
Arp9
Las17
Sup35
Dom34
Prm15
Cof1
Srv2
Cyr1
Act1
Sla2
Swa2
Sky1
Ste7
Apl1
Apl2
Sto1
Sgv1
Fus3
Rpn10
Sec23
Sly1
Sec18
Syp1
Vrp1
Sla1
Rps9a
Mtr4
Rps8a
Fun12
Rpn1
Duf1
Rad53
Art5
Ecm21
Ldb19
Yhr131c
Rsp5
Bul2
Ent1
Ent5
Apl3
Sec16
Apm4
Akl1
Rpn11
Nip1
Prt1
Rpg1
Ded1
Tif35
Cdc9
Chk1
Rad23
Ubi4
Ede1
Ent2
Vps27
Chc1
Ptp3
Stb1
Spa2
Bem2
Bud14
Rck2
Glc7
Ipl1
Cdc5
Dbf4
Pfk26
Tpk1
Gpa2
Ras2
Med6
Rpb10
Paf1
Rpo26
Bdp1
Spt15
Spt5
Rgr1
Srb7
Ssn2
Nop1
Nsr1
Nop56
Snu13
Rps31
Rad9
Rtf1
Rpo21
Npl3
Air2
Fps1
Ask10
Cdc28
Hog1
Pbs2
Ssk2
Clb2
Bud3
Bud4
Msn2
Sch9
Rim15
Yak1
Ypl247c
Vps52
Kss1
Sko1
Slt2
Msg5
Swe1
Igo2
Cdc55
Igo1
Vps53
Vps54
Rtn1
Fmp42
Cot1
Zrc1
10
20
30
40
50
t/s

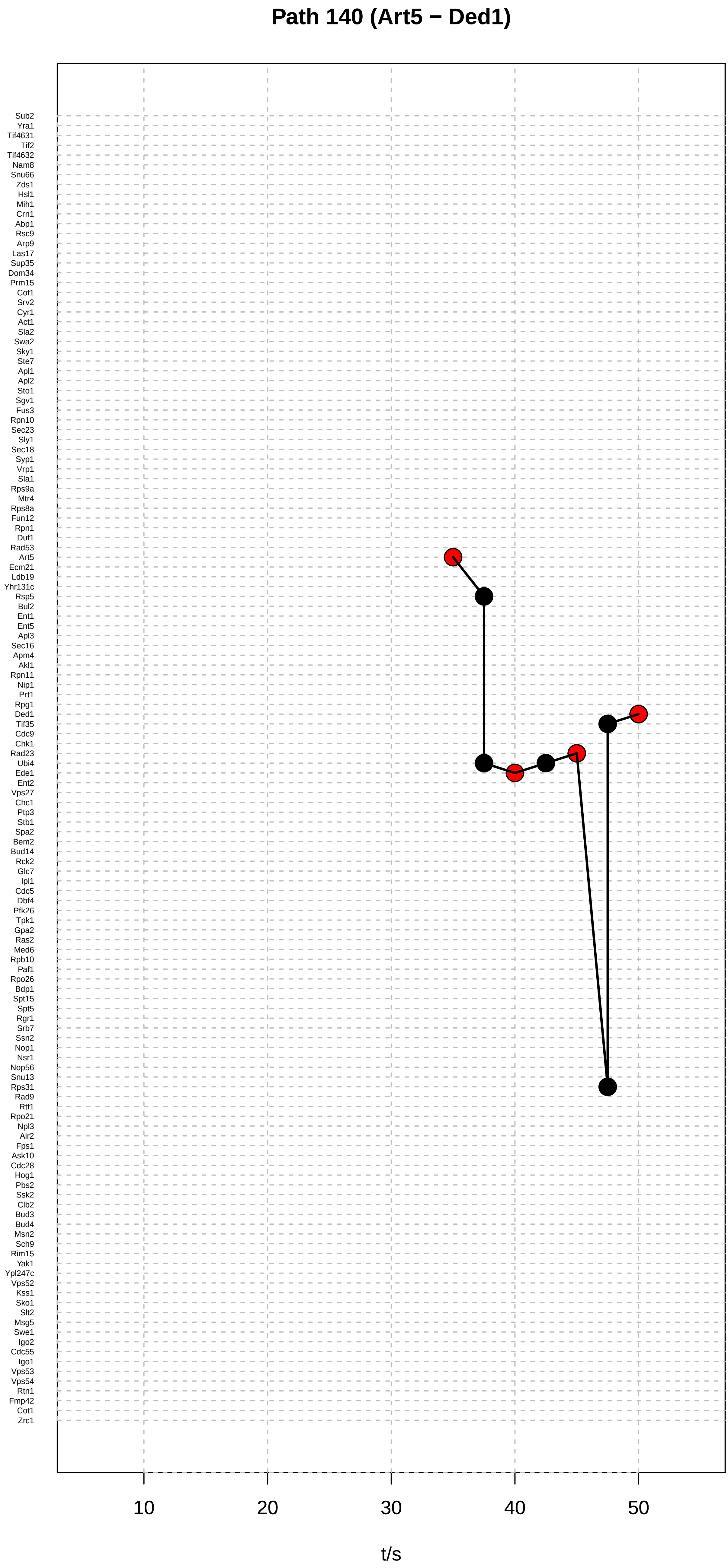

Path 140 (Art5 − Ded1)
Sub2
Yra1
Tif4631
Tif2
Tif4632
Nam8
Snu66
Zds1
Hsl1
Mih1
Crn1
Abp1
Rsc9
Arp9
Las17
Sup35
Dom34
Prm15
Cof1
Srv2
Cyr1
Act1
Sla2
Swa2
Sky1
Ste7
Apl1
Apl2
Sto1
Sgv1
Fus3
Rpn10
Sec23
Sly1
Sec18
Syp1
Vrp1
Sla1
Rps9a
Mtr4
Rps8a
Fun12
Rpn1
Duf1
Rad53
Art5
Ecm21
Ldb19
Yhr131c
Rsp5
Bul2
Ent1
Ent5
Apl3
Sec16
Apm4
Akl1
Rpn11
Nip1
Prt1
Rpg1
Ded1
Tif35
Cdc9
Chk1
Rad23
Ubi4
Ede1
Ent2
Vps27
Chc1
Ptp3
Stb1
Spa2
Bem2
Bud14
Rck2
Glc7
Ipl1
Cdc5
Dbf4
Pfk26
Tpk1
Gpa2
Ras2
Med6
Rpb10
Paf1
Rpo26
Bdp1
Spt15
Spt5
Rgr1
Srb7
Ssn2
Nop1
Nsr1
Nop56
Snu13
Rps31
Rad9
Rtf1
Rpo21
Npl3
Air2
Fps1
Ask10
Cdc28
Hog1
Pbs2
Ssk2
Clb2
Bud3
Bud4
Msn2
Sch9
Rim15
Yak1
Ypl247c
Vps52
Kss1
Sko1
Slt2
Msg5
Swe1
Igo2
Cdc55
Igo1
Vps53
Vps54
Rtn1
Fmp42
Cot1
Zrc1
10
20
30
40
50
t/s

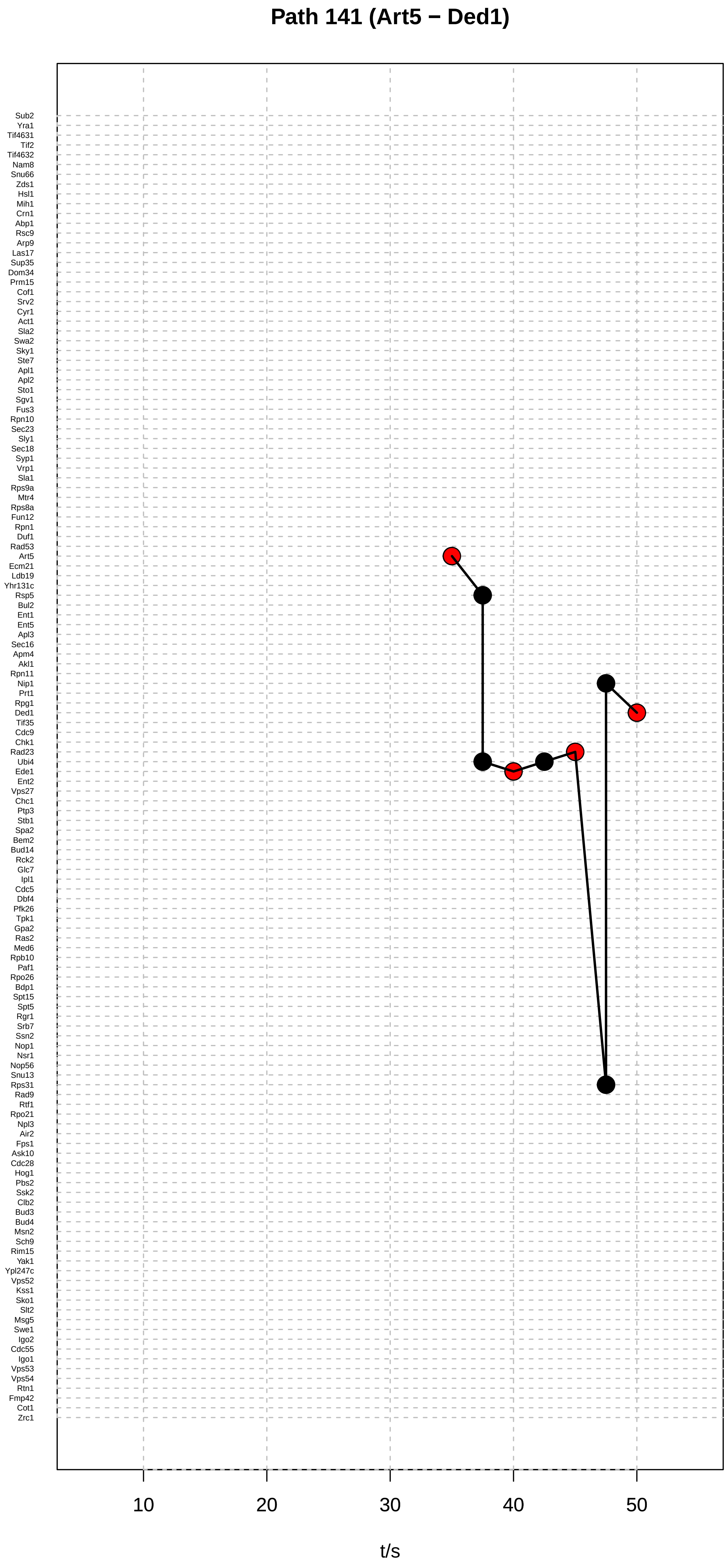
Path 141 (Art5 − Ded1)
Sub2
Yra1
Tif4631
Tif2
Tif4632
Nam8
Snu66
Zds1
Hsl1
Mih1
Crn1
Abp1
Rsc9
Arp9
Las17
Sup35
Dom34
Prm15
Cof1
Srv2
Cyr1
Act1
Sla2
Swa2
Sky1
Ste7
Apl1
Apl2
Sto1
Sgv1
Fus3
Rpn10
Sec23
Sly1
Sec18
Syp1
Vrp1
Sla1
Rps9a
Mtr4
Rps8a
Fun12
Rpn1
Duf1
Rad53
Art5
Ecm21
Ldb19
Yhr131c
Rsp5
Bul2
Ent1
Ent5
Apl3
Sec16
Apm4
Akl1
Rpn11
Nip1
Prt1
Rpg1
Ded1
Tif35
Cdc9
Chk1
Rad23
Ubi4
Ede1
Ent2
Vps27
Chc1
Ptp3
Stb1
Spa2
Bem2
Bud14
Rck2
Glc7
Ipl1
Cdc5
Dbf4
Pfk26
Tpk1
Gpa2
Ras2
Med6
Rpb10
Paf1
Rpo26
Bdp1
Spt15
Spt5
Rgr1
Srb7
Ssn2
Nop1
Nsr1
Nop56
Snu13
Rps31
Rad9
Rtf1
Rpo21
Npl3
Air2
Fps1
Ask10
Cdc28
Hog1
Pbs2
Ssk2
Clb2
Bud3
Bud4
Msn2
Sch9
Rim15
Yak1
Ypl247c
Vps52
Kss1
Sko1
Slt2
Msg5
Swe1
Igo2
Cdc55
Igo1
Vps53
Vps54
Rtn1
Fmp42
Cot1
Zrc1
10
20
30
40
50
t/s

### Path 142 (Art5 − Ded1)

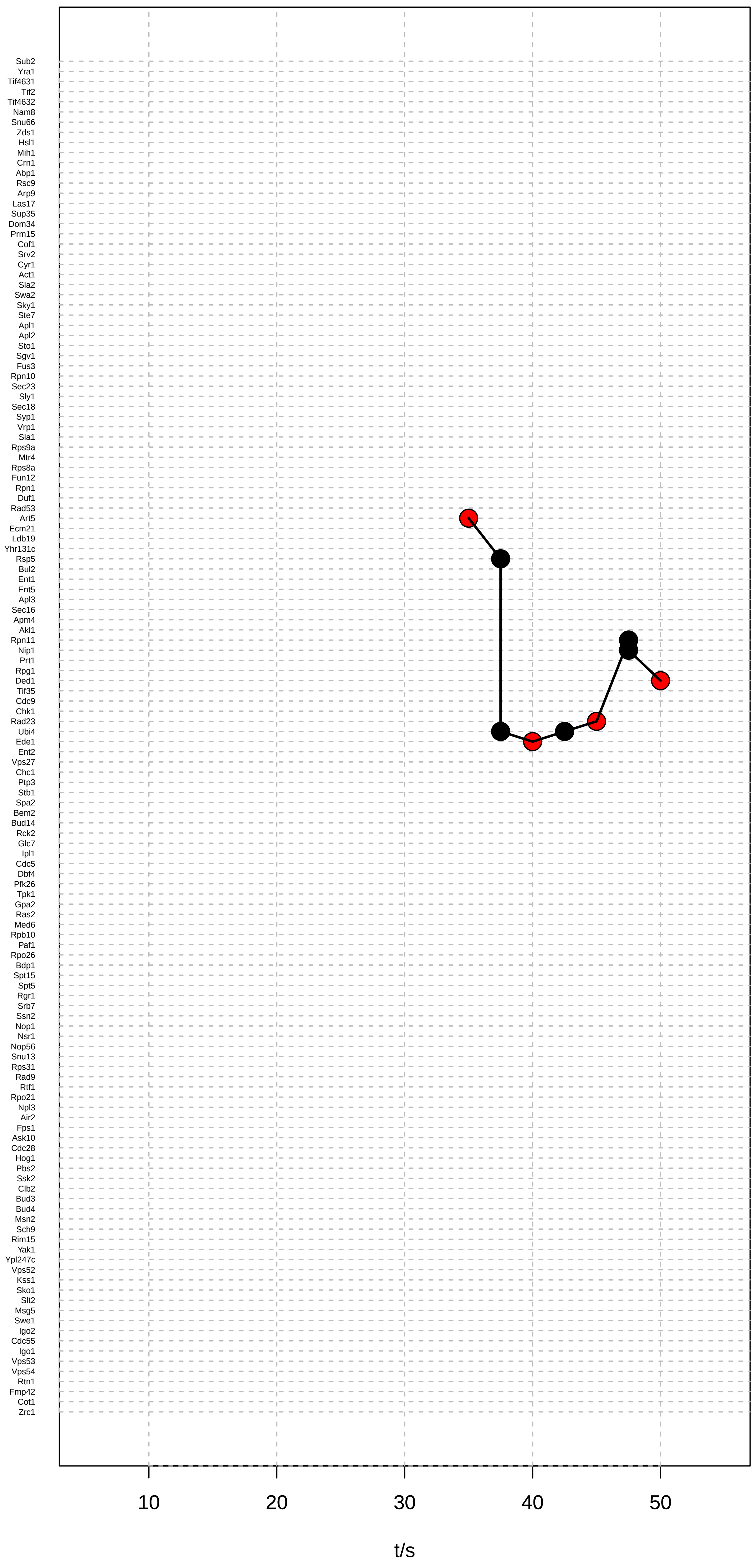

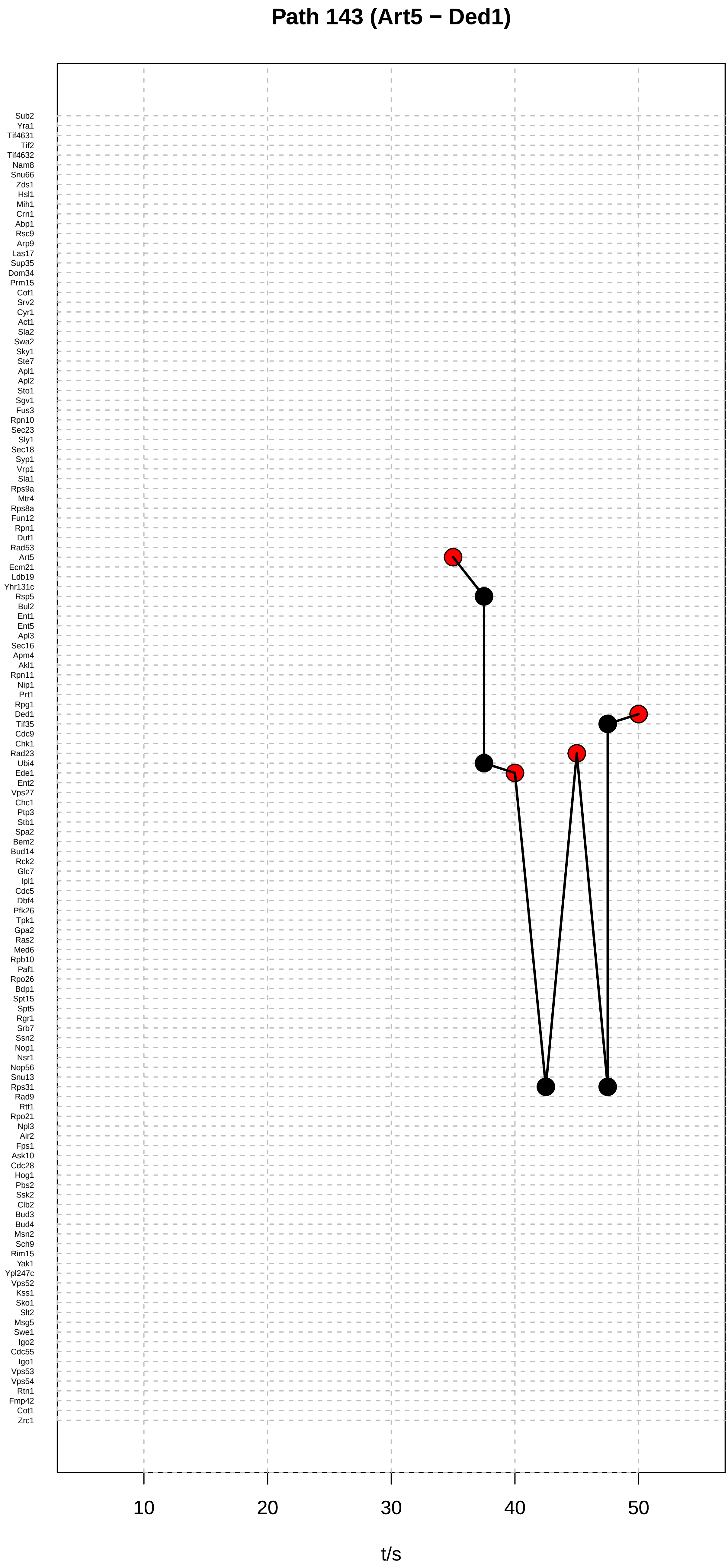
Path 143 (Art5 − Ded1)
Sub2
Yra1
Tif4631
Tif2
Tif4632
Nam8
Snu66
Zds1
Hsl1
Mih1
Crn1
Abp1
Rsc9
Arp9
Las17
Sup35
Dom34
Prm15
Cof1
Srv2
Cyr1
Act1
Sla2
Swa2
Sky1
Ste7
Apl1
Apl2
Sto1
Sgv1
Fus3
Rpn10
Sec23
Sly1
Sec18
Syp1
Vrp1
Sla1
Rps9a
Mtr4
Rps8a
Fun12
Rpn1
Duf1
Rad53
Art5
Ecm21
Ldb19
Yhr131c
Rsp5
Bul2
Ent1
Ent5
Apl3
Sec16
Apm4
Akl1
Rpn11
Nip1
Prt1
Rpg1
Ded1
Tif35
Cdc9
Chk1
Rad23
Ubi4
Ede1
Ent2
Vps27
Chc1
Ptp3
Stb1
Spa2
Bem2
Bud14
Rck2
Glc7
Ipl1
Cdc5
Dbf4
Pfk26
Tpk1
Gpa2
Ras2
Med6
Rpb10
Paf1
Rpo26
Bdp1
Spt15
Spt5
Rgr1
Srb7
Ssn2
Nop1
Nsr1
Nop56
Snu13
Rps31
Rad9
Rtf1
Rpo21
Npl3
Air2
Fps1
Ask10
Cdc28
Hog1
Pbs2
Ssk2
Clb2
Bud3
Bud4
Msn2
Sch9
Rim15
Yak1
Ypl247c
Vps52
Kss1
Sko1
Slt2
Msg5
Swe1
Igo2
Cdc55
Igo1
Vps53
Vps54
Rtn1
Fmp42
Cot1
Zrc1
10
20
30
40
50
t/s

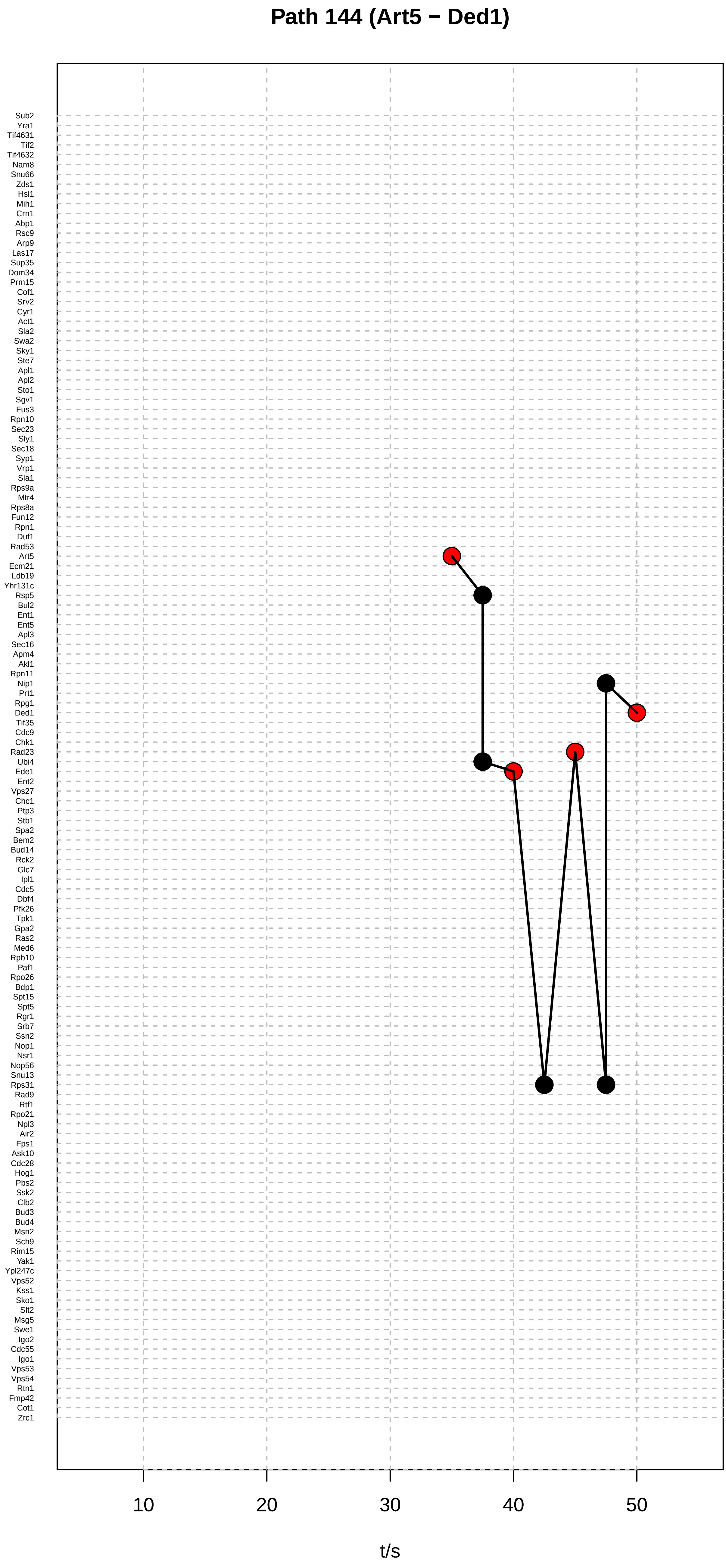

Path 144 (Art5 − Ded1)
Sub2
Yra1
Tif4631
Tif2
Tif4632
Nam8
Snu66
Zds1
Hsl1
Mih1
Crn1
Abp1
Rsc9
Arp9
Las17
Sup35
Dom34
Prm15
Cof1
Srv2
Cyr1
Act1
Sla2
Swa2
Sky1
Ste7
Apl1
Apl2
Sto1
Sgv1
Fus3
Rpn10
Sec23
Sly1
Sec18
Syp1
Vrp1
Sla1
Rps9a
Mtr4
Rps8a
Fun12
Rpn1
Duf1
Rad53
Art5
Ecm21
Ldb19
Yhr131c
Rsp5
Bul2
Ent1
Ent5
Apl3
Sec16
Apm4
Akl1
Rpn11
Nip1
Prt1
Rpg1
Ded1
Tif35
Cdc9
Chk1
Rad23
Ubi4
Ede1
Ent2
Vps27
Chc1
Ptp3
Stb1
Spa2
Bem2
Bud14
Rck2
Glc7
Ipl1
Cdc5
Dbf4
Pfk26
Tpk1
Gpa2
Ras2
Med6
Rpb10
Paf1
Rpo26
Bdp1
Spt15
Spt5
Rgr1
Srb7
Ssn2
Nop1
Nsr1
Nop56
Snu13
Rps31
Rad9
Rtf1
Rpo21
Npl3
Air2
Fps1
Ask10
Cdc28
Hog1
Pbs2
Ssk2
Clb2
Bud3
Bud4
Msn2
Sch9
Rim15
Yak1
Ypl247c
Vps52
Kss1
Sko1
Slt2
Msg5
Swe1
Igo2
Cdc55
Igo1
Vps53
Vps54
Rtn1
Fmp42
Cot1
Zrc1
10
20
30
40
50
t/s

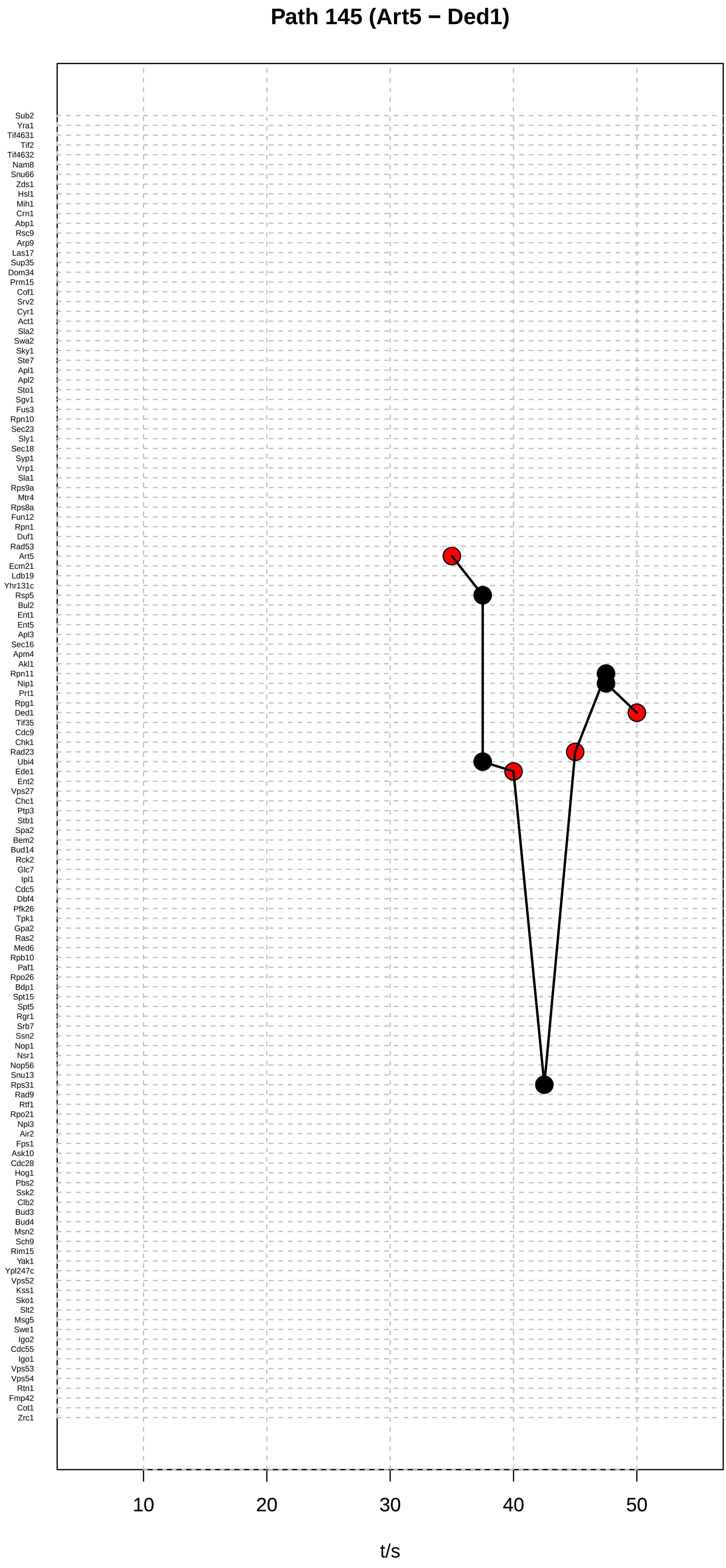
Path 145 (Art5 − Ded1)
Sub2
Yra1
Tif4631
Tif2
Tif4632
Nam8
Snu66
Zds1
Hsl1
Mih1
Crn1
Abp1
Rsc9
Arp9
Las17
Sup35
Dom34
Prm15
Cof1
Srv2
Cyr1
Act1
Sla2
Swa2
Sky1
Ste7
Apl1
Apl2
Sto1
Sgv1
Fus3
Rpn10
Sec23
Sly1
Sec18
Syp1
Vrp1
Sla1
Rps9a
Mtr4
Rps8a
Fun12
Rpn1
Duf1
Rad53
Art5
Ecm21
Ldb19
Yhr131c
Rsp5
Bul2
Ent1
Ent5
Apl3
Sec16
Apm4
Akl1
Rpn11
Nip1
Prt1
Rpg1
Ded1
Tif35
Cdc9
Chk1
Rad23
Ubi4
Ede1
Ent2
Vps27
Chc1
Ptp3
Stb1
Spa2
Bem2
Bud14
Rck2
Glc7
Ipl1
Cdc5
Dbf4
Pfk26
Tpk1
Gpa2
Ras2
Med6
Rpb10
Paf1
Rpo26
Bdp1
Spt15
Spt5
Rgr1
Srb7
Ssn2
Nop1
Nsr1
Nop56
Snu13
Rps31
Rad9
Rtf1
Rpo21
Npl3
Air2
Fps1
Ask10
Cdc28
Hog1
Pbs2
Ssk2
Clb2
Bud3
Bud4
Msn2
Sch9
Rim15
Yak1
Ypl247c
Vps52
Kss1
Sko1
Slt2
Msg5
Swe1
Igo2
Cdc55
Igo1
Vps53
Vps54
Rtn1
Fmp42
Cot1
Zrc1
10
20
30
40
50
t/s

#### Path 146 (Art5 − Ded1)

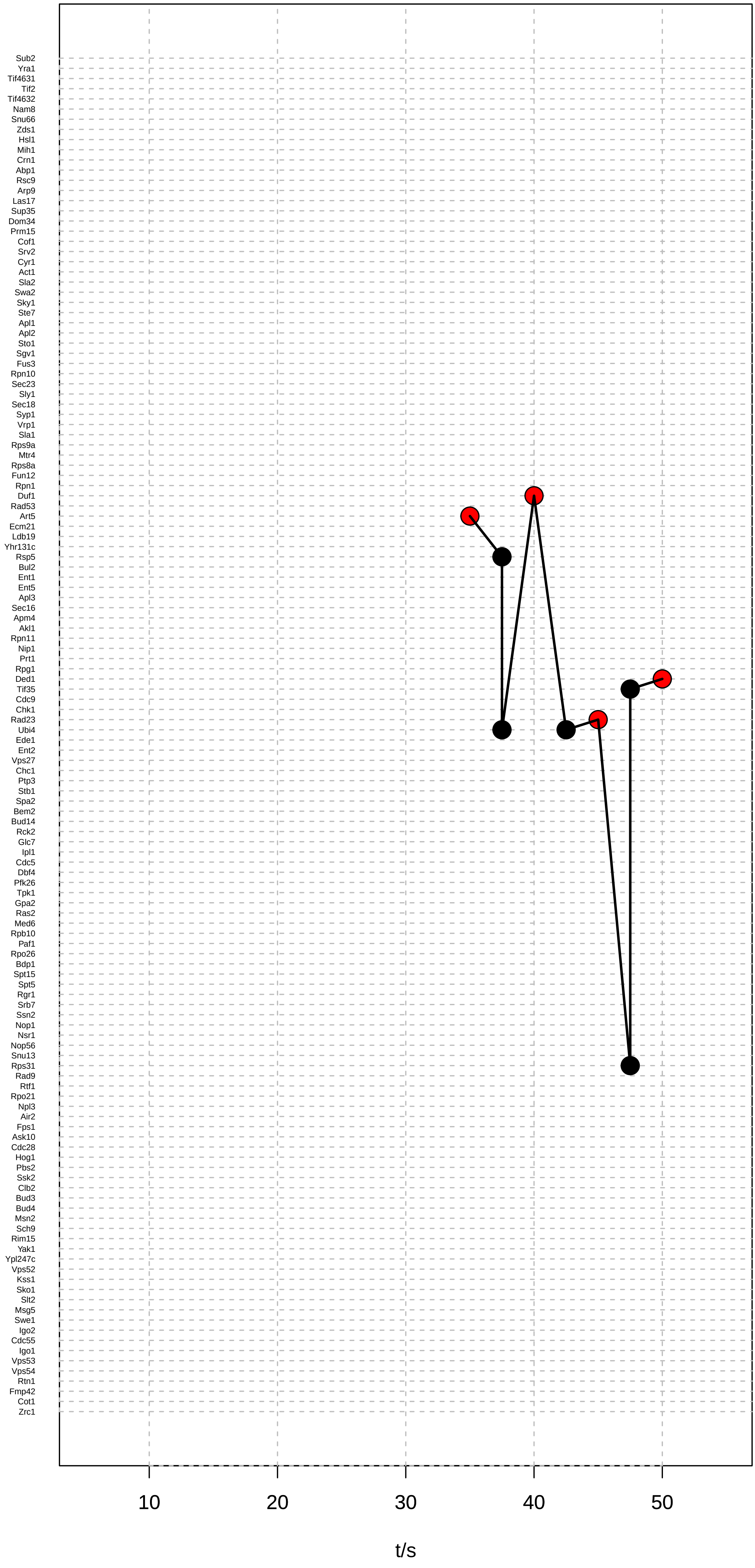

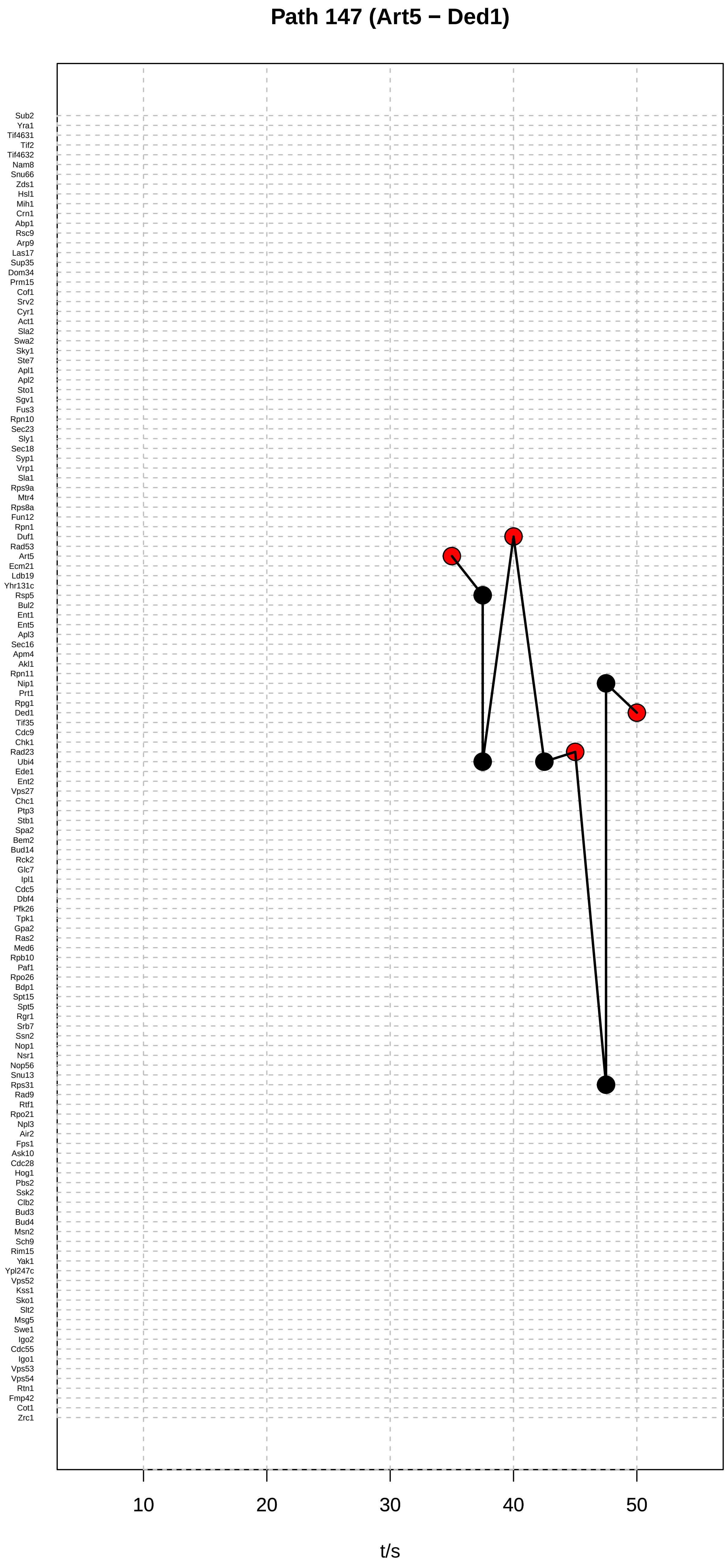

Path 147 (Art5 − Ded1)
Sub2
Yra1
Tif4631
Tif2
Tif4632
Nam8
Snu66
Zds1
Hsl1
Mih1
Crn1
Abp1
Rsc9
Arp9
Las17
Sup35
Dom34
Prm15
Cof1
Srv2
Cyr1
Act1
Sla2
Swa2
Sky1
Ste7
Apl1
Apl2
Sto1
Sgv1
Fus3
Rpn10
Sec23
Sly1
Sec18
Syp1
Vrp1
Sla1
Rps9a
Mtr4
Rps8a
Fun12
Rpn1
Duf1
Rad53
Art5
Ecm21
Ldb19
Yhr131c
Rsp5
Bul2
Ent1
Ent5
Apl3
Sec16
Apm4
Akl1
Rpn11
Nip1
Prt1
Rpg1
Ded1
Tif35
Cdc9
Chk1
Rad23
Ubi4
Ede1
Ent2
Vps27
Chc1
Ptp3
Stb1
Spa2
Bem2
Bud14
Rck2
Glc7
Ipl1
Cdc5
Dbf4
Pfk26
Tpk1
Gpa2
Ras2
Med6
Rpb10
Paf1
Rpo26
Bdp1
Spt15
Spt5
Rgr1
Srb7
Ssn2
Nop1
Nsr1
Nop56
Snu13
Rps31
Rad9
Rtf1
Rpo21
Npl3
Air2
Fps1
Ask10
Cdc28
Hog1
Pbs2
Ssk2
Clb2
Bud3
Bud4
Msn2
Sch9
Rim15
Yak1
Ypl247c
Vps52
Kss1
Sko1
Slt2
Msg5
Swe1
Igo2
Cdc55
Igo1
Vps53
Vps54
Rtn1
Fmp42
Cot1
Zrc1
10
20
30
40
50
t/s

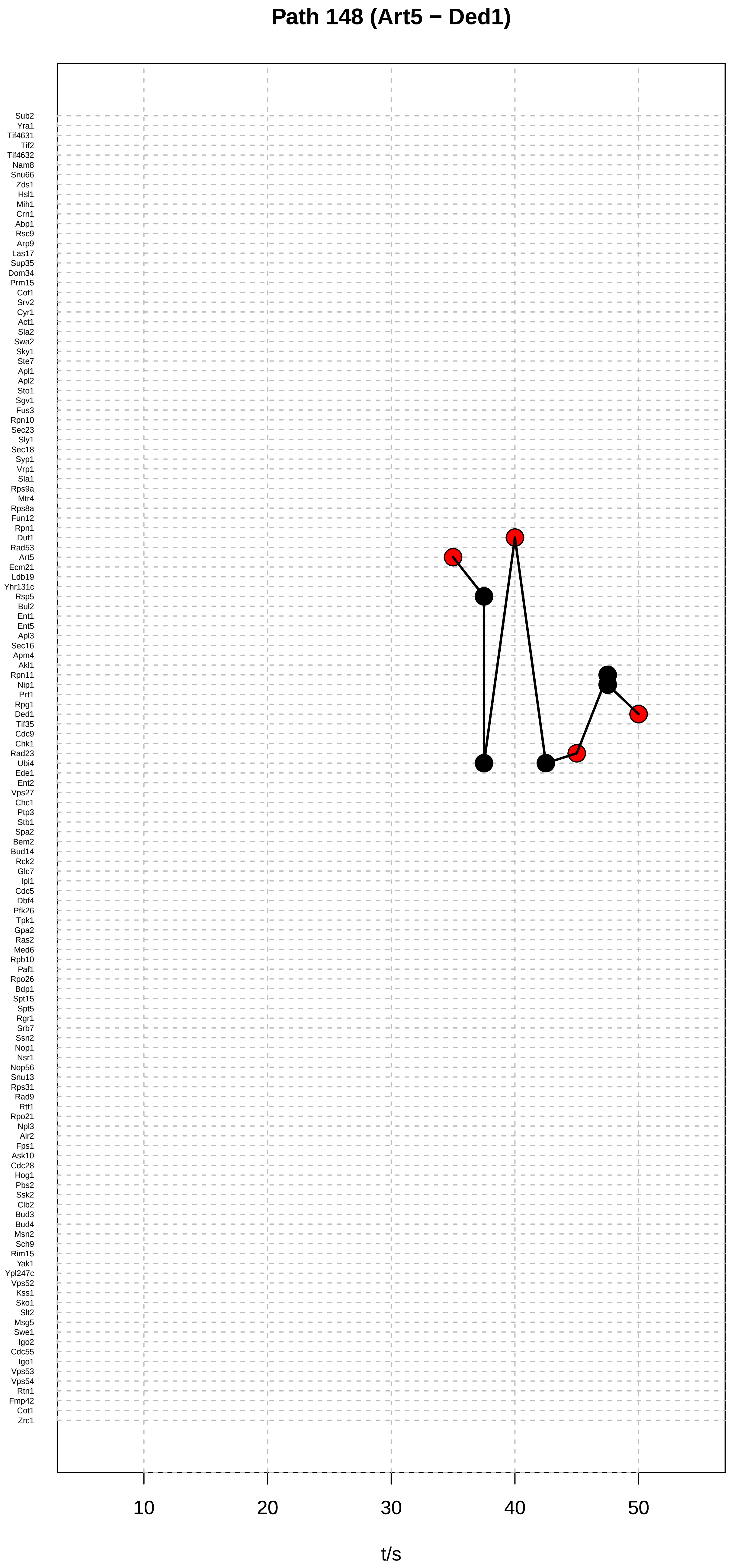
Path 148 (Art5 − Ded1)
Sub2
Yra1
Tif4631
Tif2
Tif4632
Nam8
Snu66
Zds1
Hsl1
Mih1
Crn1
Abp1
Rsc9
Arp9
Las17
Sup35
Dom34
Prm15
Cof1
Srv2
Cyr1
Act1
Sla2
Swa2
Sky1
Ste7
Apl1
Apl2
Sto1
Sgv1
Fus3
Rpn10
Sec23
Sly1
Sec18
Syp1
Vrp1
Sla1
Rps9a
Mtr4
Rps8a
Fun12
Rpn1
Duf1
Rad53
Art5
Ecm21
Ldb19
Yhr131c
Rsp5
Bul2
Ent1
Ent5
Apl3
Sec16
Apm4
Akl1
Rpn11
Nip1
Prt1
Rpg1
Ded1
Tif35
Cdc9
Chk1
Rad23
Ubi4
Ede1
Ent2
Vps27
Chc1
Ptp3
Stb1
Spa2
Bem2
Bud14
Rck2
Glc7
Ipl1
Cdc5
Dbf4
Pfk26
Tpk1
Gpa2
Ras2
Med6
Rpb10
Paf1
Rpo26
Bdp1
Spt15
Spt5
Rgr1
Srb7
Ssn2
Nop1
Nsr1
Nop56
Snu13
Rps31
Rad9
Rtf1
Rpo21
Npl3
Air2
Fps1
Ask10
Cdc28
Hog1
Pbs2
Ssk2
Clb2
Bud3
Bud4
Msn2
Sch9
Rim15
Yak1
Ypl247c
Vps52
Kss1
Sko1
Slt2
Msg5
Swe1
Igo2
Cdc55
Igo1
Vps53
Vps54
Rtn1
Fmp42
Cot1
Zrc1
10
20
30
40
50
t/s

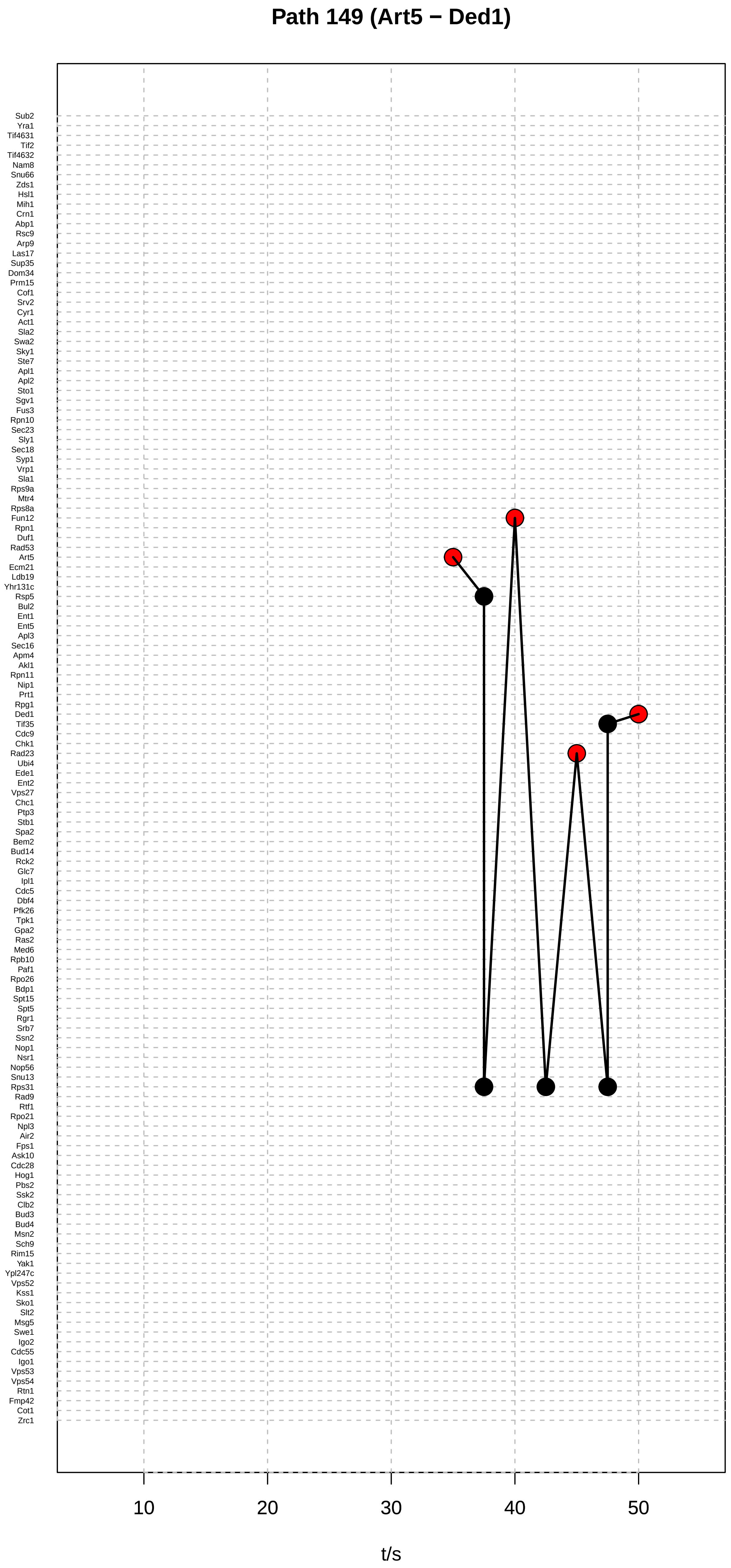

Path 149 (Art5 − Ded1)
Sub2
Yra1
Tif4631
Tif2
Tif4632
Nam8
Snu66
Zds1
Hsl1
Mih1
Crn1
Abp1
Rsc9
Arp9
Las17
Sup35
Dom34
Prm15
Cof1
Srv2
Cyr1
Act1
Sla2
Swa2
Sky1
Ste7
Apl1
Apl2
Sto1
Sgv1
Fus3
Rpn10
Sec23
Sly1
Sec18
Syp1
Vrp1
Sla1
Rps9a
Mtr4
Rps8a
Fun12
Rpn1
Duf1
Rad53
Art5
Ecm21
Ldb19
Yhr131c
Rsp5
Bul2
Ent1
Ent5
Apl3
Sec16
Apm4
Akl1
Rpn11
Nip1
Prt1
Rpg1
Ded1
Tif35
Cdc9
Chk1
Rad23
Ubi4
Ede1
Ent2
Vps27
Chc1
Ptp3
Stb1
Spa2
Bem2
Bud14
Rck2
Glc7
Ipl1
Cdc5
Dbf4
Pfk26
Tpk1
Gpa2
Ras2
Med6
Rpb10
Paf1
Rpo26
Bdp1
Spt15
Spt5
Rgr1
Srb7
Ssn2
Nop1
Nsr1
Nop56
Snu13
Rps31
Rad9
Rtf1
Rpo21
Npl3
Air2
Fps1
Ask10
Cdc28
Hog1
Pbs2
Ssk2
Clb2
Bud3
Bud4
Msn2
Sch9
Rim15
Yak1
Ypl247c
Vps52
Kss1
Sko1
Slt2
Msg5
Swe1
Igo2
Cdc55
Igo1
Vps53
Vps54
Rtn1
Fmp42
Cot1
Zrc1
10
20
30
40
50
t/s

#### Path 150 (Art5 − Ded1)

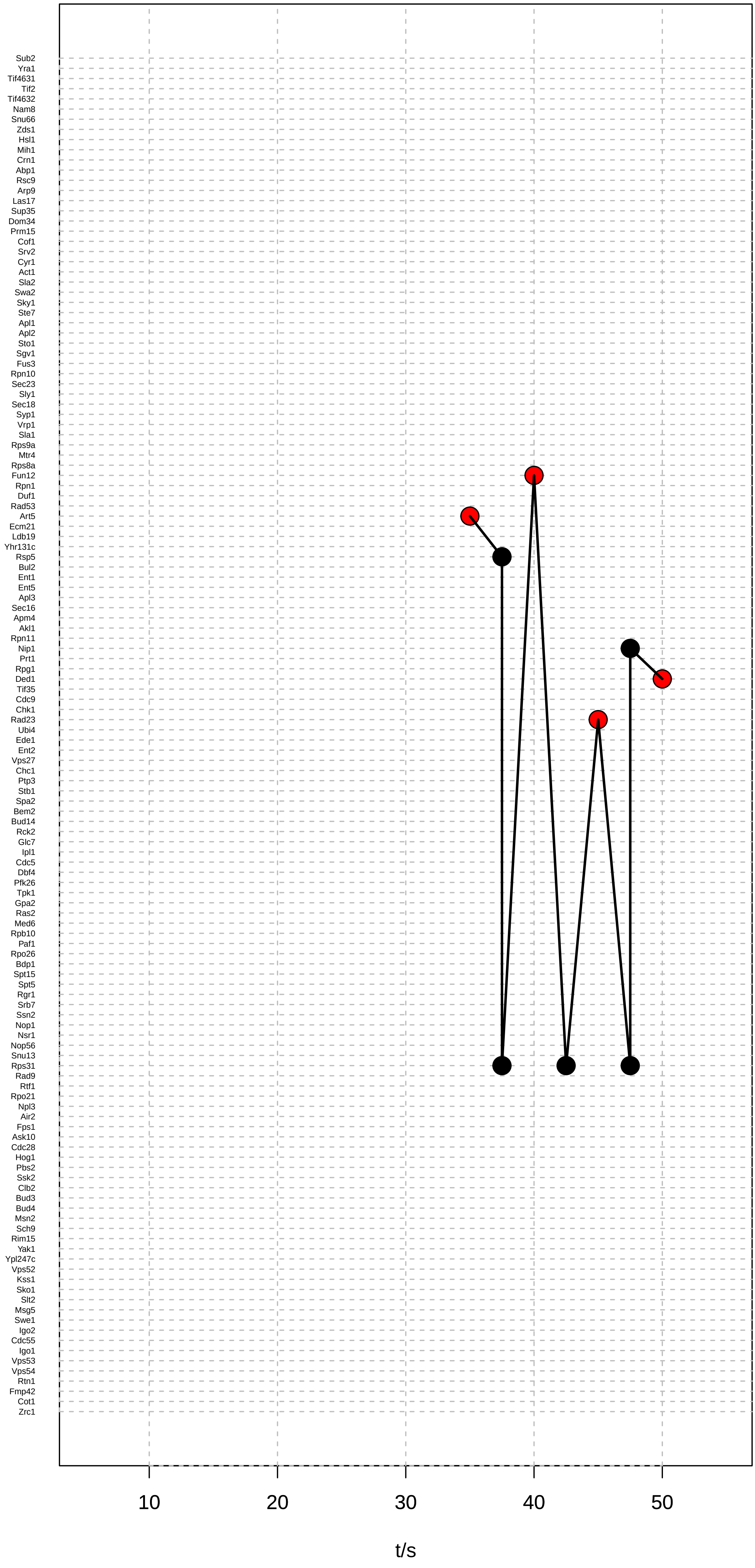

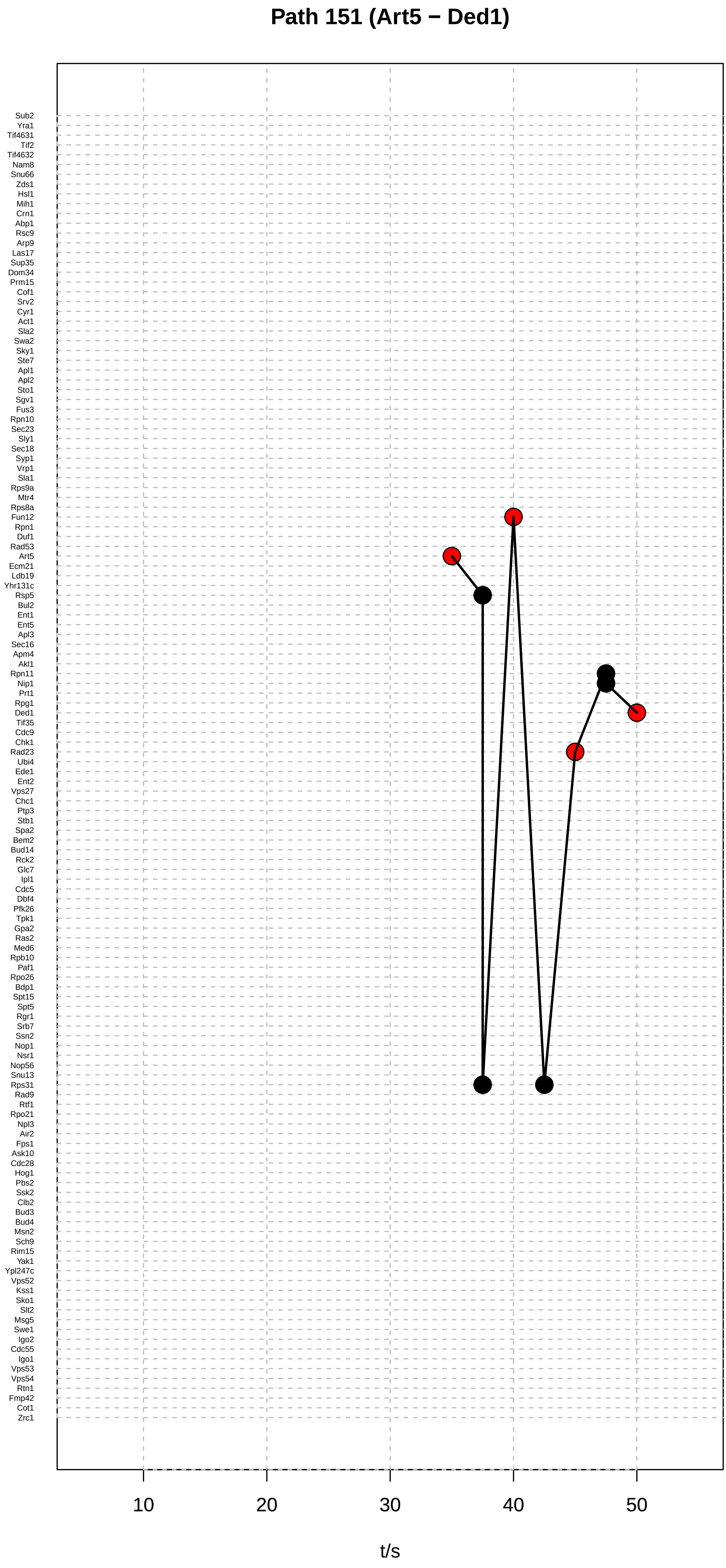

Path 151 (Art5 − Ded1)
Sub2
Yra1
Tif4631
Tif2
Tif4632
Nam8
Snu66
Zds1
Hsl1
Mih1
Crn1
Abp1
Rsc9
Arp9
Las17
Sup35
Dom34
Prm15
Cof1
Srv2
Cyr1
Act1
Sla2
Swa2
Sky1
Ste7
Apl1
Apl2
Sto1
Sgv1
Fus3
Rpn10
Sec23
Sly1
Sec18
Syp1
Vrp1
Sla1
Rps9a
Mtr4
Rps8a
Fun12
Rpn1
Duf1
Rad53
Art5
Ecm21
Ldb19
Yhr131c
Rsp5
Bul2
Ent1
Ent5
Apl3
Sec16
Apm4
Akl1
Rpn11
Nip1
Prt1
Rpg1
Ded1
Tif35
Cdc9
Chk1
Rad23
Ubi4
Ede1
Ent2
Vps27
Chc1
Ptp3
Stb1
Spa2
Bem2
Bud14
Rck2
Glc7
Ipl1
Cdc5
Dbf4
Pfk26
Tpk1
Gpa2
Ras2
Med6
Rpb10
Paf1
Rpo26
Bdp1
Spt15
Spt5
Rgr1
Srb7
Ssn2
Nop1
Nsr1
Nop56
Snu13
Rps31
Rad9
Rtf1
Rpo21
Npl3
Air2
Fps1
Ask10
Cdc28
Hog1
Pbs2
Ssk2
Clb2
Bud3
Bud4
Msn2
Sch9
Rim15
Yak1
Ypl247c
Vps52
Kss1
Sko1
Slt2
Msg5
Swe1
Igo2
Cdc55
Igo1
Vps53
Vps54
Rtn1
Fmp42
Cot1
Zrc1
10
20
30
40
50
t/s

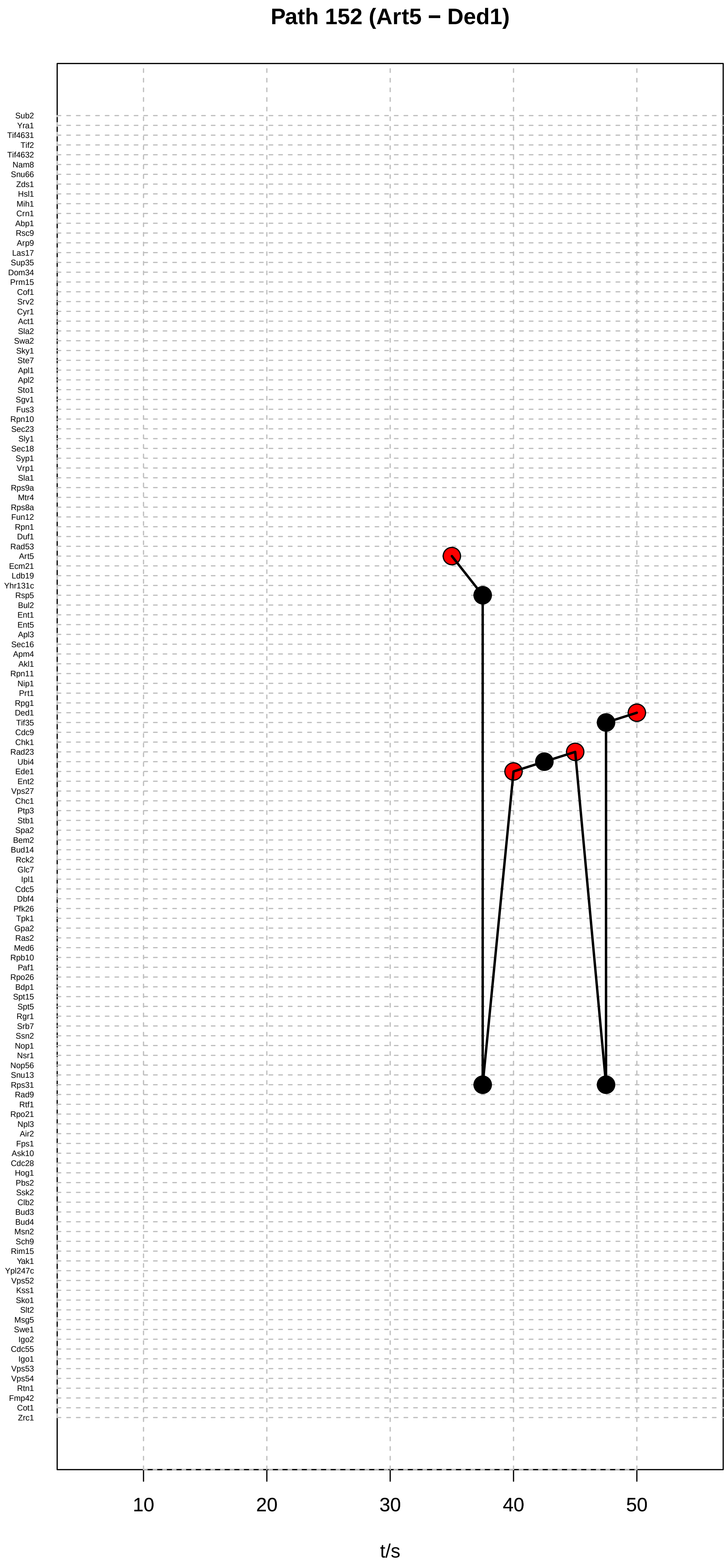
Path 152 (Art5 − Ded1)
Sub2
Yra1
Tif4631
Tif2
Tif4632
Nam8
Snu66
Zds1
Hsl1
Mih1
Crn1
Abp1
Rsc9
Arp9
Las17
Sup35
Dom34
Prm15
Cof1
Srv2
Cyr1
Act1
Sla2
Swa2
Sky1
Ste7
Apl1
Apl2
Sto1
Sgv1
Fus3
Rpn10
Sec23
Sly1
Sec18
Syp1
Vrp1
Sla1
Rps9a
Mtr4
Rps8a
Fun12
Rpn1
Duf1
Rad53
Art5
Ecm21
Ldb19
Yhr131c
Rsp5
Bul2
Ent1
Ent5
Apl3
Sec16
Apm4
Akl1
Rpn11
Nip1
Prt1
Rpg1
Ded1
Tif35
Cdc9
Chk1
Rad23
Ubi4
Ede1
Ent2
Vps27
Chc1
Ptp3
Stb1
Spa2
Bem2
Bud14
Rck2
Glc7
Ipl1
Cdc5
Dbf4
Pfk26
Tpk1
Gpa2
Ras2
Med6
Rpb10
Paf1
Rpo26
Bdp1
Spt15
Spt5
Rgr1
Srb7
Ssn2
Nop1
Nsr1
Nop56
Snu13
Rps31
Rad9
Rtf1
Rpo21
Npl3
Air2
Fps1
Ask10
Cdc28
Hog1
Pbs2
Ssk2
Clb2
Bud3
Bud4
Msn2
Sch9
Rim15
Yak1
Ypl247c
Vps52
Kss1
Sko1
Slt2
Msg5
Swe1
Igo2
Cdc55
Igo1
Vps53
Vps54
Rtn1
Fmp42
Cot1
Zrc1
10
20
30
40
50
t/s

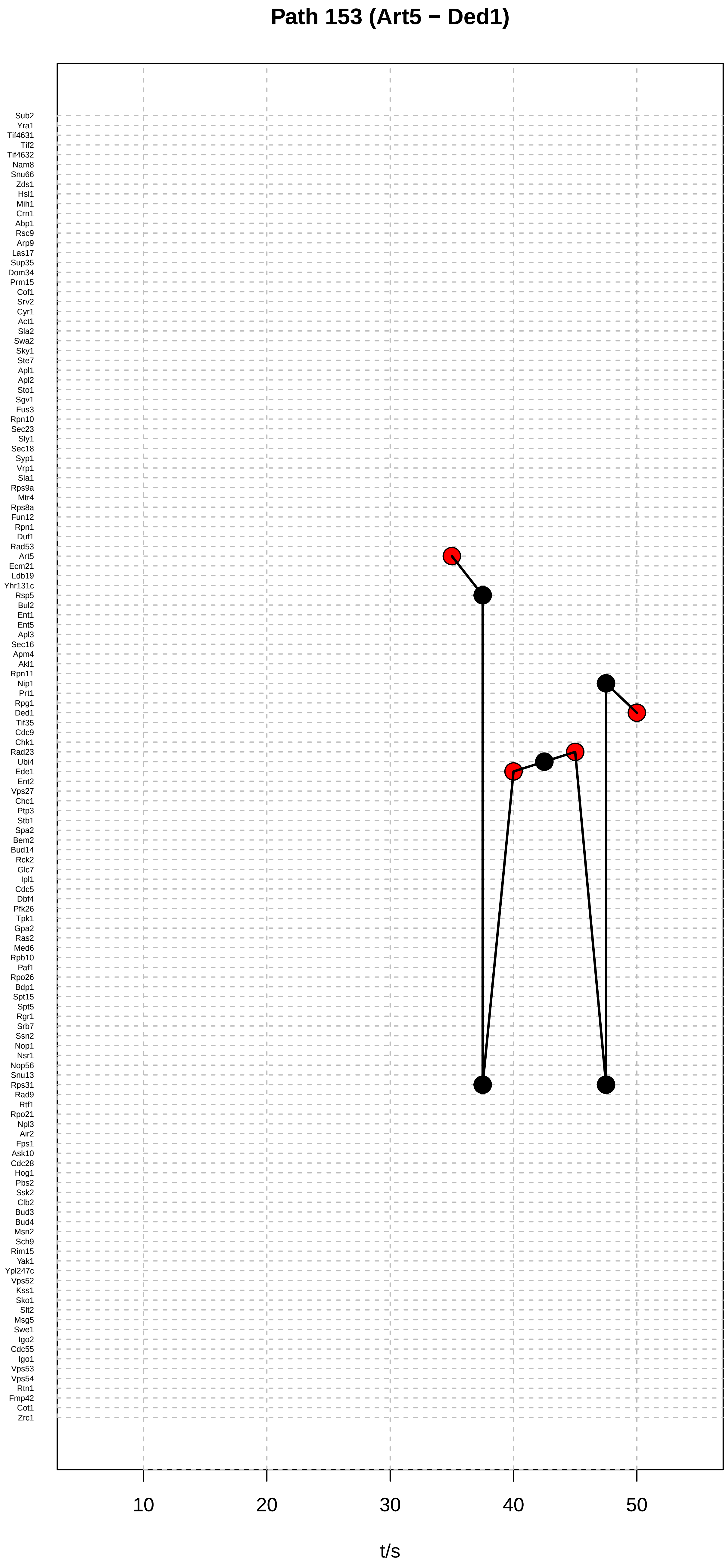

Path 153 (Art5 − Ded1)
Sub2
Yra1
Tif4631
Tif2
Tif4632
Nam8
Snu66
Zds1
Hsl1
Mih1
Crn1
Abp1
Rsc9
Arp9
Las17
Sup35
Dom34
Prm15
Cof1
Srv2
Cyr1
Act1
Sla2
Swa2
Sky1
Ste7
Apl1
Apl2
Sto1
Sgv1
Fus3
Rpn10
Sec23
Sly1
Sec18
Syp1
Vrp1
Sla1
Rps9a
Mtr4
Rps8a
Fun12
Rpn1
Duf1
Rad53
Art5
Ecm21
Ldb19
Yhr131c
Rsp5
Bul2
Ent1
Ent5
Apl3
Sec16
Apm4
Akl1
Rpn11
Nip1
Prt1
Rpg1
Ded1
Tif35
Cdc9
Chk1
Rad23
Ubi4
Ede1
Ent2
Vps27
Chc1
Ptp3
Stb1
Spa2
Bem2
Bud14
Rck2
Glc7
Ipl1
Cdc5
Dbf4
Pfk26
Tpk1
Gpa2
Ras2
Med6
Rpb10
Paf1
Rpo26
Bdp1
Spt15
Spt5
Rgr1
Srb7
Ssn2
Nop1
Nsr1
Nop56
Snu13
Rps31
Rad9
Rtf1
Rpo21
Npl3
Air2
Fps1
Ask10
Cdc28
Hog1
Pbs2
Ssk2
Clb2
Bud3
Bud4
Msn2
Sch9
Rim15
Yak1
Ypl247c
Vps52
Kss1
Sko1
Slt2
Msg5
Swe1
Igo2
Cdc55
Igo1
Vps53
Vps54
Rtn1
Fmp42
Cot1
Zrc1
10
20
30
40
50
t/s

#### Path 154 (Art5 − Ded1)

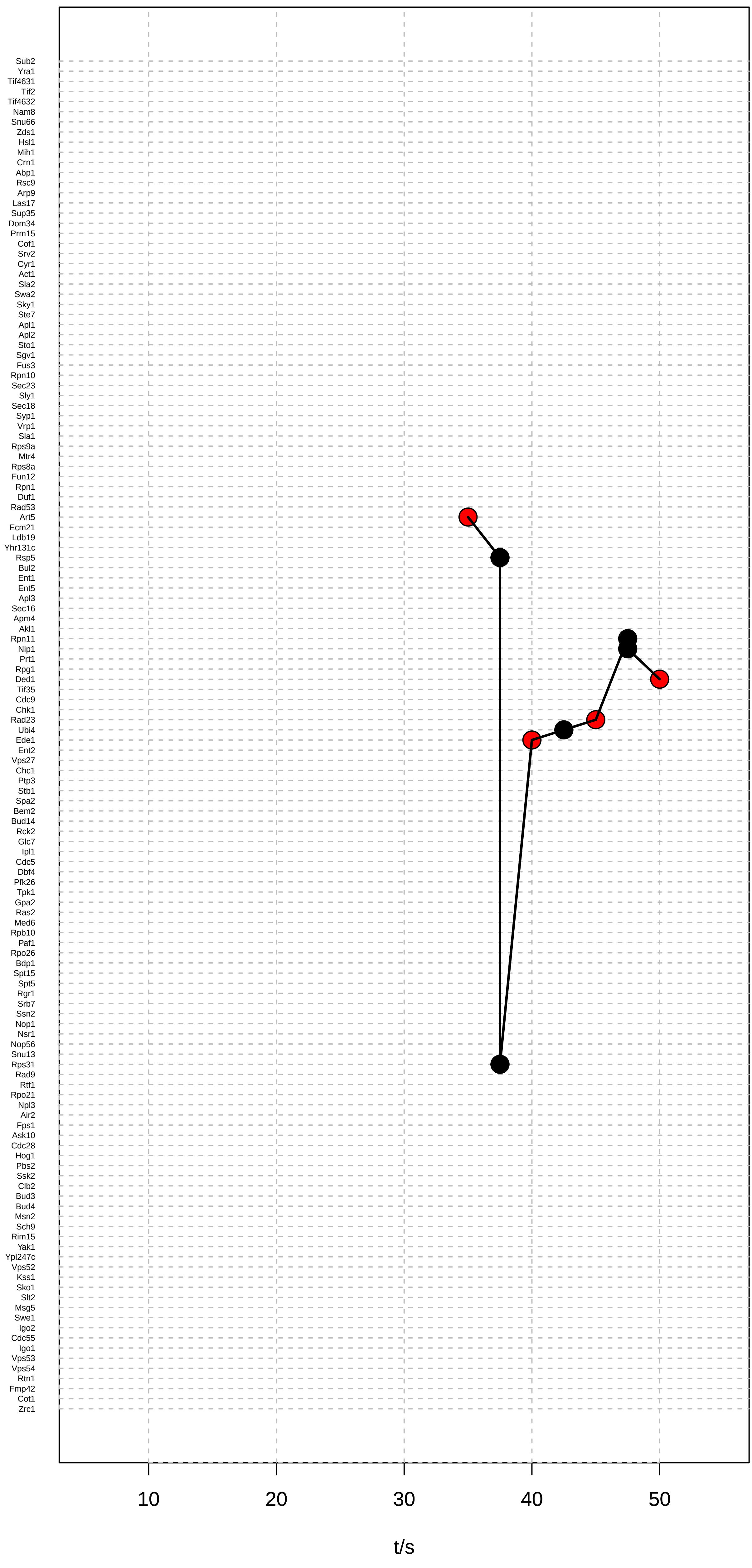

#### Path 155 (Art5 − Ded1)

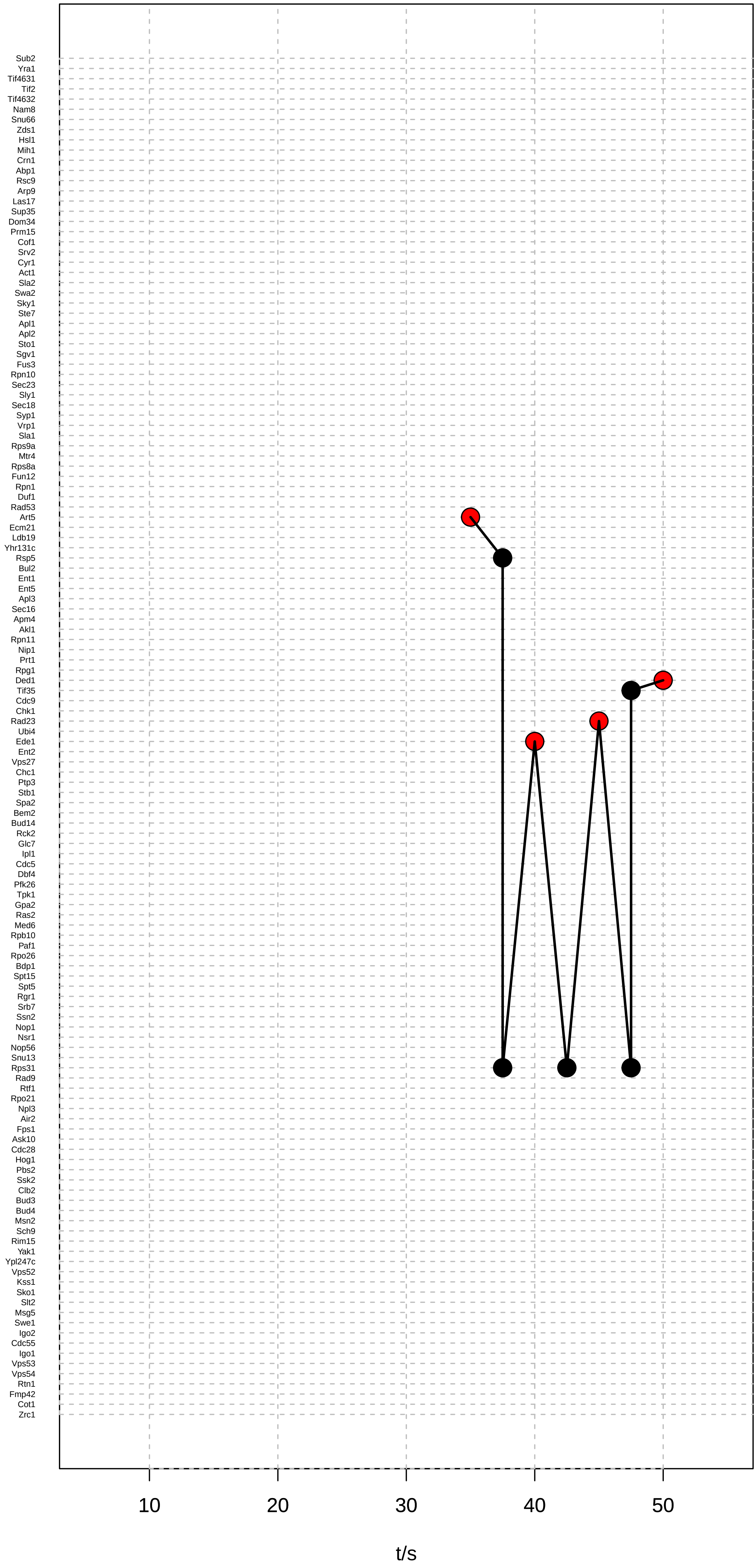

#### Path 156 (Art5 − Ded1)

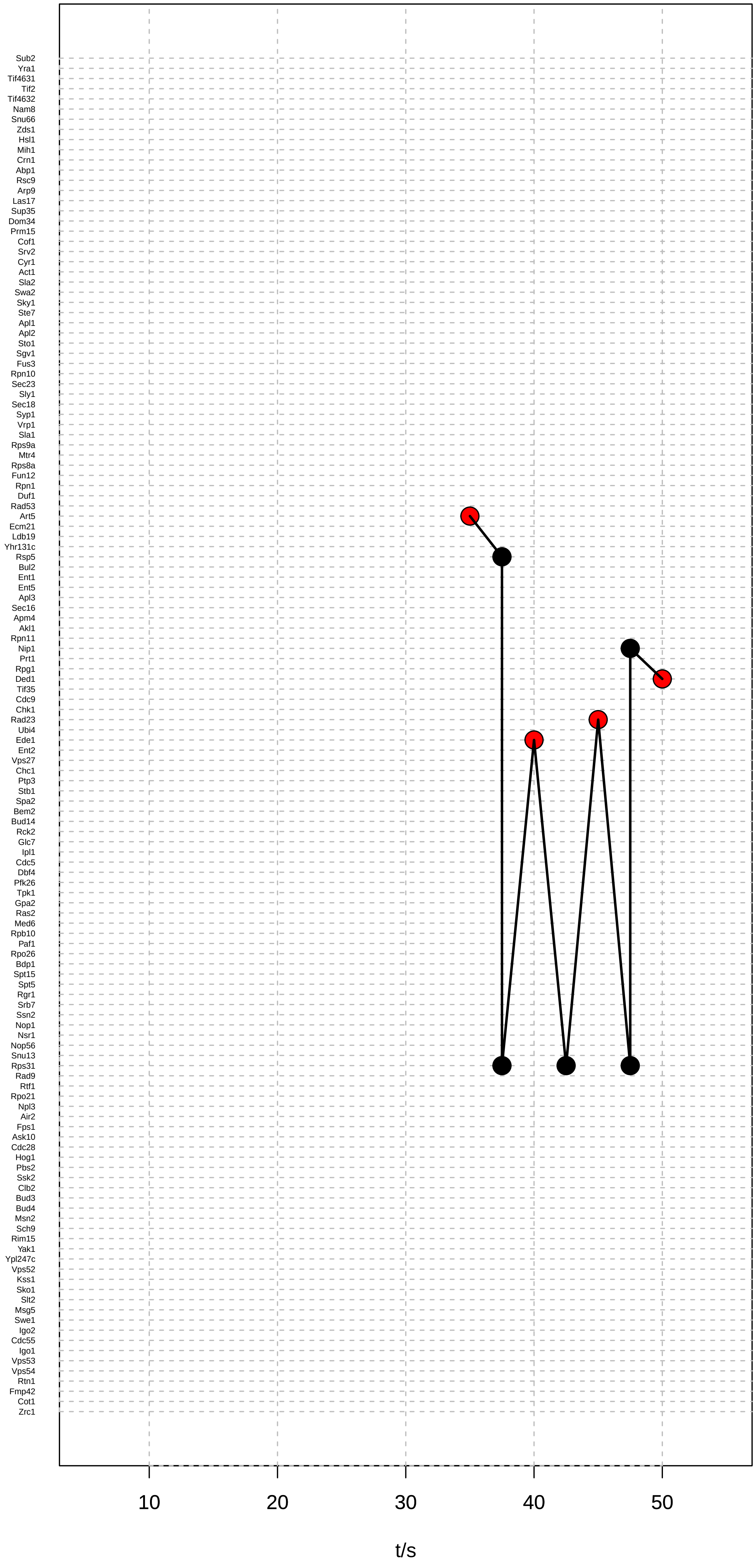

### Path 157 (Art5 − Ded1)

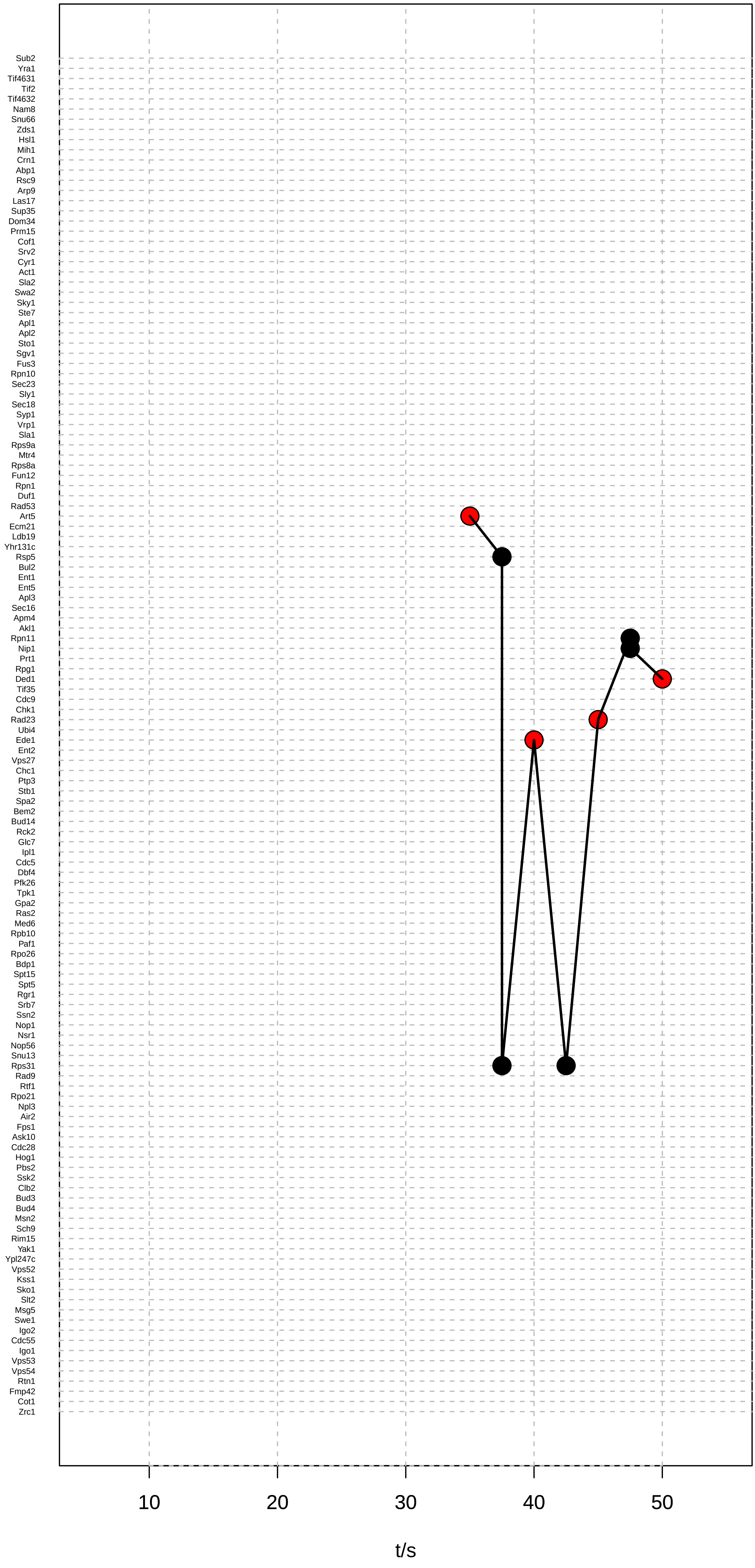

#### Path 158 (Air2 − Ded1)

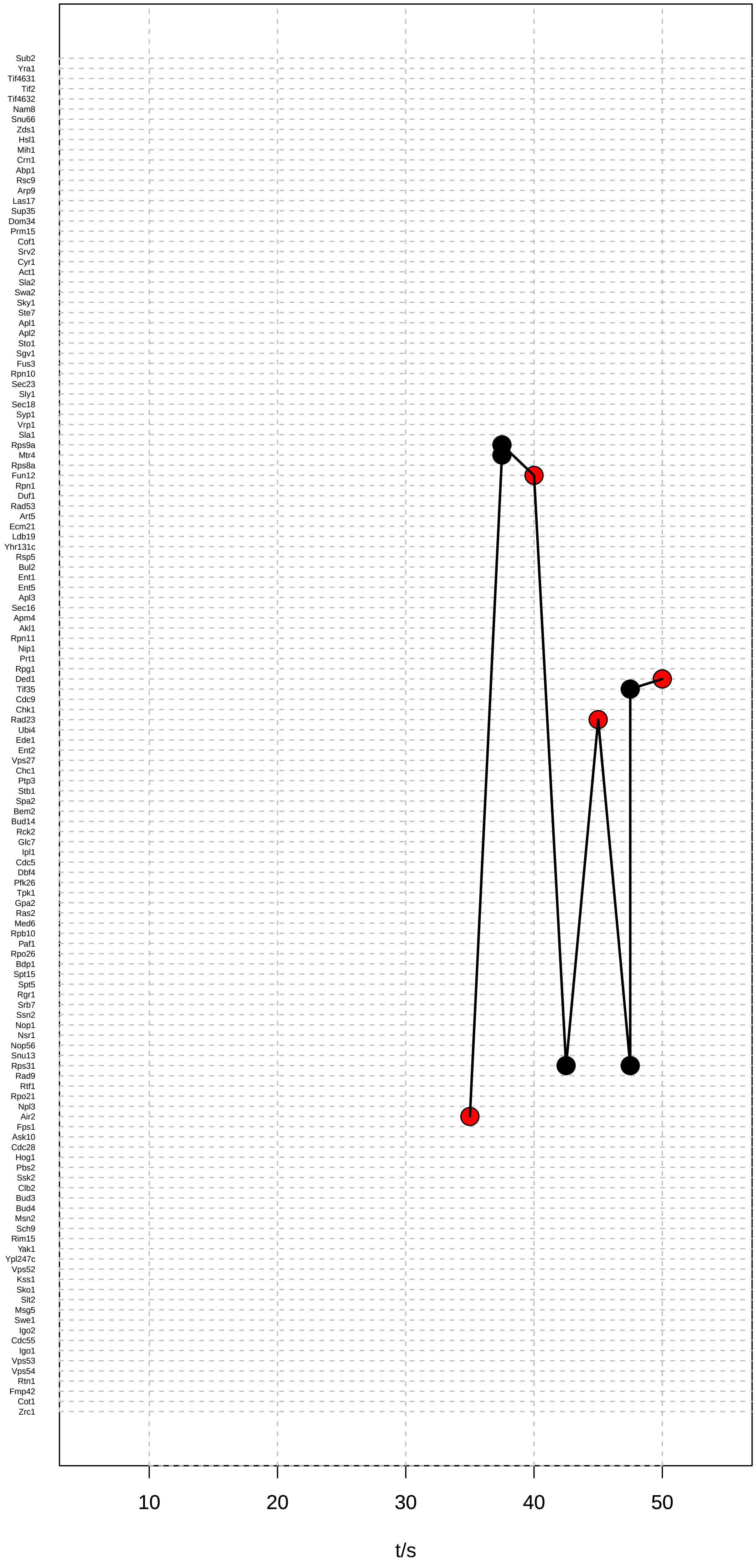

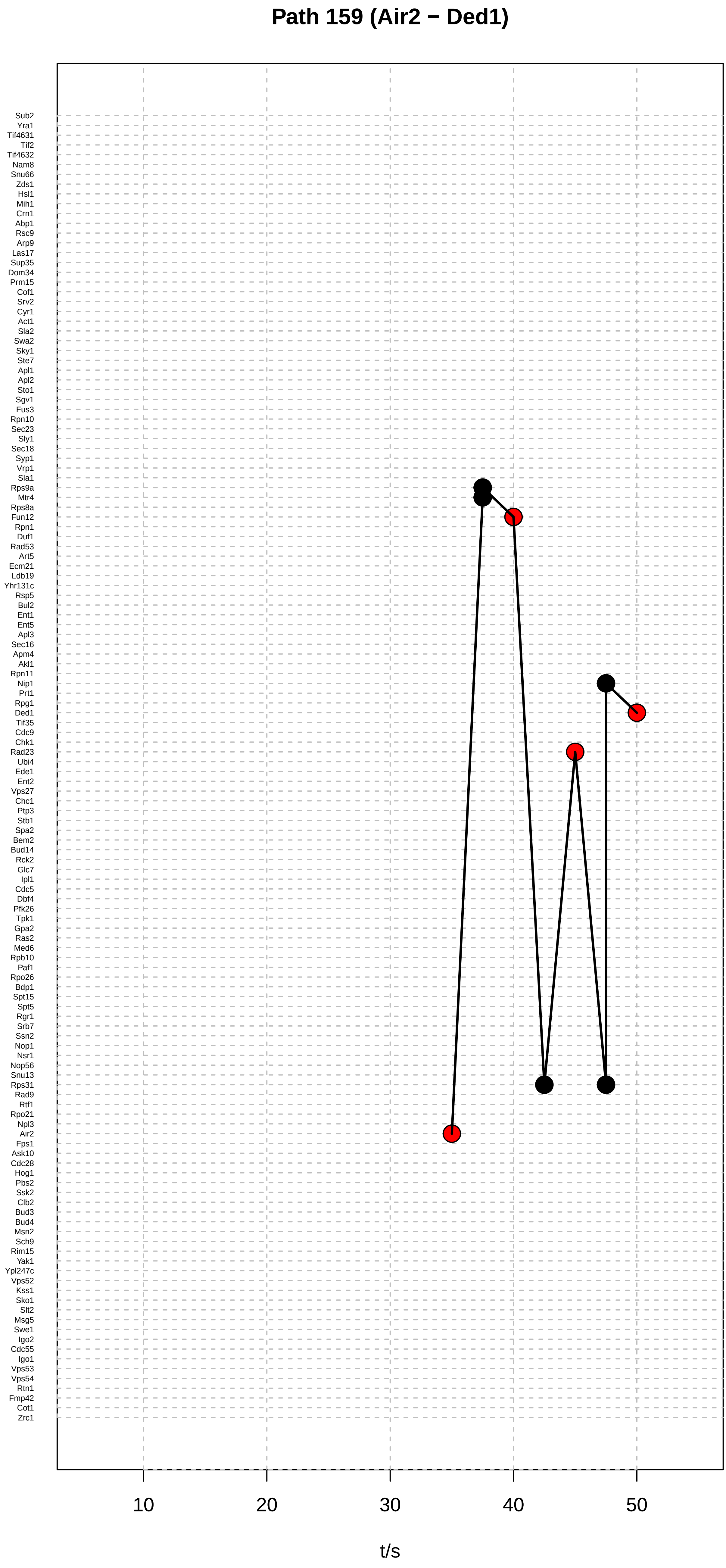

Path 159 (Air2 − Ded1)
Sub2
Yra1
Tif4631
Tif2
Tif4632
Nam8
Snu66
Zds1
Hsl1
Mih1
Crn1
Abp1
Rsc9
Arp9
Las17
Sup35
Dom34
Prm15
Cof1
Srv2
Cyr1
Act1
Sla2
Swa2
Sky1
Ste7
Apl1
Apl2
Sto1
Sgv1
Fus3
Rpn10
Sec23
Sly1
Sec18
Syp1
Vrp1
Sla1
Rps9a
Mtr4
Rps8a
Fun12
Rpn1
Duf1
Rad53
Art5
Ecm21
Ldb19
Yhr131c
Rsp5
Bul2
Ent1
Ent5
Apl3
Sec16
Apm4
Akl1
Rpn11
Nip1
Prt1
Rpg1
Ded1
Tif35
Cdc9
Chk1
Rad23
Ubi4
Ede1
Ent2
Vps27
Chc1
Ptp3
Stb1
Spa2
Bem2
Bud14
Rck2
Glc7
Ipl1
Cdc5
Dbf4
Pfk26
Tpk1
Gpa2
Ras2
Med6
Rpb10
Paf1
Rpo26
Bdp1
Spt15
Spt5
Rgr1
Srb7
Ssn2
Nop1
Nsr1
Nop56
Snu13
Rps31
Rad9
Rtf1
Rpo21
Npl3
Air2
Fps1
Ask10
Cdc28
Hog1
Pbs2
Ssk2
Clb2
Bud3
Bud4
Msn2
Sch9
Rim15
Yak1
Ypl247c
Vps52
Kss1
Sko1
Slt2
Msg5
Swe1
Igo2
Cdc55
Igo1
Vps53
Vps54
Rtn1
Fmp42
Cot1
Zrc1
10
20
30
40
50
t/s

#### Path 160 (Air2 − Ded1)

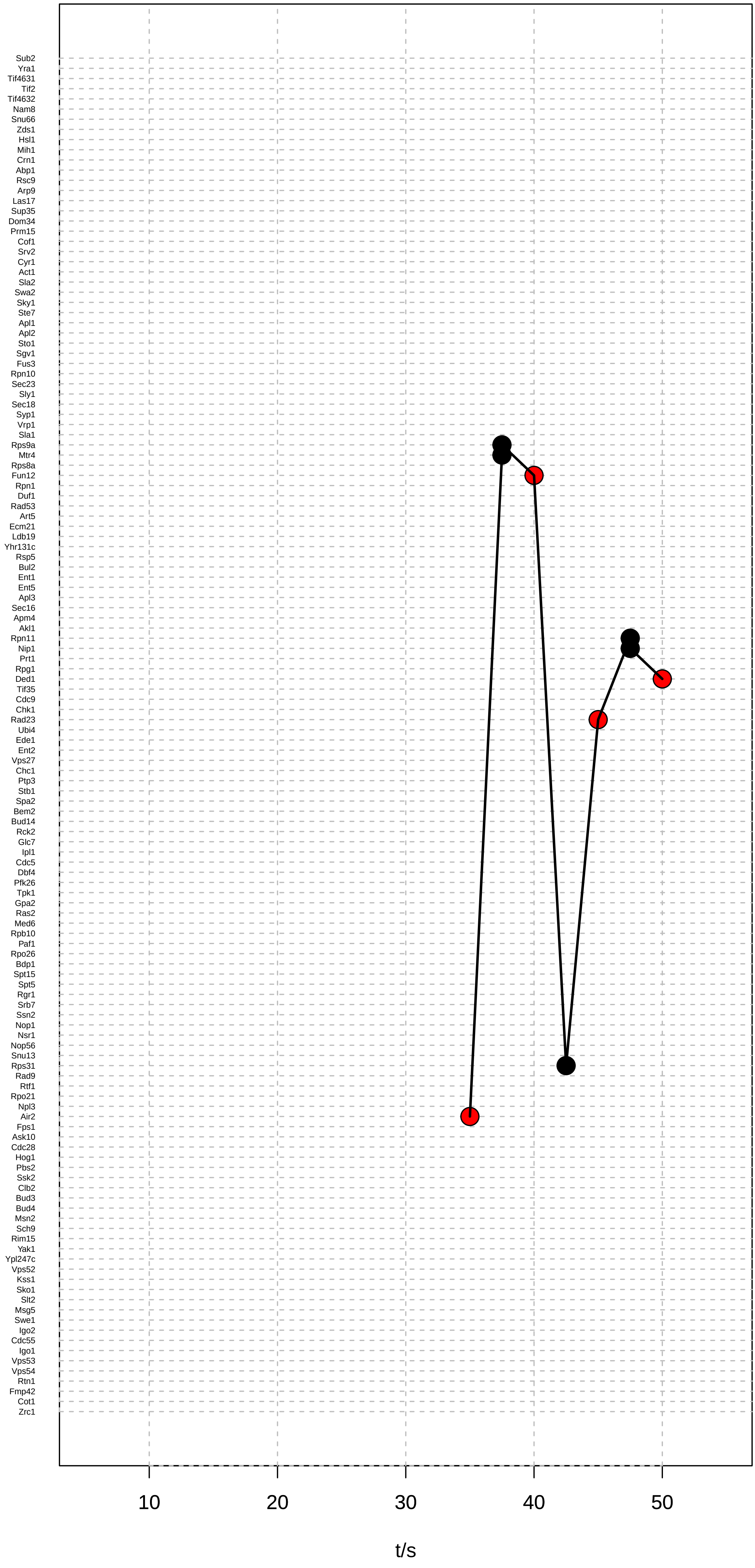

### Path 161 (Yhr131c − Ent5)

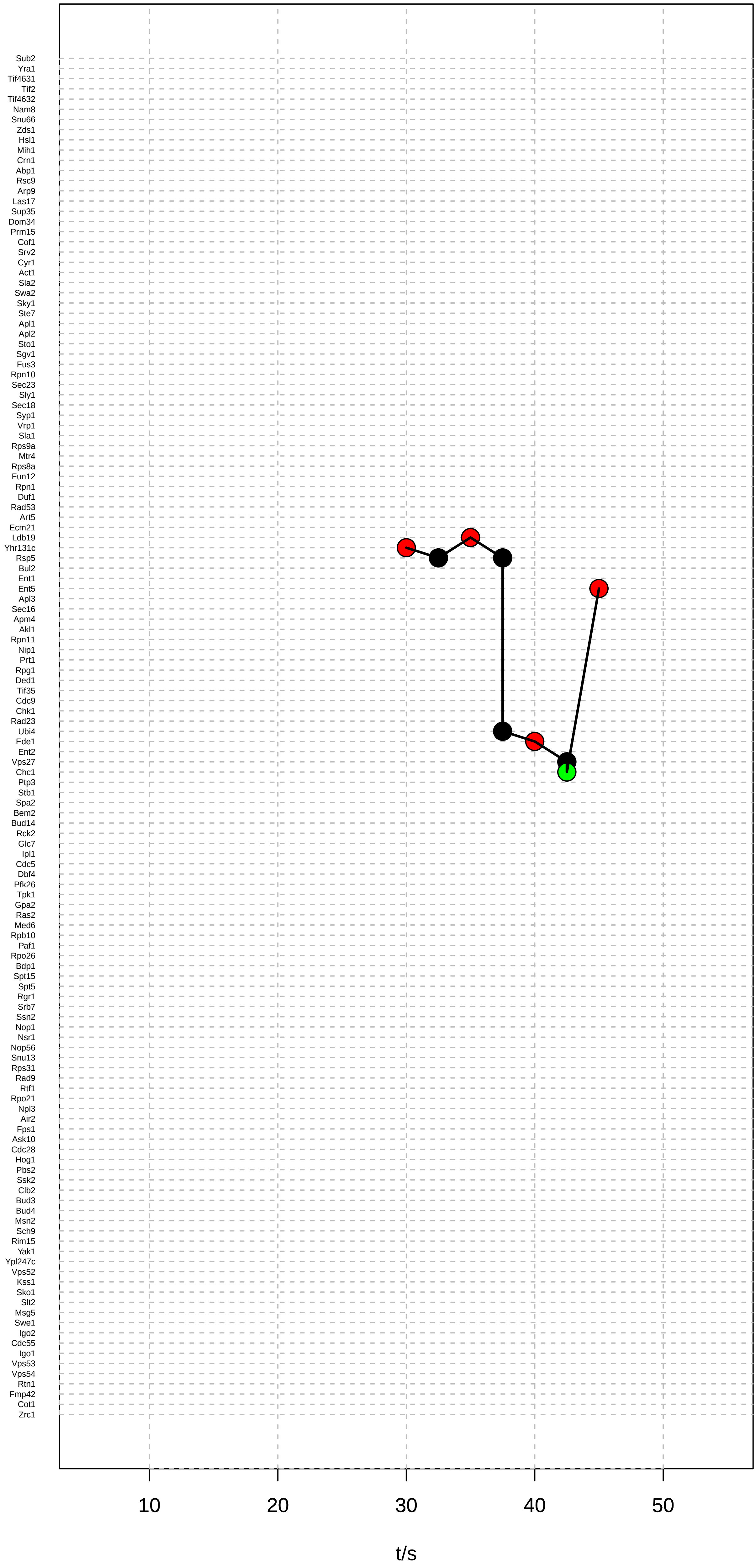

#### Path 162 (Yhr131c − Ent5)

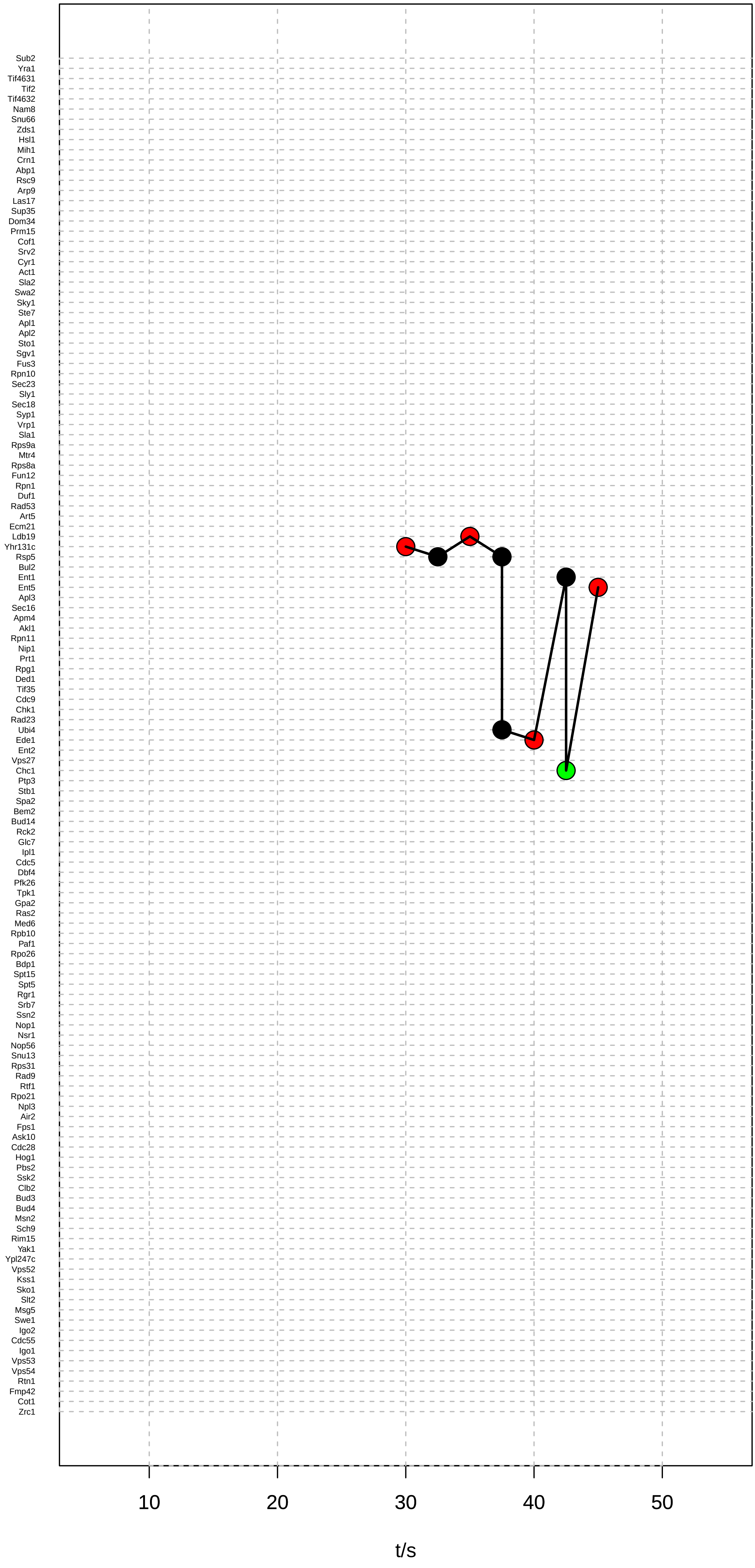

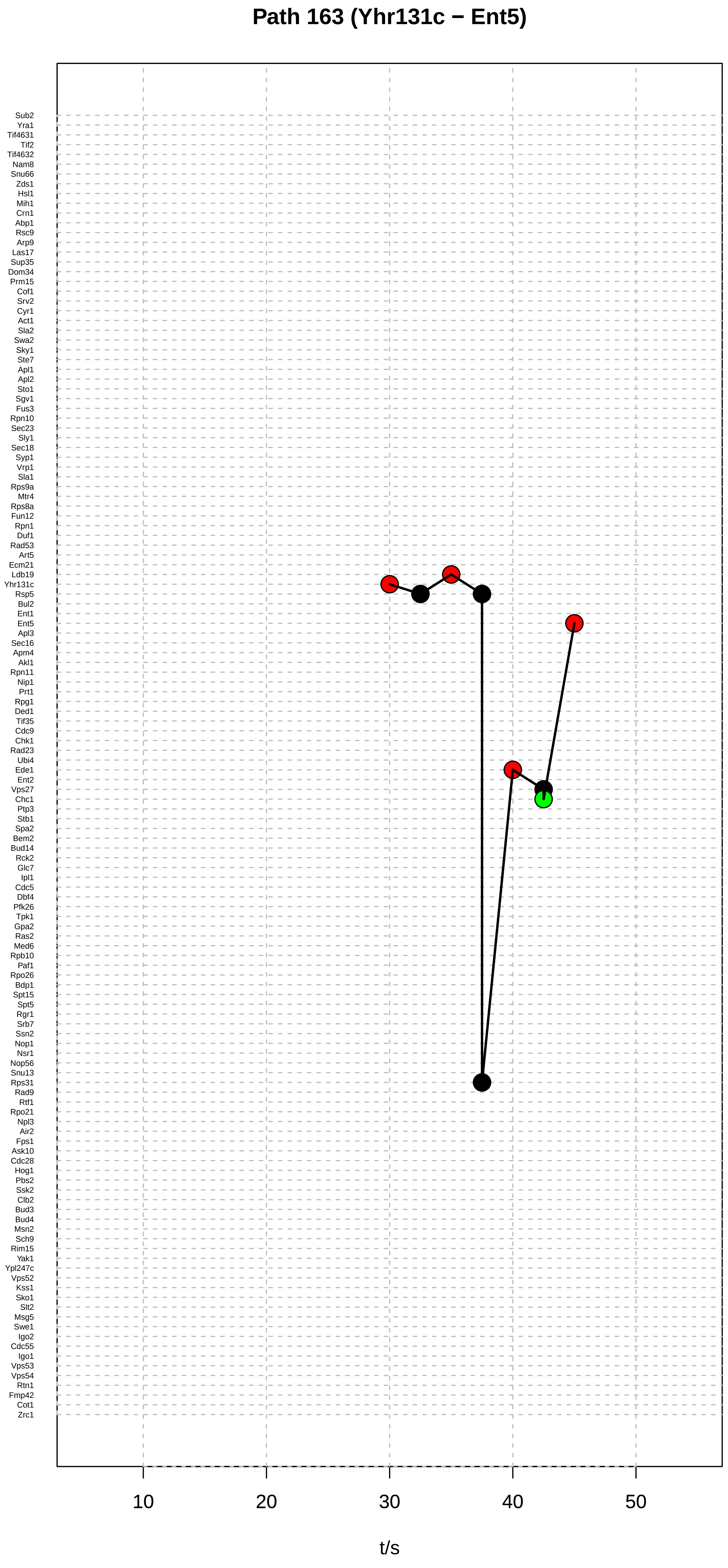

Path 163 (Yhr131c − Ent5)
Sub2
Yra1
Tif4631
Tif2
Tif4632
Nam8
Snu66
Zds1
Hsl1
Mih1
Crn1
Abp1
Rsc9
Arp9
Las17
Sup35
Dom34
Prm15
Cof1
Srv2
Cyr1
Act1
Sla2
Swa2
Sky1
Ste7
Apl1
Apl2
Sto1
Sgv1
Fus3
Rpn10
Sec23
Sly1
Sec18
Syp1
Vrp1
Sla1
Rps9a
Mtr4
Rps8a
Fun12
Rpn1
Duf1
Rad53
Art5
Ecm21
Ldb19
Yhr131c
Rsp5
Bul2
Ent1
Ent5
Apl3
Sec16
Apm4
Akl1
Rpn11
Nip1
Prt1
Rpg1
Ded1
Tif35
Cdc9
Chk1
Rad23
Ubi4
Ede1
Ent2
Vps27
Chc1
Ptp3
Stb1
Spa2
Bem2
Bud14
Rck2
Glc7
Ipl1
Cdc5
Dbf4
Pfk26
Tpk1
Gpa2
Ras2
Med6
Rpb10
Paf1
Rpo26
Bdp1
Spt15
Spt5
Rgr1
Srb7
Ssn2
Nop1
Nsr1
Nop56
Snu13
Rps31
Rad9
Rtf1
Rpo21
Npl3
Air2
Fps1
Ask10
Cdc28
Hog1
Pbs2
Ssk2
Clb2
Bud3
Bud4
Msn2
Sch9
Rim15
Yak1
Ypl247c
Vps52
Kss1
Sko1
Slt2
Msg5
Swe1
Igo2
Cdc55
Igo1
Vps53
Vps54
Rtn1
Fmp42
Cot1
Zrc1
10
20
30
40
50
t/s

### Path 164 (Yhr131c − Ent5)

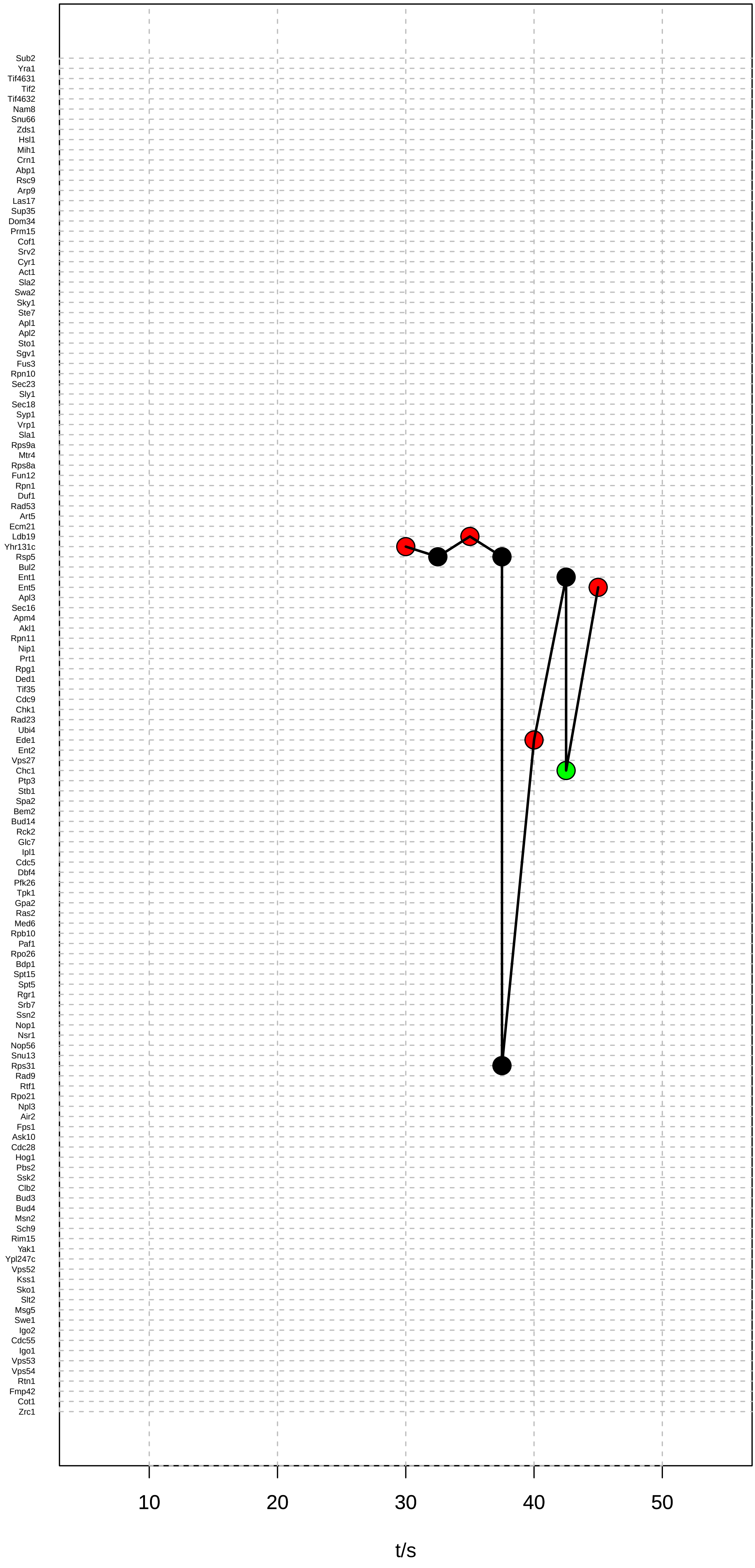

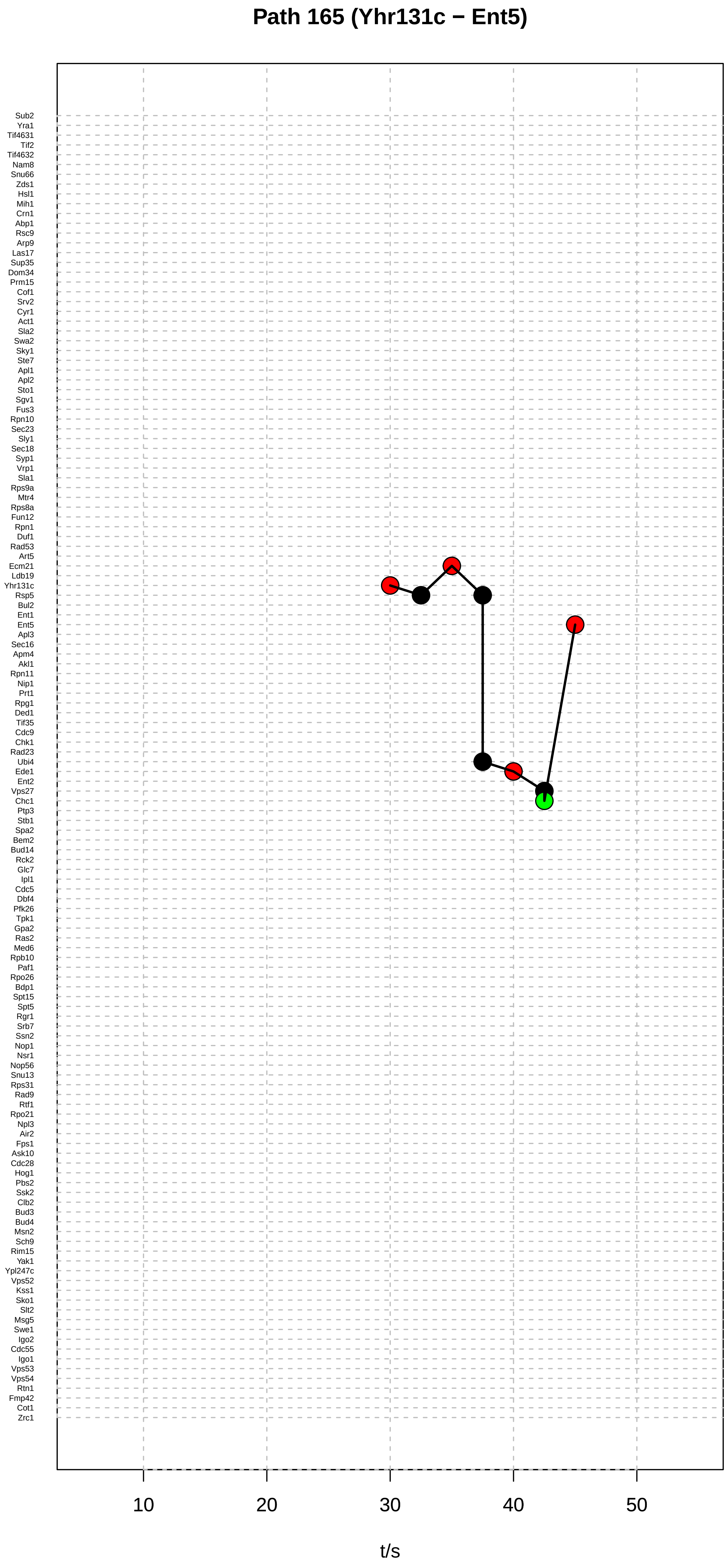
Path 165 (Yhr131c − Ent5)
Sub2
Yra1
Tif4631
Tif2
Tif4632
Nam8
Snu66
Zds1
Hsl1
Mih1
Crn1
Abp1
Rsc9
Arp9
Las17
Sup35
Dom34
Prm15
Cof1
Srv2
Cyr1
Act1
Sla2
Swa2
Sky1
Ste7
Apl1
Apl2
Sto1
Sgv1
Fus3
Rpn10
Sec23
Sly1
Sec18
Syp1
Vrp1
Sla1
Rps9a
Mtr4
Rps8a
Fun12
Rpn1
Duf1
Rad53
Art5
Ecm21
Ldb19
Yhr131c
Rsp5
Bul2
Ent1
Ent5
Apl3
Sec16
Apm4
Akl1
Rpn11
Nip1
Prt1
Rpg1
Ded1
Tif35
Cdc9
Chk1
Rad23
Ubi4
Ede1
Ent2
Vps27
Chc1
Ptp3
Stb1
Spa2
Bem2
Bud14
Rck2
Glc7
Ipl1
Cdc5
Dbf4
Pfk26
Tpk1
Gpa2
Ras2
Med6
Rpb10
Paf1
Rpo26
Bdp1
Spt15
Spt5
Rgr1
Srb7
Ssn2
Nop1
Nsr1
Nop56
Snu13
Rps31
Rad9
Rtf1
Rpo21
Npl3
Air2
Fps1
Ask10
Cdc28
Hog1
Pbs2
Ssk2
Clb2
Bud3
Bud4
Msn2
Sch9
Rim15
Yak1
Ypl247c
Vps52
Kss1
Sko1
Slt2
Msg5
Swe1
Igo2
Cdc55
Igo1
Vps53
Vps54
Rtn1
Fmp42
Cot1
Zrc1
10
20
30
40
50
t/s

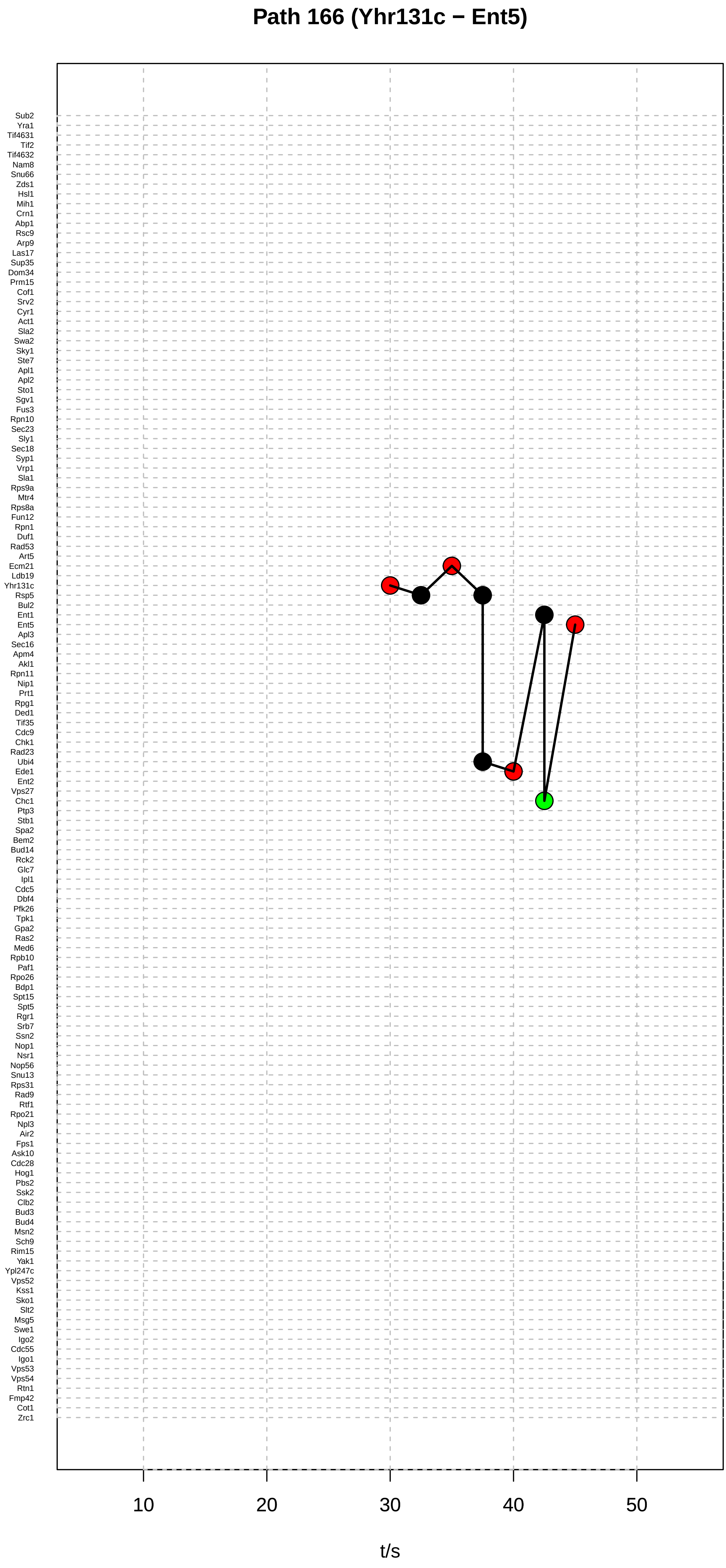
Path 166 (Yhr131c − Ent5)
Sub2
Yra1
Tif4631
Tif2
Tif4632
Nam8
Snu66
Zds1
Hsl1
Mih1
Crn1
Abp1
Rsc9
Arp9
Las17
Sup35
Dom34
Prm15
Cof1
Srv2
Cyr1
Act1
Sla2
Swa2
Sky1
Ste7
Apl1
Apl2
Sto1
Sgv1
Fus3
Rpn10
Sec23
Sly1
Sec18
Syp1
Vrp1
Sla1
Rps9a
Mtr4
Rps8a
Fun12
Rpn1
Duf1
Rad53
Art5
Ecm21
Ldb19
Yhr131c
Rsp5
Bul2
Ent1
Ent5
Apl3
Sec16
Apm4
Akl1
Rpn11
Nip1
Prt1
Rpg1
Ded1
Tif35
Cdc9
Chk1
Rad23
Ubi4
Ede1
Ent2
Vps27
Chc1
Ptp3
Stb1
Spa2
Bem2
Bud14
Rck2
Glc7
Ipl1
Cdc5
Dbf4
Pfk26
Tpk1
Gpa2
Ras2
Med6
Rpb10
Paf1
Rpo26
Bdp1
Spt15
Spt5
Rgr1
Srb7
Ssn2
Nop1
Nsr1
Nop56
Snu13
Rps31
Rad9
Rtf1
Rpo21
Npl3
Air2
Fps1
Ask10
Cdc28
Hog1
Pbs2
Ssk2
Clb2
Bud3
Bud4
Msn2
Sch9
Rim15
Yak1
Ypl247c
Vps52
Kss1
Sko1
Slt2
Msg5
Swe1
Igo2
Cdc55
Igo1
Vps53
Vps54
Rtn1
Fmp42
Cot1
Zrc1
10
20
30
40
50
t/s

### Path 167 (Yhr131c − Ent5)

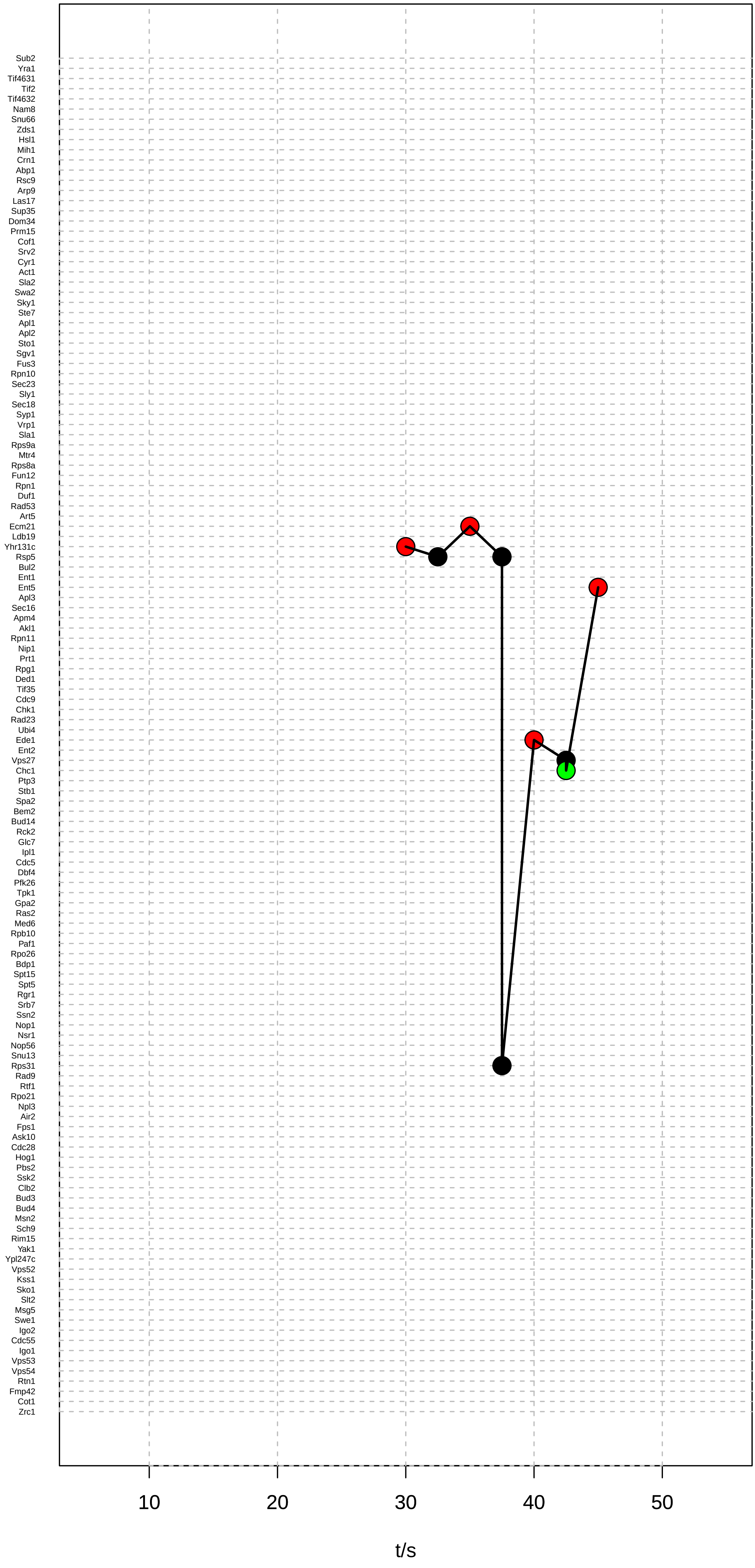

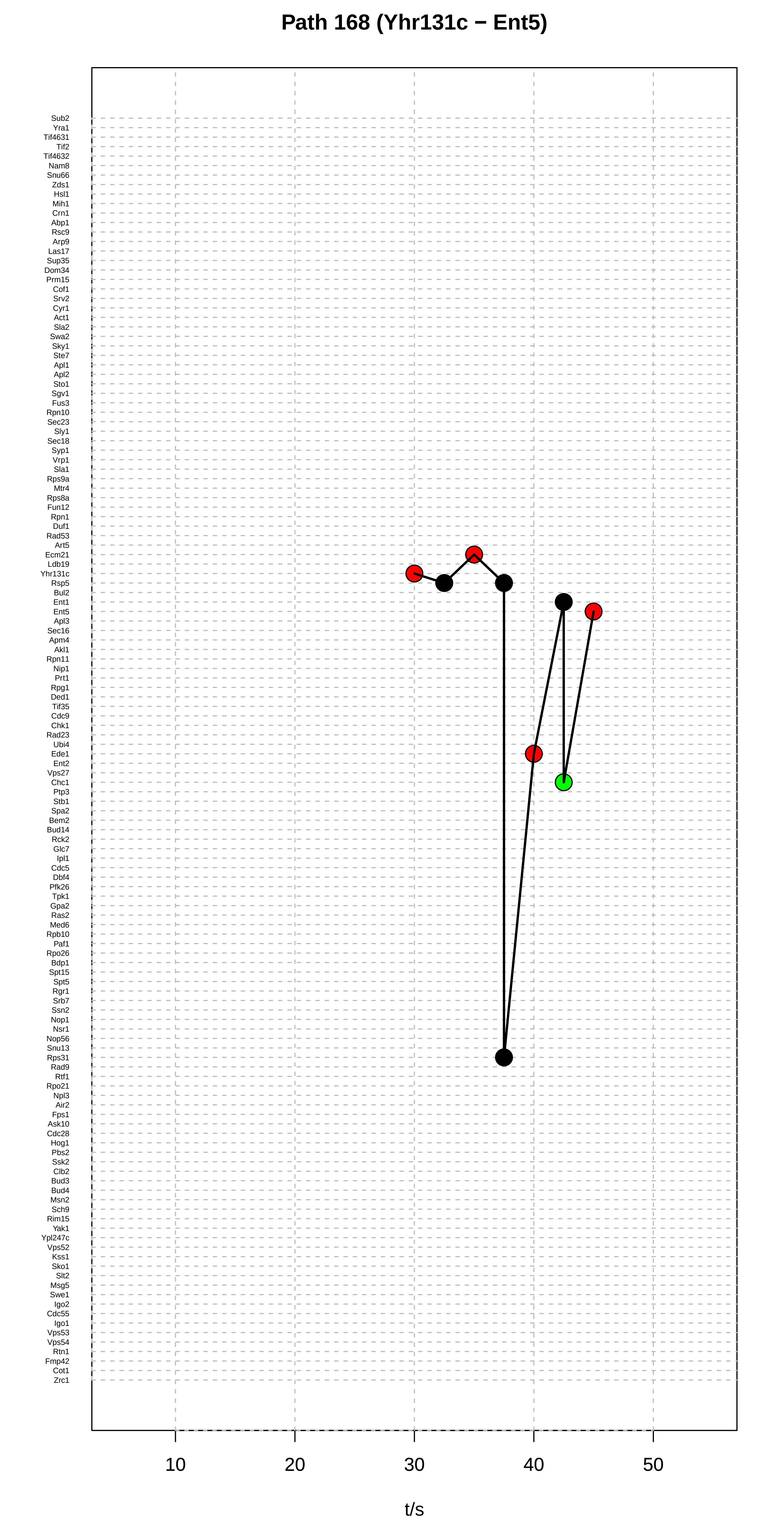

Path 168 (Yhr131c − Ent5)
Sub2
Yra1
Tif4631
Tif2
Tif4632
Nam8
Snu66
Zds1
Hsl1
Mih1
Crn1
Abp1
Rsc9
Arp9
Las17
Sup35
Dom34
Prm15
Cof1
Srv2
Cyr1
Act1
Sla2
Swa2
Sky1
Ste7
Apl1
Apl2
Sto1
Sgv1
Fus3
Rpn10
Sec23
Sly1
Sec18
Syp1
Vrp1
Sla1
Rps9a
Mtr4
Rps8a
Fun12
Rpn1
Duf1
Rad53
Art5
Ecm21
Ldb19
Yhr131c
Rsp5
Bul2
Ent1
Ent5
Apl3
Sec16
Apm4
Akl1
Rpn11
Nip1
Prt1
Rpg1
Ded1
Tif35
Cdc9
Chk1
Rad23
Ubi4
Ede1
Ent2
Vps27
Chc1
Ptp3
Stb1
Spa2
Bem2
Bud14
Rck2
Glc7
Ipl1
Cdc5
Dbf4
Pfk26
Tpk1
Gpa2
Ras2
Med6
Rpb10
Paf1
Rpo26
Bdp1
Spt15
Spt5
Rgr1
Srb7
Ssn2
Nop1
Nsr1
Nop56
Snu13
Rps31
Rad9
Rtf1
Rpo21
Npl3
Air2
Fps1
Ask10
Cdc28
Hog1
Pbs2
Ssk2
Clb2
Bud3
Bud4
Msn2
Sch9
Rim15
Yak1
Ypl247c
Vps52
Kss1
Sko1
Slt2
Msg5
Swe1
Igo2
Cdc55
Igo1
Vps53
Vps54
Rtn1
Fmp42
Cot1
Zrc1
10
20
30
40
50
t/s

### Path 169 (Yhr131c − Ent5)

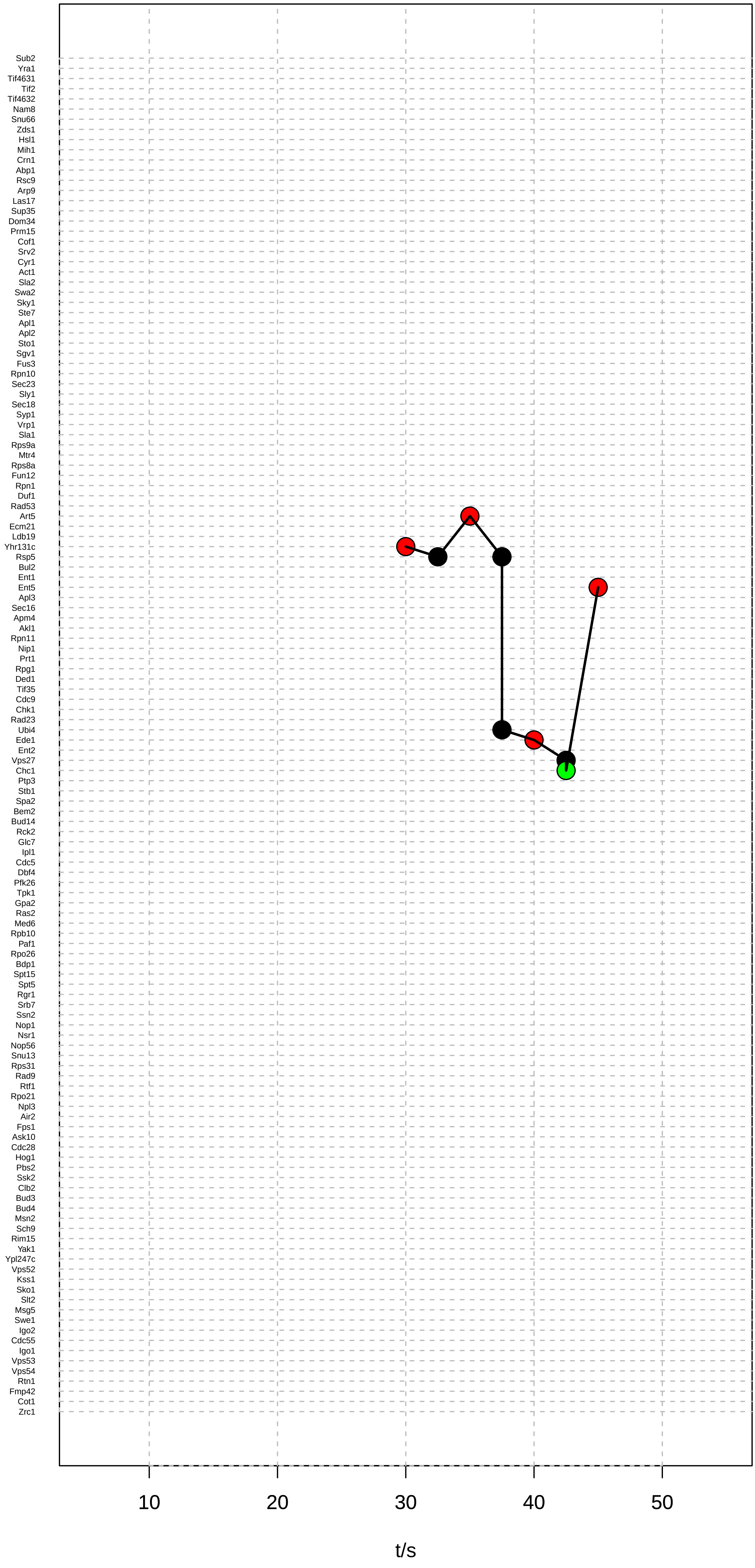

#### Path 170 (Yhr131c − Ent5)

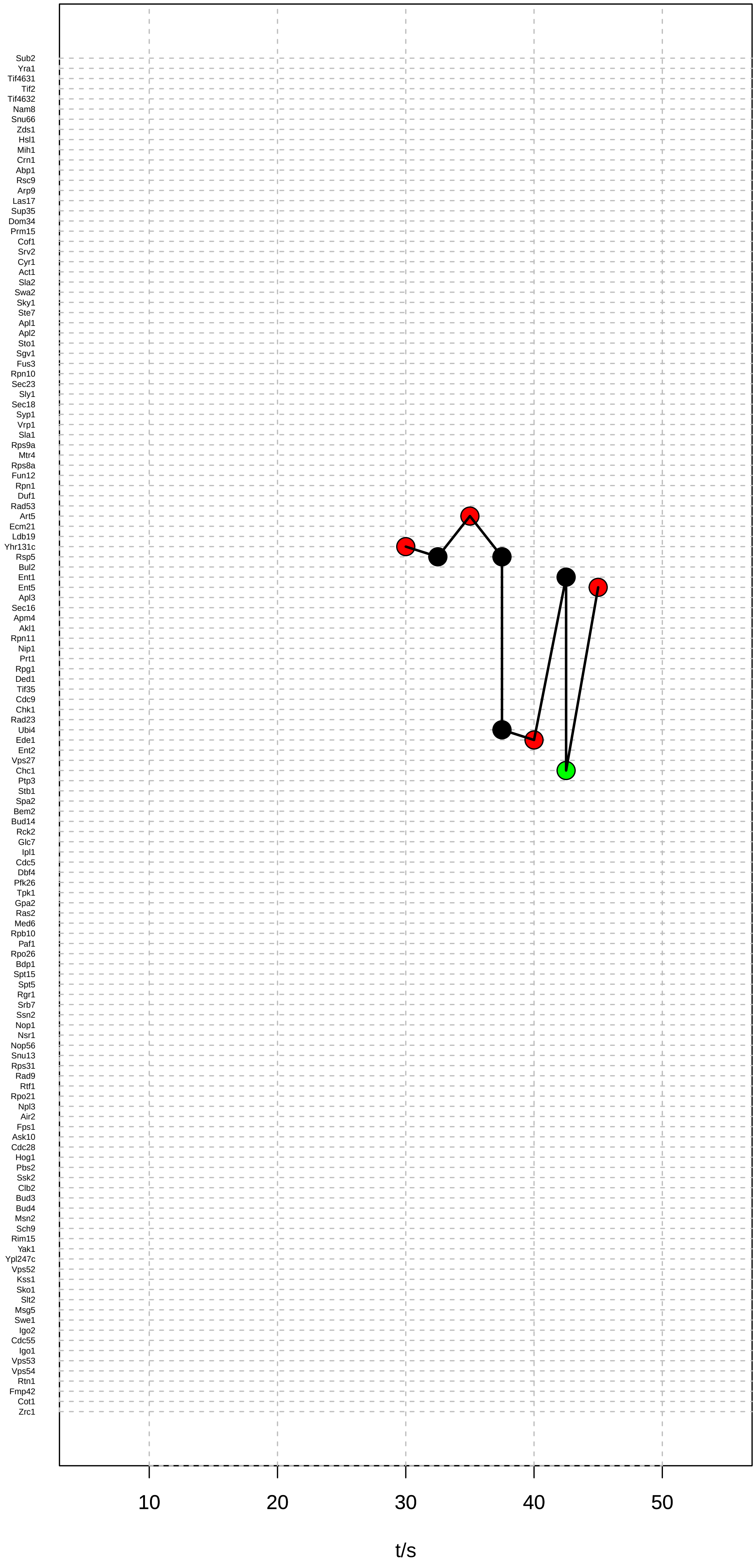

### Path 171 (Yhr131c – Ent5)

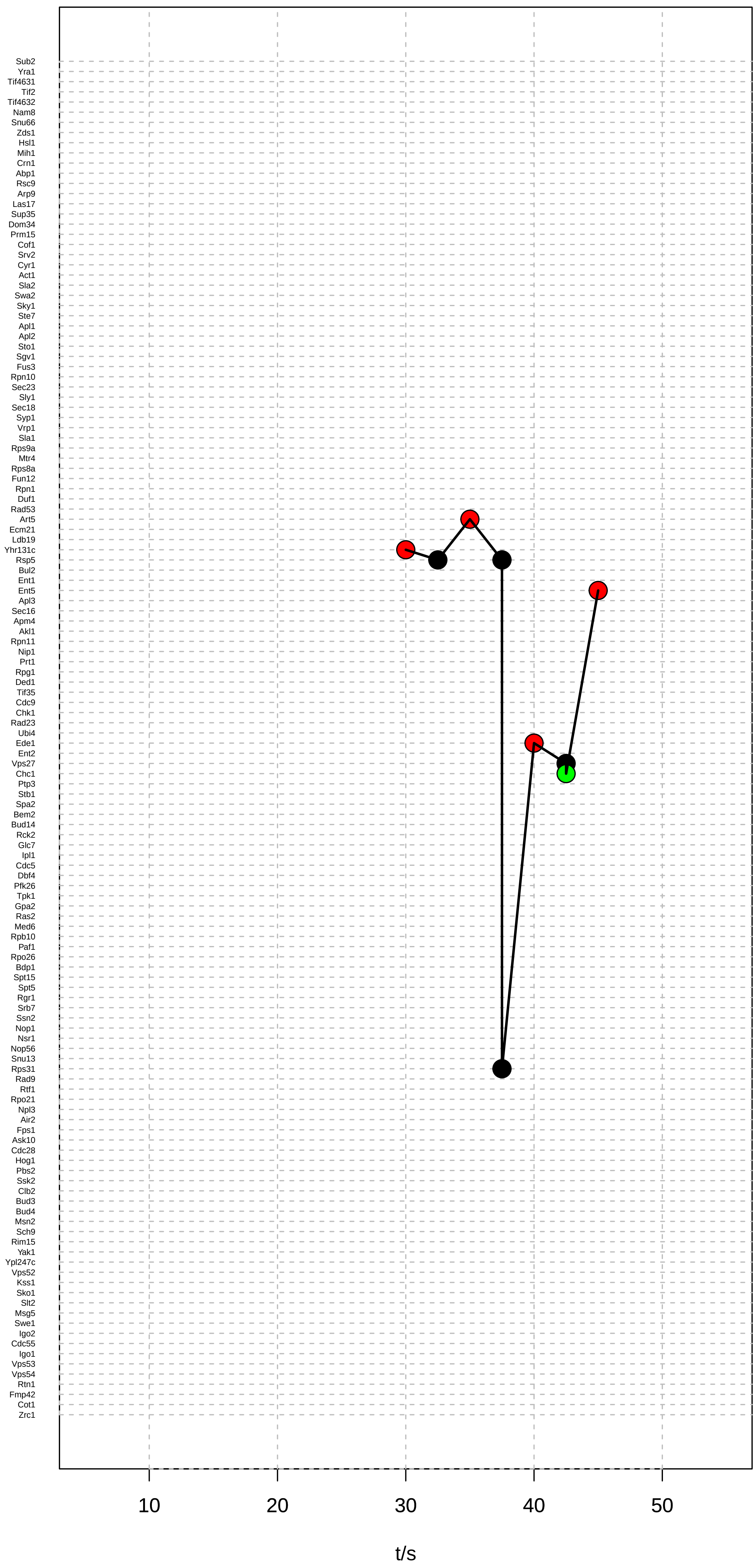

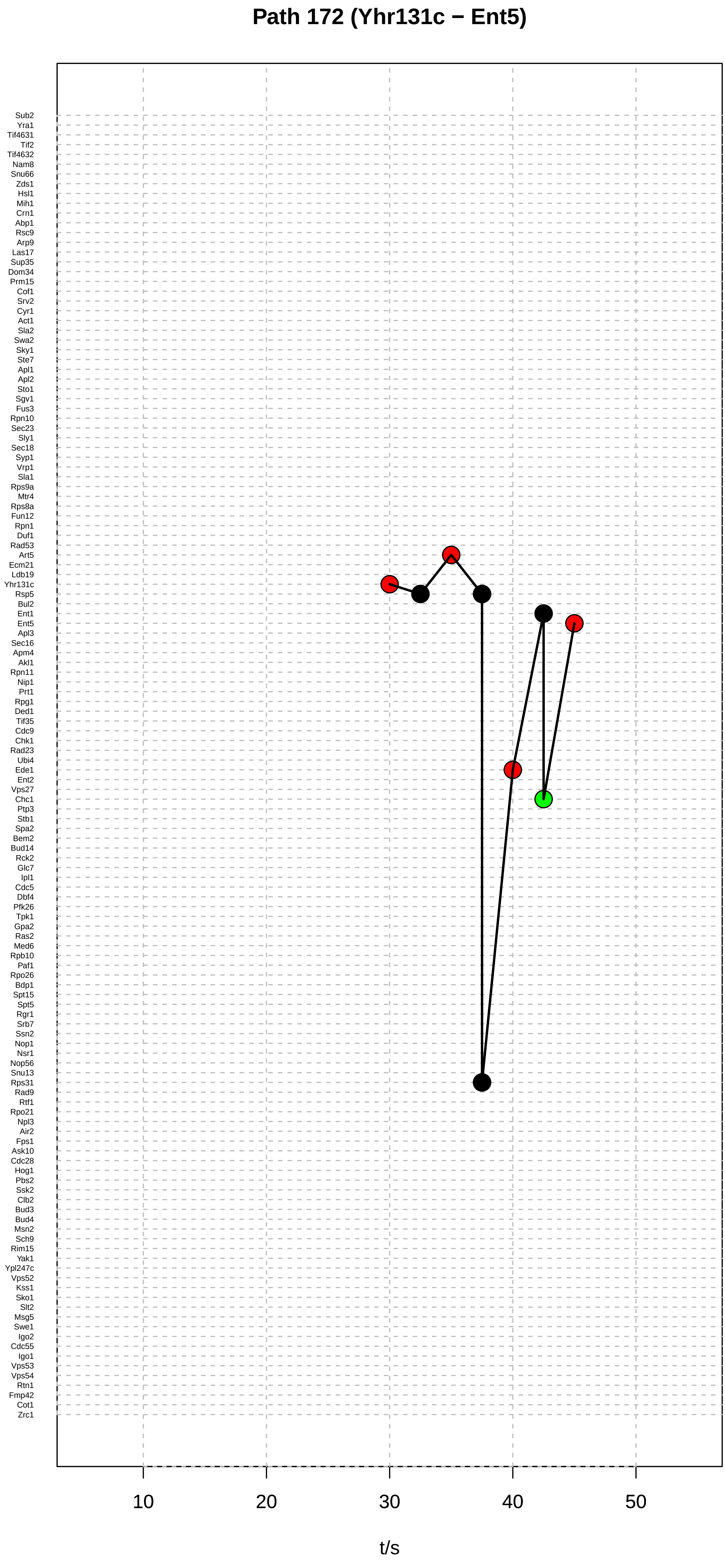

Path 172 (Yhr131c − Ent5)
Sub2
Yra1
Tif4631
Tif2
Tif4632
Nam8
Snu66
Zds1
Hsl1
Mih1
Crn1
Abp1
Rsc9
Arp9
Las17
Sup35
Dom34
Prm15
Cof1
Srv2
Cyr1
Act1
Sla2
Swa2
Sky1
Ste7
Apl1
Apl2
Sto1
Sgv1
Fus3
Rpn10
Sec23
Sly1
Sec18
Syp1
Vrp1
Sla1
Rps9a
Mtr4
Rps8a
Fun12
Rpn1
Duf1
Rad53
Art5
Ecm21
Ldb19
Yhr131c
Rsp5
Bul2
Ent1
Ent5
Apl3
Sec16
Apm4
Akl1
Rpn11
Nip1
Prt1
Rpg1
Ded1
Tif35
Cdc9
Chk1
Rad23
Ubi4
Ede1
Ent2
Vps27
Chc1
Ptp3
Stb1
Spa2
Bem2
Bud14
Rck2
Glc7
Ipl1
Cdc5
Dbf4
Pfk26
Tpk1
Gpa2
Ras2
Med6
Rpb10
Paf1
Rpo26
Bdp1
Spt15
Spt5
Rgr1
Srb7
Ssn2
Nop1
Nsr1
Nop56
Snu13
Rps31
Rad9
Rtf1
Rpo21
Npl3
Air2
Fps1
Ask10
Cdc28
Hog1
Pbs2
Ssk2
Clb2
Bud3
Bud4
Msn2
Sch9
Rim15
Yak1
Ypl247c
Vps52
Kss1
Sko1
Slt2
Msg5
Swe1
Igo2
Cdc55
Igo1
Vps53
Vps54
Rtn1
Fmp42
Cot1
Zrc1
10
20
30
40
50
t/s

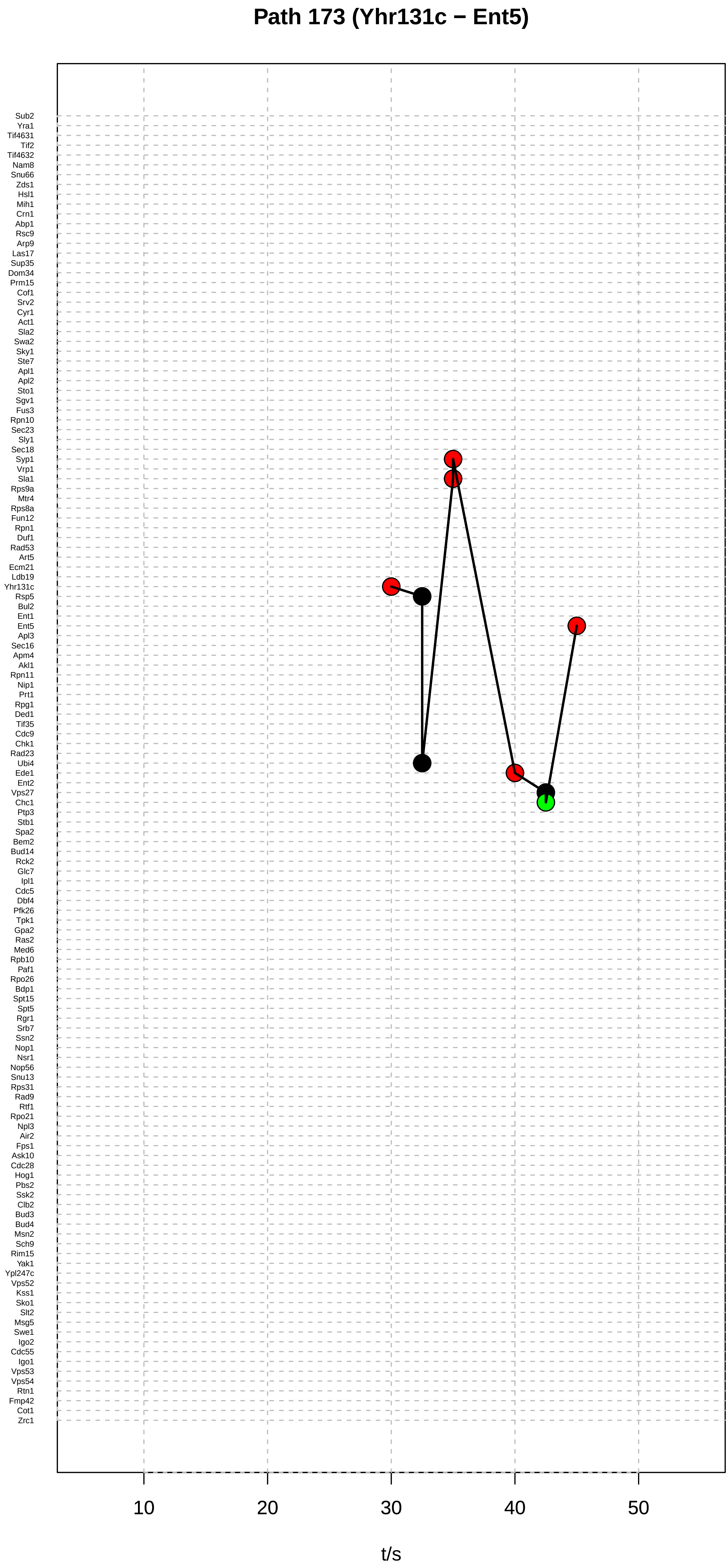

Path 173 (Yhr131c − Ent5)
Sub2
Yra1
Tif4631
Tif2
Tif4632
Nam8
Snu66
Zds1
Hsl1
Mih1
Crn1
Abp1
Rsc9
Arp9
Las17
Sup35
Dom34
Prm15
Cof1
Srv2
Cyr1
Act1
Sla2
Swa2
Sky1
Ste7
Apl1
Apl2
Sto1
Sgv1
Fus3
Rpn10
Sec23
Sly1
Sec18
Syp1
Vrp1
Sla1
Rps9a
Mtr4
Rps8a
Fun12
Rpn1
Duf1
Rad53
Art5
Ecm21
Ldb19
Yhr131c
Rsp5
Bul2
Ent1
Ent5
Apl3
Sec16
Apm4
Akl1
Rpn11
Nip1
Prt1
Rpg1
Ded1
Tif35
Cdc9
Chk1
Rad23
Ubi4
Ede1
Ent2
Vps27
Chc1
Ptp3
Stb1
Spa2
Bem2
Bud14
Rck2
Glc7
Ipl1
Cdc5
Dbf4
Pfk26
Tpk1
Gpa2
Ras2
Med6
Rpb10
Paf1
Rpo26
Bdp1
Spt15
Spt5
Rgr1
Srb7
Ssn2
Nop1
Nsr1
Nop56
Snu13
Rps31
Rad9
Rtf1
Rpo21
Npl3
Air2
Fps1
Ask10
Cdc28
Hog1
Pbs2
Ssk2
Clb2
Bud3
Bud4
Msn2
Sch9
Rim15
Yak1
Ypl247c
Vps52
Kss1
Sko1
Slt2
Msg5
Swe1
Igo2
Cdc55
Igo1
Vps53
Vps54
Rtn1
Fmp42
Cot1
Zrc1
10
20
30
40
50
t/s

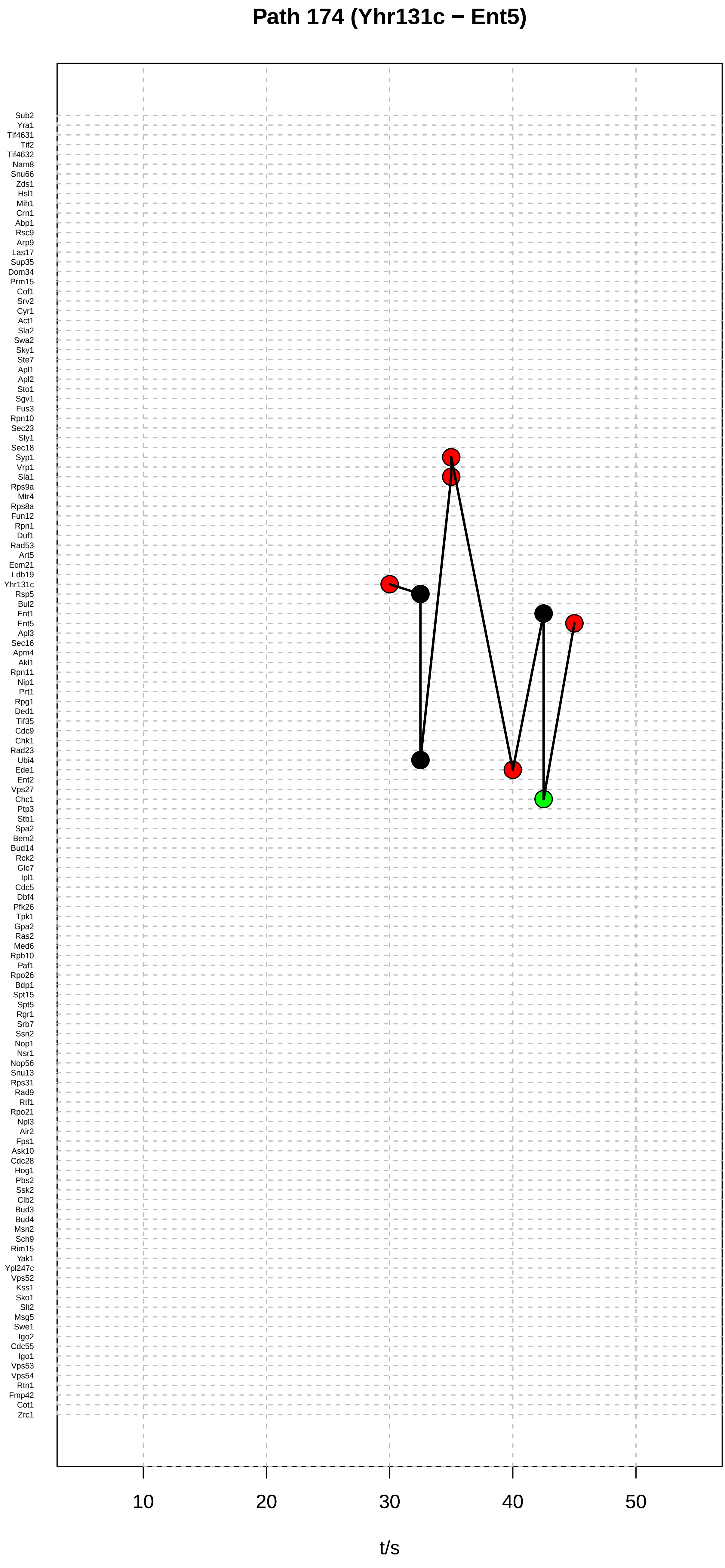
Path 174 (Yhr131c − Ent5)
Sub2
Yra1
Tif4631
Tif2
Tif4632
Nam8
Snu66
Zds1
Hsl1
Mih1
Crn1
Abp1
Rsc9
Arp9
Las17
Sup35
Dom34
Prm15
Cof1
Srv2
Cyr1
Act1
Sla2
Swa2
Sky1
Ste7
Apl1
Apl2
Sto1
Sgv1
Fus3
Rpn10
Sec23
Sly1
Sec18
Syp1
Vrp1
Sla1
Rps9a
Mtr4
Rps8a
Fun12
Rpn1
Duf1
Rad53
Art5
Ecm21
Ldb19
Yhr131c
Rsp5
Bul2
Ent1
Ent5
Apl3
Sec16
Apm4
Akl1
Rpn11
Nip1
Prt1
Rpg1
Ded1
Tif35
Cdc9
Chk1
Rad23
Ubi4
Ede1
Ent2
Vps27
Chc1
Ptp3
Stb1
Spa2
Bem2
Bud14
Rck2
Glc7
Ipl1
Cdc5
Dbf4
Pfk26
Tpk1
Gpa2
Ras2
Med6
Rpb10
Paf1
Rpo26
Bdp1
Spt15
Spt5
Rgr1
Srb7
Ssn2
Nop1
Nsr1
Nop56
Snu13
Rps31
Rad9
Rtf1
Rpo21
Npl3
Air2
Fps1
Ask10
Cdc28
Hog1
Pbs2
Ssk2
Clb2
Bud3
Bud4
Msn2
Sch9
Rim15
Yak1
Ypl247c
Vps52
Kss1
Sko1
Slt2
Msg5
Swe1
Igo2
Cdc55
Igo1
Vps53
Vps54
Rtn1
Fmp42
Cot1
Zrc1
10
20
30
40
50
t/s

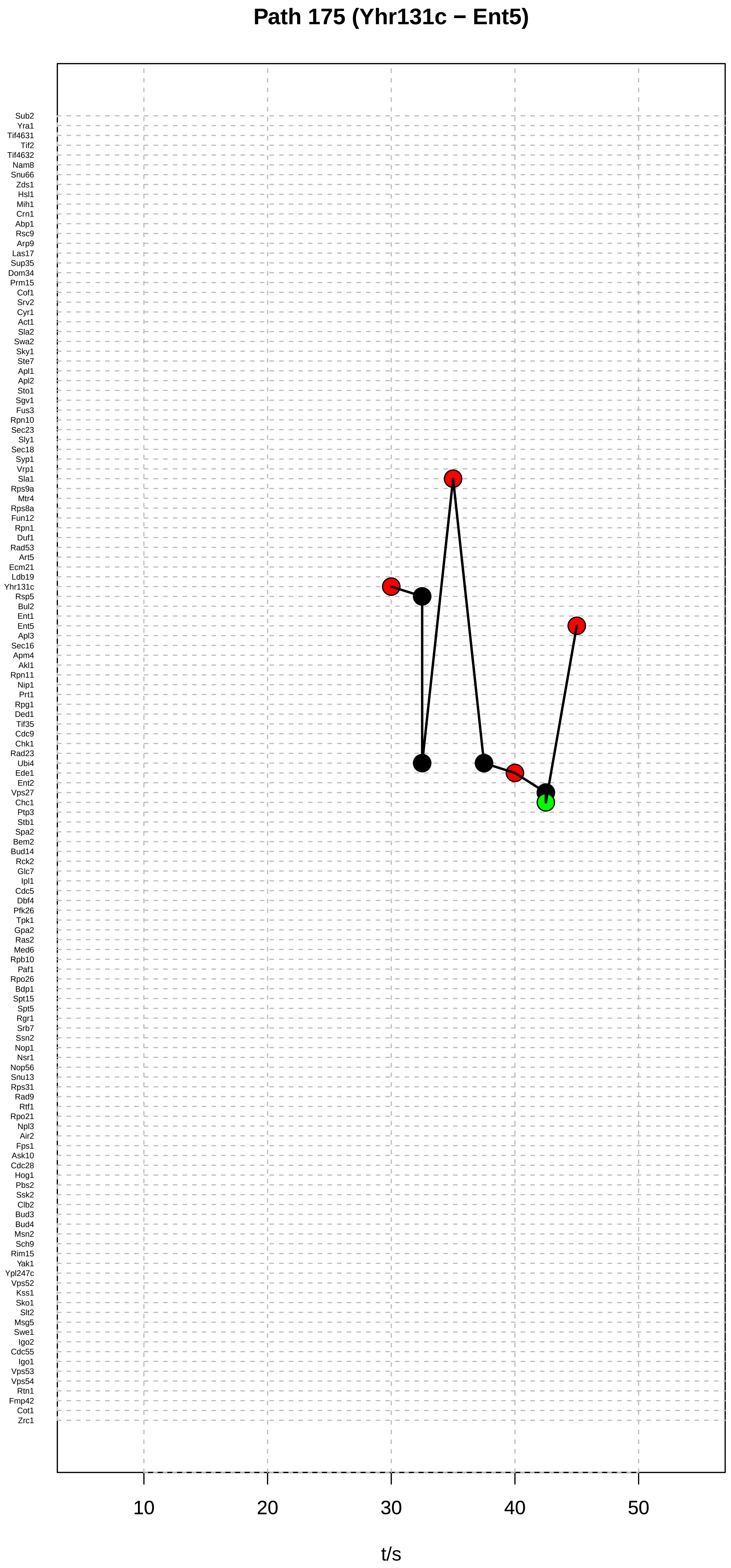
Path 175 (Yhr131c − Ent5)
Sub2
Yra1
Tif4631
Tif2
Tif4632
Nam8
Snu66
Zds1
Hsl1
Mih1
Crn1
Abp1
Rsc9
Arp9
Las17
Sup35
Dom34
Prm15
Cof1
Srv2
Cyr1
Act1
Sla2
Swa2
Sky1
Ste7
Apl1
Apl2
Sto1
Sgv1
Fus3
Rpn10
Sec23
Sly1
Sec18
Syp1
Vrp1
Sla1
Rps9a
Mtr4
Rps8a
Fun12
Rpn1
Duf1
Rad53
Art5
Ecm21
Ldb19
Yhr131c
Rsp5
Bul2
Ent1
Ent5
Apl3
Sec16
Apm4
Akl1
Rpn11
Nip1
Prt1
Rpg1
Ded1
Tif35
Cdc9
Chk1
Rad23
Ubi4
Ede1
Ent2
Vps27
Chc1
Ptp3
Stb1
Spa2
Bem2
Bud14
Rck2
Glc7
Ipl1
Cdc5
Dbf4
Pfk26
Tpk1
Gpa2
Ras2
Med6
Rpb10
Paf1
Rpo26
Bdp1
Spt15
Spt5
Rgr1
Srb7
Ssn2
Nop1
Nsr1
Nop56
Snu13
Rps31
Rad9
Rtf1
Rpo21
Npl3
Air2
Fps1
Ask10
Cdc28
Hog1
Pbs2
Ssk2
Clb2
Bud3
Bud4
Msn2
Sch9
Rim15
Yak1
Ypl247c
Vps52
Kss1
Sko1
Slt2
Msg5
Swe1
Igo2
Cdc55
Igo1
Vps53
Vps54
Rtn1
Fmp42
Cot1
Zrc1
10
20
30
40
50
t/s

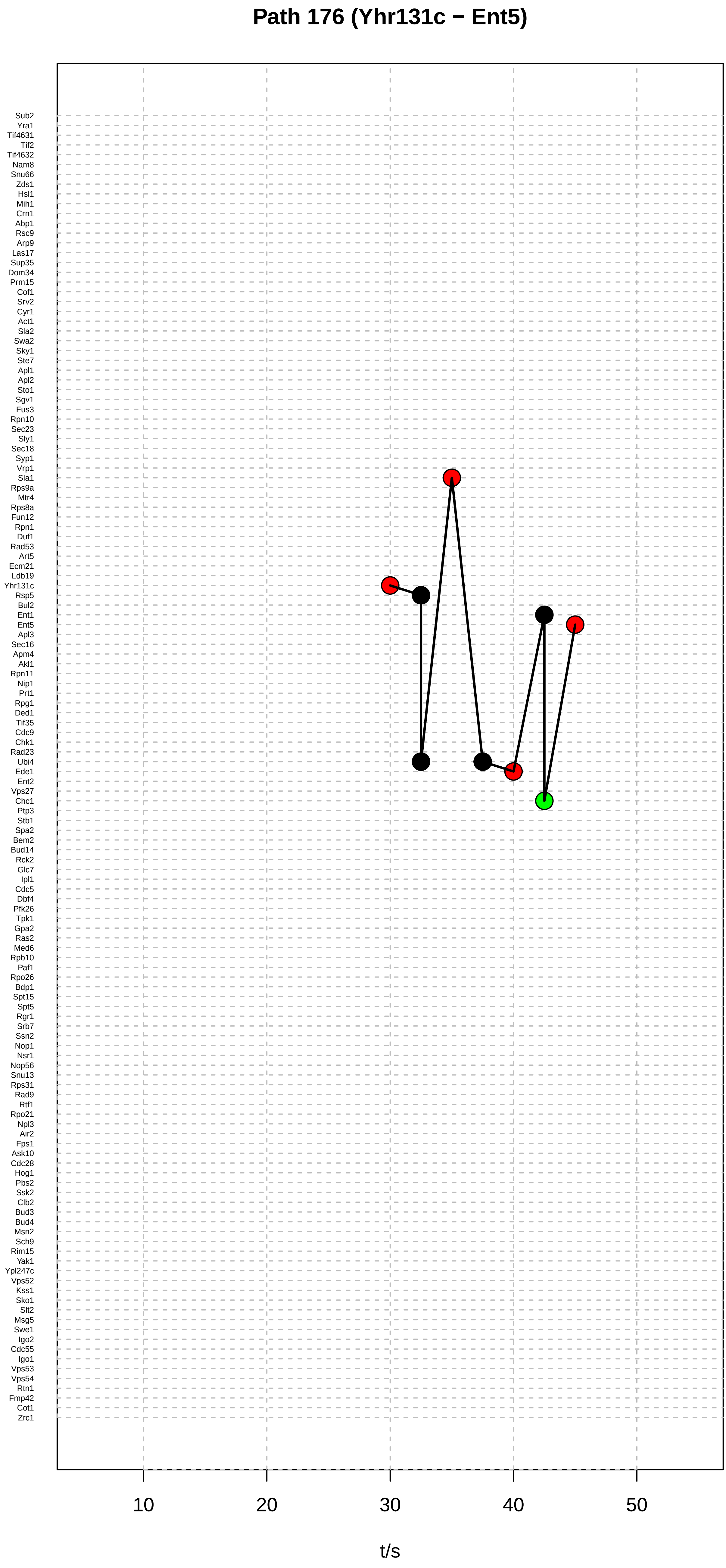

Path 176 (Yhr131c − Ent5)
Sub2
Yra1
Tif4631
Tif2
Tif4632
Nam8
Snu66
Zds1
Hsl1
Mih1
Crn1
Abp1
Rsc9
Arp9
Las17
Sup35
Dom34
Prm15
Cof1
Srv2
Cyr1
Act1
Sla2
Swa2
Sky1
Ste7
Apl1
Apl2
Sto1
Sgv1
Fus3
Rpn10
Sec23
Sly1
Sec18
Syp1
Vrp1
Sla1
Rps9a
Mtr4
Rps8a
Fun12
Rpn1
Duf1
Rad53
Art5
Ecm21
Ldb19
Yhr131c
Rsp5
Bul2
Ent1
Ent5
Apl3
Sec16
Apm4
Akl1
Rpn11
Nip1
Prt1
Rpg1
Ded1
Tif35
Cdc9
Chk1
Rad23
Ubi4
Ede1
Ent2
Vps27
Chc1
Ptp3
Stb1
Spa2
Bem2
Bud14
Rck2
Glc7
Ipl1
Cdc5
Dbf4
Pfk26
Tpk1
Gpa2
Ras2
Med6
Rpb10
Paf1
Rpo26
Bdp1
Spt15
Spt5
Rgr1
Srb7
Ssn2
Nop1
Nsr1
Nop56
Snu13
Rps31
Rad9
Rtf1
Rpo21
Npl3
Air2
Fps1
Ask10
Cdc28
Hog1
Pbs2
Ssk2
Clb2
Bud3
Bud4
Msn2
Sch9
Rim15
Yak1
Ypl247c
Vps52
Kss1
Sko1
Slt2
Msg5
Swe1
Igo2
Cdc55
Igo1
Vps53
Vps54
Rtn1
Fmp42
Cot1
Zrc1
10
20
30
40
50
t/s

### Path 177 (Yhr131c − Ent5)

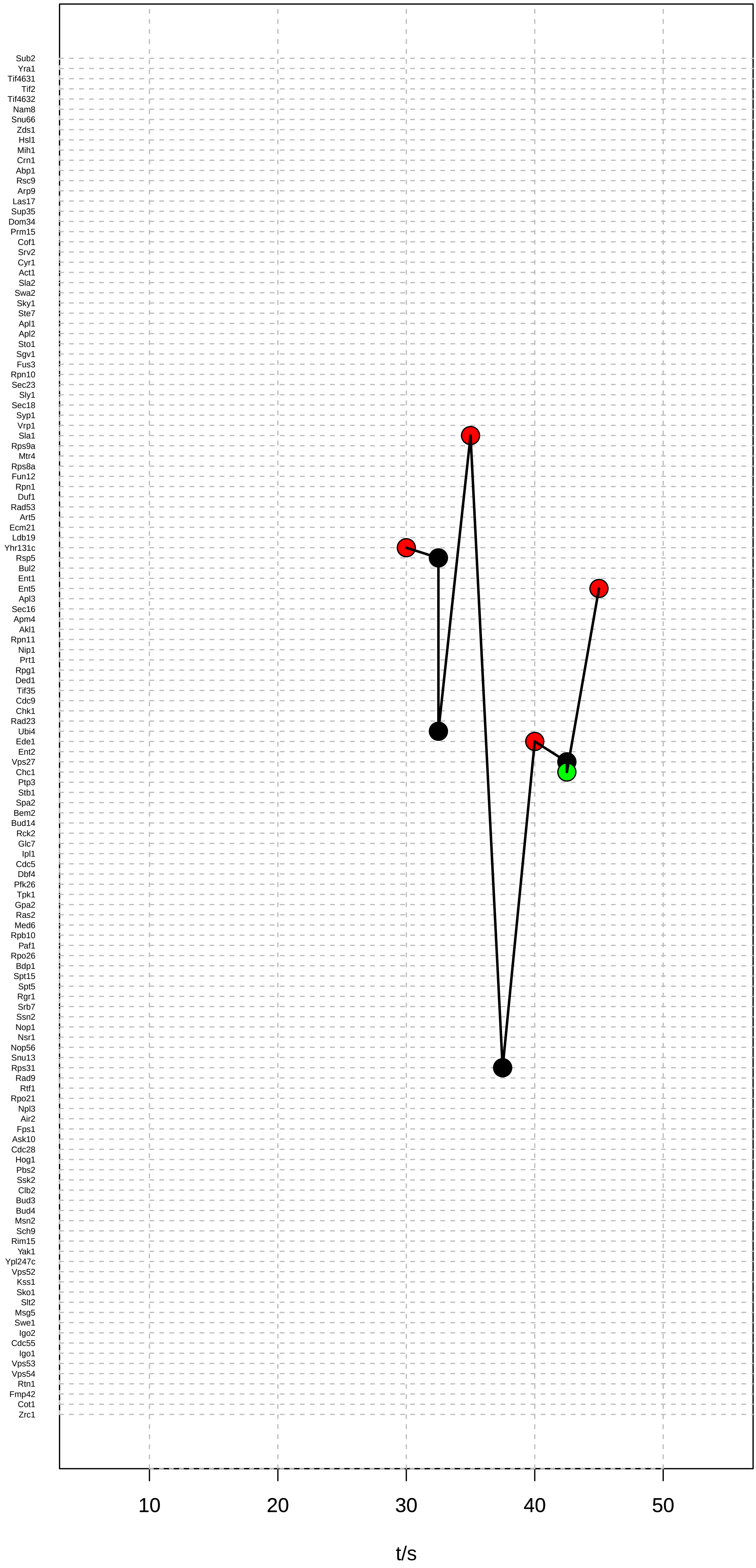

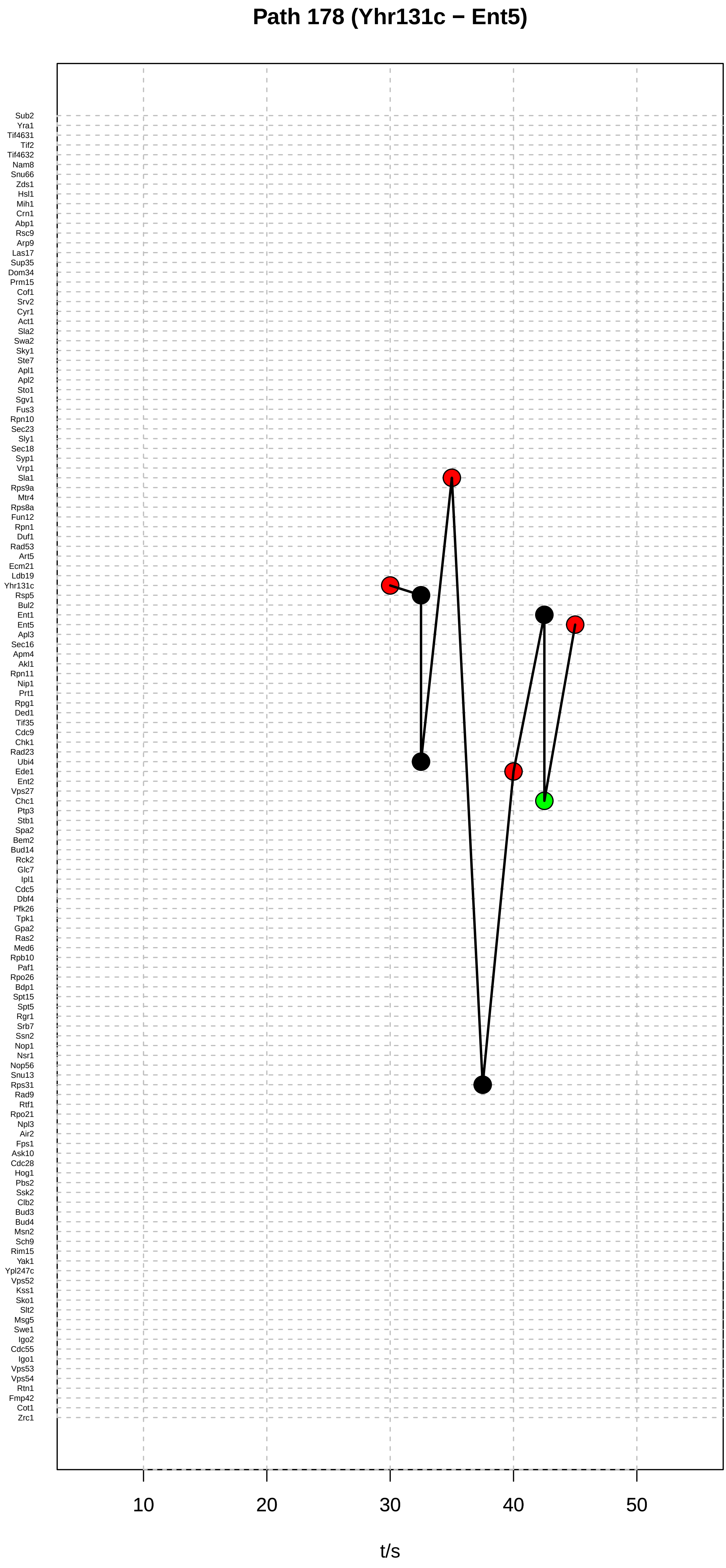
Path 178 (Yhr131c − Ent5)
Sub2
Yra1
Tif4631
Tif2
Tif4632
Nam8
Snu66
Zds1
Hsl1
Mih1
Crn1
Abp1
Rsc9
Arp9
Las17
Sup35
Dom34
Prm15
Cof1
Srv2
Cyr1
Act1
Sla2
Swa2
Sky1
Ste7
Apl1
Apl2
Sto1
Sgv1
Fus3
Rpn10
Sec23
Sly1
Sec18
Syp1
Vrp1
Sla1
Rps9a
Mtr4
Rps8a
Fun12
Rpn1
Duf1
Rad53
Art5
Ecm21
Ldb19
Yhr131c
Rsp5
Bul2
Ent1
Ent5
Apl3
Sec16
Apm4
Akl1
Rpn11
Nip1
Prt1
Rpg1
Ded1
Tif35
Cdc9
Chk1
Rad23
Ubi4
Ede1
Ent2
Vps27
Chc1
Ptp3
Stb1
Spa2
Bem2
Bud14
Rck2
Glc7
Ipl1
Cdc5
Dbf4
Pfk26
Tpk1
Gpa2
Ras2
Med6
Rpb10
Paf1
Rpo26
Bdp1
Spt15
Spt5
Rgr1
Srb7
Ssn2
Nop1
Nsr1
Nop56
Snu13
Rps31
Rad9
Rtf1
Rpo21
Npl3
Air2
Fps1
Ask10
Cdc28
Hog1
Pbs2
Ssk2
Clb2
Bud3
Bud4
Msn2
Sch9
Rim15
Yak1
Ypl247c
Vps52
Kss1
Sko1
Slt2
Msg5
Swe1
Igo2
Cdc55
Igo1
Vps53
Vps54
Rtn1
Fmp42
Cot1
Zrc1
10
20
30
40
50
t/s

### Path 179 (Sec18 − Ent5)

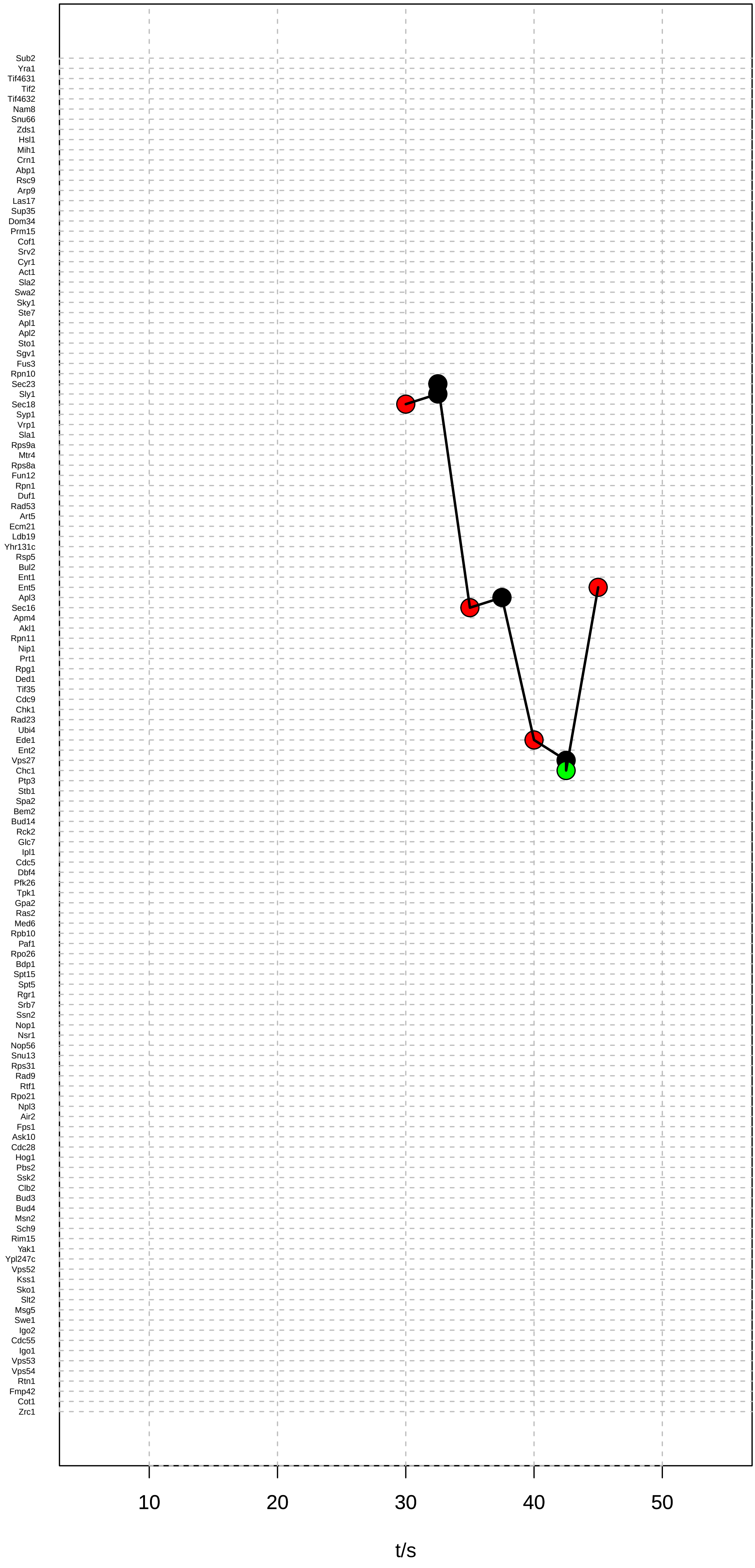

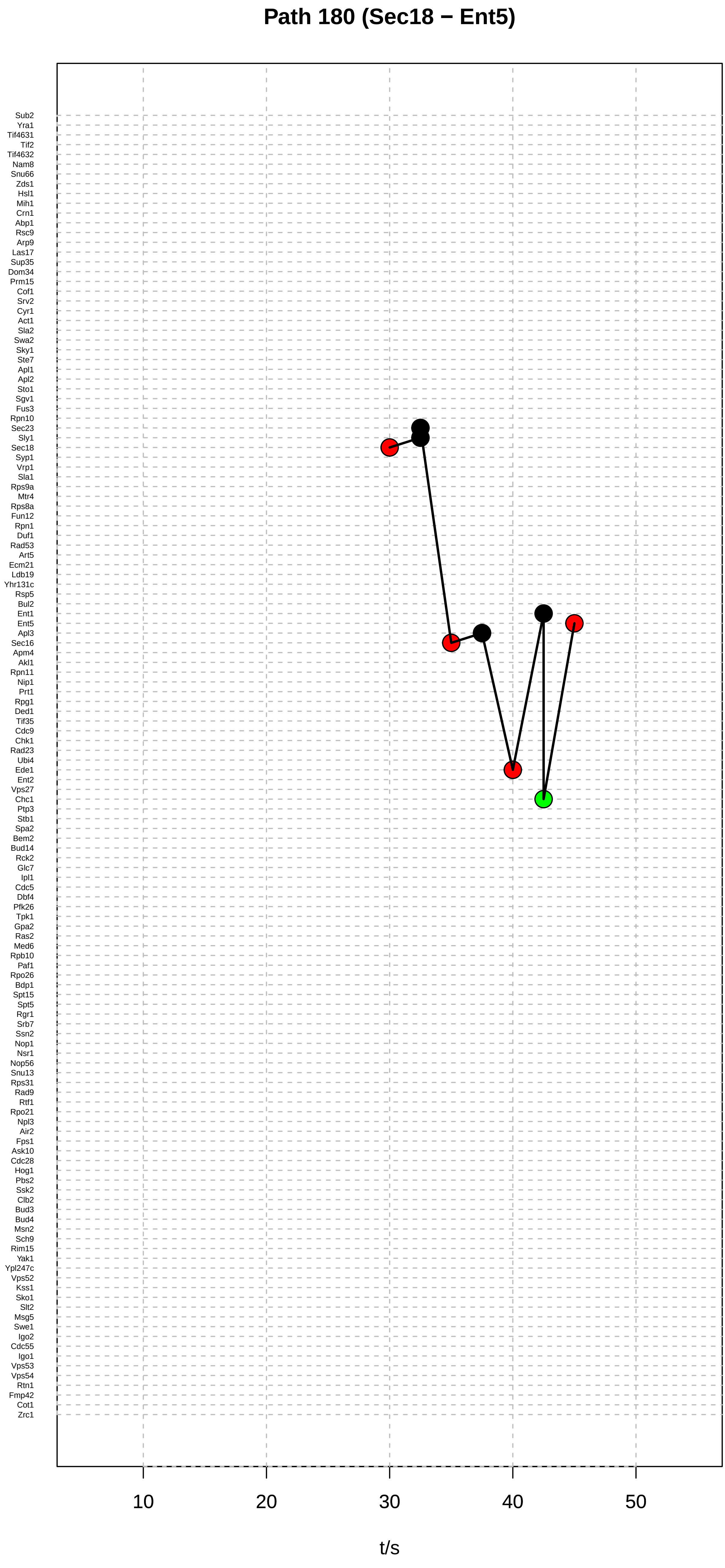

Path 180 (Sec18 − Ent5)
Sub2
Yra1
Tif4631
Tif2
Tif4632
Nam8
Snu66
Zds1
Hsl1
Mih1
Crn1
Abp1
Rsc9
Arp9
Las17
Sup35
Dom34
Prm15
Cof1
Srv2
Cyr1
Act1
Sla2
Swa2
Sky1
Ste7
Apl1
Apl2
Sto1
Sgv1
Fus3
Rpn10
Sec23
Sly1
Sec18
Syp1
Vrp1
Sla1
Rps9a
Mtr4
Rps8a
Fun12
Rpn1
Duf1
Rad53
Art5
Ecm21
Ldb19
Yhr131c
Rsp5
Bul2
Ent1
Ent5
Apl3
Sec16
Apm4
Akl1
Rpn11
Nip1
Prt1
Rpg1
Ded1
Tif35
Cdc9
Chk1
Rad23
Ubi4
Ede1
Ent2
Vps27
Chc1
Ptp3
Stb1
Spa2
Bem2
Bud14
Rck2
Glc7
Ipl1
Cdc5
Dbf4
Pfk26
Tpk1
Gpa2
Ras2
Med6
Rpb10
Paf1
Rpo26
Bdp1
Spt15
Spt5
Rgr1
Srb7
Ssn2
Nop1
Nsr1
Nop56
Snu13
Rps31
Rad9
Rtf1
Rpo21
Npl3
Air2
Fps1
Ask10
Cdc28
Hog1
Pbs2
Ssk2
Clb2
Bud3
Bud4
Msn2
Sch9
Rim15
Yak1
Ypl247c
Vps52
Kss1
Sko1
Slt2
Msg5
Swe1
Igo2
Cdc55
Igo1
Vps53
Vps54
Rtn1
Fmp42
Cot1
Zrc1
10
20
30
40
50
t/s

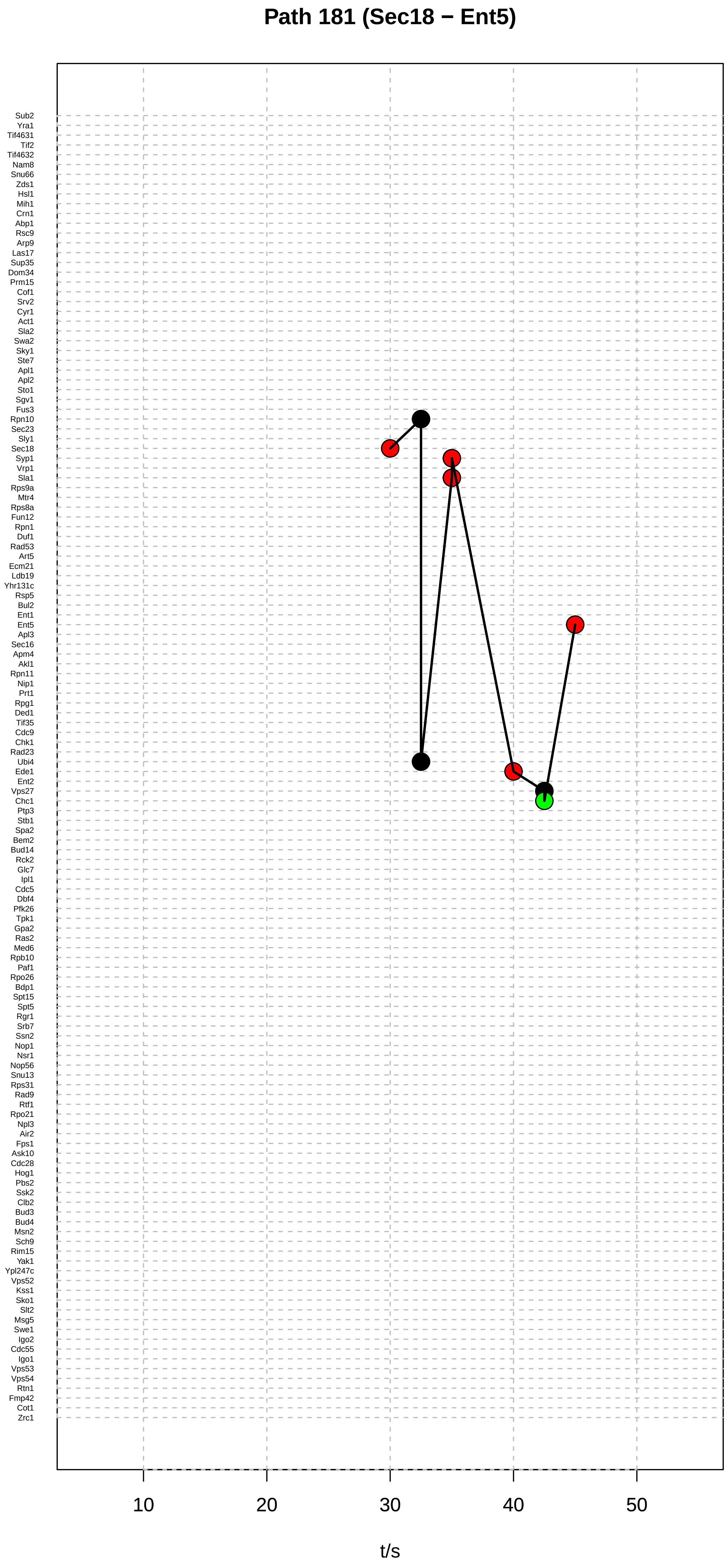

Path 181 (Sec18 − Ent5)
Sub2
Yra1
Tif4631
Tif2
Tif4632
Nam8
Snu66
Zds1
Hsl1
Mih1
Crn1
Abp1
Rsc9
Arp9
Las17
Sup35
Dom34
Prm15
Cof1
Srv2
Cyr1
Act1
Sla2
Swa2
Sky1
Ste7
Apl1
Apl2
Sto1
Sgv1
Fus3
Rpn10
Sec23
Sly1
Sec18
Syp1
Vrp1
Sla1
Rps9a
Mtr4
Rps8a
Fun12
Rpn1
Duf1
Rad53
Art5
Ecm21
Ldb19
Yhr131c
Rsp5
Bul2
Ent1
Ent5
Apl3
Sec16
Apm4
Akl1
Rpn11
Nip1
Prt1
Rpg1
Ded1
Tif35
Cdc9
Chk1
Rad23
Ubi4
Ede1
Ent2
Vps27
Chc1
Ptp3
Stb1
Spa2
Bem2
Bud14
Rck2
Glc7
Ipl1
Cdc5
Dbf4
Pfk26
Tpk1
Gpa2
Ras2
Med6
Rpb10
Paf1
Rpo26
Bdp1
Spt15
Spt5
Rgr1
Srb7
Ssn2
Nop1
Nsr1
Nop56
Snu13
Rps31
Rad9
Rtf1
Rpo21
Npl3
Air2
Fps1
Ask10
Cdc28
Hog1
Pbs2
Ssk2
Clb2
Bud3
Bud4
Msn2
Sch9
Rim15
Yak1
Ypl247c
Vps52
Kss1
Sko1
Slt2
Msg5
Swe1
Igo2
Cdc55
Igo1
Vps53
Vps54
Rtn1
Fmp42
Cot1
Zrc1
10
20
30
40
50
t/s

### Path 182 (Sec18 − Ent5)

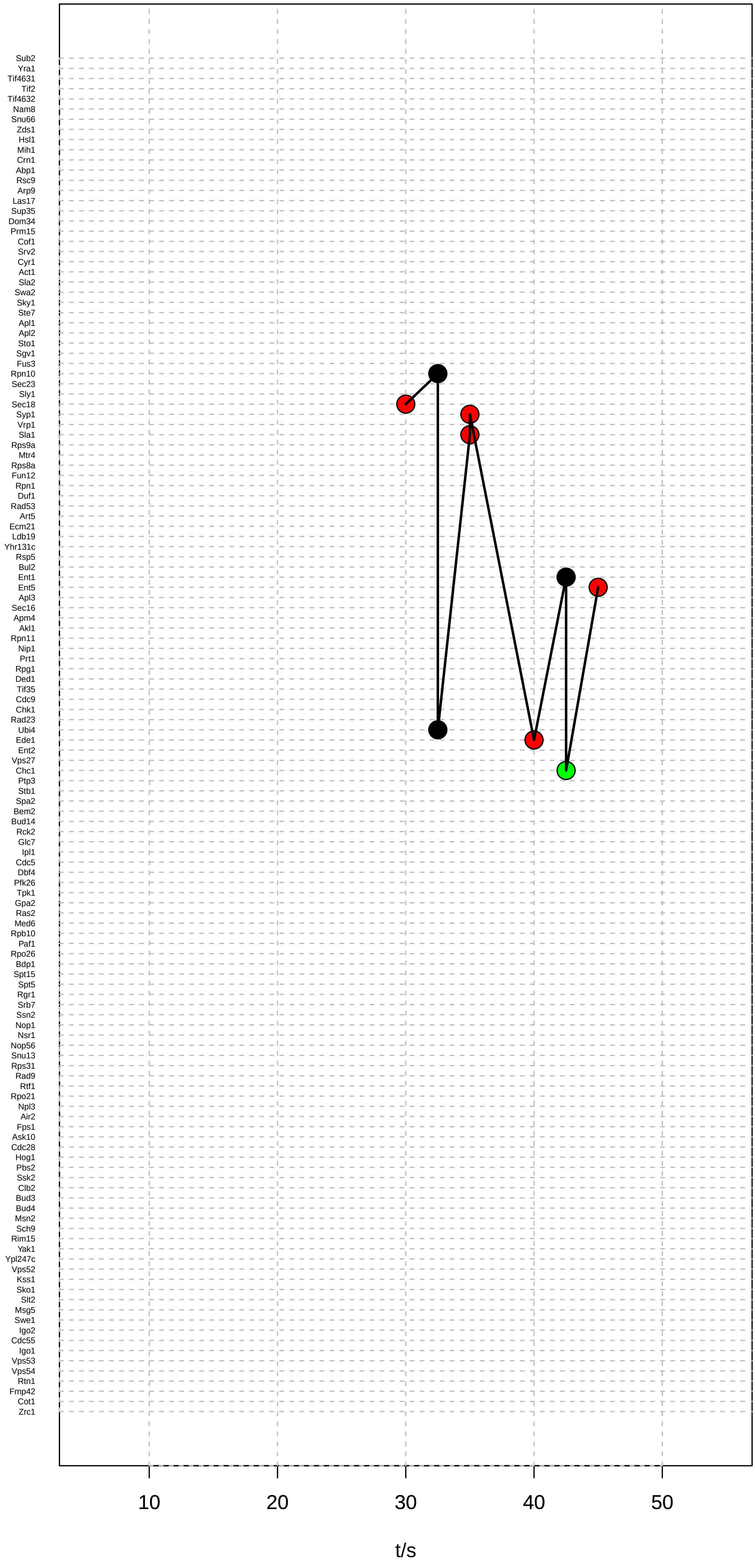

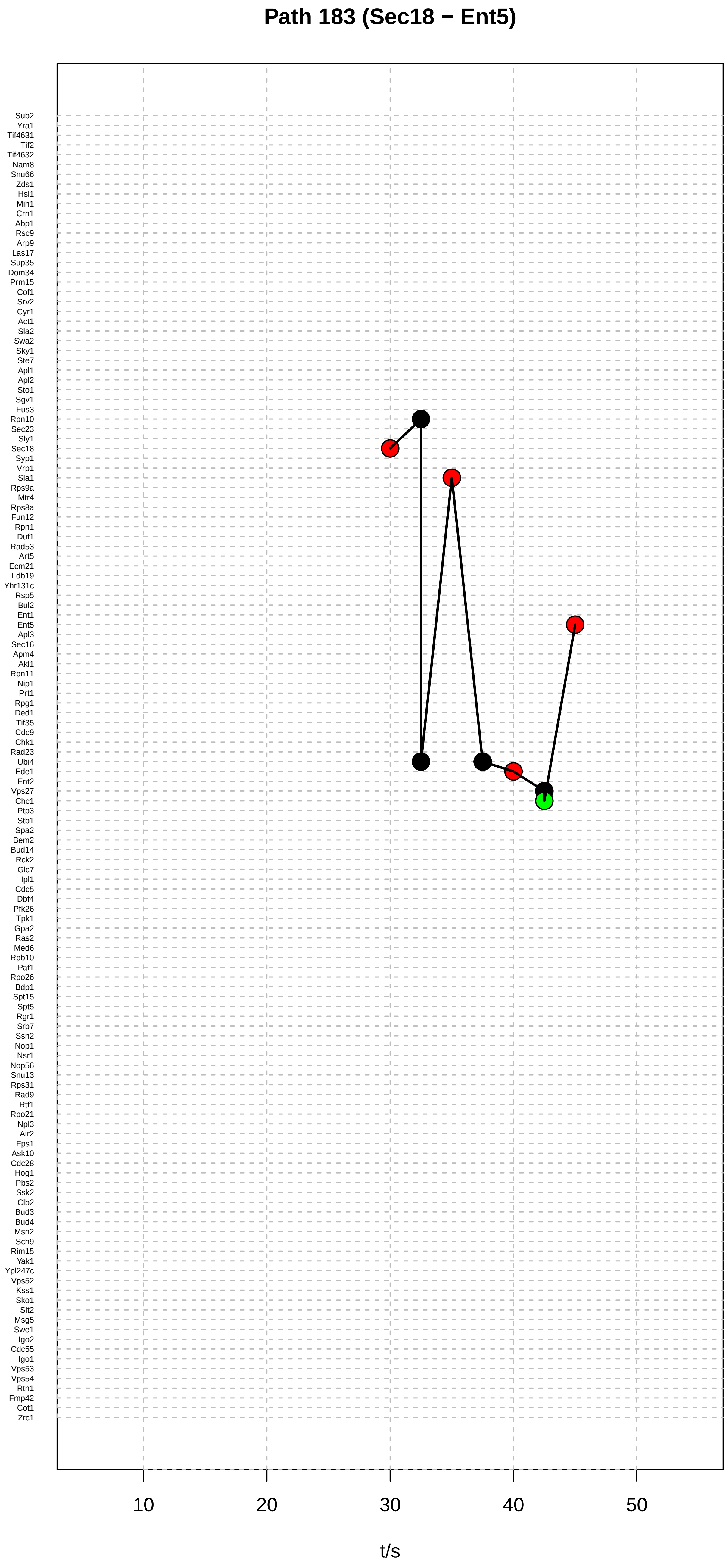
Path 183 (Sec18 − Ent5)
Sub2
Yra1
Tif4631
Tif2
Tif4632
Nam8
Snu66
Zds1
Hsl1
Mih1
Crn1
Abp1
Rsc9
Arp9
Las17
Sup35
Dom34
Prm15
Cof1
Srv2
Cyr1
Act1
Sla2
Swa2
Sky1
Ste7
Apl1
Apl2
Sto1
Sgv1
Fus3
Rpn10
Sec23
Sly1
Sec18
Syp1
Vrp1
Sla1
Rps9a
Mtr4
Rps8a
Fun12
Rpn1
Duf1
Rad53
Art5
Ecm21
Ldb19
Yhr131c
Rsp5
Bul2
Ent1
Ent5
Apl3
Sec16
Apm4
Akl1
Rpn11
Nip1
Prt1
Rpg1
Ded1
Tif35
Cdc9
Chk1
Rad23
Ubi4
Ede1
Ent2
Vps27
Chc1
Ptp3
Stb1
Spa2
Bem2
Bud14
Rck2
Glc7
Ipl1
Cdc5
Dbf4
Pfk26
Tpk1
Gpa2
Ras2
Med6
Rpb10
Paf1
Rpo26
Bdp1
Spt15
Spt5
Rgr1
Srb7
Ssn2
Nop1
Nsr1
Nop56
Snu13
Rps31
Rad9
Rtf1
Rpo21
Npl3
Air2
Fps1
Ask10
Cdc28
Hog1
Pbs2
Ssk2
Clb2
Bud3
Bud4
Msn2
Sch9
Rim15
Yak1
Ypl247c
Vps52
Kss1
Sko1
Slt2
Msg5
Swe1
Igo2
Cdc55
Igo1
Vps53
Vps54
Rtn1
Fmp42
Cot1
Zrc1
10
20
30
40
50
t/s

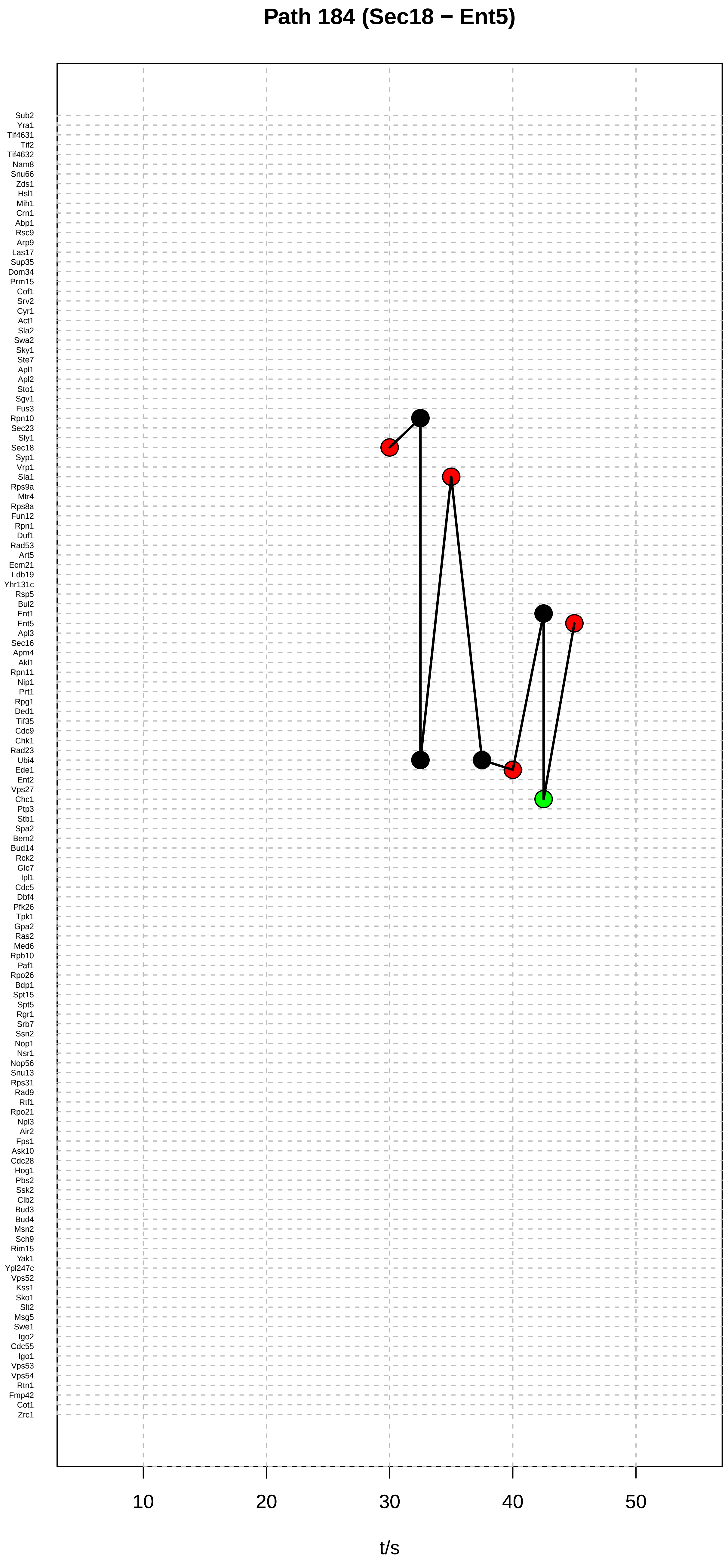
Path 184 (Sec18 − Ent5)
Sub2
Yra1
Tif4631
Tif2
Tif4632
Nam8
Snu66
Zds1
Hsl1
Mih1
Crn1
Abp1
Rsc9
Arp9
Las17
Sup35
Dom34
Prm15
Cof1
Srv2
Cyr1
Act1
Sla2
Swa2
Sky1
Ste7
Apl1
Apl2
Sto1
Sgv1
Fus3
Rpn10
Sec23
Sly1
Sec18
Syp1
Vrp1
Sla1
Rps9a
Mtr4
Rps8a
Fun12
Rpn1
Duf1
Rad53
Art5
Ecm21
Ldb19
Yhr131c
Rsp5
Bul2
Ent1
Ent5
Apl3
Sec16
Apm4
Akl1
Rpn11
Nip1
Prt1
Rpg1
Ded1
Tif35
Cdc9
Chk1
Rad23
Ubi4
Ede1
Ent2
Vps27
Chc1
Ptp3
Stb1
Spa2
Bem2
Bud14
Rck2
Glc7
Ipl1
Cdc5
Dbf4
Pfk26
Tpk1
Gpa2
Ras2
Med6
Rpb10
Paf1
Rpo26
Bdp1
Spt15
Spt5
Rgr1
Srb7
Ssn2
Nop1
Nsr1
Nop56
Snu13
Rps31
Rad9
Rtf1
Rpo21
Npl3
Air2
Fps1
Ask10
Cdc28
Hog1
Pbs2
Ssk2
Clb2
Bud3
Bud4
Msn2
Sch9
Rim15
Yak1
Ypl247c
Vps52
Kss1
Sko1
Slt2
Msg5
Swe1
Igo2
Cdc55
Igo1
Vps53
Vps54
Rtn1
Fmp42
Cot1
Zrc1
10
20
30
40
50
t/s

Path 185 (Sec18 − Ent5)

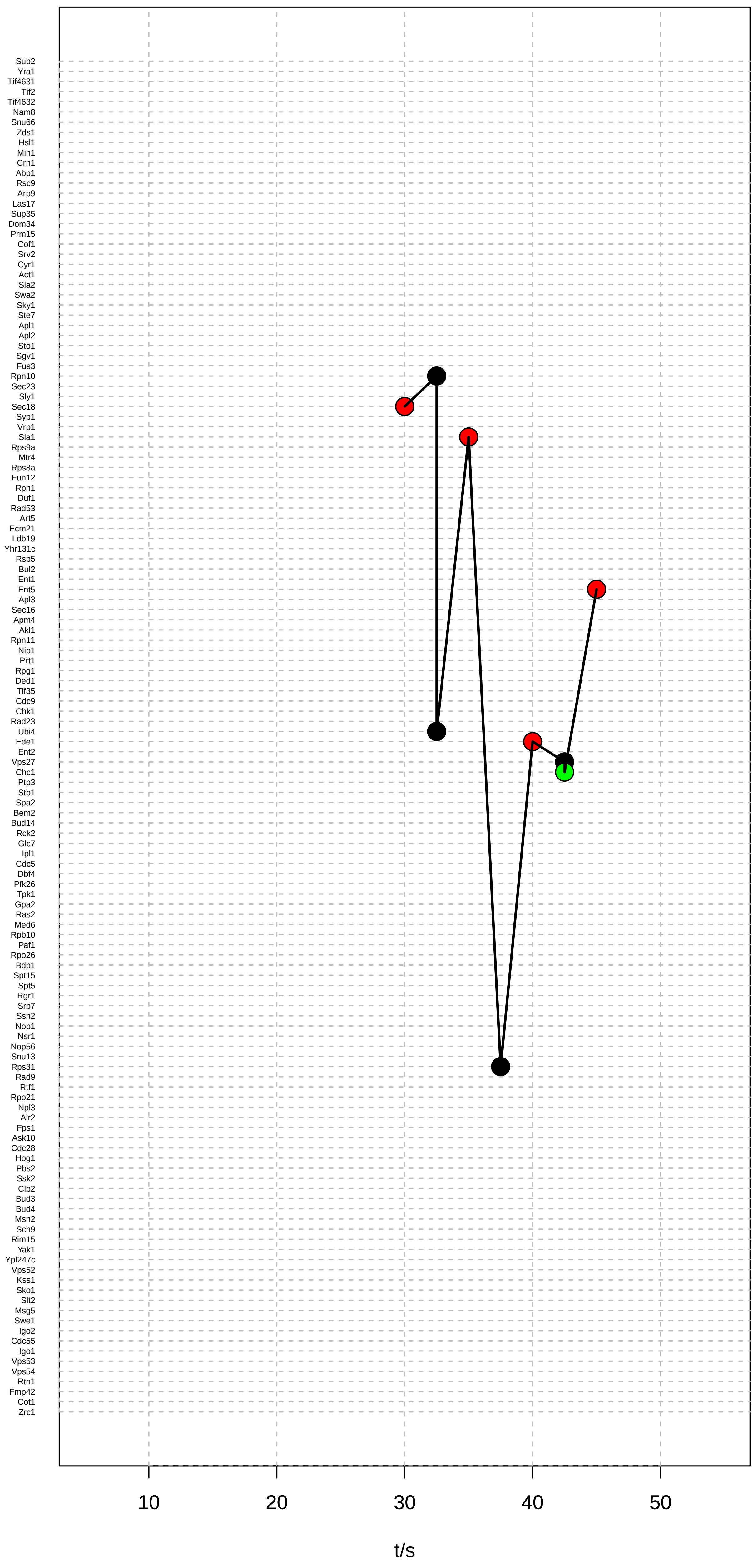

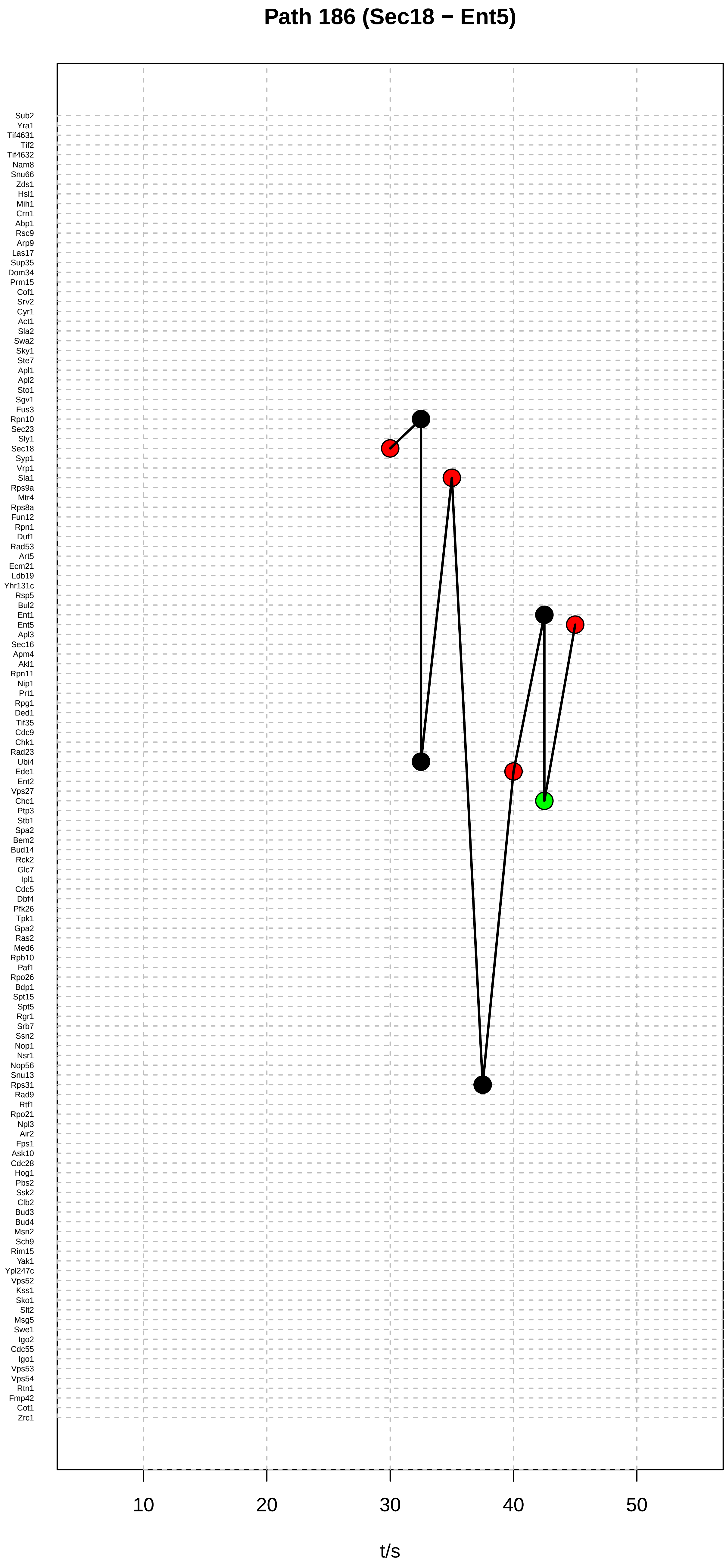

Path 186 (Sec18 − Ent5)
Sub2
Yra1
Tif4631
Tif2
Tif4632
Nam8
Snu66
Zds1
Hsl1
Mih1
Crn1
Abp1
Rsc9
Arp9
Las17
Sup35
Dom34
Prm15
Cof1
Srv2
Cyr1
Act1
Sla2
Swa2
Sky1
Ste7
Apl1
Apl2
Sto1
Sgv1
Fus3
Rpn10
Sec23
Sly1
Sec18
Syp1
Vrp1
Sla1
Rps9a
Mtr4
Rps8a
Fun12
Rpn1
Duf1
Rad53
Art5
Ecm21
Ldb19
Yhr131c
Rsp5
Bul2
Ent1
Ent5
Apl3
Sec16
Apm4
Akl1
Rpn11
Nip1
Prt1
Rpg1
Ded1
Tif35
Cdc9
Chk1
Rad23
Ubi4
Ede1
Ent2
Vps27
Chc1
Ptp3
Stb1
Spa2
Bem2
Bud14
Rck2
Glc7
Ipl1
Cdc5
Dbf4
Pfk26
Tpk1
Gpa2
Ras2
Med6
Rpb10
Paf1
Rpo26
Bdp1
Spt15
Spt5
Rgr1
Srb7
Ssn2
Nop1
Nsr1
Nop56
Snu13
Rps31
Rad9
Rtf1
Rpo21
Npl3
Air2
Fps1
Ask10
Cdc28
Hog1
Pbs2
Ssk2
Clb2
Bud3
Bud4
Msn2
Sch9
Rim15
Yak1
Ypl247c
Vps52
Kss1
Sko1
Slt2
Msg5
Swe1
Igo2
Cdc55
Igo1
Vps53
Vps54
Rtn1
Fmp42
Cot1
Zrc1
10
20
30
40
50
t/s

#### Path 187 (Akl1 − Ent5)

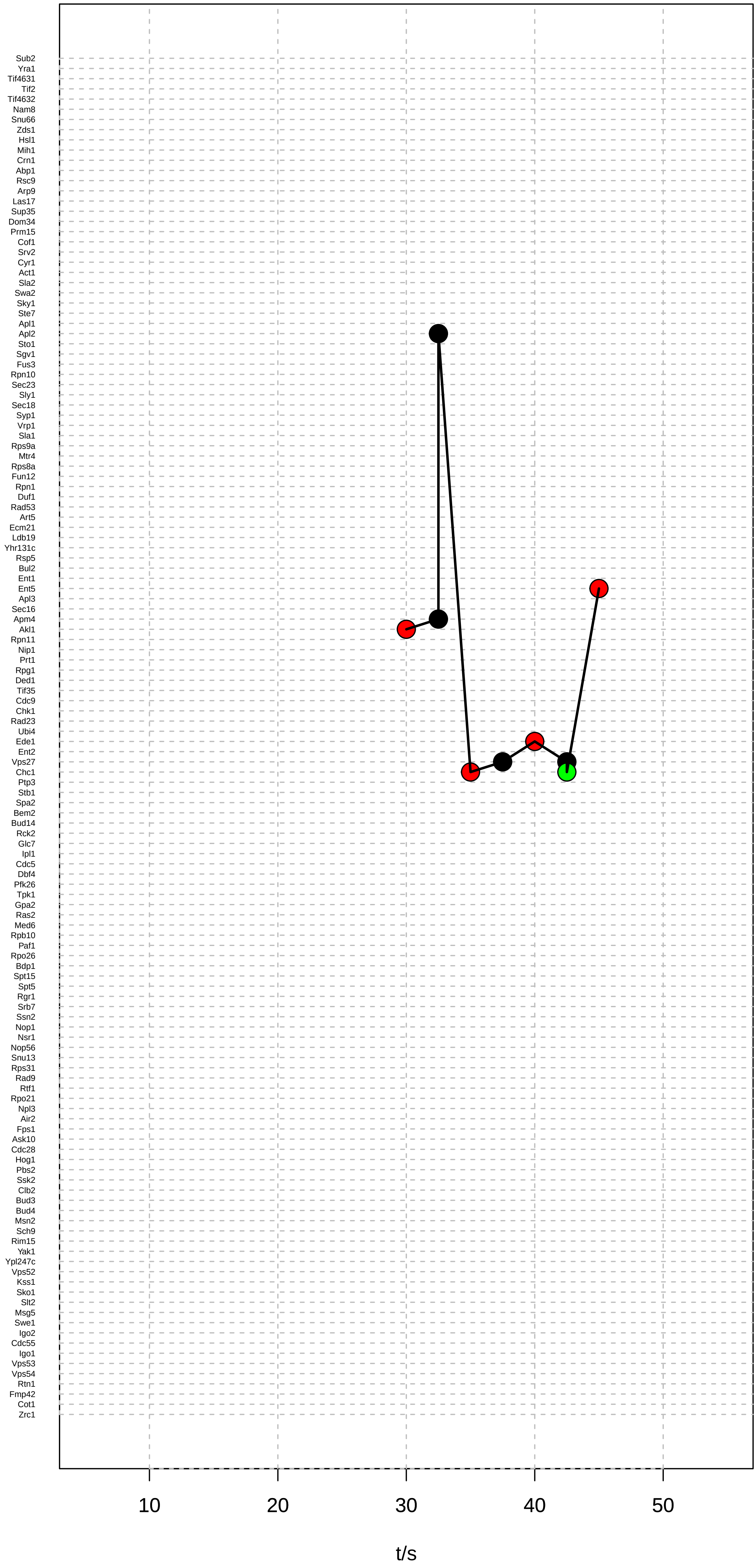

#### Path 188 (Akl1 − Ent5)

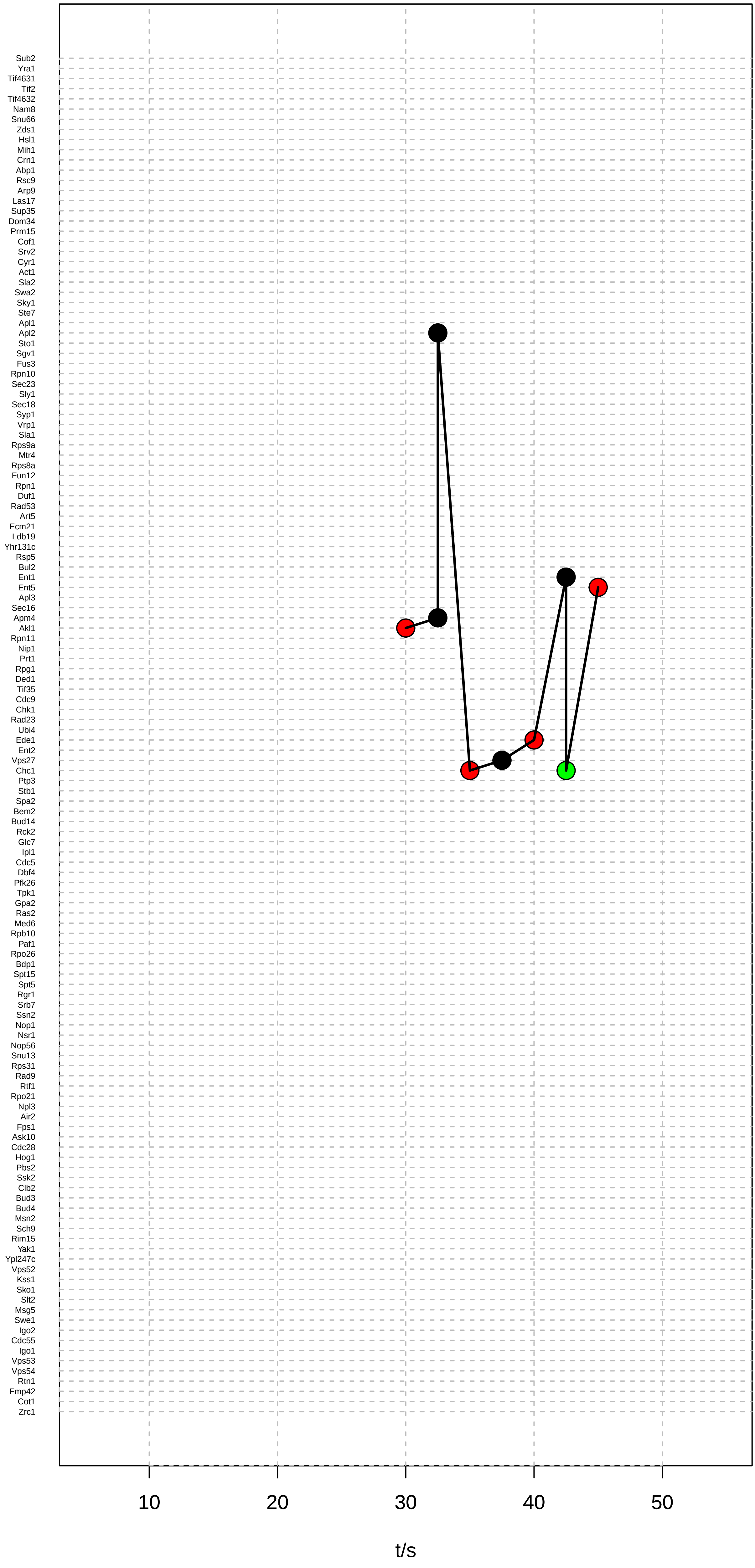

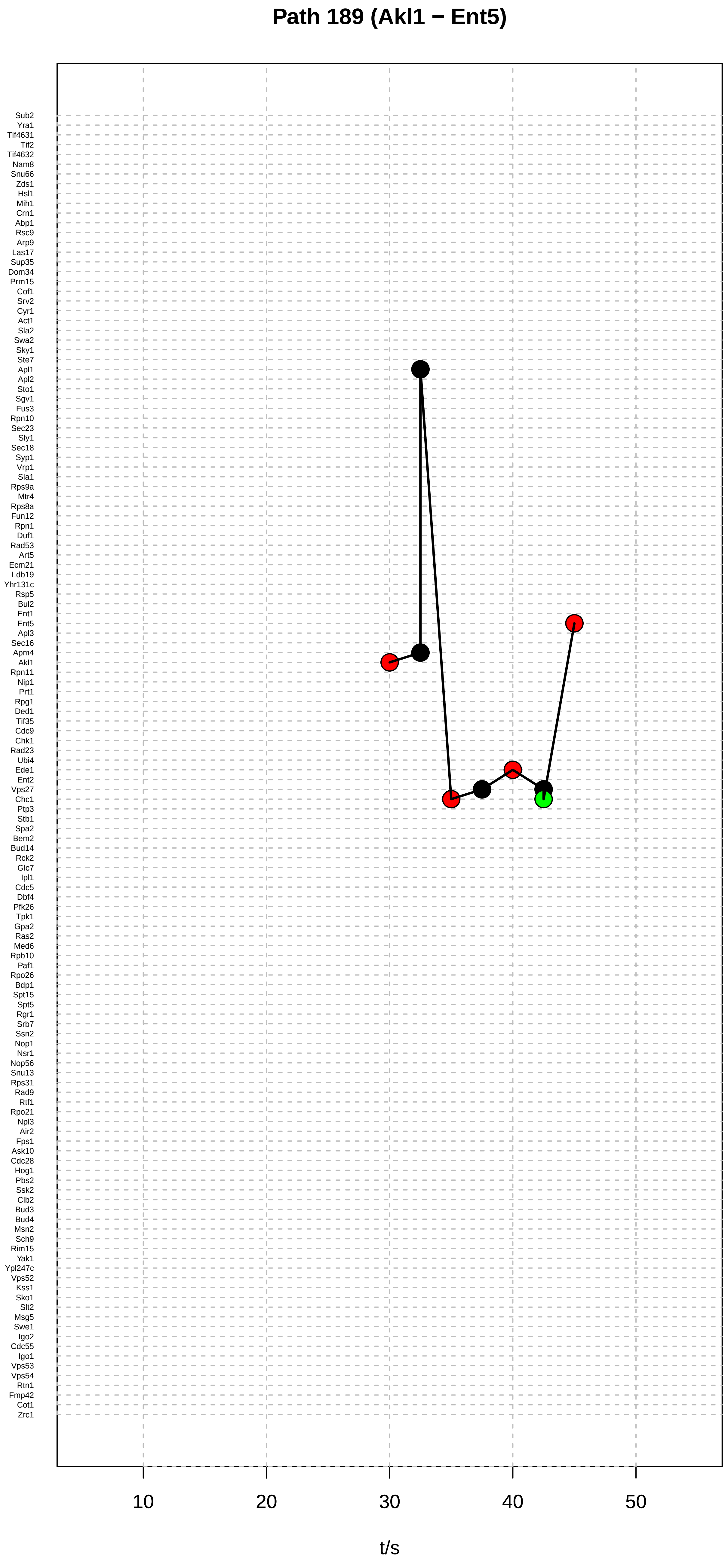
Path 189 (Akl1 − Ent5)
Sub2
Yra1
Tif4631
Tif2
Tif4632
Nam8
Snu66
Zds1
Hsl1
Mih1
Crn1
Abp1
Rsc9
Arp9
Las17
Sup35
Dom34
Prm15
Cof1
Srv2
Cyr1
Act1
Sla2
Swa2
Sky1
Ste7
Apl1
Apl2
Sto1
Sgv1
Fus3
Rpn10
Sec23
Sly1
Sec18
Syp1
Vrp1
Sla1
Rps9a
Mtr4
Rps8a
Fun12
Rpn1
Duf1
Rad53
Art5
Ecm21
Ldb19
Yhr131c
Rsp5
Bul2
Ent1
Ent5
Apl3
Sec16
Apm4
Akl1
Rpn11
Nip1
Prt1
Rpg1
Ded1
Tif35
Cdc9
Chk1
Rad23
Ubi4
Ede1
Ent2
Vps27
Chc1
Ptp3
Stb1
Spa2
Bem2
Bud14
Rck2
Glc7
Ipl1
Cdc5
Dbf4
Pfk26
Tpk1
Gpa2
Ras2
Med6
Rpb10
Paf1
Rpo26
Bdp1
Spt15
Spt5
Rgr1
Srb7
Ssn2
Nop1
Nsr1
Nop56
Snu13
Rps31
Rad9
Rtf1
Rpo21
Npl3
Air2
Fps1
Ask10
Cdc28
Hog1
Pbs2
Ssk2
Clb2
Bud3
Bud4
Msn2
Sch9
Rim15
Yak1
Ypl247c
Vps52
Kss1
Sko1
Slt2
Msg5
Swe1
Igo2
Cdc55
Igo1
Vps53
Vps54
Rtn1
Fmp42
Cot1
Zrc1
10
20
30
40
50
t/s

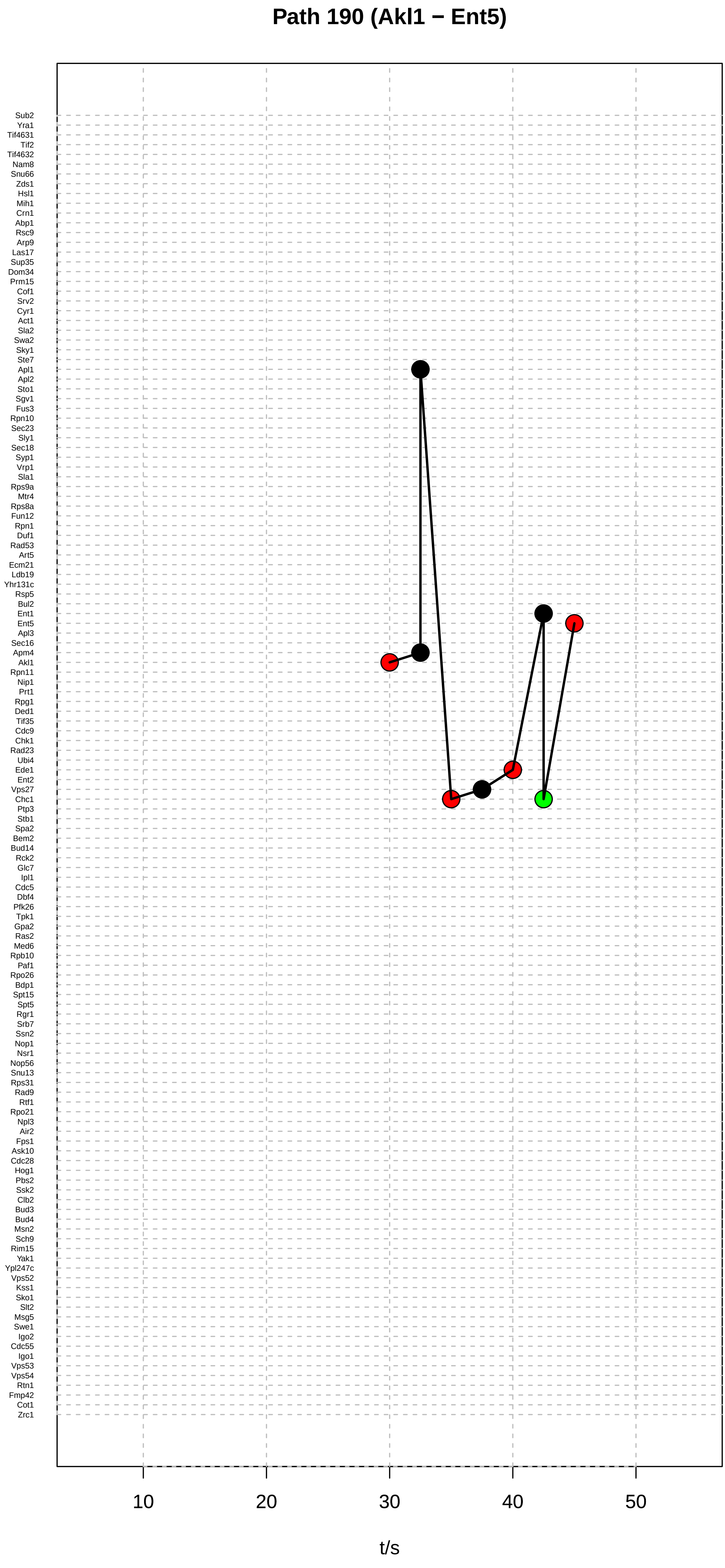
Path 190 (Akl1 − Ent5)
Sub2
Yra1
Tif4631
Tif2
Tif4632
Nam8
Snu66
Zds1
Hsl1
Mih1
Crn1
Abp1
Rsc9
Arp9
Las17
Sup35
Dom34
Prm15
Cof1
Srv2
Cyr1
Act1
Sla2
Swa2
Sky1
Ste7
Apl1
Apl2
Sto1
Sgv1
Fus3
Rpn10
Sec23
Sly1
Sec18
Syp1
Vrp1
Sla1
Rps9a
Mtr4
Rps8a
Fun12
Rpn1
Duf1
Rad53
Art5
Ecm21
Ldb19
Yhr131c
Rsp5
Bul2
Ent1
Ent5
Apl3
Sec16
Apm4
Akl1
Rpn11
Nip1
Prt1
Rpg1
Ded1
Tif35
Cdc9
Chk1
Rad23
Ubi4
Ede1
Ent2
Vps27
Chc1
Ptp3
Stb1
Spa2
Bem2
Bud14
Rck2
Glc7
Ipl1
Cdc5
Dbf4
Pfk26
Tpk1
Gpa2
Ras2
Med6
Rpb10
Paf1
Rpo26
Bdp1
Spt15
Spt5
Rgr1
Srb7
Ssn2
Nop1
Nsr1
Nop56
Snu13
Rps31
Rad9
Rtf1
Rpo21
Npl3
Air2
Fps1
Ask10
Cdc28
Hog1
Pbs2
Ssk2
Clb2
Bud3
Bud4
Msn2
Sch9
Rim15
Yak1
Ypl247c
Vps52
Kss1
Sko1
Slt2
Msg5
Swe1
Igo2
Cdc55
Igo1
Vps53
Vps54
Rtn1
Fmp42
Cot1
Zrc1
10
20
30
40
50
t/s

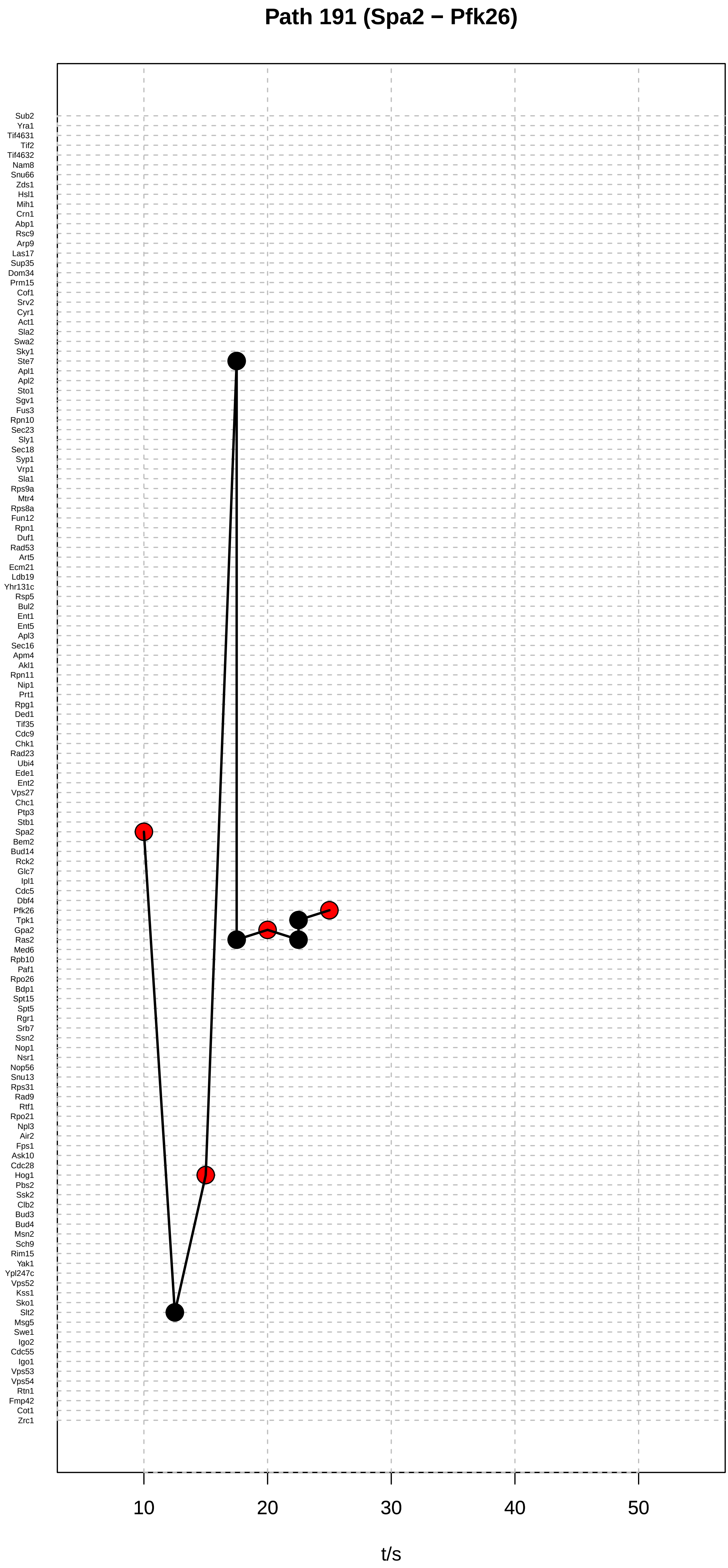

Path 191 (Spa2 − Pfk26)
Sub2
Yra1
Tif4631
Tif2
Tif4632
Nam8
Snu66
Zds1
Hsl1
Mih1
Crn1
Abp1
Rsc9
Arp9
Las17
Sup35
Dom34
Prm15
Cof1
Srv2
Cyr1
Act1
Sla2
Swa2
Sky1
Ste7
Apl1
Apl2
Sto1
Sgv1
Fus3
Rpn10
Sec23
Sly1
Sec18
Syp1
Vrp1
Sla1
Rps9a
Mtr4
Rps8a
Fun12
Rpn1
Duf1
Rad53
Art5
Ecm21
Ldb19
Yhr131c
Rsp5
Bul2
Ent1
Ent5
Apl3
Sec16
Apm4
Akl1
Rpn11
Nip1
Prt1
Rpg1
Ded1
Tif35
Cdc9
Chk1
Rad23
Ubi4
Ede1
Ent2
Vps27
Chc1
Ptp3
Stb1
Spa2
Bem2
Bud14
Rck2
Glc7
Ipl1
Cdc5
Dbf4
Pfk26
Tpk1
Gpa2
Ras2
Med6
Rpb10
Paf1
Rpo26
Bdp1
Spt15
Spt5
Rgr1
Srb7
Ssn2
Nop1
Nsr1
Nop56
Snu13
Rps31
Rad9
Rtf1
Rpo21
Npl3
Air2
Fps1
Ask10
Cdc28
Hog1
Pbs2
Ssk2
Clb2
Bud3
Bud4
Msn2
Sch9
Rim15
Yak1
Ypl247c
Vps52
Kss1
Sko1
Slt2
Msg5
Swe1
Igo2
Cdc55
Igo1
Vps53
Vps54
Rtn1
Fmp42
Cot1
Zrc1
10
20
30
40
50
t/s

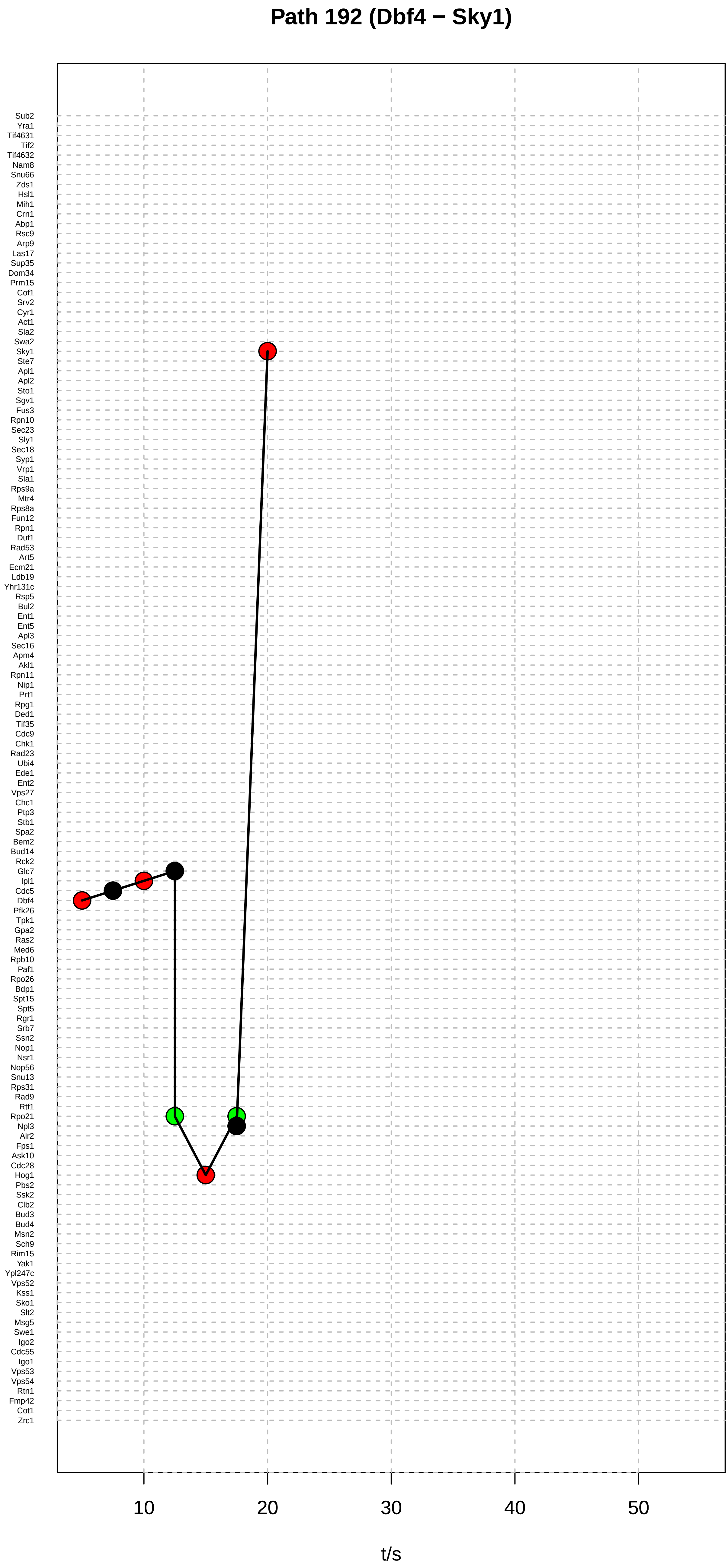
Path 192 (Dbf4 − Sky1)
Sub2
Yra1
Tif4631
Tif2
Tif4632
Nam8
Snu66
Zds1
Hsl1
Mih1
Crn1
Abp1
Rsc9
Arp9
Las17
Sup35
Dom34
Prm15
Cof1
Srv2
Cyr1
Act1
Sla2
Swa2
Sky1
Ste7
Apl1
Apl2
Sto1
Sgv1
Fus3
Rpn10
Sec23
Sly1
Sec18
Syp1
Vrp1
Sla1
Rps9a
Mtr4
Rps8a
Fun12
Rpn1
Duf1
Rad53
Art5
Ecm21
Ldb19
Yhr131c
Rsp5
Bul2
Ent1
Ent5
Apl3
Sec16
Apm4
Akl1
Rpn11
Nip1
Prt1
Rpg1
Ded1
Tif35
Cdc9
Chk1
Rad23
Ubi4
Ede1
Ent2
Vps27
Chc1
Ptp3
Stb1
Spa2
Bem2
Bud14
Rck2
Glc7
Ipl1
Cdc5
Dbf4
Pfk26
Tpk1
Gpa2
Ras2
Med6
Rpb10
Paf1
Rpo26
Bdp1
Spt15
Spt5
Rgr1
Srb7
Ssn2
Nop1
Nsr1
Nop56
Snu13
Rps31
Rad9
Rtf1
Rpo21
Npl3
Air2
Fps1
Ask10
Cdc28
Hog1
Pbs2
Ssk2
Clb2
Bud3
Bud4
Msn2
Sch9
Rim15
Yak1
Ypl247c
Vps52
Kss1
Sko1
Slt2
Msg5
Swe1
Igo2
Cdc55
Igo1
Vps53
Vps54
Rtn1
Fmp42
Cot1
Zrc1
10
20
30
40
50
t/s

#### Path 193 (Dbf4 − Bud14)

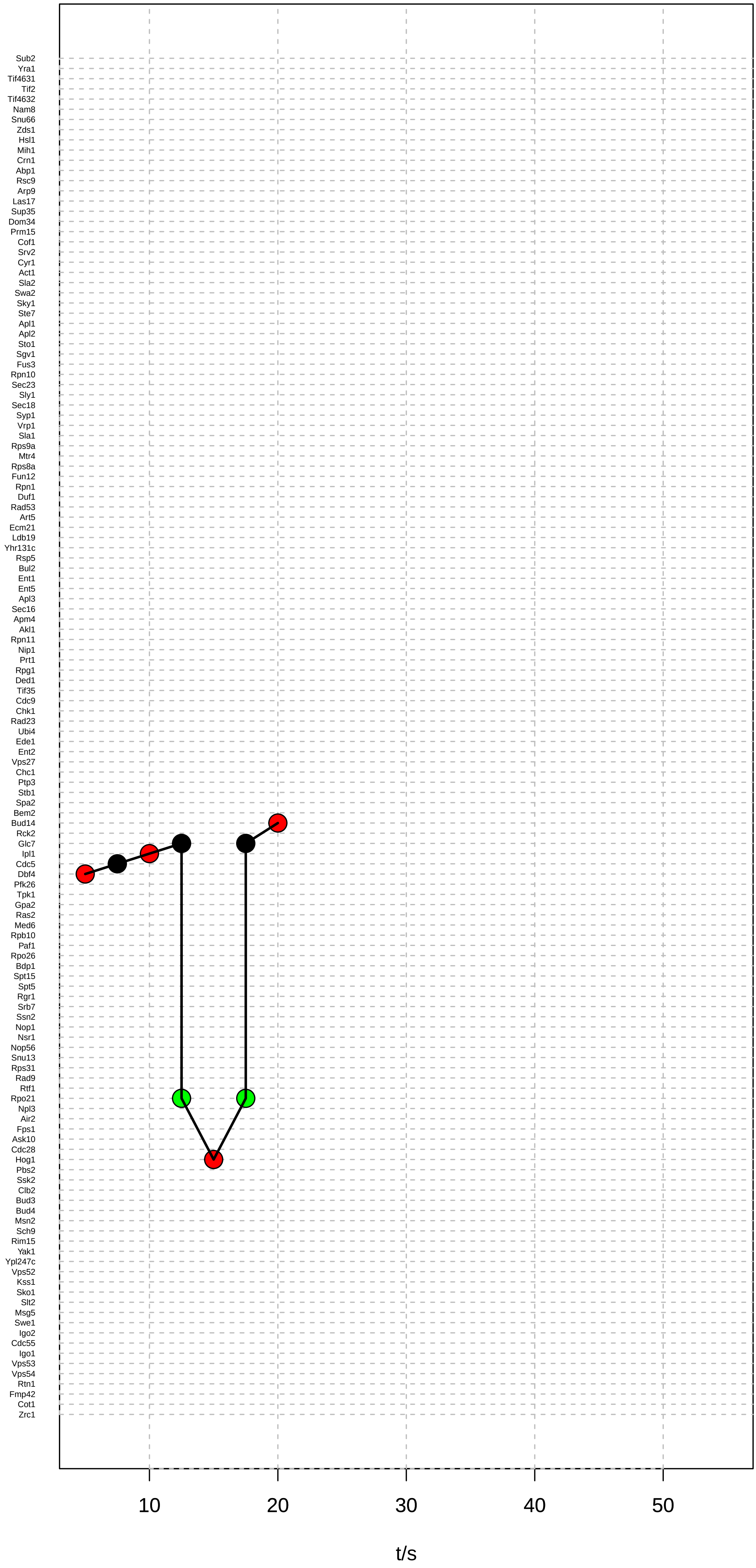

### Path 194 (Rad9 − Sky1)

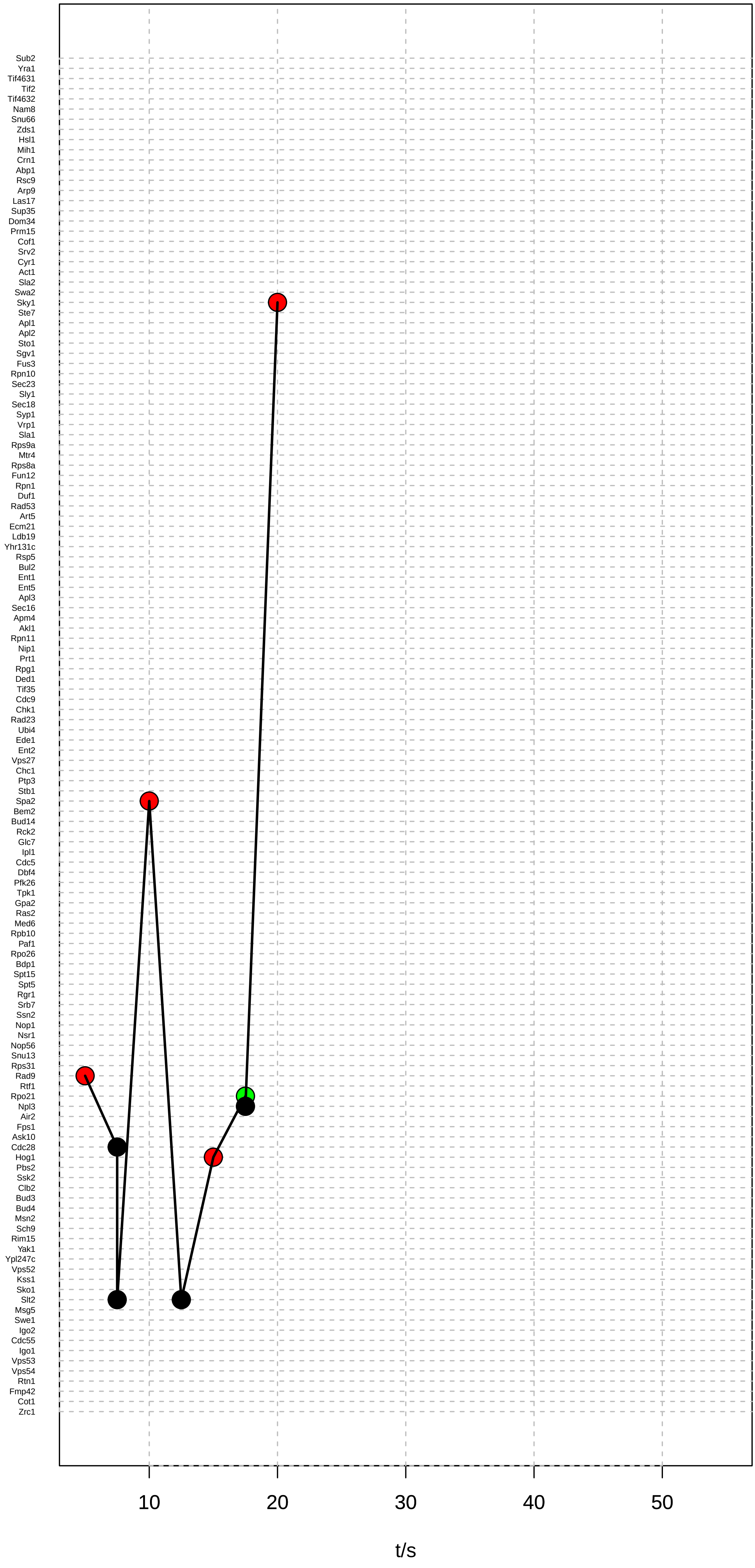

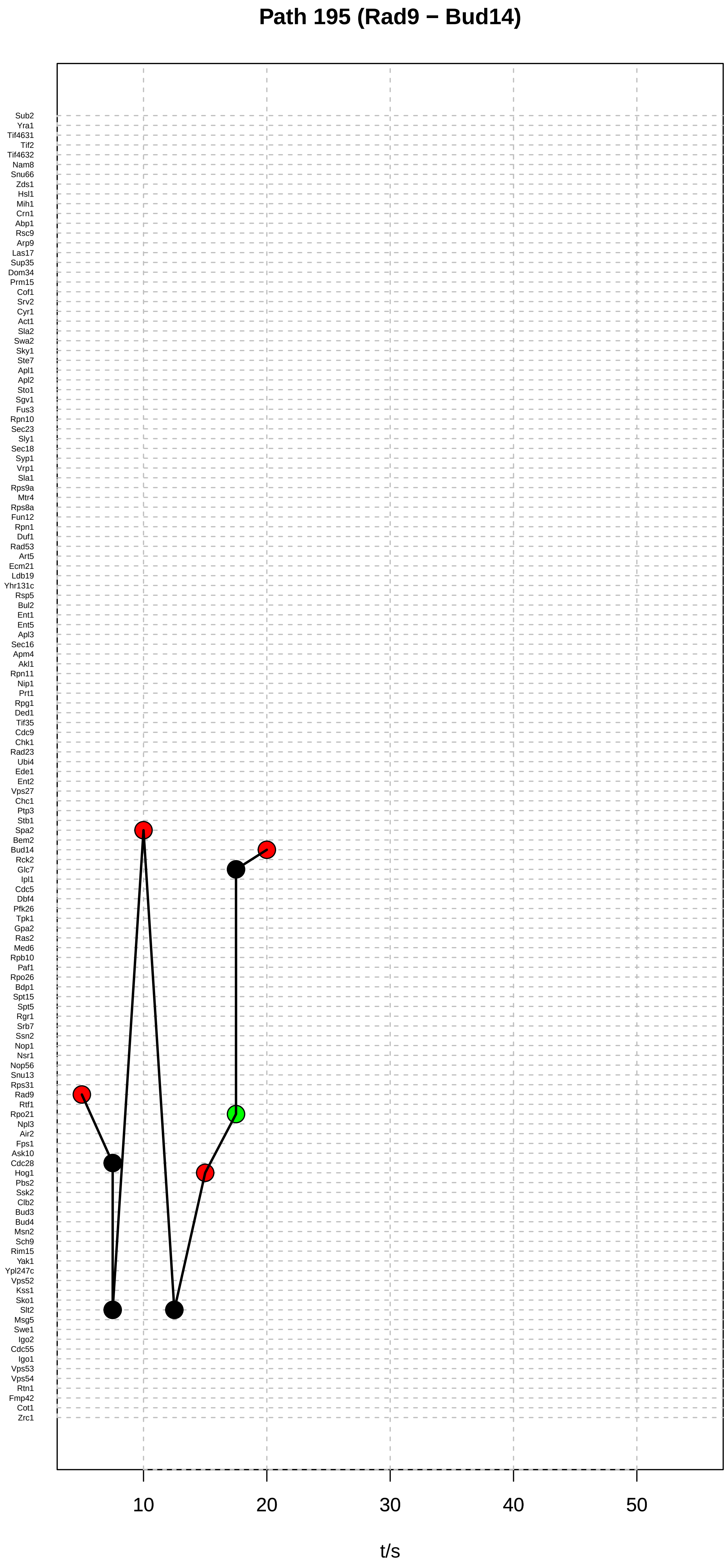

Path 195 (Rad9 − Bud14)
Sub2
Yra1
Tif4631
Tif2
Tif4632
Nam8
Snu66
Zds1
Hsl1
Mih1
Crn1
Abp1
Rsc9
Arp9
Las17
Sup35
Dom34
Prm15
Cof1
Srv2
Cyr1
Act1
Sla2
Swa2
Sky1
Ste7
Apl1
Apl2
Sto1
Sgv1
Fus3
Rpn10
Sec23
Sly1
Sec18
Syp1
Vrp1
Sla1
Rps9a
Mtr4
Rps8a
Fun12
Rpn1
Duf1
Rad53
Art5
Ecm21
Ldb19
Yhr131c
Rsp5
Bul2
Ent1
Ent5
Apl3
Sec16
Apm4
Akl1
Rpn11
Nip1
Prt1
Rpg1
Ded1
Tif35
Cdc9
Chk1
Rad23
Ubi4
Ede1
Ent2
Vps27
Chc1
Ptp3
Stb1
Spa2
Bem2
Bud14
Rck2
Glc7
Ipl1
Cdc5
Dbf4
Pfk26
Tpk1
Gpa2
Ras2
Med6
Rpb10
Paf1
Rpo26
Bdp1
Spt15
Spt5
Rgr1
Srb7
Ssn2
Nop1
Nsr1
Nop56
Snu13
Rps31
Rad9
Rtf1
Rpo21
Npl3
Air2
Fps1
Ask10
Cdc28
Hog1
Pbs2
Ssk2
Clb2
Bud3
Bud4
Msn2
Sch9
Rim15
Yak1
Ypl247c
Vps52
Kss1
Sko1
Slt2
Msg5
Swe1
Igo2
Cdc55
Igo1
Vps53
Vps54
Rtn1
Fmp42
Cot1
Zrc1
10
20
30
40
50
t/s

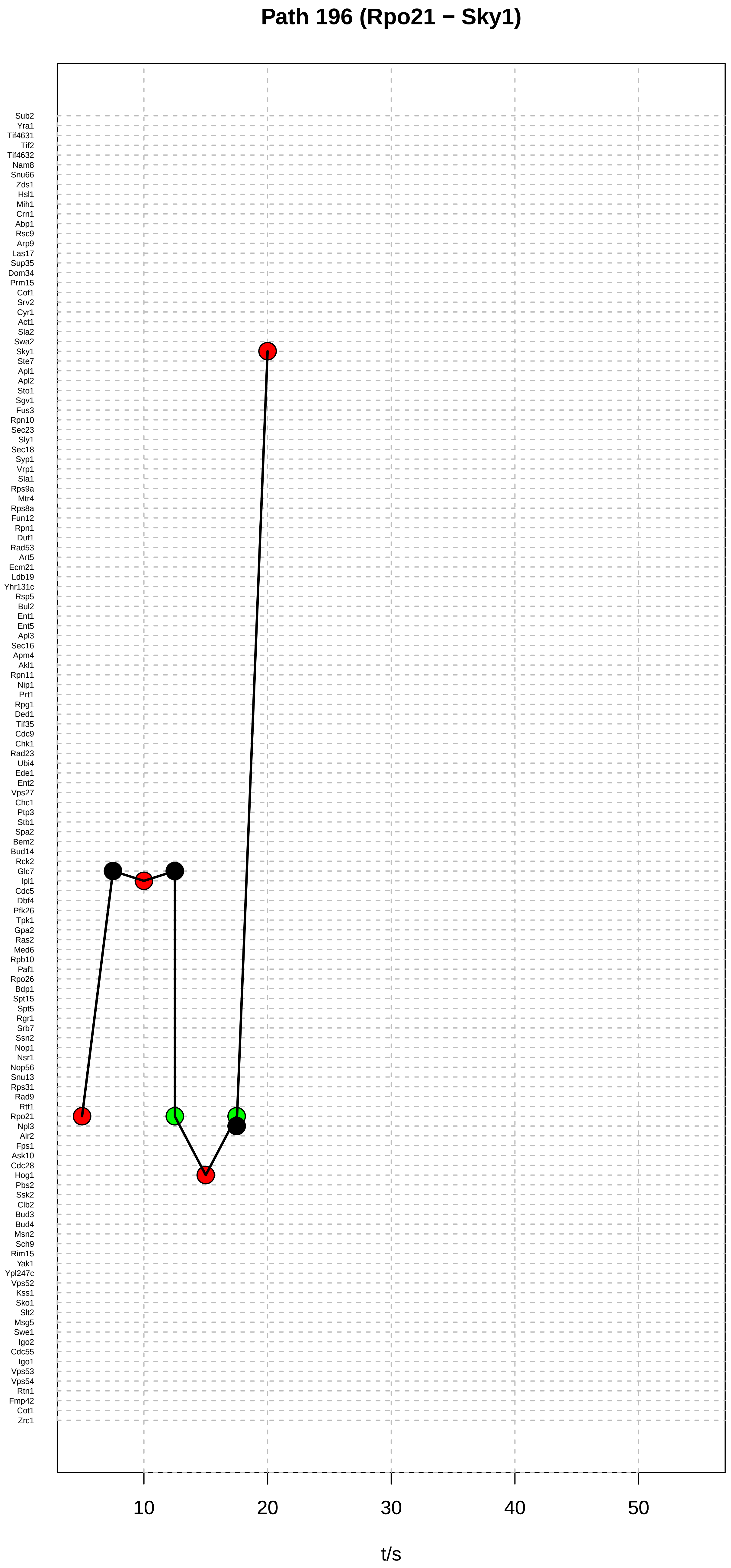

Path 196 (Rpo21 − Sky1)
Sub2
Yra1
Tif4631
Tif2
Tif4632
Nam8
Snu66
Zds1
Hsl1
Mih1
Crn1
Abp1
Rsc9
Arp9
Las17
Sup35
Dom34
Prm15
Cof1
Srv2
Cyr1
Act1
Sla2
Swa2
Sky1
Ste7
Apl1
Apl2
Sto1
Sgv1
Fus3
Rpn10
Sec23
Sly1
Sec18
Syp1
Vrp1
Sla1
Rps9a
Mtr4
Rps8a
Fun12
Rpn1
Duf1
Rad53
Art5
Ecm21
Ldb19
Yhr131c
Rsp5
Bul2
Ent1
Ent5
Apl3
Sec16
Apm4
Akl1
Rpn11
Nip1
Prt1
Rpg1
Ded1
Tif35
Cdc9
Chk1
Rad23
Ubi4
Ede1
Ent2
Vps27
Chc1
Ptp3
Stb1
Spa2
Bem2
Bud14
Rck2
Glc7
Ipl1
Cdc5
Dbf4
Pfk26
Tpk1
Gpa2
Ras2
Med6
Rpb10
Paf1
Rpo26
Bdp1
Spt15
Spt5
Rgr1
Srb7
Ssn2
Nop1
Nsr1
Nop56
Snu13
Rps31
Rad9
Rtf1
Rpo21
Npl3
Air2
Fps1
Ask10
Cdc28
Hog1
Pbs2
Ssk2
Clb2
Bud3
Bud4
Msn2
Sch9
Rim15
Yak1
Ypl247c
Vps52
Kss1
Sko1
Slt2
Msg5
Swe1
Igo2
Cdc55
Igo1
Vps53
Vps54
Rtn1
Fmp42
Cot1
Zrc1
10
20
30
40
50
t/s

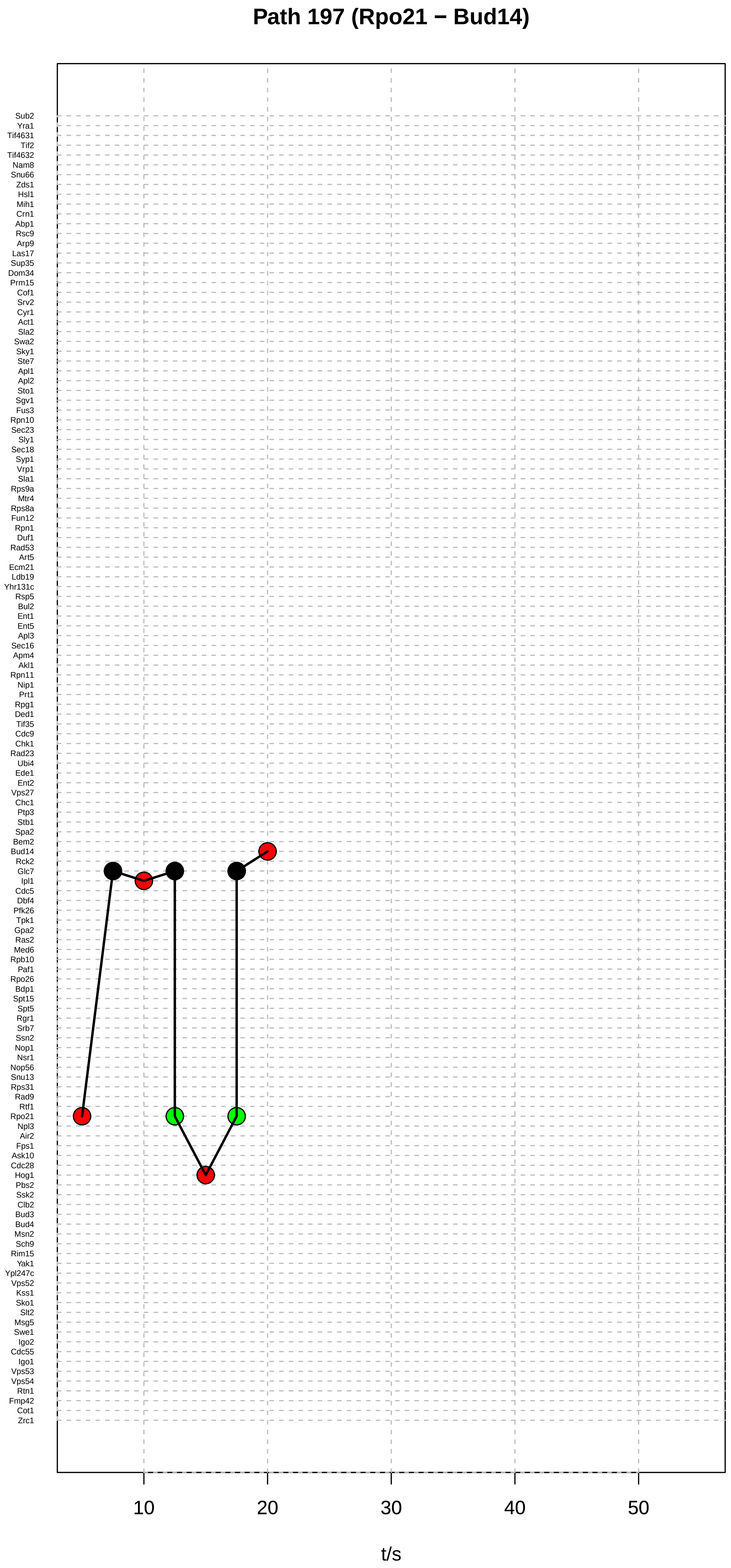

Path 197 (Rpo21 − Bud14)
Sub2
Yra1
Tif4631
Tif2
Tif4632
Nam8
Snu66
Zds1
Hsl1
Mih1
Crn1
Abp1
Rsc9
Arp9
Las17
Sup35
Dom34
Prm15
Cof1
Srv2
Cyr1
Act1
Sla2
Swa2
Sky1
Ste7
Apl1
Apl2
Sto1
Sgv1
Fus3
Rpn10
Sec23
Sly1
Sec18
Syp1
Vrp1
Sla1
Rps9a
Mtr4
Rps8a
Fun12
Rpn1
Duf1
Rad53
Art5
Ecm21
Ldb19
Yhr131c
Rsp5
Bul2
Ent1
Ent5
Apl3
Sec16
Apm4
Akl1
Rpn11
Nip1
Prt1
Rpg1
Ded1
Tif35
Cdc9
Chk1
Rad23
Ubi4
Ede1
Ent2
Vps27
Chc1
Ptp3
Stb1
Spa2
Bem2
Bud14
Rck2
Glc7
Ipl1
Cdc5
Dbf4
Pfk26
Tpk1
Gpa2
Ras2
Med6
Rpb10
Paf1
Rpo26
Bdp1
Spt15
Spt5
Rgr1
Srb7
Ssn2
Nop1
Nsr1
Nop56
Snu13
Rps31
Rad9
Rtf1
Rpo21
Npl3
Air2
Fps1
Ask10
Cdc28
Hog1
Pbs2
Ssk2
Clb2
Bud3
Bud4
Msn2
Sch9
Rim15
Yak1
Ypl247c
Vps52
Kss1
Sko1
Slt2
Msg5
Swe1
Igo2
Cdc55
Igo1
Vps53
Vps54
Rtn1
Fmp42
Cot1
Zrc1
10
20
30
40
50
t/s

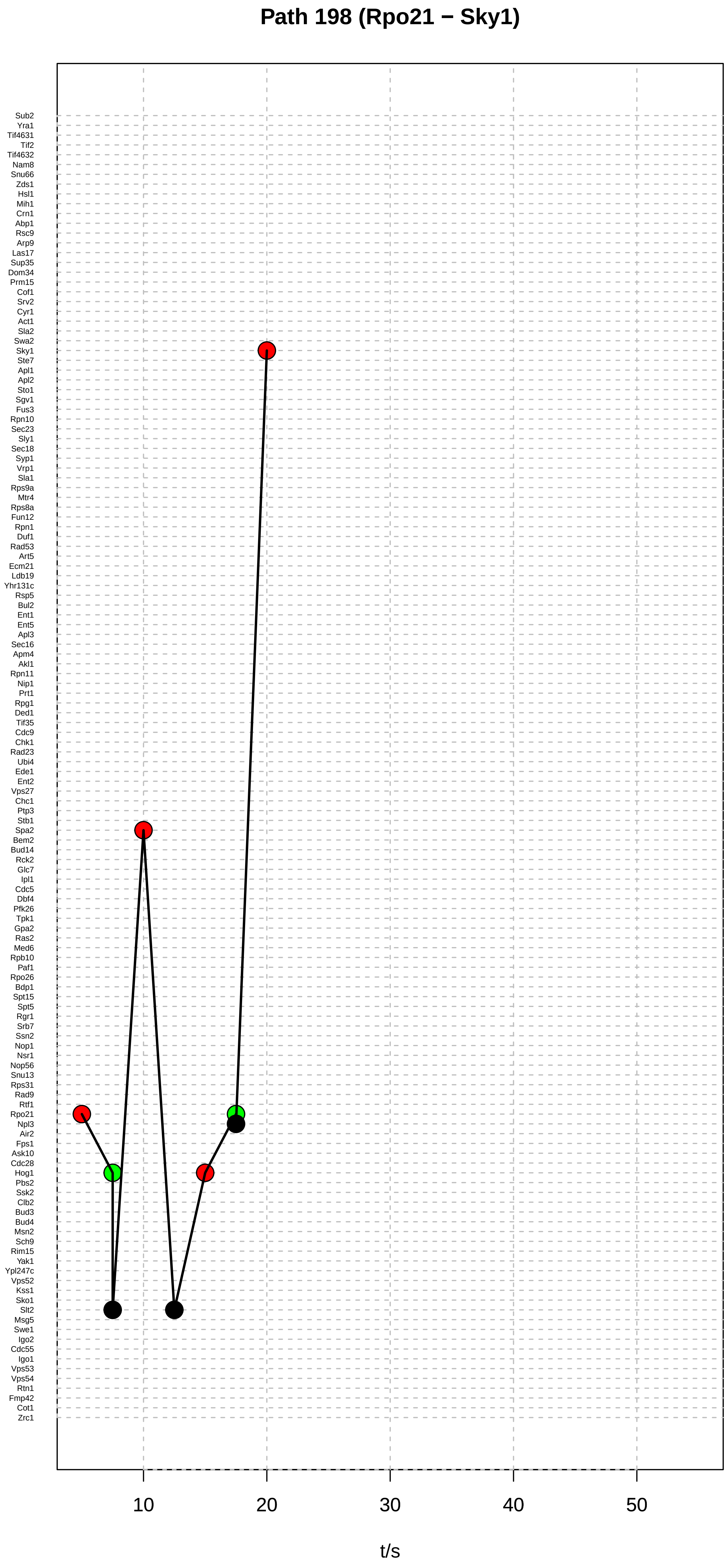
Path 198 (Rpo21 − Sky1)
Sub2
Yra1
Tif4631
Tif2
Tif4632
Nam8
Snu66
Zds1
Hsl1
Mih1
Crn1
Abp1
Rsc9
Arp9
Las17
Sup35
Dom34
Prm15
Cof1
Srv2
Cyr1
Act1
Sla2
Swa2
Sky1
Ste7
Apl1
Apl2
Sto1
Sgv1
Fus3
Rpn10
Sec23
Sly1
Sec18
Syp1
Vrp1
Sla1
Rps9a
Mtr4
Rps8a
Fun12
Rpn1
Duf1
Rad53
Art5
Ecm21
Ldb19
Yhr131c
Rsp5
Bul2
Ent1
Ent5
Apl3
Sec16
Apm4
Akl1
Rpn11
Nip1
Prt1
Rpg1
Ded1
Tif35
Cdc9
Chk1
Rad23
Ubi4
Ede1
Ent2
Vps27
Chc1
Ptp3
Stb1
Spa2
Bem2
Bud14
Rck2
Glc7
Ipl1
Cdc5
Dbf4
Pfk26
Tpk1
Gpa2
Ras2
Med6
Rpb10
Paf1
Rpo26
Bdp1
Spt15
Spt5
Rgr1
Srb7
Ssn2
Nop1
Nsr1
Nop56
Snu13
Rps31
Rad9
Rtf1
Rpo21
Npl3
Air2
Fps1
Ask10
Cdc28
Hog1
Pbs2
Ssk2
Clb2
Bud3
Bud4
Msn2
Sch9
Rim15
Yak1
Ypl247c
Vps52
Kss1
Sko1
Slt2
Msg5
Swe1
Igo2
Cdc55
Igo1
Vps53
Vps54
Rtn1
Fmp42
Cot1
Zrc1
10
20
30
40
50
t/s

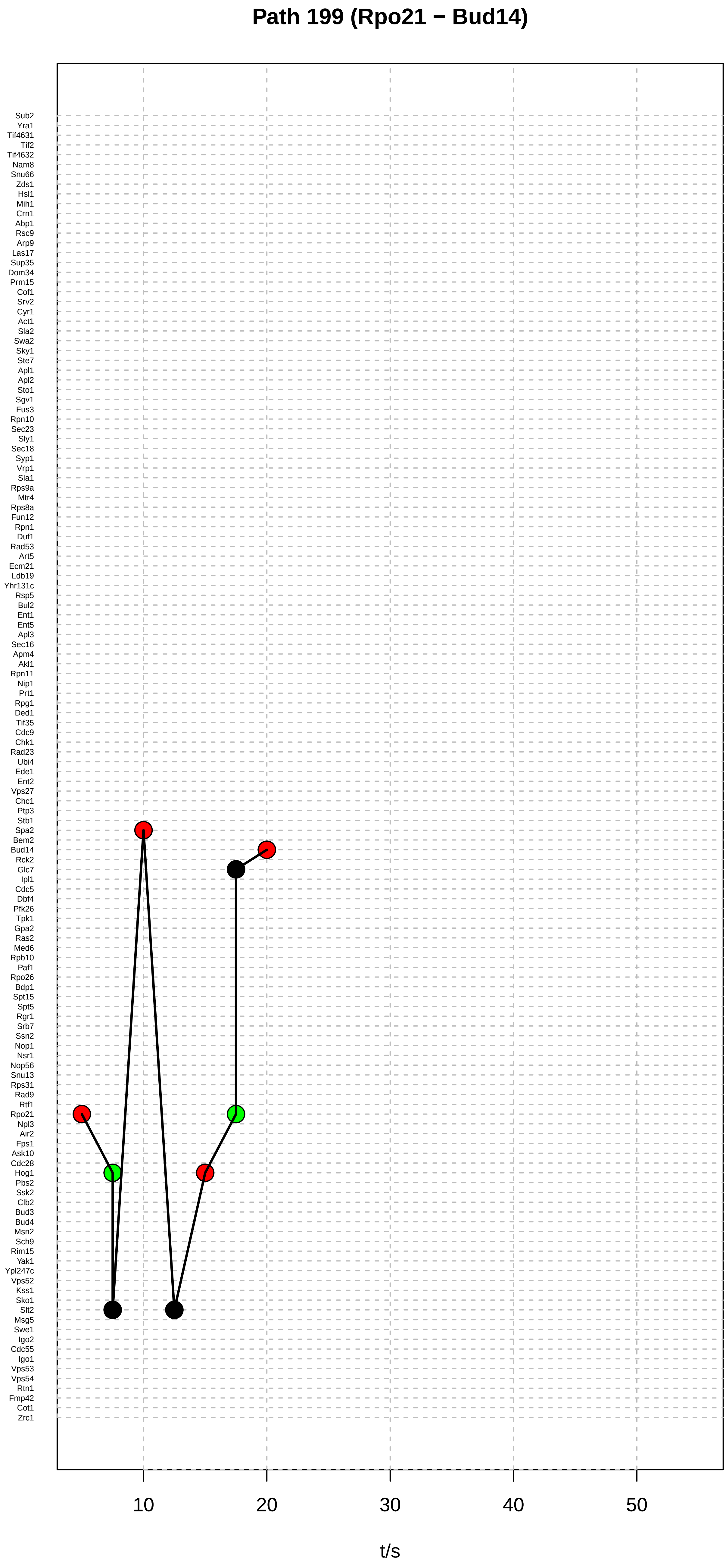

Path 199 (Rpo21 − Bud14)
Sub2
Yra1
Tif4631
Tif2
Tif4632
Nam8
Snu66
Zds1
Hsl1
Mih1
Crn1
Abp1
Rsc9
Arp9
Las17
Sup35
Dom34
Prm15
Cof1
Srv2
Cyr1
Act1
Sla2
Swa2
Sky1
Ste7
Apl1
Apl2
Sto1
Sgv1
Fus3
Rpn10
Sec23
Sly1
Sec18
Syp1
Vrp1
Sla1
Rps9a
Mtr4
Rps8a
Fun12
Rpn1
Duf1
Rad53
Art5
Ecm21
Ldb19
Yhr131c
Rsp5
Bul2
Ent1
Ent5
Apl3
Sec16
Apm4
Akl1
Rpn11
Nip1
Prt1
Rpg1
Ded1
Tif35
Cdc9
Chk1
Rad23
Ubi4
Ede1
Ent2
Vps27
Chc1
Ptp3
Stb1
Spa2
Bem2
Bud14
Rck2
Glc7
Ipl1
Cdc5
Dbf4
Pfk26
Tpk1
Gpa2
Ras2
Med6
Rpb10
Paf1
Rpo26
Bdp1
Spt15
Spt5
Rgr1
Srb7
Ssn2
Nop1
Nsr1
Nop56
Snu13
Rps31
Rad9
Rtf1
Rpo21
Npl3
Air2
Fps1
Ask10
Cdc28
Hog1
Pbs2
Ssk2
Clb2
Bud3
Bud4
Msn2
Sch9
Rim15
Yak1
Ypl247c
Vps52
Kss1
Sko1
Slt2
Msg5
Swe1
Igo2
Cdc55
Igo1
Vps53
Vps54
Rtn1
Fmp42
Cot1
Zrc1
10
20
30
40
50
t/s

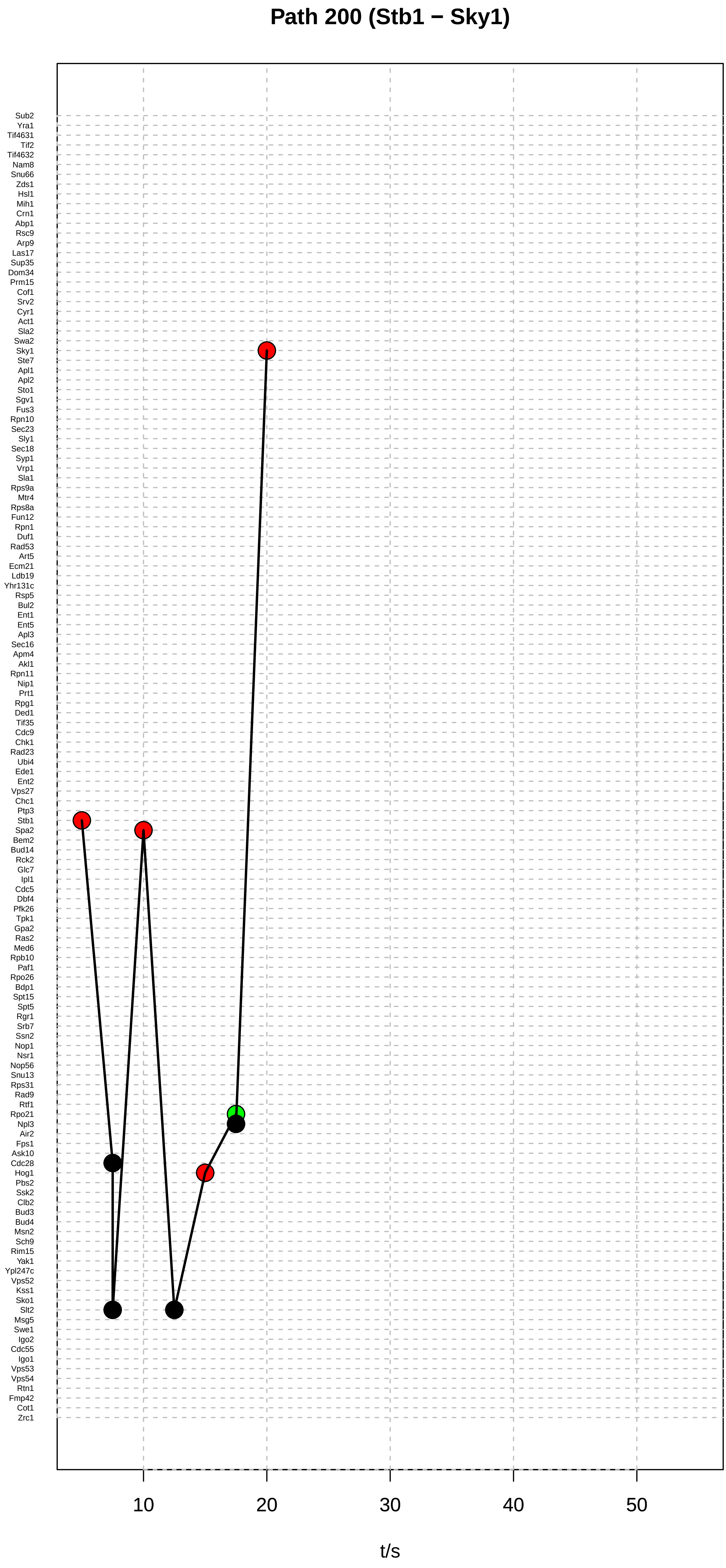

Path 200 (Stb1 − Sky1)
Sub2
Yra1
Tif4631
Tif2
Tif4632
Nam8
Snu66
Zds1
Hsl1
Mih1
Crn1
Abp1
Rsc9
Arp9
Las17
Sup35
Dom34
Prm15
Cof1
Srv2
Cyr1
Act1
Sla2
Swa2
Sky1
Ste7
Apl1
Apl2
Sto1
Sgv1
Fus3
Rpn10
Sec23
Sly1
Sec18
Syp1
Vrp1
Sla1
Rps9a
Mtr4
Rps8a
Fun12
Rpn1
Duf1
Rad53
Art5
Ecm21
Ldb19
Yhr131c
Rsp5
Bul2
Ent1
Ent5
Apl3
Sec16
Apm4
Akl1
Rpn11
Nip1
Prt1
Rpg1
Ded1
Tif35
Cdc9
Chk1
Rad23
Ubi4
Ede1
Ent2
Vps27
Chc1
Ptp3
Stb1
Spa2
Bem2
Bud14
Rck2
Glc7
Ipl1
Cdc5
Dbf4
Pfk26
Tpk1
Gpa2
Ras2
Med6
Rpb10
Paf1
Rpo26
Bdp1
Spt15
Spt5
Rgr1
Srb7
Ssn2
Nop1
Nsr1
Nop56
Snu13
Rps31
Rad9
Rtf1
Rpo21
Npl3
Air2
Fps1
Ask10
Cdc28
Hog1
Pbs2
Ssk2
Clb2
Bud3
Bud4
Msn2
Sch9
Rim15
Yak1
Ypl247c
Vps52
Kss1
Sko1
Slt2
Msg5
Swe1
Igo2
Cdc55
Igo1
Vps53
Vps54
Rtn1
Fmp42
Cot1
Zrc1
10
20
30
40
50
t/s

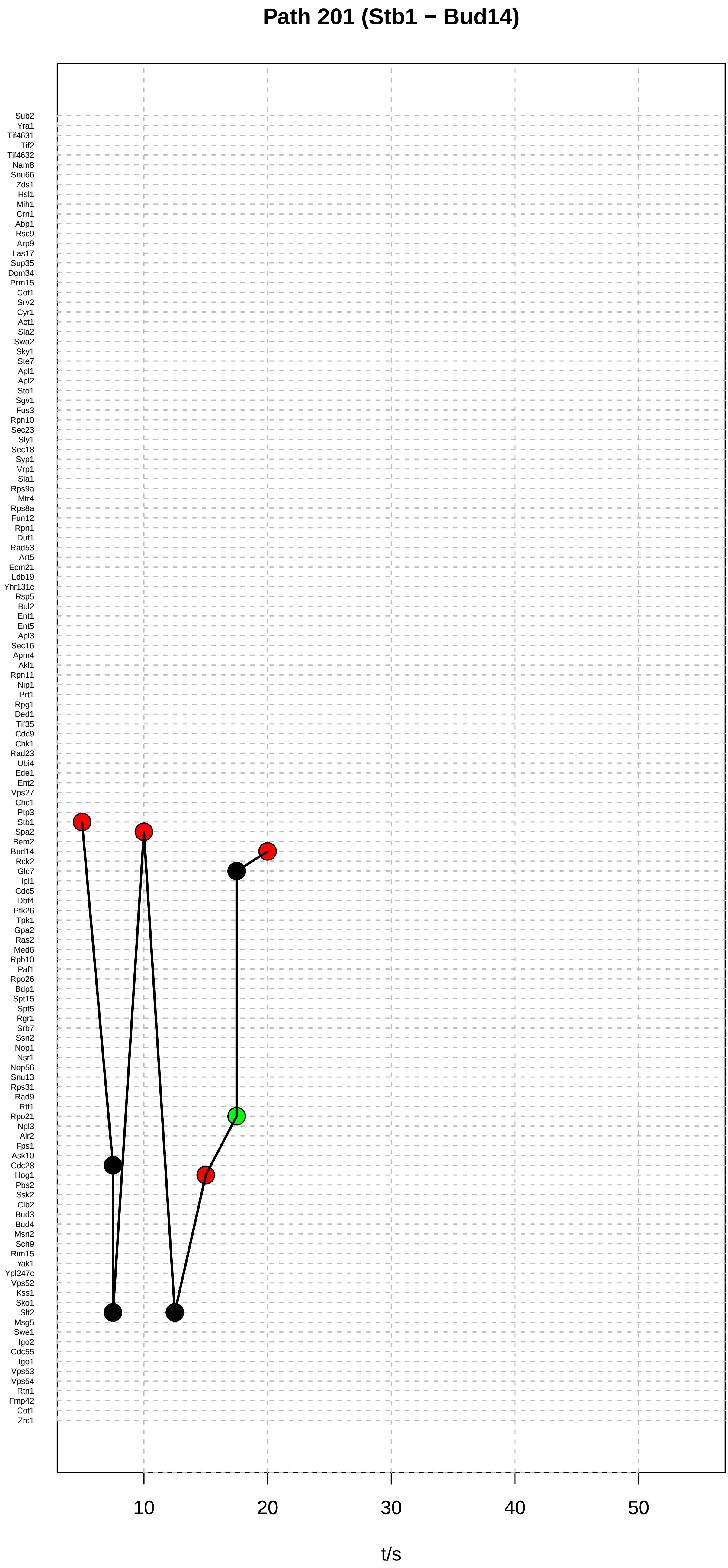

Path 201 (Stb1 − Bud14)
Sub2
Yra1
Tif4631
Tif2
Tif4632
Nam8
Snu66
Zds1
Hsl1
Mih1
Crn1
Abp1
Rsc9
Arp9
Las17
Sup35
Dom34
Prm15
Cof1
Srv2
Cyr1
Act1
Sla2
Swa2
Sky1
Ste7
Apl1
Apl2
Sto1
Sgv1
Fus3
Rpn10
Sec23
Sly1
Sec18
Syp1
Vrp1
Sla1
Rps9a
Mtr4
Rps8a
Fun12
Rpn1
Duf1
Rad53
Art5
Ecm21
Ldb19
Yhr131c
Rsp5
Bul2
Ent1
Ent5
Apl3
Sec16
Apm4
Akl1
Rpn11
Nip1
Prt1
Rpg1
Ded1
Tif35
Cdc9
Chk1
Rad23
Ubi4
Ede1
Ent2
Vps27
Chc1
Ptp3
Stb1
Spa2
Bem2
Bud14
Rck2
Glc7
Ipl1
Cdc5
Dbf4
Pfk26
Tpk1
Gpa2
Ras2
Med6
Rpb10
Paf1
Rpo26
Bdp1
Spt15
Spt5
Rgr1
Srb7
Ssn2
Nop1
Nsr1
Nop56
Snu13
Rps31
Rad9
Rtf1
Rpo21
Npl3
Air2
Fps1
Ask10
Cdc28
Hog1
Pbs2
Ssk2
Clb2
Bud3
Bud4
Msn2
Sch9
Rim15
Yak1
Ypl247c
Vps52
Kss1
Sko1
Slt2
Msg5
Swe1
Igo2
Cdc55
Igo1
Vps53
Vps54
Rtn1
Fmp42
Cot1
Zrc1
10
20
30
40
50
t/s

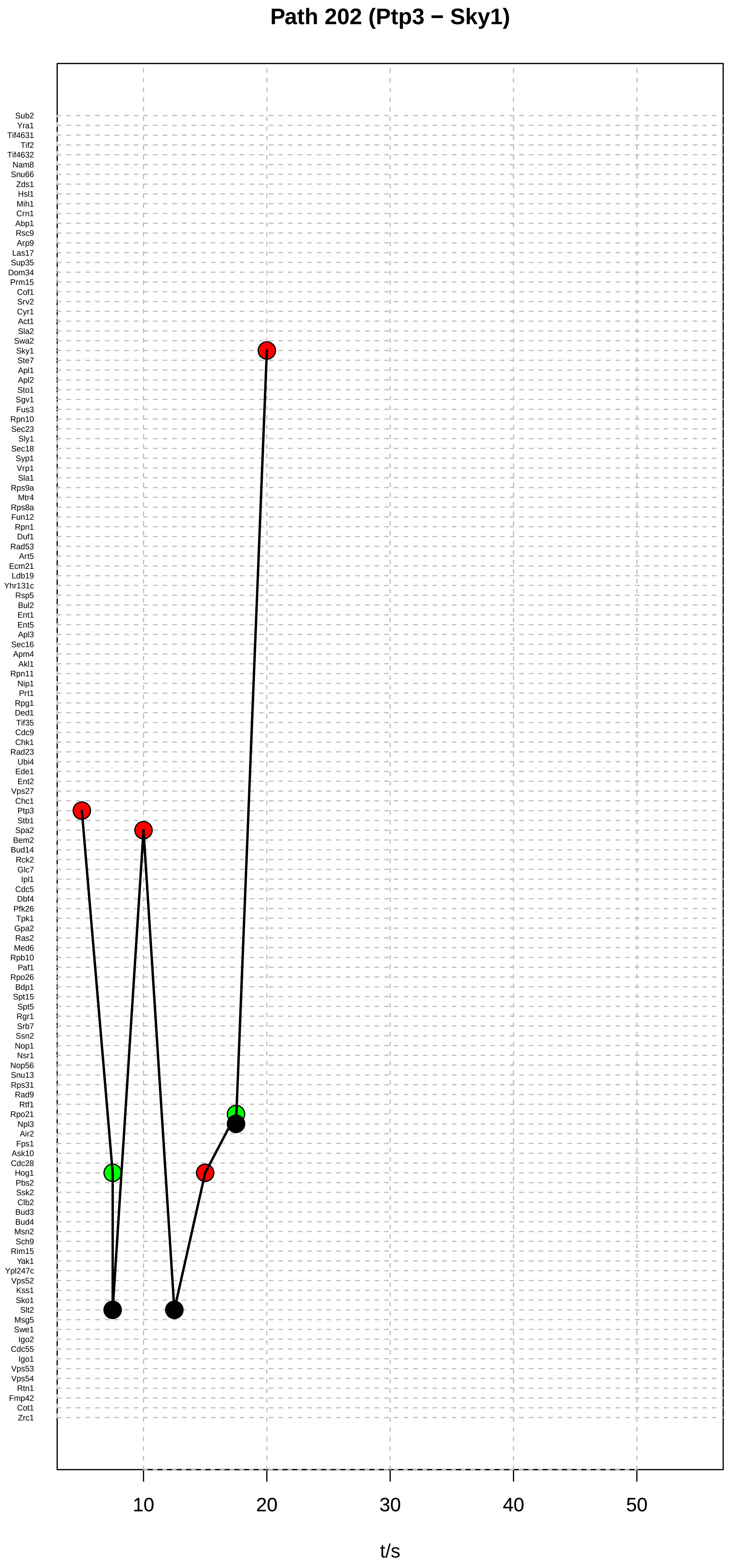

Path 202 (Ptp3 − Sky1)
Sub2
Yra1
Tif4631
Tif2
Tif4632
Nam8
Snu66
Zds1
Hsl1
Mih1
Crn1
Abp1
Rsc9
Arp9
Las17
Sup35
Dom34
Prm15
Cof1
Srv2
Cyr1
Act1
Sla2
Swa2
Sky1
Ste7
Apl1
Apl2
Sto1
Sgv1
Fus3
Rpn10
Sec23
Sly1
Sec18
Syp1
Vrp1
Sla1
Rps9a
Mtr4
Rps8a
Fun12
Rpn1
Duf1
Rad53
Art5
Ecm21
Ldb19
Yhr131c
Rsp5
Bul2
Ent1
Ent5
Apl3
Sec16
Apm4
Akl1
Rpn11
Nip1
Prt1
Rpg1
Ded1
Tif35
Cdc9
Chk1
Rad23
Ubi4
Ede1
Ent2
Vps27
Chc1
Ptp3
Stb1
Spa2
Bem2
Bud14
Rck2
Glc7
Ipl1
Cdc5
Dbf4
Pfk26
Tpk1
Gpa2
Ras2
Med6
Rpb10
Paf1
Rpo26
Bdp1
Spt15
Spt5
Rgr1
Srb7
Ssn2
Nop1
Nsr1
Nop56
Snu13
Rps31
Rad9
Rtf1
Rpo21
Npl3
Air2
Fps1
Ask10
Cdc28
Hog1
Pbs2
Ssk2
Clb2
Bud3
Bud4
Msn2
Sch9
Rim15
Yak1
Ypl247c
Vps52
Kss1
Sko1
Slt2
Msg5
Swe1
Igo2
Cdc55
Igo1
Vps53
Vps54
Rtn1
Fmp42
Cot1
Zrc1
10
20
30
40
50
t/s

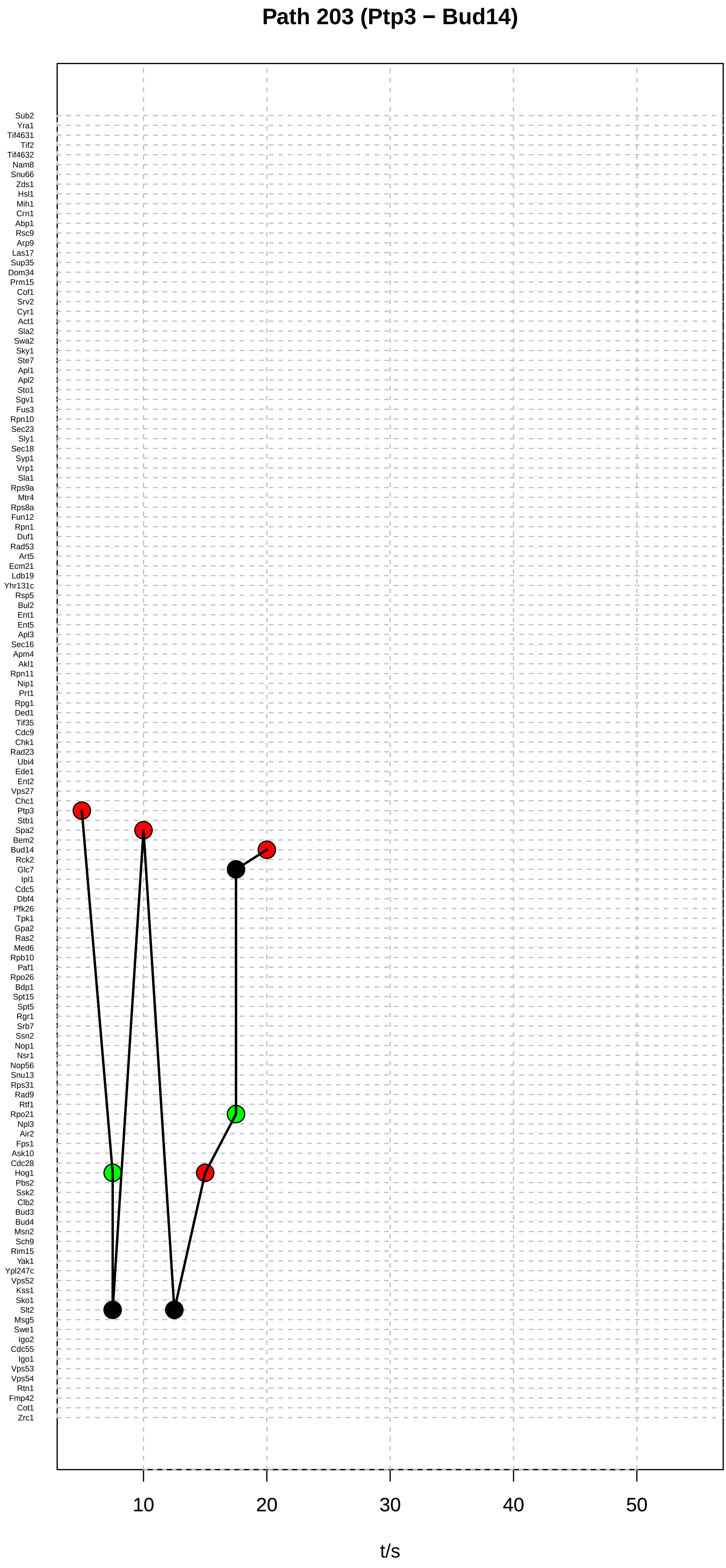
Path 203 (Ptp3 − Bud14)
Sub2
Yra1
Tif4631
Tif2
Tif4632
Nam8
Snu66
Zds1
Hsl1
Mih1
Crn1
Abp1
Rsc9
Arp9
Las17
Sup35
Dom34
Prm15
Cof1
Srv2
Cyr1
Act1
Sla2
Swa2
Sky1
Ste7
Apl1
Apl2
Sto1
Sgv1
Fus3
Rpn10
Sec23
Sly1
Sec18
Syp1
Vrp1
Sla1
Rps9a
Mtr4
Rps8a
Fun12
Rpn1
Duf1
Rad53
Art5
Ecm21
Ldb19
Yhr131c
Rsp5
Bul2
Ent1
Ent5
Apl3
Sec16
Apm4
Akl1
Rpn11
Nip1
Prt1
Rpg1
Ded1
Tif35
Cdc9
Chk1
Rad23
Ubi4
Ede1
Ent2
Vps27
Chc1
Ptp3
Stb1
Spa2
Bem2
Bud14
Rck2
Glc7
Ipl1
Cdc5
Dbf4
Pfk26
Tpk1
Gpa2
Ras2
Med6
Rpb10
Paf1
Rpo26
Bdp1
Spt15
Spt5
Rgr1
Srb7
Ssn2
Nop1
Nsr1
Nop56
Snu13
Rps31
Rad9
Rtf1
Rpo21
Npl3
Air2
Fps1
Ask10
Cdc28
Hog1
Pbs2
Ssk2
Clb2
Bud3
Bud4
Msn2
Sch9
Rim15
Yak1
Ypl247c
Vps52
Kss1
Sko1
Slt2
Msg5
Swe1
Igo2
Cdc55
Igo1
Vps53
Vps54
Rtn1
Fmp42
Cot1
Zrc1
10
20
30
40
50
t/s

### Path 204 (Syp1 − Ede1)

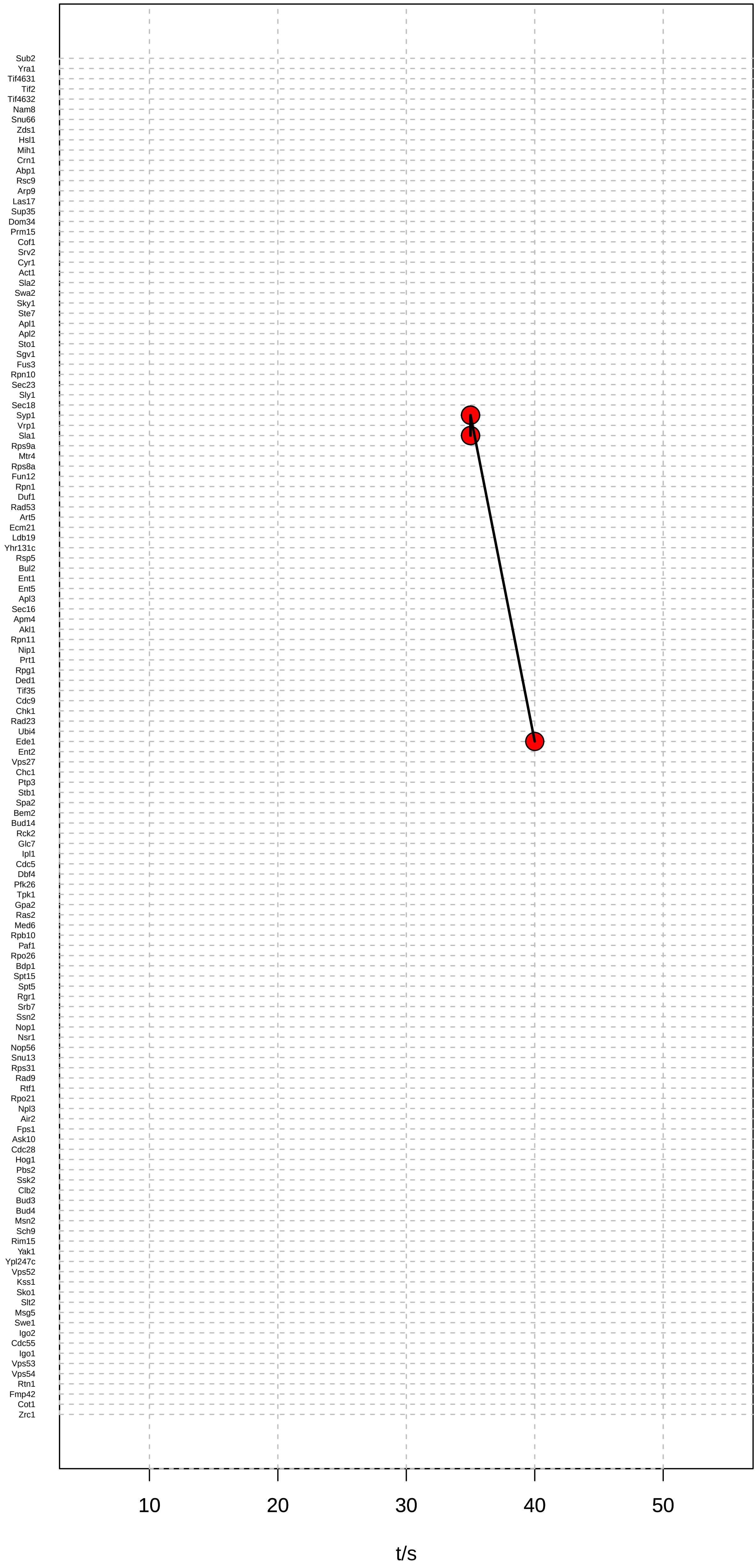

### Path 205 (Syp1 − Ede1)

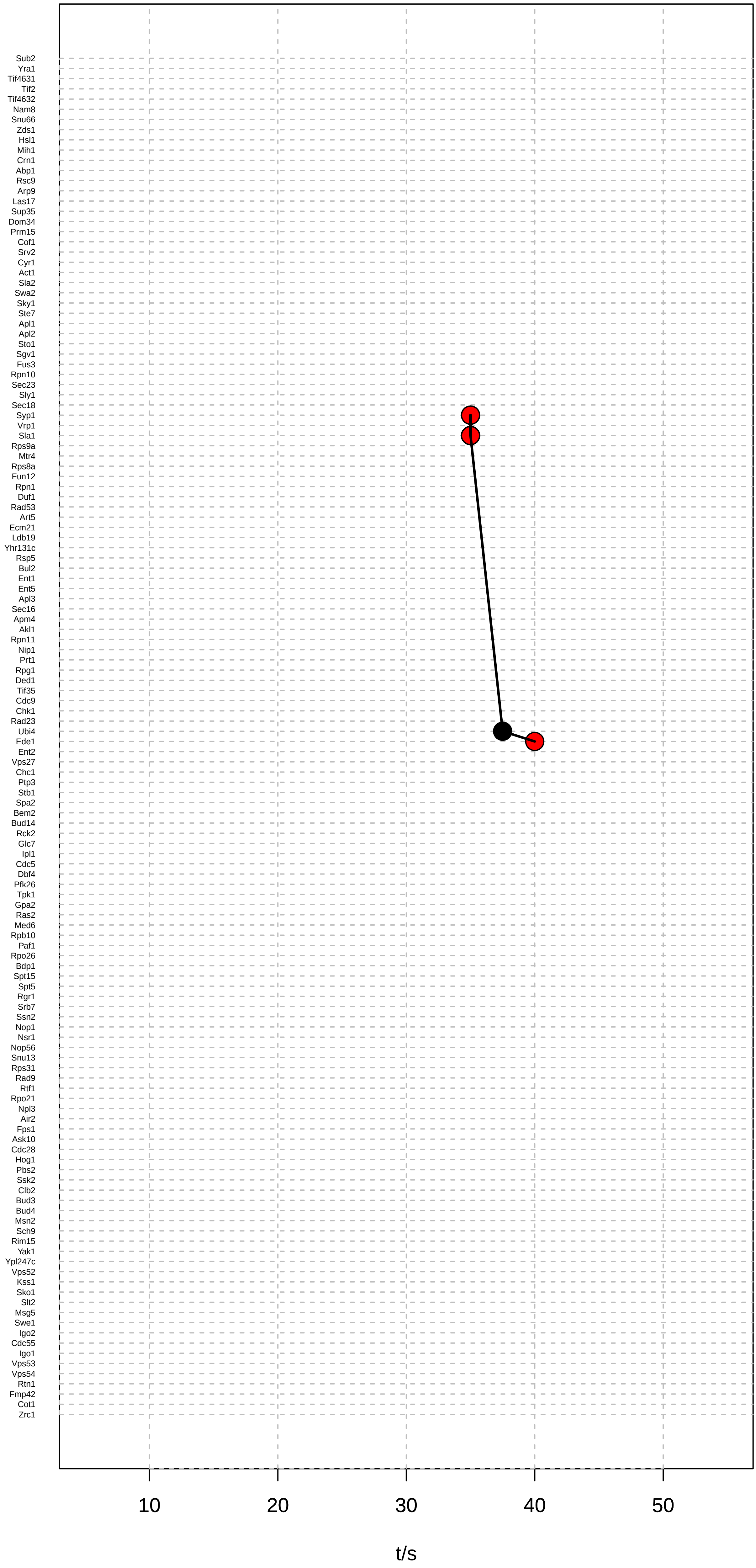

#### Path 206 (Syp1 − Duf1)

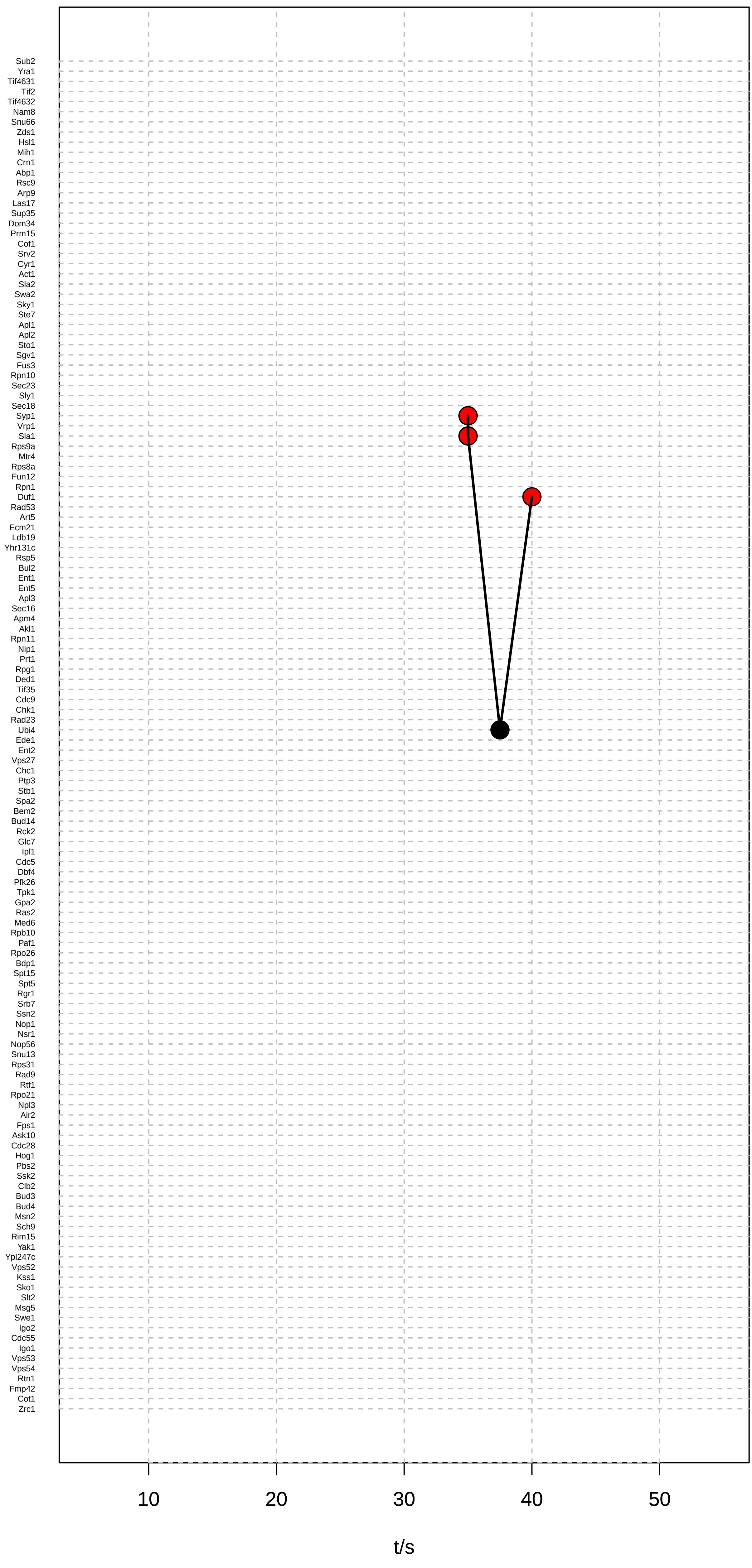

### Path 207 (Syp1 − Fun12)

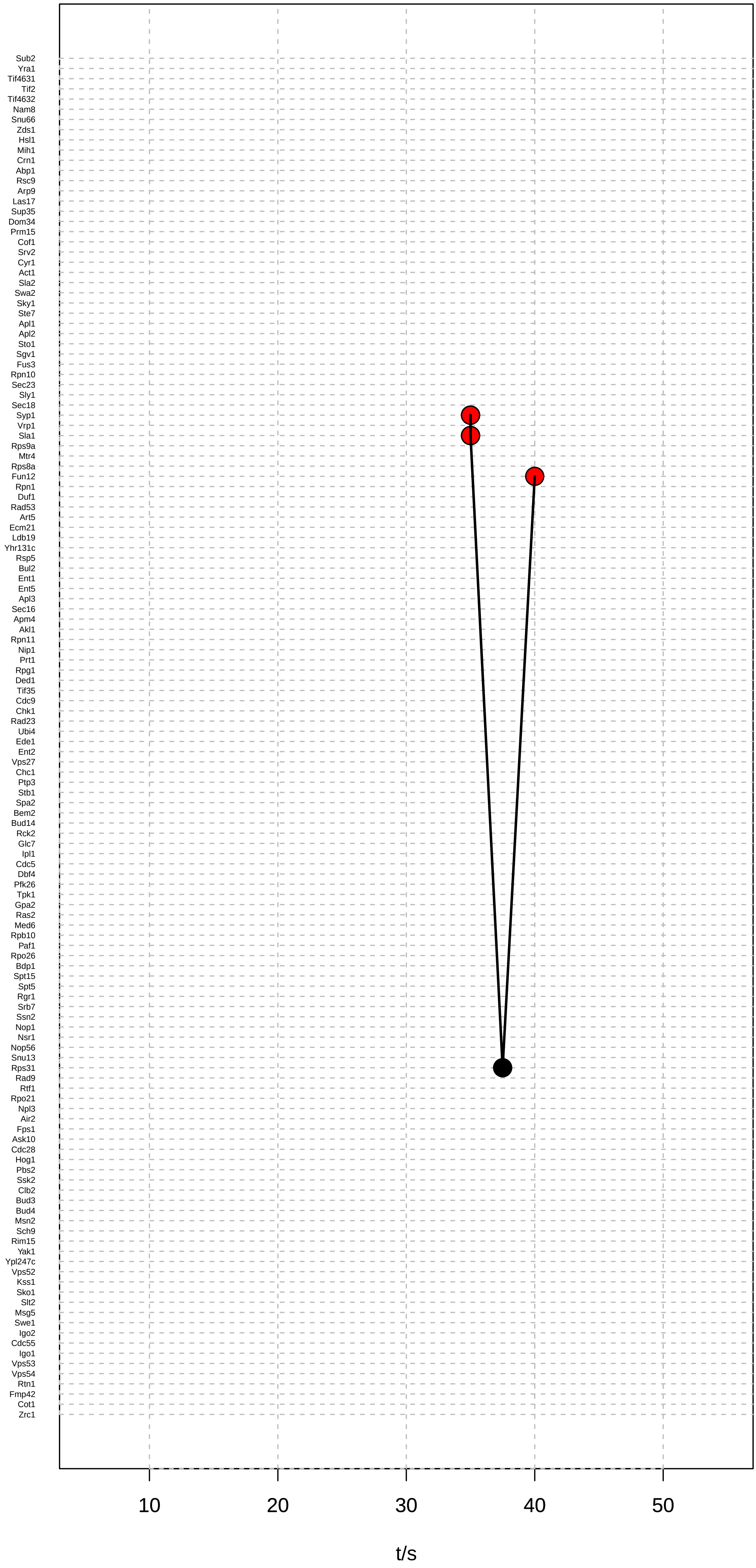

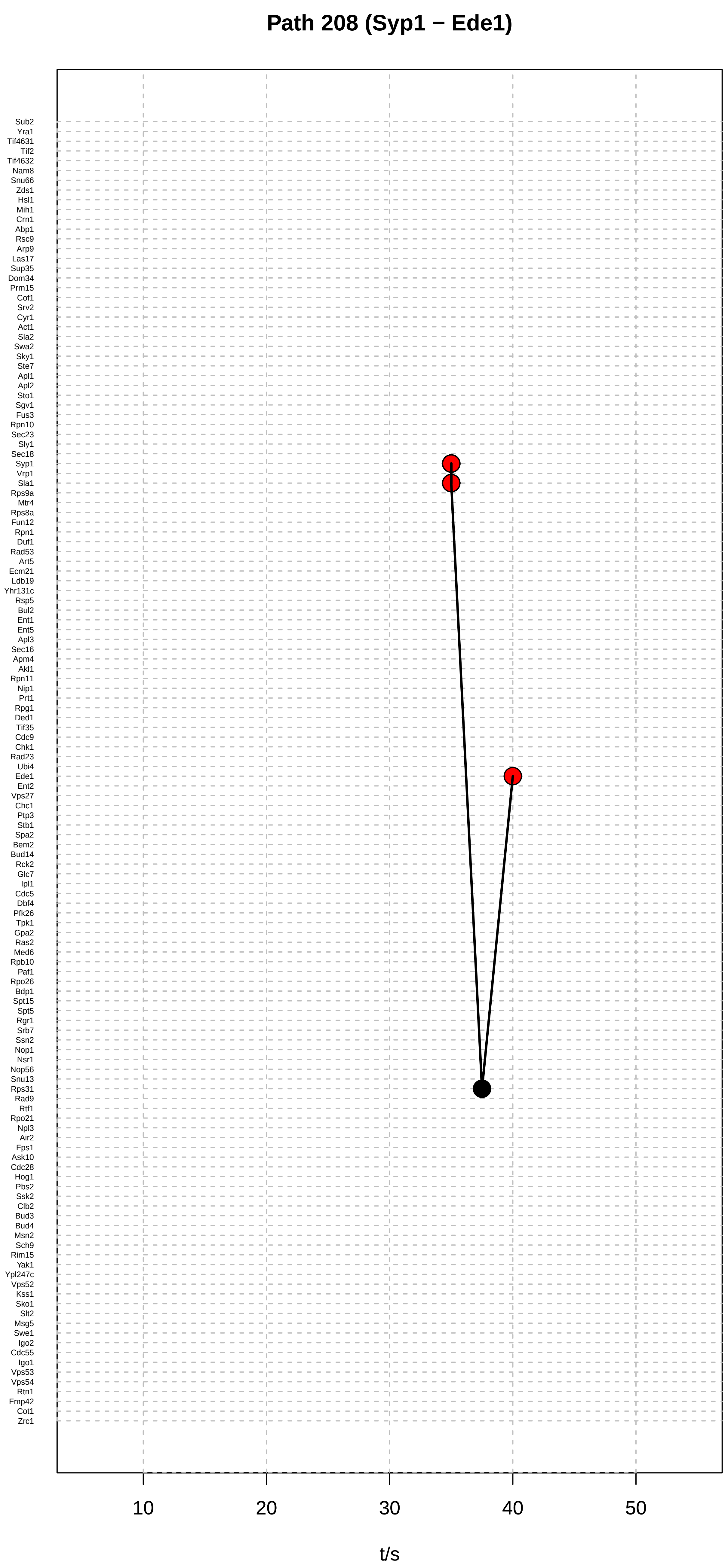

Path 208 (Syp1 − Ede1)
Sub2
Yra1
Tif4631
Tif2
Tif4632
Nam8
Snu66
Zds1
Hsl1
Mih1
Crn1
Abp1
Rsc9
Arp9
Las17
Sup35
Dom34
Prm15
Cof1
Srv2
Cyr1
Act1
Sla2
Swa2
Sky1
Ste7
Apl1
Apl2
Sto1
Sgv1
Fus3
Rpn10
Sec23
Sly1
Sec18
Syp1
Vrp1
Sla1
Rps9a
Mtr4
Rps8a
Fun12
Rpn1
Duf1
Rad53
Art5
Ecm21
Ldb19
Yhr131c
Rsp5
Bul2
Ent1
Ent5
Apl3
Sec16
Apm4
Akl1
Rpn11
Nip1
Prt1
Rpg1
Ded1
Tif35
Cdc9
Chk1
Rad23
Ubi4
Ede1
Ent2
Vps27
Chc1
Ptp3
Stb1
Spa2
Bem2
Bud14
Rck2
Glc7
Ipl1
Cdc5
Dbf4
Pfk26
Tpk1
Gpa2
Ras2
Med6
Rpb10
Paf1
Rpo26
Bdp1
Spt15
Spt5
Rgr1
Srb7
Ssn2
Nop1
Nsr1
Nop56
Snu13
Rps31
Rad9
Rtf1
Rpo21
Npl3
Air2
Fps1
Ask10
Cdc28
Hog1
Pbs2
Ssk2
Clb2
Bud3
Bud4
Msn2
Sch9
Rim15
Yak1
Ypl247c
Vps52
Kss1
Sko1
Slt2
Msg5
Swe1
Igo2
Cdc55
Igo1
Vps53
Vps54
Rtn1
Fmp42
Cot1
Zrc1
10
20
30
40
50
t/s

#### Path 209 (Syp1 − Rad23)

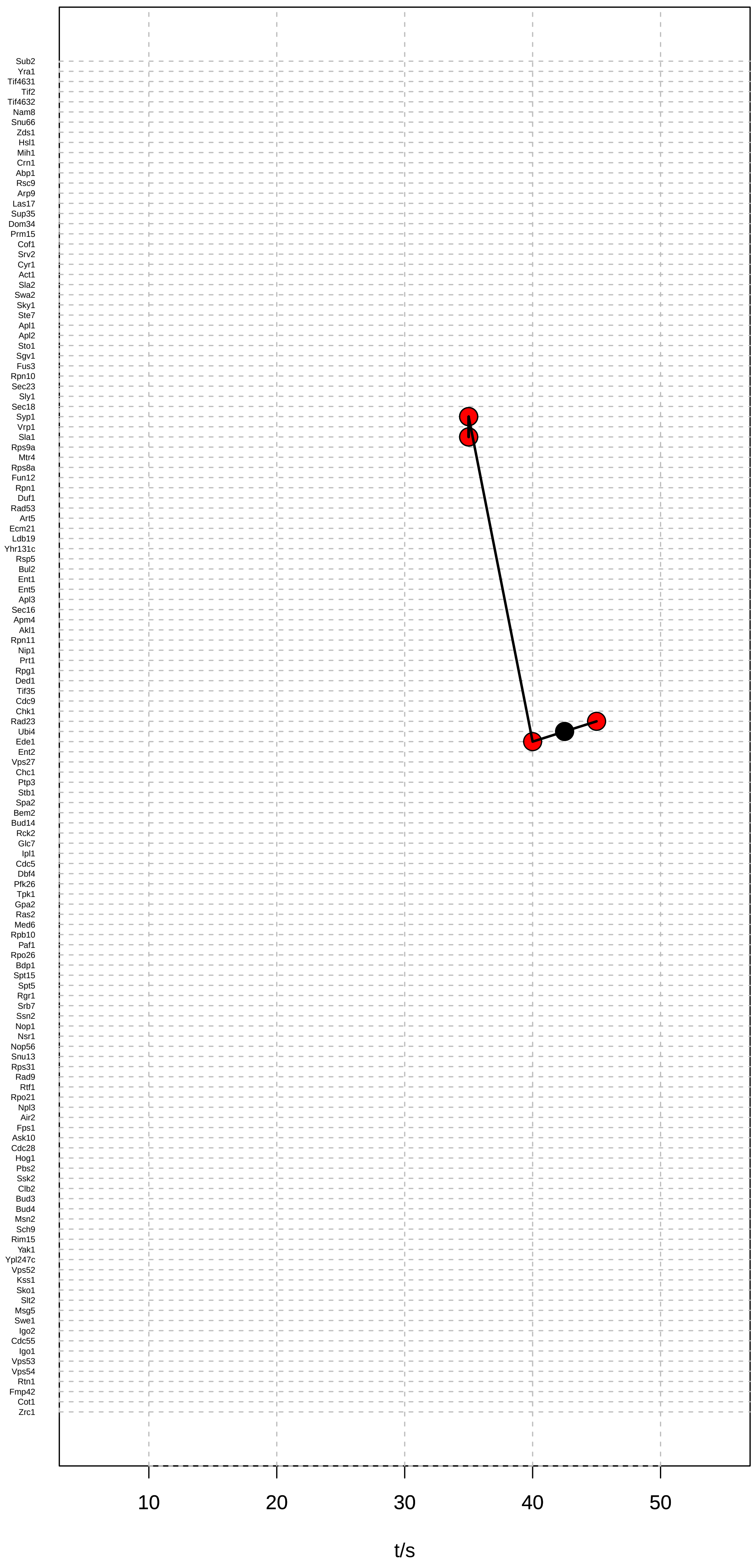

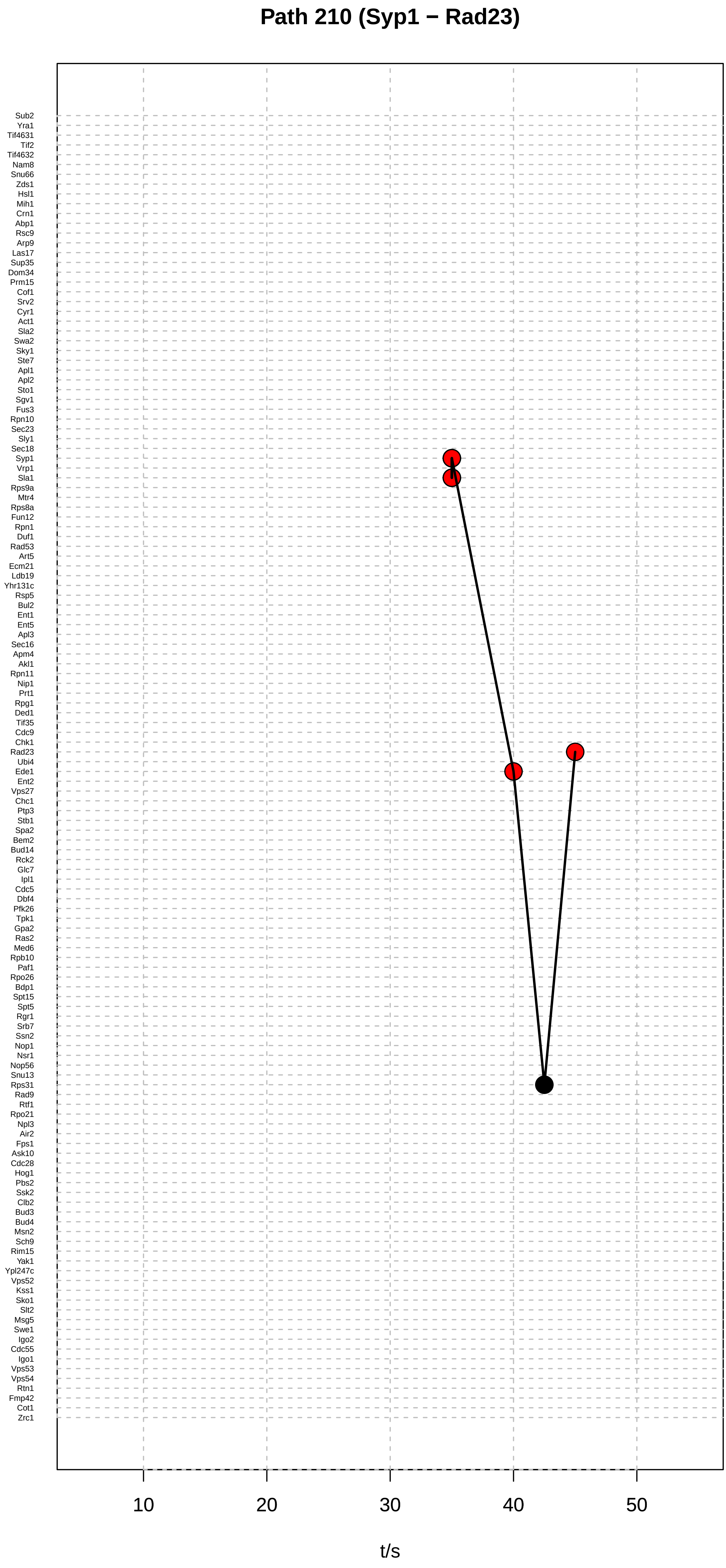
Path 210 (Syp1 − Rad23)
Sub2
Yra1
Tif4631
Tif2
Tif4632
Nam8
Snu66
Zds1
Hsl1
Mih1
Crn1
Abp1
Rsc9
Arp9
Las17
Sup35
Dom34
Prm15
Cof1
Srv2
Cyr1
Act1
Sla2
Swa2
Sky1
Ste7
Apl1
Apl2
Sto1
Sgv1
Fus3
Rpn10
Sec23
Sly1
Sec18
Syp1
Vrp1
Sla1
Rps9a
Mtr4
Rps8a
Fun12
Rpn1
Duf1
Rad53
Art5
Ecm21
Ldb19
Yhr131c
Rsp5
Bul2
Ent1
Ent5
Apl3
Sec16
Apm4
Akl1
Rpn11
Nip1
Prt1
Rpg1
Ded1
Tif35
Cdc9
Chk1
Rad23
Ubi4
Ede1
Ent2
Vps27
Chc1
Ptp3
Stb1
Spa2
Bem2
Bud14
Rck2
Glc7
Ipl1
Cdc5
Dbf4
Pfk26
Tpk1
Gpa2
Ras2
Med6
Rpb10
Paf1
Rpo26
Bdp1
Spt15
Spt5
Rgr1
Srb7
Ssn2
Nop1
Nsr1
Nop56
Snu13
Rps31
Rad9
Rtf1
Rpo21
Npl3
Air2
Fps1
Ask10
Cdc28
Hog1
Pbs2
Ssk2
Clb2
Bud3
Bud4
Msn2
Sch9
Rim15
Yak1
Ypl247c
Vps52
Kss1
Sko1
Slt2
Msg5
Swe1
Igo2
Cdc55
Igo1
Vps53
Vps54
Rtn1
Fmp42
Cot1
Zrc1
10
20
30
40
50
t/s

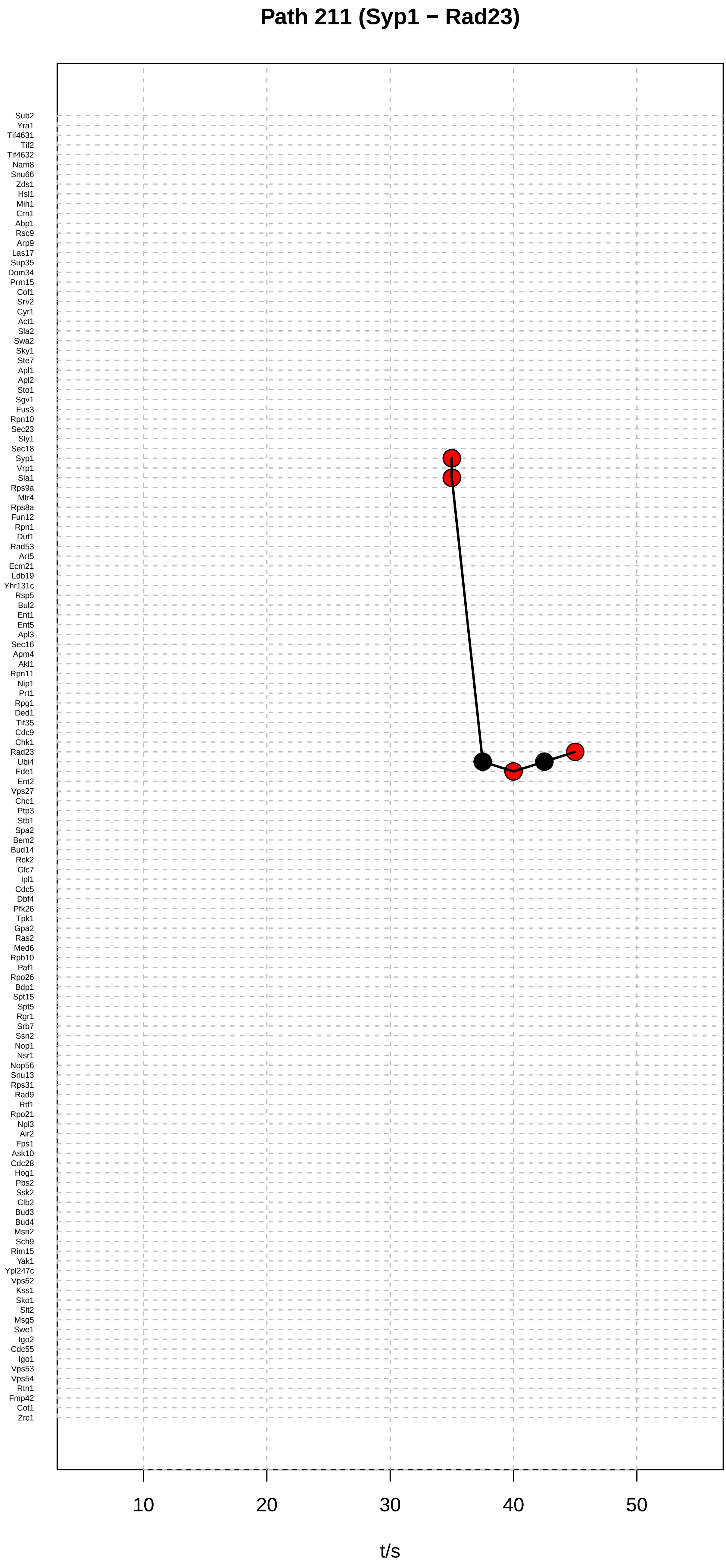
Path 211 (Syp1 − Rad23)
Sub2
Yra1
Tif4631
Tif2
Tif4632
Nam8
Snu66
Zds1
Hsl1
Mih1
Crn1
Abp1
Rsc9
Arp9
Las17
Sup35
Dom34
Prm15
Cof1
Srv2
Cyr1
Act1
Sla2
Swa2
Sky1
Ste7
Apl1
Apl2
Sto1
Sgv1
Fus3
Rpn10
Sec23
Sly1
Sec18
Syp1
Vrp1
Sla1
Rps9a
Mtr4
Rps8a
Fun12
Rpn1
Duf1
Rad53
Art5
Ecm21
Ldb19
Yhr131c
Rsp5
Bul2
Ent1
Ent5
Apl3
Sec16
Apm4
Akl1
Rpn11
Nip1
Prt1
Rpg1
Ded1
Tif35
Cdc9
Chk1
Rad23
Ubi4
Ede1
Ent2
Vps27
Chc1
Ptp3
Stb1
Spa2
Bem2
Bud14
Rck2
Glc7
Ipl1
Cdc5
Dbf4
Pfk26
Tpk1
Gpa2
Ras2
Med6
Rpb10
Paf1
Rpo26
Bdp1
Spt15
Spt5
Rgr1
Srb7
Ssn2
Nop1
Nsr1
Nop56
Snu13
Rps31
Rad9
Rtf1
Rpo21
Npl3
Air2
Fps1
Ask10
Cdc28
Hog1
Pbs2
Ssk2
Clb2
Bud3
Bud4
Msn2
Sch9
Rim15
Yak1
Ypl247c
Vps52
Kss1
Sko1
Slt2
Msg5
Swe1
Igo2
Cdc55
Igo1
Vps53
Vps54
Rtn1
Fmp42
Cot1
Zrc1
10
20
30
40
50
t/s

### Path 212 (Syp1 − Rad23)

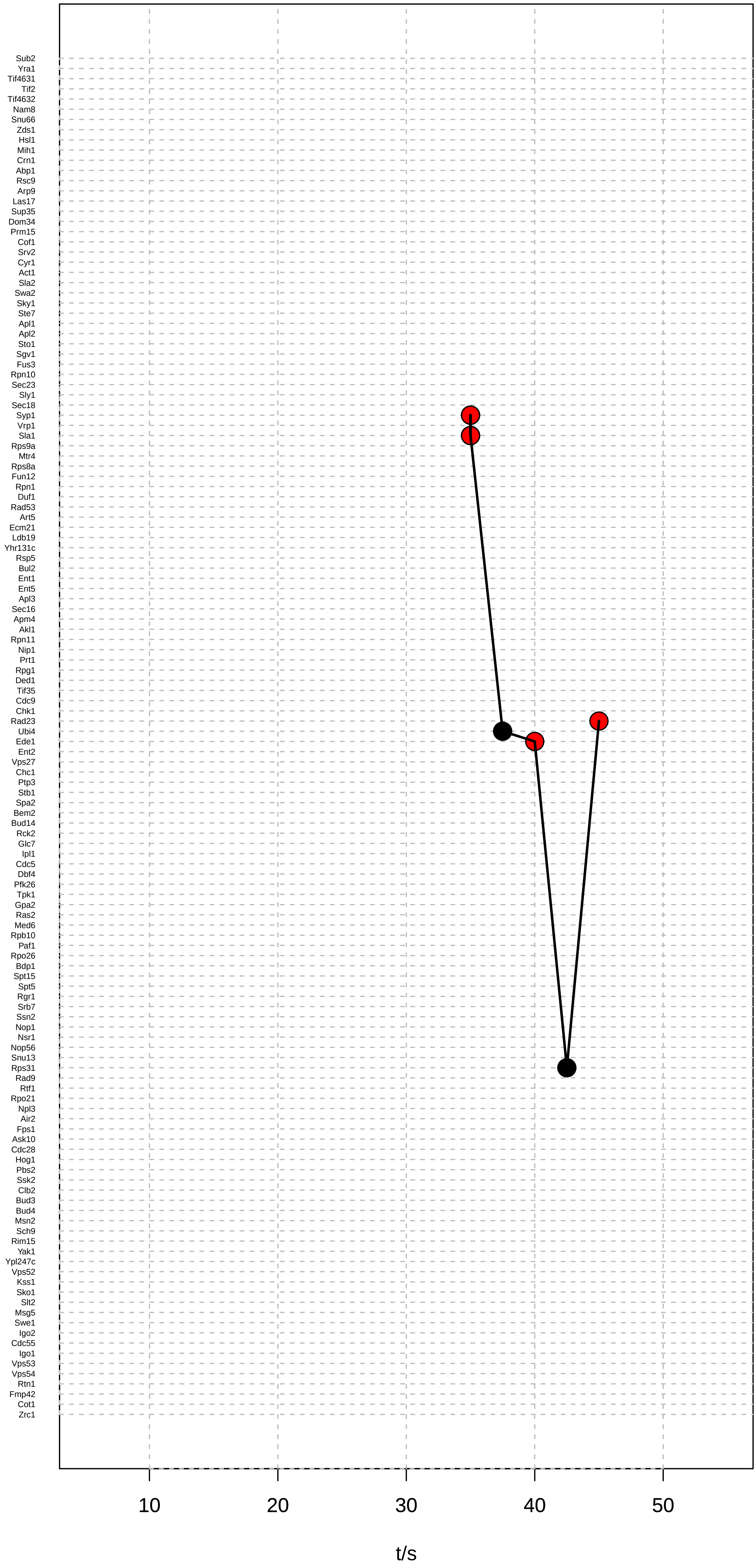

#### Path 213 (Syp1 − Rad23)

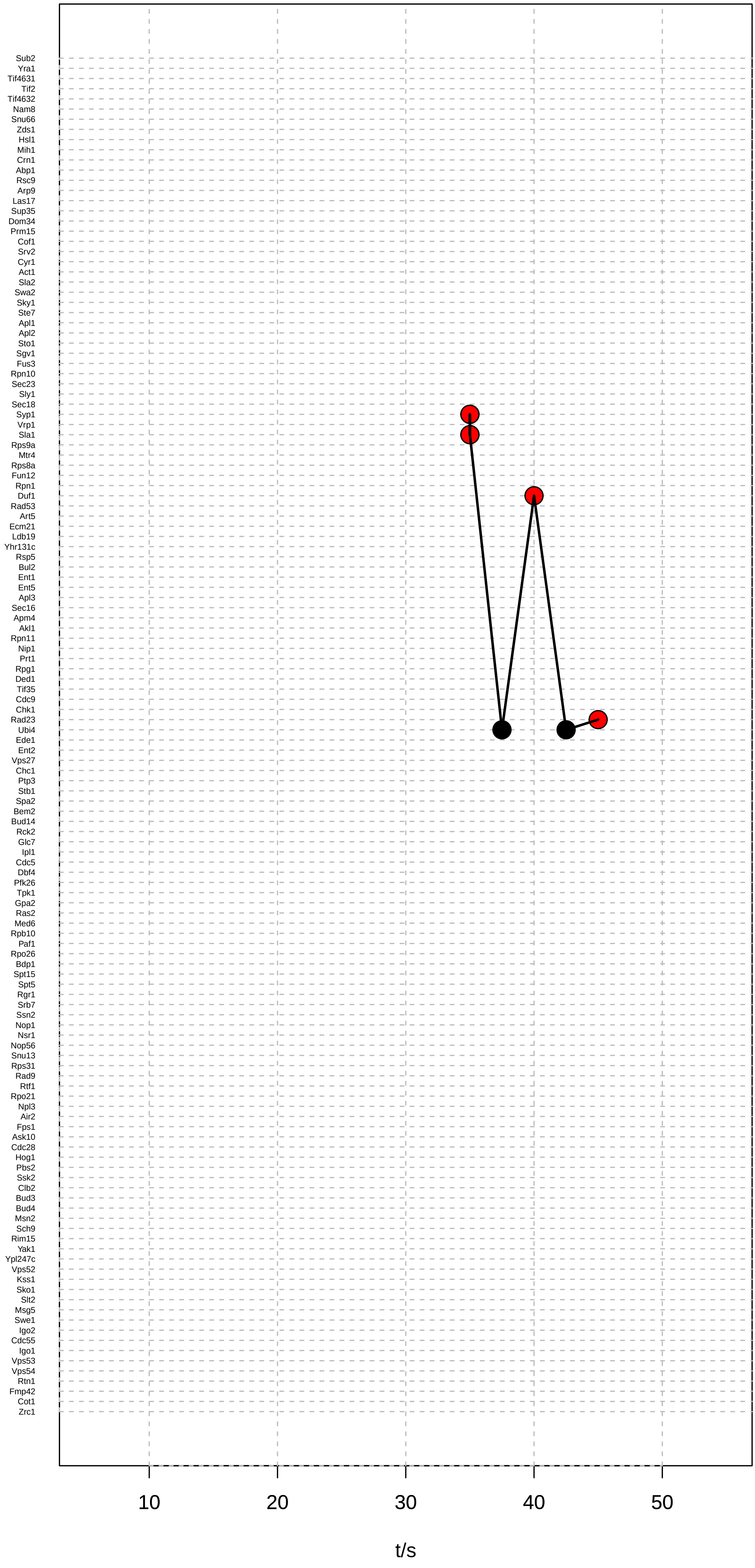

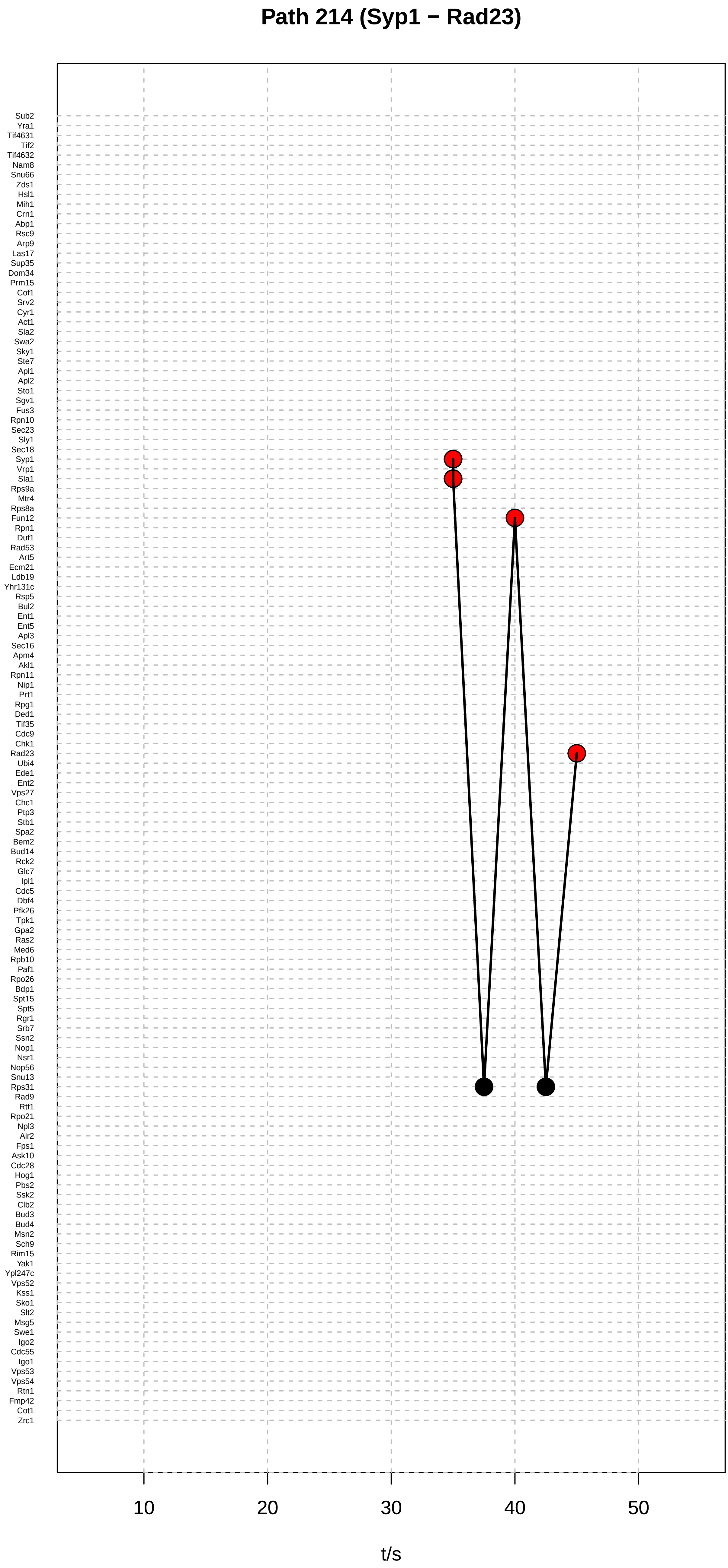
Path 214 (Syp1 − Rad23)
Sub2
Yra1
Tif4631
Tif2
Tif4632
Nam8
Snu66
Zds1
Hsl1
Mih1
Crn1
Abp1
Rsc9
Arp9
Las17
Sup35
Dom34
Prm15
Cof1
Srv2
Cyr1
Act1
Sla2
Swa2
Sky1
Ste7
Apl1
Apl2
Sto1
Sgv1
Fus3
Rpn10
Sec23
Sly1
Sec18
Syp1
Vrp1
Sla1
Rps9a
Mtr4
Rps8a
Fun12
Rpn1
Duf1
Rad53
Art5
Ecm21
Ldb19
Yhr131c
Rsp5
Bul2
Ent1
Ent5
Apl3
Sec16
Apm4
Akl1
Rpn11
Nip1
Prt1
Rpg1
Ded1
Tif35
Cdc9
Chk1
Rad23
Ubi4
Ede1
Ent2
Vps27
Chc1
Ptp3
Stb1
Spa2
Bem2
Bud14
Rck2
Glc7
Ipl1
Cdc5
Dbf4
Pfk26
Tpk1
Gpa2
Ras2
Med6
Rpb10
Paf1
Rpo26
Bdp1
Spt15
Spt5
Rgr1
Srb7
Ssn2
Nop1
Nsr1
Nop56
Snu13
Rps31
Rad9
Rtf1
Rpo21
Npl3
Air2
Fps1
Ask10
Cdc28
Hog1
Pbs2
Ssk2
Clb2
Bud3
Bud4
Msn2
Sch9
Rim15
Yak1
Ypl247c
Vps52
Kss1
Sko1
Slt2
Msg5
Swe1
Igo2
Cdc55
Igo1
Vps53
Vps54
Rtn1
Fmp42
Cot1
Zrc1
10
20
30
40
50
t/s

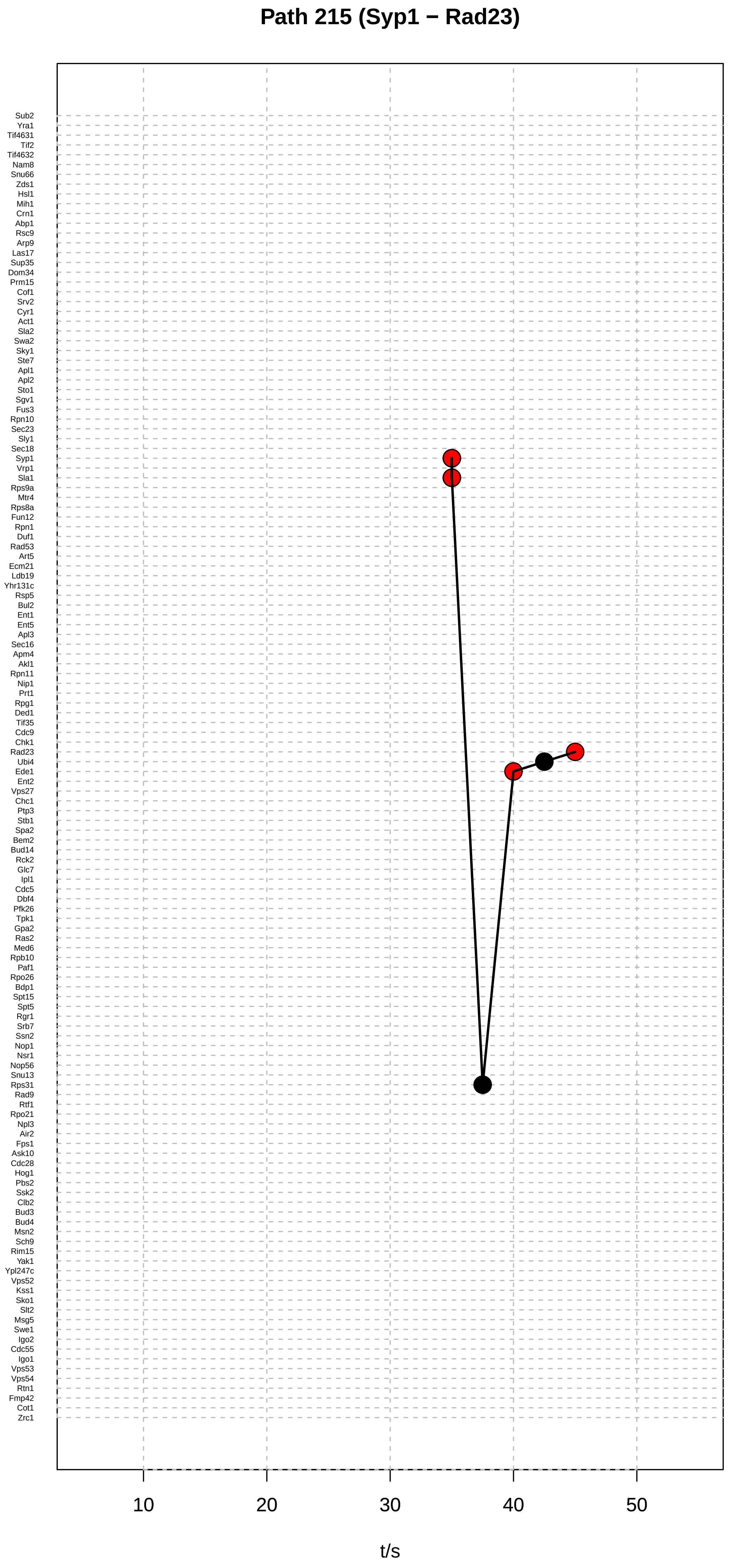

Path 215 (Syp1 − Rad23)
Sub2
Yra1
Tif4631
Tif2
Tif4632
Nam8
Snu66
Zds1
Hsl1
Mih1
Crn1
Abp1
Rsc9
Arp9
Las17
Sup35
Dom34
Prm15
Cof1
Srv2
Cyr1
Act1
Sla2
Swa2
Sky1
Ste7
Apl1
Apl2
Sto1
Sgv1
Fus3
Rpn10
Sec23
Sly1
Sec18
Syp1
Vrp1
Sla1
Rps9a
Mtr4
Rps8a
Fun12
Rpn1
Duf1
Rad53
Art5
Ecm21
Ldb19
Yhr131c
Rsp5
Bul2
Ent1
Ent5
Apl3
Sec16
Apm4
Akl1
Rpn11
Nip1
Prt1
Rpg1
Ded1
Tif35
Cdc9
Chk1
Rad23
Ubi4
Ede1
Ent2
Vps27
Chc1
Ptp3
Stb1
Spa2
Bem2
Bud14
Rck2
Glc7
Ipl1
Cdc5
Dbf4
Pfk26
Tpk1
Gpa2
Ras2
Med6
Rpb10
Paf1
Rpo26
Bdp1
Spt15
Spt5
Rgr1
Srb7
Ssn2
Nop1
Nsr1
Nop56
Snu13
Rps31
Rad9
Rtf1
Rpo21
Npl3
Air2
Fps1
Ask10
Cdc28
Hog1
Pbs2
Ssk2
Clb2
Bud3
Bud4
Msn2
Sch9
Rim15
Yak1
Ypl247c
Vps52
Kss1
Sko1
Slt2
Msg5
Swe1
Igo2
Cdc55
Igo1
Vps53
Vps54
Rtn1
Fmp42
Cot1
Zrc1
10
20
30
40
50
t/s

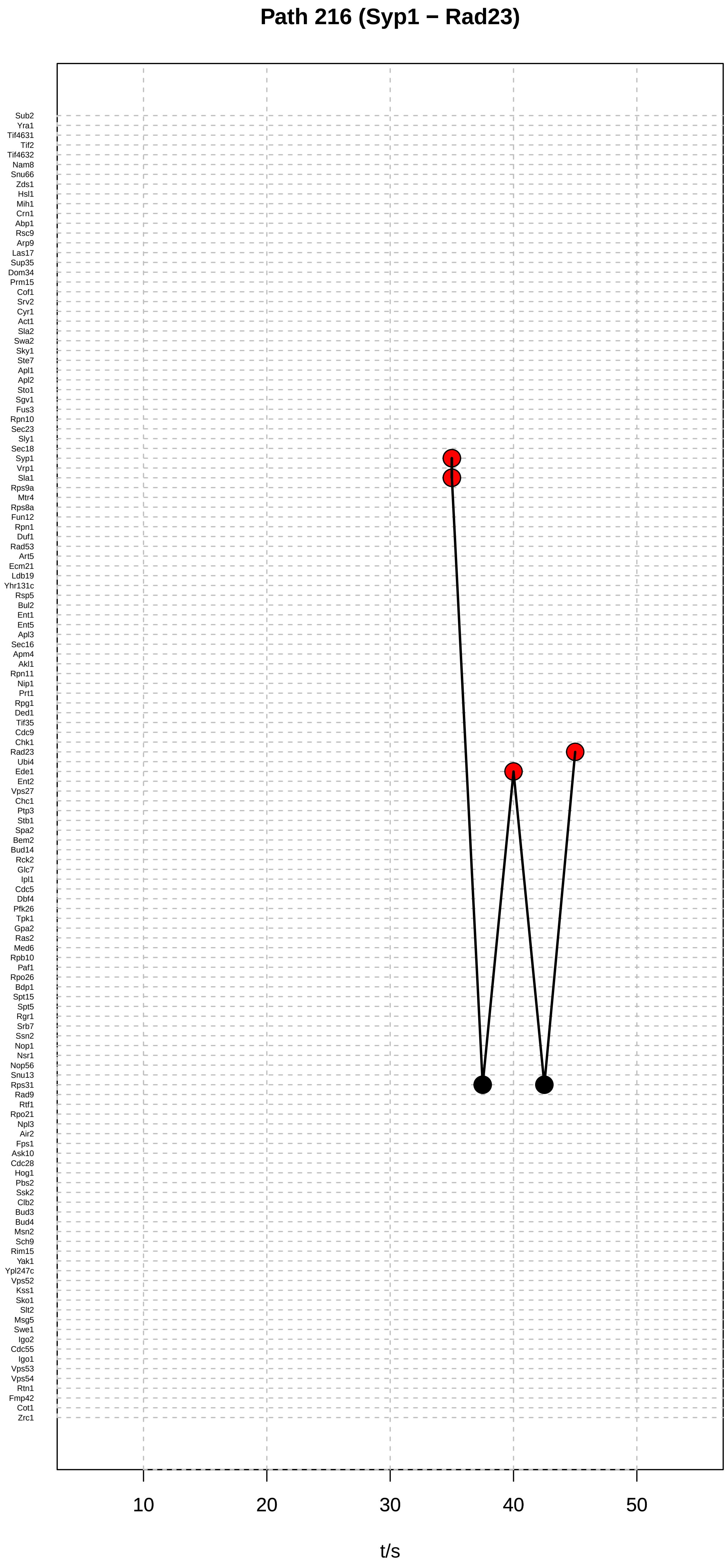
Path 216 (Syp1 − Rad23)
Sub2
Yra1
Tif4631
Tif2
Tif4632
Nam8
Snu66
Zds1
Hsl1
Mih1
Crn1
Abp1
Rsc9
Arp9
Las17
Sup35
Dom34
Prm15
Cof1
Srv2
Cyr1
Act1
Sla2
Swa2
Sky1
Ste7
Apl1
Apl2
Sto1
Sgv1
Fus3
Rpn10
Sec23
Sly1
Sec18
Syp1
Vrp1
Sla1
Rps9a
Mtr4
Rps8a
Fun12
Rpn1
Duf1
Rad53
Art5
Ecm21
Ldb19
Yhr131c
Rsp5
Bul2
Ent1
Ent5
Apl3
Sec16
Apm4
Akl1
Rpn11
Nip1
Prt1
Rpg1
Ded1
Tif35
Cdc9
Chk1
Rad23
Ubi4
Ede1
Ent2
Vps27
Chc1
Ptp3
Stb1
Spa2
Bem2
Bud14
Rck2
Glc7
Ipl1
Cdc5
Dbf4
Pfk26
Tpk1
Gpa2
Ras2
Med6
Rpb10
Paf1
Rpo26
Bdp1
Spt15
Spt5
Rgr1
Srb7
Ssn2
Nop1
Nsr1
Nop56
Snu13
Rps31
Rad9
Rtf1
Rpo21
Npl3
Air2
Fps1
Ask10
Cdc28
Hog1
Pbs2
Ssk2
Clb2
Bud3
Bud4
Msn2
Sch9
Rim15
Yak1
Ypl247c
Vps52
Kss1
Sko1
Slt2
Msg5
Swe1
Igo2
Cdc55
Igo1
Vps53
Vps54
Rtn1
Fmp42
Cot1
Zrc1
10
20
30
40
50
t/s

#### Path 217 (Syp1 − Rad23)

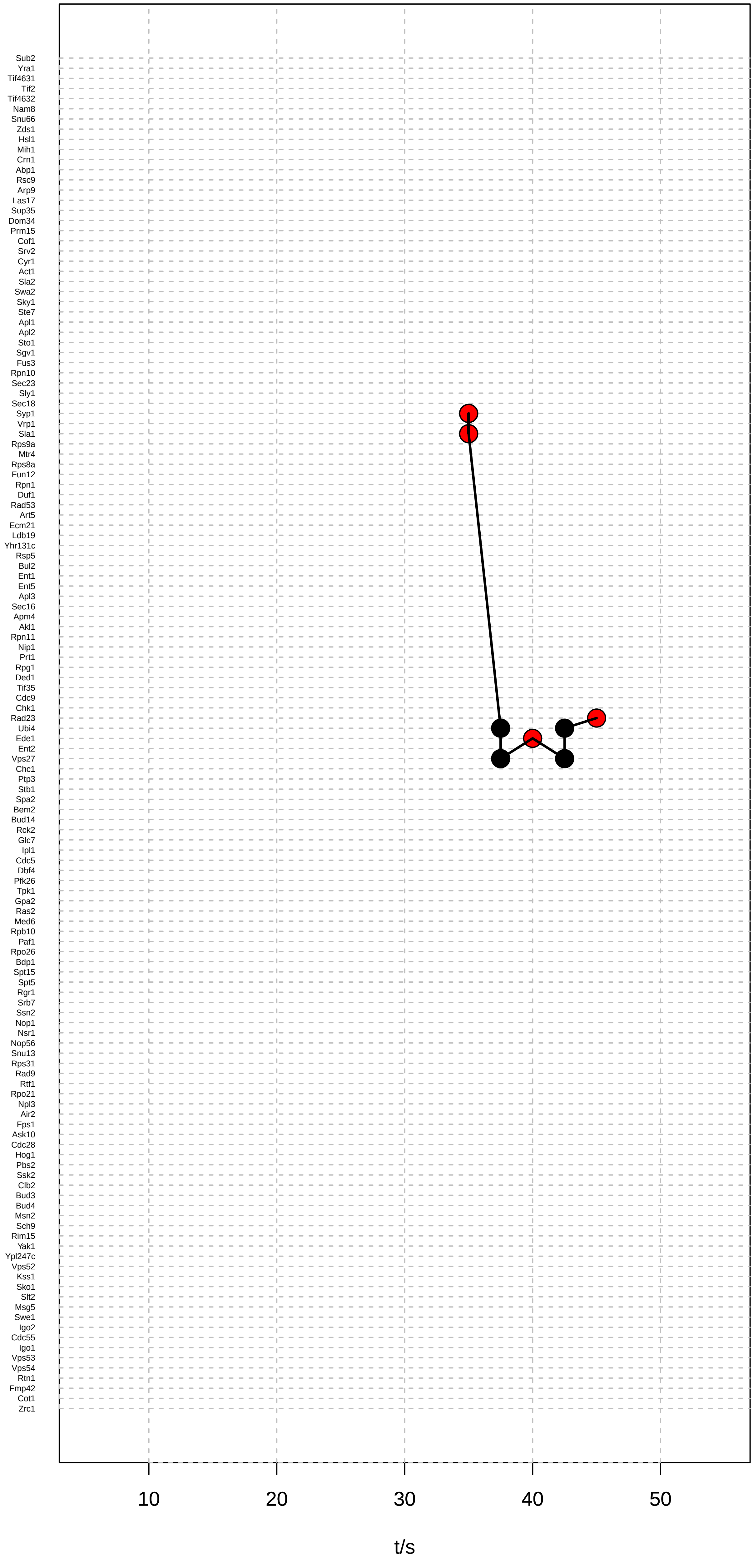

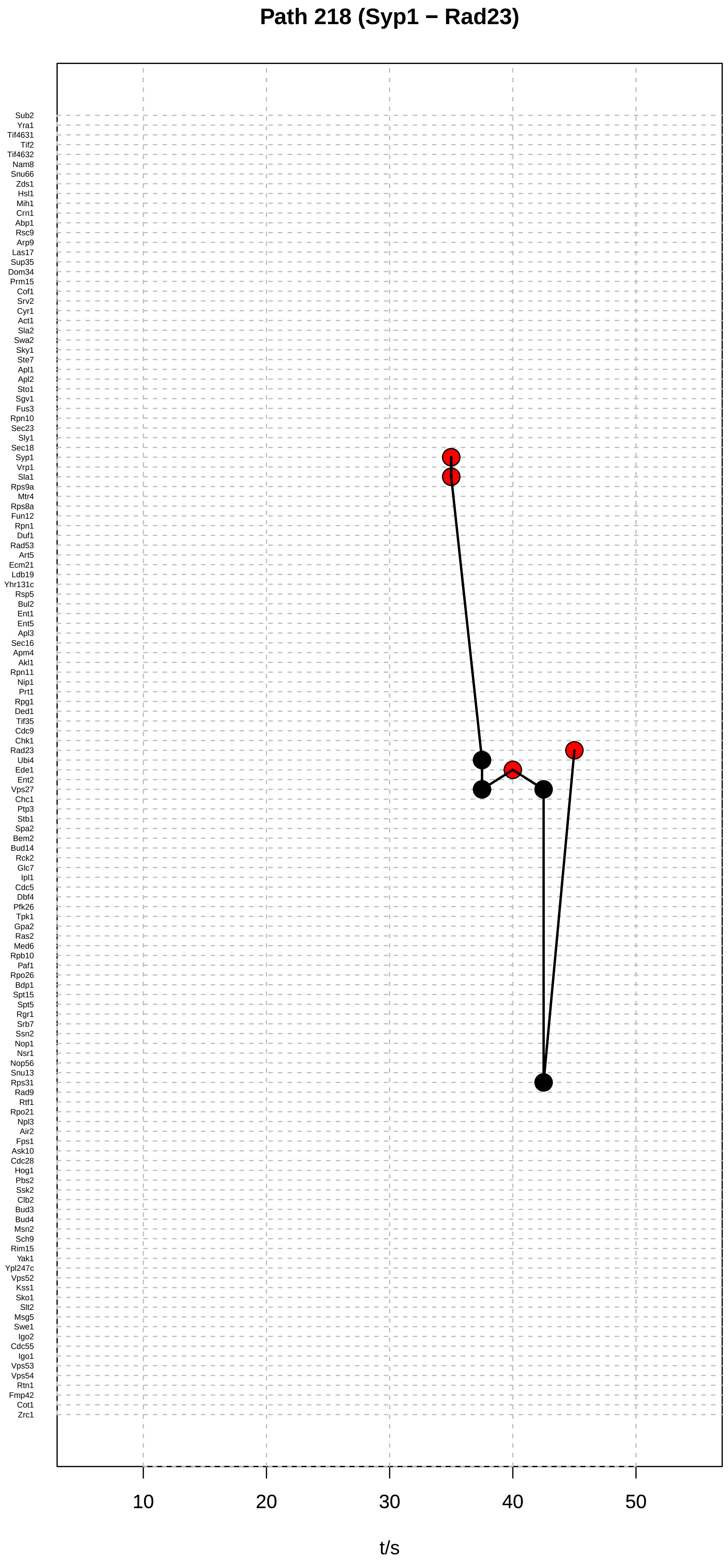
Path 218 (Syp1 − Rad23)
Sub2
Yra1
Tif4631
Tif2
Tif4632
Nam8
Snu66
Zds1
Hsl1
Mih1
Crn1
Abp1
Rsc9
Arp9
Las17
Sup35
Dom34
Prm15
Cof1
Srv2
Cyr1
Act1
Sla2
Swa2
Sky1
Ste7
Apl1
Apl2
Sto1
Sgv1
Fus3
Rpn10
Sec23
Sly1
Sec18
Syp1
Vrp1
Sla1
Rps9a
Mtr4
Rps8a
Fun12
Rpn1
Duf1
Rad53
Art5
Ecm21
Ldb19
Yhr131c
Rsp5
Bul2
Ent1
Ent5
Apl3
Sec16
Apm4
Akl1
Rpn11
Nip1
Prt1
Rpg1
Ded1
Tif35
Cdc9
Chk1
Rad23
Ubi4
Ede1
Ent2
Vps27
Chc1
Ptp3
Stb1
Spa2
Bem2
Bud14
Rck2
Glc7
Ipl1
Cdc5
Dbf4
Pfk26
Tpk1
Gpa2
Ras2
Med6
Rpb10
Paf1
Rpo26
Bdp1
Spt15
Spt5
Rgr1
Srb7
Ssn2
Nop1
Nsr1
Nop56
Snu13
Rps31
Rad9
Rtf1
Rpo21
Npl3
Air2
Fps1
Ask10
Cdc28
Hog1
Pbs2
Ssk2
Clb2
Bud3
Bud4
Msn2
Sch9
Rim15
Yak1
Ypl247c
Vps52
Kss1
Sko1
Slt2
Msg5
Swe1
Igo2
Cdc55
Igo1
Vps53
Vps54
Rtn1
Fmp42
Cot1
Zrc1
10
20
30
40
50
t/s

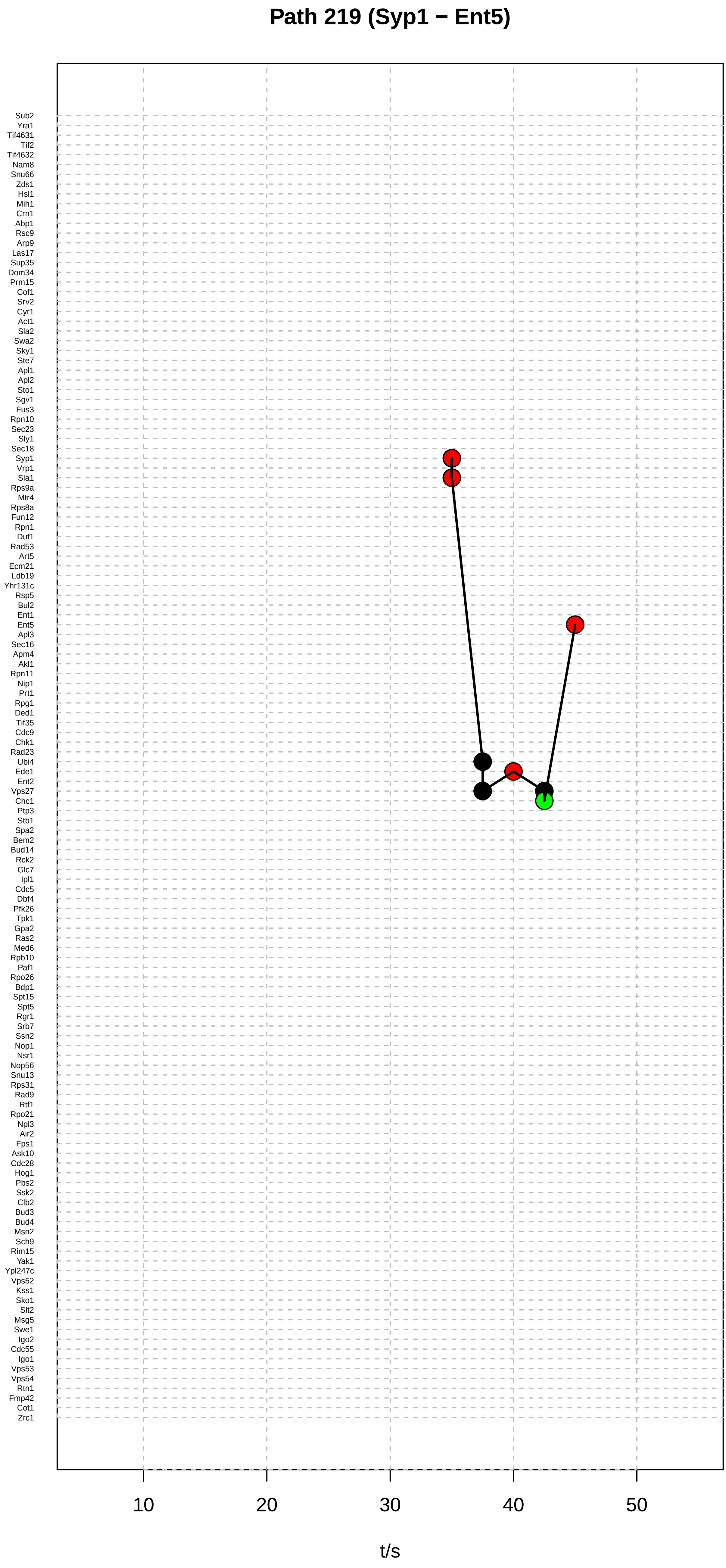

Path 219 (Syp1 − Ent5)
Sub2
Yra1
Tif4631
Tif2
Tif4632
Nam8
Snu66
Zds1
Hsl1
Mih1
Crn1
Abp1
Rsc9
Arp9
Las17
Sup35
Dom34
Prm15
Cof1
Srv2
Cyr1
Act1
Sla2
Swa2
Sky1
Ste7
Apl1
Apl2
Sto1
Sgv1
Fus3
Rpn10
Sec23
Sly1
Sec18
Syp1
Vrp1
Sla1
Rps9a
Mtr4
Rps8a
Fun12
Rpn1
Duf1
Rad53
Art5
Ecm21
Ldb19
Yhr131c
Rsp5
Bul2
Ent1
Ent5
Apl3
Sec16
Apm4
Akl1
Rpn11
Nip1
Prt1
Rpg1
Ded1
Tif35
Cdc9
Chk1
Rad23
Ubi4
Ede1
Ent2
Vps27
Chc1
Ptp3
Stb1
Spa2
Bem2
Bud14
Rck2
Glc7
Ipl1
Cdc5
Dbf4
Pfk26
Tpk1
Gpa2
Ras2
Med6
Rpb10
Paf1
Rpo26
Bdp1
Spt15
Spt5
Rgr1
Srb7
Ssn2
Nop1
Nsr1
Nop56
Snu13
Rps31
Rad9
Rtf1
Rpo21
Npl3
Air2
Fps1
Ask10
Cdc28
Hog1
Pbs2
Ssk2
Clb2
Bud3
Bud4
Msn2
Sch9
Rim15
Yak1
Ypl247c
Vps52
Kss1
Sko1
Slt2
Msg5
Swe1
Igo2
Cdc55
Igo1
Vps53
Vps54
Rtn1
Fmp42
Cot1
Zrc1
10
20
30
40
50
t/s

### Path 220 (Syp1 − Rad23)

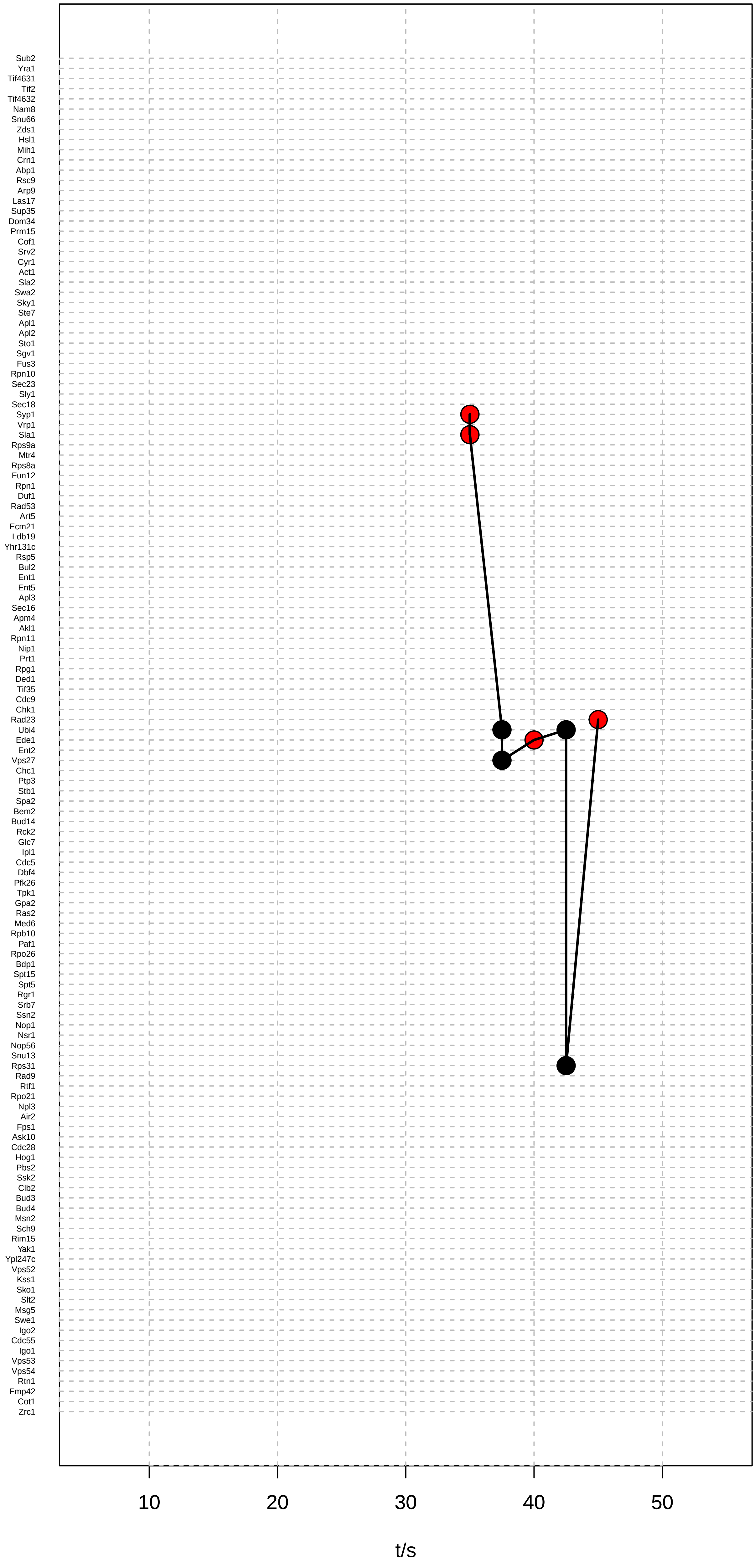

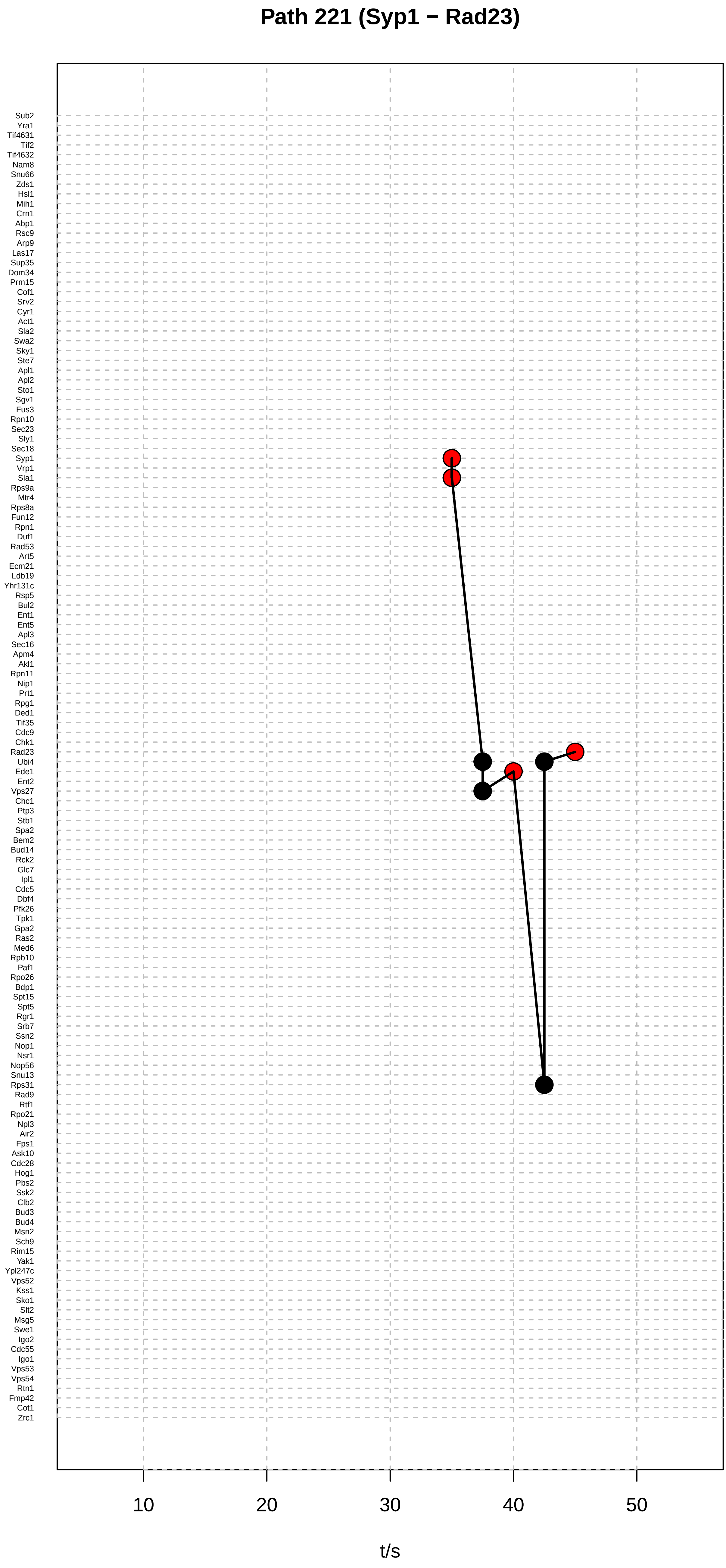
Path 221 (Syp1 − Rad23)
Sub2
Yra1
Tif4631
Tif2
Tif4632
Nam8
Snu66
Zds1
Hsl1
Mih1
Crn1
Abp1
Rsc9
Arp9
Las17
Sup35
Dom34
Prm15
Cof1
Srv2
Cyr1
Act1
Sla2
Swa2
Sky1
Ste7
Apl1
Apl2
Sto1
Sgv1
Fus3
Rpn10
Sec23
Sly1
Sec18
Syp1
Vrp1
Sla1
Rps9a
Mtr4
Rps8a
Fun12
Rpn1
Duf1
Rad53
Art5
Ecm21
Ldb19
Yhr131c
Rsp5
Bul2
Ent1
Ent5
Apl3
Sec16
Apm4
Akl1
Rpn11
Nip1
Prt1
Rpg1
Ded1
Tif35
Cdc9
Chk1
Rad23
Ubi4
Ede1
Ent2
Vps27
Chc1
Ptp3
Stb1
Spa2
Bem2
Bud14
Rck2
Glc7
Ipl1
Cdc5
Dbf4
Pfk26
Tpk1
Gpa2
Ras2
Med6
Rpb10
Paf1
Rpo26
Bdp1
Spt15
Spt5
Rgr1
Srb7
Ssn2
Nop1
Nsr1
Nop56
Snu13
Rps31
Rad9
Rtf1
Rpo21
Npl3
Air2
Fps1
Ask10
Cdc28
Hog1
Pbs2
Ssk2
Clb2
Bud3
Bud4
Msn2
Sch9
Rim15
Yak1
Ypl247c
Vps52
Kss1
Sko1
Slt2
Msg5
Swe1
Igo2
Cdc55
Igo1
Vps53
Vps54
Rtn1
Fmp42
Cot1
Zrc1
10
20
30
40
50
t/s

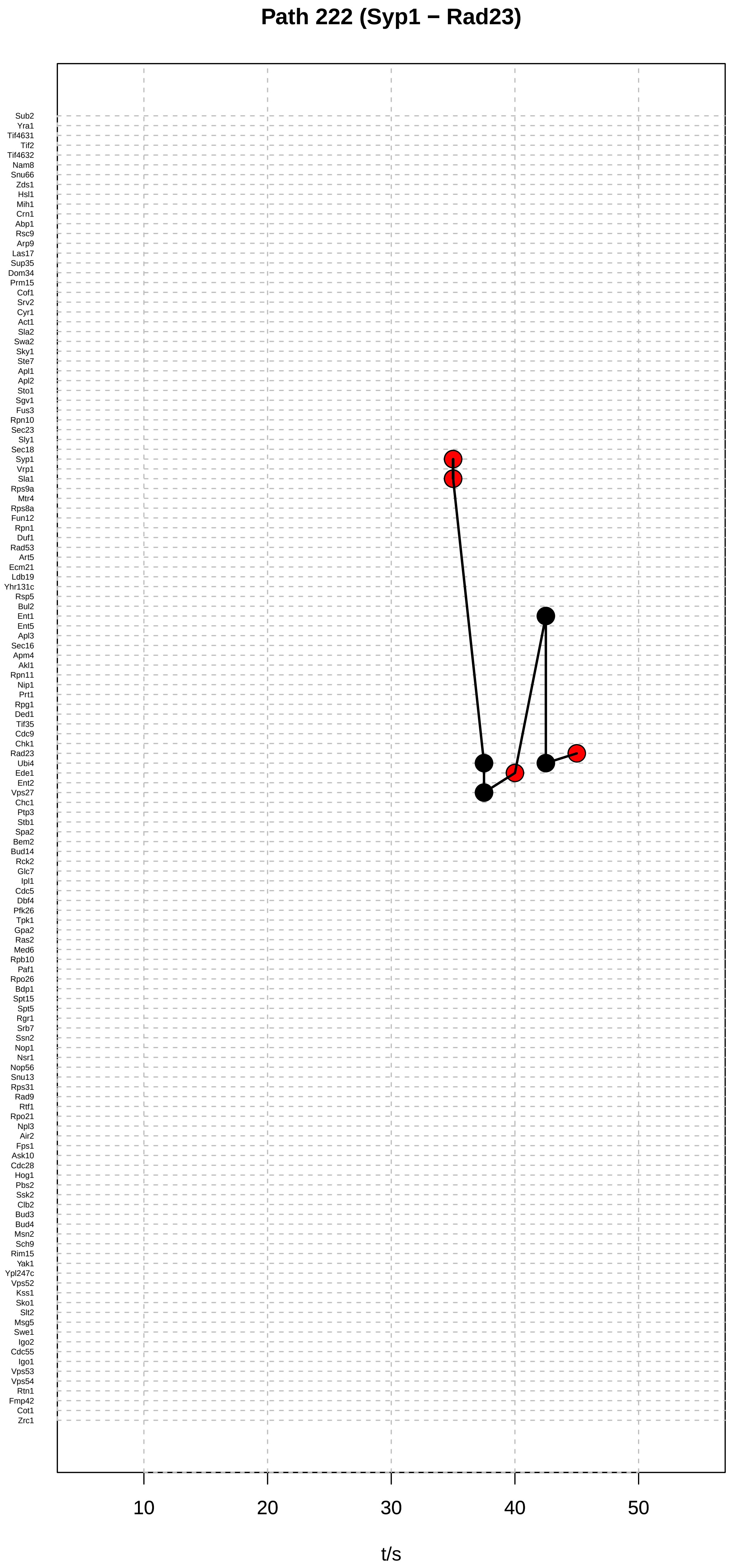

Path 222 (Syp1 − Rad23)
Sub2
Yra1
Tif4631
Tif2
Tif4632
Nam8
Snu66
Zds1
Hsl1
Mih1
Crn1
Abp1
Rsc9
Arp9
Las17
Sup35
Dom34
Prm15
Cof1
Srv2
Cyr1
Act1
Sla2
Swa2
Sky1
Ste7
Apl1
Apl2
Sto1
Sgv1
Fus3
Rpn10
Sec23
Sly1
Sec18
Syp1
Vrp1
Sla1
Rps9a
Mtr4
Rps8a
Fun12
Rpn1
Duf1
Rad53
Art5
Ecm21
Ldb19
Yhr131c
Rsp5
Bul2
Ent1
Ent5
Apl3
Sec16
Apm4
Akl1
Rpn11
Nip1
Prt1
Rpg1
Ded1
Tif35
Cdc9
Chk1
Rad23
Ubi4
Ede1
Ent2
Vps27
Chc1
Ptp3
Stb1
Spa2
Bem2
Bud14
Rck2
Glc7
Ipl1
Cdc5
Dbf4
Pfk26
Tpk1
Gpa2
Ras2
Med6
Rpb10
Paf1
Rpo26
Bdp1
Spt15
Spt5
Rgr1
Srb7
Ssn2
Nop1
Nsr1
Nop56
Snu13
Rps31
Rad9
Rtf1
Rpo21
Npl3
Air2
Fps1
Ask10
Cdc28
Hog1
Pbs2
Ssk2
Clb2
Bud3
Bud4
Msn2
Sch9
Rim15
Yak1
Ypl247c
Vps52
Kss1
Sko1
Slt2
Msg5
Swe1
Igo2
Cdc55
Igo1
Vps53
Vps54
Rtn1
Fmp42
Cot1
Zrc1
10
20
30
40
50
t/s

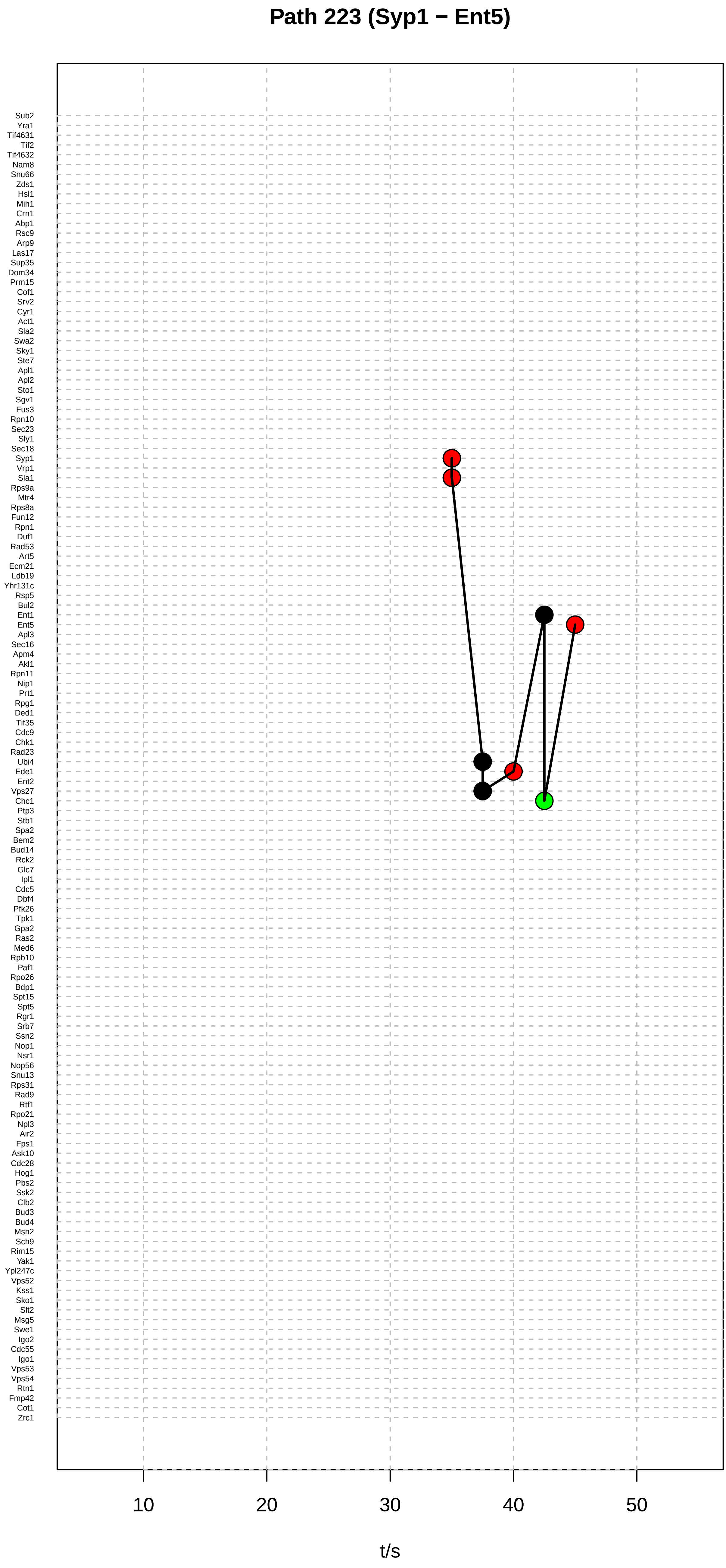
Path 223 (Syp1 − Ent5)
Sub2
Yra1
Tif4631
Tif2
Tif4632
Nam8
Snu66
Zds1
Hsl1
Mih1
Crn1
Abp1
Rsc9
Arp9
Las17
Sup35
Dom34
Prm15
Cof1
Srv2
Cyr1
Act1
Sla2
Swa2
Sky1
Ste7
Apl1
Apl2
Sto1
Sgv1
Fus3
Rpn10
Sec23
Sly1
Sec18
Syp1
Vrp1
Sla1
Rps9a
Mtr4
Rps8a
Fun12
Rpn1
Duf1
Rad53
Art5
Ecm21
Ldb19
Yhr131c
Rsp5
Bul2
Ent1
Ent5
Apl3
Sec16
Apm4
Akl1
Rpn11
Nip1
Prt1
Rpg1
Ded1
Tif35
Cdc9
Chk1
Rad23
Ubi4
Ede1
Ent2
Vps27
Chc1
Ptp3
Stb1
Spa2
Bem2
Bud14
Rck2
Glc7
Ipl1
Cdc5
Dbf4
Pfk26
Tpk1
Gpa2
Ras2
Med6
Rpb10
Paf1
Rpo26
Bdp1
Spt15
Spt5
Rgr1
Srb7
Ssn2
Nop1
Nsr1
Nop56
Snu13
Rps31
Rad9
Rtf1
Rpo21
Npl3
Air2
Fps1
Ask10
Cdc28
Hog1
Pbs2
Ssk2
Clb2
Bud3
Bud4
Msn2
Sch9
Rim15
Yak1
Ypl247c
Vps52
Kss1
Sko1
Slt2
Msg5
Swe1
Igo2
Cdc55
Igo1
Vps53
Vps54
Rtn1
Fmp42
Cot1
Zrc1
10
20
30
40
50
t/s

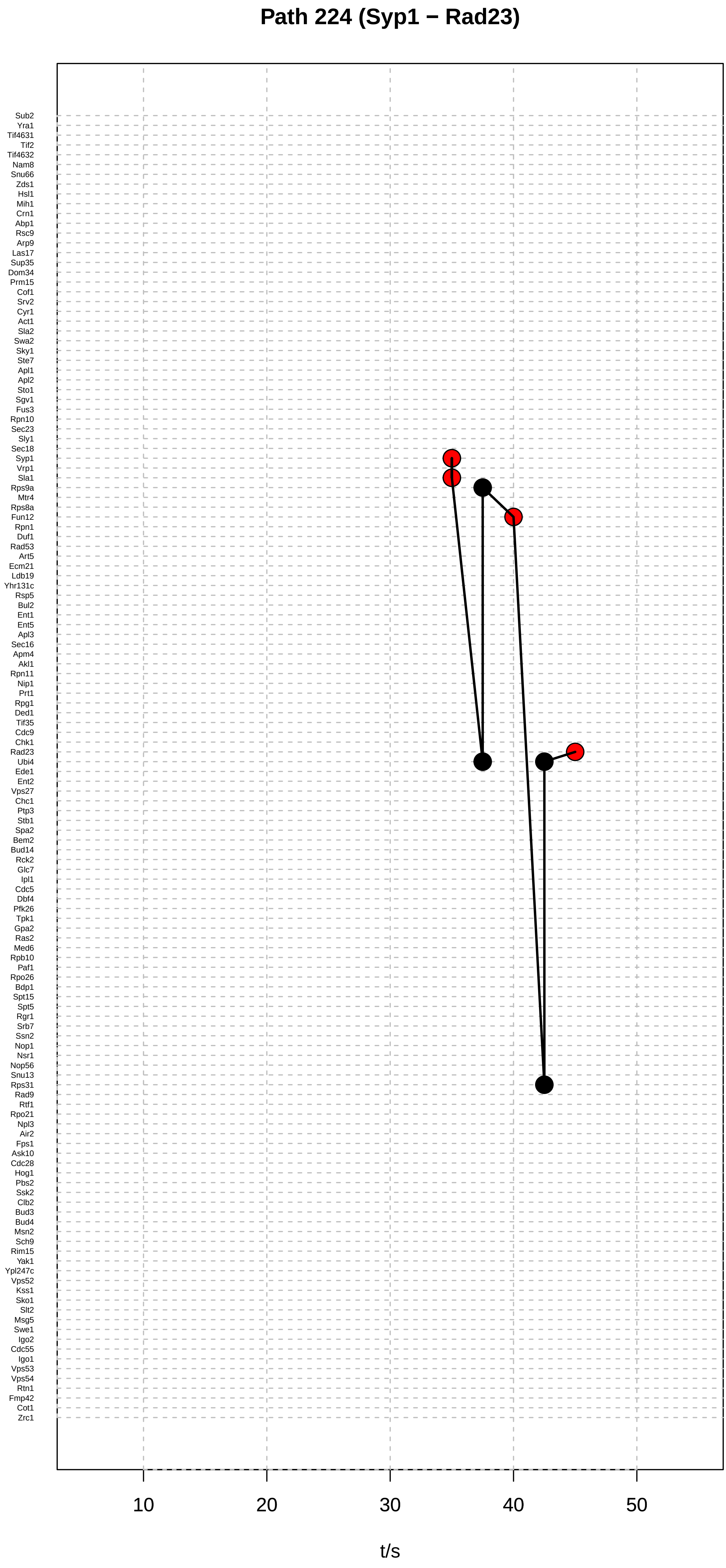
Path 224 (Syp1 − Rad23)
Sub2
Yra1
Tif4631
Tif2
Tif4632
Nam8
Snu66
Zds1
Hsl1
Mih1
Crn1
Abp1
Rsc9
Arp9
Las17
Sup35
Dom34
Prm15
Cof1
Srv2
Cyr1
Act1
Sla2
Swa2
Sky1
Ste7
Apl1
Apl2
Sto1
Sgv1
Fus3
Rpn10
Sec23
Sly1
Sec18
Syp1
Vrp1
Sla1
Rps9a
Mtr4
Rps8a
Fun12
Rpn1
Duf1
Rad53
Art5
Ecm21
Ldb19
Yhr131c
Rsp5
Bul2
Ent1
Ent5
Apl3
Sec16
Apm4
Akl1
Rpn11
Nip1
Prt1
Rpg1
Ded1
Tif35
Cdc9
Chk1
Rad23
Ubi4
Ede1
Ent2
Vps27
Chc1
Ptp3
Stb1
Spa2
Bem2
Bud14
Rck2
Glc7
Ipl1
Cdc5
Dbf4
Pfk26
Tpk1
Gpa2
Ras2
Med6
Rpb10
Paf1
Rpo26
Bdp1
Spt15
Spt5
Rgr1
Srb7
Ssn2
Nop1
Nsr1
Nop56
Snu13
Rps31
Rad9
Rtf1
Rpo21
Npl3
Air2
Fps1
Ask10
Cdc28
Hog1
Pbs2
Ssk2
Clb2
Bud3
Bud4
Msn2
Sch9
Rim15
Yak1
Ypl247c
Vps52
Kss1
Sko1
Slt2
Msg5
Swe1
Igo2
Cdc55
Igo1
Vps53
Vps54
Rtn1
Fmp42
Cot1
Zrc1
10
20
30
40
50
t/s

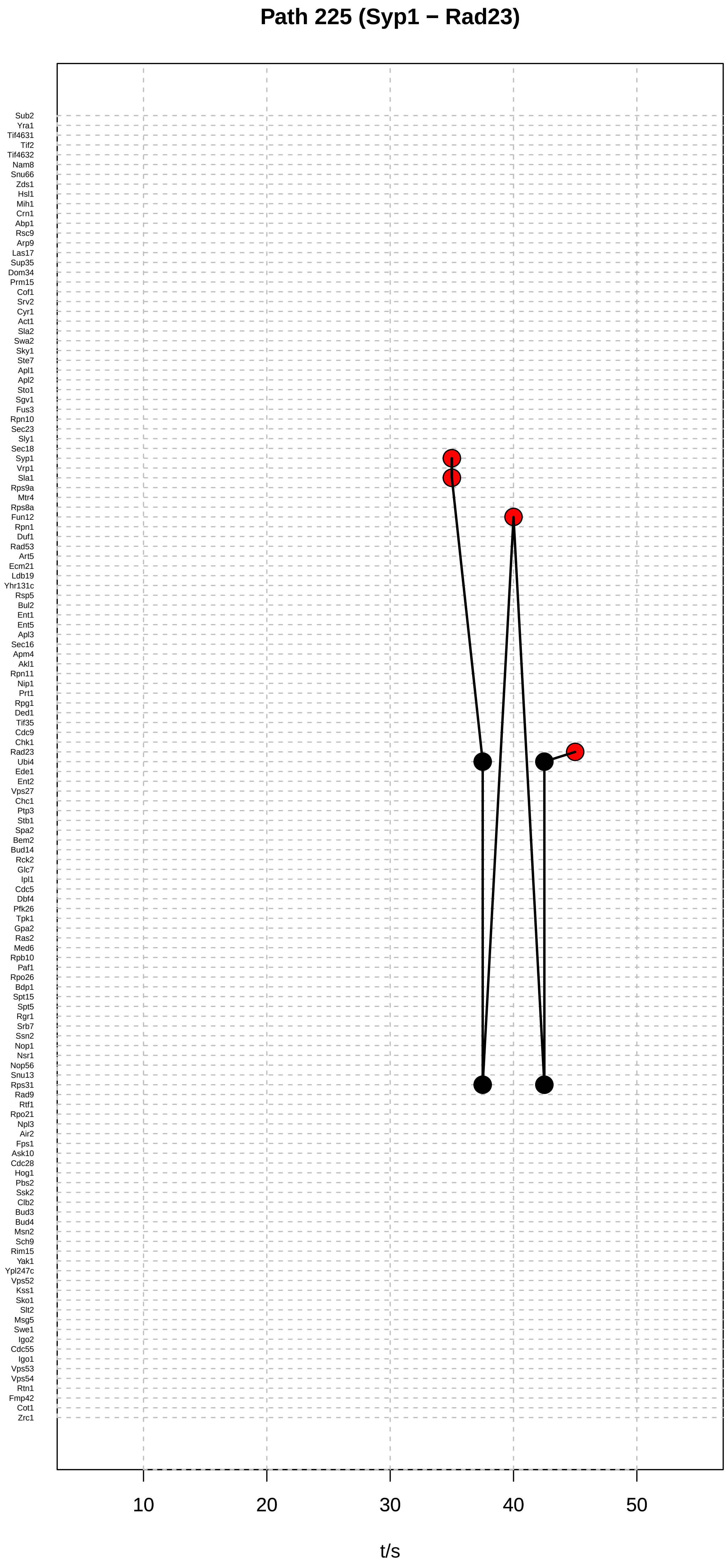
Path 225 (Syp1 − Rad23)
Sub2
Yra1
Tif4631
Tif2
Tif4632
Nam8
Snu66
Zds1
Hsl1
Mih1
Crn1
Abp1
Rsc9
Arp9
Las17
Sup35
Dom34
Prm15
Cof1
Srv2
Cyr1
Act1
Sla2
Swa2
Sky1
Ste7
Apl1
Apl2
Sto1
Sgv1
Fus3
Rpn10
Sec23
Sly1
Sec18
Syp1
Vrp1
Sla1
Rps9a
Mtr4
Rps8a
Fun12
Rpn1
Duf1
Rad53
Art5
Ecm21
Ldb19
Yhr131c
Rsp5
Bul2
Ent1
Ent5
Apl3
Sec16
Apm4
Akl1
Rpn11
Nip1
Prt1
Rpg1
Ded1
Tif35
Cdc9
Chk1
Rad23
Ubi4
Ede1
Ent2
Vps27
Chc1
Ptp3
Stb1
Spa2
Bem2
Bud14
Rck2
Glc7
Ipl1
Cdc5
Dbf4
Pfk26
Tpk1
Gpa2
Ras2
Med6
Rpb10
Paf1
Rpo26
Bdp1
Spt15
Spt5
Rgr1
Srb7
Ssn2
Nop1
Nsr1
Nop56
Snu13
Rps31
Rad9
Rtf1
Rpo21
Npl3
Air2
Fps1
Ask10
Cdc28
Hog1
Pbs2
Ssk2
Clb2
Bud3
Bud4
Msn2
Sch9
Rim15
Yak1
Ypl247c
Vps52
Kss1
Sko1
Slt2
Msg5
Swe1
Igo2
Cdc55
Igo1
Vps53
Vps54
Rtn1
Fmp42
Cot1
Zrc1
10
20
30
40
50
t/s

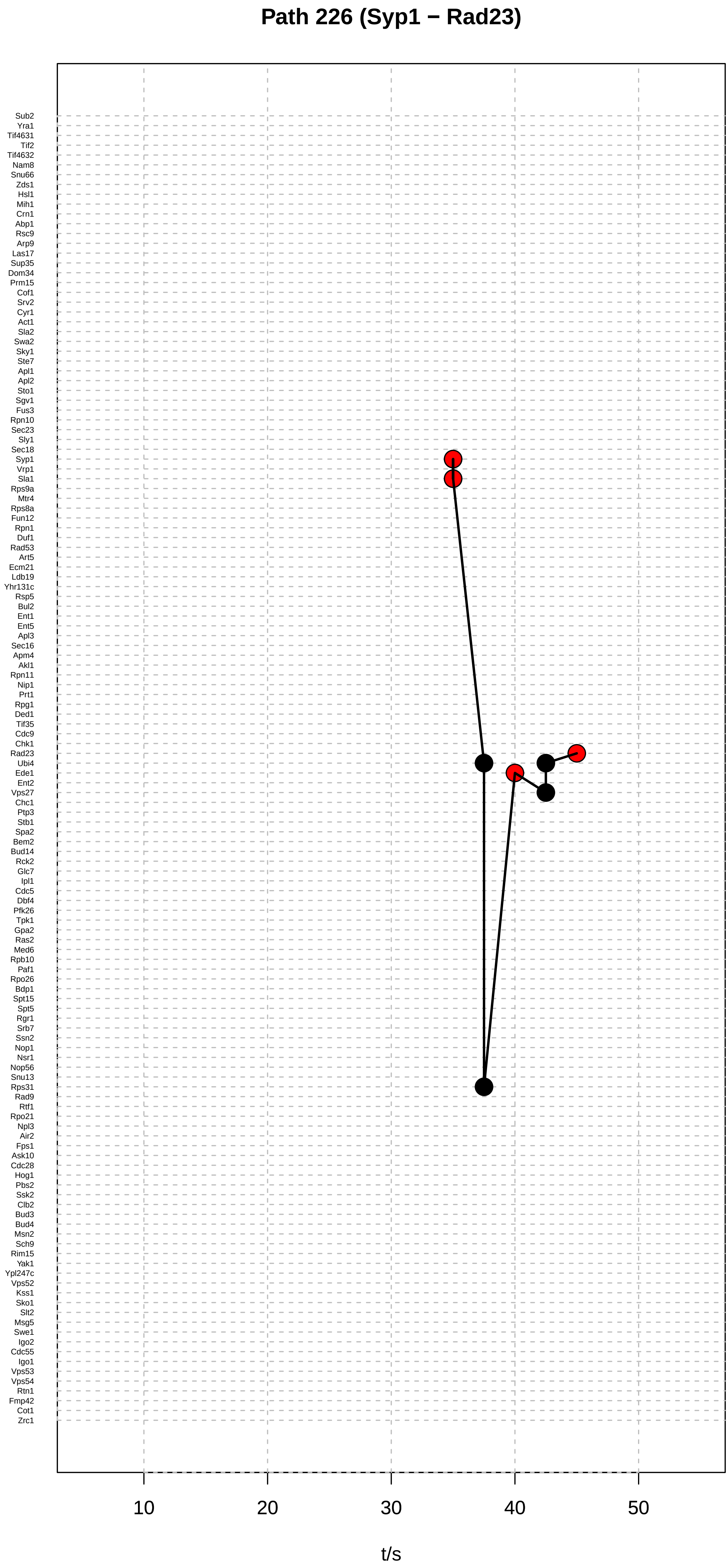

Path 226 (Syp1 − Rad23)
Sub2
Yra1
Tif4631
Tif2
Tif4632
Nam8
Snu66
Zds1
Hsl1
Mih1
Crn1
Abp1
Rsc9
Arp9
Las17
Sup35
Dom34
Prm15
Cof1
Srv2
Cyr1
Act1
Sla2
Swa2
Sky1
Ste7
Apl1
Apl2
Sto1
Sgv1
Fus3
Rpn10
Sec23
Sly1
Sec18
Syp1
Vrp1
Sla1
Rps9a
Mtr4
Rps8a
Fun12
Rpn1
Duf1
Rad53
Art5
Ecm21
Ldb19
Yhr131c
Rsp5
Bul2
Ent1
Ent5
Apl3
Sec16
Apm4
Akl1
Rpn11
Nip1
Prt1
Rpg1
Ded1
Tif35
Cdc9
Chk1
Rad23
Ubi4
Ede1
Ent2
Vps27
Chc1
Ptp3
Stb1
Spa2
Bem2
Bud14
Rck2
Glc7
Ipl1
Cdc5
Dbf4
Pfk26
Tpk1
Gpa2
Ras2
Med6
Rpb10
Paf1
Rpo26
Bdp1
Spt15
Spt5
Rgr1
Srb7
Ssn2
Nop1
Nsr1
Nop56
Snu13
Rps31
Rad9
Rtf1
Rpo21
Npl3
Air2
Fps1
Ask10
Cdc28
Hog1
Pbs2
Ssk2
Clb2
Bud3
Bud4
Msn2
Sch9
Rim15
Yak1
Ypl247c
Vps52
Kss1
Sko1
Slt2
Msg5
Swe1
Igo2
Cdc55
Igo1
Vps53
Vps54
Rtn1
Fmp42
Cot1
Zrc1
10
20
30
40
50
t/s

### Path 227 (Syp1 − Rad23)

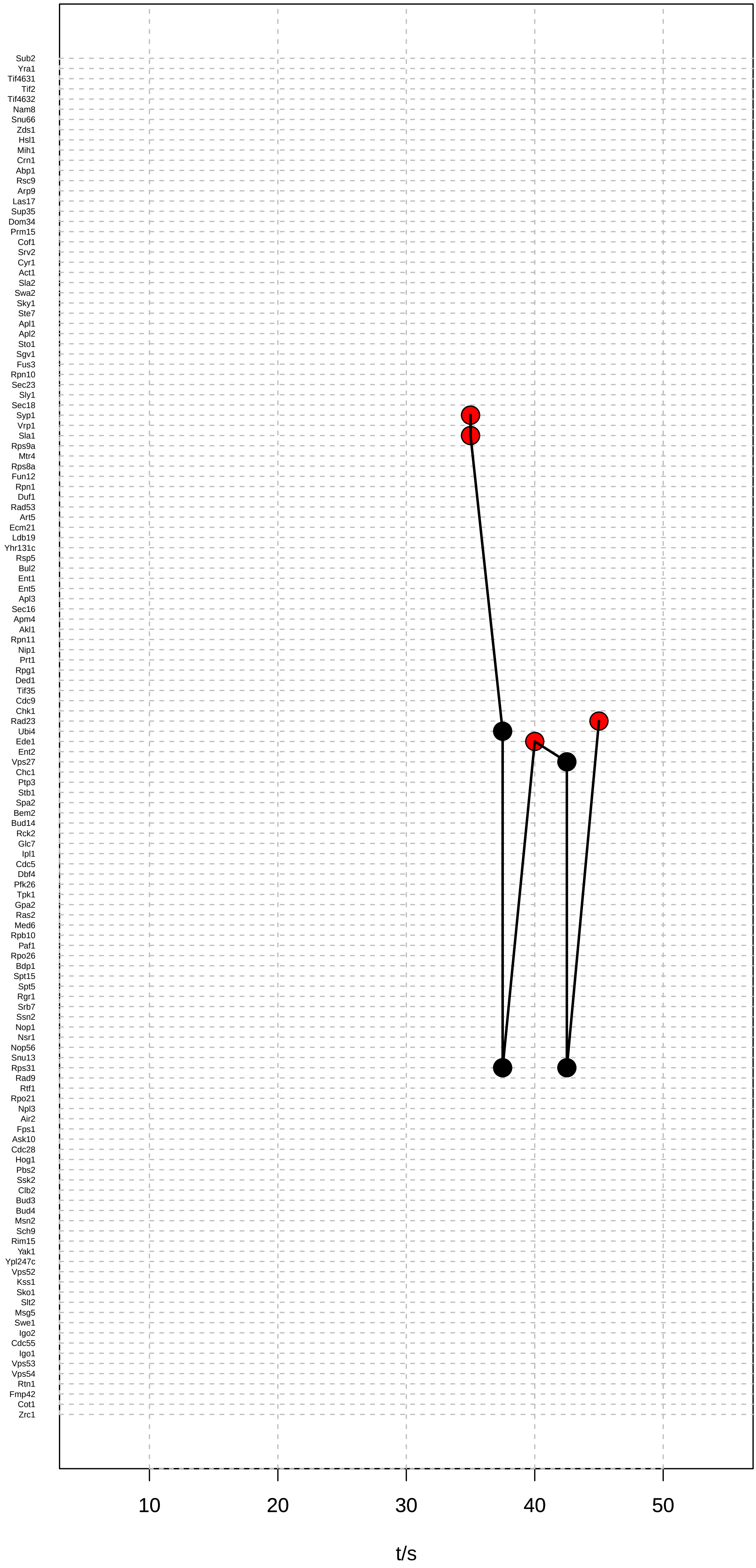

### Path 228 (Syp1 − Ent5)

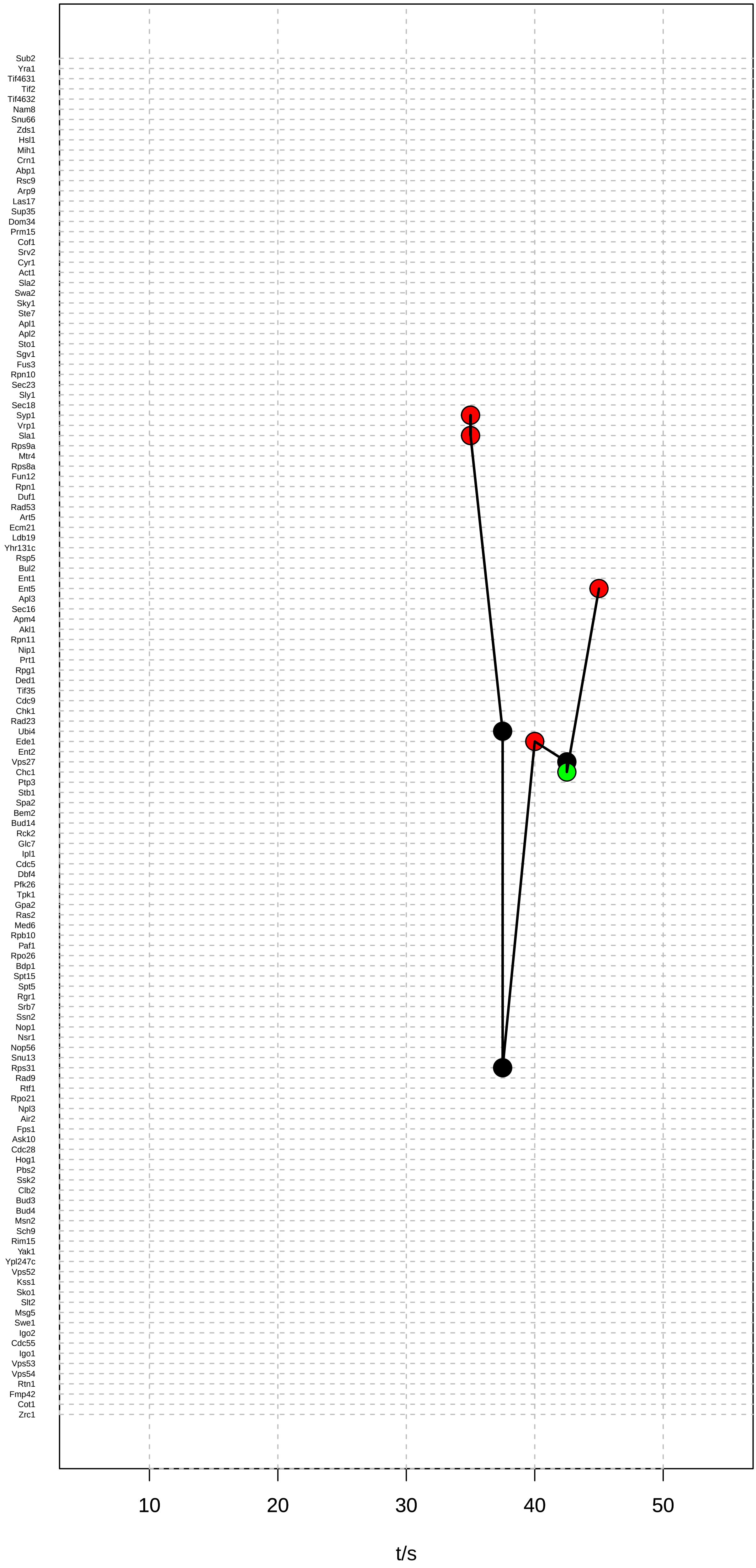

### Path 229 (Syp1 − Rad23)

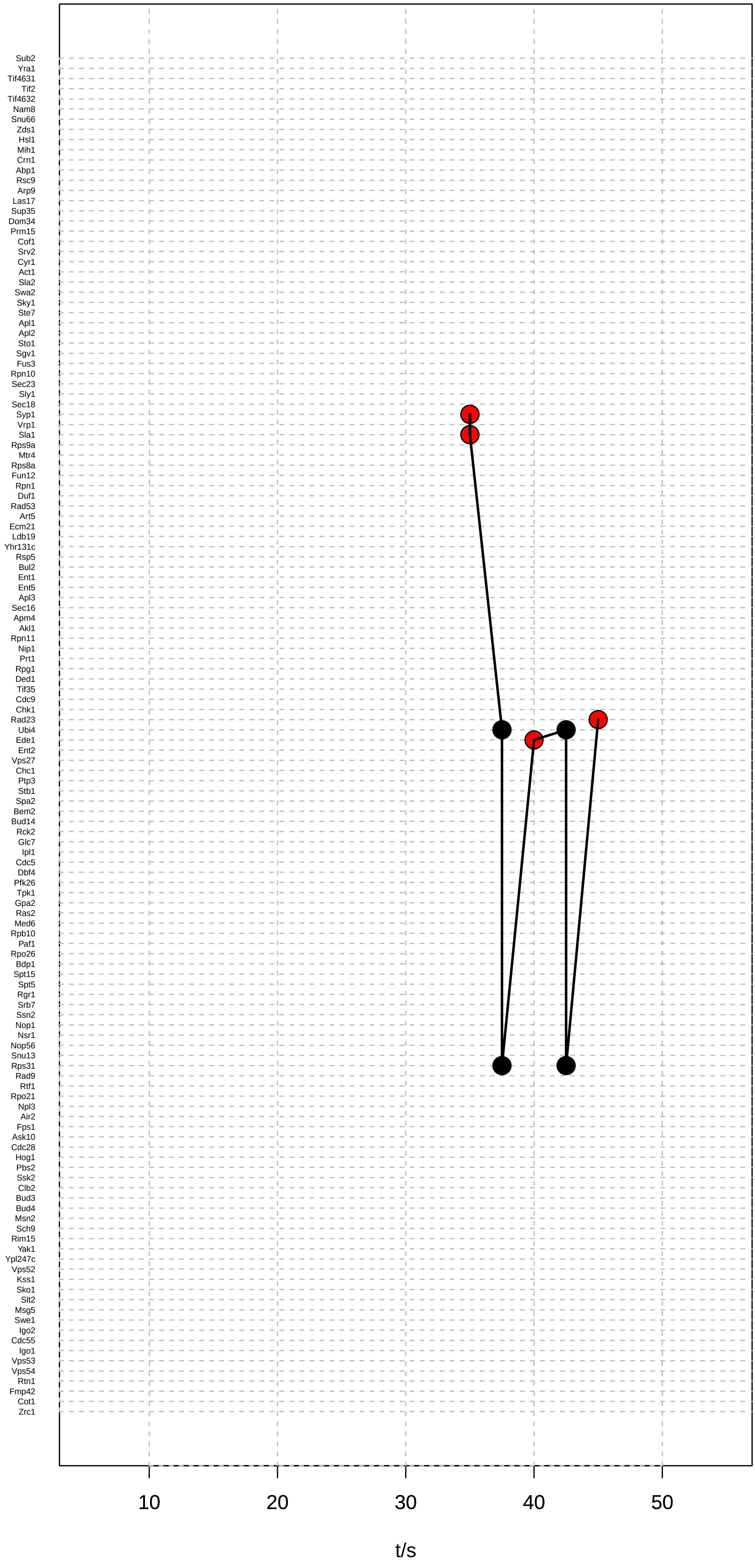

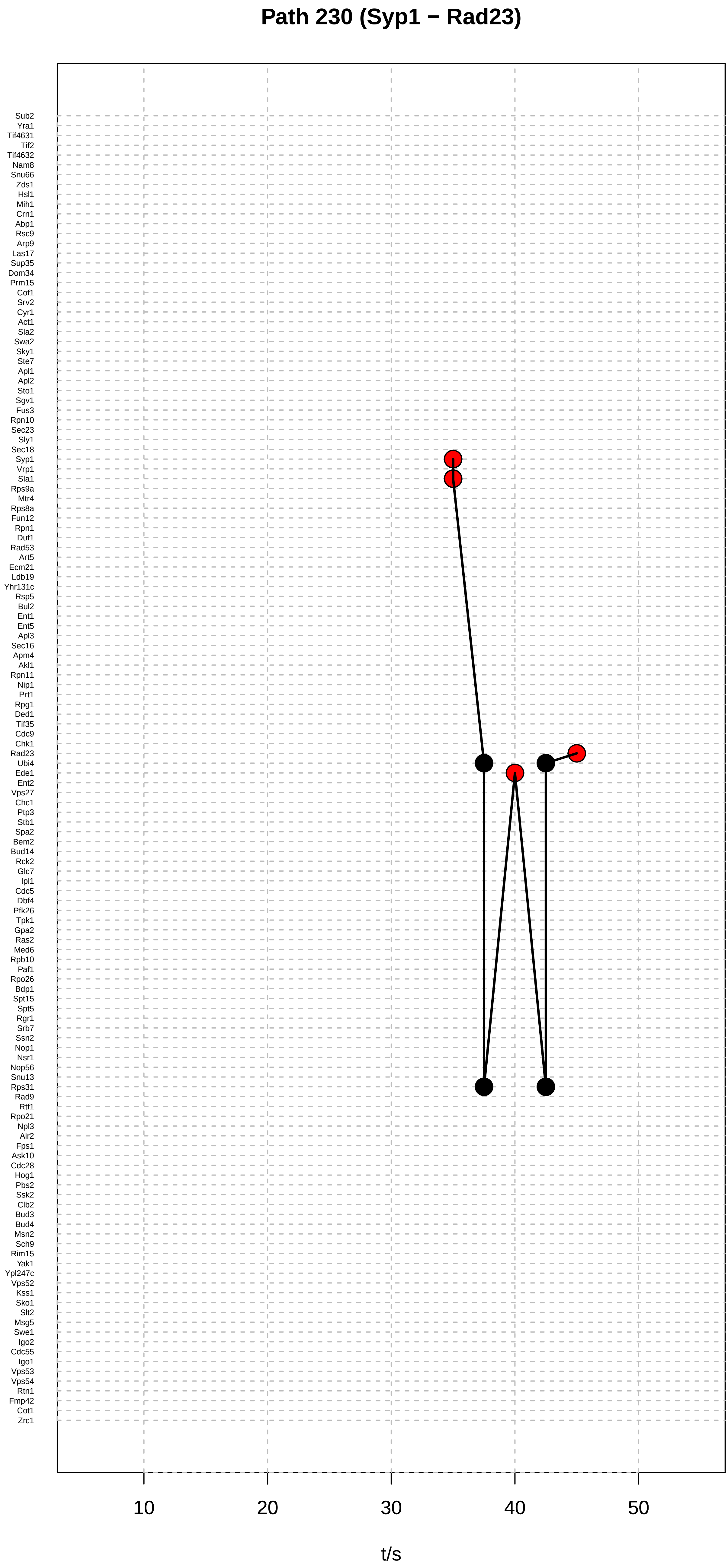
Path 230 (Syp1 − Rad23)
Sub2
Yra1
Tif4631
Tif2
Tif4632
Nam8
Snu66
Zds1
Hsl1
Mih1
Crn1
Abp1
Rsc9
Arp9
Las17
Sup35
Dom34
Prm15
Cof1
Srv2
Cyr1
Act1
Sla2
Swa2
Sky1
Ste7
Apl1
Apl2
Sto1
Sgv1
Fus3
Rpn10
Sec23
Sly1
Sec18
Syp1
Vrp1
Sla1
Rps9a
Mtr4
Rps8a
Fun12
Rpn1
Duf1
Rad53
Art5
Ecm21
Ldb19
Yhr131c
Rsp5
Bul2
Ent1
Ent5
Apl3
Sec16
Apm4
Akl1
Rpn11
Nip1
Prt1
Rpg1
Ded1
Tif35
Cdc9
Chk1
Rad23
Ubi4
Ede1
Ent2
Vps27
Chc1
Ptp3
Stb1
Spa2
Bem2
Bud14
Rck2
Glc7
Ipl1
Cdc5
Dbf4
Pfk26
Tpk1
Gpa2
Ras2
Med6
Rpb10
Paf1
Rpo26
Bdp1
Spt15
Spt5
Rgr1
Srb7
Ssn2
Nop1
Nsr1
Nop56
Snu13
Rps31
Rad9
Rtf1
Rpo21
Npl3
Air2
Fps1
Ask10
Cdc28
Hog1
Pbs2
Ssk2
Clb2
Bud3
Bud4
Msn2
Sch9
Rim15
Yak1
Ypl247c
Vps52
Kss1
Sko1
Slt2
Msg5
Swe1
Igo2
Cdc55
Igo1
Vps53
Vps54
Rtn1
Fmp42
Cot1
Zrc1
10
20
30
40
50
t/s

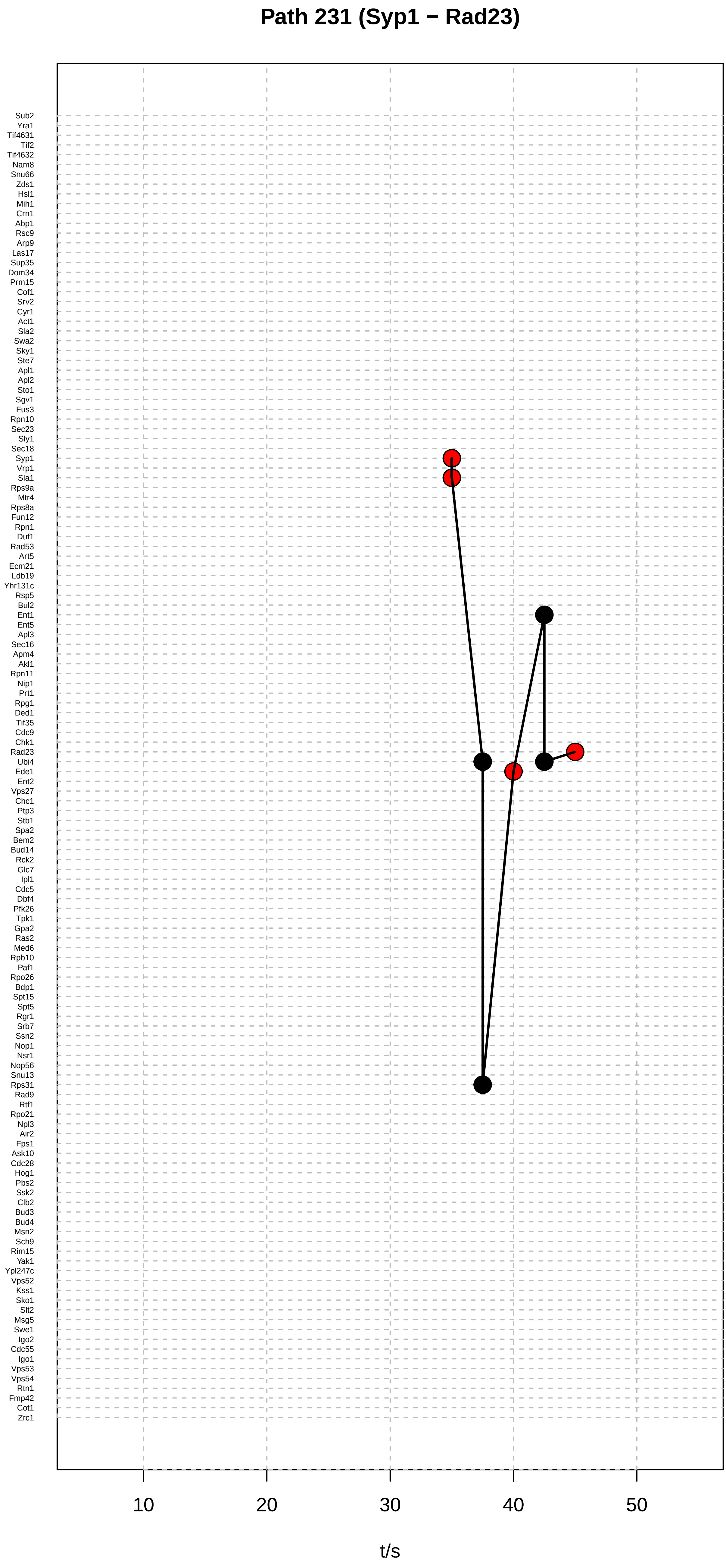

Path 231 (Syp1 − Rad23)
Sub2
Yra1
Tif4631
Tif2
Tif4632
Nam8
Snu66
Zds1
Hsl1
Mih1
Crn1
Abp1
Rsc9
Arp9
Las17
Sup35
Dom34
Prm15
Cof1
Srv2
Cyr1
Act1
Sla2
Swa2
Sky1
Ste7
Apl1
Apl2
Sto1
Sgv1
Fus3
Rpn10
Sec23
Sly1
Sec18
Syp1
Vrp1
Sla1
Rps9a
Mtr4
Rps8a
Fun12
Rpn1
Duf1
Rad53
Art5
Ecm21
Ldb19
Yhr131c
Rsp5
Bul2
Ent1
Ent5
Apl3
Sec16
Apm4
Akl1
Rpn11
Nip1
Prt1
Rpg1
Ded1
Tif35
Cdc9
Chk1
Rad23
Ubi4
Ede1
Ent2
Vps27
Chc1
Ptp3
Stb1
Spa2
Bem2
Bud14
Rck2
Glc7
Ipl1
Cdc5
Dbf4
Pfk26
Tpk1
Gpa2
Ras2
Med6
Rpb10
Paf1
Rpo26
Bdp1
Spt15
Spt5
Rgr1
Srb7
Ssn2
Nop1
Nsr1
Nop56
Snu13
Rps31
Rad9
Rtf1
Rpo21
Npl3
Air2
Fps1
Ask10
Cdc28
Hog1
Pbs2
Ssk2
Clb2
Bud3
Bud4
Msn2
Sch9
Rim15
Yak1
Ypl247c
Vps52
Kss1
Sko1
Slt2
Msg5
Swe1
Igo2
Cdc55
Igo1
Vps53
Vps54
Rtn1
Fmp42
Cot1
Zrc1
10
20
30
40
50
t/s

### Path 232 (Syp1 − Ent5)

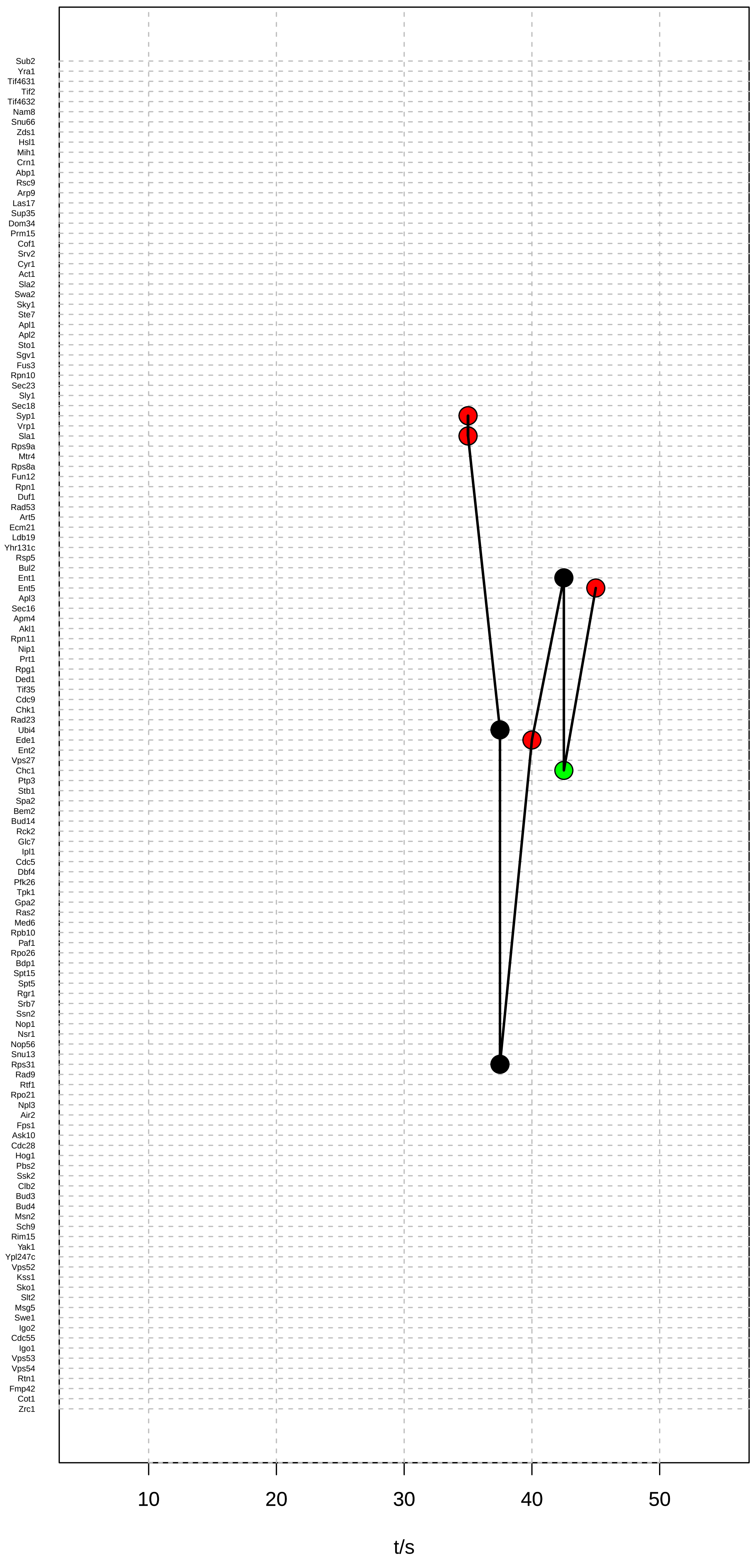

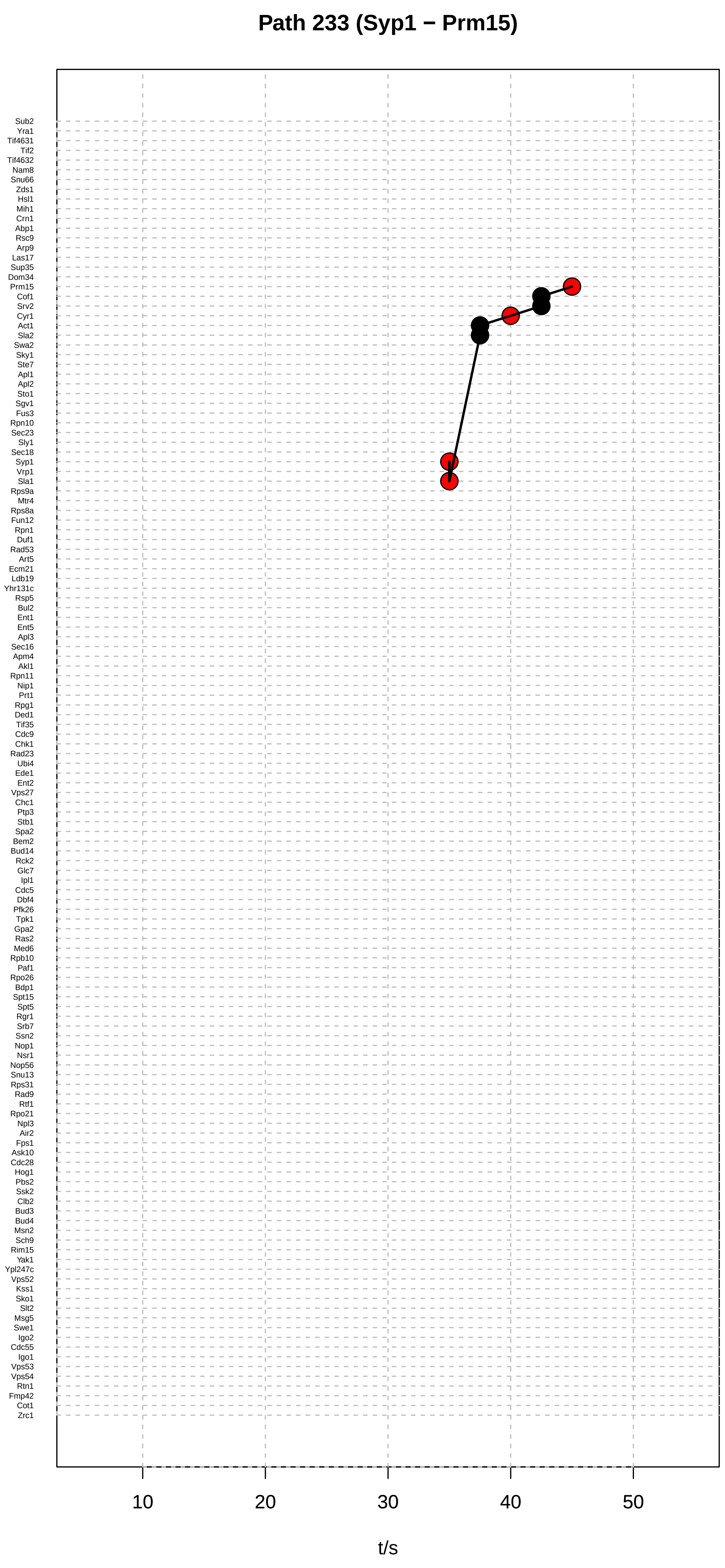
Path 233 (Syp1 − Prm15)
Sub2
Yra1
Tif4631
Tif2
Tif4632
Nam8
Snu66
Zds1
Hsl1
Mih1
Crn1
Abp1
Rsc9
Arp9
Las17
Sup35
Dom34
Prm15
Cof1
Srv2
Cyr1
Act1
Sla2
Swa2
Sky1
Ste7
Apl1
Apl2
Sto1
Sgv1
Fus3
Rpn10
Sec23
Sly1
Sec18
Syp1
Vrp1
Sla1
Rps9a
Mtr4
Rps8a
Fun12
Rpn1
Duf1
Rad53
Art5
Ecm21
Ldb19
Yhr131c
Rsp5
Bul2
Ent1
Ent5
Apl3
Sec16
Apm4
Akl1
Rpn11
Nip1
Prt1
Rpg1
Ded1
Tif35
Cdc9
Chk1
Rad23
Ubi4
Ede1
Ent2
Vps27
Chc1
Ptp3
Stb1
Spa2
Bem2
Bud14
Rck2
Glc7
Ipl1
Cdc5
Dbf4
Pfk26
Tpk1
Gpa2
Ras2
Med6
Rpb10
Paf1
Rpo26
Bdp1
Spt15
Spt5
Rgr1
Srb7
Ssn2
Nop1
Nsr1
Nop56
Snu13
Rps31
Rad9
Rtf1
Rpo21
Npl3
Air2
Fps1
Ask10
Cdc28
Hog1
Pbs2
Ssk2
Clb2
Bud3
Bud4
Msn2
Sch9
Rim15
Yak1
Ypl247c
Vps52
Kss1
Sko1
Slt2
Msg5
Swe1
Igo2
Cdc55
Igo1
Vps53
Vps54
Rtn1
Fmp42
Cot1
Zrc1
10
20
30
40
50
t/s

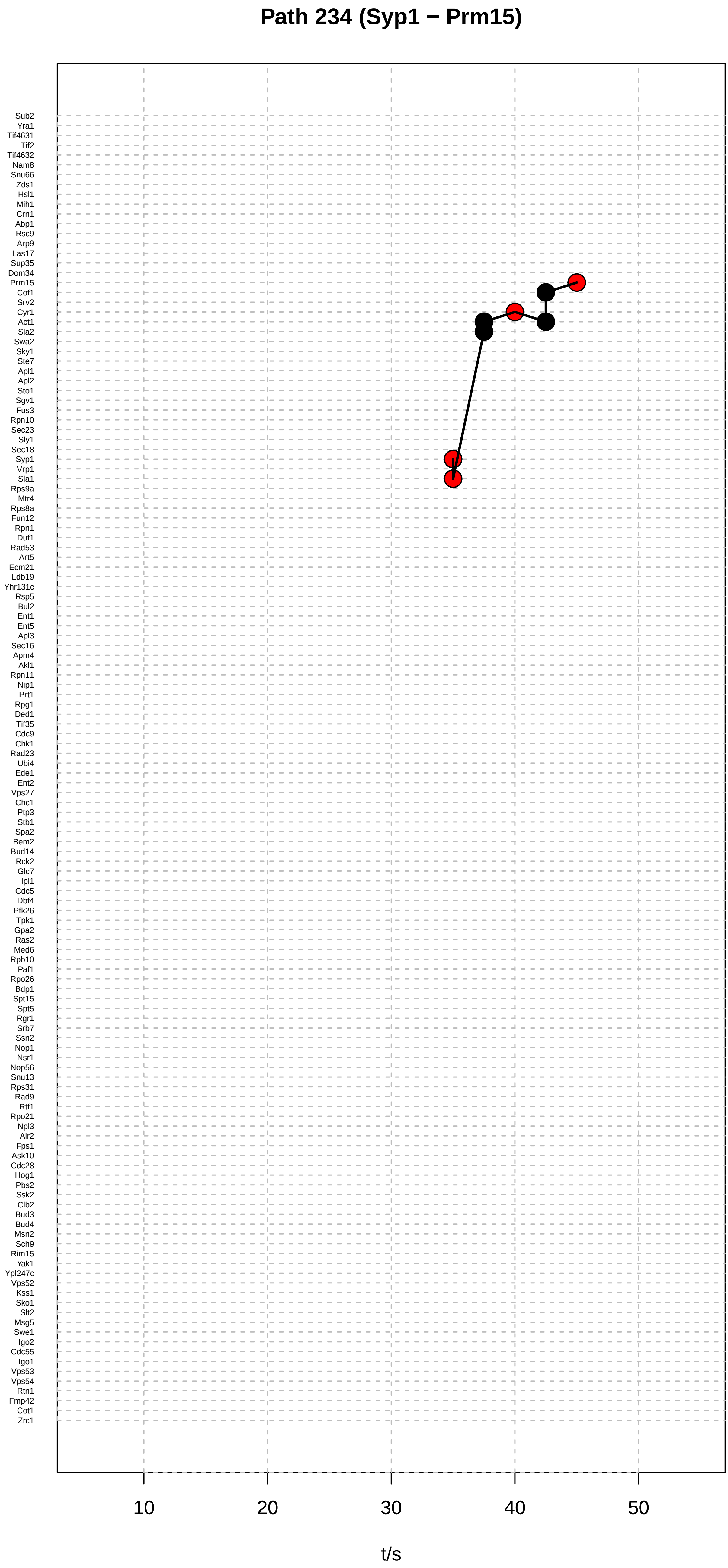
Path 234 (Syp1 − Prm15)
Sub2
Yra1
Tif4631
Tif2
Tif4632
Nam8
Snu66
Zds1
Hsl1
Mih1
Crn1
Abp1
Rsc9
Arp9
Las17
Sup35
Dom34
Prm15
Cof1
Srv2
Cyr1
Act1
Sla2
Swa2
Sky1
Ste7
Apl1
Apl2
Sto1
Sgv1
Fus3
Rpn10
Sec23
Sly1
Sec18
Syp1
Vrp1
Sla1
Rps9a
Mtr4
Rps8a
Fun12
Rpn1
Duf1
Rad53
Art5
Ecm21
Ldb19
Yhr131c
Rsp5
Bul2
Ent1
Ent5
Apl3
Sec16
Apm4
Akl1
Rpn11
Nip1
Prt1
Rpg1
Ded1
Tif35
Cdc9
Chk1
Rad23
Ubi4
Ede1
Ent2
Vps27
Chc1
Ptp3
Stb1
Spa2
Bem2
Bud14
Rck2
Glc7
Ipl1
Cdc5
Dbf4
Pfk26
Tpk1
Gpa2
Ras2
Med6
Rpb10
Paf1
Rpo26
Bdp1
Spt15
Spt5
Rgr1
Srb7
Ssn2
Nop1
Nsr1
Nop56
Snu13
Rps31
Rad9
Rtf1
Rpo21
Npl3
Air2
Fps1
Ask10
Cdc28
Hog1
Pbs2
Ssk2
Clb2
Bud3
Bud4
Msn2
Sch9
Rim15
Yak1
Ypl247c
Vps52
Kss1
Sko1
Slt2
Msg5
Swe1
Igo2
Cdc55
Igo1
Vps53
Vps54
Rtn1
Fmp42
Cot1
Zrc1
10
20
30
40
50
t/s

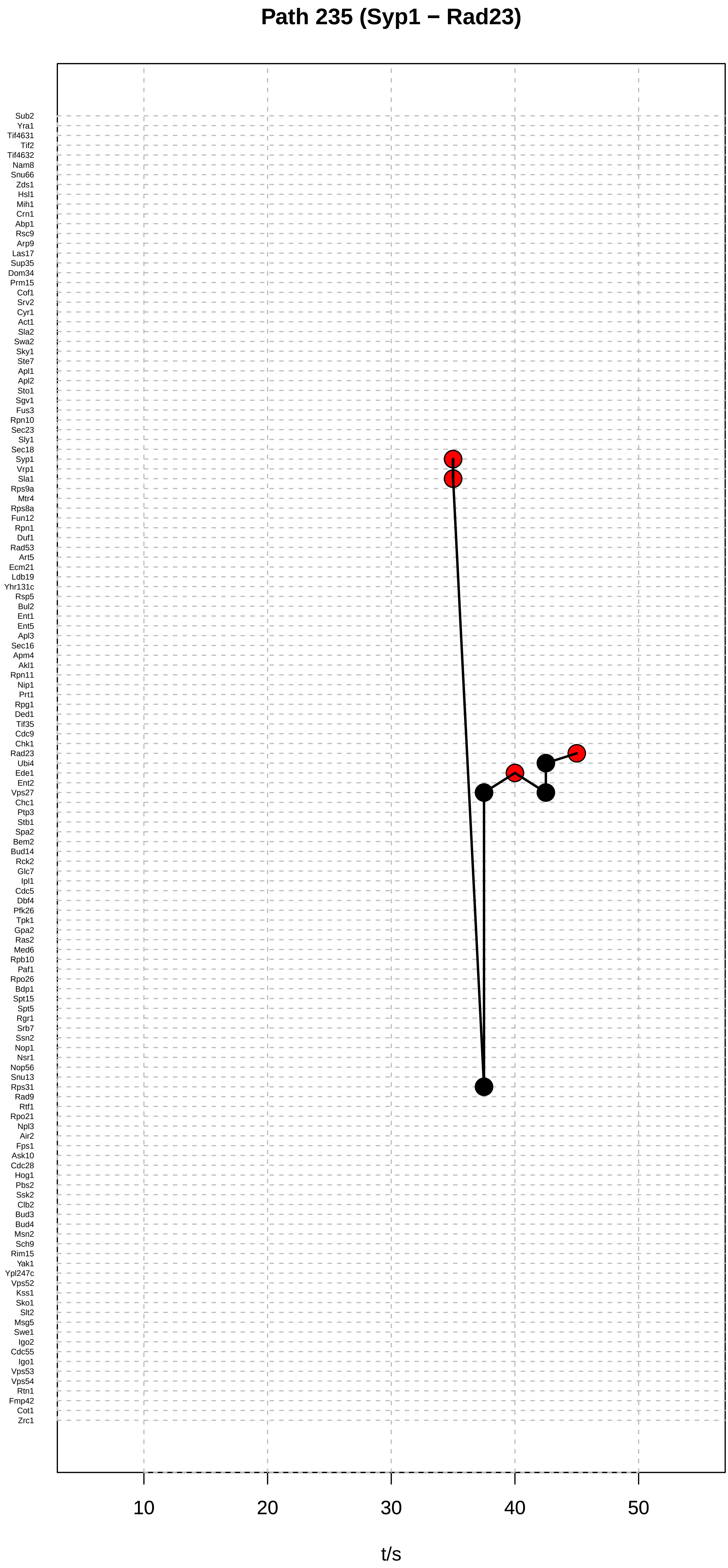

Path 235 (Syp1 − Rad23)
Sub2
Yra1
Tif4631
Tif2
Tif4632
Nam8
Snu66
Zds1
Hsl1
Mih1
Crn1
Abp1
Rsc9
Arp9
Las17
Sup35
Dom34
Prm15
Cof1
Srv2
Cyr1
Act1
Sla2
Swa2
Sky1
Ste7
Apl1
Apl2
Sto1
Sgv1
Fus3
Rpn10
Sec23
Sly1
Sec18
Syp1
Vrp1
Sla1
Rps9a
Mtr4
Rps8a
Fun12
Rpn1
Duf1
Rad53
Art5
Ecm21
Ldb19
Yhr131c
Rsp5
Bul2
Ent1
Ent5
Apl3
Sec16
Apm4
Akl1
Rpn11
Nip1
Prt1
Rpg1
Ded1
Tif35
Cdc9
Chk1
Rad23
Ubi4
Ede1
Ent2
Vps27
Chc1
Ptp3
Stb1
Spa2
Bem2
Bud14
Rck2
Glc7
Ipl1
Cdc5
Dbf4
Pfk26
Tpk1
Gpa2
Ras2
Med6
Rpb10
Paf1
Rpo26
Bdp1
Spt15
Spt5
Rgr1
Srb7
Ssn2
Nop1
Nsr1
Nop56
Snu13
Rps31
Rad9
Rtf1
Rpo21
Npl3
Air2
Fps1
Ask10
Cdc28
Hog1
Pbs2
Ssk2
Clb2
Bud3
Bud4
Msn2
Sch9
Rim15
Yak1
Ypl247c
Vps52
Kss1
Sko1
Slt2
Msg5
Swe1
Igo2
Cdc55
Igo1
Vps53
Vps54
Rtn1
Fmp42
Cot1
Zrc1
10
20
30
40
50
t/s

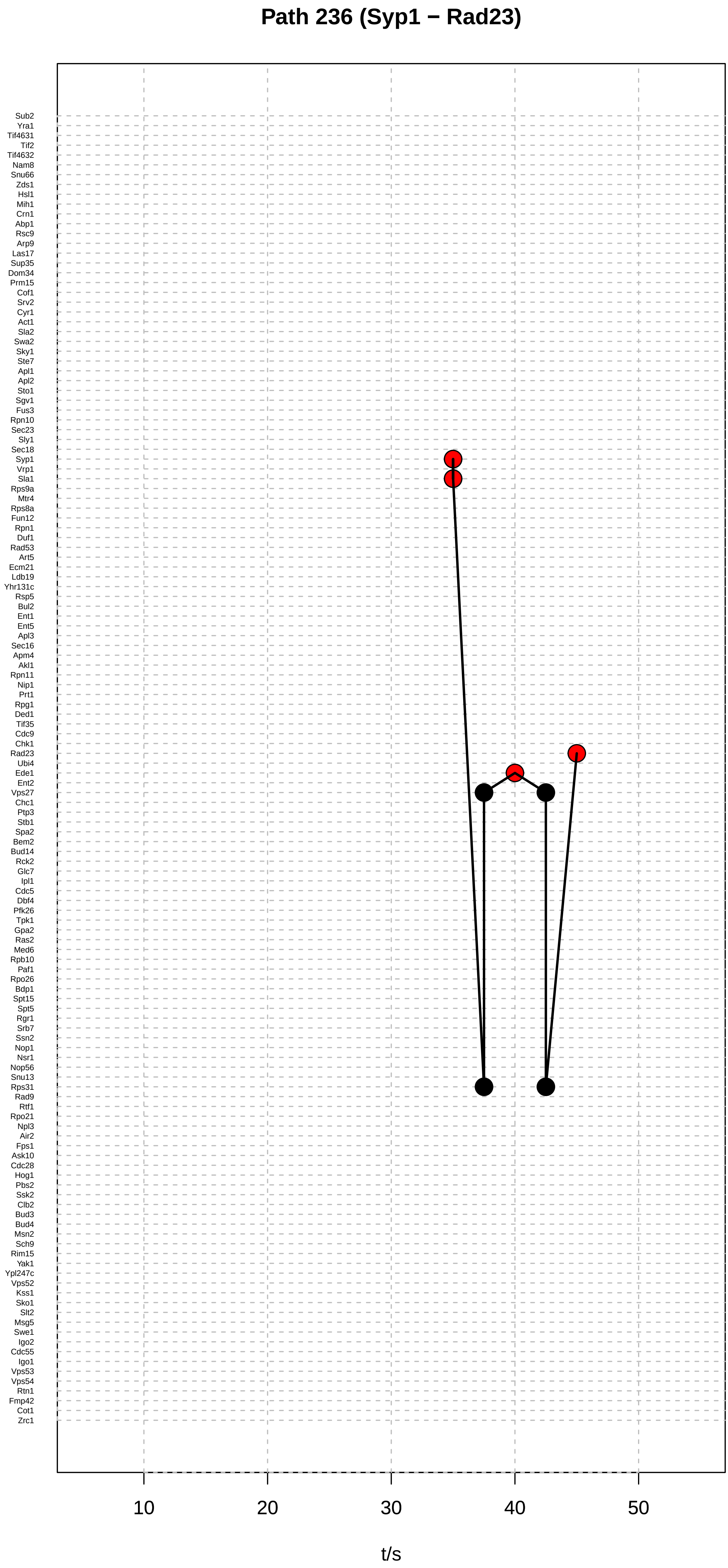

Path 236 (Syp1 − Rad23)
Sub2
Yra1
Tif4631
Tif2
Tif4632
Nam8
Snu66
Zds1
Hsl1
Mih1
Crn1
Abp1
Rsc9
Arp9
Las17
Sup35
Dom34
Prm15
Cof1
Srv2
Cyr1
Act1
Sla2
Swa2
Sky1
Ste7
Apl1
Apl2
Sto1
Sgv1
Fus3
Rpn10
Sec23
Sly1
Sec18
Syp1
Vrp1
Sla1
Rps9a
Mtr4
Rps8a
Fun12
Rpn1
Duf1
Rad53
Art5
Ecm21
Ldb19
Yhr131c
Rsp5
Bul2
Ent1
Ent5
Apl3
Sec16
Apm4
Akl1
Rpn11
Nip1
Prt1
Rpg1
Ded1
Tif35
Cdc9
Chk1
Rad23
Ubi4
Ede1
Ent2
Vps27
Chc1
Ptp3
Stb1
Spa2
Bem2
Bud14
Rck2
Glc7
Ipl1
Cdc5
Dbf4
Pfk26
Tpk1
Gpa2
Ras2
Med6
Rpb10
Paf1
Rpo26
Bdp1
Spt15
Spt5
Rgr1
Srb7
Ssn2
Nop1
Nsr1
Nop56
Snu13
Rps31
Rad9
Rtf1
Rpo21
Npl3
Air2
Fps1
Ask10
Cdc28
Hog1
Pbs2
Ssk2
Clb2
Bud3
Bud4
Msn2
Sch9
Rim15
Yak1
Ypl247c
Vps52
Kss1
Sko1
Slt2
Msg5
Swe1
Igo2
Cdc55
Igo1
Vps53
Vps54
Rtn1
Fmp42
Cot1
Zrc1
10
20
30
40
50
t/s

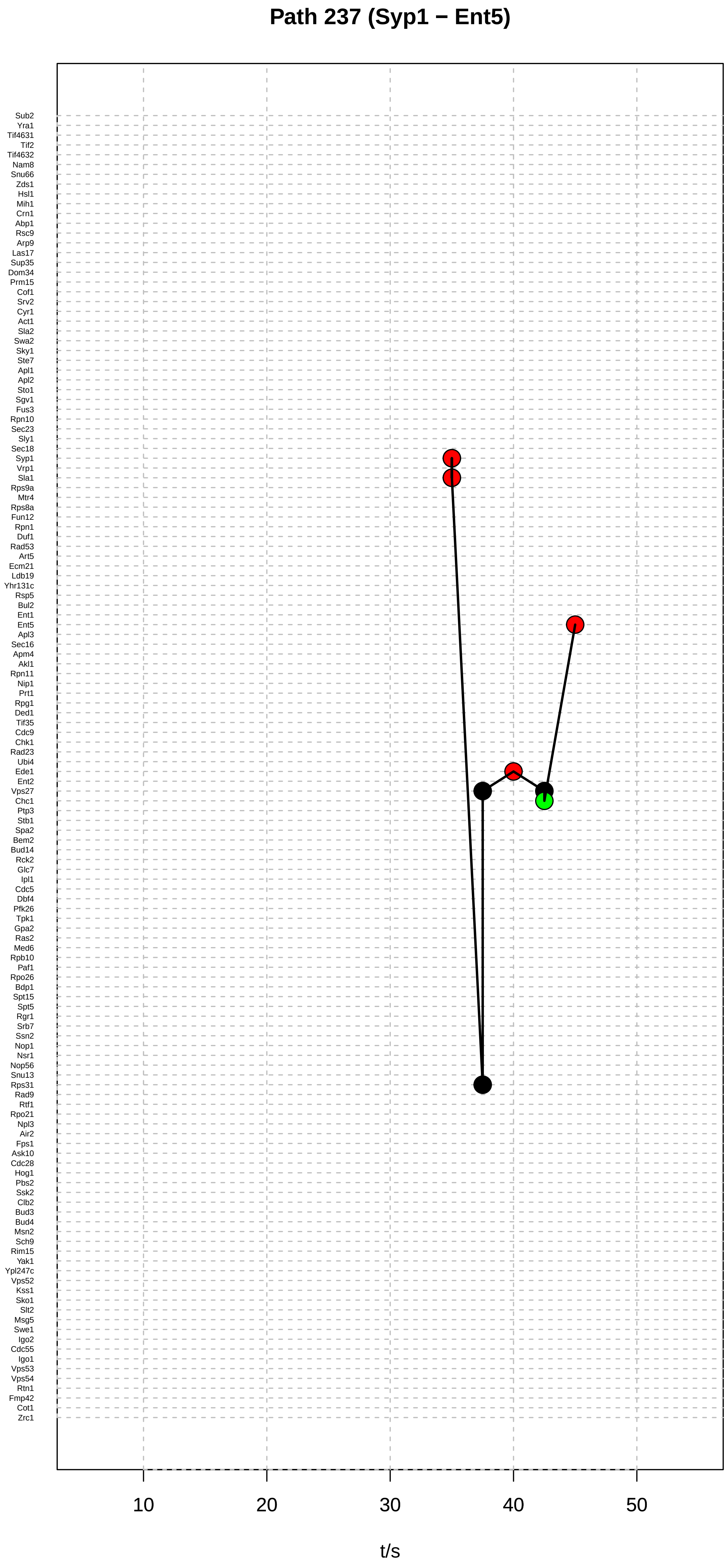
Path 237 (Syp1 − Ent5)
Sub2
Yra1
Tif4631
Tif2
Tif4632
Nam8
Snu66
Zds1
Hsl1
Mih1
Crn1
Abp1
Rsc9
Arp9
Las17
Sup35
Dom34
Prm15
Cof1
Srv2
Cyr1
Act1
Sla2
Swa2
Sky1
Ste7
Apl1
Apl2
Sto1
Sgv1
Fus3
Rpn10
Sec23
Sly1
Sec18
Syp1
Vrp1
Sla1
Rps9a
Mtr4
Rps8a
Fun12
Rpn1
Duf1
Rad53
Art5
Ecm21
Ldb19
Yhr131c
Rsp5
Bul2
Ent1
Ent5
Apl3
Sec16
Apm4
Akl1
Rpn11
Nip1
Prt1
Rpg1
Ded1
Tif35
Cdc9
Chk1
Rad23
Ubi4
Ede1
Ent2
Vps27
Chc1
Ptp3
Stb1
Spa2
Bem2
Bud14
Rck2
Glc7
Ipl1
Cdc5
Dbf4
Pfk26
Tpk1
Gpa2
Ras2
Med6
Rpb10
Paf1
Rpo26
Bdp1
Spt15
Spt5
Rgr1
Srb7
Ssn2
Nop1
Nsr1
Nop56
Snu13
Rps31
Rad9
Rtf1
Rpo21
Npl3
Air2
Fps1
Ask10
Cdc28
Hog1
Pbs2
Ssk2
Clb2
Bud3
Bud4
Msn2
Sch9
Rim15
Yak1
Ypl247c
Vps52
Kss1
Sko1
Slt2
Msg5
Swe1
Igo2
Cdc55
Igo1
Vps53
Vps54
Rtn1
Fmp42
Cot1
Zrc1
10
20
30
40
50
t/s

### Path 238 (Syp1 − Rad23)

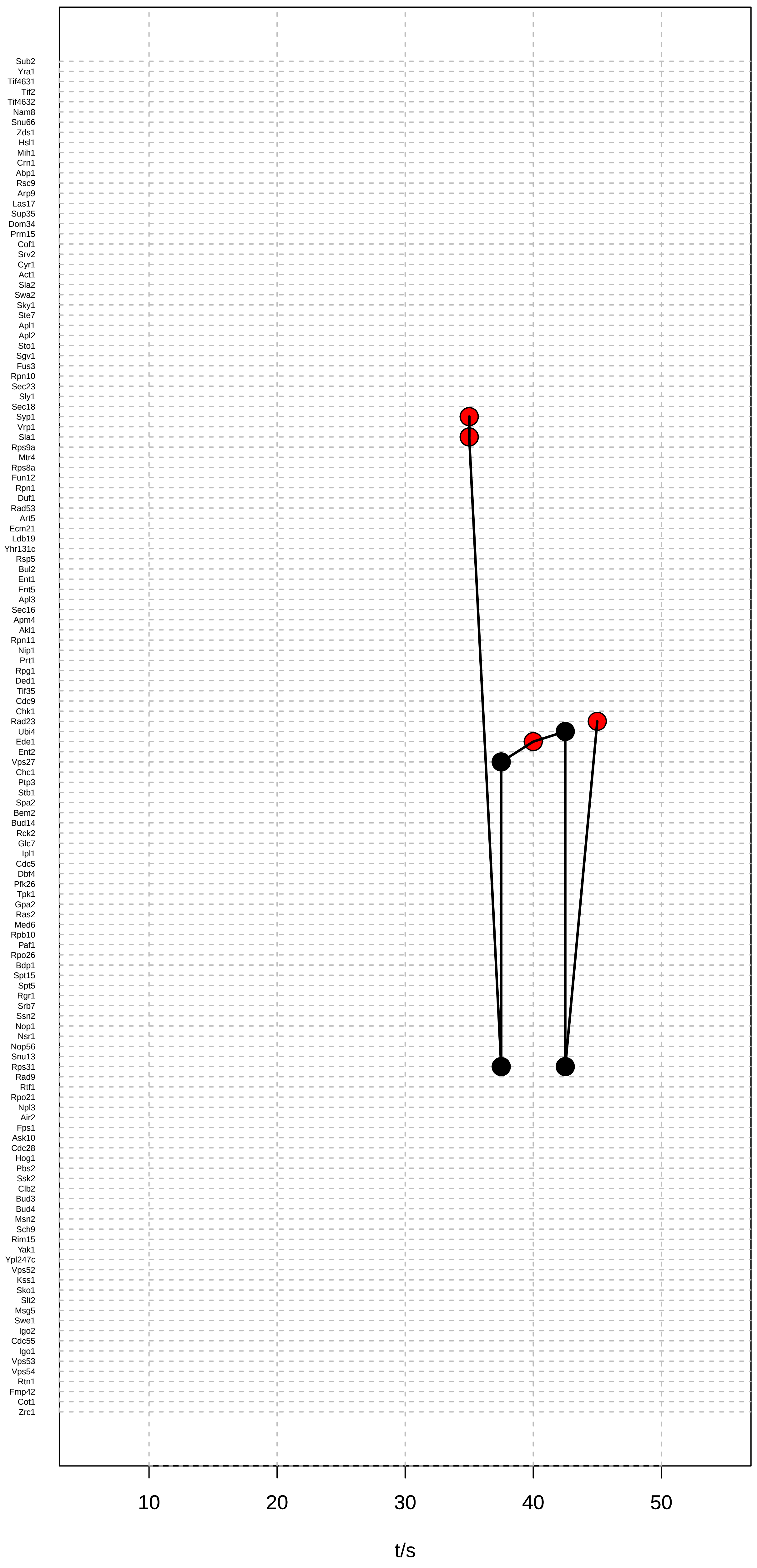

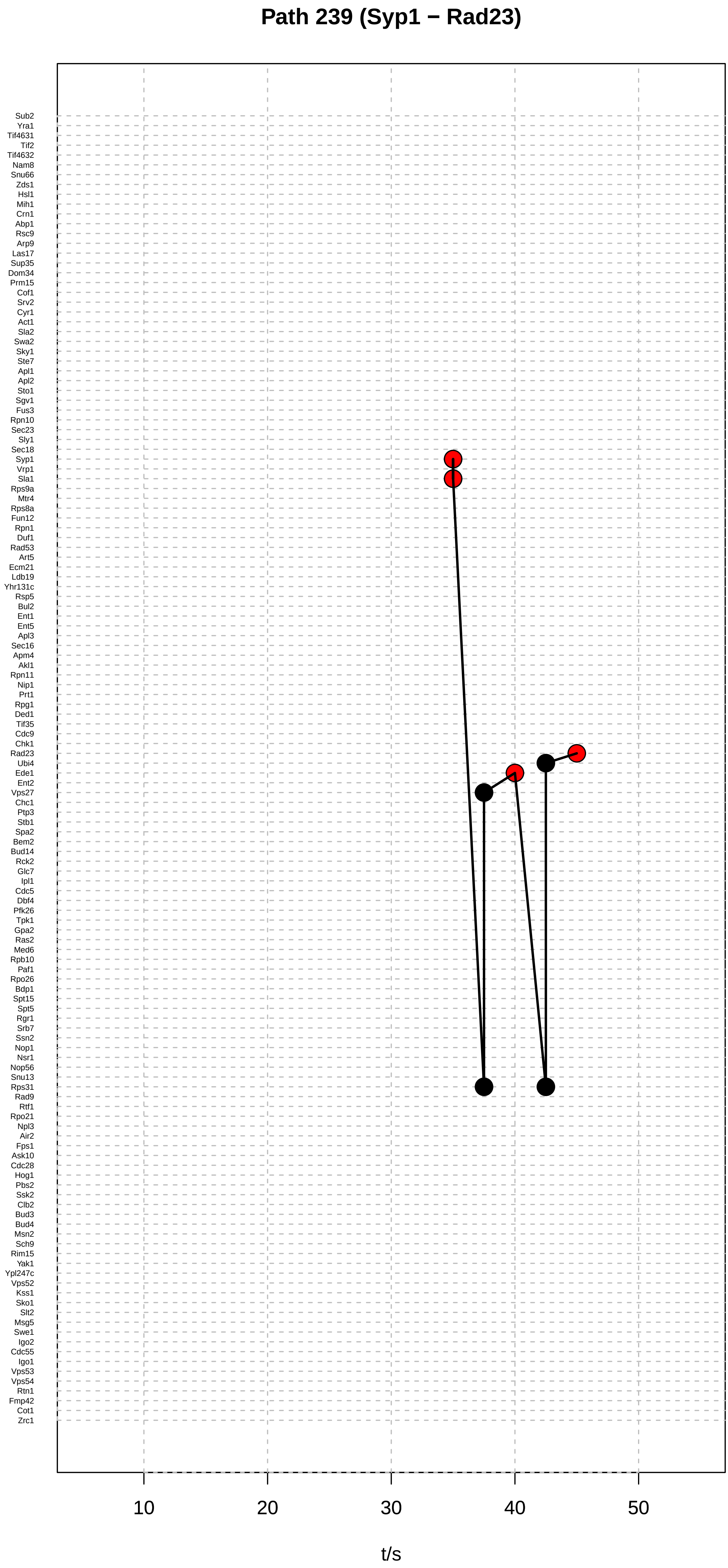
Path 239 (Syp1 − Rad23)
Sub2
Yra1
Tif4631
Tif2
Tif4632
Nam8
Snu66
Zds1
Hsl1
Mih1
Crn1
Abp1
Rsc9
Arp9
Las17
Sup35
Dom34
Prm15
Cof1
Srv2
Cyr1
Act1
Sla2
Swa2
Sky1
Ste7
Apl1
Apl2
Sto1
Sgv1
Fus3
Rpn10
Sec23
Sly1
Sec18
Syp1
Vrp1
Sla1
Rps9a
Mtr4
Rps8a
Fun12
Rpn1
Duf1
Rad53
Art5
Ecm21
Ldb19
Yhr131c
Rsp5
Bul2
Ent1
Ent5
Apl3
Sec16
Apm4
Akl1
Rpn11
Nip1
Prt1
Rpg1
Ded1
Tif35
Cdc9
Chk1
Rad23
Ubi4
Ede1
Ent2
Vps27
Chc1
Ptp3
Stb1
Spa2
Bem2
Bud14
Rck2
Glc7
Ipl1
Cdc5
Dbf4
Pfk26
Tpk1
Gpa2
Ras2
Med6
Rpb10
Paf1
Rpo26
Bdp1
Spt15
Spt5
Rgr1
Srb7
Ssn2
Nop1
Nsr1
Nop56
Snu13
Rps31
Rad9
Rtf1
Rpo21
Npl3
Air2
Fps1
Ask10
Cdc28
Hog1
Pbs2
Ssk2
Clb2
Bud3
Bud4
Msn2
Sch9
Rim15
Yak1
Ypl247c
Vps52
Kss1
Sko1
Slt2
Msg5
Swe1
Igo2
Cdc55
Igo1
Vps53
Vps54
Rtn1
Fmp42
Cot1
Zrc1
10
20
30
40
50
t/s

#### Path 240 (Syp1 − Rad23)

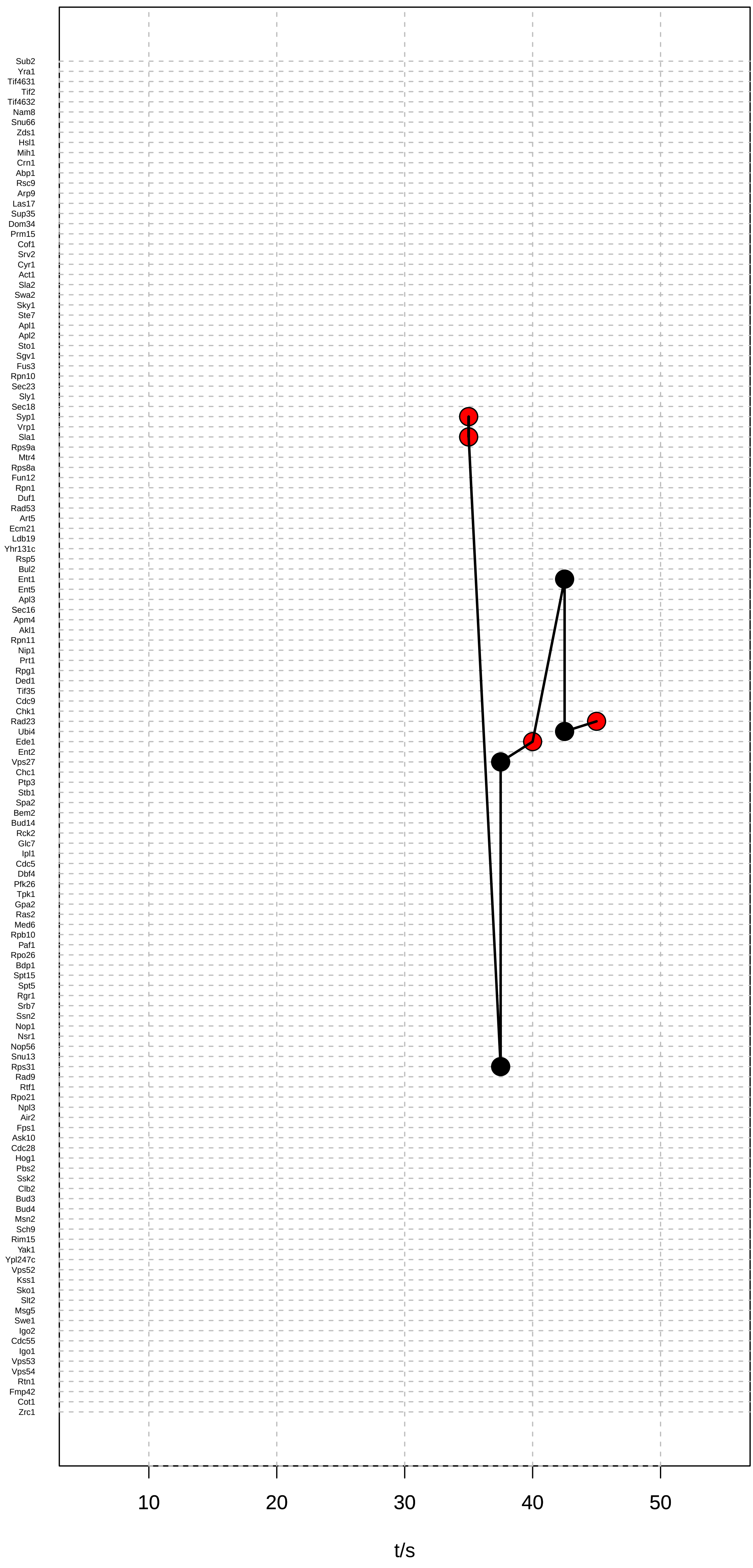

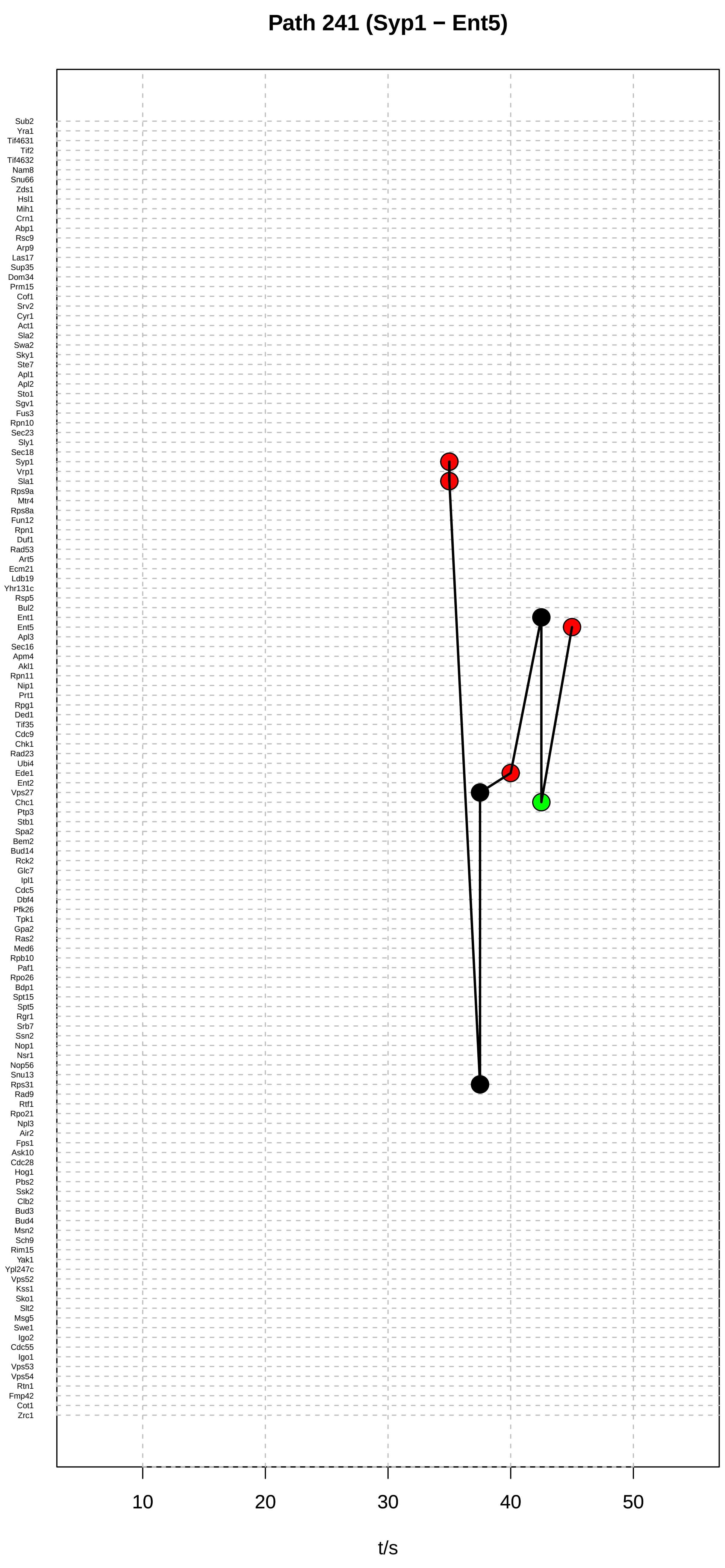

Path 241 (Syp1 − Ent5)
Sub2
Yra1
Tif4631
Tif2
Tif4632
Nam8
Snu66
Zds1
Hsl1
Mih1
Crn1
Abp1
Rsc9
Arp9
Las17
Sup35
Dom34
Prm15
Cof1
Srv2
Cyr1
Act1
Sla2
Swa2
Sky1
Ste7
Apl1
Apl2
Sto1
Sgv1
Fus3
Rpn10
Sec23
Sly1
Sec18
Syp1
Vrp1
Sla1
Rps9a
Mtr4
Rps8a
Fun12
Rpn1
Duf1
Rad53
Art5
Ecm21
Ldb19
Yhr131c
Rsp5
Bul2
Ent1
Ent5
Apl3
Sec16
Apm4
Akl1
Rpn11
Nip1
Prt1
Rpg1
Ded1
Tif35
Cdc9
Chk1
Rad23
Ubi4
Ede1
Ent2
Vps27
Chc1
Ptp3
Stb1
Spa2
Bem2
Bud14
Rck2
Glc7
Ipl1
Cdc5
Dbf4
Pfk26
Tpk1
Gpa2
Ras2
Med6
Rpb10
Paf1
Rpo26
Bdp1
Spt15
Spt5
Rgr1
Srb7
Ssn2
Nop1
Nsr1
Nop56
Snu13
Rps31
Rad9
Rtf1
Rpo21
Npl3
Air2
Fps1
Ask10
Cdc28
Hog1
Pbs2
Ssk2
Clb2
Bud3
Bud4
Msn2
Sch9
Rim15
Yak1
Ypl247c
Vps52
Kss1
Sko1
Slt2
Msg5
Swe1
Igo2
Cdc55
Igo1
Vps53
Vps54
Rtn1
Fmp42
Cot1
Zrc1
10
20
30
40
50
t/s

### Path 242 (Syp1 − Rad23)

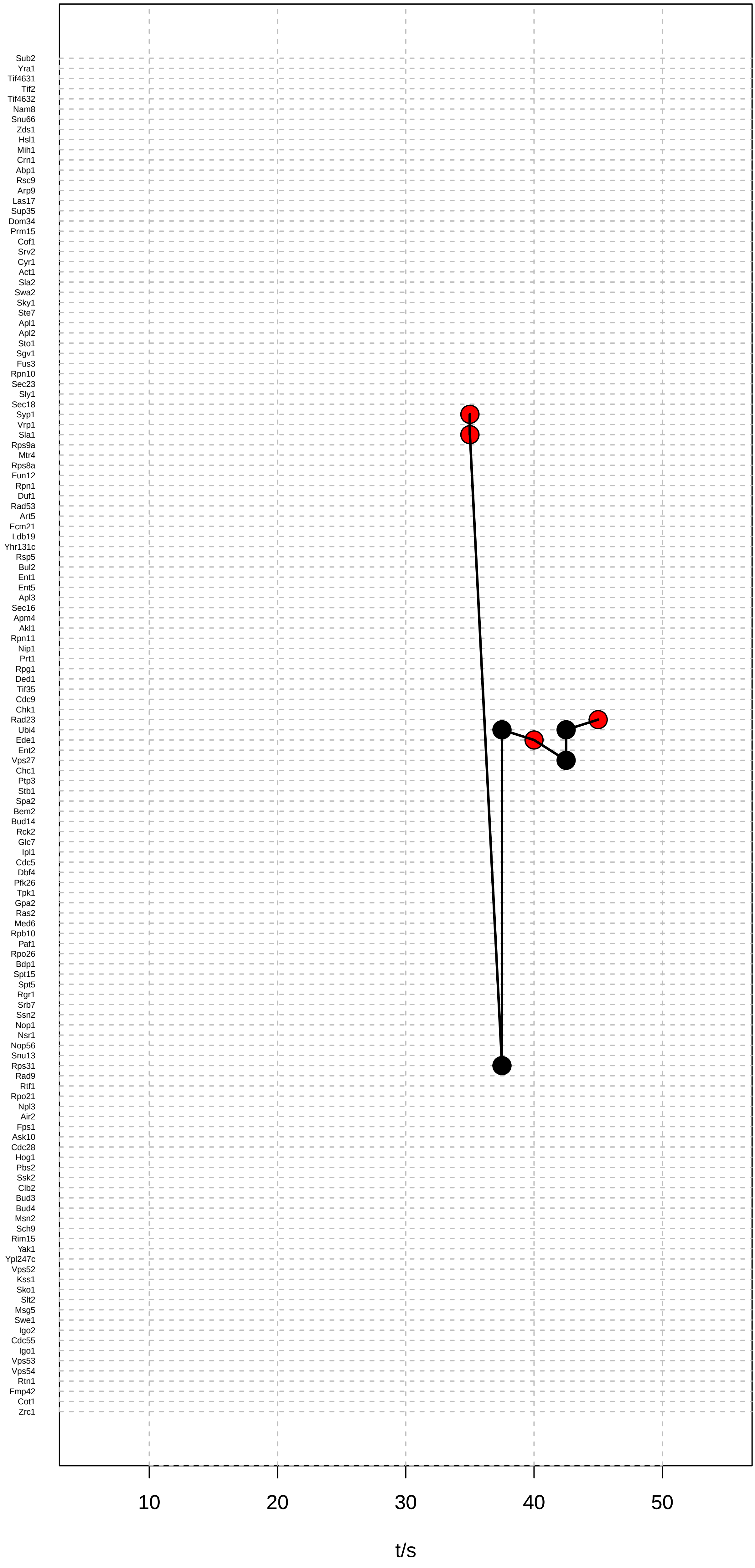

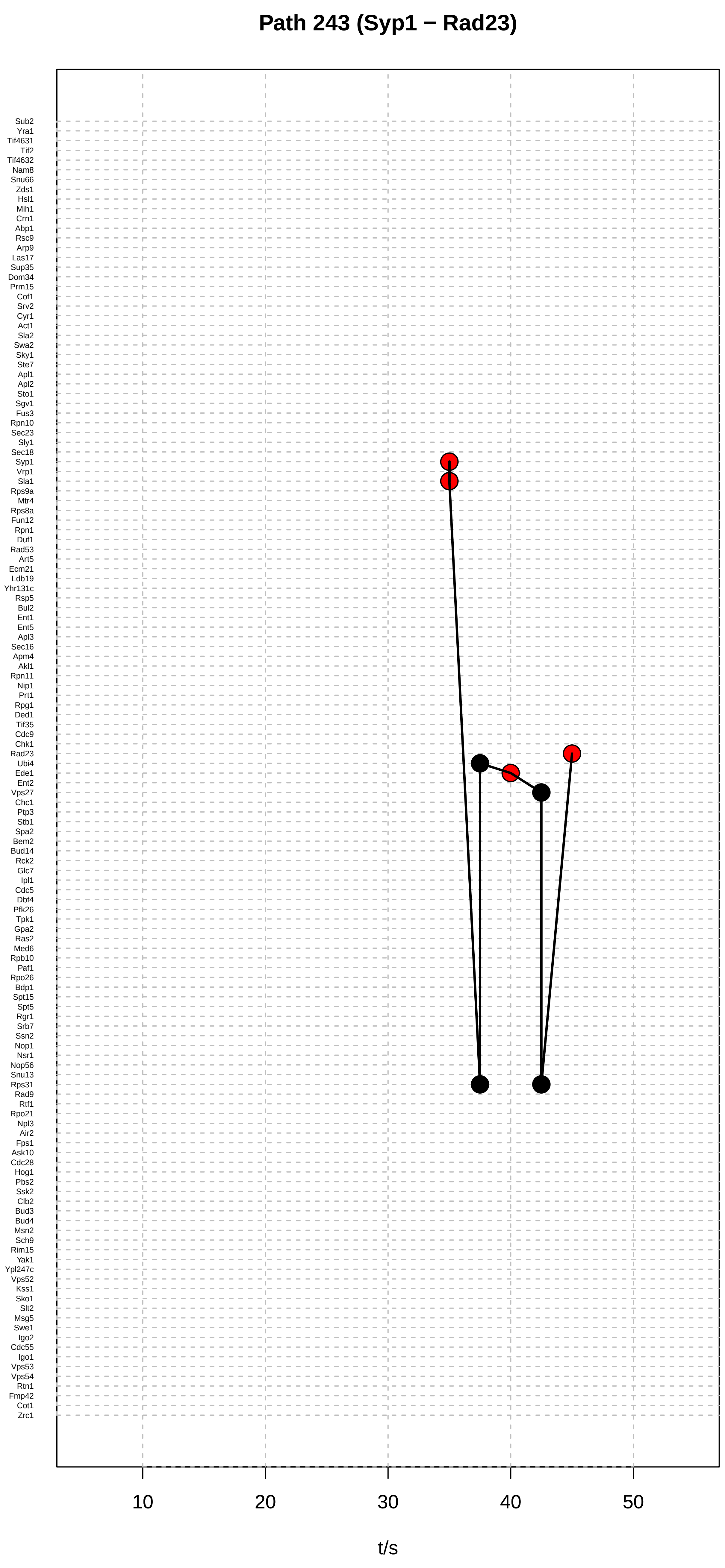

Path 243 (Syp1 − Rad23)
Sub2
Yra1
Tif4631
Tif2
Tif4632
Nam8
Snu66
Zds1
Hsl1
Mih1
Crn1
Abp1
Rsc9
Arp9
Las17
Sup35
Dom34
Prm15
Cof1
Srv2
Cyr1
Act1
Sla2
Swa2
Sky1
Ste7
Apl1
Apl2
Sto1
Sgv1
Fus3
Rpn10
Sec23
Sly1
Sec18
Syp1
Vrp1
Sla1
Rps9a
Mtr4
Rps8a
Fun12
Rpn1
Duf1
Rad53
Art5
Ecm21
Ldb19
Yhr131c
Rsp5
Bul2
Ent1
Ent5
Apl3
Sec16
Apm4
Akl1
Rpn11
Nip1
Prt1
Rpg1
Ded1
Tif35
Cdc9
Chk1
Rad23
Ubi4
Ede1
Ent2
Vps27
Chc1
Ptp3
Stb1
Spa2
Bem2
Bud14
Rck2
Glc7
Ipl1
Cdc5
Dbf4
Pfk26
Tpk1
Gpa2
Ras2
Med6
Rpb10
Paf1
Rpo26
Bdp1
Spt15
Spt5
Rgr1
Srb7
Ssn2
Nop1
Nsr1
Nop56
Snu13
Rps31
Rad9
Rtf1
Rpo21
Npl3
Air2
Fps1
Ask10
Cdc28
Hog1
Pbs2
Ssk2
Clb2
Bud3
Bud4
Msn2
Sch9
Rim15
Yak1
Ypl247c
Vps52
Kss1
Sko1
Slt2
Msg5
Swe1
Igo2
Cdc55
Igo1
Vps53
Vps54
Rtn1
Fmp42
Cot1
Zrc1
10
20
30
40
50
t/s

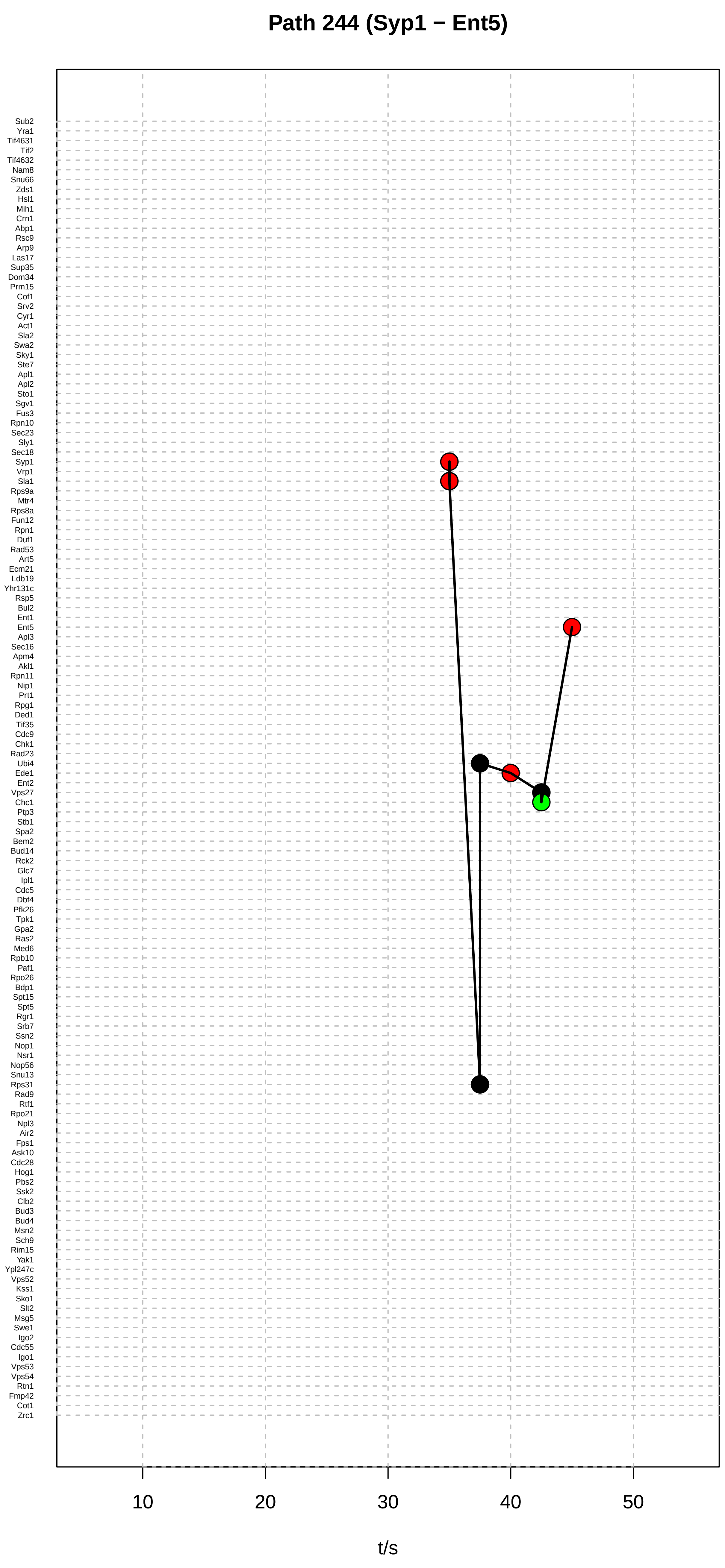
Path 244 (Syp1 − Ent5)
Sub2
Yra1
Tif4631
Tif2
Tif4632
Nam8
Snu66
Zds1
Hsl1
Mih1
Crn1
Abp1
Rsc9
Arp9
Las17
Sup35
Dom34
Prm15
Cof1
Srv2
Cyr1
Act1
Sla2
Swa2
Sky1
Ste7
Apl1
Apl2
Sto1
Sgv1
Fus3
Rpn10
Sec23
Sly1
Sec18
Syp1
Vrp1
Sla1
Rps9a
Mtr4
Rps8a
Fun12
Rpn1
Duf1
Rad53
Art5
Ecm21
Ldb19
Yhr131c
Rsp5
Bul2
Ent1
Ent5
Apl3
Sec16
Apm4
Akl1
Rpn11
Nip1
Prt1
Rpg1
Ded1
Tif35
Cdc9
Chk1
Rad23
Ubi4
Ede1
Ent2
Vps27
Chc1
Ptp3
Stb1
Spa2
Bem2
Bud14
Rck2
Glc7
Ipl1
Cdc5
Dbf4
Pfk26
Tpk1
Gpa2
Ras2
Med6
Rpb10
Paf1
Rpo26
Bdp1
Spt15
Spt5
Rgr1
Srb7
Ssn2
Nop1
Nsr1
Nop56
Snu13
Rps31
Rad9
Rtf1
Rpo21
Npl3
Air2
Fps1
Ask10
Cdc28
Hog1
Pbs2
Ssk2
Clb2
Bud3
Bud4
Msn2
Sch9
Rim15
Yak1
Ypl247c
Vps52
Kss1
Sko1
Slt2
Msg5
Swe1
Igo2
Cdc55
Igo1
Vps53
Vps54
Rtn1
Fmp42
Cot1
Zrc1
10
20
30
40
50
t/s

### Path 245 (Syp1 − Rad23)

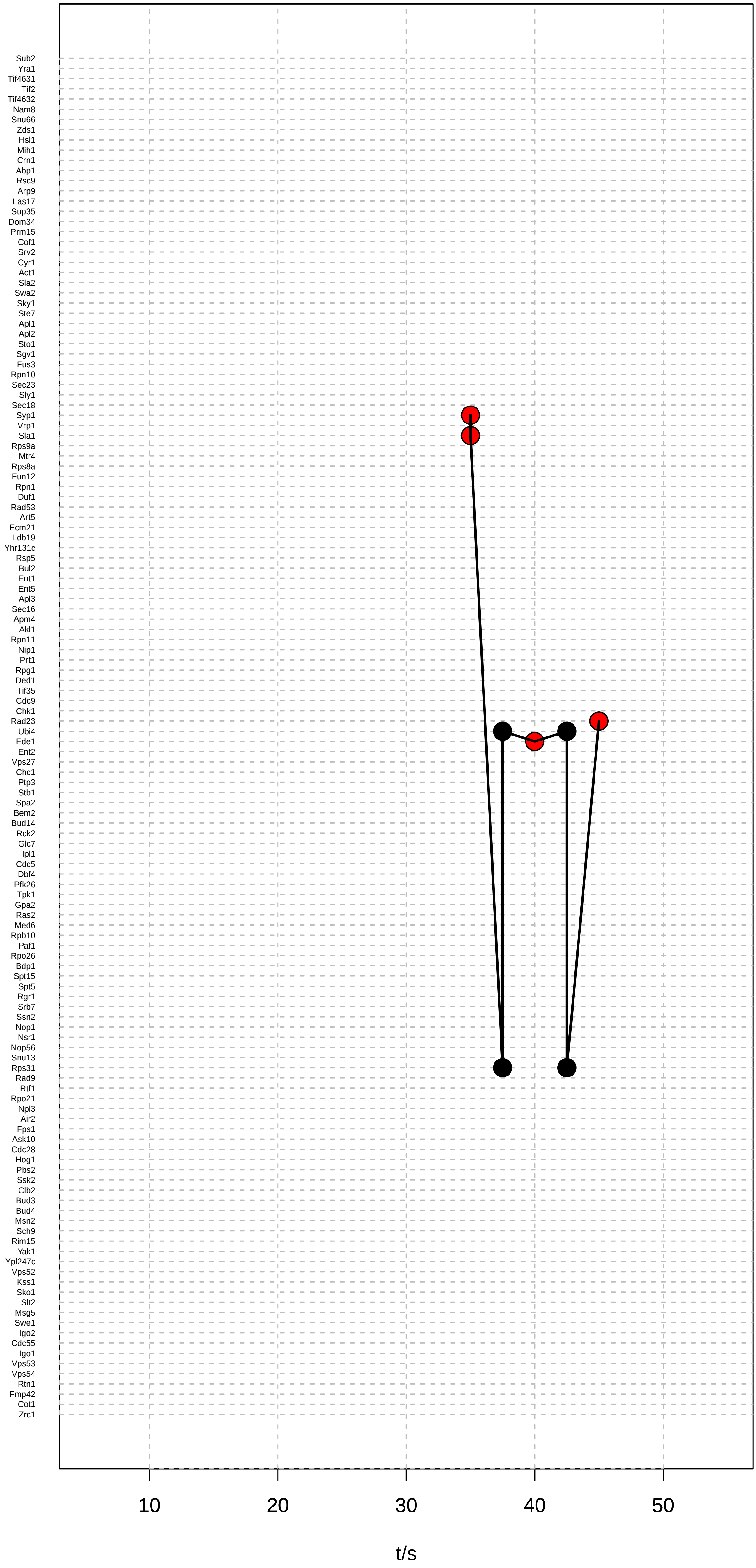

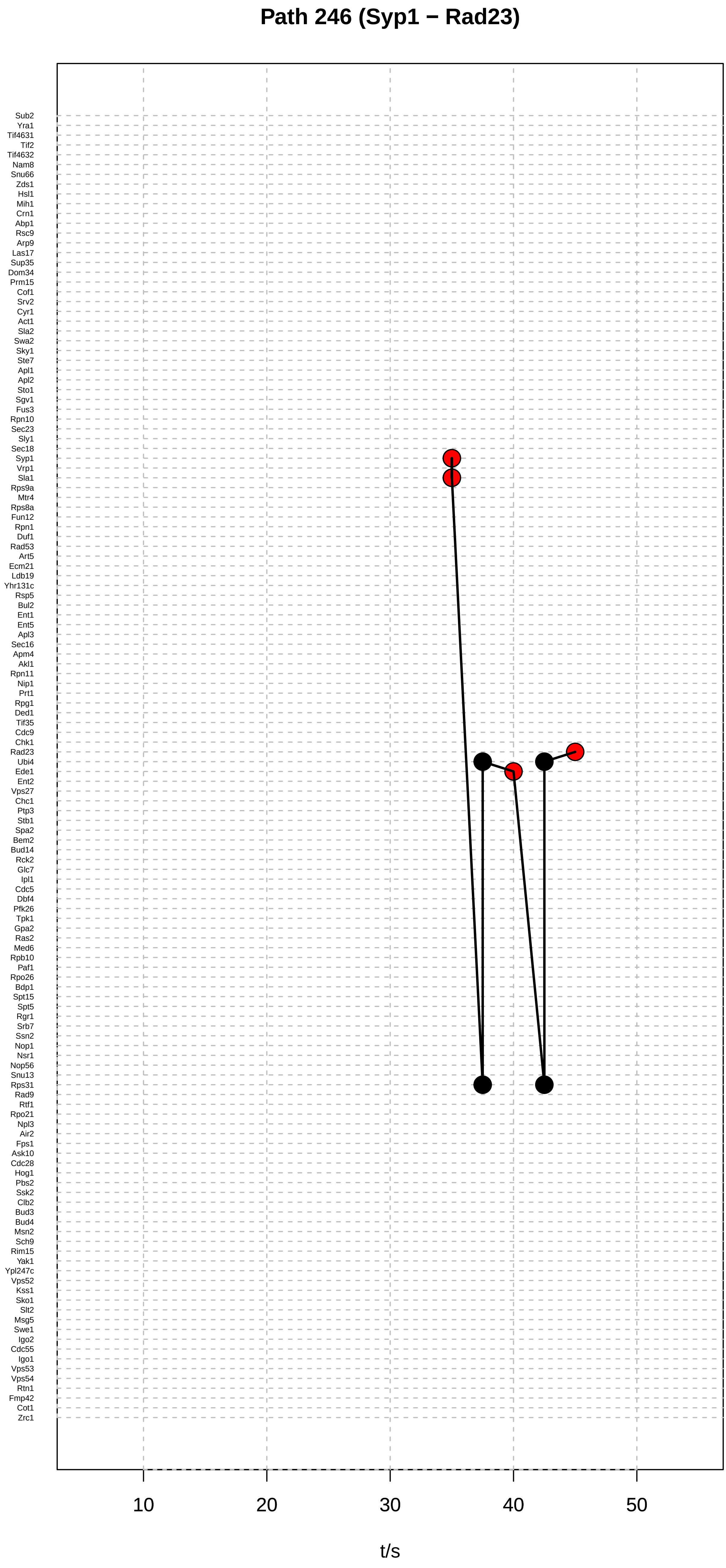
Path 246 (Syp1 − Rad23)
Sub2
Yra1
Tif4631
Tif2
Tif4632
Nam8
Snu66
Zds1
Hsl1
Mih1
Crn1
Abp1
Rsc9
Arp9
Las17
Sup35
Dom34
Prm15
Cof1
Srv2
Cyr1
Act1
Sla2
Swa2
Sky1
Ste7
Apl1
Apl2
Sto1
Sgv1
Fus3
Rpn10
Sec23
Sly1
Sec18
Syp1
Vrp1
Sla1
Rps9a
Mtr4
Rps8a
Fun12
Rpn1
Duf1
Rad53
Art5
Ecm21
Ldb19
Yhr131c
Rsp5
Bul2
Ent1
Ent5
Apl3
Sec16
Apm4
Akl1
Rpn11
Nip1
Prt1
Rpg1
Ded1
Tif35
Cdc9
Chk1
Rad23
Ubi4
Ede1
Ent2
Vps27
Chc1
Ptp3
Stb1
Spa2
Bem2
Bud14
Rck2
Glc7
Ipl1
Cdc5
Dbf4
Pfk26
Tpk1
Gpa2
Ras2
Med6
Rpb10
Paf1
Rpo26
Bdp1
Spt15
Spt5
Rgr1
Srb7
Ssn2
Nop1
Nsr1
Nop56
Snu13
Rps31
Rad9
Rtf1
Rpo21
Npl3
Air2
Fps1
Ask10
Cdc28
Hog1
Pbs2
Ssk2
Clb2
Bud3
Bud4
Msn2
Sch9
Rim15
Yak1
Ypl247c
Vps52
Kss1
Sko1
Slt2
Msg5
Swe1
Igo2
Cdc55
Igo1
Vps53
Vps54
Rtn1
Fmp42
Cot1
Zrc1
10
20
30
40
50
t/s

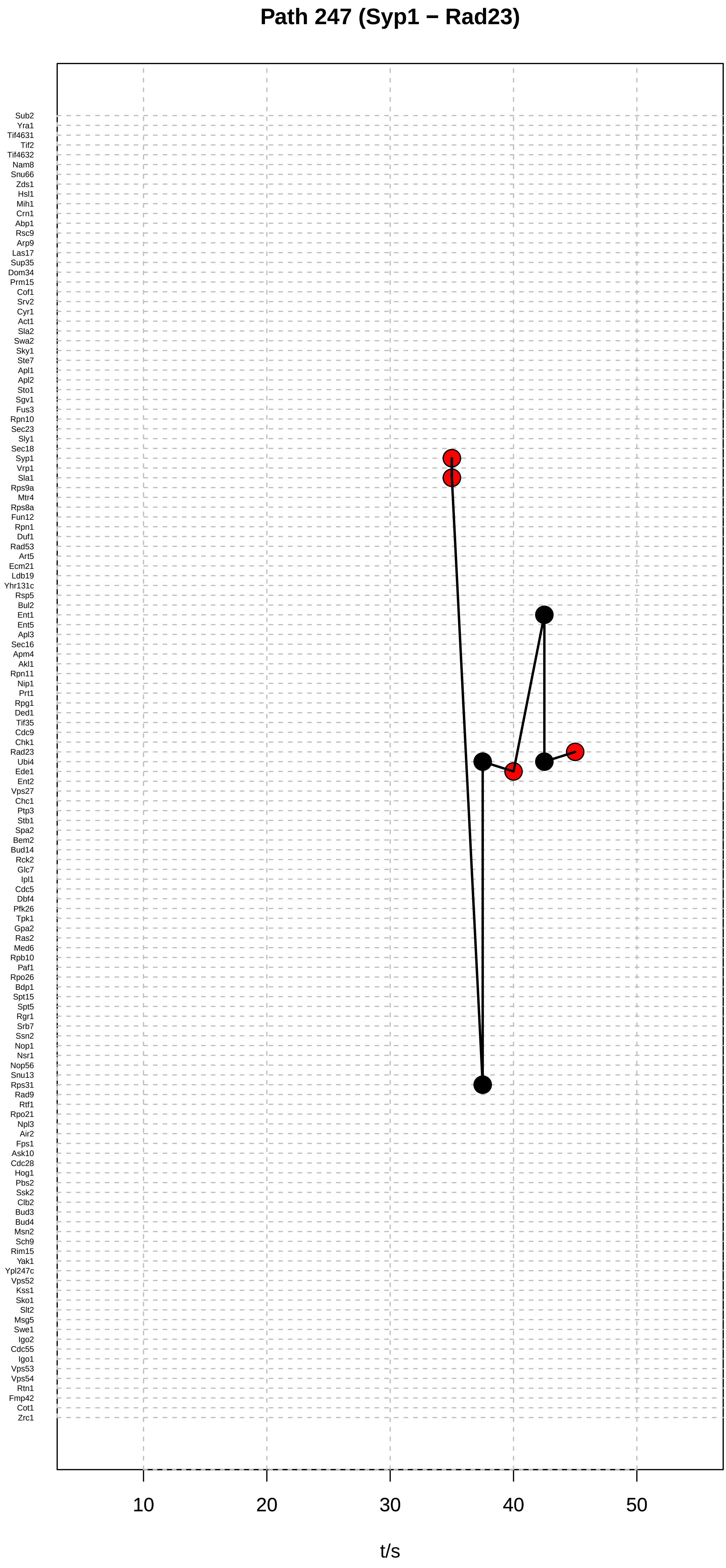
Path 247 (Syp1 − Rad23)
Sub2
Yra1
Tif4631
Tif2
Tif4632
Nam8
Snu66
Zds1
Hsl1
Mih1
Crn1
Abp1
Rsc9
Arp9
Las17
Sup35
Dom34
Prm15
Cof1
Srv2
Cyr1
Act1
Sla2
Swa2
Sky1
Ste7
Apl1
Apl2
Sto1
Sgv1
Fus3
Rpn10
Sec23
Sly1
Sec18
Syp1
Vrp1
Sla1
Rps9a
Mtr4
Rps8a
Fun12
Rpn1
Duf1
Rad53
Art5
Ecm21
Ldb19
Yhr131c
Rsp5
Bul2
Ent1
Ent5
Apl3
Sec16
Apm4
Akl1
Rpn11
Nip1
Prt1
Rpg1
Ded1
Tif35
Cdc9
Chk1
Rad23
Ubi4
Ede1
Ent2
Vps27
Chc1
Ptp3
Stb1
Spa2
Bem2
Bud14
Rck2
Glc7
Ipl1
Cdc5
Dbf4
Pfk26
Tpk1
Gpa2
Ras2
Med6
Rpb10
Paf1
Rpo26
Bdp1
Spt15
Spt5
Rgr1
Srb7
Ssn2
Nop1
Nsr1
Nop56
Snu13
Rps31
Rad9
Rtf1
Rpo21
Npl3
Air2
Fps1
Ask10
Cdc28
Hog1
Pbs2
Ssk2
Clb2
Bud3
Bud4
Msn2
Sch9
Rim15
Yak1
Ypl247c
Vps52
Kss1
Sko1
Slt2
Msg5
Swe1
Igo2
Cdc55
Igo1
Vps53
Vps54
Rtn1
Fmp42
Cot1
Zrc1
10
20
30
40
50
t/s

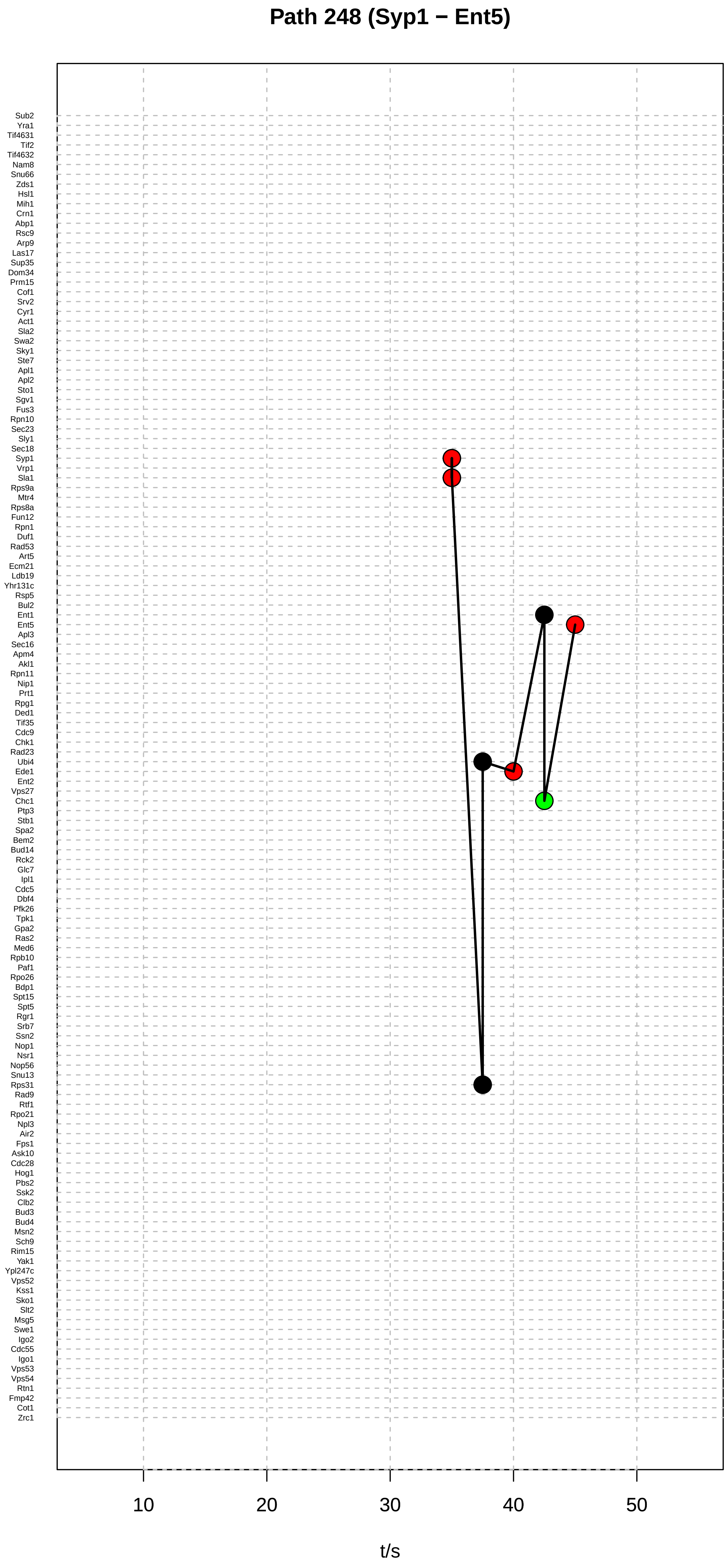
Path 248 (Syp1 − Ent5)
Sub2
Yra1
Tif4631
Tif2
Tif4632
Nam8
Snu66
Zds1
Hsl1
Mih1
Crn1
Abp1
Rsc9
Arp9
Las17
Sup35
Dom34
Prm15
Cof1
Srv2
Cyr1
Act1
Sla2
Swa2
Sky1
Ste7
Apl1
Apl2
Sto1
Sgv1
Fus3
Rpn10
Sec23
Sly1
Sec18
Syp1
Vrp1
Sla1
Rps9a
Mtr4
Rps8a
Fun12
Rpn1
Duf1
Rad53
Art5
Ecm21
Ldb19
Yhr131c
Rsp5
Bul2
Ent1
Ent5
Apl3
Sec16
Apm4
Akl1
Rpn11
Nip1
Prt1
Rpg1
Ded1
Tif35
Cdc9
Chk1
Rad23
Ubi4
Ede1
Ent2
Vps27
Chc1
Ptp3
Stb1
Spa2
Bem2
Bud14
Rck2
Glc7
Ipl1
Cdc5
Dbf4
Pfk26
Tpk1
Gpa2
Ras2
Med6
Rpb10
Paf1
Rpo26
Bdp1
Spt15
Spt5
Rgr1
Srb7
Ssn2
Nop1
Nsr1
Nop56
Snu13
Rps31
Rad9
Rtf1
Rpo21
Npl3
Air2
Fps1
Ask10
Cdc28
Hog1
Pbs2
Ssk2
Clb2
Bud3
Bud4
Msn2
Sch9
Rim15
Yak1
Ypl247c
Vps52
Kss1
Sko1
Slt2
Msg5
Swe1
Igo2
Cdc55
Igo1
Vps53
Vps54
Rtn1
Fmp42
Cot1
Zrc1
10
20
30
40
50
t/s

### Path 249 (Syp1 − Rad23)

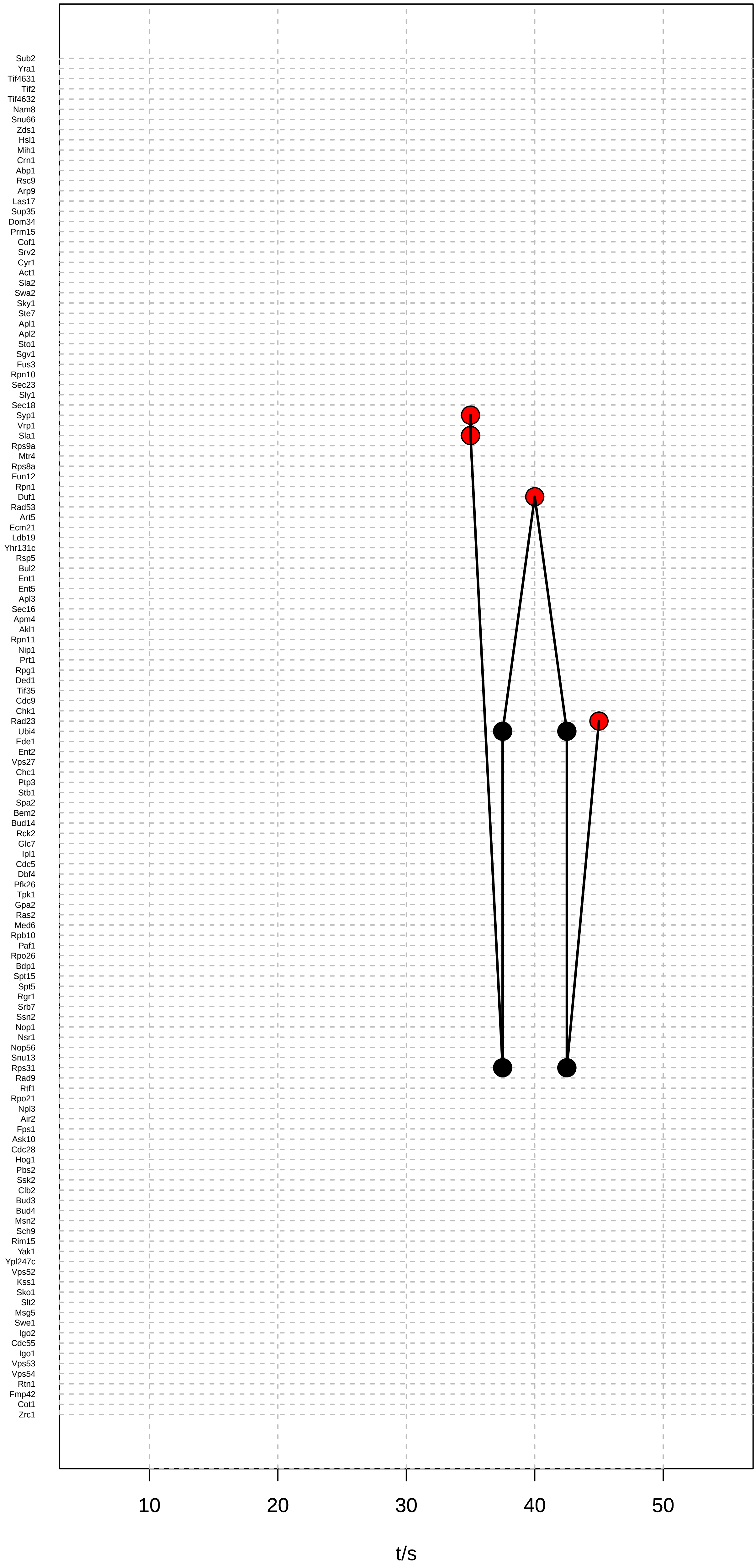

### Path 250 (Syp1 − Rad23)

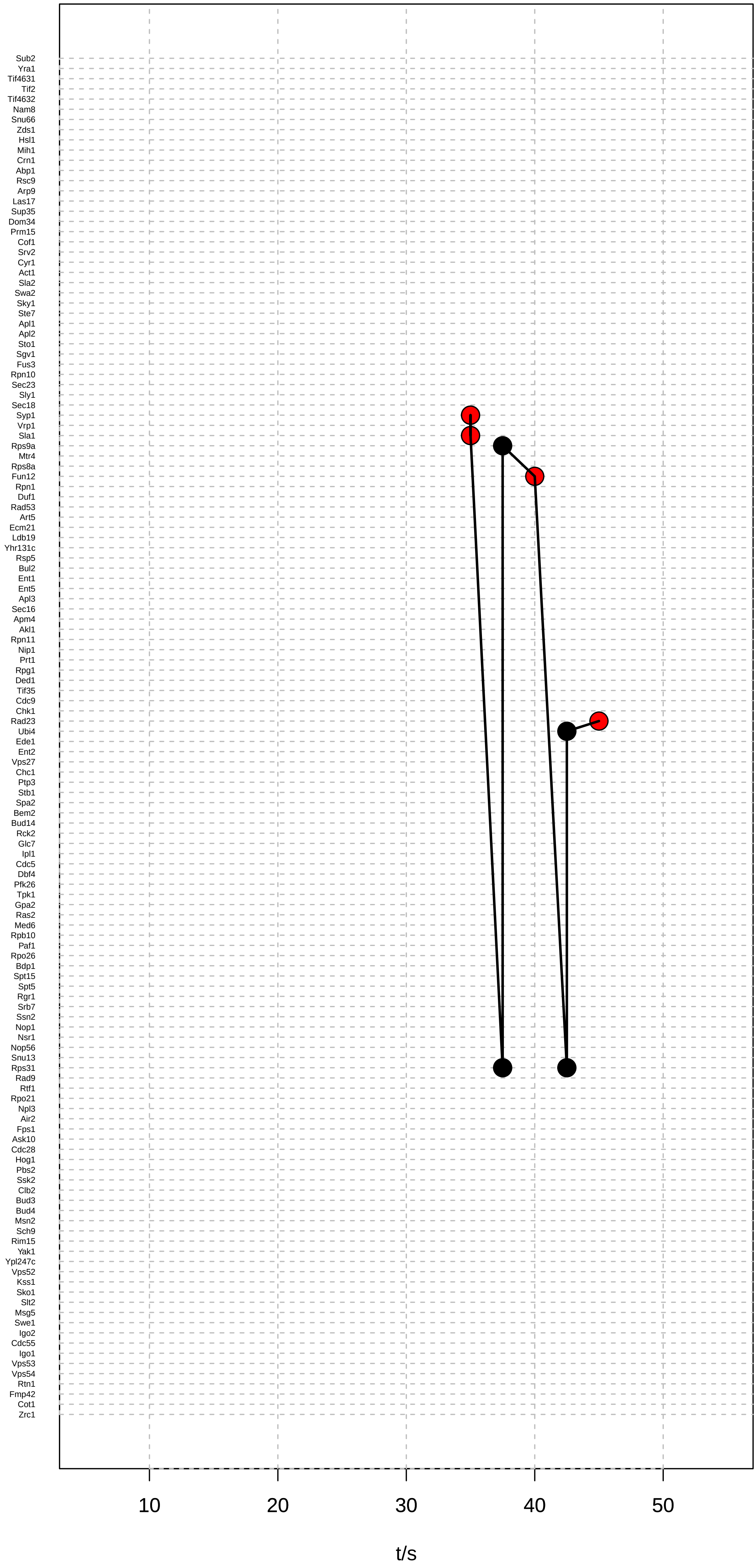

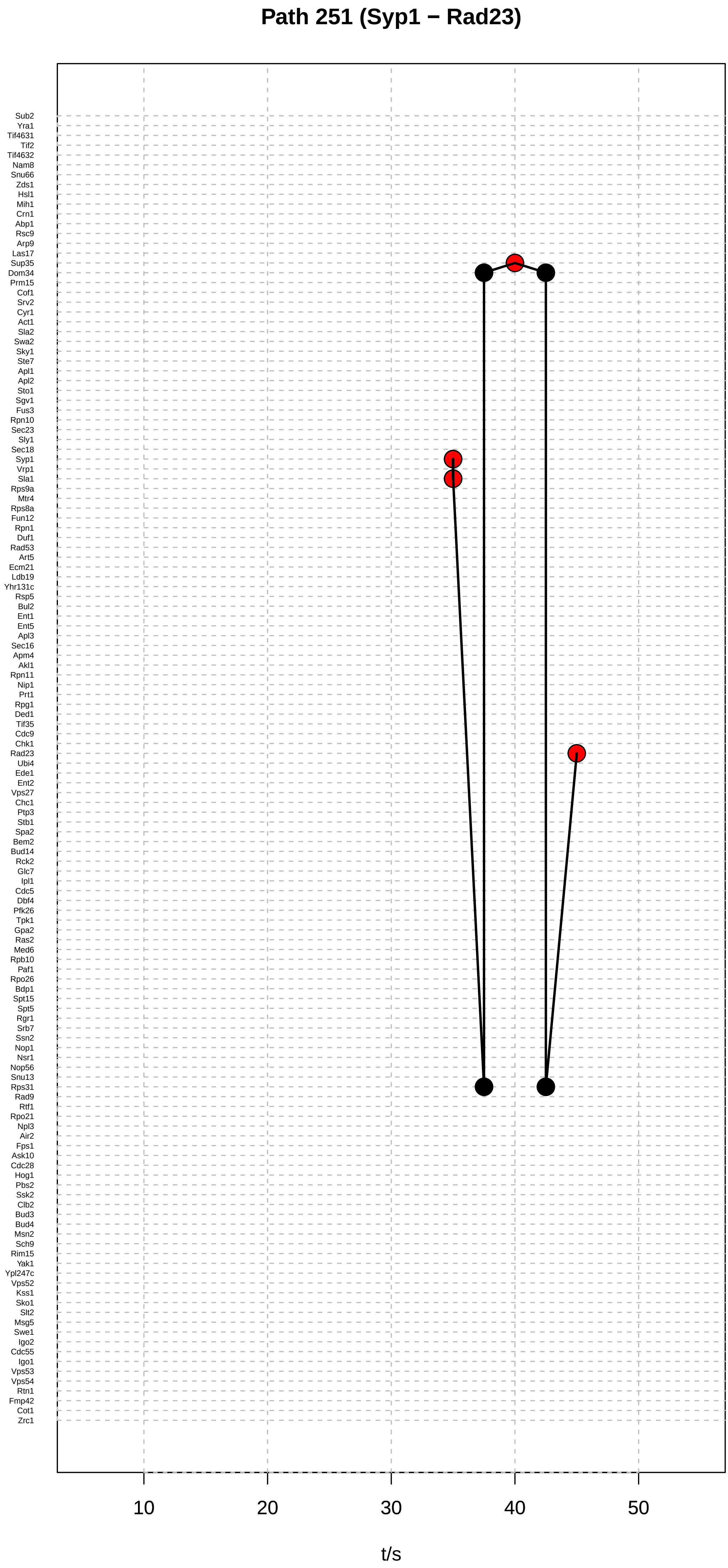

Path 251 (Syp1 − Rad23)
Sub2
Yra1
Tif4631
Tif2
Tif4632
Nam8
Snu66
Zds1
Hsl1
Mih1
Crn1
Abp1
Rsc9
Arp9
Las17
Sup35
Dom34
Prm15
Cof1
Srv2
Cyr1
Act1
Sla2
Swa2
Sky1
Ste7
Apl1
Apl2
Sto1
Sgv1
Fus3
Rpn10
Sec23
Sly1
Sec18
Syp1
Vrp1
Sla1
Rps9a
Mtr4
Rps8a
Fun12
Rpn1
Duf1
Rad53
Art5
Ecm21
Ldb19
Yhr131c
Rsp5
Bul2
Ent1
Ent5
Apl3
Sec16
Apm4
Akl1
Rpn11
Nip1
Prt1
Rpg1
Ded1
Tif35
Cdc9
Chk1
Rad23
Ubi4
Ede1
Ent2
Vps27
Chc1
Ptp3
Stb1
Spa2
Bem2
Bud14
Rck2
Glc7
Ipl1
Cdc5
Dbf4
Pfk26
Tpk1
Gpa2
Ras2
Med6
Rpb10
Paf1
Rpo26
Bdp1
Spt15
Spt5
Rgr1
Srb7
Ssn2
Nop1
Nsr1
Nop56
Snu13
Rps31
Rad9
Rtf1
Rpo21
Npl3
Air2
Fps1
Ask10
Cdc28
Hog1
Pbs2
Ssk2
Clb2
Bud3
Bud4
Msn2
Sch9
Rim15
Yak1
Ypl247c
Vps52
Kss1
Sko1
Slt2
Msg5
Swe1
Igo2
Cdc55
Igo1
Vps53
Vps54
Rtn1
Fmp42
Cot1
Zrc1
10
20
30
40
50
t/s

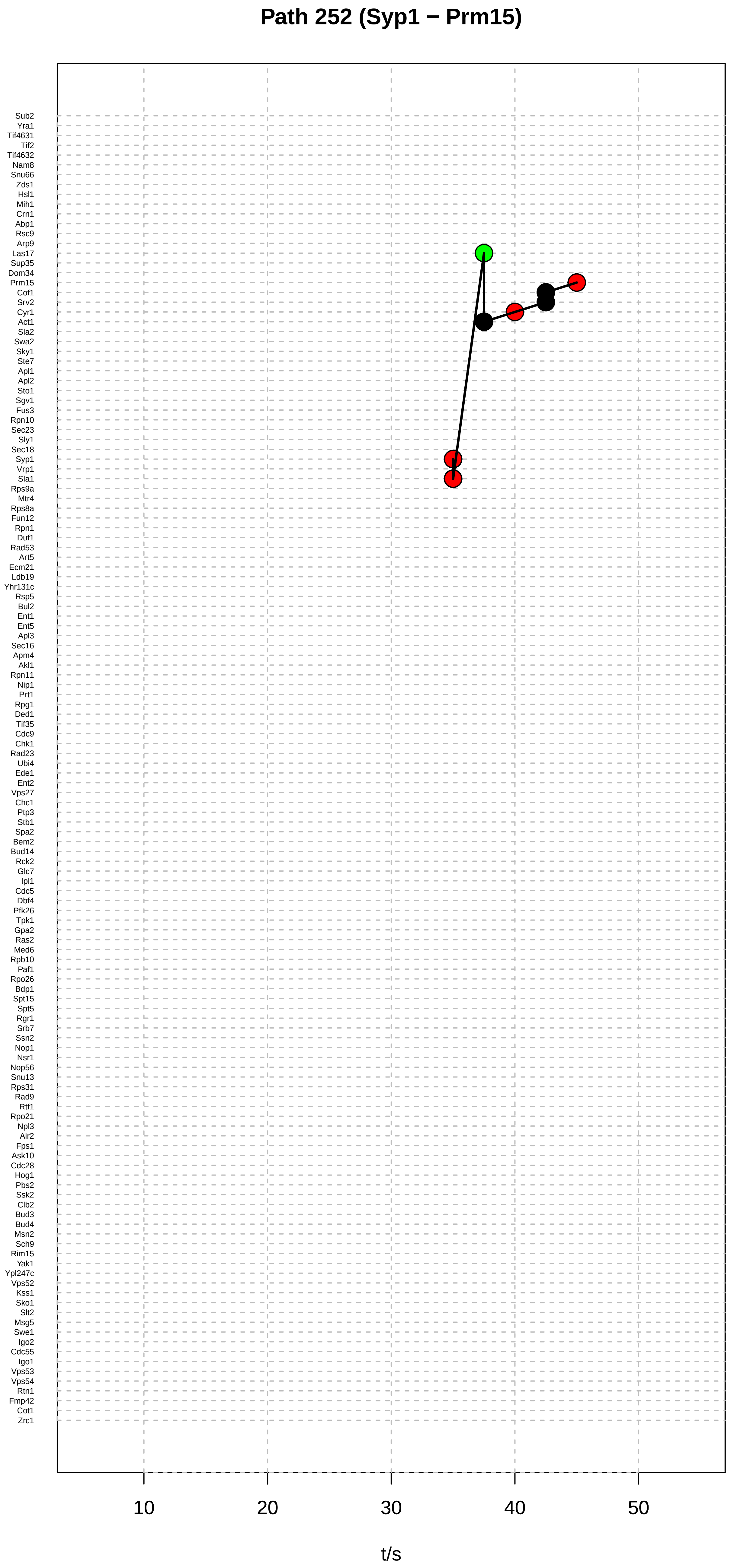

Path 252 (Syp1 − Prm15)
Sub2
Yra1
Tif4631
Tif2
Tif4632
Nam8
Snu66
Zds1
Hsl1
Mih1
Crn1
Abp1
Rsc9
Arp9
Las17
Sup35
Dom34
Prm15
Cof1
Srv2
Cyr1
Act1
Sla2
Swa2
Sky1
Ste7
Apl1
Apl2
Sto1
Sgv1
Fus3
Rpn10
Sec23
Sly1
Sec18
Syp1
Vrp1
Sla1
Rps9a
Mtr4
Rps8a
Fun12
Rpn1
Duf1
Rad53
Art5
Ecm21
Ldb19
Yhr131c
Rsp5
Bul2
Ent1
Ent5
Apl3
Sec16
Apm4
Akl1
Rpn11
Nip1
Prt1
Rpg1
Ded1
Tif35
Cdc9
Chk1
Rad23
Ubi4
Ede1
Ent2
Vps27
Chc1
Ptp3
Stb1
Spa2
Bem2
Bud14
Rck2
Glc7
Ipl1
Cdc5
Dbf4
Pfk26
Tpk1
Gpa2
Ras2
Med6
Rpb10
Paf1
Rpo26
Bdp1
Spt15
Spt5
Rgr1
Srb7
Ssn2
Nop1
Nsr1
Nop56
Snu13
Rps31
Rad9
Rtf1
Rpo21
Npl3
Air2
Fps1
Ask10
Cdc28
Hog1
Pbs2
Ssk2
Clb2
Bud3
Bud4
Msn2
Sch9
Rim15
Yak1
Ypl247c
Vps52
Kss1
Sko1
Slt2
Msg5
Swe1
Igo2
Cdc55
Igo1
Vps53
Vps54
Rtn1
Fmp42
Cot1
Zrc1
10
20
30
40
50
t/s

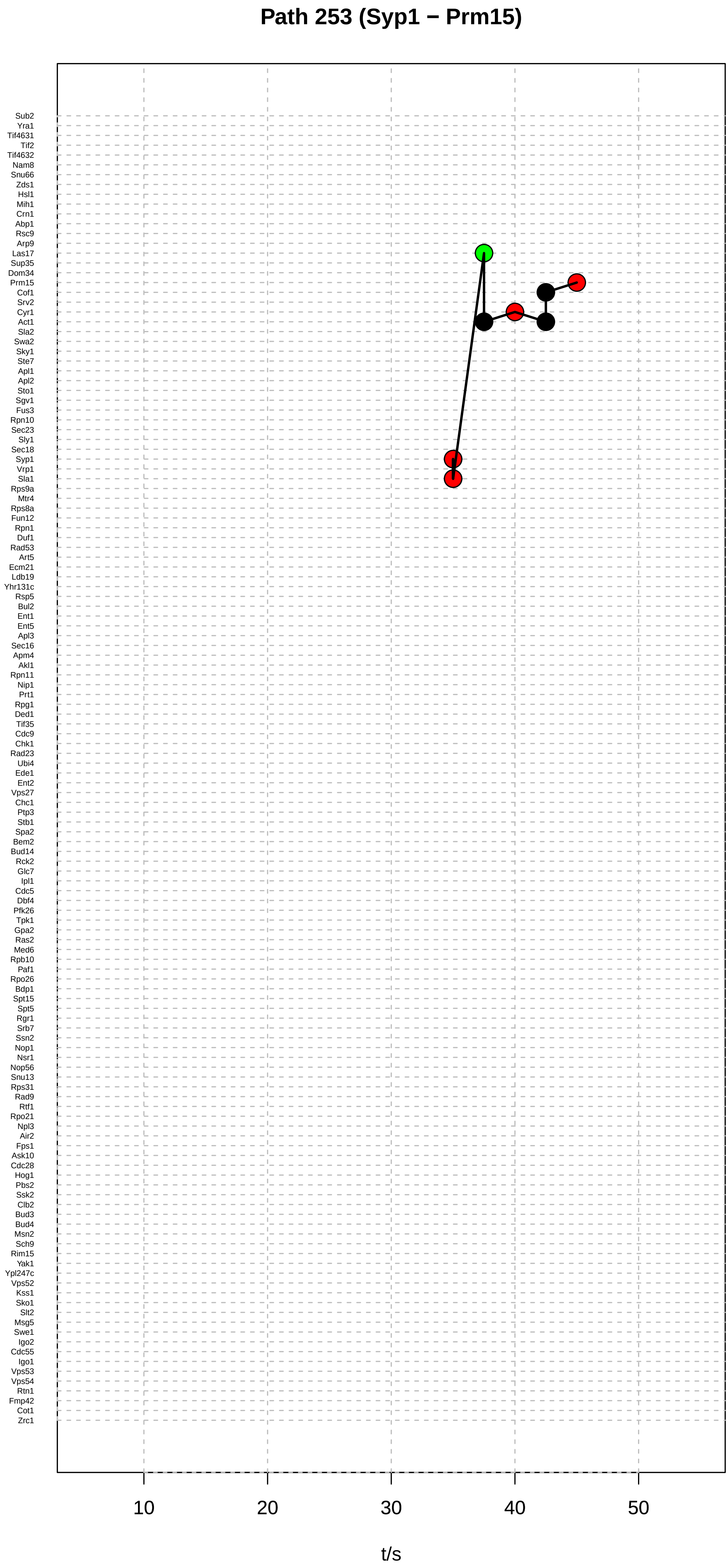

Path 253 (Syp1 − Prm15)
Sub2
Yra1
Tif4631
Tif2
Tif4632
Nam8
Snu66
Zds1
Hsl1
Mih1
Crn1
Abp1
Rsc9
Arp9
Las17
Sup35
Dom34
Prm15
Cof1
Srv2
Cyr1
Act1
Sla2
Swa2
Sky1
Ste7
Apl1
Apl2
Sto1
Sgv1
Fus3
Rpn10
Sec23
Sly1
Sec18
Syp1
Vrp1
Sla1
Rps9a
Mtr4
Rps8a
Fun12
Rpn1
Duf1
Rad53
Art5
Ecm21
Ldb19
Yhr131c
Rsp5
Bul2
Ent1
Ent5
Apl3
Sec16
Apm4
Akl1
Rpn11
Nip1
Prt1
Rpg1
Ded1
Tif35
Cdc9
Chk1
Rad23
Ubi4
Ede1
Ent2
Vps27
Chc1
Ptp3
Stb1
Spa2
Bem2
Bud14
Rck2
Glc7
Ipl1
Cdc5
Dbf4
Pfk26
Tpk1
Gpa2
Ras2
Med6
Rpb10
Paf1
Rpo26
Bdp1
Spt15
Spt5
Rgr1
Srb7
Ssn2
Nop1
Nsr1
Nop56
Snu13
Rps31
Rad9
Rtf1
Rpo21
Npl3
Air2
Fps1
Ask10
Cdc28
Hog1
Pbs2
Ssk2
Clb2
Bud3
Bud4
Msn2
Sch9
Rim15
Yak1
Ypl247c
Vps52
Kss1
Sko1
Slt2
Msg5
Swe1
Igo2
Cdc55
Igo1
Vps53
Vps54
Rtn1
Fmp42
Cot1
Zrc1
10
20
30
40
50
t/s

#### Path 254 (Syp1 − Ded1)

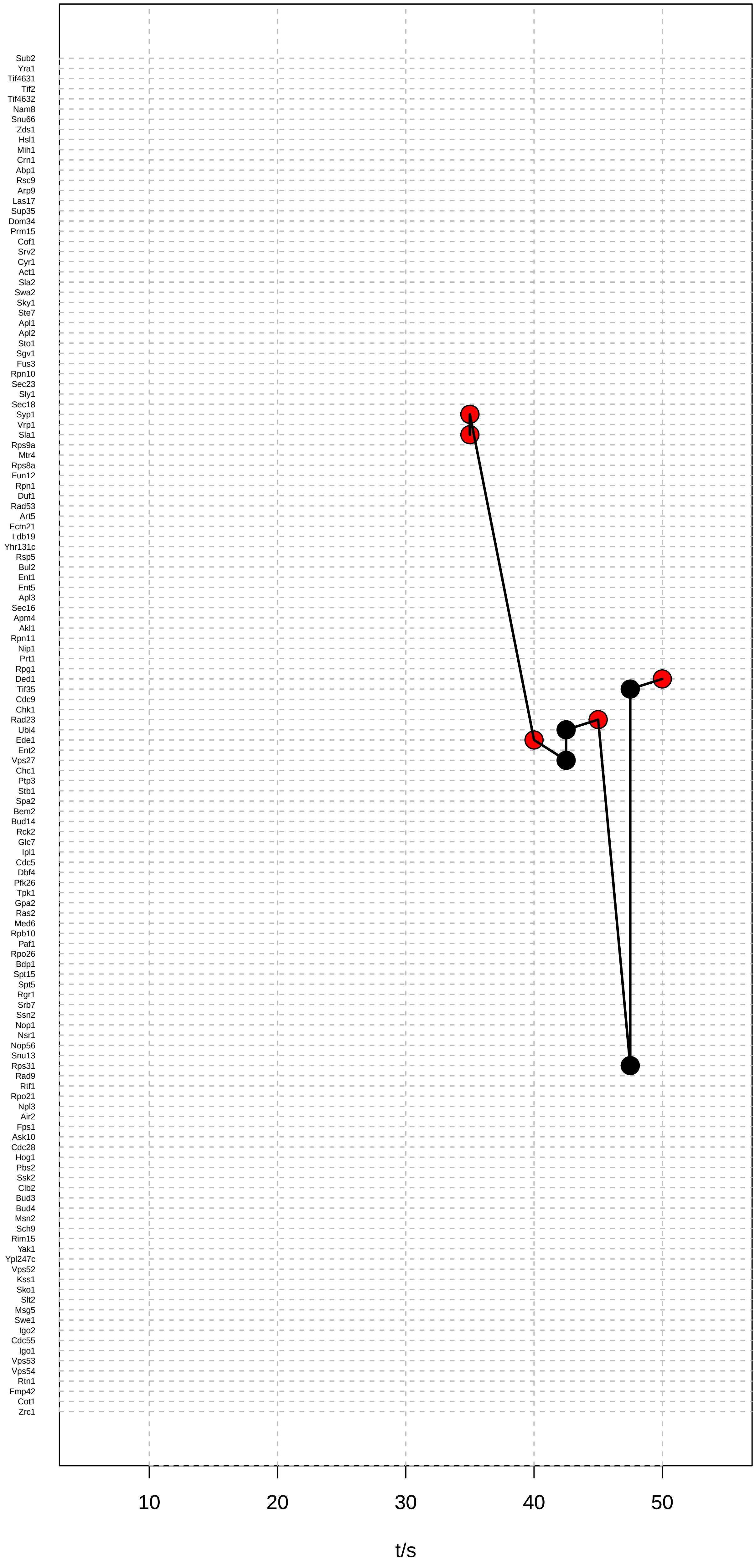

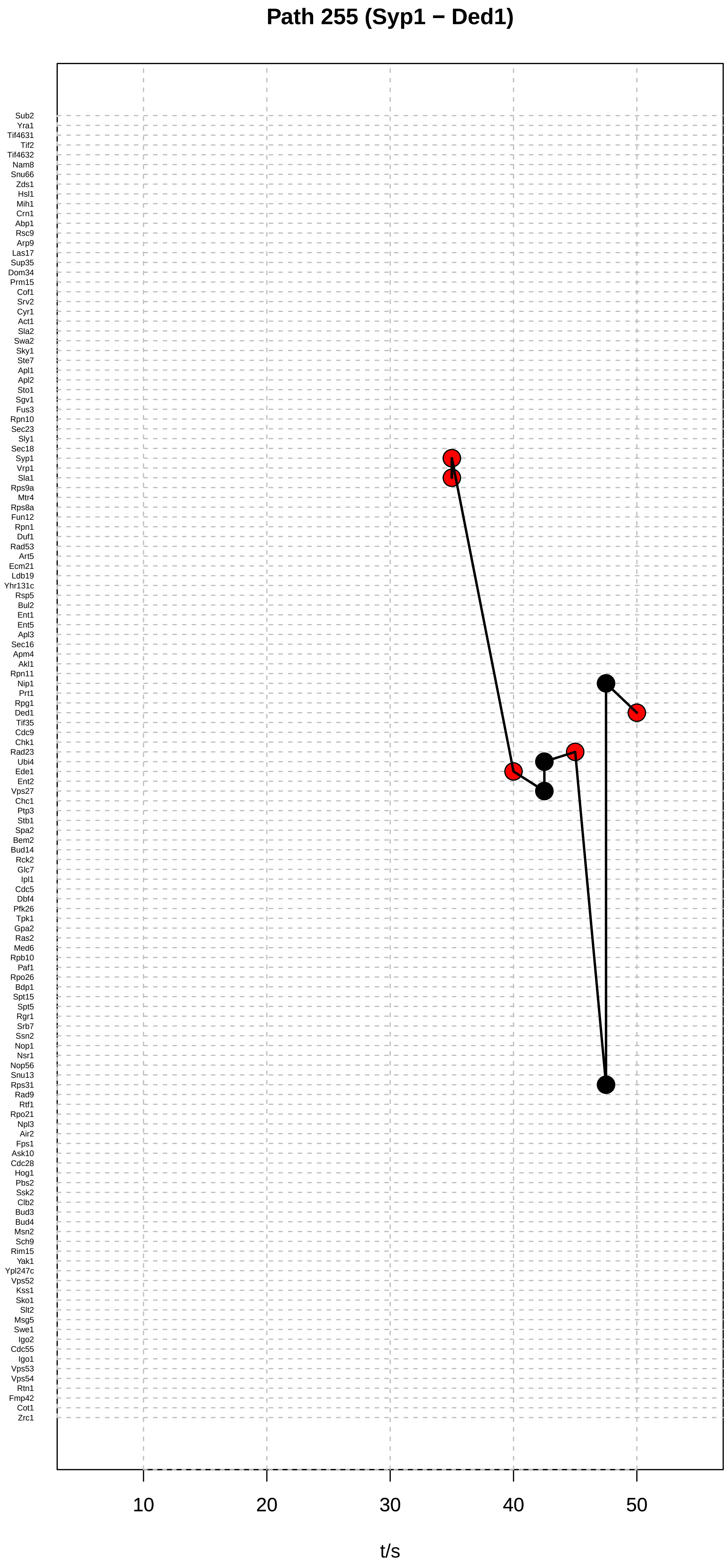

Path 255 (Syp1 − Ded1)
Sub2
Yra1
Tif4631
Tif2
Tif4632
Nam8
Snu66
Zds1
Hsl1
Mih1
Crn1
Abp1
Rsc9
Arp9
Las17
Sup35
Dom34
Prm15
Cof1
Srv2
Cyr1
Act1
Sla2
Swa2
Sky1
Ste7
Apl1
Apl2
Sto1
Sgv1
Fus3
Rpn10
Sec23
Sly1
Sec18
Syp1
Vrp1
Sla1
Rps9a
Mtr4
Rps8a
Fun12
Rpn1
Duf1
Rad53
Art5
Ecm21
Ldb19
Yhr131c
Rsp5
Bul2
Ent1
Ent5
Apl3
Sec16
Apm4
Akl1
Rpn11
Nip1
Prt1
Rpg1
Ded1
Tif35
Cdc9
Chk1
Rad23
Ubi4
Ede1
Ent2
Vps27
Chc1
Ptp3
Stb1
Spa2
Bem2
Bud14
Rck2
Glc7
Ipl1
Cdc5
Dbf4
Pfk26
Tpk1
Gpa2
Ras2
Med6
Rpb10
Paf1
Rpo26
Bdp1
Spt15
Spt5
Rgr1
Srb7
Ssn2
Nop1
Nsr1
Nop56
Snu13
Rps31
Rad9
Rtf1
Rpo21
Npl3
Air2
Fps1
Ask10
Cdc28
Hog1
Pbs2
Ssk2
Clb2
Bud3
Bud4
Msn2
Sch9
Rim15
Yak1
Ypl247c
Vps52
Kss1
Sko1
Slt2
Msg5
Swe1
Igo2
Cdc55
Igo1
Vps53
Vps54
Rtn1
Fmp42
Cot1
Zrc1
10
20
30
40
50
t/s

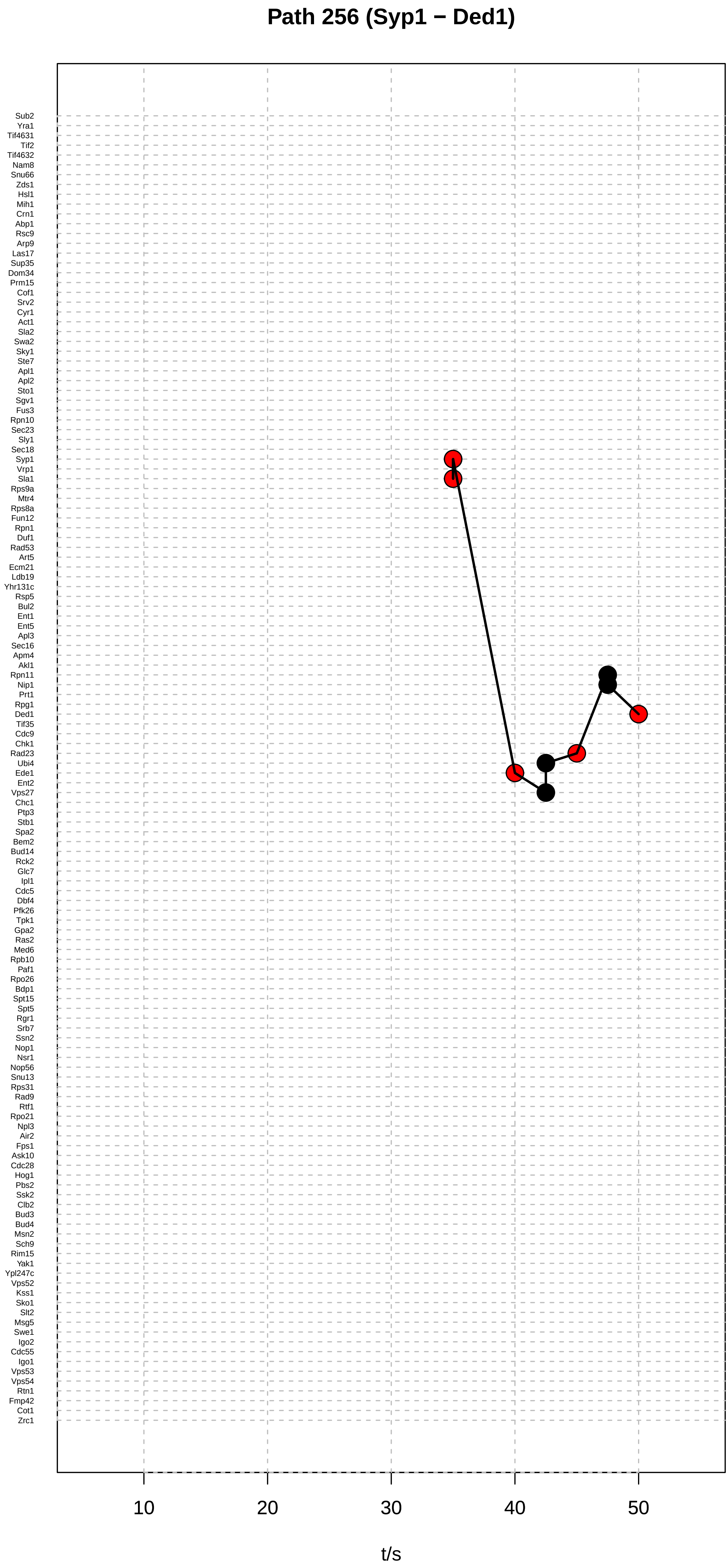

Path 256 (Syp1 − Ded1)
Sub2
Yra1
Tif4631
Tif2
Tif4632
Nam8
Snu66
Zds1
Hsl1
Mih1
Crn1
Abp1
Rsc9
Arp9
Las17
Sup35
Dom34
Prm15
Cof1
Srv2
Cyr1
Act1
Sla2
Swa2
Sky1
Ste7
Apl1
Apl2
Sto1
Sgv1
Fus3
Rpn10
Sec23
Sly1
Sec18
Syp1
Vrp1
Sla1
Rps9a
Mtr4
Rps8a
Fun12
Rpn1
Duf1
Rad53
Art5
Ecm21
Ldb19
Yhr131c
Rsp5
Bul2
Ent1
Ent5
Apl3
Sec16
Apm4
Akl1
Rpn11
Nip1
Prt1
Rpg1
Ded1
Tif35
Cdc9
Chk1
Rad23
Ubi4
Ede1
Ent2
Vps27
Chc1
Ptp3
Stb1
Spa2
Bem2
Bud14
Rck2
Glc7
Ipl1
Cdc5
Dbf4
Pfk26
Tpk1
Gpa2
Ras2
Med6
Rpb10
Paf1
Rpo26
Bdp1
Spt15
Spt5
Rgr1
Srb7
Ssn2
Nop1
Nsr1
Nop56
Snu13
Rps31
Rad9
Rtf1
Rpo21
Npl3
Air2
Fps1
Ask10
Cdc28
Hog1
Pbs2
Ssk2
Clb2
Bud3
Bud4
Msn2
Sch9
Rim15
Yak1
Ypl247c
Vps52
Kss1
Sko1
Slt2
Msg5
Swe1
Igo2
Cdc55
Igo1
Vps53
Vps54
Rtn1
Fmp42
Cot1
Zrc1
10
20
30
40
50
t/s

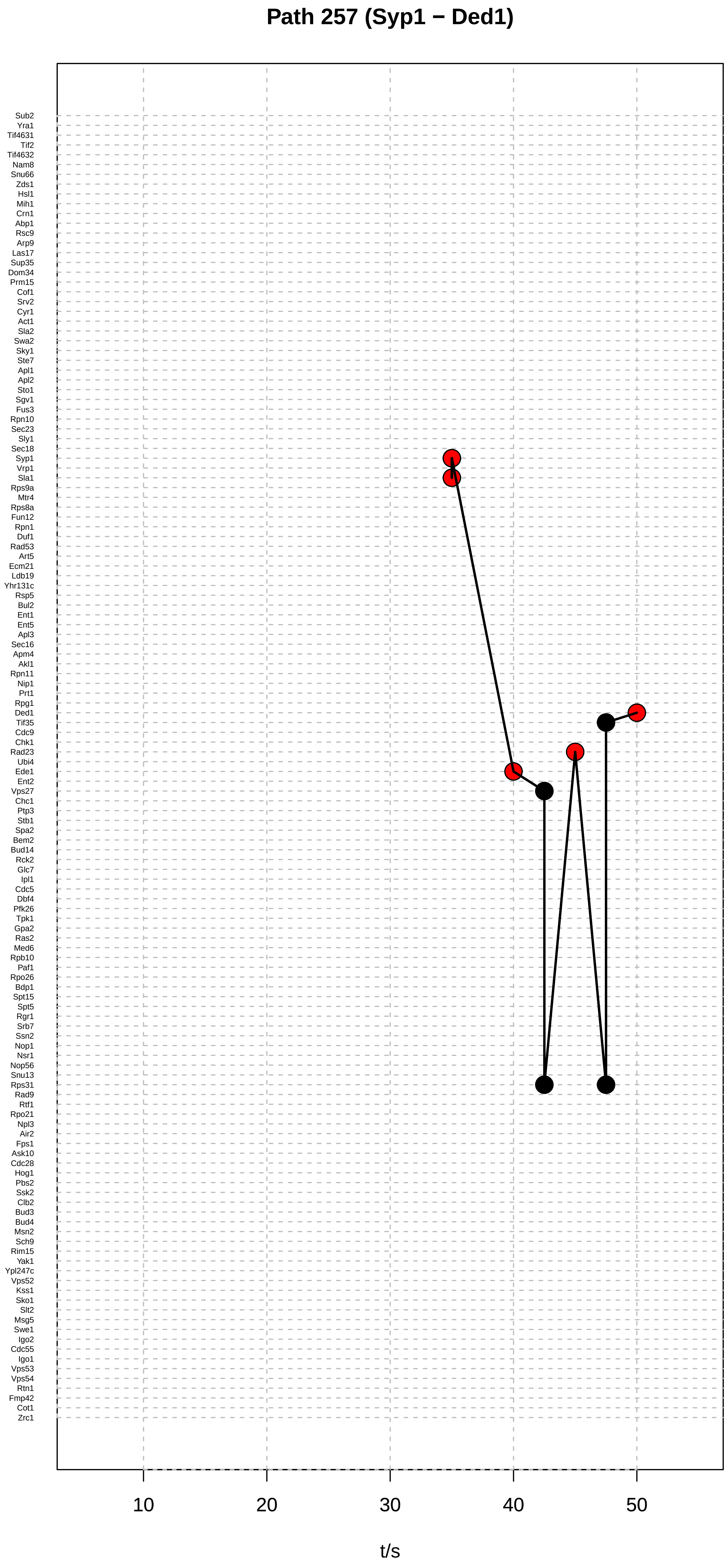
Path 257 (Syp1 − Ded1)
Sub2
Yra1
Tif4631
Tif2
Tif4632
Nam8
Snu66
Zds1
Hsl1
Mih1
Crn1
Abp1
Rsc9
Arp9
Las17
Sup35
Dom34
Prm15
Cof1
Srv2
Cyr1
Act1
Sla2
Swa2
Sky1
Ste7
Apl1
Apl2
Sto1
Sgv1
Fus3
Rpn10
Sec23
Sly1
Sec18
Syp1
Vrp1
Sla1
Rps9a
Mtr4
Rps8a
Fun12
Rpn1
Duf1
Rad53
Art5
Ecm21
Ldb19
Yhr131c
Rsp5
Bul2
Ent1
Ent5
Apl3
Sec16
Apm4
Akl1
Rpn11
Nip1
Prt1
Rpg1
Ded1
Tif35
Cdc9
Chk1
Rad23
Ubi4
Ede1
Ent2
Vps27
Chc1
Ptp3
Stb1
Spa2
Bem2
Bud14
Rck2
Glc7
Ipl1
Cdc5
Dbf4
Pfk26
Tpk1
Gpa2
Ras2
Med6
Rpb10
Paf1
Rpo26
Bdp1
Spt15
Spt5
Rgr1
Srb7
Ssn2
Nop1
Nsr1
Nop56
Snu13
Rps31
Rad9
Rtf1
Rpo21
Npl3
Air2
Fps1
Ask10
Cdc28
Hog1
Pbs2
Ssk2
Clb2
Bud3
Bud4
Msn2
Sch9
Rim15
Yak1
Ypl247c
Vps52
Kss1
Sko1
Slt2
Msg5
Swe1
Igo2
Cdc55
Igo1
Vps53
Vps54
Rtn1
Fmp42
Cot1
Zrc1
10
20
30
40
50
t/s

### Path 258 (Syp1 − Ded1)

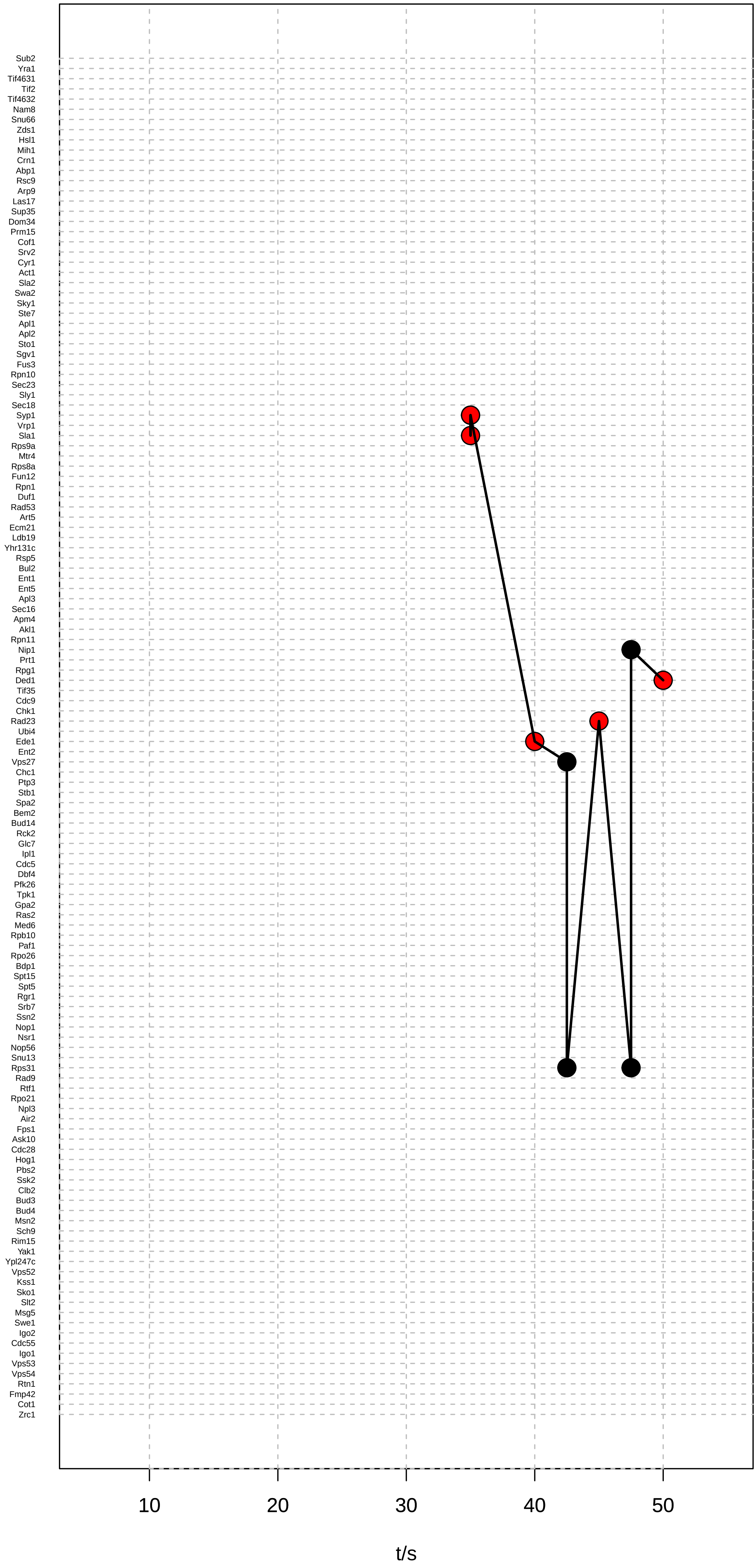

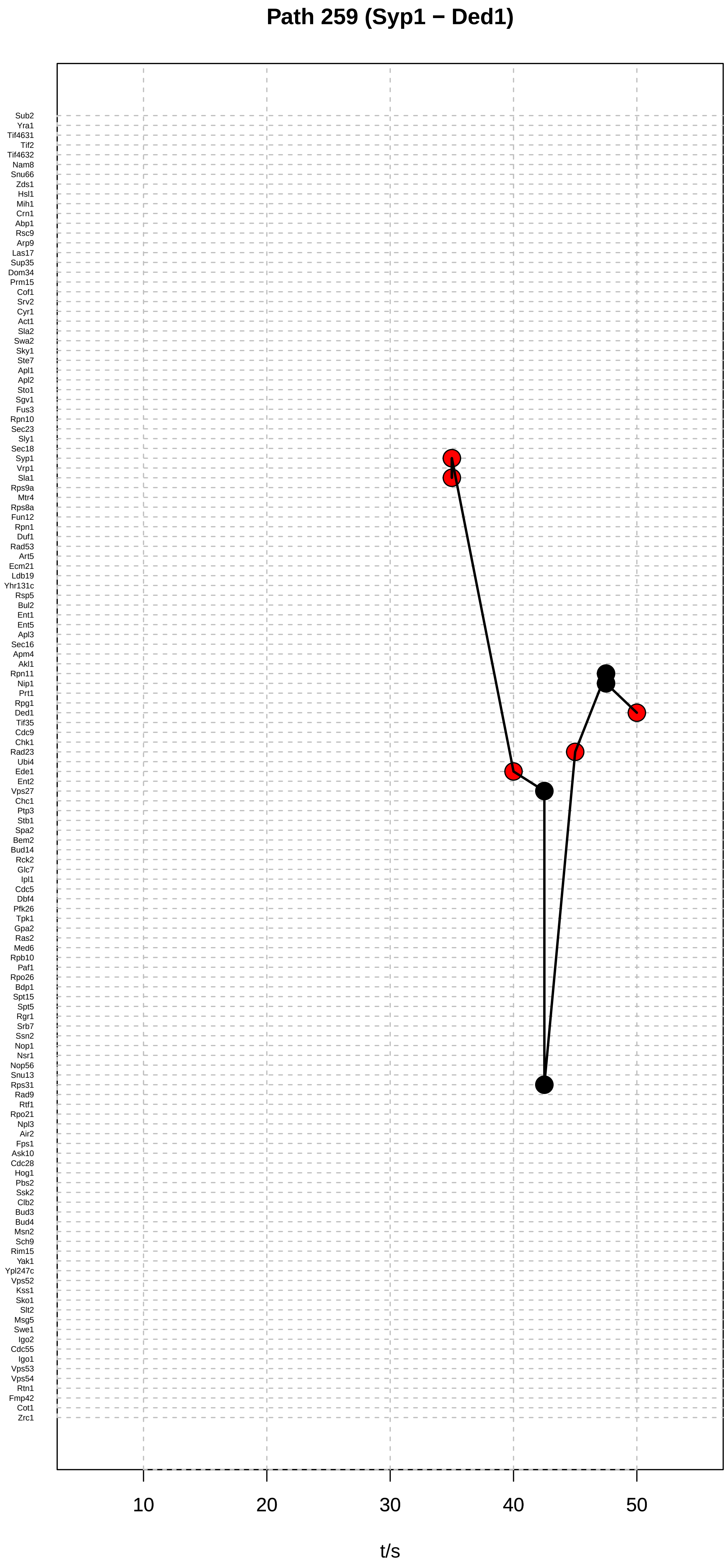
Path 259 (Syp1 − Ded1)
Sub2
Yra1
Tif4631
Tif2
Tif4632
Nam8
Snu66
Zds1
Hsl1
Mih1
Crn1
Abp1
Rsc9
Arp9
Las17
Sup35
Dom34
Prm15
Cof1
Srv2
Cyr1
Act1
Sla2
Swa2
Sky1
Ste7
Apl1
Apl2
Sto1
Sgv1
Fus3
Rpn10
Sec23
Sly1
Sec18
Syp1
Vrp1
Sla1
Rps9a
Mtr4
Rps8a
Fun12
Rpn1
Duf1
Rad53
Art5
Ecm21
Ldb19
Yhr131c
Rsp5
Bul2
Ent1
Ent5
Apl3
Sec16
Apm4
Akl1
Rpn11
Nip1
Prt1
Rpg1
Ded1
Tif35
Cdc9
Chk1
Rad23
Ubi4
Ede1
Ent2
Vps27
Chc1
Ptp3
Stb1
Spa2
Bem2
Bud14
Rck2
Glc7
Ipl1
Cdc5
Dbf4
Pfk26
Tpk1
Gpa2
Ras2
Med6
Rpb10
Paf1
Rpo26
Bdp1
Spt15
Spt5
Rgr1
Srb7
Ssn2
Nop1
Nsr1
Nop56
Snu13
Rps31
Rad9
Rtf1
Rpo21
Npl3
Air2
Fps1
Ask10
Cdc28
Hog1
Pbs2
Ssk2
Clb2
Bud3
Bud4
Msn2
Sch9
Rim15
Yak1
Ypl247c
Vps52
Kss1
Sko1
Slt2
Msg5
Swe1
Igo2
Cdc55
Igo1
Vps53
Vps54
Rtn1
Fmp42
Cot1
Zrc1
10
20
30
40
50
t/s

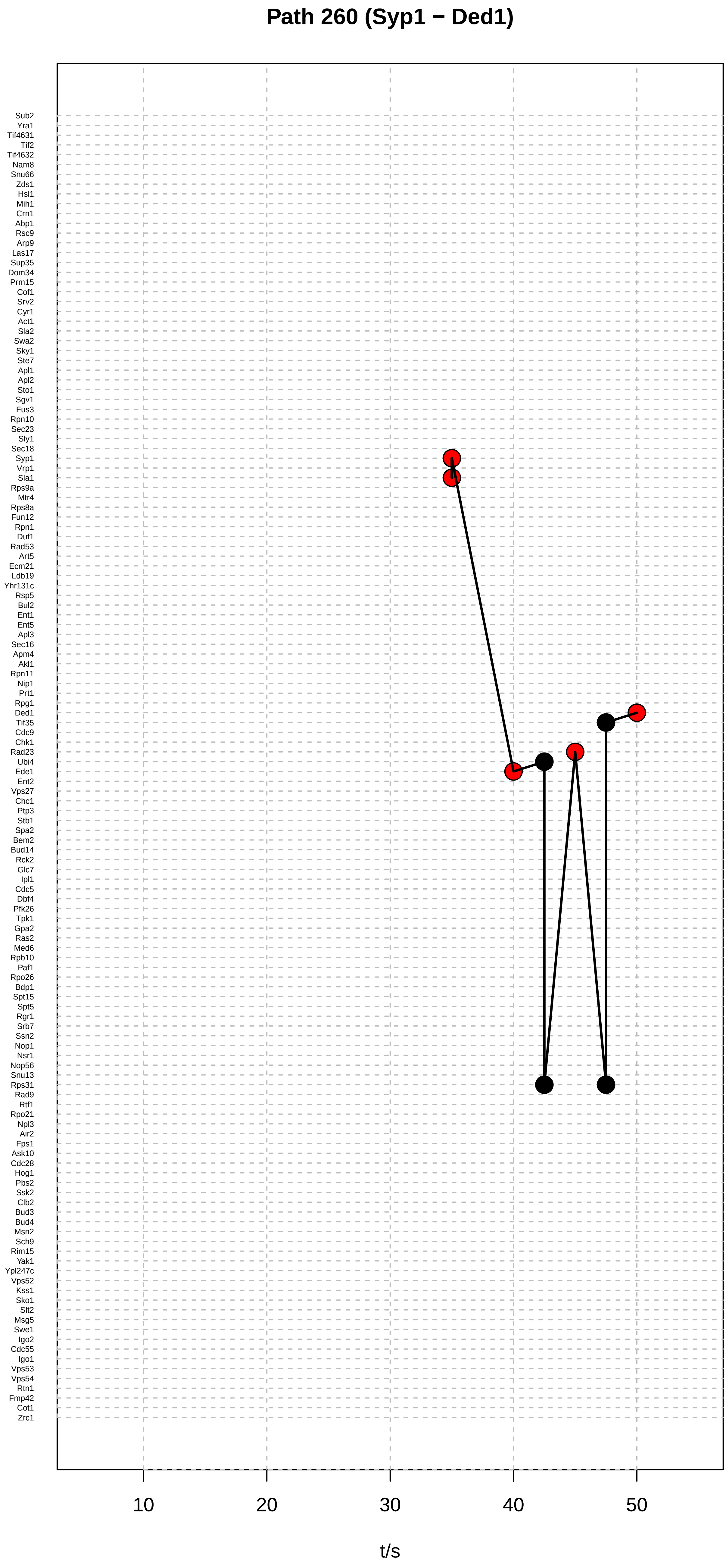

Path 260 (Syp1 − Ded1)
Sub2
Yra1
Tif4631
Tif2
Tif4632
Nam8
Snu66
Zds1
Hsl1
Mih1
Crn1
Abp1
Rsc9
Arp9
Las17
Sup35
Dom34
Prm15
Cof1
Srv2
Cyr1
Act1
Sla2
Swa2
Sky1
Ste7
Apl1
Apl2
Sto1
Sgv1
Fus3
Rpn10
Sec23
Sly1
Sec18
Syp1
Vrp1
Sla1
Rps9a
Mtr4
Rps8a
Fun12
Rpn1
Duf1
Rad53
Art5
Ecm21
Ldb19
Yhr131c
Rsp5
Bul2
Ent1
Ent5
Apl3
Sec16
Apm4
Akl1
Rpn11
Nip1
Prt1
Rpg1
Ded1
Tif35
Cdc9
Chk1
Rad23
Ubi4
Ede1
Ent2
Vps27
Chc1
Ptp3
Stb1
Spa2
Bem2
Bud14
Rck2
Glc7
Ipl1
Cdc5
Dbf4
Pfk26
Tpk1
Gpa2
Ras2
Med6
Rpb10
Paf1
Rpo26
Bdp1
Spt15
Spt5
Rgr1
Srb7
Ssn2
Nop1
Nsr1
Nop56
Snu13
Rps31
Rad9
Rtf1
Rpo21
Npl3
Air2
Fps1
Ask10
Cdc28
Hog1
Pbs2
Ssk2
Clb2
Bud3
Bud4
Msn2
Sch9
Rim15
Yak1
Ypl247c
Vps52
Kss1
Sko1
Slt2
Msg5
Swe1
Igo2
Cdc55
Igo1
Vps53
Vps54
Rtn1
Fmp42
Cot1
Zrc1
10
20
30
40
50
t/s

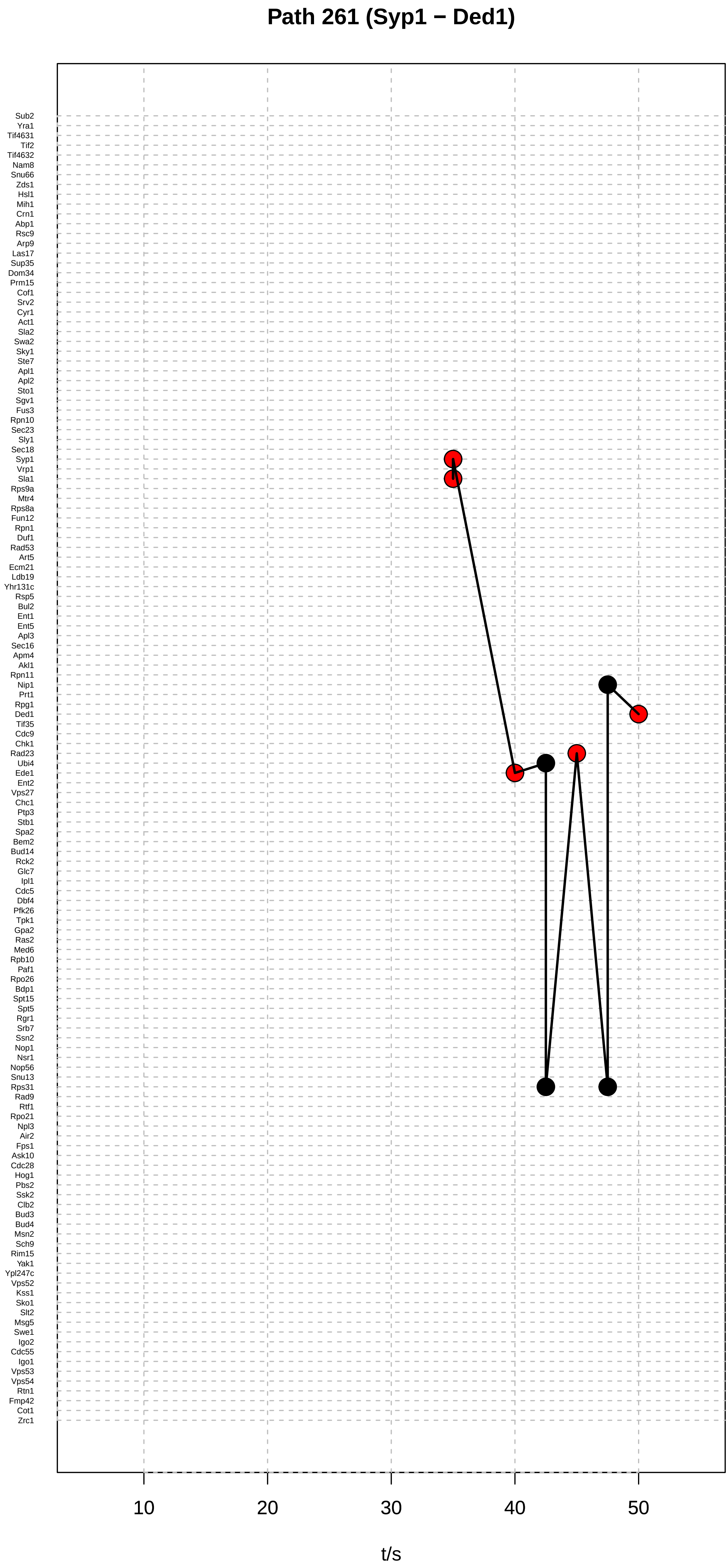

Path 261 (Syp1 − Ded1)
Sub2
Yra1
Tif4631
Tif2
Tif4632
Nam8
Snu66
Zds1
Hsl1
Mih1
Crn1
Abp1
Rsc9
Arp9
Las17
Sup35
Dom34
Prm15
Cof1
Srv2
Cyr1
Act1
Sla2
Swa2
Sky1
Ste7
Apl1
Apl2
Sto1
Sgv1
Fus3
Rpn10
Sec23
Sly1
Sec18
Syp1
Vrp1
Sla1
Rps9a
Mtr4
Rps8a
Fun12
Rpn1
Duf1
Rad53
Art5
Ecm21
Ldb19
Yhr131c
Rsp5
Bul2
Ent1
Ent5
Apl3
Sec16
Apm4
Akl1
Rpn11
Nip1
Prt1
Rpg1
Ded1
Tif35
Cdc9
Chk1
Rad23
Ubi4
Ede1
Ent2
Vps27
Chc1
Ptp3
Stb1
Spa2
Bem2
Bud14
Rck2
Glc7
Ipl1
Cdc5
Dbf4
Pfk26
Tpk1
Gpa2
Ras2
Med6
Rpb10
Paf1
Rpo26
Bdp1
Spt15
Spt5
Rgr1
Srb7
Ssn2
Nop1
Nsr1
Nop56
Snu13
Rps31
Rad9
Rtf1
Rpo21
Npl3
Air2
Fps1
Ask10
Cdc28
Hog1
Pbs2
Ssk2
Clb2
Bud3
Bud4
Msn2
Sch9
Rim15
Yak1
Ypl247c
Vps52
Kss1
Sko1
Slt2
Msg5
Swe1
Igo2
Cdc55
Igo1
Vps53
Vps54
Rtn1
Fmp42
Cot1
Zrc1
10
20
30
40
50
t/s

#### Path 262 (Syp1 − Ded1)

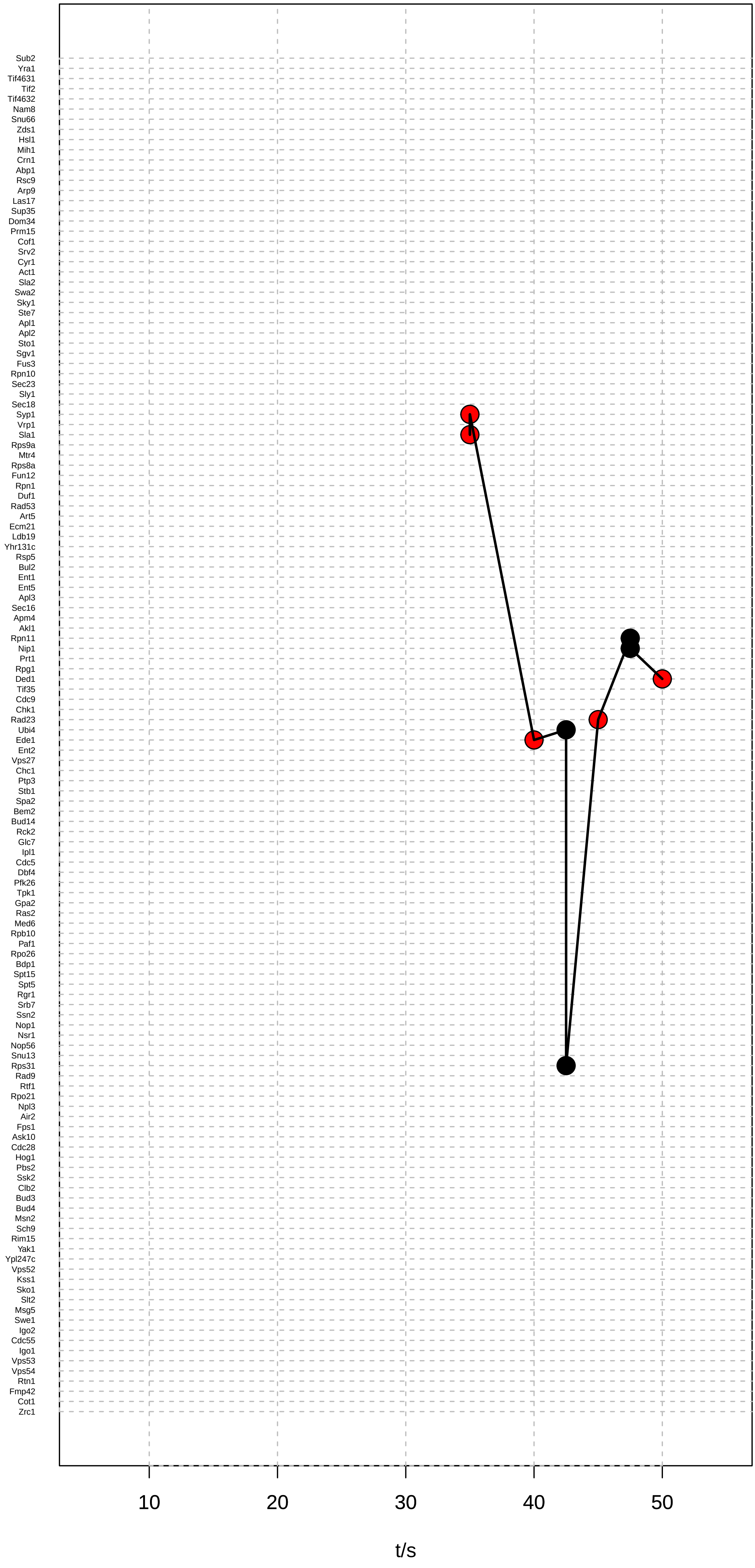

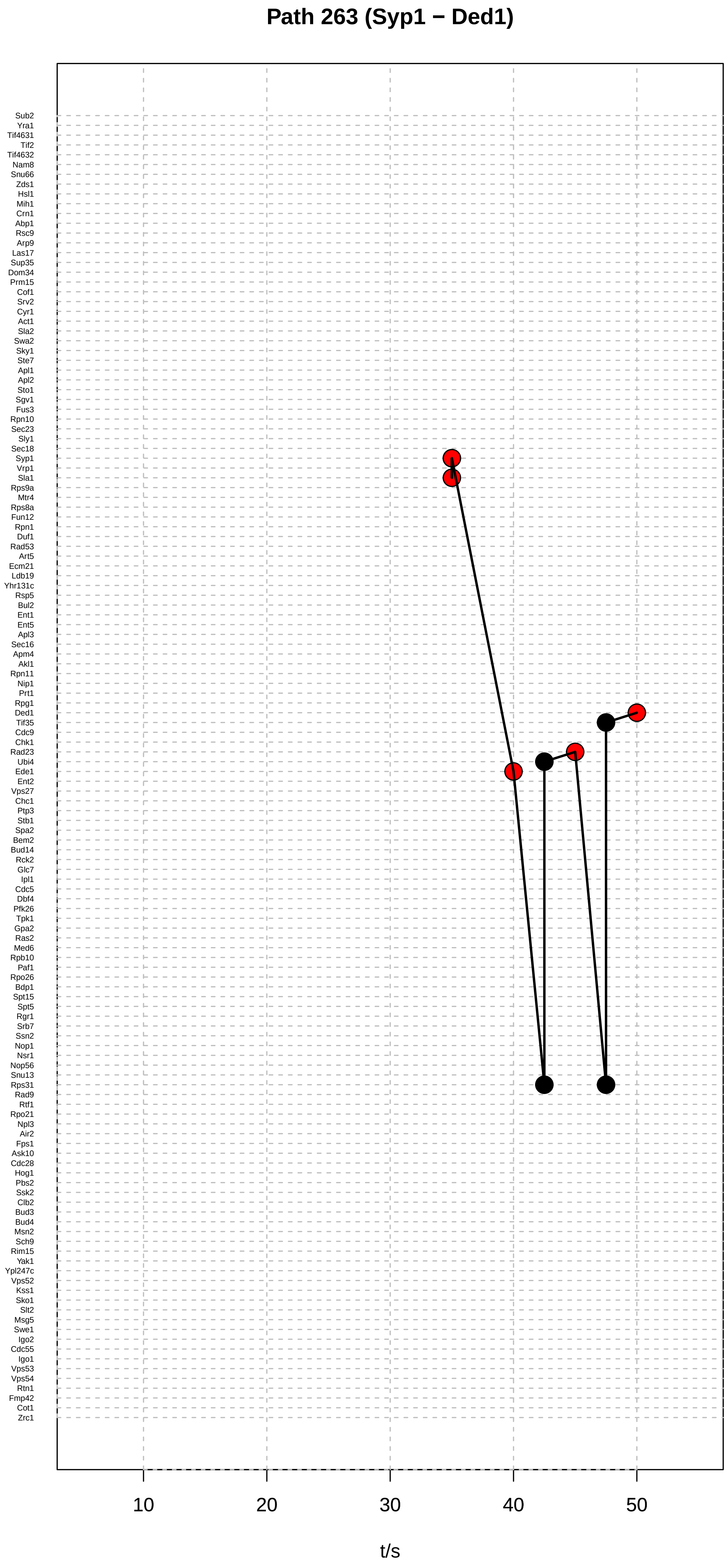
Path 263 (Syp1 − Ded1)
Sub2
Yra1
Tif4631
Tif2
Tif4632
Nam8
Snu66
Zds1
Hsl1
Mih1
Crn1
Abp1
Rsc9
Arp9
Las17
Sup35
Dom34
Prm15
Cof1
Srv2
Cyr1
Act1
Sla2
Swa2
Sky1
Ste7
Apl1
Apl2
Sto1
Sgv1
Fus3
Rpn10
Sec23
Sly1
Sec18
Syp1
Vrp1
Sla1
Rps9a
Mtr4
Rps8a
Fun12
Rpn1
Duf1
Rad53
Art5
Ecm21
Ldb19
Yhr131c
Rsp5
Bul2
Ent1
Ent5
Apl3
Sec16
Apm4
Akl1
Rpn11
Nip1
Prt1
Rpg1
Ded1
Tif35
Cdc9
Chk1
Rad23
Ubi4
Ede1
Ent2
Vps27
Chc1
Ptp3
Stb1
Spa2
Bem2
Bud14
Rck2
Glc7
Ipl1
Cdc5
Dbf4
Pfk26
Tpk1
Gpa2
Ras2
Med6
Rpb10
Paf1
Rpo26
Bdp1
Spt15
Spt5
Rgr1
Srb7
Ssn2
Nop1
Nsr1
Nop56
Snu13
Rps31
Rad9
Rtf1
Rpo21
Npl3
Air2
Fps1
Ask10
Cdc28
Hog1
Pbs2
Ssk2
Clb2
Bud3
Bud4
Msn2
Sch9
Rim15
Yak1
Ypl247c
Vps52
Kss1
Sko1
Slt2
Msg5
Swe1
Igo2
Cdc55
Igo1
Vps53
Vps54
Rtn1
Fmp42
Cot1
Zrc1
10
20
30
40
50
t/s

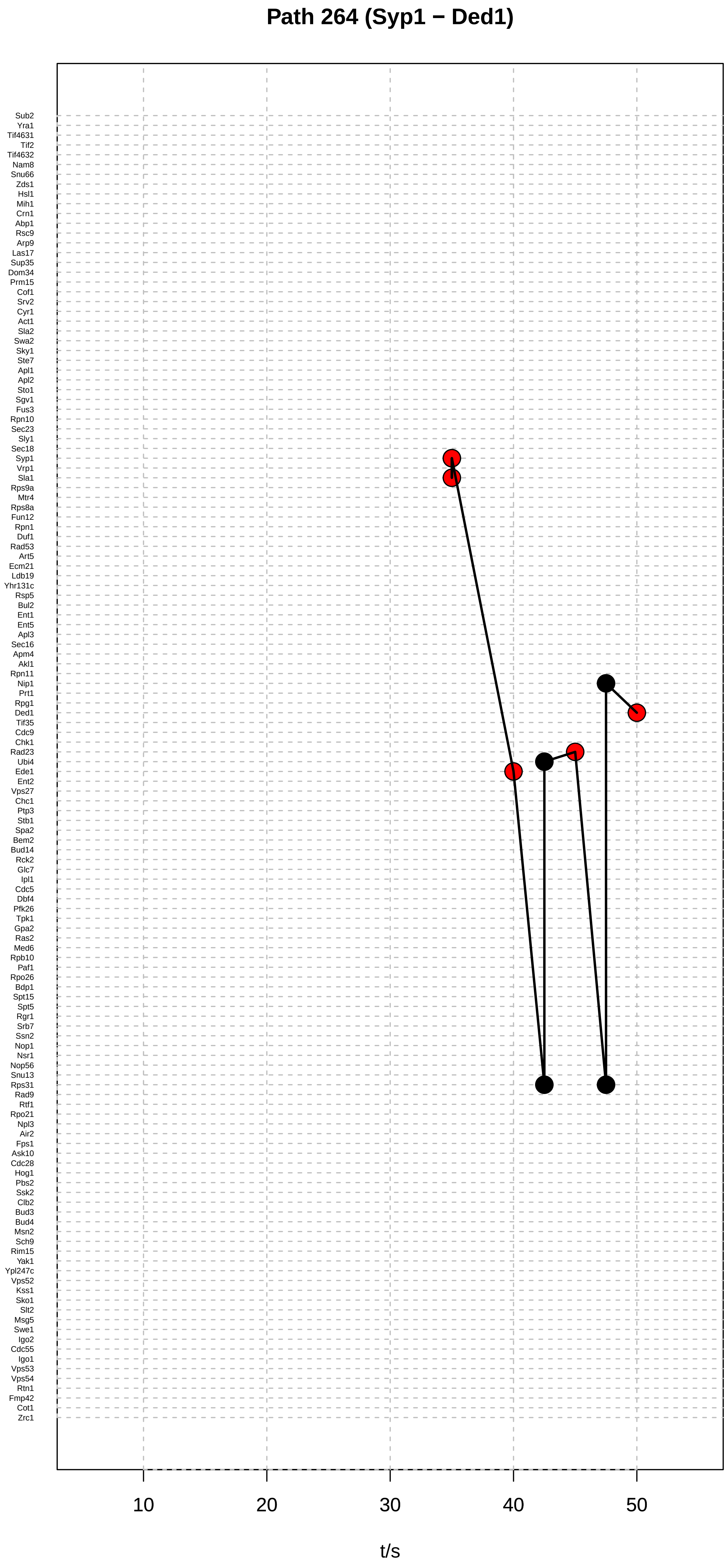
Path 264 (Syp1 − Ded1)
Sub2
Yra1
Tif4631
Tif2
Tif4632
Nam8
Snu66
Zds1
Hsl1
Mih1
Crn1
Abp1
Rsc9
Arp9
Las17
Sup35
Dom34
Prm15
Cof1
Srv2
Cyr1
Act1
Sla2
Swa2
Sky1
Ste7
Apl1
Apl2
Sto1
Sgv1
Fus3
Rpn10
Sec23
Sly1
Sec18
Syp1
Vrp1
Sla1
Rps9a
Mtr4
Rps8a
Fun12
Rpn1
Duf1
Rad53
Art5
Ecm21
Ldb19
Yhr131c
Rsp5
Bul2
Ent1
Ent5
Apl3
Sec16
Apm4
Akl1
Rpn11
Nip1
Prt1
Rpg1
Ded1
Tif35
Cdc9
Chk1
Rad23
Ubi4
Ede1
Ent2
Vps27
Chc1
Ptp3
Stb1
Spa2
Bem2
Bud14
Rck2
Glc7
Ipl1
Cdc5
Dbf4
Pfk26
Tpk1
Gpa2
Ras2
Med6
Rpb10
Paf1
Rpo26
Bdp1
Spt15
Spt5
Rgr1
Srb7
Ssn2
Nop1
Nsr1
Nop56
Snu13
Rps31
Rad9
Rtf1
Rpo21
Npl3
Air2
Fps1
Ask10
Cdc28
Hog1
Pbs2
Ssk2
Clb2
Bud3
Bud4
Msn2
Sch9
Rim15
Yak1
Ypl247c
Vps52
Kss1
Sko1
Slt2
Msg5
Swe1
Igo2
Cdc55
Igo1
Vps53
Vps54
Rtn1
Fmp42
Cot1
Zrc1
10
20
30
40
50
t/s

### Path 265 (Syp1 − Ded1)

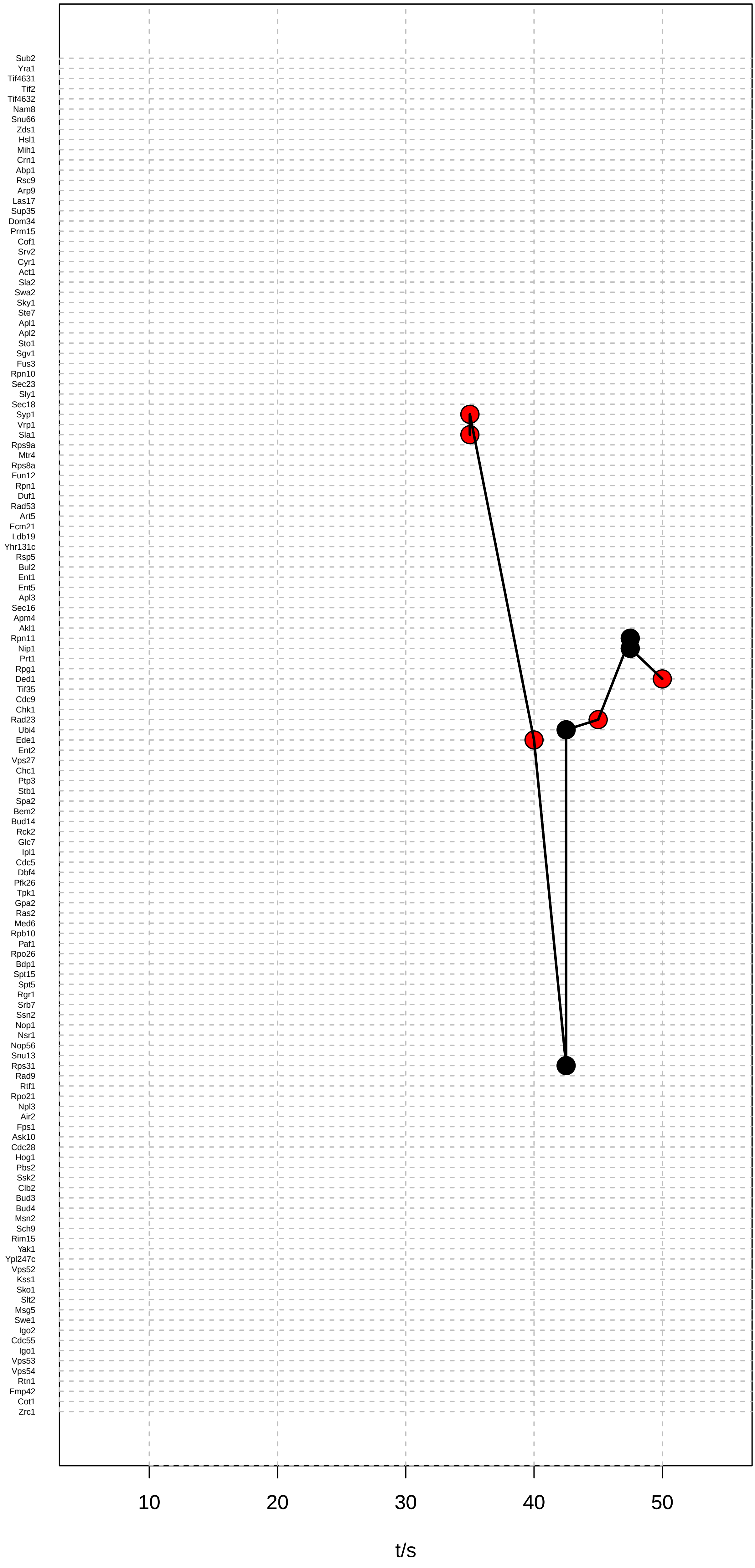

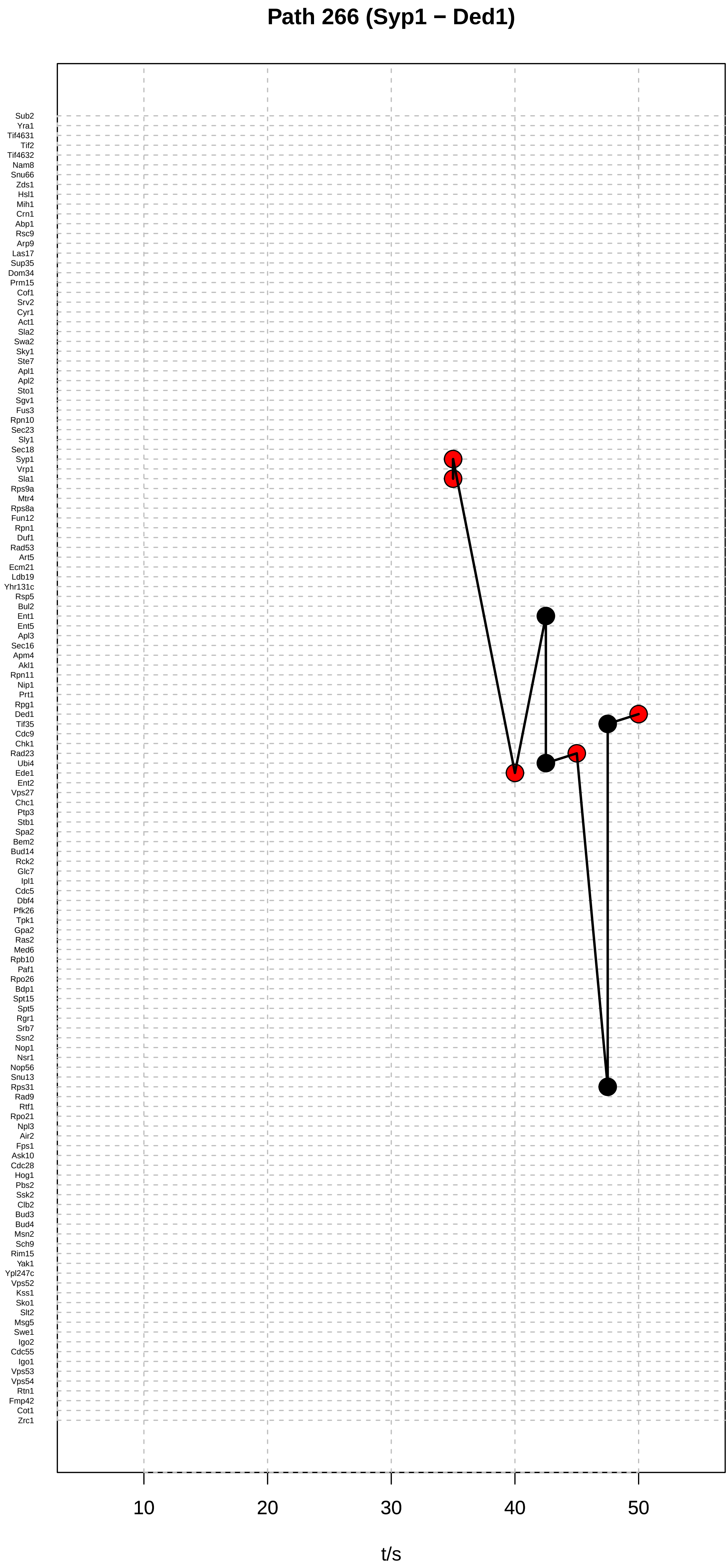

Path 266 (Syp1 − Ded1)
Sub2
Yra1
Tif4631
Tif2
Tif4632
Nam8
Snu66
Zds1
Hsl1
Mih1
Crn1
Abp1
Rsc9
Arp9
Las17
Sup35
Dom34
Prm15
Cof1
Srv2
Cyr1
Act1
Sla2
Swa2
Sky1
Ste7
Apl1
Apl2
Sto1
Sgv1
Fus3
Rpn10
Sec23
Sly1
Sec18
Syp1
Vrp1
Sla1
Rps9a
Mtr4
Rps8a
Fun12
Rpn1
Duf1
Rad53
Art5
Ecm21
Ldb19
Yhr131c
Rsp5
Bul2
Ent1
Ent5
Apl3
Sec16
Apm4
Akl1
Rpn11
Nip1
Prt1
Rpg1
Ded1
Tif35
Cdc9
Chk1
Rad23
Ubi4
Ede1
Ent2
Vps27
Chc1
Ptp3
Stb1
Spa2
Bem2
Bud14
Rck2
Glc7
Ipl1
Cdc5
Dbf4
Pfk26
Tpk1
Gpa2
Ras2
Med6
Rpb10
Paf1
Rpo26
Bdp1
Spt15
Spt5
Rgr1
Srb7
Ssn2
Nop1
Nsr1
Nop56
Snu13
Rps31
Rad9
Rtf1
Rpo21
Npl3
Air2
Fps1
Ask10
Cdc28
Hog1
Pbs2
Ssk2
Clb2
Bud3
Bud4
Msn2
Sch9
Rim15
Yak1
Ypl247c
Vps52
Kss1
Sko1
Slt2
Msg5
Swe1
Igo2
Cdc55
Igo1
Vps53
Vps54
Rtn1
Fmp42
Cot1
Zrc1
10
20
30
40
50
t/s

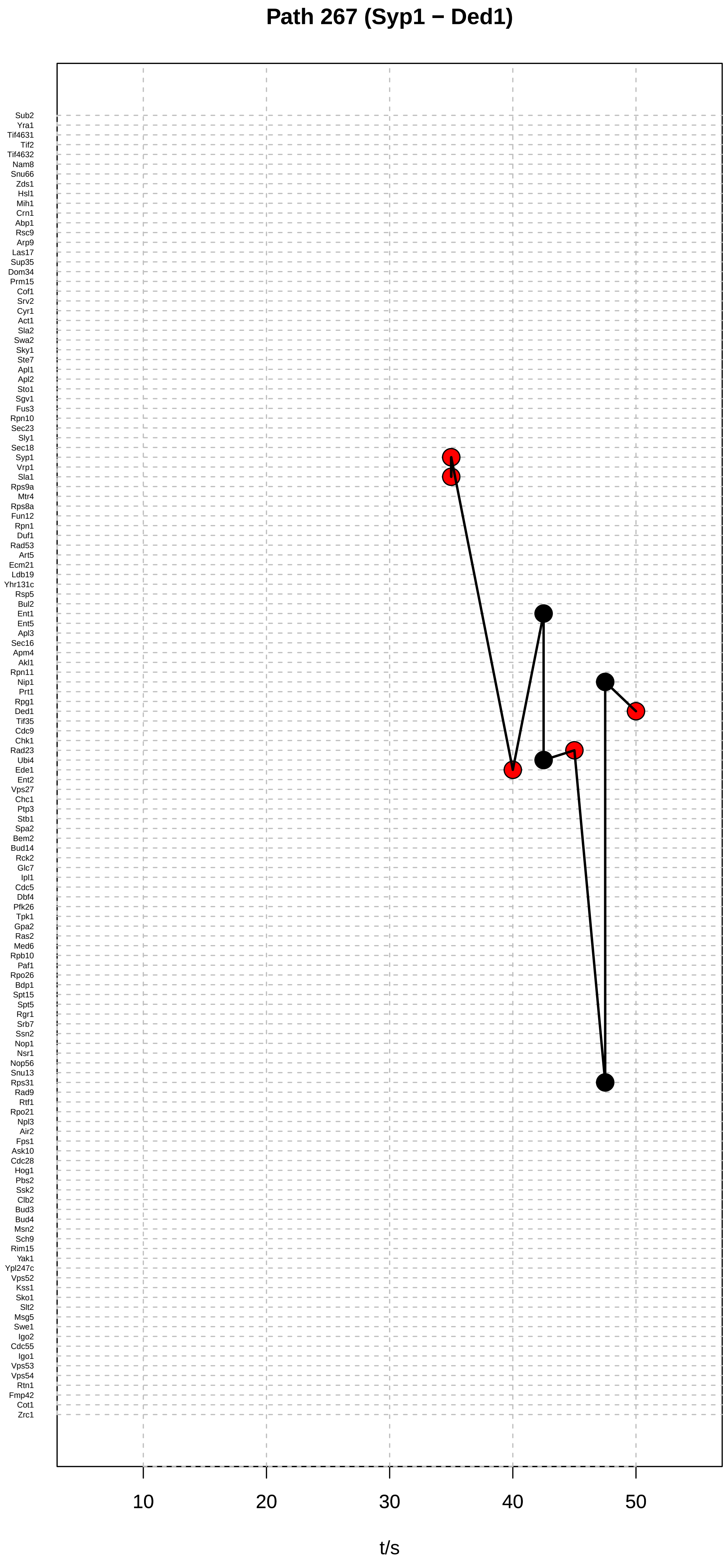

Path 267 (Syp1 − Ded1)
Sub2
Yra1
Tif4631
Tif2
Tif4632
Nam8
Snu66
Zds1
Hsl1
Mih1
Crn1
Abp1
Rsc9
Arp9
Las17
Sup35
Dom34
Prm15
Cof1
Srv2
Cyr1
Act1
Sla2
Swa2
Sky1
Ste7
Apl1
Apl2
Sto1
Sgv1
Fus3
Rpn10
Sec23
Sly1
Sec18
Syp1
Vrp1
Sla1
Rps9a
Mtr4
Rps8a
Fun12
Rpn1
Duf1
Rad53
Art5
Ecm21
Ldb19
Yhr131c
Rsp5
Bul2
Ent1
Ent5
Apl3
Sec16
Apm4
Akl1
Rpn11
Nip1
Prt1
Rpg1
Ded1
Tif35
Cdc9
Chk1
Rad23
Ubi4
Ede1
Ent2
Vps27
Chc1
Ptp3
Stb1
Spa2
Bem2
Bud14
Rck2
Glc7
Ipl1
Cdc5
Dbf4
Pfk26
Tpk1
Gpa2
Ras2
Med6
Rpb10
Paf1
Rpo26
Bdp1
Spt15
Spt5
Rgr1
Srb7
Ssn2
Nop1
Nsr1
Nop56
Snu13
Rps31
Rad9
Rtf1
Rpo21
Npl3
Air2
Fps1
Ask10
Cdc28
Hog1
Pbs2
Ssk2
Clb2
Bud3
Bud4
Msn2
Sch9
Rim15
Yak1
Ypl247c
Vps52
Kss1
Sko1
Slt2
Msg5
Swe1
Igo2
Cdc55
Igo1
Vps53
Vps54
Rtn1
Fmp42
Cot1
Zrc1
10
20
30
40
50
t/s

**Path 268 (Syp1 − Ded1)**

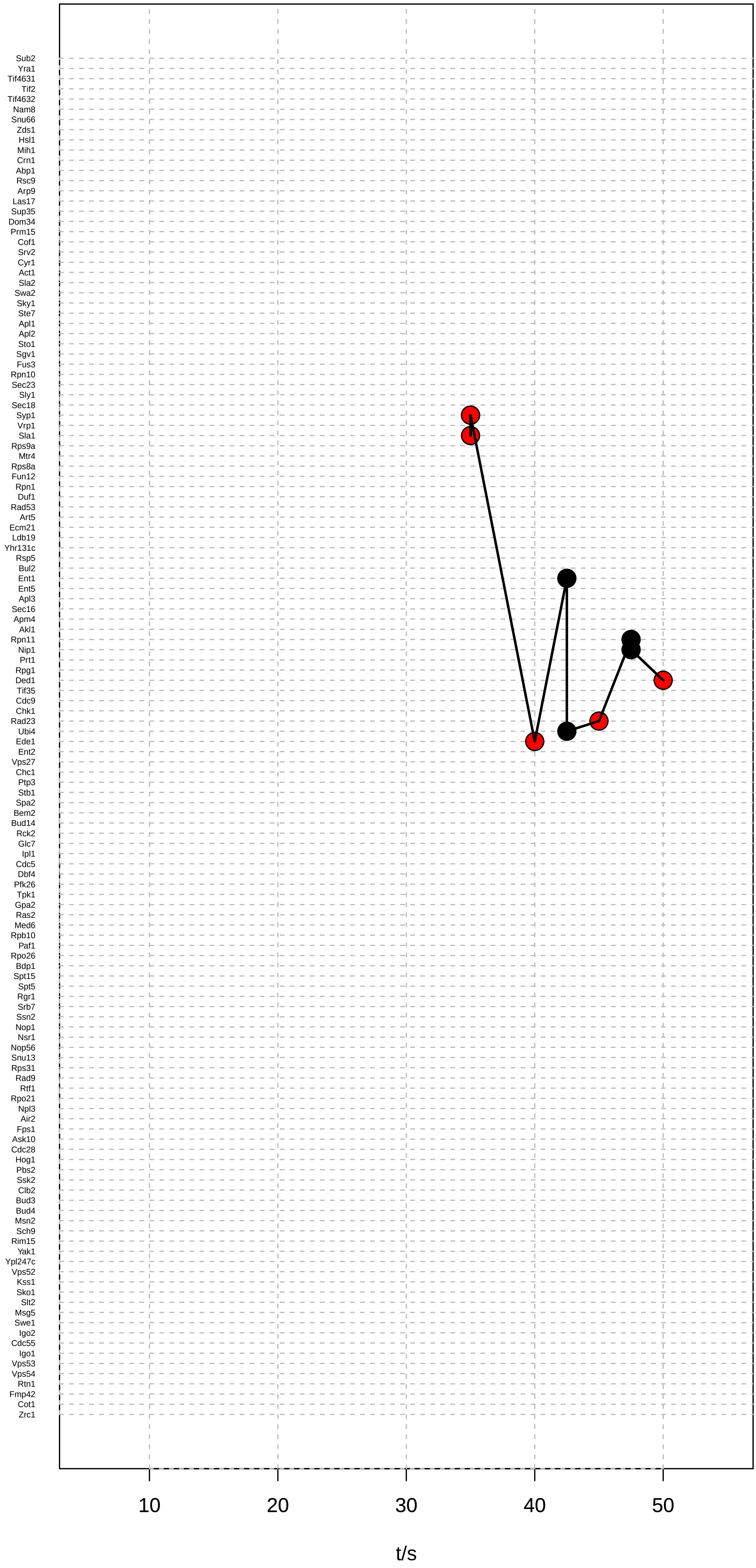

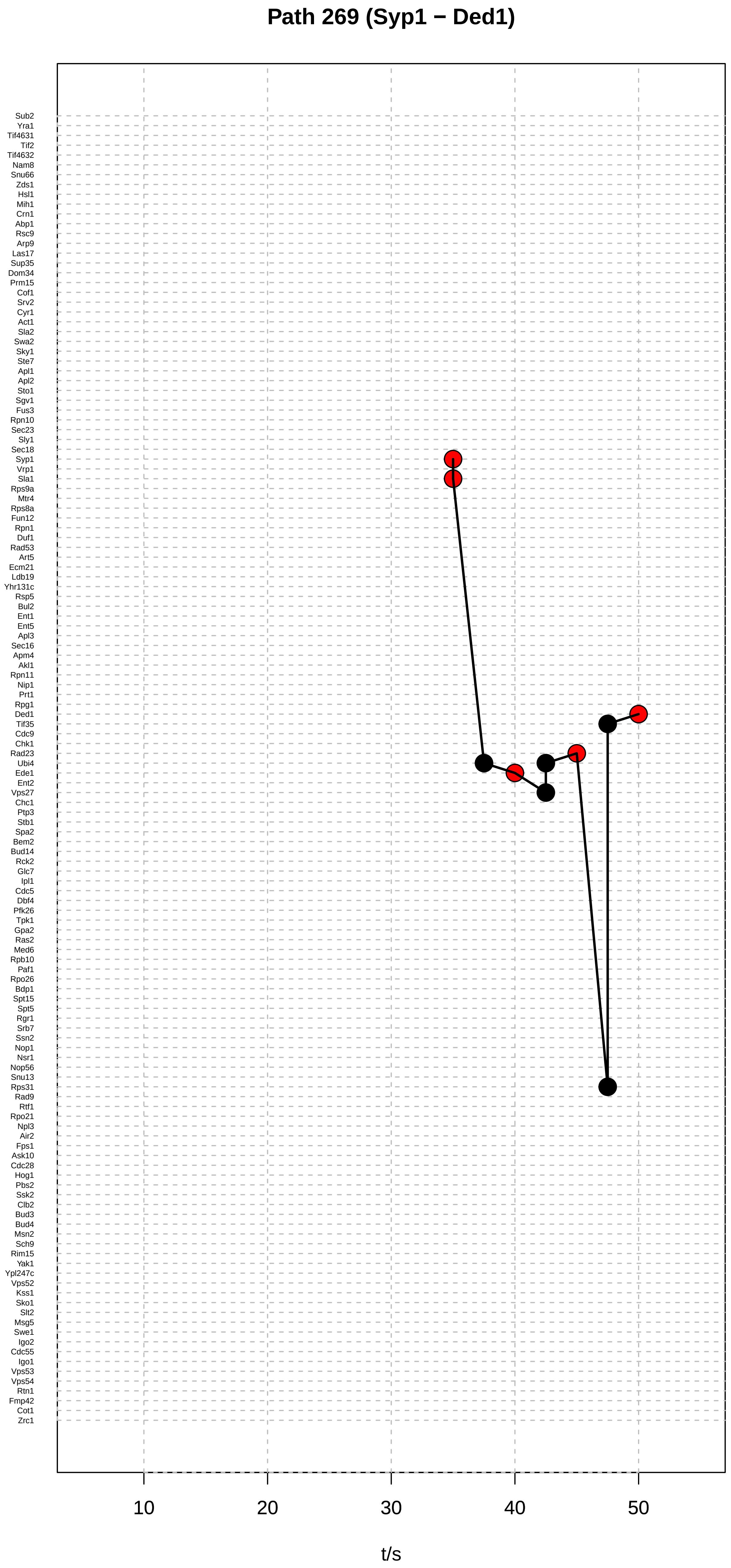

Path 269 (Syp1 − Ded1)
Sub2
Yra1
Tif4631
Tif2
Tif4632
Nam8
Snu66
Zds1
Hsl1
Mih1
Crn1
Abp1
Rsc9
Arp9
Las17
Sup35
Dom34
Prm15
Cof1
Srv2
Cyr1
Act1
Sla2
Swa2
Sky1
Ste7
Apl1
Apl2
Sto1
Sgv1
Fus3
Rpn10
Sec23
Sly1
Sec18
Syp1
Vrp1
Sla1
Rps9a
Mtr4
Rps8a
Fun12
Rpn1
Duf1
Rad53
Art5
Ecm21
Ldb19
Yhr131c
Rsp5
Bul2
Ent1
Ent5
Apl3
Sec16
Apm4
Akl1
Rpn11
Nip1
Prt1
Rpg1
Ded1
Tif35
Cdc9
Chk1
Rad23
Ubi4
Ede1
Ent2
Vps27
Chc1
Ptp3
Stb1
Spa2
Bem2
Bud14
Rck2
Glc7
Ipl1
Cdc5
Dbf4
Pfk26
Tpk1
Gpa2
Ras2
Med6
Rpb10
Paf1
Rpo26
Bdp1
Spt15
Spt5
Rgr1
Srb7
Ssn2
Nop1
Nsr1
Nop56
Snu13
Rps31
Rad9
Rtf1
Rpo21
Npl3
Air2
Fps1
Ask10
Cdc28
Hog1
Pbs2
Ssk2
Clb2
Bud3
Bud4
Msn2
Sch9
Rim15
Yak1
Ypl247c
Vps52
Kss1
Sko1
Slt2
Msg5
Swe1
Igo2
Cdc55
Igo1
Vps53
Vps54
Rtn1
Fmp42
Cot1
Zrc1
10
20
30
40
50
t/s

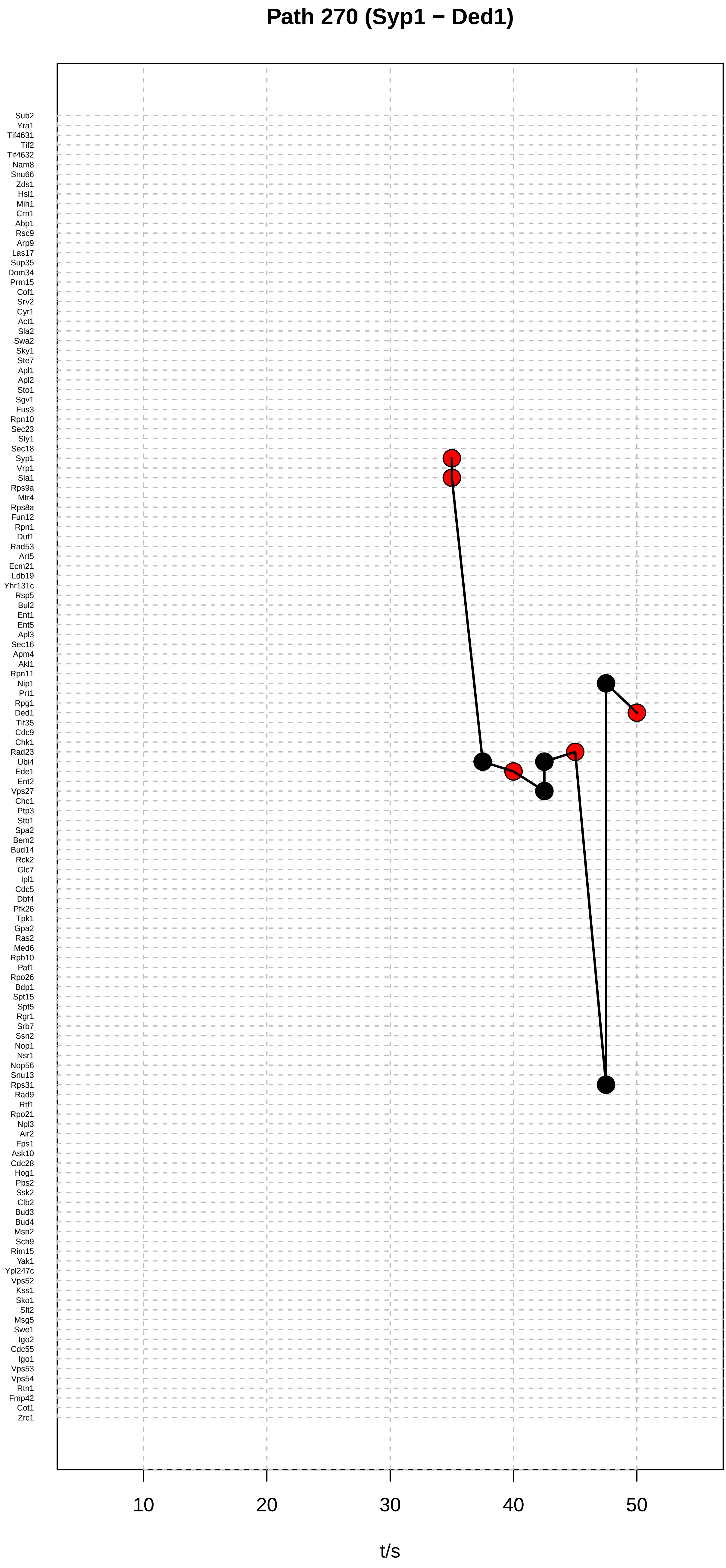

Path 270 (Syp1 − Ded1)
Sub2
Yra1
Tif4631
Tif2
Tif4632
Nam8
Snu66
Zds1
Hsl1
Mih1
Crn1
Abp1
Rsc9
Arp9
Las17
Sup35
Dom34
Prm15
Cof1
Srv2
Cyr1
Act1
Sla2
Swa2
Sky1
Ste7
Apl1
Apl2
Sto1
Sgv1
Fus3
Rpn10
Sec23
Sly1
Sec18
Syp1
Vrp1
Sla1
Rps9a
Mtr4
Rps8a
Fun12
Rpn1
Duf1
Rad53
Art5
Ecm21
Ldb19
Yhr131c
Rsp5
Bul2
Ent1
Ent5
Apl3
Sec16
Apm4
Akl1
Rpn11
Nip1
Prt1
Rpg1
Ded1
Tif35
Cdc9
Chk1
Rad23
Ubi4
Ede1
Ent2
Vps27
Chc1
Ptp3
Stb1
Spa2
Bem2
Bud14
Rck2
Glc7
Ipl1
Cdc5
Dbf4
Pfk26
Tpk1
Gpa2
Ras2
Med6
Rpb10
Paf1
Rpo26
Bdp1
Spt15
Spt5
Rgr1
Srb7
Ssn2
Nop1
Nsr1
Nop56
Snu13
Rps31
Rad9
Rtf1
Rpo21
Npl3
Air2
Fps1
Ask10
Cdc28
Hog1
Pbs2
Ssk2
Clb2
Bud3
Bud4
Msn2
Sch9
Rim15
Yak1
Ypl247c
Vps52
Kss1
Sko1
Slt2
Msg5
Swe1
Igo2
Cdc55
Igo1
Vps53
Vps54
Rtn1
Fmp42
Cot1
Zrc1
10
20
30
40
50
t/s

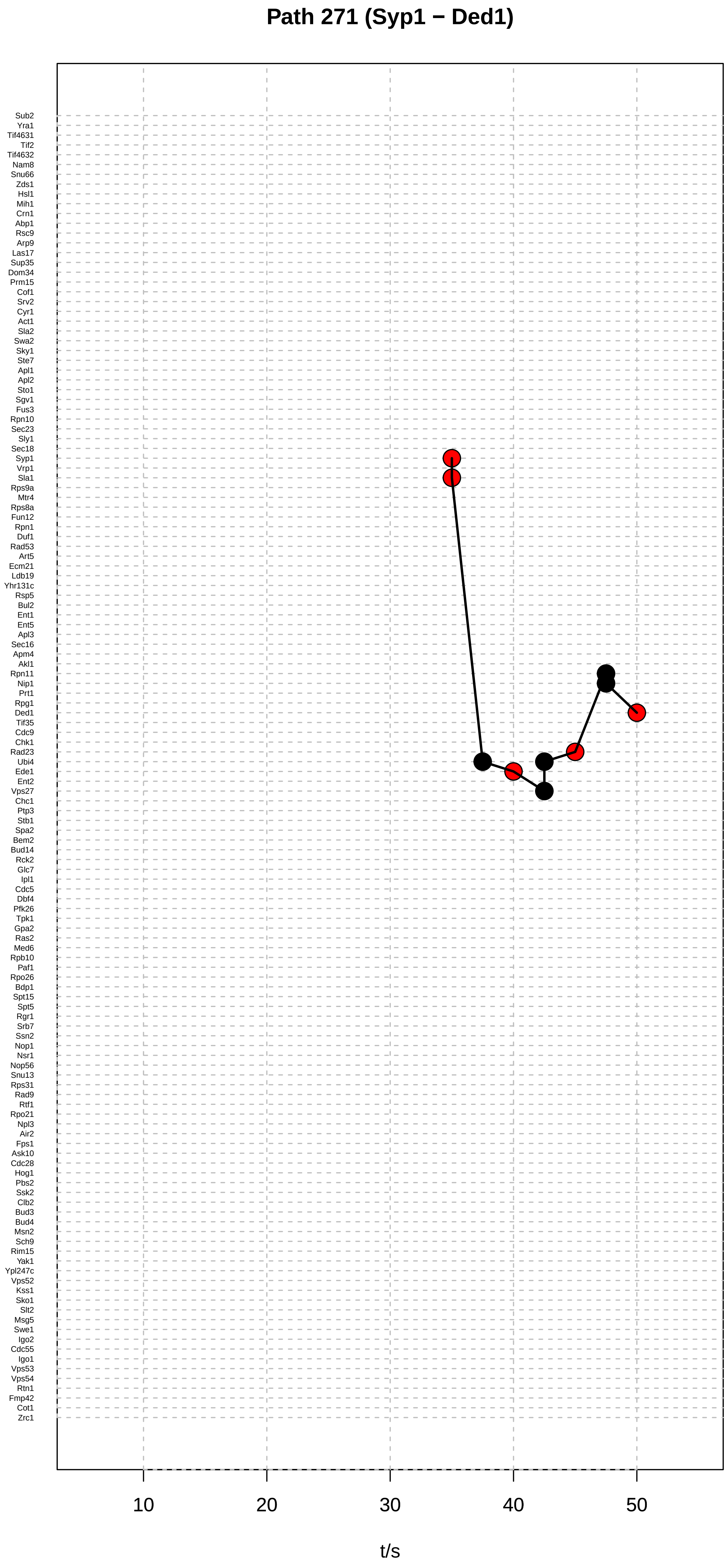

Path 271 (Syp1 − Ded1)
Sub2
Yra1
Tif4631
Tif2
Tif4632
Nam8
Snu66
Zds1
Hsl1
Mih1
Crn1
Abp1
Rsc9
Arp9
Las17
Sup35
Dom34
Prm15
Cof1
Srv2
Cyr1
Act1
Sla2
Swa2
Sky1
Ste7
Apl1
Apl2
Sto1
Sgv1
Fus3
Rpn10
Sec23
Sly1
Sec18
Syp1
Vrp1
Sla1
Rps9a
Mtr4
Rps8a
Fun12
Rpn1
Duf1
Rad53
Art5
Ecm21
Ldb19
Yhr131c
Rsp5
Bul2
Ent1
Ent5
Apl3
Sec16
Apm4
Akl1
Rpn11
Nip1
Prt1
Rpg1
Ded1
Tif35
Cdc9
Chk1
Rad23
Ubi4
Ede1
Ent2
Vps27
Chc1
Ptp3
Stb1
Spa2
Bem2
Bud14
Rck2
Glc7
Ipl1
Cdc5
Dbf4
Pfk26
Tpk1
Gpa2
Ras2
Med6
Rpb10
Paf1
Rpo26
Bdp1
Spt15
Spt5
Rgr1
Srb7
Ssn2
Nop1
Nsr1
Nop56
Snu13
Rps31
Rad9
Rtf1
Rpo21
Npl3
Air2
Fps1
Ask10
Cdc28
Hog1
Pbs2
Ssk2
Clb2
Bud3
Bud4
Msn2
Sch9
Rim15
Yak1
Ypl247c
Vps52
Kss1
Sko1
Slt2
Msg5
Swe1
Igo2
Cdc55
Igo1
Vps53
Vps54
Rtn1
Fmp42
Cot1
Zrc1
10
20
30
40
50
t/s

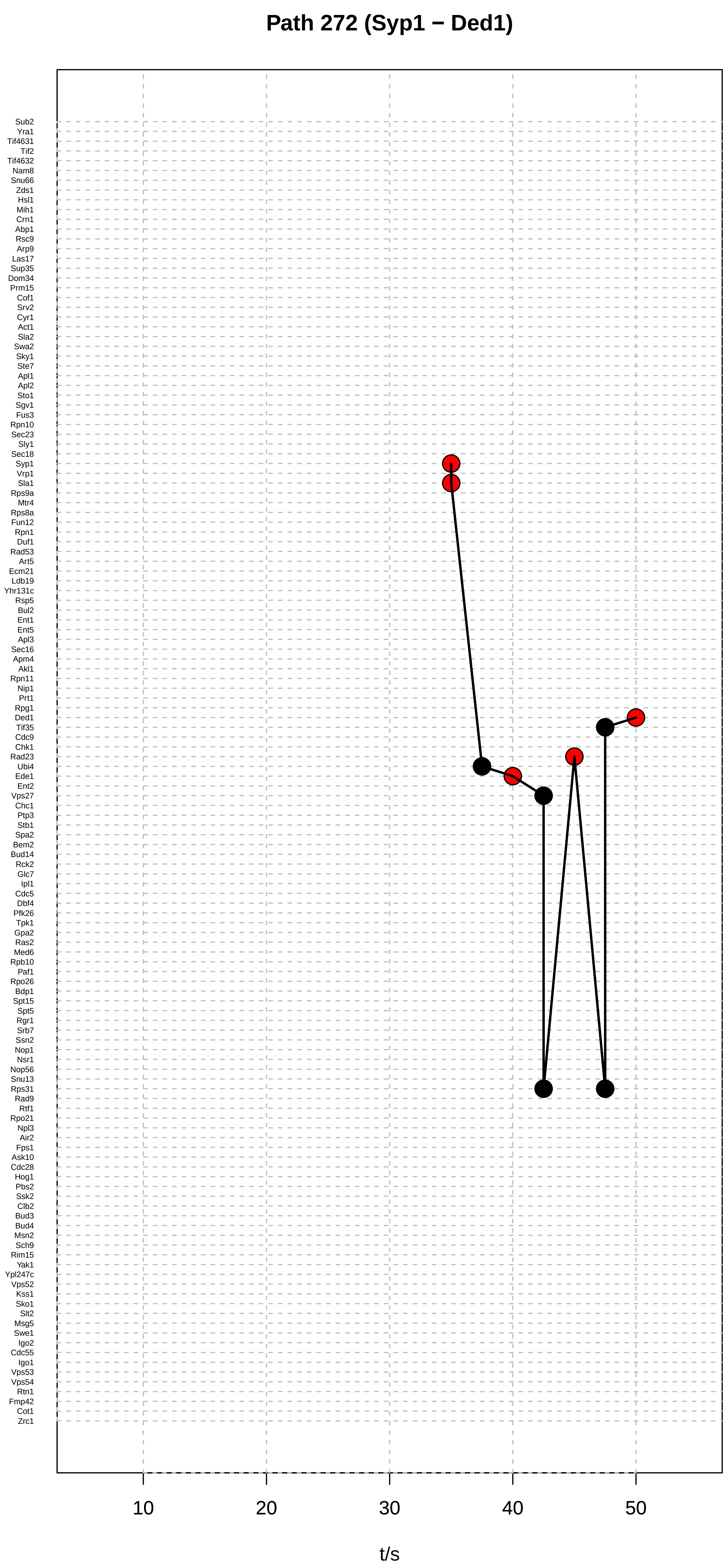

Path 272 (Syp1 − Ded1)
Sub2
Yra1
Tif4631
Tif2
Tif4632
Nam8
Snu66
Zds1
Hsl1
Mih1
Crn1
Abp1
Rsc9
Arp9
Las17
Sup35
Dom34
Prm15
Cof1
Srv2
Cyr1
Act1
Sla2
Swa2
Sky1
Ste7
Apl1
Apl2
Sto1
Sgv1
Fus3
Rpn10
Sec23
Sly1
Sec18
Syp1
Vrp1
Sla1
Rps9a
Mtr4
Rps8a
Fun12
Rpn1
Duf1
Rad53
Art5
Ecm21
Ldb19
Yhr131c
Rsp5
Bul2
Ent1
Ent5
Apl3
Sec16
Apm4
Akl1
Rpn11
Nip1
Prt1
Rpg1
Ded1
Tif35
Cdc9
Chk1
Rad23
Ubi4
Ede1
Ent2
Vps27
Chc1
Ptp3
Stb1
Spa2
Bem2
Bud14
Rck2
Glc7
Ipl1
Cdc5
Dbf4
Pfk26
Tpk1
Gpa2
Ras2
Med6
Rpb10
Paf1
Rpo26
Bdp1
Spt15
Spt5
Rgr1
Srb7
Ssn2
Nop1
Nsr1
Nop56
Snu13
Rps31
Rad9
Rtf1
Rpo21
Npl3
Air2
Fps1
Ask10
Cdc28
Hog1
Pbs2
Ssk2
Clb2
Bud3
Bud4
Msn2
Sch9
Rim15
Yak1
Ypl247c
Vps52
Kss1
Sko1
Slt2
Msg5
Swe1
Igo2
Cdc55
Igo1
Vps53
Vps54
Rtn1
Fmp42
Cot1
Zrc1
10
20
30
40
50
t/s

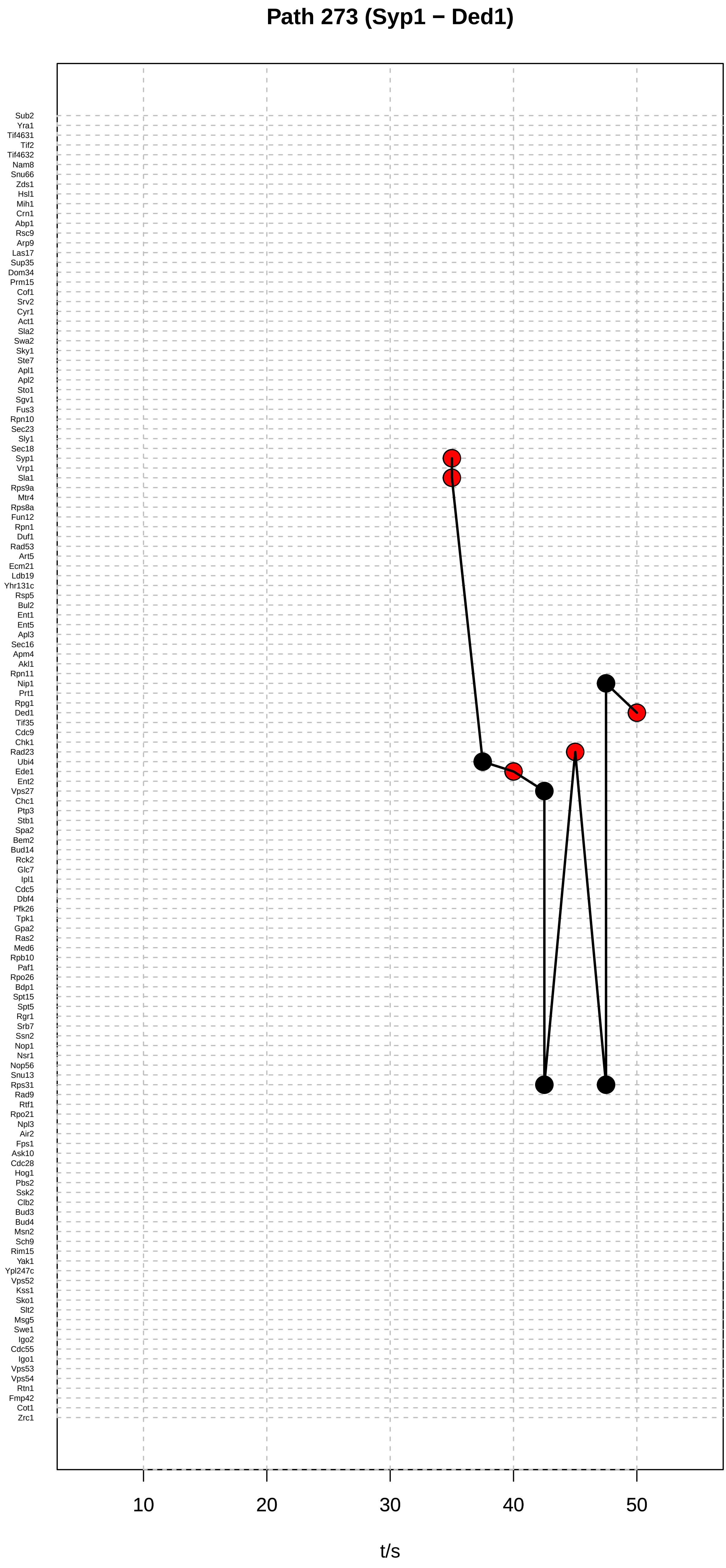
Path 273 (Syp1 − Ded1)
Sub2
Yra1
Tif4631
Tif2
Tif4632
Nam8
Snu66
Zds1
Hsl1
Mih1
Crn1
Abp1
Rsc9
Arp9
Las17
Sup35
Dom34
Prm15
Cof1
Srv2
Cyr1
Act1
Sla2
Swa2
Sky1
Ste7
Apl1
Apl2
Sto1
Sgv1
Fus3
Rpn10
Sec23
Sly1
Sec18
Syp1
Vrp1
Sla1
Rps9a
Mtr4
Rps8a
Fun12
Rpn1
Duf1
Rad53
Art5
Ecm21
Ldb19
Yhr131c
Rsp5
Bul2
Ent1
Ent5
Apl3
Sec16
Apm4
Akl1
Rpn11
Nip1
Prt1
Rpg1
Ded1
Tif35
Cdc9
Chk1
Rad23
Ubi4
Ede1
Ent2
Vps27
Chc1
Ptp3
Stb1
Spa2
Bem2
Bud14
Rck2
Glc7
Ipl1
Cdc5
Dbf4
Pfk26
Tpk1
Gpa2
Ras2
Med6
Rpb10
Paf1
Rpo26
Bdp1
Spt15
Spt5
Rgr1
Srb7
Ssn2
Nop1
Nsr1
Nop56
Snu13
Rps31
Rad9
Rtf1
Rpo21
Npl3
Air2
Fps1
Ask10
Cdc28
Hog1
Pbs2
Ssk2
Clb2
Bud3
Bud4
Msn2
Sch9
Rim15
Yak1
Ypl247c
Vps52
Kss1
Sko1
Slt2
Msg5
Swe1
Igo2
Cdc55
Igo1
Vps53
Vps54
Rtn1
Fmp42
Cot1
Zrc1
10
20
30
40
50
t/s

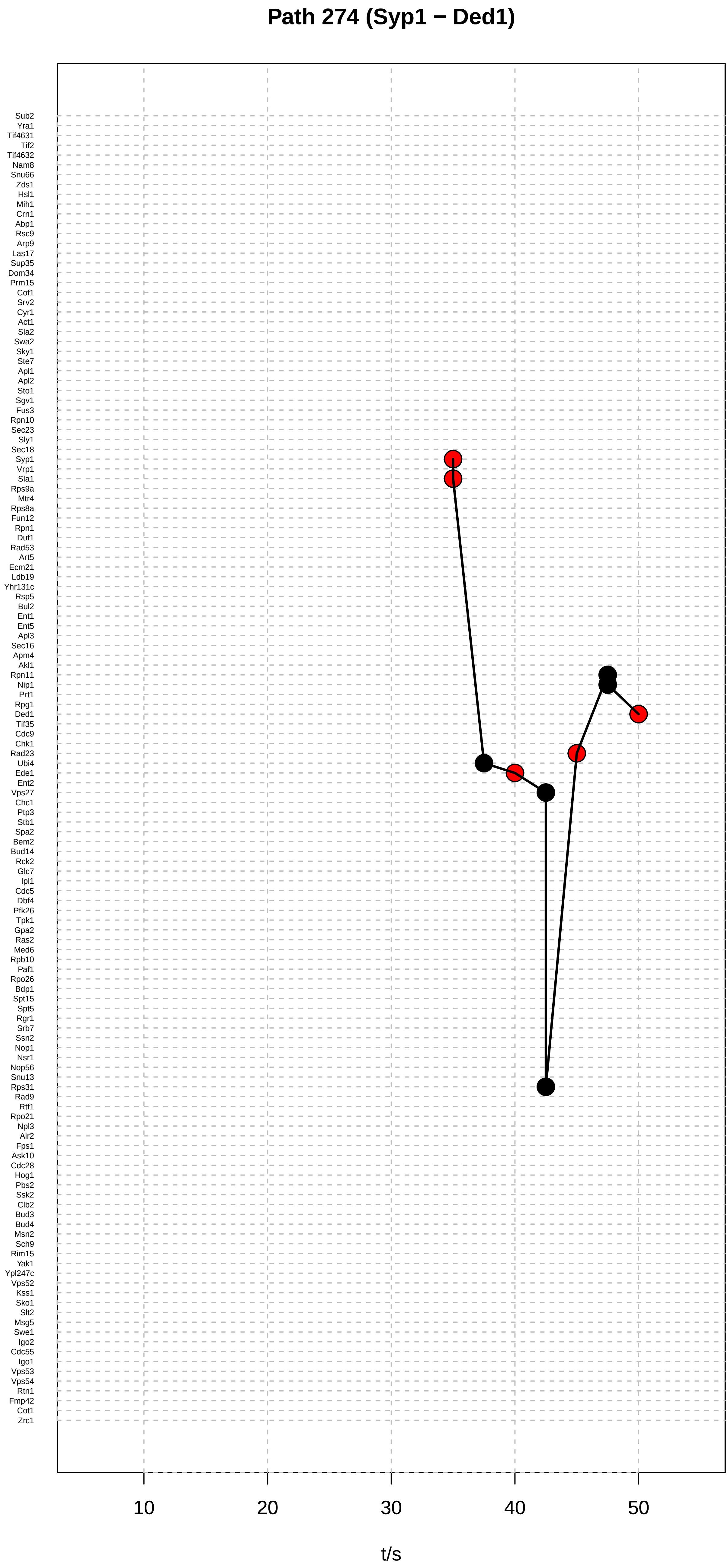
Path 274 (Syp1 − Ded1)
Sub2
Yra1
Tif4631
Tif2
Tif4632
Nam8
Snu66
Zds1
Hsl1
Mih1
Crn1
Abp1
Rsc9
Arp9
Las17
Sup35
Dom34
Prm15
Cof1
Srv2
Cyr1
Act1
Sla2
Swa2
Sky1
Ste7
Apl1
Apl2
Sto1
Sgv1
Fus3
Rpn10
Sec23
Sly1
Sec18
Syp1
Vrp1
Sla1
Rps9a
Mtr4
Rps8a
Fun12
Rpn1
Duf1
Rad53
Art5
Ecm21
Ldb19
Yhr131c
Rsp5
Bul2
Ent1
Ent5
Apl3
Sec16
Apm4
Akl1
Rpn11
Nip1
Prt1
Rpg1
Ded1
Tif35
Cdc9
Chk1
Rad23
Ubi4
Ede1
Ent2
Vps27
Chc1
Ptp3
Stb1
Spa2
Bem2
Bud14
Rck2
Glc7
Ipl1
Cdc5
Dbf4
Pfk26
Tpk1
Gpa2
Ras2
Med6
Rpb10
Paf1
Rpo26
Bdp1
Spt15
Spt5
Rgr1
Srb7
Ssn2
Nop1
Nsr1
Nop56
Snu13
Rps31
Rad9
Rtf1
Rpo21
Npl3
Air2
Fps1
Ask10
Cdc28
Hog1
Pbs2
Ssk2
Clb2
Bud3
Bud4
Msn2
Sch9
Rim15
Yak1
Ypl247c
Vps52
Kss1
Sko1
Slt2
Msg5
Swe1
Igo2
Cdc55
Igo1
Vps53
Vps54
Rtn1
Fmp42
Cot1
Zrc1
10
20
30
40
50
t/s

#### Path 275 (Syp1 − Ded1)

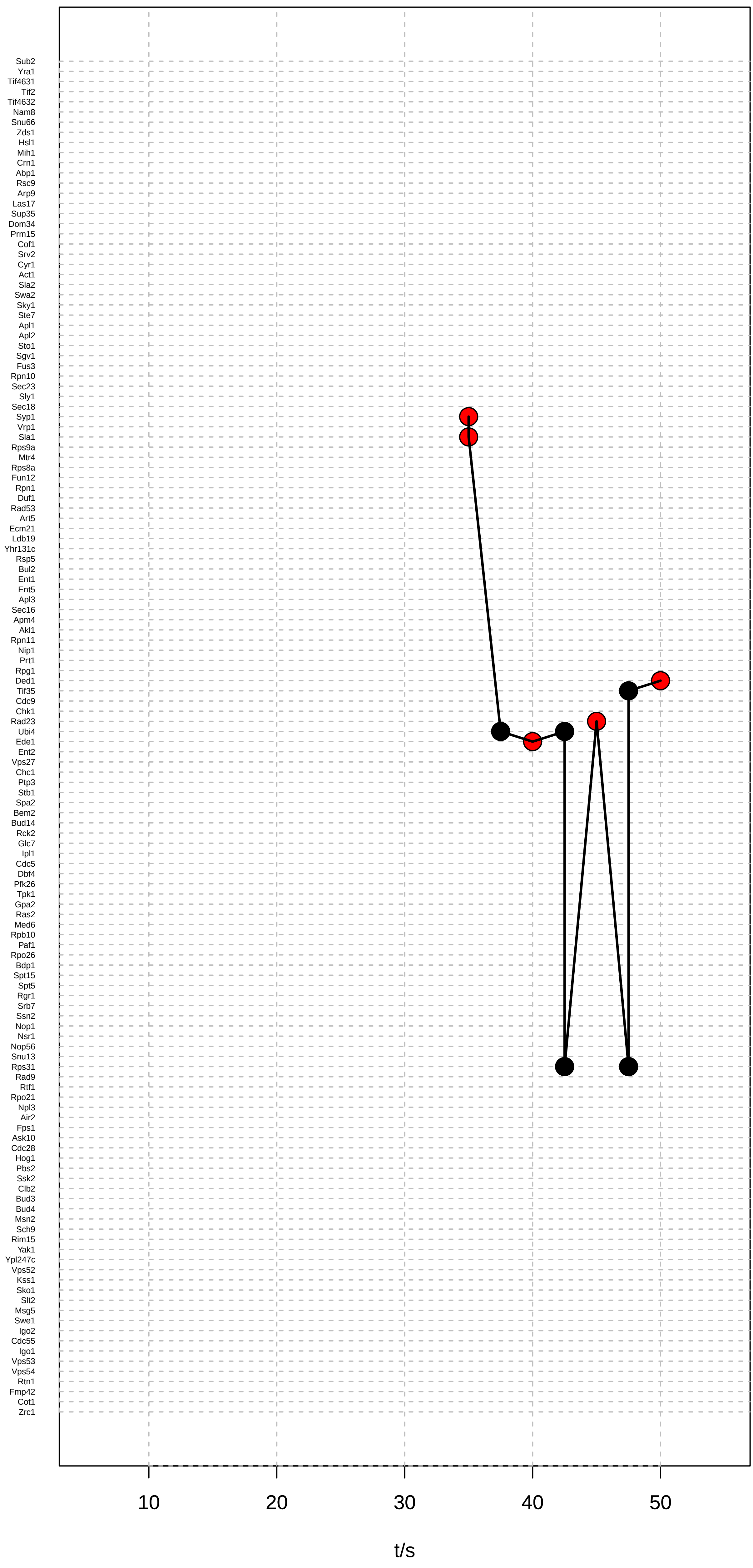

#### Path 276 (Syp1 − Ded1)

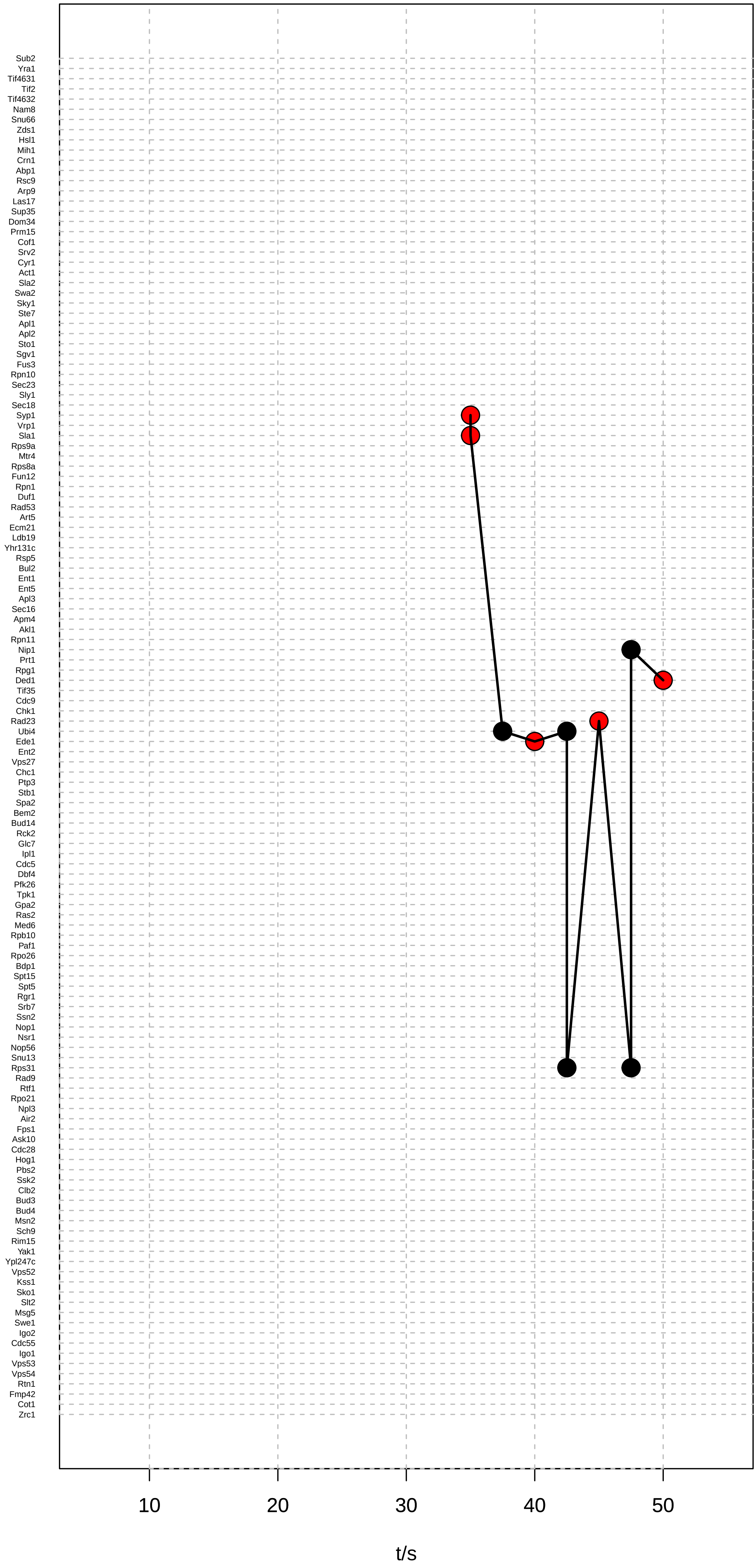

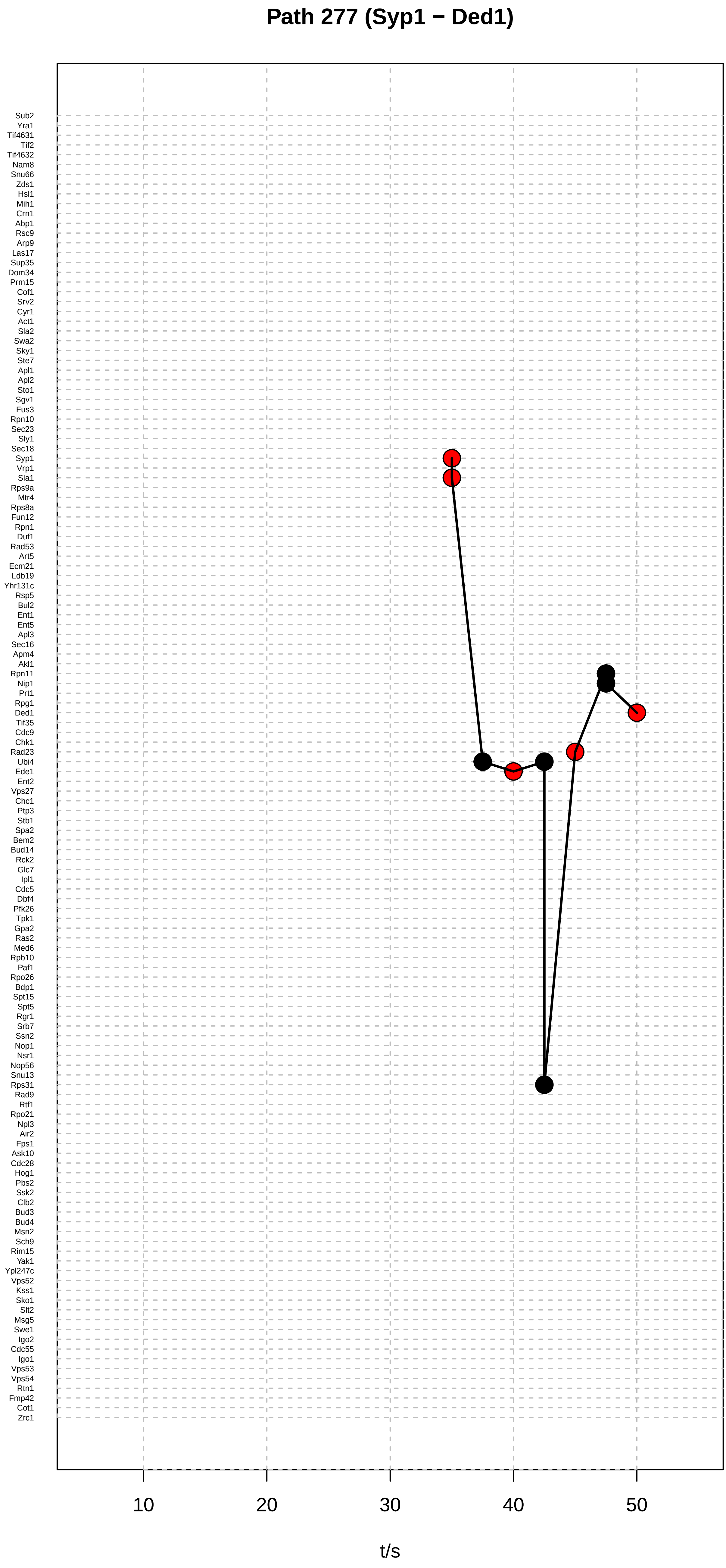

Path 277 (Syp1 − Ded1)
Sub2
Yra1
Tif4631
Tif2
Tif4632
Nam8
Snu66
Zds1
Hsl1
Mih1
Crn1
Abp1
Rsc9
Arp9
Las17
Sup35
Dom34
Prm15
Cof1
Srv2
Cyr1
Act1
Sla2
Swa2
Sky1
Ste7
Apl1
Apl2
Sto1
Sgv1
Fus3
Rpn10
Sec23
Sly1
Sec18
Syp1
Vrp1
Sla1
Rps9a
Mtr4
Rps8a
Fun12
Rpn1
Duf1
Rad53
Art5
Ecm21
Ldb19
Yhr131c
Rsp5
Bul2
Ent1
Ent5
Apl3
Sec16
Apm4
Akl1
Rpn11
Nip1
Prt1
Rpg1
Ded1
Tif35
Cdc9
Chk1
Rad23
Ubi4
Ede1
Ent2
Vps27
Chc1
Ptp3
Stb1
Spa2
Bem2
Bud14
Rck2
Glc7
Ipl1
Cdc5
Dbf4
Pfk26
Tpk1
Gpa2
Ras2
Med6
Rpb10
Paf1
Rpo26
Bdp1
Spt15
Spt5
Rgr1
Srb7
Ssn2
Nop1
Nsr1
Nop56
Snu13
Rps31
Rad9
Rtf1
Rpo21
Npl3
Air2
Fps1
Ask10
Cdc28
Hog1
Pbs2
Ssk2
Clb2
Bud3
Bud4
Msn2
Sch9
Rim15
Yak1
Ypl247c
Vps52
Kss1
Sko1
Slt2
Msg5
Swe1
Igo2
Cdc55
Igo1
Vps53
Vps54
Rtn1
Fmp42
Cot1
Zrc1
10
20
30
40
50
t/s

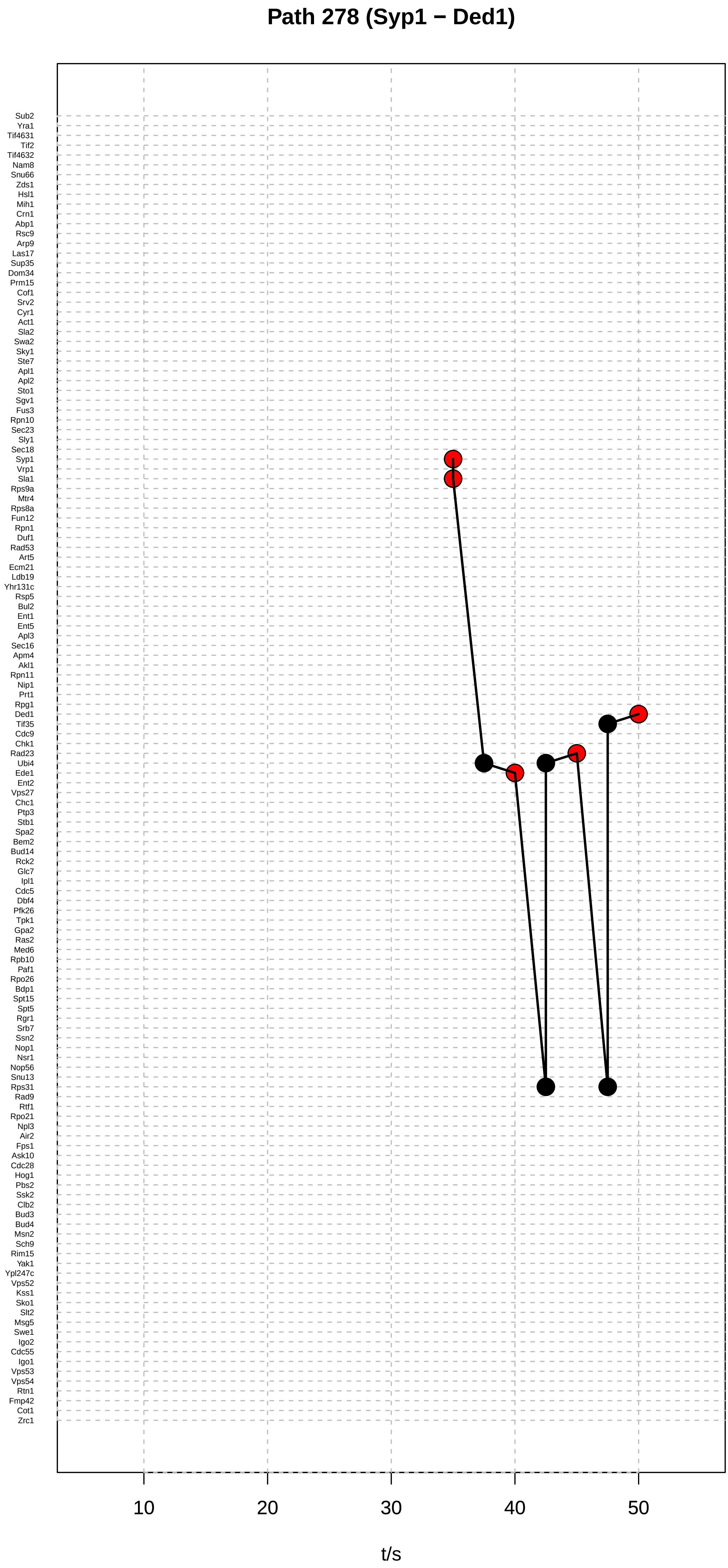

Path 278 (Syp1 − Ded1)
Sub2
Yra1
Tif4631
Tif2
Tif4632
Nam8
Snu66
Zds1
Hsl1
Mih1
Crn1
Abp1
Rsc9
Arp9
Las17
Sup35
Dom34
Prm15
Cof1
Srv2
Cyr1
Act1
Sla2
Swa2
Sky1
Ste7
Apl1
Apl2
Sto1
Sgv1
Fus3
Rpn10
Sec23
Sly1
Sec18
Syp1
Vrp1
Sla1
Rps9a
Mtr4
Rps8a
Fun12
Rpn1
Duf1
Rad53
Art5
Ecm21
Ldb19
Yhr131c
Rsp5
Bul2
Ent1
Ent5
Apl3
Sec16
Apm4
Akl1
Rpn11
Nip1
Prt1
Rpg1
Ded1
Tif35
Cdc9
Chk1
Rad23
Ubi4
Ede1
Ent2
Vps27
Chc1
Ptp3
Stb1
Spa2
Bem2
Bud14
Rck2
Glc7
Ipl1
Cdc5
Dbf4
Pfk26
Tpk1
Gpa2
Ras2
Med6
Rpb10
Paf1
Rpo26
Bdp1
Spt15
Spt5
Rgr1
Srb7
Ssn2
Nop1
Nsr1
Nop56
Snu13
Rps31
Rad9
Rtf1
Rpo21
Npl3
Air2
Fps1
Ask10
Cdc28
Hog1
Pbs2
Ssk2
Clb2
Bud3
Bud4
Msn2
Sch9
Rim15
Yak1
Ypl247c
Vps52
Kss1
Sko1
Slt2
Msg5
Swe1
Igo2
Cdc55
Igo1
Vps53
Vps54
Rtn1
Fmp42
Cot1
Zrc1
10
20
30
40
50
t/s

### Path 279 (Syp1 − Ded1)

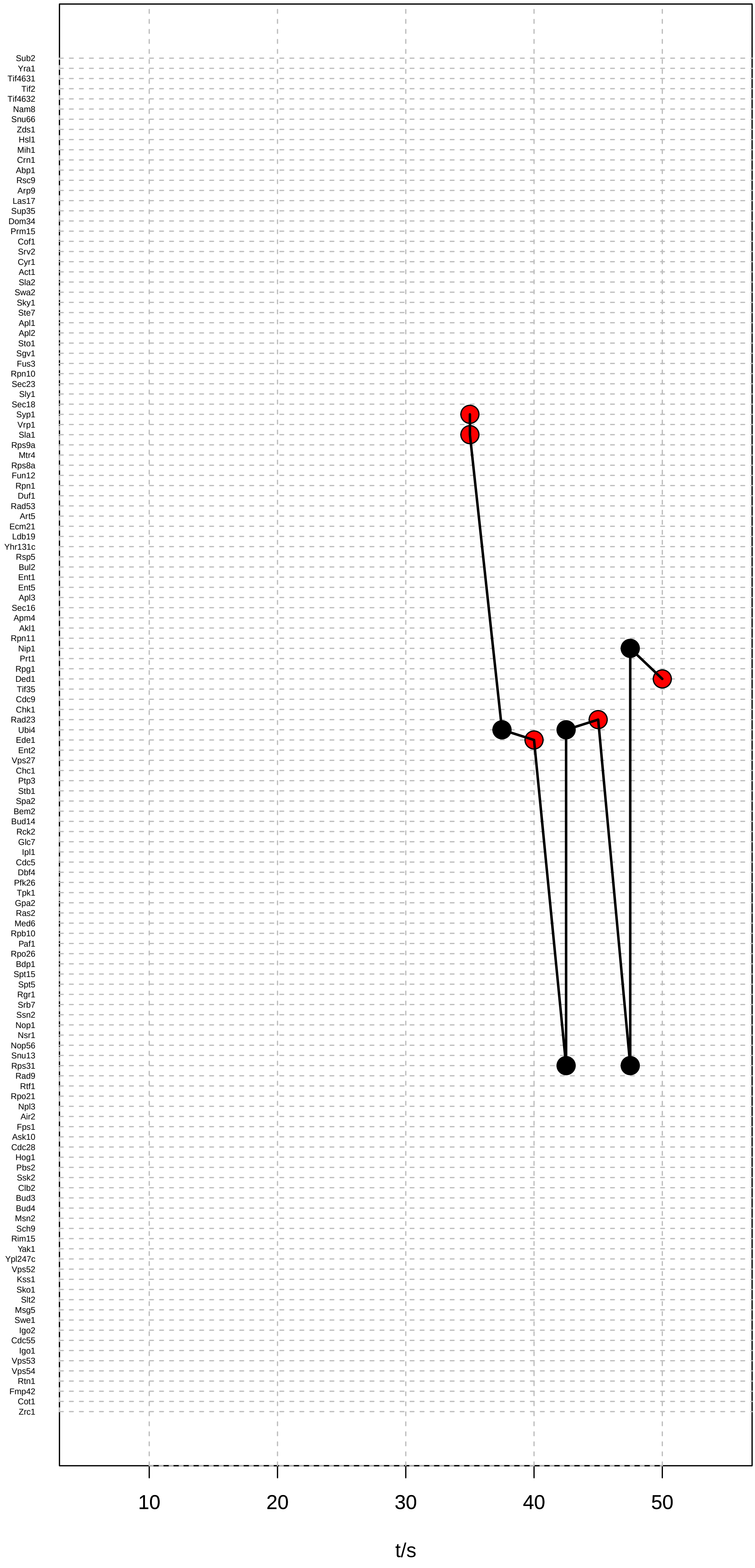

### Path 280 (Syp1 − Ded1)

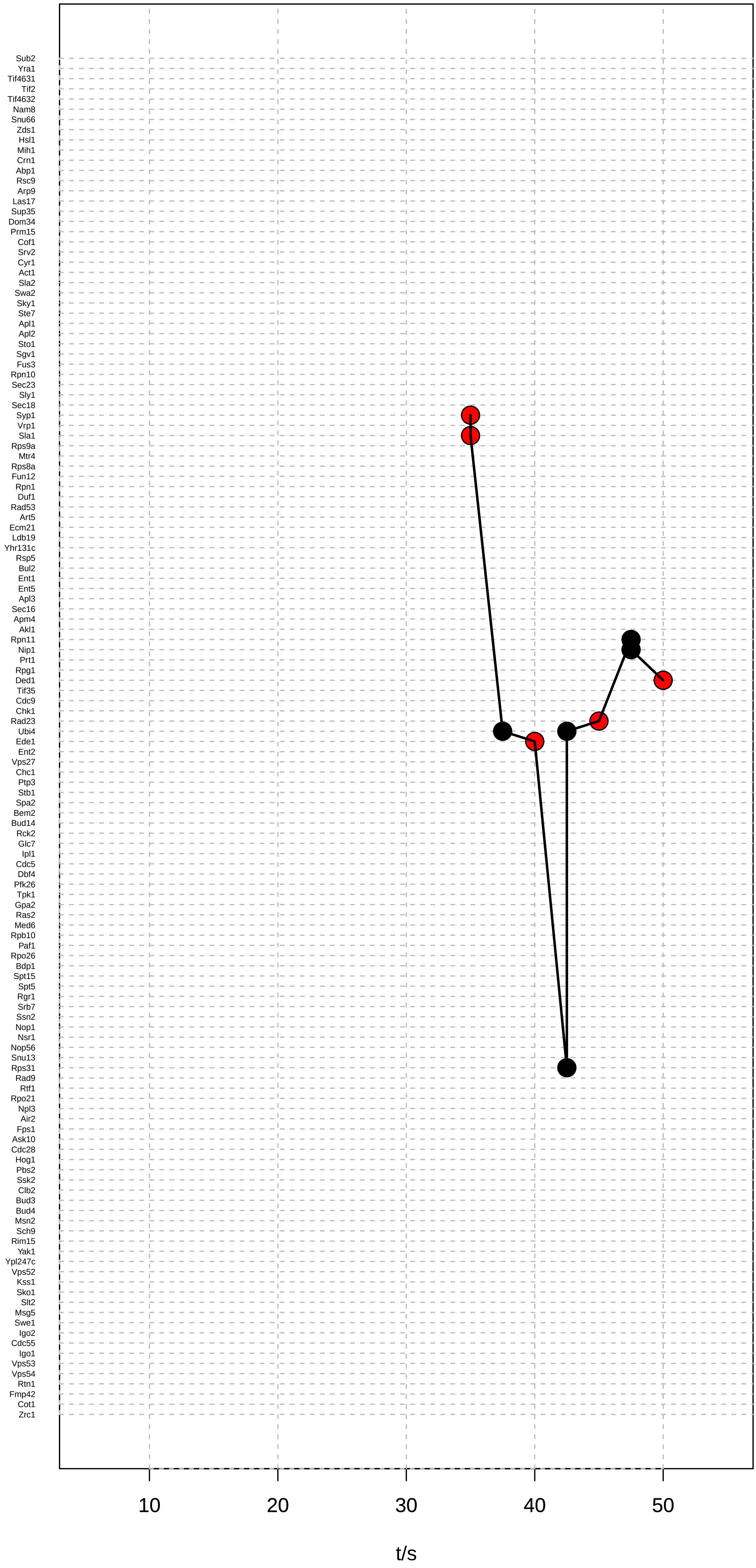

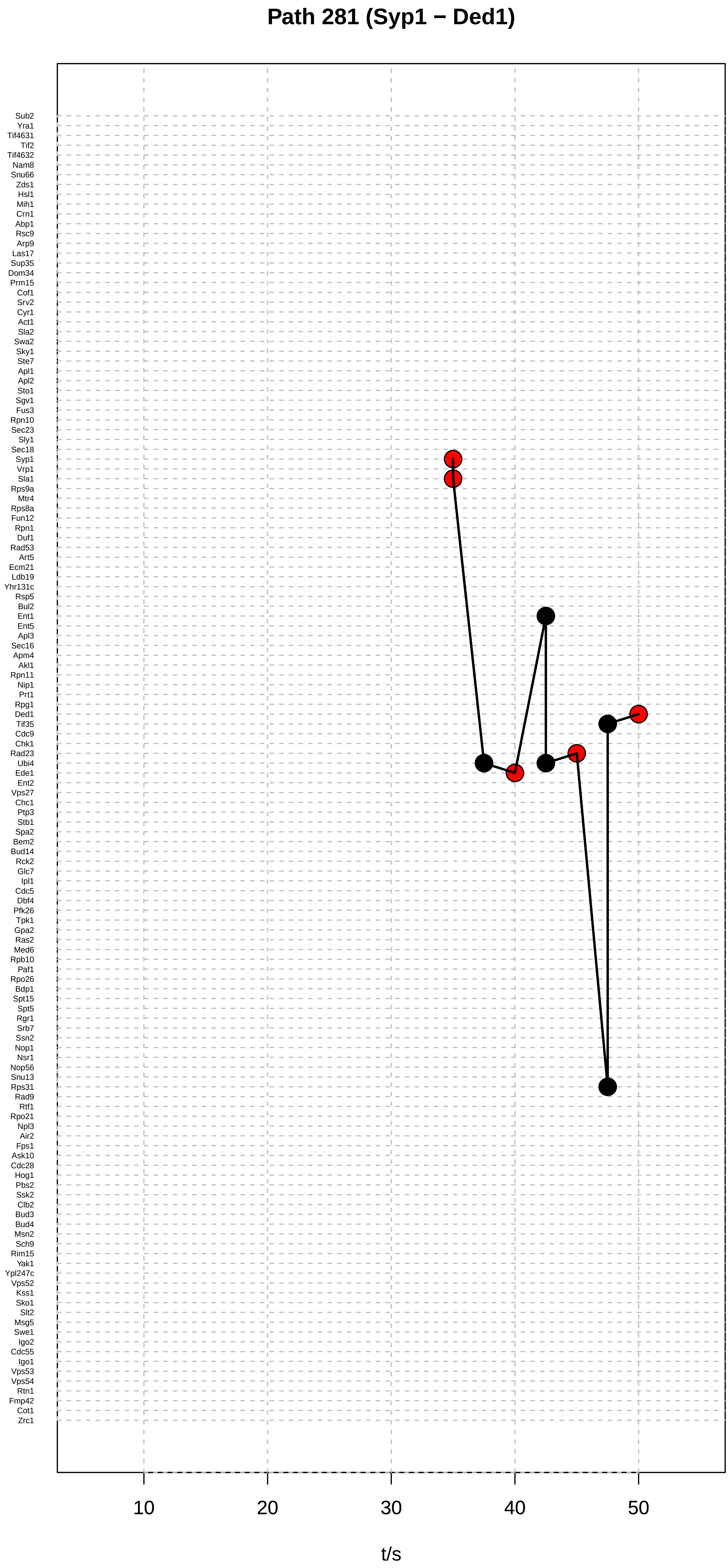
Path 281 (Syp1 − Ded1)
Sub2
Yra1
Tif4631
Tif2
Tif4632
Nam8
Snu66
Zds1
Hsl1
Mih1
Crn1
Abp1
Rsc9
Arp9
Las17
Sup35
Dom34
Prm15
Cof1
Srv2
Cyr1
Act1
Sla2
Swa2
Sky1
Ste7
Apl1
Apl2
Sto1
Sgv1
Fus3
Rpn10
Sec23
Sly1
Sec18
Syp1
Vrp1
Sla1
Rps9a
Mtr4
Rps8a
Fun12
Rpn1
Duf1
Rad53
Art5
Ecm21
Ldb19
Yhr131c
Rsp5
Bul2
Ent1
Ent5
Apl3
Sec16
Apm4
Akl1
Rpn11
Nip1
Prt1
Rpg1
Ded1
Tif35
Cdc9
Chk1
Rad23
Ubi4
Ede1
Ent2
Vps27
Chc1
Ptp3
Stb1
Spa2
Bem2
Bud14
Rck2
Glc7
Ipl1
Cdc5
Dbf4
Pfk26
Tpk1
Gpa2
Ras2
Med6
Rpb10
Paf1
Rpo26
Bdp1
Spt15
Spt5
Rgr1
Srb7
Ssn2
Nop1
Nsr1
Nop56
Snu13
Rps31
Rad9
Rtf1
Rpo21
Npl3
Air2
Fps1
Ask10
Cdc28
Hog1
Pbs2
Ssk2
Clb2
Bud3
Bud4
Msn2
Sch9
Rim15
Yak1
Ypl247c
Vps52
Kss1
Sko1
Slt2
Msg5
Swe1
Igo2
Cdc55
Igo1
Vps53
Vps54
Rtn1
Fmp42
Cot1
Zrc1
10
20
30
40
50
t/s

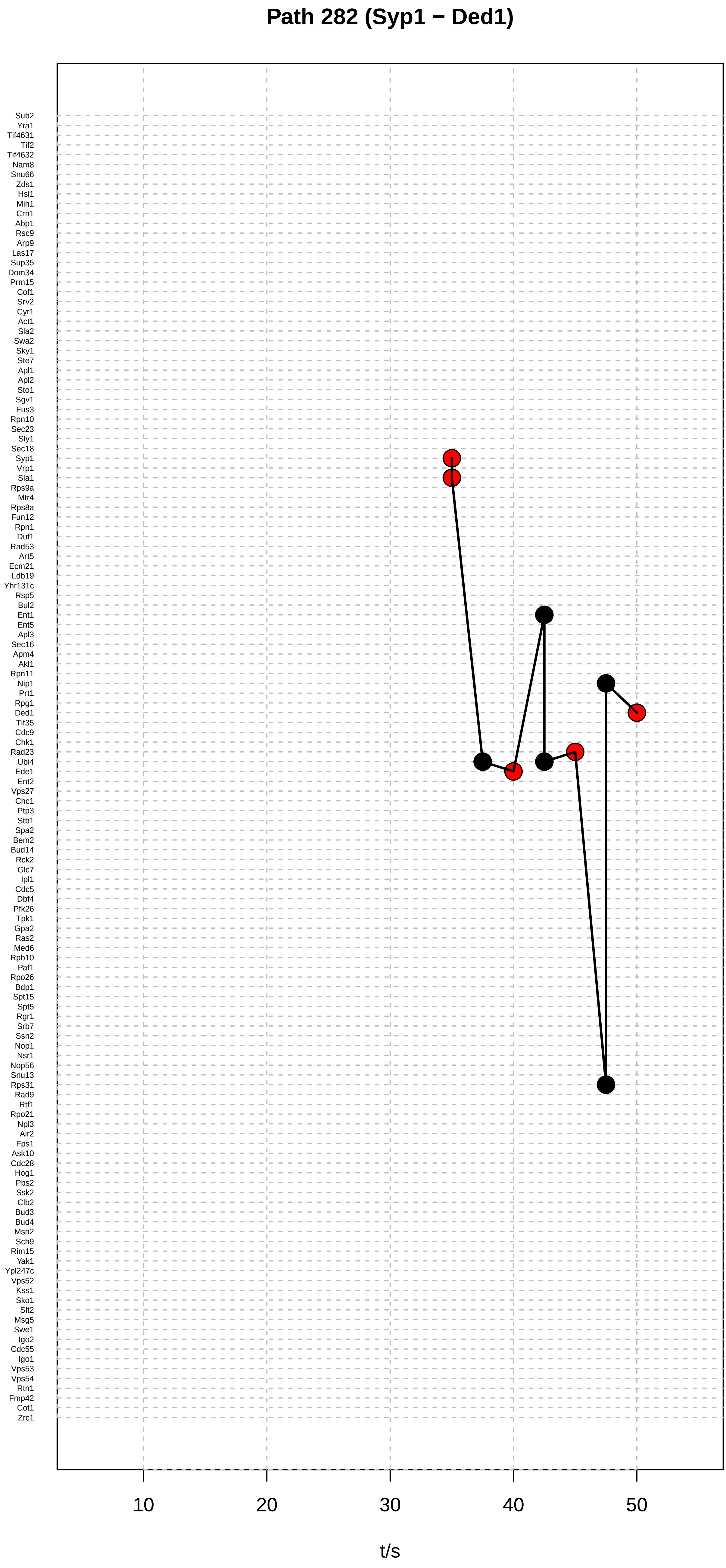

Path 282 (Syp1 − Ded1)
Sub2
Yra1
Tif4631
Tif2
Tif4632
Nam8
Snu66
Zds1
Hsl1
Mih1
Crn1
Abp1
Rsc9
Arp9
Las17
Sup35
Dom34
Prm15
Cof1
Srv2
Cyr1
Act1
Sla2
Swa2
Sky1
Ste7
Apl1
Apl2
Sto1
Sgv1
Fus3
Rpn10
Sec23
Sly1
Sec18
Syp1
Vrp1
Sla1
Rps9a
Mtr4
Rps8a
Fun12
Rpn1
Duf1
Rad53
Art5
Ecm21
Ldb19
Yhr131c
Rsp5
Bul2
Ent1
Ent5
Apl3
Sec16
Apm4
Akl1
Rpn11
Nip1
Prt1
Rpg1
Ded1
Tif35
Cdc9
Chk1
Rad23
Ubi4
Ede1
Ent2
Vps27
Chc1
Ptp3
Stb1
Spa2
Bem2
Bud14
Rck2
Glc7
Ipl1
Cdc5
Dbf4
Pfk26
Tpk1
Gpa2
Ras2
Med6
Rpb10
Paf1
Rpo26
Bdp1
Spt15
Spt5
Rgr1
Srb7
Ssn2
Nop1
Nsr1
Nop56
Snu13
Rps31
Rad9
Rtf1
Rpo21
Npl3
Air2
Fps1
Ask10
Cdc28
Hog1
Pbs2
Ssk2
Clb2
Bud3
Bud4
Msn2
Sch9
Rim15
Yak1
Ypl247c
Vps52
Kss1
Sko1
Slt2
Msg5
Swe1
Igo2
Cdc55
Igo1
Vps53
Vps54
Rtn1
Fmp42
Cot1
Zrc1
10
20
30
40
50
t/s

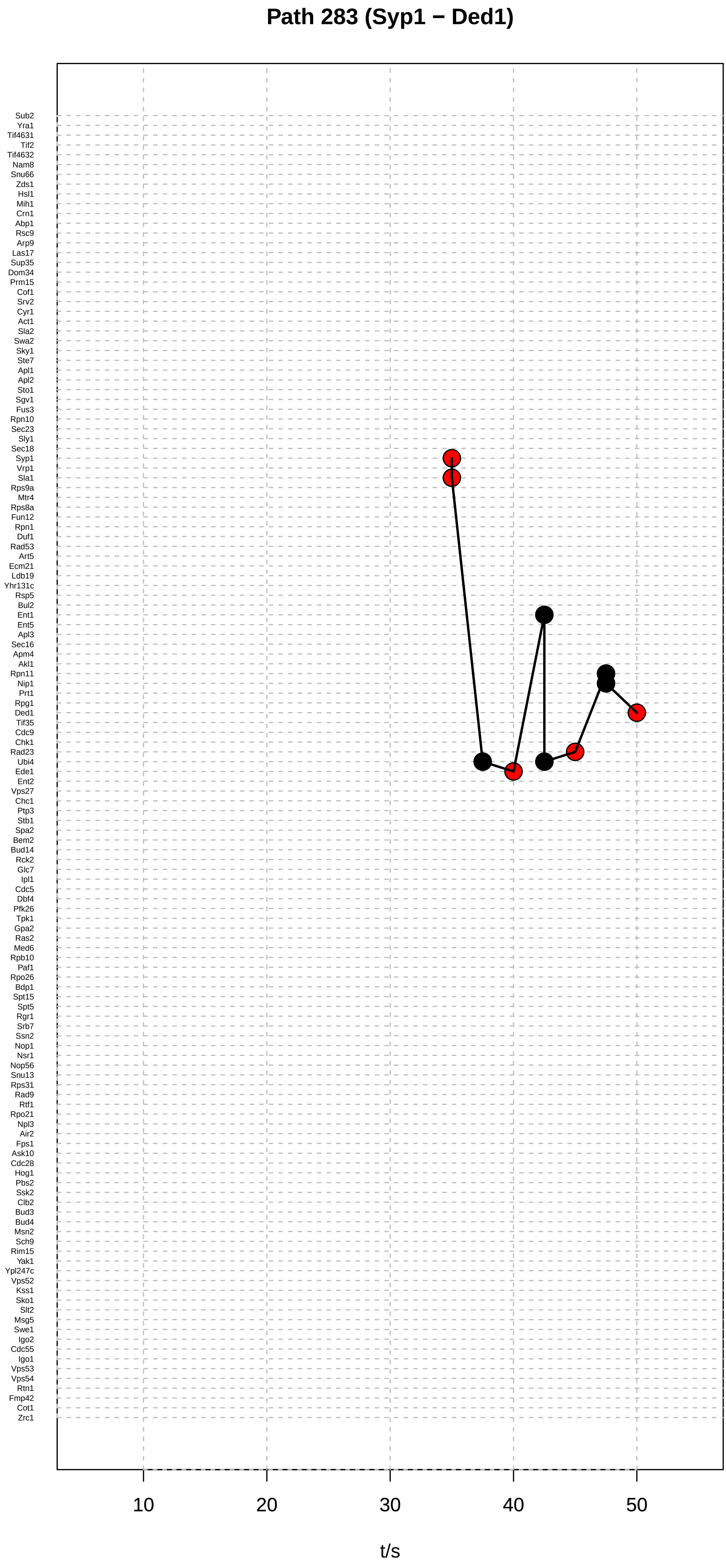

Path 283 (Syp1 − Ded1)
Sub2
Yra1
Tif4631
Tif2
Tif4632
Nam8
Snu66
Zds1
Hsl1
Mih1
Crn1
Abp1
Rsc9
Arp9
Las17
Sup35
Dom34
Prm15
Cof1
Srv2
Cyr1
Act1
Sla2
Swa2
Sky1
Ste7
Apl1
Apl2
Sto1
Sgv1
Fus3
Rpn10
Sec23
Sly1
Sec18
Syp1
Vrp1
Sla1
Rps9a
Mtr4
Rps8a
Fun12
Rpn1
Duf1
Rad53
Art5
Ecm21
Ldb19
Yhr131c
Rsp5
Bul2
Ent1
Ent5
Apl3
Sec16
Apm4
Akl1
Rpn11
Nip1
Prt1
Rpg1
Ded1
Tif35
Cdc9
Chk1
Rad23
Ubi4
Ede1
Ent2
Vps27
Chc1
Ptp3
Stb1
Spa2
Bem2
Bud14
Rck2
Glc7
Ipl1
Cdc5
Dbf4
Pfk26
Tpk1
Gpa2
Ras2
Med6
Rpb10
Paf1
Rpo26
Bdp1
Spt15
Spt5
Rgr1
Srb7
Ssn2
Nop1
Nsr1
Nop56
Snu13
Rps31
Rad9
Rtf1
Rpo21
Npl3
Air2
Fps1
Ask10
Cdc28
Hog1
Pbs2
Ssk2
Clb2
Bud3
Bud4
Msn2
Sch9
Rim15
Yak1
Ypl247c
Vps52
Kss1
Sko1
Slt2
Msg5
Swe1
Igo2
Cdc55
Igo1
Vps53
Vps54
Rtn1
Fmp42
Cot1
Zrc1
10
20
30
40
50
t/s

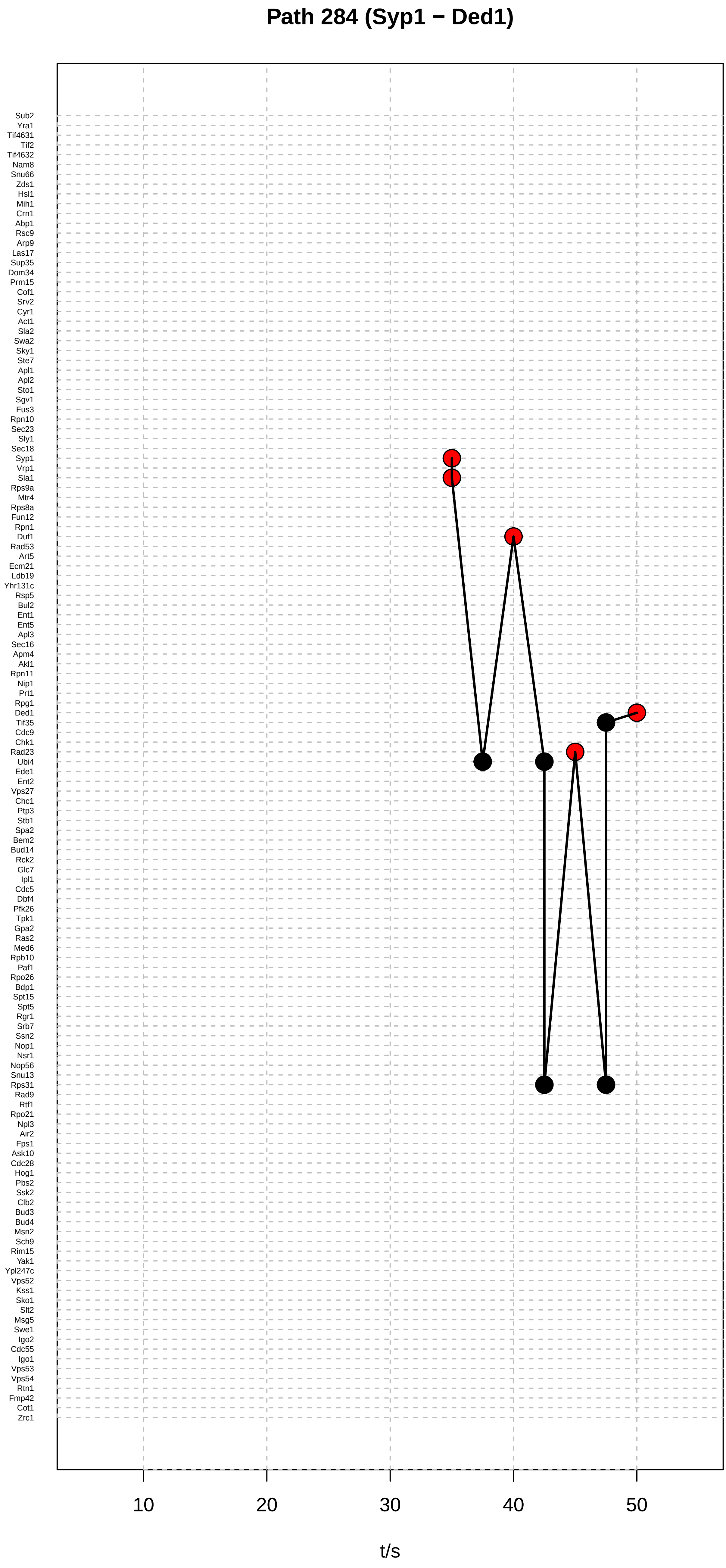

Path 284 (Syp1 − Ded1)
Sub2
Yra1
Tif4631
Tif2
Tif4632
Nam8
Snu66
Zds1
Hsl1
Mih1
Crn1
Abp1
Rsc9
Arp9
Las17
Sup35
Dom34
Prm15
Cof1
Srv2
Cyr1
Act1
Sla2
Swa2
Sky1
Ste7
Apl1
Apl2
Sto1
Sgv1
Fus3
Rpn10
Sec23
Sly1
Sec18
Syp1
Vrp1
Sla1
Rps9a
Mtr4
Rps8a
Fun12
Rpn1
Duf1
Rad53
Art5
Ecm21
Ldb19
Yhr131c
Rsp5
Bul2
Ent1
Ent5
Apl3
Sec16
Apm4
Akl1
Rpn11
Nip1
Prt1
Rpg1
Ded1
Tif35
Cdc9
Chk1
Rad23
Ubi4
Ede1
Ent2
Vps27
Chc1
Ptp3
Stb1
Spa2
Bem2
Bud14
Rck2
Glc7
Ipl1
Cdc5
Dbf4
Pfk26
Tpk1
Gpa2
Ras2
Med6
Rpb10
Paf1
Rpo26
Bdp1
Spt15
Spt5
Rgr1
Srb7
Ssn2
Nop1
Nsr1
Nop56
Snu13
Rps31
Rad9
Rtf1
Rpo21
Npl3
Air2
Fps1
Ask10
Cdc28
Hog1
Pbs2
Ssk2
Clb2
Bud3
Bud4
Msn2
Sch9
Rim15
Yak1
Ypl247c
Vps52
Kss1
Sko1
Slt2
Msg5
Swe1
Igo2
Cdc55
Igo1
Vps53
Vps54
Rtn1
Fmp42
Cot1
Zrc1
10
20
30
40
50
t/s

#### Path 285 (Syp1 − Ded1)

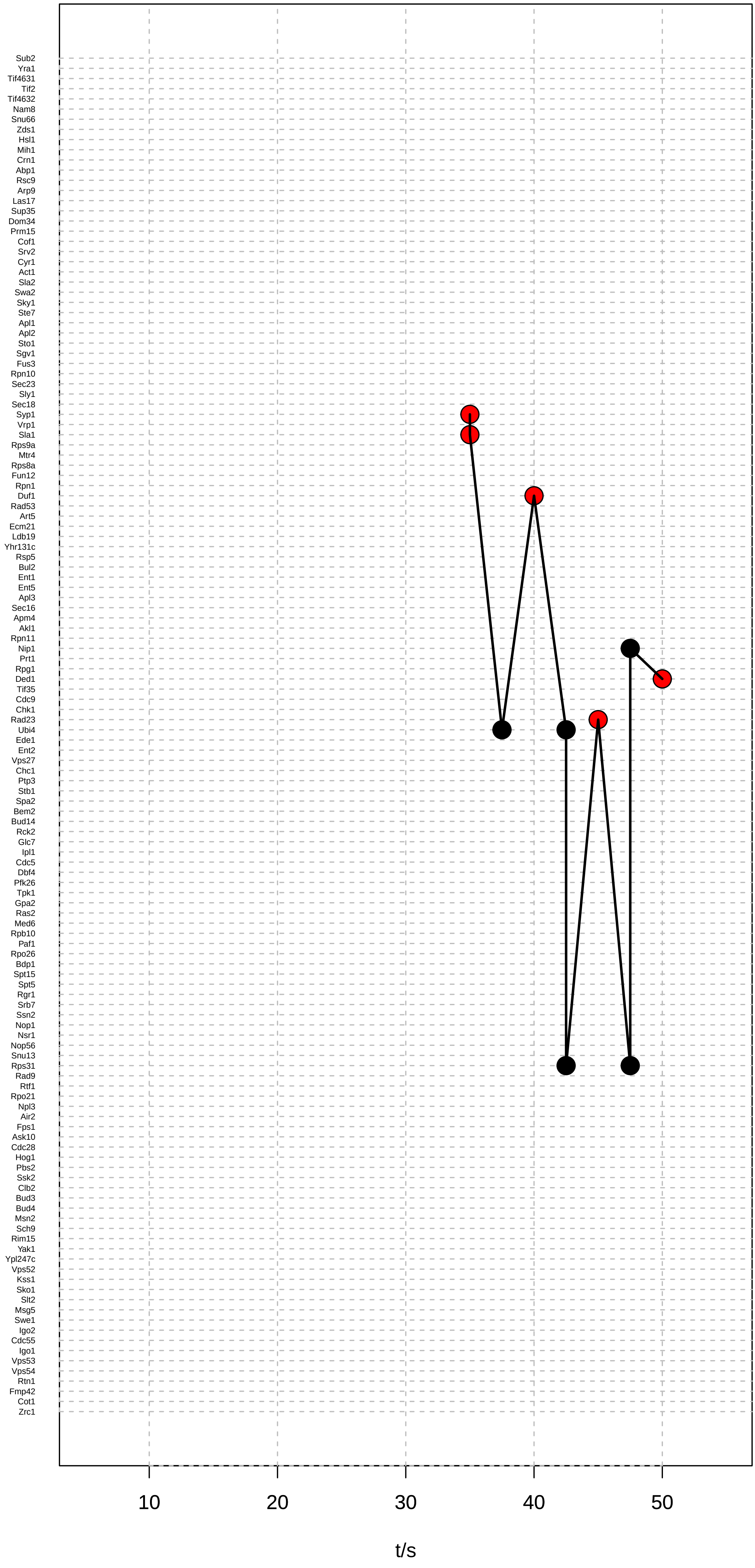

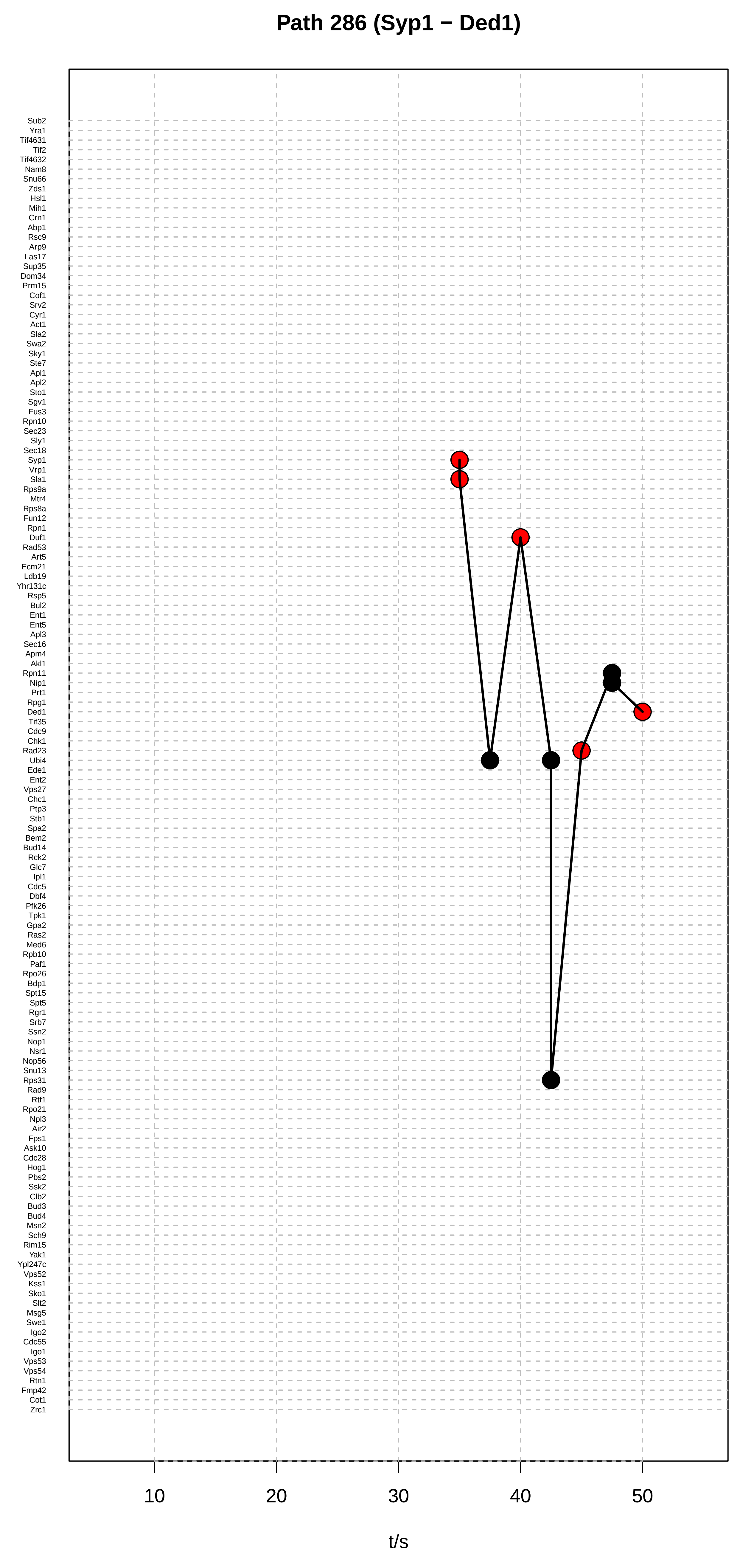

Path 286 (Syp1 − Ded1)
Sub2
Yra1
Tif4631
Tif2
Tif4632
Nam8
Snu66
Zds1
Hsl1
Mih1
Crn1
Abp1
Rsc9
Arp9
Las17
Sup35
Dom34
Prm15
Cof1
Srv2
Cyr1
Act1
Sla2
Swa2
Sky1
Ste7
Apl1
Apl2
Sto1
Sgv1
Fus3
Rpn10
Sec23
Sly1
Sec18
Syp1
Vrp1
Sla1
Rps9a
Mtr4
Rps8a
Fun12
Rpn1
Duf1
Rad53
Art5
Ecm21
Ldb19
Yhr131c
Rsp5
Bul2
Ent1
Ent5
Apl3
Sec16
Apm4
Akl1
Rpn11
Nip1
Prt1
Rpg1
Ded1
Tif35
Cdc9
Chk1
Rad23
Ubi4
Ede1
Ent2
Vps27
Chc1
Ptp3
Stb1
Spa2
Bem2
Bud14
Rck2
Glc7
Ipl1
Cdc5
Dbf4
Pfk26
Tpk1
Gpa2
Ras2
Med6
Rpb10
Paf1
Rpo26
Bdp1
Spt15
Spt5
Rgr1
Srb7
Ssn2
Nop1
Nsr1
Nop56
Snu13
Rps31
Rad9
Rtf1
Rpo21
Npl3
Air2
Fps1
Ask10
Cdc28
Hog1
Pbs2
Ssk2
Clb2
Bud3
Bud4
Msn2
Sch9
Rim15
Yak1
Ypl247c
Vps52
Kss1
Sko1
Slt2
Msg5
Swe1
Igo2
Cdc55
Igo1
Vps53
Vps54
Rtn1
Fmp42
Cot1
Zrc1
10
20
30
40
50
t/s

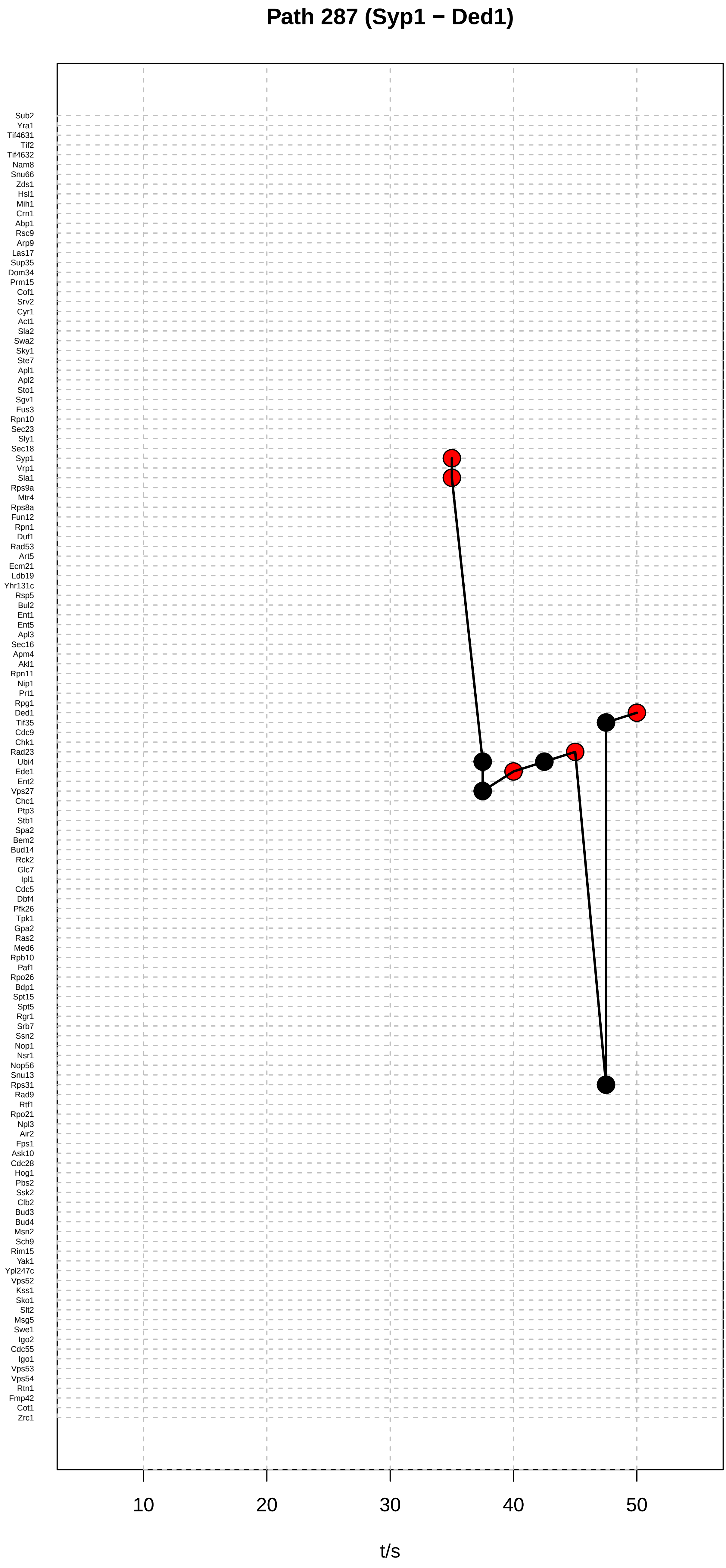

Path 287 (Syp1 − Ded1)
Sub2
Yra1
Tif4631
Tif2
Tif4632
Nam8
Snu66
Zds1
Hsl1
Mih1
Crn1
Abp1
Rsc9
Arp9
Las17
Sup35
Dom34
Prm15
Cof1
Srv2
Cyr1
Act1
Sla2
Swa2
Sky1
Ste7
Apl1
Apl2
Sto1
Sgv1
Fus3
Rpn10
Sec23
Sly1
Sec18
Syp1
Vrp1
Sla1
Rps9a
Mtr4
Rps8a
Fun12
Rpn1
Duf1
Rad53
Art5
Ecm21
Ldb19
Yhr131c
Rsp5
Bul2
Ent1
Ent5
Apl3
Sec16
Apm4
Akl1
Rpn11
Nip1
Prt1
Rpg1
Ded1
Tif35
Cdc9
Chk1
Rad23
Ubi4
Ede1
Ent2
Vps27
Chc1
Ptp3
Stb1
Spa2
Bem2
Bud14
Rck2
Glc7
Ipl1
Cdc5
Dbf4
Pfk26
Tpk1
Gpa2
Ras2
Med6
Rpb10
Paf1
Rpo26
Bdp1
Spt15
Spt5
Rgr1
Srb7
Ssn2
Nop1
Nsr1
Nop56
Snu13
Rps31
Rad9
Rtf1
Rpo21
Npl3
Air2
Fps1
Ask10
Cdc28
Hog1
Pbs2
Ssk2
Clb2
Bud3
Bud4
Msn2
Sch9
Rim15
Yak1
Ypl247c
Vps52
Kss1
Sko1
Slt2
Msg5
Swe1
Igo2
Cdc55
Igo1
Vps53
Vps54
Rtn1
Fmp42
Cot1
Zrc1
10
20
30
40
50
t/s

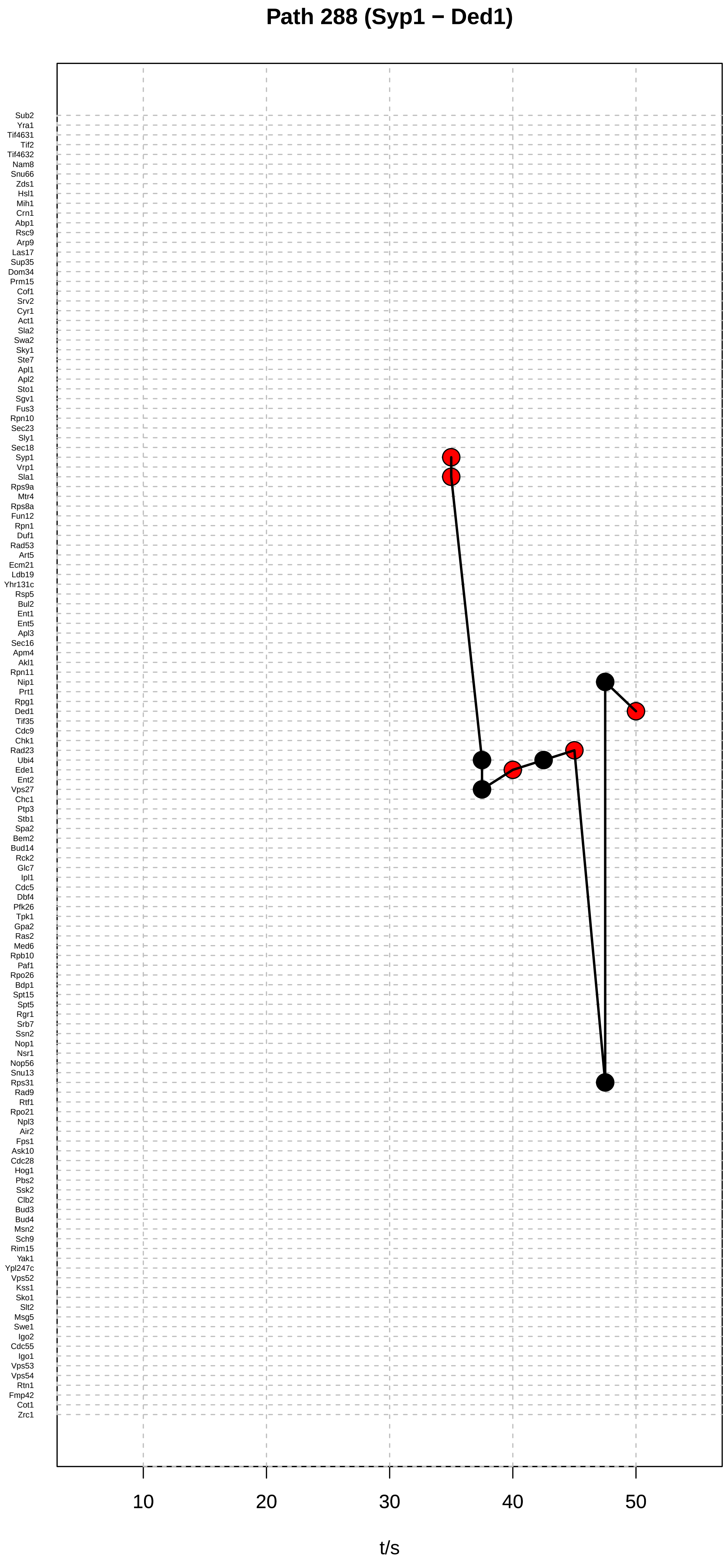
Path 288 (Syp1 − Ded1)
Sub2
Yra1
Tif4631
Tif2
Tif4632
Nam8
Snu66
Zds1
Hsl1
Mih1
Crn1
Abp1
Rsc9
Arp9
Las17
Sup35
Dom34
Prm15
Cof1
Srv2
Cyr1
Act1
Sla2
Swa2
Sky1
Ste7
Apl1
Apl2
Sto1
Sgv1
Fus3
Rpn10
Sec23
Sly1
Sec18
Syp1
Vrp1
Sla1
Rps9a
Mtr4
Rps8a
Fun12
Rpn1
Duf1
Rad53
Art5
Ecm21
Ldb19
Yhr131c
Rsp5
Bul2
Ent1
Ent5
Apl3
Sec16
Apm4
Akl1
Rpn11
Nip1
Prt1
Rpg1
Ded1
Tif35
Cdc9
Chk1
Rad23
Ubi4
Ede1
Ent2
Vps27
Chc1
Ptp3
Stb1
Spa2
Bem2
Bud14
Rck2
Glc7
Ipl1
Cdc5
Dbf4
Pfk26
Tpk1
Gpa2
Ras2
Med6
Rpb10
Paf1
Rpo26
Bdp1
Spt15
Spt5
Rgr1
Srb7
Ssn2
Nop1
Nsr1
Nop56
Snu13
Rps31
Rad9
Rtf1
Rpo21
Npl3
Air2
Fps1
Ask10
Cdc28
Hog1
Pbs2
Ssk2
Clb2
Bud3
Bud4
Msn2
Sch9
Rim15
Yak1
Ypl247c
Vps52
Kss1
Sko1
Slt2
Msg5
Swe1
Igo2
Cdc55
Igo1
Vps53
Vps54
Rtn1
Fmp42
Cot1
Zrc1
10
20
30
40
50
t/s

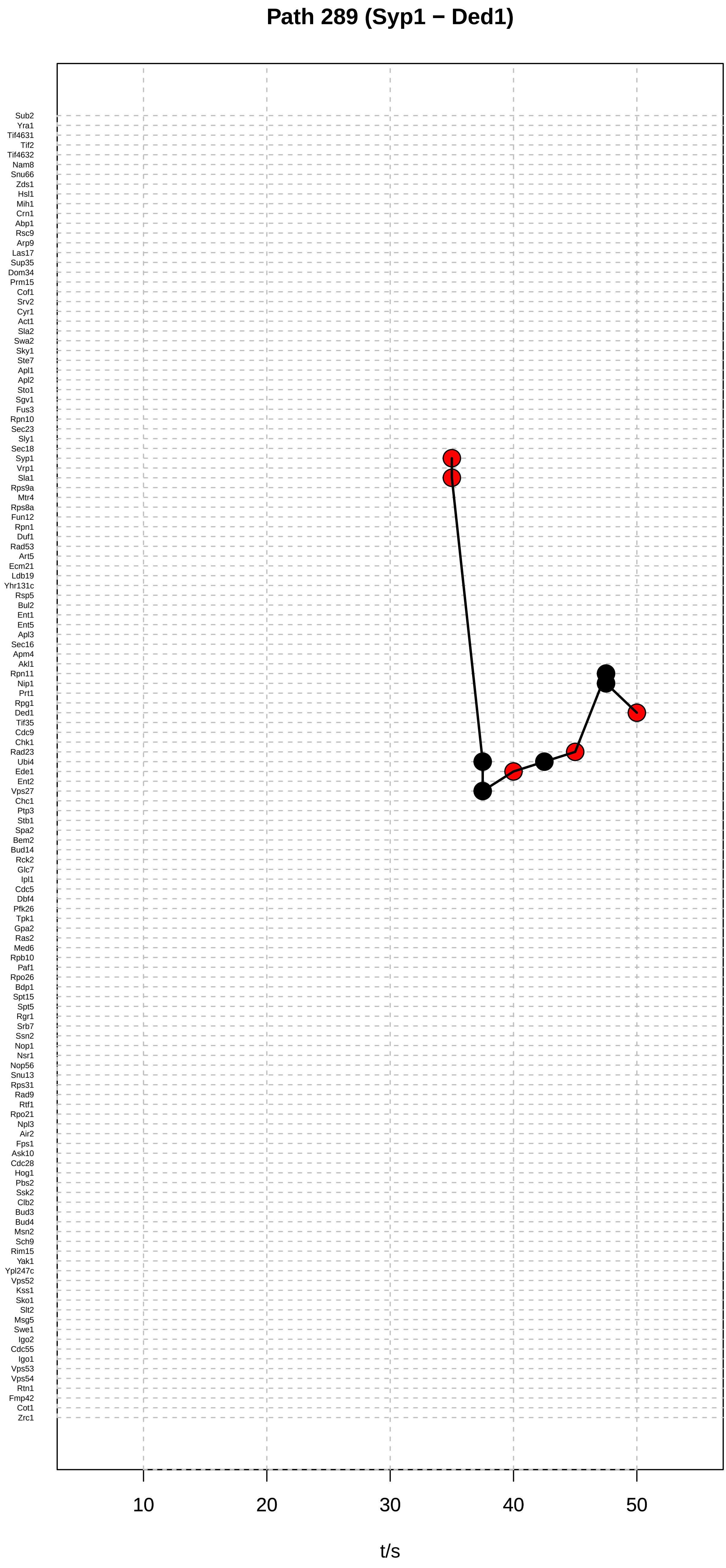

Path 289 (Syp1 − Ded1)
Sub2
Yra1
Tif4631
Tif2
Tif4632
Nam8
Snu66
Zds1
Hsl1
Mih1
Crn1
Abp1
Rsc9
Arp9
Las17
Sup35
Dom34
Prm15
Cof1
Srv2
Cyr1
Act1
Sla2
Swa2
Sky1
Ste7
Apl1
Apl2
Sto1
Sgv1
Fus3
Rpn10
Sec23
Sly1
Sec18
Syp1
Vrp1
Sla1
Rps9a
Mtr4
Rps8a
Fun12
Rpn1
Duf1
Rad53
Art5
Ecm21
Ldb19
Yhr131c
Rsp5
Bul2
Ent1
Ent5
Apl3
Sec16
Apm4
Akl1
Rpn11
Nip1
Prt1
Rpg1
Ded1
Tif35
Cdc9
Chk1
Rad23
Ubi4
Ede1
Ent2
Vps27
Chc1
Ptp3
Stb1
Spa2
Bem2
Bud14
Rck2
Glc7
Ipl1
Cdc5
Dbf4
Pfk26
Tpk1
Gpa2
Ras2
Med6
Rpb10
Paf1
Rpo26
Bdp1
Spt15
Spt5
Rgr1
Srb7
Ssn2
Nop1
Nsr1
Nop56
Snu13
Rps31
Rad9
Rtf1
Rpo21
Npl3
Air2
Fps1
Ask10
Cdc28
Hog1
Pbs2
Ssk2
Clb2
Bud3
Bud4
Msn2
Sch9
Rim15
Yak1
Ypl247c
Vps52
Kss1
Sko1
Slt2
Msg5
Swe1
Igo2
Cdc55
Igo1
Vps53
Vps54
Rtn1
Fmp42
Cot1
Zrc1
10
20
30
40
50
t/s

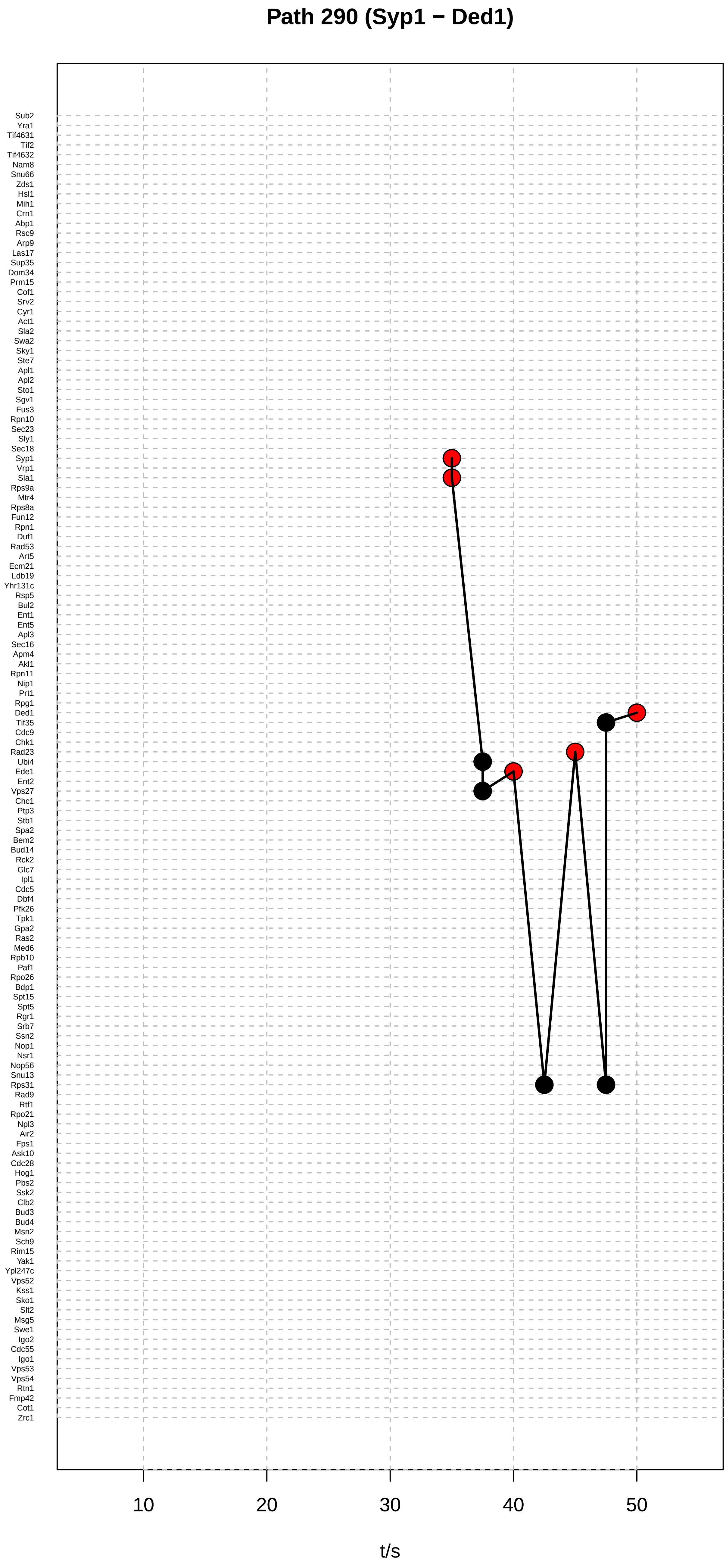
Path 290 (Syp1 − Ded1)
Sub2
Yra1
Tif4631
Tif2
Tif4632
Nam8
Snu66
Zds1
Hsl1
Mih1
Crn1
Abp1
Rsc9
Arp9
Las17
Sup35
Dom34
Prm15
Cof1
Srv2
Cyr1
Act1
Sla2
Swa2
Sky1
Ste7
Apl1
Apl2
Sto1
Sgv1
Fus3
Rpn10
Sec23
Sly1
Sec18
Syp1
Vrp1
Sla1
Rps9a
Mtr4
Rps8a
Fun12
Rpn1
Duf1
Rad53
Art5
Ecm21
Ldb19
Yhr131c
Rsp5
Bul2
Ent1
Ent5
Apl3
Sec16
Apm4
Akl1
Rpn11
Nip1
Prt1
Rpg1
Ded1
Tif35
Cdc9
Chk1
Rad23
Ubi4
Ede1
Ent2
Vps27
Chc1
Ptp3
Stb1
Spa2
Bem2
Bud14
Rck2
Glc7
Ipl1
Cdc5
Dbf4
Pfk26
Tpk1
Gpa2
Ras2
Med6
Rpb10
Paf1
Rpo26
Bdp1
Spt15
Spt5
Rgr1
Srb7
Ssn2
Nop1
Nsr1
Nop56
Snu13
Rps31
Rad9
Rtf1
Rpo21
Npl3
Air2
Fps1
Ask10
Cdc28
Hog1
Pbs2
Ssk2
Clb2
Bud3
Bud4
Msn2
Sch9
Rim15
Yak1
Ypl247c
Vps52
Kss1
Sko1
Slt2
Msg5
Swe1
Igo2
Cdc55
Igo1
Vps53
Vps54
Rtn1
Fmp42
Cot1
Zrc1
10
20
30
40
50
t/s

### Path 291 (Syp1 − Ded1)

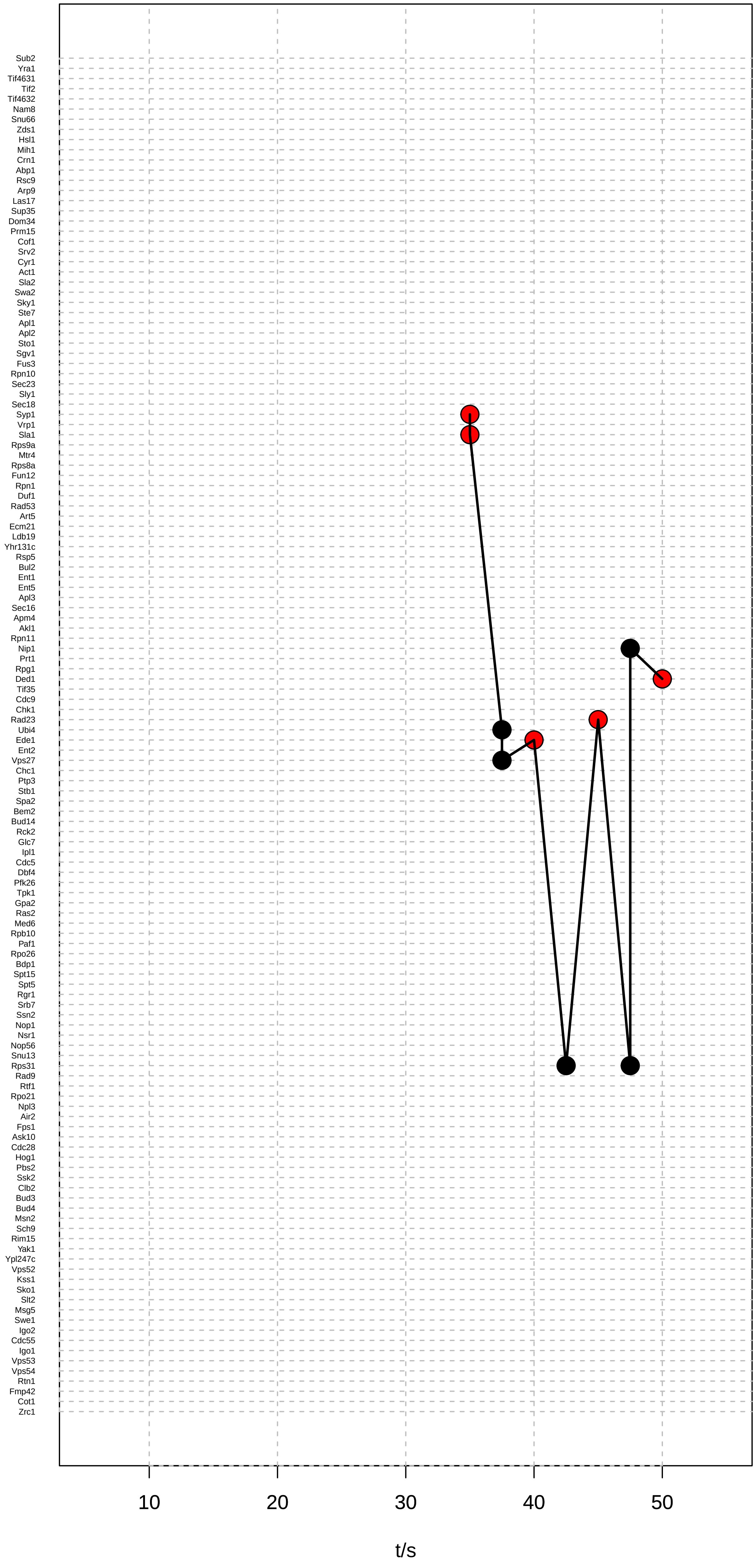

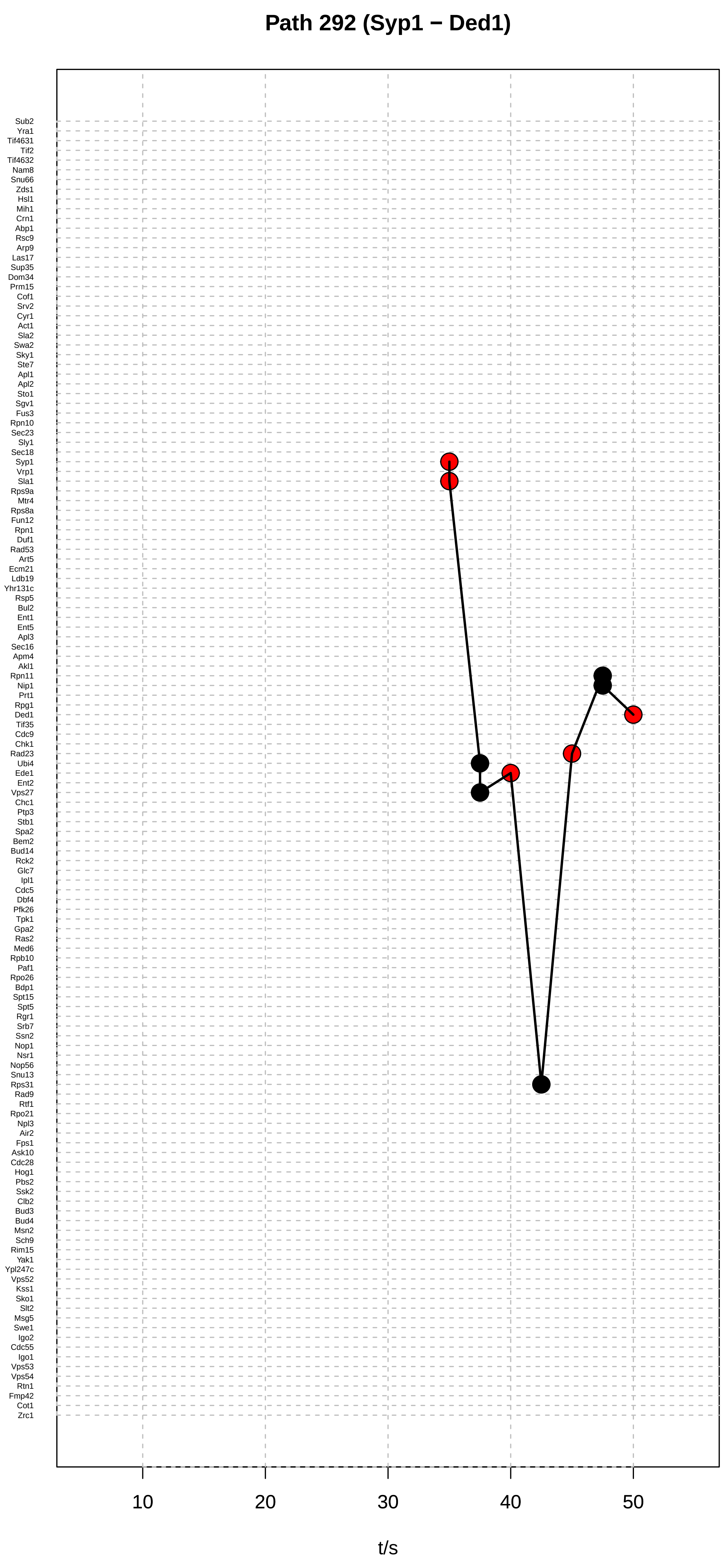

Path 292 (Syp1 − Ded1)
Sub2
Yra1
Tif4631
Tif2
Tif4632
Nam8
Snu66
Zds1
Hsl1
Mih1
Crn1
Abp1
Rsc9
Arp9
Las17
Sup35
Dom34
Prm15
Cof1
Srv2
Cyr1
Act1
Sla2
Swa2
Sky1
Ste7
Apl1
Apl2
Sto1
Sgv1
Fus3
Rpn10
Sec23
Sly1
Sec18
Syp1
Vrp1
Sla1
Rps9a
Mtr4
Rps8a
Fun12
Rpn1
Duf1
Rad53
Art5
Ecm21
Ldb19
Yhr131c
Rsp5
Bul2
Ent1
Ent5
Apl3
Sec16
Apm4
Akl1
Rpn11
Nip1
Prt1
Rpg1
Ded1
Tif35
Cdc9
Chk1
Rad23
Ubi4
Ede1
Ent2
Vps27
Chc1
Ptp3
Stb1
Spa2
Bem2
Bud14
Rck2
Glc7
Ipl1
Cdc5
Dbf4
Pfk26
Tpk1
Gpa2
Ras2
Med6
Rpb10
Paf1
Rpo26
Bdp1
Spt15
Spt5
Rgr1
Srb7
Ssn2
Nop1
Nsr1
Nop56
Snu13
Rps31
Rad9
Rtf1
Rpo21
Npl3
Air2
Fps1
Ask10
Cdc28
Hog1
Pbs2
Ssk2
Clb2
Bud3
Bud4
Msn2
Sch9
Rim15
Yak1
Ypl247c
Vps52
Kss1
Sko1
Slt2
Msg5
Swe1
Igo2
Cdc55
Igo1
Vps53
Vps54
Rtn1
Fmp42
Cot1
Zrc1
10
20
30
40
50
t/s

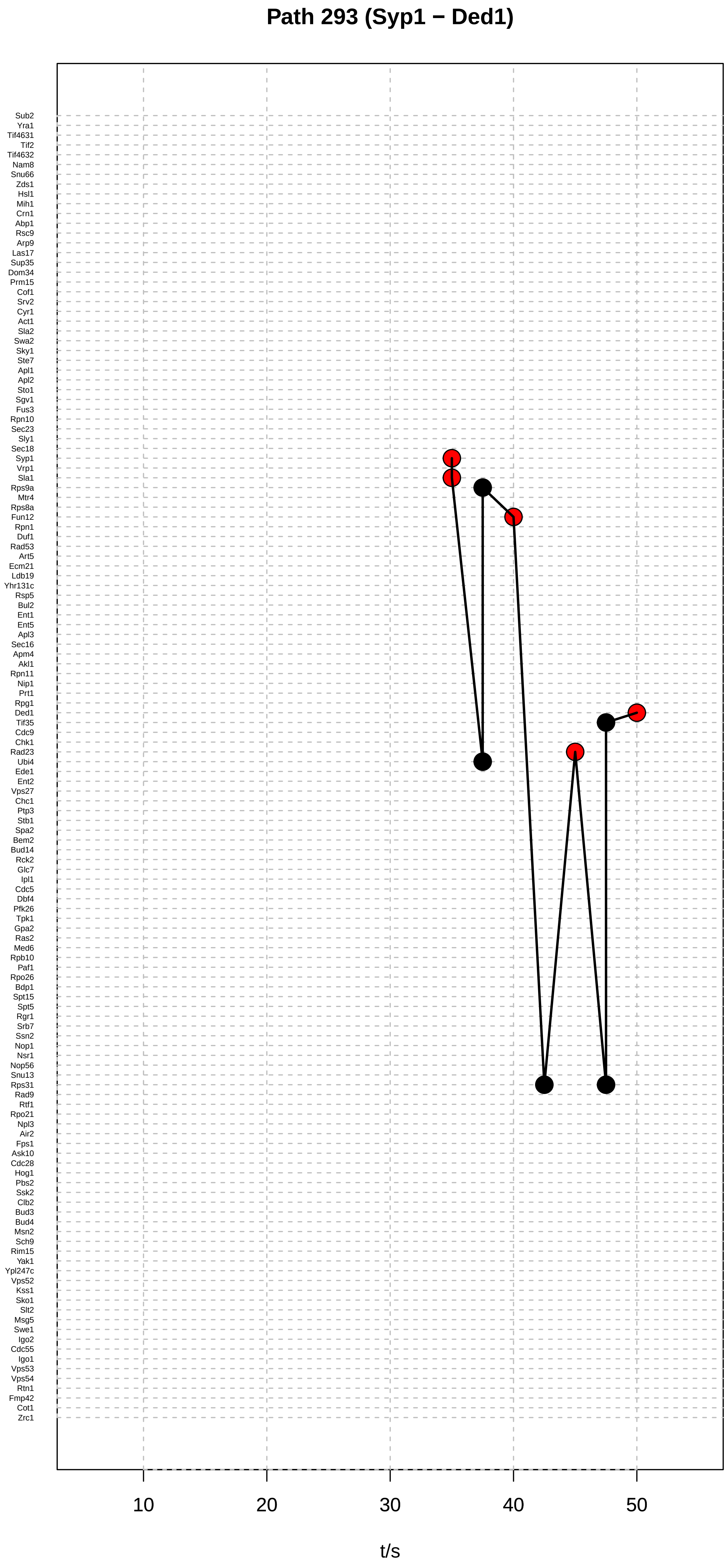

Path 293 (Syp1 − Ded1)
Sub2
Yra1
Tif4631
Tif2
Tif4632
Nam8
Snu66
Zds1
Hsl1
Mih1
Crn1
Abp1
Rsc9
Arp9
Las17
Sup35
Dom34
Prm15
Cof1
Srv2
Cyr1
Act1
Sla2
Swa2
Sky1
Ste7
Apl1
Apl2
Sto1
Sgv1
Fus3
Rpn10
Sec23
Sly1
Sec18
Syp1
Vrp1
Sla1
Rps9a
Mtr4
Rps8a
Fun12
Rpn1
Duf1
Rad53
Art5
Ecm21
Ldb19
Yhr131c
Rsp5
Bul2
Ent1
Ent5
Apl3
Sec16
Apm4
Akl1
Rpn11
Nip1
Prt1
Rpg1
Ded1
Tif35
Cdc9
Chk1
Rad23
Ubi4
Ede1
Ent2
Vps27
Chc1
Ptp3
Stb1
Spa2
Bem2
Bud14
Rck2
Glc7
Ipl1
Cdc5
Dbf4
Pfk26
Tpk1
Gpa2
Ras2
Med6
Rpb10
Paf1
Rpo26
Bdp1
Spt15
Spt5
Rgr1
Srb7
Ssn2
Nop1
Nsr1
Nop56
Snu13
Rps31
Rad9
Rtf1
Rpo21
Npl3
Air2
Fps1
Ask10
Cdc28
Hog1
Pbs2
Ssk2
Clb2
Bud3
Bud4
Msn2
Sch9
Rim15
Yak1
Ypl247c
Vps52
Kss1
Sko1
Slt2
Msg5
Swe1
Igo2
Cdc55
Igo1
Vps53
Vps54
Rtn1
Fmp42
Cot1
Zrc1
10
20
30
40
50
t/s

#### Path 294 (Syp1 − Ded1)

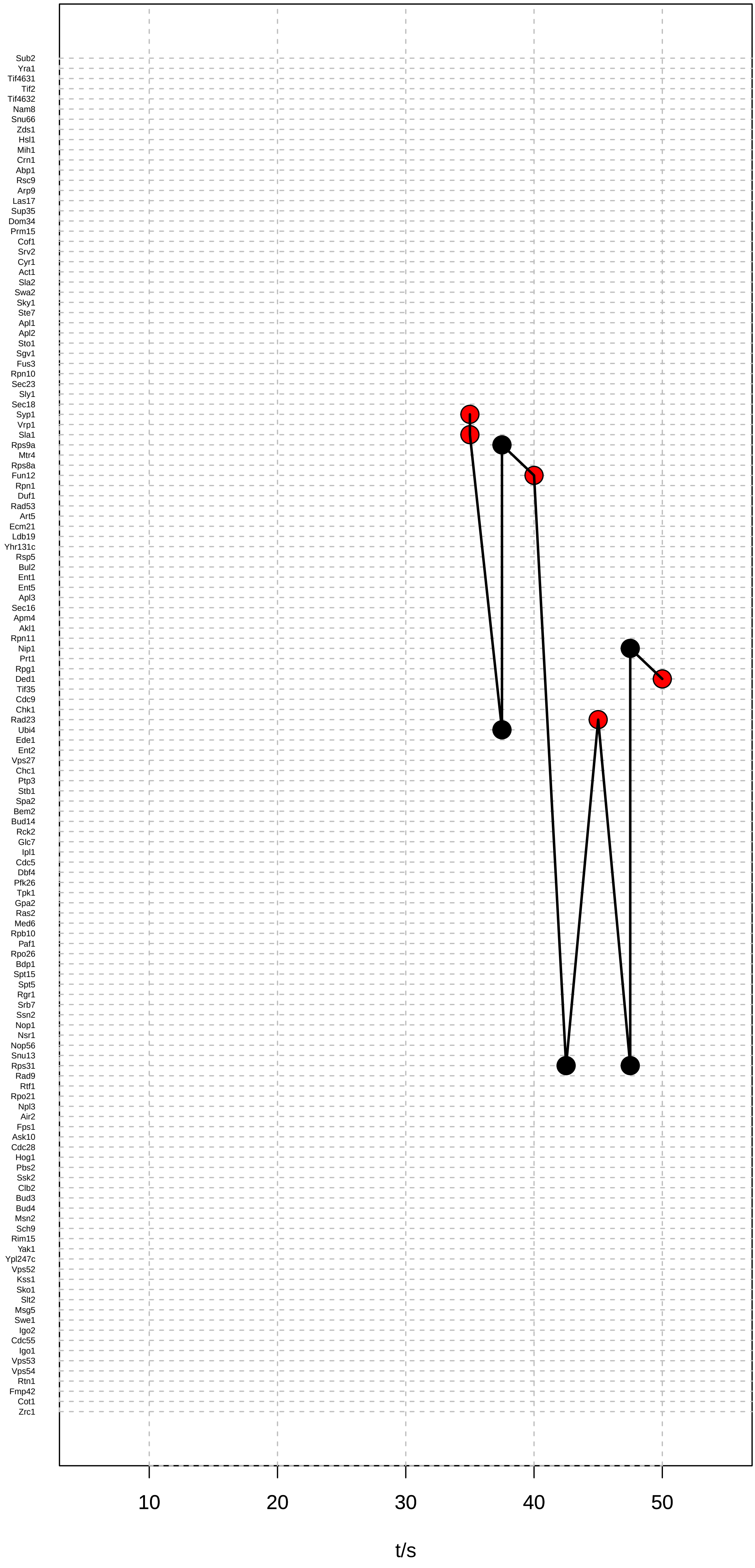

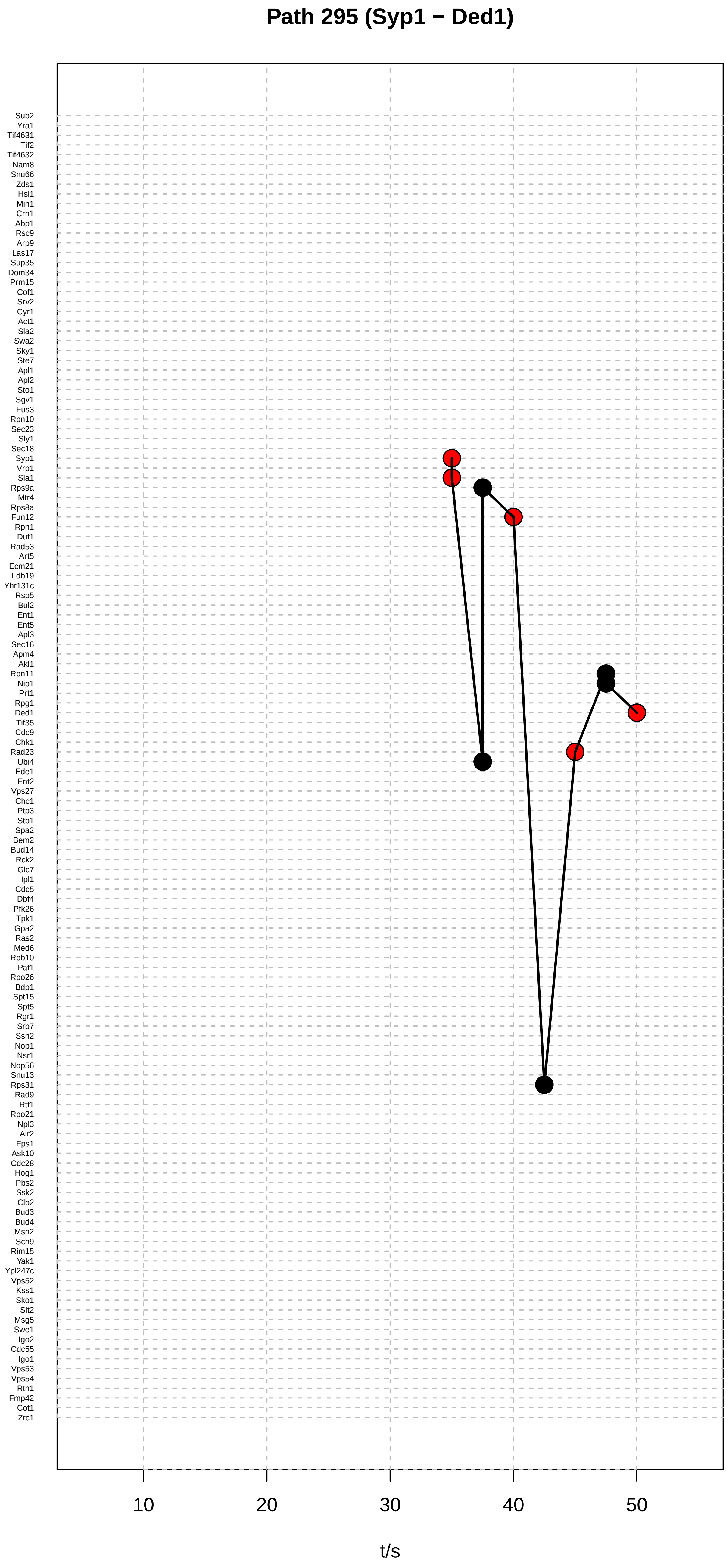

Path 295 (Syp1 − Ded1)
Sub2
Yra1
Tif4631
Tif2
Tif4632
Nam8
Snu66
Zds1
Hsl1
Mih1
Crn1
Abp1
Rsc9
Arp9
Las17
Sup35
Dom34
Prm15
Cof1
Srv2
Cyr1
Act1
Sla2
Swa2
Sky1
Ste7
Apl1
Apl2
Sto1
Sgv1
Fus3
Rpn10
Sec23
Sly1
Sec18
Syp1
Vrp1
Sla1
Rps9a
Mtr4
Rps8a
Fun12
Rpn1
Duf1
Rad53
Art5
Ecm21
Ldb19
Yhr131c
Rsp5
Bul2
Ent1
Ent5
Apl3
Sec16
Apm4
Akl1
Rpn11
Nip1
Prt1
Rpg1
Ded1
Tif35
Cdc9
Chk1
Rad23
Ubi4
Ede1
Ent2
Vps27
Chc1
Ptp3
Stb1
Spa2
Bem2
Bud14
Rck2
Glc7
Ipl1
Cdc5
Dbf4
Pfk26
Tpk1
Gpa2
Ras2
Med6
Rpb10
Paf1
Rpo26
Bdp1
Spt15
Spt5
Rgr1
Srb7
Ssn2
Nop1
Nsr1
Nop56
Snu13
Rps31
Rad9
Rtf1
Rpo21
Npl3
Air2
Fps1
Ask10
Cdc28
Hog1
Pbs2
Ssk2
Clb2
Bud3
Bud4
Msn2
Sch9
Rim15
Yak1
Ypl247c
Vps52
Kss1
Sko1
Slt2
Msg5
Swe1
Igo2
Cdc55
Igo1
Vps53
Vps54
Rtn1
Fmp42
Cot1
Zrc1
10
20
30
40
50
t/s

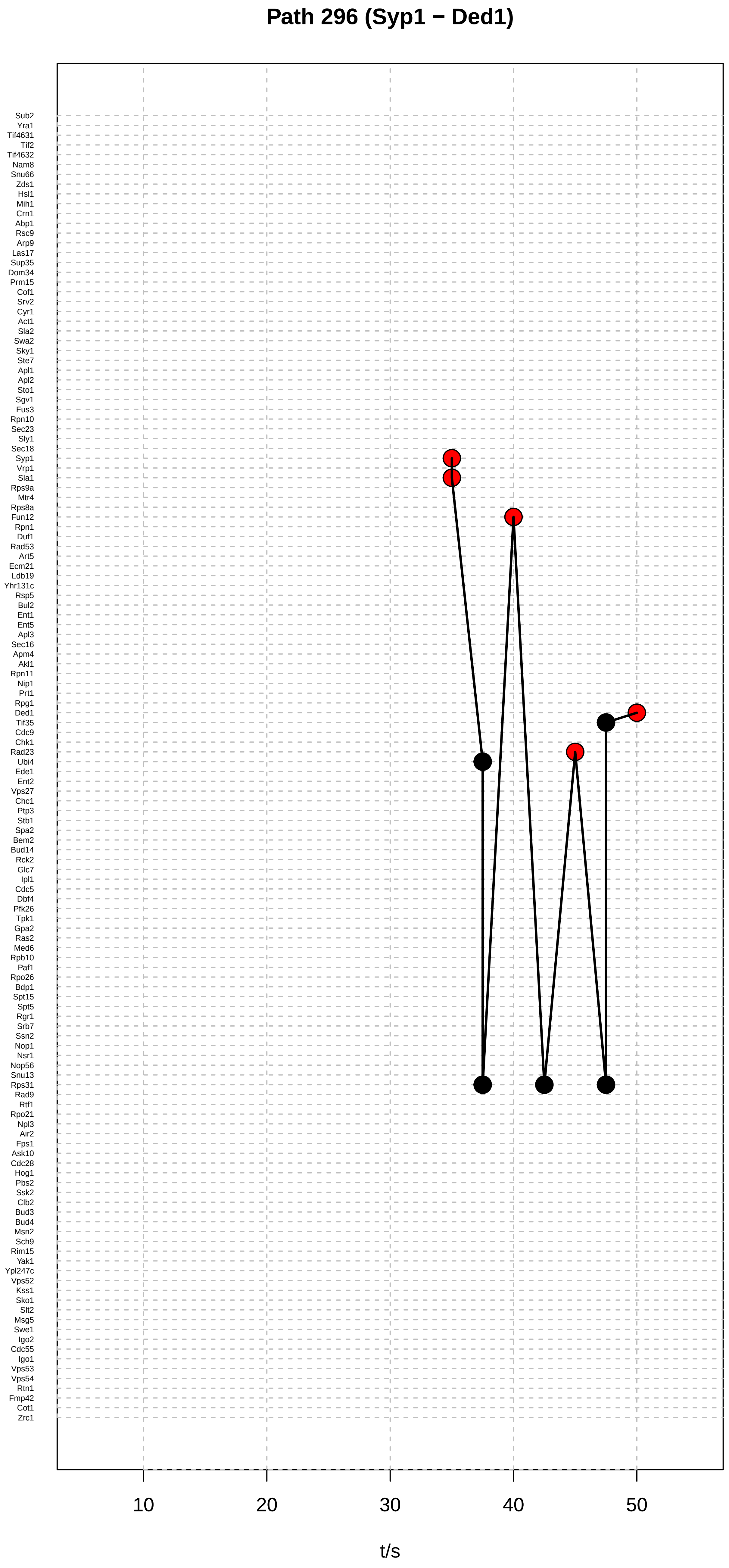
Path 296 (Syp1 − Ded1)
Sub2
Yra1
Tif4631
Tif2
Tif4632
Nam8
Snu66
Zds1
Hsl1
Mih1
Crn1
Abp1
Rsc9
Arp9
Las17
Sup35
Dom34
Prm15
Cof1
Srv2
Cyr1
Act1
Sla2
Swa2
Sky1
Ste7
Apl1
Apl2
Sto1
Sgv1
Fus3
Rpn10
Sec23
Sly1
Sec18
Syp1
Vrp1
Sla1
Rps9a
Mtr4
Rps8a
Fun12
Rpn1
Duf1
Rad53
Art5
Ecm21
Ldb19
Yhr131c
Rsp5
Bul2
Ent1
Ent5
Apl3
Sec16
Apm4
Akl1
Rpn11
Nip1
Prt1
Rpg1
Ded1
Tif35
Cdc9
Chk1
Rad23
Ubi4
Ede1
Ent2
Vps27
Chc1
Ptp3
Stb1
Spa2
Bem2
Bud14
Rck2
Glc7
Ipl1
Cdc5
Dbf4
Pfk26
Tpk1
Gpa2
Ras2
Med6
Rpb10
Paf1
Rpo26
Bdp1
Spt15
Spt5
Rgr1
Srb7
Ssn2
Nop1
Nsr1
Nop56
Snu13
Rps31
Rad9
Rtf1
Rpo21
Npl3
Air2
Fps1
Ask10
Cdc28
Hog1
Pbs2
Ssk2
Clb2
Bud3
Bud4
Msn2
Sch9
Rim15
Yak1
Ypl247c
Vps52
Kss1
Sko1
Slt2
Msg5
Swe1
Igo2
Cdc55
Igo1
Vps53
Vps54
Rtn1
Fmp42
Cot1
Zrc1
10
20
30
40
50
t/s

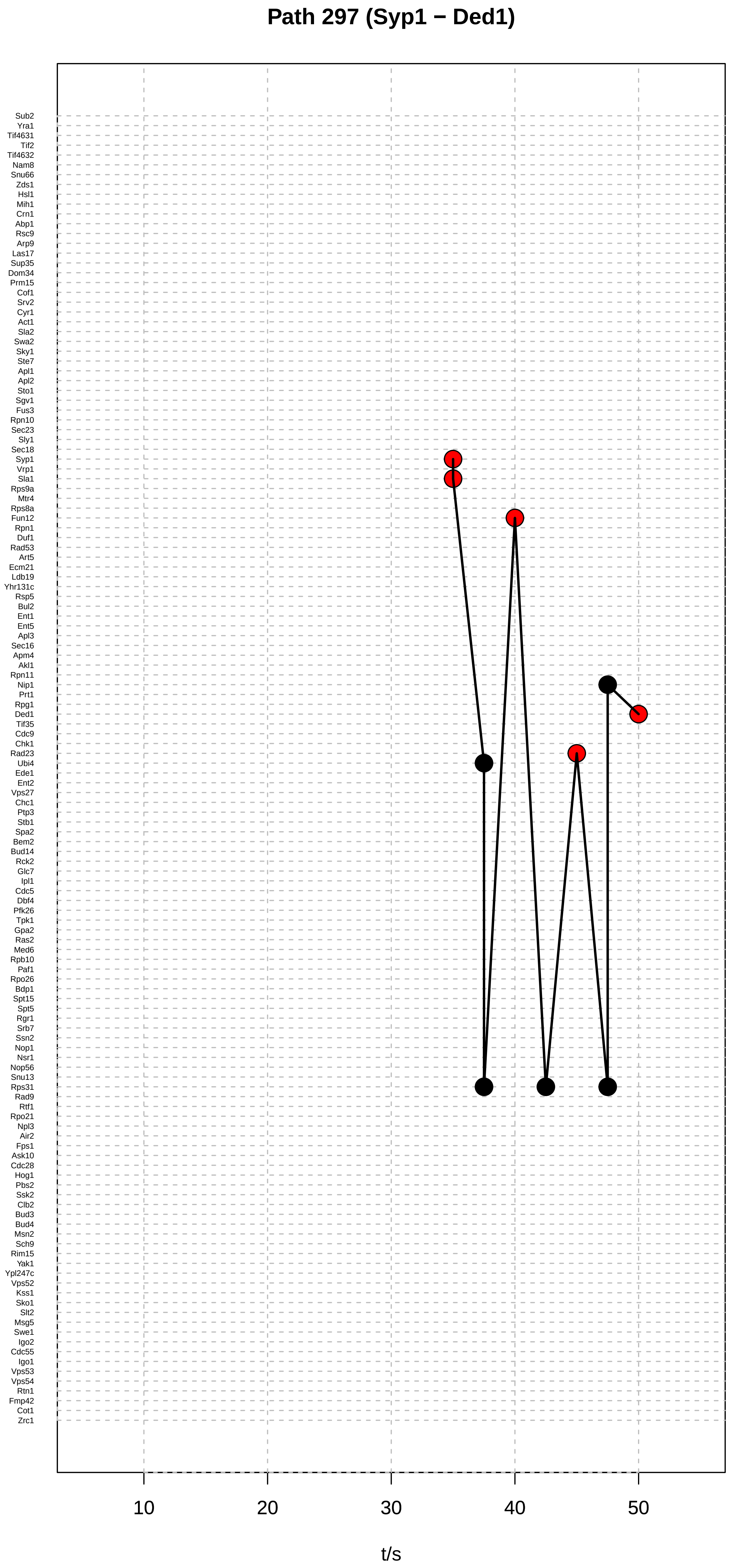

Path 297 (Syp1 − Ded1)
Sub2
Yra1
Tif4631
Tif2
Tif4632
Nam8
Snu66
Zds1
Hsl1
Mih1
Crn1
Abp1
Rsc9
Arp9
Las17
Sup35
Dom34
Prm15
Cof1
Srv2
Cyr1
Act1
Sla2
Swa2
Sky1
Ste7
Apl1
Apl2
Sto1
Sgv1
Fus3
Rpn10
Sec23
Sly1
Sec18
Syp1
Vrp1
Sla1
Rps9a
Mtr4
Rps8a
Fun12
Rpn1
Duf1
Rad53
Art5
Ecm21
Ldb19
Yhr131c
Rsp5
Bul2
Ent1
Ent5
Apl3
Sec16
Apm4
Akl1
Rpn11
Nip1
Prt1
Rpg1
Ded1
Tif35
Cdc9
Chk1
Rad23
Ubi4
Ede1
Ent2
Vps27
Chc1
Ptp3
Stb1
Spa2
Bem2
Bud14
Rck2
Glc7
Ipl1
Cdc5
Dbf4
Pfk26
Tpk1
Gpa2
Ras2
Med6
Rpb10
Paf1
Rpo26
Bdp1
Spt15
Spt5
Rgr1
Srb7
Ssn2
Nop1
Nsr1
Nop56
Snu13
Rps31
Rad9
Rtf1
Rpo21
Npl3
Air2
Fps1
Ask10
Cdc28
Hog1
Pbs2
Ssk2
Clb2
Bud3
Bud4
Msn2
Sch9
Rim15
Yak1
Ypl247c
Vps52
Kss1
Sko1
Slt2
Msg5
Swe1
Igo2
Cdc55
Igo1
Vps53
Vps54
Rtn1
Fmp42
Cot1
Zrc1
10
20
30
40
50
t/s

### Path 298 (Syp1 − Ded1)

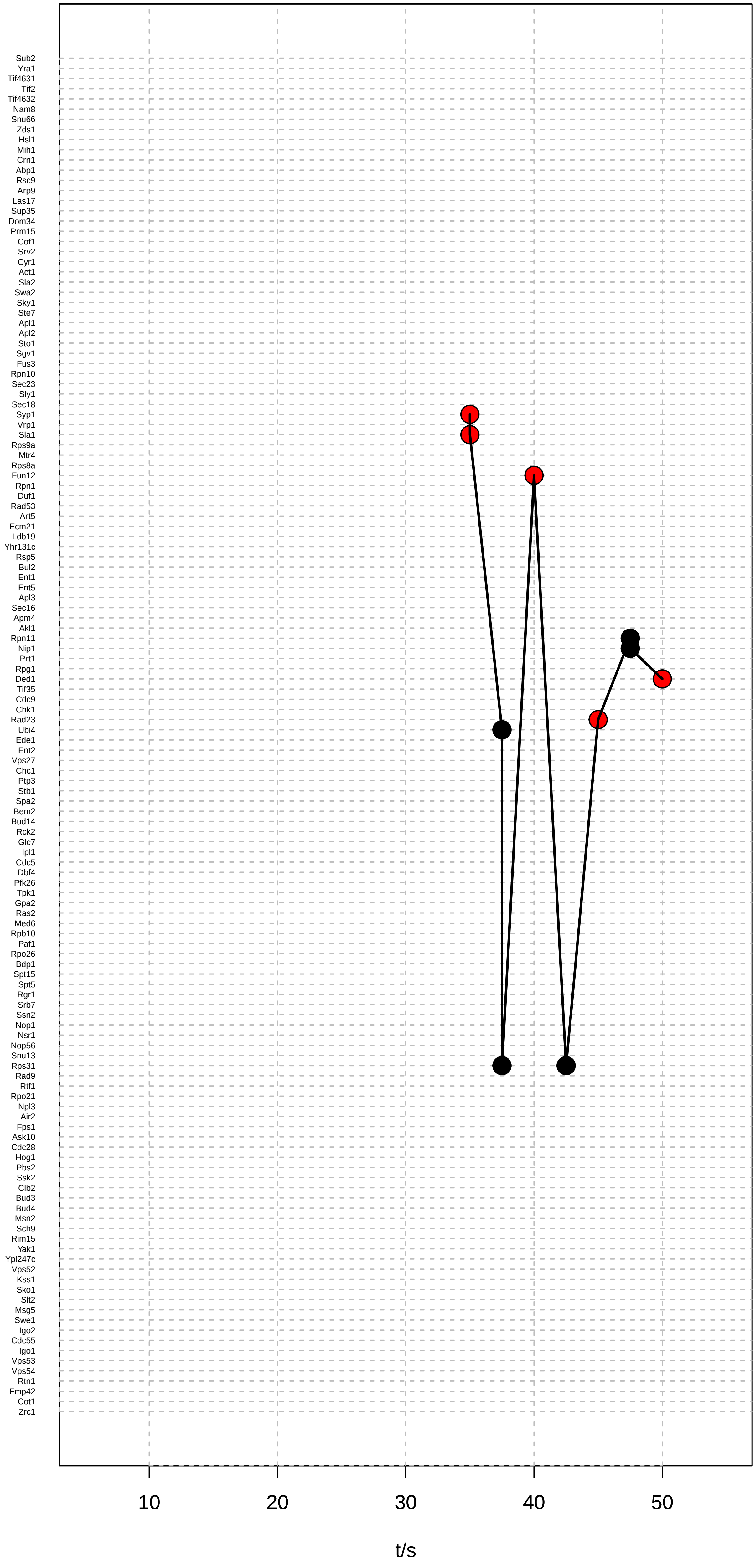

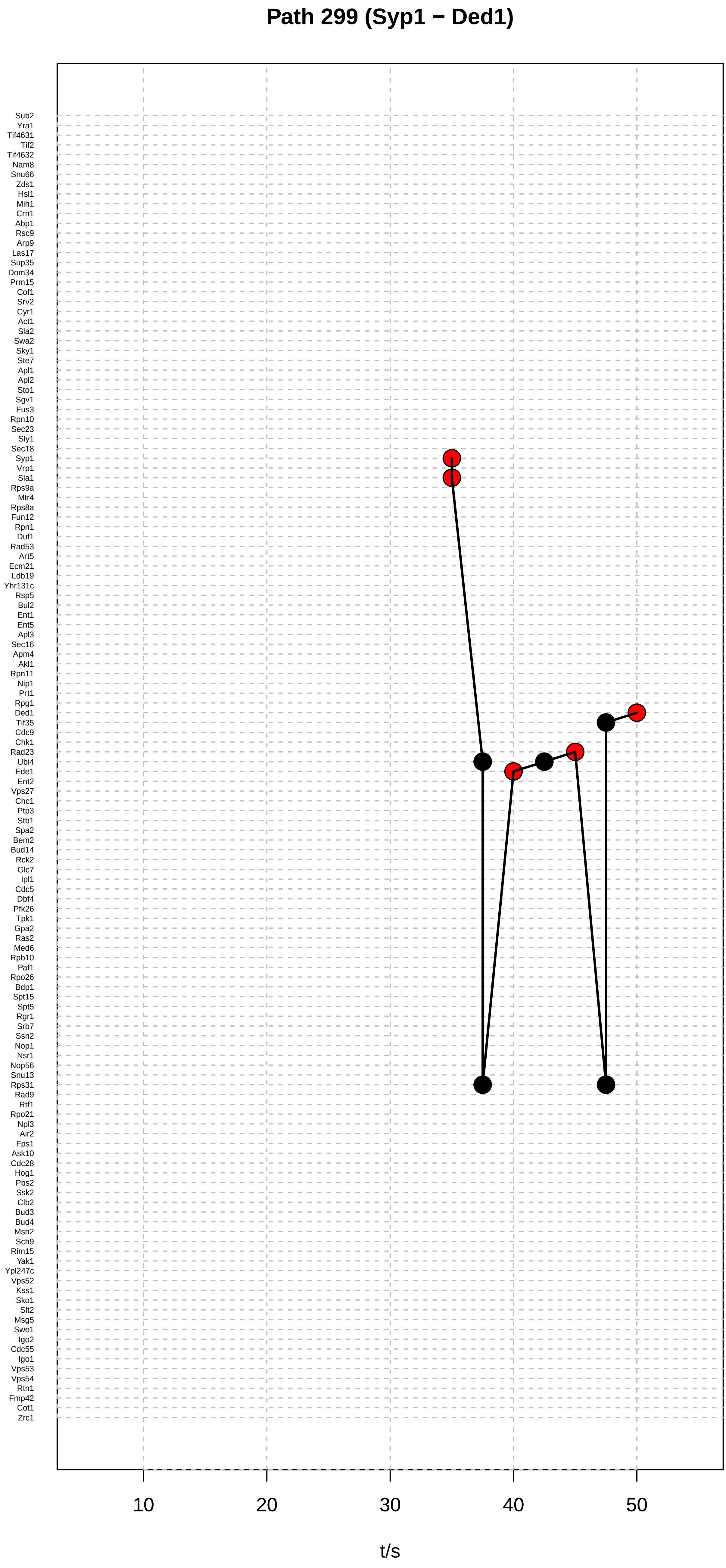

Path 299 (Syp1 − Ded1)
Sub2
Yra1
Tif4631
Tif2
Tif4632
Nam8
Snu66
Zds1
Hsl1
Mih1
Crn1
Abp1
Rsc9
Arp9
Las17
Sup35
Dom34
Prm15
Cof1
Srv2
Cyr1
Act1
Sla2
Swa2
Sky1
Ste7
Apl1
Apl2
Sto1
Sgv1
Fus3
Rpn10
Sec23
Sly1
Sec18
Syp1
Vrp1
Sla1
Rps9a
Mtr4
Rps8a
Fun12
Rpn1
Duf1
Rad53
Art5
Ecm21
Ldb19
Yhr131c
Rsp5
Bul2
Ent1
Ent5
Apl3
Sec16
Apm4
Akl1
Rpn11
Nip1
Prt1
Rpg1
Ded1
Tif35
Cdc9
Chk1
Rad23
Ubi4
Ede1
Ent2
Vps27
Chc1
Ptp3
Stb1
Spa2
Bem2
Bud14
Rck2
Glc7
Ipl1
Cdc5
Dbf4
Pfk26
Tpk1
Gpa2
Ras2
Med6
Rpb10
Paf1
Rpo26
Bdp1
Spt15
Spt5
Rgr1
Srb7
Ssn2
Nop1
Nsr1
Nop56
Snu13
Rps31
Rad9
Rtf1
Rpo21
Npl3
Air2
Fps1
Ask10
Cdc28
Hog1
Pbs2
Ssk2
Clb2
Bud3
Bud4
Msn2
Sch9
Rim15
Yak1
Ypl247c
Vps52
Kss1
Sko1
Slt2
Msg5
Swe1
Igo2
Cdc55
Igo1
Vps53
Vps54
Rtn1
Fmp42
Cot1
Zrc1
10
20
30
40
50
t/s

### Path 300 (Syp1 − Ded1)

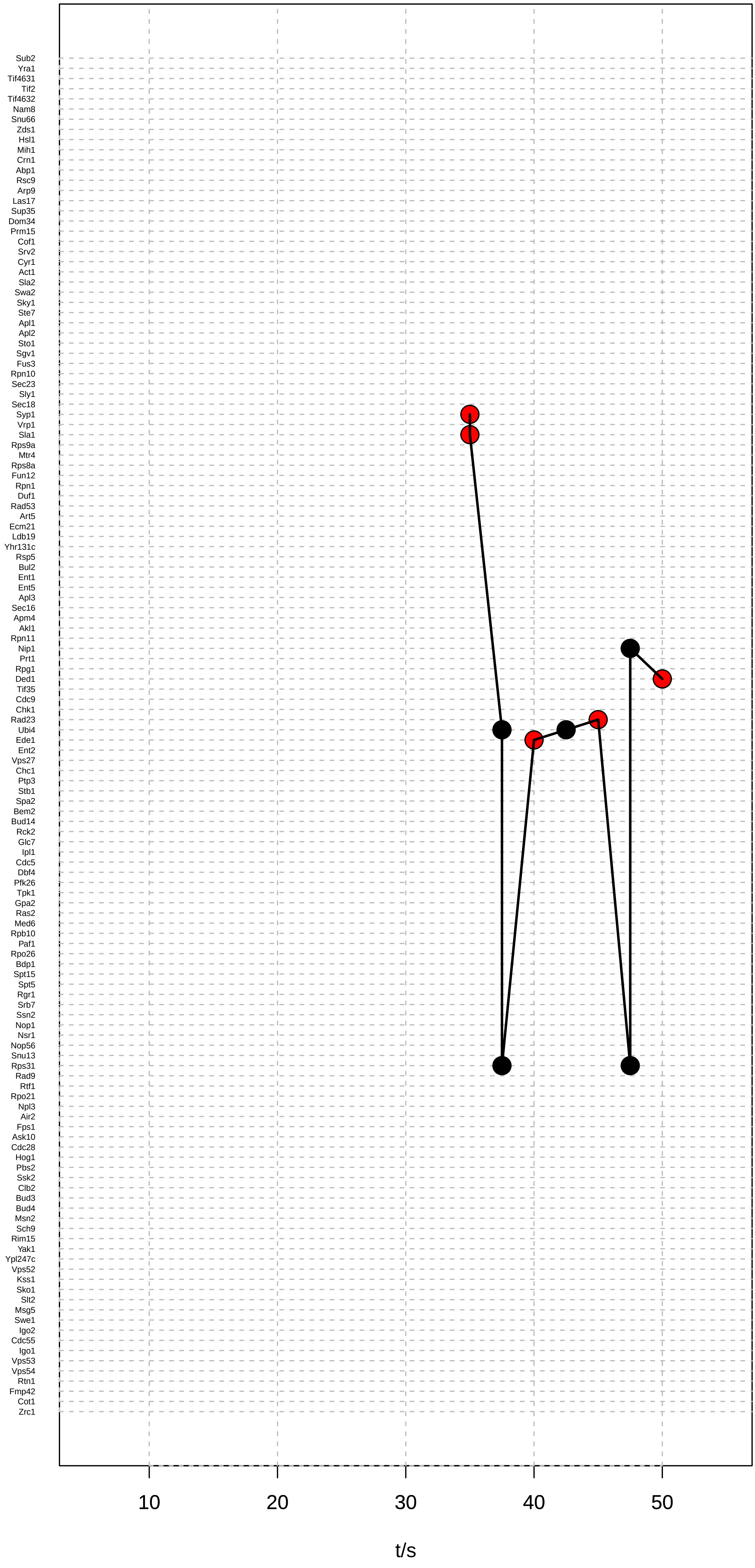

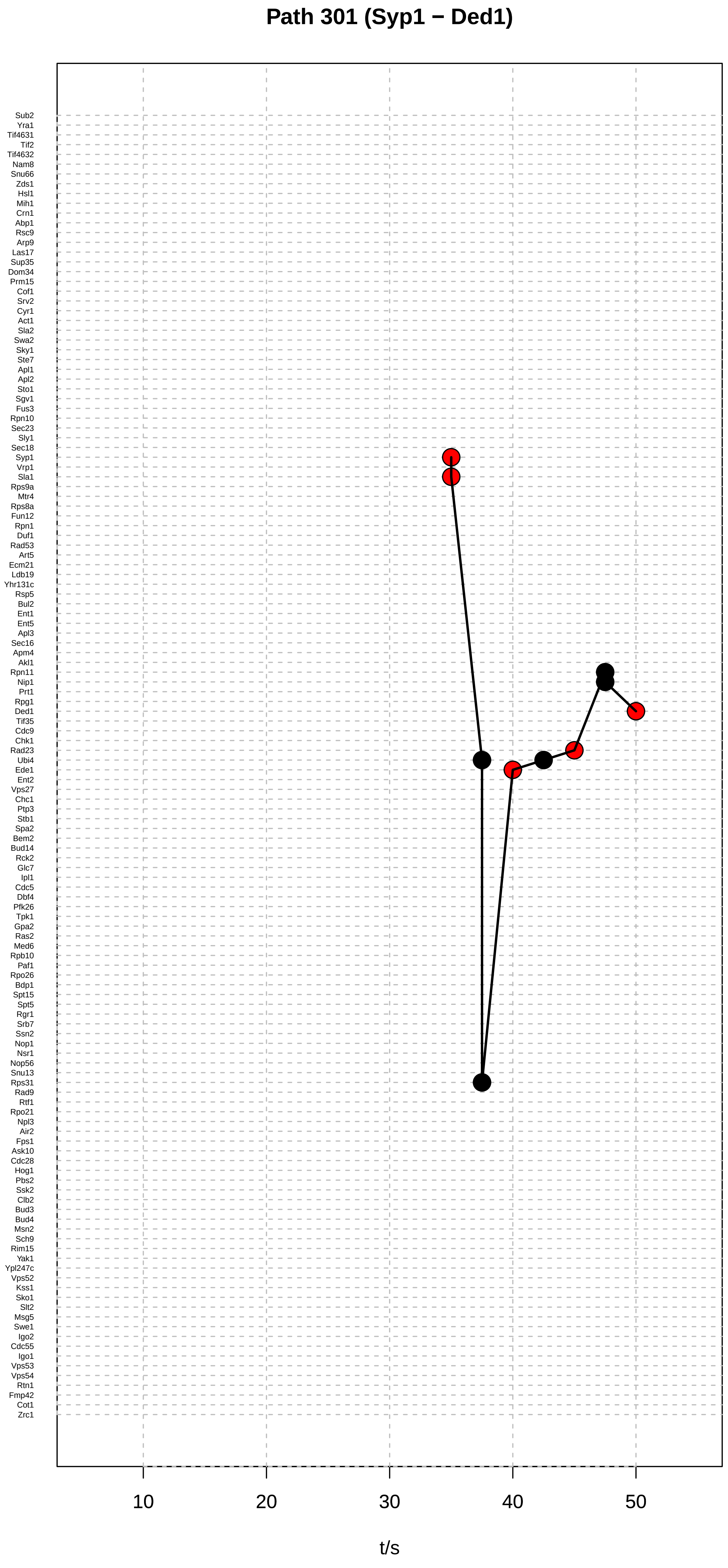

Path 301 (Syp1 − Ded1)
Sub2
Yra1
Tif4631
Tif2
Tif4632
Nam8
Snu66
Zds1
Hsl1
Mih1
Crn1
Abp1
Rsc9
Arp9
Las17
Sup35
Dom34
Prm15
Cof1
Srv2
Cyr1
Act1
Sla2
Swa2
Sky1
Ste7
Apl1
Apl2
Sto1
Sgv1
Fus3
Rpn10
Sec23
Sly1
Sec18
Syp1
Vrp1
Sla1
Rps9a
Mtr4
Rps8a
Fun12
Rpn1
Duf1
Rad53
Art5
Ecm21
Ldb19
Yhr131c
Rsp5
Bul2
Ent1
Ent5
Apl3
Sec16
Apm4
Akl1
Rpn11
Nip1
Prt1
Rpg1
Ded1
Tif35
Cdc9
Chk1
Rad23
Ubi4
Ede1
Ent2
Vps27
Chc1
Ptp3
Stb1
Spa2
Bem2
Bud14
Rck2
Glc7
Ipl1
Cdc5
Dbf4
Pfk26
Tpk1
Gpa2
Ras2
Med6
Rpb10
Paf1
Rpo26
Bdp1
Spt15
Spt5
Rgr1
Srb7
Ssn2
Nop1
Nsr1
Nop56
Snu13
Rps31
Rad9
Rtf1
Rpo21
Npl3
Air2
Fps1
Ask10
Cdc28
Hog1
Pbs2
Ssk2
Clb2
Bud3
Bud4
Msn2
Sch9
Rim15
Yak1
Ypl247c
Vps52
Kss1
Sko1
Slt2
Msg5
Swe1
Igo2
Cdc55
Igo1
Vps53
Vps54
Rtn1
Fmp42
Cot1
Zrc1
10
20
30
40
50
t/s

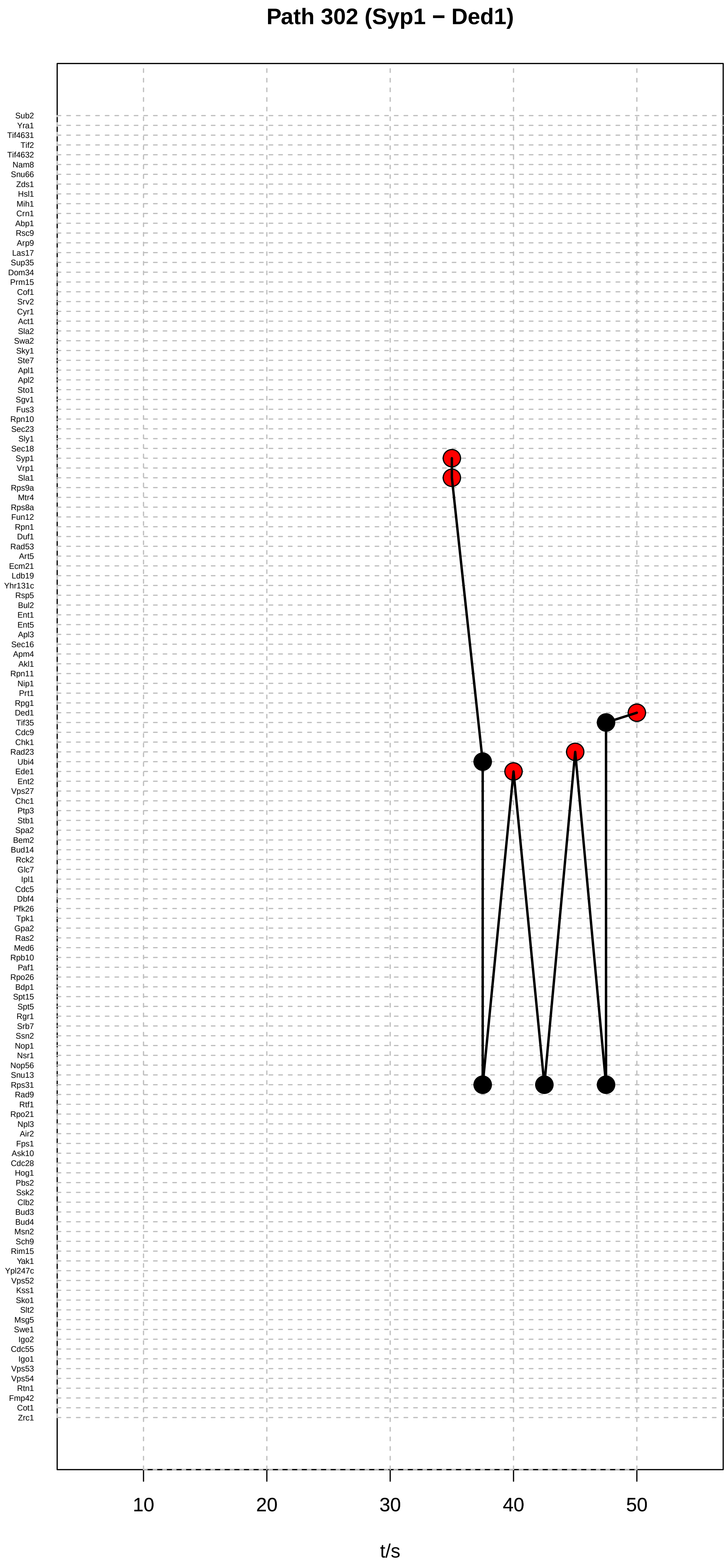

Path 302 (Syp1 − Ded1)
Sub2
Yra1
Tif4631
Tif2
Tif4632
Nam8
Snu66
Zds1
Hsl1
Mih1
Crn1
Abp1
Rsc9
Arp9
Las17
Sup35
Dom34
Prm15
Cof1
Srv2
Cyr1
Act1
Sla2
Swa2
Sky1
Ste7
Apl1
Apl2
Sto1
Sgv1
Fus3
Rpn10
Sec23
Sly1
Sec18
Syp1
Vrp1
Sla1
Rps9a
Mtr4
Rps8a
Fun12
Rpn1
Duf1
Rad53
Art5
Ecm21
Ldb19
Yhr131c
Rsp5
Bul2
Ent1
Ent5
Apl3
Sec16
Apm4
Akl1
Rpn11
Nip1
Prt1
Rpg1
Ded1
Tif35
Cdc9
Chk1
Rad23
Ubi4
Ede1
Ent2
Vps27
Chc1
Ptp3
Stb1
Spa2
Bem2
Bud14
Rck2
Glc7
Ipl1
Cdc5
Dbf4
Pfk26
Tpk1
Gpa2
Ras2
Med6
Rpb10
Paf1
Rpo26
Bdp1
Spt15
Spt5
Rgr1
Srb7
Ssn2
Nop1
Nsr1
Nop56
Snu13
Rps31
Rad9
Rtf1
Rpo21
Npl3
Air2
Fps1
Ask10
Cdc28
Hog1
Pbs2
Ssk2
Clb2
Bud3
Bud4
Msn2
Sch9
Rim15
Yak1
Ypl247c
Vps52
Kss1
Sko1
Slt2
Msg5
Swe1
Igo2
Cdc55
Igo1
Vps53
Vps54
Rtn1
Fmp42
Cot1
Zrc1
10
20
30
40
50
t/s

#### Path 303 (Syp1 − Ded1)

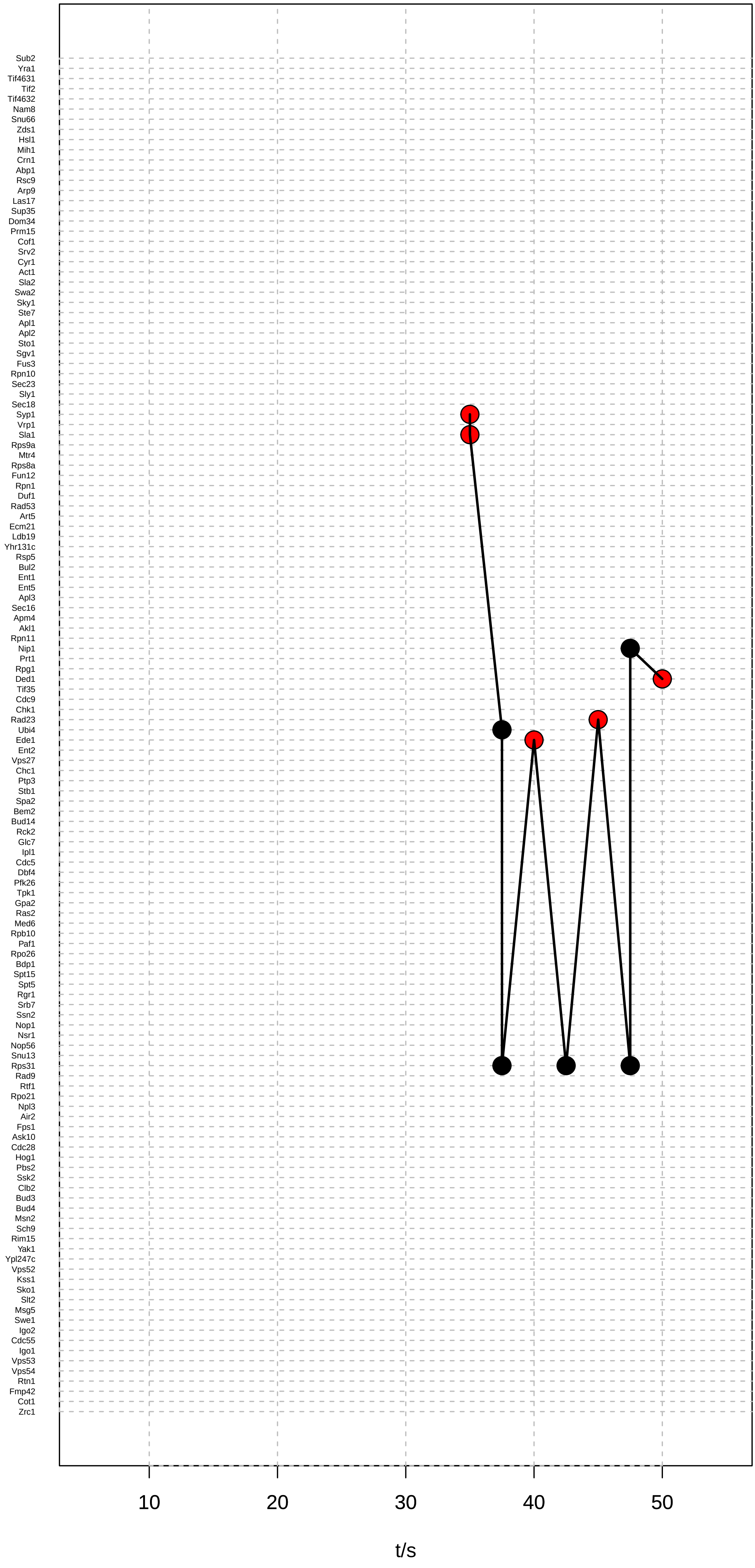

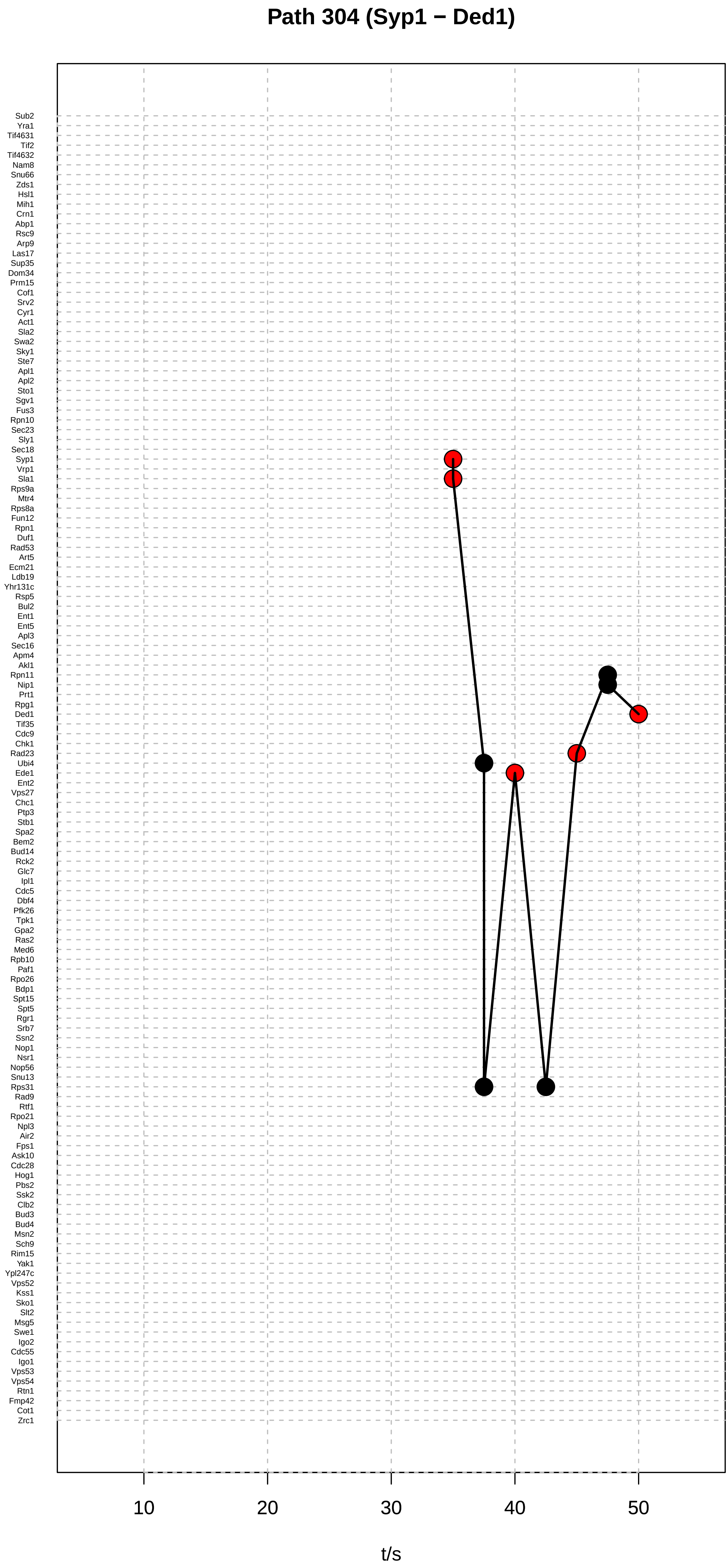

Path 304 (Syp1 − Ded1)
Sub2
Yra1
Tif4631
Tif2
Tif4632
Nam8
Snu66
Zds1
Hsl1
Mih1
Crn1
Abp1
Rsc9
Arp9
Las17
Sup35
Dom34
Prm15
Cof1
Srv2
Cyr1
Act1
Sla2
Swa2
Sky1
Ste7
Apl1
Apl2
Sto1
Sgv1
Fus3
Rpn10
Sec23
Sly1
Sec18
Syp1
Vrp1
Sla1
Rps9a
Mtr4
Rps8a
Fun12
Rpn1
Duf1
Rad53
Art5
Ecm21
Ldb19
Yhr131c
Rsp5
Bul2
Ent1
Ent5
Apl3
Sec16
Apm4
Akl1
Rpn11
Nip1
Prt1
Rpg1
Ded1
Tif35
Cdc9
Chk1
Rad23
Ubi4
Ede1
Ent2
Vps27
Chc1
Ptp3
Stb1
Spa2
Bem2
Bud14
Rck2
Glc7
Ipl1
Cdc5
Dbf4
Pfk26
Tpk1
Gpa2
Ras2
Med6
Rpb10
Paf1
Rpo26
Bdp1
Spt15
Spt5
Rgr1
Srb7
Ssn2
Nop1
Nsr1
Nop56
Snu13
Rps31
Rad9
Rtf1
Rpo21
Npl3
Air2
Fps1
Ask10
Cdc28
Hog1
Pbs2
Ssk2
Clb2
Bud3
Bud4
Msn2
Sch9
Rim15
Yak1
Ypl247c
Vps52
Kss1
Sko1
Slt2
Msg5
Swe1
Igo2
Cdc55
Igo1
Vps53
Vps54
Rtn1
Fmp42
Cot1
Zrc1
10
20
30
40
50
t/s

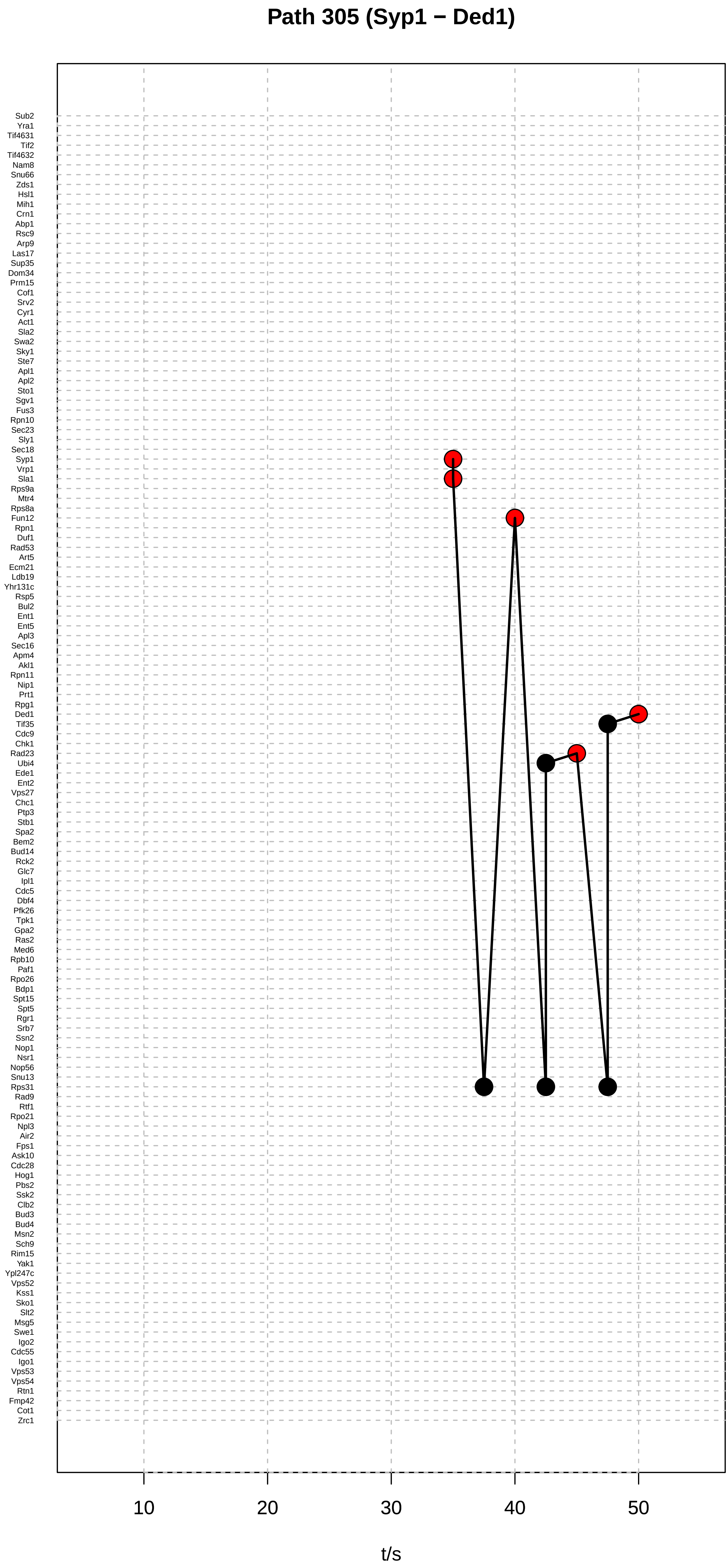
Path 305 (Syp1 − Ded1)
Sub2
Yra1
Tif4631
Tif2
Tif4632
Nam8
Snu66
Zds1
Hsl1
Mih1
Crn1
Abp1
Rsc9
Arp9
Las17
Sup35
Dom34
Prm15
Cof1
Srv2
Cyr1
Act1
Sla2
Swa2
Sky1
Ste7
Apl1
Apl2
Sto1
Sgv1
Fus3
Rpn10
Sec23
Sly1
Sec18
Syp1
Vrp1
Sla1
Rps9a
Mtr4
Rps8a
Fun12
Rpn1
Duf1
Rad53
Art5
Ecm21
Ldb19
Yhr131c
Rsp5
Bul2
Ent1
Ent5
Apl3
Sec16
Apm4
Akl1
Rpn11
Nip1
Prt1
Rpg1
Ded1
Tif35
Cdc9
Chk1
Rad23
Ubi4
Ede1
Ent2
Vps27
Chc1
Ptp3
Stb1
Spa2
Bem2
Bud14
Rck2
Glc7
Ipl1
Cdc5
Dbf4
Pfk26
Tpk1
Gpa2
Ras2
Med6
Rpb10
Paf1
Rpo26
Bdp1
Spt15
Spt5
Rgr1
Srb7
Ssn2
Nop1
Nsr1
Nop56
Snu13
Rps31
Rad9
Rtf1
Rpo21
Npl3
Air2
Fps1
Ask10
Cdc28
Hog1
Pbs2
Ssk2
Clb2
Bud3
Bud4
Msn2
Sch9
Rim15
Yak1
Ypl247c
Vps52
Kss1
Sko1
Slt2
Msg5
Swe1
Igo2
Cdc55
Igo1
Vps53
Vps54
Rtn1
Fmp42
Cot1
Zrc1
10
20
30
40
50
t/s

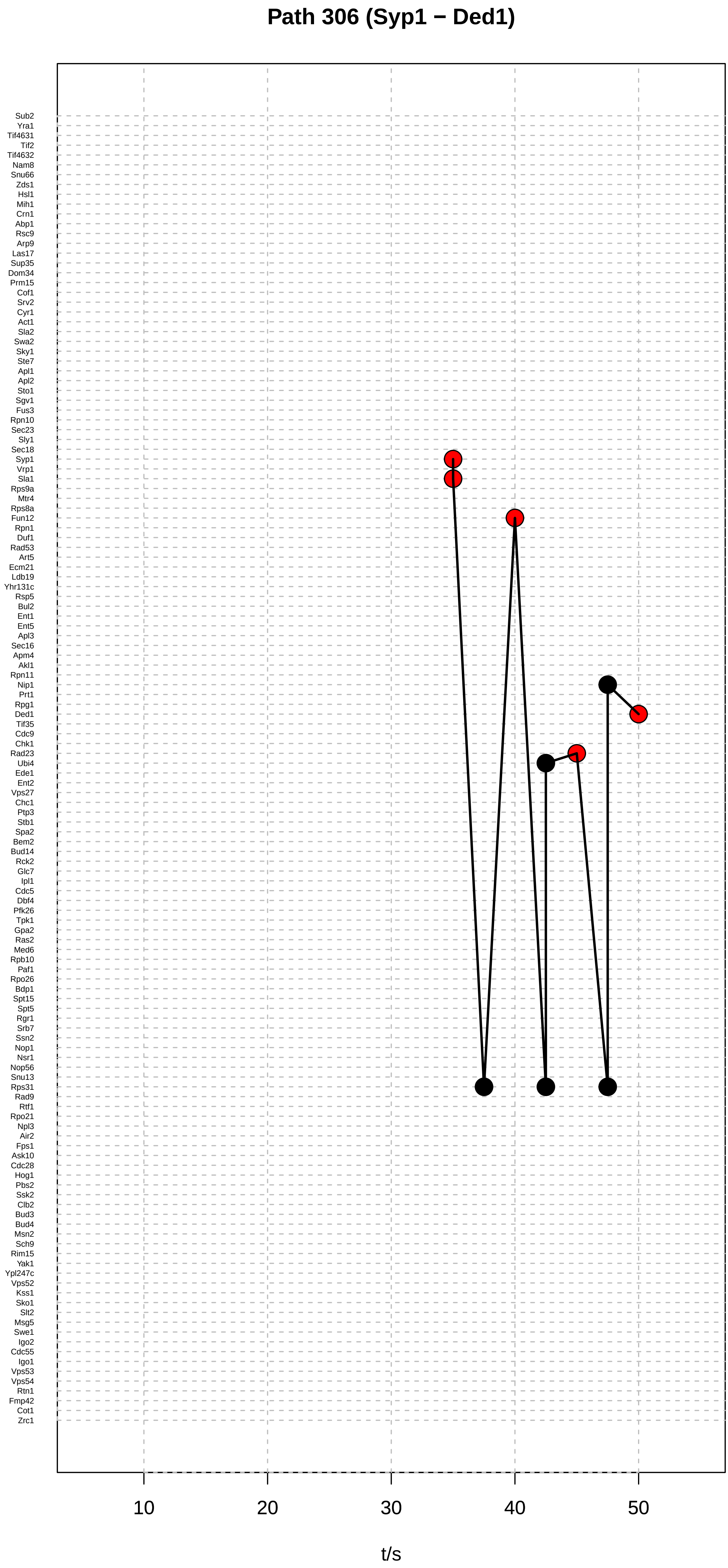
Path 306 (Syp1 − Ded1)
Sub2
Yra1
Tif4631
Tif2
Tif4632
Nam8
Snu66
Zds1
Hsl1
Mih1
Crn1
Abp1
Rsc9
Arp9
Las17
Sup35
Dom34
Prm15
Cof1
Srv2
Cyr1
Act1
Sla2
Swa2
Sky1
Ste7
Apl1
Apl2
Sto1
Sgv1
Fus3
Rpn10
Sec23
Sly1
Sec18
Syp1
Vrp1
Sla1
Rps9a
Mtr4
Rps8a
Fun12
Rpn1
Duf1
Rad53
Art5
Ecm21
Ldb19
Yhr131c
Rsp5
Bul2
Ent1
Ent5
Apl3
Sec16
Apm4
Akl1
Rpn11
Nip1
Prt1
Rpg1
Ded1
Tif35
Cdc9
Chk1
Rad23
Ubi4
Ede1
Ent2
Vps27
Chc1
Ptp3
Stb1
Spa2
Bem2
Bud14
Rck2
Glc7
Ipl1
Cdc5
Dbf4
Pfk26
Tpk1
Gpa2
Ras2
Med6
Rpb10
Paf1
Rpo26
Bdp1
Spt15
Spt5
Rgr1
Srb7
Ssn2
Nop1
Nsr1
Nop56
Snu13
Rps31
Rad9
Rtf1
Rpo21
Npl3
Air2
Fps1
Ask10
Cdc28
Hog1
Pbs2
Ssk2
Clb2
Bud3
Bud4
Msn2
Sch9
Rim15
Yak1
Ypl247c
Vps52
Kss1
Sko1
Slt2
Msg5
Swe1
Igo2
Cdc55
Igo1
Vps53
Vps54
Rtn1
Fmp42
Cot1
Zrc1
10
20
30
40
50
t/s

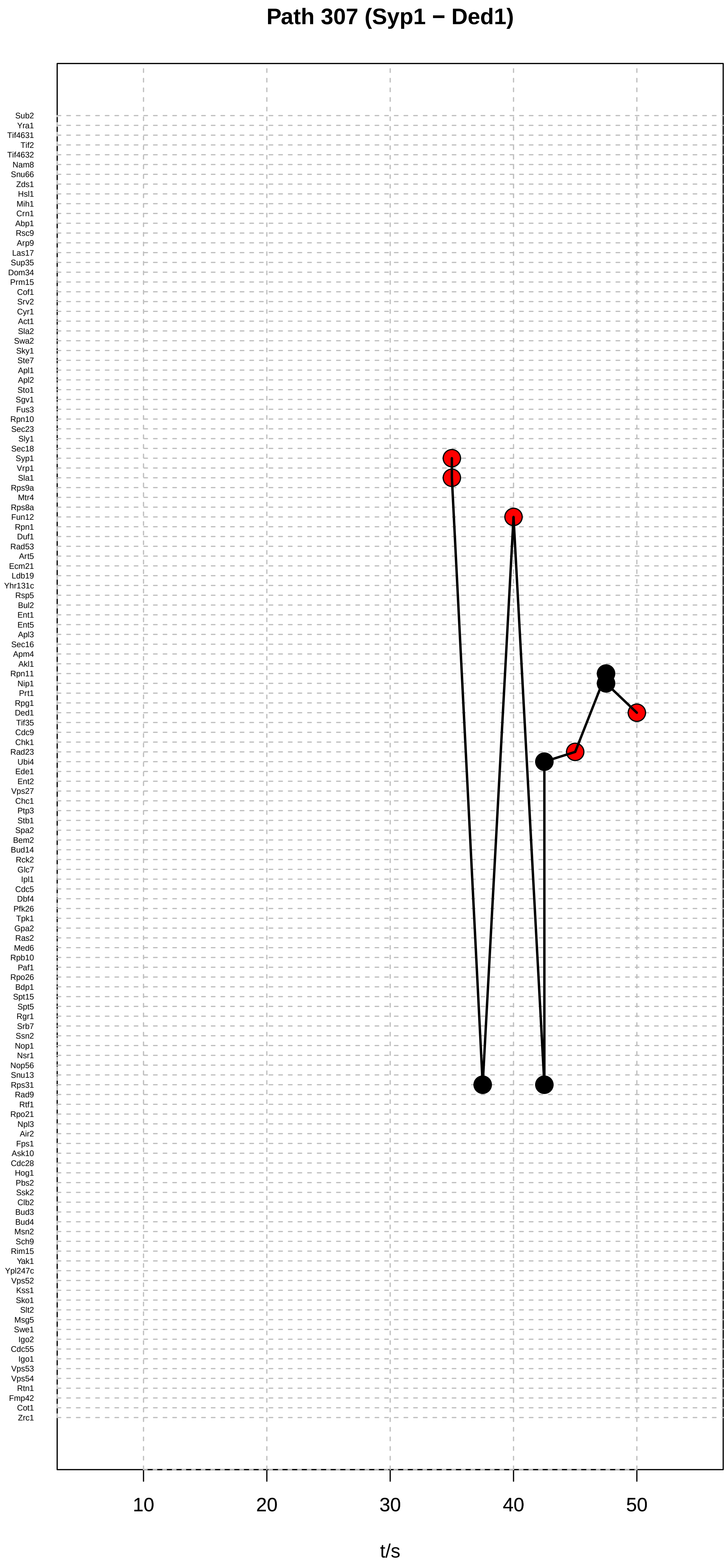

Path 307 (Syp1 − Ded1)
Sub2
Yra1
Tif4631
Tif2
Tif4632
Nam8
Snu66
Zds1
Hsl1
Mih1
Crn1
Abp1
Rsc9
Arp9
Las17
Sup35
Dom34
Prm15
Cof1
Srv2
Cyr1
Act1
Sla2
Swa2
Sky1
Ste7
Apl1
Apl2
Sto1
Sgv1
Fus3
Rpn10
Sec23
Sly1
Sec18
Syp1
Vrp1
Sla1
Rps9a
Mtr4
Rps8a
Fun12
Rpn1
Duf1
Rad53
Art5
Ecm21
Ldb19
Yhr131c
Rsp5
Bul2
Ent1
Ent5
Apl3
Sec16
Apm4
Akl1
Rpn11
Nip1
Prt1
Rpg1
Ded1
Tif35
Cdc9
Chk1
Rad23
Ubi4
Ede1
Ent2
Vps27
Chc1
Ptp3
Stb1
Spa2
Bem2
Bud14
Rck2
Glc7
Ipl1
Cdc5
Dbf4
Pfk26
Tpk1
Gpa2
Ras2
Med6
Rpb10
Paf1
Rpo26
Bdp1
Spt15
Spt5
Rgr1
Srb7
Ssn2
Nop1
Nsr1
Nop56
Snu13
Rps31
Rad9
Rtf1
Rpo21
Npl3
Air2
Fps1
Ask10
Cdc28
Hog1
Pbs2
Ssk2
Clb2
Bud3
Bud4
Msn2
Sch9
Rim15
Yak1
Ypl247c
Vps52
Kss1
Sko1
Slt2
Msg5
Swe1
Igo2
Cdc55
Igo1
Vps53
Vps54
Rtn1
Fmp42
Cot1
Zrc1
10
20
30
40
50
t/s

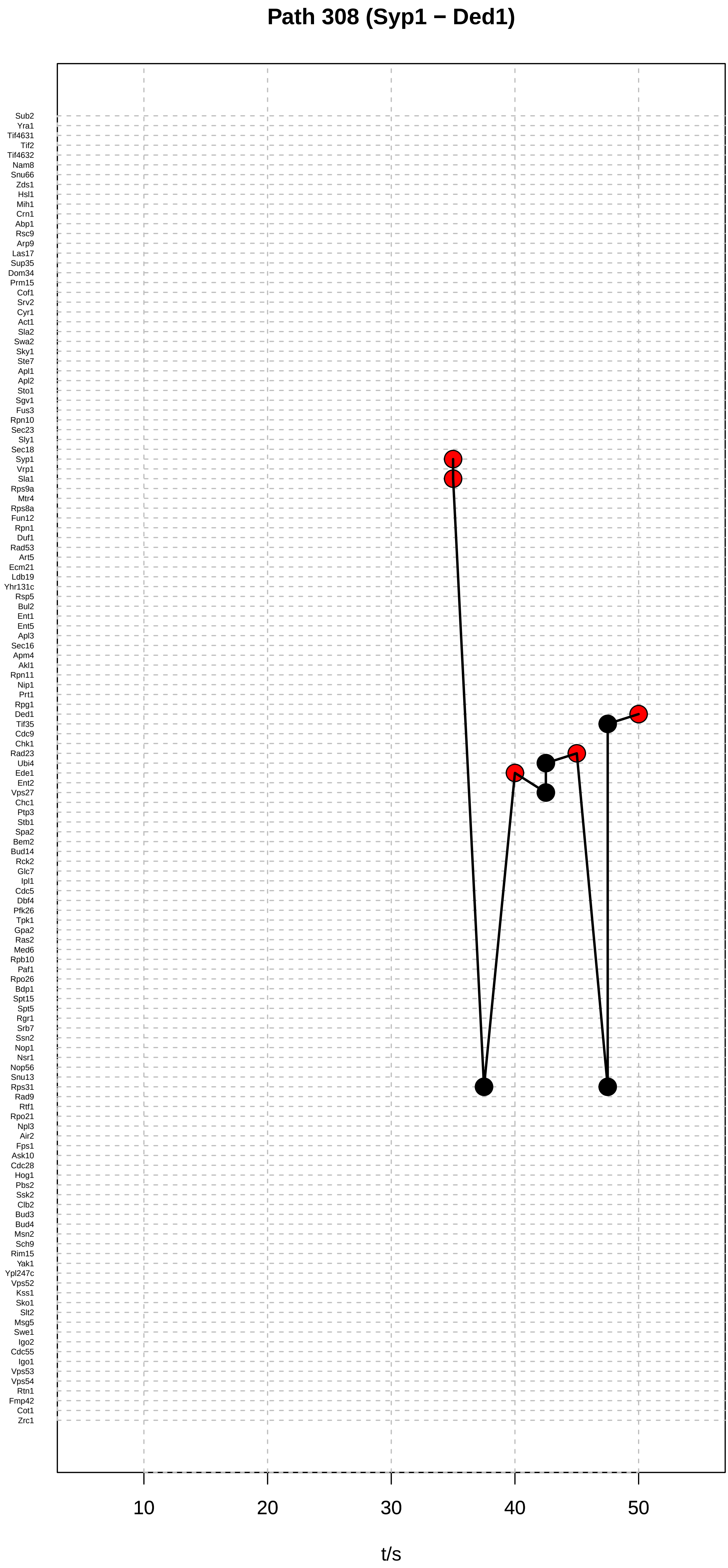

Path 308 (Syp1 − Ded1)
Sub2
Yra1
Tif4631
Tif2
Tif4632
Nam8
Snu66
Zds1
Hsl1
Mih1
Crn1
Abp1
Rsc9
Arp9
Las17
Sup35
Dom34
Prm15
Cof1
Srv2
Cyr1
Act1
Sla2
Swa2
Sky1
Ste7
Apl1
Apl2
Sto1
Sgv1
Fus3
Rpn10
Sec23
Sly1
Sec18
Syp1
Vrp1
Sla1
Rps9a
Mtr4
Rps8a
Fun12
Rpn1
Duf1
Rad53
Art5
Ecm21
Ldb19
Yhr131c
Rsp5
Bul2
Ent1
Ent5
Apl3
Sec16
Apm4
Akl1
Rpn11
Nip1
Prt1
Rpg1
Ded1
Tif35
Cdc9
Chk1
Rad23
Ubi4
Ede1
Ent2
Vps27
Chc1
Ptp3
Stb1
Spa2
Bem2
Bud14
Rck2
Glc7
Ipl1
Cdc5
Dbf4
Pfk26
Tpk1
Gpa2
Ras2
Med6
Rpb10
Paf1
Rpo26
Bdp1
Spt15
Spt5
Rgr1
Srb7
Ssn2
Nop1
Nsr1
Nop56
Snu13
Rps31
Rad9
Rtf1
Rpo21
Npl3
Air2
Fps1
Ask10
Cdc28
Hog1
Pbs2
Ssk2
Clb2
Bud3
Bud4
Msn2
Sch9
Rim15
Yak1
Ypl247c
Vps52
Kss1
Sko1
Slt2
Msg5
Swe1
Igo2
Cdc55
Igo1
Vps53
Vps54
Rtn1
Fmp42
Cot1
Zrc1
10
20
30
40
50
t/s

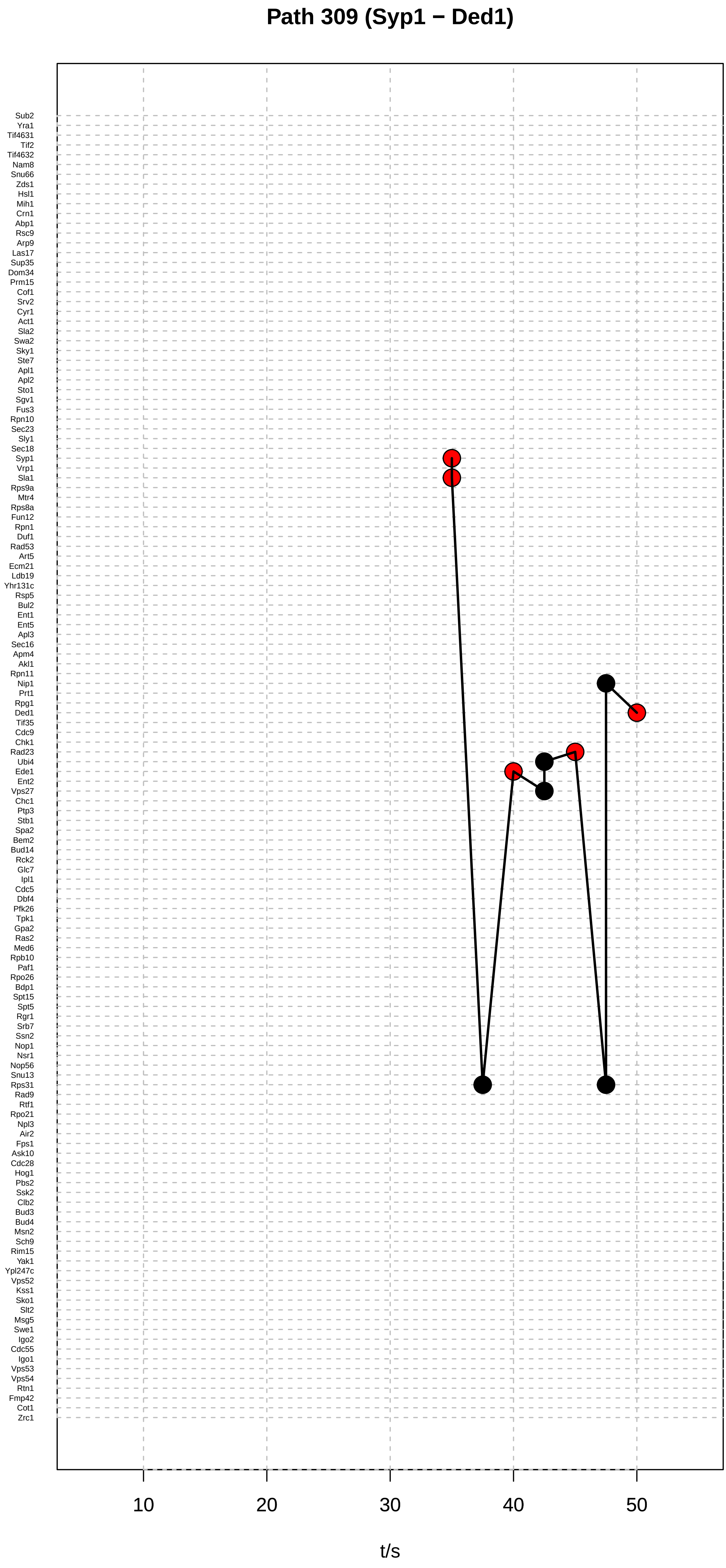

Path 309 (Syp1 − Ded1)
Sub2
Yra1
Tif4631
Tif2
Tif4632
Nam8
Snu66
Zds1
Hsl1
Mih1
Crn1
Abp1
Rsc9
Arp9
Las17
Sup35
Dom34
Prm15
Cof1
Srv2
Cyr1
Act1
Sla2
Swa2
Sky1
Ste7
Apl1
Apl2
Sto1
Sgv1
Fus3
Rpn10
Sec23
Sly1
Sec18
Syp1
Vrp1
Sla1
Rps9a
Mtr4
Rps8a
Fun12
Rpn1
Duf1
Rad53
Art5
Ecm21
Ldb19
Yhr131c
Rsp5
Bul2
Ent1
Ent5
Apl3
Sec16
Apm4
Akl1
Rpn11
Nip1
Prt1
Rpg1
Ded1
Tif35
Cdc9
Chk1
Rad23
Ubi4
Ede1
Ent2
Vps27
Chc1
Ptp3
Stb1
Spa2
Bem2
Bud14
Rck2
Glc7
Ipl1
Cdc5
Dbf4
Pfk26
Tpk1
Gpa2
Ras2
Med6
Rpb10
Paf1
Rpo26
Bdp1
Spt15
Spt5
Rgr1
Srb7
Ssn2
Nop1
Nsr1
Nop56
Snu13
Rps31
Rad9
Rtf1
Rpo21
Npl3
Air2
Fps1
Ask10
Cdc28
Hog1
Pbs2
Ssk2
Clb2
Bud3
Bud4
Msn2
Sch9
Rim15
Yak1
Ypl247c
Vps52
Kss1
Sko1
Slt2
Msg5
Swe1
Igo2
Cdc55
Igo1
Vps53
Vps54
Rtn1
Fmp42
Cot1
Zrc1
10
20
30
40
50
t/s

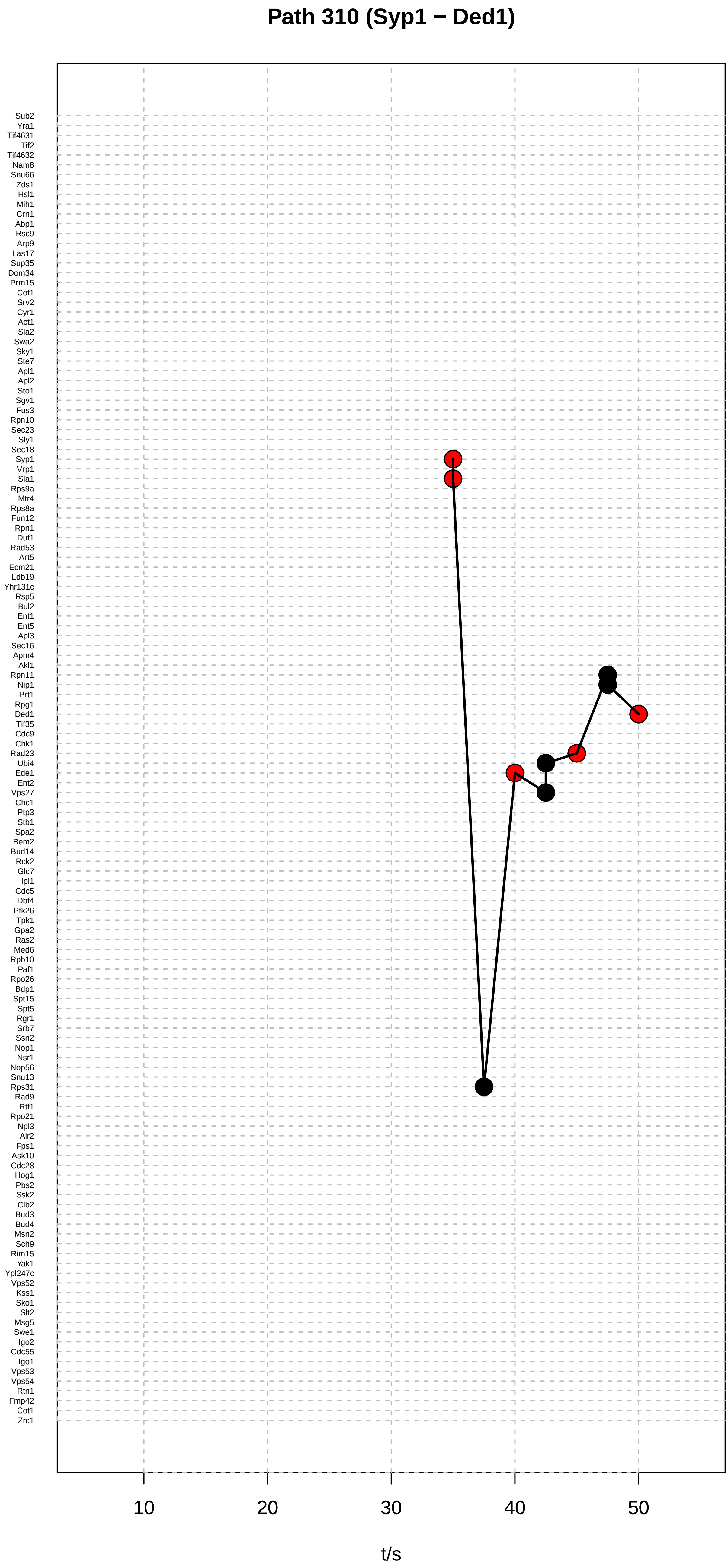
Path 310 (Syp1 − Ded1)
Sub2
Yra1
Tif4631
Tif2
Tif4632
Nam8
Snu66
Zds1
Hsl1
Mih1
Crn1
Abp1
Rsc9
Arp9
Las17
Sup35
Dom34
Prm15
Cof1
Srv2
Cyr1
Act1
Sla2
Swa2
Sky1
Ste7
Apl1
Apl2
Sto1
Sgv1
Fus3
Rpn10
Sec23
Sly1
Sec18
Syp1
Vrp1
Sla1
Rps9a
Mtr4
Rps8a
Fun12
Rpn1
Duf1
Rad53
Art5
Ecm21
Ldb19
Yhr131c
Rsp5
Bul2
Ent1
Ent5
Apl3
Sec16
Apm4
Akl1
Rpn11
Nip1
Prt1
Rpg1
Ded1
Tif35
Cdc9
Chk1
Rad23
Ubi4
Ede1
Ent2
Vps27
Chc1
Ptp3
Stb1
Spa2
Bem2
Bud14
Rck2
Glc7
Ipl1
Cdc5
Dbf4
Pfk26
Tpk1
Gpa2
Ras2
Med6
Rpb10
Paf1
Rpo26
Bdp1
Spt15
Spt5
Rgr1
Srb7
Ssn2
Nop1
Nsr1
Nop56
Snu13
Rps31
Rad9
Rtf1
Rpo21
Npl3
Air2
Fps1
Ask10
Cdc28
Hog1
Pbs2
Ssk2
Clb2
Bud3
Bud4
Msn2
Sch9
Rim15
Yak1
Ypl247c
Vps52
Kss1
Sko1
Slt2
Msg5
Swe1
Igo2
Cdc55
Igo1
Vps53
Vps54
Rtn1
Fmp42
Cot1
Zrc1
10
20
30
40
50
t/s

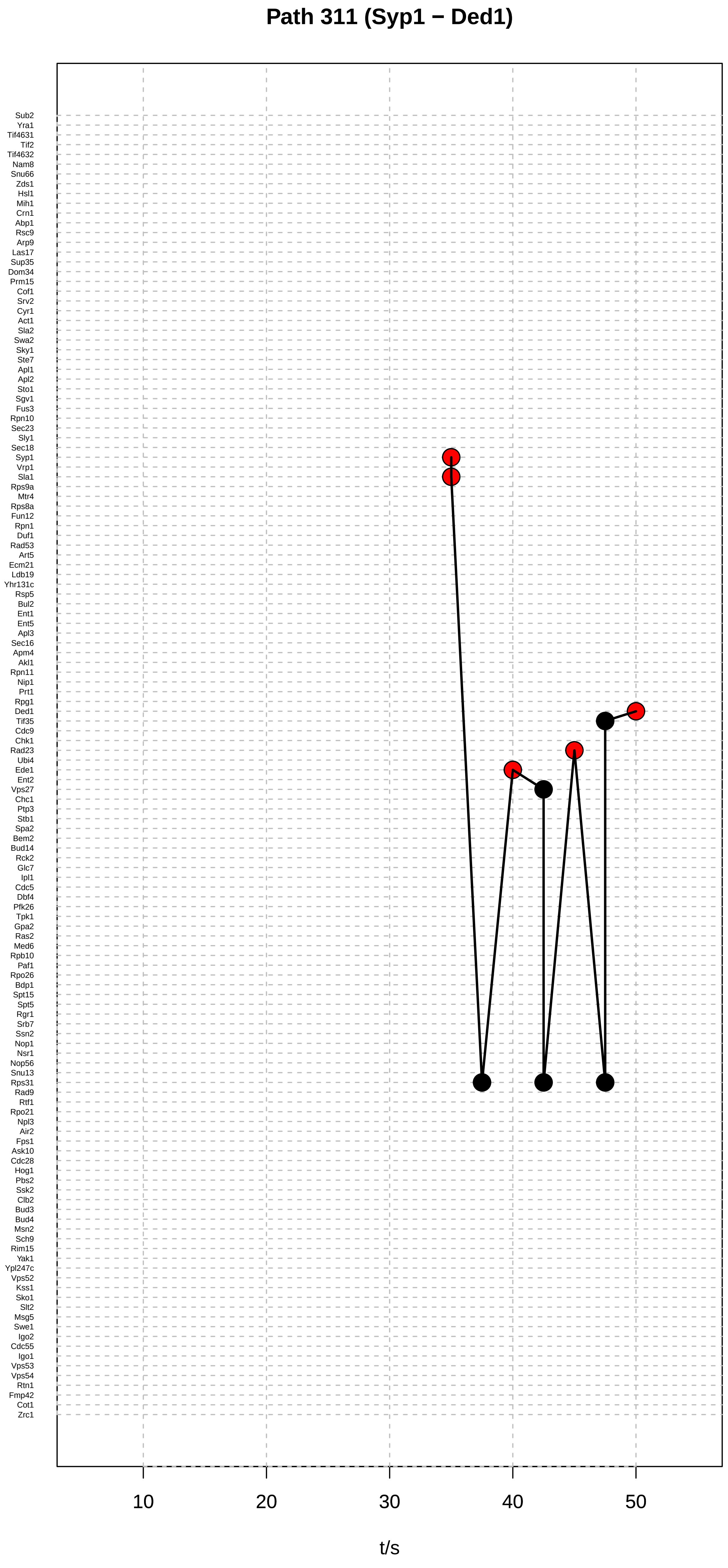

Path 311 (Syp1 − Ded1)
Sub2
Yra1
Tif4631
Tif2
Tif4632
Nam8
Snu66
Zds1
Hsl1
Mih1
Crn1
Abp1
Rsc9
Arp9
Las17
Sup35
Dom34
Prm15
Cof1
Srv2
Cyr1
Act1
Sla2
Swa2
Sky1
Ste7
Apl1
Apl2
Sto1
Sgv1
Fus3
Rpn10
Sec23
Sly1
Sec18
Syp1
Vrp1
Sla1
Rps9a
Mtr4
Rps8a
Fun12
Rpn1
Duf1
Rad53
Art5
Ecm21
Ldb19
Yhr131c
Rsp5
Bul2
Ent1
Ent5
Apl3
Sec16
Apm4
Akl1
Rpn11
Nip1
Prt1
Rpg1
Ded1
Tif35
Cdc9
Chk1
Rad23
Ubi4
Ede1
Ent2
Vps27
Chc1
Ptp3
Stb1
Spa2
Bem2
Bud14
Rck2
Glc7
Ipl1
Cdc5
Dbf4
Pfk26
Tpk1
Gpa2
Ras2
Med6
Rpb10
Paf1
Rpo26
Bdp1
Spt15
Spt5
Rgr1
Srb7
Ssn2
Nop1
Nsr1
Nop56
Snu13
Rps31
Rad9
Rtf1
Rpo21
Npl3
Air2
Fps1
Ask10
Cdc28
Hog1
Pbs2
Ssk2
Clb2
Bud3
Bud4
Msn2
Sch9
Rim15
Yak1
Ypl247c
Vps52
Kss1
Sko1
Slt2
Msg5
Swe1
Igo2
Cdc55
Igo1
Vps53
Vps54
Rtn1
Fmp42
Cot1
Zrc1
10
20
30
40
50
t/s

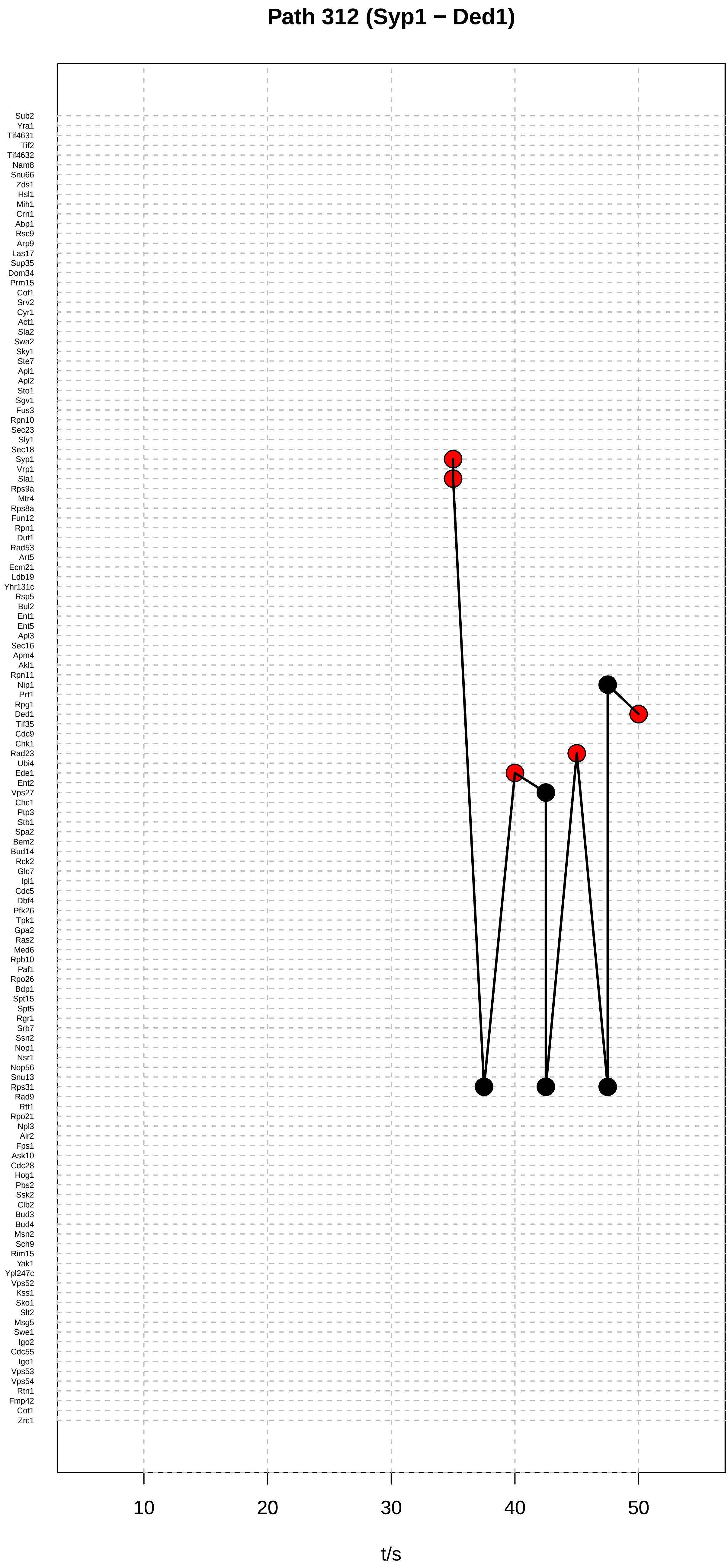

Path 312 (Syp1 − Ded1)
Sub2
Yra1
Tif4631
Tif2
Tif4632
Nam8
Snu66
Zds1
Hsl1
Mih1
Crn1
Abp1
Rsc9
Arp9
Las17
Sup35
Dom34
Prm15
Cof1
Srv2
Cyr1
Act1
Sla2
Swa2
Sky1
Ste7
Apl1
Apl2
Sto1
Sgv1
Fus3
Rpn10
Sec23
Sly1
Sec18
Syp1
Vrp1
Sla1
Rps9a
Mtr4
Rps8a
Fun12
Rpn1
Duf1
Rad53
Art5
Ecm21
Ldb19
Yhr131c
Rsp5
Bul2
Ent1
Ent5
Apl3
Sec16
Apm4
Akl1
Rpn11
Nip1
Prt1
Rpg1
Ded1
Tif35
Cdc9
Chk1
Rad23
Ubi4
Ede1
Ent2
Vps27
Chc1
Ptp3
Stb1
Spa2
Bem2
Bud14
Rck2
Glc7
Ipl1
Cdc5
Dbf4
Pfk26
Tpk1
Gpa2
Ras2
Med6
Rpb10
Paf1
Rpo26
Bdp1
Spt15
Spt5
Rgr1
Srb7
Ssn2
Nop1
Nsr1
Nop56
Snu13
Rps31
Rad9
Rtf1
Rpo21
Npl3
Air2
Fps1
Ask10
Cdc28
Hog1
Pbs2
Ssk2
Clb2
Bud3
Bud4
Msn2
Sch9
Rim15
Yak1
Ypl247c
Vps52
Kss1
Sko1
Slt2
Msg5
Swe1
Igo2
Cdc55
Igo1
Vps53
Vps54
Rtn1
Fmp42
Cot1
Zrc1
10
20
30
40
50
t/s

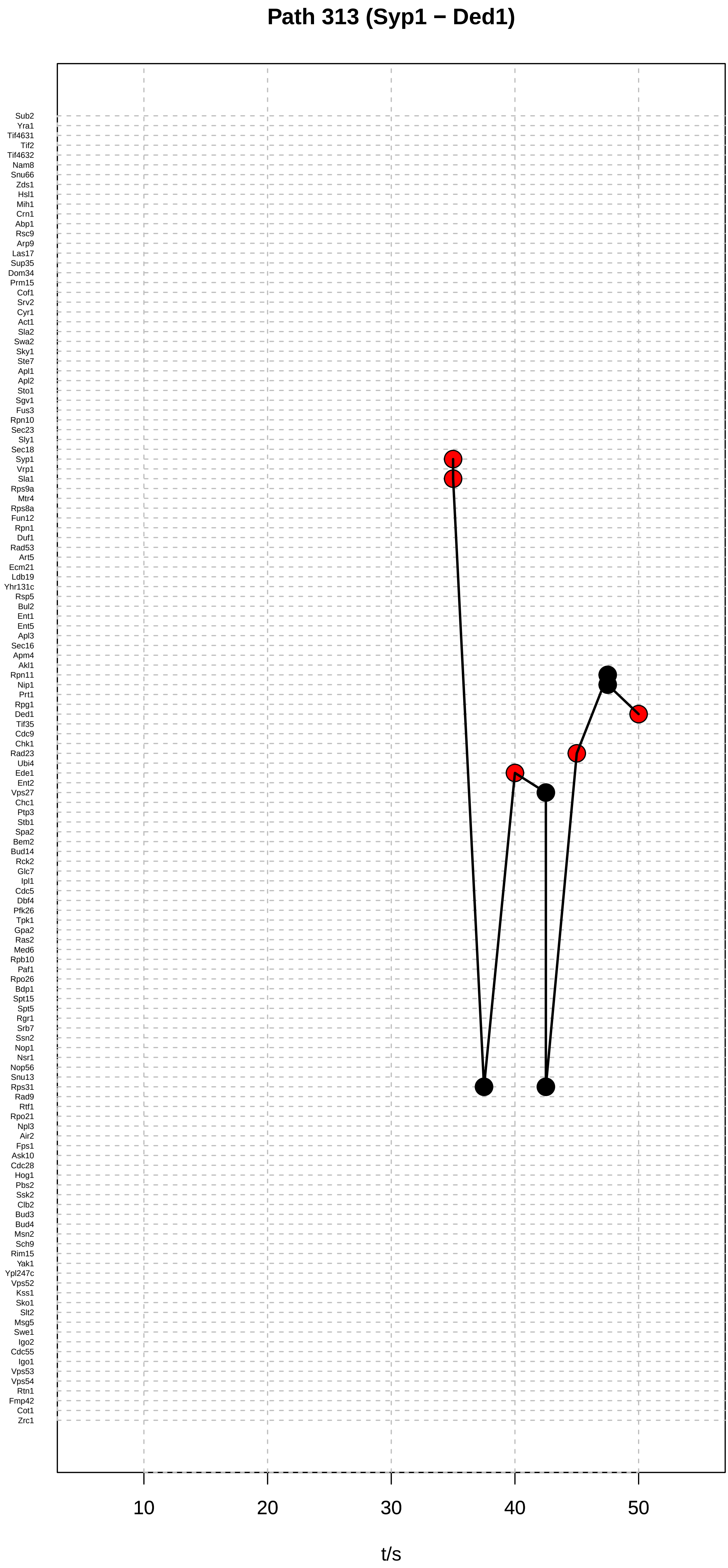
Path 313 (Syp1 − Ded1)
Sub2
Yra1
Tif4631
Tif2
Tif4632
Nam8
Snu66
Zds1
Hsl1
Mih1
Crn1
Abp1
Rsc9
Arp9
Las17
Sup35
Dom34
Prm15
Cof1
Srv2
Cyr1
Act1
Sla2
Swa2
Sky1
Ste7
Apl1
Apl2
Sto1
Sgv1
Fus3
Rpn10
Sec23
Sly1
Sec18
Syp1
Vrp1
Sla1
Rps9a
Mtr4
Rps8a
Fun12
Rpn1
Duf1
Rad53
Art5
Ecm21
Ldb19
Yhr131c
Rsp5
Bul2
Ent1
Ent5
Apl3
Sec16
Apm4
Akl1
Rpn11
Nip1
Prt1
Rpg1
Ded1
Tif35
Cdc9
Chk1
Rad23
Ubi4
Ede1
Ent2
Vps27
Chc1
Ptp3
Stb1
Spa2
Bem2
Bud14
Rck2
Glc7
Ipl1
Cdc5
Dbf4
Pfk26
Tpk1
Gpa2
Ras2
Med6
Rpb10
Paf1
Rpo26
Bdp1
Spt15
Spt5
Rgr1
Srb7
Ssn2
Nop1
Nsr1
Nop56
Snu13
Rps31
Rad9
Rtf1
Rpo21
Npl3
Air2
Fps1
Ask10
Cdc28
Hog1
Pbs2
Ssk2
Clb2
Bud3
Bud4
Msn2
Sch9
Rim15
Yak1
Ypl247c
Vps52
Kss1
Sko1
Slt2
Msg5
Swe1
Igo2
Cdc55
Igo1
Vps53
Vps54
Rtn1
Fmp42
Cot1
Zrc1
10
20
30
40
50
t/s

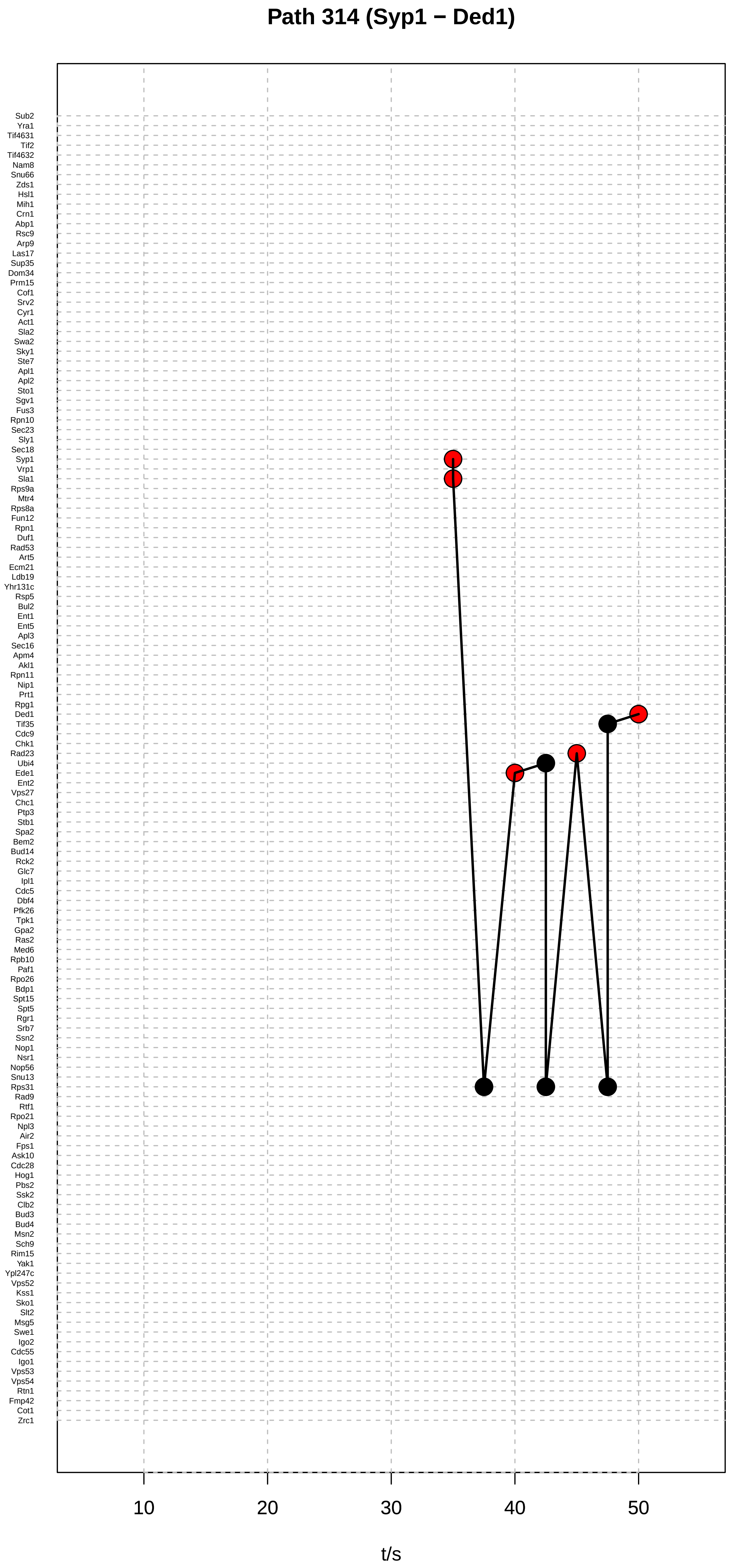

Path 314 (Syp1 − Ded1)
Sub2
Yra1
Tif4631
Tif2
Tif4632
Nam8
Snu66
Zds1
Hsl1
Mih1
Crn1
Abp1
Rsc9
Arp9
Las17
Sup35
Dom34
Prm15
Cof1
Srv2
Cyr1
Act1
Sla2
Swa2
Sky1
Ste7
Apl1
Apl2
Sto1
Sgv1
Fus3
Rpn10
Sec23
Sly1
Sec18
Syp1
Vrp1
Sla1
Rps9a
Mtr4
Rps8a
Fun12
Rpn1
Duf1
Rad53
Art5
Ecm21
Ldb19
Yhr131c
Rsp5
Bul2
Ent1
Ent5
Apl3
Sec16
Apm4
Akl1
Rpn11
Nip1
Prt1
Rpg1
Ded1
Tif35
Cdc9
Chk1
Rad23
Ubi4
Ede1
Ent2
Vps27
Chc1
Ptp3
Stb1
Spa2
Bem2
Bud14
Rck2
Glc7
Ipl1
Cdc5
Dbf4
Pfk26
Tpk1
Gpa2
Ras2
Med6
Rpb10
Paf1
Rpo26
Bdp1
Spt15
Spt5
Rgr1
Srb7
Ssn2
Nop1
Nsr1
Nop56
Snu13
Rps31
Rad9
Rtf1
Rpo21
Npl3
Air2
Fps1
Ask10
Cdc28
Hog1
Pbs2
Ssk2
Clb2
Bud3
Bud4
Msn2
Sch9
Rim15
Yak1
Ypl247c
Vps52
Kss1
Sko1
Slt2
Msg5
Swe1
Igo2
Cdc55
Igo1
Vps53
Vps54
Rtn1
Fmp42
Cot1
Zrc1
10
20
30
40
50
t/s

#### Path 315 (Syp1 − Ded1)

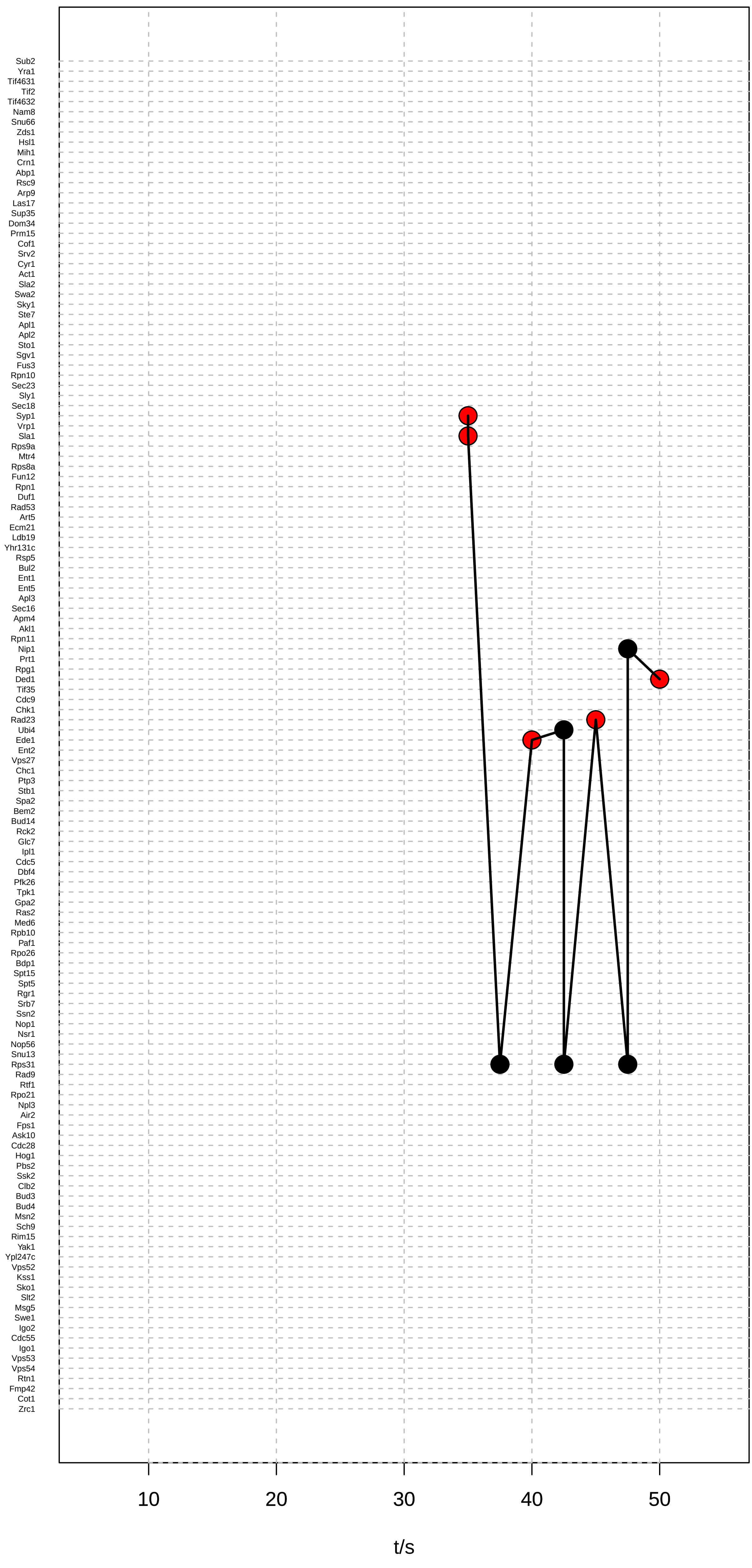

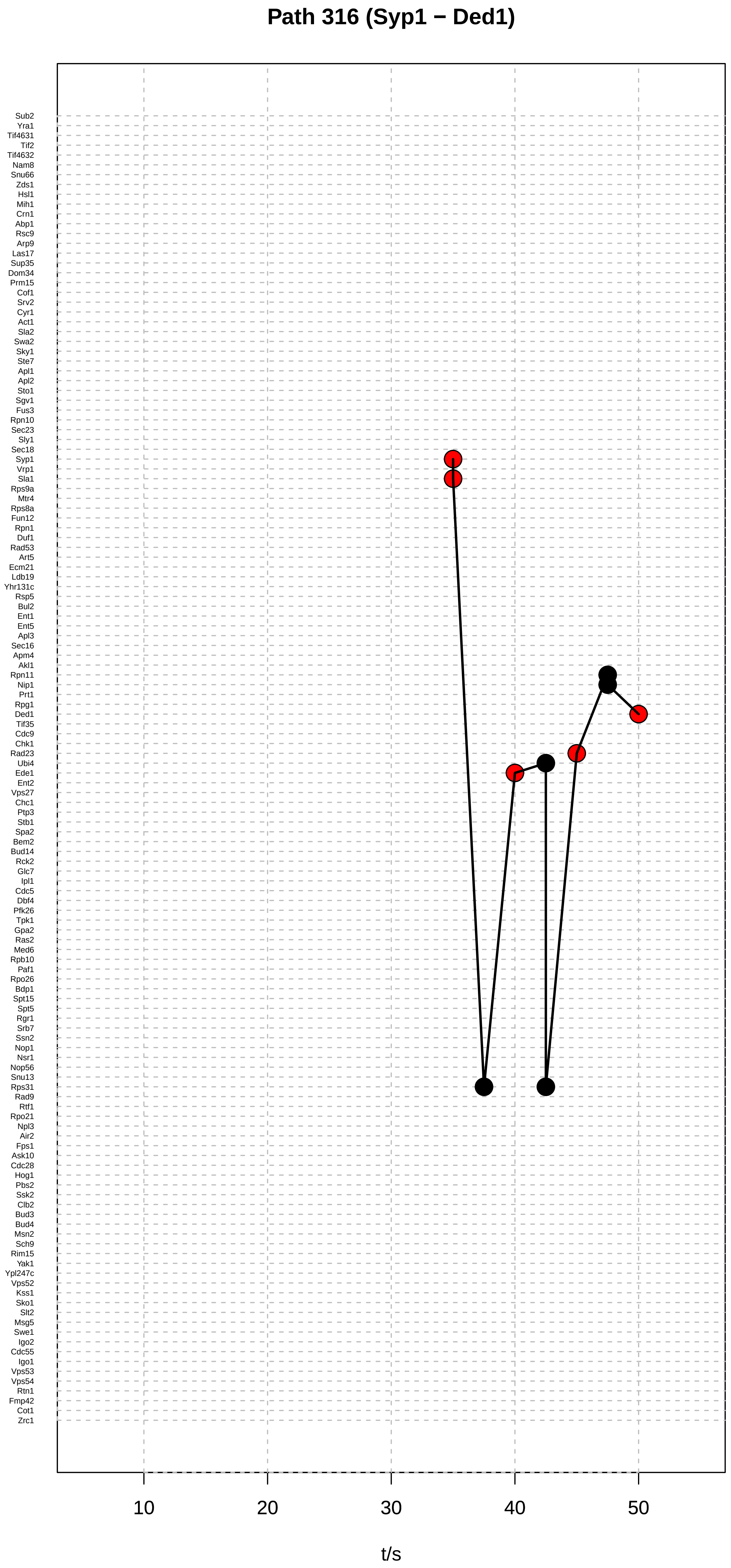
Path 316 (Syp1 − Ded1)
Sub2
Yra1
Tif4631
Tif2
Tif4632
Nam8
Snu66
Zds1
Hsl1
Mih1
Crn1
Abp1
Rsc9
Arp9
Las17
Sup35
Dom34
Prm15
Cof1
Srv2
Cyr1
Act1
Sla2
Swa2
Sky1
Ste7
Apl1
Apl2
Sto1
Sgv1
Fus3
Rpn10
Sec23
Sly1
Sec18
Syp1
Vrp1
Sla1
Rps9a
Mtr4
Rps8a
Fun12
Rpn1
Duf1
Rad53
Art5
Ecm21
Ldb19
Yhr131c
Rsp5
Bul2
Ent1
Ent5
Apl3
Sec16
Apm4
Akl1
Rpn11
Nip1
Prt1
Rpg1
Ded1
Tif35
Cdc9
Chk1
Rad23
Ubi4
Ede1
Ent2
Vps27
Chc1
Ptp3
Stb1
Spa2
Bem2
Bud14
Rck2
Glc7
Ipl1
Cdc5
Dbf4
Pfk26
Tpk1
Gpa2
Ras2
Med6
Rpb10
Paf1
Rpo26
Bdp1
Spt15
Spt5
Rgr1
Srb7
Ssn2
Nop1
Nsr1
Nop56
Snu13
Rps31
Rad9
Rtf1
Rpo21
Npl3
Air2
Fps1
Ask10
Cdc28
Hog1
Pbs2
Ssk2
Clb2
Bud3
Bud4
Msn2
Sch9
Rim15
Yak1
Ypl247c
Vps52
Kss1
Sko1
Slt2
Msg5
Swe1
Igo2
Cdc55
Igo1
Vps53
Vps54
Rtn1
Fmp42
Cot1
Zrc1
10
20
30
40
50
t/s

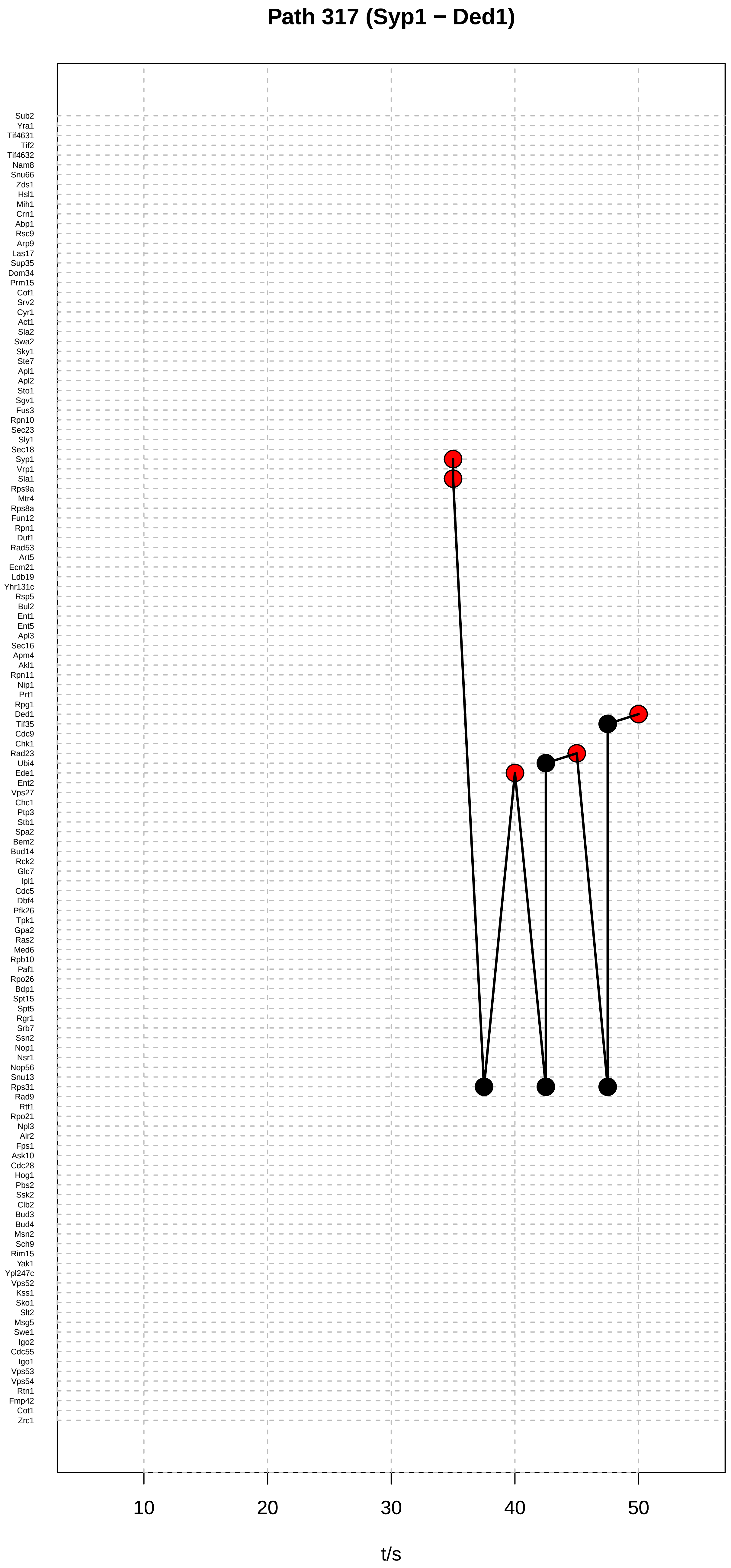
Path 317 (Syp1 − Ded1)
Sub2
Yra1
Tif4631
Tif2
Tif4632
Nam8
Snu66
Zds1
Hsl1
Mih1
Crn1
Abp1
Rsc9
Arp9
Las17
Sup35
Dom34
Prm15
Cof1
Srv2
Cyr1
Act1
Sla2
Swa2
Sky1
Ste7
Apl1
Apl2
Sto1
Sgv1
Fus3
Rpn10
Sec23
Sly1
Sec18
Syp1
Vrp1
Sla1
Rps9a
Mtr4
Rps8a
Fun12
Rpn1
Duf1
Rad53
Art5
Ecm21
Ldb19
Yhr131c
Rsp5
Bul2
Ent1
Ent5
Apl3
Sec16
Apm4
Akl1
Rpn11
Nip1
Prt1
Rpg1
Ded1
Tif35
Cdc9
Chk1
Rad23
Ubi4
Ede1
Ent2
Vps27
Chc1
Ptp3
Stb1
Spa2
Bem2
Bud14
Rck2
Glc7
Ipl1
Cdc5
Dbf4
Pfk26
Tpk1
Gpa2
Ras2
Med6
Rpb10
Paf1
Rpo26
Bdp1
Spt15
Spt5
Rgr1
Srb7
Ssn2
Nop1
Nsr1
Nop56
Snu13
Rps31
Rad9
Rtf1
Rpo21
Npl3
Air2
Fps1
Ask10
Cdc28
Hog1
Pbs2
Ssk2
Clb2
Bud3
Bud4
Msn2
Sch9
Rim15
Yak1
Ypl247c
Vps52
Kss1
Sko1
Slt2
Msg5
Swe1
Igo2
Cdc55
Igo1
Vps53
Vps54
Rtn1
Fmp42
Cot1
Zrc1
10
20
30
40
50
t/s

#### Path 318 (Syp1 − Ded1)

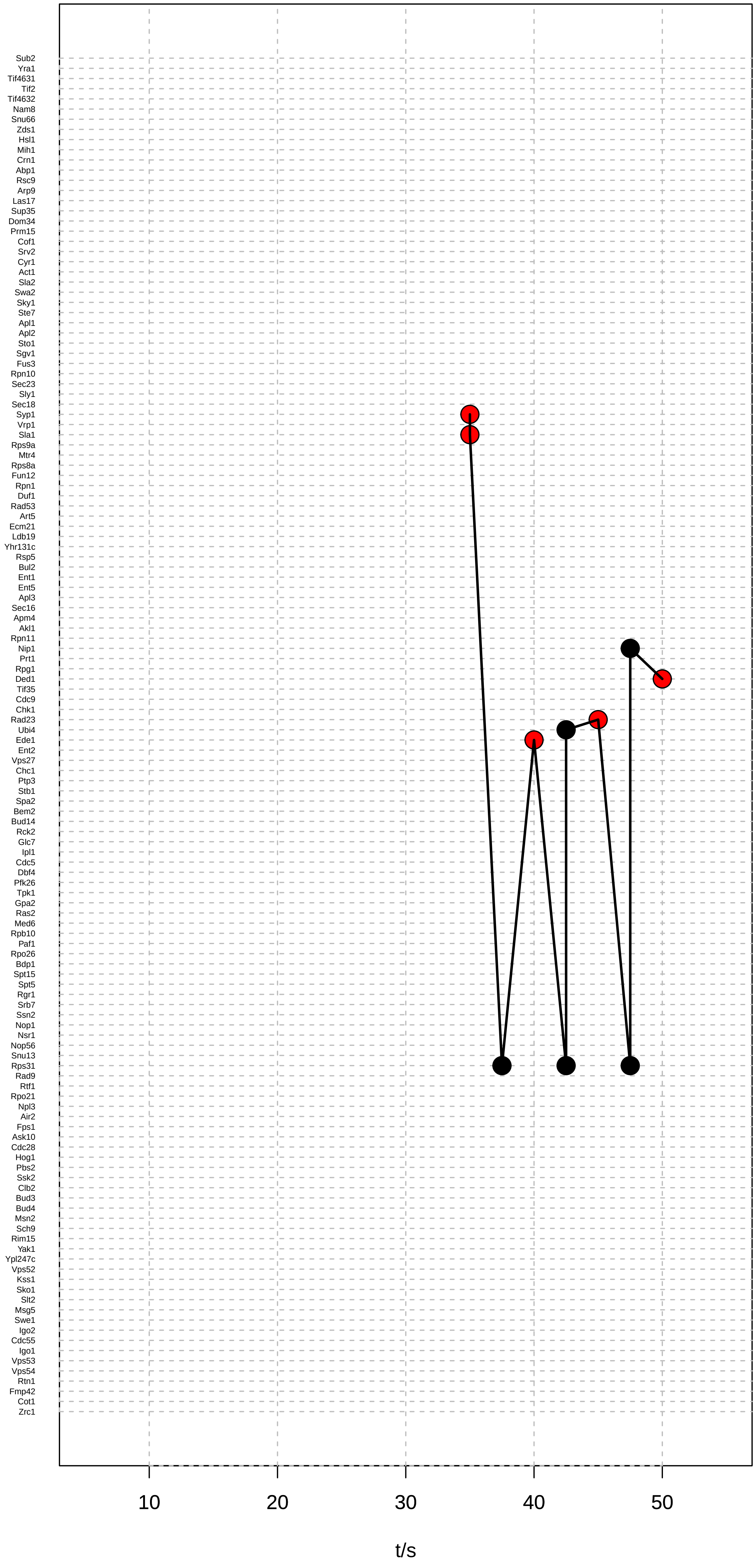

#### Path 319 (Syp1 − Ded1)

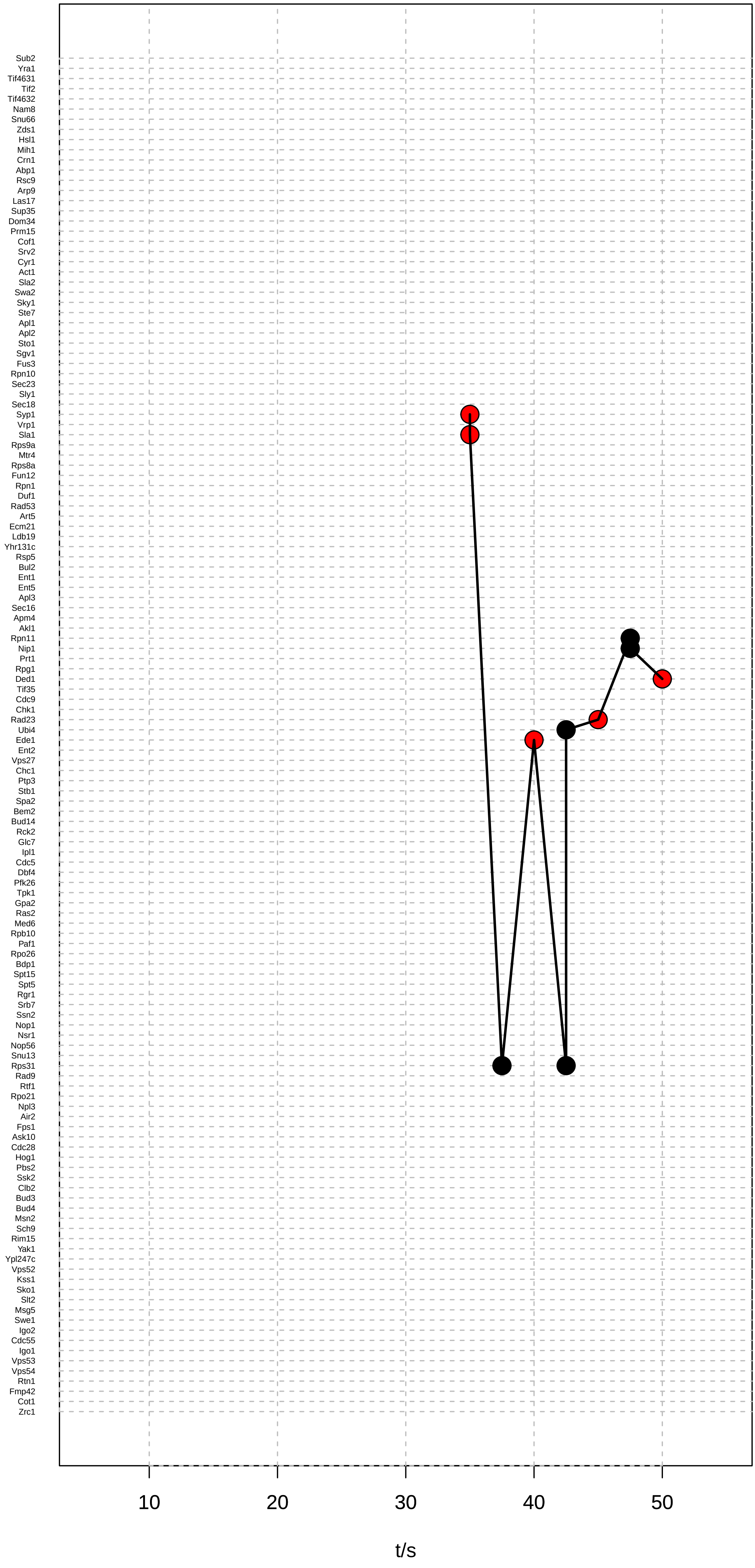

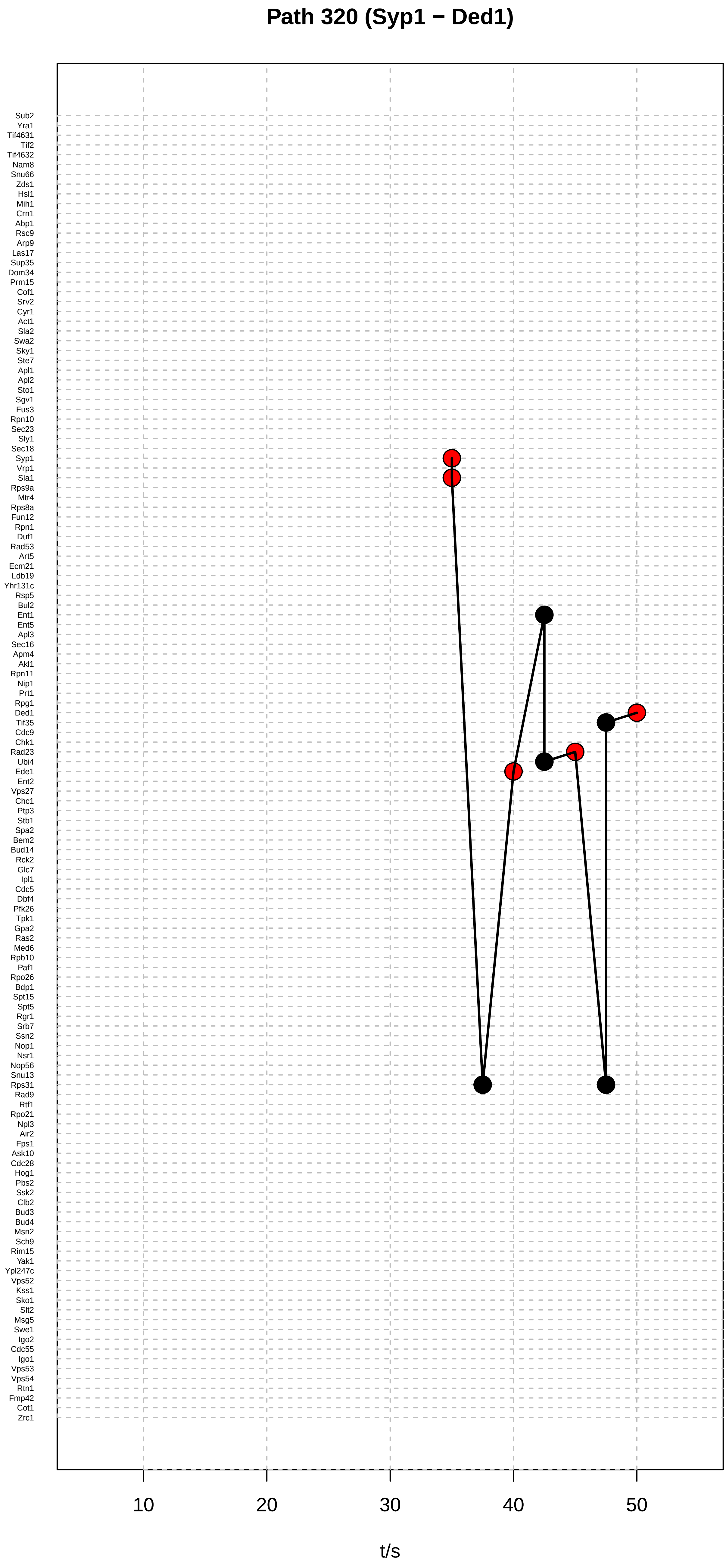

Path 320 (Syp1 − Ded1)
Sub2
Yra1
Tif4631
Tif2
Tif4632
Nam8
Snu66
Zds1
Hsl1
Mih1
Crn1
Abp1
Rsc9
Arp9
Las17
Sup35
Dom34
Prm15
Cof1
Srv2
Cyr1
Act1
Sla2
Swa2
Sky1
Ste7
Apl1
Apl2
Sto1
Sgv1
Fus3
Rpn10
Sec23
Sly1
Sec18
Syp1
Vrp1
Sla1
Rps9a
Mtr4
Rps8a
Fun12
Rpn1
Duf1
Rad53
Art5
Ecm21
Ldb19
Yhr131c
Rsp5
Bul2
Ent1
Ent5
Apl3
Sec16
Apm4
Akl1
Rpn11
Nip1
Prt1
Rpg1
Ded1
Tif35
Cdc9
Chk1
Rad23
Ubi4
Ede1
Ent2
Vps27
Chc1
Ptp3
Stb1
Spa2
Bem2
Bud14
Rck2
Glc7
Ipl1
Cdc5
Dbf4
Pfk26
Tpk1
Gpa2
Ras2
Med6
Rpb10
Paf1
Rpo26
Bdp1
Spt15
Spt5
Rgr1
Srb7
Ssn2
Nop1
Nsr1
Nop56
Snu13
Rps31
Rad9
Rtf1
Rpo21
Npl3
Air2
Fps1
Ask10
Cdc28
Hog1
Pbs2
Ssk2
Clb2
Bud3
Bud4
Msn2
Sch9
Rim15
Yak1
Ypl247c
Vps52
Kss1
Sko1
Slt2
Msg5
Swe1
Igo2
Cdc55
Igo1
Vps53
Vps54
Rtn1
Fmp42
Cot1
Zrc1
10
20
30
40
50
t/s

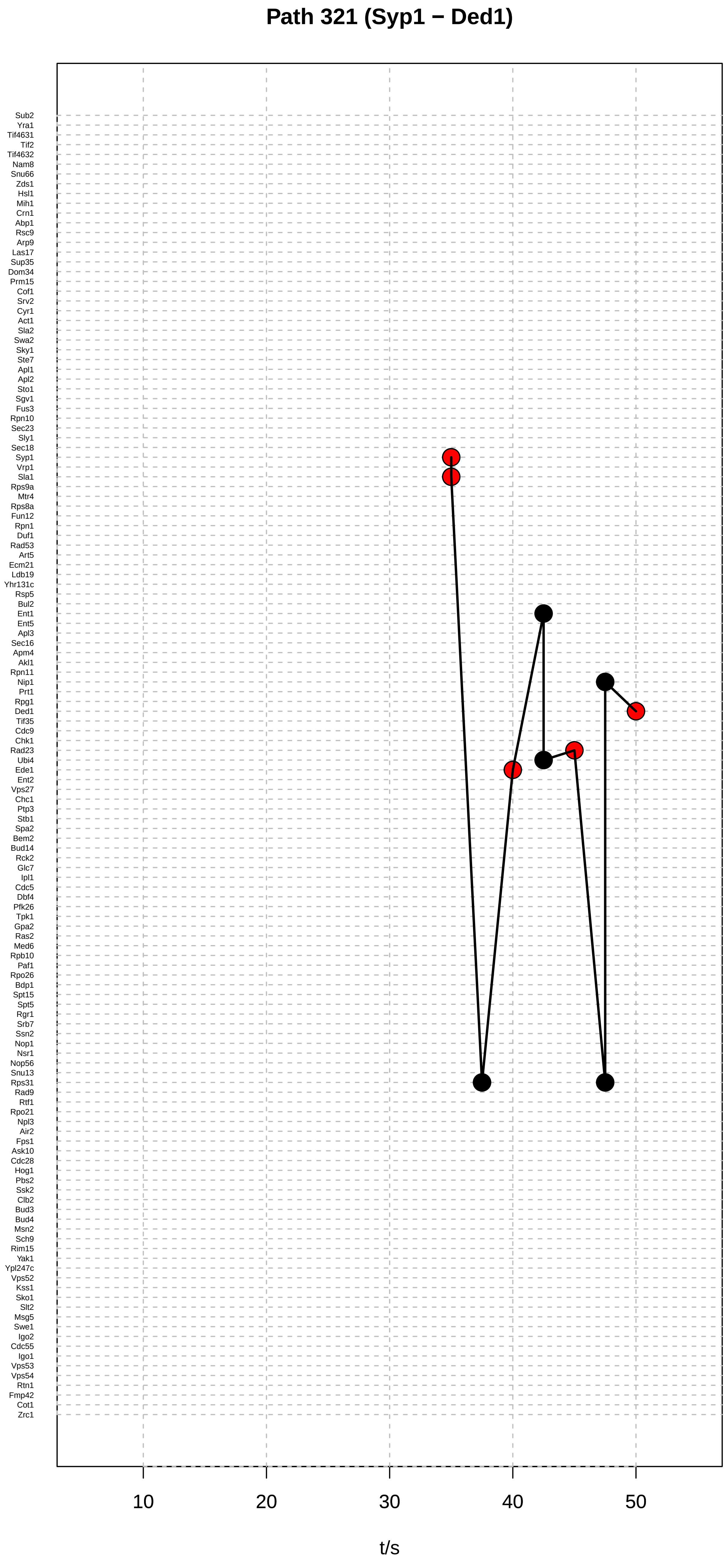

Path 321 (Syp1 − Ded1)
Sub2
Yra1
Tif4631
Tif2
Tif4632
Nam8
Snu66
Zds1
Hsl1
Mih1
Crn1
Abp1
Rsc9
Arp9
Las17
Sup35
Dom34
Prm15
Cof1
Srv2
Cyr1
Act1
Sla2
Swa2
Sky1
Ste7
Apl1
Apl2
Sto1
Sgv1
Fus3
Rpn10
Sec23
Sly1
Sec18
Syp1
Vrp1
Sla1
Rps9a
Mtr4
Rps8a
Fun12
Rpn1
Duf1
Rad53
Art5
Ecm21
Ldb19
Yhr131c
Rsp5
Bul2
Ent1
Ent5
Apl3
Sec16
Apm4
Akl1
Rpn11
Nip1
Prt1
Rpg1
Ded1
Tif35
Cdc9
Chk1
Rad23
Ubi4
Ede1
Ent2
Vps27
Chc1
Ptp3
Stb1
Spa2
Bem2
Bud14
Rck2
Glc7
Ipl1
Cdc5
Dbf4
Pfk26
Tpk1
Gpa2
Ras2
Med6
Rpb10
Paf1
Rpo26
Bdp1
Spt15
Spt5
Rgr1
Srb7
Ssn2
Nop1
Nsr1
Nop56
Snu13
Rps31
Rad9
Rtf1
Rpo21
Npl3
Air2
Fps1
Ask10
Cdc28
Hog1
Pbs2
Ssk2
Clb2
Bud3
Bud4
Msn2
Sch9
Rim15
Yak1
Ypl247c
Vps52
Kss1
Sko1
Slt2
Msg5
Swe1
Igo2
Cdc55
Igo1
Vps53
Vps54
Rtn1
Fmp42
Cot1
Zrc1
10
20
30
40
50
t/s

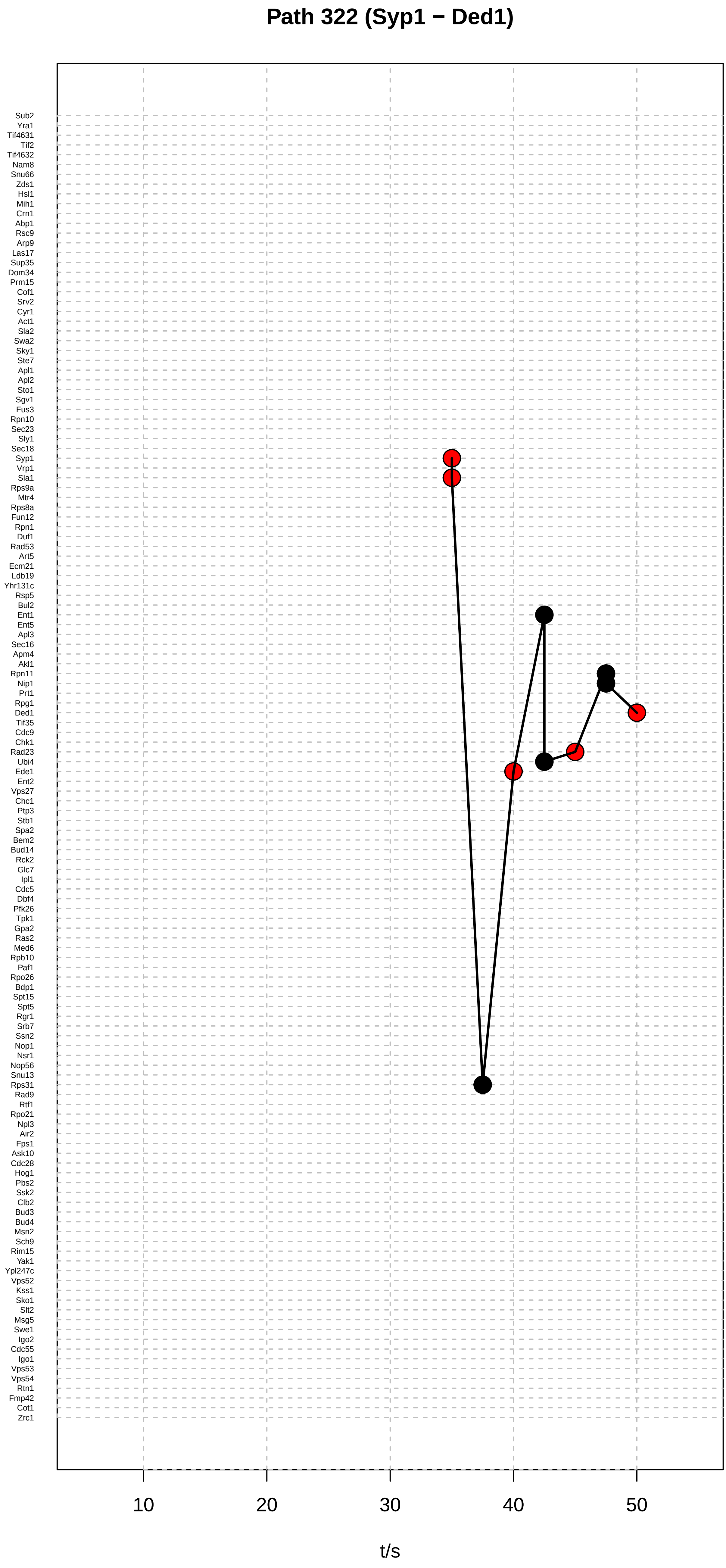
Path 322 (Syp1 − Ded1)
Sub2
Yra1
Tif4631
Tif2
Tif4632
Nam8
Snu66
Zds1
Hsl1
Mih1
Crn1
Abp1
Rsc9
Arp9
Las17
Sup35
Dom34
Prm15
Cof1
Srv2
Cyr1
Act1
Sla2
Swa2
Sky1
Ste7
Apl1
Apl2
Sto1
Sgv1
Fus3
Rpn10
Sec23
Sly1
Sec18
Syp1
Vrp1
Sla1
Rps9a
Mtr4
Rps8a
Fun12
Rpn1
Duf1
Rad53
Art5
Ecm21
Ldb19
Yhr131c
Rsp5
Bul2
Ent1
Ent5
Apl3
Sec16
Apm4
Akl1
Rpn11
Nip1
Prt1
Rpg1
Ded1
Tif35
Cdc9
Chk1
Rad23
Ubi4
Ede1
Ent2
Vps27
Chc1
Ptp3
Stb1
Spa2
Bem2
Bud14
Rck2
Glc7
Ipl1
Cdc5
Dbf4
Pfk26
Tpk1
Gpa2
Ras2
Med6
Rpb10
Paf1
Rpo26
Bdp1
Spt15
Spt5
Rgr1
Srb7
Ssn2
Nop1
Nsr1
Nop56
Snu13
Rps31
Rad9
Rtf1
Rpo21
Npl3
Air2
Fps1
Ask10
Cdc28
Hog1
Pbs2
Ssk2
Clb2
Bud3
Bud4
Msn2
Sch9
Rim15
Yak1
Ypl247c
Vps52
Kss1
Sko1
Slt2
Msg5
Swe1
Igo2
Cdc55
Igo1
Vps53
Vps54
Rtn1
Fmp42
Cot1
Zrc1
10
20
30
40
50
t/s

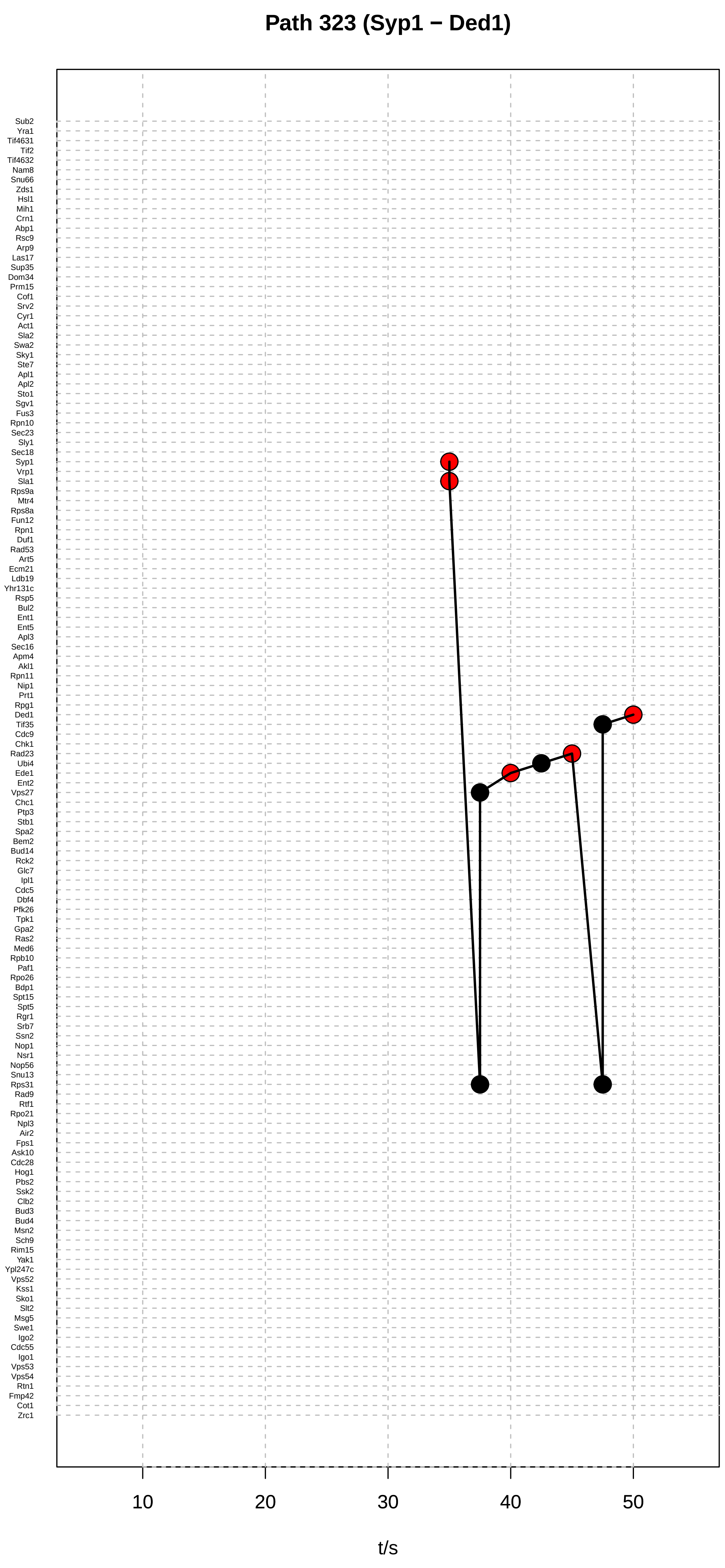
Path 323 (Syp1 − Ded1)
Sub2
Yra1
Tif4631
Tif2
Tif4632
Nam8
Snu66
Zds1
Hsl1
Mih1
Crn1
Abp1
Rsc9
Arp9
Las17
Sup35
Dom34
Prm15
Cof1
Srv2
Cyr1
Act1
Sla2
Swa2
Sky1
Ste7
Apl1
Apl2
Sto1
Sgv1
Fus3
Rpn10
Sec23
Sly1
Sec18
Syp1
Vrp1
Sla1
Rps9a
Mtr4
Rps8a
Fun12
Rpn1
Duf1
Rad53
Art5
Ecm21
Ldb19
Yhr131c
Rsp5
Bul2
Ent1
Ent5
Apl3
Sec16
Apm4
Akl1
Rpn11
Nip1
Prt1
Rpg1
Ded1
Tif35
Cdc9
Chk1
Rad23
Ubi4
Ede1
Ent2
Vps27
Chc1
Ptp3
Stb1
Spa2
Bem2
Bud14
Rck2
Glc7
Ipl1
Cdc5
Dbf4
Pfk26
Tpk1
Gpa2
Ras2
Med6
Rpb10
Paf1
Rpo26
Bdp1
Spt15
Spt5
Rgr1
Srb7
Ssn2
Nop1
Nsr1
Nop56
Snu13
Rps31
Rad9
Rtf1
Rpo21
Npl3
Air2
Fps1
Ask10
Cdc28
Hog1
Pbs2
Ssk2
Clb2
Bud3
Bud4
Msn2
Sch9
Rim15
Yak1
Ypl247c
Vps52
Kss1
Sko1
Slt2
Msg5
Swe1
Igo2
Cdc55
Igo1
Vps53
Vps54
Rtn1
Fmp42
Cot1
Zrc1
10
20
30
40
50
t/s

### Path 324 (Syp1 − Ded1)

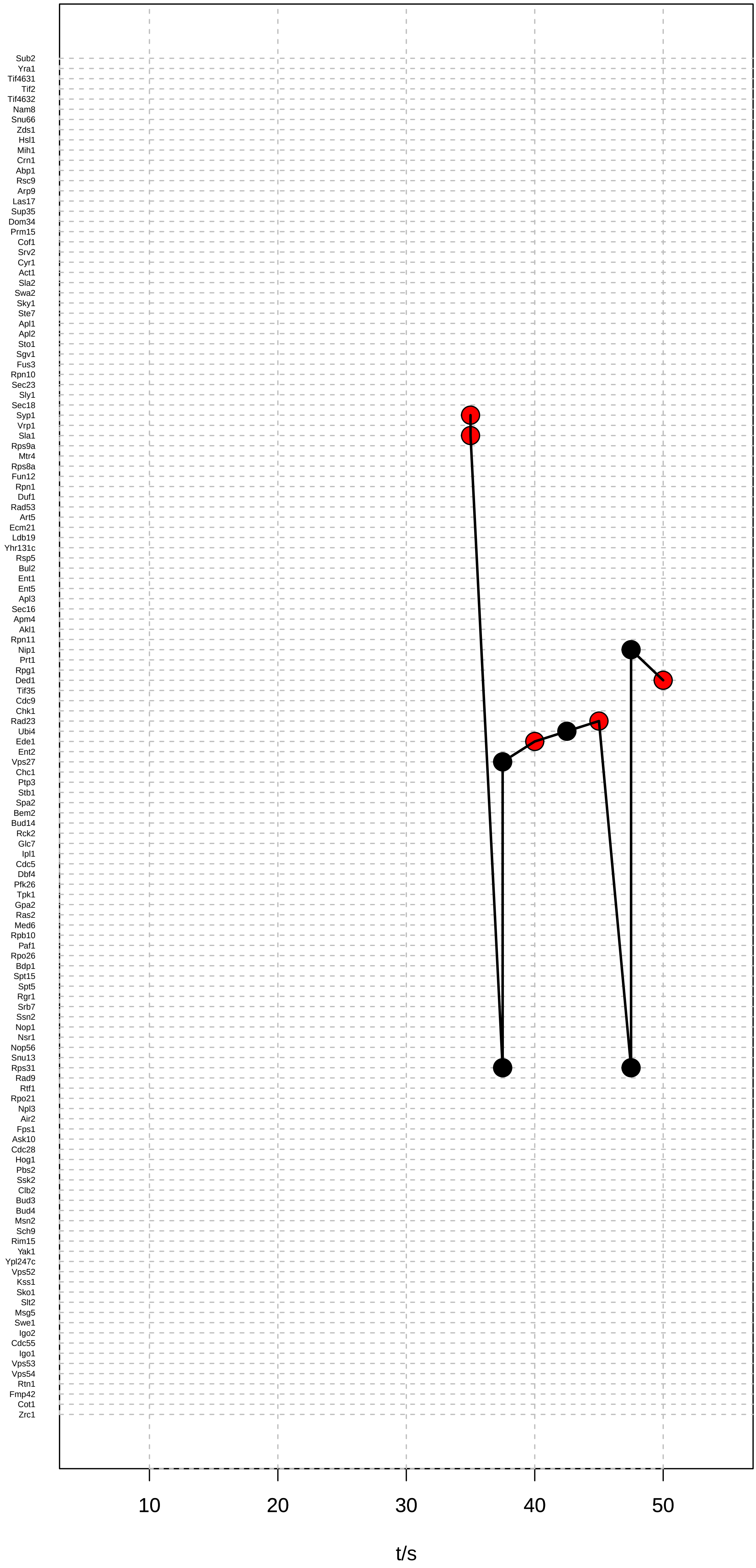

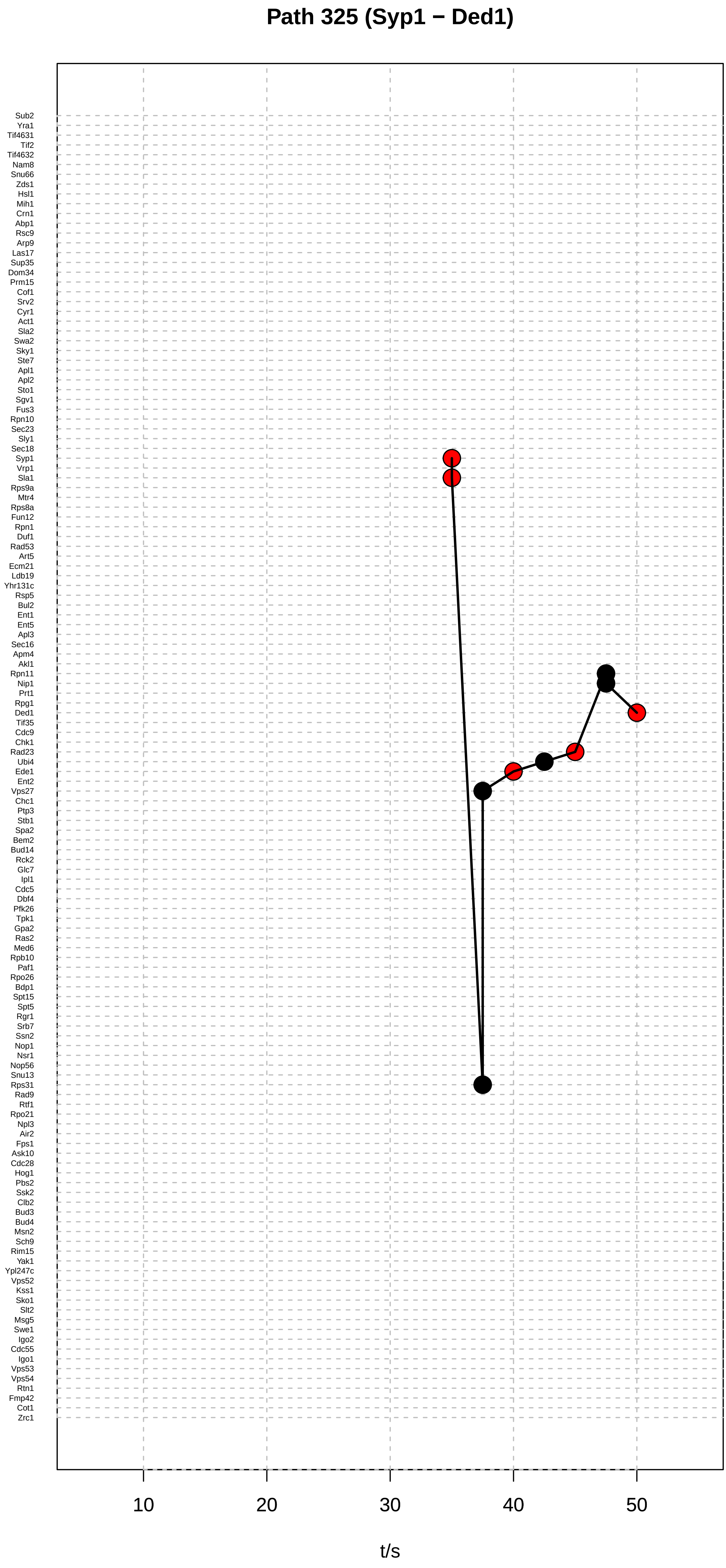
Path 325 (Syp1 − Ded1)
Sub2
Yra1
Tif4631
Tif2
Tif4632
Nam8
Snu66
Zds1
Hsl1
Mih1
Crn1
Abp1
Rsc9
Arp9
Las17
Sup35
Dom34
Prm15
Cof1
Srv2
Cyr1
Act1
Sla2
Swa2
Sky1
Ste7
Apl1
Apl2
Sto1
Sgv1
Fus3
Rpn10
Sec23
Sly1
Sec18
Syp1
Vrp1
Sla1
Rps9a
Mtr4
Rps8a
Fun12
Rpn1
Duf1
Rad53
Art5
Ecm21
Ldb19
Yhr131c
Rsp5
Bul2
Ent1
Ent5
Apl3
Sec16
Apm4
Akl1
Rpn11
Nip1
Prt1
Rpg1
Ded1
Tif35
Cdc9
Chk1
Rad23
Ubi4
Ede1
Ent2
Vps27
Chc1
Ptp3
Stb1
Spa2
Bem2
Bud14
Rck2
Glc7
Ipl1
Cdc5
Dbf4
Pfk26
Tpk1
Gpa2
Ras2
Med6
Rpb10
Paf1
Rpo26
Bdp1
Spt15
Spt5
Rgr1
Srb7
Ssn2
Nop1
Nsr1
Nop56
Snu13
Rps31
Rad9
Rtf1
Rpo21
Npl3
Air2
Fps1
Ask10
Cdc28
Hog1
Pbs2
Ssk2
Clb2
Bud3
Bud4
Msn2
Sch9
Rim15
Yak1
Ypl247c
Vps52
Kss1
Sko1
Slt2
Msg5
Swe1
Igo2
Cdc55
Igo1
Vps53
Vps54
Rtn1
Fmp42
Cot1
Zrc1
10
20
30
40
50
t/s

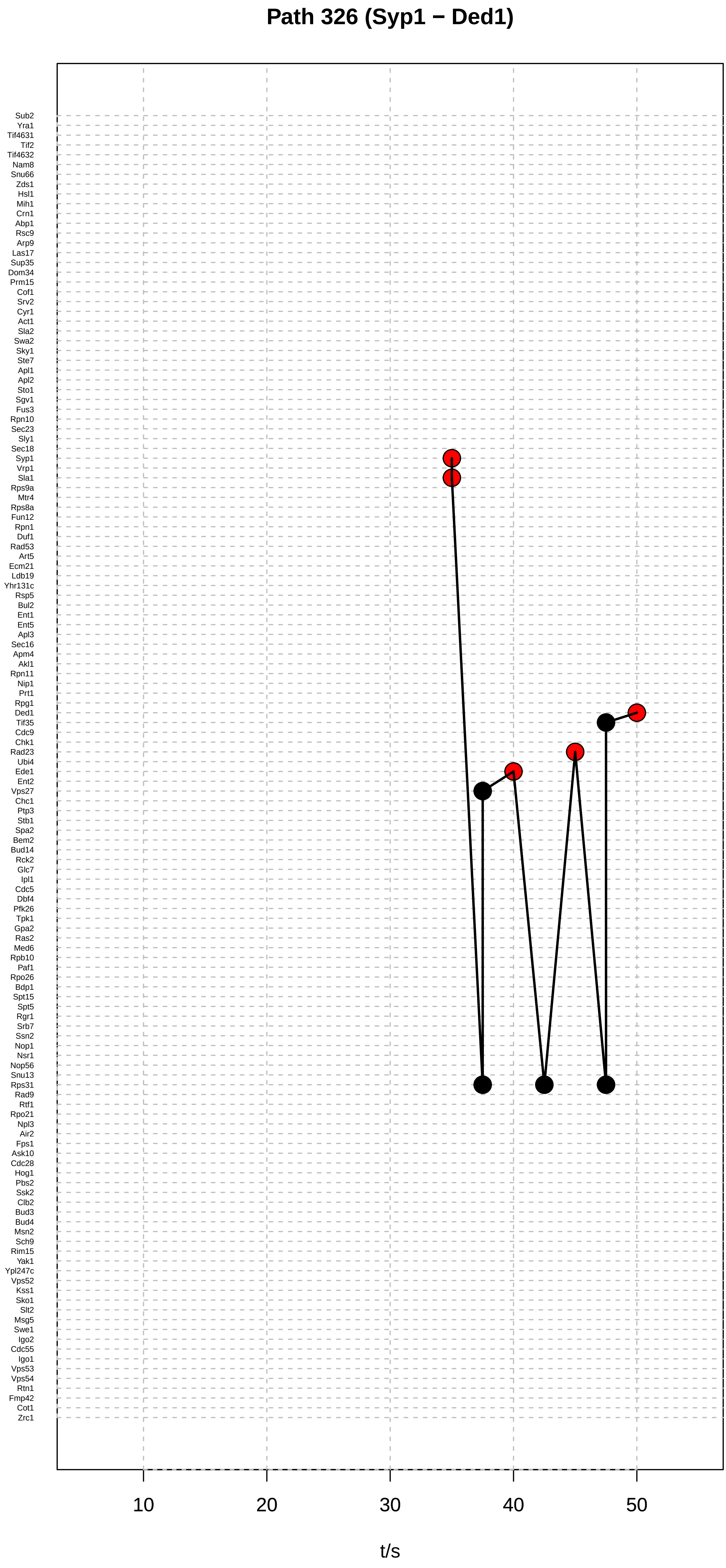

Path 326 (Syp1 − Ded1)
Sub2
Yra1
Tif4631
Tif2
Tif4632
Nam8
Snu66
Zds1
Hsl1
Mih1
Crn1
Abp1
Rsc9
Arp9
Las17
Sup35
Dom34
Prm15
Cof1
Srv2
Cyr1
Act1
Sla2
Swa2
Sky1
Ste7
Apl1
Apl2
Sto1
Sgv1
Fus3
Rpn10
Sec23
Sly1
Sec18
Syp1
Vrp1
Sla1
Rps9a
Mtr4
Rps8a
Fun12
Rpn1
Duf1
Rad53
Art5
Ecm21
Ldb19
Yhr131c
Rsp5
Bul2
Ent1
Ent5
Apl3
Sec16
Apm4
Akl1
Rpn11
Nip1
Prt1
Rpg1
Ded1
Tif35
Cdc9
Chk1
Rad23
Ubi4
Ede1
Ent2
Vps27
Chc1
Ptp3
Stb1
Spa2
Bem2
Bud14
Rck2
Glc7
Ipl1
Cdc5
Dbf4
Pfk26
Tpk1
Gpa2
Ras2
Med6
Rpb10
Paf1
Rpo26
Bdp1
Spt15
Spt5
Rgr1
Srb7
Ssn2
Nop1
Nsr1
Nop56
Snu13
Rps31
Rad9
Rtf1
Rpo21
Npl3
Air2
Fps1
Ask10
Cdc28
Hog1
Pbs2
Ssk2
Clb2
Bud3
Bud4
Msn2
Sch9
Rim15
Yak1
Ypl247c
Vps52
Kss1
Sko1
Slt2
Msg5
Swe1
Igo2
Cdc55
Igo1
Vps53
Vps54
Rtn1
Fmp42
Cot1
Zrc1
10
20
30
40
50
t/s

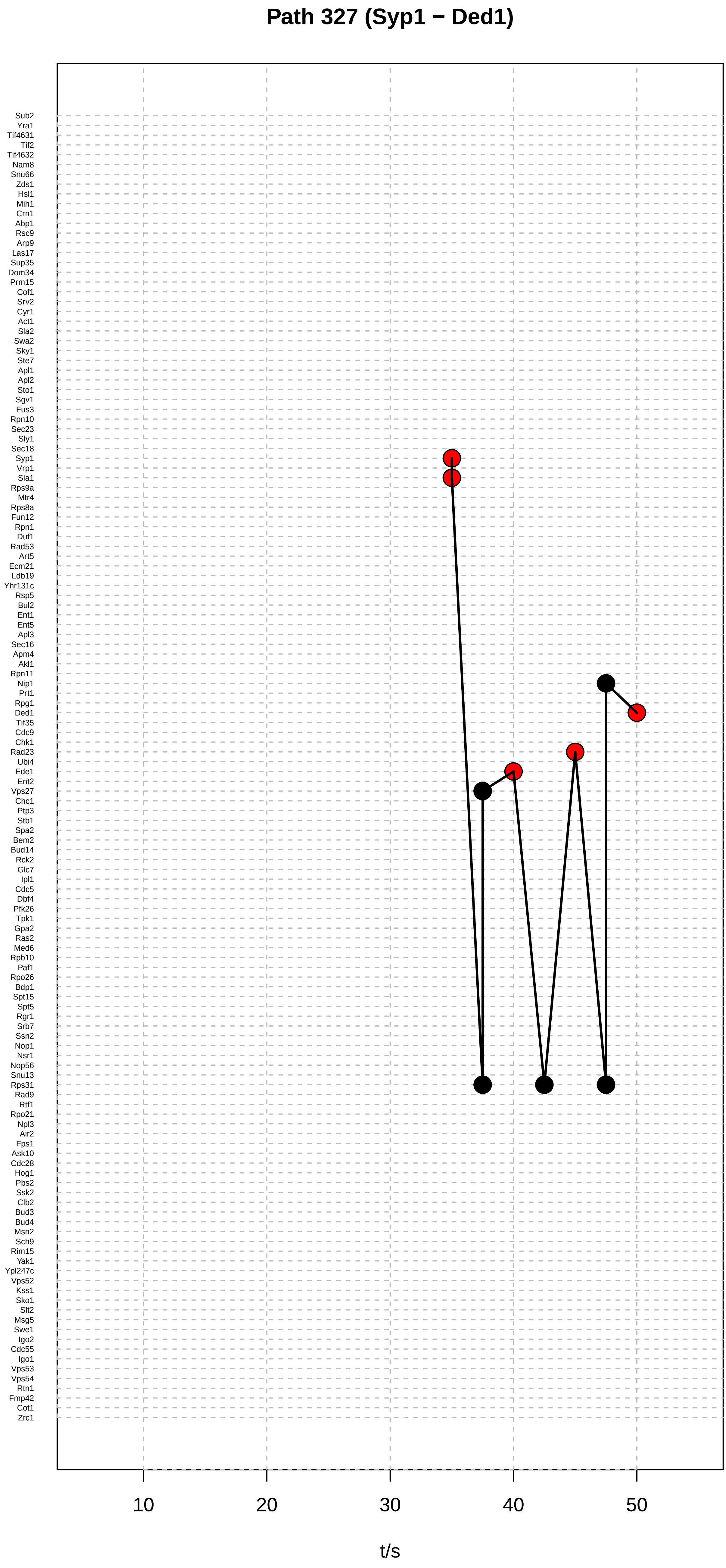

Path 327 (Syp1 − Ded1)
Sub2
Yra1
Tif4631
Tif2
Tif4632
Nam8
Snu66
Zds1
Hsl1
Mih1
Crn1
Abp1
Rsc9
Arp9
Las17
Sup35
Dom34
Prm15
Cof1
Srv2
Cyr1
Act1
Sla2
Swa2
Sky1
Ste7
Apl1
Apl2
Sto1
Sgv1
Fus3
Rpn10
Sec23
Sly1
Sec18
Syp1
Vrp1
Sla1
Rps9a
Mtr4
Rps8a
Fun12
Rpn1
Duf1
Rad53
Art5
Ecm21
Ldb19
Yhr131c
Rsp5
Bul2
Ent1
Ent5
Apl3
Sec16
Apm4
Akl1
Rpn11
Nip1
Prt1
Rpg1
Ded1
Tif35
Cdc9
Chk1
Rad23
Ubi4
Ede1
Ent2
Vps27
Chc1
Ptp3
Stb1
Spa2
Bem2
Bud14
Rck2
Glc7
Ipl1
Cdc5
Dbf4
Pfk26
Tpk1
Gpa2
Ras2
Med6
Rpb10
Paf1
Rpo26
Bdp1
Spt15
Spt5
Rgr1
Srb7
Ssn2
Nop1
Nsr1
Nop56
Snu13
Rps31
Rad9
Rtf1
Rpo21
Npl3
Air2
Fps1
Ask10
Cdc28
Hog1
Pbs2
Ssk2
Clb2
Bud3
Bud4
Msn2
Sch9
Rim15
Yak1
Ypl247c
Vps52
Kss1
Sko1
Slt2
Msg5
Swe1
Igo2
Cdc55
Igo1
Vps53
Vps54
Rtn1
Fmp42
Cot1
Zrc1
10
20
30
40
50
t/s

### Path 328 (Syp1 − Ded1)

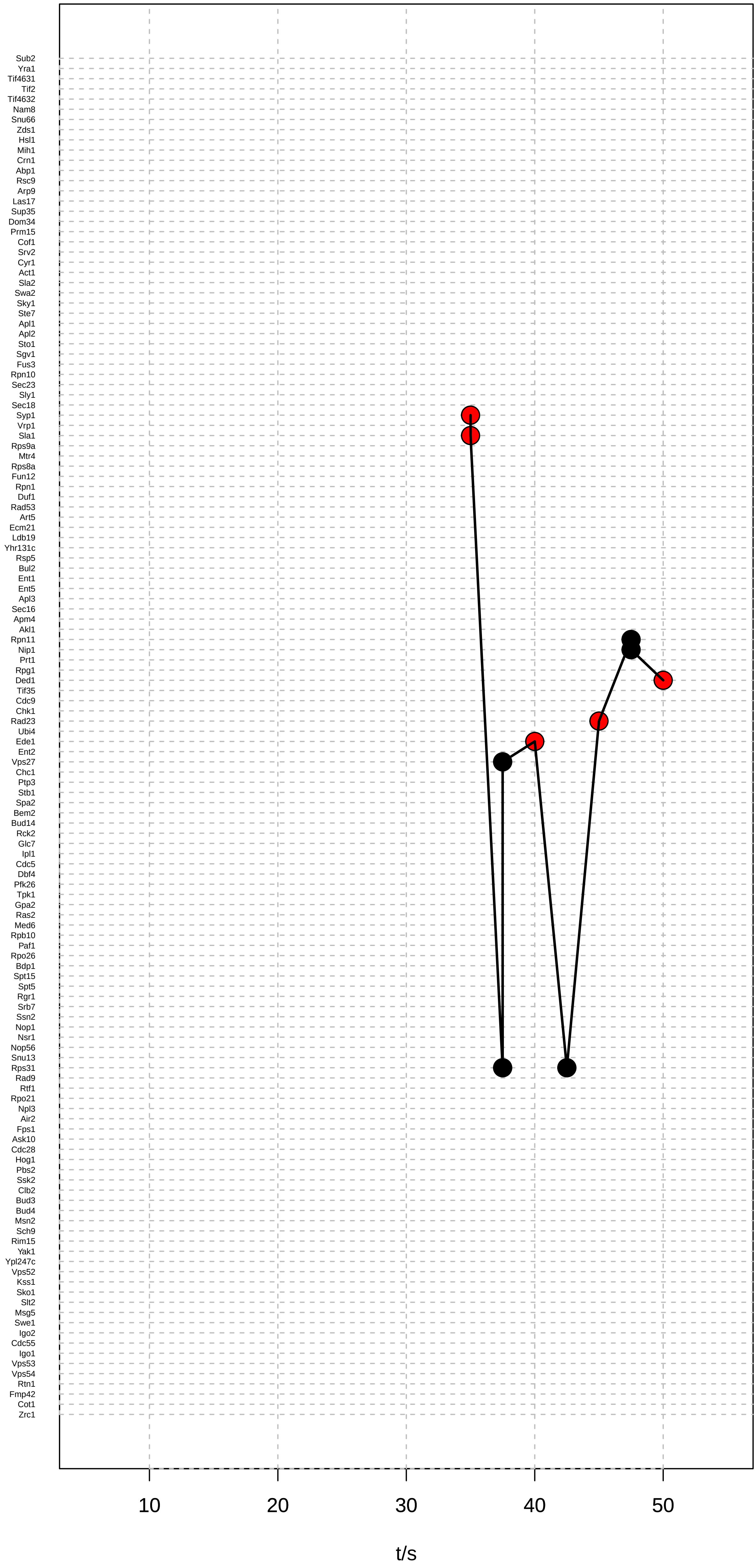

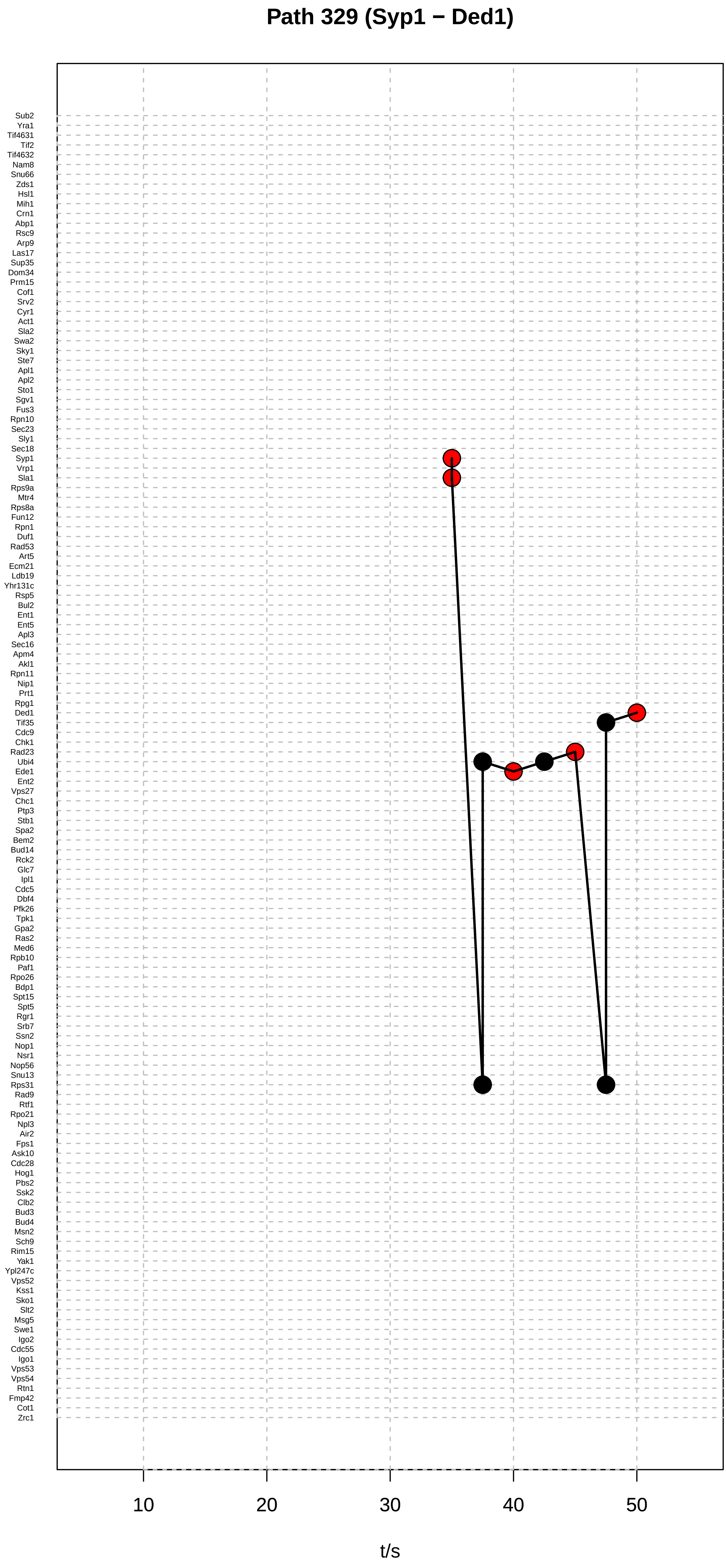

Path 329 (Syp1 − Ded1)
Sub2
Yra1
Tif4631
Tif2
Tif4632
Nam8
Snu66
Zds1
Hsl1
Mih1
Crn1
Abp1
Rsc9
Arp9
Las17
Sup35
Dom34
Prm15
Cof1
Srv2
Cyr1
Act1
Sla2
Swa2
Sky1
Ste7
Apl1
Apl2
Sto1
Sgv1
Fus3
Rpn10
Sec23
Sly1
Sec18
Syp1
Vrp1
Sla1
Rps9a
Mtr4
Rps8a
Fun12
Rpn1
Duf1
Rad53
Art5
Ecm21
Ldb19
Yhr131c
Rsp5
Bul2
Ent1
Ent5
Apl3
Sec16
Apm4
Akl1
Rpn11
Nip1
Prt1
Rpg1
Ded1
Tif35
Cdc9
Chk1
Rad23
Ubi4
Ede1
Ent2
Vps27
Chc1
Ptp3
Stb1
Spa2
Bem2
Bud14
Rck2
Glc7
Ipl1
Cdc5
Dbf4
Pfk26
Tpk1
Gpa2
Ras2
Med6
Rpb10
Paf1
Rpo26
Bdp1
Spt15
Spt5
Rgr1
Srb7
Ssn2
Nop1
Nsr1
Nop56
Snu13
Rps31
Rad9
Rtf1
Rpo21
Npl3
Air2
Fps1
Ask10
Cdc28
Hog1
Pbs2
Ssk2
Clb2
Bud3
Bud4
Msn2
Sch9
Rim15
Yak1
Ypl247c
Vps52
Kss1
Sko1
Slt2
Msg5
Swe1
Igo2
Cdc55
Igo1
Vps53
Vps54
Rtn1
Fmp42
Cot1
Zrc1
10
20
30
40
50
t/s

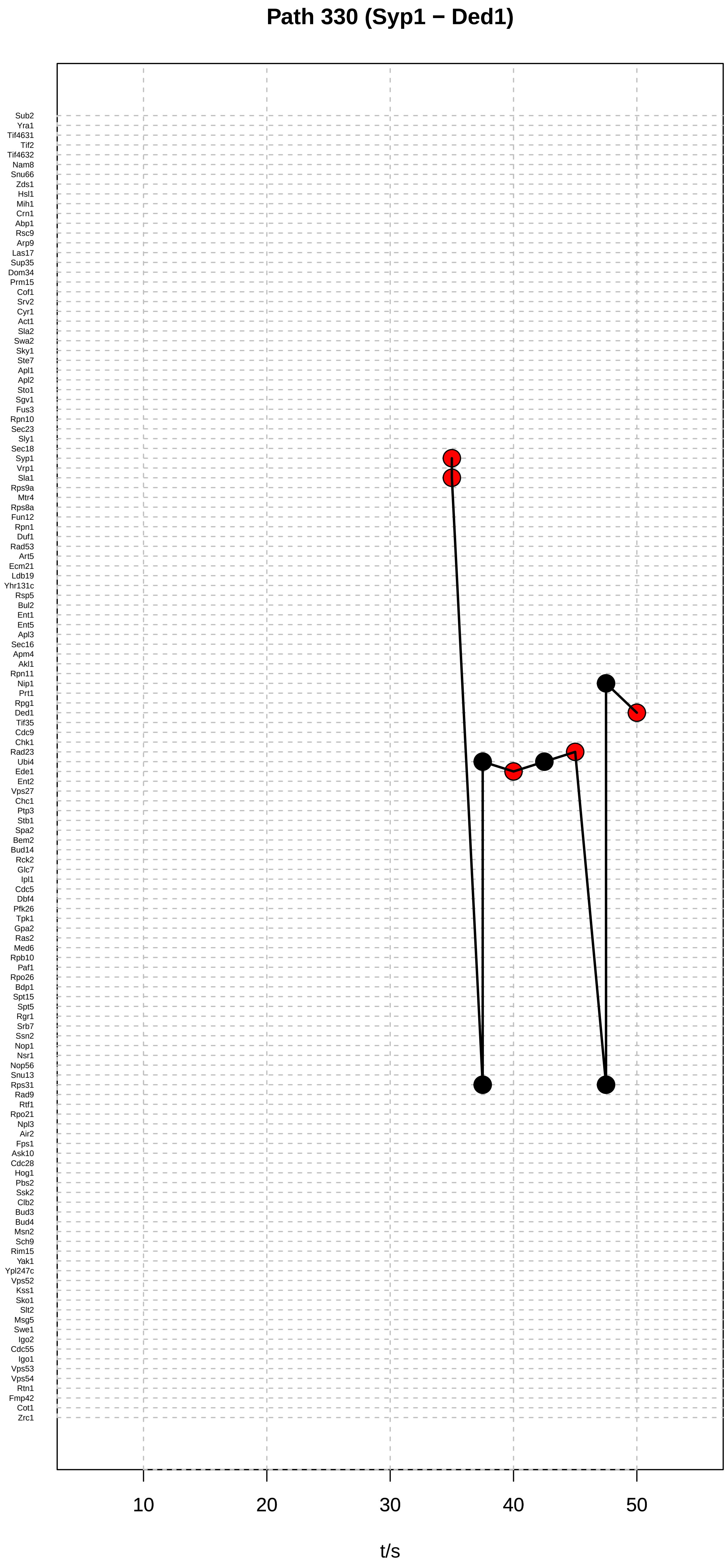

Path 330 (Syp1 − Ded1)
Sub2
Yra1
Tif4631
Tif2
Tif4632
Nam8
Snu66
Zds1
Hsl1
Mih1
Crn1
Abp1
Rsc9
Arp9
Las17
Sup35
Dom34
Prm15
Cof1
Srv2
Cyr1
Act1
Sla2
Swa2
Sky1
Ste7
Apl1
Apl2
Sto1
Sgv1
Fus3
Rpn10
Sec23
Sly1
Sec18
Syp1
Vrp1
Sla1
Rps9a
Mtr4
Rps8a
Fun12
Rpn1
Duf1
Rad53
Art5
Ecm21
Ldb19
Yhr131c
Rsp5
Bul2
Ent1
Ent5
Apl3
Sec16
Apm4
Akl1
Rpn11
Nip1
Prt1
Rpg1
Ded1
Tif35
Cdc9
Chk1
Rad23
Ubi4
Ede1
Ent2
Vps27
Chc1
Ptp3
Stb1
Spa2
Bem2
Bud14
Rck2
Glc7
Ipl1
Cdc5
Dbf4
Pfk26
Tpk1
Gpa2
Ras2
Med6
Rpb10
Paf1
Rpo26
Bdp1
Spt15
Spt5
Rgr1
Srb7
Ssn2
Nop1
Nsr1
Nop56
Snu13
Rps31
Rad9
Rtf1
Rpo21
Npl3
Air2
Fps1
Ask10
Cdc28
Hog1
Pbs2
Ssk2
Clb2
Bud3
Bud4
Msn2
Sch9
Rim15
Yak1
Ypl247c
Vps52
Kss1
Sko1
Slt2
Msg5
Swe1
Igo2
Cdc55
Igo1
Vps53
Vps54
Rtn1
Fmp42
Cot1
Zrc1
10
20
30
40
50
t/s

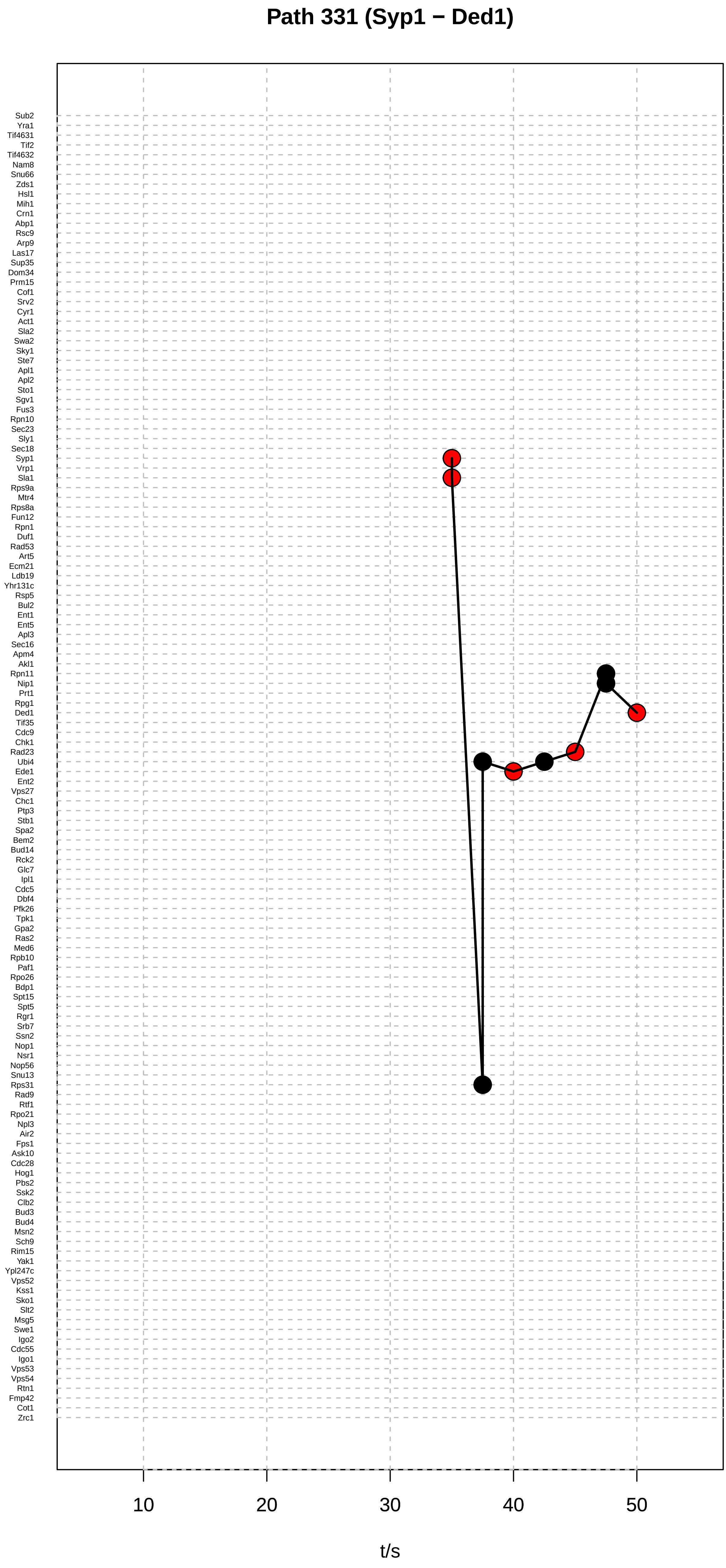

Path 331 (Syp1 − Ded1)
Sub2
Yra1
Tif4631
Tif2
Tif4632
Nam8
Snu66
Zds1
Hsl1
Mih1
Crn1
Abp1
Rsc9
Arp9
Las17
Sup35
Dom34
Prm15
Cof1
Srv2
Cyr1
Act1
Sla2
Swa2
Sky1
Ste7
Apl1
Apl2
Sto1
Sgv1
Fus3
Rpn10
Sec23
Sly1
Sec18
Syp1
Vrp1
Sla1
Rps9a
Mtr4
Rps8a
Fun12
Rpn1
Duf1
Rad53
Art5
Ecm21
Ldb19
Yhr131c
Rsp5
Bul2
Ent1
Ent5
Apl3
Sec16
Apm4
Akl1
Rpn11
Nip1
Prt1
Rpg1
Ded1
Tif35
Cdc9
Chk1
Rad23
Ubi4
Ede1
Ent2
Vps27
Chc1
Ptp3
Stb1
Spa2
Bem2
Bud14
Rck2
Glc7
Ipl1
Cdc5
Dbf4
Pfk26
Tpk1
Gpa2
Ras2
Med6
Rpb10
Paf1
Rpo26
Bdp1
Spt15
Spt5
Rgr1
Srb7
Ssn2
Nop1
Nsr1
Nop56
Snu13
Rps31
Rad9
Rtf1
Rpo21
Npl3
Air2
Fps1
Ask10
Cdc28
Hog1
Pbs2
Ssk2
Clb2
Bud3
Bud4
Msn2
Sch9
Rim15
Yak1
Ypl247c
Vps52
Kss1
Sko1
Slt2
Msg5
Swe1
Igo2
Cdc55
Igo1
Vps53
Vps54
Rtn1
Fmp42
Cot1
Zrc1
10
20
30
40
50
t/s

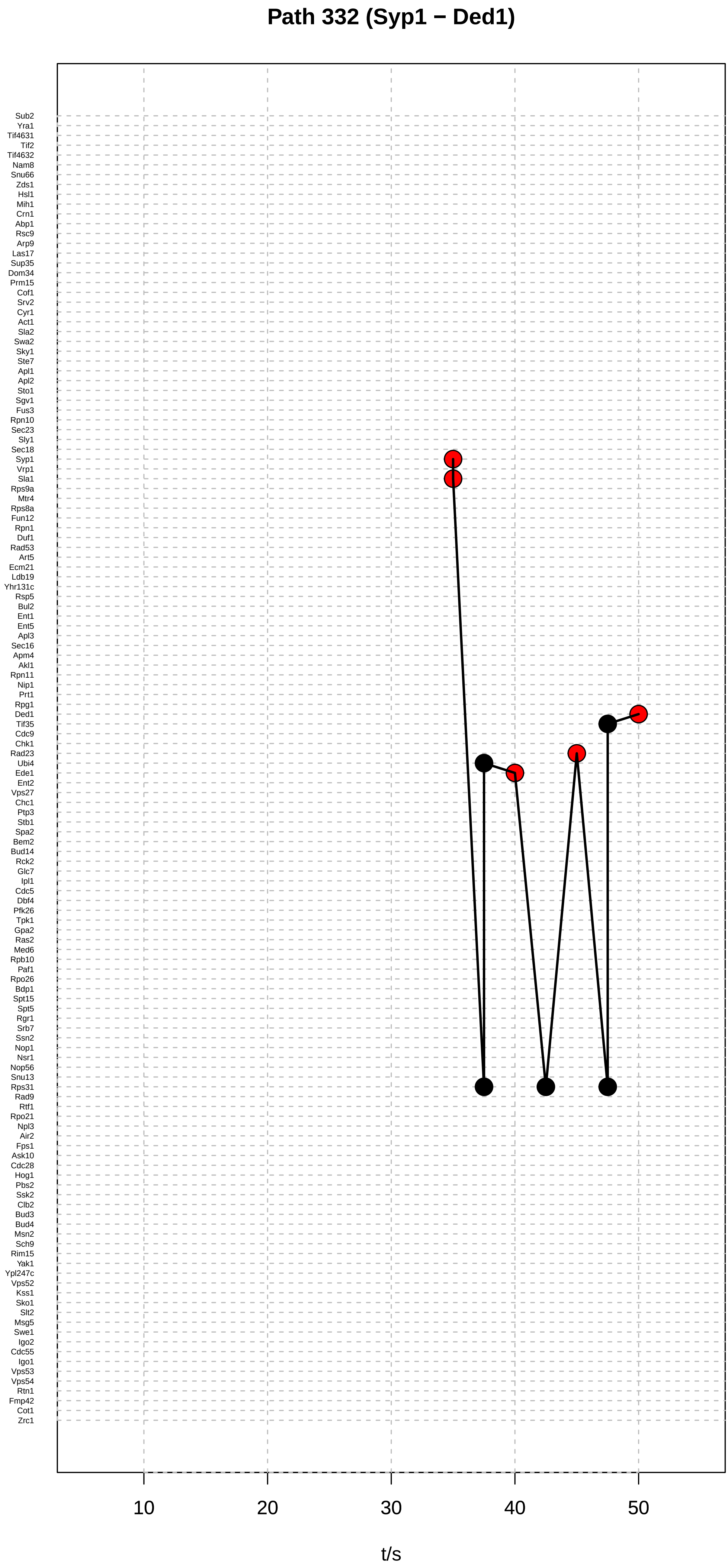

Path 332 (Syp1 − Ded1)
Sub2
Yra1
Tif4631
Tif2
Tif4632
Nam8
Snu66
Zds1
Hsl1
Mih1
Crn1
Abp1
Rsc9
Arp9
Las17
Sup35
Dom34
Prm15
Cof1
Srv2
Cyr1
Act1
Sla2
Swa2
Sky1
Ste7
Apl1
Apl2
Sto1
Sgv1
Fus3
Rpn10
Sec23
Sly1
Sec18
Syp1
Vrp1
Sla1
Rps9a
Mtr4
Rps8a
Fun12
Rpn1
Duf1
Rad53
Art5
Ecm21
Ldb19
Yhr131c
Rsp5
Bul2
Ent1
Ent5
Apl3
Sec16
Apm4
Akl1
Rpn11
Nip1
Prt1
Rpg1
Ded1
Tif35
Cdc9
Chk1
Rad23
Ubi4
Ede1
Ent2
Vps27
Chc1
Ptp3
Stb1
Spa2
Bem2
Bud14
Rck2
Glc7
Ipl1
Cdc5
Dbf4
Pfk26
Tpk1
Gpa2
Ras2
Med6
Rpb10
Paf1
Rpo26
Bdp1
Spt15
Spt5
Rgr1
Srb7
Ssn2
Nop1
Nsr1
Nop56
Snu13
Rps31
Rad9
Rtf1
Rpo21
Npl3
Air2
Fps1
Ask10
Cdc28
Hog1
Pbs2
Ssk2
Clb2
Bud3
Bud4
Msn2
Sch9
Rim15
Yak1
Ypl247c
Vps52
Kss1
Sko1
Slt2
Msg5
Swe1
Igo2
Cdc55
Igo1
Vps53
Vps54
Rtn1
Fmp42
Cot1
Zrc1
10
20
30
40
50
t/s

#### Path 333 (Syp1 − Ded1)

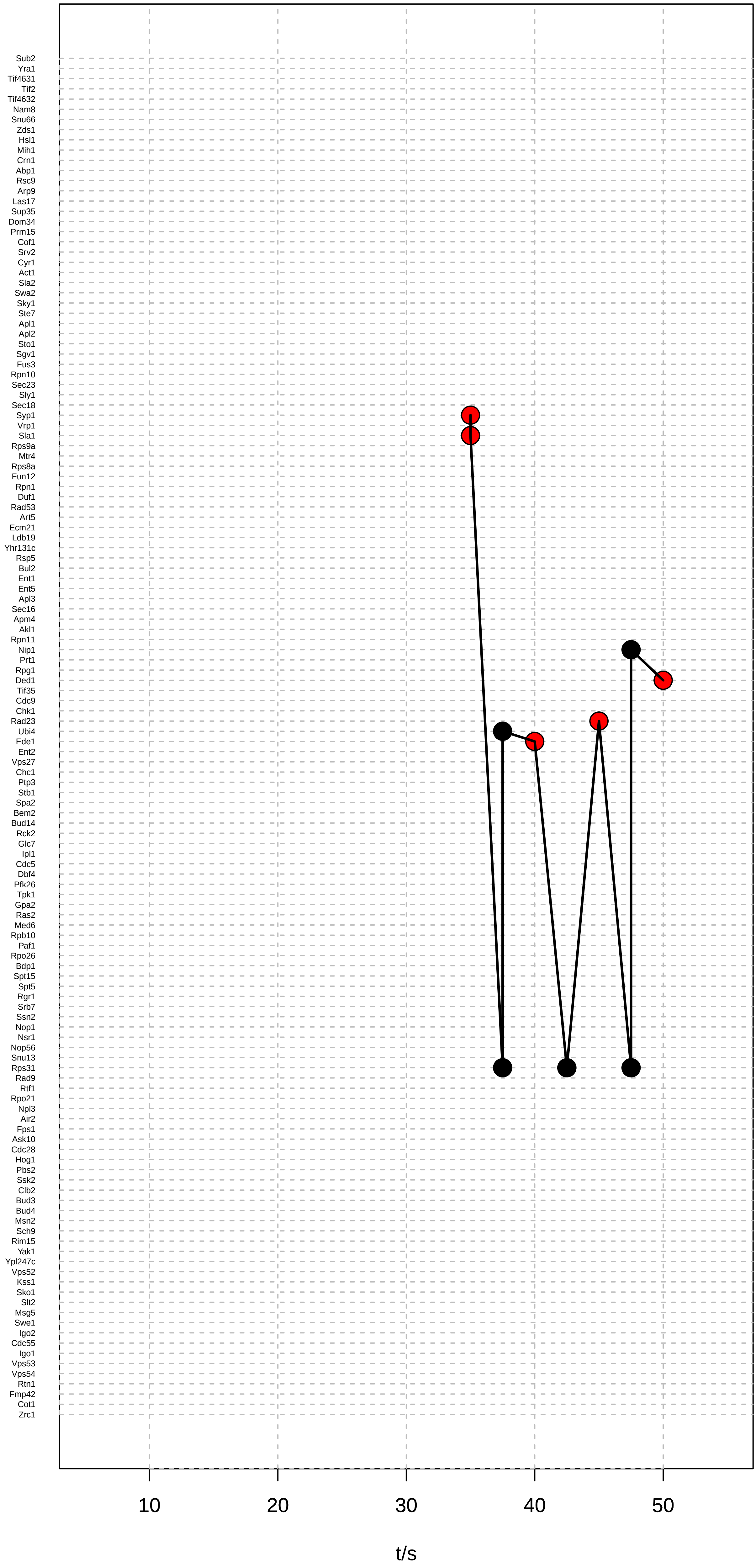

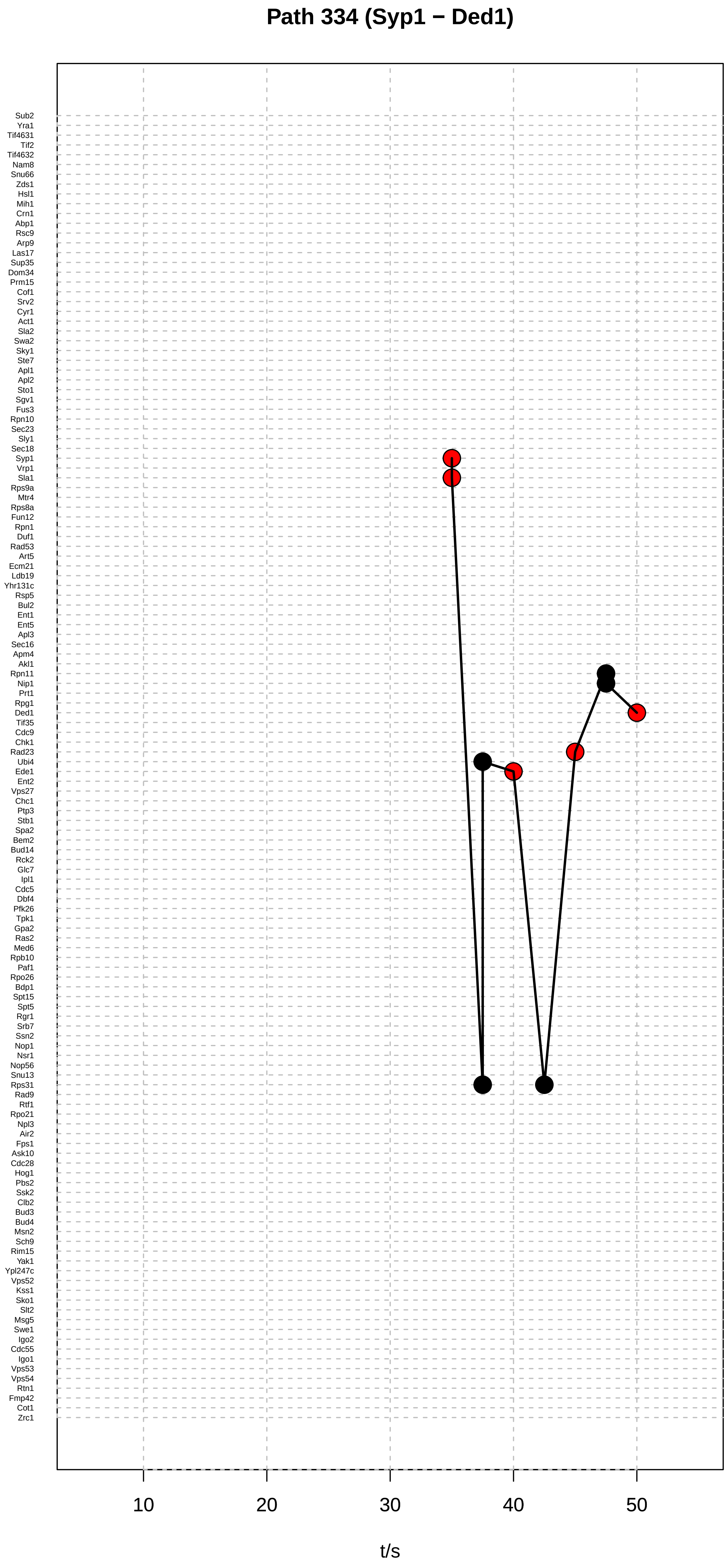

Path 334 (Syp1 − Ded1)
Sub2
Yra1
Tif4631
Tif2
Tif4632
Nam8
Snu66
Zds1
Hsl1
Mih1
Crn1
Abp1
Rsc9
Arp9
Las17
Sup35
Dom34
Prm15
Cof1
Srv2
Cyr1
Act1
Sla2
Swa2
Sky1
Ste7
Apl1
Apl2
Sto1
Sgv1
Fus3
Rpn10
Sec23
Sly1
Sec18
Syp1
Vrp1
Sla1
Rps9a
Mtr4
Rps8a
Fun12
Rpn1
Duf1
Rad53
Art5
Ecm21
Ldb19
Yhr131c
Rsp5
Bul2
Ent1
Ent5
Apl3
Sec16
Apm4
Akl1
Rpn11
Nip1
Prt1
Rpg1
Ded1
Tif35
Cdc9
Chk1
Rad23
Ubi4
Ede1
Ent2
Vps27
Chc1
Ptp3
Stb1
Spa2
Bem2
Bud14
Rck2
Glc7
Ipl1
Cdc5
Dbf4
Pfk26
Tpk1
Gpa2
Ras2
Med6
Rpb10
Paf1
Rpo26
Bdp1
Spt15
Spt5
Rgr1
Srb7
Ssn2
Nop1
Nsr1
Nop56
Snu13
Rps31
Rad9
Rtf1
Rpo21
Npl3
Air2
Fps1
Ask10
Cdc28
Hog1
Pbs2
Ssk2
Clb2
Bud3
Bud4
Msn2
Sch9
Rim15
Yak1
Ypl247c
Vps52
Kss1
Sko1
Slt2
Msg5
Swe1
Igo2
Cdc55
Igo1
Vps53
Vps54
Rtn1
Fmp42
Cot1
Zrc1
10
20
30
40
50
t/s

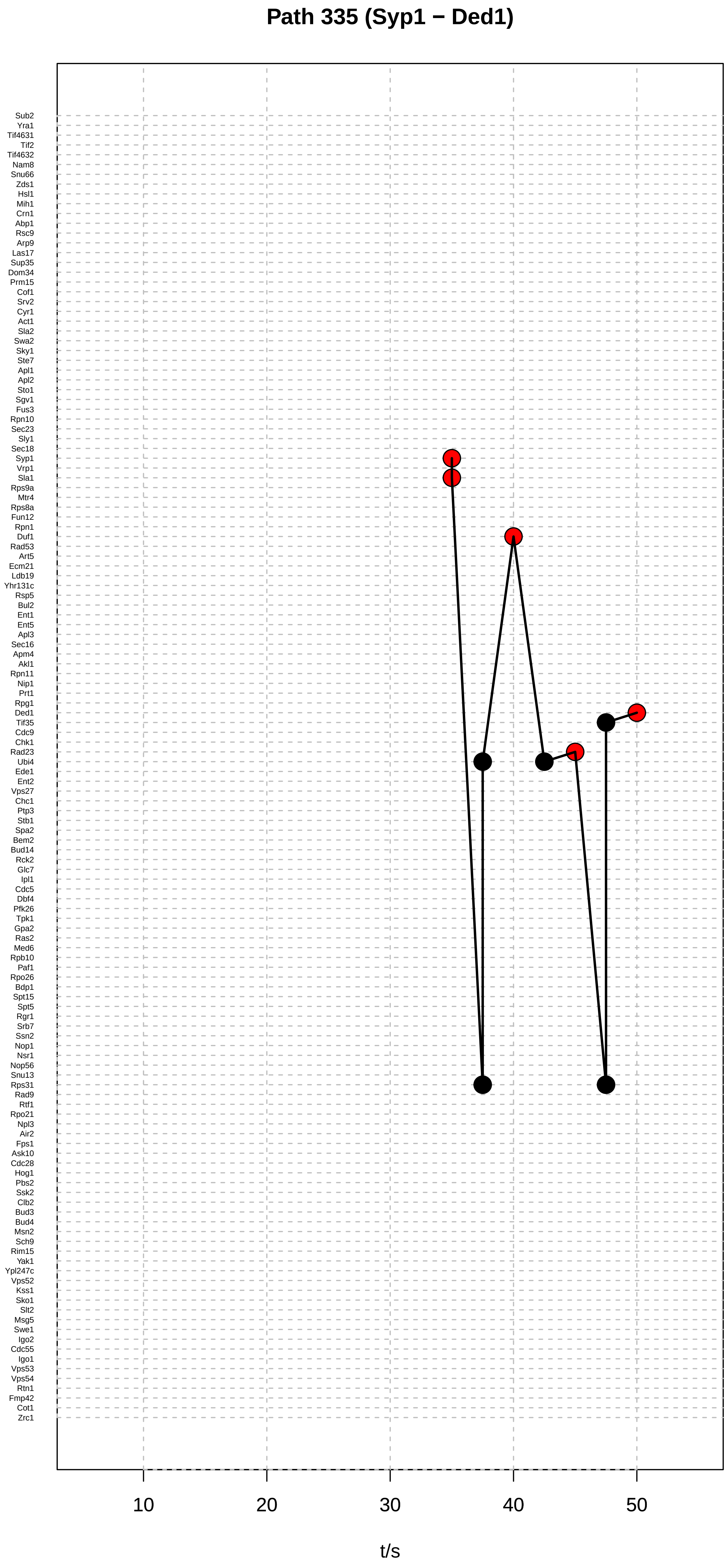

Path 335 (Syp1 − Ded1)
Sub2
Yra1
Tif4631
Tif2
Tif4632
Nam8
Snu66
Zds1
Hsl1
Mih1
Crn1
Abp1
Rsc9
Arp9
Las17
Sup35
Dom34
Prm15
Cof1
Srv2
Cyr1
Act1
Sla2
Swa2
Sky1
Ste7
Apl1
Apl2
Sto1
Sgv1
Fus3
Rpn10
Sec23
Sly1
Sec18
Syp1
Vrp1
Sla1
Rps9a
Mtr4
Rps8a
Fun12
Rpn1
Duf1
Rad53
Art5
Ecm21
Ldb19
Yhr131c
Rsp5
Bul2
Ent1
Ent5
Apl3
Sec16
Apm4
Akl1
Rpn11
Nip1
Prt1
Rpg1
Ded1
Tif35
Cdc9
Chk1
Rad23
Ubi4
Ede1
Ent2
Vps27
Chc1
Ptp3
Stb1
Spa2
Bem2
Bud14
Rck2
Glc7
Ipl1
Cdc5
Dbf4
Pfk26
Tpk1
Gpa2
Ras2
Med6
Rpb10
Paf1
Rpo26
Bdp1
Spt15
Spt5
Rgr1
Srb7
Ssn2
Nop1
Nsr1
Nop56
Snu13
Rps31
Rad9
Rtf1
Rpo21
Npl3
Air2
Fps1
Ask10
Cdc28
Hog1
Pbs2
Ssk2
Clb2
Bud3
Bud4
Msn2
Sch9
Rim15
Yak1
Ypl247c
Vps52
Kss1
Sko1
Slt2
Msg5
Swe1
Igo2
Cdc55
Igo1
Vps53
Vps54
Rtn1
Fmp42
Cot1
Zrc1
10
20
30
40
50
t/s

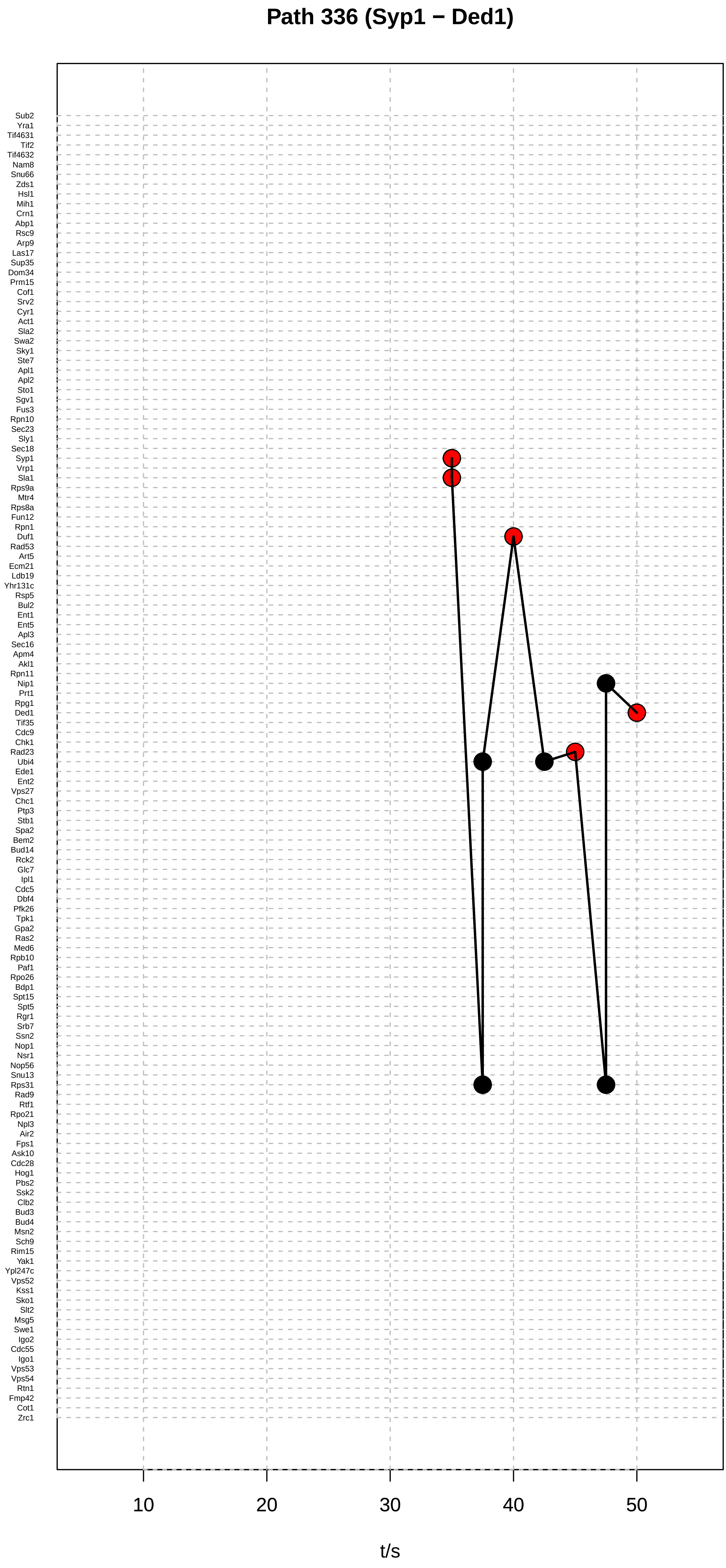
Path 336 (Syp1 − Ded1)
Sub2
Yra1
Tif4631
Tif2
Tif4632
Nam8
Snu66
Zds1
Hsl1
Mih1
Crn1
Abp1
Rsc9
Arp9
Las17
Sup35
Dom34
Prm15
Cof1
Srv2
Cyr1
Act1
Sla2
Swa2
Sky1
Ste7
Apl1
Apl2
Sto1
Sgv1
Fus3
Rpn10
Sec23
Sly1
Sec18
Syp1
Vrp1
Sla1
Rps9a
Mtr4
Rps8a
Fun12
Rpn1
Duf1
Rad53
Art5
Ecm21
Ldb19
Yhr131c
Rsp5
Bul2
Ent1
Ent5
Apl3
Sec16
Apm4
Akl1
Rpn11
Nip1
Prt1
Rpg1
Ded1
Tif35
Cdc9
Chk1
Rad23
Ubi4
Ede1
Ent2
Vps27
Chc1
Ptp3
Stb1
Spa2
Bem2
Bud14
Rck2
Glc7
Ipl1
Cdc5
Dbf4
Pfk26
Tpk1
Gpa2
Ras2
Med6
Rpb10
Paf1
Rpo26
Bdp1
Spt15
Spt5
Rgr1
Srb7
Ssn2
Nop1
Nsr1
Nop56
Snu13
Rps31
Rad9
Rtf1
Rpo21
Npl3
Air2
Fps1
Ask10
Cdc28
Hog1
Pbs2
Ssk2
Clb2
Bud3
Bud4
Msn2
Sch9
Rim15
Yak1
Ypl247c
Vps52
Kss1
Sko1
Slt2
Msg5
Swe1
Igo2
Cdc55
Igo1
Vps53
Vps54
Rtn1
Fmp42
Cot1
Zrc1
10
20
30
40
50
t/s

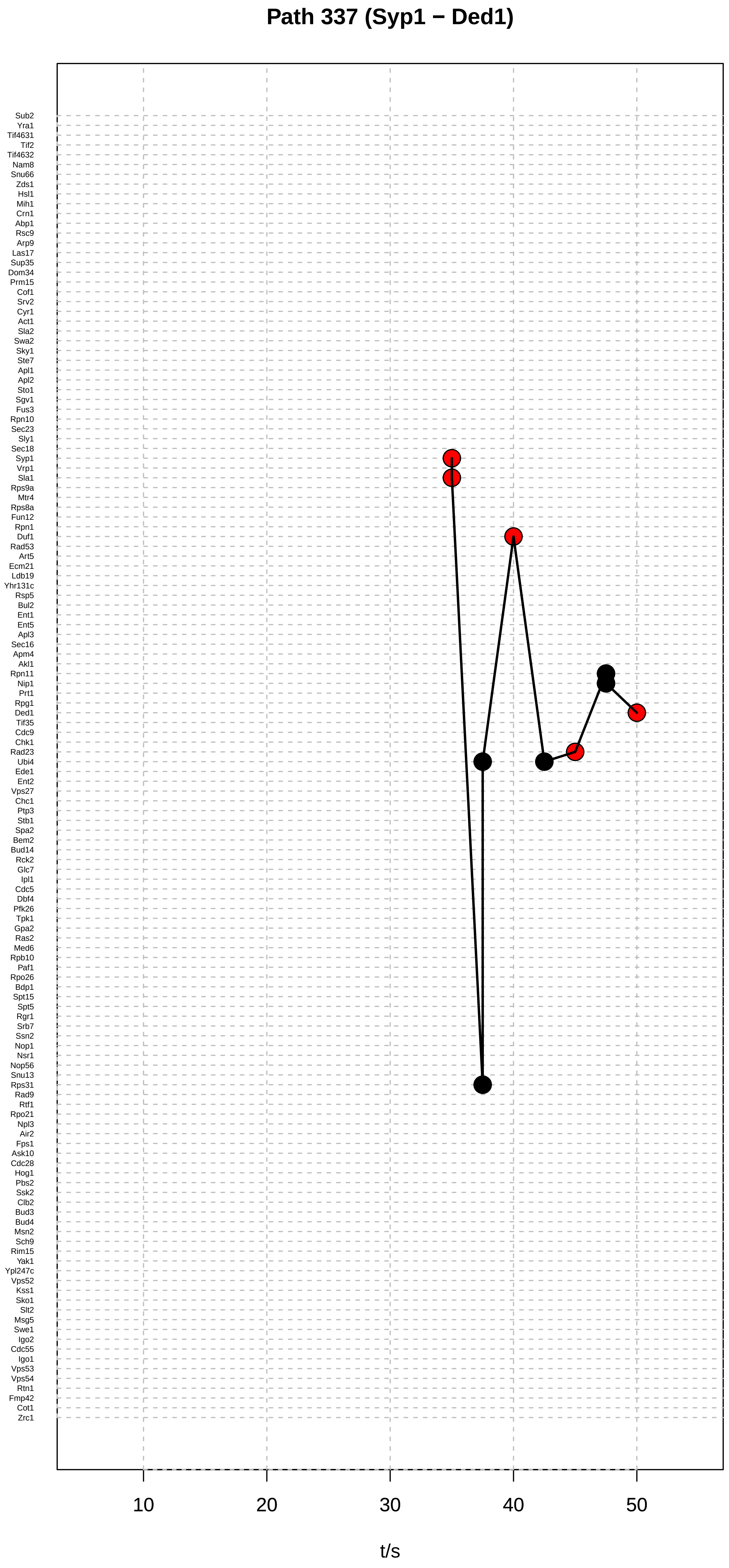
Path 337 (Syp1 − Ded1)
Sub2
Yra1
Tif4631
Tif2
Tif4632
Nam8
Snu66
Zds1
Hsl1
Mih1
Crn1
Abp1
Rsc9
Arp9
Las17
Sup35
Dom34
Prm15
Cof1
Srv2
Cyr1
Act1
Sla2
Swa2
Sky1
Ste7
Apl1
Apl2
Sto1
Sgv1
Fus3
Rpn10
Sec23
Sly1
Sec18
Syp1
Vrp1
Sla1
Rps9a
Mtr4
Rps8a
Fun12
Rpn1
Duf1
Rad53
Art5
Ecm21
Ldb19
Yhr131c
Rsp5
Bul2
Ent1
Ent5
Apl3
Sec16
Apm4
Akl1
Rpn11
Nip1
Prt1
Rpg1
Ded1
Tif35
Cdc9
Chk1
Rad23
Ubi4
Ede1
Ent2
Vps27
Chc1
Ptp3
Stb1
Spa2
Bem2
Bud14
Rck2
Glc7
Ipl1
Cdc5
Dbf4
Pfk26
Tpk1
Gpa2
Ras2
Med6
Rpb10
Paf1
Rpo26
Bdp1
Spt15
Spt5
Rgr1
Srb7
Ssn2
Nop1
Nsr1
Nop56
Snu13
Rps31
Rad9
Rtf1
Rpo21
Npl3
Air2
Fps1
Ask10
Cdc28
Hog1
Pbs2
Ssk2
Clb2
Bud3
Bud4
Msn2
Sch9
Rim15
Yak1
Ypl247c
Vps52
Kss1
Sko1
Slt2
Msg5
Swe1
Igo2
Cdc55
Igo1
Vps53
Vps54
Rtn1
Fmp42
Cot1
Zrc1
10
20
30
40
50
t/s

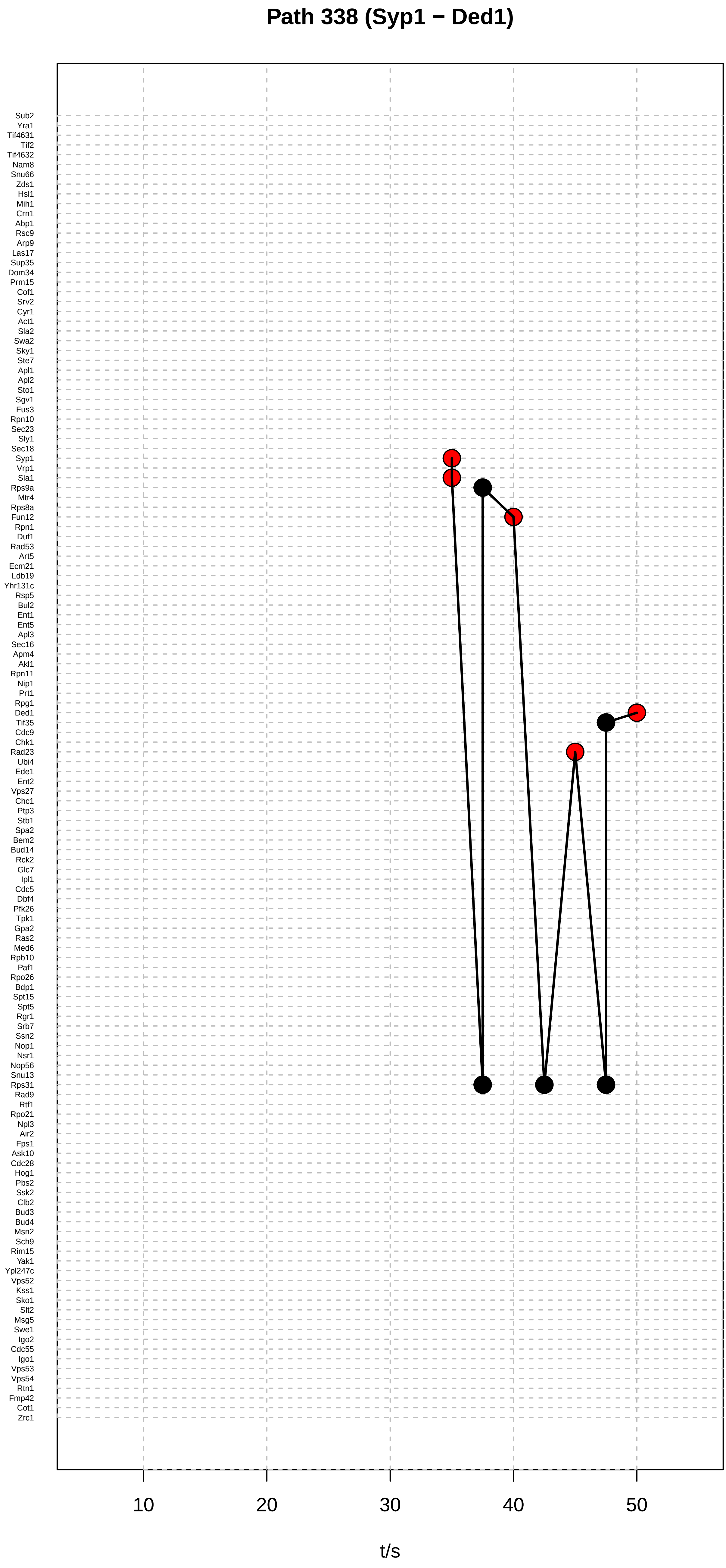

Path 338 (Syp1 − Ded1)
Sub2
Yra1
Tif4631
Tif2
Tif4632
Nam8
Snu66
Zds1
Hsl1
Mih1
Crn1
Abp1
Rsc9
Arp9
Las17
Sup35
Dom34
Prm15
Cof1
Srv2
Cyr1
Act1
Sla2
Swa2
Sky1
Ste7
Apl1
Apl2
Sto1
Sgv1
Fus3
Rpn10
Sec23
Sly1
Sec18
Syp1
Vrp1
Sla1
Rps9a
Mtr4
Rps8a
Fun12
Rpn1
Duf1
Rad53
Art5
Ecm21
Ldb19
Yhr131c
Rsp5
Bul2
Ent1
Ent5
Apl3
Sec16
Apm4
Akl1
Rpn11
Nip1
Prt1
Rpg1
Ded1
Tif35
Cdc9
Chk1
Rad23
Ubi4
Ede1
Ent2
Vps27
Chc1
Ptp3
Stb1
Spa2
Bem2
Bud14
Rck2
Glc7
Ipl1
Cdc5
Dbf4
Pfk26
Tpk1
Gpa2
Ras2
Med6
Rpb10
Paf1
Rpo26
Bdp1
Spt15
Spt5
Rgr1
Srb7
Ssn2
Nop1
Nsr1
Nop56
Snu13
Rps31
Rad9
Rtf1
Rpo21
Npl3
Air2
Fps1
Ask10
Cdc28
Hog1
Pbs2
Ssk2
Clb2
Bud3
Bud4
Msn2
Sch9
Rim15
Yak1
Ypl247c
Vps52
Kss1
Sko1
Slt2
Msg5
Swe1
Igo2
Cdc55
Igo1
Vps53
Vps54
Rtn1
Fmp42
Cot1
Zrc1
10
20
30
40
50
t/s

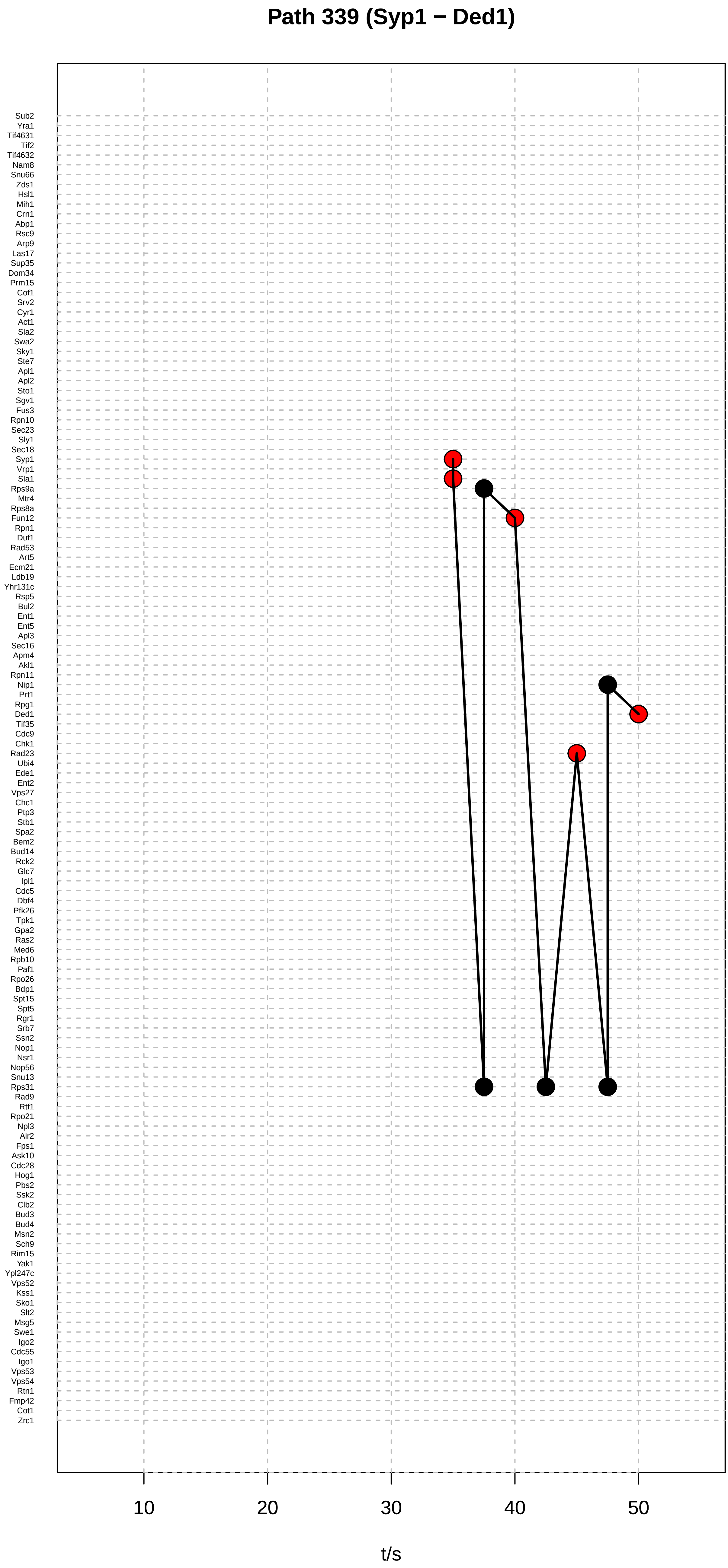

Path 339 (Syp1 − Ded1)
Sub2
Yra1
Tif4631
Tif2
Tif4632
Nam8
Snu66
Zds1
Hsl1
Mih1
Crn1
Abp1
Rsc9
Arp9
Las17
Sup35
Dom34
Prm15
Cof1
Srv2
Cyr1
Act1
Sla2
Swa2
Sky1
Ste7
Apl1
Apl2
Sto1
Sgv1
Fus3
Rpn10
Sec23
Sly1
Sec18
Syp1
Vrp1
Sla1
Rps9a
Mtr4
Rps8a
Fun12
Rpn1
Duf1
Rad53
Art5
Ecm21
Ldb19
Yhr131c
Rsp5
Bul2
Ent1
Ent5
Apl3
Sec16
Apm4
Akl1
Rpn11
Nip1
Prt1
Rpg1
Ded1
Tif35
Cdc9
Chk1
Rad23
Ubi4
Ede1
Ent2
Vps27
Chc1
Ptp3
Stb1
Spa2
Bem2
Bud14
Rck2
Glc7
Ipl1
Cdc5
Dbf4
Pfk26
Tpk1
Gpa2
Ras2
Med6
Rpb10
Paf1
Rpo26
Bdp1
Spt15
Spt5
Rgr1
Srb7
Ssn2
Nop1
Nsr1
Nop56
Snu13
Rps31
Rad9
Rtf1
Rpo21
Npl3
Air2
Fps1
Ask10
Cdc28
Hog1
Pbs2
Ssk2
Clb2
Bud3
Bud4
Msn2
Sch9
Rim15
Yak1
Ypl247c
Vps52
Kss1
Sko1
Slt2
Msg5
Swe1
Igo2
Cdc55
Igo1
Vps53
Vps54
Rtn1
Fmp42
Cot1
Zrc1
10
20
30
40
50
t/s

### Path 340 (Syp1 − Ded1)

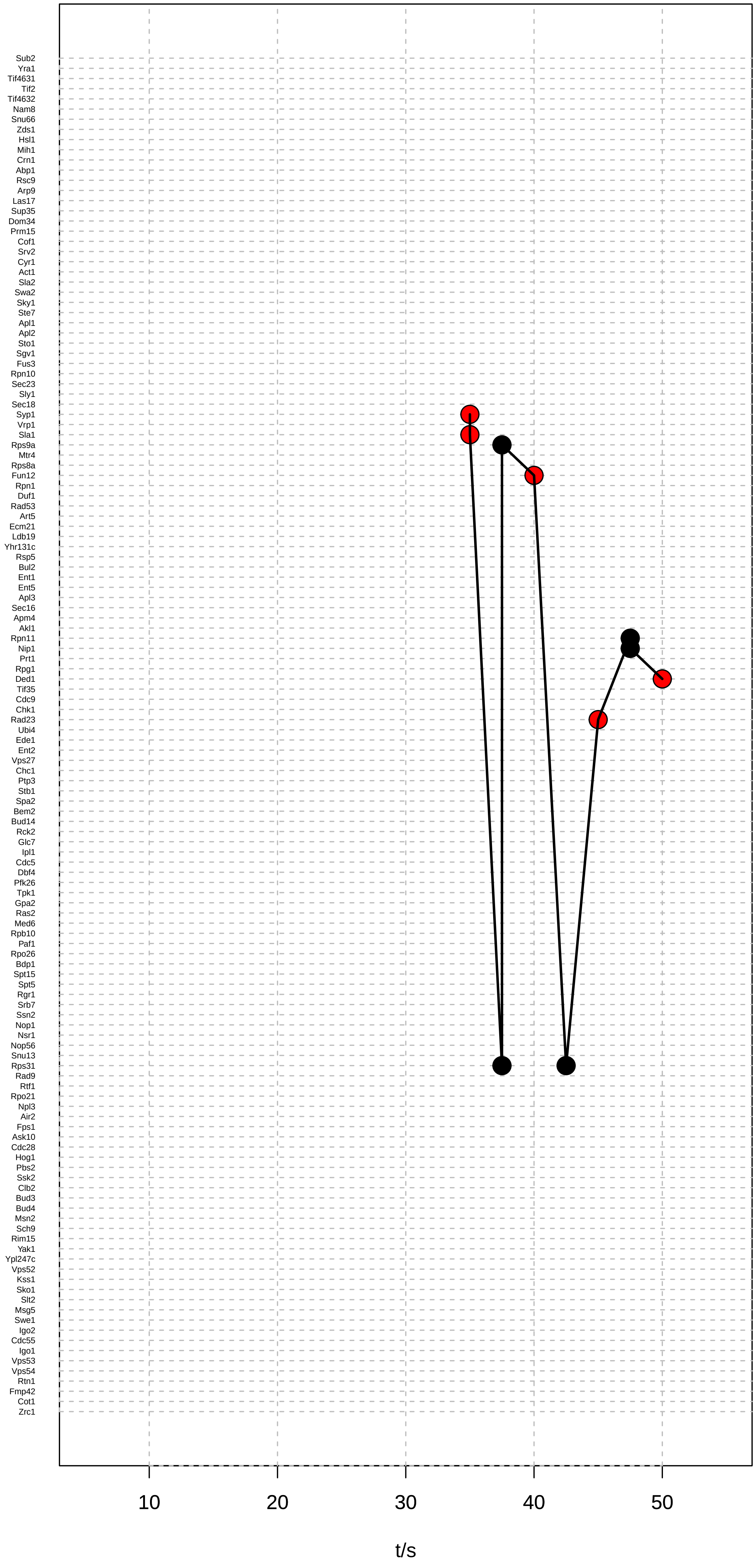

#### Path 341 (Sla1 − Ede1)

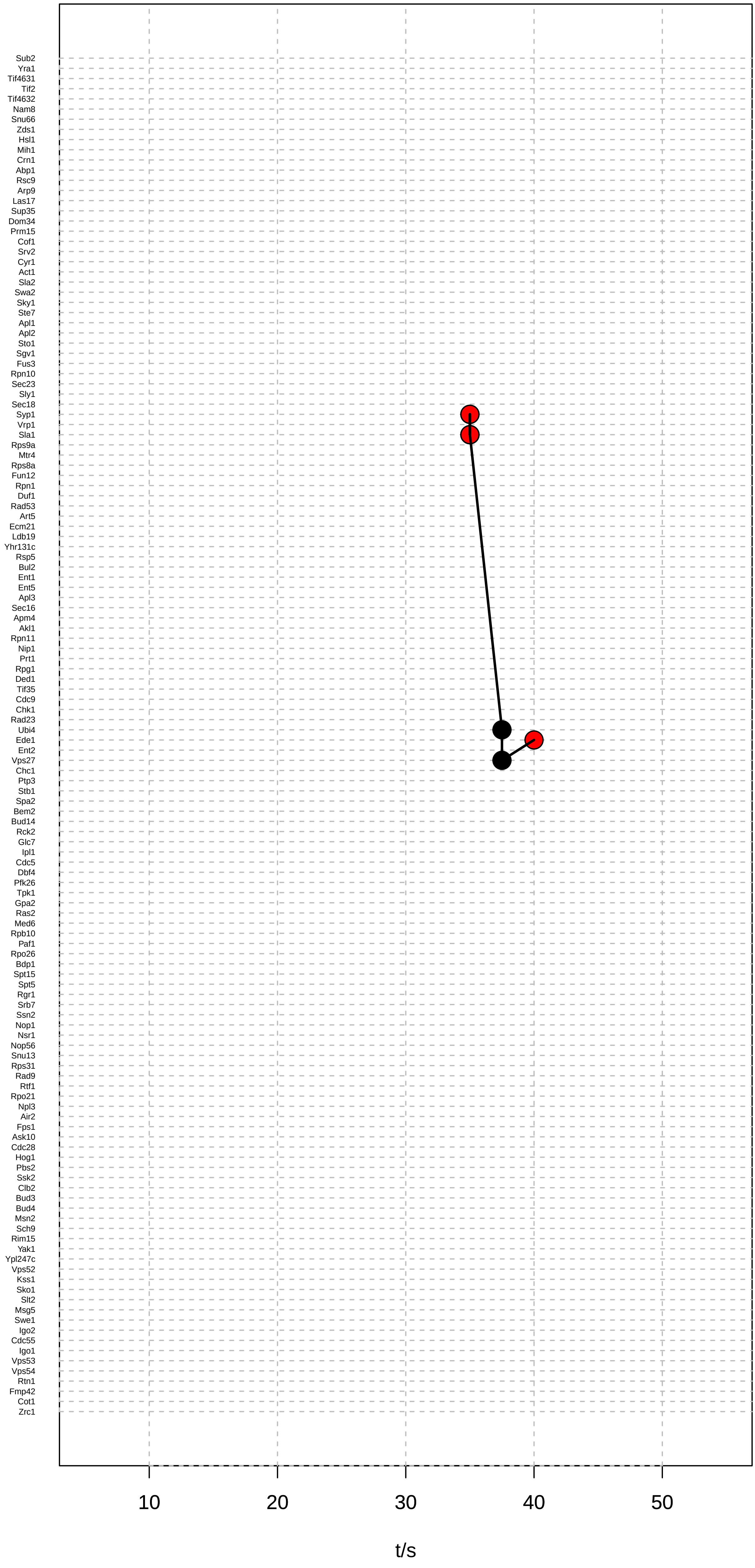

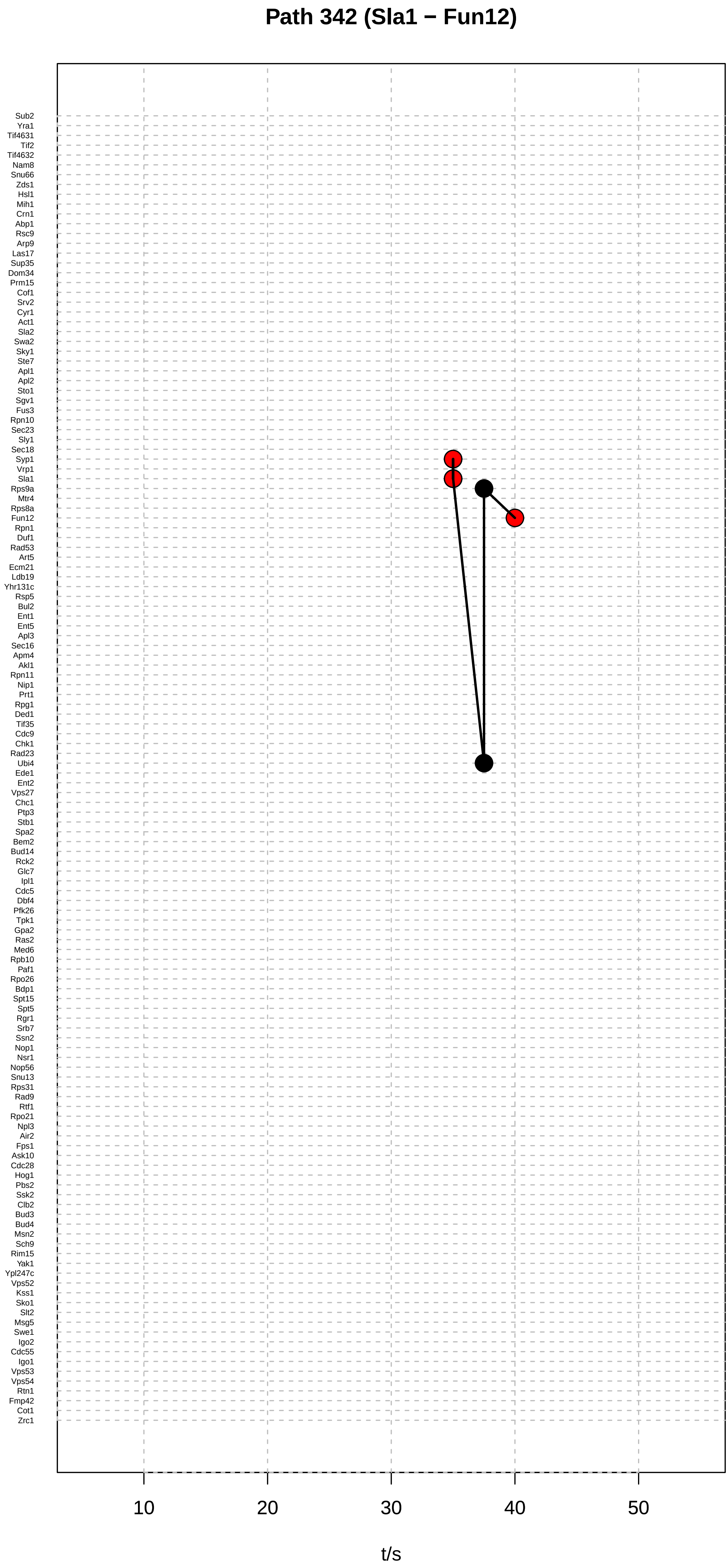

Path 342 (Sla1 − Fun12)
Sub2
Yra1
Tif4631
Tif2
Tif4632
Nam8
Snu66
Zds1
Hsl1
Mih1
Crn1
Abp1
Rsc9
Arp9
Las17
Sup35
Dom34
Prm15
Cof1
Srv2
Cyr1
Act1
Sla2
Swa2
Sky1
Ste7
Apl1
Apl2
Sto1
Sgv1
Fus3
Rpn10
Sec23
Sly1
Sec18
Syp1
Vrp1
Sla1
Rps9a
Mtr4
Rps8a
Fun12
Rpn1
Duf1
Rad53
Art5
Ecm21
Ldb19
Yhr131c
Rsp5
Bul2
Ent1
Ent5
Apl3
Sec16
Apm4
Akl1
Rpn11
Nip1
Prt1
Rpg1
Ded1
Tif35
Cdc9
Chk1
Rad23
Ubi4
Ede1
Ent2
Vps27
Chc1
Ptp3
Stb1
Spa2
Bem2
Bud14
Rck2
Glc7
Ipl1
Cdc5
Dbf4
Pfk26
Tpk1
Gpa2
Ras2
Med6
Rpb10
Paf1
Rpo26
Bdp1
Spt15
Spt5
Rgr1
Srb7
Ssn2
Nop1
Nsr1
Nop56
Snu13
Rps31
Rad9
Rtf1
Rpo21
Npl3
Air2
Fps1
Ask10
Cdc28
Hog1
Pbs2
Ssk2
Clb2
Bud3
Bud4
Msn2
Sch9
Rim15
Yak1
Ypl247c
Vps52
Kss1
Sko1
Slt2
Msg5
Swe1
Igo2
Cdc55
Igo1
Vps53
Vps54
Rtn1
Fmp42
Cot1
Zrc1
10
20
30
40
50
t/s

### Path 343 (Sla1 − Fun12)

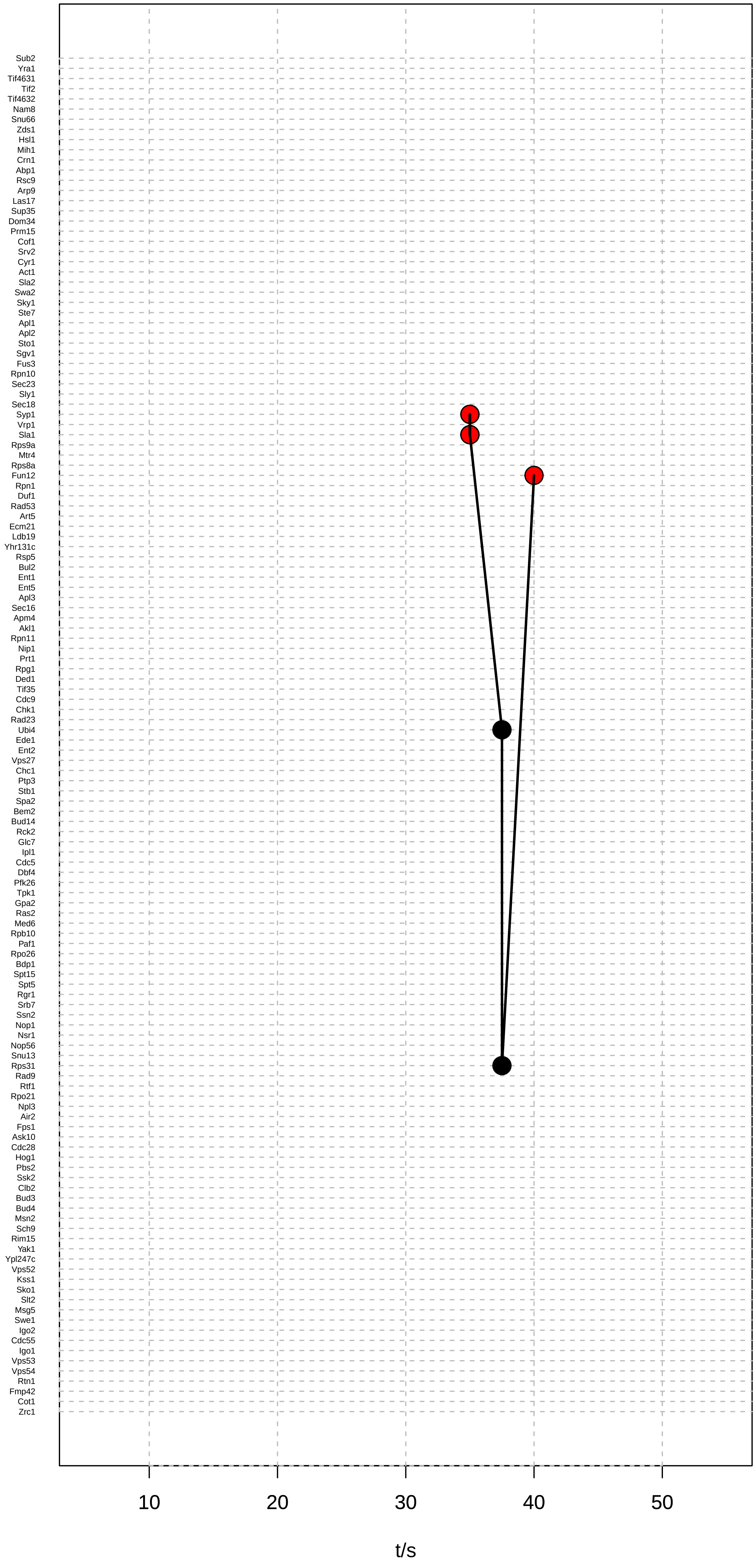

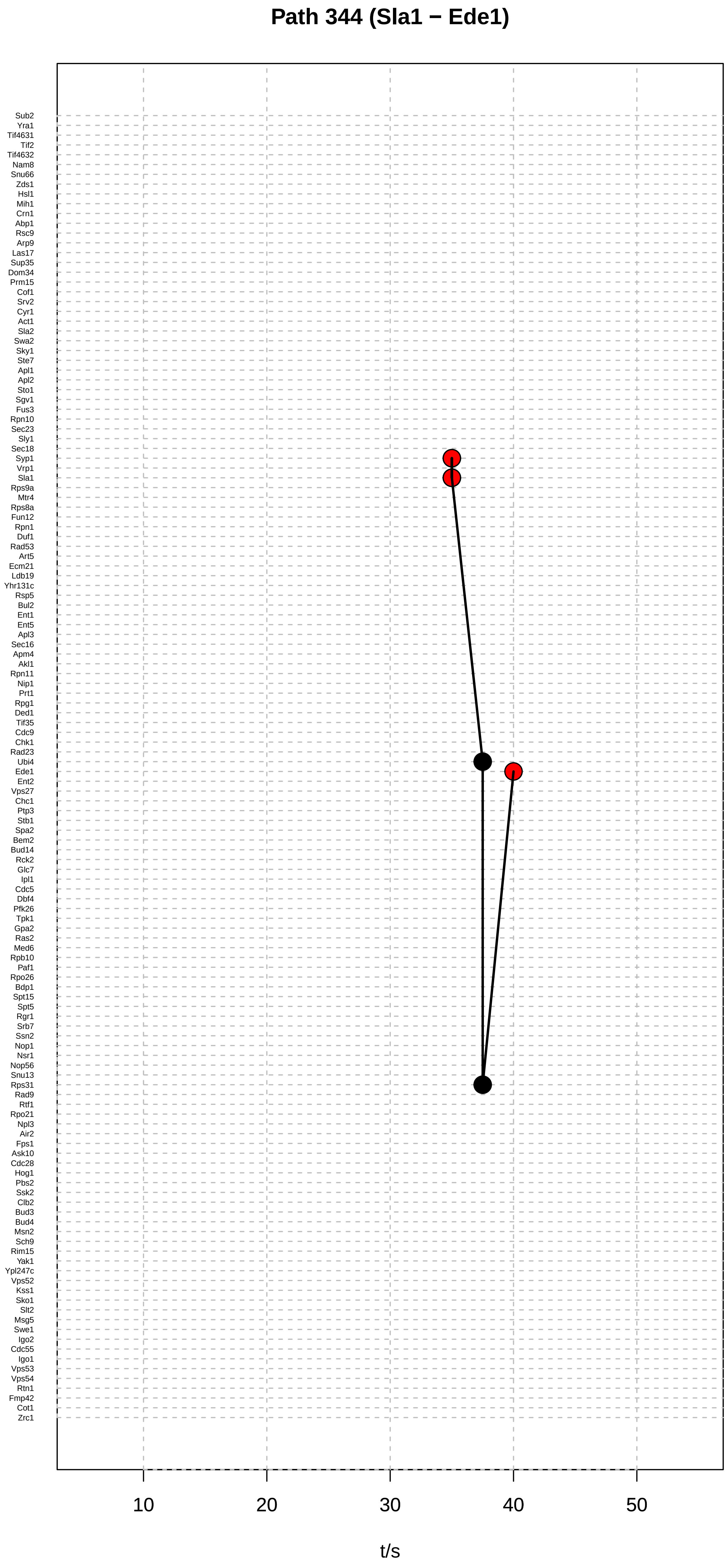

Path 344 (Sla1 − Ede1)
Sub2
Yra1
Tif4631
Tif2
Tif4632
Nam8
Snu66
Zds1
Hsl1
Mih1
Crn1
Abp1
Rsc9
Arp9
Las17
Sup35
Dom34
Prm15
Cof1
Srv2
Cyr1
Act1
Sla2
Swa2
Sky1
Ste7
Apl1
Apl2
Sto1
Sgv1
Fus3
Rpn10
Sec23
Sly1
Sec18
Syp1
Vrp1
Sla1
Rps9a
Mtr4
Rps8a
Fun12
Rpn1
Duf1
Rad53
Art5
Ecm21
Ldb19
Yhr131c
Rsp5
Bul2
Ent1
Ent5
Apl3
Sec16
Apm4
Akl1
Rpn11
Nip1
Prt1
Rpg1
Ded1
Tif35
Cdc9
Chk1
Rad23
Ubi4
Ede1
Ent2
Vps27
Chc1
Ptp3
Stb1
Spa2
Bem2
Bud14
Rck2
Glc7
Ipl1
Cdc5
Dbf4
Pfk26
Tpk1
Gpa2
Ras2
Med6
Rpb10
Paf1
Rpo26
Bdp1
Spt15
Spt5
Rgr1
Srb7
Ssn2
Nop1
Nsr1
Nop56
Snu13
Rps31
Rad9
Rtf1
Rpo21
Npl3
Air2
Fps1
Ask10
Cdc28
Hog1
Pbs2
Ssk2
Clb2
Bud3
Bud4
Msn2
Sch9
Rim15
Yak1
Ypl247c
Vps52
Kss1
Sko1
Slt2
Msg5
Swe1
Igo2
Cdc55
Igo1
Vps53
Vps54
Rtn1
Fmp42
Cot1
Zrc1
10
20
30
40
50
t/s

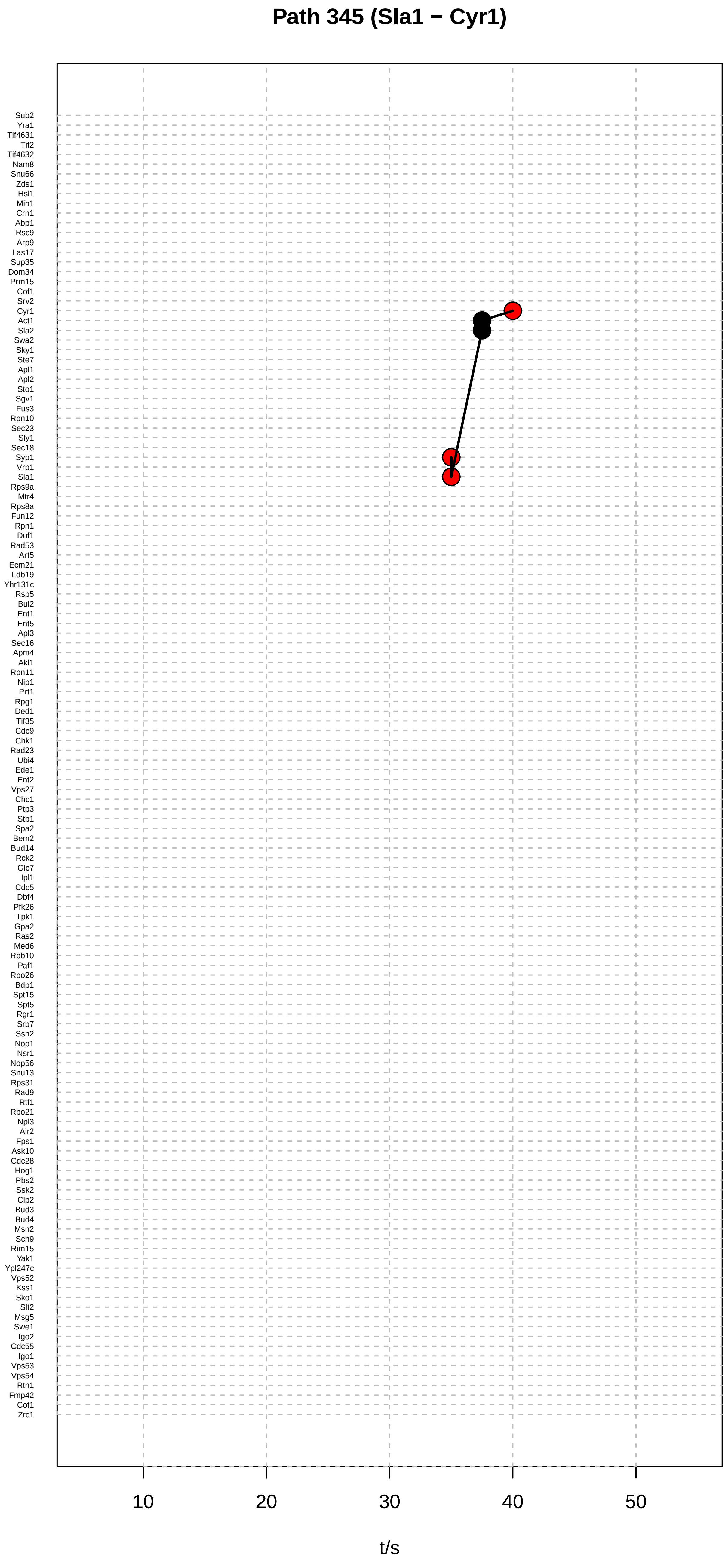

Path 345 (Sla1 − Cyr1)
Sub2
Yra1
Tif4631
Tif2
Tif4632
Nam8
Snu66
Zds1
Hsl1
Mih1
Crn1
Abp1
Rsc9
Arp9
Las17
Sup35
Dom34
Prm15
Cof1
Srv2
Cyr1
Act1
Sla2
Swa2
Sky1
Ste7
Apl1
Apl2
Sto1
Sgv1
Fus3
Rpn10
Sec23
Sly1
Sec18
Syp1
Vrp1
Sla1
Rps9a
Mtr4
Rps8a
Fun12
Rpn1
Duf1
Rad53
Art5
Ecm21
Ldb19
Yhr131c
Rsp5
Bul2
Ent1
Ent5
Apl3
Sec16
Apm4
Akl1
Rpn11
Nip1
Prt1
Rpg1
Ded1
Tif35
Cdc9
Chk1
Rad23
Ubi4
Ede1
Ent2
Vps27
Chc1
Ptp3
Stb1
Spa2
Bem2
Bud14
Rck2
Glc7
Ipl1
Cdc5
Dbf4
Pfk26
Tpk1
Gpa2
Ras2
Med6
Rpb10
Paf1
Rpo26
Bdp1
Spt15
Spt5
Rgr1
Srb7
Ssn2
Nop1
Nsr1
Nop56
Snu13
Rps31
Rad9
Rtf1
Rpo21
Npl3
Air2
Fps1
Ask10
Cdc28
Hog1
Pbs2
Ssk2
Clb2
Bud3
Bud4
Msn2
Sch9
Rim15
Yak1
Ypl247c
Vps52
Kss1
Sko1
Slt2
Msg5
Swe1
Igo2
Cdc55
Igo1
Vps53
Vps54
Rtn1
Fmp42
Cot1
Zrc1
10
20
30
40
50
t/s

### Path 346 (Sla1 − Ede1)

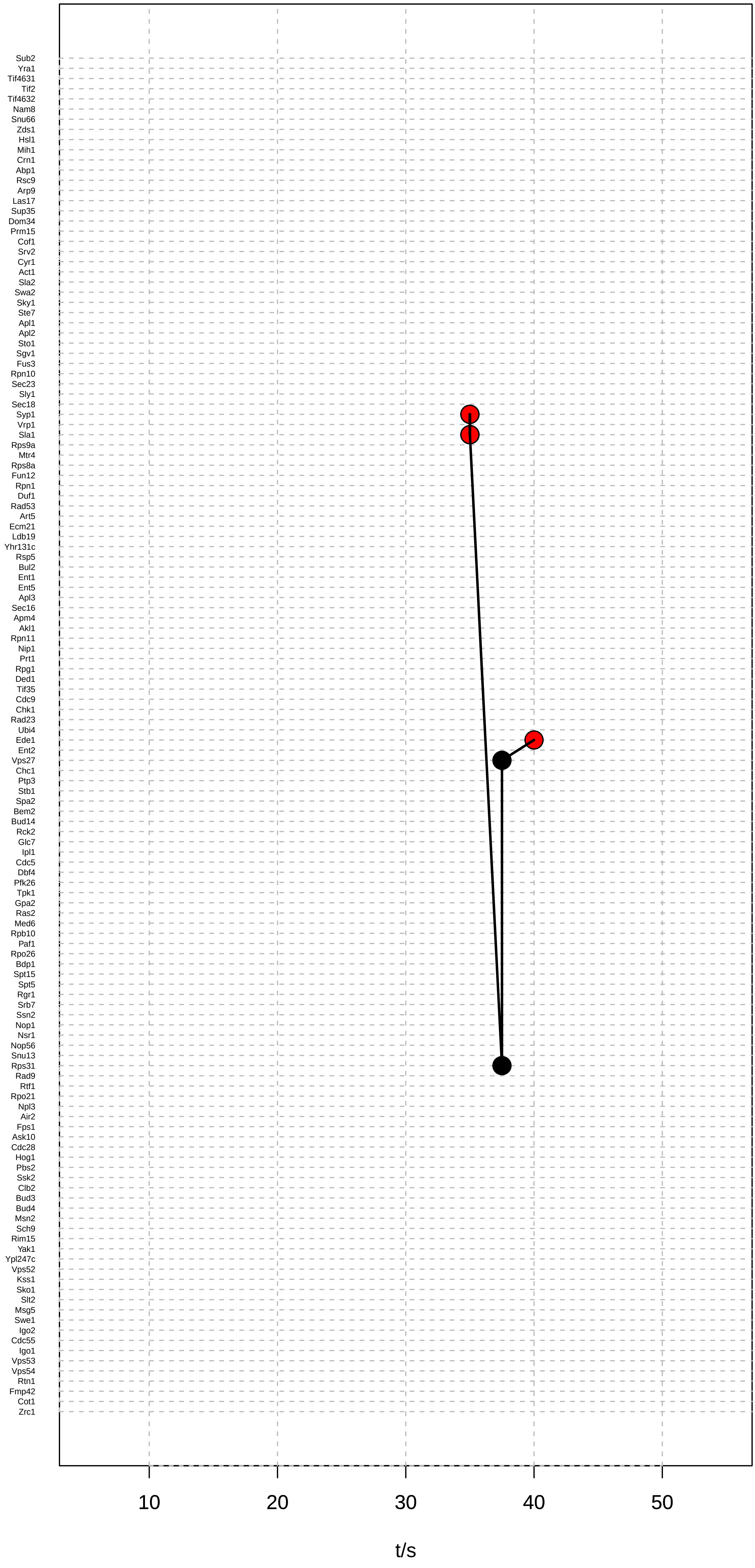

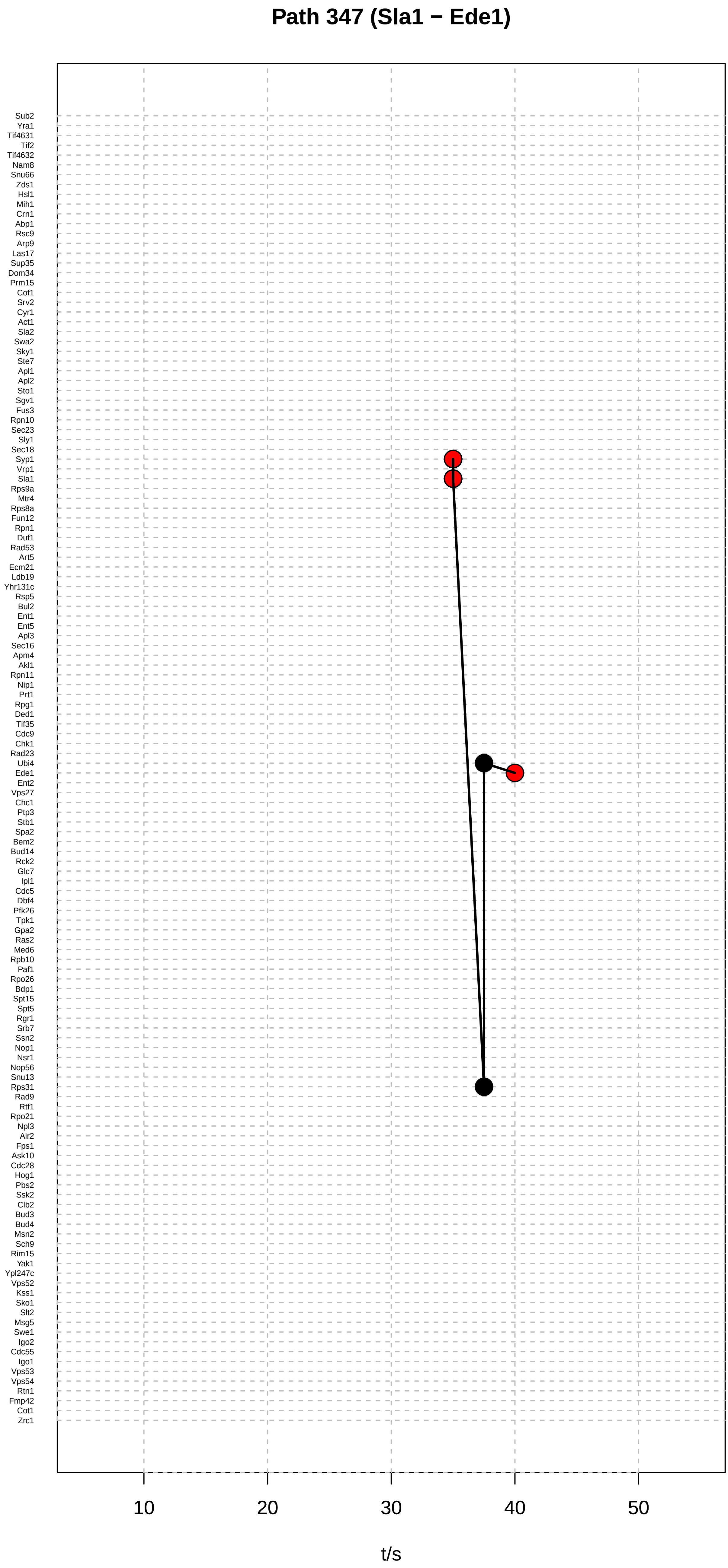

Path 347 (Sla1 − Ede1)
Sub2
Yra1
Tif4631
Tif2
Tif4632
Nam8
Snu66
Zds1
Hsl1
Mih1
Crn1
Abp1
Rsc9
Arp9
Las17
Sup35
Dom34
Prm15
Cof1
Srv2
Cyr1
Act1
Sla2
Swa2
Sky1
Ste7
Apl1
Apl2
Sto1
Sgv1
Fus3
Rpn10
Sec23
Sly1
Sec18
Syp1
Vrp1
Sla1
Rps9a
Mtr4
Rps8a
Fun12
Rpn1
Duf1
Rad53
Art5
Ecm21
Ldb19
Yhr131c
Rsp5
Bul2
Ent1
Ent5
Apl3
Sec16
Apm4
Akl1
Rpn11
Nip1
Prt1
Rpg1
Ded1
Tif35
Cdc9
Chk1
Rad23
Ubi4
Ede1
Ent2
Vps27
Chc1
Ptp3
Stb1
Spa2
Bem2
Bud14
Rck2
Glc7
Ipl1
Cdc5
Dbf4
Pfk26
Tpk1
Gpa2
Ras2
Med6
Rpb10
Paf1
Rpo26
Bdp1
Spt15
Spt5
Rgr1
Srb7
Ssn2
Nop1
Nsr1
Nop56
Snu13
Rps31
Rad9
Rtf1
Rpo21
Npl3
Air2
Fps1
Ask10
Cdc28
Hog1
Pbs2
Ssk2
Clb2
Bud3
Bud4
Msn2
Sch9
Rim15
Yak1
Ypl247c
Vps52
Kss1
Sko1
Slt2
Msg5
Swe1
Igo2
Cdc55
Igo1
Vps53
Vps54
Rtn1
Fmp42
Cot1
Zrc1
10
20
30
40
50
t/s

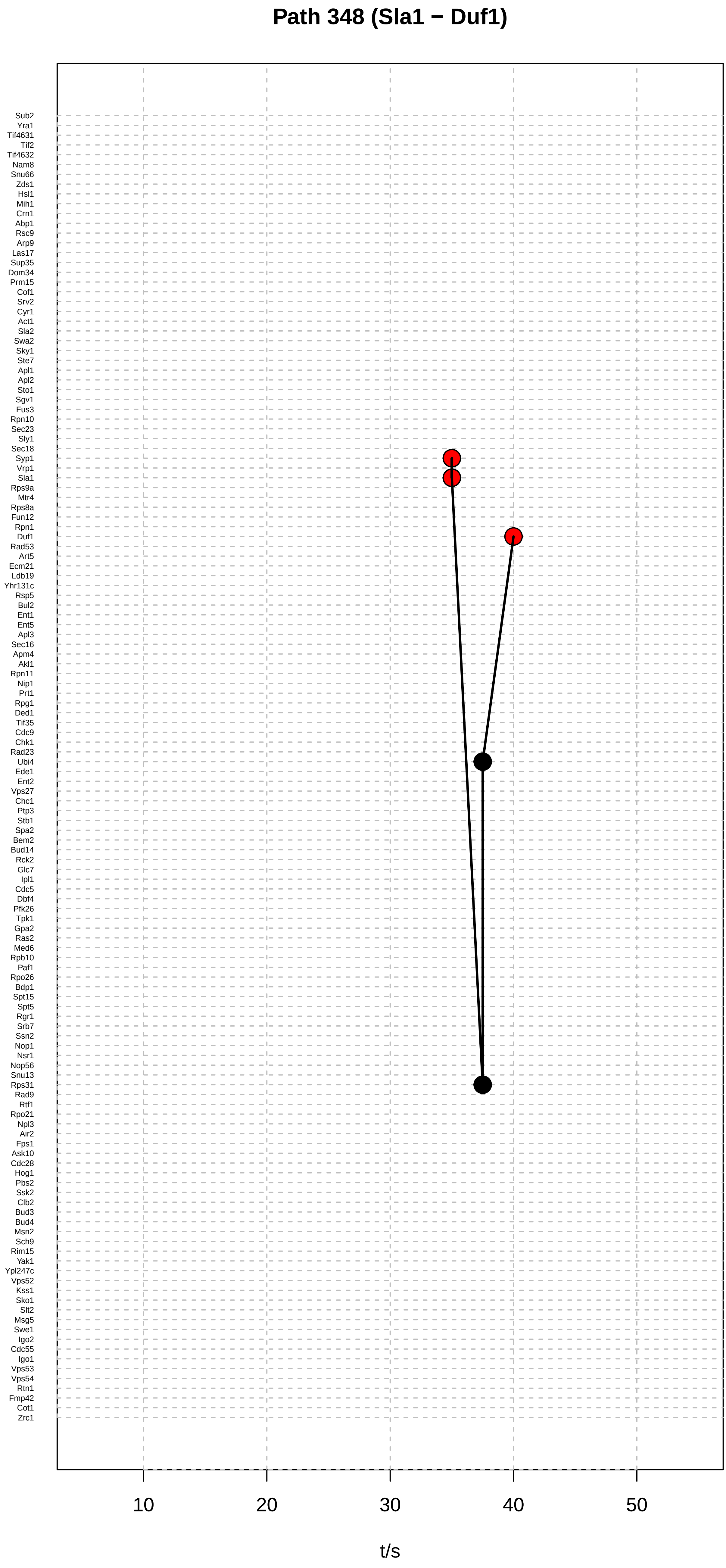

Path 348 (Sla1 − Duf1)
Sub2
Yra1
Tif4631
Tif2
Tif4632
Nam8
Snu66
Zds1
Hsl1
Mih1
Crn1
Abp1
Rsc9
Arp9
Las17
Sup35
Dom34
Prm15
Cof1
Srv2
Cyr1
Act1
Sla2
Swa2
Sky1
Ste7
Apl1
Apl2
Sto1
Sgv1
Fus3
Rpn10
Sec23
Sly1
Sec18
Syp1
Vrp1
Sla1
Rps9a
Mtr4
Rps8a
Fun12
Rpn1
Duf1
Rad53
Art5
Ecm21
Ldb19
Yhr131c
Rsp5
Bul2
Ent1
Ent5
Apl3
Sec16
Apm4
Akl1
Rpn11
Nip1
Prt1
Rpg1
Ded1
Tif35
Cdc9
Chk1
Rad23
Ubi4
Ede1
Ent2
Vps27
Chc1
Ptp3
Stb1
Spa2
Bem2
Bud14
Rck2
Glc7
Ipl1
Cdc5
Dbf4
Pfk26
Tpk1
Gpa2
Ras2
Med6
Rpb10
Paf1
Rpo26
Bdp1
Spt15
Spt5
Rgr1
Srb7
Ssn2
Nop1
Nsr1
Nop56
Snu13
Rps31
Rad9
Rtf1
Rpo21
Npl3
Air2
Fps1
Ask10
Cdc28
Hog1
Pbs2
Ssk2
Clb2
Bud3
Bud4
Msn2
Sch9
Rim15
Yak1
Ypl247c
Vps52
Kss1
Sko1
Slt2
Msg5
Swe1
Igo2
Cdc55
Igo1
Vps53
Vps54
Rtn1
Fmp42
Cot1
Zrc1
10
20
30
40
50
t/s

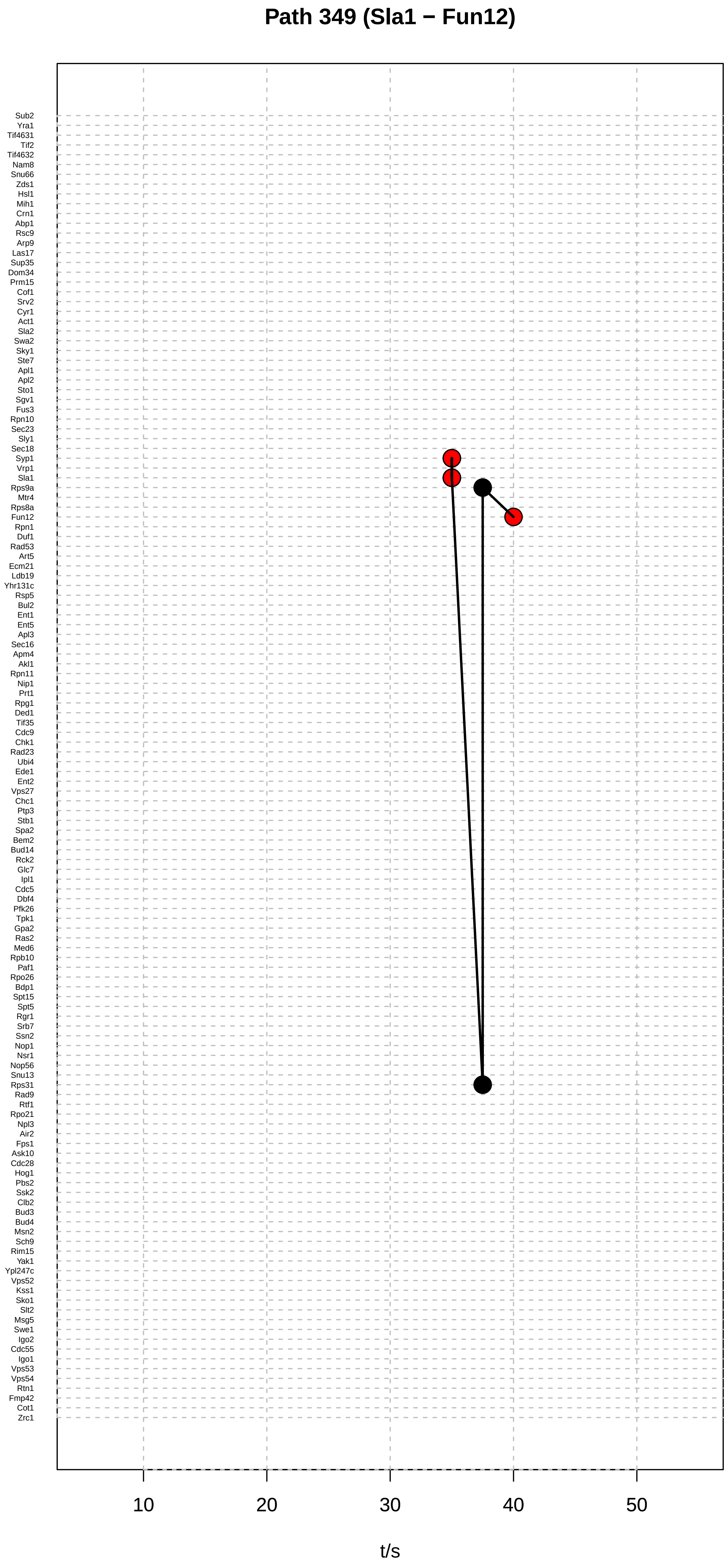

Path 349 (Sla1 − Fun12)
Sub2
Yra1
Tif4631
Tif2
Tif4632
Nam8
Snu66
Zds1
Hsl1
Mih1
Crn1
Abp1
Rsc9
Arp9
Las17
Sup35
Dom34
Prm15
Cof1
Srv2
Cyr1
Act1
Sla2
Swa2
Sky1
Ste7
Apl1
Apl2
Sto1
Sgv1
Fus3
Rpn10
Sec23
Sly1
Sec18
Syp1
Vrp1
Sla1
Rps9a
Mtr4
Rps8a
Fun12
Rpn1
Duf1
Rad53
Art5
Ecm21
Ldb19
Yhr131c
Rsp5
Bul2
Ent1
Ent5
Apl3
Sec16
Apm4
Akl1
Rpn11
Nip1
Prt1
Rpg1
Ded1
Tif35
Cdc9
Chk1
Rad23
Ubi4
Ede1
Ent2
Vps27
Chc1
Ptp3
Stb1
Spa2
Bem2
Bud14
Rck2
Glc7
Ipl1
Cdc5
Dbf4
Pfk26
Tpk1
Gpa2
Ras2
Med6
Rpb10
Paf1
Rpo26
Bdp1
Spt15
Spt5
Rgr1
Srb7
Ssn2
Nop1
Nsr1
Nop56
Snu13
Rps31
Rad9
Rtf1
Rpo21
Npl3
Air2
Fps1
Ask10
Cdc28
Hog1
Pbs2
Ssk2
Clb2
Bud3
Bud4
Msn2
Sch9
Rim15
Yak1
Ypl247c
Vps52
Kss1
Sko1
Slt2
Msg5
Swe1
Igo2
Cdc55
Igo1
Vps53
Vps54
Rtn1
Fmp42
Cot1
Zrc1
10
20
30
40
50
t/s

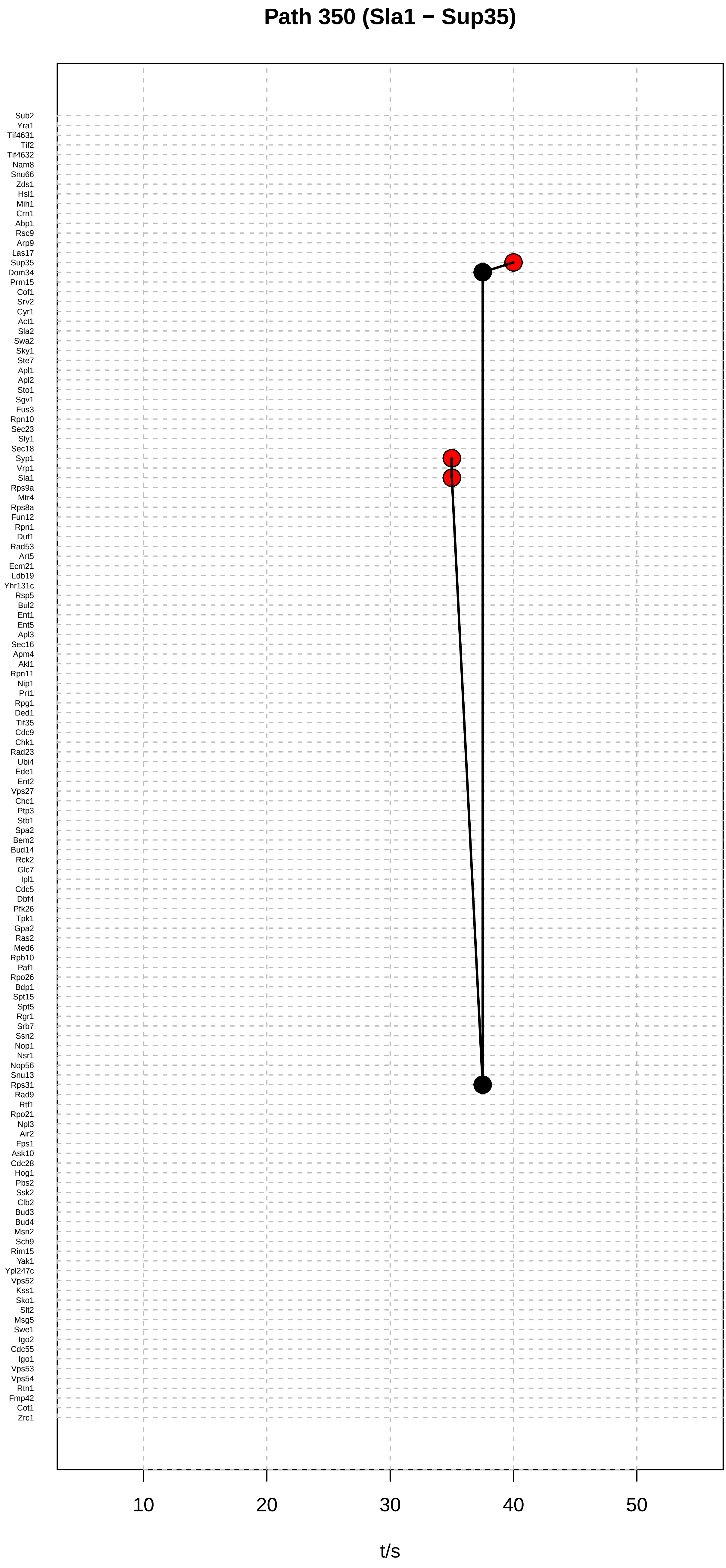
Path 350 (Sla1 − Sup35)
Sub2
Yra1
Tif4631
Tif2
Tif4632
Nam8
Snu66
Zds1
Hsl1
Mih1
Crn1
Abp1
Rsc9
Arp9
Las17
Sup35
Dom34
Prm15
Cof1
Srv2
Cyr1
Act1
Sla2
Swa2
Sky1
Ste7
Apl1
Apl2
Sto1
Sgv1
Fus3
Rpn10
Sec23
Sly1
Sec18
Syp1
Vrp1
Sla1
Rps9a
Mtr4
Rps8a
Fun12
Rpn1
Duf1
Rad53
Art5
Ecm21
Ldb19
Yhr131c
Rsp5
Bul2
Ent1
Ent5
Apl3
Sec16
Apm4
Akl1
Rpn11
Nip1
Prt1
Rpg1
Ded1
Tif35
Cdc9
Chk1
Rad23
Ubi4
Ede1
Ent2
Vps27
Chc1
Ptp3
Stb1
Spa2
Bem2
Bud14
Rck2
Glc7
Ipl1
Cdc5
Dbf4
Pfk26
Tpk1
Gpa2
Ras2
Med6
Rpb10
Paf1
Rpo26
Bdp1
Spt15
Spt5
Rgr1
Srb7
Ssn2
Nop1
Nsr1
Nop56
Snu13
Rps31
Rad9
Rtf1
Rpo21
Npl3
Air2
Fps1
Ask10
Cdc28
Hog1
Pbs2
Ssk2
Clb2
Bud3
Bud4
Msn2
Sch9
Rim15
Yak1
Ypl247c
Vps52
Kss1
Sko1
Slt2
Msg5
Swe1
Igo2
Cdc55
Igo1
Vps53
Vps54
Rtn1
Fmp42
Cot1
Zrc1
10
20
30
40
50
t/s

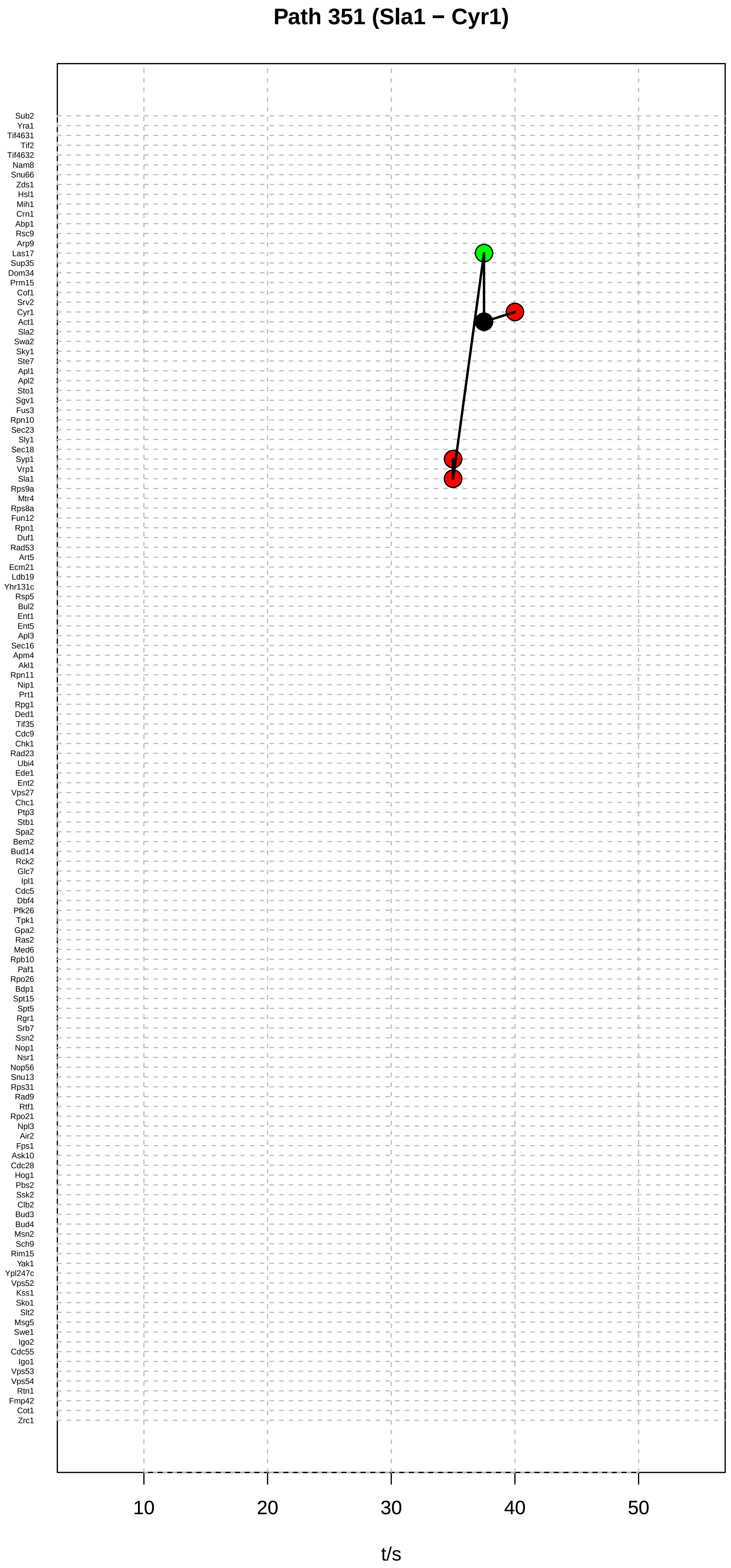
Path 351 (Sla1 − Cyr1)
Sub2
Yra1
Tif4631
Tif2
Tif4632
Nam8
Snu66
Zds1
Hsl1
Mih1
Crn1
Abp1
Rsc9
Arp9
Las17
Sup35
Dom34
Prm15
Cof1
Srv2
Cyr1
Act1
Sla2
Swa2
Sky1
Ste7
Apl1
Apl2
Sto1
Sgv1
Fus3
Rpn10
Sec23
Sly1
Sec18
Syp1
Vrp1
Sla1
Rps9a
Mtr4
Rps8a
Fun12
Rpn1
Duf1
Rad53
Art5
Ecm21
Ldb19
Yhr131c
Rsp5
Bul2
Ent1
Ent5
Apl3
Sec16
Apm4
Akl1
Rpn11
Nip1
Prt1
Rpg1
Ded1
Tif35
Cdc9
Chk1
Rad23
Ubi4
Ede1
Ent2
Vps27
Chc1
Ptp3
Stb1
Spa2
Bem2
Bud14
Rck2
Glc7
Ipl1
Cdc5
Dbf4
Pfk26
Tpk1
Gpa2
Ras2
Med6
Rpb10
Paf1
Rpo26
Bdp1
Spt15
Spt5
Rgr1
Srb7
Ssn2
Nop1
Nsr1
Nop56
Snu13
Rps31
Rad9
Rtf1
Rpo21
Npl3
Air2
Fps1
Ask10
Cdc28
Hog1
Pbs2
Ssk2
Clb2
Bud3
Bud4
Msn2
Sch9
Rim15
Yak1
Ypl247c
Vps52
Kss1
Sko1
Slt2
Msg5
Swe1
Igo2
Cdc55
Igo1
Vps53
Vps54
Rtn1
Fmp42
Cot1
Zrc1
10
20
30
40
50
t/s

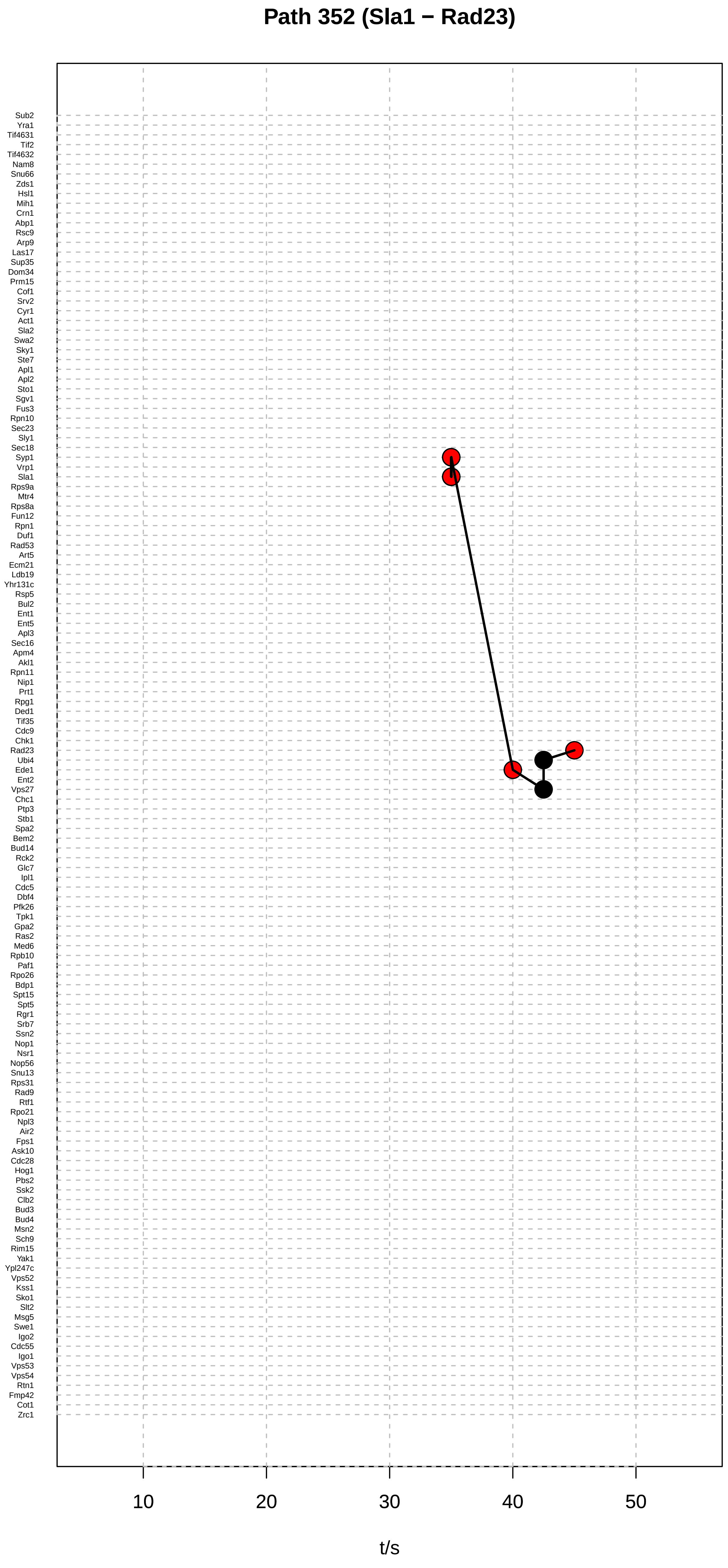

Path 352 (Sla1 − Rad23)
Sub2
Yra1
Tif4631
Tif2
Tif4632
Nam8
Snu66
Zds1
Hsl1
Mih1
Crn1
Abp1
Rsc9
Arp9
Las17
Sup35
Dom34
Prm15
Cof1
Srv2
Cyr1
Act1
Sla2
Swa2
Sky1
Ste7
Apl1
Apl2
Sto1
Sgv1
Fus3
Rpn10
Sec23
Sly1
Sec18
Syp1
Vrp1
Sla1
Rps9a
Mtr4
Rps8a
Fun12
Rpn1
Duf1
Rad53
Art5
Ecm21
Ldb19
Yhr131c
Rsp5
Bul2
Ent1
Ent5
Apl3
Sec16
Apm4
Akl1
Rpn11
Nip1
Prt1
Rpg1
Ded1
Tif35
Cdc9
Chk1
Rad23
Ubi4
Ede1
Ent2
Vps27
Chc1
Ptp3
Stb1
Spa2
Bem2
Bud14
Rck2
Glc7
Ipl1
Cdc5
Dbf4
Pfk26
Tpk1
Gpa2
Ras2
Med6
Rpb10
Paf1
Rpo26
Bdp1
Spt15
Spt5
Rgr1
Srb7
Ssn2
Nop1
Nsr1
Nop56
Snu13
Rps31
Rad9
Rtf1
Rpo21
Npl3
Air2
Fps1
Ask10
Cdc28
Hog1
Pbs2
Ssk2
Clb2
Bud3
Bud4
Msn2
Sch9
Rim15
Yak1
Ypl247c
Vps52
Kss1
Sko1
Slt2
Msg5
Swe1
Igo2
Cdc55
Igo1
Vps53
Vps54
Rtn1
Fmp42
Cot1
Zrc1
10
20
30
40
50
t/s

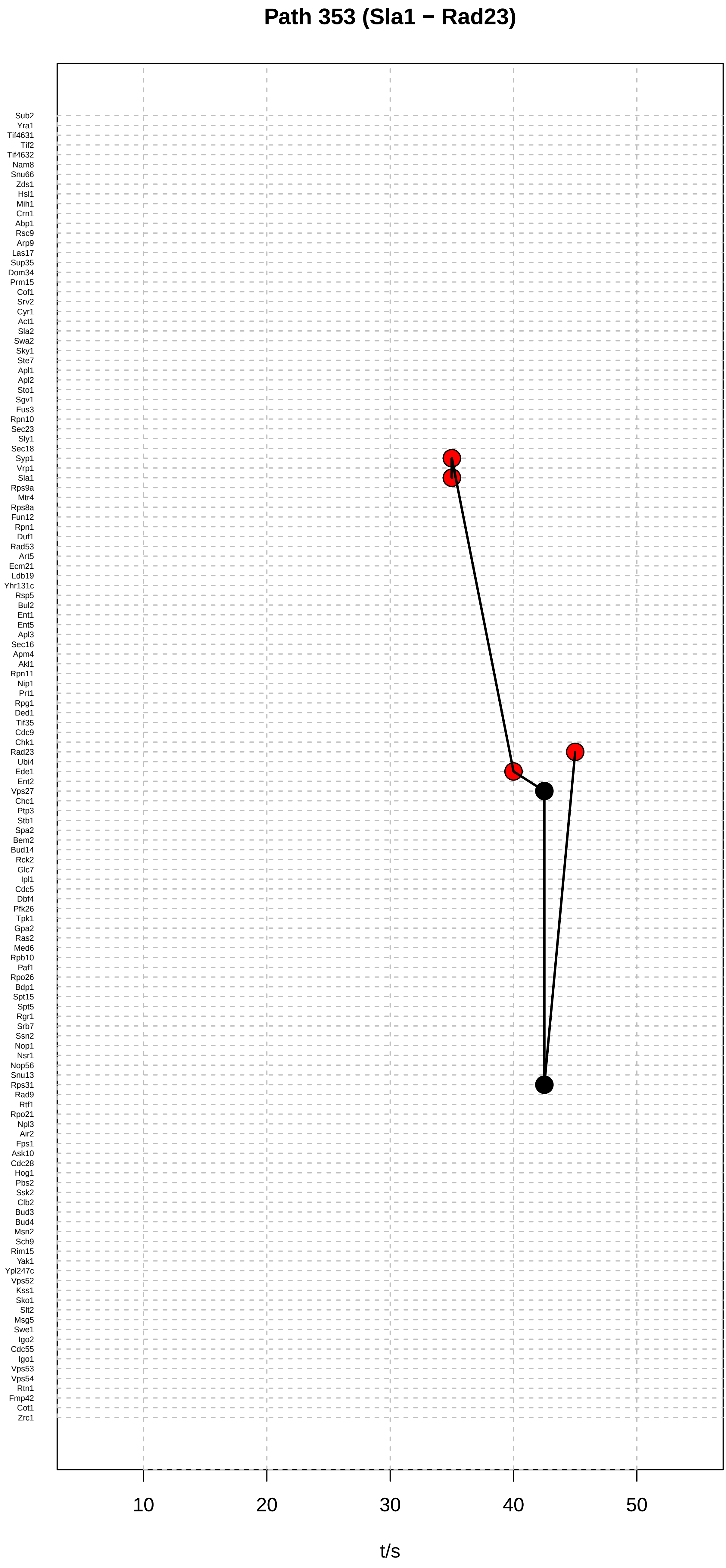
Path 353 (Sla1 − Rad23)
Sub2
Yra1
Tif4631
Tif2
Tif4632
Nam8
Snu66
Zds1
Hsl1
Mih1
Crn1
Abp1
Rsc9
Arp9
Las17
Sup35
Dom34
Prm15
Cof1
Srv2
Cyr1
Act1
Sla2
Swa2
Sky1
Ste7
Apl1
Apl2
Sto1
Sgv1
Fus3
Rpn10
Sec23
Sly1
Sec18
Syp1
Vrp1
Sla1
Rps9a
Mtr4
Rps8a
Fun12
Rpn1
Duf1
Rad53
Art5
Ecm21
Ldb19
Yhr131c
Rsp5
Bul2
Ent1
Ent5
Apl3
Sec16
Apm4
Akl1
Rpn11
Nip1
Prt1
Rpg1
Ded1
Tif35
Cdc9
Chk1
Rad23
Ubi4
Ede1
Ent2
Vps27
Chc1
Ptp3
Stb1
Spa2
Bem2
Bud14
Rck2
Glc7
Ipl1
Cdc5
Dbf4
Pfk26
Tpk1
Gpa2
Ras2
Med6
Rpb10
Paf1
Rpo26
Bdp1
Spt15
Spt5
Rgr1
Srb7
Ssn2
Nop1
Nsr1
Nop56
Snu13
Rps31
Rad9
Rtf1
Rpo21
Npl3
Air2
Fps1
Ask10
Cdc28
Hog1
Pbs2
Ssk2
Clb2
Bud3
Bud4
Msn2
Sch9
Rim15
Yak1
Ypl247c
Vps52
Kss1
Sko1
Slt2
Msg5
Swe1
Igo2
Cdc55
Igo1
Vps53
Vps54
Rtn1
Fmp42
Cot1
Zrc1
10
20
30
40
50
t/s

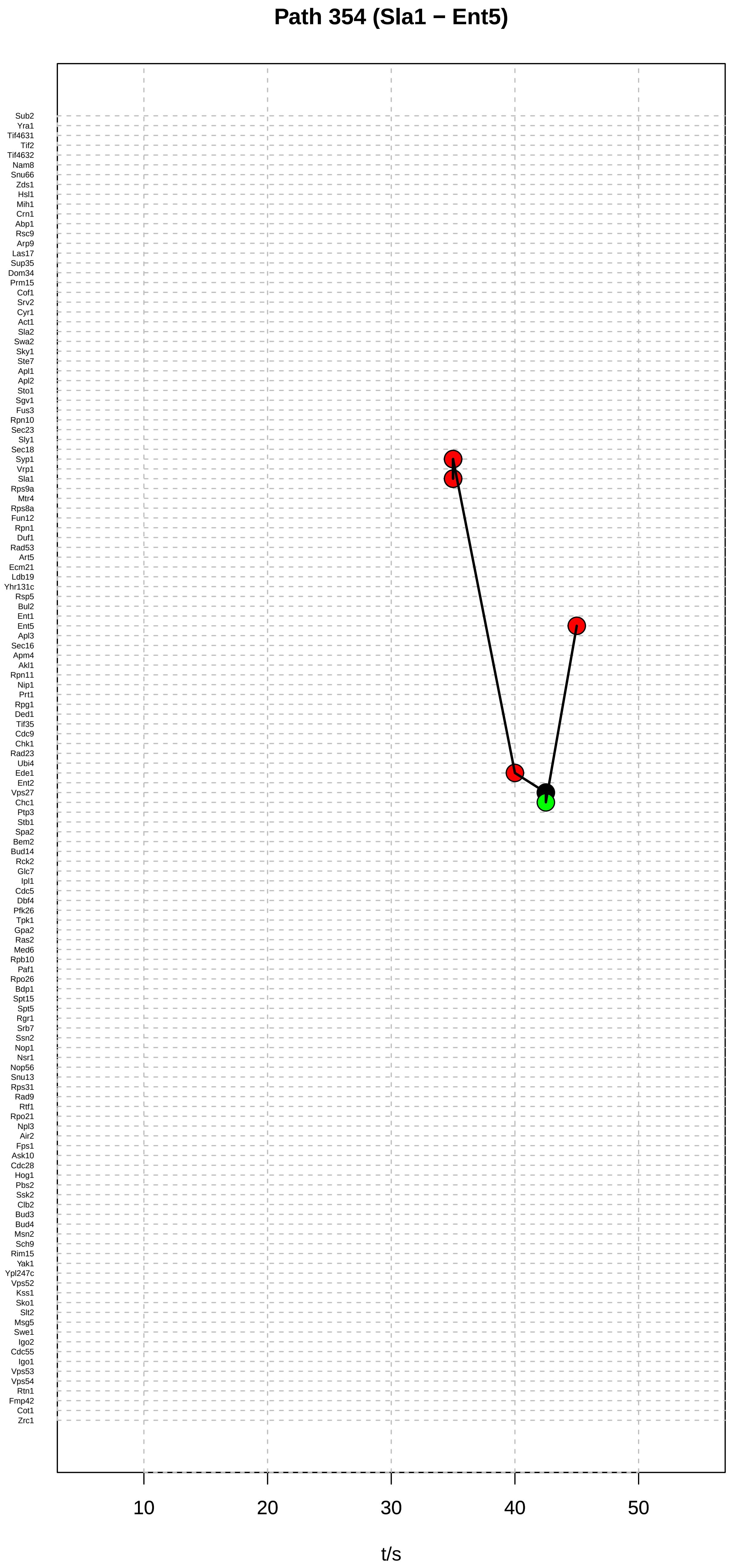

Path 354 (Sla1 − Ent5)
Sub2
Yra1
Tif4631
Tif2
Tif4632
Nam8
Snu66
Zds1
Hsl1
Mih1
Crn1
Abp1
Rsc9
Arp9
Las17
Sup35
Dom34
Prm15
Cof1
Srv2
Cyr1
Act1
Sla2
Swa2
Sky1
Ste7
Apl1
Apl2
Sto1
Sgv1
Fus3
Rpn10
Sec23
Sly1
Sec18
Syp1
Vrp1
Sla1
Rps9a
Mtr4
Rps8a
Fun12
Rpn1
Duf1
Rad53
Art5
Ecm21
Ldb19
Yhr131c
Rsp5
Bul2
Ent1
Ent5
Apl3
Sec16
Apm4
Akl1
Rpn11
Nip1
Prt1
Rpg1
Ded1
Tif35
Cdc9
Chk1
Rad23
Ubi4
Ede1
Ent2
Vps27
Chc1
Ptp3
Stb1
Spa2
Bem2
Bud14
Rck2
Glc7
Ipl1
Cdc5
Dbf4
Pfk26
Tpk1
Gpa2
Ras2
Med6
Rpb10
Paf1
Rpo26
Bdp1
Spt15
Spt5
Rgr1
Srb7
Ssn2
Nop1
Nsr1
Nop56
Snu13
Rps31
Rad9
Rtf1
Rpo21
Npl3
Air2
Fps1
Ask10
Cdc28
Hog1
Pbs2
Ssk2
Clb2
Bud3
Bud4
Msn2
Sch9
Rim15
Yak1
Ypl247c
Vps52
Kss1
Sko1
Slt2
Msg5
Swe1
Igo2
Cdc55
Igo1
Vps53
Vps54
Rtn1
Fmp42
Cot1
Zrc1
10
20
30
40
50
t/s

#### Path 355 (Sla1 − Rad23)

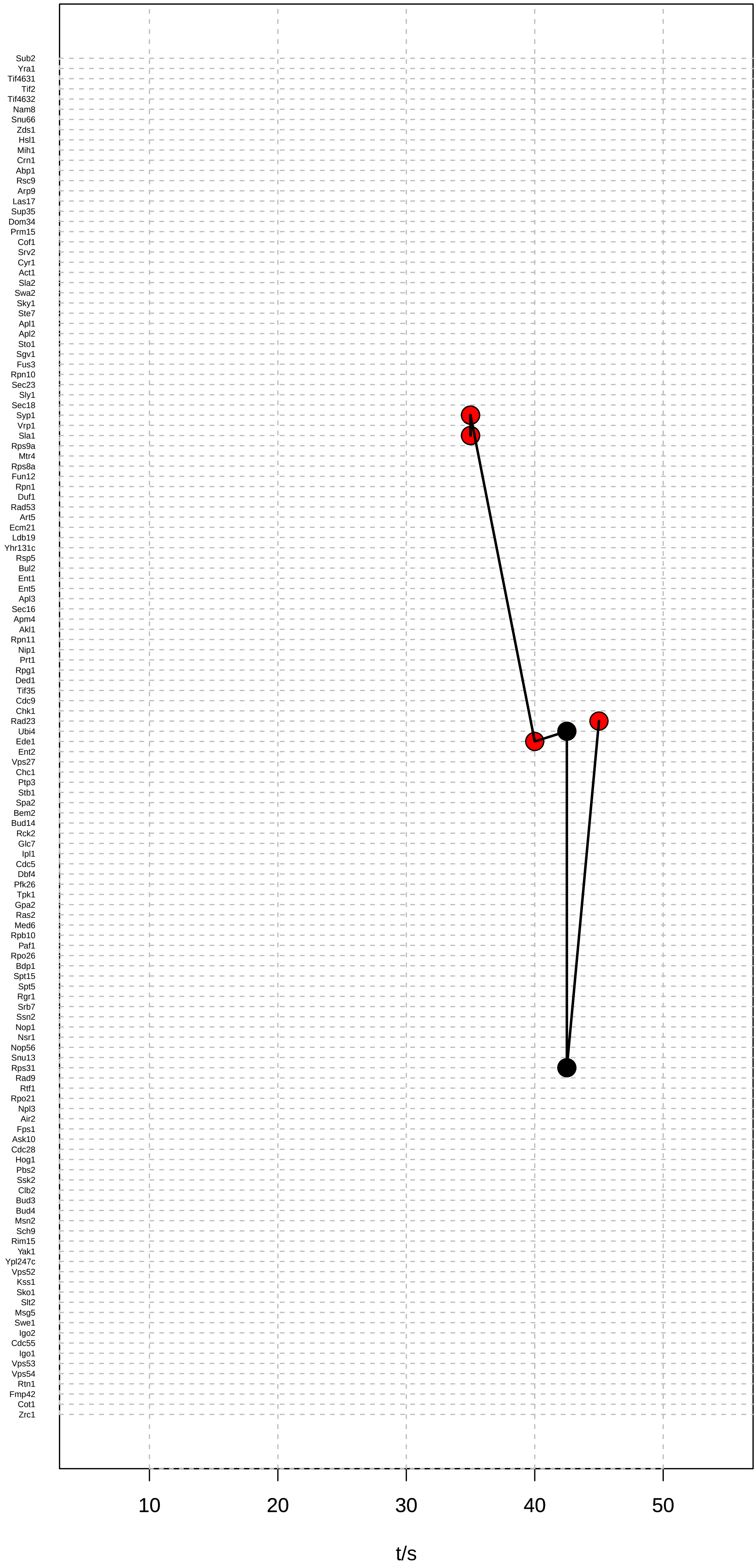

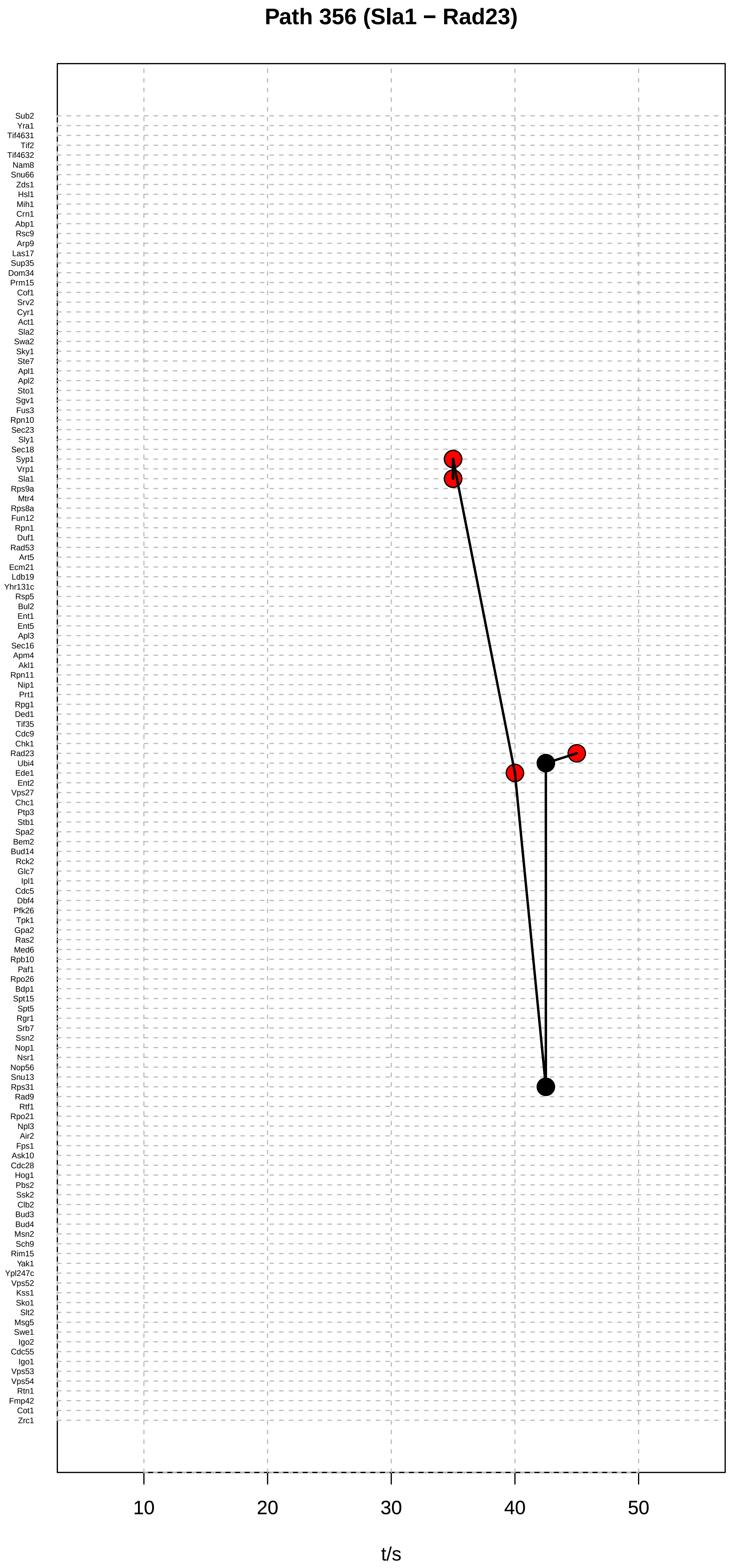
Path 356 (Sla1 − Rad23)
Sub2
Yra1
Tif4631
Tif2
Tif4632
Nam8
Snu66
Zds1
Hsl1
Mih1
Crn1
Abp1
Rsc9
Arp9
Las17
Sup35
Dom34
Prm15
Cof1
Srv2
Cyr1
Act1
Sla2
Swa2
Sky1
Ste7
Apl1
Apl2
Sto1
Sgv1
Fus3
Rpn10
Sec23
Sly1
Sec18
Syp1
Vrp1
Sla1
Rps9a
Mtr4
Rps8a
Fun12
Rpn1
Duf1
Rad53
Art5
Ecm21
Ldb19
Yhr131c
Rsp5
Bul2
Ent1
Ent5
Apl3
Sec16
Apm4
Akl1
Rpn11
Nip1
Prt1
Rpg1
Ded1
Tif35
Cdc9
Chk1
Rad23
Ubi4
Ede1
Ent2
Vps27
Chc1
Ptp3
Stb1
Spa2
Bem2
Bud14
Rck2
Glc7
Ipl1
Cdc5
Dbf4
Pfk26
Tpk1
Gpa2
Ras2
Med6
Rpb10
Paf1
Rpo26
Bdp1
Spt15
Spt5
Rgr1
Srb7
Ssn2
Nop1
Nsr1
Nop56
Snu13
Rps31
Rad9
Rtf1
Rpo21
Npl3
Air2
Fps1
Ask10
Cdc28
Hog1
Pbs2
Ssk2
Clb2
Bud3
Bud4
Msn2
Sch9
Rim15
Yak1
Ypl247c
Vps52
Kss1
Sko1
Slt2
Msg5
Swe1
Igo2
Cdc55
Igo1
Vps53
Vps54
Rtn1
Fmp42
Cot1
Zrc1
10
20
30
40
50
t/s

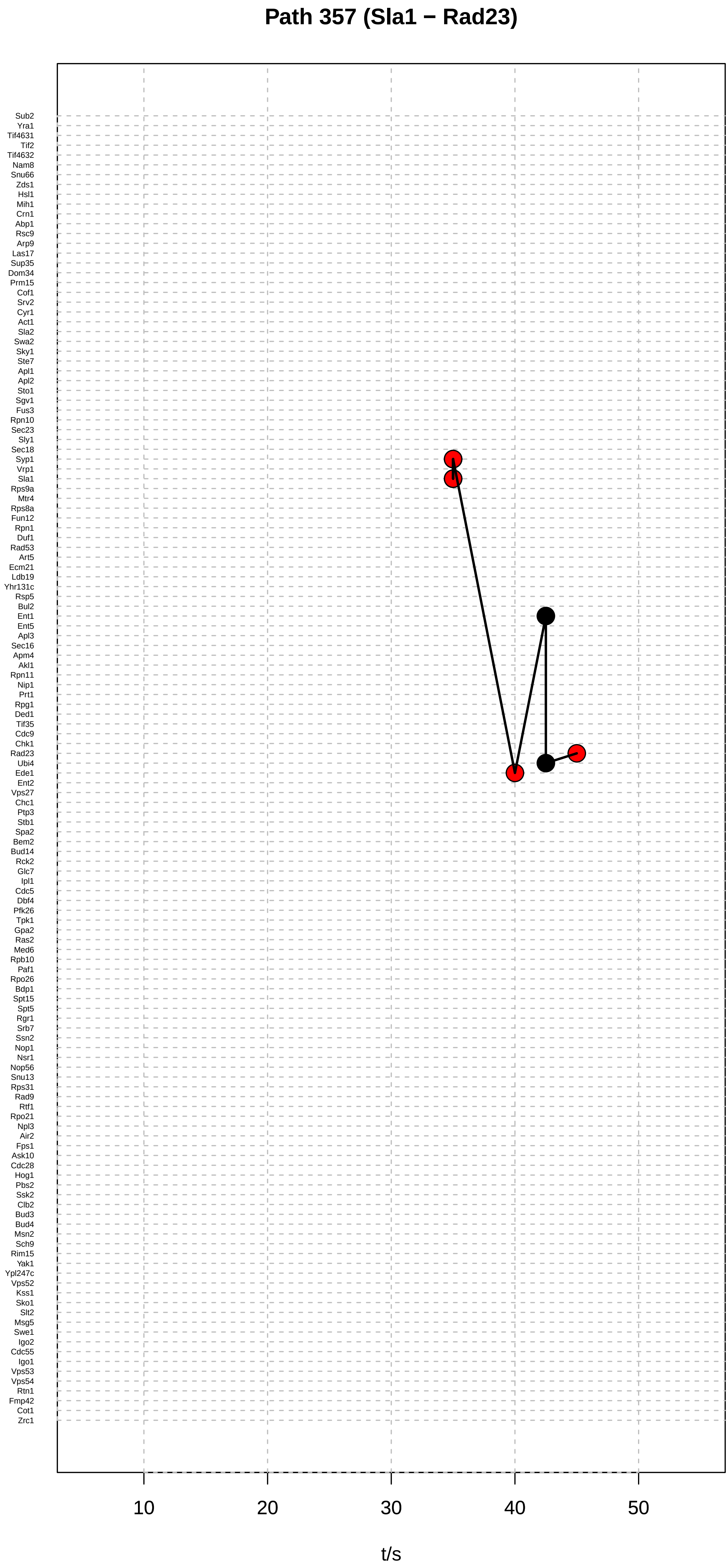
Path 357 (Sla1 − Rad23)
Sub2
Yra1
Tif4631
Tif2
Tif4632
Nam8
Snu66
Zds1
Hsl1
Mih1
Crn1
Abp1
Rsc9
Arp9
Las17
Sup35
Dom34
Prm15
Cof1
Srv2
Cyr1
Act1
Sla2
Swa2
Sky1
Ste7
Apl1
Apl2
Sto1
Sgv1
Fus3
Rpn10
Sec23
Sly1
Sec18
Syp1
Vrp1
Sla1
Rps9a
Mtr4
Rps8a
Fun12
Rpn1
Duf1
Rad53
Art5
Ecm21
Ldb19
Yhr131c
Rsp5
Bul2
Ent1
Ent5
Apl3
Sec16
Apm4
Akl1
Rpn11
Nip1
Prt1
Rpg1
Ded1
Tif35
Cdc9
Chk1
Rad23
Ubi4
Ede1
Ent2
Vps27
Chc1
Ptp3
Stb1
Spa2
Bem2
Bud14
Rck2
Glc7
Ipl1
Cdc5
Dbf4
Pfk26
Tpk1
Gpa2
Ras2
Med6
Rpb10
Paf1
Rpo26
Bdp1
Spt15
Spt5
Rgr1
Srb7
Ssn2
Nop1
Nsr1
Nop56
Snu13
Rps31
Rad9
Rtf1
Rpo21
Npl3
Air2
Fps1
Ask10
Cdc28
Hog1
Pbs2
Ssk2
Clb2
Bud3
Bud4
Msn2
Sch9
Rim15
Yak1
Ypl247c
Vps52
Kss1
Sko1
Slt2
Msg5
Swe1
Igo2
Cdc55
Igo1
Vps53
Vps54
Rtn1
Fmp42
Cot1
Zrc1
10
20
30
40
50
t/s

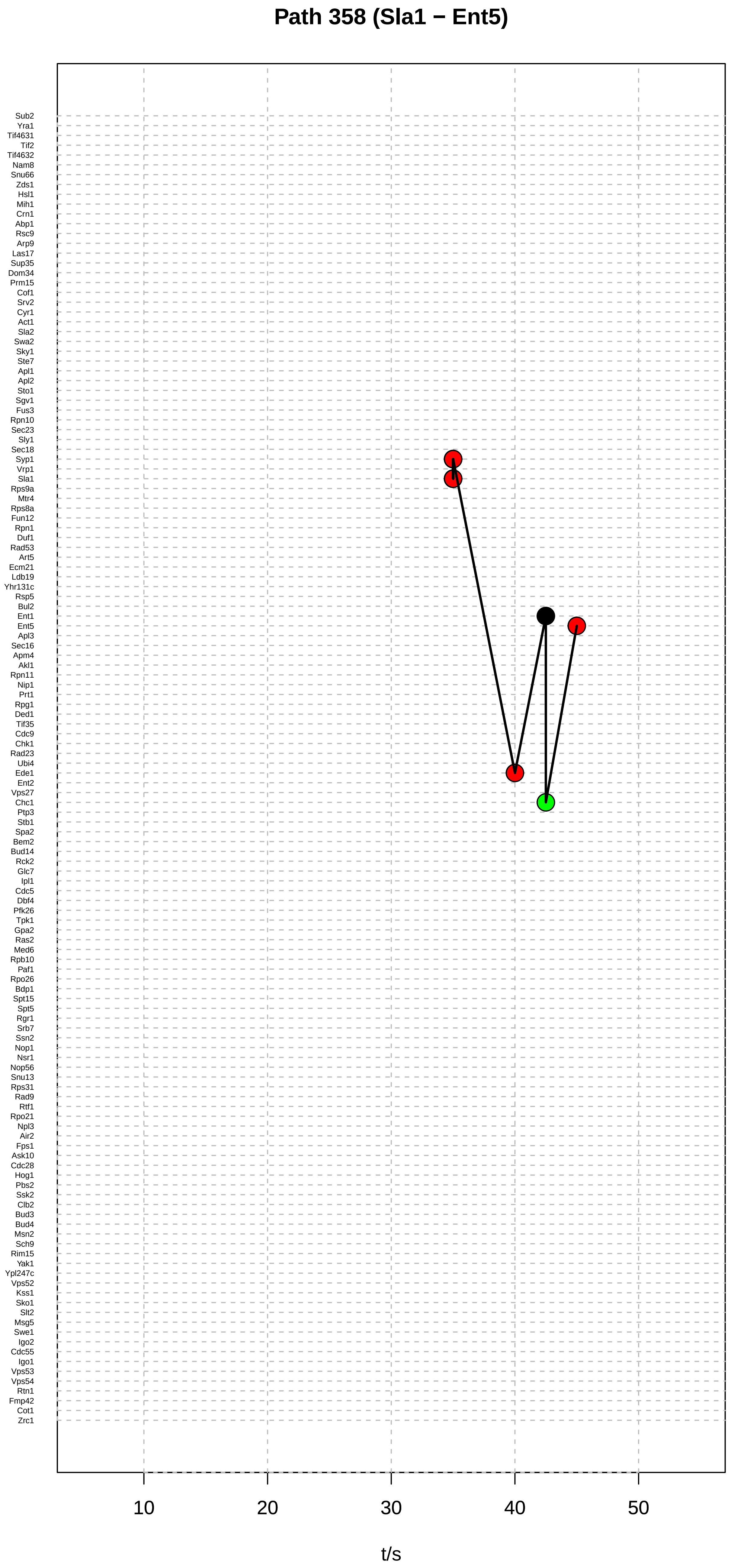
Path 358 (Sla1 − Ent5)
Sub2
Yra1
Tif4631
Tif2
Tif4632
Nam8
Snu66
Zds1
Hsl1
Mih1
Crn1
Abp1
Rsc9
Arp9
Las17
Sup35
Dom34
Prm15
Cof1
Srv2
Cyr1
Act1
Sla2
Swa2
Sky1
Ste7
Apl1
Apl2
Sto1
Sgv1
Fus3
Rpn10
Sec23
Sly1
Sec18
Syp1
Vrp1
Sla1
Rps9a
Mtr4
Rps8a
Fun12
Rpn1
Duf1
Rad53
Art5
Ecm21
Ldb19
Yhr131c
Rsp5
Bul2
Ent1
Ent5
Apl3
Sec16
Apm4
Akl1
Rpn11
Nip1
Prt1
Rpg1
Ded1
Tif35
Cdc9
Chk1
Rad23
Ubi4
Ede1
Ent2
Vps27
Chc1
Ptp3
Stb1
Spa2
Bem2
Bud14
Rck2
Glc7
Ipl1
Cdc5
Dbf4
Pfk26
Tpk1
Gpa2
Ras2
Med6
Rpb10
Paf1
Rpo26
Bdp1
Spt15
Spt5
Rgr1
Srb7
Ssn2
Nop1
Nsr1
Nop56
Snu13
Rps31
Rad9
Rtf1
Rpo21
Npl3
Air2
Fps1
Ask10
Cdc28
Hog1
Pbs2
Ssk2
Clb2
Bud3
Bud4
Msn2
Sch9
Rim15
Yak1
Ypl247c
Vps52
Kss1
Sko1
Slt2
Msg5
Swe1
Igo2
Cdc55
Igo1
Vps53
Vps54
Rtn1
Fmp42
Cot1
Zrc1
10
20
30
40
50
t/s

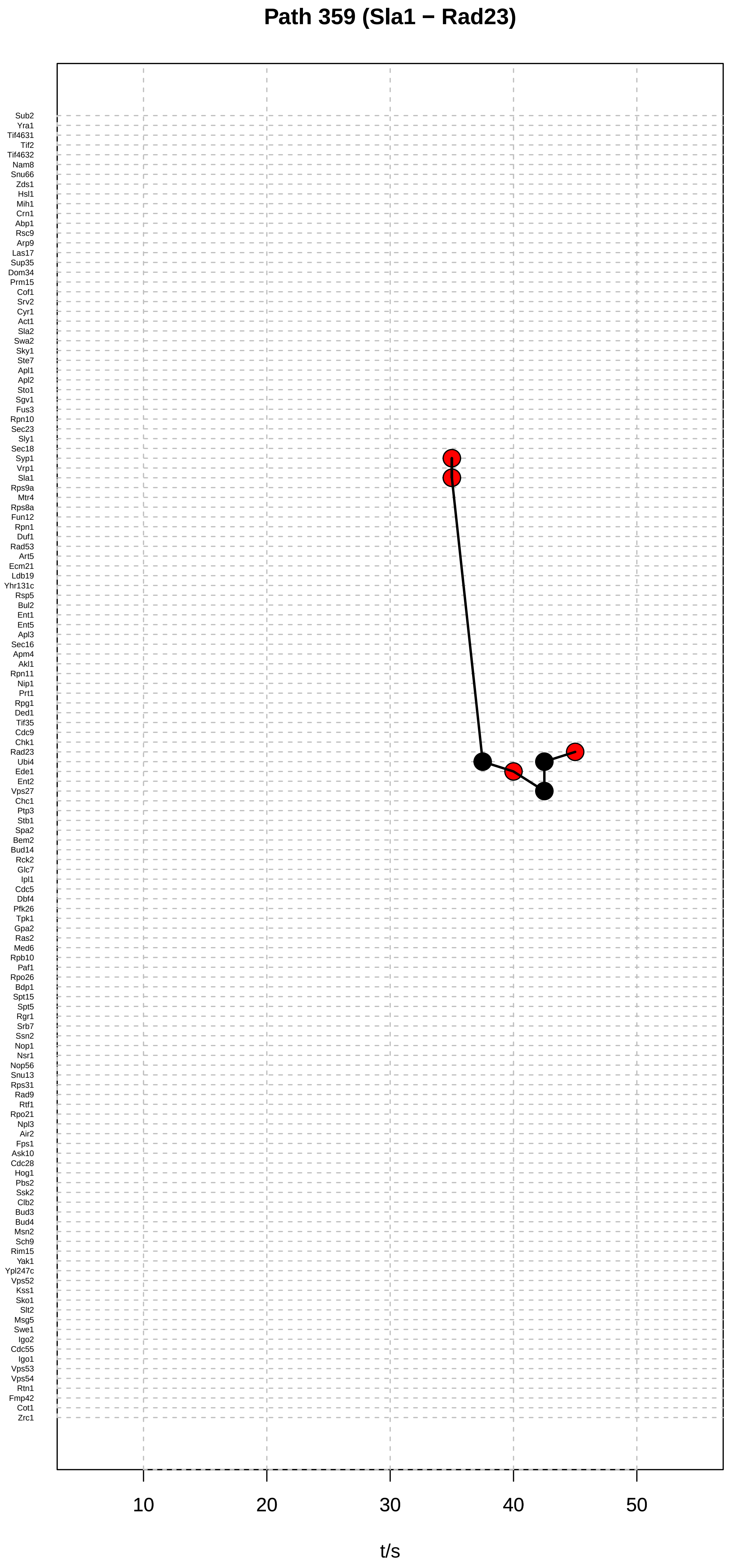
Path 359 (Sla1 − Rad23)
Sub2
Yra1
Tif4631
Tif2
Tif4632
Nam8
Snu66
Zds1
Hsl1
Mih1
Crn1
Abp1
Rsc9
Arp9
Las17
Sup35
Dom34
Prm15
Cof1
Srv2
Cyr1
Act1
Sla2
Swa2
Sky1
Ste7
Apl1
Apl2
Sto1
Sgv1
Fus3
Rpn10
Sec23
Sly1
Sec18
Syp1
Vrp1
Sla1
Rps9a
Mtr4
Rps8a
Fun12
Rpn1
Duf1
Rad53
Art5
Ecm21
Ldb19
Yhr131c
Rsp5
Bul2
Ent1
Ent5
Apl3
Sec16
Apm4
Akl1
Rpn11
Nip1
Prt1
Rpg1
Ded1
Tif35
Cdc9
Chk1
Rad23
Ubi4
Ede1
Ent2
Vps27
Chc1
Ptp3
Stb1
Spa2
Bem2
Bud14
Rck2
Glc7
Ipl1
Cdc5
Dbf4
Pfk26
Tpk1
Gpa2
Ras2
Med6
Rpb10
Paf1
Rpo26
Bdp1
Spt15
Spt5
Rgr1
Srb7
Ssn2
Nop1
Nsr1
Nop56
Snu13
Rps31
Rad9
Rtf1
Rpo21
Npl3
Air2
Fps1
Ask10
Cdc28
Hog1
Pbs2
Ssk2
Clb2
Bud3
Bud4
Msn2
Sch9
Rim15
Yak1
Ypl247c
Vps52
Kss1
Sko1
Slt2
Msg5
Swe1
Igo2
Cdc55
Igo1
Vps53
Vps54
Rtn1
Fmp42
Cot1
Zrc1
10
20
30
40
50
t/s

### Path 360 (Sla1 − Rad23)

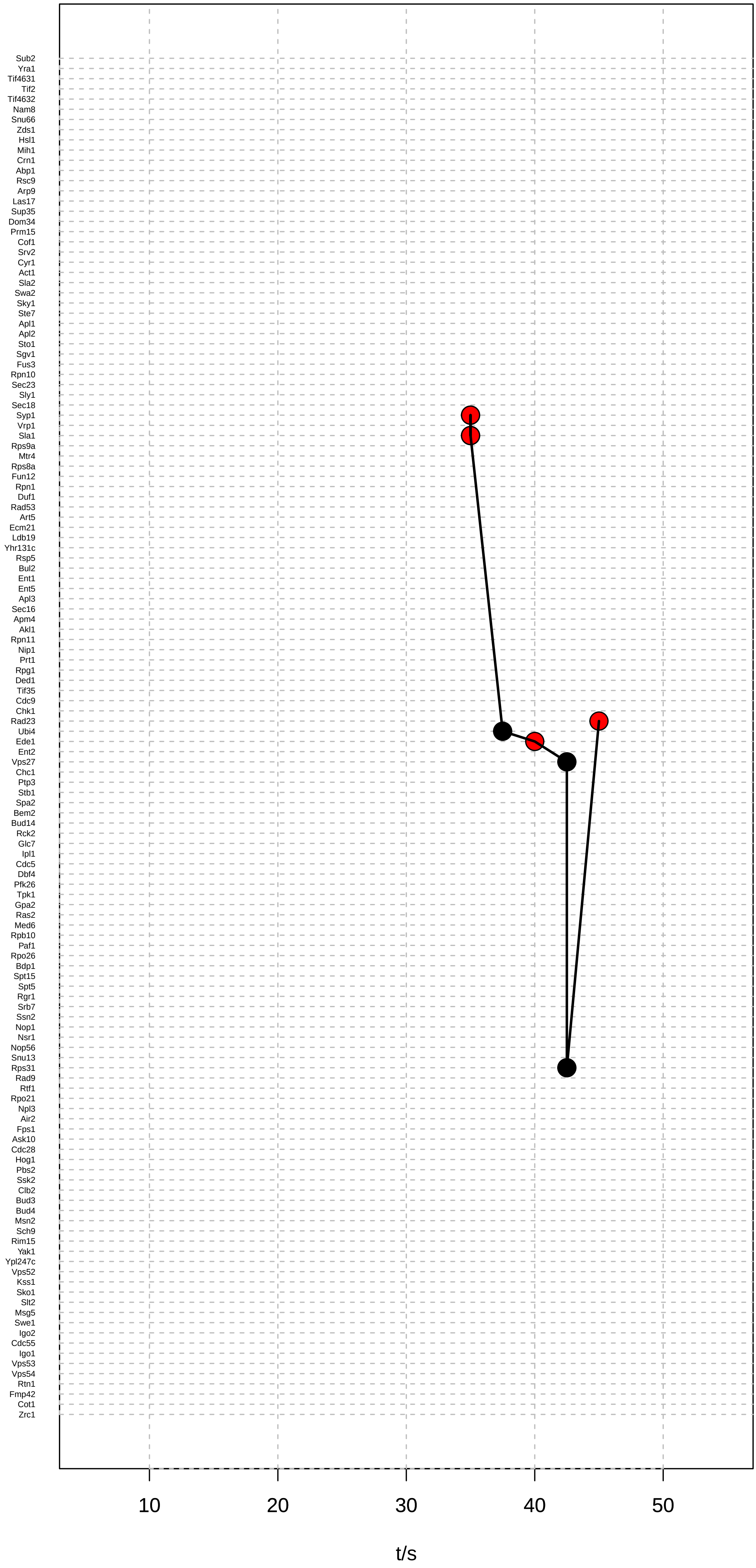

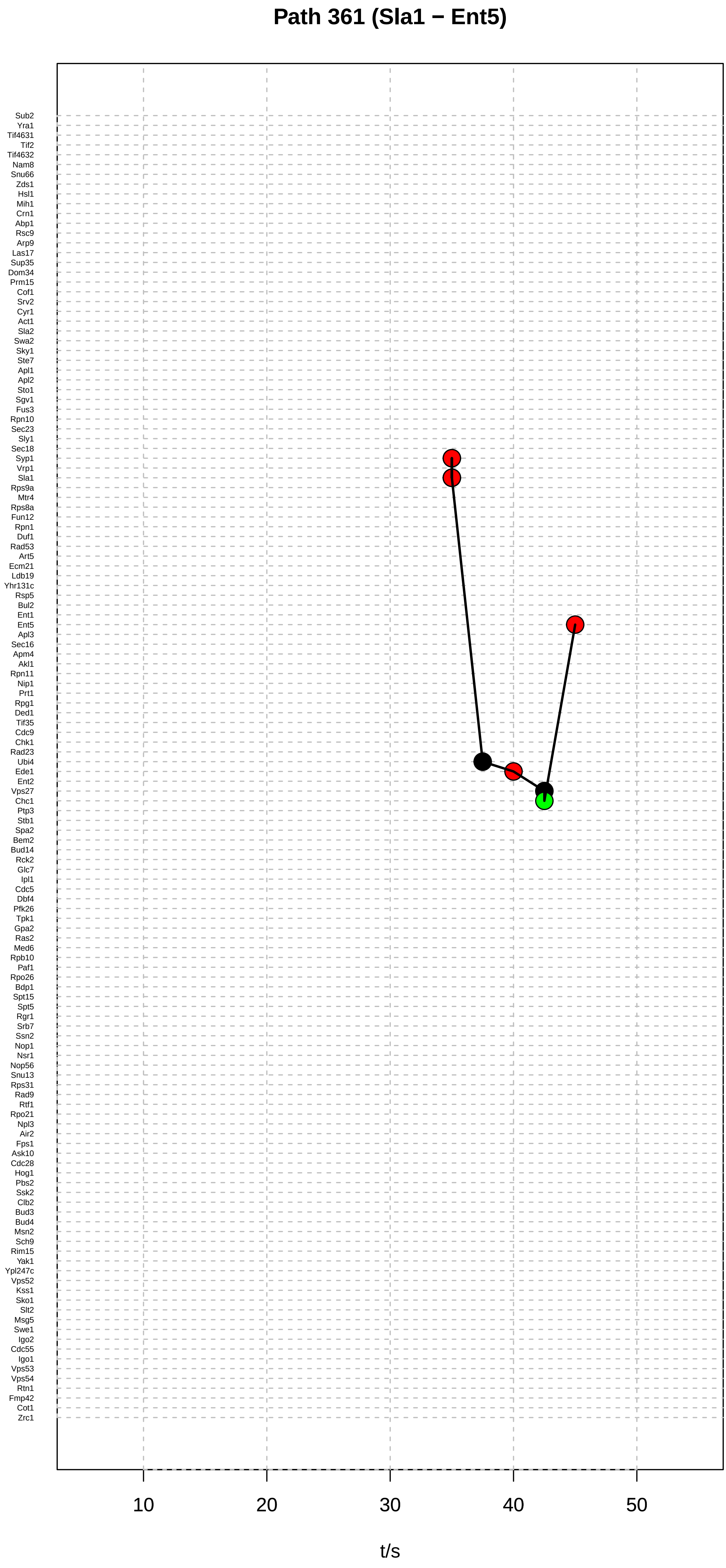

Path 361 (Sla1 − Ent5)
Sub2
Yra1
Tif4631
Tif2
Tif4632
Nam8
Snu66
Zds1
Hsl1
Mih1
Crn1
Abp1
Rsc9
Arp9
Las17
Sup35
Dom34
Prm15
Cof1
Srv2
Cyr1
Act1
Sla2
Swa2
Sky1
Ste7
Apl1
Apl2
Sto1
Sgv1
Fus3
Rpn10
Sec23
Sly1
Sec18
Syp1
Vrp1
Sla1
Rps9a
Mtr4
Rps8a
Fun12
Rpn1
Duf1
Rad53
Art5
Ecm21
Ldb19
Yhr131c
Rsp5
Bul2
Ent1
Ent5
Apl3
Sec16
Apm4
Akl1
Rpn11
Nip1
Prt1
Rpg1
Ded1
Tif35
Cdc9
Chk1
Rad23
Ubi4
Ede1
Ent2
Vps27
Chc1
Ptp3
Stb1
Spa2
Bem2
Bud14
Rck2
Glc7
Ipl1
Cdc5
Dbf4
Pfk26
Tpk1
Gpa2
Ras2
Med6
Rpb10
Paf1
Rpo26
Bdp1
Spt15
Spt5
Rgr1
Srb7
Ssn2
Nop1
Nsr1
Nop56
Snu13
Rps31
Rad9
Rtf1
Rpo21
Npl3
Air2
Fps1
Ask10
Cdc28
Hog1
Pbs2
Ssk2
Clb2
Bud3
Bud4
Msn2
Sch9
Rim15
Yak1
Ypl247c
Vps52
Kss1
Sko1
Slt2
Msg5
Swe1
Igo2
Cdc55
Igo1
Vps53
Vps54
Rtn1
Fmp42
Cot1
Zrc1
10
20
30
40
50
t/s

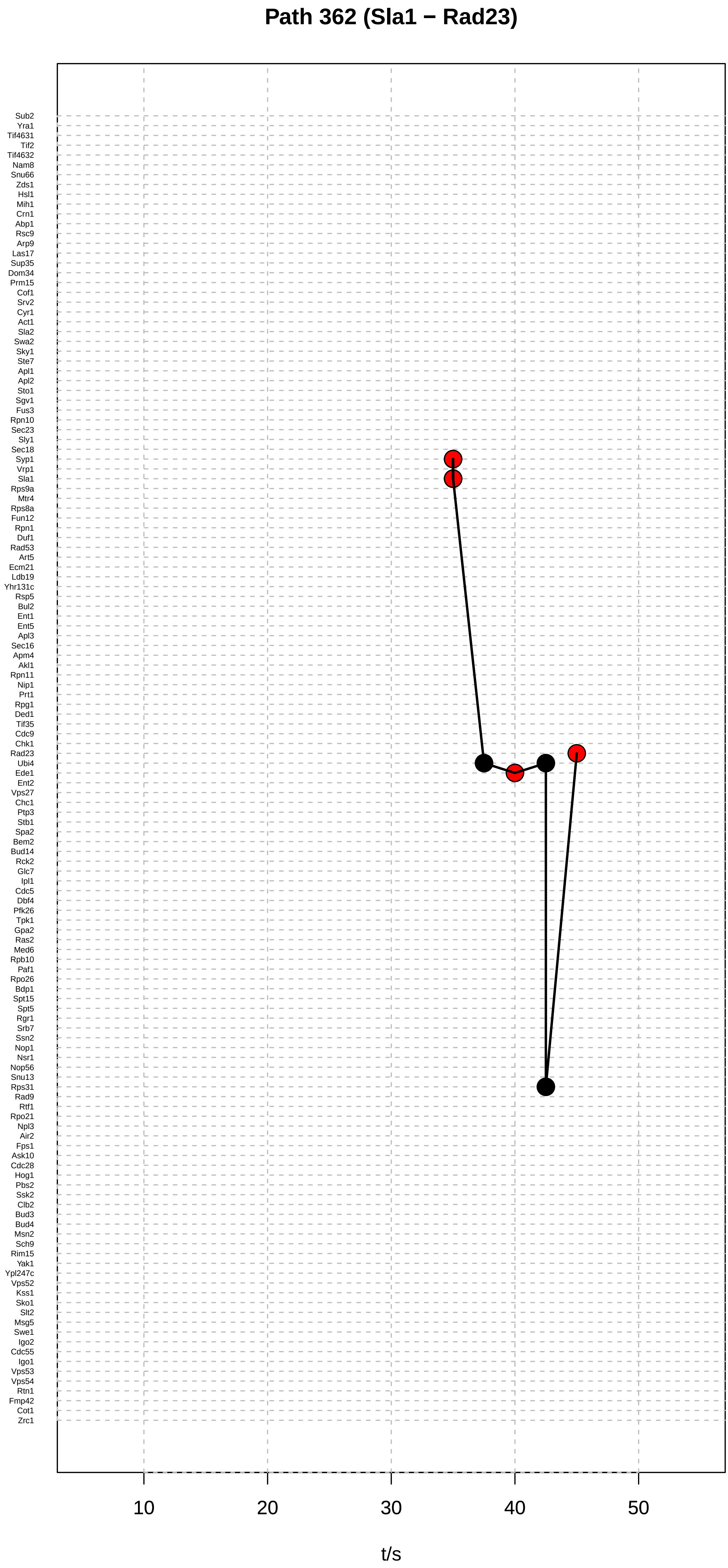

Path 362 (Sla1 − Rad23)
Sub2
Yra1
Tif4631
Tif2
Tif4632
Nam8
Snu66
Zds1
Hsl1
Mih1
Crn1
Abp1
Rsc9
Arp9
Las17
Sup35
Dom34
Prm15
Cof1
Srv2
Cyr1
Act1
Sla2
Swa2
Sky1
Ste7
Apl1
Apl2
Sto1
Sgv1
Fus3
Rpn10
Sec23
Sly1
Sec18
Syp1
Vrp1
Sla1
Rps9a
Mtr4
Rps8a
Fun12
Rpn1
Duf1
Rad53
Art5
Ecm21
Ldb19
Yhr131c
Rsp5
Bul2
Ent1
Ent5
Apl3
Sec16
Apm4
Akl1
Rpn11
Nip1
Prt1
Rpg1
Ded1
Tif35
Cdc9
Chk1
Rad23
Ubi4
Ede1
Ent2
Vps27
Chc1
Ptp3
Stb1
Spa2
Bem2
Bud14
Rck2
Glc7
Ipl1
Cdc5
Dbf4
Pfk26
Tpk1
Gpa2
Ras2
Med6
Rpb10
Paf1
Rpo26
Bdp1
Spt15
Spt5
Rgr1
Srb7
Ssn2
Nop1
Nsr1
Nop56
Snu13
Rps31
Rad9
Rtf1
Rpo21
Npl3
Air2
Fps1
Ask10
Cdc28
Hog1
Pbs2
Ssk2
Clb2
Bud3
Bud4
Msn2
Sch9
Rim15
Yak1
Ypl247c
Vps52
Kss1
Sko1
Slt2
Msg5
Swe1
Igo2
Cdc55
Igo1
Vps53
Vps54
Rtn1
Fmp42
Cot1
Zrc1
10
20
30
40
50
t/s

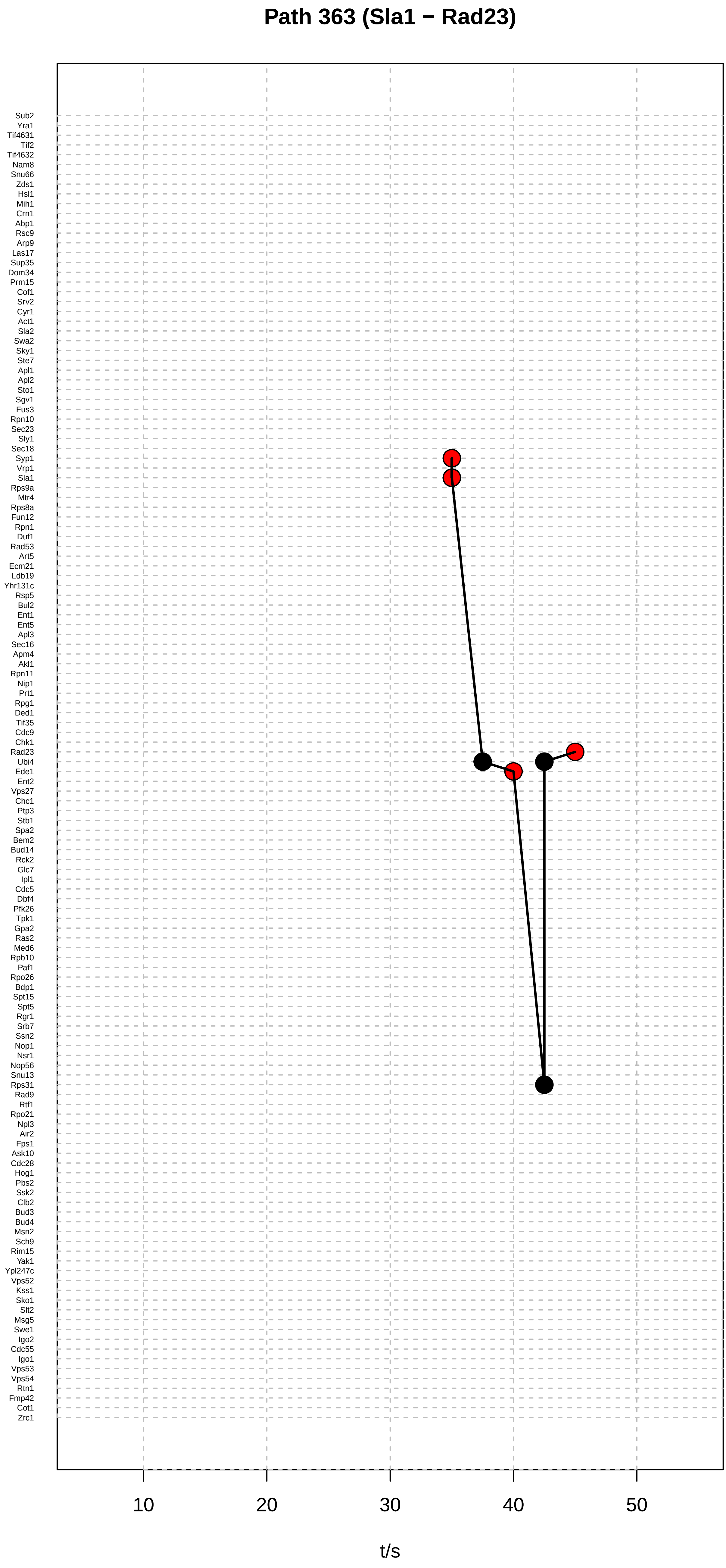

Path 363 (Sla1 − Rad23)
Sub2
Yra1
Tif4631
Tif2
Tif4632
Nam8
Snu66
Zds1
Hsl1
Mih1
Crn1
Abp1
Rsc9
Arp9
Las17
Sup35
Dom34
Prm15
Cof1
Srv2
Cyr1
Act1
Sla2
Swa2
Sky1
Ste7
Apl1
Apl2
Sto1
Sgv1
Fus3
Rpn10
Sec23
Sly1
Sec18
Syp1
Vrp1
Sla1
Rps9a
Mtr4
Rps8a
Fun12
Rpn1
Duf1
Rad53
Art5
Ecm21
Ldb19
Yhr131c
Rsp5
Bul2
Ent1
Ent5
Apl3
Sec16
Apm4
Akl1
Rpn11
Nip1
Prt1
Rpg1
Ded1
Tif35
Cdc9
Chk1
Rad23
Ubi4
Ede1
Ent2
Vps27
Chc1
Ptp3
Stb1
Spa2
Bem2
Bud14
Rck2
Glc7
Ipl1
Cdc5
Dbf4
Pfk26
Tpk1
Gpa2
Ras2
Med6
Rpb10
Paf1
Rpo26
Bdp1
Spt15
Spt5
Rgr1
Srb7
Ssn2
Nop1
Nsr1
Nop56
Snu13
Rps31
Rad9
Rtf1
Rpo21
Npl3
Air2
Fps1
Ask10
Cdc28
Hog1
Pbs2
Ssk2
Clb2
Bud3
Bud4
Msn2
Sch9
Rim15
Yak1
Ypl247c
Vps52
Kss1
Sko1
Slt2
Msg5
Swe1
Igo2
Cdc55
Igo1
Vps53
Vps54
Rtn1
Fmp42
Cot1
Zrc1
10
20
30
40
50
t/s

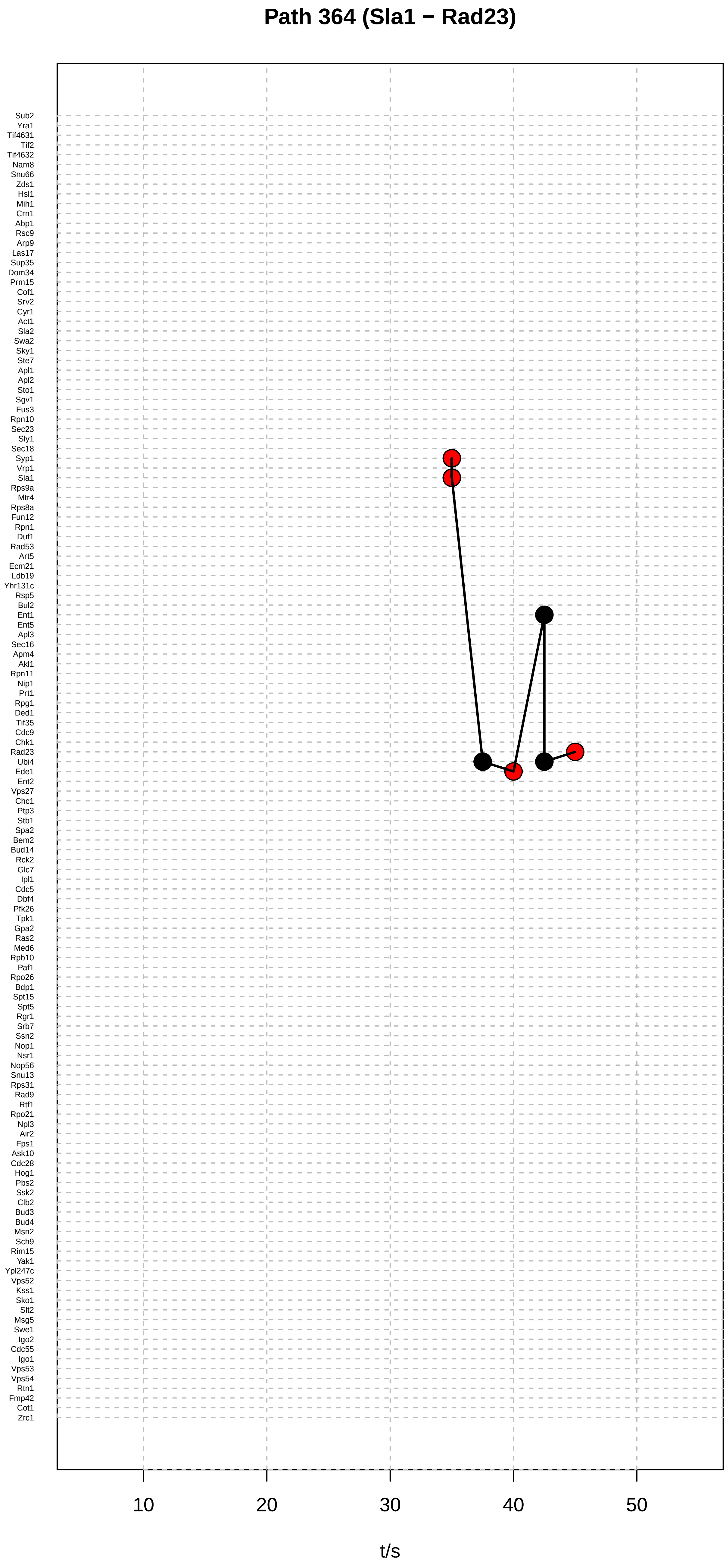

Path 364 (Sla1 − Rad23)
Sub2
Yra1
Tif4631
Tif2
Tif4632
Nam8
Snu66
Zds1
Hsl1
Mih1
Crn1
Abp1
Rsc9
Arp9
Las17
Sup35
Dom34
Prm15
Cof1
Srv2
Cyr1
Act1
Sla2
Swa2
Sky1
Ste7
Apl1
Apl2
Sto1
Sgv1
Fus3
Rpn10
Sec23
Sly1
Sec18
Syp1
Vrp1
Sla1
Rps9a
Mtr4
Rps8a
Fun12
Rpn1
Duf1
Rad53
Art5
Ecm21
Ldb19
Yhr131c
Rsp5
Bul2
Ent1
Ent5
Apl3
Sec16
Apm4
Akl1
Rpn11
Nip1
Prt1
Rpg1
Ded1
Tif35
Cdc9
Chk1
Rad23
Ubi4
Ede1
Ent2
Vps27
Chc1
Ptp3
Stb1
Spa2
Bem2
Bud14
Rck2
Glc7
Ipl1
Cdc5
Dbf4
Pfk26
Tpk1
Gpa2
Ras2
Med6
Rpb10
Paf1
Rpo26
Bdp1
Spt15
Spt5
Rgr1
Srb7
Ssn2
Nop1
Nsr1
Nop56
Snu13
Rps31
Rad9
Rtf1
Rpo21
Npl3
Air2
Fps1
Ask10
Cdc28
Hog1
Pbs2
Ssk2
Clb2
Bud3
Bud4
Msn2
Sch9
Rim15
Yak1
Ypl247c
Vps52
Kss1
Sko1
Slt2
Msg5
Swe1
Igo2
Cdc55
Igo1
Vps53
Vps54
Rtn1
Fmp42
Cot1
Zrc1
10
20
30
40
50
t/s

### Path 365 (Sla1 − Ent5)

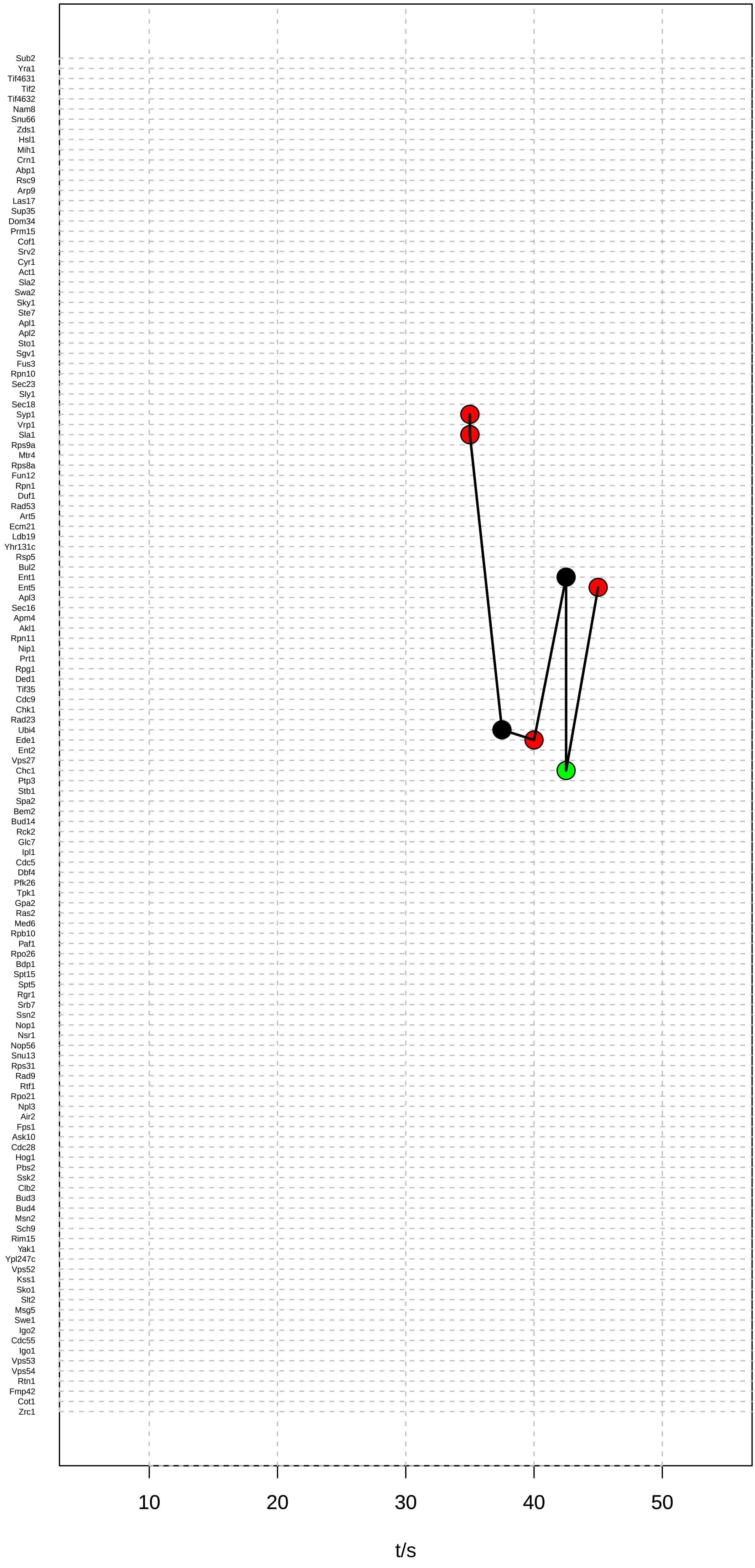

### Path 366 (Sla1 − Rad23)

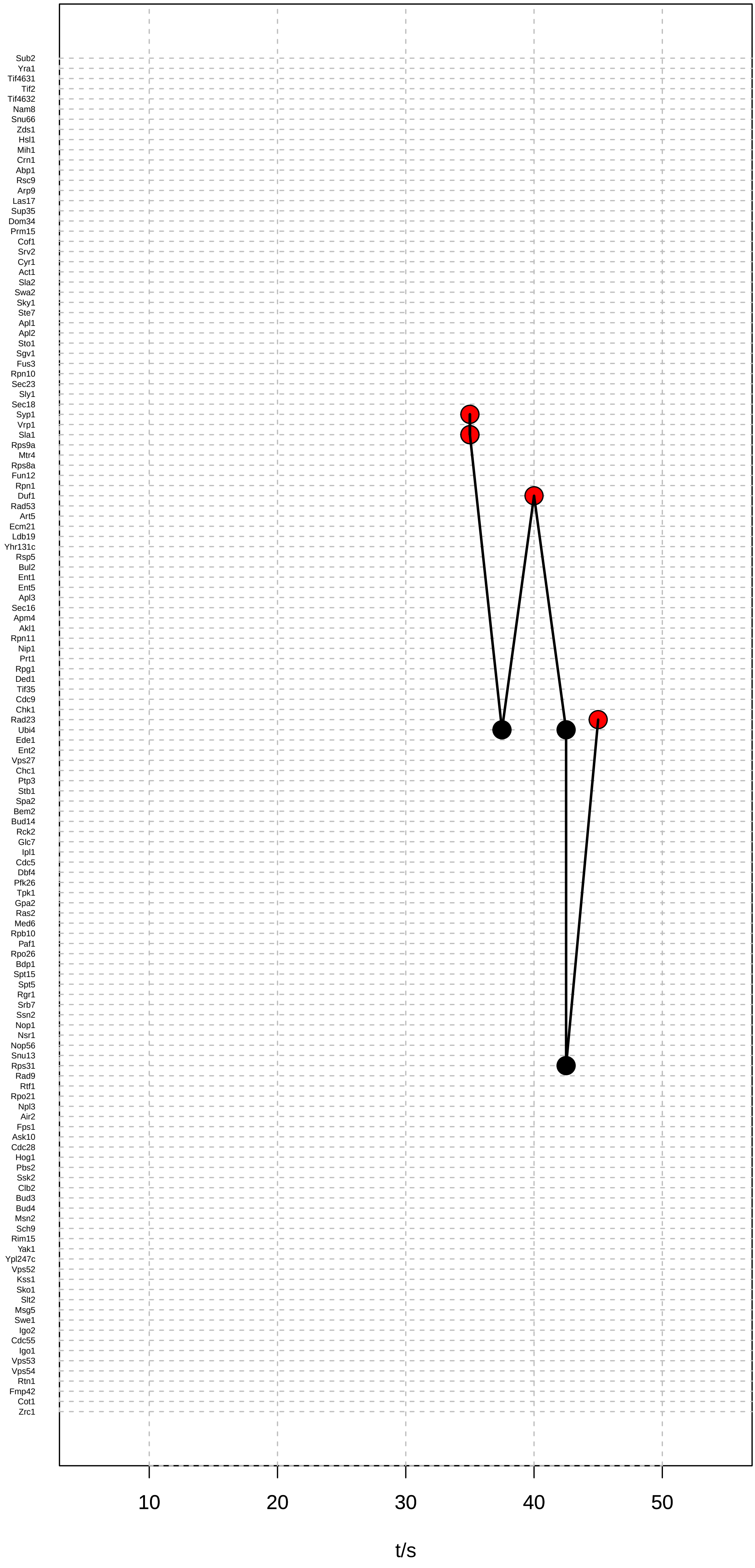

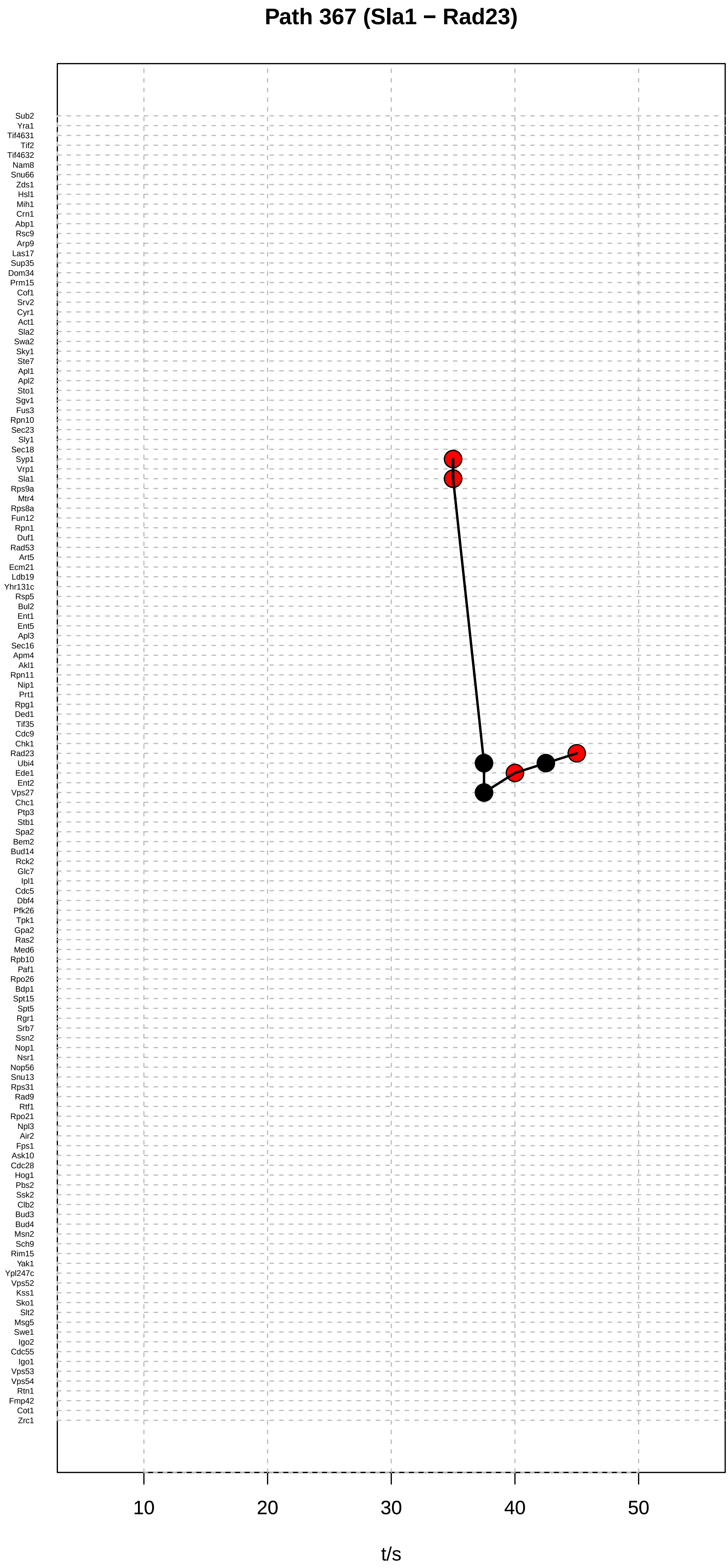
Path 367 (Sla1 − Rad23)
Sub2
Yra1
Tif4631
Tif2
Tif4632
Nam8
Snu66
Zds1
Hsl1
Mih1
Crn1
Abp1
Rsc9
Arp9
Las17
Sup35
Dom34
Prm15
Cof1
Srv2
Cyr1
Act1
Sla2
Swa2
Sky1
Ste7
Apl1
Apl2
Sto1
Sgv1
Fus3
Rpn10
Sec23
Sly1
Sec18
Syp1
Vrp1
Sla1
Rps9a
Mtr4
Rps8a
Fun12
Rpn1
Duf1
Rad53
Art5
Ecm21
Ldb19
Yhr131c
Rsp5
Bul2
Ent1
Ent5
Apl3
Sec16
Apm4
Akl1
Rpn11
Nip1
Prt1
Rpg1
Ded1
Tif35
Cdc9
Chk1
Rad23
Ubi4
Ede1
Ent2
Vps27
Chc1
Ptp3
Stb1
Spa2
Bem2
Bud14
Rck2
Glc7
Ipl1
Cdc5
Dbf4
Pfk26
Tpk1
Gpa2
Ras2
Med6
Rpb10
Paf1
Rpo26
Bdp1
Spt15
Spt5
Rgr1
Srb7
Ssn2
Nop1
Nsr1
Nop56
Snu13
Rps31
Rad9
Rtf1
Rpo21
Npl3
Air2
Fps1
Ask10
Cdc28
Hog1
Pbs2
Ssk2
Clb2
Bud3
Bud4
Msn2
Sch9
Rim15
Yak1
Ypl247c
Vps52
Kss1
Sko1
Slt2
Msg5
Swe1
Igo2
Cdc55
Igo1
Vps53
Vps54
Rtn1
Fmp42
Cot1
Zrc1
10
20
30
40
50
t/s

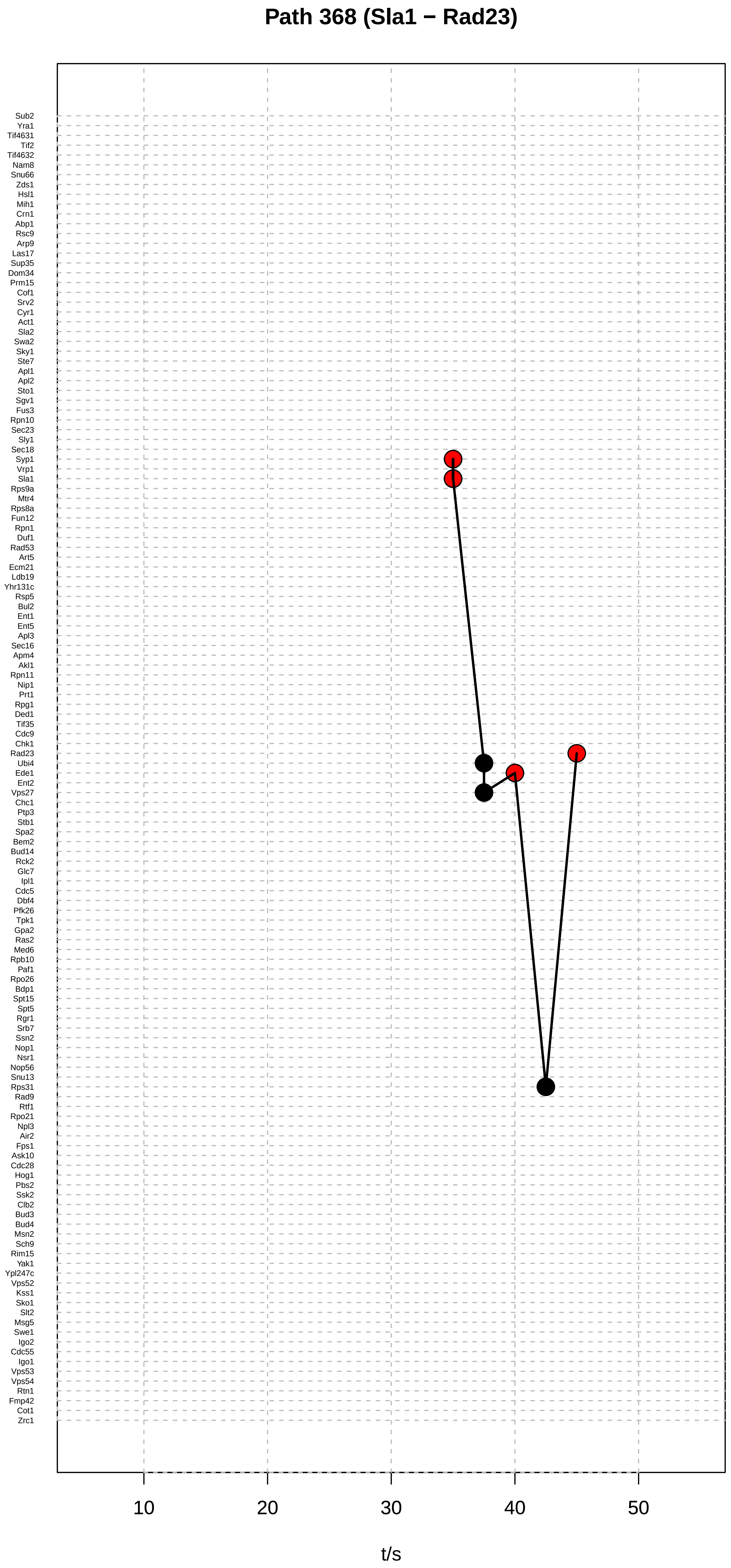
Path 368 (Sla1 − Rad23)
Sub2
Yra1
Tif4631
Tif2
Tif4632
Nam8
Snu66
Zds1
Hsl1
Mih1
Crn1
Abp1
Rsc9
Arp9
Las17
Sup35
Dom34
Prm15
Cof1
Srv2
Cyr1
Act1
Sla2
Swa2
Sky1
Ste7
Apl1
Apl2
Sto1
Sgv1
Fus3
Rpn10
Sec23
Sly1
Sec18
Syp1
Vrp1
Sla1
Rps9a
Mtr4
Rps8a
Fun12
Rpn1
Duf1
Rad53
Art5
Ecm21
Ldb19
Yhr131c
Rsp5
Bul2
Ent1
Ent5
Apl3
Sec16
Apm4
Akl1
Rpn11
Nip1
Prt1
Rpg1
Ded1
Tif35
Cdc9
Chk1
Rad23
Ubi4
Ede1
Ent2
Vps27
Chc1
Ptp3
Stb1
Spa2
Bem2
Bud14
Rck2
Glc7
Ipl1
Cdc5
Dbf4
Pfk26
Tpk1
Gpa2
Ras2
Med6
Rpb10
Paf1
Rpo26
Bdp1
Spt15
Spt5
Rgr1
Srb7
Ssn2
Nop1
Nsr1
Nop56
Snu13
Rps31
Rad9
Rtf1
Rpo21
Npl3
Air2
Fps1
Ask10
Cdc28
Hog1
Pbs2
Ssk2
Clb2
Bud3
Bud4
Msn2
Sch9
Rim15
Yak1
Ypl247c
Vps52
Kss1
Sko1
Slt2
Msg5
Swe1
Igo2
Cdc55
Igo1
Vps53
Vps54
Rtn1
Fmp42
Cot1
Zrc1
10
20
30
40
50
t/s

#### Path 369 (Sla1 − Rad23)

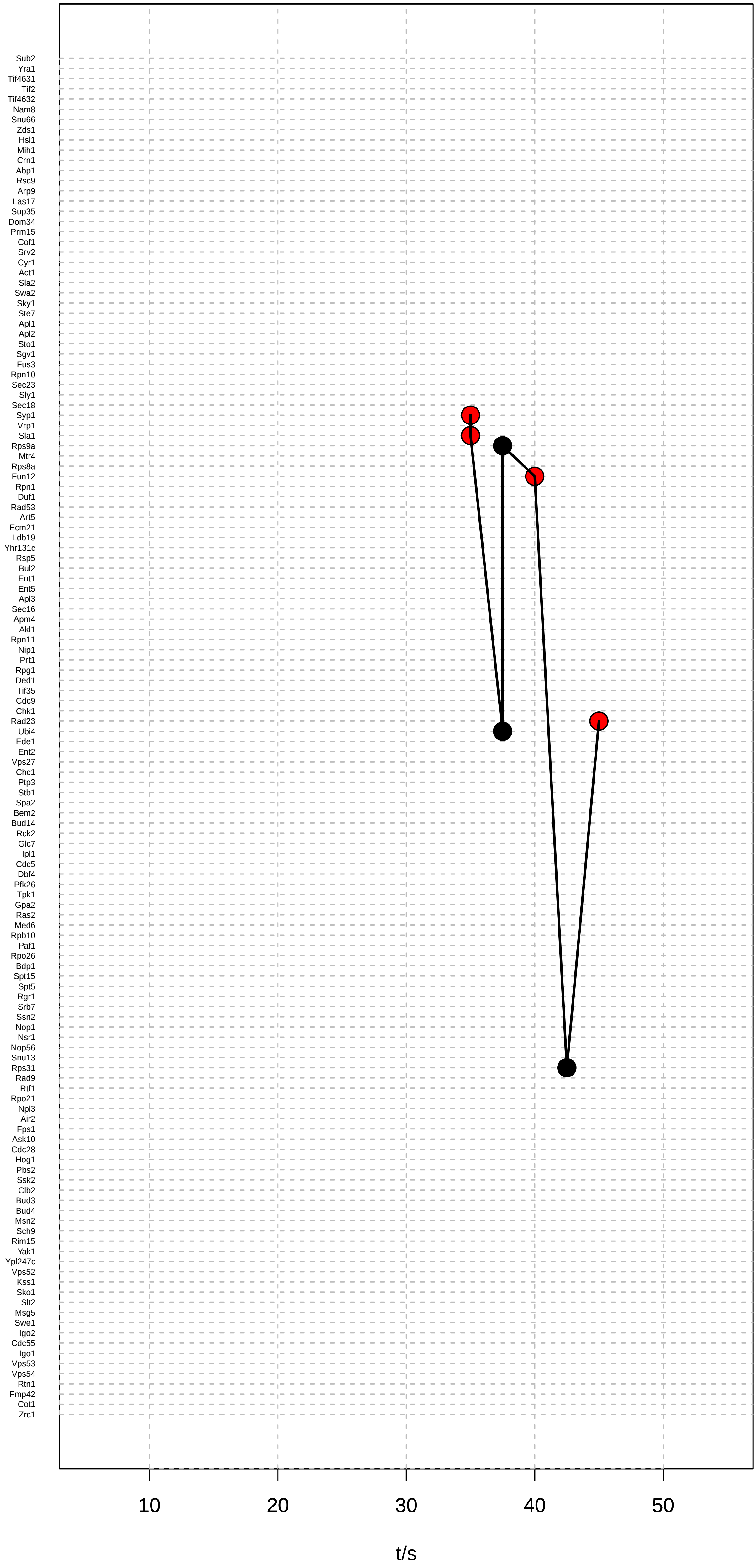

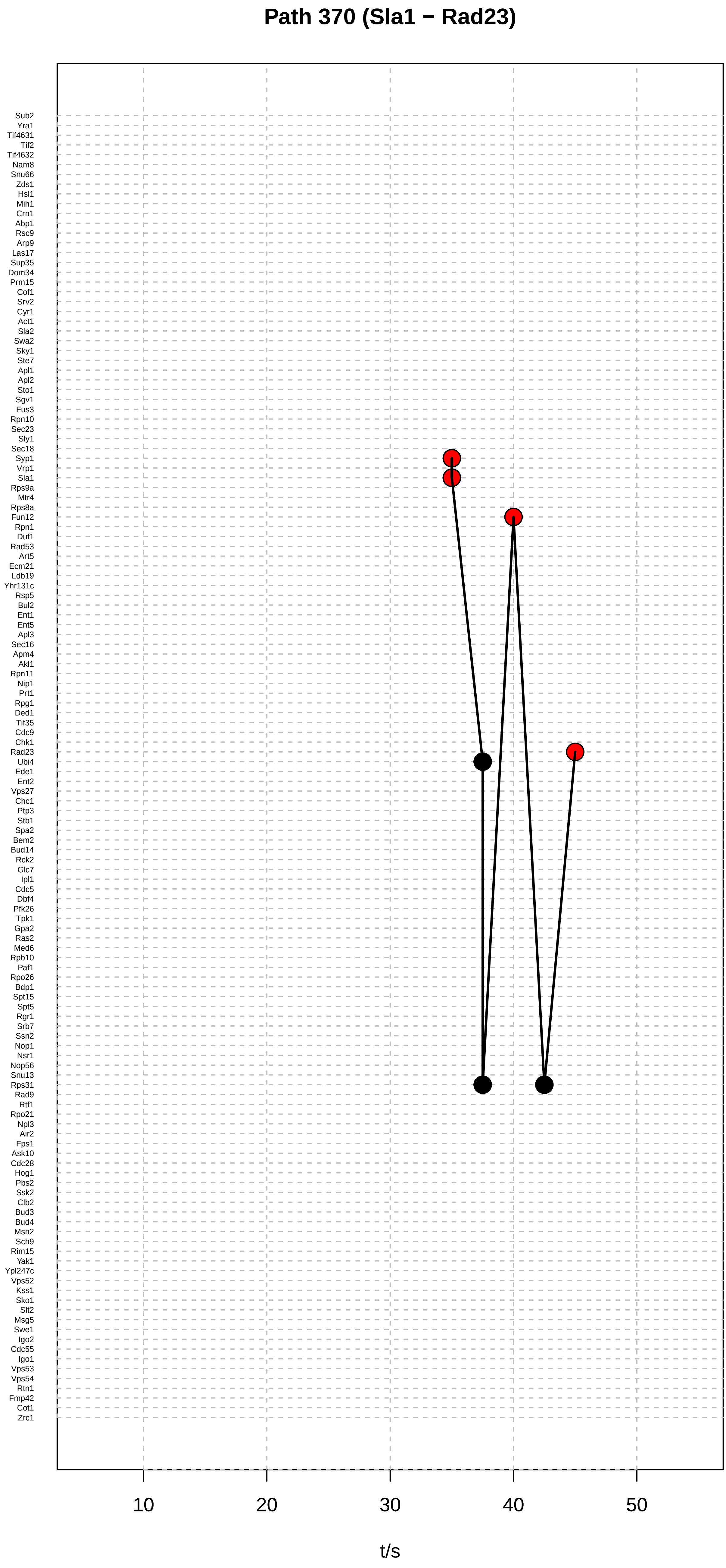

Path 370 (Sla1 − Rad23)
Sub2
Yra1
Tif4631
Tif2
Tif4632
Nam8
Snu66
Zds1
Hsl1
Mih1
Crn1
Abp1
Rsc9
Arp9
Las17
Sup35
Dom34
Prm15
Cof1
Srv2
Cyr1
Act1
Sla2
Swa2
Sky1
Ste7
Apl1
Apl2
Sto1
Sgv1
Fus3
Rpn10
Sec23
Sly1
Sec18
Syp1
Vrp1
Sla1
Rps9a
Mtr4
Rps8a
Fun12
Rpn1
Duf1
Rad53
Art5
Ecm21
Ldb19
Yhr131c
Rsp5
Bul2
Ent1
Ent5
Apl3
Sec16
Apm4
Akl1
Rpn11
Nip1
Prt1
Rpg1
Ded1
Tif35
Cdc9
Chk1
Rad23
Ubi4
Ede1
Ent2
Vps27
Chc1
Ptp3
Stb1
Spa2
Bem2
Bud14
Rck2
Glc7
Ipl1
Cdc5
Dbf4
Pfk26
Tpk1
Gpa2
Ras2
Med6
Rpb10
Paf1
Rpo26
Bdp1
Spt15
Spt5
Rgr1
Srb7
Ssn2
Nop1
Nsr1
Nop56
Snu13
Rps31
Rad9
Rtf1
Rpo21
Npl3
Air2
Fps1
Ask10
Cdc28
Hog1
Pbs2
Ssk2
Clb2
Bud3
Bud4
Msn2
Sch9
Rim15
Yak1
Ypl247c
Vps52
Kss1
Sko1
Slt2
Msg5
Swe1
Igo2
Cdc55
Igo1
Vps53
Vps54
Rtn1
Fmp42
Cot1
Zrc1
10
20
30
40
50
t/s

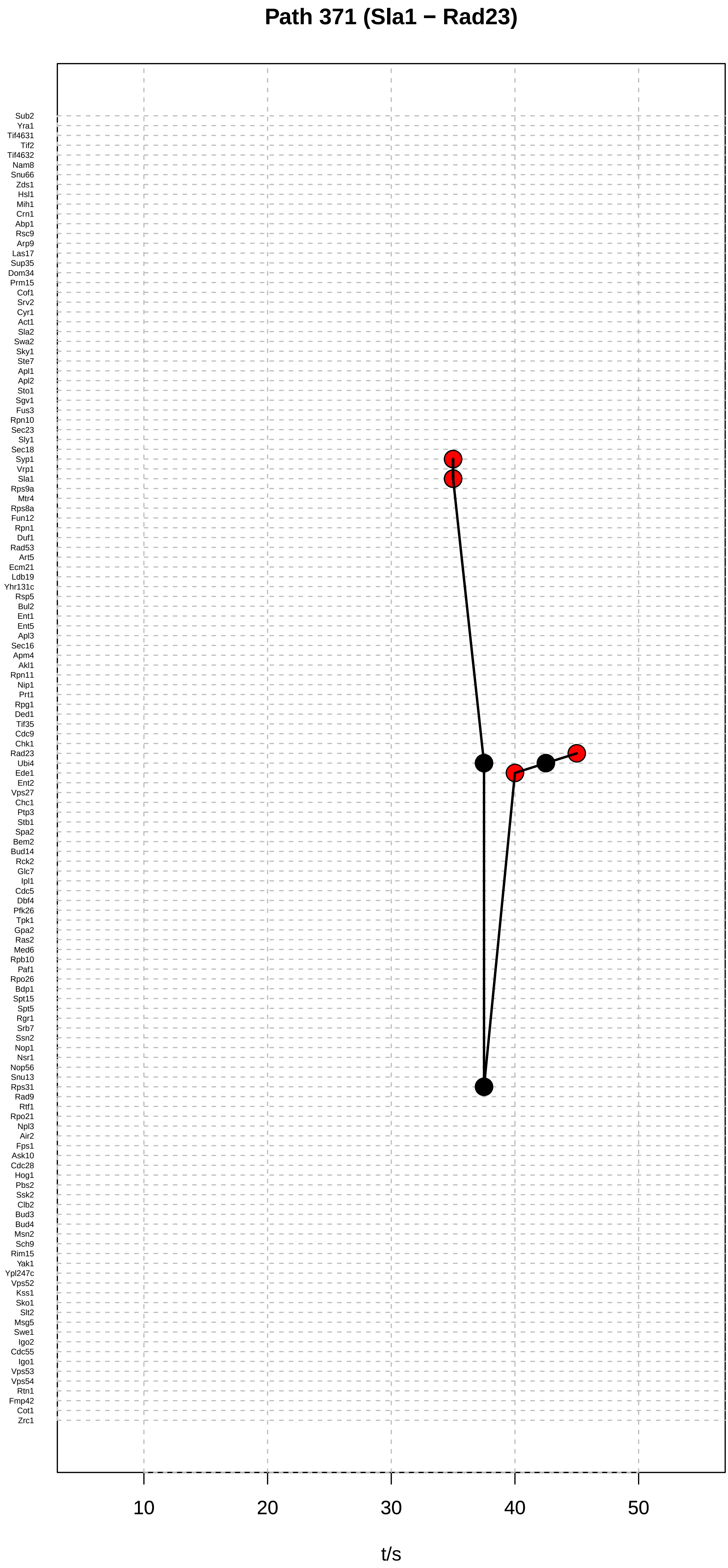
Path 371 (Sla1 − Rad23)
Sub2
Yra1
Tif4631
Tif2
Tif4632
Nam8
Snu66
Zds1
Hsl1
Mih1
Crn1
Abp1
Rsc9
Arp9
Las17
Sup35
Dom34
Prm15
Cof1
Srv2
Cyr1
Act1
Sla2
Swa2
Sky1
Ste7
Apl1
Apl2
Sto1
Sgv1
Fus3
Rpn10
Sec23
Sly1
Sec18
Syp1
Vrp1
Sla1
Rps9a
Mtr4
Rps8a
Fun12
Rpn1
Duf1
Rad53
Art5
Ecm21
Ldb19
Yhr131c
Rsp5
Bul2
Ent1
Ent5
Apl3
Sec16
Apm4
Akl1
Rpn11
Nip1
Prt1
Rpg1
Ded1
Tif35
Cdc9
Chk1
Rad23
Ubi4
Ede1
Ent2
Vps27
Chc1
Ptp3
Stb1
Spa2
Bem2
Bud14
Rck2
Glc7
Ipl1
Cdc5
Dbf4
Pfk26
Tpk1
Gpa2
Ras2
Med6
Rpb10
Paf1
Rpo26
Bdp1
Spt15
Spt5
Rgr1
Srb7
Ssn2
Nop1
Nsr1
Nop56
Snu13
Rps31
Rad9
Rtf1
Rpo21
Npl3
Air2
Fps1
Ask10
Cdc28
Hog1
Pbs2
Ssk2
Clb2
Bud3
Bud4
Msn2
Sch9
Rim15
Yak1
Ypl247c
Vps52
Kss1
Sko1
Slt2
Msg5
Swe1
Igo2
Cdc55
Igo1
Vps53
Vps54
Rtn1
Fmp42
Cot1
Zrc1
10
20
30
40
50
t/s

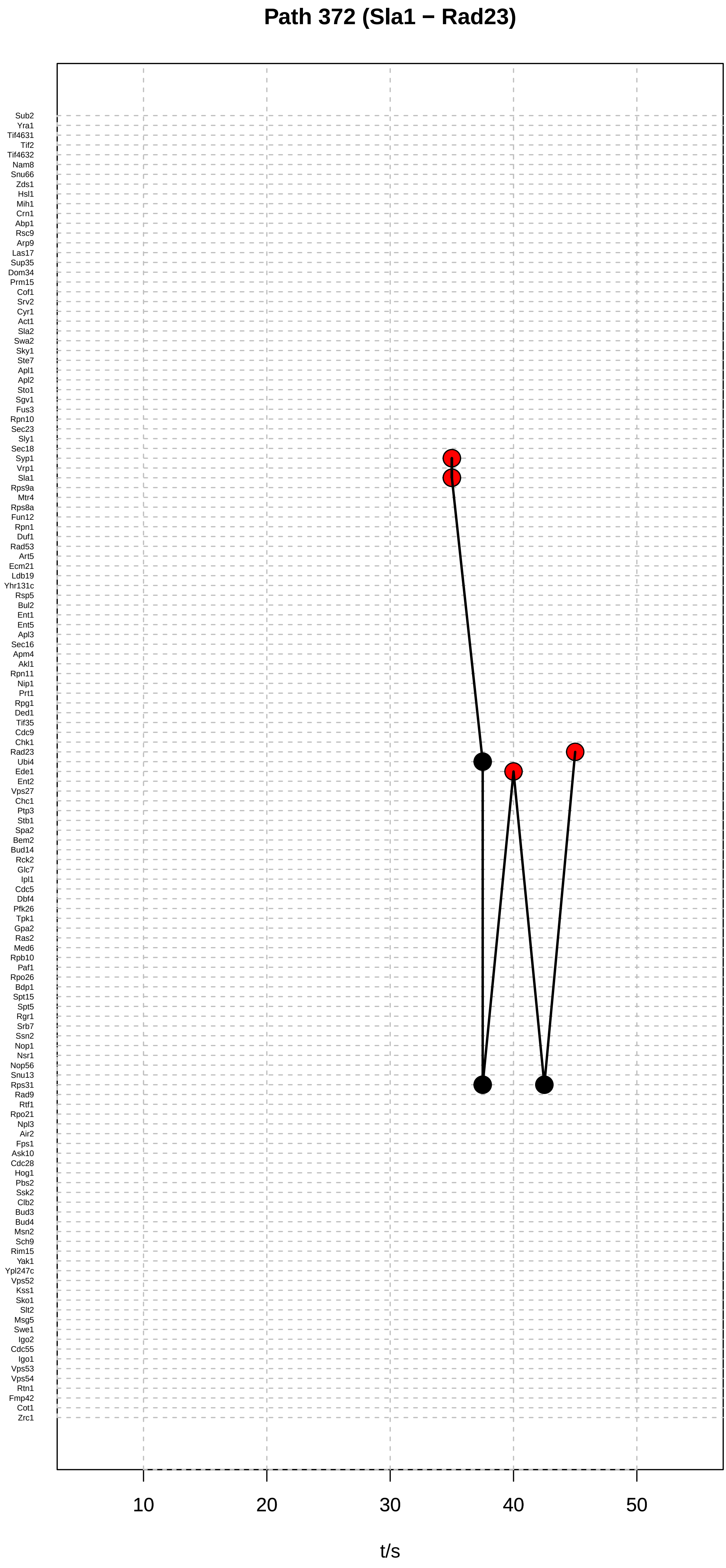
Path 372 (Sla1 − Rad23)
Sub2
Yra1
Tif4631
Tif2
Tif4632
Nam8
Snu66
Zds1
Hsl1
Mih1
Crn1
Abp1
Rsc9
Arp9
Las17
Sup35
Dom34
Prm15
Cof1
Srv2
Cyr1
Act1
Sla2
Swa2
Sky1
Ste7
Apl1
Apl2
Sto1
Sgv1
Fus3
Rpn10
Sec23
Sly1
Sec18
Syp1
Vrp1
Sla1
Rps9a
Mtr4
Rps8a
Fun12
Rpn1
Duf1
Rad53
Art5
Ecm21
Ldb19
Yhr131c
Rsp5
Bul2
Ent1
Ent5
Apl3
Sec16
Apm4
Akl1
Rpn11
Nip1
Prt1
Rpg1
Ded1
Tif35
Cdc9
Chk1
Rad23
Ubi4
Ede1
Ent2
Vps27
Chc1
Ptp3
Stb1
Spa2
Bem2
Bud14
Rck2
Glc7
Ipl1
Cdc5
Dbf4
Pfk26
Tpk1
Gpa2
Ras2
Med6
Rpb10
Paf1
Rpo26
Bdp1
Spt15
Spt5
Rgr1
Srb7
Ssn2
Nop1
Nsr1
Nop56
Snu13
Rps31
Rad9
Rtf1
Rpo21
Npl3
Air2
Fps1
Ask10
Cdc28
Hog1
Pbs2
Ssk2
Clb2
Bud3
Bud4
Msn2
Sch9
Rim15
Yak1
Ypl247c
Vps52
Kss1
Sko1
Slt2
Msg5
Swe1
Igo2
Cdc55
Igo1
Vps53
Vps54
Rtn1
Fmp42
Cot1
Zrc1
10
20
30
40
50
t/s

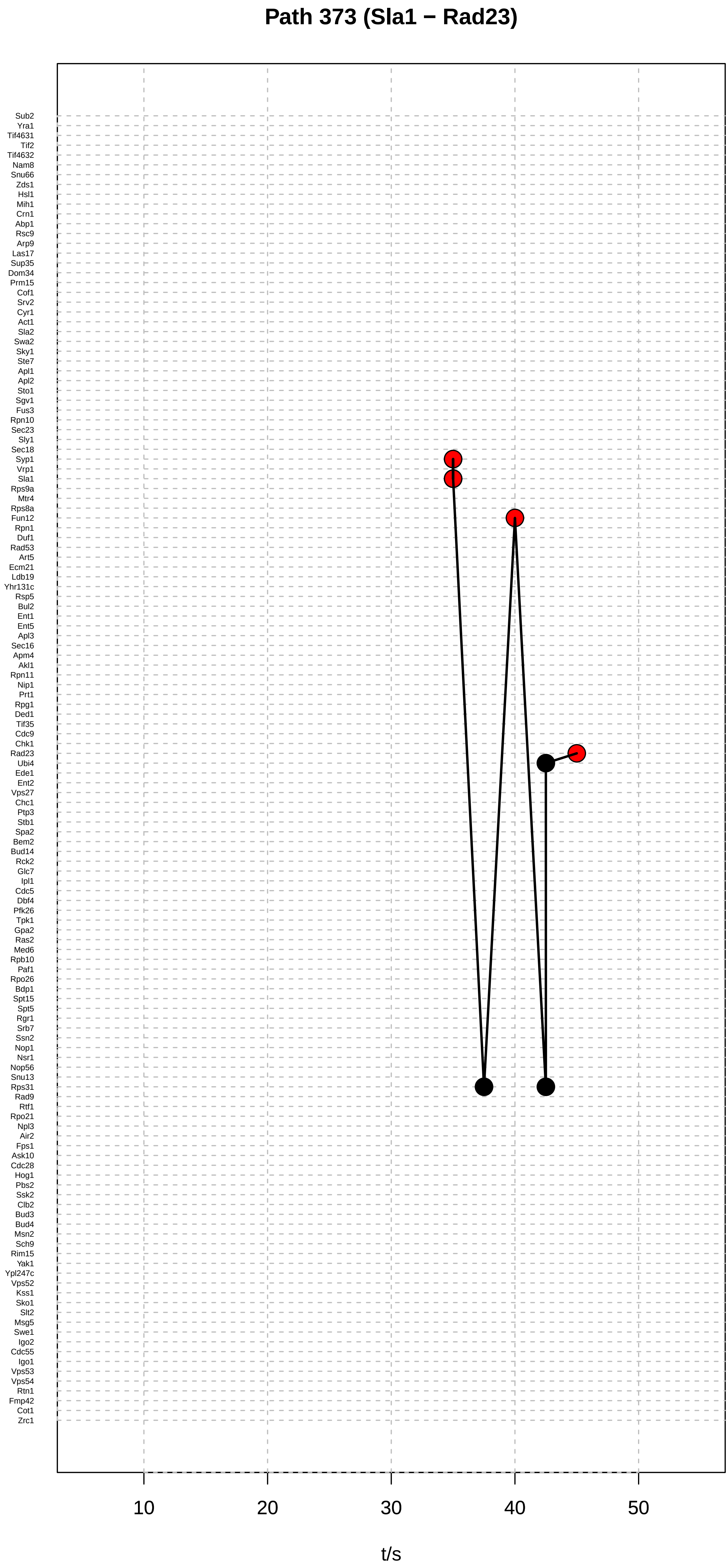
Path 373 (Sla1 − Rad23)
Sub2
Yra1
Tif4631
Tif2
Tif4632
Nam8
Snu66
Zds1
Hsl1
Mih1
Crn1
Abp1
Rsc9
Arp9
Las17
Sup35
Dom34
Prm15
Cof1
Srv2
Cyr1
Act1
Sla2
Swa2
Sky1
Ste7
Apl1
Apl2
Sto1
Sgv1
Fus3
Rpn10
Sec23
Sly1
Sec18
Syp1
Vrp1
Sla1
Rps9a
Mtr4
Rps8a
Fun12
Rpn1
Duf1
Rad53
Art5
Ecm21
Ldb19
Yhr131c
Rsp5
Bul2
Ent1
Ent5
Apl3
Sec16
Apm4
Akl1
Rpn11
Nip1
Prt1
Rpg1
Ded1
Tif35
Cdc9
Chk1
Rad23
Ubi4
Ede1
Ent2
Vps27
Chc1
Ptp3
Stb1
Spa2
Bem2
Bud14
Rck2
Glc7
Ipl1
Cdc5
Dbf4
Pfk26
Tpk1
Gpa2
Ras2
Med6
Rpb10
Paf1
Rpo26
Bdp1
Spt15
Spt5
Rgr1
Srb7
Ssn2
Nop1
Nsr1
Nop56
Snu13
Rps31
Rad9
Rtf1
Rpo21
Npl3
Air2
Fps1
Ask10
Cdc28
Hog1
Pbs2
Ssk2
Clb2
Bud3
Bud4
Msn2
Sch9
Rim15
Yak1
Ypl247c
Vps52
Kss1
Sko1
Slt2
Msg5
Swe1
Igo2
Cdc55
Igo1
Vps53
Vps54
Rtn1
Fmp42
Cot1
Zrc1
10
20
30
40
50
t/s

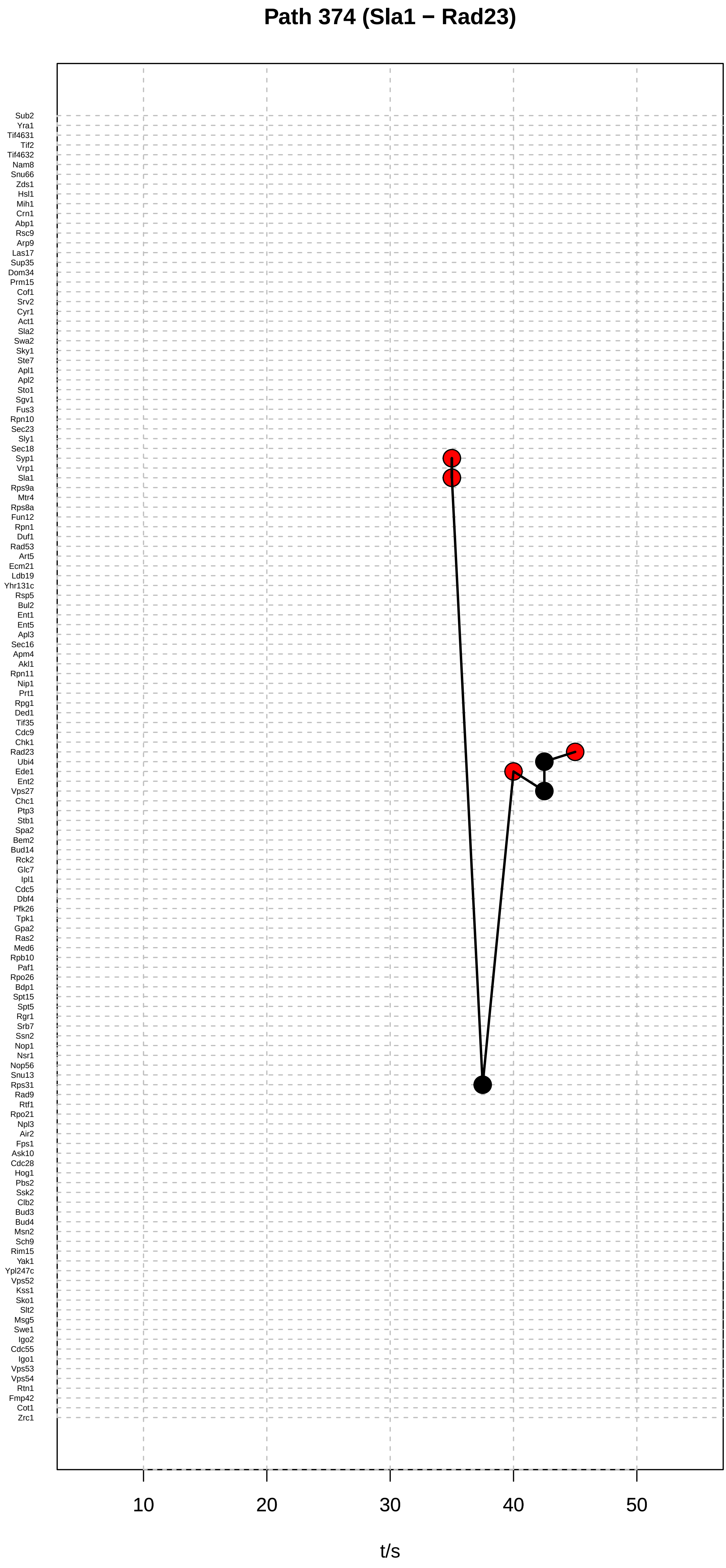
Path 374 (Sla1 − Rad23)
Sub2
Yra1
Tif4631
Tif2
Tif4632
Nam8
Snu66
Zds1
Hsl1
Mih1
Crn1
Abp1
Rsc9
Arp9
Las17
Sup35
Dom34
Prm15
Cof1
Srv2
Cyr1
Act1
Sla2
Swa2
Sky1
Ste7
Apl1
Apl2
Sto1
Sgv1
Fus3
Rpn10
Sec23
Sly1
Sec18
Syp1
Vrp1
Sla1
Rps9a
Mtr4
Rps8a
Fun12
Rpn1
Duf1
Rad53
Art5
Ecm21
Ldb19
Yhr131c
Rsp5
Bul2
Ent1
Ent5
Apl3
Sec16
Apm4
Akl1
Rpn11
Nip1
Prt1
Rpg1
Ded1
Tif35
Cdc9
Chk1
Rad23
Ubi4
Ede1
Ent2
Vps27
Chc1
Ptp3
Stb1
Spa2
Bem2
Bud14
Rck2
Glc7
Ipl1
Cdc5
Dbf4
Pfk26
Tpk1
Gpa2
Ras2
Med6
Rpb10
Paf1
Rpo26
Bdp1
Spt15
Spt5
Rgr1
Srb7
Ssn2
Nop1
Nsr1
Nop56
Snu13
Rps31
Rad9
Rtf1
Rpo21
Npl3
Air2
Fps1
Ask10
Cdc28
Hog1
Pbs2
Ssk2
Clb2
Bud3
Bud4
Msn2
Sch9
Rim15
Yak1
Ypl247c
Vps52
Kss1
Sko1
Slt2
Msg5
Swe1
Igo2
Cdc55
Igo1
Vps53
Vps54
Rtn1
Fmp42
Cot1
Zrc1
10
20
30
40
50
t/s

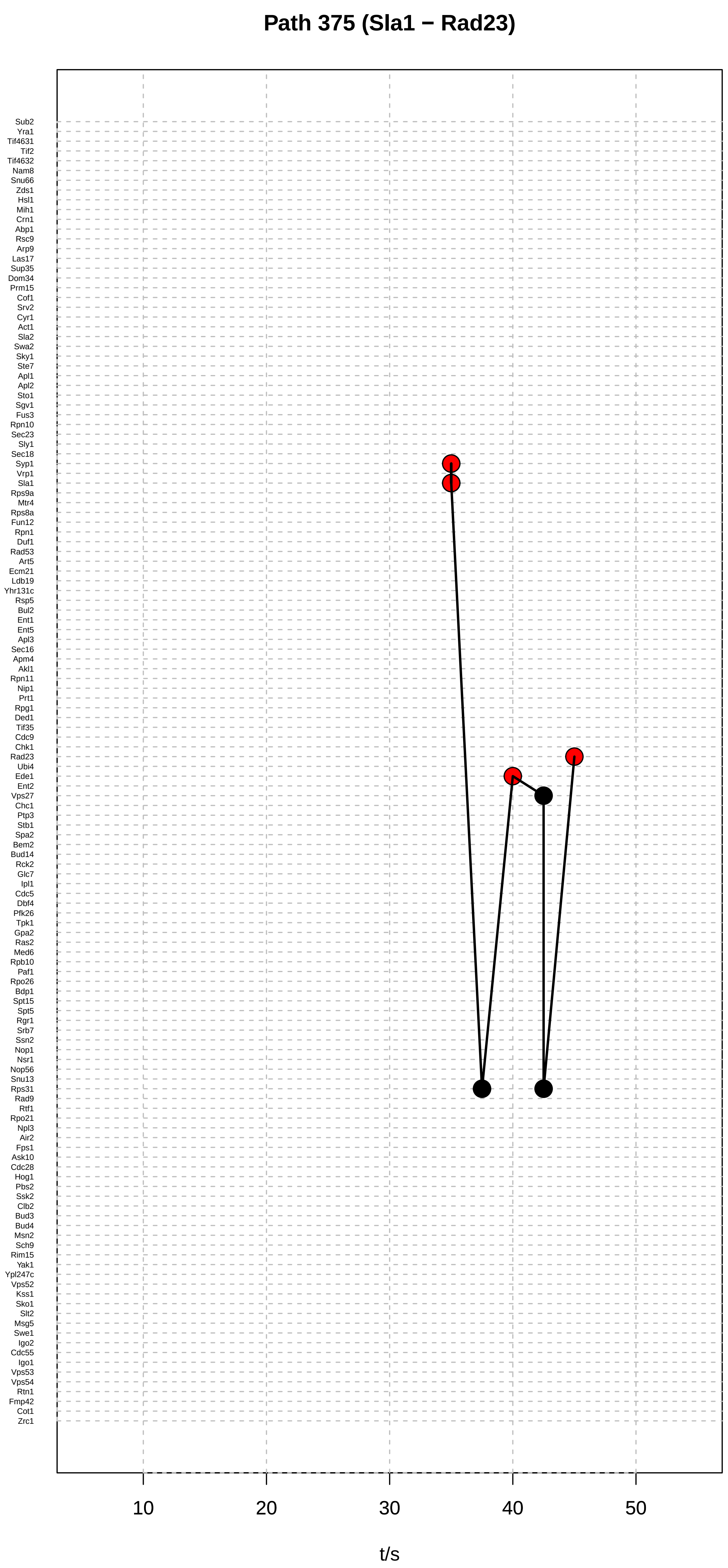

Path 375 (Sla1 − Rad23)
Sub2
Yra1
Tif4631
Tif2
Tif4632
Nam8
Snu66
Zds1
Hsl1
Mih1
Crn1
Abp1
Rsc9
Arp9
Las17
Sup35
Dom34
Prm15
Cof1
Srv2
Cyr1
Act1
Sla2
Swa2
Sky1
Ste7
Apl1
Apl2
Sto1
Sgv1
Fus3
Rpn10
Sec23
Sly1
Sec18
Syp1
Vrp1
Sla1
Rps9a
Mtr4
Rps8a
Fun12
Rpn1
Duf1
Rad53
Art5
Ecm21
Ldb19
Yhr131c
Rsp5
Bul2
Ent1
Ent5
Apl3
Sec16
Apm4
Akl1
Rpn11
Nip1
Prt1
Rpg1
Ded1
Tif35
Cdc9
Chk1
Rad23
Ubi4
Ede1
Ent2
Vps27
Chc1
Ptp3
Stb1
Spa2
Bem2
Bud14
Rck2
Glc7
Ipl1
Cdc5
Dbf4
Pfk26
Tpk1
Gpa2
Ras2
Med6
Rpb10
Paf1
Rpo26
Bdp1
Spt15
Spt5
Rgr1
Srb7
Ssn2
Nop1
Nsr1
Nop56
Snu13
Rps31
Rad9
Rtf1
Rpo21
Npl3
Air2
Fps1
Ask10
Cdc28
Hog1
Pbs2
Ssk2
Clb2
Bud3
Bud4
Msn2
Sch9
Rim15
Yak1
Ypl247c
Vps52
Kss1
Sko1
Slt2
Msg5
Swe1
Igo2
Cdc55
Igo1
Vps53
Vps54
Rtn1
Fmp42
Cot1
Zrc1
10
20
30
40
50
t/s

### Path 376 (Sla1 − Ent5)

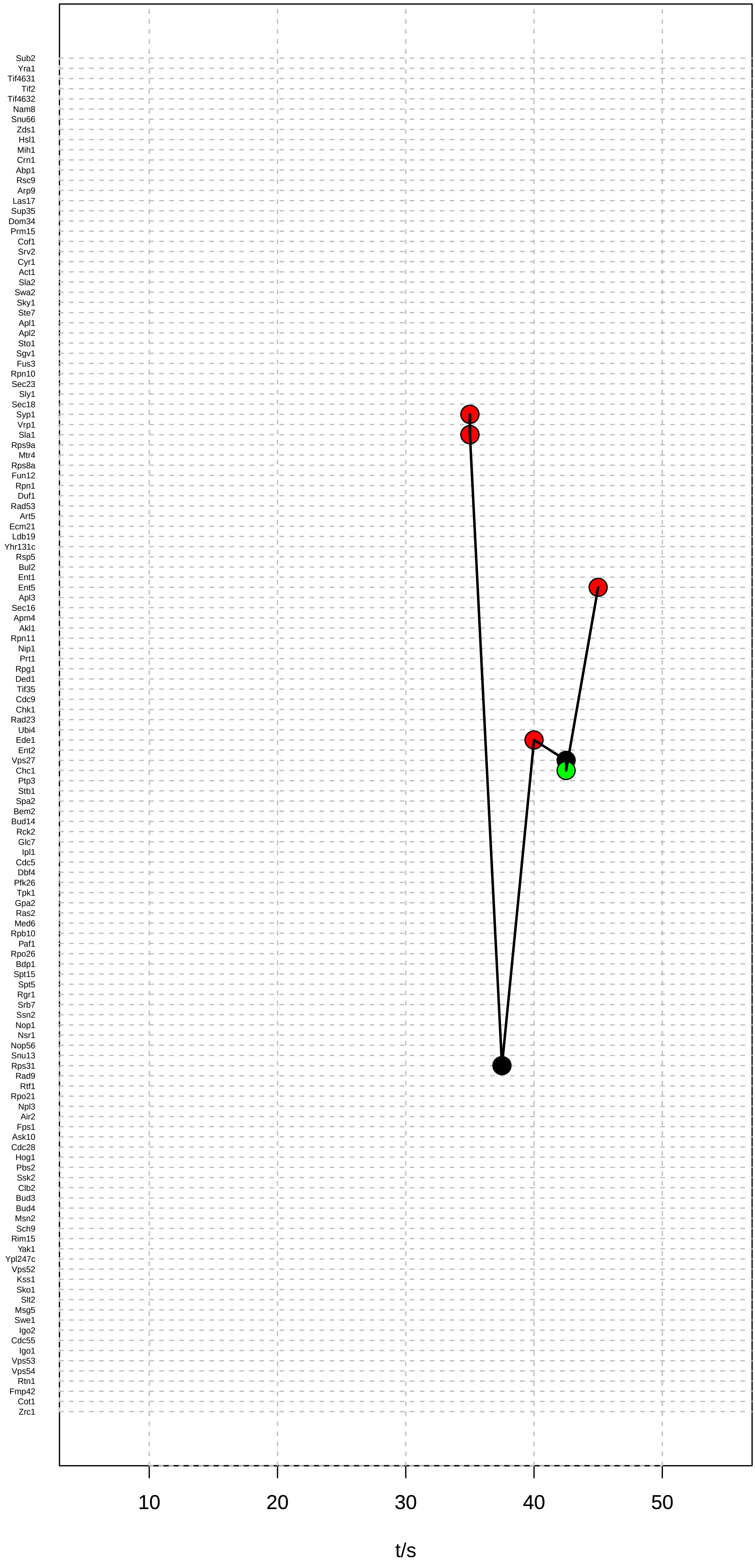

### Path 377 (Sla1 − Rad23)

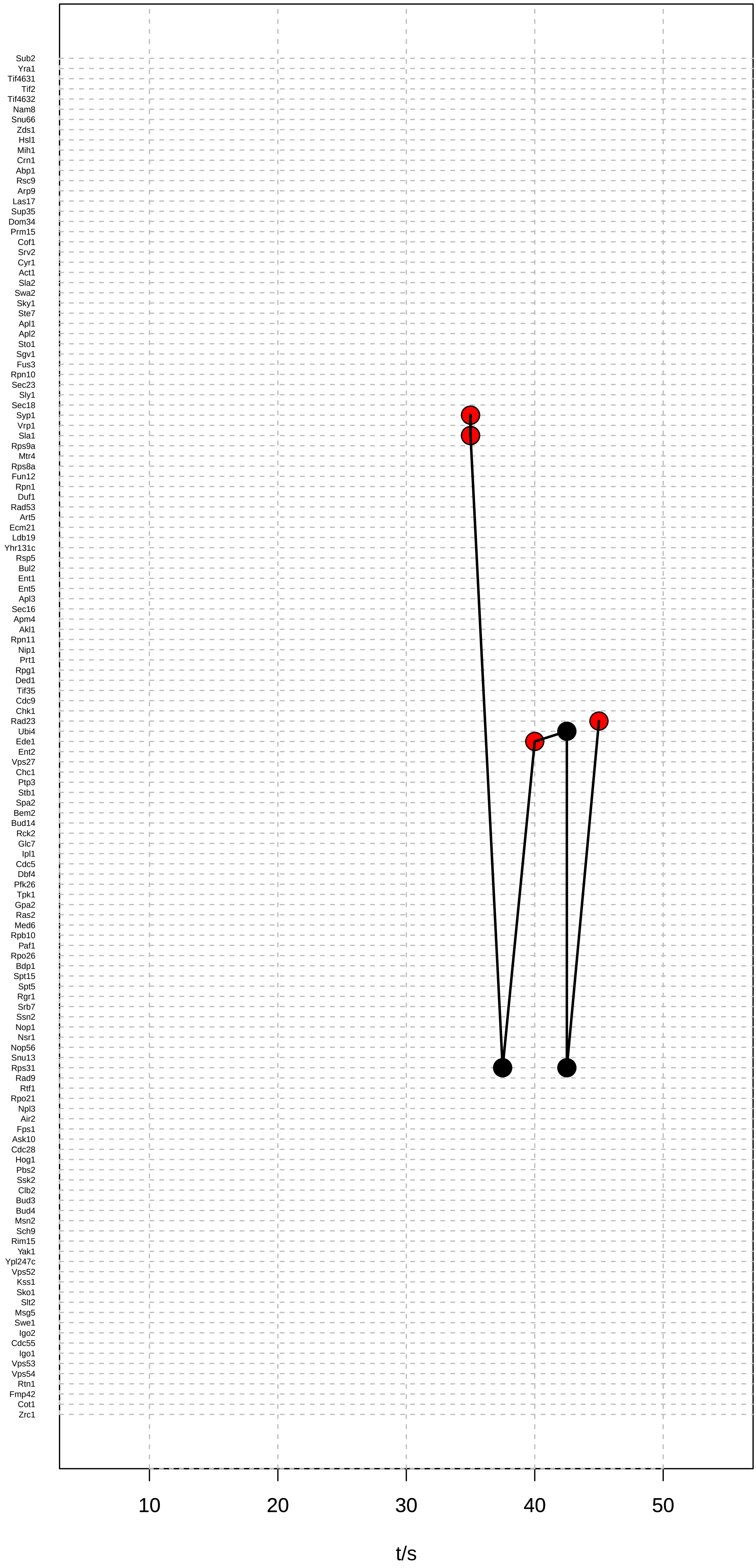

### Path 378 (Sla1 − Rad23)

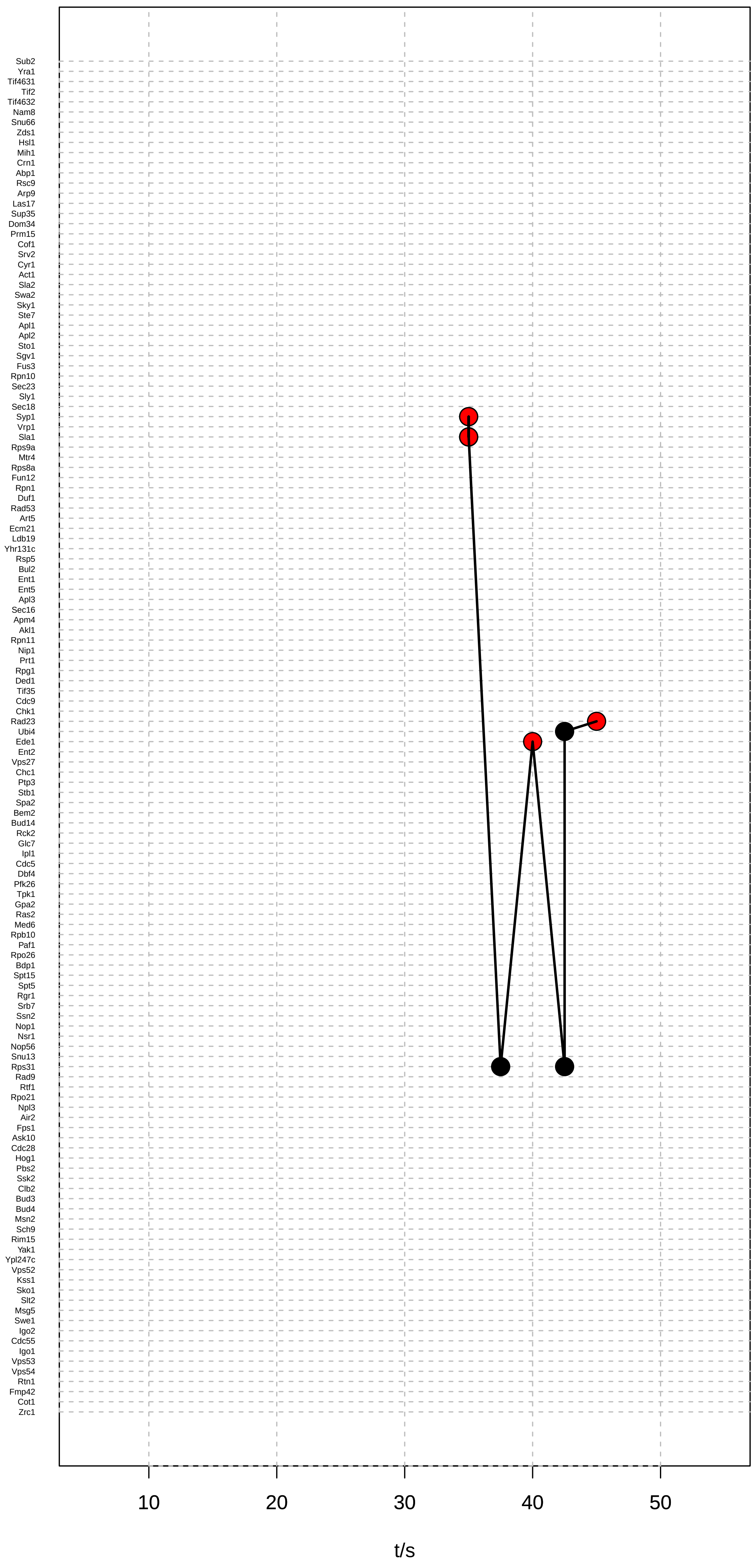

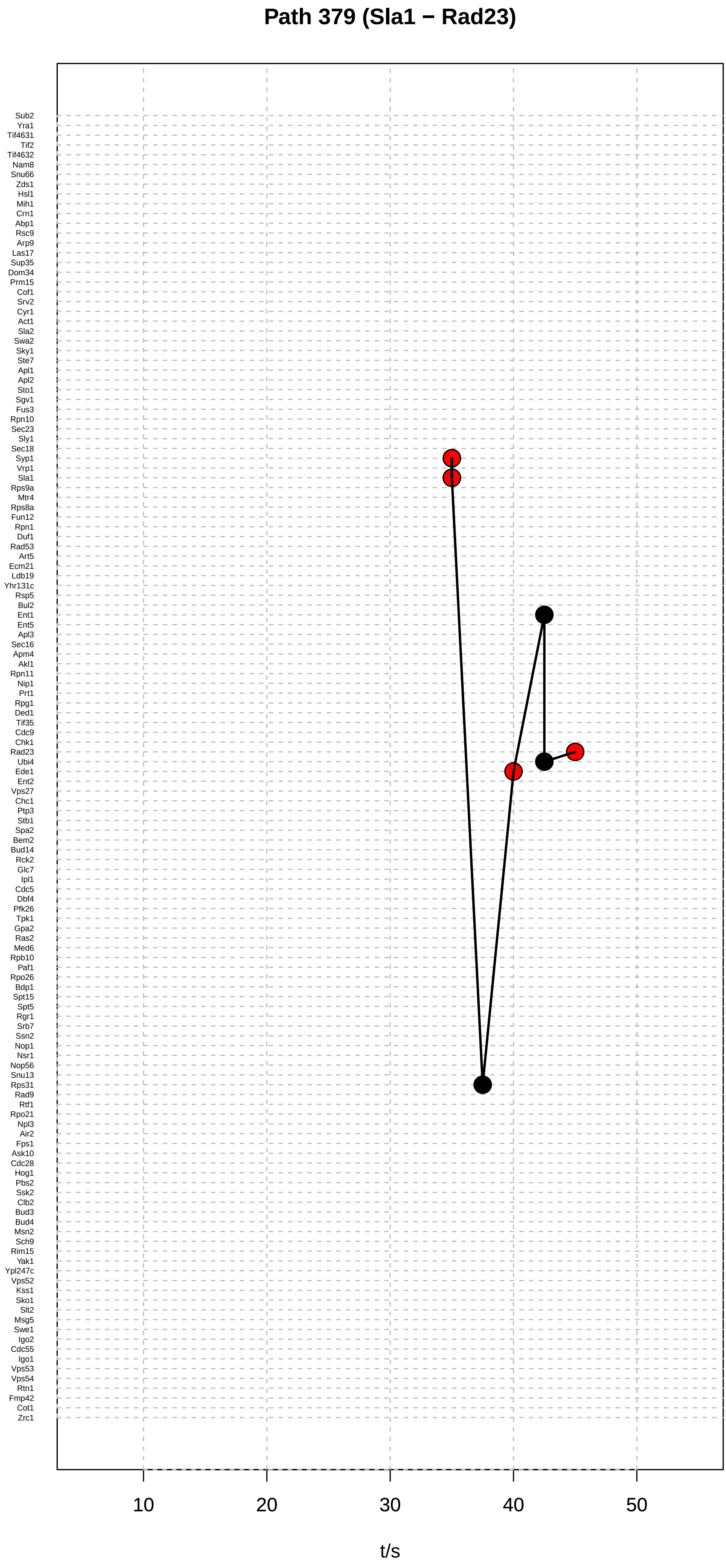
Path 379 (Sla1 − Rad23)
Sub2
Yra1
Tif4631
Tif2
Tif4632
Nam8
Snu66
Zds1
Hsl1
Mih1
Crn1
Abp1
Rsc9
Arp9
Las17
Sup35
Dom34
Prm15
Cof1
Srv2
Cyr1
Act1
Sla2
Swa2
Sky1
Ste7
Apl1
Apl2
Sto1
Sgv1
Fus3
Rpn10
Sec23
Sly1
Sec18
Syp1
Vrp1
Sla1
Rps9a
Mtr4
Rps8a
Fun12
Rpn1
Duf1
Rad53
Art5
Ecm21
Ldb19
Yhr131c
Rsp5
Bul2
Ent1
Ent5
Apl3
Sec16
Apm4
Akl1
Rpn11
Nip1
Prt1
Rpg1
Ded1
Tif35
Cdc9
Chk1
Rad23
Ubi4
Ede1
Ent2
Vps27
Chc1
Ptp3
Stb1
Spa2
Bem2
Bud14
Rck2
Glc7
Ipl1
Cdc5
Dbf4
Pfk26
Tpk1
Gpa2
Ras2
Med6
Rpb10
Paf1
Rpo26
Bdp1
Spt15
Spt5
Rgr1
Srb7
Ssn2
Nop1
Nsr1
Nop56
Snu13
Rps31
Rad9
Rtf1
Rpo21
Npl3
Air2
Fps1
Ask10
Cdc28
Hog1
Pbs2
Ssk2
Clb2
Bud3
Bud4
Msn2
Sch9
Rim15
Yak1
Ypl247c
Vps52
Kss1
Sko1
Slt2
Msg5
Swe1
Igo2
Cdc55
Igo1
Vps53
Vps54
Rtn1
Fmp42
Cot1
Zrc1
10
20
30
40
50
t/s

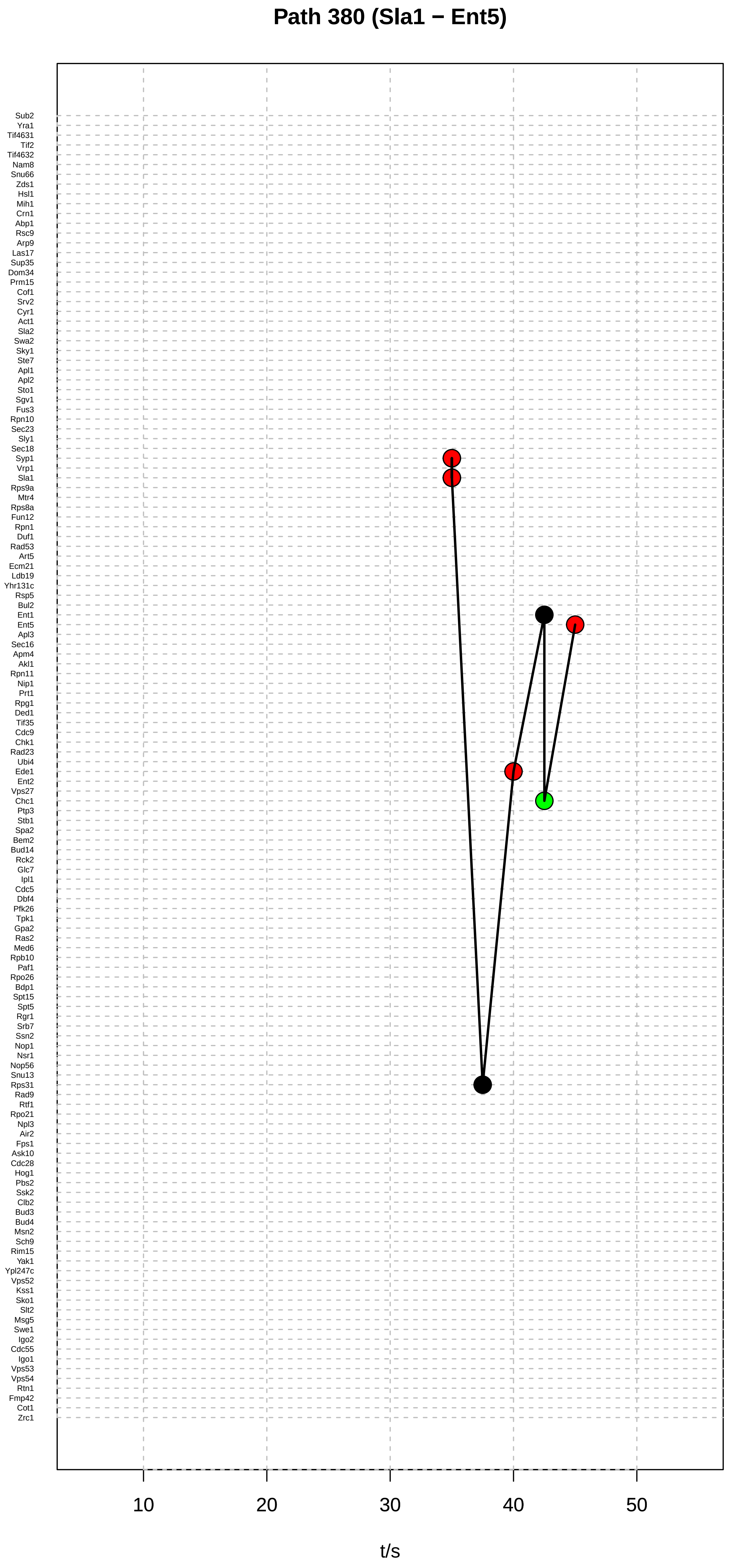
Path 380 (Sla1 − Ent5)
Sub2
Yra1
Tif4631
Tif2
Tif4632
Nam8
Snu66
Zds1
Hsl1
Mih1
Crn1
Abp1
Rsc9
Arp9
Las17
Sup35
Dom34
Prm15
Cof1
Srv2
Cyr1
Act1
Sla2
Swa2
Sky1
Ste7
Apl1
Apl2
Sto1
Sgv1
Fus3
Rpn10
Sec23
Sly1
Sec18
Syp1
Vrp1
Sla1
Rps9a
Mtr4
Rps8a
Fun12
Rpn1
Duf1
Rad53
Art5
Ecm21
Ldb19
Yhr131c
Rsp5
Bul2
Ent1
Ent5
Apl3
Sec16
Apm4
Akl1
Rpn11
Nip1
Prt1
Rpg1
Ded1
Tif35
Cdc9
Chk1
Rad23
Ubi4
Ede1
Ent2
Vps27
Chc1
Ptp3
Stb1
Spa2
Bem2
Bud14
Rck2
Glc7
Ipl1
Cdc5
Dbf4
Pfk26
Tpk1
Gpa2
Ras2
Med6
Rpb10
Paf1
Rpo26
Bdp1
Spt15
Spt5
Rgr1
Srb7
Ssn2
Nop1
Nsr1
Nop56
Snu13
Rps31
Rad9
Rtf1
Rpo21
Npl3
Air2
Fps1
Ask10
Cdc28
Hog1
Pbs2
Ssk2
Clb2
Bud3
Bud4
Msn2
Sch9
Rim15
Yak1
Ypl247c
Vps52
Kss1
Sko1
Slt2
Msg5
Swe1
Igo2
Cdc55
Igo1
Vps53
Vps54
Rtn1
Fmp42
Cot1
Zrc1
10
20
30
40
50
t/s

#### Path 381 (Sla1 − Rad23)

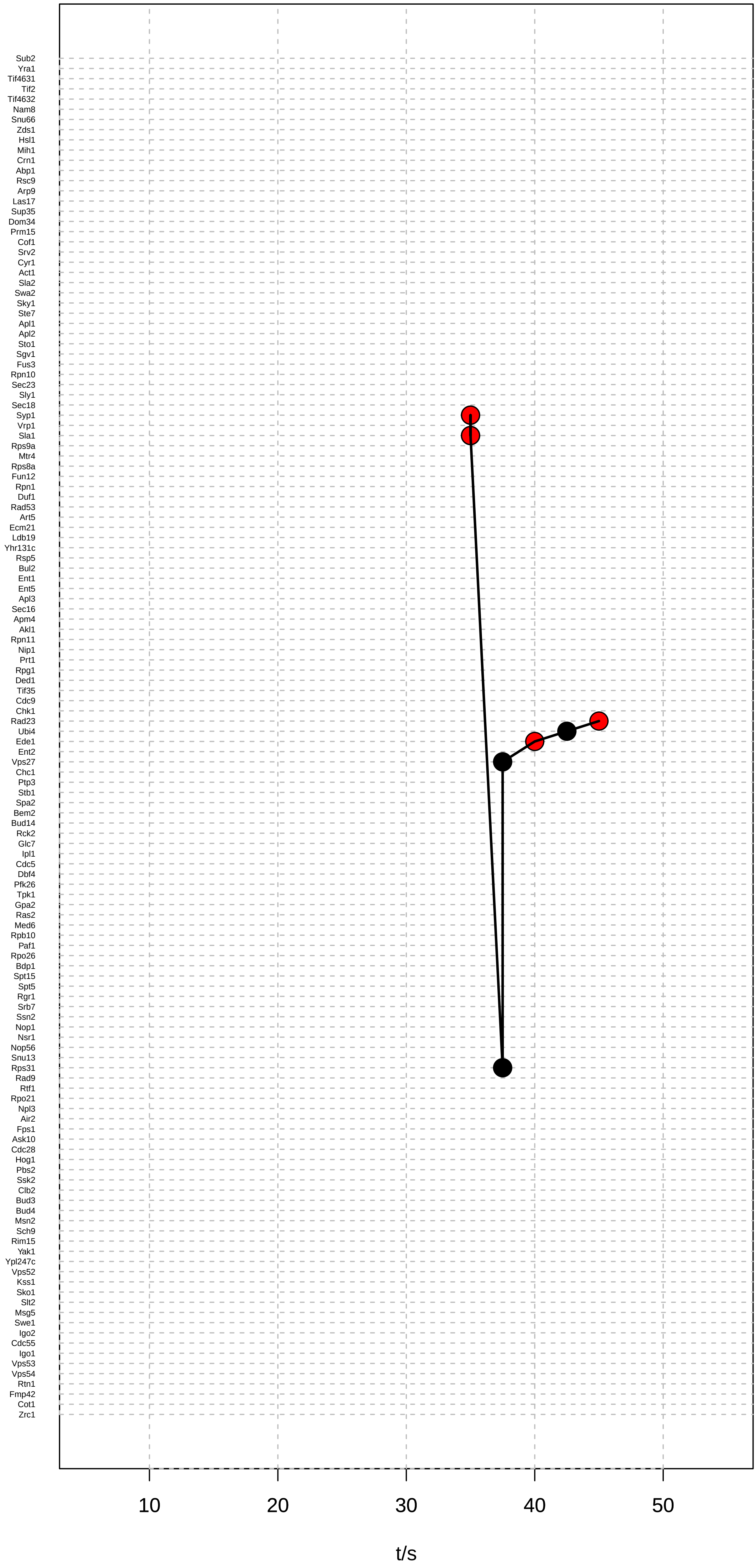

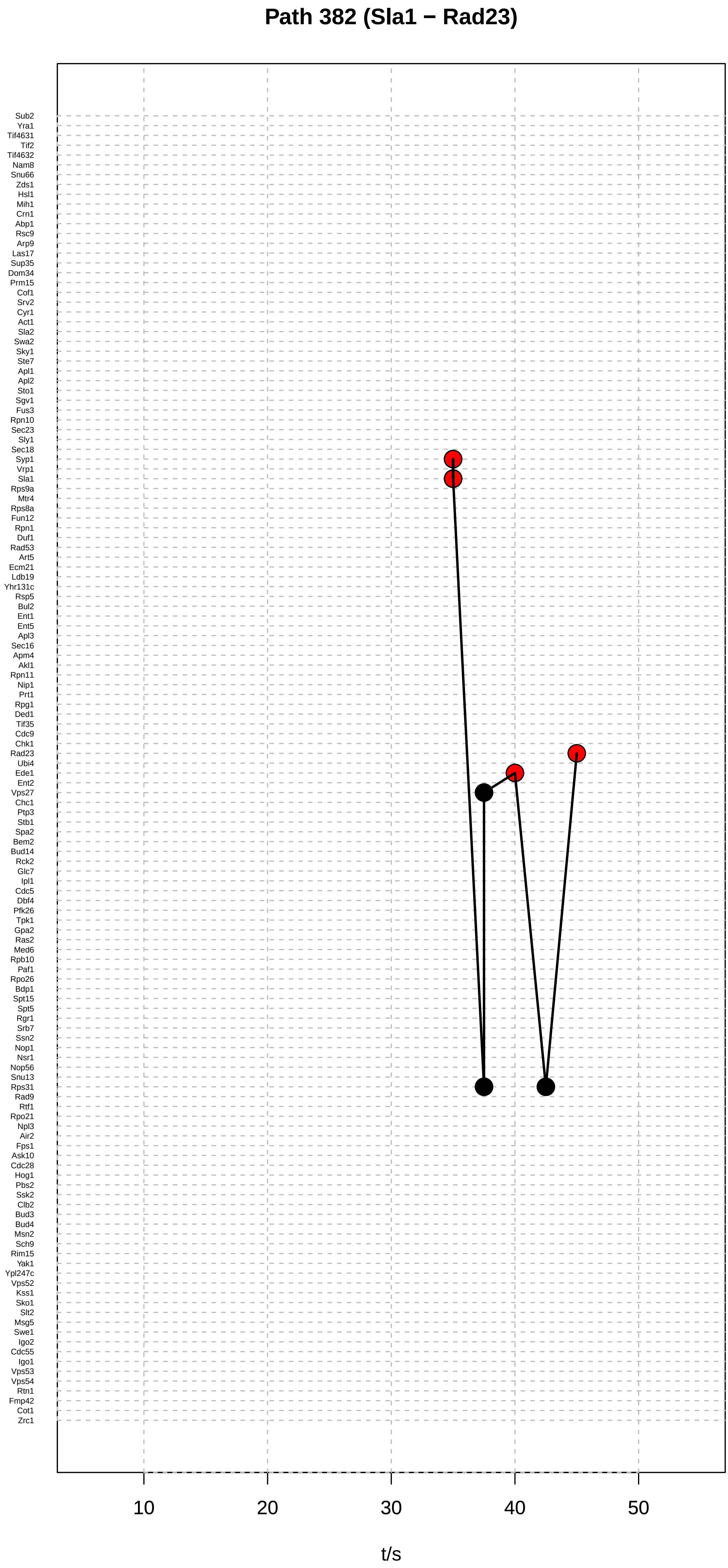

Path 382 (Sla1 − Rad23)
Sub2
Yra1
Tif4631
Tif2
Tif4632
Nam8
Snu66
Zds1
Hsl1
Mih1
Crn1
Abp1
Rsc9
Arp9
Las17
Sup35
Dom34
Prm15
Cof1
Srv2
Cyr1
Act1
Sla2
Swa2
Sky1
Ste7
Apl1
Apl2
Sto1
Sgv1
Fus3
Rpn10
Sec23
Sly1
Sec18
Syp1
Vrp1
Sla1
Rps9a
Mtr4
Rps8a
Fun12
Rpn1
Duf1
Rad53
Art5
Ecm21
Ldb19
Yhr131c
Rsp5
Bul2
Ent1
Ent5
Apl3
Sec16
Apm4
Akl1
Rpn11
Nip1
Prt1
Rpg1
Ded1
Tif35
Cdc9
Chk1
Rad23
Ubi4
Ede1
Ent2
Vps27
Chc1
Ptp3
Stb1
Spa2
Bem2
Bud14
Rck2
Glc7
Ipl1
Cdc5
Dbf4
Pfk26
Tpk1
Gpa2
Ras2
Med6
Rpb10
Paf1
Rpo26
Bdp1
Spt15
Spt5
Rgr1
Srb7
Ssn2
Nop1
Nsr1
Nop56
Snu13
Rps31
Rad9
Rtf1
Rpo21
Npl3
Air2
Fps1
Ask10
Cdc28
Hog1
Pbs2
Ssk2
Clb2
Bud3
Bud4
Msn2
Sch9
Rim15
Yak1
Ypl247c
Vps52
Kss1
Sko1
Slt2
Msg5
Swe1
Igo2
Cdc55
Igo1
Vps53
Vps54
Rtn1
Fmp42
Cot1
Zrc1
10
20
30
40
50
t/s

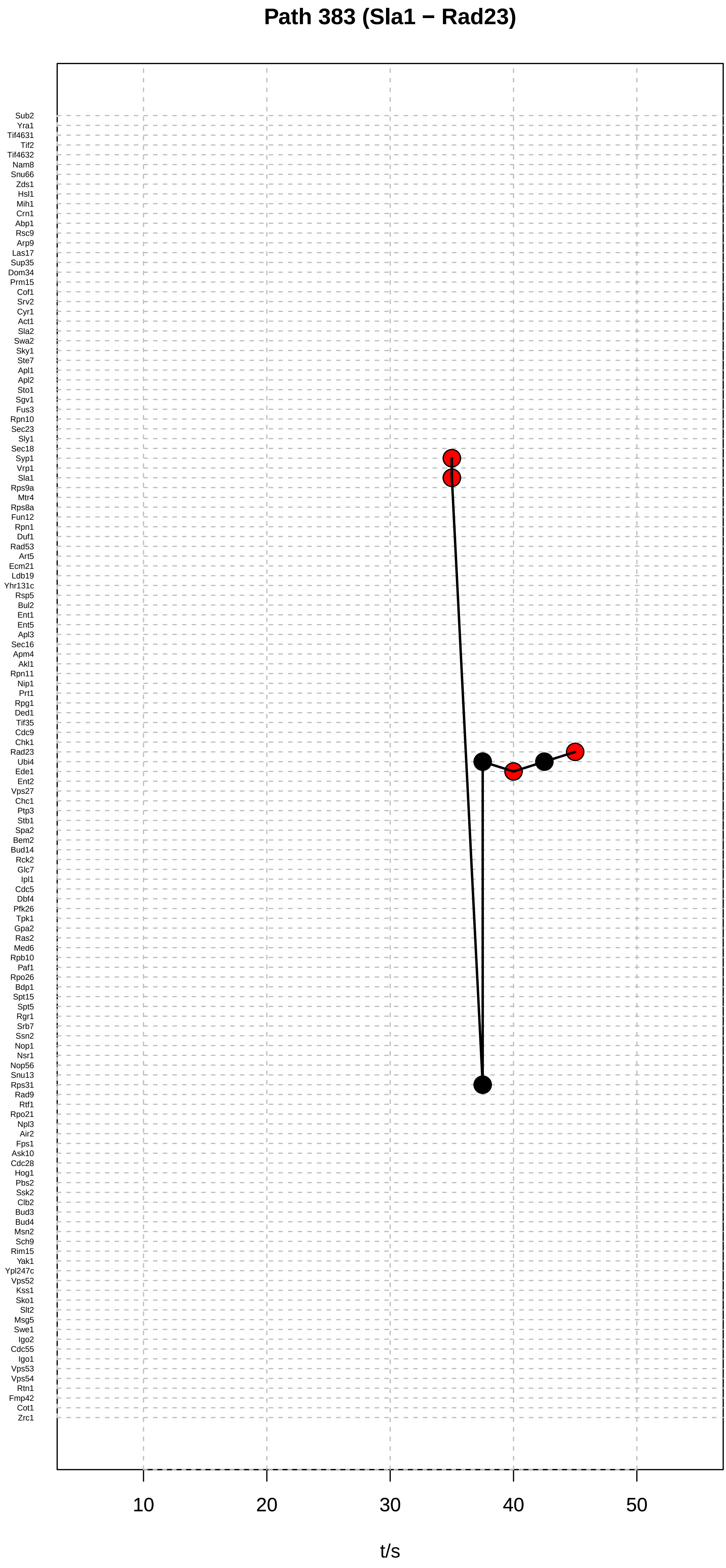
Path 383 (Sla1 − Rad23)
Sub2
Yra1
Tif4631
Tif2
Tif4632
Nam8
Snu66
Zds1
Hsl1
Mih1
Crn1
Abp1
Rsc9
Arp9
Las17
Sup35
Dom34
Prm15
Cof1
Srv2
Cyr1
Act1
Sla2
Swa2
Sky1
Ste7
Apl1
Apl2
Sto1
Sgv1
Fus3
Rpn10
Sec23
Sly1
Sec18
Syp1
Vrp1
Sla1
Rps9a
Mtr4
Rps8a
Fun12
Rpn1
Duf1
Rad53
Art5
Ecm21
Ldb19
Yhr131c
Rsp5
Bul2
Ent1
Ent5
Apl3
Sec16
Apm4
Akl1
Rpn11
Nip1
Prt1
Rpg1
Ded1
Tif35
Cdc9
Chk1
Rad23
Ubi4
Ede1
Ent2
Vps27
Chc1
Ptp3
Stb1
Spa2
Bem2
Bud14
Rck2
Glc7
Ipl1
Cdc5
Dbf4
Pfk26
Tpk1
Gpa2
Ras2
Med6
Rpb10
Paf1
Rpo26
Bdp1
Spt15
Spt5
Rgr1
Srb7
Ssn2
Nop1
Nsr1
Nop56
Snu13
Rps31
Rad9
Rtf1
Rpo21
Npl3
Air2
Fps1
Ask10
Cdc28
Hog1
Pbs2
Ssk2
Clb2
Bud3
Bud4
Msn2
Sch9
Rim15
Yak1
Ypl247c
Vps52
Kss1
Sko1
Slt2
Msg5
Swe1
Igo2
Cdc55
Igo1
Vps53
Vps54
Rtn1
Fmp42
Cot1
Zrc1
10
20
30
40
50
t/s

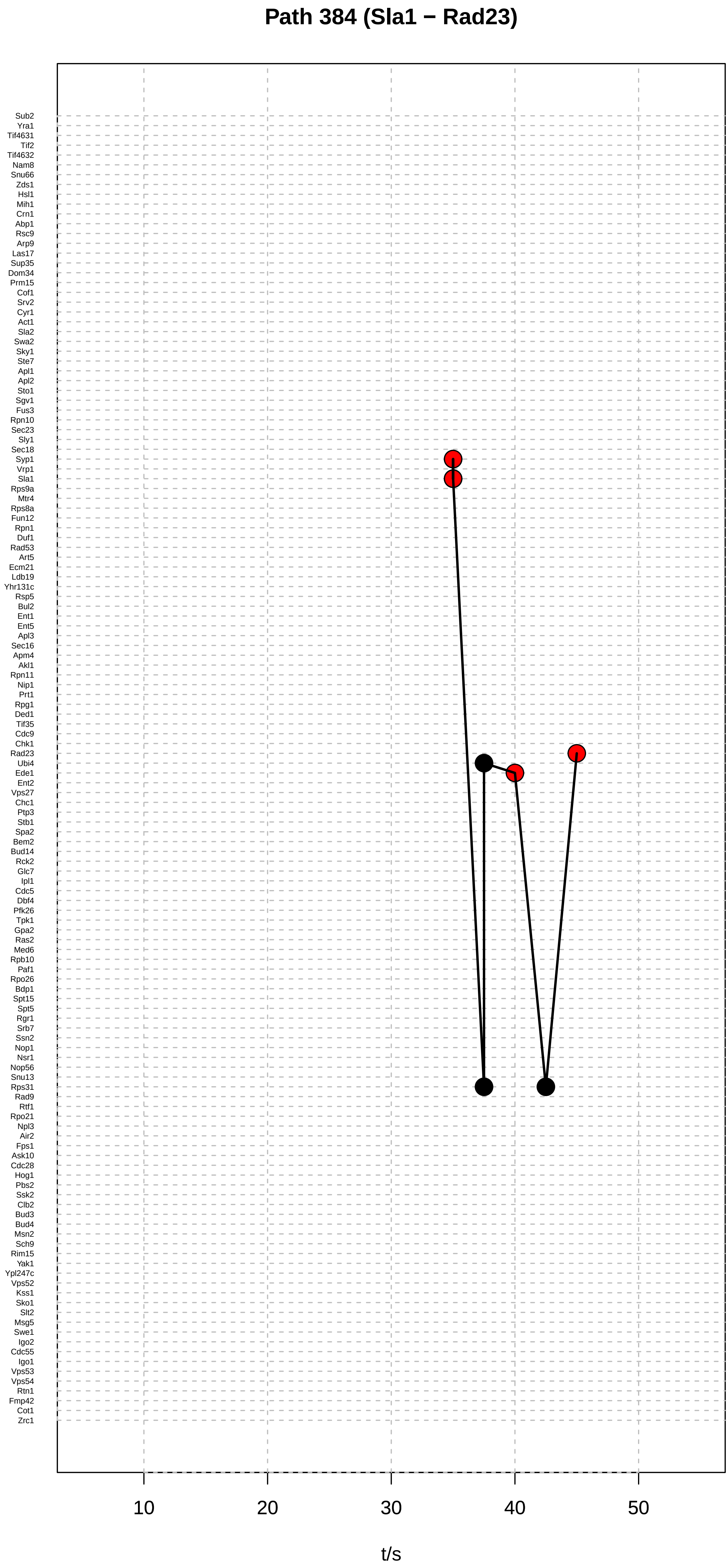
Path 384 (Sla1 − Rad23)
Sub2
Yra1
Tif4631
Tif2
Tif4632
Nam8
Snu66
Zds1
Hsl1
Mih1
Crn1
Abp1
Rsc9
Arp9
Las17
Sup35
Dom34
Prm15
Cof1
Srv2
Cyr1
Act1
Sla2
Swa2
Sky1
Ste7
Apl1
Apl2
Sto1
Sgv1
Fus3
Rpn10
Sec23
Sly1
Sec18
Syp1
Vrp1
Sla1
Rps9a
Mtr4
Rps8a
Fun12
Rpn1
Duf1
Rad53
Art5
Ecm21
Ldb19
Yhr131c
Rsp5
Bul2
Ent1
Ent5
Apl3
Sec16
Apm4
Akl1
Rpn11
Nip1
Prt1
Rpg1
Ded1
Tif35
Cdc9
Chk1
Rad23
Ubi4
Ede1
Ent2
Vps27
Chc1
Ptp3
Stb1
Spa2
Bem2
Bud14
Rck2
Glc7
Ipl1
Cdc5
Dbf4
Pfk26
Tpk1
Gpa2
Ras2
Med6
Rpb10
Paf1
Rpo26
Bdp1
Spt15
Spt5
Rgr1
Srb7
Ssn2
Nop1
Nsr1
Nop56
Snu13
Rps31
Rad9
Rtf1
Rpo21
Npl3
Air2
Fps1
Ask10
Cdc28
Hog1
Pbs2
Ssk2
Clb2
Bud3
Bud4
Msn2
Sch9
Rim15
Yak1
Ypl247c
Vps52
Kss1
Sko1
Slt2
Msg5
Swe1
Igo2
Cdc55
Igo1
Vps53
Vps54
Rtn1
Fmp42
Cot1
Zrc1
10
20
30
40
50
t/s

### Path 385 (Sla1 − Rad23)

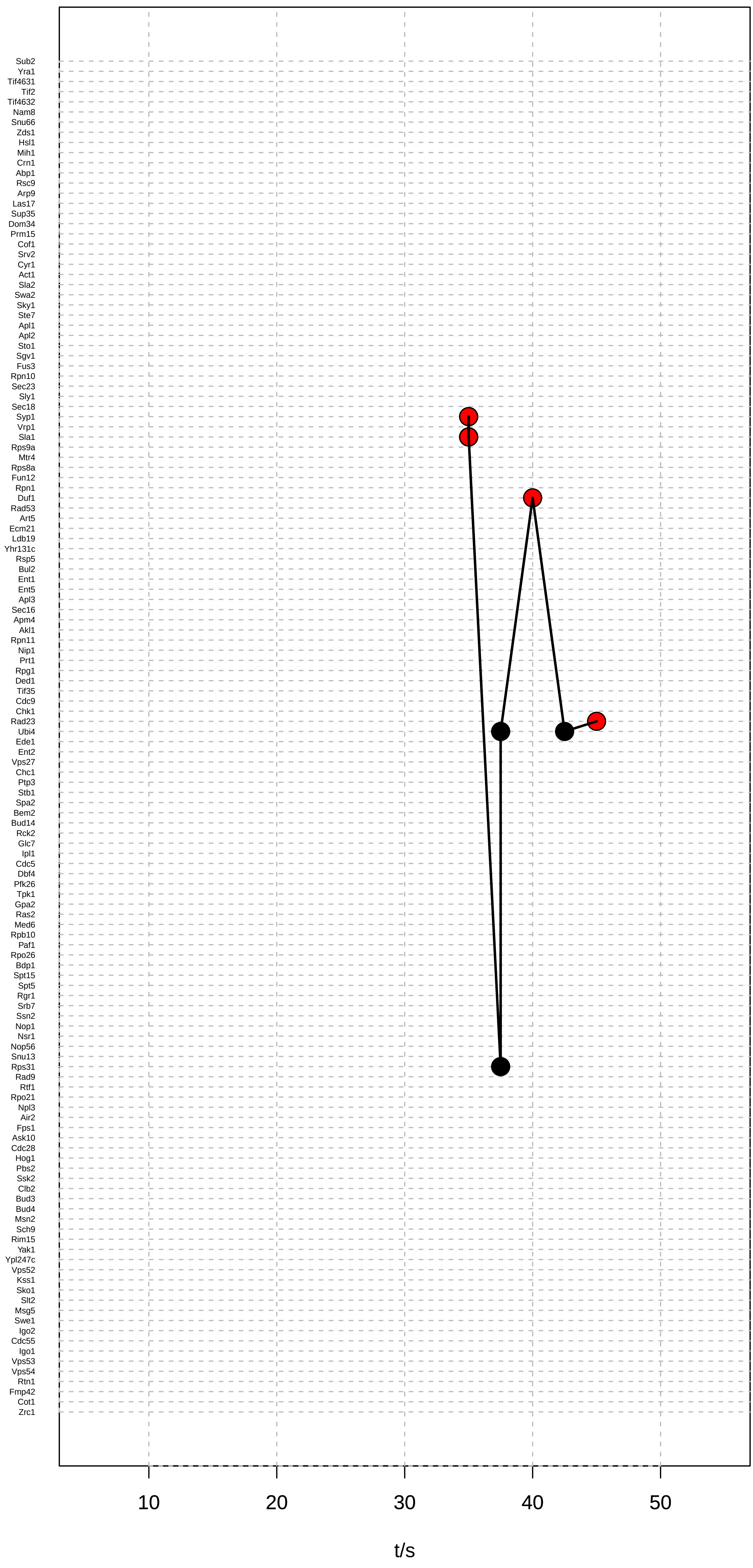

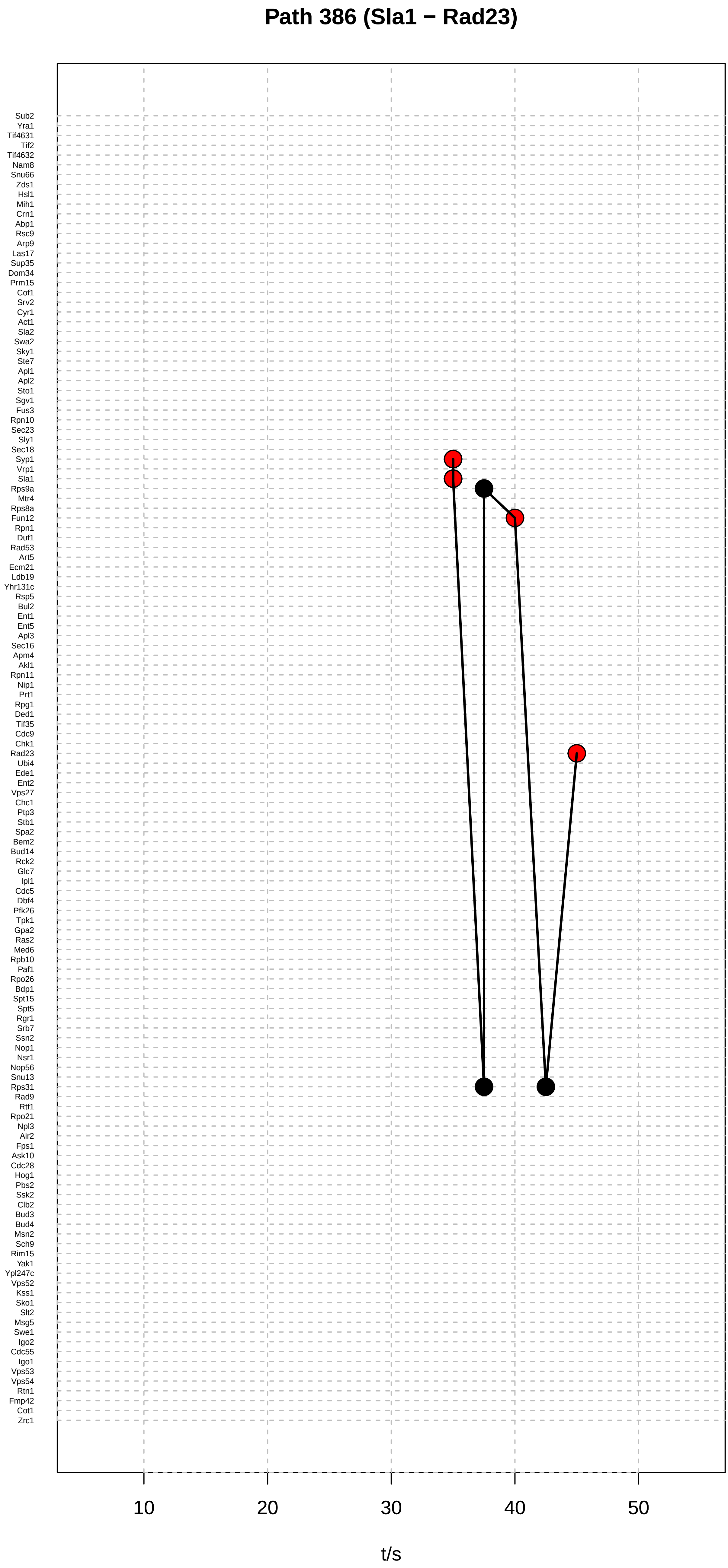

Path 386 (Sla1 − Rad23)
Sub2
Yra1
Tif4631
Tif2
Tif4632
Nam8
Snu66
Zds1
Hsl1
Mih1
Crn1
Abp1
Rsc9
Arp9
Las17
Sup35
Dom34
Prm15
Cof1
Srv2
Cyr1
Act1
Sla2
Swa2
Sky1
Ste7
Apl1
Apl2
Sto1
Sgv1
Fus3
Rpn10
Sec23
Sly1
Sec18
Syp1
Vrp1
Sla1
Rps9a
Mtr4
Rps8a
Fun12
Rpn1
Duf1
Rad53
Art5
Ecm21
Ldb19
Yhr131c
Rsp5
Bul2
Ent1
Ent5
Apl3
Sec16
Apm4
Akl1
Rpn11
Nip1
Prt1
Rpg1
Ded1
Tif35
Cdc9
Chk1
Rad23
Ubi4
Ede1
Ent2
Vps27
Chc1
Ptp3
Stb1
Spa2
Bem2
Bud14
Rck2
Glc7
Ipl1
Cdc5
Dbf4
Pfk26
Tpk1
Gpa2
Ras2
Med6
Rpb10
Paf1
Rpo26
Bdp1
Spt15
Spt5
Rgr1
Srb7
Ssn2
Nop1
Nsr1
Nop56
Snu13
Rps31
Rad9
Rtf1
Rpo21
Npl3
Air2
Fps1
Ask10
Cdc28
Hog1
Pbs2
Ssk2
Clb2
Bud3
Bud4
Msn2
Sch9
Rim15
Yak1
Ypl247c
Vps52
Kss1
Sko1
Slt2
Msg5
Swe1
Igo2
Cdc55
Igo1
Vps53
Vps54
Rtn1
Fmp42
Cot1
Zrc1
10
20
30
40
50
t/s

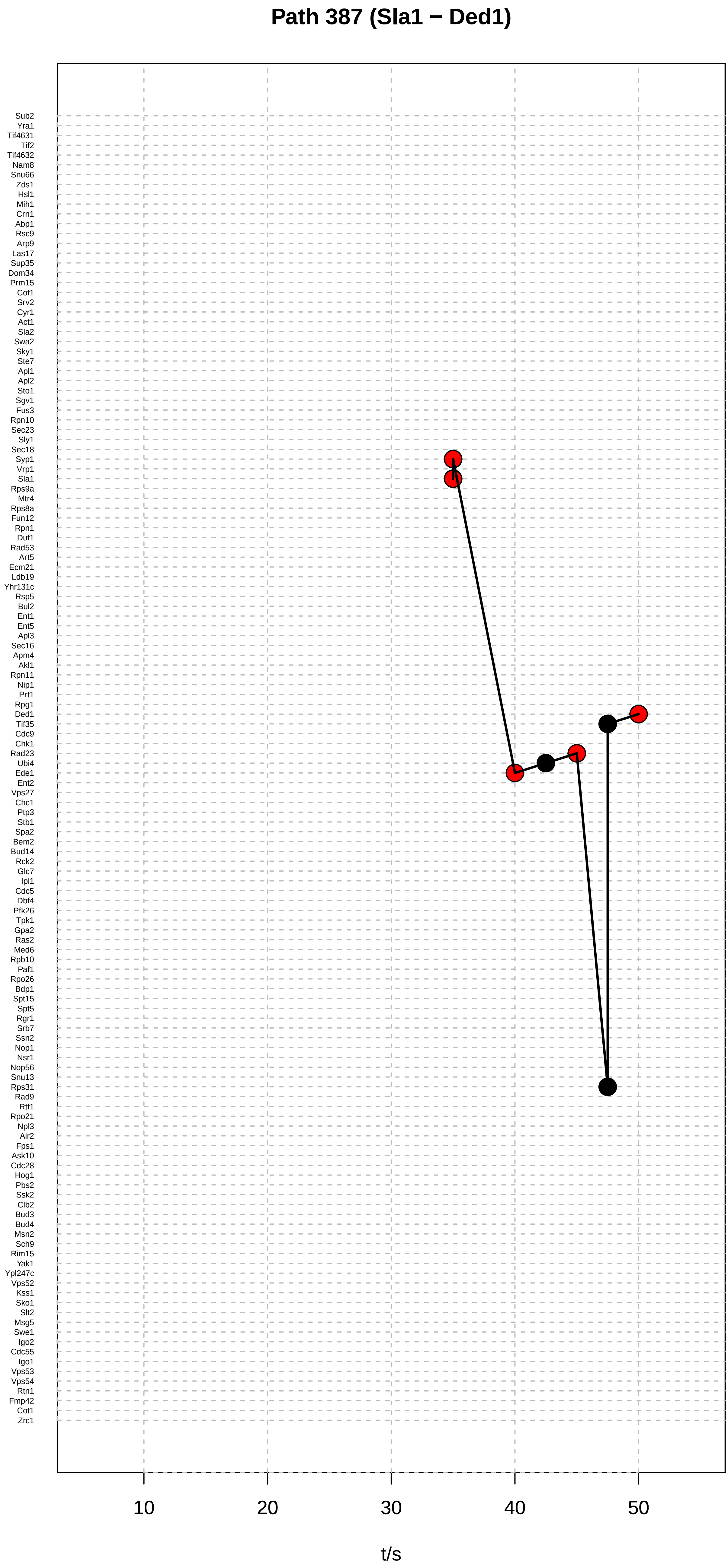

Path 387 (Sla1 − Ded1)
Sub2
Yra1
Tif4631
Tif2
Tif4632
Nam8
Snu66
Zds1
Hsl1
Mih1
Crn1
Abp1
Rsc9
Arp9
Las17
Sup35
Dom34
Prm15
Cof1
Srv2
Cyr1
Act1
Sla2
Swa2
Sky1
Ste7
Apl1
Apl2
Sto1
Sgv1
Fus3
Rpn10
Sec23
Sly1
Sec18
Syp1
Vrp1
Sla1
Rps9a
Mtr4
Rps8a
Fun12
Rpn1
Duf1
Rad53
Art5
Ecm21
Ldb19
Yhr131c
Rsp5
Bul2
Ent1
Ent5
Apl3
Sec16
Apm4
Akl1
Rpn11
Nip1
Prt1
Rpg1
Ded1
Tif35
Cdc9
Chk1
Rad23
Ubi4
Ede1
Ent2
Vps27
Chc1
Ptp3
Stb1
Spa2
Bem2
Bud14
Rck2
Glc7
Ipl1
Cdc5
Dbf4
Pfk26
Tpk1
Gpa2
Ras2
Med6
Rpb10
Paf1
Rpo26
Bdp1
Spt15
Spt5
Rgr1
Srb7
Ssn2
Nop1
Nsr1
Nop56
Snu13
Rps31
Rad9
Rtf1
Rpo21
Npl3
Air2
Fps1
Ask10
Cdc28
Hog1
Pbs2
Ssk2
Clb2
Bud3
Bud4
Msn2
Sch9
Rim15
Yak1
Ypl247c
Vps52
Kss1
Sko1
Slt2
Msg5
Swe1
Igo2
Cdc55
Igo1
Vps53
Vps54
Rtn1
Fmp42
Cot1
Zrc1
10
20
30
40
50
t/s

### Path 388 (Sla1 − Ded1)

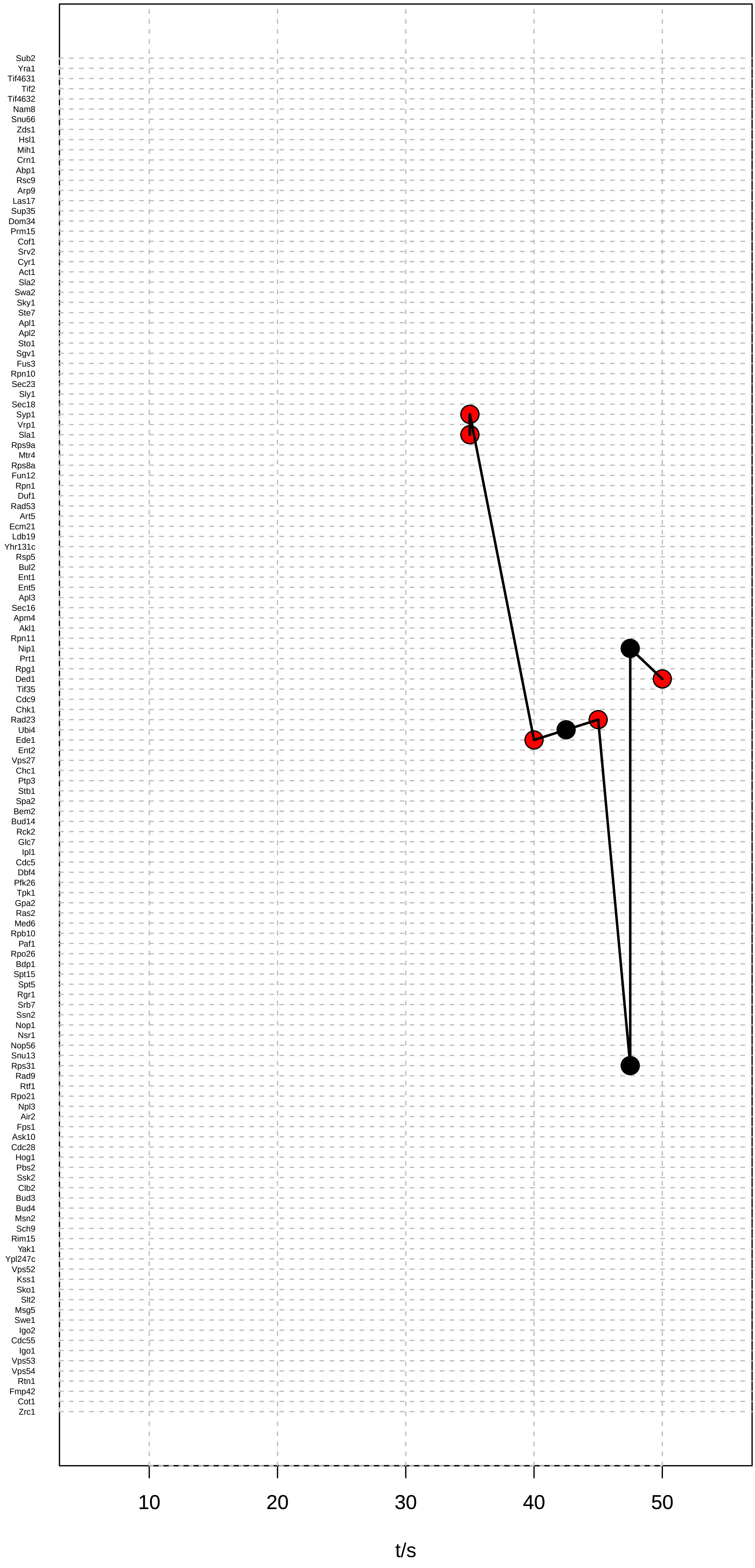

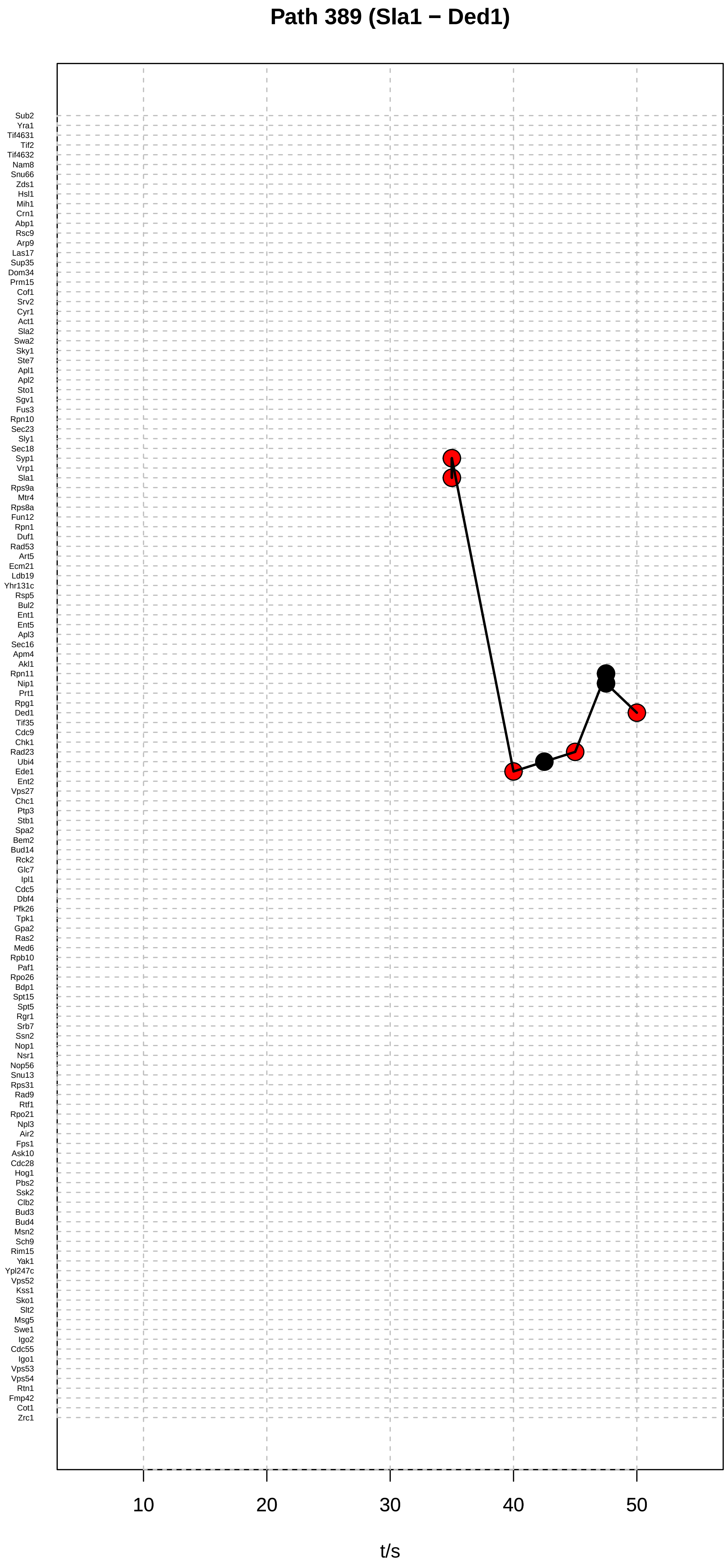

Path 389 (Sla1 − Ded1)
Sub2
Yra1
Tif4631
Tif2
Tif4632
Nam8
Snu66
Zds1
Hsl1
Mih1
Crn1
Abp1
Rsc9
Arp9
Las17
Sup35
Dom34
Prm15
Cof1
Srv2
Cyr1
Act1
Sla2
Swa2
Sky1
Ste7
Apl1
Apl2
Sto1
Sgv1
Fus3
Rpn10
Sec23
Sly1
Sec18
Syp1
Vrp1
Sla1
Rps9a
Mtr4
Rps8a
Fun12
Rpn1
Duf1
Rad53
Art5
Ecm21
Ldb19
Yhr131c
Rsp5
Bul2
Ent1
Ent5
Apl3
Sec16
Apm4
Akl1
Rpn11
Nip1
Prt1
Rpg1
Ded1
Tif35
Cdc9
Chk1
Rad23
Ubi4
Ede1
Ent2
Vps27
Chc1
Ptp3
Stb1
Spa2
Bem2
Bud14
Rck2
Glc7
Ipl1
Cdc5
Dbf4
Pfk26
Tpk1
Gpa2
Ras2
Med6
Rpb10
Paf1
Rpo26
Bdp1
Spt15
Spt5
Rgr1
Srb7
Ssn2
Nop1
Nsr1
Nop56
Snu13
Rps31
Rad9
Rtf1
Rpo21
Npl3
Air2
Fps1
Ask10
Cdc28
Hog1
Pbs2
Ssk2
Clb2
Bud3
Bud4
Msn2
Sch9
Rim15
Yak1
Ypl247c
Vps52
Kss1
Sko1
Slt2
Msg5
Swe1
Igo2
Cdc55
Igo1
Vps53
Vps54
Rtn1
Fmp42
Cot1
Zrc1
10
20
30
40
50
t/s

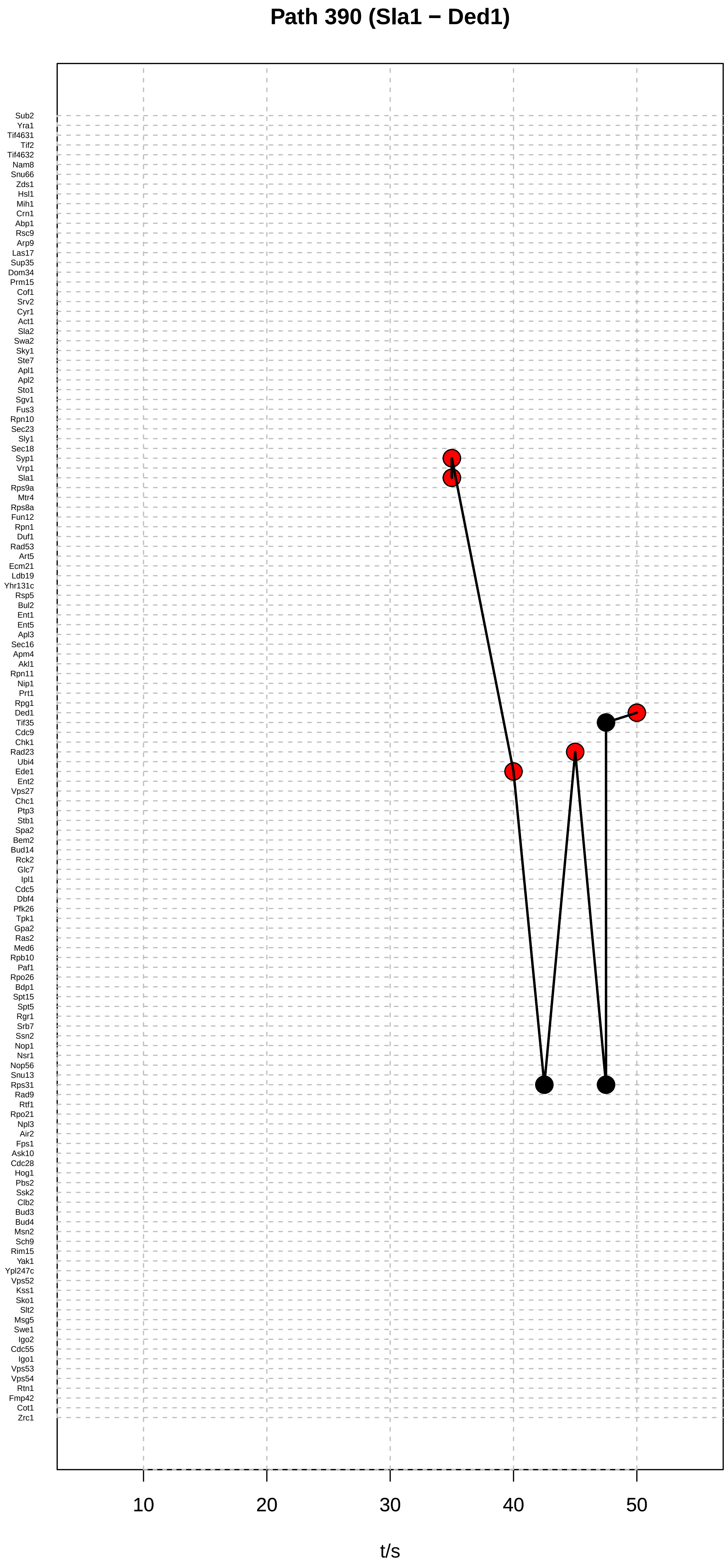
Path 390 (Sla1 − Ded1)
Sub2
Yra1
Tif4631
Tif2
Tif4632
Nam8
Snu66
Zds1
Hsl1
Mih1
Crn1
Abp1
Rsc9
Arp9
Las17
Sup35
Dom34
Prm15
Cof1
Srv2
Cyr1
Act1
Sla2
Swa2
Sky1
Ste7
Apl1
Apl2
Sto1
Sgv1
Fus3
Rpn10
Sec23
Sly1
Sec18
Syp1
Vrp1
Sla1
Rps9a
Mtr4
Rps8a
Fun12
Rpn1
Duf1
Rad53
Art5
Ecm21
Ldb19
Yhr131c
Rsp5
Bul2
Ent1
Ent5
Apl3
Sec16
Apm4
Akl1
Rpn11
Nip1
Prt1
Rpg1
Ded1
Tif35
Cdc9
Chk1
Rad23
Ubi4
Ede1
Ent2
Vps27
Chc1
Ptp3
Stb1
Spa2
Bem2
Bud14
Rck2
Glc7
Ipl1
Cdc5
Dbf4
Pfk26
Tpk1
Gpa2
Ras2
Med6
Rpb10
Paf1
Rpo26
Bdp1
Spt15
Spt5
Rgr1
Srb7
Ssn2
Nop1
Nsr1
Nop56
Snu13
Rps31
Rad9
Rtf1
Rpo21
Npl3
Air2
Fps1
Ask10
Cdc28
Hog1
Pbs2
Ssk2
Clb2
Bud3
Bud4
Msn2
Sch9
Rim15
Yak1
Ypl247c
Vps52
Kss1
Sko1
Slt2
Msg5
Swe1
Igo2
Cdc55
Igo1
Vps53
Vps54
Rtn1
Fmp42
Cot1
Zrc1
10
20
30
40
50
t/s

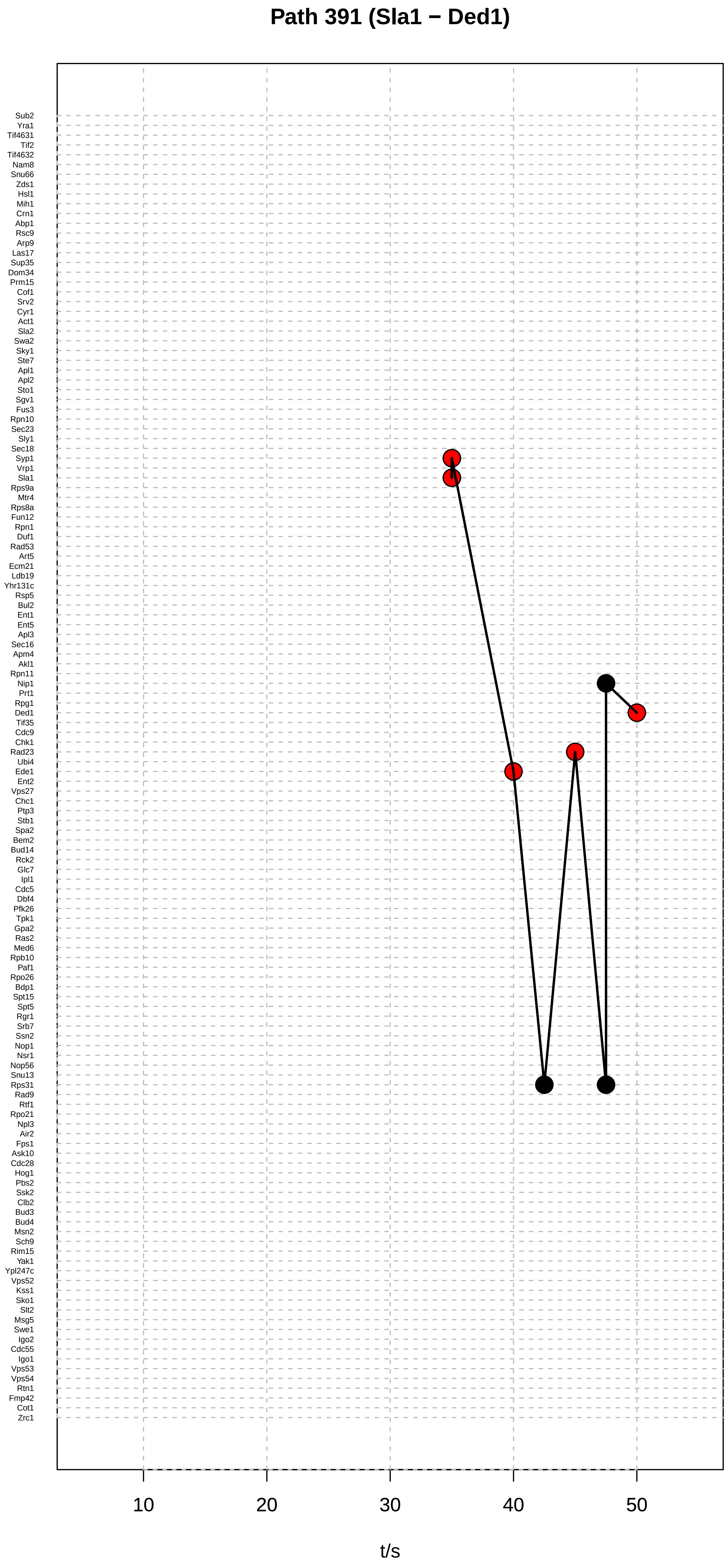
Path 391 (Sla1 − Ded1)
Sub2
Yra1
Tif4631
Tif2
Tif4632
Nam8
Snu66
Zds1
Hsl1
Mih1
Crn1
Abp1
Rsc9
Arp9
Las17
Sup35
Dom34
Prm15
Cof1
Srv2
Cyr1
Act1
Sla2
Swa2
Sky1
Ste7
Apl1
Apl2
Sto1
Sgv1
Fus3
Rpn10
Sec23
Sly1
Sec18
Syp1
Vrp1
Sla1
Rps9a
Mtr4
Rps8a
Fun12
Rpn1
Duf1
Rad53
Art5
Ecm21
Ldb19
Yhr131c
Rsp5
Bul2
Ent1
Ent5
Apl3
Sec16
Apm4
Akl1
Rpn11
Nip1
Prt1
Rpg1
Ded1
Tif35
Cdc9
Chk1
Rad23
Ubi4
Ede1
Ent2
Vps27
Chc1
Ptp3
Stb1
Spa2
Bem2
Bud14
Rck2
Glc7
Ipl1
Cdc5
Dbf4
Pfk26
Tpk1
Gpa2
Ras2
Med6
Rpb10
Paf1
Rpo26
Bdp1
Spt15
Spt5
Rgr1
Srb7
Ssn2
Nop1
Nsr1
Nop56
Snu13
Rps31
Rad9
Rtf1
Rpo21
Npl3
Air2
Fps1
Ask10
Cdc28
Hog1
Pbs2
Ssk2
Clb2
Bud3
Bud4
Msn2
Sch9
Rim15
Yak1
Ypl247c
Vps52
Kss1
Sko1
Slt2
Msg5
Swe1
Igo2
Cdc55
Igo1
Vps53
Vps54
Rtn1
Fmp42
Cot1
Zrc1
10
20
30
40
50
t/s

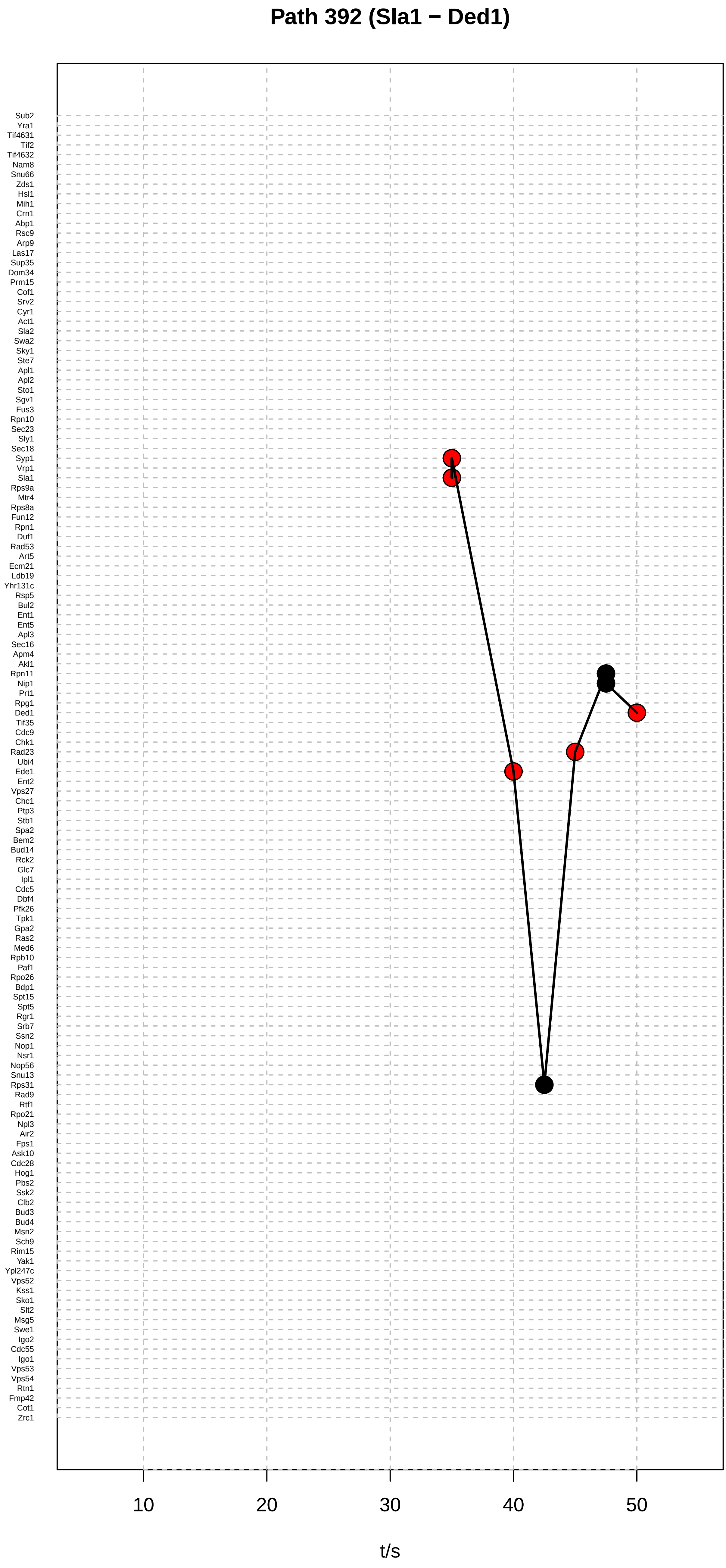

Path 392 (Sla1 − Ded1)
Sub2
Yra1
Tif4631
Tif2
Tif4632
Nam8
Snu66
Zds1
Hsl1
Mih1
Crn1
Abp1
Rsc9
Arp9
Las17
Sup35
Dom34
Prm15
Cof1
Srv2
Cyr1
Act1
Sla2
Swa2
Sky1
Ste7
Apl1
Apl2
Sto1
Sgv1
Fus3
Rpn10
Sec23
Sly1
Sec18
Syp1
Vrp1
Sla1
Rps9a
Mtr4
Rps8a
Fun12
Rpn1
Duf1
Rad53
Art5
Ecm21
Ldb19
Yhr131c
Rsp5
Bul2
Ent1
Ent5
Apl3
Sec16
Apm4
Akl1
Rpn11
Nip1
Prt1
Rpg1
Ded1
Tif35
Cdc9
Chk1
Rad23
Ubi4
Ede1
Ent2
Vps27
Chc1
Ptp3
Stb1
Spa2
Bem2
Bud14
Rck2
Glc7
Ipl1
Cdc5
Dbf4
Pfk26
Tpk1
Gpa2
Ras2
Med6
Rpb10
Paf1
Rpo26
Bdp1
Spt15
Spt5
Rgr1
Srb7
Ssn2
Nop1
Nsr1
Nop56
Snu13
Rps31
Rad9
Rtf1
Rpo21
Npl3
Air2
Fps1
Ask10
Cdc28
Hog1
Pbs2
Ssk2
Clb2
Bud3
Bud4
Msn2
Sch9
Rim15
Yak1
Ypl247c
Vps52
Kss1
Sko1
Slt2
Msg5
Swe1
Igo2
Cdc55
Igo1
Vps53
Vps54
Rtn1
Fmp42
Cot1
Zrc1
10
20
30
40
50
t/s

#### Path 393 (Sla1 − Ded1)

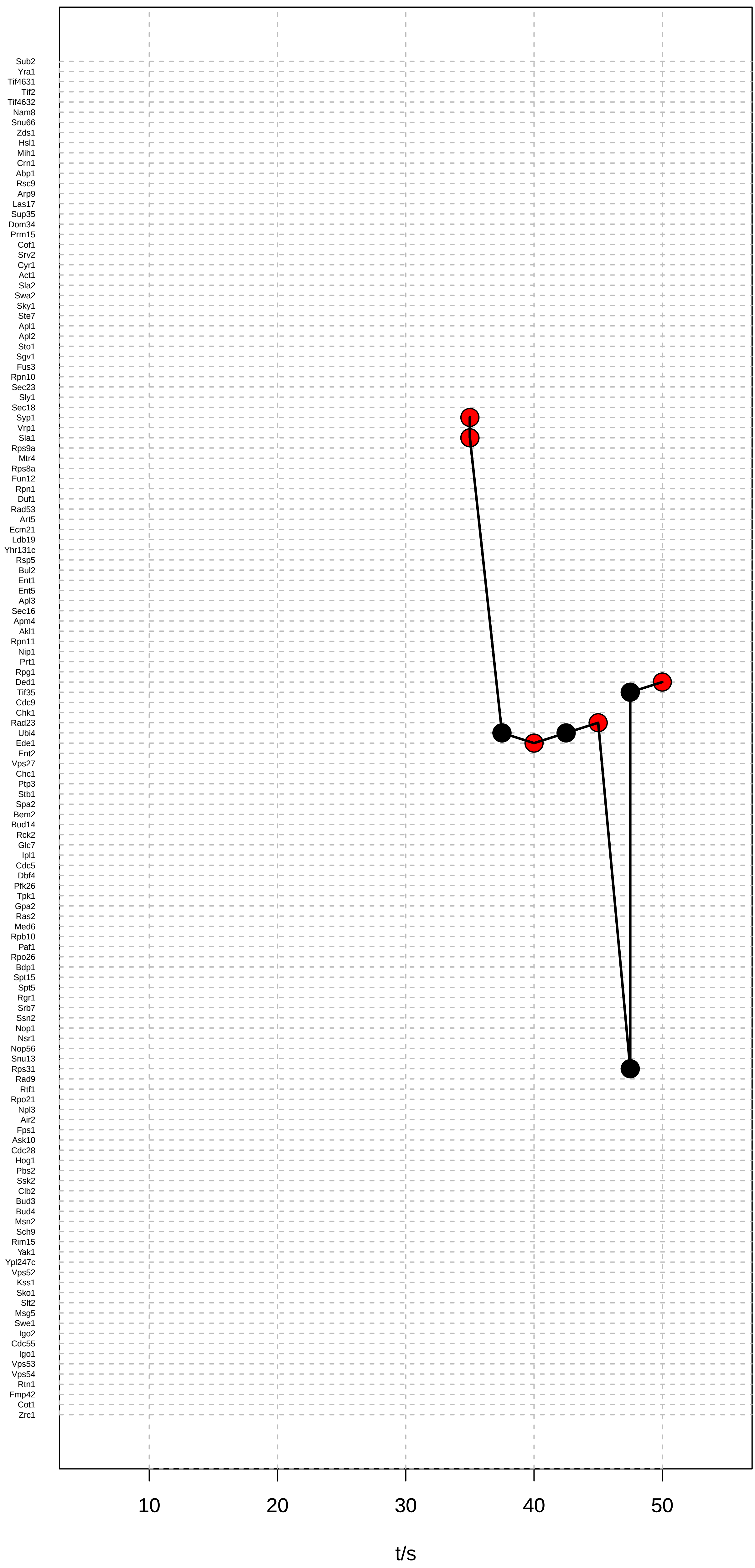

Path 394 (Sla1 − Ded1)

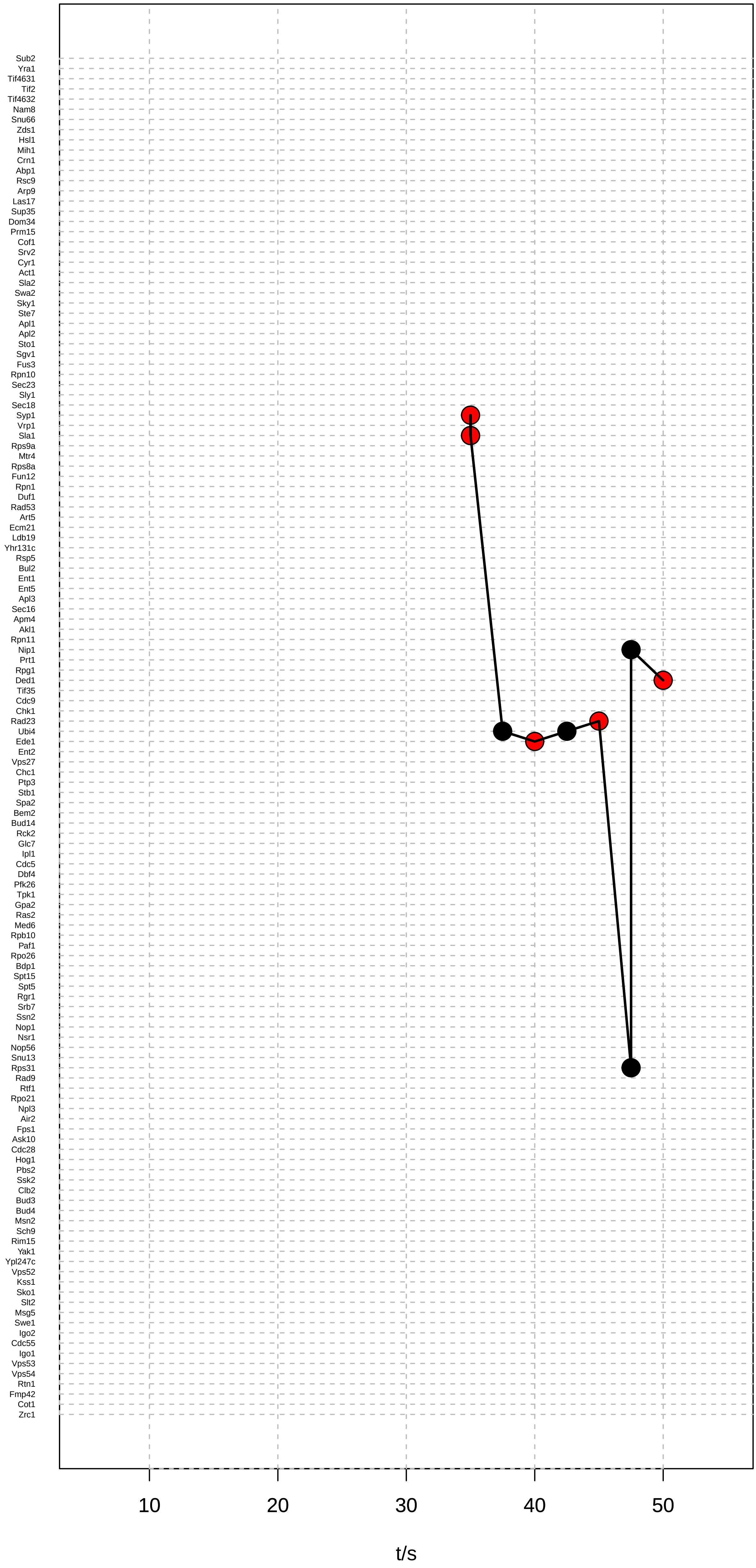

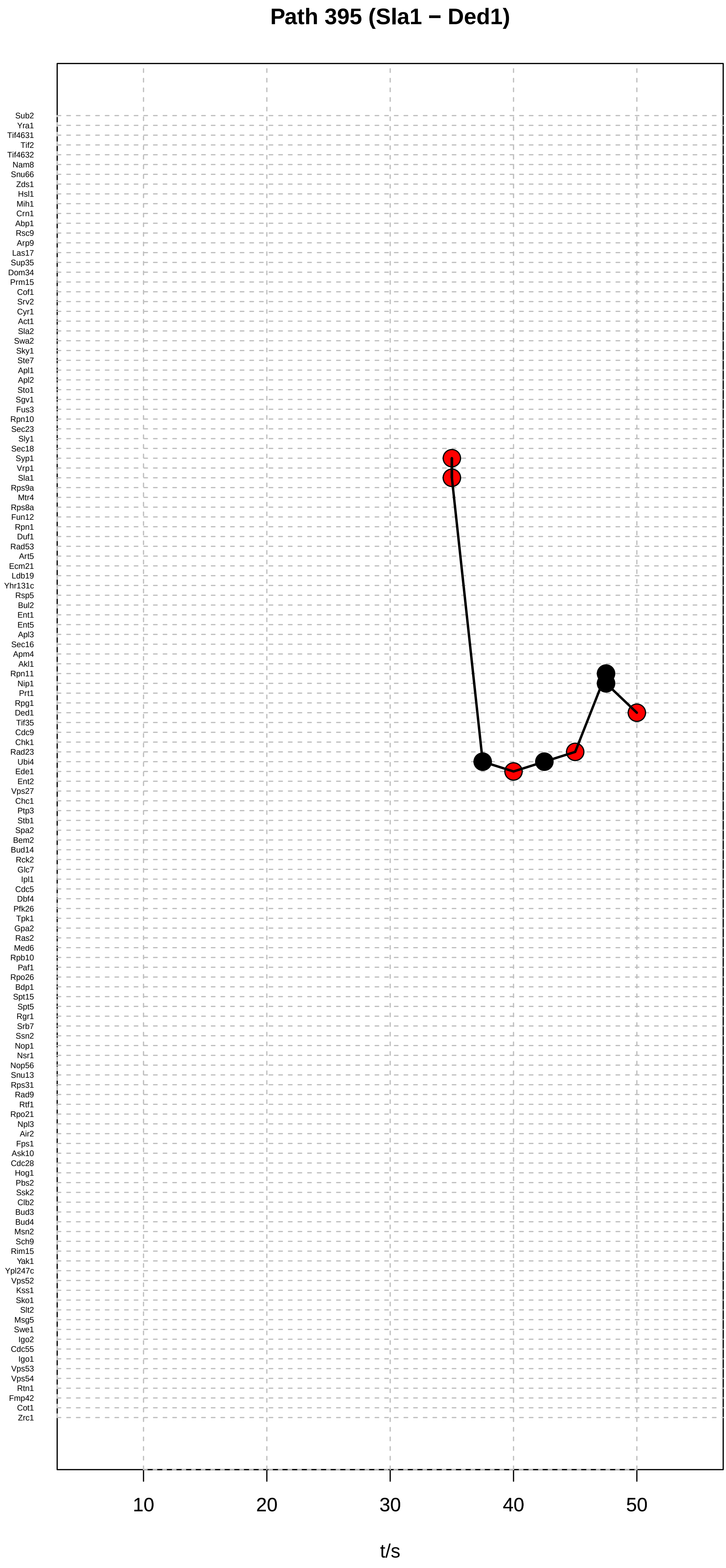
Path 395 (Sla1 − Ded1)
Sub2
Yra1
Tif4631
Tif2
Tif4632
Nam8
Snu66
Zds1
Hsl1
Mih1
Crn1
Abp1
Rsc9
Arp9
Las17
Sup35
Dom34
Prm15
Cof1
Srv2
Cyr1
Act1
Sla2
Swa2
Sky1
Ste7
Apl1
Apl2
Sto1
Sgv1
Fus3
Rpn10
Sec23
Sly1
Sec18
Syp1
Vrp1
Sla1
Rps9a
Mtr4
Rps8a
Fun12
Rpn1
Duf1
Rad53
Art5
Ecm21
Ldb19
Yhr131c
Rsp5
Bul2
Ent1
Ent5
Apl3
Sec16
Apm4
Akl1
Rpn11
Nip1
Prt1
Rpg1
Ded1
Tif35
Cdc9
Chk1
Rad23
Ubi4
Ede1
Ent2
Vps27
Chc1
Ptp3
Stb1
Spa2
Bem2
Bud14
Rck2
Glc7
Ipl1
Cdc5
Dbf4
Pfk26
Tpk1
Gpa2
Ras2
Med6
Rpb10
Paf1
Rpo26
Bdp1
Spt15
Spt5
Rgr1
Srb7
Ssn2
Nop1
Nsr1
Nop56
Snu13
Rps31
Rad9
Rtf1
Rpo21
Npl3
Air2
Fps1
Ask10
Cdc28
Hog1
Pbs2
Ssk2
Clb2
Bud3
Bud4
Msn2
Sch9
Rim15
Yak1
Ypl247c
Vps52
Kss1
Sko1
Slt2
Msg5
Swe1
Igo2
Cdc55
Igo1
Vps53
Vps54
Rtn1
Fmp42
Cot1
Zrc1
10
20
30
40
50
t/s

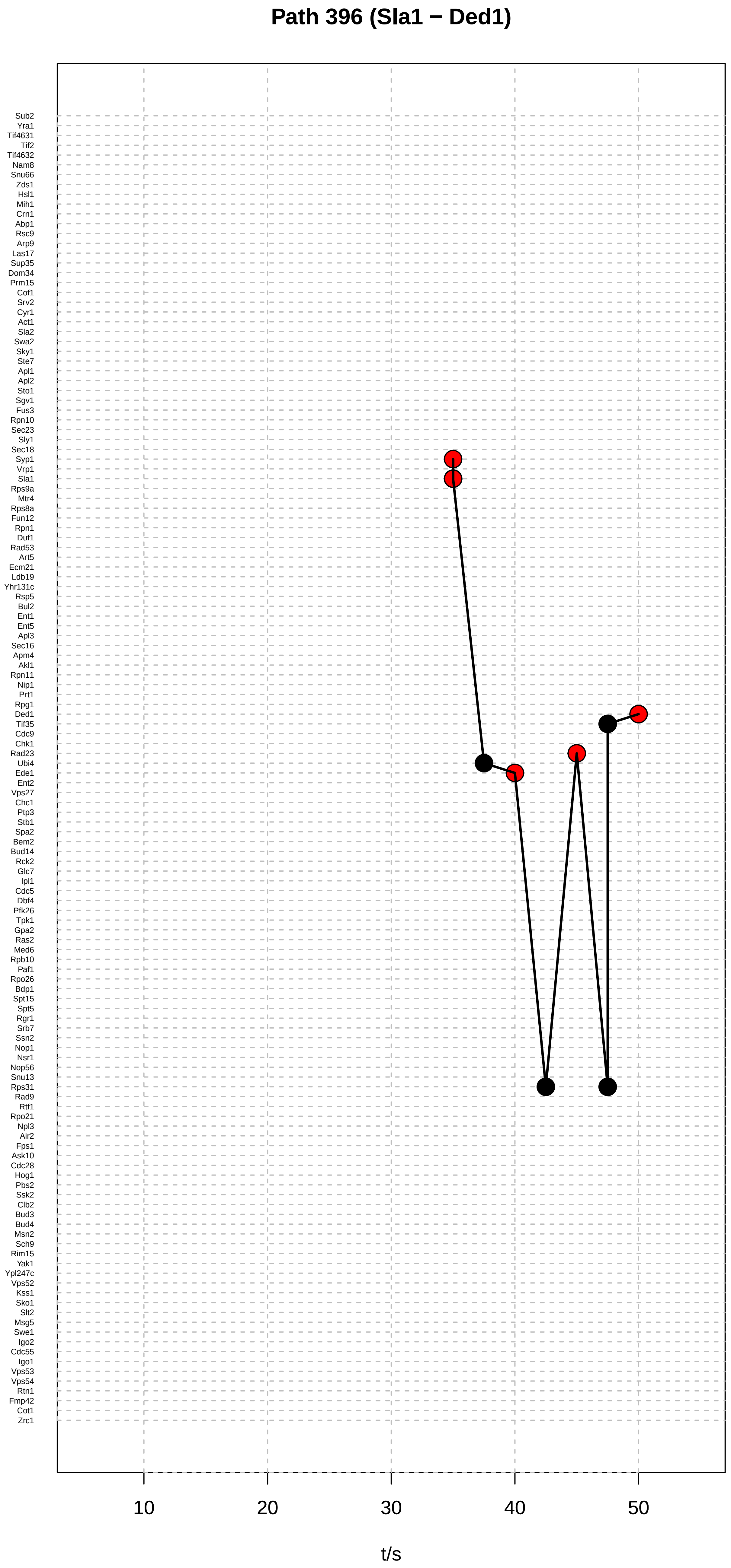

Path 396 (Sla1 − Ded1)
Sub2
Yra1
Tif4631
Tif2
Tif4632
Nam8
Snu66
Zds1
Hsl1
Mih1
Crn1
Abp1
Rsc9
Arp9
Las17
Sup35
Dom34
Prm15
Cof1
Srv2
Cyr1
Act1
Sla2
Swa2
Sky1
Ste7
Apl1
Apl2
Sto1
Sgv1
Fus3
Rpn10
Sec23
Sly1
Sec18
Syp1
Vrp1
Sla1
Rps9a
Mtr4
Rps8a
Fun12
Rpn1
Duf1
Rad53
Art5
Ecm21
Ldb19
Yhr131c
Rsp5
Bul2
Ent1
Ent5
Apl3
Sec16
Apm4
Akl1
Rpn11
Nip1
Prt1
Rpg1
Ded1
Tif35
Cdc9
Chk1
Rad23
Ubi4
Ede1
Ent2
Vps27
Chc1
Ptp3
Stb1
Spa2
Bem2
Bud14
Rck2
Glc7
Ipl1
Cdc5
Dbf4
Pfk26
Tpk1
Gpa2
Ras2
Med6
Rpb10
Paf1
Rpo26
Bdp1
Spt15
Spt5
Rgr1
Srb7
Ssn2
Nop1
Nsr1
Nop56
Snu13
Rps31
Rad9
Rtf1
Rpo21
Npl3
Air2
Fps1
Ask10
Cdc28
Hog1
Pbs2
Ssk2
Clb2
Bud3
Bud4
Msn2
Sch9
Rim15
Yak1
Ypl247c
Vps52
Kss1
Sko1
Slt2
Msg5
Swe1
Igo2
Cdc55
Igo1
Vps53
Vps54
Rtn1
Fmp42
Cot1
Zrc1
10
20
30
40
50
t/s

#### Path 397 (Sla1 − Ded1)

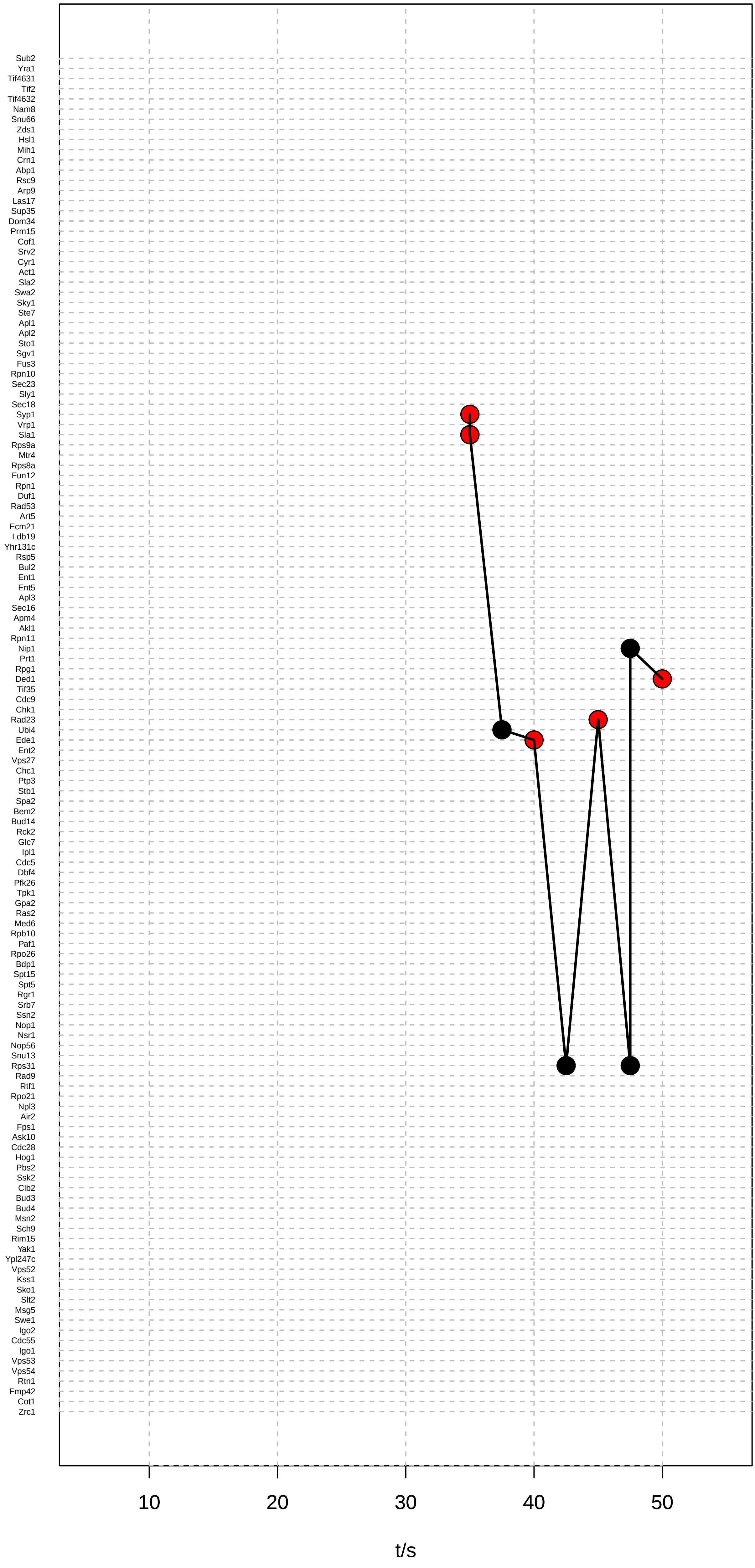

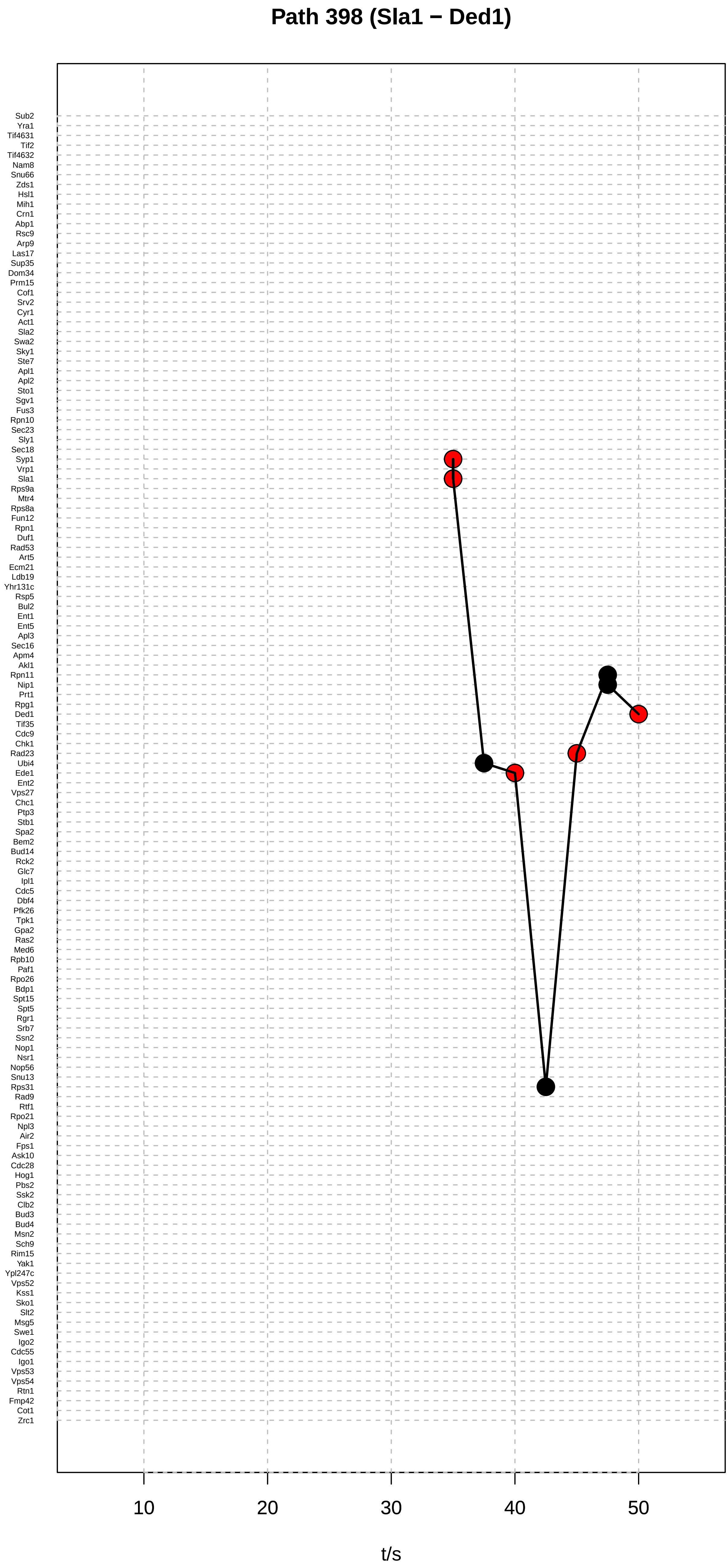
Path 398 (Sla1 − Ded1)
Sub2
Yra1
Tif4631
Tif2
Tif4632
Nam8
Snu66
Zds1
Hsl1
Mih1
Crn1
Abp1
Rsc9
Arp9
Las17
Sup35
Dom34
Prm15
Cof1
Srv2
Cyr1
Act1
Sla2
Swa2
Sky1
Ste7
Apl1
Apl2
Sto1
Sgv1
Fus3
Rpn10
Sec23
Sly1
Sec18
Syp1
Vrp1
Sla1
Rps9a
Mtr4
Rps8a
Fun12
Rpn1
Duf1
Rad53
Art5
Ecm21
Ldb19
Yhr131c
Rsp5
Bul2
Ent1
Ent5
Apl3
Sec16
Apm4
Akl1
Rpn11
Nip1
Prt1
Rpg1
Ded1
Tif35
Cdc9
Chk1
Rad23
Ubi4
Ede1
Ent2
Vps27
Chc1
Ptp3
Stb1
Spa2
Bem2
Bud14
Rck2
Glc7
Ipl1
Cdc5
Dbf4
Pfk26
Tpk1
Gpa2
Ras2
Med6
Rpb10
Paf1
Rpo26
Bdp1
Spt15
Spt5
Rgr1
Srb7
Ssn2
Nop1
Nsr1
Nop56
Snu13
Rps31
Rad9
Rtf1
Rpo21
Npl3
Air2
Fps1
Ask10
Cdc28
Hog1
Pbs2
Ssk2
Clb2
Bud3
Bud4
Msn2
Sch9
Rim15
Yak1
Ypl247c
Vps52
Kss1
Sko1
Slt2
Msg5
Swe1
Igo2
Cdc55
Igo1
Vps53
Vps54
Rtn1
Fmp42
Cot1
Zrc1
10
20
30
40
50
t/s

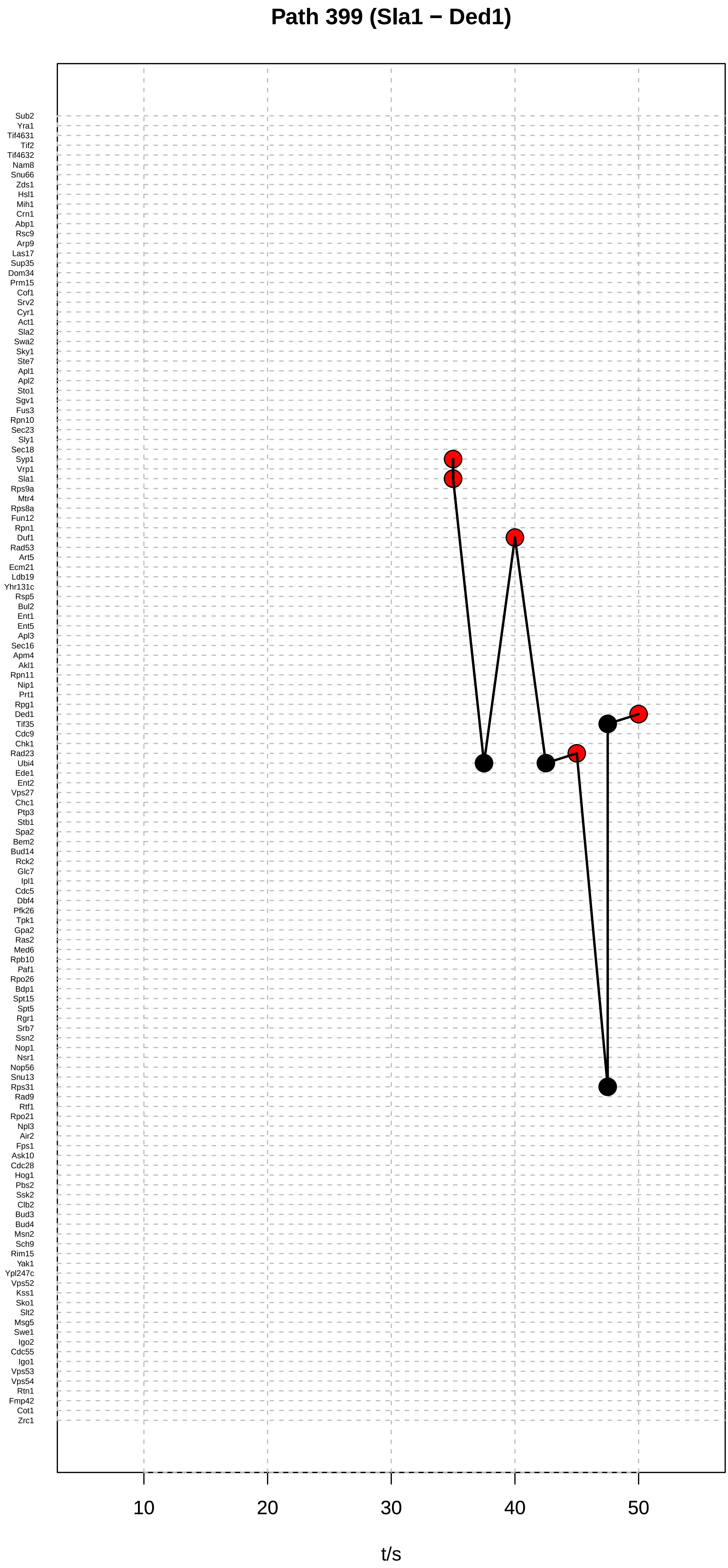

Path 399 (Sla1 − Ded1)
Sub2
Yra1
Tif4631
Tif2
Tif4632
Nam8
Snu66
Zds1
Hsl1
Mih1
Crn1
Abp1
Rsc9
Arp9
Las17
Sup35
Dom34
Prm15
Cof1
Srv2
Cyr1
Act1
Sla2
Swa2
Sky1
Ste7
Apl1
Apl2
Sto1
Sgv1
Fus3
Rpn10
Sec23
Sly1
Sec18
Syp1
Vrp1
Sla1
Rps9a
Mtr4
Rps8a
Fun12
Rpn1
Duf1
Rad53
Art5
Ecm21
Ldb19
Yhr131c
Rsp5
Bul2
Ent1
Ent5
Apl3
Sec16
Apm4
Akl1
Rpn11
Nip1
Prt1
Rpg1
Ded1
Tif35
Cdc9
Chk1
Rad23
Ubi4
Ede1
Ent2
Vps27
Chc1
Ptp3
Stb1
Spa2
Bem2
Bud14
Rck2
Glc7
Ipl1
Cdc5
Dbf4
Pfk26
Tpk1
Gpa2
Ras2
Med6
Rpb10
Paf1
Rpo26
Bdp1
Spt15
Spt5
Rgr1
Srb7
Ssn2
Nop1
Nsr1
Nop56
Snu13
Rps31
Rad9
Rtf1
Rpo21
Npl3
Air2
Fps1
Ask10
Cdc28
Hog1
Pbs2
Ssk2
Clb2
Bud3
Bud4
Msn2
Sch9
Rim15
Yak1
Ypl247c
Vps52
Kss1
Sko1
Slt2
Msg5
Swe1
Igo2
Cdc55
Igo1
Vps53
Vps54
Rtn1
Fmp42
Cot1
Zrc1
10
20
30
40
50
t/s

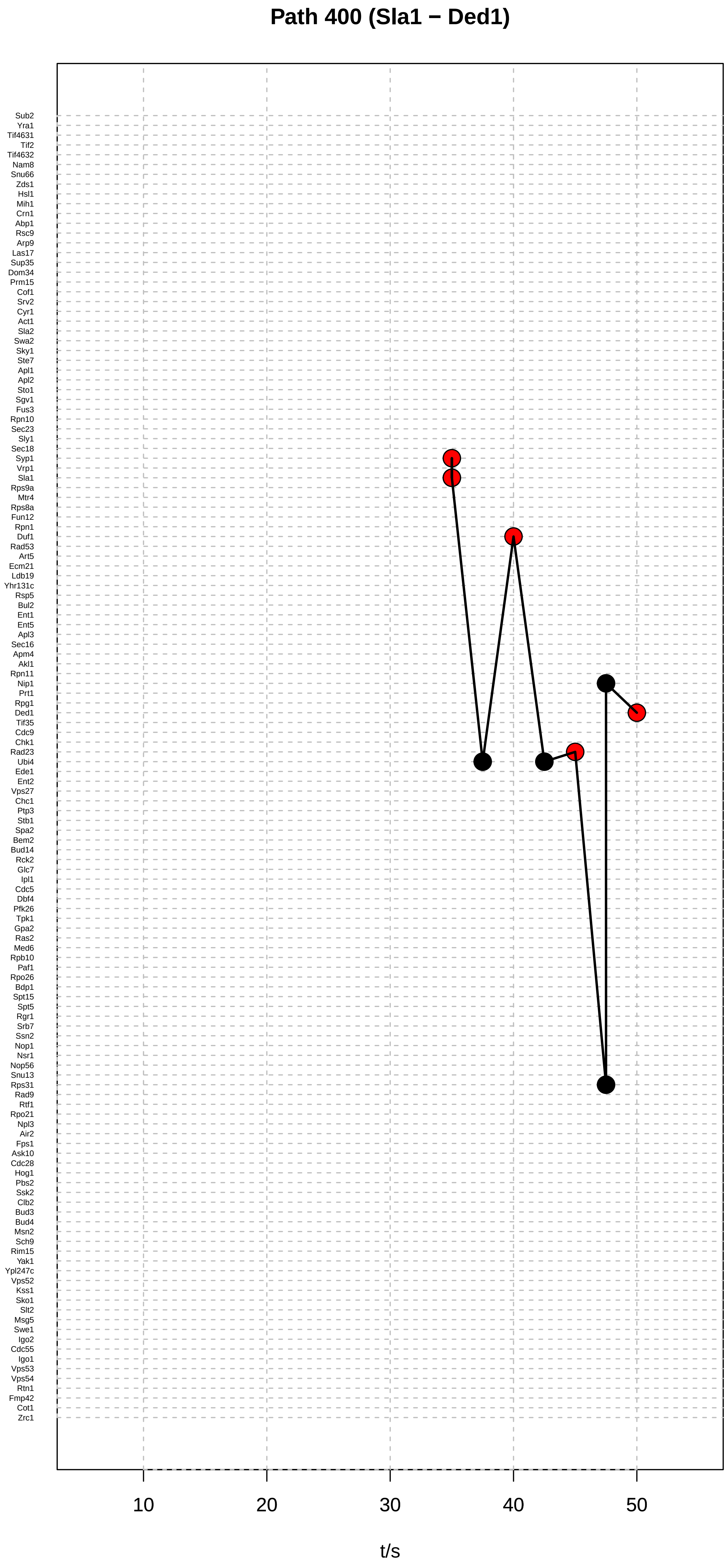

Path 400 (Sla1 − Ded1)
Sub2
Yra1
Tif4631
Tif2
Tif4632
Nam8
Snu66
Zds1
Hsl1
Mih1
Crn1
Abp1
Rsc9
Arp9
Las17
Sup35
Dom34
Prm15
Cof1
Srv2
Cyr1
Act1
Sla2
Swa2
Sky1
Ste7
Apl1
Apl2
Sto1
Sgv1
Fus3
Rpn10
Sec23
Sly1
Sec18
Syp1
Vrp1
Sla1
Rps9a
Mtr4
Rps8a
Fun12
Rpn1
Duf1
Rad53
Art5
Ecm21
Ldb19
Yhr131c
Rsp5
Bul2
Ent1
Ent5
Apl3
Sec16
Apm4
Akl1
Rpn11
Nip1
Prt1
Rpg1
Ded1
Tif35
Cdc9
Chk1
Rad23
Ubi4
Ede1
Ent2
Vps27
Chc1
Ptp3
Stb1
Spa2
Bem2
Bud14
Rck2
Glc7
Ipl1
Cdc5
Dbf4
Pfk26
Tpk1
Gpa2
Ras2
Med6
Rpb10
Paf1
Rpo26
Bdp1
Spt15
Spt5
Rgr1
Srb7
Ssn2
Nop1
Nsr1
Nop56
Snu13
Rps31
Rad9
Rtf1
Rpo21
Npl3
Air2
Fps1
Ask10
Cdc28
Hog1
Pbs2
Ssk2
Clb2
Bud3
Bud4
Msn2
Sch9
Rim15
Yak1
Ypl247c
Vps52
Kss1
Sko1
Slt2
Msg5
Swe1
Igo2
Cdc55
Igo1
Vps53
Vps54
Rtn1
Fmp42
Cot1
Zrc1
10
20
30
40
50
t/s

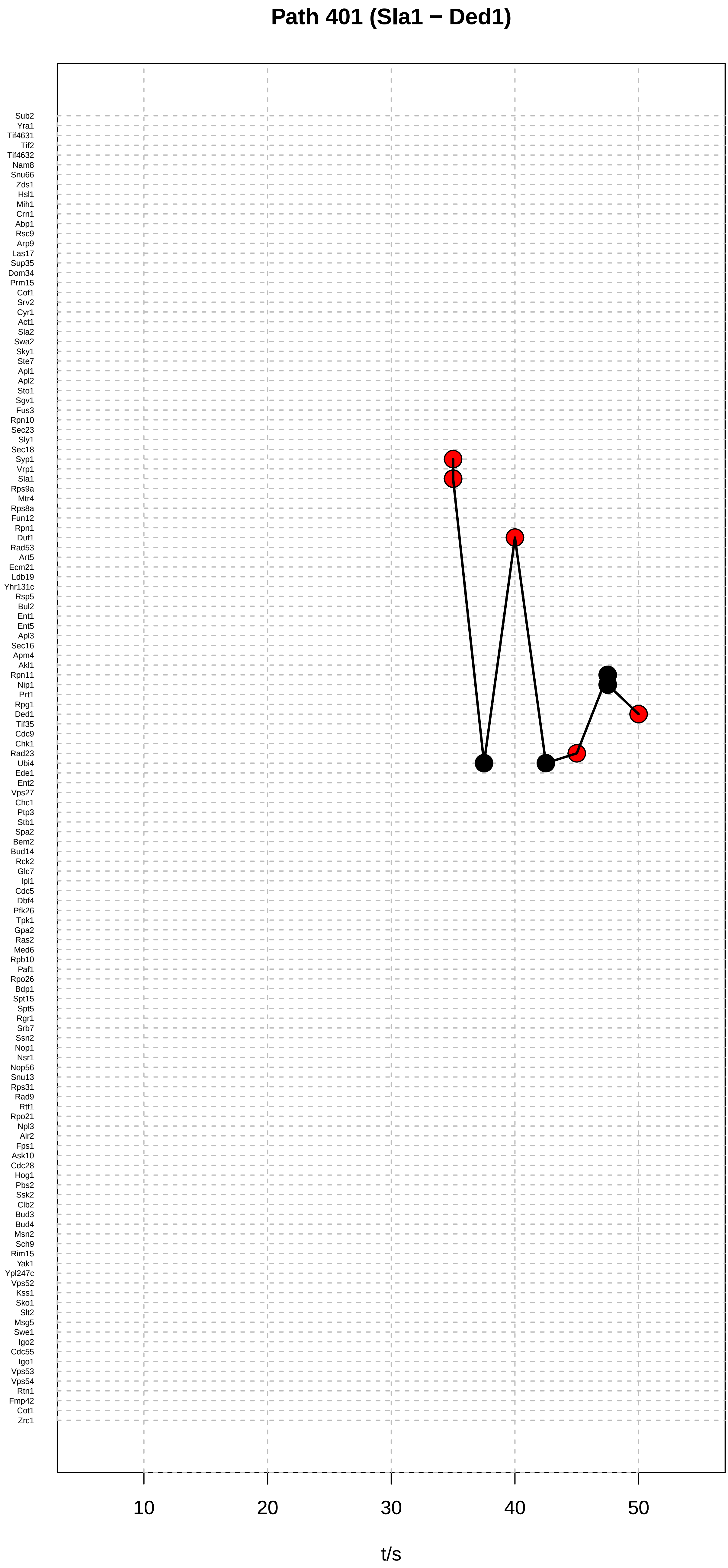

Path 401 (Sla1 − Ded1)
Sub2
Yra1
Tif4631
Tif2
Tif4632
Nam8
Snu66
Zds1
Hsl1
Mih1
Crn1
Abp1
Rsc9
Arp9
Las17
Sup35
Dom34
Prm15
Cof1
Srv2
Cyr1
Act1
Sla2
Swa2
Sky1
Ste7
Apl1
Apl2
Sto1
Sgv1
Fus3
Rpn10
Sec23
Sly1
Sec18
Syp1
Vrp1
Sla1
Rps9a
Mtr4
Rps8a
Fun12
Rpn1
Duf1
Rad53
Art5
Ecm21
Ldb19
Yhr131c
Rsp5
Bul2
Ent1
Ent5
Apl3
Sec16
Apm4
Akl1
Rpn11
Nip1
Prt1
Rpg1
Ded1
Tif35
Cdc9
Chk1
Rad23
Ubi4
Ede1
Ent2
Vps27
Chc1
Ptp3
Stb1
Spa2
Bem2
Bud14
Rck2
Glc7
Ipl1
Cdc5
Dbf4
Pfk26
Tpk1
Gpa2
Ras2
Med6
Rpb10
Paf1
Rpo26
Bdp1
Spt15
Spt5
Rgr1
Srb7
Ssn2
Nop1
Nsr1
Nop56
Snu13
Rps31
Rad9
Rtf1
Rpo21
Npl3
Air2
Fps1
Ask10
Cdc28
Hog1
Pbs2
Ssk2
Clb2
Bud3
Bud4
Msn2
Sch9
Rim15
Yak1
Ypl247c
Vps52
Kss1
Sko1
Slt2
Msg5
Swe1
Igo2
Cdc55
Igo1
Vps53
Vps54
Rtn1
Fmp42
Cot1
Zrc1
10
20
30
40
50
t/s

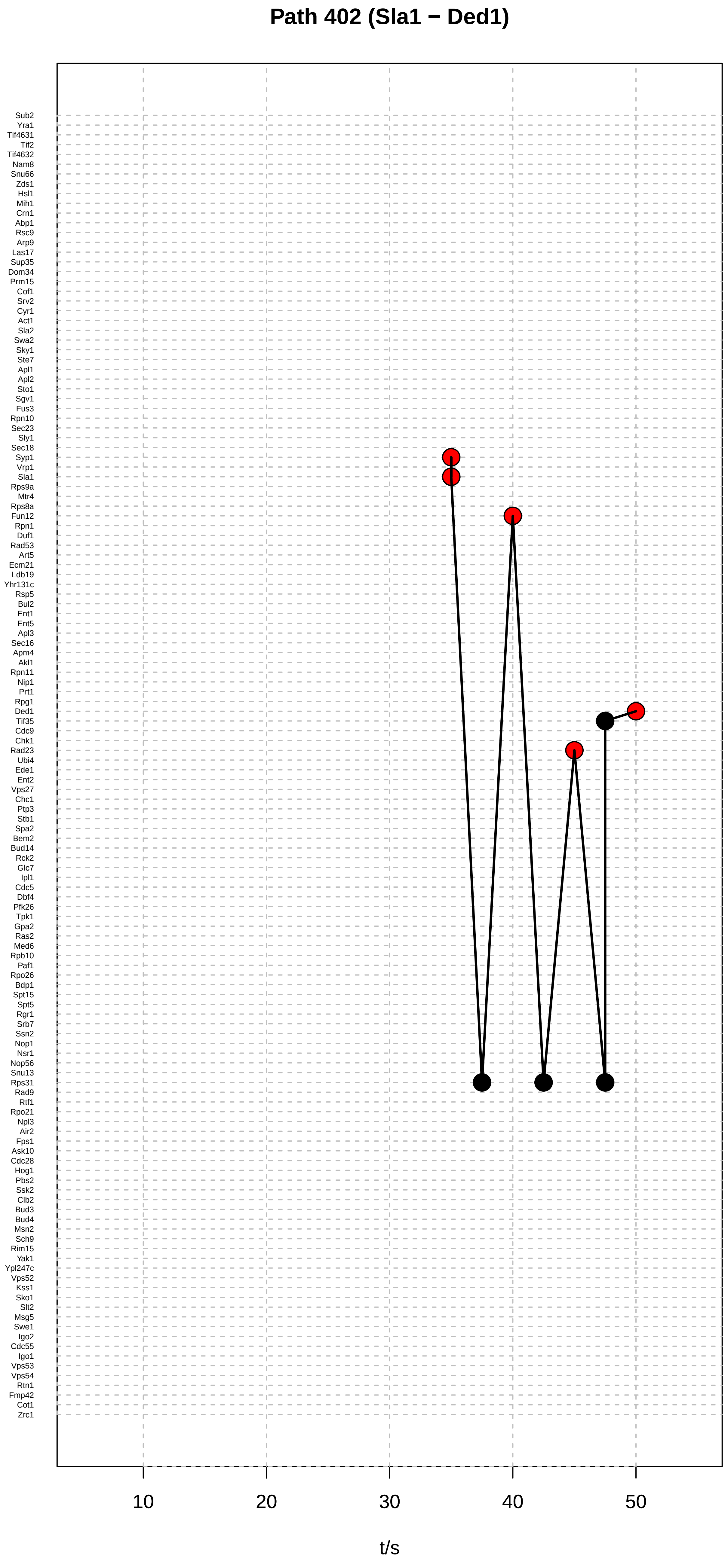

Path 402 (Sla1 − Ded1)
Sub2
Yra1
Tif4631
Tif2
Tif4632
Nam8
Snu66
Zds1
Hsl1
Mih1
Crn1
Abp1
Rsc9
Arp9
Las17
Sup35
Dom34
Prm15
Cof1
Srv2
Cyr1
Act1
Sla2
Swa2
Sky1
Ste7
Apl1
Apl2
Sto1
Sgv1
Fus3
Rpn10
Sec23
Sly1
Sec18
Syp1
Vrp1
Sla1
Rps9a
Mtr4
Rps8a
Fun12
Rpn1
Duf1
Rad53
Art5
Ecm21
Ldb19
Yhr131c
Rsp5
Bul2
Ent1
Ent5
Apl3
Sec16
Apm4
Akl1
Rpn11
Nip1
Prt1
Rpg1
Ded1
Tif35
Cdc9
Chk1
Rad23
Ubi4
Ede1
Ent2
Vps27
Chc1
Ptp3
Stb1
Spa2
Bem2
Bud14
Rck2
Glc7
Ipl1
Cdc5
Dbf4
Pfk26
Tpk1
Gpa2
Ras2
Med6
Rpb10
Paf1
Rpo26
Bdp1
Spt15
Spt5
Rgr1
Srb7
Ssn2
Nop1
Nsr1
Nop56
Snu13
Rps31
Rad9
Rtf1
Rpo21
Npl3
Air2
Fps1
Ask10
Cdc28
Hog1
Pbs2
Ssk2
Clb2
Bud3
Bud4
Msn2
Sch9
Rim15
Yak1
Ypl247c
Vps52
Kss1
Sko1
Slt2
Msg5
Swe1
Igo2
Cdc55
Igo1
Vps53
Vps54
Rtn1
Fmp42
Cot1
Zrc1
10
20
30
40
50
t/s

#### Path 403 (Sla1 − Ded1)

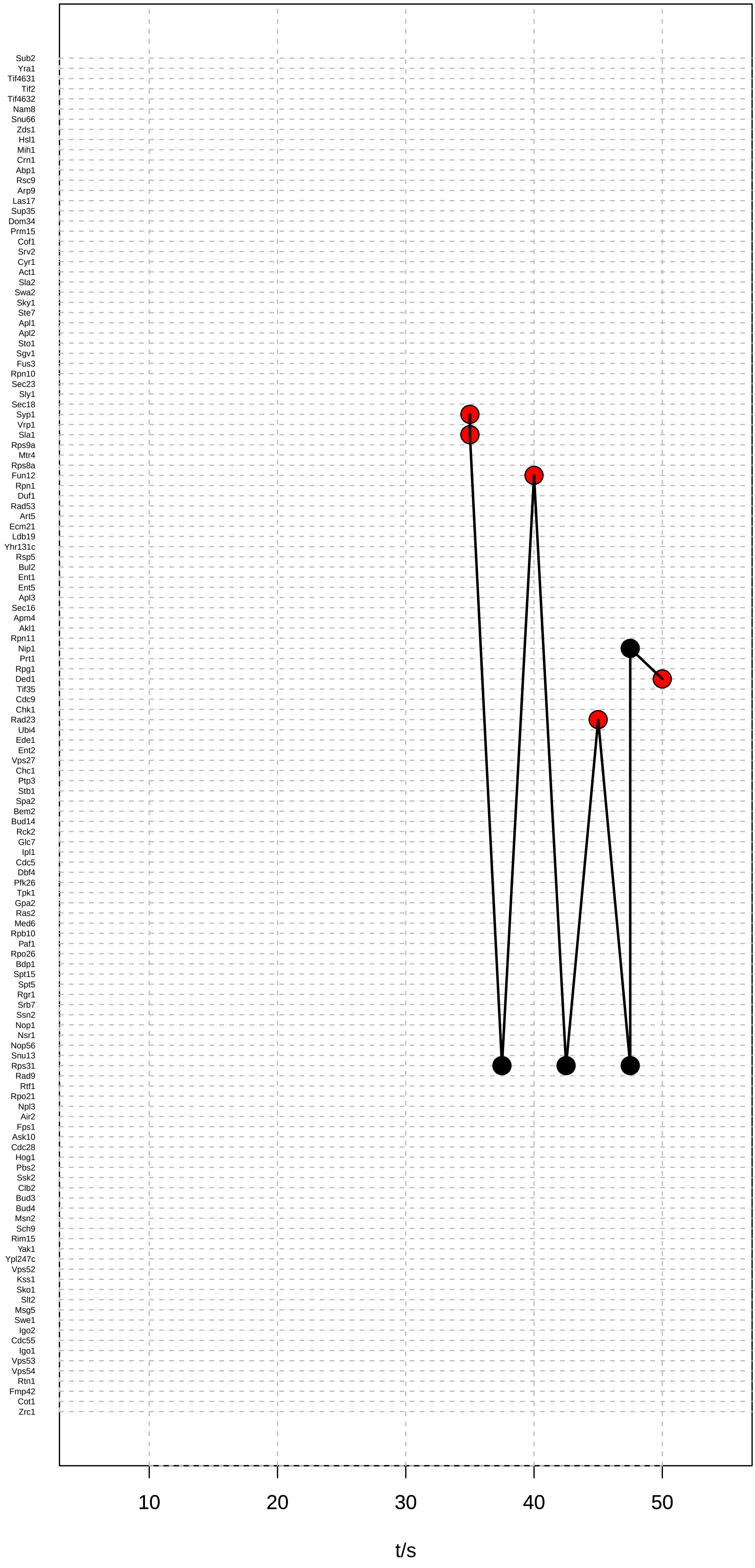

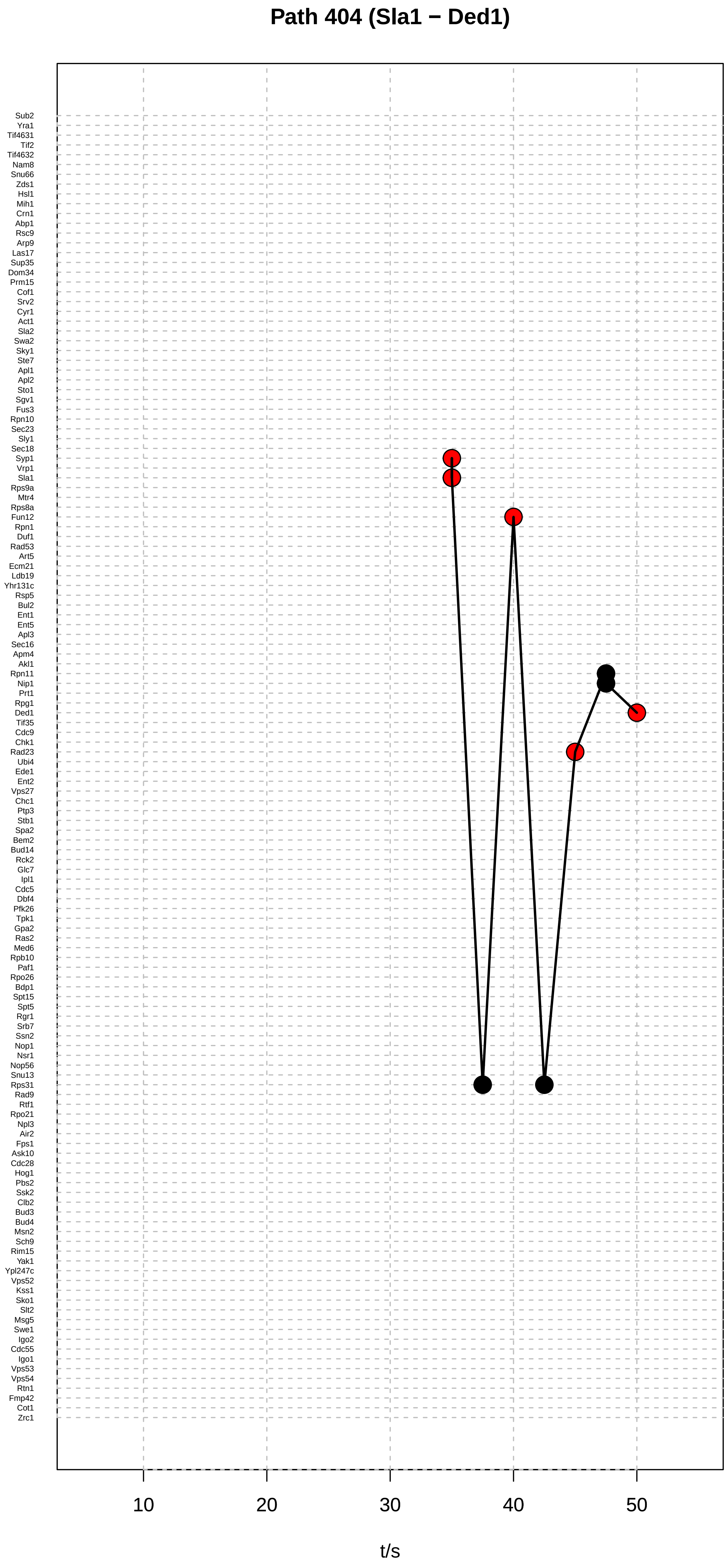
Path 404 (Sla1 − Ded1)
Sub2
Yra1
Tif4631
Tif2
Tif4632
Nam8
Snu66
Zds1
Hsl1
Mih1
Crn1
Abp1
Rsc9
Arp9
Las17
Sup35
Dom34
Prm15
Cof1
Srv2
Cyr1
Act1
Sla2
Swa2
Sky1
Ste7
Apl1
Apl2
Sto1
Sgv1
Fus3
Rpn10
Sec23
Sly1
Sec18
Syp1
Vrp1
Sla1
Rps9a
Mtr4
Rps8a
Fun12
Rpn1
Duf1
Rad53
Art5
Ecm21
Ldb19
Yhr131c
Rsp5
Bul2
Ent1
Ent5
Apl3
Sec16
Apm4
Akl1
Rpn11
Nip1
Prt1
Rpg1
Ded1
Tif35
Cdc9
Chk1
Rad23
Ubi4
Ede1
Ent2
Vps27
Chc1
Ptp3
Stb1
Spa2
Bem2
Bud14
Rck2
Glc7
Ipl1
Cdc5
Dbf4
Pfk26
Tpk1
Gpa2
Ras2
Med6
Rpb10
Paf1
Rpo26
Bdp1
Spt15
Spt5
Rgr1
Srb7
Ssn2
Nop1
Nsr1
Nop56
Snu13
Rps31
Rad9
Rtf1
Rpo21
Npl3
Air2
Fps1
Ask10
Cdc28
Hog1
Pbs2
Ssk2
Clb2
Bud3
Bud4
Msn2
Sch9
Rim15
Yak1
Ypl247c
Vps52
Kss1
Sko1
Slt2
Msg5
Swe1
Igo2
Cdc55
Igo1
Vps53
Vps54
Rtn1
Fmp42
Cot1
Zrc1
10
20
30
40
50
t/s

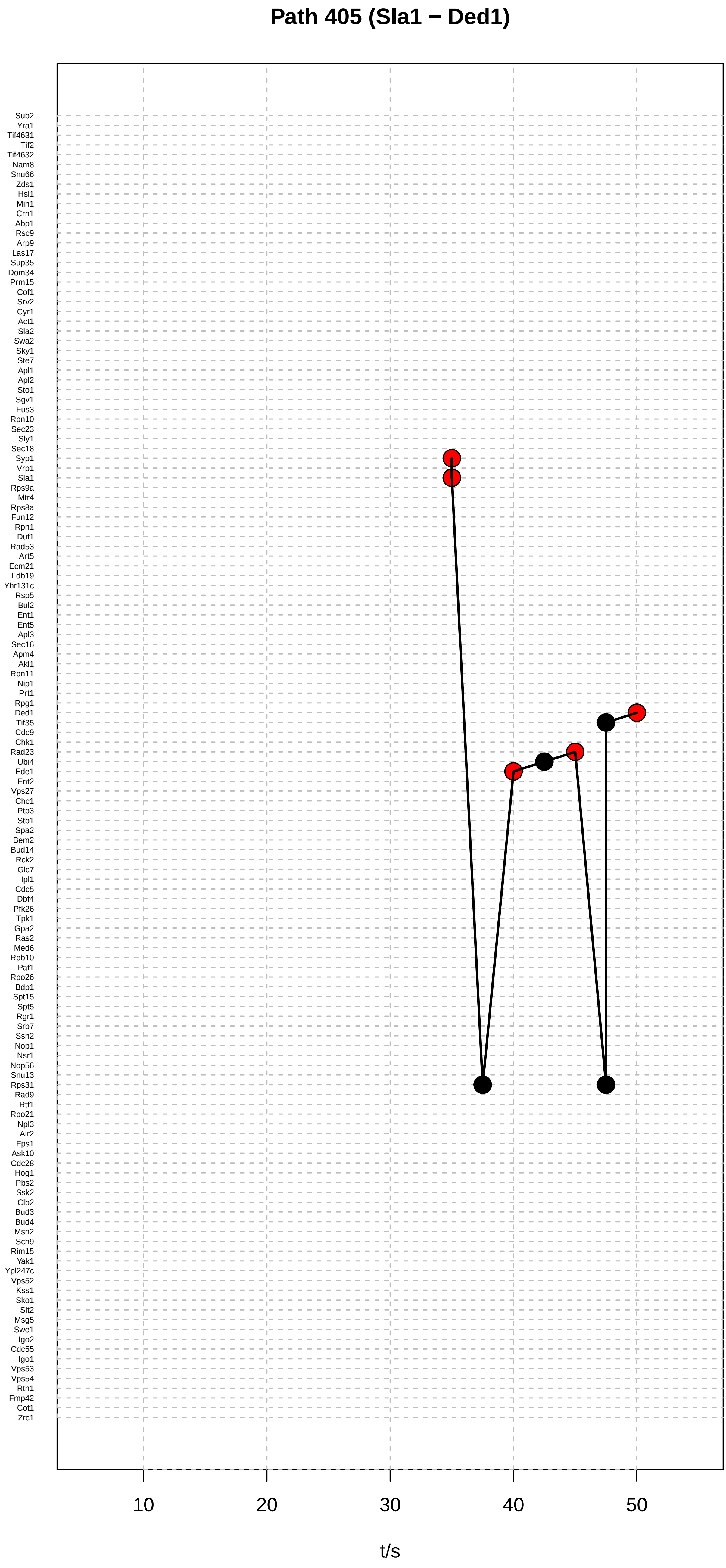

Path 405 (Sla1 − Ded1)
Sub2
Yra1
Tif4631
Tif2
Tif4632
Nam8
Snu66
Zds1
Hsl1
Mih1
Crn1
Abp1
Rsc9
Arp9
Las17
Sup35
Dom34
Prm15
Cof1
Srv2
Cyr1
Act1
Sla2
Swa2
Sky1
Ste7
Apl1
Apl2
Sto1
Sgv1
Fus3
Rpn10
Sec23
Sly1
Sec18
Syp1
Vrp1
Sla1
Rps9a
Mtr4
Rps8a
Fun12
Rpn1
Duf1
Rad53
Art5
Ecm21
Ldb19
Yhr131c
Rsp5
Bul2
Ent1
Ent5
Apl3
Sec16
Apm4
Akl1
Rpn11
Nip1
Prt1
Rpg1
Ded1
Tif35
Cdc9
Chk1
Rad23
Ubi4
Ede1
Ent2
Vps27
Chc1
Ptp3
Stb1
Spa2
Bem2
Bud14
Rck2
Glc7
Ipl1
Cdc5
Dbf4
Pfk26
Tpk1
Gpa2
Ras2
Med6
Rpb10
Paf1
Rpo26
Bdp1
Spt15
Spt5
Rgr1
Srb7
Ssn2
Nop1
Nsr1
Nop56
Snu13
Rps31
Rad9
Rtf1
Rpo21
Npl3
Air2
Fps1
Ask10
Cdc28
Hog1
Pbs2
Ssk2
Clb2
Bud3
Bud4
Msn2
Sch9
Rim15
Yak1
Ypl247c
Vps52
Kss1
Sko1
Slt2
Msg5
Swe1
Igo2
Cdc55
Igo1
Vps53
Vps54
Rtn1
Fmp42
Cot1
Zrc1
10
20
30
40
50
t/s

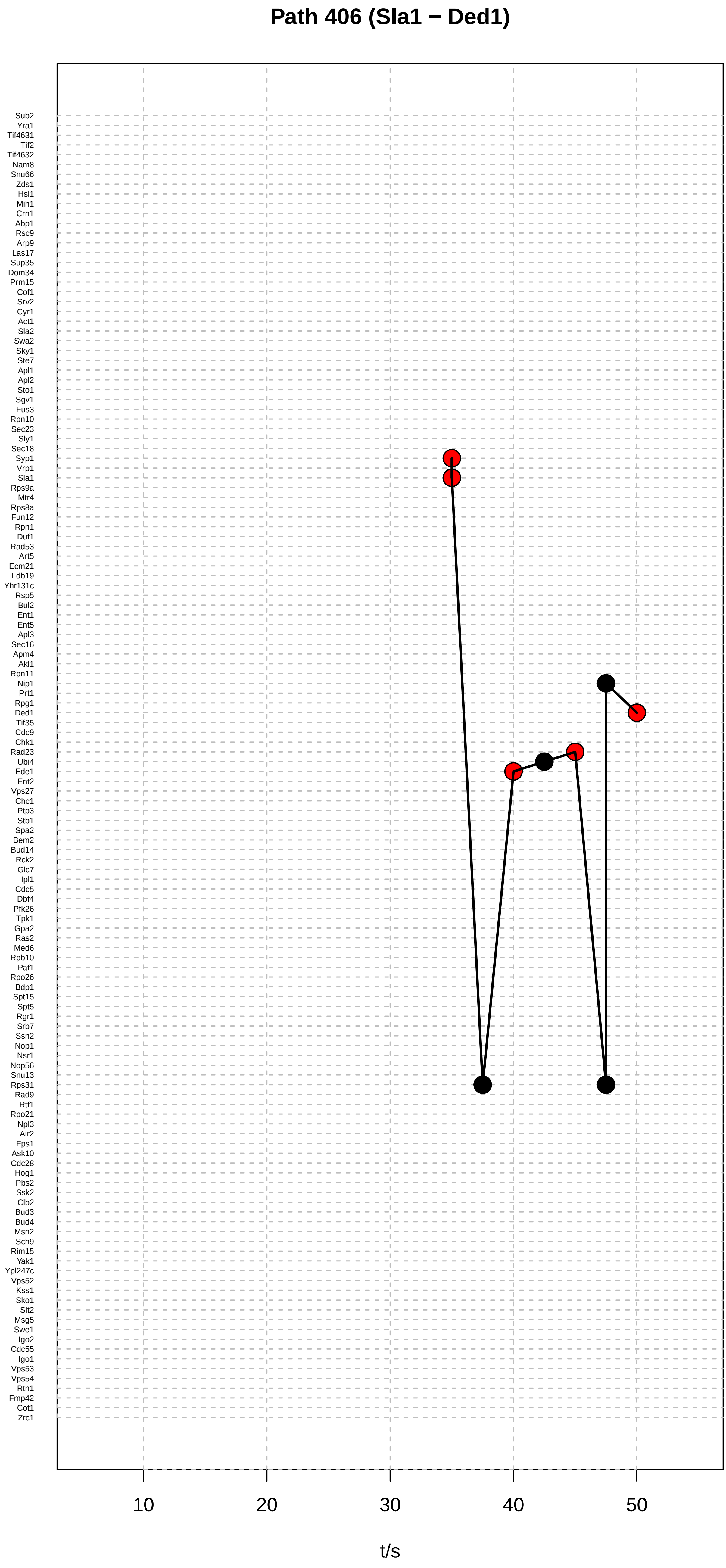

Path 406 (Sla1 − Ded1)
Sub2
Yra1
Tif4631
Tif2
Tif4632
Nam8
Snu66
Zds1
Hsl1
Mih1
Crn1
Abp1
Rsc9
Arp9
Las17
Sup35
Dom34
Prm15
Cof1
Srv2
Cyr1
Act1
Sla2
Swa2
Sky1
Ste7
Apl1
Apl2
Sto1
Sgv1
Fus3
Rpn10
Sec23
Sly1
Sec18
Syp1
Vrp1
Sla1
Rps9a
Mtr4
Rps8a
Fun12
Rpn1
Duf1
Rad53
Art5
Ecm21
Ldb19
Yhr131c
Rsp5
Bul2
Ent1
Ent5
Apl3
Sec16
Apm4
Akl1
Rpn11
Nip1
Prt1
Rpg1
Ded1
Tif35
Cdc9
Chk1
Rad23
Ubi4
Ede1
Ent2
Vps27
Chc1
Ptp3
Stb1
Spa2
Bem2
Bud14
Rck2
Glc7
Ipl1
Cdc5
Dbf4
Pfk26
Tpk1
Gpa2
Ras2
Med6
Rpb10
Paf1
Rpo26
Bdp1
Spt15
Spt5
Rgr1
Srb7
Ssn2
Nop1
Nsr1
Nop56
Snu13
Rps31
Rad9
Rtf1
Rpo21
Npl3
Air2
Fps1
Ask10
Cdc28
Hog1
Pbs2
Ssk2
Clb2
Bud3
Bud4
Msn2
Sch9
Rim15
Yak1
Ypl247c
Vps52
Kss1
Sko1
Slt2
Msg5
Swe1
Igo2
Cdc55
Igo1
Vps53
Vps54
Rtn1
Fmp42
Cot1
Zrc1
10
20
30
40
50
t/s

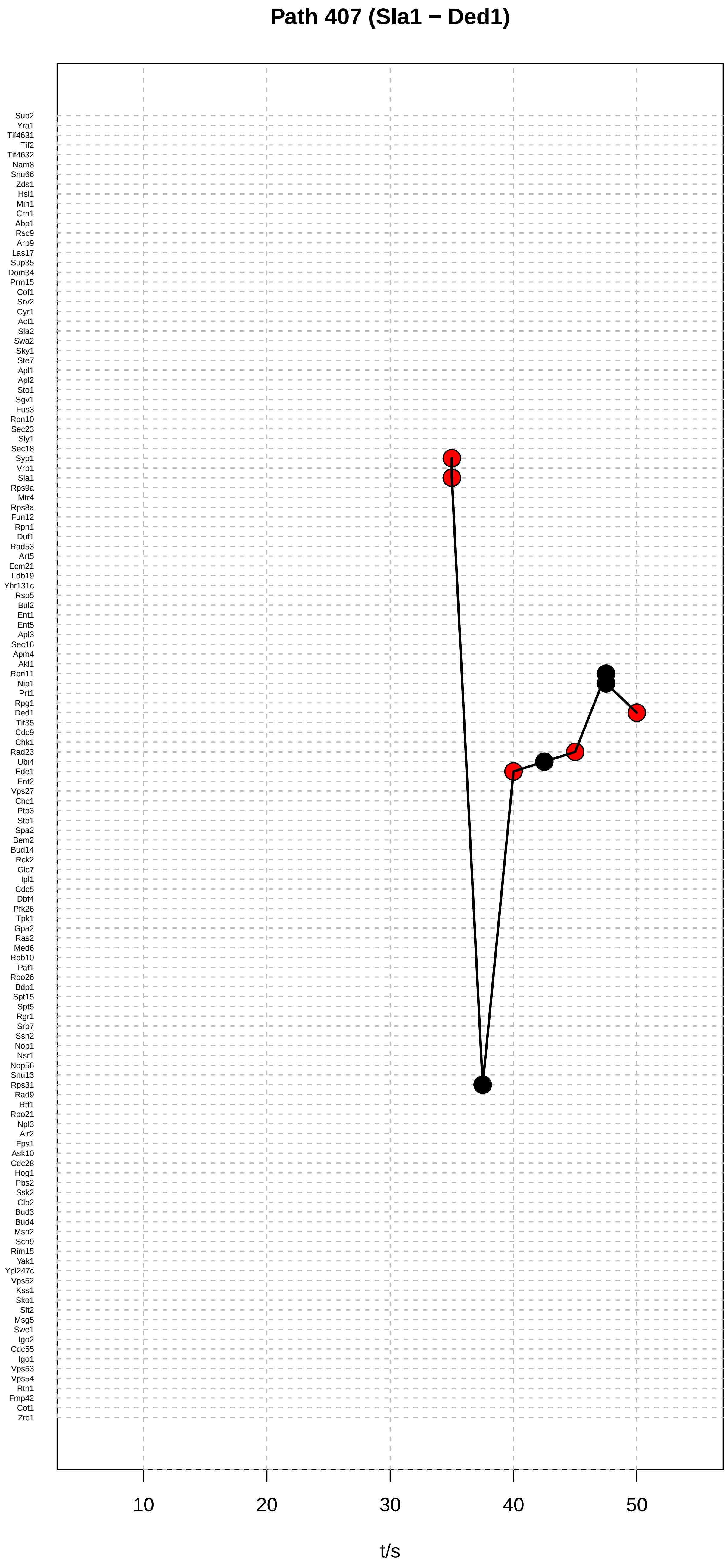
Path 407 (Sla1 − Ded1)
Sub2
Yra1
Tif4631
Tif2
Tif4632
Nam8
Snu66
Zds1
Hsl1
Mih1
Crn1
Abp1
Rsc9
Arp9
Las17
Sup35
Dom34
Prm15
Cof1
Srv2
Cyr1
Act1
Sla2
Swa2
Sky1
Ste7
Apl1
Apl2
Sto1
Sgv1
Fus3
Rpn10
Sec23
Sly1
Sec18
Syp1
Vrp1
Sla1
Rps9a
Mtr4
Rps8a
Fun12
Rpn1
Duf1
Rad53
Art5
Ecm21
Ldb19
Yhr131c
Rsp5
Bul2
Ent1
Ent5
Apl3
Sec16
Apm4
Akl1
Rpn11
Nip1
Prt1
Rpg1
Ded1
Tif35
Cdc9
Chk1
Rad23
Ubi4
Ede1
Ent2
Vps27
Chc1
Ptp3
Stb1
Spa2
Bem2
Bud14
Rck2
Glc7
Ipl1
Cdc5
Dbf4
Pfk26
Tpk1
Gpa2
Ras2
Med6
Rpb10
Paf1
Rpo26
Bdp1
Spt15
Spt5
Rgr1
Srb7
Ssn2
Nop1
Nsr1
Nop56
Snu13
Rps31
Rad9
Rtf1
Rpo21
Npl3
Air2
Fps1
Ask10
Cdc28
Hog1
Pbs2
Ssk2
Clb2
Bud3
Bud4
Msn2
Sch9
Rim15
Yak1
Ypl247c
Vps52
Kss1
Sko1
Slt2
Msg5
Swe1
Igo2
Cdc55
Igo1
Vps53
Vps54
Rtn1
Fmp42
Cot1
Zrc1
10
20
30
40
50
t/s

#### Path 408 (Sla1 − Ded1)

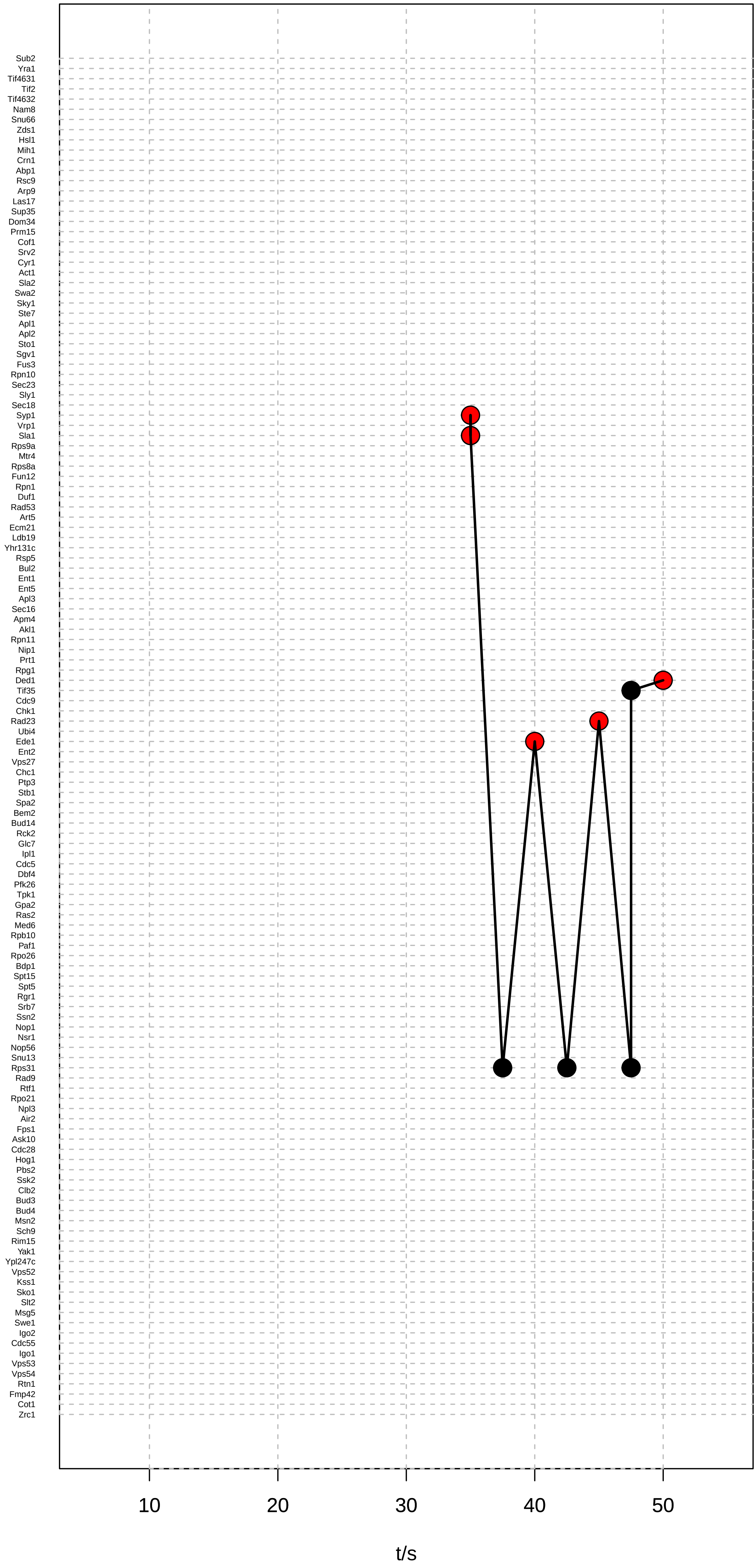

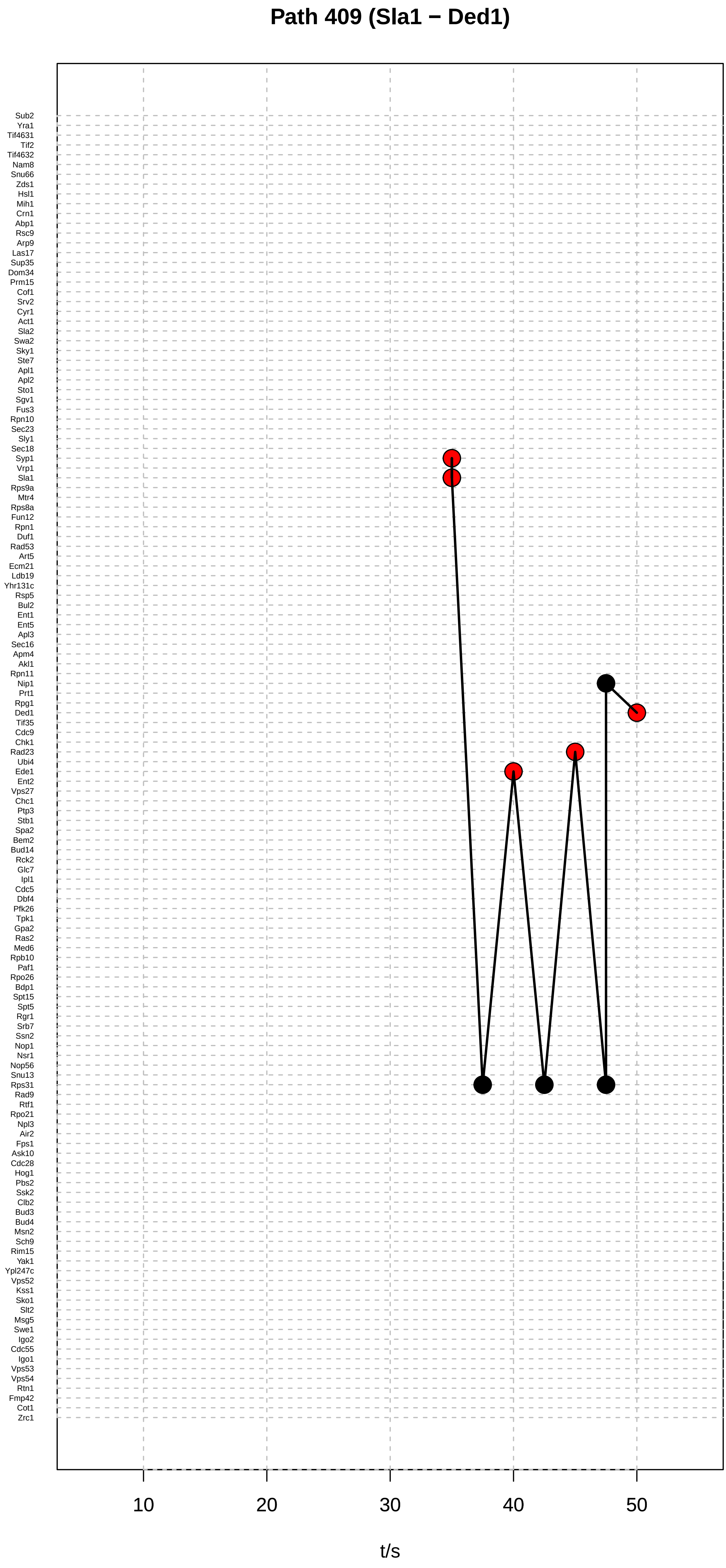

Path 409 (Sla1 − Ded1)
Sub2
Yra1
Tif4631
Tif2
Tif4632
Nam8
Snu66
Zds1
Hsl1
Mih1
Crn1
Abp1
Rsc9
Arp9
Las17
Sup35
Dom34
Prm15
Cof1
Srv2
Cyr1
Act1
Sla2
Swa2
Sky1
Ste7
Apl1
Apl2
Sto1
Sgv1
Fus3
Rpn10
Sec23
Sly1
Sec18
Syp1
Vrp1
Sla1
Rps9a
Mtr4
Rps8a
Fun12
Rpn1
Duf1
Rad53
Art5
Ecm21
Ldb19
Yhr131c
Rsp5
Bul2
Ent1
Ent5
Apl3
Sec16
Apm4
Akl1
Rpn11
Nip1
Prt1
Rpg1
Ded1
Tif35
Cdc9
Chk1
Rad23
Ubi4
Ede1
Ent2
Vps27
Chc1
Ptp3
Stb1
Spa2
Bem2
Bud14
Rck2
Glc7
Ipl1
Cdc5
Dbf4
Pfk26
Tpk1
Gpa2
Ras2
Med6
Rpb10
Paf1
Rpo26
Bdp1
Spt15
Spt5
Rgr1
Srb7
Ssn2
Nop1
Nsr1
Nop56
Snu13
Rps31
Rad9
Rtf1
Rpo21
Npl3
Air2
Fps1
Ask10
Cdc28
Hog1
Pbs2
Ssk2
Clb2
Bud3
Bud4
Msn2
Sch9
Rim15
Yak1
Ypl247c
Vps52
Kss1
Sko1
Slt2
Msg5
Swe1
Igo2
Cdc55
Igo1
Vps53
Vps54
Rtn1
Fmp42
Cot1
Zrc1
10
20
30
40
50
t/s

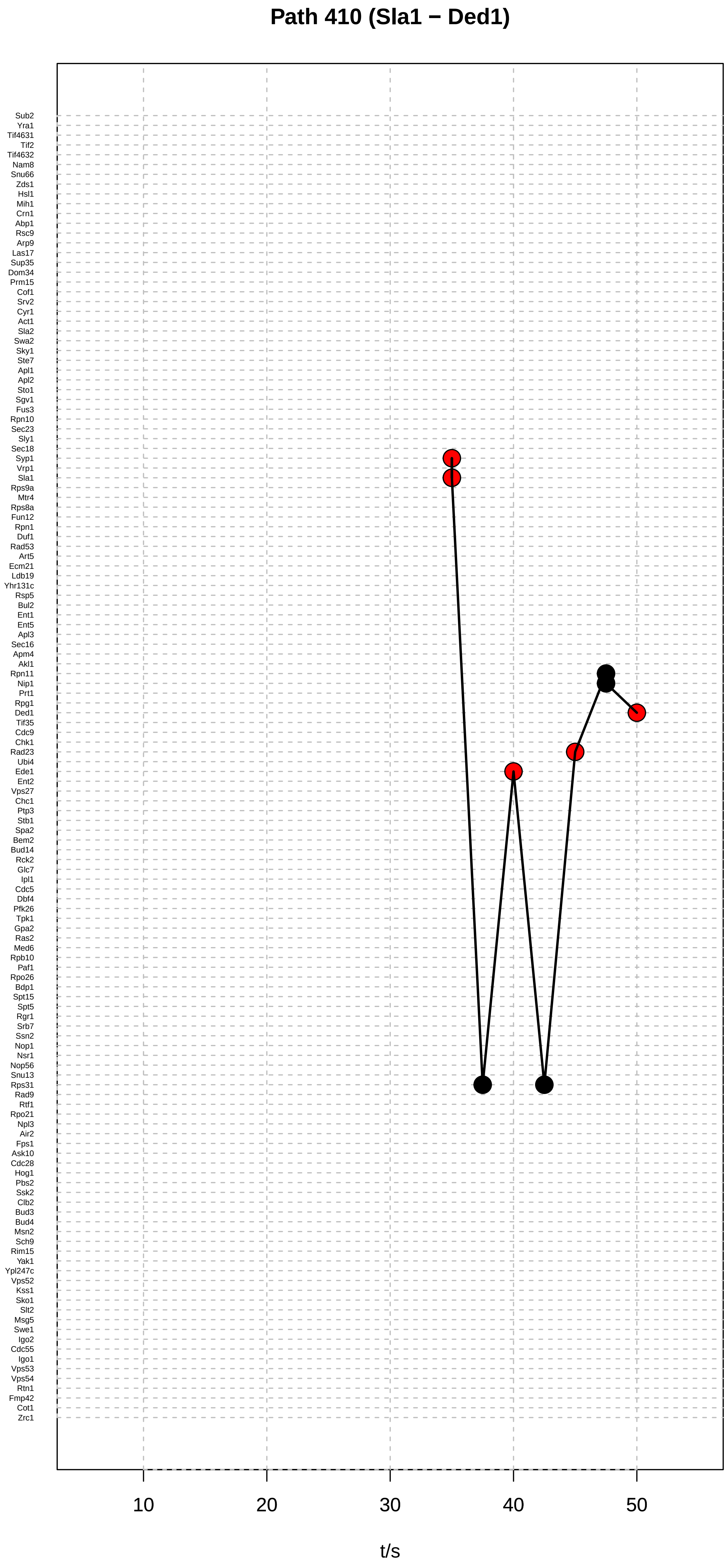
Path 410 (Sla1 − Ded1)
Sub2
Yra1
Tif4631
Tif2
Tif4632
Nam8
Snu66
Zds1
Hsl1
Mih1
Crn1
Abp1
Rsc9
Arp9
Las17
Sup35
Dom34
Prm15
Cof1
Srv2
Cyr1
Act1
Sla2
Swa2
Sky1
Ste7
Apl1
Apl2
Sto1
Sgv1
Fus3
Rpn10
Sec23
Sly1
Sec18
Syp1
Vrp1
Sla1
Rps9a
Mtr4
Rps8a
Fun12
Rpn1
Duf1
Rad53
Art5
Ecm21
Ldb19
Yhr131c
Rsp5
Bul2
Ent1
Ent5
Apl3
Sec16
Apm4
Akl1
Rpn11
Nip1
Prt1
Rpg1
Ded1
Tif35
Cdc9
Chk1
Rad23
Ubi4
Ede1
Ent2
Vps27
Chc1
Ptp3
Stb1
Spa2
Bem2
Bud14
Rck2
Glc7
Ipl1
Cdc5
Dbf4
Pfk26
Tpk1
Gpa2
Ras2
Med6
Rpb10
Paf1
Rpo26
Bdp1
Spt15
Spt5
Rgr1
Srb7
Ssn2
Nop1
Nsr1
Nop56
Snu13
Rps31
Rad9
Rtf1
Rpo21
Npl3
Air2
Fps1
Ask10
Cdc28
Hog1
Pbs2
Ssk2
Clb2
Bud3
Bud4
Msn2
Sch9
Rim15
Yak1
Ypl247c
Vps52
Kss1
Sko1
Slt2
Msg5
Swe1
Igo2
Cdc55
Igo1
Vps53
Vps54
Rtn1
Fmp42
Cot1
Zrc1
10
20
30
40
50
t/s

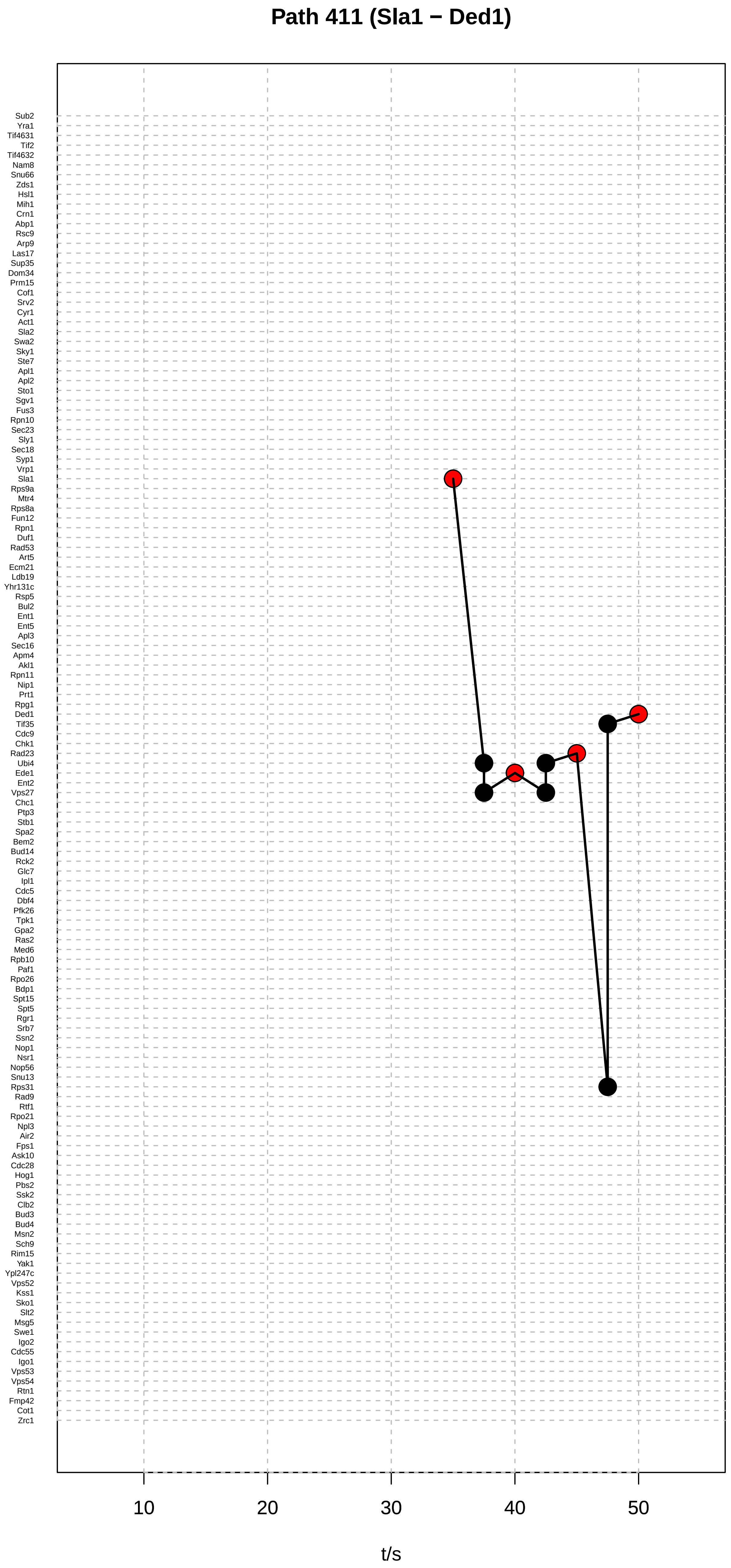

Path 411 (Sla1 − Ded1)
Sub2
Yra1
Tif4631
Tif2
Tif4632
Nam8
Snu66
Zds1
Hsl1
Mih1
Crn1
Abp1
Rsc9
Arp9
Las17
Sup35
Dom34
Prm15
Cof1
Srv2
Cyr1
Act1
Sla2
Swa2
Sky1
Ste7
Apl1
Apl2
Sto1
Sgv1
Fus3
Rpn10
Sec23
Sly1
Sec18
Syp1
Vrp1
Sla1
Rps9a
Mtr4
Rps8a
Fun12
Rpn1
Duf1
Rad53
Art5
Ecm21
Ldb19
Yhr131c
Rsp5
Bul2
Ent1
Ent5
Apl3
Sec16
Apm4
Akl1
Rpn11
Nip1
Prt1
Rpg1
Ded1
Tif35
Cdc9
Chk1
Rad23
Ubi4
Ede1
Ent2
Vps27
Chc1
Ptp3
Stb1
Spa2
Bem2
Bud14
Rck2
Glc7
Ipl1
Cdc5
Dbf4
Pfk26
Tpk1
Gpa2
Ras2
Med6
Rpb10
Paf1
Rpo26
Bdp1
Spt15
Spt5
Rgr1
Srb7
Ssn2
Nop1
Nsr1
Nop56
Snu13
Rps31
Rad9
Rtf1
Rpo21
Npl3
Air2
Fps1
Ask10
Cdc28
Hog1
Pbs2
Ssk2
Clb2
Bud3
Bud4
Msn2
Sch9
Rim15
Yak1
Ypl247c
Vps52
Kss1
Sko1
Slt2
Msg5
Swe1
Igo2
Cdc55
Igo1
Vps53
Vps54
Rtn1
Fmp42
Cot1
Zrc1
10
20
30
40
50
t/s

### Path 412 (Sla1 − Ded1)

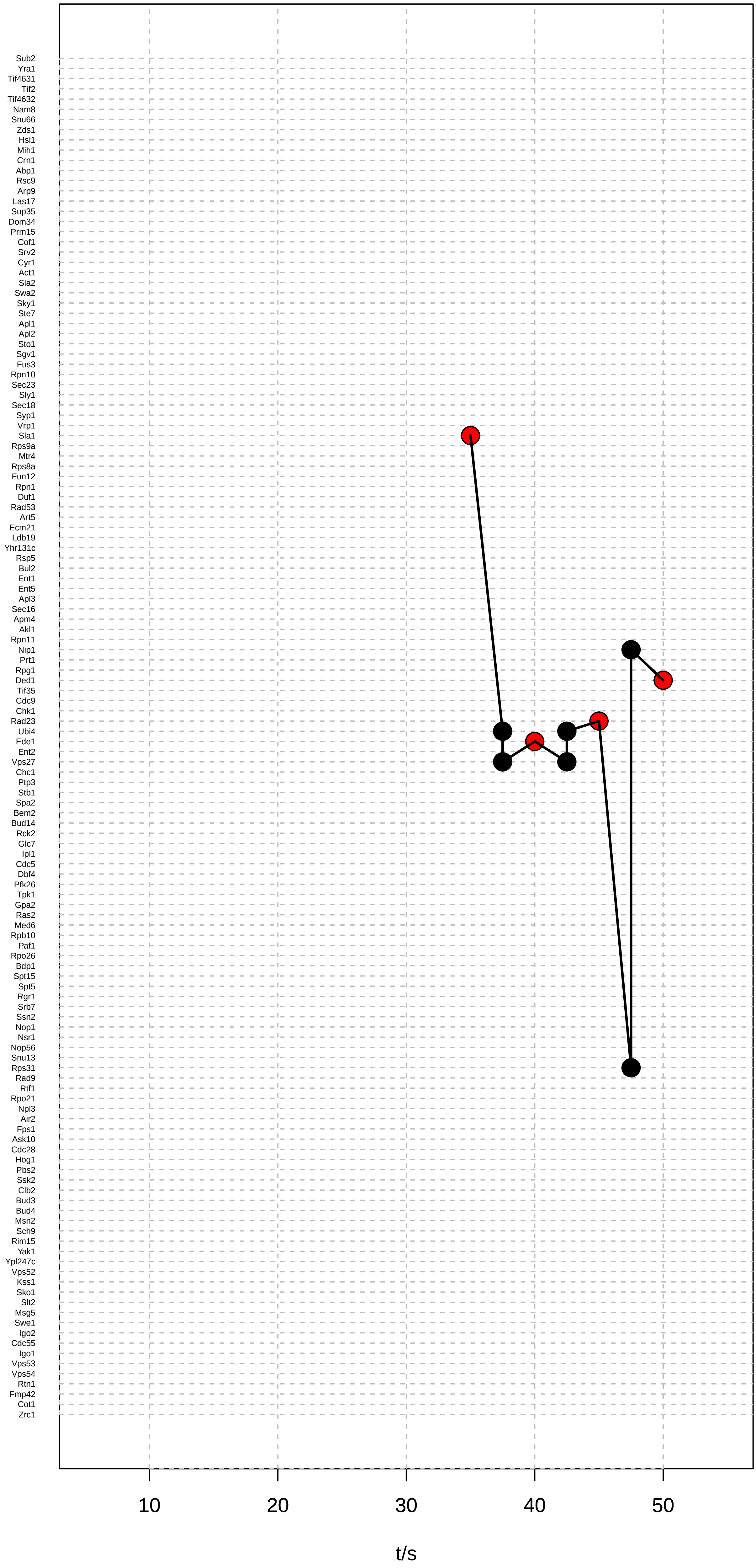

#### Path 413 (Sla1 − Ded1)

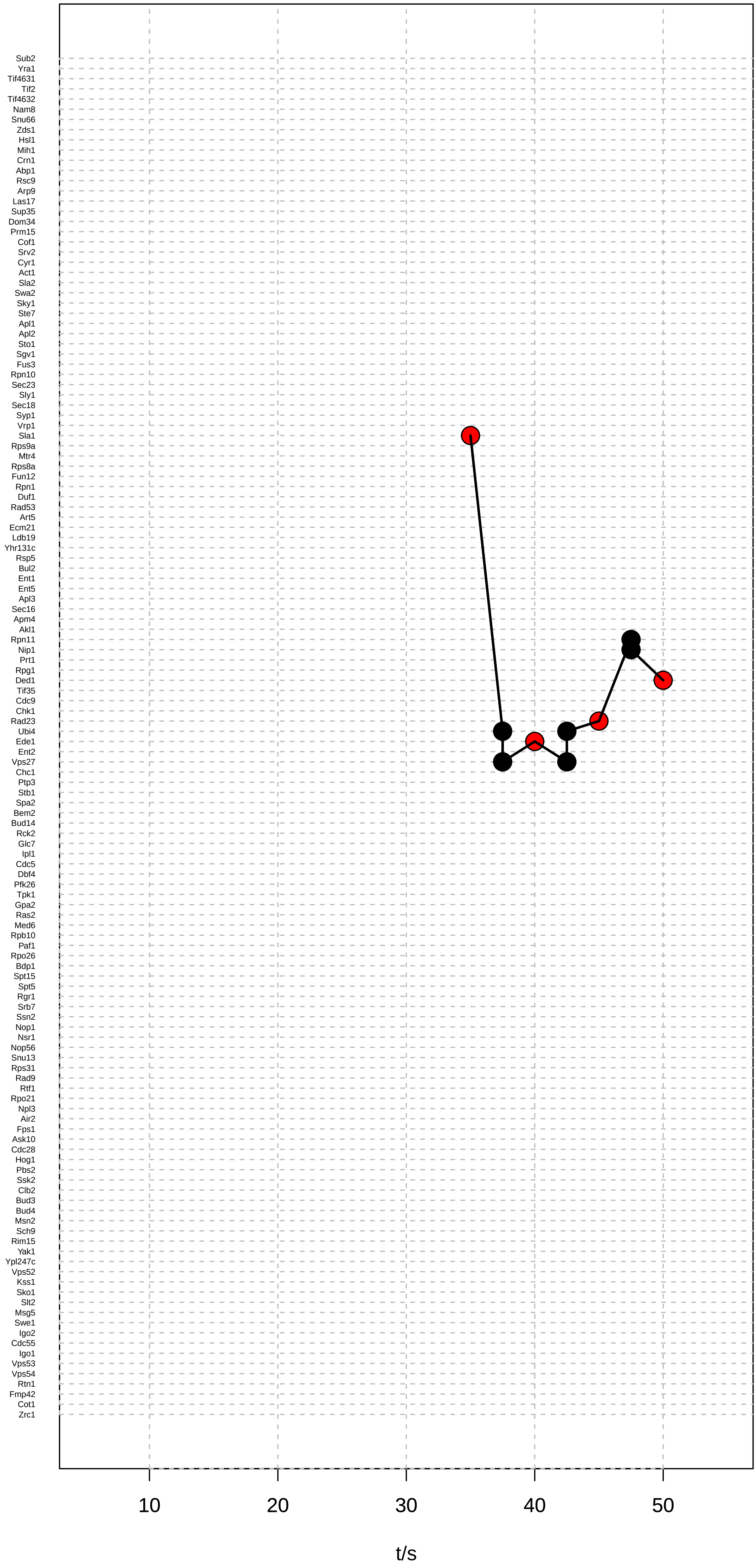

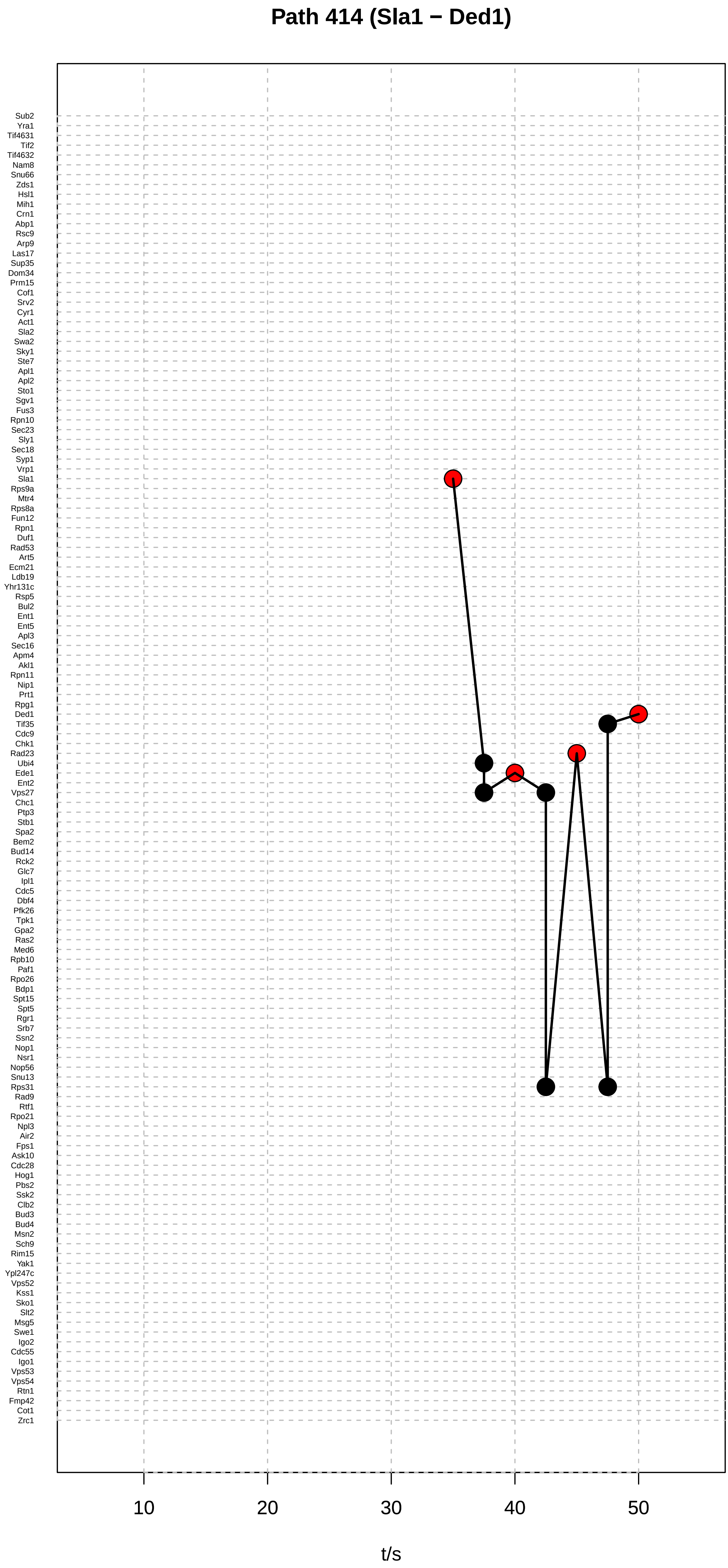

Path 414 (Sla1 − Ded1)
Sub2
Yra1
Tif4631
Tif2
Tif4632
Nam8
Snu66
Zds1
Hsl1
Mih1
Crn1
Abp1
Rsc9
Arp9
Las17
Sup35
Dom34
Prm15
Cof1
Srv2
Cyr1
Act1
Sla2
Swa2
Sky1
Ste7
Apl1
Apl2
Sto1
Sgv1
Fus3
Rpn10
Sec23
Sly1
Sec18
Syp1
Vrp1
Sla1
Rps9a
Mtr4
Rps8a
Fun12
Rpn1
Duf1
Rad53
Art5
Ecm21
Ldb19
Yhr131c
Rsp5
Bul2
Ent1
Ent5
Apl3
Sec16
Apm4
Akl1
Rpn11
Nip1
Prt1
Rpg1
Ded1
Tif35
Cdc9
Chk1
Rad23
Ubi4
Ede1
Ent2
Vps27
Chc1
Ptp3
Stb1
Spa2
Bem2
Bud14
Rck2
Glc7
Ipl1
Cdc5
Dbf4
Pfk26
Tpk1
Gpa2
Ras2
Med6
Rpb10
Paf1
Rpo26
Bdp1
Spt15
Spt5
Rgr1
Srb7
Ssn2
Nop1
Nsr1
Nop56
Snu13
Rps31
Rad9
Rtf1
Rpo21
Npl3
Air2
Fps1
Ask10
Cdc28
Hog1
Pbs2
Ssk2
Clb2
Bud3
Bud4
Msn2
Sch9
Rim15
Yak1
Ypl247c
Vps52
Kss1
Sko1
Slt2
Msg5
Swe1
Igo2
Cdc55
Igo1
Vps53
Vps54
Rtn1
Fmp42
Cot1
Zrc1
10
20
30
40
50
t/s

### Path 415 (Sla1 − Ded1)

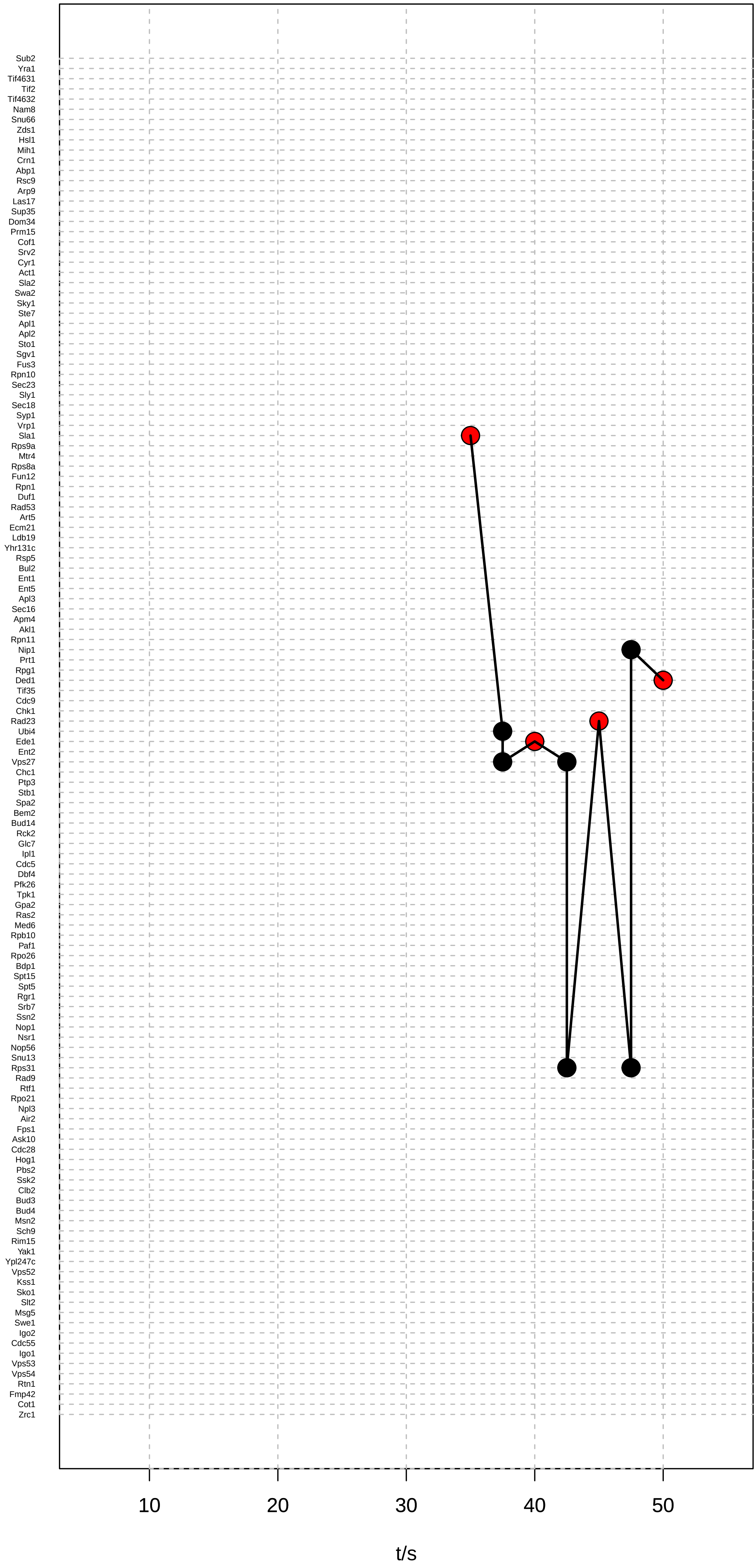

#### Path 416 (Sla1 − Ded1)

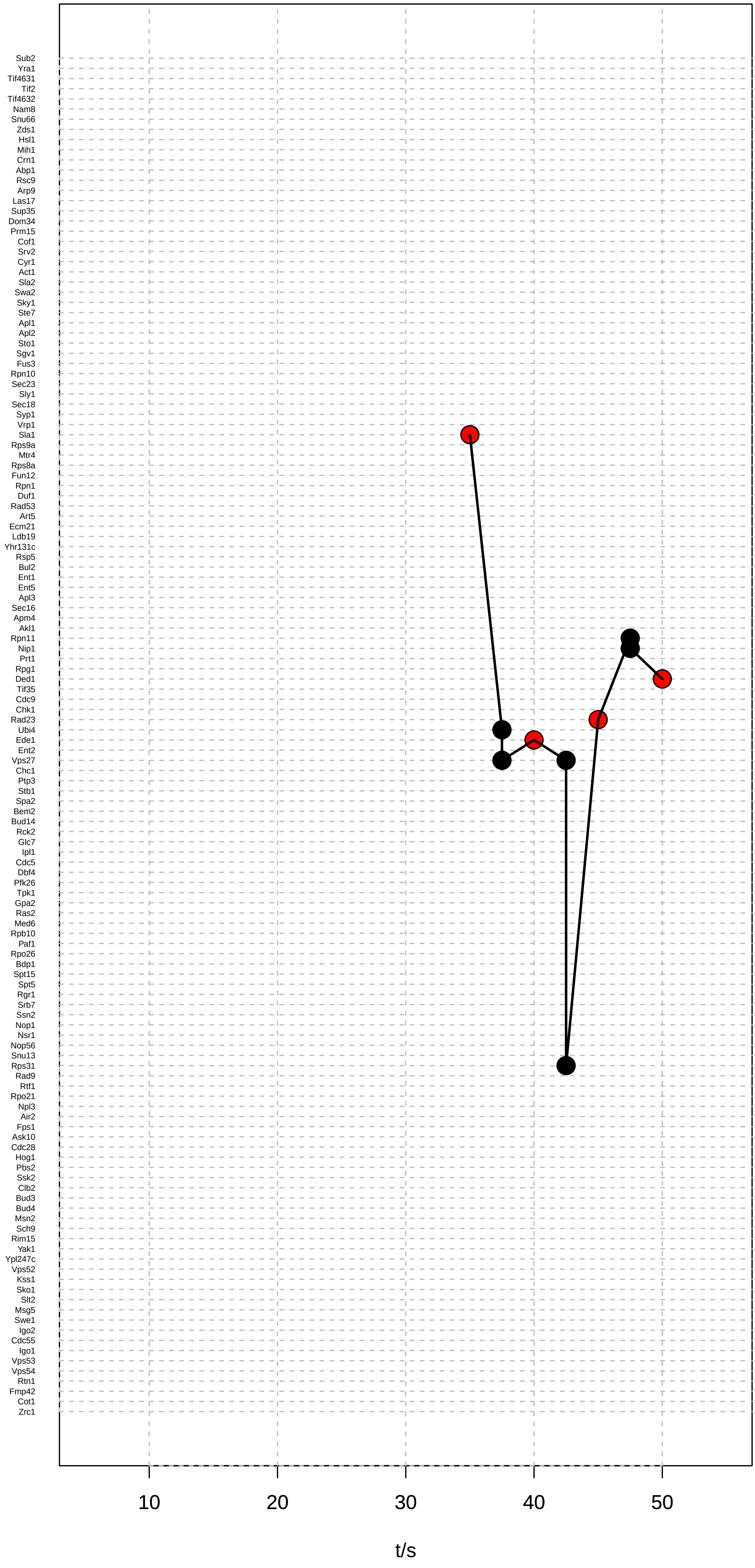

### Path 417 (Sla1 − Ded1)

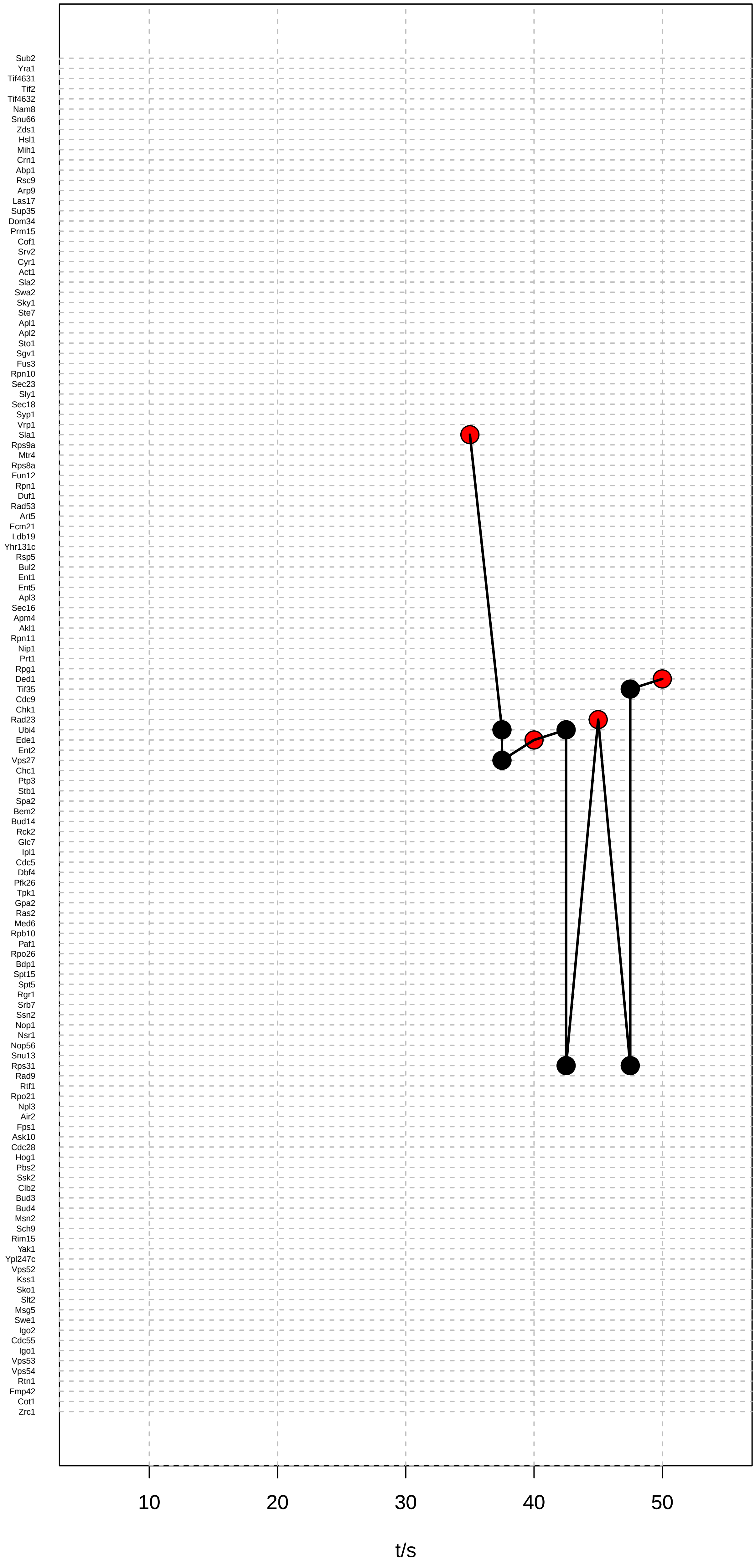

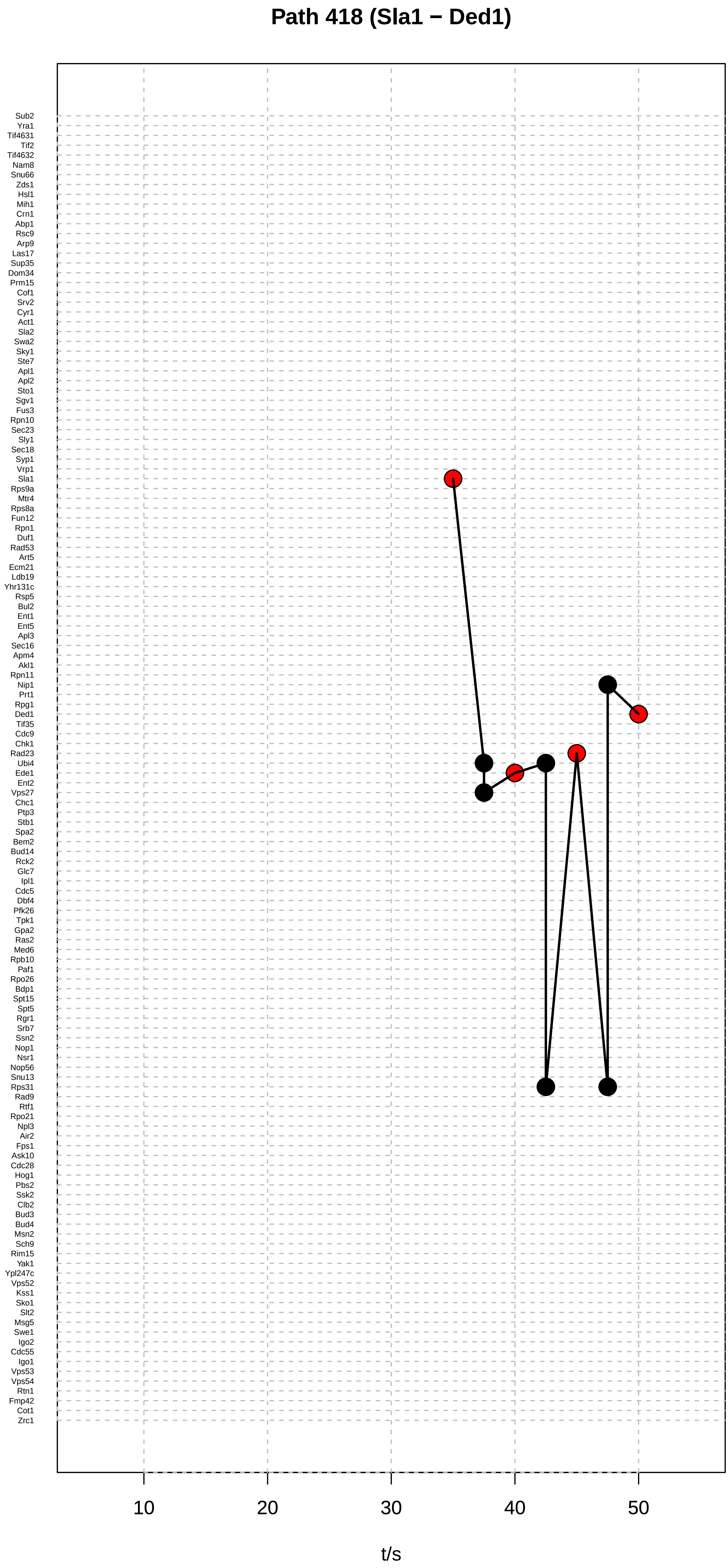

Path 418 (Sla1 − Ded1)
Sub2
Yra1
Tif4631
Tif2
Tif4632
Nam8
Snu66
Zds1
Hsl1
Mih1
Crn1
Abp1
Rsc9
Arp9
Las17
Sup35
Dom34
Prm15
Cof1
Srv2
Cyr1
Act1
Sla2
Swa2
Sky1
Ste7
Apl1
Apl2
Sto1
Sgv1
Fus3
Rpn10
Sec23
Sly1
Sec18
Syp1
Vrp1
Sla1
Rps9a
Mtr4
Rps8a
Fun12
Rpn1
Duf1
Rad53
Art5
Ecm21
Ldb19
Yhr131c
Rsp5
Bul2
Ent1
Ent5
Apl3
Sec16
Apm4
Akl1
Rpn11
Nip1
Prt1
Rpg1
Ded1
Tif35
Cdc9
Chk1
Rad23
Ubi4
Ede1
Ent2
Vps27
Chc1
Ptp3
Stb1
Spa2
Bem2
Bud14
Rck2
Glc7
Ipl1
Cdc5
Dbf4
Pfk26
Tpk1
Gpa2
Ras2
Med6
Rpb10
Paf1
Rpo26
Bdp1
Spt15
Spt5
Rgr1
Srb7
Ssn2
Nop1
Nsr1
Nop56
Snu13
Rps31
Rad9
Rtf1
Rpo21
Npl3
Air2
Fps1
Ask10
Cdc28
Hog1
Pbs2
Ssk2
Clb2
Bud3
Bud4
Msn2
Sch9
Rim15
Yak1
Ypl247c
Vps52
Kss1
Sko1
Slt2
Msg5
Swe1
Igo2
Cdc55
Igo1
Vps53
Vps54
Rtn1
Fmp42
Cot1
Zrc1
10
20
30
40
50
t/s

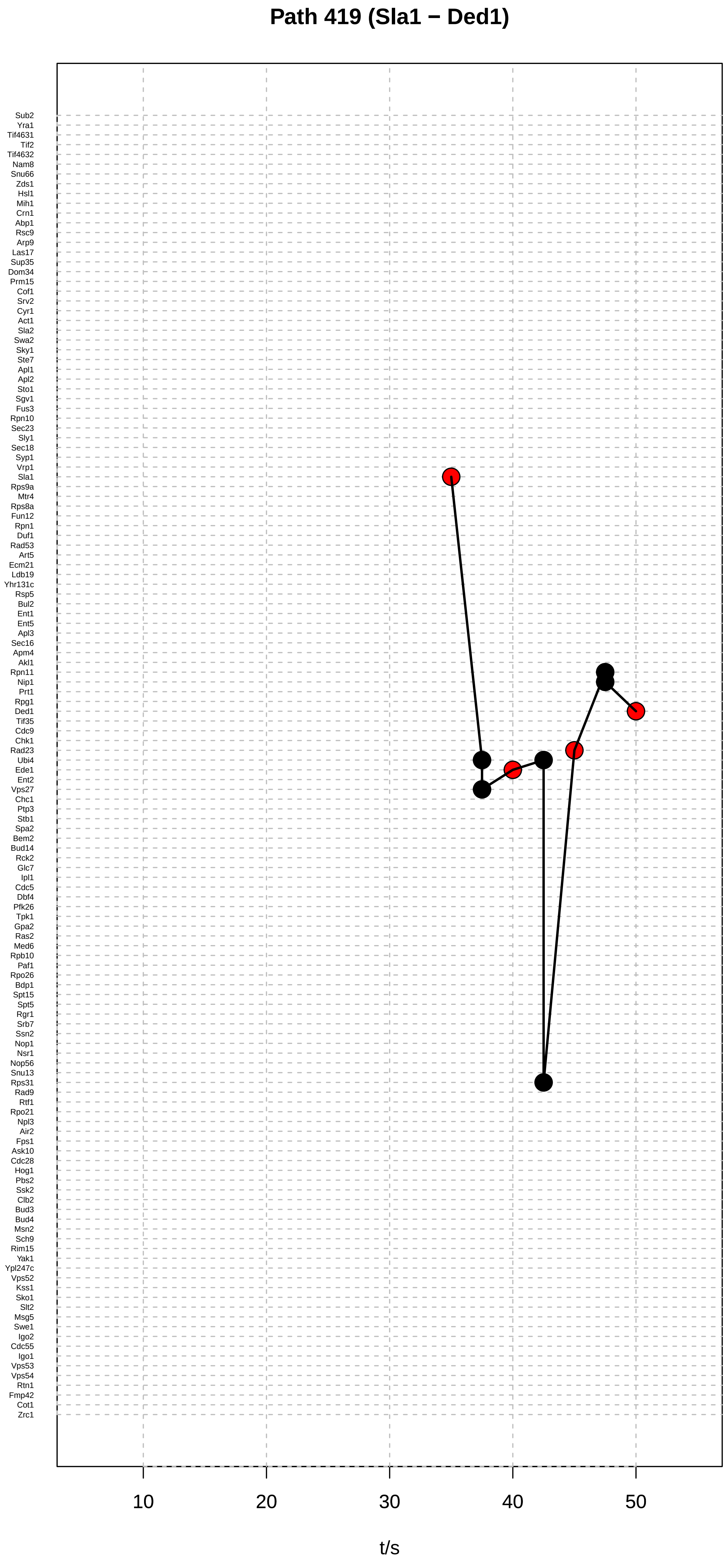

Path 419 (Sla1 − Ded1)
Sub2
Yra1
Tif4631
Tif2
Tif4632
Nam8
Snu66
Zds1
Hsl1
Mih1
Crn1
Abp1
Rsc9
Arp9
Las17
Sup35
Dom34
Prm15
Cof1
Srv2
Cyr1
Act1
Sla2
Swa2
Sky1
Ste7
Apl1
Apl2
Sto1
Sgv1
Fus3
Rpn10
Sec23
Sly1
Sec18
Syp1
Vrp1
Sla1
Rps9a
Mtr4
Rps8a
Fun12
Rpn1
Duf1
Rad53
Art5
Ecm21
Ldb19
Yhr131c
Rsp5
Bul2
Ent1
Ent5
Apl3
Sec16
Apm4
Akl1
Rpn11
Nip1
Prt1
Rpg1
Ded1
Tif35
Cdc9
Chk1
Rad23
Ubi4
Ede1
Ent2
Vps27
Chc1
Ptp3
Stb1
Spa2
Bem2
Bud14
Rck2
Glc7
Ipl1
Cdc5
Dbf4
Pfk26
Tpk1
Gpa2
Ras2
Med6
Rpb10
Paf1
Rpo26
Bdp1
Spt15
Spt5
Rgr1
Srb7
Ssn2
Nop1
Nsr1
Nop56
Snu13
Rps31
Rad9
Rtf1
Rpo21
Npl3
Air2
Fps1
Ask10
Cdc28
Hog1
Pbs2
Ssk2
Clb2
Bud3
Bud4
Msn2
Sch9
Rim15
Yak1
Ypl247c
Vps52
Kss1
Sko1
Slt2
Msg5
Swe1
Igo2
Cdc55
Igo1
Vps53
Vps54
Rtn1
Fmp42
Cot1
Zrc1
10
20
30
40
50
t/s

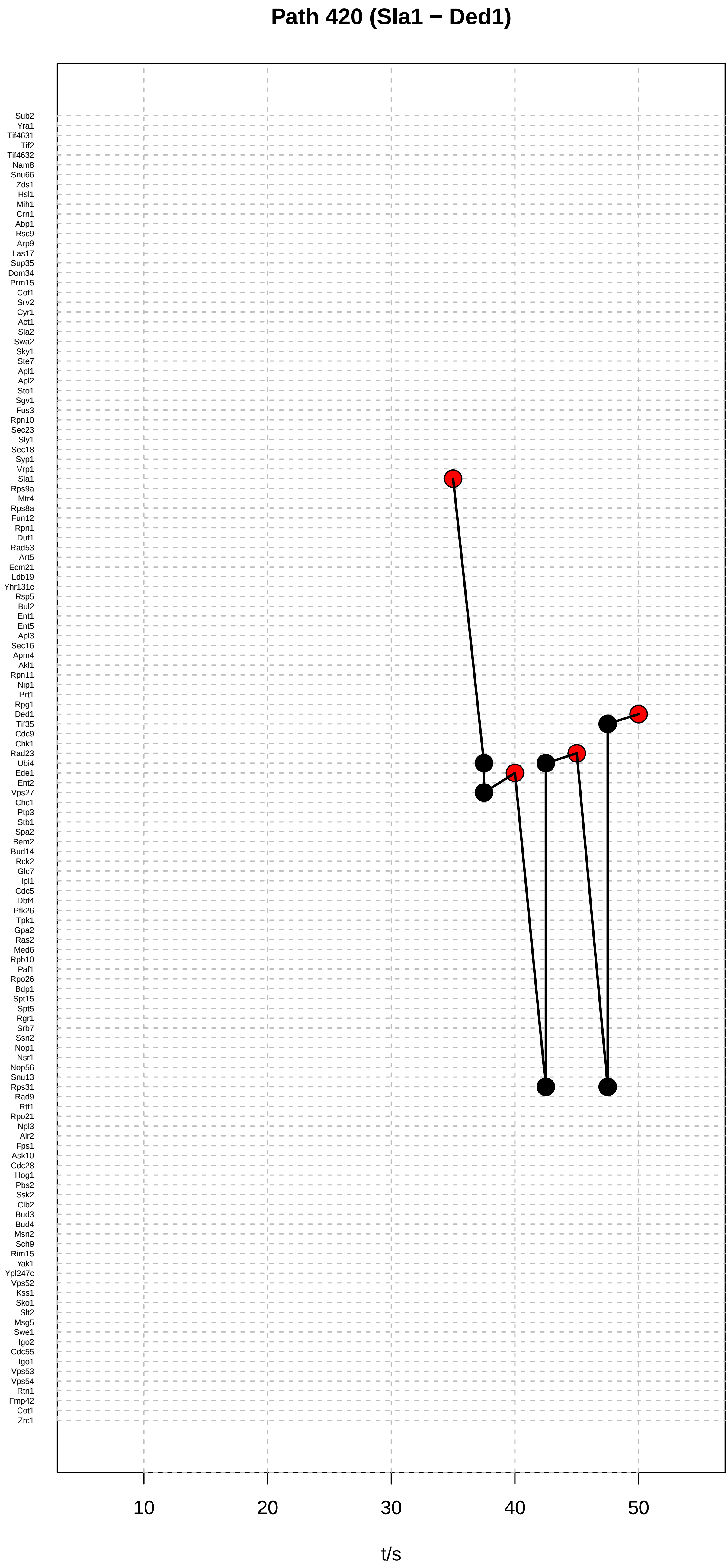

Path 420 (Sla1 − Ded1)
Sub2
Yra1
Tif4631
Tif2
Tif4632
Nam8
Snu66
Zds1
Hsl1
Mih1
Crn1
Abp1
Rsc9
Arp9
Las17
Sup35
Dom34
Prm15
Cof1
Srv2
Cyr1
Act1
Sla2
Swa2
Sky1
Ste7
Apl1
Apl2
Sto1
Sgv1
Fus3
Rpn10
Sec23
Sly1
Sec18
Syp1
Vrp1
Sla1
Rps9a
Mtr4
Rps8a
Fun12
Rpn1
Duf1
Rad53
Art5
Ecm21
Ldb19
Yhr131c
Rsp5
Bul2
Ent1
Ent5
Apl3
Sec16
Apm4
Akl1
Rpn11
Nip1
Prt1
Rpg1
Ded1
Tif35
Cdc9
Chk1
Rad23
Ubi4
Ede1
Ent2
Vps27
Chc1
Ptp3
Stb1
Spa2
Bem2
Bud14
Rck2
Glc7
Ipl1
Cdc5
Dbf4
Pfk26
Tpk1
Gpa2
Ras2
Med6
Rpb10
Paf1
Rpo26
Bdp1
Spt15
Spt5
Rgr1
Srb7
Ssn2
Nop1
Nsr1
Nop56
Snu13
Rps31
Rad9
Rtf1
Rpo21
Npl3
Air2
Fps1
Ask10
Cdc28
Hog1
Pbs2
Ssk2
Clb2
Bud3
Bud4
Msn2
Sch9
Rim15
Yak1
Ypl247c
Vps52
Kss1
Sko1
Slt2
Msg5
Swe1
Igo2
Cdc55
Igo1
Vps53
Vps54
Rtn1
Fmp42
Cot1
Zrc1
10
20
30
40
50
t/s

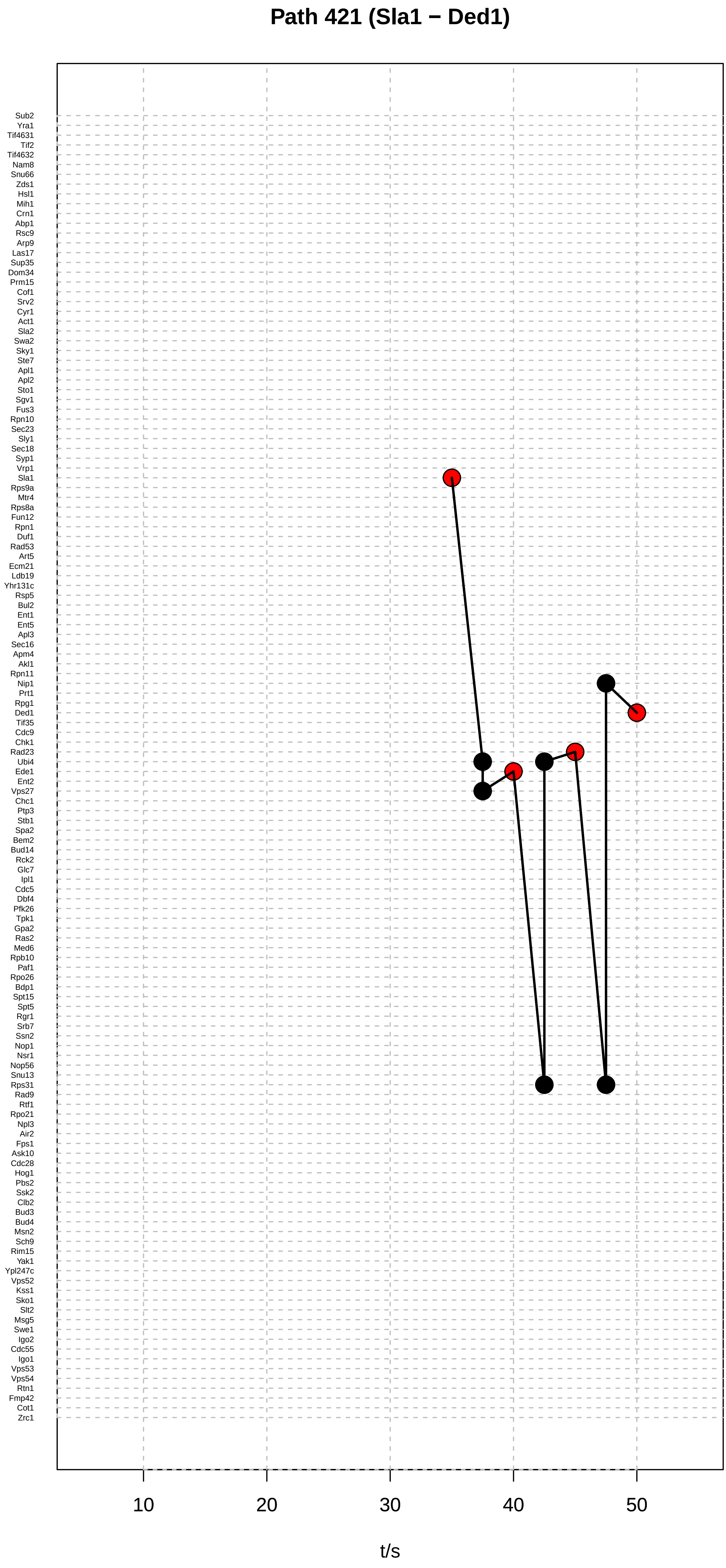

Path 421 (Sla1 − Ded1)
Sub2
Yra1
Tif4631
Tif2
Tif4632
Nam8
Snu66
Zds1
Hsl1
Mih1
Crn1
Abp1
Rsc9
Arp9
Las17
Sup35
Dom34
Prm15
Cof1
Srv2
Cyr1
Act1
Sla2
Swa2
Sky1
Ste7
Apl1
Apl2
Sto1
Sgv1
Fus3
Rpn10
Sec23
Sly1
Sec18
Syp1
Vrp1
Sla1
Rps9a
Mtr4
Rps8a
Fun12
Rpn1
Duf1
Rad53
Art5
Ecm21
Ldb19
Yhr131c
Rsp5
Bul2
Ent1
Ent5
Apl3
Sec16
Apm4
Akl1
Rpn11
Nip1
Prt1
Rpg1
Ded1
Tif35
Cdc9
Chk1
Rad23
Ubi4
Ede1
Ent2
Vps27
Chc1
Ptp3
Stb1
Spa2
Bem2
Bud14
Rck2
Glc7
Ipl1
Cdc5
Dbf4
Pfk26
Tpk1
Gpa2
Ras2
Med6
Rpb10
Paf1
Rpo26
Bdp1
Spt15
Spt5
Rgr1
Srb7
Ssn2
Nop1
Nsr1
Nop56
Snu13
Rps31
Rad9
Rtf1
Rpo21
Npl3
Air2
Fps1
Ask10
Cdc28
Hog1
Pbs2
Ssk2
Clb2
Bud3
Bud4
Msn2
Sch9
Rim15
Yak1
Ypl247c
Vps52
Kss1
Sko1
Slt2
Msg5
Swe1
Igo2
Cdc55
Igo1
Vps53
Vps54
Rtn1
Fmp42
Cot1
Zrc1
10
20
30
40
50
t/s

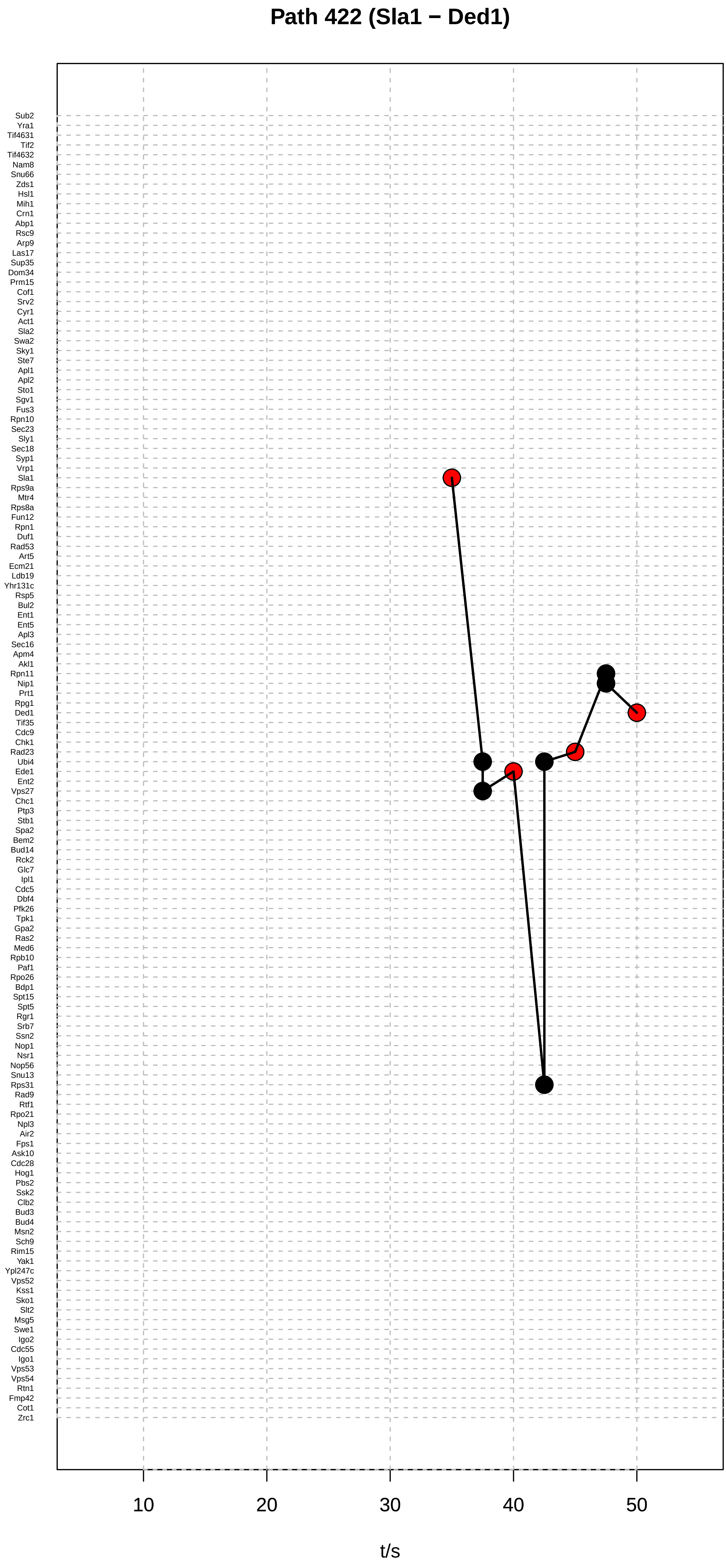

Path 422 (Sla1 − Ded1)
Sub2
Yra1
Tif4631
Tif2
Tif4632
Nam8
Snu66
Zds1
Hsl1
Mih1
Crn1
Abp1
Rsc9
Arp9
Las17
Sup35
Dom34
Prm15
Cof1
Srv2
Cyr1
Act1
Sla2
Swa2
Sky1
Ste7
Apl1
Apl2
Sto1
Sgv1
Fus3
Rpn10
Sec23
Sly1
Sec18
Syp1
Vrp1
Sla1
Rps9a
Mtr4
Rps8a
Fun12
Rpn1
Duf1
Rad53
Art5
Ecm21
Ldb19
Yhr131c
Rsp5
Bul2
Ent1
Ent5
Apl3
Sec16
Apm4
Akl1
Rpn11
Nip1
Prt1
Rpg1
Ded1
Tif35
Cdc9
Chk1
Rad23
Ubi4
Ede1
Ent2
Vps27
Chc1
Ptp3
Stb1
Spa2
Bem2
Bud14
Rck2
Glc7
Ipl1
Cdc5
Dbf4
Pfk26
Tpk1
Gpa2
Ras2
Med6
Rpb10
Paf1
Rpo26
Bdp1
Spt15
Spt5
Rgr1
Srb7
Ssn2
Nop1
Nsr1
Nop56
Snu13
Rps31
Rad9
Rtf1
Rpo21
Npl3
Air2
Fps1
Ask10
Cdc28
Hog1
Pbs2
Ssk2
Clb2
Bud3
Bud4
Msn2
Sch9
Rim15
Yak1
Ypl247c
Vps52
Kss1
Sko1
Slt2
Msg5
Swe1
Igo2
Cdc55
Igo1
Vps53
Vps54
Rtn1
Fmp42
Cot1
Zrc1
10
20
30
40
50
t/s

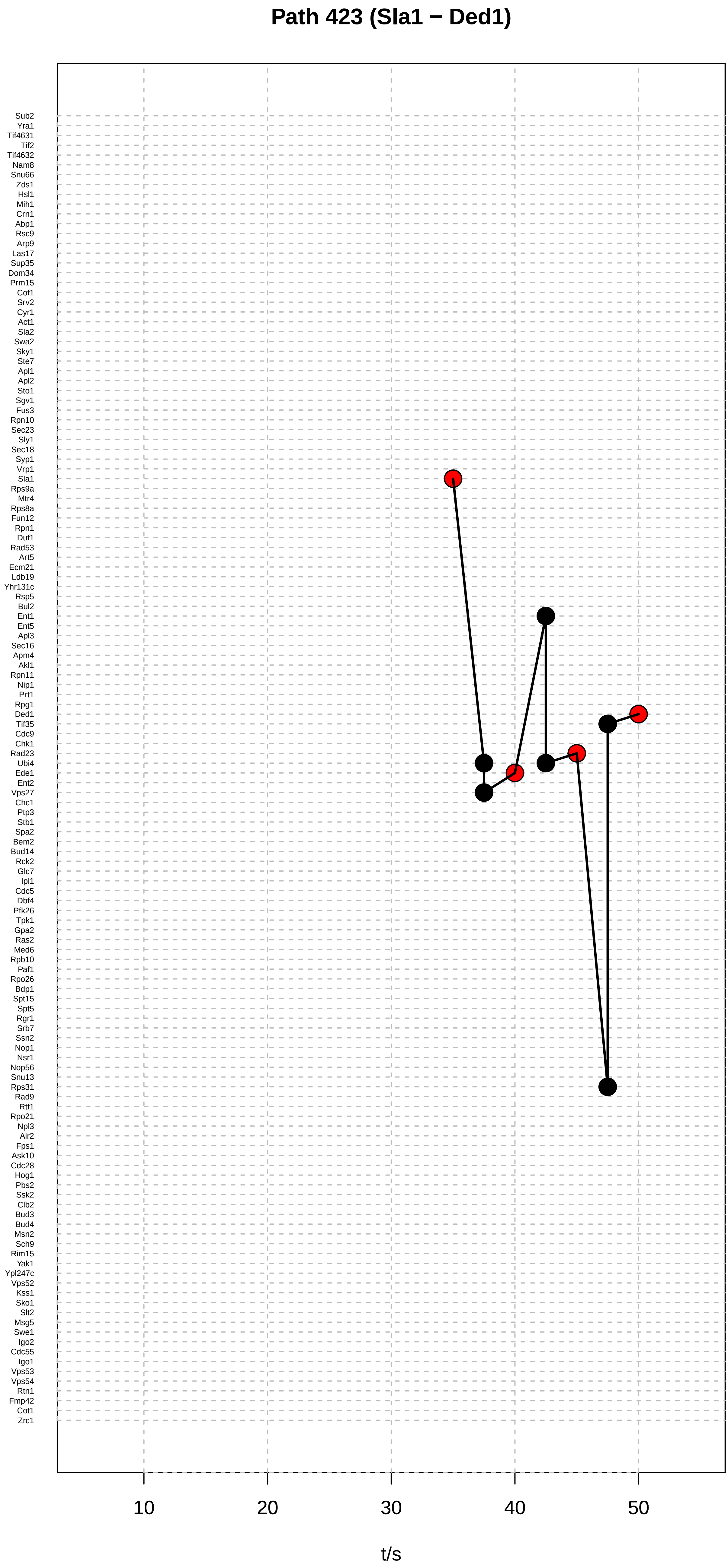

Path 423 (Sla1 − Ded1)
Sub2
Yra1
Tif4631
Tif2
Tif4632
Nam8
Snu66
Zds1
Hsl1
Mih1
Crn1
Abp1
Rsc9
Arp9
Las17
Sup35
Dom34
Prm15
Cof1
Srv2
Cyr1
Act1
Sla2
Swa2
Sky1
Ste7
Apl1
Apl2
Sto1
Sgv1
Fus3
Rpn10
Sec23
Sly1
Sec18
Syp1
Vrp1
Sla1
Rps9a
Mtr4
Rps8a
Fun12
Rpn1
Duf1
Rad53
Art5
Ecm21
Ldb19
Yhr131c
Rsp5
Bul2
Ent1
Ent5
Apl3
Sec16
Apm4
Akl1
Rpn11
Nip1
Prt1
Rpg1
Ded1
Tif35
Cdc9
Chk1
Rad23
Ubi4
Ede1
Ent2
Vps27
Chc1
Ptp3
Stb1
Spa2
Bem2
Bud14
Rck2
Glc7
Ipl1
Cdc5
Dbf4
Pfk26
Tpk1
Gpa2
Ras2
Med6
Rpb10
Paf1
Rpo26
Bdp1
Spt15
Spt5
Rgr1
Srb7
Ssn2
Nop1
Nsr1
Nop56
Snu13
Rps31
Rad9
Rtf1
Rpo21
Npl3
Air2
Fps1
Ask10
Cdc28
Hog1
Pbs2
Ssk2
Clb2
Bud3
Bud4
Msn2
Sch9
Rim15
Yak1
Ypl247c
Vps52
Kss1
Sko1
Slt2
Msg5
Swe1
Igo2
Cdc55
Igo1
Vps53
Vps54
Rtn1
Fmp42
Cot1
Zrc1
10
20
30
40
50
t/s

### Path 424 (Sla1 − Ded1)

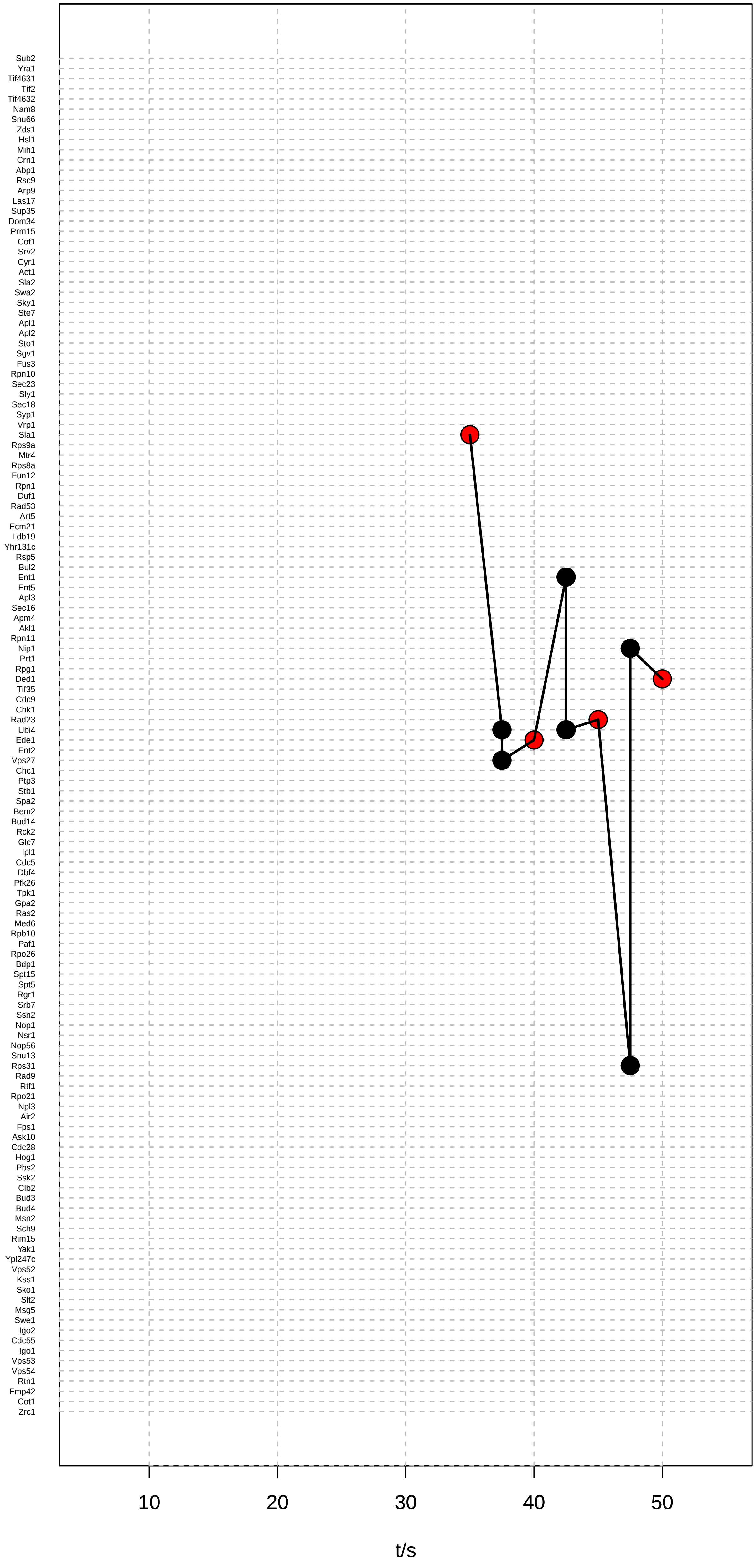

Path 425 (Sla1 − Ded1)

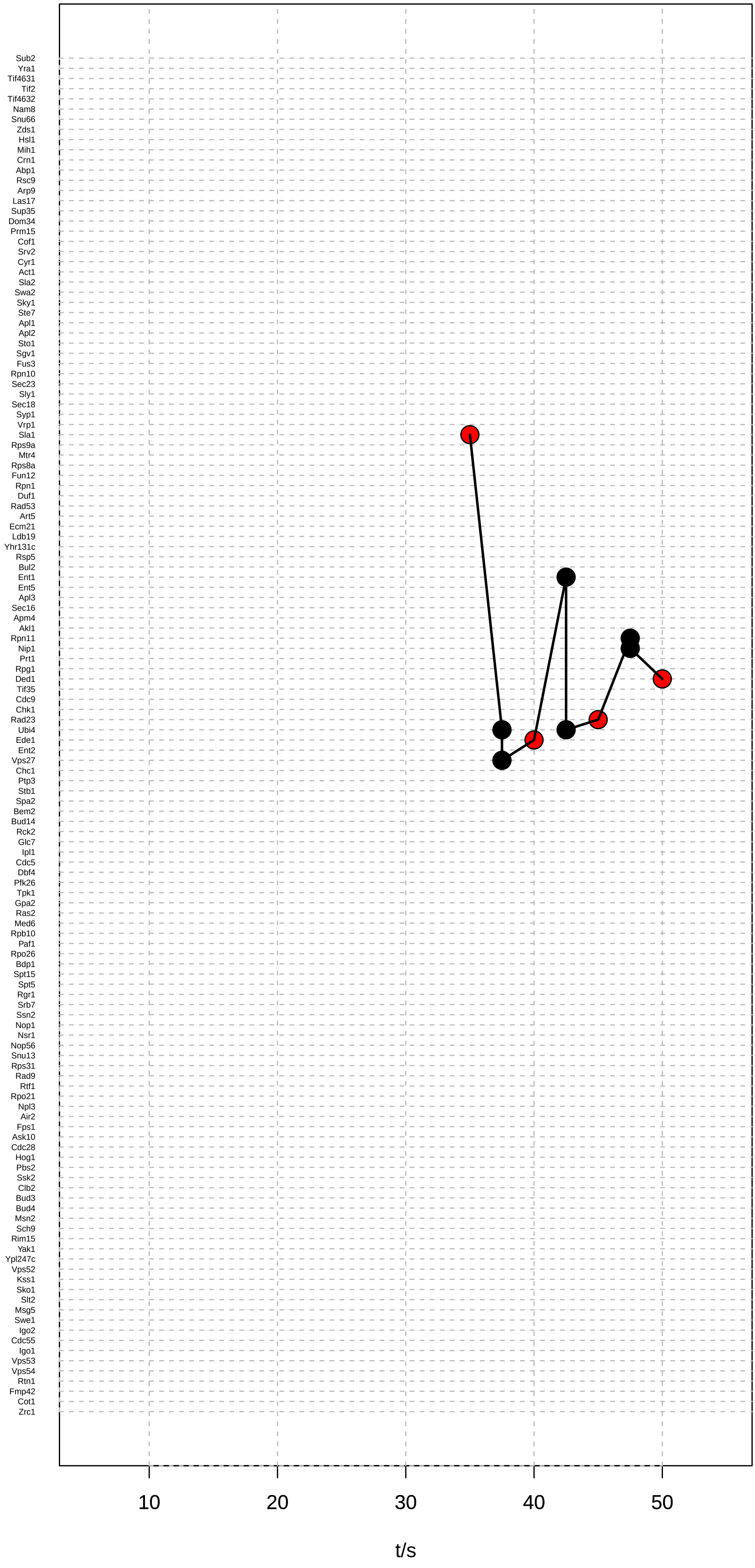

### Path 426 (Sla1 − Ded1)

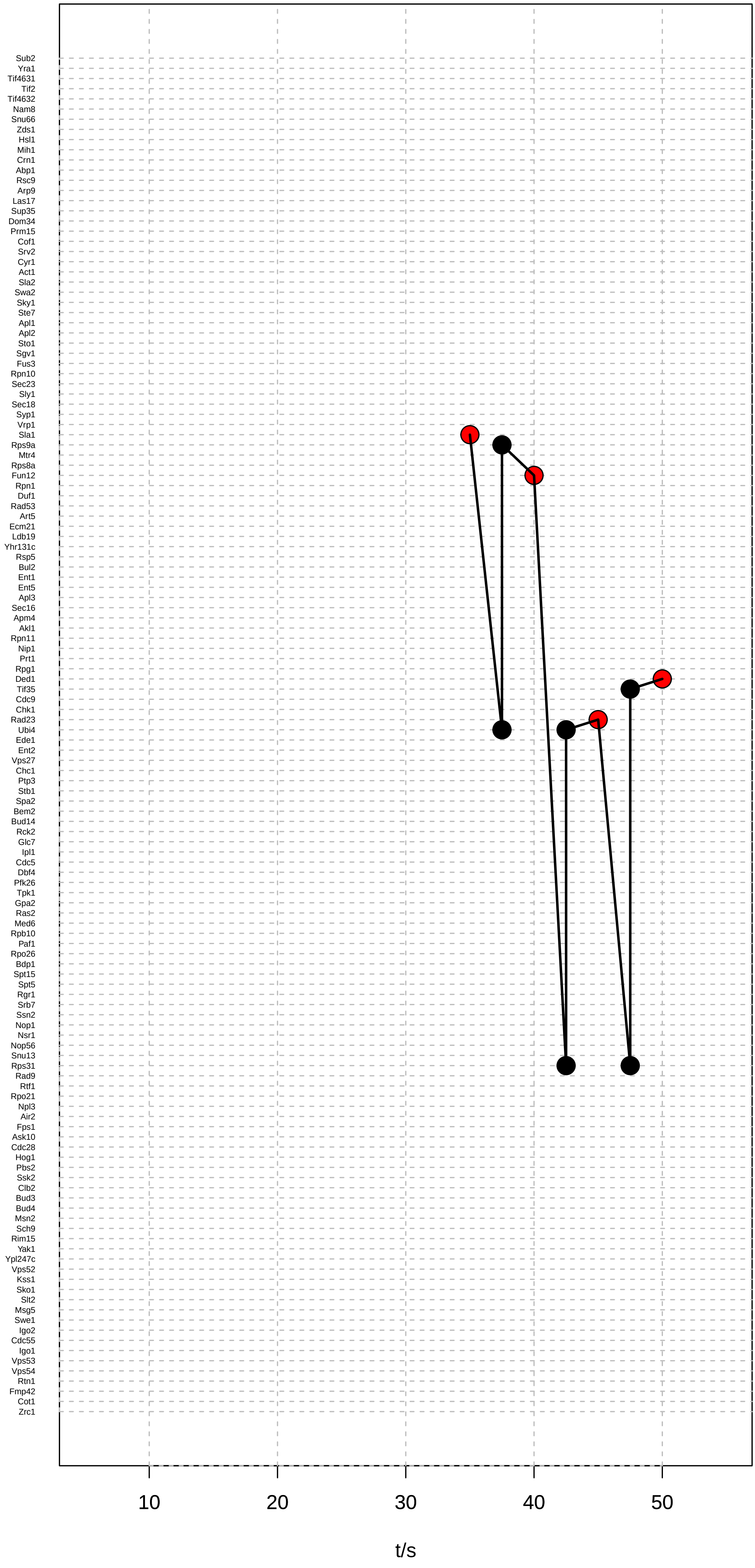

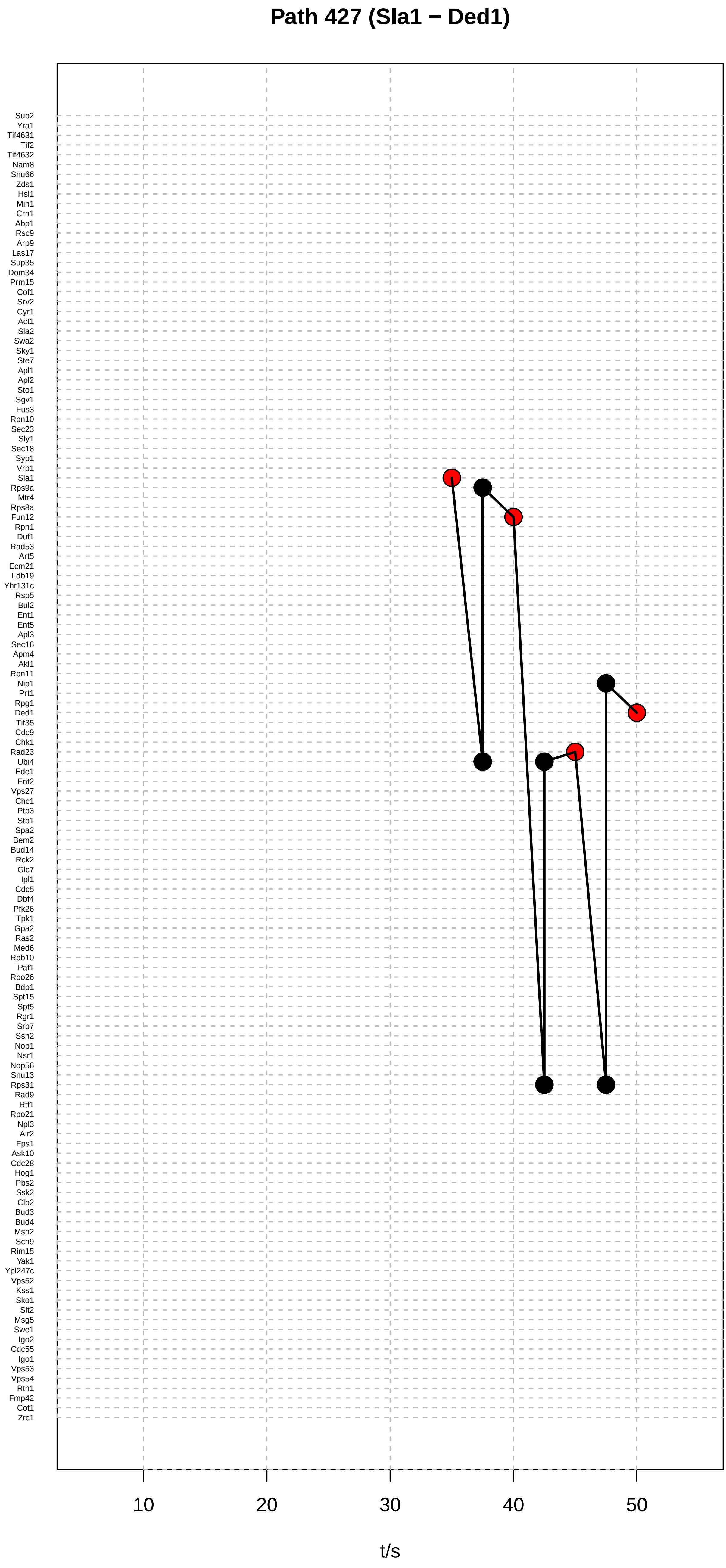
Path 427 (Sla1 − Ded1)
Sub2
Yra1
Tif4631
Tif2
Tif4632
Nam8
Snu66
Zds1
Hsl1
Mih1
Crn1
Abp1
Rsc9
Arp9
Las17
Sup35
Dom34
Prm15
Cof1
Srv2
Cyr1
Act1
Sla2
Swa2
Sky1
Ste7
Apl1
Apl2
Sto1
Sgv1
Fus3
Rpn10
Sec23
Sly1
Sec18
Syp1
Vrp1
Sla1
Rps9a
Mtr4
Rps8a
Fun12
Rpn1
Duf1
Rad53
Art5
Ecm21
Ldb19
Yhr131c
Rsp5
Bul2
Ent1
Ent5
Apl3
Sec16
Apm4
Akl1
Rpn11
Nip1
Prt1
Rpg1
Ded1
Tif35
Cdc9
Chk1
Rad23
Ubi4
Ede1
Ent2
Vps27
Chc1
Ptp3
Stb1
Spa2
Bem2
Bud14
Rck2
Glc7
Ipl1
Cdc5
Dbf4
Pfk26
Tpk1
Gpa2
Ras2
Med6
Rpb10
Paf1
Rpo26
Bdp1
Spt15
Spt5
Rgr1
Srb7
Ssn2
Nop1
Nsr1
Nop56
Snu13
Rps31
Rad9
Rtf1
Rpo21
Npl3
Air2
Fps1
Ask10
Cdc28
Hog1
Pbs2
Ssk2
Clb2
Bud3
Bud4
Msn2
Sch9
Rim15
Yak1
Ypl247c
Vps52
Kss1
Sko1
Slt2
Msg5
Swe1
Igo2
Cdc55
Igo1
Vps53
Vps54
Rtn1
Fmp42
Cot1
Zrc1
10
20
30
40
50
t/s

### Path 428 (Sla1 − Ded1)

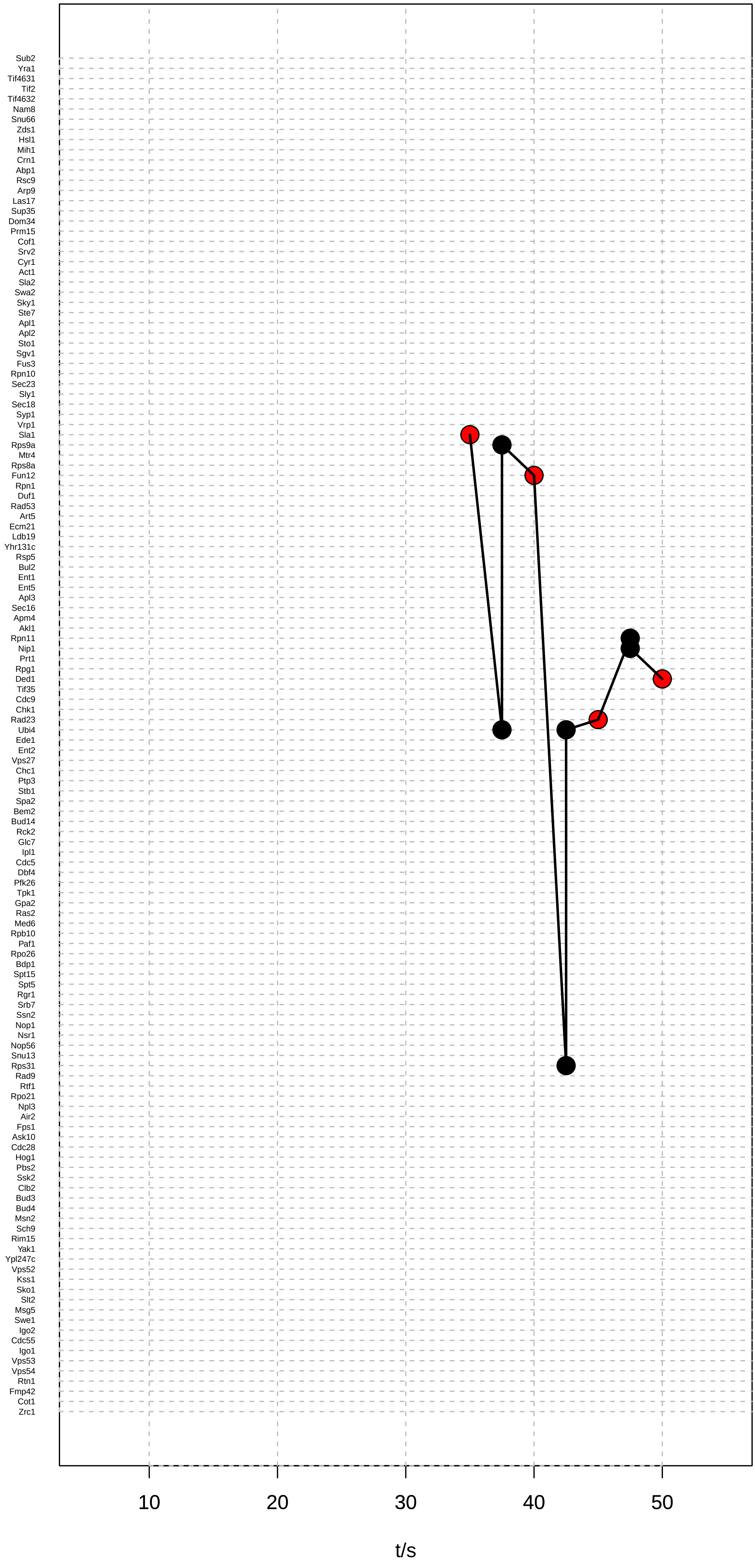

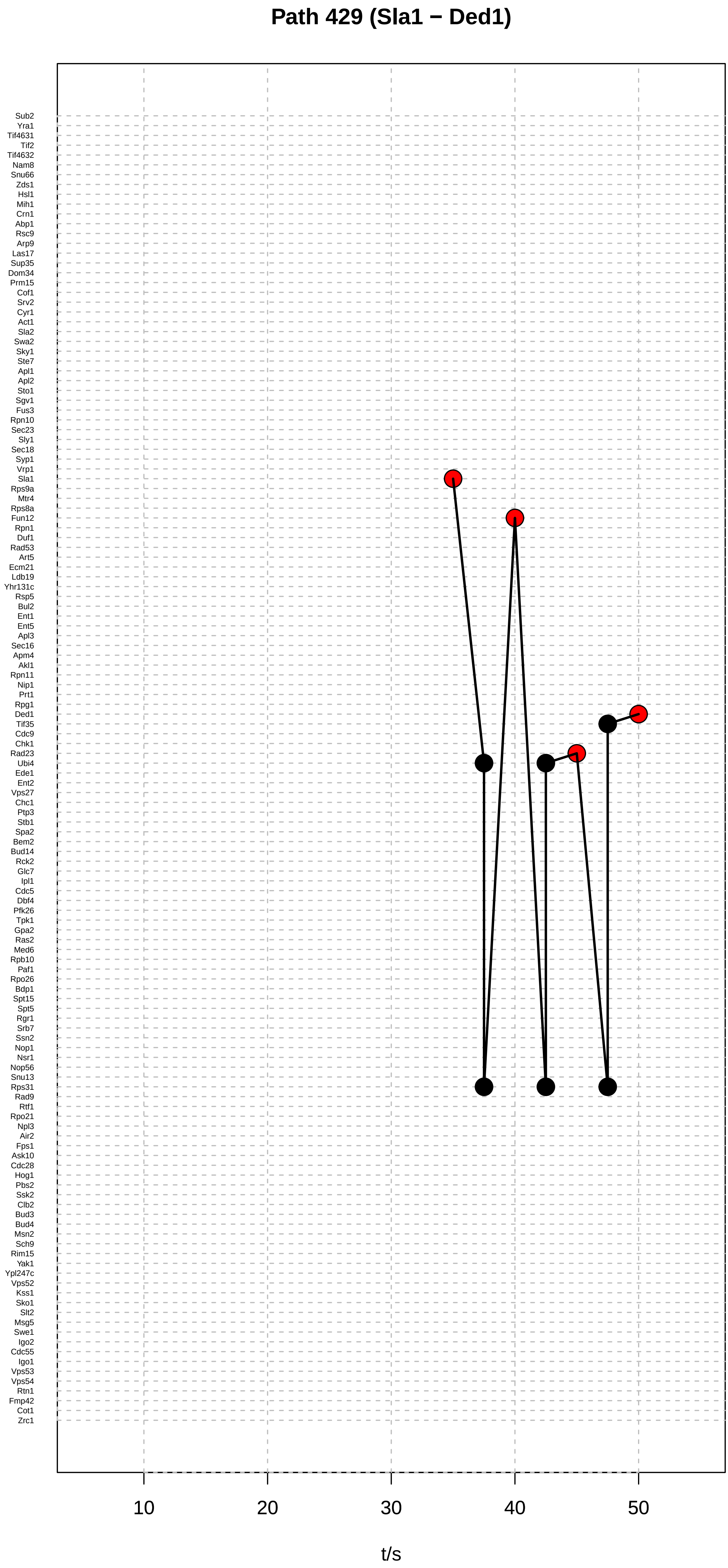

Path 429 (Sla1 − Ded1)
Sub2
Yra1
Tif4631
Tif2
Tif4632
Nam8
Snu66
Zds1
Hsl1
Mih1
Crn1
Abp1
Rsc9
Arp9
Las17
Sup35
Dom34
Prm15
Cof1
Srv2
Cyr1
Act1
Sla2
Swa2
Sky1
Ste7
Apl1
Apl2
Sto1
Sgv1
Fus3
Rpn10
Sec23
Sly1
Sec18
Syp1
Vrp1
Sla1
Rps9a
Mtr4
Rps8a
Fun12
Rpn1
Duf1
Rad53
Art5
Ecm21
Ldb19
Yhr131c
Rsp5
Bul2
Ent1
Ent5
Apl3
Sec16
Apm4
Akl1
Rpn11
Nip1
Prt1
Rpg1
Ded1
Tif35
Cdc9
Chk1
Rad23
Ubi4
Ede1
Ent2
Vps27
Chc1
Ptp3
Stb1
Spa2
Bem2
Bud14
Rck2
Glc7
Ipl1
Cdc5
Dbf4
Pfk26
Tpk1
Gpa2
Ras2
Med6
Rpb10
Paf1
Rpo26
Bdp1
Spt15
Spt5
Rgr1
Srb7
Ssn2
Nop1
Nsr1
Nop56
Snu13
Rps31
Rad9
Rtf1
Rpo21
Npl3
Air2
Fps1
Ask10
Cdc28
Hog1
Pbs2
Ssk2
Clb2
Bud3
Bud4
Msn2
Sch9
Rim15
Yak1
Ypl247c
Vps52
Kss1
Sko1
Slt2
Msg5
Swe1
Igo2
Cdc55
Igo1
Vps53
Vps54
Rtn1
Fmp42
Cot1
Zrc1
10
20
30
40
50
t/s

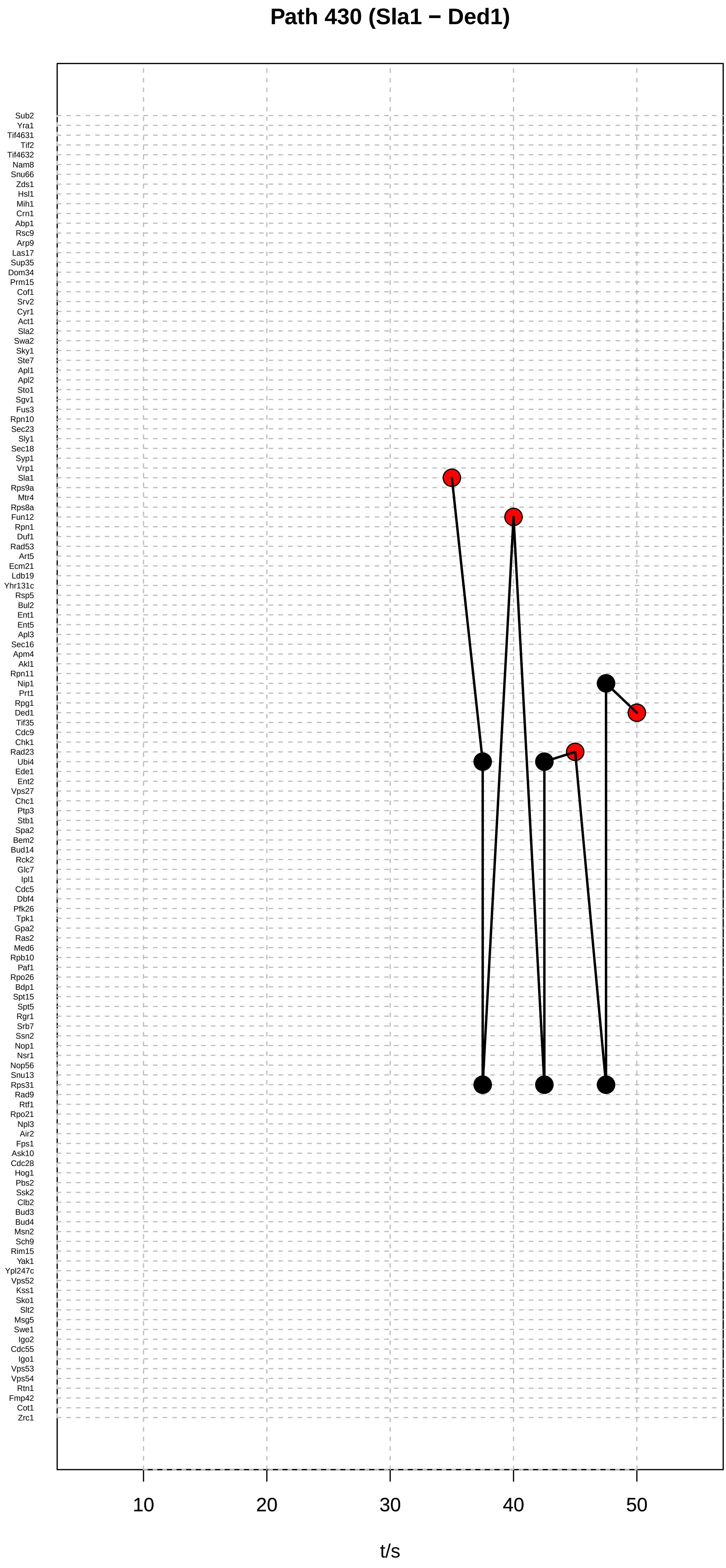
Path 430 (Sla1 − Ded1)
Sub2
Yra1
Tif4631
Tif2
Tif4632
Nam8
Snu66
Zds1
Hsl1
Mih1
Crn1
Abp1
Rsc9
Arp9
Las17
Sup35
Dom34
Prm15
Cof1
Srv2
Cyr1
Act1
Sla2
Swa2
Sky1
Ste7
Apl1
Apl2
Sto1
Sgv1
Fus3
Rpn10
Sec23
Sly1
Sec18
Syp1
Vrp1
Sla1
Rps9a
Mtr4
Rps8a
Fun12
Rpn1
Duf1
Rad53
Art5
Ecm21
Ldb19
Yhr131c
Rsp5
Bul2
Ent1
Ent5
Apl3
Sec16
Apm4
Akl1
Rpn11
Nip1
Prt1
Rpg1
Ded1
Tif35
Cdc9
Chk1
Rad23
Ubi4
Ede1
Ent2
Vps27
Chc1
Ptp3
Stb1
Spa2
Bem2
Bud14
Rck2
Glc7
Ipl1
Cdc5
Dbf4
Pfk26
Tpk1
Gpa2
Ras2
Med6
Rpb10
Paf1
Rpo26
Bdp1
Spt15
Spt5
Rgr1
Srb7
Ssn2
Nop1
Nsr1
Nop56
Snu13
Rps31
Rad9
Rtf1
Rpo21
Npl3
Air2
Fps1
Ask10
Cdc28
Hog1
Pbs2
Ssk2
Clb2
Bud3
Bud4
Msn2
Sch9
Rim15
Yak1
Ypl247c
Vps52
Kss1
Sko1
Slt2
Msg5
Swe1
Igo2
Cdc55
Igo1
Vps53
Vps54
Rtn1
Fmp42
Cot1
Zrc1
10
20
30
40
50
t/s

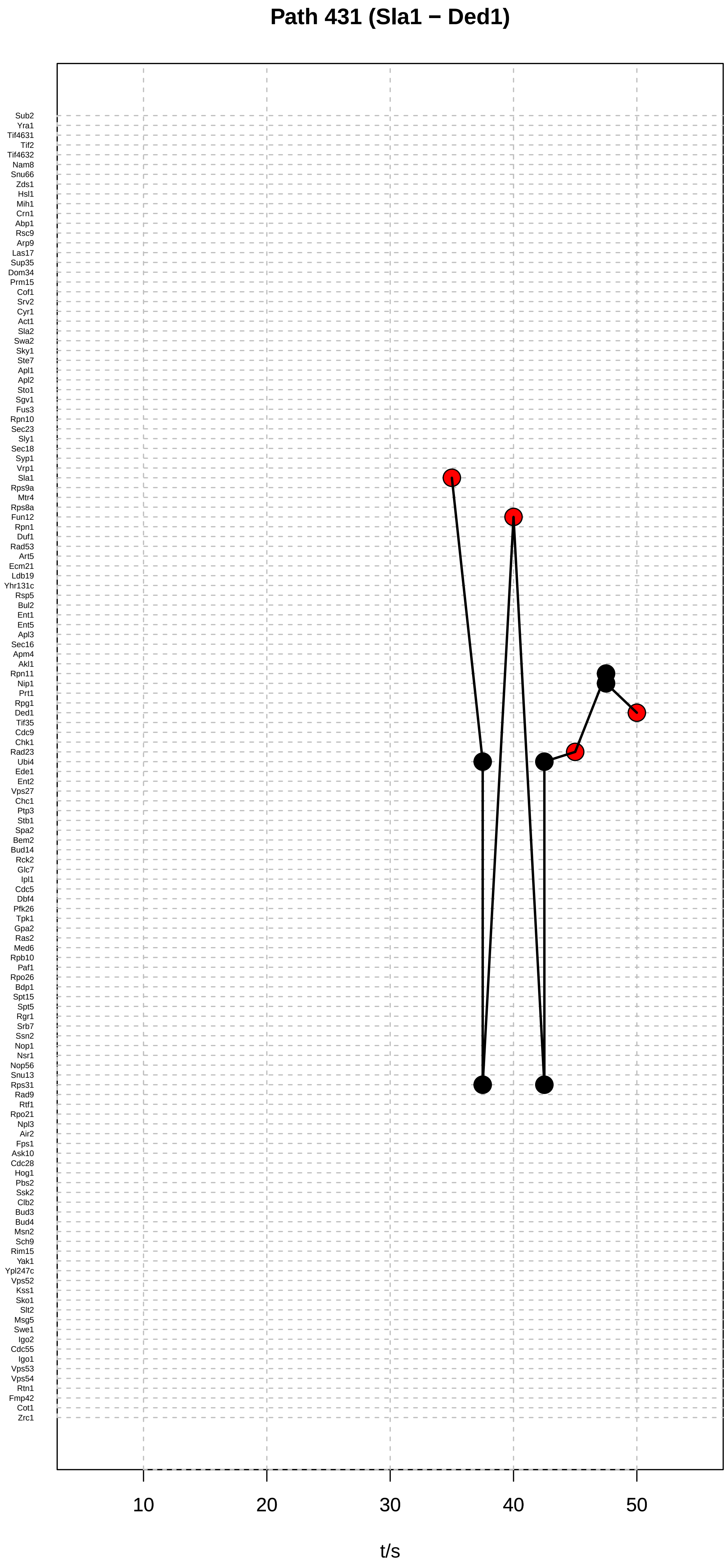

Path 431 (Sla1 − Ded1)
Sub2
Yra1
Tif4631
Tif2
Tif4632
Nam8
Snu66
Zds1
Hsl1
Mih1
Crn1
Abp1
Rsc9
Arp9
Las17
Sup35
Dom34
Prm15
Cof1
Srv2
Cyr1
Act1
Sla2
Swa2
Sky1
Ste7
Apl1
Apl2
Sto1
Sgv1
Fus3
Rpn10
Sec23
Sly1
Sec18
Syp1
Vrp1
Sla1
Rps9a
Mtr4
Rps8a
Fun12
Rpn1
Duf1
Rad53
Art5
Ecm21
Ldb19
Yhr131c
Rsp5
Bul2
Ent1
Ent5
Apl3
Sec16
Apm4
Akl1
Rpn11
Nip1
Prt1
Rpg1
Ded1
Tif35
Cdc9
Chk1
Rad23
Ubi4
Ede1
Ent2
Vps27
Chc1
Ptp3
Stb1
Spa2
Bem2
Bud14
Rck2
Glc7
Ipl1
Cdc5
Dbf4
Pfk26
Tpk1
Gpa2
Ras2
Med6
Rpb10
Paf1
Rpo26
Bdp1
Spt15
Spt5
Rgr1
Srb7
Ssn2
Nop1
Nsr1
Nop56
Snu13
Rps31
Rad9
Rtf1
Rpo21
Npl3
Air2
Fps1
Ask10
Cdc28
Hog1
Pbs2
Ssk2
Clb2
Bud3
Bud4
Msn2
Sch9
Rim15
Yak1
Ypl247c
Vps52
Kss1
Sko1
Slt2
Msg5
Swe1
Igo2
Cdc55
Igo1
Vps53
Vps54
Rtn1
Fmp42
Cot1
Zrc1
10
20
30
40
50
t/s

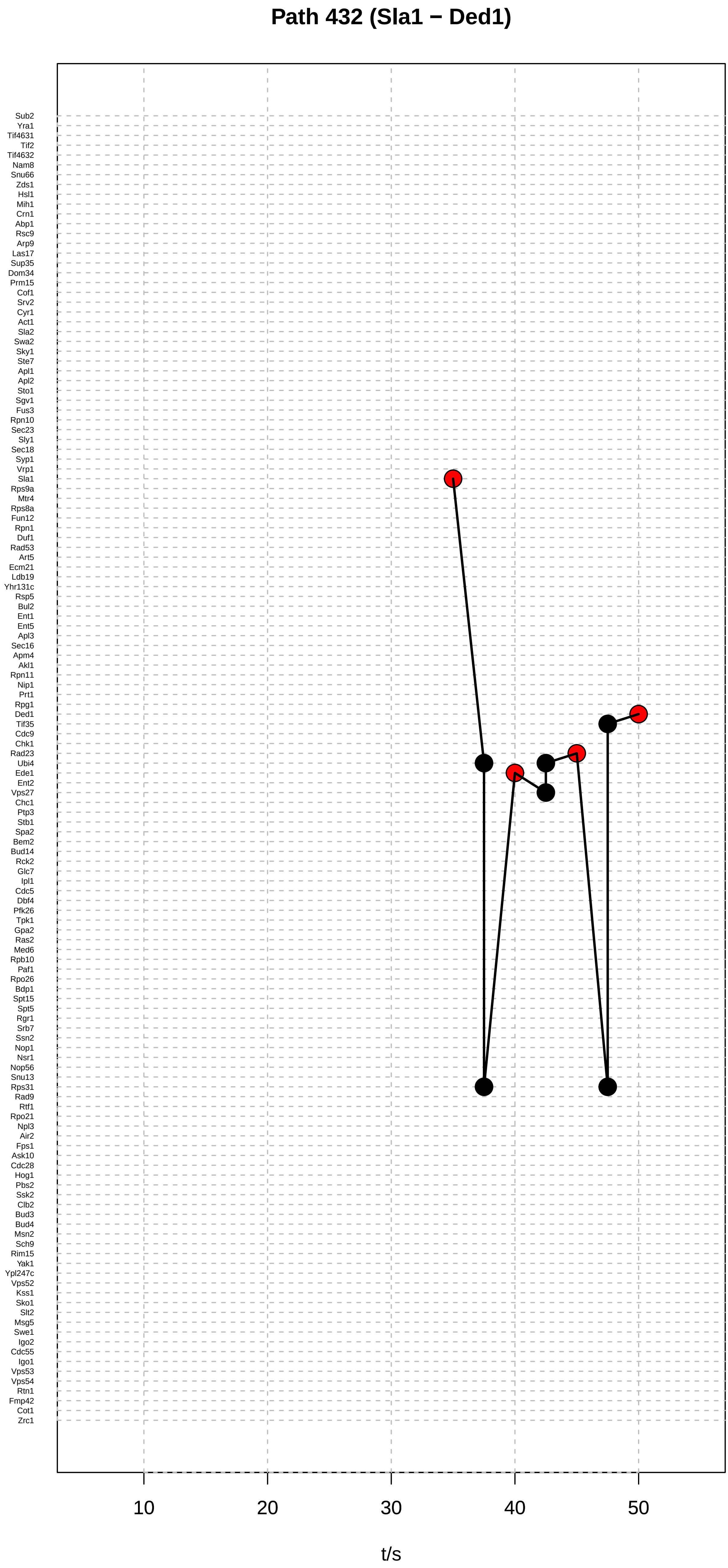

Path 432 (Sla1 − Ded1)
Sub2
Yra1
Tif4631
Tif2
Tif4632
Nam8
Snu66
Zds1
Hsl1
Mih1
Crn1
Abp1
Rsc9
Arp9
Las17
Sup35
Dom34
Prm15
Cof1
Srv2
Cyr1
Act1
Sla2
Swa2
Sky1
Ste7
Apl1
Apl2
Sto1
Sgv1
Fus3
Rpn10
Sec23
Sly1
Sec18
Syp1
Vrp1
Sla1
Rps9a
Mtr4
Rps8a
Fun12
Rpn1
Duf1
Rad53
Art5
Ecm21
Ldb19
Yhr131c
Rsp5
Bul2
Ent1
Ent5
Apl3
Sec16
Apm4
Akl1
Rpn11
Nip1
Prt1
Rpg1
Ded1
Tif35
Cdc9
Chk1
Rad23
Ubi4
Ede1
Ent2
Vps27
Chc1
Ptp3
Stb1
Spa2
Bem2
Bud14
Rck2
Glc7
Ipl1
Cdc5
Dbf4
Pfk26
Tpk1
Gpa2
Ras2
Med6
Rpb10
Paf1
Rpo26
Bdp1
Spt15
Spt5
Rgr1
Srb7
Ssn2
Nop1
Nsr1
Nop56
Snu13
Rps31
Rad9
Rtf1
Rpo21
Npl3
Air2
Fps1
Ask10
Cdc28
Hog1
Pbs2
Ssk2
Clb2
Bud3
Bud4
Msn2
Sch9
Rim15
Yak1
Ypl247c
Vps52
Kss1
Sko1
Slt2
Msg5
Swe1
Igo2
Cdc55
Igo1
Vps53
Vps54
Rtn1
Fmp42
Cot1
Zrc1
10
20
30
40
50
t/s

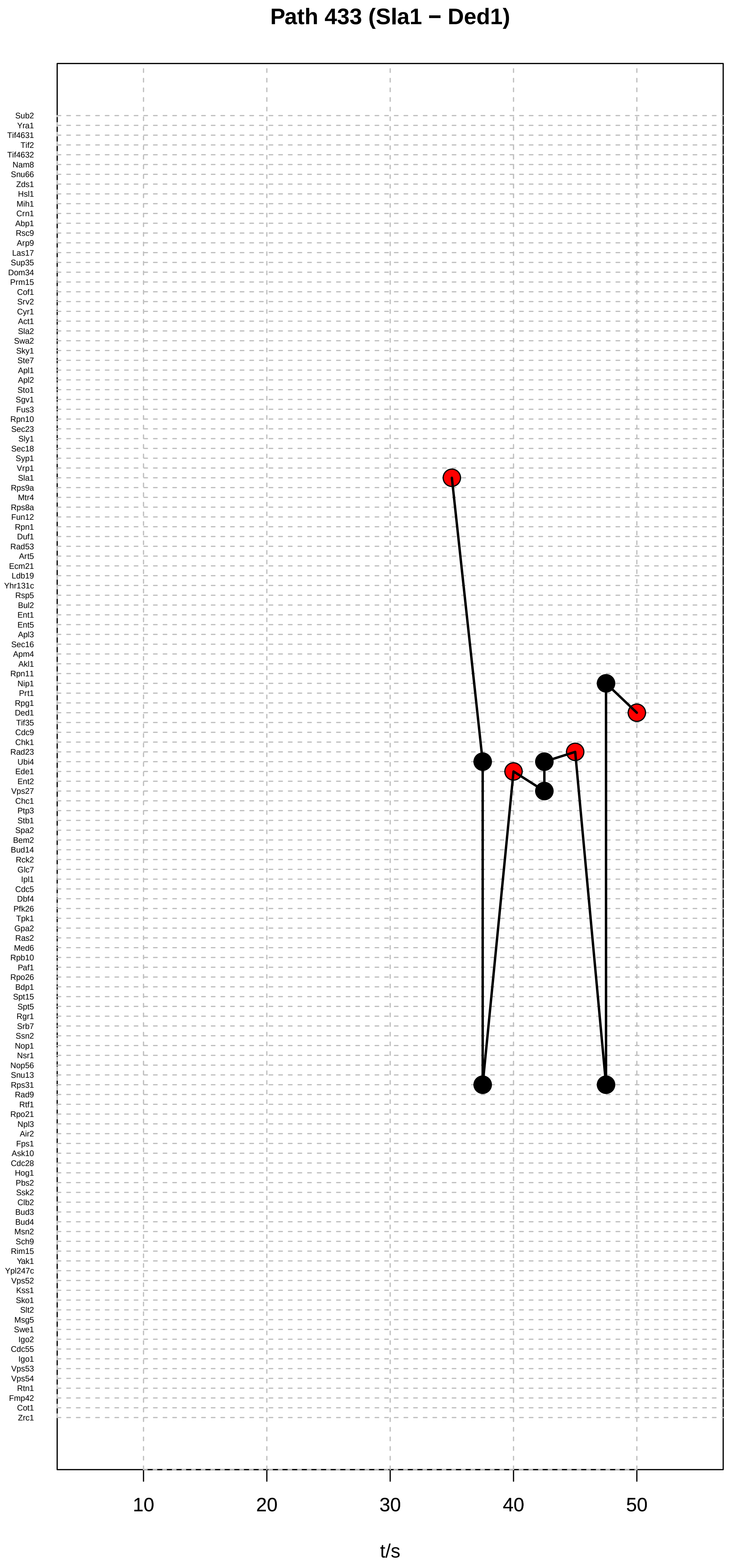
Path 433 (Sla1 − Ded1)
Sub2
Yra1
Tif4631
Tif2
Tif4632
Nam8
Snu66
Zds1
Hsl1
Mih1
Crn1
Abp1
Rsc9
Arp9
Las17
Sup35
Dom34
Prm15
Cof1
Srv2
Cyr1
Act1
Sla2
Swa2
Sky1
Ste7
Apl1
Apl2
Sto1
Sgv1
Fus3
Rpn10
Sec23
Sly1
Sec18
Syp1
Vrp1
Sla1
Rps9a
Mtr4
Rps8a
Fun12
Rpn1
Duf1
Rad53
Art5
Ecm21
Ldb19
Yhr131c
Rsp5
Bul2
Ent1
Ent5
Apl3
Sec16
Apm4
Akl1
Rpn11
Nip1
Prt1
Rpg1
Ded1
Tif35
Cdc9
Chk1
Rad23
Ubi4
Ede1
Ent2
Vps27
Chc1
Ptp3
Stb1
Spa2
Bem2
Bud14
Rck2
Glc7
Ipl1
Cdc5
Dbf4
Pfk26
Tpk1
Gpa2
Ras2
Med6
Rpb10
Paf1
Rpo26
Bdp1
Spt15
Spt5
Rgr1
Srb7
Ssn2
Nop1
Nsr1
Nop56
Snu13
Rps31
Rad9
Rtf1
Rpo21
Npl3
Air2
Fps1
Ask10
Cdc28
Hog1
Pbs2
Ssk2
Clb2
Bud3
Bud4
Msn2
Sch9
Rim15
Yak1
Ypl247c
Vps52
Kss1
Sko1
Slt2
Msg5
Swe1
Igo2
Cdc55
Igo1
Vps53
Vps54
Rtn1
Fmp42
Cot1
Zrc1
10
20
30
40
50
t/s

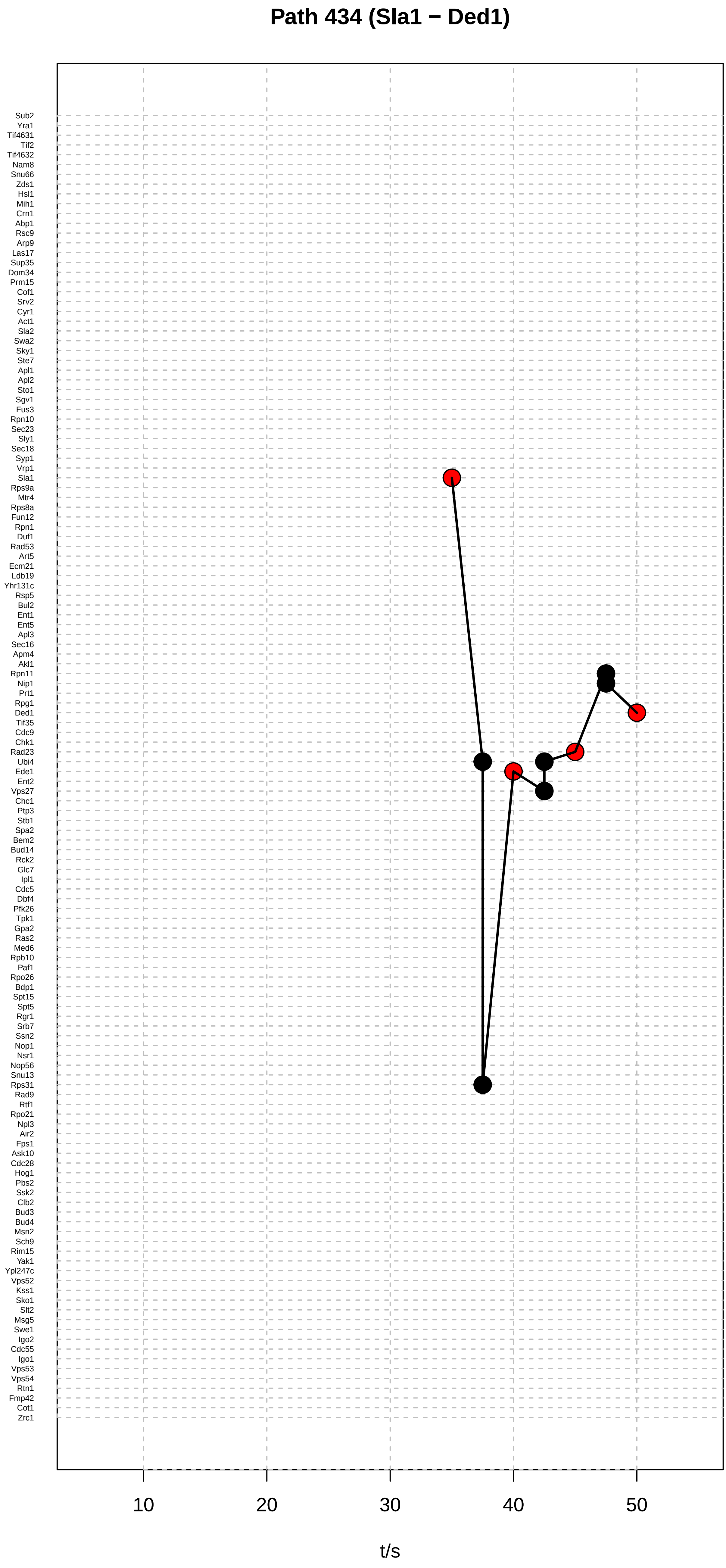
Path 434 (Sla1 − Ded1)
Sub2
Yra1
Tif4631
Tif2
Tif4632
Nam8
Snu66
Zds1
Hsl1
Mih1
Crn1
Abp1
Rsc9
Arp9
Las17
Sup35
Dom34
Prm15
Cof1
Srv2
Cyr1
Act1
Sla2
Swa2
Sky1
Ste7
Apl1
Apl2
Sto1
Sgv1
Fus3
Rpn10
Sec23
Sly1
Sec18
Syp1
Vrp1
Sla1
Rps9a
Mtr4
Rps8a
Fun12
Rpn1
Duf1
Rad53
Art5
Ecm21
Ldb19
Yhr131c
Rsp5
Bul2
Ent1
Ent5
Apl3
Sec16
Apm4
Akl1
Rpn11
Nip1
Prt1
Rpg1
Ded1
Tif35
Cdc9
Chk1
Rad23
Ubi4
Ede1
Ent2
Vps27
Chc1
Ptp3
Stb1
Spa2
Bem2
Bud14
Rck2
Glc7
Ipl1
Cdc5
Dbf4
Pfk26
Tpk1
Gpa2
Ras2
Med6
Rpb10
Paf1
Rpo26
Bdp1
Spt15
Spt5
Rgr1
Srb7
Ssn2
Nop1
Nsr1
Nop56
Snu13
Rps31
Rad9
Rtf1
Rpo21
Npl3
Air2
Fps1
Ask10
Cdc28
Hog1
Pbs2
Ssk2
Clb2
Bud3
Bud4
Msn2
Sch9
Rim15
Yak1
Ypl247c
Vps52
Kss1
Sko1
Slt2
Msg5
Swe1
Igo2
Cdc55
Igo1
Vps53
Vps54
Rtn1
Fmp42
Cot1
Zrc1
10
20
30
40
50
t/s

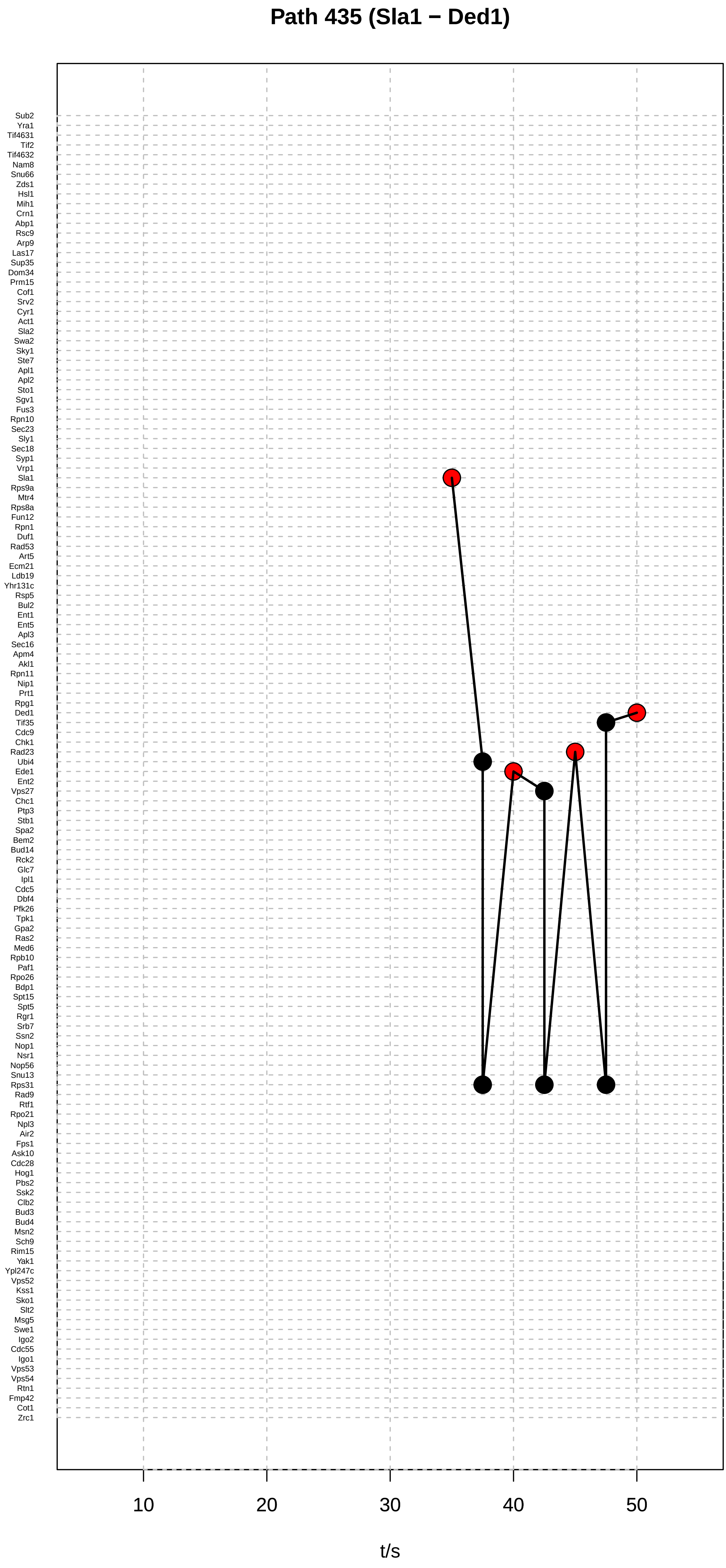
Path 435 (Sla1 − Ded1)
Sub2
Yra1
Tif4631
Tif2
Tif4632
Nam8
Snu66
Zds1
Hsl1
Mih1
Crn1
Abp1
Rsc9
Arp9
Las17
Sup35
Dom34
Prm15
Cof1
Srv2
Cyr1
Act1
Sla2
Swa2
Sky1
Ste7
Apl1
Apl2
Sto1
Sgv1
Fus3
Rpn10
Sec23
Sly1
Sec18
Syp1
Vrp1
Sla1
Rps9a
Mtr4
Rps8a
Fun12
Rpn1
Duf1
Rad53
Art5
Ecm21
Ldb19
Yhr131c
Rsp5
Bul2
Ent1
Ent5
Apl3
Sec16
Apm4
Akl1
Rpn11
Nip1
Prt1
Rpg1
Ded1
Tif35
Cdc9
Chk1
Rad23
Ubi4
Ede1
Ent2
Vps27
Chc1
Ptp3
Stb1
Spa2
Bem2
Bud14
Rck2
Glc7
Ipl1
Cdc5
Dbf4
Pfk26
Tpk1
Gpa2
Ras2
Med6
Rpb10
Paf1
Rpo26
Bdp1
Spt15
Spt5
Rgr1
Srb7
Ssn2
Nop1
Nsr1
Nop56
Snu13
Rps31
Rad9
Rtf1
Rpo21
Npl3
Air2
Fps1
Ask10
Cdc28
Hog1
Pbs2
Ssk2
Clb2
Bud3
Bud4
Msn2
Sch9
Rim15
Yak1
Ypl247c
Vps52
Kss1
Sko1
Slt2
Msg5
Swe1
Igo2
Cdc55
Igo1
Vps53
Vps54
Rtn1
Fmp42
Cot1
Zrc1
10
20
30
40
50
t/s

### Path 436 (Sla1 − Ded1)

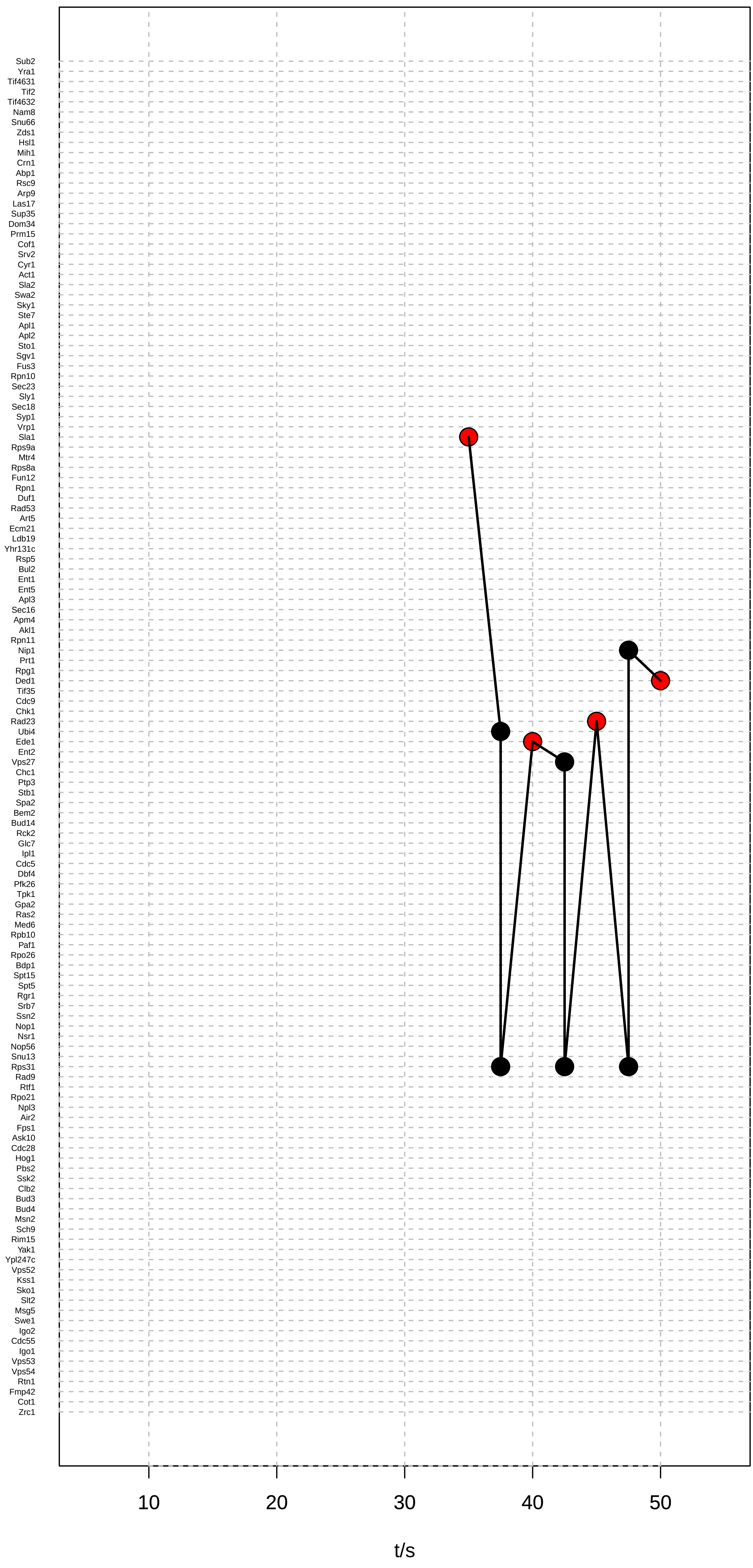

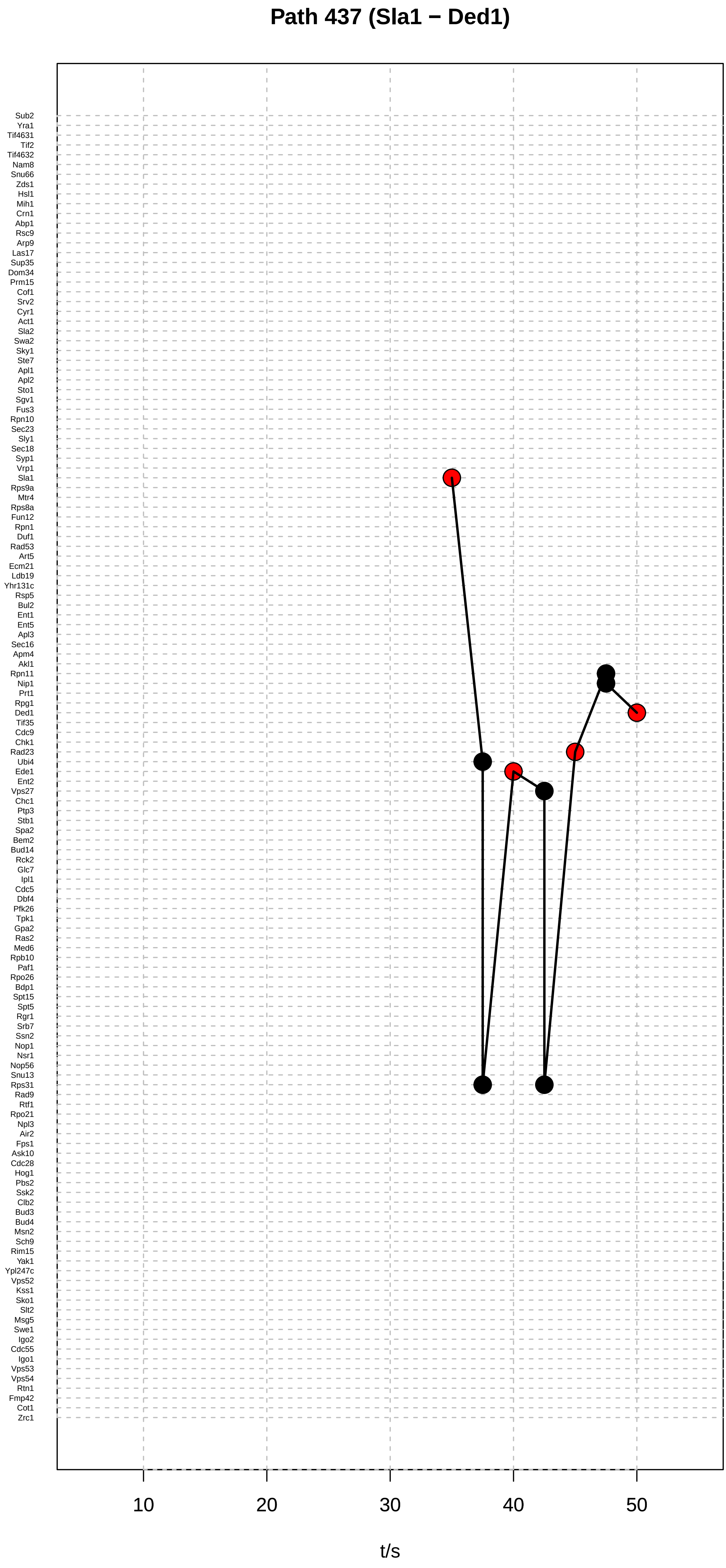
Path 437 (Sla1 − Ded1)
Sub2
Yra1
Tif4631
Tif2
Tif4632
Nam8
Snu66
Zds1
Hsl1
Mih1
Crn1
Abp1
Rsc9
Arp9
Las17
Sup35
Dom34
Prm15
Cof1
Srv2
Cyr1
Act1
Sla2
Swa2
Sky1
Ste7
Apl1
Apl2
Sto1
Sgv1
Fus3
Rpn10
Sec23
Sly1
Sec18
Syp1
Vrp1
Sla1
Rps9a
Mtr4
Rps8a
Fun12
Rpn1
Duf1
Rad53
Art5
Ecm21
Ldb19
Yhr131c
Rsp5
Bul2
Ent1
Ent5
Apl3
Sec16
Apm4
Akl1
Rpn11
Nip1
Prt1
Rpg1
Ded1
Tif35
Cdc9
Chk1
Rad23
Ubi4
Ede1
Ent2
Vps27
Chc1
Ptp3
Stb1
Spa2
Bem2
Bud14
Rck2
Glc7
Ipl1
Cdc5
Dbf4
Pfk26
Tpk1
Gpa2
Ras2
Med6
Rpb10
Paf1
Rpo26
Bdp1
Spt15
Spt5
Rgr1
Srb7
Ssn2
Nop1
Nsr1
Nop56
Snu13
Rps31
Rad9
Rtf1
Rpo21
Npl3
Air2
Fps1
Ask10
Cdc28
Hog1
Pbs2
Ssk2
Clb2
Bud3
Bud4
Msn2
Sch9
Rim15
Yak1
Ypl247c
Vps52
Kss1
Sko1
Slt2
Msg5
Swe1
Igo2
Cdc55
Igo1
Vps53
Vps54
Rtn1
Fmp42
Cot1
Zrc1
10
20
30
40
50
t/s

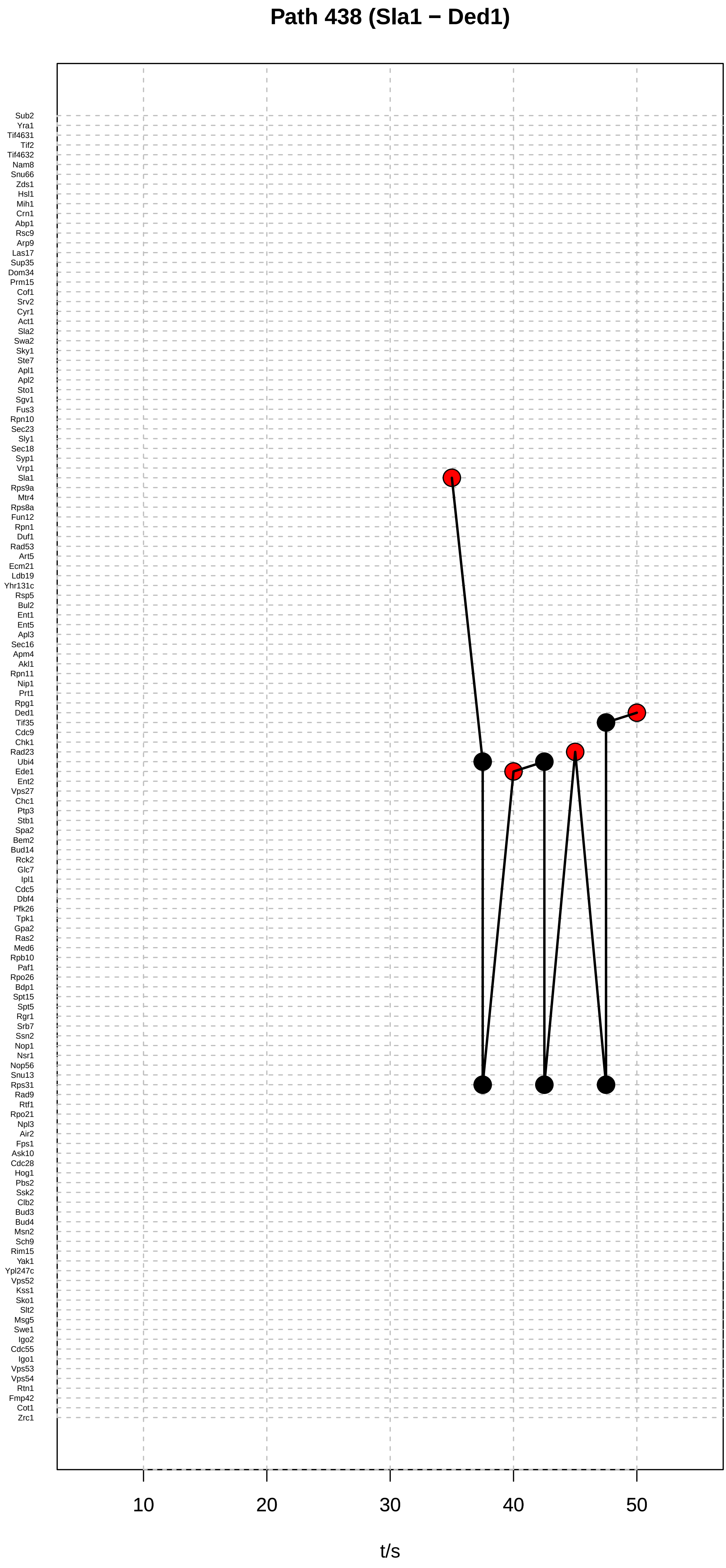

Path 438 (Sla1 − Ded1)
Sub2
Yra1
Tif4631
Tif2
Tif4632
Nam8
Snu66
Zds1
Hsl1
Mih1
Crn1
Abp1
Rsc9
Arp9
Las17
Sup35
Dom34
Prm15
Cof1
Srv2
Cyr1
Act1
Sla2
Swa2
Sky1
Ste7
Apl1
Apl2
Sto1
Sgv1
Fus3
Rpn10
Sec23
Sly1
Sec18
Syp1
Vrp1
Sla1
Rps9a
Mtr4
Rps8a
Fun12
Rpn1
Duf1
Rad53
Art5
Ecm21
Ldb19
Yhr131c
Rsp5
Bul2
Ent1
Ent5
Apl3
Sec16
Apm4
Akl1
Rpn11
Nip1
Prt1
Rpg1
Ded1
Tif35
Cdc9
Chk1
Rad23
Ubi4
Ede1
Ent2
Vps27
Chc1
Ptp3
Stb1
Spa2
Bem2
Bud14
Rck2
Glc7
Ipl1
Cdc5
Dbf4
Pfk26
Tpk1
Gpa2
Ras2
Med6
Rpb10
Paf1
Rpo26
Bdp1
Spt15
Spt5
Rgr1
Srb7
Ssn2
Nop1
Nsr1
Nop56
Snu13
Rps31
Rad9
Rtf1
Rpo21
Npl3
Air2
Fps1
Ask10
Cdc28
Hog1
Pbs2
Ssk2
Clb2
Bud3
Bud4
Msn2
Sch9
Rim15
Yak1
Ypl247c
Vps52
Kss1
Sko1
Slt2
Msg5
Swe1
Igo2
Cdc55
Igo1
Vps53
Vps54
Rtn1
Fmp42
Cot1
Zrc1
10
20
30
40
50
t/s

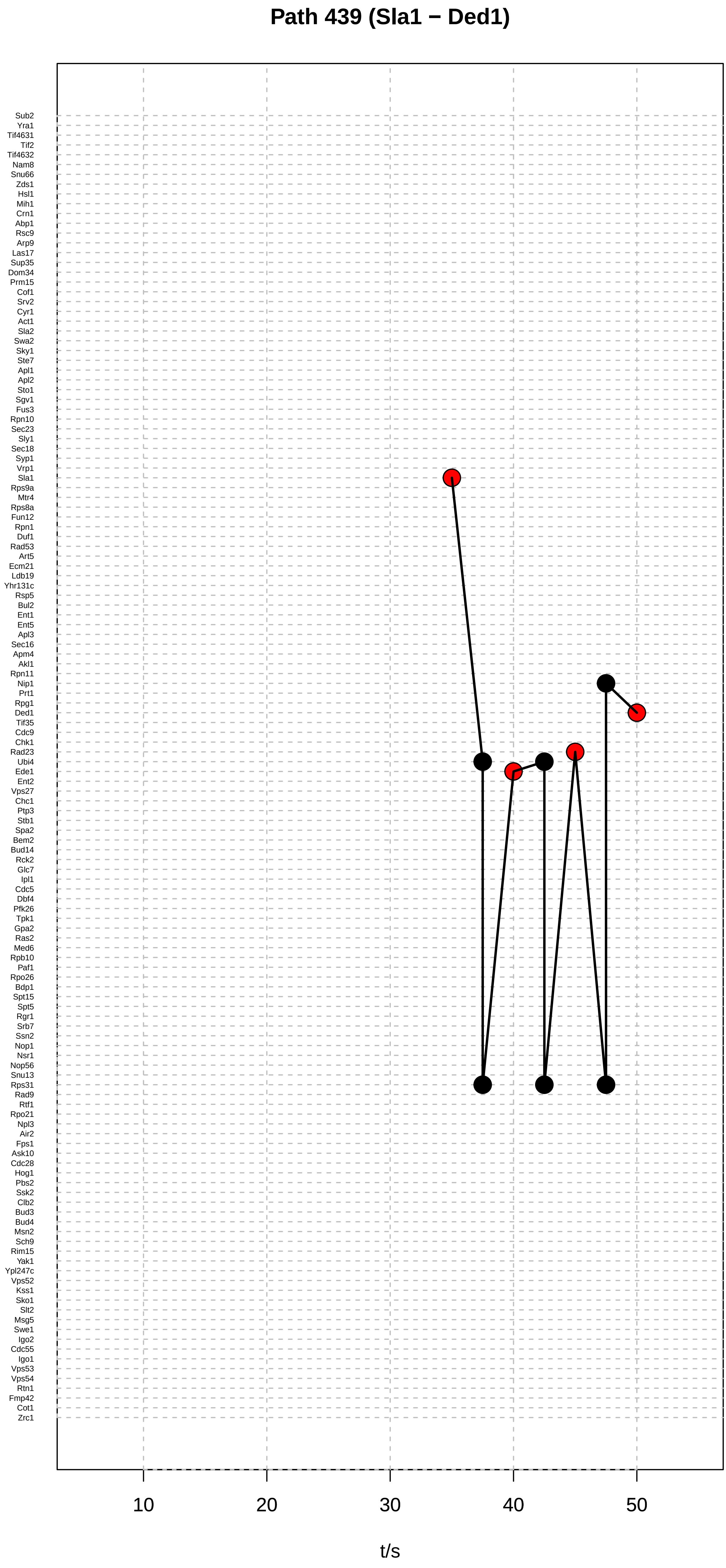

Path 439 (Sla1 − Ded1)
Sub2
Yra1
Tif4631
Tif2
Tif4632
Nam8
Snu66
Zds1
Hsl1
Mih1
Crn1
Abp1
Rsc9
Arp9
Las17
Sup35
Dom34
Prm15
Cof1
Srv2
Cyr1
Act1
Sla2
Swa2
Sky1
Ste7
Apl1
Apl2
Sto1
Sgv1
Fus3
Rpn10
Sec23
Sly1
Sec18
Syp1
Vrp1
Sla1
Rps9a
Mtr4
Rps8a
Fun12
Rpn1
Duf1
Rad53
Art5
Ecm21
Ldb19
Yhr131c
Rsp5
Bul2
Ent1
Ent5
Apl3
Sec16
Apm4
Akl1
Rpn11
Nip1
Prt1
Rpg1
Ded1
Tif35
Cdc9
Chk1
Rad23
Ubi4
Ede1
Ent2
Vps27
Chc1
Ptp3
Stb1
Spa2
Bem2
Bud14
Rck2
Glc7
Ipl1
Cdc5
Dbf4
Pfk26
Tpk1
Gpa2
Ras2
Med6
Rpb10
Paf1
Rpo26
Bdp1
Spt15
Spt5
Rgr1
Srb7
Ssn2
Nop1
Nsr1
Nop56
Snu13
Rps31
Rad9
Rtf1
Rpo21
Npl3
Air2
Fps1
Ask10
Cdc28
Hog1
Pbs2
Ssk2
Clb2
Bud3
Bud4
Msn2
Sch9
Rim15
Yak1
Ypl247c
Vps52
Kss1
Sko1
Slt2
Msg5
Swe1
Igo2
Cdc55
Igo1
Vps53
Vps54
Rtn1
Fmp42
Cot1
Zrc1
10
20
30
40
50
t/s

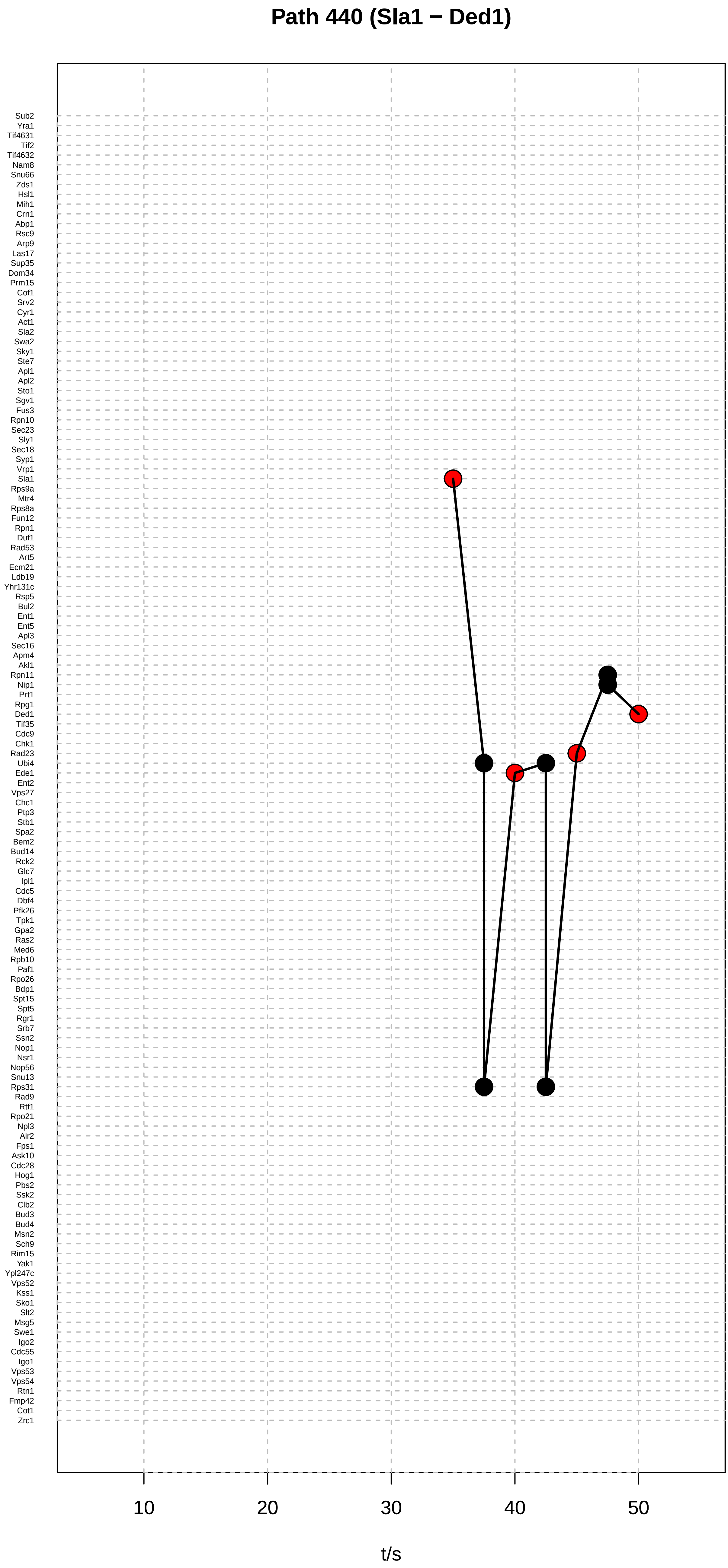

Path 440 (Sla1 − Ded1)
Sub2
Yra1
Tif4631
Tif2
Tif4632
Nam8
Snu66
Zds1
Hsl1
Mih1
Crn1
Abp1
Rsc9
Arp9
Las17
Sup35
Dom34
Prm15
Cof1
Srv2
Cyr1
Act1
Sla2
Swa2
Sky1
Ste7
Apl1
Apl2
Sto1
Sgv1
Fus3
Rpn10
Sec23
Sly1
Sec18
Syp1
Vrp1
Sla1
Rps9a
Mtr4
Rps8a
Fun12
Rpn1
Duf1
Rad53
Art5
Ecm21
Ldb19
Yhr131c
Rsp5
Bul2
Ent1
Ent5
Apl3
Sec16
Apm4
Akl1
Rpn11
Nip1
Prt1
Rpg1
Ded1
Tif35
Cdc9
Chk1
Rad23
Ubi4
Ede1
Ent2
Vps27
Chc1
Ptp3
Stb1
Spa2
Bem2
Bud14
Rck2
Glc7
Ipl1
Cdc5
Dbf4
Pfk26
Tpk1
Gpa2
Ras2
Med6
Rpb10
Paf1
Rpo26
Bdp1
Spt15
Spt5
Rgr1
Srb7
Ssn2
Nop1
Nsr1
Nop56
Snu13
Rps31
Rad9
Rtf1
Rpo21
Npl3
Air2
Fps1
Ask10
Cdc28
Hog1
Pbs2
Ssk2
Clb2
Bud3
Bud4
Msn2
Sch9
Rim15
Yak1
Ypl247c
Vps52
Kss1
Sko1
Slt2
Msg5
Swe1
Igo2
Cdc55
Igo1
Vps53
Vps54
Rtn1
Fmp42
Cot1
Zrc1
10
20
30
40
50
t/s

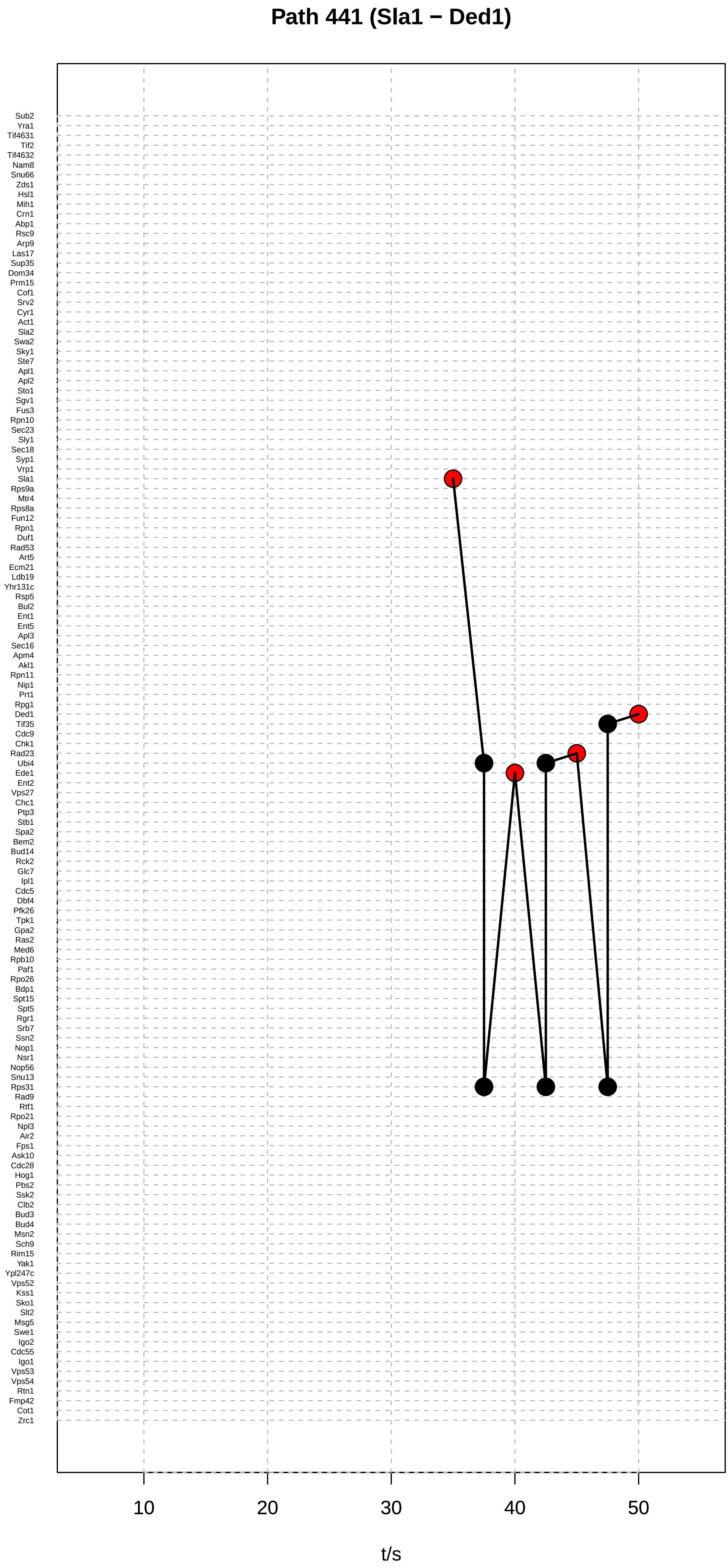
Path 441 (Sla1 − Ded1)
Sub2
Yra1
Tif4631
Tif2
Tif4632
Nam8
Snu66
Zds1
Hsl1
Mih1
Crn1
Abp1
Rsc9
Arp9
Las17
Sup35
Dom34
Prm15
Cof1
Srv2
Cyr1
Act1
Sla2
Swa2
Sky1
Ste7
Apl1
Apl2
Sto1
Sgv1
Fus3
Rpn10
Sec23
Sly1
Sec18
Syp1
Vrp1
Sla1
Rps9a
Mtr4
Rps8a
Fun12
Rpn1
Duf1
Rad53
Art5
Ecm21
Ldb19
Yhr131c
Rsp5
Bul2
Ent1
Ent5
Apl3
Sec16
Apm4
Akl1
Rpn11
Nip1
Prt1
Rpg1
Ded1
Tif35
Cdc9
Chk1
Rad23
Ubi4
Ede1
Ent2
Vps27
Chc1
Ptp3
Stb1
Spa2
Bem2
Bud14
Rck2
Glc7
Ipl1
Cdc5
Dbf4
Pfk26
Tpk1
Gpa2
Ras2
Med6
Rpb10
Paf1
Rpo26
Bdp1
Spt15
Spt5
Rgr1
Srb7
Ssn2
Nop1
Nsr1
Nop56
Snu13
Rps31
Rad9
Rtf1
Rpo21
Npl3
Air2
Fps1
Ask10
Cdc28
Hog1
Pbs2
Ssk2
Clb2
Bud3
Bud4
Msn2
Sch9
Rim15
Yak1
Ypl247c
Vps52
Kss1
Sko1
Slt2
Msg5
Swe1
Igo2
Cdc55
Igo1
Vps53
Vps54
Rtn1
Fmp42
Cot1
Zrc1
10
20
30
40
50
t/s

#### Path 442 (Sla1 − Ded1)

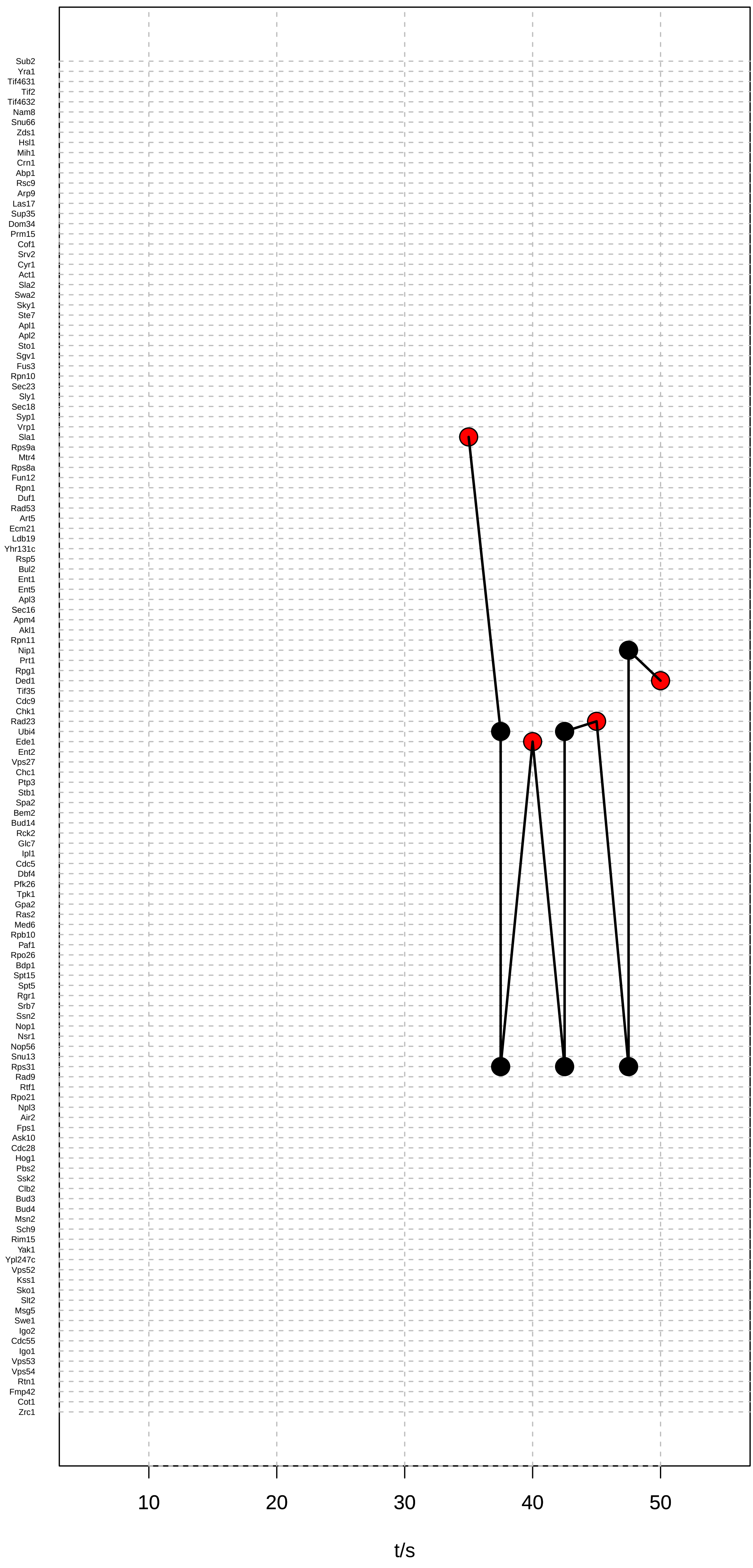

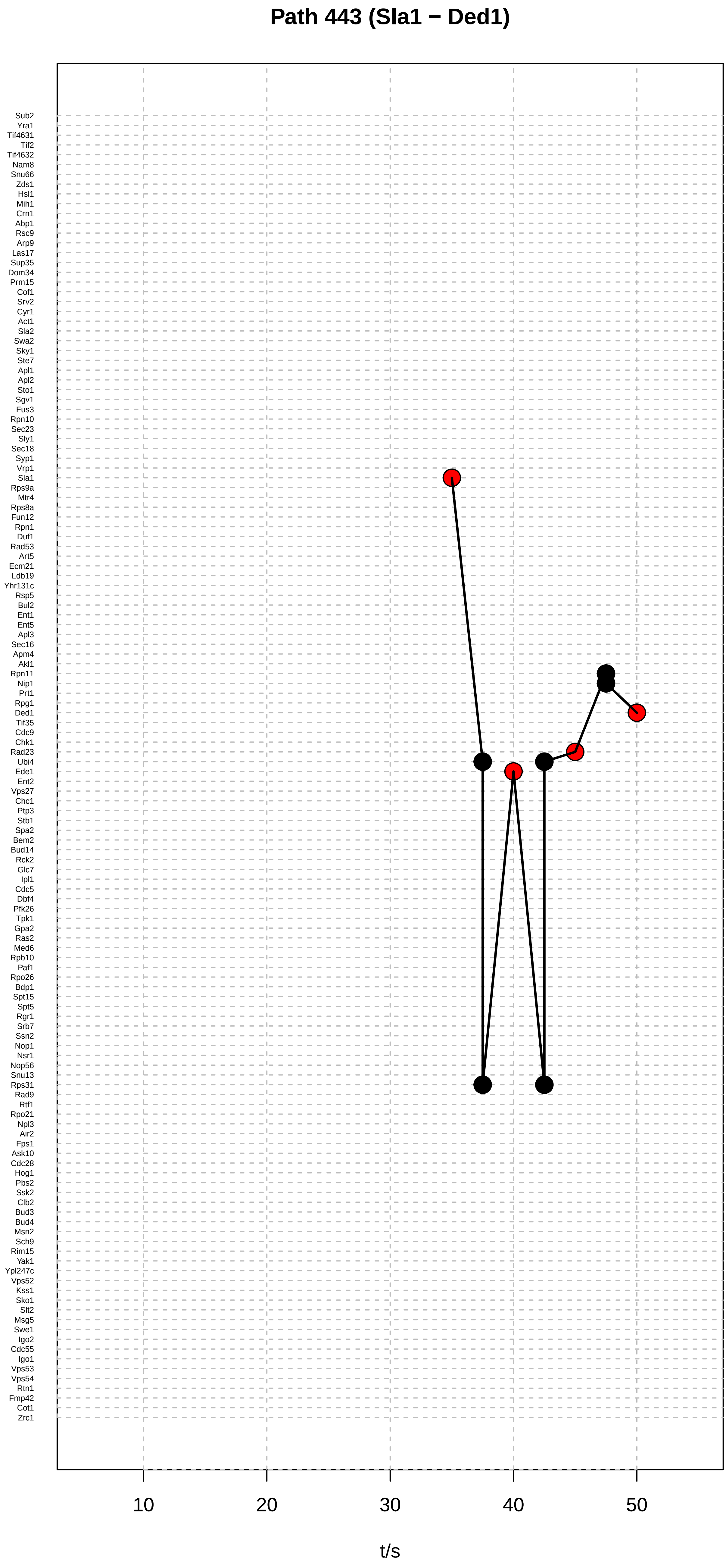

Path 443 (Sla1 − Ded1)
Sub2
Yra1
Tif4631
Tif2
Tif4632
Nam8
Snu66
Zds1
Hsl1
Mih1
Crn1
Abp1
Rsc9
Arp9
Las17
Sup35
Dom34
Prm15
Cof1
Srv2
Cyr1
Act1
Sla2
Swa2
Sky1
Ste7
Apl1
Apl2
Sto1
Sgv1
Fus3
Rpn10
Sec23
Sly1
Sec18
Syp1
Vrp1
Sla1
Rps9a
Mtr4
Rps8a
Fun12
Rpn1
Duf1
Rad53
Art5
Ecm21
Ldb19
Yhr131c
Rsp5
Bul2
Ent1
Ent5
Apl3
Sec16
Apm4
Akl1
Rpn11
Nip1
Prt1
Rpg1
Ded1
Tif35
Cdc9
Chk1
Rad23
Ubi4
Ede1
Ent2
Vps27
Chc1
Ptp3
Stb1
Spa2
Bem2
Bud14
Rck2
Glc7
Ipl1
Cdc5
Dbf4
Pfk26
Tpk1
Gpa2
Ras2
Med6
Rpb10
Paf1
Rpo26
Bdp1
Spt15
Spt5
Rgr1
Srb7
Ssn2
Nop1
Nsr1
Nop56
Snu13
Rps31
Rad9
Rtf1
Rpo21
Npl3
Air2
Fps1
Ask10
Cdc28
Hog1
Pbs2
Ssk2
Clb2
Bud3
Bud4
Msn2
Sch9
Rim15
Yak1
Ypl247c
Vps52
Kss1
Sko1
Slt2
Msg5
Swe1
Igo2
Cdc55
Igo1
Vps53
Vps54
Rtn1
Fmp42
Cot1
Zrc1
10
20
30
40
50
t/s

#### Path 444 (Sla1 − Ded1)

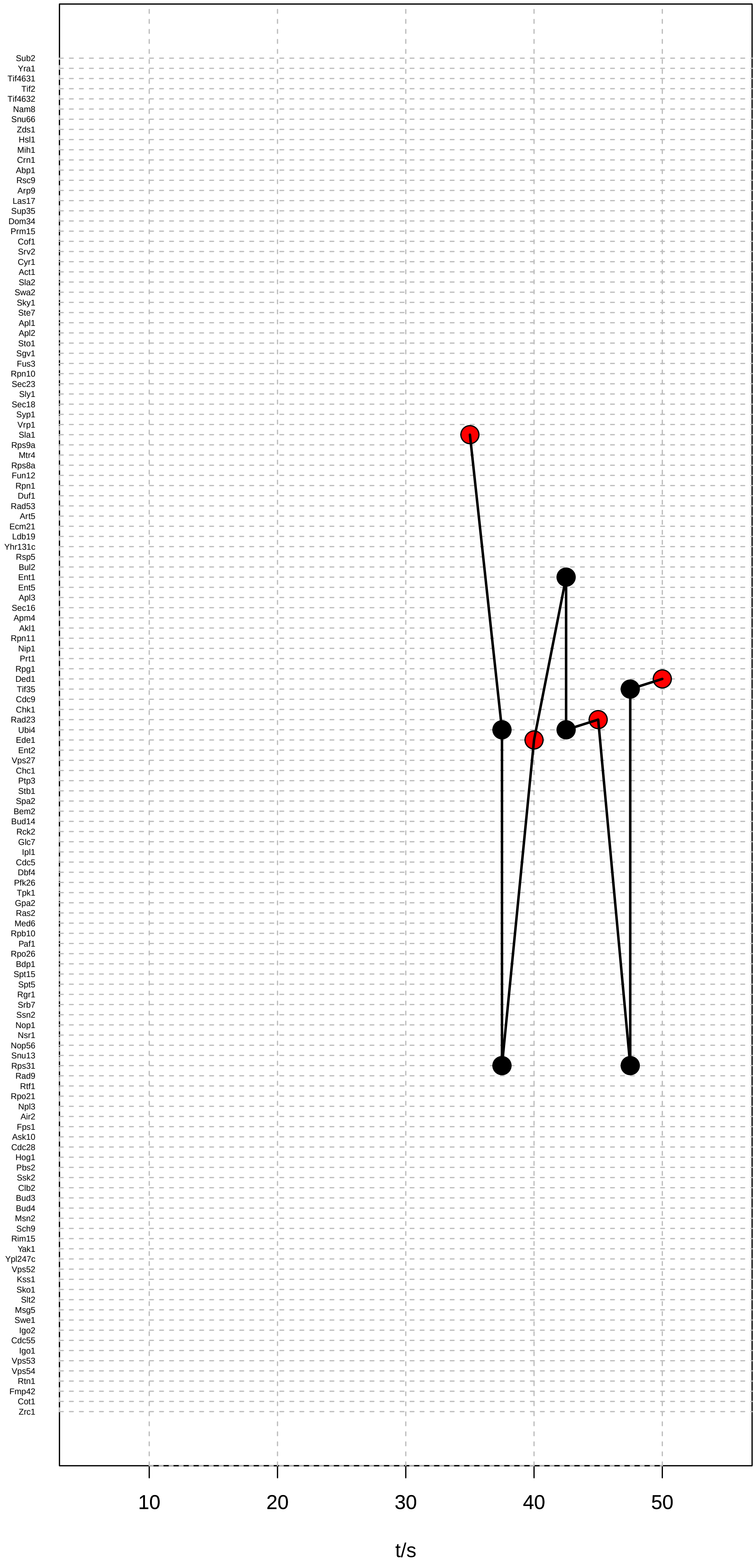

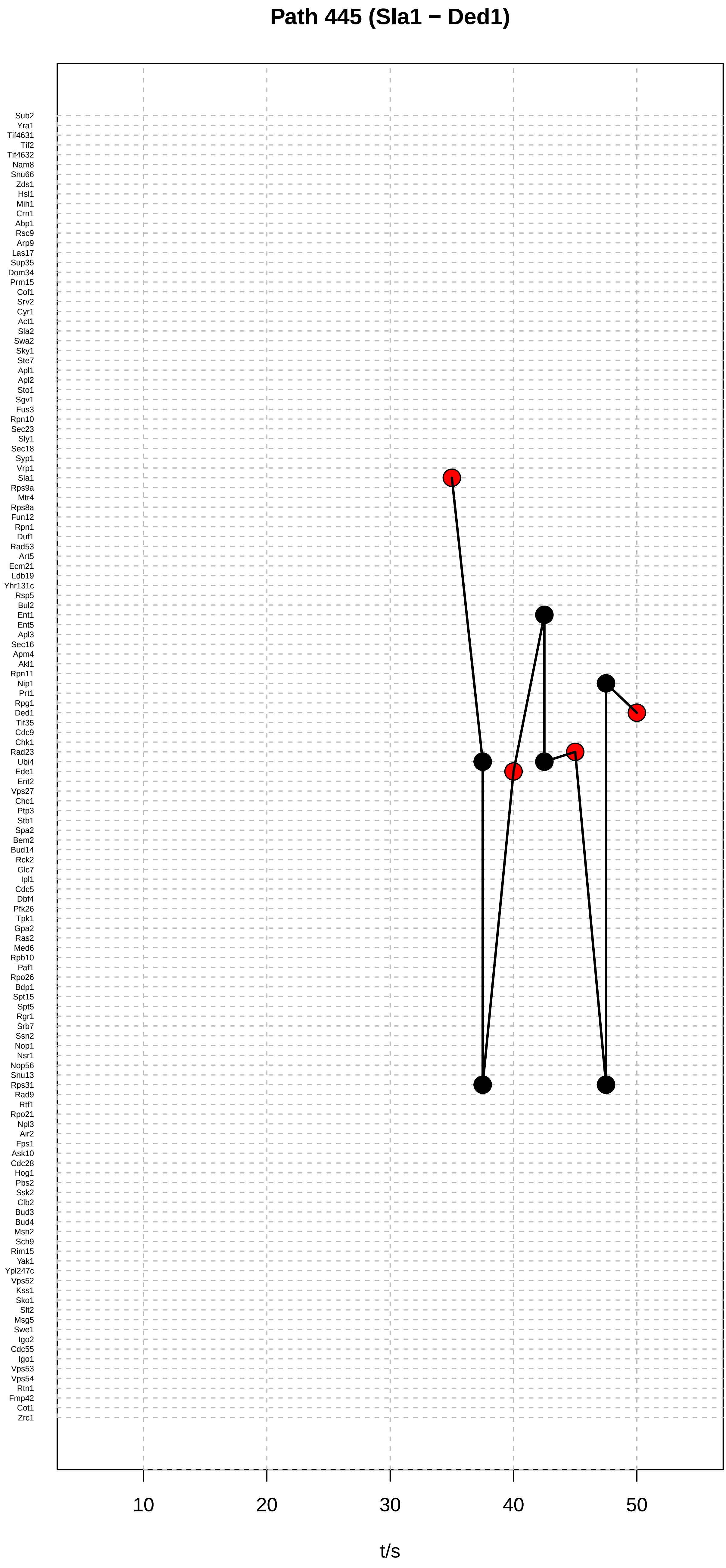

Path 445 (Sla1 − Ded1)
Sub2
Yra1
Tif4631
Tif2
Tif4632
Nam8
Snu66
Zds1
Hsl1
Mih1
Crn1
Abp1
Rsc9
Arp9
Las17
Sup35
Dom34
Prm15
Cof1
Srv2
Cyr1
Act1
Sla2
Swa2
Sky1
Ste7
Apl1
Apl2
Sto1
Sgv1
Fus3
Rpn10
Sec23
Sly1
Sec18
Syp1
Vrp1
Sla1
Rps9a
Mtr4
Rps8a
Fun12
Rpn1
Duf1
Rad53
Art5
Ecm21
Ldb19
Yhr131c
Rsp5
Bul2
Ent1
Ent5
Apl3
Sec16
Apm4
Akl1
Rpn11
Nip1
Prt1
Rpg1
Ded1
Tif35
Cdc9
Chk1
Rad23
Ubi4
Ede1
Ent2
Vps27
Chc1
Ptp3
Stb1
Spa2
Bem2
Bud14
Rck2
Glc7
Ipl1
Cdc5
Dbf4
Pfk26
Tpk1
Gpa2
Ras2
Med6
Rpb10
Paf1
Rpo26
Bdp1
Spt15
Spt5
Rgr1
Srb7
Ssn2
Nop1
Nsr1
Nop56
Snu13
Rps31
Rad9
Rtf1
Rpo21
Npl3
Air2
Fps1
Ask10
Cdc28
Hog1
Pbs2
Ssk2
Clb2
Bud3
Bud4
Msn2
Sch9
Rim15
Yak1
Ypl247c
Vps52
Kss1
Sko1
Slt2
Msg5
Swe1
Igo2
Cdc55
Igo1
Vps53
Vps54
Rtn1
Fmp42
Cot1
Zrc1
10
20
30
40
50
t/s

### Path 446 (Sla1 − Ded1)

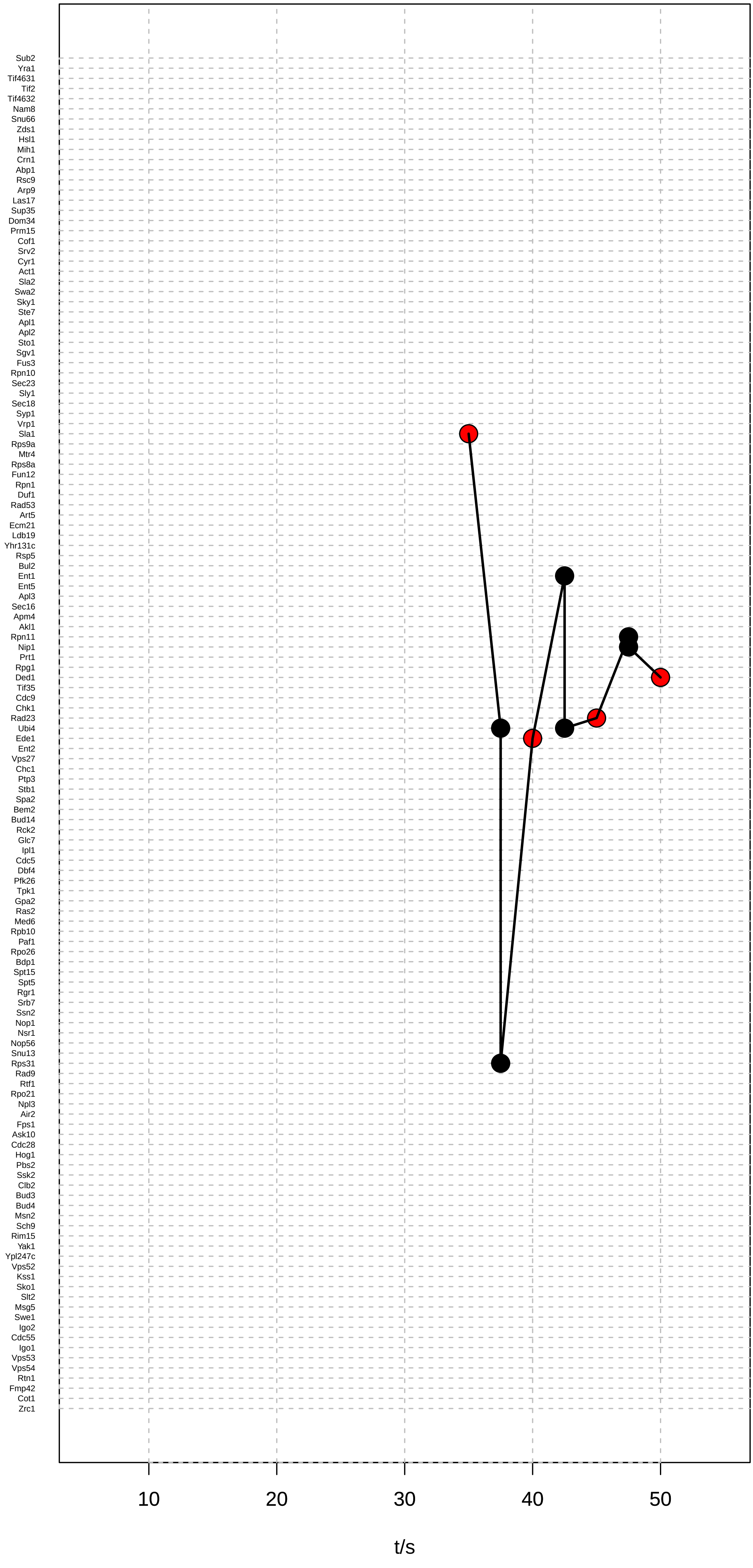

### Path 447 (Sla1 − Rsc9)

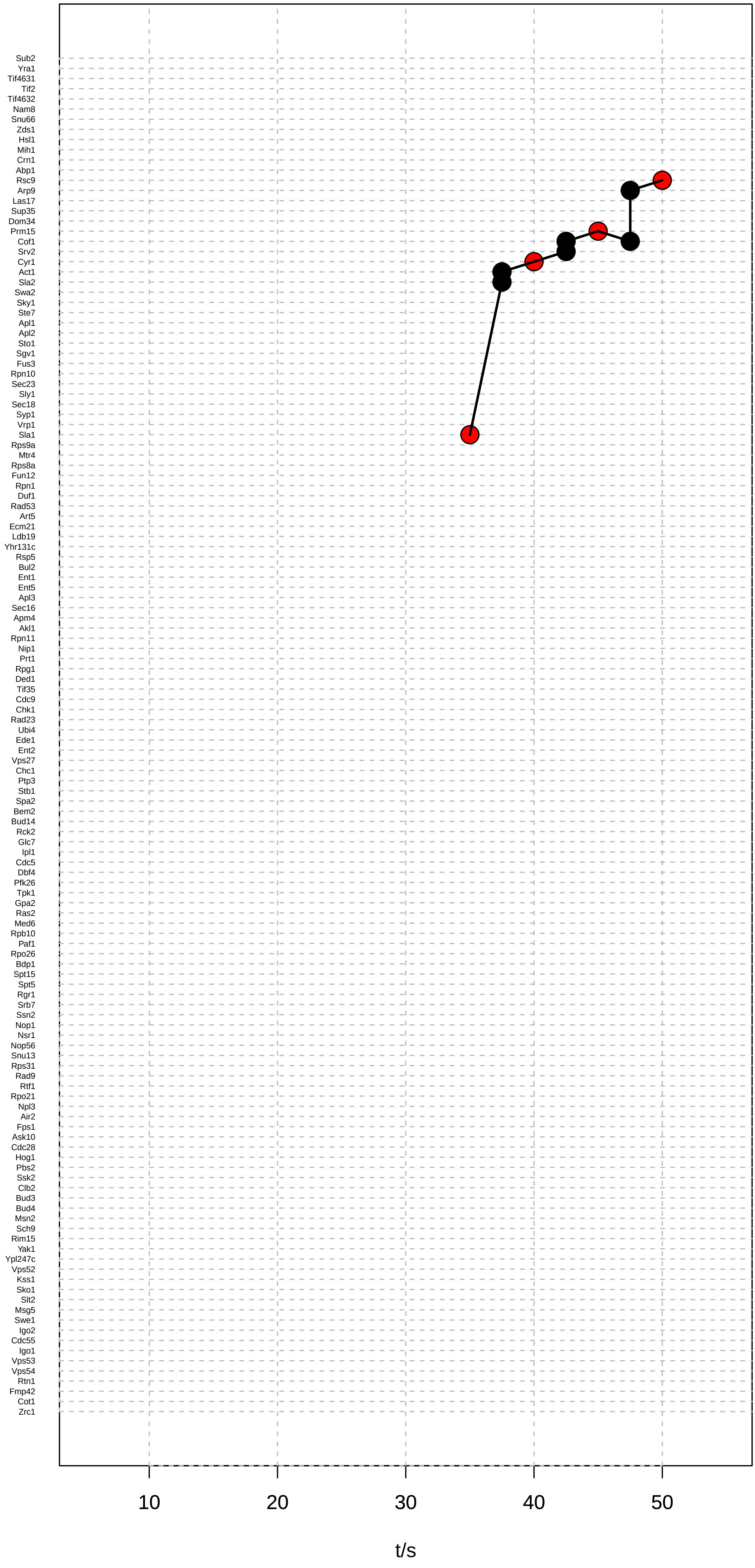

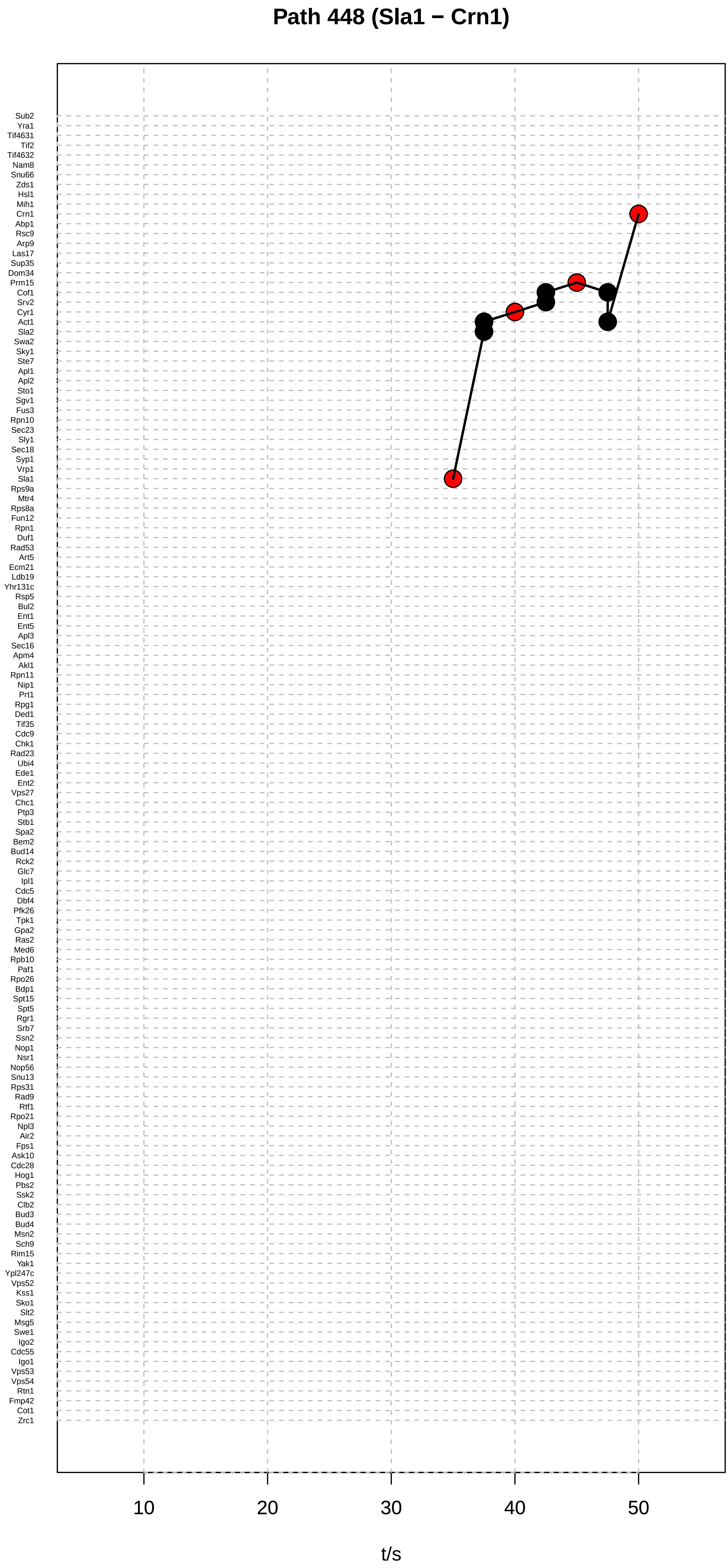
Path 448 (Sla1 − Crn1)
Sub2
Yra1
Tif4631
Tif2
Tif4632
Nam8
Snu66
Zds1
Hsl1
Mih1
Crn1
Abp1
Rsc9
Arp9
Las17
Sup35
Dom34
Prm15
Cof1
Srv2
Cyr1
Act1
Sla2
Swa2
Sky1
Ste7
Apl1
Apl2
Sto1
Sgv1
Fus3
Rpn10
Sec23
Sly1
Sec18
Syp1
Vrp1
Sla1
Rps9a
Mtr4
Rps8a
Fun12
Rpn1
Duf1
Rad53
Art5
Ecm21
Ldb19
Yhr131c
Rsp5
Bul2
Ent1
Ent5
Apl3
Sec16
Apm4
Akl1
Rpn11
Nip1
Prt1
Rpg1
Ded1
Tif35
Cdc9
Chk1
Rad23
Ubi4
Ede1
Ent2
Vps27
Chc1
Ptp3
Stb1
Spa2
Bem2
Bud14
Rck2
Glc7
Ipl1
Cdc5
Dbf4
Pfk26
Tpk1
Gpa2
Ras2
Med6
Rpb10
Paf1
Rpo26
Bdp1
Spt15
Spt5
Rgr1
Srb7
Ssn2
Nop1
Nsr1
Nop56
Snu13
Rps31
Rad9
Rtf1
Rpo21
Npl3
Air2
Fps1
Ask10
Cdc28
Hog1
Pbs2
Ssk2
Clb2
Bud3
Bud4
Msn2
Sch9
Rim15
Yak1
Ypl247c
Vps52
Kss1
Sko1
Slt2
Msg5
Swe1
Igo2
Cdc55
Igo1
Vps53
Vps54
Rtn1
Fmp42
Cot1
Zrc1
10
20
30
40
50
t/s

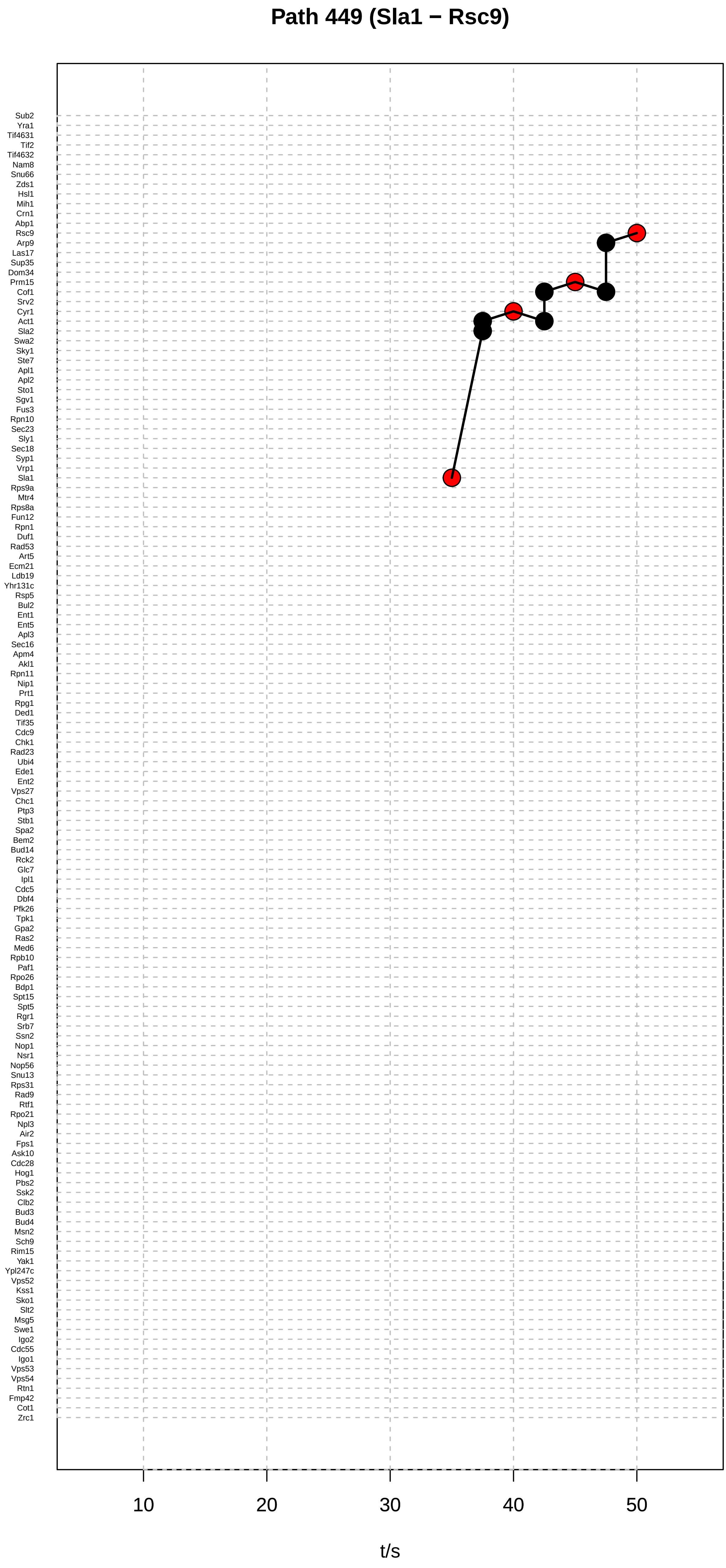
Path 449 (Sla1 − Rsc9)
Sub2
Yra1
Tif4631
Tif2
Tif4632
Nam8
Snu66
Zds1
Hsl1
Mih1
Crn1
Abp1
Rsc9
Arp9
Las17
Sup35
Dom34
Prm15
Cof1
Srv2
Cyr1
Act1
Sla2
Swa2
Sky1
Ste7
Apl1
Apl2
Sto1
Sgv1
Fus3
Rpn10
Sec23
Sly1
Sec18
Syp1
Vrp1
Sla1
Rps9a
Mtr4
Rps8a
Fun12
Rpn1
Duf1
Rad53
Art5
Ecm21
Ldb19
Yhr131c
Rsp5
Bul2
Ent1
Ent5
Apl3
Sec16
Apm4
Akl1
Rpn11
Nip1
Prt1
Rpg1
Ded1
Tif35
Cdc9
Chk1
Rad23
Ubi4
Ede1
Ent2
Vps27
Chc1
Ptp3
Stb1
Spa2
Bem2
Bud14
Rck2
Glc7
Ipl1
Cdc5
Dbf4
Pfk26
Tpk1
Gpa2
Ras2
Med6
Rpb10
Paf1
Rpo26
Bdp1
Spt15
Spt5
Rgr1
Srb7
Ssn2
Nop1
Nsr1
Nop56
Snu13
Rps31
Rad9
Rtf1
Rpo21
Npl3
Air2
Fps1
Ask10
Cdc28
Hog1
Pbs2
Ssk2
Clb2
Bud3
Bud4
Msn2
Sch9
Rim15
Yak1
Ypl247c
Vps52
Kss1
Sko1
Slt2
Msg5
Swe1
Igo2
Cdc55
Igo1
Vps53
Vps54
Rtn1
Fmp42
Cot1
Zrc1
10
20
30
40
50
t/s

### Path 450 (Sla1 − Crn1)

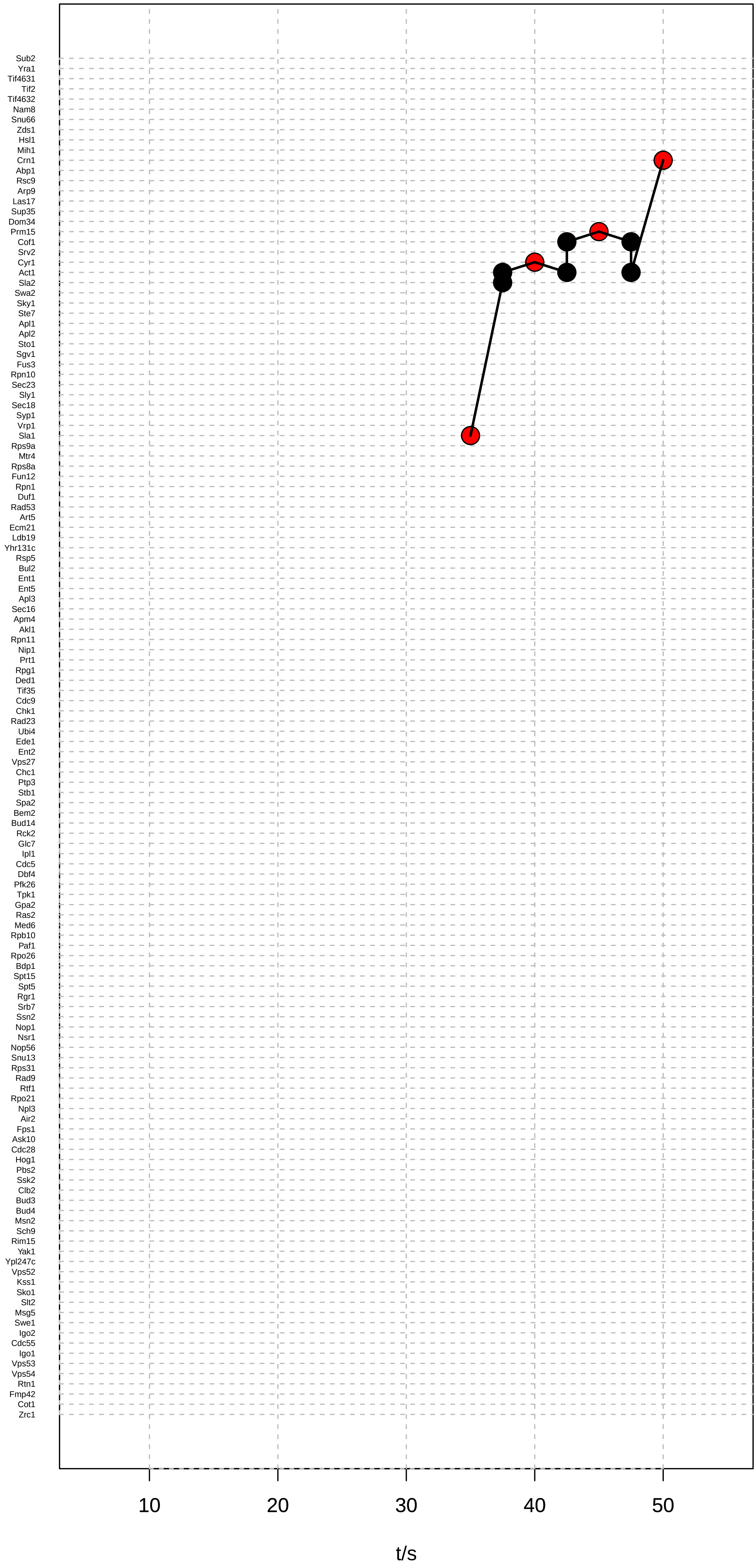

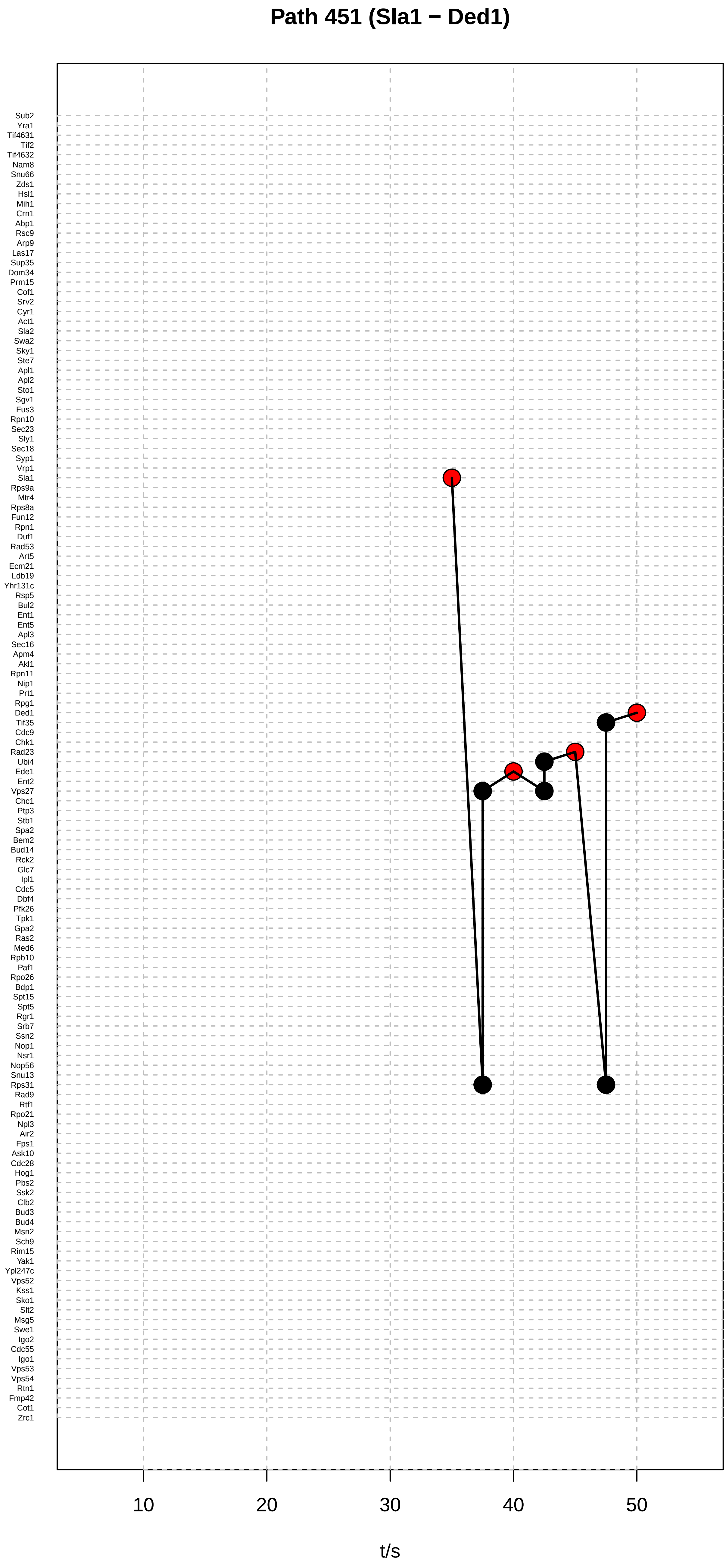

Path 451 (Sla1 − Ded1)
Sub2
Yra1
Tif4631
Tif2
Tif4632
Nam8
Snu66
Zds1
Hsl1
Mih1
Crn1
Abp1
Rsc9
Arp9
Las17
Sup35
Dom34
Prm15
Cof1
Srv2
Cyr1
Act1
Sla2
Swa2
Sky1
Ste7
Apl1
Apl2
Sto1
Sgv1
Fus3
Rpn10
Sec23
Sly1
Sec18
Syp1
Vrp1
Sla1
Rps9a
Mtr4
Rps8a
Fun12
Rpn1
Duf1
Rad53
Art5
Ecm21
Ldb19
Yhr131c
Rsp5
Bul2
Ent1
Ent5
Apl3
Sec16
Apm4
Akl1
Rpn11
Nip1
Prt1
Rpg1
Ded1
Tif35
Cdc9
Chk1
Rad23
Ubi4
Ede1
Ent2
Vps27
Chc1
Ptp3
Stb1
Spa2
Bem2
Bud14
Rck2
Glc7
Ipl1
Cdc5
Dbf4
Pfk26
Tpk1
Gpa2
Ras2
Med6
Rpb10
Paf1
Rpo26
Bdp1
Spt15
Spt5
Rgr1
Srb7
Ssn2
Nop1
Nsr1
Nop56
Snu13
Rps31
Rad9
Rtf1
Rpo21
Npl3
Air2
Fps1
Ask10
Cdc28
Hog1
Pbs2
Ssk2
Clb2
Bud3
Bud4
Msn2
Sch9
Rim15
Yak1
Ypl247c
Vps52
Kss1
Sko1
Slt2
Msg5
Swe1
Igo2
Cdc55
Igo1
Vps53
Vps54
Rtn1
Fmp42
Cot1
Zrc1
10
20
30
40
50
t/s

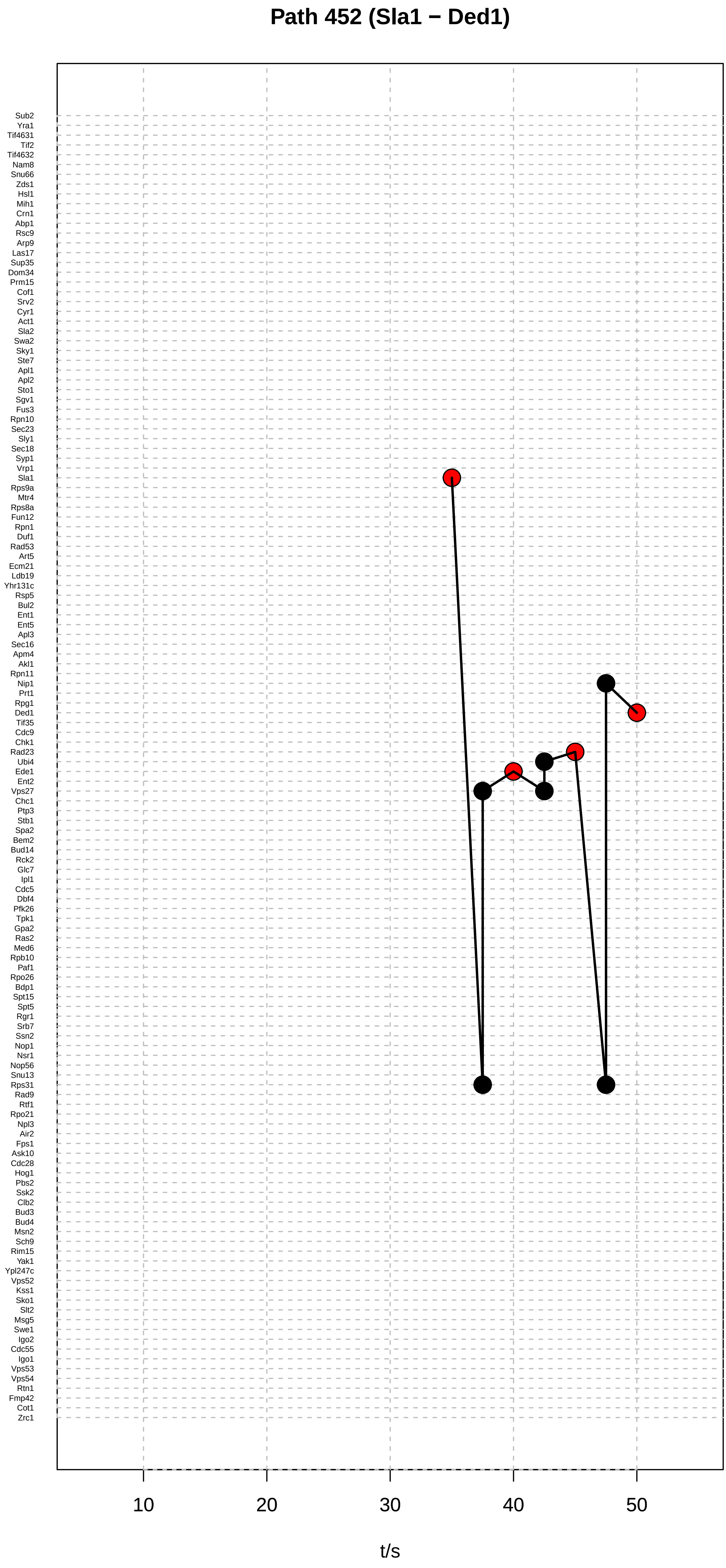

Path 452 (Sla1 − Ded1)
Sub2
Yra1
Tif4631
Tif2
Tif4632
Nam8
Snu66
Zds1
Hsl1
Mih1
Crn1
Abp1
Rsc9
Arp9
Las17
Sup35
Dom34
Prm15
Cof1
Srv2
Cyr1
Act1
Sla2
Swa2
Sky1
Ste7
Apl1
Apl2
Sto1
Sgv1
Fus3
Rpn10
Sec23
Sly1
Sec18
Syp1
Vrp1
Sla1
Rps9a
Mtr4
Rps8a
Fun12
Rpn1
Duf1
Rad53
Art5
Ecm21
Ldb19
Yhr131c
Rsp5
Bul2
Ent1
Ent5
Apl3
Sec16
Apm4
Akl1
Rpn11
Nip1
Prt1
Rpg1
Ded1
Tif35
Cdc9
Chk1
Rad23
Ubi4
Ede1
Ent2
Vps27
Chc1
Ptp3
Stb1
Spa2
Bem2
Bud14
Rck2
Glc7
Ipl1
Cdc5
Dbf4
Pfk26
Tpk1
Gpa2
Ras2
Med6
Rpb10
Paf1
Rpo26
Bdp1
Spt15
Spt5
Rgr1
Srb7
Ssn2
Nop1
Nsr1
Nop56
Snu13
Rps31
Rad9
Rtf1
Rpo21
Npl3
Air2
Fps1
Ask10
Cdc28
Hog1
Pbs2
Ssk2
Clb2
Bud3
Bud4
Msn2
Sch9
Rim15
Yak1
Ypl247c
Vps52
Kss1
Sko1
Slt2
Msg5
Swe1
Igo2
Cdc55
Igo1
Vps53
Vps54
Rtn1
Fmp42
Cot1
Zrc1
10
20
30
40
50
t/s

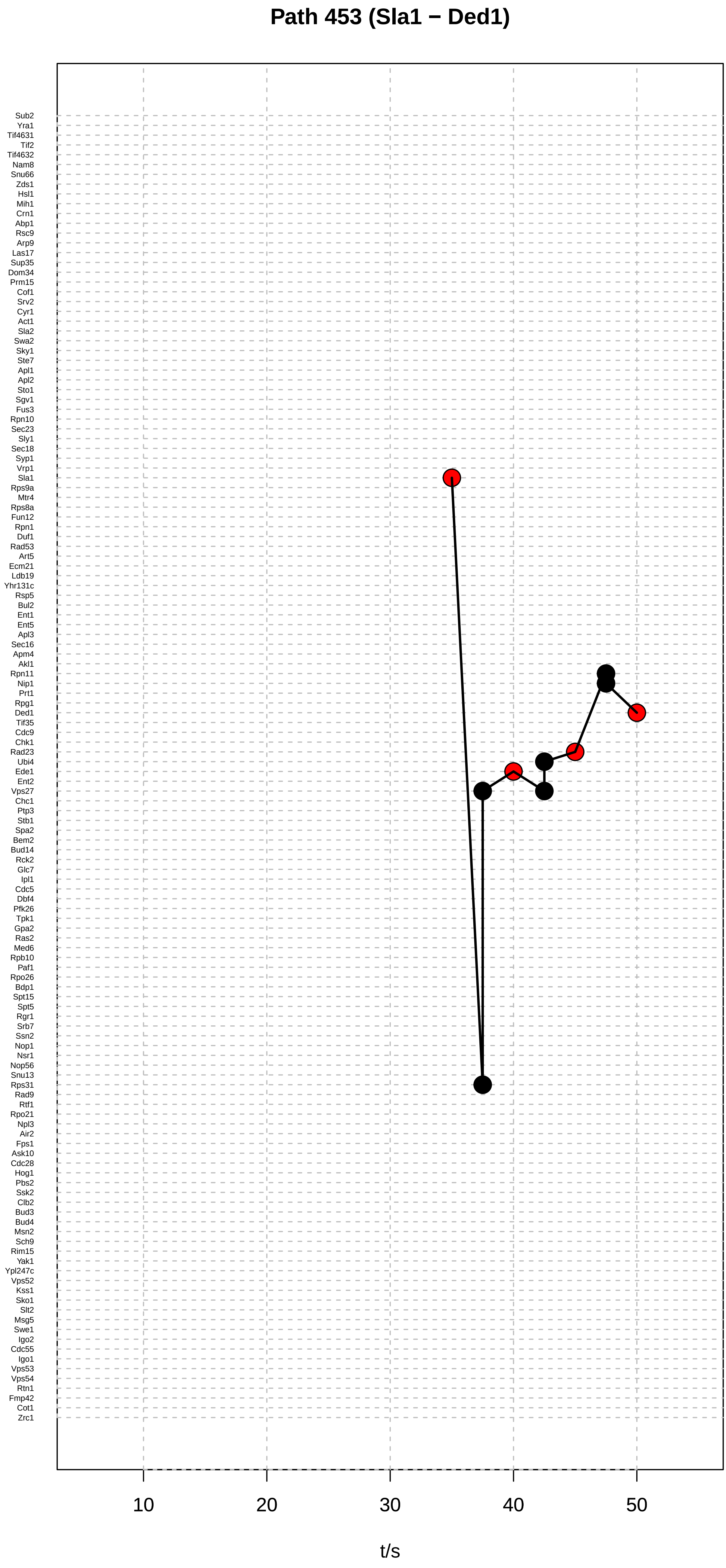
Path 453 (Sla1 − Ded1)
Sub2
Yra1
Tif4631
Tif2
Tif4632
Nam8
Snu66
Zds1
Hsl1
Mih1
Crn1
Abp1
Rsc9
Arp9
Las17
Sup35
Dom34
Prm15
Cof1
Srv2
Cyr1
Act1
Sla2
Swa2
Sky1
Ste7
Apl1
Apl2
Sto1
Sgv1
Fus3
Rpn10
Sec23
Sly1
Sec18
Syp1
Vrp1
Sla1
Rps9a
Mtr4
Rps8a
Fun12
Rpn1
Duf1
Rad53
Art5
Ecm21
Ldb19
Yhr131c
Rsp5
Bul2
Ent1
Ent5
Apl3
Sec16
Apm4
Akl1
Rpn11
Nip1
Prt1
Rpg1
Ded1
Tif35
Cdc9
Chk1
Rad23
Ubi4
Ede1
Ent2
Vps27
Chc1
Ptp3
Stb1
Spa2
Bem2
Bud14
Rck2
Glc7
Ipl1
Cdc5
Dbf4
Pfk26
Tpk1
Gpa2
Ras2
Med6
Rpb10
Paf1
Rpo26
Bdp1
Spt15
Spt5
Rgr1
Srb7
Ssn2
Nop1
Nsr1
Nop56
Snu13
Rps31
Rad9
Rtf1
Rpo21
Npl3
Air2
Fps1
Ask10
Cdc28
Hog1
Pbs2
Ssk2
Clb2
Bud3
Bud4
Msn2
Sch9
Rim15
Yak1
Ypl247c
Vps52
Kss1
Sko1
Slt2
Msg5
Swe1
Igo2
Cdc55
Igo1
Vps53
Vps54
Rtn1
Fmp42
Cot1
Zrc1
10
20
30
40
50
t/s

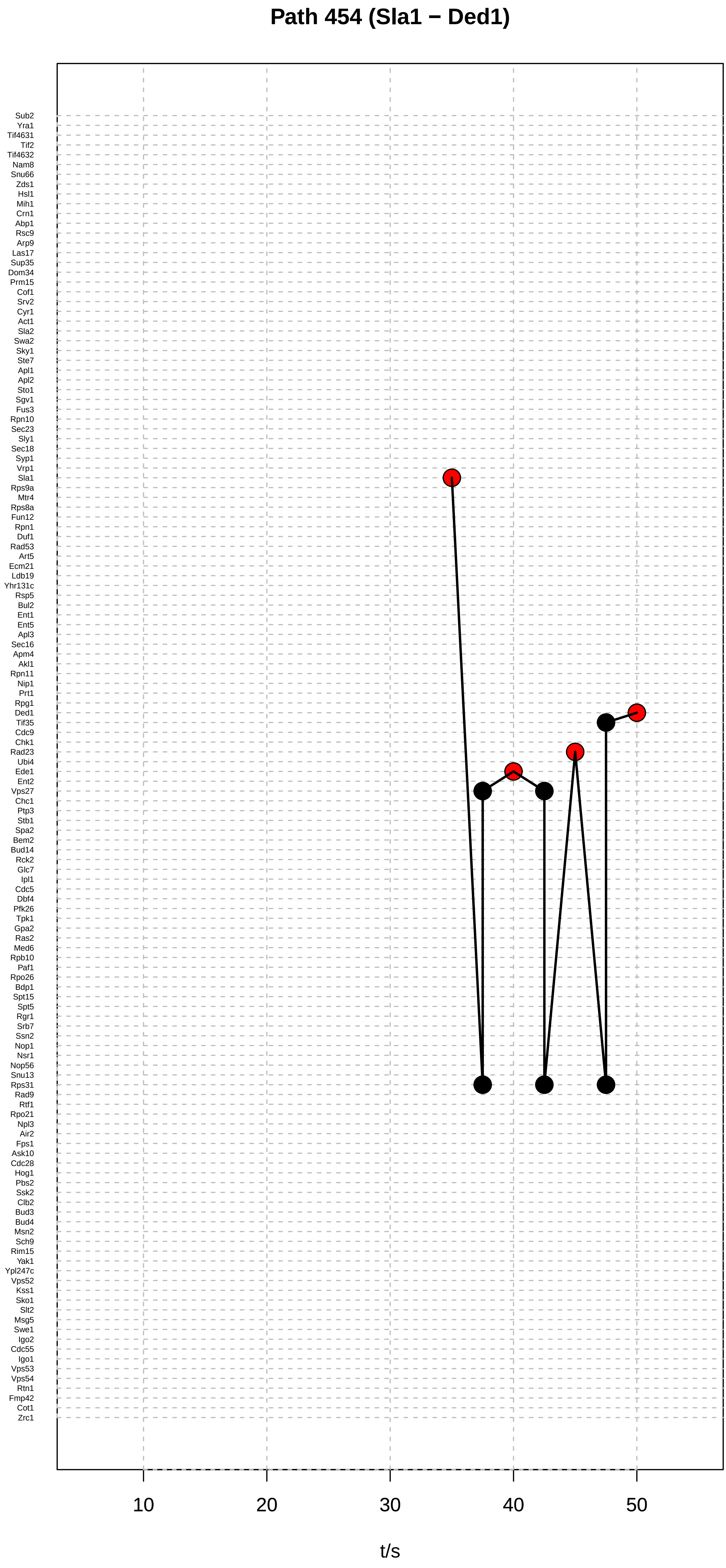

Path 454 (Sla1 − Ded1)
Sub2
Yra1
Tif4631
Tif2
Tif4632
Nam8
Snu66
Zds1
Hsl1
Mih1
Crn1
Abp1
Rsc9
Arp9
Las17
Sup35
Dom34
Prm15
Cof1
Srv2
Cyr1
Act1
Sla2
Swa2
Sky1
Ste7
Apl1
Apl2
Sto1
Sgv1
Fus3
Rpn10
Sec23
Sly1
Sec18
Syp1
Vrp1
Sla1
Rps9a
Mtr4
Rps8a
Fun12
Rpn1
Duf1
Rad53
Art5
Ecm21
Ldb19
Yhr131c
Rsp5
Bul2
Ent1
Ent5
Apl3
Sec16
Apm4
Akl1
Rpn11
Nip1
Prt1
Rpg1
Ded1
Tif35
Cdc9
Chk1
Rad23
Ubi4
Ede1
Ent2
Vps27
Chc1
Ptp3
Stb1
Spa2
Bem2
Bud14
Rck2
Glc7
Ipl1
Cdc5
Dbf4
Pfk26
Tpk1
Gpa2
Ras2
Med6
Rpb10
Paf1
Rpo26
Bdp1
Spt15
Spt5
Rgr1
Srb7
Ssn2
Nop1
Nsr1
Nop56
Snu13
Rps31
Rad9
Rtf1
Rpo21
Npl3
Air2
Fps1
Ask10
Cdc28
Hog1
Pbs2
Ssk2
Clb2
Bud3
Bud4
Msn2
Sch9
Rim15
Yak1
Ypl247c
Vps52
Kss1
Sko1
Slt2
Msg5
Swe1
Igo2
Cdc55
Igo1
Vps53
Vps54
Rtn1
Fmp42
Cot1
Zrc1
10
20
30
40
50
t/s

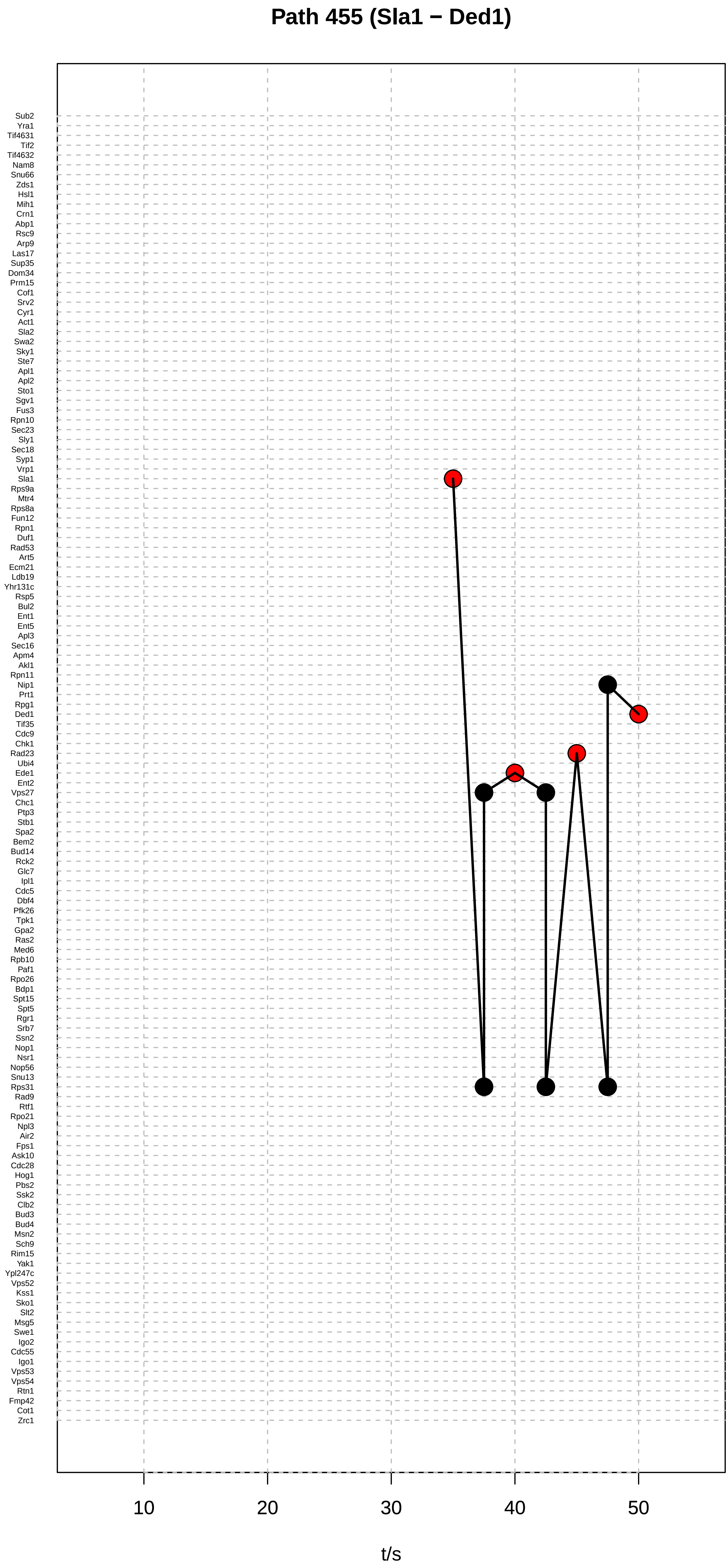
Path 455 (Sla1 − Ded1)
Sub2
Yra1
Tif4631
Tif2
Tif4632
Nam8
Snu66
Zds1
Hsl1
Mih1
Crn1
Abp1
Rsc9
Arp9
Las17
Sup35
Dom34
Prm15
Cof1
Srv2
Cyr1
Act1
Sla2
Swa2
Sky1
Ste7
Apl1
Apl2
Sto1
Sgv1
Fus3
Rpn10
Sec23
Sly1
Sec18
Syp1
Vrp1
Sla1
Rps9a
Mtr4
Rps8a
Fun12
Rpn1
Duf1
Rad53
Art5
Ecm21
Ldb19
Yhr131c
Rsp5
Bul2
Ent1
Ent5
Apl3
Sec16
Apm4
Akl1
Rpn11
Nip1
Prt1
Rpg1
Ded1
Tif35
Cdc9
Chk1
Rad23
Ubi4
Ede1
Ent2
Vps27
Chc1
Ptp3
Stb1
Spa2
Bem2
Bud14
Rck2
Glc7
Ipl1
Cdc5
Dbf4
Pfk26
Tpk1
Gpa2
Ras2
Med6
Rpb10
Paf1
Rpo26
Bdp1
Spt15
Spt5
Rgr1
Srb7
Ssn2
Nop1
Nsr1
Nop56
Snu13
Rps31
Rad9
Rtf1
Rpo21
Npl3
Air2
Fps1
Ask10
Cdc28
Hog1
Pbs2
Ssk2
Clb2
Bud3
Bud4
Msn2
Sch9
Rim15
Yak1
Ypl247c
Vps52
Kss1
Sko1
Slt2
Msg5
Swe1
Igo2
Cdc55
Igo1
Vps53
Vps54
Rtn1
Fmp42
Cot1
Zrc1
10
20
30
40
50
t/s

### Path 456 (Sla1 − Ded1)

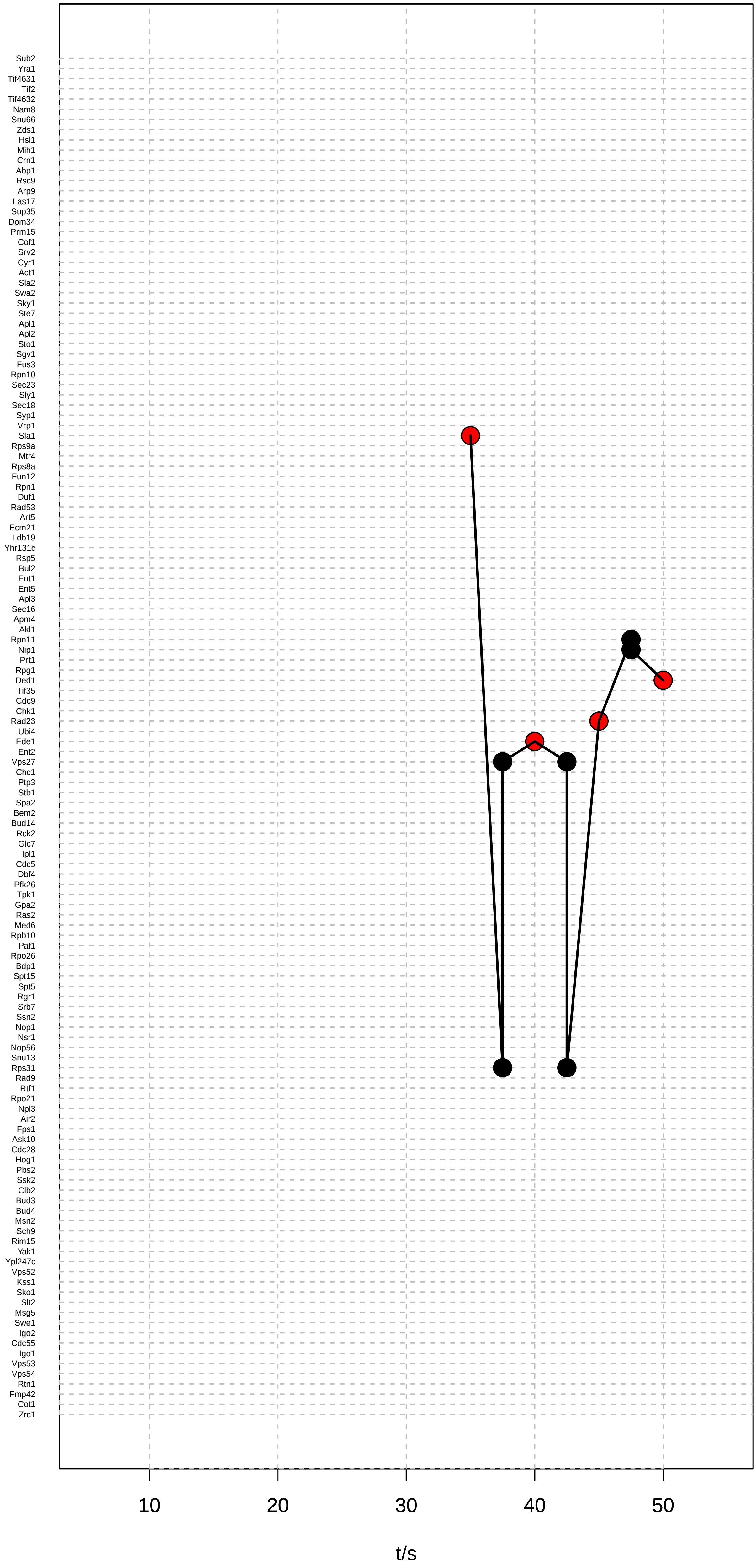

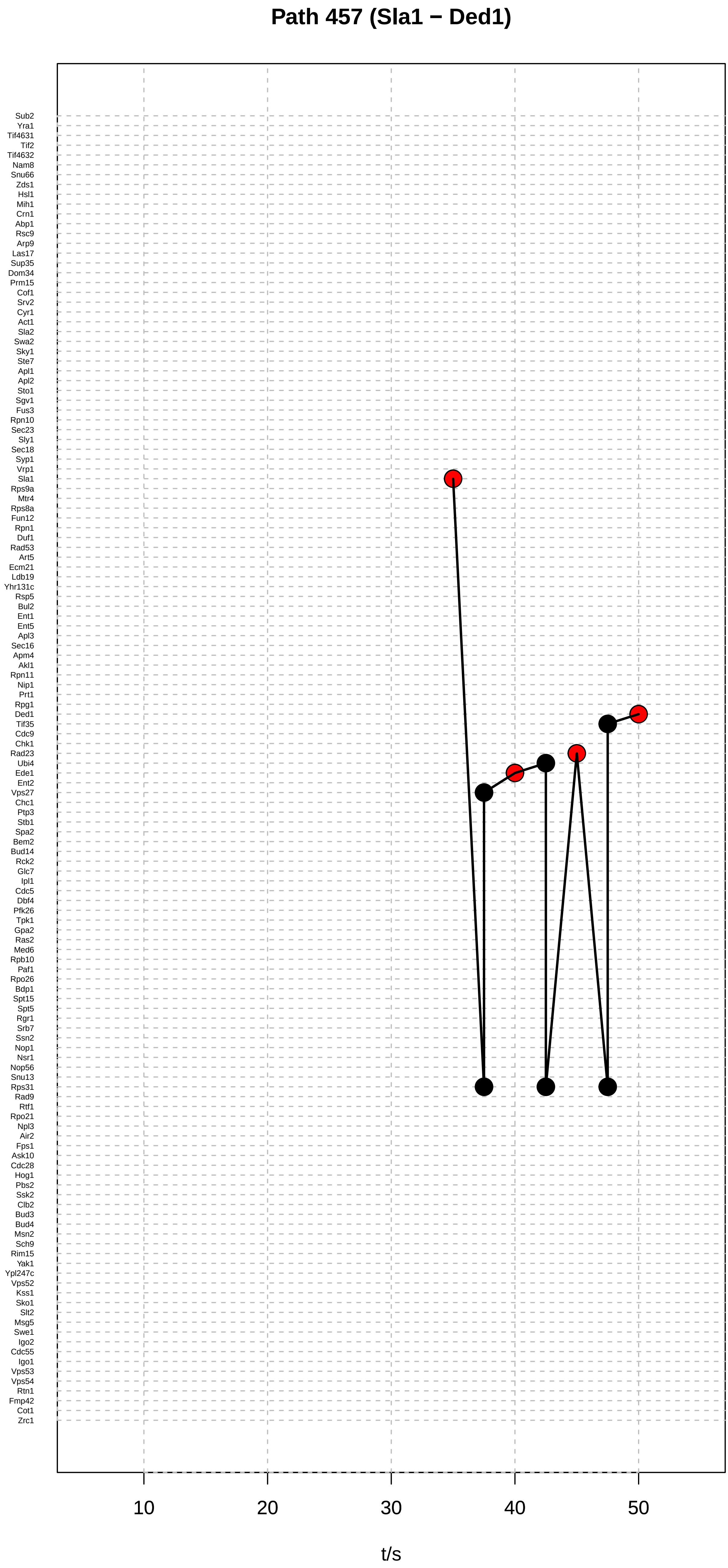
Path 457 (Sla1 − Ded1)
Sub2
Yra1
Tif4631
Tif2
Tif4632
Nam8
Snu66
Zds1
Hsl1
Mih1
Crn1
Abp1
Rsc9
Arp9
Las17
Sup35
Dom34
Prm15
Cof1
Srv2
Cyr1
Act1
Sla2
Swa2
Sky1
Ste7
Apl1
Apl2
Sto1
Sgv1
Fus3
Rpn10
Sec23
Sly1
Sec18
Syp1
Vrp1
Sla1
Rps9a
Mtr4
Rps8a
Fun12
Rpn1
Duf1
Rad53
Art5
Ecm21
Ldb19
Yhr131c
Rsp5
Bul2
Ent1
Ent5
Apl3
Sec16
Apm4
Akl1
Rpn11
Nip1
Prt1
Rpg1
Ded1
Tif35
Cdc9
Chk1
Rad23
Ubi4
Ede1
Ent2
Vps27
Chc1
Ptp3
Stb1
Spa2
Bem2
Bud14
Rck2
Glc7
Ipl1
Cdc5
Dbf4
Pfk26
Tpk1
Gpa2
Ras2
Med6
Rpb10
Paf1
Rpo26
Bdp1
Spt15
Spt5
Rgr1
Srb7
Ssn2
Nop1
Nsr1
Nop56
Snu13
Rps31
Rad9
Rtf1
Rpo21
Npl3
Air2
Fps1
Ask10
Cdc28
Hog1
Pbs2
Ssk2
Clb2
Bud3
Bud4
Msn2
Sch9
Rim15
Yak1
Ypl247c
Vps52
Kss1
Sko1
Slt2
Msg5
Swe1
Igo2
Cdc55
Igo1
Vps53
Vps54
Rtn1
Fmp42
Cot1
Zrc1
10
20
30
40
50
t/s

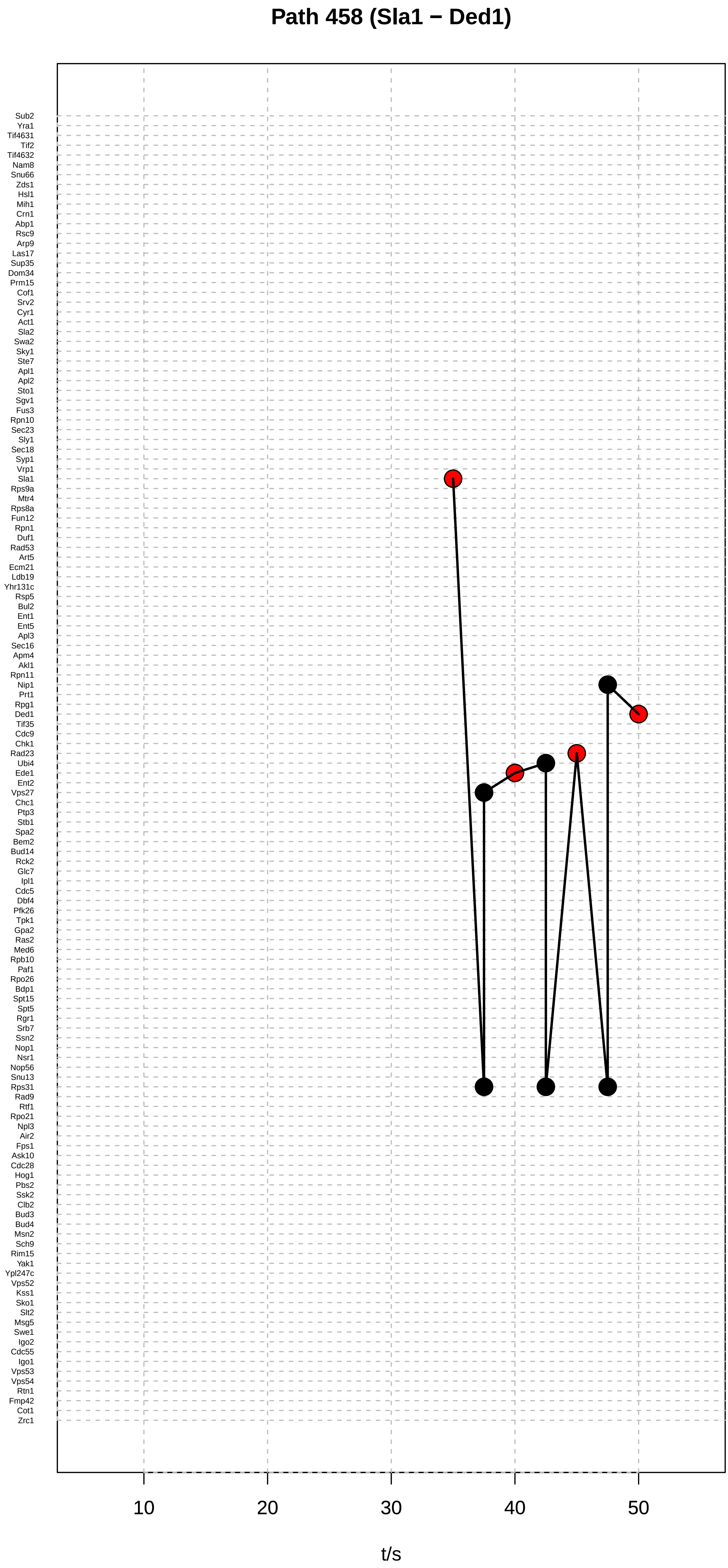

Path 458 (Sla1 − Ded1)
Sub2
Yra1
Tif4631
Tif2
Tif4632
Nam8
Snu66
Zds1
Hsl1
Mih1
Crn1
Abp1
Rsc9
Arp9
Las17
Sup35
Dom34
Prm15
Cof1
Srv2
Cyr1
Act1
Sla2
Swa2
Sky1
Ste7
Apl1
Apl2
Sto1
Sgv1
Fus3
Rpn10
Sec23
Sly1
Sec18
Syp1
Vrp1
Sla1
Rps9a
Mtr4
Rps8a
Fun12
Rpn1
Duf1
Rad53
Art5
Ecm21
Ldb19
Yhr131c
Rsp5
Bul2
Ent1
Ent5
Apl3
Sec16
Apm4
Akl1
Rpn11
Nip1
Prt1
Rpg1
Ded1
Tif35
Cdc9
Chk1
Rad23
Ubi4
Ede1
Ent2
Vps27
Chc1
Ptp3
Stb1
Spa2
Bem2
Bud14
Rck2
Glc7
Ipl1
Cdc5
Dbf4
Pfk26
Tpk1
Gpa2
Ras2
Med6
Rpb10
Paf1
Rpo26
Bdp1
Spt15
Spt5
Rgr1
Srb7
Ssn2
Nop1
Nsr1
Nop56
Snu13
Rps31
Rad9
Rtf1
Rpo21
Npl3
Air2
Fps1
Ask10
Cdc28
Hog1
Pbs2
Ssk2
Clb2
Bud3
Bud4
Msn2
Sch9
Rim15
Yak1
Ypl247c
Vps52
Kss1
Sko1
Slt2
Msg5
Swe1
Igo2
Cdc55
Igo1
Vps53
Vps54
Rtn1
Fmp42
Cot1
Zrc1
10
20
30
40
50
t/s

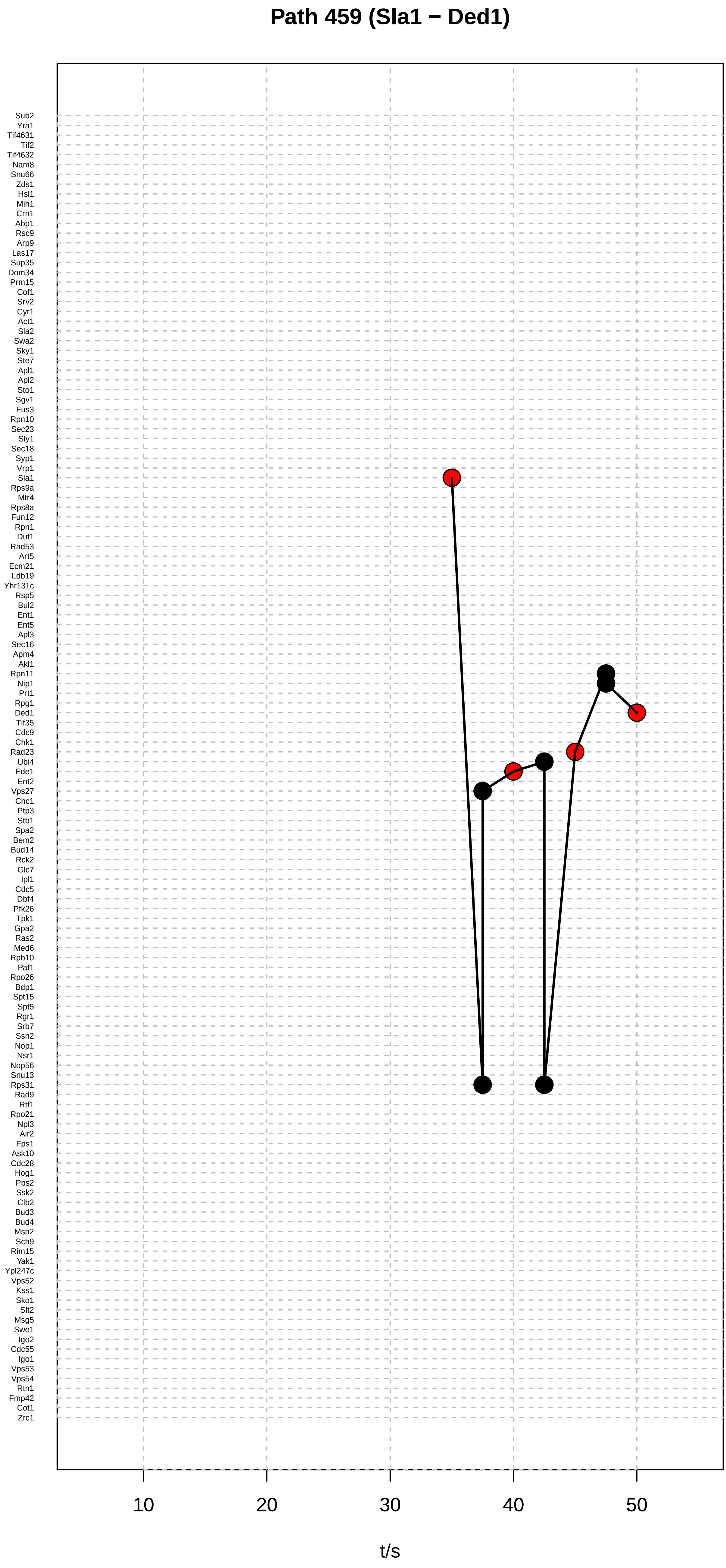

Path 459 (Sla1 − Ded1)
Sub2
Yra1
Tif4631
Tif2
Tif4632
Nam8
Snu66
Zds1
Hsl1
Mih1
Crn1
Abp1
Rsc9
Arp9
Las17
Sup35
Dom34
Prm15
Cof1
Srv2
Cyr1
Act1
Sla2
Swa2
Sky1
Ste7
Apl1
Apl2
Sto1
Sgv1
Fus3
Rpn10
Sec23
Sly1
Sec18
Syp1
Vrp1
Sla1
Rps9a
Mtr4
Rps8a
Fun12
Rpn1
Duf1
Rad53
Art5
Ecm21
Ldb19
Yhr131c
Rsp5
Bul2
Ent1
Ent5
Apl3
Sec16
Apm4
Akl1
Rpn11
Nip1
Prt1
Rpg1
Ded1
Tif35
Cdc9
Chk1
Rad23
Ubi4
Ede1
Ent2
Vps27
Chc1
Ptp3
Stb1
Spa2
Bem2
Bud14
Rck2
Glc7
Ipl1
Cdc5
Dbf4
Pfk26
Tpk1
Gpa2
Ras2
Med6
Rpb10
Paf1
Rpo26
Bdp1
Spt15
Spt5
Rgr1
Srb7
Ssn2
Nop1
Nsr1
Nop56
Snu13
Rps31
Rad9
Rtf1
Rpo21
Npl3
Air2
Fps1
Ask10
Cdc28
Hog1
Pbs2
Ssk2
Clb2
Bud3
Bud4
Msn2
Sch9
Rim15
Yak1
Ypl247c
Vps52
Kss1
Sko1
Slt2
Msg5
Swe1
Igo2
Cdc55
Igo1
Vps53
Vps54
Rtn1
Fmp42
Cot1
Zrc1
10
20
30
40
50
t/s

#### Path 460 (Sla1 − Ded1)

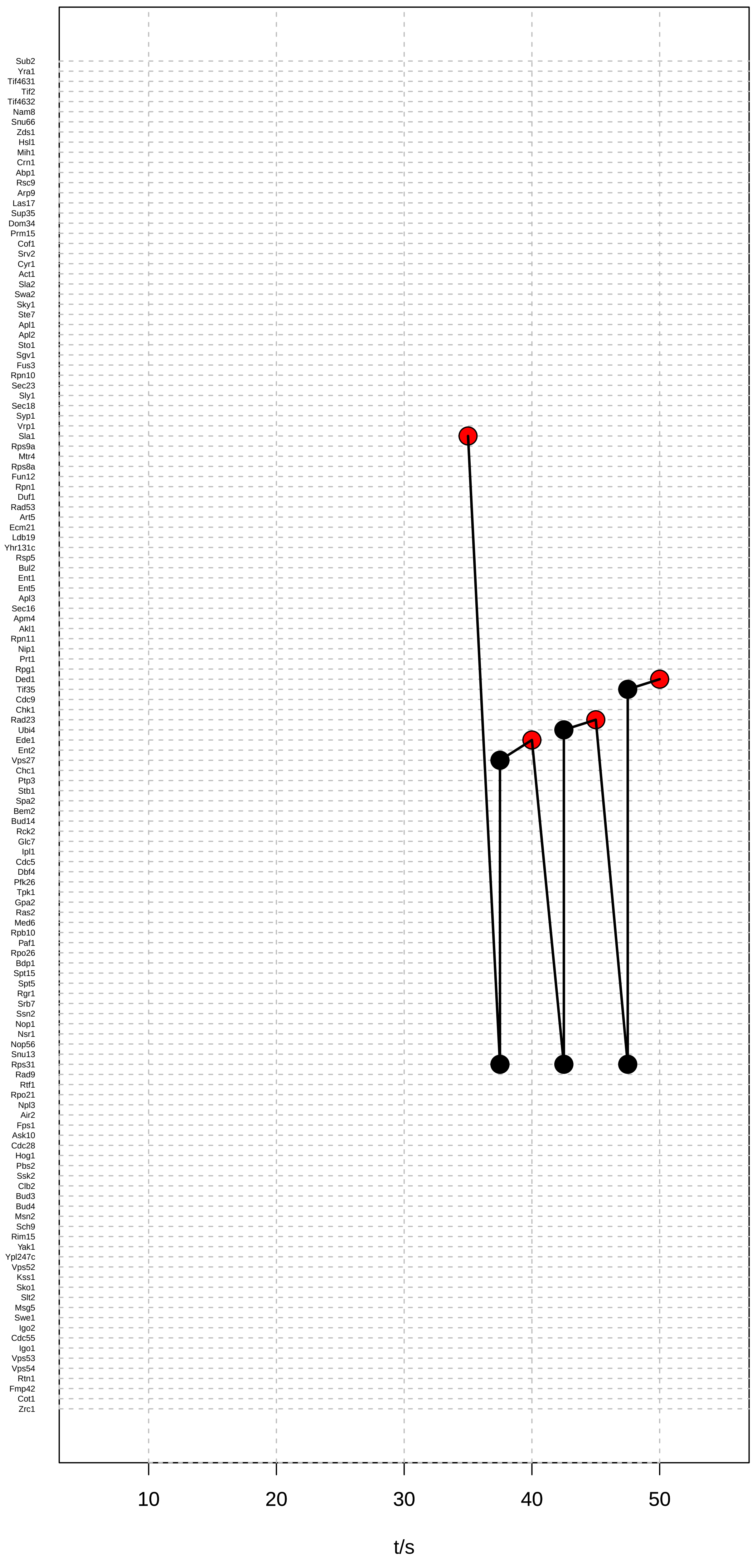

### Path 461 (Sla1 − Ded1)

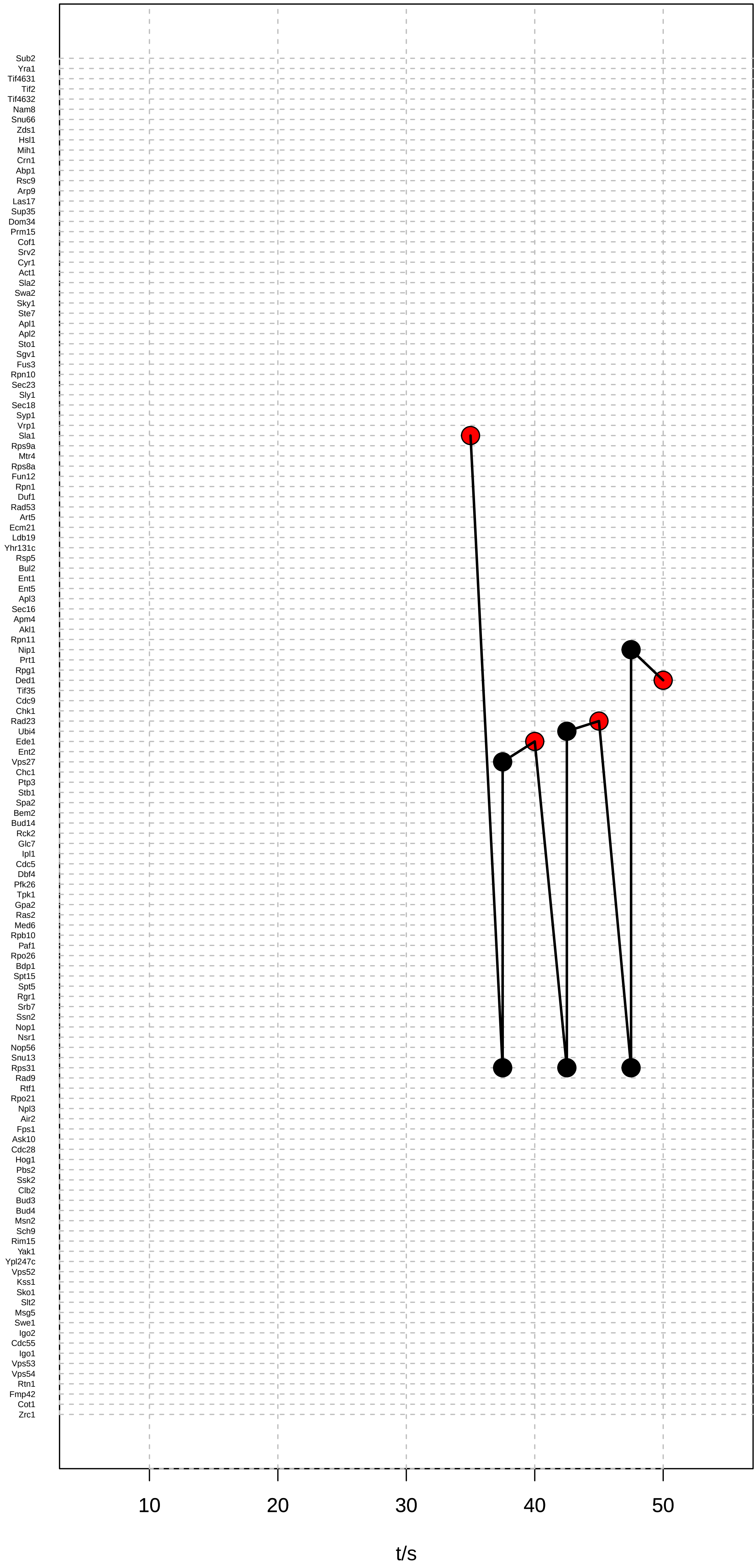

#### Path 462 (Sla1 − Ded1)

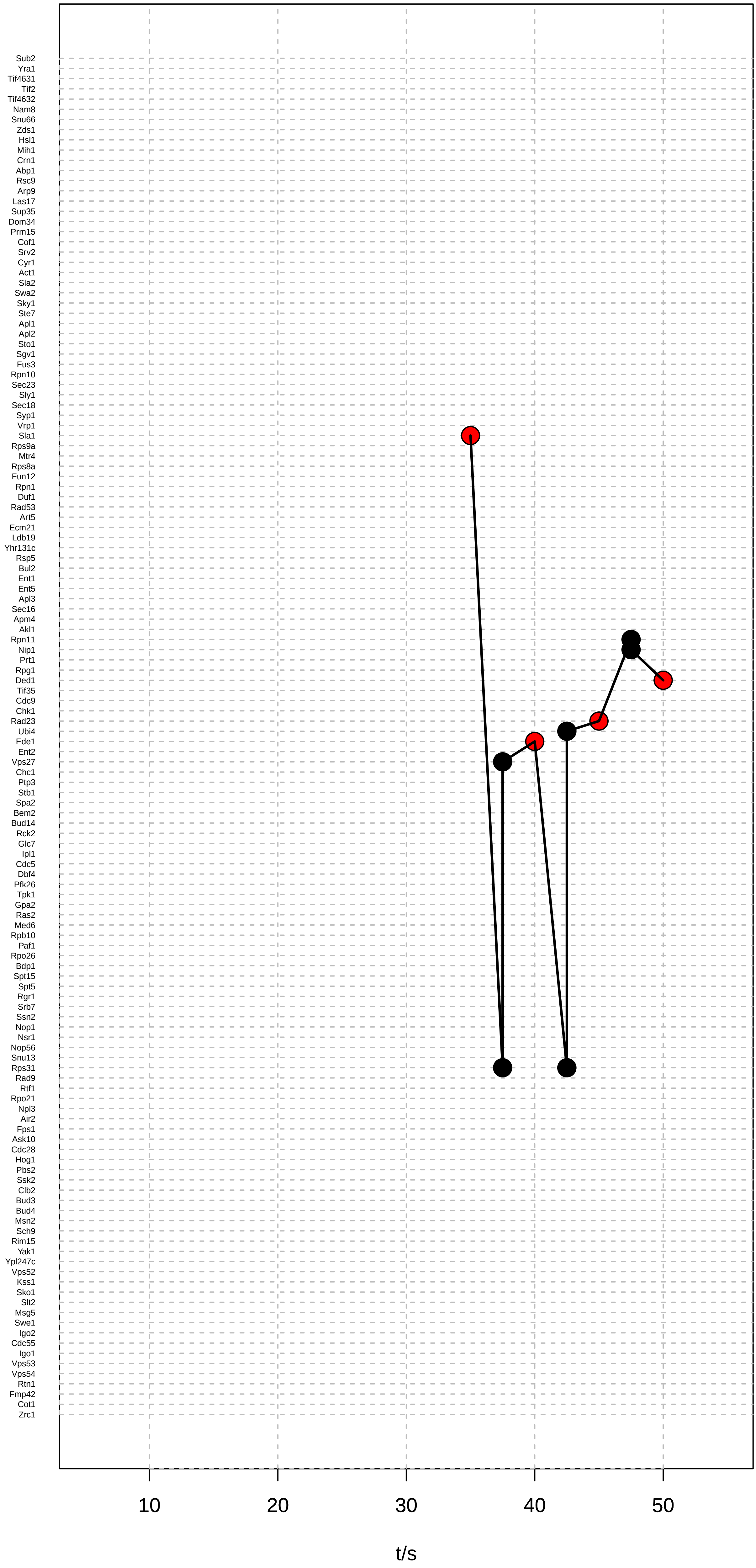

#### Path 463 (Sla1 − Ded1)

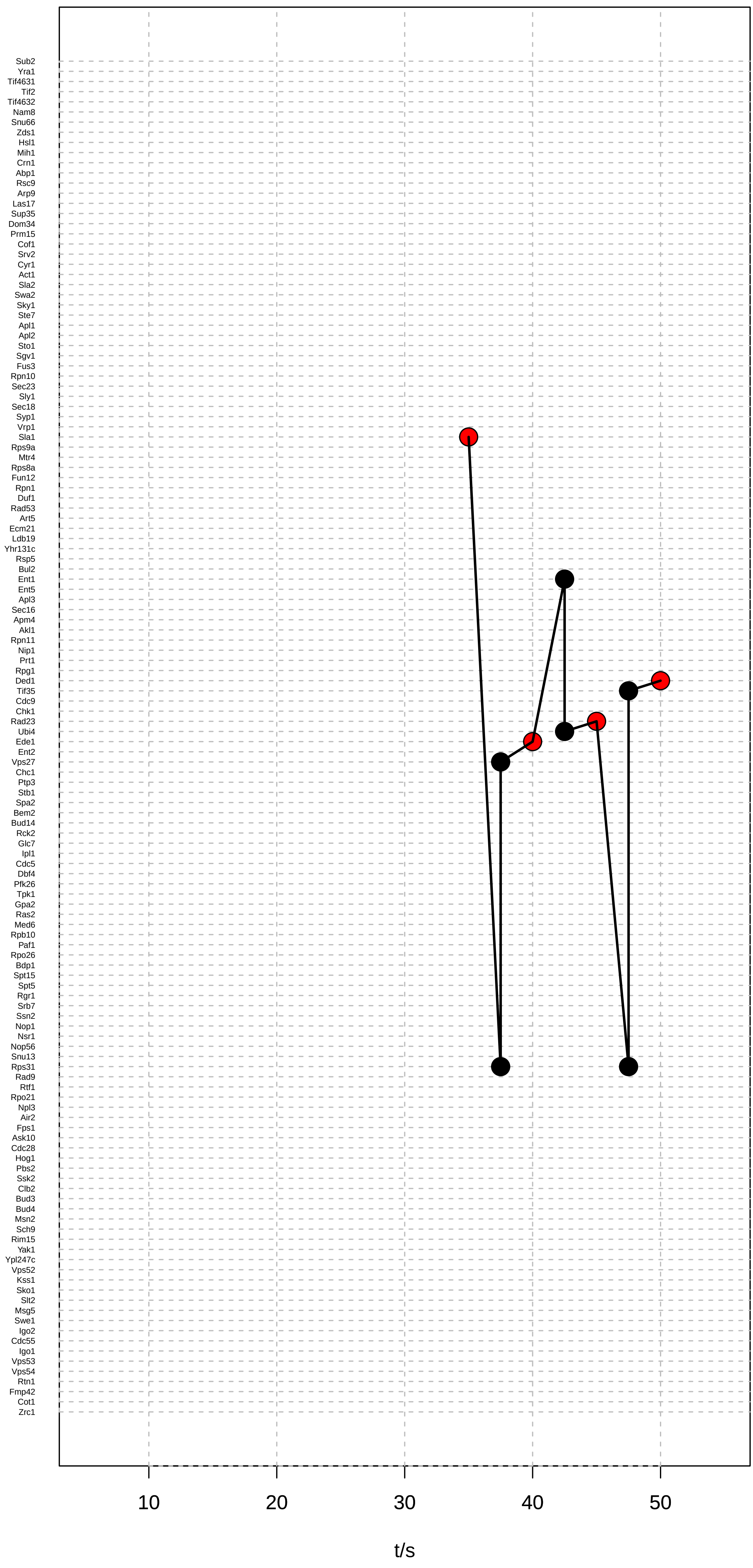

#### Path 464 (Sla1 − Ded1)

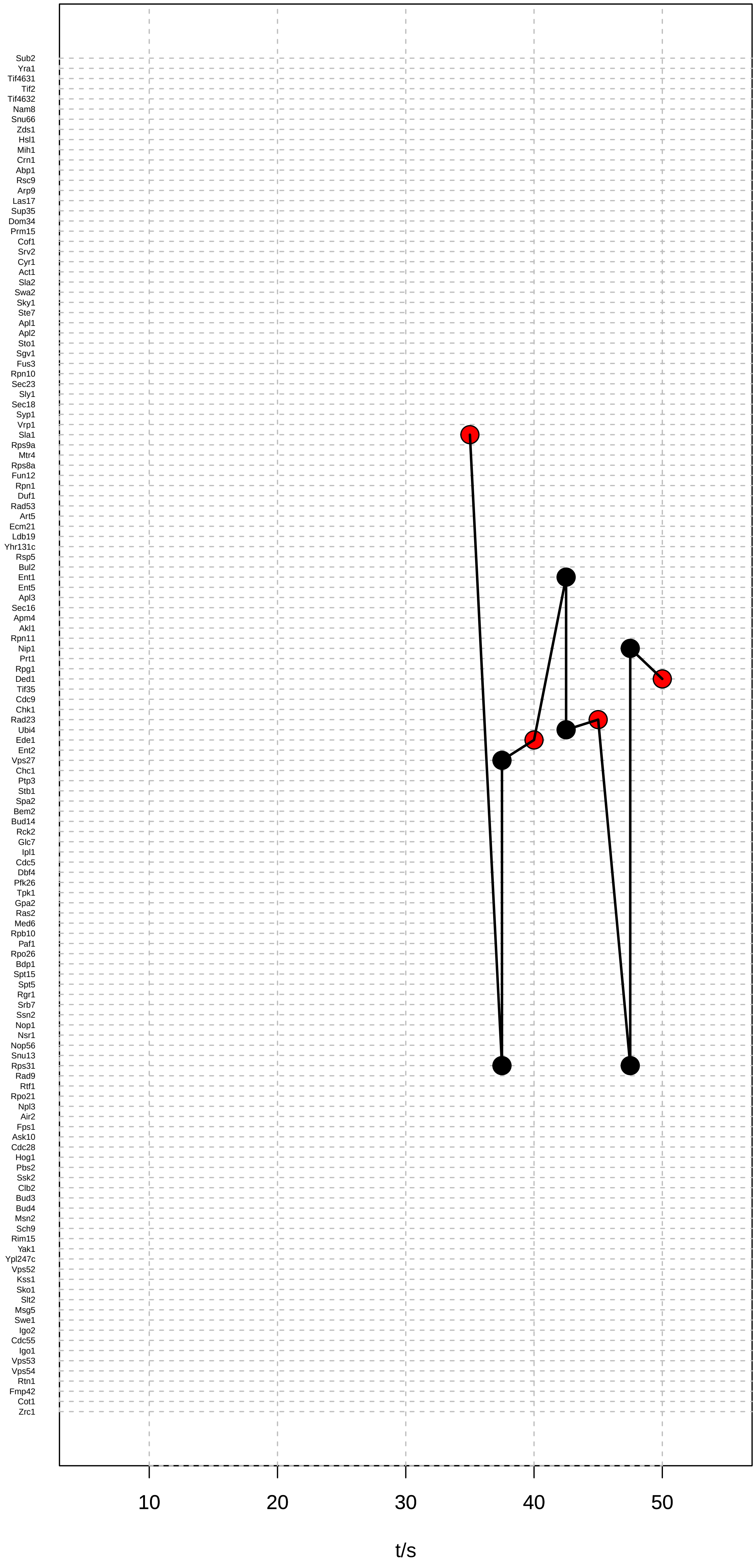

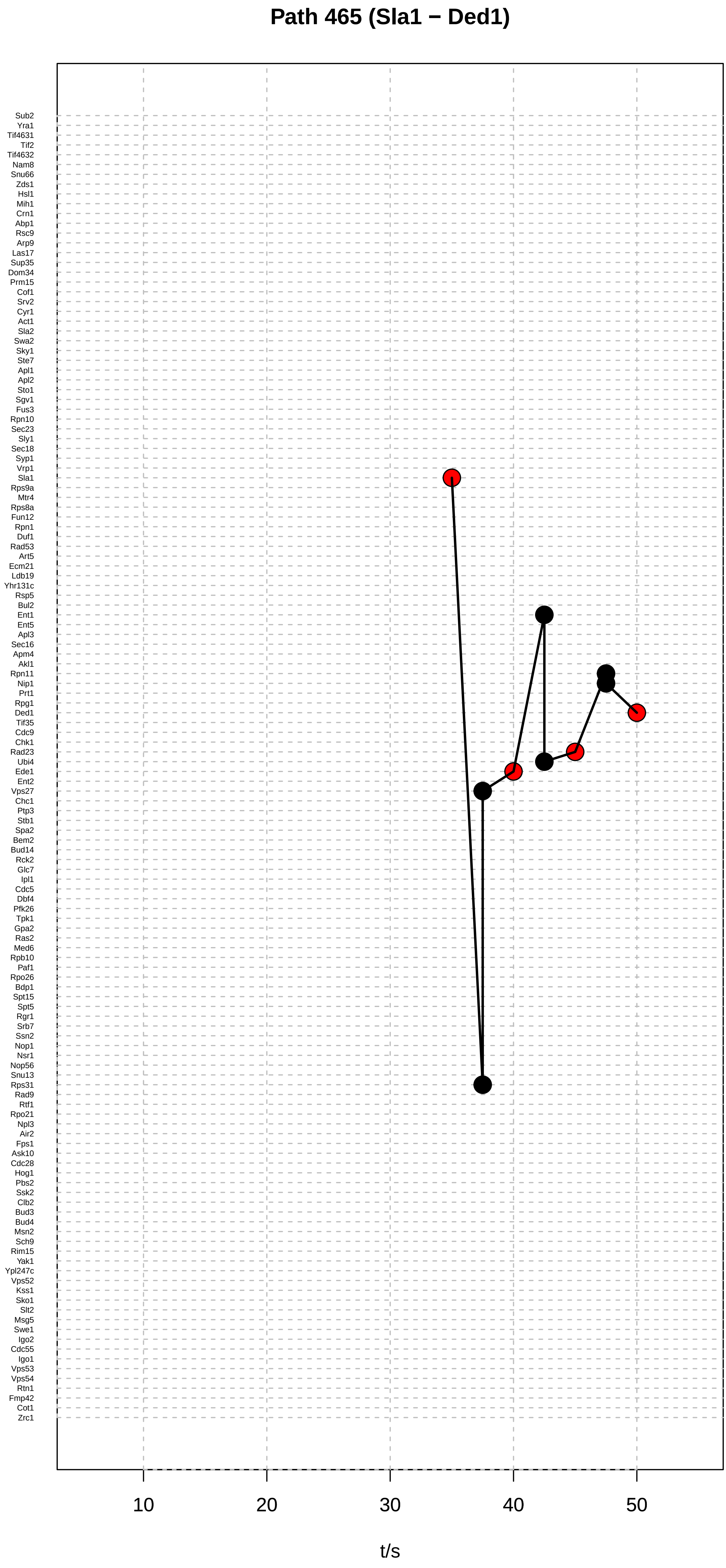

Path 465 (Sla1 − Ded1)
Sub2
Yra1
Tif4631
Tif2
Tif4632
Nam8
Snu66
Zds1
Hsl1
Mih1
Crn1
Abp1
Rsc9
Arp9
Las17
Sup35
Dom34
Prm15
Cof1
Srv2
Cyr1
Act1
Sla2
Swa2
Sky1
Ste7
Apl1
Apl2
Sto1
Sgv1
Fus3
Rpn10
Sec23
Sly1
Sec18
Syp1
Vrp1
Sla1
Rps9a
Mtr4
Rps8a
Fun12
Rpn1
Duf1
Rad53
Art5
Ecm21
Ldb19
Yhr131c
Rsp5
Bul2
Ent1
Ent5
Apl3
Sec16
Apm4
Akl1
Rpn11
Nip1
Prt1
Rpg1
Ded1
Tif35
Cdc9
Chk1
Rad23
Ubi4
Ede1
Ent2
Vps27
Chc1
Ptp3
Stb1
Spa2
Bem2
Bud14
Rck2
Glc7
Ipl1
Cdc5
Dbf4
Pfk26
Tpk1
Gpa2
Ras2
Med6
Rpb10
Paf1
Rpo26
Bdp1
Spt15
Spt5
Rgr1
Srb7
Ssn2
Nop1
Nsr1
Nop56
Snu13
Rps31
Rad9
Rtf1
Rpo21
Npl3
Air2
Fps1
Ask10
Cdc28
Hog1
Pbs2
Ssk2
Clb2
Bud3
Bud4
Msn2
Sch9
Rim15
Yak1
Ypl247c
Vps52
Kss1
Sko1
Slt2
Msg5
Swe1
Igo2
Cdc55
Igo1
Vps53
Vps54
Rtn1
Fmp42
Cot1
Zrc1
10
20
30
40
50
t/s

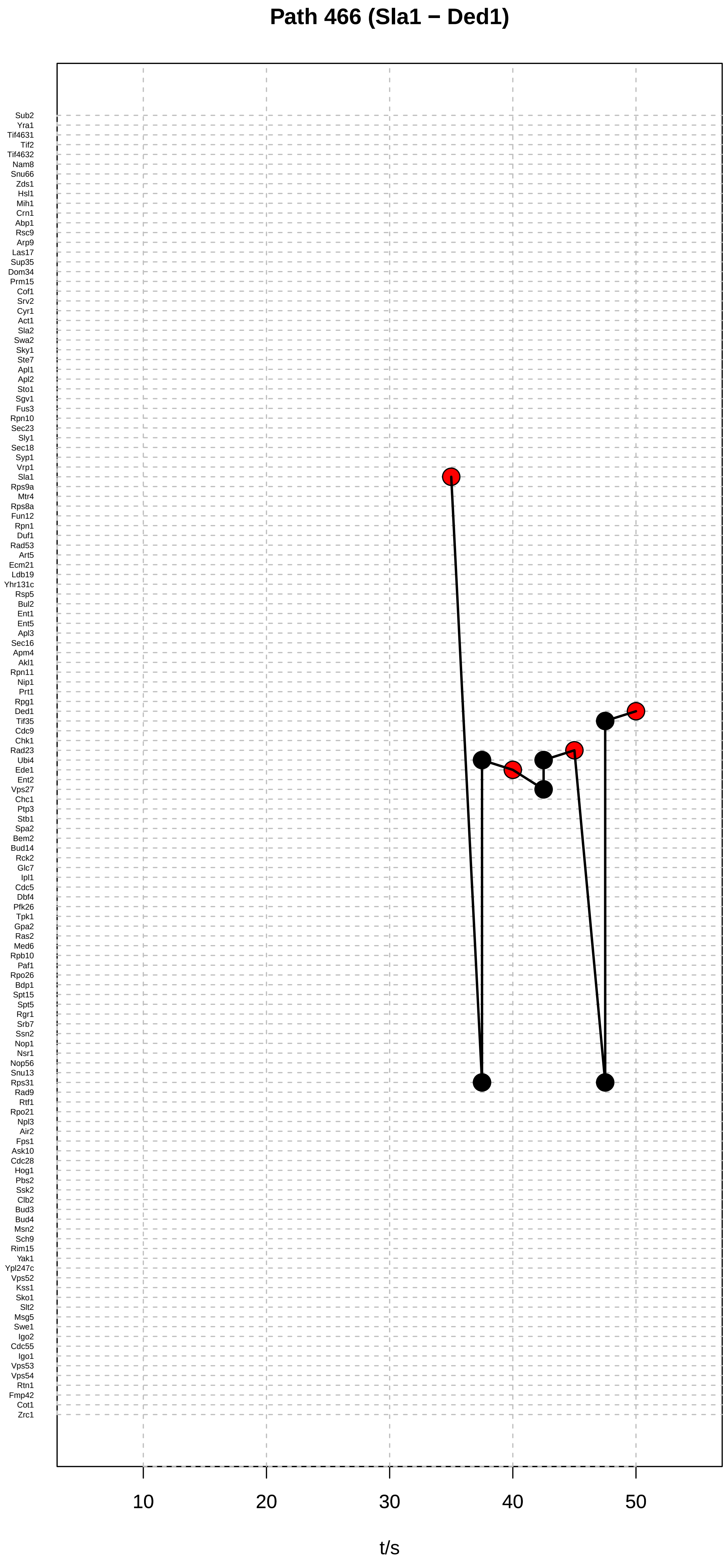

Path 466 (Sla1 − Ded1)
Sub2
Yra1
Tif4631
Tif2
Tif4632
Nam8
Snu66
Zds1
Hsl1
Mih1
Crn1
Abp1
Rsc9
Arp9
Las17
Sup35
Dom34
Prm15
Cof1
Srv2
Cyr1
Act1
Sla2
Swa2
Sky1
Ste7
Apl1
Apl2
Sto1
Sgv1
Fus3
Rpn10
Sec23
Sly1
Sec18
Syp1
Vrp1
Sla1
Rps9a
Mtr4
Rps8a
Fun12
Rpn1
Duf1
Rad53
Art5
Ecm21
Ldb19
Yhr131c
Rsp5
Bul2
Ent1
Ent5
Apl3
Sec16
Apm4
Akl1
Rpn11
Nip1
Prt1
Rpg1
Ded1
Tif35
Cdc9
Chk1
Rad23
Ubi4
Ede1
Ent2
Vps27
Chc1
Ptp3
Stb1
Spa2
Bem2
Bud14
Rck2
Glc7
Ipl1
Cdc5
Dbf4
Pfk26
Tpk1
Gpa2
Ras2
Med6
Rpb10
Paf1
Rpo26
Bdp1
Spt15
Spt5
Rgr1
Srb7
Ssn2
Nop1
Nsr1
Nop56
Snu13
Rps31
Rad9
Rtf1
Rpo21
Npl3
Air2
Fps1
Ask10
Cdc28
Hog1
Pbs2
Ssk2
Clb2
Bud3
Bud4
Msn2
Sch9
Rim15
Yak1
Ypl247c
Vps52
Kss1
Sko1
Slt2
Msg5
Swe1
Igo2
Cdc55
Igo1
Vps53
Vps54
Rtn1
Fmp42
Cot1
Zrc1
10
20
30
40
50
t/s

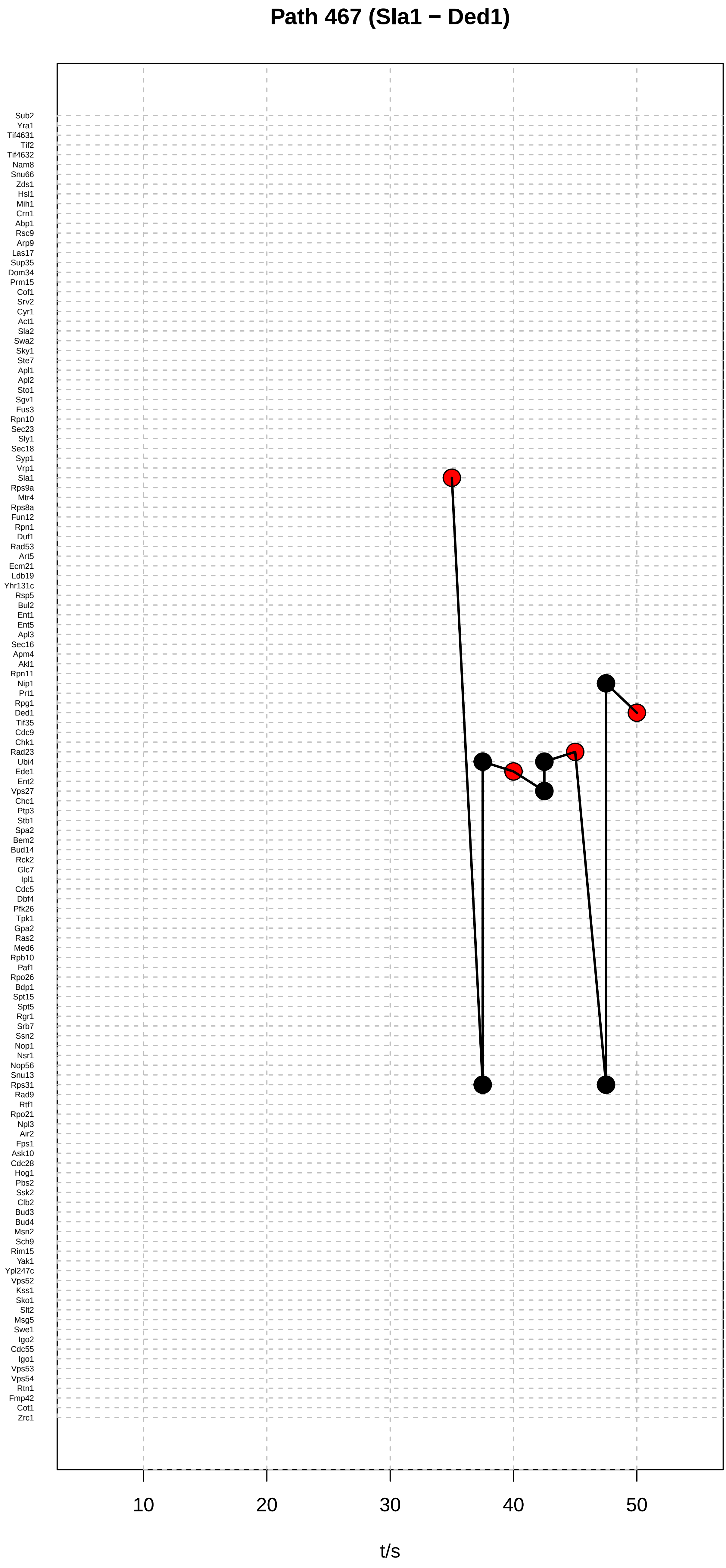

Path 467 (Sla1 − Ded1)
Sub2
Yra1
Tif4631
Tif2
Tif4632
Nam8
Snu66
Zds1
Hsl1
Mih1
Crn1
Abp1
Rsc9
Arp9
Las17
Sup35
Dom34
Prm15
Cof1
Srv2
Cyr1
Act1
Sla2
Swa2
Sky1
Ste7
Apl1
Apl2
Sto1
Sgv1
Fus3
Rpn10
Sec23
Sly1
Sec18
Syp1
Vrp1
Sla1
Rps9a
Mtr4
Rps8a
Fun12
Rpn1
Duf1
Rad53
Art5
Ecm21
Ldb19
Yhr131c
Rsp5
Bul2
Ent1
Ent5
Apl3
Sec16
Apm4
Akl1
Rpn11
Nip1
Prt1
Rpg1
Ded1
Tif35
Cdc9
Chk1
Rad23
Ubi4
Ede1
Ent2
Vps27
Chc1
Ptp3
Stb1
Spa2
Bem2
Bud14
Rck2
Glc7
Ipl1
Cdc5
Dbf4
Pfk26
Tpk1
Gpa2
Ras2
Med6
Rpb10
Paf1
Rpo26
Bdp1
Spt15
Spt5
Rgr1
Srb7
Ssn2
Nop1
Nsr1
Nop56
Snu13
Rps31
Rad9
Rtf1
Rpo21
Npl3
Air2
Fps1
Ask10
Cdc28
Hog1
Pbs2
Ssk2
Clb2
Bud3
Bud4
Msn2
Sch9
Rim15
Yak1
Ypl247c
Vps52
Kss1
Sko1
Slt2
Msg5
Swe1
Igo2
Cdc55
Igo1
Vps53
Vps54
Rtn1
Fmp42
Cot1
Zrc1
10
20
30
40
50
t/s

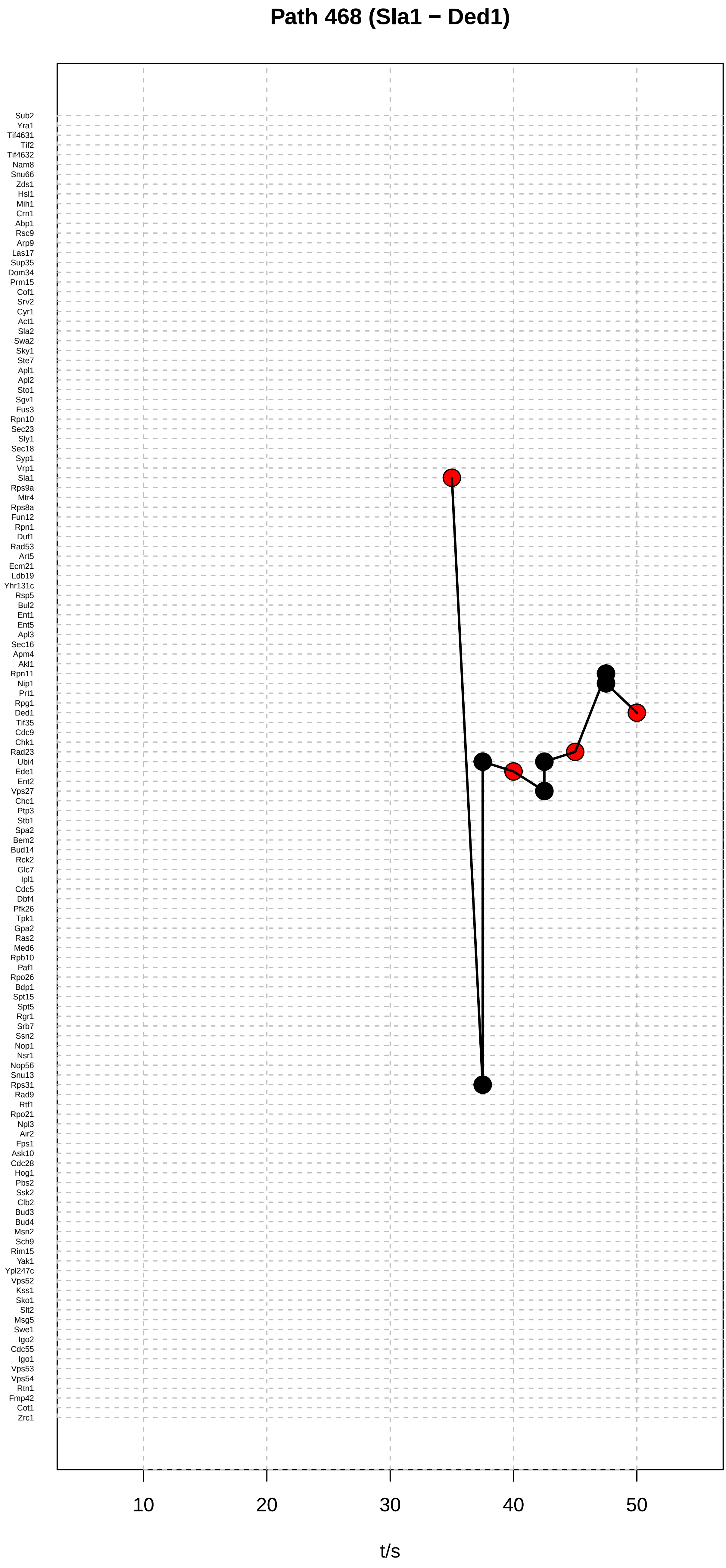

Path 468 (Sla1 − Ded1)
Sub2
Yra1
Tif4631
Tif2
Tif4632
Nam8
Snu66
Zds1
Hsl1
Mih1
Crn1
Abp1
Rsc9
Arp9
Las17
Sup35
Dom34
Prm15
Cof1
Srv2
Cyr1
Act1
Sla2
Swa2
Sky1
Ste7
Apl1
Apl2
Sto1
Sgv1
Fus3
Rpn10
Sec23
Sly1
Sec18
Syp1
Vrp1
Sla1
Rps9a
Mtr4
Rps8a
Fun12
Rpn1
Duf1
Rad53
Art5
Ecm21
Ldb19
Yhr131c
Rsp5
Bul2
Ent1
Ent5
Apl3
Sec16
Apm4
Akl1
Rpn11
Nip1
Prt1
Rpg1
Ded1
Tif35
Cdc9
Chk1
Rad23
Ubi4
Ede1
Ent2
Vps27
Chc1
Ptp3
Stb1
Spa2
Bem2
Bud14
Rck2
Glc7
Ipl1
Cdc5
Dbf4
Pfk26
Tpk1
Gpa2
Ras2
Med6
Rpb10
Paf1
Rpo26
Bdp1
Spt15
Spt5
Rgr1
Srb7
Ssn2
Nop1
Nsr1
Nop56
Snu13
Rps31
Rad9
Rtf1
Rpo21
Npl3
Air2
Fps1
Ask10
Cdc28
Hog1
Pbs2
Ssk2
Clb2
Bud3
Bud4
Msn2
Sch9
Rim15
Yak1
Ypl247c
Vps52
Kss1
Sko1
Slt2
Msg5
Swe1
Igo2
Cdc55
Igo1
Vps53
Vps54
Rtn1
Fmp42
Cot1
Zrc1
10
20
30
40
50
t/s

#### Path 469 (Sla1 − Ded1)

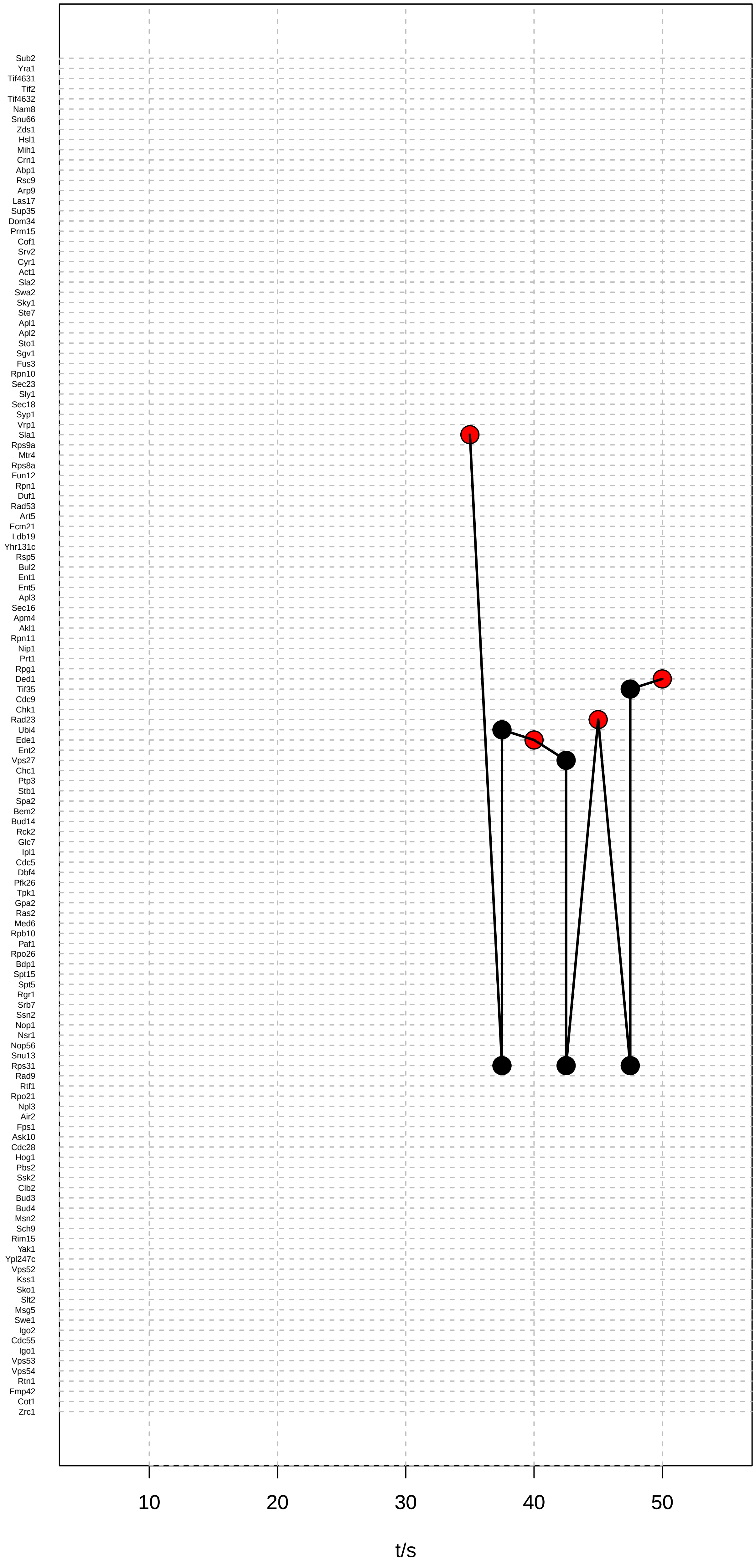

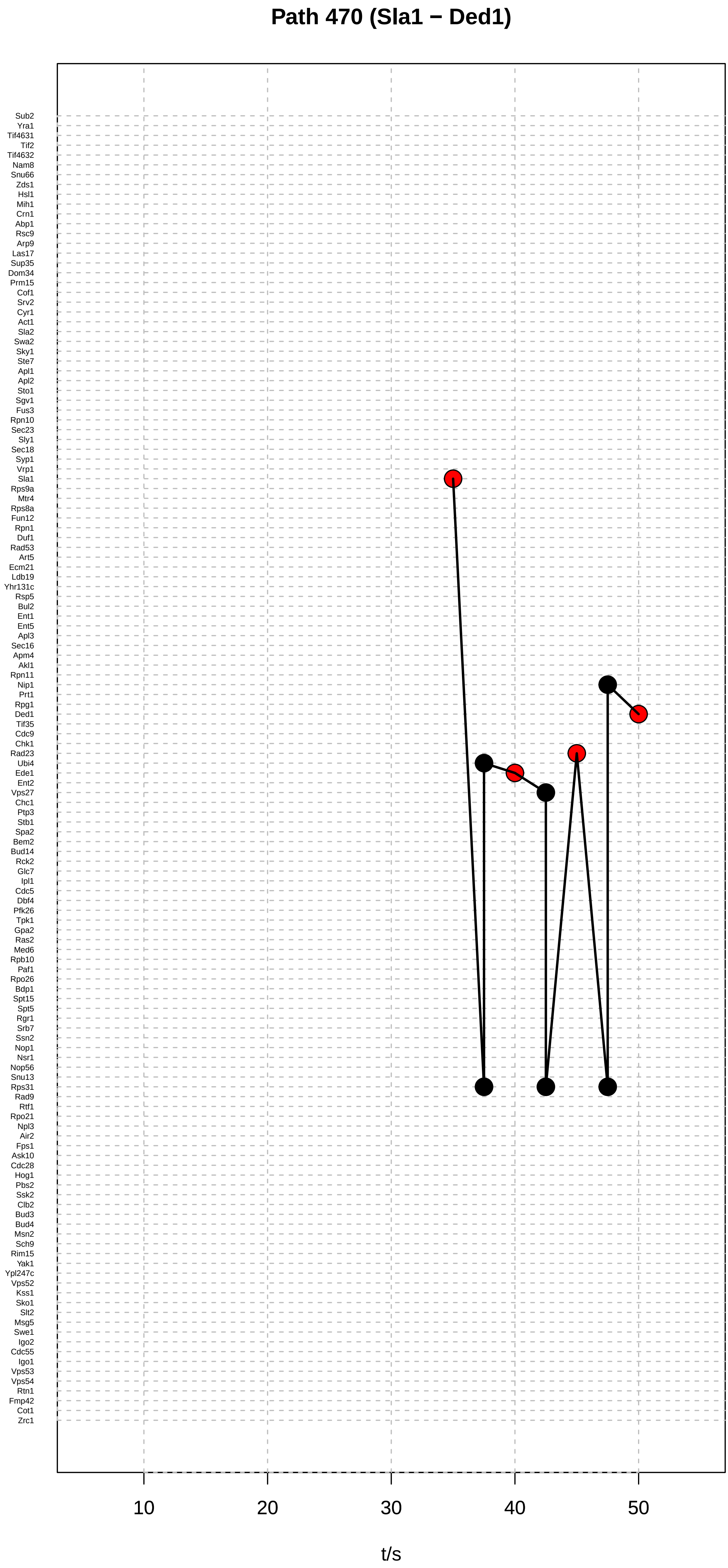

Path 470 (Sla1 − Ded1)
Sub2
Yra1
Tif4631
Tif2
Tif4632
Nam8
Snu66
Zds1
Hsl1
Mih1
Crn1
Abp1
Rsc9
Arp9
Las17
Sup35
Dom34
Prm15
Cof1
Srv2
Cyr1
Act1
Sla2
Swa2
Sky1
Ste7
Apl1
Apl2
Sto1
Sgv1
Fus3
Rpn10
Sec23
Sly1
Sec18
Syp1
Vrp1
Sla1
Rps9a
Mtr4
Rps8a
Fun12
Rpn1
Duf1
Rad53
Art5
Ecm21
Ldb19
Yhr131c
Rsp5
Bul2
Ent1
Ent5
Apl3
Sec16
Apm4
Akl1
Rpn11
Nip1
Prt1
Rpg1
Ded1
Tif35
Cdc9
Chk1
Rad23
Ubi4
Ede1
Ent2
Vps27
Chc1
Ptp3
Stb1
Spa2
Bem2
Bud14
Rck2
Glc7
Ipl1
Cdc5
Dbf4
Pfk26
Tpk1
Gpa2
Ras2
Med6
Rpb10
Paf1
Rpo26
Bdp1
Spt15
Spt5
Rgr1
Srb7
Ssn2
Nop1
Nsr1
Nop56
Snu13
Rps31
Rad9
Rtf1
Rpo21
Npl3
Air2
Fps1
Ask10
Cdc28
Hog1
Pbs2
Ssk2
Clb2
Bud3
Bud4
Msn2
Sch9
Rim15
Yak1
Ypl247c
Vps52
Kss1
Sko1
Slt2
Msg5
Swe1
Igo2
Cdc55
Igo1
Vps53
Vps54
Rtn1
Fmp42
Cot1
Zrc1
10
20
30
40
50
t/s

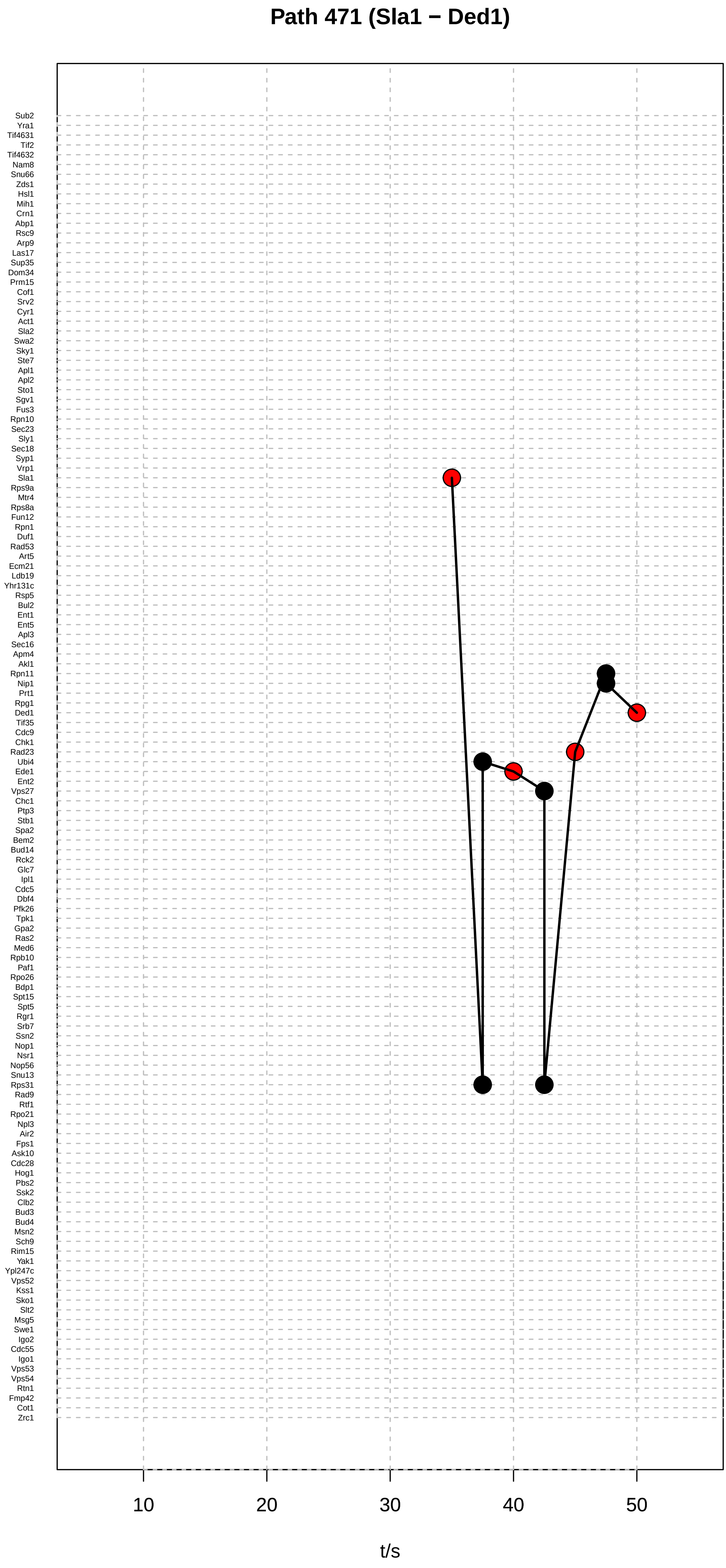

Path 471 (Sla1 − Ded1)
Sub2
Yra1
Tif4631
Tif2
Tif4632
Nam8
Snu66
Zds1
Hsl1
Mih1
Crn1
Abp1
Rsc9
Arp9
Las17
Sup35
Dom34
Prm15
Cof1
Srv2
Cyr1
Act1
Sla2
Swa2
Sky1
Ste7
Apl1
Apl2
Sto1
Sgv1
Fus3
Rpn10
Sec23
Sly1
Sec18
Syp1
Vrp1
Sla1
Rps9a
Mtr4
Rps8a
Fun12
Rpn1
Duf1
Rad53
Art5
Ecm21
Ldb19
Yhr131c
Rsp5
Bul2
Ent1
Ent5
Apl3
Sec16
Apm4
Akl1
Rpn11
Nip1
Prt1
Rpg1
Ded1
Tif35
Cdc9
Chk1
Rad23
Ubi4
Ede1
Ent2
Vps27
Chc1
Ptp3
Stb1
Spa2
Bem2
Bud14
Rck2
Glc7
Ipl1
Cdc5
Dbf4
Pfk26
Tpk1
Gpa2
Ras2
Med6
Rpb10
Paf1
Rpo26
Bdp1
Spt15
Spt5
Rgr1
Srb7
Ssn2
Nop1
Nsr1
Nop56
Snu13
Rps31
Rad9
Rtf1
Rpo21
Npl3
Air2
Fps1
Ask10
Cdc28
Hog1
Pbs2
Ssk2
Clb2
Bud3
Bud4
Msn2
Sch9
Rim15
Yak1
Ypl247c
Vps52
Kss1
Sko1
Slt2
Msg5
Swe1
Igo2
Cdc55
Igo1
Vps53
Vps54
Rtn1
Fmp42
Cot1
Zrc1
10
20
30
40
50
t/s

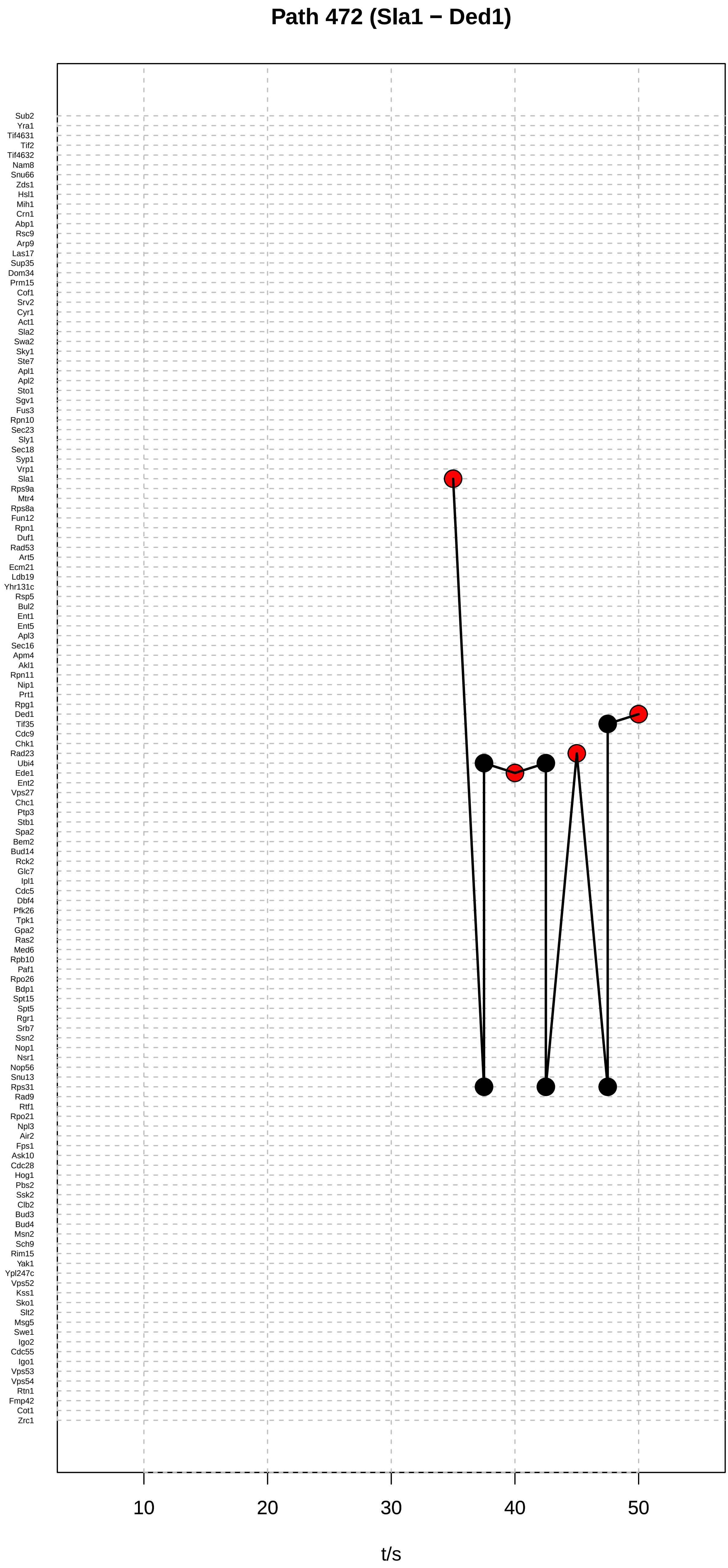

Path 472 (Sla1 − Ded1)
Sub2
Yra1
Tif4631
Tif2
Tif4632
Nam8
Snu66
Zds1
Hsl1
Mih1
Crn1
Abp1
Rsc9
Arp9
Las17
Sup35
Dom34
Prm15
Cof1
Srv2
Cyr1
Act1
Sla2
Swa2
Sky1
Ste7
Apl1
Apl2
Sto1
Sgv1
Fus3
Rpn10
Sec23
Sly1
Sec18
Syp1
Vrp1
Sla1
Rps9a
Mtr4
Rps8a
Fun12
Rpn1
Duf1
Rad53
Art5
Ecm21
Ldb19
Yhr131c
Rsp5
Bul2
Ent1
Ent5
Apl3
Sec16
Apm4
Akl1
Rpn11
Nip1
Prt1
Rpg1
Ded1
Tif35
Cdc9
Chk1
Rad23
Ubi4
Ede1
Ent2
Vps27
Chc1
Ptp3
Stb1
Spa2
Bem2
Bud14
Rck2
Glc7
Ipl1
Cdc5
Dbf4
Pfk26
Tpk1
Gpa2
Ras2
Med6
Rpb10
Paf1
Rpo26
Bdp1
Spt15
Spt5
Rgr1
Srb7
Ssn2
Nop1
Nsr1
Nop56
Snu13
Rps31
Rad9
Rtf1
Rpo21
Npl3
Air2
Fps1
Ask10
Cdc28
Hog1
Pbs2
Ssk2
Clb2
Bud3
Bud4
Msn2
Sch9
Rim15
Yak1
Ypl247c
Vps52
Kss1
Sko1
Slt2
Msg5
Swe1
Igo2
Cdc55
Igo1
Vps53
Vps54
Rtn1
Fmp42
Cot1
Zrc1
10
20
30
40
50
t/s

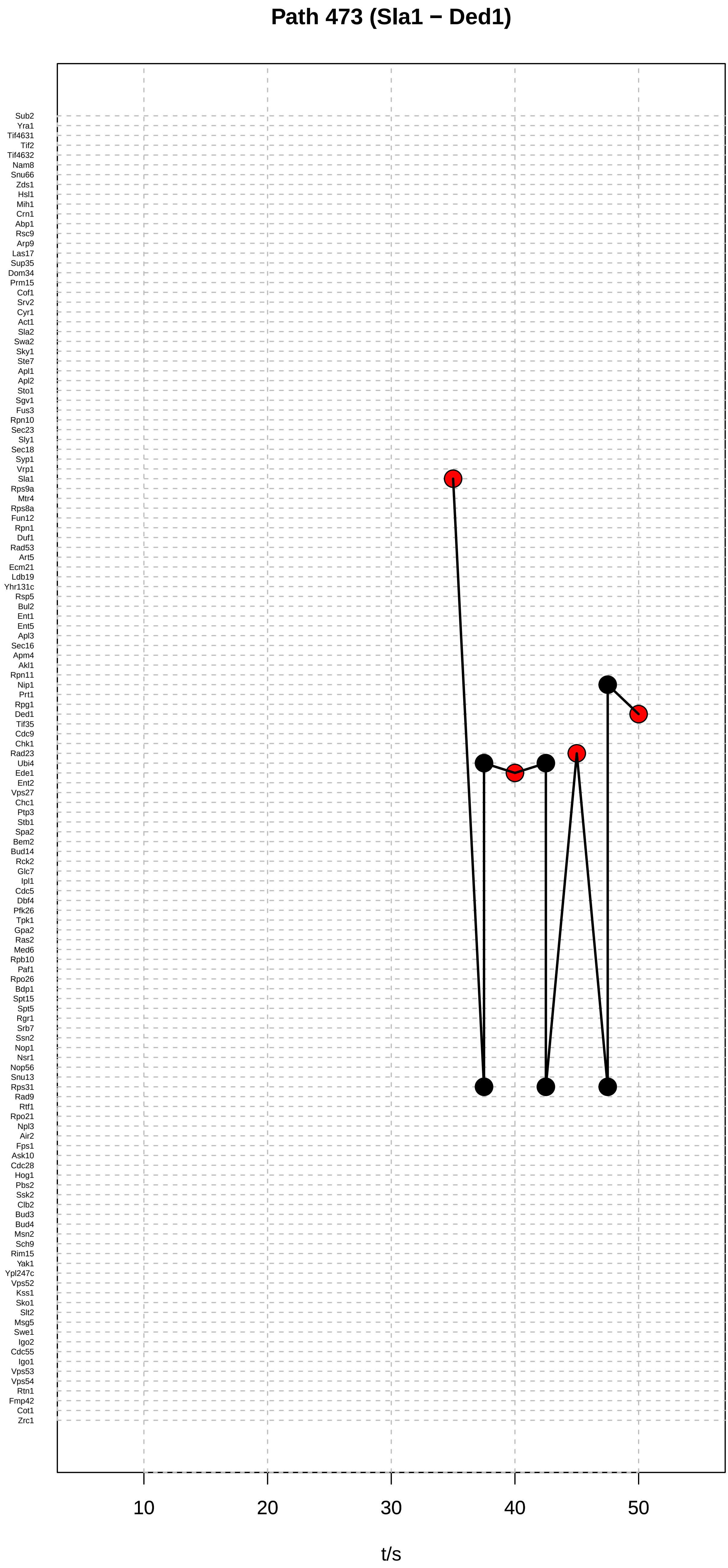

Path 473 (Sla1 − Ded1)
Sub2
Yra1
Tif4631
Tif2
Tif4632
Nam8
Snu66
Zds1
Hsl1
Mih1
Crn1
Abp1
Rsc9
Arp9
Las17
Sup35
Dom34
Prm15
Cof1
Srv2
Cyr1
Act1
Sla2
Swa2
Sky1
Ste7
Apl1
Apl2
Sto1
Sgv1
Fus3
Rpn10
Sec23
Sly1
Sec18
Syp1
Vrp1
Sla1
Rps9a
Mtr4
Rps8a
Fun12
Rpn1
Duf1
Rad53
Art5
Ecm21
Ldb19
Yhr131c
Rsp5
Bul2
Ent1
Ent5
Apl3
Sec16
Apm4
Akl1
Rpn11
Nip1
Prt1
Rpg1
Ded1
Tif35
Cdc9
Chk1
Rad23
Ubi4
Ede1
Ent2
Vps27
Chc1
Ptp3
Stb1
Spa2
Bem2
Bud14
Rck2
Glc7
Ipl1
Cdc5
Dbf4
Pfk26
Tpk1
Gpa2
Ras2
Med6
Rpb10
Paf1
Rpo26
Bdp1
Spt15
Spt5
Rgr1
Srb7
Ssn2
Nop1
Nsr1
Nop56
Snu13
Rps31
Rad9
Rtf1
Rpo21
Npl3
Air2
Fps1
Ask10
Cdc28
Hog1
Pbs2
Ssk2
Clb2
Bud3
Bud4
Msn2
Sch9
Rim15
Yak1
Ypl247c
Vps52
Kss1
Sko1
Slt2
Msg5
Swe1
Igo2
Cdc55
Igo1
Vps53
Vps54
Rtn1
Fmp42
Cot1
Zrc1
10
20
30
40
50
t/s

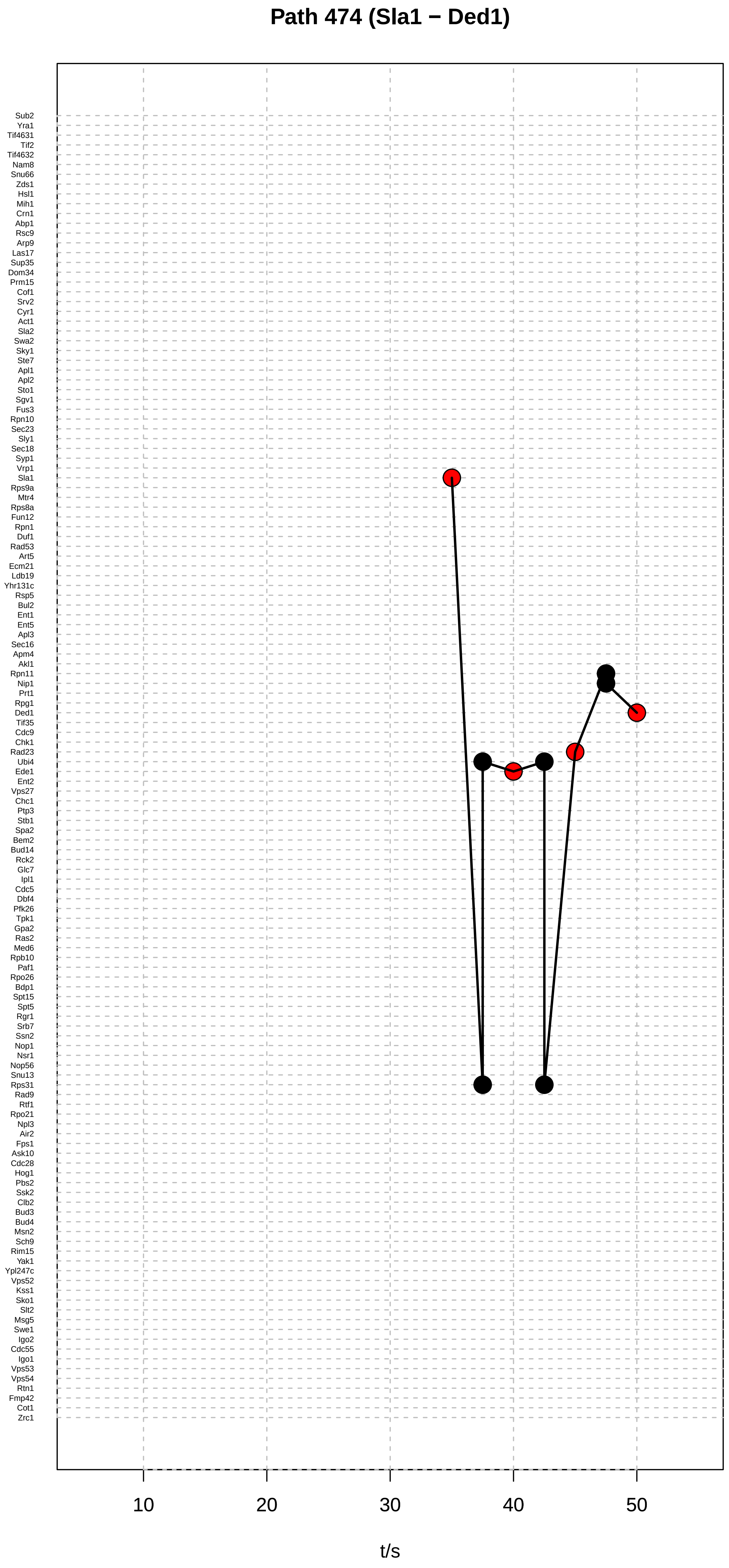
Path 474 (Sla1 − Ded1)
Sub2
Yra1
Tif4631
Tif2
Tif4632
Nam8
Snu66
Zds1
Hsl1
Mih1
Crn1
Abp1
Rsc9
Arp9
Las17
Sup35
Dom34
Prm15
Cof1
Srv2
Cyr1
Act1
Sla2
Swa2
Sky1
Ste7
Apl1
Apl2
Sto1
Sgv1
Fus3
Rpn10
Sec23
Sly1
Sec18
Syp1
Vrp1
Sla1
Rps9a
Mtr4
Rps8a
Fun12
Rpn1
Duf1
Rad53
Art5
Ecm21
Ldb19
Yhr131c
Rsp5
Bul2
Ent1
Ent5
Apl3
Sec16
Apm4
Akl1
Rpn11
Nip1
Prt1
Rpg1
Ded1
Tif35
Cdc9
Chk1
Rad23
Ubi4
Ede1
Ent2
Vps27
Chc1
Ptp3
Stb1
Spa2
Bem2
Bud14
Rck2
Glc7
Ipl1
Cdc5
Dbf4
Pfk26
Tpk1
Gpa2
Ras2
Med6
Rpb10
Paf1
Rpo26
Bdp1
Spt15
Spt5
Rgr1
Srb7
Ssn2
Nop1
Nsr1
Nop56
Snu13
Rps31
Rad9
Rtf1
Rpo21
Npl3
Air2
Fps1
Ask10
Cdc28
Hog1
Pbs2
Ssk2
Clb2
Bud3
Bud4
Msn2
Sch9
Rim15
Yak1
Ypl247c
Vps52
Kss1
Sko1
Slt2
Msg5
Swe1
Igo2
Cdc55
Igo1
Vps53
Vps54
Rtn1
Fmp42
Cot1
Zrc1
10
20
30
40
50
t/s

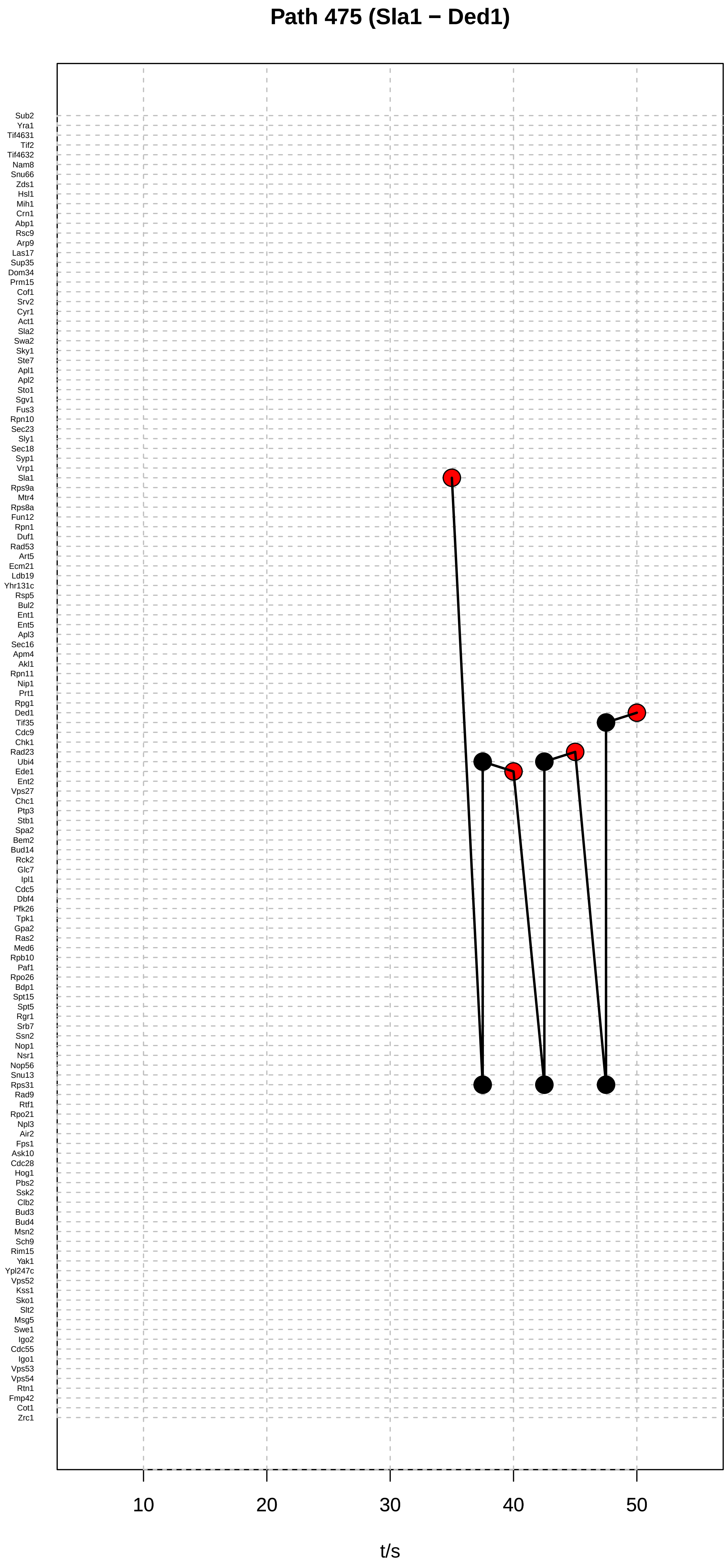

Path 475 (Sla1 − Ded1)
Sub2
Yra1
Tif4631
Tif2
Tif4632
Nam8
Snu66
Zds1
Hsl1
Mih1
Crn1
Abp1
Rsc9
Arp9
Las17
Sup35
Dom34
Prm15
Cof1
Srv2
Cyr1
Act1
Sla2
Swa2
Sky1
Ste7
Apl1
Apl2
Sto1
Sgv1
Fus3
Rpn10
Sec23
Sly1
Sec18
Syp1
Vrp1
Sla1
Rps9a
Mtr4
Rps8a
Fun12
Rpn1
Duf1
Rad53
Art5
Ecm21
Ldb19
Yhr131c
Rsp5
Bul2
Ent1
Ent5
Apl3
Sec16
Apm4
Akl1
Rpn11
Nip1
Prt1
Rpg1
Ded1
Tif35
Cdc9
Chk1
Rad23
Ubi4
Ede1
Ent2
Vps27
Chc1
Ptp3
Stb1
Spa2
Bem2
Bud14
Rck2
Glc7
Ipl1
Cdc5
Dbf4
Pfk26
Tpk1
Gpa2
Ras2
Med6
Rpb10
Paf1
Rpo26
Bdp1
Spt15
Spt5
Rgr1
Srb7
Ssn2
Nop1
Nsr1
Nop56
Snu13
Rps31
Rad9
Rtf1
Rpo21
Npl3
Air2
Fps1
Ask10
Cdc28
Hog1
Pbs2
Ssk2
Clb2
Bud3
Bud4
Msn2
Sch9
Rim15
Yak1
Ypl247c
Vps52
Kss1
Sko1
Slt2
Msg5
Swe1
Igo2
Cdc55
Igo1
Vps53
Vps54
Rtn1
Fmp42
Cot1
Zrc1
10
20
30
40
50
t/s

#### Path 476 (Sla1 − Ded1)

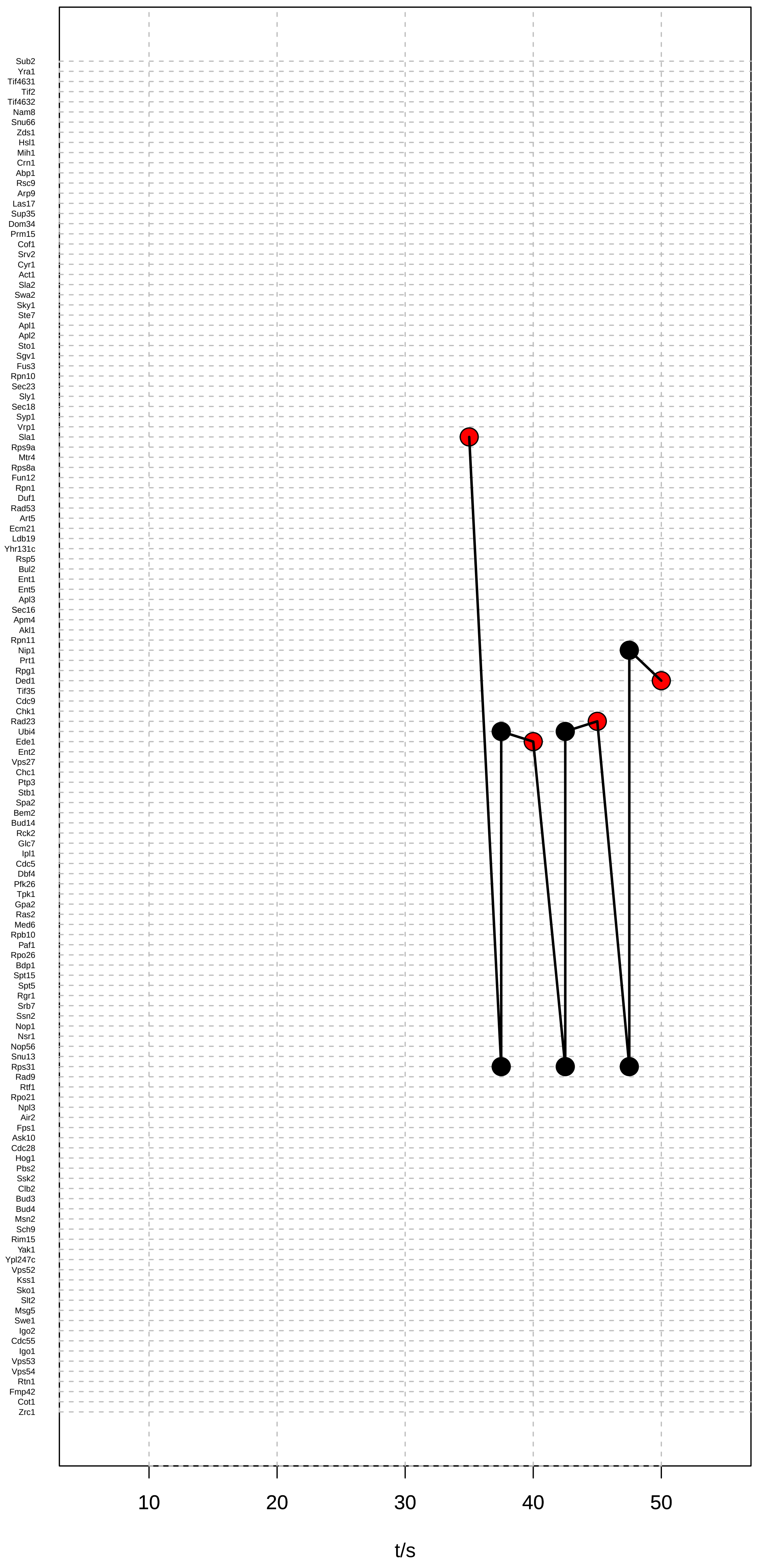

### Path 477 (Sla1 − Ded1)

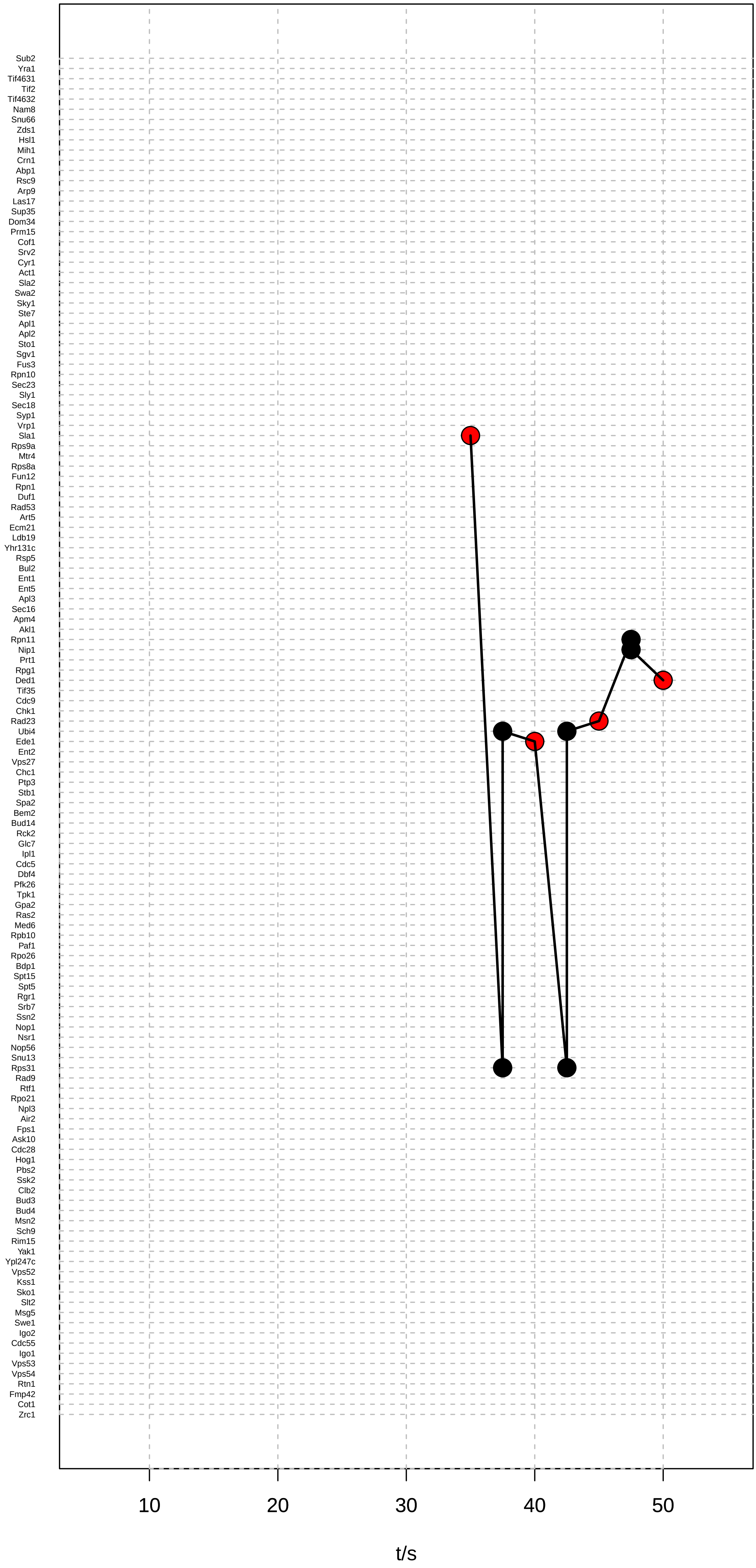

#### Path 478 (Sla1 − Ded1)

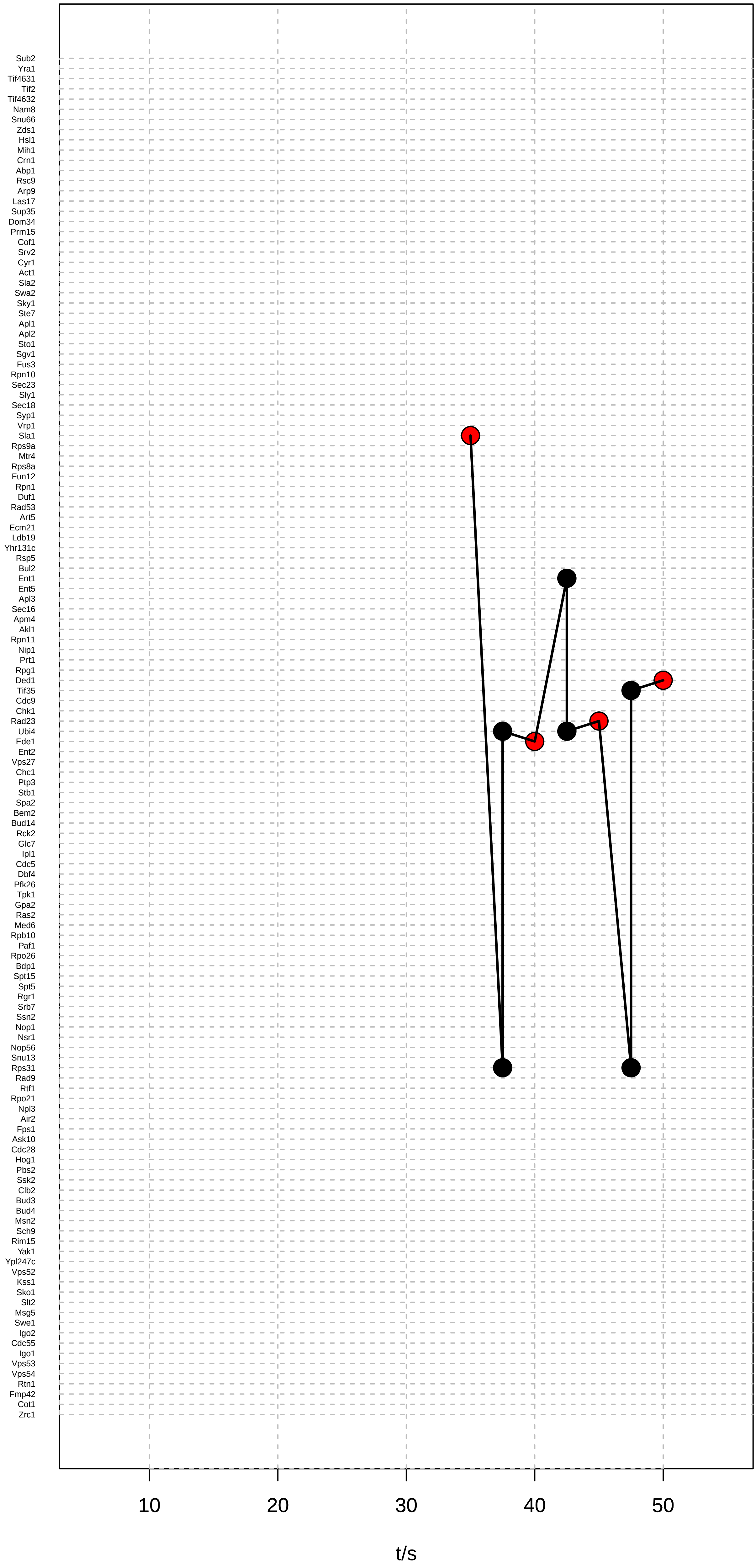

### Path 479 (Sla1 − Ded1)

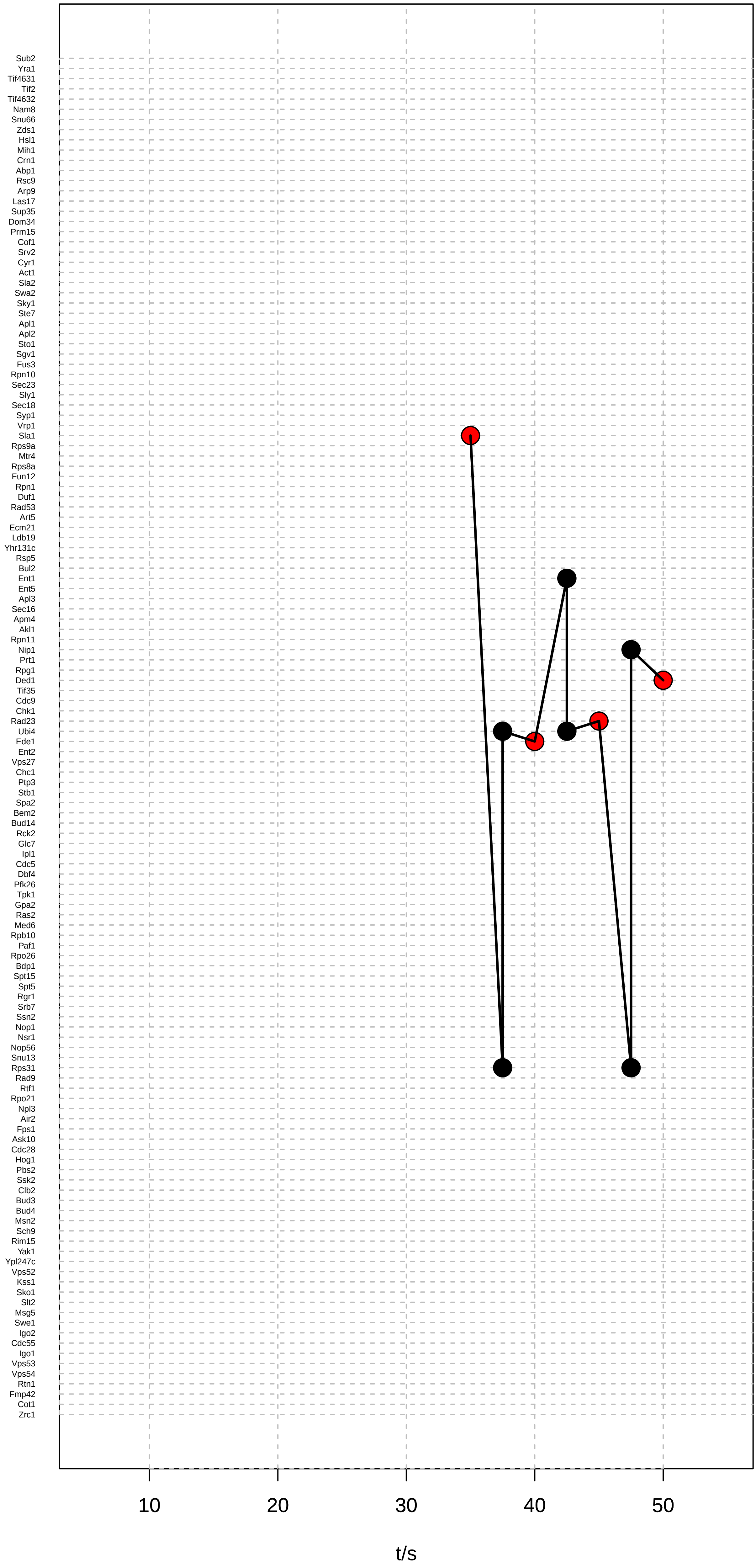

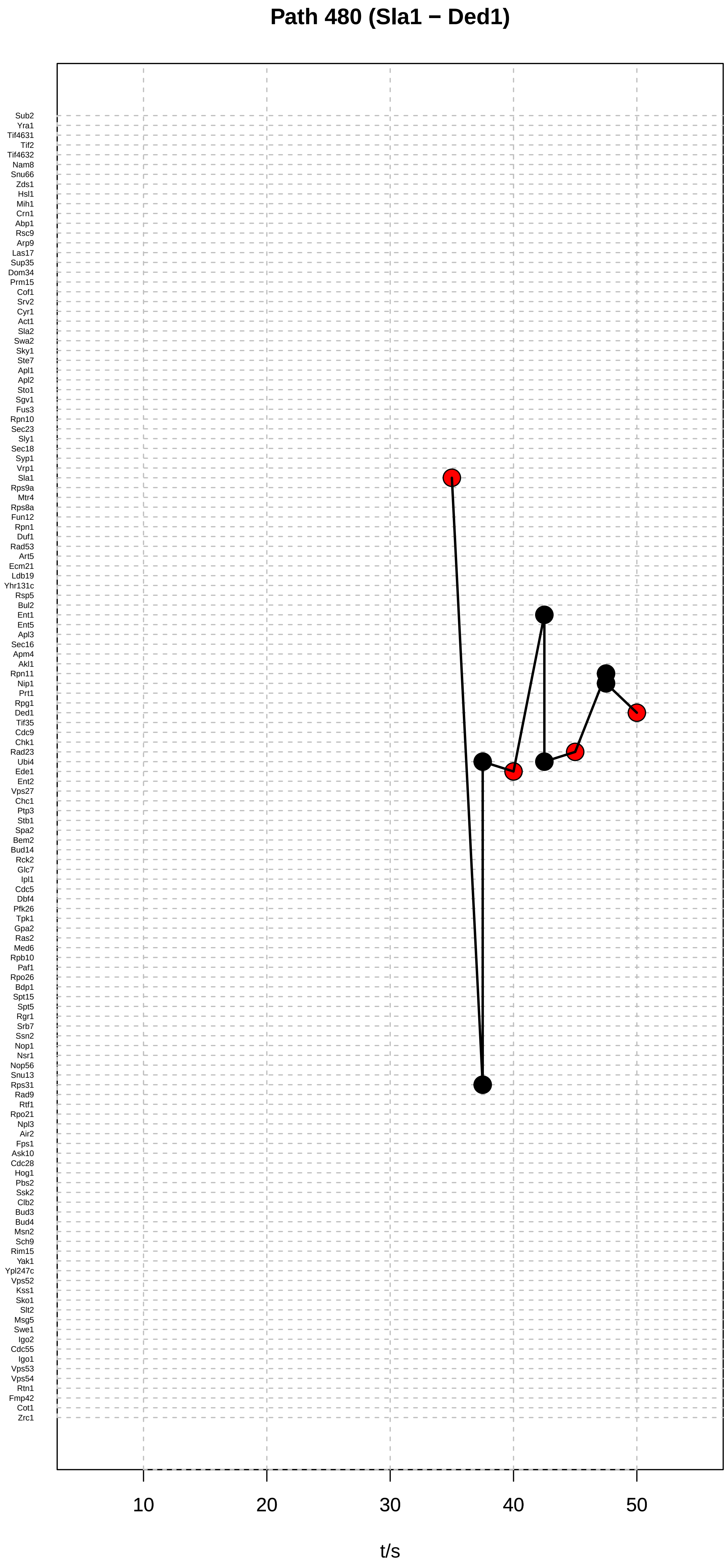

Path 480 (Sla1 − Ded1)
Sub2
Yra1
Tif4631
Tif2
Tif4632
Nam8
Snu66
Zds1
Hsl1
Mih1
Crn1
Abp1
Rsc9
Arp9
Las17
Sup35
Dom34
Prm15
Cof1
Srv2
Cyr1
Act1
Sla2
Swa2
Sky1
Ste7
Apl1
Apl2
Sto1
Sgv1
Fus3
Rpn10
Sec23
Sly1
Sec18
Syp1
Vrp1
Sla1
Rps9a
Mtr4
Rps8a
Fun12
Rpn1
Duf1
Rad53
Art5
Ecm21
Ldb19
Yhr131c
Rsp5
Bul2
Ent1
Ent5
Apl3
Sec16
Apm4
Akl1
Rpn11
Nip1
Prt1
Rpg1
Ded1
Tif35
Cdc9
Chk1
Rad23
Ubi4
Ede1
Ent2
Vps27
Chc1
Ptp3
Stb1
Spa2
Bem2
Bud14
Rck2
Glc7
Ipl1
Cdc5
Dbf4
Pfk26
Tpk1
Gpa2
Ras2
Med6
Rpb10
Paf1
Rpo26
Bdp1
Spt15
Spt5
Rgr1
Srb7
Ssn2
Nop1
Nsr1
Nop56
Snu13
Rps31
Rad9
Rtf1
Rpo21
Npl3
Air2
Fps1
Ask10
Cdc28
Hog1
Pbs2
Ssk2
Clb2
Bud3
Bud4
Msn2
Sch9
Rim15
Yak1
Ypl247c
Vps52
Kss1
Sko1
Slt2
Msg5
Swe1
Igo2
Cdc55
Igo1
Vps53
Vps54
Rtn1
Fmp42
Cot1
Zrc1
10
20
30
40
50
t/s

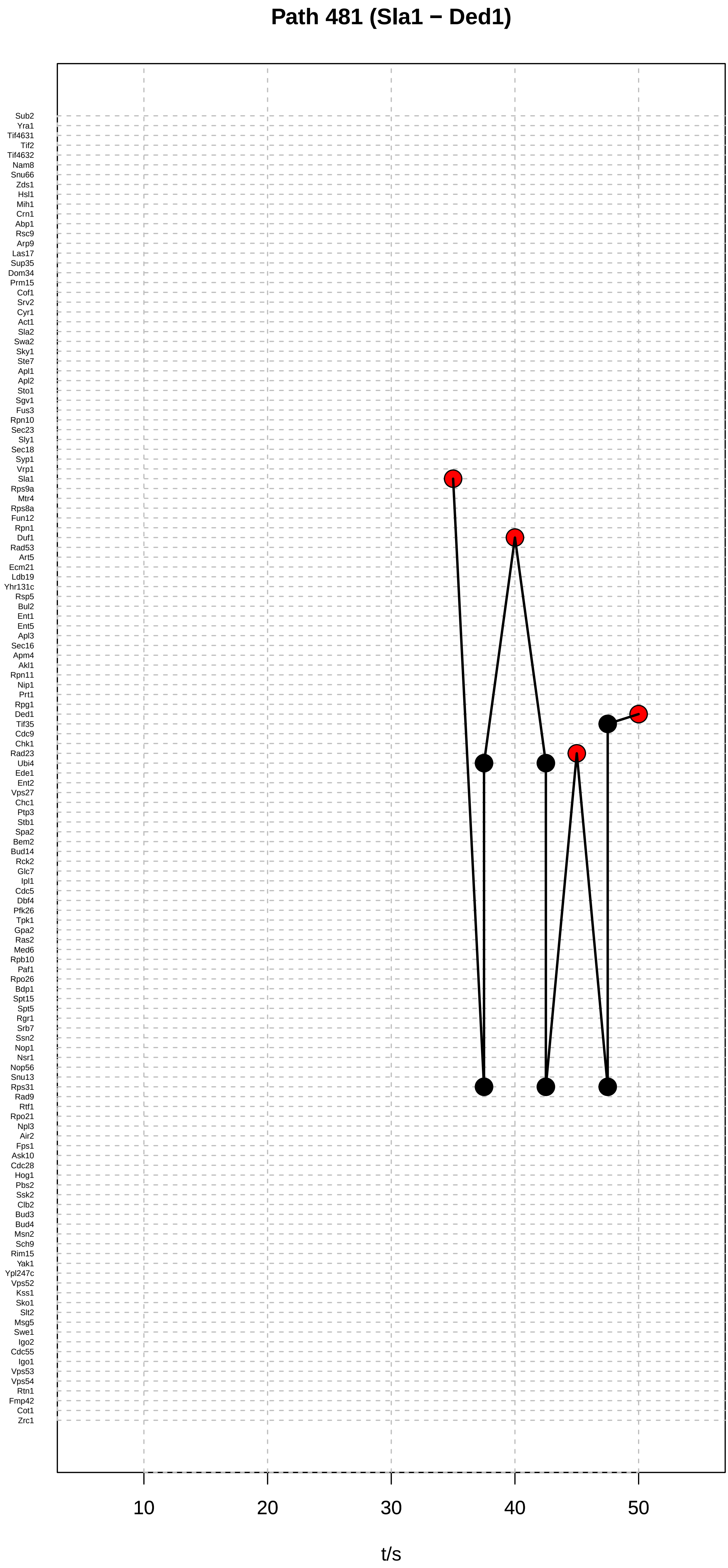

Path 481 (Sla1 − Ded1)
Sub2
Yra1
Tif4631
Tif2
Tif4632
Nam8
Snu66
Zds1
Hsl1
Mih1
Crn1
Abp1
Rsc9
Arp9
Las17
Sup35
Dom34
Prm15
Cof1
Srv2
Cyr1
Act1
Sla2
Swa2
Sky1
Ste7
Apl1
Apl2
Sto1
Sgv1
Fus3
Rpn10
Sec23
Sly1
Sec18
Syp1
Vrp1
Sla1
Rps9a
Mtr4
Rps8a
Fun12
Rpn1
Duf1
Rad53
Art5
Ecm21
Ldb19
Yhr131c
Rsp5
Bul2
Ent1
Ent5
Apl3
Sec16
Apm4
Akl1
Rpn11
Nip1
Prt1
Rpg1
Ded1
Tif35
Cdc9
Chk1
Rad23
Ubi4
Ede1
Ent2
Vps27
Chc1
Ptp3
Stb1
Spa2
Bem2
Bud14
Rck2
Glc7
Ipl1
Cdc5
Dbf4
Pfk26
Tpk1
Gpa2
Ras2
Med6
Rpb10
Paf1
Rpo26
Bdp1
Spt15
Spt5
Rgr1
Srb7
Ssn2
Nop1
Nsr1
Nop56
Snu13
Rps31
Rad9
Rtf1
Rpo21
Npl3
Air2
Fps1
Ask10
Cdc28
Hog1
Pbs2
Ssk2
Clb2
Bud3
Bud4
Msn2
Sch9
Rim15
Yak1
Ypl247c
Vps52
Kss1
Sko1
Slt2
Msg5
Swe1
Igo2
Cdc55
Igo1
Vps53
Vps54
Rtn1
Fmp42
Cot1
Zrc1
10
20
30
40
50
t/s

### Path 482 (Sla1 − Ded1)

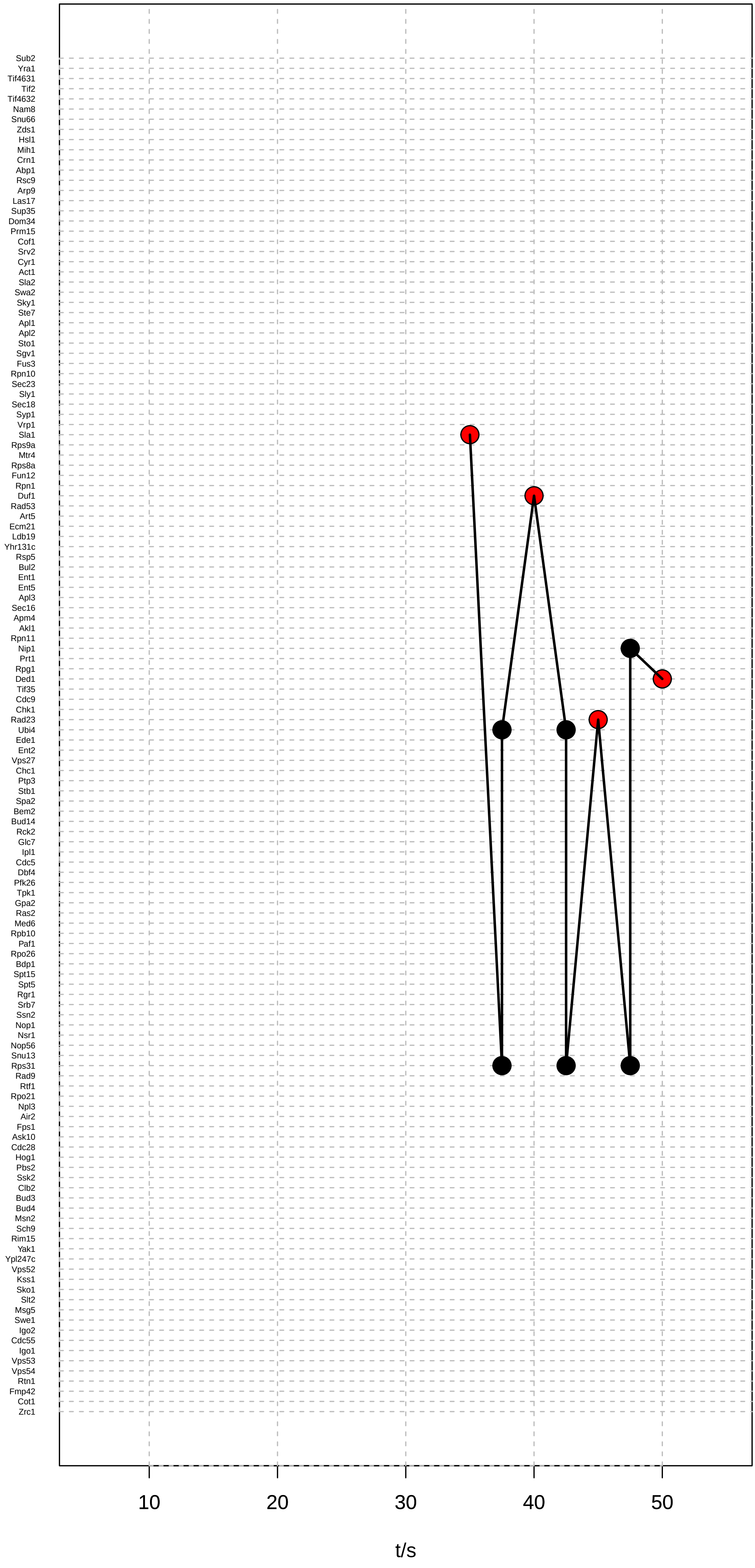

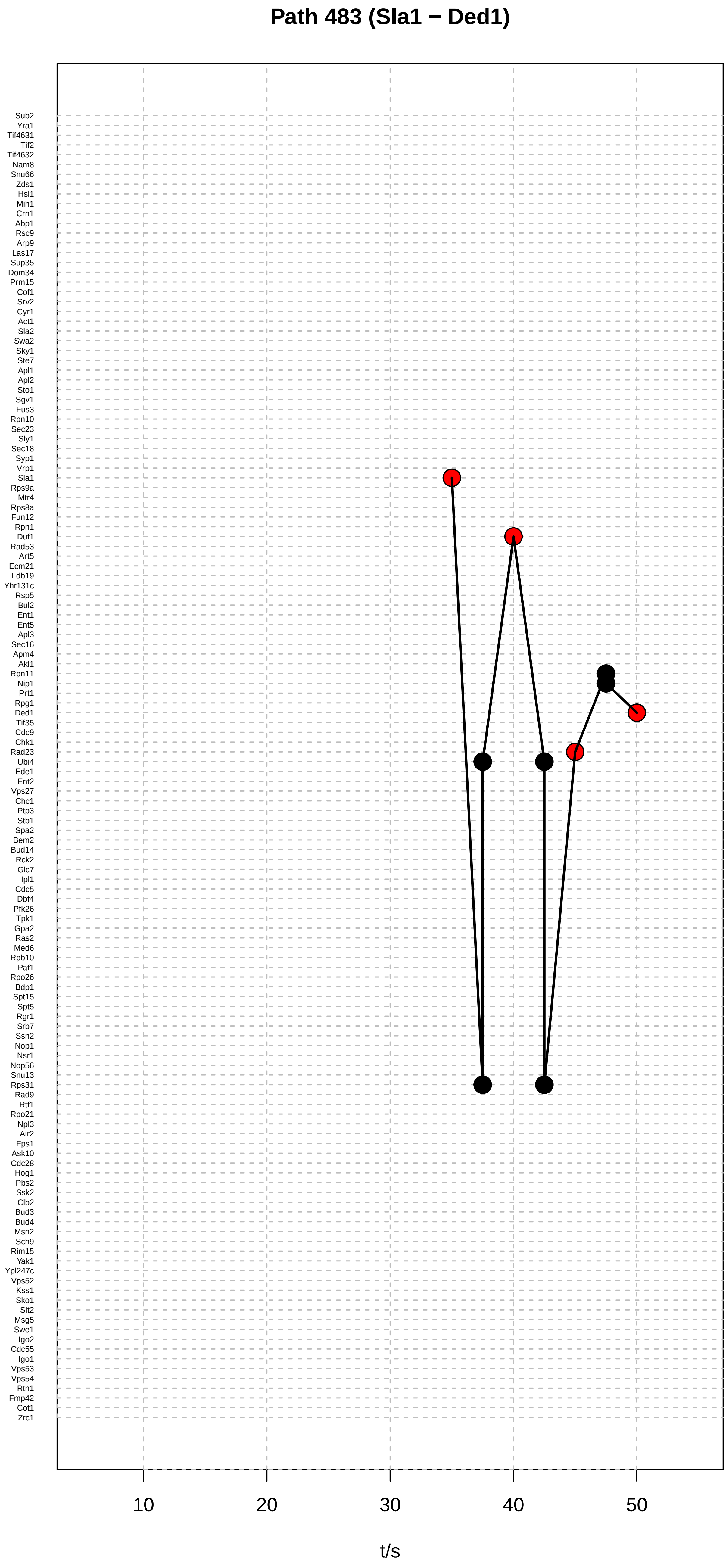

Path 483 (Sla1 − Ded1)
Sub2
Yra1
Tif4631
Tif2
Tif4632
Nam8
Snu66
Zds1
Hsl1
Mih1
Crn1
Abp1
Rsc9
Arp9
Las17
Sup35
Dom34
Prm15
Cof1
Srv2
Cyr1
Act1
Sla2
Swa2
Sky1
Ste7
Apl1
Apl2
Sto1
Sgv1
Fus3
Rpn10
Sec23
Sly1
Sec18
Syp1
Vrp1
Sla1
Rps9a
Mtr4
Rps8a
Fun12
Rpn1
Duf1
Rad53
Art5
Ecm21
Ldb19
Yhr131c
Rsp5
Bul2
Ent1
Ent5
Apl3
Sec16
Apm4
Akl1
Rpn11
Nip1
Prt1
Rpg1
Ded1
Tif35
Cdc9
Chk1
Rad23
Ubi4
Ede1
Ent2
Vps27
Chc1
Ptp3
Stb1
Spa2
Bem2
Bud14
Rck2
Glc7
Ipl1
Cdc5
Dbf4
Pfk26
Tpk1
Gpa2
Ras2
Med6
Rpb10
Paf1
Rpo26
Bdp1
Spt15
Spt5
Rgr1
Srb7
Ssn2
Nop1
Nsr1
Nop56
Snu13
Rps31
Rad9
Rtf1
Rpo21
Npl3
Air2
Fps1
Ask10
Cdc28
Hog1
Pbs2
Ssk2
Clb2
Bud3
Bud4
Msn2
Sch9
Rim15
Yak1
Ypl247c
Vps52
Kss1
Sko1
Slt2
Msg5
Swe1
Igo2
Cdc55
Igo1
Vps53
Vps54
Rtn1
Fmp42
Cot1
Zrc1
10
20
30
40
50
t/s

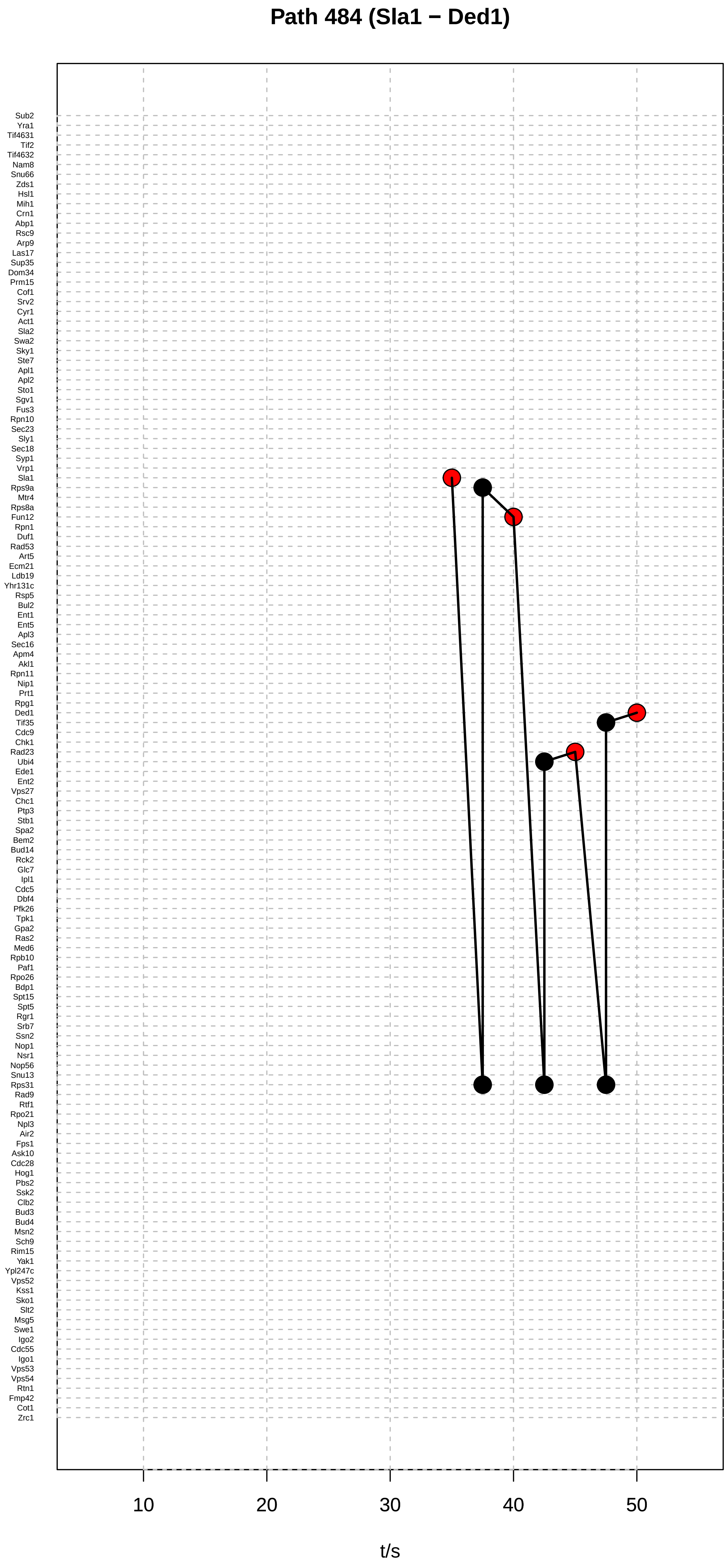

Path 484 (Sla1 − Ded1)
Sub2
Yra1
Tif4631
Tif2
Tif4632
Nam8
Snu66
Zds1
Hsl1
Mih1
Crn1
Abp1
Rsc9
Arp9
Las17
Sup35
Dom34
Prm15
Cof1
Srv2
Cyr1
Act1
Sla2
Swa2
Sky1
Ste7
Apl1
Apl2
Sto1
Sgv1
Fus3
Rpn10
Sec23
Sly1
Sec18
Syp1
Vrp1
Sla1
Rps9a
Mtr4
Rps8a
Fun12
Rpn1
Duf1
Rad53
Art5
Ecm21
Ldb19
Yhr131c
Rsp5
Bul2
Ent1
Ent5
Apl3
Sec16
Apm4
Akl1
Rpn11
Nip1
Prt1
Rpg1
Ded1
Tif35
Cdc9
Chk1
Rad23
Ubi4
Ede1
Ent2
Vps27
Chc1
Ptp3
Stb1
Spa2
Bem2
Bud14
Rck2
Glc7
Ipl1
Cdc5
Dbf4
Pfk26
Tpk1
Gpa2
Ras2
Med6
Rpb10
Paf1
Rpo26
Bdp1
Spt15
Spt5
Rgr1
Srb7
Ssn2
Nop1
Nsr1
Nop56
Snu13
Rps31
Rad9
Rtf1
Rpo21
Npl3
Air2
Fps1
Ask10
Cdc28
Hog1
Pbs2
Ssk2
Clb2
Bud3
Bud4
Msn2
Sch9
Rim15
Yak1
Ypl247c
Vps52
Kss1
Sko1
Slt2
Msg5
Swe1
Igo2
Cdc55
Igo1
Vps53
Vps54
Rtn1
Fmp42
Cot1
Zrc1
10
20
30
40
50
t/s

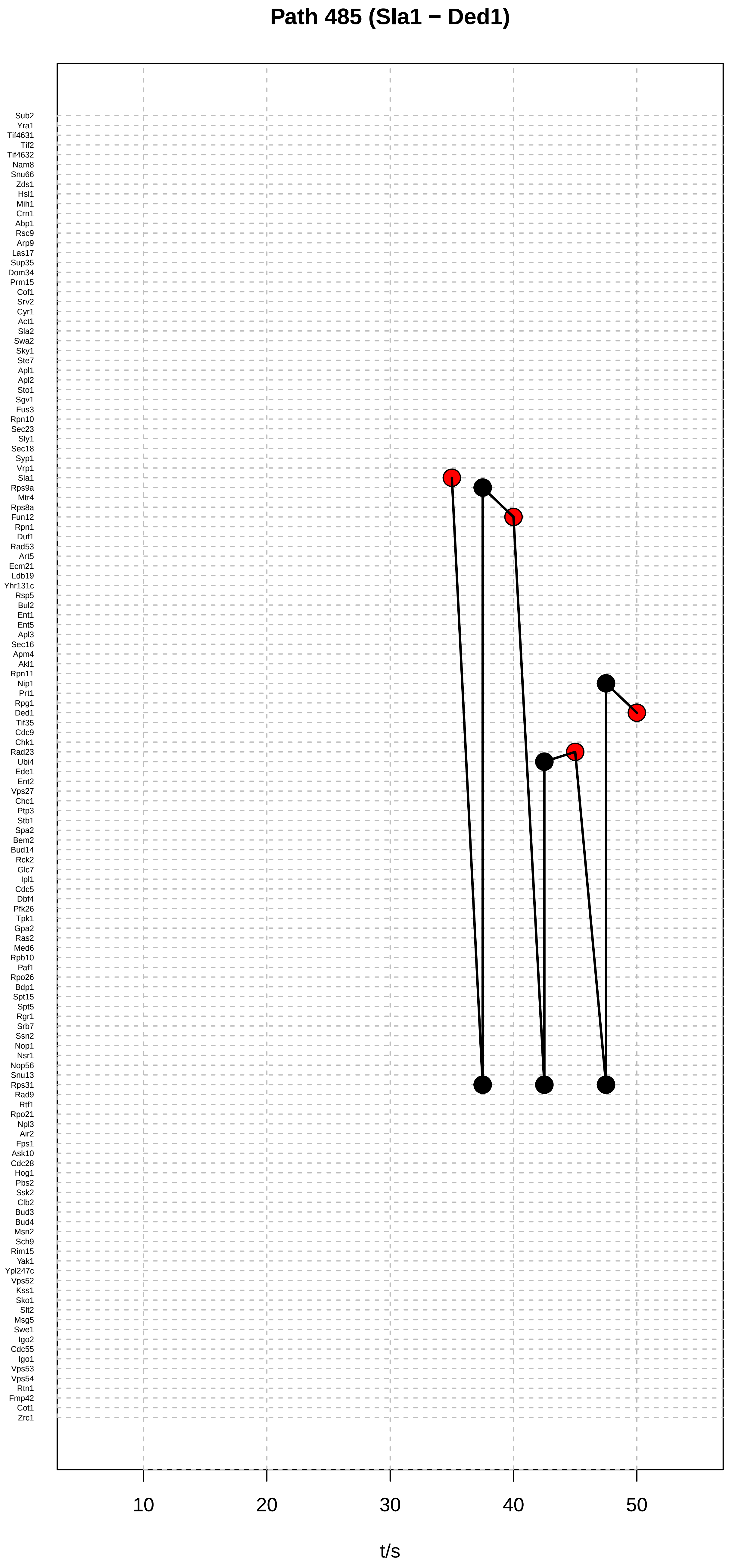

Path 485 (Sla1 − Ded1)
Sub2
Yra1
Tif4631
Tif2
Tif4632
Nam8
Snu66
Zds1
Hsl1
Mih1
Crn1
Abp1
Rsc9
Arp9
Las17
Sup35
Dom34
Prm15
Cof1
Srv2
Cyr1
Act1
Sla2
Swa2
Sky1
Ste7
Apl1
Apl2
Sto1
Sgv1
Fus3
Rpn10
Sec23
Sly1
Sec18
Syp1
Vrp1
Sla1
Rps9a
Mtr4
Rps8a
Fun12
Rpn1
Duf1
Rad53
Art5
Ecm21
Ldb19
Yhr131c
Rsp5
Bul2
Ent1
Ent5
Apl3
Sec16
Apm4
Akl1
Rpn11
Nip1
Prt1
Rpg1
Ded1
Tif35
Cdc9
Chk1
Rad23
Ubi4
Ede1
Ent2
Vps27
Chc1
Ptp3
Stb1
Spa2
Bem2
Bud14
Rck2
Glc7
Ipl1
Cdc5
Dbf4
Pfk26
Tpk1
Gpa2
Ras2
Med6
Rpb10
Paf1
Rpo26
Bdp1
Spt15
Spt5
Rgr1
Srb7
Ssn2
Nop1
Nsr1
Nop56
Snu13
Rps31
Rad9
Rtf1
Rpo21
Npl3
Air2
Fps1
Ask10
Cdc28
Hog1
Pbs2
Ssk2
Clb2
Bud3
Bud4
Msn2
Sch9
Rim15
Yak1
Ypl247c
Vps52
Kss1
Sko1
Slt2
Msg5
Swe1
Igo2
Cdc55
Igo1
Vps53
Vps54
Rtn1
Fmp42
Cot1
Zrc1
10
20
30
40
50
t/s

#### Path 486 (Sla1 − Ded1)

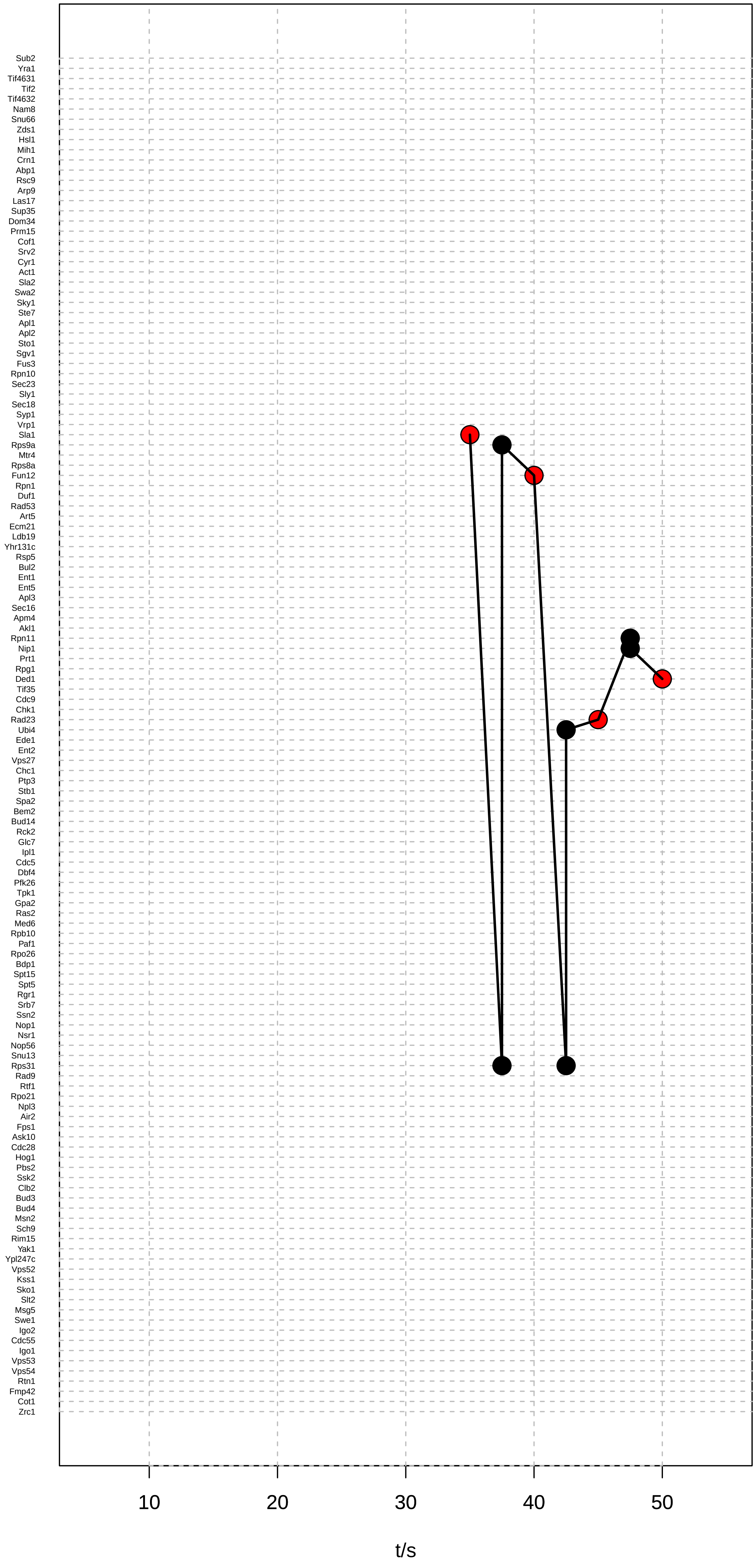

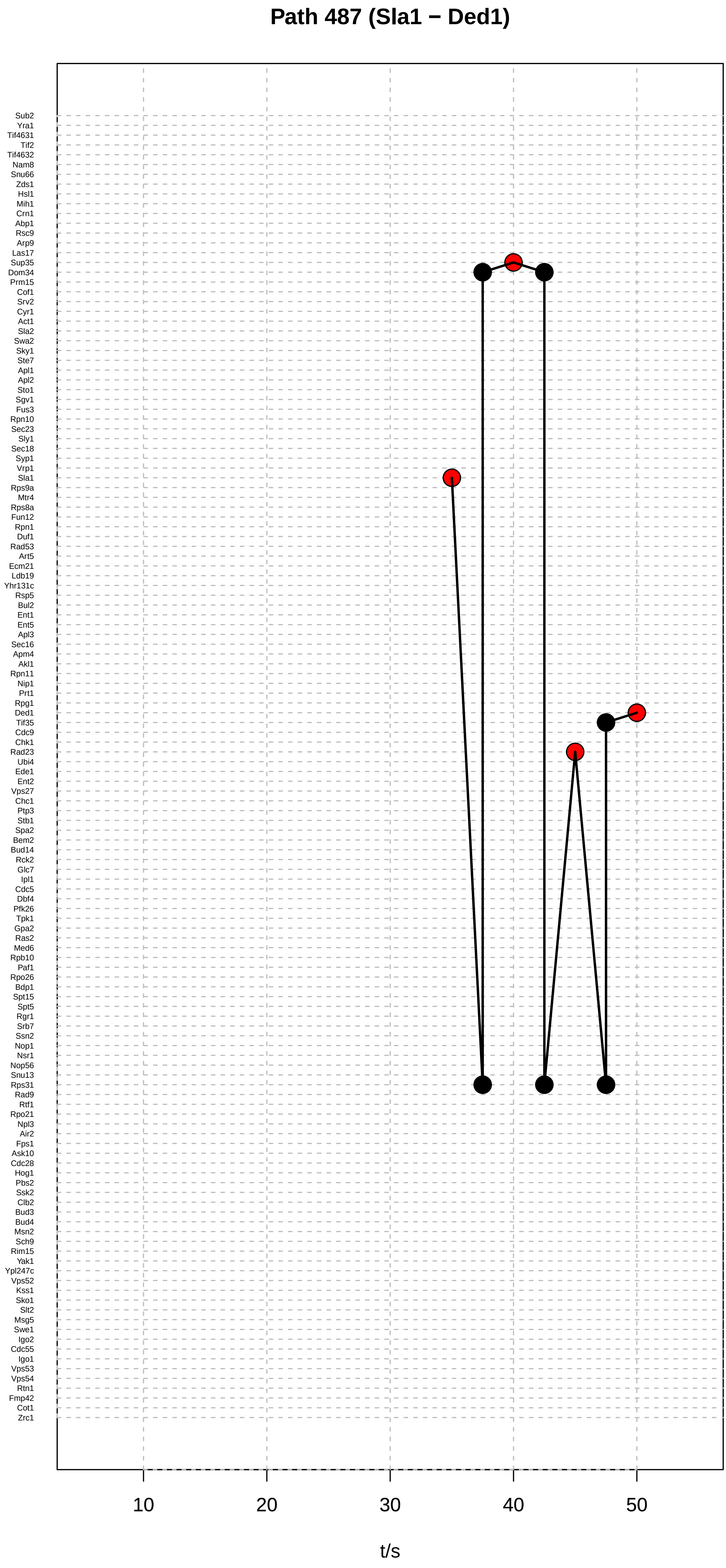
Path 487 (Sla1 − Ded1)
Sub2
Yra1
Tif4631
Tif2
Tif4632
Nam8
Snu66
Zds1
Hsl1
Mih1
Crn1
Abp1
Rsc9
Arp9
Las17
Sup35
Dom34
Prm15
Cof1
Srv2
Cyr1
Act1
Sla2
Swa2
Sky1
Ste7
Apl1
Apl2
Sto1
Sgv1
Fus3
Rpn10
Sec23
Sly1
Sec18
Syp1
Vrp1
Sla1
Rps9a
Mtr4
Rps8a
Fun12
Rpn1
Duf1
Rad53
Art5
Ecm21
Ldb19
Yhr131c
Rsp5
Bul2
Ent1
Ent5
Apl3
Sec16
Apm4
Akl1
Rpn11
Nip1
Prt1
Rpg1
Ded1
Tif35
Cdc9
Chk1
Rad23
Ubi4
Ede1
Ent2
Vps27
Chc1
Ptp3
Stb1
Spa2
Bem2
Bud14
Rck2
Glc7
Ipl1
Cdc5
Dbf4
Pfk26
Tpk1
Gpa2
Ras2
Med6
Rpb10
Paf1
Rpo26
Bdp1
Spt15
Spt5
Rgr1
Srb7
Ssn2
Nop1
Nsr1
Nop56
Snu13
Rps31
Rad9
Rtf1
Rpo21
Npl3
Air2
Fps1
Ask10
Cdc28
Hog1
Pbs2
Ssk2
Clb2
Bud3
Bud4
Msn2
Sch9
Rim15
Yak1
Ypl247c
Vps52
Kss1
Sko1
Slt2
Msg5
Swe1
Igo2
Cdc55
Igo1
Vps53
Vps54
Rtn1
Fmp42
Cot1
Zrc1
10
20
30
40
50
t/s

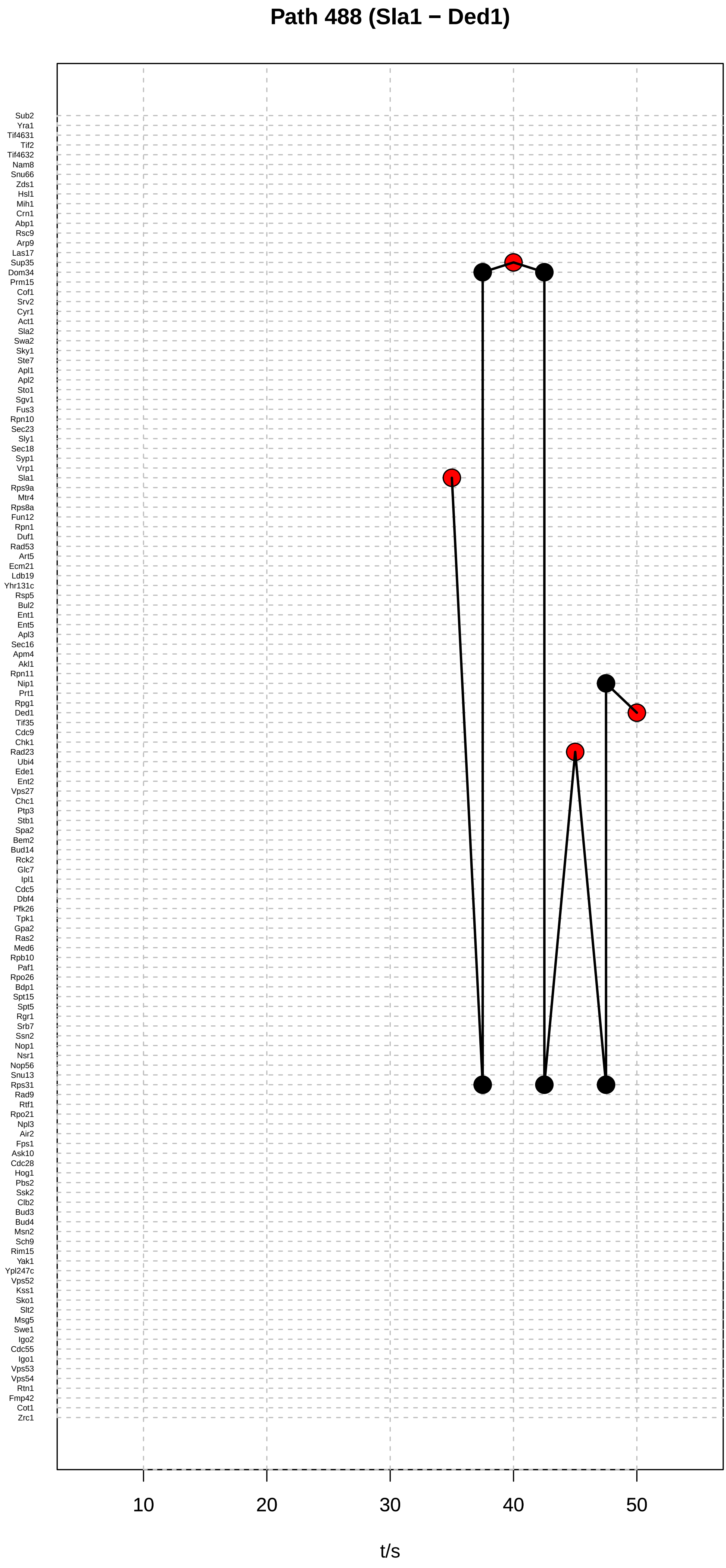

Path 488 (Sla1 − Ded1)
Sub2
Yra1
Tif4631
Tif2
Tif4632
Nam8
Snu66
Zds1
Hsl1
Mih1
Crn1
Abp1
Rsc9
Arp9
Las17
Sup35
Dom34
Prm15
Cof1
Srv2
Cyr1
Act1
Sla2
Swa2
Sky1
Ste7
Apl1
Apl2
Sto1
Sgv1
Fus3
Rpn10
Sec23
Sly1
Sec18
Syp1
Vrp1
Sla1
Rps9a
Mtr4
Rps8a
Fun12
Rpn1
Duf1
Rad53
Art5
Ecm21
Ldb19
Yhr131c
Rsp5
Bul2
Ent1
Ent5
Apl3
Sec16
Apm4
Akl1
Rpn11
Nip1
Prt1
Rpg1
Ded1
Tif35
Cdc9
Chk1
Rad23
Ubi4
Ede1
Ent2
Vps27
Chc1
Ptp3
Stb1
Spa2
Bem2
Bud14
Rck2
Glc7
Ipl1
Cdc5
Dbf4
Pfk26
Tpk1
Gpa2
Ras2
Med6
Rpb10
Paf1
Rpo26
Bdp1
Spt15
Spt5
Rgr1
Srb7
Ssn2
Nop1
Nsr1
Nop56
Snu13
Rps31
Rad9
Rtf1
Rpo21
Npl3
Air2
Fps1
Ask10
Cdc28
Hog1
Pbs2
Ssk2
Clb2
Bud3
Bud4
Msn2
Sch9
Rim15
Yak1
Ypl247c
Vps52
Kss1
Sko1
Slt2
Msg5
Swe1
Igo2
Cdc55
Igo1
Vps53
Vps54
Rtn1
Fmp42
Cot1
Zrc1
10
20
30
40
50
t/s

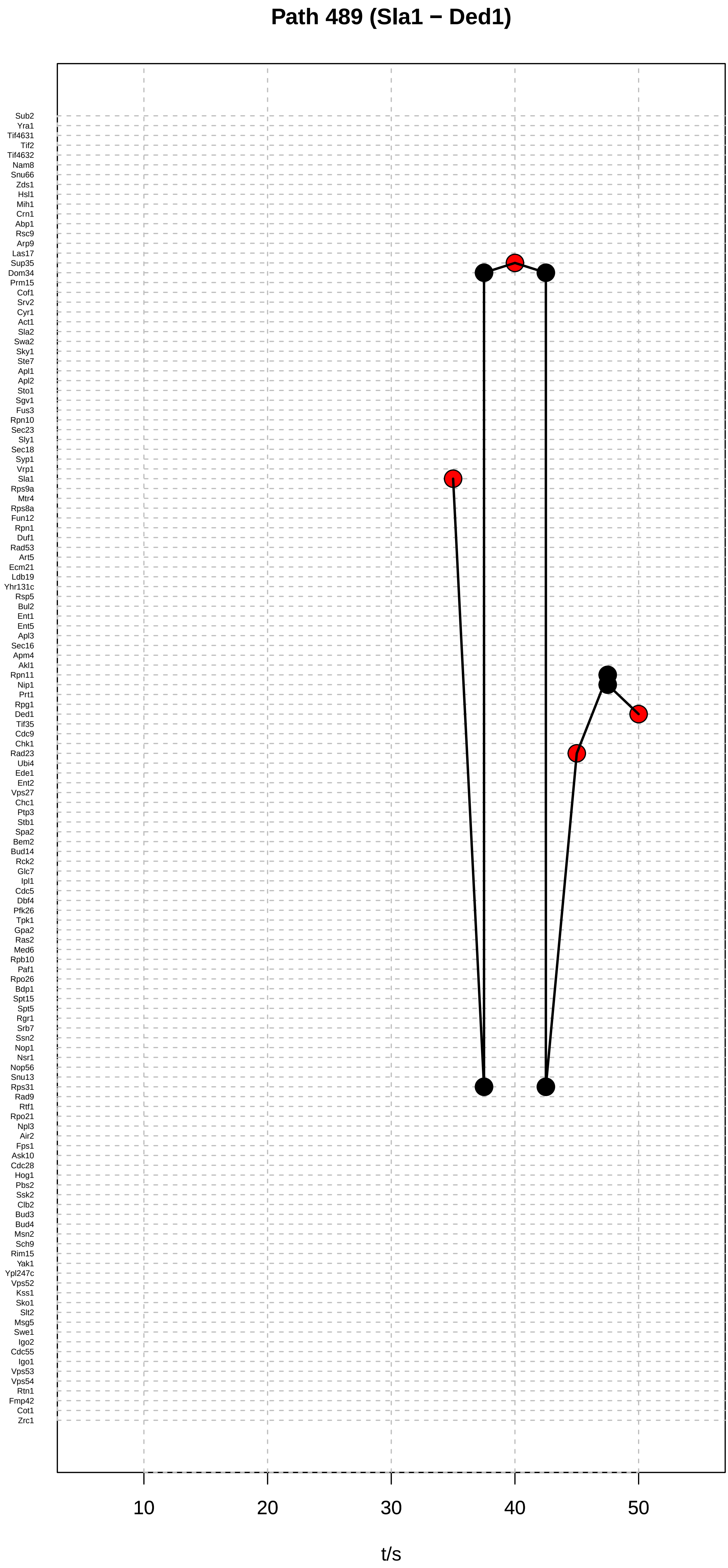

Path 489 (Sla1 − Ded1)
Sub2
Yra1
Tif4631
Tif2
Tif4632
Nam8
Snu66
Zds1
Hsl1
Mih1
Crn1
Abp1
Rsc9
Arp9
Las17
Sup35
Dom34
Prm15
Cof1
Srv2
Cyr1
Act1
Sla2
Swa2
Sky1
Ste7
Apl1
Apl2
Sto1
Sgv1
Fus3
Rpn10
Sec23
Sly1
Sec18
Syp1
Vrp1
Sla1
Rps9a
Mtr4
Rps8a
Fun12
Rpn1
Duf1
Rad53
Art5
Ecm21
Ldb19
Yhr131c
Rsp5
Bul2
Ent1
Ent5
Apl3
Sec16
Apm4
Akl1
Rpn11
Nip1
Prt1
Rpg1
Ded1
Tif35
Cdc9
Chk1
Rad23
Ubi4
Ede1
Ent2
Vps27
Chc1
Ptp3
Stb1
Spa2
Bem2
Bud14
Rck2
Glc7
Ipl1
Cdc5
Dbf4
Pfk26
Tpk1
Gpa2
Ras2
Med6
Rpb10
Paf1
Rpo26
Bdp1
Spt15
Spt5
Rgr1
Srb7
Ssn2
Nop1
Nsr1
Nop56
Snu13
Rps31
Rad9
Rtf1
Rpo21
Npl3
Air2
Fps1
Ask10
Cdc28
Hog1
Pbs2
Ssk2
Clb2
Bud3
Bud4
Msn2
Sch9
Rim15
Yak1
Ypl247c
Vps52
Kss1
Sko1
Slt2
Msg5
Swe1
Igo2
Cdc55
Igo1
Vps53
Vps54
Rtn1
Fmp42
Cot1
Zrc1
10
20
30
40
50
t/s

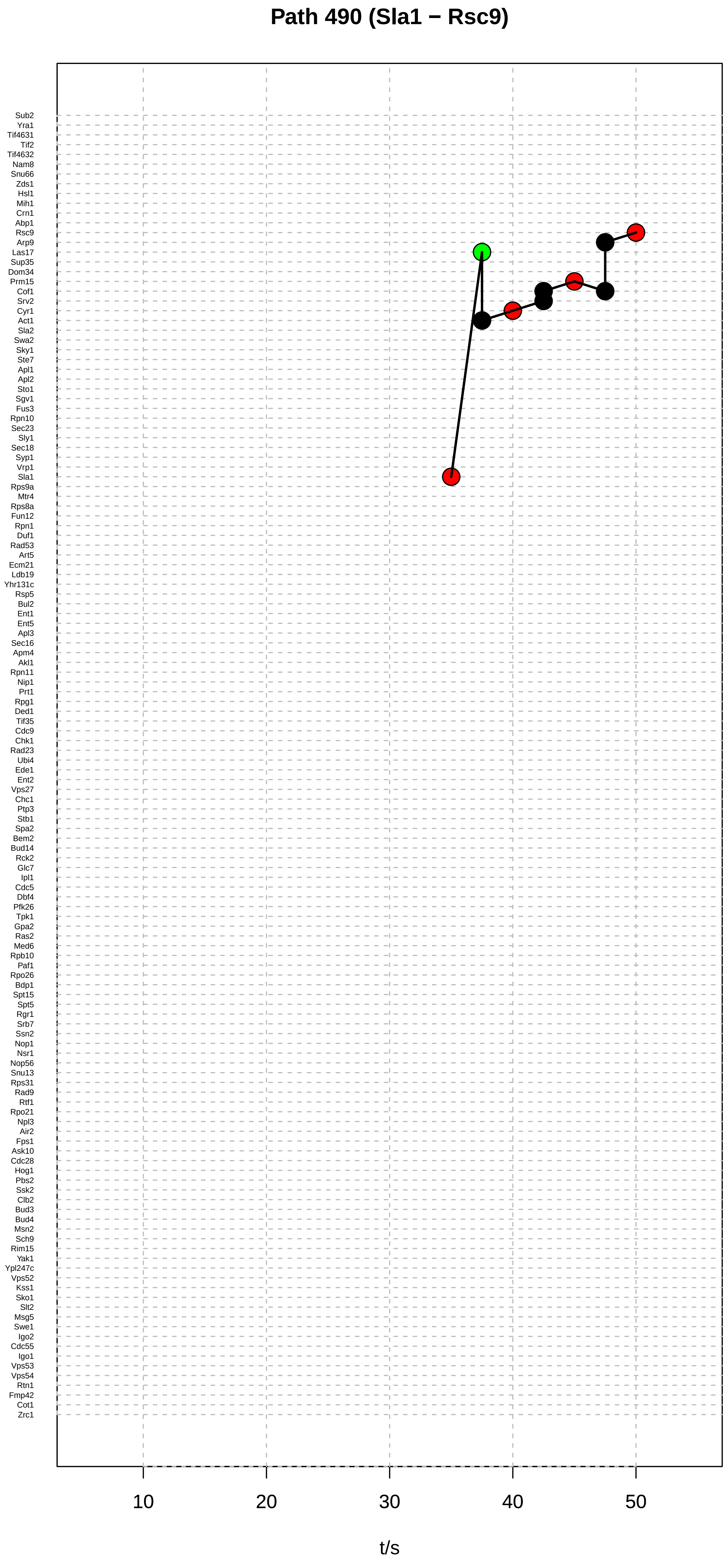
Path 490 (Sla1 − Rsc9)
Sub2
Yra1
Tif4631
Tif2
Tif4632
Nam8
Snu66
Zds1
Hsl1
Mih1
Crn1
Abp1
Rsc9
Arp9
Las17
Sup35
Dom34
Prm15
Cof1
Srv2
Cyr1
Act1
Sla2
Swa2
Sky1
Ste7
Apl1
Apl2
Sto1
Sgv1
Fus3
Rpn10
Sec23
Sly1
Sec18
Syp1
Vrp1
Sla1
Rps9a
Mtr4
Rps8a
Fun12
Rpn1
Duf1
Rad53
Art5
Ecm21
Ldb19
Yhr131c
Rsp5
Bul2
Ent1
Ent5
Apl3
Sec16
Apm4
Akl1
Rpn11
Nip1
Prt1
Rpg1
Ded1
Tif35
Cdc9
Chk1
Rad23
Ubi4
Ede1
Ent2
Vps27
Chc1
Ptp3
Stb1
Spa2
Bem2
Bud14
Rck2
Glc7
Ipl1
Cdc5
Dbf4
Pfk26
Tpk1
Gpa2
Ras2
Med6
Rpb10
Paf1
Rpo26
Bdp1
Spt15
Spt5
Rgr1
Srb7
Ssn2
Nop1
Nsr1
Nop56
Snu13
Rps31
Rad9
Rtf1
Rpo21
Npl3
Air2
Fps1
Ask10
Cdc28
Hog1
Pbs2
Ssk2
Clb2
Bud3
Bud4
Msn2
Sch9
Rim15
Yak1
Ypl247c
Vps52
Kss1
Sko1
Slt2
Msg5
Swe1
Igo2
Cdc55
Igo1
Vps53
Vps54
Rtn1
Fmp42
Cot1
Zrc1
10
20
30
40
50
t/s

### Path 491 (Sla1 − Crn1)

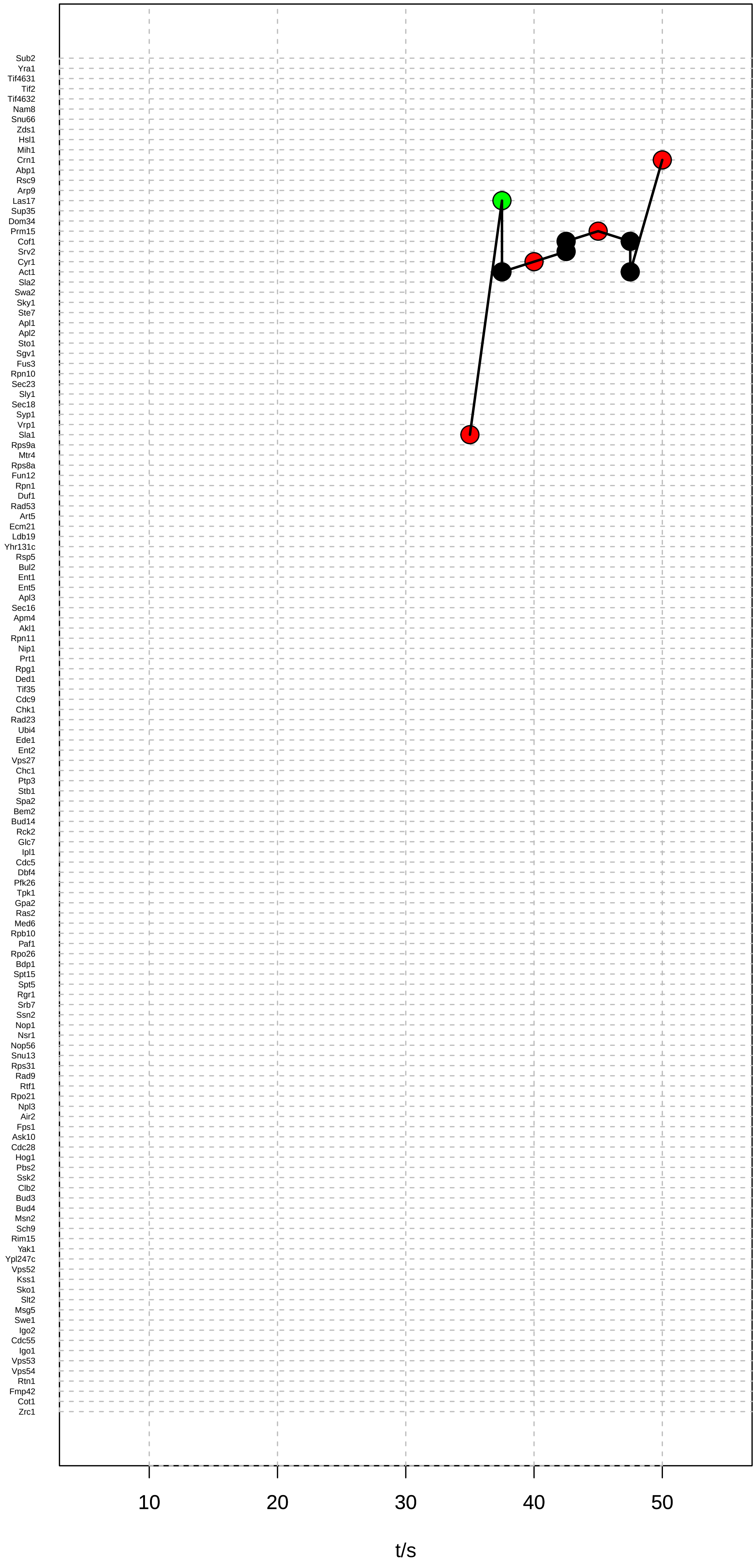

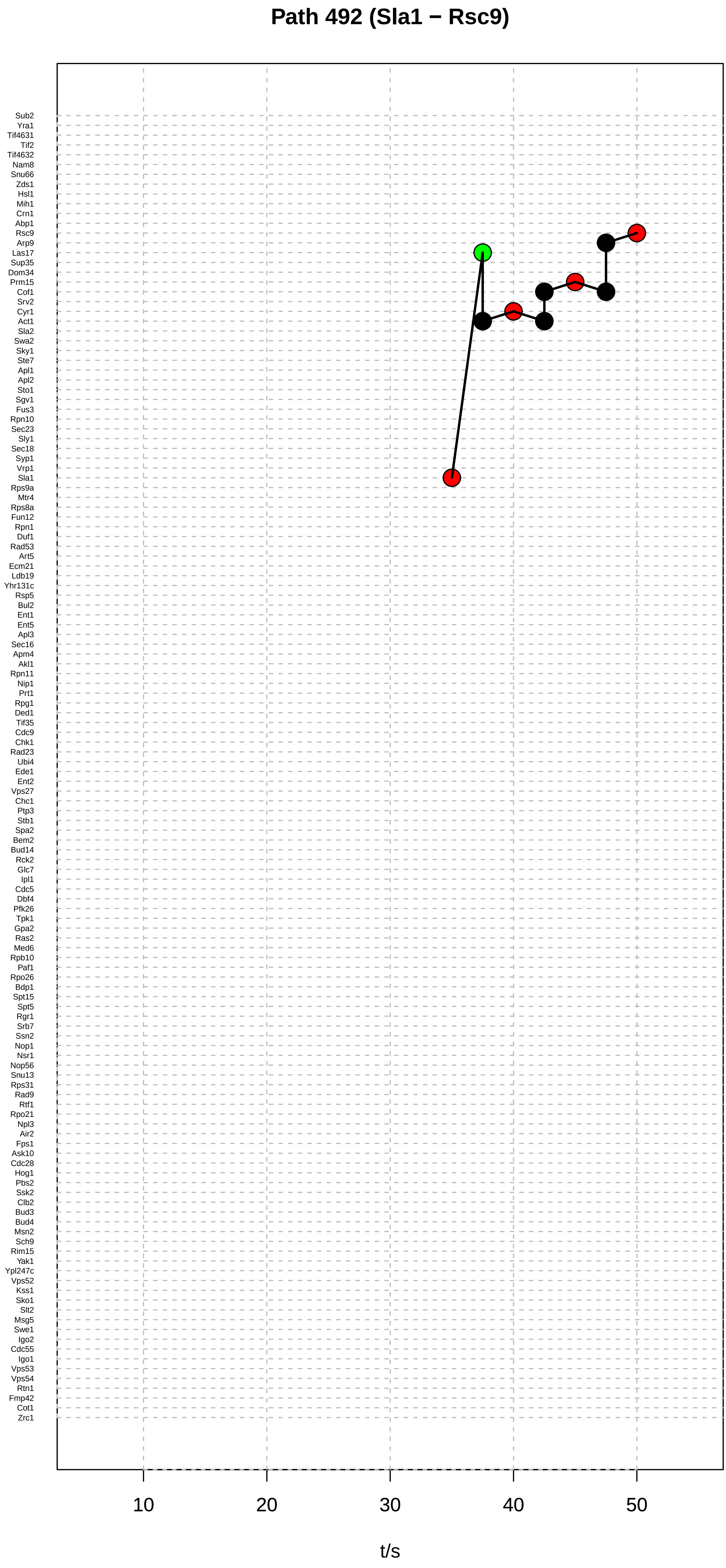

Path 492 (Sla1 − Rsc9)
Sub2
Yra1
Tif4631
Tif2
Tif4632
Nam8
Snu66
Zds1
Hsl1
Mih1
Crn1
Abp1
Rsc9
Arp9
Las17
Sup35
Dom34
Prm15
Cof1
Srv2
Cyr1
Act1
Sla2
Swa2
Sky1
Ste7
Apl1
Apl2
Sto1
Sgv1
Fus3
Rpn10
Sec23
Sly1
Sec18
Syp1
Vrp1
Sla1
Rps9a
Mtr4
Rps8a
Fun12
Rpn1
Duf1
Rad53
Art5
Ecm21
Ldb19
Yhr131c
Rsp5
Bul2
Ent1
Ent5
Apl3
Sec16
Apm4
Akl1
Rpn11
Nip1
Prt1
Rpg1
Ded1
Tif35
Cdc9
Chk1
Rad23
Ubi4
Ede1
Ent2
Vps27
Chc1
Ptp3
Stb1
Spa2
Bem2
Bud14
Rck2
Glc7
Ipl1
Cdc5
Dbf4
Pfk26
Tpk1
Gpa2
Ras2
Med6
Rpb10
Paf1
Rpo26
Bdp1
Spt15
Spt5
Rgr1
Srb7
Ssn2
Nop1
Nsr1
Nop56
Snu13
Rps31
Rad9
Rtf1
Rpo21
Npl3
Air2
Fps1
Ask10
Cdc28
Hog1
Pbs2
Ssk2
Clb2
Bud3
Bud4
Msn2
Sch9
Rim15
Yak1
Ypl247c
Vps52
Kss1
Sko1
Slt2
Msg5
Swe1
Igo2
Cdc55
Igo1
Vps53
Vps54
Rtn1
Fmp42
Cot1
Zrc1
10
20
30
40
50
t/s

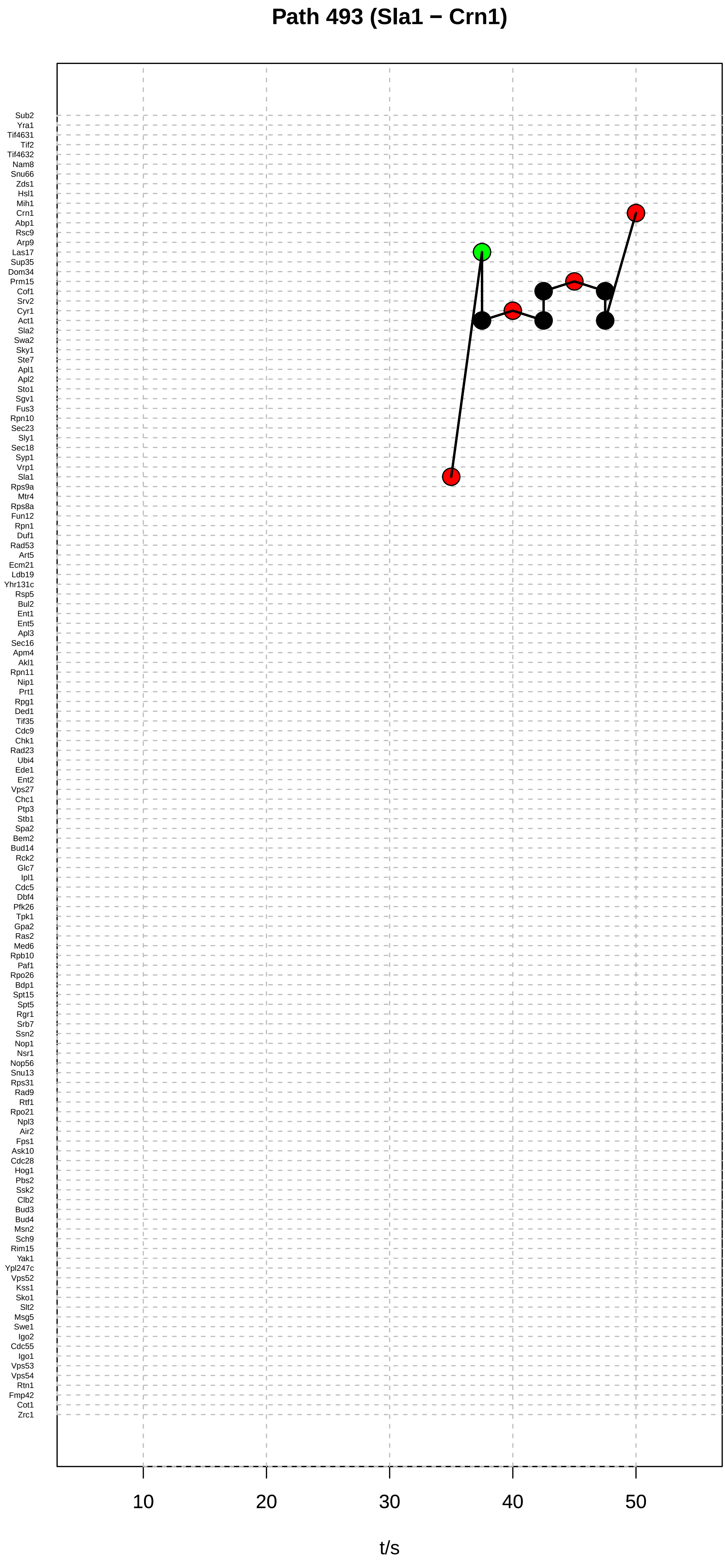
Path 493 (Sla1 − Crn1)
Sub2
Yra1
Tif4631
Tif2
Tif4632
Nam8
Snu66
Zds1
Hsl1
Mih1
Crn1
Abp1
Rsc9
Arp9
Las17
Sup35
Dom34
Prm15
Cof1
Srv2
Cyr1
Act1
Sla2
Swa2
Sky1
Ste7
Apl1
Apl2
Sto1
Sgv1
Fus3
Rpn10
Sec23
Sly1
Sec18
Syp1
Vrp1
Sla1
Rps9a
Mtr4
Rps8a
Fun12
Rpn1
Duf1
Rad53
Art5
Ecm21
Ldb19
Yhr131c
Rsp5
Bul2
Ent1
Ent5
Apl3
Sec16
Apm4
Akl1
Rpn11
Nip1
Prt1
Rpg1
Ded1
Tif35
Cdc9
Chk1
Rad23
Ubi4
Ede1
Ent2
Vps27
Chc1
Ptp3
Stb1
Spa2
Bem2
Bud14
Rck2
Glc7
Ipl1
Cdc5
Dbf4
Pfk26
Tpk1
Gpa2
Ras2
Med6
Rpb10
Paf1
Rpo26
Bdp1
Spt15
Spt5
Rgr1
Srb7
Ssn2
Nop1
Nsr1
Nop56
Snu13
Rps31
Rad9
Rtf1
Rpo21
Npl3
Air2
Fps1
Ask10
Cdc28
Hog1
Pbs2
Ssk2
Clb2
Bud3
Bud4
Msn2
Sch9
Rim15
Yak1
Ypl247c
Vps52
Kss1
Sko1
Slt2
Msg5
Swe1
Igo2
Cdc55
Igo1
Vps53
Vps54
Rtn1
Fmp42
Cot1
Zrc1
10
20
30
40
50
t/s

#### Path 494 (Ldb19 − Ded1)

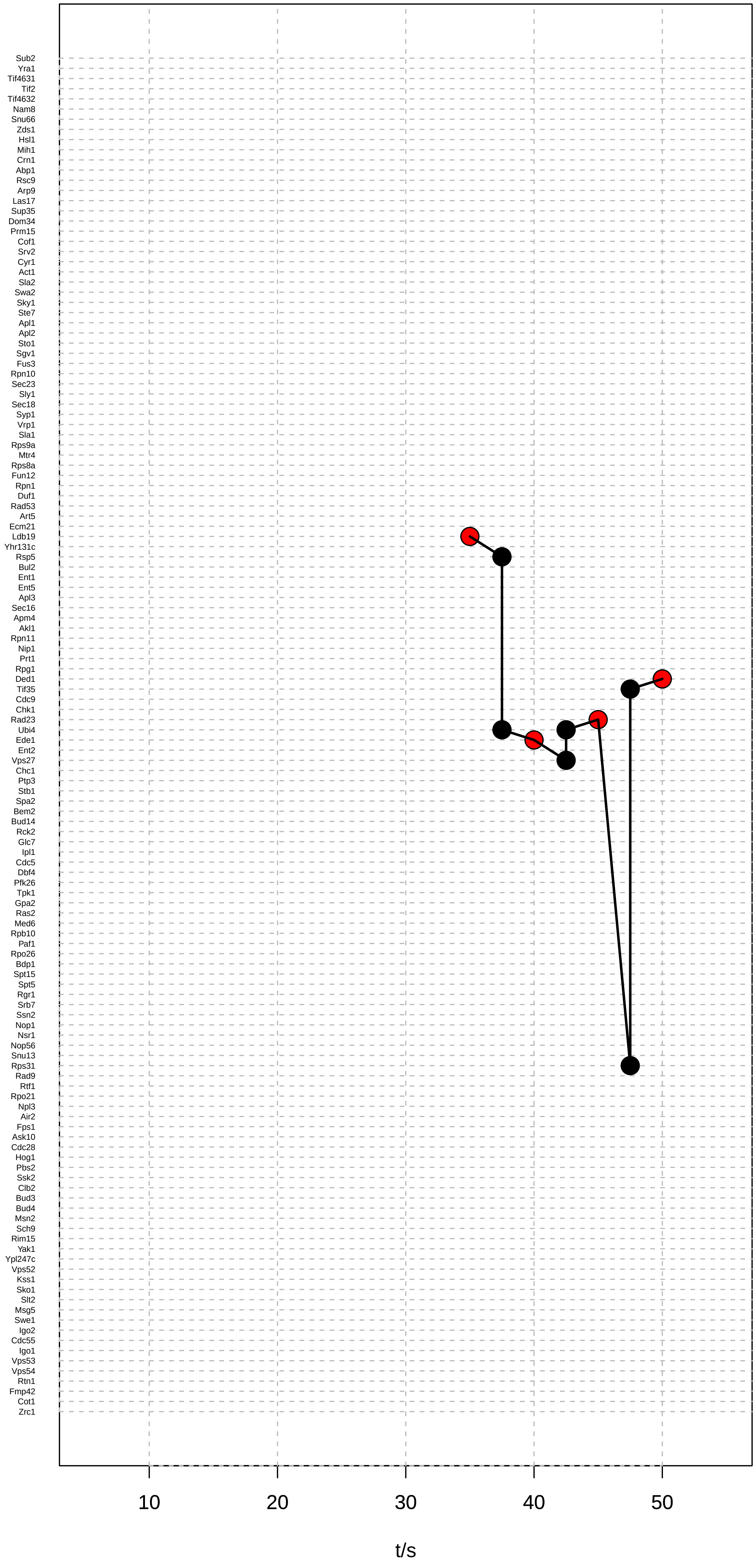

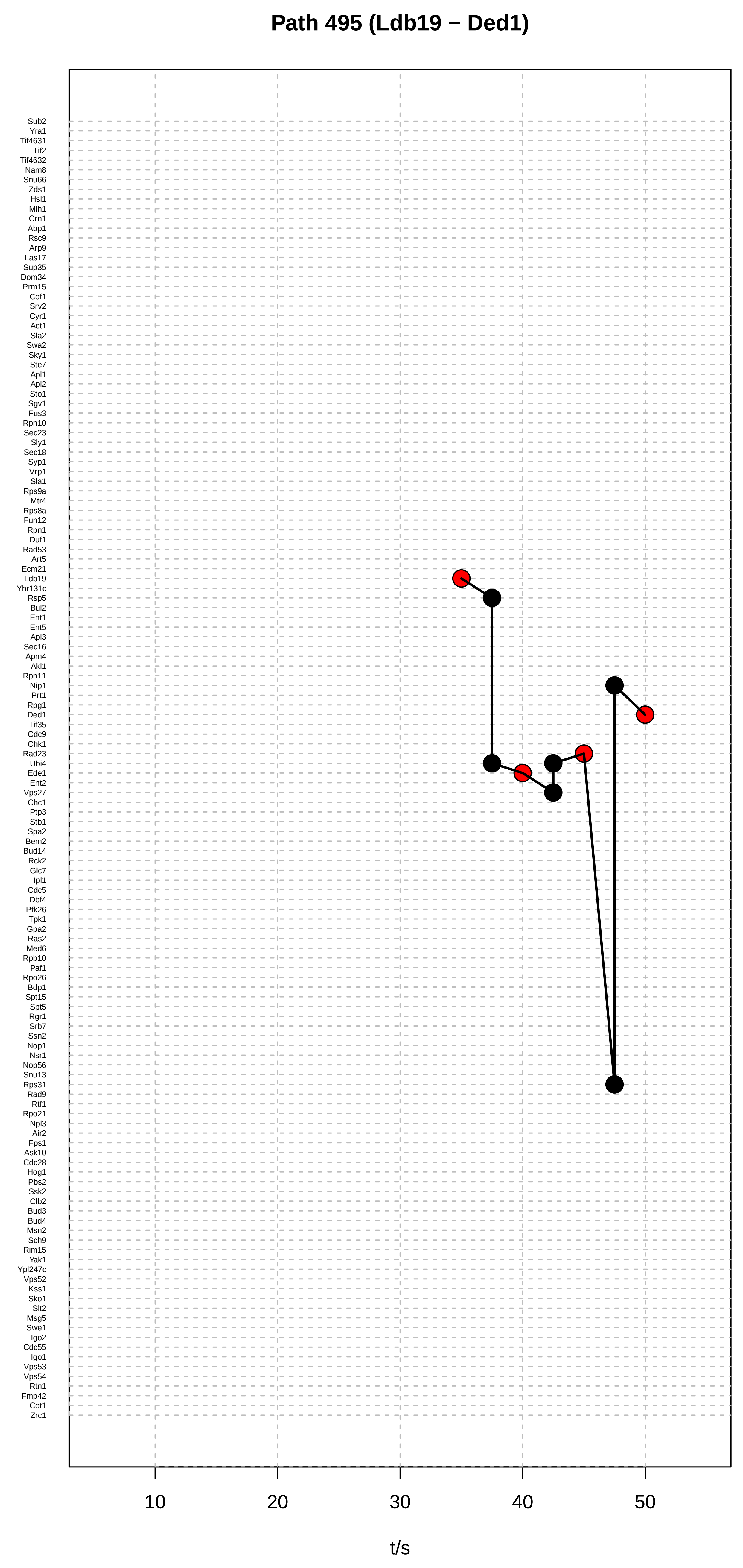

Path 495 (Ldb19 − Ded1)
Sub2
Yra1
Tif4631
Tif2
Tif4632
Nam8
Snu66
Zds1
Hsl1
Mih1
Crn1
Abp1
Rsc9
Arp9
Las17
Sup35
Dom34
Prm15
Cof1
Srv2
Cyr1
Act1
Sla2
Swa2
Sky1
Ste7
Apl1
Apl2
Sto1
Sgv1
Fus3
Rpn10
Sec23
Sly1
Sec18
Syp1
Vrp1
Sla1
Rps9a
Mtr4
Rps8a
Fun12
Rpn1
Duf1
Rad53
Art5
Ecm21
Ldb19
Yhr131c
Rsp5
Bul2
Ent1
Ent5
Apl3
Sec16
Apm4
Akl1
Rpn11
Nip1
Prt1
Rpg1
Ded1
Tif35
Cdc9
Chk1
Rad23
Ubi4
Ede1
Ent2
Vps27
Chc1
Ptp3
Stb1
Spa2
Bem2
Bud14
Rck2
Glc7
Ipl1
Cdc5
Dbf4
Pfk26
Tpk1
Gpa2
Ras2
Med6
Rpb10
Paf1
Rpo26
Bdp1
Spt15
Spt5
Rgr1
Srb7
Ssn2
Nop1
Nsr1
Nop56
Snu13
Rps31
Rad9
Rtf1
Rpo21
Npl3
Air2
Fps1
Ask10
Cdc28
Hog1
Pbs2
Ssk2
Clb2
Bud3
Bud4
Msn2
Sch9
Rim15
Yak1
Ypl247c
Vps52
Kss1
Sko1
Slt2
Msg5
Swe1
Igo2
Cdc55
Igo1
Vps53
Vps54
Rtn1
Fmp42
Cot1
Zrc1
10
20
30
40
50
t/s

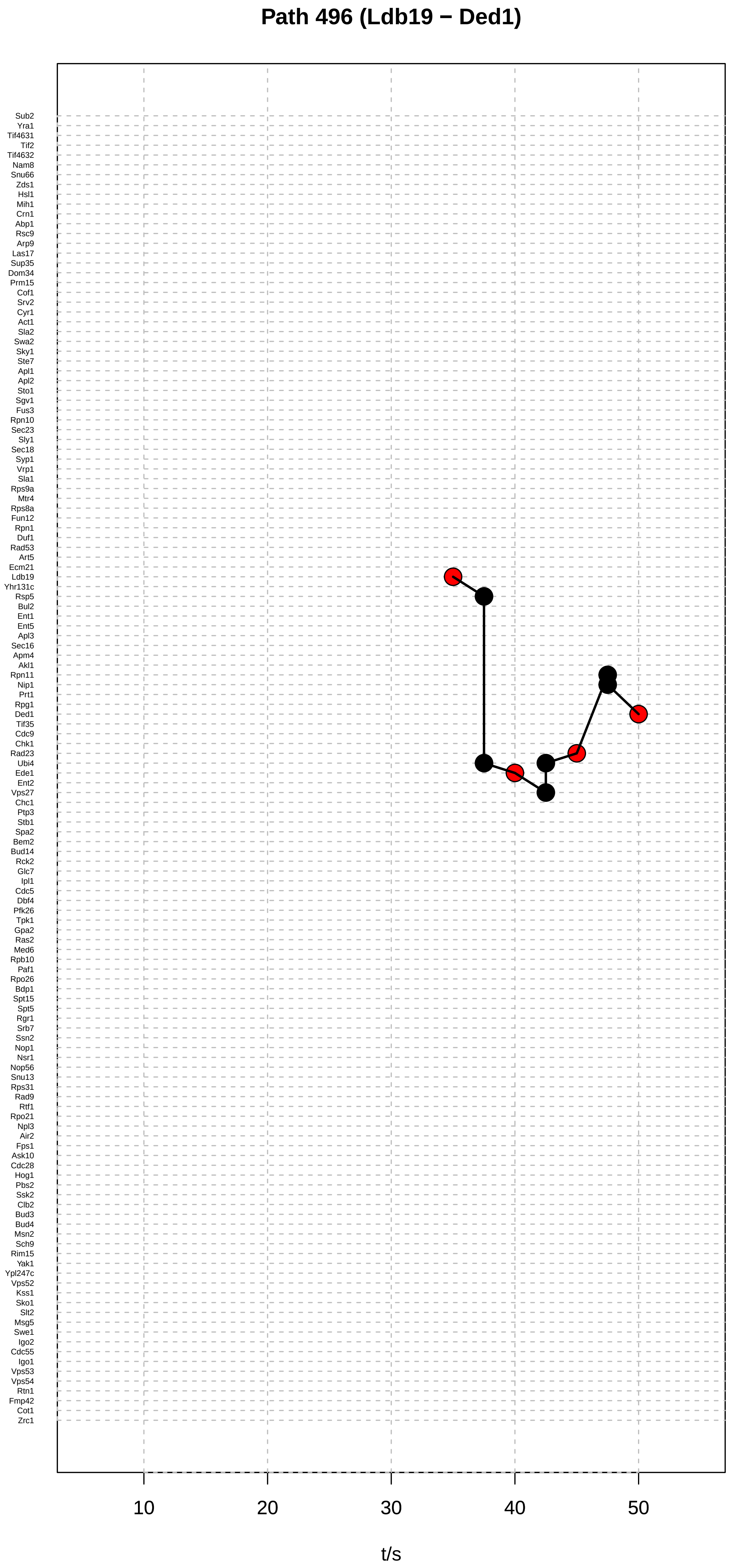

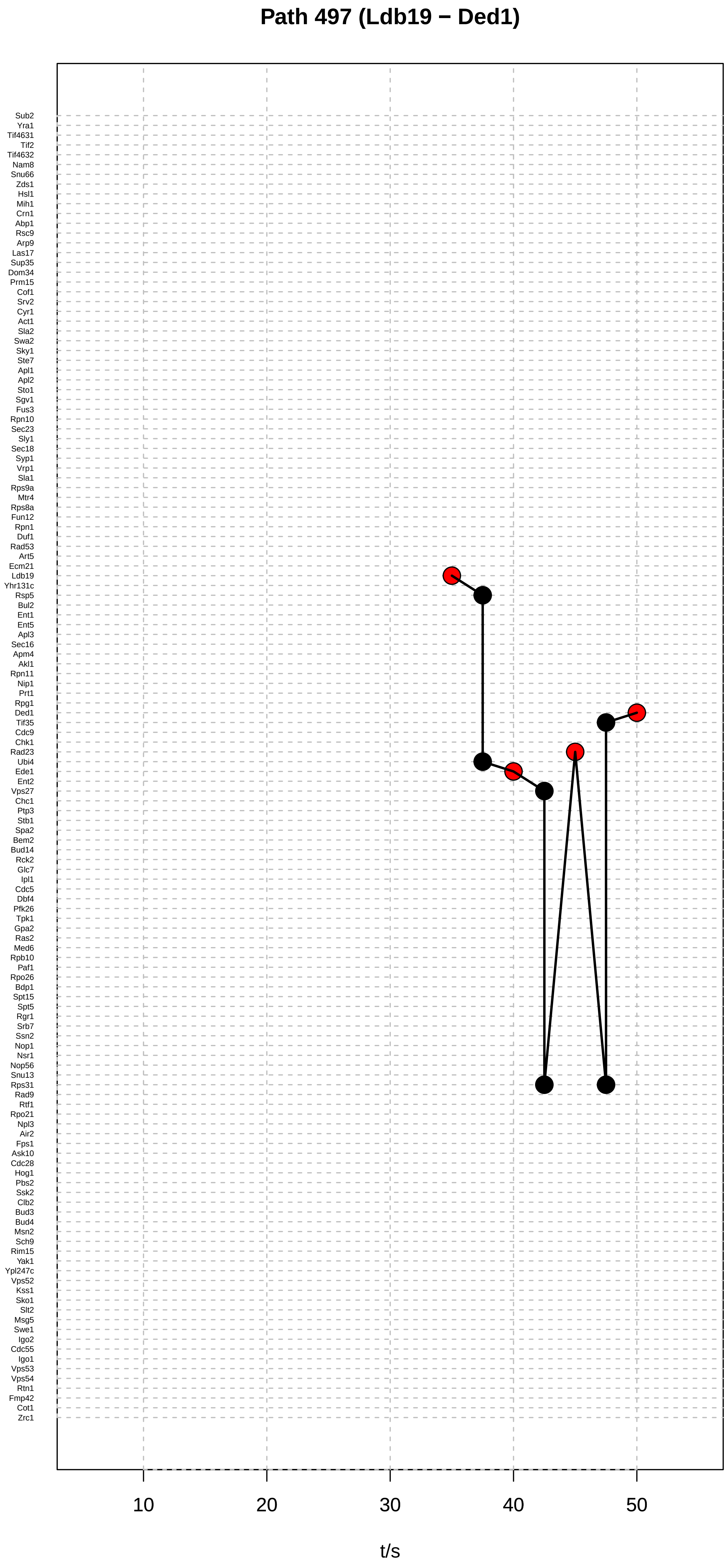
Path 497 (Ldb19 − Ded1)
Sub2
Yra1
Tif4631
Tif2
Tif4632
Nam8
Snu66
Zds1
Hsl1
Mih1
Crn1
Abp1
Rsc9
Arp9
Las17
Sup35
Dom34
Prm15
Cof1
Srv2
Cyr1
Act1
Sla2
Swa2
Sky1
Ste7
Apl1
Apl2
Sto1
Sgv1
Fus3
Rpn10
Sec23
Sly1
Sec18
Syp1
Vrp1
Sla1
Rps9a
Mtr4
Rps8a
Fun12
Rpn1
Duf1
Rad53
Art5
Ecm21
Ldb19
Yhr131c
Rsp5
Bul2
Ent1
Ent5
Apl3
Sec16
Apm4
Akl1
Rpn11
Nip1
Prt1
Rpg1
Ded1
Tif35
Cdc9
Chk1
Rad23
Ubi4
Ede1
Ent2
Vps27
Chc1
Ptp3
Stb1
Spa2
Bem2
Bud14
Rck2
Glc7
Ipl1
Cdc5
Dbf4
Pfk26
Tpk1
Gpa2
Ras2
Med6
Rpb10
Paf1
Rpo26
Bdp1
Spt15
Spt5
Rgr1
Srb7
Ssn2
Nop1
Nsr1
Nop56
Snu13
Rps31
Rad9
Rtf1
Rpo21
Npl3
Air2
Fps1
Ask10
Cdc28
Hog1
Pbs2
Ssk2
Clb2
Bud3
Bud4
Msn2
Sch9
Rim15
Yak1
Ypl247c
Vps52
Kss1
Sko1
Slt2
Msg5
Swe1
Igo2
Cdc55
Igo1
Vps53
Vps54
Rtn1
Fmp42
Cot1
Zrc1
10
20
30
40
50
t/s

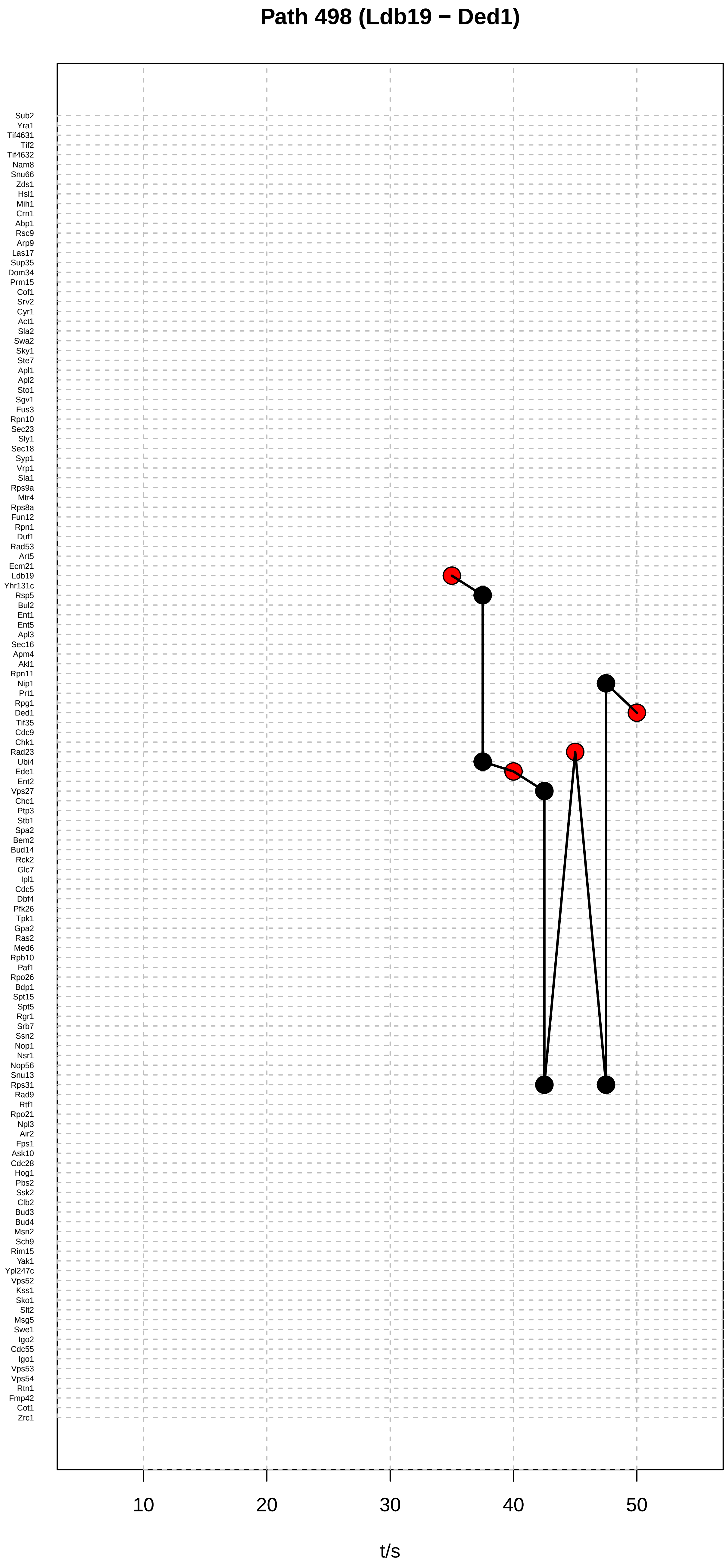

Path 498 (Ldb19 − Ded1)
Sub2
Yra1
Tif4631
Tif2
Tif4632
Nam8
Snu66
Zds1
Hsl1
Mih1
Crn1
Abp1
Rsc9
Arp9
Las17
Sup35
Dom34
Prm15
Cof1
Srv2
Cyr1
Act1
Sla2
Swa2
Sky1
Ste7
Apl1
Apl2
Sto1
Sgv1
Fus3
Rpn10
Sec23
Sly1
Sec18
Syp1
Vrp1
Sla1
Rps9a
Mtr4
Rps8a
Fun12
Rpn1
Duf1
Rad53
Art5
Ecm21
Ldb19
Yhr131c
Rsp5
Bul2
Ent1
Ent5
Apl3
Sec16
Apm4
Akl1
Rpn11
Nip1
Prt1
Rpg1
Ded1
Tif35
Cdc9
Chk1
Rad23
Ubi4
Ede1
Ent2
Vps27
Chc1
Ptp3
Stb1
Spa2
Bem2
Bud14
Rck2
Glc7
Ipl1
Cdc5
Dbf4
Pfk26
Tpk1
Gpa2
Ras2
Med6
Rpb10
Paf1
Rpo26
Bdp1
Spt15
Spt5
Rgr1
Srb7
Ssn2
Nop1
Nsr1
Nop56
Snu13
Rps31
Rad9
Rtf1
Rpo21
Npl3
Air2
Fps1
Ask10
Cdc28
Hog1
Pbs2
Ssk2
Clb2
Bud3
Bud4
Msn2
Sch9
Rim15
Yak1
Ypl247c
Vps52
Kss1
Sko1
Slt2
Msg5
Swe1
Igo2
Cdc55
Igo1
Vps53
Vps54
Rtn1
Fmp42
Cot1
Zrc1
10
20
30
40
50
t/s

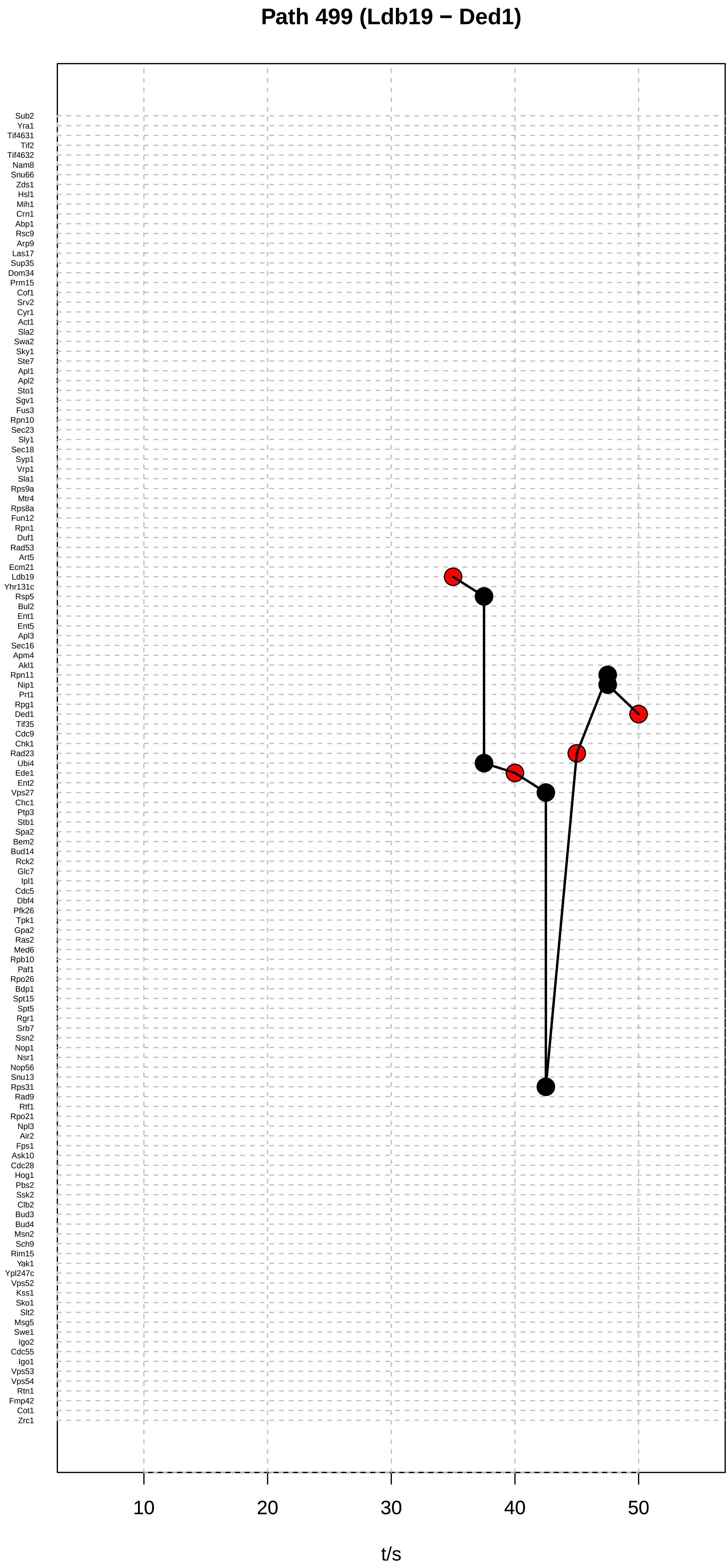

Path 499 (Ldb19 − Ded1)
Sub2
Yra1
Tif4631
Tif2
Tif4632
Nam8
Snu66
Zds1
Hsl1
Mih1
Crn1
Abp1
Rsc9
Arp9
Las17
Sup35
Dom34
Prm15
Cof1
Srv2
Cyr1
Act1
Sla2
Swa2
Sky1
Ste7
Apl1
Apl2
Sto1
Sgv1
Fus3
Rpn10
Sec23
Sly1
Sec18
Syp1
Vrp1
Sla1
Rps9a
Mtr4
Rps8a
Fun12
Rpn1
Duf1
Rad53
Art5
Ecm21
Ldb19
Yhr131c
Rsp5
Bul2
Ent1
Ent5
Apl3
Sec16
Apm4
Akl1
Rpn11
Nip1
Prt1
Rpg1
Ded1
Tif35
Cdc9
Chk1
Rad23
Ubi4
Ede1
Ent2
Vps27
Chc1
Ptp3
Stb1
Spa2
Bem2
Bud14
Rck2
Glc7
Ipl1
Cdc5
Dbf4
Pfk26
Tpk1
Gpa2
Ras2
Med6
Rpb10
Paf1
Rpo26
Bdp1
Spt15
Spt5
Rgr1
Srb7
Ssn2
Nop1
Nsr1
Nop56
Snu13
Rps31
Rad9
Rtf1
Rpo21
Npl3
Air2
Fps1
Ask10
Cdc28
Hog1
Pbs2
Ssk2
Clb2
Bud3
Bud4
Msn2
Sch9
Rim15
Yak1
Ypl247c
Vps52
Kss1
Sko1
Slt2
Msg5
Swe1
Igo2
Cdc55
Igo1
Vps53
Vps54
Rtn1
Fmp42
Cot1
Zrc1
10
20
30
40
50
t/s

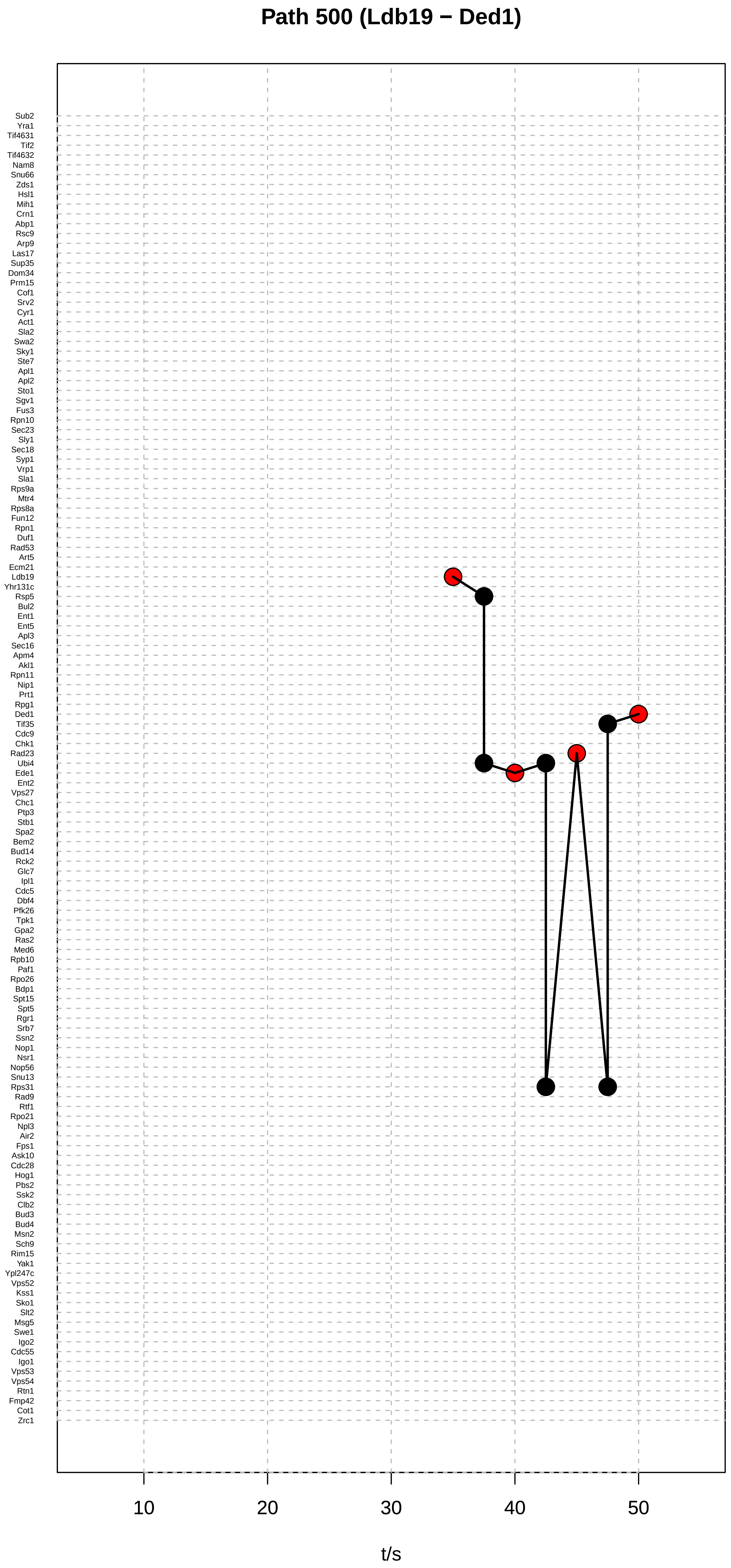

Path 500 (Ldb19 − Ded1)
Sub2
Yra1
Tif4631
Tif2
Tif4632
Nam8
Snu66
Zds1
Hsl1
Mih1
Crn1
Abp1
Rsc9
Arp9
Las17
Sup35
Dom34
Prm15
Cof1
Srv2
Cyr1
Act1
Sla2
Swa2
Sky1
Ste7
Apl1
Apl2
Sto1
Sgv1
Fus3
Rpn10
Sec23
Sly1
Sec18
Syp1
Vrp1
Sla1
Rps9a
Mtr4
Rps8a
Fun12
Rpn1
Duf1
Rad53
Art5
Ecm21
Ldb19
Yhr131c
Rsp5
Bul2
Ent1
Ent5
Apl3
Sec16
Apm4
Akl1
Rpn11
Nip1
Prt1
Rpg1
Ded1
Tif35
Cdc9
Chk1
Rad23
Ubi4
Ede1
Ent2
Vps27
Chc1
Ptp3
Stb1
Spa2
Bem2
Bud14
Rck2
Glc7
Ipl1
Cdc5
Dbf4
Pfk26
Tpk1
Gpa2
Ras2
Med6
Rpb10
Paf1
Rpo26
Bdp1
Spt15
Spt5
Rgr1
Srb7
Ssn2
Nop1
Nsr1
Nop56
Snu13
Rps31
Rad9
Rtf1
Rpo21
Npl3
Air2
Fps1
Ask10
Cdc28
Hog1
Pbs2
Ssk2
Clb2
Bud3
Bud4
Msn2
Sch9
Rim15
Yak1
Ypl247c
Vps52
Kss1
Sko1
Slt2
Msg5
Swe1
Igo2
Cdc55
Igo1
Vps53
Vps54
Rtn1
Fmp42
Cot1
Zrc1
10
20
30
40
50
t/s

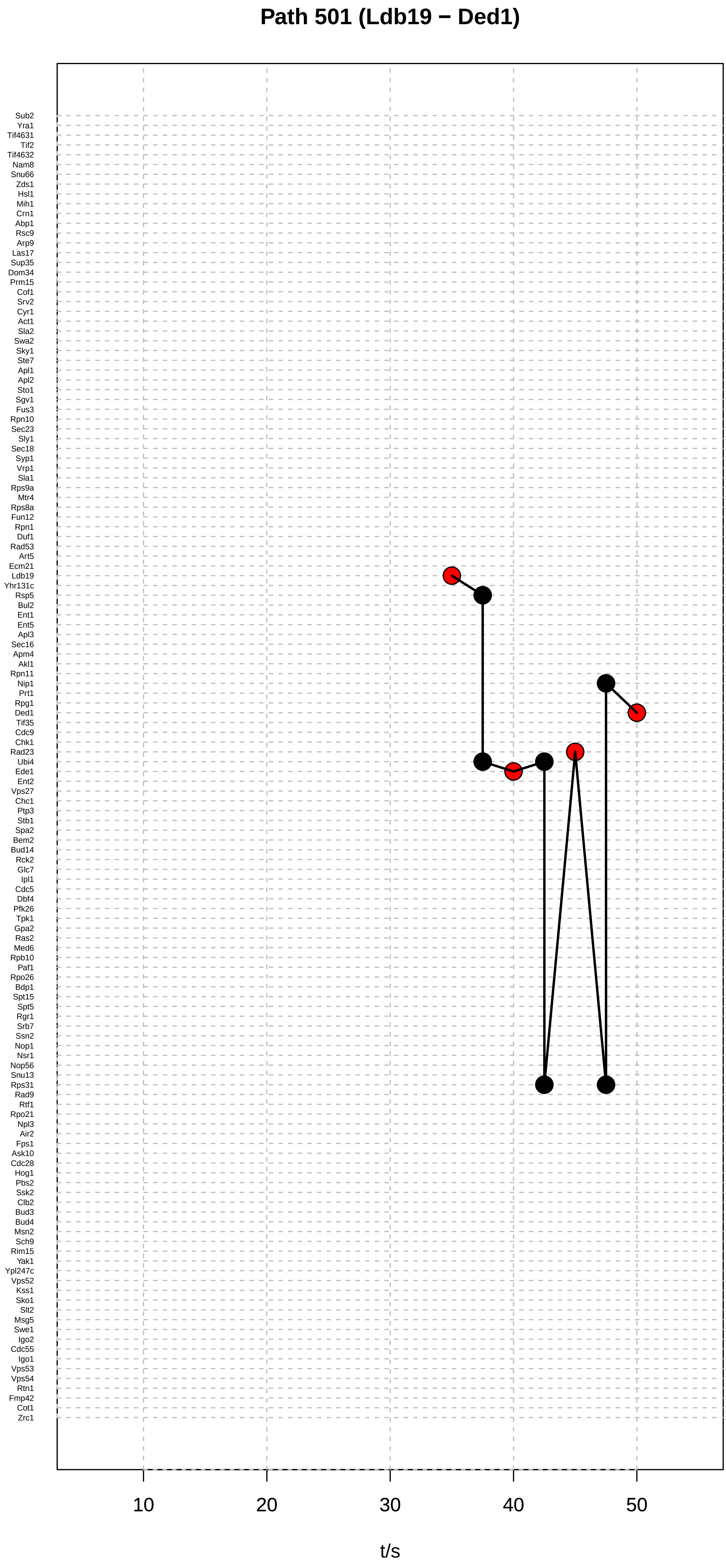
Path 501 (Ldb19 − Ded1)
Sub2
Yra1
Tif4631
Tif2
Tif4632
Nam8
Snu66
Zds1
Hsl1
Mih1
Crn1
Abp1
Rsc9
Arp9
Las17
Sup35
Dom34
Prm15
Cof1
Srv2
Cyr1
Act1
Sla2
Swa2
Sky1
Ste7
Apl1
Apl2
Sto1
Sgv1
Fus3
Rpn10
Sec23
Sly1
Sec18
Syp1
Vrp1
Sla1
Rps9a
Mtr4
Rps8a
Fun12
Rpn1
Duf1
Rad53
Art5
Ecm21
Ldb19
Yhr131c
Rsp5
Bul2
Ent1
Ent5
Apl3
Sec16
Apm4
Akl1
Rpn11
Nip1
Prt1
Rpg1
Ded1
Tif35
Cdc9
Chk1
Rad23
Ubi4
Ede1
Ent2
Vps27
Chc1
Ptp3
Stb1
Spa2
Bem2
Bud14
Rck2
Glc7
Ipl1
Cdc5
Dbf4
Pfk26
Tpk1
Gpa2
Ras2
Med6
Rpb10
Paf1
Rpo26
Bdp1
Spt15
Spt5
Rgr1
Srb7
Ssn2
Nop1
Nsr1
Nop56
Snu13
Rps31
Rad9
Rtf1
Rpo21
Npl3
Air2
Fps1
Ask10
Cdc28
Hog1
Pbs2
Ssk2
Clb2
Bud3
Bud4
Msn2
Sch9
Rim15
Yak1
Ypl247c
Vps52
Kss1
Sko1
Slt2
Msg5
Swe1
Igo2
Cdc55
Igo1
Vps53
Vps54
Rtn1
Fmp42
Cot1
Zrc1
10
20
30
40
50
t/s

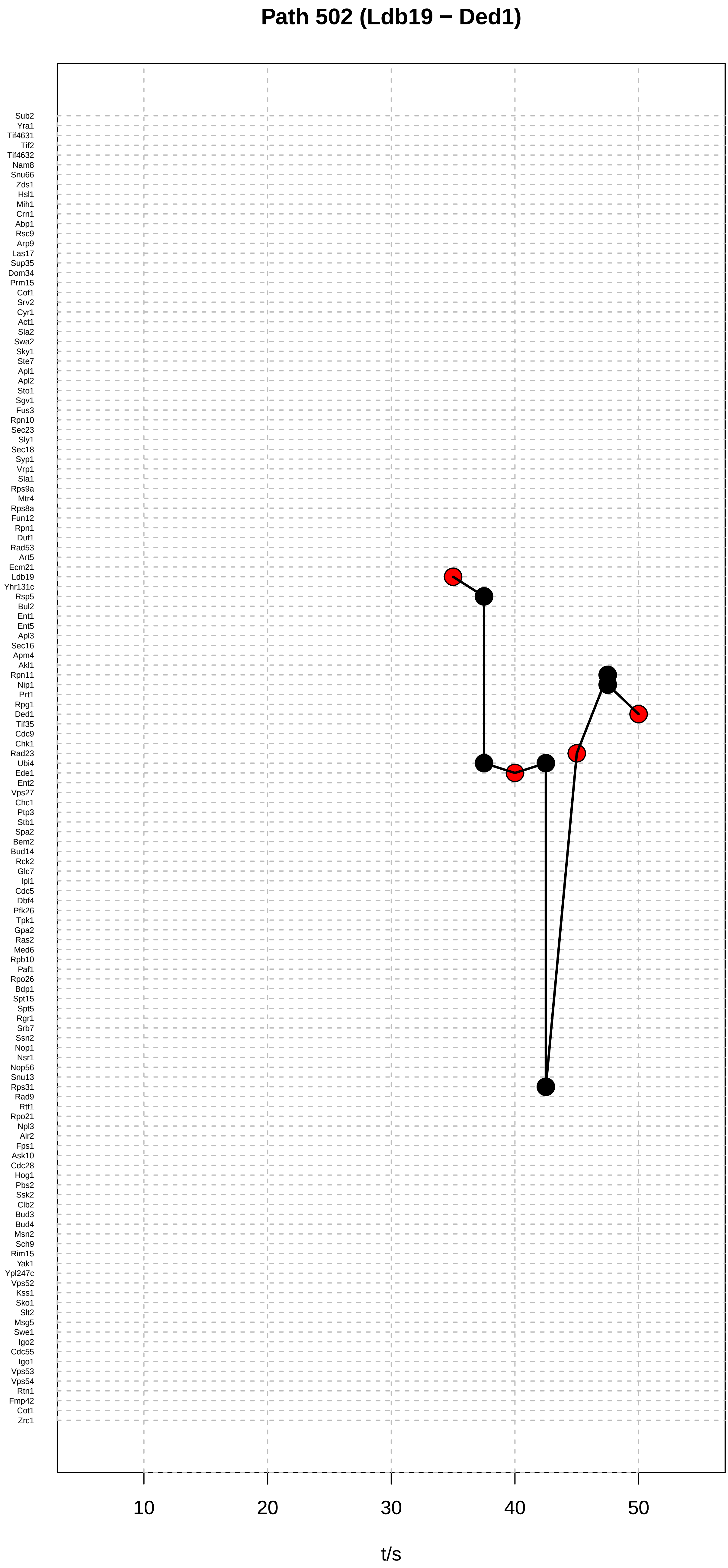

Path 502 (Ldb19 − Ded1)
Sub2
Yra1
Tif4631
Tif2
Tif4632
Nam8
Snu66
Zds1
Hsl1
Mih1
Crn1
Abp1
Rsc9
Arp9
Las17
Sup35
Dom34
Prm15
Cof1
Srv2
Cyr1
Act1
Sla2
Swa2
Sky1
Ste7
Apl1
Apl2
Sto1
Sgv1
Fus3
Rpn10
Sec23
Sly1
Sec18
Syp1
Vrp1
Sla1
Rps9a
Mtr4
Rps8a
Fun12
Rpn1
Duf1
Rad53
Art5
Ecm21
Ldb19
Yhr131c
Rsp5
Bul2
Ent1
Ent5
Apl3
Sec16
Apm4
Akl1
Rpn11
Nip1
Prt1
Rpg1
Ded1
Tif35
Cdc9
Chk1
Rad23
Ubi4
Ede1
Ent2
Vps27
Chc1
Ptp3
Stb1
Spa2
Bem2
Bud14
Rck2
Glc7
Ipl1
Cdc5
Dbf4
Pfk26
Tpk1
Gpa2
Ras2
Med6
Rpb10
Paf1
Rpo26
Bdp1
Spt15
Spt5
Rgr1
Srb7
Ssn2
Nop1
Nsr1
Nop56
Snu13
Rps31
Rad9
Rtf1
Rpo21
Npl3
Air2
Fps1
Ask10
Cdc28
Hog1
Pbs2
Ssk2
Clb2
Bud3
Bud4
Msn2
Sch9
Rim15
Yak1
Ypl247c
Vps52
Kss1
Sko1
Slt2
Msg5
Swe1
Igo2
Cdc55
Igo1
Vps53
Vps54
Rtn1
Fmp42
Cot1
Zrc1
10
20
30
40
50
t/s

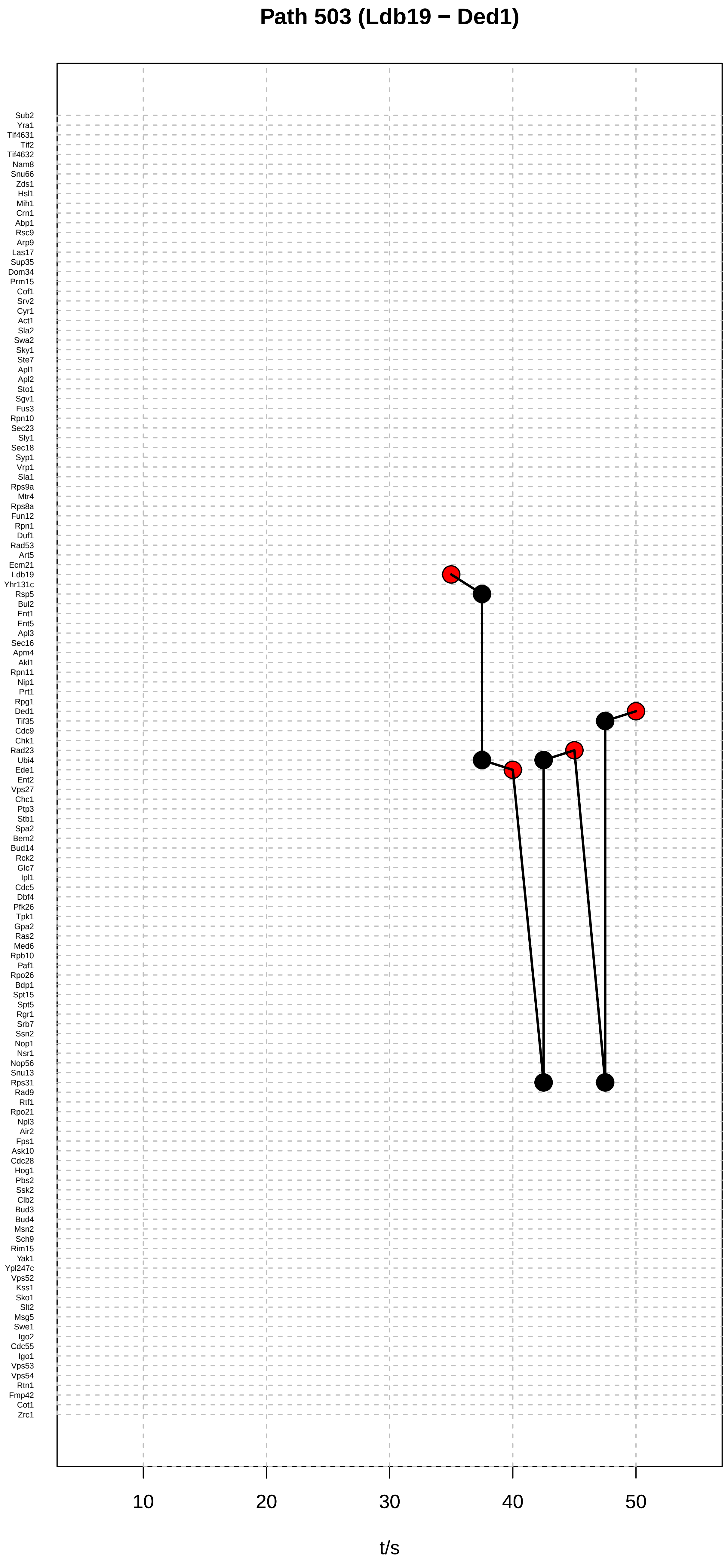

Path 503 (Ldb19 − Ded1)
Sub2
Yra1
Tif4631
Tif2
Tif4632
Nam8
Snu66
Zds1
Hsl1
Mih1
Crn1
Abp1
Rsc9
Arp9
Las17
Sup35
Dom34
Prm15
Cof1
Srv2
Cyr1
Act1
Sla2
Swa2
Sky1
Ste7
Apl1
Apl2
Sto1
Sgv1
Fus3
Rpn10
Sec23
Sly1
Sec18
Syp1
Vrp1
Sla1
Rps9a
Mtr4
Rps8a
Fun12
Rpn1
Duf1
Rad53
Art5
Ecm21
Ldb19
Yhr131c
Rsp5
Bul2
Ent1
Ent5
Apl3
Sec16
Apm4
Akl1
Rpn11
Nip1
Prt1
Rpg1
Ded1
Tif35
Cdc9
Chk1
Rad23
Ubi4
Ede1
Ent2
Vps27
Chc1
Ptp3
Stb1
Spa2
Bem2
Bud14
Rck2
Glc7
Ipl1
Cdc5
Dbf4
Pfk26
Tpk1
Gpa2
Ras2
Med6
Rpb10
Paf1
Rpo26
Bdp1
Spt15
Spt5
Rgr1
Srb7
Ssn2
Nop1
Nsr1
Nop56
Snu13
Rps31
Rad9
Rtf1
Rpo21
Npl3
Air2
Fps1
Ask10
Cdc28
Hog1
Pbs2
Ssk2
Clb2
Bud3
Bud4
Msn2
Sch9
Rim15
Yak1
Ypl247c
Vps52
Kss1
Sko1
Slt2
Msg5
Swe1
Igo2
Cdc55
Igo1
Vps53
Vps54
Rtn1
Fmp42
Cot1
Zrc1
10
20
30
40
50
t/s

### Path 504 (Ldb19 − Ded1)

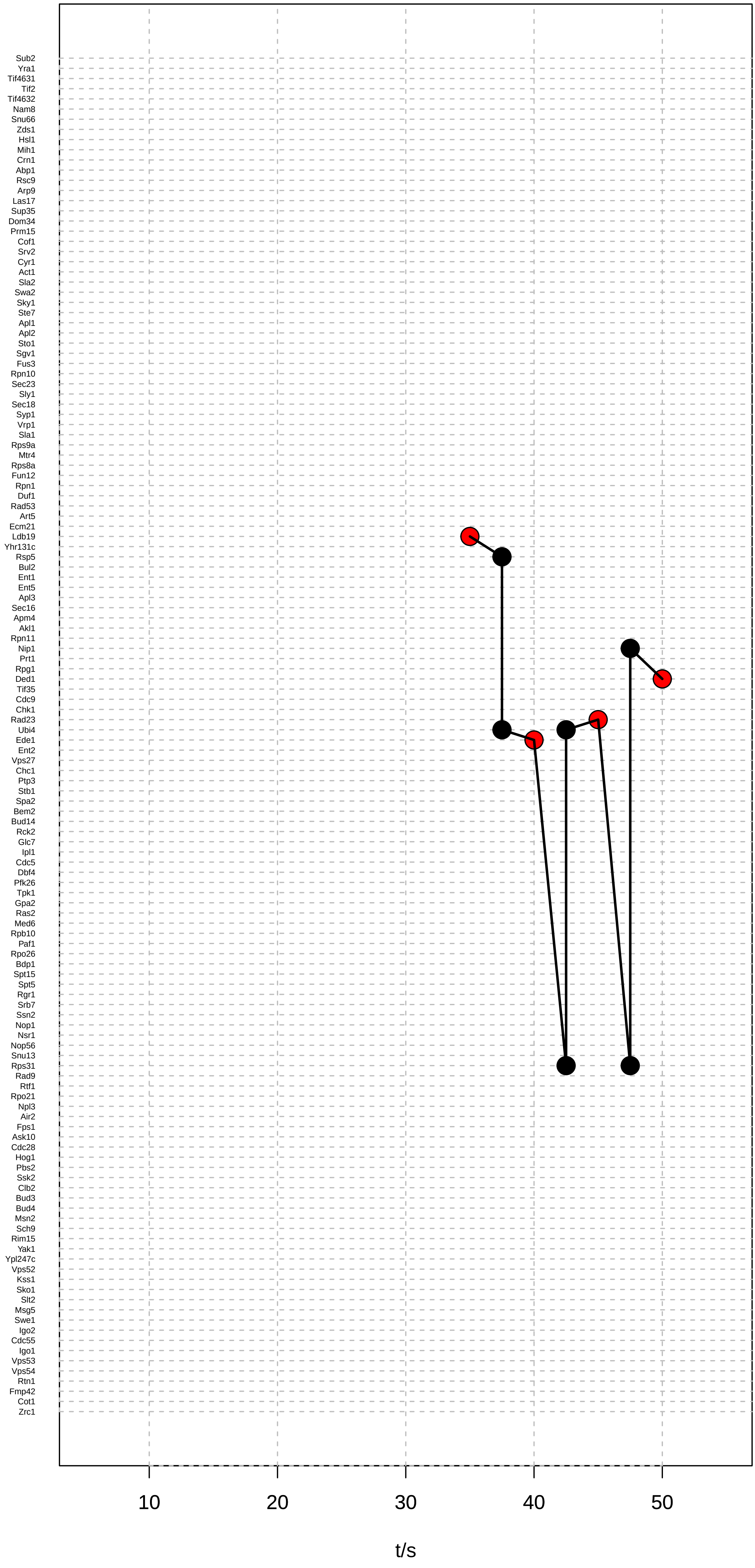

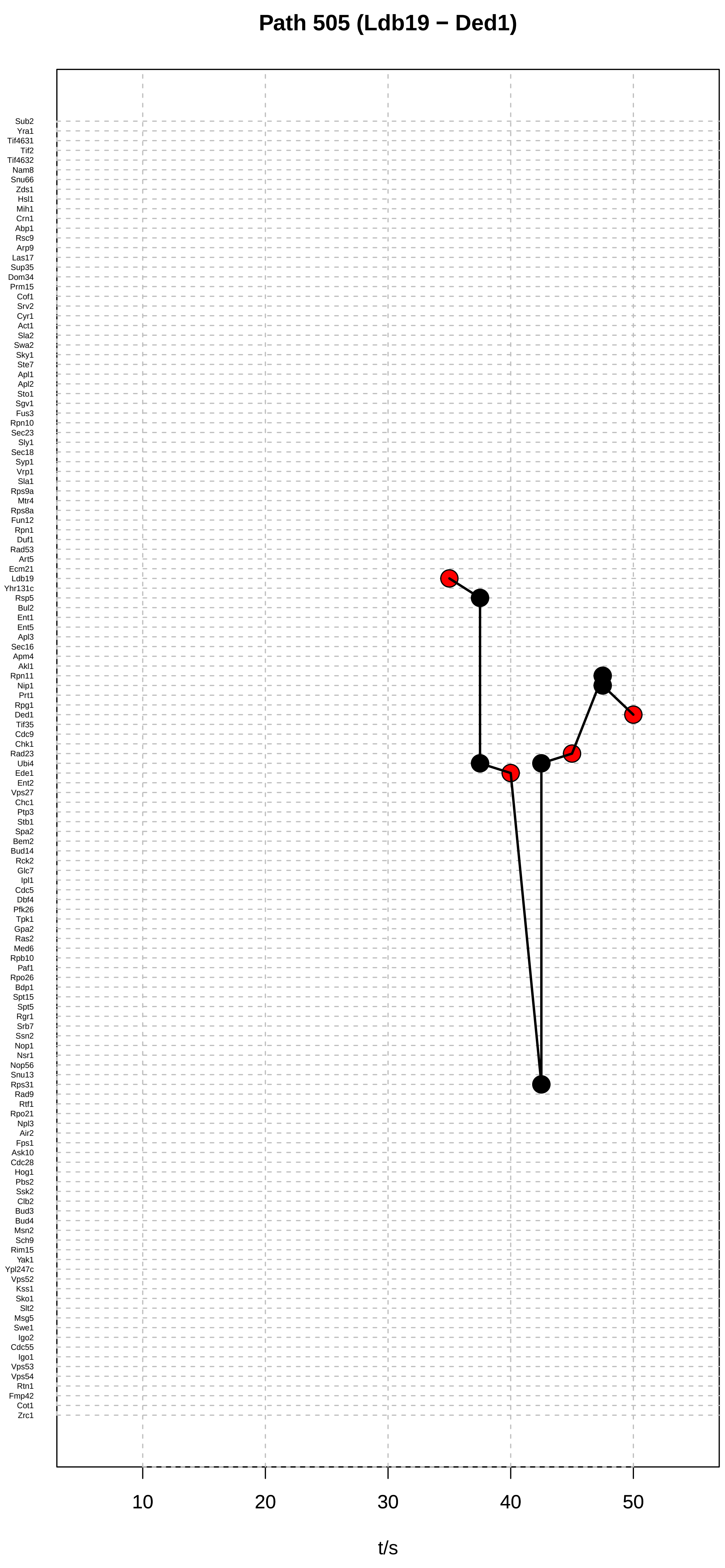

Path 505 (Ldb19 − Ded1)
Sub2
Yra1
Tif4631
Tif2
Tif4632
Nam8
Snu66
Zds1
Hsl1
Mih1
Crn1
Abp1
Rsc9
Arp9
Las17
Sup35
Dom34
Prm15
Cof1
Srv2
Cyr1
Act1
Sla2
Swa2
Sky1
Ste7
Apl1
Apl2
Sto1
Sgv1
Fus3
Rpn10
Sec23
Sly1
Sec18
Syp1
Vrp1
Sla1
Rps9a
Mtr4
Rps8a
Fun12
Rpn1
Duf1
Rad53
Art5
Ecm21
Ldb19
Yhr131c
Rsp5
Bul2
Ent1
Ent5
Apl3
Sec16
Apm4
Akl1
Rpn11
Nip1
Prt1
Rpg1
Ded1
Tif35
Cdc9
Chk1
Rad23
Ubi4
Ede1
Ent2
Vps27
Chc1
Ptp3
Stb1
Spa2
Bem2
Bud14
Rck2
Glc7
Ipl1
Cdc5
Dbf4
Pfk26
Tpk1
Gpa2
Ras2
Med6
Rpb10
Paf1
Rpo26
Bdp1
Spt15
Spt5
Rgr1
Srb7
Ssn2
Nop1
Nsr1
Nop56
Snu13
Rps31
Rad9
Rtf1
Rpo21
Npl3
Air2
Fps1
Ask10
Cdc28
Hog1
Pbs2
Ssk2
Clb2
Bud3
Bud4
Msn2
Sch9
Rim15
Yak1
Ypl247c
Vps52
Kss1
Sko1
Slt2
Msg5
Swe1
Igo2
Cdc55
Igo1
Vps53
Vps54
Rtn1
Fmp42
Cot1
Zrc1
10
20
30
40
50
t/s

### Path 506 (Ldb19 − Ded1)

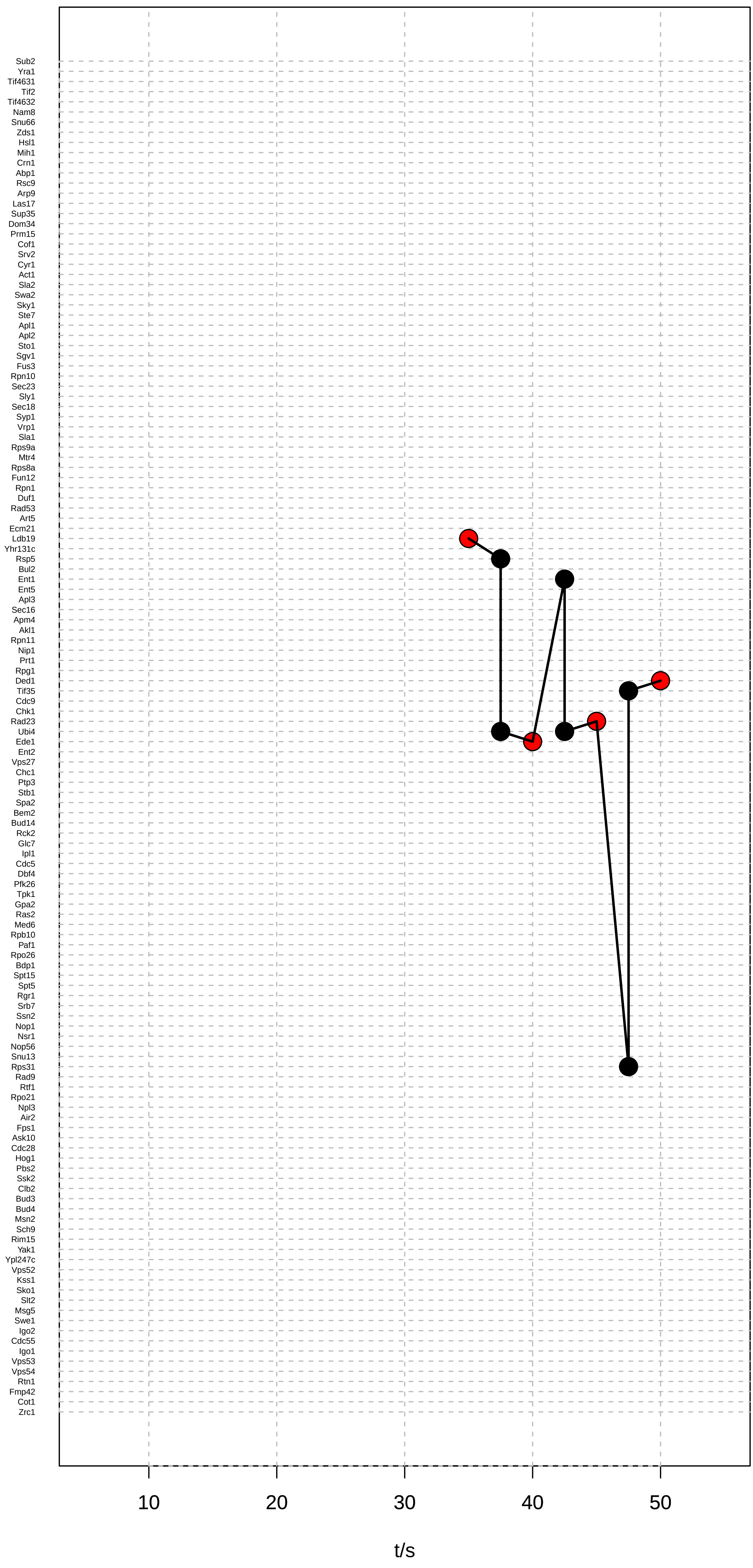

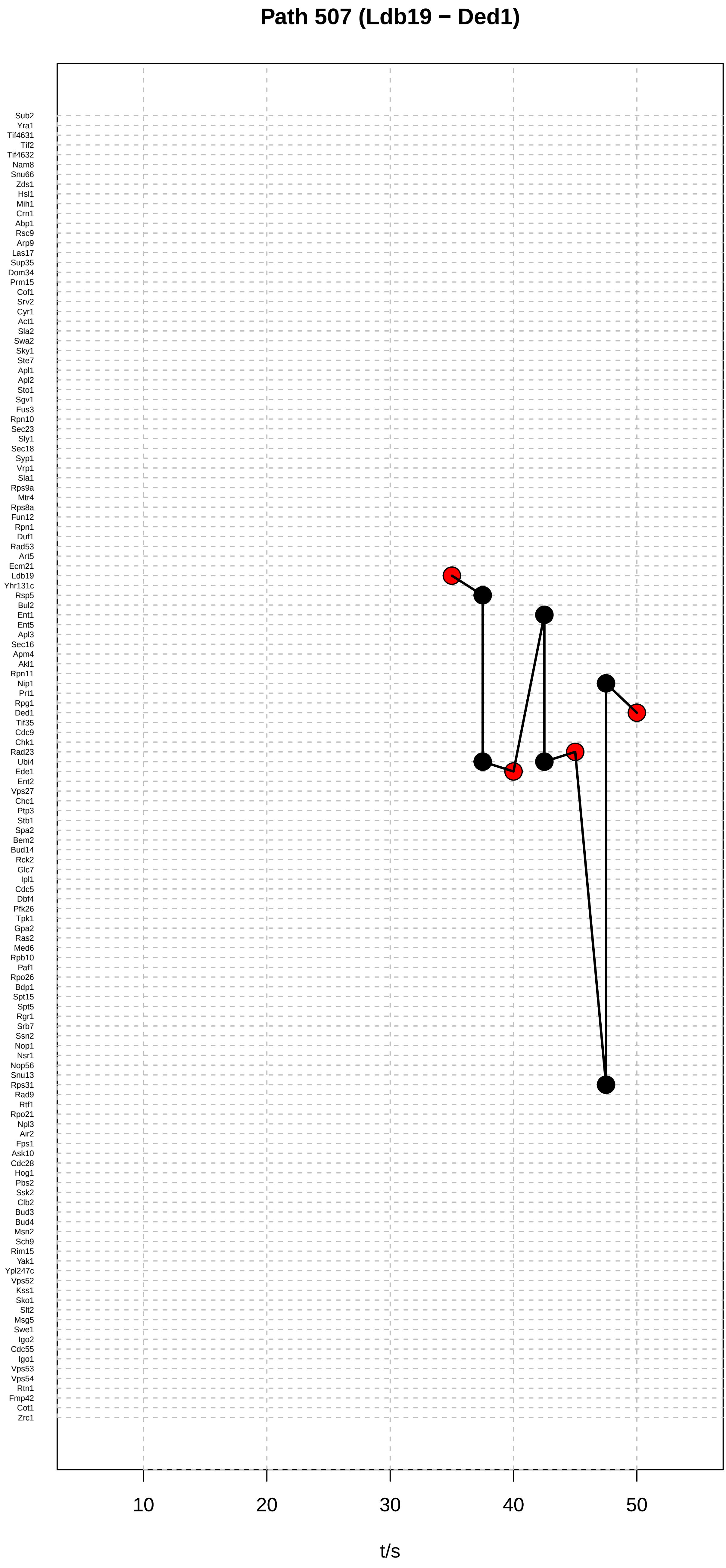
Path 507 (Ldb19 − Ded1)
Sub2
Yra1
Tif4631
Tif2
Tif4632
Nam8
Snu66
Zds1
Hsl1
Mih1
Crn1
Abp1
Rsc9
Arp9
Las17
Sup35
Dom34
Prm15
Cof1
Srv2
Cyr1
Act1
Sla2
Swa2
Sky1
Ste7
Apl1
Apl2
Sto1
Sgv1
Fus3
Rpn10
Sec23
Sly1
Sec18
Syp1
Vrp1
Sla1
Rps9a
Mtr4
Rps8a
Fun12
Rpn1
Duf1
Rad53
Art5
Ecm21
Ldb19
Yhr131c
Rsp5
Bul2
Ent1
Ent5
Apl3
Sec16
Apm4
Akl1
Rpn11
Nip1
Prt1
Rpg1
Ded1
Tif35
Cdc9
Chk1
Rad23
Ubi4
Ede1
Ent2
Vps27
Chc1
Ptp3
Stb1
Spa2
Bem2
Bud14
Rck2
Glc7
Ipl1
Cdc5
Dbf4
Pfk26
Tpk1
Gpa2
Ras2
Med6
Rpb10
Paf1
Rpo26
Bdp1
Spt15
Spt5
Rgr1
Srb7
Ssn2
Nop1
Nsr1
Nop56
Snu13
Rps31
Rad9
Rtf1
Rpo21
Npl3
Air2
Fps1
Ask10
Cdc28
Hog1
Pbs2
Ssk2
Clb2
Bud3
Bud4
Msn2
Sch9
Rim15
Yak1
Ypl247c
Vps52
Kss1
Sko1
Slt2
Msg5
Swe1
Igo2
Cdc55
Igo1
Vps53
Vps54
Rtn1
Fmp42
Cot1
Zrc1
10
20
30
40
50
t/s

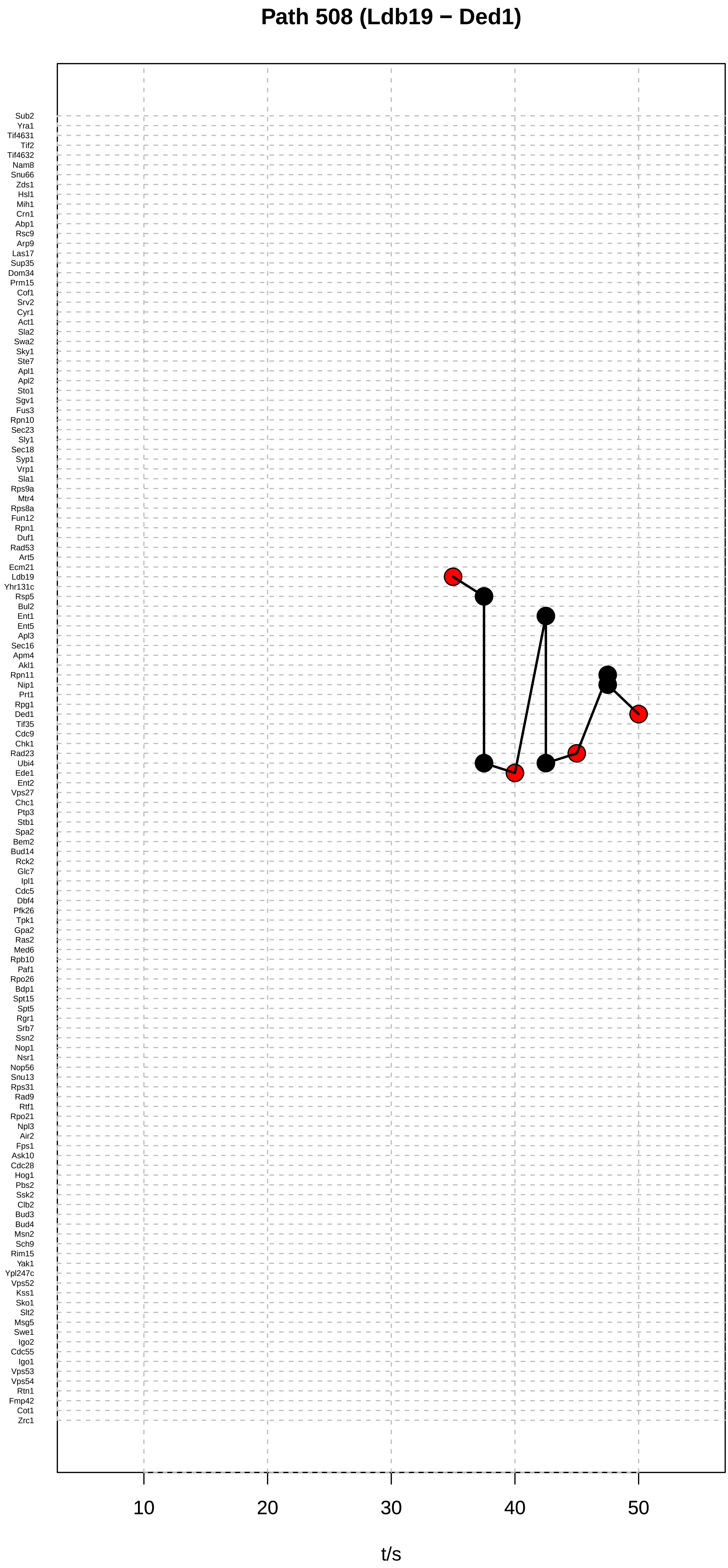

Path 508 (Ldb19 − Ded1)
Sub2
Yra1
Tif4631
Tif2
Tif4632
Nam8
Snu66
Zds1
Hsl1
Mih1
Crn1
Abp1
Rsc9
Arp9
Las17
Sup35
Dom34
Prm15
Cof1
Srv2
Cyr1
Act1
Sla2
Swa2
Sky1
Ste7
Apl1
Apl2
Sto1
Sgv1
Fus3
Rpn10
Sec23
Sly1
Sec18
Syp1
Vrp1
Sla1
Rps9a
Mtr4
Rps8a
Fun12
Rpn1
Duf1
Rad53
Art5
Ecm21
Ldb19
Yhr131c
Rsp5
Bul2
Ent1
Ent5
Apl3
Sec16
Apm4
Akl1
Rpn11
Nip1
Prt1
Rpg1
Ded1
Tif35
Cdc9
Chk1
Rad23
Ubi4
Ede1
Ent2
Vps27
Chc1
Ptp3
Stb1
Spa2
Bem2
Bud14
Rck2
Glc7
Ipl1
Cdc5
Dbf4
Pfk26
Tpk1
Gpa2
Ras2
Med6
Rpb10
Paf1
Rpo26
Bdp1
Spt15
Spt5
Rgr1
Srb7
Ssn2
Nop1
Nsr1
Nop56
Snu13
Rps31
Rad9
Rtf1
Rpo21
Npl3
Air2
Fps1
Ask10
Cdc28
Hog1
Pbs2
Ssk2
Clb2
Bud3
Bud4
Msn2
Sch9
Rim15
Yak1
Ypl247c
Vps52
Kss1
Sko1
Slt2
Msg5
Swe1
Igo2
Cdc55
Igo1
Vps53
Vps54
Rtn1
Fmp42
Cot1
Zrc1
10
20
30
40
50
t/s

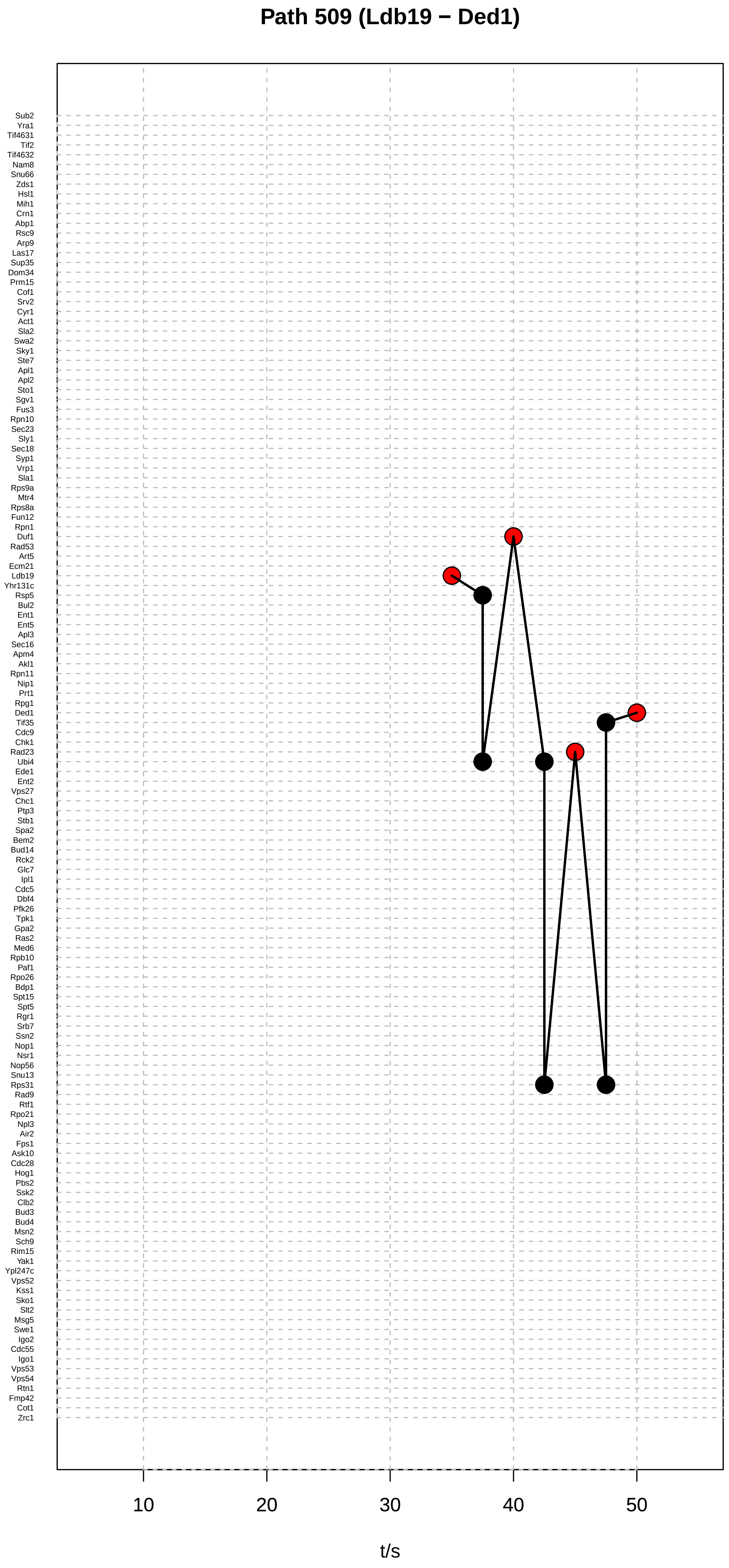
Path 509 (Ldb19 − Ded1)
Sub2
Yra1
Tif4631
Tif2
Tif4632
Nam8
Snu66
Zds1
Hsl1
Mih1
Crn1
Abp1
Rsc9
Arp9
Las17
Sup35
Dom34
Prm15
Cof1
Srv2
Cyr1
Act1
Sla2
Swa2
Sky1
Ste7
Apl1
Apl2
Sto1
Sgv1
Fus3
Rpn10
Sec23
Sly1
Sec18
Syp1
Vrp1
Sla1
Rps9a
Mtr4
Rps8a
Fun12
Rpn1
Duf1
Rad53
Art5
Ecm21
Ldb19
Yhr131c
Rsp5
Bul2
Ent1
Ent5
Apl3
Sec16
Apm4
Akl1
Rpn11
Nip1
Prt1
Rpg1
Ded1
Tif35
Cdc9
Chk1
Rad23
Ubi4
Ede1
Ent2
Vps27
Chc1
Ptp3
Stb1
Spa2
Bem2
Bud14
Rck2
Glc7
Ipl1
Cdc5
Dbf4
Pfk26
Tpk1
Gpa2
Ras2
Med6
Rpb10
Paf1
Rpo26
Bdp1
Spt15
Spt5
Rgr1
Srb7
Ssn2
Nop1
Nsr1
Nop56
Snu13
Rps31
Rad9
Rtf1
Rpo21
Npl3
Air2
Fps1
Ask10
Cdc28
Hog1
Pbs2
Ssk2
Clb2
Bud3
Bud4
Msn2
Sch9
Rim15
Yak1
Ypl247c
Vps52
Kss1
Sko1
Slt2
Msg5
Swe1
Igo2
Cdc55
Igo1
Vps53
Vps54
Rtn1
Fmp42
Cot1
Zrc1
10
20
30
40
50
t/s

#### Path 510 (Ldb19 − Ded1)

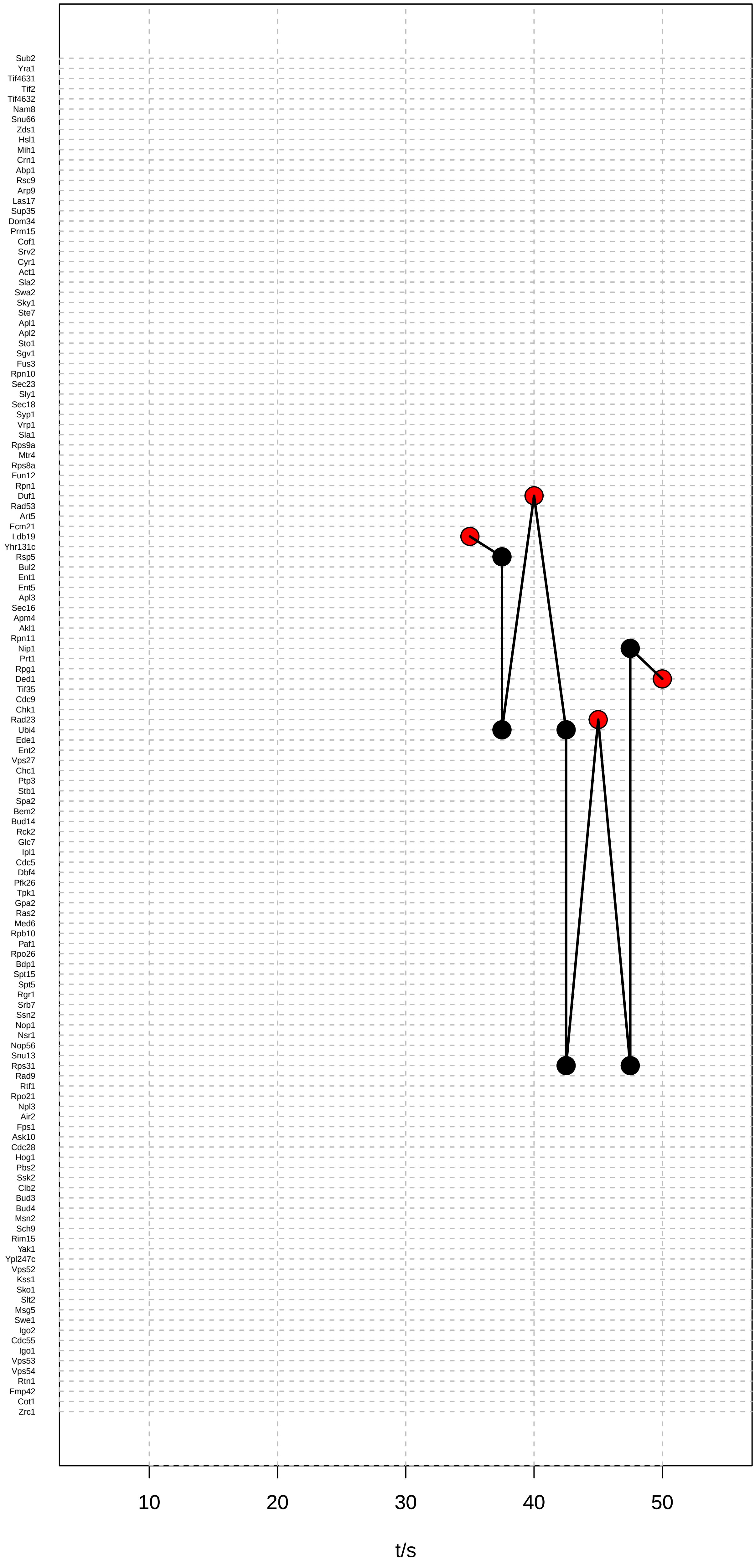

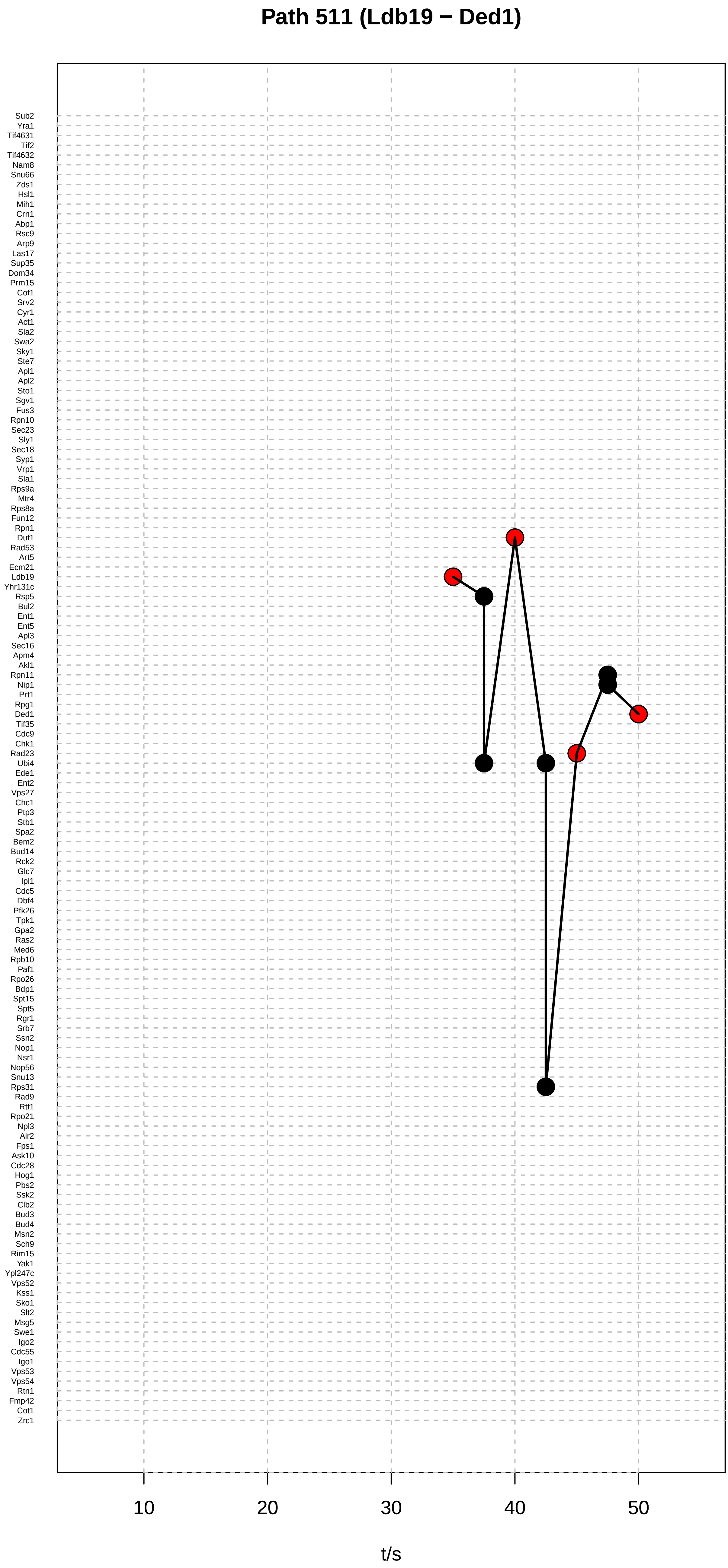

Path 511 (Ldb19 − Ded1)
Sub2
Yra1
Tif4631
Tif2
Tif4632
Nam8
Snu66
Zds1
Hsl1
Mih1
Crn1
Abp1
Rsc9
Arp9
Las17
Sup35
Dom34
Prm15
Cof1
Srv2
Cyr1
Act1
Sla2
Swa2
Sky1
Ste7
Apl1
Apl2
Sto1
Sgv1
Fus3
Rpn10
Sec23
Sly1
Sec18
Syp1
Vrp1
Sla1
Rps9a
Mtr4
Rps8a
Fun12
Rpn1
Duf1
Rad53
Art5
Ecm21
Ldb19
Yhr131c
Rsp5
Bul2
Ent1
Ent5
Apl3
Sec16
Apm4
Akl1
Rpn11
Nip1
Prt1
Rpg1
Ded1
Tif35
Cdc9
Chk1
Rad23
Ubi4
Ede1
Ent2
Vps27
Chc1
Ptp3
Stb1
Spa2
Bem2
Bud14
Rck2
Glc7
Ipl1
Cdc5
Dbf4
Pfk26
Tpk1
Gpa2
Ras2
Med6
Rpb10
Paf1
Rpo26
Bdp1
Spt15
Spt5
Rgr1
Srb7
Ssn2
Nop1
Nsr1
Nop56
Snu13
Rps31
Rad9
Rtf1
Rpo21
Npl3
Air2
Fps1
Ask10
Cdc28
Hog1
Pbs2
Ssk2
Clb2
Bud3
Bud4
Msn2
Sch9
Rim15
Yak1
Ypl247c
Vps52
Kss1
Sko1
Slt2
Msg5
Swe1
Igo2
Cdc55
Igo1
Vps53
Vps54
Rtn1
Fmp42
Cot1
Zrc1
10
20
30
40
50
t/s

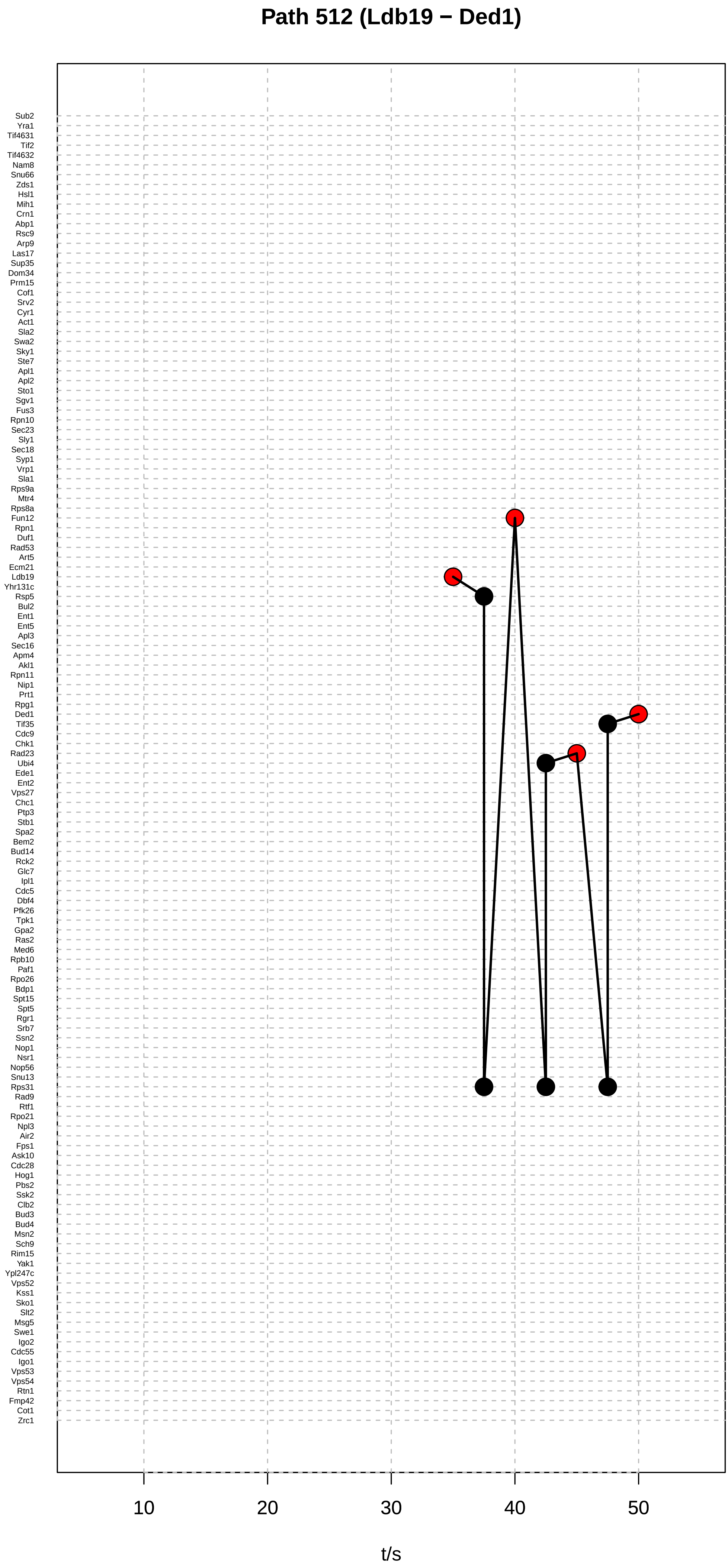

Path 512 (Ldb19 − Ded1)
Sub2
Yra1
Tif4631
Tif2
Tif4632
Nam8
Snu66
Zds1
Hsl1
Mih1
Crn1
Abp1
Rsc9
Arp9
Las17
Sup35
Dom34
Prm15
Cof1
Srv2
Cyr1
Act1
Sla2
Swa2
Sky1
Ste7
Apl1
Apl2
Sto1
Sgv1
Fus3
Rpn10
Sec23
Sly1
Sec18
Syp1
Vrp1
Sla1
Rps9a
Mtr4
Rps8a
Fun12
Rpn1
Duf1
Rad53
Art5
Ecm21
Ldb19
Yhr131c
Rsp5
Bul2
Ent1
Ent5
Apl3
Sec16
Apm4
Akl1
Rpn11
Nip1
Prt1
Rpg1
Ded1
Tif35
Cdc9
Chk1
Rad23
Ubi4
Ede1
Ent2
Vps27
Chc1
Ptp3
Stb1
Spa2
Bem2
Bud14
Rck2
Glc7
Ipl1
Cdc5
Dbf4
Pfk26
Tpk1
Gpa2
Ras2
Med6
Rpb10
Paf1
Rpo26
Bdp1
Spt15
Spt5
Rgr1
Srb7
Ssn2
Nop1
Nsr1
Nop56
Snu13
Rps31
Rad9
Rtf1
Rpo21
Npl3
Air2
Fps1
Ask10
Cdc28
Hog1
Pbs2
Ssk2
Clb2
Bud3
Bud4
Msn2
Sch9
Rim15
Yak1
Ypl247c
Vps52
Kss1
Sko1
Slt2
Msg5
Swe1
Igo2
Cdc55
Igo1
Vps53
Vps54
Rtn1
Fmp42
Cot1
Zrc1
10
20
30
40
50
t/s

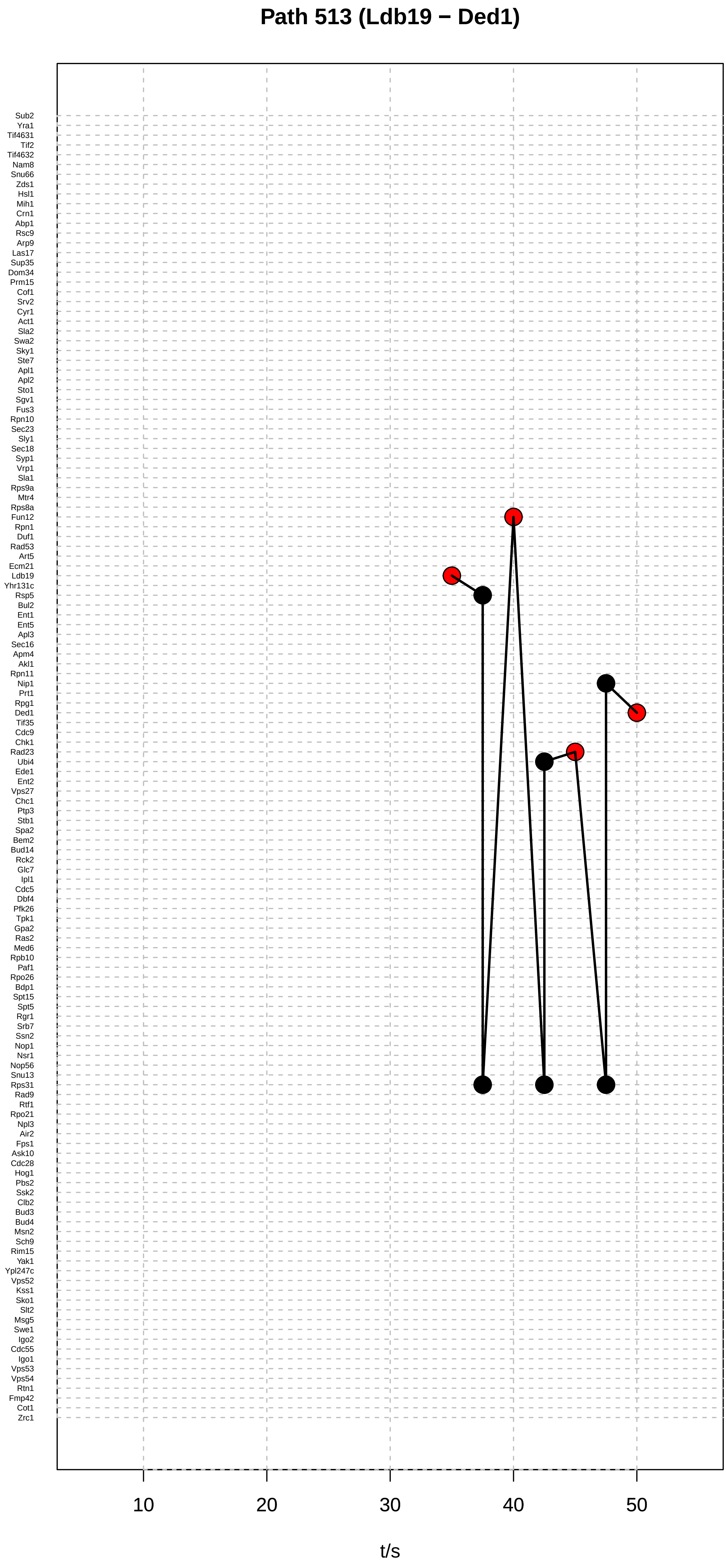

Path 513 (Ldb19 − Ded1)
Sub2
Yra1
Tif4631
Tif2
Tif4632
Nam8
Snu66
Zds1
Hsl1
Mih1
Crn1
Abp1
Rsc9
Arp9
Las17
Sup35
Dom34
Prm15
Cof1
Srv2
Cyr1
Act1
Sla2
Swa2
Sky1
Ste7
Apl1
Apl2
Sto1
Sgv1
Fus3
Rpn10
Sec23
Sly1
Sec18
Syp1
Vrp1
Sla1
Rps9a
Mtr4
Rps8a
Fun12
Rpn1
Duf1
Rad53
Art5
Ecm21
Ldb19
Yhr131c
Rsp5
Bul2
Ent1
Ent5
Apl3
Sec16
Apm4
Akl1
Rpn11
Nip1
Prt1
Rpg1
Ded1
Tif35
Cdc9
Chk1
Rad23
Ubi4
Ede1
Ent2
Vps27
Chc1
Ptp3
Stb1
Spa2
Bem2
Bud14
Rck2
Glc7
Ipl1
Cdc5
Dbf4
Pfk26
Tpk1
Gpa2
Ras2
Med6
Rpb10
Paf1
Rpo26
Bdp1
Spt15
Spt5
Rgr1
Srb7
Ssn2
Nop1
Nsr1
Nop56
Snu13
Rps31
Rad9
Rtf1
Rpo21
Npl3
Air2
Fps1
Ask10
Cdc28
Hog1
Pbs2
Ssk2
Clb2
Bud3
Bud4
Msn2
Sch9
Rim15
Yak1
Ypl247c
Vps52
Kss1
Sko1
Slt2
Msg5
Swe1
Igo2
Cdc55
Igo1
Vps53
Vps54
Rtn1
Fmp42
Cot1
Zrc1
10
20
30
40
50
t/s

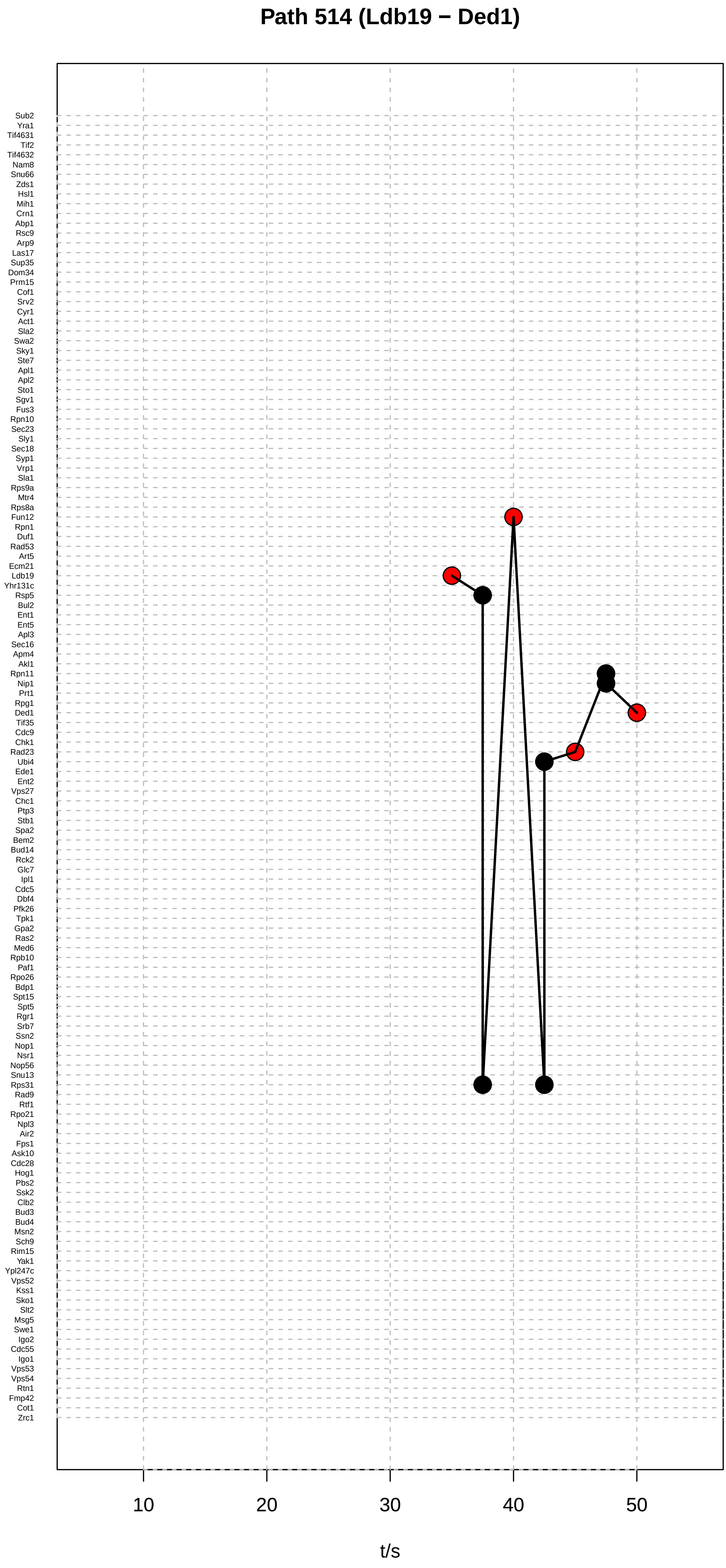
Path 514 (Ldb19 − Ded1)
Sub2
Yra1
Tif4631
Tif2
Tif4632
Nam8
Snu66
Zds1
Hsl1
Mih1
Crn1
Abp1
Rsc9
Arp9
Las17
Sup35
Dom34
Prm15
Cof1
Srv2
Cyr1
Act1
Sla2
Swa2
Sky1
Ste7
Apl1
Apl2
Sto1
Sgv1
Fus3
Rpn10
Sec23
Sly1
Sec18
Syp1
Vrp1
Sla1
Rps9a
Mtr4
Rps8a
Fun12
Rpn1
Duf1
Rad53
Art5
Ecm21
Ldb19
Yhr131c
Rsp5
Bul2
Ent1
Ent5
Apl3
Sec16
Apm4
Akl1
Rpn11
Nip1
Prt1
Rpg1
Ded1
Tif35
Cdc9
Chk1
Rad23
Ubi4
Ede1
Ent2
Vps27
Chc1
Ptp3
Stb1
Spa2
Bem2
Bud14
Rck2
Glc7
Ipl1
Cdc5
Dbf4
Pfk26
Tpk1
Gpa2
Ras2
Med6
Rpb10
Paf1
Rpo26
Bdp1
Spt15
Spt5
Rgr1
Srb7
Ssn2
Nop1
Nsr1
Nop56
Snu13
Rps31
Rad9
Rtf1
Rpo21
Npl3
Air2
Fps1
Ask10
Cdc28
Hog1
Pbs2
Ssk2
Clb2
Bud3
Bud4
Msn2
Sch9
Rim15
Yak1
Ypl247c
Vps52
Kss1
Sko1
Slt2
Msg5
Swe1
Igo2
Cdc55
Igo1
Vps53
Vps54
Rtn1
Fmp42
Cot1
Zrc1
10
20
30
40
50
t/s

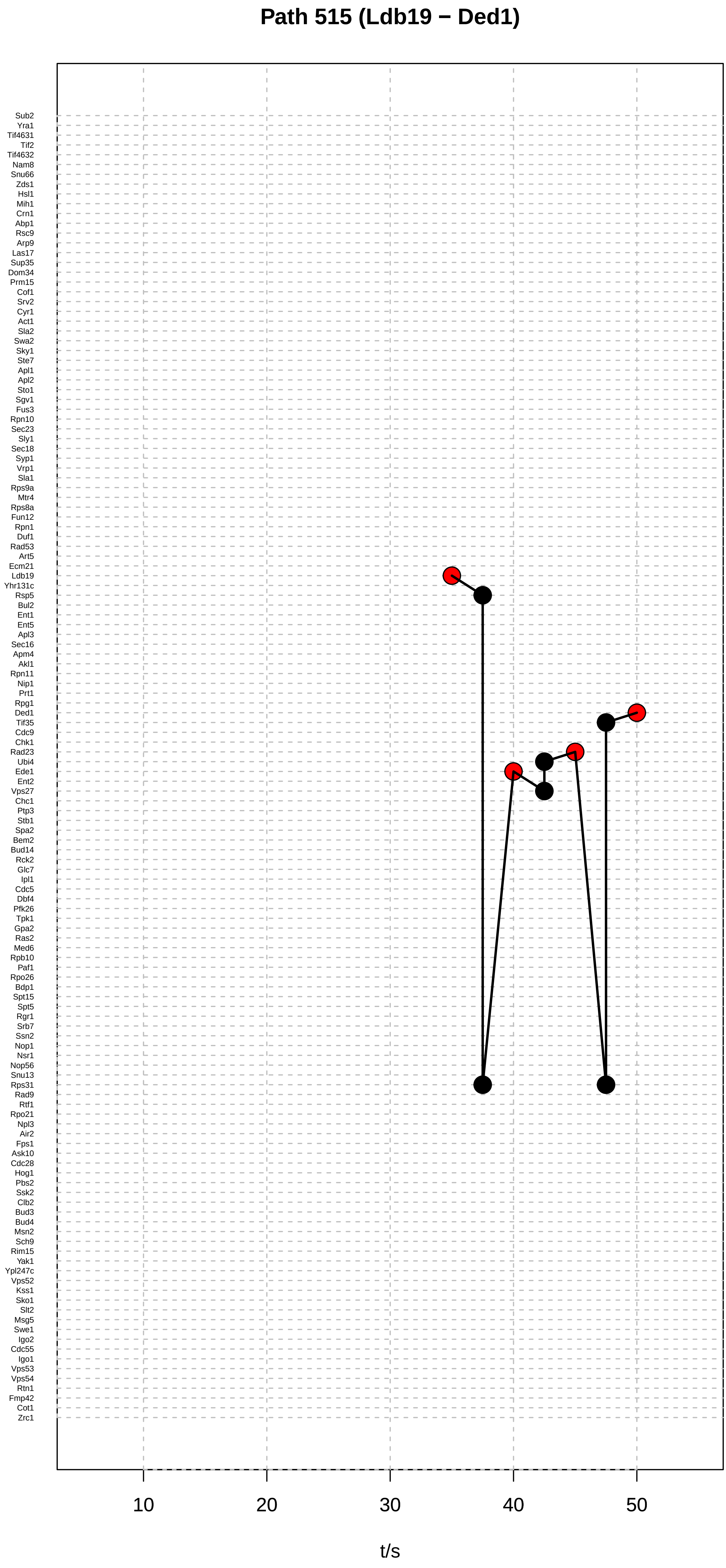

Path 515 (Ldb19 − Ded1)
Sub2
Yra1
Tif4631
Tif2
Tif4632
Nam8
Snu66
Zds1
Hsl1
Mih1
Crn1
Abp1
Rsc9
Arp9
Las17
Sup35
Dom34
Prm15
Cof1
Srv2
Cyr1
Act1
Sla2
Swa2
Sky1
Ste7
Apl1
Apl2
Sto1
Sgv1
Fus3
Rpn10
Sec23
Sly1
Sec18
Syp1
Vrp1
Sla1
Rps9a
Mtr4
Rps8a
Fun12
Rpn1
Duf1
Rad53
Art5
Ecm21
Ldb19
Yhr131c
Rsp5
Bul2
Ent1
Ent5
Apl3
Sec16
Apm4
Akl1
Rpn11
Nip1
Prt1
Rpg1
Ded1
Tif35
Cdc9
Chk1
Rad23
Ubi4
Ede1
Ent2
Vps27
Chc1
Ptp3
Stb1
Spa2
Bem2
Bud14
Rck2
Glc7
Ipl1
Cdc5
Dbf4
Pfk26
Tpk1
Gpa2
Ras2
Med6
Rpb10
Paf1
Rpo26
Bdp1
Spt15
Spt5
Rgr1
Srb7
Ssn2
Nop1
Nsr1
Nop56
Snu13
Rps31
Rad9
Rtf1
Rpo21
Npl3
Air2
Fps1
Ask10
Cdc28
Hog1
Pbs2
Ssk2
Clb2
Bud3
Bud4
Msn2
Sch9
Rim15
Yak1
Ypl247c
Vps52
Kss1
Sko1
Slt2
Msg5
Swe1
Igo2
Cdc55
Igo1
Vps53
Vps54
Rtn1
Fmp42
Cot1
Zrc1
10
20
30
40
50
t/s

#### Path 516 (Ldb19 − Ded1)

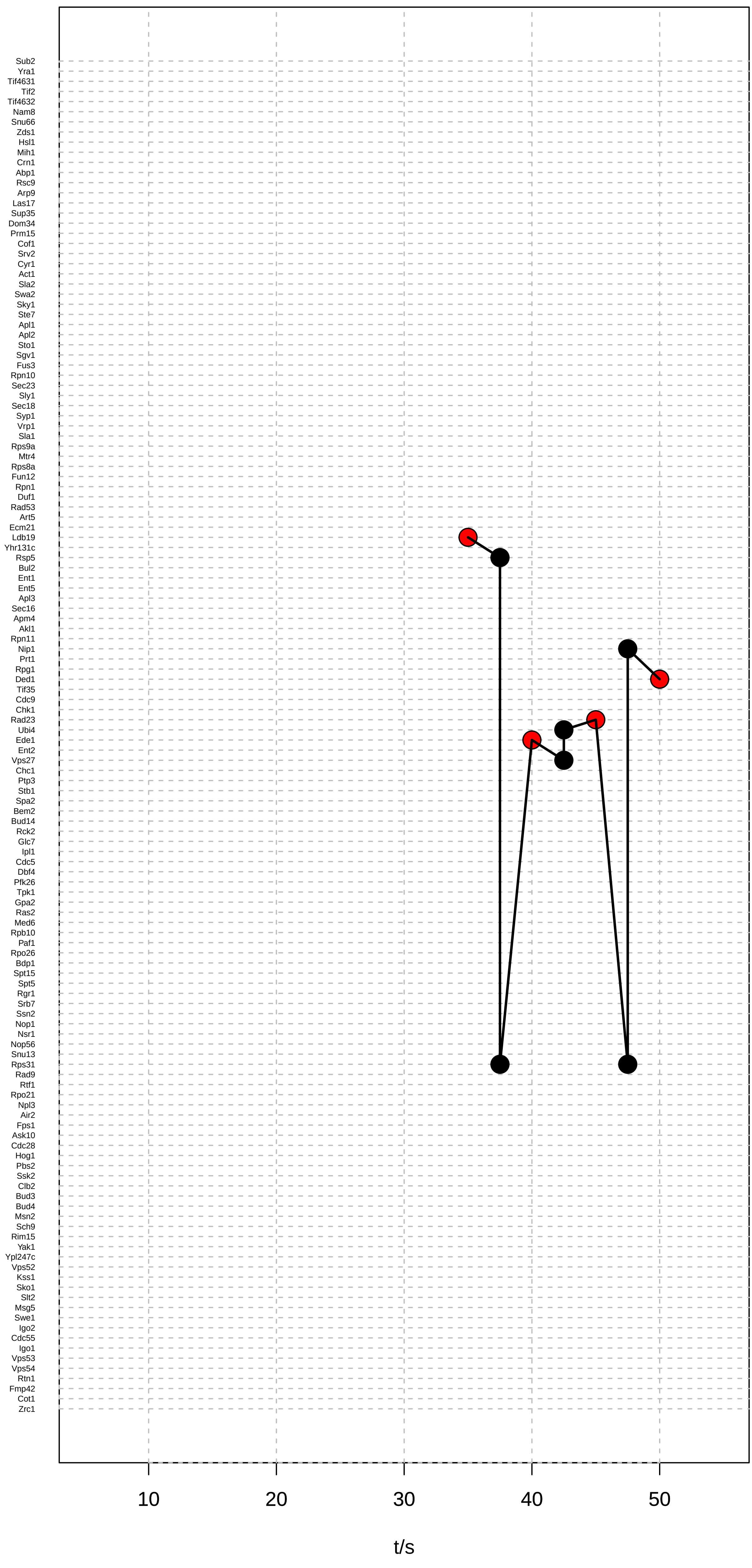

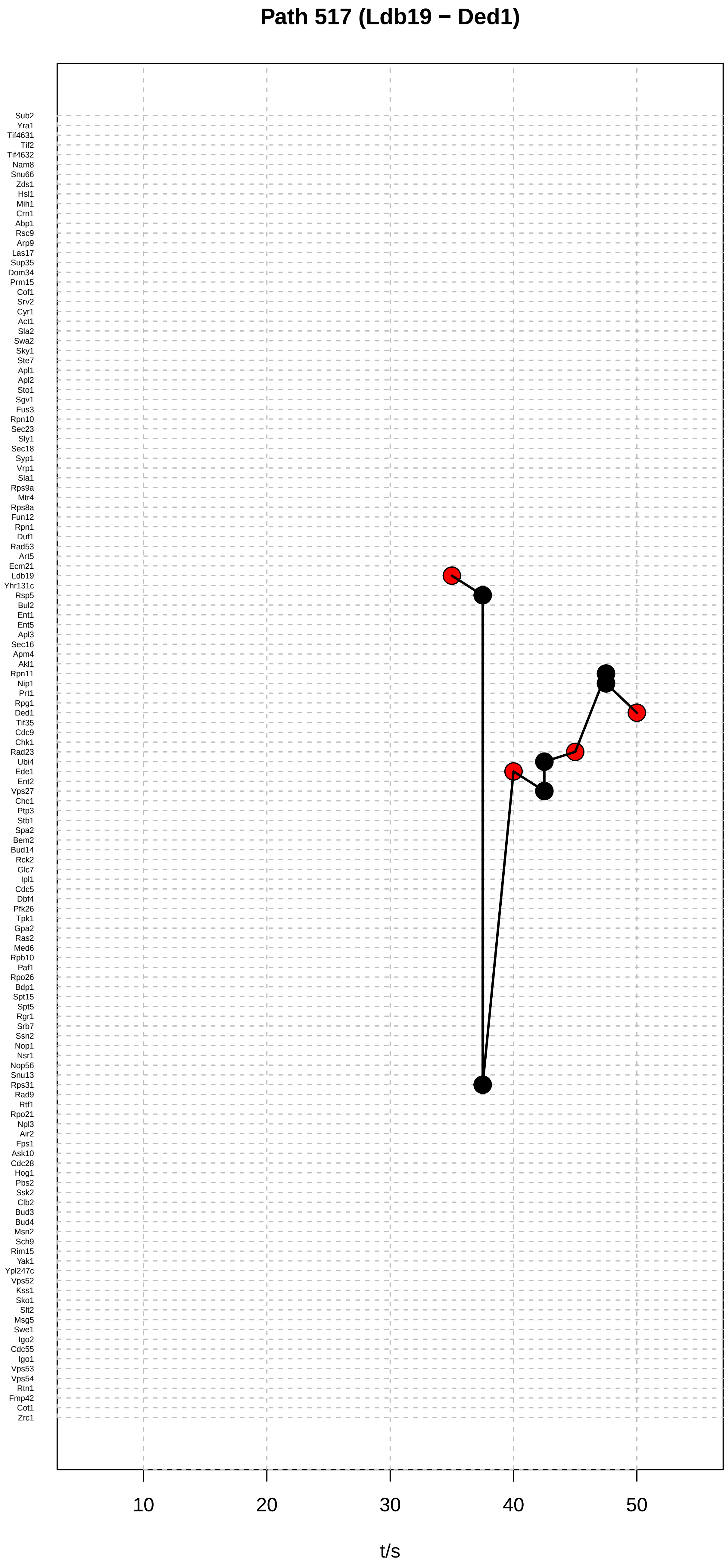

Path 517 (Ldb19 − Ded1)
Sub2
Yra1
Tif4631
Tif2
Tif4632
Nam8
Snu66
Zds1
Hsl1
Mih1
Crn1
Abp1
Rsc9
Arp9
Las17
Sup35
Dom34
Prm15
Cof1
Srv2
Cyr1
Act1
Sla2
Swa2
Sky1
Ste7
Apl1
Apl2
Sto1
Sgv1
Fus3
Rpn10
Sec23
Sly1
Sec18
Syp1
Vrp1
Sla1
Rps9a
Mtr4
Rps8a
Fun12
Rpn1
Duf1
Rad53
Art5
Ecm21
Ldb19
Yhr131c
Rsp5
Bul2
Ent1
Ent5
Apl3
Sec16
Apm4
Akl1
Rpn11
Nip1
Prt1
Rpg1
Ded1
Tif35
Cdc9
Chk1
Rad23
Ubi4
Ede1
Ent2
Vps27
Chc1
Ptp3
Stb1
Spa2
Bem2
Bud14
Rck2
Glc7
Ipl1
Cdc5
Dbf4
Pfk26
Tpk1
Gpa2
Ras2
Med6
Rpb10
Paf1
Rpo26
Bdp1
Spt15
Spt5
Rgr1
Srb7
Ssn2
Nop1
Nsr1
Nop56
Snu13
Rps31
Rad9
Rtf1
Rpo21
Npl3
Air2
Fps1
Ask10
Cdc28
Hog1
Pbs2
Ssk2
Clb2
Bud3
Bud4
Msn2
Sch9
Rim15
Yak1
Ypl247c
Vps52
Kss1
Sko1
Slt2
Msg5
Swe1
Igo2
Cdc55
Igo1
Vps53
Vps54
Rtn1
Fmp42
Cot1
Zrc1
10
20
30
40
50
t/s

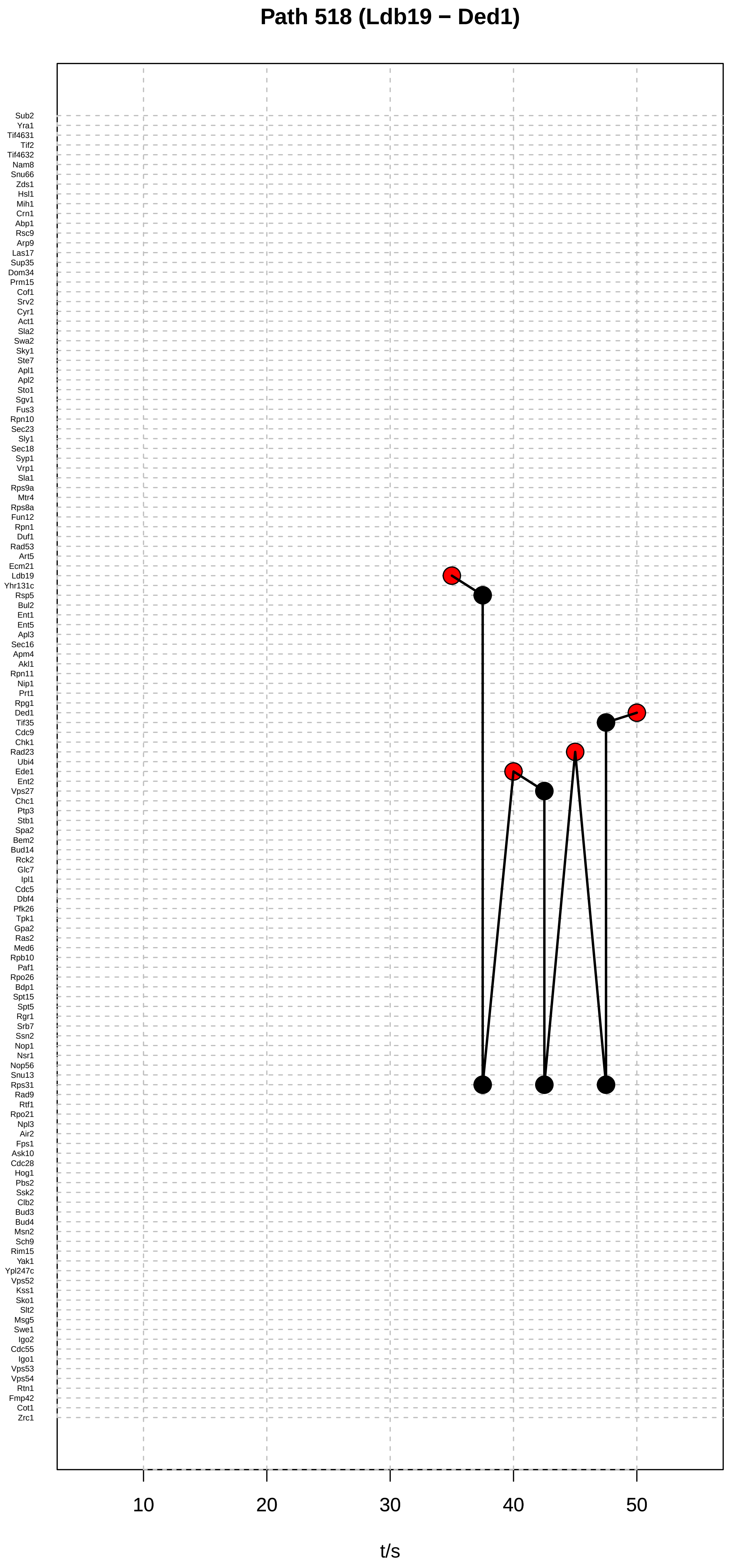
Path 518 (Ldb19 − Ded1)
Sub2
Yra1
Tif4631
Tif2
Tif4632
Nam8
Snu66
Zds1
Hsl1
Mih1
Crn1
Abp1
Rsc9
Arp9
Las17
Sup35
Dom34
Prm15
Cof1
Srv2
Cyr1
Act1
Sla2
Swa2
Sky1
Ste7
Apl1
Apl2
Sto1
Sgv1
Fus3
Rpn10
Sec23
Sly1
Sec18
Syp1
Vrp1
Sla1
Rps9a
Mtr4
Rps8a
Fun12
Rpn1
Duf1
Rad53
Art5
Ecm21
Ldb19
Yhr131c
Rsp5
Bul2
Ent1
Ent5
Apl3
Sec16
Apm4
Akl1
Rpn11
Nip1
Prt1
Rpg1
Ded1
Tif35
Cdc9
Chk1
Rad23
Ubi4
Ede1
Ent2
Vps27
Chc1
Ptp3
Stb1
Spa2
Bem2
Bud14
Rck2
Glc7
Ipl1
Cdc5
Dbf4
Pfk26
Tpk1
Gpa2
Ras2
Med6
Rpb10
Paf1
Rpo26
Bdp1
Spt15
Spt5
Rgr1
Srb7
Ssn2
Nop1
Nsr1
Nop56
Snu13
Rps31
Rad9
Rtf1
Rpo21
Npl3
Air2
Fps1
Ask10
Cdc28
Hog1
Pbs2
Ssk2
Clb2
Bud3
Bud4
Msn2
Sch9
Rim15
Yak1
Ypl247c
Vps52
Kss1
Sko1
Slt2
Msg5
Swe1
Igo2
Cdc55
Igo1
Vps53
Vps54
Rtn1
Fmp42
Cot1
Zrc1
10
20
30
40
50
t/s

### Path 519 (Ldb19 − Ded1)

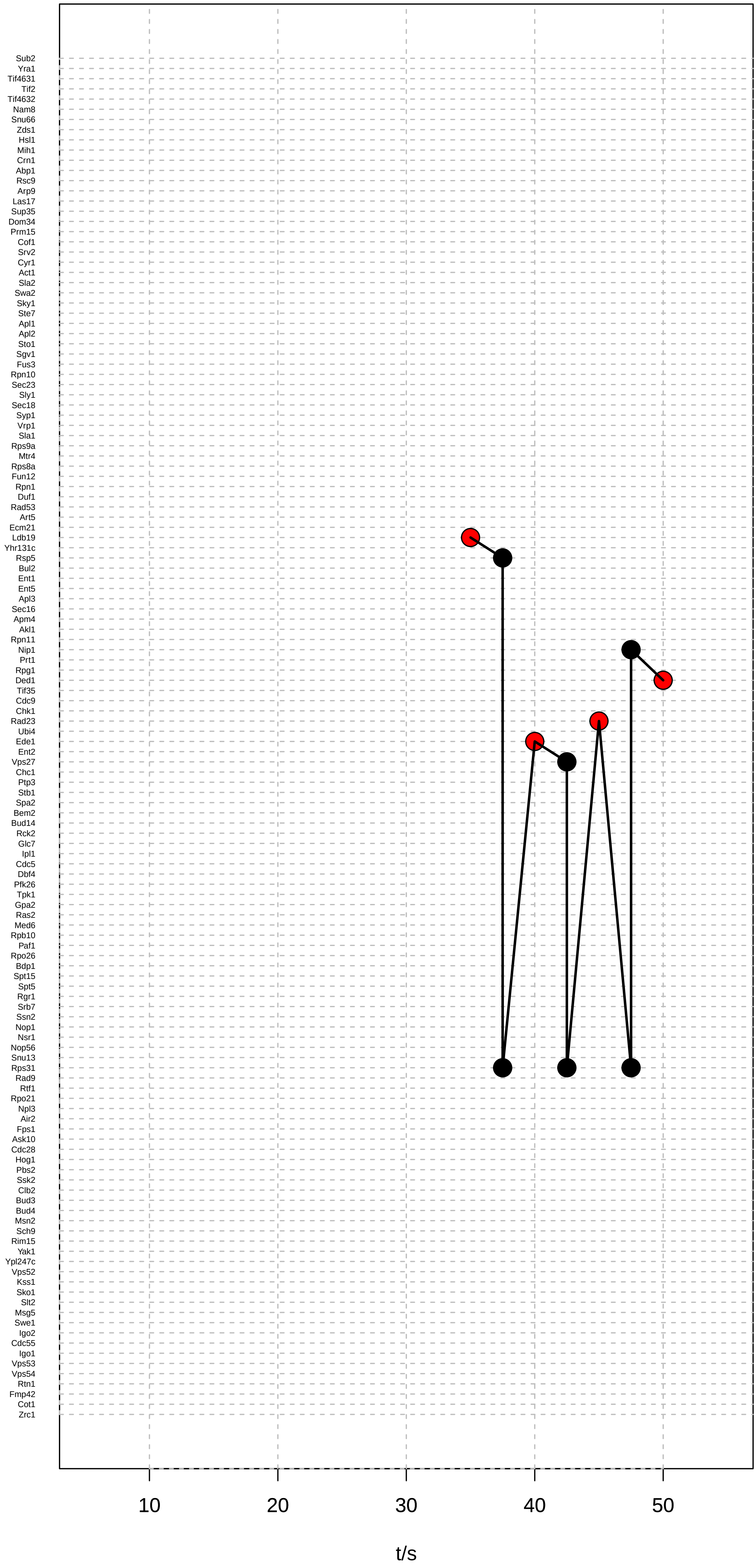

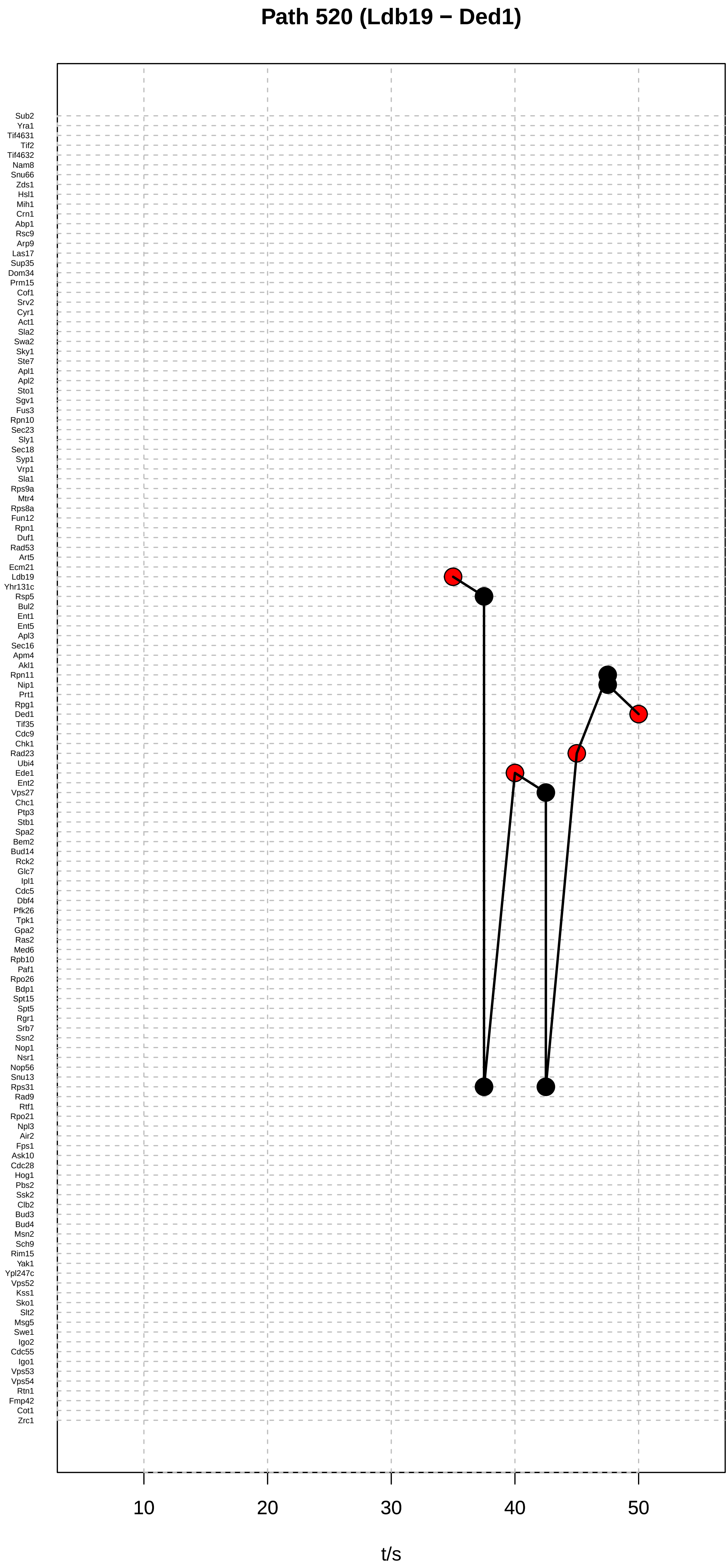

Path 520 (Ldb19 − Ded1)
Sub2
Yra1
Tif4631
Tif2
Tif4632
Nam8
Snu66
Zds1
Hsl1
Mih1
Crn1
Abp1
Rsc9
Arp9
Las17
Sup35
Dom34
Prm15
Cof1
Srv2
Cyr1
Act1
Sla2
Swa2
Sky1
Ste7
Apl1
Apl2
Sto1
Sgv1
Fus3
Rpn10
Sec23
Sly1
Sec18
Syp1
Vrp1
Sla1
Rps9a
Mtr4
Rps8a
Fun12
Rpn1
Duf1
Rad53
Art5
Ecm21
Ldb19
Yhr131c
Rsp5
Bul2
Ent1
Ent5
Apl3
Sec16
Apm4
Akl1
Rpn11
Nip1
Prt1
Rpg1
Ded1
Tif35
Cdc9
Chk1
Rad23
Ubi4
Ede1
Ent2
Vps27
Chc1
Ptp3
Stb1
Spa2
Bem2
Bud14
Rck2
Glc7
Ipl1
Cdc5
Dbf4
Pfk26
Tpk1
Gpa2
Ras2
Med6
Rpb10
Paf1
Rpo26
Bdp1
Spt15
Spt5
Rgr1
Srb7
Ssn2
Nop1
Nsr1
Nop56
Snu13
Rps31
Rad9
Rtf1
Rpo21
Npl3
Air2
Fps1
Ask10
Cdc28
Hog1
Pbs2
Ssk2
Clb2
Bud3
Bud4
Msn2
Sch9
Rim15
Yak1
Ypl247c
Vps52
Kss1
Sko1
Slt2
Msg5
Swe1
Igo2
Cdc55
Igo1
Vps53
Vps54
Rtn1
Fmp42
Cot1
Zrc1
10
20
30
40
50
t/s

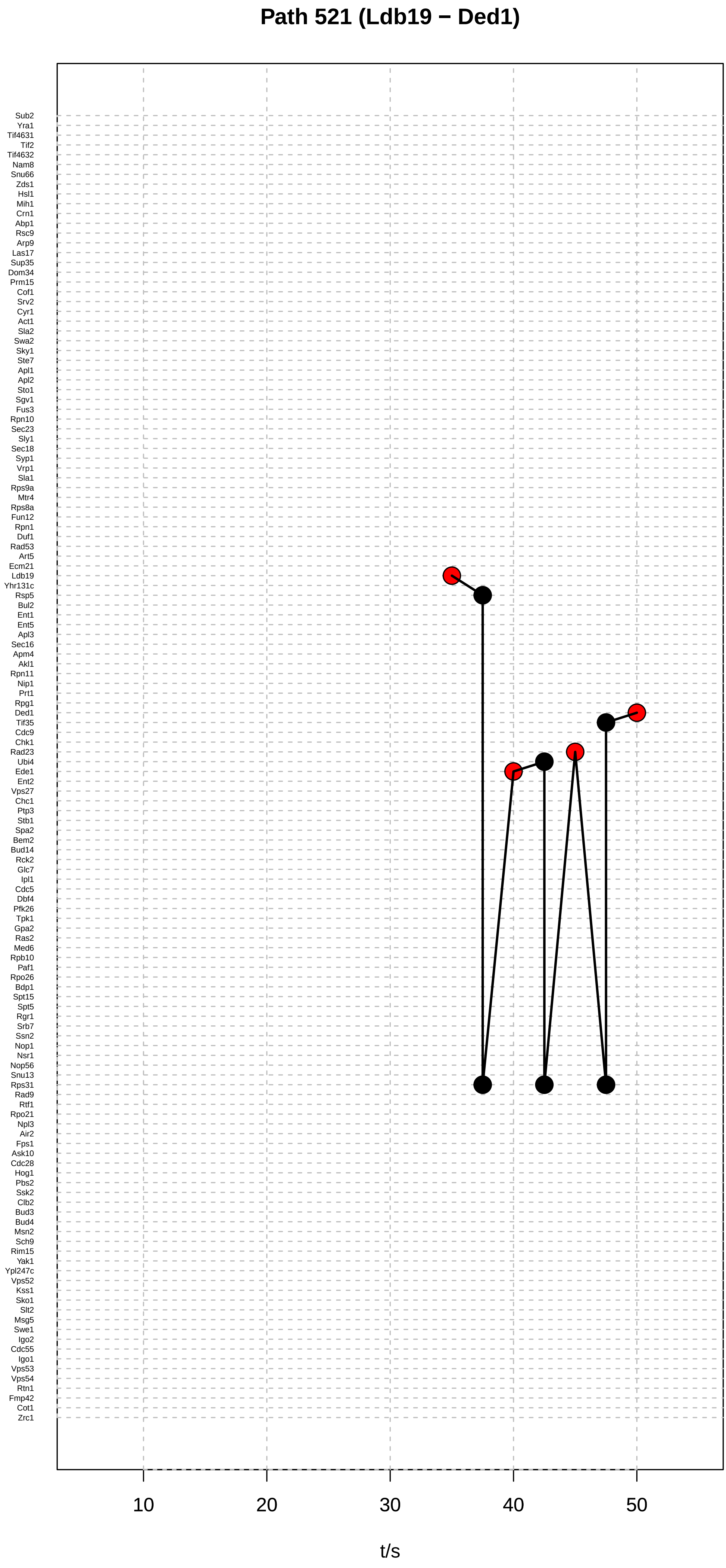

Path 521 (Ldb19 − Ded1)
Sub2
Yra1
Tif4631
Tif2
Tif4632
Nam8
Snu66
Zds1
Hsl1
Mih1
Crn1
Abp1
Rsc9
Arp9
Las17
Sup35
Dom34
Prm15
Cof1
Srv2
Cyr1
Act1
Sla2
Swa2
Sky1
Ste7
Apl1
Apl2
Sto1
Sgv1
Fus3
Rpn10
Sec23
Sly1
Sec18
Syp1
Vrp1
Sla1
Rps9a
Mtr4
Rps8a
Fun12
Rpn1
Duf1
Rad53
Art5
Ecm21
Ldb19
Yhr131c
Rsp5
Bul2
Ent1
Ent5
Apl3
Sec16
Apm4
Akl1
Rpn11
Nip1
Prt1
Rpg1
Ded1
Tif35
Cdc9
Chk1
Rad23
Ubi4
Ede1
Ent2
Vps27
Chc1
Ptp3
Stb1
Spa2
Bem2
Bud14
Rck2
Glc7
Ipl1
Cdc5
Dbf4
Pfk26
Tpk1
Gpa2
Ras2
Med6
Rpb10
Paf1
Rpo26
Bdp1
Spt15
Spt5
Rgr1
Srb7
Ssn2
Nop1
Nsr1
Nop56
Snu13
Rps31
Rad9
Rtf1
Rpo21
Npl3
Air2
Fps1
Ask10
Cdc28
Hog1
Pbs2
Ssk2
Clb2
Bud3
Bud4
Msn2
Sch9
Rim15
Yak1
Ypl247c
Vps52
Kss1
Sko1
Slt2
Msg5
Swe1
Igo2
Cdc55
Igo1
Vps53
Vps54
Rtn1
Fmp42
Cot1
Zrc1
10
20
30
40
50
t/s

#### Path 522 (Ldb19 − Ded1)

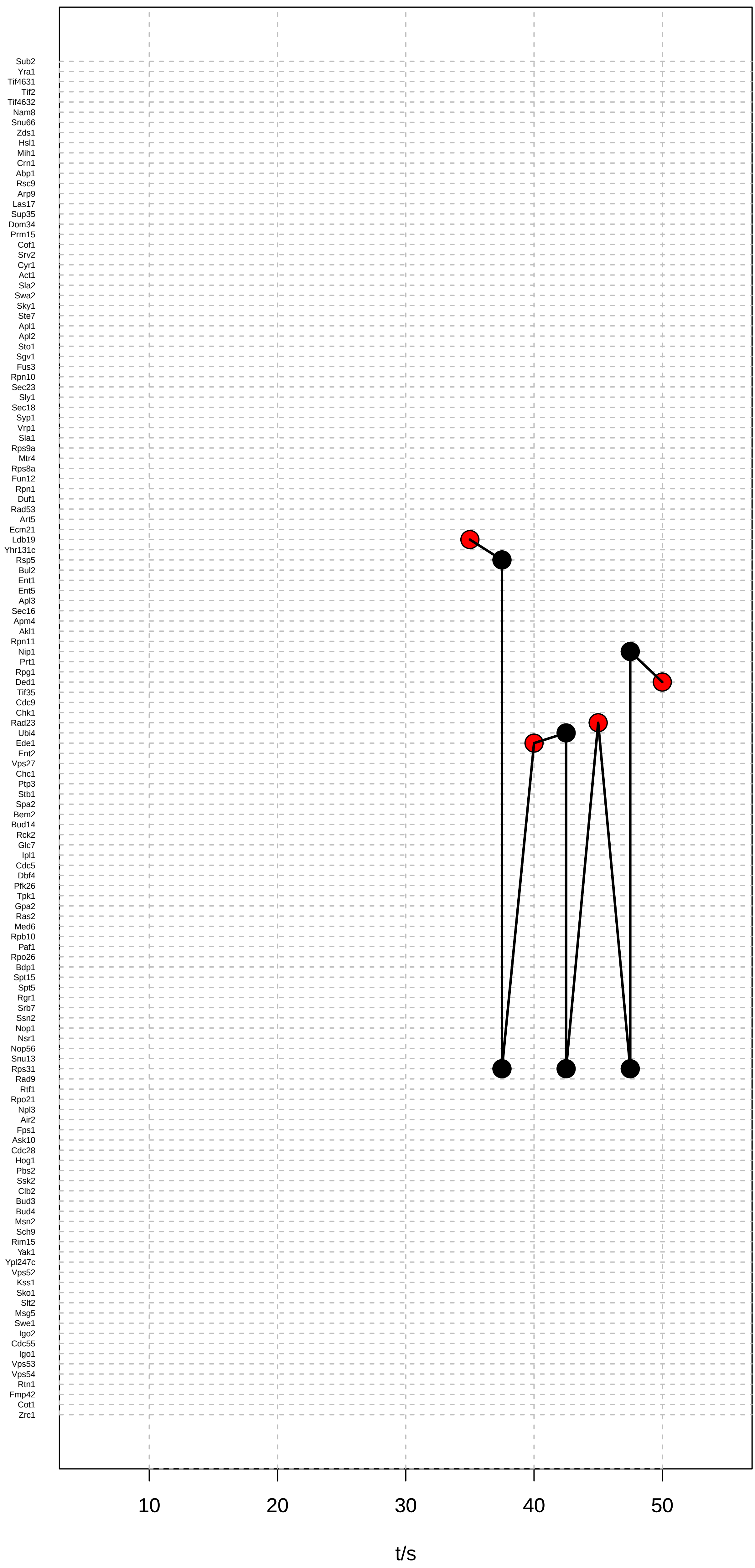

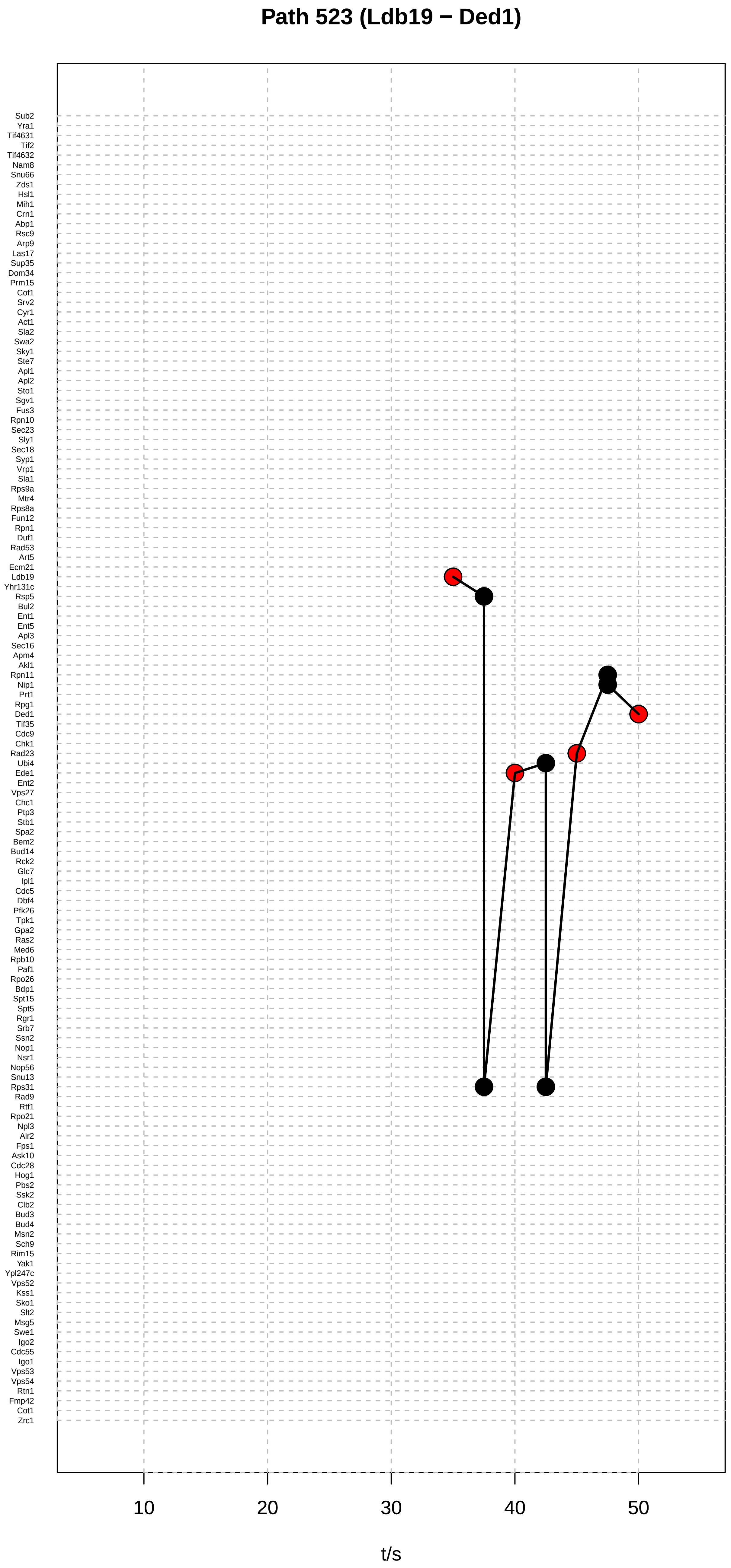
Path 523 (Ldb19 − Ded1)
Sub2
Yra1
Tif4631
Tif2
Tif4632
Nam8
Snu66
Zds1
Hsl1
Mih1
Crn1
Abp1
Rsc9
Arp9
Las17
Sup35
Dom34
Prm15
Cof1
Srv2
Cyr1
Act1
Sla2
Swa2
Sky1
Ste7
Apl1
Apl2
Sto1
Sgv1
Fus3
Rpn10
Sec23
Sly1
Sec18
Syp1
Vrp1
Sla1
Rps9a
Mtr4
Rps8a
Fun12
Rpn1
Duf1
Rad53
Art5
Ecm21
Ldb19
Yhr131c
Rsp5
Bul2
Ent1
Ent5
Apl3
Sec16
Apm4
Akl1
Rpn11
Nip1
Prt1
Rpg1
Ded1
Tif35
Cdc9
Chk1
Rad23
Ubi4
Ede1
Ent2
Vps27
Chc1
Ptp3
Stb1
Spa2
Bem2
Bud14
Rck2
Glc7
Ipl1
Cdc5
Dbf4
Pfk26
Tpk1
Gpa2
Ras2
Med6
Rpb10
Paf1
Rpo26
Bdp1
Spt15
Spt5
Rgr1
Srb7
Ssn2
Nop1
Nsr1
Nop56
Snu13
Rps31
Rad9
Rtf1
Rpo21
Npl3
Air2
Fps1
Ask10
Cdc28
Hog1
Pbs2
Ssk2
Clb2
Bud3
Bud4
Msn2
Sch9
Rim15
Yak1
Ypl247c
Vps52
Kss1
Sko1
Slt2
Msg5
Swe1
Igo2
Cdc55
Igo1
Vps53
Vps54
Rtn1
Fmp42
Cot1
Zrc1
10
20
30
40
50
t/s

### Path 524 (Ldb19 − Ded1)

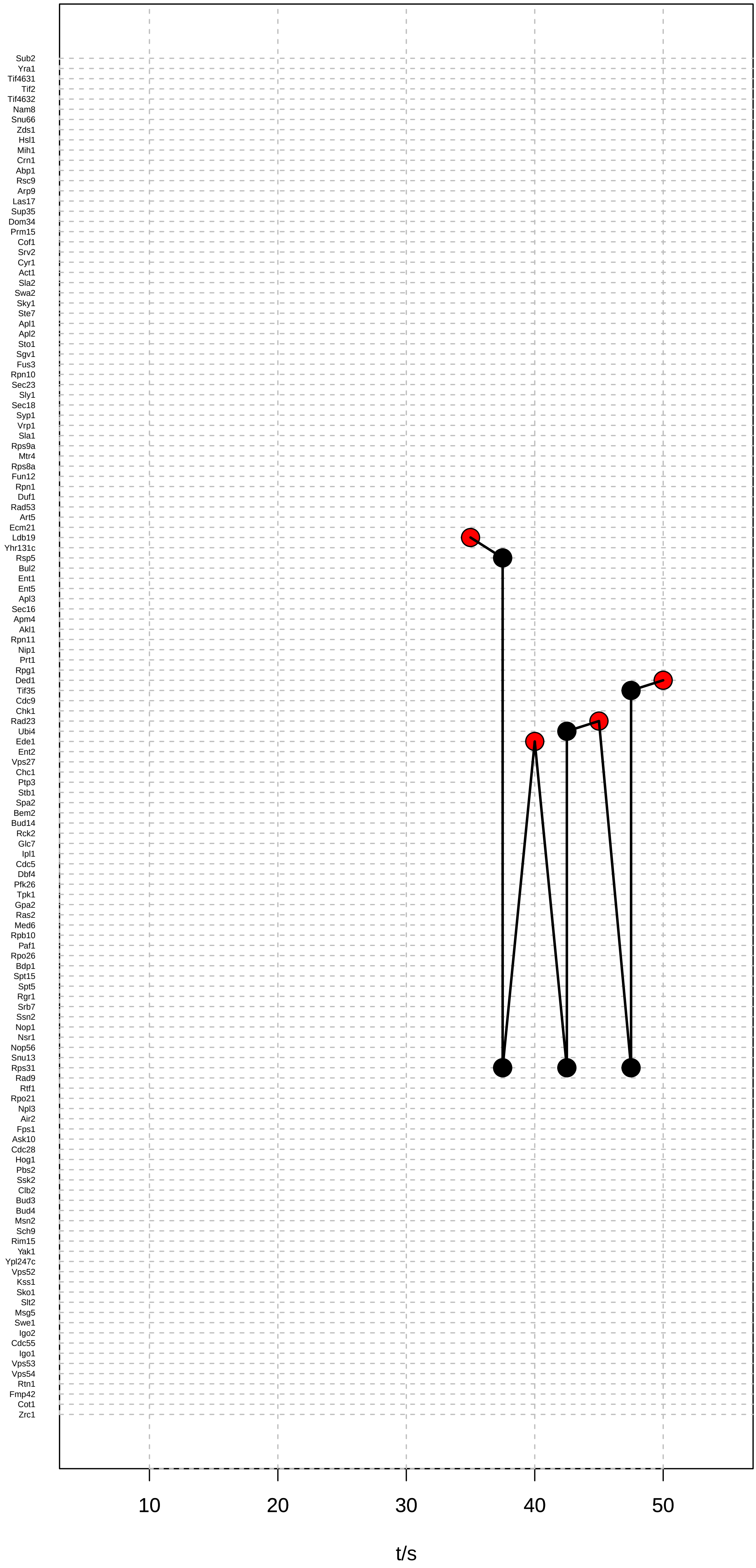

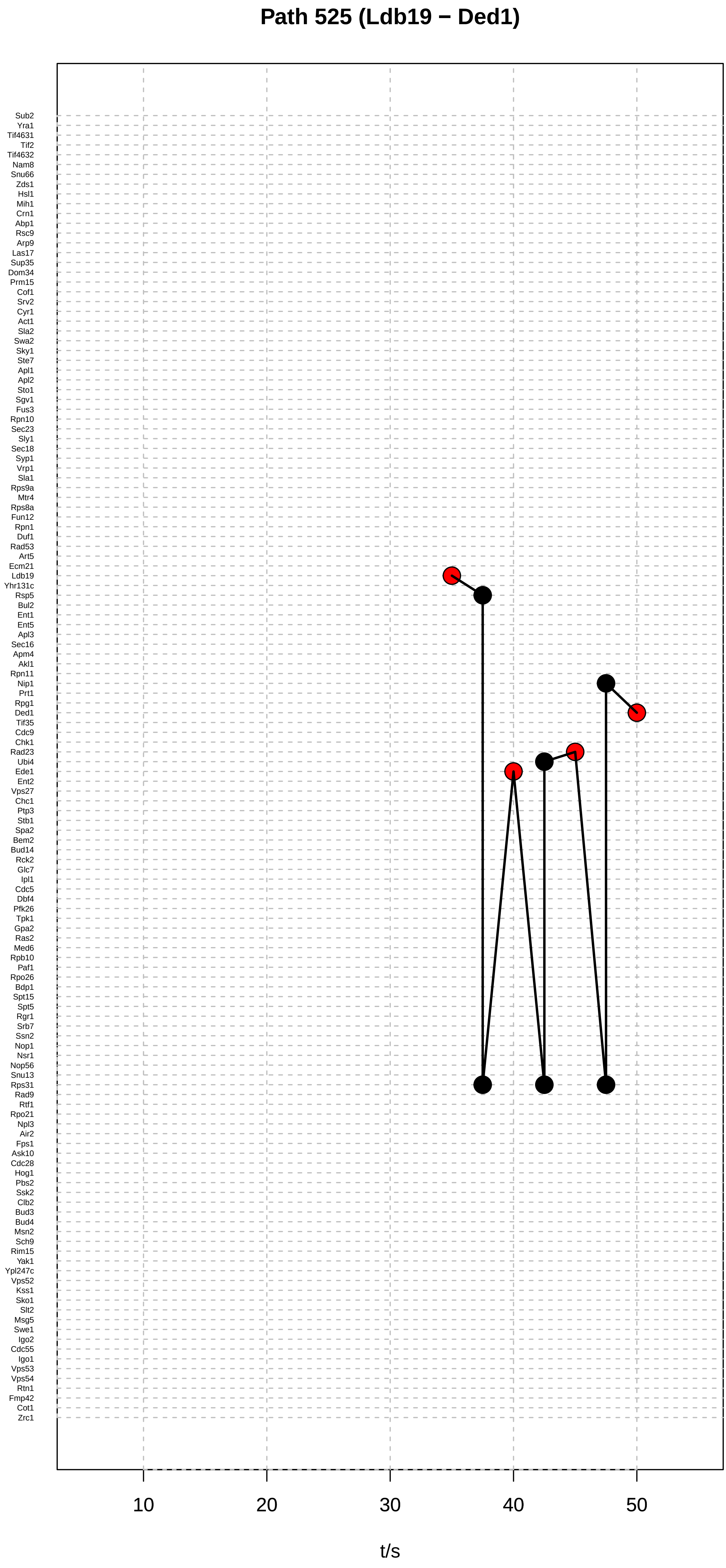

Path 525 (Ldb19 − Ded1)
Sub2
Yra1
Tif4631
Tif2
Tif4632
Nam8
Snu66
Zds1
Hsl1
Mih1
Crn1
Abp1
Rsc9
Arp9
Las17
Sup35
Dom34
Prm15
Cof1
Srv2
Cyr1
Act1
Sla2
Swa2
Sky1
Ste7
Apl1
Apl2
Sto1
Sgv1
Fus3
Rpn10
Sec23
Sly1
Sec18
Syp1
Vrp1
Sla1
Rps9a
Mtr4
Rps8a
Fun12
Rpn1
Duf1
Rad53
Art5
Ecm21
Ldb19
Yhr131c
Rsp5
Bul2
Ent1
Ent5
Apl3
Sec16
Apm4
Akl1
Rpn11
Nip1
Prt1
Rpg1
Ded1
Tif35
Cdc9
Chk1
Rad23
Ubi4
Ede1
Ent2
Vps27
Chc1
Ptp3
Stb1
Spa2
Bem2
Bud14
Rck2
Glc7
Ipl1
Cdc5
Dbf4
Pfk26
Tpk1
Gpa2
Ras2
Med6
Rpb10
Paf1
Rpo26
Bdp1
Spt15
Spt5
Rgr1
Srb7
Ssn2
Nop1
Nsr1
Nop56
Snu13
Rps31
Rad9
Rtf1
Rpo21
Npl3
Air2
Fps1
Ask10
Cdc28
Hog1
Pbs2
Ssk2
Clb2
Bud3
Bud4
Msn2
Sch9
Rim15
Yak1
Ypl247c
Vps52
Kss1
Sko1
Slt2
Msg5
Swe1
Igo2
Cdc55
Igo1
Vps53
Vps54
Rtn1
Fmp42
Cot1
Zrc1
10
20
30
40
50
t/s

### Path 526 (Ldb19 − Ded1)

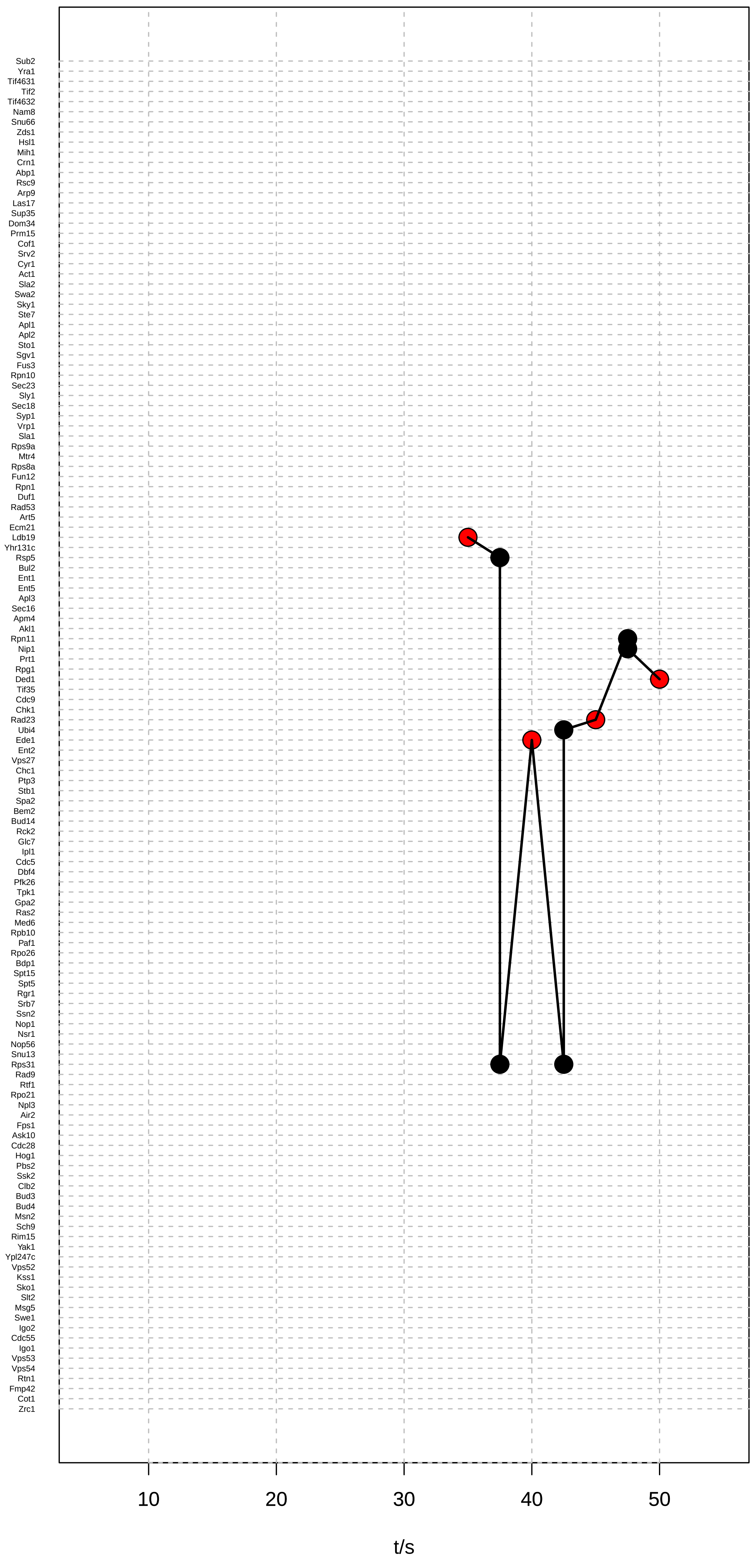

### Path 527 (Ldb19 − Ded1)

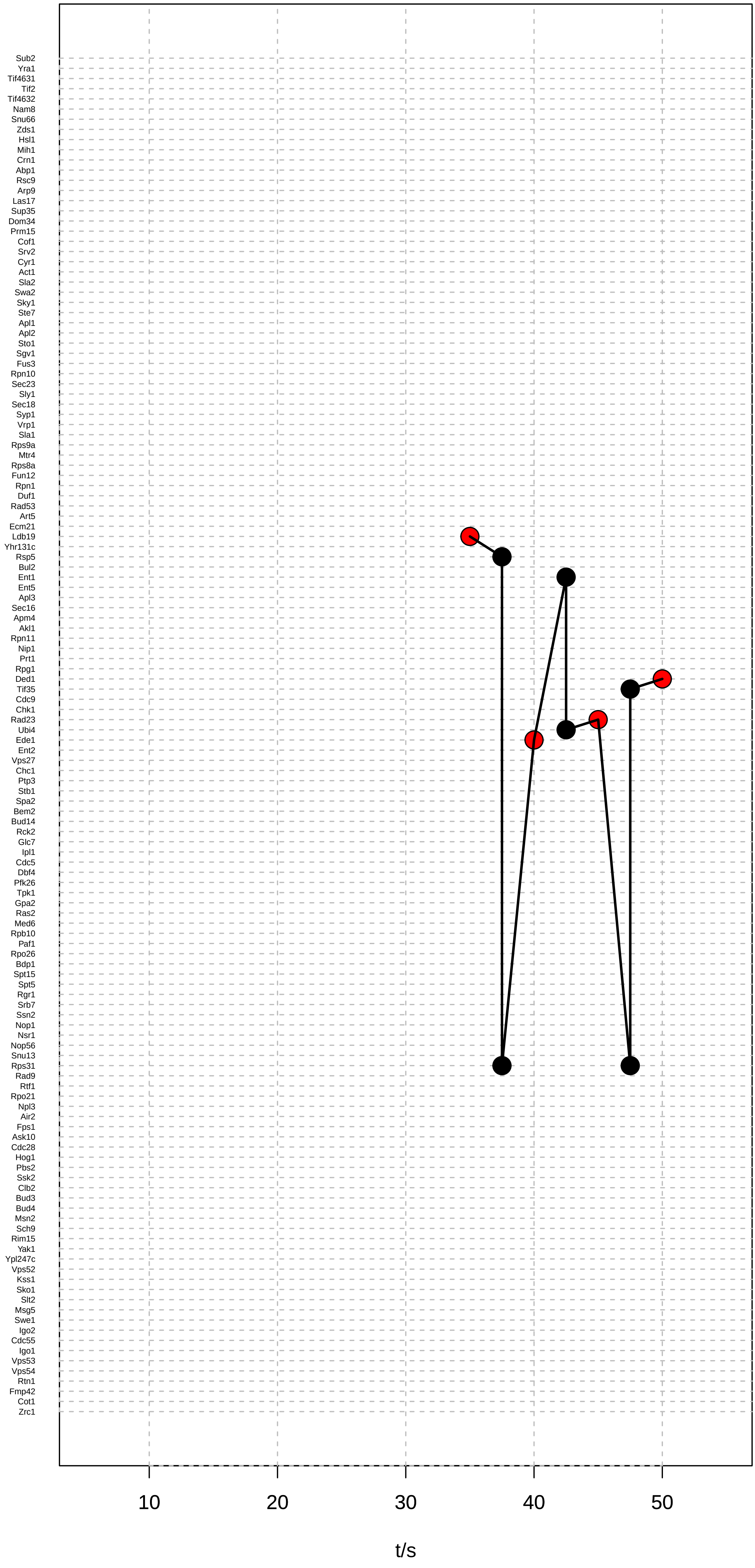

#### Path 528 (Ldb19 − Ded1)

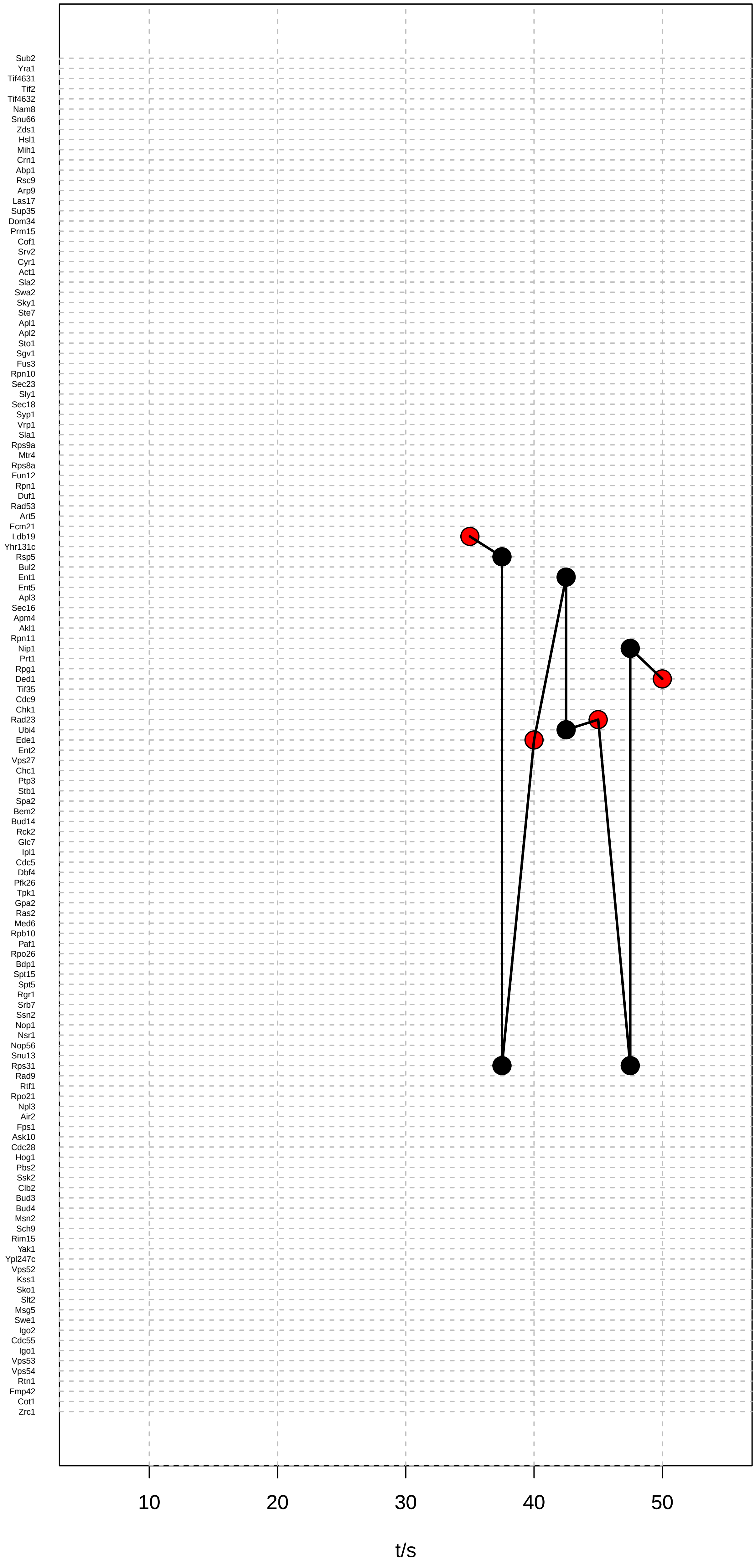

#### Path 529 (Ldb19 − Ded1)

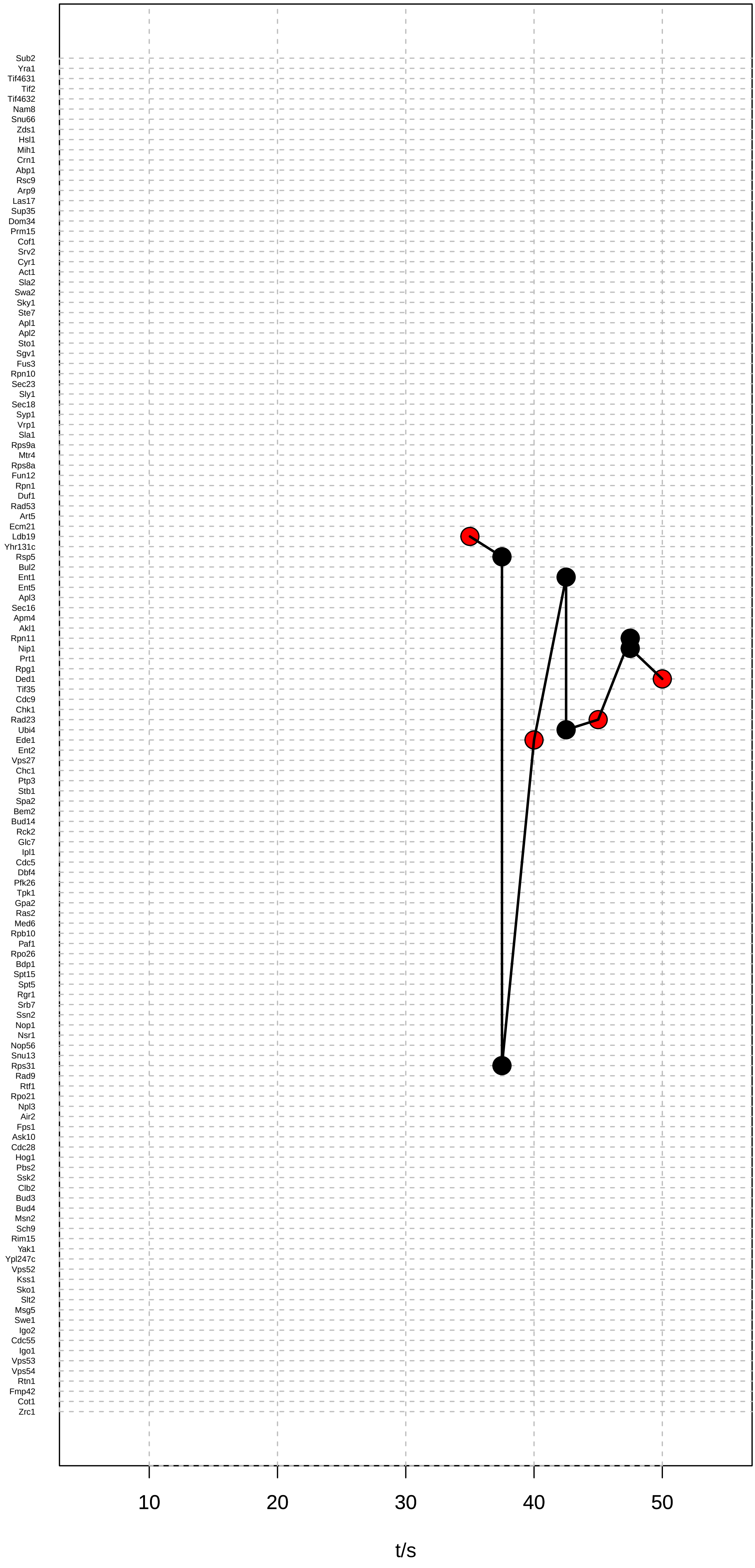

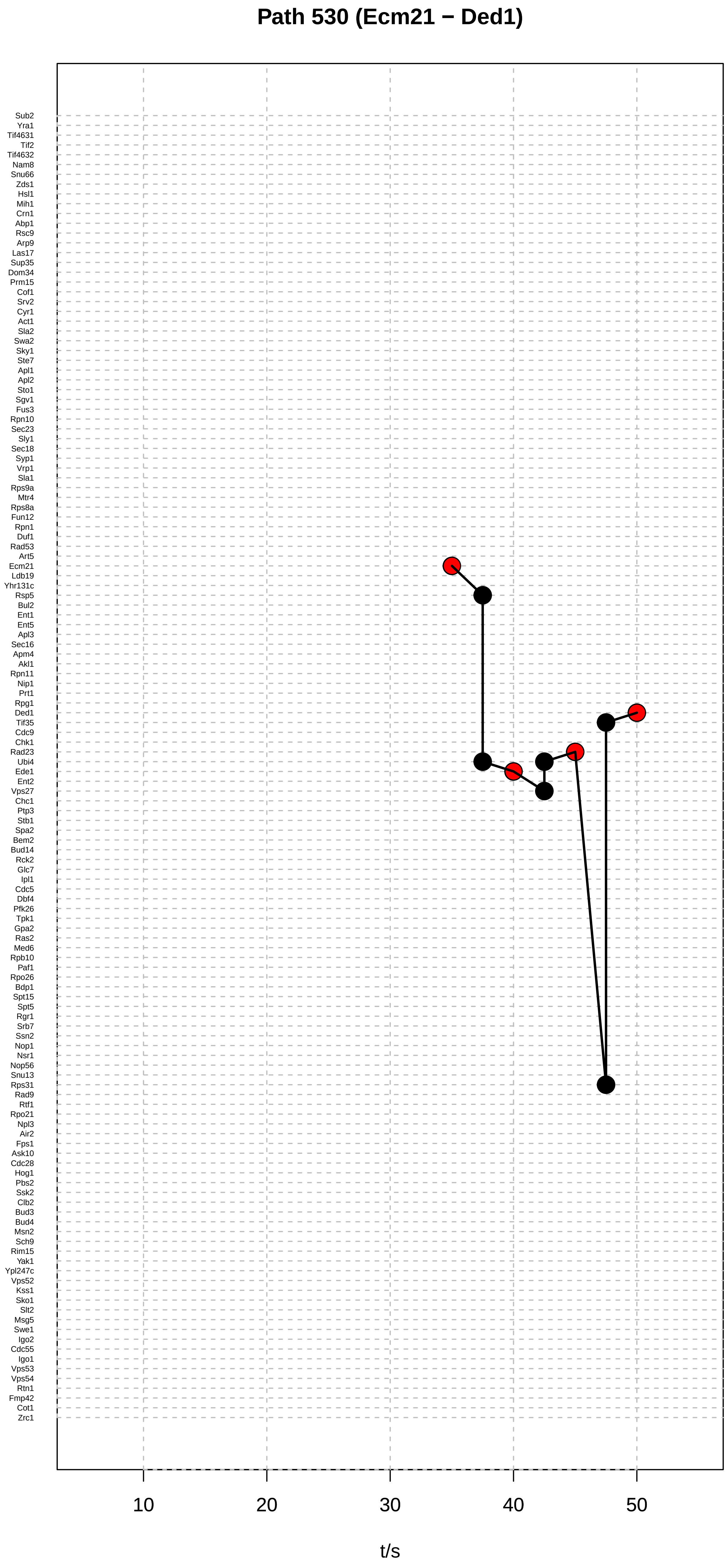

Path 530 (Ecm21 − Ded1)
Sub2
Yra1
Tif4631
Tif2
Tif4632
Nam8
Snu66
Zds1
Hsl1
Mih1
Crn1
Abp1
Rsc9
Arp9
Las17
Sup35
Dom34
Prm15
Cof1
Srv2
Cyr1
Act1
Sla2
Swa2
Sky1
Ste7
Apl1
Apl2
Sto1
Sgv1
Fus3
Rpn10
Sec23
Sly1
Sec18
Syp1
Vrp1
Sla1
Rps9a
Mtr4
Rps8a
Fun12
Rpn1
Duf1
Rad53
Art5
Ecm21
Ldb19
Yhr131c
Rsp5
Bul2
Ent1
Ent5
Apl3
Sec16
Apm4
Akl1
Rpn11
Nip1
Prt1
Rpg1
Ded1
Tif35
Cdc9
Chk1
Rad23
Ubi4
Ede1
Ent2
Vps27
Chc1
Ptp3
Stb1
Spa2
Bem2
Bud14
Rck2
Glc7
Ipl1
Cdc5
Dbf4
Pfk26
Tpk1
Gpa2
Ras2
Med6
Rpb10
Paf1
Rpo26
Bdp1
Spt15
Spt5
Rgr1
Srb7
Ssn2
Nop1
Nsr1
Nop56
Snu13
Rps31
Rad9
Rtf1
Rpo21
Npl3
Air2
Fps1
Ask10
Cdc28
Hog1
Pbs2
Ssk2
Clb2
Bud3
Bud4
Msn2
Sch9
Rim15
Yak1
Ypl247c
Vps52
Kss1
Sko1
Slt2
Msg5
Swe1
Igo2
Cdc55
Igo1
Vps53
Vps54
Rtn1
Fmp42
Cot1
Zrc1
10
20
30
40
50
t/s

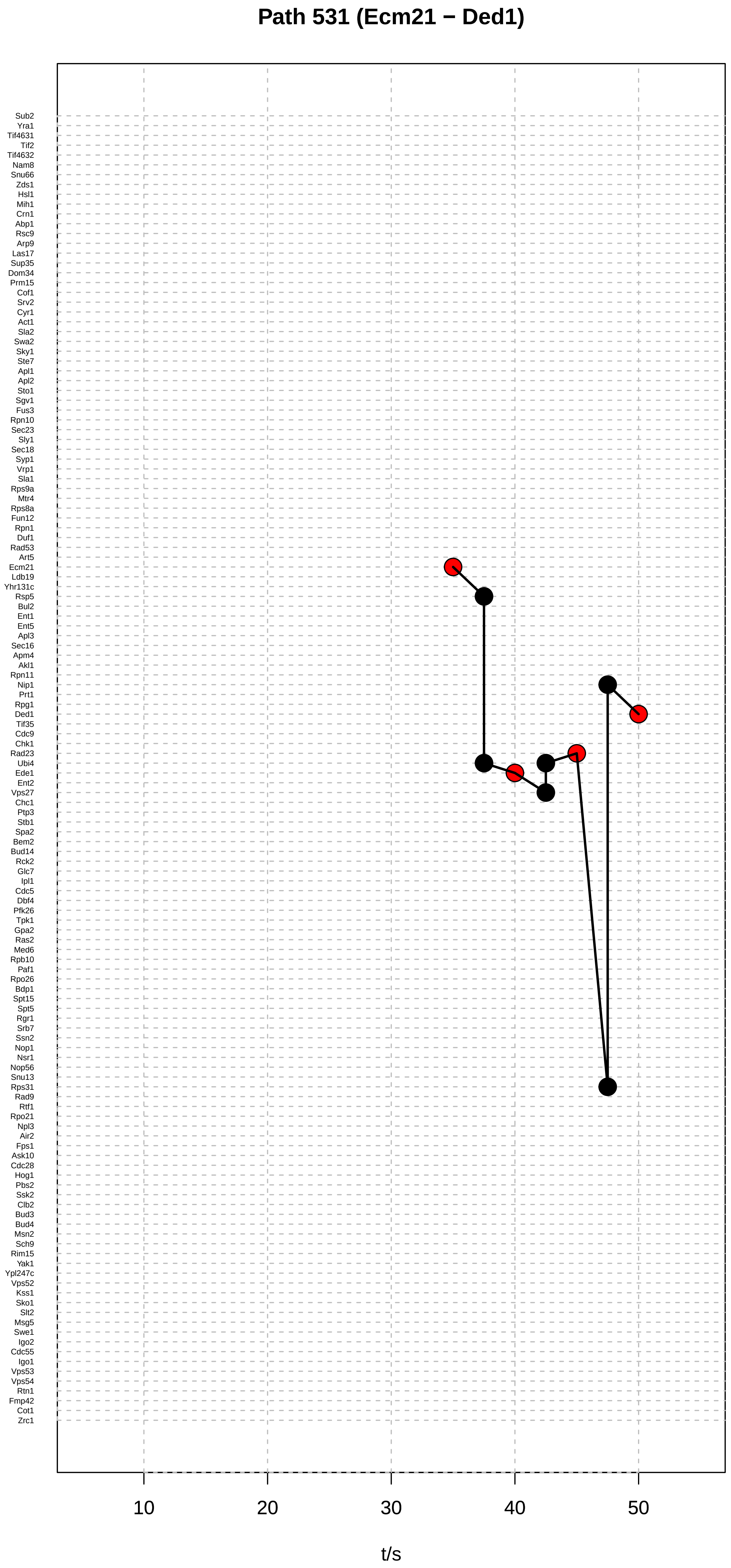

Path 531 (Ecm21 − Ded1)
Sub2
Yra1
Tif4631
Tif2
Tif4632
Nam8
Snu66
Zds1
Hsl1
Mih1
Crn1
Abp1
Rsc9
Arp9
Las17
Sup35
Dom34
Prm15
Cof1
Srv2
Cyr1
Act1
Sla2
Swa2
Sky1
Ste7
Apl1
Apl2
Sto1
Sgv1
Fus3
Rpn10
Sec23
Sly1
Sec18
Syp1
Vrp1
Sla1
Rps9a
Mtr4
Rps8a
Fun12
Rpn1
Duf1
Rad53
Art5
Ecm21
Ldb19
Yhr131c
Rsp5
Bul2
Ent1
Ent5
Apl3
Sec16
Apm4
Akl1
Rpn11
Nip1
Prt1
Rpg1
Ded1
Tif35
Cdc9
Chk1
Rad23
Ubi4
Ede1
Ent2
Vps27
Chc1
Ptp3
Stb1
Spa2
Bem2
Bud14
Rck2
Glc7
Ipl1
Cdc5
Dbf4
Pfk26
Tpk1
Gpa2
Ras2
Med6
Rpb10
Paf1
Rpo26
Bdp1
Spt15
Spt5
Rgr1
Srb7
Ssn2
Nop1
Nsr1
Nop56
Snu13
Rps31
Rad9
Rtf1
Rpo21
Npl3
Air2
Fps1
Ask10
Cdc28
Hog1
Pbs2
Ssk2
Clb2
Bud3
Bud4
Msn2
Sch9
Rim15
Yak1
Ypl247c
Vps52
Kss1
Sko1
Slt2
Msg5
Swe1
Igo2
Cdc55
Igo1
Vps53
Vps54
Rtn1
Fmp42
Cot1
Zrc1
10
20
30
40
50
t/s

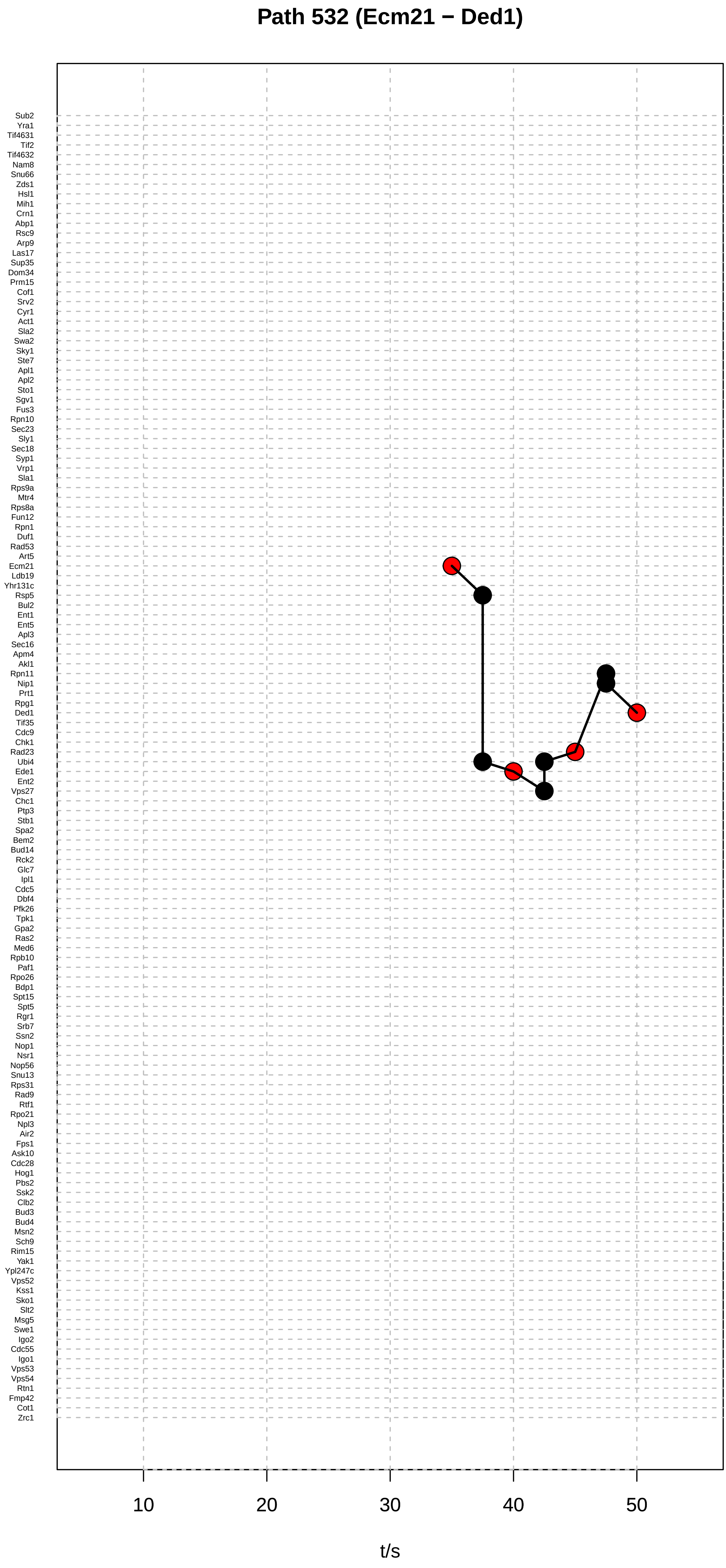

Path 532 (Ecm21 − Ded1)
Sub2
Yra1
Tif4631
Tif2
Tif4632
Nam8
Snu66
Zds1
Hsl1
Mih1
Crn1
Abp1
Rsc9
Arp9
Las17
Sup35
Dom34
Prm15
Cof1
Srv2
Cyr1
Act1
Sla2
Swa2
Sky1
Ste7
Apl1
Apl2
Sto1
Sgv1
Fus3
Rpn10
Sec23
Sly1
Sec18
Syp1
Vrp1
Sla1
Rps9a
Mtr4
Rps8a
Fun12
Rpn1
Duf1
Rad53
Art5
Ecm21
Ldb19
Yhr131c
Rsp5
Bul2
Ent1
Ent5
Apl3
Sec16
Apm4
Akl1
Rpn11
Nip1
Prt1
Rpg1
Ded1
Tif35
Cdc9
Chk1
Rad23
Ubi4
Ede1
Ent2
Vps27
Chc1
Ptp3
Stb1
Spa2
Bem2
Bud14
Rck2
Glc7
Ipl1
Cdc5
Dbf4
Pfk26
Tpk1
Gpa2
Ras2
Med6
Rpb10
Paf1
Rpo26
Bdp1
Spt15
Spt5
Rgr1
Srb7
Ssn2
Nop1
Nsr1
Nop56
Snu13
Rps31
Rad9
Rtf1
Rpo21
Npl3
Air2
Fps1
Ask10
Cdc28
Hog1
Pbs2
Ssk2
Clb2
Bud3
Bud4
Msn2
Sch9
Rim15
Yak1
Ypl247c
Vps52
Kss1
Sko1
Slt2
Msg5
Swe1
Igo2
Cdc55
Igo1
Vps53
Vps54
Rtn1
Fmp42
Cot1
Zrc1
10
20
30
40
50
t/s

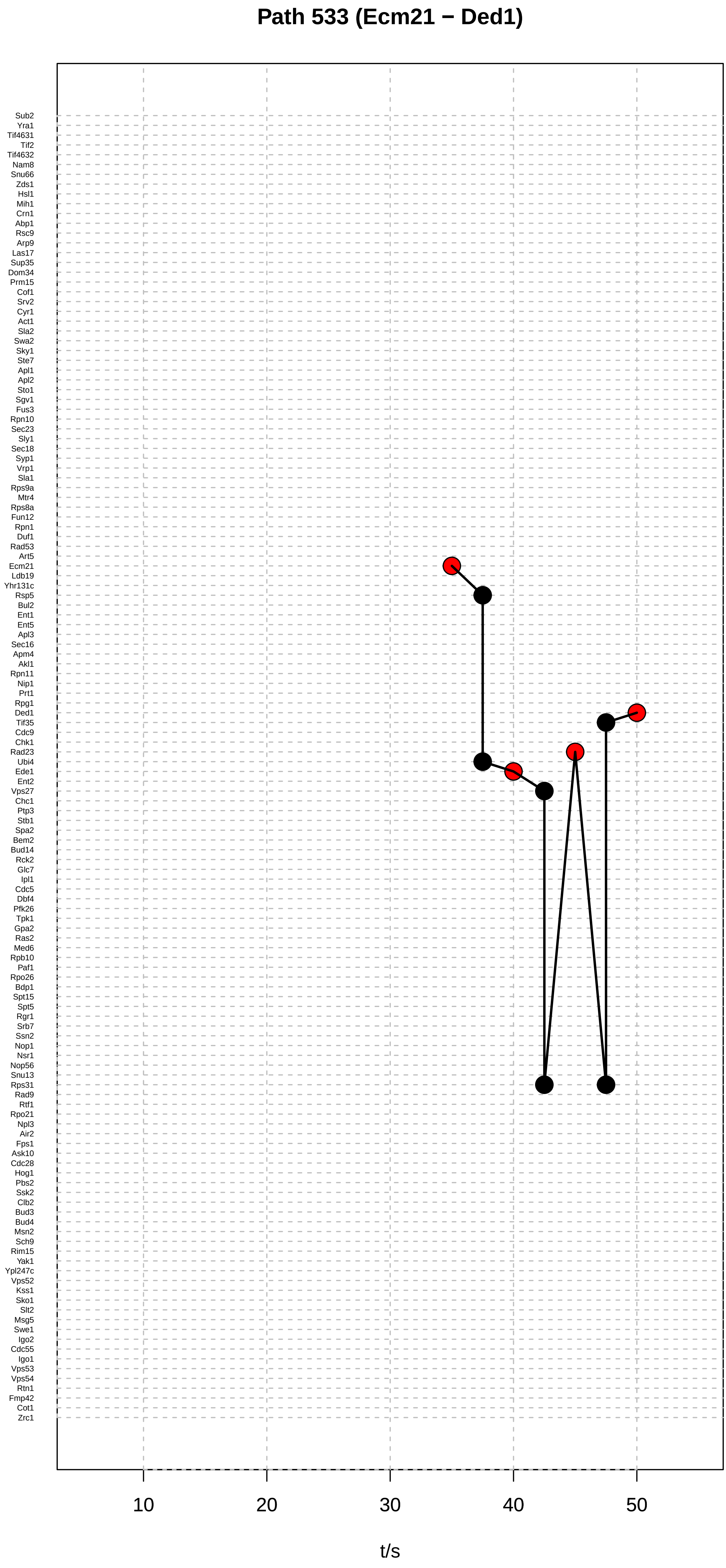

Path 533 (Ecm21 − Ded1)
Sub2
Yra1
Tif4631
Tif2
Tif4632
Nam8
Snu66
Zds1
Hsl1
Mih1
Crn1
Abp1
Rsc9
Arp9
Las17
Sup35
Dom34
Prm15
Cof1
Srv2
Cyr1
Act1
Sla2
Swa2
Sky1
Ste7
Apl1
Apl2
Sto1
Sgv1
Fus3
Rpn10
Sec23
Sly1
Sec18
Syp1
Vrp1
Sla1
Rps9a
Mtr4
Rps8a
Fun12
Rpn1
Duf1
Rad53
Art5
Ecm21
Ldb19
Yhr131c
Rsp5
Bul2
Ent1
Ent5
Apl3
Sec16
Apm4
Akl1
Rpn11
Nip1
Prt1
Rpg1
Ded1
Tif35
Cdc9
Chk1
Rad23
Ubi4
Ede1
Ent2
Vps27
Chc1
Ptp3
Stb1
Spa2
Bem2
Bud14
Rck2
Glc7
Ipl1
Cdc5
Dbf4
Pfk26
Tpk1
Gpa2
Ras2
Med6
Rpb10
Paf1
Rpo26
Bdp1
Spt15
Spt5
Rgr1
Srb7
Ssn2
Nop1
Nsr1
Nop56
Snu13
Rps31
Rad9
Rtf1
Rpo21
Npl3
Air2
Fps1
Ask10
Cdc28
Hog1
Pbs2
Ssk2
Clb2
Bud3
Bud4
Msn2
Sch9
Rim15
Yak1
Ypl247c
Vps52
Kss1
Sko1
Slt2
Msg5
Swe1
Igo2
Cdc55
Igo1
Vps53
Vps54
Rtn1
Fmp42
Cot1
Zrc1
10
20
30
40
50
t/s

#### Path 534 (Ecm21 − Ded1)

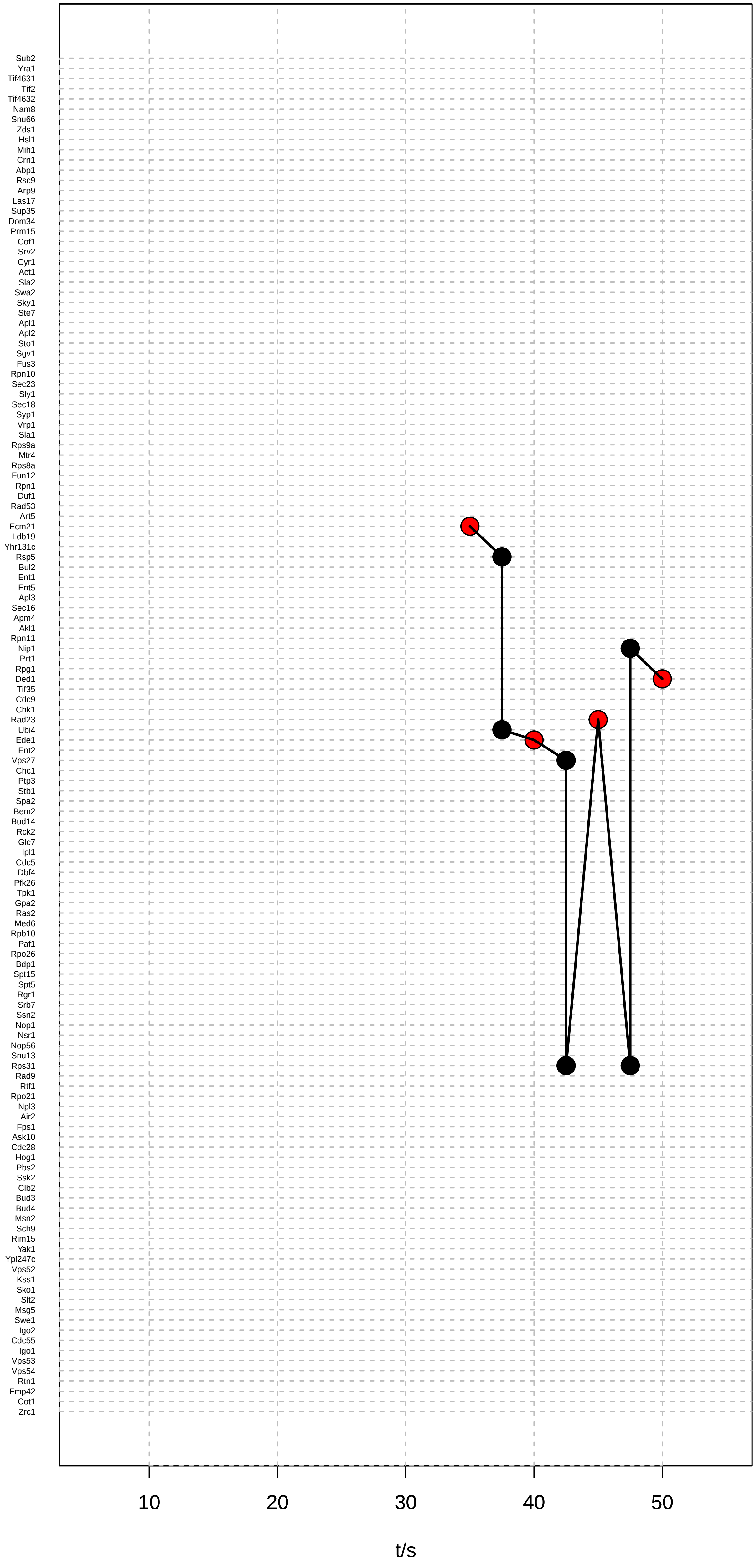

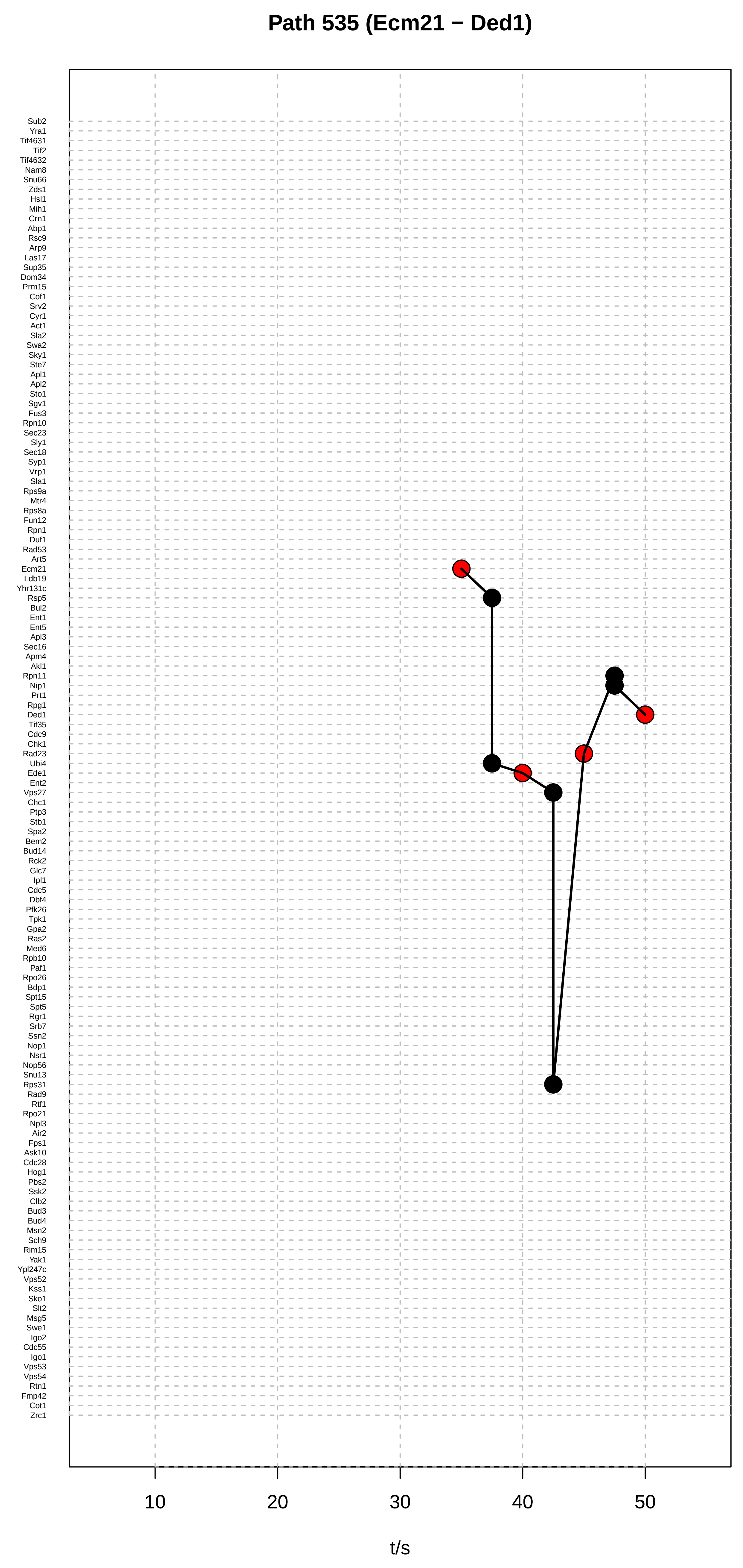
Path 535 (Ecm21 − Ded1)
Sub2
Yra1
Tif4631
Tif2
Tif4632
Nam8
Snu66
Zds1
Hsl1
Mih1
Crn1
Abp1
Rsc9
Arp9
Las17
Sup35
Dom34
Prm15
Cof1
Srv2
Cyr1
Act1
Sla2
Swa2
Sky1
Ste7
Apl1
Apl2
Sto1
Sgv1
Fus3
Rpn10
Sec23
Sly1
Sec18
Syp1
Vrp1
Sla1
Rps9a
Mtr4
Rps8a
Fun12
Rpn1
Duf1
Rad53
Art5
Ecm21
Ldb19
Yhr131c
Rsp5
Bul2
Ent1
Ent5
Apl3
Sec16
Apm4
Akl1
Rpn11
Nip1
Prt1
Rpg1
Ded1
Tif35
Cdc9
Chk1
Rad23
Ubi4
Ede1
Ent2
Vps27
Chc1
Ptp3
Stb1
Spa2
Bem2
Bud14
Rck2
Glc7
Ipl1
Cdc5
Dbf4
Pfk26
Tpk1
Gpa2
Ras2
Med6
Rpb10
Paf1
Rpo26
Bdp1
Spt15
Spt5
Rgr1
Srb7
Ssn2
Nop1
Nsr1
Nop56
Snu13
Rps31
Rad9
Rtf1
Rpo21
Npl3
Air2
Fps1
Ask10
Cdc28
Hog1
Pbs2
Ssk2
Clb2
Bud3
Bud4
Msn2
Sch9
Rim15
Yak1
Ypl247c
Vps52
Kss1
Sko1
Slt2
Msg5
Swe1
Igo2
Cdc55
Igo1
Vps53
Vps54
Rtn1
Fmp42
Cot1
Zrc1
10
20
30
40
50
t/s

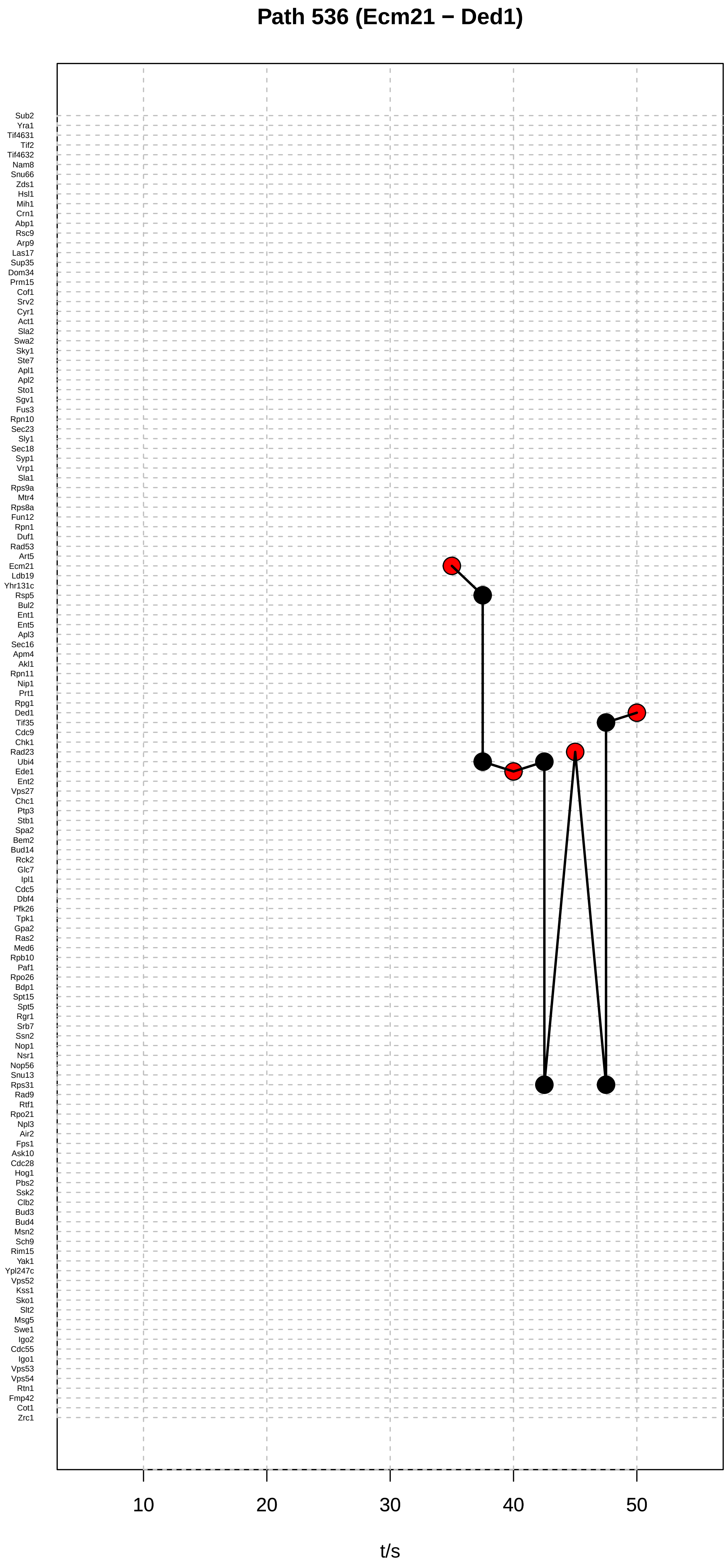

Path 536 (Ecm21 − Ded1)
Sub2
Yra1
Tif4631
Tif2
Tif4632
Nam8
Snu66
Zds1
Hsl1
Mih1
Crn1
Abp1
Rsc9
Arp9
Las17
Sup35
Dom34
Prm15
Cof1
Srv2
Cyr1
Act1
Sla2
Swa2
Sky1
Ste7
Apl1
Apl2
Sto1
Sgv1
Fus3
Rpn10
Sec23
Sly1
Sec18
Syp1
Vrp1
Sla1
Rps9a
Mtr4
Rps8a
Fun12
Rpn1
Duf1
Rad53
Art5
Ecm21
Ldb19
Yhr131c
Rsp5
Bul2
Ent1
Ent5
Apl3
Sec16
Apm4
Akl1
Rpn11
Nip1
Prt1
Rpg1
Ded1
Tif35
Cdc9
Chk1
Rad23
Ubi4
Ede1
Ent2
Vps27
Chc1
Ptp3
Stb1
Spa2
Bem2
Bud14
Rck2
Glc7
Ipl1
Cdc5
Dbf4
Pfk26
Tpk1
Gpa2
Ras2
Med6
Rpb10
Paf1
Rpo26
Bdp1
Spt15
Spt5
Rgr1
Srb7
Ssn2
Nop1
Nsr1
Nop56
Snu13
Rps31
Rad9
Rtf1
Rpo21
Npl3
Air2
Fps1
Ask10
Cdc28
Hog1
Pbs2
Ssk2
Clb2
Bud3
Bud4
Msn2
Sch9
Rim15
Yak1
Ypl247c
Vps52
Kss1
Sko1
Slt2
Msg5
Swe1
Igo2
Cdc55
Igo1
Vps53
Vps54
Rtn1
Fmp42
Cot1
Zrc1
10
20
30
40
50
t/s

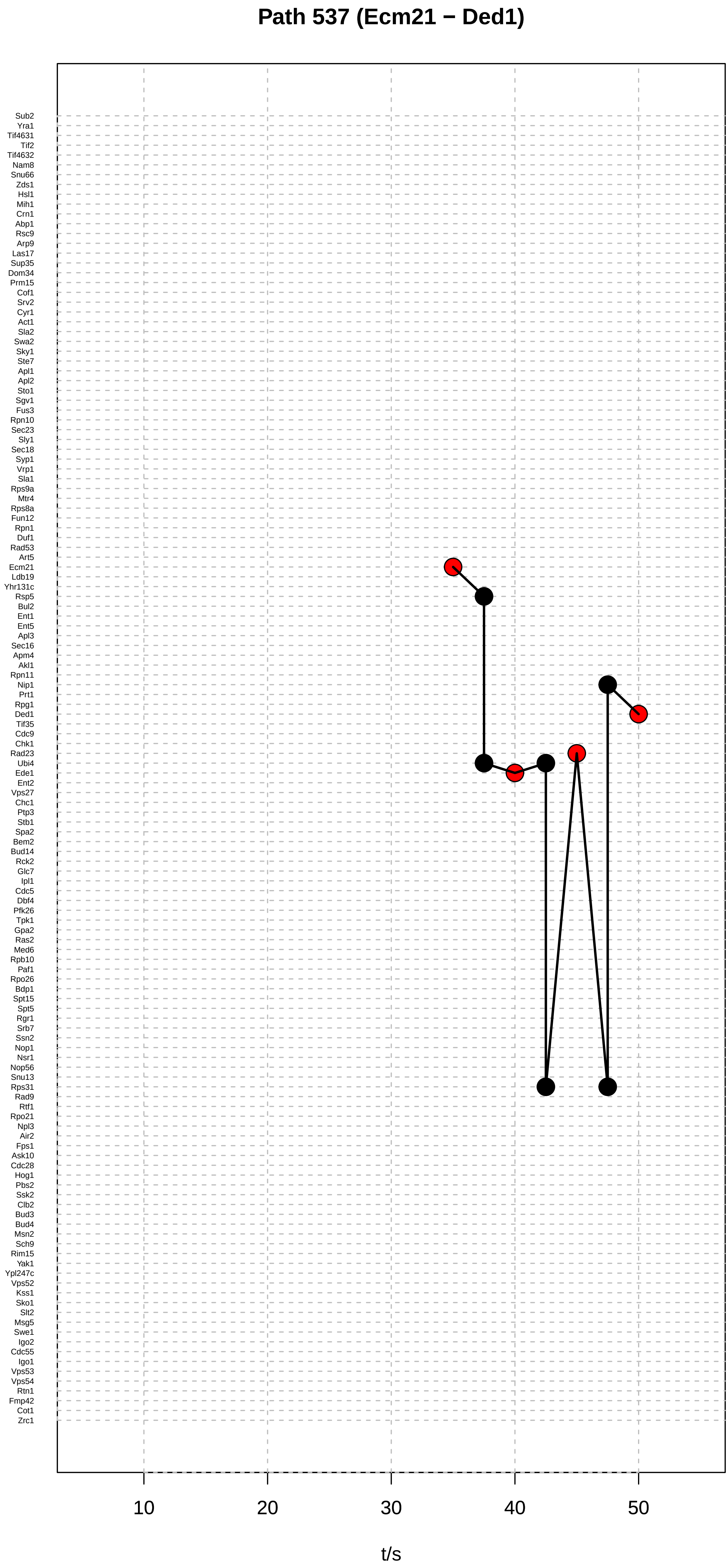

Path 537 (Ecm21 − Ded1)
Sub2
Yra1
Tif4631
Tif2
Tif4632
Nam8
Snu66
Zds1
Hsl1
Mih1
Crn1
Abp1
Rsc9
Arp9
Las17
Sup35
Dom34
Prm15
Cof1
Srv2
Cyr1
Act1
Sla2
Swa2
Sky1
Ste7
Apl1
Apl2
Sto1
Sgv1
Fus3
Rpn10
Sec23
Sly1
Sec18
Syp1
Vrp1
Sla1
Rps9a
Mtr4
Rps8a
Fun12
Rpn1
Duf1
Rad53
Art5
Ecm21
Ldb19
Yhr131c
Rsp5
Bul2
Ent1
Ent5
Apl3
Sec16
Apm4
Akl1
Rpn11
Nip1
Prt1
Rpg1
Ded1
Tif35
Cdc9
Chk1
Rad23
Ubi4
Ede1
Ent2
Vps27
Chc1
Ptp3
Stb1
Spa2
Bem2
Bud14
Rck2
Glc7
Ipl1
Cdc5
Dbf4
Pfk26
Tpk1
Gpa2
Ras2
Med6
Rpb10
Paf1
Rpo26
Bdp1
Spt15
Spt5
Rgr1
Srb7
Ssn2
Nop1
Nsr1
Nop56
Snu13
Rps31
Rad9
Rtf1
Rpo21
Npl3
Air2
Fps1
Ask10
Cdc28
Hog1
Pbs2
Ssk2
Clb2
Bud3
Bud4
Msn2
Sch9
Rim15
Yak1
Ypl247c
Vps52
Kss1
Sko1
Slt2
Msg5
Swe1
Igo2
Cdc55
Igo1
Vps53
Vps54
Rtn1
Fmp42
Cot1
Zrc1
10
20
30
40
50
t/s

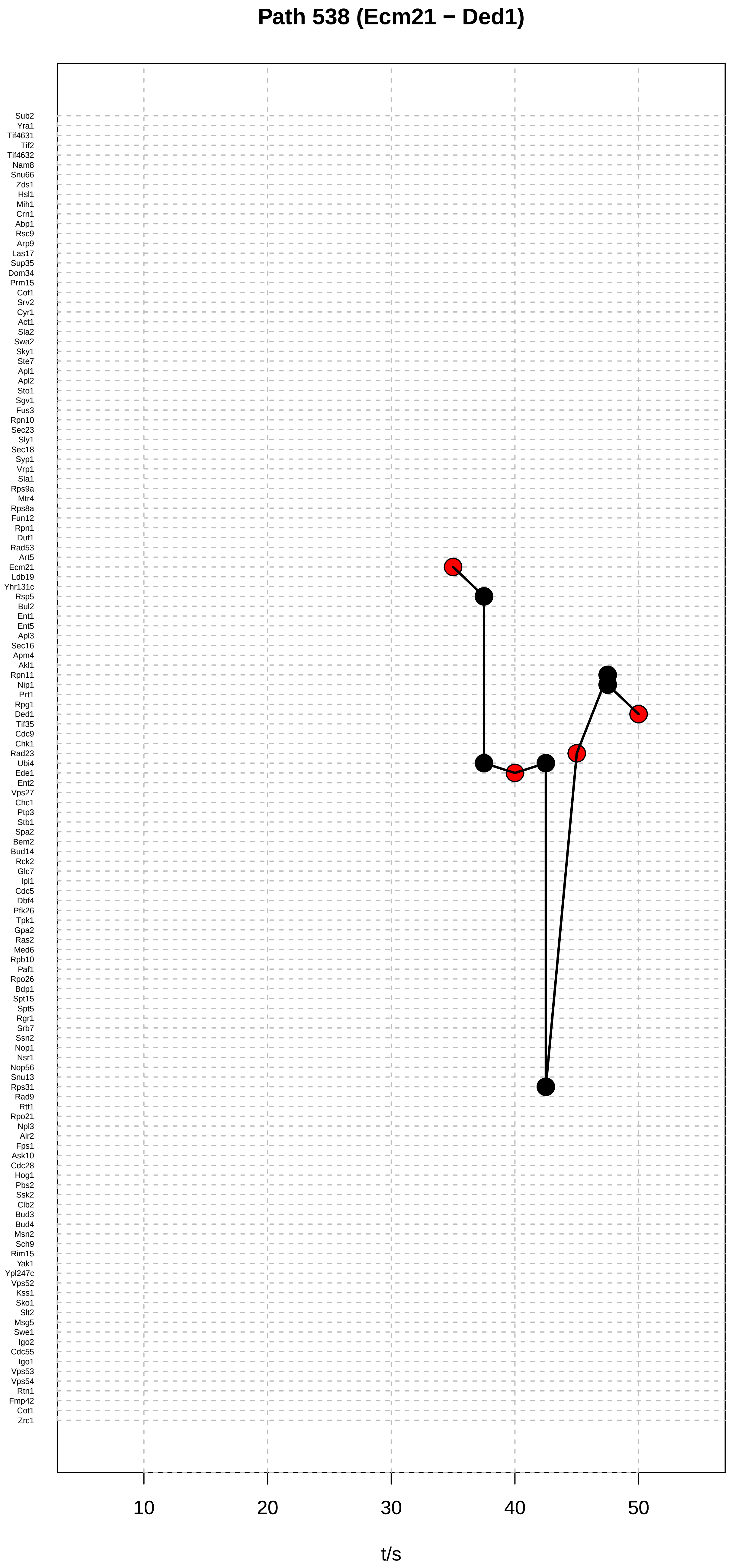
Path 538 (Ecm21 − Ded1)
Sub2
Yra1
Tif4631
Tif2
Tif4632
Nam8
Snu66
Zds1
Hsl1
Mih1
Crn1
Abp1
Rsc9
Arp9
Las17
Sup35
Dom34
Prm15
Cof1
Srv2
Cyr1
Act1
Sla2
Swa2
Sky1
Ste7
Apl1
Apl2
Sto1
Sgv1
Fus3
Rpn10
Sec23
Sly1
Sec18
Syp1
Vrp1
Sla1
Rps9a
Mtr4
Rps8a
Fun12
Rpn1
Duf1
Rad53
Art5
Ecm21
Ldb19
Yhr131c
Rsp5
Bul2
Ent1
Ent5
Apl3
Sec16
Apm4
Akl1
Rpn11
Nip1
Prt1
Rpg1
Ded1
Tif35
Cdc9
Chk1
Rad23
Ubi4
Ede1
Ent2
Vps27
Chc1
Ptp3
Stb1
Spa2
Bem2
Bud14
Rck2
Glc7
Ipl1
Cdc5
Dbf4
Pfk26
Tpk1
Gpa2
Ras2
Med6
Rpb10
Paf1
Rpo26
Bdp1
Spt15
Spt5
Rgr1
Srb7
Ssn2
Nop1
Nsr1
Nop56
Snu13
Rps31
Rad9
Rtf1
Rpo21
Npl3
Air2
Fps1
Ask10
Cdc28
Hog1
Pbs2
Ssk2
Clb2
Bud3
Bud4
Msn2
Sch9
Rim15
Yak1
Ypl247c
Vps52
Kss1
Sko1
Slt2
Msg5
Swe1
Igo2
Cdc55
Igo1
Vps53
Vps54
Rtn1
Fmp42
Cot1
Zrc1
10
20
30
40
50
t/s

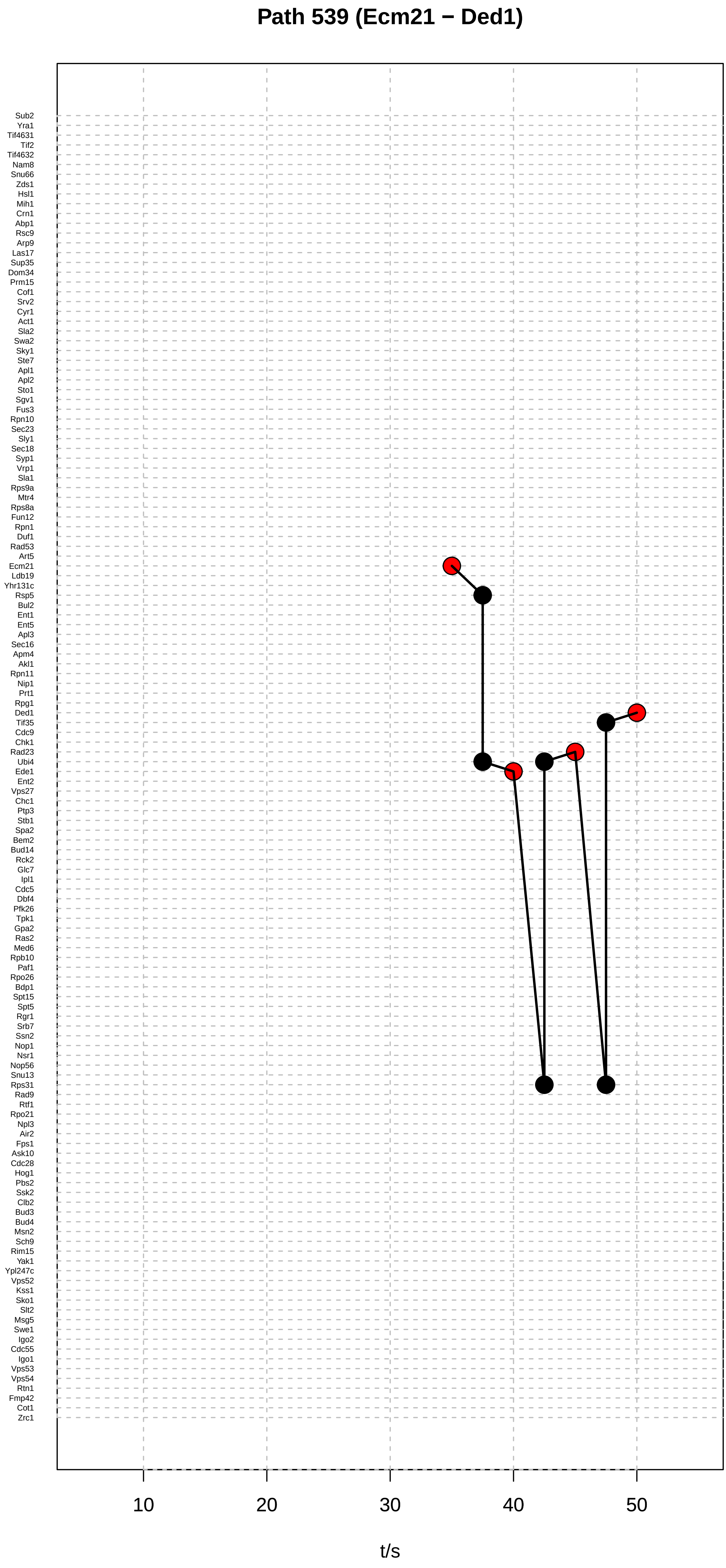

Path 539 (Ecm21 − Ded1)
Sub2
Yra1
Tif4631
Tif2
Tif4632
Nam8
Snu66
Zds1
Hsl1
Mih1
Crn1
Abp1
Rsc9
Arp9
Las17
Sup35
Dom34
Prm15
Cof1
Srv2
Cyr1
Act1
Sla2
Swa2
Sky1
Ste7
Apl1
Apl2
Sto1
Sgv1
Fus3
Rpn10
Sec23
Sly1
Sec18
Syp1
Vrp1
Sla1
Rps9a
Mtr4
Rps8a
Fun12
Rpn1
Duf1
Rad53
Art5
Ecm21
Ldb19
Yhr131c
Rsp5
Bul2
Ent1
Ent5
Apl3
Sec16
Apm4
Akl1
Rpn11
Nip1
Prt1
Rpg1
Ded1
Tif35
Cdc9
Chk1
Rad23
Ubi4
Ede1
Ent2
Vps27
Chc1
Ptp3
Stb1
Spa2
Bem2
Bud14
Rck2
Glc7
Ipl1
Cdc5
Dbf4
Pfk26
Tpk1
Gpa2
Ras2
Med6
Rpb10
Paf1
Rpo26
Bdp1
Spt15
Spt5
Rgr1
Srb7
Ssn2
Nop1
Nsr1
Nop56
Snu13
Rps31
Rad9
Rtf1
Rpo21
Npl3
Air2
Fps1
Ask10
Cdc28
Hog1
Pbs2
Ssk2
Clb2
Bud3
Bud4
Msn2
Sch9
Rim15
Yak1
Ypl247c
Vps52
Kss1
Sko1
Slt2
Msg5
Swe1
Igo2
Cdc55
Igo1
Vps53
Vps54
Rtn1
Fmp42
Cot1
Zrc1
10
20
30
40
50
t/s

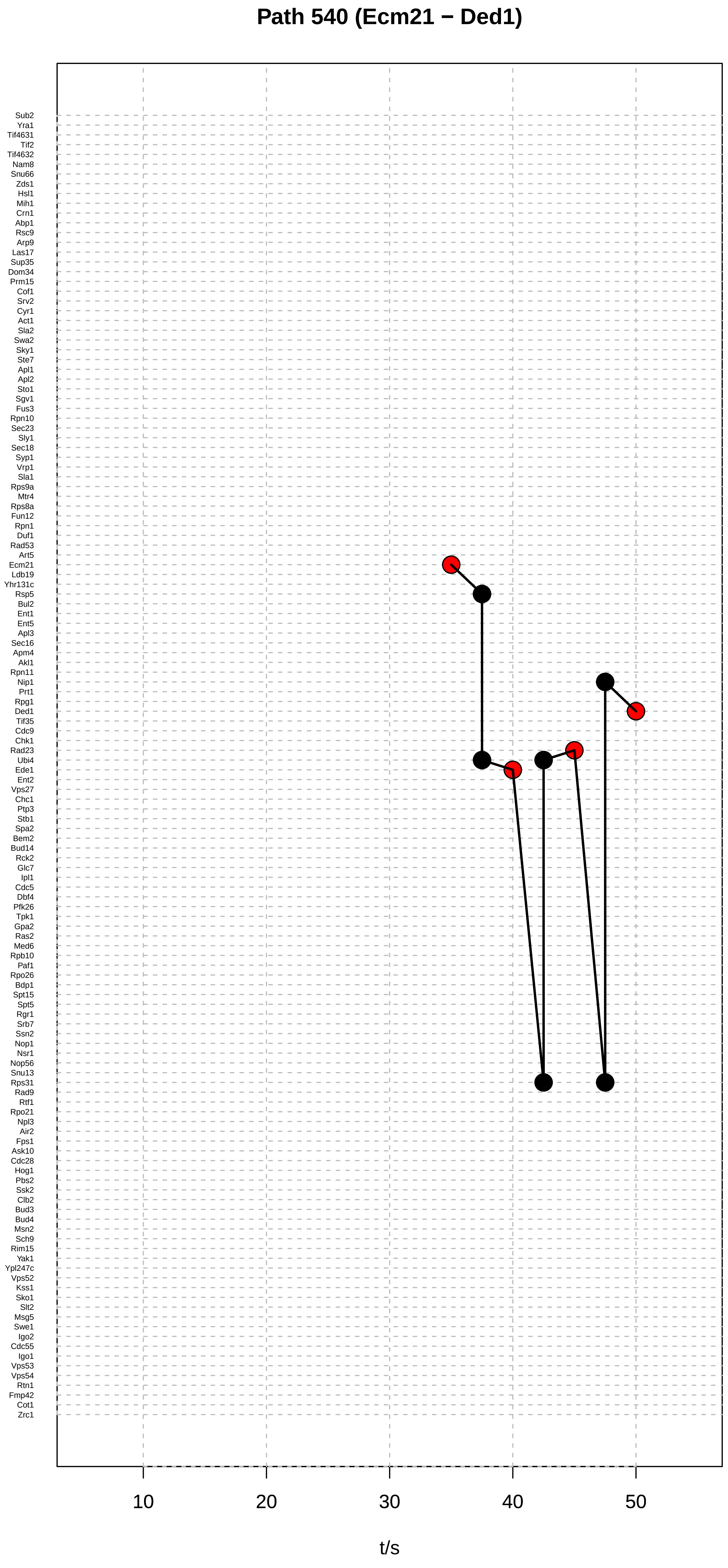

Path 540 (Ecm21 − Ded1)
Sub2
Yra1
Tif4631
Tif2
Tif4632
Nam8
Snu66
Zds1
Hsl1
Mih1
Crn1
Abp1
Rsc9
Arp9
Las17
Sup35
Dom34
Prm15
Cof1
Srv2
Cyr1
Act1
Sla2
Swa2
Sky1
Ste7
Apl1
Apl2
Sto1
Sgv1
Fus3
Rpn10
Sec23
Sly1
Sec18
Syp1
Vrp1
Sla1
Rps9a
Mtr4
Rps8a
Fun12
Rpn1
Duf1
Rad53
Art5
Ecm21
Ldb19
Yhr131c
Rsp5
Bul2
Ent1
Ent5
Apl3
Sec16
Apm4
Akl1
Rpn11
Nip1
Prt1
Rpg1
Ded1
Tif35
Cdc9
Chk1
Rad23
Ubi4
Ede1
Ent2
Vps27
Chc1
Ptp3
Stb1
Spa2
Bem2
Bud14
Rck2
Glc7
Ipl1
Cdc5
Dbf4
Pfk26
Tpk1
Gpa2
Ras2
Med6
Rpb10
Paf1
Rpo26
Bdp1
Spt15
Spt5
Rgr1
Srb7
Ssn2
Nop1
Nsr1
Nop56
Snu13
Rps31
Rad9
Rtf1
Rpo21
Npl3
Air2
Fps1
Ask10
Cdc28
Hog1
Pbs2
Ssk2
Clb2
Bud3
Bud4
Msn2
Sch9
Rim15
Yak1
Ypl247c
Vps52
Kss1
Sko1
Slt2
Msg5
Swe1
Igo2
Cdc55
Igo1
Vps53
Vps54
Rtn1
Fmp42
Cot1
Zrc1
10
20
30
40
50
t/s

#### Path 541 (Ecm21 − Ded1)

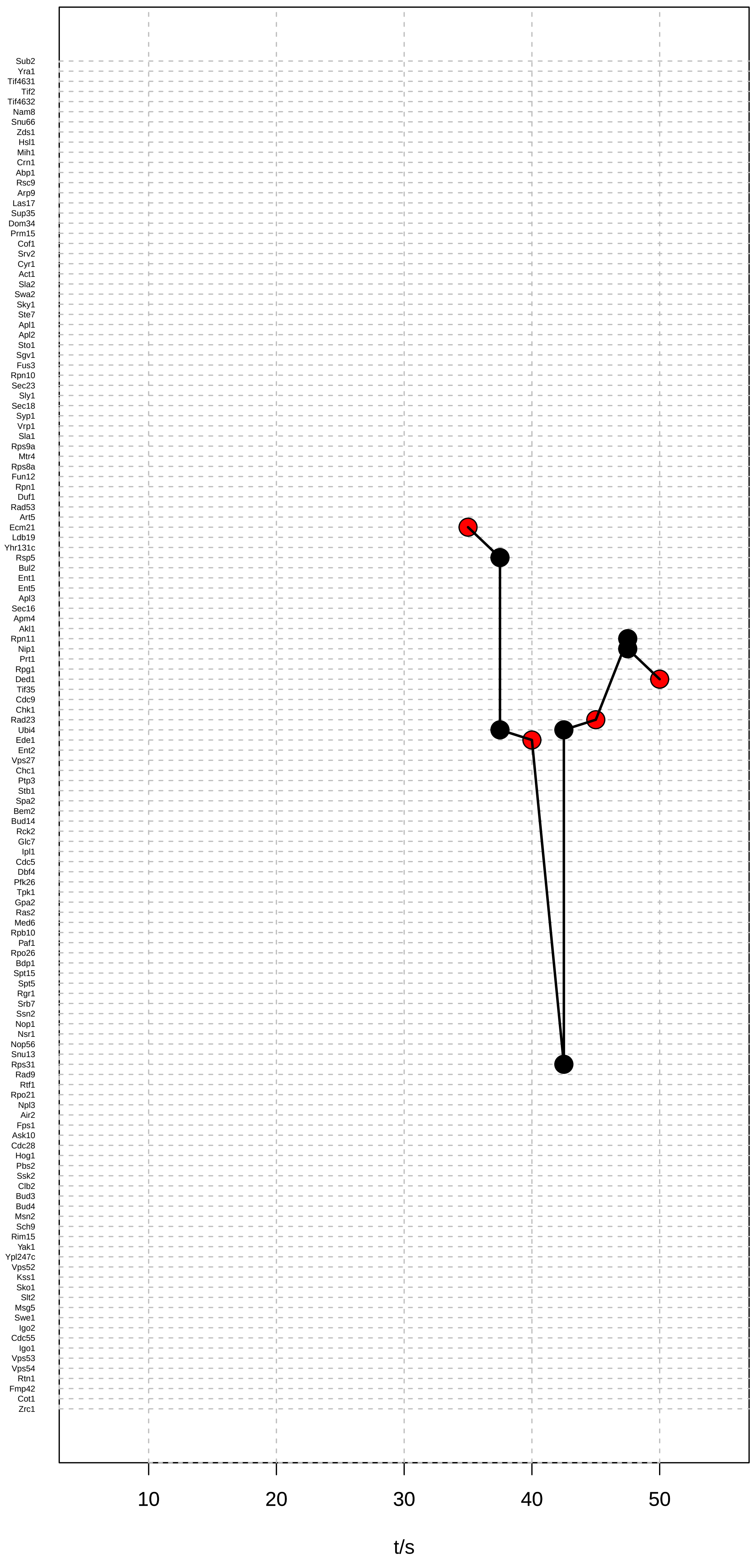

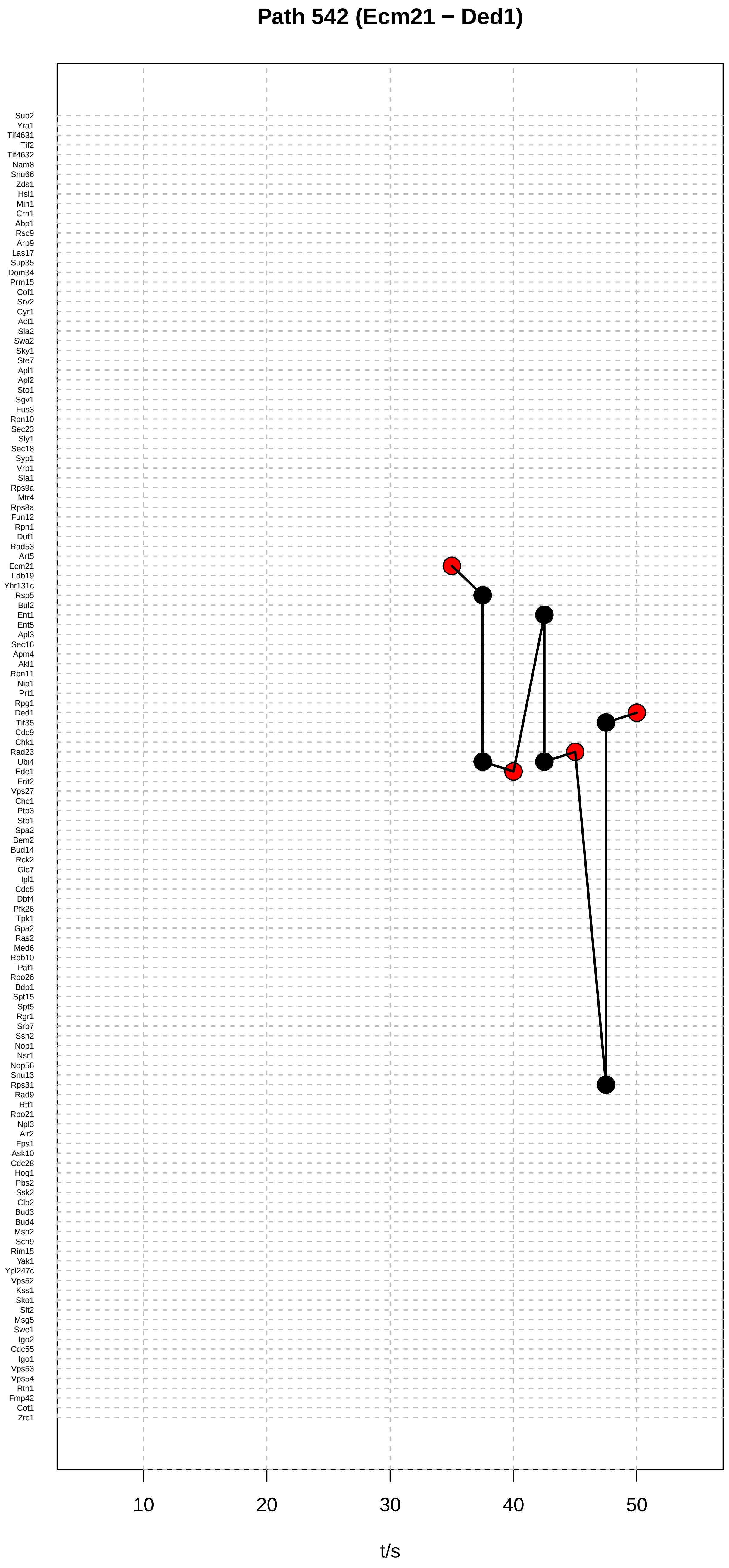

Path 542 (Ecm21 − Ded1)
Sub2
Yra1
Tif4631
Tif2
Tif4632
Nam8
Snu66
Zds1
Hsl1
Mih1
Crn1
Abp1
Rsc9
Arp9
Las17
Sup35
Dom34
Prm15
Cof1
Srv2
Cyr1
Act1
Sla2
Swa2
Sky1
Ste7
Apl1
Apl2
Sto1
Sgv1
Fus3
Rpn10
Sec23
Sly1
Sec18
Syp1
Vrp1
Sla1
Rps9a
Mtr4
Rps8a
Fun12
Rpn1
Duf1
Rad53
Art5
Ecm21
Ldb19
Yhr131c
Rsp5
Bul2
Ent1
Ent5
Apl3
Sec16
Apm4
Akl1
Rpn11
Nip1
Prt1
Rpg1
Ded1
Tif35
Cdc9
Chk1
Rad23
Ubi4
Ede1
Ent2
Vps27
Chc1
Ptp3
Stb1
Spa2
Bem2
Bud14
Rck2
Glc7
Ipl1
Cdc5
Dbf4
Pfk26
Tpk1
Gpa2
Ras2
Med6
Rpb10
Paf1
Rpo26
Bdp1
Spt15
Spt5
Rgr1
Srb7
Ssn2
Nop1
Nsr1
Nop56
Snu13
Rps31
Rad9
Rtf1
Rpo21
Npl3
Air2
Fps1
Ask10
Cdc28
Hog1
Pbs2
Ssk2
Clb2
Bud3
Bud4
Msn2
Sch9
Rim15
Yak1
Ypl247c
Vps52
Kss1
Sko1
Slt2
Msg5
Swe1
Igo2
Cdc55
Igo1
Vps53
Vps54
Rtn1
Fmp42
Cot1
Zrc1
10
20
30
40
50
t/s

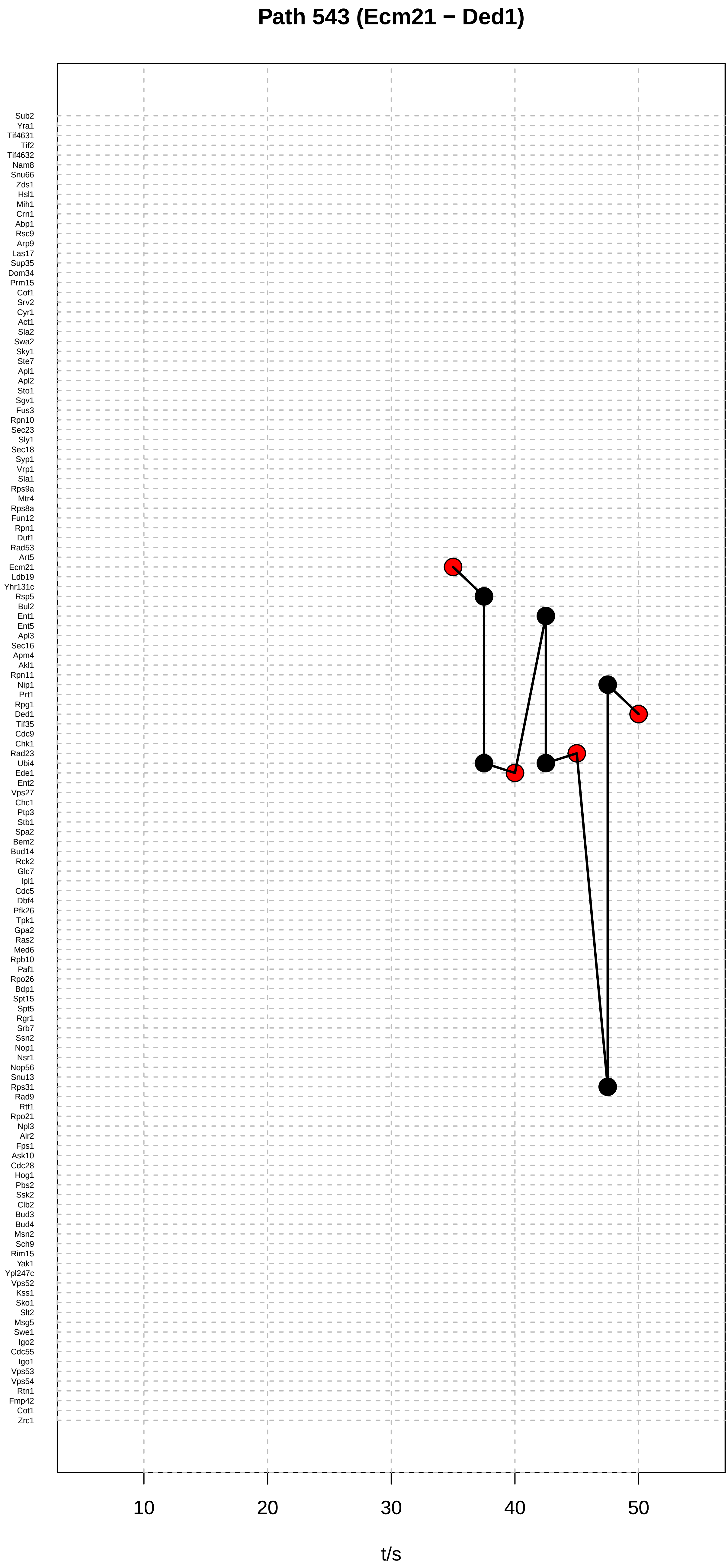
Path 543 (Ecm21 − Ded1)
Sub2
Yra1
Tif4631
Tif2
Tif4632
Nam8
Snu66
Zds1
Hsl1
Mih1
Crn1
Abp1
Rsc9
Arp9
Las17
Sup35
Dom34
Prm15
Cof1
Srv2
Cyr1
Act1
Sla2
Swa2
Sky1
Ste7
Apl1
Apl2
Sto1
Sgv1
Fus3
Rpn10
Sec23
Sly1
Sec18
Syp1
Vrp1
Sla1
Rps9a
Mtr4
Rps8a
Fun12
Rpn1
Duf1
Rad53
Art5
Ecm21
Ldb19
Yhr131c
Rsp5
Bul2
Ent1
Ent5
Apl3
Sec16
Apm4
Akl1
Rpn11
Nip1
Prt1
Rpg1
Ded1
Tif35
Cdc9
Chk1
Rad23
Ubi4
Ede1
Ent2
Vps27
Chc1
Ptp3
Stb1
Spa2
Bem2
Bud14
Rck2
Glc7
Ipl1
Cdc5
Dbf4
Pfk26
Tpk1
Gpa2
Ras2
Med6
Rpb10
Paf1
Rpo26
Bdp1
Spt15
Spt5
Rgr1
Srb7
Ssn2
Nop1
Nsr1
Nop56
Snu13
Rps31
Rad9
Rtf1
Rpo21
Npl3
Air2
Fps1
Ask10
Cdc28
Hog1
Pbs2
Ssk2
Clb2
Bud3
Bud4
Msn2
Sch9
Rim15
Yak1
Ypl247c
Vps52
Kss1
Sko1
Slt2
Msg5
Swe1
Igo2
Cdc55
Igo1
Vps53
Vps54
Rtn1
Fmp42
Cot1
Zrc1
10
20
30
40
50
t/s

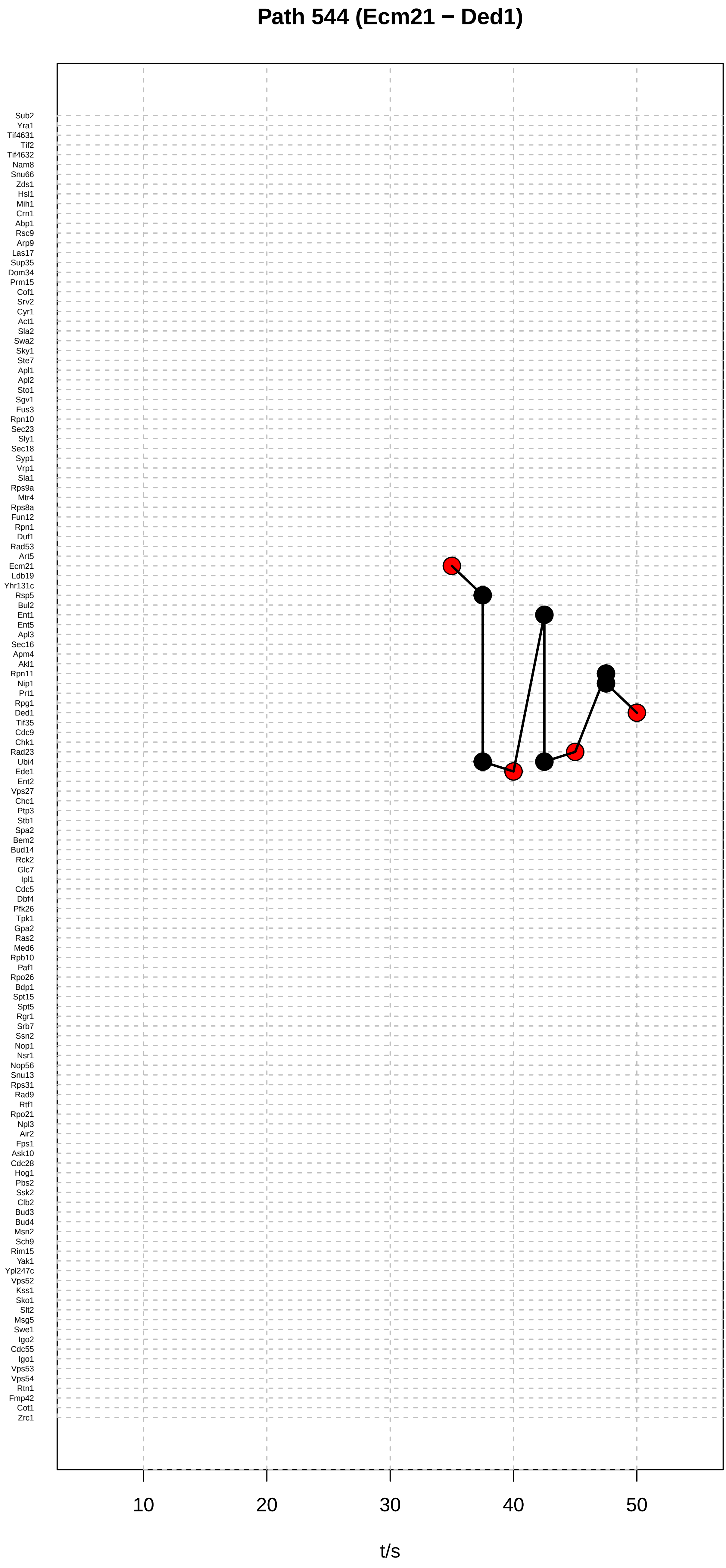
Path 544 (Ecm21 − Ded1)
Sub2
Yra1
Tif4631
Tif2
Tif4632
Nam8
Snu66
Zds1
Hsl1
Mih1
Crn1
Abp1
Rsc9
Arp9
Las17
Sup35
Dom34
Prm15
Cof1
Srv2
Cyr1
Act1
Sla2
Swa2
Sky1
Ste7
Apl1
Apl2
Sto1
Sgv1
Fus3
Rpn10
Sec23
Sly1
Sec18
Syp1
Vrp1
Sla1
Rps9a
Mtr4
Rps8a
Fun12
Rpn1
Duf1
Rad53
Art5
Ecm21
Ldb19
Yhr131c
Rsp5
Bul2
Ent1
Ent5
Apl3
Sec16
Apm4
Akl1
Rpn11
Nip1
Prt1
Rpg1
Ded1
Tif35
Cdc9
Chk1
Rad23
Ubi4
Ede1
Ent2
Vps27
Chc1
Ptp3
Stb1
Spa2
Bem2
Bud14
Rck2
Glc7
Ipl1
Cdc5
Dbf4
Pfk26
Tpk1
Gpa2
Ras2
Med6
Rpb10
Paf1
Rpo26
Bdp1
Spt15
Spt5
Rgr1
Srb7
Ssn2
Nop1
Nsr1
Nop56
Snu13
Rps31
Rad9
Rtf1
Rpo21
Npl3
Air2
Fps1
Ask10
Cdc28
Hog1
Pbs2
Ssk2
Clb2
Bud3
Bud4
Msn2
Sch9
Rim15
Yak1
Ypl247c
Vps52
Kss1
Sko1
Slt2
Msg5
Swe1
Igo2
Cdc55
Igo1
Vps53
Vps54
Rtn1
Fmp42
Cot1
Zrc1
10
20
30
40
50
t/s

#### Path 545 (Ecm21 − Ded1)

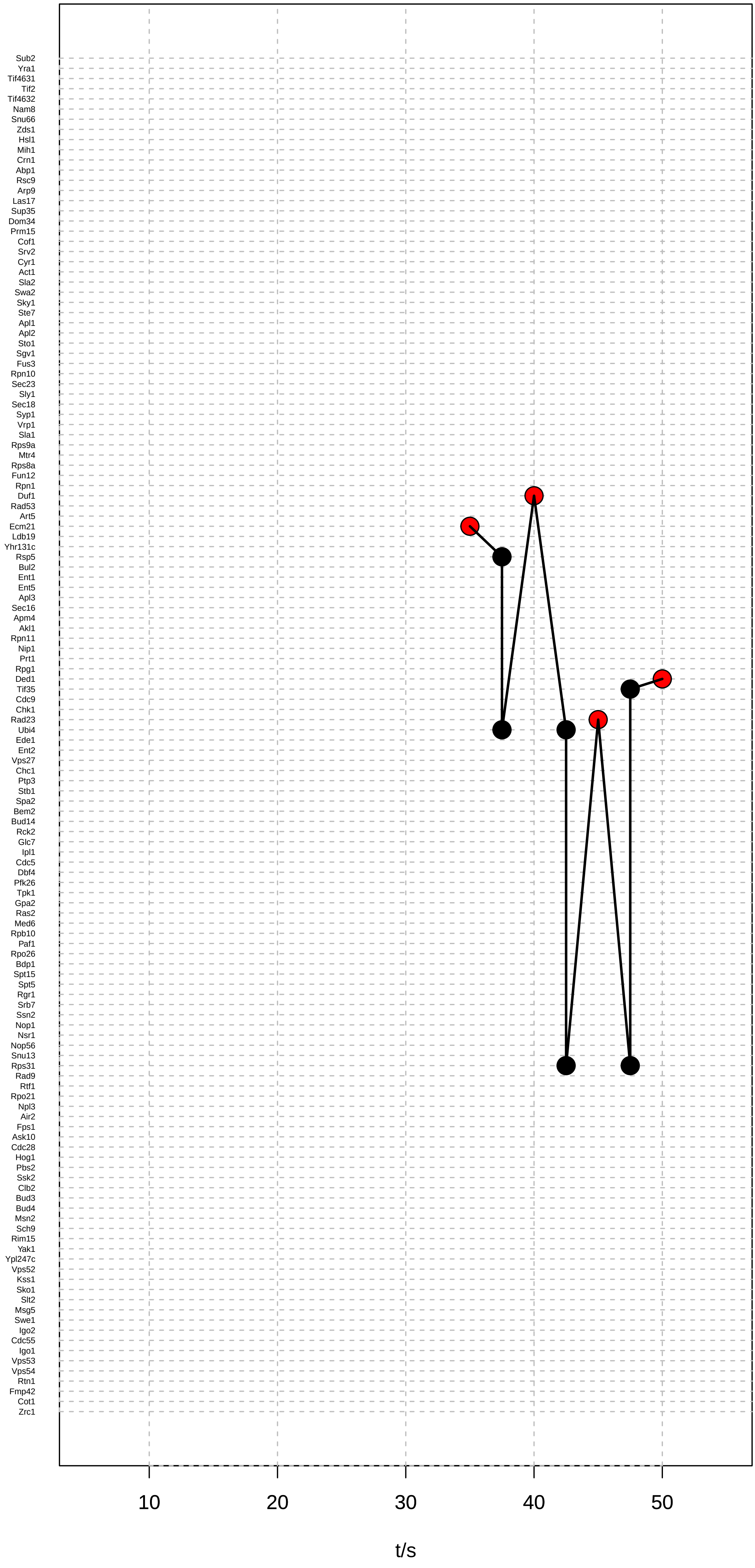

#### Path 546 (Ecm21 − Ded1)

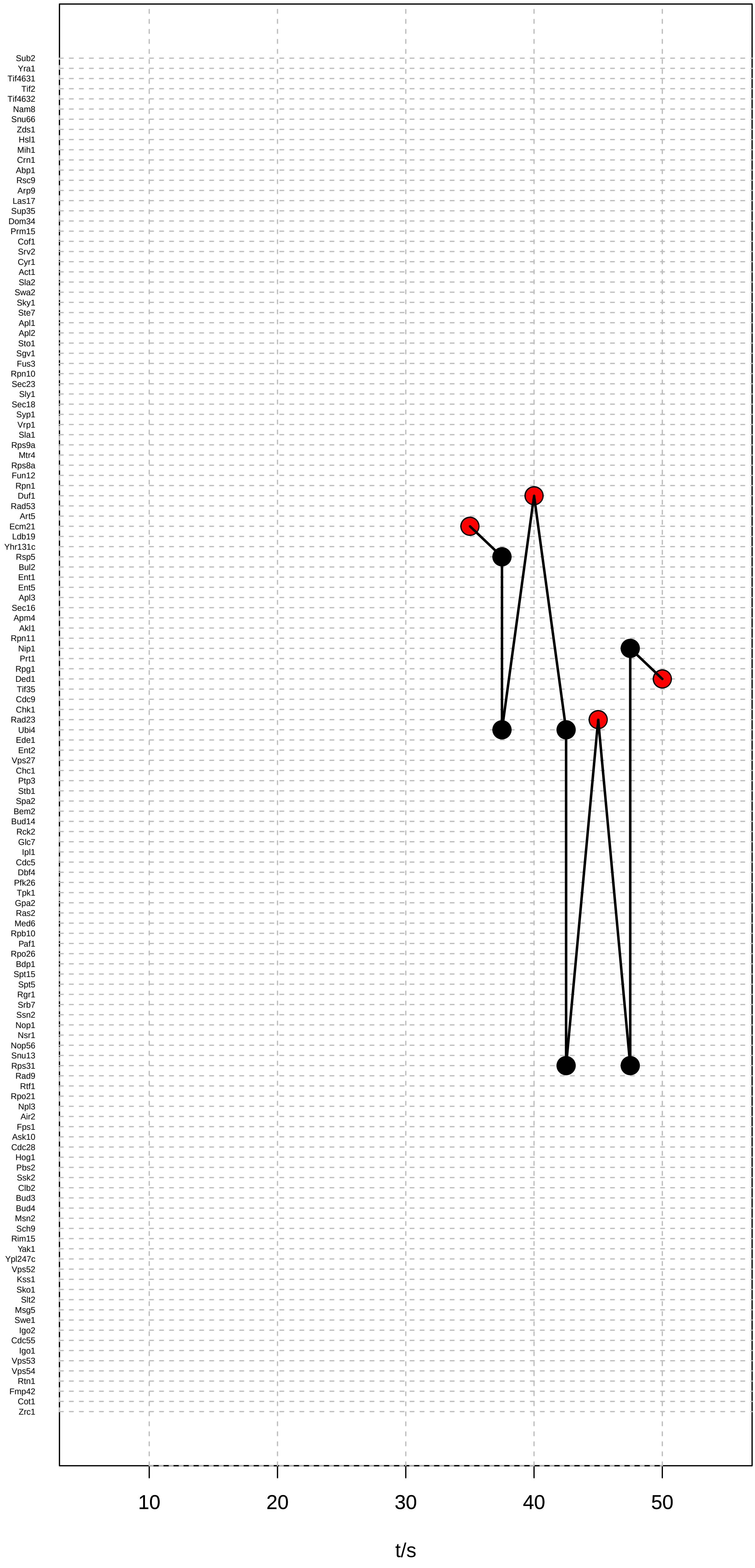

### Path 547 (Ecm21 − Ded1)

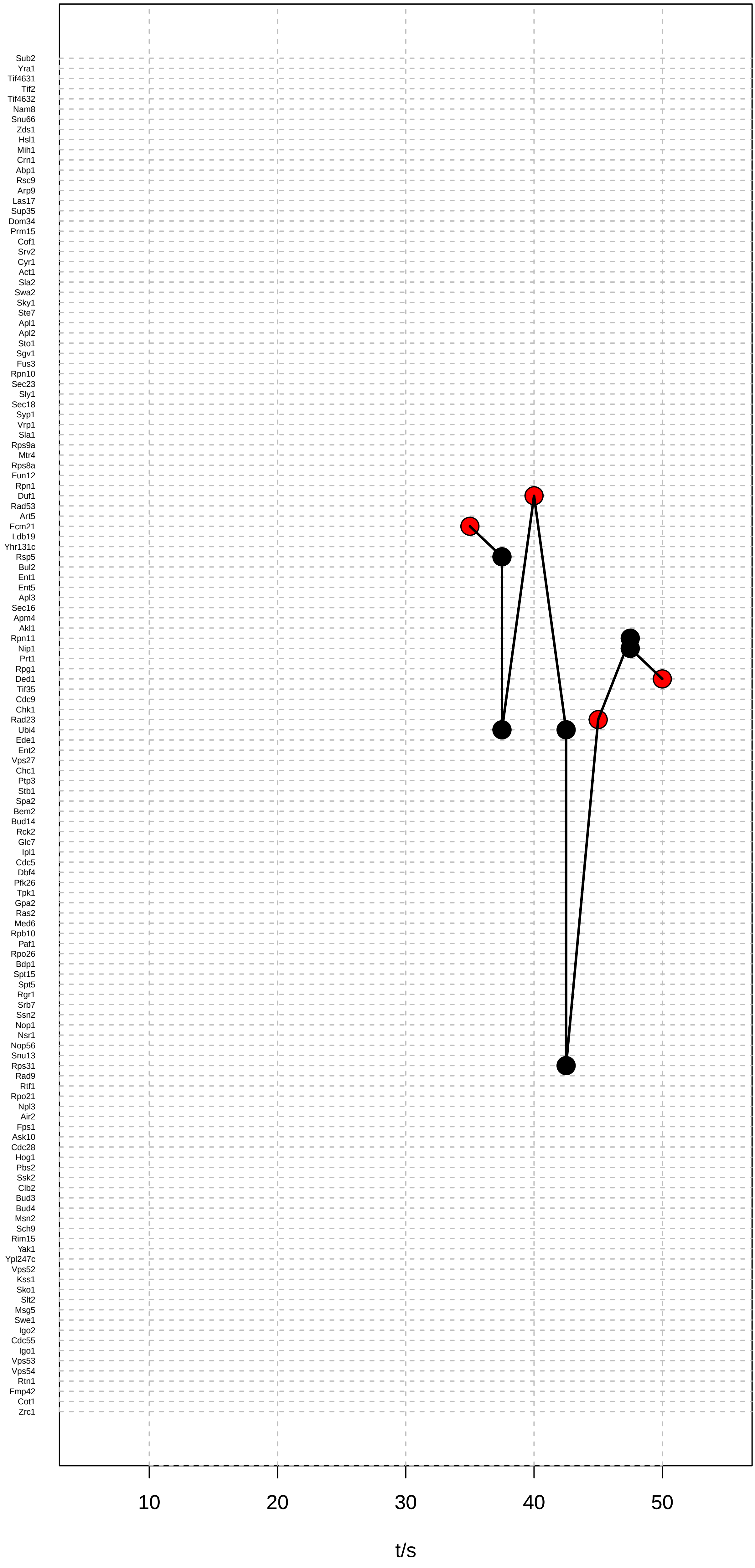

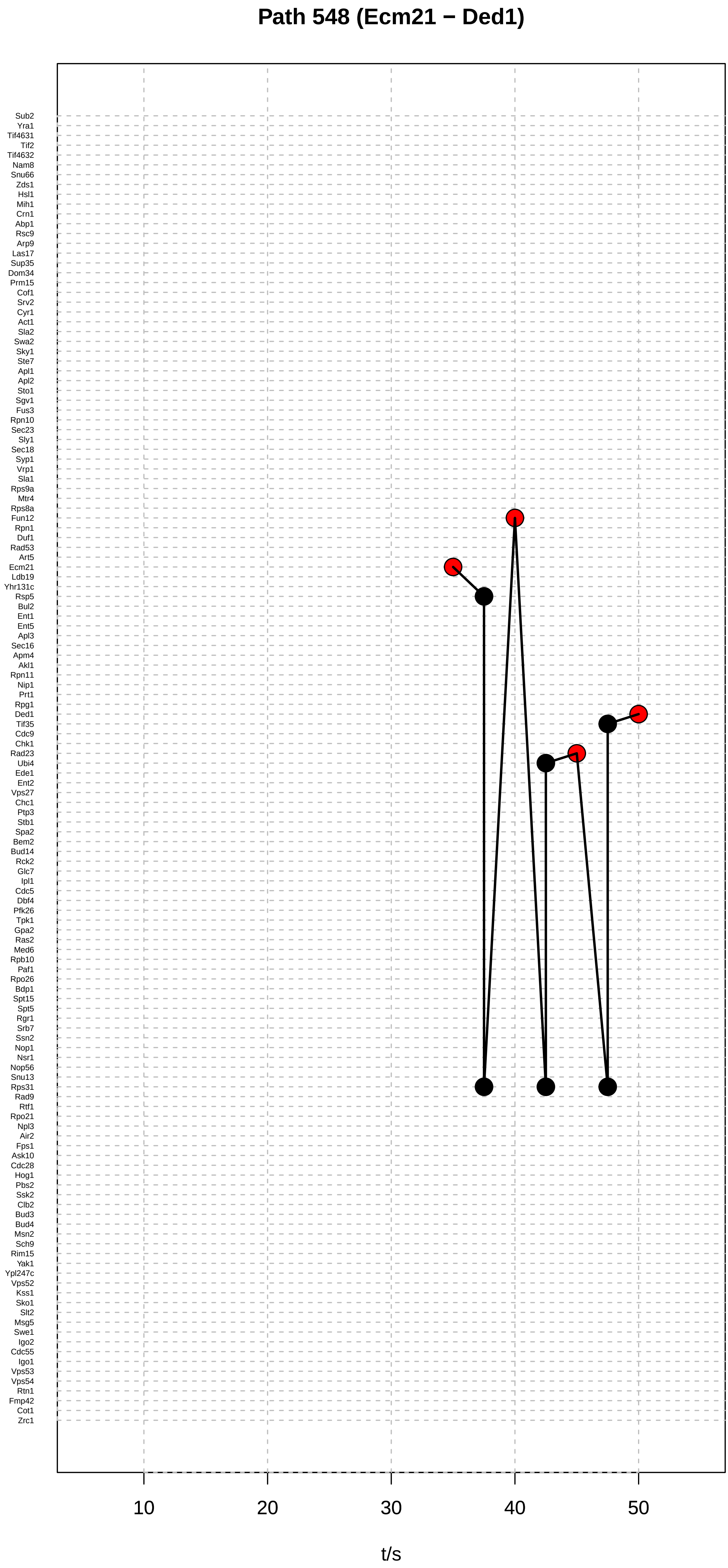
Path 548 (Ecm21 − Ded1)
Sub2
Yra1
Tif4631
Tif2
Tif4632
Nam8
Snu66
Zds1
Hsl1
Mih1
Crn1
Abp1
Rsc9
Arp9
Las17
Sup35
Dom34
Prm15
Cof1
Srv2
Cyr1
Act1
Sla2
Swa2
Sky1
Ste7
Apl1
Apl2
Sto1
Sgv1
Fus3
Rpn10
Sec23
Sly1
Sec18
Syp1
Vrp1
Sla1
Rps9a
Mtr4
Rps8a
Fun12
Rpn1
Duf1
Rad53
Art5
Ecm21
Ldb19
Yhr131c
Rsp5
Bul2
Ent1
Ent5
Apl3
Sec16
Apm4
Akl1
Rpn11
Nip1
Prt1
Rpg1
Ded1
Tif35
Cdc9
Chk1
Rad23
Ubi4
Ede1
Ent2
Vps27
Chc1
Ptp3
Stb1
Spa2
Bem2
Bud14
Rck2
Glc7
Ipl1
Cdc5
Dbf4
Pfk26
Tpk1
Gpa2
Ras2
Med6
Rpb10
Paf1
Rpo26
Bdp1
Spt15
Spt5
Rgr1
Srb7
Ssn2
Nop1
Nsr1
Nop56
Snu13
Rps31
Rad9
Rtf1
Rpo21
Npl3
Air2
Fps1
Ask10
Cdc28
Hog1
Pbs2
Ssk2
Clb2
Bud3
Bud4
Msn2
Sch9
Rim15
Yak1
Ypl247c
Vps52
Kss1
Sko1
Slt2
Msg5
Swe1
Igo2
Cdc55
Igo1
Vps53
Vps54
Rtn1
Fmp42
Cot1
Zrc1
10
20
30
40
50
t/s

#### Path 549 (Ecm21 − Ded1)

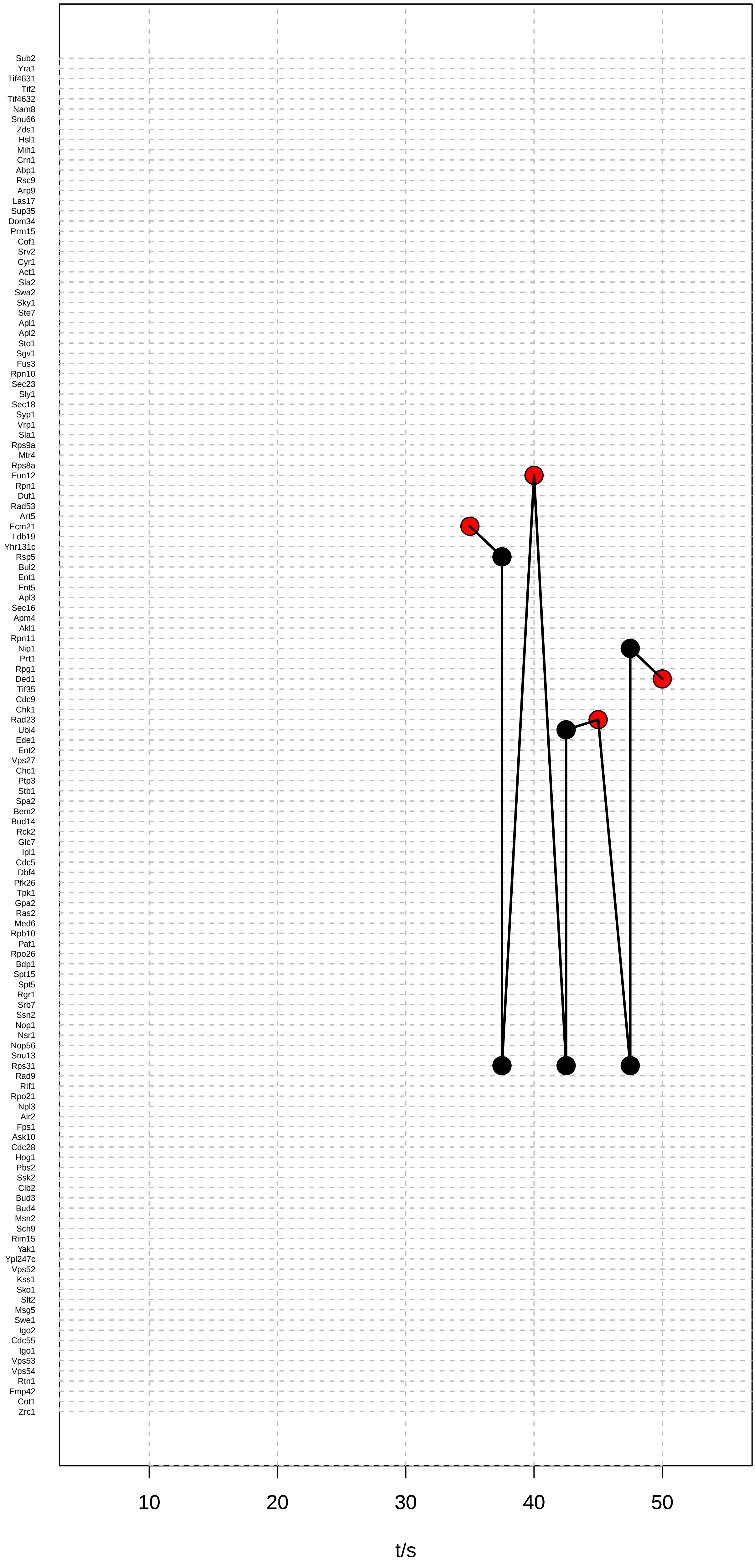

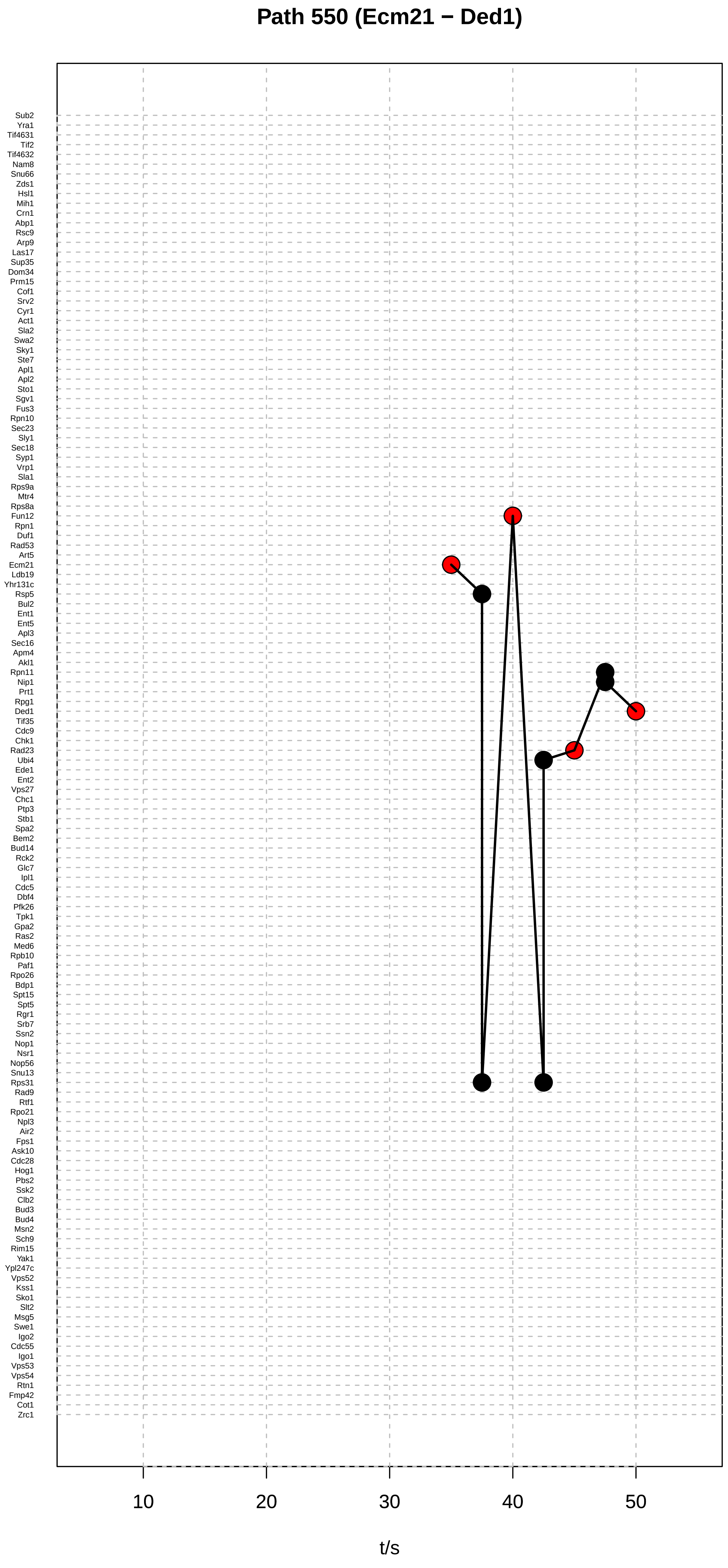

Path 550 (Ecm21 − Ded1)
Sub2
Yra1
Tif4631
Tif2
Tif4632
Nam8
Snu66
Zds1
Hsl1
Mih1
Crn1
Abp1
Rsc9
Arp9
Las17
Sup35
Dom34
Prm15
Cof1
Srv2
Cyr1
Act1
Sla2
Swa2
Sky1
Ste7
Apl1
Apl2
Sto1
Sgv1
Fus3
Rpn10
Sec23
Sly1
Sec18
Syp1
Vrp1
Sla1
Rps9a
Mtr4
Rps8a
Fun12
Rpn1
Duf1
Rad53
Art5
Ecm21
Ldb19
Yhr131c
Rsp5
Bul2
Ent1
Ent5
Apl3
Sec16
Apm4
Akl1
Rpn11
Nip1
Prt1
Rpg1
Ded1
Tif35
Cdc9
Chk1
Rad23
Ubi4
Ede1
Ent2
Vps27
Chc1
Ptp3
Stb1
Spa2
Bem2
Bud14
Rck2
Glc7
Ipl1
Cdc5
Dbf4
Pfk26
Tpk1
Gpa2
Ras2
Med6
Rpb10
Paf1
Rpo26
Bdp1
Spt15
Spt5
Rgr1
Srb7
Ssn2
Nop1
Nsr1
Nop56
Snu13
Rps31
Rad9
Rtf1
Rpo21
Npl3
Air2
Fps1
Ask10
Cdc28
Hog1
Pbs2
Ssk2
Clb2
Bud3
Bud4
Msn2
Sch9
Rim15
Yak1
Ypl247c
Vps52
Kss1
Sko1
Slt2
Msg5
Swe1
Igo2
Cdc55
Igo1
Vps53
Vps54
Rtn1
Fmp42
Cot1
Zrc1
10
20
30
40
50
t/s

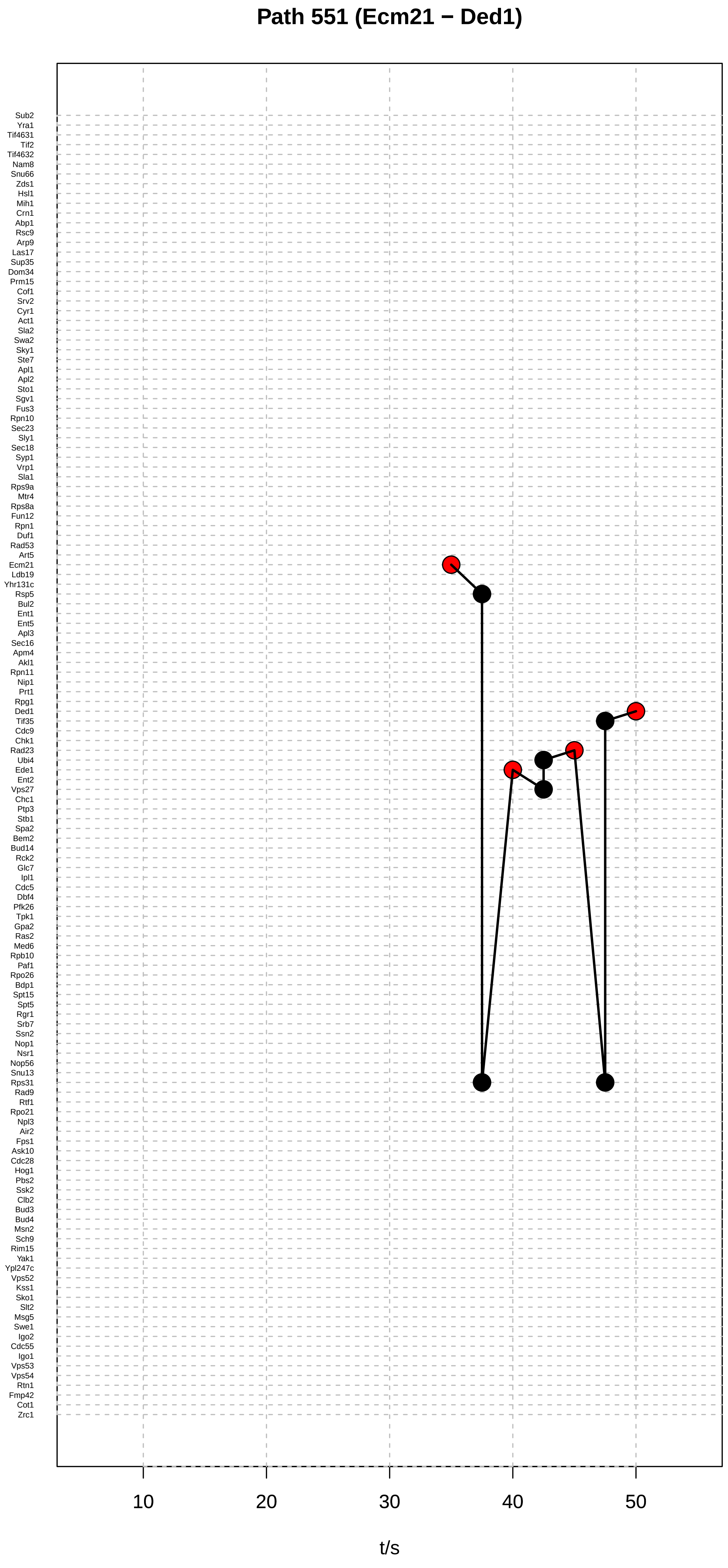

Path 551 (Ecm21 − Ded1)
Sub2
Yra1
Tif4631
Tif2
Tif4632
Nam8
Snu66
Zds1
Hsl1
Mih1
Crn1
Abp1
Rsc9
Arp9
Las17
Sup35
Dom34
Prm15
Cof1
Srv2
Cyr1
Act1
Sla2
Swa2
Sky1
Ste7
Apl1
Apl2
Sto1
Sgv1
Fus3
Rpn10
Sec23
Sly1
Sec18
Syp1
Vrp1
Sla1
Rps9a
Mtr4
Rps8a
Fun12
Rpn1
Duf1
Rad53
Art5
Ecm21
Ldb19
Yhr131c
Rsp5
Bul2
Ent1
Ent5
Apl3
Sec16
Apm4
Akl1
Rpn11
Nip1
Prt1
Rpg1
Ded1
Tif35
Cdc9
Chk1
Rad23
Ubi4
Ede1
Ent2
Vps27
Chc1
Ptp3
Stb1
Spa2
Bem2
Bud14
Rck2
Glc7
Ipl1
Cdc5
Dbf4
Pfk26
Tpk1
Gpa2
Ras2
Med6
Rpb10
Paf1
Rpo26
Bdp1
Spt15
Spt5
Rgr1
Srb7
Ssn2
Nop1
Nsr1
Nop56
Snu13
Rps31
Rad9
Rtf1
Rpo21
Npl3
Air2
Fps1
Ask10
Cdc28
Hog1
Pbs2
Ssk2
Clb2
Bud3
Bud4
Msn2
Sch9
Rim15
Yak1
Ypl247c
Vps52
Kss1
Sko1
Slt2
Msg5
Swe1
Igo2
Cdc55
Igo1
Vps53
Vps54
Rtn1
Fmp42
Cot1
Zrc1
10
20
30
40
50
t/s

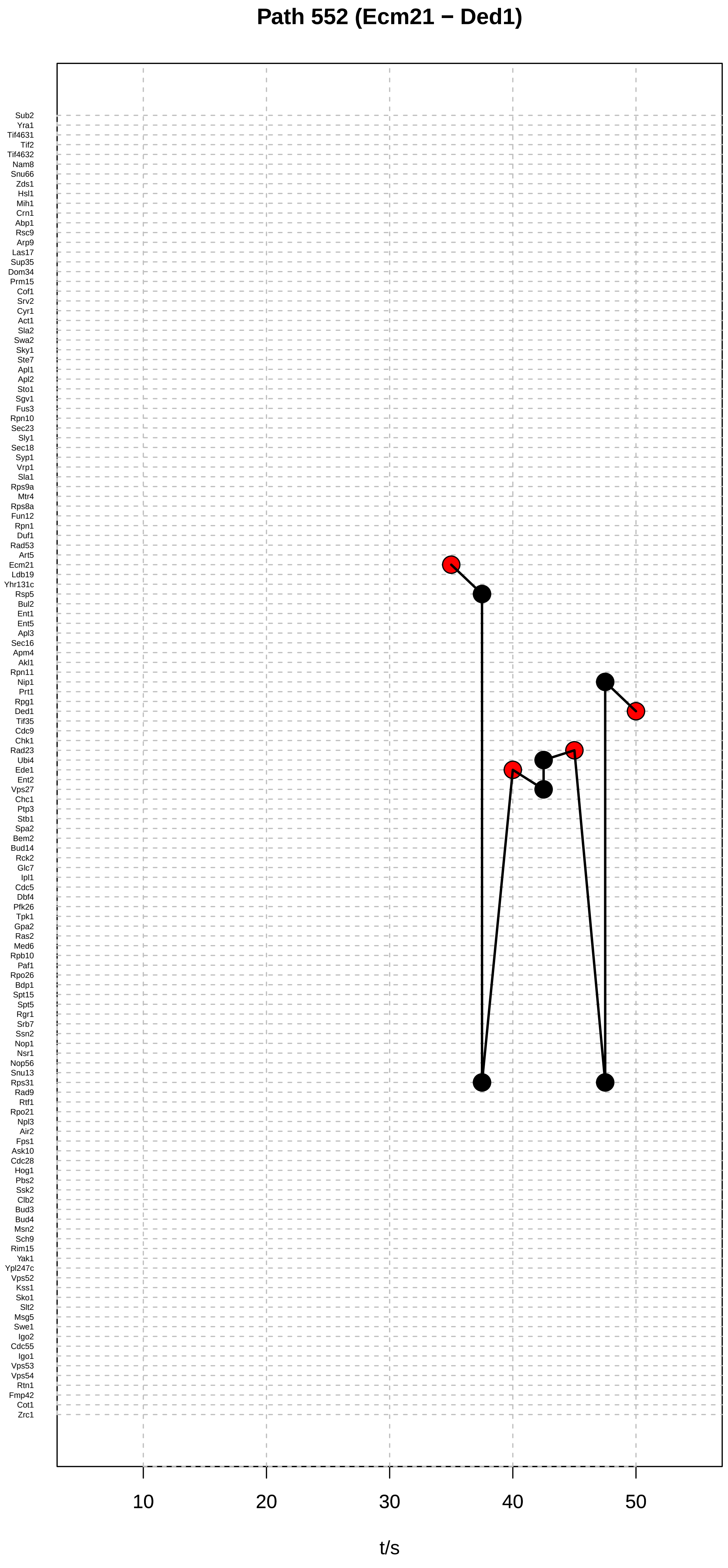

Path 552 (Ecm21 − Ded1)
Sub2
Yra1
Tif4631
Tif2
Tif4632
Nam8
Snu66
Zds1
Hsl1
Mih1
Crn1
Abp1
Rsc9
Arp9
Las17
Sup35
Dom34
Prm15
Cof1
Srv2
Cyr1
Act1
Sla2
Swa2
Sky1
Ste7
Apl1
Apl2
Sto1
Sgv1
Fus3
Rpn10
Sec23
Sly1
Sec18
Syp1
Vrp1
Sla1
Rps9a
Mtr4
Rps8a
Fun12
Rpn1
Duf1
Rad53
Art5
Ecm21
Ldb19
Yhr131c
Rsp5
Bul2
Ent1
Ent5
Apl3
Sec16
Apm4
Akl1
Rpn11
Nip1
Prt1
Rpg1
Ded1
Tif35
Cdc9
Chk1
Rad23
Ubi4
Ede1
Ent2
Vps27
Chc1
Ptp3
Stb1
Spa2
Bem2
Bud14
Rck2
Glc7
Ipl1
Cdc5
Dbf4
Pfk26
Tpk1
Gpa2
Ras2
Med6
Rpb10
Paf1
Rpo26
Bdp1
Spt15
Spt5
Rgr1
Srb7
Ssn2
Nop1
Nsr1
Nop56
Snu13
Rps31
Rad9
Rtf1
Rpo21
Npl3
Air2
Fps1
Ask10
Cdc28
Hog1
Pbs2
Ssk2
Clb2
Bud3
Bud4
Msn2
Sch9
Rim15
Yak1
Ypl247c
Vps52
Kss1
Sko1
Slt2
Msg5
Swe1
Igo2
Cdc55
Igo1
Vps53
Vps54
Rtn1
Fmp42
Cot1
Zrc1
10
20
30
40
50
t/s

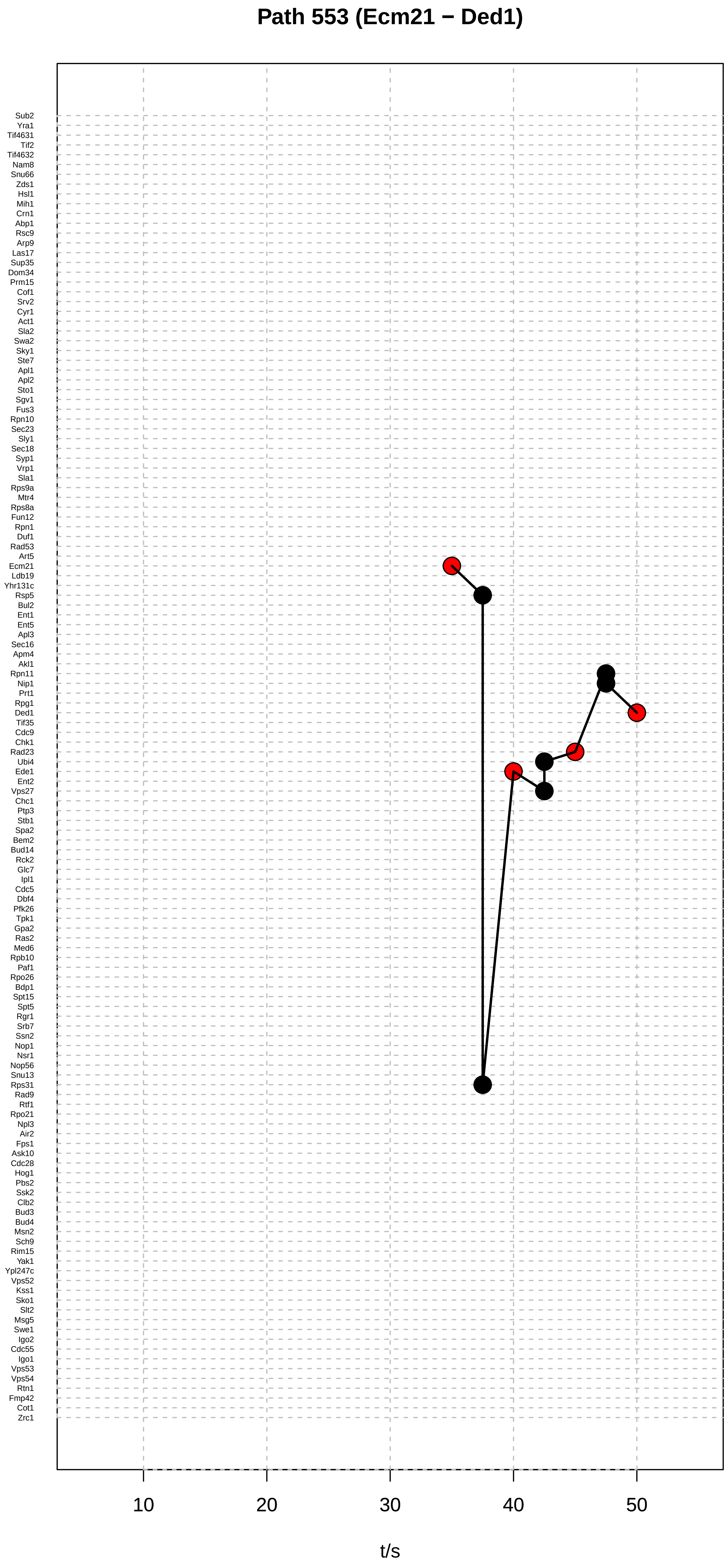

Path 553 (Ecm21 − Ded1)
Sub2
Yra1
Tif4631
Tif2
Tif4632
Nam8
Snu66
Zds1
Hsl1
Mih1
Crn1
Abp1
Rsc9
Arp9
Las17
Sup35
Dom34
Prm15
Cof1
Srv2
Cyr1
Act1
Sla2
Swa2
Sky1
Ste7
Apl1
Apl2
Sto1
Sgv1
Fus3
Rpn10
Sec23
Sly1
Sec18
Syp1
Vrp1
Sla1
Rps9a
Mtr4
Rps8a
Fun12
Rpn1
Duf1
Rad53
Art5
Ecm21
Ldb19
Yhr131c
Rsp5
Bul2
Ent1
Ent5
Apl3
Sec16
Apm4
Akl1
Rpn11
Nip1
Prt1
Rpg1
Ded1
Tif35
Cdc9
Chk1
Rad23
Ubi4
Ede1
Ent2
Vps27
Chc1
Ptp3
Stb1
Spa2
Bem2
Bud14
Rck2
Glc7
Ipl1
Cdc5
Dbf4
Pfk26
Tpk1
Gpa2
Ras2
Med6
Rpb10
Paf1
Rpo26
Bdp1
Spt15
Spt5
Rgr1
Srb7
Ssn2
Nop1
Nsr1
Nop56
Snu13
Rps31
Rad9
Rtf1
Rpo21
Npl3
Air2
Fps1
Ask10
Cdc28
Hog1
Pbs2
Ssk2
Clb2
Bud3
Bud4
Msn2
Sch9
Rim15
Yak1
Ypl247c
Vps52
Kss1
Sko1
Slt2
Msg5
Swe1
Igo2
Cdc55
Igo1
Vps53
Vps54
Rtn1
Fmp42
Cot1
Zrc1
10
20
30
40
50
t/s

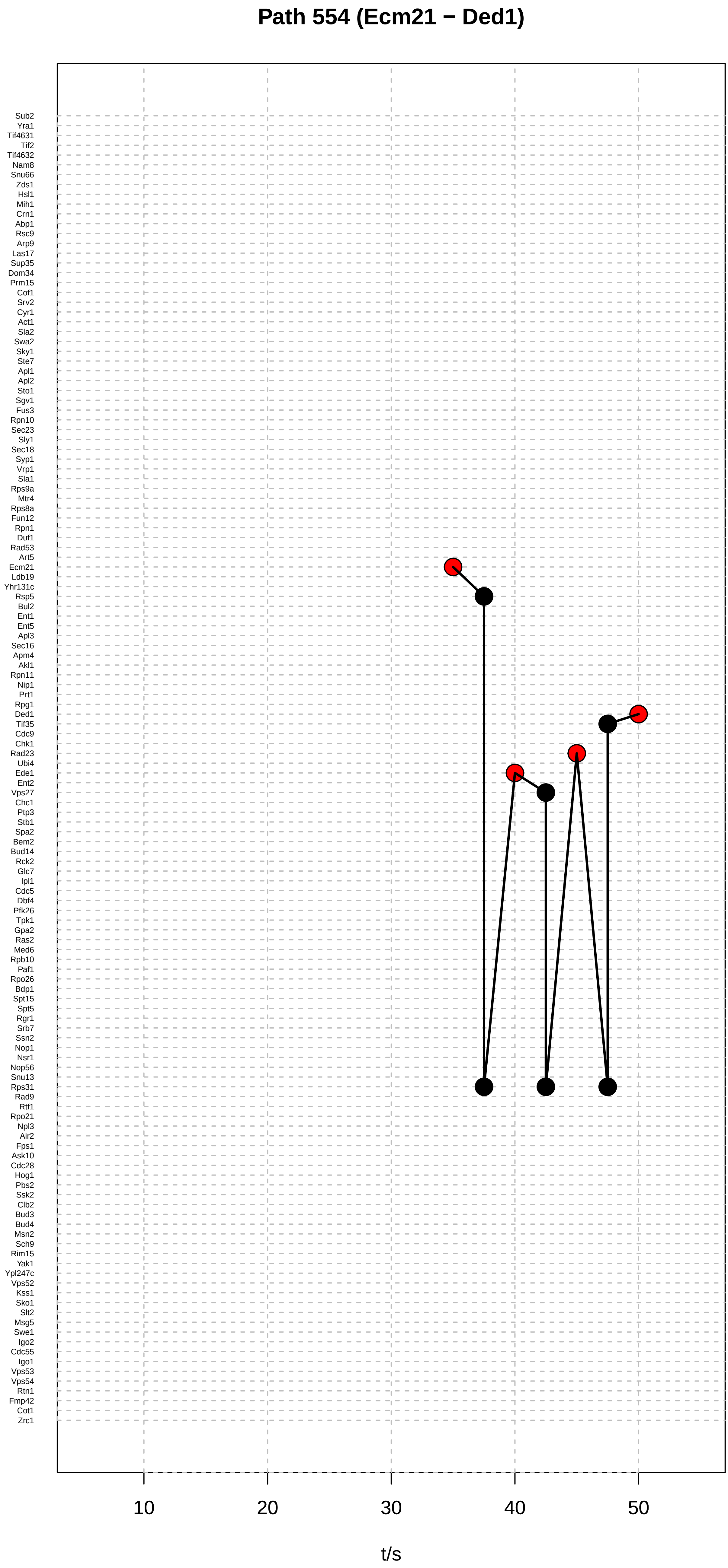
Path 554 (Ecm21 − Ded1)
Sub2
Yra1
Tif4631
Tif2
Tif4632
Nam8
Snu66
Zds1
Hsl1
Mih1
Crn1
Abp1
Rsc9
Arp9
Las17
Sup35
Dom34
Prm15
Cof1
Srv2
Cyr1
Act1
Sla2
Swa2
Sky1
Ste7
Apl1
Apl2
Sto1
Sgv1
Fus3
Rpn10
Sec23
Sly1
Sec18
Syp1
Vrp1
Sla1
Rps9a
Mtr4
Rps8a
Fun12
Rpn1
Duf1
Rad53
Art5
Ecm21
Ldb19
Yhr131c
Rsp5
Bul2
Ent1
Ent5
Apl3
Sec16
Apm4
Akl1
Rpn11
Nip1
Prt1
Rpg1
Ded1
Tif35
Cdc9
Chk1
Rad23
Ubi4
Ede1
Ent2
Vps27
Chc1
Ptp3
Stb1
Spa2
Bem2
Bud14
Rck2
Glc7
Ipl1
Cdc5
Dbf4
Pfk26
Tpk1
Gpa2
Ras2
Med6
Rpb10
Paf1
Rpo26
Bdp1
Spt15
Spt5
Rgr1
Srb7
Ssn2
Nop1
Nsr1
Nop56
Snu13
Rps31
Rad9
Rtf1
Rpo21
Npl3
Air2
Fps1
Ask10
Cdc28
Hog1
Pbs2
Ssk2
Clb2
Bud3
Bud4
Msn2
Sch9
Rim15
Yak1
Ypl247c
Vps52
Kss1
Sko1
Slt2
Msg5
Swe1
Igo2
Cdc55
Igo1
Vps53
Vps54
Rtn1
Fmp42
Cot1
Zrc1
10
20
30
40
50
t/s

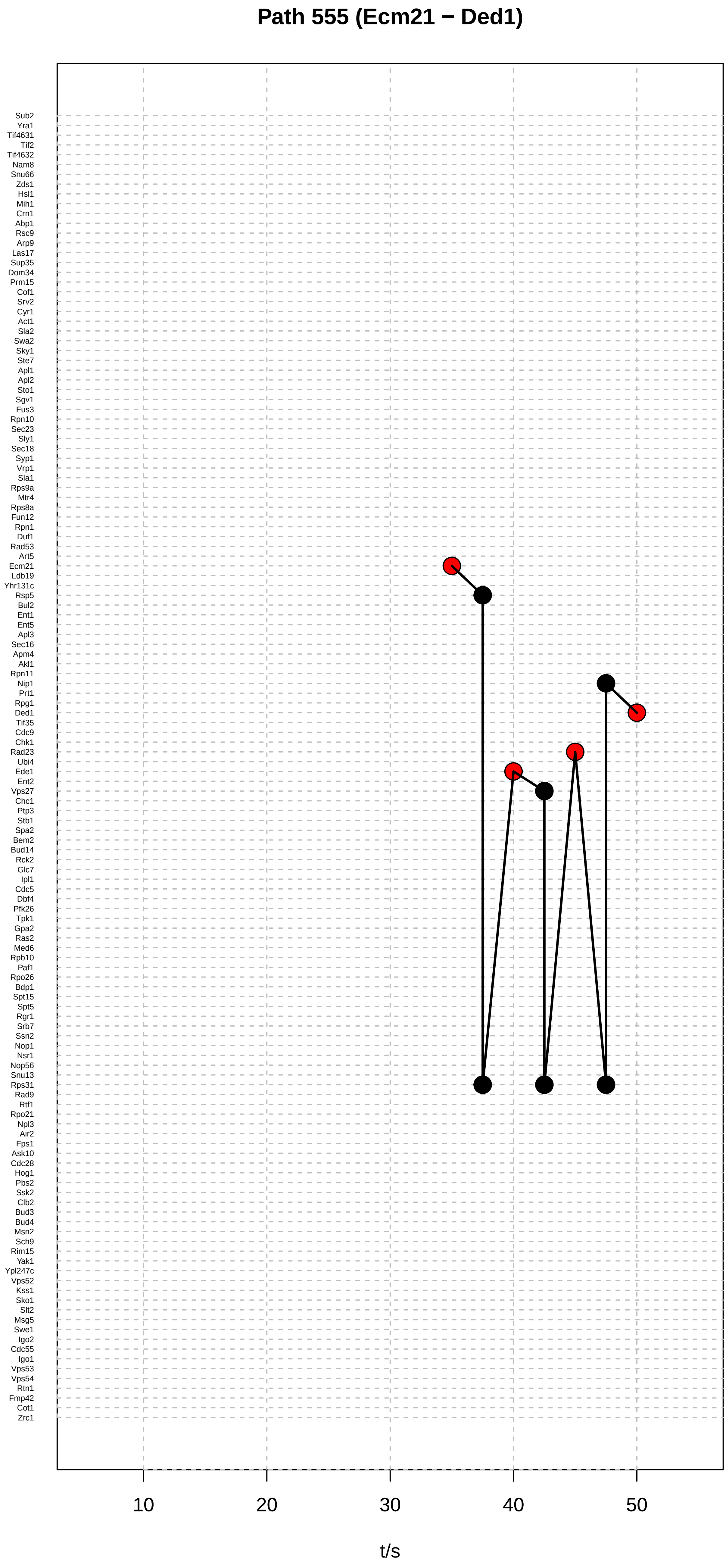

Path 555 (Ecm21 − Ded1)
Sub2
Yra1
Tif4631
Tif2
Tif4632
Nam8
Snu66
Zds1
Hsl1
Mih1
Crn1
Abp1
Rsc9
Arp9
Las17
Sup35
Dom34
Prm15
Cof1
Srv2
Cyr1
Act1
Sla2
Swa2
Sky1
Ste7
Apl1
Apl2
Sto1
Sgv1
Fus3
Rpn10
Sec23
Sly1
Sec18
Syp1
Vrp1
Sla1
Rps9a
Mtr4
Rps8a
Fun12
Rpn1
Duf1
Rad53
Art5
Ecm21
Ldb19
Yhr131c
Rsp5
Bul2
Ent1
Ent5
Apl3
Sec16
Apm4
Akl1
Rpn11
Nip1
Prt1
Rpg1
Ded1
Tif35
Cdc9
Chk1
Rad23
Ubi4
Ede1
Ent2
Vps27
Chc1
Ptp3
Stb1
Spa2
Bem2
Bud14
Rck2
Glc7
Ipl1
Cdc5
Dbf4
Pfk26
Tpk1
Gpa2
Ras2
Med6
Rpb10
Paf1
Rpo26
Bdp1
Spt15
Spt5
Rgr1
Srb7
Ssn2
Nop1
Nsr1
Nop56
Snu13
Rps31
Rad9
Rtf1
Rpo21
Npl3
Air2
Fps1
Ask10
Cdc28
Hog1
Pbs2
Ssk2
Clb2
Bud3
Bud4
Msn2
Sch9
Rim15
Yak1
Ypl247c
Vps52
Kss1
Sko1
Slt2
Msg5
Swe1
Igo2
Cdc55
Igo1
Vps53
Vps54
Rtn1
Fmp42
Cot1
Zrc1
10
20
30
40
50
t/s

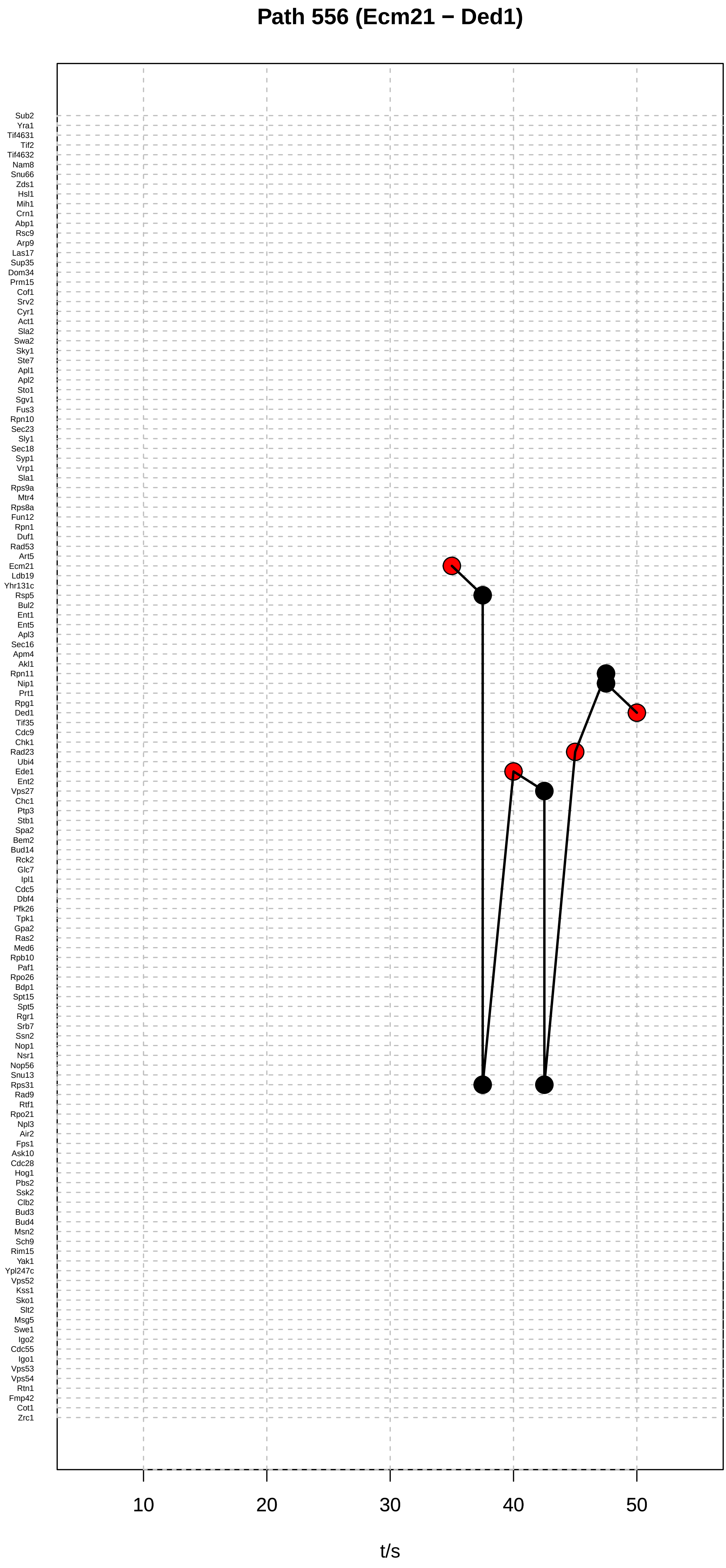

Path 556 (Ecm21 − Ded1)
Sub2
Yra1
Tif4631
Tif2
Tif4632
Nam8
Snu66
Zds1
Hsl1
Mih1
Crn1
Abp1
Rsc9
Arp9
Las17
Sup35
Dom34
Prm15
Cof1
Srv2
Cyr1
Act1
Sla2
Swa2
Sky1
Ste7
Apl1
Apl2
Sto1
Sgv1
Fus3
Rpn10
Sec23
Sly1
Sec18
Syp1
Vrp1
Sla1
Rps9a
Mtr4
Rps8a
Fun12
Rpn1
Duf1
Rad53
Art5
Ecm21
Ldb19
Yhr131c
Rsp5
Bul2
Ent1
Ent5
Apl3
Sec16
Apm4
Akl1
Rpn11
Nip1
Prt1
Rpg1
Ded1
Tif35
Cdc9
Chk1
Rad23
Ubi4
Ede1
Ent2
Vps27
Chc1
Ptp3
Stb1
Spa2
Bem2
Bud14
Rck2
Glc7
Ipl1
Cdc5
Dbf4
Pfk26
Tpk1
Gpa2
Ras2
Med6
Rpb10
Paf1
Rpo26
Bdp1
Spt15
Spt5
Rgr1
Srb7
Ssn2
Nop1
Nsr1
Nop56
Snu13
Rps31
Rad9
Rtf1
Rpo21
Npl3
Air2
Fps1
Ask10
Cdc28
Hog1
Pbs2
Ssk2
Clb2
Bud3
Bud4
Msn2
Sch9
Rim15
Yak1
Ypl247c
Vps52
Kss1
Sko1
Slt2
Msg5
Swe1
Igo2
Cdc55
Igo1
Vps53
Vps54
Rtn1
Fmp42
Cot1
Zrc1
10
20
30
40
50
t/s

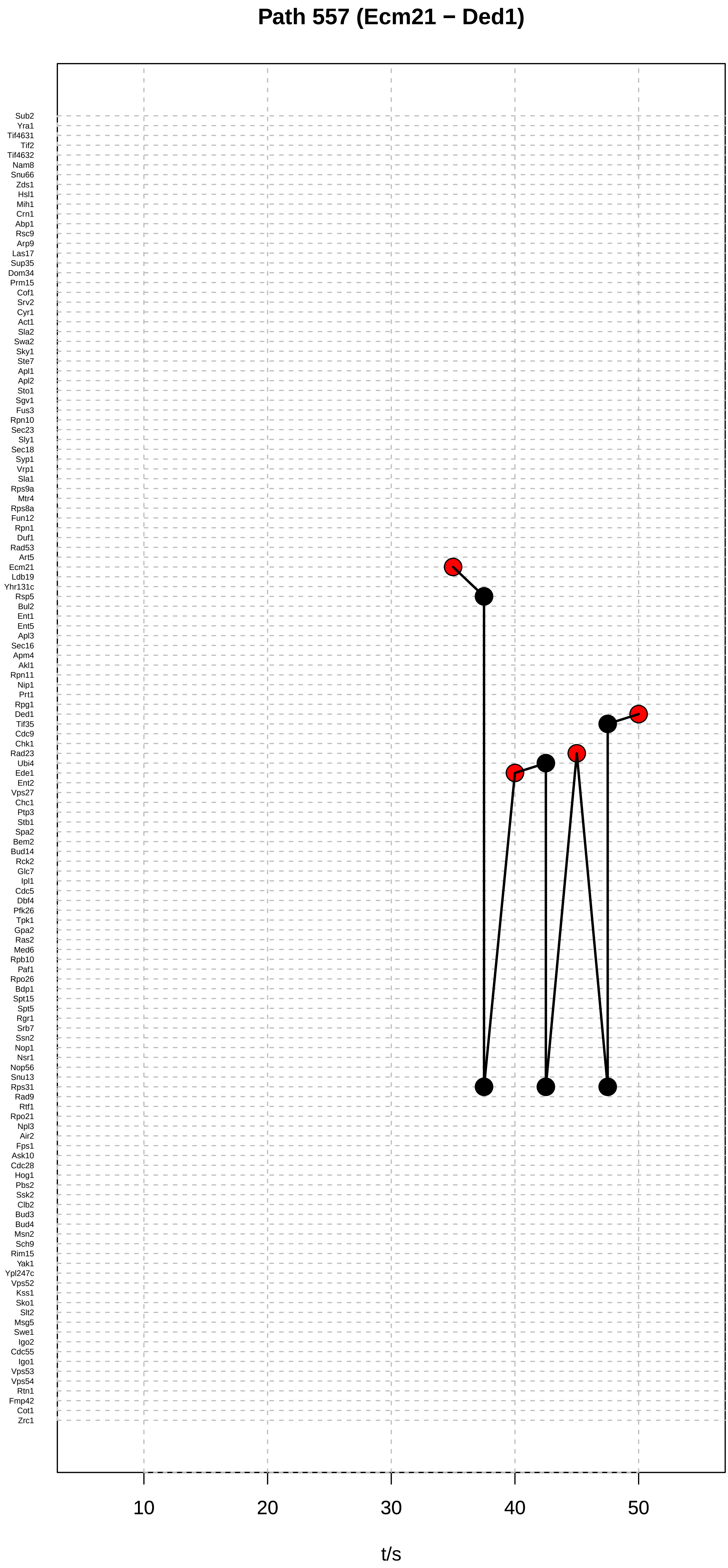

Path 557 (Ecm21 − Ded1)
Sub2
Yra1
Tif4631
Tif2
Tif4632
Nam8
Snu66
Zds1
Hsl1
Mih1
Crn1
Abp1
Rsc9
Arp9
Las17
Sup35
Dom34
Prm15
Cof1
Srv2
Cyr1
Act1
Sla2
Swa2
Sky1
Ste7
Apl1
Apl2
Sto1
Sgv1
Fus3
Rpn10
Sec23
Sly1
Sec18
Syp1
Vrp1
Sla1
Rps9a
Mtr4
Rps8a
Fun12
Rpn1
Duf1
Rad53
Art5
Ecm21
Ldb19
Yhr131c
Rsp5
Bul2
Ent1
Ent5
Apl3
Sec16
Apm4
Akl1
Rpn11
Nip1
Prt1
Rpg1
Ded1
Tif35
Cdc9
Chk1
Rad23
Ubi4
Ede1
Ent2
Vps27
Chc1
Ptp3
Stb1
Spa2
Bem2
Bud14
Rck2
Glc7
Ipl1
Cdc5
Dbf4
Pfk26
Tpk1
Gpa2
Ras2
Med6
Rpb10
Paf1
Rpo26
Bdp1
Spt15
Spt5
Rgr1
Srb7
Ssn2
Nop1
Nsr1
Nop56
Snu13
Rps31
Rad9
Rtf1
Rpo21
Npl3
Air2
Fps1
Ask10
Cdc28
Hog1
Pbs2
Ssk2
Clb2
Bud3
Bud4
Msn2
Sch9
Rim15
Yak1
Ypl247c
Vps52
Kss1
Sko1
Slt2
Msg5
Swe1
Igo2
Cdc55
Igo1
Vps53
Vps54
Rtn1
Fmp42
Cot1
Zrc1
10
20
30
40
50
t/s

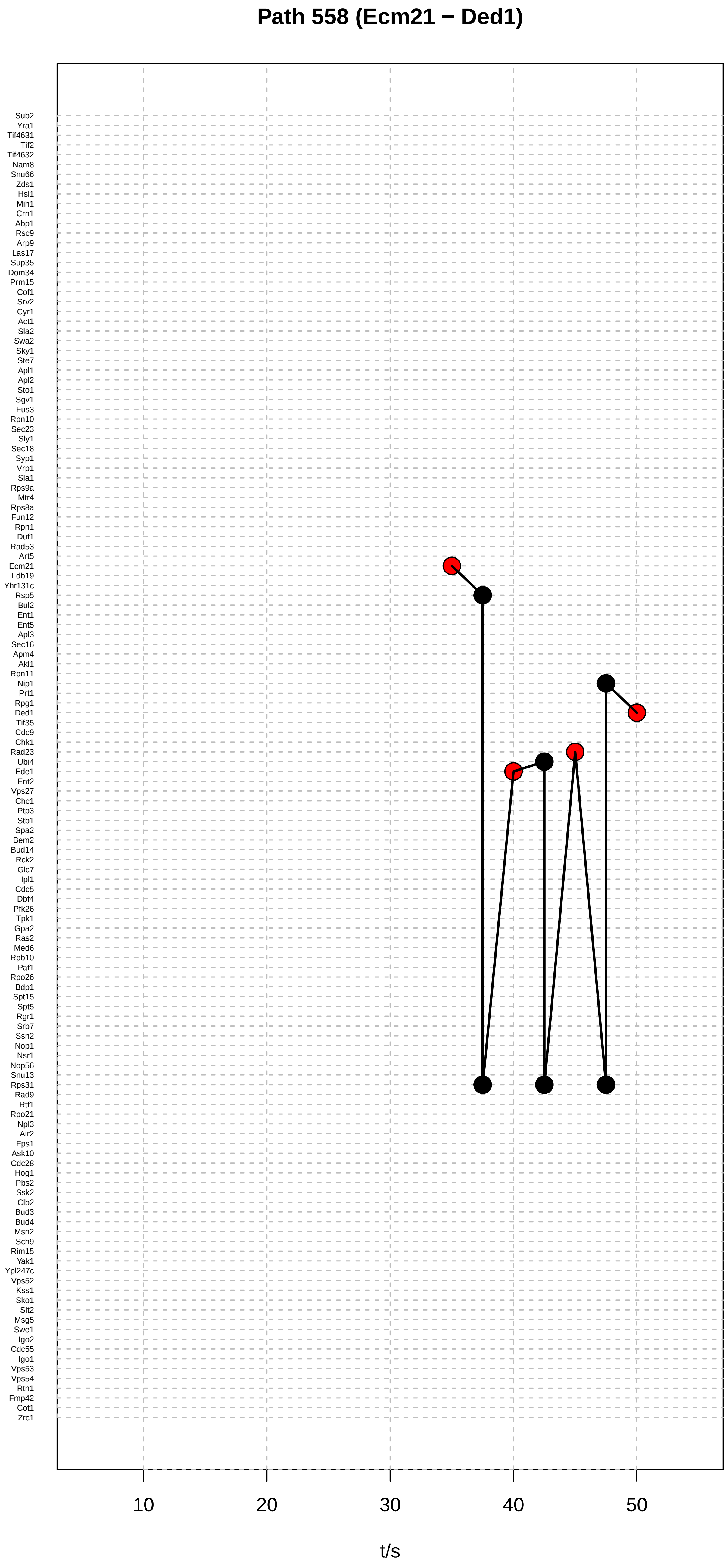

Path 558 (Ecm21 − Ded1)
Sub2
Yra1
Tif4631
Tif2
Tif4632
Nam8
Snu66
Zds1
Hsl1
Mih1
Crn1
Abp1
Rsc9
Arp9
Las17
Sup35
Dom34
Prm15
Cof1
Srv2
Cyr1
Act1
Sla2
Swa2
Sky1
Ste7
Apl1
Apl2
Sto1
Sgv1
Fus3
Rpn10
Sec23
Sly1
Sec18
Syp1
Vrp1
Sla1
Rps9a
Mtr4
Rps8a
Fun12
Rpn1
Duf1
Rad53
Art5
Ecm21
Ldb19
Yhr131c
Rsp5
Bul2
Ent1
Ent5
Apl3
Sec16
Apm4
Akl1
Rpn11
Nip1
Prt1
Rpg1
Ded1
Tif35
Cdc9
Chk1
Rad23
Ubi4
Ede1
Ent2
Vps27
Chc1
Ptp3
Stb1
Spa2
Bem2
Bud14
Rck2
Glc7
Ipl1
Cdc5
Dbf4
Pfk26
Tpk1
Gpa2
Ras2
Med6
Rpb10
Paf1
Rpo26
Bdp1
Spt15
Spt5
Rgr1
Srb7
Ssn2
Nop1
Nsr1
Nop56
Snu13
Rps31
Rad9
Rtf1
Rpo21
Npl3
Air2
Fps1
Ask10
Cdc28
Hog1
Pbs2
Ssk2
Clb2
Bud3
Bud4
Msn2
Sch9
Rim15
Yak1
Ypl247c
Vps52
Kss1
Sko1
Slt2
Msg5
Swe1
Igo2
Cdc55
Igo1
Vps53
Vps54
Rtn1
Fmp42
Cot1
Zrc1
10
20
30
40
50
t/s

### Path 559 (Ecm21 − Ded1)

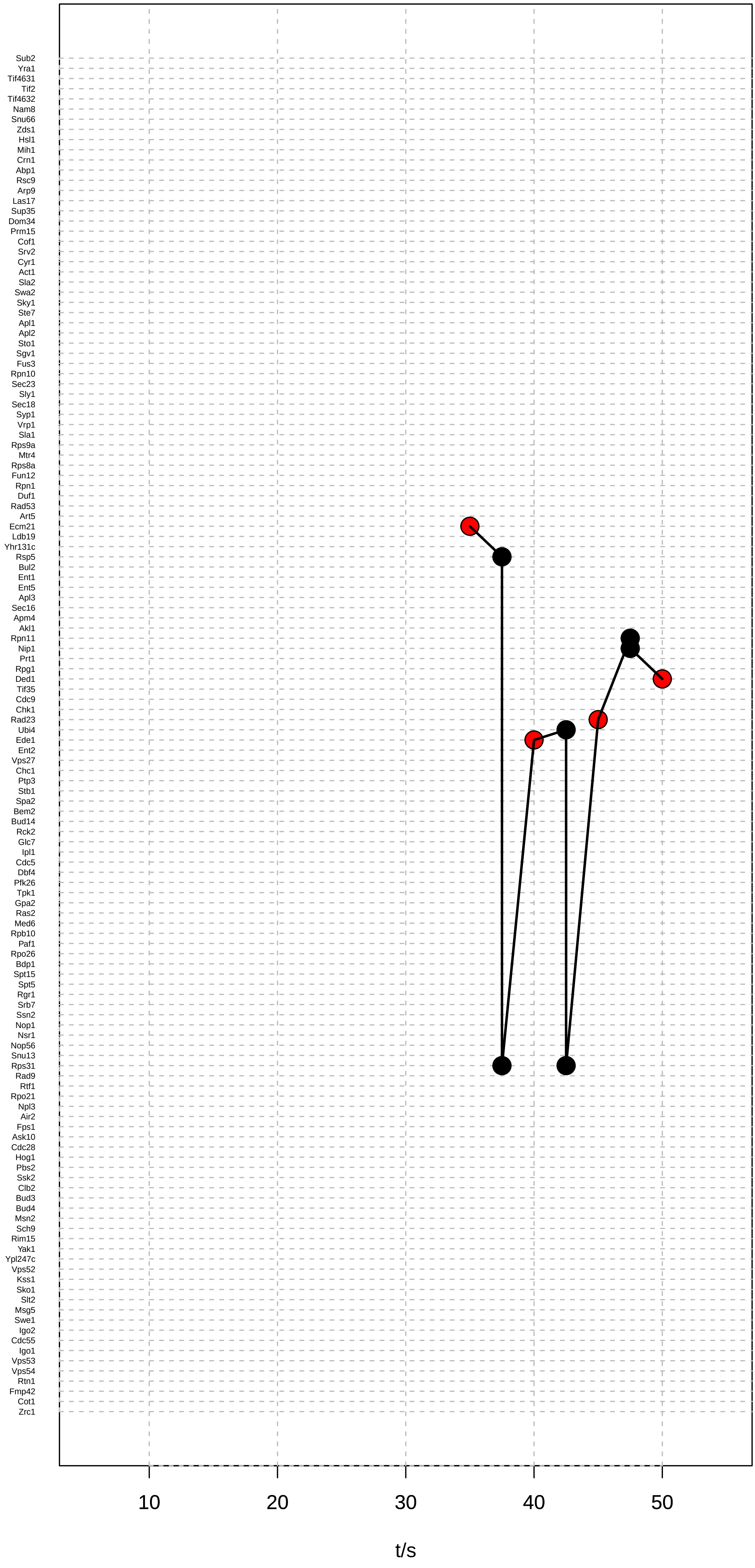

#### Path 560 (Ecm21 − Ded1)

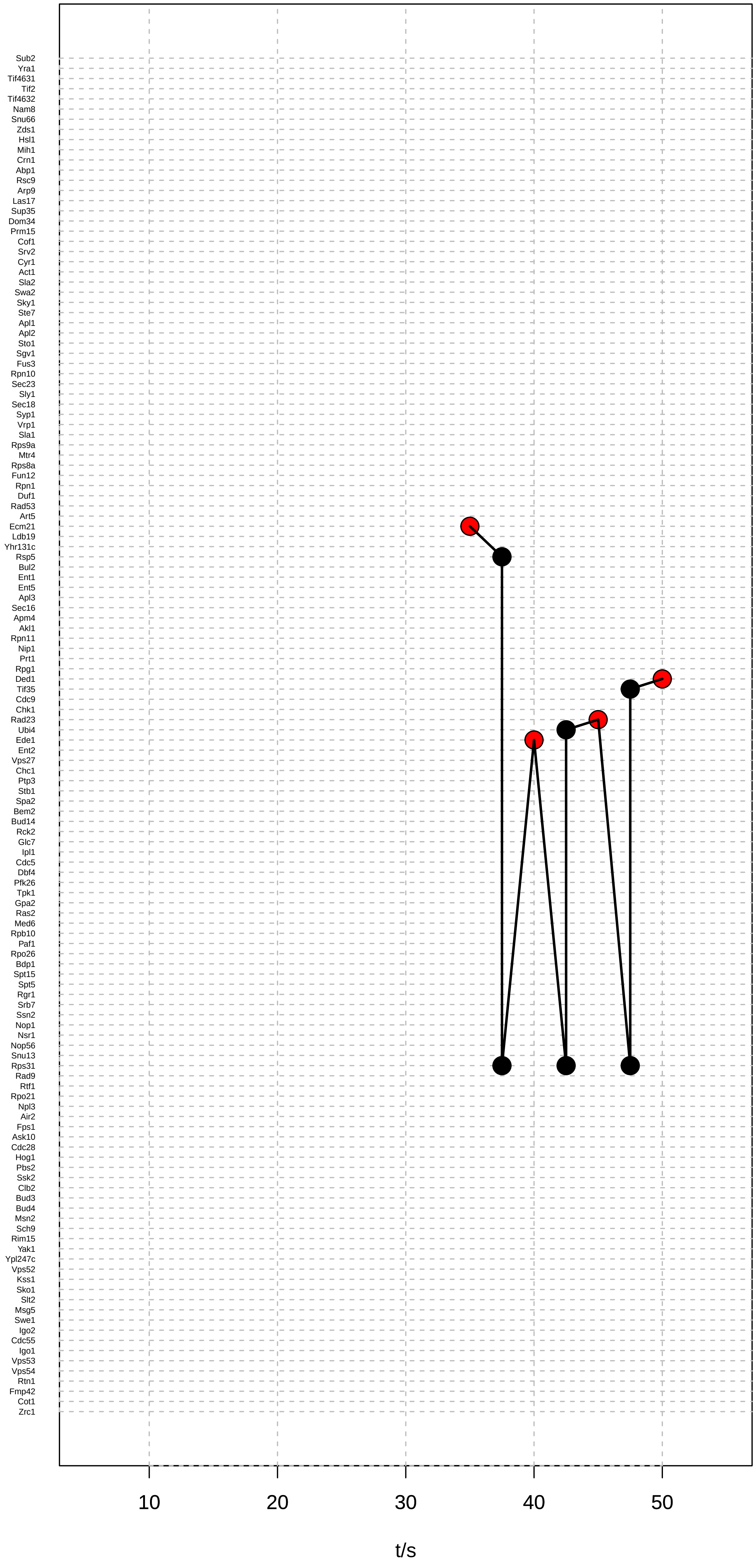

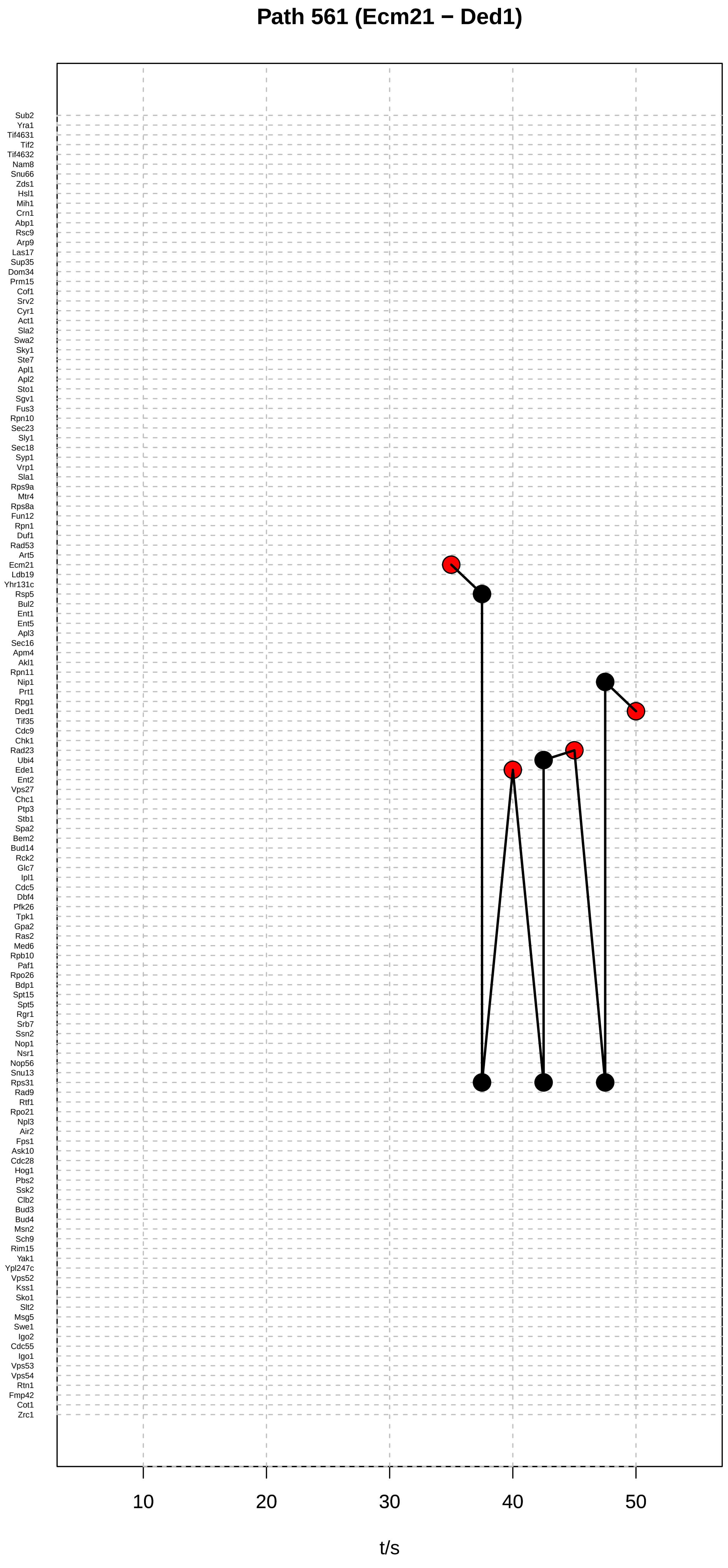

Path 561 (Ecm21 − Ded1)
Sub2
Yra1
Tif4631
Tif2
Tif4632
Nam8
Snu66
Zds1
Hsl1
Mih1
Crn1
Abp1
Rsc9
Arp9
Las17
Sup35
Dom34
Prm15
Cof1
Srv2
Cyr1
Act1
Sla2
Swa2
Sky1
Ste7
Apl1
Apl2
Sto1
Sgv1
Fus3
Rpn10
Sec23
Sly1
Sec18
Syp1
Vrp1
Sla1
Rps9a
Mtr4
Rps8a
Fun12
Rpn1
Duf1
Rad53
Art5
Ecm21
Ldb19
Yhr131c
Rsp5
Bul2
Ent1
Ent5
Apl3
Sec16
Apm4
Akl1
Rpn11
Nip1
Prt1
Rpg1
Ded1
Tif35
Cdc9
Chk1
Rad23
Ubi4
Ede1
Ent2
Vps27
Chc1
Ptp3
Stb1
Spa2
Bem2
Bud14
Rck2
Glc7
Ipl1
Cdc5
Dbf4
Pfk26
Tpk1
Gpa2
Ras2
Med6
Rpb10
Paf1
Rpo26
Bdp1
Spt15
Spt5
Rgr1
Srb7
Ssn2
Nop1
Nsr1
Nop56
Snu13
Rps31
Rad9
Rtf1
Rpo21
Npl3
Air2
Fps1
Ask10
Cdc28
Hog1
Pbs2
Ssk2
Clb2
Bud3
Bud4
Msn2
Sch9
Rim15
Yak1
Ypl247c
Vps52
Kss1
Sko1
Slt2
Msg5
Swe1
Igo2
Cdc55
Igo1
Vps53
Vps54
Rtn1
Fmp42
Cot1
Zrc1
10
20
30
40
50
t/s

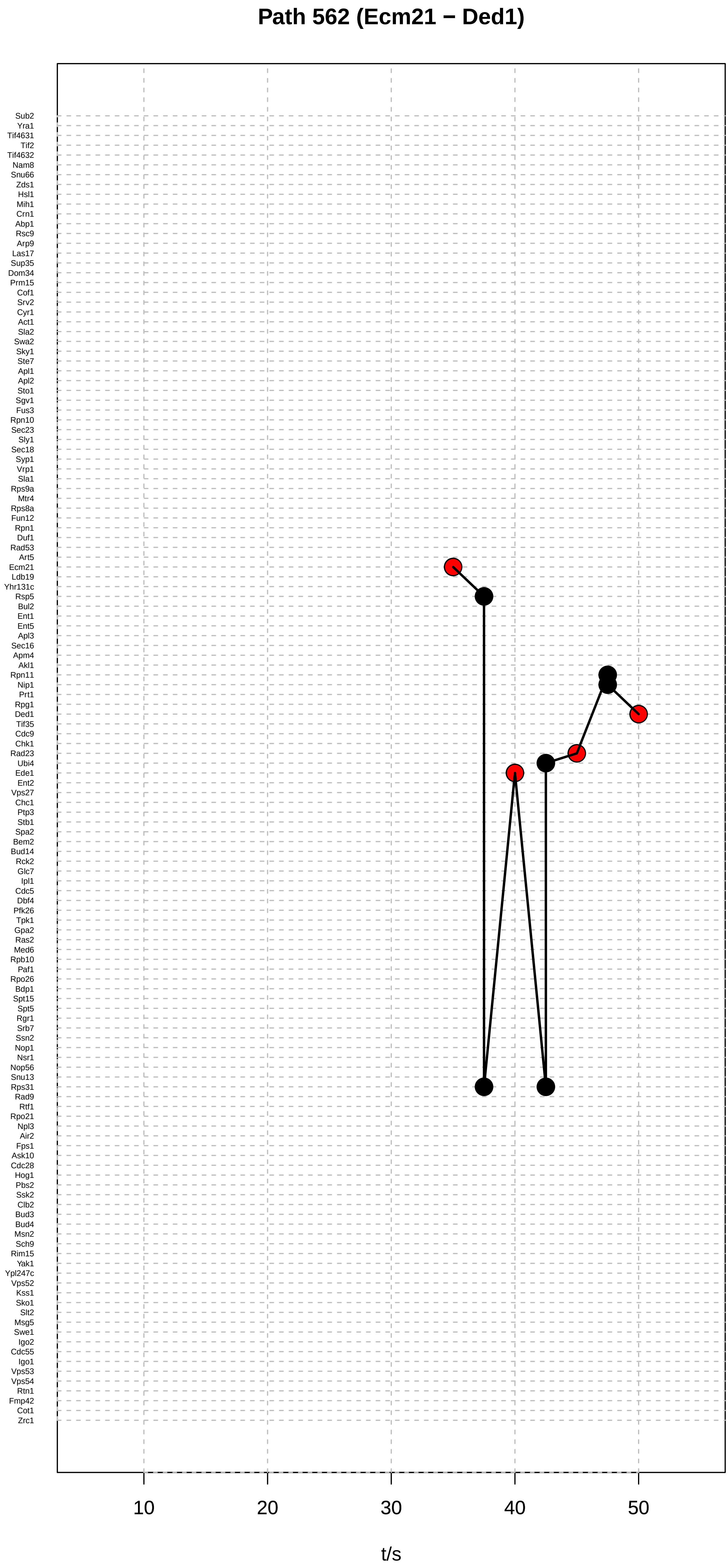

Path 562 (Ecm21 − Ded1)
Sub2
Yra1
Tif4631
Tif2
Tif4632
Nam8
Snu66
Zds1
Hsl1
Mih1
Crn1
Abp1
Rsc9
Arp9
Las17
Sup35
Dom34
Prm15
Cof1
Srv2
Cyr1
Act1
Sla2
Swa2
Sky1
Ste7
Apl1
Apl2
Sto1
Sgv1
Fus3
Rpn10
Sec23
Sly1
Sec18
Syp1
Vrp1
Sla1
Rps9a
Mtr4
Rps8a
Fun12
Rpn1
Duf1
Rad53
Art5
Ecm21
Ldb19
Yhr131c
Rsp5
Bul2
Ent1
Ent5
Apl3
Sec16
Apm4
Akl1
Rpn11
Nip1
Prt1
Rpg1
Ded1
Tif35
Cdc9
Chk1
Rad23
Ubi4
Ede1
Ent2
Vps27
Chc1
Ptp3
Stb1
Spa2
Bem2
Bud14
Rck2
Glc7
Ipl1
Cdc5
Dbf4
Pfk26
Tpk1
Gpa2
Ras2
Med6
Rpb10
Paf1
Rpo26
Bdp1
Spt15
Spt5
Rgr1
Srb7
Ssn2
Nop1
Nsr1
Nop56
Snu13
Rps31
Rad9
Rtf1
Rpo21
Npl3
Air2
Fps1
Ask10
Cdc28
Hog1
Pbs2
Ssk2
Clb2
Bud3
Bud4
Msn2
Sch9
Rim15
Yak1
Ypl247c
Vps52
Kss1
Sko1
Slt2
Msg5
Swe1
Igo2
Cdc55
Igo1
Vps53
Vps54
Rtn1
Fmp42
Cot1
Zrc1
10
20
30
40
50
t/s

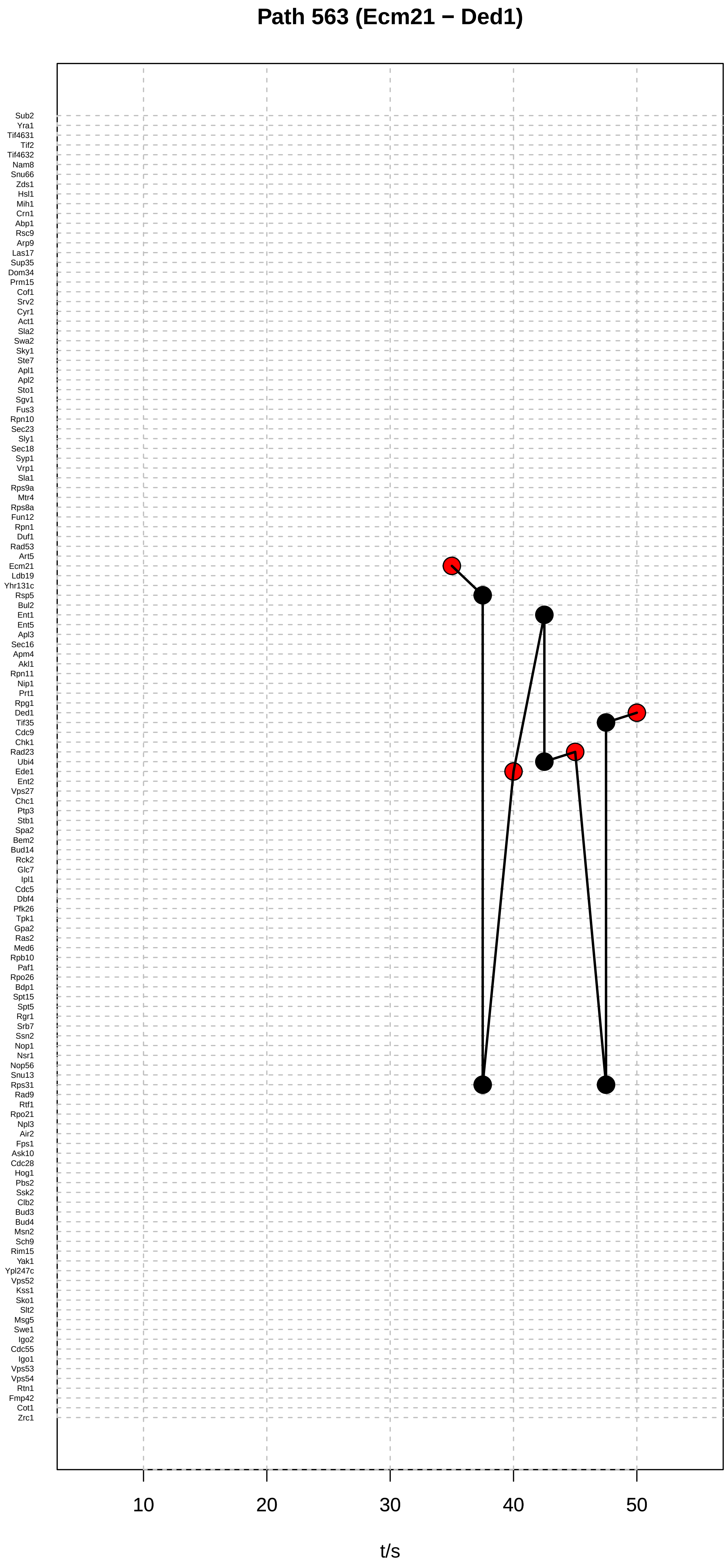
Path 563 (Ecm21 − Ded1)
Sub2
Yra1
Tif4631
Tif2
Tif4632
Nam8
Snu66
Zds1
Hsl1
Mih1
Crn1
Abp1
Rsc9
Arp9
Las17
Sup35
Dom34
Prm15
Cof1
Srv2
Cyr1
Act1
Sla2
Swa2
Sky1
Ste7
Apl1
Apl2
Sto1
Sgv1
Fus3
Rpn10
Sec23
Sly1
Sec18
Syp1
Vrp1
Sla1
Rps9a
Mtr4
Rps8a
Fun12
Rpn1
Duf1
Rad53
Art5
Ecm21
Ldb19
Yhr131c
Rsp5
Bul2
Ent1
Ent5
Apl3
Sec16
Apm4
Akl1
Rpn11
Nip1
Prt1
Rpg1
Ded1
Tif35
Cdc9
Chk1
Rad23
Ubi4
Ede1
Ent2
Vps27
Chc1
Ptp3
Stb1
Spa2
Bem2
Bud14
Rck2
Glc7
Ipl1
Cdc5
Dbf4
Pfk26
Tpk1
Gpa2
Ras2
Med6
Rpb10
Paf1
Rpo26
Bdp1
Spt15
Spt5
Rgr1
Srb7
Ssn2
Nop1
Nsr1
Nop56
Snu13
Rps31
Rad9
Rtf1
Rpo21
Npl3
Air2
Fps1
Ask10
Cdc28
Hog1
Pbs2
Ssk2
Clb2
Bud3
Bud4
Msn2
Sch9
Rim15
Yak1
Ypl247c
Vps52
Kss1
Sko1
Slt2
Msg5
Swe1
Igo2
Cdc55
Igo1
Vps53
Vps54
Rtn1
Fmp42
Cot1
Zrc1
10
20
30
40
50
t/s

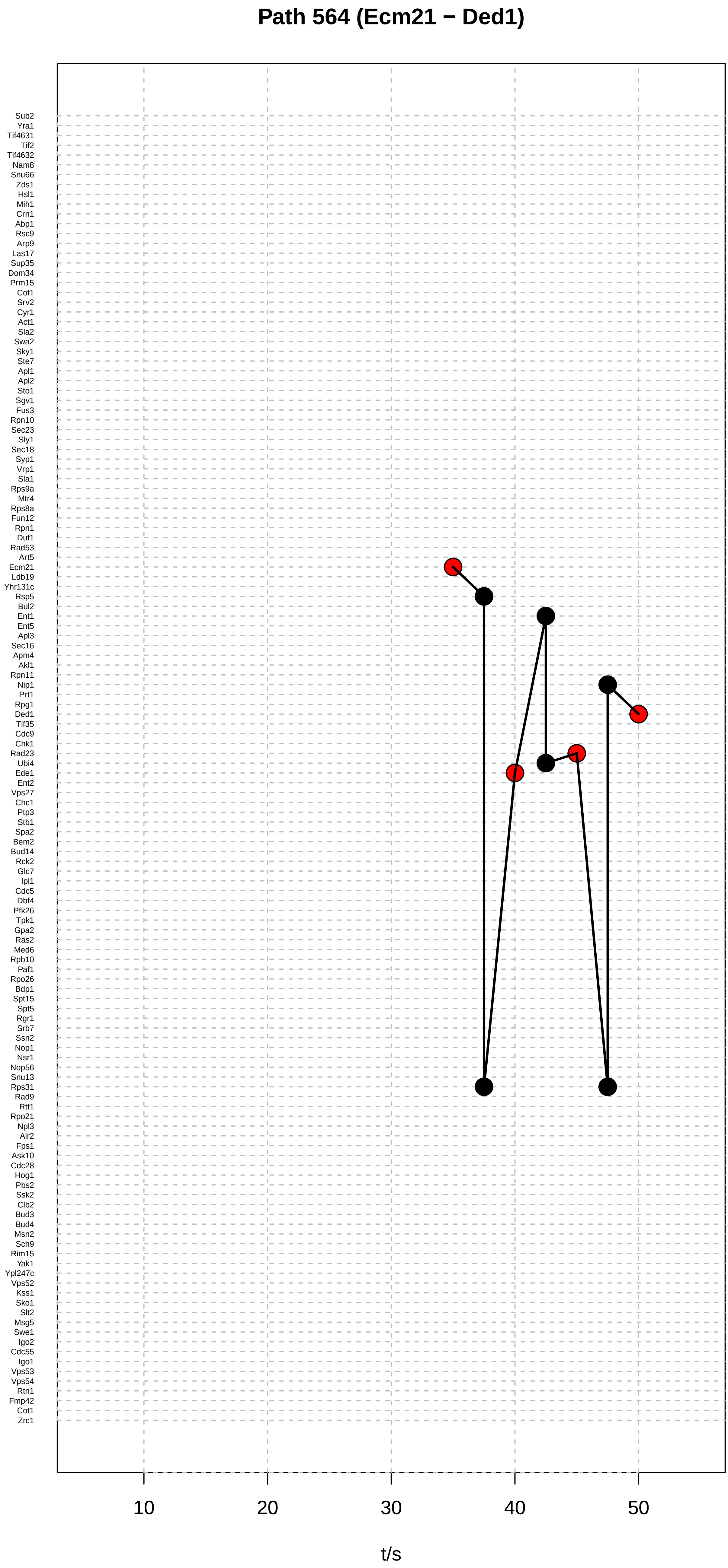

Path 564 (Ecm21 − Ded1)
Sub2
Yra1
Tif4631
Tif2
Tif4632
Nam8
Snu66
Zds1
Hsl1
Mih1
Crn1
Abp1
Rsc9
Arp9
Las17
Sup35
Dom34
Prm15
Cof1
Srv2
Cyr1
Act1
Sla2
Swa2
Sky1
Ste7
Apl1
Apl2
Sto1
Sgv1
Fus3
Rpn10
Sec23
Sly1
Sec18
Syp1
Vrp1
Sla1
Rps9a
Mtr4
Rps8a
Fun12
Rpn1
Duf1
Rad53
Art5
Ecm21
Ldb19
Yhr131c
Rsp5
Bul2
Ent1
Ent5
Apl3
Sec16
Apm4
Akl1
Rpn11
Nip1
Prt1
Rpg1
Ded1
Tif35
Cdc9
Chk1
Rad23
Ubi4
Ede1
Ent2
Vps27
Chc1
Ptp3
Stb1
Spa2
Bem2
Bud14
Rck2
Glc7
Ipl1
Cdc5
Dbf4
Pfk26
Tpk1
Gpa2
Ras2
Med6
Rpb10
Paf1
Rpo26
Bdp1
Spt15
Spt5
Rgr1
Srb7
Ssn2
Nop1
Nsr1
Nop56
Snu13
Rps31
Rad9
Rtf1
Rpo21
Npl3
Air2
Fps1
Ask10
Cdc28
Hog1
Pbs2
Ssk2
Clb2
Bud3
Bud4
Msn2
Sch9
Rim15
Yak1
Ypl247c
Vps52
Kss1
Sko1
Slt2
Msg5
Swe1
Igo2
Cdc55
Igo1
Vps53
Vps54
Rtn1
Fmp42
Cot1
Zrc1
10
20
30
40
50
t/s

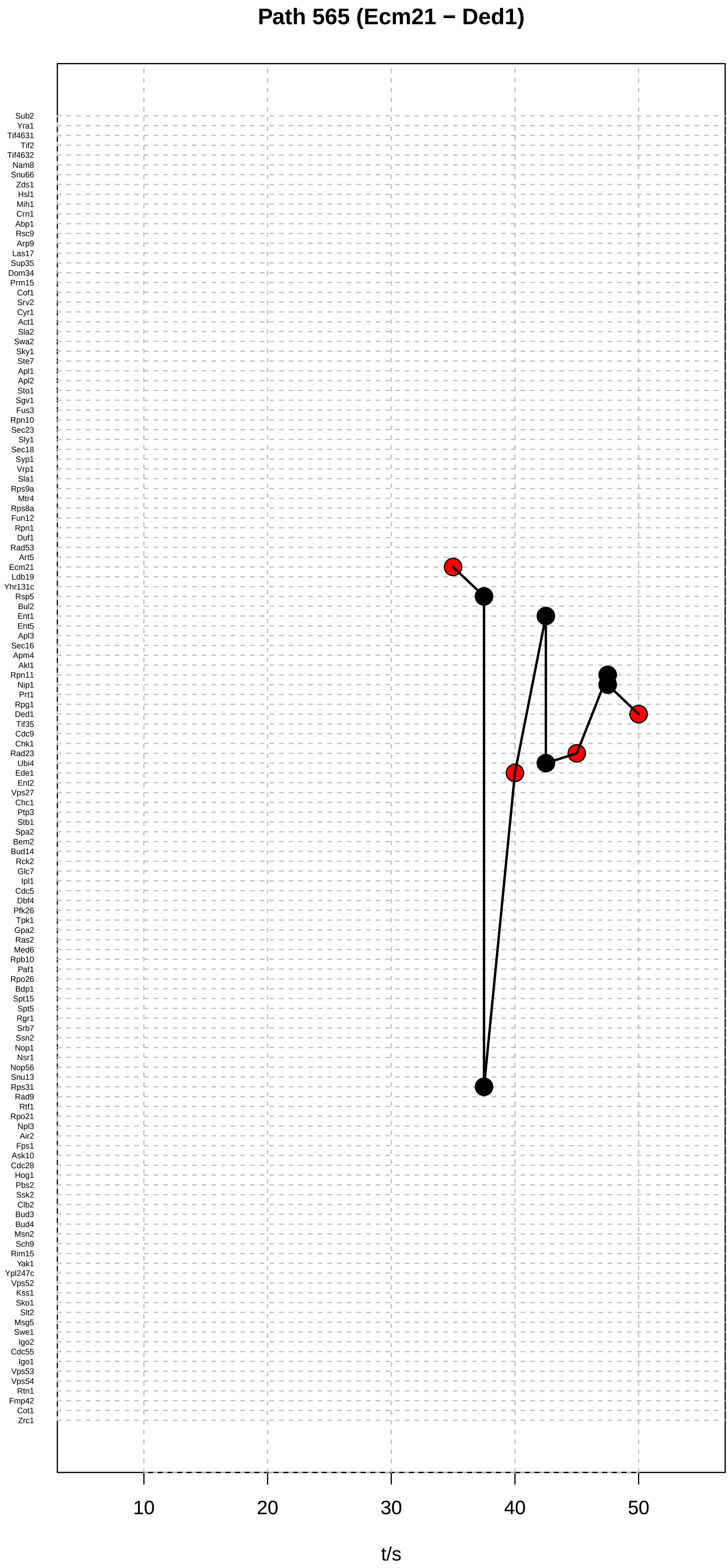
Path 565 (Ecm21 − Ded1)
Sub2
Yra1
Tif4631
Tif2
Tif4632
Nam8
Snu66
Zds1
Hsl1
Mih1
Crn1
Abp1
Rsc9
Arp9
Las17
Sup35
Dom34
Prm15
Cof1
Srv2
Cyr1
Act1
Sla2
Swa2
Sky1
Ste7
Apl1
Apl2
Sto1
Sgv1
Fus3
Rpn10
Sec23
Sly1
Sec18
Syp1
Vrp1
Sla1
Rps9a
Mtr4
Rps8a
Fun12
Rpn1
Duf1
Rad53
Art5
Ecm21
Ldb19
Yhr131c
Rsp5
Bul2
Ent1
Ent5
Apl3
Sec16
Apm4
Akl1
Rpn11
Nip1
Prt1
Rpg1
Ded1
Tif35
Cdc9
Chk1
Rad23
Ubi4
Ede1
Ent2
Vps27
Chc1
Ptp3
Stb1
Spa2
Bem2
Bud14
Rck2
Glc7
Ipl1
Cdc5
Dbf4
Pfk26
Tpk1
Gpa2
Ras2
Med6
Rpb10
Paf1
Rpo26
Bdp1
Spt15
Spt5
Rgr1
Srb7
Ssn2
Nop1
Nsr1
Nop56
Snu13
Rps31
Rad9
Rtf1
Rpo21
Npl3
Air2
Fps1
Ask10
Cdc28
Hog1
Pbs2
Ssk2
Clb2
Bud3
Bud4
Msn2
Sch9
Rim15
Yak1
Ypl247c
Vps52
Kss1
Sko1
Slt2
Msg5
Swe1
Igo2
Cdc55
Igo1
Vps53
Vps54
Rtn1
Fmp42
Cot1
Zrc1
10
20
30
40
50
t/s

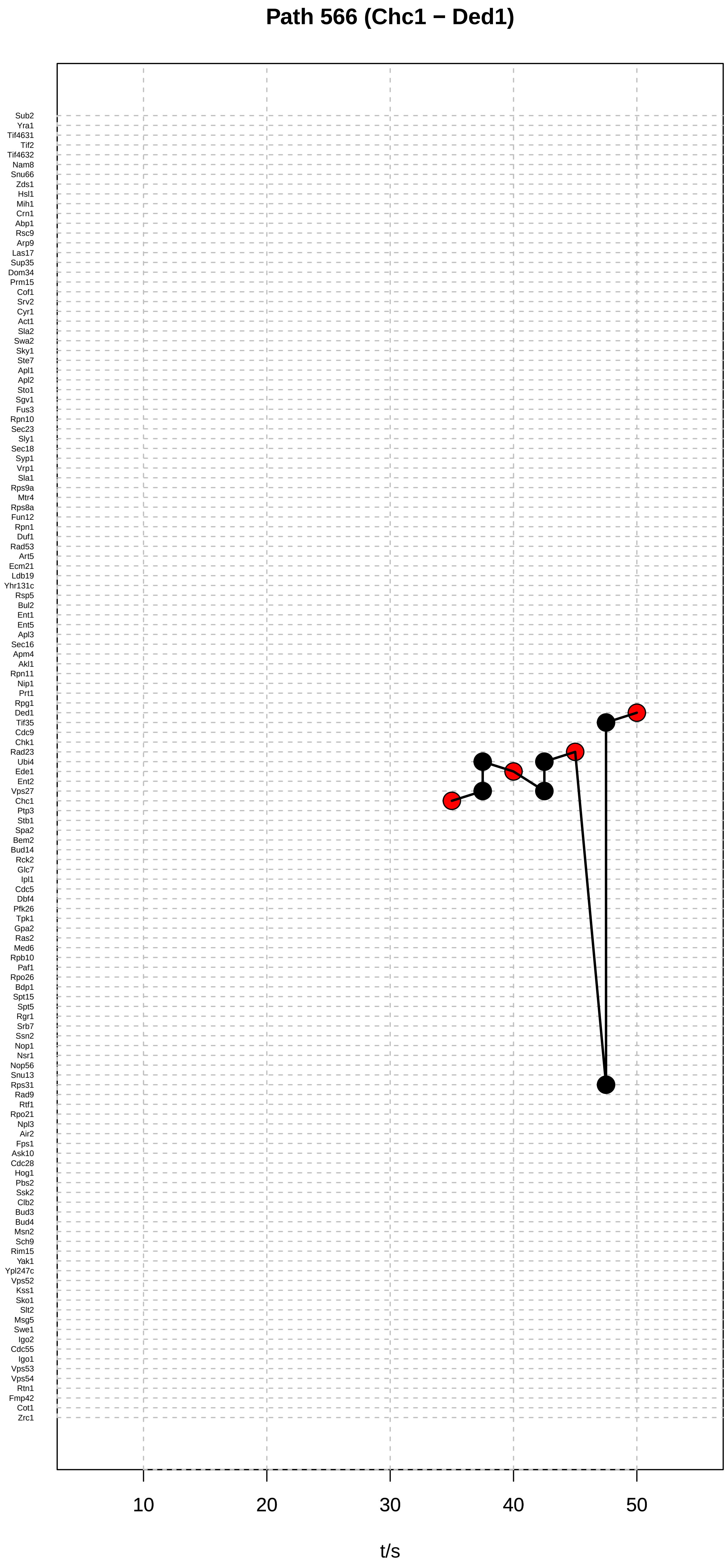

Path 566 (Chc1 − Ded1)
Sub2
Yra1
Tif4631
Tif2
Tif4632
Nam8
Snu66
Zds1
Hsl1
Mih1
Crn1
Abp1
Rsc9
Arp9
Las17
Sup35
Dom34
Prm15
Cof1
Srv2
Cyr1
Act1
Sla2
Swa2
Sky1
Ste7
Apl1
Apl2
Sto1
Sgv1
Fus3
Rpn10
Sec23
Sly1
Sec18
Syp1
Vrp1
Sla1
Rps9a
Mtr4
Rps8a
Fun12
Rpn1
Duf1
Rad53
Art5
Ecm21
Ldb19
Yhr131c
Rsp5
Bul2
Ent1
Ent5
Apl3
Sec16
Apm4
Akl1
Rpn11
Nip1
Prt1
Rpg1
Ded1
Tif35
Cdc9
Chk1
Rad23
Ubi4
Ede1
Ent2
Vps27
Chc1
Ptp3
Stb1
Spa2
Bem2
Bud14
Rck2
Glc7
Ipl1
Cdc5
Dbf4
Pfk26
Tpk1
Gpa2
Ras2
Med6
Rpb10
Paf1
Rpo26
Bdp1
Spt15
Spt5
Rgr1
Srb7
Ssn2
Nop1
Nsr1
Nop56
Snu13
Rps31
Rad9
Rtf1
Rpo21
Npl3
Air2
Fps1
Ask10
Cdc28
Hog1
Pbs2
Ssk2
Clb2
Bud3
Bud4
Msn2
Sch9
Rim15
Yak1
Ypl247c
Vps52
Kss1
Sko1
Slt2
Msg5
Swe1
Igo2
Cdc55
Igo1
Vps53
Vps54
Rtn1
Fmp42
Cot1
Zrc1
10
20
30
40
50
t/s

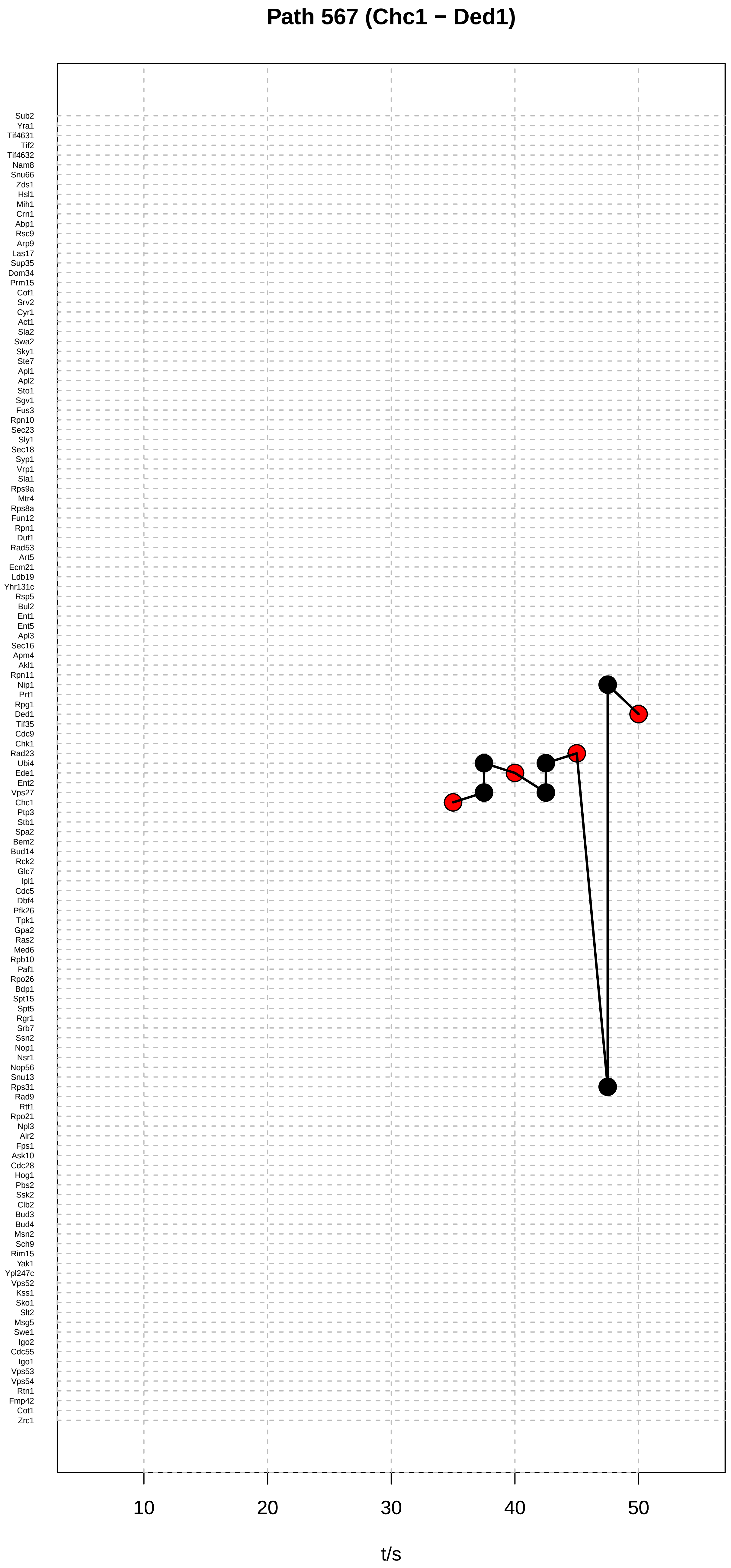

Path 567 (Chc1 − Ded1)
Sub2
Yra1
Tif4631
Tif2
Tif4632
Nam8
Snu66
Zds1
Hsl1
Mih1
Crn1
Abp1
Rsc9
Arp9
Las17
Sup35
Dom34
Prm15
Cof1
Srv2
Cyr1
Act1
Sla2
Swa2
Sky1
Ste7
Apl1
Apl2
Sto1
Sgv1
Fus3
Rpn10
Sec23
Sly1
Sec18
Syp1
Vrp1
Sla1
Rps9a
Mtr4
Rps8a
Fun12
Rpn1
Duf1
Rad53
Art5
Ecm21
Ldb19
Yhr131c
Rsp5
Bul2
Ent1
Ent5
Apl3
Sec16
Apm4
Akl1
Rpn11
Nip1
Prt1
Rpg1
Ded1
Tif35
Cdc9
Chk1
Rad23
Ubi4
Ede1
Ent2
Vps27
Chc1
Ptp3
Stb1
Spa2
Bem2
Bud14
Rck2
Glc7
Ipl1
Cdc5
Dbf4
Pfk26
Tpk1
Gpa2
Ras2
Med6
Rpb10
Paf1
Rpo26
Bdp1
Spt15
Spt5
Rgr1
Srb7
Ssn2
Nop1
Nsr1
Nop56
Snu13
Rps31
Rad9
Rtf1
Rpo21
Npl3
Air2
Fps1
Ask10
Cdc28
Hog1
Pbs2
Ssk2
Clb2
Bud3
Bud4
Msn2
Sch9
Rim15
Yak1
Ypl247c
Vps52
Kss1
Sko1
Slt2
Msg5
Swe1
Igo2
Cdc55
Igo1
Vps53
Vps54
Rtn1
Fmp42
Cot1
Zrc1
10
20
30
40
50
t/s

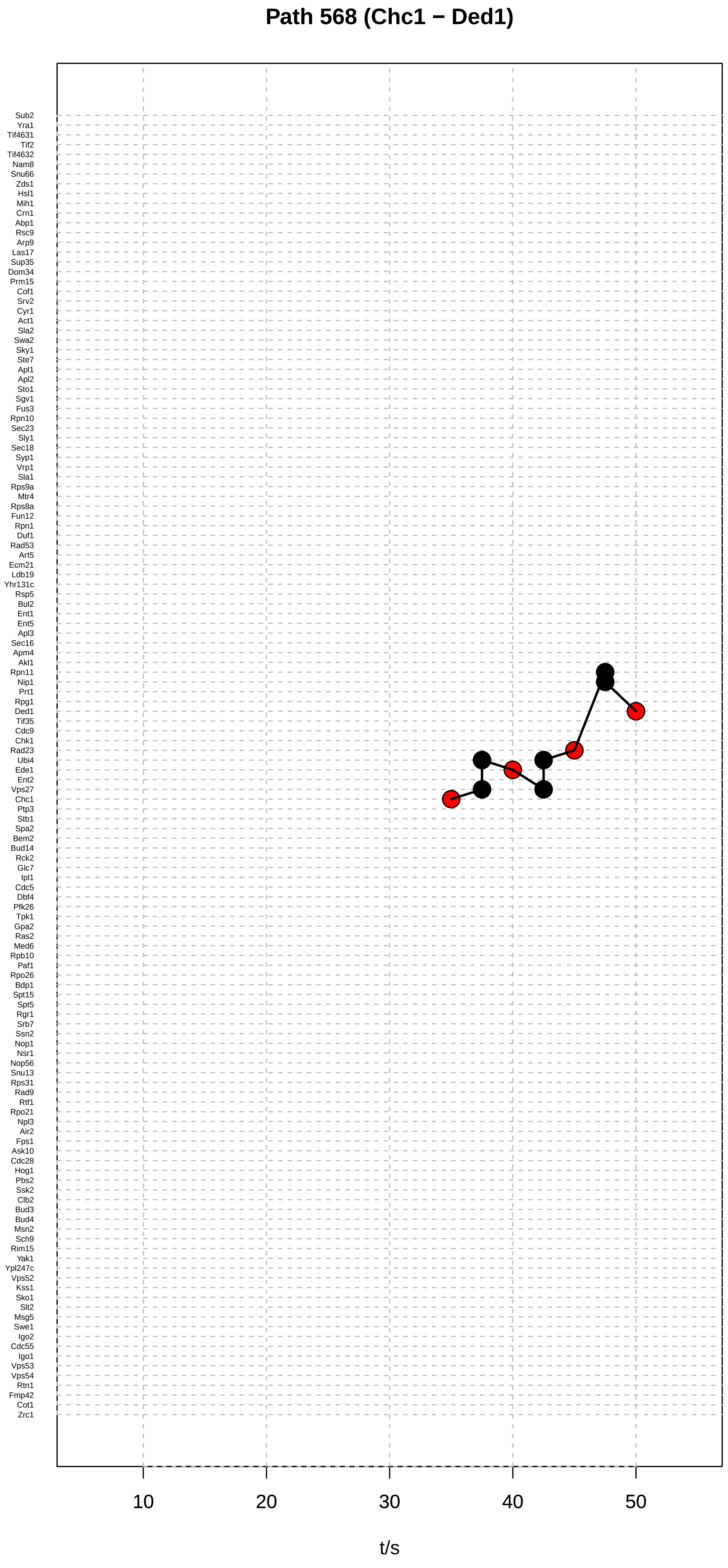
Path 568 (Chc1 − Ded1)
Sub2
Yra1
Tif4631
Tif2
Tif4632
Nam8
Snu66
Zds1
Hsl1
Mih1
Crn1
Abp1
Rsc9
Arp9
Las17
Sup35
Dom34
Prm15
Cof1
Srv2
Cyr1
Act1
Sla2
Swa2
Sky1
Ste7
Apl1
Apl2
Sto1
Sgv1
Fus3
Rpn10
Sec23
Sly1
Sec18
Syp1
Vrp1
Sla1
Rps9a
Mtr4
Rps8a
Fun12
Rpn1
Duf1
Rad53
Art5
Ecm21
Ldb19
Yhr131c
Rsp5
Bul2
Ent1
Ent5
Apl3
Sec16
Apm4
Akl1
Rpn11
Nip1
Prt1
Rpg1
Ded1
Tif35
Cdc9
Chk1
Rad23
Ubi4
Ede1
Ent2
Vps27
Chc1
Ptp3
Stb1
Spa2
Bem2
Bud14
Rck2
Glc7
Ipl1
Cdc5
Dbf4
Pfk26
Tpk1
Gpa2
Ras2
Med6
Rpb10
Paf1
Rpo26
Bdp1
Spt15
Spt5
Rgr1
Srb7
Ssn2
Nop1
Nsr1
Nop56
Snu13
Rps31
Rad9
Rtf1
Rpo21
Npl3
Air2
Fps1
Ask10
Cdc28
Hog1
Pbs2
Ssk2
Clb2
Bud3
Bud4
Msn2
Sch9
Rim15
Yak1
Ypl247c
Vps52
Kss1
Sko1
Slt2
Msg5
Swe1
Igo2
Cdc55
Igo1
Vps53
Vps54
Rtn1
Fmp42
Cot1
Zrc1
10
20
30
40
50
t/s

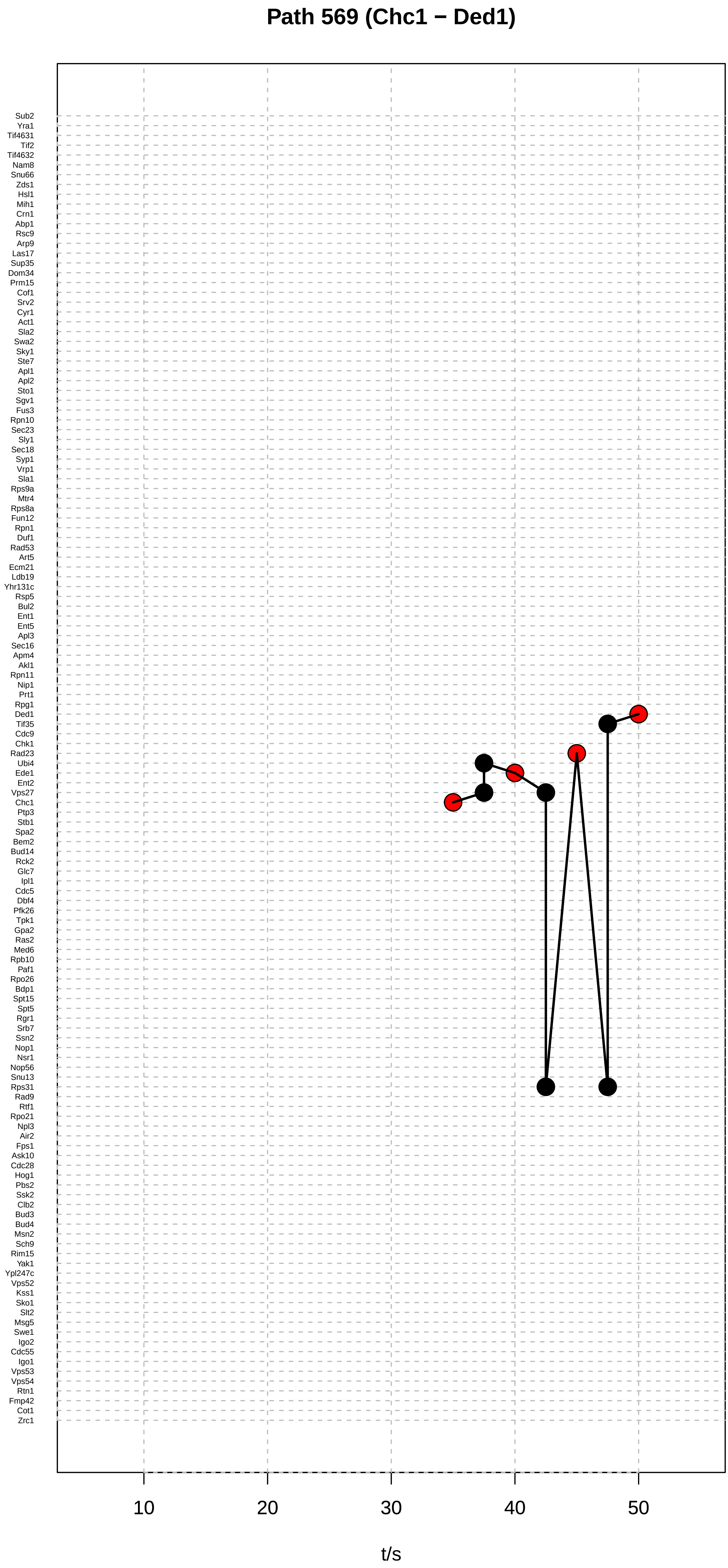

Path 569 (Chc1 − Ded1)
Sub2
Yra1
Tif4631
Tif2
Tif4632
Nam8
Snu66
Zds1
Hsl1
Mih1
Crn1
Abp1
Rsc9
Arp9
Las17
Sup35
Dom34
Prm15
Cof1
Srv2
Cyr1
Act1
Sla2
Swa2
Sky1
Ste7
Apl1
Apl2
Sto1
Sgv1
Fus3
Rpn10
Sec23
Sly1
Sec18
Syp1
Vrp1
Sla1
Rps9a
Mtr4
Rps8a
Fun12
Rpn1
Duf1
Rad53
Art5
Ecm21
Ldb19
Yhr131c
Rsp5
Bul2
Ent1
Ent5
Apl3
Sec16
Apm4
Akl1
Rpn11
Nip1
Prt1
Rpg1
Ded1
Tif35
Cdc9
Chk1
Rad23
Ubi4
Ede1
Ent2
Vps27
Chc1
Ptp3
Stb1
Spa2
Bem2
Bud14
Rck2
Glc7
Ipl1
Cdc5
Dbf4
Pfk26
Tpk1
Gpa2
Ras2
Med6
Rpb10
Paf1
Rpo26
Bdp1
Spt15
Spt5
Rgr1
Srb7
Ssn2
Nop1
Nsr1
Nop56
Snu13
Rps31
Rad9
Rtf1
Rpo21
Npl3
Air2
Fps1
Ask10
Cdc28
Hog1
Pbs2
Ssk2
Clb2
Bud3
Bud4
Msn2
Sch9
Rim15
Yak1
Ypl247c
Vps52
Kss1
Sko1
Slt2
Msg5
Swe1
Igo2
Cdc55
Igo1
Vps53
Vps54
Rtn1
Fmp42
Cot1
Zrc1
10
20
30
40
50
t/s

### Path 570 (Chc1 − Ded1)

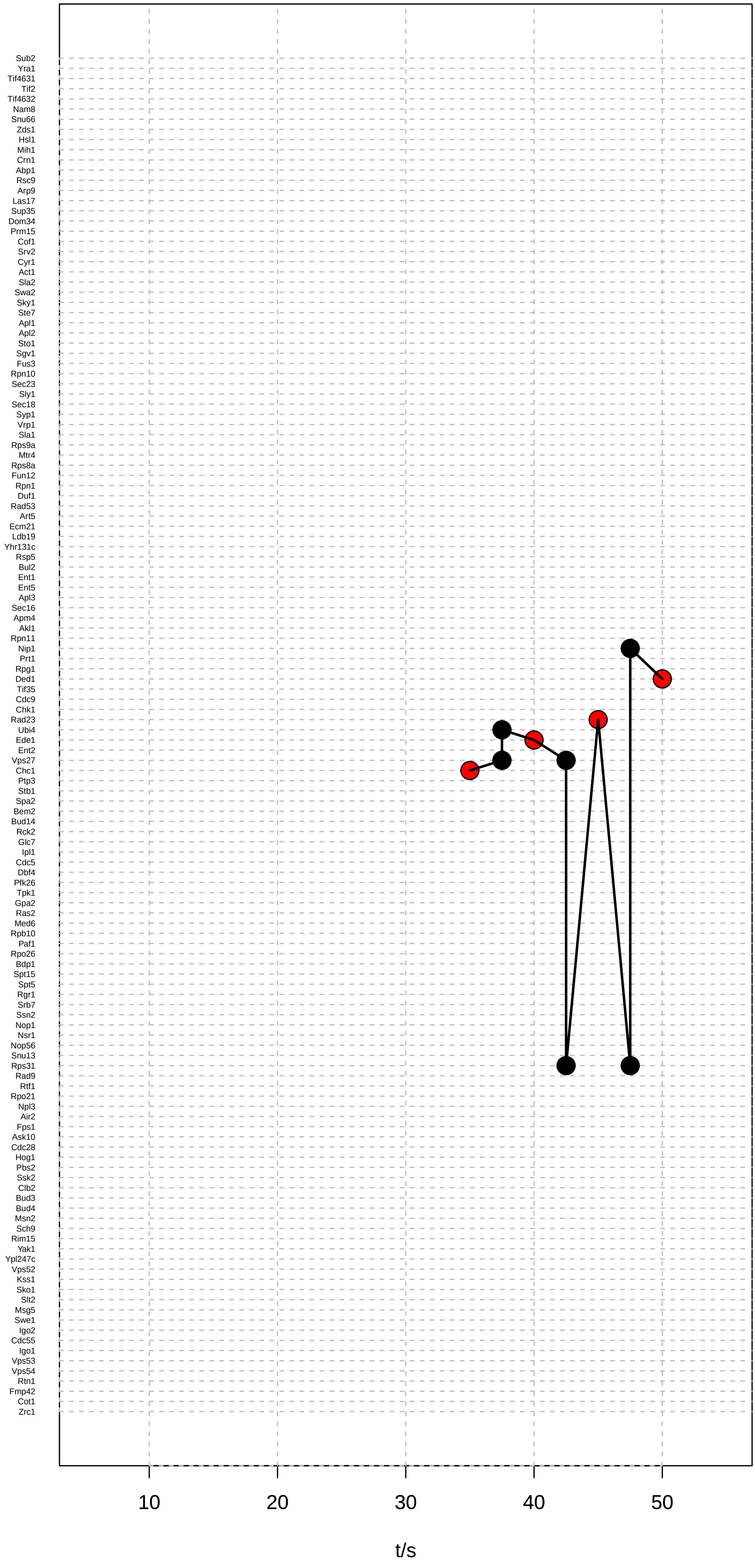

#### Path 571 (Chc1 − Ded1)

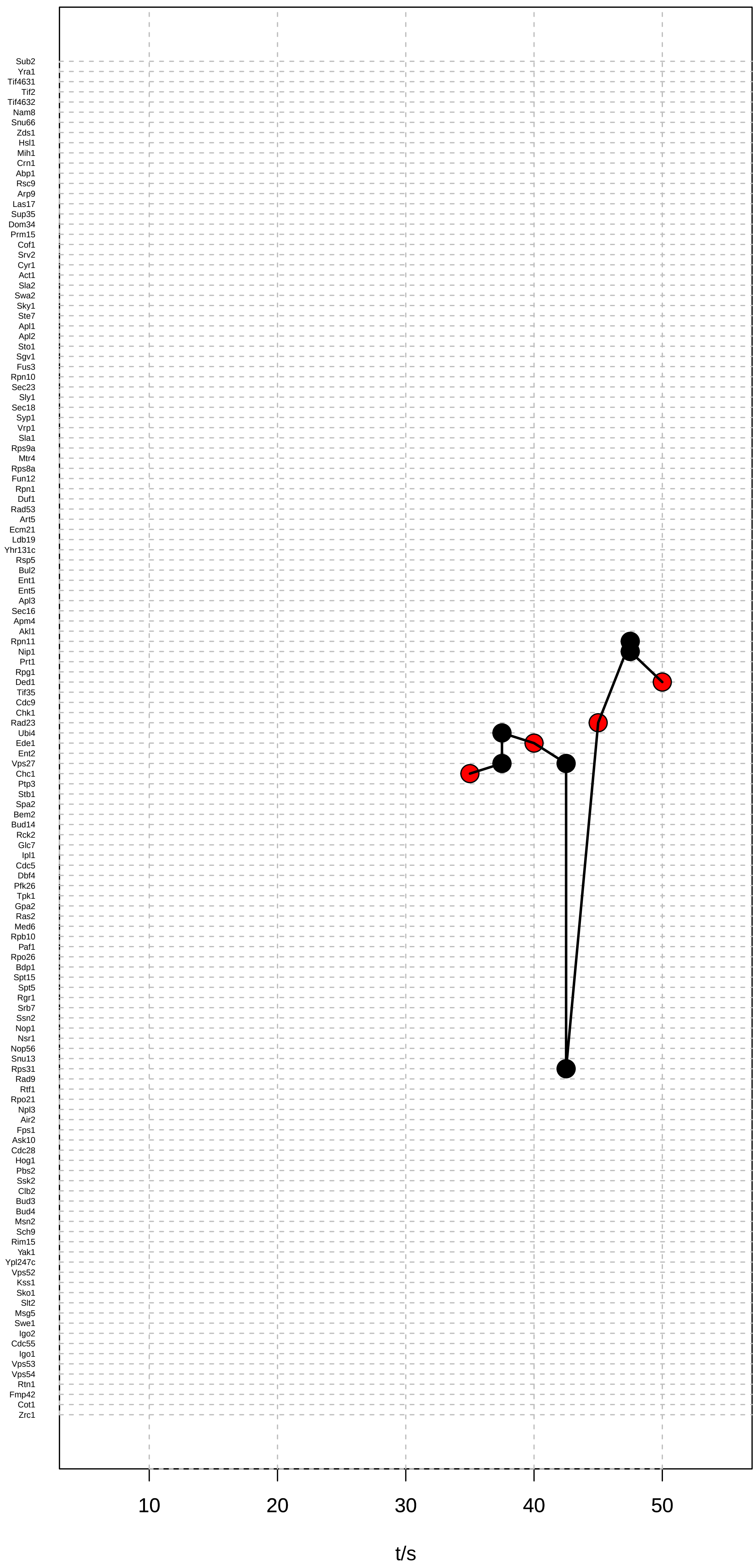

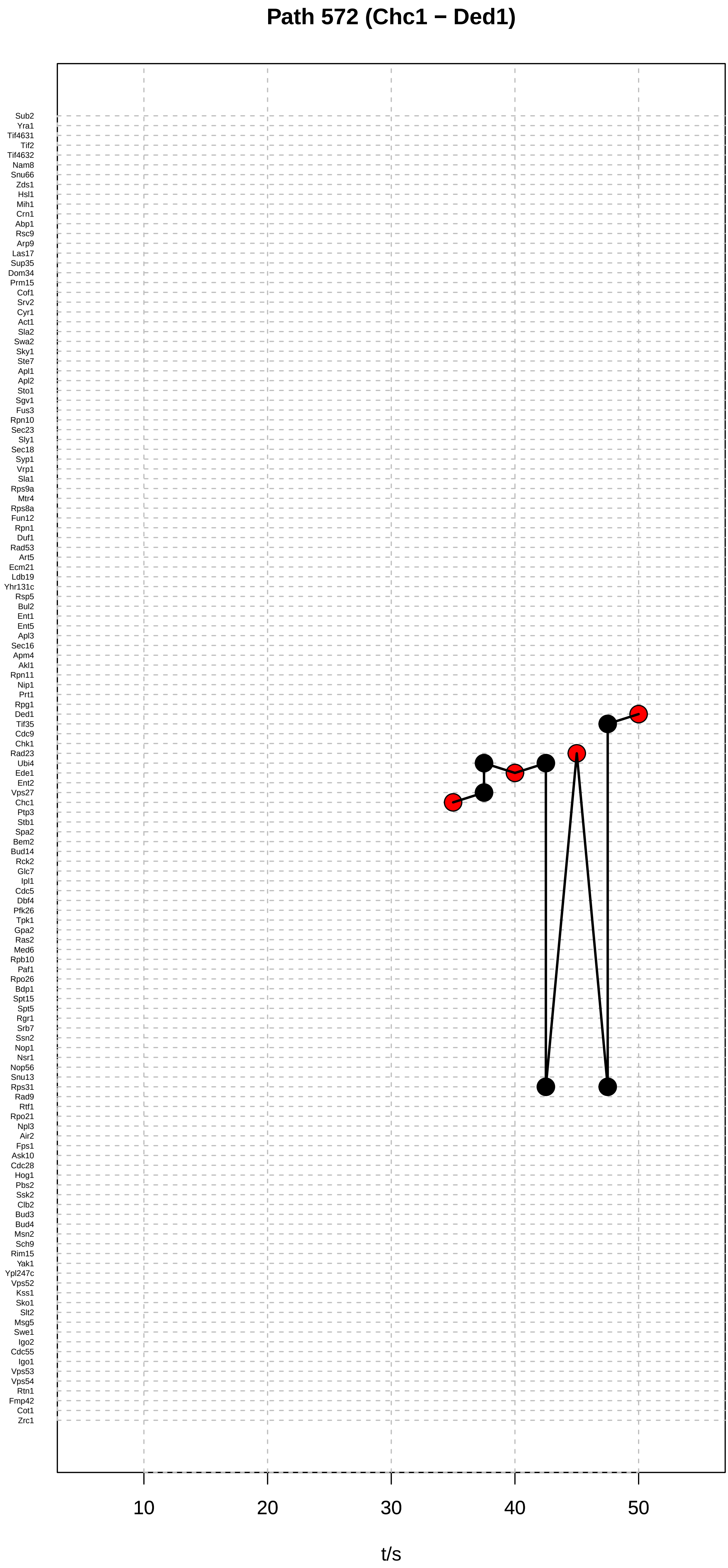

Path 572 (Chc1 − Ded1)
Sub2
Yra1
Tif4631
Tif2
Tif4632
Nam8
Snu66
Zds1
Hsl1
Mih1
Crn1
Abp1
Rsc9
Arp9
Las17
Sup35
Dom34
Prm15
Cof1
Srv2
Cyr1
Act1
Sla2
Swa2
Sky1
Ste7
Apl1
Apl2
Sto1
Sgv1
Fus3
Rpn10
Sec23
Sly1
Sec18
Syp1
Vrp1
Sla1
Rps9a
Mtr4
Rps8a
Fun12
Rpn1
Duf1
Rad53
Art5
Ecm21
Ldb19
Yhr131c
Rsp5
Bul2
Ent1
Ent5
Apl3
Sec16
Apm4
Akl1
Rpn11
Nip1
Prt1
Rpg1
Ded1
Tif35
Cdc9
Chk1
Rad23
Ubi4
Ede1
Ent2
Vps27
Chc1
Ptp3
Stb1
Spa2
Bem2
Bud14
Rck2
Glc7
Ipl1
Cdc5
Dbf4
Pfk26
Tpk1
Gpa2
Ras2
Med6
Rpb10
Paf1
Rpo26
Bdp1
Spt15
Spt5
Rgr1
Srb7
Ssn2
Nop1
Nsr1
Nop56
Snu13
Rps31
Rad9
Rtf1
Rpo21
Npl3
Air2
Fps1
Ask10
Cdc28
Hog1
Pbs2
Ssk2
Clb2
Bud3
Bud4
Msn2
Sch9
Rim15
Yak1
Ypl247c
Vps52
Kss1
Sko1
Slt2
Msg5
Swe1
Igo2
Cdc55
Igo1
Vps53
Vps54
Rtn1
Fmp42
Cot1
Zrc1
10
20
30
40
50
t/s

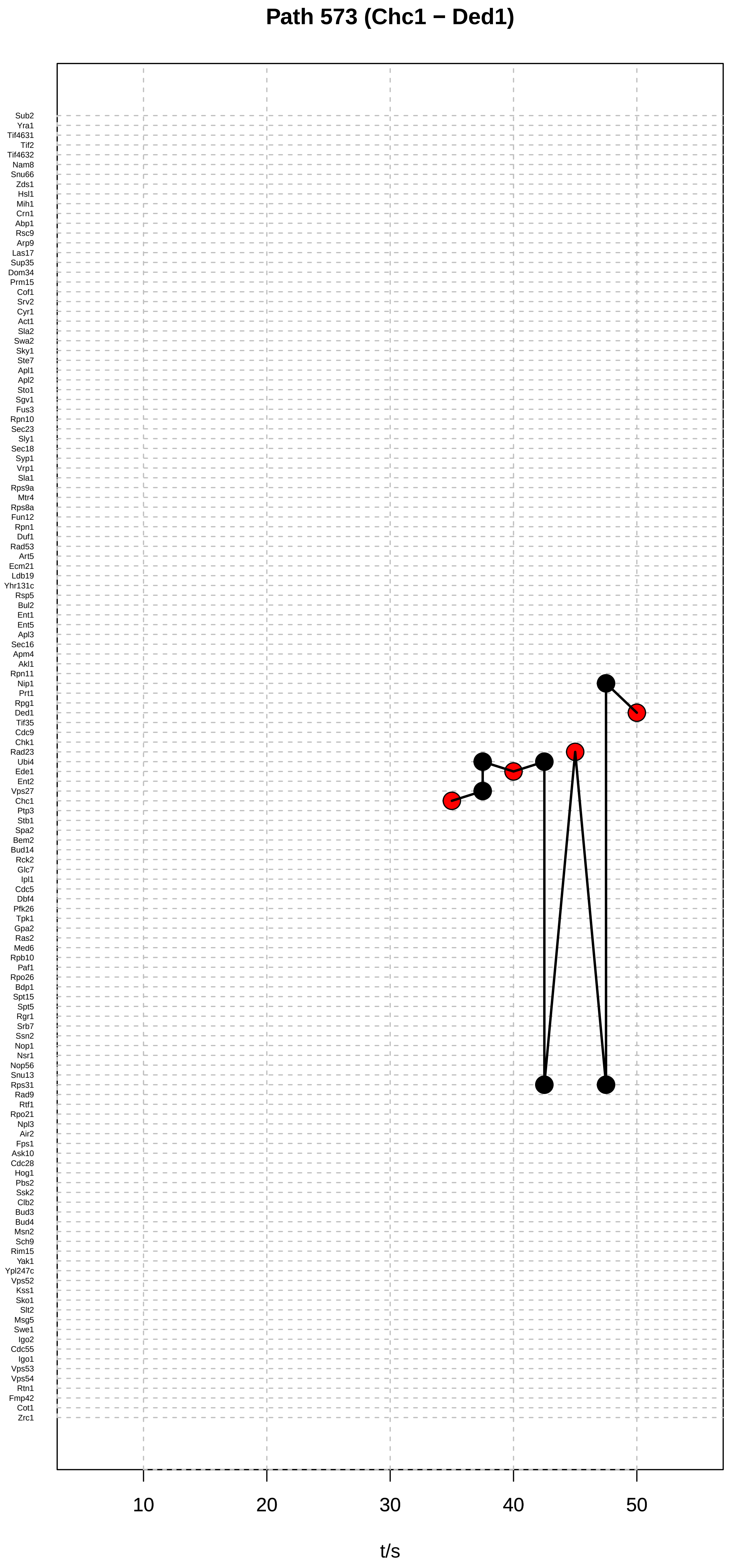

Path 573 (Chc1 − Ded1)
Sub2
Yra1
Tif4631
Tif2
Tif4632
Nam8
Snu66
Zds1
Hsl1
Mih1
Crn1
Abp1
Rsc9
Arp9
Las17
Sup35
Dom34
Prm15
Cof1
Srv2
Cyr1
Act1
Sla2
Swa2
Sky1
Ste7
Apl1
Apl2
Sto1
Sgv1
Fus3
Rpn10
Sec23
Sly1
Sec18
Syp1
Vrp1
Sla1
Rps9a
Mtr4
Rps8a
Fun12
Rpn1
Duf1
Rad53
Art5
Ecm21
Ldb19
Yhr131c
Rsp5
Bul2
Ent1
Ent5
Apl3
Sec16
Apm4
Akl1
Rpn11
Nip1
Prt1
Rpg1
Ded1
Tif35
Cdc9
Chk1
Rad23
Ubi4
Ede1
Ent2
Vps27
Chc1
Ptp3
Stb1
Spa2
Bem2
Bud14
Rck2
Glc7
Ipl1
Cdc5
Dbf4
Pfk26
Tpk1
Gpa2
Ras2
Med6
Rpb10
Paf1
Rpo26
Bdp1
Spt15
Spt5
Rgr1
Srb7
Ssn2
Nop1
Nsr1
Nop56
Snu13
Rps31
Rad9
Rtf1
Rpo21
Npl3
Air2
Fps1
Ask10
Cdc28
Hog1
Pbs2
Ssk2
Clb2
Bud3
Bud4
Msn2
Sch9
Rim15
Yak1
Ypl247c
Vps52
Kss1
Sko1
Slt2
Msg5
Swe1
Igo2
Cdc55
Igo1
Vps53
Vps54
Rtn1
Fmp42
Cot1
Zrc1
10
20
30
40
50
t/s

### Path 574 (Chc1 − Ded1)

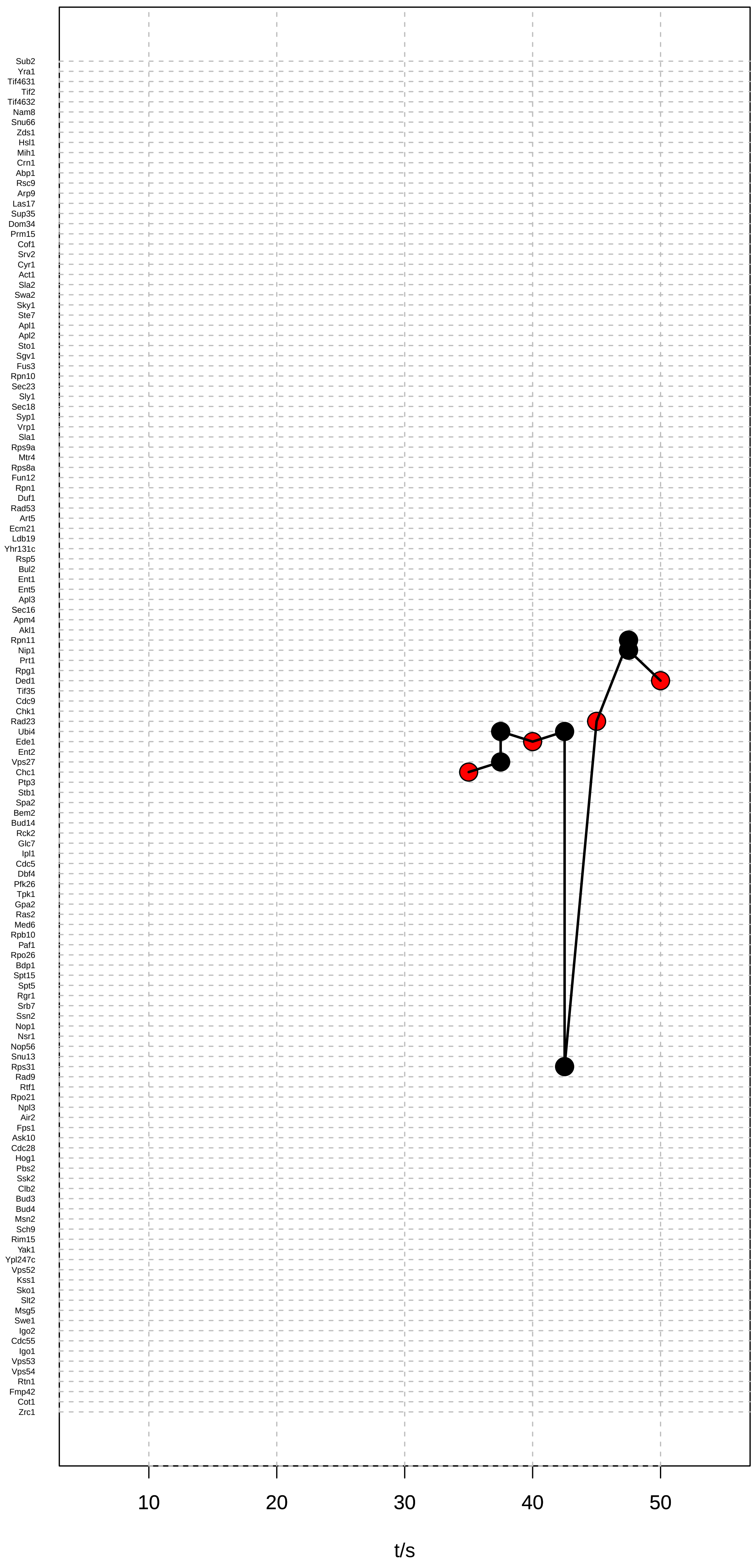

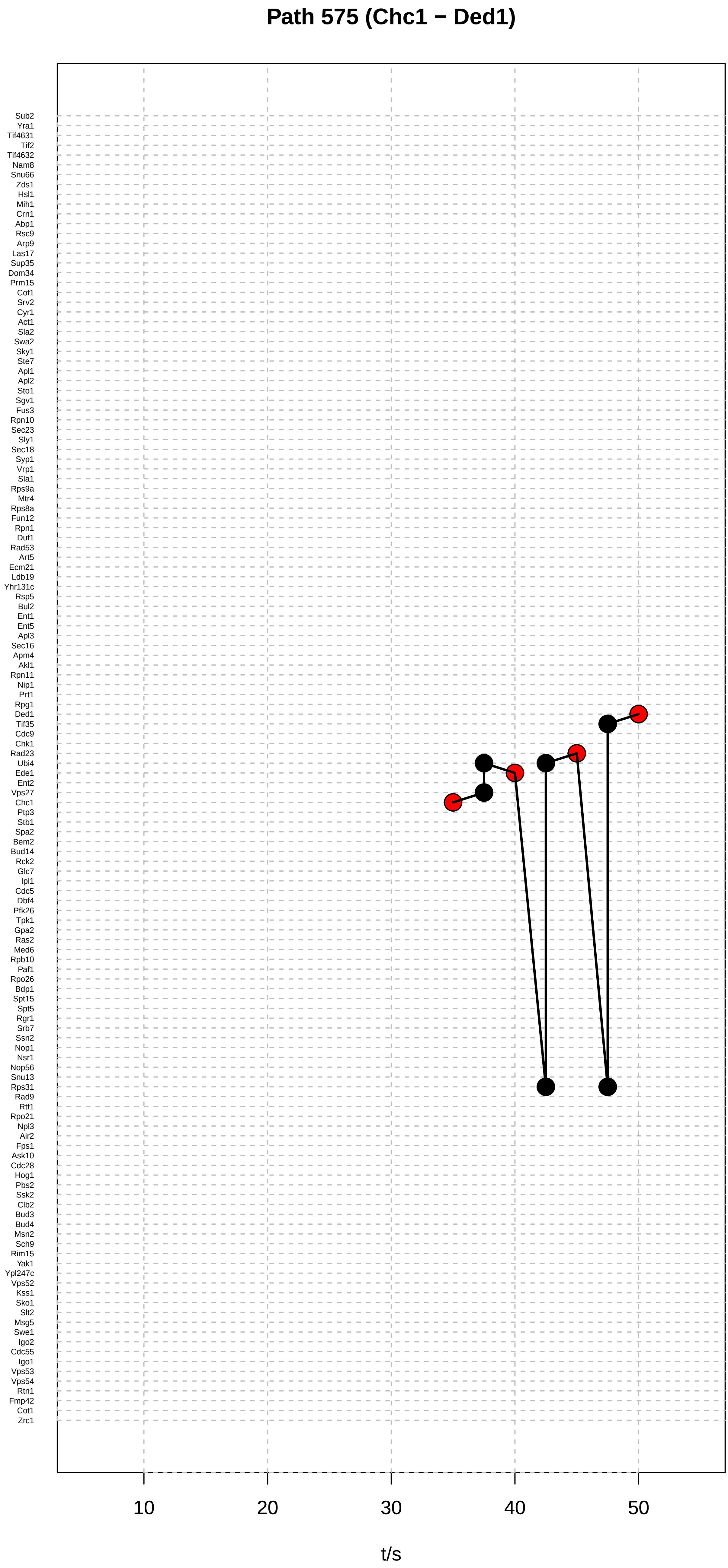
Path 575 (Chc1 − Ded1)
Sub2
Yra1
Tif4631
Tif2
Tif4632
Nam8
Snu66
Zds1
Hsl1
Mih1
Crn1
Abp1
Rsc9
Arp9
Las17
Sup35
Dom34
Prm15
Cof1
Srv2
Cyr1
Act1
Sla2
Swa2
Sky1
Ste7
Apl1
Apl2
Sto1
Sgv1
Fus3
Rpn10
Sec23
Sly1
Sec18
Syp1
Vrp1
Sla1
Rps9a
Mtr4
Rps8a
Fun12
Rpn1
Duf1
Rad53
Art5
Ecm21
Ldb19
Yhr131c
Rsp5
Bul2
Ent1
Ent5
Apl3
Sec16
Apm4
Akl1
Rpn11
Nip1
Prt1
Rpg1
Ded1
Tif35
Cdc9
Chk1
Rad23
Ubi4
Ede1
Ent2
Vps27
Chc1
Ptp3
Stb1
Spa2
Bem2
Bud14
Rck2
Glc7
Ipl1
Cdc5
Dbf4
Pfk26
Tpk1
Gpa2
Ras2
Med6
Rpb10
Paf1
Rpo26
Bdp1
Spt15
Spt5
Rgr1
Srb7
Ssn2
Nop1
Nsr1
Nop56
Snu13
Rps31
Rad9
Rtf1
Rpo21
Npl3
Air2
Fps1
Ask10
Cdc28
Hog1
Pbs2
Ssk2
Clb2
Bud3
Bud4
Msn2
Sch9
Rim15
Yak1
Ypl247c
Vps52
Kss1
Sko1
Slt2
Msg5
Swe1
Igo2
Cdc55
Igo1
Vps53
Vps54
Rtn1
Fmp42
Cot1
Zrc1
10
20
30
40
50
t/s

### Path 576 (Chc1 − Ded1)

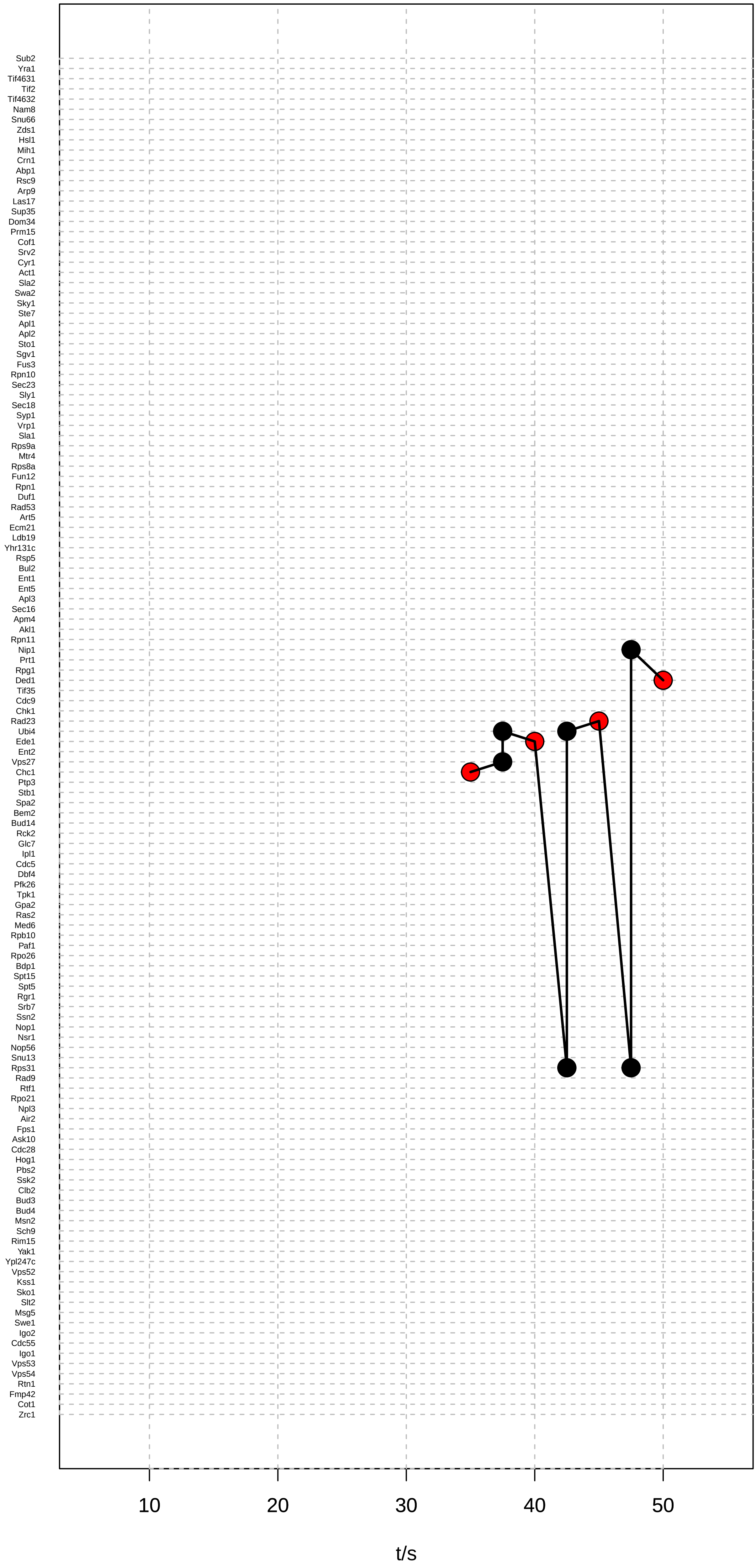

### Path 577 (Chc1 − Ded1)

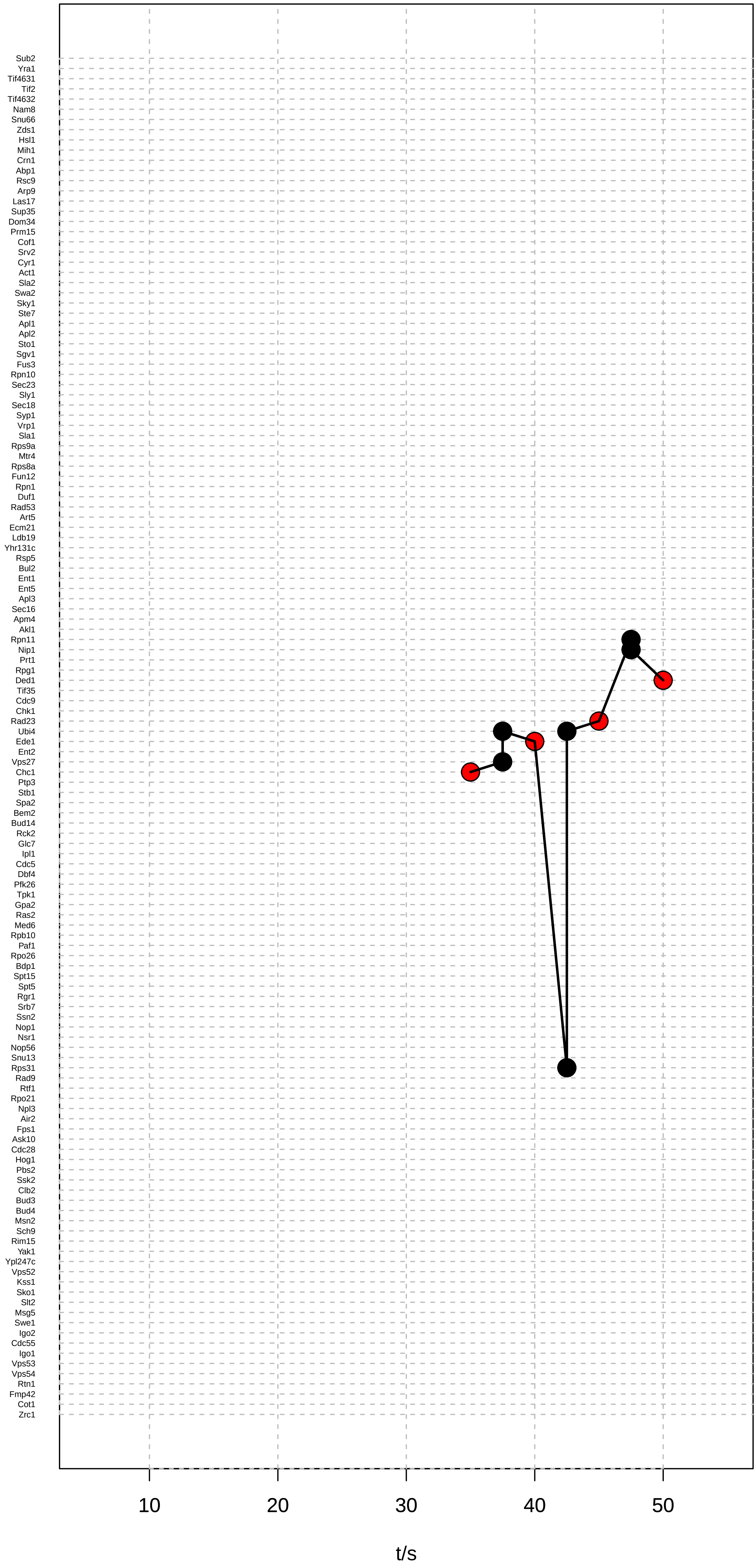

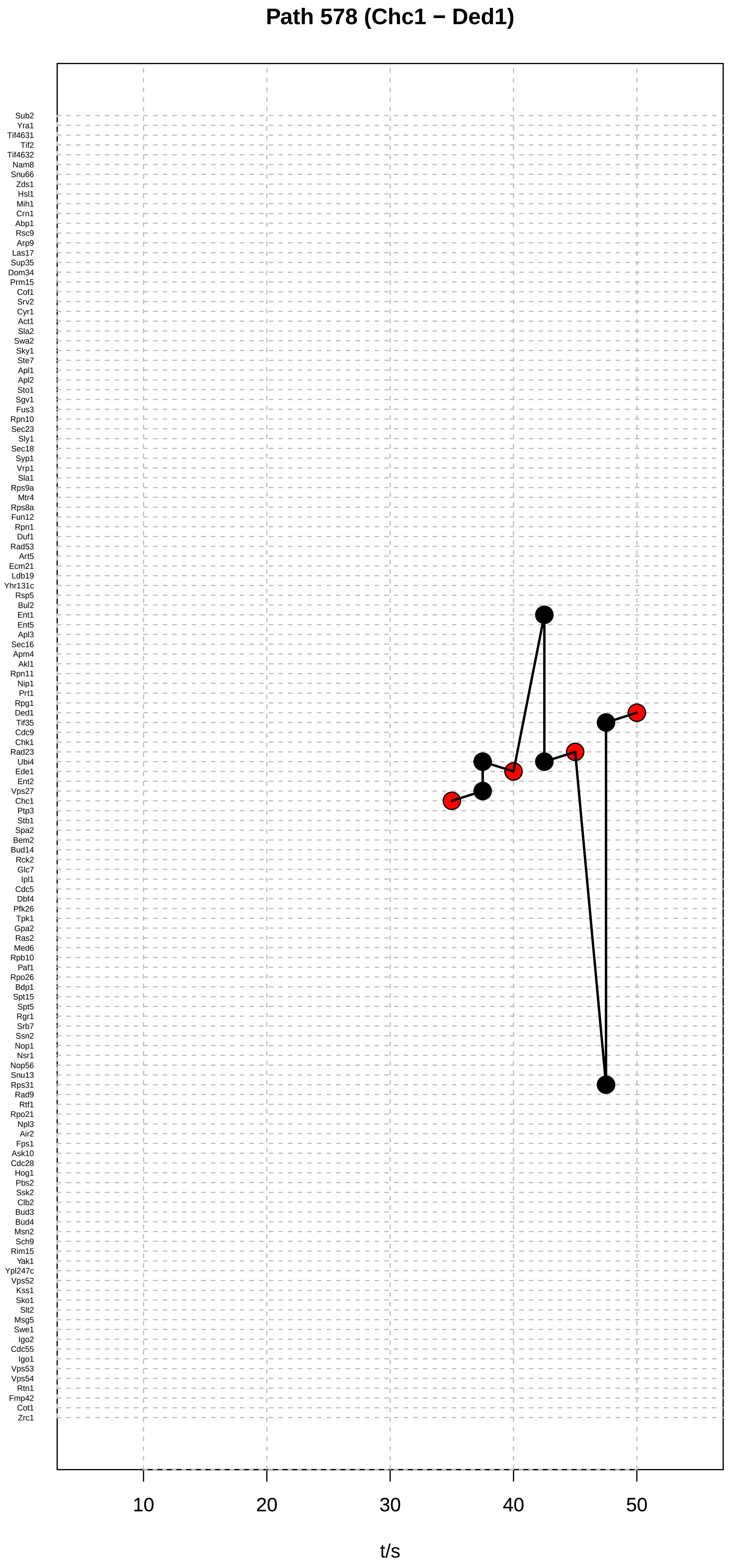

Path 578 (Chc1 − Ded1)
Sub2
Yra1
Tif4631
Tif2
Tif4632
Nam8
Snu66
Zds1
Hsl1
Mih1
Crn1
Abp1
Rsc9
Arp9
Las17
Sup35
Dom34
Prm15
Cof1
Srv2
Cyr1
Act1
Sla2
Swa2
Sky1
Ste7
Apl1
Apl2
Sto1
Sgv1
Fus3
Rpn10
Sec23
Sly1
Sec18
Syp1
Vrp1
Sla1
Rps9a
Mtr4
Rps8a
Fun12
Rpn1
Duf1
Rad53
Art5
Ecm21
Ldb19
Yhr131c
Rsp5
Bul2
Ent1
Ent5
Apl3
Sec16
Apm4
Akl1
Rpn11
Nip1
Prt1
Rpg1
Ded1
Tif35
Cdc9
Chk1
Rad23
Ubi4
Ede1
Ent2
Vps27
Chc1
Ptp3
Stb1
Spa2
Bem2
Bud14
Rck2
Glc7
Ipl1
Cdc5
Dbf4
Pfk26
Tpk1
Gpa2
Ras2
Med6
Rpb10
Paf1
Rpo26
Bdp1
Spt15
Spt5
Rgr1
Srb7
Ssn2
Nop1
Nsr1
Nop56
Snu13
Rps31
Rad9
Rtf1
Rpo21
Npl3
Air2
Fps1
Ask10
Cdc28
Hog1
Pbs2
Ssk2
Clb2
Bud3
Bud4
Msn2
Sch9
Rim15
Yak1
Ypl247c
Vps52
Kss1
Sko1
Slt2
Msg5
Swe1
Igo2
Cdc55
Igo1
Vps53
Vps54
Rtn1
Fmp42
Cot1
Zrc1
10
20
30
40
50
t/s

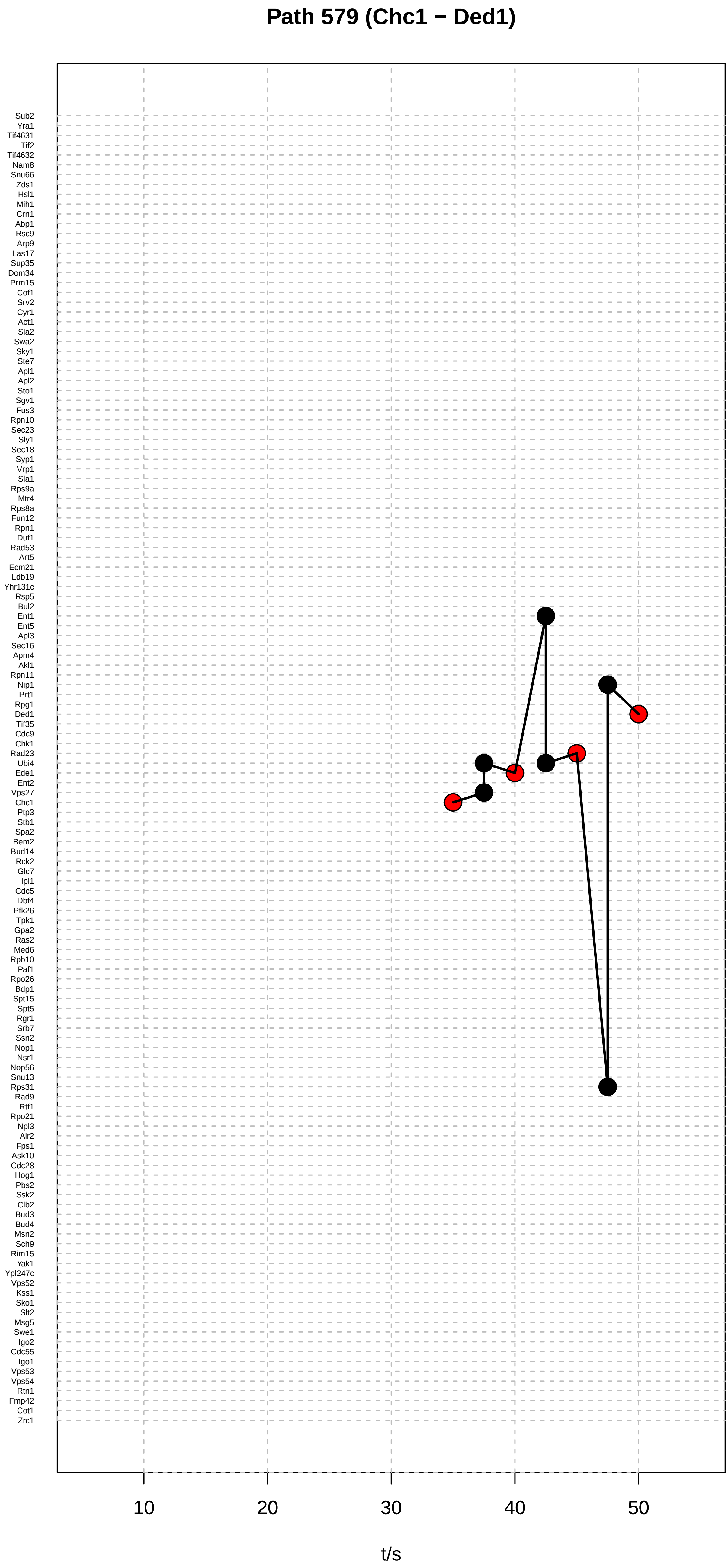
Path 579 (Chc1 − Ded1)
Sub2
Yra1
Tif4631
Tif2
Tif4632
Nam8
Snu66
Zds1
Hsl1
Mih1
Crn1
Abp1
Rsc9
Arp9
Las17
Sup35
Dom34
Prm15
Cof1
Srv2
Cyr1
Act1
Sla2
Swa2
Sky1
Ste7
Apl1
Apl2
Sto1
Sgv1
Fus3
Rpn10
Sec23
Sly1
Sec18
Syp1
Vrp1
Sla1
Rps9a
Mtr4
Rps8a
Fun12
Rpn1
Duf1
Rad53
Art5
Ecm21
Ldb19
Yhr131c
Rsp5
Bul2
Ent1
Ent5
Apl3
Sec16
Apm4
Akl1
Rpn11
Nip1
Prt1
Rpg1
Ded1
Tif35
Cdc9
Chk1
Rad23
Ubi4
Ede1
Ent2
Vps27
Chc1
Ptp3
Stb1
Spa2
Bem2
Bud14
Rck2
Glc7
Ipl1
Cdc5
Dbf4
Pfk26
Tpk1
Gpa2
Ras2
Med6
Rpb10
Paf1
Rpo26
Bdp1
Spt15
Spt5
Rgr1
Srb7
Ssn2
Nop1
Nsr1
Nop56
Snu13
Rps31
Rad9
Rtf1
Rpo21
Npl3
Air2
Fps1
Ask10
Cdc28
Hog1
Pbs2
Ssk2
Clb2
Bud3
Bud4
Msn2
Sch9
Rim15
Yak1
Ypl247c
Vps52
Kss1
Sko1
Slt2
Msg5
Swe1
Igo2
Cdc55
Igo1
Vps53
Vps54
Rtn1
Fmp42
Cot1
Zrc1
10
20
30
40
50
t/s

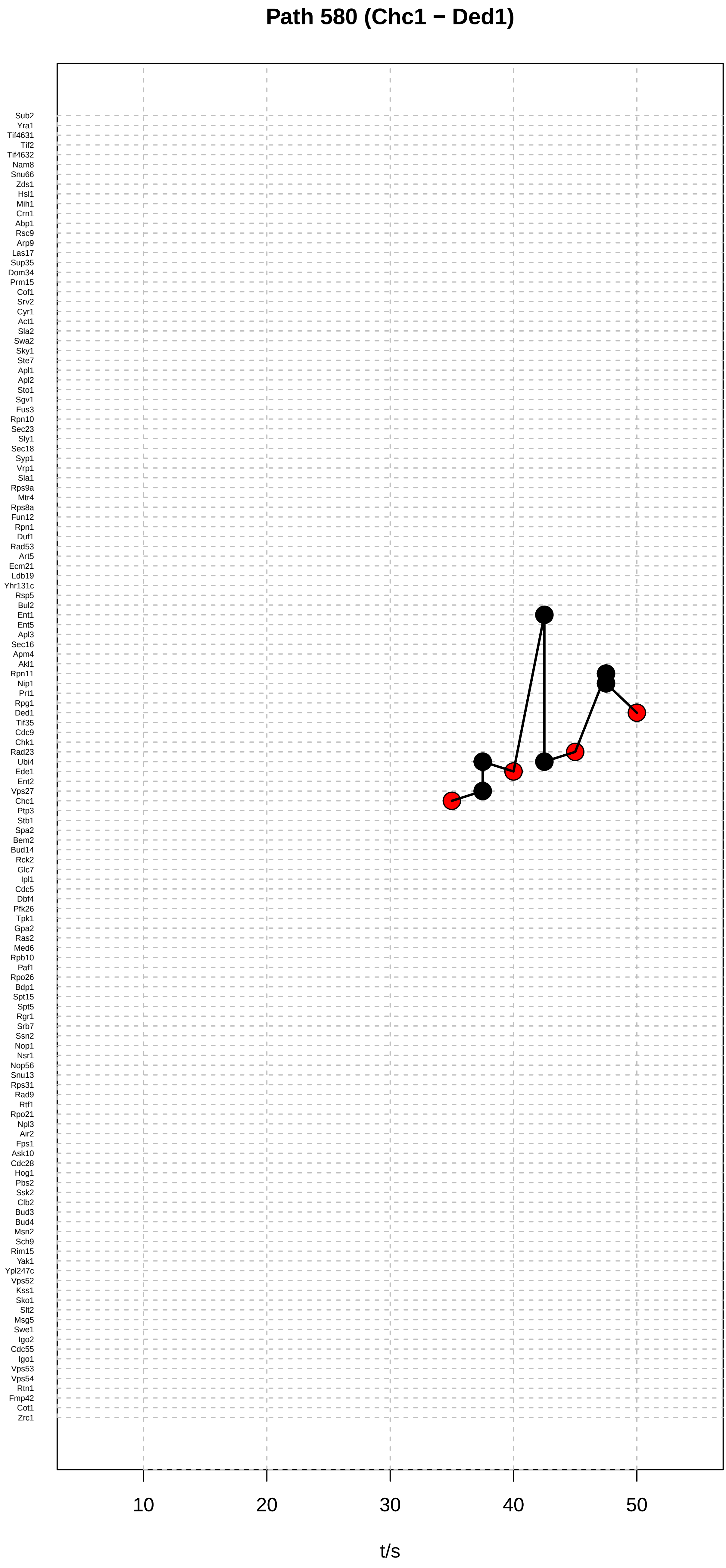

Path 580 (Chc1 − Ded1)
Sub2
Yra1
Tif4631
Tif2
Tif4632
Nam8
Snu66
Zds1
Hsl1
Mih1
Crn1
Abp1
Rsc9
Arp9
Las17
Sup35
Dom34
Prm15
Cof1
Srv2
Cyr1
Act1
Sla2
Swa2
Sky1
Ste7
Apl1
Apl2
Sto1
Sgv1
Fus3
Rpn10
Sec23
Sly1
Sec18
Syp1
Vrp1
Sla1
Rps9a
Mtr4
Rps8a
Fun12
Rpn1
Duf1
Rad53
Art5
Ecm21
Ldb19
Yhr131c
Rsp5
Bul2
Ent1
Ent5
Apl3
Sec16
Apm4
Akl1
Rpn11
Nip1
Prt1
Rpg1
Ded1
Tif35
Cdc9
Chk1
Rad23
Ubi4
Ede1
Ent2
Vps27
Chc1
Ptp3
Stb1
Spa2
Bem2
Bud14
Rck2
Glc7
Ipl1
Cdc5
Dbf4
Pfk26
Tpk1
Gpa2
Ras2
Med6
Rpb10
Paf1
Rpo26
Bdp1
Spt15
Spt5
Rgr1
Srb7
Ssn2
Nop1
Nsr1
Nop56
Snu13
Rps31
Rad9
Rtf1
Rpo21
Npl3
Air2
Fps1
Ask10
Cdc28
Hog1
Pbs2
Ssk2
Clb2
Bud3
Bud4
Msn2
Sch9
Rim15
Yak1
Ypl247c
Vps52
Kss1
Sko1
Slt2
Msg5
Swe1
Igo2
Cdc55
Igo1
Vps53
Vps54
Rtn1
Fmp42
Cot1
Zrc1
10
20
30
40
50
t/s

#### Path 581 (Chc1 − Ded1)

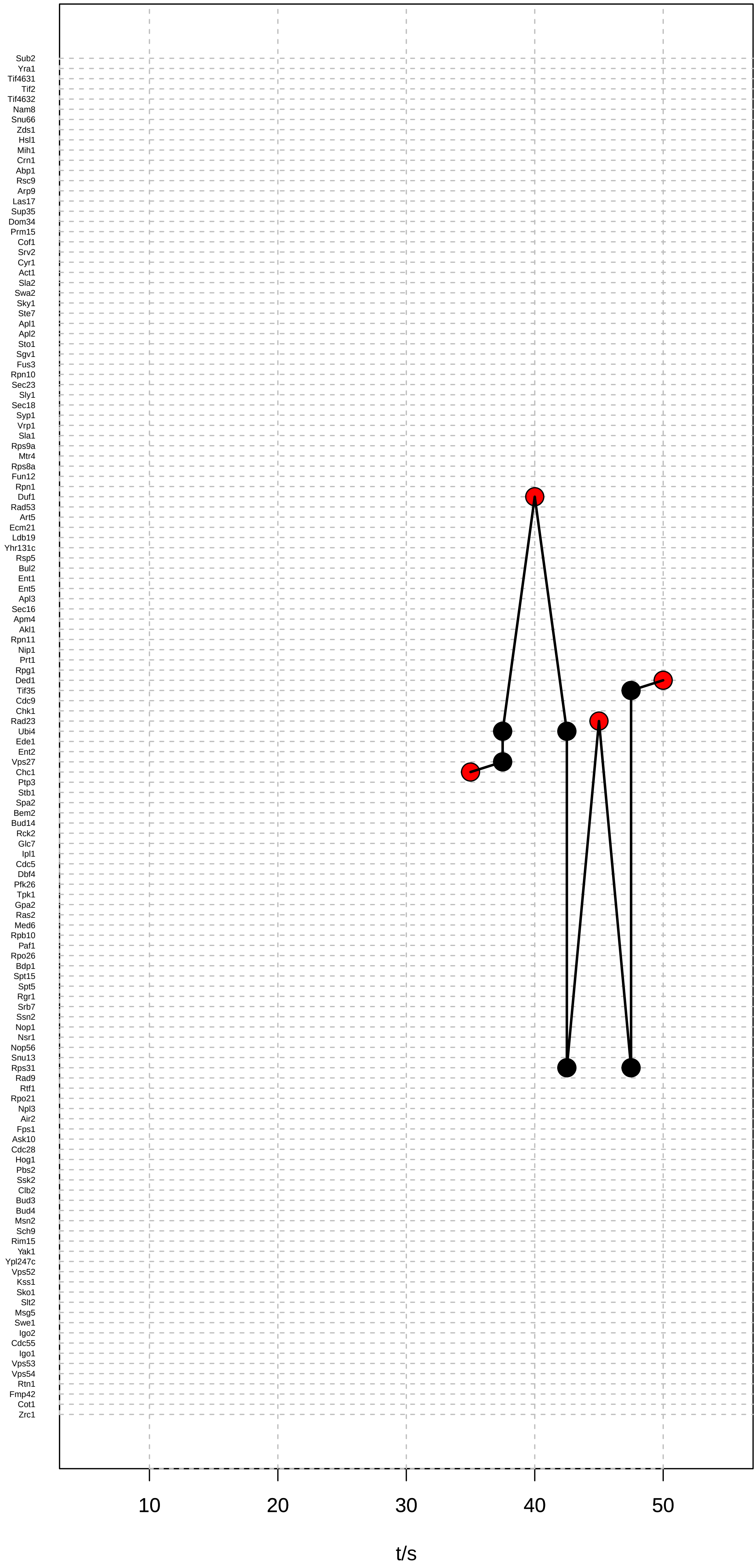

Path 582 (Chc1 − Ded1)

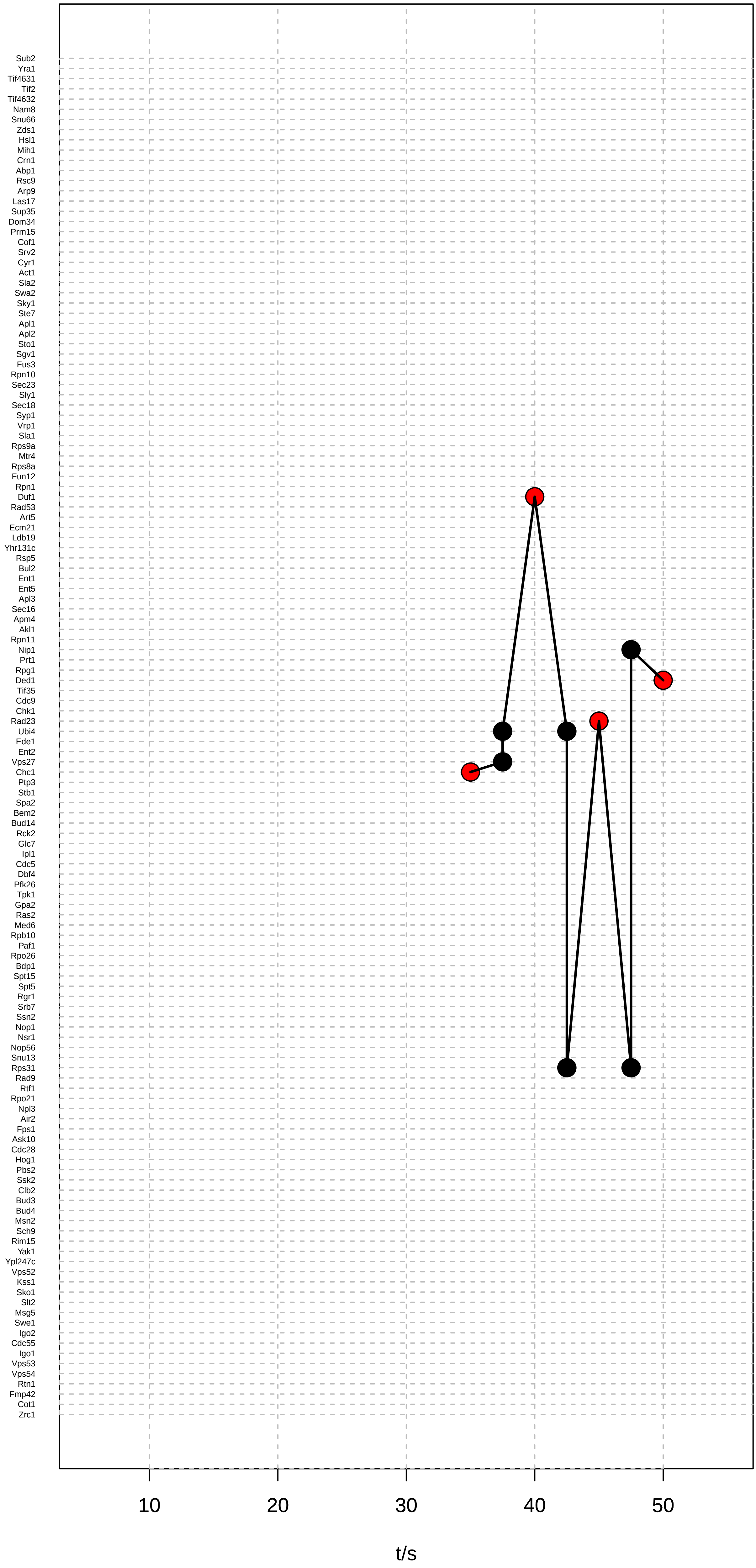

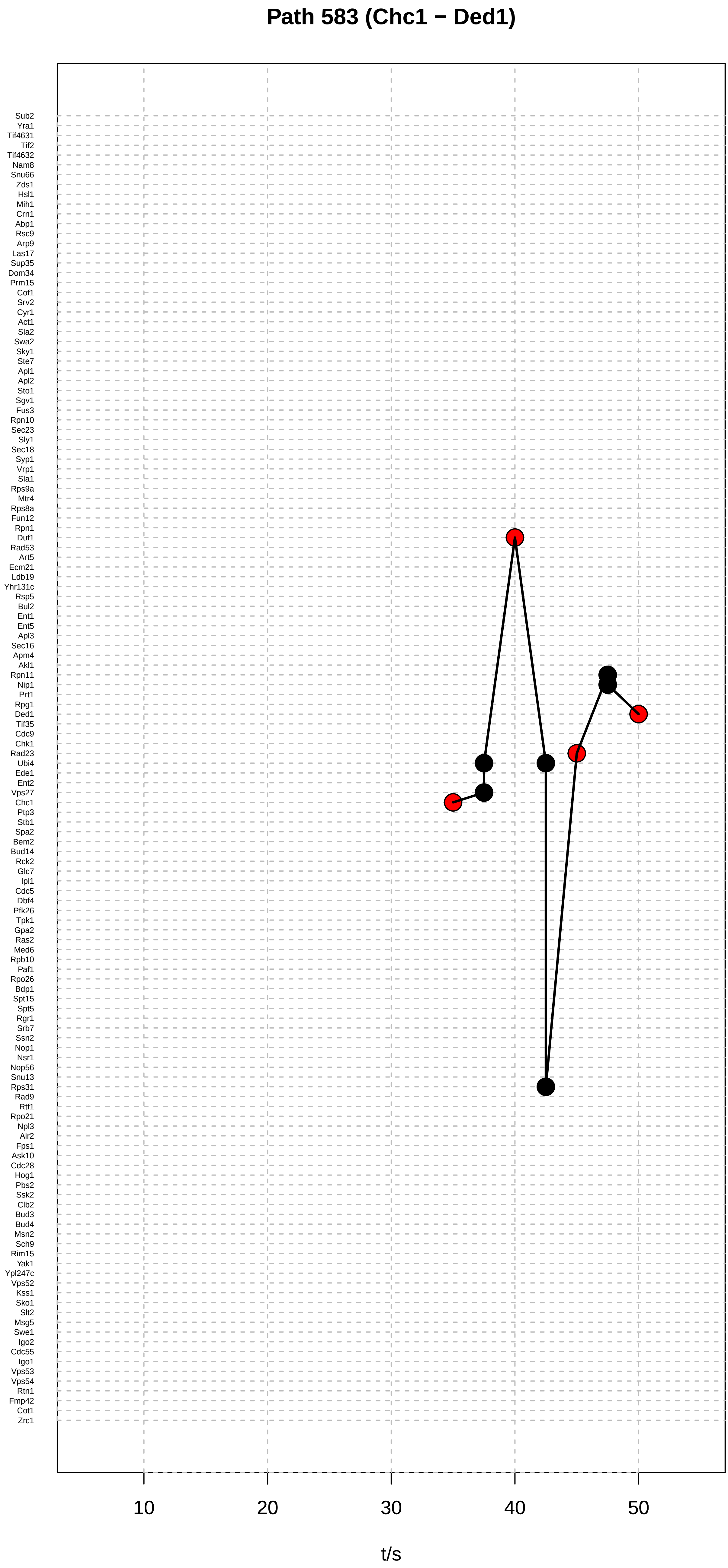

Path 583 (Chc1 − Ded1)
Sub2
Yra1
Tif4631
Tif2
Tif4632
Nam8
Snu66
Zds1
Hsl1
Mih1
Crn1
Abp1
Rsc9
Arp9
Las17
Sup35
Dom34
Prm15
Cof1
Srv2
Cyr1
Act1
Sla2
Swa2
Sky1
Ste7
Apl1
Apl2
Sto1
Sgv1
Fus3
Rpn10
Sec23
Sly1
Sec18
Syp1
Vrp1
Sla1
Rps9a
Mtr4
Rps8a
Fun12
Rpn1
Duf1
Rad53
Art5
Ecm21
Ldb19
Yhr131c
Rsp5
Bul2
Ent1
Ent5
Apl3
Sec16
Apm4
Akl1
Rpn11
Nip1
Prt1
Rpg1
Ded1
Tif35
Cdc9
Chk1
Rad23
Ubi4
Ede1
Ent2
Vps27
Chc1
Ptp3
Stb1
Spa2
Bem2
Bud14
Rck2
Glc7
Ipl1
Cdc5
Dbf4
Pfk26
Tpk1
Gpa2
Ras2
Med6
Rpb10
Paf1
Rpo26
Bdp1
Spt15
Spt5
Rgr1
Srb7
Ssn2
Nop1
Nsr1
Nop56
Snu13
Rps31
Rad9
Rtf1
Rpo21
Npl3
Air2
Fps1
Ask10
Cdc28
Hog1
Pbs2
Ssk2
Clb2
Bud3
Bud4
Msn2
Sch9
Rim15
Yak1
Ypl247c
Vps52
Kss1
Sko1
Slt2
Msg5
Swe1
Igo2
Cdc55
Igo1
Vps53
Vps54
Rtn1
Fmp42
Cot1
Zrc1
10
20
30
40
50
t/s

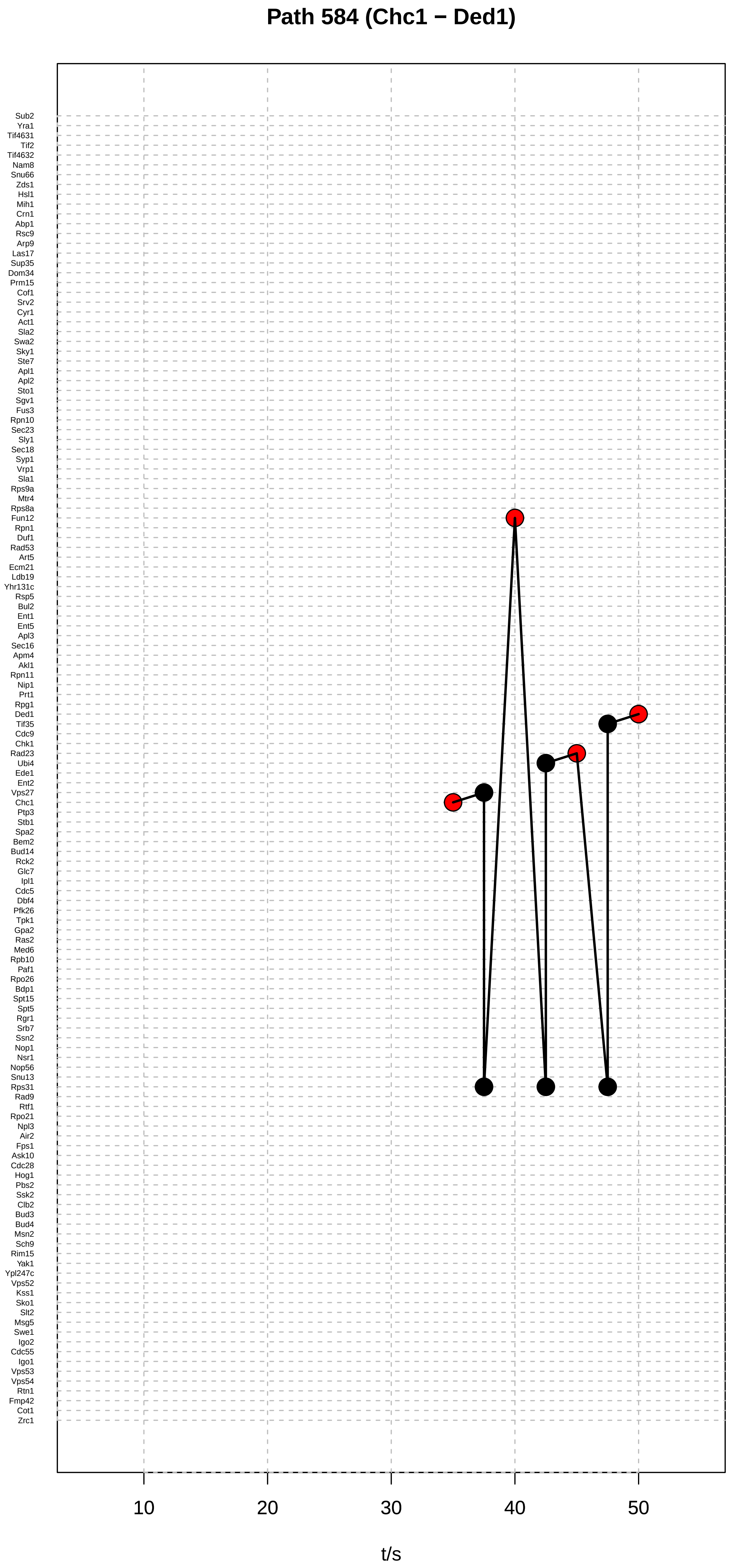
Path 584 (Chc1 − Ded1)
Sub2
Yra1
Tif4631
Tif2
Tif4632
Nam8
Snu66
Zds1
Hsl1
Mih1
Crn1
Abp1
Rsc9
Arp9
Las17
Sup35
Dom34
Prm15
Cof1
Srv2
Cyr1
Act1
Sla2
Swa2
Sky1
Ste7
Apl1
Apl2
Sto1
Sgv1
Fus3
Rpn10
Sec23
Sly1
Sec18
Syp1
Vrp1
Sla1
Rps9a
Mtr4
Rps8a
Fun12
Rpn1
Duf1
Rad53
Art5
Ecm21
Ldb19
Yhr131c
Rsp5
Bul2
Ent1
Ent5
Apl3
Sec16
Apm4
Akl1
Rpn11
Nip1
Prt1
Rpg1
Ded1
Tif35
Cdc9
Chk1
Rad23
Ubi4
Ede1
Ent2
Vps27
Chc1
Ptp3
Stb1
Spa2
Bem2
Bud14
Rck2
Glc7
Ipl1
Cdc5
Dbf4
Pfk26
Tpk1
Gpa2
Ras2
Med6
Rpb10
Paf1
Rpo26
Bdp1
Spt15
Spt5
Rgr1
Srb7
Ssn2
Nop1
Nsr1
Nop56
Snu13
Rps31
Rad9
Rtf1
Rpo21
Npl3
Air2
Fps1
Ask10
Cdc28
Hog1
Pbs2
Ssk2
Clb2
Bud3
Bud4
Msn2
Sch9
Rim15
Yak1
Ypl247c
Vps52
Kss1
Sko1
Slt2
Msg5
Swe1
Igo2
Cdc55
Igo1
Vps53
Vps54
Rtn1
Fmp42
Cot1
Zrc1
10
20
30
40
50
t/s

#### Path 585 (Chc1 − Ded1)

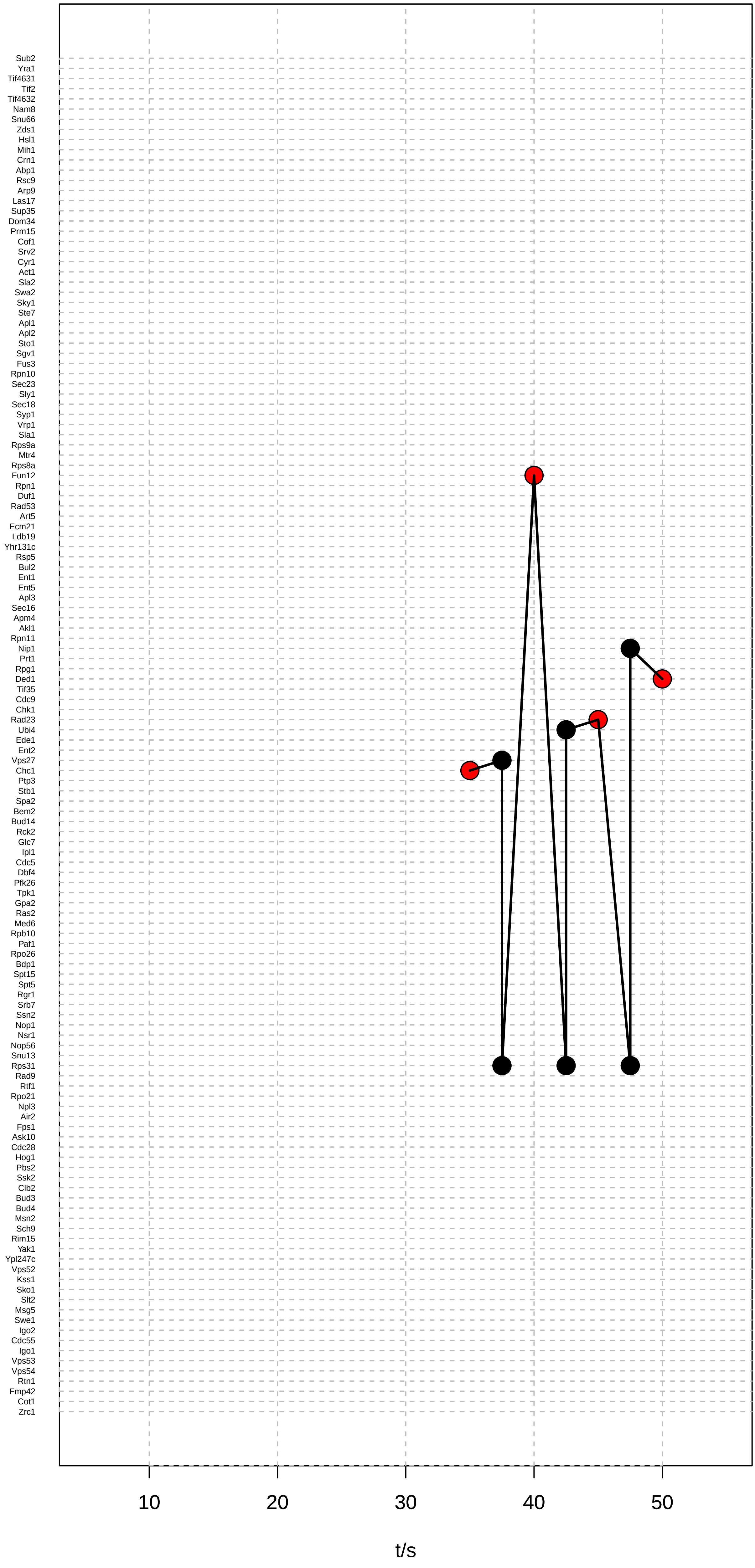

### Path 586 (Chc1 − Ded1)

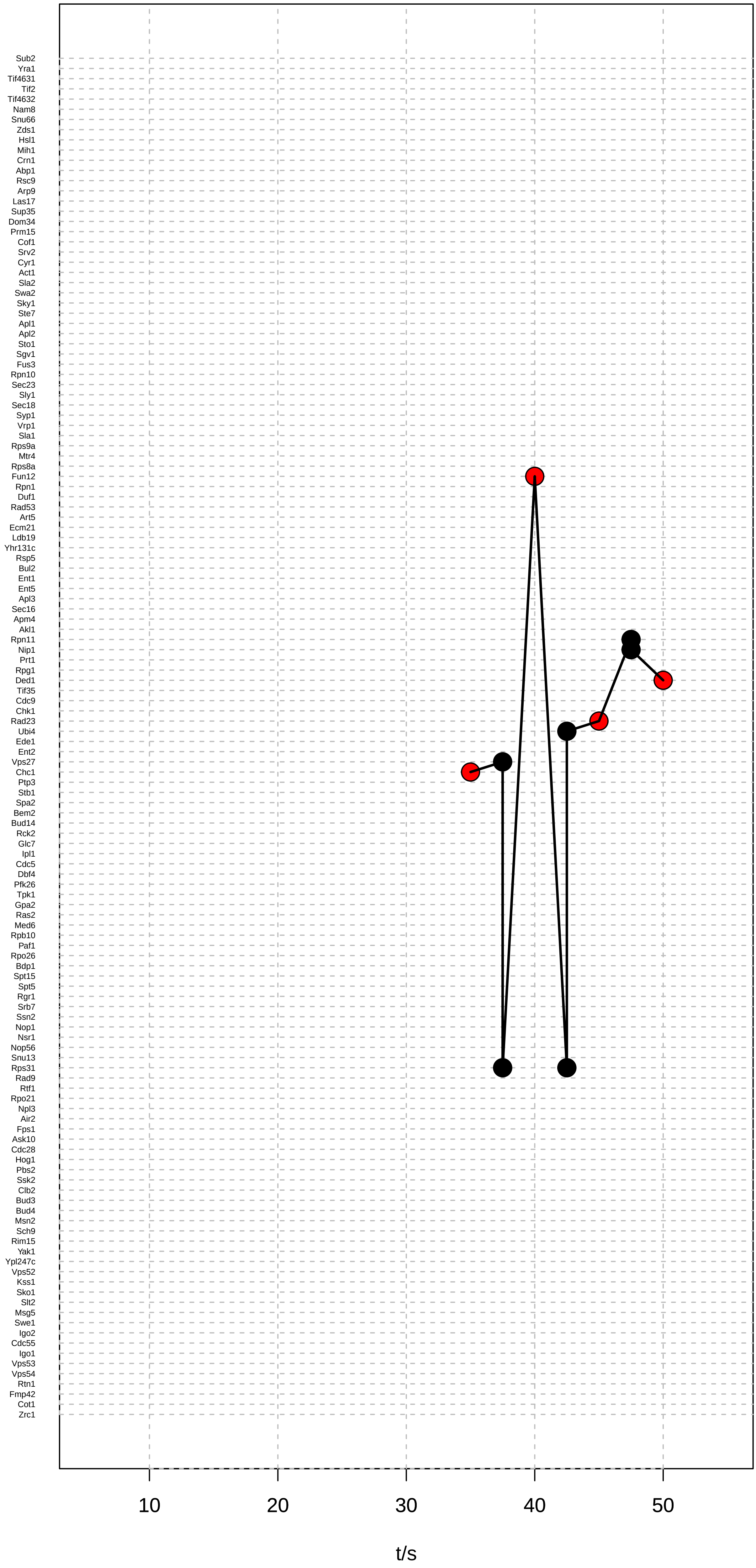

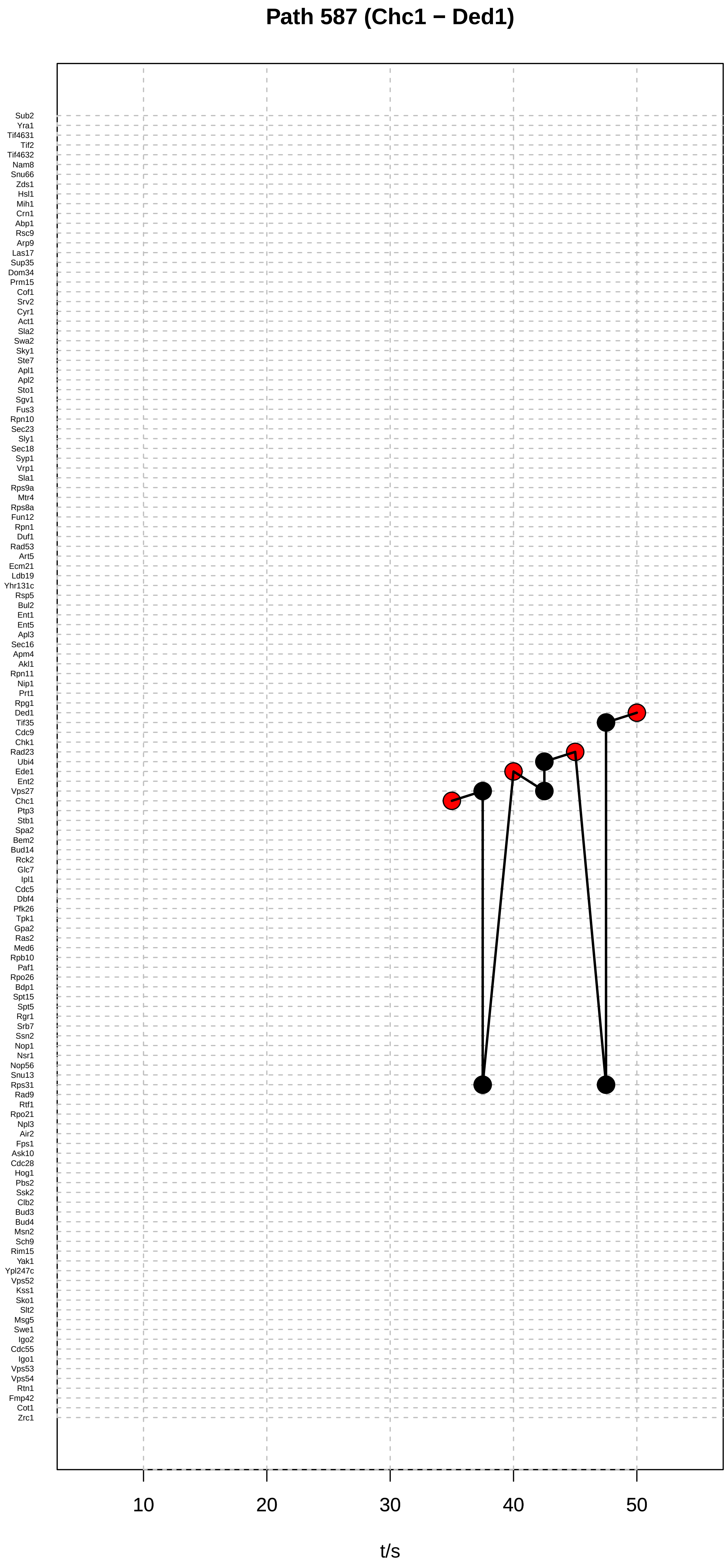
Path 587 (Chc1 − Ded1)
Sub2
Yra1
Tif4631
Tif2
Tif4632
Nam8
Snu66
Zds1
Hsl1
Mih1
Crn1
Abp1
Rsc9
Arp9
Las17
Sup35
Dom34
Prm15
Cof1
Srv2
Cyr1
Act1
Sla2
Swa2
Sky1
Ste7
Apl1
Apl2
Sto1
Sgv1
Fus3
Rpn10
Sec23
Sly1
Sec18
Syp1
Vrp1
Sla1
Rps9a
Mtr4
Rps8a
Fun12
Rpn1
Duf1
Rad53
Art5
Ecm21
Ldb19
Yhr131c
Rsp5
Bul2
Ent1
Ent5
Apl3
Sec16
Apm4
Akl1
Rpn11
Nip1
Prt1
Rpg1
Ded1
Tif35
Cdc9
Chk1
Rad23
Ubi4
Ede1
Ent2
Vps27
Chc1
Ptp3
Stb1
Spa2
Bem2
Bud14
Rck2
Glc7
Ipl1
Cdc5
Dbf4
Pfk26
Tpk1
Gpa2
Ras2
Med6
Rpb10
Paf1
Rpo26
Bdp1
Spt15
Spt5
Rgr1
Srb7
Ssn2
Nop1
Nsr1
Nop56
Snu13
Rps31
Rad9
Rtf1
Rpo21
Npl3
Air2
Fps1
Ask10
Cdc28
Hog1
Pbs2
Ssk2
Clb2
Bud3
Bud4
Msn2
Sch9
Rim15
Yak1
Ypl247c
Vps52
Kss1
Sko1
Slt2
Msg5
Swe1
Igo2
Cdc55
Igo1
Vps53
Vps54
Rtn1
Fmp42
Cot1
Zrc1
10
20
30
40
50
t/s

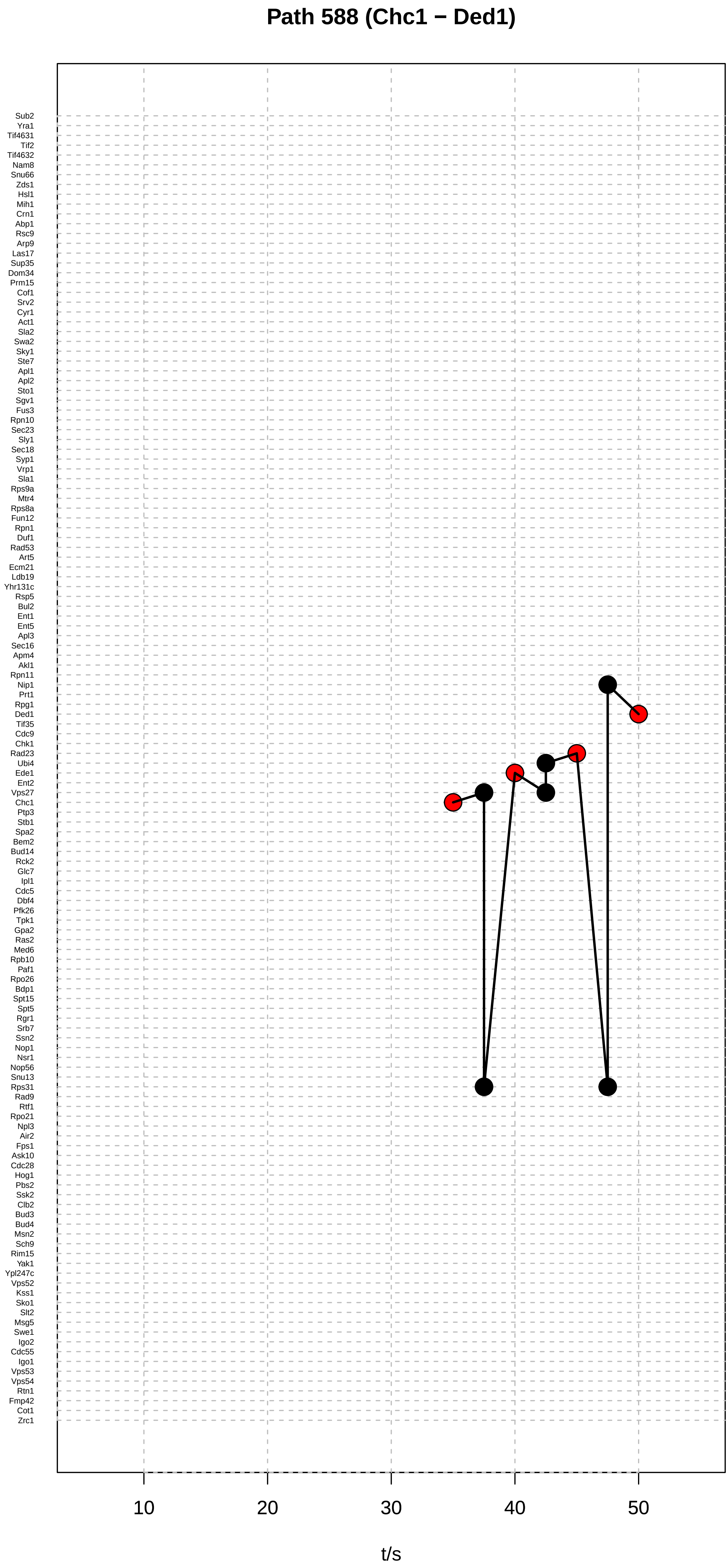
Path 588 (Chc1 − Ded1)
Sub2
Yra1
Tif4631
Tif2
Tif4632
Nam8
Snu66
Zds1
Hsl1
Mih1
Crn1
Abp1
Rsc9
Arp9
Las17
Sup35
Dom34
Prm15
Cof1
Srv2
Cyr1
Act1
Sla2
Swa2
Sky1
Ste7
Apl1
Apl2
Sto1
Sgv1
Fus3
Rpn10
Sec23
Sly1
Sec18
Syp1
Vrp1
Sla1
Rps9a
Mtr4
Rps8a
Fun12
Rpn1
Duf1
Rad53
Art5
Ecm21
Ldb19
Yhr131c
Rsp5
Bul2
Ent1
Ent5
Apl3
Sec16
Apm4
Akl1
Rpn11
Nip1
Prt1
Rpg1
Ded1
Tif35
Cdc9
Chk1
Rad23
Ubi4
Ede1
Ent2
Vps27
Chc1
Ptp3
Stb1
Spa2
Bem2
Bud14
Rck2
Glc7
Ipl1
Cdc5
Dbf4
Pfk26
Tpk1
Gpa2
Ras2
Med6
Rpb10
Paf1
Rpo26
Bdp1
Spt15
Spt5
Rgr1
Srb7
Ssn2
Nop1
Nsr1
Nop56
Snu13
Rps31
Rad9
Rtf1
Rpo21
Npl3
Air2
Fps1
Ask10
Cdc28
Hog1
Pbs2
Ssk2
Clb2
Bud3
Bud4
Msn2
Sch9
Rim15
Yak1
Ypl247c
Vps52
Kss1
Sko1
Slt2
Msg5
Swe1
Igo2
Cdc55
Igo1
Vps53
Vps54
Rtn1
Fmp42
Cot1
Zrc1
10
20
30
40
50
t/s

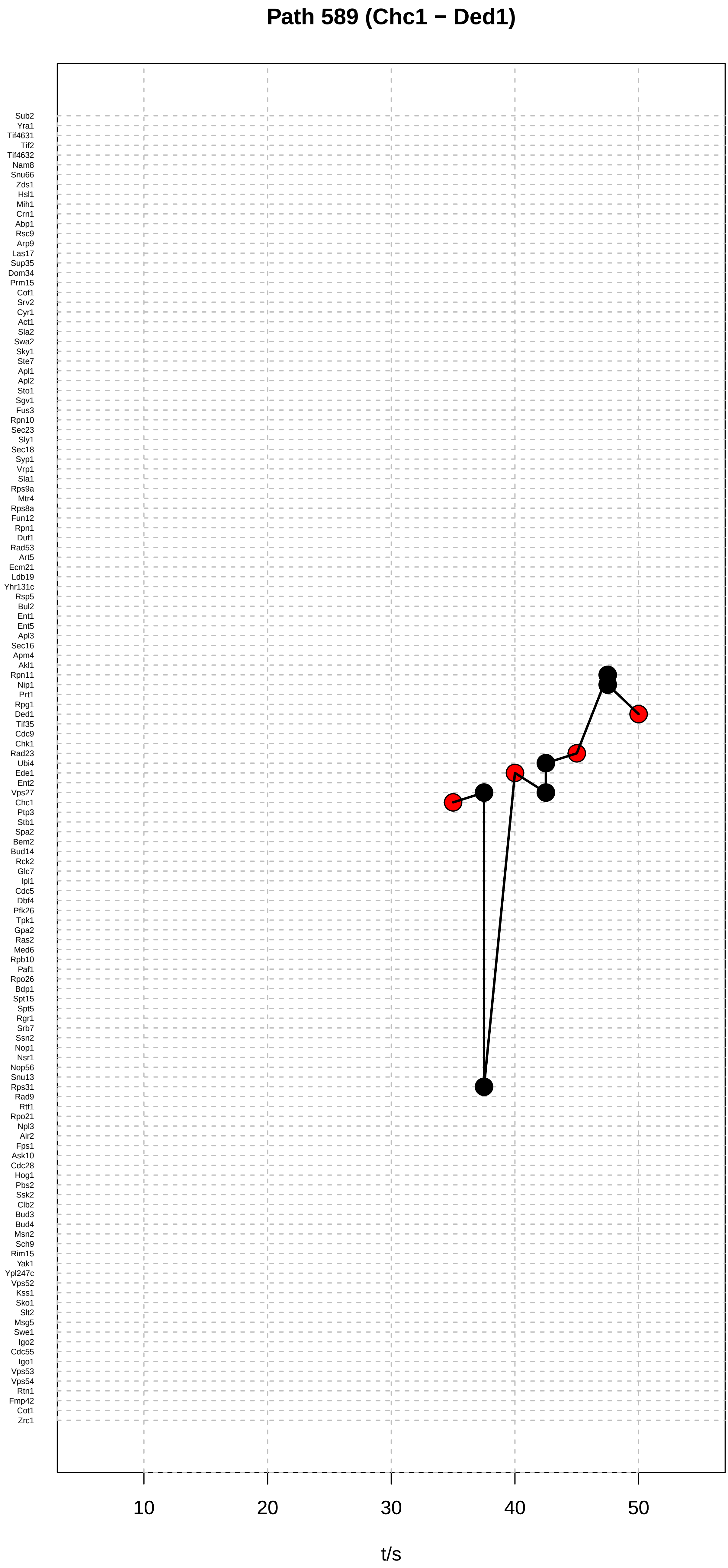
Path 589 (Chc1 − Ded1)
Sub2
Yra1
Tif4631
Tif2
Tif4632
Nam8
Snu66
Zds1
Hsl1
Mih1
Crn1
Abp1
Rsc9
Arp9
Las17
Sup35
Dom34
Prm15
Cof1
Srv2
Cyr1
Act1
Sla2
Swa2
Sky1
Ste7
Apl1
Apl2
Sto1
Sgv1
Fus3
Rpn10
Sec23
Sly1
Sec18
Syp1
Vrp1
Sla1
Rps9a
Mtr4
Rps8a
Fun12
Rpn1
Duf1
Rad53
Art5
Ecm21
Ldb19
Yhr131c
Rsp5
Bul2
Ent1
Ent5
Apl3
Sec16
Apm4
Akl1
Rpn11
Nip1
Prt1
Rpg1
Ded1
Tif35
Cdc9
Chk1
Rad23
Ubi4
Ede1
Ent2
Vps27
Chc1
Ptp3
Stb1
Spa2
Bem2
Bud14
Rck2
Glc7
Ipl1
Cdc5
Dbf4
Pfk26
Tpk1
Gpa2
Ras2
Med6
Rpb10
Paf1
Rpo26
Bdp1
Spt15
Spt5
Rgr1
Srb7
Ssn2
Nop1
Nsr1
Nop56
Snu13
Rps31
Rad9
Rtf1
Rpo21
Npl3
Air2
Fps1
Ask10
Cdc28
Hog1
Pbs2
Ssk2
Clb2
Bud3
Bud4
Msn2
Sch9
Rim15
Yak1
Ypl247c
Vps52
Kss1
Sko1
Slt2
Msg5
Swe1
Igo2
Cdc55
Igo1
Vps53
Vps54
Rtn1
Fmp42
Cot1
Zrc1
10
20
30
40
50
t/s

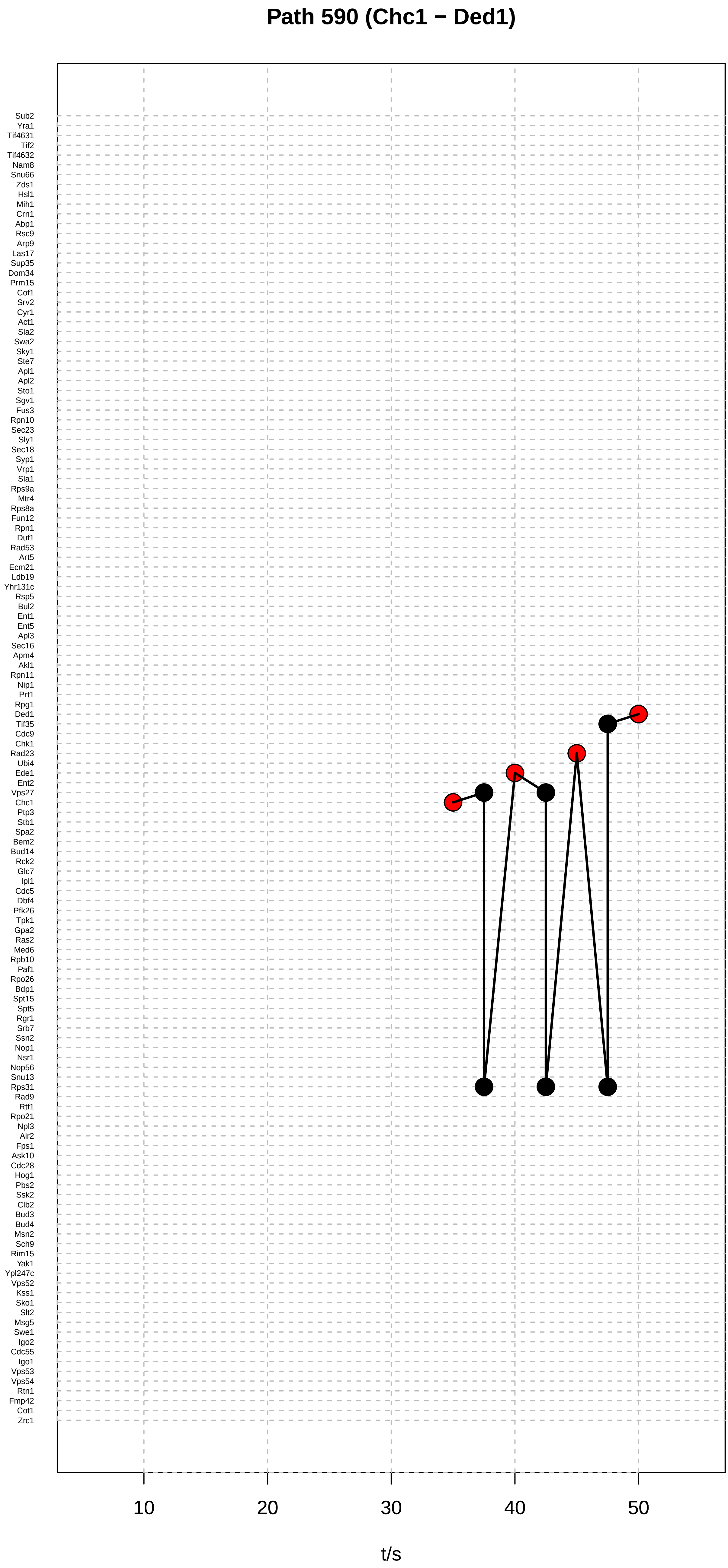
Path 590 (Chc1 − Ded1)
Sub2
Yra1
Tif4631
Tif2
Tif4632
Nam8
Snu66
Zds1
Hsl1
Mih1
Crn1
Abp1
Rsc9
Arp9
Las17
Sup35
Dom34
Prm15
Cof1
Srv2
Cyr1
Act1
Sla2
Swa2
Sky1
Ste7
Apl1
Apl2
Sto1
Sgv1
Fus3
Rpn10
Sec23
Sly1
Sec18
Syp1
Vrp1
Sla1
Rps9a
Mtr4
Rps8a
Fun12
Rpn1
Duf1
Rad53
Art5
Ecm21
Ldb19
Yhr131c
Rsp5
Bul2
Ent1
Ent5
Apl3
Sec16
Apm4
Akl1
Rpn11
Nip1
Prt1
Rpg1
Ded1
Tif35
Cdc9
Chk1
Rad23
Ubi4
Ede1
Ent2
Vps27
Chc1
Ptp3
Stb1
Spa2
Bem2
Bud14
Rck2
Glc7
Ipl1
Cdc5
Dbf4
Pfk26
Tpk1
Gpa2
Ras2
Med6
Rpb10
Paf1
Rpo26
Bdp1
Spt15
Spt5
Rgr1
Srb7
Ssn2
Nop1
Nsr1
Nop56
Snu13
Rps31
Rad9
Rtf1
Rpo21
Npl3
Air2
Fps1
Ask10
Cdc28
Hog1
Pbs2
Ssk2
Clb2
Bud3
Bud4
Msn2
Sch9
Rim15
Yak1
Ypl247c
Vps52
Kss1
Sko1
Slt2
Msg5
Swe1
Igo2
Cdc55
Igo1
Vps53
Vps54
Rtn1
Fmp42
Cot1
Zrc1
10
20
30
40
50
t/s

#### Path 591 (Chc1 − Ded1)

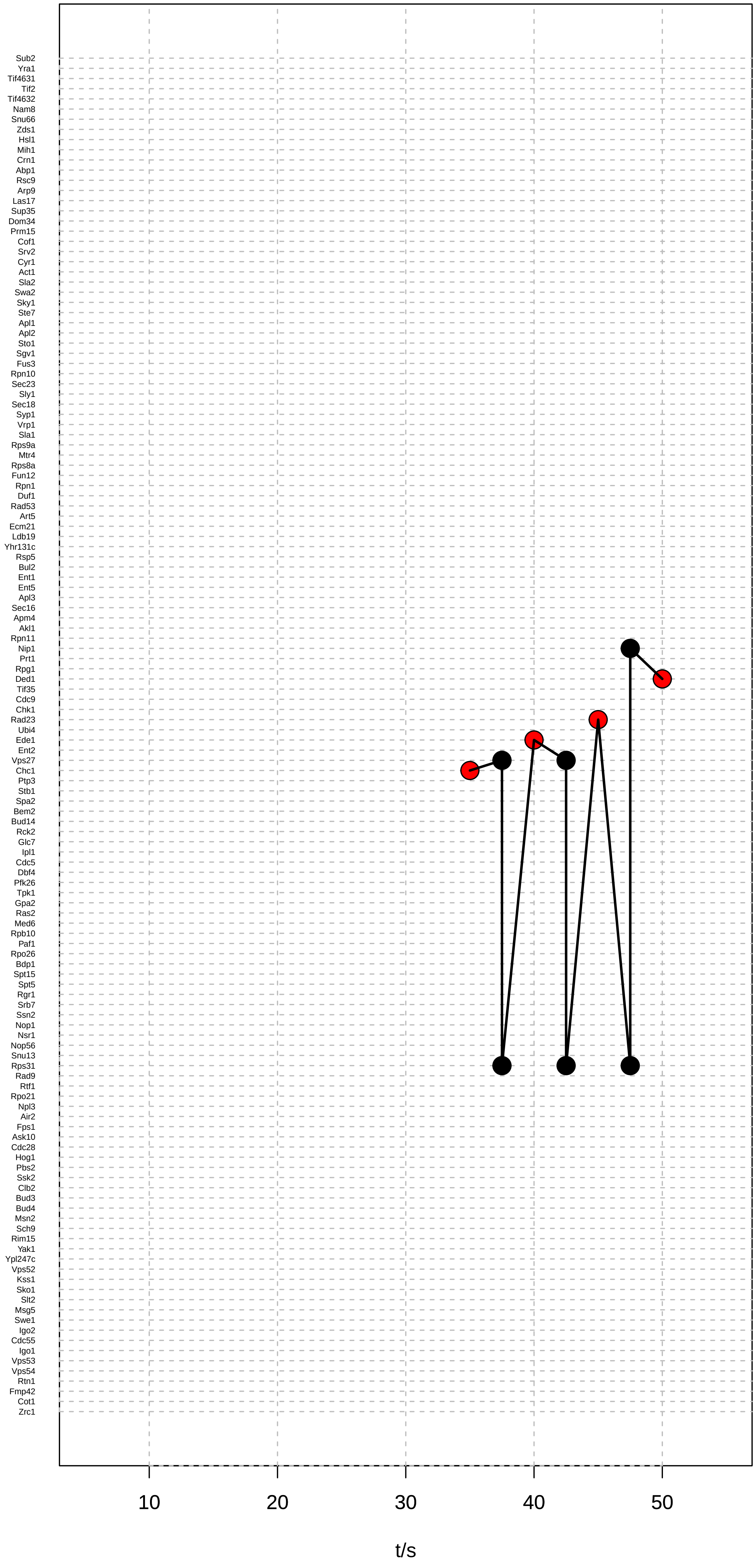

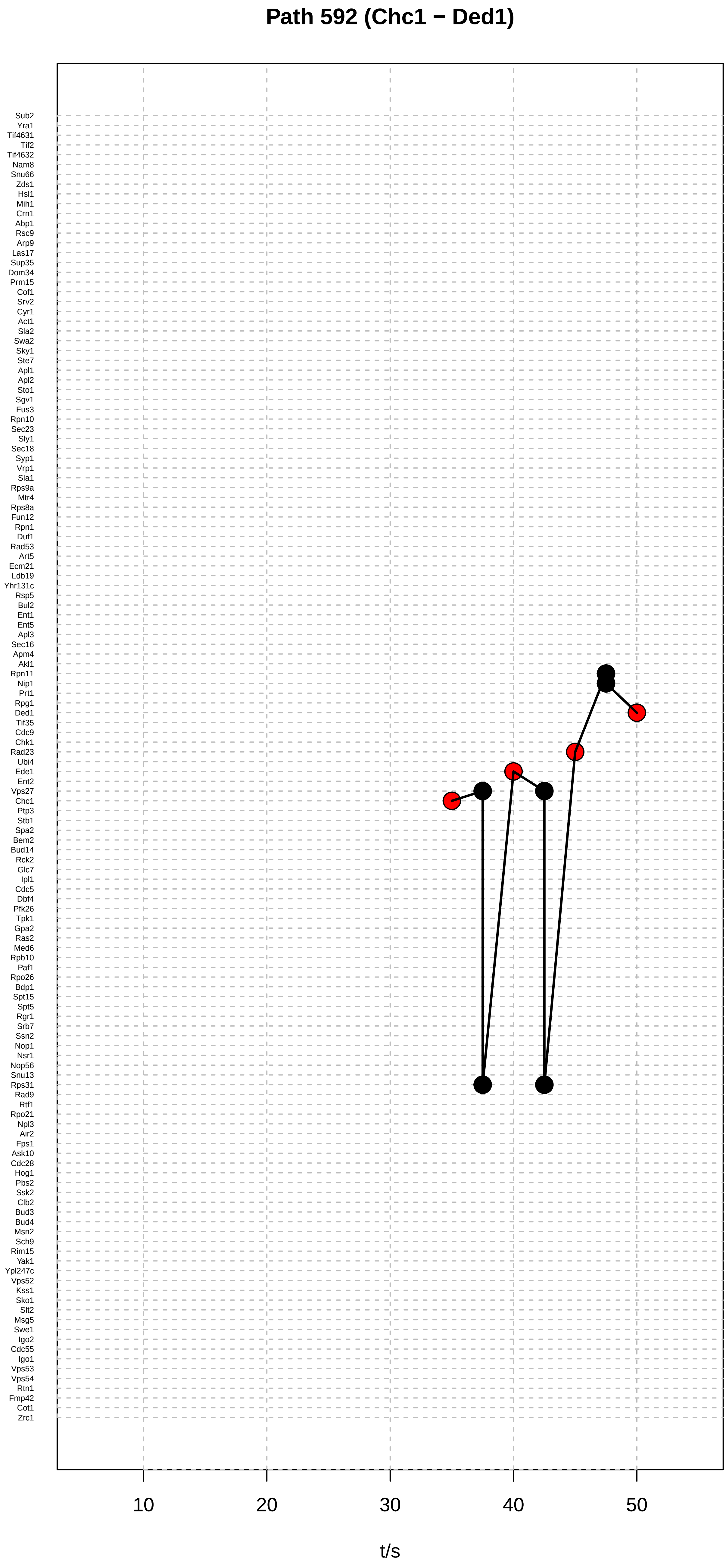

Path 592 (Chc1 − Ded1)
Sub2
Yra1
Tif4631
Tif2
Tif4632
Nam8
Snu66
Zds1
Hsl1
Mih1
Crn1
Abp1
Rsc9
Arp9
Las17
Sup35
Dom34
Prm15
Cof1
Srv2
Cyr1
Act1
Sla2
Swa2
Sky1
Ste7
Apl1
Apl2
Sto1
Sgv1
Fus3
Rpn10
Sec23
Sly1
Sec18
Syp1
Vrp1
Sla1
Rps9a
Mtr4
Rps8a
Fun12
Rpn1
Duf1
Rad53
Art5
Ecm21
Ldb19
Yhr131c
Rsp5
Bul2
Ent1
Ent5
Apl3
Sec16
Apm4
Akl1
Rpn11
Nip1
Prt1
Rpg1
Ded1
Tif35
Cdc9
Chk1
Rad23
Ubi4
Ede1
Ent2
Vps27
Chc1
Ptp3
Stb1
Spa2
Bem2
Bud14
Rck2
Glc7
Ipl1
Cdc5
Dbf4
Pfk26
Tpk1
Gpa2
Ras2
Med6
Rpb10
Paf1
Rpo26
Bdp1
Spt15
Spt5
Rgr1
Srb7
Ssn2
Nop1
Nsr1
Nop56
Snu13
Rps31
Rad9
Rtf1
Rpo21
Npl3
Air2
Fps1
Ask10
Cdc28
Hog1
Pbs2
Ssk2
Clb2
Bud3
Bud4
Msn2
Sch9
Rim15
Yak1
Ypl247c
Vps52
Kss1
Sko1
Slt2
Msg5
Swe1
Igo2
Cdc55
Igo1
Vps53
Vps54
Rtn1
Fmp42
Cot1
Zrc1
10
20
30
40
50
t/s

#### Path 593 (Chc1 − Ded1)

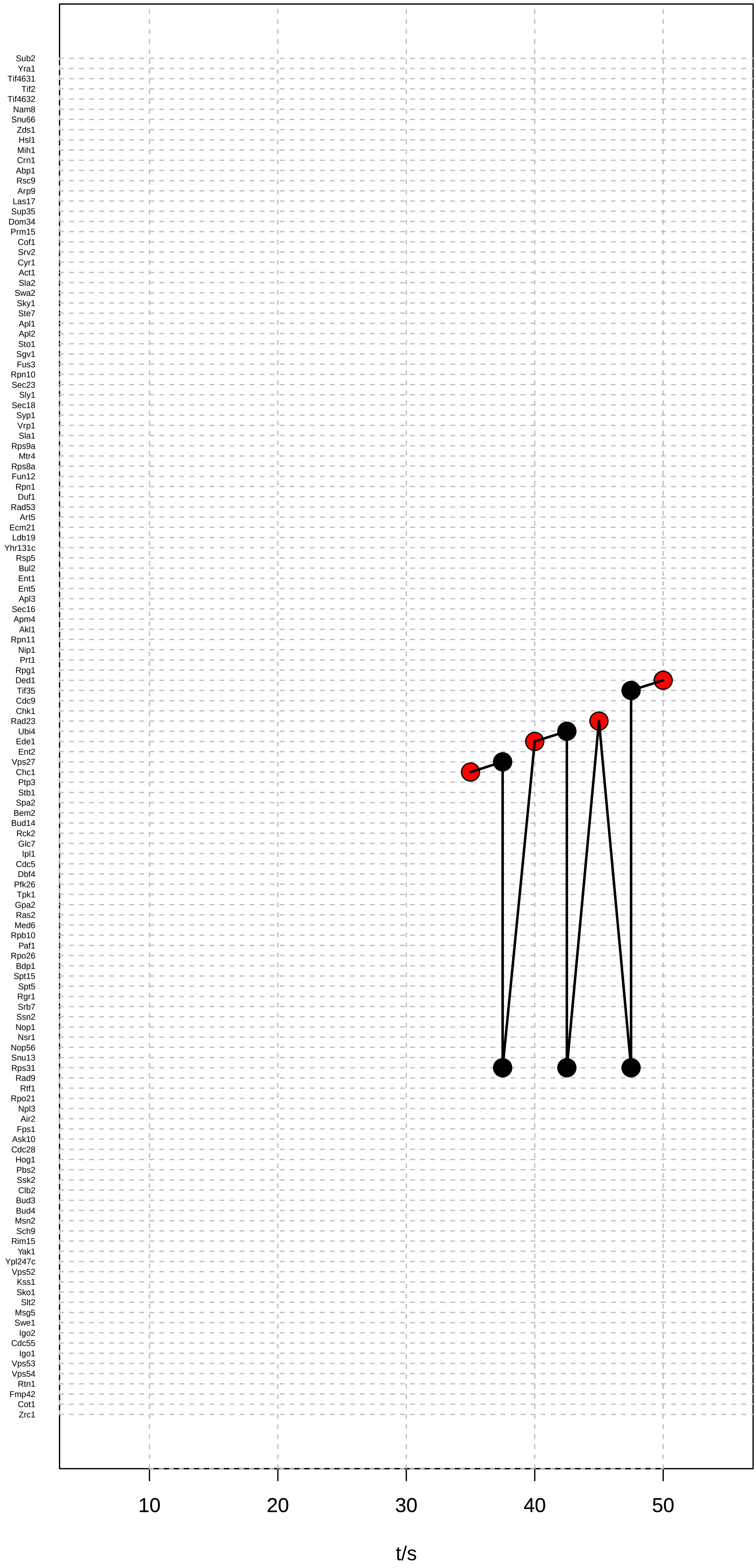

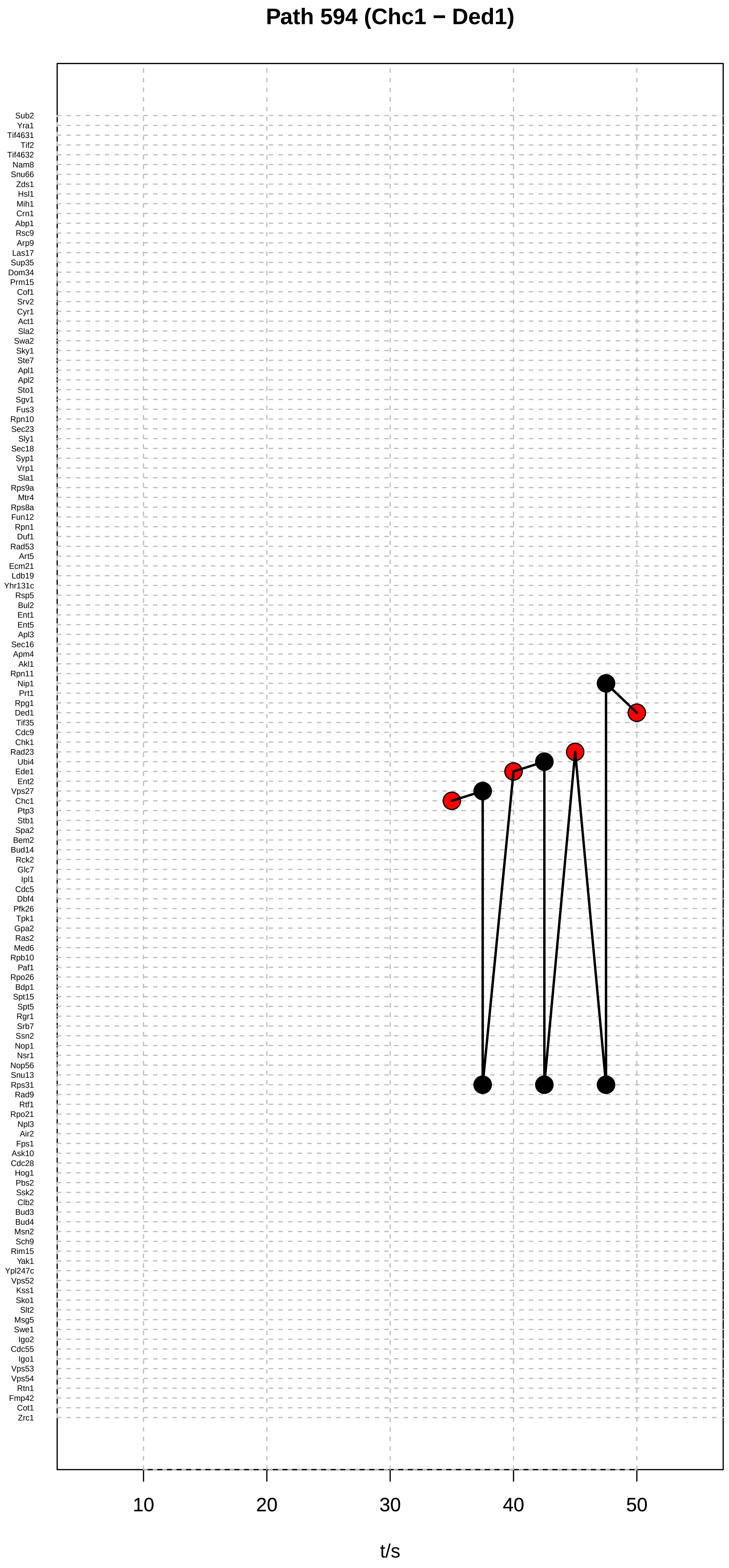

Path 594 (Chc1 − Ded1)
Sub2
Yra1
Tif4631
Tif2
Tif4632
Nam8
Snu66
Zds1
Hsl1
Mih1
Crn1
Abp1
Rsc9
Arp9
Las17
Sup35
Dom34
Prm15
Cof1
Srv2
Cyr1
Act1
Sla2
Swa2
Sky1
Ste7
Apl1
Apl2
Sto1
Sgv1
Fus3
Rpn10
Sec23
Sly1
Sec18
Syp1
Vrp1
Sla1
Rps9a
Mtr4
Rps8a
Fun12
Rpn1
Duf1
Rad53
Art5
Ecm21
Ldb19
Yhr131c
Rsp5
Bul2
Ent1
Ent5
Apl3
Sec16
Apm4
Akl1
Rpn11
Nip1
Prt1
Rpg1
Ded1
Tif35
Cdc9
Chk1
Rad23
Ubi4
Ede1
Ent2
Vps27
Chc1
Ptp3
Stb1
Spa2
Bem2
Bud14
Rck2
Glc7
Ipl1
Cdc5
Dbf4
Pfk26
Tpk1
Gpa2
Ras2
Med6
Rpb10
Paf1
Rpo26
Bdp1
Spt15
Spt5
Rgr1
Srb7
Ssn2
Nop1
Nsr1
Nop56
Snu13
Rps31
Rad9
Rtf1
Rpo21
Npl3
Air2
Fps1
Ask10
Cdc28
Hog1
Pbs2
Ssk2
Clb2
Bud3
Bud4
Msn2
Sch9
Rim15
Yak1
Ypl247c
Vps52
Kss1
Sko1
Slt2
Msg5
Swe1
Igo2
Cdc55
Igo1
Vps53
Vps54
Rtn1
Fmp42
Cot1
Zrc1
10
20
30
40
50
t/s

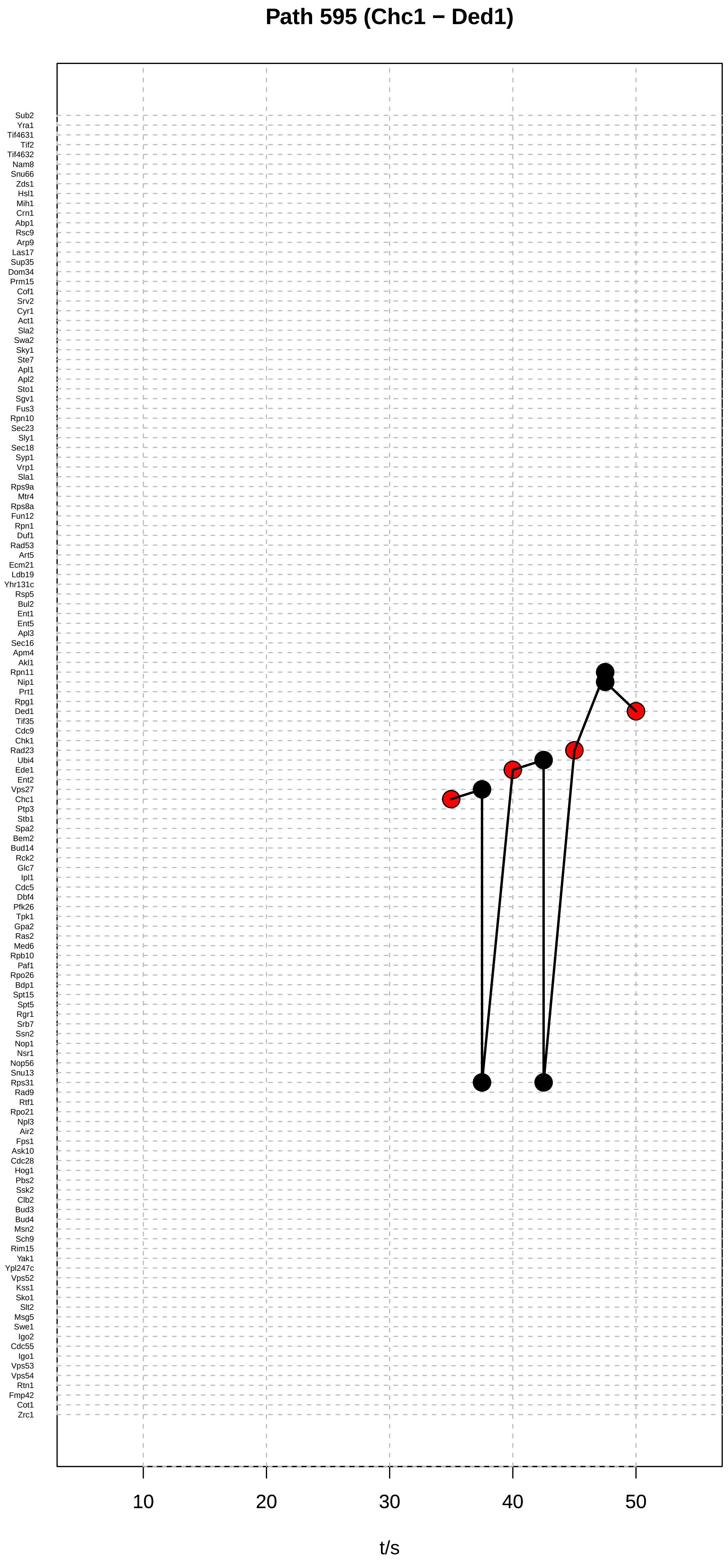

Path 595 (Chc1 − Ded1)
Sub2
Yra1
Tif4631
Tif2
Tif4632
Nam8
Snu66
Zds1
Hsl1
Mih1
Crn1
Abp1
Rsc9
Arp9
Las17
Sup35
Dom34
Prm15
Cof1
Srv2
Cyr1
Act1
Sla2
Swa2
Sky1
Ste7
Apl1
Apl2
Sto1
Sgv1
Fus3
Rpn10
Sec23
Sly1
Sec18
Syp1
Vrp1
Sla1
Rps9a
Mtr4
Rps8a
Fun12
Rpn1
Duf1
Rad53
Art5
Ecm21
Ldb19
Yhr131c
Rsp5
Bul2
Ent1
Ent5
Apl3
Sec16
Apm4
Akl1
Rpn11
Nip1
Prt1
Rpg1
Ded1
Tif35
Cdc9
Chk1
Rad23
Ubi4
Ede1
Ent2
Vps27
Chc1
Ptp3
Stb1
Spa2
Bem2
Bud14
Rck2
Glc7
Ipl1
Cdc5
Dbf4
Pfk26
Tpk1
Gpa2
Ras2
Med6
Rpb10
Paf1
Rpo26
Bdp1
Spt15
Spt5
Rgr1
Srb7
Ssn2
Nop1
Nsr1
Nop56
Snu13
Rps31
Rad9
Rtf1
Rpo21
Npl3
Air2
Fps1
Ask10
Cdc28
Hog1
Pbs2
Ssk2
Clb2
Bud3
Bud4
Msn2
Sch9
Rim15
Yak1
Ypl247c
Vps52
Kss1
Sko1
Slt2
Msg5
Swe1
Igo2
Cdc55
Igo1
Vps53
Vps54
Rtn1
Fmp42
Cot1
Zrc1
10
20
30
40
50
t/s

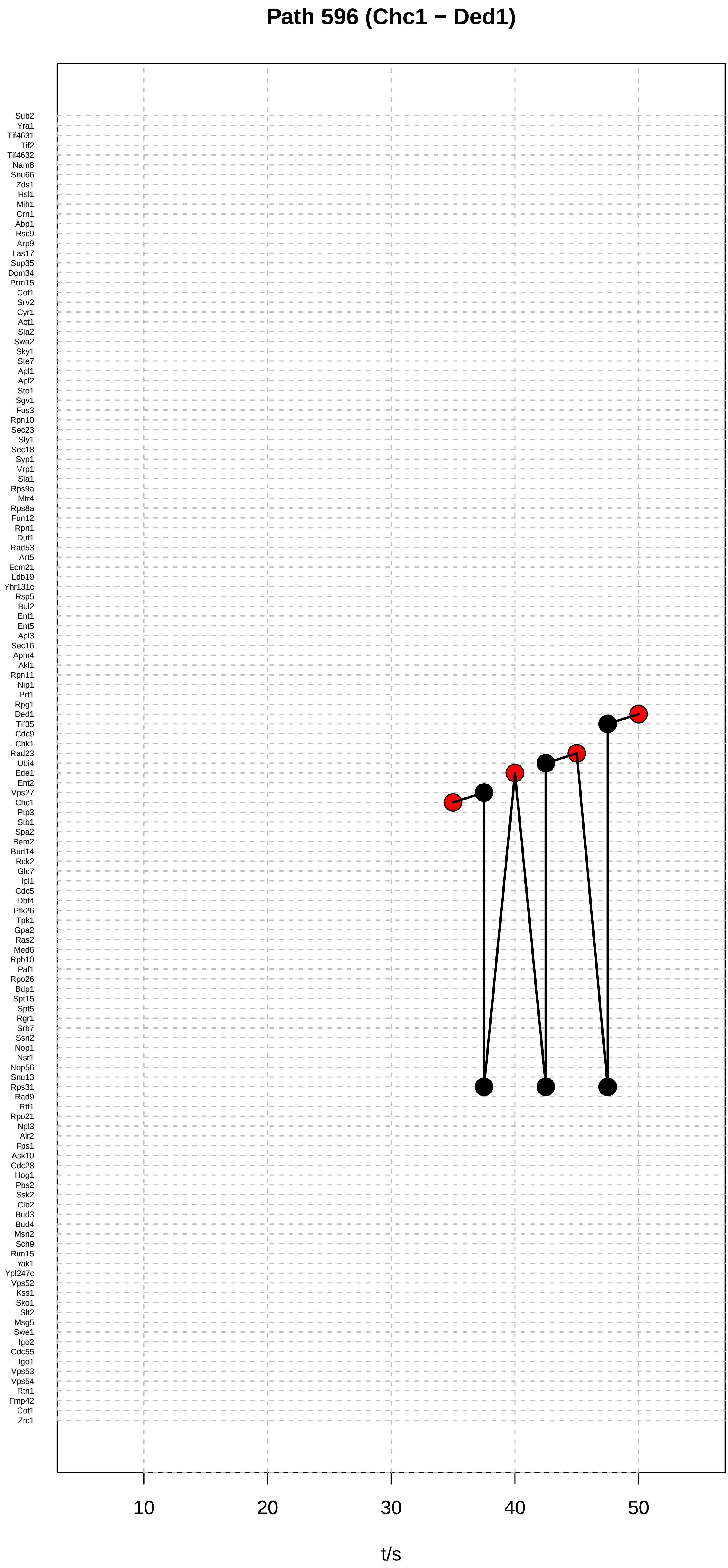
Path 596 (Chc1 − Ded1)
Sub2
Yra1
Tif4631
Tif2
Tif4632
Nam8
Snu66
Zds1
Hsl1
Mih1
Crn1
Abp1
Rsc9
Arp9
Las17
Sup35
Dom34
Prm15
Cof1
Srv2
Cyr1
Act1
Sla2
Swa2
Sky1
Ste7
Apl1
Apl2
Sto1
Sgv1
Fus3
Rpn10
Sec23
Sly1
Sec18
Syp1
Vrp1
Sla1
Rps9a
Mtr4
Rps8a
Fun12
Rpn1
Duf1
Rad53
Art5
Ecm21
Ldb19
Yhr131c
Rsp5
Bul2
Ent1
Ent5
Apl3
Sec16
Apm4
Akl1
Rpn11
Nip1
Prt1
Rpg1
Ded1
Tif35
Cdc9
Chk1
Rad23
Ubi4
Ede1
Ent2
Vps27
Chc1
Ptp3
Stb1
Spa2
Bem2
Bud14
Rck2
Glc7
Ipl1
Cdc5
Dbf4
Pfk26
Tpk1
Gpa2
Ras2
Med6
Rpb10
Paf1
Rpo26
Bdp1
Spt15
Spt5
Rgr1
Srb7
Ssn2
Nop1
Nsr1
Nop56
Snu13
Rps31
Rad9
Rtf1
Rpo21
Npl3
Air2
Fps1
Ask10
Cdc28
Hog1
Pbs2
Ssk2
Clb2
Bud3
Bud4
Msn2
Sch9
Rim15
Yak1
Ypl247c
Vps52
Kss1
Sko1
Slt2
Msg5
Swe1
Igo2
Cdc55
Igo1
Vps53
Vps54
Rtn1
Fmp42
Cot1
Zrc1
10
20
30
40
50
t/s

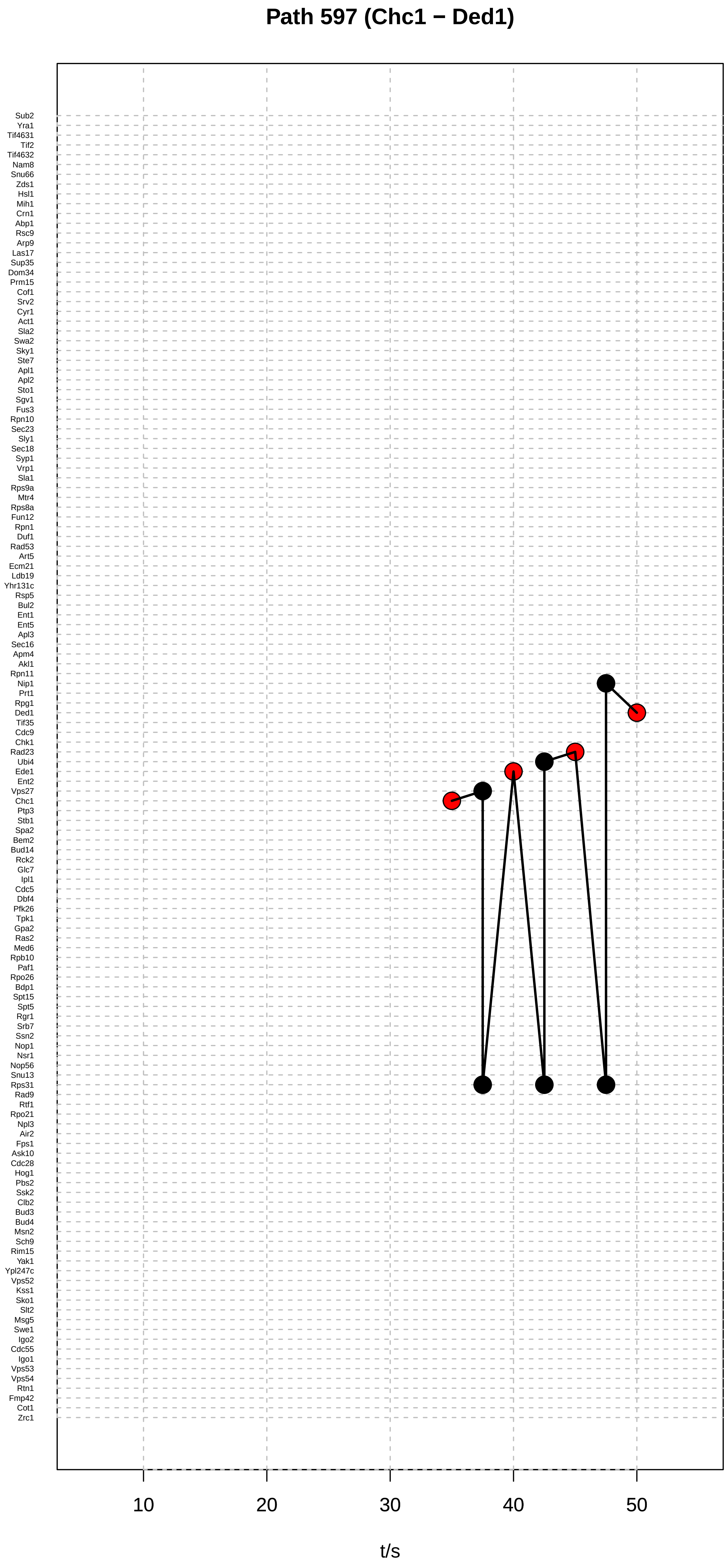

Path 597 (Chc1 − Ded1)
Sub2
Yra1
Tif4631
Tif2
Tif4632
Nam8
Snu66
Zds1
Hsl1
Mih1
Crn1
Abp1
Rsc9
Arp9
Las17
Sup35
Dom34
Prm15
Cof1
Srv2
Cyr1
Act1
Sla2
Swa2
Sky1
Ste7
Apl1
Apl2
Sto1
Sgv1
Fus3
Rpn10
Sec23
Sly1
Sec18
Syp1
Vrp1
Sla1
Rps9a
Mtr4
Rps8a
Fun12
Rpn1
Duf1
Rad53
Art5
Ecm21
Ldb19
Yhr131c
Rsp5
Bul2
Ent1
Ent5
Apl3
Sec16
Apm4
Akl1
Rpn11
Nip1
Prt1
Rpg1
Ded1
Tif35
Cdc9
Chk1
Rad23
Ubi4
Ede1
Ent2
Vps27
Chc1
Ptp3
Stb1
Spa2
Bem2
Bud14
Rck2
Glc7
Ipl1
Cdc5
Dbf4
Pfk26
Tpk1
Gpa2
Ras2
Med6
Rpb10
Paf1
Rpo26
Bdp1
Spt15
Spt5
Rgr1
Srb7
Ssn2
Nop1
Nsr1
Nop56
Snu13
Rps31
Rad9
Rtf1
Rpo21
Npl3
Air2
Fps1
Ask10
Cdc28
Hog1
Pbs2
Ssk2
Clb2
Bud3
Bud4
Msn2
Sch9
Rim15
Yak1
Ypl247c
Vps52
Kss1
Sko1
Slt2
Msg5
Swe1
Igo2
Cdc55
Igo1
Vps53
Vps54
Rtn1
Fmp42
Cot1
Zrc1
10
20
30
40
50
t/s

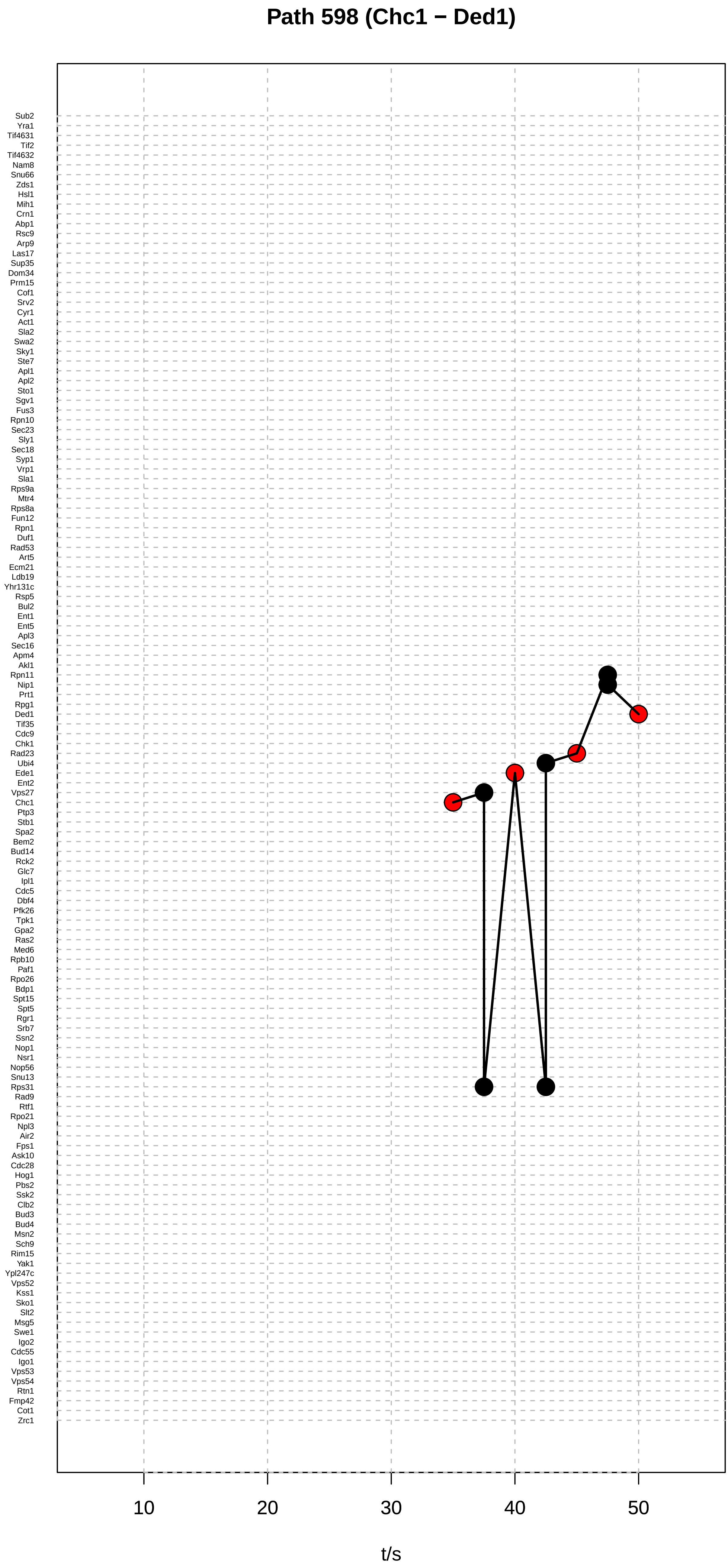

Path 598 (Chc1 − Ded1)
Sub2
Yra1
Tif4631
Tif2
Tif4632
Nam8
Snu66
Zds1
Hsl1
Mih1
Crn1
Abp1
Rsc9
Arp9
Las17
Sup35
Dom34
Prm15
Cof1
Srv2
Cyr1
Act1
Sla2
Swa2
Sky1
Ste7
Apl1
Apl2
Sto1
Sgv1
Fus3
Rpn10
Sec23
Sly1
Sec18
Syp1
Vrp1
Sla1
Rps9a
Mtr4
Rps8a
Fun12
Rpn1
Duf1
Rad53
Art5
Ecm21
Ldb19
Yhr131c
Rsp5
Bul2
Ent1
Ent5
Apl3
Sec16
Apm4
Akl1
Rpn11
Nip1
Prt1
Rpg1
Ded1
Tif35
Cdc9
Chk1
Rad23
Ubi4
Ede1
Ent2
Vps27
Chc1
Ptp3
Stb1
Spa2
Bem2
Bud14
Rck2
Glc7
Ipl1
Cdc5
Dbf4
Pfk26
Tpk1
Gpa2
Ras2
Med6
Rpb10
Paf1
Rpo26
Bdp1
Spt15
Spt5
Rgr1
Srb7
Ssn2
Nop1
Nsr1
Nop56
Snu13
Rps31
Rad9
Rtf1
Rpo21
Npl3
Air2
Fps1
Ask10
Cdc28
Hog1
Pbs2
Ssk2
Clb2
Bud3
Bud4
Msn2
Sch9
Rim15
Yak1
Ypl247c
Vps52
Kss1
Sko1
Slt2
Msg5
Swe1
Igo2
Cdc55
Igo1
Vps53
Vps54
Rtn1
Fmp42
Cot1
Zrc1
10
20
30
40
50
t/s

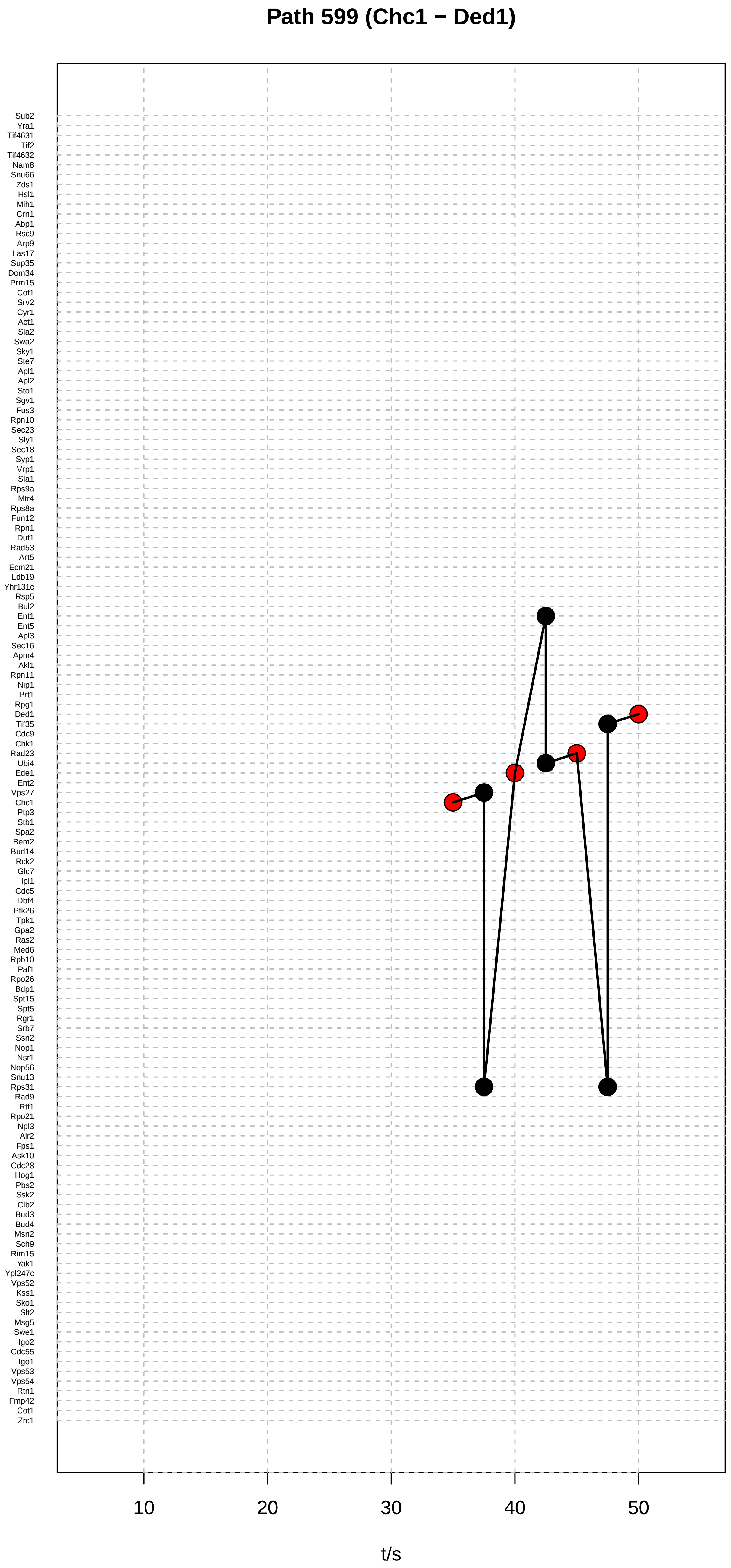

Path 599 (Chc1 − Ded1)
Sub2
Yra1
Tif4631
Tif2
Tif4632
Nam8
Snu66
Zds1
Hsl1
Mih1
Crn1
Abp1
Rsc9
Arp9
Las17
Sup35
Dom34
Prm15
Cof1
Srv2
Cyr1
Act1
Sla2
Swa2
Sky1
Ste7
Apl1
Apl2
Sto1
Sgv1
Fus3
Rpn10
Sec23
Sly1
Sec18
Syp1
Vrp1
Sla1
Rps9a
Mtr4
Rps8a
Fun12
Rpn1
Duf1
Rad53
Art5
Ecm21
Ldb19
Yhr131c
Rsp5
Bul2
Ent1
Ent5
Apl3
Sec16
Apm4
Akl1
Rpn11
Nip1
Prt1
Rpg1
Ded1
Tif35
Cdc9
Chk1
Rad23
Ubi4
Ede1
Ent2
Vps27
Chc1
Ptp3
Stb1
Spa2
Bem2
Bud14
Rck2
Glc7
Ipl1
Cdc5
Dbf4
Pfk26
Tpk1
Gpa2
Ras2
Med6
Rpb10
Paf1
Rpo26
Bdp1
Spt15
Spt5
Rgr1
Srb7
Ssn2
Nop1
Nsr1
Nop56
Snu13
Rps31
Rad9
Rtf1
Rpo21
Npl3
Air2
Fps1
Ask10
Cdc28
Hog1
Pbs2
Ssk2
Clb2
Bud3
Bud4
Msn2
Sch9
Rim15
Yak1
Ypl247c
Vps52
Kss1
Sko1
Slt2
Msg5
Swe1
Igo2
Cdc55
Igo1
Vps53
Vps54
Rtn1
Fmp42
Cot1
Zrc1
10
20
30
40
50
t/s

#### Path 600 (Chc1 − Ded1)

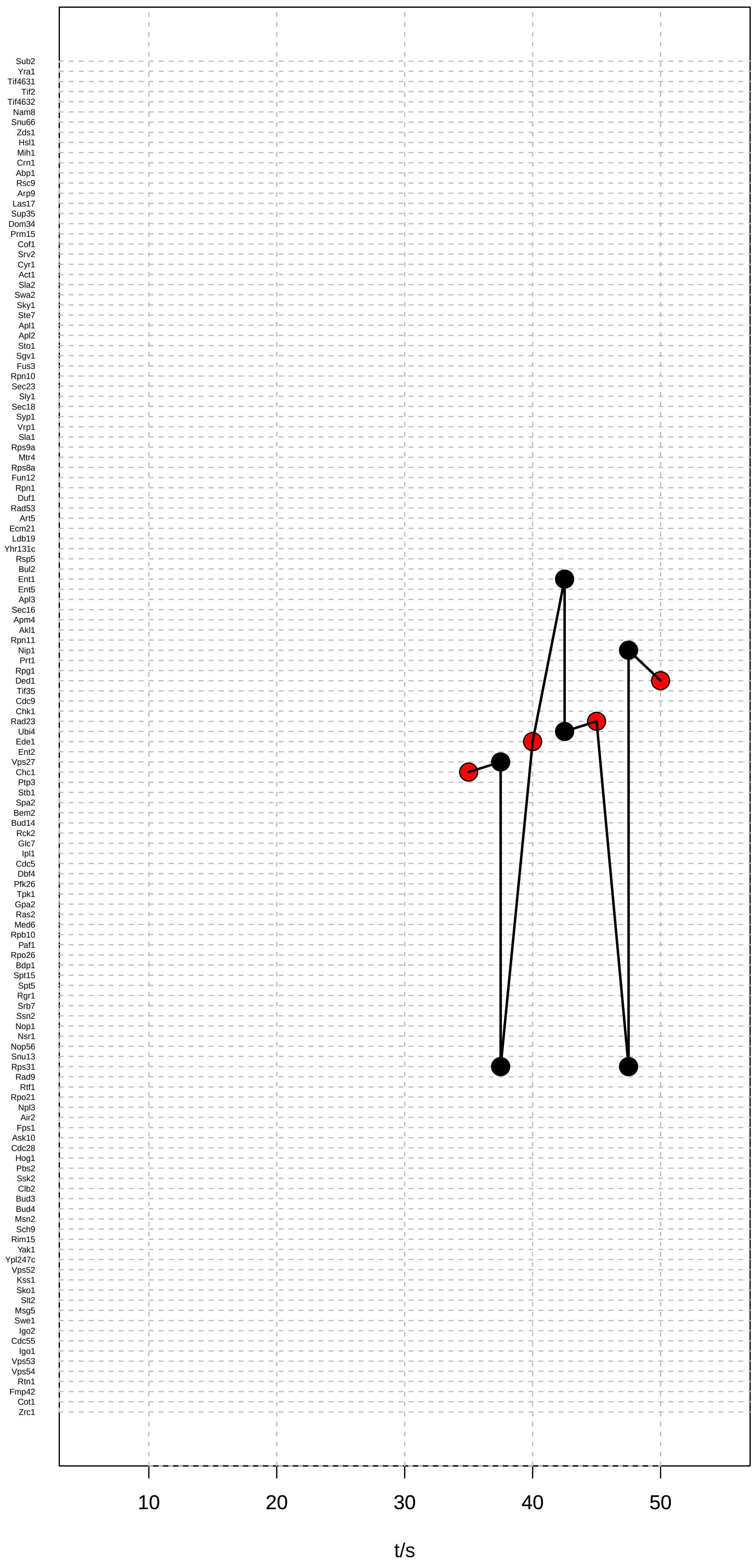

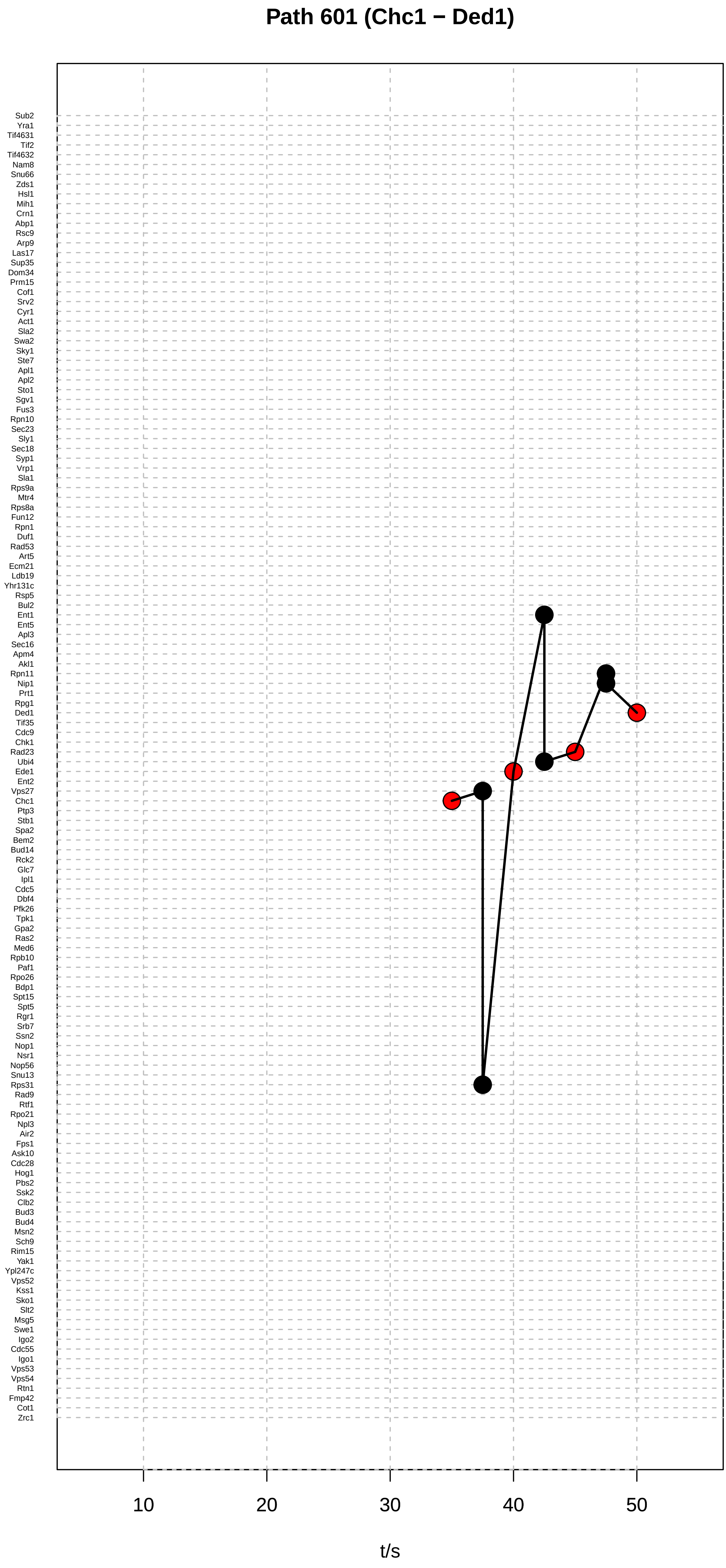

Path 601 (Chc1 − Ded1)
Sub2
Yra1
Tif4631
Tif2
Tif4632
Nam8
Snu66
Zds1
Hsl1
Mih1
Crn1
Abp1
Rsc9
Arp9
Las17
Sup35
Dom34
Prm15
Cof1
Srv2
Cyr1
Act1
Sla2
Swa2
Sky1
Ste7
Apl1
Apl2
Sto1
Sgv1
Fus3
Rpn10
Sec23
Sly1
Sec18
Syp1
Vrp1
Sla1
Rps9a
Mtr4
Rps8a
Fun12
Rpn1
Duf1
Rad53
Art5
Ecm21
Ldb19
Yhr131c
Rsp5
Bul2
Ent1
Ent5
Apl3
Sec16
Apm4
Akl1
Rpn11
Nip1
Prt1
Rpg1
Ded1
Tif35
Cdc9
Chk1
Rad23
Ubi4
Ede1
Ent2
Vps27
Chc1
Ptp3
Stb1
Spa2
Bem2
Bud14
Rck2
Glc7
Ipl1
Cdc5
Dbf4
Pfk26
Tpk1
Gpa2
Ras2
Med6
Rpb10
Paf1
Rpo26
Bdp1
Spt15
Spt5
Rgr1
Srb7
Ssn2
Nop1
Nsr1
Nop56
Snu13
Rps31
Rad9
Rtf1
Rpo21
Npl3
Air2
Fps1
Ask10
Cdc28
Hog1
Pbs2
Ssk2
Clb2
Bud3
Bud4
Msn2
Sch9
Rim15
Yak1
Ypl247c
Vps52
Kss1
Sko1
Slt2
Msg5
Swe1
Igo2
Cdc55
Igo1
Vps53
Vps54
Rtn1
Fmp42
Cot1
Zrc1
10
20
30
40
50
t/s

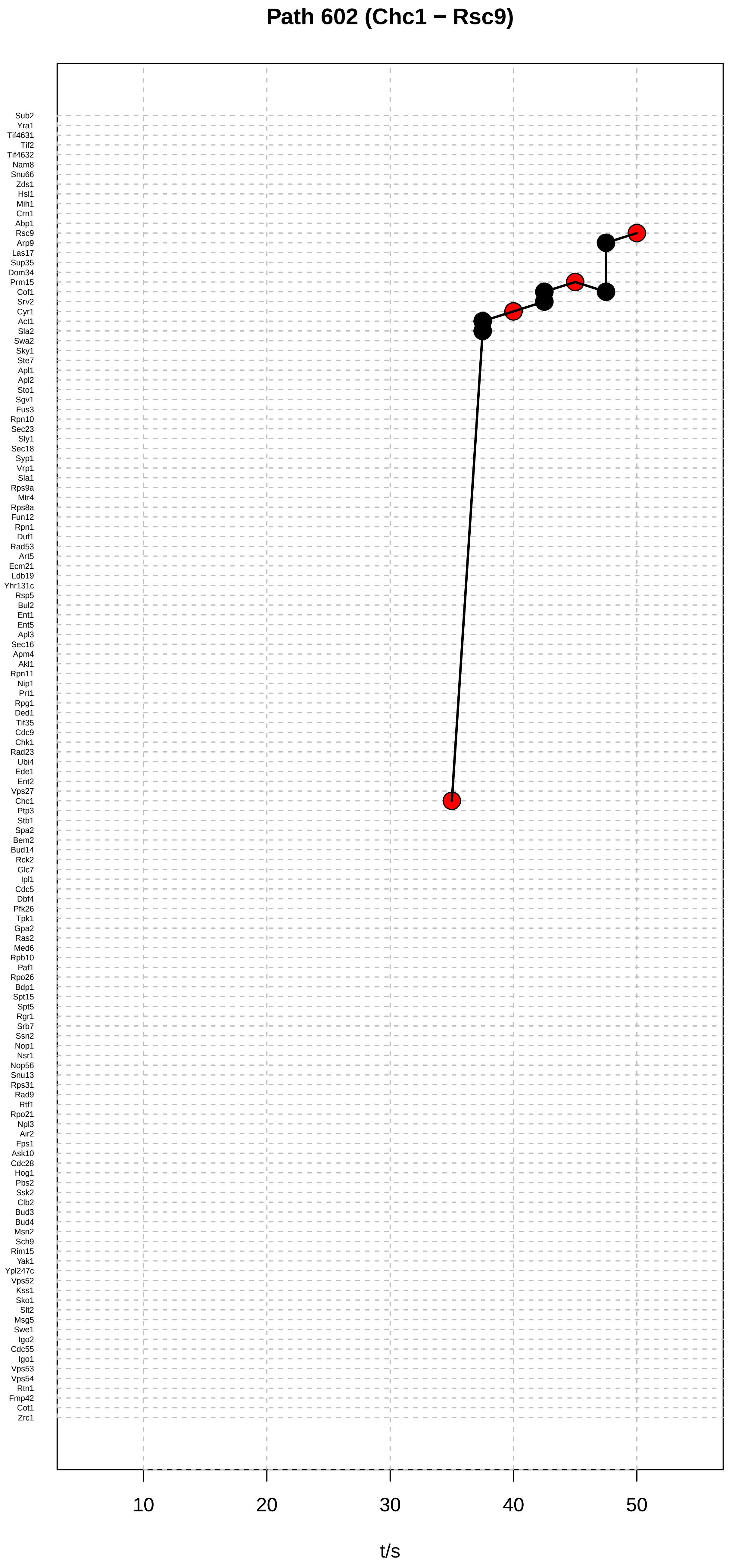

Path 602 (Chc1 − Rsc9)
Sub2
Yra1
Tif4631
Tif2
Tif4632
Nam8
Snu66
Zds1
Hsl1
Mih1
Crn1
Abp1
Rsc9
Arp9
Las17
Sup35
Dom34
Prm15
Cof1
Srv2
Cyr1
Act1
Sla2
Swa2
Sky1
Ste7
Apl1
Apl2
Sto1
Sgv1
Fus3
Rpn10
Sec23
Sly1
Sec18
Syp1
Vrp1
Sla1
Rps9a
Mtr4
Rps8a
Fun12
Rpn1
Duf1
Rad53
Art5
Ecm21
Ldb19
Yhr131c
Rsp5
Bul2
Ent1
Ent5
Apl3
Sec16
Apm4
Akl1
Rpn11
Nip1
Prt1
Rpg1
Ded1
Tif35
Cdc9
Chk1
Rad23
Ubi4
Ede1
Ent2
Vps27
Chc1
Ptp3
Stb1
Spa2
Bem2
Bud14
Rck2
Glc7
Ipl1
Cdc5
Dbf4
Pfk26
Tpk1
Gpa2
Ras2
Med6
Rpb10
Paf1
Rpo26
Bdp1
Spt15
Spt5
Rgr1
Srb7
Ssn2
Nop1
Nsr1
Nop56
Snu13
Rps31
Rad9
Rtf1
Rpo21
Npl3
Air2
Fps1
Ask10
Cdc28
Hog1
Pbs2
Ssk2
Clb2
Bud3
Bud4
Msn2
Sch9
Rim15
Yak1
Ypl247c
Vps52
Kss1
Sko1
Slt2
Msg5
Swe1
Igo2
Cdc55
Igo1
Vps53
Vps54
Rtn1
Fmp42
Cot1
Zrc1
10
20
30
40
50
t/s

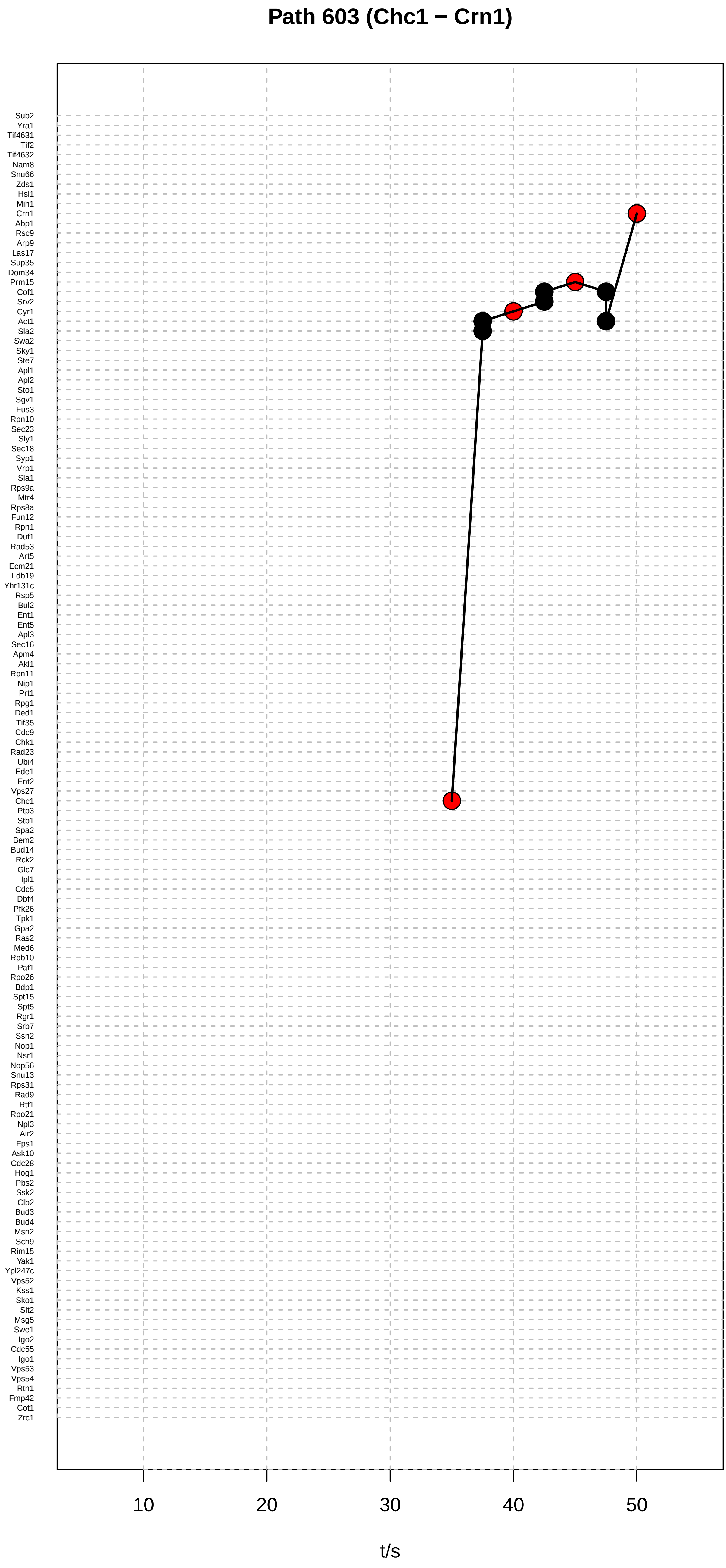

Path 603 (Chc1 − Crn1)
Sub2
Yra1
Tif4631
Tif2
Tif4632
Nam8
Snu66
Zds1
Hsl1
Mih1
Crn1
Abp1
Rsc9
Arp9
Las17
Sup35
Dom34
Prm15
Cof1
Srv2
Cyr1
Act1
Sla2
Swa2
Sky1
Ste7
Apl1
Apl2
Sto1
Sgv1
Fus3
Rpn10
Sec23
Sly1
Sec18
Syp1
Vrp1
Sla1
Rps9a
Mtr4
Rps8a
Fun12
Rpn1
Duf1
Rad53
Art5
Ecm21
Ldb19
Yhr131c
Rsp5
Bul2
Ent1
Ent5
Apl3
Sec16
Apm4
Akl1
Rpn11
Nip1
Prt1
Rpg1
Ded1
Tif35
Cdc9
Chk1
Rad23
Ubi4
Ede1
Ent2
Vps27
Chc1
Ptp3
Stb1
Spa2
Bem2
Bud14
Rck2
Glc7
Ipl1
Cdc5
Dbf4
Pfk26
Tpk1
Gpa2
Ras2
Med6
Rpb10
Paf1
Rpo26
Bdp1
Spt15
Spt5
Rgr1
Srb7
Ssn2
Nop1
Nsr1
Nop56
Snu13
Rps31
Rad9
Rtf1
Rpo21
Npl3
Air2
Fps1
Ask10
Cdc28
Hog1
Pbs2
Ssk2
Clb2
Bud3
Bud4
Msn2
Sch9
Rim15
Yak1
Ypl247c
Vps52
Kss1
Sko1
Slt2
Msg5
Swe1
Igo2
Cdc55
Igo1
Vps53
Vps54
Rtn1
Fmp42
Cot1
Zrc1
10
20
30
40
50
t/s

#### Path 604 (Chc1 − Rsc9)

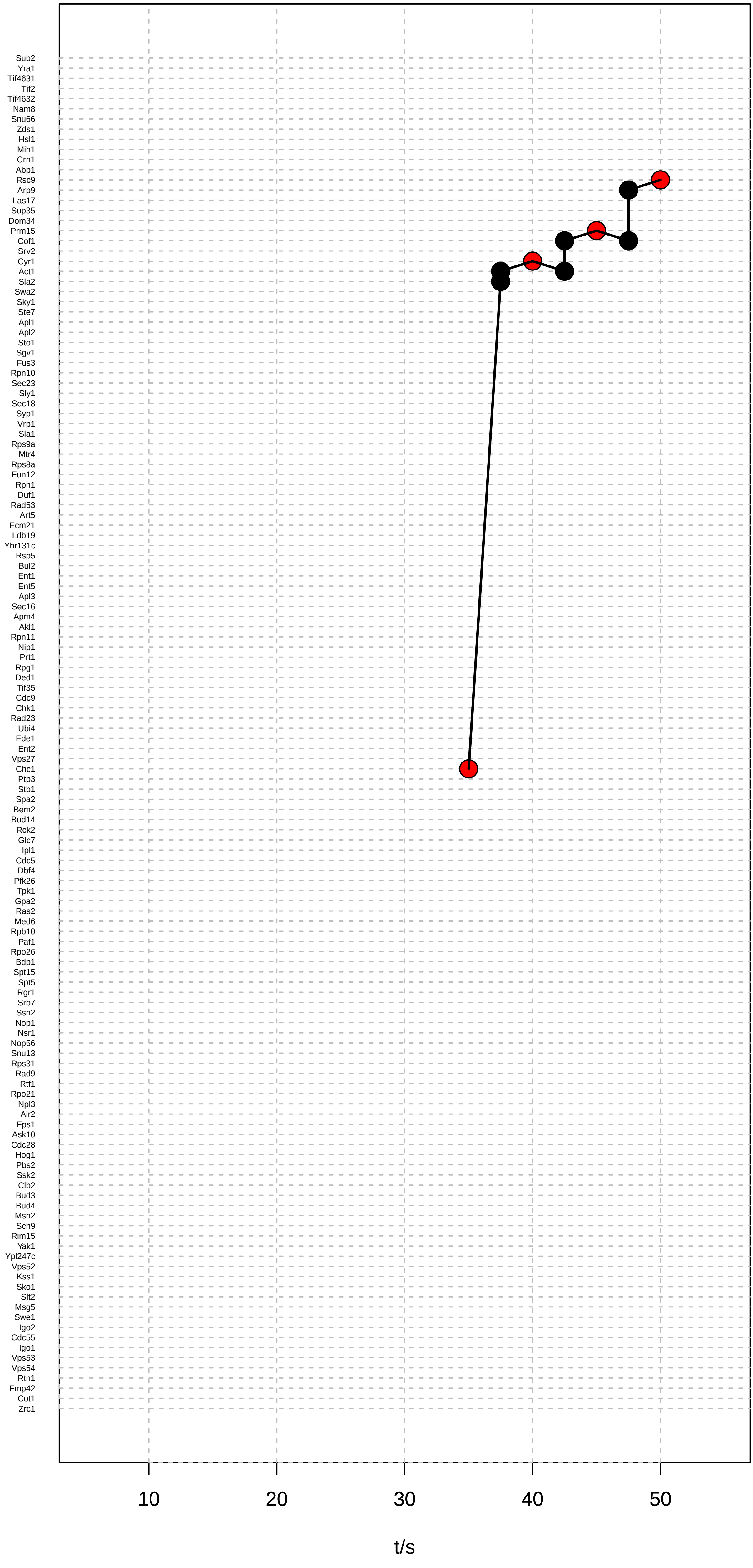

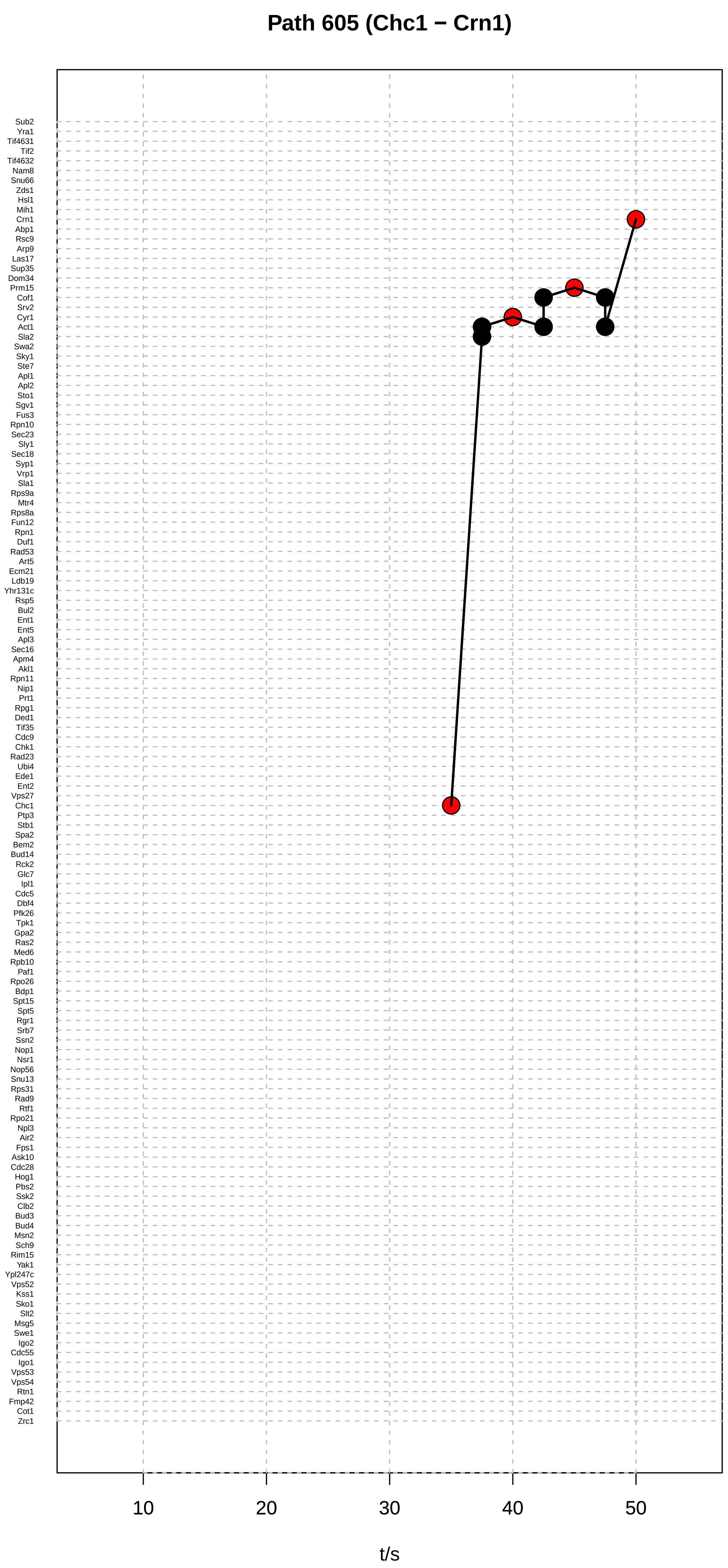
Path 605 (Chc1 − Crn1)
Sub2
Yra1
Tif4631
Tif2
Tif4632
Nam8
Snu66
Zds1
Hsl1
Mih1
Crn1
Abp1
Rsc9
Arp9
Las17
Sup35
Dom34
Prm15
Cof1
Srv2
Cyr1
Act1
Sla2
Swa2
Sky1
Ste7
Apl1
Apl2
Sto1
Sgv1
Fus3
Rpn10
Sec23
Sly1
Sec18
Syp1
Vrp1
Sla1
Rps9a
Mtr4
Rps8a
Fun12
Rpn1
Duf1
Rad53
Art5
Ecm21
Ldb19
Yhr131c
Rsp5
Bul2
Ent1
Ent5
Apl3
Sec16
Apm4
Akl1
Rpn11
Nip1
Prt1
Rpg1
Ded1
Tif35
Cdc9
Chk1
Rad23
Ubi4
Ede1
Ent2
Vps27
Chc1
Ptp3
Stb1
Spa2
Bem2
Bud14
Rck2
Glc7
Ipl1
Cdc5
Dbf4
Pfk26
Tpk1
Gpa2
Ras2
Med6
Rpb10
Paf1
Rpo26
Bdp1
Spt15
Spt5
Rgr1
Srb7
Ssn2
Nop1
Nsr1
Nop56
Snu13
Rps31
Rad9
Rtf1
Rpo21
Npl3
Air2
Fps1
Ask10
Cdc28
Hog1
Pbs2
Ssk2
Clb2
Bud3
Bud4
Msn2
Sch9
Rim15
Yak1
Ypl247c
Vps52
Kss1
Sko1
Slt2
Msg5
Swe1
Igo2
Cdc55
Igo1
Vps53
Vps54
Rtn1
Fmp42
Cot1
Zrc1
10
20
30
40
50
t/s

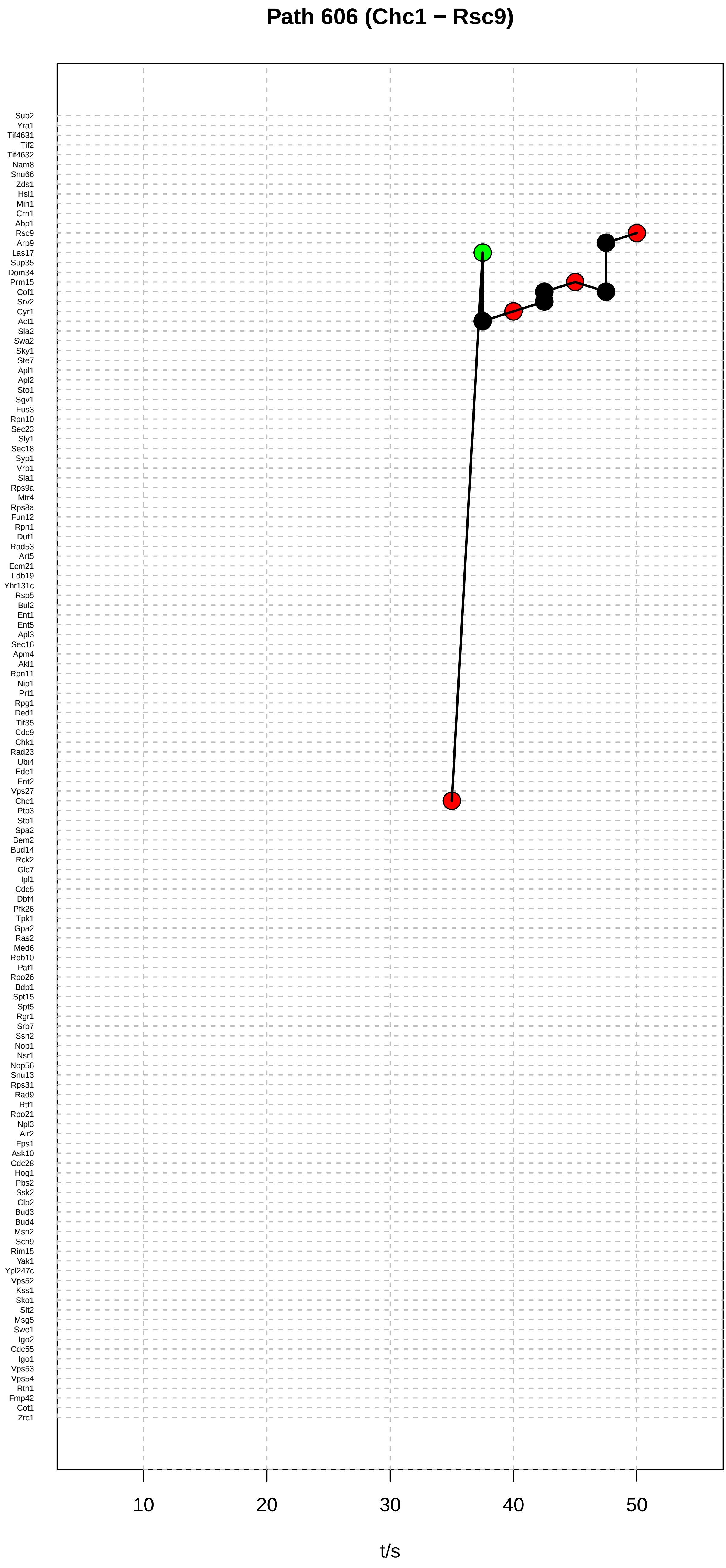

Path 606 (Chc1 − Rsc9)
Sub2
Yra1
Tif4631
Tif2
Tif4632
Nam8
Snu66
Zds1
Hsl1
Mih1
Crn1
Abp1
Rsc9
Arp9
Las17
Sup35
Dom34
Prm15
Cof1
Srv2
Cyr1
Act1
Sla2
Swa2
Sky1
Ste7
Apl1
Apl2
Sto1
Sgv1
Fus3
Rpn10
Sec23
Sly1
Sec18
Syp1
Vrp1
Sla1
Rps9a
Mtr4
Rps8a
Fun12
Rpn1
Duf1
Rad53
Art5
Ecm21
Ldb19
Yhr131c
Rsp5
Bul2
Ent1
Ent5
Apl3
Sec16
Apm4
Akl1
Rpn11
Nip1
Prt1
Rpg1
Ded1
Tif35
Cdc9
Chk1
Rad23
Ubi4
Ede1
Ent2
Vps27
Chc1
Ptp3
Stb1
Spa2
Bem2
Bud14
Rck2
Glc7
Ipl1
Cdc5
Dbf4
Pfk26
Tpk1
Gpa2
Ras2
Med6
Rpb10
Paf1
Rpo26
Bdp1
Spt15
Spt5
Rgr1
Srb7
Ssn2
Nop1
Nsr1
Nop56
Snu13
Rps31
Rad9
Rtf1
Rpo21
Npl3
Air2
Fps1
Ask10
Cdc28
Hog1
Pbs2
Ssk2
Clb2
Bud3
Bud4
Msn2
Sch9
Rim15
Yak1
Ypl247c
Vps52
Kss1
Sko1
Slt2
Msg5
Swe1
Igo2
Cdc55
Igo1
Vps53
Vps54
Rtn1
Fmp42
Cot1
Zrc1
10
20
30
40
50
t/s

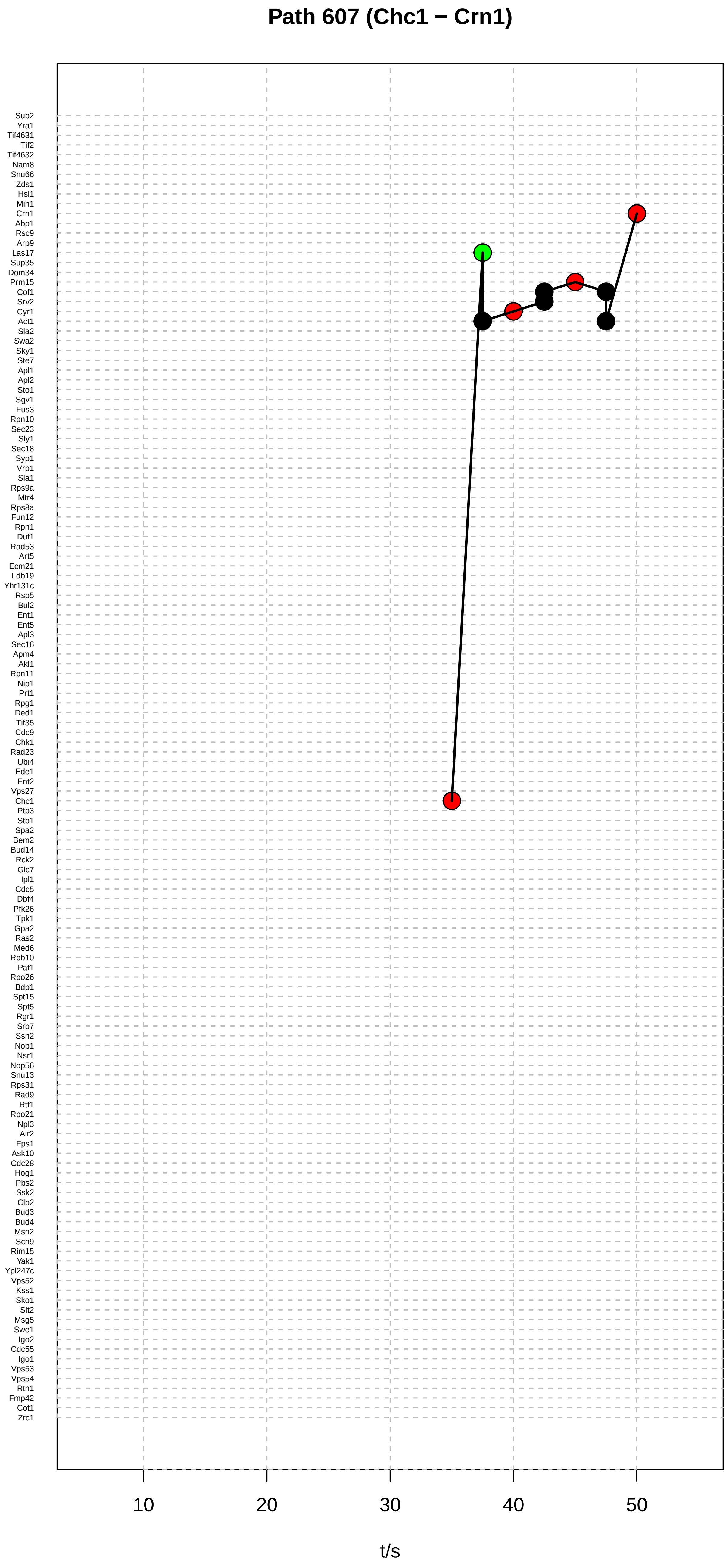
Path 607 (Chc1 − Crn1)
Sub2
Yra1
Tif4631
Tif2
Tif4632
Nam8
Snu66
Zds1
Hsl1
Mih1
Crn1
Abp1
Rsc9
Arp9
Las17
Sup35
Dom34
Prm15
Cof1
Srv2
Cyr1
Act1
Sla2
Swa2
Sky1
Ste7
Apl1
Apl2
Sto1
Sgv1
Fus3
Rpn10
Sec23
Sly1
Sec18
Syp1
Vrp1
Sla1
Rps9a
Mtr4
Rps8a
Fun12
Rpn1
Duf1
Rad53
Art5
Ecm21
Ldb19
Yhr131c
Rsp5
Bul2
Ent1
Ent5
Apl3
Sec16
Apm4
Akl1
Rpn11
Nip1
Prt1
Rpg1
Ded1
Tif35
Cdc9
Chk1
Rad23
Ubi4
Ede1
Ent2
Vps27
Chc1
Ptp3
Stb1
Spa2
Bem2
Bud14
Rck2
Glc7
Ipl1
Cdc5
Dbf4
Pfk26
Tpk1
Gpa2
Ras2
Med6
Rpb10
Paf1
Rpo26
Bdp1
Spt15
Spt5
Rgr1
Srb7
Ssn2
Nop1
Nsr1
Nop56
Snu13
Rps31
Rad9
Rtf1
Rpo21
Npl3
Air2
Fps1
Ask10
Cdc28
Hog1
Pbs2
Ssk2
Clb2
Bud3
Bud4
Msn2
Sch9
Rim15
Yak1
Ypl247c
Vps52
Kss1
Sko1
Slt2
Msg5
Swe1
Igo2
Cdc55
Igo1
Vps53
Vps54
Rtn1
Fmp42
Cot1
Zrc1
10
20
30
40
50
t/s

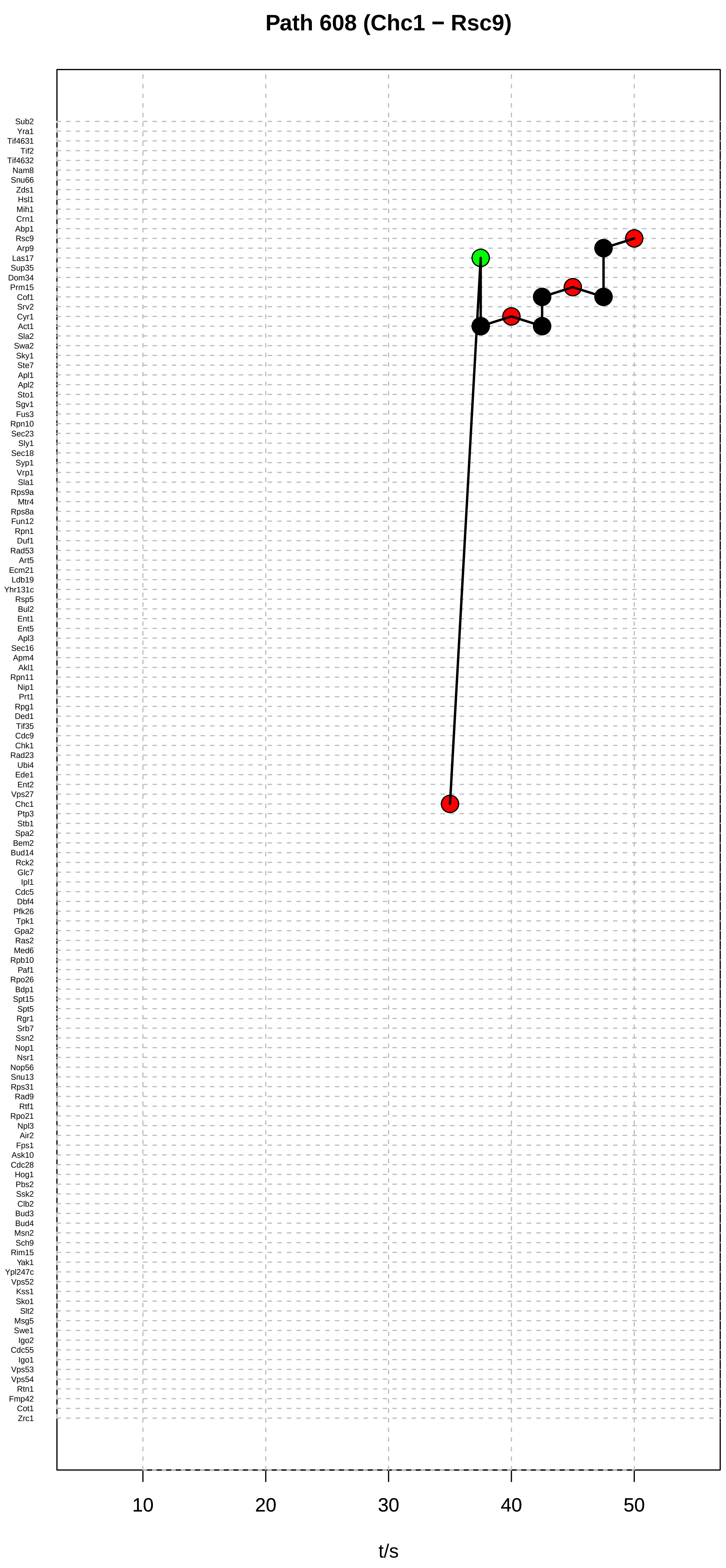
Path 608 (Chc1 − Rsc9)
Sub2
Yra1
Tif4631
Tif2
Tif4632
Nam8
Snu66
Zds1
Hsl1
Mih1
Crn1
Abp1
Rsc9
Arp9
Las17
Sup35
Dom34
Prm15
Cof1
Srv2
Cyr1
Act1
Sla2
Swa2
Sky1
Ste7
Apl1
Apl2
Sto1
Sgv1
Fus3
Rpn10
Sec23
Sly1
Sec18
Syp1
Vrp1
Sla1
Rps9a
Mtr4
Rps8a
Fun12
Rpn1
Duf1
Rad53
Art5
Ecm21
Ldb19
Yhr131c
Rsp5
Bul2
Ent1
Ent5
Apl3
Sec16
Apm4
Akl1
Rpn11
Nip1
Prt1
Rpg1
Ded1
Tif35
Cdc9
Chk1
Rad23
Ubi4
Ede1
Ent2
Vps27
Chc1
Ptp3
Stb1
Spa2
Bem2
Bud14
Rck2
Glc7
Ipl1
Cdc5
Dbf4
Pfk26
Tpk1
Gpa2
Ras2
Med6
Rpb10
Paf1
Rpo26
Bdp1
Spt15
Spt5
Rgr1
Srb7
Ssn2
Nop1
Nsr1
Nop56
Snu13
Rps31
Rad9
Rtf1
Rpo21
Npl3
Air2
Fps1
Ask10
Cdc28
Hog1
Pbs2
Ssk2
Clb2
Bud3
Bud4
Msn2
Sch9
Rim15
Yak1
Ypl247c
Vps52
Kss1
Sko1
Slt2
Msg5
Swe1
Igo2
Cdc55
Igo1
Vps53
Vps54
Rtn1
Fmp42
Cot1
Zrc1
10
20
30
40
50
t/s

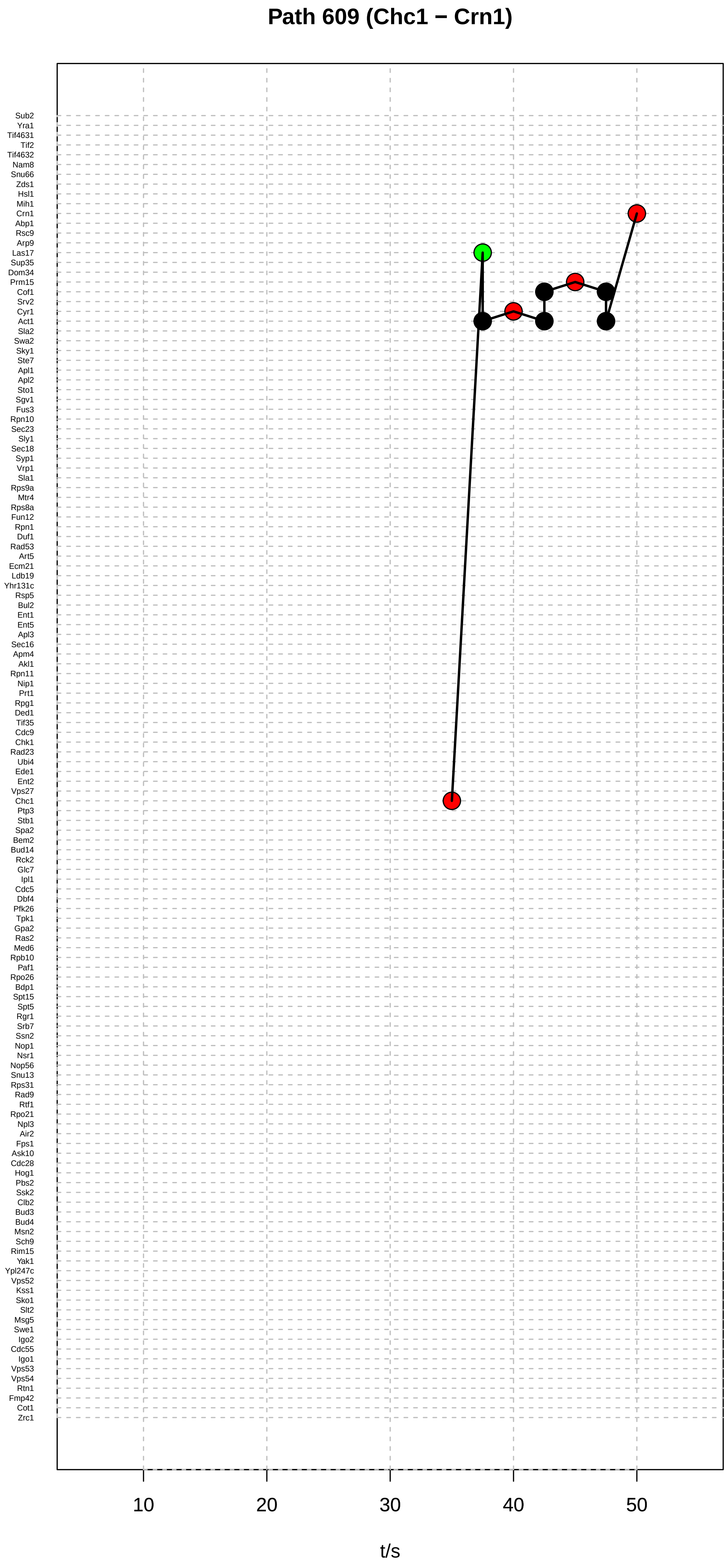
Path 609 (Chc1 − Crn1)
Sub2
Yra1
Tif4631
Tif2
Tif4632
Nam8
Snu66
Zds1
Hsl1
Mih1
Crn1
Abp1
Rsc9
Arp9
Las17
Sup35
Dom34
Prm15
Cof1
Srv2
Cyr1
Act1
Sla2
Swa2
Sky1
Ste7
Apl1
Apl2
Sto1
Sgv1
Fus3
Rpn10
Sec23
Sly1
Sec18
Syp1
Vrp1
Sla1
Rps9a
Mtr4
Rps8a
Fun12
Rpn1
Duf1
Rad53
Art5
Ecm21
Ldb19
Yhr131c
Rsp5
Bul2
Ent1
Ent5
Apl3
Sec16
Apm4
Akl1
Rpn11
Nip1
Prt1
Rpg1
Ded1
Tif35
Cdc9
Chk1
Rad23
Ubi4
Ede1
Ent2
Vps27
Chc1
Ptp3
Stb1
Spa2
Bem2
Bud14
Rck2
Glc7
Ipl1
Cdc5
Dbf4
Pfk26
Tpk1
Gpa2
Ras2
Med6
Rpb10
Paf1
Rpo26
Bdp1
Spt15
Spt5
Rgr1
Srb7
Ssn2
Nop1
Nsr1
Nop56
Snu13
Rps31
Rad9
Rtf1
Rpo21
Npl3
Air2
Fps1
Ask10
Cdc28
Hog1
Pbs2
Ssk2
Clb2
Bud3
Bud4
Msn2
Sch9
Rim15
Yak1
Ypl247c
Vps52
Kss1
Sko1
Slt2
Msg5
Swe1
Igo2
Cdc55
Igo1
Vps53
Vps54
Rtn1
Fmp42
Cot1
Zrc1
10
20
30
40
50
t/s

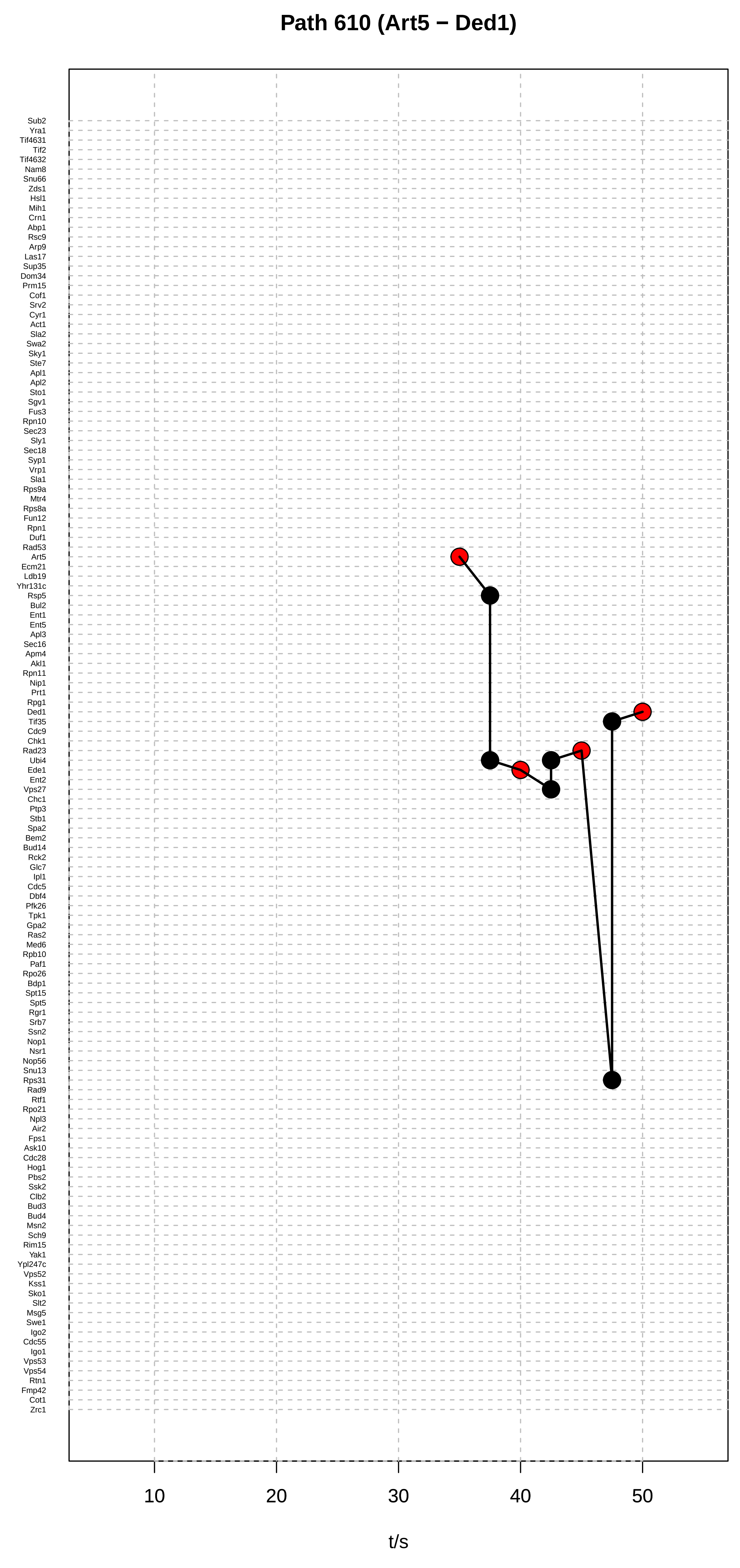
Path 610 (Art5 − Ded1)
Sub2
Yra1
Tif4631
Tif2
Tif4632
Nam8
Snu66
Zds1
Hsl1
Mih1
Crn1
Abp1
Rsc9
Arp9
Las17
Sup35
Dom34
Prm15
Cof1
Srv2
Cyr1
Act1
Sla2
Swa2
Sky1
Ste7
Apl1
Apl2
Sto1
Sgv1
Fus3
Rpn10
Sec23
Sly1
Sec18
Syp1
Vrp1
Sla1
Rps9a
Mtr4
Rps8a
Fun12
Rpn1
Duf1
Rad53
Art5
Ecm21
Ldb19
Yhr131c
Rsp5
Bul2
Ent1
Ent5
Apl3
Sec16
Apm4
Akl1
Rpn11
Nip1
Prt1
Rpg1
Ded1
Tif35
Cdc9
Chk1
Rad23
Ubi4
Ede1
Ent2
Vps27
Chc1
Ptp3
Stb1
Spa2
Bem2
Bud14
Rck2
Glc7
Ipl1
Cdc5
Dbf4
Pfk26
Tpk1
Gpa2
Ras2
Med6
Rpb10
Paf1
Rpo26
Bdp1
Spt15
Spt5
Rgr1
Srb7
Ssn2
Nop1
Nsr1
Nop56
Snu13
Rps31
Rad9
Rtf1
Rpo21
Npl3
Air2
Fps1
Ask10
Cdc28
Hog1
Pbs2
Ssk2
Clb2
Bud3
Bud4
Msn2
Sch9
Rim15
Yak1
Ypl247c
Vps52
Kss1
Sko1
Slt2
Msg5
Swe1
Igo2
Cdc55
Igo1
Vps53
Vps54
Rtn1
Fmp42
Cot1
Zrc1
10
20
30
40
50
t/s

#### Path 611 (Art5 − Ded1)

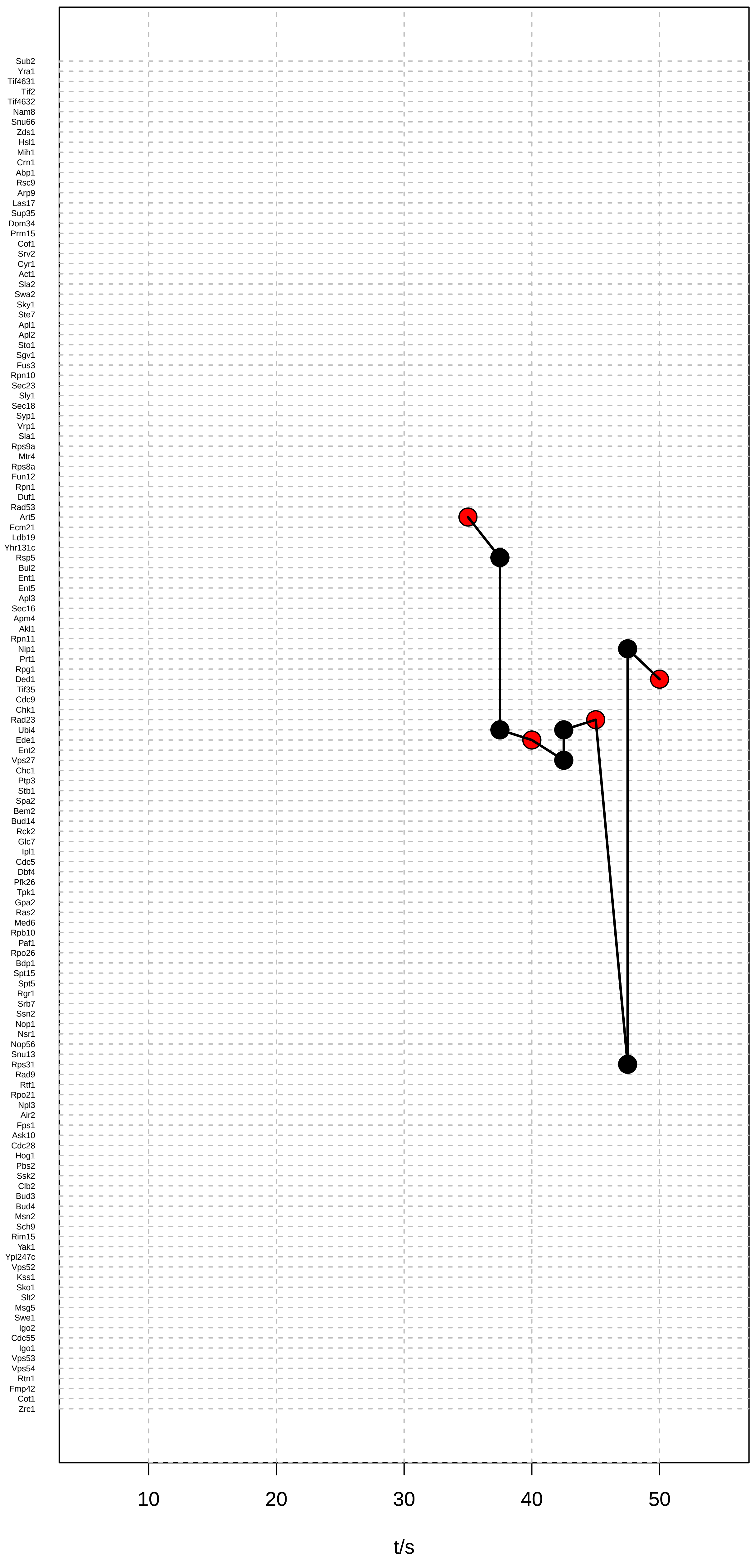

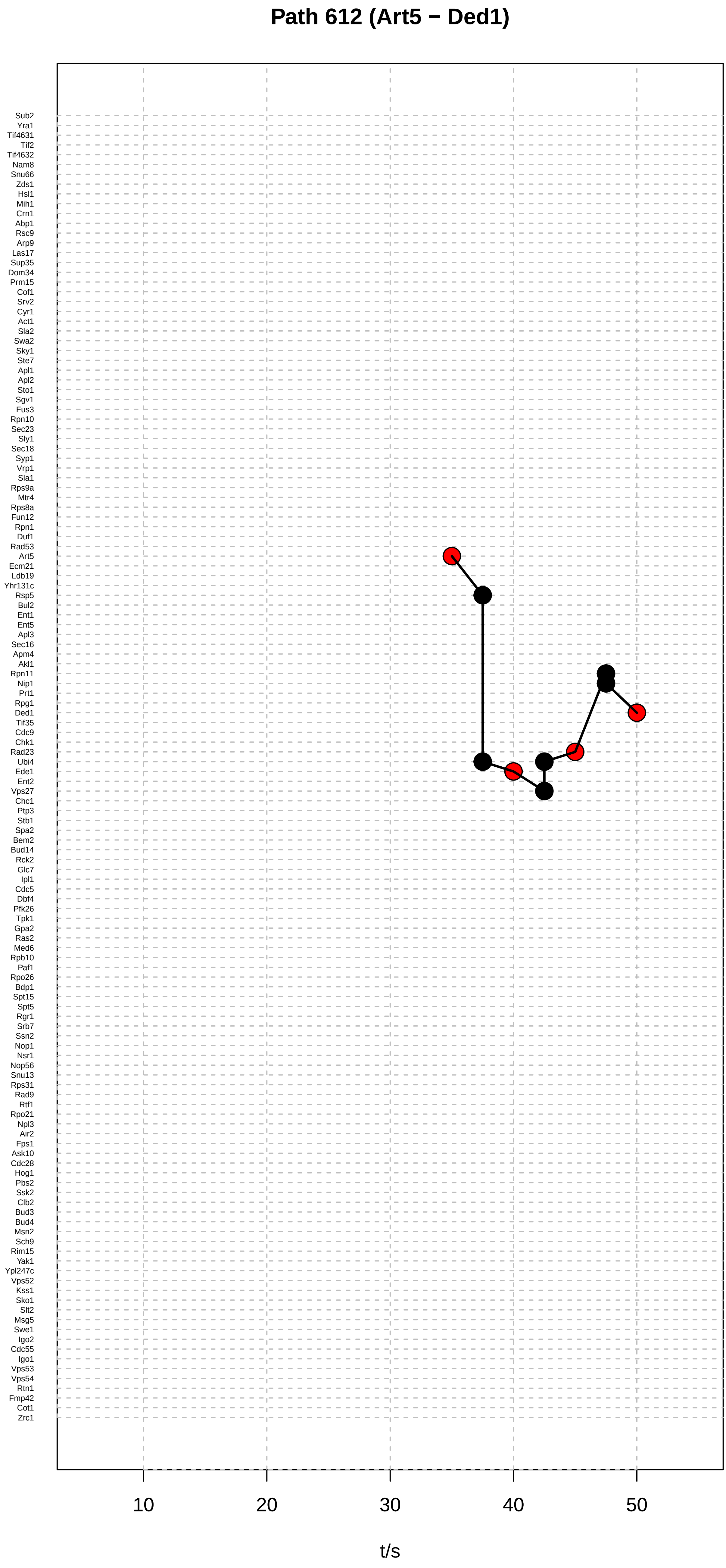
Path 612 (Art5 − Ded1)
Sub2
Yra1
Tif4631
Tif2
Tif4632
Nam8
Snu66
Zds1
Hsl1
Mih1
Crn1
Abp1
Rsc9
Arp9
Las17
Sup35
Dom34
Prm15
Cof1
Srv2
Cyr1
Act1
Sla2
Swa2
Sky1
Ste7
Apl1
Apl2
Sto1
Sgv1
Fus3
Rpn10
Sec23
Sly1
Sec18
Syp1
Vrp1
Sla1
Rps9a
Mtr4
Rps8a
Fun12
Rpn1
Duf1
Rad53
Art5
Ecm21
Ldb19
Yhr131c
Rsp5
Bul2
Ent1
Ent5
Apl3
Sec16
Apm4
Akl1
Rpn11
Nip1
Prt1
Rpg1
Ded1
Tif35
Cdc9
Chk1
Rad23
Ubi4
Ede1
Ent2
Vps27
Chc1
Ptp3
Stb1
Spa2
Bem2
Bud14
Rck2
Glc7
Ipl1
Cdc5
Dbf4
Pfk26
Tpk1
Gpa2
Ras2
Med6
Rpb10
Paf1
Rpo26
Bdp1
Spt15
Spt5
Rgr1
Srb7
Ssn2
Nop1
Nsr1
Nop56
Snu13
Rps31
Rad9
Rtf1
Rpo21
Npl3
Air2
Fps1
Ask10
Cdc28
Hog1
Pbs2
Ssk2
Clb2
Bud3
Bud4
Msn2
Sch9
Rim15
Yak1
Ypl247c
Vps52
Kss1
Sko1
Slt2
Msg5
Swe1
Igo2
Cdc55
Igo1
Vps53
Vps54
Rtn1
Fmp42
Cot1
Zrc1
10
20
30
40
50
t/s

### Path 613 (Art5 − Ded1)

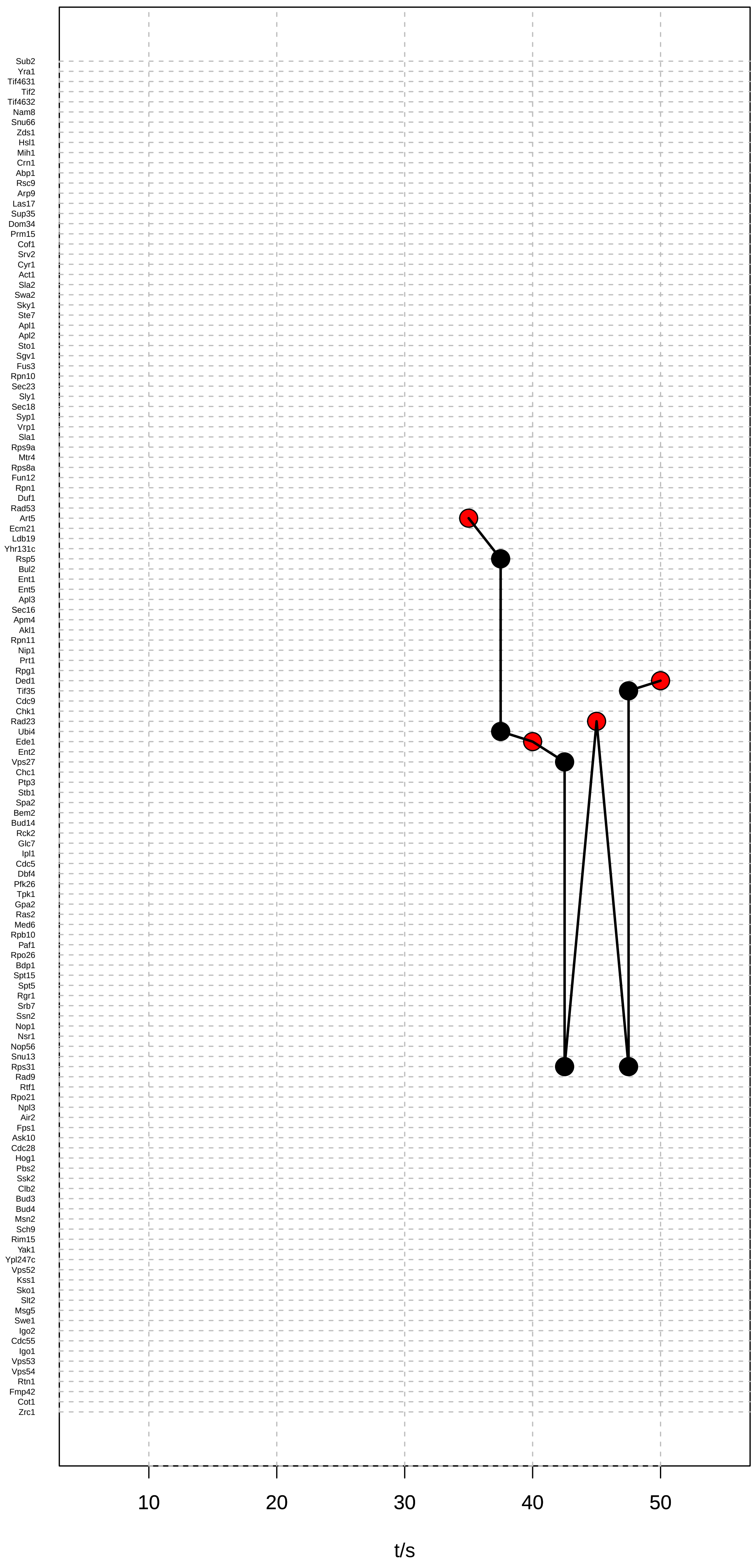

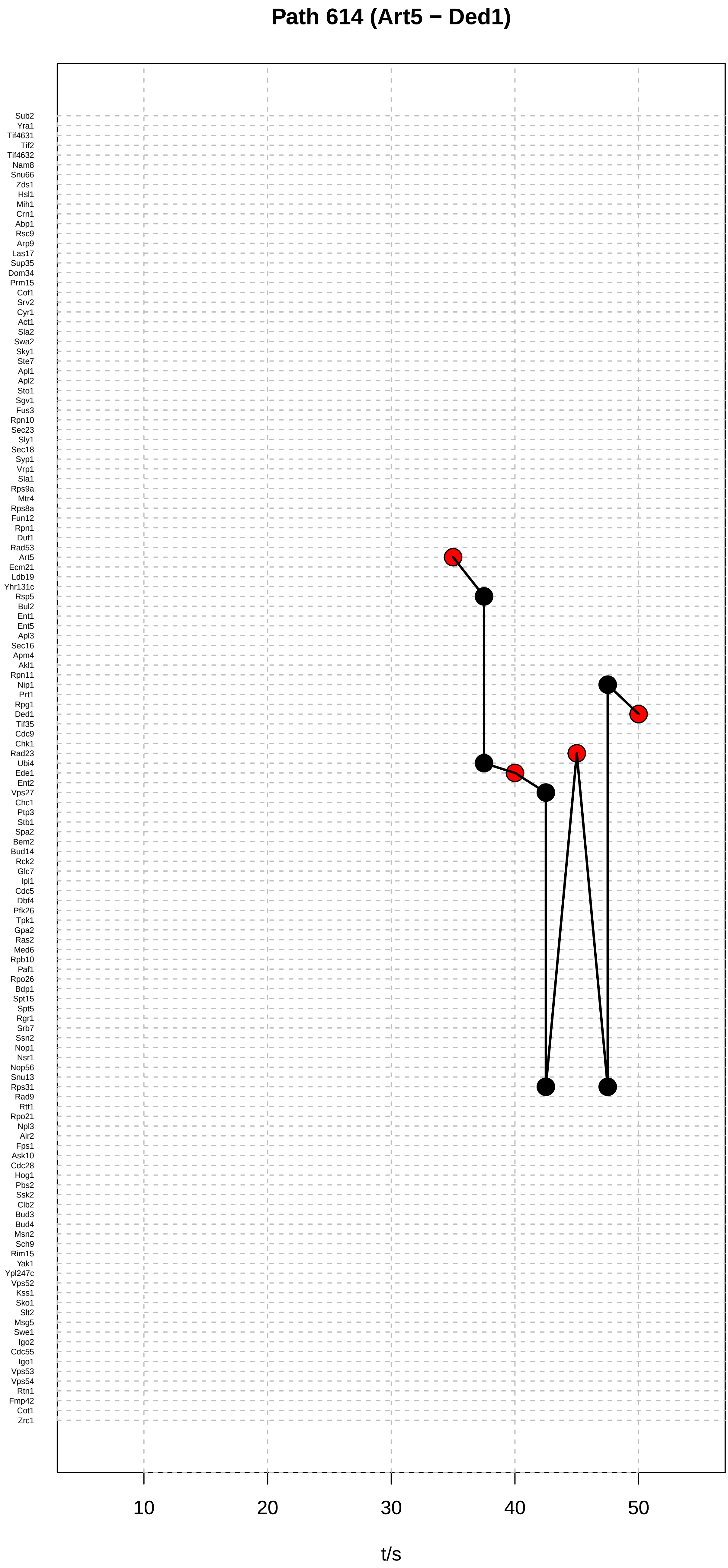
Path 614 (Art5 − Ded1)
Sub2
Yra1
Tif4631
Tif2
Tif4632
Nam8
Snu66
Zds1
Hsl1
Mih1
Crn1
Abp1
Rsc9
Arp9
Las17
Sup35
Dom34
Prm15
Cof1
Srv2
Cyr1
Act1
Sla2
Swa2
Sky1
Ste7
Apl1
Apl2
Sto1
Sgv1
Fus3
Rpn10
Sec23
Sly1
Sec18
Syp1
Vrp1
Sla1
Rps9a
Mtr4
Rps8a
Fun12
Rpn1
Duf1
Rad53
Art5
Ecm21
Ldb19
Yhr131c
Rsp5
Bul2
Ent1
Ent5
Apl3
Sec16
Apm4
Akl1
Rpn11
Nip1
Prt1
Rpg1
Ded1
Tif35
Cdc9
Chk1
Rad23
Ubi4
Ede1
Ent2
Vps27
Chc1
Ptp3
Stb1
Spa2
Bem2
Bud14
Rck2
Glc7
Ipl1
Cdc5
Dbf4
Pfk26
Tpk1
Gpa2
Ras2
Med6
Rpb10
Paf1
Rpo26
Bdp1
Spt15
Spt5
Rgr1
Srb7
Ssn2
Nop1
Nsr1
Nop56
Snu13
Rps31
Rad9
Rtf1
Rpo21
Npl3
Air2
Fps1
Ask10
Cdc28
Hog1
Pbs2
Ssk2
Clb2
Bud3
Bud4
Msn2
Sch9
Rim15
Yak1
Ypl247c
Vps52
Kss1
Sko1
Slt2
Msg5
Swe1
Igo2
Cdc55
Igo1
Vps53
Vps54
Rtn1
Fmp42
Cot1
Zrc1
10
20
30
40
50
t/s

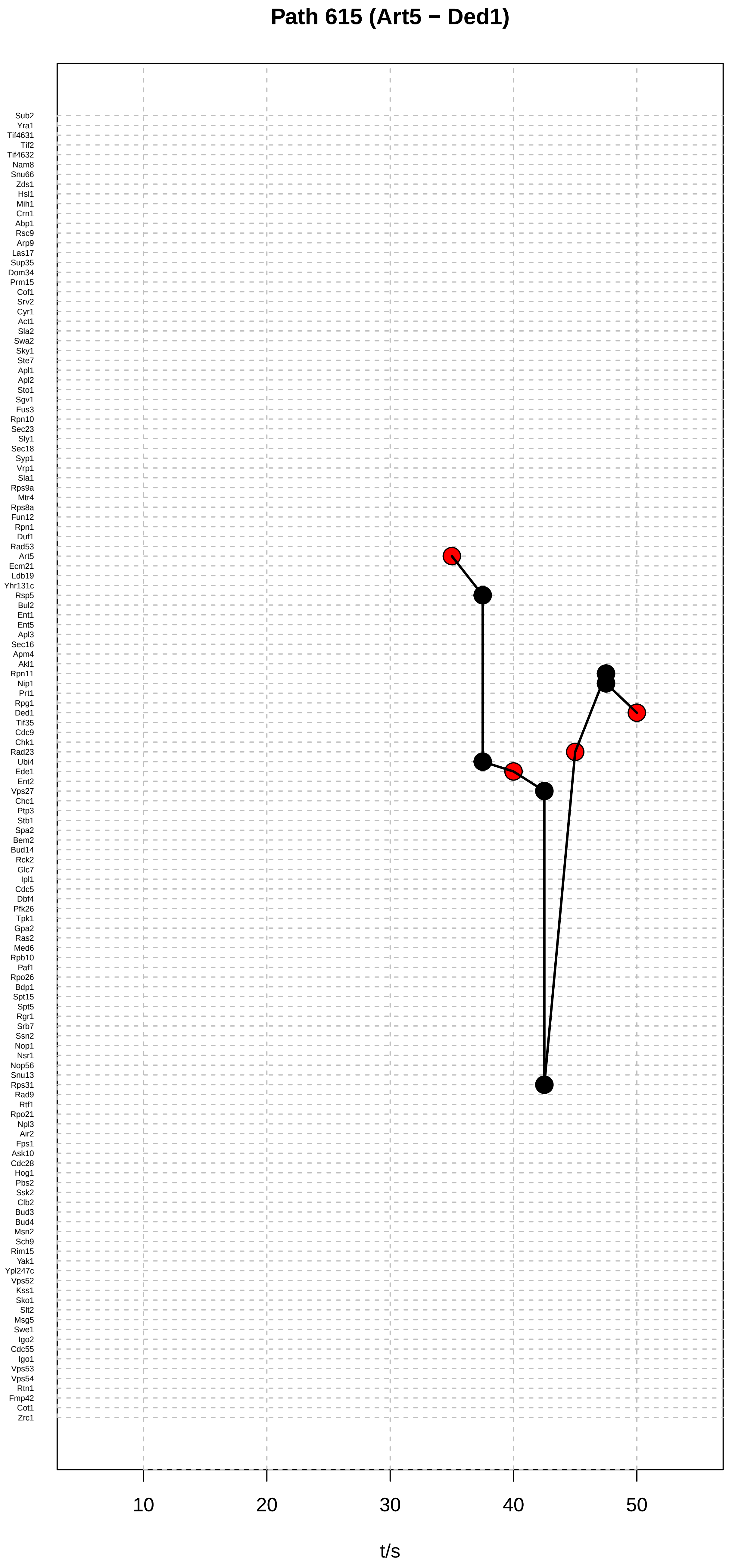
Path 615 (Art5 − Ded1)
Sub2
Yra1
Tif4631
Tif2
Tif4632
Nam8
Snu66
Zds1
Hsl1
Mih1
Crn1
Abp1
Rsc9
Arp9
Las17
Sup35
Dom34
Prm15
Cof1
Srv2
Cyr1
Act1
Sla2
Swa2
Sky1
Ste7
Apl1
Apl2
Sto1
Sgv1
Fus3
Rpn10
Sec23
Sly1
Sec18
Syp1
Vrp1
Sla1
Rps9a
Mtr4
Rps8a
Fun12
Rpn1
Duf1
Rad53
Art5
Ecm21
Ldb19
Yhr131c
Rsp5
Bul2
Ent1
Ent5
Apl3
Sec16
Apm4
Akl1
Rpn11
Nip1
Prt1
Rpg1
Ded1
Tif35
Cdc9
Chk1
Rad23
Ubi4
Ede1
Ent2
Vps27
Chc1
Ptp3
Stb1
Spa2
Bem2
Bud14
Rck2
Glc7
Ipl1
Cdc5
Dbf4
Pfk26
Tpk1
Gpa2
Ras2
Med6
Rpb10
Paf1
Rpo26
Bdp1
Spt15
Spt5
Rgr1
Srb7
Ssn2
Nop1
Nsr1
Nop56
Snu13
Rps31
Rad9
Rtf1
Rpo21
Npl3
Air2
Fps1
Ask10
Cdc28
Hog1
Pbs2
Ssk2
Clb2
Bud3
Bud4
Msn2
Sch9
Rim15
Yak1
Ypl247c
Vps52
Kss1
Sko1
Slt2
Msg5
Swe1
Igo2
Cdc55
Igo1
Vps53
Vps54
Rtn1
Fmp42
Cot1
Zrc1
10
20
30
40
50
t/s

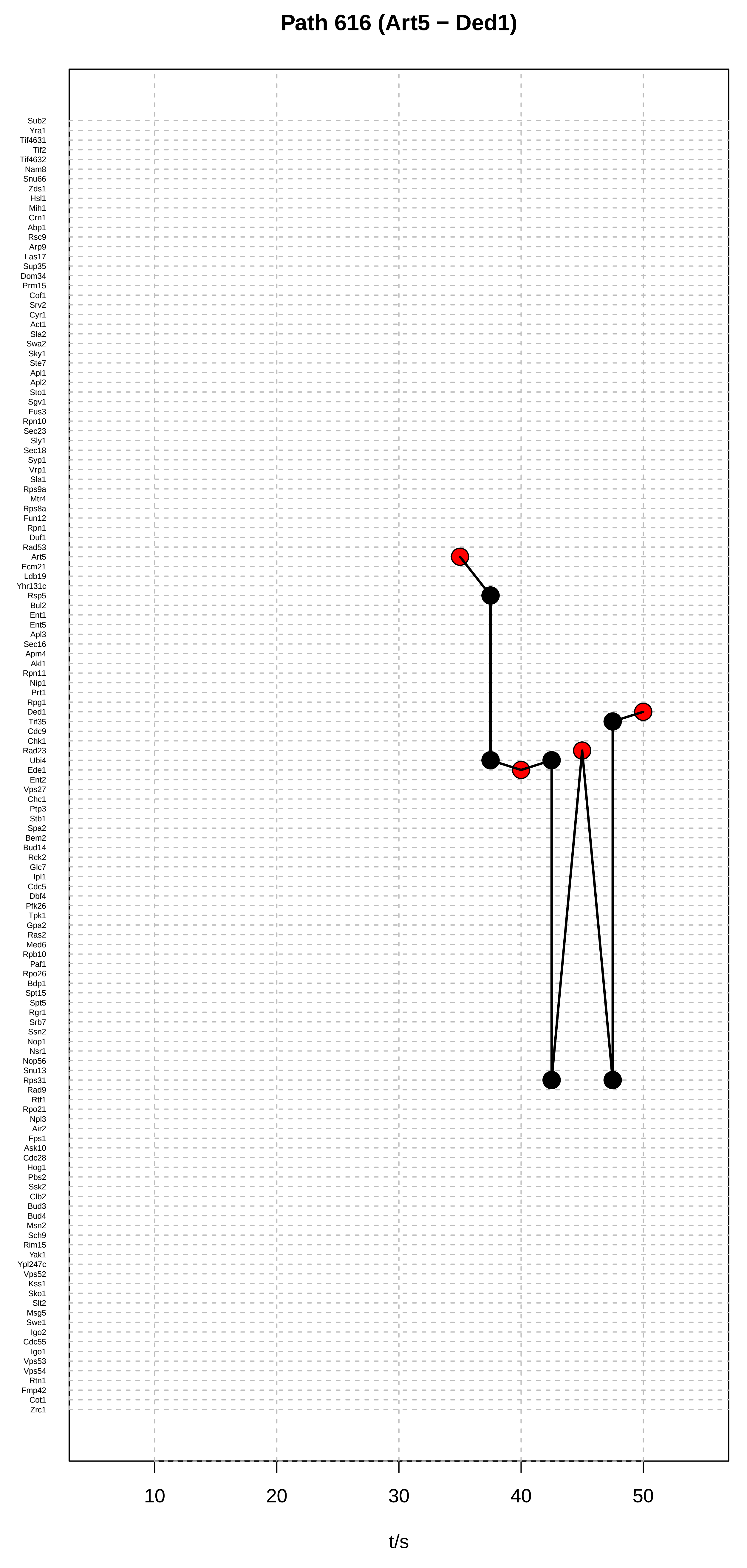

Path 616 (Art5 − Ded1)
Sub2
Yra1
Tif4631
Tif2
Tif4632
Nam8
Snu66
Zds1
Hsl1
Mih1
Crn1
Abp1
Rsc9
Arp9
Las17
Sup35
Dom34
Prm15
Cof1
Srv2
Cyr1
Act1
Sla2
Swa2
Sky1
Ste7
Apl1
Apl2
Sto1
Sgv1
Fus3
Rpn10
Sec23
Sly1
Sec18
Syp1
Vrp1
Sla1
Rps9a
Mtr4
Rps8a
Fun12
Rpn1
Duf1
Rad53
Art5
Ecm21
Ldb19
Yhr131c
Rsp5
Bul2
Ent1
Ent5
Apl3
Sec16
Apm4
Akl1
Rpn11
Nip1
Prt1
Rpg1
Ded1
Tif35
Cdc9
Chk1
Rad23
Ubi4
Ede1
Ent2
Vps27
Chc1
Ptp3
Stb1
Spa2
Bem2
Bud14
Rck2
Glc7
Ipl1
Cdc5
Dbf4
Pfk26
Tpk1
Gpa2
Ras2
Med6
Rpb10
Paf1
Rpo26
Bdp1
Spt15
Spt5
Rgr1
Srb7
Ssn2
Nop1
Nsr1
Nop56
Snu13
Rps31
Rad9
Rtf1
Rpo21
Npl3
Air2
Fps1
Ask10
Cdc28
Hog1
Pbs2
Ssk2
Clb2
Bud3
Bud4
Msn2
Sch9
Rim15
Yak1
Ypl247c
Vps52
Kss1
Sko1
Slt2
Msg5
Swe1
Igo2
Cdc55
Igo1
Vps53
Vps54
Rtn1
Fmp42
Cot1
Zrc1
10
20
30
40
50
t/s

### Path 617 (Art5 − Ded1)

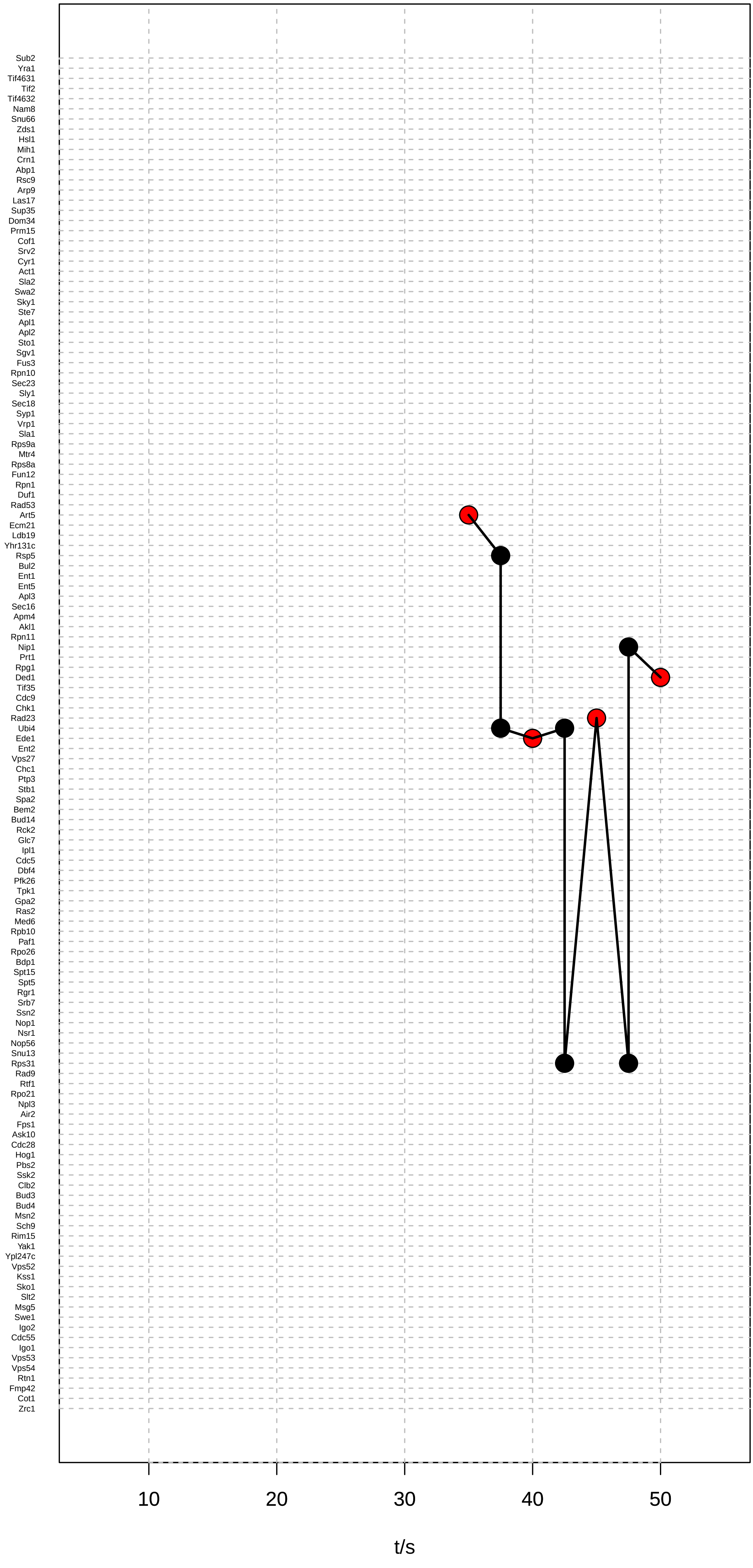

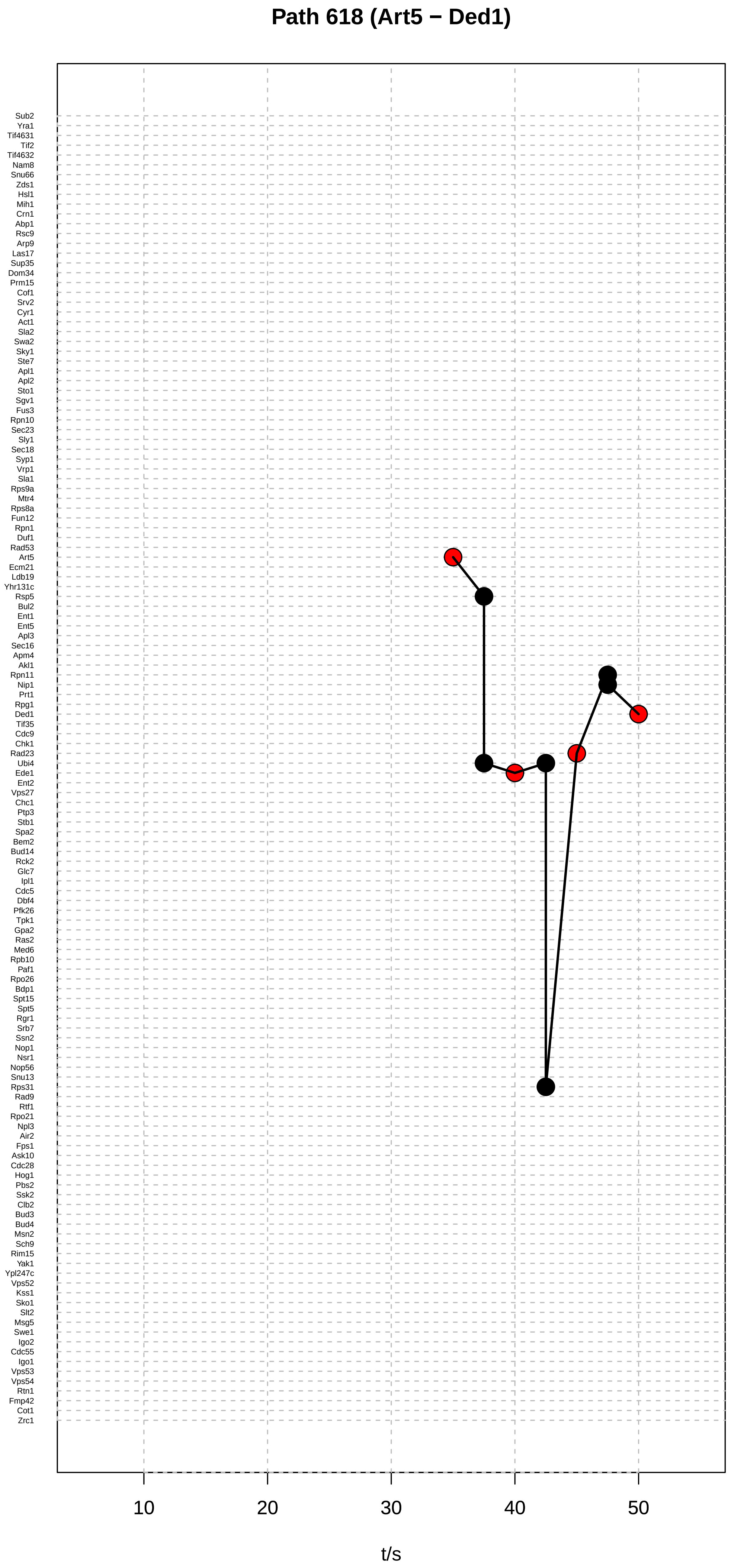
Path 618 (Art5 − Ded1)
Sub2
Yra1
Tif4631
Tif2
Tif4632
Nam8
Snu66
Zds1
Hsl1
Mih1
Crn1
Abp1
Rsc9
Arp9
Las17
Sup35
Dom34
Prm15
Cof1
Srv2
Cyr1
Act1
Sla2
Swa2
Sky1
Ste7
Apl1
Apl2
Sto1
Sgv1
Fus3
Rpn10
Sec23
Sly1
Sec18
Syp1
Vrp1
Sla1
Rps9a
Mtr4
Rps8a
Fun12
Rpn1
Duf1
Rad53
Art5
Ecm21
Ldb19
Yhr131c
Rsp5
Bul2
Ent1
Ent5
Apl3
Sec16
Apm4
Akl1
Rpn11
Nip1
Prt1
Rpg1
Ded1
Tif35
Cdc9
Chk1
Rad23
Ubi4
Ede1
Ent2
Vps27
Chc1
Ptp3
Stb1
Spa2
Bem2
Bud14
Rck2
Glc7
Ipl1
Cdc5
Dbf4
Pfk26
Tpk1
Gpa2
Ras2
Med6
Rpb10
Paf1
Rpo26
Bdp1
Spt15
Spt5
Rgr1
Srb7
Ssn2
Nop1
Nsr1
Nop56
Snu13
Rps31
Rad9
Rtf1
Rpo21
Npl3
Air2
Fps1
Ask10
Cdc28
Hog1
Pbs2
Ssk2
Clb2
Bud3
Bud4
Msn2
Sch9
Rim15
Yak1
Ypl247c
Vps52
Kss1
Sko1
Slt2
Msg5
Swe1
Igo2
Cdc55
Igo1
Vps53
Vps54
Rtn1
Fmp42
Cot1
Zrc1
10
20
30
40
50
t/s

### Path 619 (Art5 − Ded1)

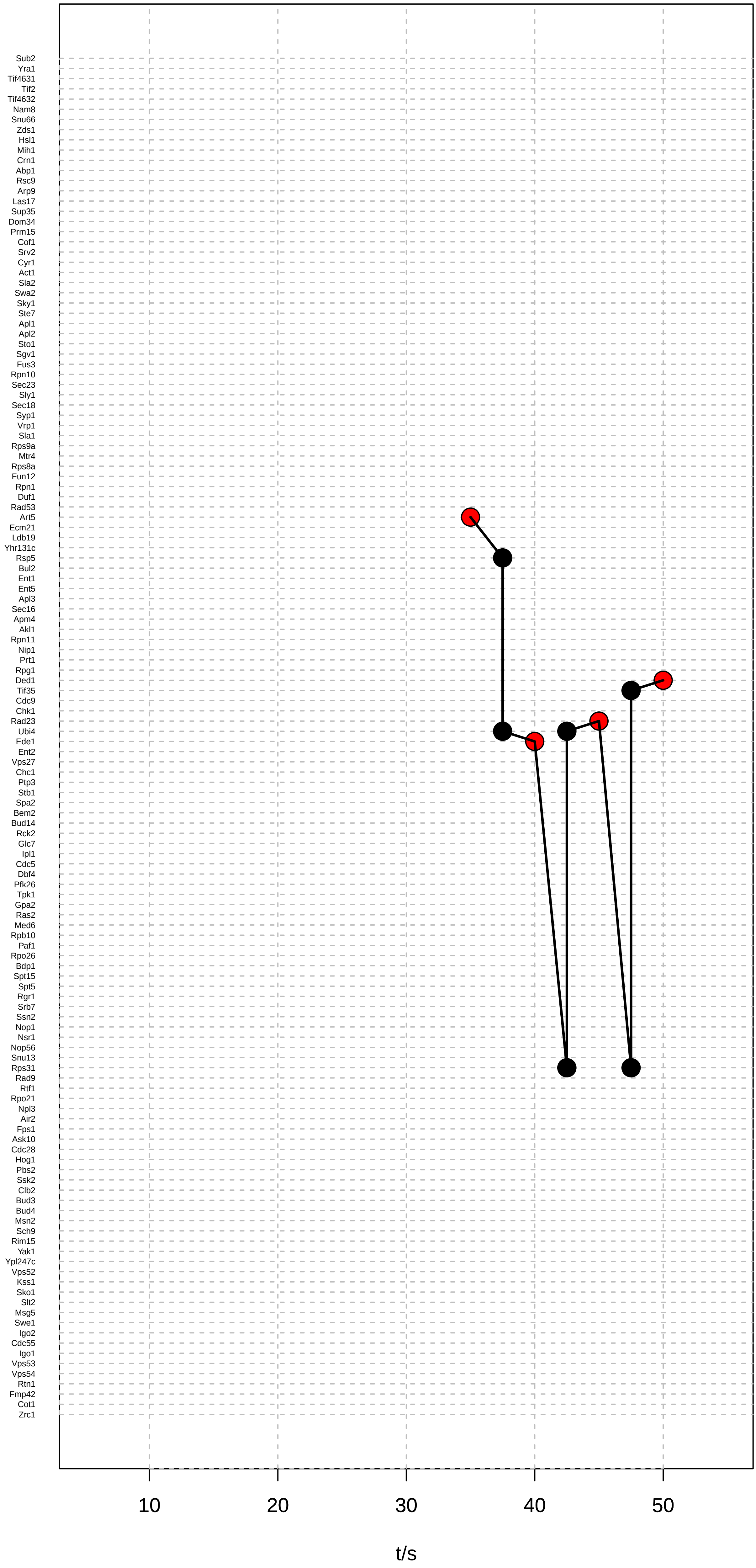

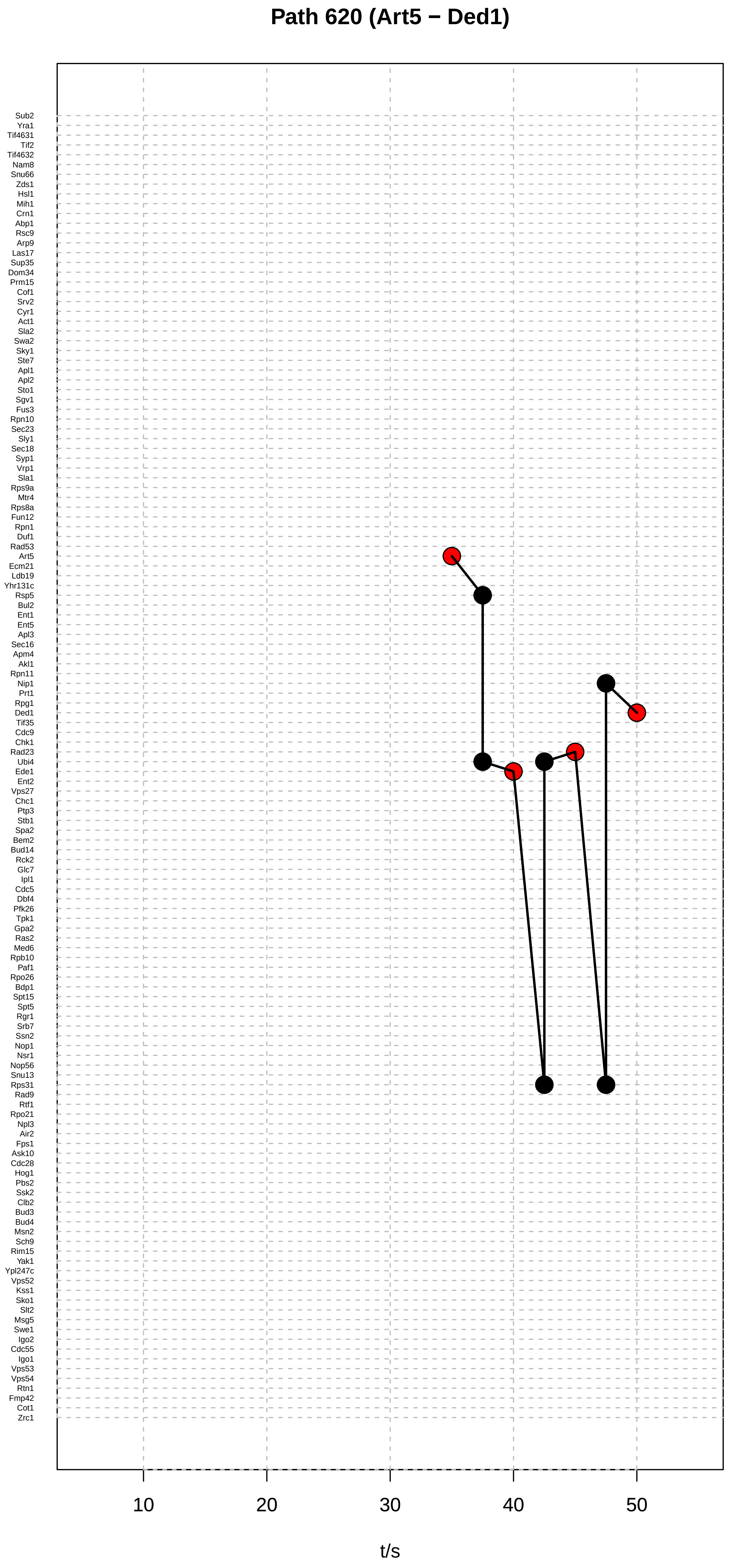

Path 620 (Art5 − Ded1)
Sub2
Yra1
Tif4631
Tif2
Tif4632
Nam8
Snu66
Zds1
Hsl1
Mih1
Crn1
Abp1
Rsc9
Arp9
Las17
Sup35
Dom34
Prm15
Cof1
Srv2
Cyr1
Act1
Sla2
Swa2
Sky1
Ste7
Apl1
Apl2
Sto1
Sgv1
Fus3
Rpn10
Sec23
Sly1
Sec18
Syp1
Vrp1
Sla1
Rps9a
Mtr4
Rps8a
Fun12
Rpn1
Duf1
Rad53
Art5
Ecm21
Ldb19
Yhr131c
Rsp5
Bul2
Ent1
Ent5
Apl3
Sec16
Apm4
Akl1
Rpn11
Nip1
Prt1
Rpg1
Ded1
Tif35
Cdc9
Chk1
Rad23
Ubi4
Ede1
Ent2
Vps27
Chc1
Ptp3
Stb1
Spa2
Bem2
Bud14
Rck2
Glc7
Ipl1
Cdc5
Dbf4
Pfk26
Tpk1
Gpa2
Ras2
Med6
Rpb10
Paf1
Rpo26
Bdp1
Spt15
Spt5
Rgr1
Srb7
Ssn2
Nop1
Nsr1
Nop56
Snu13
Rps31
Rad9
Rtf1
Rpo21
Npl3
Air2
Fps1
Ask10
Cdc28
Hog1
Pbs2
Ssk2
Clb2
Bud3
Bud4
Msn2
Sch9
Rim15
Yak1
Ypl247c
Vps52
Kss1
Sko1
Slt2
Msg5
Swe1
Igo2
Cdc55
Igo1
Vps53
Vps54
Rtn1
Fmp42
Cot1
Zrc1
10
20
30
40
50
t/s

### Path 621 (Art5 − Ded1)

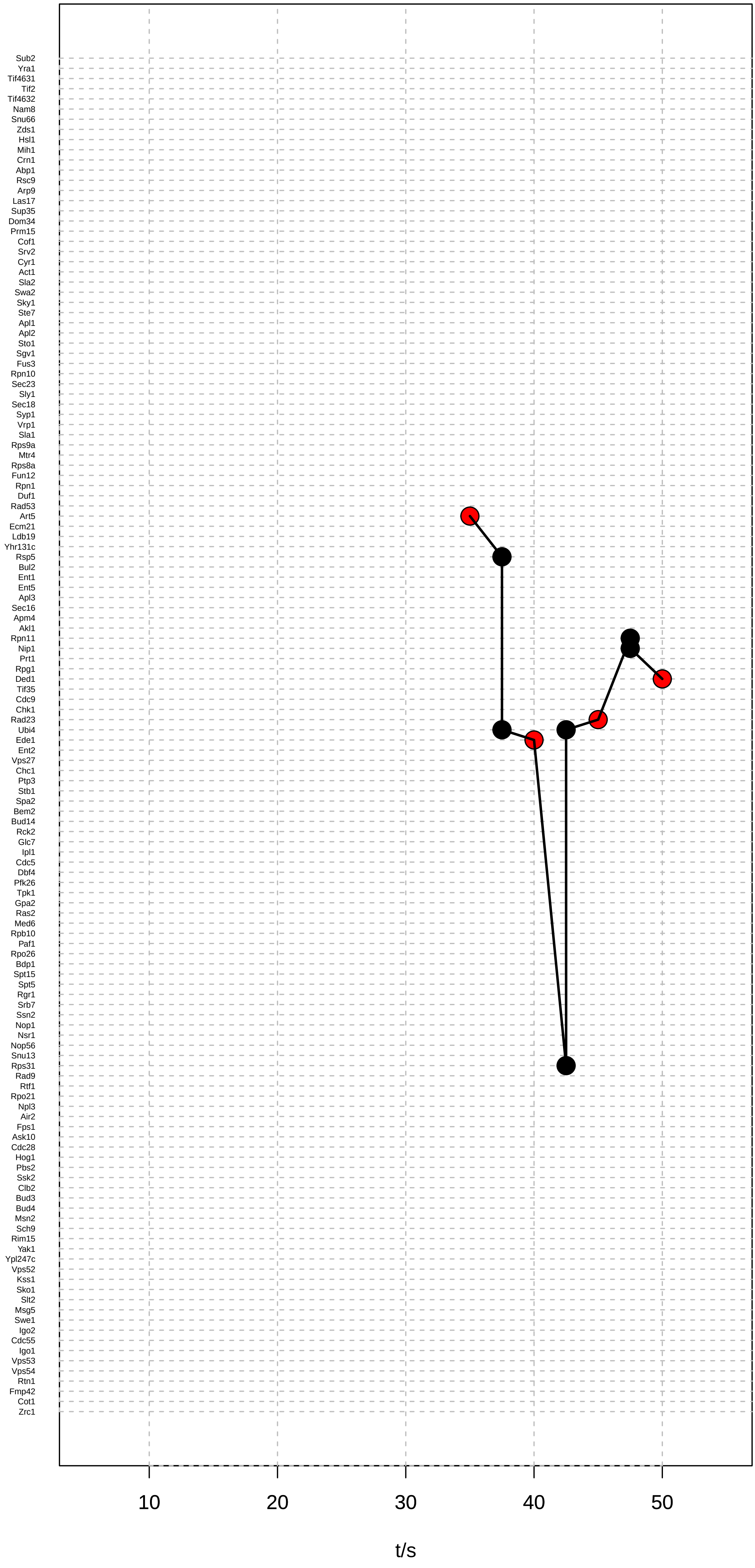

### Path 622 (Art5 − Ded1)

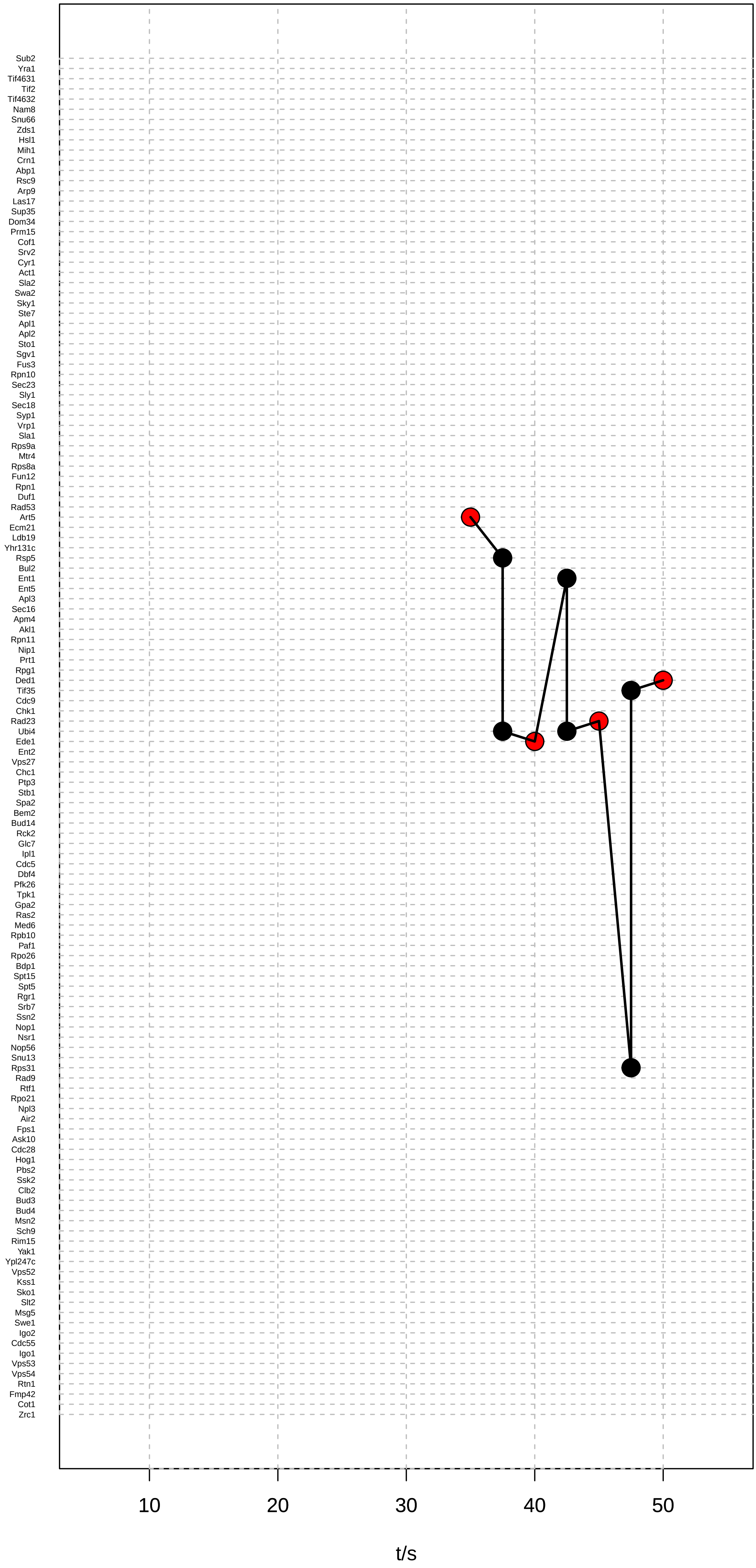

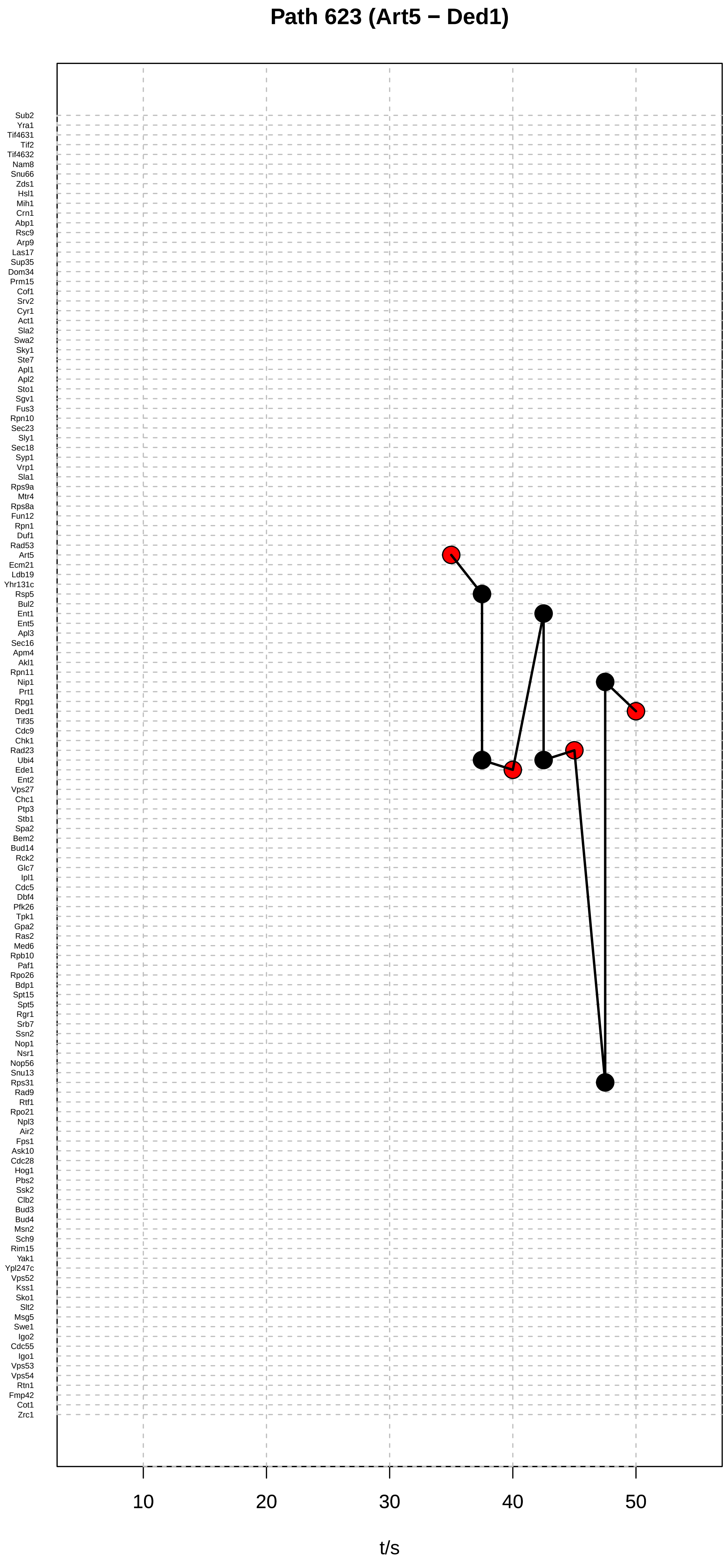
Path 623 (Art5 − Ded1)
Sub2
Yra1
Tif4631
Tif2
Tif4632
Nam8
Snu66
Zds1
Hsl1
Mih1
Crn1
Abp1
Rsc9
Arp9
Las17
Sup35
Dom34
Prm15
Cof1
Srv2
Cyr1
Act1
Sla2
Swa2
Sky1
Ste7
Apl1
Apl2
Sto1
Sgv1
Fus3
Rpn10
Sec23
Sly1
Sec18
Syp1
Vrp1
Sla1
Rps9a
Mtr4
Rps8a
Fun12
Rpn1
Duf1
Rad53
Art5
Ecm21
Ldb19
Yhr131c
Rsp5
Bul2
Ent1
Ent5
Apl3
Sec16
Apm4
Akl1
Rpn11
Nip1
Prt1
Rpg1
Ded1
Tif35
Cdc9
Chk1
Rad23
Ubi4
Ede1
Ent2
Vps27
Chc1
Ptp3
Stb1
Spa2
Bem2
Bud14
Rck2
Glc7
Ipl1
Cdc5
Dbf4
Pfk26
Tpk1
Gpa2
Ras2
Med6
Rpb10
Paf1
Rpo26
Bdp1
Spt15
Spt5
Rgr1
Srb7
Ssn2
Nop1
Nsr1
Nop56
Snu13
Rps31
Rad9
Rtf1
Rpo21
Npl3
Air2
Fps1
Ask10
Cdc28
Hog1
Pbs2
Ssk2
Clb2
Bud3
Bud4
Msn2
Sch9
Rim15
Yak1
Ypl247c
Vps52
Kss1
Sko1
Slt2
Msg5
Swe1
Igo2
Cdc55
Igo1
Vps53
Vps54
Rtn1
Fmp42
Cot1
Zrc1
10
20
30
40
50
t/s

### Path 624 (Art5 − Ded1)

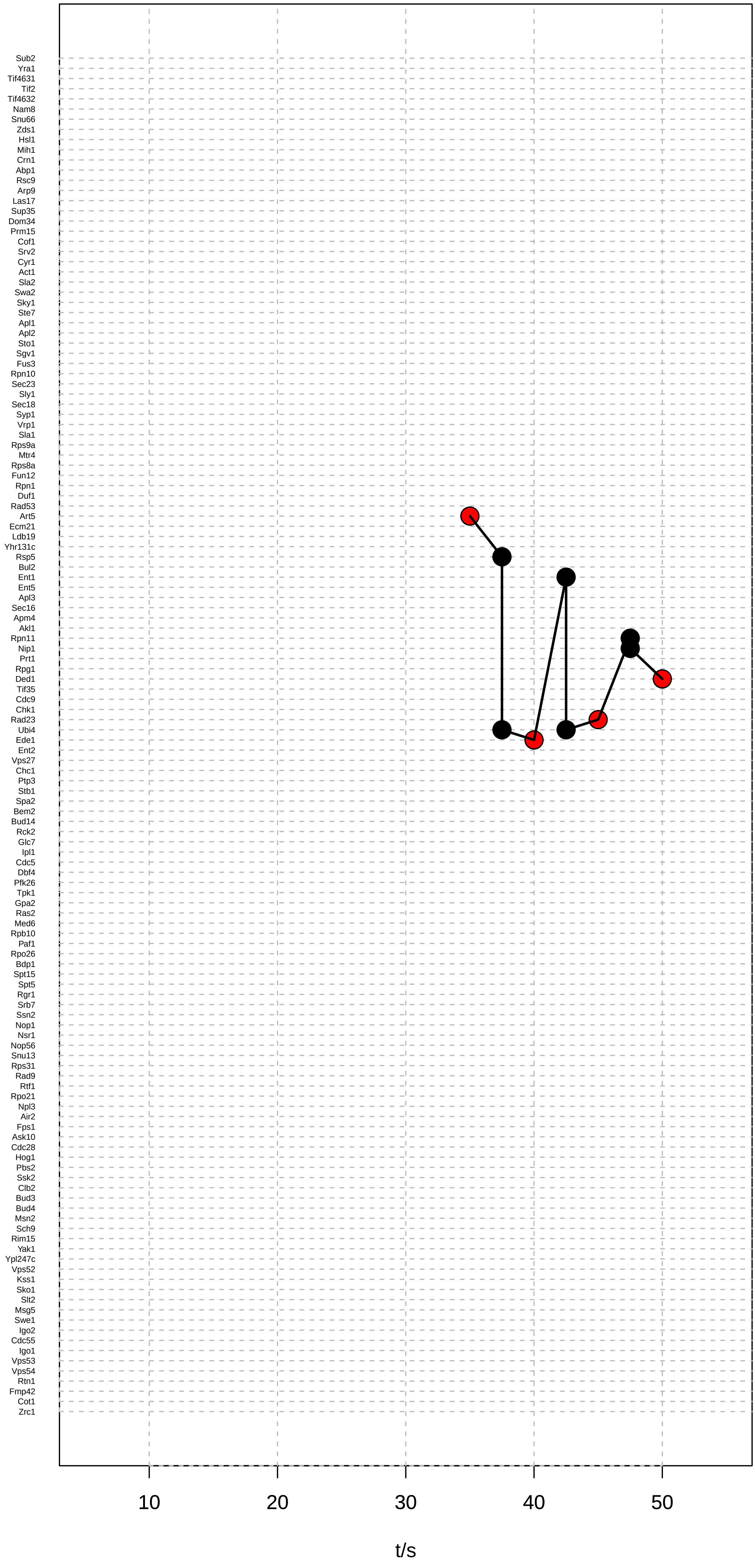

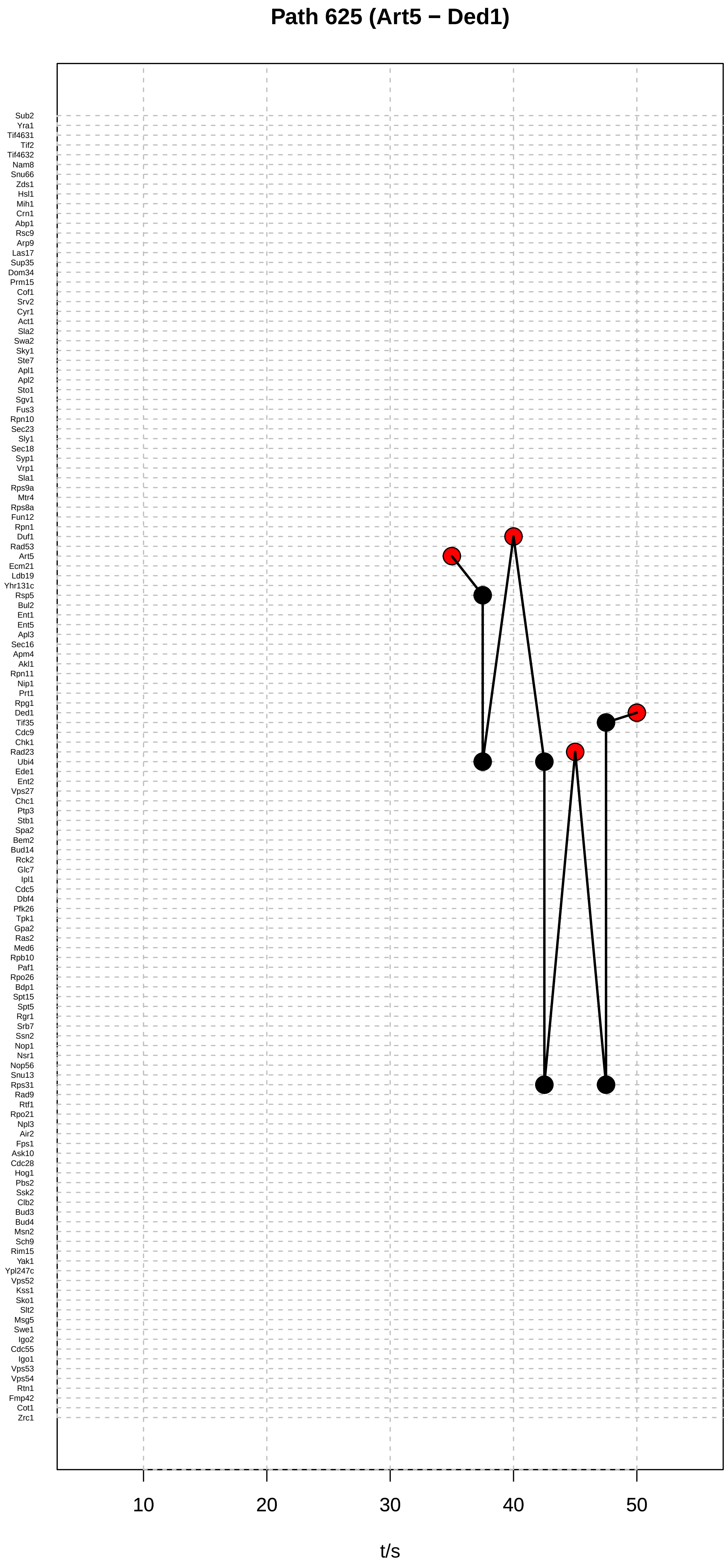

Path 625 (Art5 − Ded1)
Sub2
Yra1
Tif4631
Tif2
Tif4632
Nam8
Snu66
Zds1
Hsl1
Mih1
Crn1
Abp1
Rsc9
Arp9
Las17
Sup35
Dom34
Prm15
Cof1
Srv2
Cyr1
Act1
Sla2
Swa2
Sky1
Ste7
Apl1
Apl2
Sto1
Sgv1
Fus3
Rpn10
Sec23
Sly1
Sec18
Syp1
Vrp1
Sla1
Rps9a
Mtr4
Rps8a
Fun12
Rpn1
Duf1
Rad53
Art5
Ecm21
Ldb19
Yhr131c
Rsp5
Bul2
Ent1
Ent5
Apl3
Sec16
Apm4
Akl1
Rpn11
Nip1
Prt1
Rpg1
Ded1
Tif35
Cdc9
Chk1
Rad23
Ubi4
Ede1
Ent2
Vps27
Chc1
Ptp3
Stb1
Spa2
Bem2
Bud14
Rck2
Glc7
Ipl1
Cdc5
Dbf4
Pfk26
Tpk1
Gpa2
Ras2
Med6
Rpb10
Paf1
Rpo26
Bdp1
Spt15
Spt5
Rgr1
Srb7
Ssn2
Nop1
Nsr1
Nop56
Snu13
Rps31
Rad9
Rtf1
Rpo21
Npl3
Air2
Fps1
Ask10
Cdc28
Hog1
Pbs2
Ssk2
Clb2
Bud3
Bud4
Msn2
Sch9
Rim15
Yak1
Ypl247c
Vps52
Kss1
Sko1
Slt2
Msg5
Swe1
Igo2
Cdc55
Igo1
Vps53
Vps54
Rtn1
Fmp42
Cot1
Zrc1
10
20
30
40
50
t/s

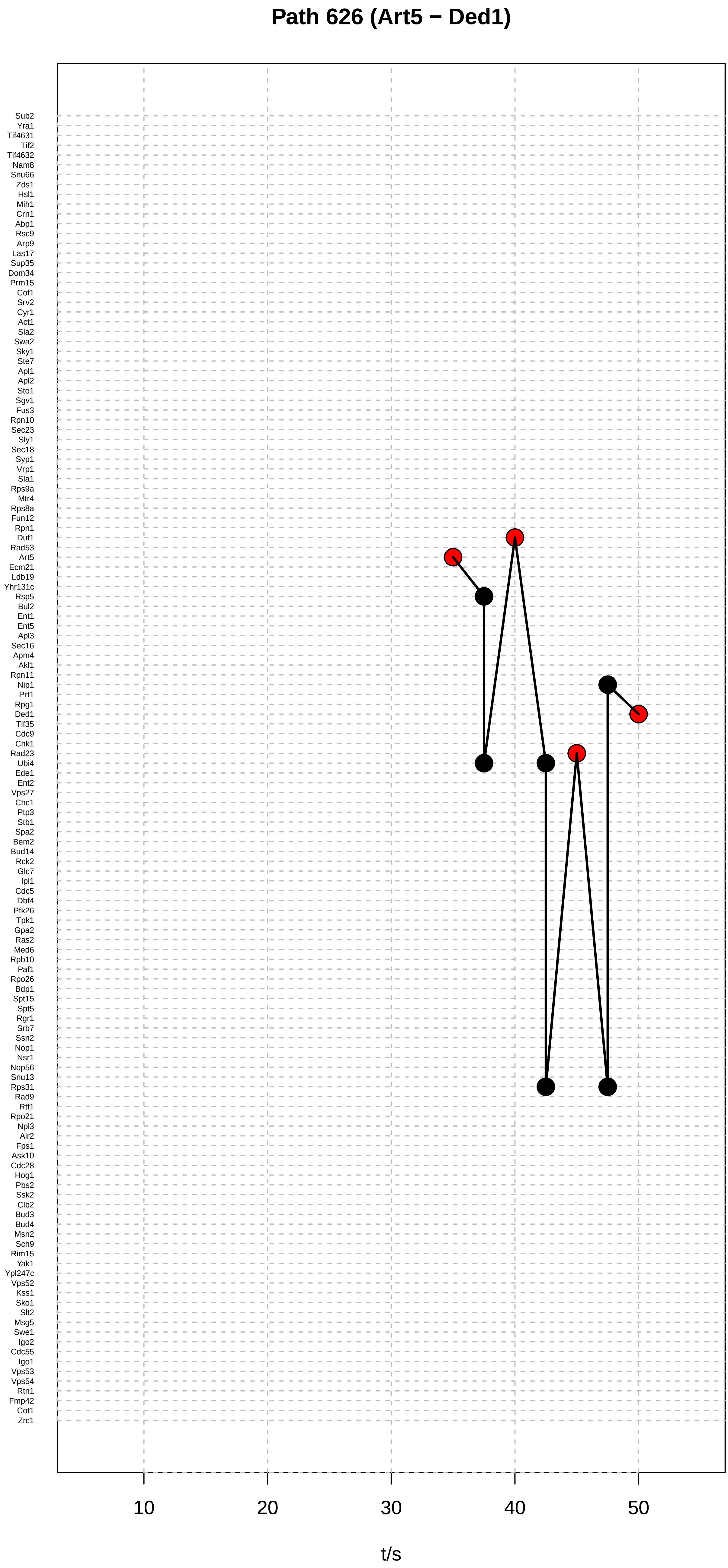

Path 626 (Art5 − Ded1)
Sub2
Yra1
Tif4631
Tif2
Tif4632
Nam8
Snu66
Zds1
Hsl1
Mih1
Crn1
Abp1
Rsc9
Arp9
Las17
Sup35
Dom34
Prm15
Cof1
Srv2
Cyr1
Act1
Sla2
Swa2
Sky1
Ste7
Apl1
Apl2
Sto1
Sgv1
Fus3
Rpn10
Sec23
Sly1
Sec18
Syp1
Vrp1
Sla1
Rps9a
Mtr4
Rps8a
Fun12
Rpn1
Duf1
Rad53
Art5
Ecm21
Ldb19
Yhr131c
Rsp5
Bul2
Ent1
Ent5
Apl3
Sec16
Apm4
Akl1
Rpn11
Nip1
Prt1
Rpg1
Ded1
Tif35
Cdc9
Chk1
Rad23
Ubi4
Ede1
Ent2
Vps27
Chc1
Ptp3
Stb1
Spa2
Bem2
Bud14
Rck2
Glc7
Ipl1
Cdc5
Dbf4
Pfk26
Tpk1
Gpa2
Ras2
Med6
Rpb10
Paf1
Rpo26
Bdp1
Spt15
Spt5
Rgr1
Srb7
Ssn2
Nop1
Nsr1
Nop56
Snu13
Rps31
Rad9
Rtf1
Rpo21
Npl3
Air2
Fps1
Ask10
Cdc28
Hog1
Pbs2
Ssk2
Clb2
Bud3
Bud4
Msn2
Sch9
Rim15
Yak1
Ypl247c
Vps52
Kss1
Sko1
Slt2
Msg5
Swe1
Igo2
Cdc55
Igo1
Vps53
Vps54
Rtn1
Fmp42
Cot1
Zrc1
10
20
30
40
50
t/s

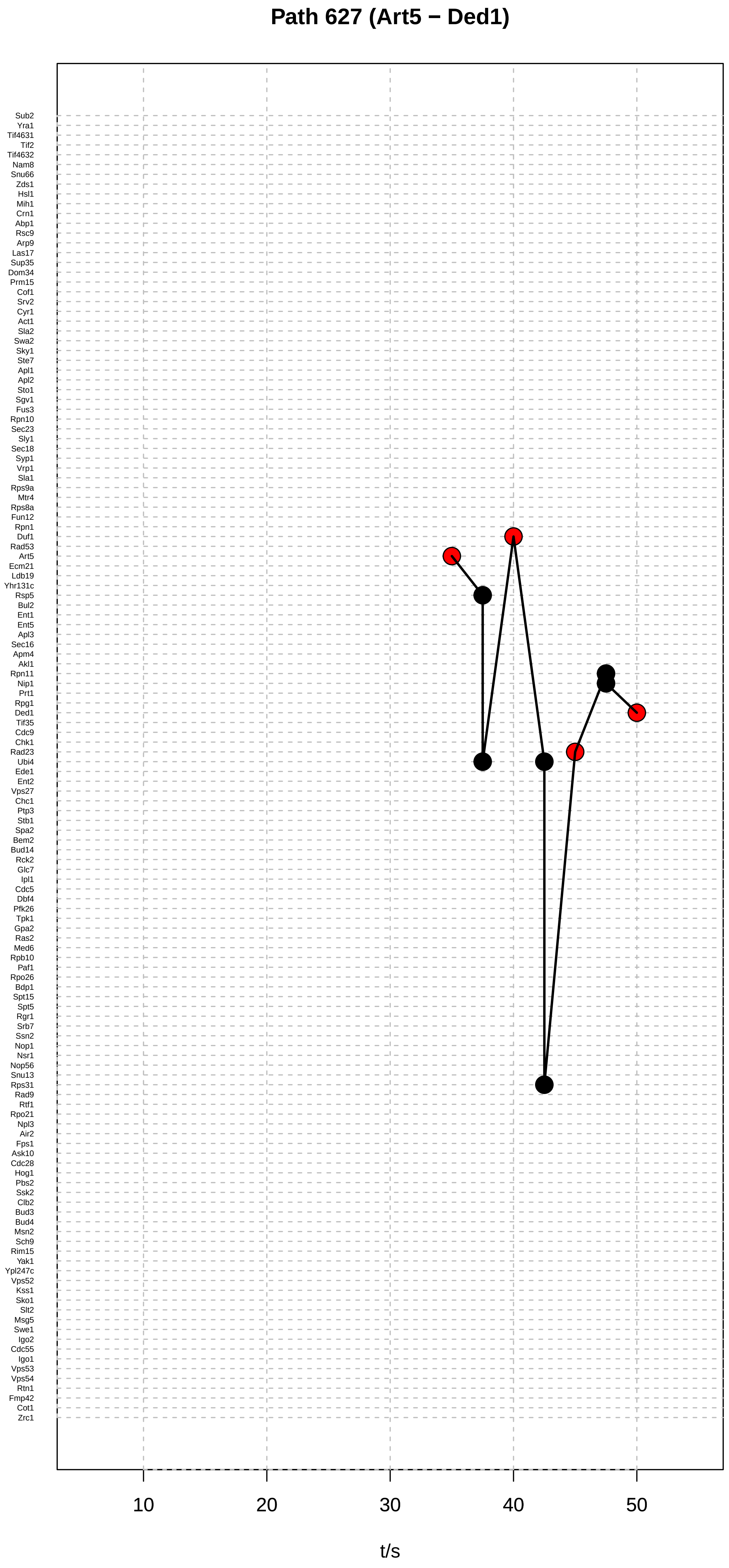

Path 627 (Art5 − Ded1)
Sub2
Yra1
Tif4631
Tif2
Tif4632
Nam8
Snu66
Zds1
Hsl1
Mih1
Crn1
Abp1
Rsc9
Arp9
Las17
Sup35
Dom34
Prm15
Cof1
Srv2
Cyr1
Act1
Sla2
Swa2
Sky1
Ste7
Apl1
Apl2
Sto1
Sgv1
Fus3
Rpn10
Sec23
Sly1
Sec18
Syp1
Vrp1
Sla1
Rps9a
Mtr4
Rps8a
Fun12
Rpn1
Duf1
Rad53
Art5
Ecm21
Ldb19
Yhr131c
Rsp5
Bul2
Ent1
Ent5
Apl3
Sec16
Apm4
Akl1
Rpn11
Nip1
Prt1
Rpg1
Ded1
Tif35
Cdc9
Chk1
Rad23
Ubi4
Ede1
Ent2
Vps27
Chc1
Ptp3
Stb1
Spa2
Bem2
Bud14
Rck2
Glc7
Ipl1
Cdc5
Dbf4
Pfk26
Tpk1
Gpa2
Ras2
Med6
Rpb10
Paf1
Rpo26
Bdp1
Spt15
Spt5
Rgr1
Srb7
Ssn2
Nop1
Nsr1
Nop56
Snu13
Rps31
Rad9
Rtf1
Rpo21
Npl3
Air2
Fps1
Ask10
Cdc28
Hog1
Pbs2
Ssk2
Clb2
Bud3
Bud4
Msn2
Sch9
Rim15
Yak1
Ypl247c
Vps52
Kss1
Sko1
Slt2
Msg5
Swe1
Igo2
Cdc55
Igo1
Vps53
Vps54
Rtn1
Fmp42
Cot1
Zrc1
10
20
30
40
50
t/s

### Path 628 (Art5 − Ded1)

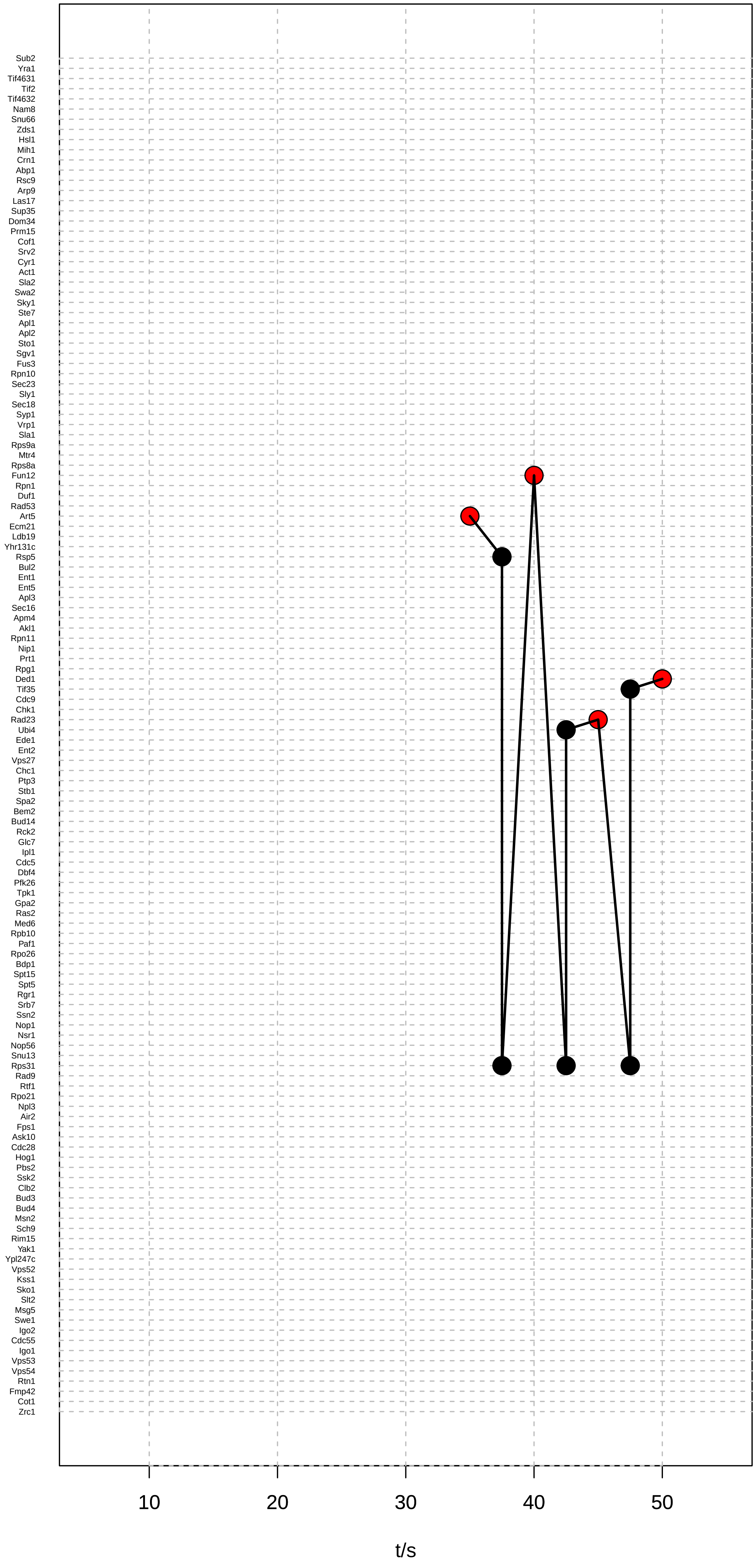

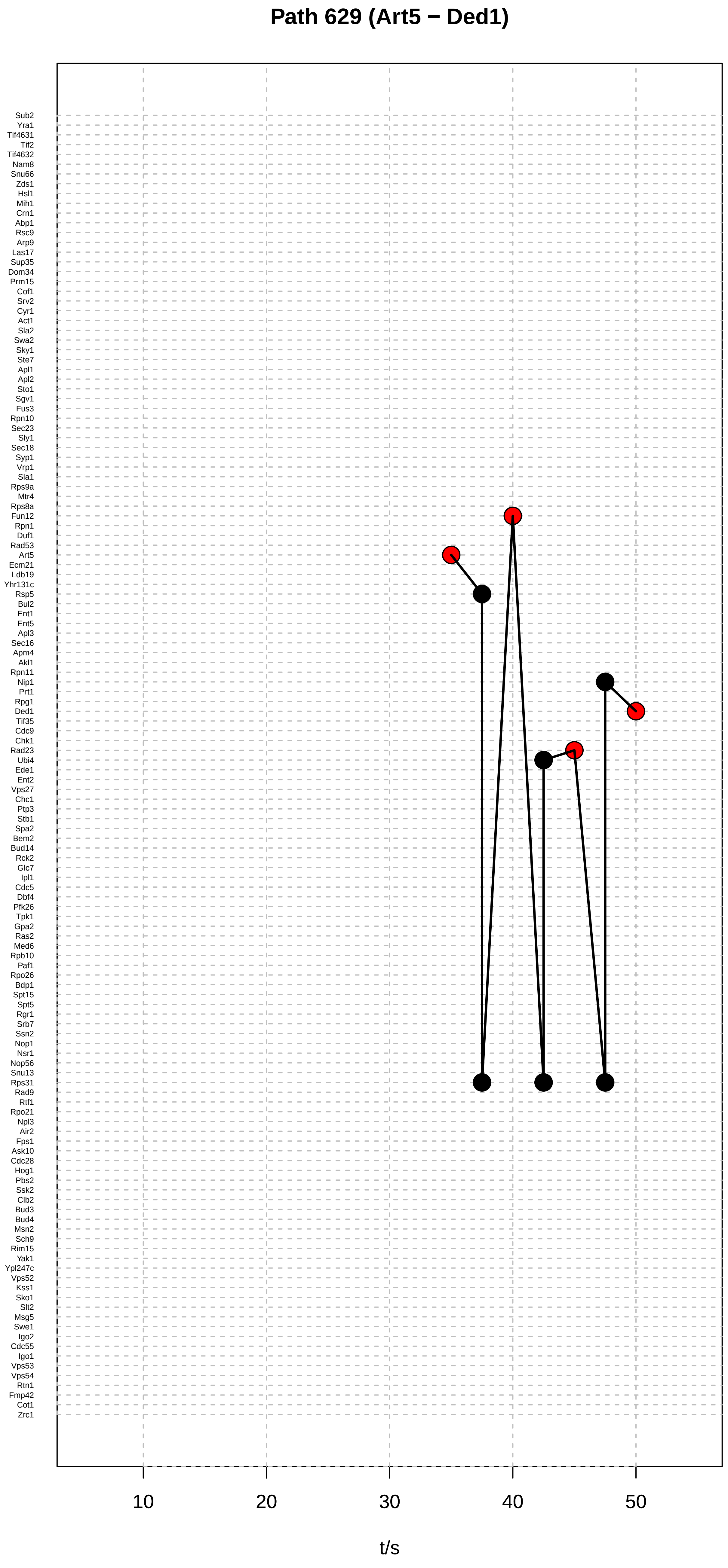

Path 629 (Art5 − Ded1)
Sub2
Yra1
Tif4631
Tif2
Tif4632
Nam8
Snu66
Zds1
Hsl1
Mih1
Crn1
Abp1
Rsc9
Arp9
Las17
Sup35
Dom34
Prm15
Cof1
Srv2
Cyr1
Act1
Sla2
Swa2
Sky1
Ste7
Apl1
Apl2
Sto1
Sgv1
Fus3
Rpn10
Sec23
Sly1
Sec18
Syp1
Vrp1
Sla1
Rps9a
Mtr4
Rps8a
Fun12
Rpn1
Duf1
Rad53
Art5
Ecm21
Ldb19
Yhr131c
Rsp5
Bul2
Ent1
Ent5
Apl3
Sec16
Apm4
Akl1
Rpn11
Nip1
Prt1
Rpg1
Ded1
Tif35
Cdc9
Chk1
Rad23
Ubi4
Ede1
Ent2
Vps27
Chc1
Ptp3
Stb1
Spa2
Bem2
Bud14
Rck2
Glc7
Ipl1
Cdc5
Dbf4
Pfk26
Tpk1
Gpa2
Ras2
Med6
Rpb10
Paf1
Rpo26
Bdp1
Spt15
Spt5
Rgr1
Srb7
Ssn2
Nop1
Nsr1
Nop56
Snu13
Rps31
Rad9
Rtf1
Rpo21
Npl3
Air2
Fps1
Ask10
Cdc28
Hog1
Pbs2
Ssk2
Clb2
Bud3
Bud4
Msn2
Sch9
Rim15
Yak1
Ypl247c
Vps52
Kss1
Sko1
Slt2
Msg5
Swe1
Igo2
Cdc55
Igo1
Vps53
Vps54
Rtn1
Fmp42
Cot1
Zrc1
10
20
30
40
50
t/s

### Path 630 (Art5 − Ded1)

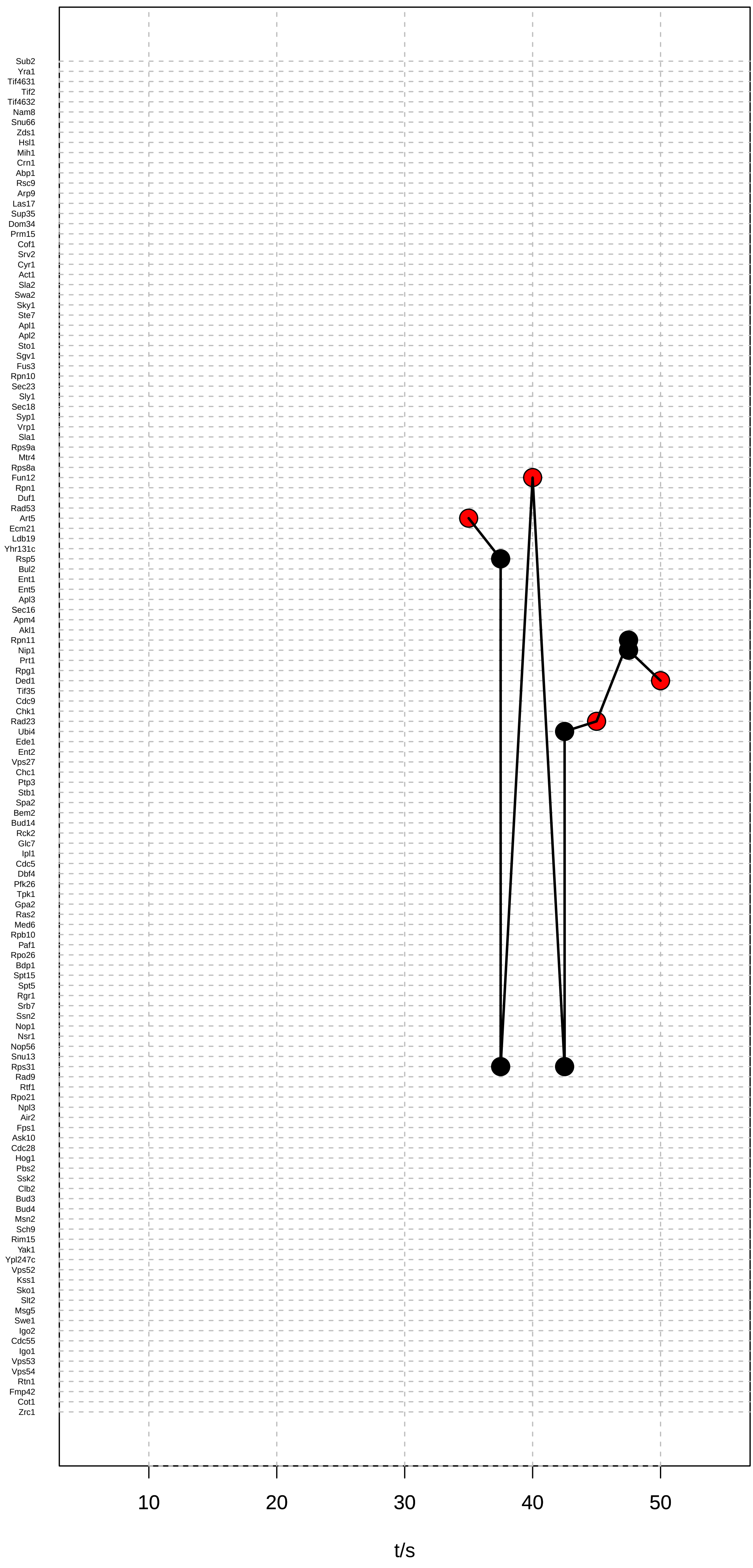

### Path 631 (Art5 − Ded1)

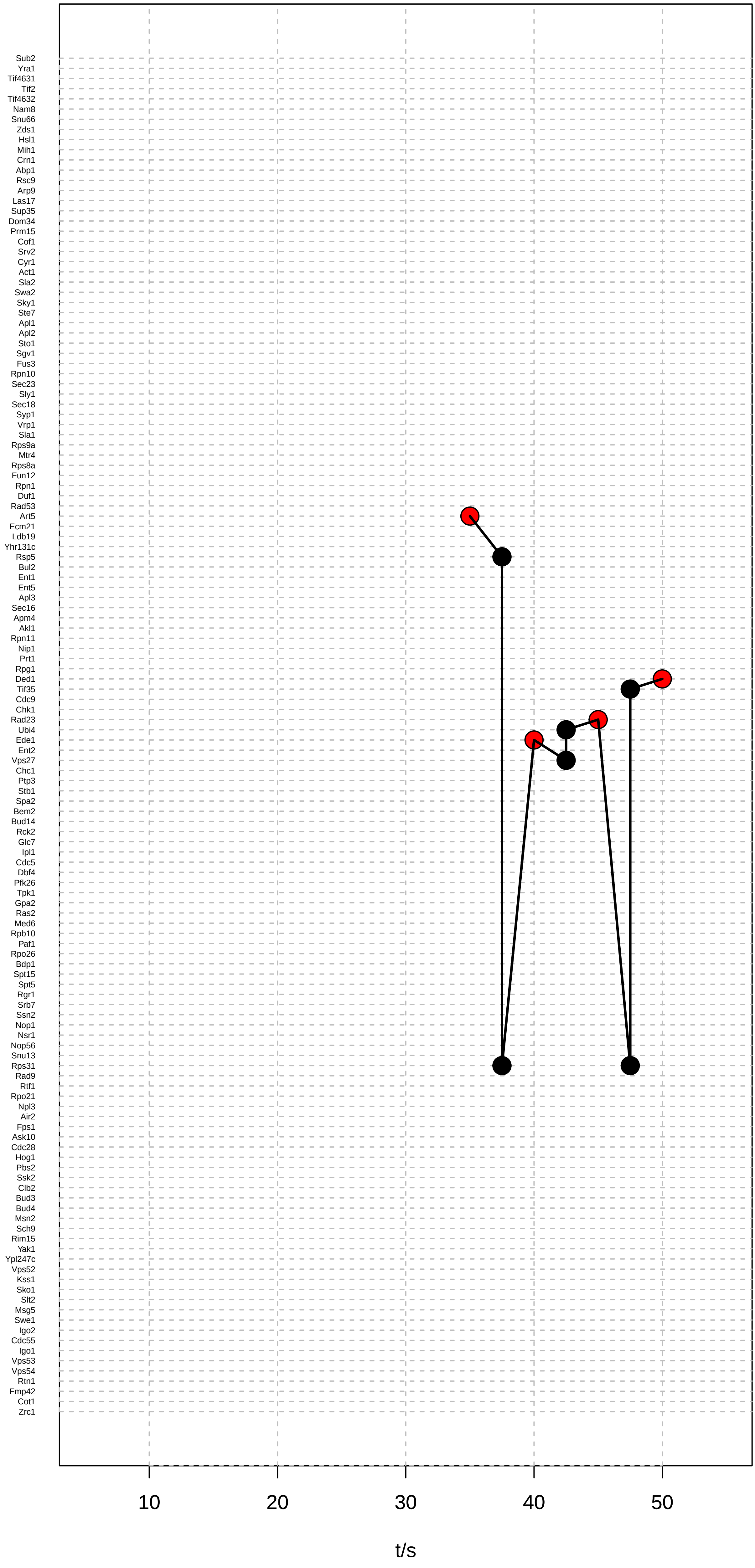

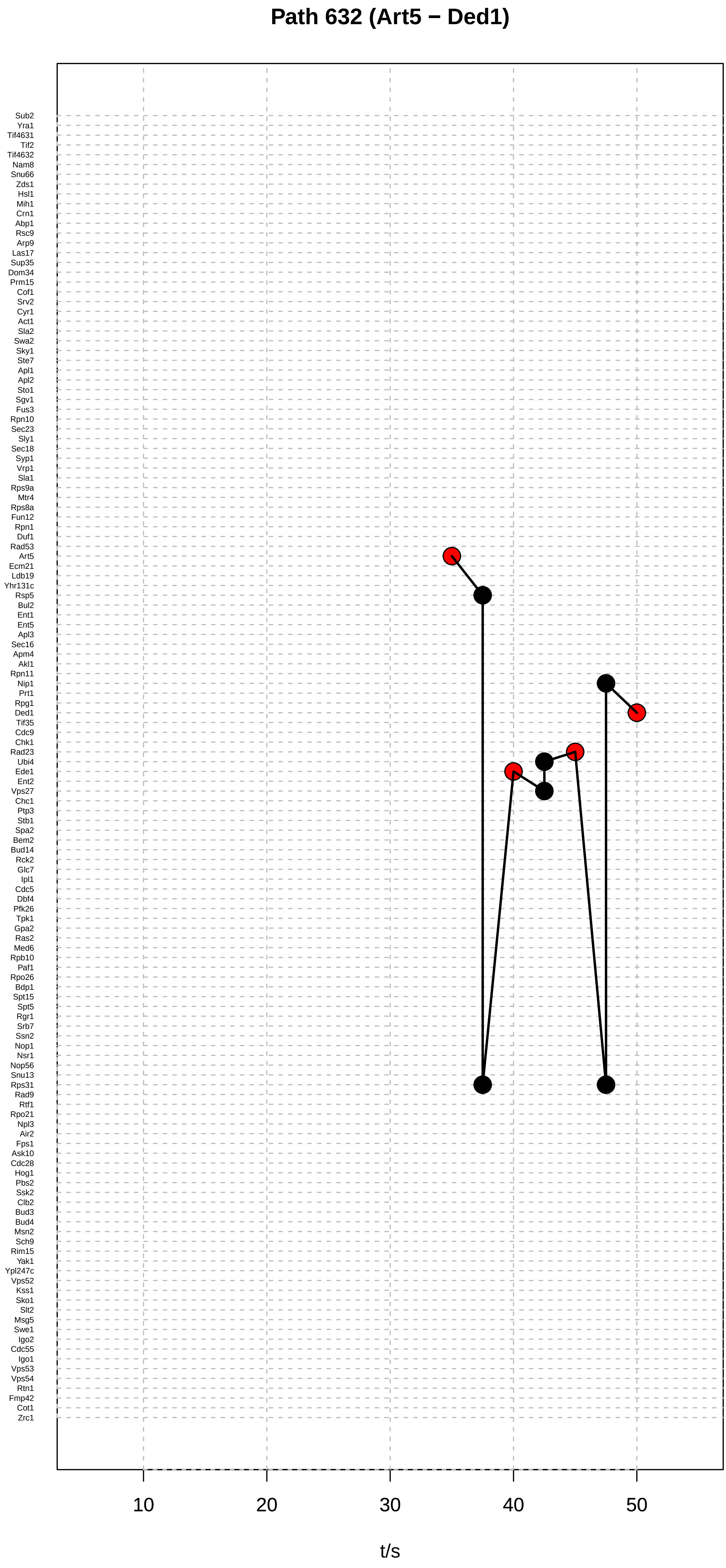

Path 632 (Art5 − Ded1)
Sub2
Yra1
Tif4631
Tif2
Tif4632
Nam8
Snu66
Zds1
Hsl1
Mih1
Crn1
Abp1
Rsc9
Arp9
Las17
Sup35
Dom34
Prm15
Cof1
Srv2
Cyr1
Act1
Sla2
Swa2
Sky1
Ste7
Apl1
Apl2
Sto1
Sgv1
Fus3
Rpn10
Sec23
Sly1
Sec18
Syp1
Vrp1
Sla1
Rps9a
Mtr4
Rps8a
Fun12
Rpn1
Duf1
Rad53
Art5
Ecm21
Ldb19
Yhr131c
Rsp5
Bul2
Ent1
Ent5
Apl3
Sec16
Apm4
Akl1
Rpn11
Nip1
Prt1
Rpg1
Ded1
Tif35
Cdc9
Chk1
Rad23
Ubi4
Ede1
Ent2
Vps27
Chc1
Ptp3
Stb1
Spa2
Bem2
Bud14
Rck2
Glc7
Ipl1
Cdc5
Dbf4
Pfk26
Tpk1
Gpa2
Ras2
Med6
Rpb10
Paf1
Rpo26
Bdp1
Spt15
Spt5
Rgr1
Srb7
Ssn2
Nop1
Nsr1
Nop56
Snu13
Rps31
Rad9
Rtf1
Rpo21
Npl3
Air2
Fps1
Ask10
Cdc28
Hog1
Pbs2
Ssk2
Clb2
Bud3
Bud4
Msn2
Sch9
Rim15
Yak1
Ypl247c
Vps52
Kss1
Sko1
Slt2
Msg5
Swe1
Igo2
Cdc55
Igo1
Vps53
Vps54
Rtn1
Fmp42
Cot1
Zrc1
10
20
30
40
50
t/s

### Path 633 (Art5 − Ded1)

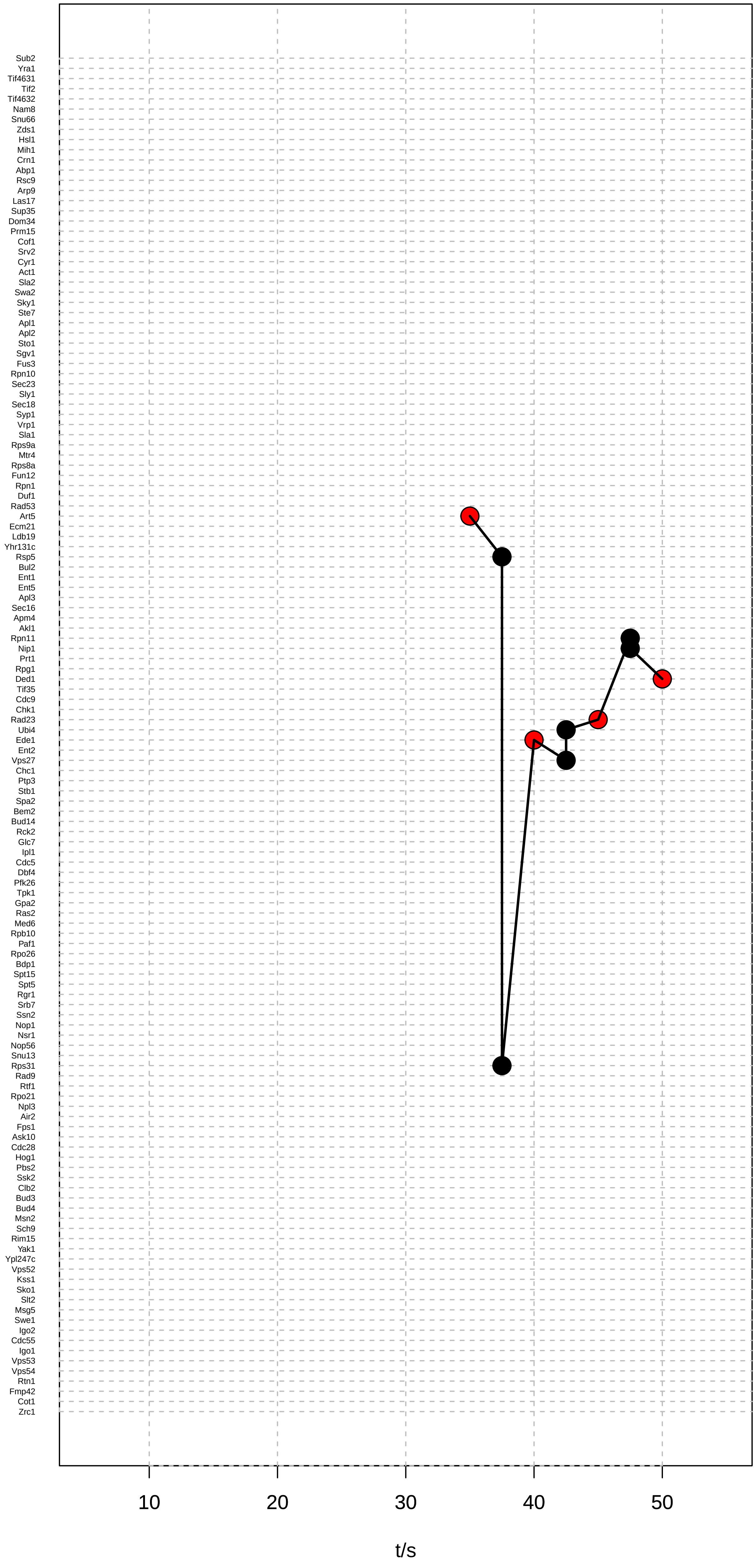

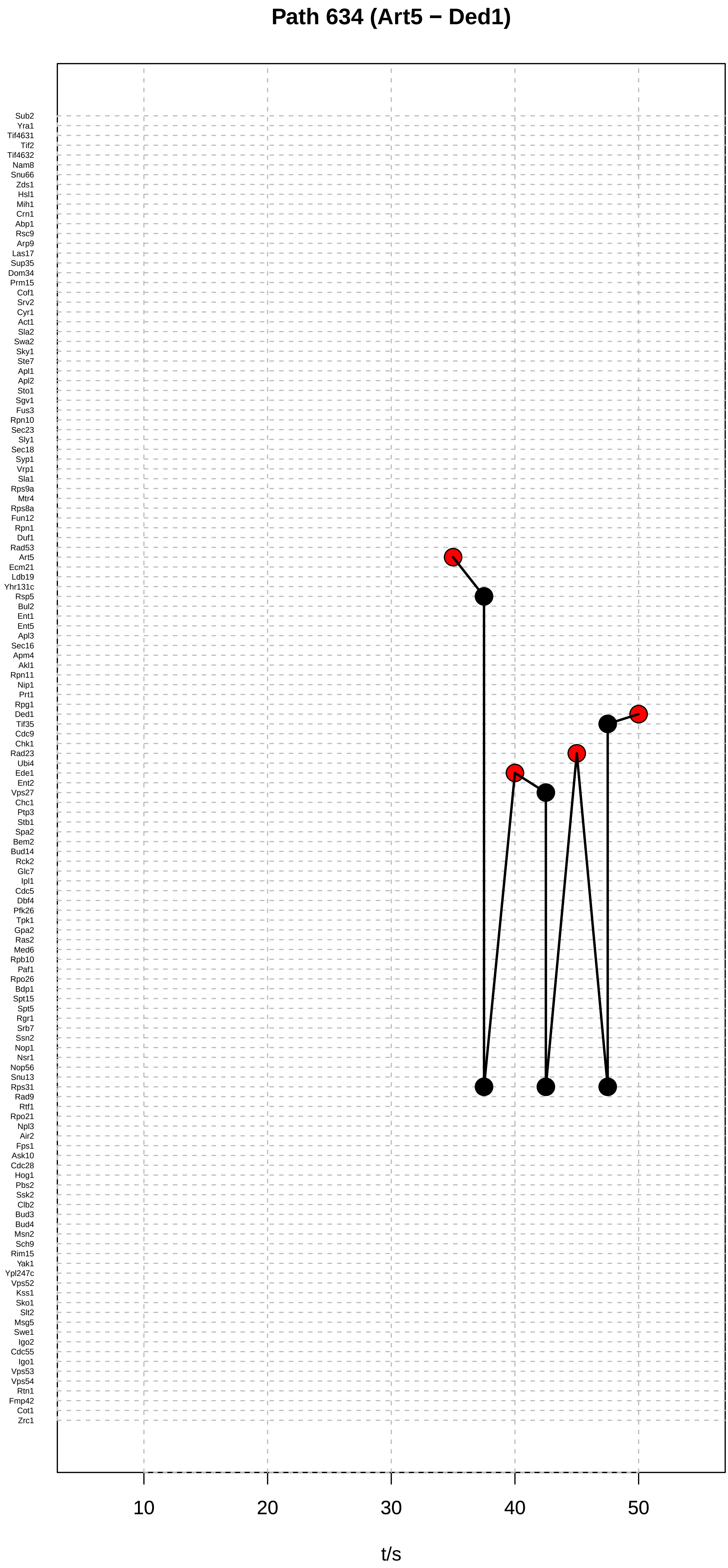

Path 634 (Art5 − Ded1)
Sub2
Yra1
Tif4631
Tif2
Tif4632
Nam8
Snu66
Zds1
Hsl1
Mih1
Crn1
Abp1
Rsc9
Arp9
Las17
Sup35
Dom34
Prm15
Cof1
Srv2
Cyr1
Act1
Sla2
Swa2
Sky1
Ste7
Apl1
Apl2
Sto1
Sgv1
Fus3
Rpn10
Sec23
Sly1
Sec18
Syp1
Vrp1
Sla1
Rps9a
Mtr4
Rps8a
Fun12
Rpn1
Duf1
Rad53
Art5
Ecm21
Ldb19
Yhr131c
Rsp5
Bul2
Ent1
Ent5
Apl3
Sec16
Apm4
Akl1
Rpn11
Nip1
Prt1
Rpg1
Ded1
Tif35
Cdc9
Chk1
Rad23
Ubi4
Ede1
Ent2
Vps27
Chc1
Ptp3
Stb1
Spa2
Bem2
Bud14
Rck2
Glc7
Ipl1
Cdc5
Dbf4
Pfk26
Tpk1
Gpa2
Ras2
Med6
Rpb10
Paf1
Rpo26
Bdp1
Spt15
Spt5
Rgr1
Srb7
Ssn2
Nop1
Nsr1
Nop56
Snu13
Rps31
Rad9
Rtf1
Rpo21
Npl3
Air2
Fps1
Ask10
Cdc28
Hog1
Pbs2
Ssk2
Clb2
Bud3
Bud4
Msn2
Sch9
Rim15
Yak1
Ypl247c
Vps52
Kss1
Sko1
Slt2
Msg5
Swe1
Igo2
Cdc55
Igo1
Vps53
Vps54
Rtn1
Fmp42
Cot1
Zrc1
10
20
30
40
50
t/s

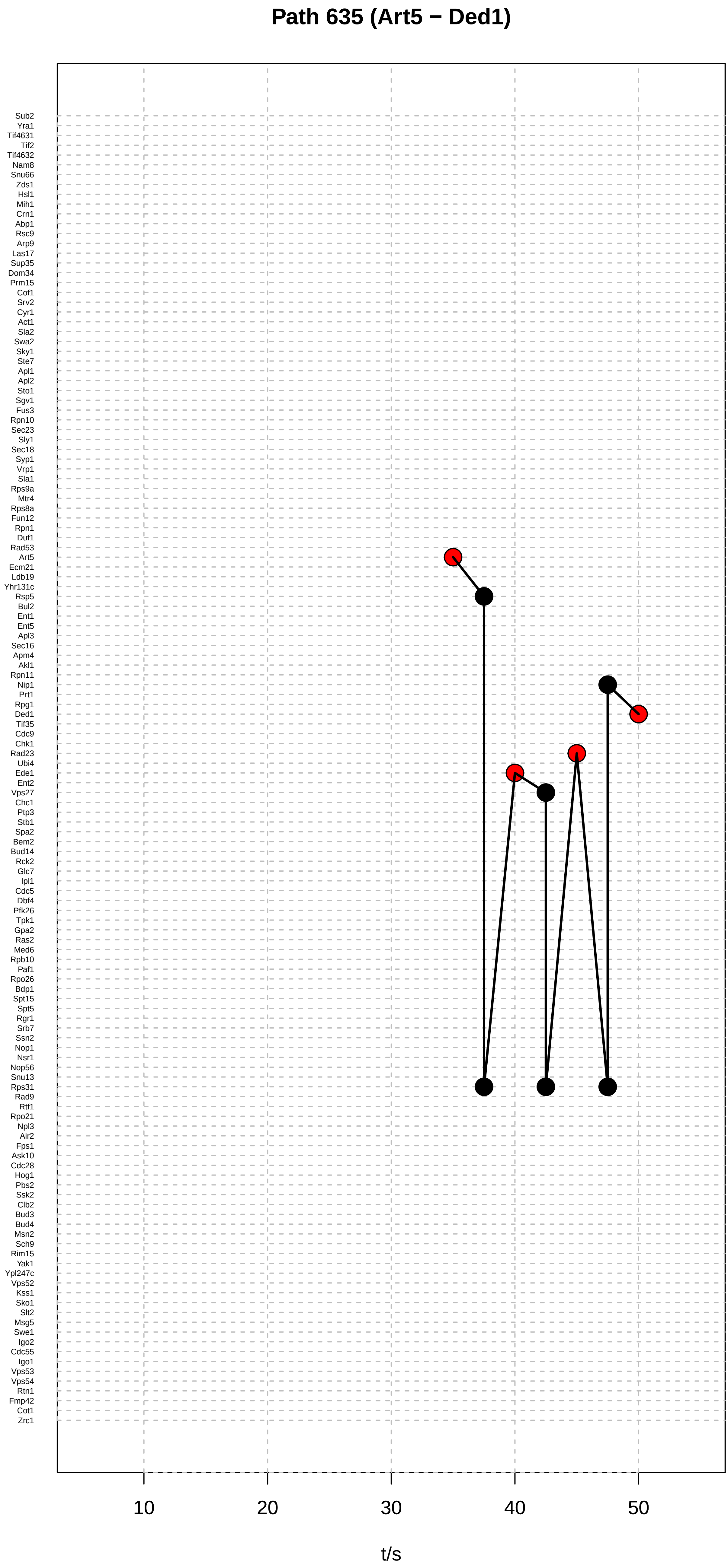

Path 635 (Art5 − Ded1)
Sub2
Yra1
Tif4631
Tif2
Tif4632
Nam8
Snu66
Zds1
Hsl1
Mih1
Crn1
Abp1
Rsc9
Arp9
Las17
Sup35
Dom34
Prm15
Cof1
Srv2
Cyr1
Act1
Sla2
Swa2
Sky1
Ste7
Apl1
Apl2
Sto1
Sgv1
Fus3
Rpn10
Sec23
Sly1
Sec18
Syp1
Vrp1
Sla1
Rps9a
Mtr4
Rps8a
Fun12
Rpn1
Duf1
Rad53
Art5
Ecm21
Ldb19
Yhr131c
Rsp5
Bul2
Ent1
Ent5
Apl3
Sec16
Apm4
Akl1
Rpn11
Nip1
Prt1
Rpg1
Ded1
Tif35
Cdc9
Chk1
Rad23
Ubi4
Ede1
Ent2
Vps27
Chc1
Ptp3
Stb1
Spa2
Bem2
Bud14
Rck2
Glc7
Ipl1
Cdc5
Dbf4
Pfk26
Tpk1
Gpa2
Ras2
Med6
Rpb10
Paf1
Rpo26
Bdp1
Spt15
Spt5
Rgr1
Srb7
Ssn2
Nop1
Nsr1
Nop56
Snu13
Rps31
Rad9
Rtf1
Rpo21
Npl3
Air2
Fps1
Ask10
Cdc28
Hog1
Pbs2
Ssk2
Clb2
Bud3
Bud4
Msn2
Sch9
Rim15
Yak1
Ypl247c
Vps52
Kss1
Sko1
Slt2
Msg5
Swe1
Igo2
Cdc55
Igo1
Vps53
Vps54
Rtn1
Fmp42
Cot1
Zrc1
10
20
30
40
50
t/s

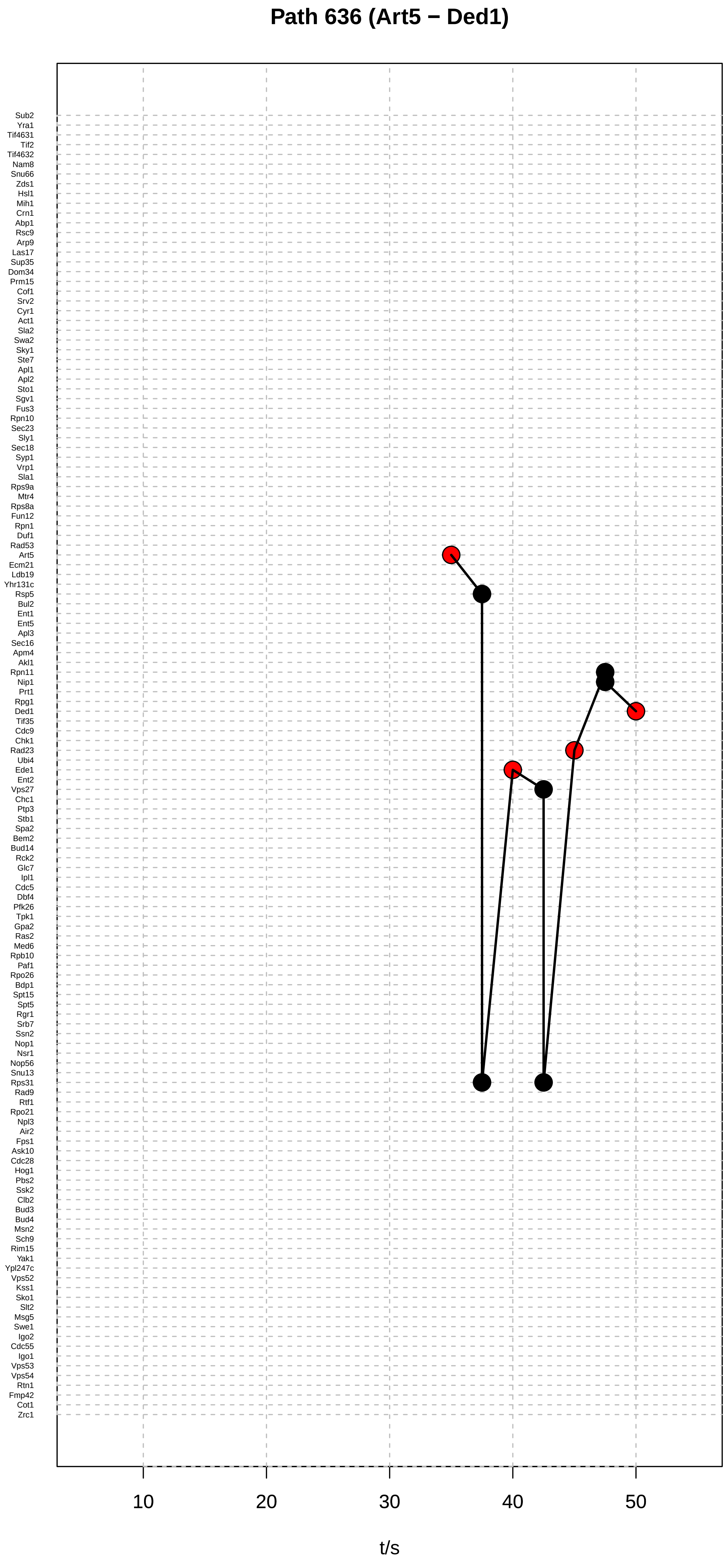

Path 636 (Art5 − Ded1)
Sub2
Yra1
Tif4631
Tif2
Tif4632
Nam8
Snu66
Zds1
Hsl1
Mih1
Crn1
Abp1
Rsc9
Arp9
Las17
Sup35
Dom34
Prm15
Cof1
Srv2
Cyr1
Act1
Sla2
Swa2
Sky1
Ste7
Apl1
Apl2
Sto1
Sgv1
Fus3
Rpn10
Sec23
Sly1
Sec18
Syp1
Vrp1
Sla1
Rps9a
Mtr4
Rps8a
Fun12
Rpn1
Duf1
Rad53
Art5
Ecm21
Ldb19
Yhr131c
Rsp5
Bul2
Ent1
Ent5
Apl3
Sec16
Apm4
Akl1
Rpn11
Nip1
Prt1
Rpg1
Ded1
Tif35
Cdc9
Chk1
Rad23
Ubi4
Ede1
Ent2
Vps27
Chc1
Ptp3
Stb1
Spa2
Bem2
Bud14
Rck2
Glc7
Ipl1
Cdc5
Dbf4
Pfk26
Tpk1
Gpa2
Ras2
Med6
Rpb10
Paf1
Rpo26
Bdp1
Spt15
Spt5
Rgr1
Srb7
Ssn2
Nop1
Nsr1
Nop56
Snu13
Rps31
Rad9
Rtf1
Rpo21
Npl3
Air2
Fps1
Ask10
Cdc28
Hog1
Pbs2
Ssk2
Clb2
Bud3
Bud4
Msn2
Sch9
Rim15
Yak1
Ypl247c
Vps52
Kss1
Sko1
Slt2
Msg5
Swe1
Igo2
Cdc55
Igo1
Vps53
Vps54
Rtn1
Fmp42
Cot1
Zrc1
10
20
30
40
50
t/s

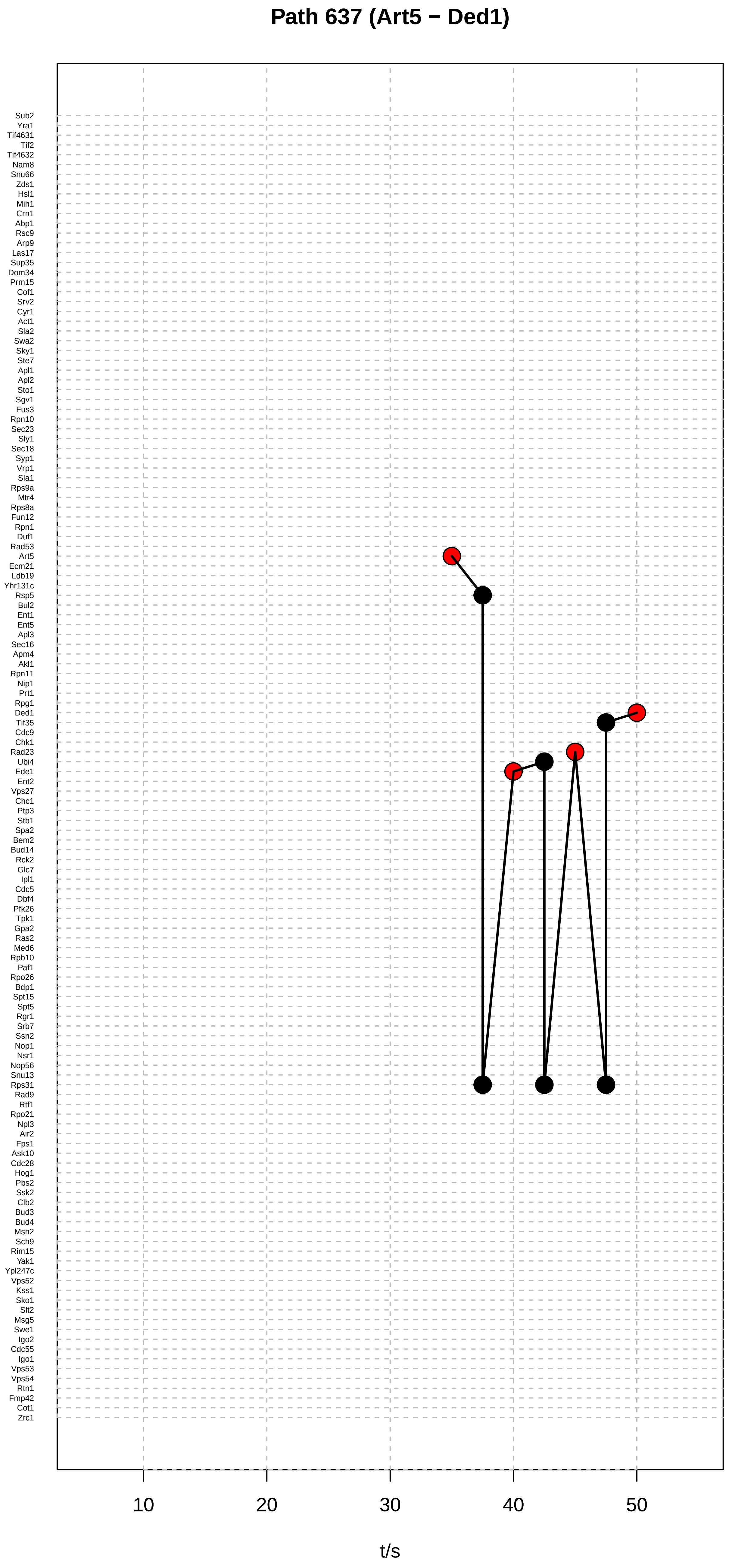
Path 637 (Art5 − Ded1)
Sub2
Yra1
Tif4631
Tif2
Tif4632
Nam8
Snu66
Zds1
Hsl1
Mih1
Crn1
Abp1
Rsc9
Arp9
Las17
Sup35
Dom34
Prm15
Cof1
Srv2
Cyr1
Act1
Sla2
Swa2
Sky1
Ste7
Apl1
Apl2
Sto1
Sgv1
Fus3
Rpn10
Sec23
Sly1
Sec18
Syp1
Vrp1
Sla1
Rps9a
Mtr4
Rps8a
Fun12
Rpn1
Duf1
Rad53
Art5
Ecm21
Ldb19
Yhr131c
Rsp5
Bul2
Ent1
Ent5
Apl3
Sec16
Apm4
Akl1
Rpn11
Nip1
Prt1
Rpg1
Ded1
Tif35
Cdc9
Chk1
Rad23
Ubi4
Ede1
Ent2
Vps27
Chc1
Ptp3
Stb1
Spa2
Bem2
Bud14
Rck2
Glc7
Ipl1
Cdc5
Dbf4
Pfk26
Tpk1
Gpa2
Ras2
Med6
Rpb10
Paf1
Rpo26
Bdp1
Spt15
Spt5
Rgr1
Srb7
Ssn2
Nop1
Nsr1
Nop56
Snu13
Rps31
Rad9
Rtf1
Rpo21
Npl3
Air2
Fps1
Ask10
Cdc28
Hog1
Pbs2
Ssk2
Clb2
Bud3
Bud4
Msn2
Sch9
Rim15
Yak1
Ypl247c
Vps52
Kss1
Sko1
Slt2
Msg5
Swe1
Igo2
Cdc55
Igo1
Vps53
Vps54
Rtn1
Fmp42
Cot1
Zrc1
10
20
30
40
50
t/s

#### Path 638 (Art5 − Ded1)

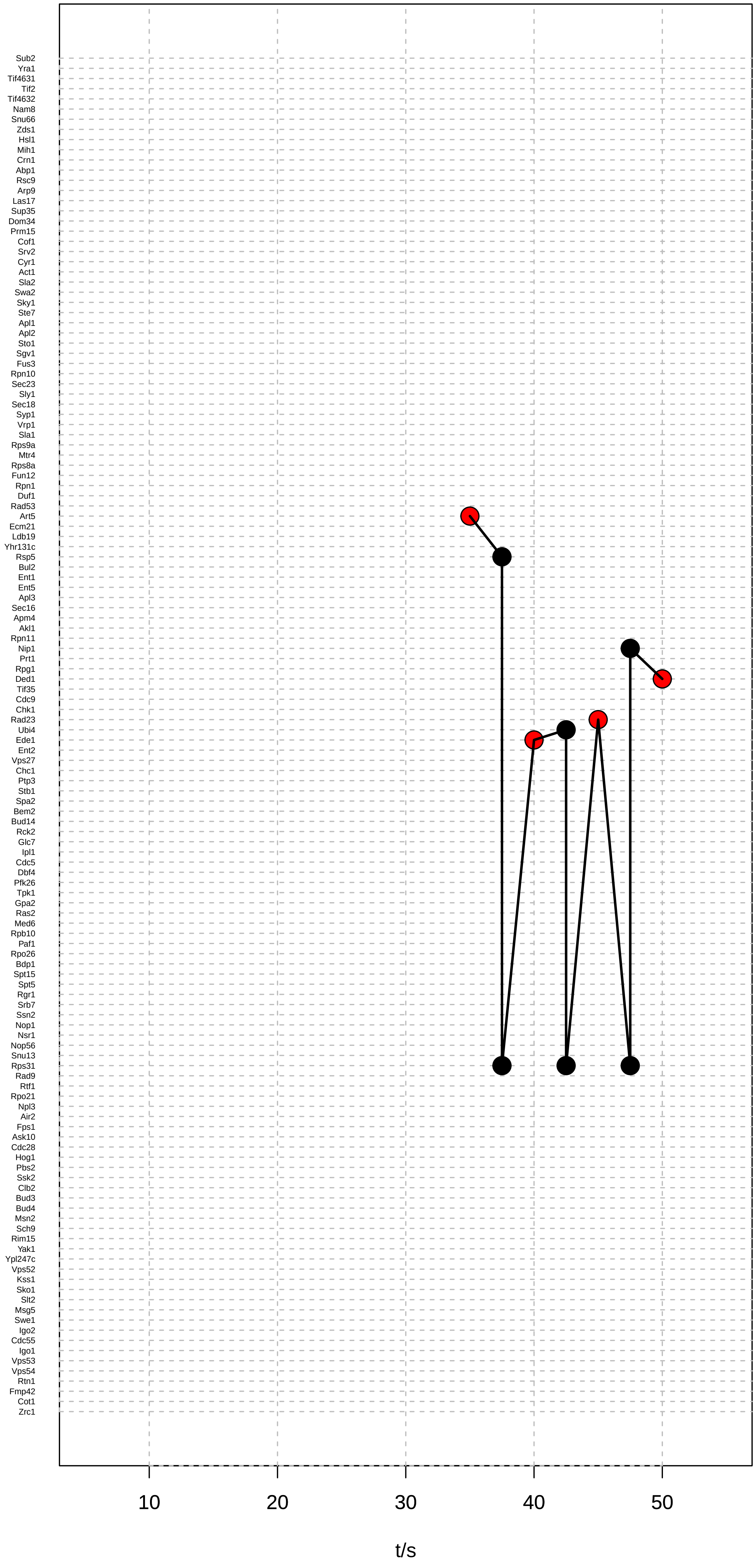

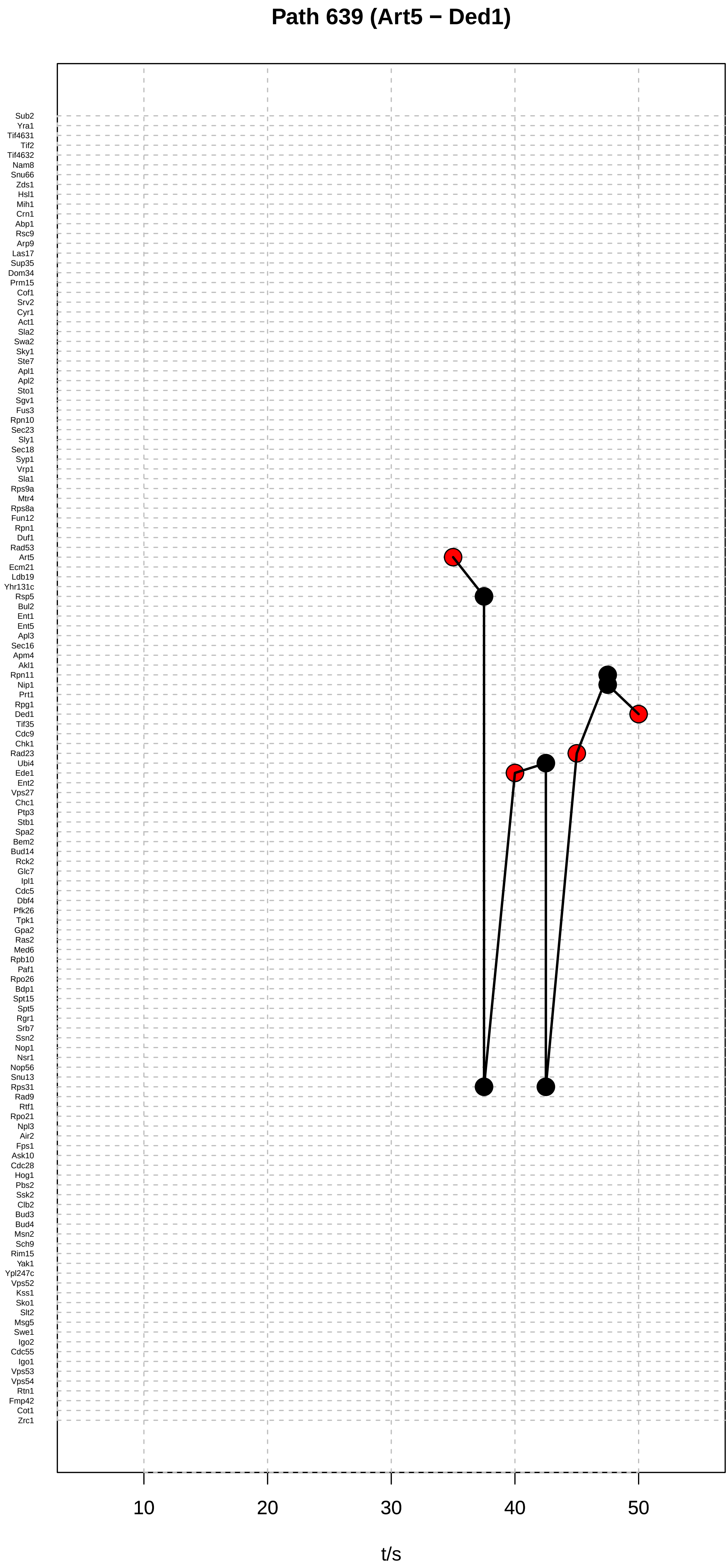

Path 639 (Art5 − Ded1)
Sub2
Yra1
Tif4631
Tif2
Tif4632
Nam8
Snu66
Zds1
Hsl1
Mih1
Crn1
Abp1
Rsc9
Arp9
Las17
Sup35
Dom34
Prm15
Cof1
Srv2
Cyr1
Act1
Sla2
Swa2
Sky1
Ste7
Apl1
Apl2
Sto1
Sgv1
Fus3
Rpn10
Sec23
Sly1
Sec18
Syp1
Vrp1
Sla1
Rps9a
Mtr4
Rps8a
Fun12
Rpn1
Duf1
Rad53
Art5
Ecm21
Ldb19
Yhr131c
Rsp5
Bul2
Ent1
Ent5
Apl3
Sec16
Apm4
Akl1
Rpn11
Nip1
Prt1
Rpg1
Ded1
Tif35
Cdc9
Chk1
Rad23
Ubi4
Ede1
Ent2
Vps27
Chc1
Ptp3
Stb1
Spa2
Bem2
Bud14
Rck2
Glc7
Ipl1
Cdc5
Dbf4
Pfk26
Tpk1
Gpa2
Ras2
Med6
Rpb10
Paf1
Rpo26
Bdp1
Spt15
Spt5
Rgr1
Srb7
Ssn2
Nop1
Nsr1
Nop56
Snu13
Rps31
Rad9
Rtf1
Rpo21
Npl3
Air2
Fps1
Ask10
Cdc28
Hog1
Pbs2
Ssk2
Clb2
Bud3
Bud4
Msn2
Sch9
Rim15
Yak1
Ypl247c
Vps52
Kss1
Sko1
Slt2
Msg5
Swe1
Igo2
Cdc55
Igo1
Vps53
Vps54
Rtn1
Fmp42
Cot1
Zrc1
10
20
30
40
50
t/s

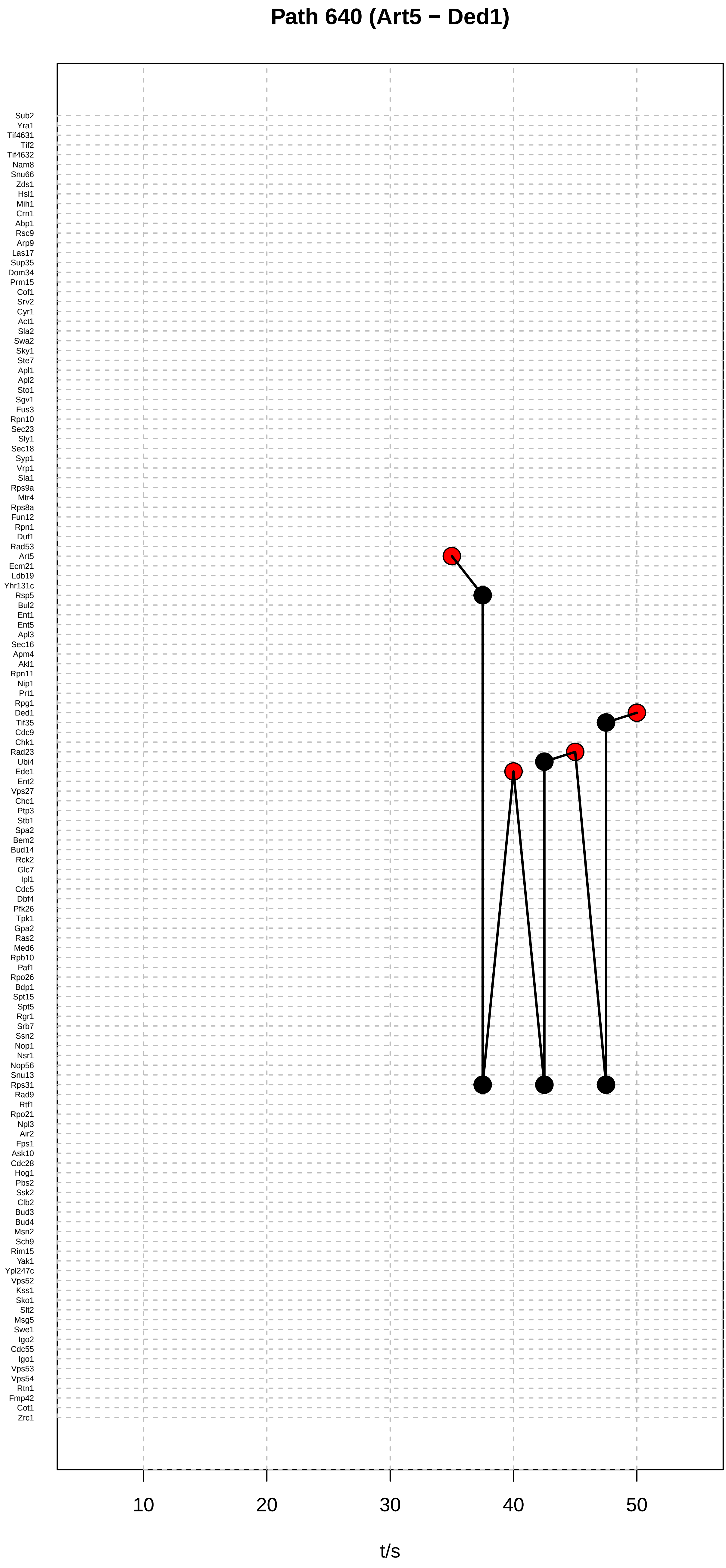
Path 640 (Art5 − Ded1)
Sub2
Yra1
Tif4631
Tif2
Tif4632
Nam8
Snu66
Zds1
Hsl1
Mih1
Crn1
Abp1
Rsc9
Arp9
Las17
Sup35
Dom34
Prm15
Cof1
Srv2
Cyr1
Act1
Sla2
Swa2
Sky1
Ste7
Apl1
Apl2
Sto1
Sgv1
Fus3
Rpn10
Sec23
Sly1
Sec18
Syp1
Vrp1
Sla1
Rps9a
Mtr4
Rps8a
Fun12
Rpn1
Duf1
Rad53
Art5
Ecm21
Ldb19
Yhr131c
Rsp5
Bul2
Ent1
Ent5
Apl3
Sec16
Apm4
Akl1
Rpn11
Nip1
Prt1
Rpg1
Ded1
Tif35
Cdc9
Chk1
Rad23
Ubi4
Ede1
Ent2
Vps27
Chc1
Ptp3
Stb1
Spa2
Bem2
Bud14
Rck2
Glc7
Ipl1
Cdc5
Dbf4
Pfk26
Tpk1
Gpa2
Ras2
Med6
Rpb10
Paf1
Rpo26
Bdp1
Spt15
Spt5
Rgr1
Srb7
Ssn2
Nop1
Nsr1
Nop56
Snu13
Rps31
Rad9
Rtf1
Rpo21
Npl3
Air2
Fps1
Ask10
Cdc28
Hog1
Pbs2
Ssk2
Clb2
Bud3
Bud4
Msn2
Sch9
Rim15
Yak1
Ypl247c
Vps52
Kss1
Sko1
Slt2
Msg5
Swe1
Igo2
Cdc55
Igo1
Vps53
Vps54
Rtn1
Fmp42
Cot1
Zrc1
10
20
30
40
50
t/s

### Path 641 (Art5 − Ded1)

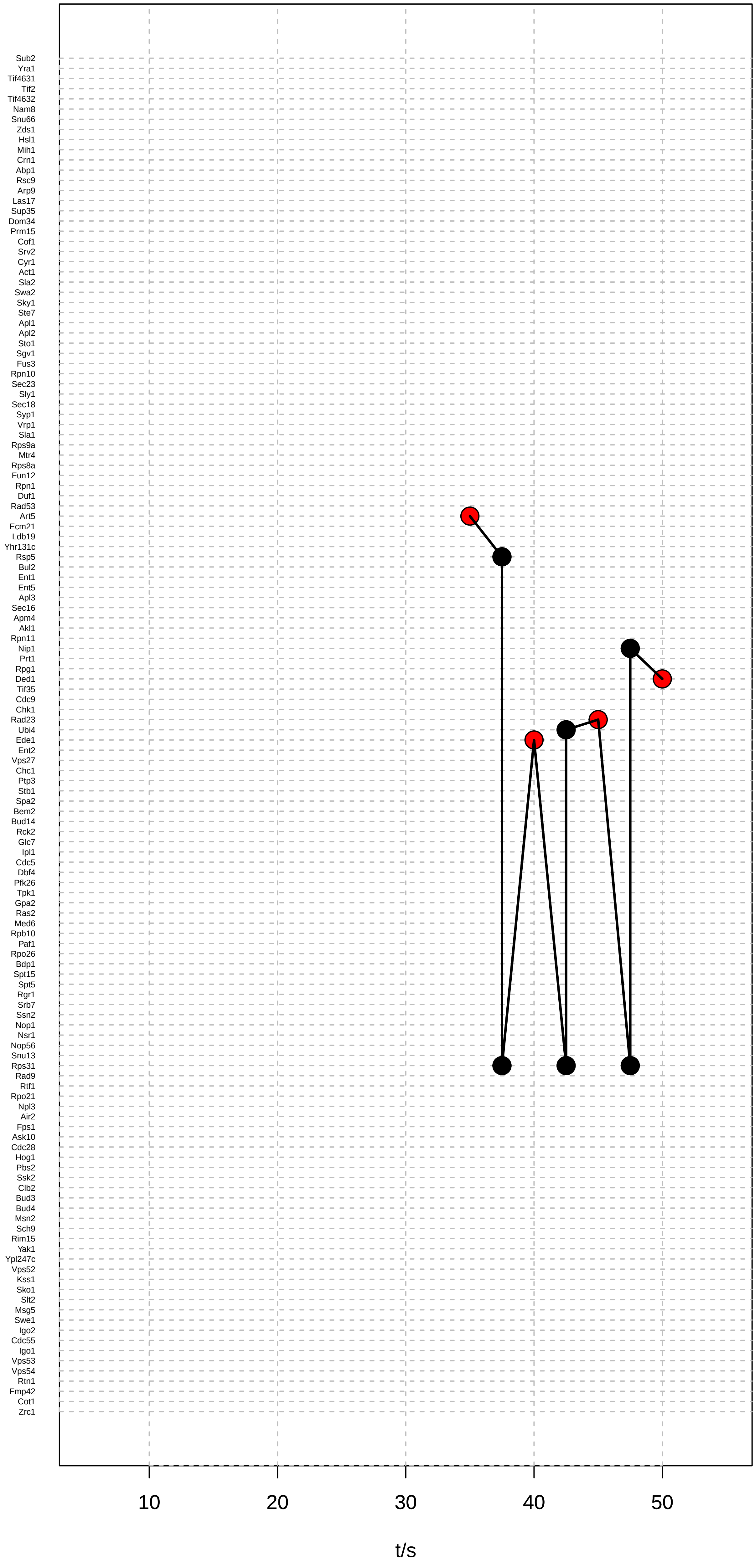

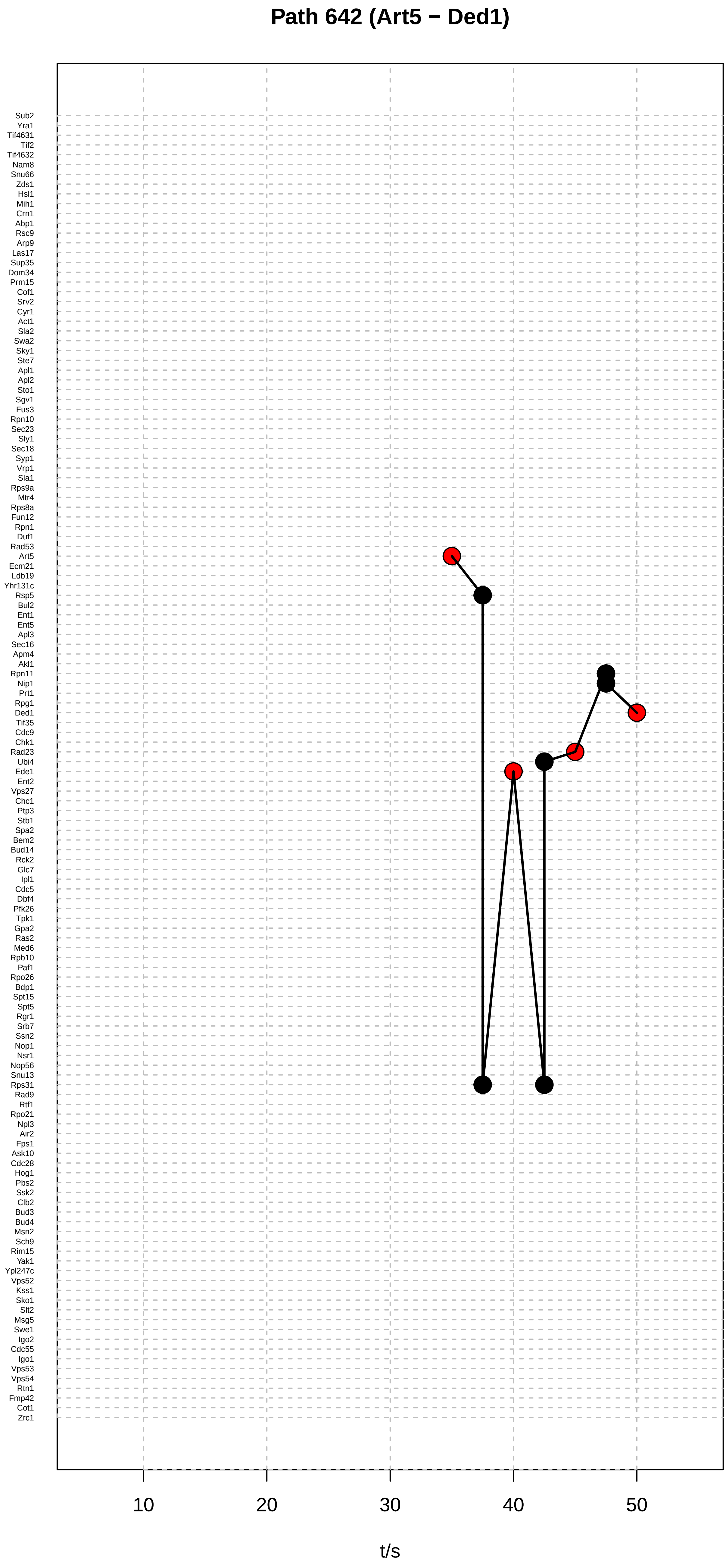
Path 642 (Art5 − Ded1)
Sub2
Yra1
Tif4631
Tif2
Tif4632
Nam8
Snu66
Zds1
Hsl1
Mih1
Crn1
Abp1
Rsc9
Arp9
Las17
Sup35
Dom34
Prm15
Cof1
Srv2
Cyr1
Act1
Sla2
Swa2
Sky1
Ste7
Apl1
Apl2
Sto1
Sgv1
Fus3
Rpn10
Sec23
Sly1
Sec18
Syp1
Vrp1
Sla1
Rps9a
Mtr4
Rps8a
Fun12
Rpn1
Duf1
Rad53
Art5
Ecm21
Ldb19
Yhr131c
Rsp5
Bul2
Ent1
Ent5
Apl3
Sec16
Apm4
Akl1
Rpn11
Nip1
Prt1
Rpg1
Ded1
Tif35
Cdc9
Chk1
Rad23
Ubi4
Ede1
Ent2
Vps27
Chc1
Ptp3
Stb1
Spa2
Bem2
Bud14
Rck2
Glc7
Ipl1
Cdc5
Dbf4
Pfk26
Tpk1
Gpa2
Ras2
Med6
Rpb10
Paf1
Rpo26
Bdp1
Spt15
Spt5
Rgr1
Srb7
Ssn2
Nop1
Nsr1
Nop56
Snu13
Rps31
Rad9
Rtf1
Rpo21
Npl3
Air2
Fps1
Ask10
Cdc28
Hog1
Pbs2
Ssk2
Clb2
Bud3
Bud4
Msn2
Sch9
Rim15
Yak1
Ypl247c
Vps52
Kss1
Sko1
Slt2
Msg5
Swe1
Igo2
Cdc55
Igo1
Vps53
Vps54
Rtn1
Fmp42
Cot1
Zrc1
10
20
30
40
50
t/s

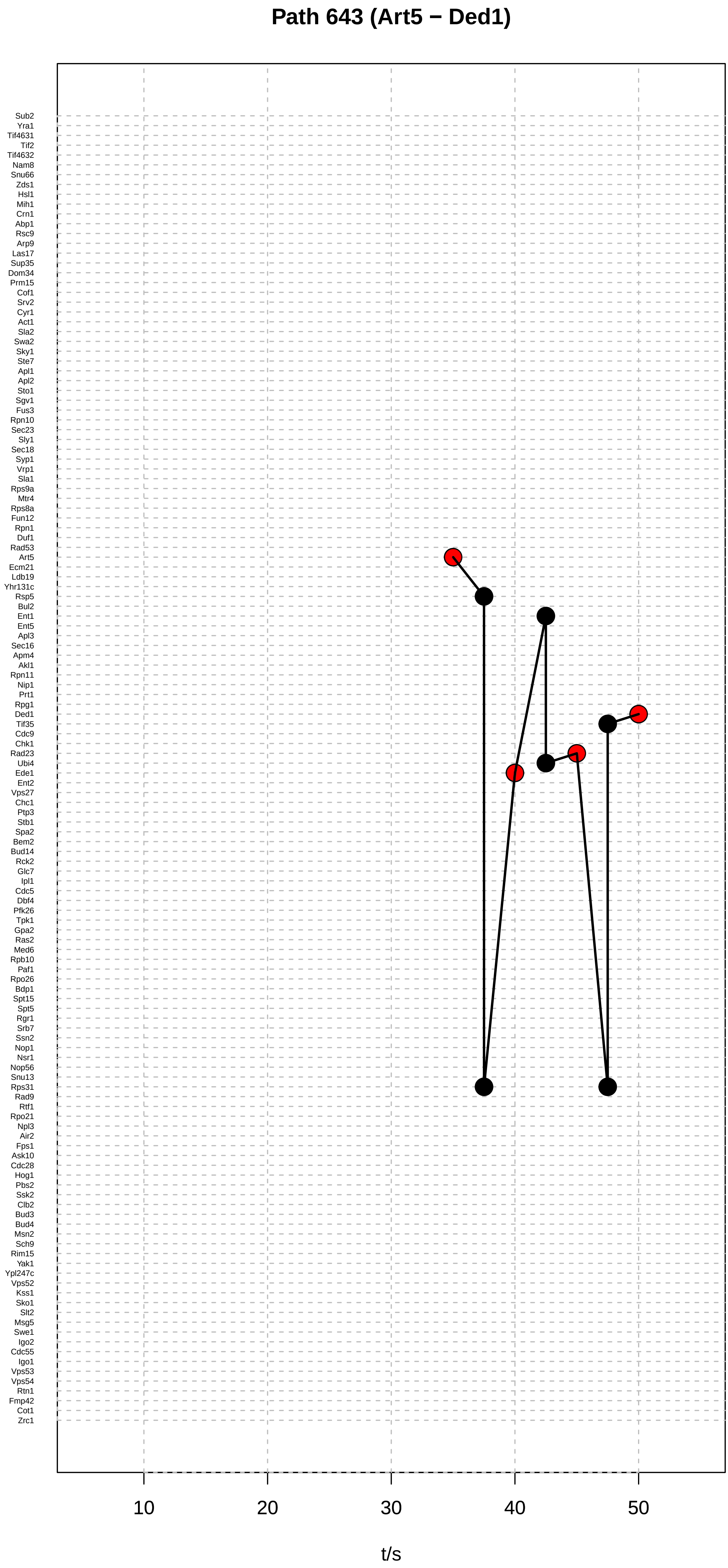
Path 643 (Art5 − Ded1)
Sub2
Yra1
Tif4631
Tif2
Tif4632
Nam8
Snu66
Zds1
Hsl1
Mih1
Crn1
Abp1
Rsc9
Arp9
Las17
Sup35
Dom34
Prm15
Cof1
Srv2
Cyr1
Act1
Sla2
Swa2
Sky1
Ste7
Apl1
Apl2
Sto1
Sgv1
Fus3
Rpn10
Sec23
Sly1
Sec18
Syp1
Vrp1
Sla1
Rps9a
Mtr4
Rps8a
Fun12
Rpn1
Duf1
Rad53
Art5
Ecm21
Ldb19
Yhr131c
Rsp5
Bul2
Ent1
Ent5
Apl3
Sec16
Apm4
Akl1
Rpn11
Nip1
Prt1
Rpg1
Ded1
Tif35
Cdc9
Chk1
Rad23
Ubi4
Ede1
Ent2
Vps27
Chc1
Ptp3
Stb1
Spa2
Bem2
Bud14
Rck2
Glc7
Ipl1
Cdc5
Dbf4
Pfk26
Tpk1
Gpa2
Ras2
Med6
Rpb10
Paf1
Rpo26
Bdp1
Spt15
Spt5
Rgr1
Srb7
Ssn2
Nop1
Nsr1
Nop56
Snu13
Rps31
Rad9
Rtf1
Rpo21
Npl3
Air2
Fps1
Ask10
Cdc28
Hog1
Pbs2
Ssk2
Clb2
Bud3
Bud4
Msn2
Sch9
Rim15
Yak1
Ypl247c
Vps52
Kss1
Sko1
Slt2
Msg5
Swe1
Igo2
Cdc55
Igo1
Vps53
Vps54
Rtn1
Fmp42
Cot1
Zrc1
10
20
30
40
50
t/s

### Path 644 (Art5 − Ded1)

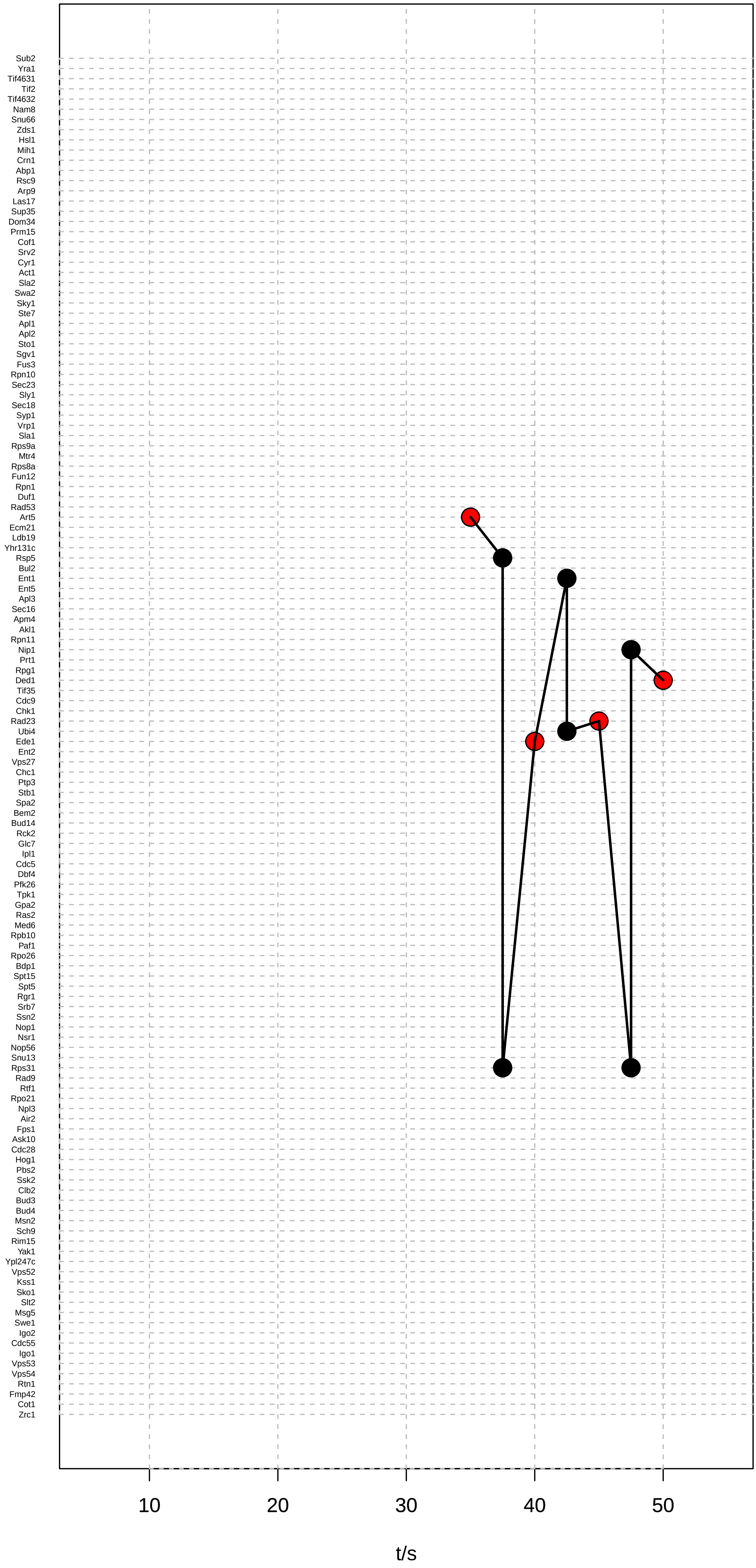

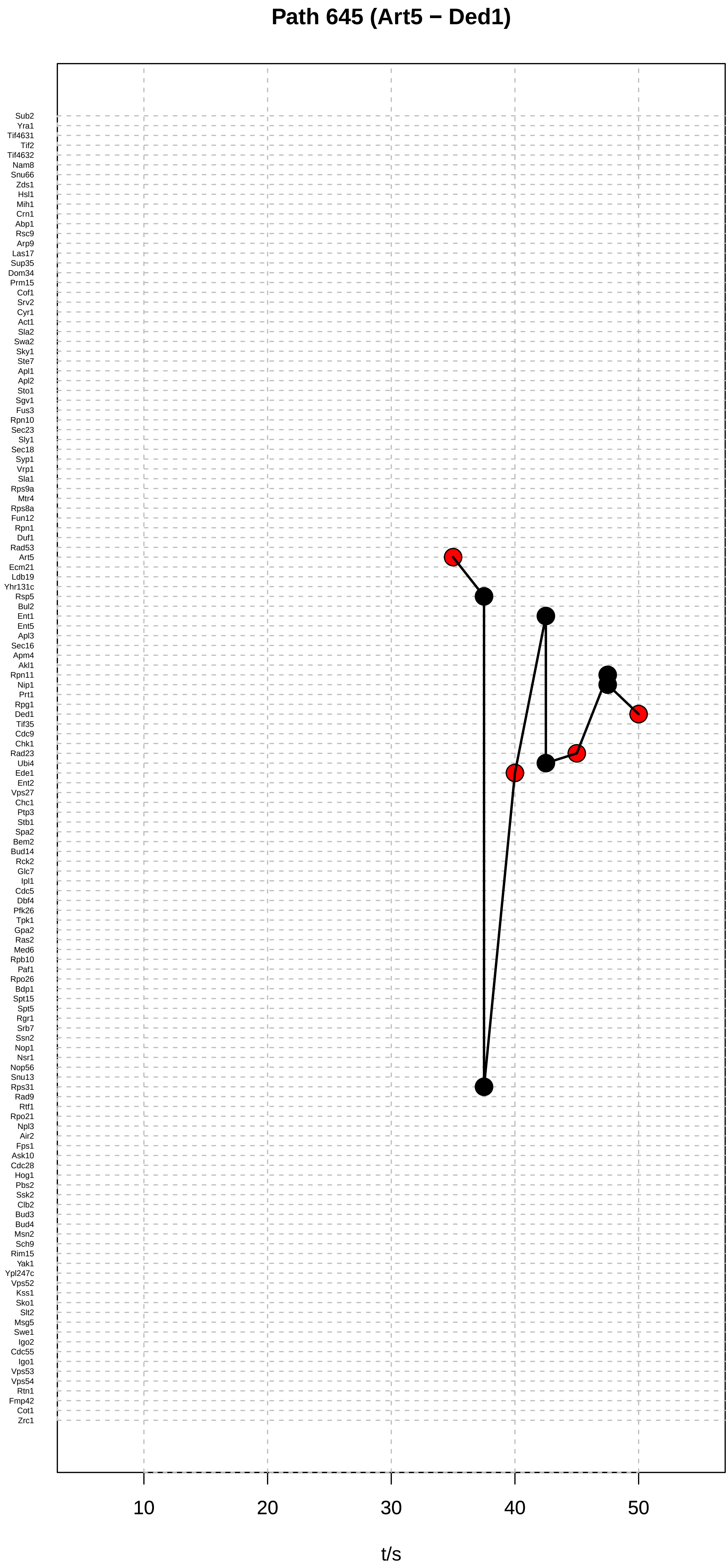
Path 645 (Art5 − Ded1)
Sub2
Yra1
Tif4631
Tif2
Tif4632
Nam8
Snu66
Zds1
Hsl1
Mih1
Crn1
Abp1
Rsc9
Arp9
Las17
Sup35
Dom34
Prm15
Cof1
Srv2
Cyr1
Act1
Sla2
Swa2
Sky1
Ste7
Apl1
Apl2
Sto1
Sgv1
Fus3
Rpn10
Sec23
Sly1
Sec18
Syp1
Vrp1
Sla1
Rps9a
Mtr4
Rps8a
Fun12
Rpn1
Duf1
Rad53
Art5
Ecm21
Ldb19
Yhr131c
Rsp5
Bul2
Ent1
Ent5
Apl3
Sec16
Apm4
Akl1
Rpn11
Nip1
Prt1
Rpg1
Ded1
Tif35
Cdc9
Chk1
Rad23
Ubi4
Ede1
Ent2
Vps27
Chc1
Ptp3
Stb1
Spa2
Bem2
Bud14
Rck2
Glc7
Ipl1
Cdc5
Dbf4
Pfk26
Tpk1
Gpa2
Ras2
Med6
Rpb10
Paf1
Rpo26
Bdp1
Spt15
Spt5
Rgr1
Srb7
Ssn2
Nop1
Nsr1
Nop56
Snu13
Rps31
Rad9
Rtf1
Rpo21
Npl3
Air2
Fps1
Ask10
Cdc28
Hog1
Pbs2
Ssk2
Clb2
Bud3
Bud4
Msn2
Sch9
Rim15
Yak1
Ypl247c
Vps52
Kss1
Sko1
Slt2
Msg5
Swe1
Igo2
Cdc55
Igo1
Vps53
Vps54
Rtn1
Fmp42
Cot1
Zrc1
10
20
30
40
50
t/s

### Path 646 (Air2 − Ded1)

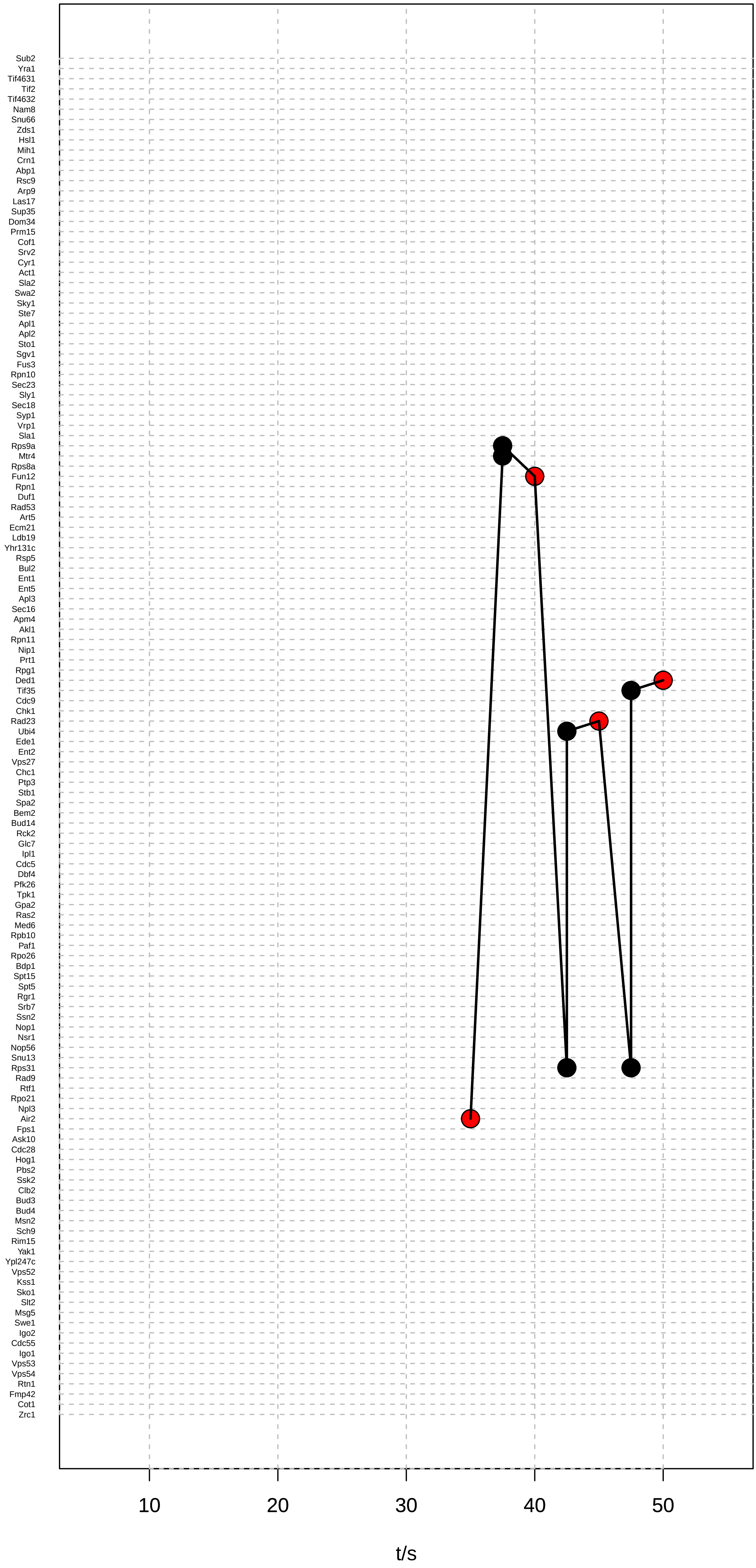

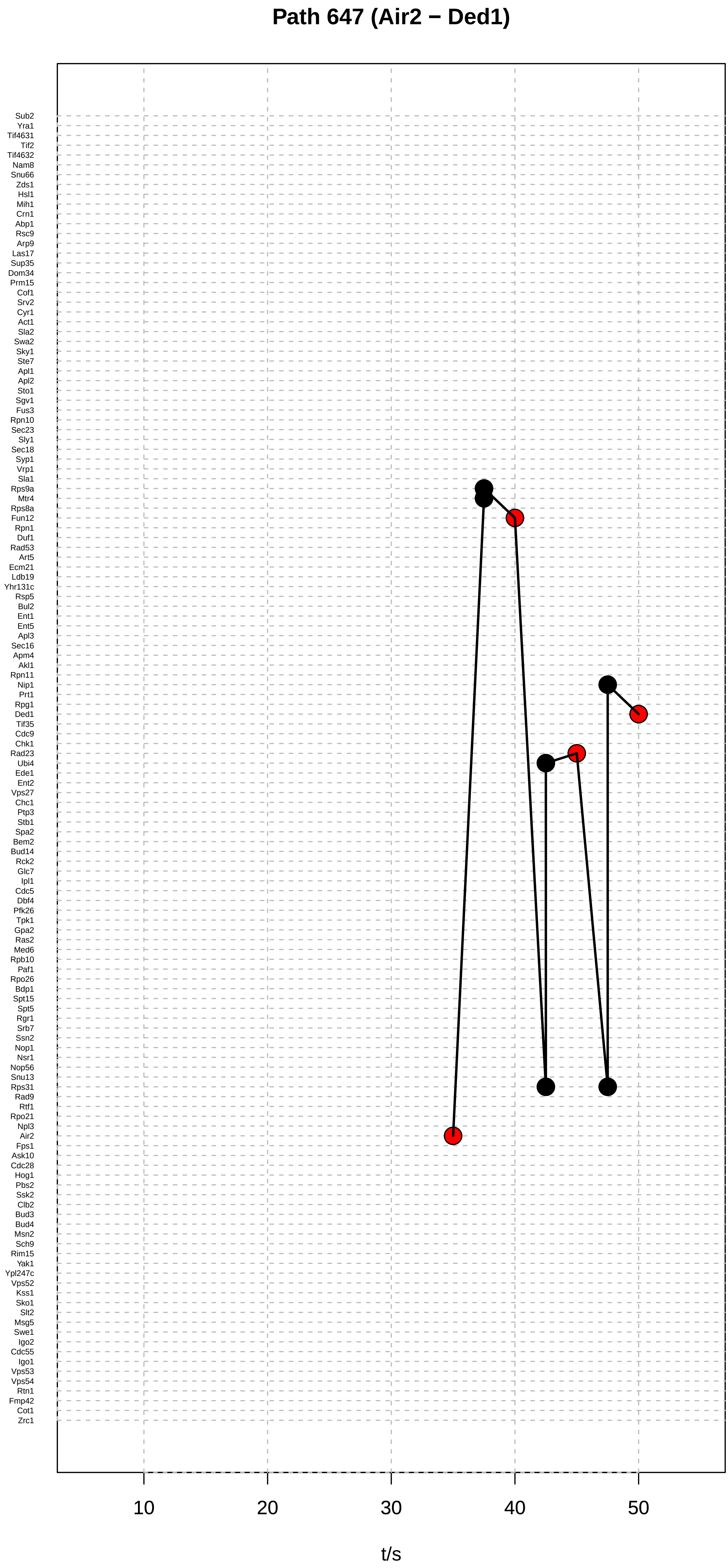

Path 647 (Air2 − Ded1)
Sub2
Yra1
Tif4631
Tif2
Tif4632
Nam8
Snu66
Zds1
Hsl1
Mih1
Crn1
Abp1
Rsc9
Arp9
Las17
Sup35
Dom34
Prm15
Cof1
Srv2
Cyr1
Act1
Sla2
Swa2
Sky1
Ste7
Apl1
Apl2
Sto1
Sgv1
Fus3
Rpn10
Sec23
Sly1
Sec18
Syp1
Vrp1
Sla1
Rps9a
Mtr4
Rps8a
Fun12
Rpn1
Duf1
Rad53
Art5
Ecm21
Ldb19
Yhr131c
Rsp5
Bul2
Ent1
Ent5
Apl3
Sec16
Apm4
Akl1
Rpn11
Nip1
Prt1
Rpg1
Ded1
Tif35
Cdc9
Chk1
Rad23
Ubi4
Ede1
Ent2
Vps27
Chc1
Ptp3
Stb1
Spa2
Bem2
Bud14
Rck2
Glc7
Ipl1
Cdc5
Dbf4
Pfk26
Tpk1
Gpa2
Ras2
Med6
Rpb10
Paf1
Rpo26
Bdp1
Spt15
Spt5
Rgr1
Srb7
Ssn2
Nop1
Nsr1
Nop56
Snu13
Rps31
Rad9
Rtf1
Rpo21
Npl3
Air2
Fps1
Ask10
Cdc28
Hog1
Pbs2
Ssk2
Clb2
Bud3
Bud4
Msn2
Sch9
Rim15
Yak1
Ypl247c
Vps52
Kss1
Sko1
Slt2
Msg5
Swe1
Igo2
Cdc55
Igo1
Vps53
Vps54
Rtn1
Fmp42
Cot1
Zrc1
10
20
30
40
50
t/s

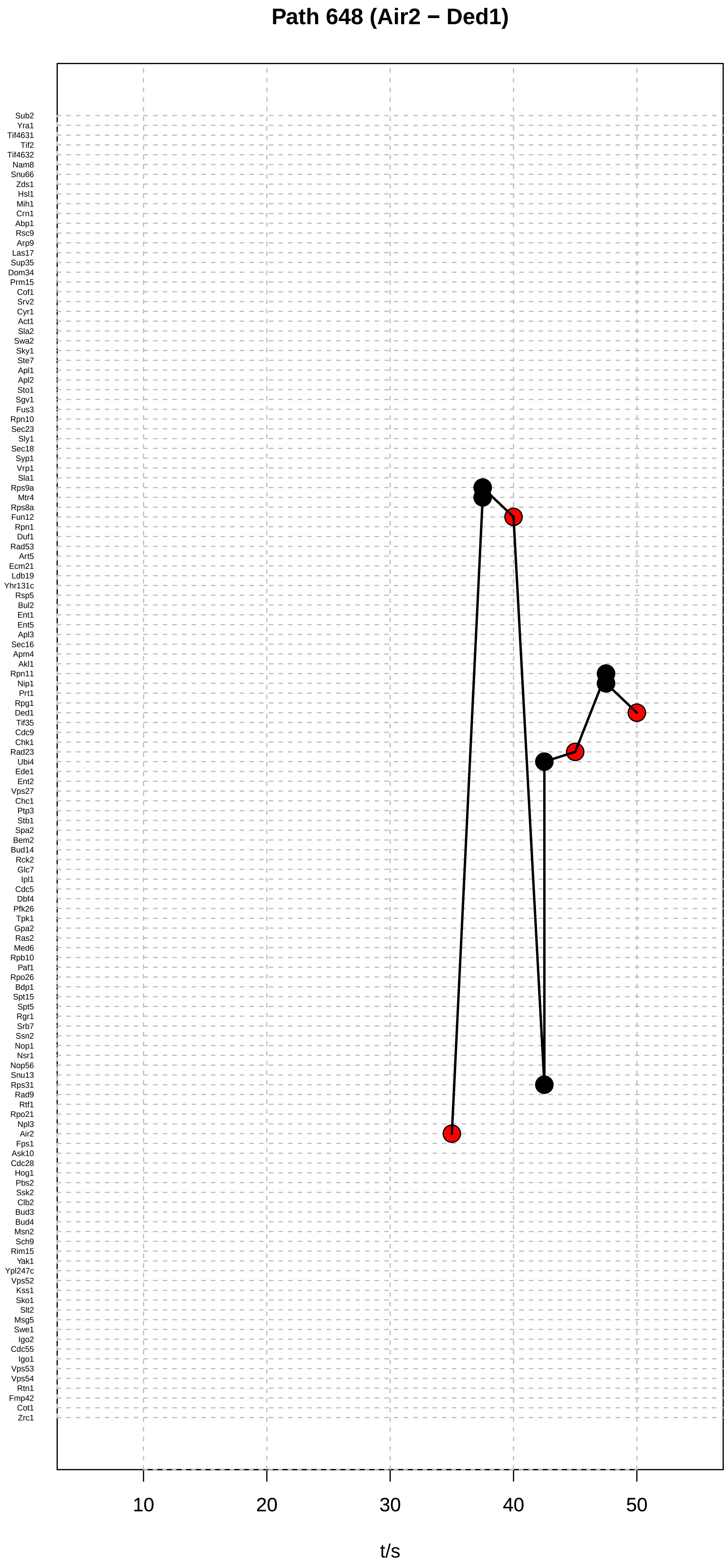
Path 648 (Air2 − Ded1)
Sub2
Yra1
Tif4631
Tif2
Tif4632
Nam8
Snu66
Zds1
Hsl1
Mih1
Crn1
Abp1
Rsc9
Arp9
Las17
Sup35
Dom34
Prm15
Cof1
Srv2
Cyr1
Act1
Sla2
Swa2
Sky1
Ste7
Apl1
Apl2
Sto1
Sgv1
Fus3
Rpn10
Sec23
Sly1
Sec18
Syp1
Vrp1
Sla1
Rps9a
Mtr4
Rps8a
Fun12
Rpn1
Duf1
Rad53
Art5
Ecm21
Ldb19
Yhr131c
Rsp5
Bul2
Ent1
Ent5
Apl3
Sec16
Apm4
Akl1
Rpn11
Nip1
Prt1
Rpg1
Ded1
Tif35
Cdc9
Chk1
Rad23
Ubi4
Ede1
Ent2
Vps27
Chc1
Ptp3
Stb1
Spa2
Bem2
Bud14
Rck2
Glc7
Ipl1
Cdc5
Dbf4
Pfk26
Tpk1
Gpa2
Ras2
Med6
Rpb10
Paf1
Rpo26
Bdp1
Spt15
Spt5
Rgr1
Srb7
Ssn2
Nop1
Nsr1
Nop56
Snu13
Rps31
Rad9
Rtf1
Rpo21
Npl3
Air2
Fps1
Ask10
Cdc28
Hog1
Pbs2
Ssk2
Clb2
Bud3
Bud4
Msn2
Sch9
Rim15
Yak1
Ypl247c
Vps52
Kss1
Sko1
Slt2
Msg5
Swe1
Igo2
Cdc55
Igo1
Vps53
Vps54
Rtn1
Fmp42
Cot1
Zrc1
10
20
30
40
50
t/s

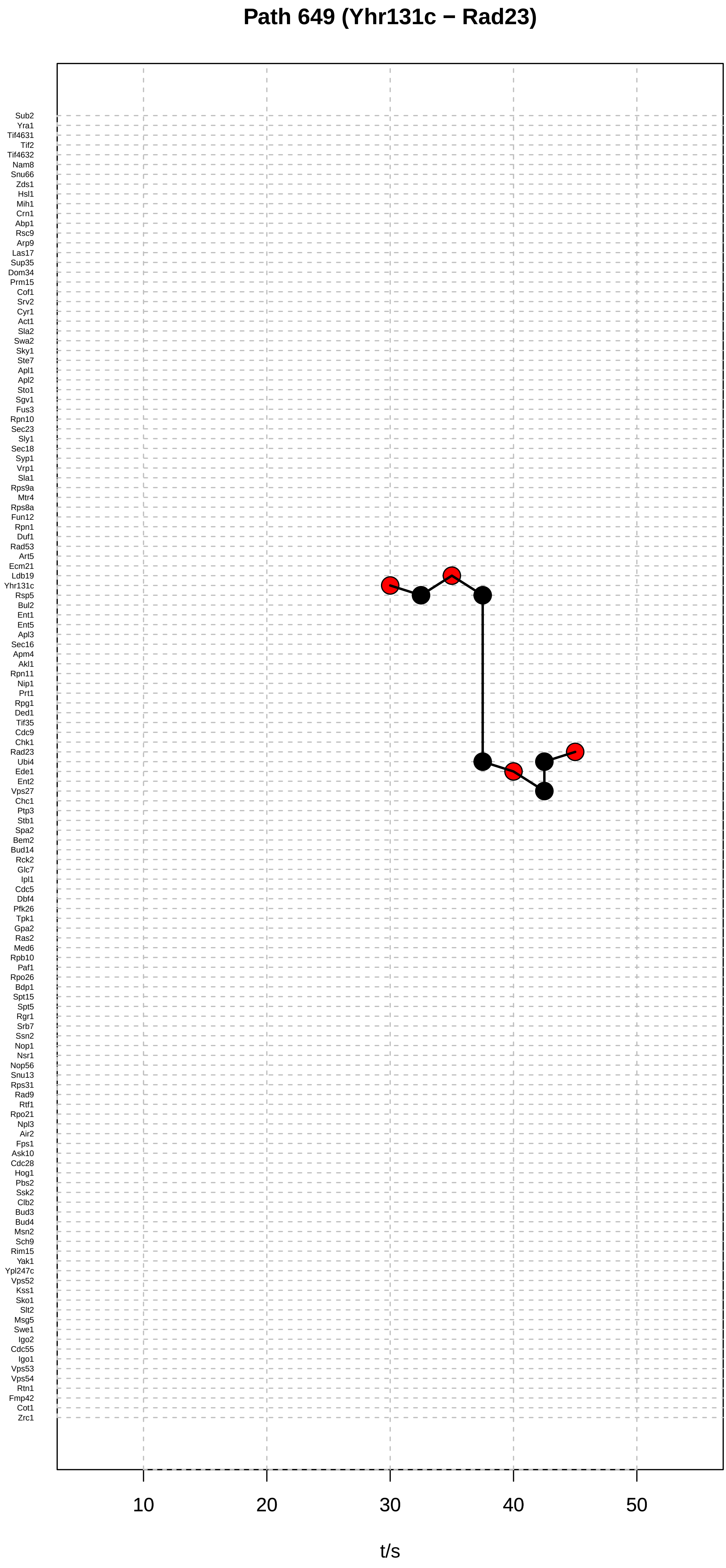

Path 649 (Yhr131c − Rad23)
Sub2
Yra1
Tif4631
Tif2
Tif4632
Nam8
Snu66
Zds1
Hsl1
Mih1
Crn1
Abp1
Rsc9
Arp9
Las17
Sup35
Dom34
Prm15
Cof1
Srv2
Cyr1
Act1
Sla2
Swa2
Sky1
Ste7
Apl1
Apl2
Sto1
Sgv1
Fus3
Rpn10
Sec23
Sly1
Sec18
Syp1
Vrp1
Sla1
Rps9a
Mtr4
Rps8a
Fun12
Rpn1
Duf1
Rad53
Art5
Ecm21
Ldb19
Yhr131c
Rsp5
Bul2
Ent1
Ent5
Apl3
Sec16
Apm4
Akl1
Rpn11
Nip1
Prt1
Rpg1
Ded1
Tif35
Cdc9
Chk1
Rad23
Ubi4
Ede1
Ent2
Vps27
Chc1
Ptp3
Stb1
Spa2
Bem2
Bud14
Rck2
Glc7
Ipl1
Cdc5
Dbf4
Pfk26
Tpk1
Gpa2
Ras2
Med6
Rpb10
Paf1
Rpo26
Bdp1
Spt15
Spt5
Rgr1
Srb7
Ssn2
Nop1
Nsr1
Nop56
Snu13
Rps31
Rad9
Rtf1
Rpo21
Npl3
Air2
Fps1
Ask10
Cdc28
Hog1
Pbs2
Ssk2
Clb2
Bud3
Bud4
Msn2
Sch9
Rim15
Yak1
Ypl247c
Vps52
Kss1
Sko1
Slt2
Msg5
Swe1
Igo2
Cdc55
Igo1
Vps53
Vps54
Rtn1
Fmp42
Cot1
Zrc1
10
20
30
40
50
t/s

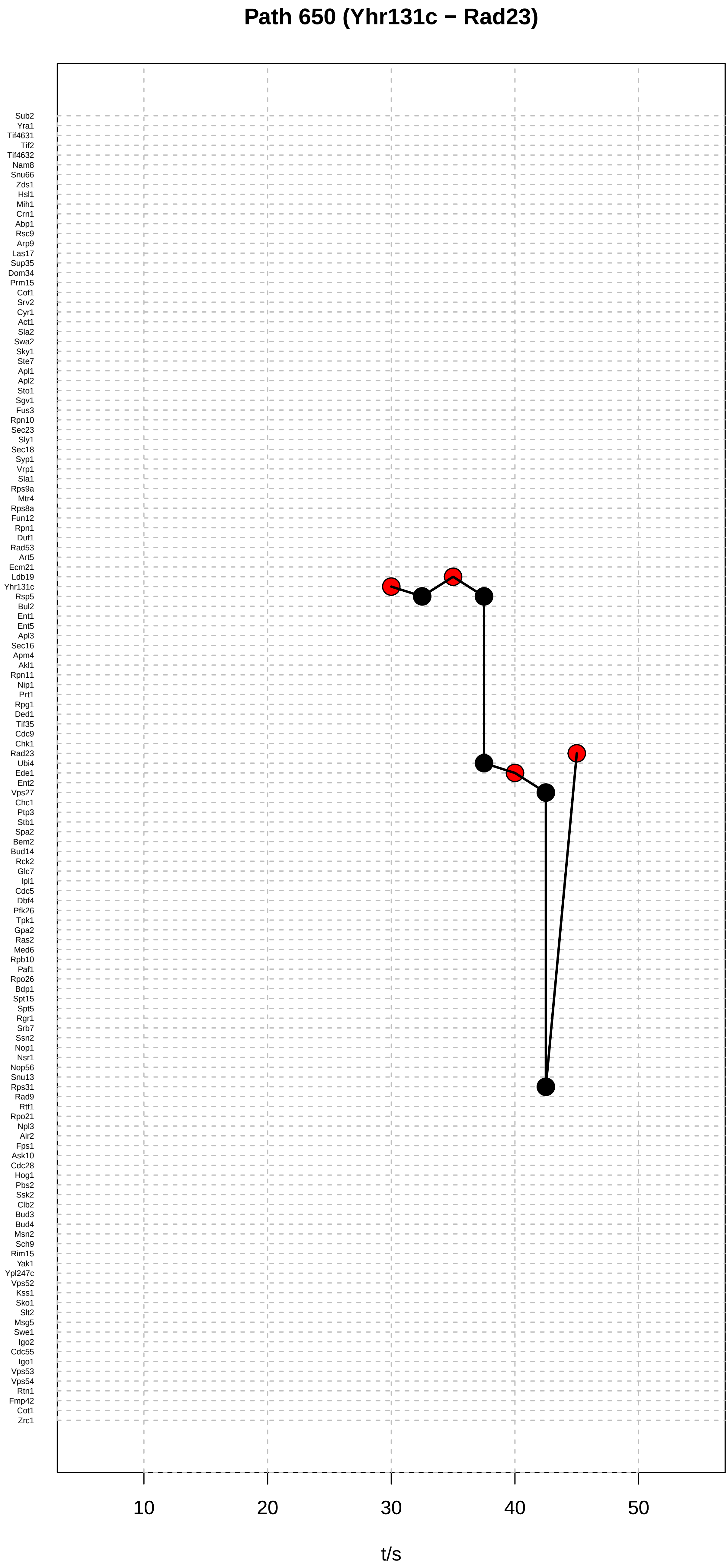

Path 650 (Yhr131c − Rad23)
Sub2
Yra1
Tif4631
Tif2
Tif4632
Nam8
Snu66
Zds1
Hsl1
Mih1
Crn1
Abp1
Rsc9
Arp9
Las17
Sup35
Dom34
Prm15
Cof1
Srv2
Cyr1
Act1
Sla2
Swa2
Sky1
Ste7
Apl1
Apl2
Sto1
Sgv1
Fus3
Rpn10
Sec23
Sly1
Sec18
Syp1
Vrp1
Sla1
Rps9a
Mtr4
Rps8a
Fun12
Rpn1
Duf1
Rad53
Art5
Ecm21
Ldb19
Yhr131c
Rsp5
Bul2
Ent1
Ent5
Apl3
Sec16
Apm4
Akl1
Rpn11
Nip1
Prt1
Rpg1
Ded1
Tif35
Cdc9
Chk1
Rad23
Ubi4
Ede1
Ent2
Vps27
Chc1
Ptp3
Stb1
Spa2
Bem2
Bud14
Rck2
Glc7
Ipl1
Cdc5
Dbf4
Pfk26
Tpk1
Gpa2
Ras2
Med6
Rpb10
Paf1
Rpo26
Bdp1
Spt15
Spt5
Rgr1
Srb7
Ssn2
Nop1
Nsr1
Nop56
Snu13
Rps31
Rad9
Rtf1
Rpo21
Npl3
Air2
Fps1
Ask10
Cdc28
Hog1
Pbs2
Ssk2
Clb2
Bud3
Bud4
Msn2
Sch9
Rim15
Yak1
Ypl247c
Vps52
Kss1
Sko1
Slt2
Msg5
Swe1
Igo2
Cdc55
Igo1
Vps53
Vps54
Rtn1
Fmp42
Cot1
Zrc1
10
20
30
40
50
t/s

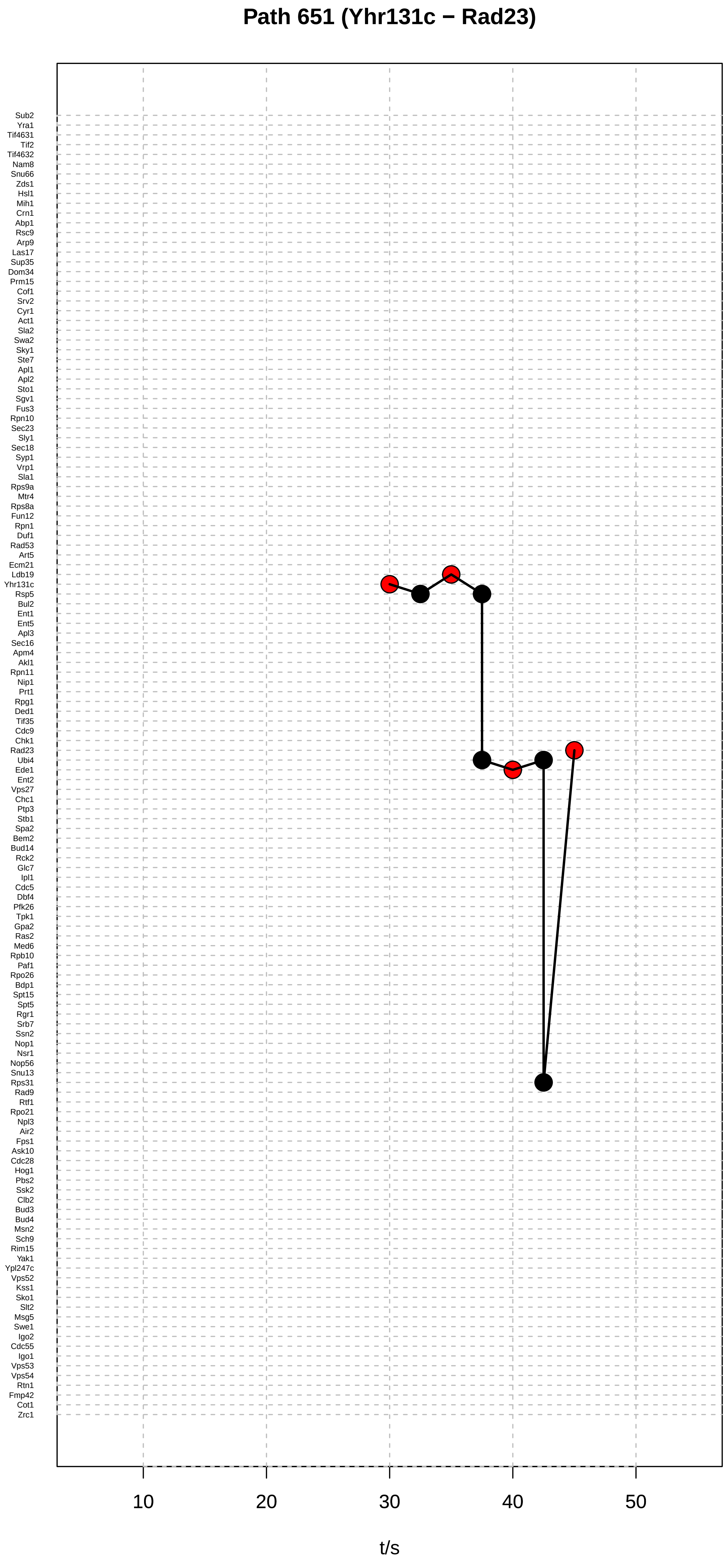
Path 651 (Yhr131c − Rad23)
Sub2
Yra1
Tif4631
Tif2
Tif4632
Nam8
Snu66
Zds1
Hsl1
Mih1
Crn1
Abp1
Rsc9
Arp9
Las17
Sup35
Dom34
Prm15
Cof1
Srv2
Cyr1
Act1
Sla2
Swa2
Sky1
Ste7
Apl1
Apl2
Sto1
Sgv1
Fus3
Rpn10
Sec23
Sly1
Sec18
Syp1
Vrp1
Sla1
Rps9a
Mtr4
Rps8a
Fun12
Rpn1
Duf1
Rad53
Art5
Ecm21
Ldb19
Yhr131c
Rsp5
Bul2
Ent1
Ent5
Apl3
Sec16
Apm4
Akl1
Rpn11
Nip1
Prt1
Rpg1
Ded1
Tif35
Cdc9
Chk1
Rad23
Ubi4
Ede1
Ent2
Vps27
Chc1
Ptp3
Stb1
Spa2
Bem2
Bud14
Rck2
Glc7
Ipl1
Cdc5
Dbf4
Pfk26
Tpk1
Gpa2
Ras2
Med6
Rpb10
Paf1
Rpo26
Bdp1
Spt15
Spt5
Rgr1
Srb7
Ssn2
Nop1
Nsr1
Nop56
Snu13
Rps31
Rad9
Rtf1
Rpo21
Npl3
Air2
Fps1
Ask10
Cdc28
Hog1
Pbs2
Ssk2
Clb2
Bud3
Bud4
Msn2
Sch9
Rim15
Yak1
Ypl247c
Vps52
Kss1
Sko1
Slt2
Msg5
Swe1
Igo2
Cdc55
Igo1
Vps53
Vps54
Rtn1
Fmp42
Cot1
Zrc1
10
20
30
40
50
t/s

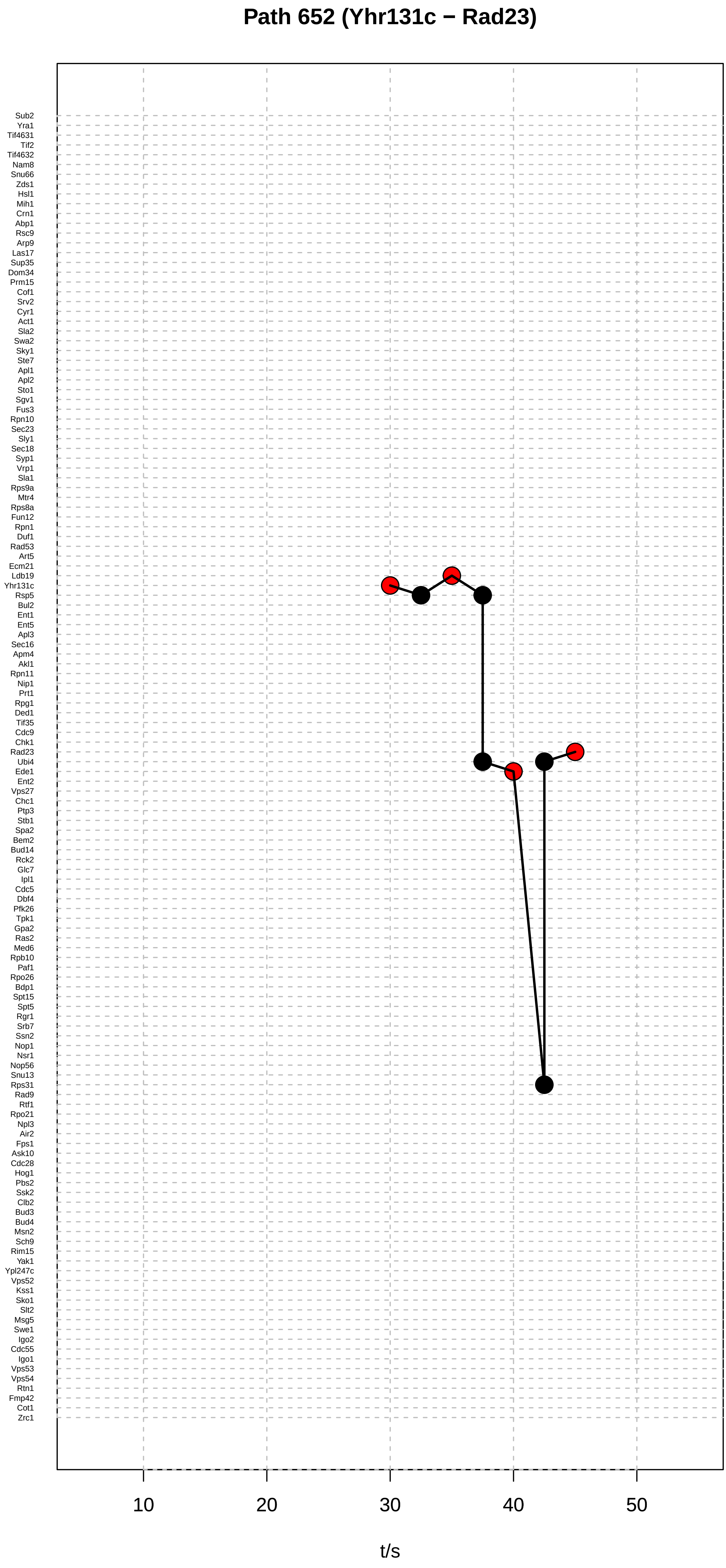

Path 652 (Yhr131c − Rad23)
Sub2
Yra1
Tif4631
Tif2
Tif4632
Nam8
Snu66
Zds1
Hsl1
Mih1
Crn1
Abp1
Rsc9
Arp9
Las17
Sup35
Dom34
Prm15
Cof1
Srv2
Cyr1
Act1
Sla2
Swa2
Sky1
Ste7
Apl1
Apl2
Sto1
Sgv1
Fus3
Rpn10
Sec23
Sly1
Sec18
Syp1
Vrp1
Sla1
Rps9a
Mtr4
Rps8a
Fun12
Rpn1
Duf1
Rad53
Art5
Ecm21
Ldb19
Yhr131c
Rsp5
Bul2
Ent1
Ent5
Apl3
Sec16
Apm4
Akl1
Rpn11
Nip1
Prt1
Rpg1
Ded1
Tif35
Cdc9
Chk1
Rad23
Ubi4
Ede1
Ent2
Vps27
Chc1
Ptp3
Stb1
Spa2
Bem2
Bud14
Rck2
Glc7
Ipl1
Cdc5
Dbf4
Pfk26
Tpk1
Gpa2
Ras2
Med6
Rpb10
Paf1
Rpo26
Bdp1
Spt15
Spt5
Rgr1
Srb7
Ssn2
Nop1
Nsr1
Nop56
Snu13
Rps31
Rad9
Rtf1
Rpo21
Npl3
Air2
Fps1
Ask10
Cdc28
Hog1
Pbs2
Ssk2
Clb2
Bud3
Bud4
Msn2
Sch9
Rim15
Yak1
Ypl247c
Vps52
Kss1
Sko1
Slt2
Msg5
Swe1
Igo2
Cdc55
Igo1
Vps53
Vps54
Rtn1
Fmp42
Cot1
Zrc1
10
20
30
40
50
t/s

#### Path 653 (Yhr131c − Rad23)

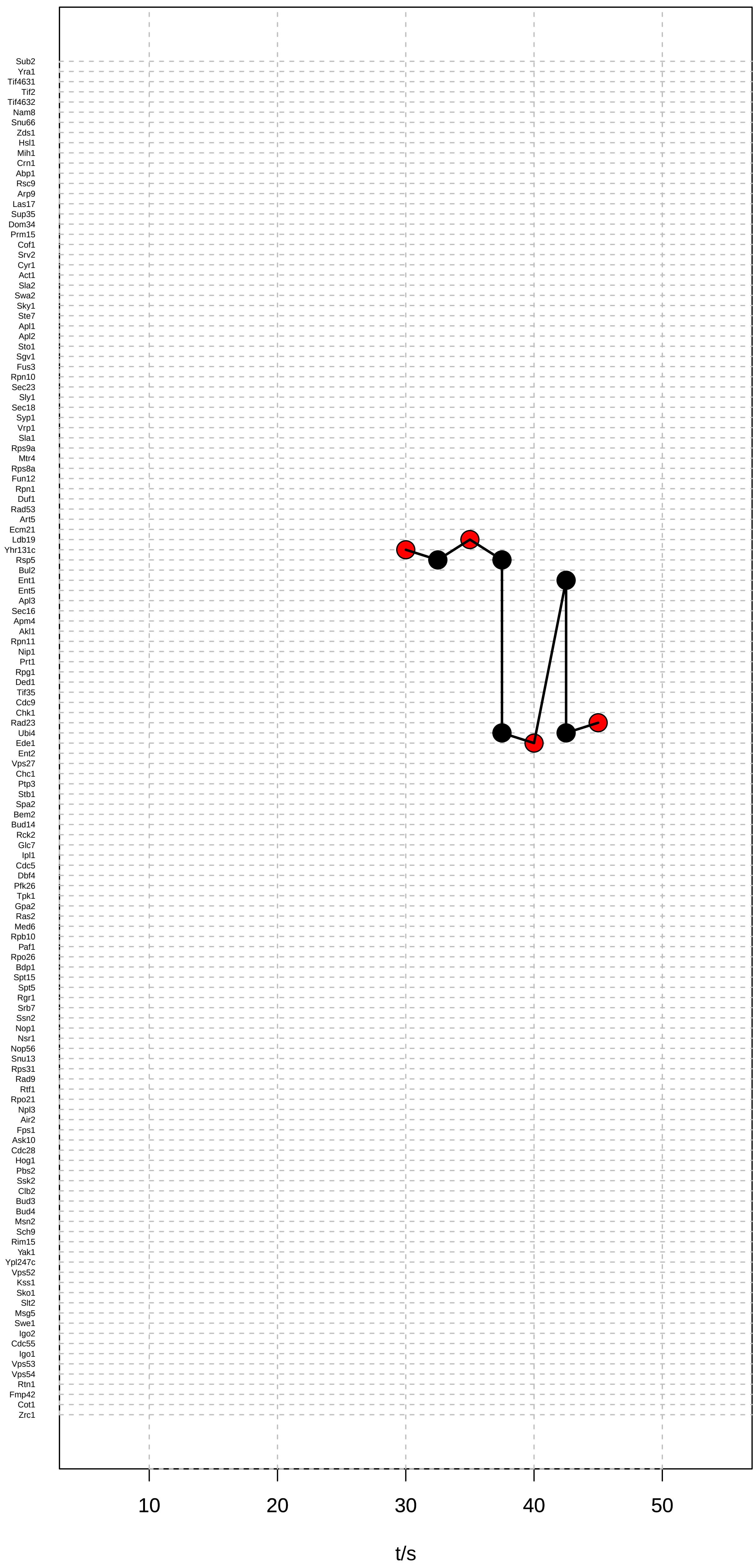

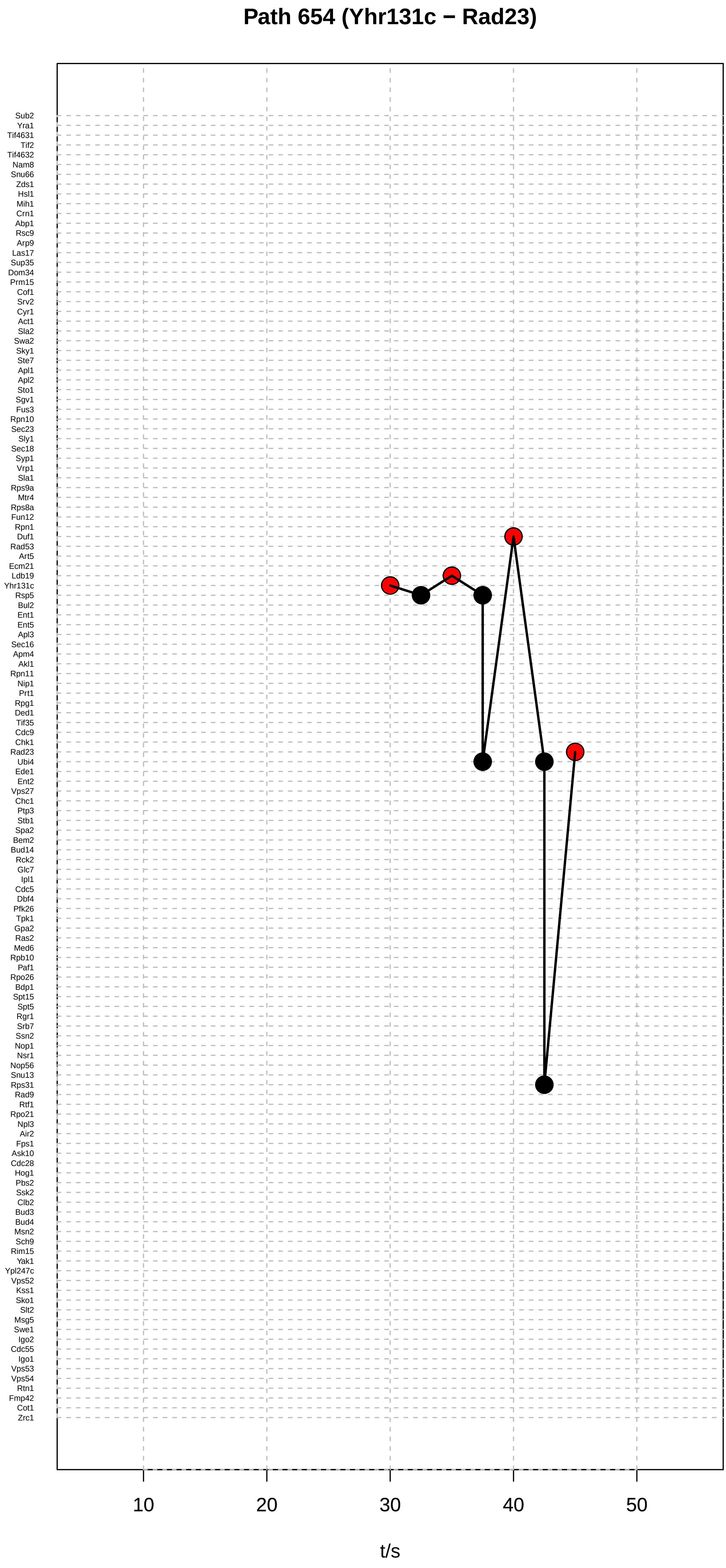

Path 654 (Yhr131c − Rad23)
Sub2
Yra1
Tif4631
Tif2
Tif4632
Nam8
Snu66
Zds1
Hsl1
Mih1
Crn1
Abp1
Rsc9
Arp9
Las17
Sup35
Dom34
Prm15
Cof1
Srv2
Cyr1
Act1
Sla2
Swa2
Sky1
Ste7
Apl1
Apl2
Sto1
Sgv1
Fus3
Rpn10
Sec23
Sly1
Sec18
Syp1
Vrp1
Sla1
Rps9a
Mtr4
Rps8a
Fun12
Rpn1
Duf1
Rad53
Art5
Ecm21
Ldb19
Yhr131c
Rsp5
Bul2
Ent1
Ent5
Apl3
Sec16
Apm4
Akl1
Rpn11
Nip1
Prt1
Rpg1
Ded1
Tif35
Cdc9
Chk1
Rad23
Ubi4
Ede1
Ent2
Vps27
Chc1
Ptp3
Stb1
Spa2
Bem2
Bud14
Rck2
Glc7
Ipl1
Cdc5
Dbf4
Pfk26
Tpk1
Gpa2
Ras2
Med6
Rpb10
Paf1
Rpo26
Bdp1
Spt15
Spt5
Rgr1
Srb7
Ssn2
Nop1
Nsr1
Nop56
Snu13
Rps31
Rad9
Rtf1
Rpo21
Npl3
Air2
Fps1
Ask10
Cdc28
Hog1
Pbs2
Ssk2
Clb2
Bud3
Bud4
Msn2
Sch9
Rim15
Yak1
Ypl247c
Vps52
Kss1
Sko1
Slt2
Msg5
Swe1
Igo2
Cdc55
Igo1
Vps53
Vps54
Rtn1
Fmp42
Cot1
Zrc1
10
20
30
40
50
t/s

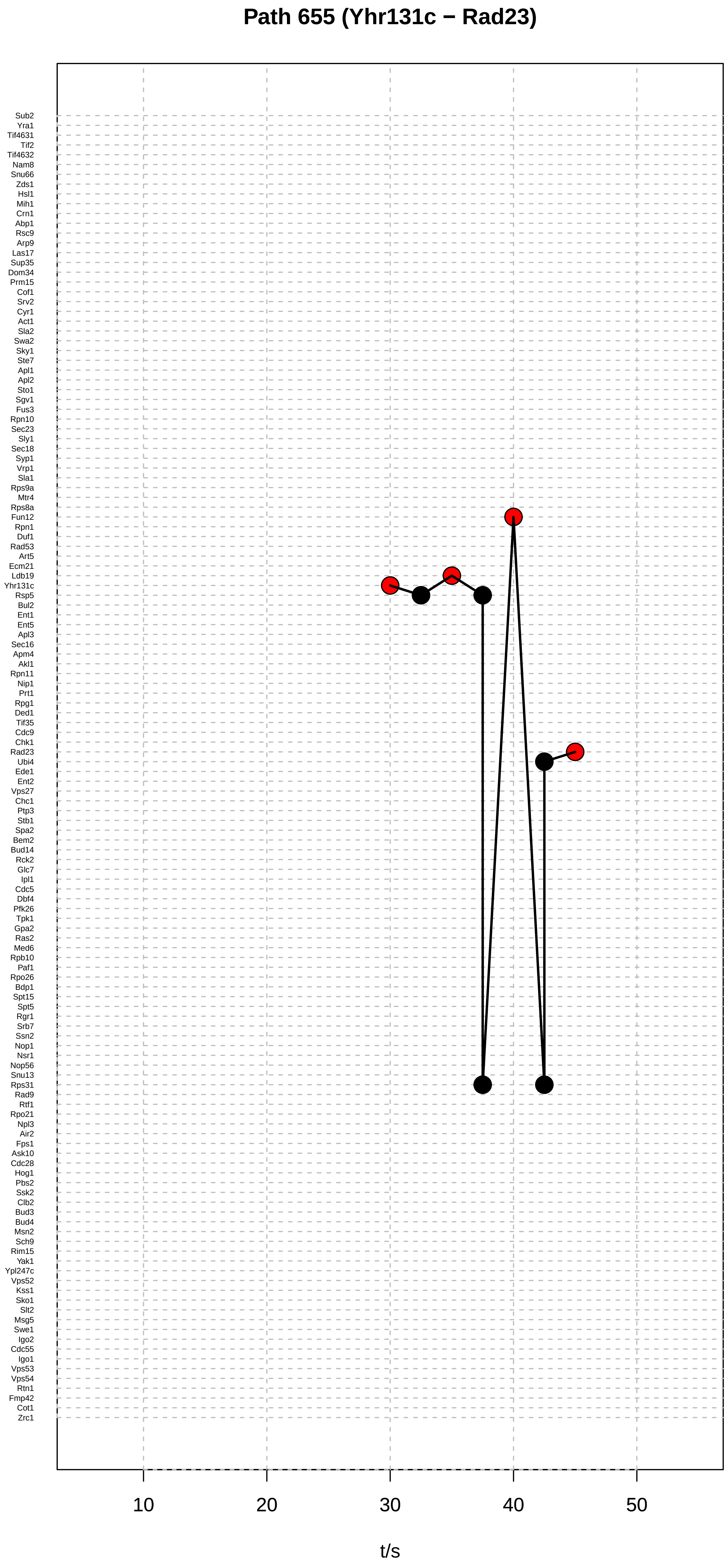

Path 655 (Yhr131c − Rad23)
Sub2
Yra1
Tif4631
Tif2
Tif4632
Nam8
Snu66
Zds1
Hsl1
Mih1
Crn1
Abp1
Rsc9
Arp9
Las17
Sup35
Dom34
Prm15
Cof1
Srv2
Cyr1
Act1
Sla2
Swa2
Sky1
Ste7
Apl1
Apl2
Sto1
Sgv1
Fus3
Rpn10
Sec23
Sly1
Sec18
Syp1
Vrp1
Sla1
Rps9a
Mtr4
Rps8a
Fun12
Rpn1
Duf1
Rad53
Art5
Ecm21
Ldb19
Yhr131c
Rsp5
Bul2
Ent1
Ent5
Apl3
Sec16
Apm4
Akl1
Rpn11
Nip1
Prt1
Rpg1
Ded1
Tif35
Cdc9
Chk1
Rad23
Ubi4
Ede1
Ent2
Vps27
Chc1
Ptp3
Stb1
Spa2
Bem2
Bud14
Rck2
Glc7
Ipl1
Cdc5
Dbf4
Pfk26
Tpk1
Gpa2
Ras2
Med6
Rpb10
Paf1
Rpo26
Bdp1
Spt15
Spt5
Rgr1
Srb7
Ssn2
Nop1
Nsr1
Nop56
Snu13
Rps31
Rad9
Rtf1
Rpo21
Npl3
Air2
Fps1
Ask10
Cdc28
Hog1
Pbs2
Ssk2
Clb2
Bud3
Bud4
Msn2
Sch9
Rim15
Yak1
Ypl247c
Vps52
Kss1
Sko1
Slt2
Msg5
Swe1
Igo2
Cdc55
Igo1
Vps53
Vps54
Rtn1
Fmp42
Cot1
Zrc1
10
20
30
40
50
t/s

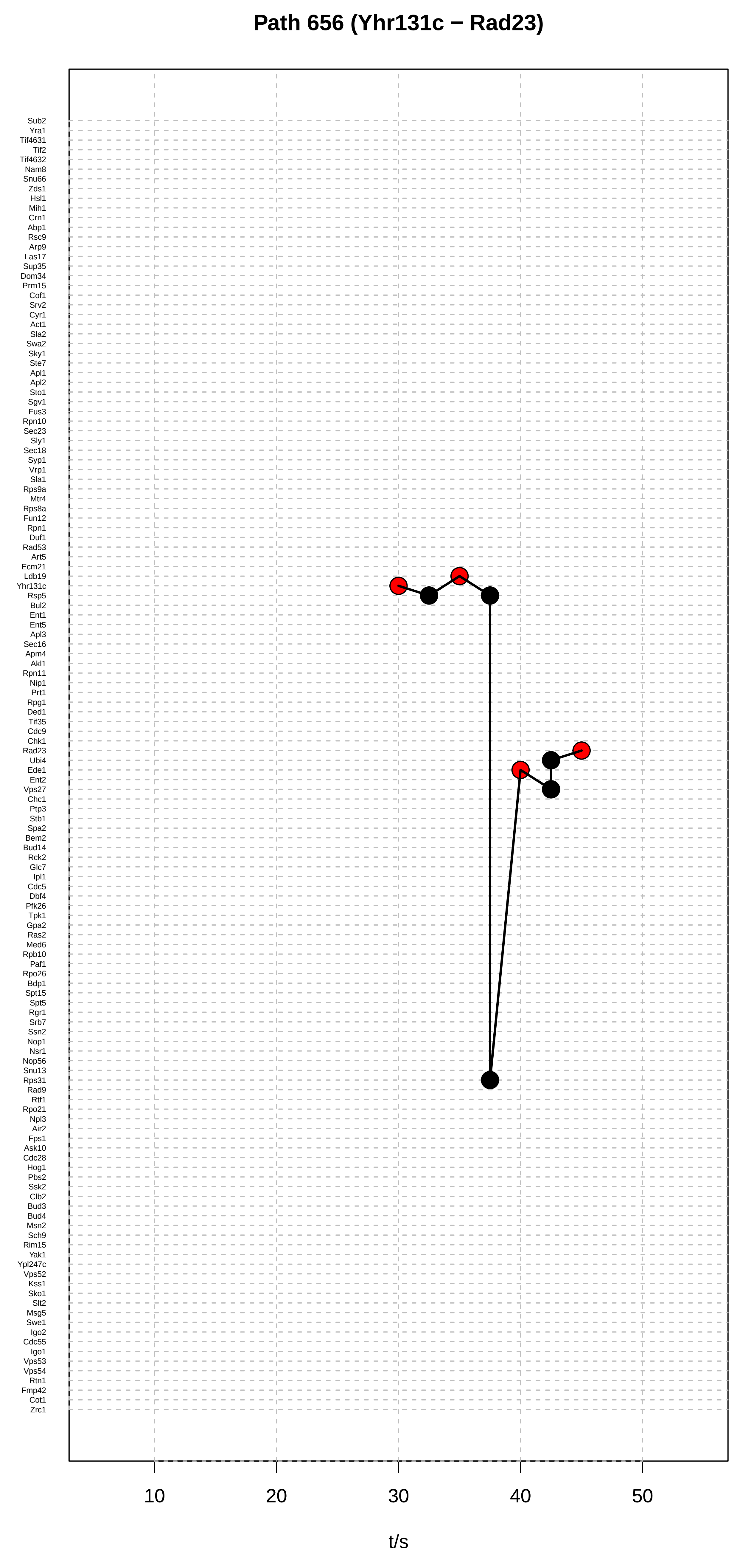

Path 656 (Yhr131c − Rad23)
Sub2
Yra1
Tif4631
Tif2
Tif4632
Nam8
Snu66
Zds1
Hsl1
Mih1
Crn1
Abp1
Rsc9
Arp9
Las17
Sup35
Dom34
Prm15
Cof1
Srv2
Cyr1
Act1
Sla2
Swa2
Sky1
Ste7
Apl1
Apl2
Sto1
Sgv1
Fus3
Rpn10
Sec23
Sly1
Sec18
Syp1
Vrp1
Sla1
Rps9a
Mtr4
Rps8a
Fun12
Rpn1
Duf1
Rad53
Art5
Ecm21
Ldb19
Yhr131c
Rsp5
Bul2
Ent1
Ent5
Apl3
Sec16
Apm4
Akl1
Rpn11
Nip1
Prt1
Rpg1
Ded1
Tif35
Cdc9
Chk1
Rad23
Ubi4
Ede1
Ent2
Vps27
Chc1
Ptp3
Stb1
Spa2
Bem2
Bud14
Rck2
Glc7
Ipl1
Cdc5
Dbf4
Pfk26
Tpk1
Gpa2
Ras2
Med6
Rpb10
Paf1
Rpo26
Bdp1
Spt15
Spt5
Rgr1
Srb7
Ssn2
Nop1
Nsr1
Nop56
Snu13
Rps31
Rad9
Rtf1
Rpo21
Npl3
Air2
Fps1
Ask10
Cdc28
Hog1
Pbs2
Ssk2
Clb2
Bud3
Bud4
Msn2
Sch9
Rim15
Yak1
Ypl247c
Vps52
Kss1
Sko1
Slt2
Msg5
Swe1
Igo2
Cdc55
Igo1
Vps53
Vps54
Rtn1
Fmp42
Cot1
Zrc1
10
20
30
40
50
t/s

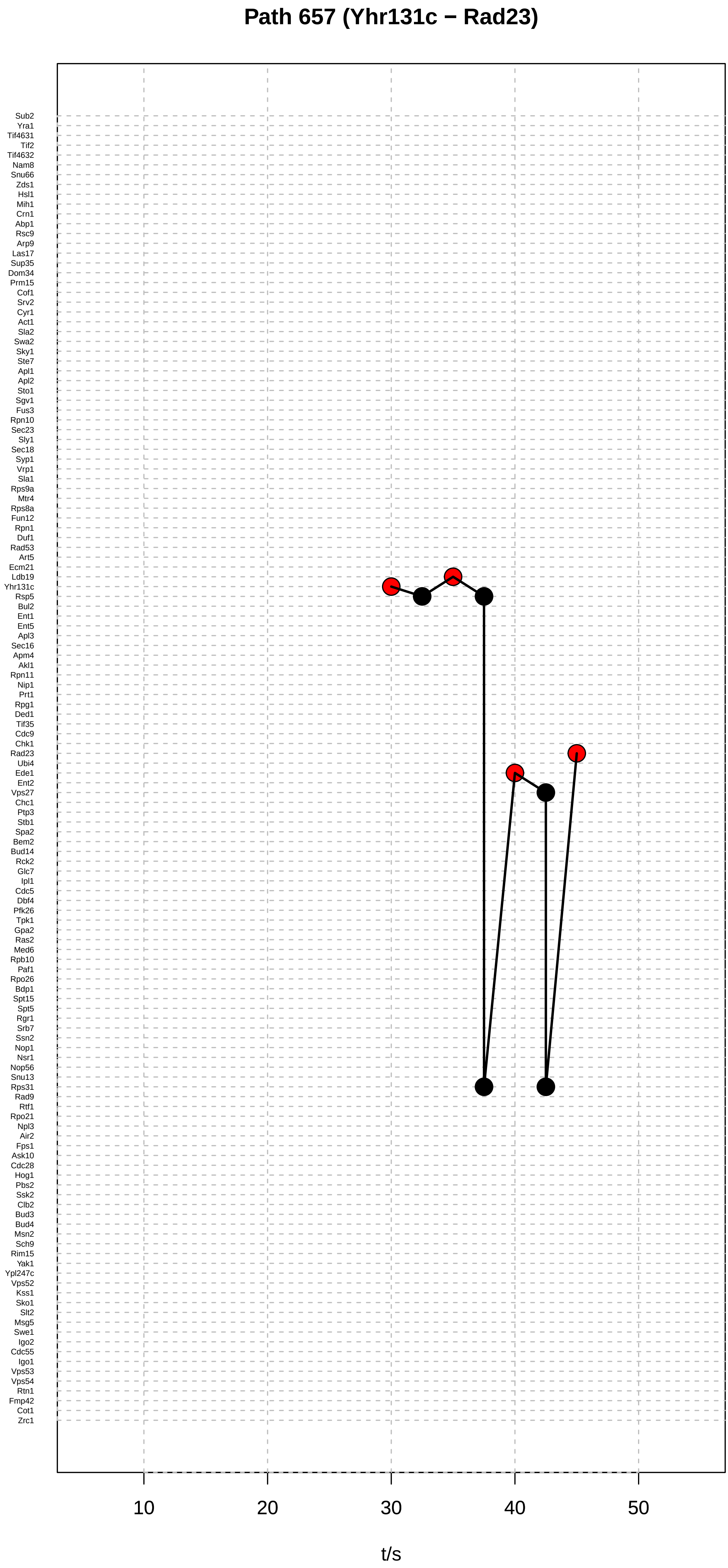

Path 657 (Yhr131c − Rad23)
Sub2
Yra1
Tif4631
Tif2
Tif4632
Nam8
Snu66
Zds1
Hsl1
Mih1
Crn1
Abp1
Rsc9
Arp9
Las17
Sup35
Dom34
Prm15
Cof1
Srv2
Cyr1
Act1
Sla2
Swa2
Sky1
Ste7
Apl1
Apl2
Sto1
Sgv1
Fus3
Rpn10
Sec23
Sly1
Sec18
Syp1
Vrp1
Sla1
Rps9a
Mtr4
Rps8a
Fun12
Rpn1
Duf1
Rad53
Art5
Ecm21
Ldb19
Yhr131c
Rsp5
Bul2
Ent1
Ent5
Apl3
Sec16
Apm4
Akl1
Rpn11
Nip1
Prt1
Rpg1
Ded1
Tif35
Cdc9
Chk1
Rad23
Ubi4
Ede1
Ent2
Vps27
Chc1
Ptp3
Stb1
Spa2
Bem2
Bud14
Rck2
Glc7
Ipl1
Cdc5
Dbf4
Pfk26
Tpk1
Gpa2
Ras2
Med6
Rpb10
Paf1
Rpo26
Bdp1
Spt15
Spt5
Rgr1
Srb7
Ssn2
Nop1
Nsr1
Nop56
Snu13
Rps31
Rad9
Rtf1
Rpo21
Npl3
Air2
Fps1
Ask10
Cdc28
Hog1
Pbs2
Ssk2
Clb2
Bud3
Bud4
Msn2
Sch9
Rim15
Yak1
Ypl247c
Vps52
Kss1
Sko1
Slt2
Msg5
Swe1
Igo2
Cdc55
Igo1
Vps53
Vps54
Rtn1
Fmp42
Cot1
Zrc1
10
20
30
40
50
t/s

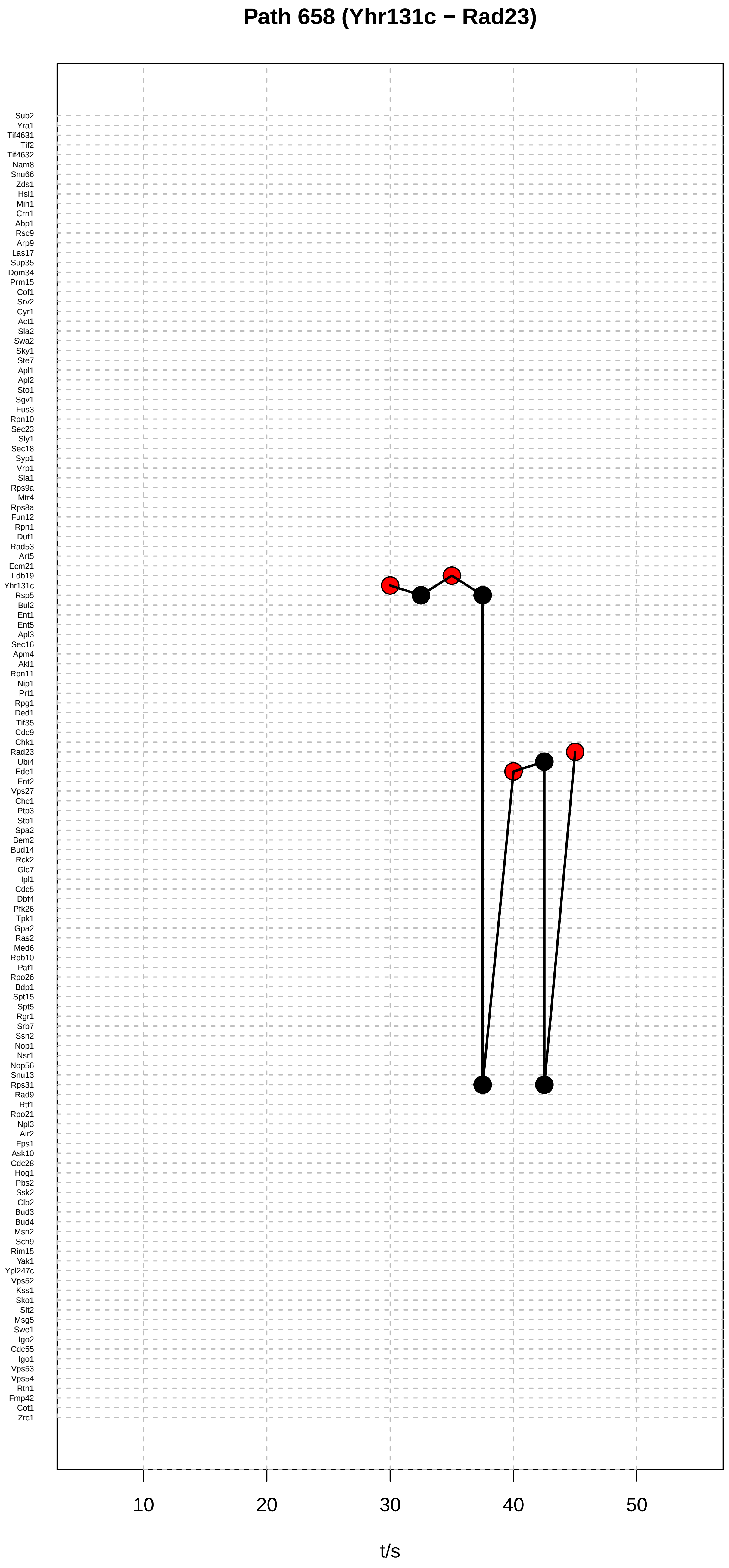
Path 658 (Yhr131c − Rad23)
Sub2
Yra1
Tif4631
Tif2
Tif4632
Nam8
Snu66
Zds1
Hsl1
Mih1
Crn1
Abp1
Rsc9
Arp9
Las17
Sup35
Dom34
Prm15
Cof1
Srv2
Cyr1
Act1
Sla2
Swa2
Sky1
Ste7
Apl1
Apl2
Sto1
Sgv1
Fus3
Rpn10
Sec23
Sly1
Sec18
Syp1
Vrp1
Sla1
Rps9a
Mtr4
Rps8a
Fun12
Rpn1
Duf1
Rad53
Art5
Ecm21
Ldb19
Yhr131c
Rsp5
Bul2
Ent1
Ent5
Apl3
Sec16
Apm4
Akl1
Rpn11
Nip1
Prt1
Rpg1
Ded1
Tif35
Cdc9
Chk1
Rad23
Ubi4
Ede1
Ent2
Vps27
Chc1
Ptp3
Stb1
Spa2
Bem2
Bud14
Rck2
Glc7
Ipl1
Cdc5
Dbf4
Pfk26
Tpk1
Gpa2
Ras2
Med6
Rpb10
Paf1
Rpo26
Bdp1
Spt15
Spt5
Rgr1
Srb7
Ssn2
Nop1
Nsr1
Nop56
Snu13
Rps31
Rad9
Rtf1
Rpo21
Npl3
Air2
Fps1
Ask10
Cdc28
Hog1
Pbs2
Ssk2
Clb2
Bud3
Bud4
Msn2
Sch9
Rim15
Yak1
Ypl247c
Vps52
Kss1
Sko1
Slt2
Msg5
Swe1
Igo2
Cdc55
Igo1
Vps53
Vps54
Rtn1
Fmp42
Cot1
Zrc1
10
20
30
40
50
t/s

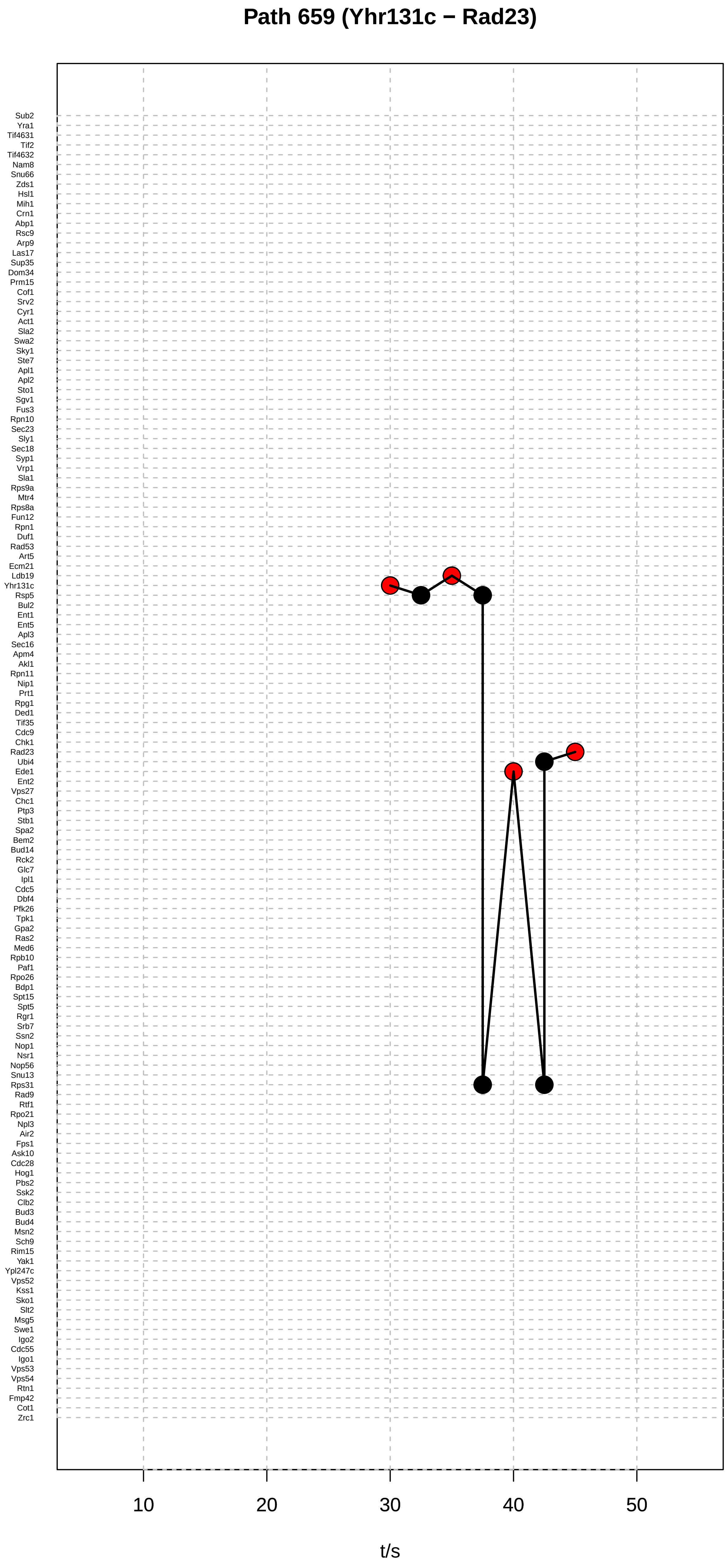

Path 659 (Yhr131c − Rad23)
Sub2
Yra1
Tif4631
Tif2
Tif4632
Nam8
Snu66
Zds1
Hsl1
Mih1
Crn1
Abp1
Rsc9
Arp9
Las17
Sup35
Dom34
Prm15
Cof1
Srv2
Cyr1
Act1
Sla2
Swa2
Sky1
Ste7
Apl1
Apl2
Sto1
Sgv1
Fus3
Rpn10
Sec23
Sly1
Sec18
Syp1
Vrp1
Sla1
Rps9a
Mtr4
Rps8a
Fun12
Rpn1
Duf1
Rad53
Art5
Ecm21
Ldb19
Yhr131c
Rsp5
Bul2
Ent1
Ent5
Apl3
Sec16
Apm4
Akl1
Rpn11
Nip1
Prt1
Rpg1
Ded1
Tif35
Cdc9
Chk1
Rad23
Ubi4
Ede1
Ent2
Vps27
Chc1
Ptp3
Stb1
Spa2
Bem2
Bud14
Rck2
Glc7
Ipl1
Cdc5
Dbf4
Pfk26
Tpk1
Gpa2
Ras2
Med6
Rpb10
Paf1
Rpo26
Bdp1
Spt15
Spt5
Rgr1
Srb7
Ssn2
Nop1
Nsr1
Nop56
Snu13
Rps31
Rad9
Rtf1
Rpo21
Npl3
Air2
Fps1
Ask10
Cdc28
Hog1
Pbs2
Ssk2
Clb2
Bud3
Bud4
Msn2
Sch9
Rim15
Yak1
Ypl247c
Vps52
Kss1
Sko1
Slt2
Msg5
Swe1
Igo2
Cdc55
Igo1
Vps53
Vps54
Rtn1
Fmp42
Cot1
Zrc1
10
20
30
40
50
t/s

**Path 660 (Yhr131c − Rad23)**

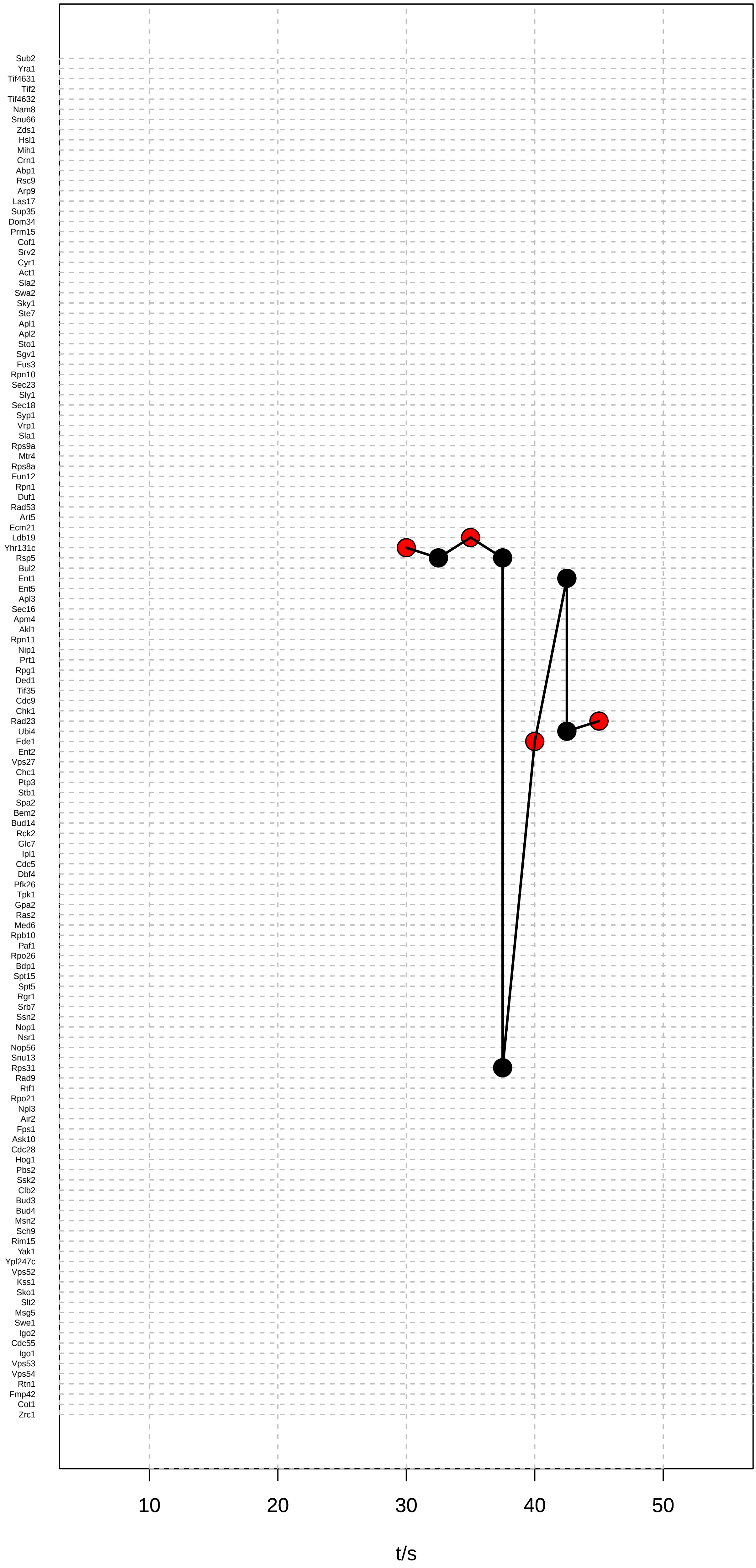

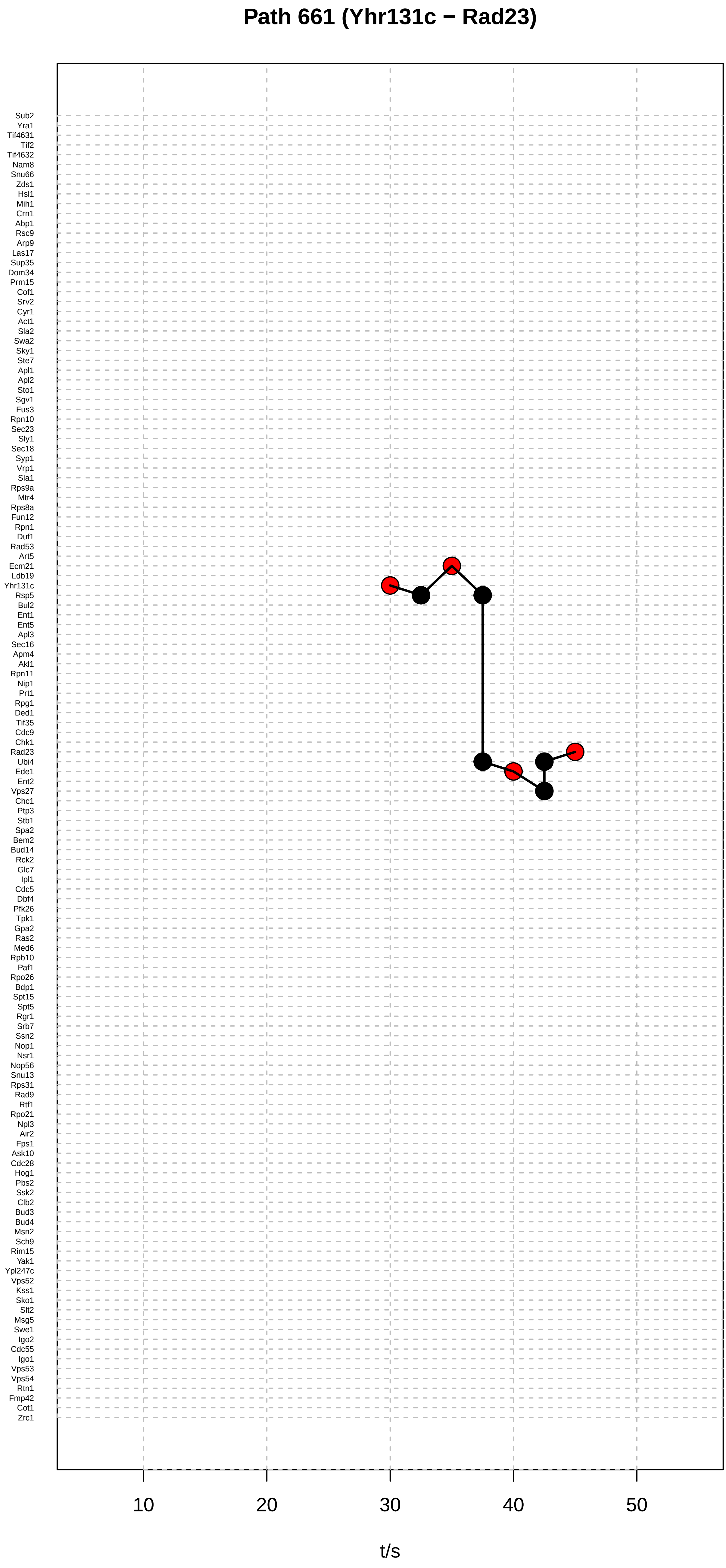

Path 661 (Yhr131c − Rad23)
Sub2
Yra1
Tif4631
Tif2
Tif4632
Nam8
Snu66
Zds1
Hsl1
Mih1
Crn1
Abp1
Rsc9
Arp9
Las17
Sup35
Dom34
Prm15
Cof1
Srv2
Cyr1
Act1
Sla2
Swa2
Sky1
Ste7
Apl1
Apl2
Sto1
Sgv1
Fus3
Rpn10
Sec23
Sly1
Sec18
Syp1
Vrp1
Sla1
Rps9a
Mtr4
Rps8a
Fun12
Rpn1
Duf1
Rad53
Art5
Ecm21
Ldb19
Yhr131c
Rsp5
Bul2
Ent1
Ent5
Apl3
Sec16
Apm4
Akl1
Rpn11
Nip1
Prt1
Rpg1
Ded1
Tif35
Cdc9
Chk1
Rad23
Ubi4
Ede1
Ent2
Vps27
Chc1
Ptp3
Stb1
Spa2
Bem2
Bud14
Rck2
Glc7
Ipl1
Cdc5
Dbf4
Pfk26
Tpk1
Gpa2
Ras2
Med6
Rpb10
Paf1
Rpo26
Bdp1
Spt15
Spt5
Rgr1
Srb7
Ssn2
Nop1
Nsr1
Nop56
Snu13
Rps31
Rad9
Rtf1
Rpo21
Npl3
Air2
Fps1
Ask10
Cdc28
Hog1
Pbs2
Ssk2
Clb2
Bud3
Bud4
Msn2
Sch9
Rim15
Yak1
Ypl247c
Vps52
Kss1
Sko1
Slt2
Msg5
Swe1
Igo2
Cdc55
Igo1
Vps53
Vps54
Rtn1
Fmp42
Cot1
Zrc1
10
20
30
40
50
t/s

### Path 662 (Yhr131c − Rad23)

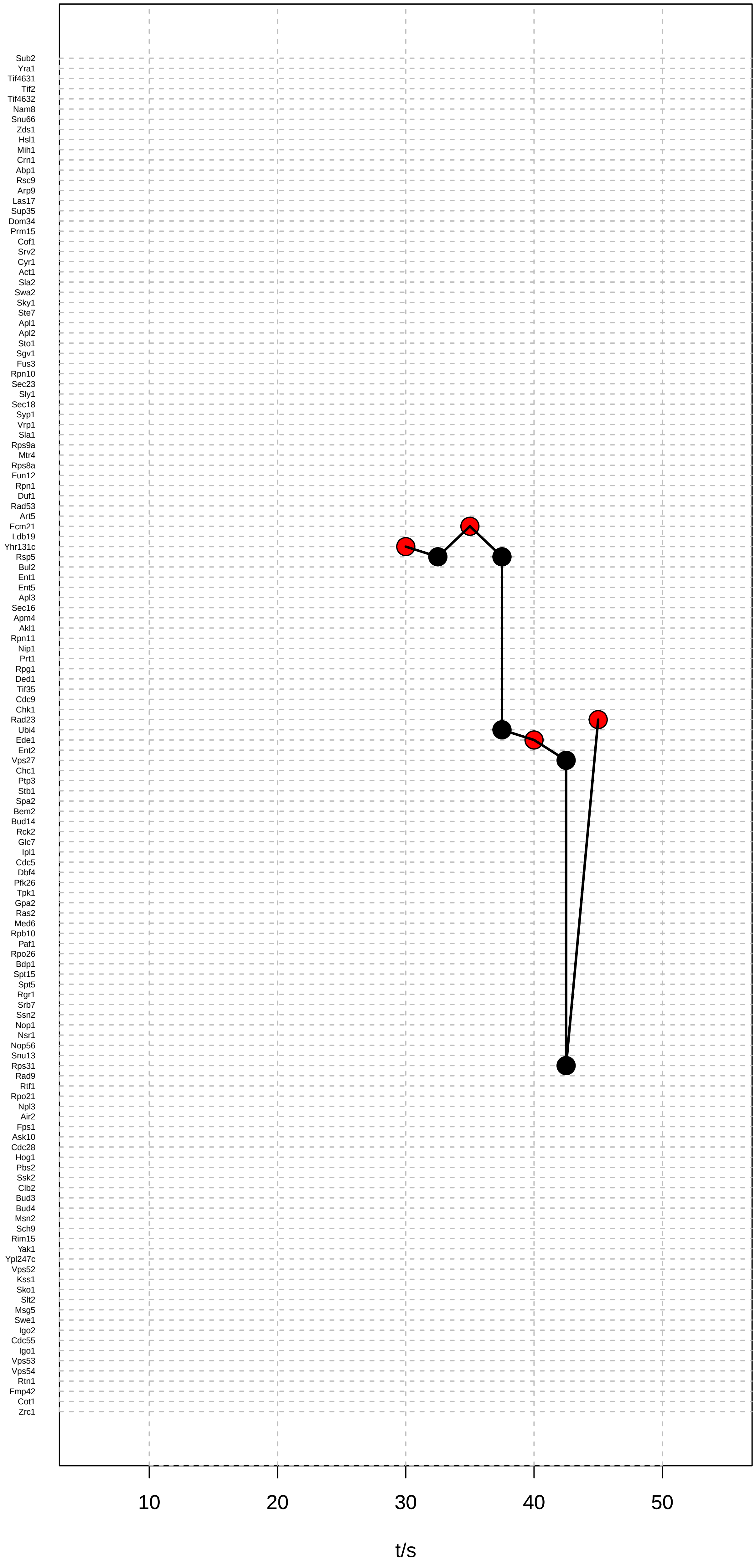

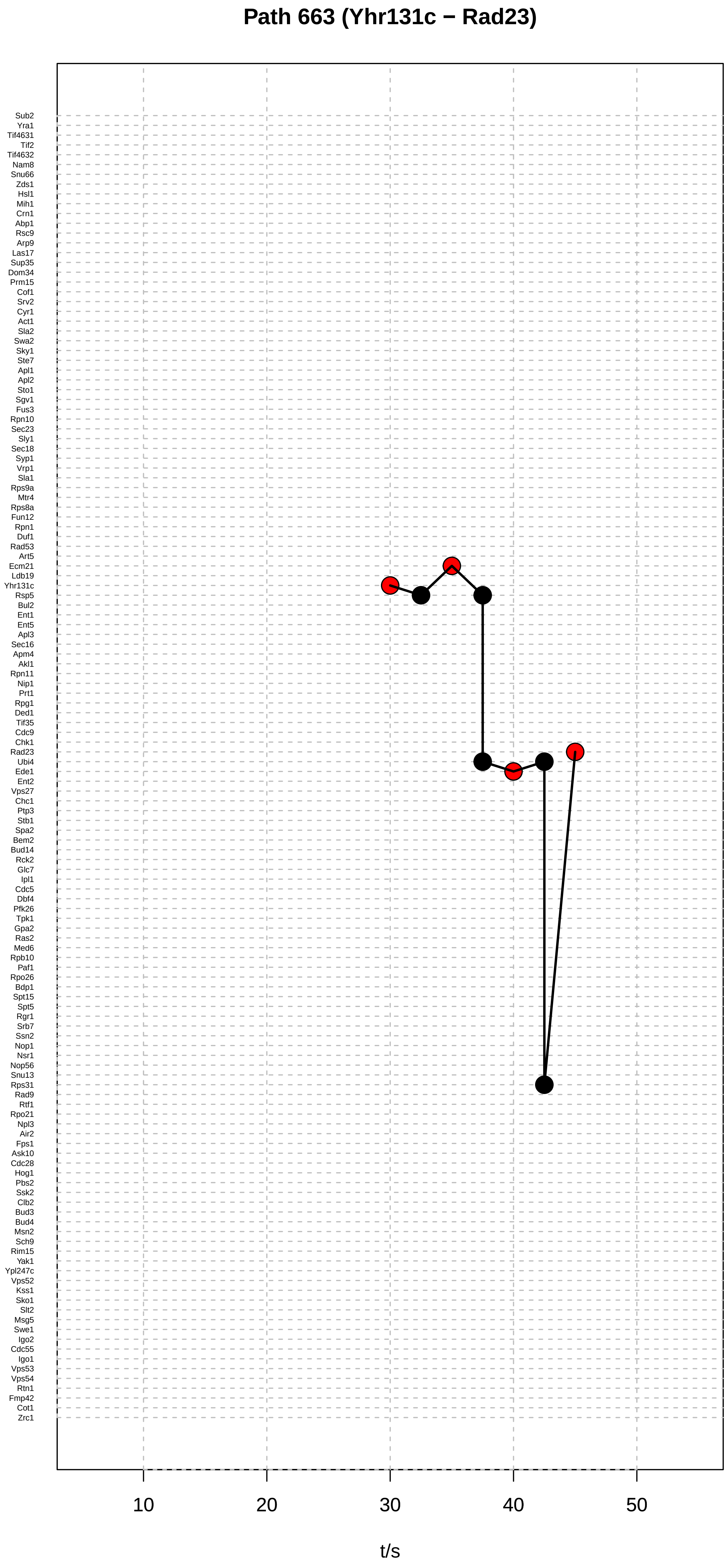

Path 663 (Yhr131c − Rad23)
Sub2
Yra1
Tif4631
Tif2
Tif4632
Nam8
Snu66
Zds1
Hsl1
Mih1
Crn1
Abp1
Rsc9
Arp9
Las17
Sup35
Dom34
Prm15
Cof1
Srv2
Cyr1
Act1
Sla2
Swa2
Sky1
Ste7
Apl1
Apl2
Sto1
Sgv1
Fus3
Rpn10
Sec23
Sly1
Sec18
Syp1
Vrp1
Sla1
Rps9a
Mtr4
Rps8a
Fun12
Rpn1
Duf1
Rad53
Art5
Ecm21
Ldb19
Yhr131c
Rsp5
Bul2
Ent1
Ent5
Apl3
Sec16
Apm4
Akl1
Rpn11
Nip1
Prt1
Rpg1
Ded1
Tif35
Cdc9
Chk1
Rad23
Ubi4
Ede1
Ent2
Vps27
Chc1
Ptp3
Stb1
Spa2
Bem2
Bud14
Rck2
Glc7
Ipl1
Cdc5
Dbf4
Pfk26
Tpk1
Gpa2
Ras2
Med6
Rpb10
Paf1
Rpo26
Bdp1
Spt15
Spt5
Rgr1
Srb7
Ssn2
Nop1
Nsr1
Nop56
Snu13
Rps31
Rad9
Rtf1
Rpo21
Npl3
Air2
Fps1
Ask10
Cdc28
Hog1
Pbs2
Ssk2
Clb2
Bud3
Bud4
Msn2
Sch9
Rim15
Yak1
Ypl247c
Vps52
Kss1
Sko1
Slt2
Msg5
Swe1
Igo2
Cdc55
Igo1
Vps53
Vps54
Rtn1
Fmp42
Cot1
Zrc1
10
20
30
40
50
t/s

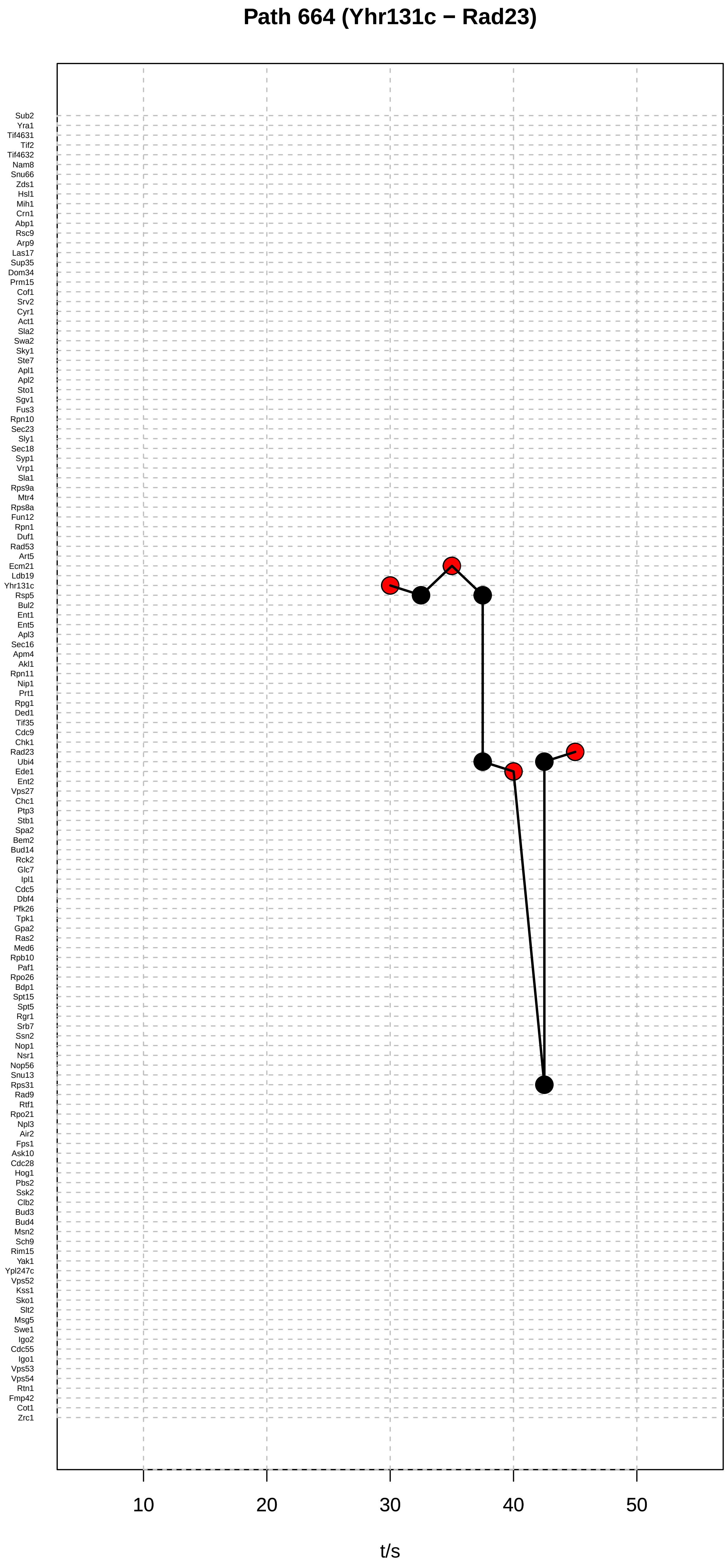

Path 664 (Yhr131c − Rad23)
Sub2
Yra1
Tif4631
Tif2
Tif4632
Nam8
Snu66
Zds1
Hsl1
Mih1
Crn1
Abp1
Rsc9
Arp9
Las17
Sup35
Dom34
Prm15
Cof1
Srv2
Cyr1
Act1
Sla2
Swa2
Sky1
Ste7
Apl1
Apl2
Sto1
Sgv1
Fus3
Rpn10
Sec23
Sly1
Sec18
Syp1
Vrp1
Sla1
Rps9a
Mtr4
Rps8a
Fun12
Rpn1
Duf1
Rad53
Art5
Ecm21
Ldb19
Yhr131c
Rsp5
Bul2
Ent1
Ent5
Apl3
Sec16
Apm4
Akl1
Rpn11
Nip1
Prt1
Rpg1
Ded1
Tif35
Cdc9
Chk1
Rad23
Ubi4
Ede1
Ent2
Vps27
Chc1
Ptp3
Stb1
Spa2
Bem2
Bud14
Rck2
Glc7
Ipl1
Cdc5
Dbf4
Pfk26
Tpk1
Gpa2
Ras2
Med6
Rpb10
Paf1
Rpo26
Bdp1
Spt15
Spt5
Rgr1
Srb7
Ssn2
Nop1
Nsr1
Nop56
Snu13
Rps31
Rad9
Rtf1
Rpo21
Npl3
Air2
Fps1
Ask10
Cdc28
Hog1
Pbs2
Ssk2
Clb2
Bud3
Bud4
Msn2
Sch9
Rim15
Yak1
Ypl247c
Vps52
Kss1
Sko1
Slt2
Msg5
Swe1
Igo2
Cdc55
Igo1
Vps53
Vps54
Rtn1
Fmp42
Cot1
Zrc1
10
20
30
40
50
t/s

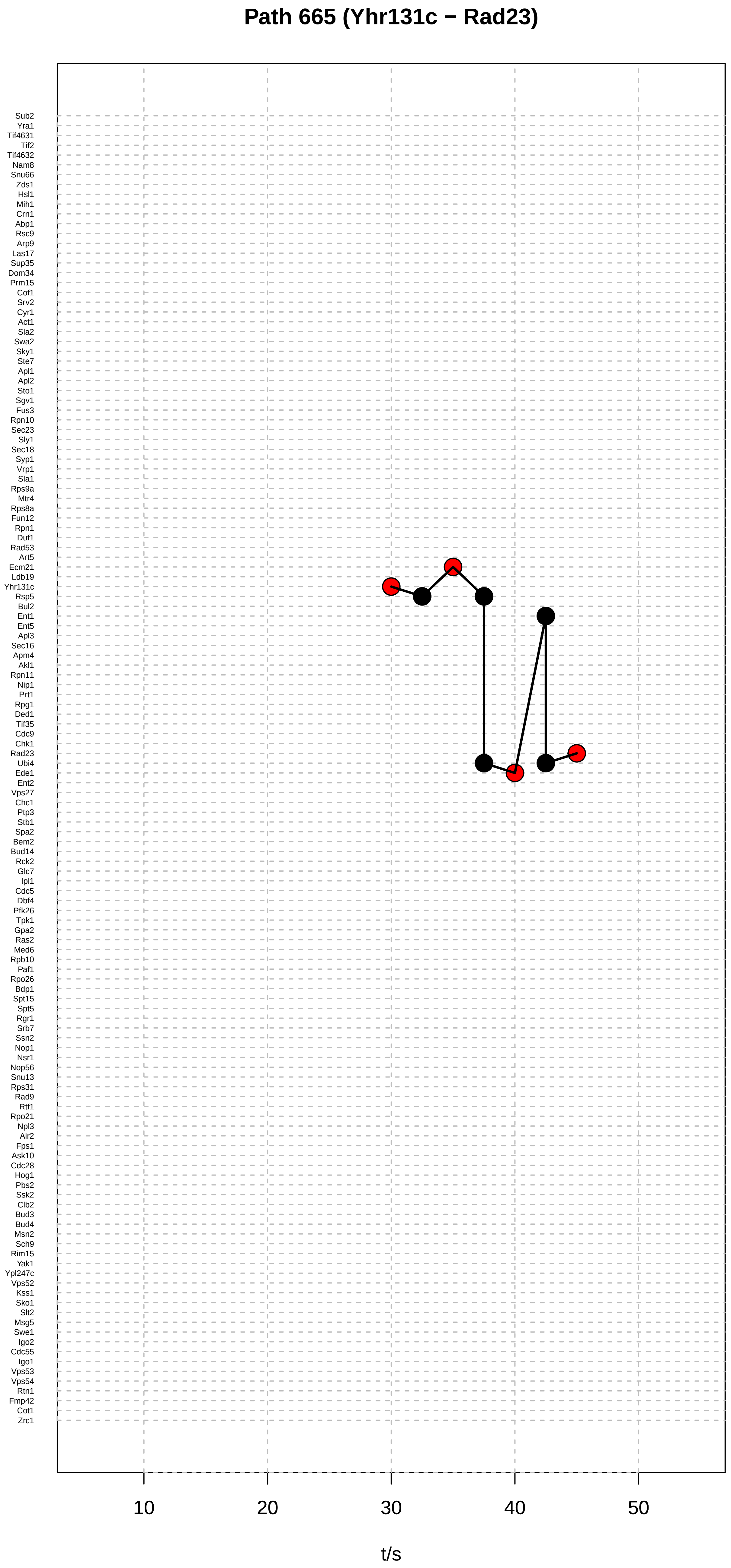
Path 665 (Yhr131c − Rad23)
Sub2
Yra1
Tif4631
Tif2
Tif4632
Nam8
Snu66
Zds1
Hsl1
Mih1
Crn1
Abp1
Rsc9
Arp9
Las17
Sup35
Dom34
Prm15
Cof1
Srv2
Cyr1
Act1
Sla2
Swa2
Sky1
Ste7
Apl1
Apl2
Sto1
Sgv1
Fus3
Rpn10
Sec23
Sly1
Sec18
Syp1
Vrp1
Sla1
Rps9a
Mtr4
Rps8a
Fun12
Rpn1
Duf1
Rad53
Art5
Ecm21
Ldb19
Yhr131c
Rsp5
Bul2
Ent1
Ent5
Apl3
Sec16
Apm4
Akl1
Rpn11
Nip1
Prt1
Rpg1
Ded1
Tif35
Cdc9
Chk1
Rad23
Ubi4
Ede1
Ent2
Vps27
Chc1
Ptp3
Stb1
Spa2
Bem2
Bud14
Rck2
Glc7
Ipl1
Cdc5
Dbf4
Pfk26
Tpk1
Gpa2
Ras2
Med6
Rpb10
Paf1
Rpo26
Bdp1
Spt15
Spt5
Rgr1
Srb7
Ssn2
Nop1
Nsr1
Nop56
Snu13
Rps31
Rad9
Rtf1
Rpo21
Npl3
Air2
Fps1
Ask10
Cdc28
Hog1
Pbs2
Ssk2
Clb2
Bud3
Bud4
Msn2
Sch9
Rim15
Yak1
Ypl247c
Vps52
Kss1
Sko1
Slt2
Msg5
Swe1
Igo2
Cdc55
Igo1
Vps53
Vps54
Rtn1
Fmp42
Cot1
Zrc1
10
20
30
40
50
t/s

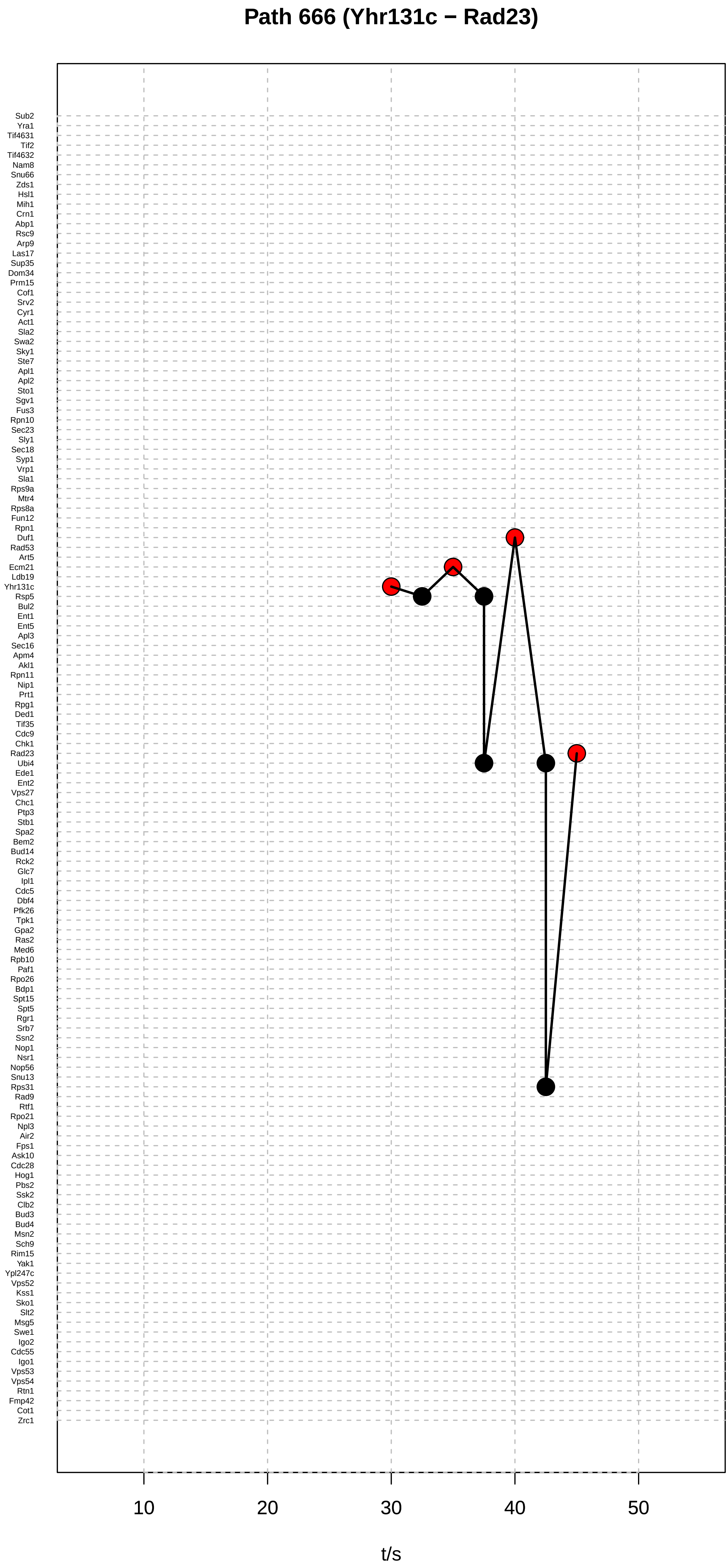

Path 666 (Yhr131c − Rad23)
Sub2
Yra1
Tif4631
Tif2
Tif4632
Nam8
Snu66
Zds1
Hsl1
Mih1
Crn1
Abp1
Rsc9
Arp9
Las17
Sup35
Dom34
Prm15
Cof1
Srv2
Cyr1
Act1
Sla2
Swa2
Sky1
Ste7
Apl1
Apl2
Sto1
Sgv1
Fus3
Rpn10
Sec23
Sly1
Sec18
Syp1
Vrp1
Sla1
Rps9a
Mtr4
Rps8a
Fun12
Rpn1
Duf1
Rad53
Art5
Ecm21
Ldb19
Yhr131c
Rsp5
Bul2
Ent1
Ent5
Apl3
Sec16
Apm4
Akl1
Rpn11
Nip1
Prt1
Rpg1
Ded1
Tif35
Cdc9
Chk1
Rad23
Ubi4
Ede1
Ent2
Vps27
Chc1
Ptp3
Stb1
Spa2
Bem2
Bud14
Rck2
Glc7
Ipl1
Cdc5
Dbf4
Pfk26
Tpk1
Gpa2
Ras2
Med6
Rpb10
Paf1
Rpo26
Bdp1
Spt15
Spt5
Rgr1
Srb7
Ssn2
Nop1
Nsr1
Nop56
Snu13
Rps31
Rad9
Rtf1
Rpo21
Npl3
Air2
Fps1
Ask10
Cdc28
Hog1
Pbs2
Ssk2
Clb2
Bud3
Bud4
Msn2
Sch9
Rim15
Yak1
Ypl247c
Vps52
Kss1
Sko1
Slt2
Msg5
Swe1
Igo2
Cdc55
Igo1
Vps53
Vps54
Rtn1
Fmp42
Cot1
Zrc1
10
20
30
40
50
t/s

#### Path 667 (Yhr131c − Rad23)

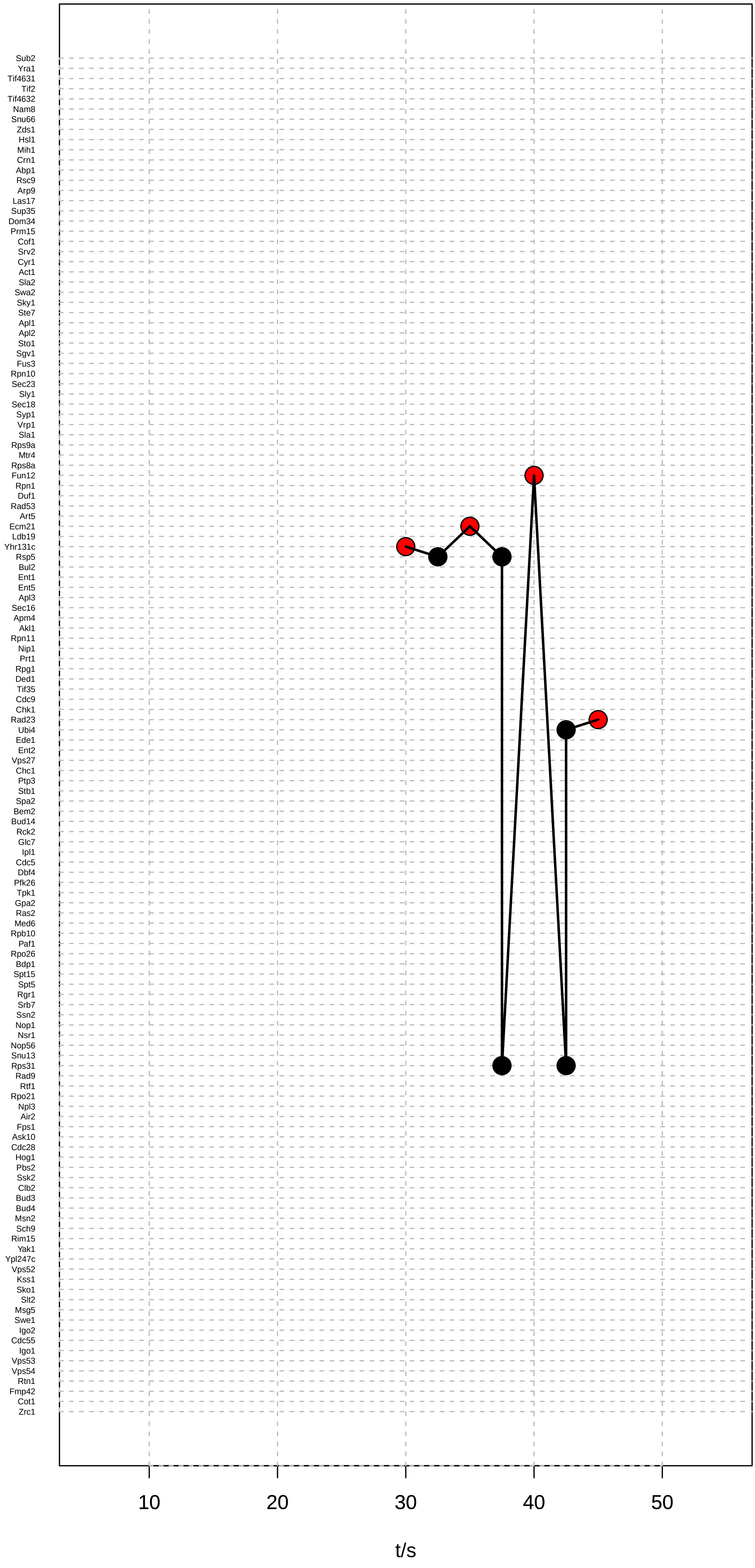

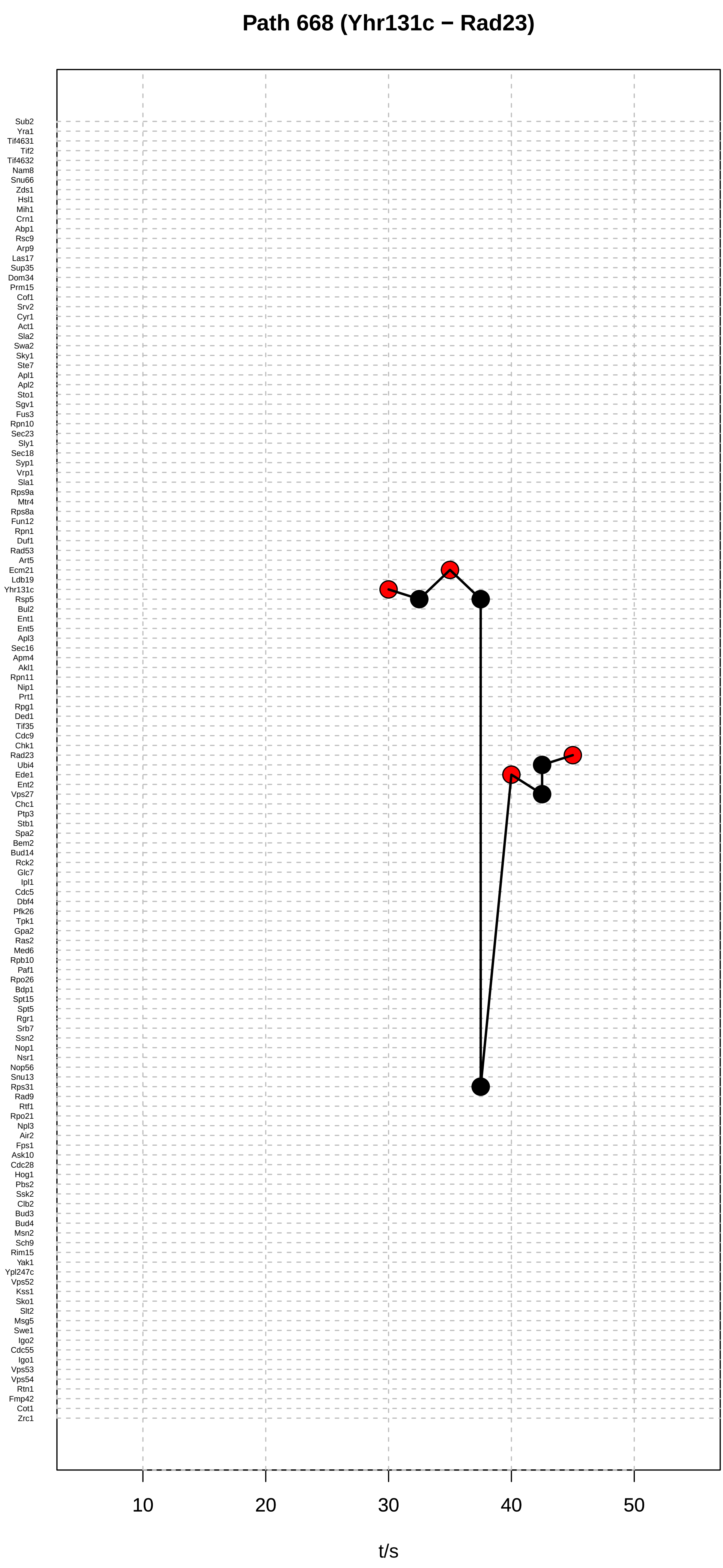
Path 668 (Yhr131c − Rad23)
Sub2
Yra1
Tif4631
Tif2
Tif4632
Nam8
Snu66
Zds1
Hsl1
Mih1
Crn1
Abp1
Rsc9
Arp9
Las17
Sup35
Dom34
Prm15
Cof1
Srv2
Cyr1
Act1
Sla2
Swa2
Sky1
Ste7
Apl1
Apl2
Sto1
Sgv1
Fus3
Rpn10
Sec23
Sly1
Sec18
Syp1
Vrp1
Sla1
Rps9a
Mtr4
Rps8a
Fun12
Rpn1
Duf1
Rad53
Art5
Ecm21
Ldb19
Yhr131c
Rsp5
Bul2
Ent1
Ent5
Apl3
Sec16
Apm4
Akl1
Rpn11
Nip1
Prt1
Rpg1
Ded1
Tif35
Cdc9
Chk1
Rad23
Ubi4
Ede1
Ent2
Vps27
Chc1
Ptp3
Stb1
Spa2
Bem2
Bud14
Rck2
Glc7
Ipl1
Cdc5
Dbf4
Pfk26
Tpk1
Gpa2
Ras2
Med6
Rpb10
Paf1
Rpo26
Bdp1
Spt15
Spt5
Rgr1
Srb7
Ssn2
Nop1
Nsr1
Nop56
Snu13
Rps31
Rad9
Rtf1
Rpo21
Npl3
Air2
Fps1
Ask10
Cdc28
Hog1
Pbs2
Ssk2
Clb2
Bud3
Bud4
Msn2
Sch9
Rim15
Yak1
Ypl247c
Vps52
Kss1
Sko1
Slt2
Msg5
Swe1
Igo2
Cdc55
Igo1
Vps53
Vps54
Rtn1
Fmp42
Cot1
Zrc1
10
20
30
40
50
t/s

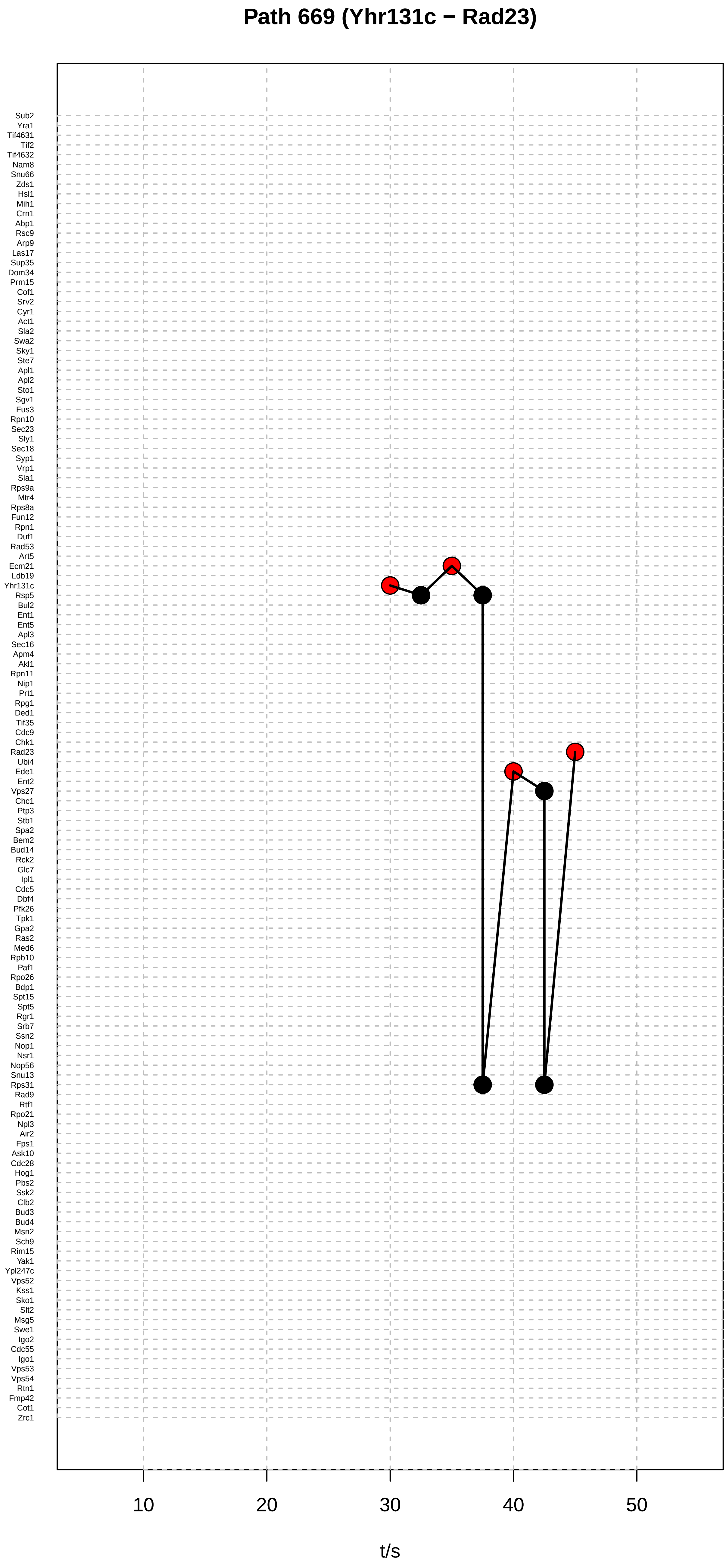

Path 669 (Yhr131c − Rad23)
Sub2
Yra1
Tif4631
Tif2
Tif4632
Nam8
Snu66
Zds1
Hsl1
Mih1
Crn1
Abp1
Rsc9
Arp9
Las17
Sup35
Dom34
Prm15
Cof1
Srv2
Cyr1
Act1
Sla2
Swa2
Sky1
Ste7
Apl1
Apl2
Sto1
Sgv1
Fus3
Rpn10
Sec23
Sly1
Sec18
Syp1
Vrp1
Sla1
Rps9a
Mtr4
Rps8a
Fun12
Rpn1
Duf1
Rad53
Art5
Ecm21
Ldb19
Yhr131c
Rsp5
Bul2
Ent1
Ent5
Apl3
Sec16
Apm4
Akl1
Rpn11
Nip1
Prt1
Rpg1
Ded1
Tif35
Cdc9
Chk1
Rad23
Ubi4
Ede1
Ent2
Vps27
Chc1
Ptp3
Stb1
Spa2
Bem2
Bud14
Rck2
Glc7
Ipl1
Cdc5
Dbf4
Pfk26
Tpk1
Gpa2
Ras2
Med6
Rpb10
Paf1
Rpo26
Bdp1
Spt15
Spt5
Rgr1
Srb7
Ssn2
Nop1
Nsr1
Nop56
Snu13
Rps31
Rad9
Rtf1
Rpo21
Npl3
Air2
Fps1
Ask10
Cdc28
Hog1
Pbs2
Ssk2
Clb2
Bud3
Bud4
Msn2
Sch9
Rim15
Yak1
Ypl247c
Vps52
Kss1
Sko1
Slt2
Msg5
Swe1
Igo2
Cdc55
Igo1
Vps53
Vps54
Rtn1
Fmp42
Cot1
Zrc1
10
20
30
40
50
t/s

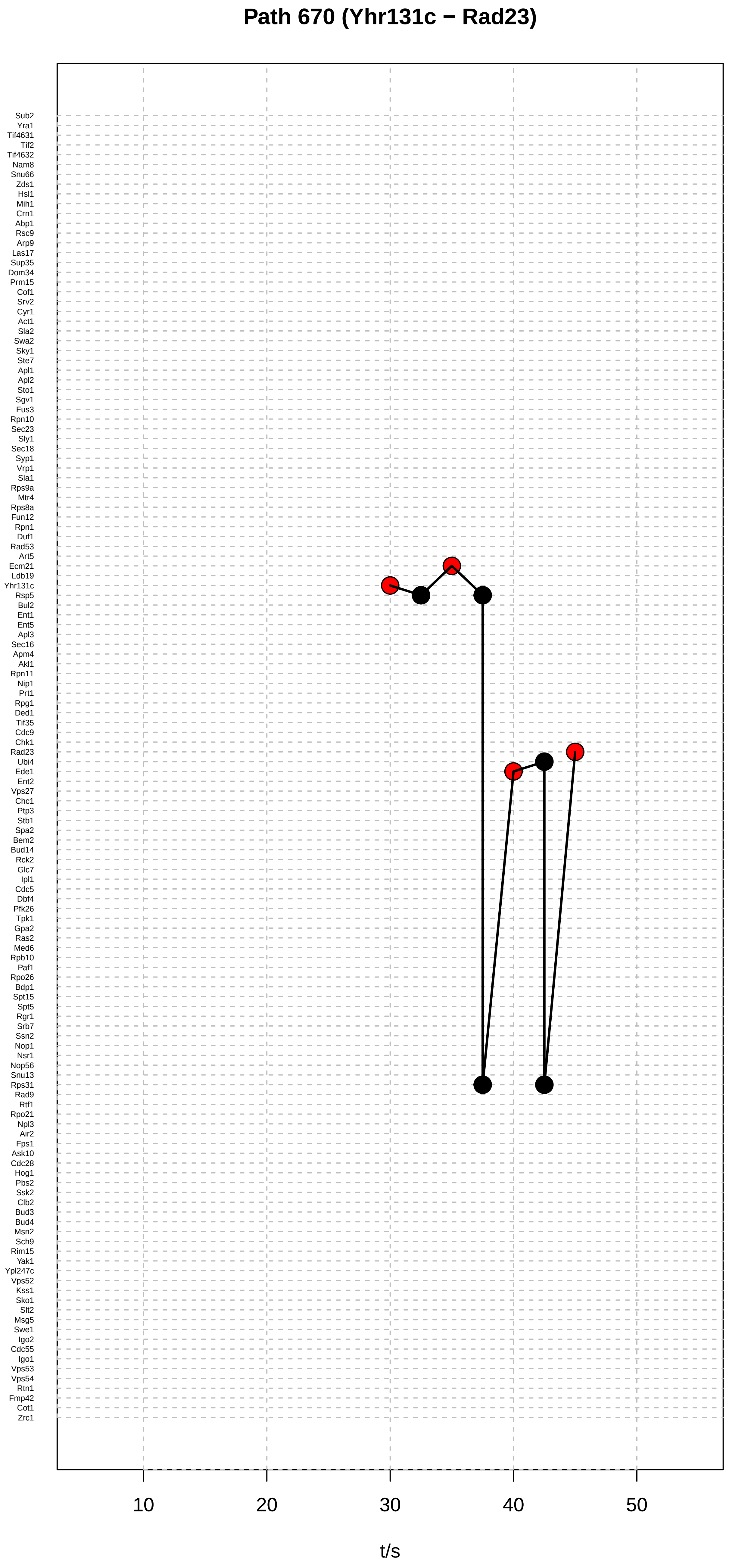

Path 670 (Yhr131c − Rad23)
Sub2
Yra1
Tif4631
Tif2
Tif4632
Nam8
Snu66
Zds1
Hsl1
Mih1
Crn1
Abp1
Rsc9
Arp9
Las17
Sup35
Dom34
Prm15
Cof1
Srv2
Cyr1
Act1
Sla2
Swa2
Sky1
Ste7
Apl1
Apl2
Sto1
Sgv1
Fus3
Rpn10
Sec23
Sly1
Sec18
Syp1
Vrp1
Sla1
Rps9a
Mtr4
Rps8a
Fun12
Rpn1
Duf1
Rad53
Art5
Ecm21
Ldb19
Yhr131c
Rsp5
Bul2
Ent1
Ent5
Apl3
Sec16
Apm4
Akl1
Rpn11
Nip1
Prt1
Rpg1
Ded1
Tif35
Cdc9
Chk1
Rad23
Ubi4
Ede1
Ent2
Vps27
Chc1
Ptp3
Stb1
Spa2
Bem2
Bud14
Rck2
Glc7
Ipl1
Cdc5
Dbf4
Pfk26
Tpk1
Gpa2
Ras2
Med6
Rpb10
Paf1
Rpo26
Bdp1
Spt15
Spt5
Rgr1
Srb7
Ssn2
Nop1
Nsr1
Nop56
Snu13
Rps31
Rad9
Rtf1
Rpo21
Npl3
Air2
Fps1
Ask10
Cdc28
Hog1
Pbs2
Ssk2
Clb2
Bud3
Bud4
Msn2
Sch9
Rim15
Yak1
Ypl247c
Vps52
Kss1
Sko1
Slt2
Msg5
Swe1
Igo2
Cdc55
Igo1
Vps53
Vps54
Rtn1
Fmp42
Cot1
Zrc1
10
20
30
40
50
t/s

Path 671 (Yhr131c − Rad23)

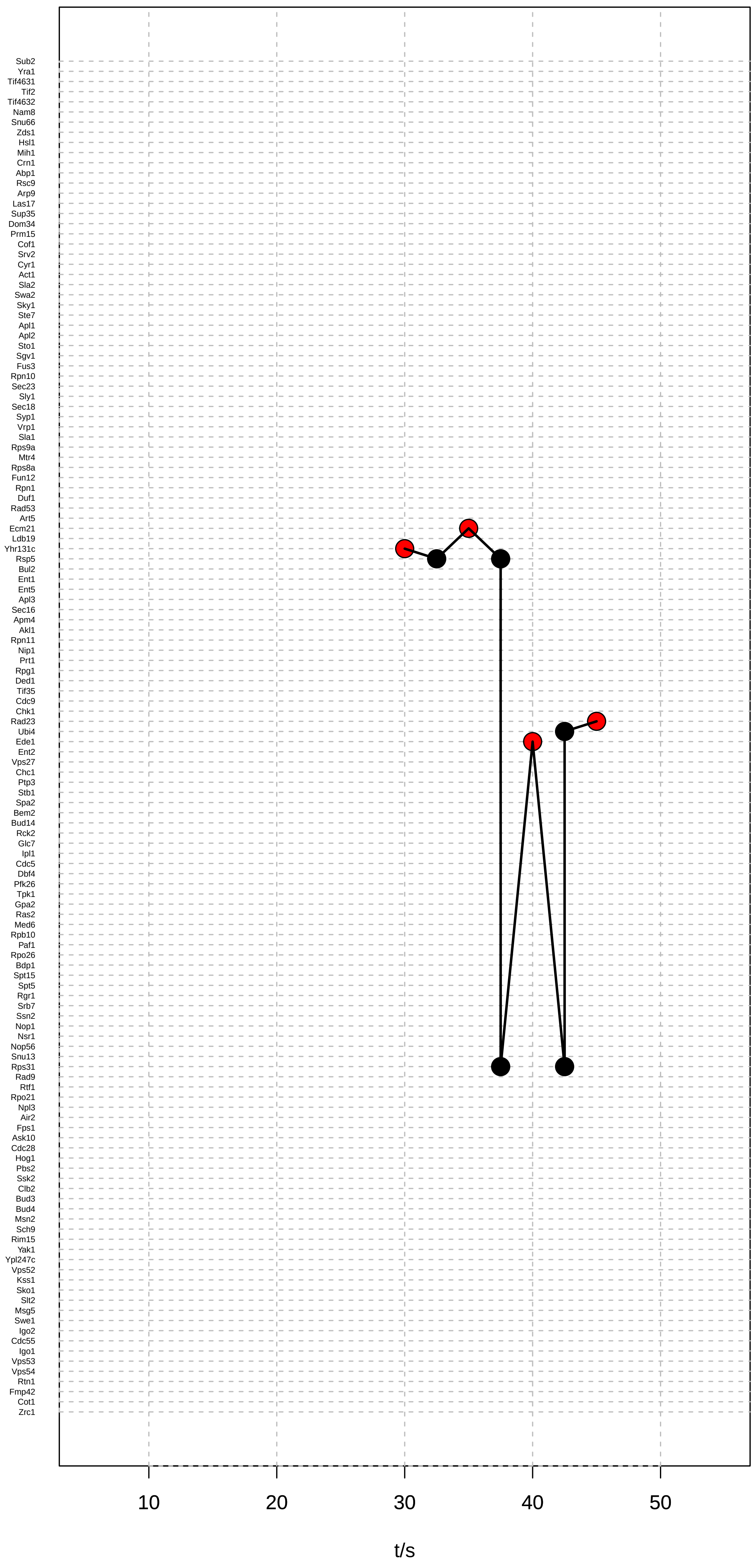

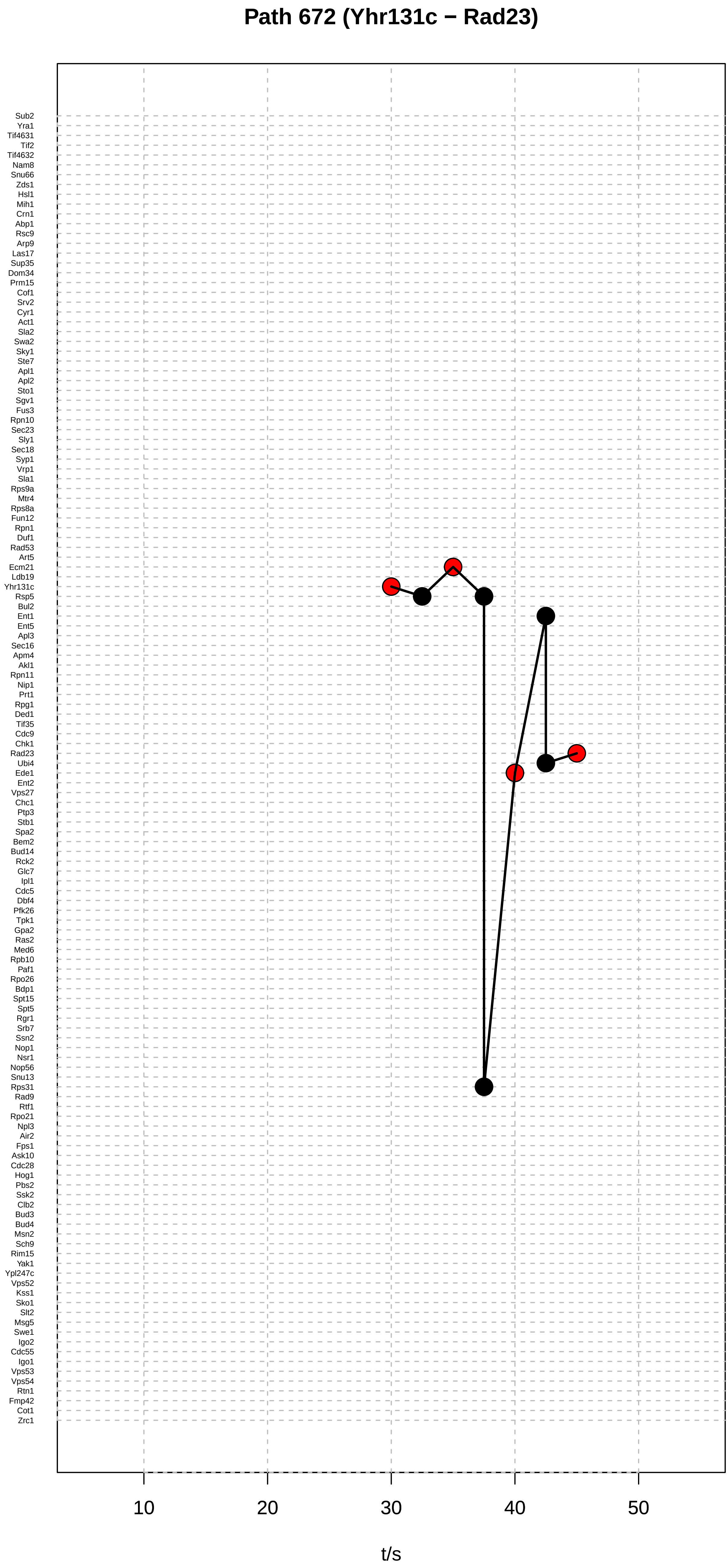

Path 672 (Yhr131c − Rad23)
Sub2
Yra1
Tif4631
Tif2
Tif4632
Nam8
Snu66
Zds1
Hsl1
Mih1
Crn1
Abp1
Rsc9
Arp9
Las17
Sup35
Dom34
Prm15
Cof1
Srv2
Cyr1
Act1
Sla2
Swa2
Sky1
Ste7
Apl1
Apl2
Sto1
Sgv1
Fus3
Rpn10
Sec23
Sly1
Sec18
Syp1
Vrp1
Sla1
Rps9a
Mtr4
Rps8a
Fun12
Rpn1
Duf1
Rad53
Art5
Ecm21
Ldb19
Yhr131c
Rsp5
Bul2
Ent1
Ent5
Apl3
Sec16
Apm4
Akl1
Rpn11
Nip1
Prt1
Rpg1
Ded1
Tif35
Cdc9
Chk1
Rad23
Ubi4
Ede1
Ent2
Vps27
Chc1
Ptp3
Stb1
Spa2
Bem2
Bud14
Rck2
Glc7
Ipl1
Cdc5
Dbf4
Pfk26
Tpk1
Gpa2
Ras2
Med6
Rpb10
Paf1
Rpo26
Bdp1
Spt15
Spt5
Rgr1
Srb7
Ssn2
Nop1
Nsr1
Nop56
Snu13
Rps31
Rad9
Rtf1
Rpo21
Npl3
Air2
Fps1
Ask10
Cdc28
Hog1
Pbs2
Ssk2
Clb2
Bud3
Bud4
Msn2
Sch9
Rim15
Yak1
Ypl247c
Vps52
Kss1
Sko1
Slt2
Msg5
Swe1
Igo2
Cdc55
Igo1
Vps53
Vps54
Rtn1
Fmp42
Cot1
Zrc1
10
20
30
40
50
t/s

### Path 673 (Yhr131c − Rad23)

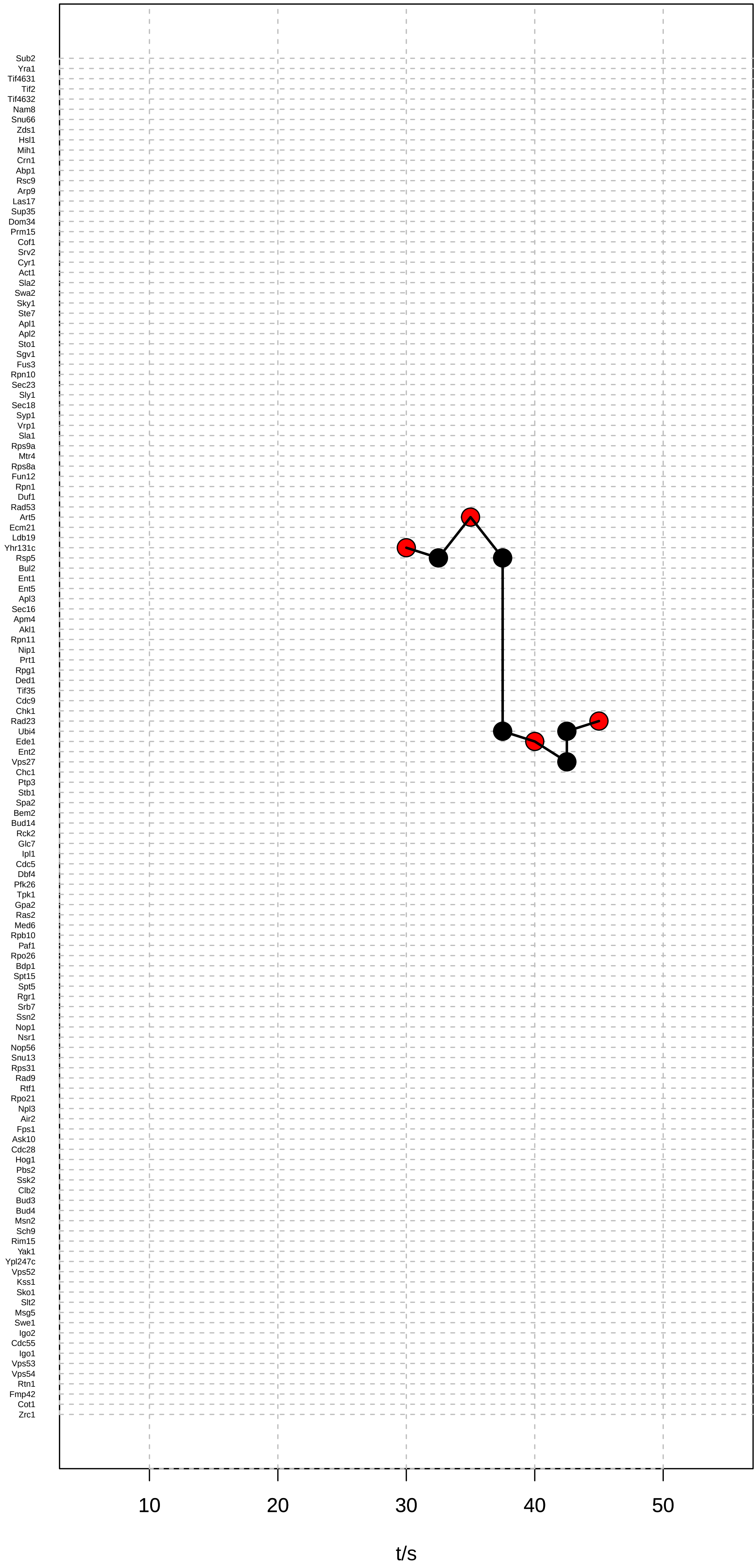

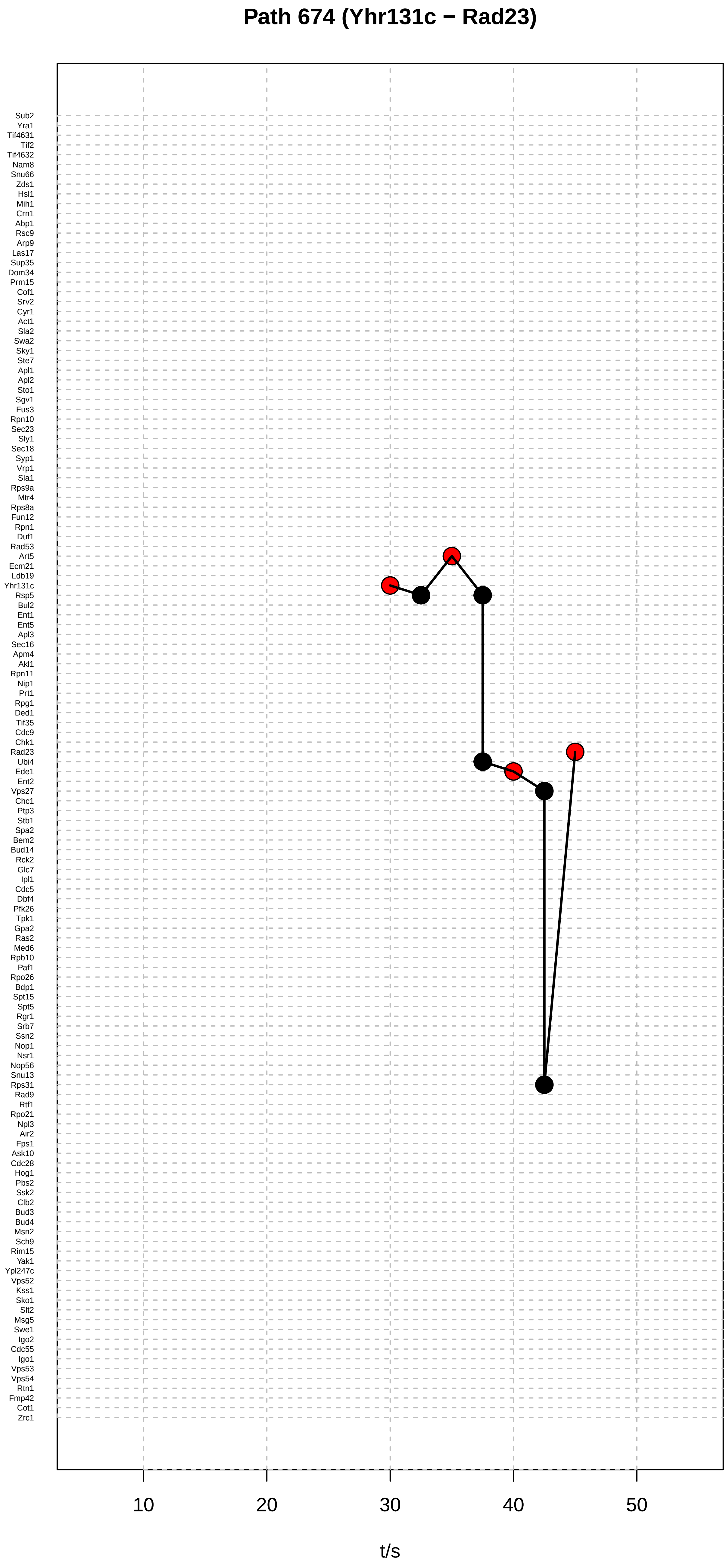

Path 674 (Yhr131c − Rad23)
Sub2
Yra1
Tif4631
Tif2
Tif4632
Nam8
Snu66
Zds1
Hsl1
Mih1
Crn1
Abp1
Rsc9
Arp9
Las17
Sup35
Dom34
Prm15
Cof1
Srv2
Cyr1
Act1
Sla2
Swa2
Sky1
Ste7
Apl1
Apl2
Sto1
Sgv1
Fus3
Rpn10
Sec23
Sly1
Sec18
Syp1
Vrp1
Sla1
Rps9a
Mtr4
Rps8a
Fun12
Rpn1
Duf1
Rad53
Art5
Ecm21
Ldb19
Yhr131c
Rsp5
Bul2
Ent1
Ent5
Apl3
Sec16
Apm4
Akl1
Rpn11
Nip1
Prt1
Rpg1
Ded1
Tif35
Cdc9
Chk1
Rad23
Ubi4
Ede1
Ent2
Vps27
Chc1
Ptp3
Stb1
Spa2
Bem2
Bud14
Rck2
Glc7
Ipl1
Cdc5
Dbf4
Pfk26
Tpk1
Gpa2
Ras2
Med6
Rpb10
Paf1
Rpo26
Bdp1
Spt15
Spt5
Rgr1
Srb7
Ssn2
Nop1
Nsr1
Nop56
Snu13
Rps31
Rad9
Rtf1
Rpo21
Npl3
Air2
Fps1
Ask10
Cdc28
Hog1
Pbs2
Ssk2
Clb2
Bud3
Bud4
Msn2
Sch9
Rim15
Yak1
Ypl247c
Vps52
Kss1
Sko1
Slt2
Msg5
Swe1
Igo2
Cdc55
Igo1
Vps53
Vps54
Rtn1
Fmp42
Cot1
Zrc1
10
20
30
40
50
t/s

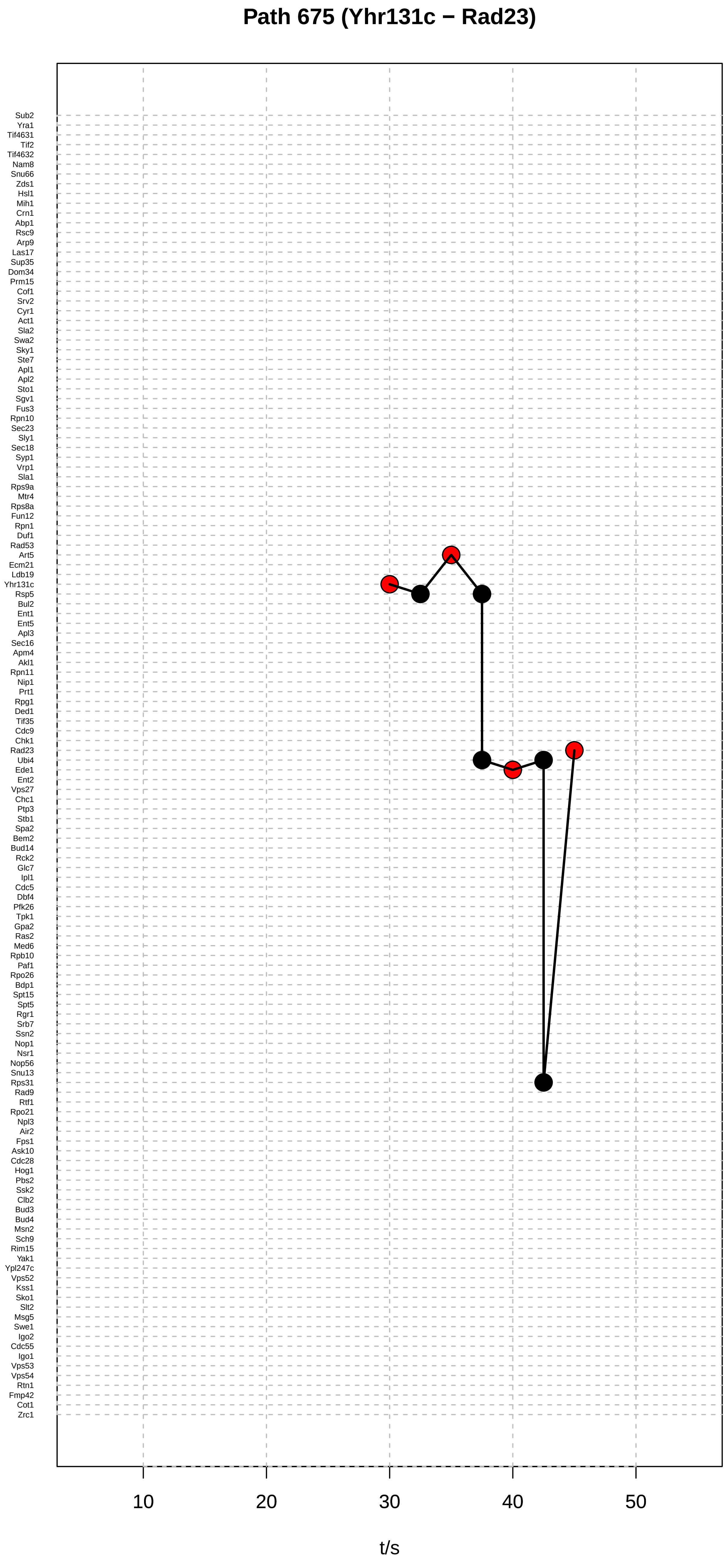

Path 675 (Yhr131c − Rad23)
Sub2
Yra1
Tif4631
Tif2
Tif4632
Nam8
Snu66
Zds1
Hsl1
Mih1
Crn1
Abp1
Rsc9
Arp9
Las17
Sup35
Dom34
Prm15
Cof1
Srv2
Cyr1
Act1
Sla2
Swa2
Sky1
Ste7
Apl1
Apl2
Sto1
Sgv1
Fus3
Rpn10
Sec23
Sly1
Sec18
Syp1
Vrp1
Sla1
Rps9a
Mtr4
Rps8a
Fun12
Rpn1
Duf1
Rad53
Art5
Ecm21
Ldb19
Yhr131c
Rsp5
Bul2
Ent1
Ent5
Apl3
Sec16
Apm4
Akl1
Rpn11
Nip1
Prt1
Rpg1
Ded1
Tif35
Cdc9
Chk1
Rad23
Ubi4
Ede1
Ent2
Vps27
Chc1
Ptp3
Stb1
Spa2
Bem2
Bud14
Rck2
Glc7
Ipl1
Cdc5
Dbf4
Pfk26
Tpk1
Gpa2
Ras2
Med6
Rpb10
Paf1
Rpo26
Bdp1
Spt15
Spt5
Rgr1
Srb7
Ssn2
Nop1
Nsr1
Nop56
Snu13
Rps31
Rad9
Rtf1
Rpo21
Npl3
Air2
Fps1
Ask10
Cdc28
Hog1
Pbs2
Ssk2
Clb2
Bud3
Bud4
Msn2
Sch9
Rim15
Yak1
Ypl247c
Vps52
Kss1
Sko1
Slt2
Msg5
Swe1
Igo2
Cdc55
Igo1
Vps53
Vps54
Rtn1
Fmp42
Cot1
Zrc1
10
20
30
40
50
t/s

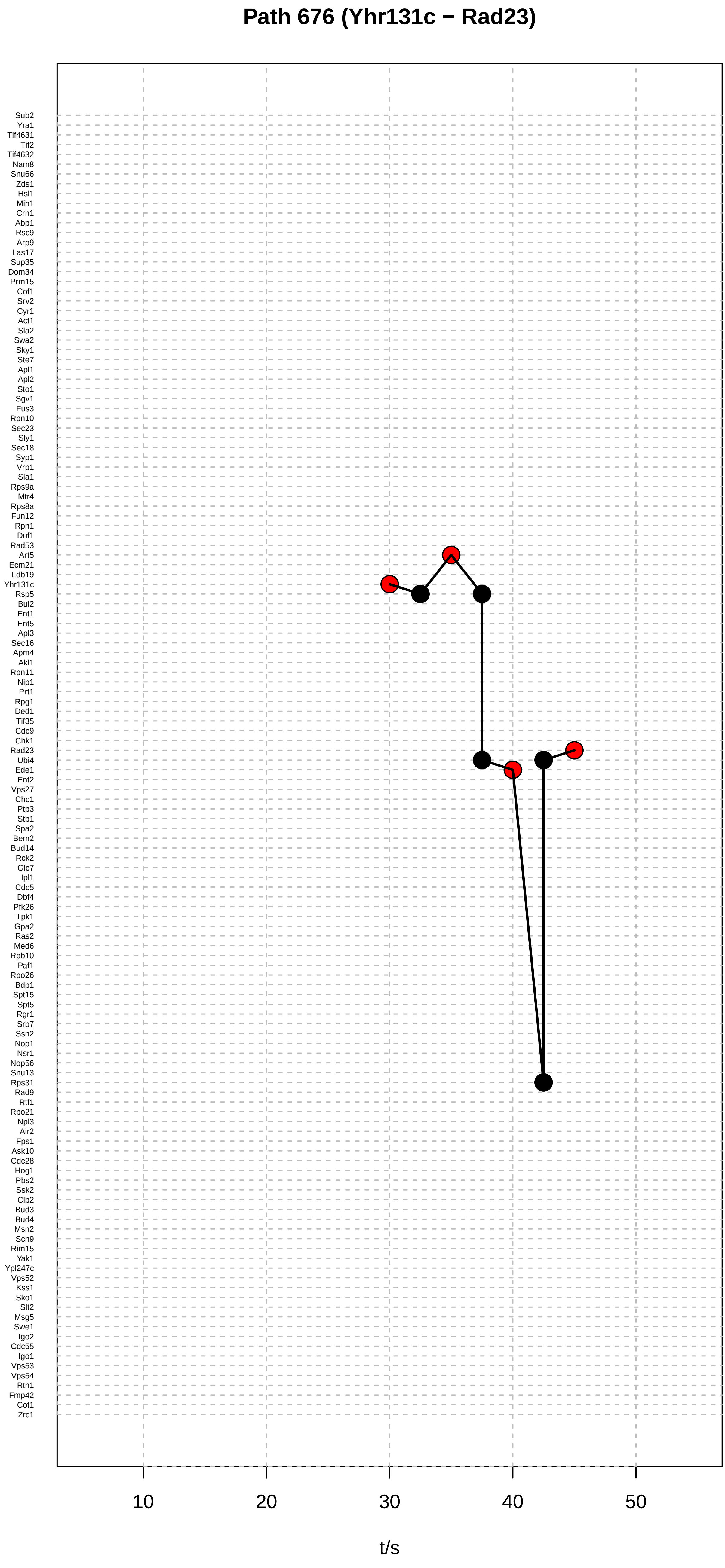

Path 676 (Yhr131c − Rad23)
Sub2
Yra1
Tif4631
Tif2
Tif4632
Nam8
Snu66
Zds1
Hsl1
Mih1
Crn1
Abp1
Rsc9
Arp9
Las17
Sup35
Dom34
Prm15
Cof1
Srv2
Cyr1
Act1
Sla2
Swa2
Sky1
Ste7
Apl1
Apl2
Sto1
Sgv1
Fus3
Rpn10
Sec23
Sly1
Sec18
Syp1
Vrp1
Sla1
Rps9a
Mtr4
Rps8a
Fun12
Rpn1
Duf1
Rad53
Art5
Ecm21
Ldb19
Yhr131c
Rsp5
Bul2
Ent1
Ent5
Apl3
Sec16
Apm4
Akl1
Rpn11
Nip1
Prt1
Rpg1
Ded1
Tif35
Cdc9
Chk1
Rad23
Ubi4
Ede1
Ent2
Vps27
Chc1
Ptp3
Stb1
Spa2
Bem2
Bud14
Rck2
Glc7
Ipl1
Cdc5
Dbf4
Pfk26
Tpk1
Gpa2
Ras2
Med6
Rpb10
Paf1
Rpo26
Bdp1
Spt15
Spt5
Rgr1
Srb7
Ssn2
Nop1
Nsr1
Nop56
Snu13
Rps31
Rad9
Rtf1
Rpo21
Npl3
Air2
Fps1
Ask10
Cdc28
Hog1
Pbs2
Ssk2
Clb2
Bud3
Bud4
Msn2
Sch9
Rim15
Yak1
Ypl247c
Vps52
Kss1
Sko1
Slt2
Msg5
Swe1
Igo2
Cdc55
Igo1
Vps53
Vps54
Rtn1
Fmp42
Cot1
Zrc1
10
20
30
40
50
t/s

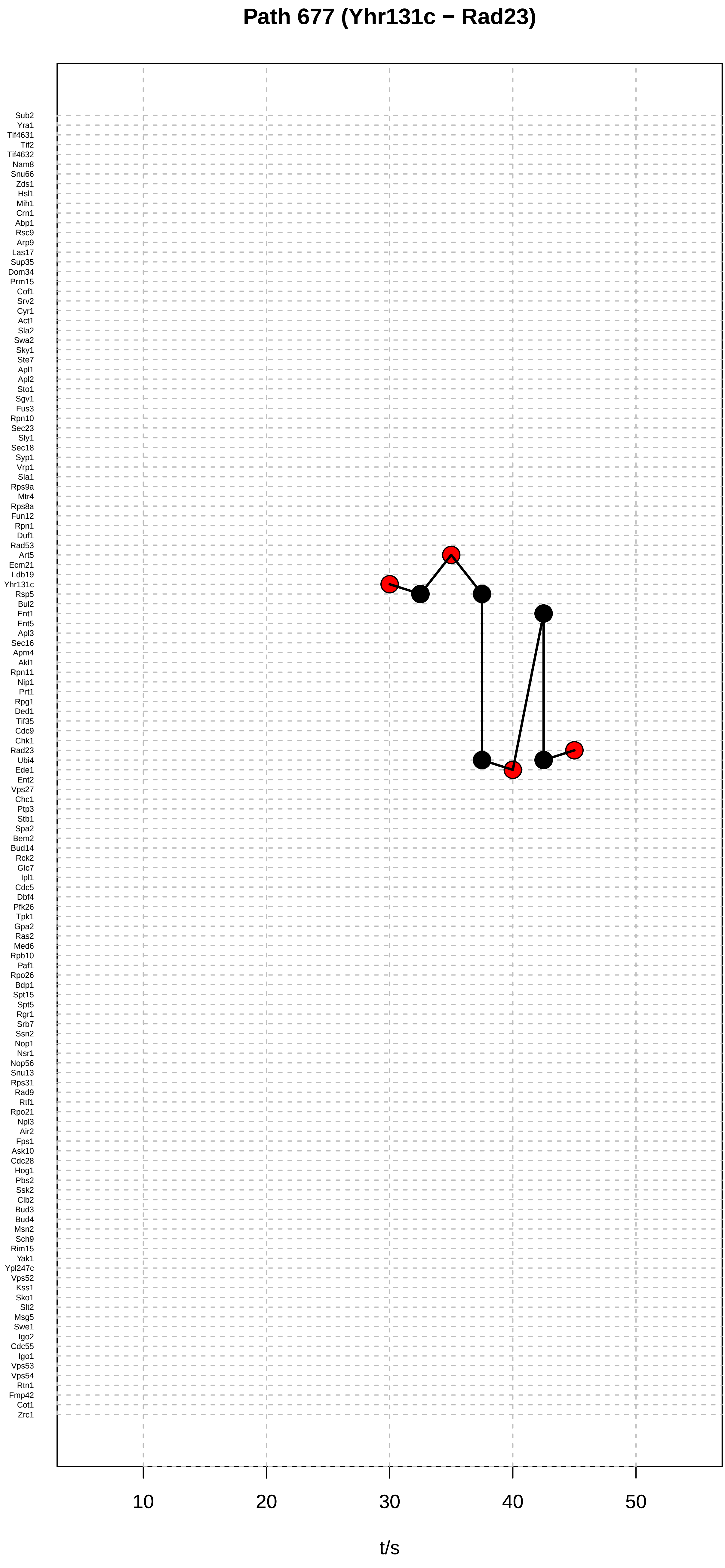
Path 677 (Yhr131c − Rad23)
Sub2
Yra1
Tif4631
Tif2
Tif4632
Nam8
Snu66
Zds1
Hsl1
Mih1
Crn1
Abp1
Rsc9
Arp9
Las17
Sup35
Dom34
Prm15
Cof1
Srv2
Cyr1
Act1
Sla2
Swa2
Sky1
Ste7
Apl1
Apl2
Sto1
Sgv1
Fus3
Rpn10
Sec23
Sly1
Sec18
Syp1
Vrp1
Sla1
Rps9a
Mtr4
Rps8a
Fun12
Rpn1
Duf1
Rad53
Art5
Ecm21
Ldb19
Yhr131c
Rsp5
Bul2
Ent1
Ent5
Apl3
Sec16
Apm4
Akl1
Rpn11
Nip1
Prt1
Rpg1
Ded1
Tif35
Cdc9
Chk1
Rad23
Ubi4
Ede1
Ent2
Vps27
Chc1
Ptp3
Stb1
Spa2
Bem2
Bud14
Rck2
Glc7
Ipl1
Cdc5
Dbf4
Pfk26
Tpk1
Gpa2
Ras2
Med6
Rpb10
Paf1
Rpo26
Bdp1
Spt15
Spt5
Rgr1
Srb7
Ssn2
Nop1
Nsr1
Nop56
Snu13
Rps31
Rad9
Rtf1
Rpo21
Npl3
Air2
Fps1
Ask10
Cdc28
Hog1
Pbs2
Ssk2
Clb2
Bud3
Bud4
Msn2
Sch9
Rim15
Yak1
Ypl247c
Vps52
Kss1
Sko1
Slt2
Msg5
Swe1
Igo2
Cdc55
Igo1
Vps53
Vps54
Rtn1
Fmp42
Cot1
Zrc1
10
20
30
40
50
t/s

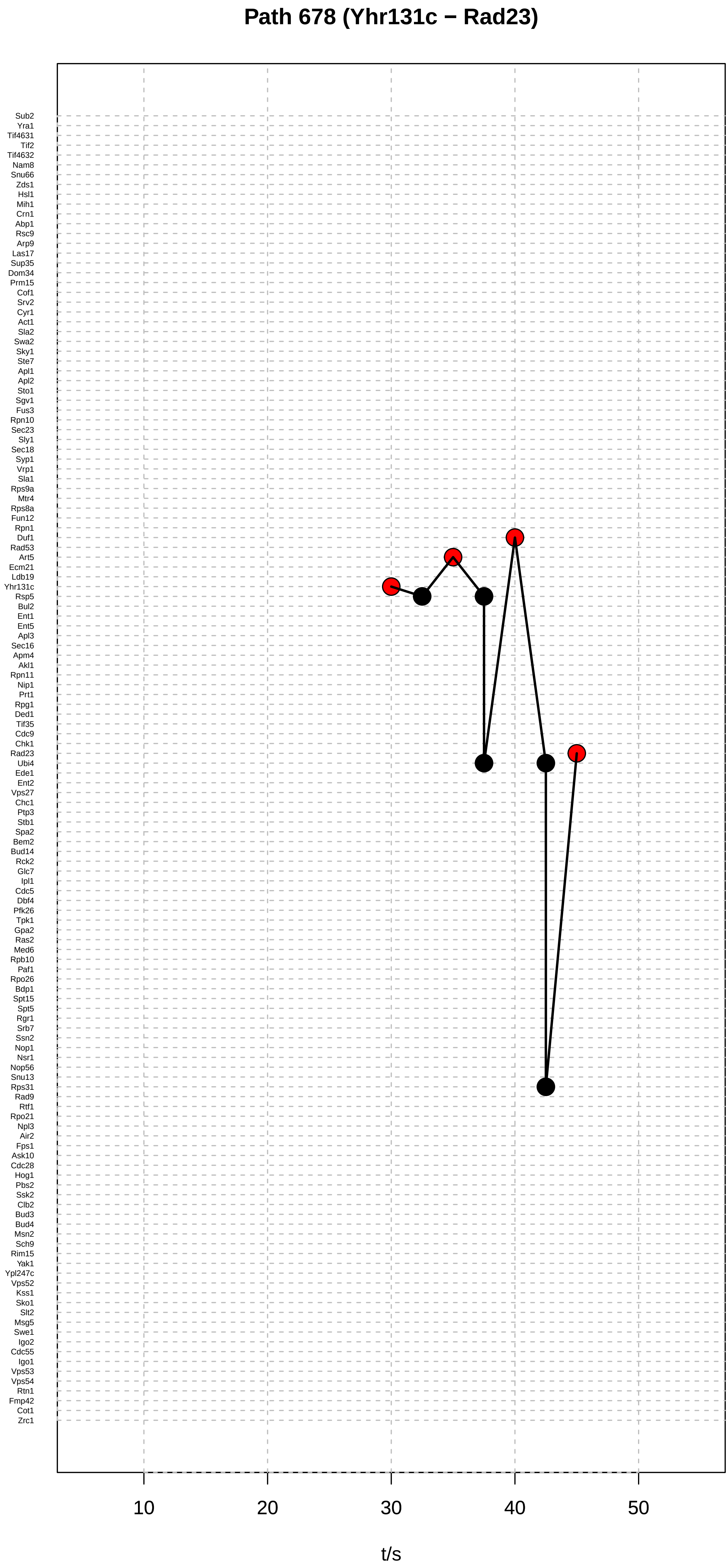

Path 678 (Yhr131c − Rad23)
Sub2
Yra1
Tif4631
Tif2
Tif4632
Nam8
Snu66
Zds1
Hsl1
Mih1
Crn1
Abp1
Rsc9
Arp9
Las17
Sup35
Dom34
Prm15
Cof1
Srv2
Cyr1
Act1
Sla2
Swa2
Sky1
Ste7
Apl1
Apl2
Sto1
Sgv1
Fus3
Rpn10
Sec23
Sly1
Sec18
Syp1
Vrp1
Sla1
Rps9a
Mtr4
Rps8a
Fun12
Rpn1
Duf1
Rad53
Art5
Ecm21
Ldb19
Yhr131c
Rsp5
Bul2
Ent1
Ent5
Apl3
Sec16
Apm4
Akl1
Rpn11
Nip1
Prt1
Rpg1
Ded1
Tif35
Cdc9
Chk1
Rad23
Ubi4
Ede1
Ent2
Vps27
Chc1
Ptp3
Stb1
Spa2
Bem2
Bud14
Rck2
Glc7
Ipl1
Cdc5
Dbf4
Pfk26
Tpk1
Gpa2
Ras2
Med6
Rpb10
Paf1
Rpo26
Bdp1
Spt15
Spt5
Rgr1
Srb7
Ssn2
Nop1
Nsr1
Nop56
Snu13
Rps31
Rad9
Rtf1
Rpo21
Npl3
Air2
Fps1
Ask10
Cdc28
Hog1
Pbs2
Ssk2
Clb2
Bud3
Bud4
Msn2
Sch9
Rim15
Yak1
Ypl247c
Vps52
Kss1
Sko1
Slt2
Msg5
Swe1
Igo2
Cdc55
Igo1
Vps53
Vps54
Rtn1
Fmp42
Cot1
Zrc1
10
20
30
40
50
t/s

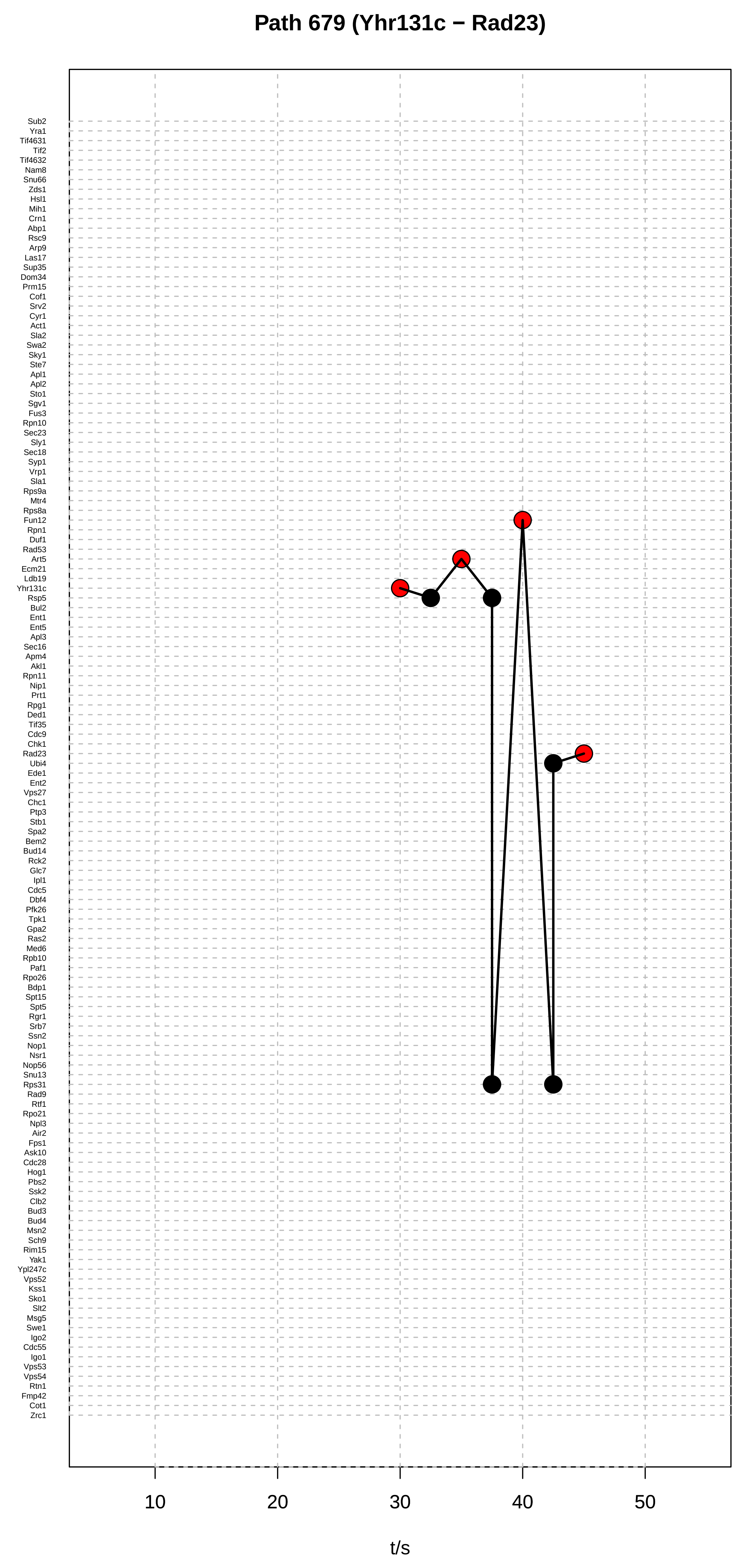
Path 679 (Yhr131c − Rad23)
Sub2
Yra1
Tif4631
Tif2
Tif4632
Nam8
Snu66
Zds1
Hsl1
Mih1
Crn1
Abp1
Rsc9
Arp9
Las17
Sup35
Dom34
Prm15
Cof1
Srv2
Cyr1
Act1
Sla2
Swa2
Sky1
Ste7
Apl1
Apl2
Sto1
Sgv1
Fus3
Rpn10
Sec23
Sly1
Sec18
Syp1
Vrp1
Sla1
Rps9a
Mtr4
Rps8a
Fun12
Rpn1
Duf1
Rad53
Art5
Ecm21
Ldb19
Yhr131c
Rsp5
Bul2
Ent1
Ent5
Apl3
Sec16
Apm4
Akl1
Rpn11
Nip1
Prt1
Rpg1
Ded1
Tif35
Cdc9
Chk1
Rad23
Ubi4
Ede1
Ent2
Vps27
Chc1
Ptp3
Stb1
Spa2
Bem2
Bud14
Rck2
Glc7
Ipl1
Cdc5
Dbf4
Pfk26
Tpk1
Gpa2
Ras2
Med6
Rpb10
Paf1
Rpo26
Bdp1
Spt15
Spt5
Rgr1
Srb7
Ssn2
Nop1
Nsr1
Nop56
Snu13
Rps31
Rad9
Rtf1
Rpo21
Npl3
Air2
Fps1
Ask10
Cdc28
Hog1
Pbs2
Ssk2
Clb2
Bud3
Bud4
Msn2
Sch9
Rim15
Yak1
Ypl247c
Vps52
Kss1
Sko1
Slt2
Msg5
Swe1
Igo2
Cdc55
Igo1
Vps53
Vps54
Rtn1
Fmp42
Cot1
Zrc1
10
20
30
40
50
t/s

#### Path 680 (Yhr131c − Rad23)

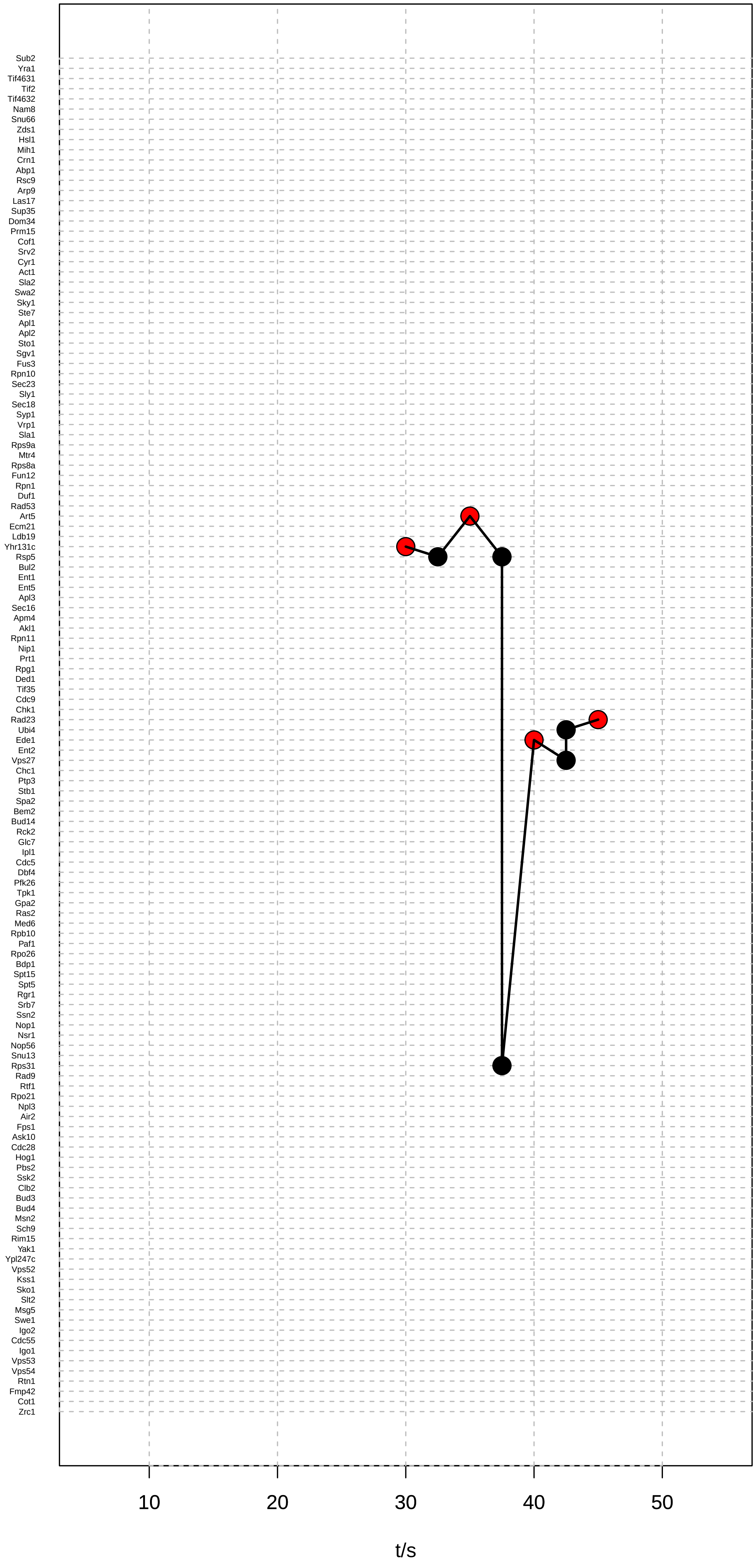

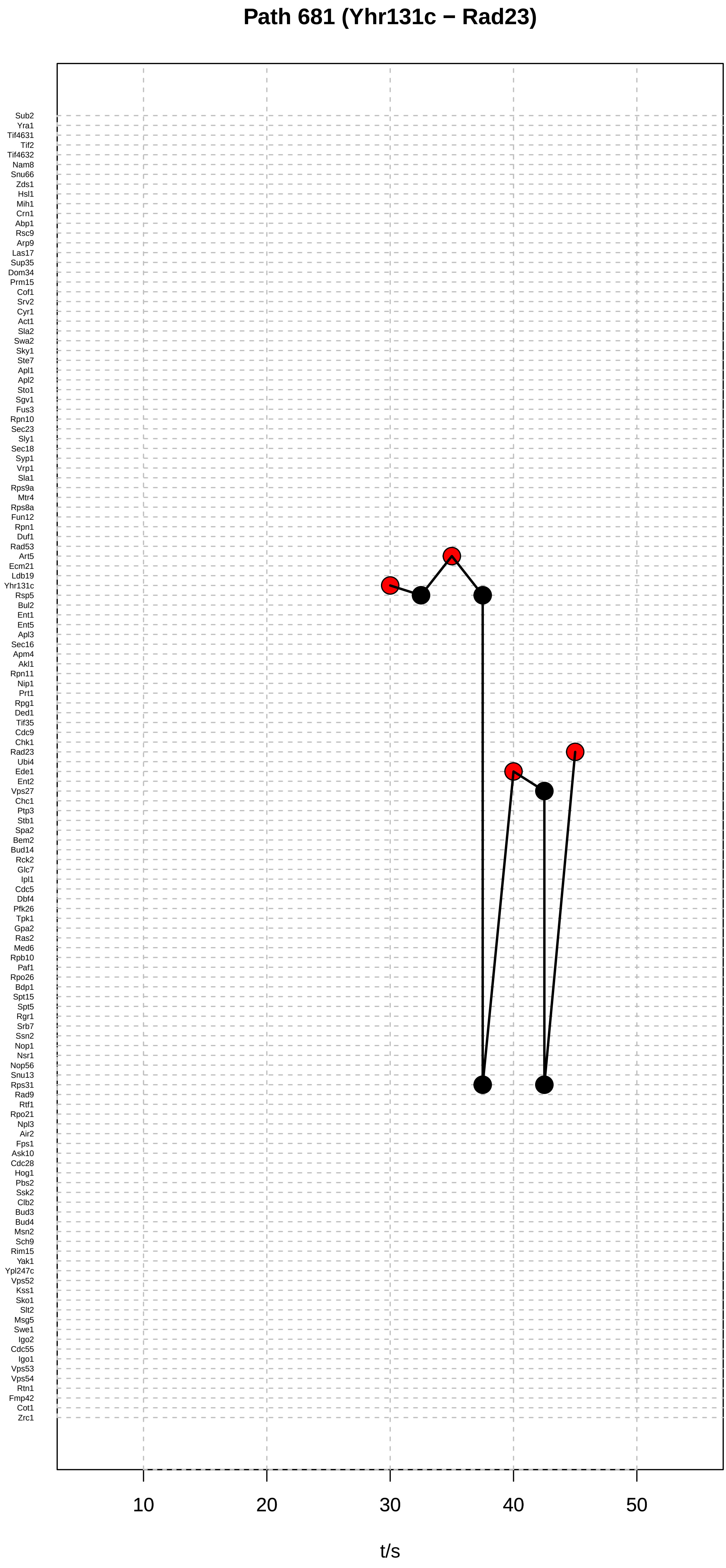
Path 681 (Yhr131c − Rad23)
Sub2
Yra1
Tif4631
Tif2
Tif4632
Nam8
Snu66
Zds1
Hsl1
Mih1
Crn1
Abp1
Rsc9
Arp9
Las17
Sup35
Dom34
Prm15
Cof1
Srv2
Cyr1
Act1
Sla2
Swa2
Sky1
Ste7
Apl1
Apl2
Sto1
Sgv1
Fus3
Rpn10
Sec23
Sly1
Sec18
Syp1
Vrp1
Sla1
Rps9a
Mtr4
Rps8a
Fun12
Rpn1
Duf1
Rad53
Art5
Ecm21
Ldb19
Yhr131c
Rsp5
Bul2
Ent1
Ent5
Apl3
Sec16
Apm4
Akl1
Rpn11
Nip1
Prt1
Rpg1
Ded1
Tif35
Cdc9
Chk1
Rad23
Ubi4
Ede1
Ent2
Vps27
Chc1
Ptp3
Stb1
Spa2
Bem2
Bud14
Rck2
Glc7
Ipl1
Cdc5
Dbf4
Pfk26
Tpk1
Gpa2
Ras2
Med6
Rpb10
Paf1
Rpo26
Bdp1
Spt15
Spt5
Rgr1
Srb7
Ssn2
Nop1
Nsr1
Nop56
Snu13
Rps31
Rad9
Rtf1
Rpo21
Npl3
Air2
Fps1
Ask10
Cdc28
Hog1
Pbs2
Ssk2
Clb2
Bud3
Bud4
Msn2
Sch9
Rim15
Yak1
Ypl247c
Vps52
Kss1
Sko1
Slt2
Msg5
Swe1
Igo2
Cdc55
Igo1
Vps53
Vps54
Rtn1
Fmp42
Cot1
Zrc1
10
20
30
40
50
t/s

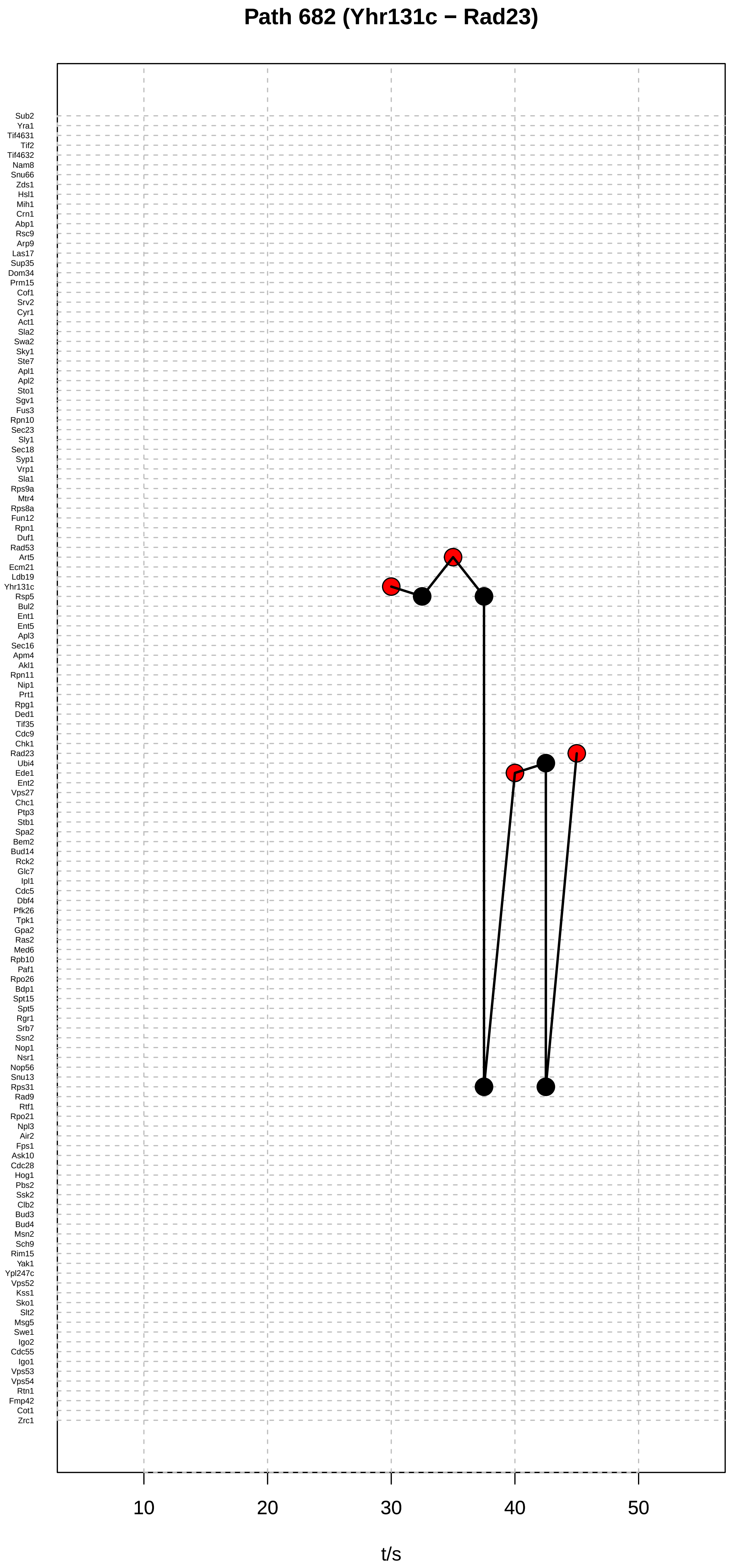

Path 682 (Yhr131c − Rad23)
Sub2
Yra1
Tif4631
Tif2
Tif4632
Nam8
Snu66
Zds1
Hsl1
Mih1
Crn1
Abp1
Rsc9
Arp9
Las17
Sup35
Dom34
Prm15
Cof1
Srv2
Cyr1
Act1
Sla2
Swa2
Sky1
Ste7
Apl1
Apl2
Sto1
Sgv1
Fus3
Rpn10
Sec23
Sly1
Sec18
Syp1
Vrp1
Sla1
Rps9a
Mtr4
Rps8a
Fun12
Rpn1
Duf1
Rad53
Art5
Ecm21
Ldb19
Yhr131c
Rsp5
Bul2
Ent1
Ent5
Apl3
Sec16
Apm4
Akl1
Rpn11
Nip1
Prt1
Rpg1
Ded1
Tif35
Cdc9
Chk1
Rad23
Ubi4
Ede1
Ent2
Vps27
Chc1
Ptp3
Stb1
Spa2
Bem2
Bud14
Rck2
Glc7
Ipl1
Cdc5
Dbf4
Pfk26
Tpk1
Gpa2
Ras2
Med6
Rpb10
Paf1
Rpo26
Bdp1
Spt15
Spt5
Rgr1
Srb7
Ssn2
Nop1
Nsr1
Nop56
Snu13
Rps31
Rad9
Rtf1
Rpo21
Npl3
Air2
Fps1
Ask10
Cdc28
Hog1
Pbs2
Ssk2
Clb2
Bud3
Bud4
Msn2
Sch9
Rim15
Yak1
Ypl247c
Vps52
Kss1
Sko1
Slt2
Msg5
Swe1
Igo2
Cdc55
Igo1
Vps53
Vps54
Rtn1
Fmp42
Cot1
Zrc1
10
20
30
40
50
t/s

#### Path 683 (Yhr131c − Rad23)

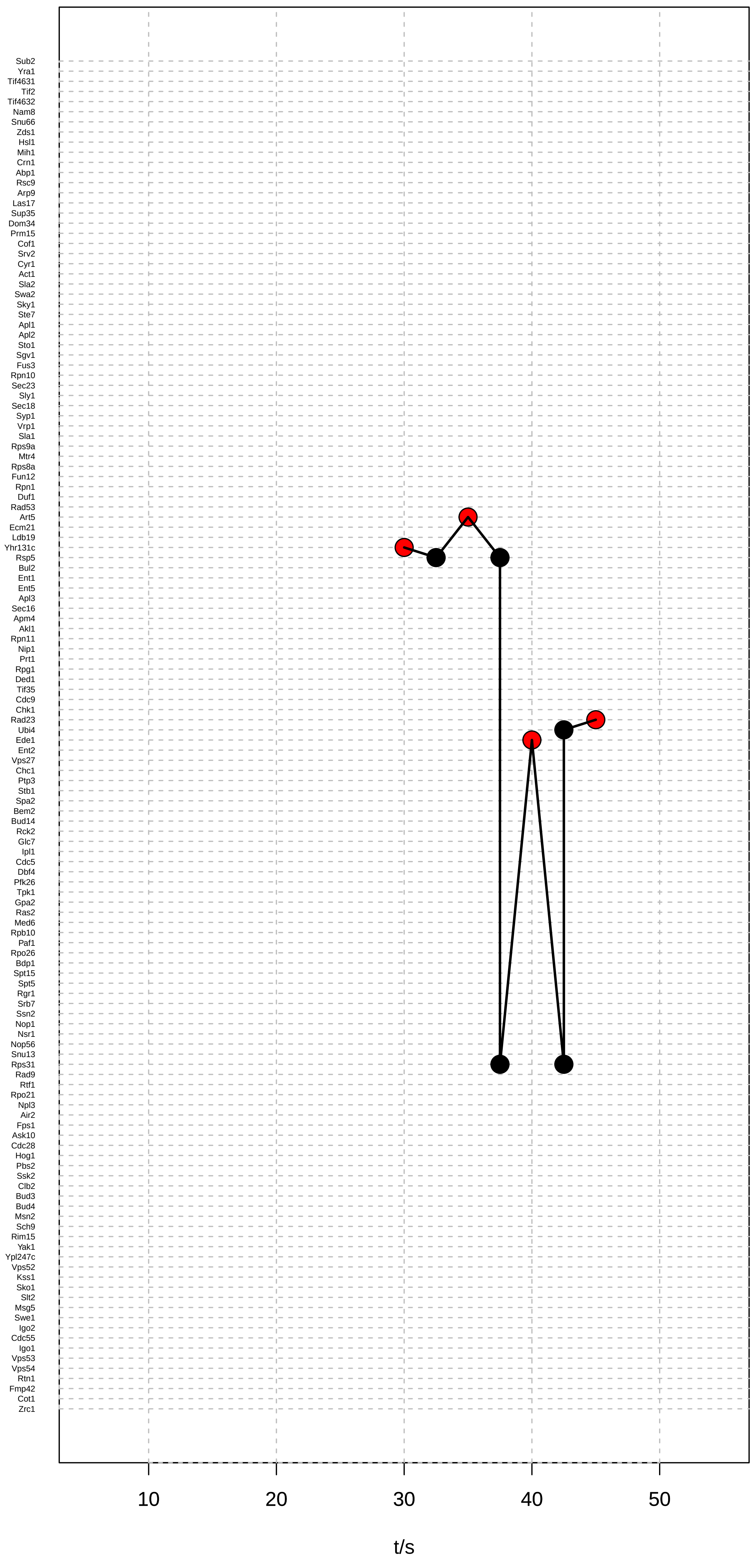

#### Path 684 (Yhr131c − Rad23)

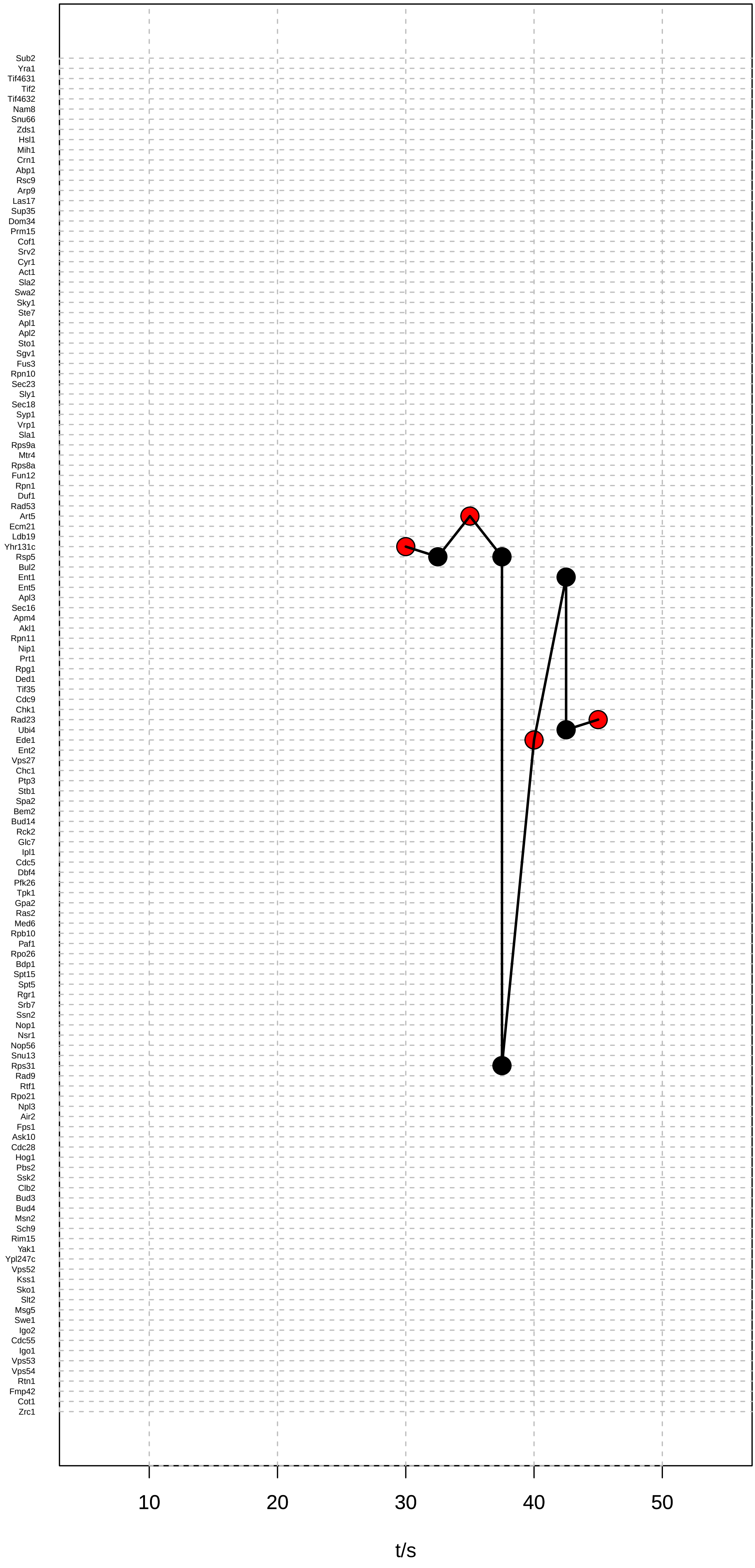

#### Path 685 (Yhr131c − Ede1)

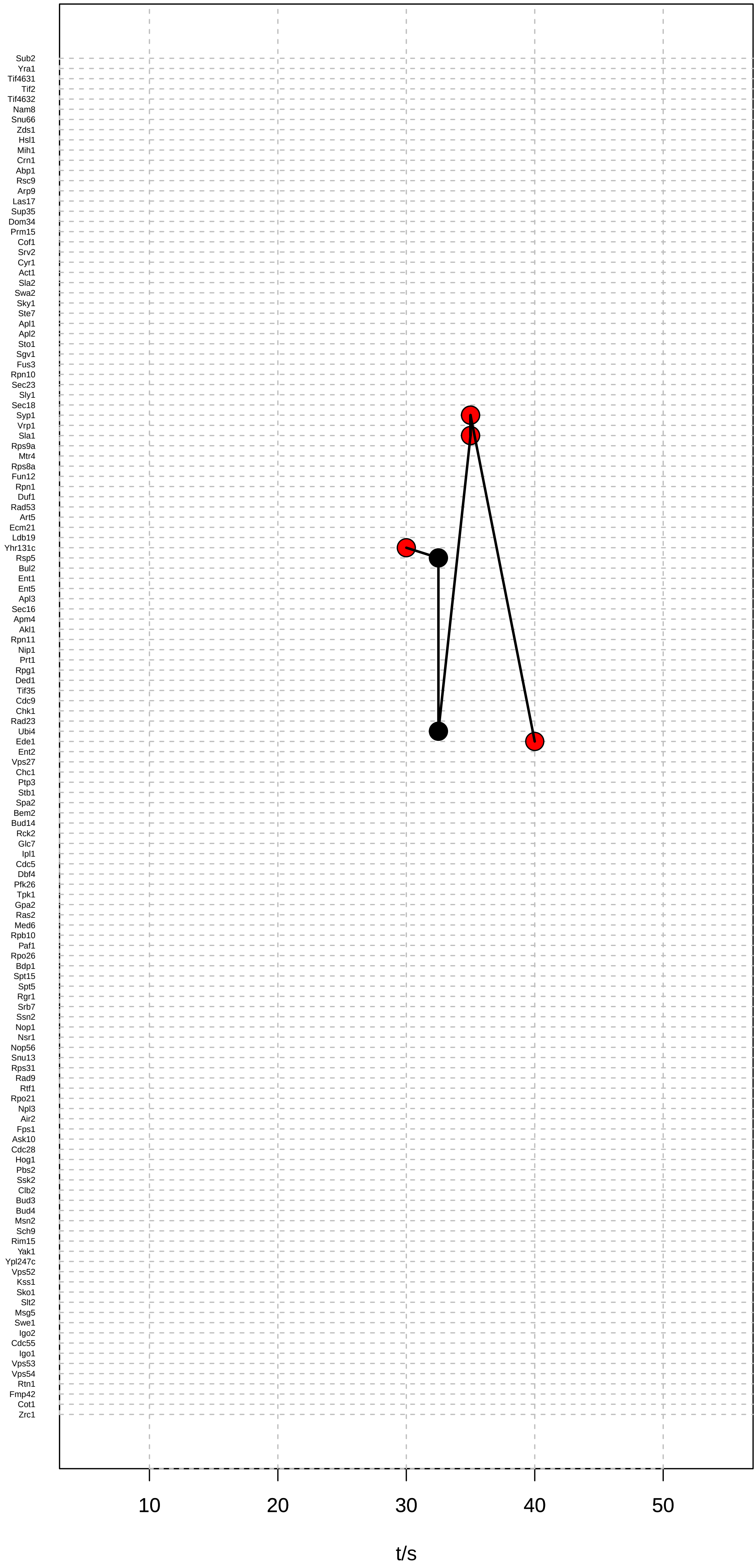

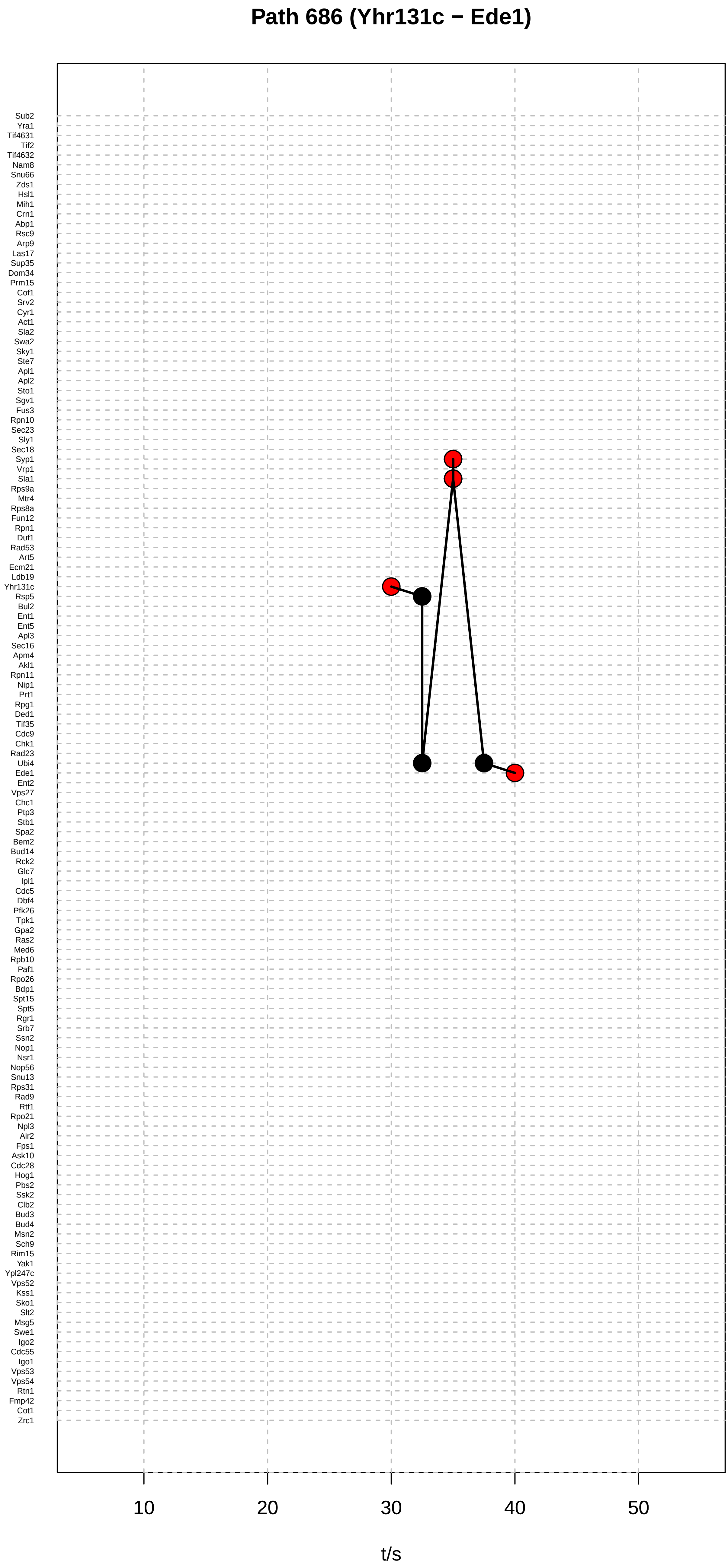

Path 686 (Yhr131c − Ede1)
Sub2
Yra1
Tif4631
Tif2
Tif4632
Nam8
Snu66
Zds1
Hsl1
Mih1
Crn1
Abp1
Rsc9
Arp9
Las17
Sup35
Dom34
Prm15
Cof1
Srv2
Cyr1
Act1
Sla2
Swa2
Sky1
Ste7
Apl1
Apl2
Sto1
Sgv1
Fus3
Rpn10
Sec23
Sly1
Sec18
Syp1
Vrp1
Sla1
Rps9a
Mtr4
Rps8a
Fun12
Rpn1
Duf1
Rad53
Art5
Ecm21
Ldb19
Yhr131c
Rsp5
Bul2
Ent1
Ent5
Apl3
Sec16
Apm4
Akl1
Rpn11
Nip1
Prt1
Rpg1
Ded1
Tif35
Cdc9
Chk1
Rad23
Ubi4
Ede1
Ent2
Vps27
Chc1
Ptp3
Stb1
Spa2
Bem2
Bud14
Rck2
Glc7
Ipl1
Cdc5
Dbf4
Pfk26
Tpk1
Gpa2
Ras2
Med6
Rpb10
Paf1
Rpo26
Bdp1
Spt15
Spt5
Rgr1
Srb7
Ssn2
Nop1
Nsr1
Nop56
Snu13
Rps31
Rad9
Rtf1
Rpo21
Npl3
Air2
Fps1
Ask10
Cdc28
Hog1
Pbs2
Ssk2
Clb2
Bud3
Bud4
Msn2
Sch9
Rim15
Yak1
Ypl247c
Vps52
Kss1
Sko1
Slt2
Msg5
Swe1
Igo2
Cdc55
Igo1
Vps53
Vps54
Rtn1
Fmp42
Cot1
Zrc1
10
20
30
40
50
t/s

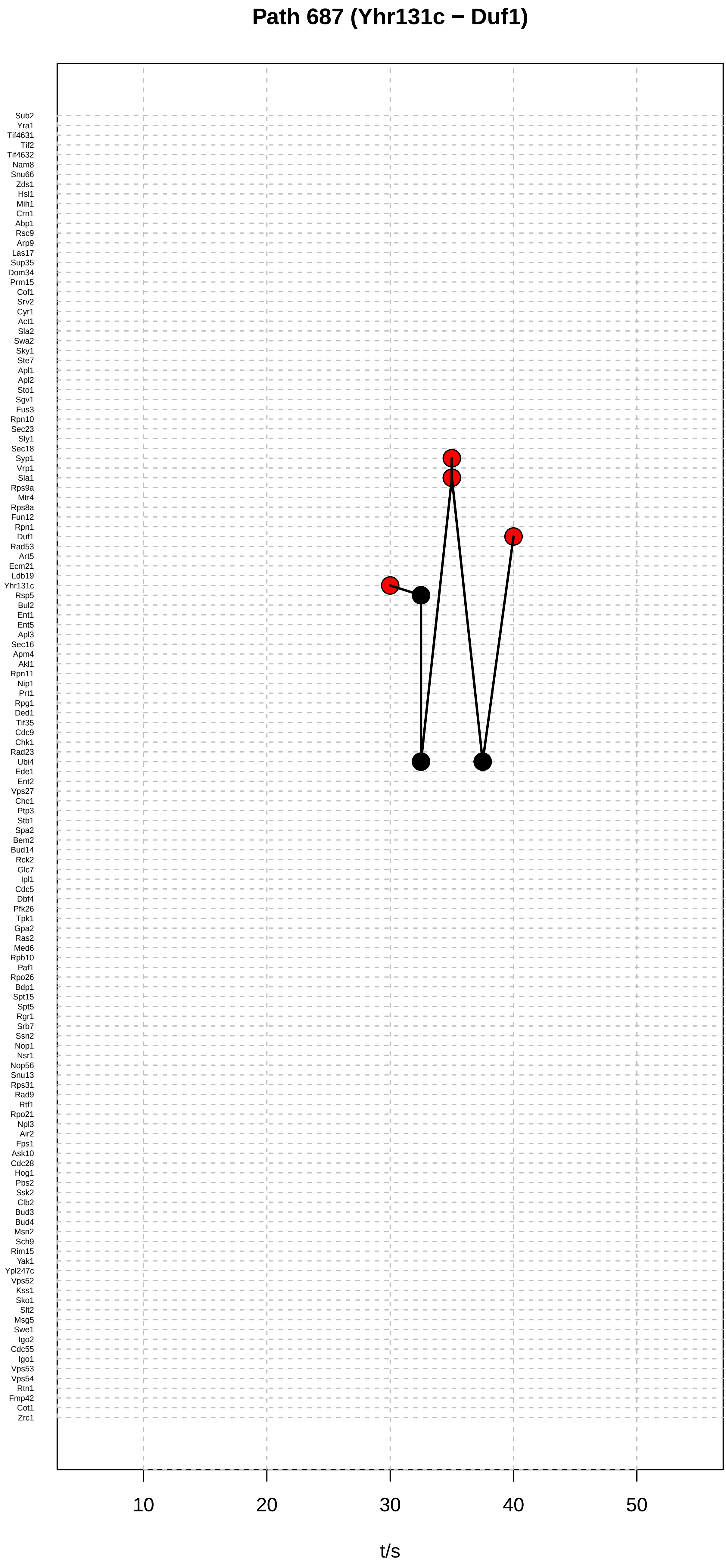
Path 687 (Yhr131c − Duf1)
Sub2
Yra1
Tif4631
Tif2
Tif4632
Nam8
Snu66
Zds1
Hsl1
Mih1
Crn1
Abp1
Rsc9
Arp9
Las17
Sup35
Dom34
Prm15
Cof1
Srv2
Cyr1
Act1
Sla2
Swa2
Sky1
Ste7
Apl1
Apl2
Sto1
Sgv1
Fus3
Rpn10
Sec23
Sly1
Sec18
Syp1
Vrp1
Sla1
Rps9a
Mtr4
Rps8a
Fun12
Rpn1
Duf1
Rad53
Art5
Ecm21
Ldb19
Yhr131c
Rsp5
Bul2
Ent1
Ent5
Apl3
Sec16
Apm4
Akl1
Rpn11
Nip1
Prt1
Rpg1
Ded1
Tif35
Cdc9
Chk1
Rad23
Ubi4
Ede1
Ent2
Vps27
Chc1
Ptp3
Stb1
Spa2
Bem2
Bud14
Rck2
Glc7
Ipl1
Cdc5
Dbf4
Pfk26
Tpk1
Gpa2
Ras2
Med6
Rpb10
Paf1
Rpo26
Bdp1
Spt15
Spt5
Rgr1
Srb7
Ssn2
Nop1
Nsr1
Nop56
Snu13
Rps31
Rad9
Rtf1
Rpo21
Npl3
Air2
Fps1
Ask10
Cdc28
Hog1
Pbs2
Ssk2
Clb2
Bud3
Bud4
Msn2
Sch9
Rim15
Yak1
Ypl247c
Vps52
Kss1
Sko1
Slt2
Msg5
Swe1
Igo2
Cdc55
Igo1
Vps53
Vps54
Rtn1
Fmp42
Cot1
Zrc1
10
20
30
40
50
t/s

Path 688 (Yhr131c − Fun12)

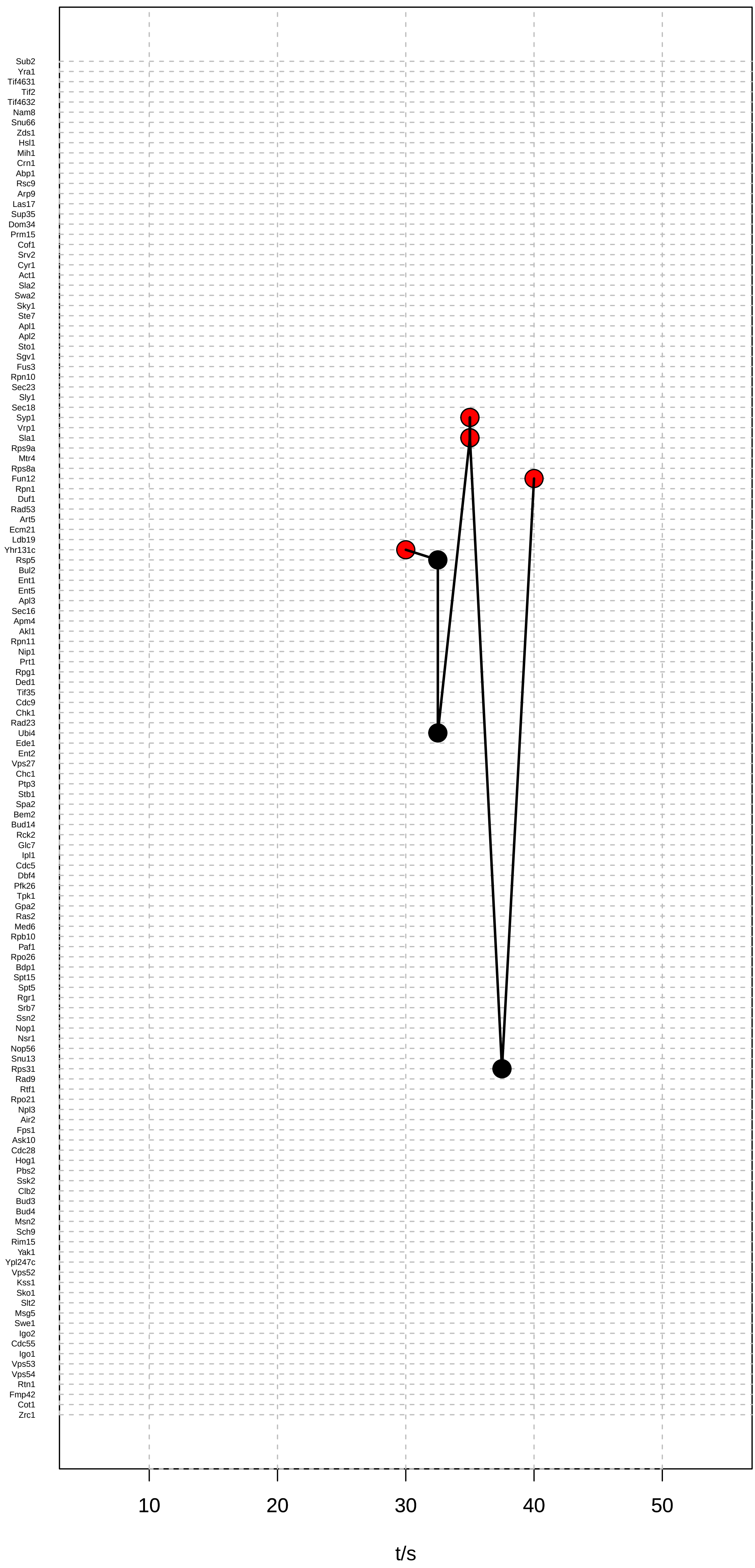

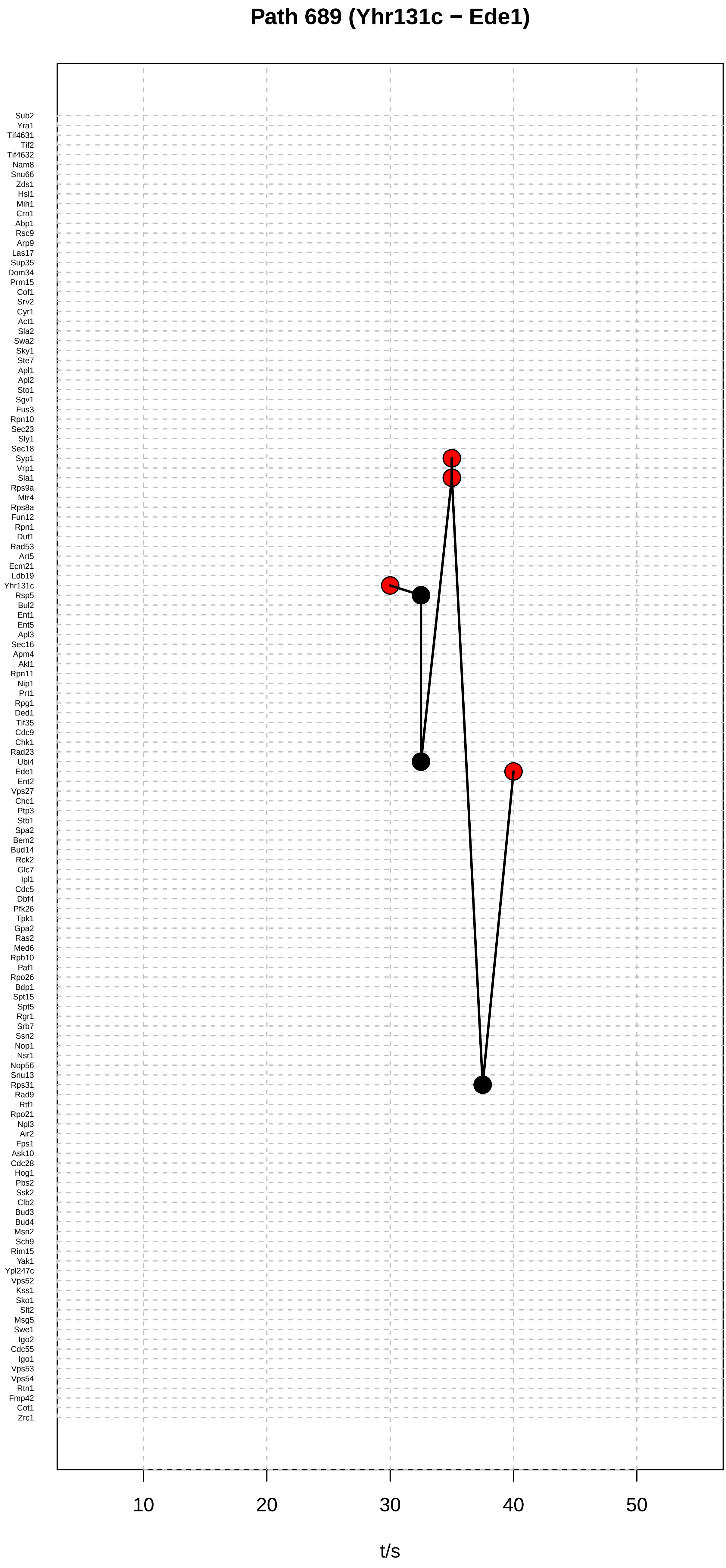

Path 689 (Yhr131c − Ede1)
Sub2
Yra1
Tif4631
Tif2
Tif4632
Nam8
Snu66
Zds1
Hsl1
Mih1
Crn1
Abp1
Rsc9
Arp9
Las17
Sup35
Dom34
Prm15
Cof1
Srv2
Cyr1
Act1
Sla2
Swa2
Sky1
Ste7
Apl1
Apl2
Sto1
Sgv1
Fus3
Rpn10
Sec23
Sly1
Sec18
Syp1
Vrp1
Sla1
Rps9a
Mtr4
Rps8a
Fun12
Rpn1
Duf1
Rad53
Art5
Ecm21
Ldb19
Yhr131c
Rsp5
Bul2
Ent1
Ent5
Apl3
Sec16
Apm4
Akl1
Rpn11
Nip1
Prt1
Rpg1
Ded1
Tif35
Cdc9
Chk1
Rad23
Ubi4
Ede1
Ent2
Vps27
Chc1
Ptp3
Stb1
Spa2
Bem2
Bud14
Rck2
Glc7
Ipl1
Cdc5
Dbf4
Pfk26
Tpk1
Gpa2
Ras2
Med6
Rpb10
Paf1
Rpo26
Bdp1
Spt15
Spt5
Rgr1
Srb7
Ssn2
Nop1
Nsr1
Nop56
Snu13
Rps31
Rad9
Rtf1
Rpo21
Npl3
Air2
Fps1
Ask10
Cdc28
Hog1
Pbs2
Ssk2
Clb2
Bud3
Bud4
Msn2
Sch9
Rim15
Yak1
Ypl247c
Vps52
Kss1
Sko1
Slt2
Msg5
Swe1
Igo2
Cdc55
Igo1
Vps53
Vps54
Rtn1
Fmp42
Cot1
Zrc1
10
20
30
40
50
t/s

### Path 690 (Yhr131c − Rad23)

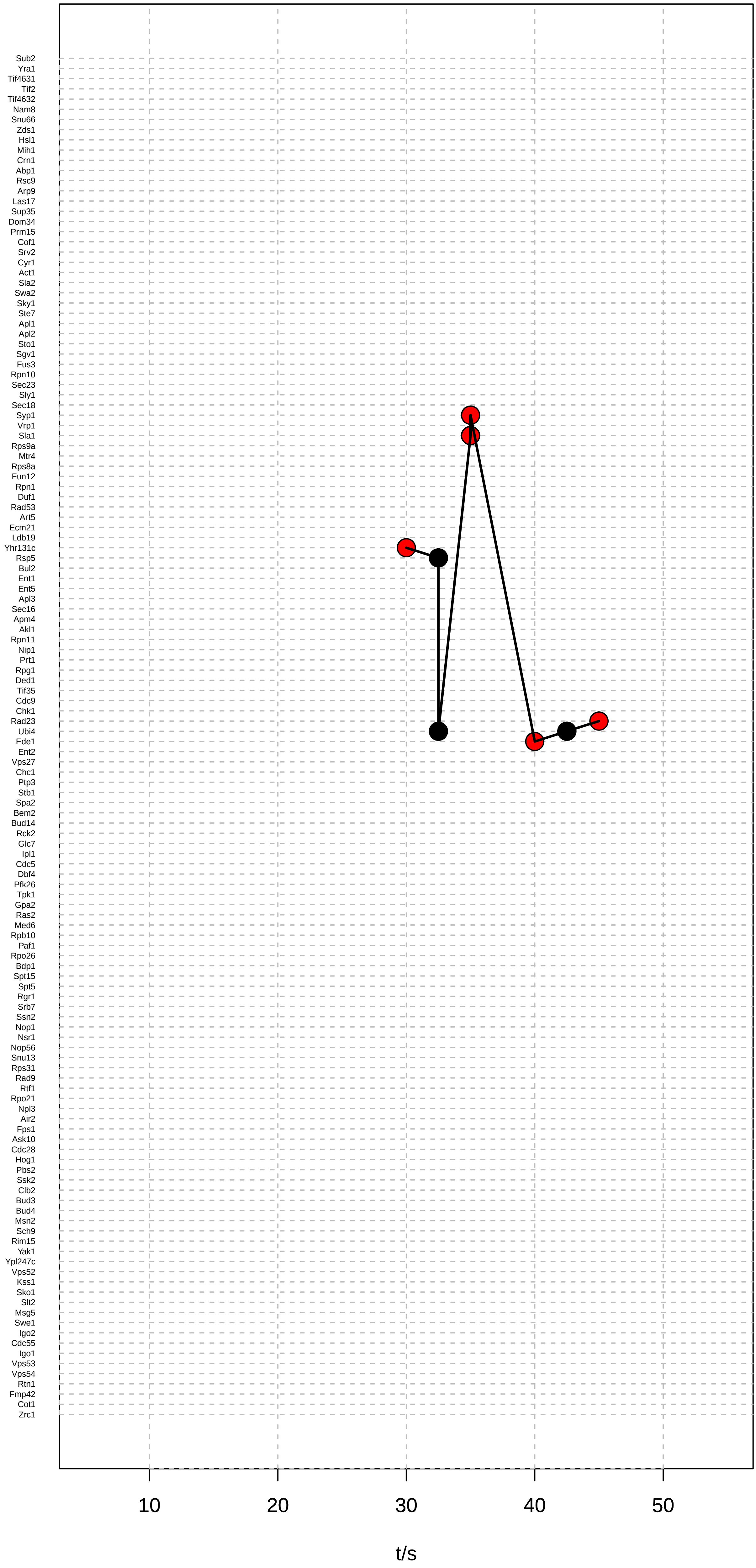

### Path 691 (Yhr131c − Rad23)

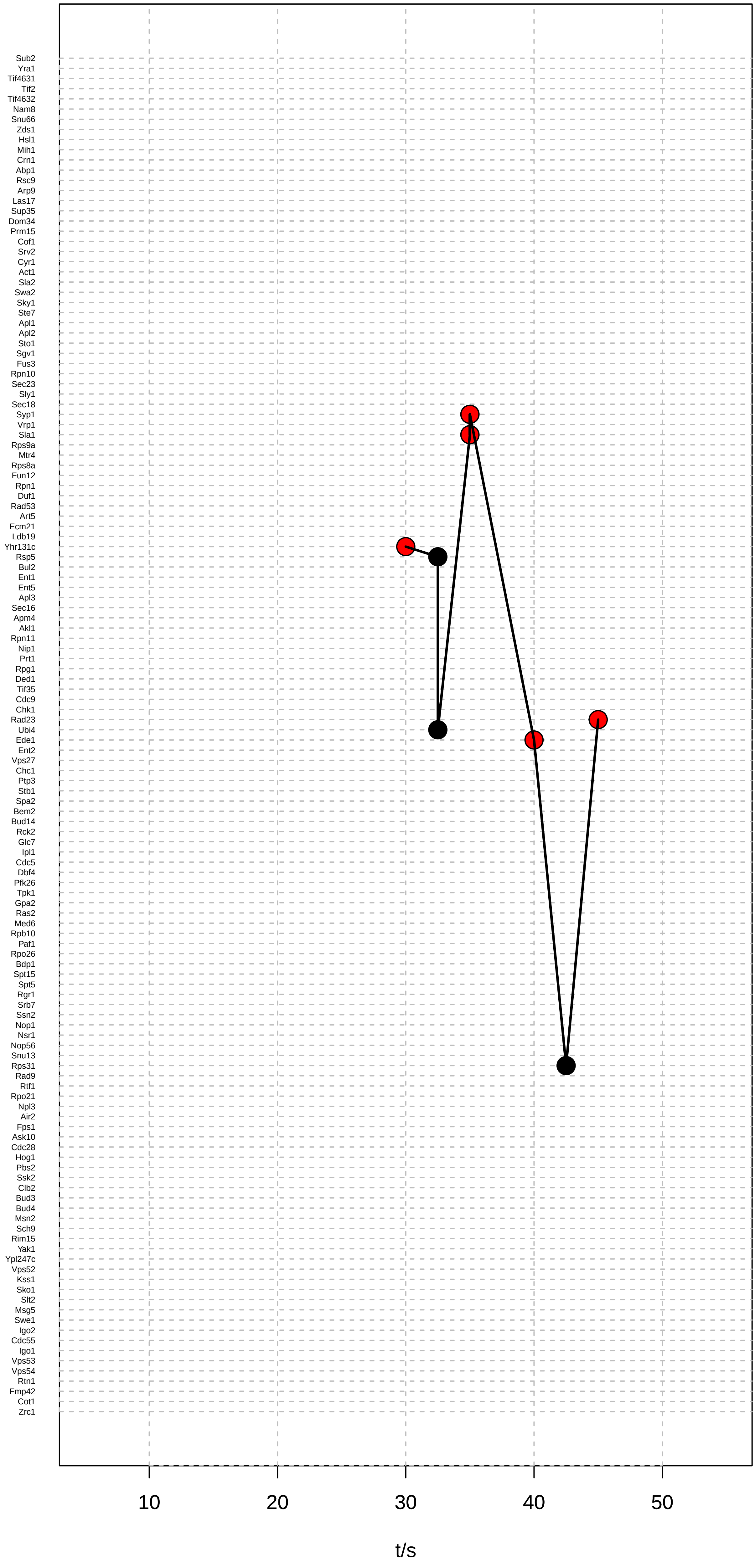

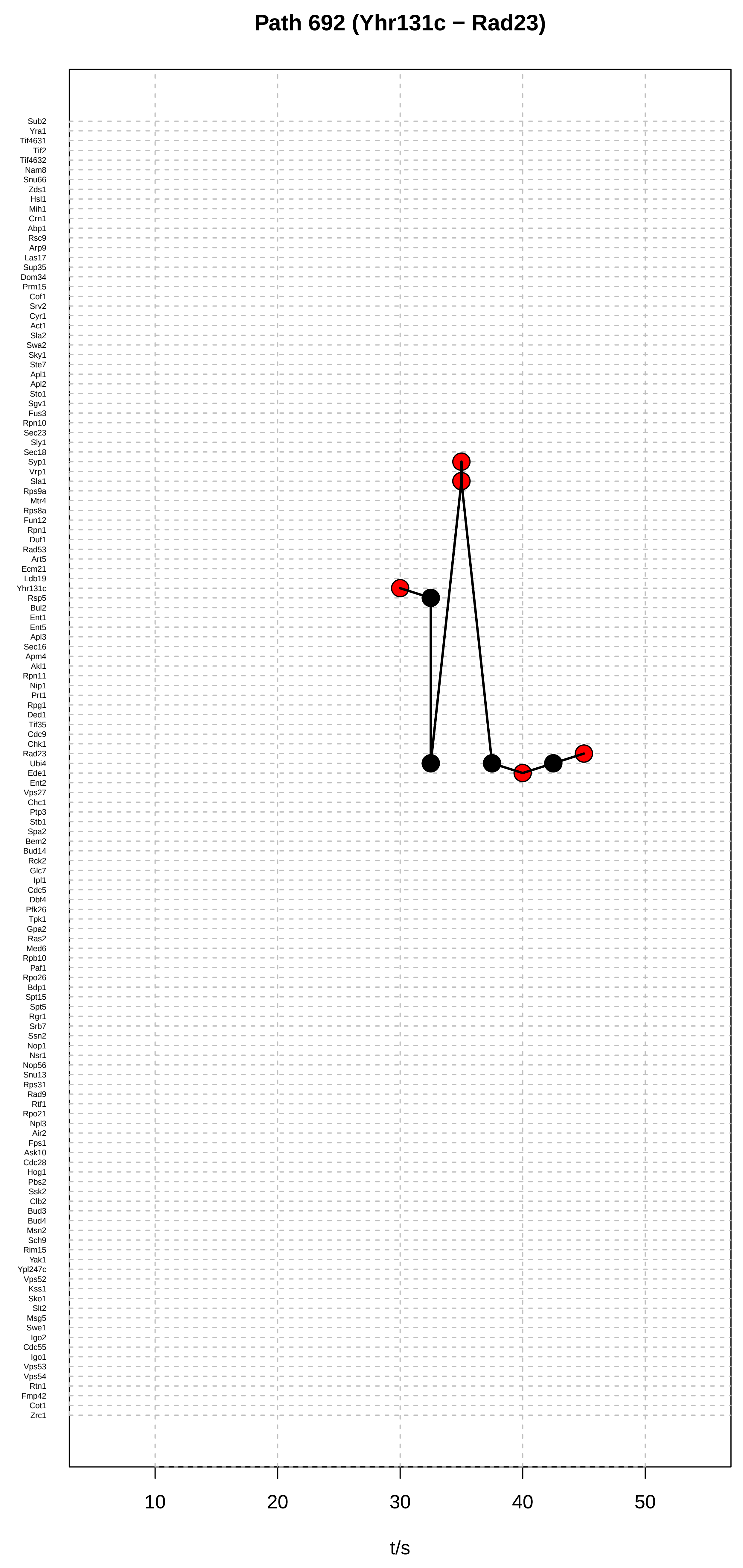
Path 692 (Yhr131c − Rad23)
Sub2
Yra1
Tif4631
Tif2
Tif4632
Nam8
Snu66
Zds1
Hsl1
Mih1
Crn1
Abp1
Rsc9
Arp9
Las17
Sup35
Dom34
Prm15
Cof1
Srv2
Cyr1
Act1
Sla2
Swa2
Sky1
Ste7
Apl1
Apl2
Sto1
Sgv1
Fus3
Rpn10
Sec23
Sly1
Sec18
Syp1
Vrp1
Sla1
Rps9a
Mtr4
Rps8a
Fun12
Rpn1
Duf1
Rad53
Art5
Ecm21
Ldb19
Yhr131c
Rsp5
Bul2
Ent1
Ent5
Apl3
Sec16
Apm4
Akl1
Rpn11
Nip1
Prt1
Rpg1
Ded1
Tif35
Cdc9
Chk1
Rad23
Ubi4
Ede1
Ent2
Vps27
Chc1
Ptp3
Stb1
Spa2
Bem2
Bud14
Rck2
Glc7
Ipl1
Cdc5
Dbf4
Pfk26
Tpk1
Gpa2
Ras2
Med6
Rpb10
Paf1
Rpo26
Bdp1
Spt15
Spt5
Rgr1
Srb7
Ssn2
Nop1
Nsr1
Nop56
Snu13
Rps31
Rad9
Rtf1
Rpo21
Npl3
Air2
Fps1
Ask10
Cdc28
Hog1
Pbs2
Ssk2
Clb2
Bud3
Bud4
Msn2
Sch9
Rim15
Yak1
Ypl247c
Vps52
Kss1
Sko1
Slt2
Msg5
Swe1
Igo2
Cdc55
Igo1
Vps53
Vps54
Rtn1
Fmp42
Cot1
Zrc1
10
20
30
40
50
t/s

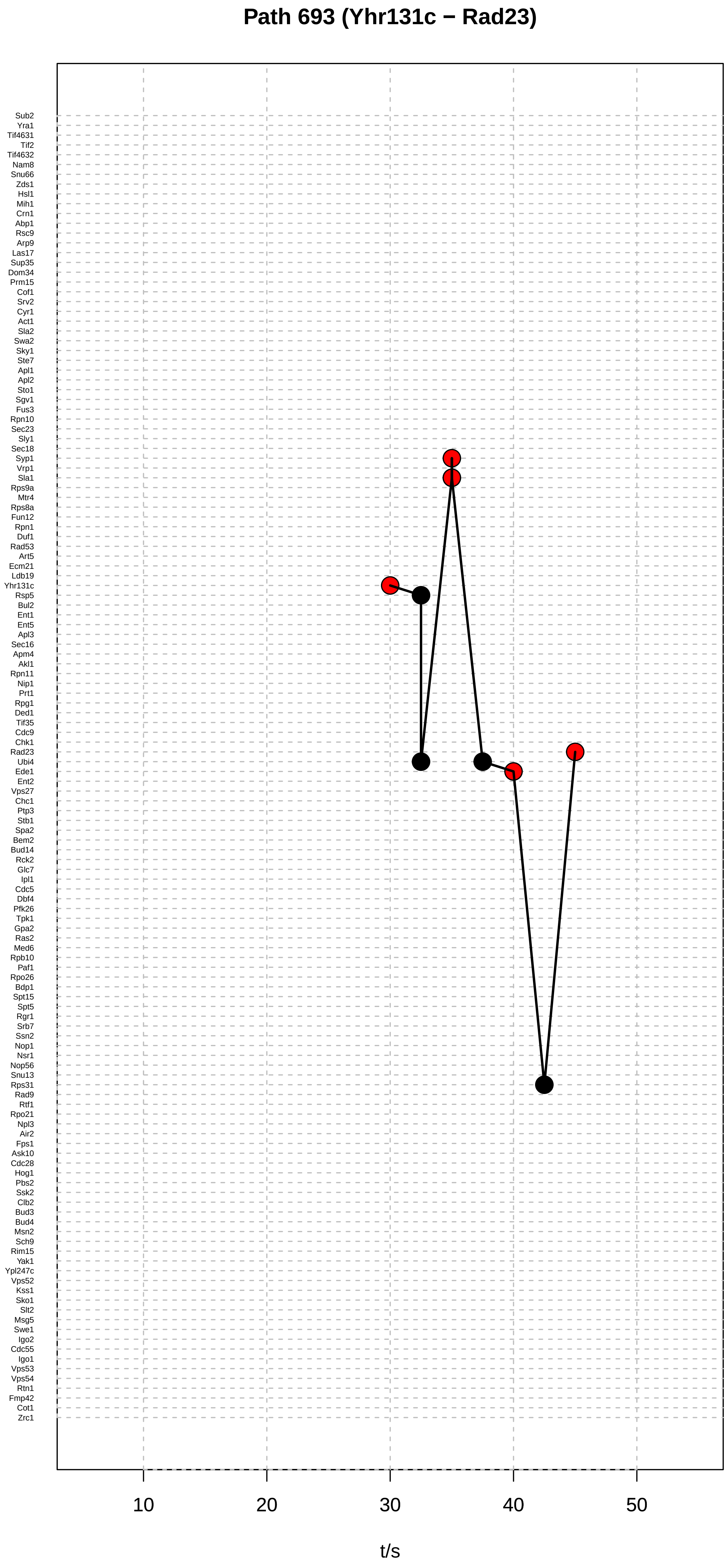

Path 693 (Yhr131c − Rad23)
Sub2
Yra1
Tif4631
Tif2
Tif4632
Nam8
Snu66
Zds1
Hsl1
Mih1
Crn1
Abp1
Rsc9
Arp9
Las17
Sup35
Dom34
Prm15
Cof1
Srv2
Cyr1
Act1
Sla2
Swa2
Sky1
Ste7
Apl1
Apl2
Sto1
Sgv1
Fus3
Rpn10
Sec23
Sly1
Sec18
Syp1
Vrp1
Sla1
Rps9a
Mtr4
Rps8a
Fun12
Rpn1
Duf1
Rad53
Art5
Ecm21
Ldb19
Yhr131c
Rsp5
Bul2
Ent1
Ent5
Apl3
Sec16
Apm4
Akl1
Rpn11
Nip1
Prt1
Rpg1
Ded1
Tif35
Cdc9
Chk1
Rad23
Ubi4
Ede1
Ent2
Vps27
Chc1
Ptp3
Stb1
Spa2
Bem2
Bud14
Rck2
Glc7
Ipl1
Cdc5
Dbf4
Pfk26
Tpk1
Gpa2
Ras2
Med6
Rpb10
Paf1
Rpo26
Bdp1
Spt15
Spt5
Rgr1
Srb7
Ssn2
Nop1
Nsr1
Nop56
Snu13
Rps31
Rad9
Rtf1
Rpo21
Npl3
Air2
Fps1
Ask10
Cdc28
Hog1
Pbs2
Ssk2
Clb2
Bud3
Bud4
Msn2
Sch9
Rim15
Yak1
Ypl247c
Vps52
Kss1
Sko1
Slt2
Msg5
Swe1
Igo2
Cdc55
Igo1
Vps53
Vps54
Rtn1
Fmp42
Cot1
Zrc1
10
20
30
40
50
t/s

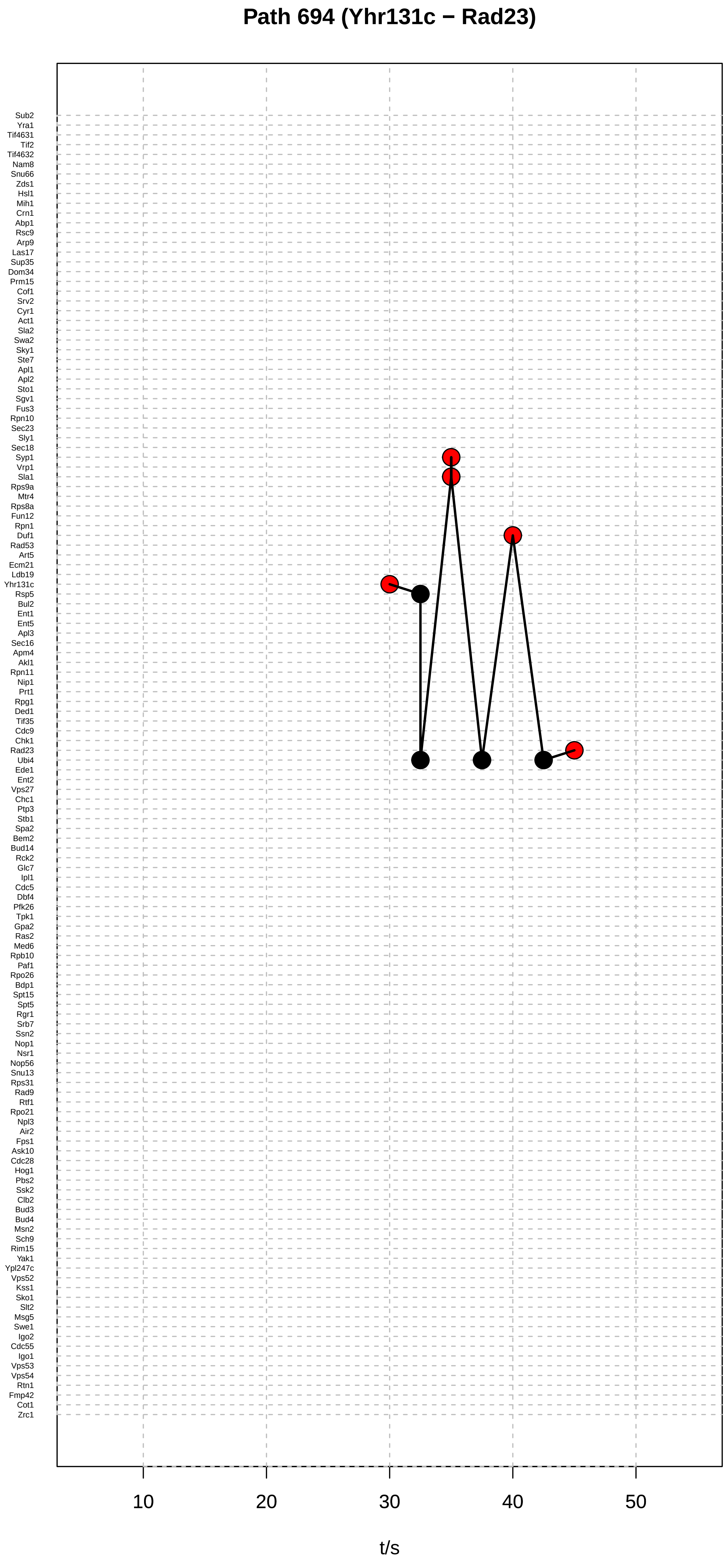

Path 694 (Yhr131c − Rad23)
Sub2
Yra1
Tif4631
Tif2
Tif4632
Nam8
Snu66
Zds1
Hsl1
Mih1
Crn1
Abp1
Rsc9
Arp9
Las17
Sup35
Dom34
Prm15
Cof1
Srv2
Cyr1
Act1
Sla2
Swa2
Sky1
Ste7
Apl1
Apl2
Sto1
Sgv1
Fus3
Rpn10
Sec23
Sly1
Sec18
Syp1
Vrp1
Sla1
Rps9a
Mtr4
Rps8a
Fun12
Rpn1
Duf1
Rad53
Art5
Ecm21
Ldb19
Yhr131c
Rsp5
Bul2
Ent1
Ent5
Apl3
Sec16
Apm4
Akl1
Rpn11
Nip1
Prt1
Rpg1
Ded1
Tif35
Cdc9
Chk1
Rad23
Ubi4
Ede1
Ent2
Vps27
Chc1
Ptp3
Stb1
Spa2
Bem2
Bud14
Rck2
Glc7
Ipl1
Cdc5
Dbf4
Pfk26
Tpk1
Gpa2
Ras2
Med6
Rpb10
Paf1
Rpo26
Bdp1
Spt15
Spt5
Rgr1
Srb7
Ssn2
Nop1
Nsr1
Nop56
Snu13
Rps31
Rad9
Rtf1
Rpo21
Npl3
Air2
Fps1
Ask10
Cdc28
Hog1
Pbs2
Ssk2
Clb2
Bud3
Bud4
Msn2
Sch9
Rim15
Yak1
Ypl247c
Vps52
Kss1
Sko1
Slt2
Msg5
Swe1
Igo2
Cdc55
Igo1
Vps53
Vps54
Rtn1
Fmp42
Cot1
Zrc1
10
20
30
40
50
t/s

### Path 695 (Yhr131c − Ede1)

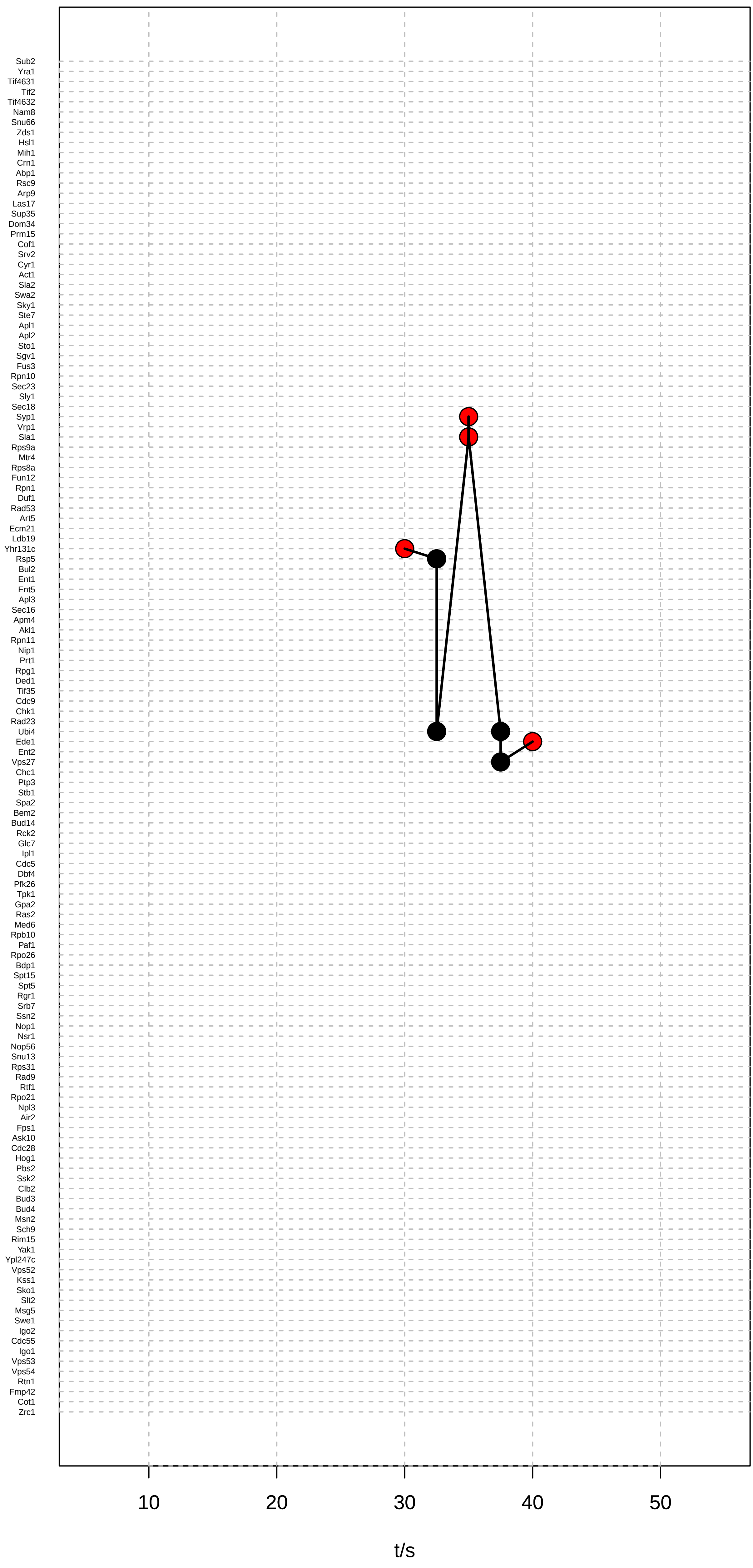

#### Path 696 (Yhr131c − Fun12)

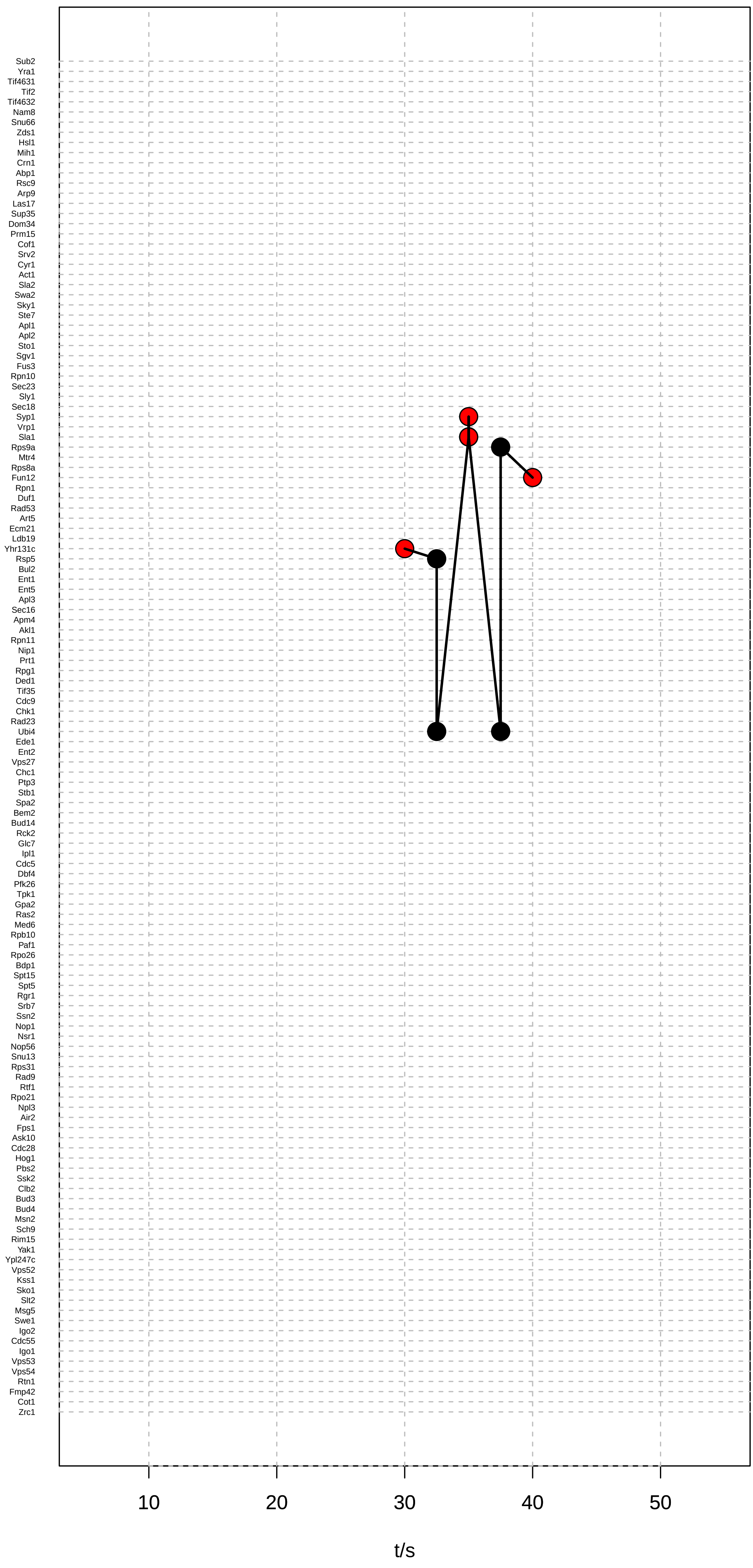

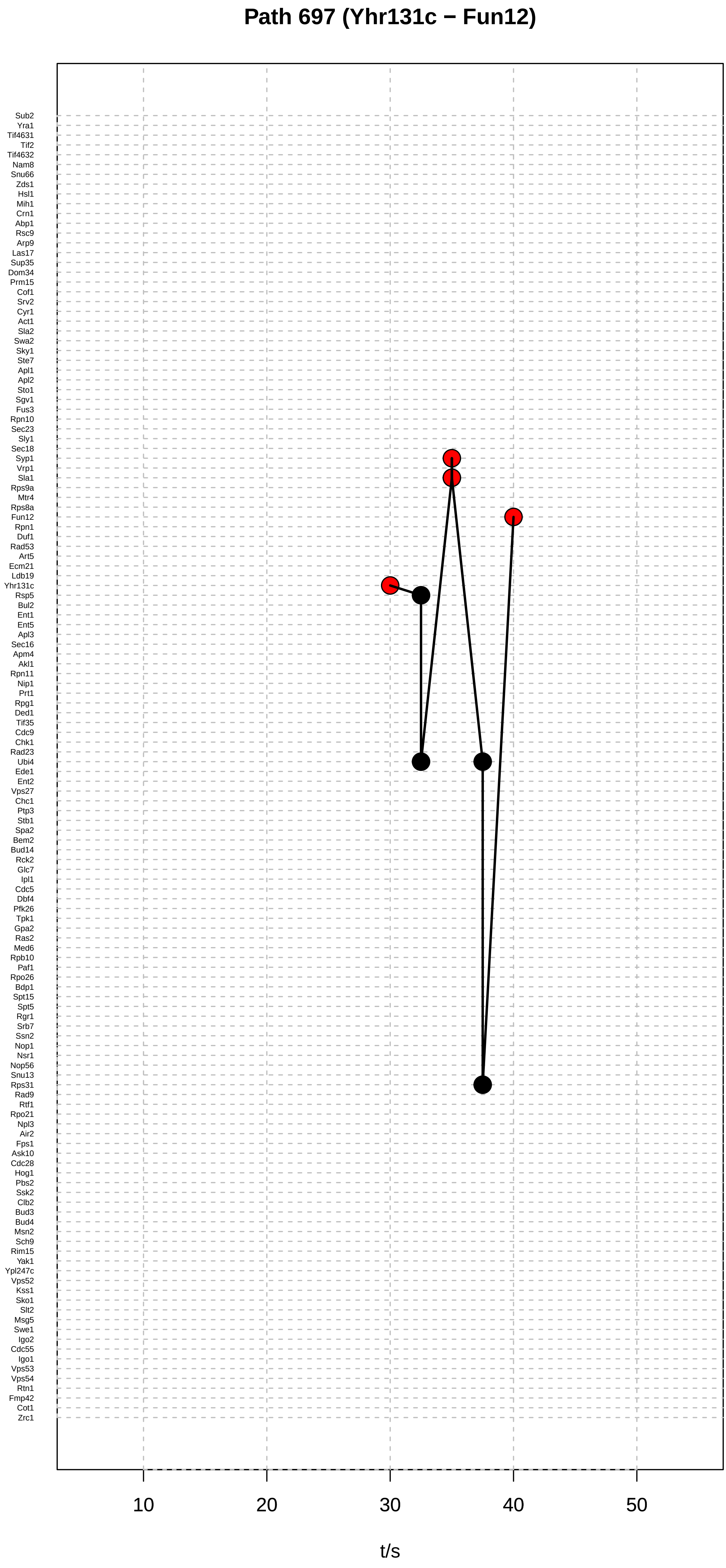

Path 697 (Yhr131c − Fun12)
Sub2
Yra1
Tif4631
Tif2
Tif4632
Nam8
Snu66
Zds1
Hsl1
Mih1
Crn1
Abp1
Rsc9
Arp9
Las17
Sup35
Dom34
Prm15
Cof1
Srv2
Cyr1
Act1
Sla2
Swa2
Sky1
Ste7
Apl1
Apl2
Sto1
Sgv1
Fus3
Rpn10
Sec23
Sly1
Sec18
Syp1
Vrp1
Sla1
Rps9a
Mtr4
Rps8a
Fun12
Rpn1
Duf1
Rad53
Art5
Ecm21
Ldb19
Yhr131c
Rsp5
Bul2
Ent1
Ent5
Apl3
Sec16
Apm4
Akl1
Rpn11
Nip1
Prt1
Rpg1
Ded1
Tif35
Cdc9
Chk1
Rad23
Ubi4
Ede1
Ent2
Vps27
Chc1
Ptp3
Stb1
Spa2
Bem2
Bud14
Rck2
Glc7
Ipl1
Cdc5
Dbf4
Pfk26
Tpk1
Gpa2
Ras2
Med6
Rpb10
Paf1
Rpo26
Bdp1
Spt15
Spt5
Rgr1
Srb7
Ssn2
Nop1
Nsr1
Nop56
Snu13
Rps31
Rad9
Rtf1
Rpo21
Npl3
Air2
Fps1
Ask10
Cdc28
Hog1
Pbs2
Ssk2
Clb2
Bud3
Bud4
Msn2
Sch9
Rim15
Yak1
Ypl247c
Vps52
Kss1
Sko1
Slt2
Msg5
Swe1
Igo2
Cdc55
Igo1
Vps53
Vps54
Rtn1
Fmp42
Cot1
Zrc1
10
20
30
40
50
t/s

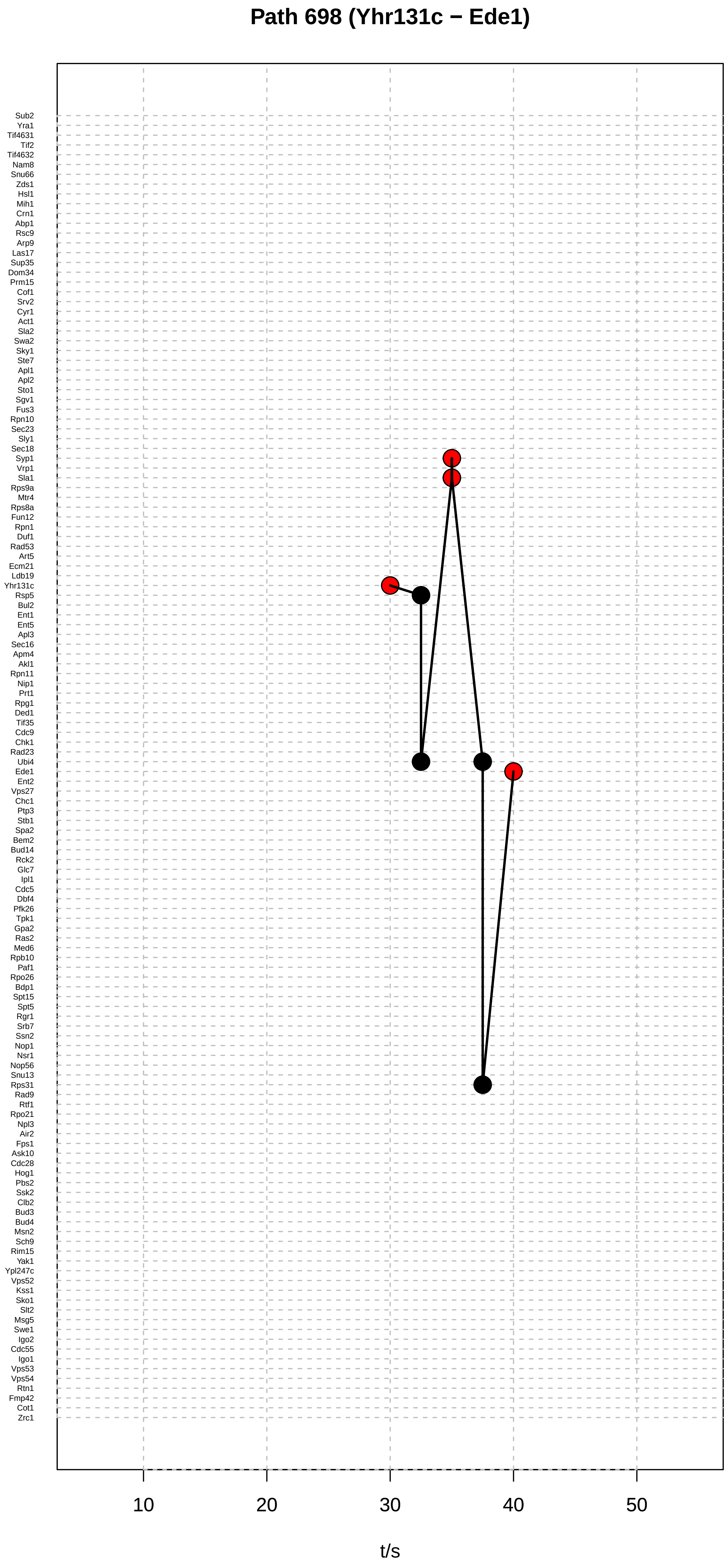

Path 698 (Yhr131c − Ede1)
Sub2
Yra1
Tif4631
Tif2
Tif4632
Nam8
Snu66
Zds1
Hsl1
Mih1
Crn1
Abp1
Rsc9
Arp9
Las17
Sup35
Dom34
Prm15
Cof1
Srv2
Cyr1
Act1
Sla2
Swa2
Sky1
Ste7
Apl1
Apl2
Sto1
Sgv1
Fus3
Rpn10
Sec23
Sly1
Sec18
Syp1
Vrp1
Sla1
Rps9a
Mtr4
Rps8a
Fun12
Rpn1
Duf1
Rad53
Art5
Ecm21
Ldb19
Yhr131c
Rsp5
Bul2
Ent1
Ent5
Apl3
Sec16
Apm4
Akl1
Rpn11
Nip1
Prt1
Rpg1
Ded1
Tif35
Cdc9
Chk1
Rad23
Ubi4
Ede1
Ent2
Vps27
Chc1
Ptp3
Stb1
Spa2
Bem2
Bud14
Rck2
Glc7
Ipl1
Cdc5
Dbf4
Pfk26
Tpk1
Gpa2
Ras2
Med6
Rpb10
Paf1
Rpo26
Bdp1
Spt15
Spt5
Rgr1
Srb7
Ssn2
Nop1
Nsr1
Nop56
Snu13
Rps31
Rad9
Rtf1
Rpo21
Npl3
Air2
Fps1
Ask10
Cdc28
Hog1
Pbs2
Ssk2
Clb2
Bud3
Bud4
Msn2
Sch9
Rim15
Yak1
Ypl247c
Vps52
Kss1
Sko1
Slt2
Msg5
Swe1
Igo2
Cdc55
Igo1
Vps53
Vps54
Rtn1
Fmp42
Cot1
Zrc1
10
20
30
40
50
t/s

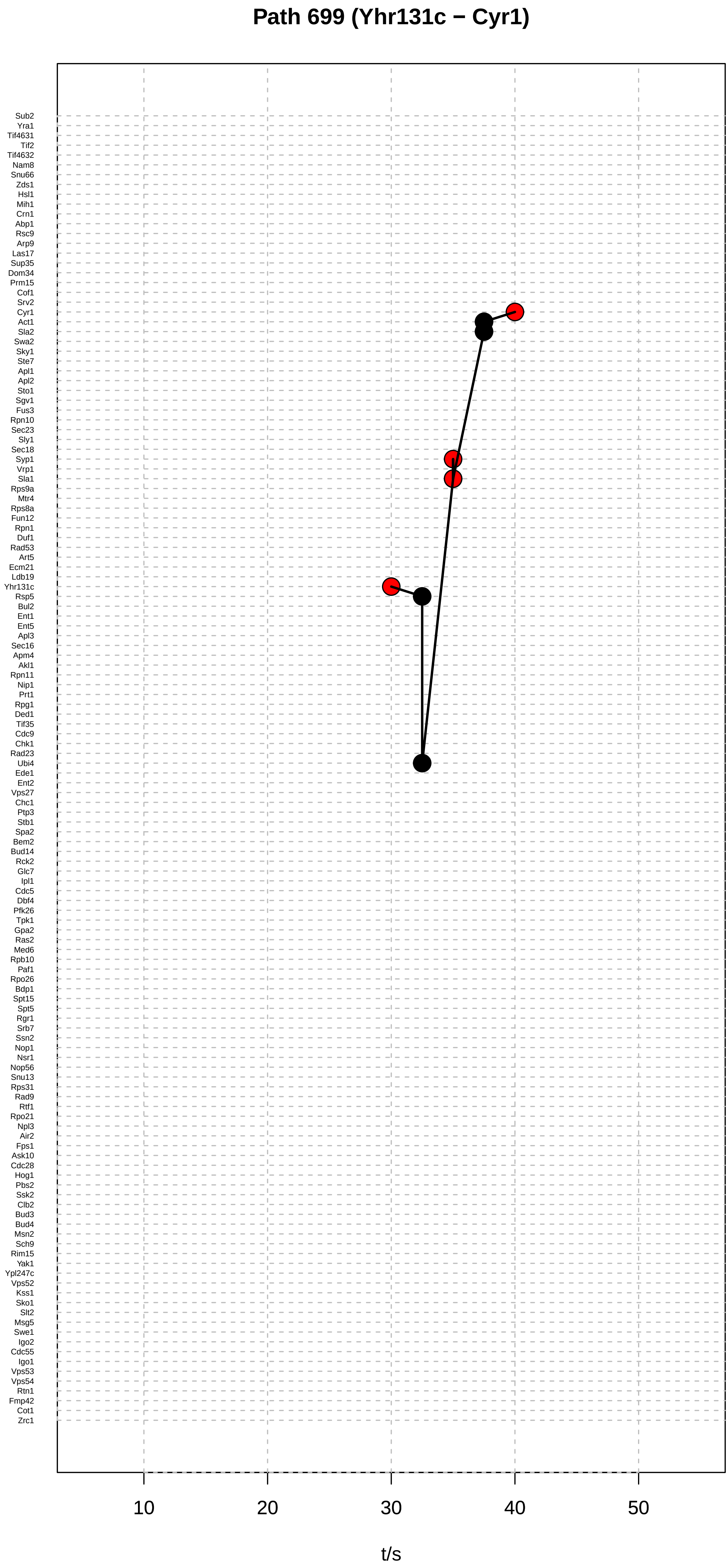

Path 699 (Yhr131c − Cyr1)
Sub2
Yra1
Tif4631
Tif2
Tif4632
Nam8
Snu66
Zds1
Hsl1
Mih1
Crn1
Abp1
Rsc9
Arp9
Las17
Sup35
Dom34
Prm15
Cof1
Srv2
Cyr1
Act1
Sla2
Swa2
Sky1
Ste7
Apl1
Apl2
Sto1
Sgv1
Fus3
Rpn10
Sec23
Sly1
Sec18
Syp1
Vrp1
Sla1
Rps9a
Mtr4
Rps8a
Fun12
Rpn1
Duf1
Rad53
Art5
Ecm21
Ldb19
Yhr131c
Rsp5
Bul2
Ent1
Ent5
Apl3
Sec16
Apm4
Akl1
Rpn11
Nip1
Prt1
Rpg1
Ded1
Tif35
Cdc9
Chk1
Rad23
Ubi4
Ede1
Ent2
Vps27
Chc1
Ptp3
Stb1
Spa2
Bem2
Bud14
Rck2
Glc7
Ipl1
Cdc5
Dbf4
Pfk26
Tpk1
Gpa2
Ras2
Med6
Rpb10
Paf1
Rpo26
Bdp1
Spt15
Spt5
Rgr1
Srb7
Ssn2
Nop1
Nsr1
Nop56
Snu13
Rps31
Rad9
Rtf1
Rpo21
Npl3
Air2
Fps1
Ask10
Cdc28
Hog1
Pbs2
Ssk2
Clb2
Bud3
Bud4
Msn2
Sch9
Rim15
Yak1
Ypl247c
Vps52
Kss1
Sko1
Slt2
Msg5
Swe1
Igo2
Cdc55
Igo1
Vps53
Vps54
Rtn1
Fmp42
Cot1
Zrc1
10
20
30
40
50
t/s

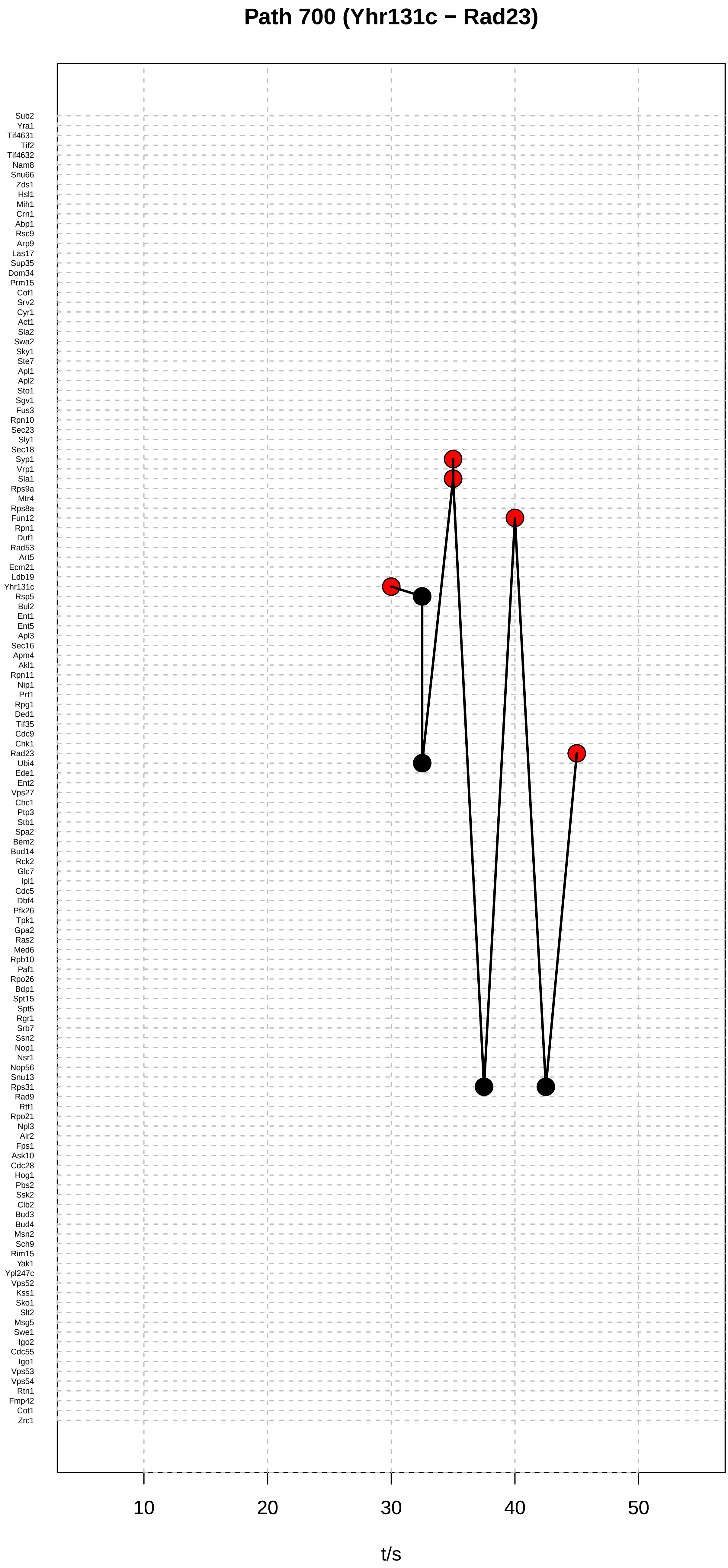

Path 700 (Yhr131c − Rad23)
Sub2
Yra1
Tif4631
Tif2
Tif4632
Nam8
Snu66
Zds1
Hsl1
Mih1
Crn1
Abp1
Rsc9
Arp9
Las17
Sup35
Dom34
Prm15
Cof1
Srv2
Cyr1
Act1
Sla2
Swa2
Sky1
Ste7
Apl1
Apl2
Sto1
Sgv1
Fus3
Rpn10
Sec23
Sly1
Sec18
Syp1
Vrp1
Sla1
Rps9a
Mtr4
Rps8a
Fun12
Rpn1
Duf1
Rad53
Art5
Ecm21
Ldb19
Yhr131c
Rsp5
Bul2
Ent1
Ent5
Apl3
Sec16
Apm4
Akl1
Rpn11
Nip1
Prt1
Rpg1
Ded1
Tif35
Cdc9
Chk1
Rad23
Ubi4
Ede1
Ent2
Vps27
Chc1
Ptp3
Stb1
Spa2
Bem2
Bud14
Rck2
Glc7
Ipl1
Cdc5
Dbf4
Pfk26
Tpk1
Gpa2
Ras2
Med6
Rpb10
Paf1
Rpo26
Bdp1
Spt15
Spt5
Rgr1
Srb7
Ssn2
Nop1
Nsr1
Nop56
Snu13
Rps31
Rad9
Rtf1
Rpo21
Npl3
Air2
Fps1
Ask10
Cdc28
Hog1
Pbs2
Ssk2
Clb2
Bud3
Bud4
Msn2
Sch9
Rim15
Yak1
Ypl247c
Vps52
Kss1
Sko1
Slt2
Msg5
Swe1
Igo2
Cdc55
Igo1
Vps53
Vps54
Rtn1
Fmp42
Cot1
Zrc1
10
20
30
40
50
t/s

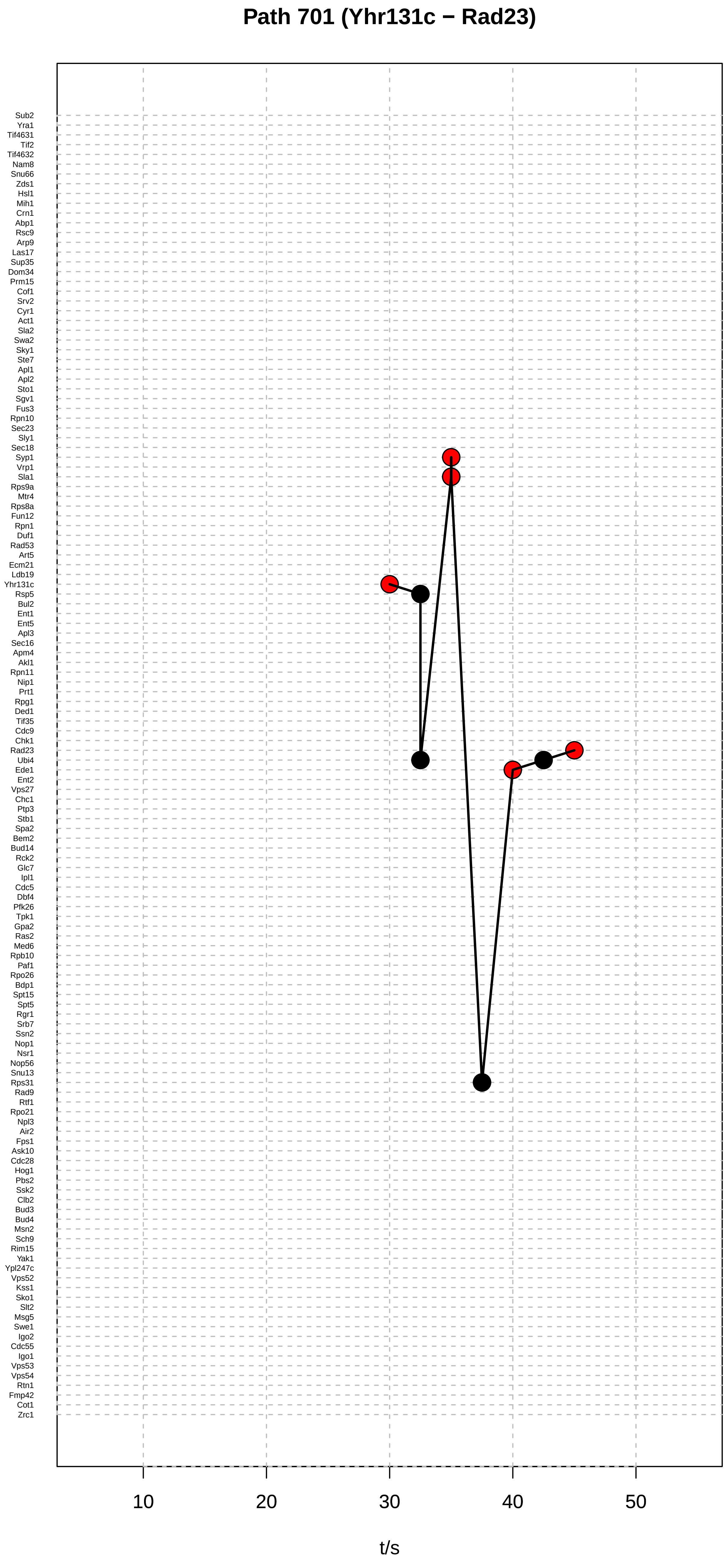

Path 701 (Yhr131c − Rad23)
Sub2
Yra1
Tif4631
Tif2
Tif4632
Nam8
Snu66
Zds1
Hsl1
Mih1
Crn1
Abp1
Rsc9
Arp9
Las17
Sup35
Dom34
Prm15
Cof1
Srv2
Cyr1
Act1
Sla2
Swa2
Sky1
Ste7
Apl1
Apl2
Sto1
Sgv1
Fus3
Rpn10
Sec23
Sly1
Sec18
Syp1
Vrp1
Sla1
Rps9a
Mtr4
Rps8a
Fun12
Rpn1
Duf1
Rad53
Art5
Ecm21
Ldb19
Yhr131c
Rsp5
Bul2
Ent1
Ent5
Apl3
Sec16
Apm4
Akl1
Rpn11
Nip1
Prt1
Rpg1
Ded1
Tif35
Cdc9
Chk1
Rad23
Ubi4
Ede1
Ent2
Vps27
Chc1
Ptp3
Stb1
Spa2
Bem2
Bud14
Rck2
Glc7
Ipl1
Cdc5
Dbf4
Pfk26
Tpk1
Gpa2
Ras2
Med6
Rpb10
Paf1
Rpo26
Bdp1
Spt15
Spt5
Rgr1
Srb7
Ssn2
Nop1
Nsr1
Nop56
Snu13
Rps31
Rad9
Rtf1
Rpo21
Npl3
Air2
Fps1
Ask10
Cdc28
Hog1
Pbs2
Ssk2
Clb2
Bud3
Bud4
Msn2
Sch9
Rim15
Yak1
Ypl247c
Vps52
Kss1
Sko1
Slt2
Msg5
Swe1
Igo2
Cdc55
Igo1
Vps53
Vps54
Rtn1
Fmp42
Cot1
Zrc1
10
20
30
40
50
t/s

### Path 702 (Yhr131c − Rad23)

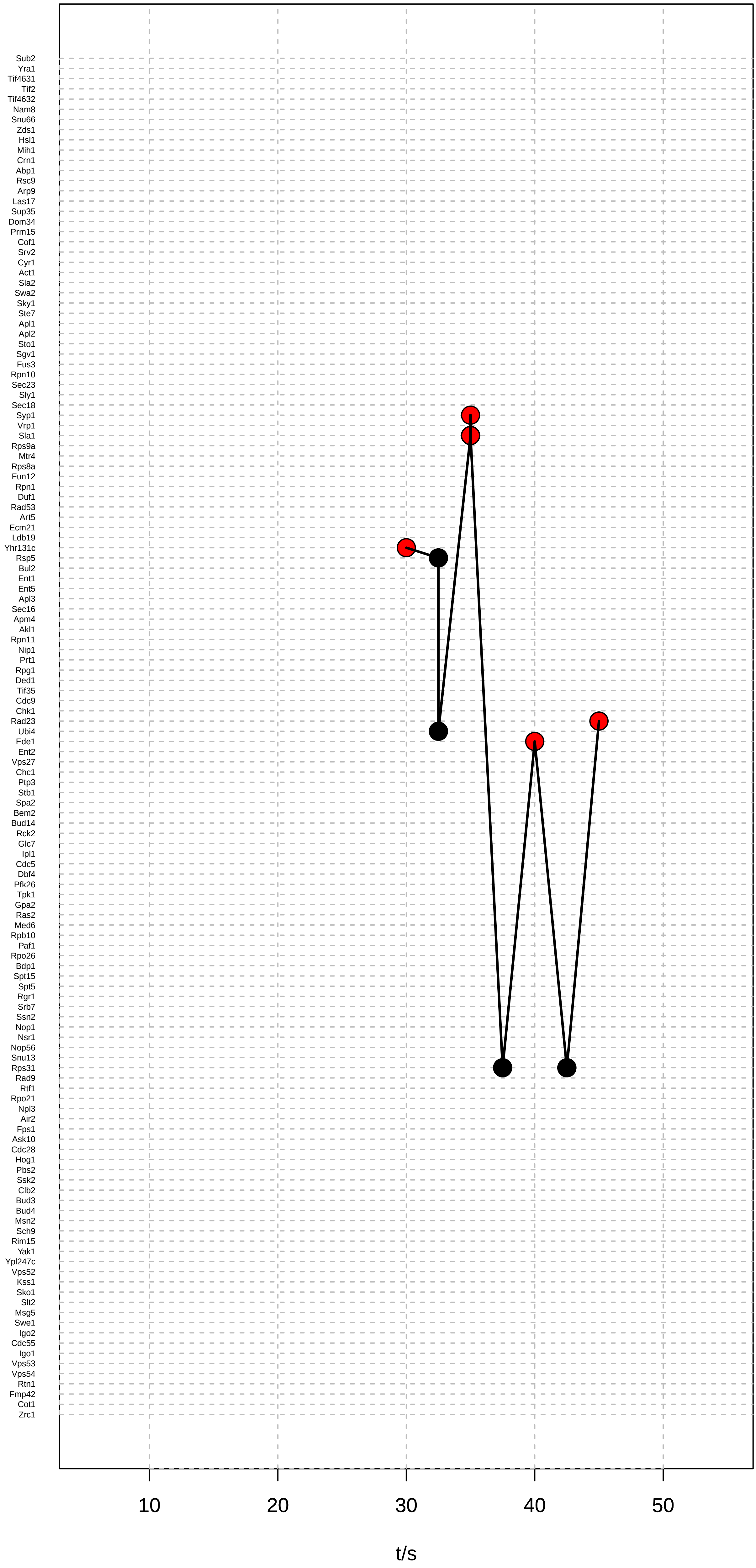

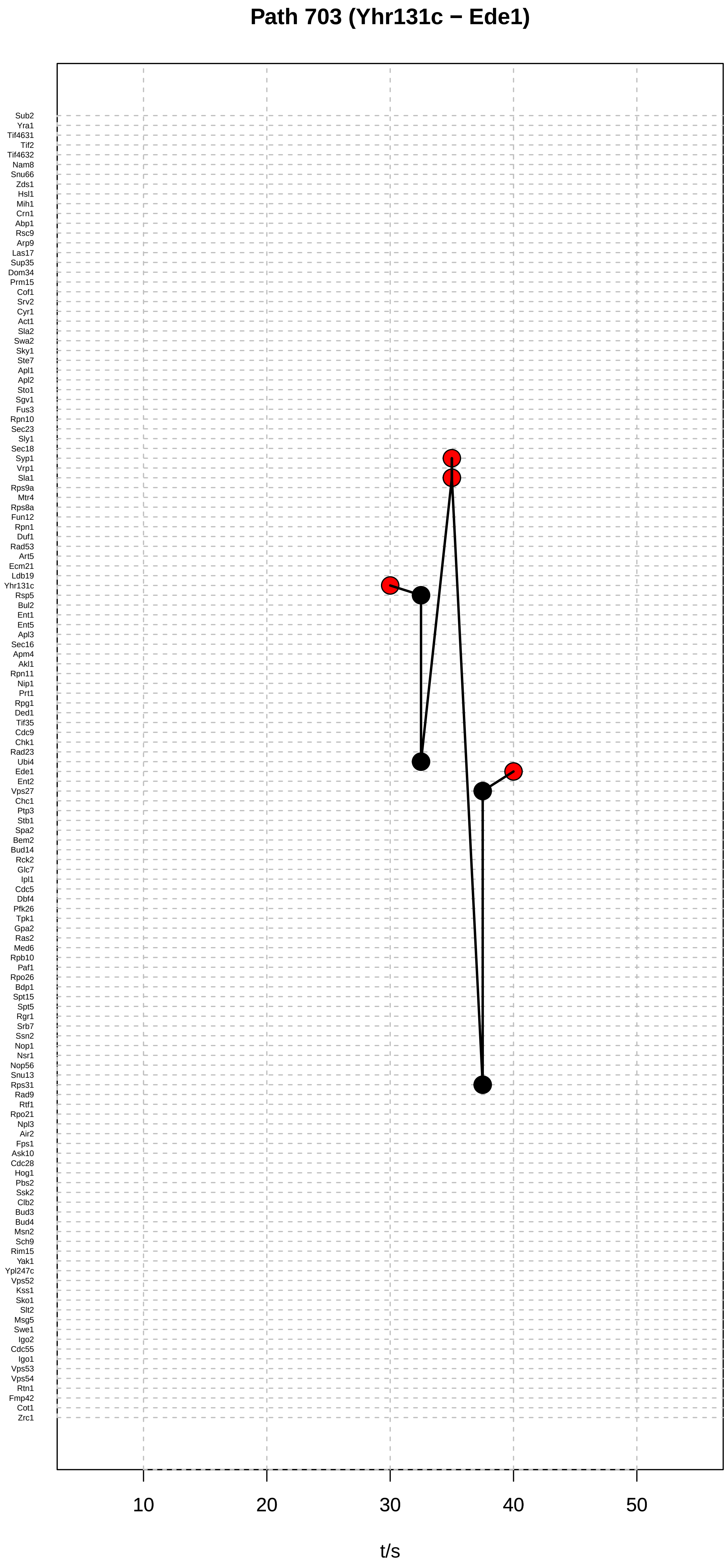

Path 703 (Yhr131c − Ede1)
Sub2
Yra1
Tif4631
Tif2
Tif4632
Nam8
Snu66
Zds1
Hsl1
Mih1
Crn1
Abp1
Rsc9
Arp9
Las17
Sup35
Dom34
Prm15
Cof1
Srv2
Cyr1
Act1
Sla2
Swa2
Sky1
Ste7
Apl1
Apl2
Sto1
Sgv1
Fus3
Rpn10
Sec23
Sly1
Sec18
Syp1
Vrp1
Sla1
Rps9a
Mtr4
Rps8a
Fun12
Rpn1
Duf1
Rad53
Art5
Ecm21
Ldb19
Yhr131c
Rsp5
Bul2
Ent1
Ent5
Apl3
Sec16
Apm4
Akl1
Rpn11
Nip1
Prt1
Rpg1
Ded1
Tif35
Cdc9
Chk1
Rad23
Ubi4
Ede1
Ent2
Vps27
Chc1
Ptp3
Stb1
Spa2
Bem2
Bud14
Rck2
Glc7
Ipl1
Cdc5
Dbf4
Pfk26
Tpk1
Gpa2
Ras2
Med6
Rpb10
Paf1
Rpo26
Bdp1
Spt15
Spt5
Rgr1
Srb7
Ssn2
Nop1
Nsr1
Nop56
Snu13
Rps31
Rad9
Rtf1
Rpo21
Npl3
Air2
Fps1
Ask10
Cdc28
Hog1
Pbs2
Ssk2
Clb2
Bud3
Bud4
Msn2
Sch9
Rim15
Yak1
Ypl247c
Vps52
Kss1
Sko1
Slt2
Msg5
Swe1
Igo2
Cdc55
Igo1
Vps53
Vps54
Rtn1
Fmp42
Cot1
Zrc1
10
20
30
40
50
t/s

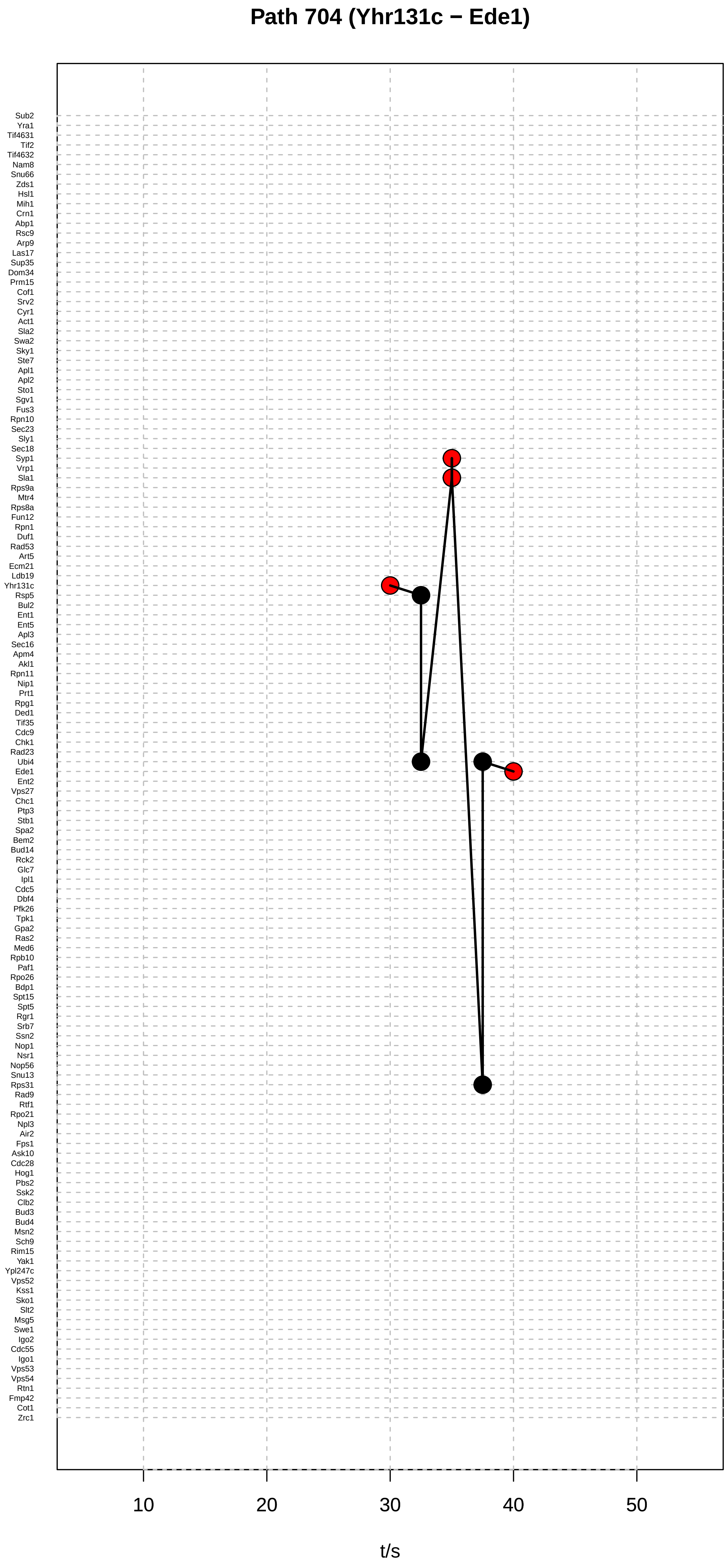

Path 704 (Yhr131c − Ede1)
Sub2
Yra1
Tif4631
Tif2
Tif4632
Nam8
Snu66
Zds1
Hsl1
Mih1
Crn1
Abp1
Rsc9
Arp9
Las17
Sup35
Dom34
Prm15
Cof1
Srv2
Cyr1
Act1
Sla2
Swa2
Sky1
Ste7
Apl1
Apl2
Sto1
Sgv1
Fus3
Rpn10
Sec23
Sly1
Sec18
Syp1
Vrp1
Sla1
Rps9a
Mtr4
Rps8a
Fun12
Rpn1
Duf1
Rad53
Art5
Ecm21
Ldb19
Yhr131c
Rsp5
Bul2
Ent1
Ent5
Apl3
Sec16
Apm4
Akl1
Rpn11
Nip1
Prt1
Rpg1
Ded1
Tif35
Cdc9
Chk1
Rad23
Ubi4
Ede1
Ent2
Vps27
Chc1
Ptp3
Stb1
Spa2
Bem2
Bud14
Rck2
Glc7
Ipl1
Cdc5
Dbf4
Pfk26
Tpk1
Gpa2
Ras2
Med6
Rpb10
Paf1
Rpo26
Bdp1
Spt15
Spt5
Rgr1
Srb7
Ssn2
Nop1
Nsr1
Nop56
Snu13
Rps31
Rad9
Rtf1
Rpo21
Npl3
Air2
Fps1
Ask10
Cdc28
Hog1
Pbs2
Ssk2
Clb2
Bud3
Bud4
Msn2
Sch9
Rim15
Yak1
Ypl247c
Vps52
Kss1
Sko1
Slt2
Msg5
Swe1
Igo2
Cdc55
Igo1
Vps53
Vps54
Rtn1
Fmp42
Cot1
Zrc1
10
20
30
40
50
t/s

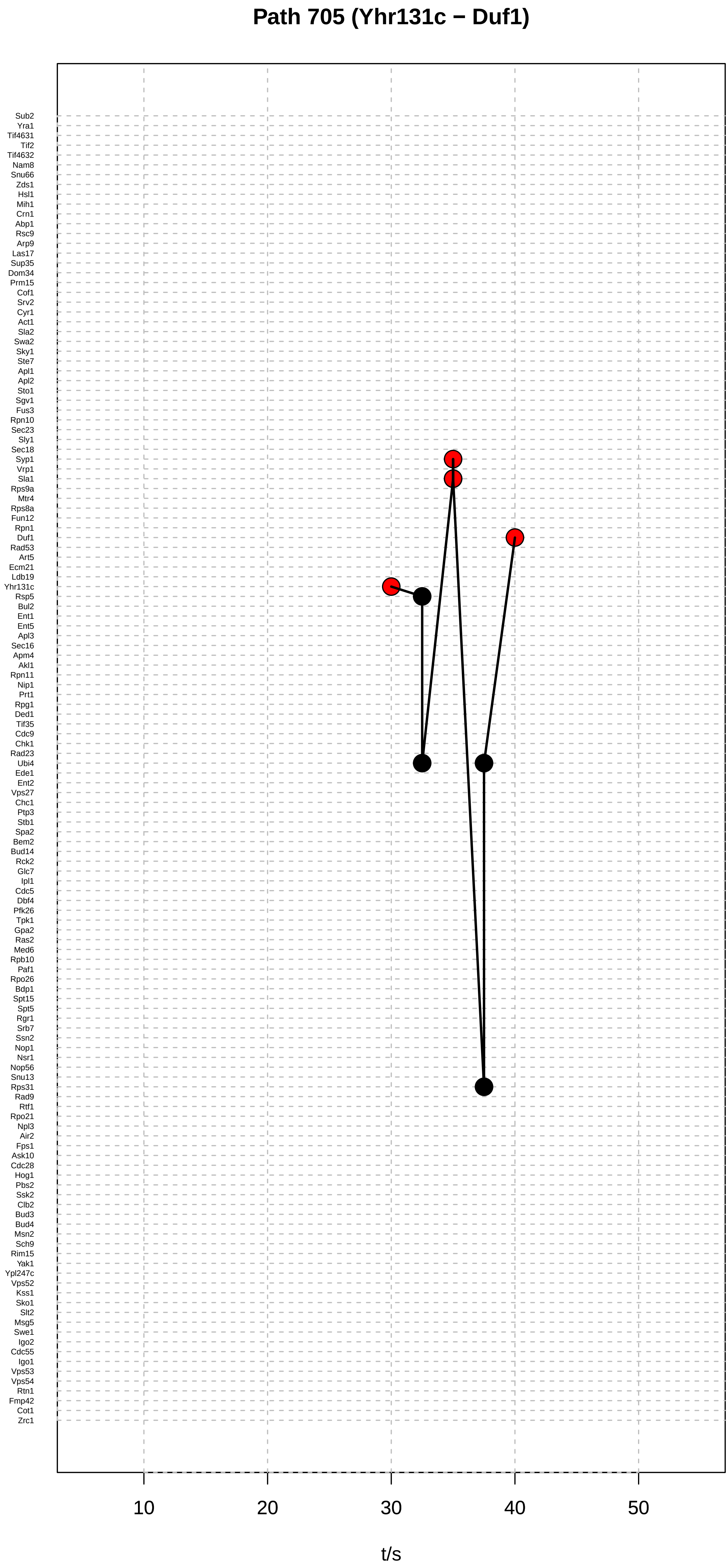

Path 705 (Yhr131c − Duf1)
Sub2
Yra1
Tif4631
Tif2
Tif4632
Nam8
Snu66
Zds1
Hsl1
Mih1
Crn1
Abp1
Rsc9
Arp9
Las17
Sup35
Dom34
Prm15
Cof1
Srv2
Cyr1
Act1
Sla2
Swa2
Sky1
Ste7
Apl1
Apl2
Sto1
Sgv1
Fus3
Rpn10
Sec23
Sly1
Sec18
Syp1
Vrp1
Sla1
Rps9a
Mtr4
Rps8a
Fun12
Rpn1
Duf1
Rad53
Art5
Ecm21
Ldb19
Yhr131c
Rsp5
Bul2
Ent1
Ent5
Apl3
Sec16
Apm4
Akl1
Rpn11
Nip1
Prt1
Rpg1
Ded1
Tif35
Cdc9
Chk1
Rad23
Ubi4
Ede1
Ent2
Vps27
Chc1
Ptp3
Stb1
Spa2
Bem2
Bud14
Rck2
Glc7
Ipl1
Cdc5
Dbf4
Pfk26
Tpk1
Gpa2
Ras2
Med6
Rpb10
Paf1
Rpo26
Bdp1
Spt15
Spt5
Rgr1
Srb7
Ssn2
Nop1
Nsr1
Nop56
Snu13
Rps31
Rad9
Rtf1
Rpo21
Npl3
Air2
Fps1
Ask10
Cdc28
Hog1
Pbs2
Ssk2
Clb2
Bud3
Bud4
Msn2
Sch9
Rim15
Yak1
Ypl247c
Vps52
Kss1
Sko1
Slt2
Msg5
Swe1
Igo2
Cdc55
Igo1
Vps53
Vps54
Rtn1
Fmp42
Cot1
Zrc1
10
20
30
40
50
t/s

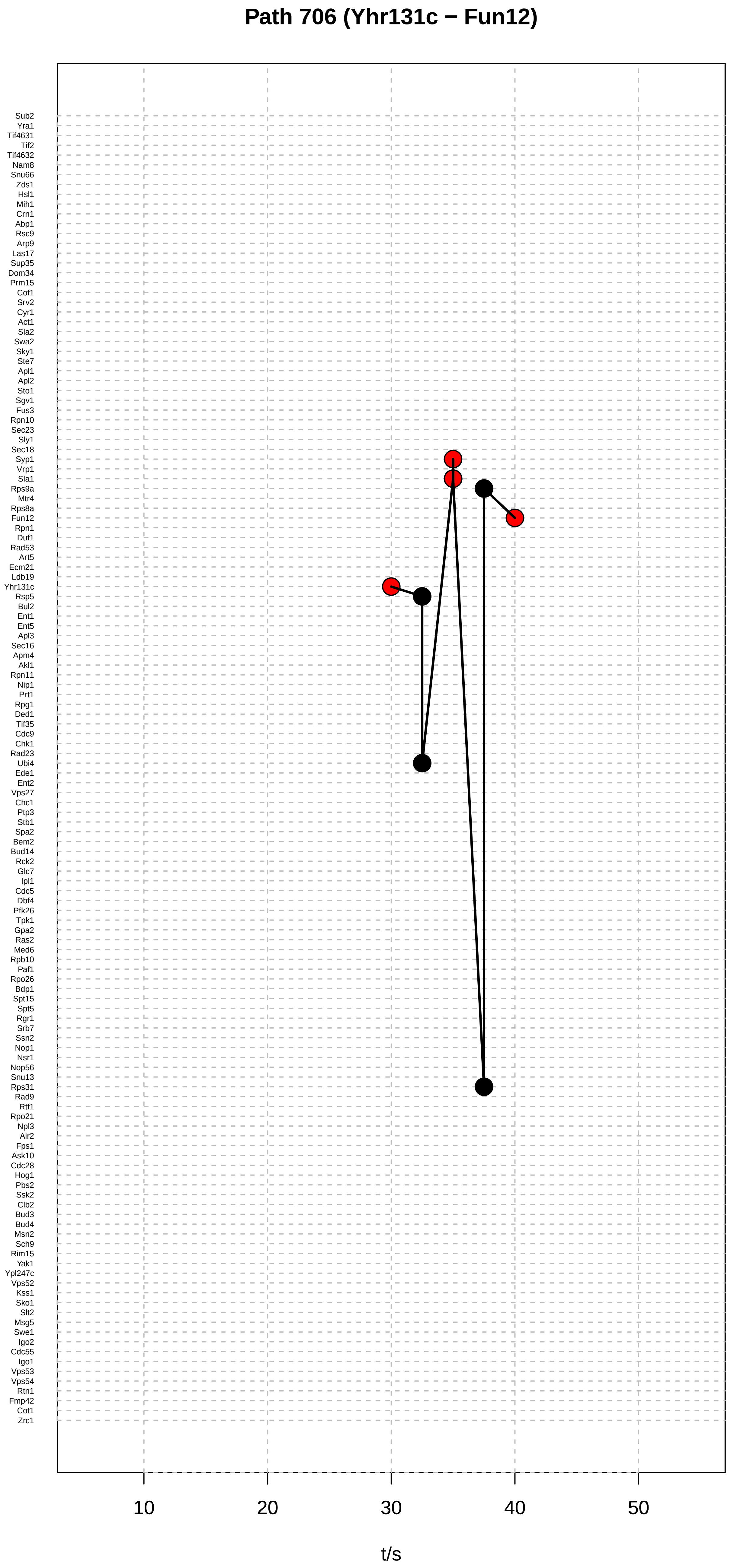

Path 706 (Yhr131c − Fun12)
Sub2
Yra1
Tif4631
Tif2
Tif4632
Nam8
Snu66
Zds1
Hsl1
Mih1
Crn1
Abp1
Rsc9
Arp9
Las17
Sup35
Dom34
Prm15
Cof1
Srv2
Cyr1
Act1
Sla2
Swa2
Sky1
Ste7
Apl1
Apl2
Sto1
Sgv1
Fus3
Rpn10
Sec23
Sly1
Sec18
Syp1
Vrp1
Sla1
Rps9a
Mtr4
Rps8a
Fun12
Rpn1
Duf1
Rad53
Art5
Ecm21
Ldb19
Yhr131c
Rsp5
Bul2
Ent1
Ent5
Apl3
Sec16
Apm4
Akl1
Rpn11
Nip1
Prt1
Rpg1
Ded1
Tif35
Cdc9
Chk1
Rad23
Ubi4
Ede1
Ent2
Vps27
Chc1
Ptp3
Stb1
Spa2
Bem2
Bud14
Rck2
Glc7
Ipl1
Cdc5
Dbf4
Pfk26
Tpk1
Gpa2
Ras2
Med6
Rpb10
Paf1
Rpo26
Bdp1
Spt15
Spt5
Rgr1
Srb7
Ssn2
Nop1
Nsr1
Nop56
Snu13
Rps31
Rad9
Rtf1
Rpo21
Npl3
Air2
Fps1
Ask10
Cdc28
Hog1
Pbs2
Ssk2
Clb2
Bud3
Bud4
Msn2
Sch9
Rim15
Yak1
Ypl247c
Vps52
Kss1
Sko1
Slt2
Msg5
Swe1
Igo2
Cdc55
Igo1
Vps53
Vps54
Rtn1
Fmp42
Cot1
Zrc1
10
20
30
40
50
t/s

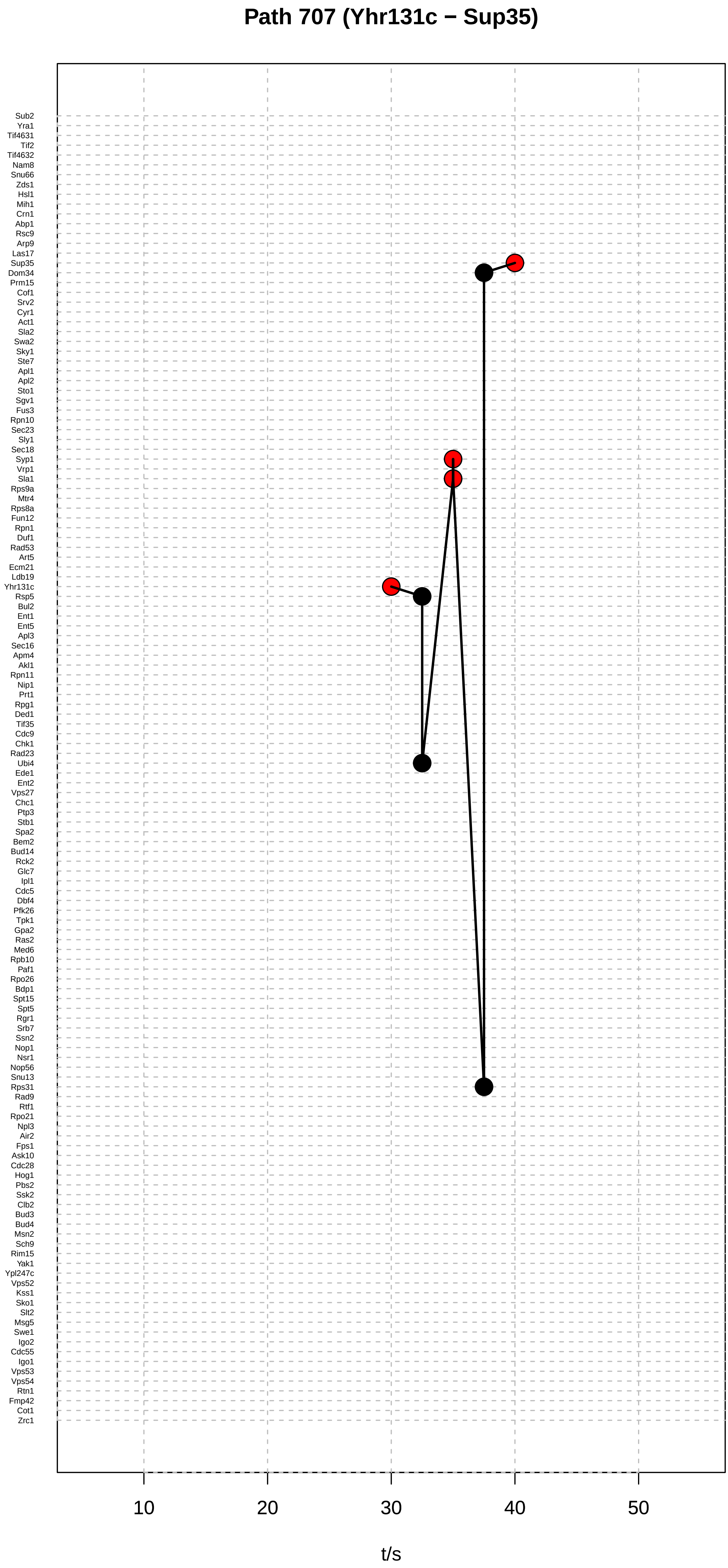
Path 707 (Yhr131c − Sup35)
Sub2
Yra1
Tif4631
Tif2
Tif4632
Nam8
Snu66
Zds1
Hsl1
Mih1
Crn1
Abp1
Rsc9
Arp9
Las17
Sup35
Dom34
Prm15
Cof1
Srv2
Cyr1
Act1
Sla2
Swa2
Sky1
Ste7
Apl1
Apl2
Sto1
Sgv1
Fus3
Rpn10
Sec23
Sly1
Sec18
Syp1
Vrp1
Sla1
Rps9a
Mtr4
Rps8a
Fun12
Rpn1
Duf1
Rad53
Art5
Ecm21
Ldb19
Yhr131c
Rsp5
Bul2
Ent1
Ent5
Apl3
Sec16
Apm4
Akl1
Rpn11
Nip1
Prt1
Rpg1
Ded1
Tif35
Cdc9
Chk1
Rad23
Ubi4
Ede1
Ent2
Vps27
Chc1
Ptp3
Stb1
Spa2
Bem2
Bud14
Rck2
Glc7
Ipl1
Cdc5
Dbf4
Pfk26
Tpk1
Gpa2
Ras2
Med6
Rpb10
Paf1
Rpo26
Bdp1
Spt15
Spt5
Rgr1
Srb7
Ssn2
Nop1
Nsr1
Nop56
Snu13
Rps31
Rad9
Rtf1
Rpo21
Npl3
Air2
Fps1
Ask10
Cdc28
Hog1
Pbs2
Ssk2
Clb2
Bud3
Bud4
Msn2
Sch9
Rim15
Yak1
Ypl247c
Vps52
Kss1
Sko1
Slt2
Msg5
Swe1
Igo2
Cdc55
Igo1
Vps53
Vps54
Rtn1
Fmp42
Cot1
Zrc1
10
20
30
40
50
t/s

### Path 708 (Yhr131c − Cyr1)

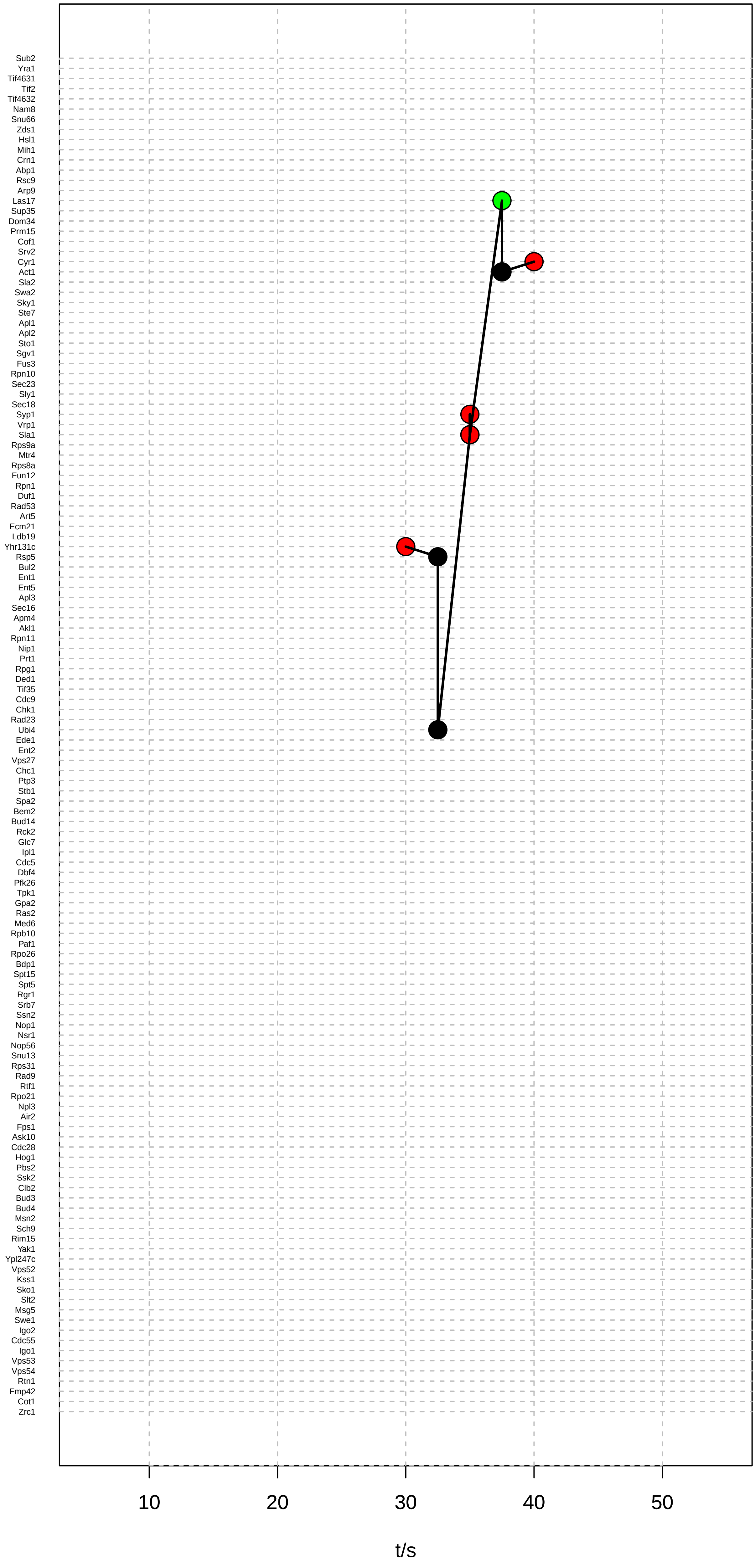

#### Path 709 (Yhr131c − Rad23)

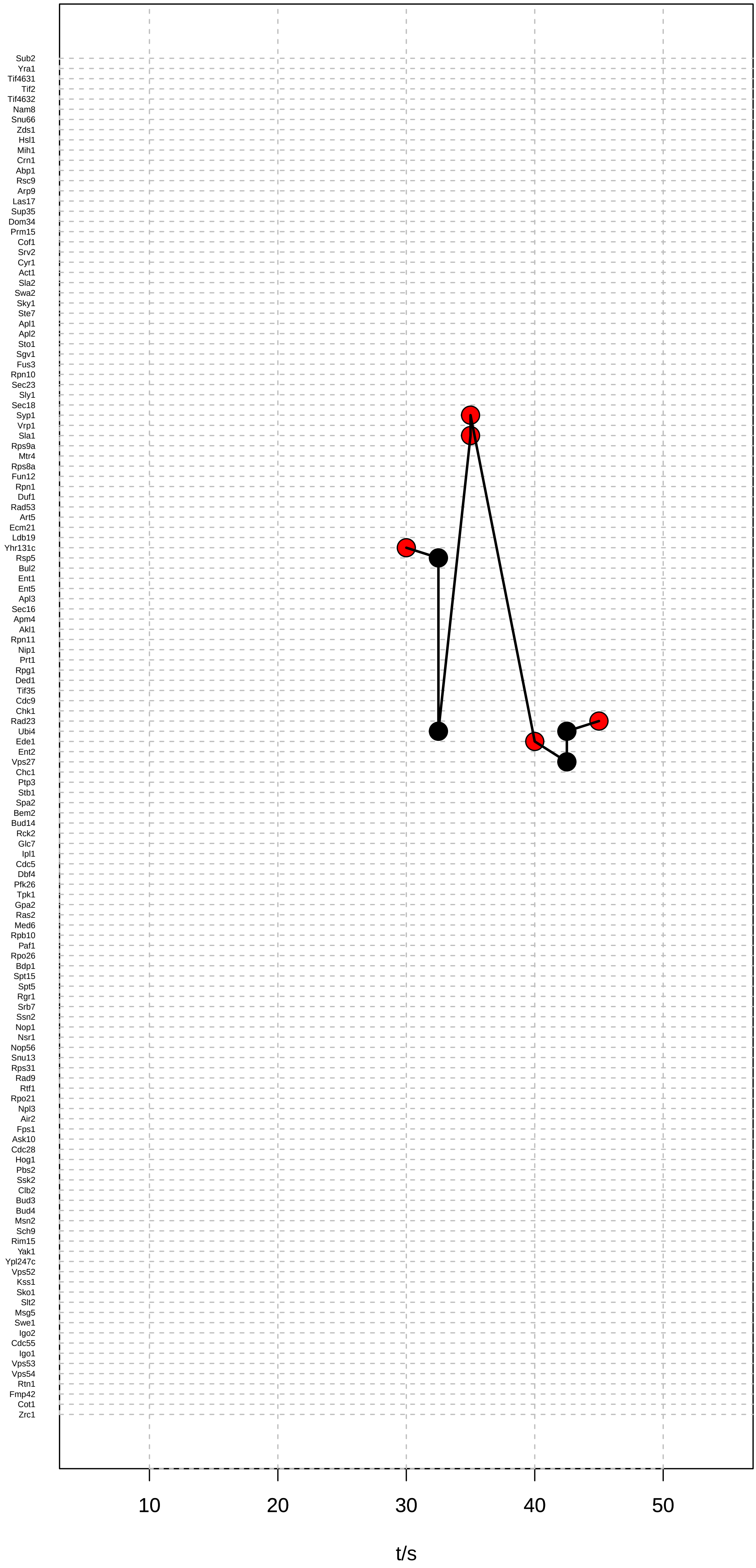

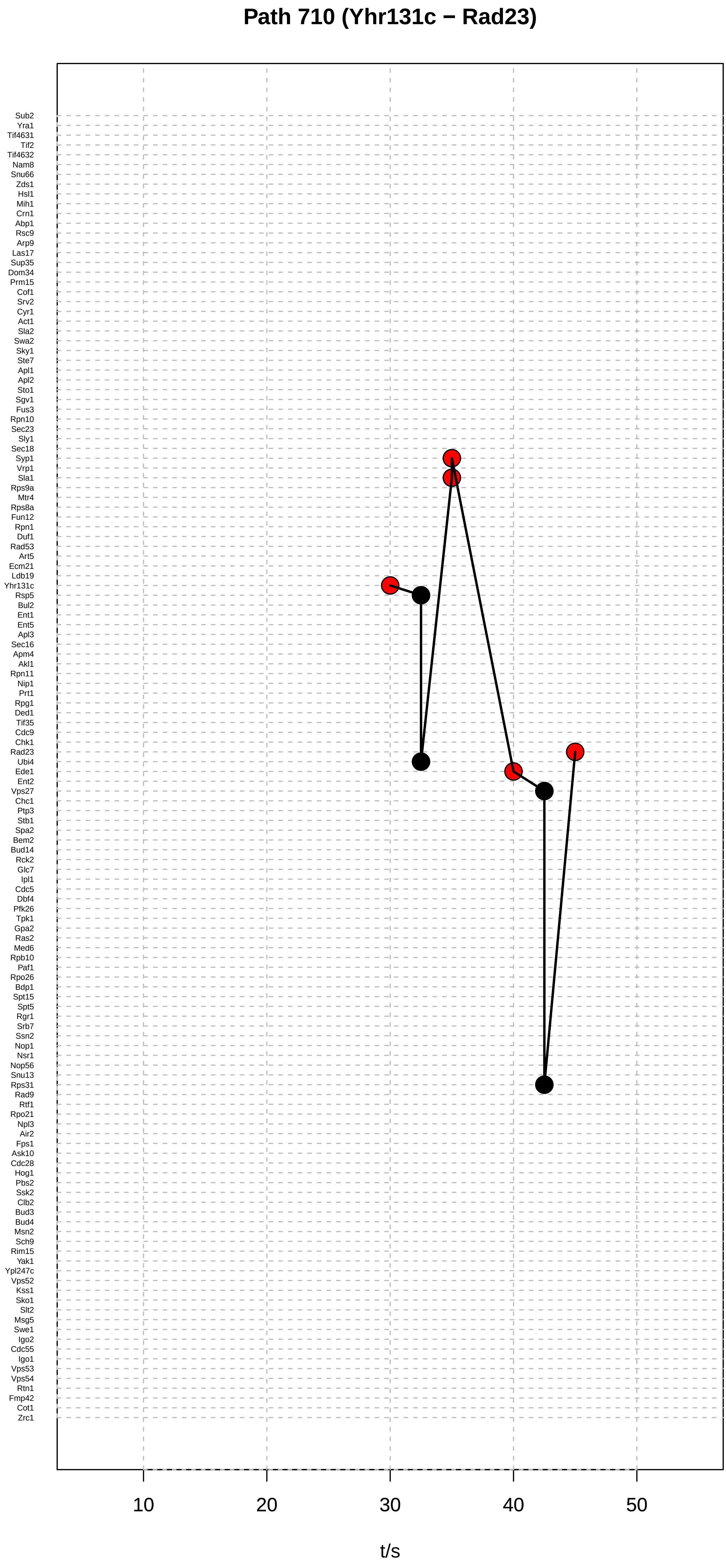
Path 710 (Yhr131c − Rad23)
Sub2
Yra1
Tif4631
Tif2
Tif4632
Nam8
Snu66
Zds1
Hsl1
Mih1
Crn1
Abp1
Rsc9
Arp9
Las17
Sup35
Dom34
Prm15
Cof1
Srv2
Cyr1
Act1
Sla2
Swa2
Sky1
Ste7
Apl1
Apl2
Sto1
Sgv1
Fus3
Rpn10
Sec23
Sly1
Sec18
Syp1
Vrp1
Sla1
Rps9a
Mtr4
Rps8a
Fun12
Rpn1
Duf1
Rad53
Art5
Ecm21
Ldb19
Yhr131c
Rsp5
Bul2
Ent1
Ent5
Apl3
Sec16
Apm4
Akl1
Rpn11
Nip1
Prt1
Rpg1
Ded1
Tif35
Cdc9
Chk1
Rad23
Ubi4
Ede1
Ent2
Vps27
Chc1
Ptp3
Stb1
Spa2
Bem2
Bud14
Rck2
Glc7
Ipl1
Cdc5
Dbf4
Pfk26
Tpk1
Gpa2
Ras2
Med6
Rpb10
Paf1
Rpo26
Bdp1
Spt15
Spt5
Rgr1
Srb7
Ssn2
Nop1
Nsr1
Nop56
Snu13
Rps31
Rad9
Rtf1
Rpo21
Npl3
Air2
Fps1
Ask10
Cdc28
Hog1
Pbs2
Ssk2
Clb2
Bud3
Bud4
Msn2
Sch9
Rim15
Yak1
Ypl247c
Vps52
Kss1
Sko1
Slt2
Msg5
Swe1
Igo2
Cdc55
Igo1
Vps53
Vps54
Rtn1
Fmp42
Cot1
Zrc1
10
20
30
40
50
t/s

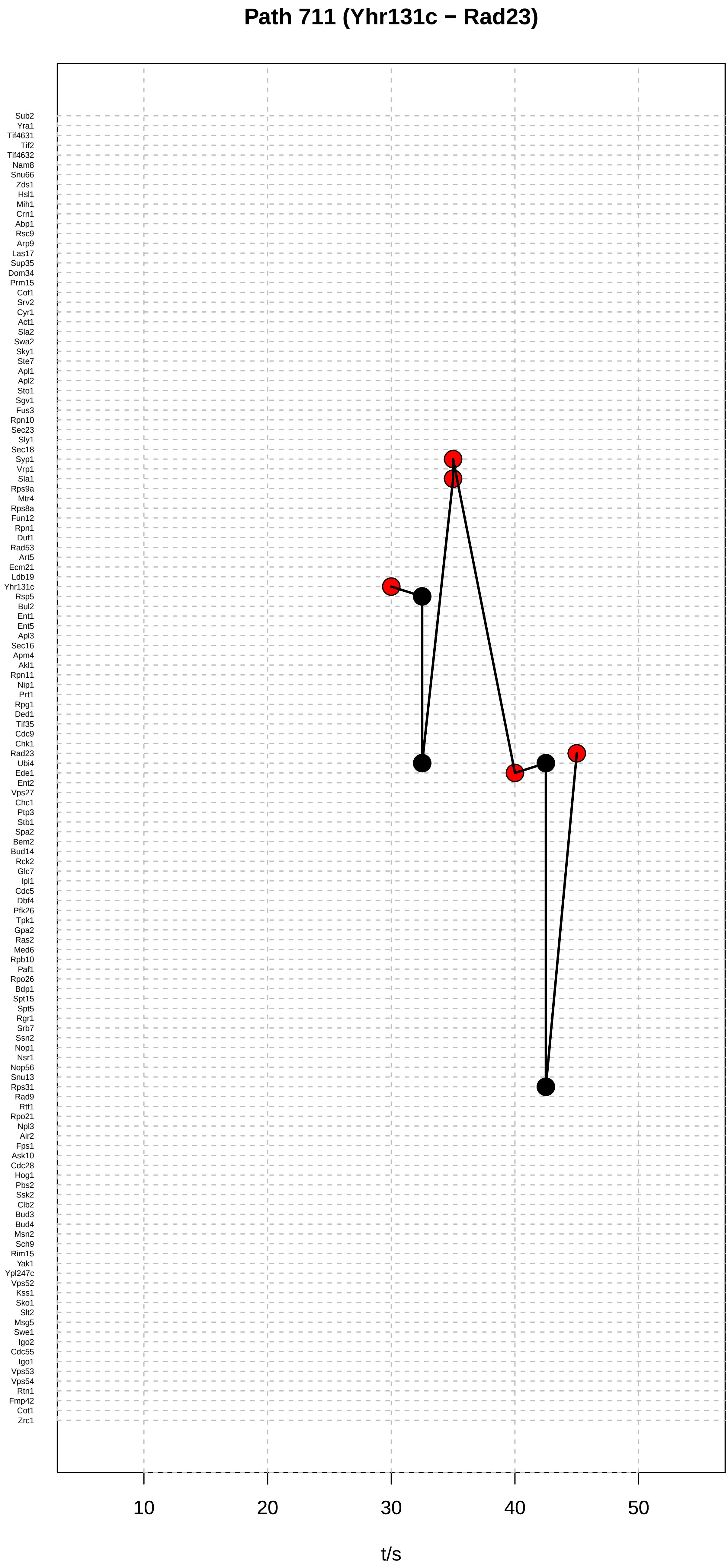

Path 711 (Yhr131c − Rad23)
Sub2
Yra1
Tif4631
Tif2
Tif4632
Nam8
Snu66
Zds1
Hsl1
Mih1
Crn1
Abp1
Rsc9
Arp9
Las17
Sup35
Dom34
Prm15
Cof1
Srv2
Cyr1
Act1
Sla2
Swa2
Sky1
Ste7
Apl1
Apl2
Sto1
Sgv1
Fus3
Rpn10
Sec23
Sly1
Sec18
Syp1
Vrp1
Sla1
Rps9a
Mtr4
Rps8a
Fun12
Rpn1
Duf1
Rad53
Art5
Ecm21
Ldb19
Yhr131c
Rsp5
Bul2
Ent1
Ent5
Apl3
Sec16
Apm4
Akl1
Rpn11
Nip1
Prt1
Rpg1
Ded1
Tif35
Cdc9
Chk1
Rad23
Ubi4
Ede1
Ent2
Vps27
Chc1
Ptp3
Stb1
Spa2
Bem2
Bud14
Rck2
Glc7
Ipl1
Cdc5
Dbf4
Pfk26
Tpk1
Gpa2
Ras2
Med6
Rpb10
Paf1
Rpo26
Bdp1
Spt15
Spt5
Rgr1
Srb7
Ssn2
Nop1
Nsr1
Nop56
Snu13
Rps31
Rad9
Rtf1
Rpo21
Npl3
Air2
Fps1
Ask10
Cdc28
Hog1
Pbs2
Ssk2
Clb2
Bud3
Bud4
Msn2
Sch9
Rim15
Yak1
Ypl247c
Vps52
Kss1
Sko1
Slt2
Msg5
Swe1
Igo2
Cdc55
Igo1
Vps53
Vps54
Rtn1
Fmp42
Cot1
Zrc1
10
20
30
40
50
t/s

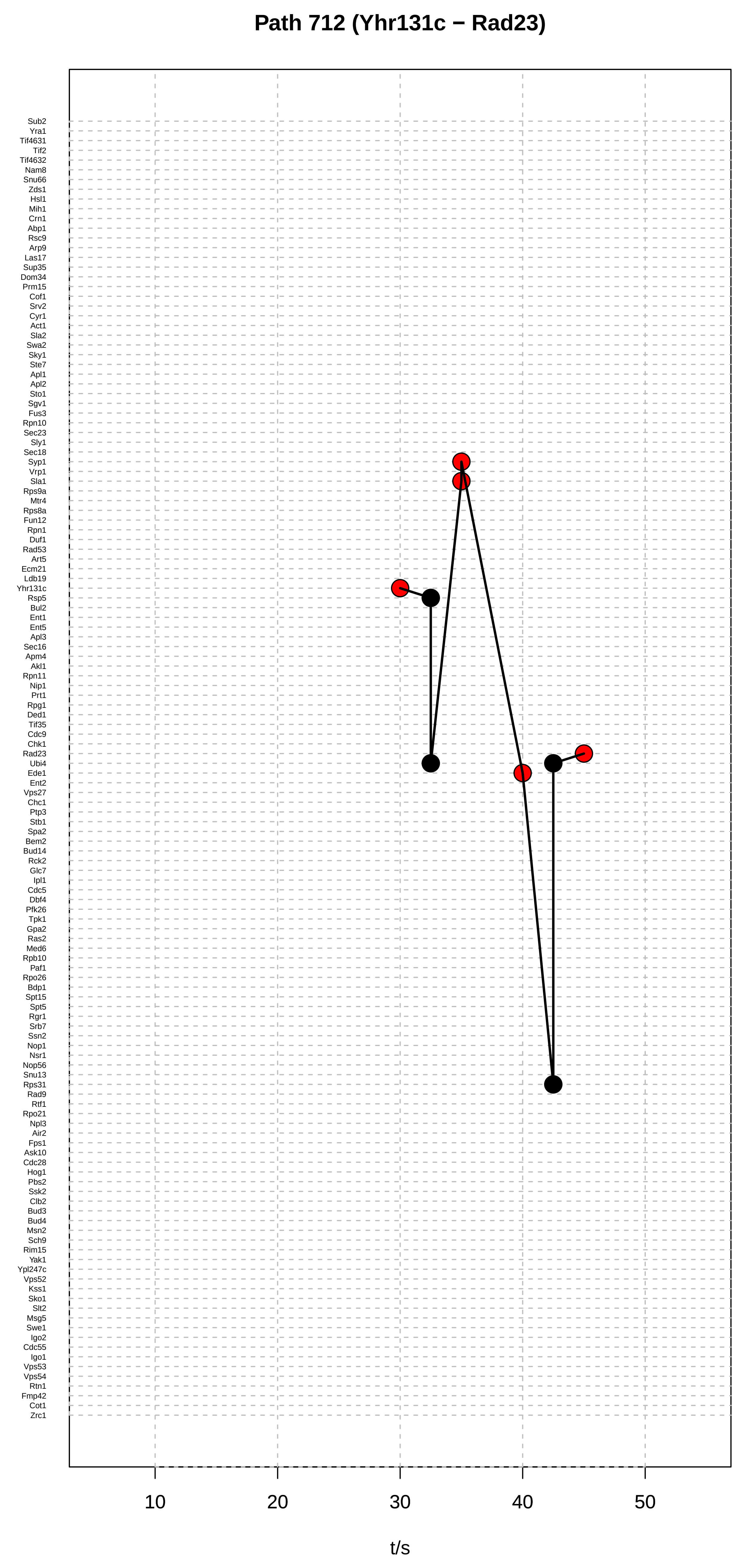
Path 712 (Yhr131c − Rad23)
Sub2
Yra1
Tif4631
Tif2
Tif4632
Nam8
Snu66
Zds1
Hsl1
Mih1
Crn1
Abp1
Rsc9
Arp9
Las17
Sup35
Dom34
Prm15
Cof1
Srv2
Cyr1
Act1
Sla2
Swa2
Sky1
Ste7
Apl1
Apl2
Sto1
Sgv1
Fus3
Rpn10
Sec23
Sly1
Sec18
Syp1
Vrp1
Sla1
Rps9a
Mtr4
Rps8a
Fun12
Rpn1
Duf1
Rad53
Art5
Ecm21
Ldb19
Yhr131c
Rsp5
Bul2
Ent1
Ent5
Apl3
Sec16
Apm4
Akl1
Rpn11
Nip1
Prt1
Rpg1
Ded1
Tif35
Cdc9
Chk1
Rad23
Ubi4
Ede1
Ent2
Vps27
Chc1
Ptp3
Stb1
Spa2
Bem2
Bud14
Rck2
Glc7
Ipl1
Cdc5
Dbf4
Pfk26
Tpk1
Gpa2
Ras2
Med6
Rpb10
Paf1
Rpo26
Bdp1
Spt15
Spt5
Rgr1
Srb7
Ssn2
Nop1
Nsr1
Nop56
Snu13
Rps31
Rad9
Rtf1
Rpo21
Npl3
Air2
Fps1
Ask10
Cdc28
Hog1
Pbs2
Ssk2
Clb2
Bud3
Bud4
Msn2
Sch9
Rim15
Yak1
Ypl247c
Vps52
Kss1
Sko1
Slt2
Msg5
Swe1
Igo2
Cdc55
Igo1
Vps53
Vps54
Rtn1
Fmp42
Cot1
Zrc1
10
20
30
40
50
t/s

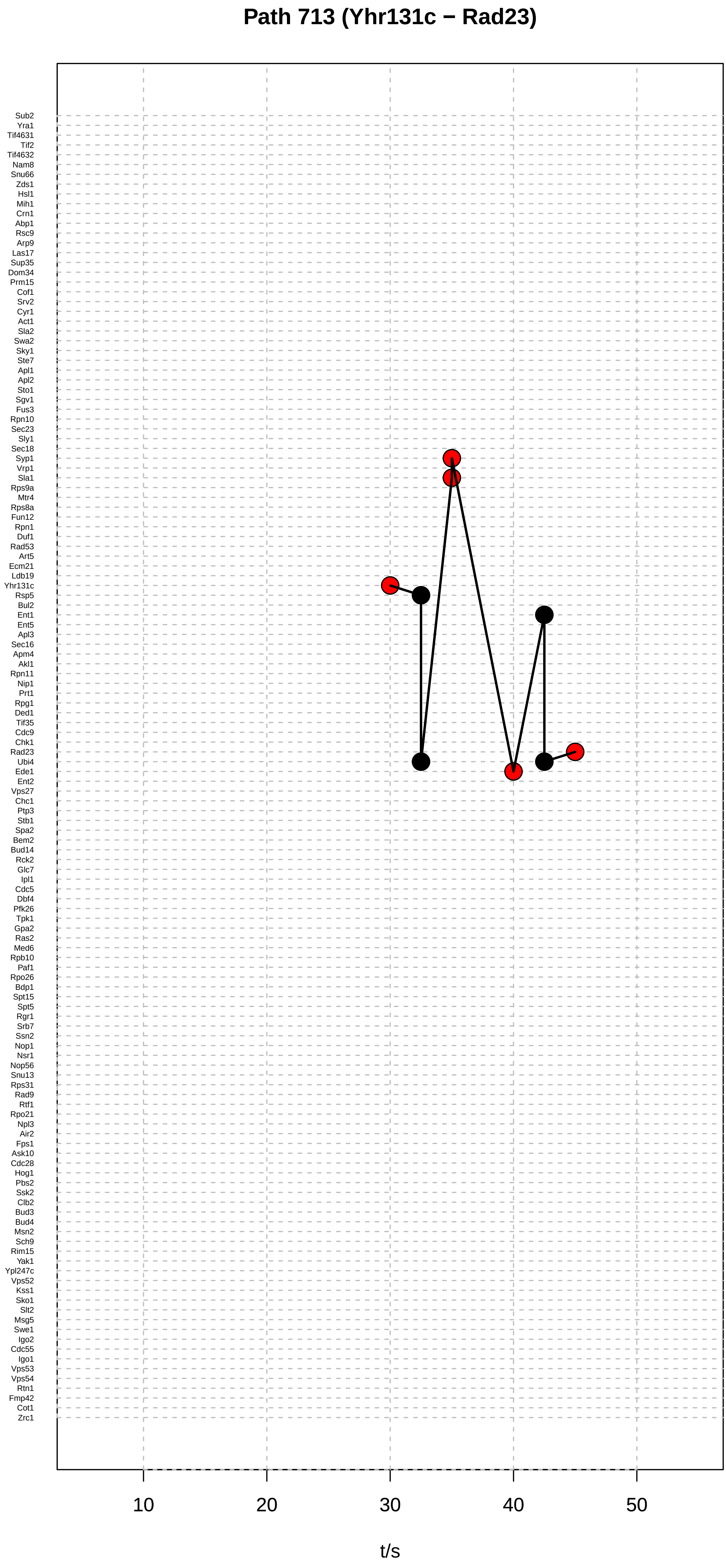

Path 713 (Yhr131c − Rad23)
Sub2
Yra1
Tif4631
Tif2
Tif4632
Nam8
Snu66
Zds1
Hsl1
Mih1
Crn1
Abp1
Rsc9
Arp9
Las17
Sup35
Dom34
Prm15
Cof1
Srv2
Cyr1
Act1
Sla2
Swa2
Sky1
Ste7
Apl1
Apl2
Sto1
Sgv1
Fus3
Rpn10
Sec23
Sly1
Sec18
Syp1
Vrp1
Sla1
Rps9a
Mtr4
Rps8a
Fun12
Rpn1
Duf1
Rad53
Art5
Ecm21
Ldb19
Yhr131c
Rsp5
Bul2
Ent1
Ent5
Apl3
Sec16
Apm4
Akl1
Rpn11
Nip1
Prt1
Rpg1
Ded1
Tif35
Cdc9
Chk1
Rad23
Ubi4
Ede1
Ent2
Vps27
Chc1
Ptp3
Stb1
Spa2
Bem2
Bud14
Rck2
Glc7
Ipl1
Cdc5
Dbf4
Pfk26
Tpk1
Gpa2
Ras2
Med6
Rpb10
Paf1
Rpo26
Bdp1
Spt15
Spt5
Rgr1
Srb7
Ssn2
Nop1
Nsr1
Nop56
Snu13
Rps31
Rad9
Rtf1
Rpo21
Npl3
Air2
Fps1
Ask10
Cdc28
Hog1
Pbs2
Ssk2
Clb2
Bud3
Bud4
Msn2
Sch9
Rim15
Yak1
Ypl247c
Vps52
Kss1
Sko1
Slt2
Msg5
Swe1
Igo2
Cdc55
Igo1
Vps53
Vps54
Rtn1
Fmp42
Cot1
Zrc1
10
20
30
40
50
t/s

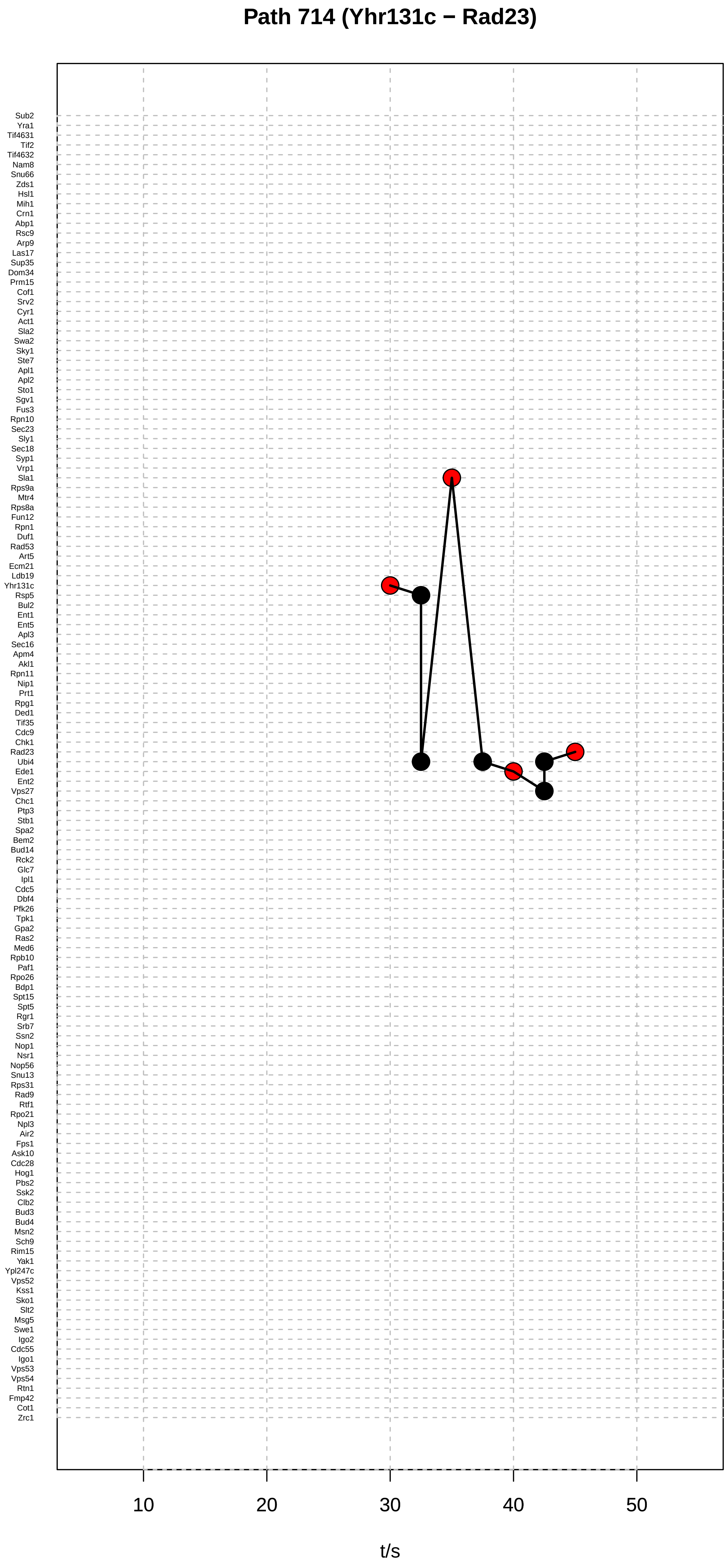

Path 714 (Yhr131c − Rad23)
Sub2
Yra1
Tif4631
Tif2
Tif4632
Nam8
Snu66
Zds1
Hsl1
Mih1
Crn1
Abp1
Rsc9
Arp9
Las17
Sup35
Dom34
Prm15
Cof1
Srv2
Cyr1
Act1
Sla2
Swa2
Sky1
Ste7
Apl1
Apl2
Sto1
Sgv1
Fus3
Rpn10
Sec23
Sly1
Sec18
Syp1
Vrp1
Sla1
Rps9a
Mtr4
Rps8a
Fun12
Rpn1
Duf1
Rad53
Art5
Ecm21
Ldb19
Yhr131c
Rsp5
Bul2
Ent1
Ent5
Apl3
Sec16
Apm4
Akl1
Rpn11
Nip1
Prt1
Rpg1
Ded1
Tif35
Cdc9
Chk1
Rad23
Ubi4
Ede1
Ent2
Vps27
Chc1
Ptp3
Stb1
Spa2
Bem2
Bud14
Rck2
Glc7
Ipl1
Cdc5
Dbf4
Pfk26
Tpk1
Gpa2
Ras2
Med6
Rpb10
Paf1
Rpo26
Bdp1
Spt15
Spt5
Rgr1
Srb7
Ssn2
Nop1
Nsr1
Nop56
Snu13
Rps31
Rad9
Rtf1
Rpo21
Npl3
Air2
Fps1
Ask10
Cdc28
Hog1
Pbs2
Ssk2
Clb2
Bud3
Bud4
Msn2
Sch9
Rim15
Yak1
Ypl247c
Vps52
Kss1
Sko1
Slt2
Msg5
Swe1
Igo2
Cdc55
Igo1
Vps53
Vps54
Rtn1
Fmp42
Cot1
Zrc1
10
20
30
40
50
t/s

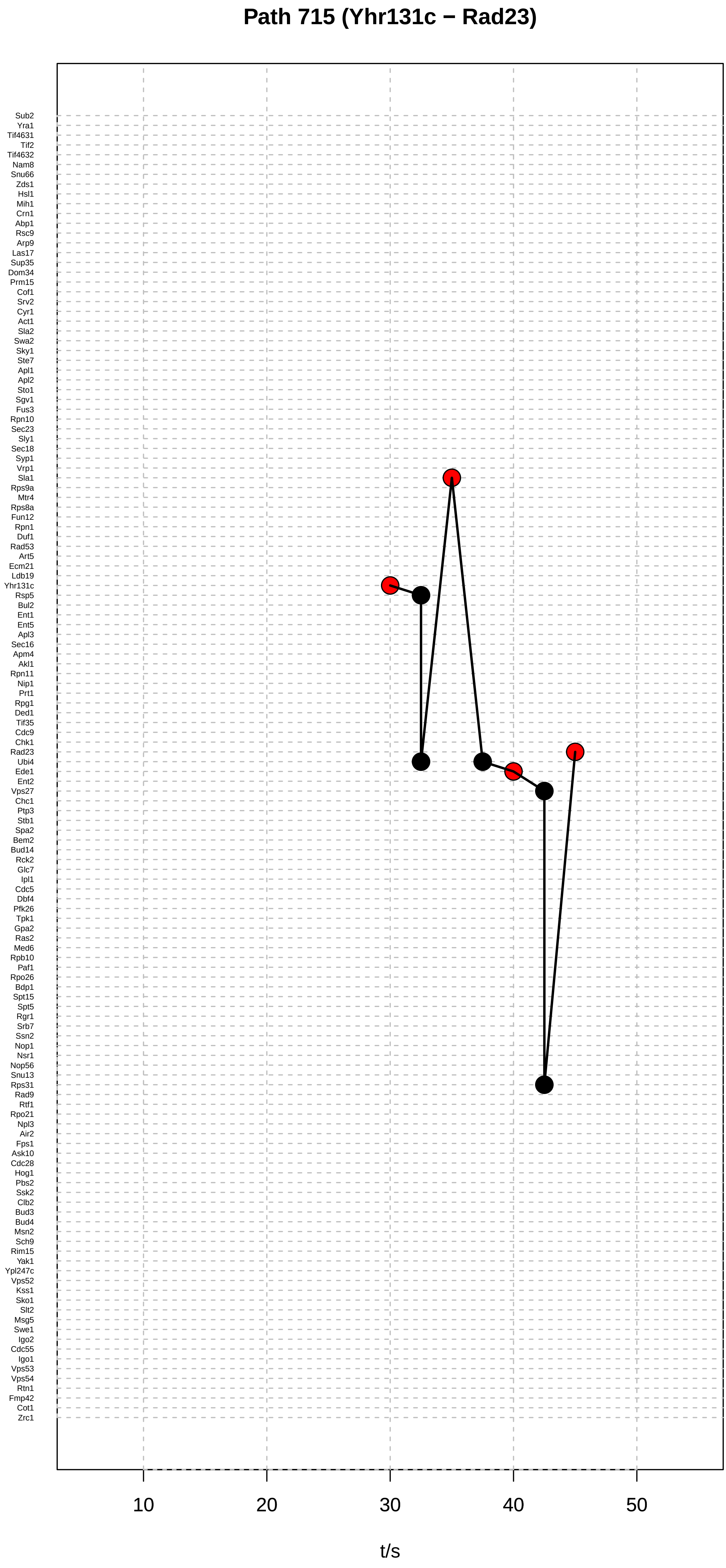
Path 715 (Yhr131c − Rad23)
t/s
10
20
30
40
50
Yhr131c
Rsp5
Sla1
Rad23
Ubi4
Ede1
Vps27
Rps31

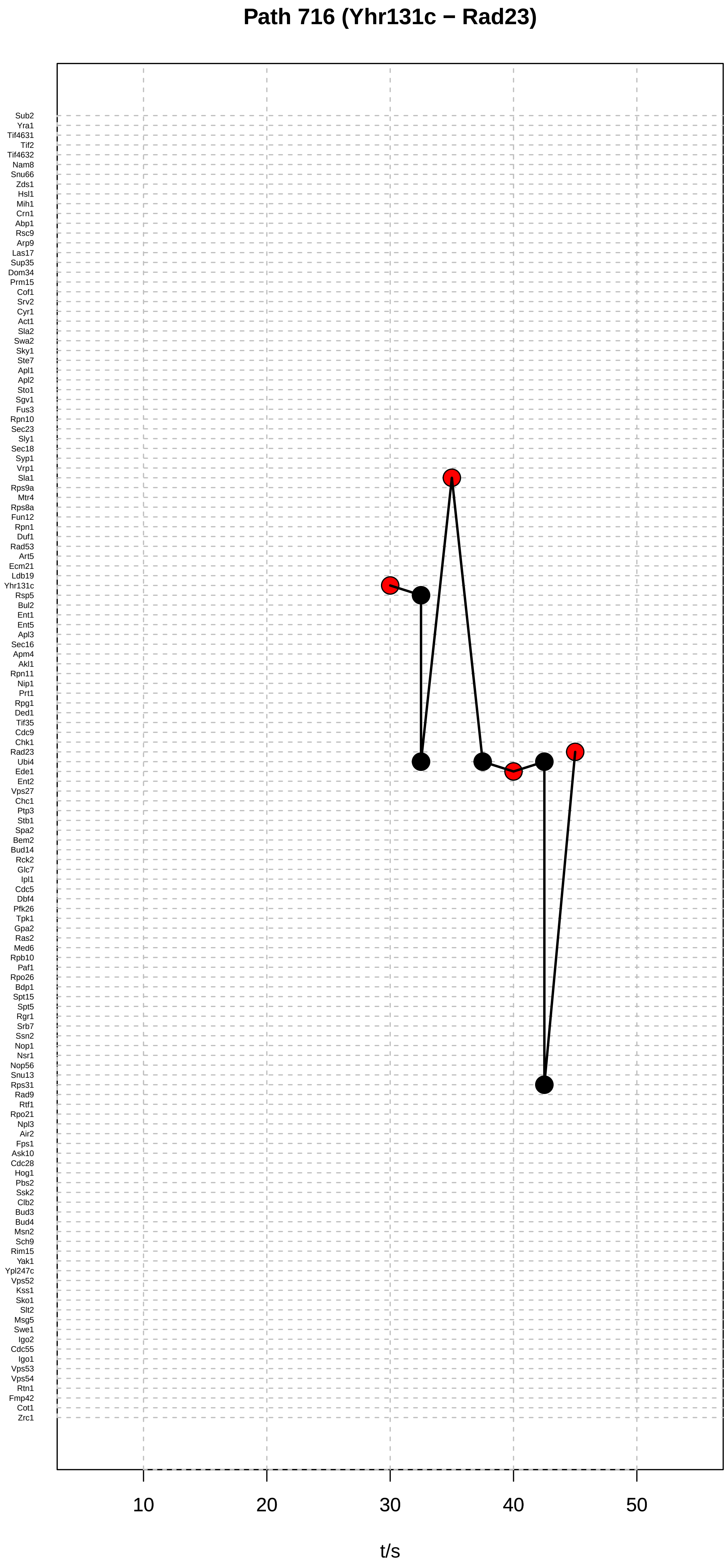

Path 716 (Yhr131c − Rad23)
Sub2
Yra1
Tif4631
Tif2
Tif4632
Nam8
Snu66
Zds1
Hsl1
Mih1
Crn1
Abp1
Rsc9
Arp9
Las17
Sup35
Dom34
Prm15
Cof1
Srv2
Cyr1
Act1
Sla2
Swa2
Sky1
Ste7
Apl1
Apl2
Sto1
Sgv1
Fus3
Rpn10
Sec23
Sly1
Sec18
Syp1
Vrp1
Sla1
Rps9a
Mtr4
Rps8a
Fun12
Rpn1
Duf1
Rad53
Art5
Ecm21
Ldb19
Yhr131c
Rsp5
Bul2
Ent1
Ent5
Apl3
Sec16
Apm4
Akl1
Rpn11
Nip1
Prt1
Rpg1
Ded1
Tif35
Cdc9
Chk1
Rad23
Ubi4
Ede1
Ent2
Vps27
Chc1
Ptp3
Stb1
Spa2
Bem2
Bud14
Rck2
Glc7
Ipl1
Cdc5
Dbf4
Pfk26
Tpk1
Gpa2
Ras2
Med6
Rpb10
Paf1
Rpo26
Bdp1
Spt15
Spt5
Rgr1
Srb7
Ssn2
Nop1
Nsr1
Nop56
Snu13
Rps31
Rad9
Rtf1
Rpo21
Npl3
Air2
Fps1
Ask10
Cdc28
Hog1
Pbs2
Ssk2
Clb2
Bud3
Bud4
Msn2
Sch9
Rim15
Yak1
Ypl247c
Vps52
Kss1
Sko1
Slt2
Msg5
Swe1
Igo2
Cdc55
Igo1
Vps53
Vps54
Rtn1
Fmp42
Cot1
Zrc1
10
20
30
40
50
t/s

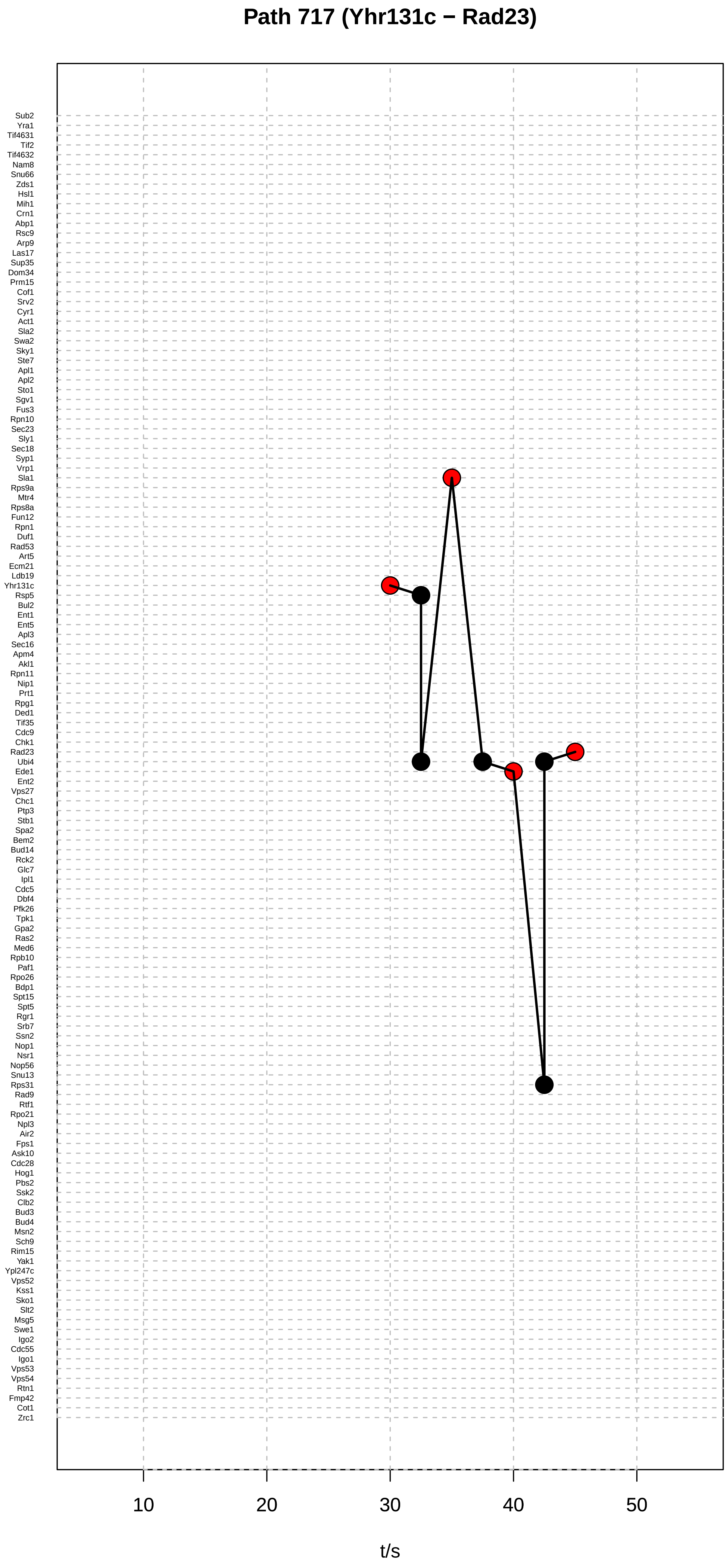

Path 717 (Yhr131c − Rad23)
Sub2
Yra1
Tif4631
Tif2
Tif4632
Nam8
Snu66
Zds1
Hsl1
Mih1
Crn1
Abp1
Rsc9
Arp9
Las17
Sup35
Dom34
Prm15
Cof1
Srv2
Cyr1
Act1
Sla2
Swa2
Sky1
Ste7
Apl1
Apl2
Sto1
Sgv1
Fus3
Rpn10
Sec23
Sly1
Sec18
Syp1
Vrp1
Sla1
Rps9a
Mtr4
Rps8a
Fun12
Rpn1
Duf1
Rad53
Art5
Ecm21
Ldb19
Yhr131c
Rsp5
Bul2
Ent1
Ent5
Apl3
Sec16
Apm4
Akl1
Rpn11
Nip1
Prt1
Rpg1
Ded1
Tif35
Cdc9
Chk1
Rad23
Ubi4
Ede1
Ent2
Vps27
Chc1
Ptp3
Stb1
Spa2
Bem2
Bud14
Rck2
Glc7
Ipl1
Cdc5
Dbf4
Pfk26
Tpk1
Gpa2
Ras2
Med6
Rpb10
Paf1
Rpo26
Bdp1
Spt15
Spt5
Rgr1
Srb7
Ssn2
Nop1
Nsr1
Nop56
Snu13
Rps31
Rad9
Rtf1
Rpo21
Npl3
Air2
Fps1
Ask10
Cdc28
Hog1
Pbs2
Ssk2
Clb2
Bud3
Bud4
Msn2
Sch9
Rim15
Yak1
Ypl247c
Vps52
Kss1
Sko1
Slt2
Msg5
Swe1
Igo2
Cdc55
Igo1
Vps53
Vps54
Rtn1
Fmp42
Cot1
Zrc1
10
20
30
40
50
t/s

#### Path 718 (Yhr131c − Rad23)

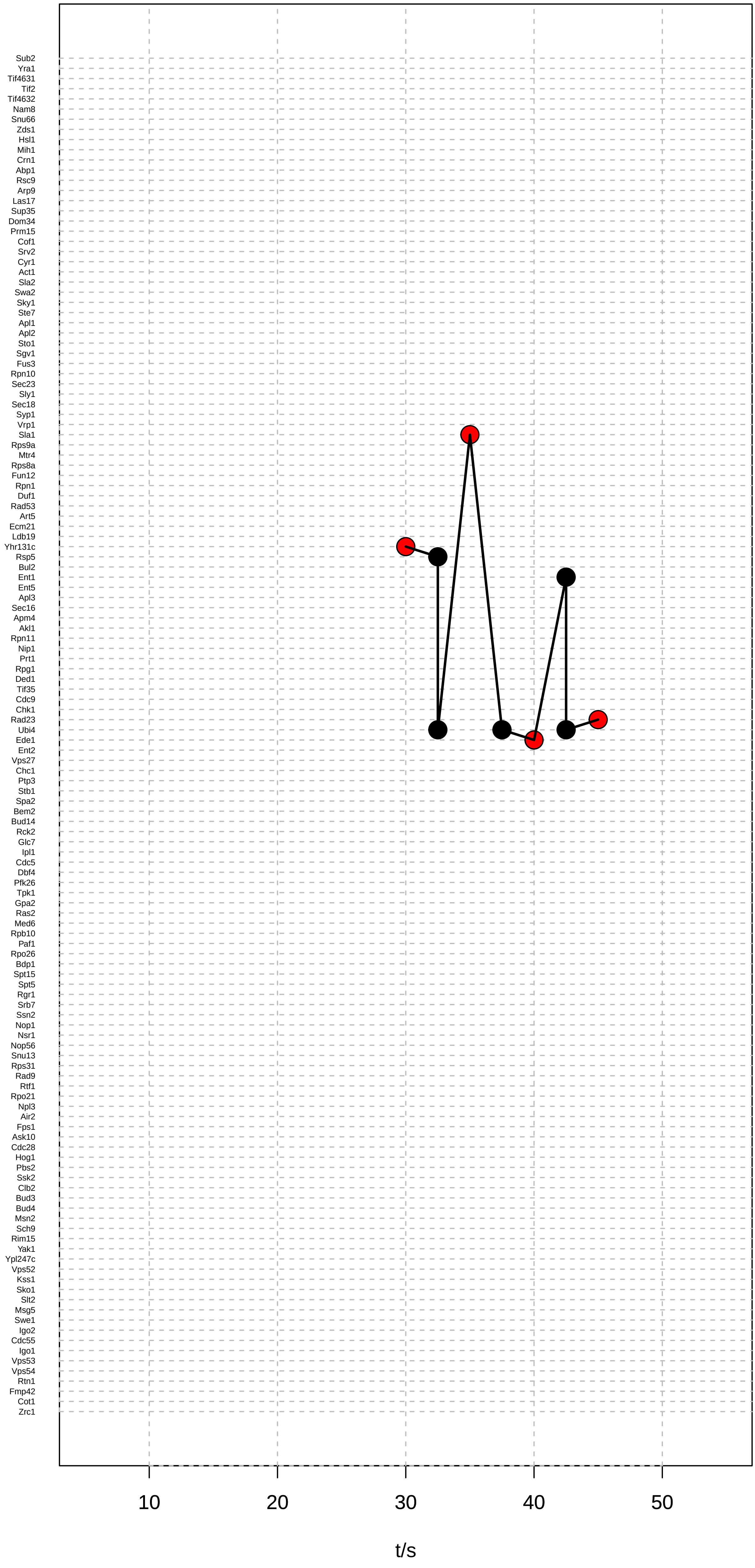

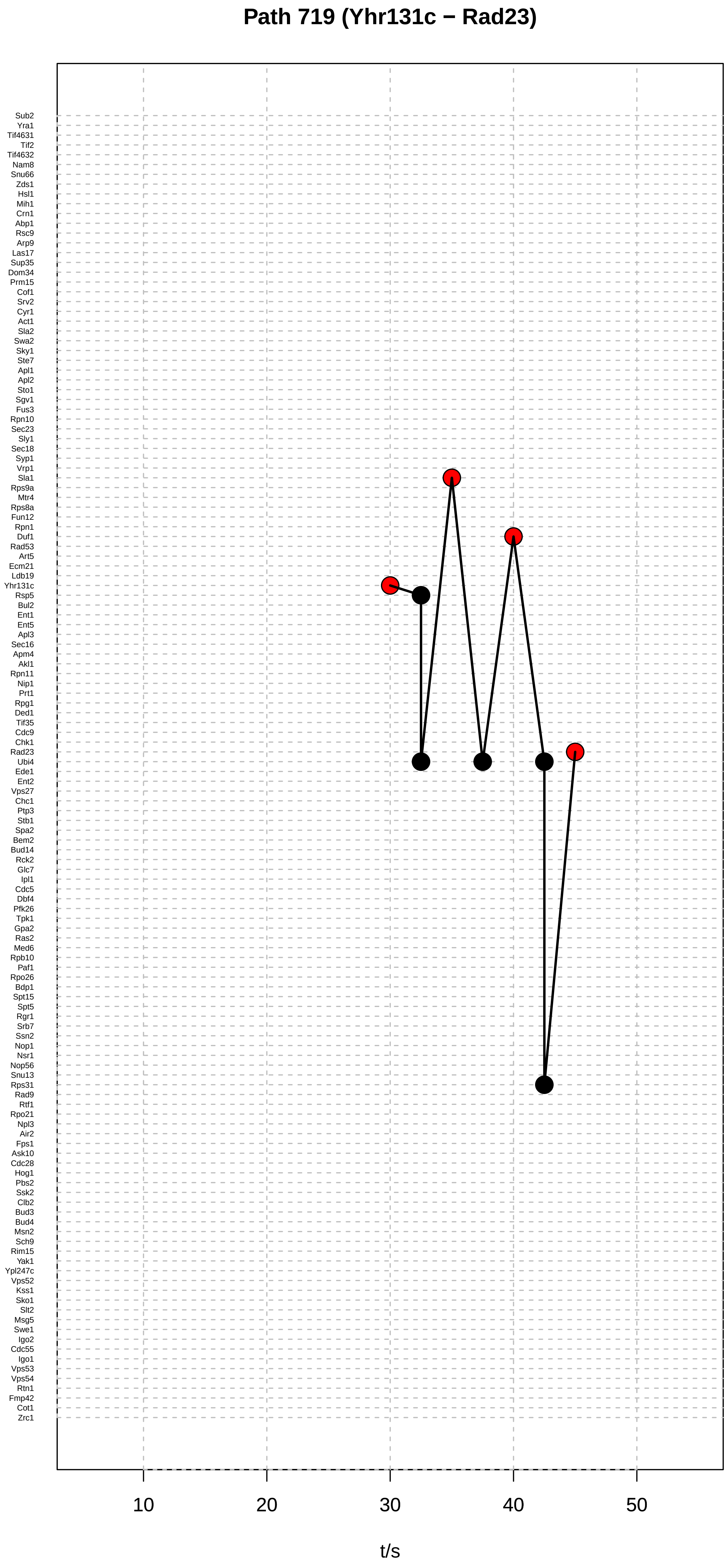

Path 719 (Yhr131c − Rad23)
Sub2
Yra1
Tif4631
Tif2
Tif4632
Nam8
Snu66
Zds1
Hsl1
Mih1
Crn1
Abp1
Rsc9
Arp9
Las17
Sup35
Dom34
Prm15
Cof1
Srv2
Cyr1
Act1
Sla2
Swa2
Sky1
Ste7
Apl1
Apl2
Sto1
Sgv1
Fus3
Rpn10
Sec23
Sly1
Sec18
Syp1
Vrp1
Sla1
Rps9a
Mtr4
Rps8a
Fun12
Rpn1
Duf1
Rad53
Art5
Ecm21
Ldb19
Yhr131c
Rsp5
Bul2
Ent1
Ent5
Apl3
Sec16
Apm4
Akl1
Rpn11
Nip1
Prt1
Rpg1
Ded1
Tif35
Cdc9
Chk1
Rad23
Ubi4
Ede1
Ent2
Vps27
Chc1
Ptp3
Stb1
Spa2
Bem2
Bud14
Rck2
Glc7
Ipl1
Cdc5
Dbf4
Pfk26
Tpk1
Gpa2
Ras2
Med6
Rpb10
Paf1
Rpo26
Bdp1
Spt15
Spt5
Rgr1
Srb7
Ssn2
Nop1
Nsr1
Nop56
Snu13
Rps31
Rad9
Rtf1
Rpo21
Npl3
Air2
Fps1
Ask10
Cdc28
Hog1
Pbs2
Ssk2
Clb2
Bud3
Bud4
Msn2
Sch9
Rim15
Yak1
Ypl247c
Vps52
Kss1
Sko1
Slt2
Msg5
Swe1
Igo2
Cdc55
Igo1
Vps53
Vps54
Rtn1
Fmp42
Cot1
Zrc1
10
20
30
40
50
t/s

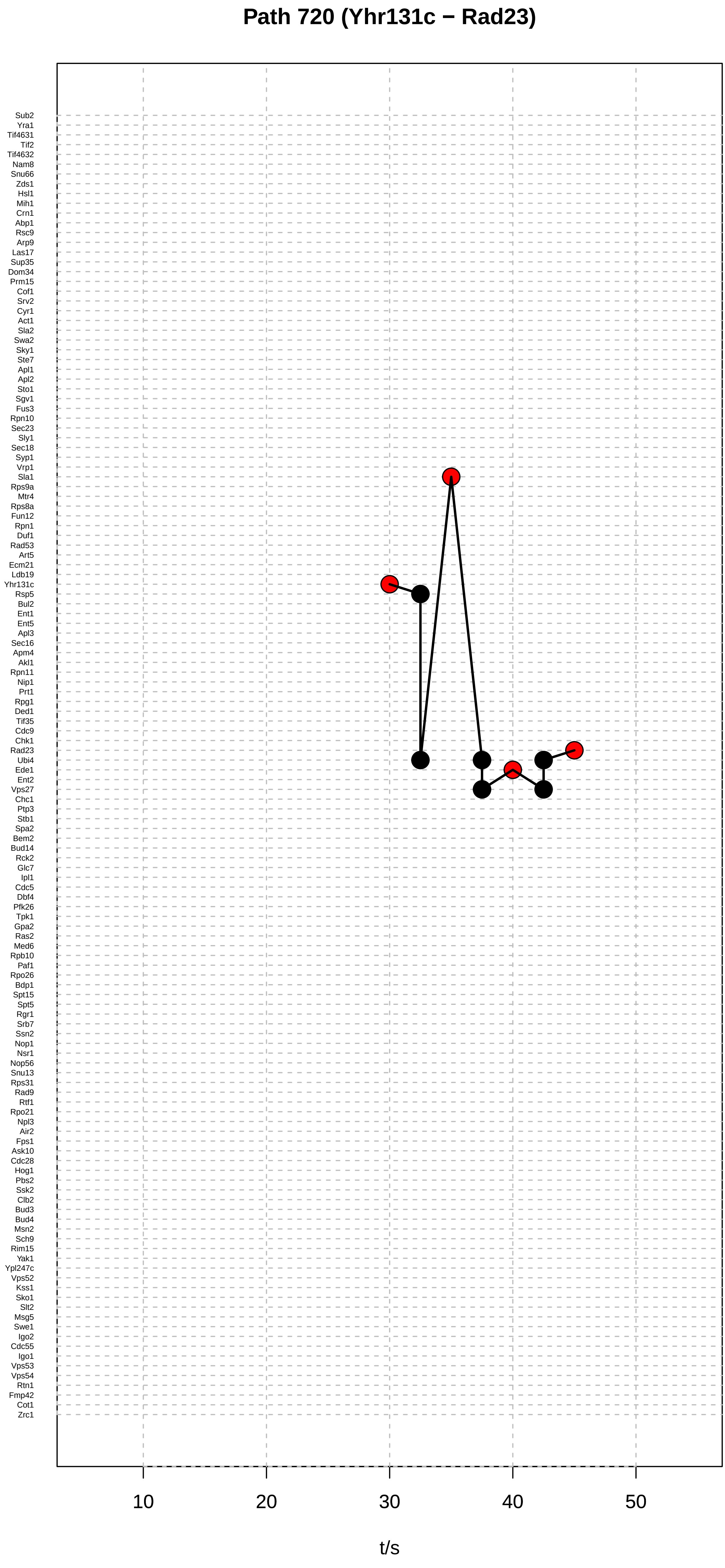

Path 720 (Yhr131c − Rad23)
Sub2
Yra1
Tif4631
Tif2
Tif4632
Nam8
Snu66
Zds1
Hsl1
Mih1
Crn1
Abp1
Rsc9
Arp9
Las17
Sup35
Dom34
Prm15
Cof1
Srv2
Cyr1
Act1
Sla2
Swa2
Sky1
Ste7
Apl1
Apl2
Sto1
Sgv1
Fus3
Rpn10
Sec23
Sly1
Sec18
Syp1
Vrp1
Sla1
Rps9a
Mtr4
Rps8a
Fun12
Rpn1
Duf1
Rad53
Art5
Ecm21
Ldb19
Yhr131c
Rsp5
Bul2
Ent1
Ent5
Apl3
Sec16
Apm4
Akl1
Rpn11
Nip1
Prt1
Rpg1
Ded1
Tif35
Cdc9
Chk1
Rad23
Ubi4
Ede1
Ent2
Vps27
Chc1
Ptp3
Stb1
Spa2
Bem2
Bud14
Rck2
Glc7
Ipl1
Cdc5
Dbf4
Pfk26
Tpk1
Gpa2
Ras2
Med6
Rpb10
Paf1
Rpo26
Bdp1
Spt15
Spt5
Rgr1
Srb7
Ssn2
Nop1
Nsr1
Nop56
Snu13
Rps31
Rad9
Rtf1
Rpo21
Npl3
Air2
Fps1
Ask10
Cdc28
Hog1
Pbs2
Ssk2
Clb2
Bud3
Bud4
Msn2
Sch9
Rim15
Yak1
Ypl247c
Vps52
Kss1
Sko1
Slt2
Msg5
Swe1
Igo2
Cdc55
Igo1
Vps53
Vps54
Rtn1
Fmp42
Cot1
Zrc1
10
20
30
40
50
t/s

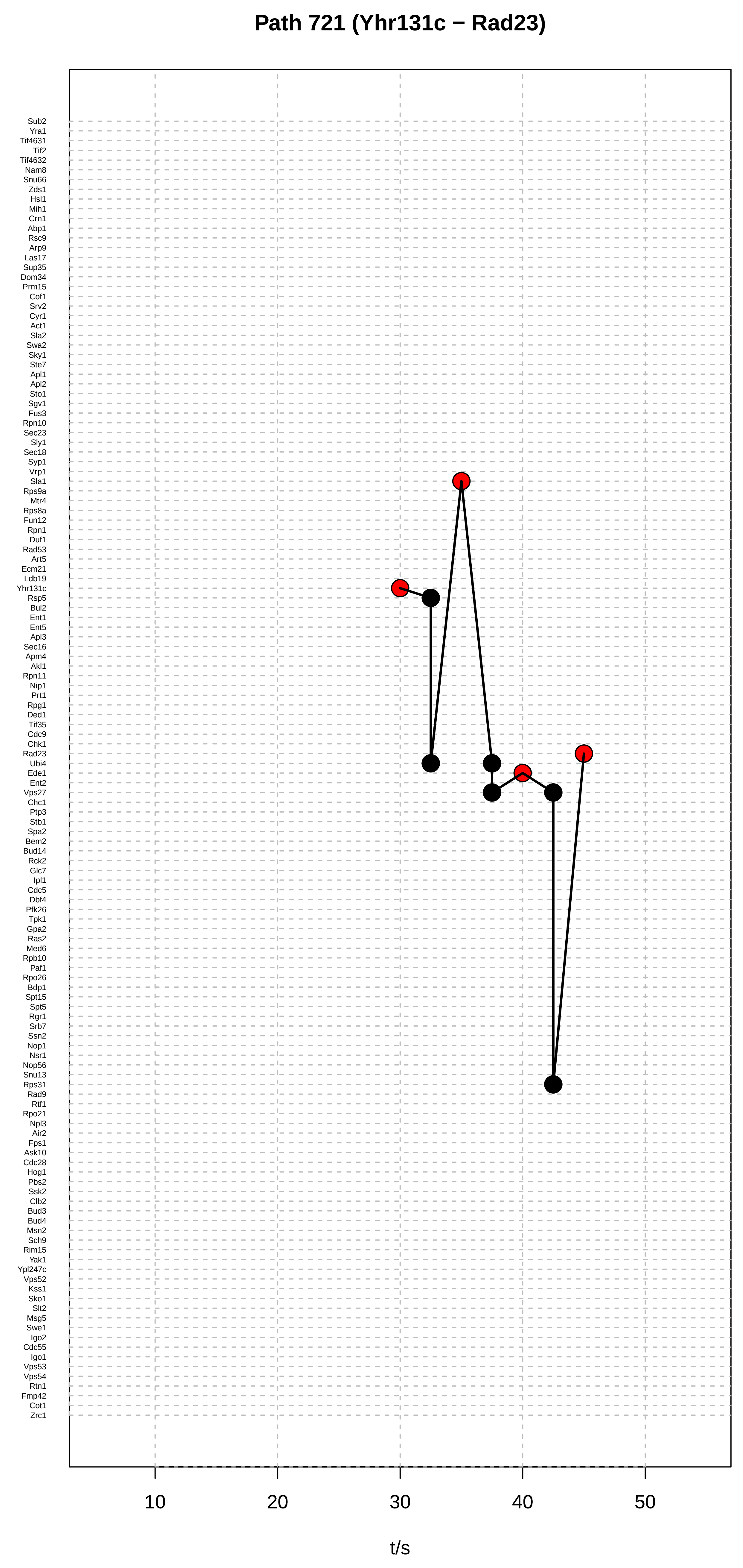

Path 721 (Yhr131c − Rad23)
Sub2
Yra1
Tif4631
Tif2
Tif4632
Nam8
Snu66
Zds1
Hsl1
Mih1
Crn1
Abp1
Rsc9
Arp9
Las17
Sup35
Dom34
Prm15
Cof1
Srv2
Cyr1
Act1
Sla2
Swa2
Sky1
Ste7
Apl1
Apl2
Sto1
Sgv1
Fus3
Rpn10
Sec23
Sly1
Sec18
Syp1
Vrp1
Sla1
Rps9a
Mtr4
Rps8a
Fun12
Rpn1
Duf1
Rad53
Art5
Ecm21
Ldb19
Yhr131c
Rsp5
Bul2
Ent1
Ent5
Apl3
Sec16
Apm4
Akl1
Rpn11
Nip1
Prt1
Rpg1
Ded1
Tif35
Cdc9
Chk1
Rad23
Ubi4
Ede1
Ent2
Vps27
Chc1
Ptp3
Stb1
Spa2
Bem2
Bud14
Rck2
Glc7
Ipl1
Cdc5
Dbf4
Pfk26
Tpk1
Gpa2
Ras2
Med6
Rpb10
Paf1
Rpo26
Bdp1
Spt15
Spt5
Rgr1
Srb7
Ssn2
Nop1
Nsr1
Nop56
Snu13
Rps31
Rad9
Rtf1
Rpo21
Npl3
Air2
Fps1
Ask10
Cdc28
Hog1
Pbs2
Ssk2
Clb2
Bud3
Bud4
Msn2
Sch9
Rim15
Yak1
Ypl247c
Vps52
Kss1
Sko1
Slt2
Msg5
Swe1
Igo2
Cdc55
Igo1
Vps53
Vps54
Rtn1
Fmp42
Cot1
Zrc1
10
20
30
40
50
t/s

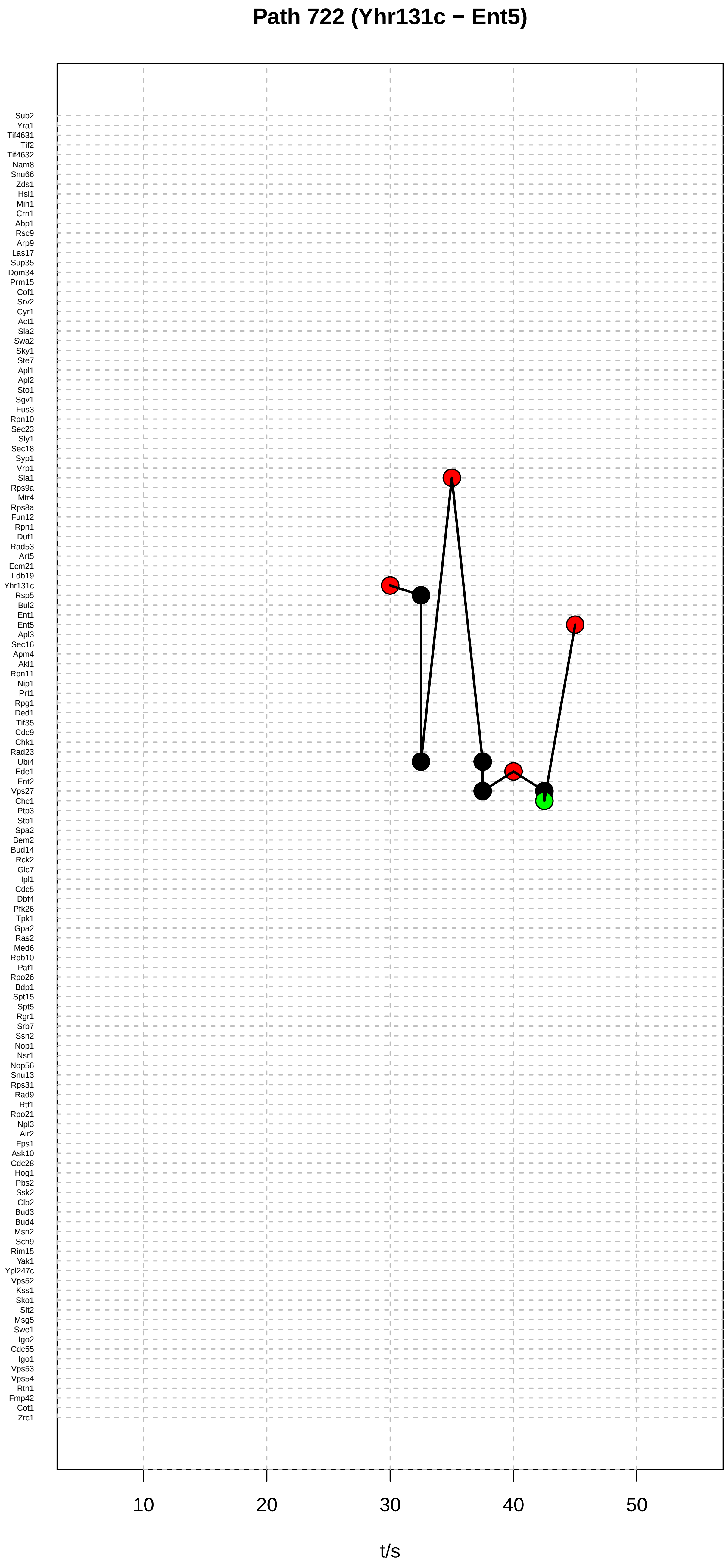

Path 722 (Yhr131c − Ent5)
Sub2
Yra1
Tif4631
Tif2
Tif4632
Nam8
Snu66
Zds1
Hsl1
Mih1
Crn1
Abp1
Rsc9
Arp9
Las17
Sup35
Dom34
Prm15
Cof1
Srv2
Cyr1
Act1
Sla2
Swa2
Sky1
Ste7
Apl1
Apl2
Sto1
Sgv1
Fus3
Rpn10
Sec23
Sly1
Sec18
Syp1
Vrp1
Sla1
Rps9a
Mtr4
Rps8a
Fun12
Rpn1
Duf1
Rad53
Art5
Ecm21
Ldb19
Yhr131c
Rsp5
Bul2
Ent1
Ent5
Apl3
Sec16
Apm4
Akl1
Rpn11
Nip1
Prt1
Rpg1
Ded1
Tif35
Cdc9
Chk1
Rad23
Ubi4
Ede1
Ent2
Vps27
Chc1
Ptp3
Stb1
Spa2
Bem2
Bud14
Rck2
Glc7
Ipl1
Cdc5
Dbf4
Pfk26
Tpk1
Gpa2
Ras2
Med6
Rpb10
Paf1
Rpo26
Bdp1
Spt15
Spt5
Rgr1
Srb7
Ssn2
Nop1
Nsr1
Nop56
Snu13
Rps31
Rad9
Rtf1
Rpo21
Npl3
Air2
Fps1
Ask10
Cdc28
Hog1
Pbs2
Ssk2
Clb2
Bud3
Bud4
Msn2
Sch9
Rim15
Yak1
Ypl247c
Vps52
Kss1
Sko1
Slt2
Msg5
Swe1
Igo2
Cdc55
Igo1
Vps53
Vps54
Rtn1
Fmp42
Cot1
Zrc1
10
20
30
40
50
t/s

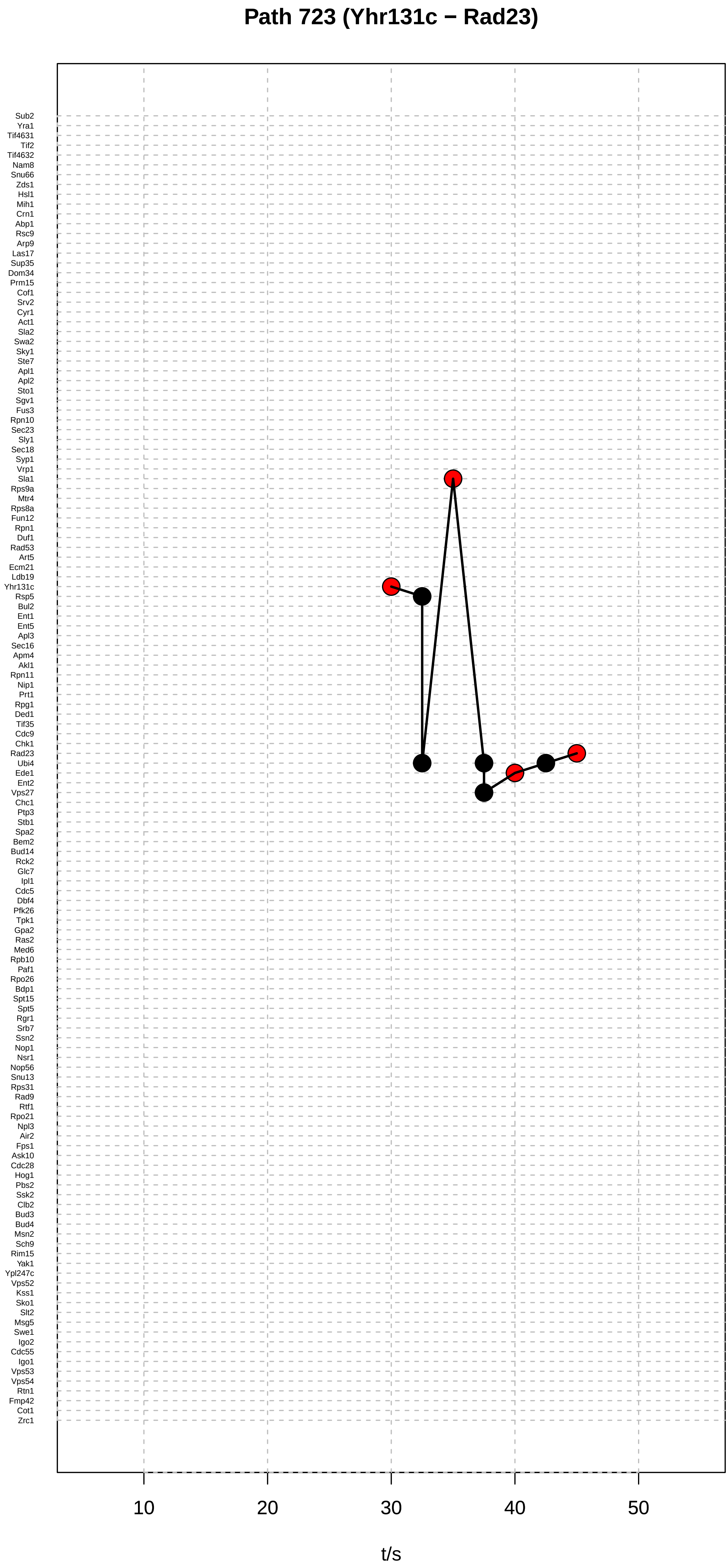

Path 723 (Yhr131c − Rad23)
Sub2
Yra1
Tif4631
Tif2
Tif4632
Nam8
Snu66
Zds1
Hsl1
Mih1
Crn1
Abp1
Rsc9
Arp9
Las17
Sup35
Dom34
Prm15
Cof1
Srv2
Cyr1
Act1
Sla2
Swa2
Sky1
Ste7
Apl1
Apl2
Sto1
Sgv1
Fus3
Rpn10
Sec23
Sly1
Sec18
Syp1
Vrp1
Sla1
Rps9a
Mtr4
Rps8a
Fun12
Rpn1
Duf1
Rad53
Art5
Ecm21
Ldb19
Yhr131c
Rsp5
Bul2
Ent1
Ent5
Apl3
Sec16
Apm4
Akl1
Rpn11
Nip1
Prt1
Rpg1
Ded1
Tif35
Cdc9
Chk1
Rad23
Ubi4
Ede1
Ent2
Vps27
Chc1
Ptp3
Stb1
Spa2
Bem2
Bud14
Rck2
Glc7
Ipl1
Cdc5
Dbf4
Pfk26
Tpk1
Gpa2
Ras2
Med6
Rpb10
Paf1
Rpo26
Bdp1
Spt15
Spt5
Rgr1
Srb7
Ssn2
Nop1
Nsr1
Nop56
Snu13
Rps31
Rad9
Rtf1
Rpo21
Npl3
Air2
Fps1
Ask10
Cdc28
Hog1
Pbs2
Ssk2
Clb2
Bud3
Bud4
Msn2
Sch9
Rim15
Yak1
Ypl247c
Vps52
Kss1
Sko1
Slt2
Msg5
Swe1
Igo2
Cdc55
Igo1
Vps53
Vps54
Rtn1
Fmp42
Cot1
Zrc1
10
20
30
40
50
t/s

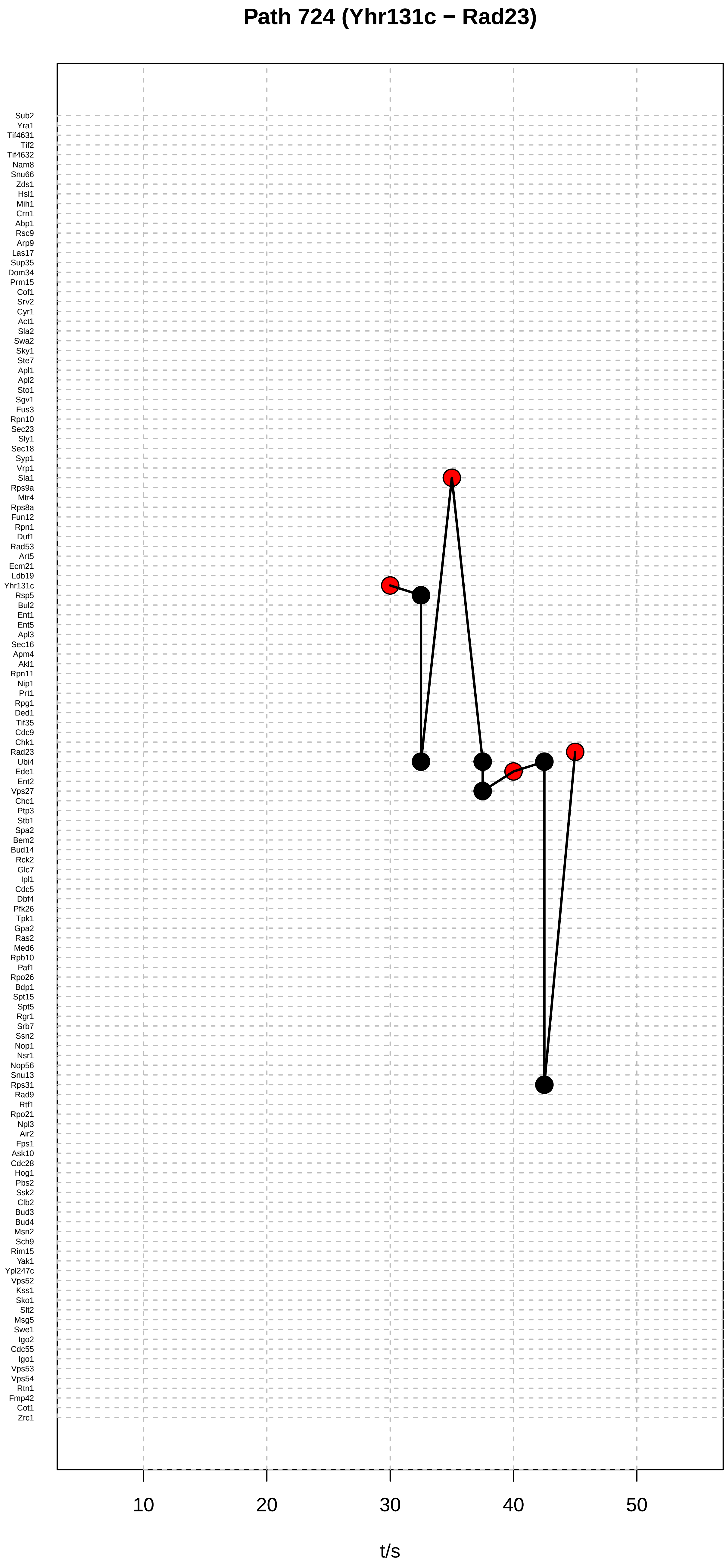
Path 724 (Yhr131c − Rad23)
Sub2
Yra1
Tif4631
Tif2
Tif4632
Nam8
Snu66
Zds1
Hsl1
Mih1
Crn1
Abp1
Rsc9
Arp9
Las17
Sup35
Dom34
Prm15
Cof1
Srv2
Cyr1
Act1
Sla2
Swa2
Sky1
Ste7
Apl1
Apl2
Sto1
Sgv1
Fus3
Rpn10
Sec23
Sly1
Sec18
Syp1
Vrp1
Sla1
Rps9a
Mtr4
Rps8a
Fun12
Rpn1
Duf1
Rad53
Art5
Ecm21
Ldb19
Yhr131c
Rsp5
Bul2
Ent1
Ent5
Apl3
Sec16
Apm4
Akl1
Rpn11
Nip1
Prt1
Rpg1
Ded1
Tif35
Cdc9
Chk1
Rad23
Ubi4
Ede1
Ent2
Vps27
Chc1
Ptp3
Stb1
Spa2
Bem2
Bud14
Rck2
Glc7
Ipl1
Cdc5
Dbf4
Pfk26
Tpk1
Gpa2
Ras2
Med6
Rpb10
Paf1
Rpo26
Bdp1
Spt15
Spt5
Rgr1
Srb7
Ssn2
Nop1
Nsr1
Nop56
Snu13
Rps31
Rad9
Rtf1
Rpo21
Npl3
Air2
Fps1
Ask10
Cdc28
Hog1
Pbs2
Ssk2
Clb2
Bud3
Bud4
Msn2
Sch9
Rim15
Yak1
Ypl247c
Vps52
Kss1
Sko1
Slt2
Msg5
Swe1
Igo2
Cdc55
Igo1
Vps53
Vps54
Rtn1
Fmp42
Cot1
Zrc1
10
20
30
40
50
t/s

### Path 725 (Yhr131c − Rad23)

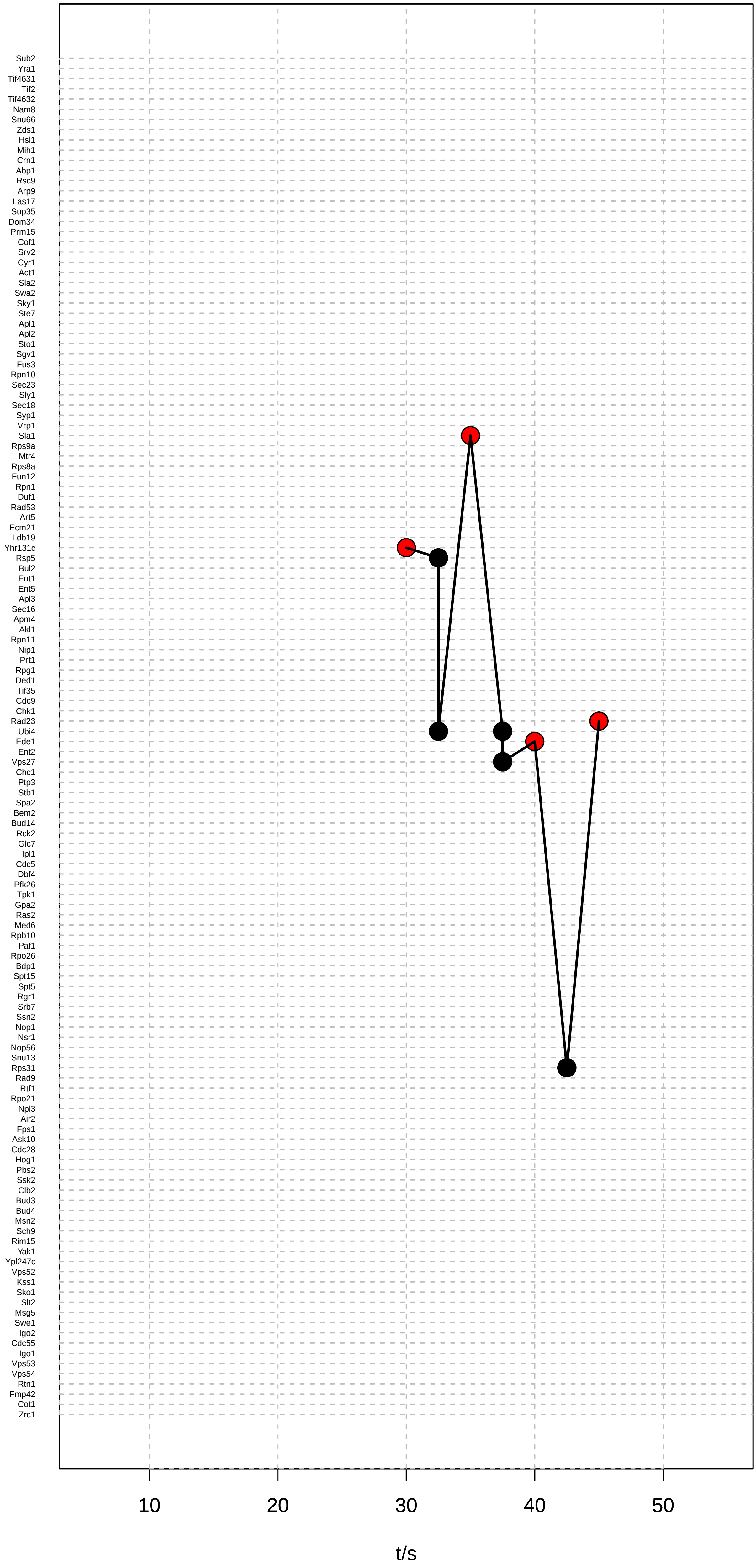

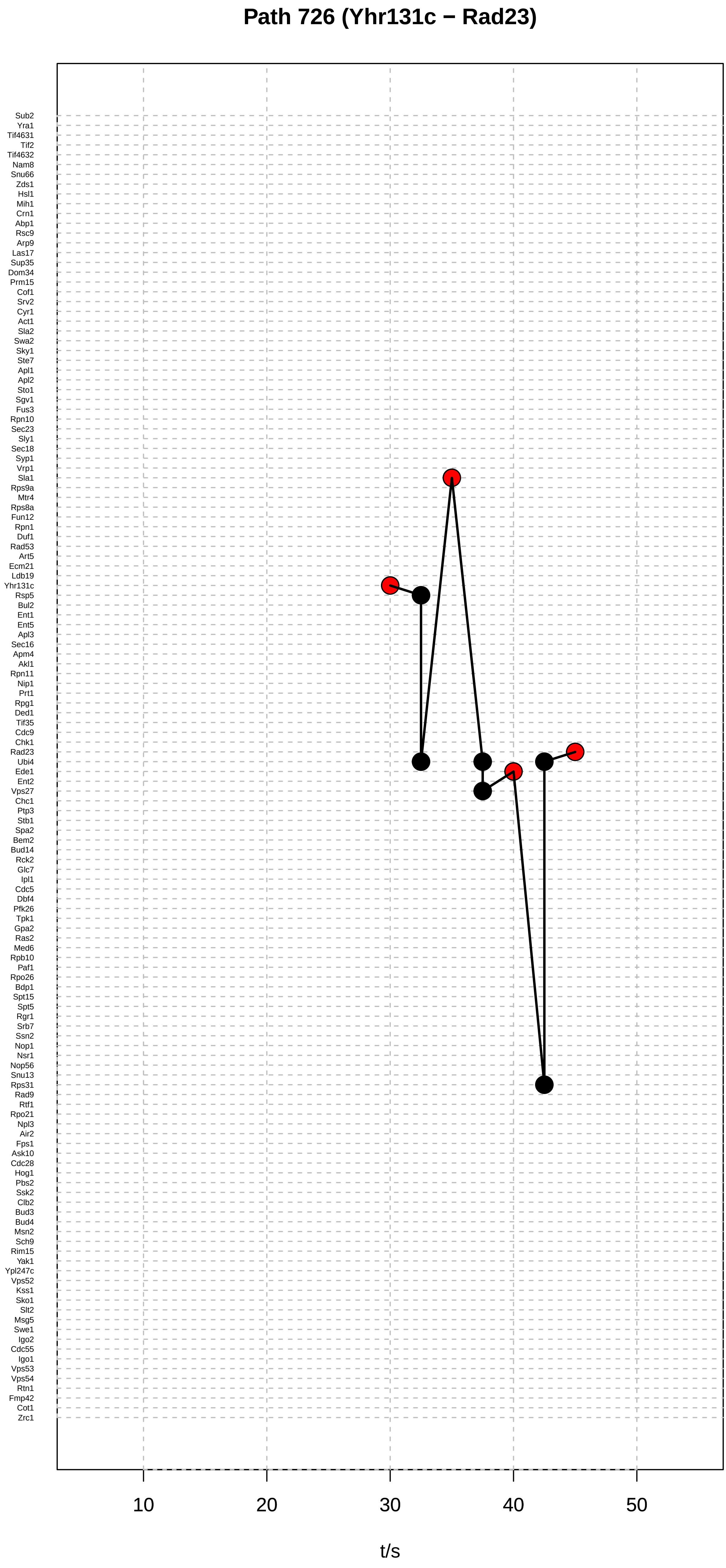
Path 726 (Yhr131c − Rad23)
Sub2
Yra1
Tif4631
Tif2
Tif4632
Nam8
Snu66
Zds1
Hsl1
Mih1
Crn1
Abp1
Rsc9
Arp9
Las17
Sup35
Dom34
Prm15
Cof1
Srv2
Cyr1
Act1
Sla2
Swa2
Sky1
Ste7
Apl1
Apl2
Sto1
Sgv1
Fus3
Rpn10
Sec23
Sly1
Sec18
Syp1
Vrp1
Sla1
Rps9a
Mtr4
Rps8a
Fun12
Rpn1
Duf1
Rad53
Art5
Ecm21
Ldb19
Yhr131c
Rsp5
Bul2
Ent1
Ent5
Apl3
Sec16
Apm4
Akl1
Rpn11
Nip1
Prt1
Rpg1
Ded1
Tif35
Cdc9
Chk1
Rad23
Ubi4
Ede1
Ent2
Vps27
Chc1
Ptp3
Stb1
Spa2
Bem2
Bud14
Rck2
Glc7
Ipl1
Cdc5
Dbf4
Pfk26
Tpk1
Gpa2
Ras2
Med6
Rpb10
Paf1
Rpo26
Bdp1
Spt15
Spt5
Rgr1
Srb7
Ssn2
Nop1
Nsr1
Nop56
Snu13
Rps31
Rad9
Rtf1
Rpo21
Npl3
Air2
Fps1
Ask10
Cdc28
Hog1
Pbs2
Ssk2
Clb2
Bud3
Bud4
Msn2
Sch9
Rim15
Yak1
Ypl247c
Vps52
Kss1
Sko1
Slt2
Msg5
Swe1
Igo2
Cdc55
Igo1
Vps53
Vps54
Rtn1
Fmp42
Cot1
Zrc1
10
20
30
40
50
t/s

### Path 727 (Yhr131c − Rad23)

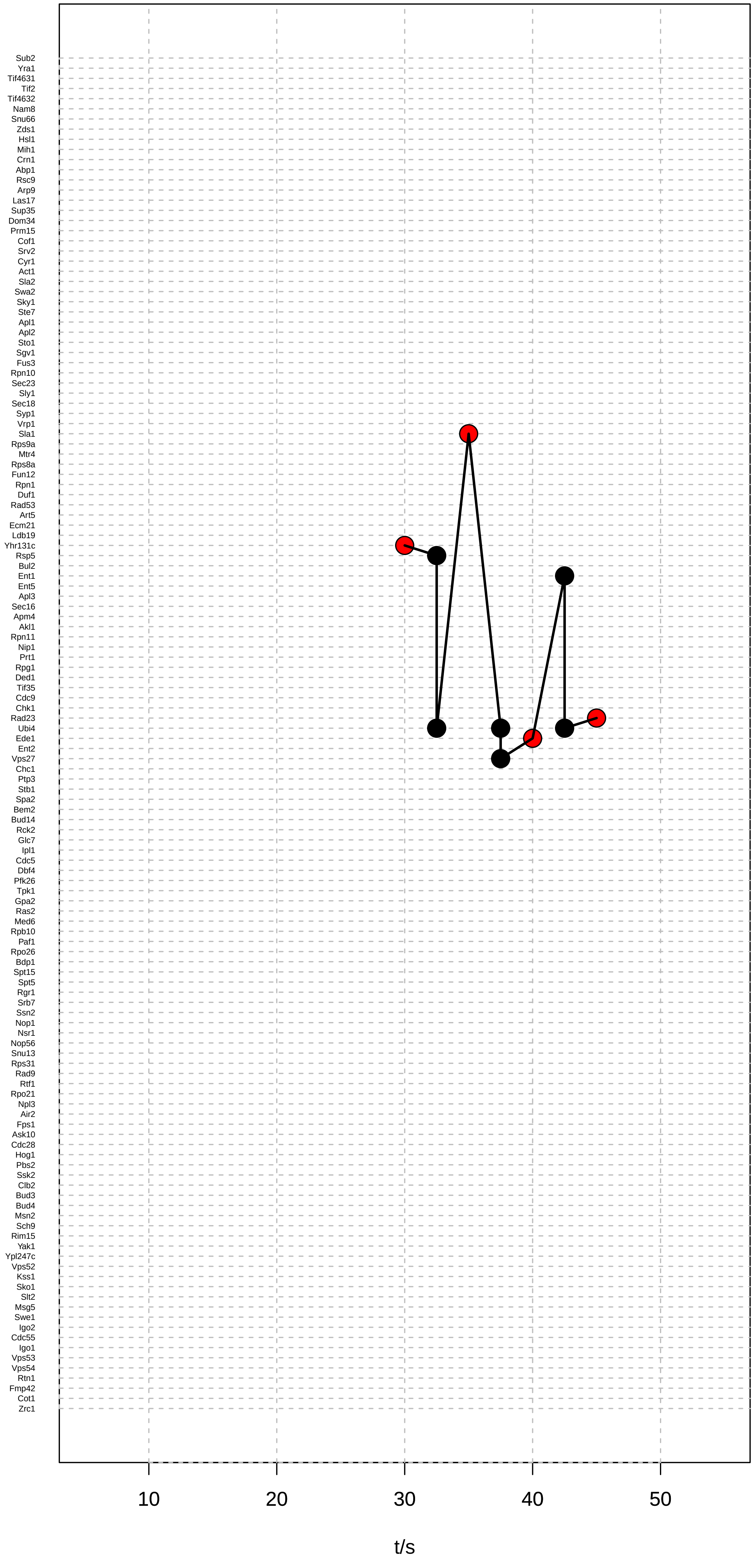

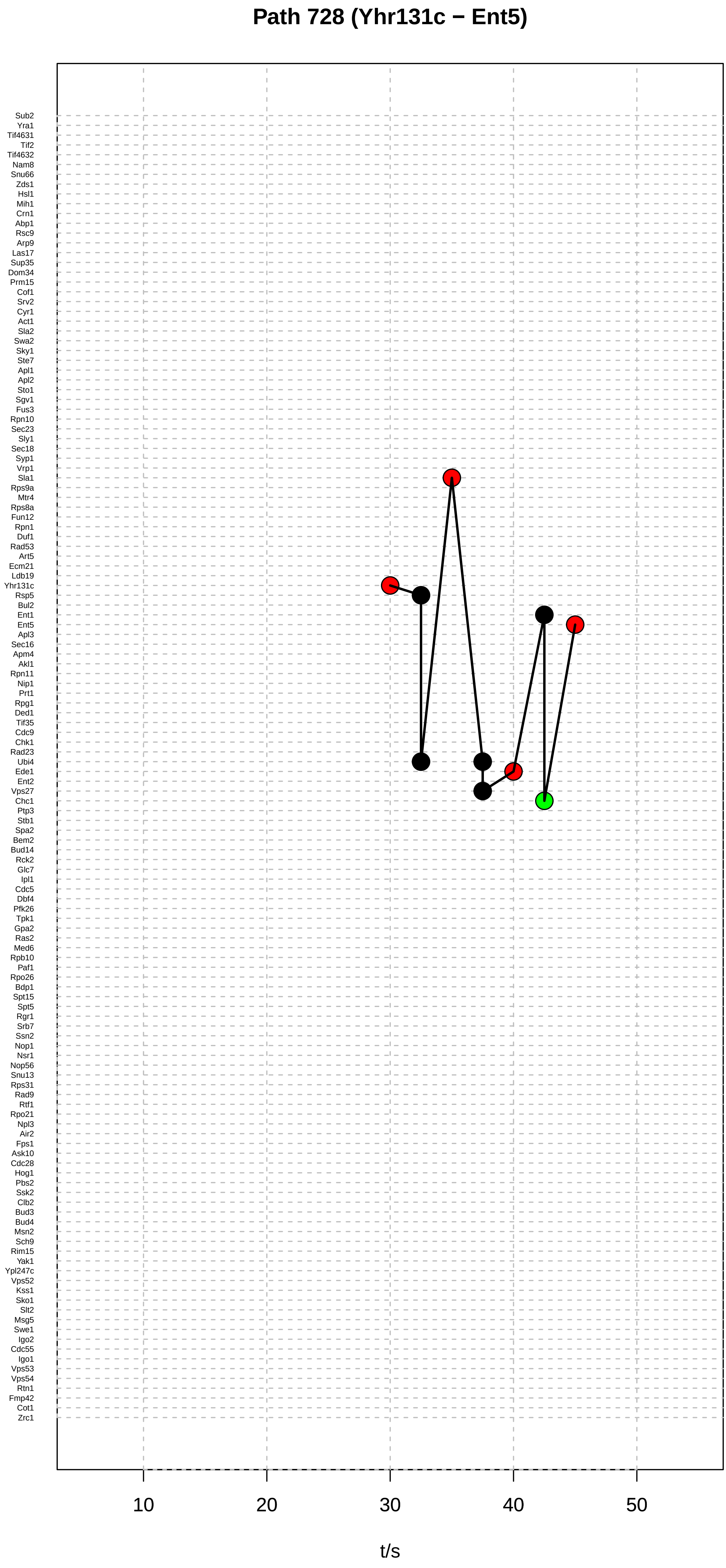
Path 728 (Yhr131c − Ent5)
Sub2
Yra1
Tif4631
Tif2
Tif4632
Nam8
Snu66
Zds1
Hsl1
Mih1
Crn1
Abp1
Rsc9
Arp9
Las17
Sup35
Dom34
Prm15
Cof1
Srv2
Cyr1
Act1
Sla2
Swa2
Sky1
Ste7
Apl1
Apl2
Sto1
Sgv1
Fus3
Rpn10
Sec23
Sly1
Sec18
Syp1
Vrp1
Sla1
Rps9a
Mtr4
Rps8a
Fun12
Rpn1
Duf1
Rad53
Art5
Ecm21
Ldb19
Yhr131c
Rsp5
Bul2
Ent1
Ent5
Apl3
Sec16
Apm4
Akl1
Rpn11
Nip1
Prt1
Rpg1
Ded1
Tif35
Cdc9
Chk1
Rad23
Ubi4
Ede1
Ent2
Vps27
Chc1
Ptp3
Stb1
Spa2
Bem2
Bud14
Rck2
Glc7
Ipl1
Cdc5
Dbf4
Pfk26
Tpk1
Gpa2
Ras2
Med6
Rpb10
Paf1
Rpo26
Bdp1
Spt15
Spt5
Rgr1
Srb7
Ssn2
Nop1
Nsr1
Nop56
Snu13
Rps31
Rad9
Rtf1
Rpo21
Npl3
Air2
Fps1
Ask10
Cdc28
Hog1
Pbs2
Ssk2
Clb2
Bud3
Bud4
Msn2
Sch9
Rim15
Yak1
Ypl247c
Vps52
Kss1
Sko1
Slt2
Msg5
Swe1
Igo2
Cdc55
Igo1
Vps53
Vps54
Rtn1
Fmp42
Cot1
Zrc1
10
20
30
40
50
t/s

### Path 729 (Yhr131c − Rad23)

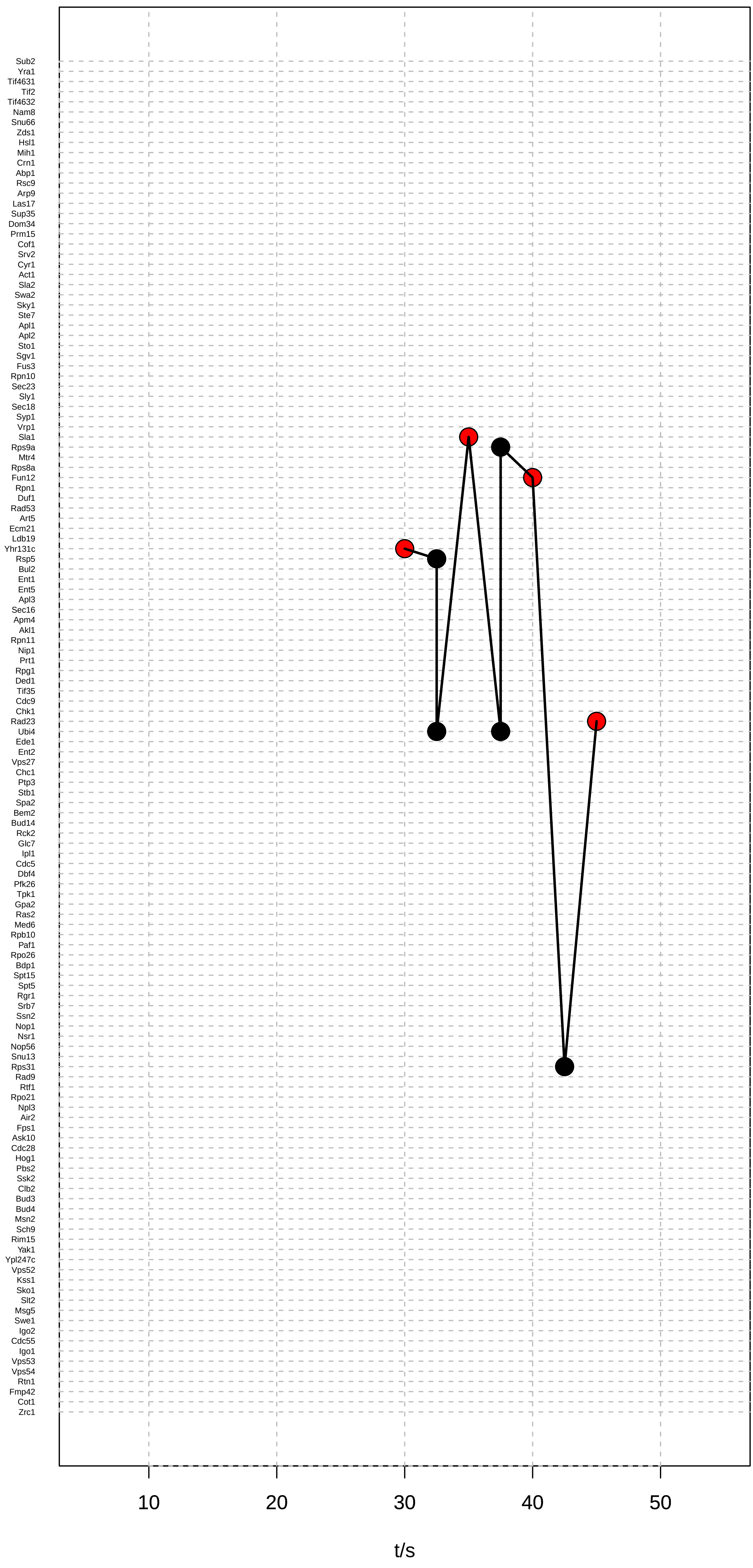

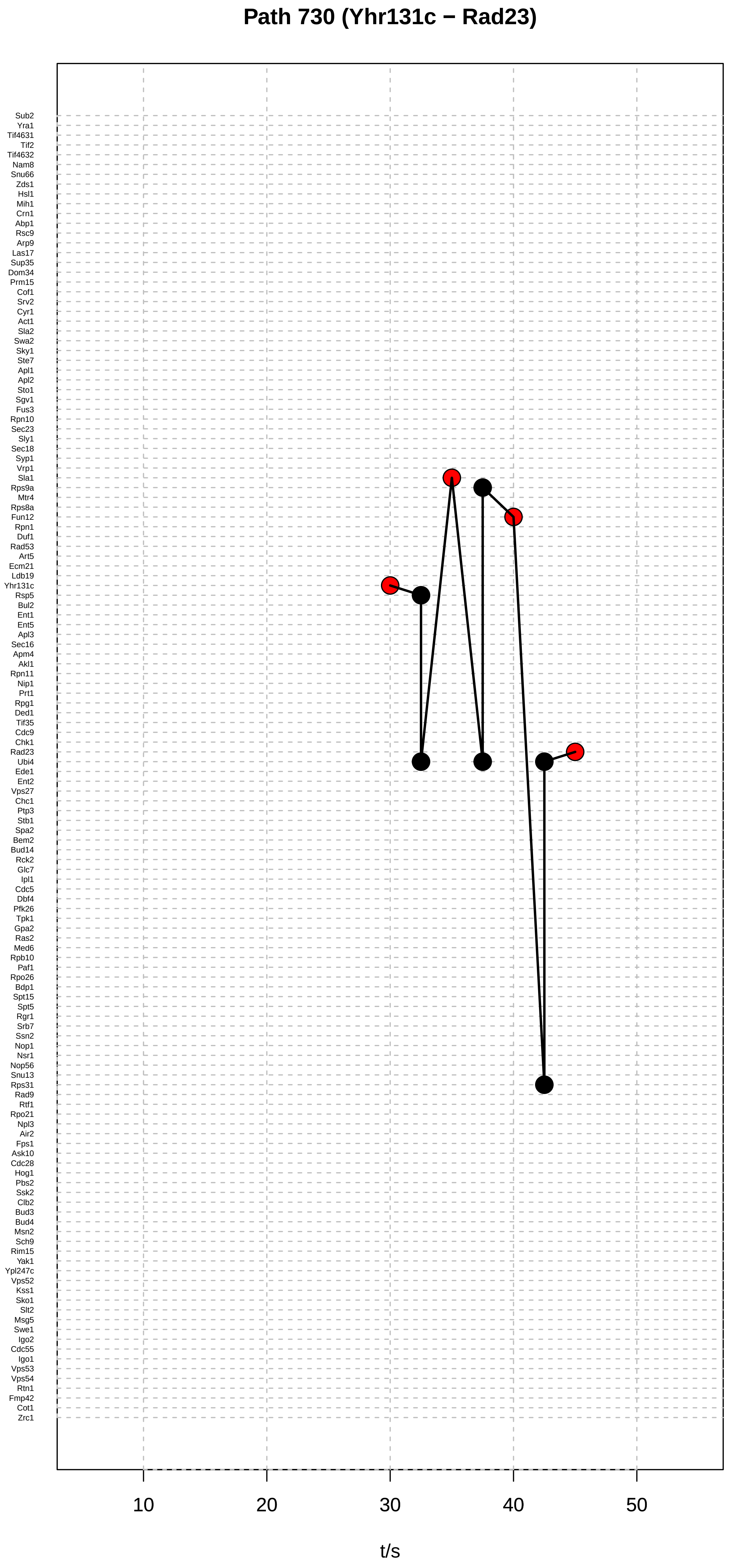

Path 730 (Yhr131c − Rad23)
Sub2
Yra1
Tif4631
Tif2
Tif4632
Nam8
Snu66
Zds1
Hsl1
Mih1
Crn1
Abp1
Rsc9
Arp9
Las17
Sup35
Dom34
Prm15
Cof1
Srv2
Cyr1
Act1
Sla2
Swa2
Sky1
Ste7
Apl1
Apl2
Sto1
Sgv1
Fus3
Rpn10
Sec23
Sly1
Sec18
Syp1
Vrp1
Sla1
Rps9a
Mtr4
Rps8a
Fun12
Rpn1
Duf1
Rad53
Art5
Ecm21
Ldb19
Yhr131c
Rsp5
Bul2
Ent1
Ent5
Apl3
Sec16
Apm4
Akl1
Rpn11
Nip1
Prt1
Rpg1
Ded1
Tif35
Cdc9
Chk1
Rad23
Ubi4
Ede1
Ent2
Vps27
Chc1
Ptp3
Stb1
Spa2
Bem2
Bud14
Rck2
Glc7
Ipl1
Cdc5
Dbf4
Pfk26
Tpk1
Gpa2
Ras2
Med6
Rpb10
Paf1
Rpo26
Bdp1
Spt15
Spt5
Rgr1
Srb7
Ssn2
Nop1
Nsr1
Nop56
Snu13
Rps31
Rad9
Rtf1
Rpo21
Npl3
Air2
Fps1
Ask10
Cdc28
Hog1
Pbs2
Ssk2
Clb2
Bud3
Bud4
Msn2
Sch9
Rim15
Yak1
Ypl247c
Vps52
Kss1
Sko1
Slt2
Msg5
Swe1
Igo2
Cdc55
Igo1
Vps53
Vps54
Rtn1
Fmp42
Cot1
Zrc1
10
20
30
40
50
t/s

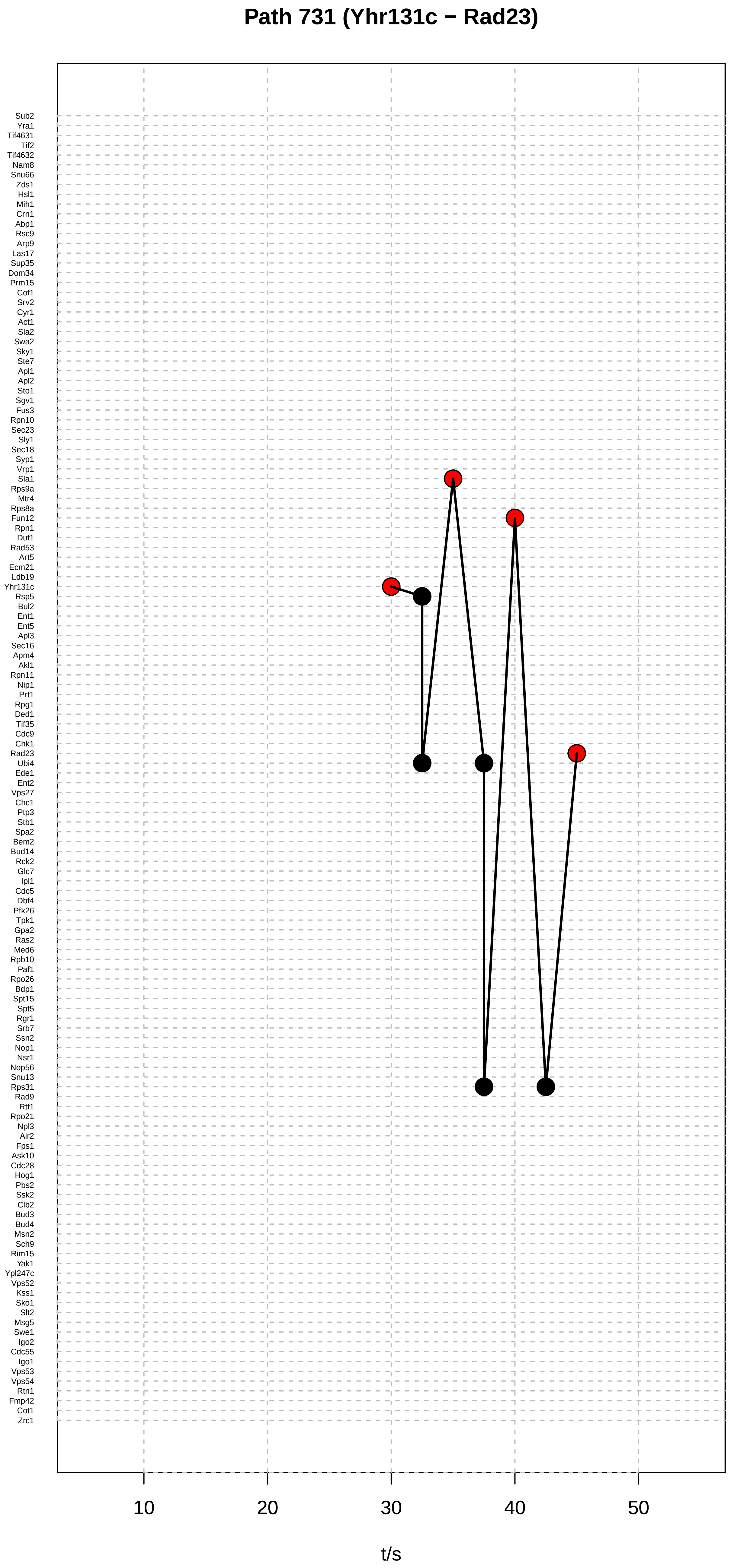

Path 731 (Yhr131c − Rad23)
Sub2
Yra1
Tif4631
Tif2
Tif4632
Nam8
Snu66
Zds1
Hsl1
Mih1
Crn1
Abp1
Rsc9
Arp9
Las17
Sup35
Dom34
Prm15
Cof1
Srv2
Cyr1
Act1
Sla2
Swa2
Sky1
Ste7
Apl1
Apl2
Sto1
Sgv1
Fus3
Rpn10
Sec23
Sly1
Sec18
Syp1
Vrp1
Sla1
Rps9a
Mtr4
Rps8a
Fun12
Rpn1
Duf1
Rad53
Art5
Ecm21
Ldb19
Yhr131c
Rsp5
Bul2
Ent1
Ent5
Apl3
Sec16
Apm4
Akl1
Rpn11
Nip1
Prt1
Rpg1
Ded1
Tif35
Cdc9
Chk1
Rad23
Ubi4
Ede1
Ent2
Vps27
Chc1
Ptp3
Stb1
Spa2
Bem2
Bud14
Rck2
Glc7
Ipl1
Cdc5
Dbf4
Pfk26
Tpk1
Gpa2
Ras2
Med6
Rpb10
Paf1
Rpo26
Bdp1
Spt15
Spt5
Rgr1
Srb7
Ssn2
Nop1
Nsr1
Nop56
Snu13
Rps31
Rad9
Rtf1
Rpo21
Npl3
Air2
Fps1
Ask10
Cdc28
Hog1
Pbs2
Ssk2
Clb2
Bud3
Bud4
Msn2
Sch9
Rim15
Yak1
Ypl247c
Vps52
Kss1
Sko1
Slt2
Msg5
Swe1
Igo2
Cdc55
Igo1
Vps53
Vps54
Rtn1
Fmp42
Cot1
Zrc1
10
20
30
40
50
t/s

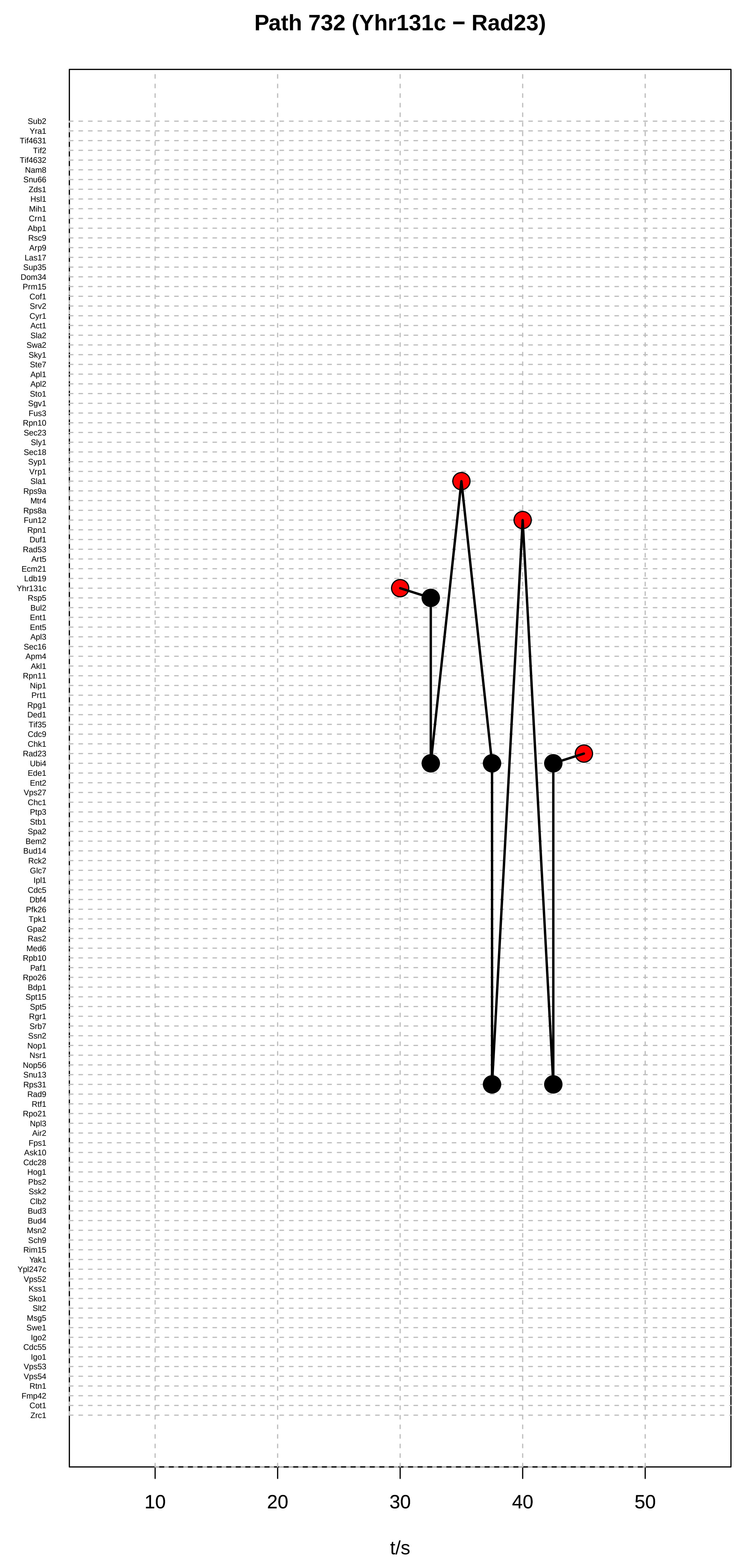
Path 732 (Yhr131c − Rad23)
Sub2
Yra1
Tif4631
Tif2
Tif4632
Nam8
Snu66
Zds1
Hsl1
Mih1
Crn1
Abp1
Rsc9
Arp9
Las17
Sup35
Dom34
Prm15
Cof1
Srv2
Cyr1
Act1
Sla2
Swa2
Sky1
Ste7
Apl1
Apl2
Sto1
Sgv1
Fus3
Rpn10
Sec23
Sly1
Sec18
Syp1
Vrp1
Sla1
Rps9a
Mtr4
Rps8a
Fun12
Rpn1
Duf1
Rad53
Art5
Ecm21
Ldb19
Yhr131c
Rsp5
Bul2
Ent1
Ent5
Apl3
Sec16
Apm4
Akl1
Rpn11
Nip1
Prt1
Rpg1
Ded1
Tif35
Cdc9
Chk1
Rad23
Ubi4
Ede1
Ent2
Vps27
Chc1
Ptp3
Stb1
Spa2
Bem2
Bud14
Rck2
Glc7
Ipl1
Cdc5
Dbf4
Pfk26
Tpk1
Gpa2
Ras2
Med6
Rpb10
Paf1
Rpo26
Bdp1
Spt15
Spt5
Rgr1
Srb7
Ssn2
Nop1
Nsr1
Nop56
Snu13
Rps31
Rad9
Rtf1
Rpo21
Npl3
Air2
Fps1
Ask10
Cdc28
Hog1
Pbs2
Ssk2
Clb2
Bud3
Bud4
Msn2
Sch9
Rim15
Yak1
Ypl247c
Vps52
Kss1
Sko1
Slt2
Msg5
Swe1
Igo2
Cdc55
Igo1
Vps53
Vps54
Rtn1
Fmp42
Cot1
Zrc1
10
20
30
40
50
t/s

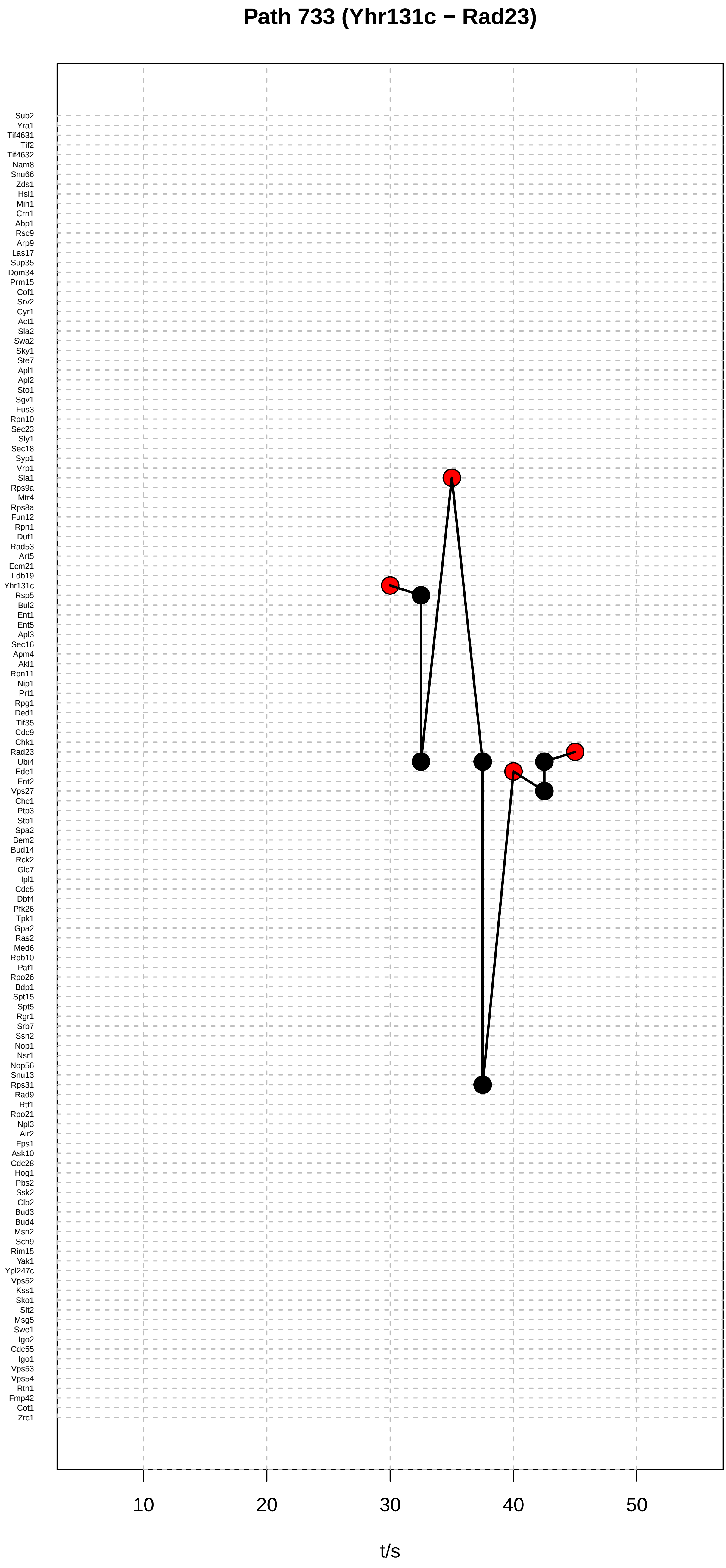

Path 733 (Yhr131c − Rad23)
Sub2
Yra1
Tif4631
Tif2
Tif4632
Nam8
Snu66
Zds1
Hsl1
Mih1
Crn1
Abp1
Rsc9
Arp9
Las17
Sup35
Dom34
Prm15
Cof1
Srv2
Cyr1
Act1
Sla2
Swa2
Sky1
Ste7
Apl1
Apl2
Sto1
Sgv1
Fus3
Rpn10
Sec23
Sly1
Sec18
Syp1
Vrp1
Sla1
Rps9a
Mtr4
Rps8a
Fun12
Rpn1
Duf1
Rad53
Art5
Ecm21
Ldb19
Yhr131c
Rsp5
Bul2
Ent1
Ent5
Apl3
Sec16
Apm4
Akl1
Rpn11
Nip1
Prt1
Rpg1
Ded1
Tif35
Cdc9
Chk1
Rad23
Ubi4
Ede1
Ent2
Vps27
Chc1
Ptp3
Stb1
Spa2
Bem2
Bud14
Rck2
Glc7
Ipl1
Cdc5
Dbf4
Pfk26
Tpk1
Gpa2
Ras2
Med6
Rpb10
Paf1
Rpo26
Bdp1
Spt15
Spt5
Rgr1
Srb7
Ssn2
Nop1
Nsr1
Nop56
Snu13
Rps31
Rad9
Rtf1
Rpo21
Npl3
Air2
Fps1
Ask10
Cdc28
Hog1
Pbs2
Ssk2
Clb2
Bud3
Bud4
Msn2
Sch9
Rim15
Yak1
Ypl247c
Vps52
Kss1
Sko1
Slt2
Msg5
Swe1
Igo2
Cdc55
Igo1
Vps53
Vps54
Rtn1
Fmp42
Cot1
Zrc1
10
20
30
40
50
t/s

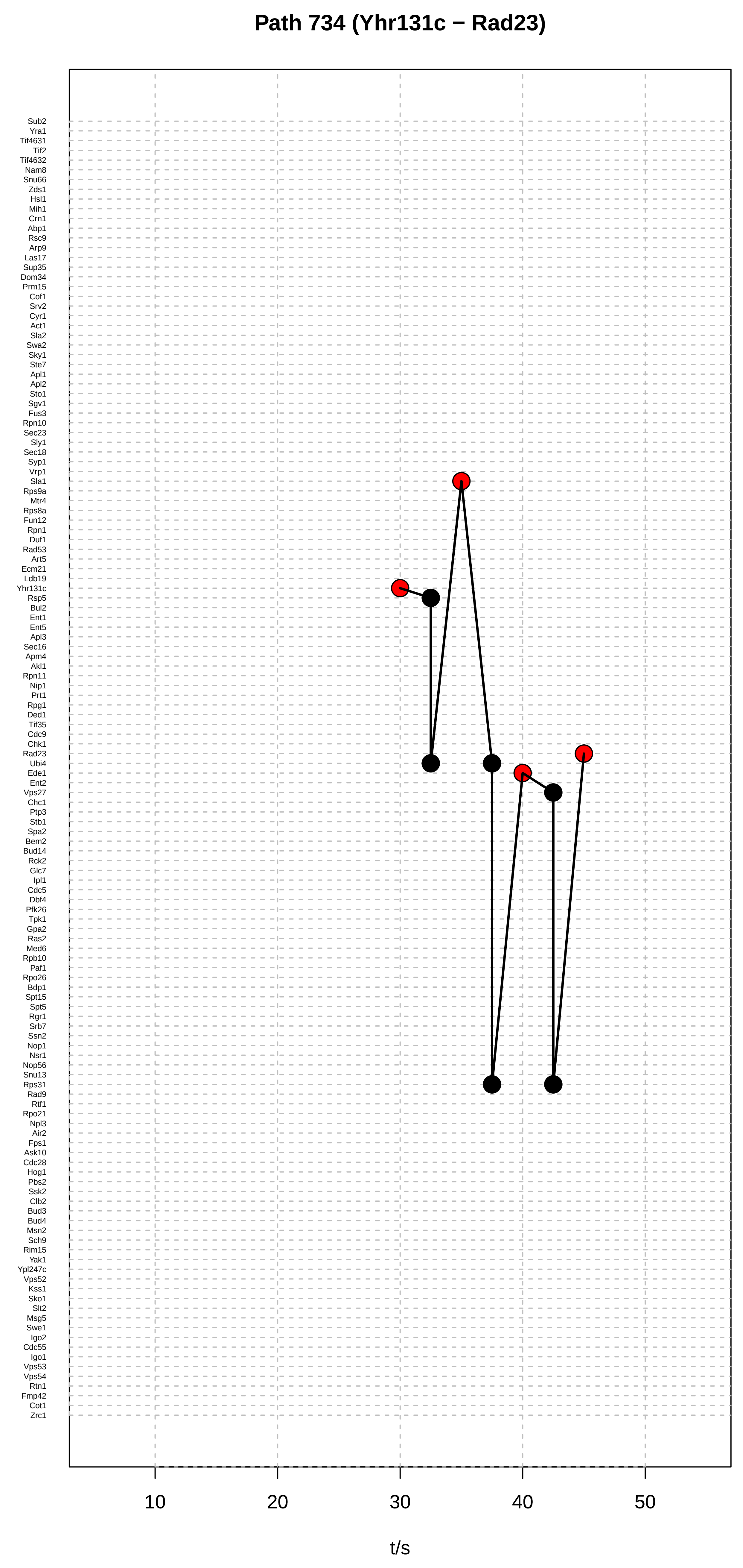

Path 734 (Yhr131c − Rad23)
Sub2
Yra1
Tif4631
Tif2
Tif4632
Nam8
Snu66
Zds1
Hsl1
Mih1
Crn1
Abp1
Rsc9
Arp9
Las17
Sup35
Dom34
Prm15
Cof1
Srv2
Cyr1
Act1
Sla2
Swa2
Sky1
Ste7
Apl1
Apl2
Sto1
Sgv1
Fus3
Rpn10
Sec23
Sly1
Sec18
Syp1
Vrp1
Sla1
Rps9a
Mtr4
Rps8a
Fun12
Rpn1
Duf1
Rad53
Art5
Ecm21
Ldb19
Yhr131c
Rsp5
Bul2
Ent1
Ent5
Apl3
Sec16
Apm4
Akl1
Rpn11
Nip1
Prt1
Rpg1
Ded1
Tif35
Cdc9
Chk1
Rad23
Ubi4
Ede1
Ent2
Vps27
Chc1
Ptp3
Stb1
Spa2
Bem2
Bud14
Rck2
Glc7
Ipl1
Cdc5
Dbf4
Pfk26
Tpk1
Gpa2
Ras2
Med6
Rpb10
Paf1
Rpo26
Bdp1
Spt15
Spt5
Rgr1
Srb7
Ssn2
Nop1
Nsr1
Nop56
Snu13
Rps31
Rad9
Rtf1
Rpo21
Npl3
Air2
Fps1
Ask10
Cdc28
Hog1
Pbs2
Ssk2
Clb2
Bud3
Bud4
Msn2
Sch9
Rim15
Yak1
Ypl247c
Vps52
Kss1
Sko1
Slt2
Msg5
Swe1
Igo2
Cdc55
Igo1
Vps53
Vps54
Rtn1
Fmp42
Cot1
Zrc1
10
20
30
40
50
t/s

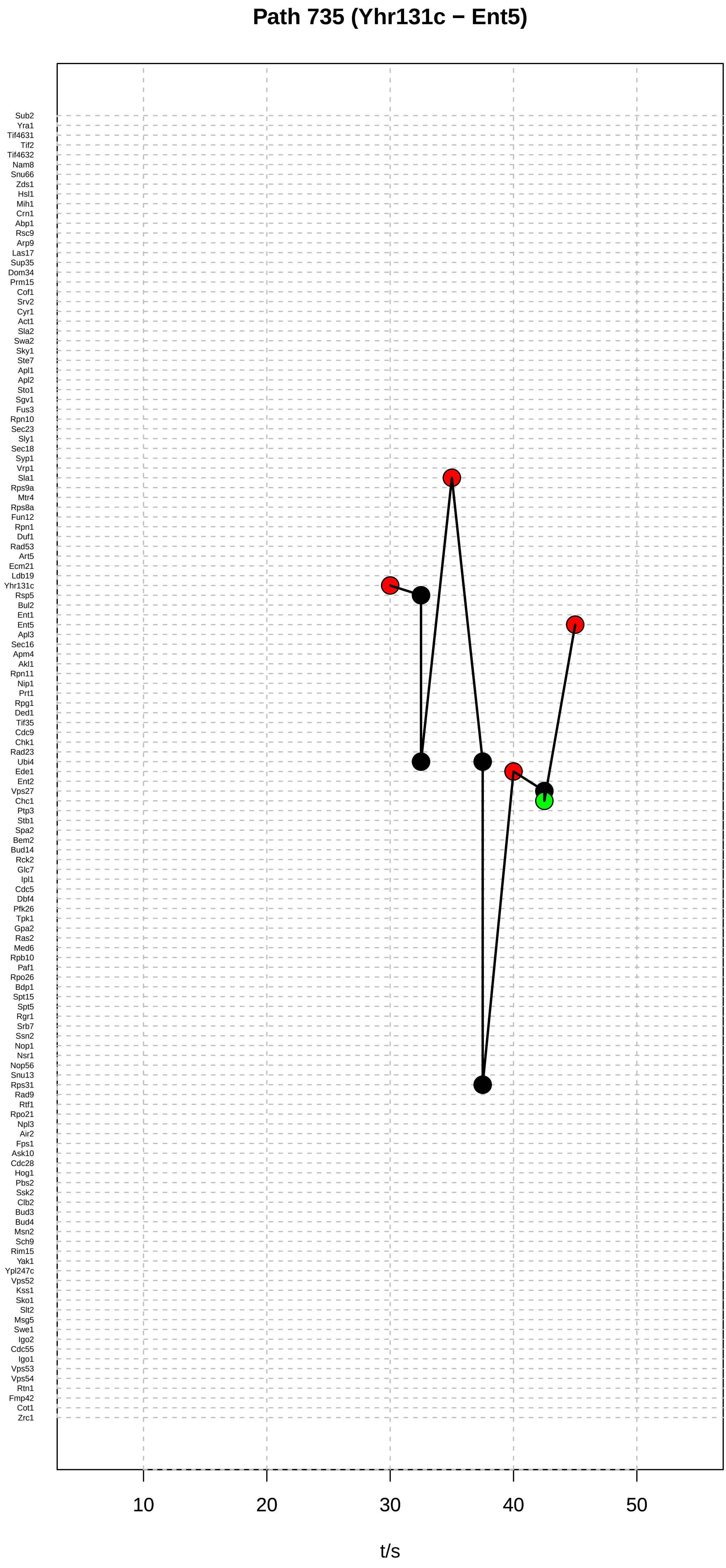
Path 735 (Yhr131c − Ent5)
Sub2
Yra1
Tif4631
Tif2
Tif4632
Nam8
Snu66
Zds1
Hsl1
Mih1
Crn1
Abp1
Rsc9
Arp9
Las17
Sup35
Dom34
Prm15
Cof1
Srv2
Cyr1
Act1
Sla2
Swa2
Sky1
Ste7
Apl1
Apl2
Sto1
Sgv1
Fus3
Rpn10
Sec23
Sly1
Sec18
Syp1
Vrp1
Sla1
Rps9a
Mtr4
Rps8a
Fun12
Rpn1
Duf1
Rad53
Art5
Ecm21
Ldb19
Yhr131c
Rsp5
Bul2
Ent1
Ent5
Apl3
Sec16
Apm4
Akl1
Rpn11
Nip1
Prt1
Rpg1
Ded1
Tif35
Cdc9
Chk1
Rad23
Ubi4
Ede1
Ent2
Vps27
Chc1
Ptp3
Stb1
Spa2
Bem2
Bud14
Rck2
Glc7
Ipl1
Cdc5
Dbf4
Pfk26
Tpk1
Gpa2
Ras2
Med6
Rpb10
Paf1
Rpo26
Bdp1
Spt15
Spt5
Rgr1
Srb7
Ssn2
Nop1
Nsr1
Nop56
Snu13
Rps31
Rad9
Rtf1
Rpo21
Npl3
Air2
Fps1
Ask10
Cdc28
Hog1
Pbs2
Ssk2
Clb2
Bud3
Bud4
Msn2
Sch9
Rim15
Yak1
Ypl247c
Vps52
Kss1
Sko1
Slt2
Msg5
Swe1
Igo2
Cdc55
Igo1
Vps53
Vps54
Rtn1
Fmp42
Cot1
Zrc1
10
20
30
40
50
t/s

#### Path 736 (Yhr131c − Rad23)

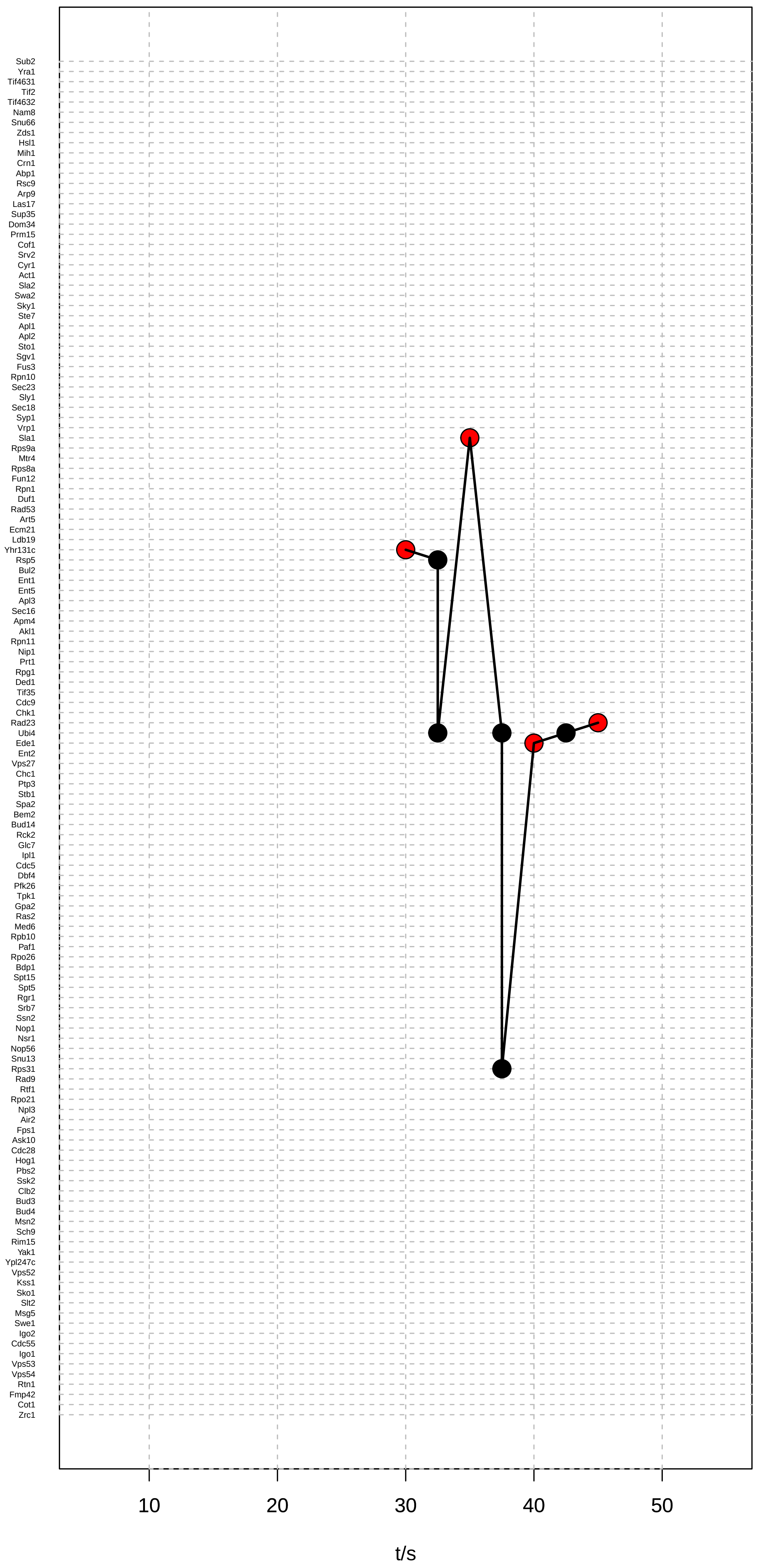

#### Path 737 (Yhr131c − Rad23)

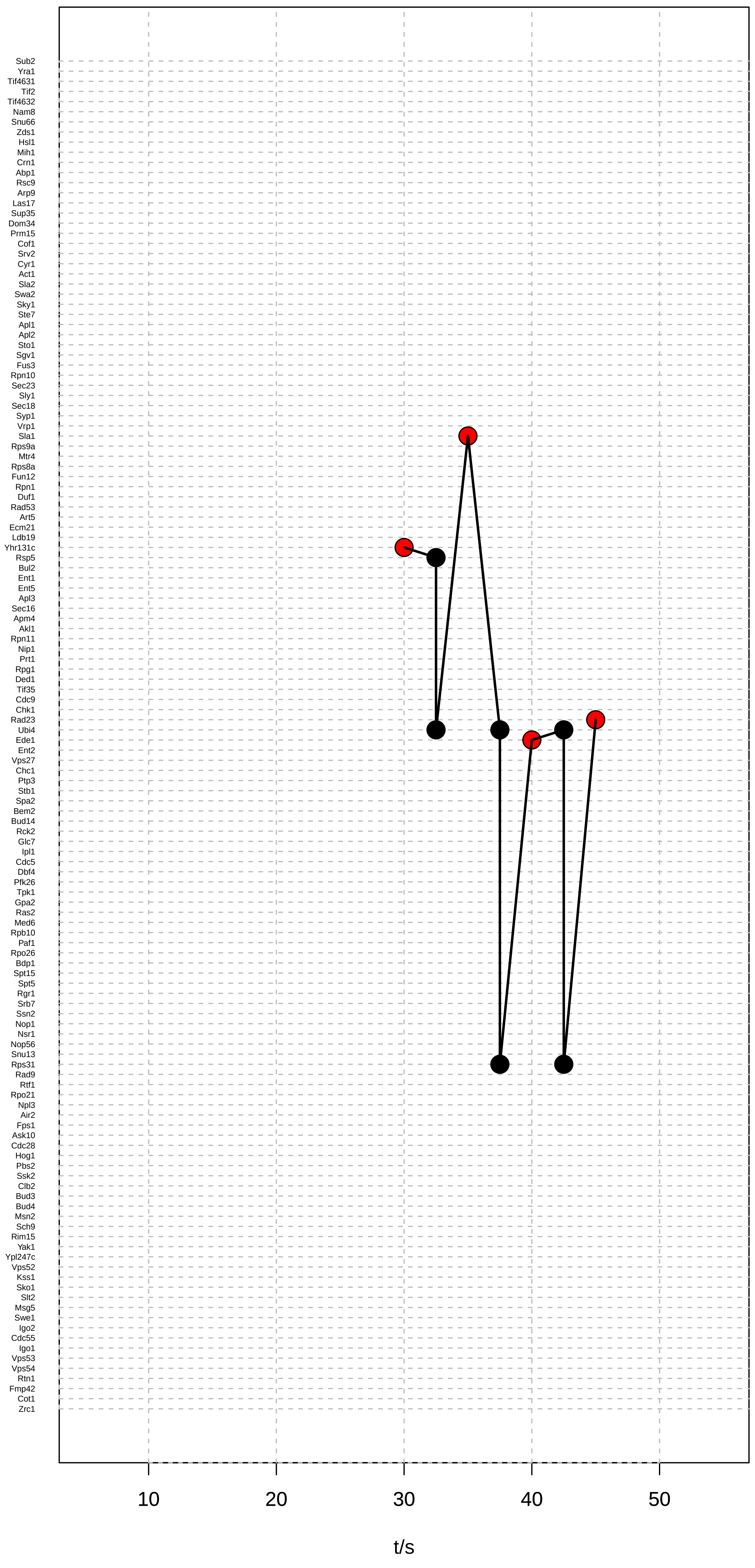

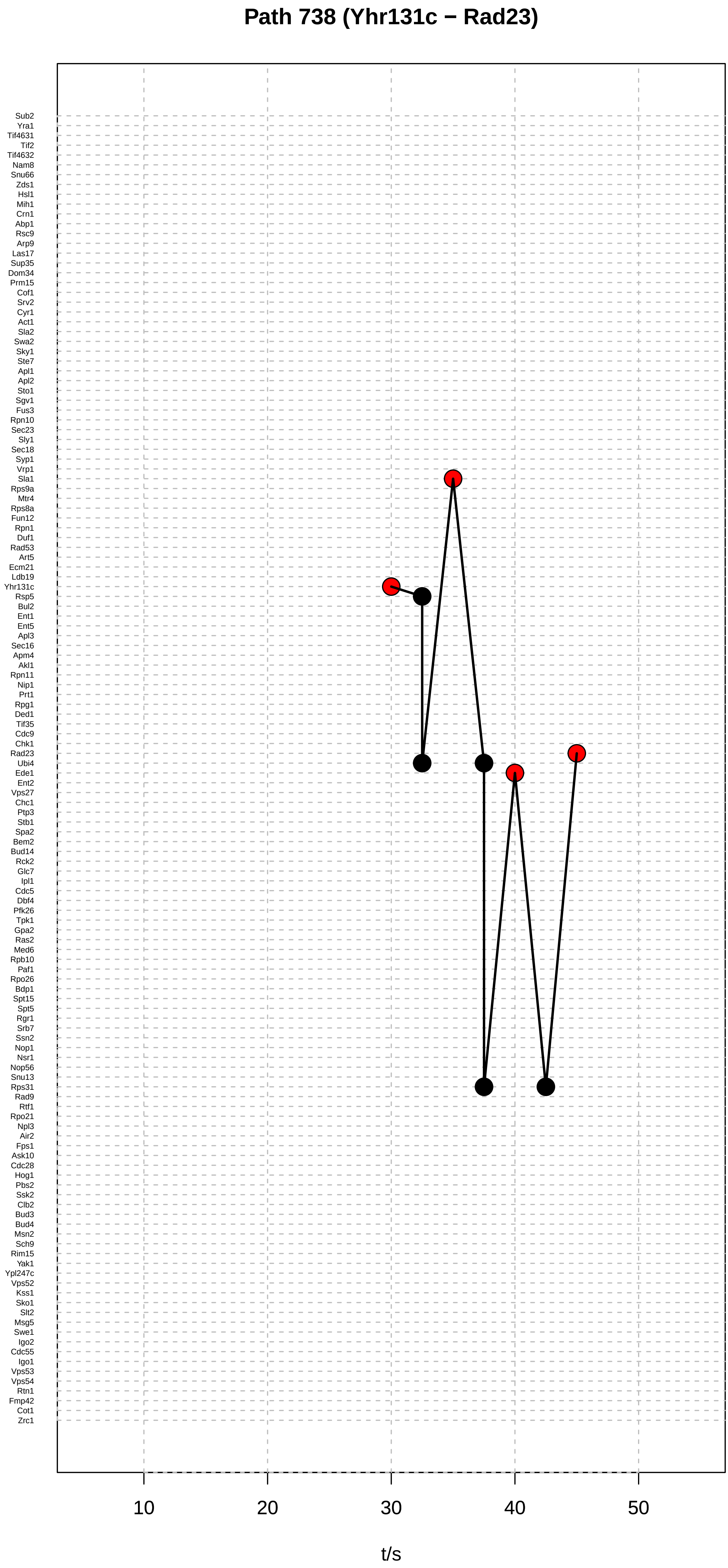
Path 738 (Yhr131c − Rad23)
Sub2
Yra1
Tif4631
Tif2
Tif4632
Nam8
Snu66
Zds1
Hsl1
Mih1
Crn1
Abp1
Rsc9
Arp9
Las17
Sup35
Dom34
Prm15
Cof1
Srv2
Cyr1
Act1
Sla2
Swa2
Sky1
Ste7
Apl1
Apl2
Sto1
Sgv1
Fus3
Rpn10
Sec23
Sly1
Sec18
Syp1
Vrp1
Sla1
Rps9a
Mtr4
Rps8a
Fun12
Rpn1
Duf1
Rad53
Art5
Ecm21
Ldb19
Yhr131c
Rsp5
Bul2
Ent1
Ent5
Apl3
Sec16
Apm4
Akl1
Rpn11
Nip1
Prt1
Rpg1
Ded1
Tif35
Cdc9
Chk1
Rad23
Ubi4
Ede1
Ent2
Vps27
Chc1
Ptp3
Stb1
Spa2
Bem2
Bud14
Rck2
Glc7
Ipl1
Cdc5
Dbf4
Pfk26
Tpk1
Gpa2
Ras2
Med6
Rpb10
Paf1
Rpo26
Bdp1
Spt15
Spt5
Rgr1
Srb7
Ssn2
Nop1
Nsr1
Nop56
Snu13
Rps31
Rad9
Rtf1
Rpo21
Npl3
Air2
Fps1
Ask10
Cdc28
Hog1
Pbs2
Ssk2
Clb2
Bud3
Bud4
Msn2
Sch9
Rim15
Yak1
Ypl247c
Vps52
Kss1
Sko1
Slt2
Msg5
Swe1
Igo2
Cdc55
Igo1
Vps53
Vps54
Rtn1
Fmp42
Cot1
Zrc1
10
20
30
40
50
t/s

### Path 739 (Yhr131c − Rad23)

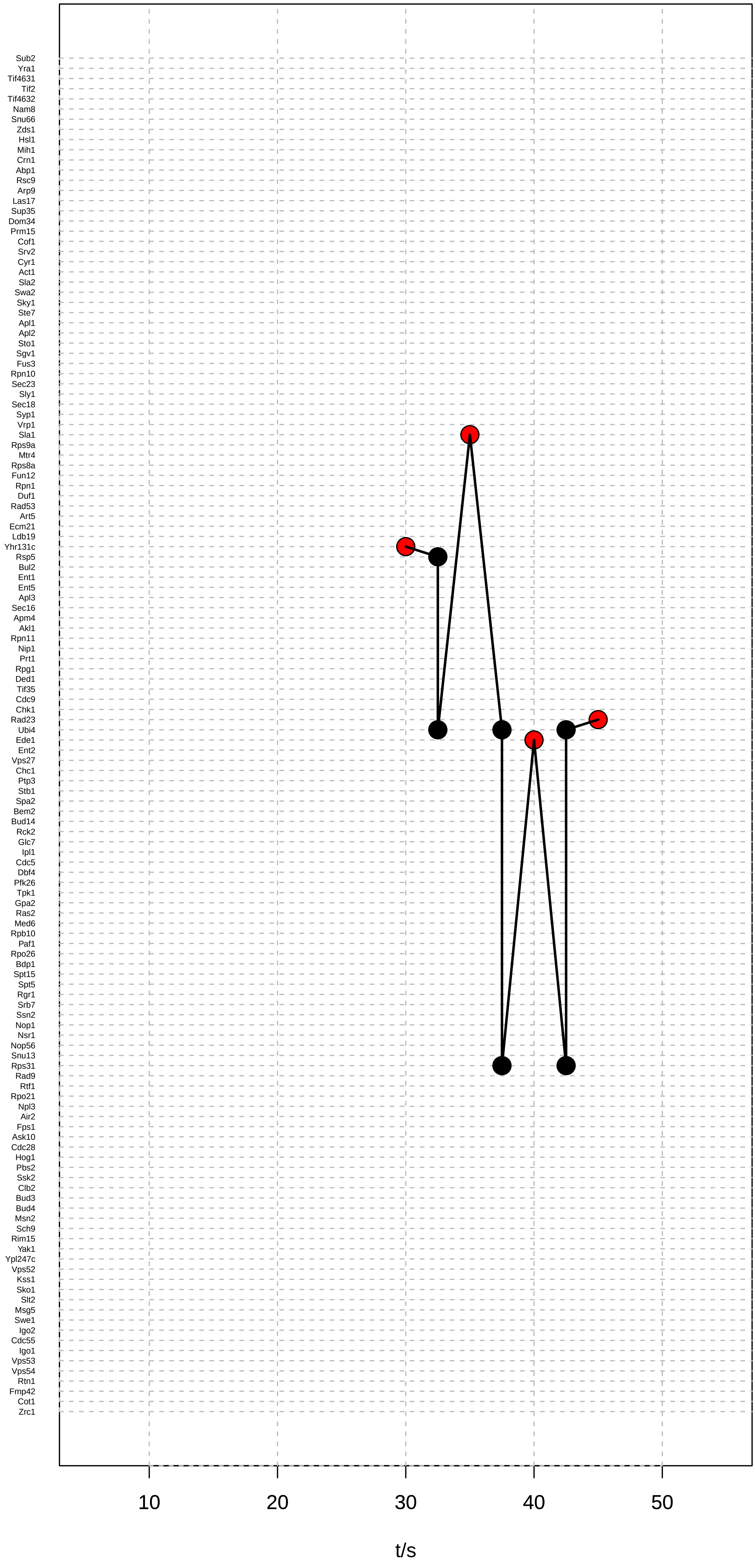

### Path 740 (Yhr131c − Rad23)

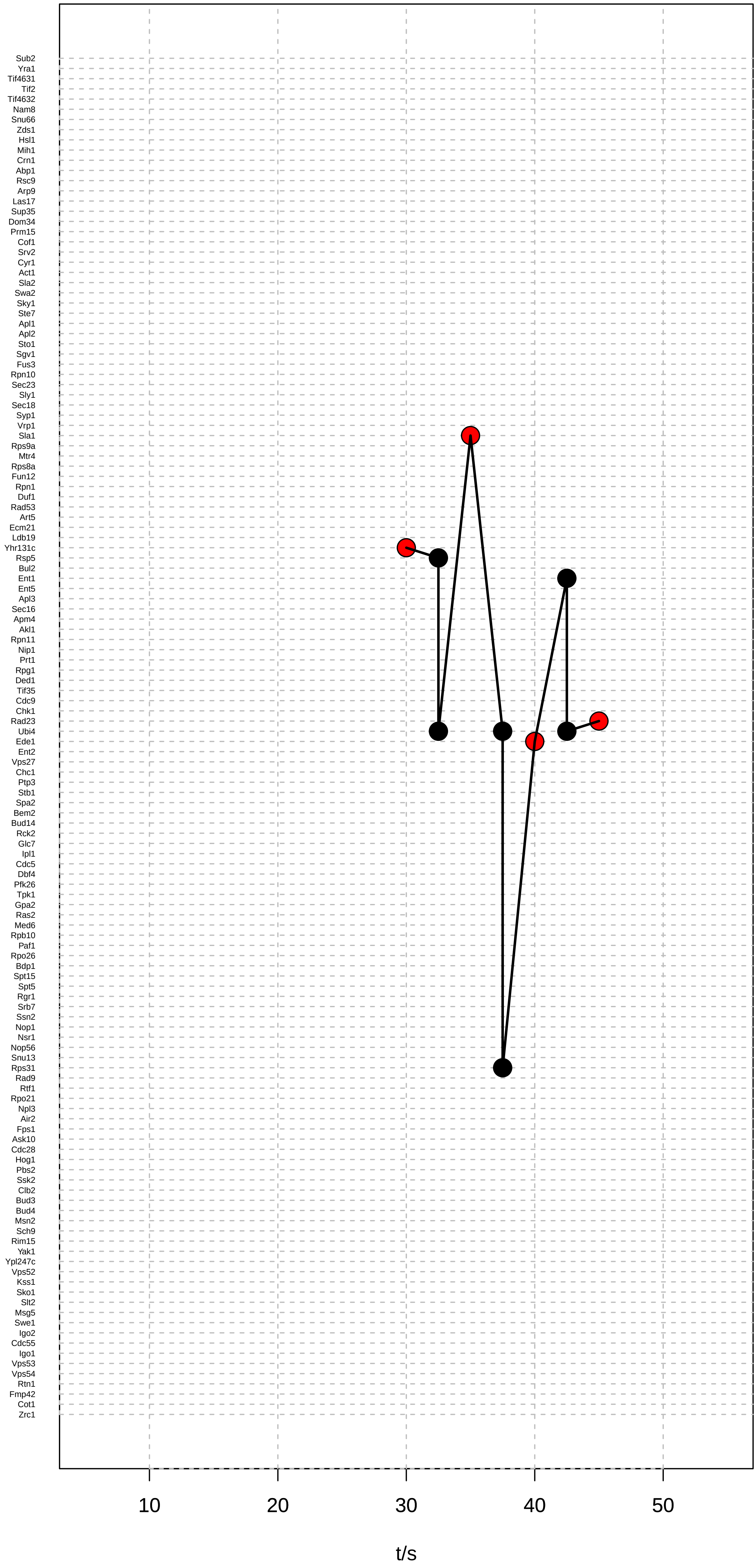

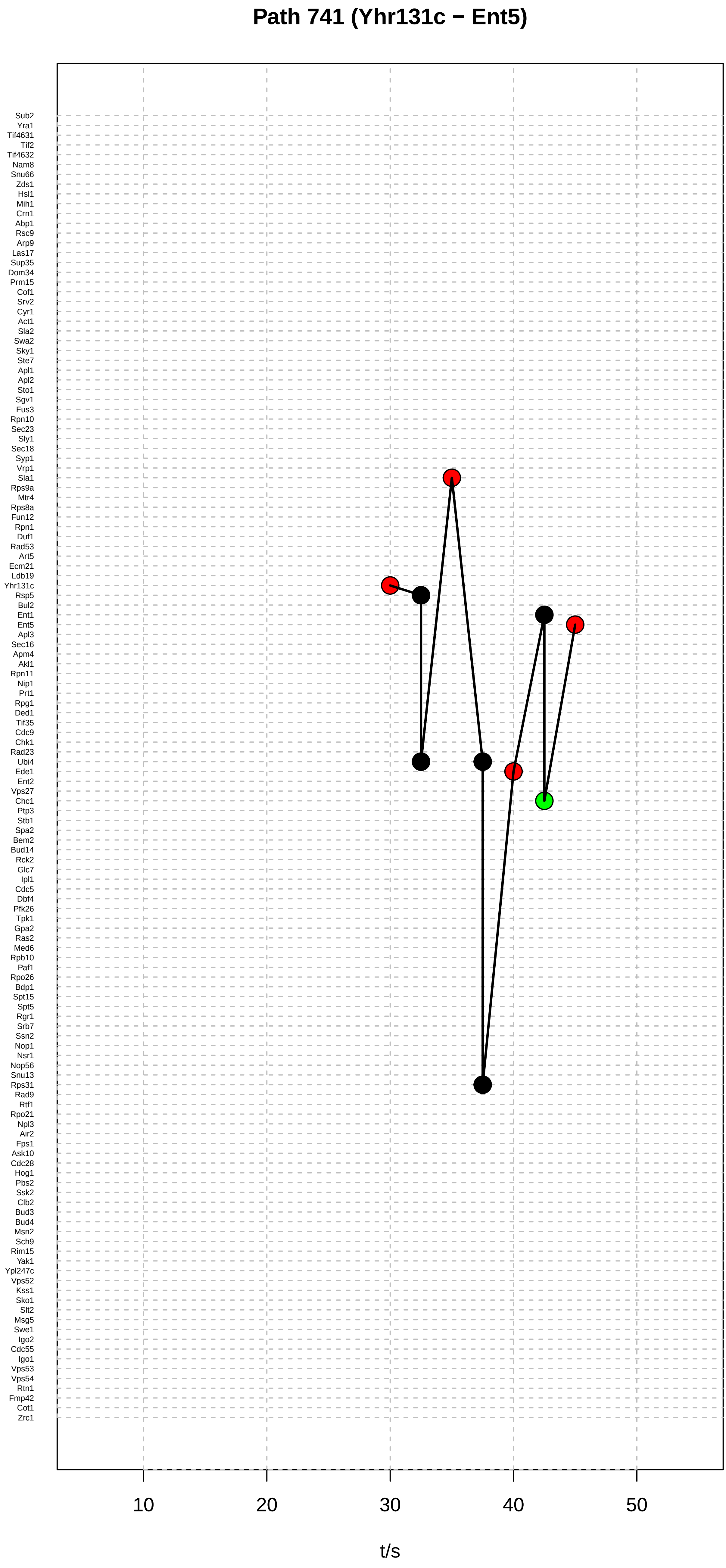

Path 741 (Yhr131c − Ent5)
Sub2
Yra1
Tif4631
Tif2
Tif4632
Nam8
Snu66
Zds1
Hsl1
Mih1
Crn1
Abp1
Rsc9
Arp9
Las17
Sup35
Dom34
Prm15
Cof1
Srv2
Cyr1
Act1
Sla2
Swa2
Sky1
Ste7
Apl1
Apl2
Sto1
Sgv1
Fus3
Rpn10
Sec23
Sly1
Sec18
Syp1
Vrp1
Sla1
Rps9a
Mtr4
Rps8a
Fun12
Rpn1
Duf1
Rad53
Art5
Ecm21
Ldb19
Yhr131c
Rsp5
Bul2
Ent1
Ent5
Apl3
Sec16
Apm4
Akl1
Rpn11
Nip1
Prt1
Rpg1
Ded1
Tif35
Cdc9
Chk1
Rad23
Ubi4
Ede1
Ent2
Vps27
Chc1
Ptp3
Stb1
Spa2
Bem2
Bud14
Rck2
Glc7
Ipl1
Cdc5
Dbf4
Pfk26
Tpk1
Gpa2
Ras2
Med6
Rpb10
Paf1
Rpo26
Bdp1
Spt15
Spt5
Rgr1
Srb7
Ssn2
Nop1
Nsr1
Nop56
Snu13
Rps31
Rad9
Rtf1
Rpo21
Npl3
Air2
Fps1
Ask10
Cdc28
Hog1
Pbs2
Ssk2
Clb2
Bud3
Bud4
Msn2
Sch9
Rim15
Yak1
Ypl247c
Vps52
Kss1
Sko1
Slt2
Msg5
Swe1
Igo2
Cdc55
Igo1
Vps53
Vps54
Rtn1
Fmp42
Cot1
Zrc1
10
20
30
40
50
t/s

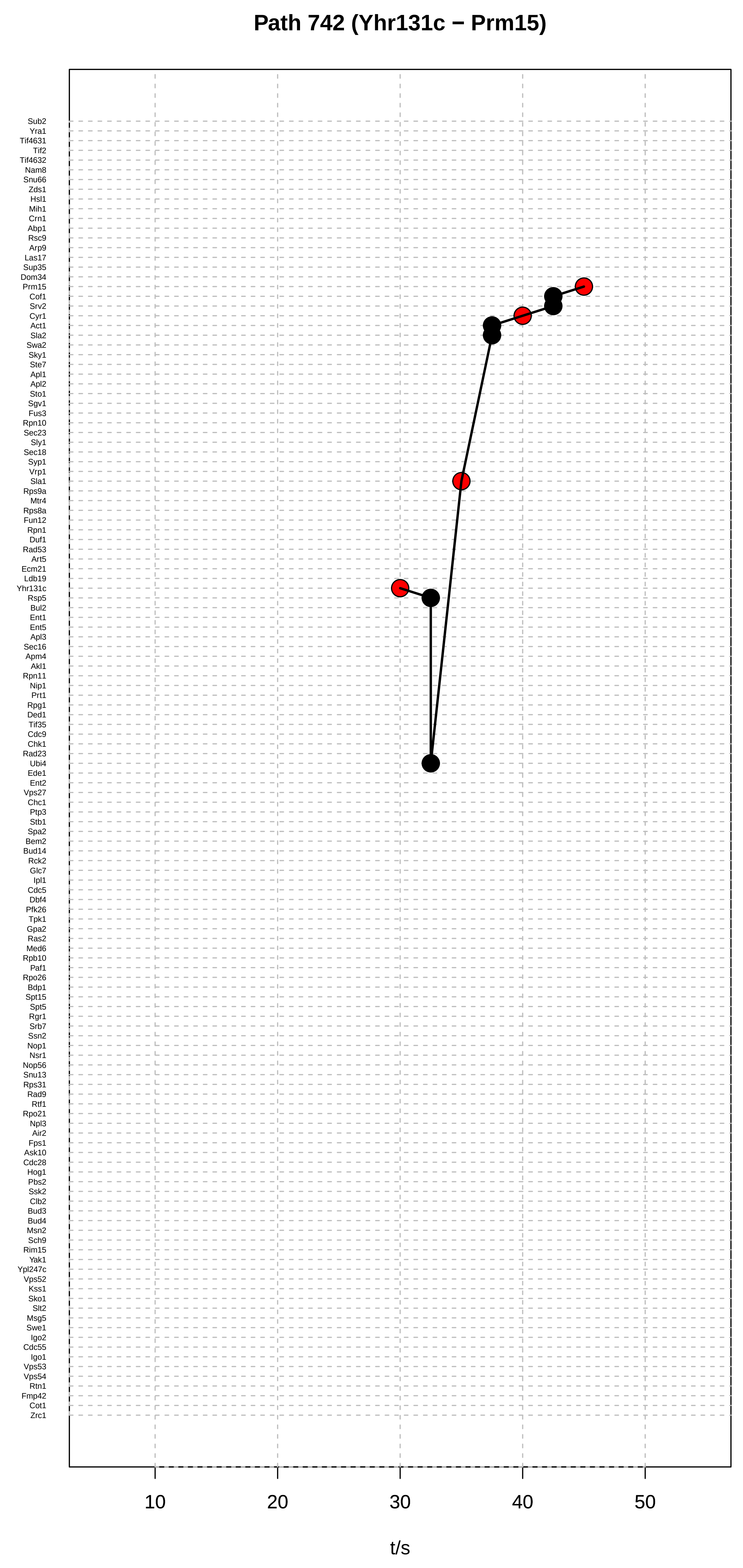

Path 742 (Yhr131c − Prm15)
Sub2
Yra1
Tif4631
Tif2
Tif4632
Nam8
Snu66
Zds1
Hsl1
Mih1
Crn1
Abp1
Rsc9
Arp9
Las17
Sup35
Dom34
Prm15
Cof1
Srv2
Cyr1
Act1
Sla2
Swa2
Sky1
Ste7
Apl1
Apl2
Sto1
Sgv1
Fus3
Rpn10
Sec23
Sly1
Sec18
Syp1
Vrp1
Sla1
Rps9a
Mtr4
Rps8a
Fun12
Rpn1
Duf1
Rad53
Art5
Ecm21
Ldb19
Yhr131c
Rsp5
Bul2
Ent1
Ent5
Apl3
Sec16
Apm4
Akl1
Rpn11
Nip1
Prt1
Rpg1
Ded1
Tif35
Cdc9
Chk1
Rad23
Ubi4
Ede1
Ent2
Vps27
Chc1
Ptp3
Stb1
Spa2
Bem2
Bud14
Rck2
Glc7
Ipl1
Cdc5
Dbf4
Pfk26
Tpk1
Gpa2
Ras2
Med6
Rpb10
Paf1
Rpo26
Bdp1
Spt15
Spt5
Rgr1
Srb7
Ssn2
Nop1
Nsr1
Nop56
Snu13
Rps31
Rad9
Rtf1
Rpo21
Npl3
Air2
Fps1
Ask10
Cdc28
Hog1
Pbs2
Ssk2
Clb2
Bud3
Bud4
Msn2
Sch9
Rim15
Yak1
Ypl247c
Vps52
Kss1
Sko1
Slt2
Msg5
Swe1
Igo2
Cdc55
Igo1
Vps53
Vps54
Rtn1
Fmp42
Cot1
Zrc1
10
20
30
40
50
t/s

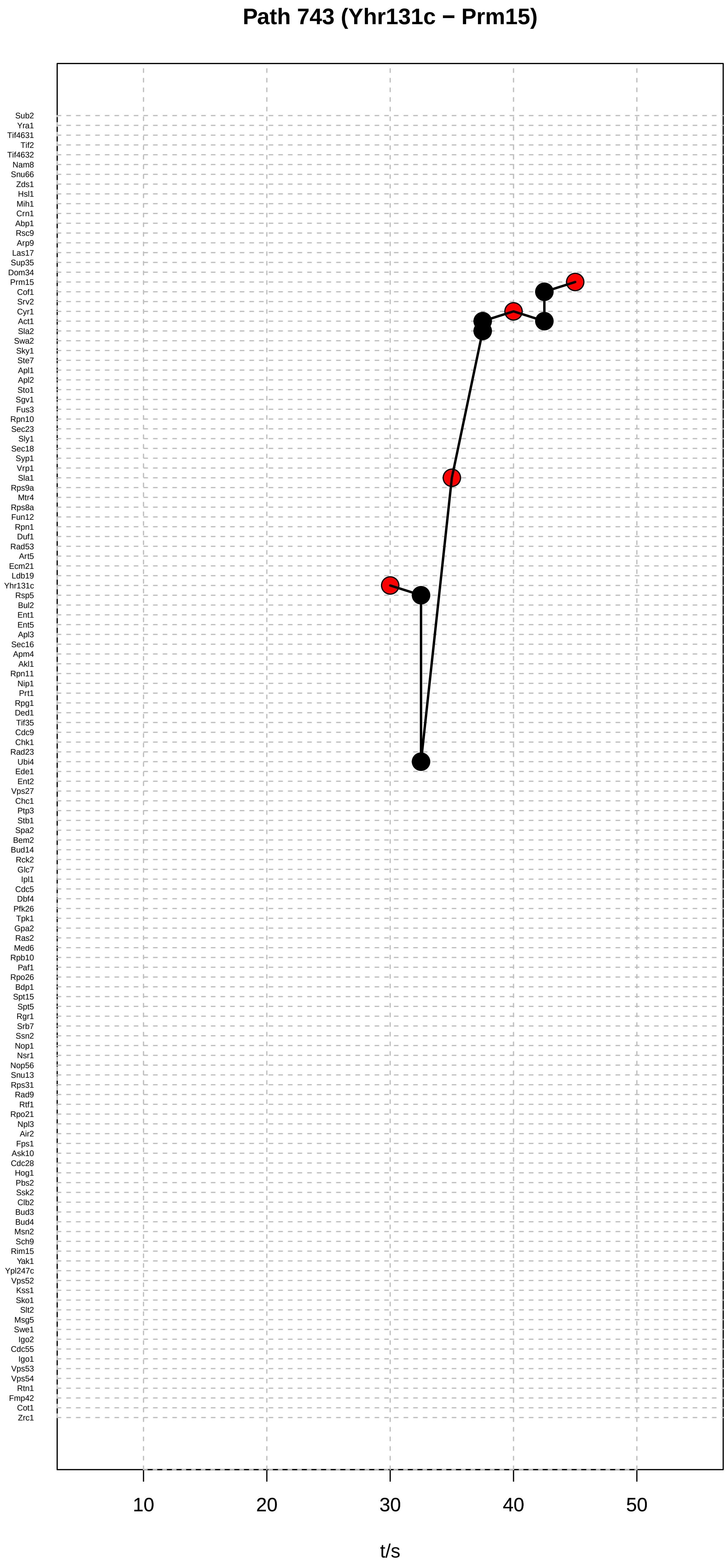
Path 743 (Yhr131c − Prm15)
Sub2
Yra1
Tif4631
Tif2
Tif4632
Nam8
Snu66
Zds1
Hsl1
Mih1
Crn1
Abp1
Rsc9
Arp9
Las17
Sup35
Dom34
Prm15
Cof1
Srv2
Cyr1
Act1
Sla2
Swa2
Sky1
Ste7
Apl1
Apl2
Sto1
Sgv1
Fus3
Rpn10
Sec23
Sly1
Sec18
Syp1
Vrp1
Sla1
Rps9a
Mtr4
Rps8a
Fun12
Rpn1
Duf1
Rad53
Art5
Ecm21
Ldb19
Yhr131c
Rsp5
Bul2
Ent1
Ent5
Apl3
Sec16
Apm4
Akl1
Rpn11
Nip1
Prt1
Rpg1
Ded1
Tif35
Cdc9
Chk1
Rad23
Ubi4
Ede1
Ent2
Vps27
Chc1
Ptp3
Stb1
Spa2
Bem2
Bud14
Rck2
Glc7
Ipl1
Cdc5
Dbf4
Pfk26
Tpk1
Gpa2
Ras2
Med6
Rpb10
Paf1
Rpo26
Bdp1
Spt15
Spt5
Rgr1
Srb7
Ssn2
Nop1
Nsr1
Nop56
Snu13
Rps31
Rad9
Rtf1
Rpo21
Npl3
Air2
Fps1
Ask10
Cdc28
Hog1
Pbs2
Ssk2
Clb2
Bud3
Bud4
Msn2
Sch9
Rim15
Yak1
Ypl247c
Vps52
Kss1
Sko1
Slt2
Msg5
Swe1
Igo2
Cdc55
Igo1
Vps53
Vps54
Rtn1
Fmp42
Cot1
Zrc1
10
20
30
40
50
t/s

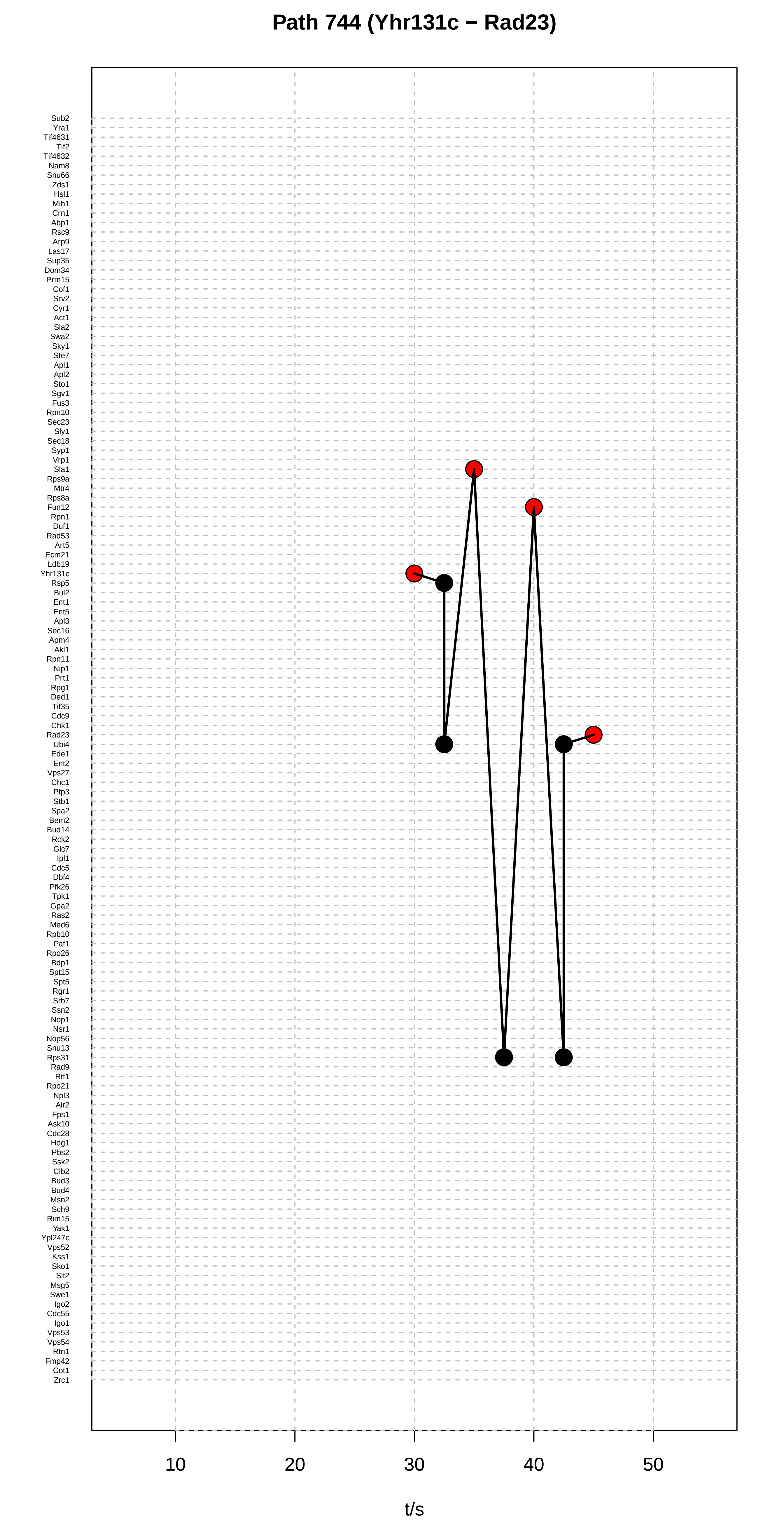

Path 744 (Yhr131c − Rad23)
Sub2
Yra1
Tif4631
Tif2
Tif4632
Nam8
Snu66
Zds1
Hsl1
Mih1
Crn1
Abp1
Rsc9
Arp9
Las17
Sup35
Dom34
Prm15
Cof1
Srv2
Cyr1
Act1
Sla2
Swa2
Sky1
Ste7
Apl1
Apl2
Sto1
Sgv1
Fus3
Rpn10
Sec23
Sly1
Sec18
Syp1
Vrp1
Sla1
Rps9a
Mtr4
Rps8a
Fun12
Rpn1
Duf1
Rad53
Art5
Ecm21
Ldb19
Yhr131c
Rsp5
Bul2
Ent1
Ent5
Apl3
Sec16
Apm4
Akl1
Rpn11
Nip1
Prt1
Rpg1
Ded1
Tif35
Cdc9
Chk1
Rad23
Ubi4
Ede1
Ent2
Vps27
Chc1
Ptp3
Stb1
Spa2
Bem2
Bud14
Rck2
Glc7
Ipl1
Cdc5
Dbf4
Pfk26
Tpk1
Gpa2
Ras2
Med6
Rpb10
Paf1
Rpo26
Bdp1
Spt15
Spt5
Rgr1
Srb7
Ssn2
Nop1
Nsr1
Nop56
Snu13
Rps31
Rad9
Rtf1
Rpo21
Npl3
Air2
Fps1
Ask10
Cdc28
Hog1
Pbs2
Ssk2
Clb2
Bud3
Bud4
Msn2
Sch9
Rim15
Yak1
Ypl247c
Vps52
Kss1
Sko1
Slt2
Msg5
Swe1
Igo2
Cdc55
Igo1
Vps53
Vps54
Rtn1
Fmp42
Cot1
Zrc1
10
20
30
40
50
t/s

#### Path 745 (Yhr131c − Rad23)

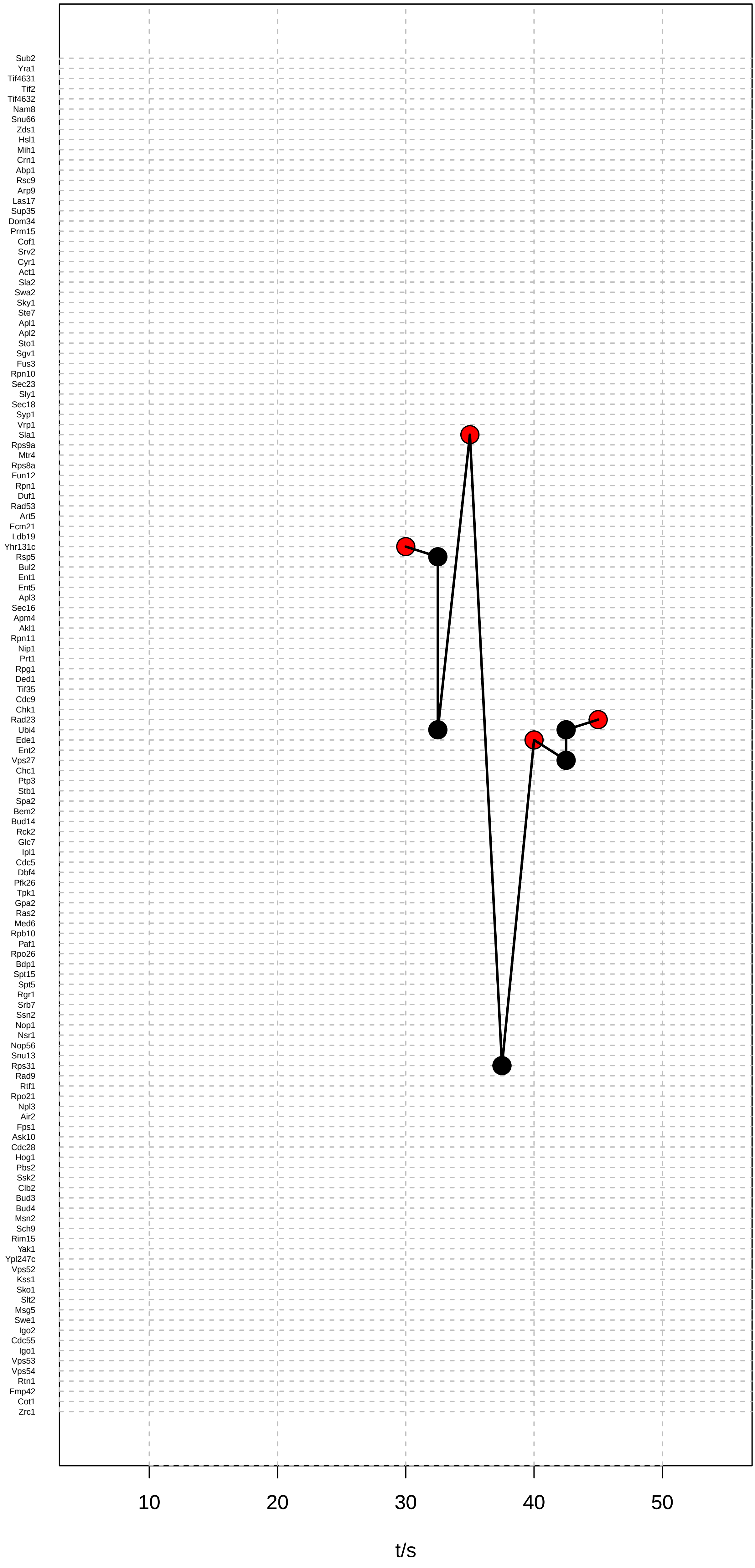

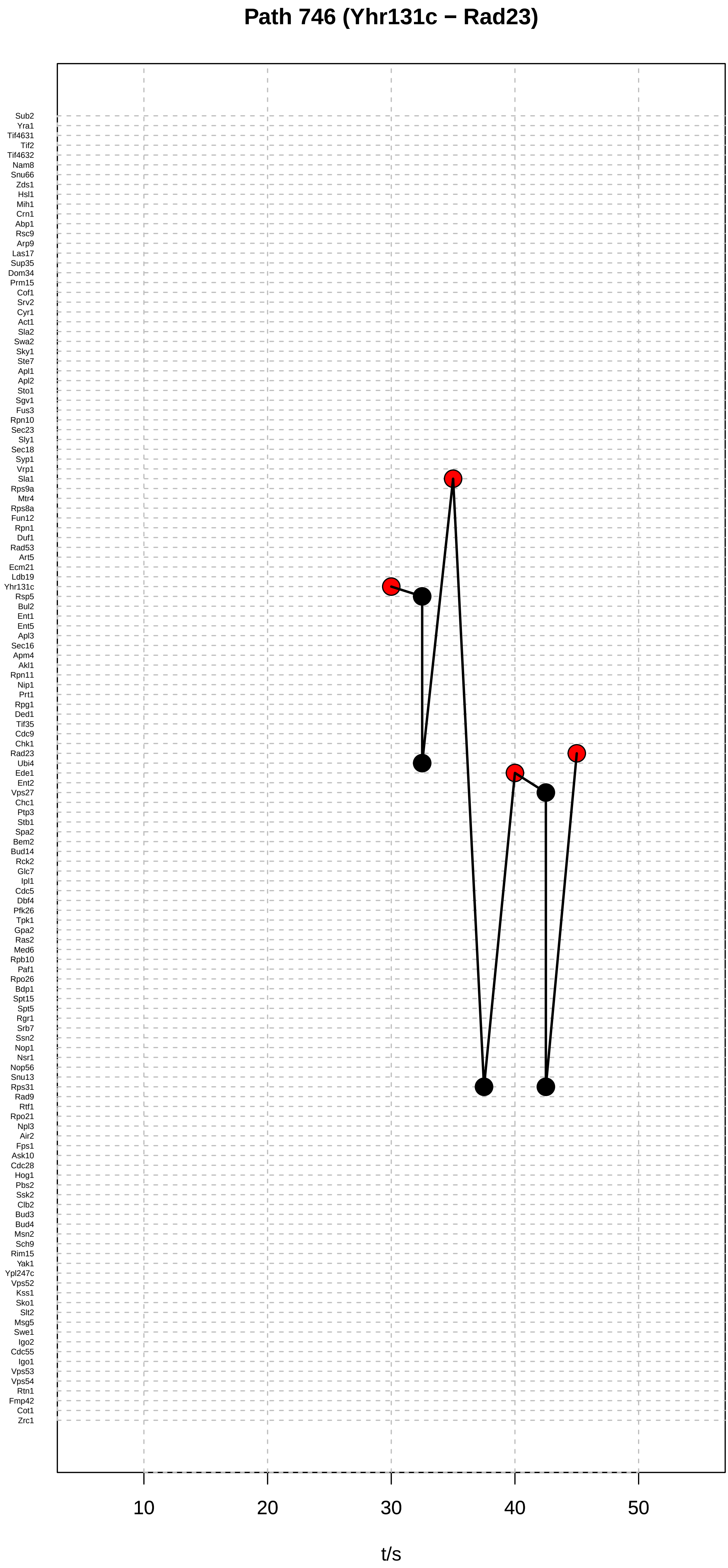

Path 746 (Yhr131c − Rad23)
Sub2
Yra1
Tif4631
Tif2
Tif4632
Nam8
Snu66
Zds1
Hsl1
Mih1
Crn1
Abp1
Rsc9
Arp9
Las17
Sup35
Dom34
Prm15
Cof1
Srv2
Cyr1
Act1
Sla2
Swa2
Sky1
Ste7
Apl1
Apl2
Sto1
Sgv1
Fus3
Rpn10
Sec23
Sly1
Sec18
Syp1
Vrp1
Sla1
Rps9a
Mtr4
Rps8a
Fun12
Rpn1
Duf1
Rad53
Art5
Ecm21
Ldb19
Yhr131c
Rsp5
Bul2
Ent1
Ent5
Apl3
Sec16
Apm4
Akl1
Rpn11
Nip1
Prt1
Rpg1
Ded1
Tif35
Cdc9
Chk1
Rad23
Ubi4
Ede1
Ent2
Vps27
Chc1
Ptp3
Stb1
Spa2
Bem2
Bud14
Rck2
Glc7
Ipl1
Cdc5
Dbf4
Pfk26
Tpk1
Gpa2
Ras2
Med6
Rpb10
Paf1
Rpo26
Bdp1
Spt15
Spt5
Rgr1
Srb7
Ssn2
Nop1
Nsr1
Nop56
Snu13
Rps31
Rad9
Rtf1
Rpo21
Npl3
Air2
Fps1
Ask10
Cdc28
Hog1
Pbs2
Ssk2
Clb2
Bud3
Bud4
Msn2
Sch9
Rim15
Yak1
Ypl247c
Vps52
Kss1
Sko1
Slt2
Msg5
Swe1
Igo2
Cdc55
Igo1
Vps53
Vps54
Rtn1
Fmp42
Cot1
Zrc1
10
20
30
40
50
t/s

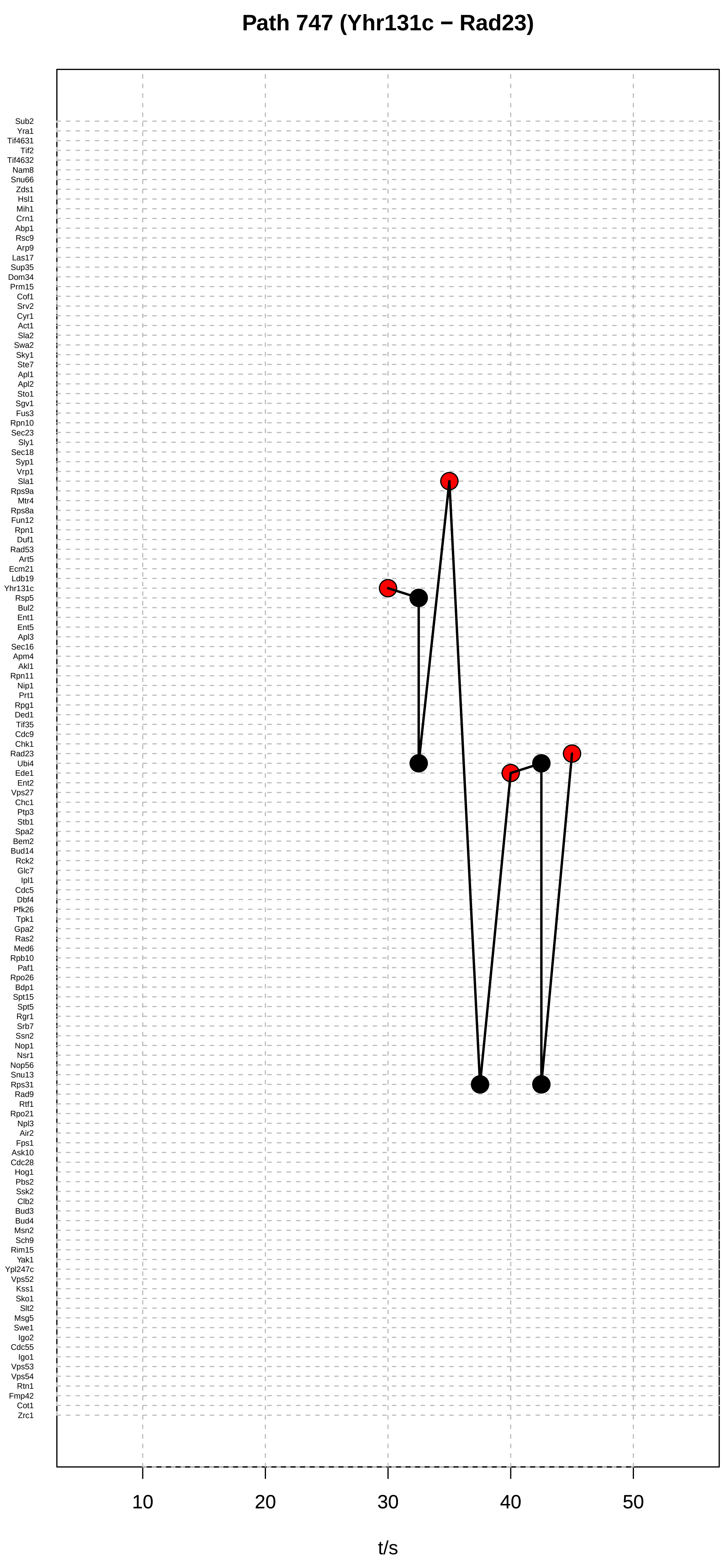
Path 747 (Yhr131c − Rad23)
Sub2
Yra1
Tif4631
Tif2
Tif4632
Nam8
Snu66
Zds1
Hsl1
Mih1
Crn1
Abp1
Rsc9
Arp9
Las17
Sup35
Dom34
Prm15
Cof1
Srv2
Cyr1
Act1
Sla2
Swa2
Sky1
Ste7
Apl1
Apl2
Sto1
Sgv1
Fus3
Rpn10
Sec23
Sly1
Sec18
Syp1
Vrp1
Sla1
Rps9a
Mtr4
Rps8a
Fun12
Rpn1
Duf1
Rad53
Art5
Ecm21
Ldb19
Yhr131c
Rsp5
Bul2
Ent1
Ent5
Apl3
Sec16
Apm4
Akl1
Rpn11
Nip1
Prt1
Rpg1
Ded1
Tif35
Cdc9
Chk1
Rad23
Ubi4
Ede1
Ent2
Vps27
Chc1
Ptp3
Stb1
Spa2
Bem2
Bud14
Rck2
Glc7
Ipl1
Cdc5
Dbf4
Pfk26
Tpk1
Gpa2
Ras2
Med6
Rpb10
Paf1
Rpo26
Bdp1
Spt15
Spt5
Rgr1
Srb7
Ssn2
Nop1
Nsr1
Nop56
Snu13
Rps31
Rad9
Rtf1
Rpo21
Npl3
Air2
Fps1
Ask10
Cdc28
Hog1
Pbs2
Ssk2
Clb2
Bud3
Bud4
Msn2
Sch9
Rim15
Yak1
Ypl247c
Vps52
Kss1
Sko1
Slt2
Msg5
Swe1
Igo2
Cdc55
Igo1
Vps53
Vps54
Rtn1
Fmp42
Cot1
Zrc1
10
20
30
40
50
t/s

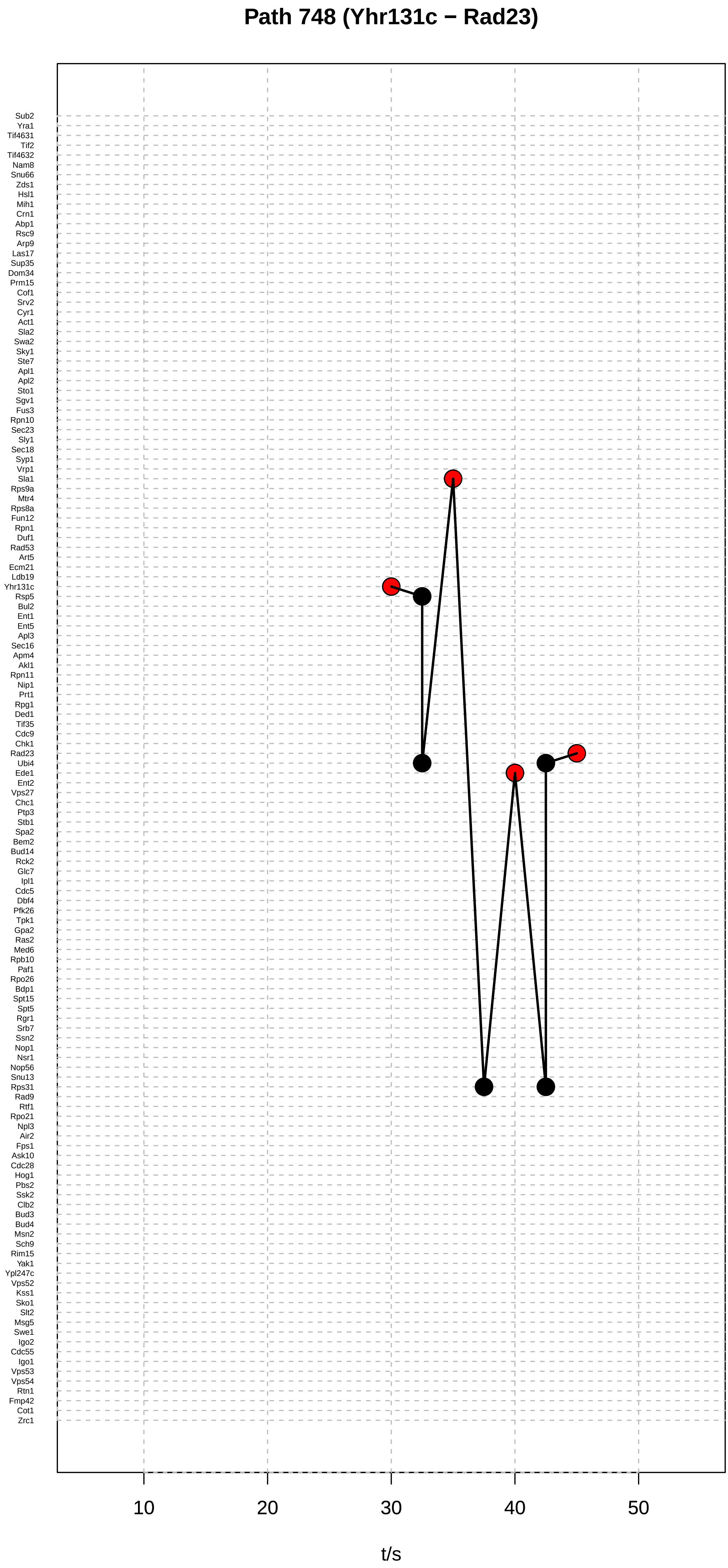

Path 748 (Yhr131c − Rad23)
Sub2
Yra1
Tif4631
Tif2
Tif4632
Nam8
Snu66
Zds1
Hsl1
Mih1
Crn1
Abp1
Rsc9
Arp9
Las17
Sup35
Dom34
Prm15
Cof1
Srv2
Cyr1
Act1
Sla2
Swa2
Sky1
Ste7
Apl1
Apl2
Sto1
Sgv1
Fus3
Rpn10
Sec23
Sly1
Sec18
Syp1
Vrp1
Sla1
Rps9a
Mtr4
Rps8a
Fun12
Rpn1
Duf1
Rad53
Art5
Ecm21
Ldb19
Yhr131c
Rsp5
Bul2
Ent1
Ent5
Apl3
Sec16
Apm4
Akl1
Rpn11
Nip1
Prt1
Rpg1
Ded1
Tif35
Cdc9
Chk1
Rad23
Ubi4
Ede1
Ent2
Vps27
Chc1
Ptp3
Stb1
Spa2
Bem2
Bud14
Rck2
Glc7
Ipl1
Cdc5
Dbf4
Pfk26
Tpk1
Gpa2
Ras2
Med6
Rpb10
Paf1
Rpo26
Bdp1
Spt15
Spt5
Rgr1
Srb7
Ssn2
Nop1
Nsr1
Nop56
Snu13
Rps31
Rad9
Rtf1
Rpo21
Npl3
Air2
Fps1
Ask10
Cdc28
Hog1
Pbs2
Ssk2
Clb2
Bud3
Bud4
Msn2
Sch9
Rim15
Yak1
Ypl247c
Vps52
Kss1
Sko1
Slt2
Msg5
Swe1
Igo2
Cdc55
Igo1
Vps53
Vps54
Rtn1
Fmp42
Cot1
Zrc1
10
20
30
40
50
t/s

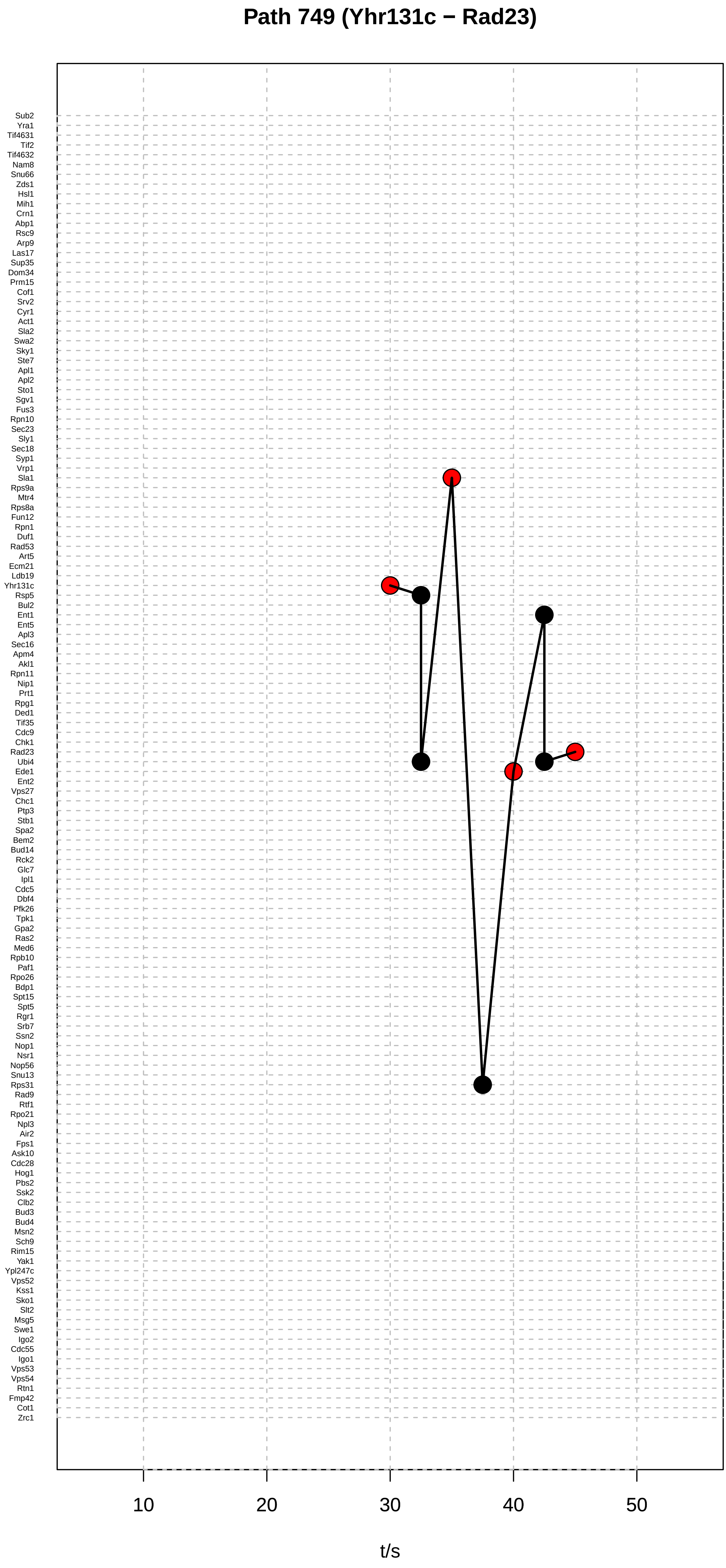
Path 749 (Yhr131c − Rad23)
Sub2
Yra1
Tif4631
Tif2
Tif4632
Nam8
Snu66
Zds1
Hsl1
Mih1
Crn1
Abp1
Rsc9
Arp9
Las17
Sup35
Dom34
Prm15
Cof1
Srv2
Cyr1
Act1
Sla2
Swa2
Sky1
Ste7
Apl1
Apl2
Sto1
Sgv1
Fus3
Rpn10
Sec23
Sly1
Sec18
Syp1
Vrp1
Sla1
Rps9a
Mtr4
Rps8a
Fun12
Rpn1
Duf1
Rad53
Art5
Ecm21
Ldb19
Yhr131c
Rsp5
Bul2
Ent1
Ent5
Apl3
Sec16
Apm4
Akl1
Rpn11
Nip1
Prt1
Rpg1
Ded1
Tif35
Cdc9
Chk1
Rad23
Ubi4
Ede1
Ent2
Vps27
Chc1
Ptp3
Stb1
Spa2
Bem2
Bud14
Rck2
Glc7
Ipl1
Cdc5
Dbf4
Pfk26
Tpk1
Gpa2
Ras2
Med6
Rpb10
Paf1
Rpo26
Bdp1
Spt15
Spt5
Rgr1
Srb7
Ssn2
Nop1
Nsr1
Nop56
Snu13
Rps31
Rad9
Rtf1
Rpo21
Npl3
Air2
Fps1
Ask10
Cdc28
Hog1
Pbs2
Ssk2
Clb2
Bud3
Bud4
Msn2
Sch9
Rim15
Yak1
Ypl247c
Vps52
Kss1
Sko1
Slt2
Msg5
Swe1
Igo2
Cdc55
Igo1
Vps53
Vps54
Rtn1
Fmp42
Cot1
Zrc1
10
20
30
40
50
t/s

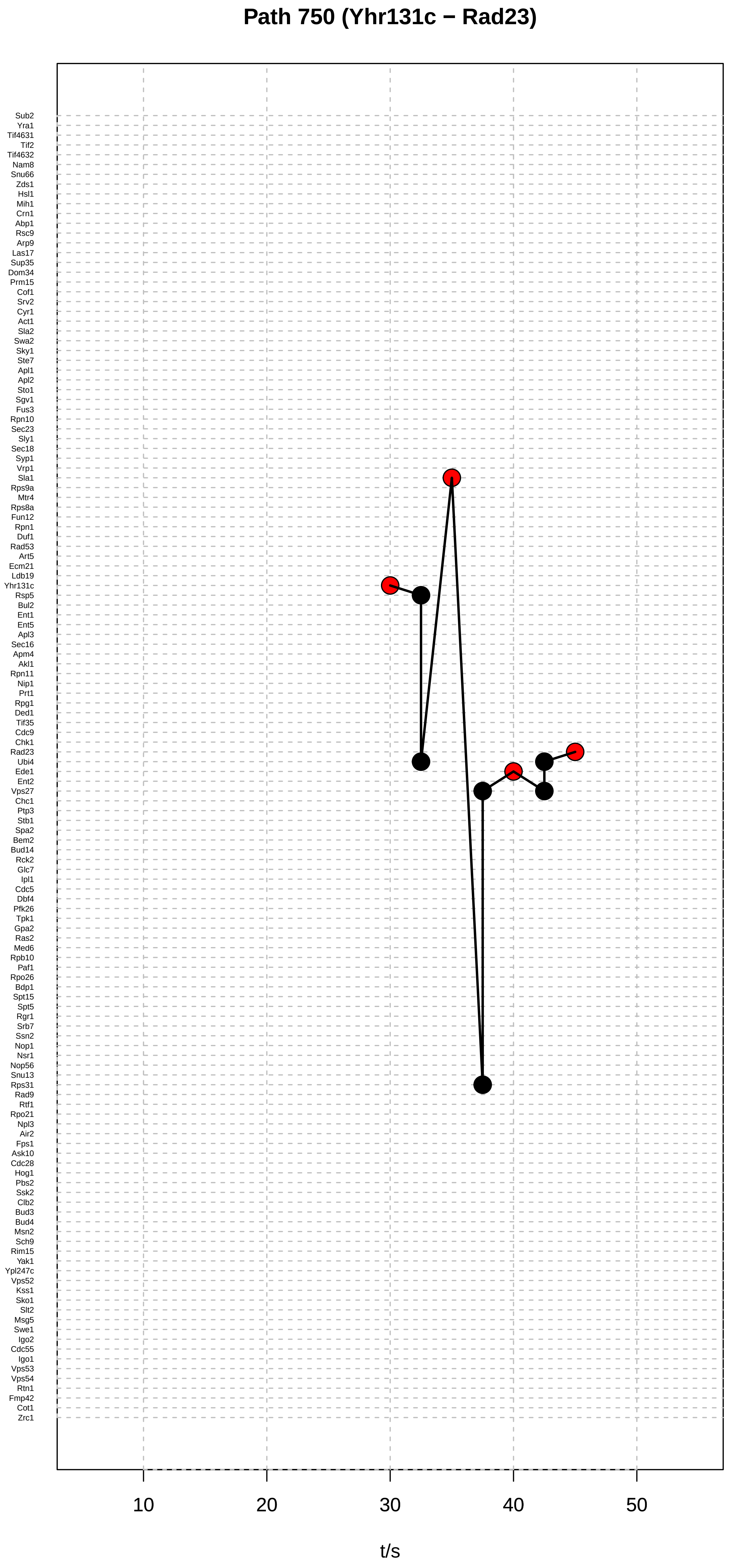

Path 750 (Yhr131c − Rad23)
Sub2
Yra1
Tif4631
Tif2
Tif4632
Nam8
Snu66
Zds1
Hsl1
Mih1
Crn1
Abp1
Rsc9
Arp9
Las17
Sup35
Dom34
Prm15
Cof1
Srv2
Cyr1
Act1
Sla2
Swa2
Sky1
Ste7
Apl1
Apl2
Sto1
Sgv1
Fus3
Rpn10
Sec23
Sly1
Sec18
Syp1
Vrp1
Sla1
Rps9a
Mtr4
Rps8a
Fun12
Rpn1
Duf1
Rad53
Art5
Ecm21
Ldb19
Yhr131c
Rsp5
Bul2
Ent1
Ent5
Apl3
Sec16
Apm4
Akl1
Rpn11
Nip1
Prt1
Rpg1
Ded1
Tif35
Cdc9
Chk1
Rad23
Ubi4
Ede1
Ent2
Vps27
Chc1
Ptp3
Stb1
Spa2
Bem2
Bud14
Rck2
Glc7
Ipl1
Cdc5
Dbf4
Pfk26
Tpk1
Gpa2
Ras2
Med6
Rpb10
Paf1
Rpo26
Bdp1
Spt15
Spt5
Rgr1
Srb7
Ssn2
Nop1
Nsr1
Nop56
Snu13
Rps31
Rad9
Rtf1
Rpo21
Npl3
Air2
Fps1
Ask10
Cdc28
Hog1
Pbs2
Ssk2
Clb2
Bud3
Bud4
Msn2
Sch9
Rim15
Yak1
Ypl247c
Vps52
Kss1
Sko1
Slt2
Msg5
Swe1
Igo2
Cdc55
Igo1
Vps53
Vps54
Rtn1
Fmp42
Cot1
Zrc1
10
20
30
40
50
t/s

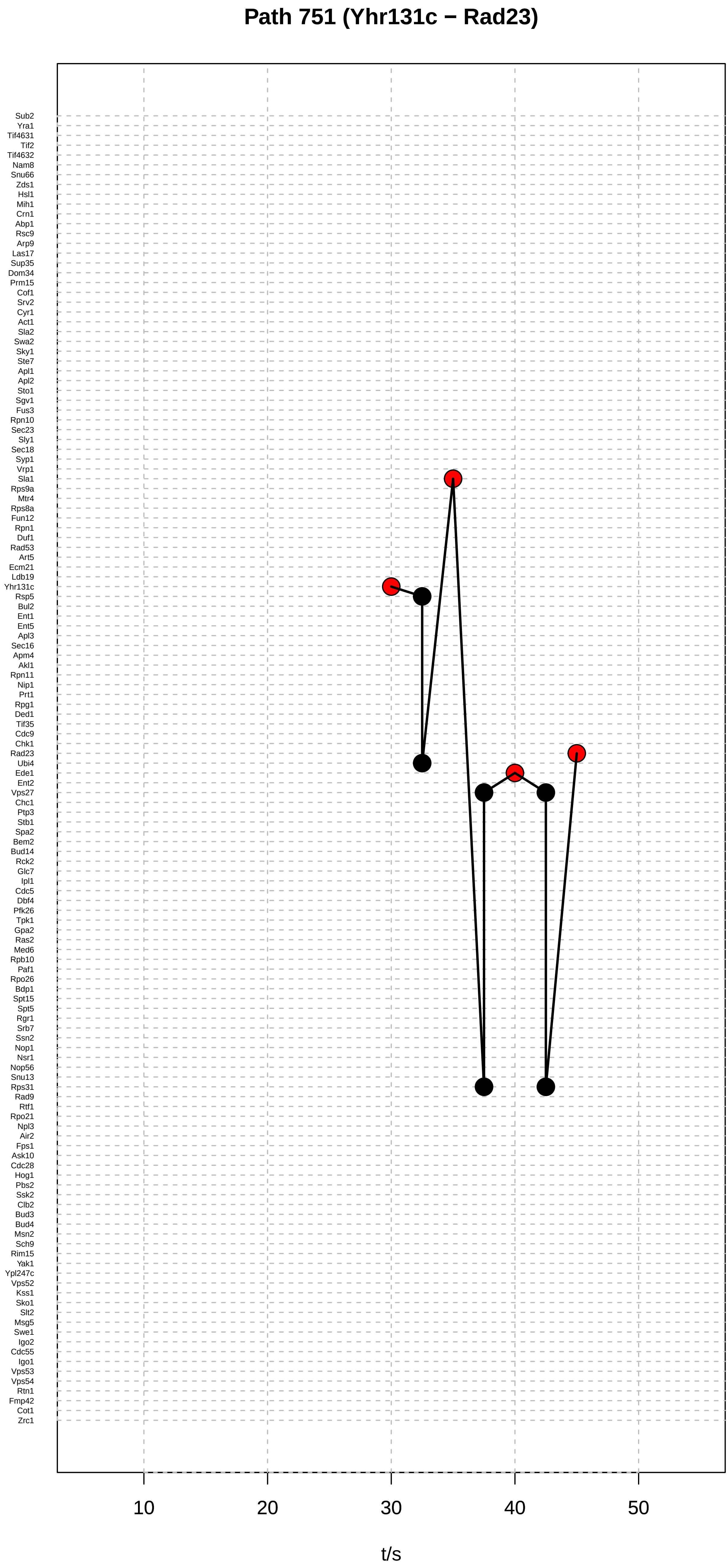

Path 751 (Yhr131c − Rad23)
Sub2
Yra1
Tif4631
Tif2
Tif4632
Nam8
Snu66
Zds1
Hsl1
Mih1
Crn1
Abp1
Rsc9
Arp9
Las17
Sup35
Dom34
Prm15
Cof1
Srv2
Cyr1
Act1
Sla2
Swa2
Sky1
Ste7
Apl1
Apl2
Sto1
Sgv1
Fus3
Rpn10
Sec23
Sly1
Sec18
Syp1
Vrp1
Sla1
Rps9a
Mtr4
Rps8a
Fun12
Rpn1
Duf1
Rad53
Art5
Ecm21
Ldb19
Yhr131c
Rsp5
Bul2
Ent1
Ent5
Apl3
Sec16
Apm4
Akl1
Rpn11
Nip1
Prt1
Rpg1
Ded1
Tif35
Cdc9
Chk1
Rad23
Ubi4
Ede1
Ent2
Vps27
Chc1
Ptp3
Stb1
Spa2
Bem2
Bud14
Rck2
Glc7
Ipl1
Cdc5
Dbf4
Pfk26
Tpk1
Gpa2
Ras2
Med6
Rpb10
Paf1
Rpo26
Bdp1
Spt15
Spt5
Rgr1
Srb7
Ssn2
Nop1
Nsr1
Nop56
Snu13
Rps31
Rad9
Rtf1
Rpo21
Npl3
Air2
Fps1
Ask10
Cdc28
Hog1
Pbs2
Ssk2
Clb2
Bud3
Bud4
Msn2
Sch9
Rim15
Yak1
Ypl247c
Vps52
Kss1
Sko1
Slt2
Msg5
Swe1
Igo2
Cdc55
Igo1
Vps53
Vps54
Rtn1
Fmp42
Cot1
Zrc1
10
20
30
40
50
t/s

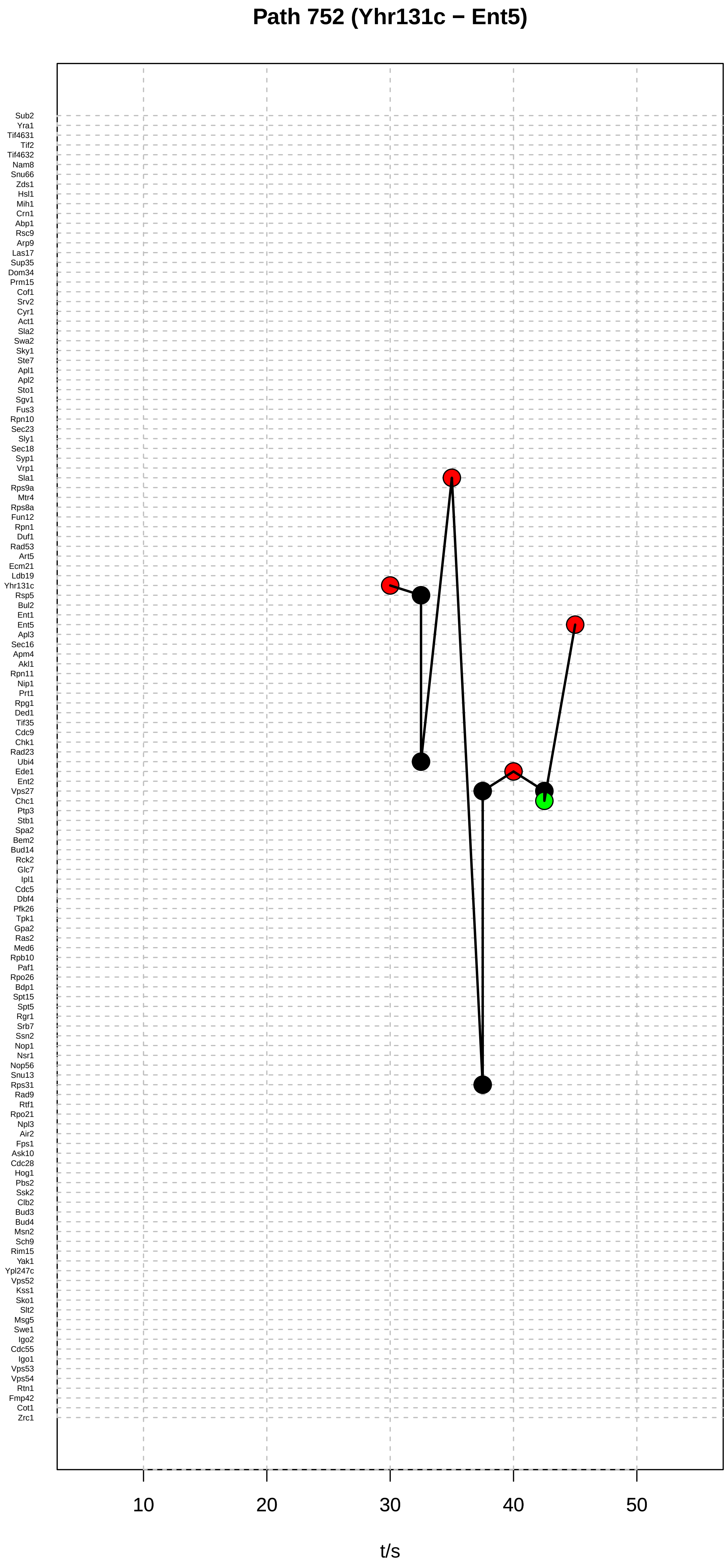

Path 752 (Yhr131c − Ent5)
Sub2
Yra1
Tif4631
Tif2
Tif4632
Nam8
Snu66
Zds1
Hsl1
Mih1
Crn1
Abp1
Rsc9
Arp9
Las17
Sup35
Dom34
Prm15
Cof1
Srv2
Cyr1
Act1
Sla2
Swa2
Sky1
Ste7
Apl1
Apl2
Sto1
Sgv1
Fus3
Rpn10
Sec23
Sly1
Sec18
Syp1
Vrp1
Sla1
Rps9a
Mtr4
Rps8a
Fun12
Rpn1
Duf1
Rad53
Art5
Ecm21
Ldb19
Yhr131c
Rsp5
Bul2
Ent1
Ent5
Apl3
Sec16
Apm4
Akl1
Rpn11
Nip1
Prt1
Rpg1
Ded1
Tif35
Cdc9
Chk1
Rad23
Ubi4
Ede1
Ent2
Vps27
Chc1
Ptp3
Stb1
Spa2
Bem2
Bud14
Rck2
Glc7
Ipl1
Cdc5
Dbf4
Pfk26
Tpk1
Gpa2
Ras2
Med6
Rpb10
Paf1
Rpo26
Bdp1
Spt15
Spt5
Rgr1
Srb7
Ssn2
Nop1
Nsr1
Nop56
Snu13
Rps31
Rad9
Rtf1
Rpo21
Npl3
Air2
Fps1
Ask10
Cdc28
Hog1
Pbs2
Ssk2
Clb2
Bud3
Bud4
Msn2
Sch9
Rim15
Yak1
Ypl247c
Vps52
Kss1
Sko1
Slt2
Msg5
Swe1
Igo2
Cdc55
Igo1
Vps53
Vps54
Rtn1
Fmp42
Cot1
Zrc1
10
20
30
40
50
t/s

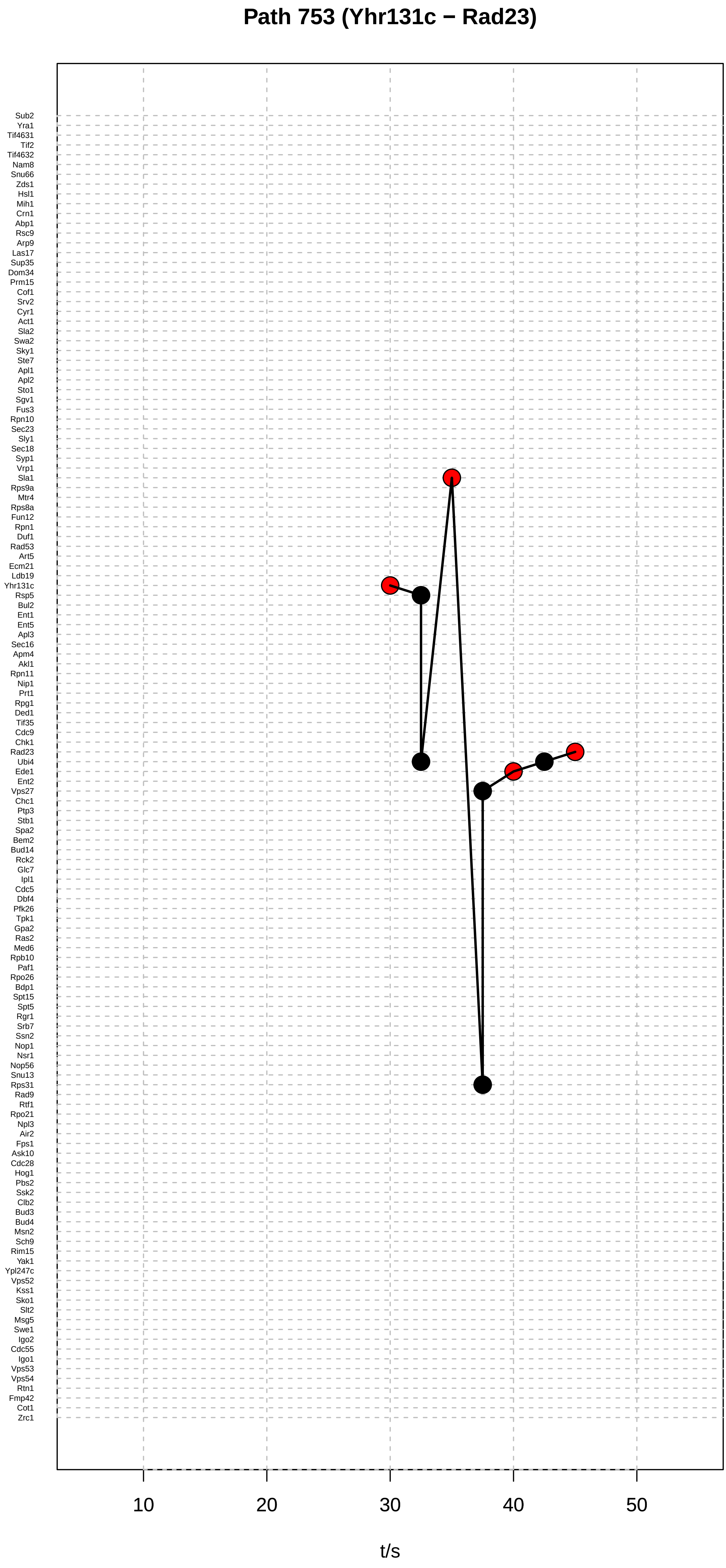
Path 753 (Yhr131c − Rad23)
Sub2
Yra1
Tif4631
Tif2
Tif4632
Nam8
Snu66
Zds1
Hsl1
Mih1
Crn1
Abp1
Rsc9
Arp9
Las17
Sup35
Dom34
Prm15
Cof1
Srv2
Cyr1
Act1
Sla2
Swa2
Sky1
Ste7
Apl1
Apl2
Sto1
Sgv1
Fus3
Rpn10
Sec23
Sly1
Sec18
Syp1
Vrp1
Sla1
Rps9a
Mtr4
Rps8a
Fun12
Rpn1
Duf1
Rad53
Art5
Ecm21
Ldb19
Yhr131c
Rsp5
Bul2
Ent1
Ent5
Apl3
Sec16
Apm4
Akl1
Rpn11
Nip1
Prt1
Rpg1
Ded1
Tif35
Cdc9
Chk1
Rad23
Ubi4
Ede1
Ent2
Vps27
Chc1
Ptp3
Stb1
Spa2
Bem2
Bud14
Rck2
Glc7
Ipl1
Cdc5
Dbf4
Pfk26
Tpk1
Gpa2
Ras2
Med6
Rpb10
Paf1
Rpo26
Bdp1
Spt15
Spt5
Rgr1
Srb7
Ssn2
Nop1
Nsr1
Nop56
Snu13
Rps31
Rad9
Rtf1
Rpo21
Npl3
Air2
Fps1
Ask10
Cdc28
Hog1
Pbs2
Ssk2
Clb2
Bud3
Bud4
Msn2
Sch9
Rim15
Yak1
Ypl247c
Vps52
Kss1
Sko1
Slt2
Msg5
Swe1
Igo2
Cdc55
Igo1
Vps53
Vps54
Rtn1
Fmp42
Cot1
Zrc1
10
20
30
40
50
t/s

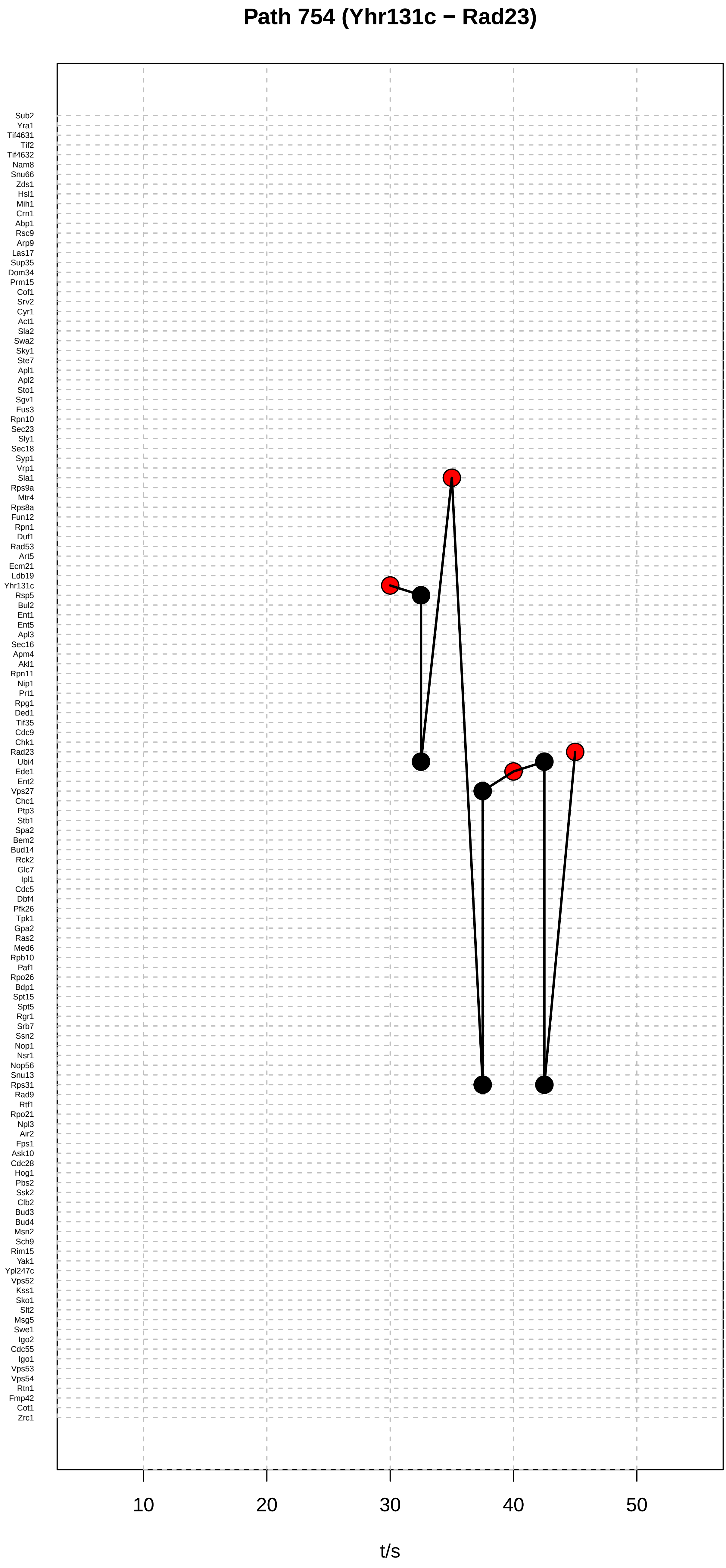

Path 754 (Yhr131c − Rad23)
Sub2
Yra1
Tif4631
Tif2
Tif4632
Nam8
Snu66
Zds1
Hsl1
Mih1
Crn1
Abp1
Rsc9
Arp9
Las17
Sup35
Dom34
Prm15
Cof1
Srv2
Cyr1
Act1
Sla2
Swa2
Sky1
Ste7
Apl1
Apl2
Sto1
Sgv1
Fus3
Rpn10
Sec23
Sly1
Sec18
Syp1
Vrp1
Sla1
Rps9a
Mtr4
Rps8a
Fun12
Rpn1
Duf1
Rad53
Art5
Ecm21
Ldb19
Yhr131c
Rsp5
Bul2
Ent1
Ent5
Apl3
Sec16
Apm4
Akl1
Rpn11
Nip1
Prt1
Rpg1
Ded1
Tif35
Cdc9
Chk1
Rad23
Ubi4
Ede1
Ent2
Vps27
Chc1
Ptp3
Stb1
Spa2
Bem2
Bud14
Rck2
Glc7
Ipl1
Cdc5
Dbf4
Pfk26
Tpk1
Gpa2
Ras2
Med6
Rpb10
Paf1
Rpo26
Bdp1
Spt15
Spt5
Rgr1
Srb7
Ssn2
Nop1
Nsr1
Nop56
Snu13
Rps31
Rad9
Rtf1
Rpo21
Npl3
Air2
Fps1
Ask10
Cdc28
Hog1
Pbs2
Ssk2
Clb2
Bud3
Bud4
Msn2
Sch9
Rim15
Yak1
Ypl247c
Vps52
Kss1
Sko1
Slt2
Msg5
Swe1
Igo2
Cdc55
Igo1
Vps53
Vps54
Rtn1
Fmp42
Cot1
Zrc1
10
20
30
40
50
t/s

### Path 755 (Yhr131c − Rad23)

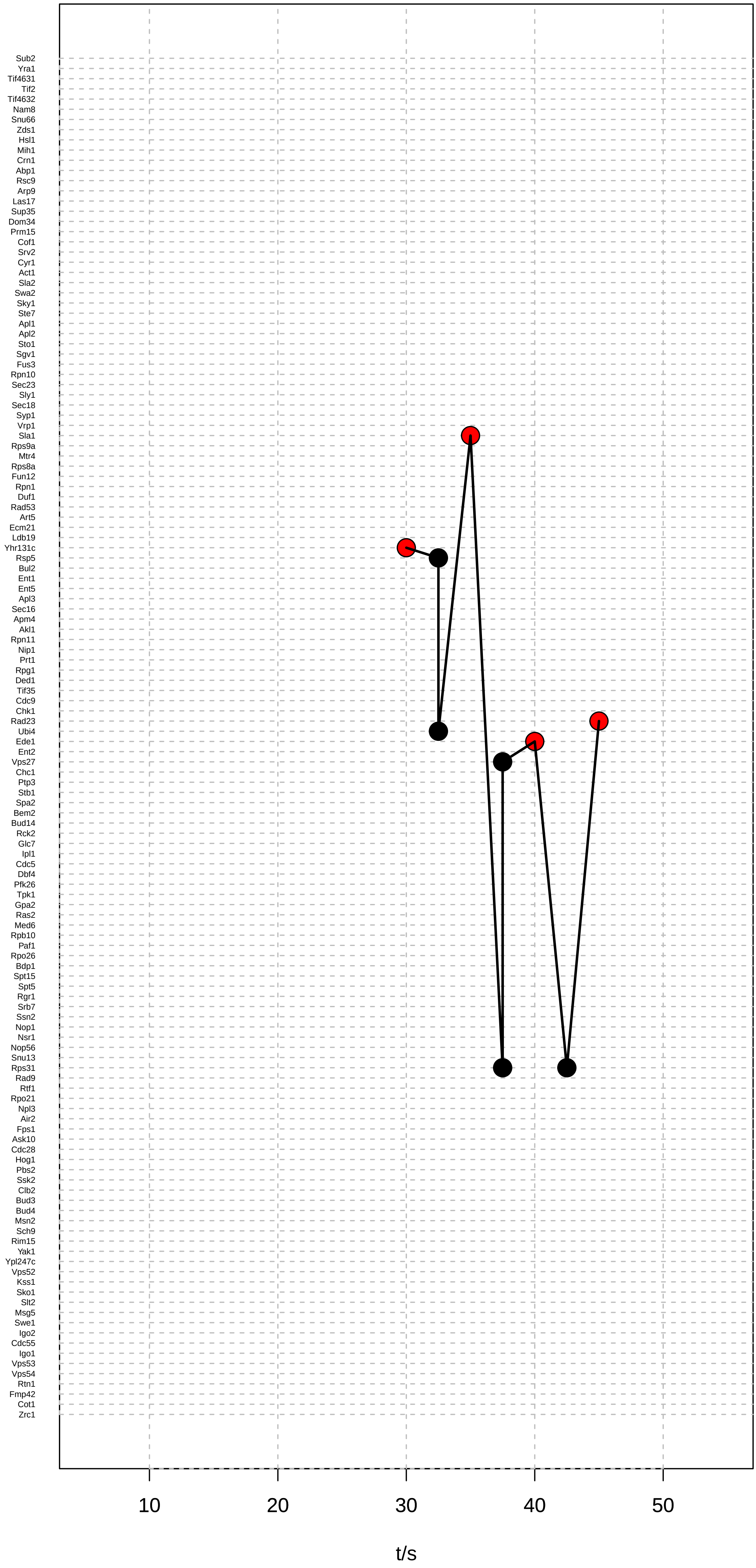

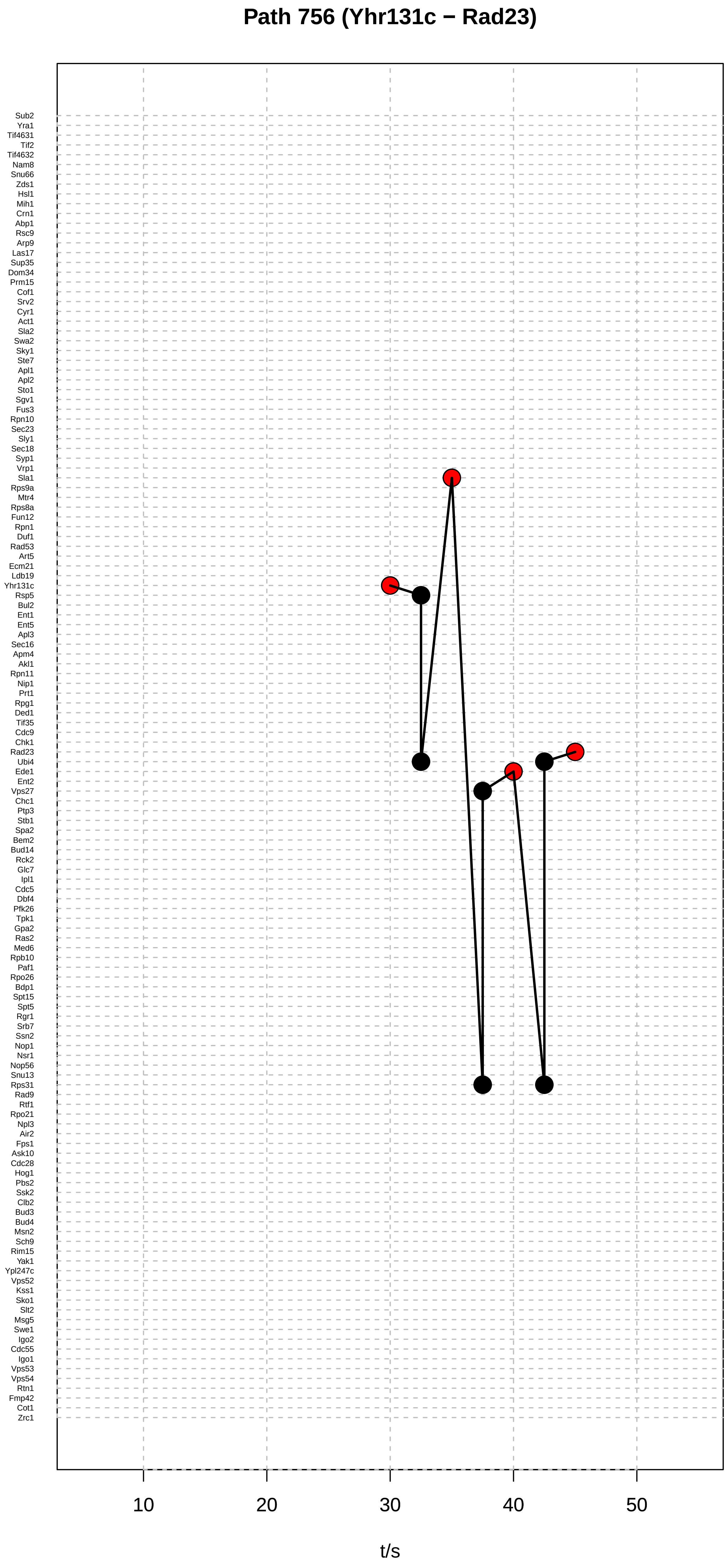

Path 756 (Yhr131c − Rad23)
Sub2
Yra1
Tif4631
Tif2
Tif4632
Nam8
Snu66
Zds1
Hsl1
Mih1
Crn1
Abp1
Rsc9
Arp9
Las17
Sup35
Dom34
Prm15
Cof1
Srv2
Cyr1
Act1
Sla2
Swa2
Sky1
Ste7
Apl1
Apl2
Sto1
Sgv1
Fus3
Rpn10
Sec23
Sly1
Sec18
Syp1
Vrp1
Sla1
Rps9a
Mtr4
Rps8a
Fun12
Rpn1
Duf1
Rad53
Art5
Ecm21
Ldb19
Yhr131c
Rsp5
Bul2
Ent1
Ent5
Apl3
Sec16
Apm4
Akl1
Rpn11
Nip1
Prt1
Rpg1
Ded1
Tif35
Cdc9
Chk1
Rad23
Ubi4
Ede1
Ent2
Vps27
Chc1
Ptp3
Stb1
Spa2
Bem2
Bud14
Rck2
Glc7
Ipl1
Cdc5
Dbf4
Pfk26
Tpk1
Gpa2
Ras2
Med6
Rpb10
Paf1
Rpo26
Bdp1
Spt15
Spt5
Rgr1
Srb7
Ssn2
Nop1
Nsr1
Nop56
Snu13
Rps31
Rad9
Rtf1
Rpo21
Npl3
Air2
Fps1
Ask10
Cdc28
Hog1
Pbs2
Ssk2
Clb2
Bud3
Bud4
Msn2
Sch9
Rim15
Yak1
Ypl247c
Vps52
Kss1
Sko1
Slt2
Msg5
Swe1
Igo2
Cdc55
Igo1
Vps53
Vps54
Rtn1
Fmp42
Cot1
Zrc1
10
20
30
40
50
t/s

**Path 757 (Yhr131c − Rad23)**

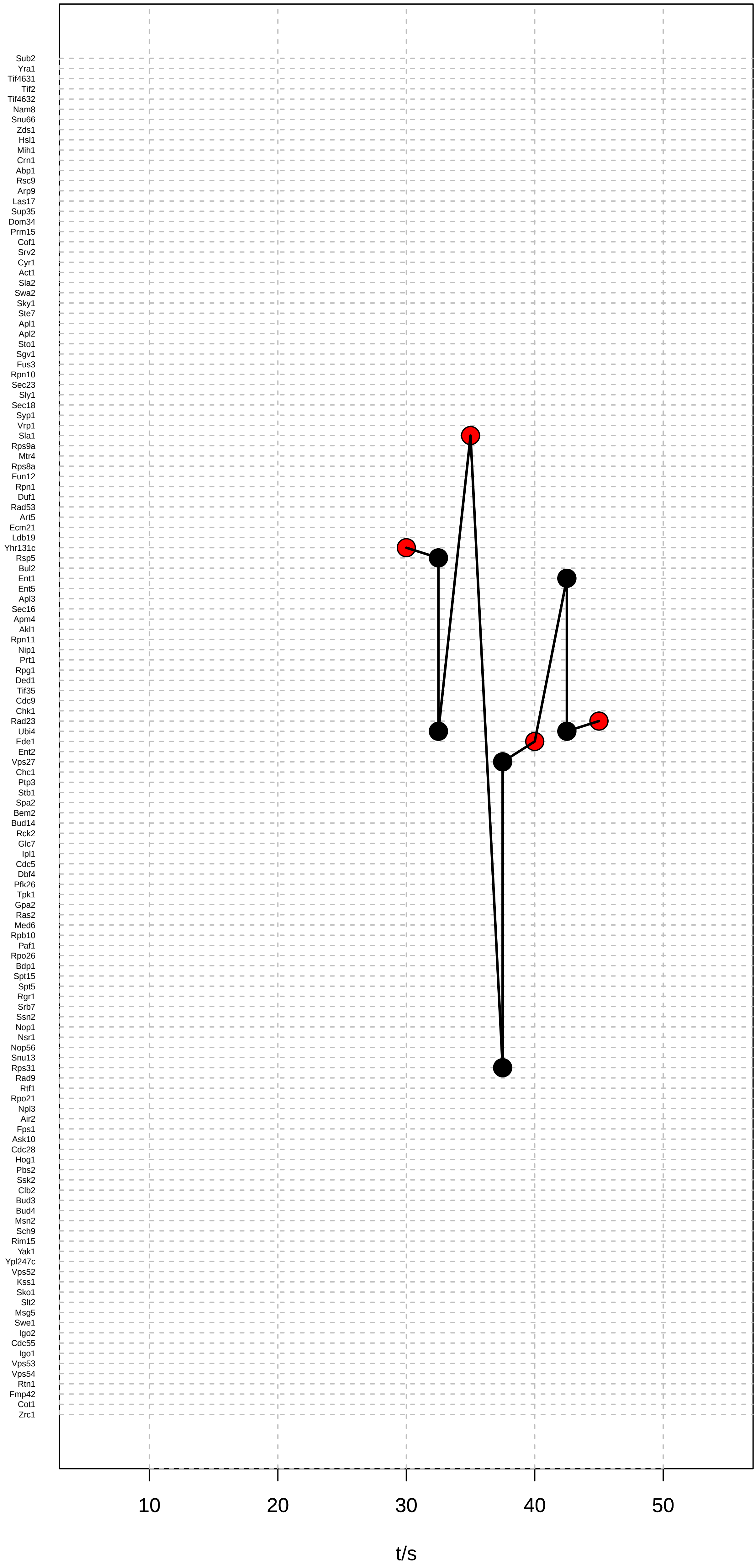

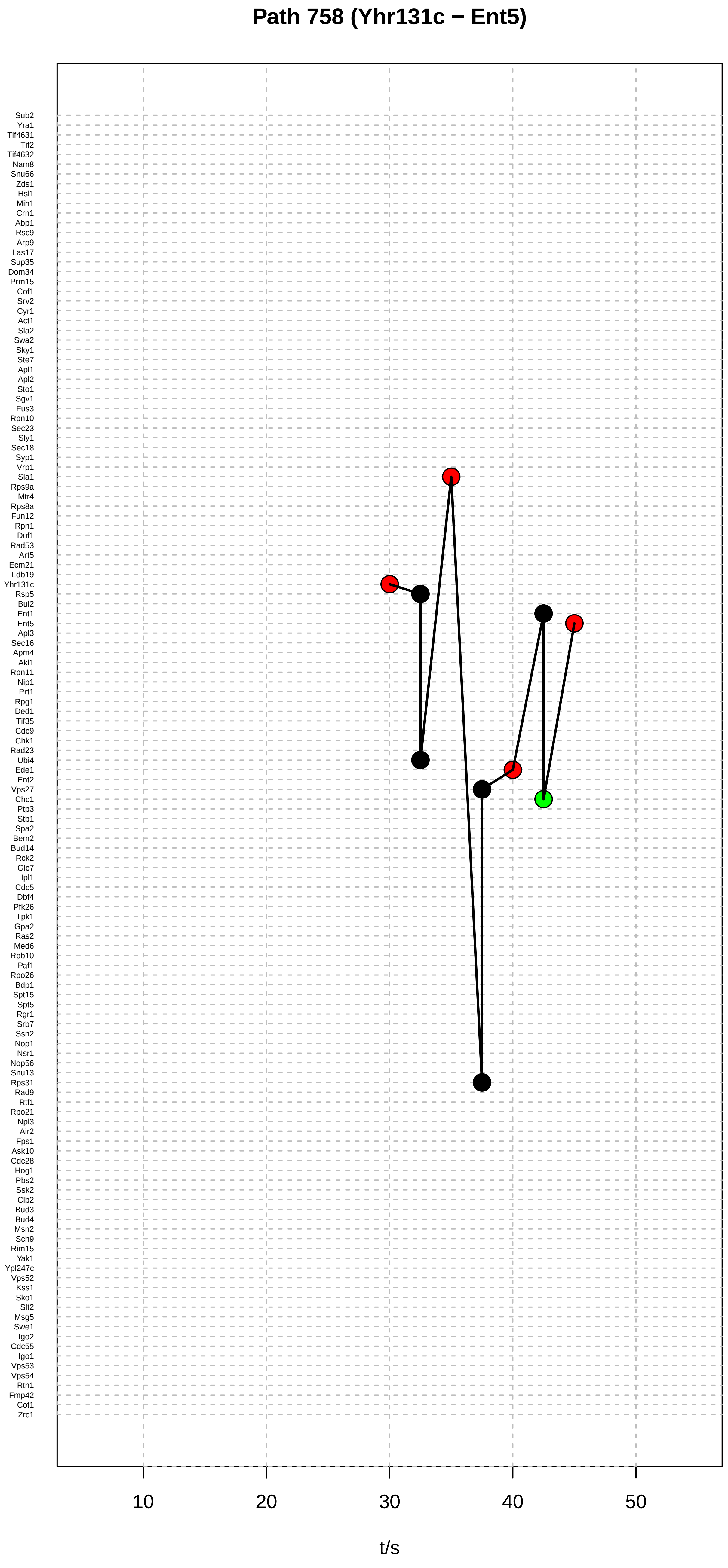

Path 758 (Yhr131c − Ent5)
Sub2
Yra1
Tif4631
Tif2
Tif4632
Nam8
Snu66
Zds1
Hsl1
Mih1
Crn1
Abp1
Rsc9
Arp9
Las17
Sup35
Dom34
Prm15
Cof1
Srv2
Cyr1
Act1
Sla2
Swa2
Sky1
Ste7
Apl1
Apl2
Sto1
Sgv1
Fus3
Rpn10
Sec23
Sly1
Sec18
Syp1
Vrp1
Sla1
Rps9a
Mtr4
Rps8a
Fun12
Rpn1
Duf1
Rad53
Art5
Ecm21
Ldb19
Yhr131c
Rsp5
Bul2
Ent1
Ent5
Apl3
Sec16
Apm4
Akl1
Rpn11
Nip1
Prt1
Rpg1
Ded1
Tif35
Cdc9
Chk1
Rad23
Ubi4
Ede1
Ent2
Vps27
Chc1
Ptp3
Stb1
Spa2
Bem2
Bud14
Rck2
Glc7
Ipl1
Cdc5
Dbf4
Pfk26
Tpk1
Gpa2
Ras2
Med6
Rpb10
Paf1
Rpo26
Bdp1
Spt15
Spt5
Rgr1
Srb7
Ssn2
Nop1
Nsr1
Nop56
Snu13
Rps31
Rad9
Rtf1
Rpo21
Npl3
Air2
Fps1
Ask10
Cdc28
Hog1
Pbs2
Ssk2
Clb2
Bud3
Bud4
Msn2
Sch9
Rim15
Yak1
Ypl247c
Vps52
Kss1
Sko1
Slt2
Msg5
Swe1
Igo2
Cdc55
Igo1
Vps53
Vps54
Rtn1
Fmp42
Cot1
Zrc1
10
20
30
40
50
t/s

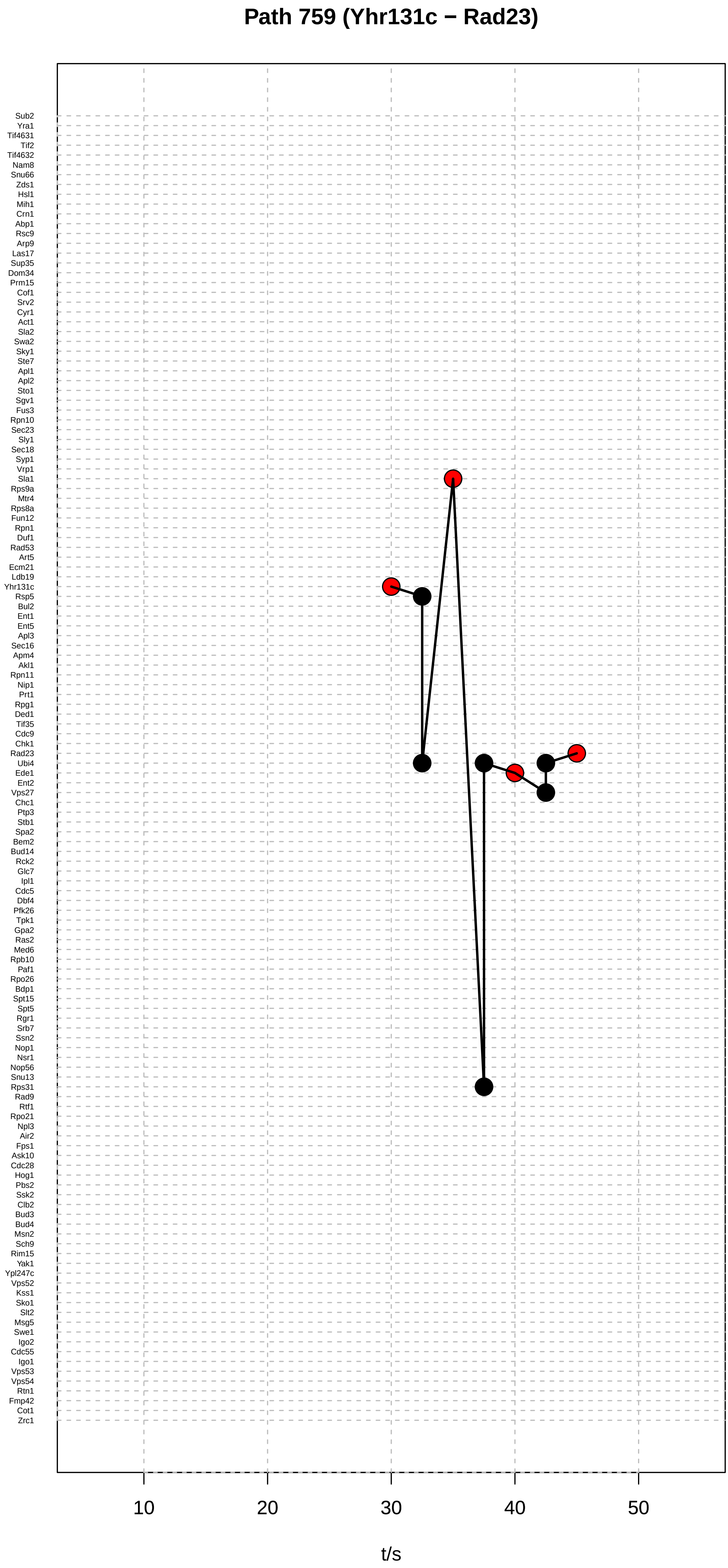

Path 759 (Yhr131c − Rad23)
Sub2
Yra1
Tif4631
Tif2
Tif4632
Nam8
Snu66
Zds1
Hsl1
Mih1
Crn1
Abp1
Rsc9
Arp9
Las17
Sup35
Dom34
Prm15
Cof1
Srv2
Cyr1
Act1
Sla2
Swa2
Sky1
Ste7
Apl1
Apl2
Sto1
Sgv1
Fus3
Rpn10
Sec23
Sly1
Sec18
Syp1
Vrp1
Sla1
Rps9a
Mtr4
Rps8a
Fun12
Rpn1
Duf1
Rad53
Art5
Ecm21
Ldb19
Yhr131c
Rsp5
Bul2
Ent1
Ent5
Apl3
Sec16
Apm4
Akl1
Rpn11
Nip1
Prt1
Rpg1
Ded1
Tif35
Cdc9
Chk1
Rad23
Ubi4
Ede1
Ent2
Vps27
Chc1
Ptp3
Stb1
Spa2
Bem2
Bud14
Rck2
Glc7
Ipl1
Cdc5
Dbf4
Pfk26
Tpk1
Gpa2
Ras2
Med6
Rpb10
Paf1
Rpo26
Bdp1
Spt15
Spt5
Rgr1
Srb7
Ssn2
Nop1
Nsr1
Nop56
Snu13
Rps31
Rad9
Rtf1
Rpo21
Npl3
Air2
Fps1
Ask10
Cdc28
Hog1
Pbs2
Ssk2
Clb2
Bud3
Bud4
Msn2
Sch9
Rim15
Yak1
Ypl247c
Vps52
Kss1
Sko1
Slt2
Msg5
Swe1
Igo2
Cdc55
Igo1
Vps53
Vps54
Rtn1
Fmp42
Cot1
Zrc1
10
20
30
40
50
t/s

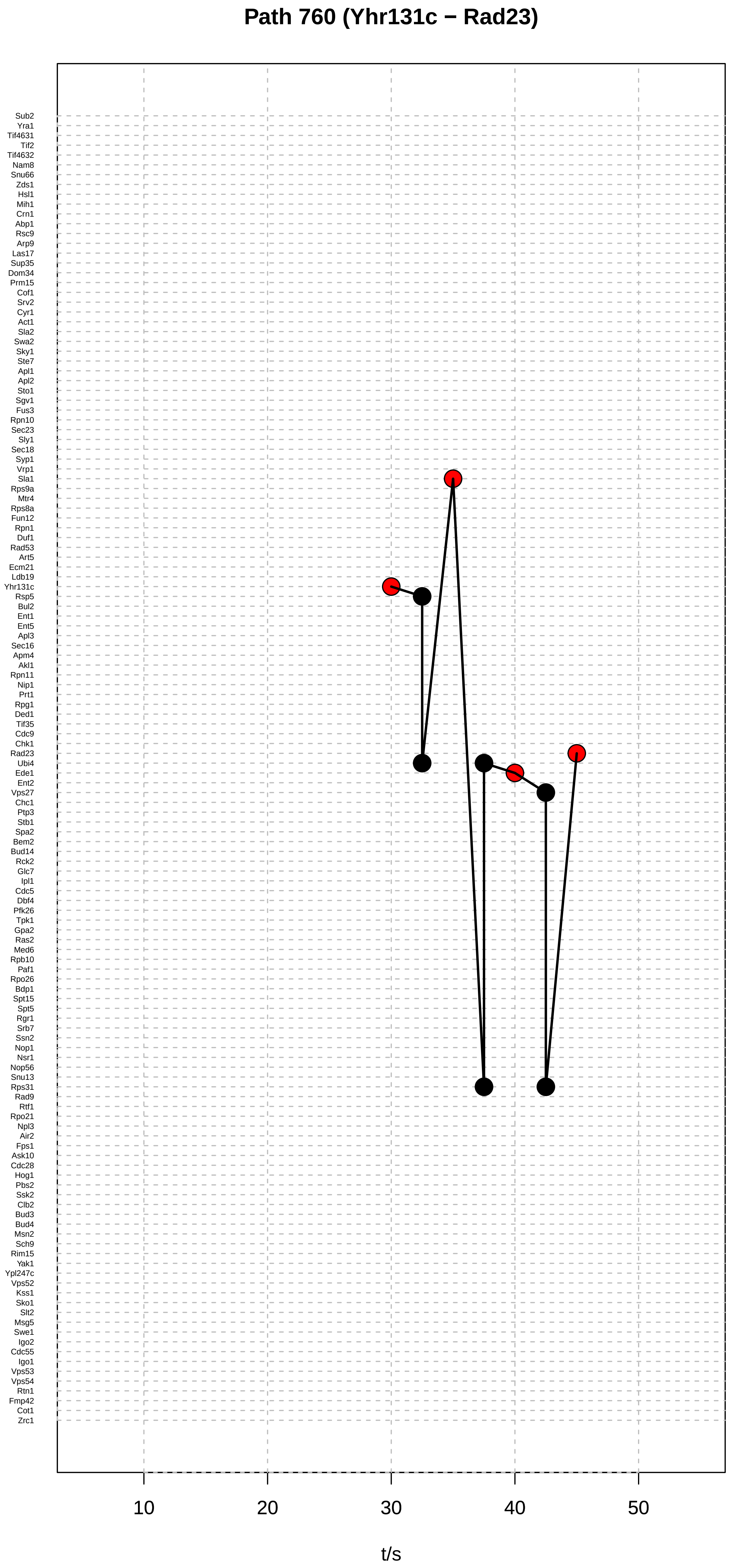

Path 760 (Yhr131c − Rad23)
Sub2
Yra1
Tif4631
Tif2
Tif4632
Nam8
Snu66
Zds1
Hsl1
Mih1
Crn1
Abp1
Rsc9
Arp9
Las17
Sup35
Dom34
Prm15
Cof1
Srv2
Cyr1
Act1
Sla2
Swa2
Sky1
Ste7
Apl1
Apl2
Sto1
Sgv1
Fus3
Rpn10
Sec23
Sly1
Sec18
Syp1
Vrp1
Sla1
Rps9a
Mtr4
Rps8a
Fun12
Rpn1
Duf1
Rad53
Art5
Ecm21
Ldb19
Yhr131c
Rsp5
Bul2
Ent1
Ent5
Apl3
Sec16
Apm4
Akl1
Rpn11
Nip1
Prt1
Rpg1
Ded1
Tif35
Cdc9
Chk1
Rad23
Ubi4
Ede1
Ent2
Vps27
Chc1
Ptp3
Stb1
Spa2
Bem2
Bud14
Rck2
Glc7
Ipl1
Cdc5
Dbf4
Pfk26
Tpk1
Gpa2
Ras2
Med6
Rpb10
Paf1
Rpo26
Bdp1
Spt15
Spt5
Rgr1
Srb7
Ssn2
Nop1
Nsr1
Nop56
Snu13
Rps31
Rad9
Rtf1
Rpo21
Npl3
Air2
Fps1
Ask10
Cdc28
Hog1
Pbs2
Ssk2
Clb2
Bud3
Bud4
Msn2
Sch9
Rim15
Yak1
Ypl247c
Vps52
Kss1
Sko1
Slt2
Msg5
Swe1
Igo2
Cdc55
Igo1
Vps53
Vps54
Rtn1
Fmp42
Cot1
Zrc1
10
20
30
40
50
t/s

#### Path 761 (Yhr131c − Ent5)

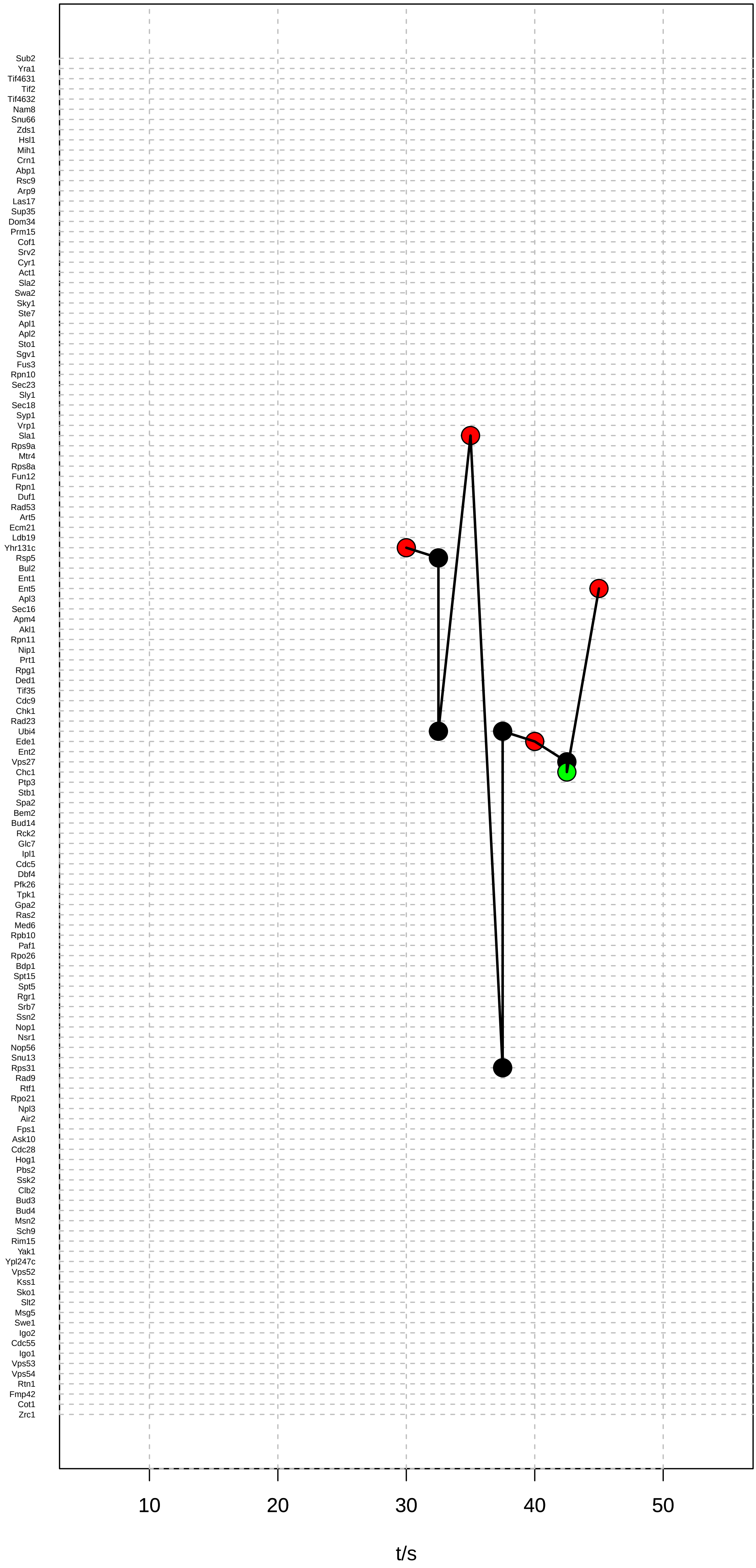

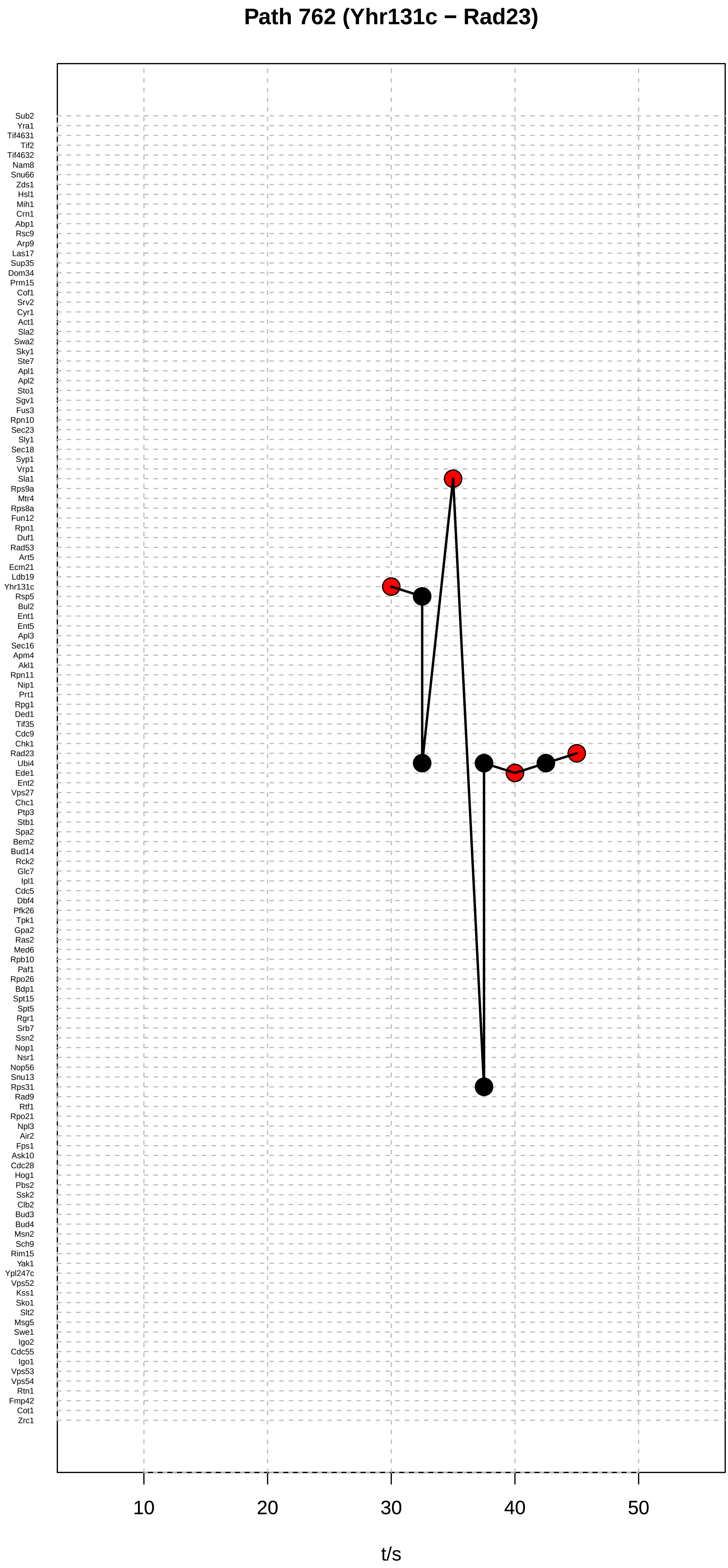

Path 762 (Yhr131c − Rad23)
Sub2
Yra1
Tif4631
Tif2
Tif4632
Nam8
Snu66
Zds1
Hsl1
Mih1
Crn1
Abp1
Rsc9
Arp9
Las17
Sup35
Dom34
Prm15
Cof1
Srv2
Cyr1
Act1
Sla2
Swa2
Sky1
Ste7
Apl1
Apl2
Sto1
Sgv1
Fus3
Rpn10
Sec23
Sly1
Sec18
Syp1
Vrp1
Sla1
Rps9a
Mtr4
Rps8a
Fun12
Rpn1
Duf1
Rad53
Art5
Ecm21
Ldb19
Yhr131c
Rsp5
Bul2
Ent1
Ent5
Apl3
Sec16
Apm4
Akl1
Rpn11
Nip1
Prt1
Rpg1
Ded1
Tif35
Cdc9
Chk1
Rad23
Ubi4
Ede1
Ent2
Vps27
Chc1
Ptp3
Stb1
Spa2
Bem2
Bud14
Rck2
Glc7
Ipl1
Cdc5
Dbf4
Pfk26
Tpk1
Gpa2
Ras2
Med6
Rpb10
Paf1
Rpo26
Bdp1
Spt15
Spt5
Rgr1
Srb7
Ssn2
Nop1
Nsr1
Nop56
Snu13
Rps31
Rad9
Rtf1
Rpo21
Npl3
Air2
Fps1
Ask10
Cdc28
Hog1
Pbs2
Ssk2
Clb2
Bud3
Bud4
Msn2
Sch9
Rim15
Yak1
Ypl247c
Vps52
Kss1
Sko1
Slt2
Msg5
Swe1
Igo2
Cdc55
Igo1
Vps53
Vps54
Rtn1
Fmp42
Cot1
Zrc1
10
20
30
40
50
t/s

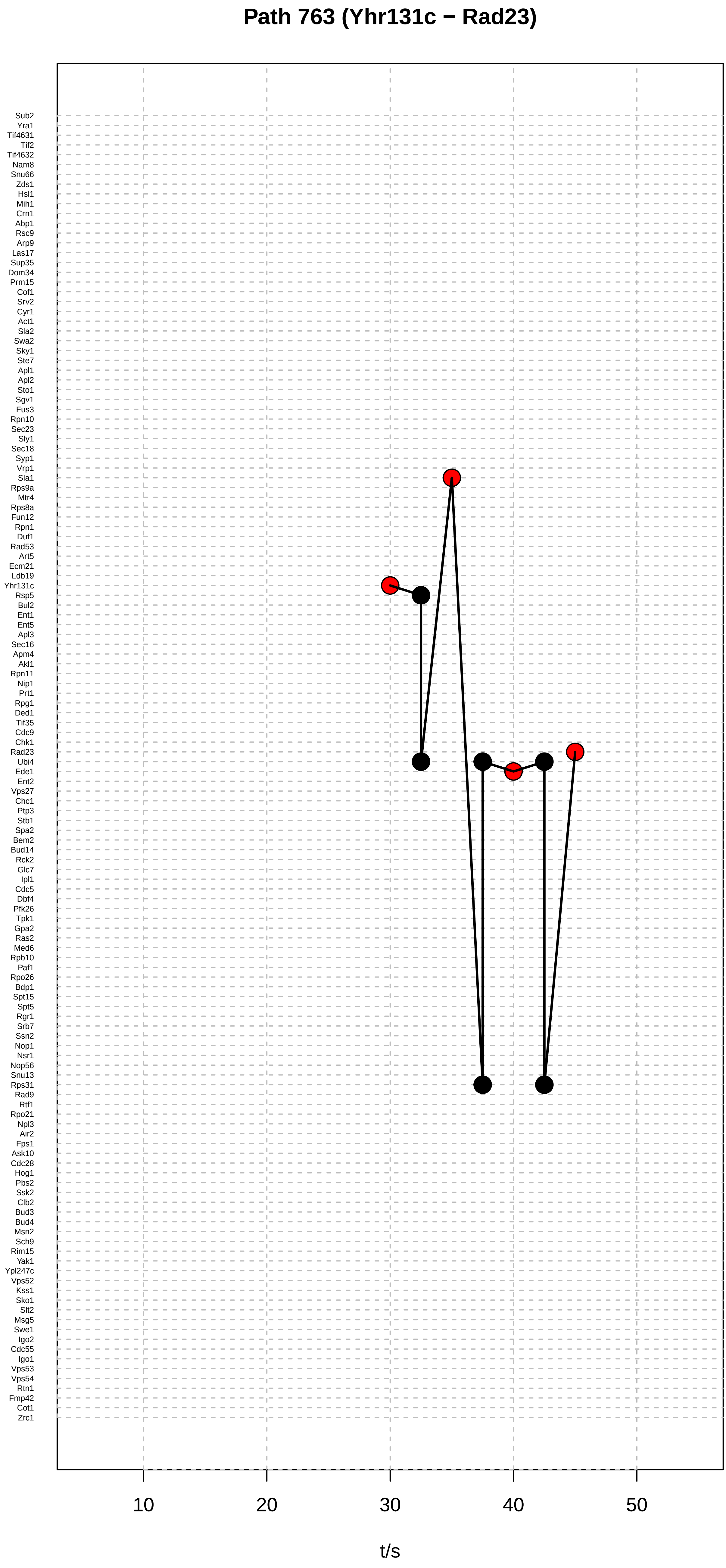
Path 763 (Yhr131c − Rad23)
Sub2
Yra1
Tif4631
Tif2
Tif4632
Nam8
Snu66
Zds1
Hsl1
Mih1
Crn1
Abp1
Rsc9
Arp9
Las17
Sup35
Dom34
Prm15
Cof1
Srv2
Cyr1
Act1
Sla2
Swa2
Sky1
Ste7
Apl1
Apl2
Sto1
Sgv1
Fus3
Rpn10
Sec23
Sly1
Sec18
Syp1
Vrp1
Sla1
Rps9a
Mtr4
Rps8a
Fun12
Rpn1
Duf1
Rad53
Art5
Ecm21
Ldb19
Yhr131c
Rsp5
Bul2
Ent1
Ent5
Apl3
Sec16
Apm4
Akl1
Rpn11
Nip1
Prt1
Rpg1
Ded1
Tif35
Cdc9
Chk1
Rad23
Ubi4
Ede1
Ent2
Vps27
Chc1
Ptp3
Stb1
Spa2
Bem2
Bud14
Rck2
Glc7
Ipl1
Cdc5
Dbf4
Pfk26
Tpk1
Gpa2
Ras2
Med6
Rpb10
Paf1
Rpo26
Bdp1
Spt15
Spt5
Rgr1
Srb7
Ssn2
Nop1
Nsr1
Nop56
Snu13
Rps31
Rad9
Rtf1
Rpo21
Npl3
Air2
Fps1
Ask10
Cdc28
Hog1
Pbs2
Ssk2
Clb2
Bud3
Bud4
Msn2
Sch9
Rim15
Yak1
Ypl247c
Vps52
Kss1
Sko1
Slt2
Msg5
Swe1
Igo2
Cdc55
Igo1
Vps53
Vps54
Rtn1
Fmp42
Cot1
Zrc1
10
20
30
40
50
t/s

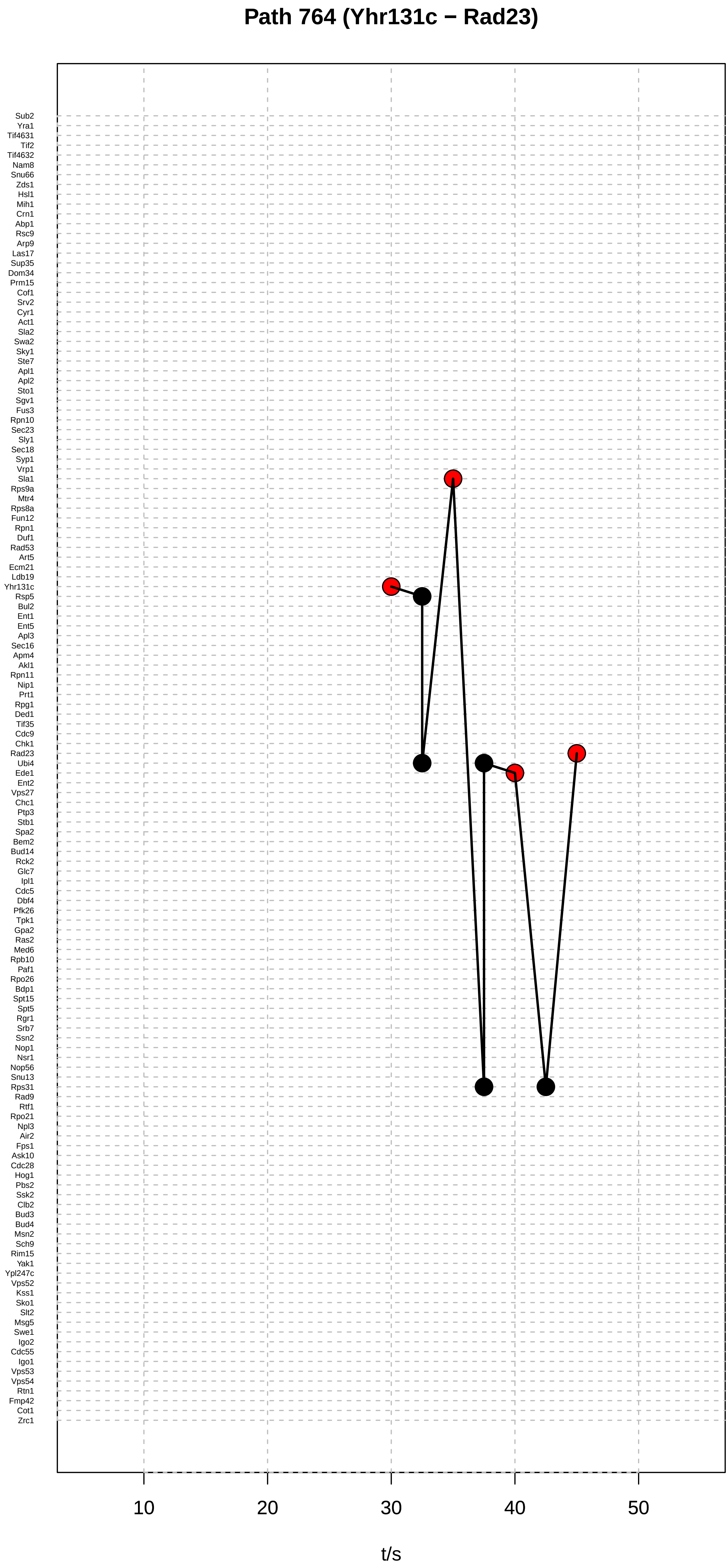

Path 764 (Yhr131c − Rad23)
Sub2
Yra1
Tif4631
Tif2
Tif4632
Nam8
Snu66
Zds1
Hsl1
Mih1
Crn1
Abp1
Rsc9
Arp9
Las17
Sup35
Dom34
Prm15
Cof1
Srv2
Cyr1
Act1
Sla2
Swa2
Sky1
Ste7
Apl1
Apl2
Sto1
Sgv1
Fus3
Rpn10
Sec23
Sly1
Sec18
Syp1
Vrp1
Sla1
Rps9a
Mtr4
Rps8a
Fun12
Rpn1
Duf1
Rad53
Art5
Ecm21
Ldb19
Yhr131c
Rsp5
Bul2
Ent1
Ent5
Apl3
Sec16
Apm4
Akl1
Rpn11
Nip1
Prt1
Rpg1
Ded1
Tif35
Cdc9
Chk1
Rad23
Ubi4
Ede1
Ent2
Vps27
Chc1
Ptp3
Stb1
Spa2
Bem2
Bud14
Rck2
Glc7
Ipl1
Cdc5
Dbf4
Pfk26
Tpk1
Gpa2
Ras2
Med6
Rpb10
Paf1
Rpo26
Bdp1
Spt15
Spt5
Rgr1
Srb7
Ssn2
Nop1
Nsr1
Nop56
Snu13
Rps31
Rad9
Rtf1
Rpo21
Npl3
Air2
Fps1
Ask10
Cdc28
Hog1
Pbs2
Ssk2
Clb2
Bud3
Bud4
Msn2
Sch9
Rim15
Yak1
Ypl247c
Vps52
Kss1
Sko1
Slt2
Msg5
Swe1
Igo2
Cdc55
Igo1
Vps53
Vps54
Rtn1
Fmp42
Cot1
Zrc1
10
20
30
40
50
t/s

### Path 765 (Yhr131c − Rad23)

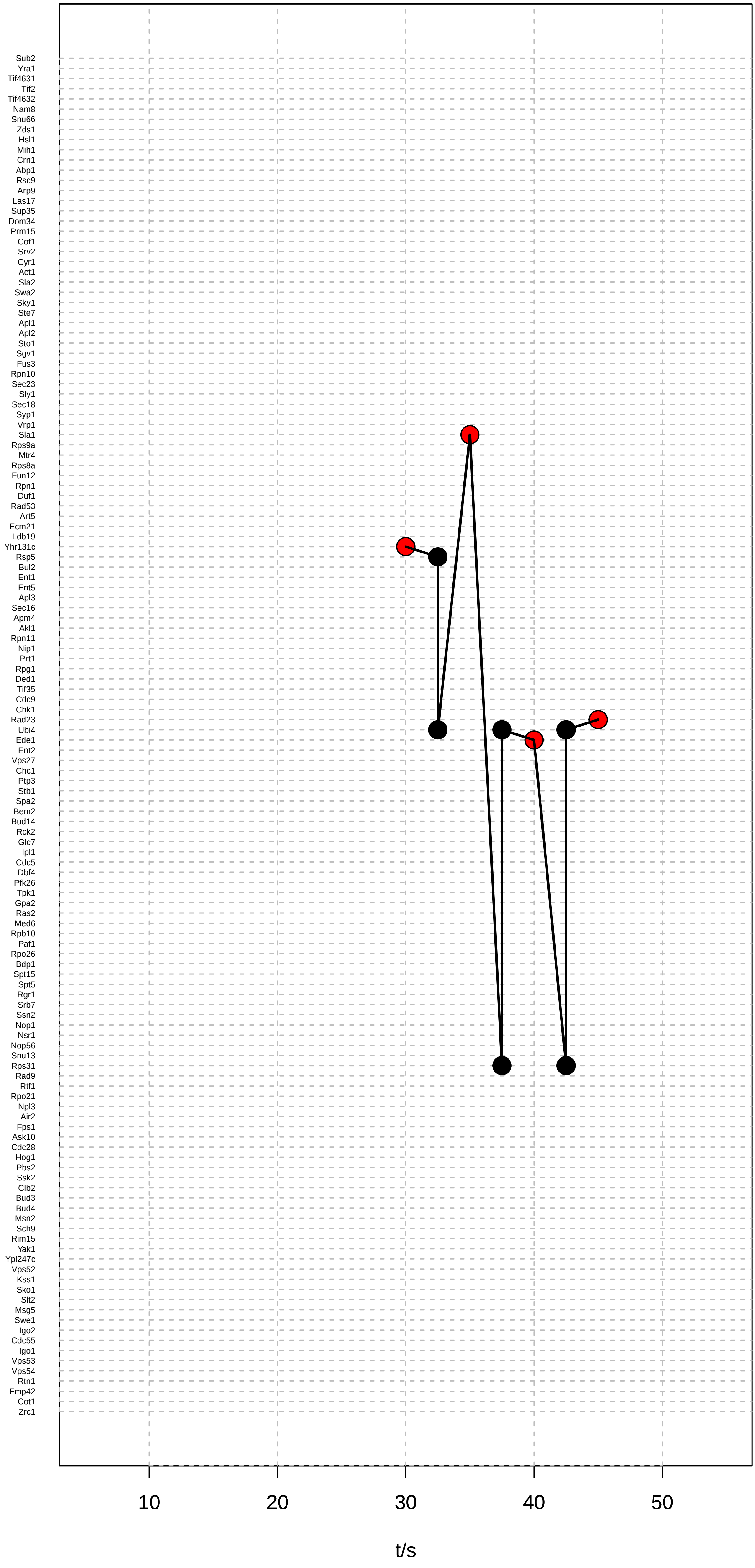

### Path 766 (Yhr131c − Rad23)

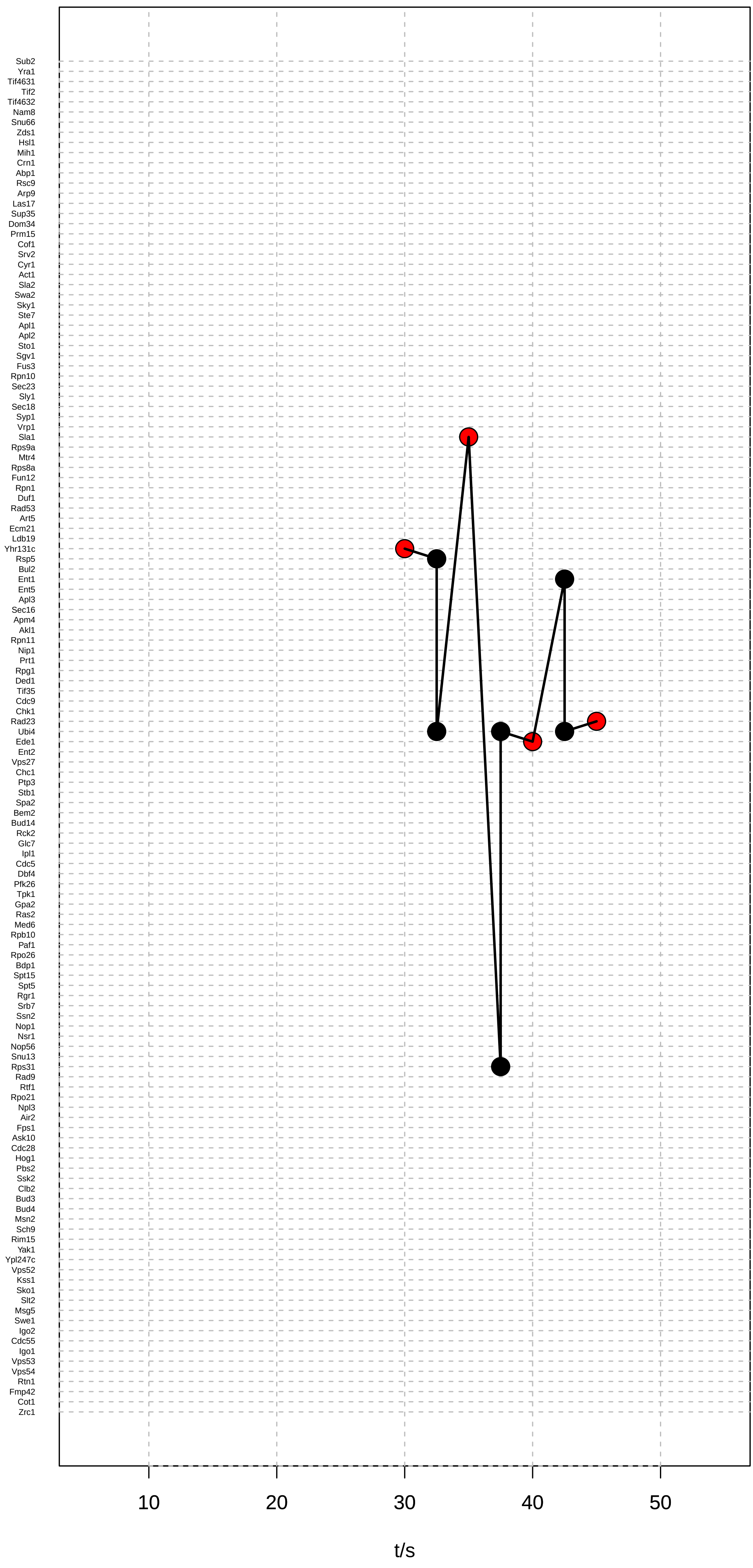

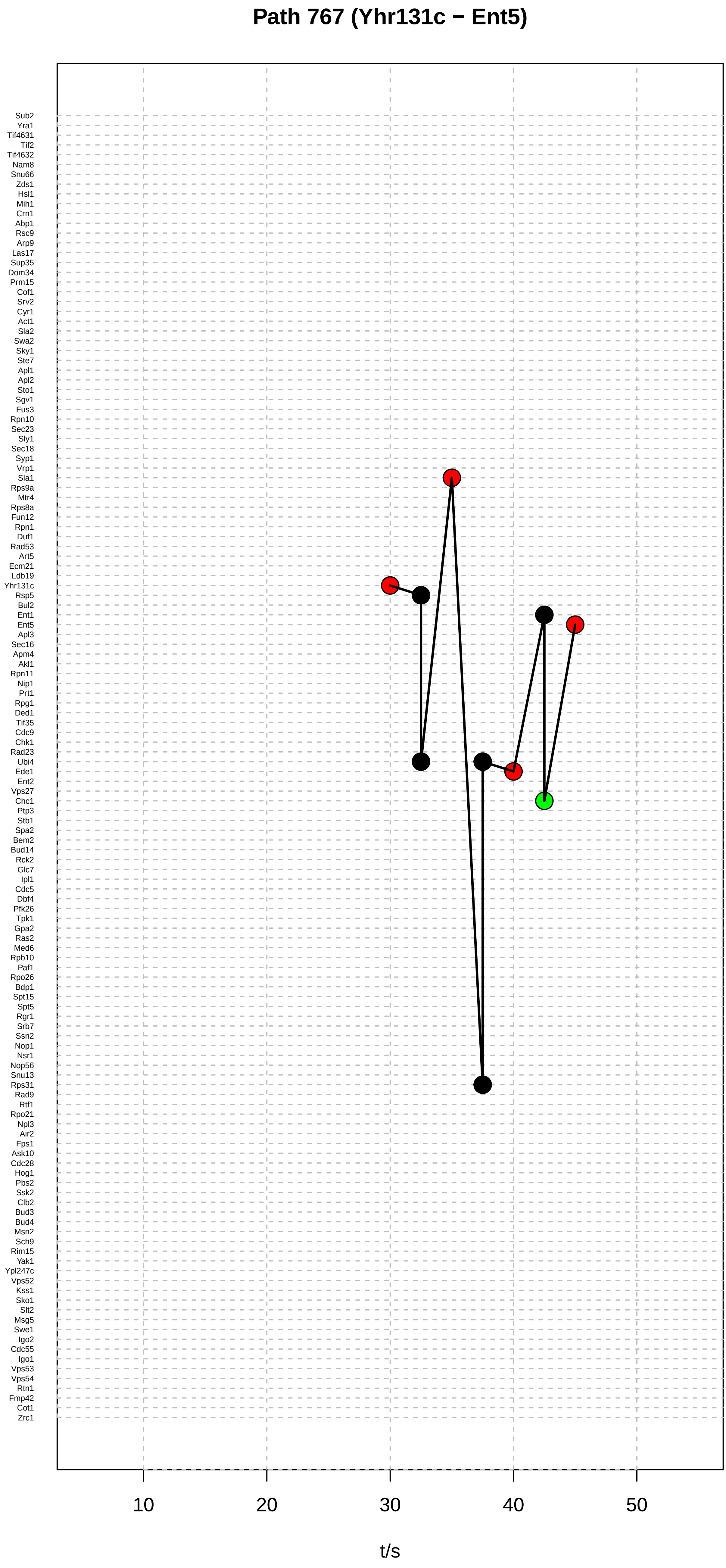
Path 767 (Yhr131c – Ent5)
Sub2
Yra1
Tif4631
Tif2
Tif4632
Nam8
Snu66
Zds1
Hsl1
Mih1
Crn1
Abp1
Rsc9
Arp9
Las17
Sup35
Dom34
Prm15
Cof1
Srv2
Cyr1
Act1
Sla2
Swa2
Sky1
Ste7
Apl1
Apl2
Sto1
Sgv1
Fus3
Rpn10
Sec23
Sly1
Sec18
Syp1
Vrp1
Sla1
Rps9a
Mtr4
Rps8a
Fun12
Rpn1
Duf1
Rad53
Art5
Ecm21
Ldb19
Yhr131c
Rsp5
Bul2
Ent1
Ent5
Apl3
Sec16
Apm4
Akl1
Rpn11
Nip1
Prt1
Rpg1
Ded1
Tif35
Cdc9
Chk1
Rad23
Ubi4
Ede1
Ent2
Vps27
Chc1
Ptp3
Stb1
Spa2
Bem2
Bud14
Rck2
Glc7
Ipl1
Cdc5
Dbf4
Pfk26
Tpk1
Gpa2
Ras2
Med6
Rpb10
Paf1
Rpo26
Bdp1
Spt15
Spt5
Rgr1
Srb7
Ssn2
Nop1
Nsr1
Nop56
Snu13
Rps31
Rad9
Rtf1
Rpo21
Npl3
Air2
Fps1
Ask10
Cdc28
Hog1
Pbs2
Ssk2
Clb2
Bud3
Bud4
Msn2
Sch9
Rim15
Yak1
Ypl247c
Vps52
Kss1
Sko1
Slt2
Msg5
Swe1
Igo2
Cdc55
Igo1
Vps53
Vps54
Rtn1
Fmp42
Cot1
Zrc1
10
20
30
40
50
t/s

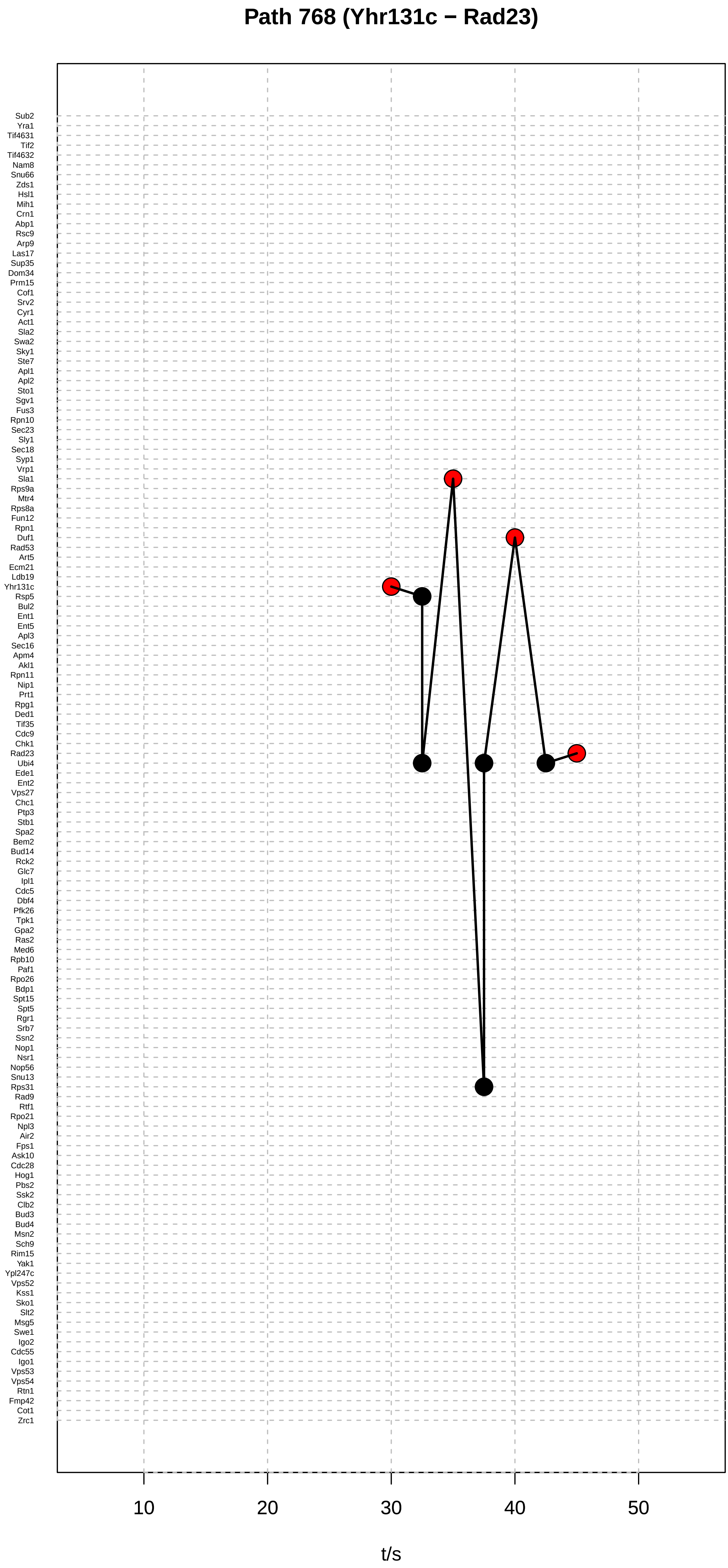
Path 768 (Yhr131c − Rad23)
Sub2
Yra1
Tif4631
Tif2
Tif4632
Nam8
Snu66
Zds1
Hsl1
Mih1
Crn1
Abp1
Rsc9
Arp9
Las17
Sup35
Dom34
Prm15
Cof1
Srv2
Cyr1
Act1
Sla2
Swa2
Sky1
Ste7
Apl1
Apl2
Sto1
Sgv1
Fus3
Rpn10
Sec23
Sly1
Sec18
Syp1
Vrp1
Sla1
Rps9a
Mtr4
Rps8a
Fun12
Rpn1
Duf1
Rad53
Art5
Ecm21
Ldb19
Yhr131c
Rsp5
Bul2
Ent1
Ent5
Apl3
Sec16
Apm4
Akl1
Rpn11
Nip1
Prt1
Rpg1
Ded1
Tif35
Cdc9
Chk1
Rad23
Ubi4
Ede1
Ent2
Vps27
Chc1
Ptp3
Stb1
Spa2
Bem2
Bud14
Rck2
Glc7
Ipl1
Cdc5
Dbf4
Pfk26
Tpk1
Gpa2
Ras2
Med6
Rpb10
Paf1
Rpo26
Bdp1
Spt15
Spt5
Rgr1
Srb7
Ssn2
Nop1
Nsr1
Nop56
Snu13
Rps31
Rad9
Rtf1
Rpo21
Npl3
Air2
Fps1
Ask10
Cdc28
Hog1
Pbs2
Ssk2
Clb2
Bud3
Bud4
Msn2
Sch9
Rim15
Yak1
Ypl247c
Vps52
Kss1
Sko1
Slt2
Msg5
Swe1
Igo2
Cdc55
Igo1
Vps53
Vps54
Rtn1
Fmp42
Cot1
Zrc1
10
20
30
40
50
t/s

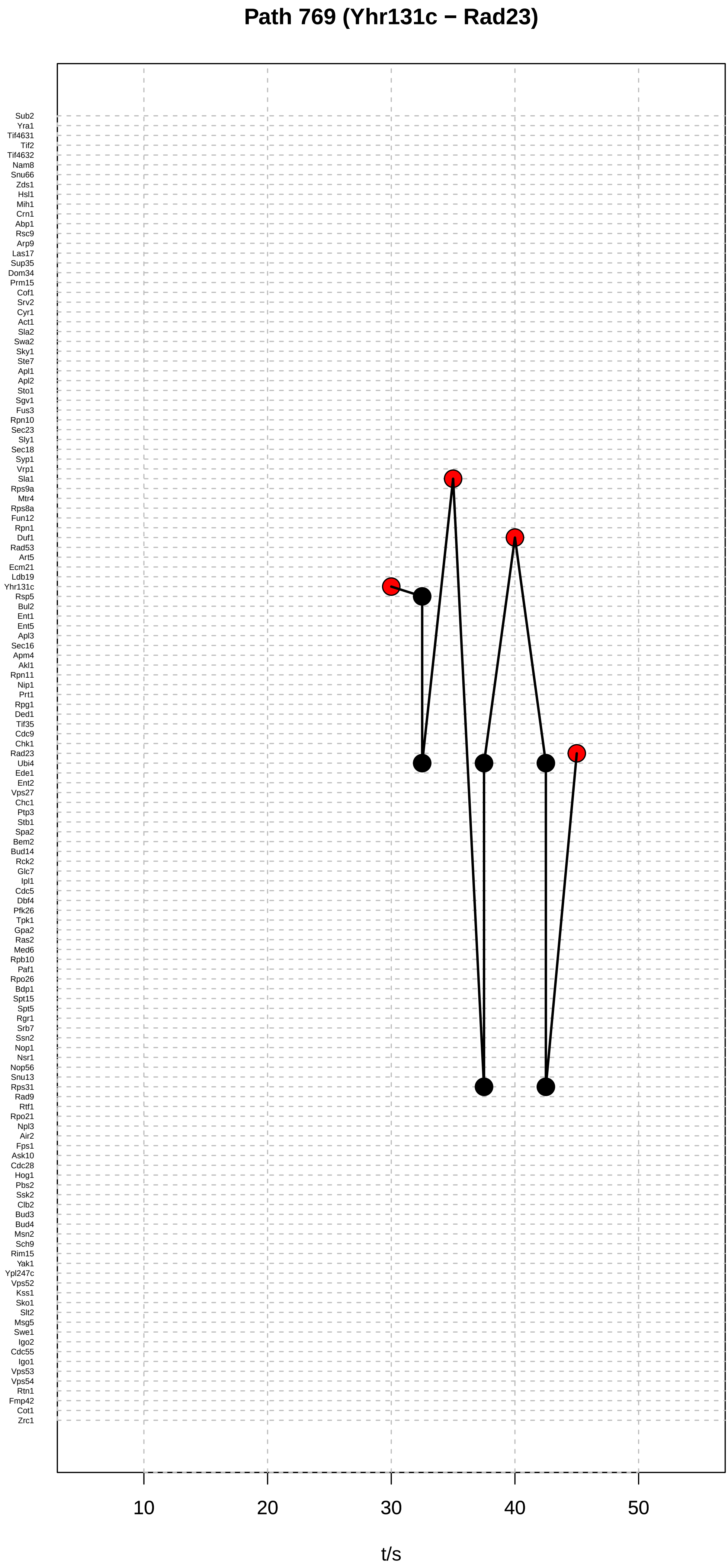

Path 769 (Yhr131c − Rad23)
Sub2
Yra1
Tif4631
Tif2
Tif4632
Nam8
Snu66
Zds1
Hsl1
Mih1
Crn1
Abp1
Rsc9
Arp9
Las17
Sup35
Dom34
Prm15
Cof1
Srv2
Cyr1
Act1
Sla2
Swa2
Sky1
Ste7
Apl1
Apl2
Sto1
Sgv1
Fus3
Rpn10
Sec23
Sly1
Sec18
Syp1
Vrp1
Sla1
Rps9a
Mtr4
Rps8a
Fun12
Rpn1
Duf1
Rad53
Art5
Ecm21
Ldb19
Yhr131c
Rsp5
Bul2
Ent1
Ent5
Apl3
Sec16
Apm4
Akl1
Rpn11
Nip1
Prt1
Rpg1
Ded1
Tif35
Cdc9
Chk1
Rad23
Ubi4
Ede1
Ent2
Vps27
Chc1
Ptp3
Stb1
Spa2
Bem2
Bud14
Rck2
Glc7
Ipl1
Cdc5
Dbf4
Pfk26
Tpk1
Gpa2
Ras2
Med6
Rpb10
Paf1
Rpo26
Bdp1
Spt15
Spt5
Rgr1
Srb7
Ssn2
Nop1
Nsr1
Nop56
Snu13
Rps31
Rad9
Rtf1
Rpo21
Npl3
Air2
Fps1
Ask10
Cdc28
Hog1
Pbs2
Ssk2
Clb2
Bud3
Bud4
Msn2
Sch9
Rim15
Yak1
Ypl247c
Vps52
Kss1
Sko1
Slt2
Msg5
Swe1
Igo2
Cdc55
Igo1
Vps53
Vps54
Rtn1
Fmp42
Cot1
Zrc1
10
20
30
40
50
t/s

### Path 770 (Yhr131c − Rad23)

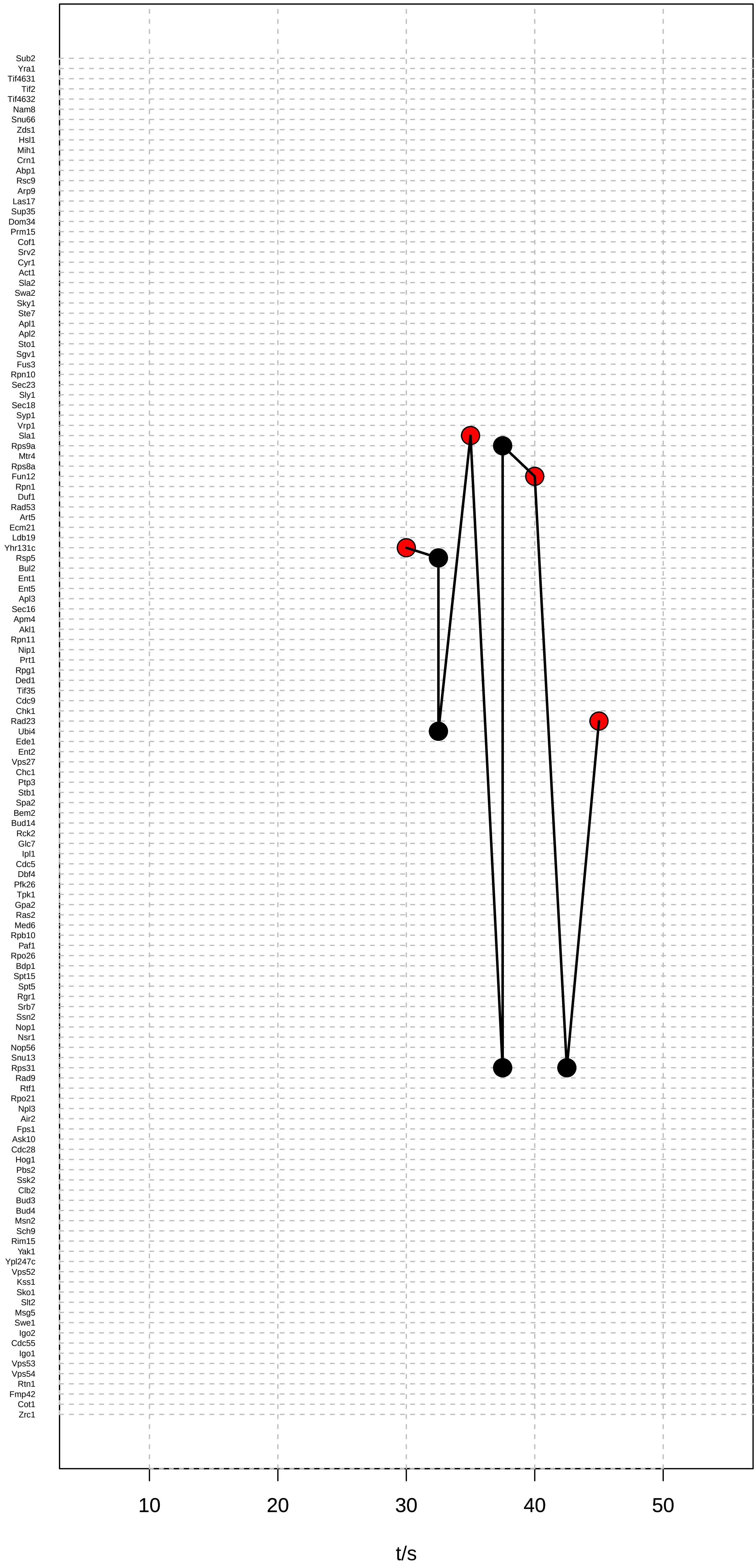

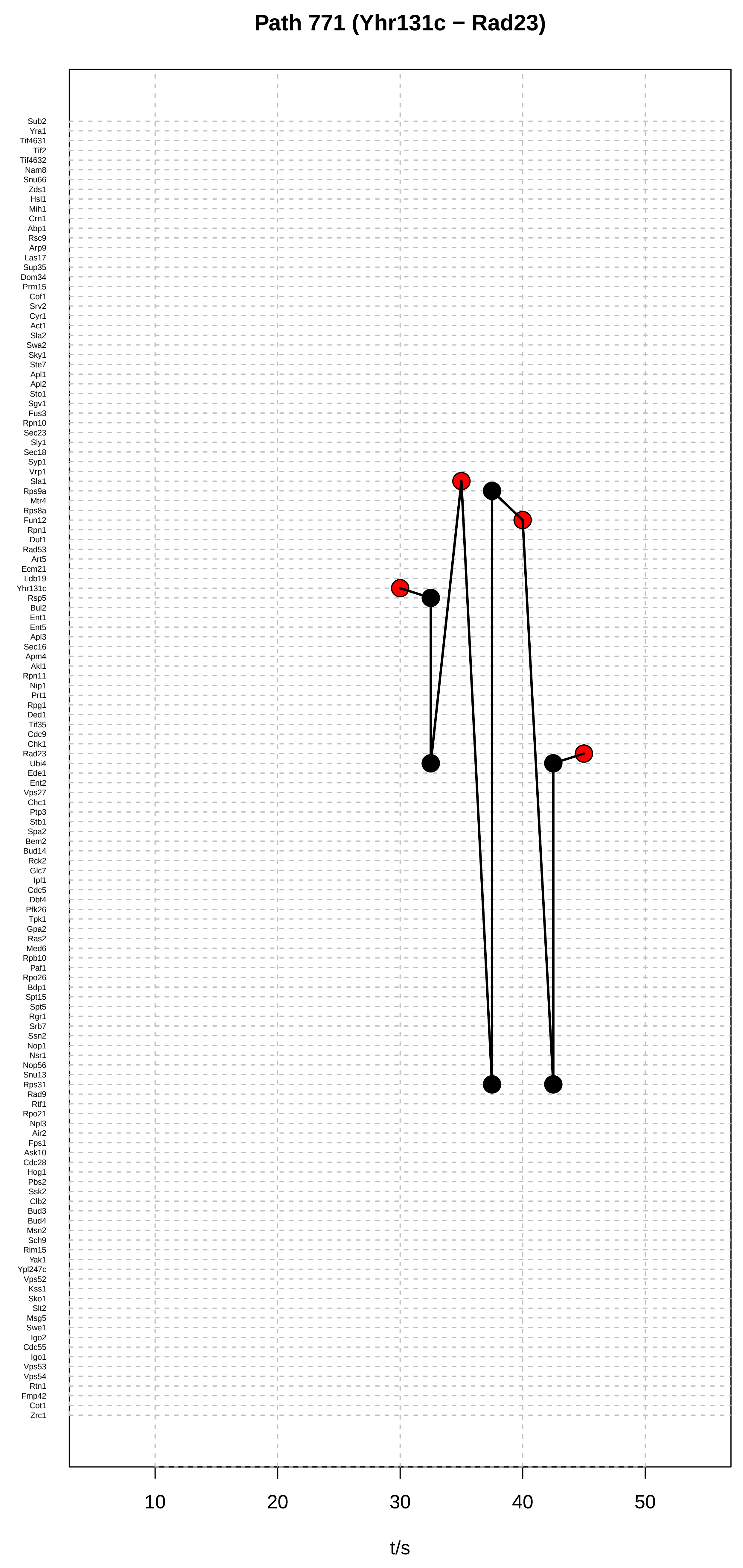
Path 771 (Yhr131c − Rad23)
Sub2
Yra1
Tif4631
Tif2
Tif4632
Nam8
Snu66
Zds1
Hsl1
Mih1
Crn1
Abp1
Rsc9
Arp9
Las17
Sup35
Dom34
Prm15
Cof1
Srv2
Cyr1
Act1
Sla2
Swa2
Sky1
Ste7
Apl1
Apl2
Sto1
Sgv1
Fus3
Rpn10
Sec23
Sly1
Sec18
Syp1
Vrp1
Sla1
Rps9a
Mtr4
Rps8a
Fun12
Rpn1
Duf1
Rad53
Art5
Ecm21
Ldb19
Yhr131c
Rsp5
Bul2
Ent1
Ent5
Apl3
Sec16
Apm4
Akl1
Rpn11
Nip1
Prt1
Rpg1
Ded1
Tif35
Cdc9
Chk1
Rad23
Ubi4
Ede1
Ent2
Vps27
Chc1
Ptp3
Stb1
Spa2
Bem2
Bud14
Rck2
Glc7
Ipl1
Cdc5
Dbf4
Pfk26
Tpk1
Gpa2
Ras2
Med6
Rpb10
Paf1
Rpo26
Bdp1
Spt15
Spt5
Rgr1
Srb7
Ssn2
Nop1
Nsr1
Nop56
Snu13
Rps31
Rad9
Rtf1
Rpo21
Npl3
Air2
Fps1
Ask10
Cdc28
Hog1
Pbs2
Ssk2
Clb2
Bud3
Bud4
Msn2
Sch9
Rim15
Yak1
Ypl247c
Vps52
Kss1
Sko1
Slt2
Msg5
Swe1
Igo2
Cdc55
Igo1
Vps53
Vps54
Rtn1
Fmp42
Cot1
Zrc1
10
20
30
40
50
t/s

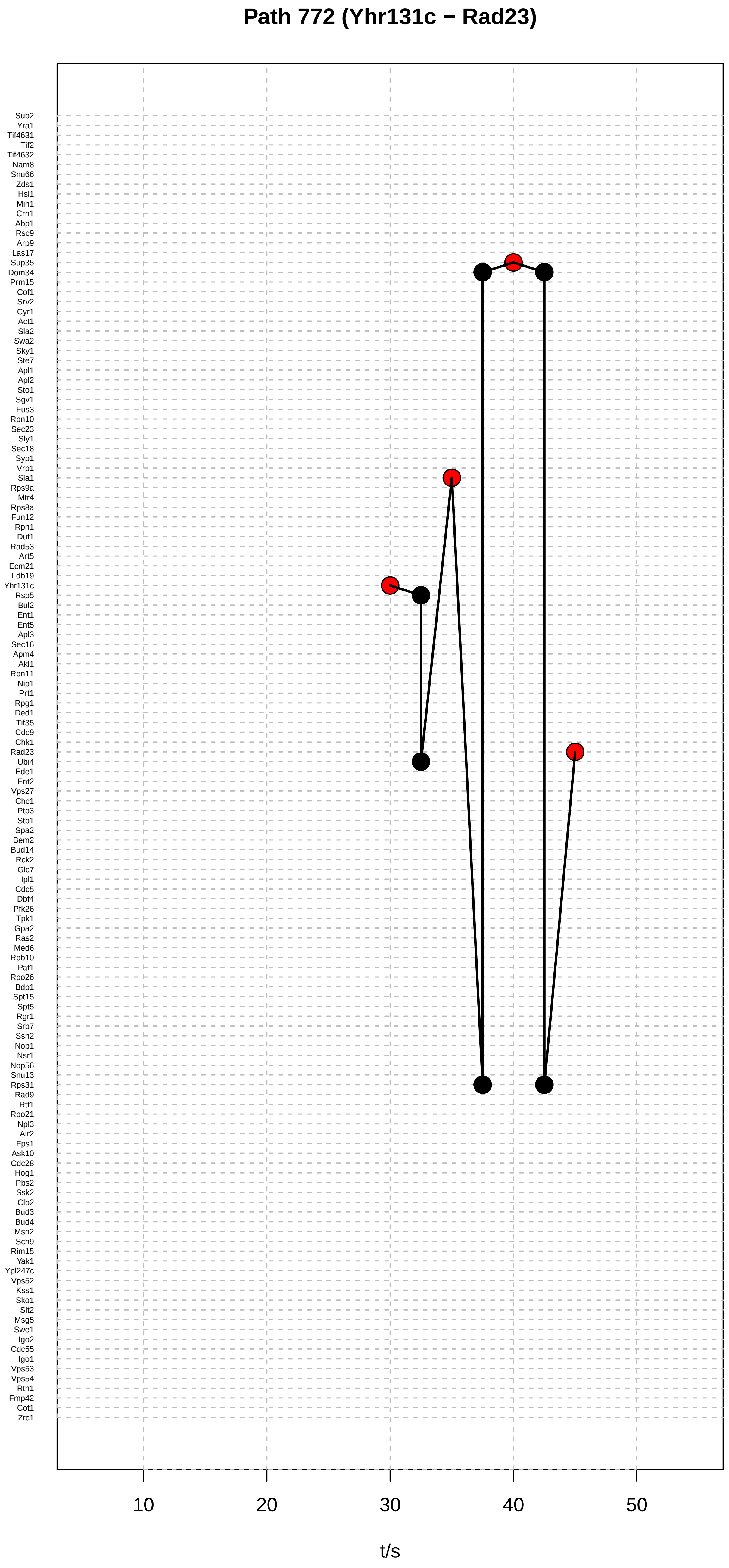
Path 772 (Yhr131c − Rad23)
Sub2
Yra1
Tif4631
Tif2
Tif4632
Nam8
Snu66
Zds1
Hsl1
Mih1
Crn1
Abp1
Rsc9
Arp9
Las17
Sup35
Dom34
Prm15
Cof1
Srv2
Cyr1
Act1
Sla2
Swa2
Sky1
Ste7
Apl1
Apl2
Sto1
Sgv1
Fus3
Rpn10
Sec23
Sly1
Sec18
Syp1
Vrp1
Sla1
Rps9a
Mtr4
Rps8a
Fun12
Rpn1
Duf1
Rad53
Art5
Ecm21
Ldb19
Yhr131c
Rsp5
Bul2
Ent1
Ent5
Apl3
Sec16
Apm4
Akl1
Rpn11
Nip1
Prt1
Rpg1
Ded1
Tif35
Cdc9
Chk1
Rad23
Ubi4
Ede1
Ent2
Vps27
Chc1
Ptp3
Stb1
Spa2
Bem2
Bud14
Rck2
Glc7
Ipl1
Cdc5
Dbf4
Pfk26
Tpk1
Gpa2
Ras2
Med6
Rpb10
Paf1
Rpo26
Bdp1
Spt15
Spt5
Rgr1
Srb7
Ssn2
Nop1
Nsr1
Nop56
Snu13
Rps31
Rad9
Rtf1
Rpo21
Npl3
Air2
Fps1
Ask10
Cdc28
Hog1
Pbs2
Ssk2
Clb2
Bud3
Bud4
Msn2
Sch9
Rim15
Yak1
Ypl247c
Vps52
Kss1
Sko1
Slt2
Msg5
Swe1
Igo2
Cdc55
Igo1
Vps53
Vps54
Rtn1
Fmp42
Cot1
Zrc1
10
20
30
40
50
t/s

### Path 773 (Yhr131c − Prm15)

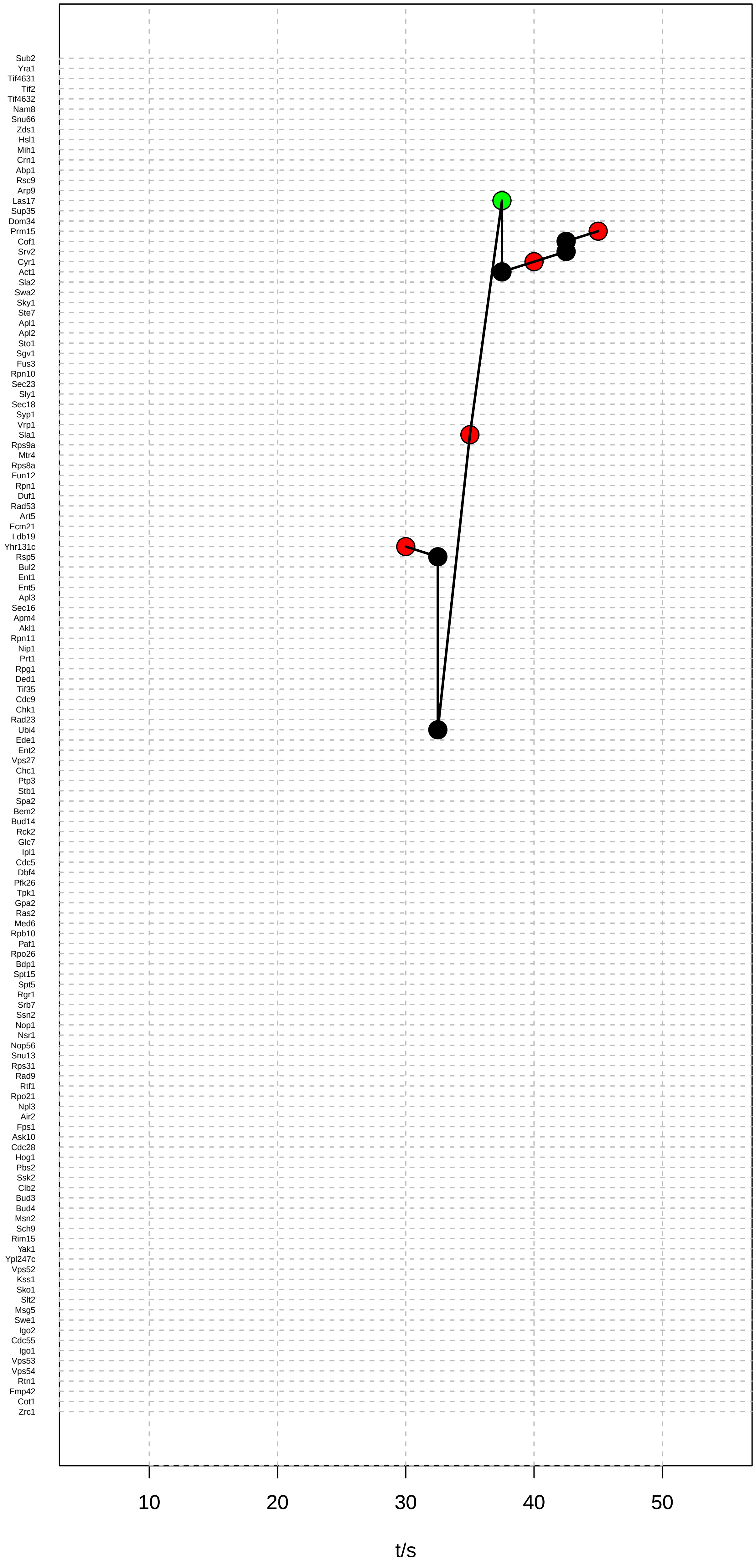

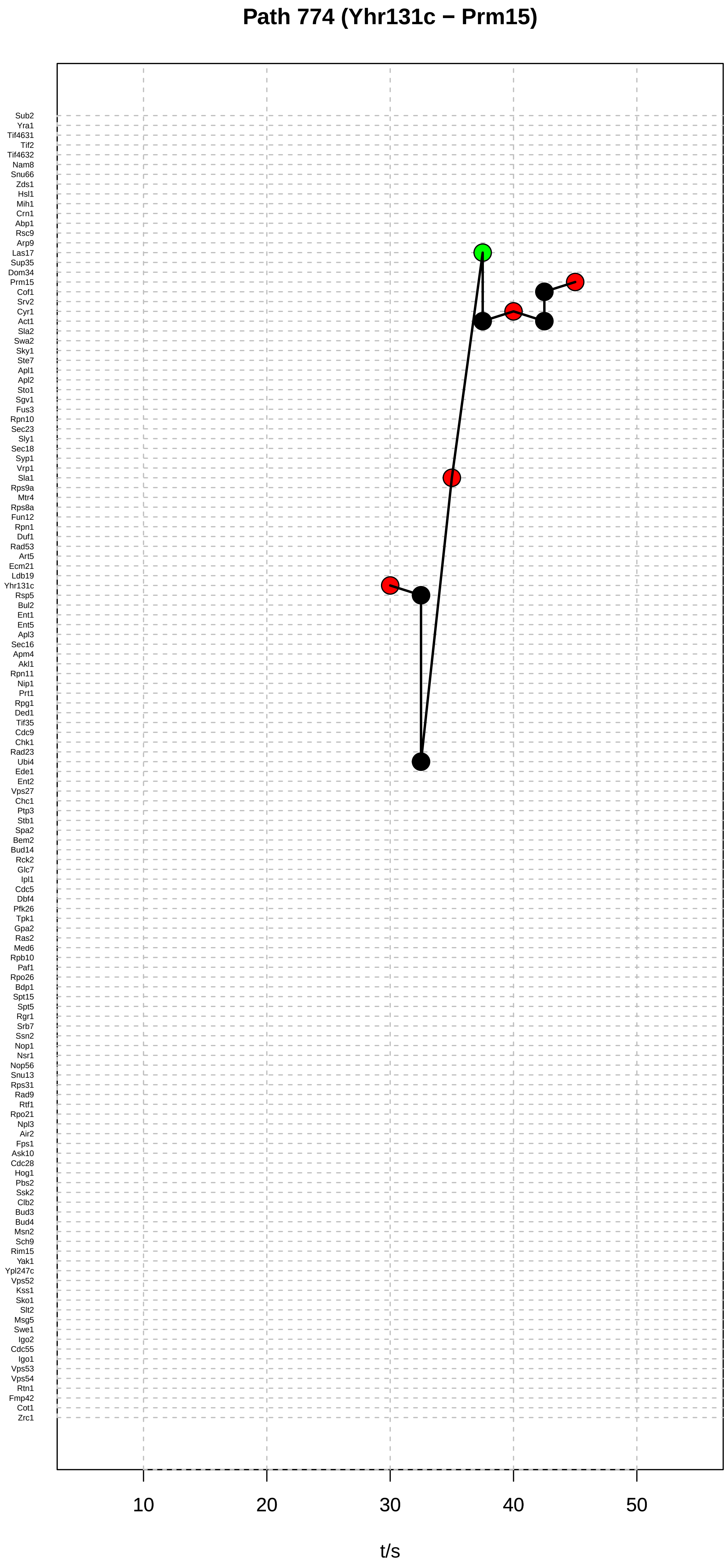

Path 774 (Yhr131c − Prm15)
Sub2
Yra1
Tif4631
Tif2
Tif4632
Nam8
Snu66
Zds1
Hsl1
Mih1
Crn1
Abp1
Rsc9
Arp9
Las17
Sup35
Dom34
Prm15
Cof1
Srv2
Cyr1
Act1
Sla2
Swa2
Sky1
Ste7
Apl1
Apl2
Sto1
Sgv1
Fus3
Rpn10
Sec23
Sly1
Sec18
Syp1
Vrp1
Sla1
Rps9a
Mtr4
Rps8a
Fun12
Rpn1
Duf1
Rad53
Art5
Ecm21
Ldb19
Yhr131c
Rsp5
Bul2
Ent1
Ent5
Apl3
Sec16
Apm4
Akl1
Rpn11
Nip1
Prt1
Rpg1
Ded1
Tif35
Cdc9
Chk1
Rad23
Ubi4
Ede1
Ent2
Vps27
Chc1
Ptp3
Stb1
Spa2
Bem2
Bud14
Rck2
Glc7
Ipl1
Cdc5
Dbf4
Pfk26
Tpk1
Gpa2
Ras2
Med6
Rpb10
Paf1
Rpo26
Bdp1
Spt15
Spt5
Rgr1
Srb7
Ssn2
Nop1
Nsr1
Nop56
Snu13
Rps31
Rad9
Rtf1
Rpo21
Npl3
Air2
Fps1
Ask10
Cdc28
Hog1
Pbs2
Ssk2
Clb2
Bud3
Bud4
Msn2
Sch9
Rim15
Yak1
Ypl247c
Vps52
Kss1
Sko1
Slt2
Msg5
Swe1
Igo2
Cdc55
Igo1
Vps53
Vps54
Rtn1
Fmp42
Cot1
Zrc1
10
20
30
40
50
t/s

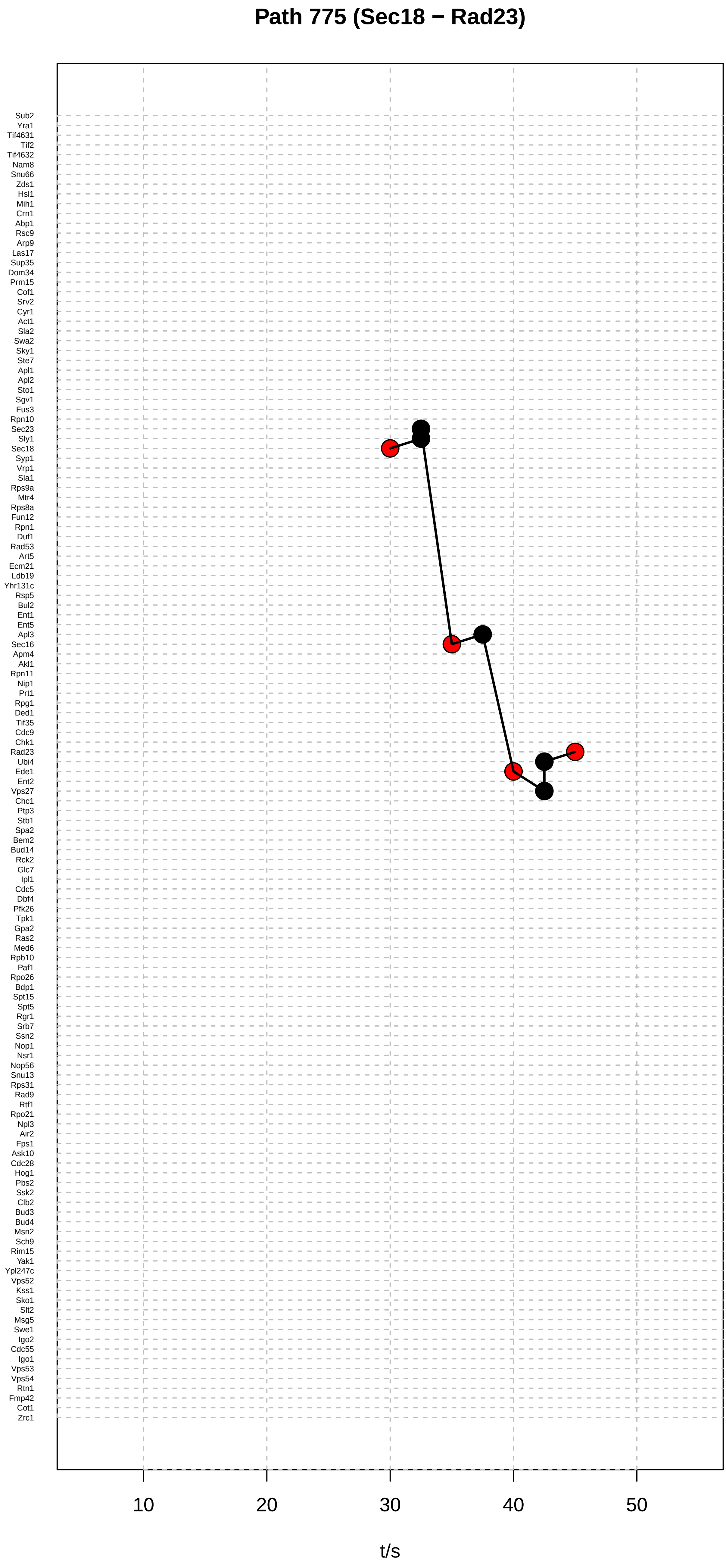

Path 775 (Sec18 − Rad23)
Sub2
Yra1
Tif4631
Tif2
Tif4632
Nam8
Snu66
Zds1
Hsl1
Mih1
Crn1
Abp1
Rsc9
Arp9
Las17
Sup35
Dom34
Prm15
Cof1
Srv2
Cyr1
Act1
Sla2
Swa2
Sky1
Ste7
Apl1
Apl2
Sto1
Sgv1
Fus3
Rpn10
Sec23
Sly1
Sec18
Syp1
Vrp1
Sla1
Rps9a
Mtr4
Rps8a
Fun12
Rpn1
Duf1
Rad53
Art5
Ecm21
Ldb19
Yhr131c
Rsp5
Bul2
Ent1
Ent5
Apl3
Sec16
Apm4
Akl1
Rpn11
Nip1
Prt1
Rpg1
Ded1
Tif35
Cdc9
Chk1
Rad23
Ubi4
Ede1
Ent2
Vps27
Chc1
Ptp3
Stb1
Spa2
Bem2
Bud14
Rck2
Glc7
Ipl1
Cdc5
Dbf4
Pfk26
Tpk1
Gpa2
Ras2
Med6
Rpb10
Paf1
Rpo26
Bdp1
Spt15
Spt5
Rgr1
Srb7
Ssn2
Nop1
Nsr1
Nop56
Snu13
Rps31
Rad9
Rtf1
Rpo21
Npl3
Air2
Fps1
Ask10
Cdc28
Hog1
Pbs2
Ssk2
Clb2
Bud3
Bud4
Msn2
Sch9
Rim15
Yak1
Ypl247c
Vps52
Kss1
Sko1
Slt2
Msg5
Swe1
Igo2
Cdc55
Igo1
Vps53
Vps54
Rtn1
Fmp42
Cot1
Zrc1
10
20
30
40
50
t/s

Path 776 (Sec18 − Rad23)

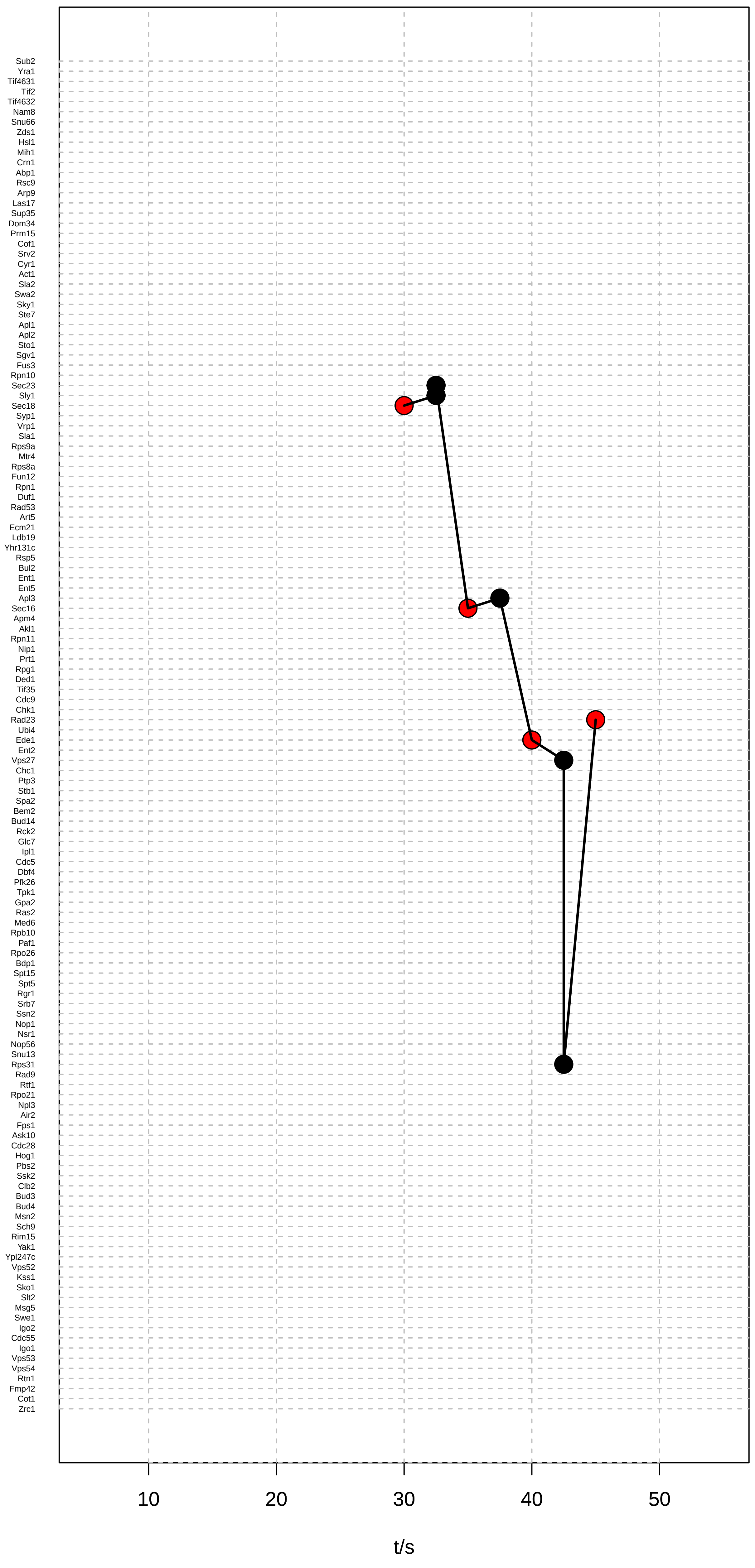

### Path 777 (Sec18 − Rad23)

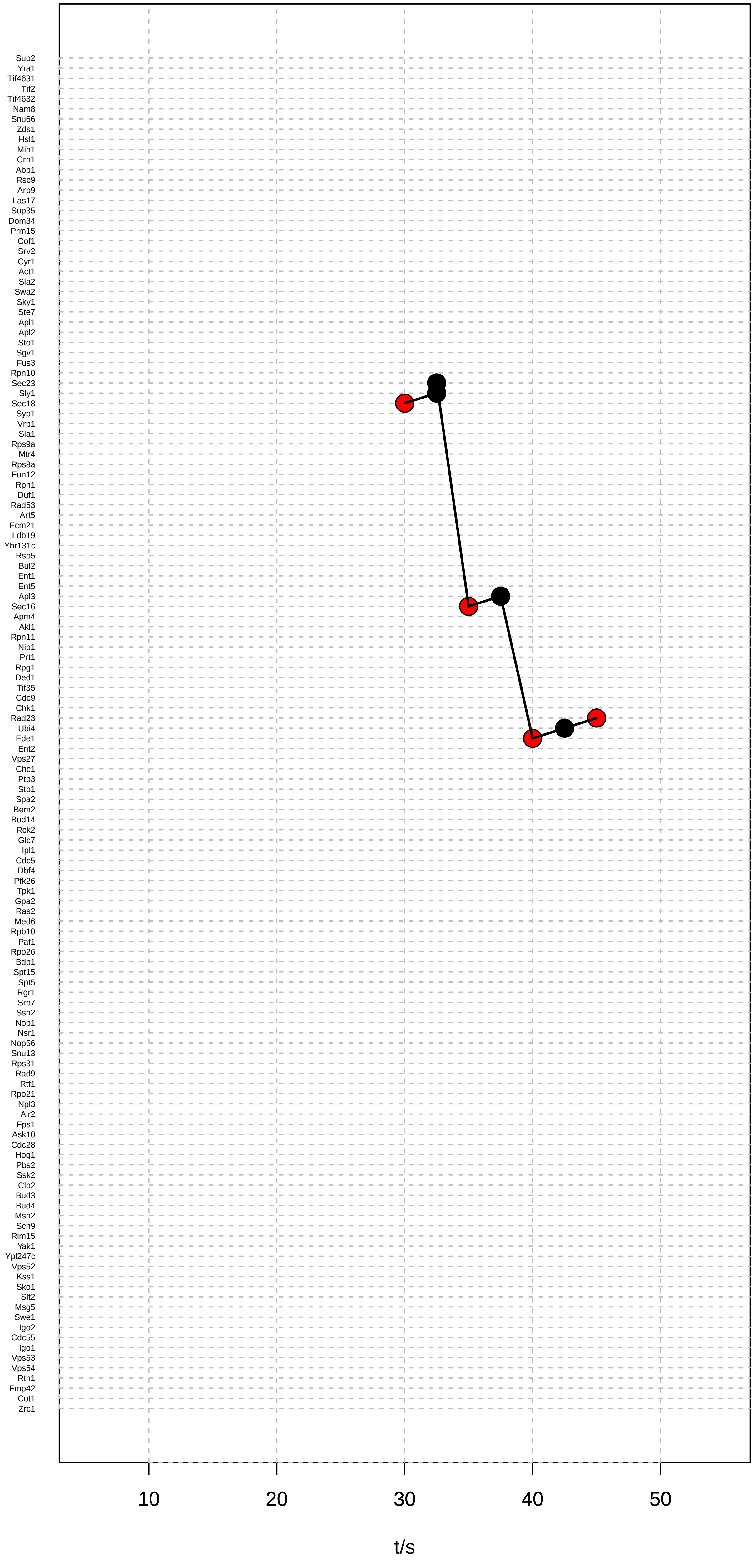

#### Path 778 (Sec18 − Rad23)

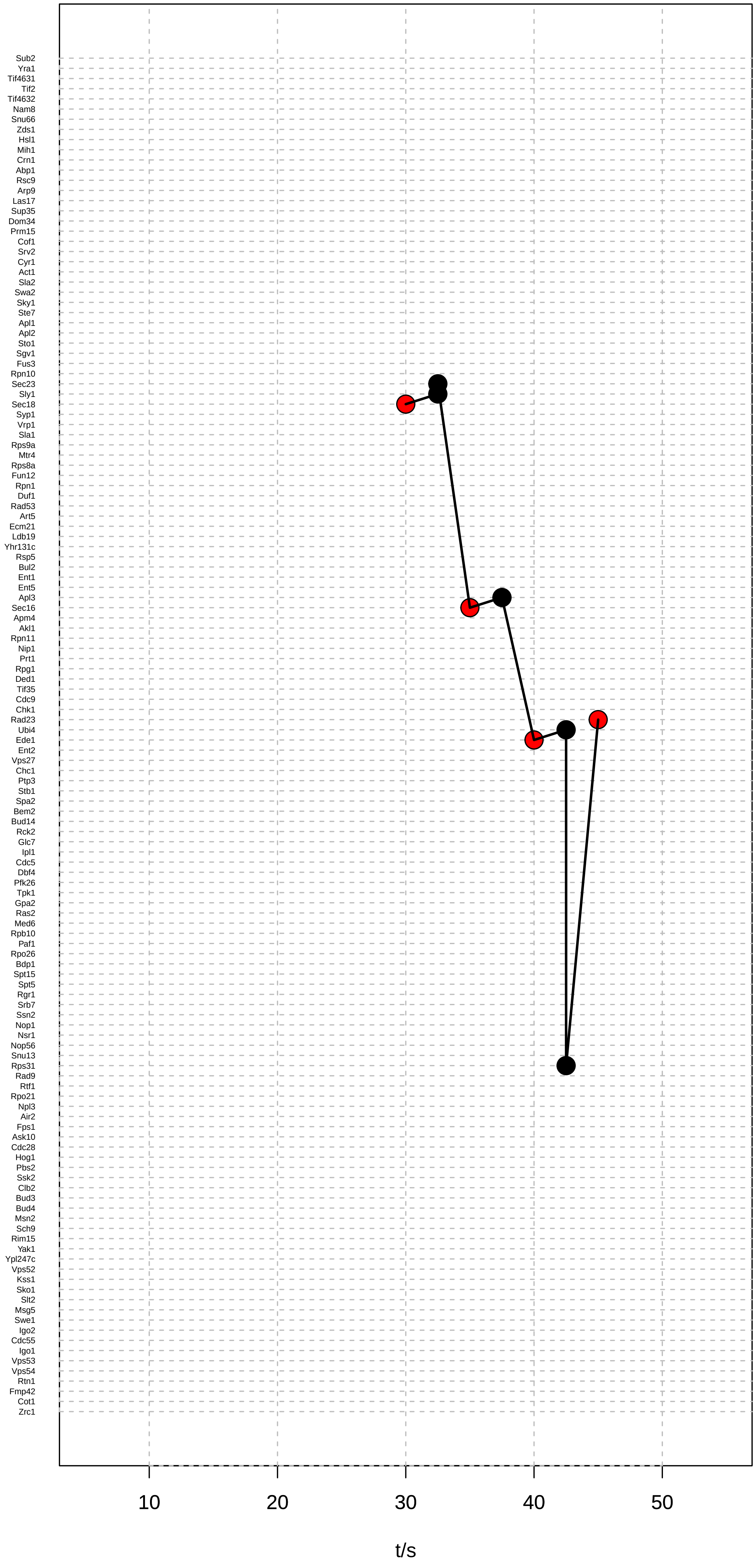

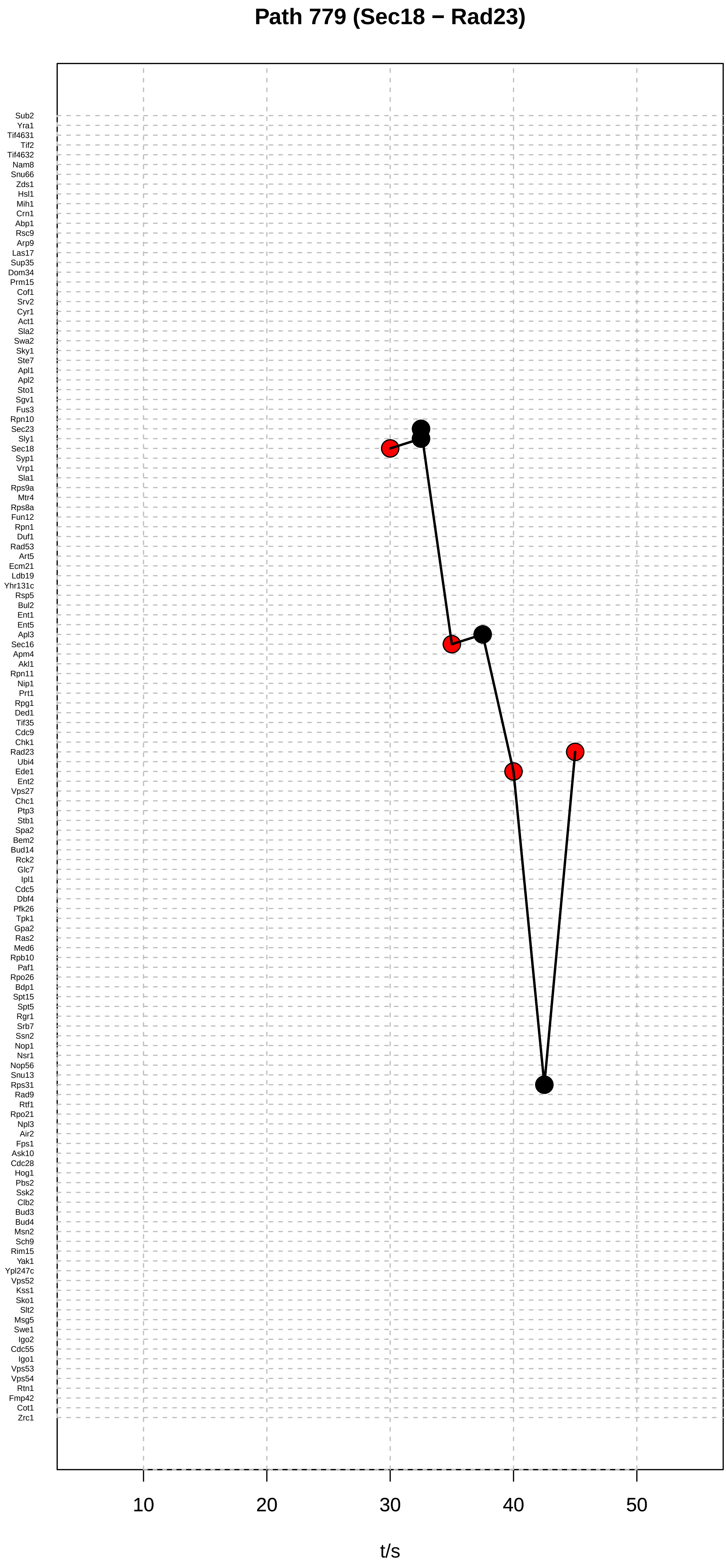

Path 779 (Sec18 − Rad23)
Sub2
Yra1
Tif4631
Tif2
Tif4632
Nam8
Snu66
Zds1
Hsl1
Mih1
Crn1
Abp1
Rsc9
Arp9
Las17
Sup35
Dom34
Prm15
Cof1
Srv2
Cyr1
Act1
Sla2
Swa2
Sky1
Ste7
Apl1
Apl2
Sto1
Sgv1
Fus3
Rpn10
Sec23
Sly1
Sec18
Syp1
Vrp1
Sla1
Rps9a
Mtr4
Rps8a
Fun12
Rpn1
Duf1
Rad53
Art5
Ecm21
Ldb19
Yhr131c
Rsp5
Bul2
Ent1
Ent5
Apl3
Sec16
Apm4
Akl1
Rpn11
Nip1
Prt1
Rpg1
Ded1
Tif35
Cdc9
Chk1
Rad23
Ubi4
Ede1
Ent2
Vps27
Chc1
Ptp3
Stb1
Spa2
Bem2
Bud14
Rck2
Glc7
Ipl1
Cdc5
Dbf4
Pfk26
Tpk1
Gpa2
Ras2
Med6
Rpb10
Paf1
Rpo26
Bdp1
Spt15
Spt5
Rgr1
Srb7
Ssn2
Nop1
Nsr1
Nop56
Snu13
Rps31
Rad9
Rtf1
Rpo21
Npl3
Air2
Fps1
Ask10
Cdc28
Hog1
Pbs2
Ssk2
Clb2
Bud3
Bud4
Msn2
Sch9
Rim15
Yak1
Ypl247c
Vps52
Kss1
Sko1
Slt2
Msg5
Swe1
Igo2
Cdc55
Igo1
Vps53
Vps54
Rtn1
Fmp42
Cot1
Zrc1
10
20
30
40
50
t/s

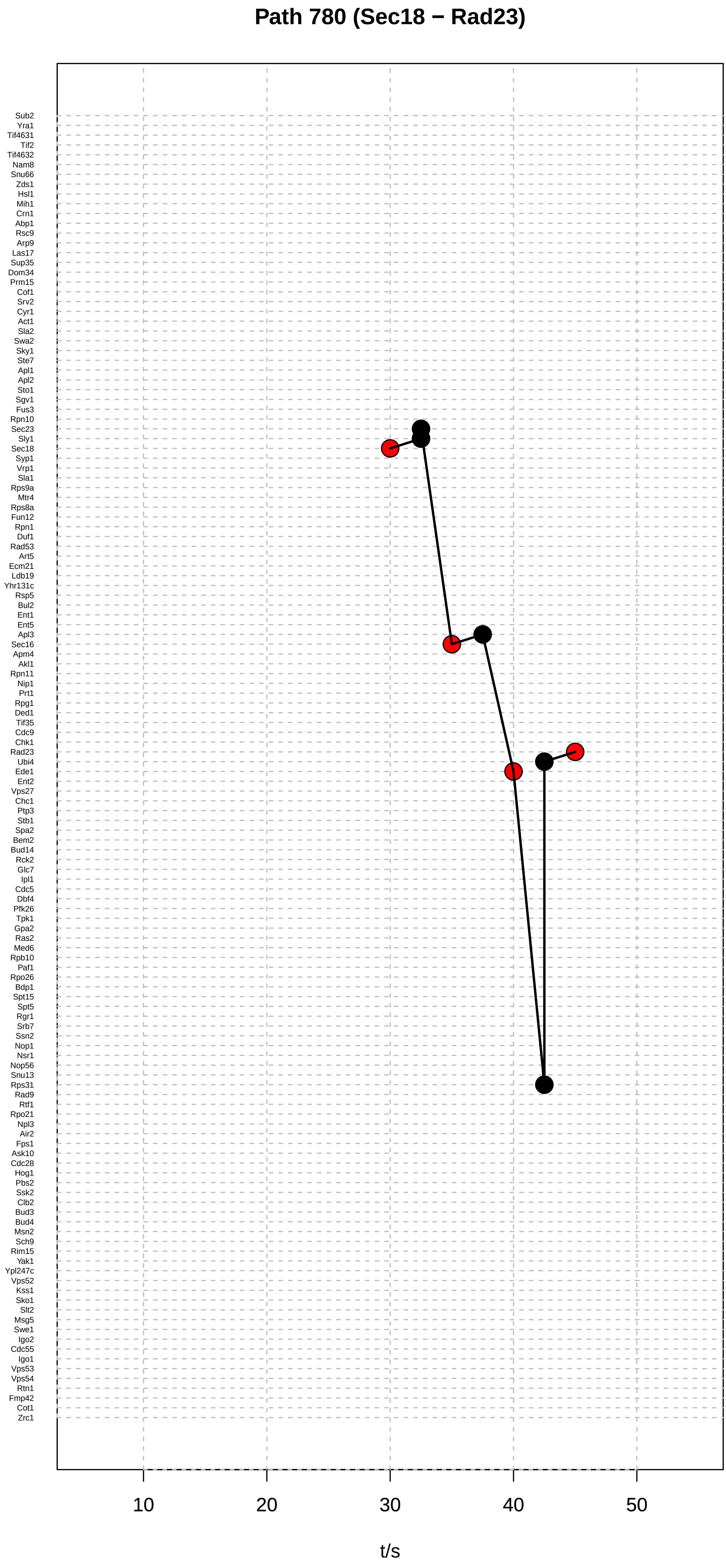

Path 780 (Sec18 − Rad23)
Sub2
Yra1
Tif4631
Tif2
Tif4632
Nam8
Snu66
Zds1
Hsl1
Mih1
Crn1
Abp1
Rsc9
Arp9
Las17
Sup35
Dom34
Prm15
Cof1
Srv2
Cyr1
Act1
Sla2
Swa2
Sky1
Ste7
Apl1
Apl2
Sto1
Sgv1
Fus3
Rpn10
Sec23
Sly1
Sec18
Syp1
Vrp1
Sla1
Rps9a
Mtr4
Rps8a
Fun12
Rpn1
Duf1
Rad53
Art5
Ecm21
Ldb19
Yhr131c
Rsp5
Bul2
Ent1
Ent5
Apl3
Sec16
Apm4
Akl1
Rpn11
Nip1
Prt1
Rpg1
Ded1
Tif35
Cdc9
Chk1
Rad23
Ubi4
Ede1
Ent2
Vps27
Chc1
Ptp3
Stb1
Spa2
Bem2
Bud14
Rck2
Glc7
Ipl1
Cdc5
Dbf4
Pfk26
Tpk1
Gpa2
Ras2
Med6
Rpb10
Paf1
Rpo26
Bdp1
Spt15
Spt5
Rgr1
Srb7
Ssn2
Nop1
Nsr1
Nop56
Snu13
Rps31
Rad9
Rtf1
Rpo21
Npl3
Air2
Fps1
Ask10
Cdc28
Hog1
Pbs2
Ssk2
Clb2
Bud3
Bud4
Msn2
Sch9
Rim15
Yak1
Ypl247c
Vps52
Kss1
Sko1
Slt2
Msg5
Swe1
Igo2
Cdc55
Igo1
Vps53
Vps54
Rtn1
Fmp42
Cot1
Zrc1
10
20
30
40
50
t/s

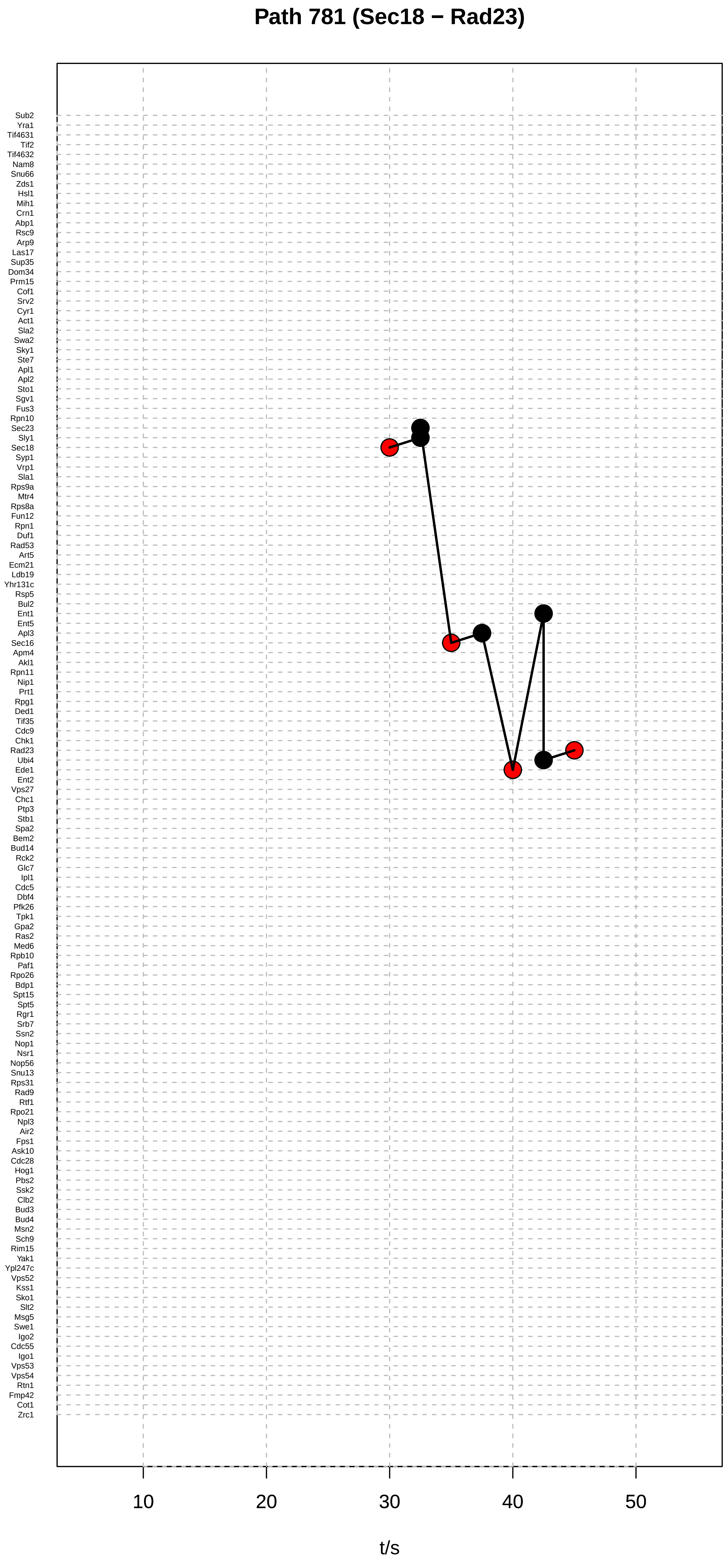

Path 781 (Sec18 − Rad23)
Sub2
Yra1
Tif4631
Tif2
Tif4632
Nam8
Snu66
Zds1
Hsl1
Mih1
Crn1
Abp1
Rsc9
Arp9
Las17
Sup35
Dom34
Prm15
Cof1
Srv2
Cyr1
Act1
Sla2
Swa2
Sky1
Ste7
Apl1
Apl2
Sto1
Sgv1
Fus3
Rpn10
Sec23
Sly1
Sec18
Syp1
Vrp1
Sla1
Rps9a
Mtr4
Rps8a
Fun12
Rpn1
Duf1
Rad53
Art5
Ecm21
Ldb19
Yhr131c
Rsp5
Bul2
Ent1
Ent5
Apl3
Sec16
Apm4
Akl1
Rpn11
Nip1
Prt1
Rpg1
Ded1
Tif35
Cdc9
Chk1
Rad23
Ubi4
Ede1
Ent2
Vps27
Chc1
Ptp3
Stb1
Spa2
Bem2
Bud14
Rck2
Glc7
Ipl1
Cdc5
Dbf4
Pfk26
Tpk1
Gpa2
Ras2
Med6
Rpb10
Paf1
Rpo26
Bdp1
Spt15
Spt5
Rgr1
Srb7
Ssn2
Nop1
Nsr1
Nop56
Snu13
Rps31
Rad9
Rtf1
Rpo21
Npl3
Air2
Fps1
Ask10
Cdc28
Hog1
Pbs2
Ssk2
Clb2
Bud3
Bud4
Msn2
Sch9
Rim15
Yak1
Ypl247c
Vps52
Kss1
Sko1
Slt2
Msg5
Swe1
Igo2
Cdc55
Igo1
Vps53
Vps54
Rtn1
Fmp42
Cot1
Zrc1
10
20
30
40
50
t/s

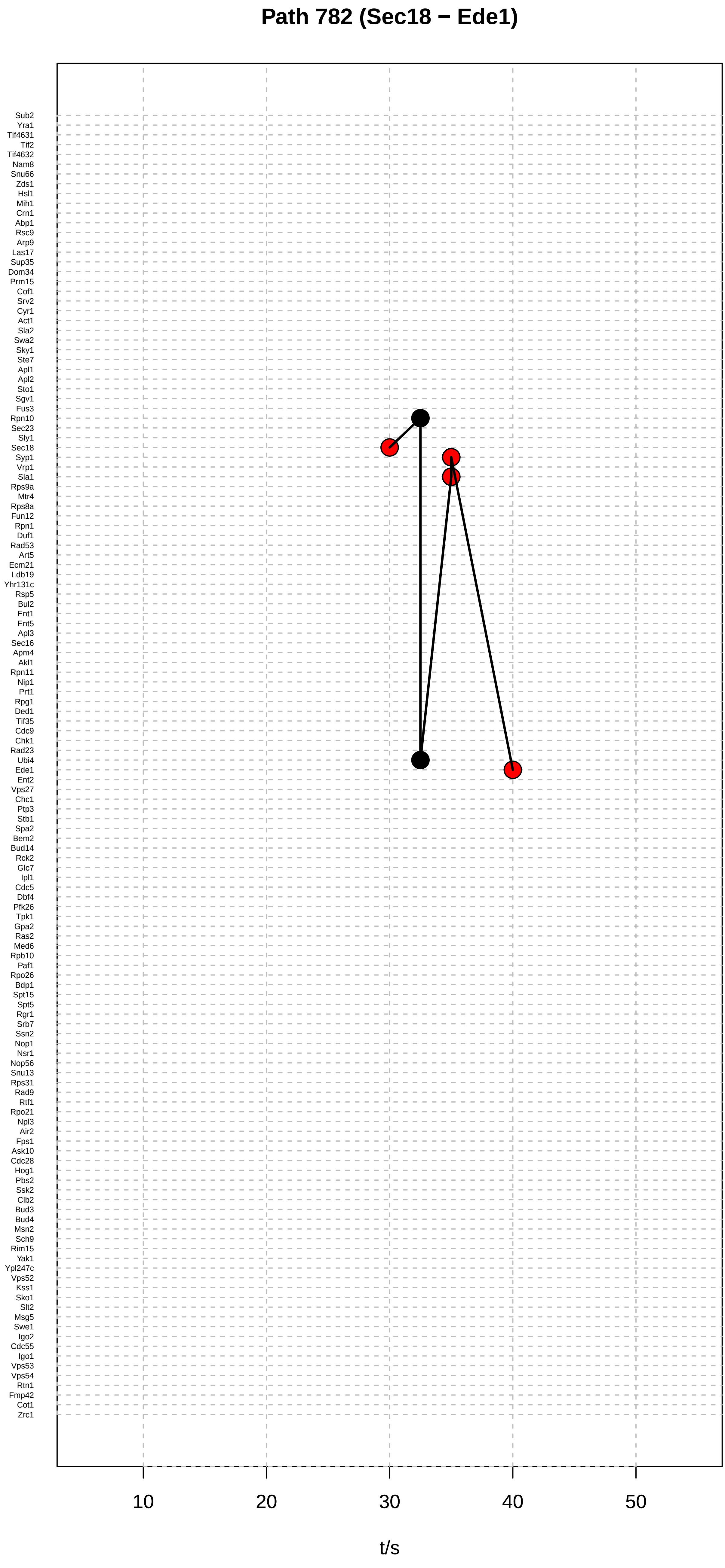

Path 782 (Sec18 − Ede1)
Sub2
Yra1
Tif4631
Tif2
Tif4632
Nam8
Snu66
Zds1
Hsl1
Mih1
Crn1
Abp1
Rsc9
Arp9
Las17
Sup35
Dom34
Prm15
Cof1
Srv2
Cyr1
Act1
Sla2
Swa2
Sky1
Ste7
Apl1
Apl2
Sto1
Sgv1
Fus3
Rpn10
Sec23
Sly1
Sec18
Syp1
Vrp1
Sla1
Rps9a
Mtr4
Rps8a
Fun12
Rpn1
Duf1
Rad53
Art5
Ecm21
Ldb19
Yhr131c
Rsp5
Bul2
Ent1
Ent5
Apl3
Sec16
Apm4
Akl1
Rpn11
Nip1
Prt1
Rpg1
Ded1
Tif35
Cdc9
Chk1
Rad23
Ubi4
Ede1
Ent2
Vps27
Chc1
Ptp3
Stb1
Spa2
Bem2
Bud14
Rck2
Glc7
Ipl1
Cdc5
Dbf4
Pfk26
Tpk1
Gpa2
Ras2
Med6
Rpb10
Paf1
Rpo26
Bdp1
Spt15
Spt5
Rgr1
Srb7
Ssn2
Nop1
Nsr1
Nop56
Snu13
Rps31
Rad9
Rtf1
Rpo21
Npl3
Air2
Fps1
Ask10
Cdc28
Hog1
Pbs2
Ssk2
Clb2
Bud3
Bud4
Msn2
Sch9
Rim15
Yak1
Ypl247c
Vps52
Kss1
Sko1
Slt2
Msg5
Swe1
Igo2
Cdc55
Igo1
Vps53
Vps54
Rtn1
Fmp42
Cot1
Zrc1
10
20
30
40
50
t/s

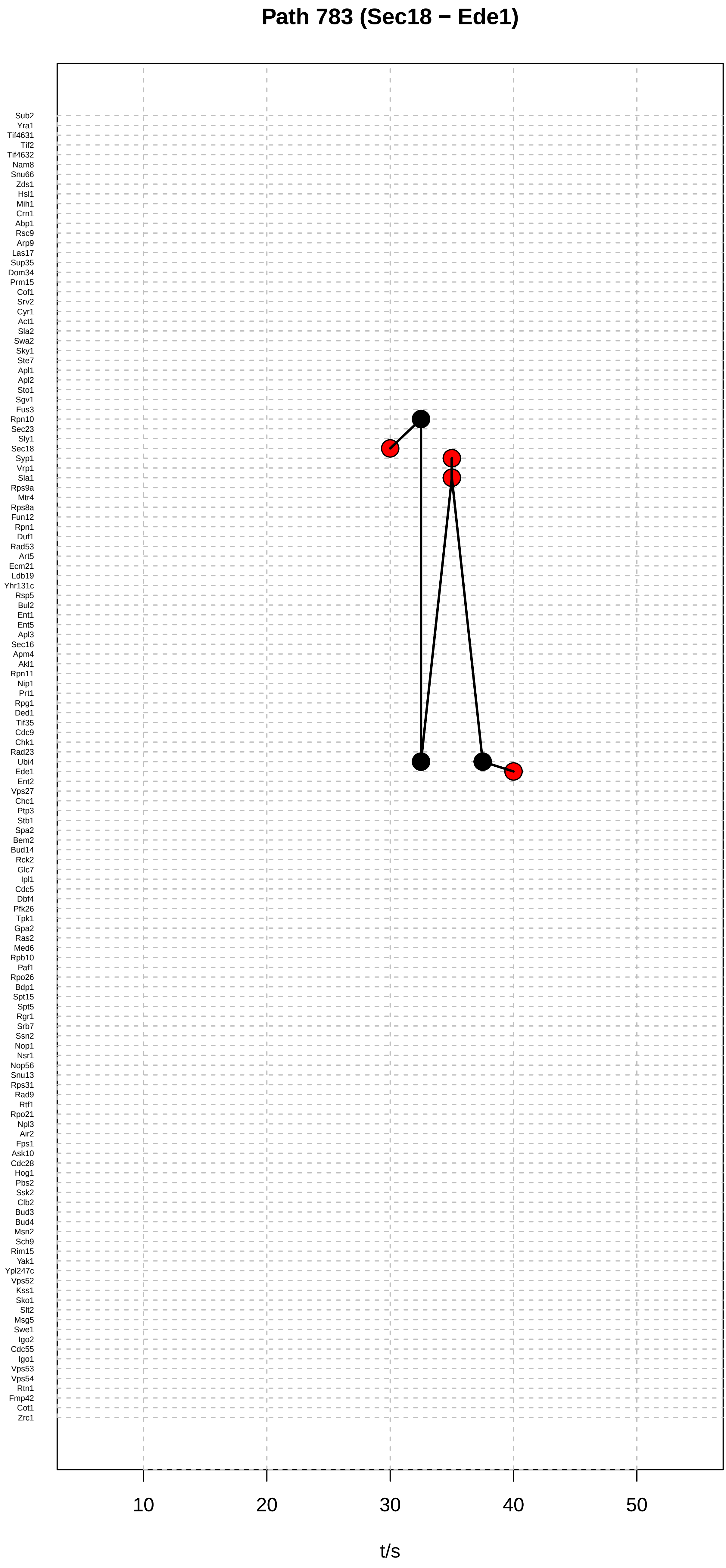
Path 783 (Sec18 − Ede1)
Sub2
Yra1
Tif4631
Tif2
Tif4632
Nam8
Snu66
Zds1
Hsl1
Mih1
Crn1
Abp1
Rsc9
Arp9
Las17
Sup35
Dom34
Prm15
Cof1
Srv2
Cyr1
Act1
Sla2
Swa2
Sky1
Ste7
Apl1
Apl2
Sto1
Sgv1
Fus3
Rpn10
Sec23
Sly1
Sec18
Syp1
Vrp1
Sla1
Rps9a
Mtr4
Rps8a
Fun12
Rpn1
Duf1
Rad53
Art5
Ecm21
Ldb19
Yhr131c
Rsp5
Bul2
Ent1
Ent5
Apl3
Sec16
Apm4
Akl1
Rpn11
Nip1
Prt1
Rpg1
Ded1
Tif35
Cdc9
Chk1
Rad23
Ubi4
Ede1
Ent2
Vps27
Chc1
Ptp3
Stb1
Spa2
Bem2
Bud14
Rck2
Glc7
Ipl1
Cdc5
Dbf4
Pfk26
Tpk1
Gpa2
Ras2
Med6
Rpb10
Paf1
Rpo26
Bdp1
Spt15
Spt5
Rgr1
Srb7
Ssn2
Nop1
Nsr1
Nop56
Snu13
Rps31
Rad9
Rtf1
Rpo21
Npl3
Air2
Fps1
Ask10
Cdc28
Hog1
Pbs2
Ssk2
Clb2
Bud3
Bud4
Msn2
Sch9
Rim15
Yak1
Ypl247c
Vps52
Kss1
Sko1
Slt2
Msg5
Swe1
Igo2
Cdc55
Igo1
Vps53
Vps54
Rtn1
Fmp42
Cot1
Zrc1
10
20
30
40
50
t/s

### Path 784 (Sec18 − Duf1)

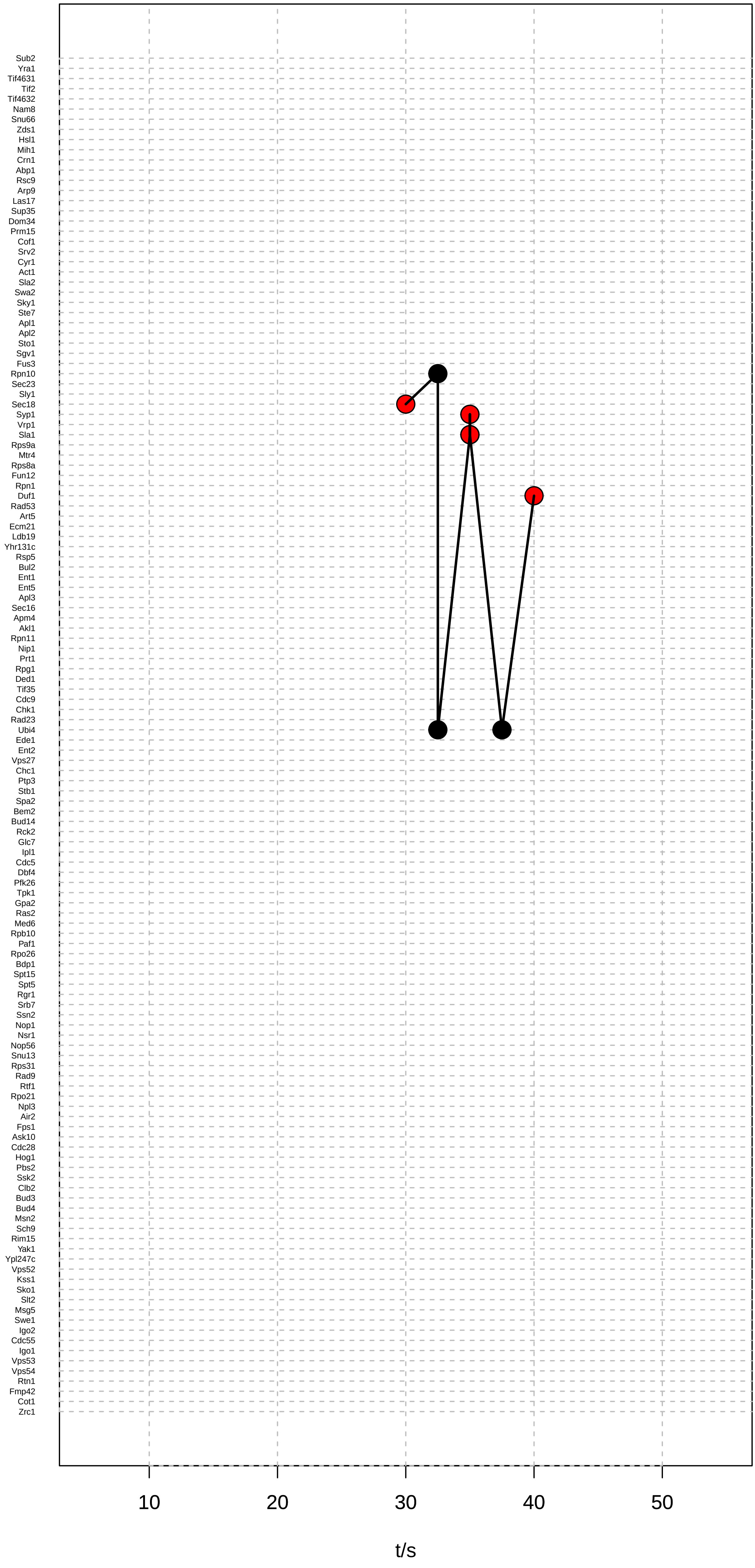

### Path 785 (Sec18 − Fun12)

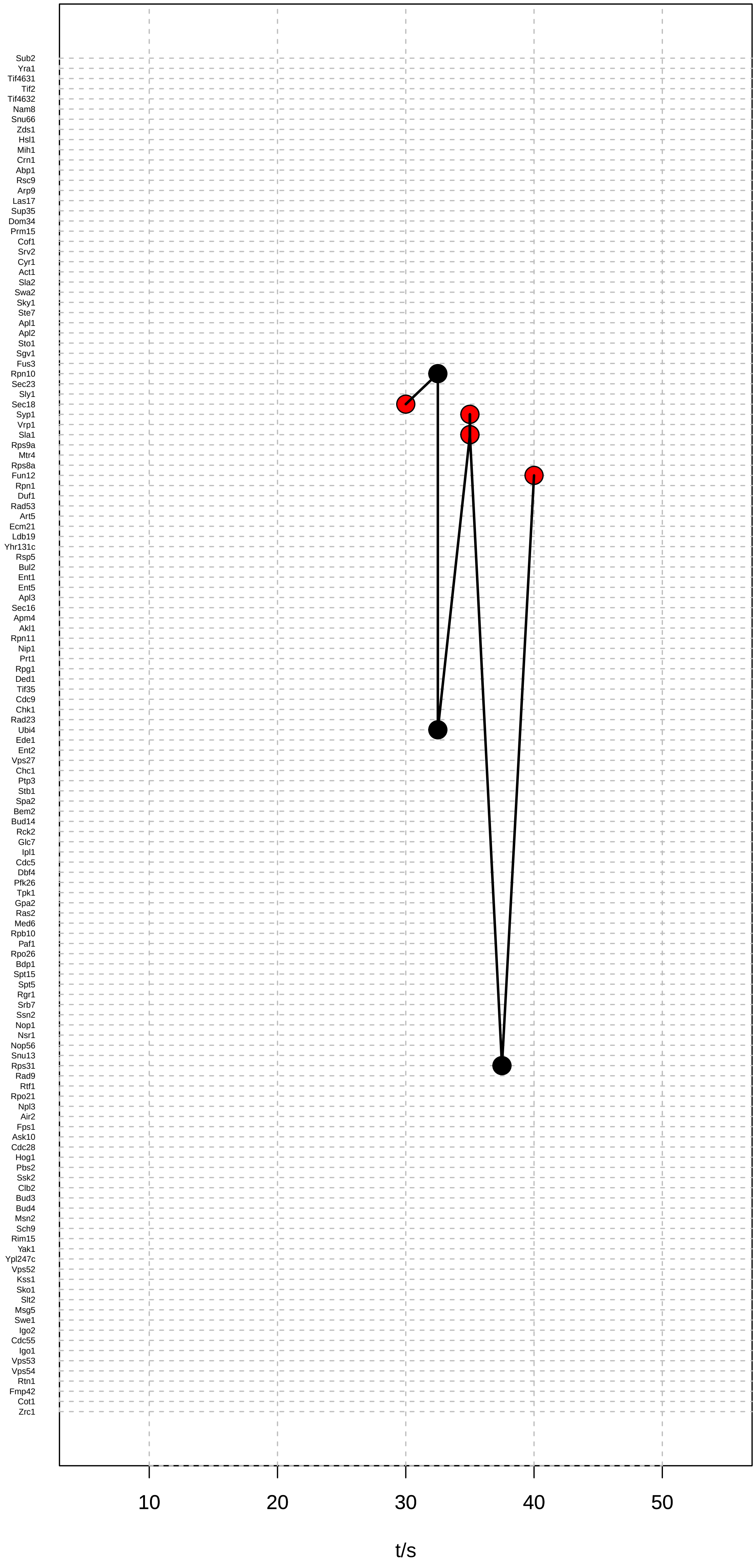

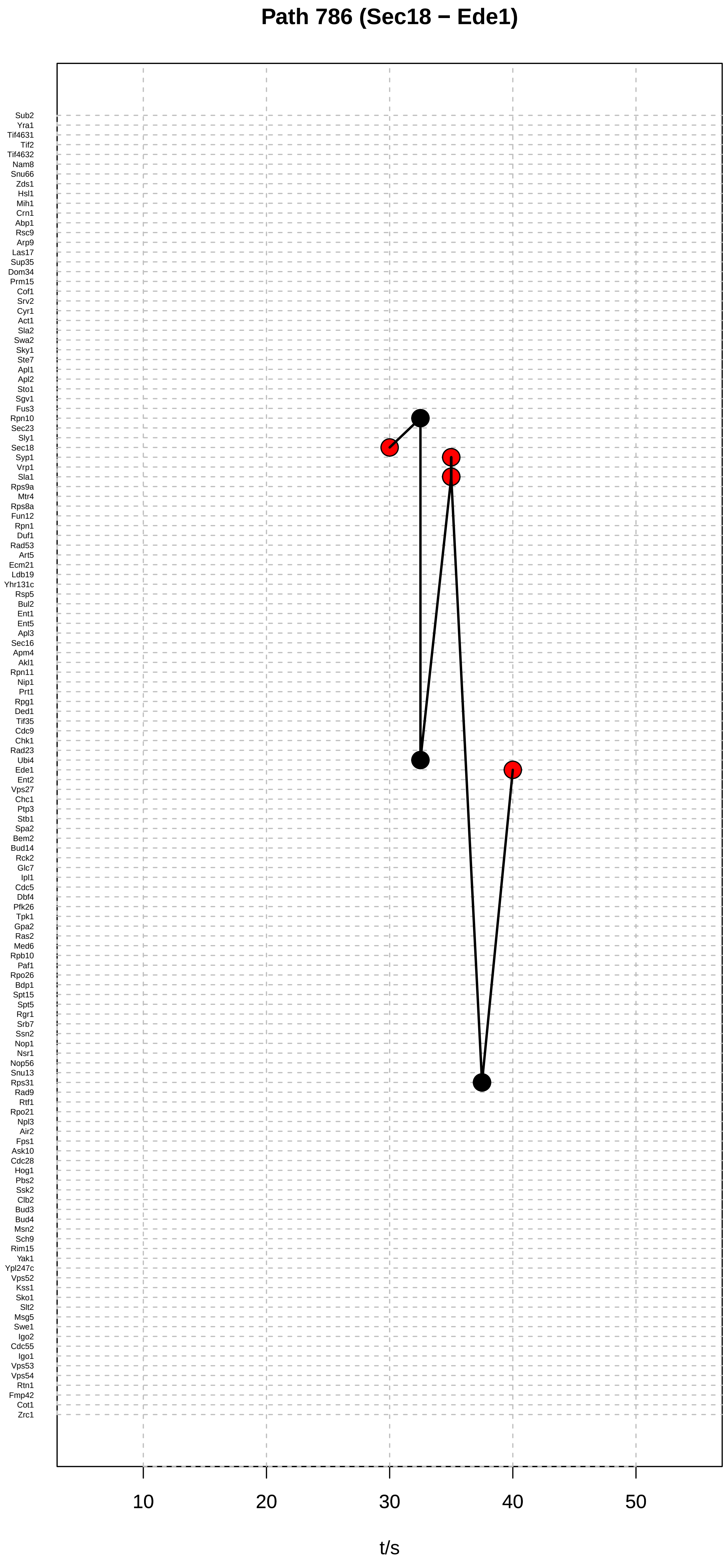

Path 786 (Sec18 − Ede1)
Sub2
Yra1
Tif4631
Tif2
Tif4632
Nam8
Snu66
Zds1
Hsl1
Mih1
Crn1
Abp1
Rsc9
Arp9
Las17
Sup35
Dom34
Prm15
Cof1
Srv2
Cyr1
Act1
Sla2
Swa2
Sky1
Ste7
Apl1
Apl2
Sto1
Sgv1
Fus3
Rpn10
Sec23
Sly1
Sec18
Syp1
Vrp1
Sla1
Rps9a
Mtr4
Rps8a
Fun12
Rpn1
Duf1
Rad53
Art5
Ecm21
Ldb19
Yhr131c
Rsp5
Bul2
Ent1
Ent5
Apl3
Sec16
Apm4
Akl1
Rpn11
Nip1
Prt1
Rpg1
Ded1
Tif35
Cdc9
Chk1
Rad23
Ubi4
Ede1
Ent2
Vps27
Chc1
Ptp3
Stb1
Spa2
Bem2
Bud14
Rck2
Glc7
Ipl1
Cdc5
Dbf4
Pfk26
Tpk1
Gpa2
Ras2
Med6
Rpb10
Paf1
Rpo26
Bdp1
Spt15
Spt5
Rgr1
Srb7
Ssn2
Nop1
Nsr1
Nop56
Snu13
Rps31
Rad9
Rtf1
Rpo21
Npl3
Air2
Fps1
Ask10
Cdc28
Hog1
Pbs2
Ssk2
Clb2
Bud3
Bud4
Msn2
Sch9
Rim15
Yak1
Ypl247c
Vps52
Kss1
Sko1
Slt2
Msg5
Swe1
Igo2
Cdc55
Igo1
Vps53
Vps54
Rtn1
Fmp42
Cot1
Zrc1
10
20
30
40
50
t/s

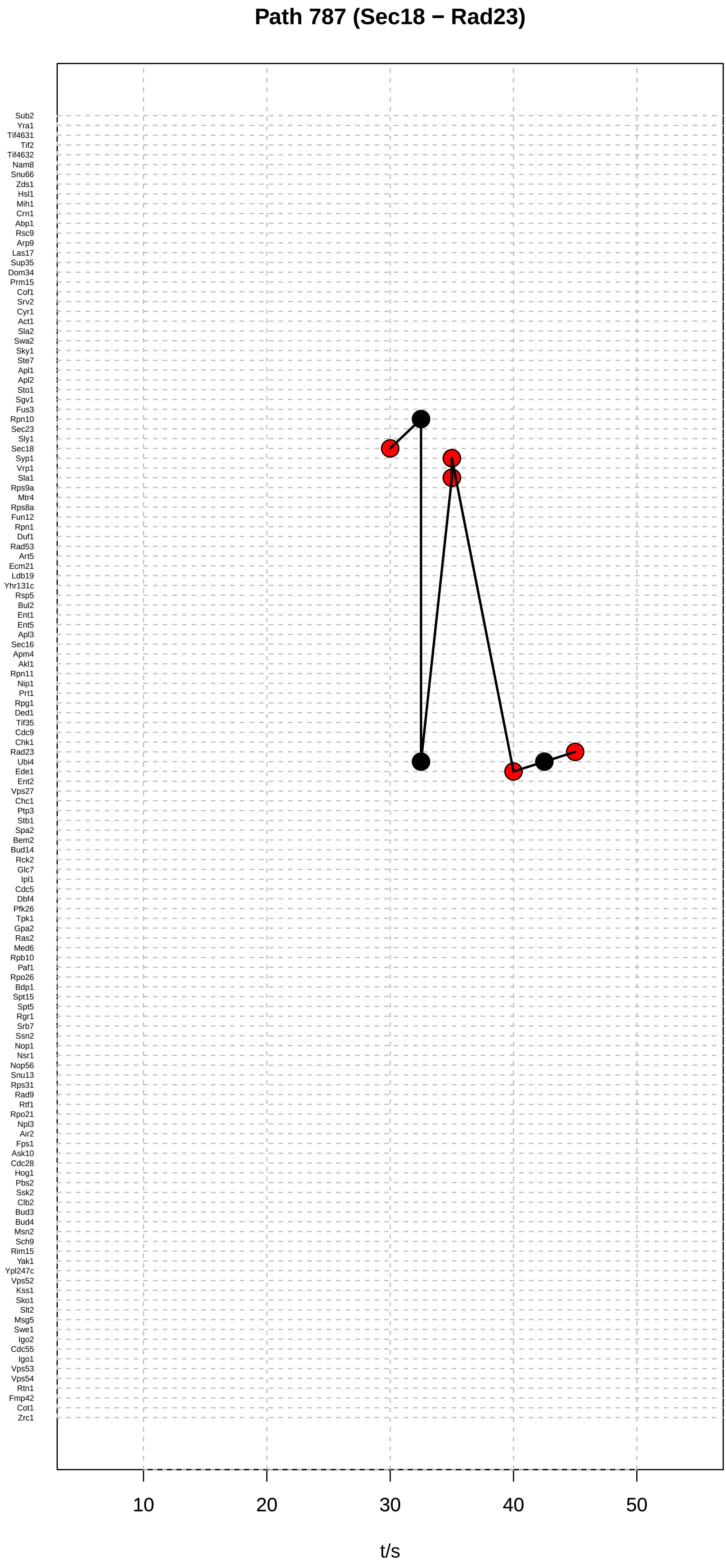

Path 787 (Sec18 − Rad23)
Sub2
Yra1
Tif4631
Tif2
Tif4632
Nam8
Snu66
Zds1
Hsl1
Mih1
Crn1
Abp1
Rsc9
Arp9
Las17
Sup35
Dom34
Prm15
Cof1
Srv2
Cyr1
Act1
Sla2
Swa2
Sky1
Ste7
Apl1
Apl2
Sto1
Sgv1
Fus3
Rpn10
Sec23
Sly1
Sec18
Syp1
Vrp1
Sla1
Rps9a
Mtr4
Rps8a
Fun12
Rpn1
Duf1
Rad53
Art5
Ecm21
Ldb19
Yhr131c
Rsp5
Bul2
Ent1
Ent5
Apl3
Sec16
Apm4
Akl1
Rpn11
Nip1
Prt1
Rpg1
Ded1
Tif35
Cdc9
Chk1
Rad23
Ubi4
Ede1
Ent2
Vps27
Chc1
Ptp3
Stb1
Spa2
Bem2
Bud14
Rck2
Glc7
Ipl1
Cdc5
Dbf4
Pfk26
Tpk1
Gpa2
Ras2
Med6
Rpb10
Paf1
Rpo26
Bdp1
Spt15
Spt5
Rgr1
Srb7
Ssn2
Nop1
Nsr1
Nop56
Snu13
Rps31
Rad9
Rtf1
Rpo21
Npl3
Air2
Fps1
Ask10
Cdc28
Hog1
Pbs2
Ssk2
Clb2
Bud3
Bud4
Msn2
Sch9
Rim15
Yak1
Ypl247c
Vps52
Kss1
Sko1
Slt2
Msg5
Swe1
Igo2
Cdc55
Igo1
Vps53
Vps54
Rtn1
Fmp42
Cot1
Zrc1
10
20
30
40
50
t/s

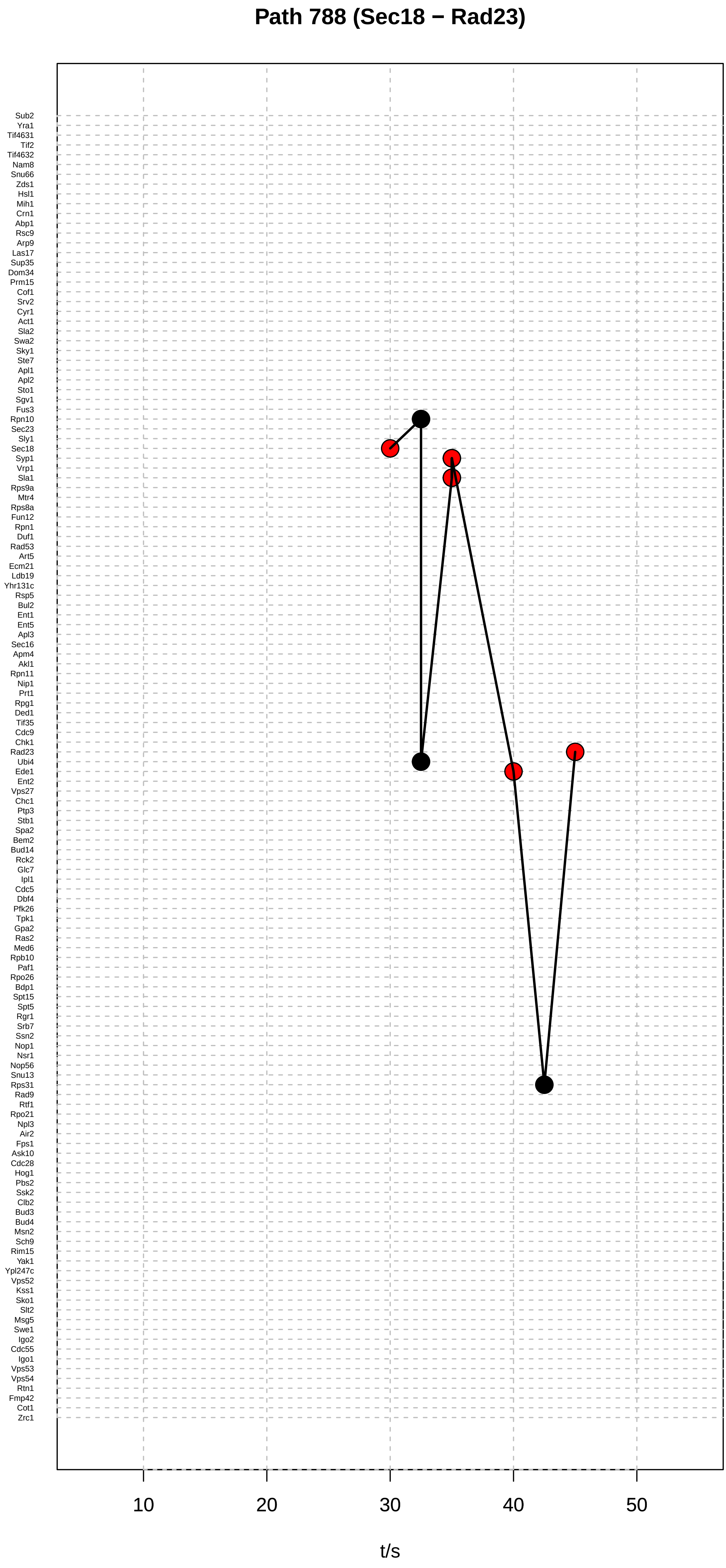
Path 788 (Sec18 − Rad23)
Sub2
Yra1
Tif4631
Tif2
Tif4632
Nam8
Snu66
Zds1
Hsl1
Mih1
Crn1
Abp1
Rsc9
Arp9
Las17
Sup35
Dom34
Prm15
Cof1
Srv2
Cyr1
Act1
Sla2
Swa2
Sky1
Ste7
Apl1
Apl2
Sto1
Sgv1
Fus3
Rpn10
Sec23
Sly1
Sec18
Syp1
Vrp1
Sla1
Rps9a
Mtr4
Rps8a
Fun12
Rpn1
Duf1
Rad53
Art5
Ecm21
Ldb19
Yhr131c
Rsp5
Bul2
Ent1
Ent5
Apl3
Sec16
Apm4
Akl1
Rpn11
Nip1
Prt1
Rpg1
Ded1
Tif35
Cdc9
Chk1
Rad23
Ubi4
Ede1
Ent2
Vps27
Chc1
Ptp3
Stb1
Spa2
Bem2
Bud14
Rck2
Glc7
Ipl1
Cdc5
Dbf4
Pfk26
Tpk1
Gpa2
Ras2
Med6
Rpb10
Paf1
Rpo26
Bdp1
Spt15
Spt5
Rgr1
Srb7
Ssn2
Nop1
Nsr1
Nop56
Snu13
Rps31
Rad9
Rtf1
Rpo21
Npl3
Air2
Fps1
Ask10
Cdc28
Hog1
Pbs2
Ssk2
Clb2
Bud3
Bud4
Msn2
Sch9
Rim15
Yak1
Ypl247c
Vps52
Kss1
Sko1
Slt2
Msg5
Swe1
Igo2
Cdc55
Igo1
Vps53
Vps54
Rtn1
Fmp42
Cot1
Zrc1
10
20
30
40
50
t/s

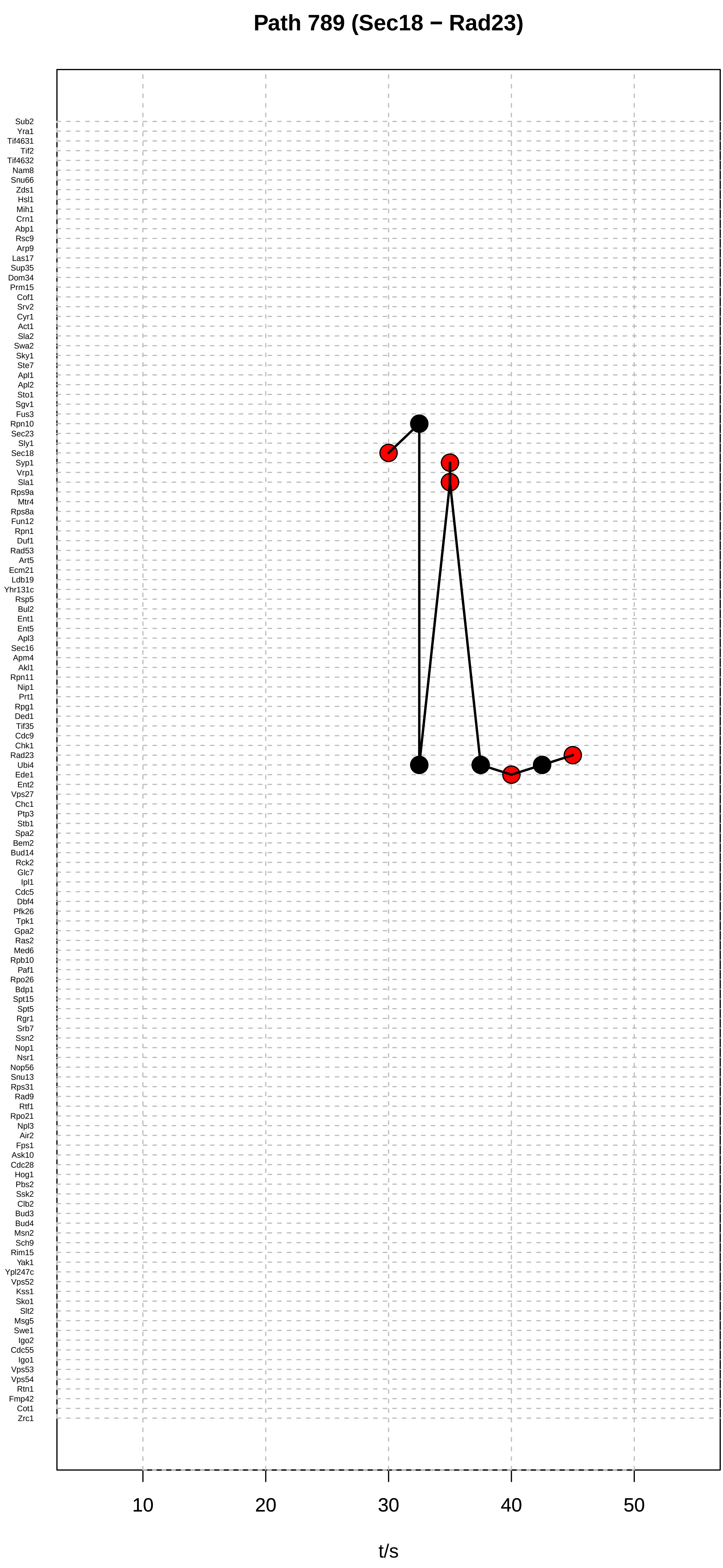
Path 789 (Sec18 − Rad23)
Sub2
Yra1
Tif4631
Tif2
Tif4632
Nam8
Snu66
Zds1
Hsl1
Mih1
Crn1
Abp1
Rsc9
Arp9
Las17
Sup35
Dom34
Prm15
Cof1
Srv2
Cyr1
Act1
Sla2
Swa2
Sky1
Ste7
Apl1
Apl2
Sto1
Sgv1
Fus3
Rpn10
Sec23
Sly1
Sec18
Syp1
Vrp1
Sla1
Rps9a
Mtr4
Rps8a
Fun12
Rpn1
Duf1
Rad53
Art5
Ecm21
Ldb19
Yhr131c
Rsp5
Bul2
Ent1
Ent5
Apl3
Sec16
Apm4
Akl1
Rpn11
Nip1
Prt1
Rpg1
Ded1
Tif35
Cdc9
Chk1
Rad23
Ubi4
Ede1
Ent2
Vps27
Chc1
Ptp3
Stb1
Spa2
Bem2
Bud14
Rck2
Glc7
Ipl1
Cdc5
Dbf4
Pfk26
Tpk1
Gpa2
Ras2
Med6
Rpb10
Paf1
Rpo26
Bdp1
Spt15
Spt5
Rgr1
Srb7
Ssn2
Nop1
Nsr1
Nop56
Snu13
Rps31
Rad9
Rtf1
Rpo21
Npl3
Air2
Fps1
Ask10
Cdc28
Hog1
Pbs2
Ssk2
Clb2
Bud3
Bud4
Msn2
Sch9
Rim15
Yak1
Ypl247c
Vps52
Kss1
Sko1
Slt2
Msg5
Swe1
Igo2
Cdc55
Igo1
Vps53
Vps54
Rtn1
Fmp42
Cot1
Zrc1
10
20
30
40
50
t/s

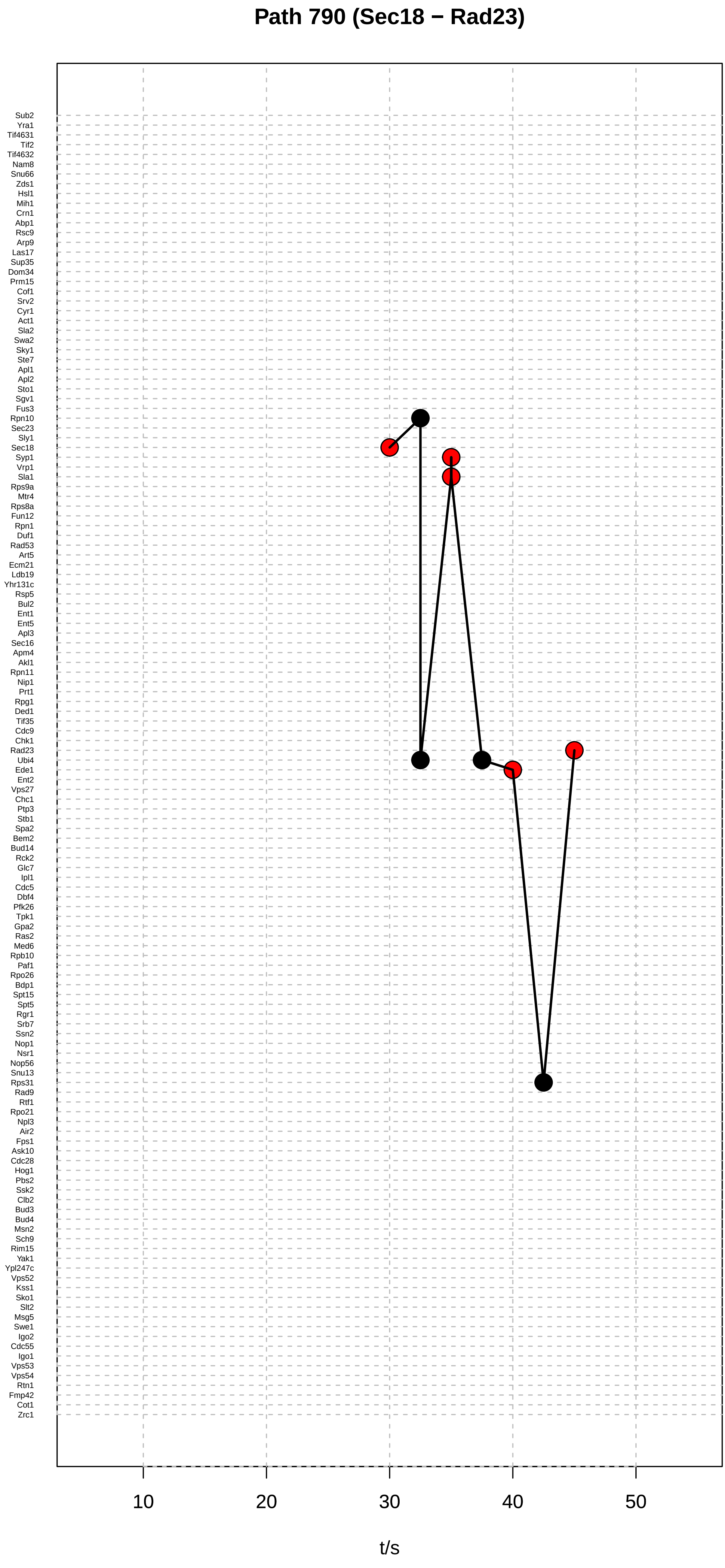
Path 790 (Sec18 − Rad23)
Sub2
Yra1
Tif4631
Tif2
Tif4632
Nam8
Snu66
Zds1
Hsl1
Mih1
Crn1
Abp1
Rsc9
Arp9
Las17
Sup35
Dom34
Prm15
Cof1
Srv2
Cyr1
Act1
Sla2
Swa2
Sky1
Ste7
Apl1
Apl2
Sto1
Sgv1
Fus3
Rpn10
Sec23
Sly1
Sec18
Syp1
Vrp1
Sla1
Rps9a
Mtr4
Rps8a
Fun12
Rpn1
Duf1
Rad53
Art5
Ecm21
Ldb19
Yhr131c
Rsp5
Bul2
Ent1
Ent5
Apl3
Sec16
Apm4
Akl1
Rpn11
Nip1
Prt1
Rpg1
Ded1
Tif35
Cdc9
Chk1
Rad23
Ubi4
Ede1
Ent2
Vps27
Chc1
Ptp3
Stb1
Spa2
Bem2
Bud14
Rck2
Glc7
Ipl1
Cdc5
Dbf4
Pfk26
Tpk1
Gpa2
Ras2
Med6
Rpb10
Paf1
Rpo26
Bdp1
Spt15
Spt5
Rgr1
Srb7
Ssn2
Nop1
Nsr1
Nop56
Snu13
Rps31
Rad9
Rtf1
Rpo21
Npl3
Air2
Fps1
Ask10
Cdc28
Hog1
Pbs2
Ssk2
Clb2
Bud3
Bud4
Msn2
Sch9
Rim15
Yak1
Ypl247c
Vps52
Kss1
Sko1
Slt2
Msg5
Swe1
Igo2
Cdc55
Igo1
Vps53
Vps54
Rtn1
Fmp42
Cot1
Zrc1
10
20
30
40
50
t/s

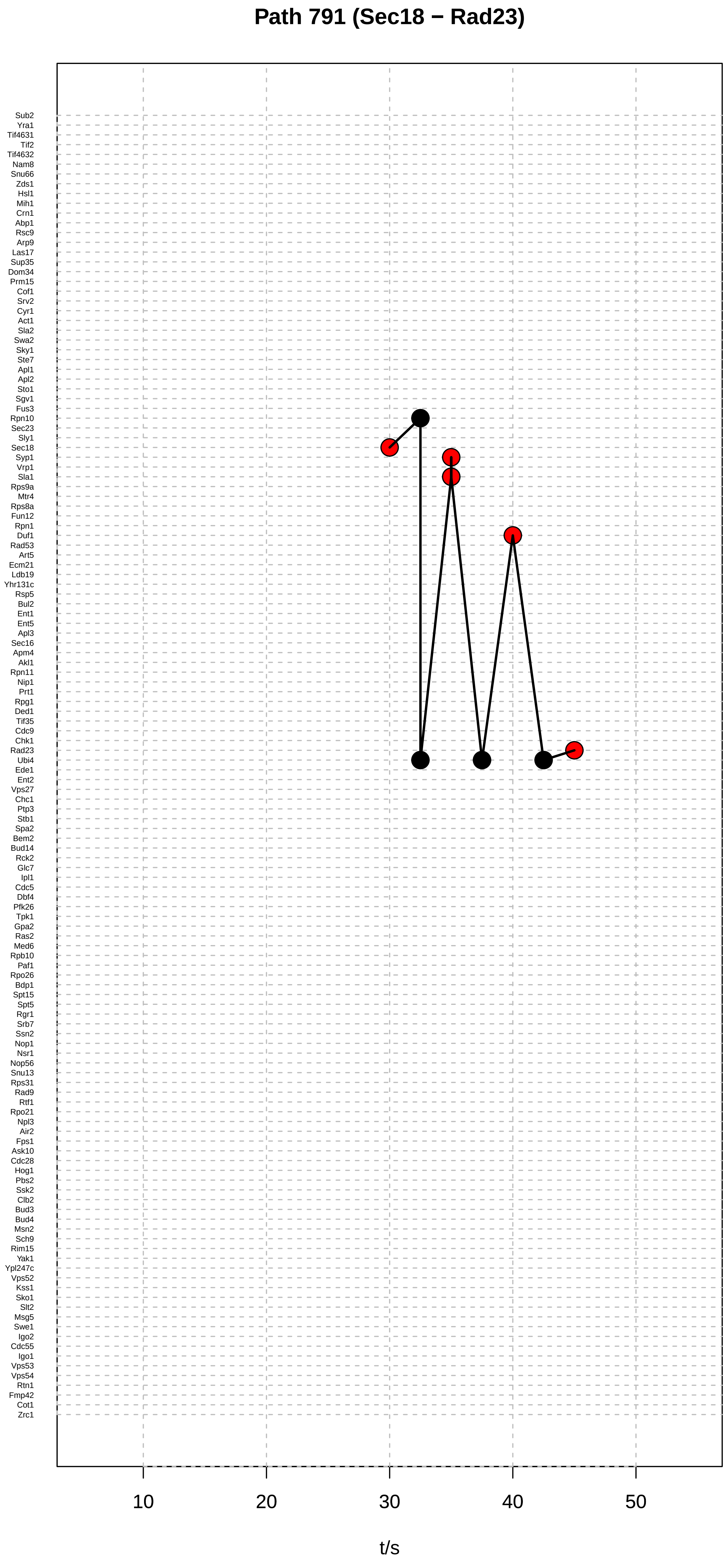

Path 791 (Sec18 − Rad23)
Sub2
Yra1
Tif4631
Tif2
Tif4632
Nam8
Snu66
Zds1
Hsl1
Mih1
Crn1
Abp1
Rsc9
Arp9
Las17
Sup35
Dom34
Prm15
Cof1
Srv2
Cyr1
Act1
Sla2
Swa2
Sky1
Ste7
Apl1
Apl2
Sto1
Sgv1
Fus3
Rpn10
Sec23
Sly1
Sec18
Syp1
Vrp1
Sla1
Rps9a
Mtr4
Rps8a
Fun12
Rpn1
Duf1
Rad53
Art5
Ecm21
Ldb19
Yhr131c
Rsp5
Bul2
Ent1
Ent5
Apl3
Sec16
Apm4
Akl1
Rpn11
Nip1
Prt1
Rpg1
Ded1
Tif35
Cdc9
Chk1
Rad23
Ubi4
Ede1
Ent2
Vps27
Chc1
Ptp3
Stb1
Spa2
Bem2
Bud14
Rck2
Glc7
Ipl1
Cdc5
Dbf4
Pfk26
Tpk1
Gpa2
Ras2
Med6
Rpb10
Paf1
Rpo26
Bdp1
Spt15
Spt5
Rgr1
Srb7
Ssn2
Nop1
Nsr1
Nop56
Snu13
Rps31
Rad9
Rtf1
Rpo21
Npl3
Air2
Fps1
Ask10
Cdc28
Hog1
Pbs2
Ssk2
Clb2
Bud3
Bud4
Msn2
Sch9
Rim15
Yak1
Ypl247c
Vps52
Kss1
Sko1
Slt2
Msg5
Swe1
Igo2
Cdc55
Igo1
Vps53
Vps54
Rtn1
Fmp42
Cot1
Zrc1
10
20
30
40
50
t/s

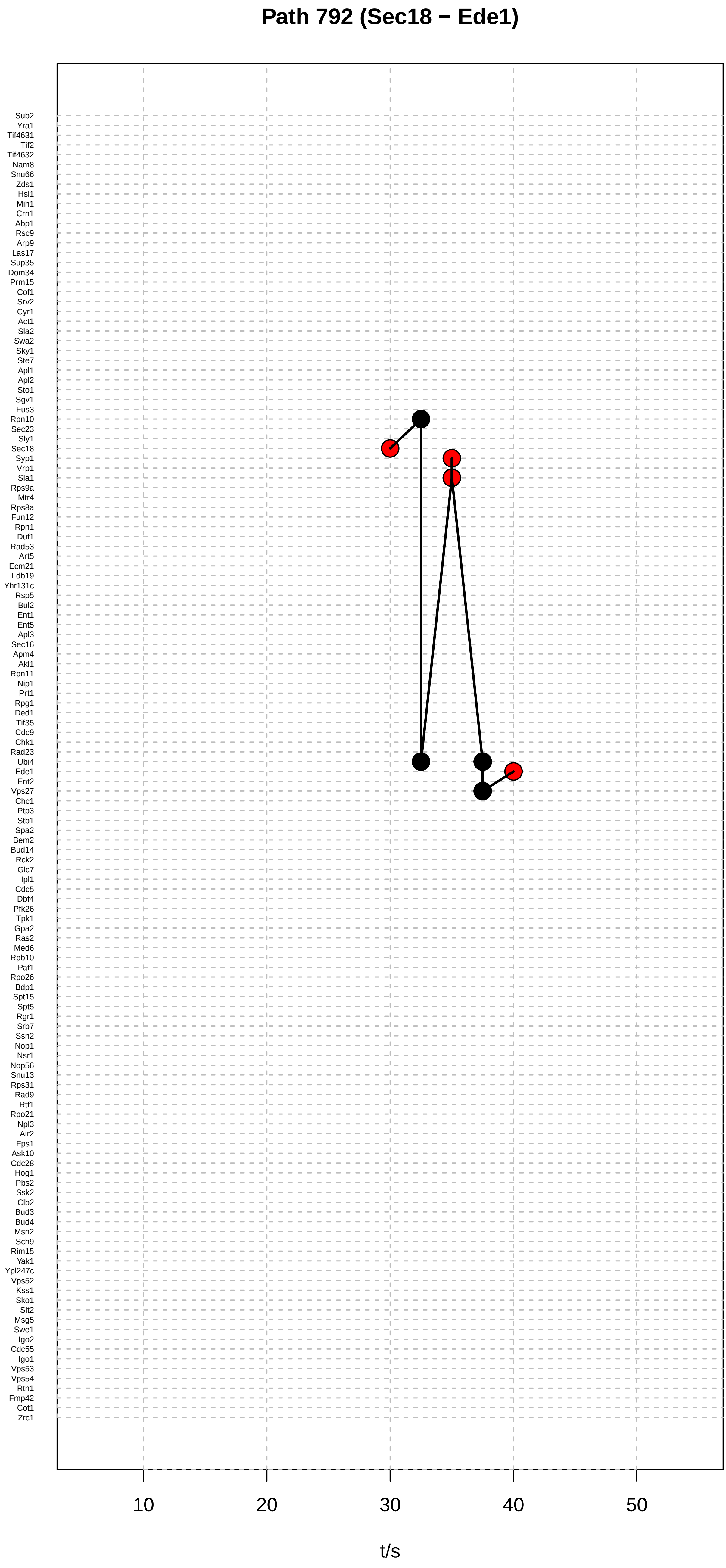
Path 792 (Sec18 − Ede1)
Sub2
Yra1
Tif4631
Tif2
Tif4632
Nam8
Snu66
Zds1
Hsl1
Mih1
Crn1
Abp1
Rsc9
Arp9
Las17
Sup35
Dom34
Prm15
Cof1
Srv2
Cyr1
Act1
Sla2
Swa2
Sky1
Ste7
Apl1
Apl2
Sto1
Sgv1
Fus3
Rpn10
Sec23
Sly1
Sec18
Syp1
Vrp1
Sla1
Rps9a
Mtr4
Rps8a
Fun12
Rpn1
Duf1
Rad53
Art5
Ecm21
Ldb19
Yhr131c
Rsp5
Bul2
Ent1
Ent5
Apl3
Sec16
Apm4
Akl1
Rpn11
Nip1
Prt1
Rpg1
Ded1
Tif35
Cdc9
Chk1
Rad23
Ubi4
Ede1
Ent2
Vps27
Chc1
Ptp3
Stb1
Spa2
Bem2
Bud14
Rck2
Glc7
Ipl1
Cdc5
Dbf4
Pfk26
Tpk1
Gpa2
Ras2
Med6
Rpb10
Paf1
Rpo26
Bdp1
Spt15
Spt5
Rgr1
Srb7
Ssn2
Nop1
Nsr1
Nop56
Snu13
Rps31
Rad9
Rtf1
Rpo21
Npl3
Air2
Fps1
Ask10
Cdc28
Hog1
Pbs2
Ssk2
Clb2
Bud3
Bud4
Msn2
Sch9
Rim15
Yak1
Ypl247c
Vps52
Kss1
Sko1
Slt2
Msg5
Swe1
Igo2
Cdc55
Igo1
Vps53
Vps54
Rtn1
Fmp42
Cot1
Zrc1
10
20
30
40
50
t/s

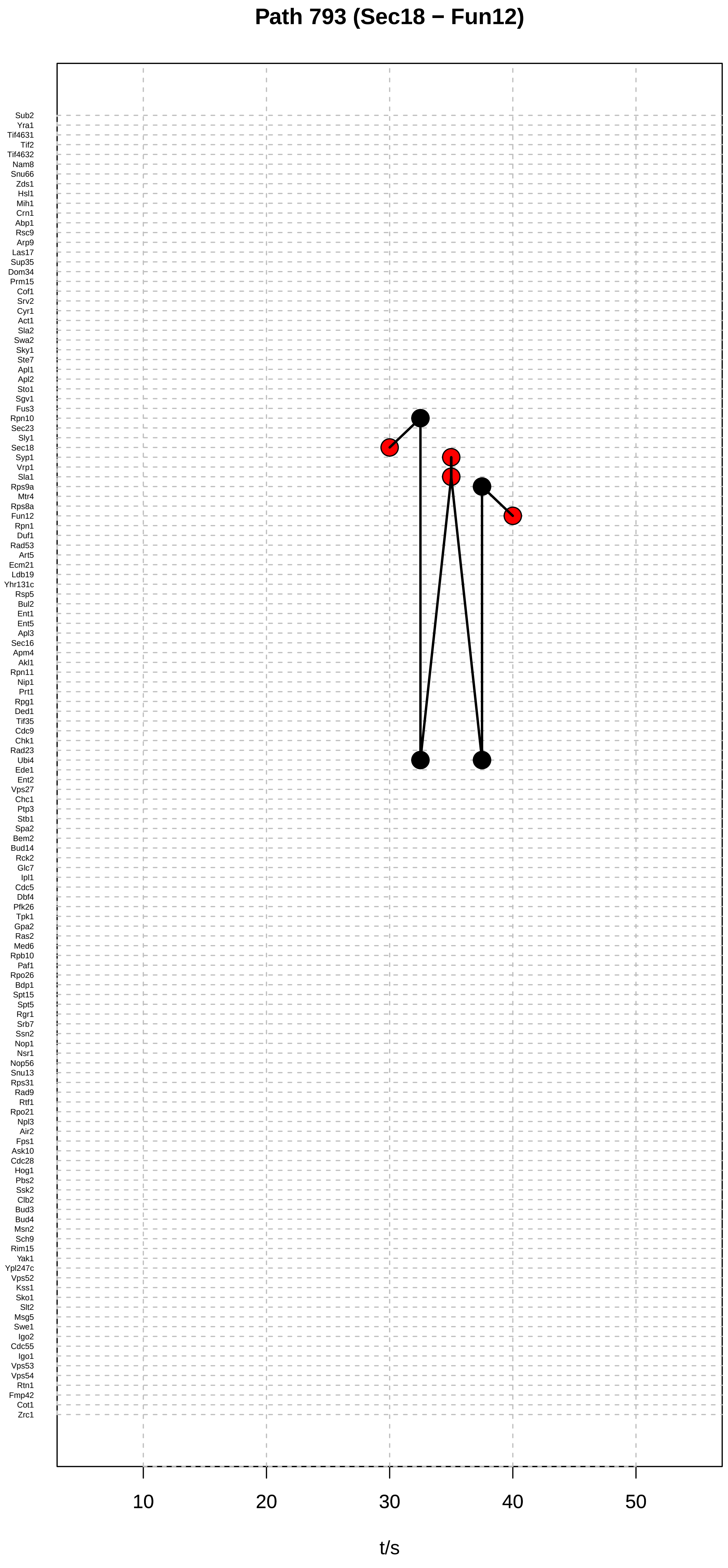
Path 793 (Sec18 − Fun12)
Sub2
Yra1
Tif4631
Tif2
Tif4632
Nam8
Snu66
Zds1
Hsl1
Mih1
Crn1
Abp1
Rsc9
Arp9
Las17
Sup35
Dom34
Prm15
Cof1
Srv2
Cyr1
Act1
Sla2
Swa2
Sky1
Ste7
Apl1
Apl2
Sto1
Sgv1
Fus3
Rpn10
Sec23
Sly1
Sec18
Syp1
Vrp1
Sla1
Rps9a
Mtr4
Rps8a
Fun12
Rpn1
Duf1
Rad53
Art5
Ecm21
Ldb19
Yhr131c
Rsp5
Bul2
Ent1
Ent5
Apl3
Sec16
Apm4
Akl1
Rpn11
Nip1
Prt1
Rpg1
Ded1
Tif35
Cdc9
Chk1
Rad23
Ubi4
Ede1
Ent2
Vps27
Chc1
Ptp3
Stb1
Spa2
Bem2
Bud14
Rck2
Glc7
Ipl1
Cdc5
Dbf4
Pfk26
Tpk1
Gpa2
Ras2
Med6
Rpb10
Paf1
Rpo26
Bdp1
Spt15
Spt5
Rgr1
Srb7
Ssn2
Nop1
Nsr1
Nop56
Snu13
Rps31
Rad9
Rtf1
Rpo21
Npl3
Air2
Fps1
Ask10
Cdc28
Hog1
Pbs2
Ssk2
Clb2
Bud3
Bud4
Msn2
Sch9
Rim15
Yak1
Ypl247c
Vps52
Kss1
Sko1
Slt2
Msg5
Swe1
Igo2
Cdc55
Igo1
Vps53
Vps54
Rtn1
Fmp42
Cot1
Zrc1
10
20
30
40
50
t/s

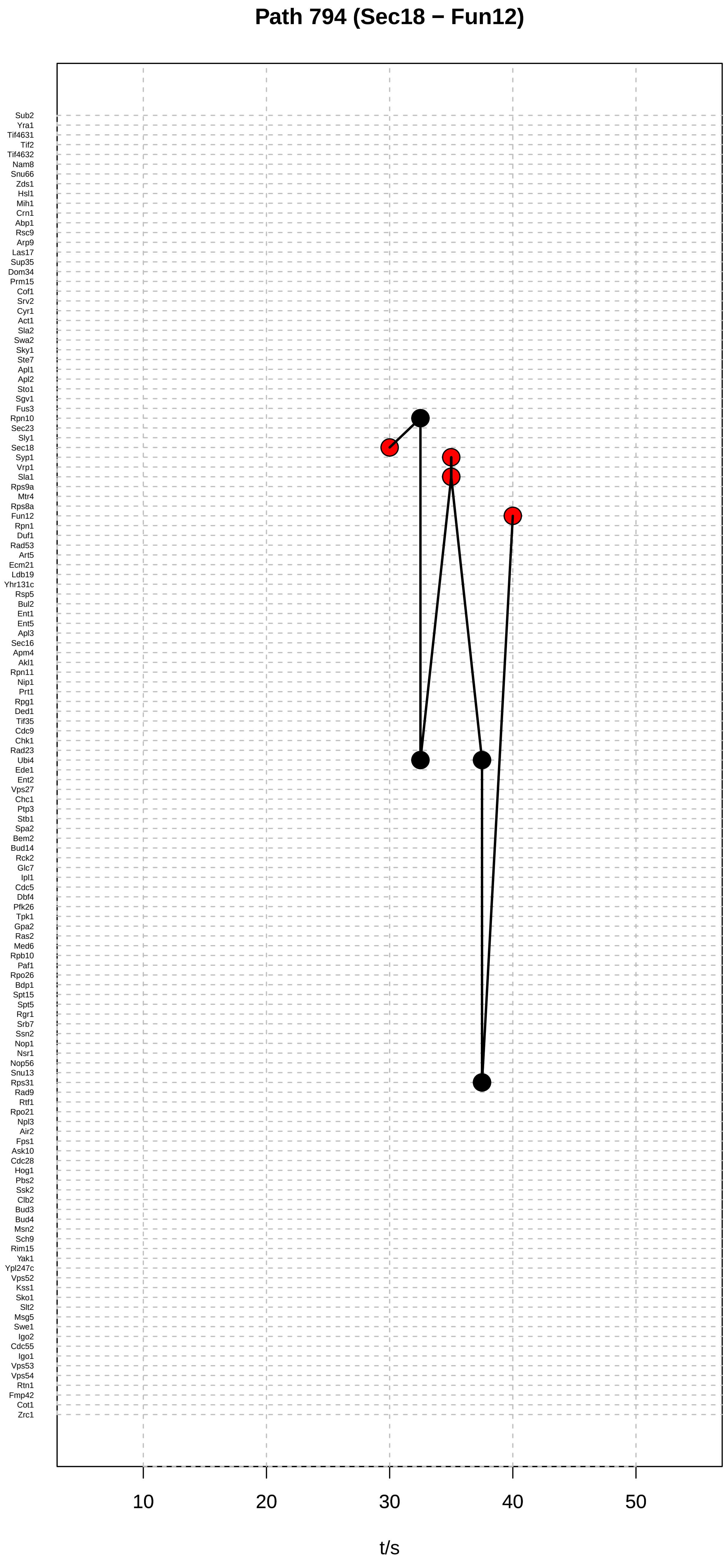
Path 794 (Sec18 − Fun12)
Sub2
Yra1
Tif4631
Tif2
Tif4632
Nam8
Snu66
Zds1
Hsl1
Mih1
Crn1
Abp1
Rsc9
Arp9
Las17
Sup35
Dom34
Prm15
Cof1
Srv2
Cyr1
Act1
Sla2
Swa2
Sky1
Ste7
Apl1
Apl2
Sto1
Sgv1
Fus3
Rpn10
Sec23
Sly1
Sec18
Syp1
Vrp1
Sla1
Rps9a
Mtr4
Rps8a
Fun12
Rpn1
Duf1
Rad53
Art5
Ecm21
Ldb19
Yhr131c
Rsp5
Bul2
Ent1
Ent5
Apl3
Sec16
Apm4
Akl1
Rpn11
Nip1
Prt1
Rpg1
Ded1
Tif35
Cdc9
Chk1
Rad23
Ubi4
Ede1
Ent2
Vps27
Chc1
Ptp3
Stb1
Spa2
Bem2
Bud14
Rck2
Glc7
Ipl1
Cdc5
Dbf4
Pfk26
Tpk1
Gpa2
Ras2
Med6
Rpb10
Paf1
Rpo26
Bdp1
Spt15
Spt5
Rgr1
Srb7
Ssn2
Nop1
Nsr1
Nop56
Snu13
Rps31
Rad9
Rtf1
Rpo21
Npl3
Air2
Fps1
Ask10
Cdc28
Hog1
Pbs2
Ssk2
Clb2
Bud3
Bud4
Msn2
Sch9
Rim15
Yak1
Ypl247c
Vps52
Kss1
Sko1
Slt2
Msg5
Swe1
Igo2
Cdc55
Igo1
Vps53
Vps54
Rtn1
Fmp42
Cot1
Zrc1
10
20
30
40
50
t/s

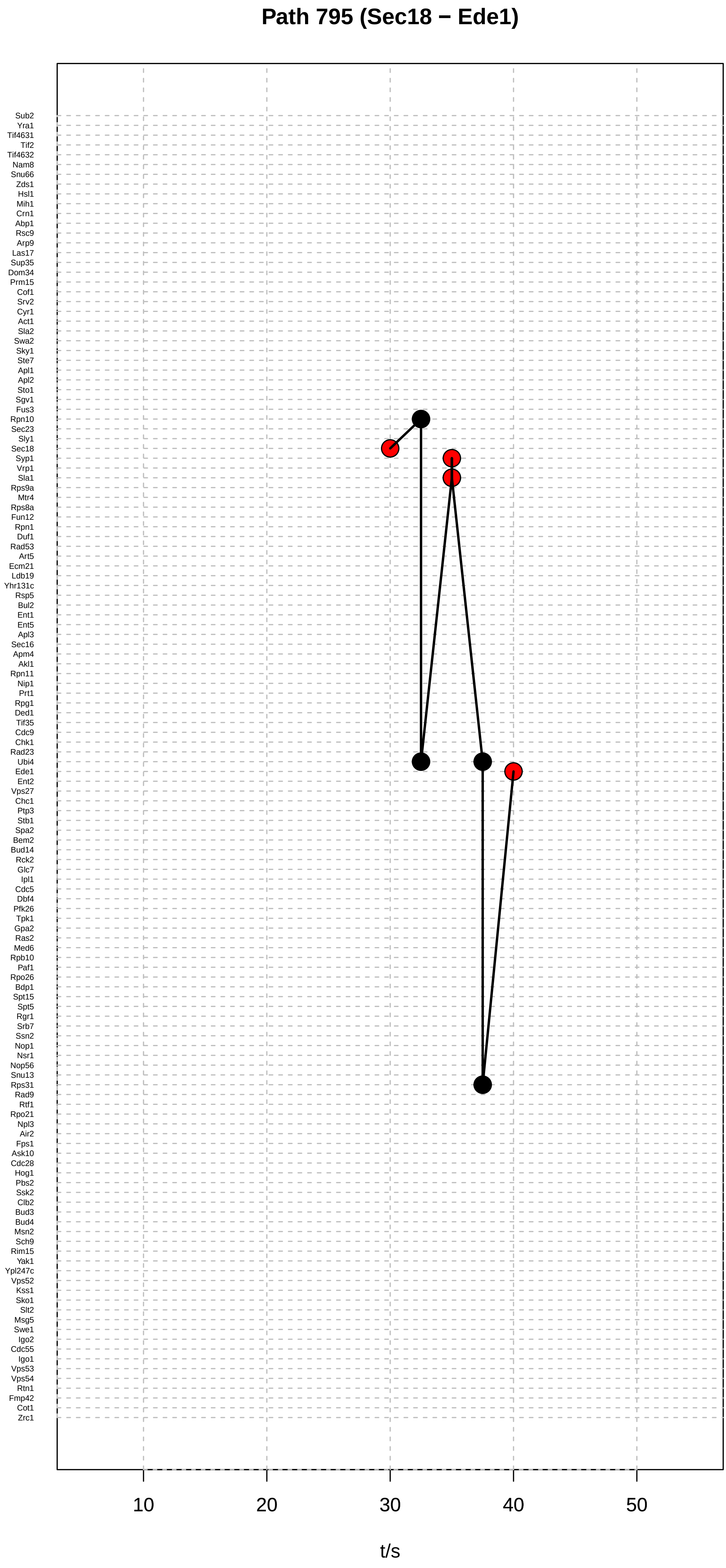

Path 795 (Sec18 − Ede1)
Sub2
Yra1
Tif4631
Tif2
Tif4632
Nam8
Snu66
Zds1
Hsl1
Mih1
Crn1
Abp1
Rsc9
Arp9
Las17
Sup35
Dom34
Prm15
Cof1
Srv2
Cyr1
Act1
Sla2
Swa2
Sky1
Ste7
Apl1
Apl2
Sto1
Sgv1
Fus3
Rpn10
Sec23
Sly1
Sec18
Syp1
Vrp1
Sla1
Rps9a
Mtr4
Rps8a
Fun12
Rpn1
Duf1
Rad53
Art5
Ecm21
Ldb19
Yhr131c
Rsp5
Bul2
Ent1
Ent5
Apl3
Sec16
Apm4
Akl1
Rpn11
Nip1
Prt1
Rpg1
Ded1
Tif35
Cdc9
Chk1
Rad23
Ubi4
Ede1
Ent2
Vps27
Chc1
Ptp3
Stb1
Spa2
Bem2
Bud14
Rck2
Glc7
Ipl1
Cdc5
Dbf4
Pfk26
Tpk1
Gpa2
Ras2
Med6
Rpb10
Paf1
Rpo26
Bdp1
Spt15
Spt5
Rgr1
Srb7
Ssn2
Nop1
Nsr1
Nop56
Snu13
Rps31
Rad9
Rtf1
Rpo21
Npl3
Air2
Fps1
Ask10
Cdc28
Hog1
Pbs2
Ssk2
Clb2
Bud3
Bud4
Msn2
Sch9
Rim15
Yak1
Ypl247c
Vps52
Kss1
Sko1
Slt2
Msg5
Swe1
Igo2
Cdc55
Igo1
Vps53
Vps54
Rtn1
Fmp42
Cot1
Zrc1
10
20
30
40
50
t/s

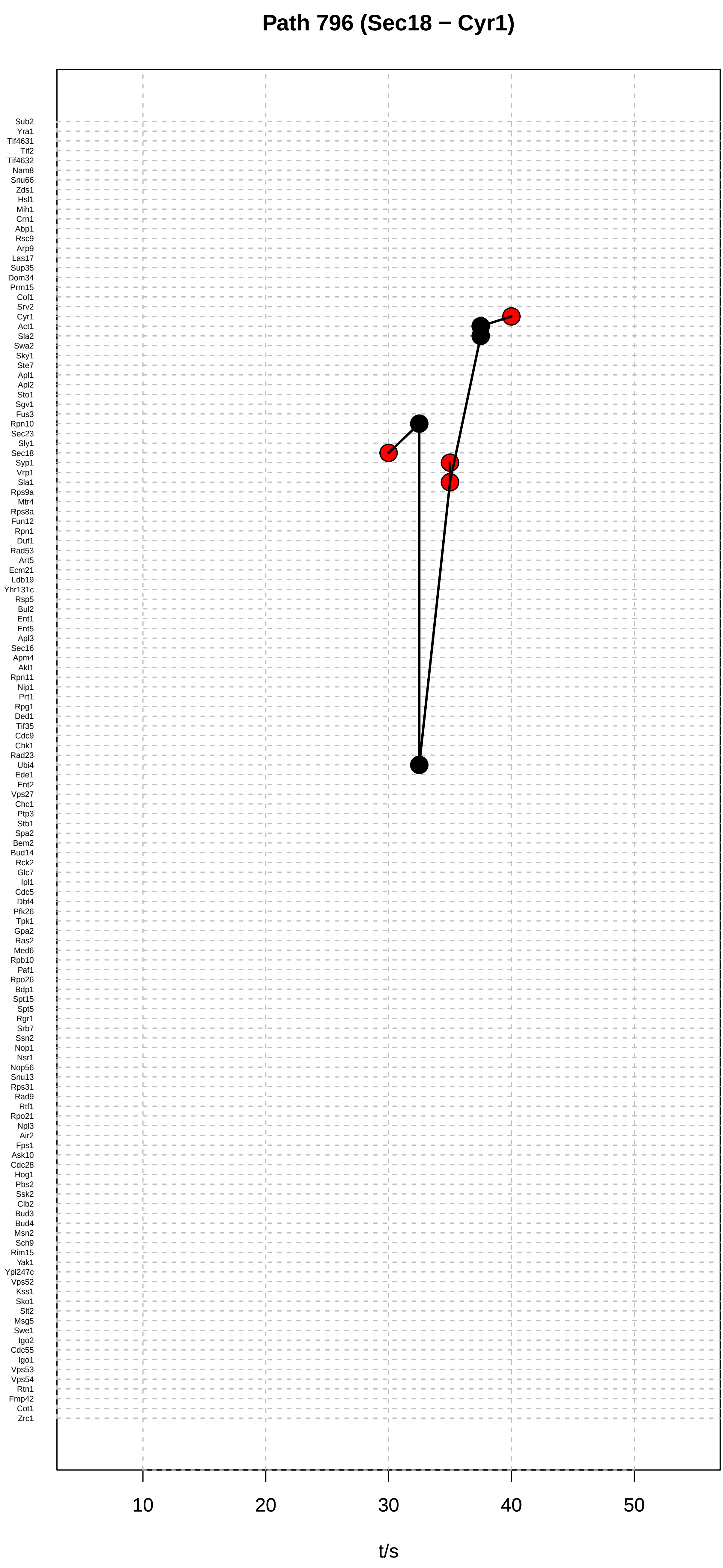

Path 796 (Sec18 − Cyr1)
Sub2
Yra1
Tif4631
Tif2
Tif4632
Nam8
Snu66
Zds1
Hsl1
Mih1
Crn1
Abp1
Rsc9
Arp9
Las17
Sup35
Dom34
Prm15
Cof1
Srv2
Cyr1
Act1
Sla2
Swa2
Sky1
Ste7
Apl1
Apl2
Sto1
Sgv1
Fus3
Rpn10
Sec23
Sly1
Sec18
Syp1
Vrp1
Sla1
Rps9a
Mtr4
Rps8a
Fun12
Rpn1
Duf1
Rad53
Art5
Ecm21
Ldb19
Yhr131c
Rsp5
Bul2
Ent1
Ent5
Apl3
Sec16
Apm4
Akl1
Rpn11
Nip1
Prt1
Rpg1
Ded1
Tif35
Cdc9
Chk1
Rad23
Ubi4
Ede1
Ent2
Vps27
Chc1
Ptp3
Stb1
Spa2
Bem2
Bud14
Rck2
Glc7
Ipl1
Cdc5
Dbf4
Pfk26
Tpk1
Gpa2
Ras2
Med6
Rpb10
Paf1
Rpo26
Bdp1
Spt15
Spt5
Rgr1
Srb7
Ssn2
Nop1
Nsr1
Nop56
Snu13
Rps31
Rad9
Rtf1
Rpo21
Npl3
Air2
Fps1
Ask10
Cdc28
Hog1
Pbs2
Ssk2
Clb2
Bud3
Bud4
Msn2
Sch9
Rim15
Yak1
Ypl247c
Vps52
Kss1
Sko1
Slt2
Msg5
Swe1
Igo2
Cdc55
Igo1
Vps53
Vps54
Rtn1
Fmp42
Cot1
Zrc1
10
20
30
40
50
t/s

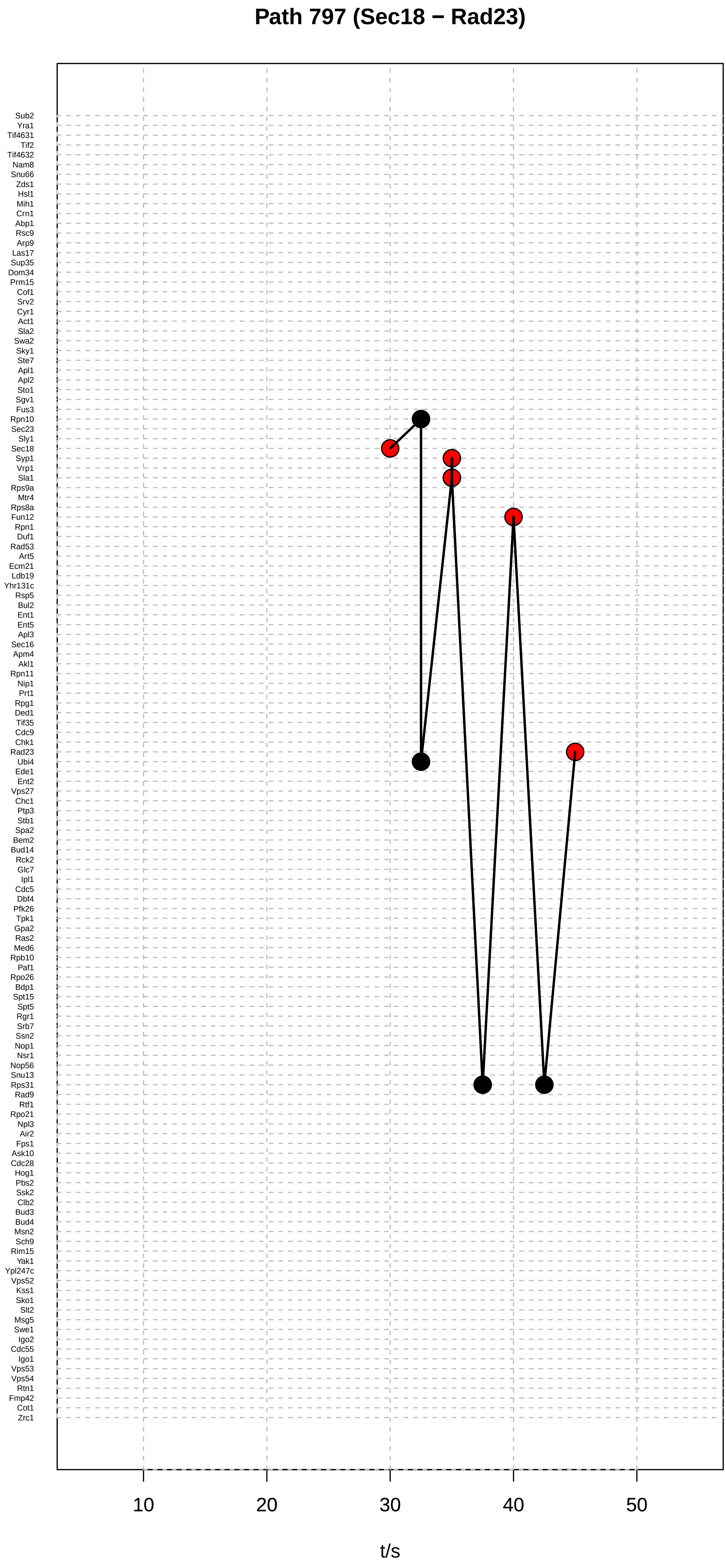

Path 797 (Sec18 − Rad23)
Sub2
Yra1
Tif4631
Tif2
Tif4632
Nam8
Snu66
Zds1
Hsl1
Mih1
Crn1
Abp1
Rsc9
Arp9
Las17
Sup35
Dom34
Prm15
Cof1
Srv2
Cyr1
Act1
Sla2
Swa2
Sky1
Ste7
Apl1
Apl2
Sto1
Sgv1
Fus3
Rpn10
Sec23
Sly1
Sec18
Syp1
Vrp1
Sla1
Rps9a
Mtr4
Rps8a
Fun12
Rpn1
Duf1
Rad53
Art5
Ecm21
Ldb19
Yhr131c
Rsp5
Bul2
Ent1
Ent5
Apl3
Sec16
Apm4
Akl1
Rpn11
Nip1
Prt1
Rpg1
Ded1
Tif35
Cdc9
Chk1
Rad23
Ubi4
Ede1
Ent2
Vps27
Chc1
Ptp3
Stb1
Spa2
Bem2
Bud14
Rck2
Glc7
Ipl1
Cdc5
Dbf4
Pfk26
Tpk1
Gpa2
Ras2
Med6
Rpb10
Paf1
Rpo26
Bdp1
Spt15
Spt5
Rgr1
Srb7
Ssn2
Nop1
Nsr1
Nop56
Snu13
Rps31
Rad9
Rtf1
Rpo21
Npl3
Air2
Fps1
Ask10
Cdc28
Hog1
Pbs2
Ssk2
Clb2
Bud3
Bud4
Msn2
Sch9
Rim15
Yak1
Ypl247c
Vps52
Kss1
Sko1
Slt2
Msg5
Swe1
Igo2
Cdc55
Igo1
Vps53
Vps54
Rtn1
Fmp42
Cot1
Zrc1
10
20
30
40
50
t/s

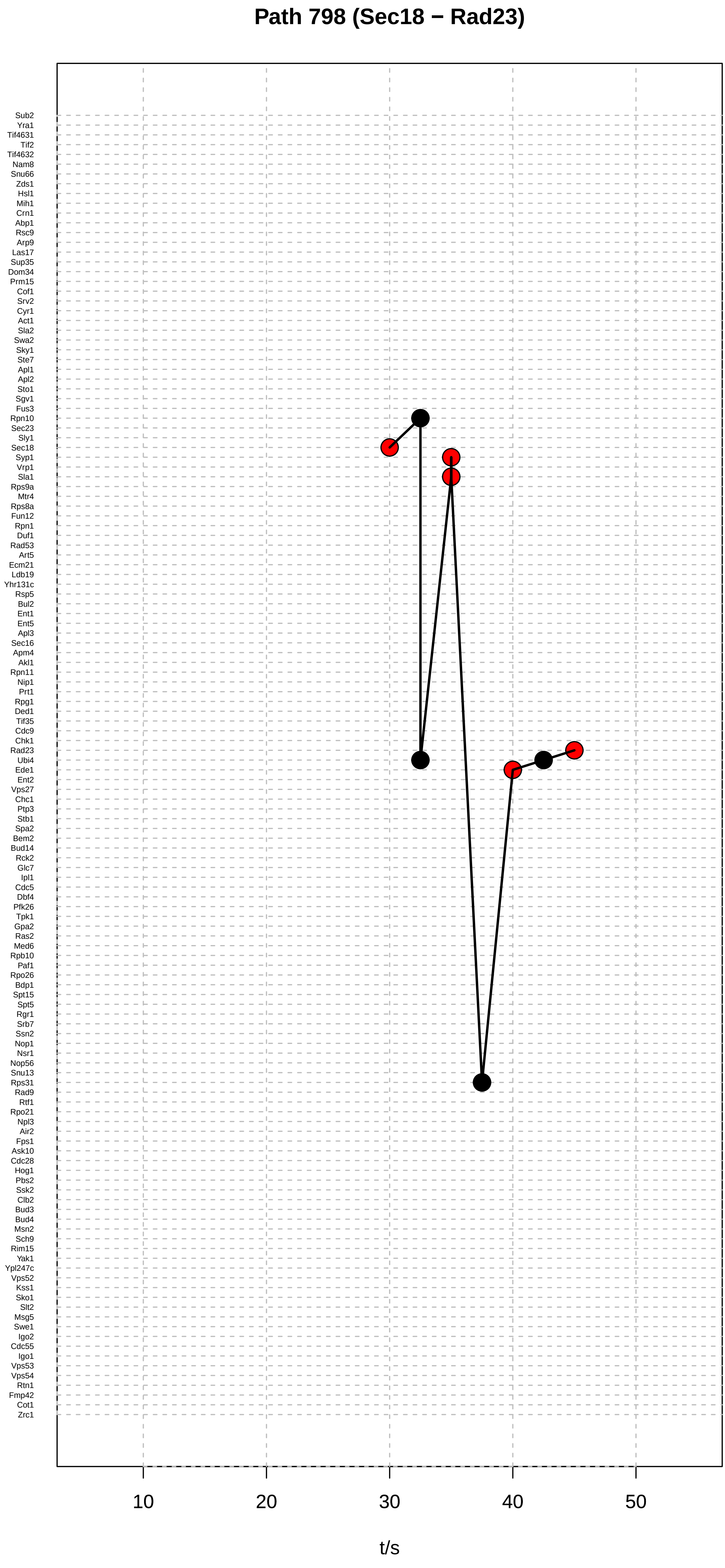

Path 798 (Sec18 − Rad23)
Sub2
Yra1
Tif4631
Tif2
Tif4632
Nam8
Snu66
Zds1
Hsl1
Mih1
Crn1
Abp1
Rsc9
Arp9
Las17
Sup35
Dom34
Prm15
Cof1
Srv2
Cyr1
Act1
Sla2
Swa2
Sky1
Ste7
Apl1
Apl2
Sto1
Sgv1
Fus3
Rpn10
Sec23
Sly1
Sec18
Syp1
Vrp1
Sla1
Rps9a
Mtr4
Rps8a
Fun12
Rpn1
Duf1
Rad53
Art5
Ecm21
Ldb19
Yhr131c
Rsp5
Bul2
Ent1
Ent5
Apl3
Sec16
Apm4
Akl1
Rpn11
Nip1
Prt1
Rpg1
Ded1
Tif35
Cdc9
Chk1
Rad23
Ubi4
Ede1
Ent2
Vps27
Chc1
Ptp3
Stb1
Spa2
Bem2
Bud14
Rck2
Glc7
Ipl1
Cdc5
Dbf4
Pfk26
Tpk1
Gpa2
Ras2
Med6
Rpb10
Paf1
Rpo26
Bdp1
Spt15
Spt5
Rgr1
Srb7
Ssn2
Nop1
Nsr1
Nop56
Snu13
Rps31
Rad9
Rtf1
Rpo21
Npl3
Air2
Fps1
Ask10
Cdc28
Hog1
Pbs2
Ssk2
Clb2
Bud3
Bud4
Msn2
Sch9
Rim15
Yak1
Ypl247c
Vps52
Kss1
Sko1
Slt2
Msg5
Swe1
Igo2
Cdc55
Igo1
Vps53
Vps54
Rtn1
Fmp42
Cot1
Zrc1
10
20
30
40
50
t/s

#### Path 799 (Sec18 − Rad23)

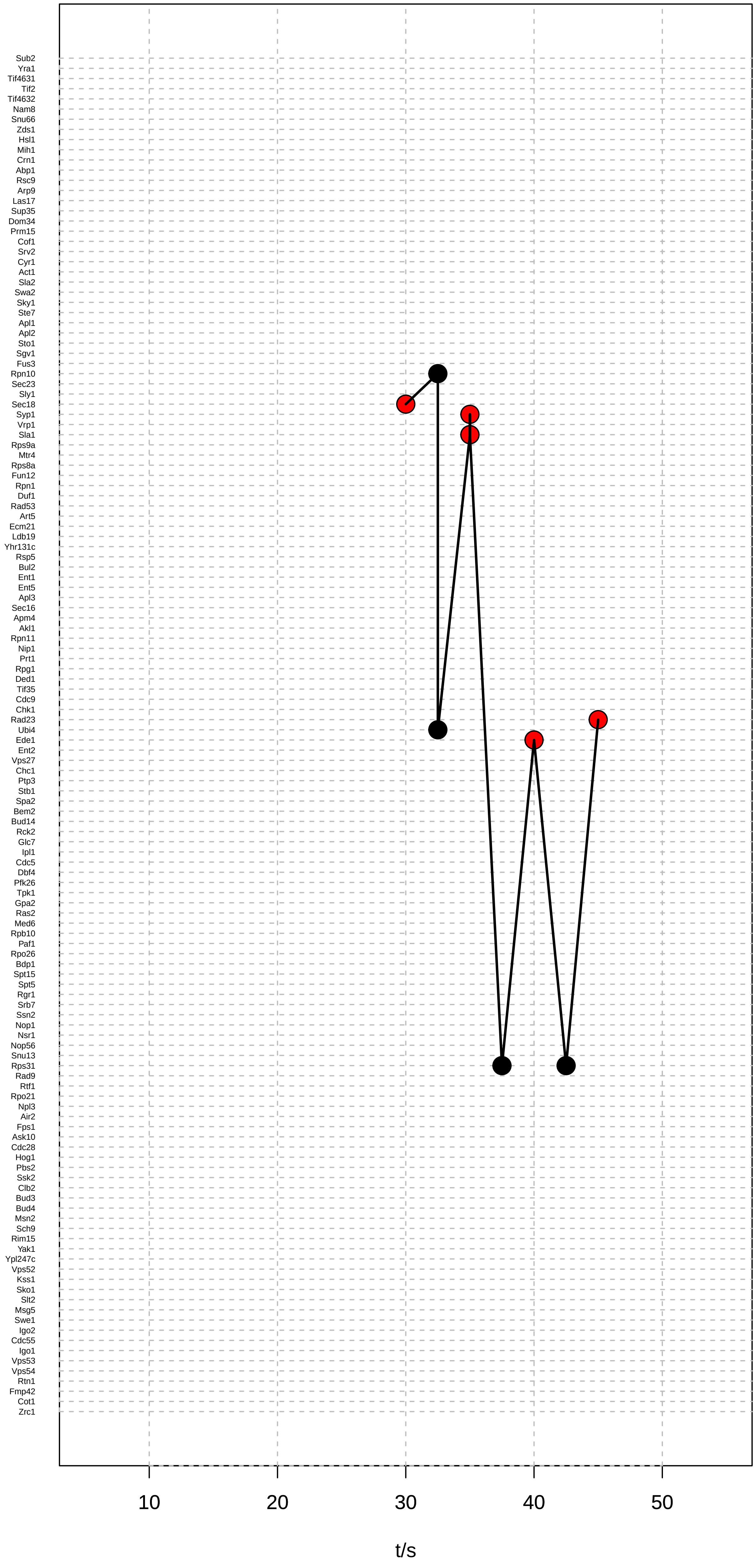

### Path 800 (Sec18 − Ede1)

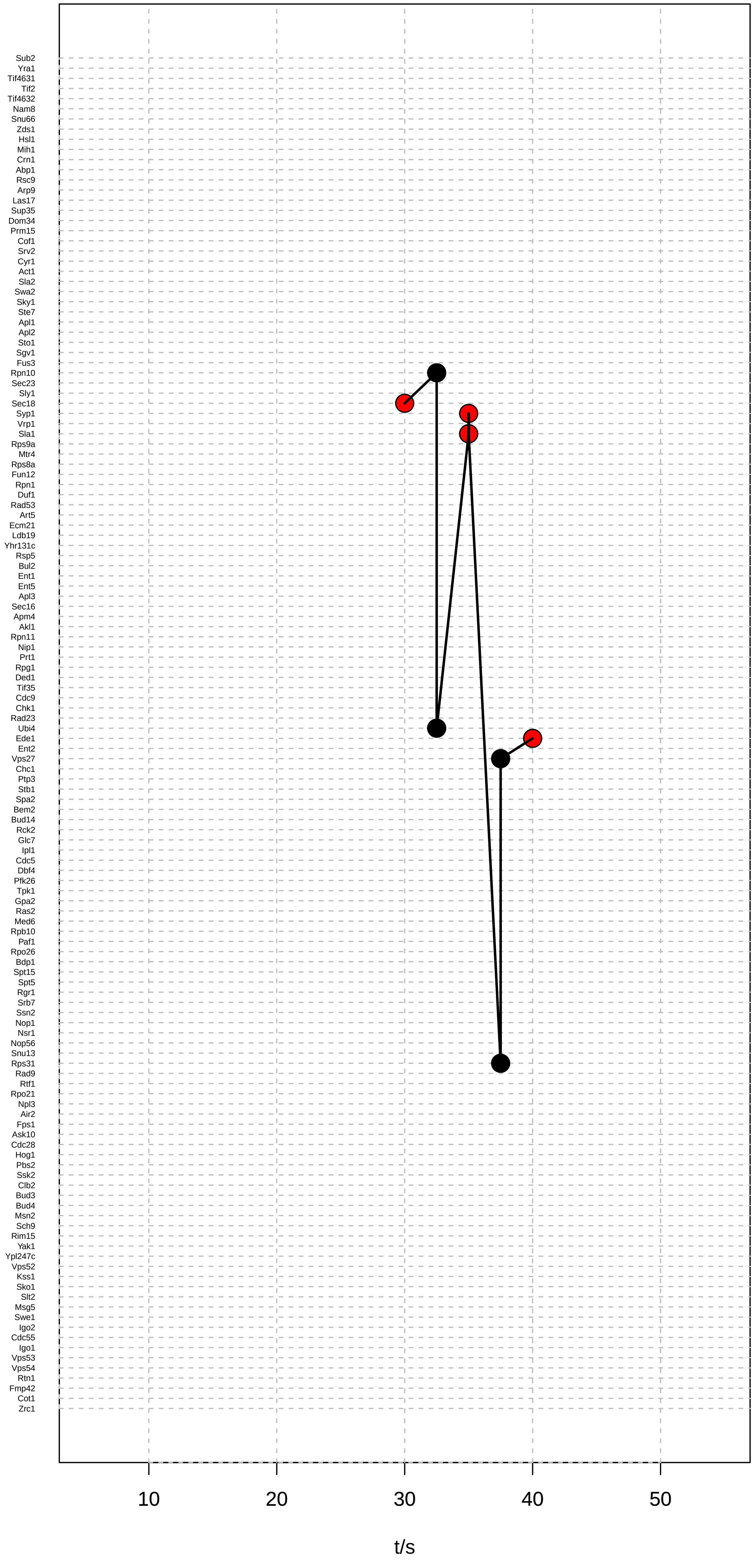

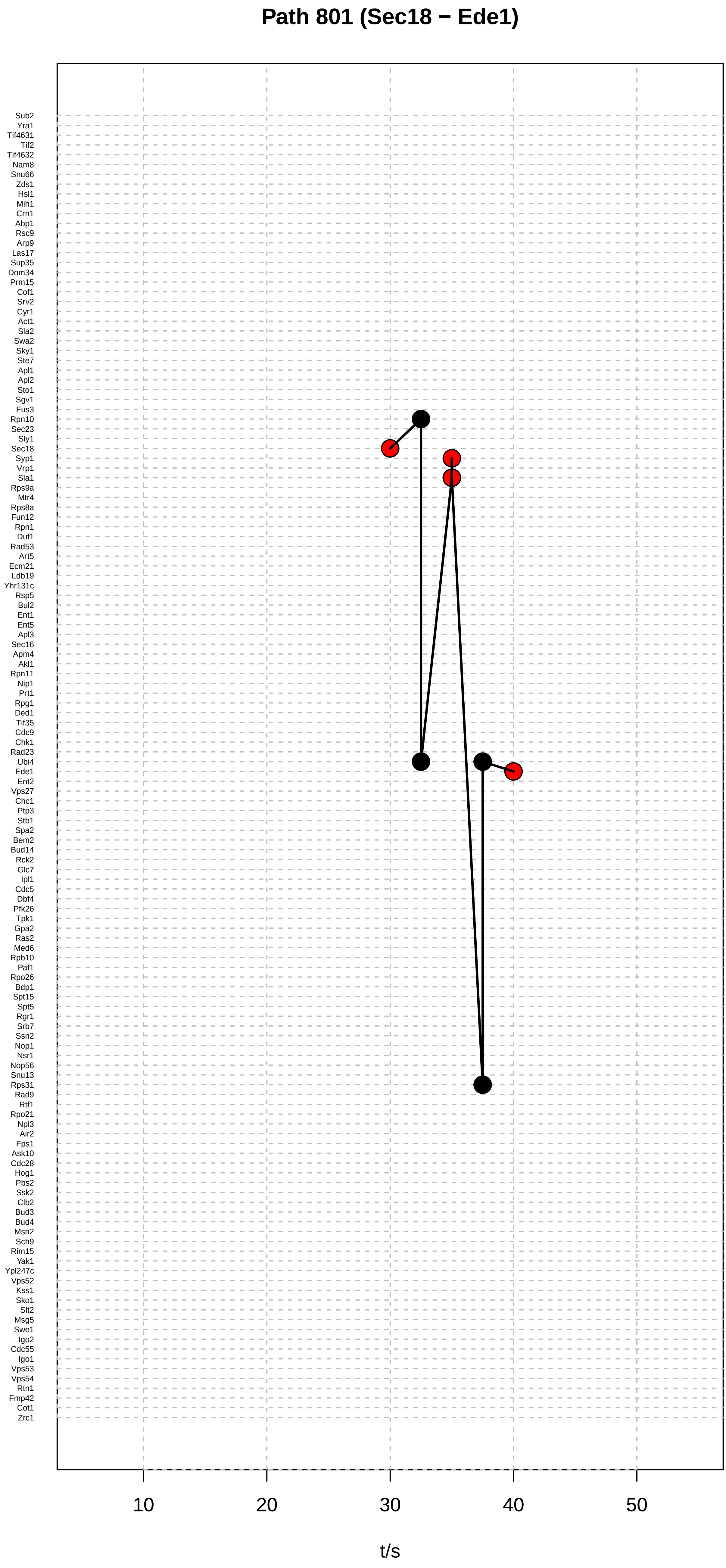

Path 801 (Sec18 − Ede1)
Sub2
Yra1
Tif4631
Tif2
Tif4632
Nam8
Snu66
Zds1
Hsl1
Mih1
Crn1
Abp1
Rsc9
Arp9
Las17
Sup35
Dom34
Prm15
Cof1
Srv2
Cyr1
Act1
Sla2
Swa2
Sky1
Ste7
Apl1
Apl2
Sto1
Sgv1
Fus3
Rpn10
Sec23
Sly1
Sec18
Syp1
Vrp1
Sla1
Rps9a
Mtr4
Rps8a
Fun12
Rpn1
Duf1
Rad53
Art5
Ecm21
Ldb19
Yhr131c
Rsp5
Bul2
Ent1
Ent5
Apl3
Sec16
Apm4
Akl1
Rpn11
Nip1
Prt1
Rpg1
Ded1
Tif35
Cdc9
Chk1
Rad23
Ubi4
Ede1
Ent2
Vps27
Chc1
Ptp3
Stb1
Spa2
Bem2
Bud14
Rck2
Glc7
Ipl1
Cdc5
Dbf4
Pfk26
Tpk1
Gpa2
Ras2
Med6
Rpb10
Paf1
Rpo26
Bdp1
Spt15
Spt5
Rgr1
Srb7
Ssn2
Nop1
Nsr1
Nop56
Snu13
Rps31
Rad9
Rtf1
Rpo21
Npl3
Air2
Fps1
Ask10
Cdc28
Hog1
Pbs2
Ssk2
Clb2
Bud3
Bud4
Msn2
Sch9
Rim15
Yak1
Ypl247c
Vps52
Kss1
Sko1
Slt2
Msg5
Swe1
Igo2
Cdc55
Igo1
Vps53
Vps54
Rtn1
Fmp42
Cot1
Zrc1
10
20
30
40
50
t/s

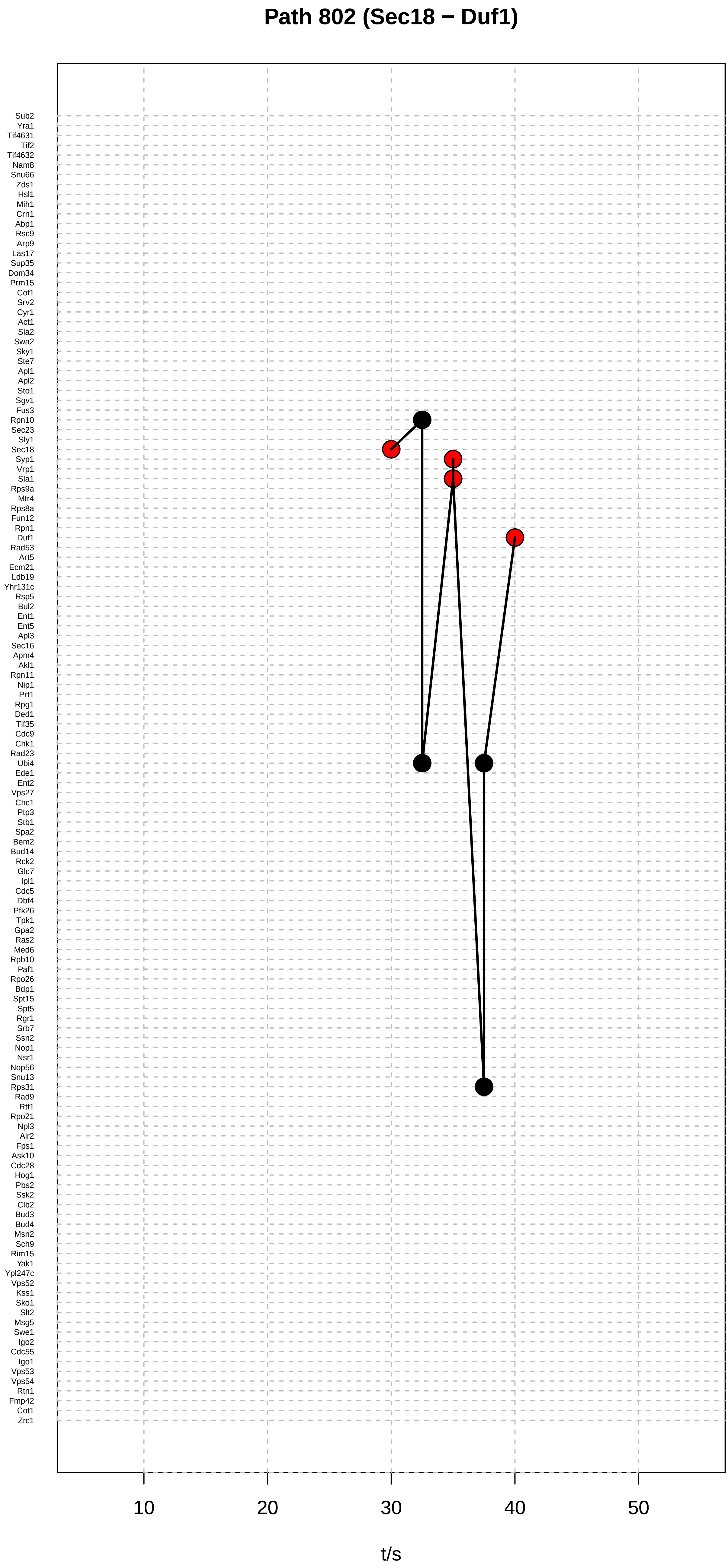

Path 802 (Sec18 − Duf1)
Sub2
Yra1
Tif4631
Tif2
Tif4632
Nam8
Snu66
Zds1
Hsl1
Mih1
Crn1
Abp1
Rsc9
Arp9
Las17
Sup35
Dom34
Prm15
Cof1
Srv2
Cyr1
Act1
Sla2
Swa2
Sky1
Ste7
Apl1
Apl2
Sto1
Sgv1
Fus3
Rpn10
Sec23
Sly1
Sec18
Syp1
Vrp1
Sla1
Rps9a
Mtr4
Rps8a
Fun12
Rpn1
Duf1
Rad53
Art5
Ecm21
Ldb19
Yhr131c
Rsp5
Bul2
Ent1
Ent5
Apl3
Sec16
Apm4
Akl1
Rpn11
Nip1
Prt1
Rpg1
Ded1
Tif35
Cdc9
Chk1
Rad23
Ubi4
Ede1
Ent2
Vps27
Chc1
Ptp3
Stb1
Spa2
Bem2
Bud14
Rck2
Glc7
Ipl1
Cdc5
Dbf4
Pfk26
Tpk1
Gpa2
Ras2
Med6
Rpb10
Paf1
Rpo26
Bdp1
Spt15
Spt5
Rgr1
Srb7
Ssn2
Nop1
Nsr1
Nop56
Snu13
Rps31
Rad9
Rtf1
Rpo21
Npl3
Air2
Fps1
Ask10
Cdc28
Hog1
Pbs2
Ssk2
Clb2
Bud3
Bud4
Msn2
Sch9
Rim15
Yak1
Ypl247c
Vps52
Kss1
Sko1
Slt2
Msg5
Swe1
Igo2
Cdc55
Igo1
Vps53
Vps54
Rtn1
Fmp42
Cot1
Zrc1
10
20
30
40
50
t/s

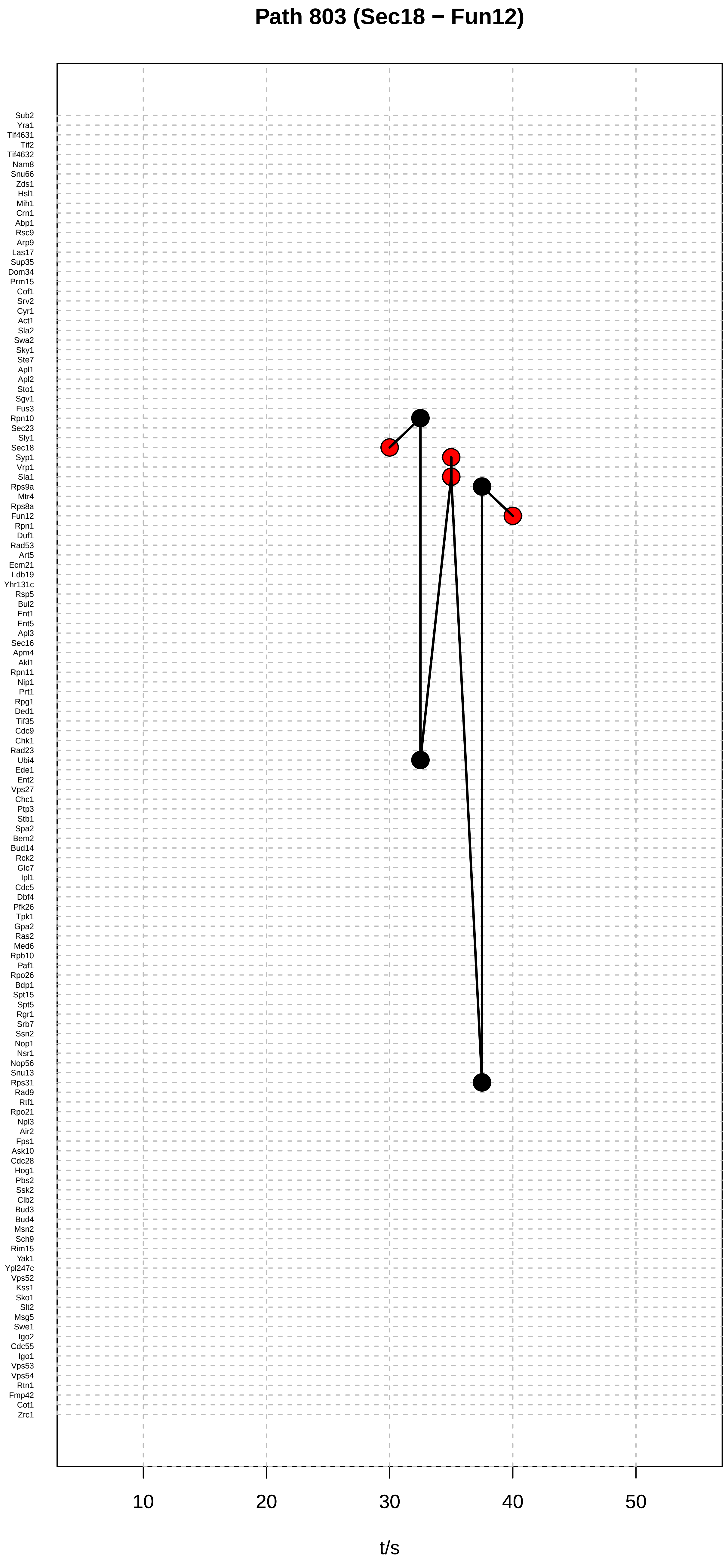

Path 803 (Sec18 − Fun12)
Sub2
Yra1
Tif4631
Tif2
Tif4632
Nam8
Snu66
Zds1
Hsl1
Mih1
Crn1
Abp1
Rsc9
Arp9
Las17
Sup35
Dom34
Prm15
Cof1
Srv2
Cyr1
Act1
Sla2
Swa2
Sky1
Ste7
Apl1
Apl2
Sto1
Sgv1
Fus3
Rpn10
Sec23
Sly1
Sec18
Syp1
Vrp1
Sla1
Rps9a
Mtr4
Rps8a
Fun12
Rpn1
Duf1
Rad53
Art5
Ecm21
Ldb19
Yhr131c
Rsp5
Bul2
Ent1
Ent5
Apl3
Sec16
Apm4
Akl1
Rpn11
Nip1
Prt1
Rpg1
Ded1
Tif35
Cdc9
Chk1
Rad23
Ubi4
Ede1
Ent2
Vps27
Chc1
Ptp3
Stb1
Spa2
Bem2
Bud14
Rck2
Glc7
Ipl1
Cdc5
Dbf4
Pfk26
Tpk1
Gpa2
Ras2
Med6
Rpb10
Paf1
Rpo26
Bdp1
Spt15
Spt5
Rgr1
Srb7
Ssn2
Nop1
Nsr1
Nop56
Snu13
Rps31
Rad9
Rtf1
Rpo21
Npl3
Air2
Fps1
Ask10
Cdc28
Hog1
Pbs2
Ssk2
Clb2
Bud3
Bud4
Msn2
Sch9
Rim15
Yak1
Ypl247c
Vps52
Kss1
Sko1
Slt2
Msg5
Swe1
Igo2
Cdc55
Igo1
Vps53
Vps54
Rtn1
Fmp42
Cot1
Zrc1
10
20
30
40
50
t/s

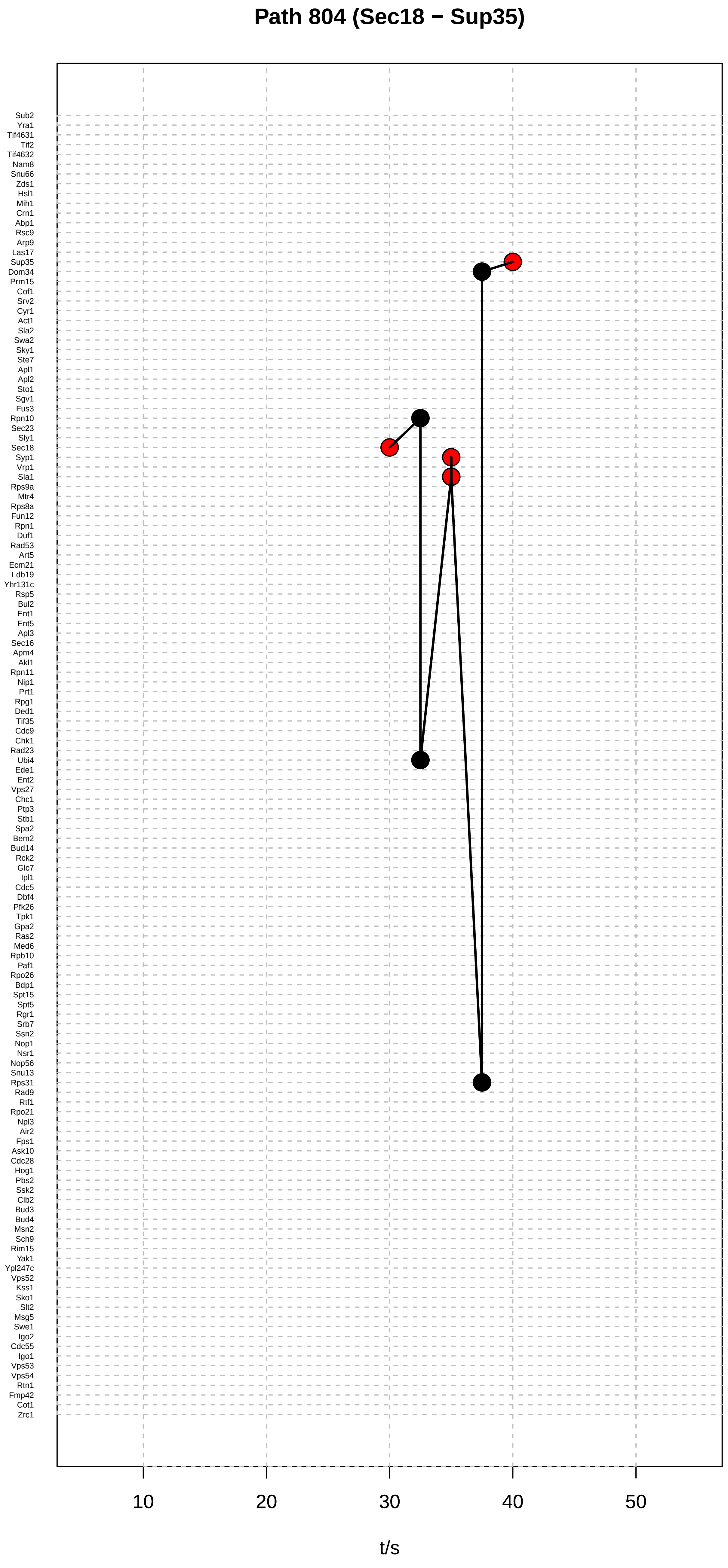
Path 804 (Sec18 − Sup35)
Sub2
Yra1
Tif4631
Tif2
Tif4632
Nam8
Snu66
Zds1
Hsl1
Mih1
Crn1
Abp1
Rsc9
Arp9
Las17
Sup35
Dom34
Prm15
Cof1
Srv2
Cyr1
Act1
Sla2
Swa2
Sky1
Ste7
Apl1
Apl2
Sto1
Sgv1
Fus3
Rpn10
Sec23
Sly1
Sec18
Syp1
Vrp1
Sla1
Rps9a
Mtr4
Rps8a
Fun12
Rpn1
Duf1
Rad53
Art5
Ecm21
Ldb19
Yhr131c
Rsp5
Bul2
Ent1
Ent5
Apl3
Sec16
Apm4
Akl1
Rpn11
Nip1
Prt1
Rpg1
Ded1
Tif35
Cdc9
Chk1
Rad23
Ubi4
Ede1
Ent2
Vps27
Chc1
Ptp3
Stb1
Spa2
Bem2
Bud14
Rck2
Glc7
Ipl1
Cdc5
Dbf4
Pfk26
Tpk1
Gpa2
Ras2
Med6
Rpb10
Paf1
Rpo26
Bdp1
Spt15
Spt5
Rgr1
Srb7
Ssn2
Nop1
Nsr1
Nop56
Snu13
Rps31
Rad9
Rtf1
Rpo21
Npl3
Air2
Fps1
Ask10
Cdc28
Hog1
Pbs2
Ssk2
Clb2
Bud3
Bud4
Msn2
Sch9
Rim15
Yak1
Ypl247c
Vps52
Kss1
Sko1
Slt2
Msg5
Swe1
Igo2
Cdc55
Igo1
Vps53
Vps54
Rtn1
Fmp42
Cot1
Zrc1
10
20
30
40
50
t/s

### Path 805 (Sec18 − Cyr1)

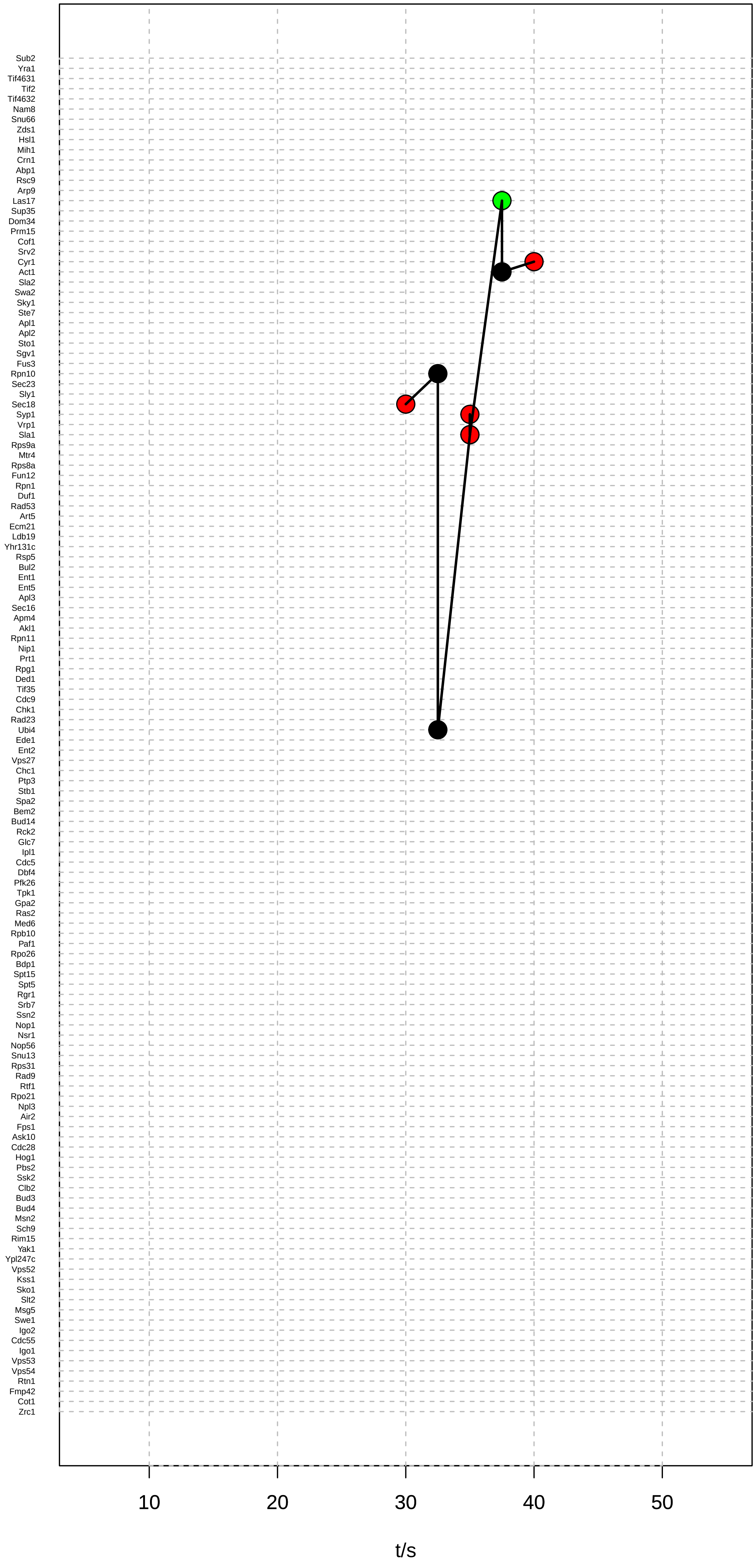

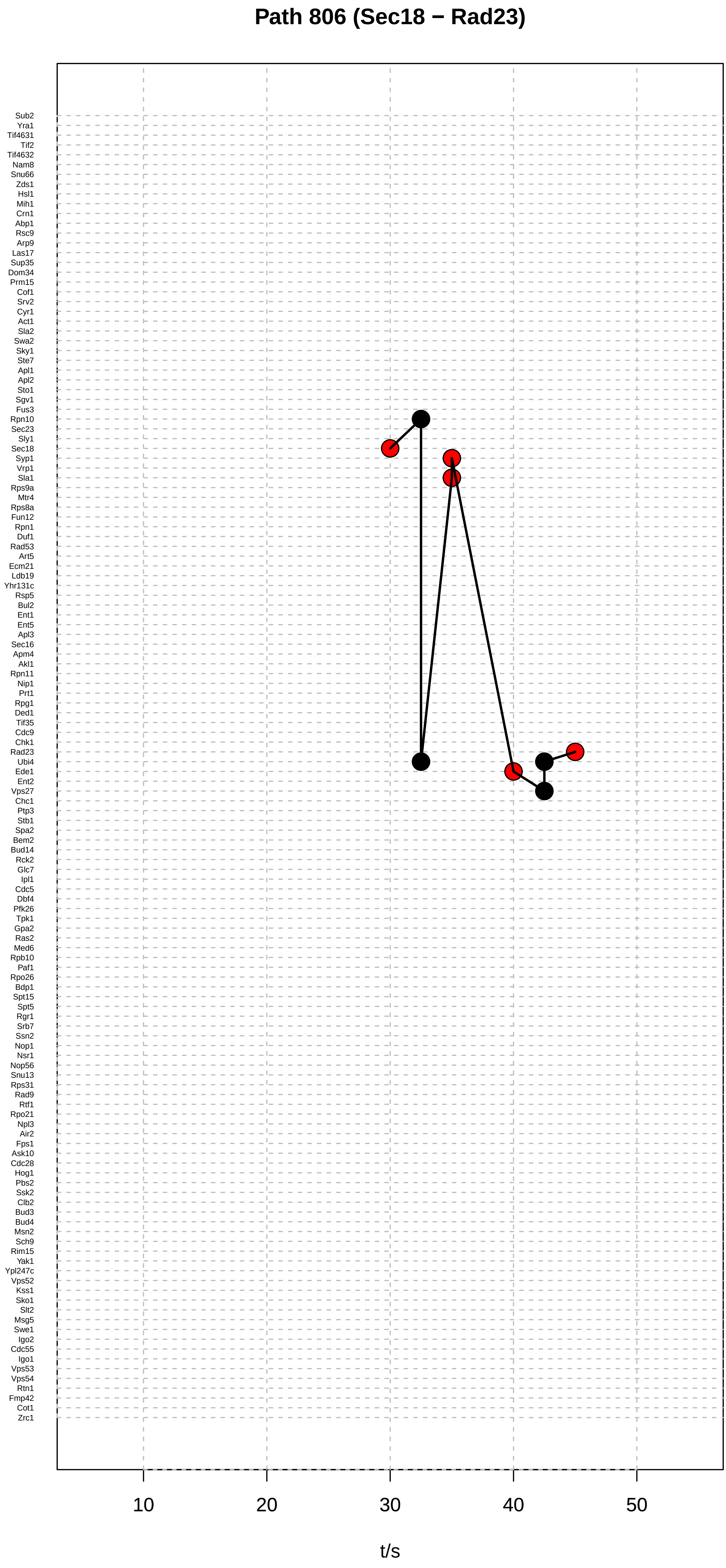
Path 806 (Sec18 − Rad23)
Sub2
Yra1
Tif4631
Tif2
Tif4632
Nam8
Snu66
Zds1
Hsl1
Mih1
Crn1
Abp1
Rsc9
Arp9
Las17
Sup35
Dom34
Prm15
Cof1
Srv2
Cyr1
Act1
Sla2
Swa2
Sky1
Ste7
Apl1
Apl2
Sto1
Sgv1
Fus3
Rpn10
Sec23
Sly1
Sec18
Syp1
Vrp1
Sla1
Rps9a
Mtr4
Rps8a
Fun12
Rpn1
Duf1
Rad53
Art5
Ecm21
Ldb19
Yhr131c
Rsp5
Bul2
Ent1
Ent5
Apl3
Sec16
Apm4
Akl1
Rpn11
Nip1
Prt1
Rpg1
Ded1
Tif35
Cdc9
Chk1
Rad23
Ubi4
Ede1
Ent2
Vps27
Chc1
Ptp3
Stb1
Spa2
Bem2
Bud14
Rck2
Glc7
Ipl1
Cdc5
Dbf4
Pfk26
Tpk1
Gpa2
Ras2
Med6
Rpb10
Paf1
Rpo26
Bdp1
Spt15
Spt5
Rgr1
Srb7
Ssn2
Nop1
Nsr1
Nop56
Snu13
Rps31
Rad9
Rtf1
Rpo21
Npl3
Air2
Fps1
Ask10
Cdc28
Hog1
Pbs2
Ssk2
Clb2
Bud3
Bud4
Msn2
Sch9
Rim15
Yak1
Ypl247c
Vps52
Kss1
Sko1
Slt2
Msg5
Swe1
Igo2
Cdc55
Igo1
Vps53
Vps54
Rtn1
Fmp42
Cot1
Zrc1
10
20
30
40
50
t/s

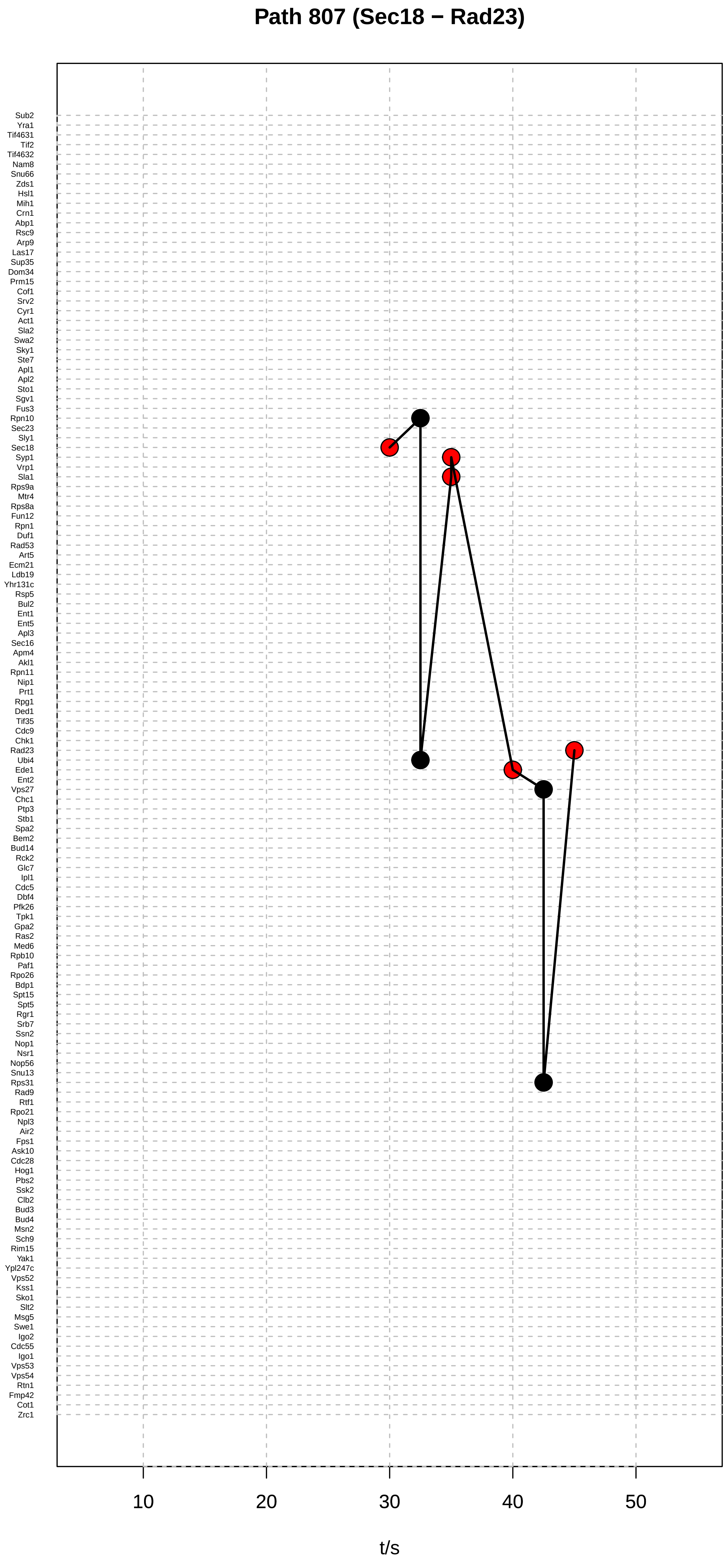

Path 807 (Sec18 − Rad23)
Sub2
Yra1
Tif4631
Tif2
Tif4632
Nam8
Snu66
Zds1
Hsl1
Mih1
Crn1
Abp1
Rsc9
Arp9
Las17
Sup35
Dom34
Prm15
Cof1
Srv2
Cyr1
Act1
Sla2
Swa2
Sky1
Ste7
Apl1
Apl2
Sto1
Sgv1
Fus3
Rpn10
Sec23
Sly1
Sec18
Syp1
Vrp1
Sla1
Rps9a
Mtr4
Rps8a
Fun12
Rpn1
Duf1
Rad53
Art5
Ecm21
Ldb19
Yhr131c
Rsp5
Bul2
Ent1
Ent5
Apl3
Sec16
Apm4
Akl1
Rpn11
Nip1
Prt1
Rpg1
Ded1
Tif35
Cdc9
Chk1
Rad23
Ubi4
Ede1
Ent2
Vps27
Chc1
Ptp3
Stb1
Spa2
Bem2
Bud14
Rck2
Glc7
Ipl1
Cdc5
Dbf4
Pfk26
Tpk1
Gpa2
Ras2
Med6
Rpb10
Paf1
Rpo26
Bdp1
Spt15
Spt5
Rgr1
Srb7
Ssn2
Nop1
Nsr1
Nop56
Snu13
Rps31
Rad9
Rtf1
Rpo21
Npl3
Air2
Fps1
Ask10
Cdc28
Hog1
Pbs2
Ssk2
Clb2
Bud3
Bud4
Msn2
Sch9
Rim15
Yak1
Ypl247c
Vps52
Kss1
Sko1
Slt2
Msg5
Swe1
Igo2
Cdc55
Igo1
Vps53
Vps54
Rtn1
Fmp42
Cot1
Zrc1
10
20
30
40
50
t/s

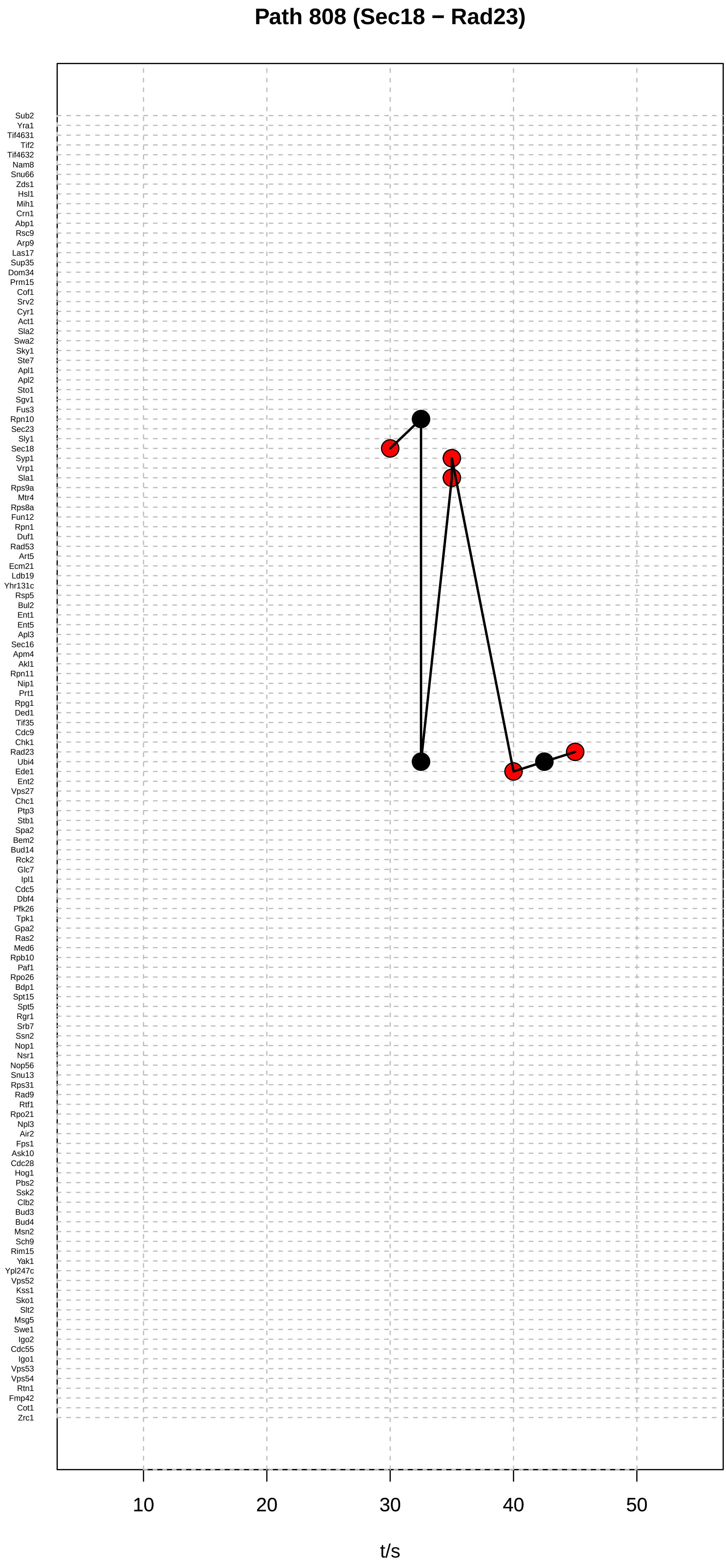

Path 808 (Sec18 − Rad23)
Sub2
Yra1
Tif4631
Tif2
Tif4632
Nam8
Snu66
Zds1
Hsl1
Mih1
Crn1
Abp1
Rsc9
Arp9
Las17
Sup35
Dom34
Prm15
Cof1
Srv2
Cyr1
Act1
Sla2
Swa2
Sky1
Ste7
Apl1
Apl2
Sto1
Sgv1
Fus3
Rpn10
Sec23
Sly1
Sec18
Syp1
Vrp1
Sla1
Rps9a
Mtr4
Rps8a
Fun12
Rpn1
Duf1
Rad53
Art5
Ecm21
Ldb19
Yhr131c
Rsp5
Bul2
Ent1
Ent5
Apl3
Sec16
Apm4
Akl1
Rpn11
Nip1
Prt1
Rpg1
Ded1
Tif35
Cdc9
Chk1
Rad23
Ubi4
Ede1
Ent2
Vps27
Chc1
Ptp3
Stb1
Spa2
Bem2
Bud14
Rck2
Glc7
Ipl1
Cdc5
Dbf4
Pfk26
Tpk1
Gpa2
Ras2
Med6
Rpb10
Paf1
Rpo26
Bdp1
Spt15
Spt5
Rgr1
Srb7
Ssn2
Nop1
Nsr1
Nop56
Snu13
Rps31
Rad9
Rtf1
Rpo21
Npl3
Air2
Fps1
Ask10
Cdc28
Hog1
Pbs2
Ssk2
Clb2
Bud3
Bud4
Msn2
Sch9
Rim15
Yak1
Ypl247c
Vps52
Kss1
Sko1
Slt2
Msg5
Swe1
Igo2
Cdc55
Igo1
Vps53
Vps54
Rtn1
Fmp42
Cot1
Zrc1
10
20
30
40
50
t/s

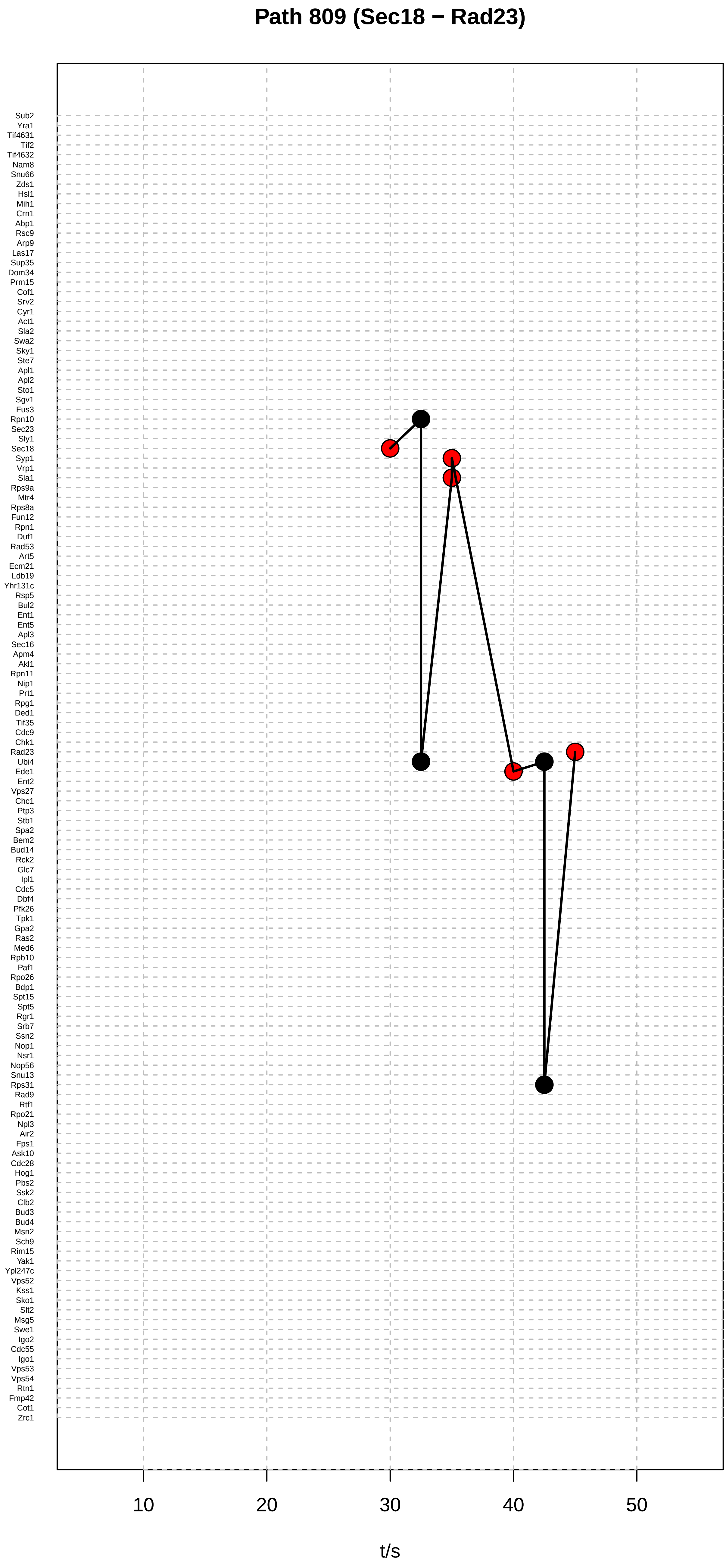
Path 809 (Sec18 − Rad23)
Sub2
Yra1
Tif4631
Tif2
Tif4632
Nam8
Snu66
Zds1
Hsl1
Mih1
Crn1
Abp1
Rsc9
Arp9
Las17
Sup35
Dom34
Prm15
Cof1
Srv2
Cyr1
Act1
Sla2
Swa2
Sky1
Ste7
Apl1
Apl2
Sto1
Sgv1
Fus3
Rpn10
Sec23
Sly1
Sec18
Syp1
Vrp1
Sla1
Rps9a
Mtr4
Rps8a
Fun12
Rpn1
Duf1
Rad53
Art5
Ecm21
Ldb19
Yhr131c
Rsp5
Bul2
Ent1
Ent5
Apl3
Sec16
Apm4
Akl1
Rpn11
Nip1
Prt1
Rpg1
Ded1
Tif35
Cdc9
Chk1
Rad23
Ubi4
Ede1
Ent2
Vps27
Chc1
Ptp3
Stb1
Spa2
Bem2
Bud14
Rck2
Glc7
Ipl1
Cdc5
Dbf4
Pfk26
Tpk1
Gpa2
Ras2
Med6
Rpb10
Paf1
Rpo26
Bdp1
Spt15
Spt5
Rgr1
Srb7
Ssn2
Nop1
Nsr1
Nop56
Snu13
Rps31
Rad9
Rtf1
Rpo21
Npl3
Air2
Fps1
Ask10
Cdc28
Hog1
Pbs2
Ssk2
Clb2
Bud3
Bud4
Msn2
Sch9
Rim15
Yak1
Ypl247c
Vps52
Kss1
Sko1
Slt2
Msg5
Swe1
Igo2
Cdc55
Igo1
Vps53
Vps54
Rtn1
Fmp42
Cot1
Zrc1
10
20
30
40
50
t/s

#### Path 810 (Sec18 − Rad23)

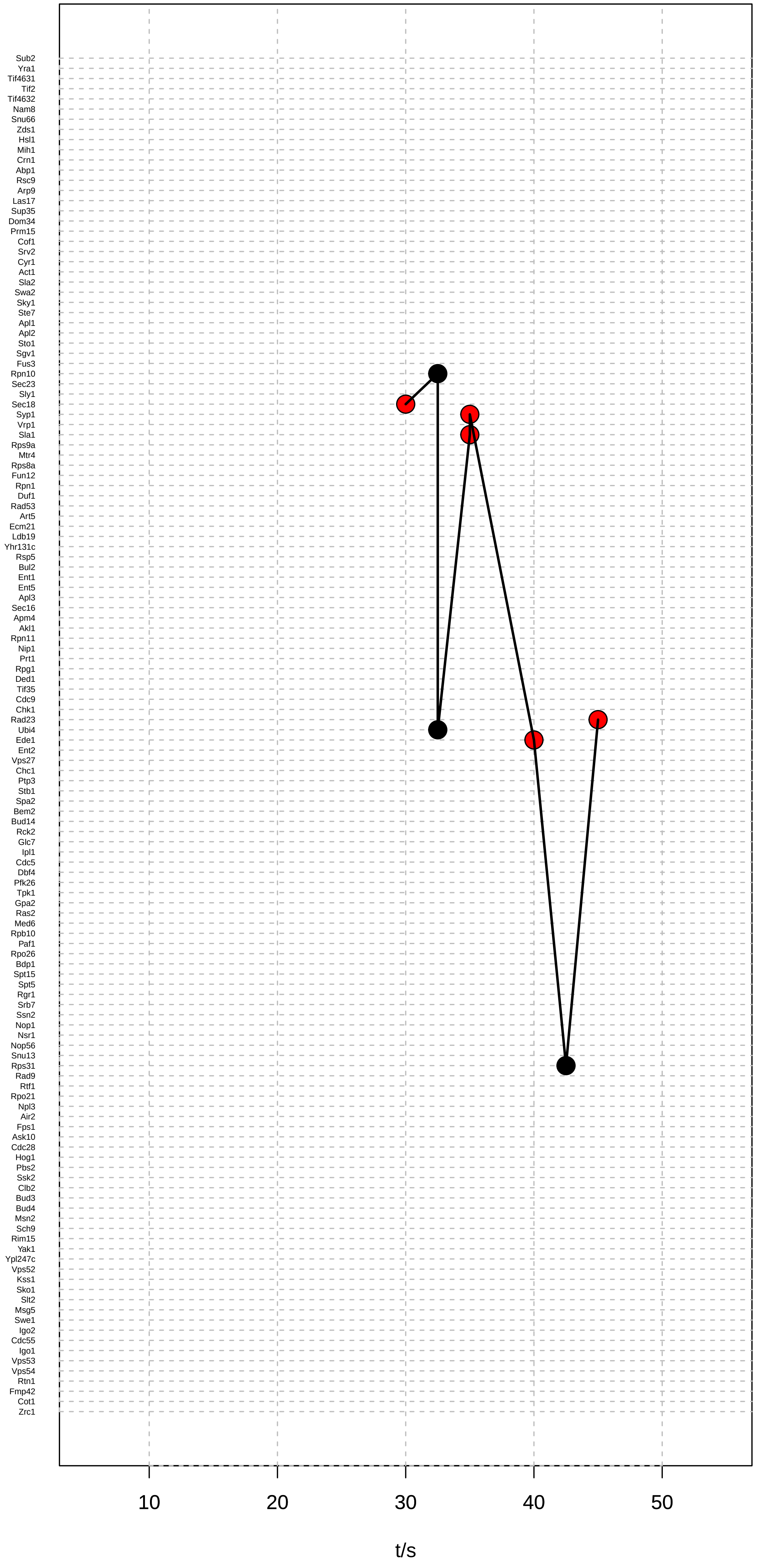

#### Path 811 (Sec18 − Rad23)

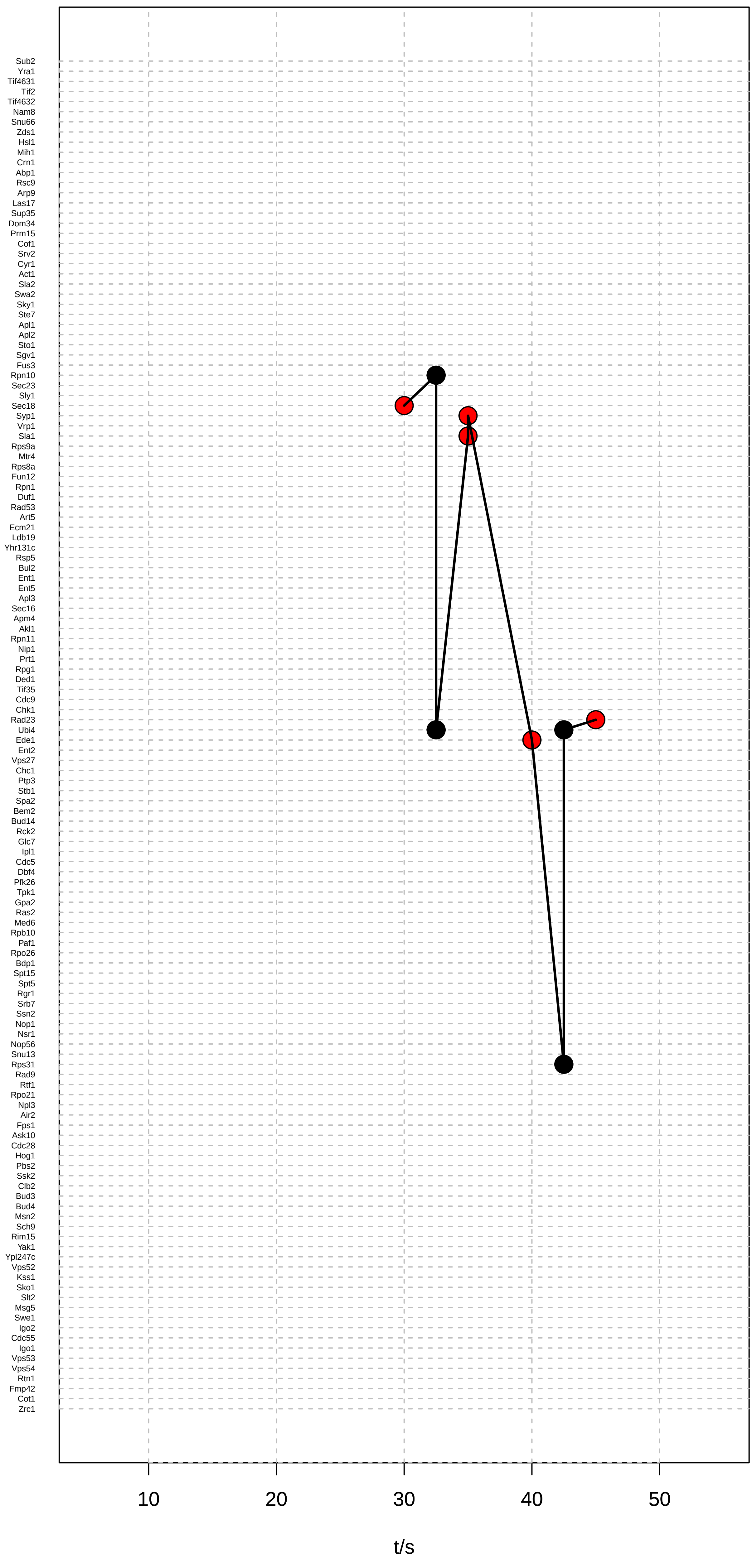

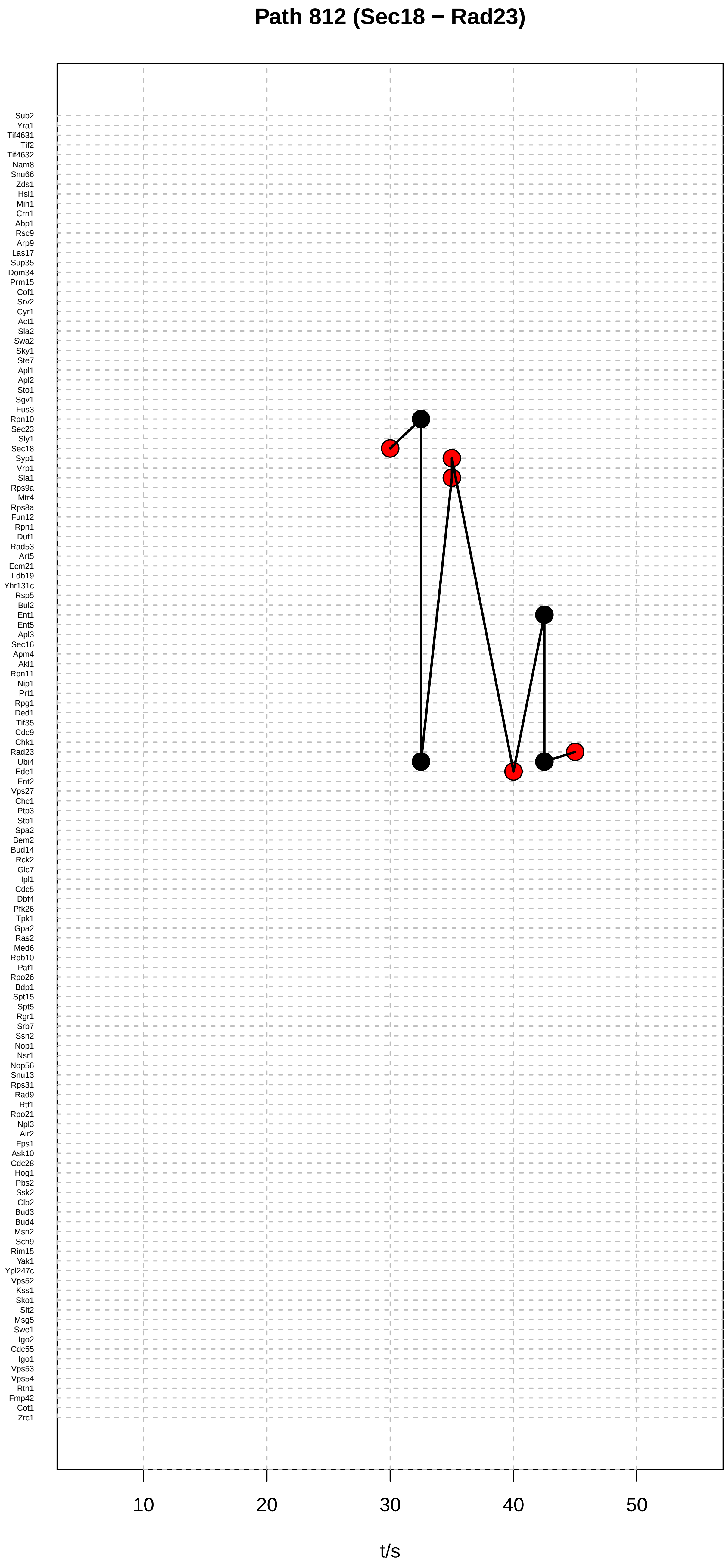

Path 812 (Sec18 − Rad23)
Sub2
Yra1
Tif4631
Tif2
Tif4632
Nam8
Snu66
Zds1
Hsl1
Mih1
Crn1
Abp1
Rsc9
Arp9
Las17
Sup35
Dom34
Prm15
Cof1
Srv2
Cyr1
Act1
Sla2
Swa2
Sky1
Ste7
Apl1
Apl2
Sto1
Sgv1
Fus3
Rpn10
Sec23
Sly1
Sec18
Syp1
Vrp1
Sla1
Rps9a
Mtr4
Rps8a
Fun12
Rpn1
Duf1
Rad53
Art5
Ecm21
Ldb19
Yhr131c
Rsp5
Bul2
Ent1
Ent5
Apl3
Sec16
Apm4
Akl1
Rpn11
Nip1
Prt1
Rpg1
Ded1
Tif35
Cdc9
Chk1
Rad23
Ubi4
Ede1
Ent2
Vps27
Chc1
Ptp3
Stb1
Spa2
Bem2
Bud14
Rck2
Glc7
Ipl1
Cdc5
Dbf4
Pfk26
Tpk1
Gpa2
Ras2
Med6
Rpb10
Paf1
Rpo26
Bdp1
Spt15
Spt5
Rgr1
Srb7
Ssn2
Nop1
Nsr1
Nop56
Snu13
Rps31
Rad9
Rtf1
Rpo21
Npl3
Air2
Fps1
Ask10
Cdc28
Hog1
Pbs2
Ssk2
Clb2
Bud3
Bud4
Msn2
Sch9
Rim15
Yak1
Ypl247c
Vps52
Kss1
Sko1
Slt2
Msg5
Swe1
Igo2
Cdc55
Igo1
Vps53
Vps54
Rtn1
Fmp42
Cot1
Zrc1
10
20
30
40
50
t/s

#### Path 813 (Sec18 − Rad23)

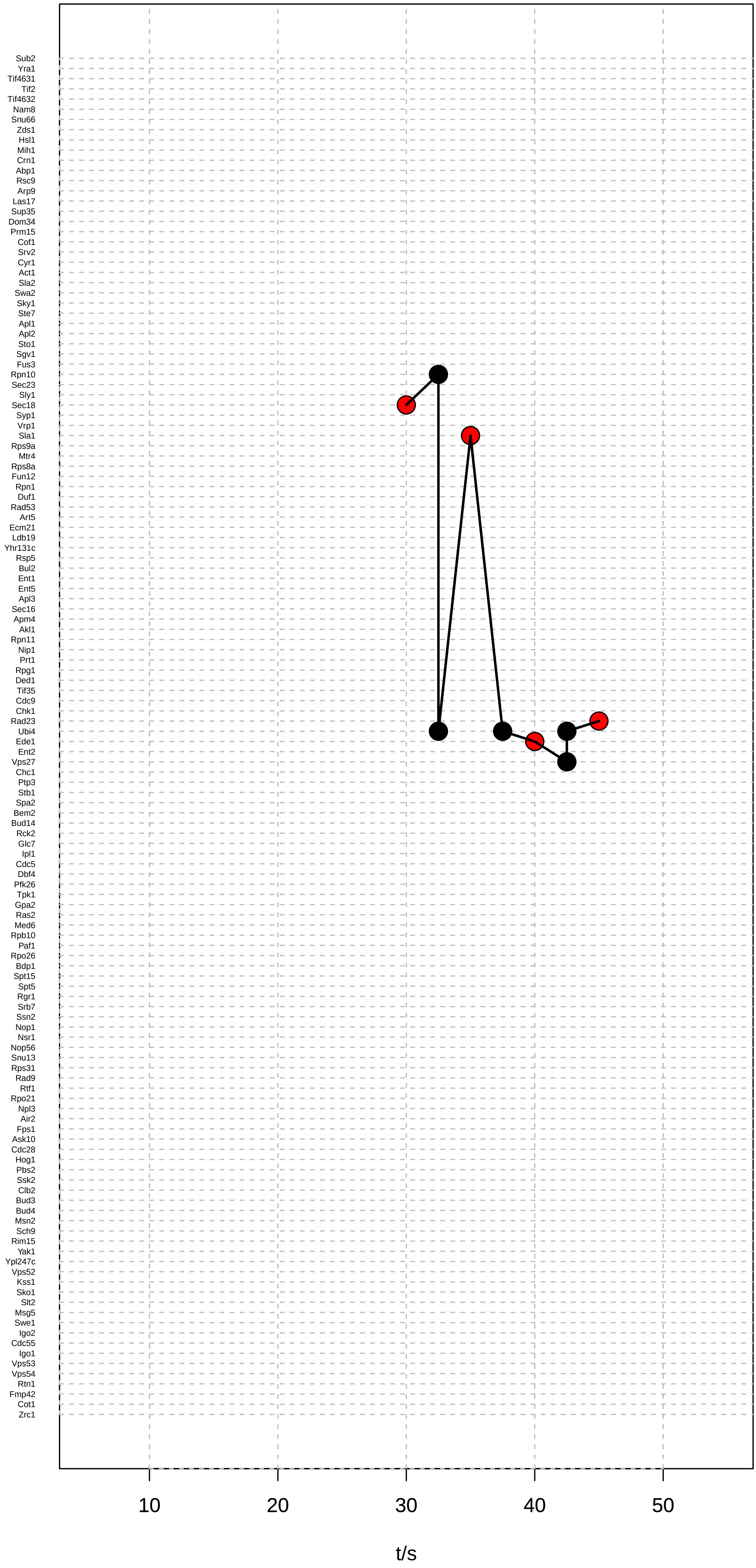

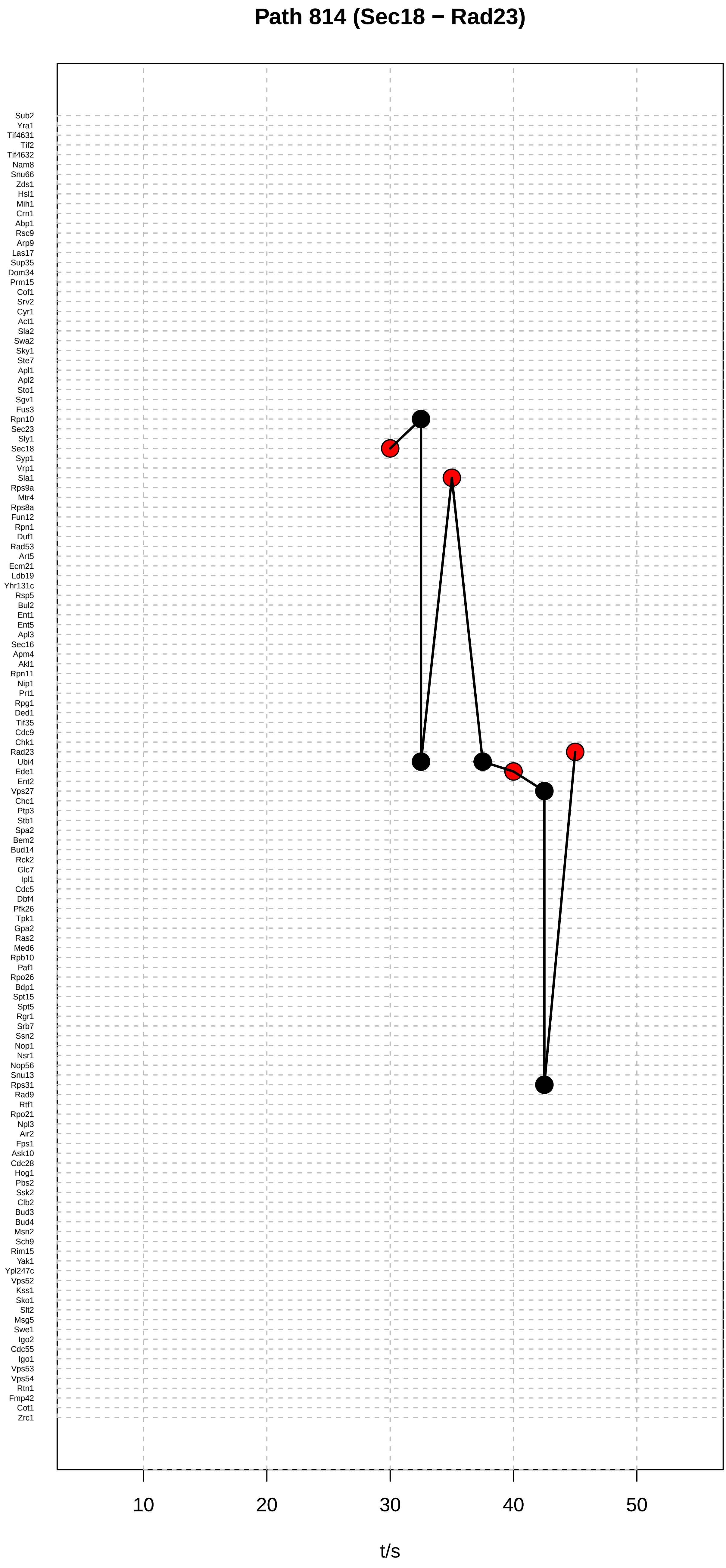
Path 814 (Sec18 − Rad23)
Sub2
Yra1
Tif4631
Tif2
Tif4632
Nam8
Snu66
Zds1
Hsl1
Mih1
Crn1
Abp1
Rsc9
Arp9
Las17
Sup35
Dom34
Prm15
Cof1
Srv2
Cyr1
Act1
Sla2
Swa2
Sky1
Ste7
Apl1
Apl2
Sto1
Sgv1
Fus3
Rpn10
Sec23
Sly1
Sec18
Syp1
Vrp1
Sla1
Rps9a
Mtr4
Rps8a
Fun12
Rpn1
Duf1
Rad53
Art5
Ecm21
Ldb19
Yhr131c
Rsp5
Bul2
Ent1
Ent5
Apl3
Sec16
Apm4
Akl1
Rpn11
Nip1
Prt1
Rpg1
Ded1
Tif35
Cdc9
Chk1
Rad23
Ubi4
Ede1
Ent2
Vps27
Chc1
Ptp3
Stb1
Spa2
Bem2
Bud14
Rck2
Glc7
Ipl1
Cdc5
Dbf4
Pfk26
Tpk1
Gpa2
Ras2
Med6
Rpb10
Paf1
Rpo26
Bdp1
Spt15
Spt5
Rgr1
Srb7
Ssn2
Nop1
Nsr1
Nop56
Snu13
Rps31
Rad9
Rtf1
Rpo21
Npl3
Air2
Fps1
Ask10
Cdc28
Hog1
Pbs2
Ssk2
Clb2
Bud3
Bud4
Msn2
Sch9
Rim15
Yak1
Ypl247c
Vps52
Kss1
Sko1
Slt2
Msg5
Swe1
Igo2
Cdc55
Igo1
Vps53
Vps54
Rtn1
Fmp42
Cot1
Zrc1
10
20
30
40
50
t/s

### Path 815 (Sec18 − Rad23)

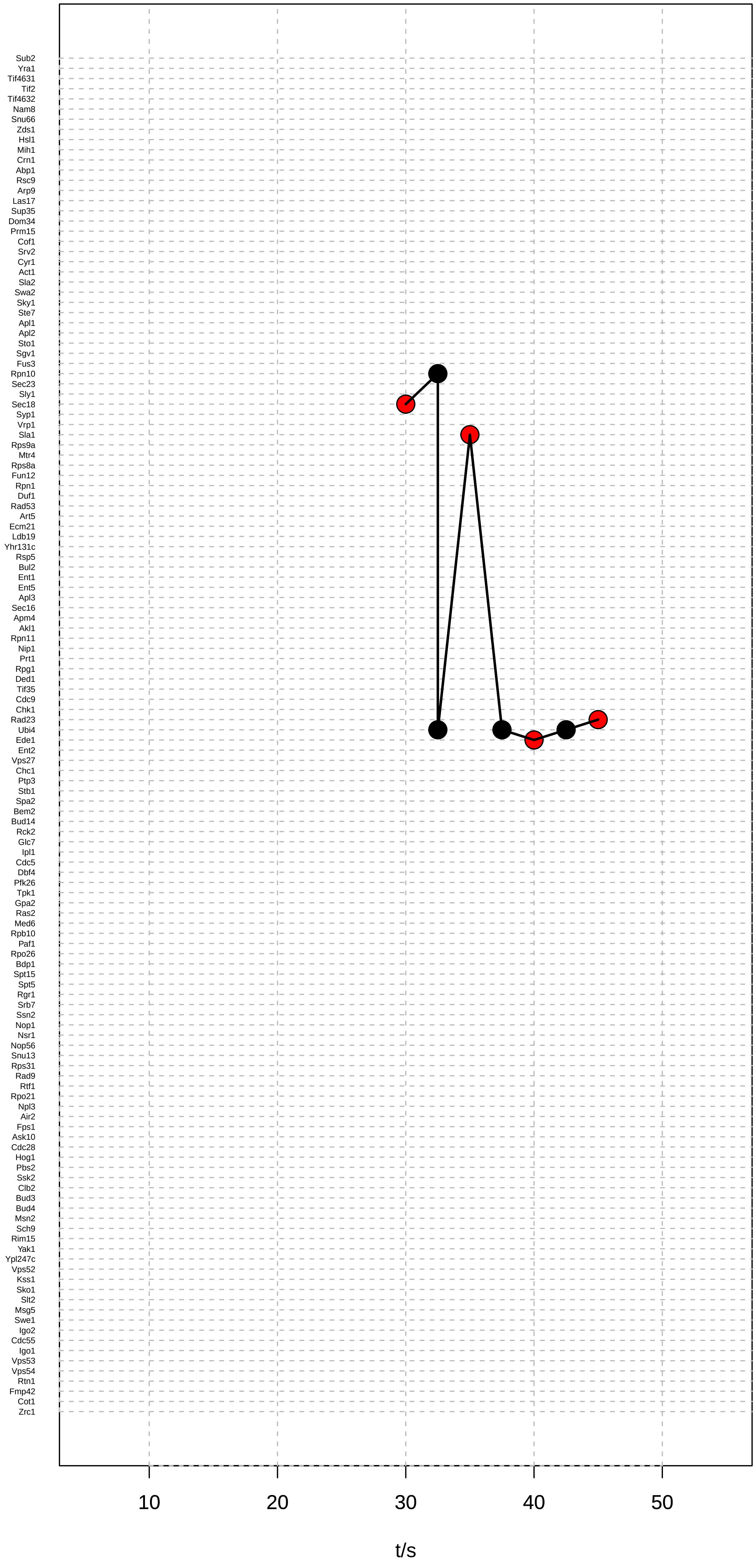

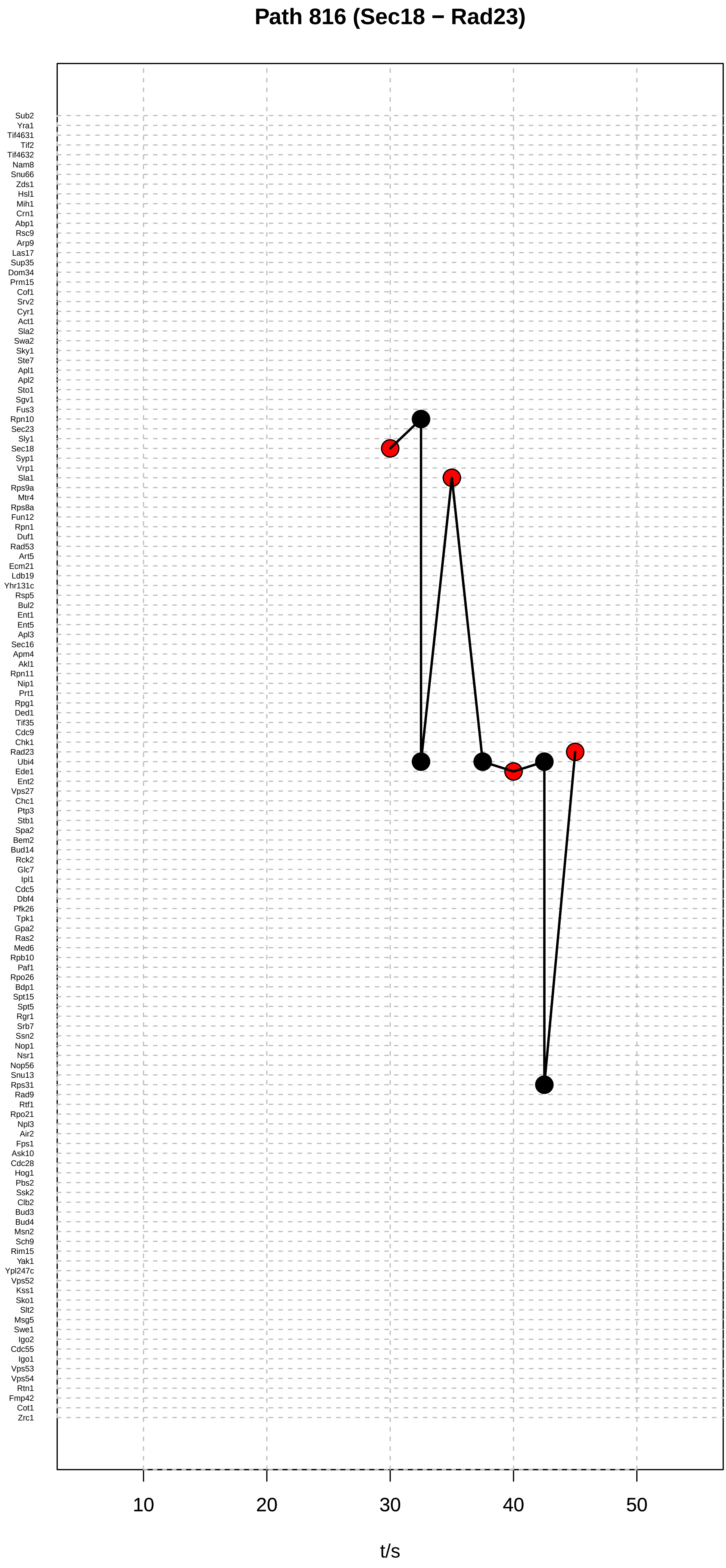

Path 816 (Sec18 − Rad23)
Sub2
Yra1
Tif4631
Tif2
Tif4632
Nam8
Snu66
Zds1
Hsl1
Mih1
Crn1
Abp1
Rsc9
Arp9
Las17
Sup35
Dom34
Prm15
Cof1
Srv2
Cyr1
Act1
Sla2
Swa2
Sky1
Ste7
Apl1
Apl2
Sto1
Sgv1
Fus3
Rpn10
Sec23
Sly1
Sec18
Syp1
Vrp1
Sla1
Rps9a
Mtr4
Rps8a
Fun12
Rpn1
Duf1
Rad53
Art5
Ecm21
Ldb19
Yhr131c
Rsp5
Bul2
Ent1
Ent5
Apl3
Sec16
Apm4
Akl1
Rpn11
Nip1
Prt1
Rpg1
Ded1
Tif35
Cdc9
Chk1
Rad23
Ubi4
Ede1
Ent2
Vps27
Chc1
Ptp3
Stb1
Spa2
Bem2
Bud14
Rck2
Glc7
Ipl1
Cdc5
Dbf4
Pfk26
Tpk1
Gpa2
Ras2
Med6
Rpb10
Paf1
Rpo26
Bdp1
Spt15
Spt5
Rgr1
Srb7
Ssn2
Nop1
Nsr1
Nop56
Snu13
Rps31
Rad9
Rtf1
Rpo21
Npl3
Air2
Fps1
Ask10
Cdc28
Hog1
Pbs2
Ssk2
Clb2
Bud3
Bud4
Msn2
Sch9
Rim15
Yak1
Ypl247c
Vps52
Kss1
Sko1
Slt2
Msg5
Swe1
Igo2
Cdc55
Igo1
Vps53
Vps54
Rtn1
Fmp42
Cot1
Zrc1
10
20
30
40
50
t/s

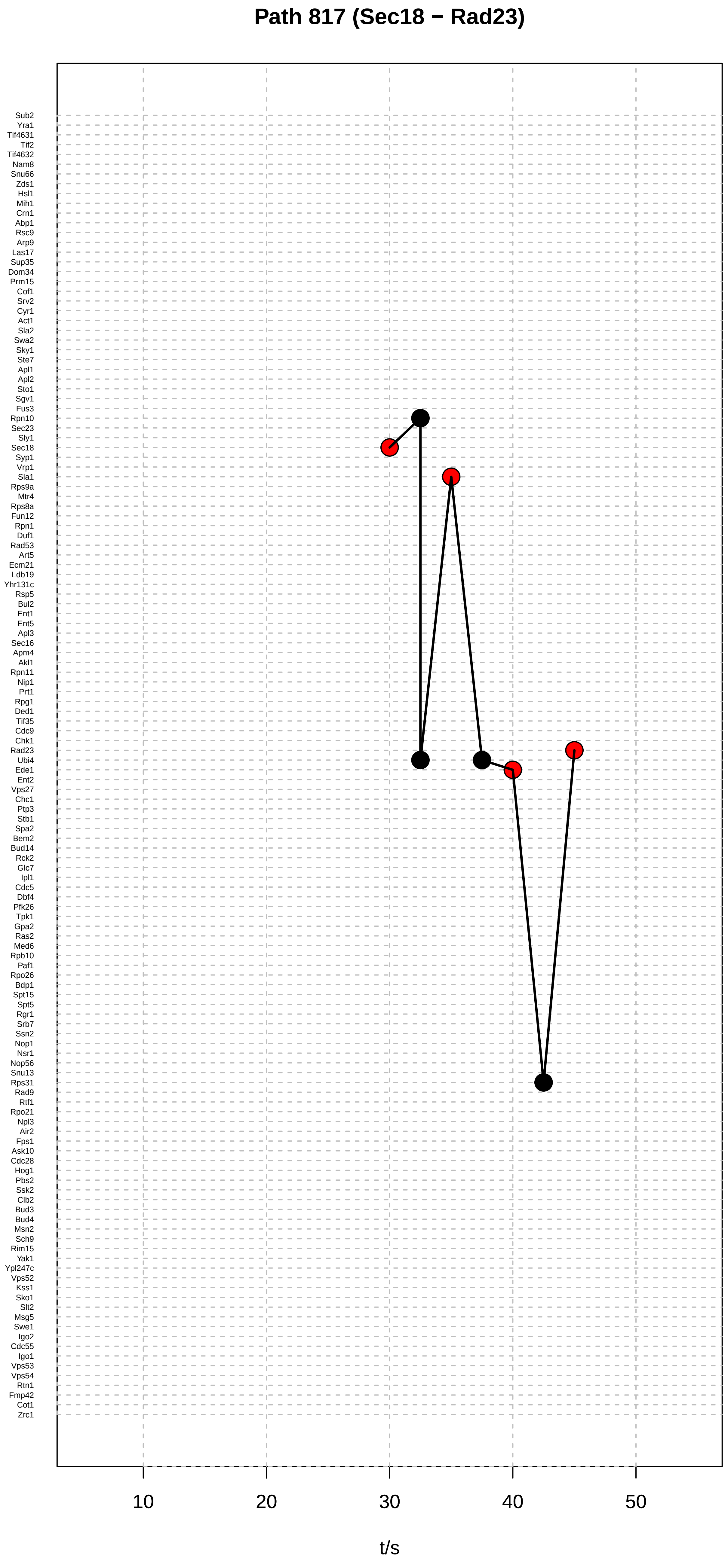

Path 817 (Sec18 − Rad23)
Sub2
Yra1
Tif4631
Tif2
Tif4632
Nam8
Snu66
Zds1
Hsl1
Mih1
Crn1
Abp1
Rsc9
Arp9
Las17
Sup35
Dom34
Prm15
Cof1
Srv2
Cyr1
Act1
Sla2
Swa2
Sky1
Ste7
Apl1
Apl2
Sto1
Sgv1
Fus3
Rpn10
Sec23
Sly1
Sec18
Syp1
Vrp1
Sla1
Rps9a
Mtr4
Rps8a
Fun12
Rpn1
Duf1
Rad53
Art5
Ecm21
Ldb19
Yhr131c
Rsp5
Bul2
Ent1
Ent5
Apl3
Sec16
Apm4
Akl1
Rpn11
Nip1
Prt1
Rpg1
Ded1
Tif35
Cdc9
Chk1
Rad23
Ubi4
Ede1
Ent2
Vps27
Chc1
Ptp3
Stb1
Spa2
Bem2
Bud14
Rck2
Glc7
Ipl1
Cdc5
Dbf4
Pfk26
Tpk1
Gpa2
Ras2
Med6
Rpb10
Paf1
Rpo26
Bdp1
Spt15
Spt5
Rgr1
Srb7
Ssn2
Nop1
Nsr1
Nop56
Snu13
Rps31
Rad9
Rtf1
Rpo21
Npl3
Air2
Fps1
Ask10
Cdc28
Hog1
Pbs2
Ssk2
Clb2
Bud3
Bud4
Msn2
Sch9
Rim15
Yak1
Ypl247c
Vps52
Kss1
Sko1
Slt2
Msg5
Swe1
Igo2
Cdc55
Igo1
Vps53
Vps54
Rtn1
Fmp42
Cot1
Zrc1
10
20
30
40
50
t/s

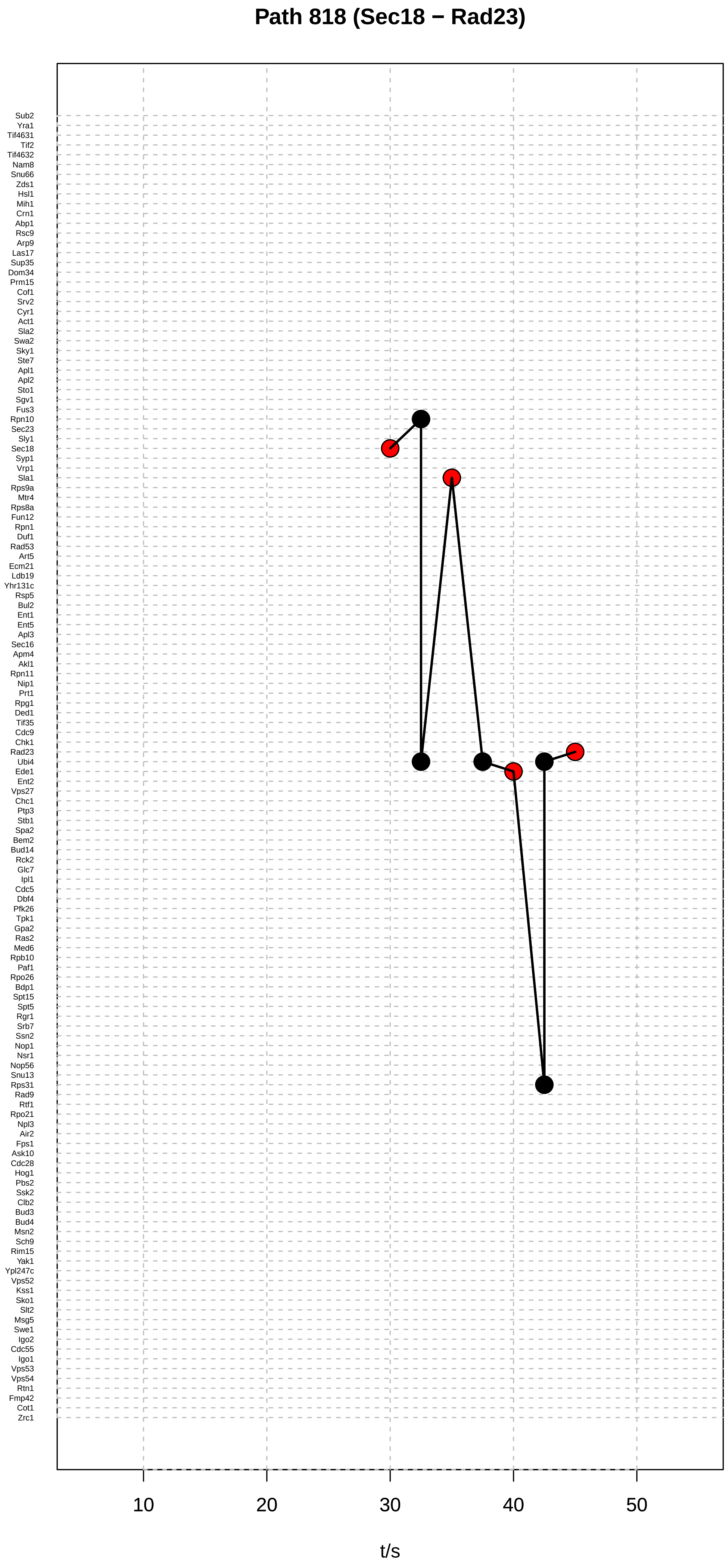

Path 818 (Sec18 − Rad23)
Sub2
Yra1
Tif4631
Tif2
Tif4632
Nam8
Snu66
Zds1
Hsl1
Mih1
Crn1
Abp1
Rsc9
Arp9
Las17
Sup35
Dom34
Prm15
Cof1
Srv2
Cyr1
Act1
Sla2
Swa2
Sky1
Ste7
Apl1
Apl2
Sto1
Sgv1
Fus3
Rpn10
Sec23
Sly1
Sec18
Syp1
Vrp1
Sla1
Rps9a
Mtr4
Rps8a
Fun12
Rpn1
Duf1
Rad53
Art5
Ecm21
Ldb19
Yhr131c
Rsp5
Bul2
Ent1
Ent5
Apl3
Sec16
Apm4
Akl1
Rpn11
Nip1
Prt1
Rpg1
Ded1
Tif35
Cdc9
Chk1
Rad23
Ubi4
Ede1
Ent2
Vps27
Chc1
Ptp3
Stb1
Spa2
Bem2
Bud14
Rck2
Glc7
Ipl1
Cdc5
Dbf4
Pfk26
Tpk1
Gpa2
Ras2
Med6
Rpb10
Paf1
Rpo26
Bdp1
Spt15
Spt5
Rgr1
Srb7
Ssn2
Nop1
Nsr1
Nop56
Snu13
Rps31
Rad9
Rtf1
Rpo21
Npl3
Air2
Fps1
Ask10
Cdc28
Hog1
Pbs2
Ssk2
Clb2
Bud3
Bud4
Msn2
Sch9
Rim15
Yak1
Ypl247c
Vps52
Kss1
Sko1
Slt2
Msg5
Swe1
Igo2
Cdc55
Igo1
Vps53
Vps54
Rtn1
Fmp42
Cot1
Zrc1
10
20
30
40
50
t/s

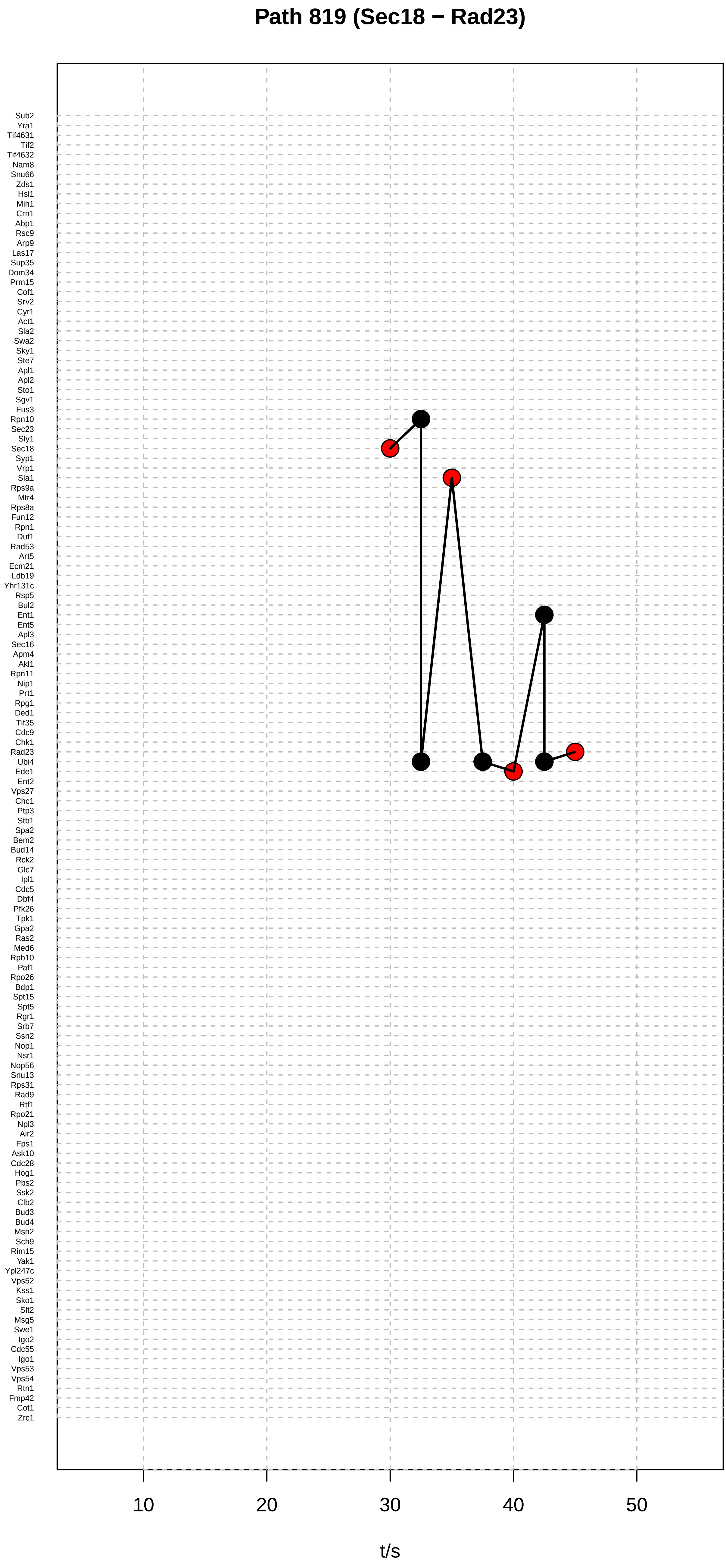
Path 819 (Sec18 − Rad23)
Sub2
Yra1
Tif4631
Tif2
Tif4632
Nam8
Snu66
Zds1
Hsl1
Mih1
Crn1
Abp1
Rsc9
Arp9
Las17
Sup35
Dom34
Prm15
Cof1
Srv2
Cyr1
Act1
Sla2
Swa2
Sky1
Ste7
Apl1
Apl2
Sto1
Sgv1
Fus3
Rpn10
Sec23
Sly1
Sec18
Syp1
Vrp1
Sla1
Rps9a
Mtr4
Rps8a
Fun12
Rpn1
Duf1
Rad53
Art5
Ecm21
Ldb19
Yhr131c
Rsp5
Bul2
Ent1
Ent5
Apl3
Sec16
Apm4
Akl1
Rpn11
Nip1
Prt1
Rpg1
Ded1
Tif35
Cdc9
Chk1
Rad23
Ubi4
Ede1
Ent2
Vps27
Chc1
Ptp3
Stb1
Spa2
Bem2
Bud14
Rck2
Glc7
Ipl1
Cdc5
Dbf4
Pfk26
Tpk1
Gpa2
Ras2
Med6
Rpb10
Paf1
Rpo26
Bdp1
Spt15
Spt5
Rgr1
Srb7
Ssn2
Nop1
Nsr1
Nop56
Snu13
Rps31
Rad9
Rtf1
Rpo21
Npl3
Air2
Fps1
Ask10
Cdc28
Hog1
Pbs2
Ssk2
Clb2
Bud3
Bud4
Msn2
Sch9
Rim15
Yak1
Ypl247c
Vps52
Kss1
Sko1
Slt2
Msg5
Swe1
Igo2
Cdc55
Igo1
Vps53
Vps54
Rtn1
Fmp42
Cot1
Zrc1
10
20
30
40
50
t/s

#### Path 820 (Sec18 − Rad23)

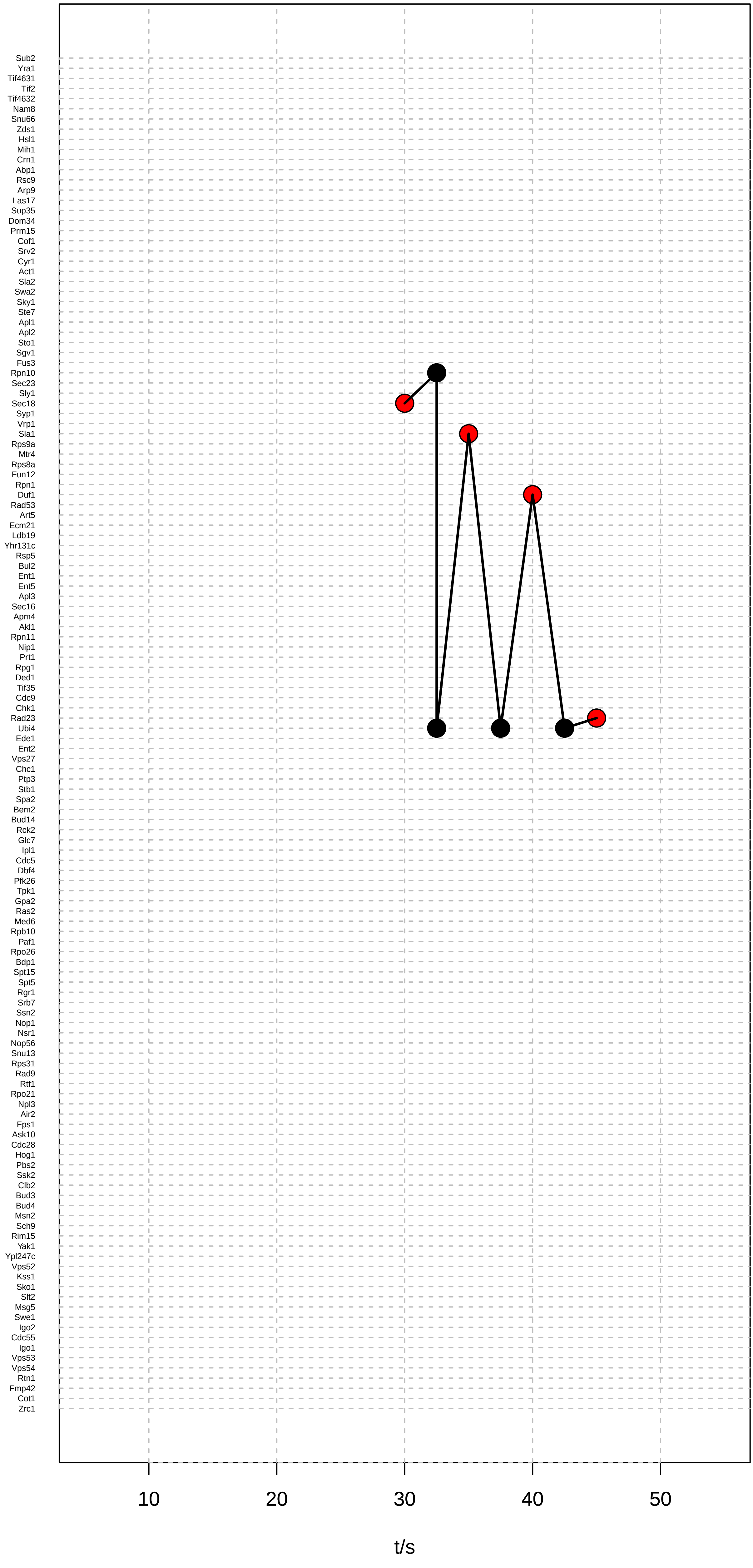

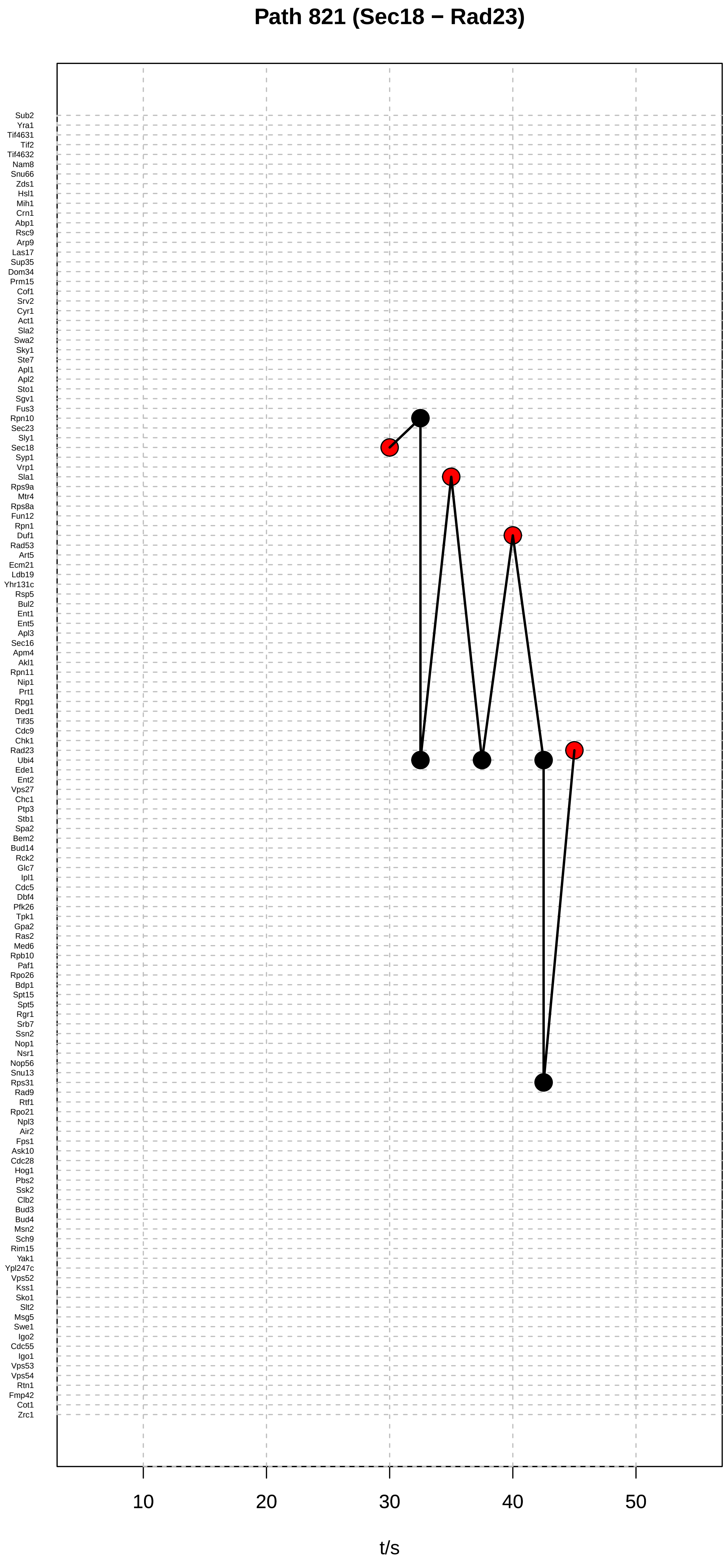
Path 821 (Sec18 − Rad23)
Sub2
Yra1
Tif4631
Tif2
Tif4632
Nam8
Snu66
Zds1
Hsl1
Mih1
Crn1
Abp1
Rsc9
Arp9
Las17
Sup35
Dom34
Prm15
Cof1
Srv2
Cyr1
Act1
Sla2
Swa2
Sky1
Ste7
Apl1
Apl2
Sto1
Sgv1
Fus3
Rpn10
Sec23
Sly1
Sec18
Syp1
Vrp1
Sla1
Rps9a
Mtr4
Rps8a
Fun12
Rpn1
Duf1
Rad53
Art5
Ecm21
Ldb19
Yhr131c
Rsp5
Bul2
Ent1
Ent5
Apl3
Sec16
Apm4
Akl1
Rpn11
Nip1
Prt1
Rpg1
Ded1
Tif35
Cdc9
Chk1
Rad23
Ubi4
Ede1
Ent2
Vps27
Chc1
Ptp3
Stb1
Spa2
Bem2
Bud14
Rck2
Glc7
Ipl1
Cdc5
Dbf4
Pfk26
Tpk1
Gpa2
Ras2
Med6
Rpb10
Paf1
Rpo26
Bdp1
Spt15
Spt5
Rgr1
Srb7
Ssn2
Nop1
Nsr1
Nop56
Snu13
Rps31
Rad9
Rtf1
Rpo21
Npl3
Air2
Fps1
Ask10
Cdc28
Hog1
Pbs2
Ssk2
Clb2
Bud3
Bud4
Msn2
Sch9
Rim15
Yak1
Ypl247c
Vps52
Kss1
Sko1
Slt2
Msg5
Swe1
Igo2
Cdc55
Igo1
Vps53
Vps54
Rtn1
Fmp42
Cot1
Zrc1
10
20
30
40
50
t/s

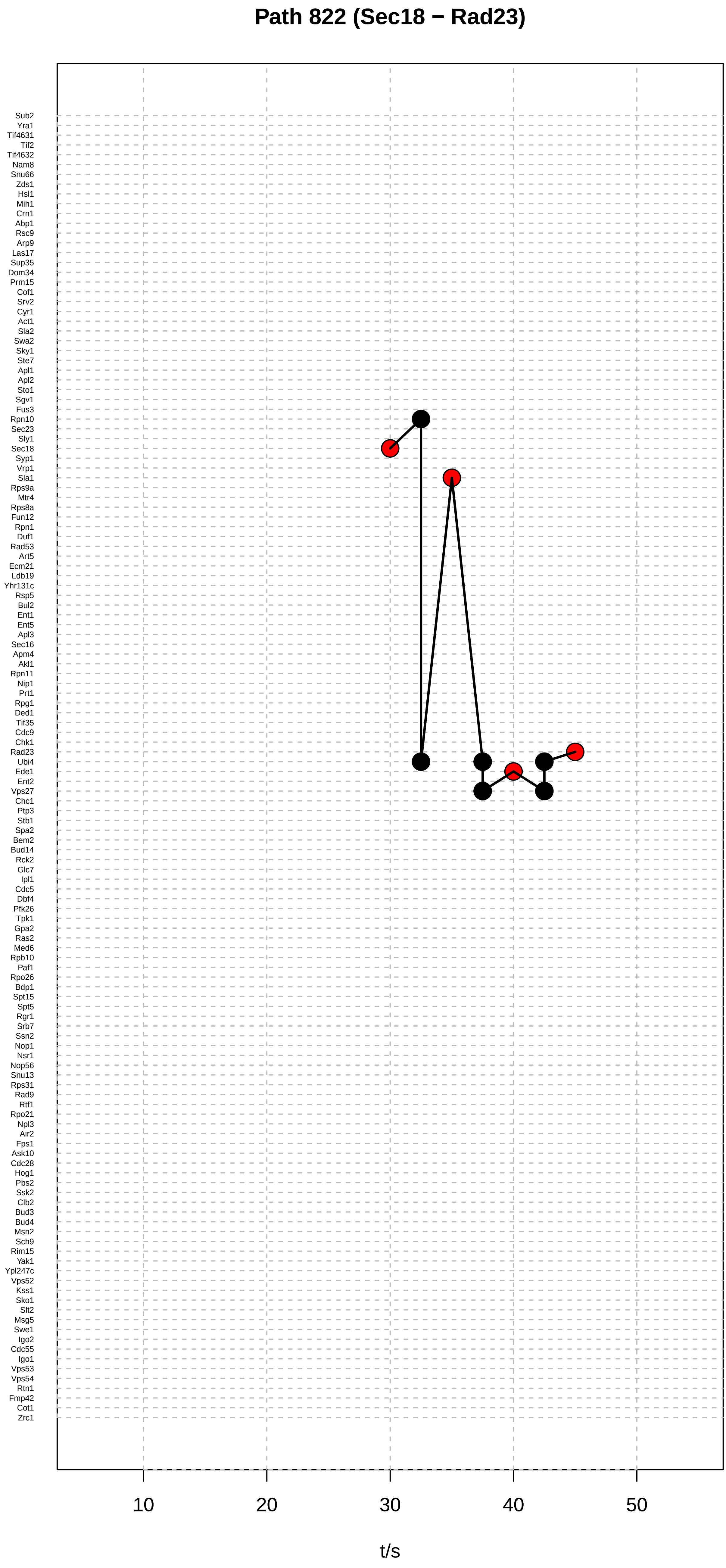

Path 822 (Sec18 − Rad23)
Sub2
Yra1
Tif4631
Tif2
Tif4632
Nam8
Snu66
Zds1
Hsl1
Mih1
Crn1
Abp1
Rsc9
Arp9
Las17
Sup35
Dom34
Prm15
Cof1
Srv2
Cyr1
Act1
Sla2
Swa2
Sky1
Ste7
Apl1
Apl2
Sto1
Sgv1
Fus3
Rpn10
Sec23
Sly1
Sec18
Syp1
Vrp1
Sla1
Rps9a
Mtr4
Rps8a
Fun12
Rpn1
Duf1
Rad53
Art5
Ecm21
Ldb19
Yhr131c
Rsp5
Bul2
Ent1
Ent5
Apl3
Sec16
Apm4
Akl1
Rpn11
Nip1
Prt1
Rpg1
Ded1
Tif35
Cdc9
Chk1
Rad23
Ubi4
Ede1
Ent2
Vps27
Chc1
Ptp3
Stb1
Spa2
Bem2
Bud14
Rck2
Glc7
Ipl1
Cdc5
Dbf4
Pfk26
Tpk1
Gpa2
Ras2
Med6
Rpb10
Paf1
Rpo26
Bdp1
Spt15
Spt5
Rgr1
Srb7
Ssn2
Nop1
Nsr1
Nop56
Snu13
Rps31
Rad9
Rtf1
Rpo21
Npl3
Air2
Fps1
Ask10
Cdc28
Hog1
Pbs2
Ssk2
Clb2
Bud3
Bud4
Msn2
Sch9
Rim15
Yak1
Ypl247c
Vps52
Kss1
Sko1
Slt2
Msg5
Swe1
Igo2
Cdc55
Igo1
Vps53
Vps54
Rtn1
Fmp42
Cot1
Zrc1
10
20
30
40
50
t/s

### Path 823 (Sec18 − Rad23)

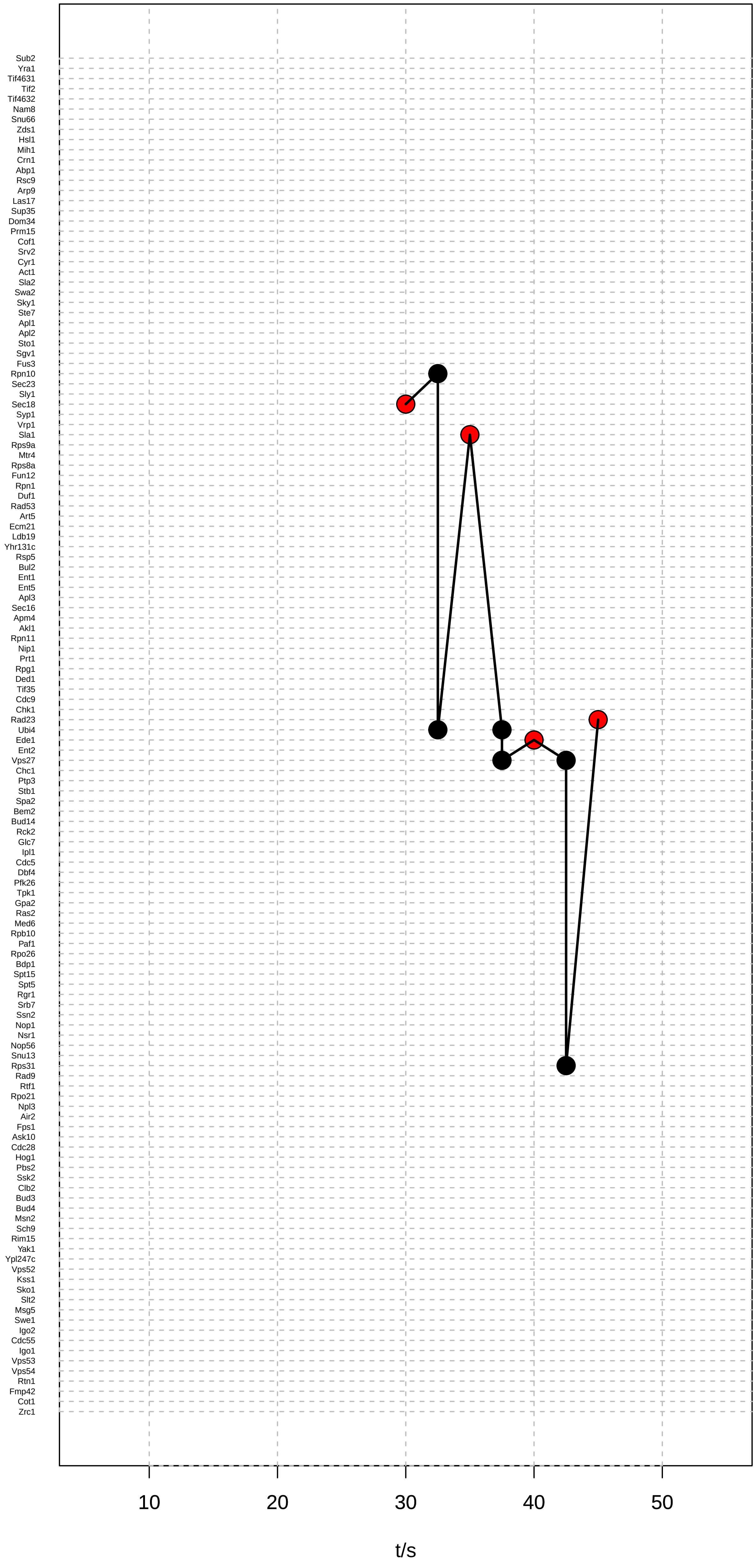

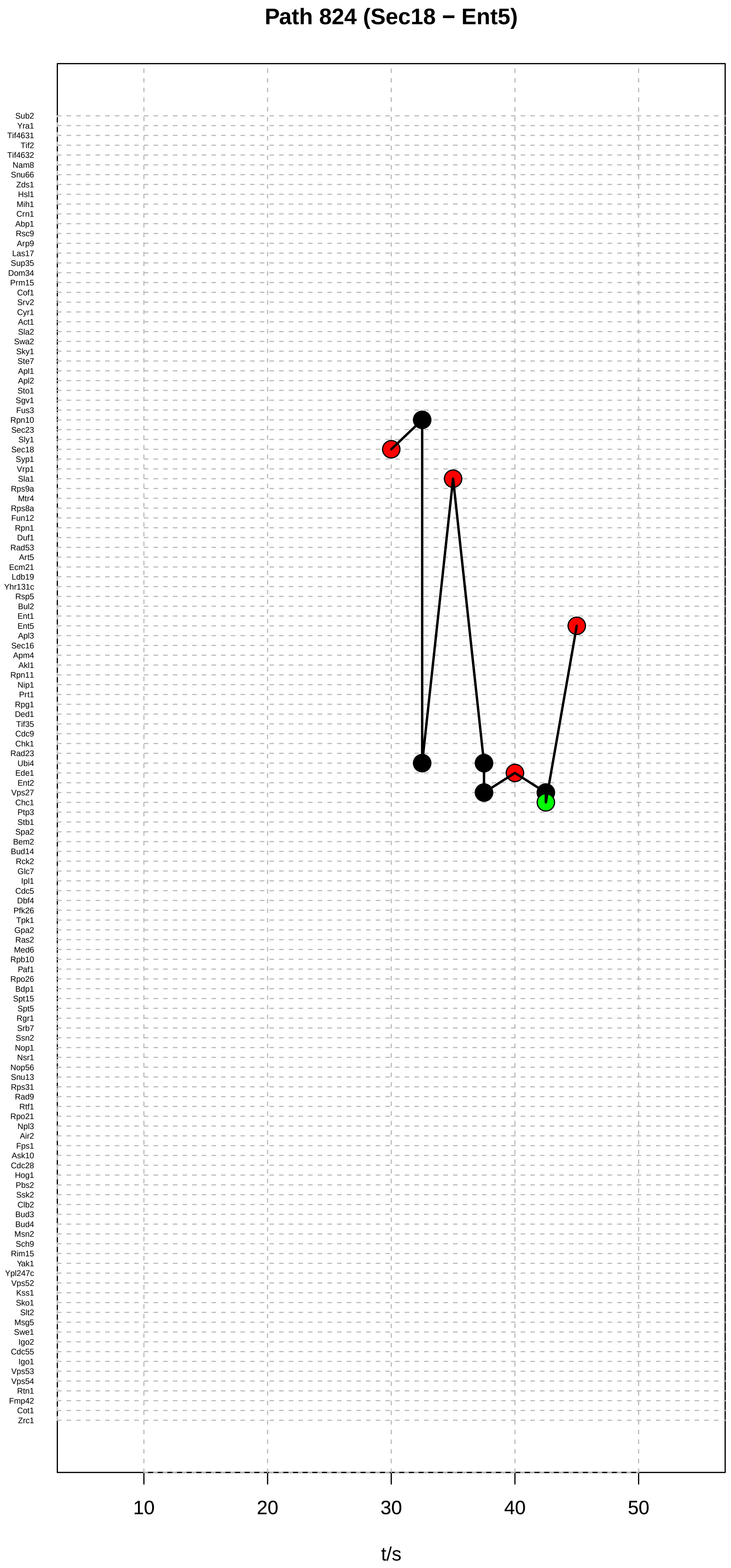

Path 824 (Sec18 − Ent5)
Sub2
Yra1
Tif4631
Tif2
Tif4632
Nam8
Snu66
Zds1
Hsl1
Mih1
Crn1
Abp1
Rsc9
Arp9
Las17
Sup35
Dom34
Prm15
Cof1
Srv2
Cyr1
Act1
Sla2
Swa2
Sky1
Ste7
Apl1
Apl2
Sto1
Sgv1
Fus3
Rpn10
Sec23
Sly1
Sec18
Syp1
Vrp1
Sla1
Rps9a
Mtr4
Rps8a
Fun12
Rpn1
Duf1
Rad53
Art5
Ecm21
Ldb19
Yhr131c
Rsp5
Bul2
Ent1
Ent5
Apl3
Sec16
Apm4
Akl1
Rpn11
Nip1
Prt1
Rpg1
Ded1
Tif35
Cdc9
Chk1
Rad23
Ubi4
Ede1
Ent2
Vps27
Chc1
Ptp3
Stb1
Spa2
Bem2
Bud14
Rck2
Glc7
Ipl1
Cdc5
Dbf4
Pfk26
Tpk1
Gpa2
Ras2
Med6
Rpb10
Paf1
Rpo26
Bdp1
Spt15
Spt5
Rgr1
Srb7
Ssn2
Nop1
Nsr1
Nop56
Snu13
Rps31
Rad9
Rtf1
Rpo21
Npl3
Air2
Fps1
Ask10
Cdc28
Hog1
Pbs2
Ssk2
Clb2
Bud3
Bud4
Msn2
Sch9
Rim15
Yak1
Ypl247c
Vps52
Kss1
Sko1
Slt2
Msg5
Swe1
Igo2
Cdc55
Igo1
Vps53
Vps54
Rtn1
Fmp42
Cot1
Zrc1
10
20
30
40
50
t/s

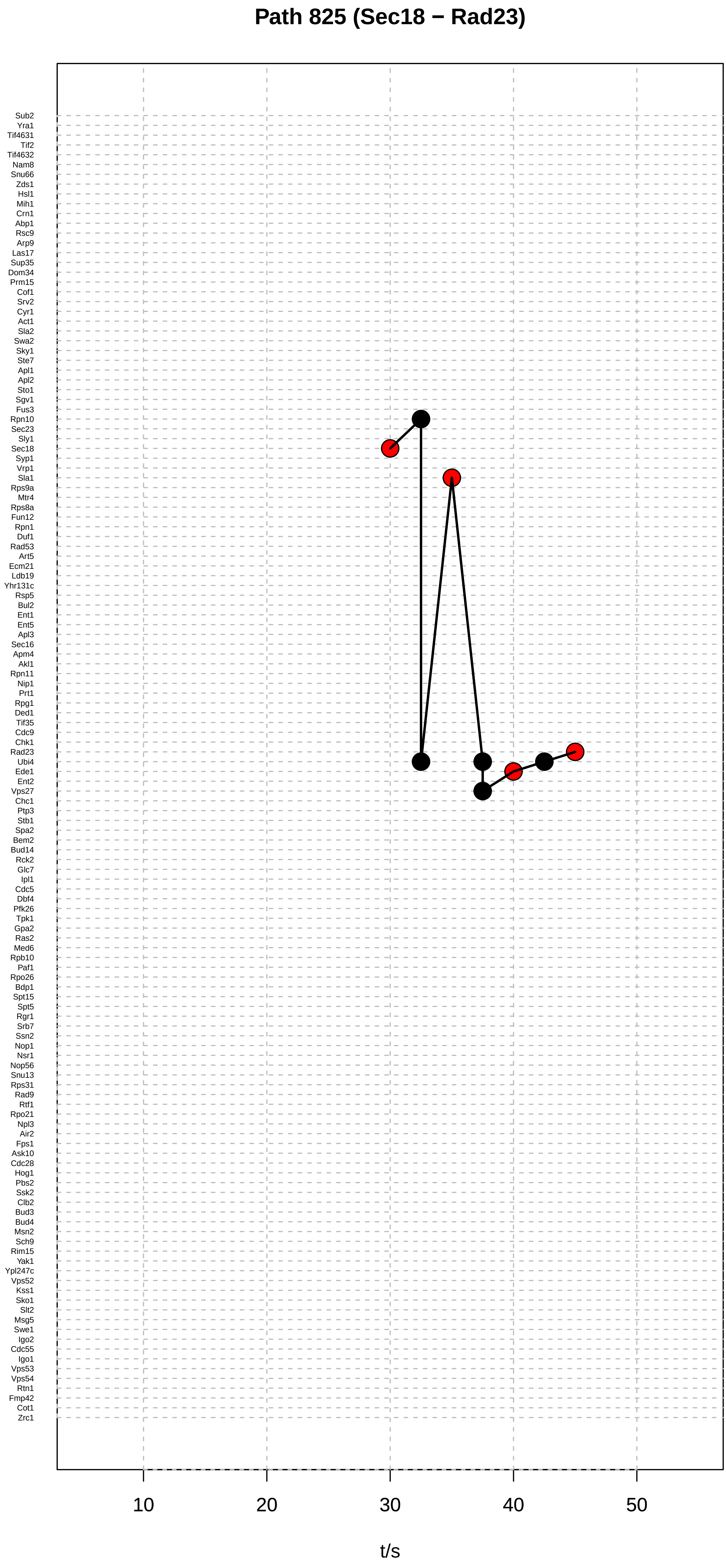
Path 825 (Sec18 − Rad23)
Sub2
Yra1
Tif4631
Tif2
Tif4632
Nam8
Snu66
Zds1
Hsl1
Mih1
Crn1
Abp1
Rsc9
Arp9
Las17
Sup35
Dom34
Prm15
Cof1
Srv2
Cyr1
Act1
Sla2
Swa2
Sky1
Ste7
Apl1
Apl2
Sto1
Sgv1
Fus3
Rpn10
Sec23
Sly1
Sec18
Syp1
Vrp1
Sla1
Rps9a
Mtr4
Rps8a
Fun12
Rpn1
Duf1
Rad53
Art5
Ecm21
Ldb19
Yhr131c
Rsp5
Bul2
Ent1
Ent5
Apl3
Sec16
Apm4
Akl1
Rpn11
Nip1
Prt1
Rpg1
Ded1
Tif35
Cdc9
Chk1
Rad23
Ubi4
Ede1
Ent2
Vps27
Chc1
Ptp3
Stb1
Spa2
Bem2
Bud14
Rck2
Glc7
Ipl1
Cdc5
Dbf4
Pfk26
Tpk1
Gpa2
Ras2
Med6
Rpb10
Paf1
Rpo26
Bdp1
Spt15
Spt5
Rgr1
Srb7
Ssn2
Nop1
Nsr1
Nop56
Snu13
Rps31
Rad9
Rtf1
Rpo21
Npl3
Air2
Fps1
Ask10
Cdc28
Hog1
Pbs2
Ssk2
Clb2
Bud3
Bud4
Msn2
Sch9
Rim15
Yak1
Ypl247c
Vps52
Kss1
Sko1
Slt2
Msg5
Swe1
Igo2
Cdc55
Igo1
Vps53
Vps54
Rtn1
Fmp42
Cot1
Zrc1
10
20
30
40
50
t/s

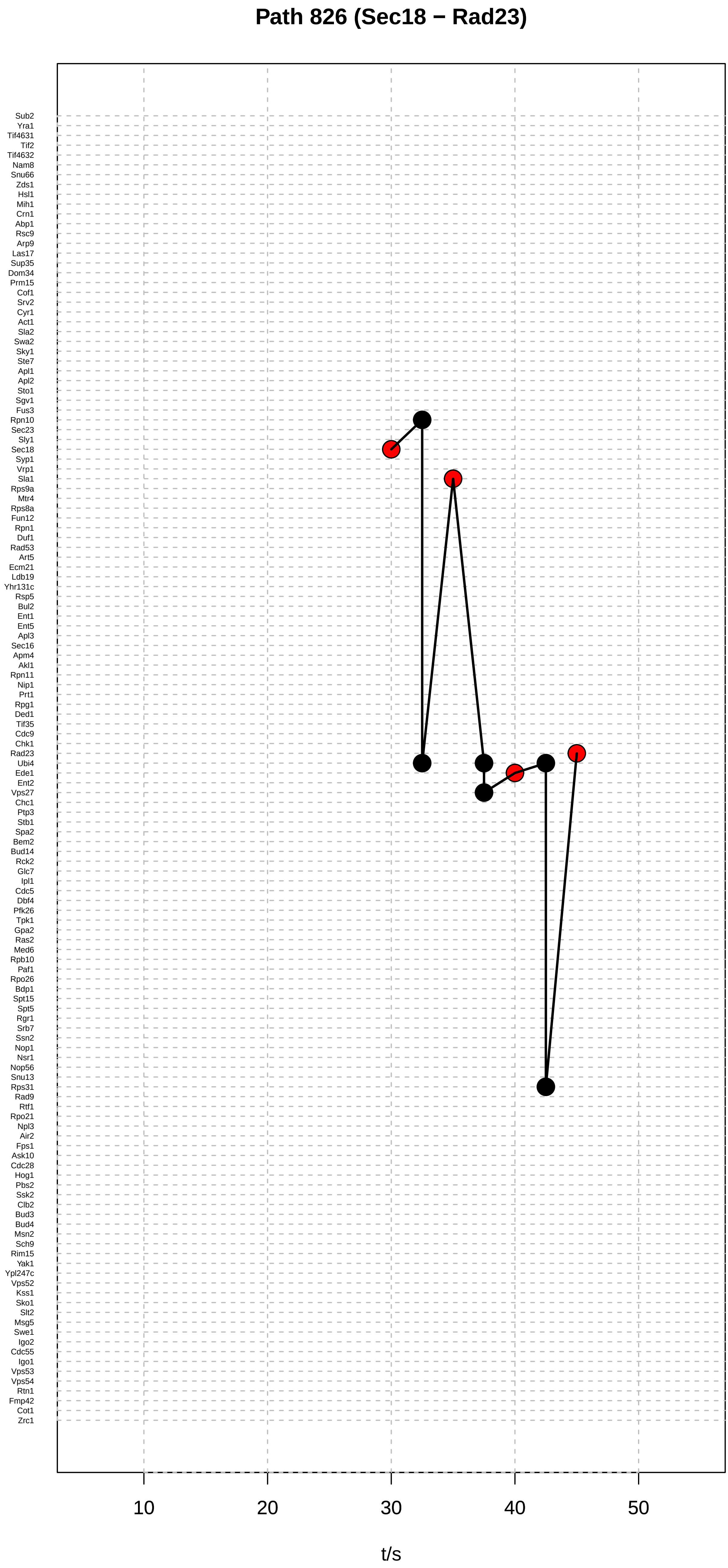

Path 826 (Sec18 − Rad23)
Sub2
Yra1
Tif4631
Tif2
Tif4632
Nam8
Snu66
Zds1
Hsl1
Mih1
Crn1
Abp1
Rsc9
Arp9
Las17
Sup35
Dom34
Prm15
Cof1
Srv2
Cyr1
Act1
Sla2
Swa2
Sky1
Ste7
Apl1
Apl2
Sto1
Sgv1
Fus3
Rpn10
Sec23
Sly1
Sec18
Syp1
Vrp1
Sla1
Rps9a
Mtr4
Rps8a
Fun12
Rpn1
Duf1
Rad53
Art5
Ecm21
Ldb19
Yhr131c
Rsp5
Bul2
Ent1
Ent5
Apl3
Sec16
Apm4
Akl1
Rpn11
Nip1
Prt1
Rpg1
Ded1
Tif35
Cdc9
Chk1
Rad23
Ubi4
Ede1
Ent2
Vps27
Chc1
Ptp3
Stb1
Spa2
Bem2
Bud14
Rck2
Glc7
Ipl1
Cdc5
Dbf4
Pfk26
Tpk1
Gpa2
Ras2
Med6
Rpb10
Paf1
Rpo26
Bdp1
Spt15
Spt5
Rgr1
Srb7
Ssn2
Nop1
Nsr1
Nop56
Snu13
Rps31
Rad9
Rtf1
Rpo21
Npl3
Air2
Fps1
Ask10
Cdc28
Hog1
Pbs2
Ssk2
Clb2
Bud3
Bud4
Msn2
Sch9
Rim15
Yak1
Ypl247c
Vps52
Kss1
Sko1
Slt2
Msg5
Swe1
Igo2
Cdc55
Igo1
Vps53
Vps54
Rtn1
Fmp42
Cot1
Zrc1
10
20
30
40
50
t/s

### Path 827 (Sec18 − Rad23)

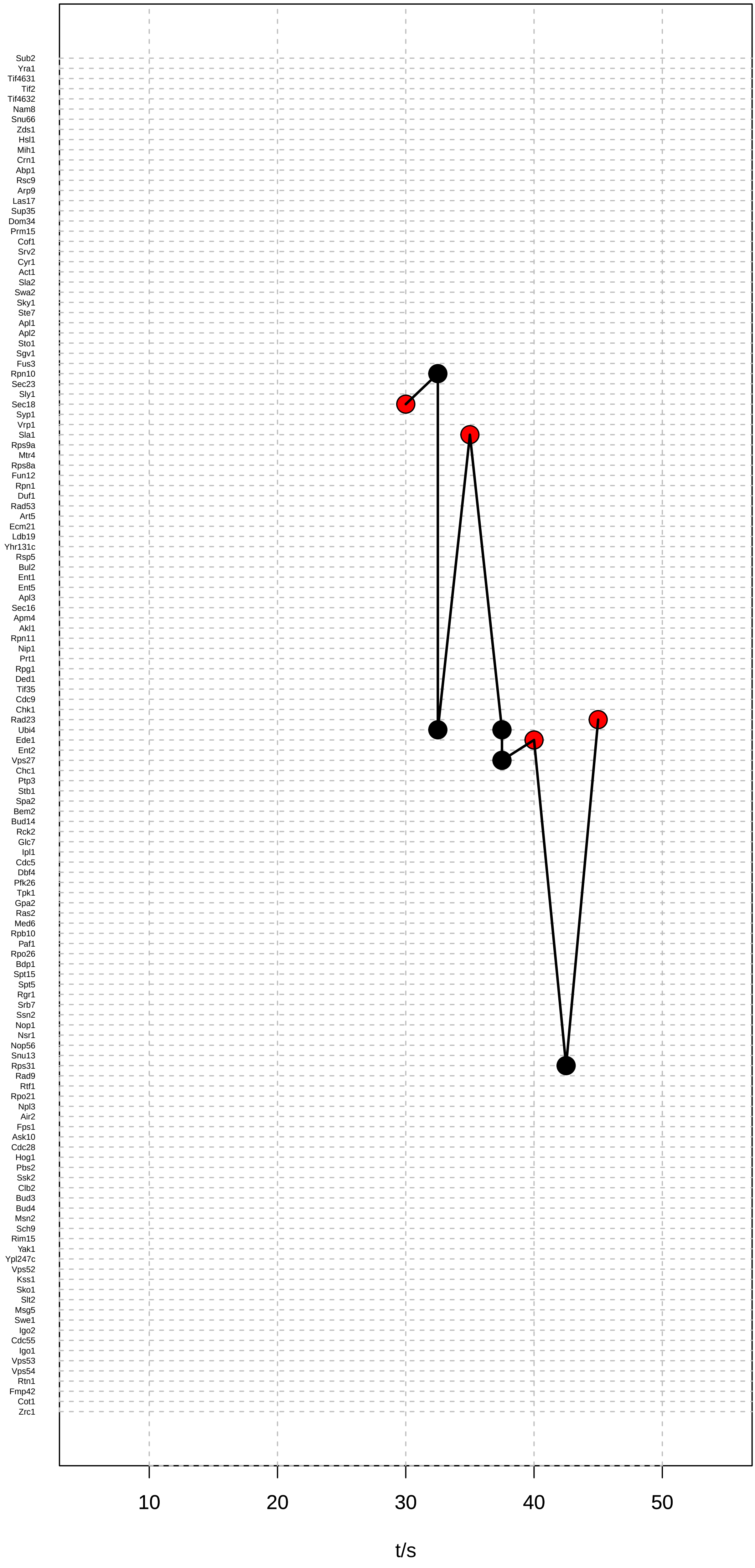

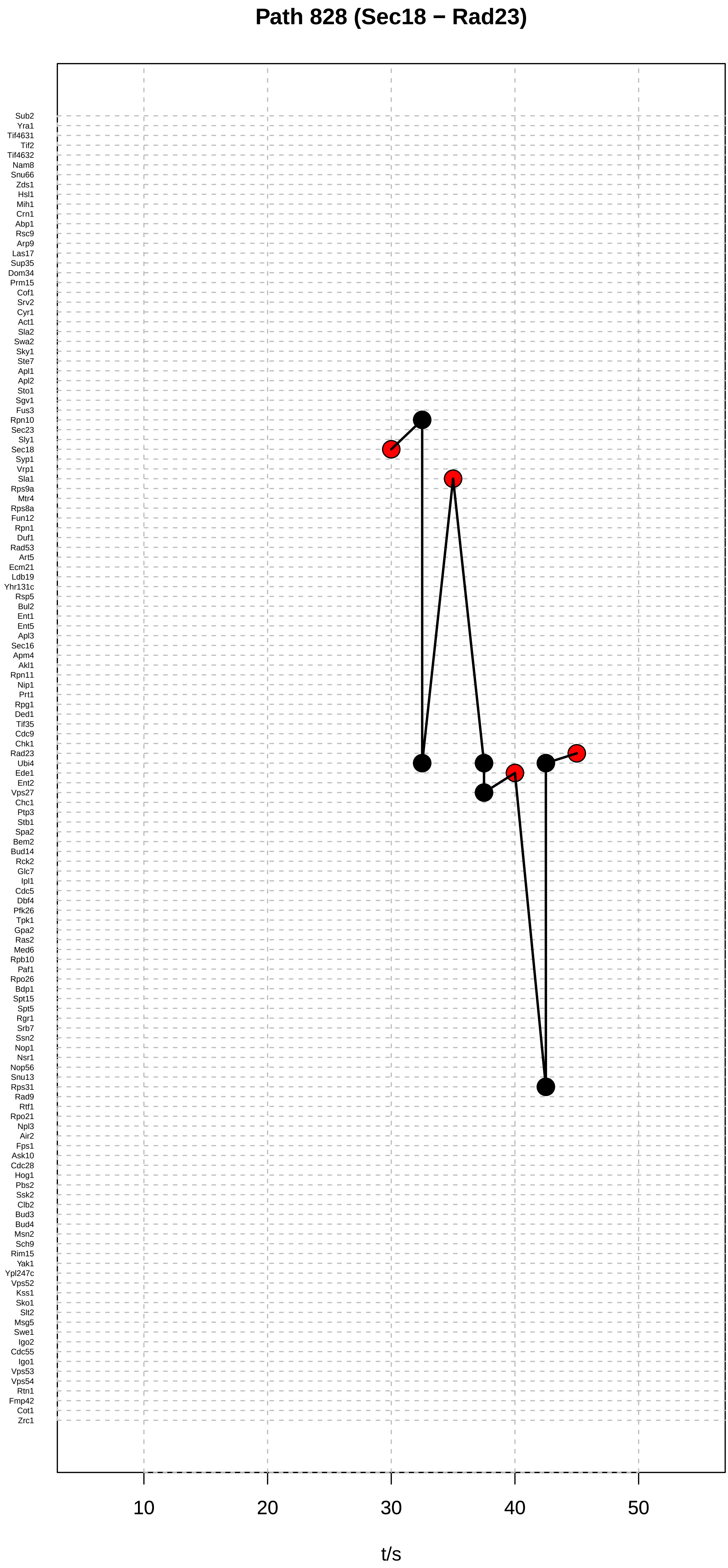

Path 828 (Sec18 − Rad23)
Sub2
Yra1
Tif4631
Tif2
Tif4632
Nam8
Snu66
Zds1
Hsl1
Mih1
Crn1
Abp1
Rsc9
Arp9
Las17
Sup35
Dom34
Prm15
Cof1
Srv2
Cyr1
Act1
Sla2
Swa2
Sky1
Ste7
Apl1
Apl2
Sto1
Sgv1
Fus3
Rpn10
Sec23
Sly1
Sec18
Syp1
Vrp1
Sla1
Rps9a
Mtr4
Rps8a
Fun12
Rpn1
Duf1
Rad53
Art5
Ecm21
Ldb19
Yhr131c
Rsp5
Bul2
Ent1
Ent5
Apl3
Sec16
Apm4
Akl1
Rpn11
Nip1
Prt1
Rpg1
Ded1
Tif35
Cdc9
Chk1
Rad23
Ubi4
Ede1
Ent2
Vps27
Chc1
Ptp3
Stb1
Spa2
Bem2
Bud14
Rck2
Glc7
Ipl1
Cdc5
Dbf4
Pfk26
Tpk1
Gpa2
Ras2
Med6
Rpb10
Paf1
Rpo26
Bdp1
Spt15
Spt5
Rgr1
Srb7
Ssn2
Nop1
Nsr1
Nop56
Snu13
Rps31
Rad9
Rtf1
Rpo21
Npl3
Air2
Fps1
Ask10
Cdc28
Hog1
Pbs2
Ssk2
Clb2
Bud3
Bud4
Msn2
Sch9
Rim15
Yak1
Ypl247c
Vps52
Kss1
Sko1
Slt2
Msg5
Swe1
Igo2
Cdc55
Igo1
Vps53
Vps54
Rtn1
Fmp42
Cot1
Zrc1
10
20
30
40
50
t/s

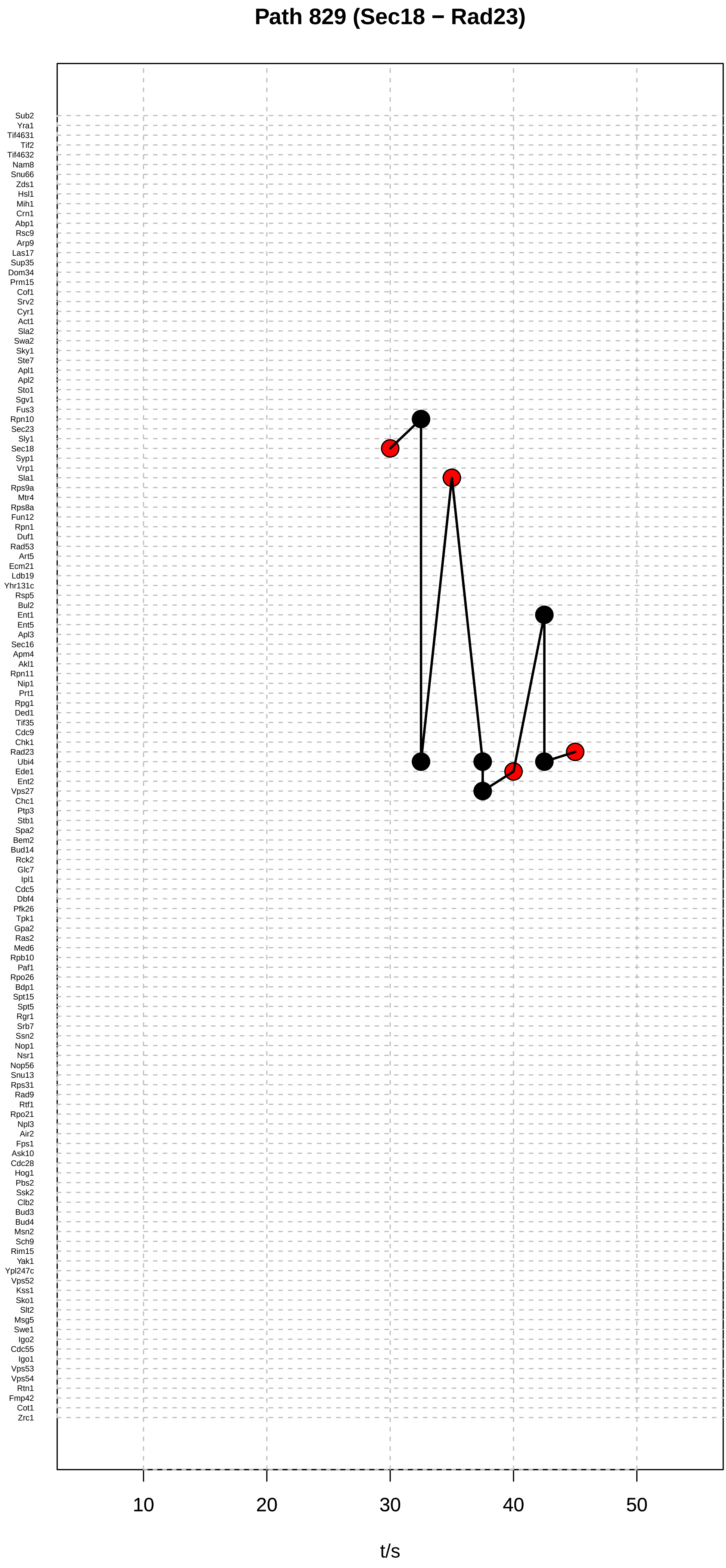

Path 829 (Sec18 − Rad23)
Sub2
Yra1
Tif4631
Tif2
Tif4632
Nam8
Snu66
Zds1
Hsl1
Mih1
Crn1
Abp1
Rsc9
Arp9
Las17
Sup35
Dom34
Prm15
Cof1
Srv2
Cyr1
Act1
Sla2
Swa2
Sky1
Ste7
Apl1
Apl2
Sto1
Sgv1
Fus3
Rpn10
Sec23
Sly1
Sec18
Syp1
Vrp1
Sla1
Rps9a
Mtr4
Rps8a
Fun12
Rpn1
Duf1
Rad53
Art5
Ecm21
Ldb19
Yhr131c
Rsp5
Bul2
Ent1
Ent5
Apl3
Sec16
Apm4
Akl1
Rpn11
Nip1
Prt1
Rpg1
Ded1
Tif35
Cdc9
Chk1
Rad23
Ubi4
Ede1
Ent2
Vps27
Chc1
Ptp3
Stb1
Spa2
Bem2
Bud14
Rck2
Glc7
Ipl1
Cdc5
Dbf4
Pfk26
Tpk1
Gpa2
Ras2
Med6
Rpb10
Paf1
Rpo26
Bdp1
Spt15
Spt5
Rgr1
Srb7
Ssn2
Nop1
Nsr1
Nop56
Snu13
Rps31
Rad9
Rtf1
Rpo21
Npl3
Air2
Fps1
Ask10
Cdc28
Hog1
Pbs2
Ssk2
Clb2
Bud3
Bud4
Msn2
Sch9
Rim15
Yak1
Ypl247c
Vps52
Kss1
Sko1
Slt2
Msg5
Swe1
Igo2
Cdc55
Igo1
Vps53
Vps54
Rtn1
Fmp42
Cot1
Zrc1
10
20
30
40
50
t/s

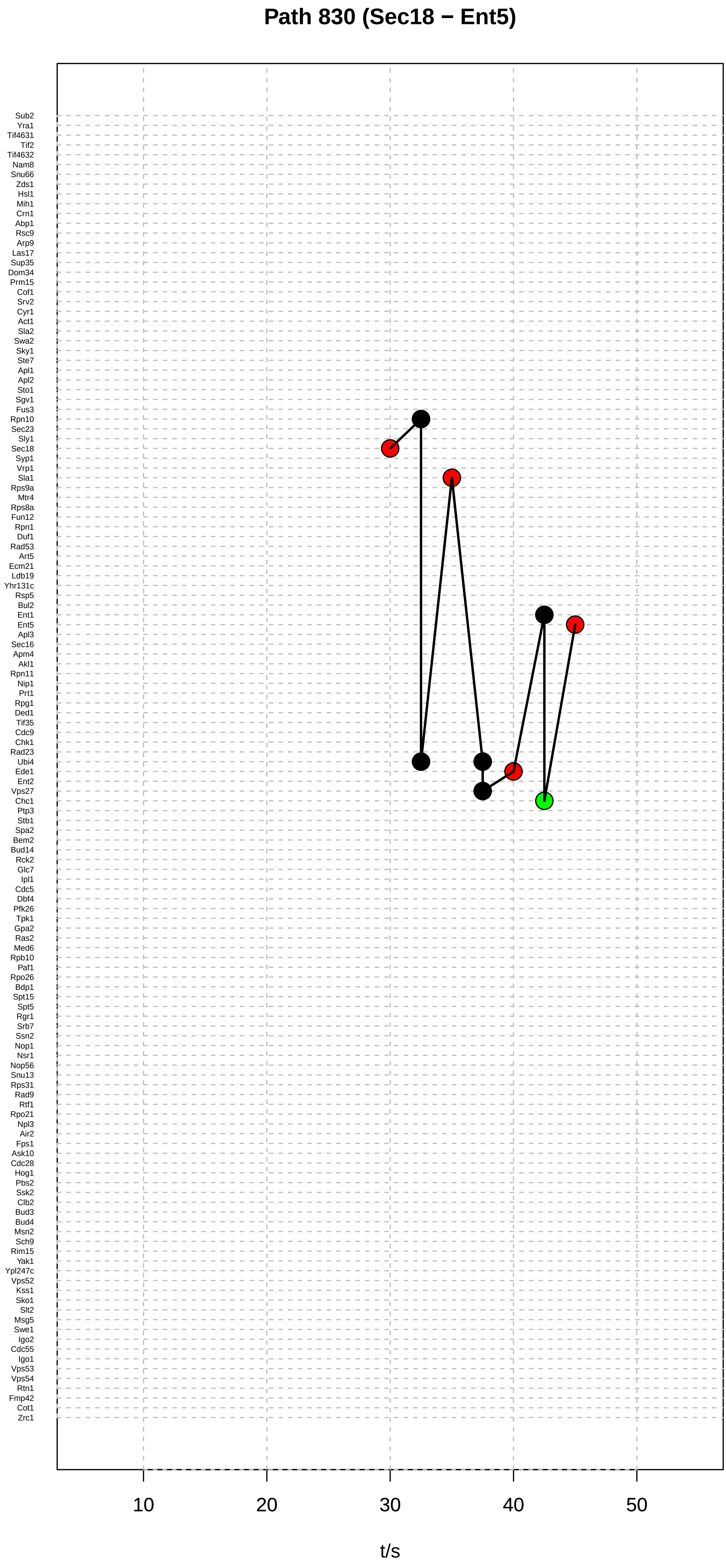

Path 830 (Sec18 − Ent5)
Sub2
Yra1
Tif4631
Tif2
Tif4632
Nam8
Snu66
Zds1
Hsl1
Mih1
Crn1
Abp1
Rsc9
Arp9
Las17
Sup35
Dom34
Prm15
Cof1
Srv2
Cyr1
Act1
Sla2
Swa2
Sky1
Ste7
Apl1
Apl2
Sto1
Sgv1
Fus3
Rpn10
Sec23
Sly1
Sec18
Syp1
Vrp1
Sla1
Rps9a
Mtr4
Rps8a
Fun12
Rpn1
Duf1
Rad53
Art5
Ecm21
Ldb19
Yhr131c
Rsp5
Bul2
Ent1
Ent5
Apl3
Sec16
Apm4
Akl1
Rpn11
Nip1
Prt1
Rpg1
Ded1
Tif35
Cdc9
Chk1
Rad23
Ubi4
Ede1
Ent2
Vps27
Chc1
Ptp3
Stb1
Spa2
Bem2
Bud14
Rck2
Glc7
Ipl1
Cdc5
Dbf4
Pfk26
Tpk1
Gpa2
Ras2
Med6
Rpb10
Paf1
Rpo26
Bdp1
Spt15
Spt5
Rgr1
Srb7
Ssn2
Nop1
Nsr1
Nop56
Snu13
Rps31
Rad9
Rtf1
Rpo21
Npl3
Air2
Fps1
Ask10
Cdc28
Hog1
Pbs2
Ssk2
Clb2
Bud3
Bud4
Msn2
Sch9
Rim15
Yak1
Ypl247c
Vps52
Kss1
Sko1
Slt2
Msg5
Swe1
Igo2
Cdc55
Igo1
Vps53
Vps54
Rtn1
Fmp42
Cot1
Zrc1
10
20
30
40
50
t/s

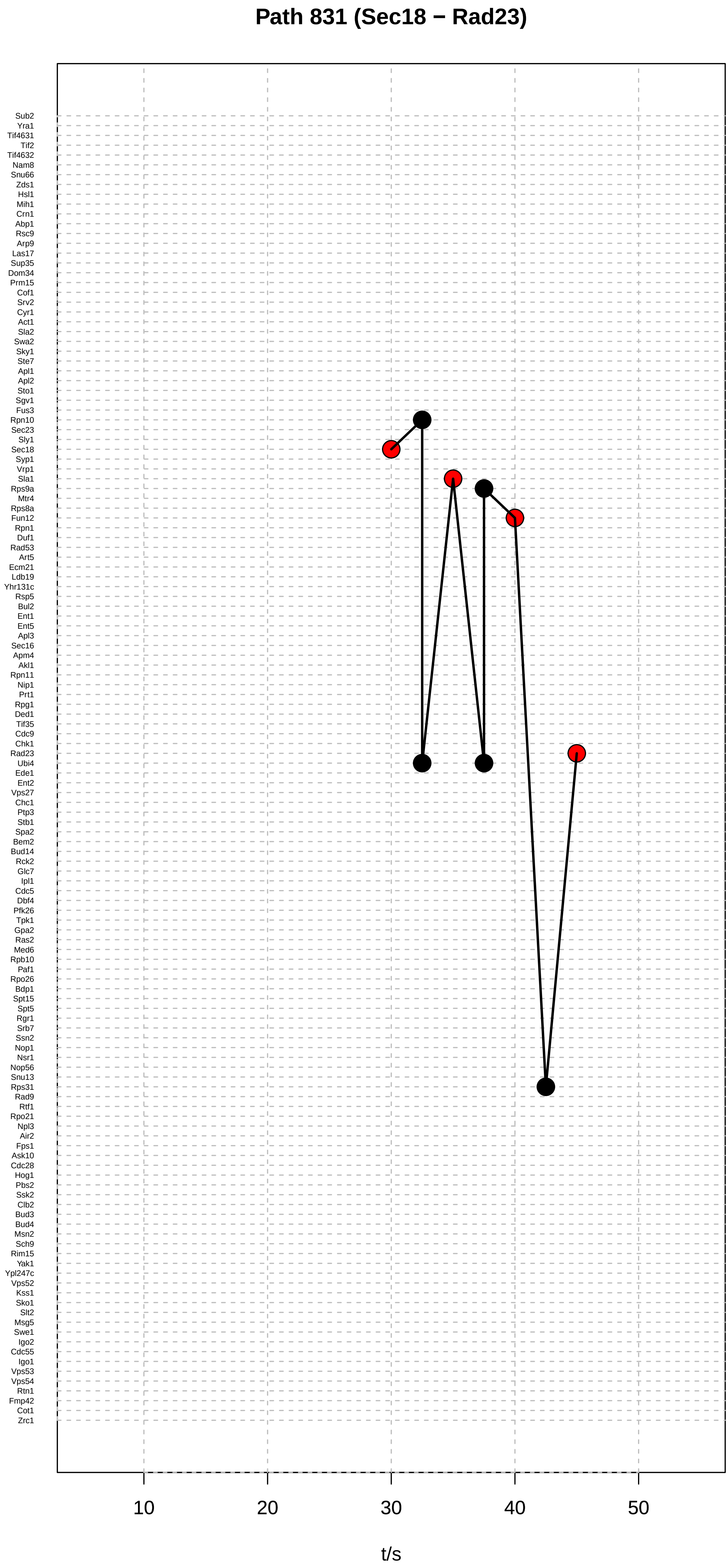
Path 831 (Sec18 − Rad23)
Sub2
Yra1
Tif4631
Tif2
Tif4632
Nam8
Snu66
Zds1
Hsl1
Mih1
Crn1
Abp1
Rsc9
Arp9
Las17
Sup35
Dom34
Prm15
Cof1
Srv2
Cyr1
Act1
Sla2
Swa2
Sky1
Ste7
Apl1
Apl2
Sto1
Sgv1
Fus3
Rpn10
Sec23
Sly1
Sec18
Syp1
Vrp1
Sla1
Rps9a
Mtr4
Rps8a
Fun12
Rpn1
Duf1
Rad53
Art5
Ecm21
Ldb19
Yhr131c
Rsp5
Bul2
Ent1
Ent5
Apl3
Sec16
Apm4
Akl1
Rpn11
Nip1
Prt1
Rpg1
Ded1
Tif35
Cdc9
Chk1
Rad23
Ubi4
Ede1
Ent2
Vps27
Chc1
Ptp3
Stb1
Spa2
Bem2
Bud14
Rck2
Glc7
Ipl1
Cdc5
Dbf4
Pfk26
Tpk1
Gpa2
Ras2
Med6
Rpb10
Paf1
Rpo26
Bdp1
Spt15
Spt5
Rgr1
Srb7
Ssn2
Nop1
Nsr1
Nop56
Snu13
Rps31
Rad9
Rtf1
Rpo21
Npl3
Air2
Fps1
Ask10
Cdc28
Hog1
Pbs2
Ssk2
Clb2
Bud3
Bud4
Msn2
Sch9
Rim15
Yak1
Ypl247c
Vps52
Kss1
Sko1
Slt2
Msg5
Swe1
Igo2
Cdc55
Igo1
Vps53
Vps54
Rtn1
Fmp42
Cot1
Zrc1
10
20
30
40
50
t/s

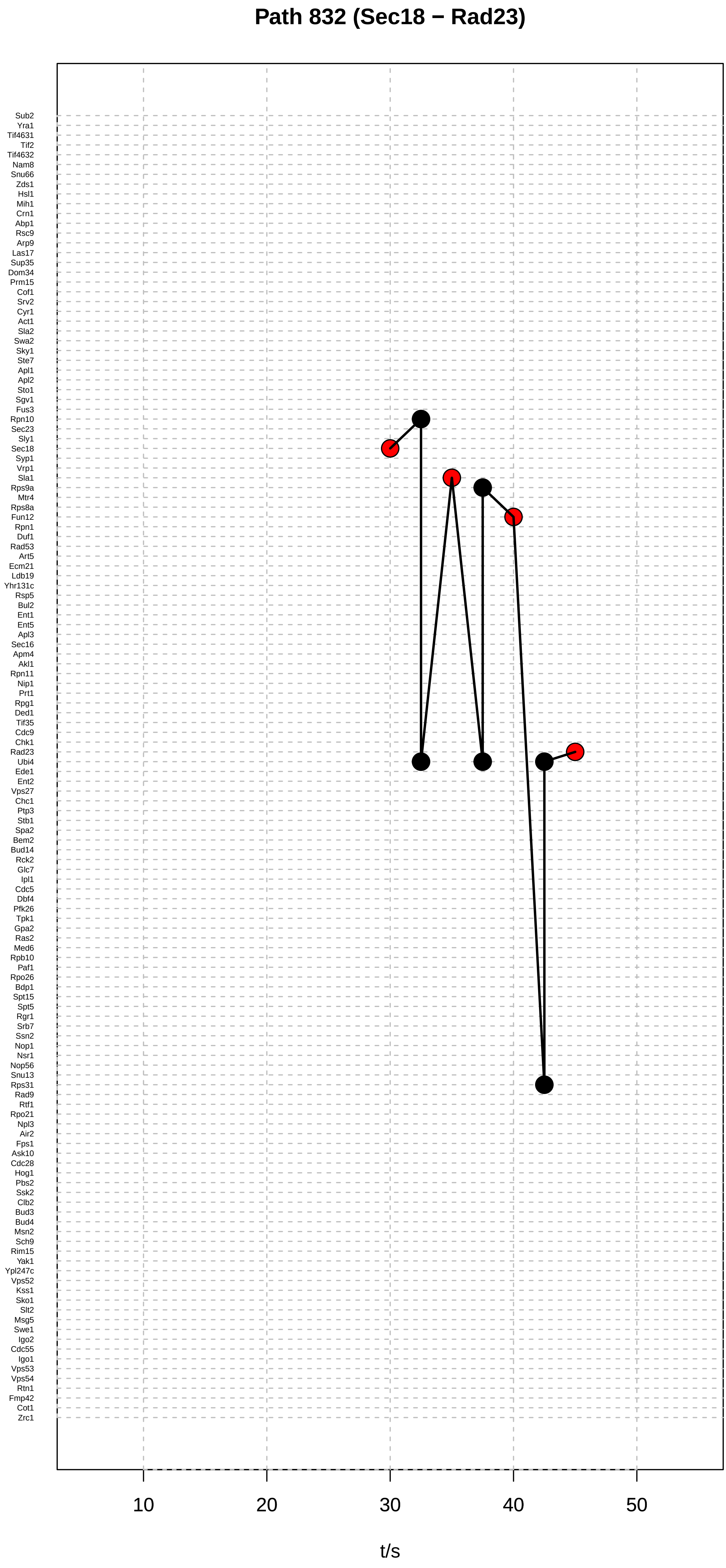
Path 832 (Sec18 − Rad23)
Sub2
Yra1
Tif4631
Tif2
Tif4632
Nam8
Snu66
Zds1
Hsl1
Mih1
Crn1
Abp1
Rsc9
Arp9
Las17
Sup35
Dom34
Prm15
Cof1
Srv2
Cyr1
Act1
Sla2
Swa2
Sky1
Ste7
Apl1
Apl2
Sto1
Sgv1
Fus3
Rpn10
Sec23
Sly1
Sec18
Syp1
Vrp1
Sla1
Rps9a
Mtr4
Rps8a
Fun12
Rpn1
Duf1
Rad53
Art5
Ecm21
Ldb19
Yhr131c
Rsp5
Bul2
Ent1
Ent5
Apl3
Sec16
Apm4
Akl1
Rpn11
Nip1
Prt1
Rpg1
Ded1
Tif35
Cdc9
Chk1
Rad23
Ubi4
Ede1
Ent2
Vps27
Chc1
Ptp3
Stb1
Spa2
Bem2
Bud14
Rck2
Glc7
Ipl1
Cdc5
Dbf4
Pfk26
Tpk1
Gpa2
Ras2
Med6
Rpb10
Paf1
Rpo26
Bdp1
Spt15
Spt5
Rgr1
Srb7
Ssn2
Nop1
Nsr1
Nop56
Snu13
Rps31
Rad9
Rtf1
Rpo21
Npl3
Air2
Fps1
Ask10
Cdc28
Hog1
Pbs2
Ssk2
Clb2
Bud3
Bud4
Msn2
Sch9
Rim15
Yak1
Ypl247c
Vps52
Kss1
Sko1
Slt2
Msg5
Swe1
Igo2
Cdc55
Igo1
Vps53
Vps54
Rtn1
Fmp42
Cot1
Zrc1
10
20
30
40
50
t/s

**Path 833 (Sec18 − Rad23)**

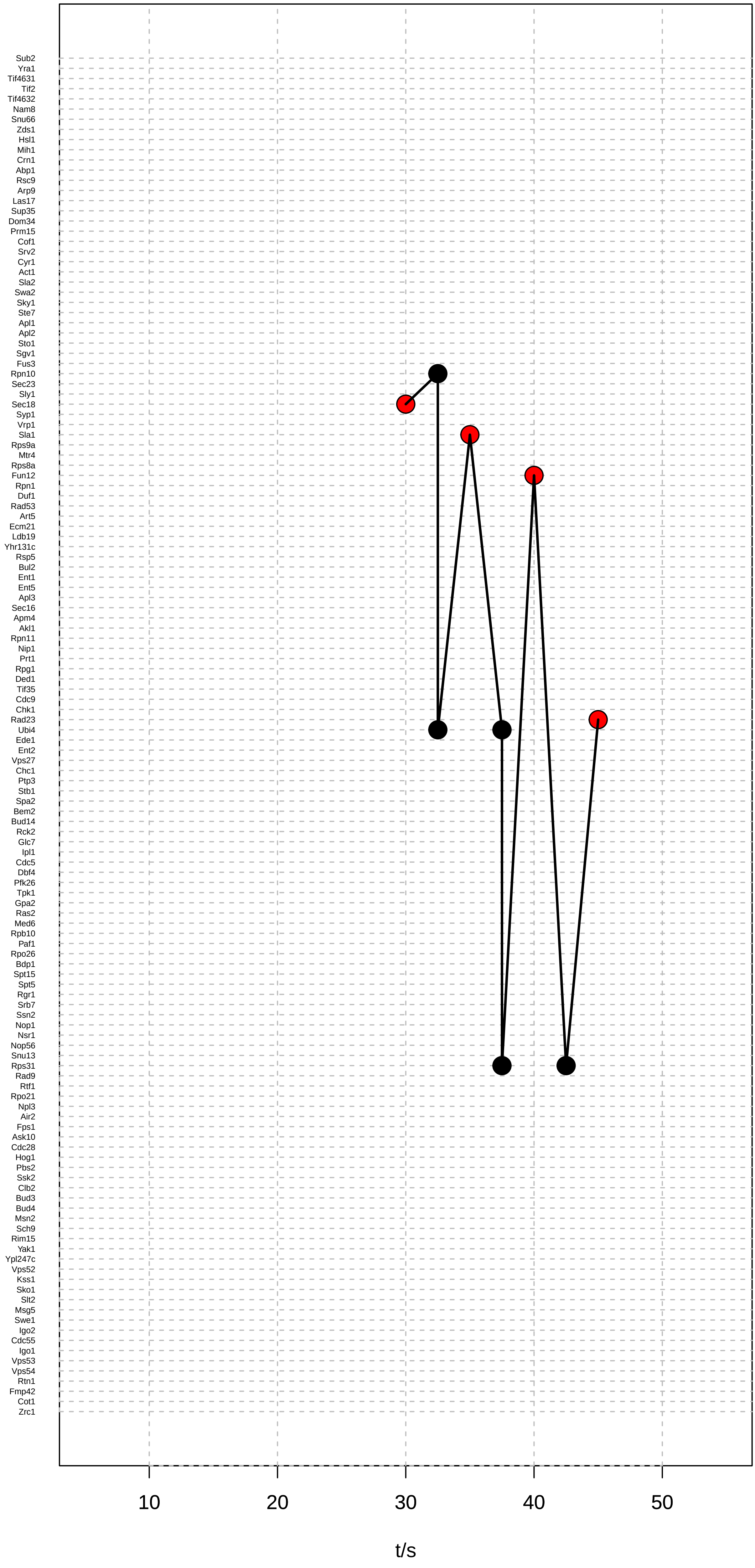

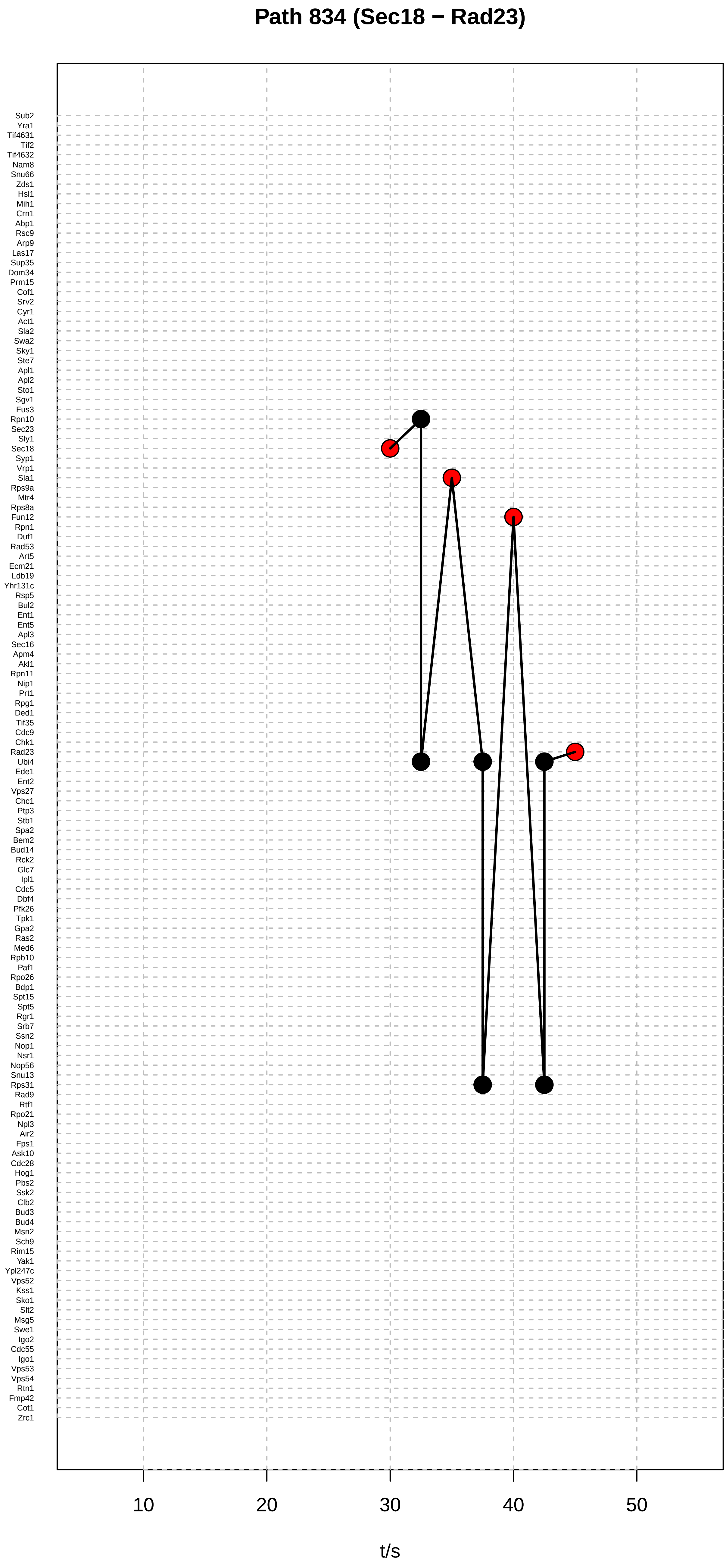
Path 834 (Sec18 − Rad23)
Sub2
Yra1
Tif4631
Tif2
Tif4632
Nam8
Snu66
Zds1
Hsl1
Mih1
Crn1
Abp1
Rsc9
Arp9
Las17
Sup35
Dom34
Prm15
Cof1
Srv2
Cyr1
Act1
Sla2
Swa2
Sky1
Ste7
Apl1
Apl2
Sto1
Sgv1
Fus3
Rpn10
Sec23
Sly1
Sec18
Syp1
Vrp1
Sla1
Rps9a
Mtr4
Rps8a
Fun12
Rpn1
Duf1
Rad53
Art5
Ecm21
Ldb19
Yhr131c
Rsp5
Bul2
Ent1
Ent5
Apl3
Sec16
Apm4
Akl1
Rpn11
Nip1
Prt1
Rpg1
Ded1
Tif35
Cdc9
Chk1
Rad23
Ubi4
Ede1
Ent2
Vps27
Chc1
Ptp3
Stb1
Spa2
Bem2
Bud14
Rck2
Glc7
Ipl1
Cdc5
Dbf4
Pfk26
Tpk1
Gpa2
Ras2
Med6
Rpb10
Paf1
Rpo26
Bdp1
Spt15
Spt5
Rgr1
Srb7
Ssn2
Nop1
Nsr1
Nop56
Snu13
Rps31
Rad9
Rtf1
Rpo21
Npl3
Air2
Fps1
Ask10
Cdc28
Hog1
Pbs2
Ssk2
Clb2
Bud3
Bud4
Msn2
Sch9
Rim15
Yak1
Ypl247c
Vps52
Kss1
Sko1
Slt2
Msg5
Swe1
Igo2
Cdc55
Igo1
Vps53
Vps54
Rtn1
Fmp42
Cot1
Zrc1
10
20
30
40
50
t/s

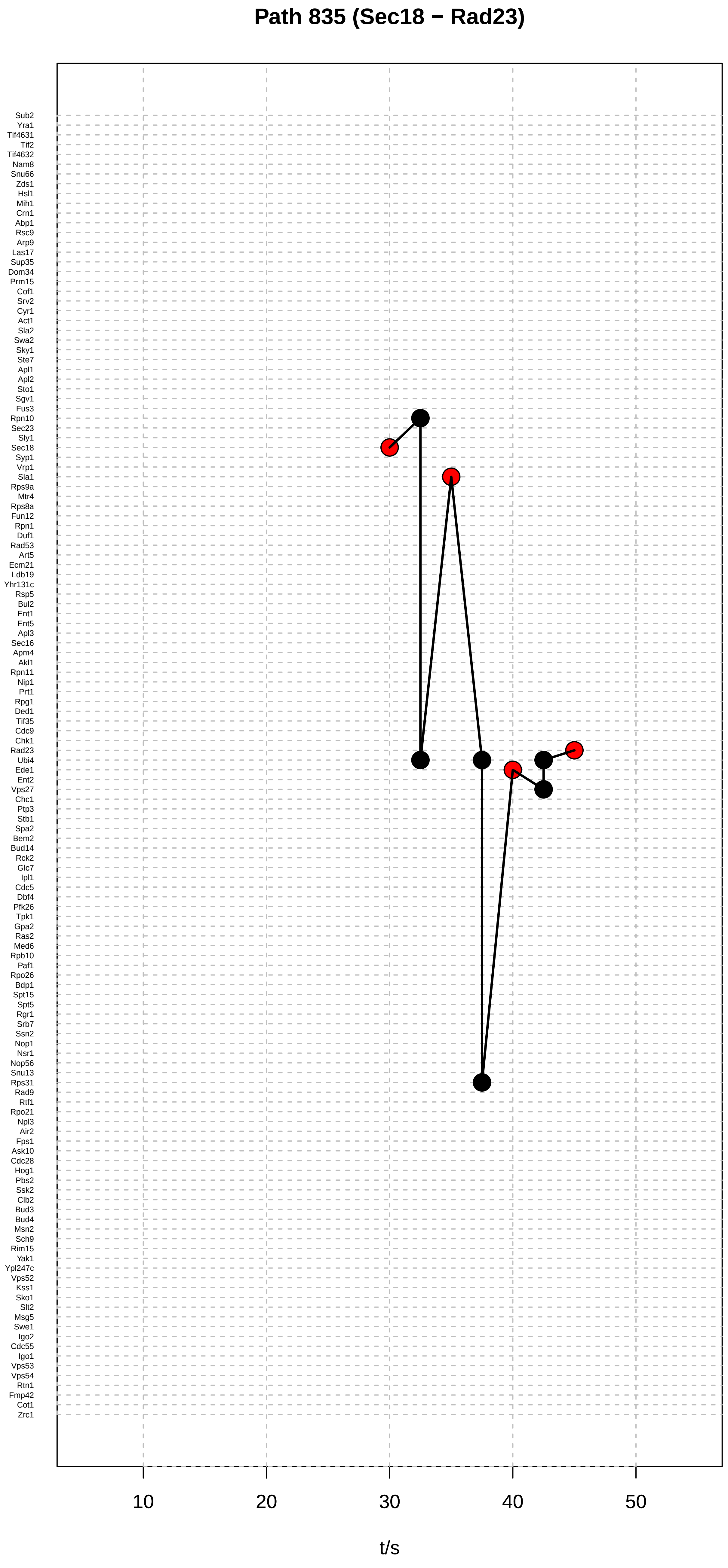
Path 835 (Sec18 − Rad23)
Sub2
Yra1
Tif4631
Tif2
Tif4632
Nam8
Snu66
Zds1
Hsl1
Mih1
Crn1
Abp1
Rsc9
Arp9
Las17
Sup35
Dom34
Prm15
Cof1
Srv2
Cyr1
Act1
Sla2
Swa2
Sky1
Ste7
Apl1
Apl2
Sto1
Sgv1
Fus3
Rpn10
Sec23
Sly1
Sec18
Syp1
Vrp1
Sla1
Rps9a
Mtr4
Rps8a
Fun12
Rpn1
Duf1
Rad53
Art5
Ecm21
Ldb19
Yhr131c
Rsp5
Bul2
Ent1
Ent5
Apl3
Sec16
Apm4
Akl1
Rpn11
Nip1
Prt1
Rpg1
Ded1
Tif35
Cdc9
Chk1
Rad23
Ubi4
Ede1
Ent2
Vps27
Chc1
Ptp3
Stb1
Spa2
Bem2
Bud14
Rck2
Glc7
Ipl1
Cdc5
Dbf4
Pfk26
Tpk1
Gpa2
Ras2
Med6
Rpb10
Paf1
Rpo26
Bdp1
Spt15
Spt5
Rgr1
Srb7
Ssn2
Nop1
Nsr1
Nop56
Snu13
Rps31
Rad9
Rtf1
Rpo21
Npl3
Air2
Fps1
Ask10
Cdc28
Hog1
Pbs2
Ssk2
Clb2
Bud3
Bud4
Msn2
Sch9
Rim15
Yak1
Ypl247c
Vps52
Kss1
Sko1
Slt2
Msg5
Swe1
Igo2
Cdc55
Igo1
Vps53
Vps54
Rtn1
Fmp42
Cot1
Zrc1
10
20
30
40
50
t/s

### Path 836 (Sec18 − Rad23)

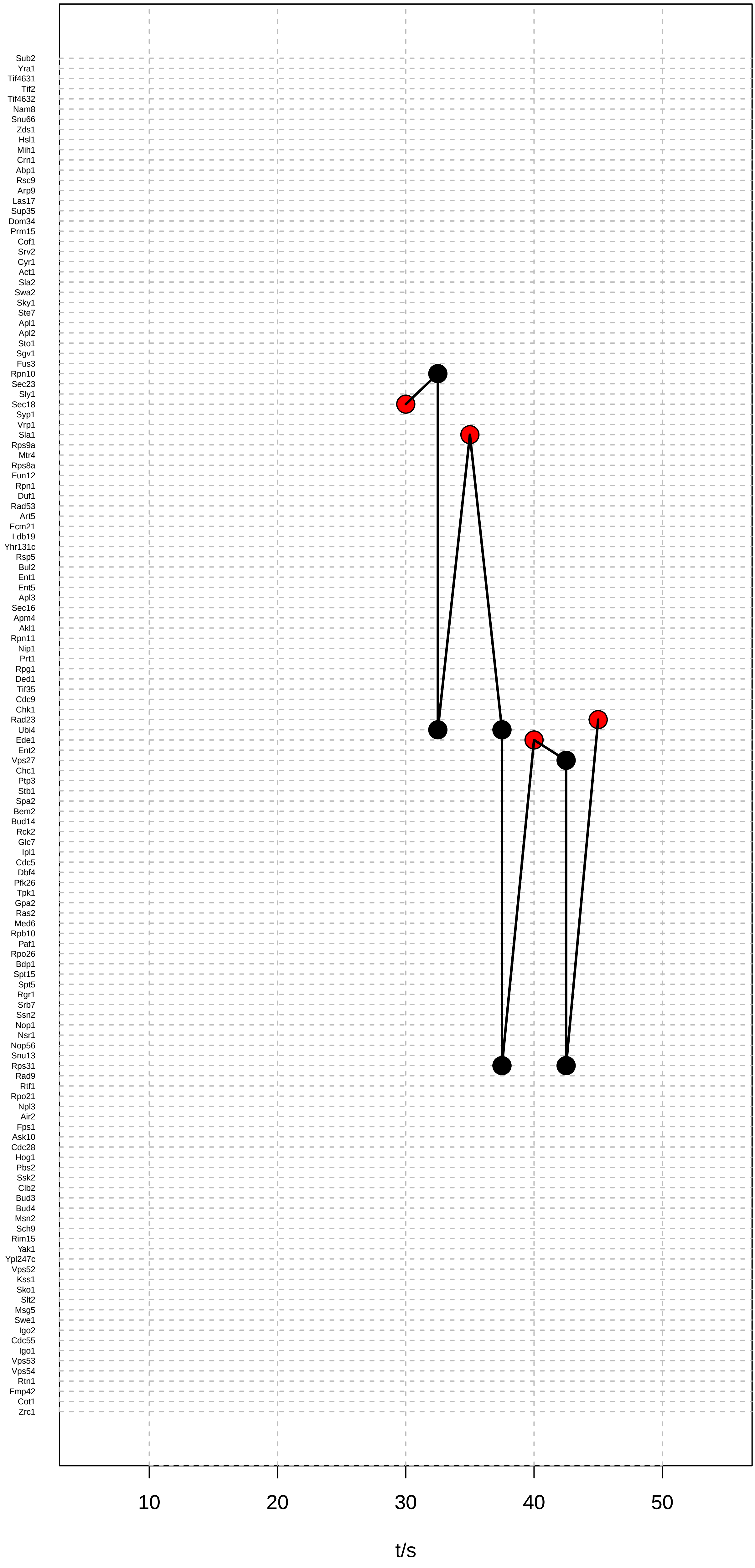

### Path 837 (Sec18 − Ent5)

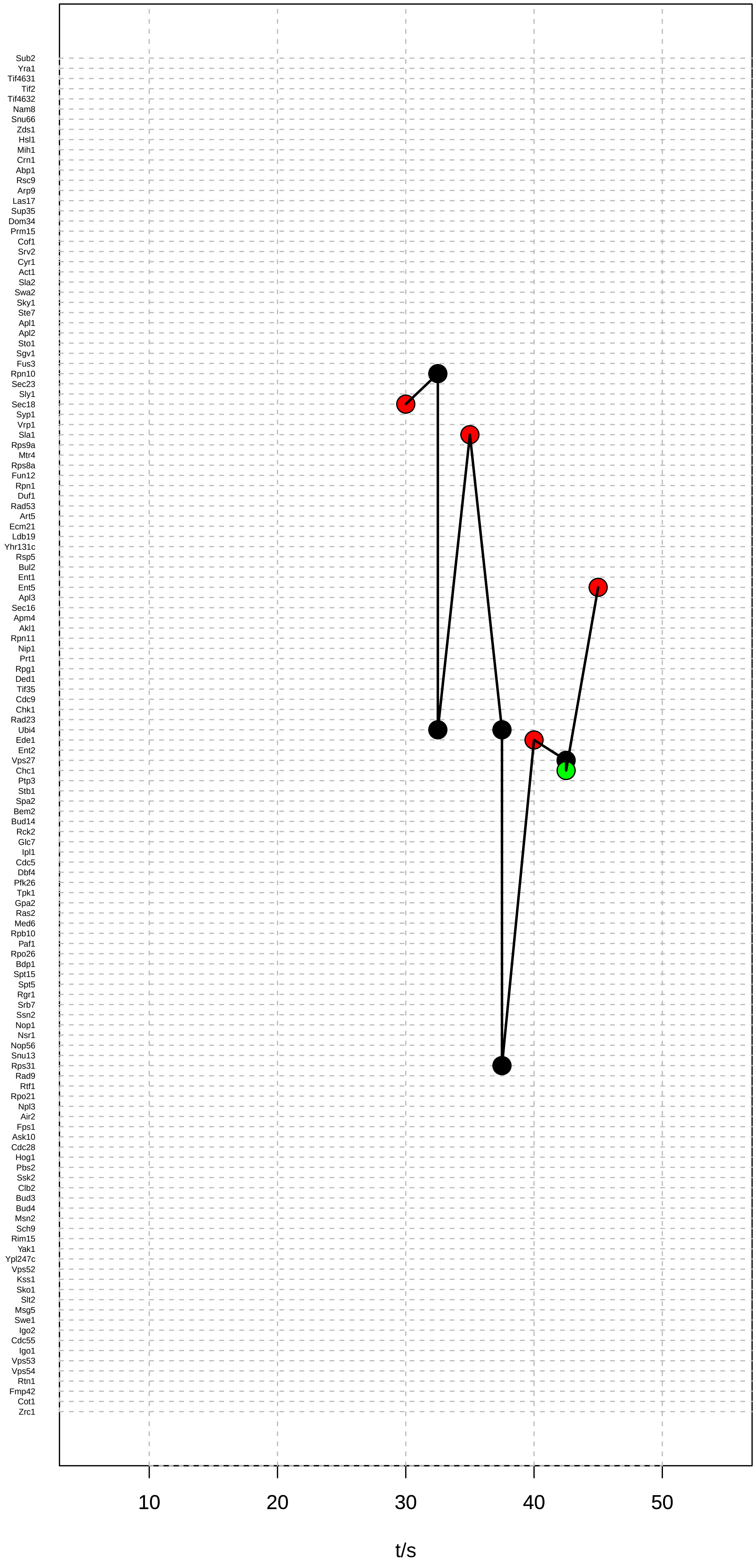

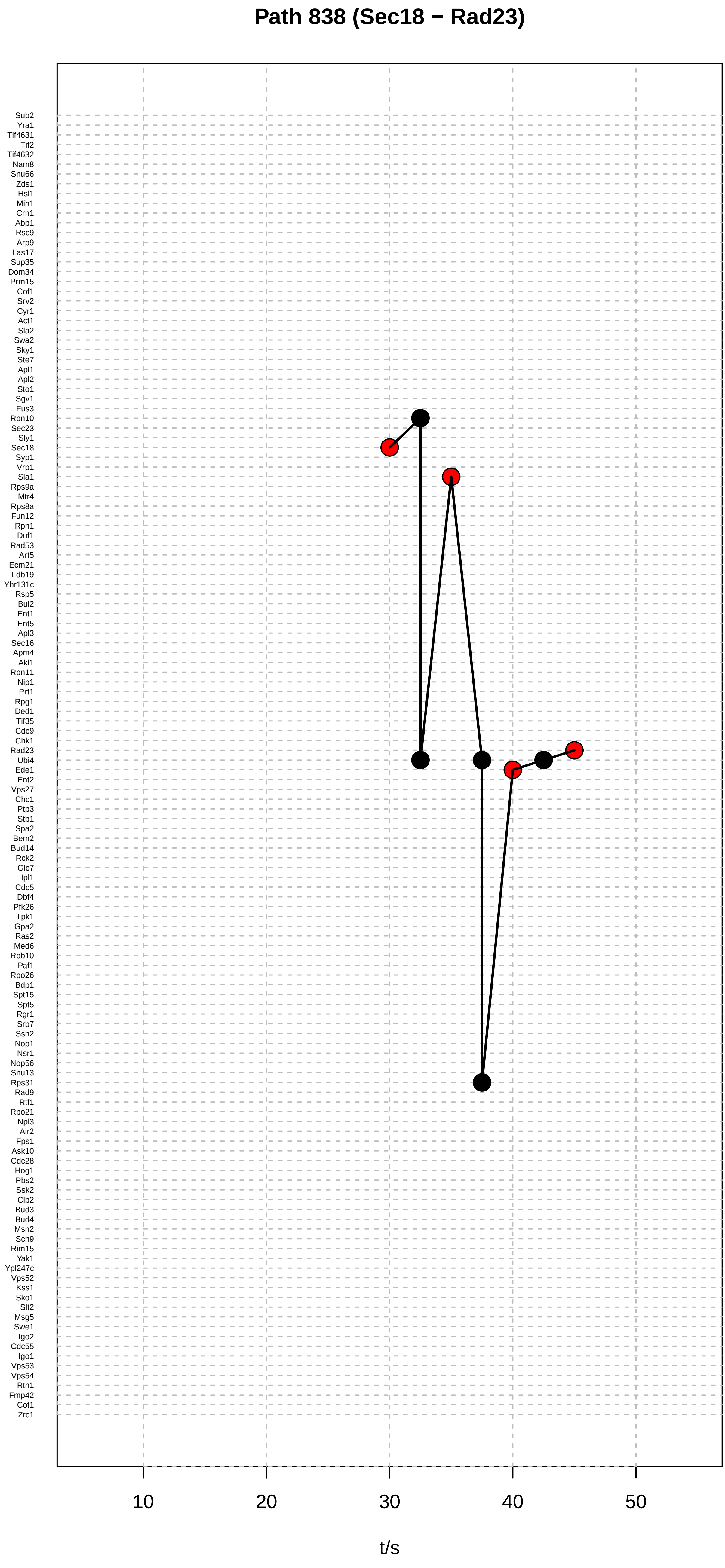

Path 838 (Sec18 − Rad23)
Sub2
Yra1
Tif4631
Tif2
Tif4632
Nam8
Snu66
Zds1
Hsl1
Mih1
Crn1
Abp1
Rsc9
Arp9
Las17
Sup35
Dom34
Prm15
Cof1
Srv2
Cyr1
Act1
Sla2
Swa2
Sky1
Ste7
Apl1
Apl2
Sto1
Sgv1
Fus3
Rpn10
Sec23
Sly1
Sec18
Syp1
Vrp1
Sla1
Rps9a
Mtr4
Rps8a
Fun12
Rpn1
Duf1
Rad53
Art5
Ecm21
Ldb19
Yhr131c
Rsp5
Bul2
Ent1
Ent5
Apl3
Sec16
Apm4
Akl1
Rpn11
Nip1
Prt1
Rpg1
Ded1
Tif35
Cdc9
Chk1
Rad23
Ubi4
Ede1
Ent2
Vps27
Chc1
Ptp3
Stb1
Spa2
Bem2
Bud14
Rck2
Glc7
Ipl1
Cdc5
Dbf4
Pfk26
Tpk1
Gpa2
Ras2
Med6
Rpb10
Paf1
Rpo26
Bdp1
Spt15
Spt5
Rgr1
Srb7
Ssn2
Nop1
Nsr1
Nop56
Snu13
Rps31
Rad9
Rtf1
Rpo21
Npl3
Air2
Fps1
Ask10
Cdc28
Hog1
Pbs2
Ssk2
Clb2
Bud3
Bud4
Msn2
Sch9
Rim15
Yak1
Ypl247c
Vps52
Kss1
Sko1
Slt2
Msg5
Swe1
Igo2
Cdc55
Igo1
Vps53
Vps54
Rtn1
Fmp42
Cot1
Zrc1
10
20
30
40
50
t/s

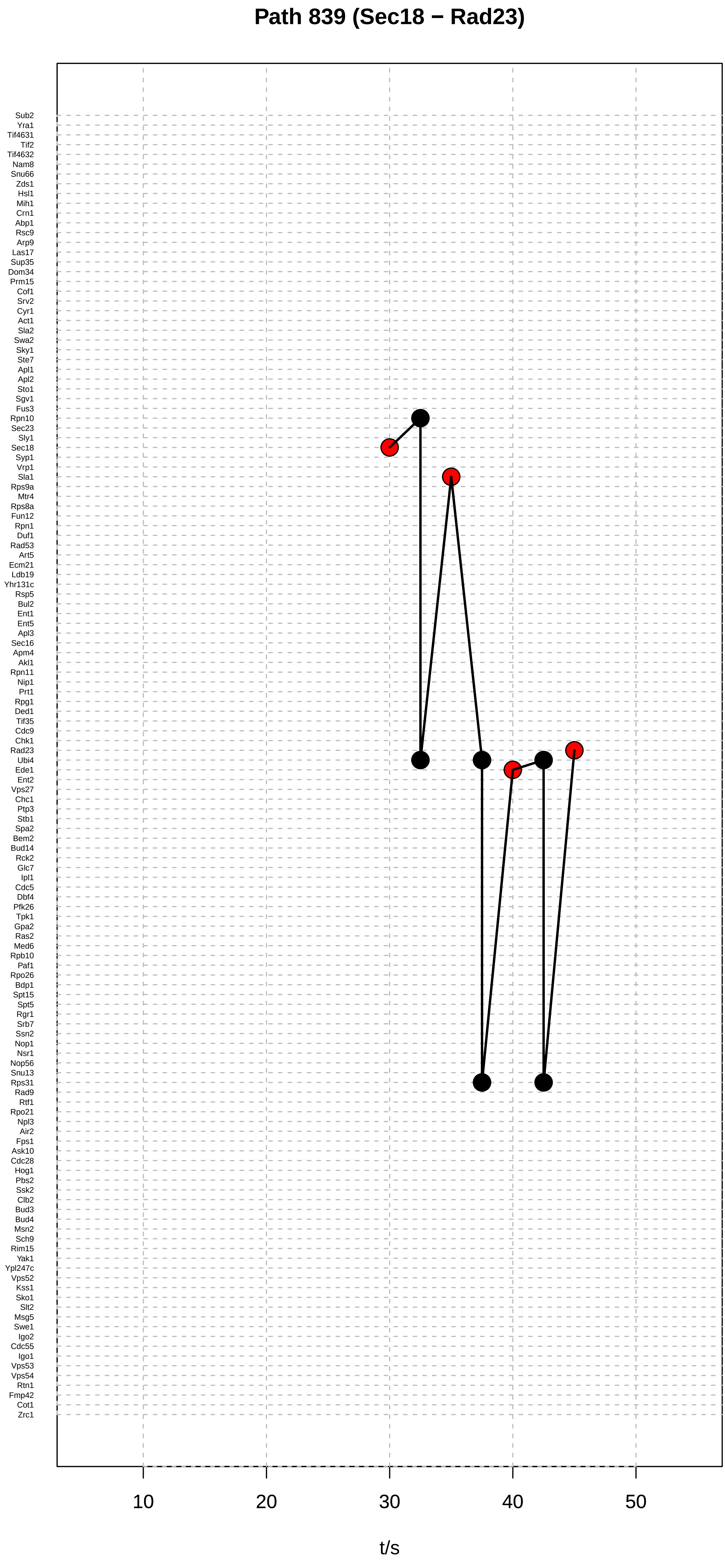
Path 839 (Sec18 − Rad23)
Sub2
Yra1
Tif4631
Tif2
Tif4632
Nam8
Snu66
Zds1
Hsl1
Mih1
Crn1
Abp1
Rsc9
Arp9
Las17
Sup35
Dom34
Prm15
Cof1
Srv2
Cyr1
Act1
Sla2
Swa2
Sky1
Ste7
Apl1
Apl2
Sto1
Sgv1
Fus3
Rpn10
Sec23
Sly1
Sec18
Syp1
Vrp1
Sla1
Rps9a
Mtr4
Rps8a
Fun12
Rpn1
Duf1
Rad53
Art5
Ecm21
Ldb19
Yhr131c
Rsp5
Bul2
Ent1
Ent5
Apl3
Sec16
Apm4
Akl1
Rpn11
Nip1
Prt1
Rpg1
Ded1
Tif35
Cdc9
Chk1
Rad23
Ubi4
Ede1
Ent2
Vps27
Chc1
Ptp3
Stb1
Spa2
Bem2
Bud14
Rck2
Glc7
Ipl1
Cdc5
Dbf4
Pfk26
Tpk1
Gpa2
Ras2
Med6
Rpb10
Paf1
Rpo26
Bdp1
Spt15
Spt5
Rgr1
Srb7
Ssn2
Nop1
Nsr1
Nop56
Snu13
Rps31
Rad9
Rtf1
Rpo21
Npl3
Air2
Fps1
Ask10
Cdc28
Hog1
Pbs2
Ssk2
Clb2
Bud3
Bud4
Msn2
Sch9
Rim15
Yak1
Ypl247c
Vps52
Kss1
Sko1
Slt2
Msg5
Swe1
Igo2
Cdc55
Igo1
Vps53
Vps54
Rtn1
Fmp42
Cot1
Zrc1
10
20
30
40
50
t/s

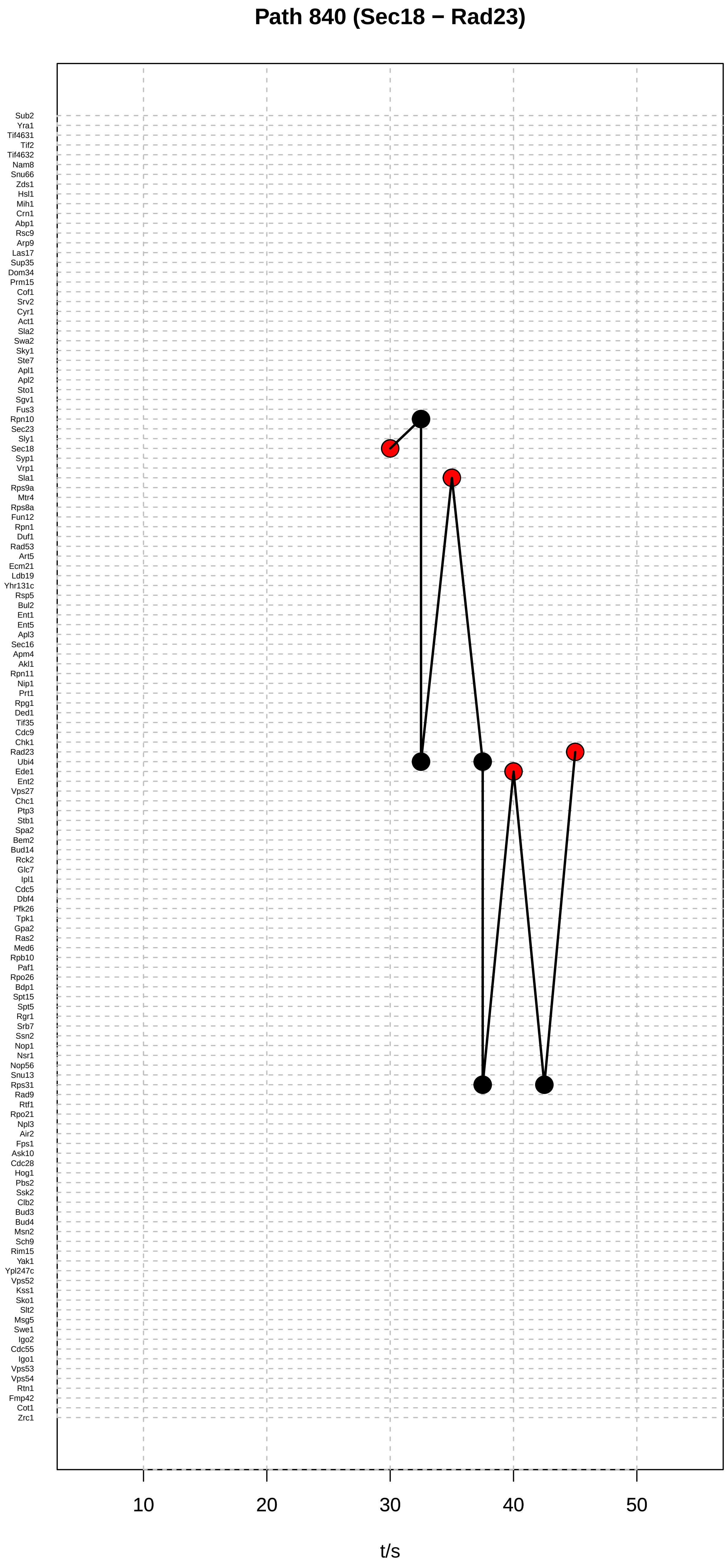

Path 840 (Sec18 − Rad23)
Sub2
Yra1
Tif4631
Tif2
Tif4632
Nam8
Snu66
Zds1
Hsl1
Mih1
Crn1
Abp1
Rsc9
Arp9
Las17
Sup35
Dom34
Prm15
Cof1
Srv2
Cyr1
Act1
Sla2
Swa2
Sky1
Ste7
Apl1
Apl2
Sto1
Sgv1
Fus3
Rpn10
Sec23
Sly1
Sec18
Syp1
Vrp1
Sla1
Rps9a
Mtr4
Rps8a
Fun12
Rpn1
Duf1
Rad53
Art5
Ecm21
Ldb19
Yhr131c
Rsp5
Bul2
Ent1
Ent5
Apl3
Sec16
Apm4
Akl1
Rpn11
Nip1
Prt1
Rpg1
Ded1
Tif35
Cdc9
Chk1
Rad23
Ubi4
Ede1
Ent2
Vps27
Chc1
Ptp3
Stb1
Spa2
Bem2
Bud14
Rck2
Glc7
Ipl1
Cdc5
Dbf4
Pfk26
Tpk1
Gpa2
Ras2
Med6
Rpb10
Paf1
Rpo26
Bdp1
Spt15
Spt5
Rgr1
Srb7
Ssn2
Nop1
Nsr1
Nop56
Snu13
Rps31
Rad9
Rtf1
Rpo21
Npl3
Air2
Fps1
Ask10
Cdc28
Hog1
Pbs2
Ssk2
Clb2
Bud3
Bud4
Msn2
Sch9
Rim15
Yak1
Ypl247c
Vps52
Kss1
Sko1
Slt2
Msg5
Swe1
Igo2
Cdc55
Igo1
Vps53
Vps54
Rtn1
Fmp42
Cot1
Zrc1
10
20
30
40
50
t/s

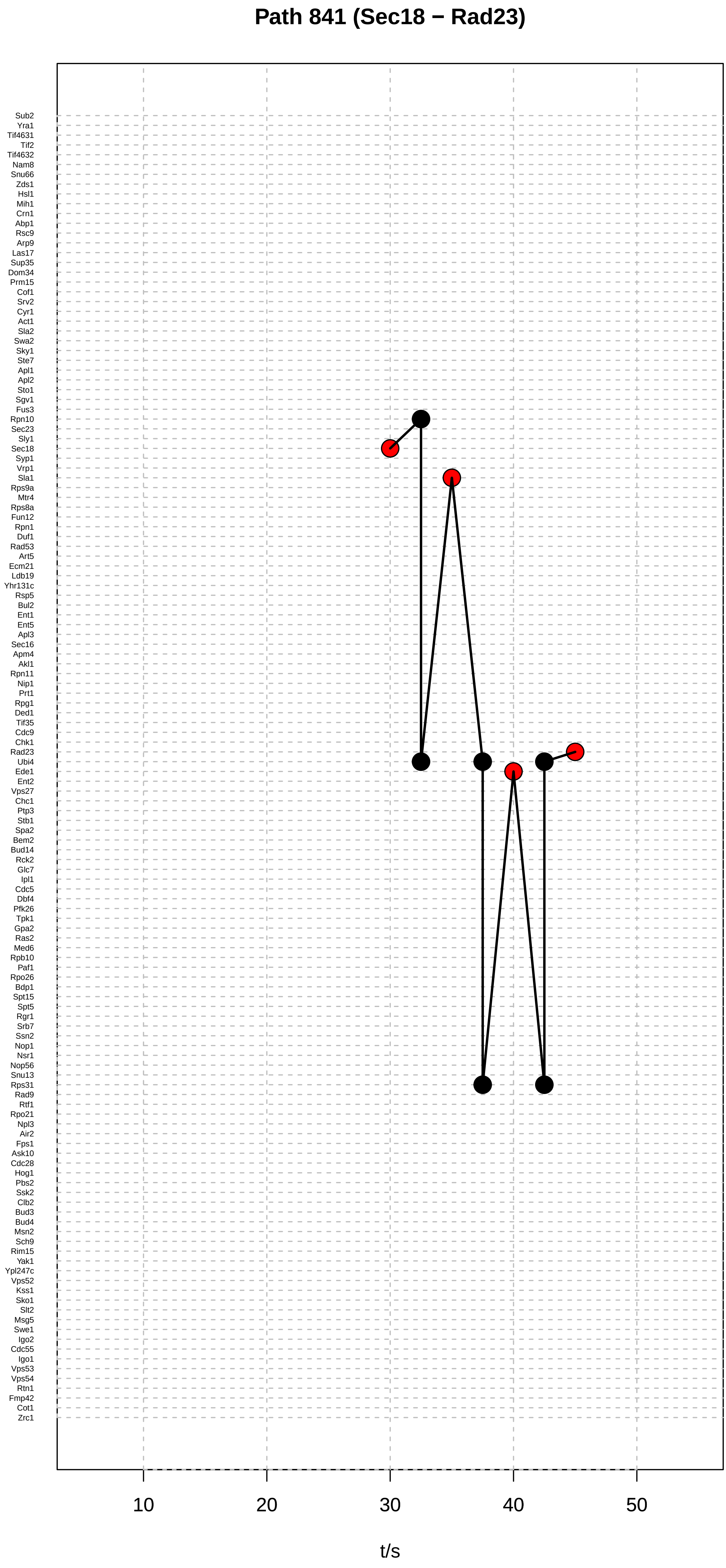

Path 841 (Sec18 − Rad23)
Sub2
Yra1
Tif4631
Tif2
Tif4632
Nam8
Snu66
Zds1
Hsl1
Mih1
Crn1
Abp1
Rsc9
Arp9
Las17
Sup35
Dom34
Prm15
Cof1
Srv2
Cyr1
Act1
Sla2
Swa2
Sky1
Ste7
Apl1
Apl2
Sto1
Sgv1
Fus3
Rpn10
Sec23
Sly1
Sec18
Syp1
Vrp1
Sla1
Rps9a
Mtr4
Rps8a
Fun12
Rpn1
Duf1
Rad53
Art5
Ecm21
Ldb19
Yhr131c
Rsp5
Bul2
Ent1
Ent5
Apl3
Sec16
Apm4
Akl1
Rpn11
Nip1
Prt1
Rpg1
Ded1
Tif35
Cdc9
Chk1
Rad23
Ubi4
Ede1
Ent2
Vps27
Chc1
Ptp3
Stb1
Spa2
Bem2
Bud14
Rck2
Glc7
Ipl1
Cdc5
Dbf4
Pfk26
Tpk1
Gpa2
Ras2
Med6
Rpb10
Paf1
Rpo26
Bdp1
Spt15
Spt5
Rgr1
Srb7
Ssn2
Nop1
Nsr1
Nop56
Snu13
Rps31
Rad9
Rtf1
Rpo21
Npl3
Air2
Fps1
Ask10
Cdc28
Hog1
Pbs2
Ssk2
Clb2
Bud3
Bud4
Msn2
Sch9
Rim15
Yak1
Ypl247c
Vps52
Kss1
Sko1
Slt2
Msg5
Swe1
Igo2
Cdc55
Igo1
Vps53
Vps54
Rtn1
Fmp42
Cot1
Zrc1
10
20
30
40
50
t/s

### Path 842 (Sec18 − Rad23)

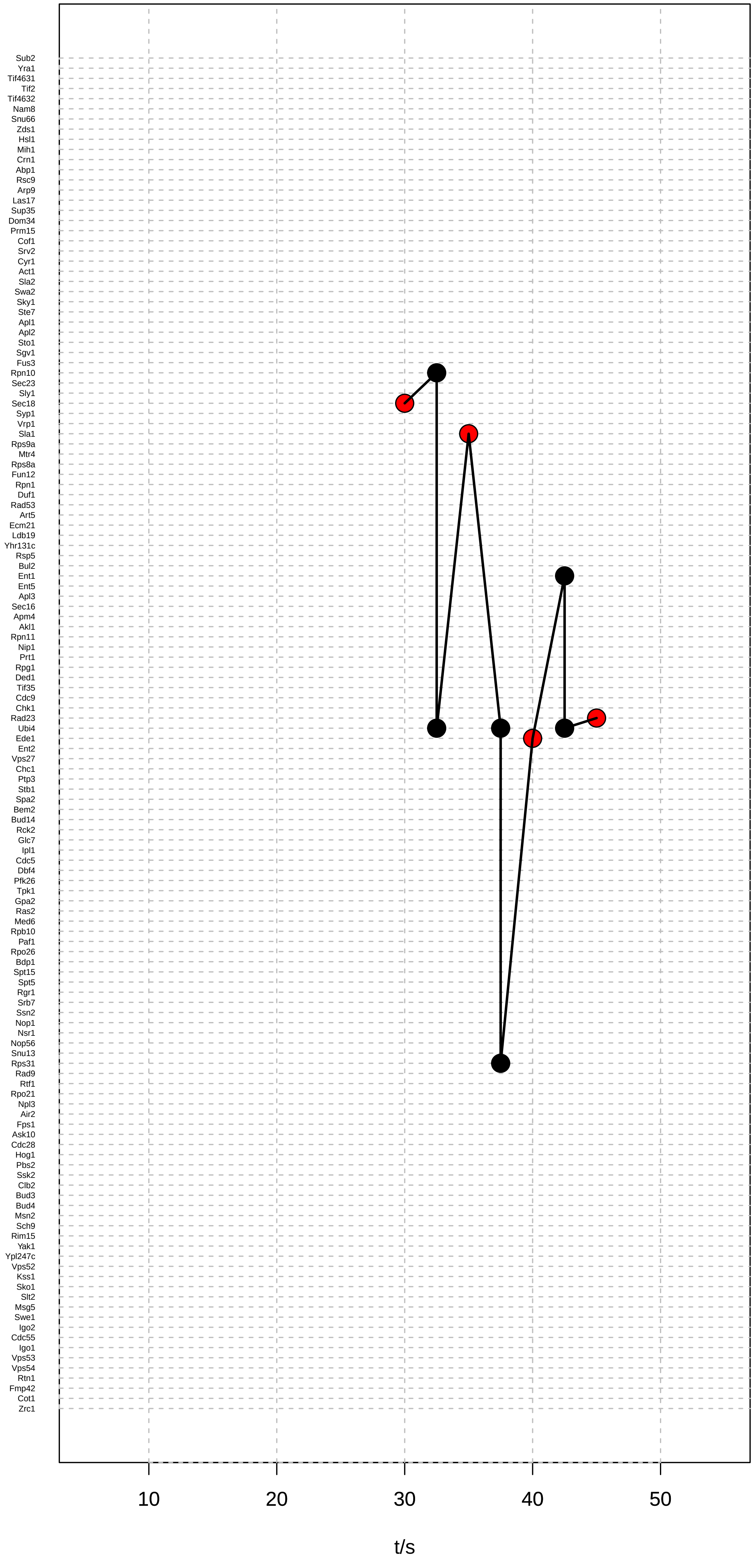

### Path 843 (Sec18 − Ent5)

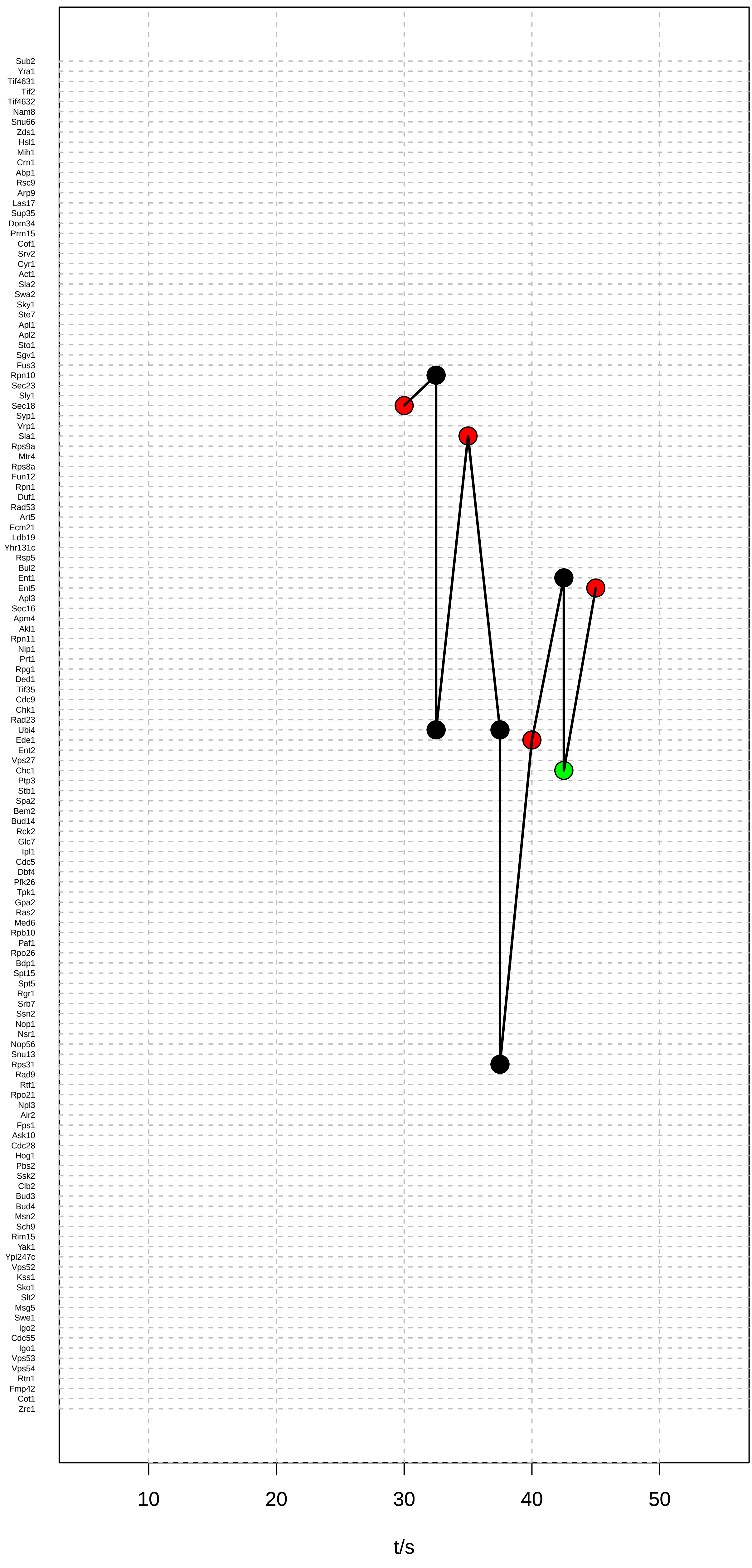

#### Path 844 (Sec18 − Prm15)

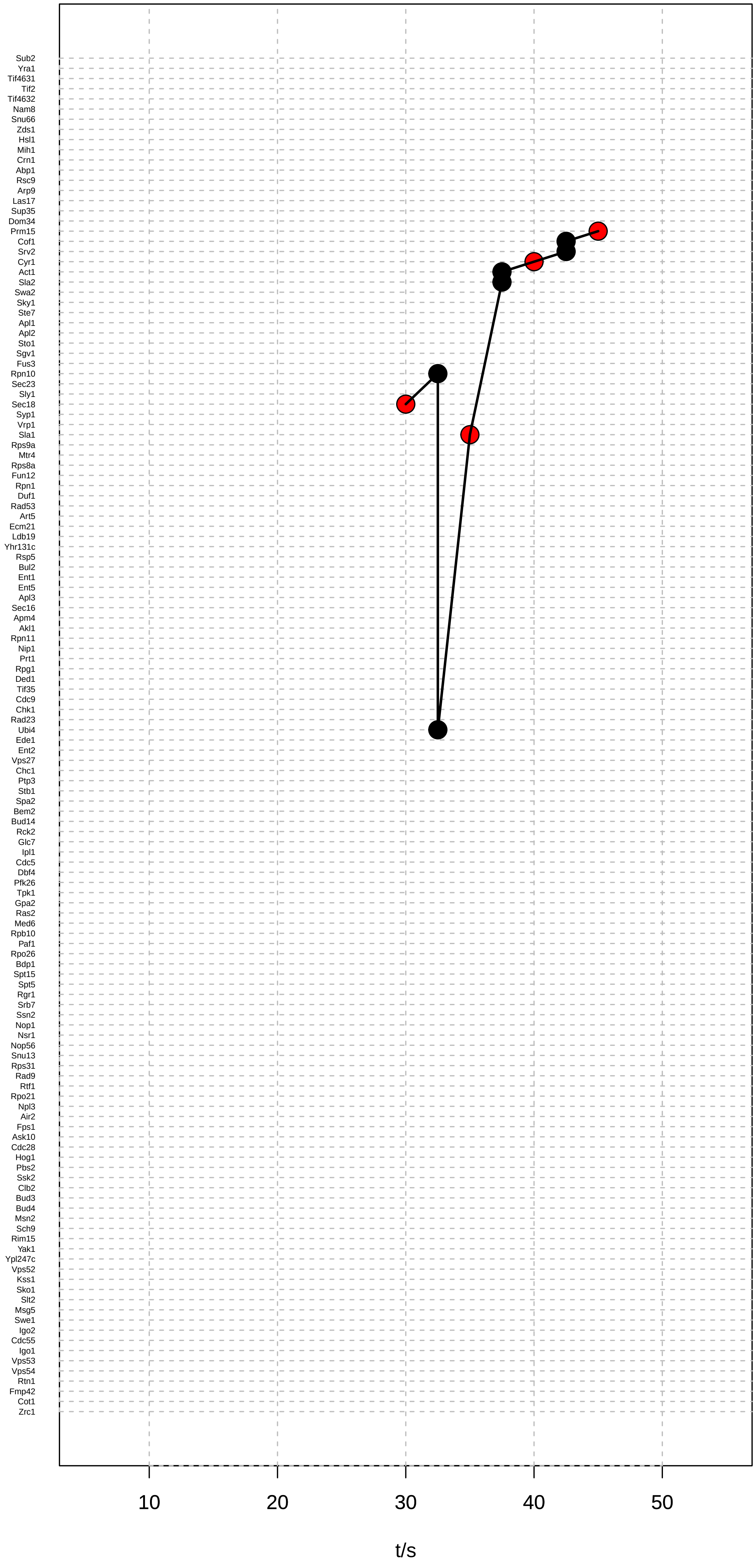

### Path 845 (Sec18 − Prm15)

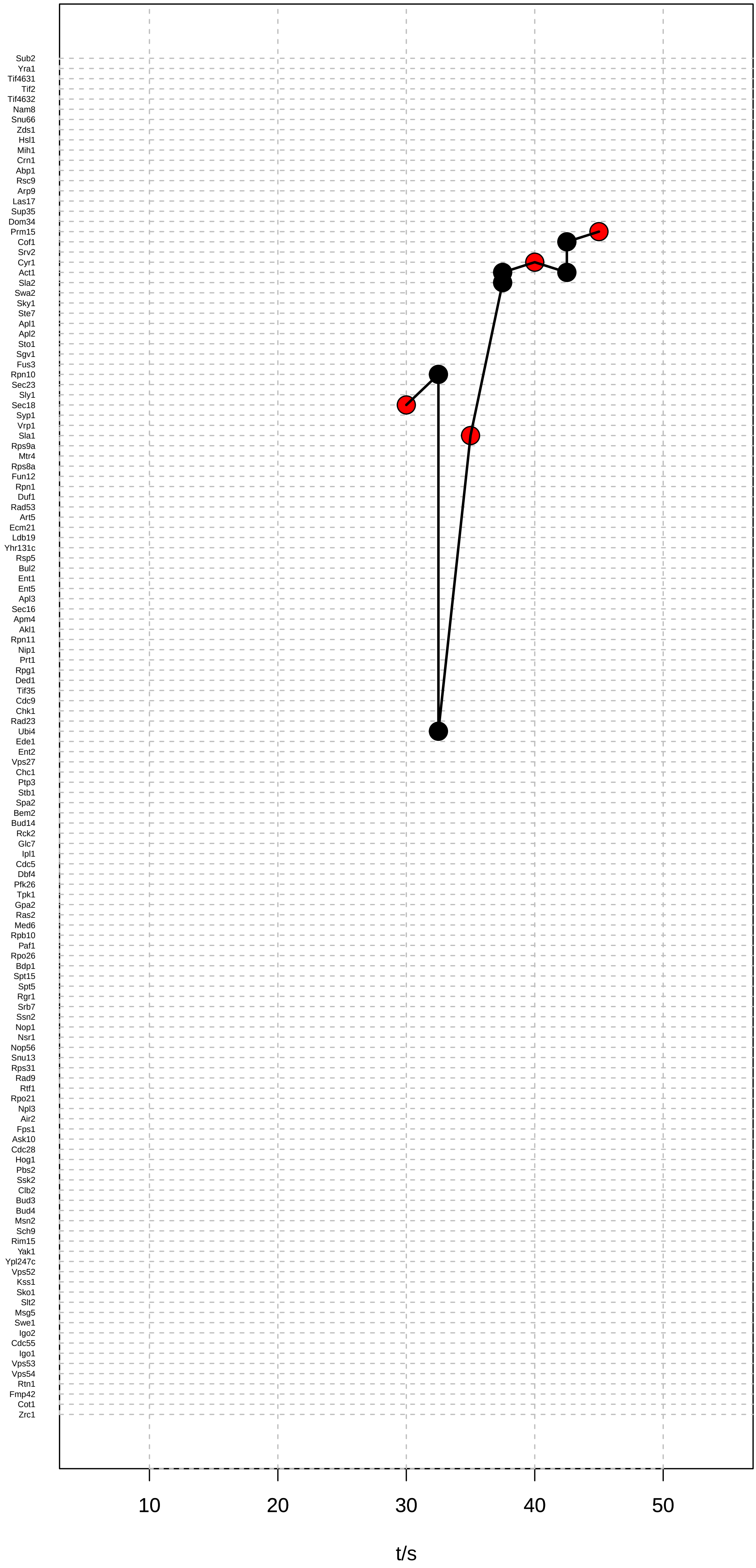

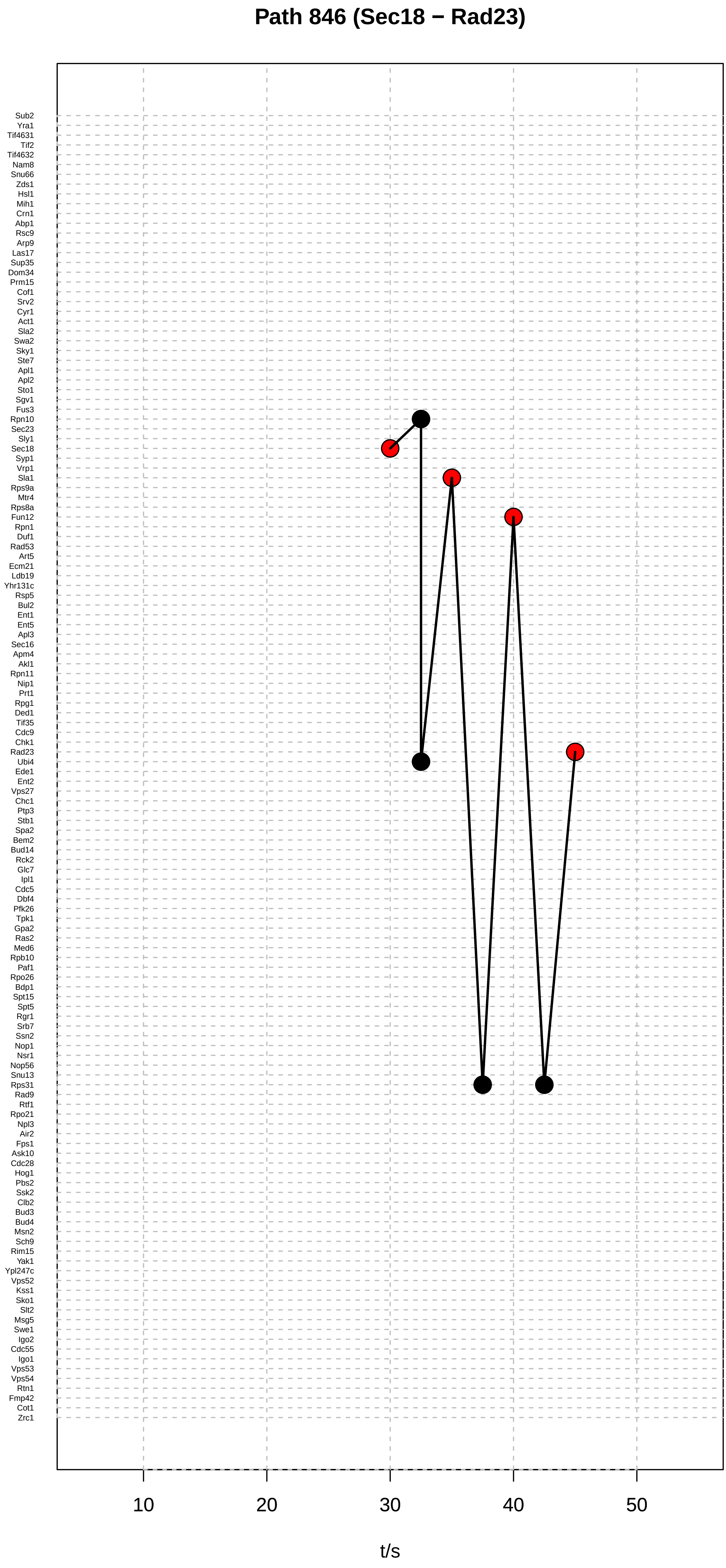

Path 846 (Sec18 − Rad23)
Sub2
Yra1
Tif4631
Tif2
Tif4632
Nam8
Snu66
Zds1
Hsl1
Mih1
Crn1
Abp1
Rsc9
Arp9
Las17
Sup35
Dom34
Prm15
Cof1
Srv2
Cyr1
Act1
Sla2
Swa2
Sky1
Ste7
Apl1
Apl2
Sto1
Sgv1
Fus3
Rpn10
Sec23
Sly1
Sec18
Syp1
Vrp1
Sla1
Rps9a
Mtr4
Rps8a
Fun12
Rpn1
Duf1
Rad53
Art5
Ecm21
Ldb19
Yhr131c
Rsp5
Bul2
Ent1
Ent5
Apl3
Sec16
Apm4
Akl1
Rpn11
Nip1
Prt1
Rpg1
Ded1
Tif35
Cdc9
Chk1
Rad23
Ubi4
Ede1
Ent2
Vps27
Chc1
Ptp3
Stb1
Spa2
Bem2
Bud14
Rck2
Glc7
Ipl1
Cdc5
Dbf4
Pfk26
Tpk1
Gpa2
Ras2
Med6
Rpb10
Paf1
Rpo26
Bdp1
Spt15
Spt5
Rgr1
Srb7
Ssn2
Nop1
Nsr1
Nop56
Snu13
Rps31
Rad9
Rtf1
Rpo21
Npl3
Air2
Fps1
Ask10
Cdc28
Hog1
Pbs2
Ssk2
Clb2
Bud3
Bud4
Msn2
Sch9
Rim15
Yak1
Ypl247c
Vps52
Kss1
Sko1
Slt2
Msg5
Swe1
Igo2
Cdc55
Igo1
Vps53
Vps54
Rtn1
Fmp42
Cot1
Zrc1
10
20
30
40
50
t/s

### Path 847 (Sec18 − Rad23)

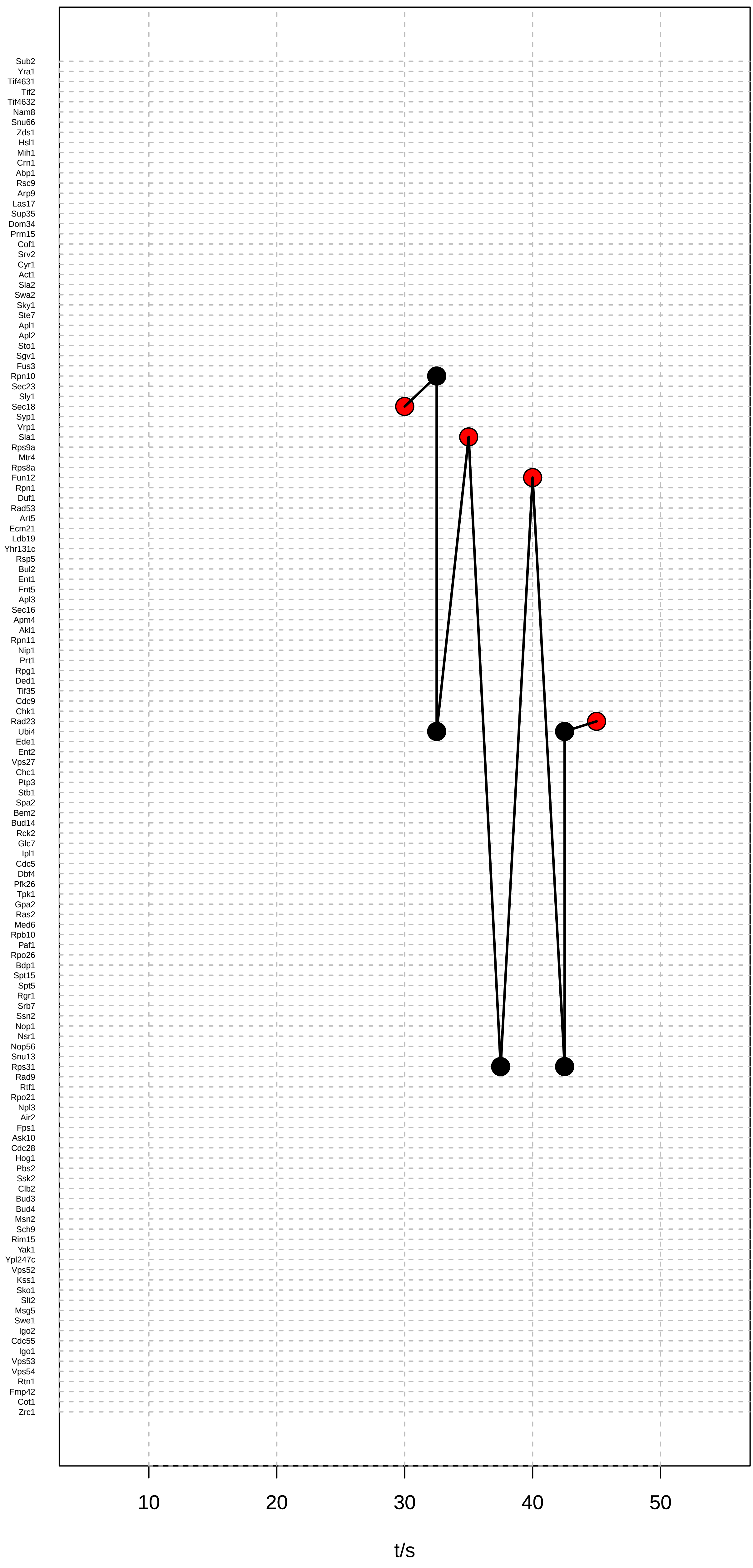

#### Path 848 (Sec18 − Rad23)

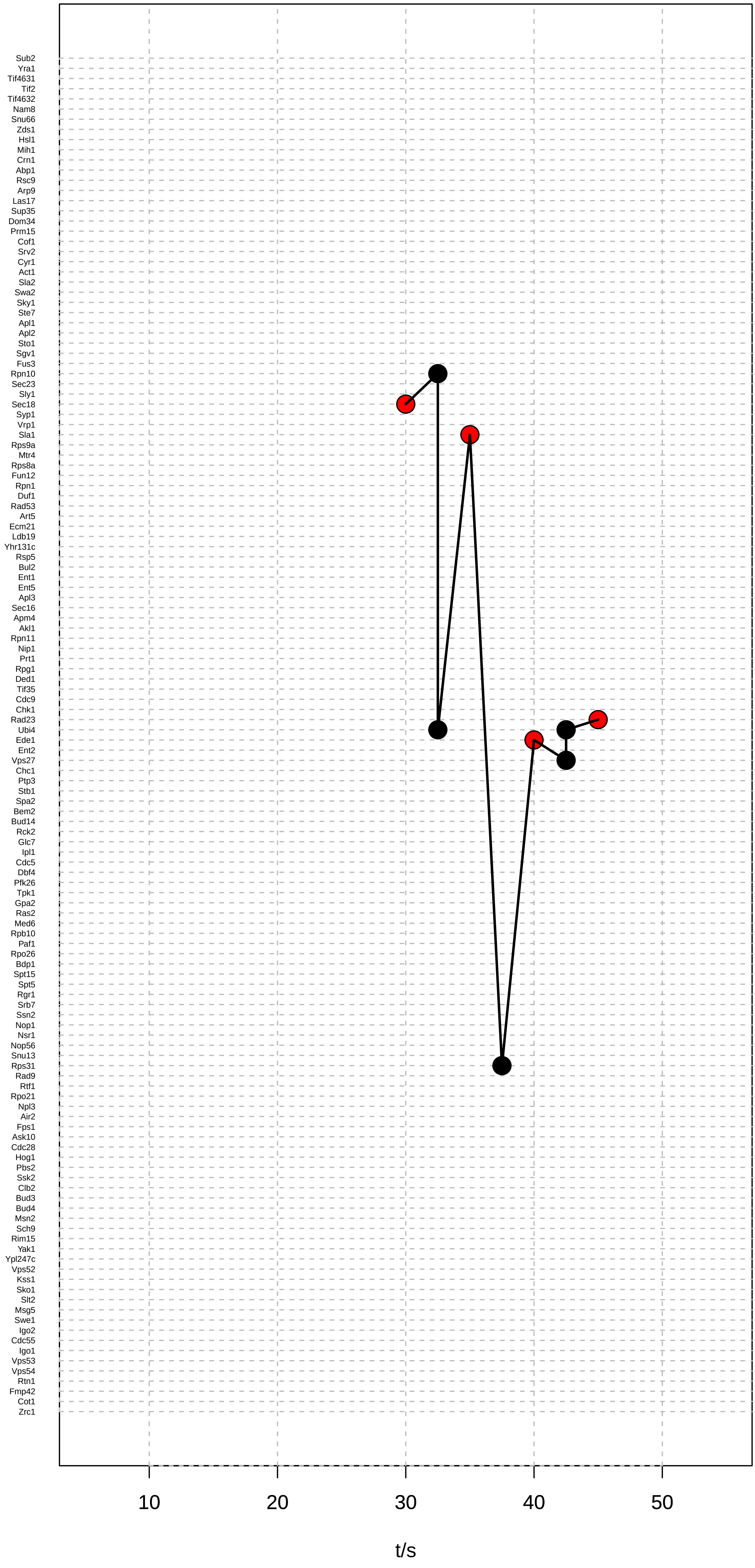

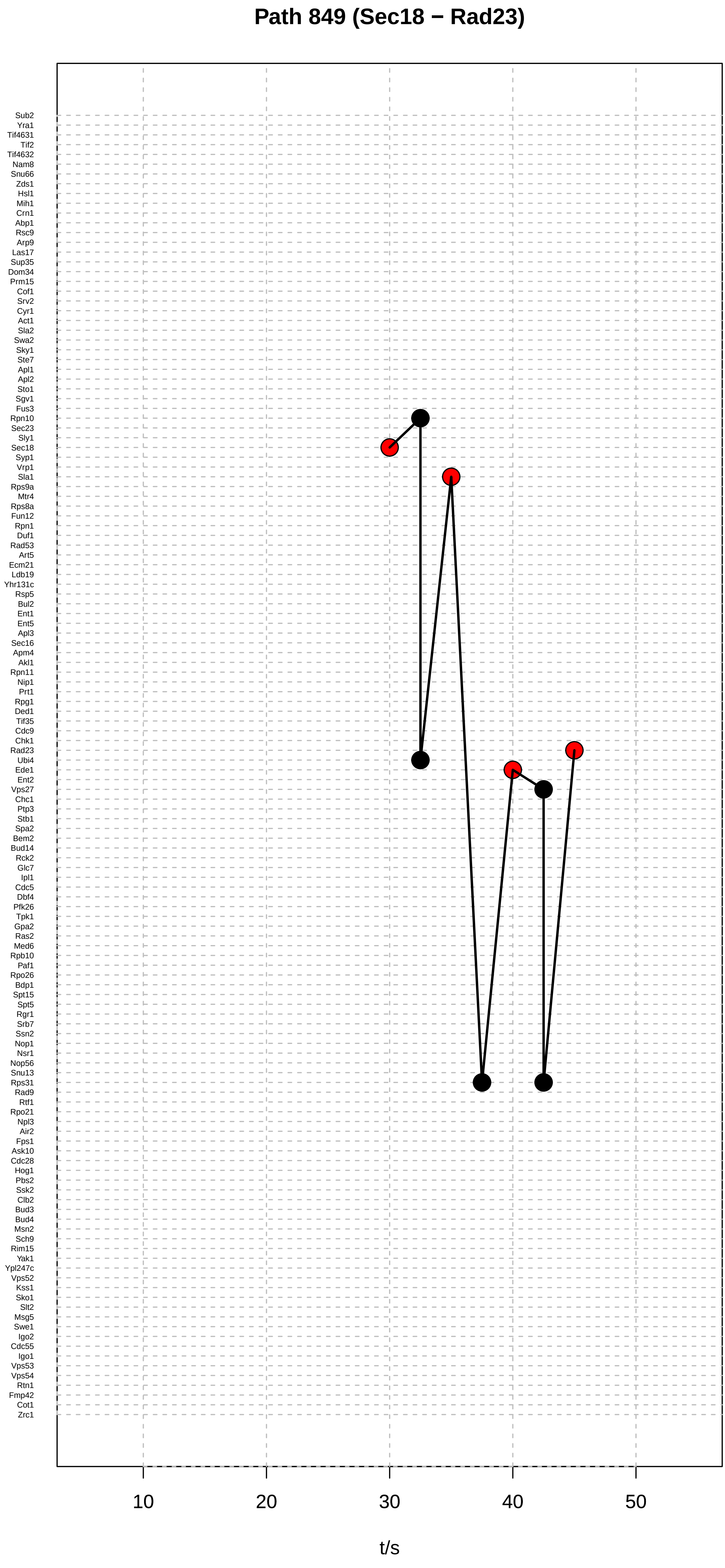

Path 849 (Sec18 − Rad23)
Sub2
Yra1
Tif4631
Tif2
Tif4632
Nam8
Snu66
Zds1
Hsl1
Mih1
Crn1
Abp1
Rsc9
Arp9
Las17
Sup35
Dom34
Prm15
Cof1
Srv2
Cyr1
Act1
Sla2
Swa2
Sky1
Ste7
Apl1
Apl2
Sto1
Sgv1
Fus3
Rpn10
Sec23
Sly1
Sec18
Syp1
Vrp1
Sla1
Rps9a
Mtr4
Rps8a
Fun12
Rpn1
Duf1
Rad53
Art5
Ecm21
Ldb19
Yhr131c
Rsp5
Bul2
Ent1
Ent5
Apl3
Sec16
Apm4
Akl1
Rpn11
Nip1
Prt1
Rpg1
Ded1
Tif35
Cdc9
Chk1
Rad23
Ubi4
Ede1
Ent2
Vps27
Chc1
Ptp3
Stb1
Spa2
Bem2
Bud14
Rck2
Glc7
Ipl1
Cdc5
Dbf4
Pfk26
Tpk1
Gpa2
Ras2
Med6
Rpb10
Paf1
Rpo26
Bdp1
Spt15
Spt5
Rgr1
Srb7
Ssn2
Nop1
Nsr1
Nop56
Snu13
Rps31
Rad9
Rtf1
Rpo21
Npl3
Air2
Fps1
Ask10
Cdc28
Hog1
Pbs2
Ssk2
Clb2
Bud3
Bud4
Msn2
Sch9
Rim15
Yak1
Ypl247c
Vps52
Kss1
Sko1
Slt2
Msg5
Swe1
Igo2
Cdc55
Igo1
Vps53
Vps54
Rtn1
Fmp42
Cot1
Zrc1
10
20
30
40
50
t/s

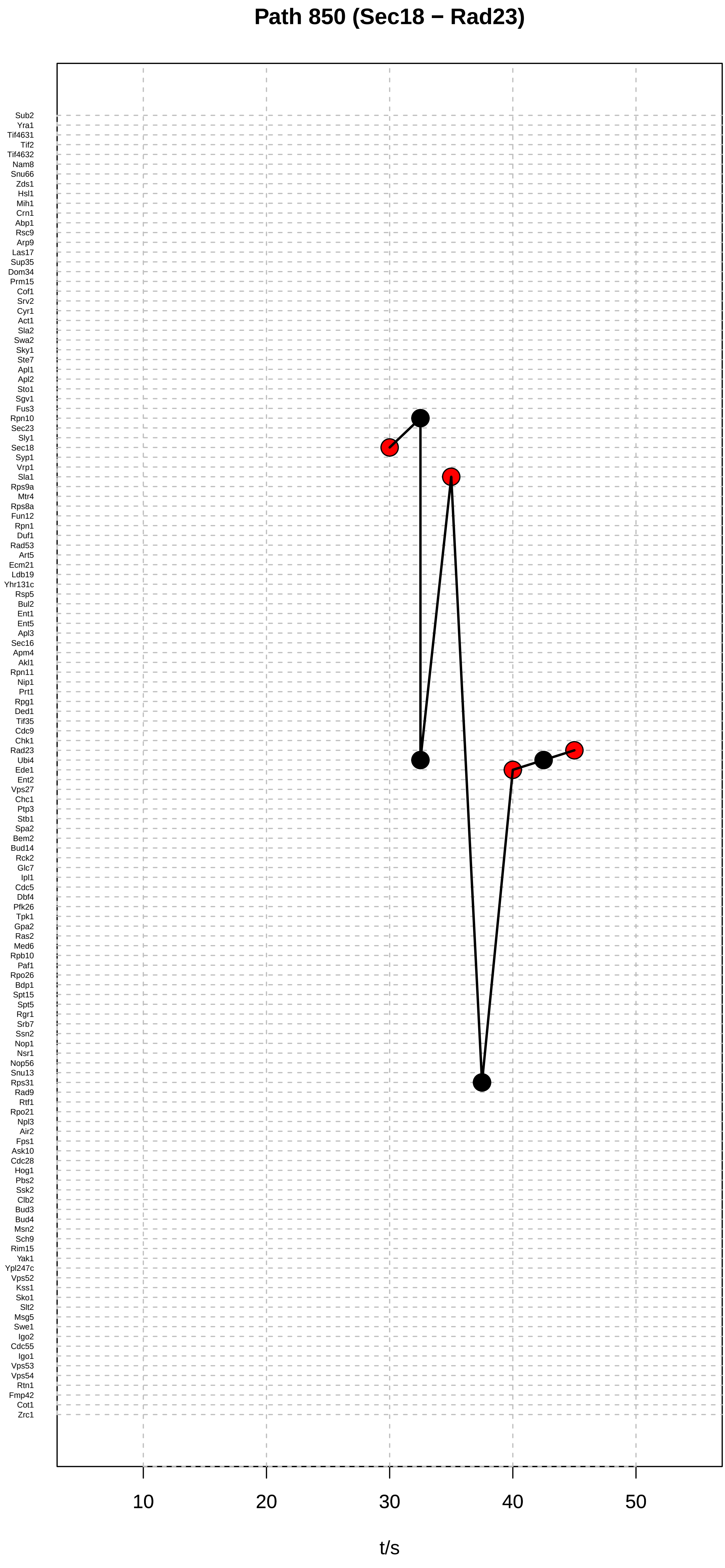
Path 850 (Sec18 − Rad23)
Sub2
Yra1
Tif4631
Tif2
Tif4632
Nam8
Snu66
Zds1
Hsl1
Mih1
Crn1
Abp1
Rsc9
Arp9
Las17
Sup35
Dom34
Prm15
Cof1
Srv2
Cyr1
Act1
Sla2
Swa2
Sky1
Ste7
Apl1
Apl2
Sto1
Sgv1
Fus3
Rpn10
Sec23
Sly1
Sec18
Syp1
Vrp1
Sla1
Rps9a
Mtr4
Rps8a
Fun12
Rpn1
Duf1
Rad53
Art5
Ecm21
Ldb19
Yhr131c
Rsp5
Bul2
Ent1
Ent5
Apl3
Sec16
Apm4
Akl1
Rpn11
Nip1
Prt1
Rpg1
Ded1
Tif35
Cdc9
Chk1
Rad23
Ubi4
Ede1
Ent2
Vps27
Chc1
Ptp3
Stb1
Spa2
Bem2
Bud14
Rck2
Glc7
Ipl1
Cdc5
Dbf4
Pfk26
Tpk1
Gpa2
Ras2
Med6
Rpb10
Paf1
Rpo26
Bdp1
Spt15
Spt5
Rgr1
Srb7
Ssn2
Nop1
Nsr1
Nop56
Snu13
Rps31
Rad9
Rtf1
Rpo21
Npl3
Air2
Fps1
Ask10
Cdc28
Hog1
Pbs2
Ssk2
Clb2
Bud3
Bud4
Msn2
Sch9
Rim15
Yak1
Ypl247c
Vps52
Kss1
Sko1
Slt2
Msg5
Swe1
Igo2
Cdc55
Igo1
Vps53
Vps54
Rtn1
Fmp42
Cot1
Zrc1
10
20
30
40
50
t/s

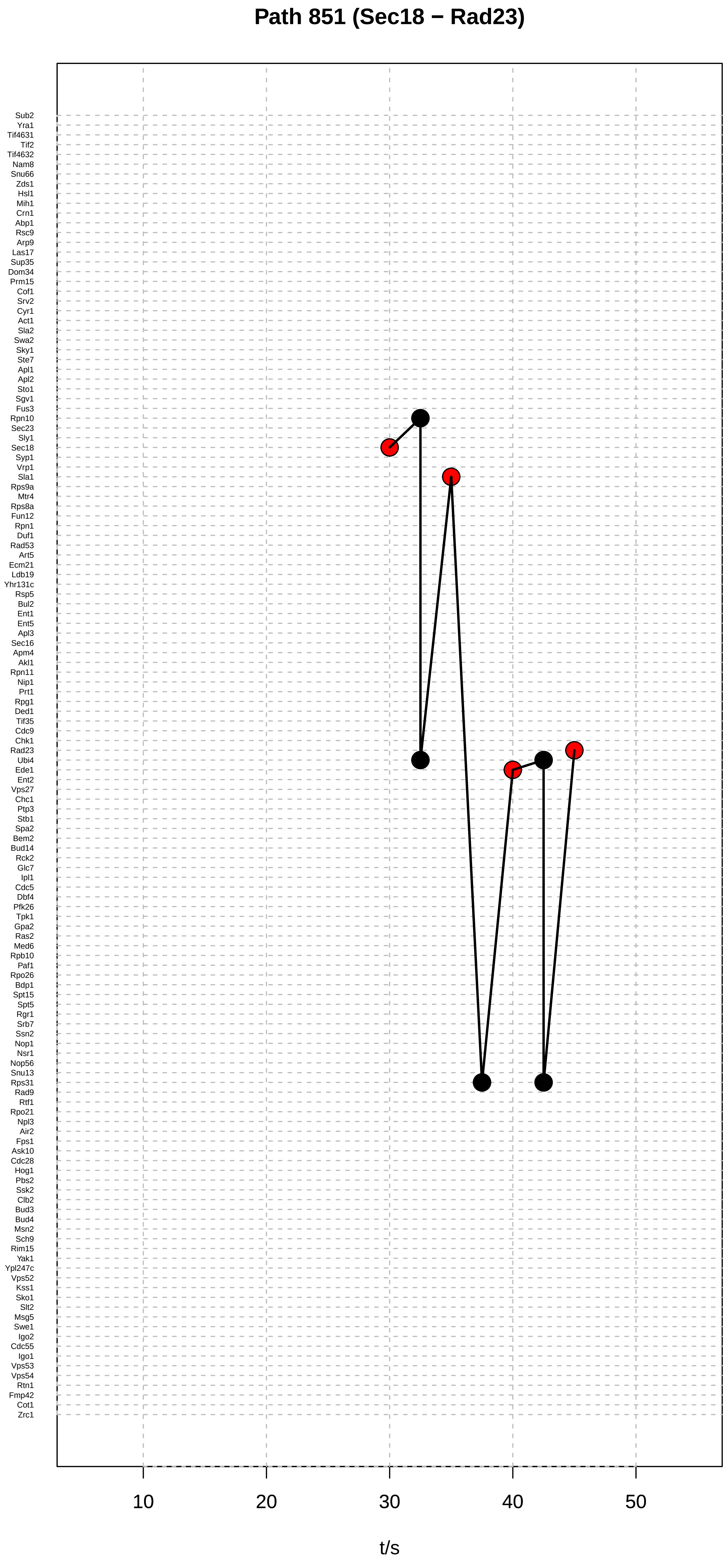
Path 851 (Sec18 − Rad23)
Sub2
Yra1
Tif4631
Tif2
Tif4632
Nam8
Snu66
Zds1
Hsl1
Mih1
Crn1
Abp1
Rsc9
Arp9
Las17
Sup35
Dom34
Prm15
Cof1
Srv2
Cyr1
Act1
Sla2
Swa2
Sky1
Ste7
Apl1
Apl2
Sto1
Sgv1
Fus3
Rpn10
Sec23
Sly1
Sec18
Syp1
Vrp1
Sla1
Rps9a
Mtr4
Rps8a
Fun12
Rpn1
Duf1
Rad53
Art5
Ecm21
Ldb19
Yhr131c
Rsp5
Bul2
Ent1
Ent5
Apl3
Sec16
Apm4
Akl1
Rpn11
Nip1
Prt1
Rpg1
Ded1
Tif35
Cdc9
Chk1
Rad23
Ubi4
Ede1
Ent2
Vps27
Chc1
Ptp3
Stb1
Spa2
Bem2
Bud14
Rck2
Glc7
Ipl1
Cdc5
Dbf4
Pfk26
Tpk1
Gpa2
Ras2
Med6
Rpb10
Paf1
Rpo26
Bdp1
Spt15
Spt5
Rgr1
Srb7
Ssn2
Nop1
Nsr1
Nop56
Snu13
Rps31
Rad9
Rtf1
Rpo21
Npl3
Air2
Fps1
Ask10
Cdc28
Hog1
Pbs2
Ssk2
Clb2
Bud3
Bud4
Msn2
Sch9
Rim15
Yak1
Ypl247c
Vps52
Kss1
Sko1
Slt2
Msg5
Swe1
Igo2
Cdc55
Igo1
Vps53
Vps54
Rtn1
Fmp42
Cot1
Zrc1
10
20
30
40
50
t/s

#### Path 852 (Sec18 − Rad23)

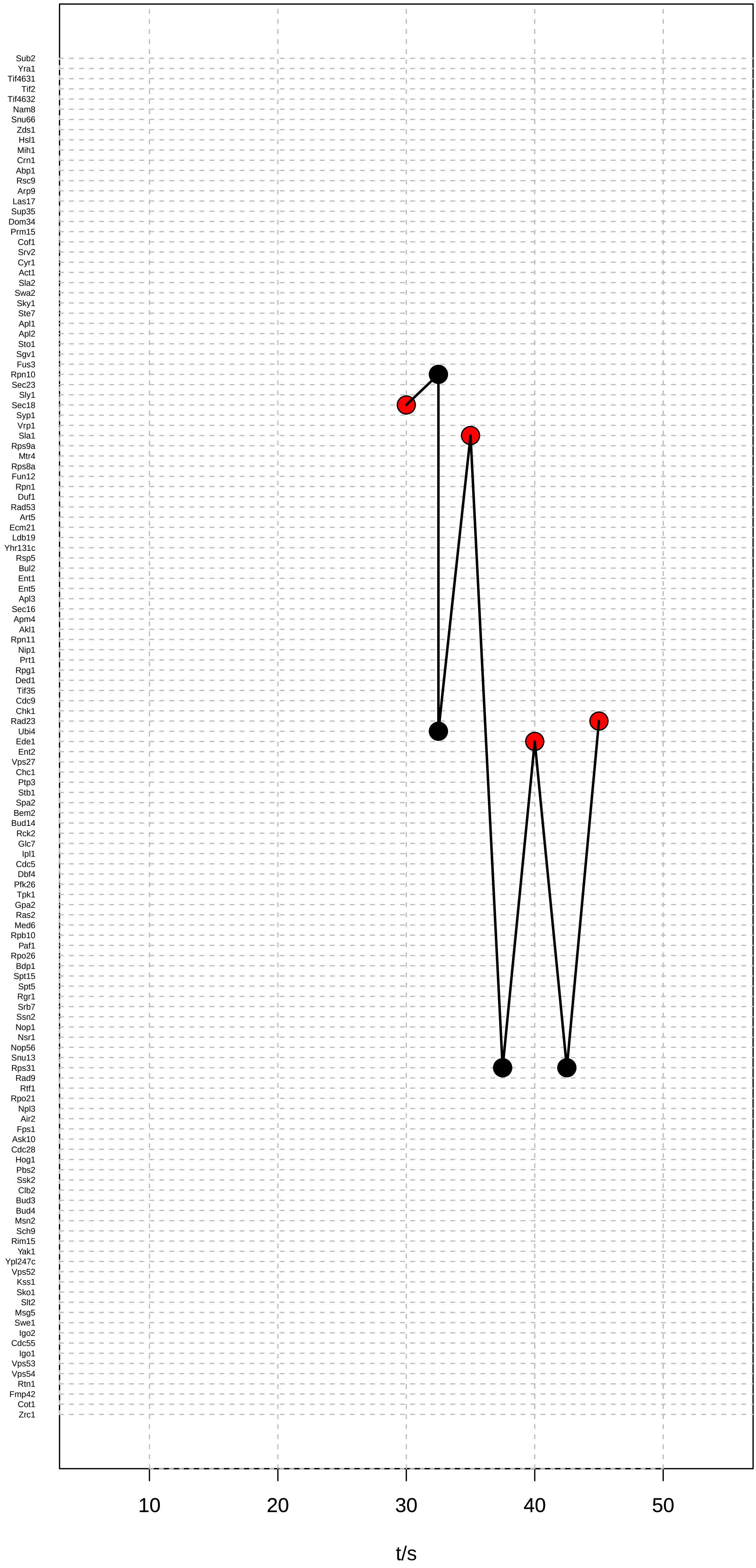

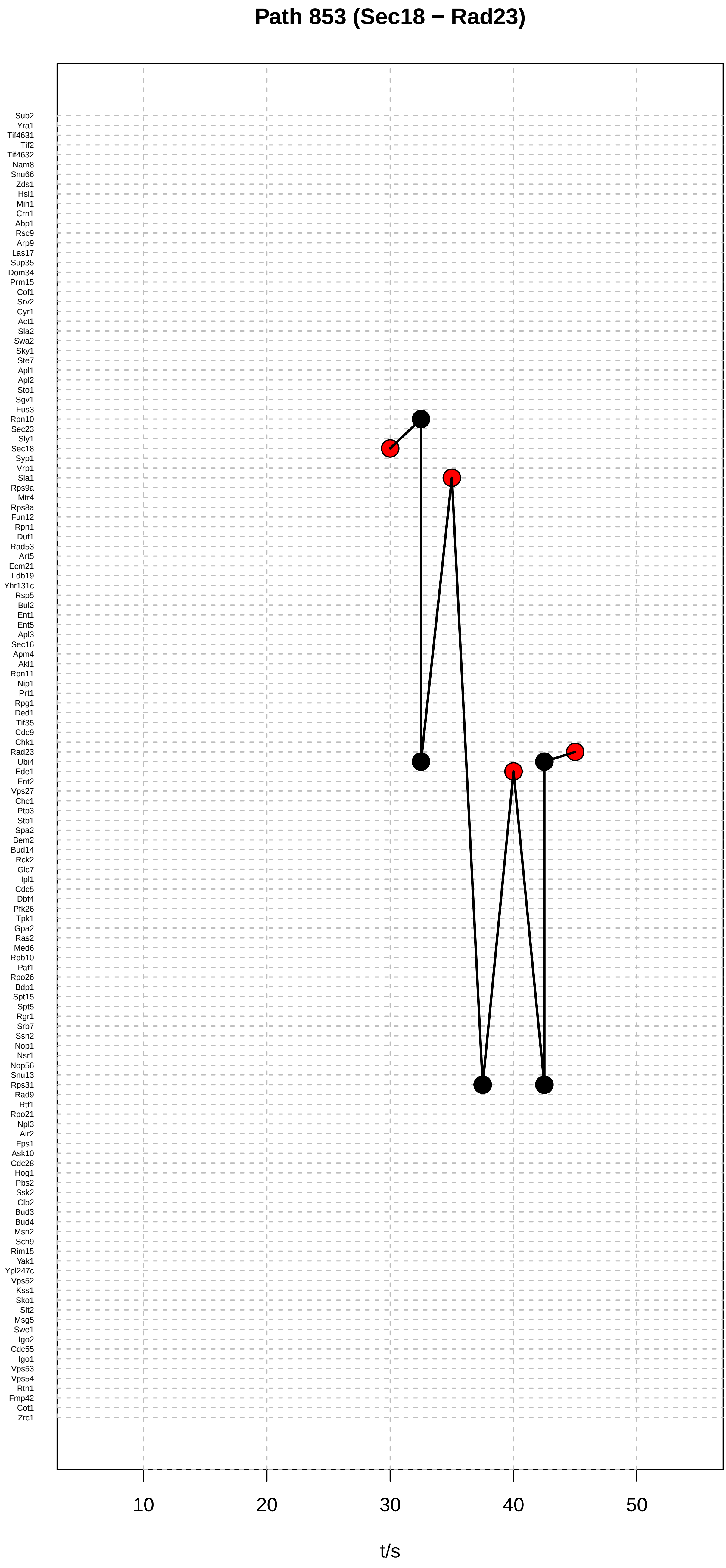

Path 853 (Sec18 − Rad23)
Sub2
Yra1
Tif4631
Tif2
Tif4632
Nam8
Snu66
Zds1
Hsl1
Mih1
Crn1
Abp1
Rsc9
Arp9
Las17
Sup35
Dom34
Prm15
Cof1
Srv2
Cyr1
Act1
Sla2
Swa2
Sky1
Ste7
Apl1
Apl2
Sto1
Sgv1
Fus3
Rpn10
Sec23
Sly1
Sec18
Syp1
Vrp1
Sla1
Rps9a
Mtr4
Rps8a
Fun12
Rpn1
Duf1
Rad53
Art5
Ecm21
Ldb19
Yhr131c
Rsp5
Bul2
Ent1
Ent5
Apl3
Sec16
Apm4
Akl1
Rpn11
Nip1
Prt1
Rpg1
Ded1
Tif35
Cdc9
Chk1
Rad23
Ubi4
Ede1
Ent2
Vps27
Chc1
Ptp3
Stb1
Spa2
Bem2
Bud14
Rck2
Glc7
Ipl1
Cdc5
Dbf4
Pfk26
Tpk1
Gpa2
Ras2
Med6
Rpb10
Paf1
Rpo26
Bdp1
Spt15
Spt5
Rgr1
Srb7
Ssn2
Nop1
Nsr1
Nop56
Snu13
Rps31
Rad9
Rtf1
Rpo21
Npl3
Air2
Fps1
Ask10
Cdc28
Hog1
Pbs2
Ssk2
Clb2
Bud3
Bud4
Msn2
Sch9
Rim15
Yak1
Ypl247c
Vps52
Kss1
Sko1
Slt2
Msg5
Swe1
Igo2
Cdc55
Igo1
Vps53
Vps54
Rtn1
Fmp42
Cot1
Zrc1
10
20
30
40
50
t/s

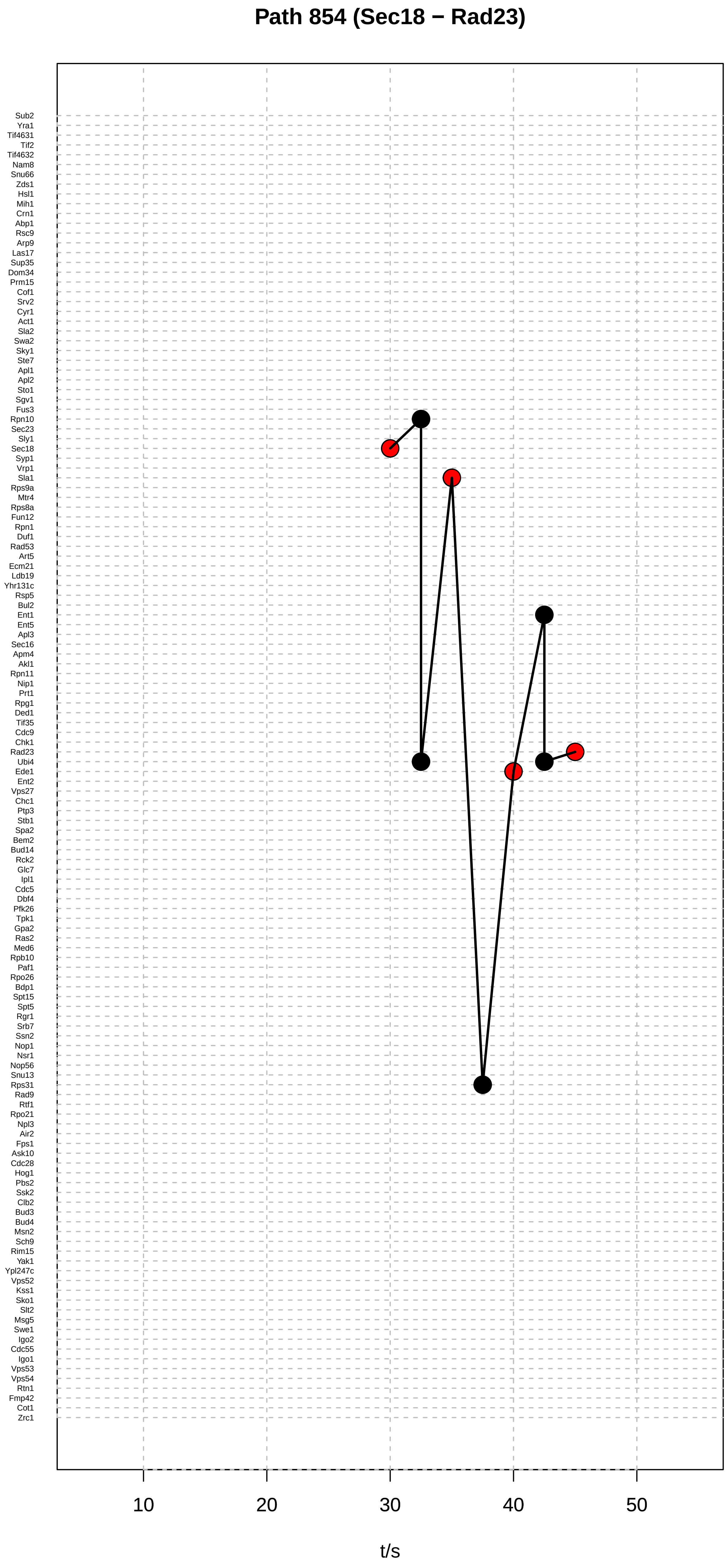
Path 854 (Sec18 − Rad23)
Sub2
Yra1
Tif4631
Tif2
Tif4632
Nam8
Snu66
Zds1
Hsl1
Mih1
Crn1
Abp1
Rsc9
Arp9
Las17
Sup35
Dom34
Prm15
Cof1
Srv2
Cyr1
Act1
Sla2
Swa2
Sky1
Ste7
Apl1
Apl2
Sto1
Sgv1
Fus3
Rpn10
Sec23
Sly1
Sec18
Syp1
Vrp1
Sla1
Rps9a
Mtr4
Rps8a
Fun12
Rpn1
Duf1
Rad53
Art5
Ecm21
Ldb19
Yhr131c
Rsp5
Bul2
Ent1
Ent5
Apl3
Sec16
Apm4
Akl1
Rpn11
Nip1
Prt1
Rpg1
Ded1
Tif35
Cdc9
Chk1
Rad23
Ubi4
Ede1
Ent2
Vps27
Chc1
Ptp3
Stb1
Spa2
Bem2
Bud14
Rck2
Glc7
Ipl1
Cdc5
Dbf4
Pfk26
Tpk1
Gpa2
Ras2
Med6
Rpb10
Paf1
Rpo26
Bdp1
Spt15
Spt5
Rgr1
Srb7
Ssn2
Nop1
Nsr1
Nop56
Snu13
Rps31
Rad9
Rtf1
Rpo21
Npl3
Air2
Fps1
Ask10
Cdc28
Hog1
Pbs2
Ssk2
Clb2
Bud3
Bud4
Msn2
Sch9
Rim15
Yak1
Ypl247c
Vps52
Kss1
Sko1
Slt2
Msg5
Swe1
Igo2
Cdc55
Igo1
Vps53
Vps54
Rtn1
Fmp42
Cot1
Zrc1
10
20
30
40
50
t/s

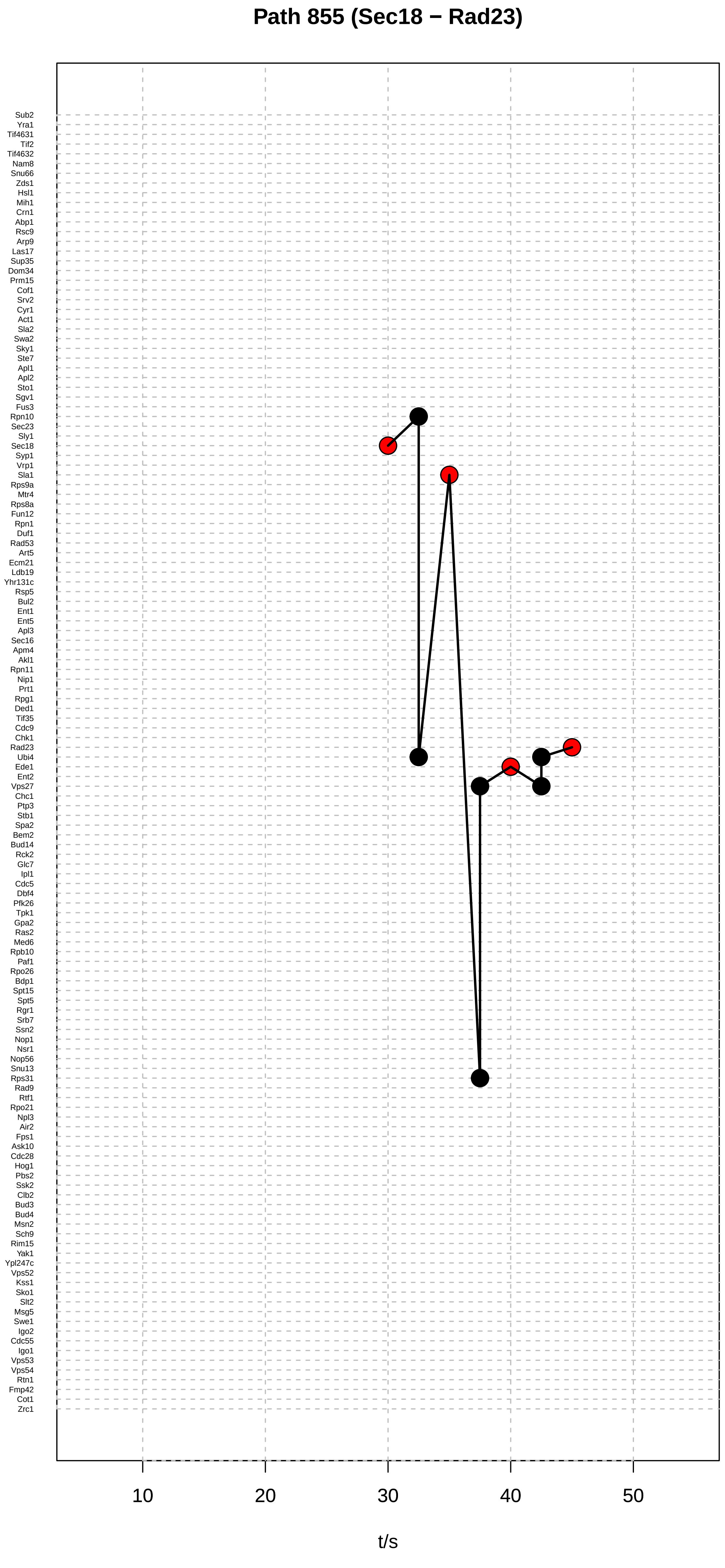

Path 855 (Sec18 − Rad23)
Sub2
Yra1
Tif4631
Tif2
Tif4632
Nam8
Snu66
Zds1
Hsl1
Mih1
Crn1
Abp1
Rsc9
Arp9
Las17
Sup35
Dom34
Prm15
Cof1
Srv2
Cyr1
Act1
Sla2
Swa2
Sky1
Ste7
Apl1
Apl2
Sto1
Sgv1
Fus3
Rpn10
Sec23
Sly1
Sec18
Syp1
Vrp1
Sla1
Rps9a
Mtr4
Rps8a
Fun12
Rpn1
Duf1
Rad53
Art5
Ecm21
Ldb19
Yhr131c
Rsp5
Bul2
Ent1
Ent5
Apl3
Sec16
Apm4
Akl1
Rpn11
Nip1
Prt1
Rpg1
Ded1
Tif35
Cdc9
Chk1
Rad23
Ubi4
Ede1
Ent2
Vps27
Chc1
Ptp3
Stb1
Spa2
Bem2
Bud14
Rck2
Glc7
Ipl1
Cdc5
Dbf4
Pfk26
Tpk1
Gpa2
Ras2
Med6
Rpb10
Paf1
Rpo26
Bdp1
Spt15
Spt5
Rgr1
Srb7
Ssn2
Nop1
Nsr1
Nop56
Snu13
Rps31
Rad9
Rtf1
Rpo21
Npl3
Air2
Fps1
Ask10
Cdc28
Hog1
Pbs2
Ssk2
Clb2
Bud3
Bud4
Msn2
Sch9
Rim15
Yak1
Ypl247c
Vps52
Kss1
Sko1
Slt2
Msg5
Swe1
Igo2
Cdc55
Igo1
Vps53
Vps54
Rtn1
Fmp42
Cot1
Zrc1
10
20
30
40
50
t/s

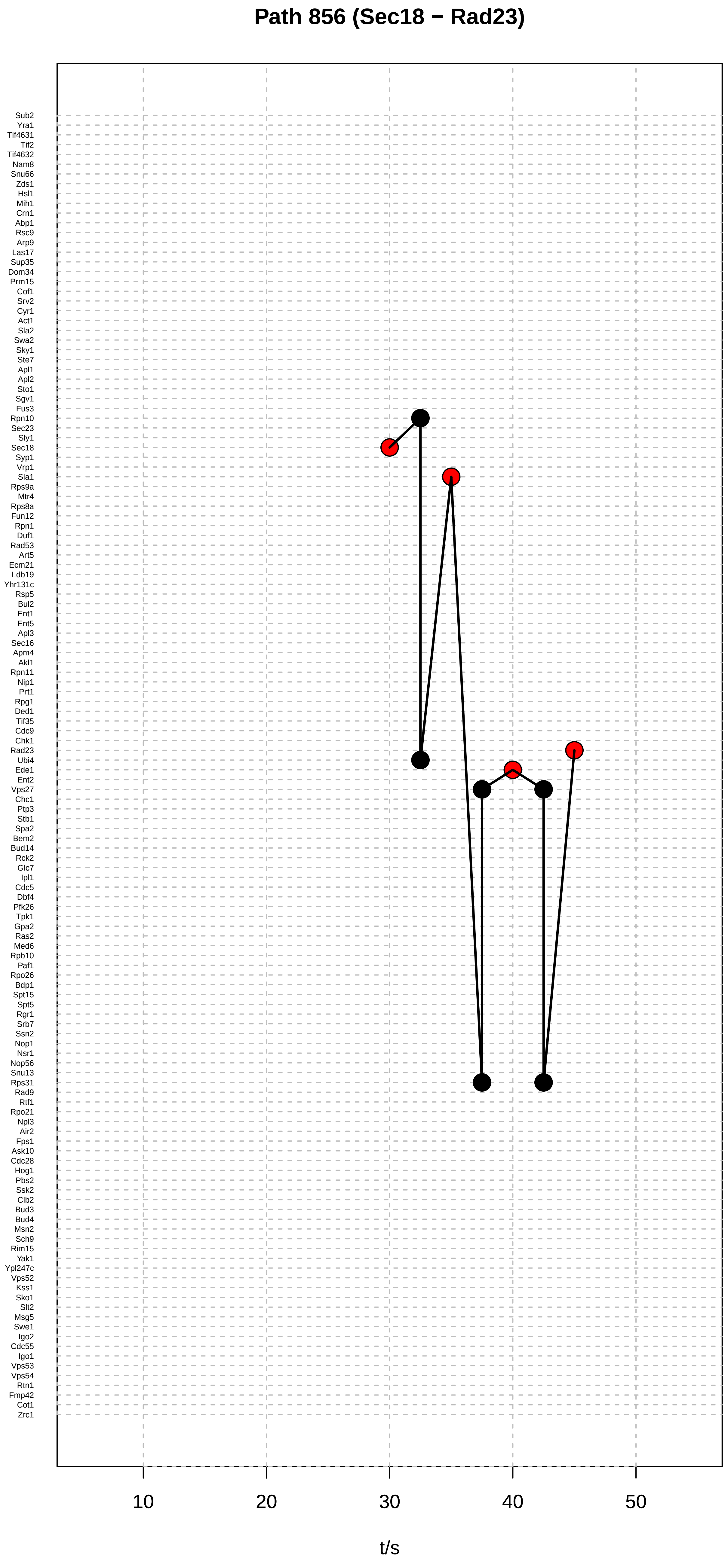
Path 856 (Sec18 − Rad23)
Sub2
Yra1
Tif4631
Tif2
Tif4632
Nam8
Snu66
Zds1
Hsl1
Mih1
Crn1
Abp1
Rsc9
Arp9
Las17
Sup35
Dom34
Prm15
Cof1
Srv2
Cyr1
Act1
Sla2
Swa2
Sky1
Ste7
Apl1
Apl2
Sto1
Sgv1
Fus3
Rpn10
Sec23
Sly1
Sec18
Syp1
Vrp1
Sla1
Rps9a
Mtr4
Rps8a
Fun12
Rpn1
Duf1
Rad53
Art5
Ecm21
Ldb19
Yhr131c
Rsp5
Bul2
Ent1
Ent5
Apl3
Sec16
Apm4
Akl1
Rpn11
Nip1
Prt1
Rpg1
Ded1
Tif35
Cdc9
Chk1
Rad23
Ubi4
Ede1
Ent2
Vps27
Chc1
Ptp3
Stb1
Spa2
Bem2
Bud14
Rck2
Glc7
Ipl1
Cdc5
Dbf4
Pfk26
Tpk1
Gpa2
Ras2
Med6
Rpb10
Paf1
Rpo26
Bdp1
Spt15
Spt5
Rgr1
Srb7
Ssn2
Nop1
Nsr1
Nop56
Snu13
Rps31
Rad9
Rtf1
Rpo21
Npl3
Air2
Fps1
Ask10
Cdc28
Hog1
Pbs2
Ssk2
Clb2
Bud3
Bud4
Msn2
Sch9
Rim15
Yak1
Ypl247c
Vps52
Kss1
Sko1
Slt2
Msg5
Swe1
Igo2
Cdc55
Igo1
Vps53
Vps54
Rtn1
Fmp42
Cot1
Zrc1
10
20
30
40
50
t/s

### Path 857 (Sec18 − Ent5)

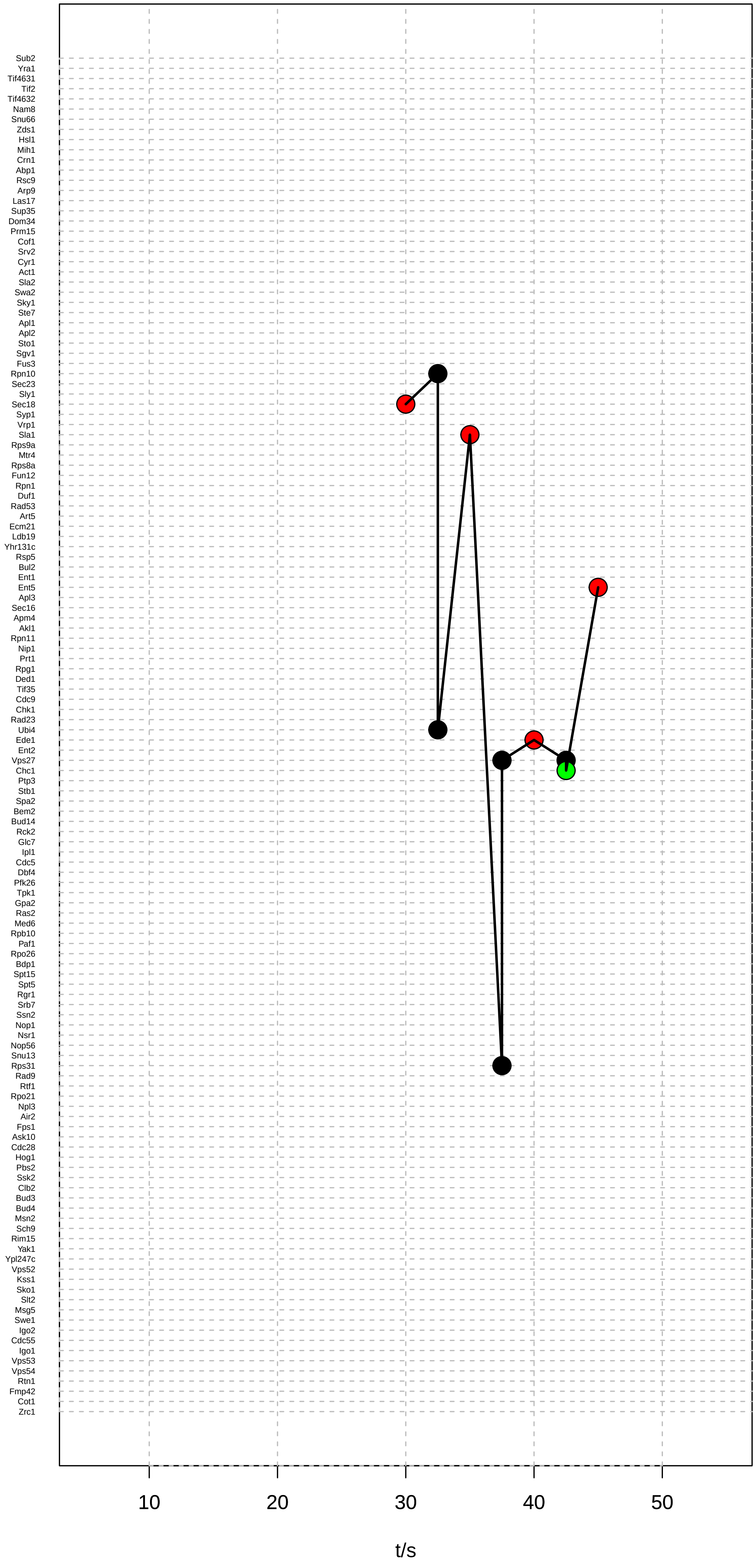

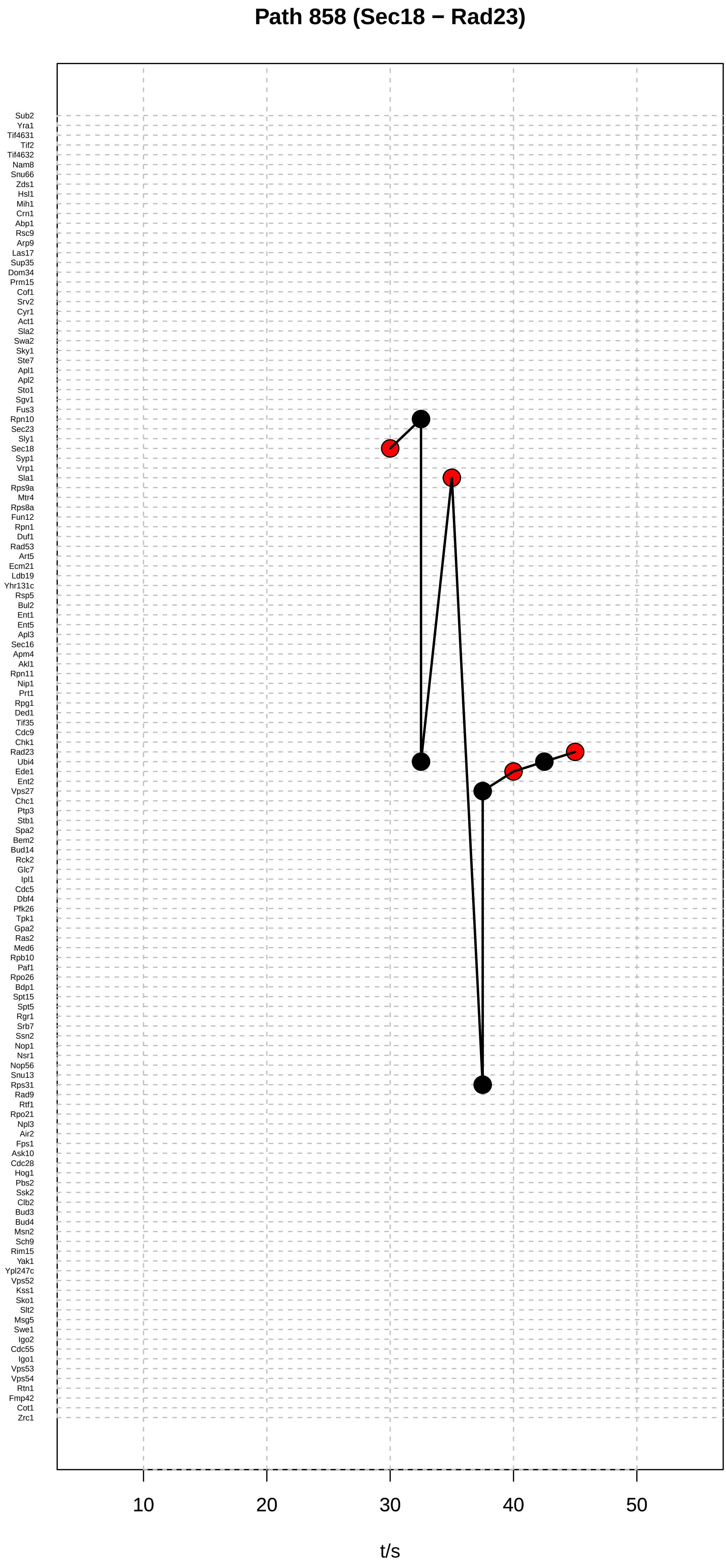

Path 858 (Sec18 − Rad23)
Sub2
Yra1
Tif4631
Tif2
Tif4632
Nam8
Snu66
Zds1
Hsl1
Mih1
Crn1
Abp1
Rsc9
Arp9
Las17
Sup35
Dom34
Prm15
Cof1
Srv2
Cyr1
Act1
Sla2
Swa2
Sky1
Ste7
Apl1
Apl2
Sto1
Sgv1
Fus3
Rpn10
Sec23
Sly1
Sec18
Syp1
Vrp1
Sla1
Rps9a
Mtr4
Rps8a
Fun12
Rpn1
Duf1
Rad53
Art5
Ecm21
Ldb19
Yhr131c
Rsp5
Bul2
Ent1
Ent5
Apl3
Sec16
Apm4
Akl1
Rpn11
Nip1
Prt1
Rpg1
Ded1
Tif35
Cdc9
Chk1
Rad23
Ubi4
Ede1
Ent2
Vps27
Chc1
Ptp3
Stb1
Spa2
Bem2
Bud14
Rck2
Glc7
Ipl1
Cdc5
Dbf4
Pfk26
Tpk1
Gpa2
Ras2
Med6
Rpb10
Paf1
Rpo26
Bdp1
Spt15
Spt5
Rgr1
Srb7
Ssn2
Nop1
Nsr1
Nop56
Snu13
Rps31
Rad9
Rtf1
Rpo21
Npl3
Air2
Fps1
Ask10
Cdc28
Hog1
Pbs2
Ssk2
Clb2
Bud3
Bud4
Msn2
Sch9
Rim15
Yak1
Ypl247c
Vps52
Kss1
Sko1
Slt2
Msg5
Swe1
Igo2
Cdc55
Igo1
Vps53
Vps54
Rtn1
Fmp42
Cot1
Zrc1
10
20
30
40
50
t/s

Path 859 (Sec18 − Rad23)

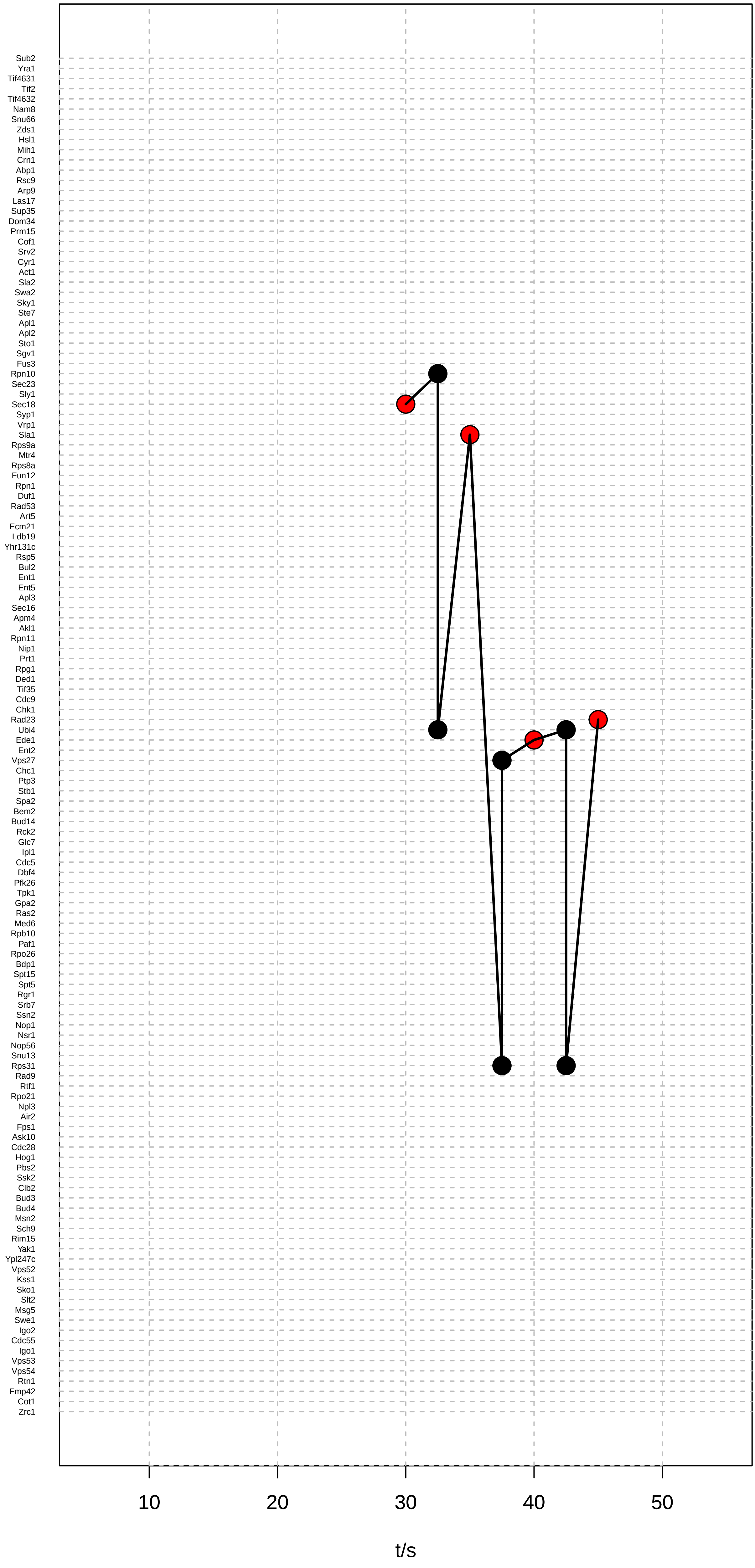

#### Path 860 (Sec18 − Rad23)

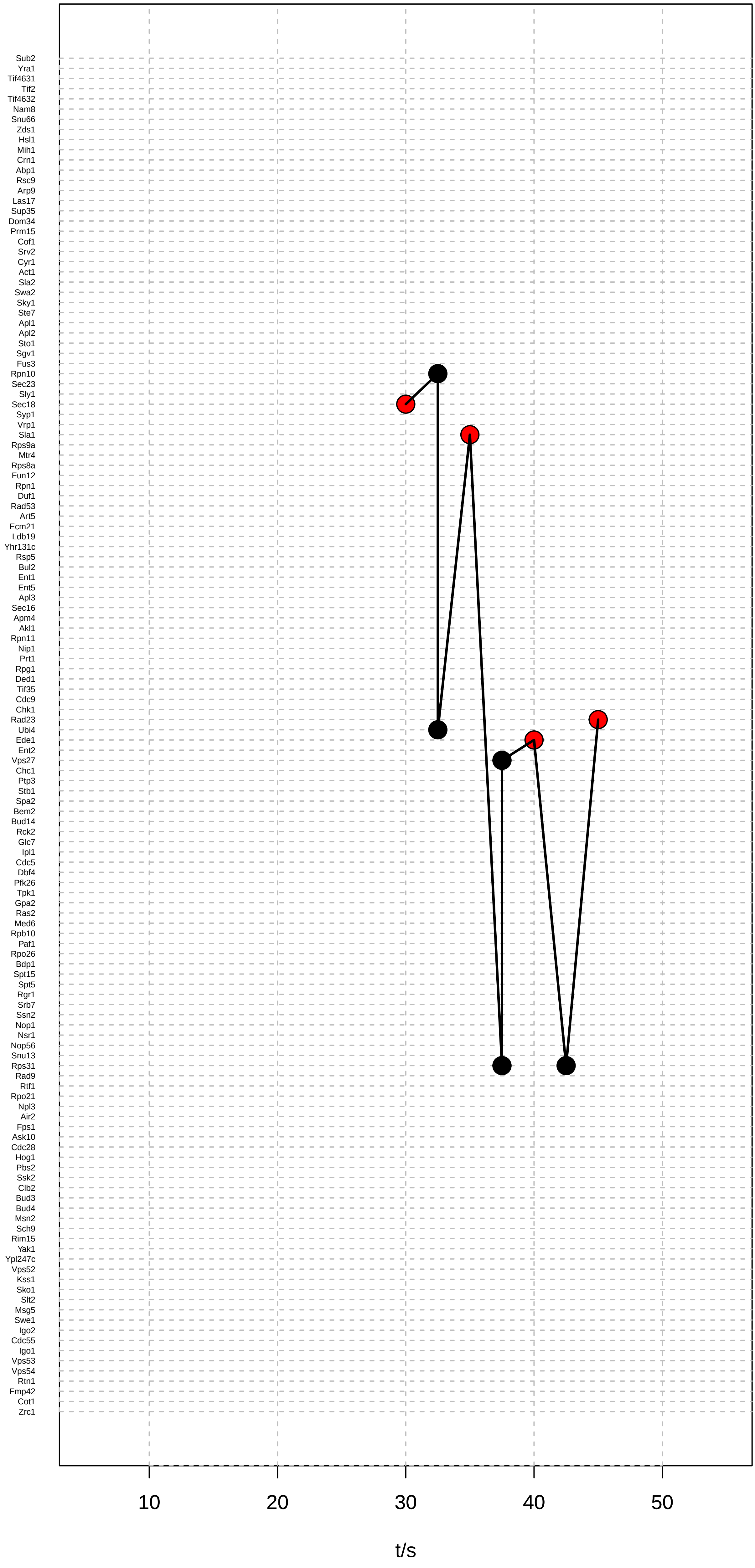

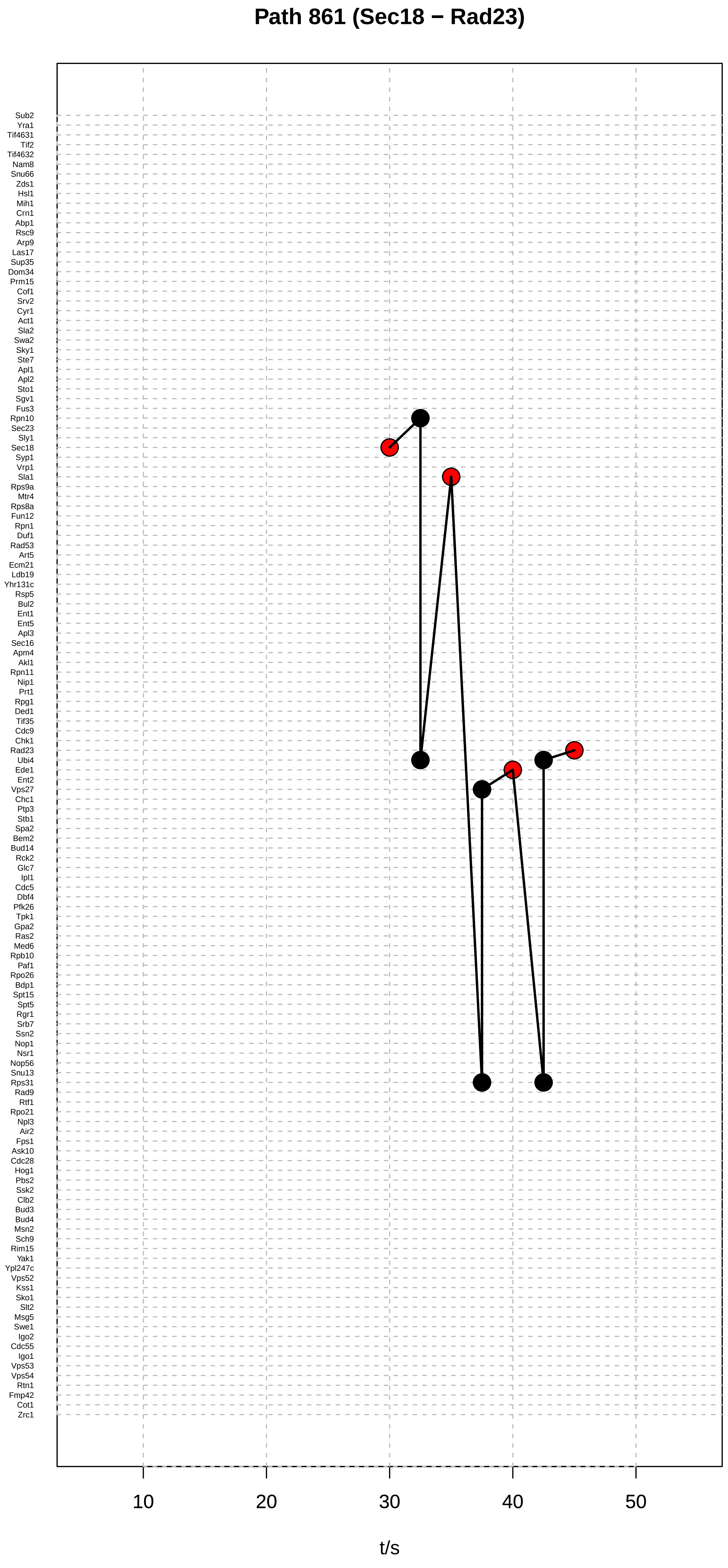

Path 861 (Sec18 − Rad23)
Sub2
Yra1
Tif4631
Tif2
Tif4632
Nam8
Snu66
Zds1
Hsl1
Mih1
Crn1
Abp1
Rsc9
Arp9
Las17
Sup35
Dom34
Prm15
Cof1
Srv2
Cyr1
Act1
Sla2
Swa2
Sky1
Ste7
Apl1
Apl2
Sto1
Sgv1
Fus3
Rpn10
Sec23
Sly1
Sec18
Syp1
Vrp1
Sla1
Rps9a
Mtr4
Rps8a
Fun12
Rpn1
Duf1
Rad53
Art5
Ecm21
Ldb19
Yhr131c
Rsp5
Bul2
Ent1
Ent5
Apl3
Sec16
Apm4
Akl1
Rpn11
Nip1
Prt1
Rpg1
Ded1
Tif35
Cdc9
Chk1
Rad23
Ubi4
Ede1
Ent2
Vps27
Chc1
Ptp3
Stb1
Spa2
Bem2
Bud14
Rck2
Glc7
Ipl1
Cdc5
Dbf4
Pfk26
Tpk1
Gpa2
Ras2
Med6
Rpb10
Paf1
Rpo26
Bdp1
Spt15
Spt5
Rgr1
Srb7
Ssn2
Nop1
Nsr1
Nop56
Snu13
Rps31
Rad9
Rtf1
Rpo21
Npl3
Air2
Fps1
Ask10
Cdc28
Hog1
Pbs2
Ssk2
Clb2
Bud3
Bud4
Msn2
Sch9
Rim15
Yak1
Ypl247c
Vps52
Kss1
Sko1
Slt2
Msg5
Swe1
Igo2
Cdc55
Igo1
Vps53
Vps54
Rtn1
Fmp42
Cot1
Zrc1
10
20
30
40
50
t/s

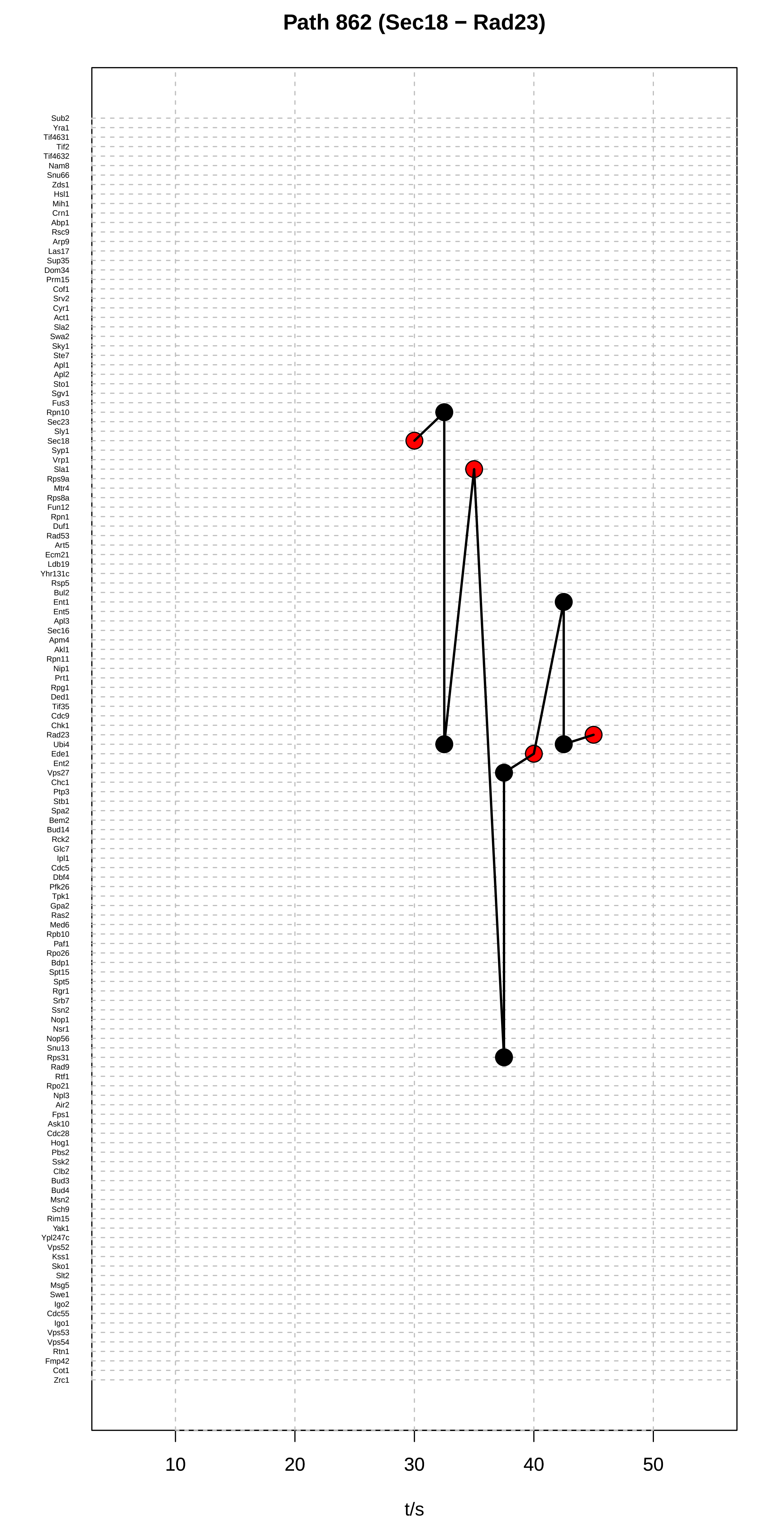

Path 862 (Sec18 − Rad23)
Sub2
Yra1
Tif4631
Tif2
Tif4632
Nam8
Snu66
Zds1
Hsl1
Mih1
Crn1
Abp1
Rsc9
Arp9
Las17
Sup35
Dom34
Prm15
Cof1
Srv2
Cyr1
Act1
Sla2
Swa2
Sky1
Ste7
Apl1
Apl2
Sto1
Sgv1
Fus3
Rpn10
Sec23
Sly1
Sec18
Syp1
Vrp1
Sla1
Rps9a
Mtr4
Rps8a
Fun12
Rpn1
Duf1
Rad53
Art5
Ecm21
Ldb19
Yhr131c
Rsp5
Bul2
Ent1
Ent5
Apl3
Sec16
Apm4
Akl1
Rpn11
Nip1
Prt1
Rpg1
Ded1
Tif35
Cdc9
Chk1
Rad23
Ubi4
Ede1
Ent2
Vps27
Chc1
Ptp3
Stb1
Spa2
Bem2
Bud14
Rck2
Glc7
Ipl1
Cdc5
Dbf4
Pfk26
Tpk1
Gpa2
Ras2
Med6
Rpb10
Paf1
Rpo26
Bdp1
Spt15
Spt5
Rgr1
Srb7
Ssn2
Nop1
Nsr1
Nop56
Snu13
Rps31
Rad9
Rtf1
Rpo21
Npl3
Air2
Fps1
Ask10
Cdc28
Hog1
Pbs2
Ssk2
Clb2
Bud3
Bud4
Msn2
Sch9
Rim15
Yak1
Ypl247c
Vps52
Kss1
Sko1
Slt2
Msg5
Swe1
Igo2
Cdc55
Igo1
Vps53
Vps54
Rtn1
Fmp42
Cot1
Zrc1
10
20
30
40
50
t/s

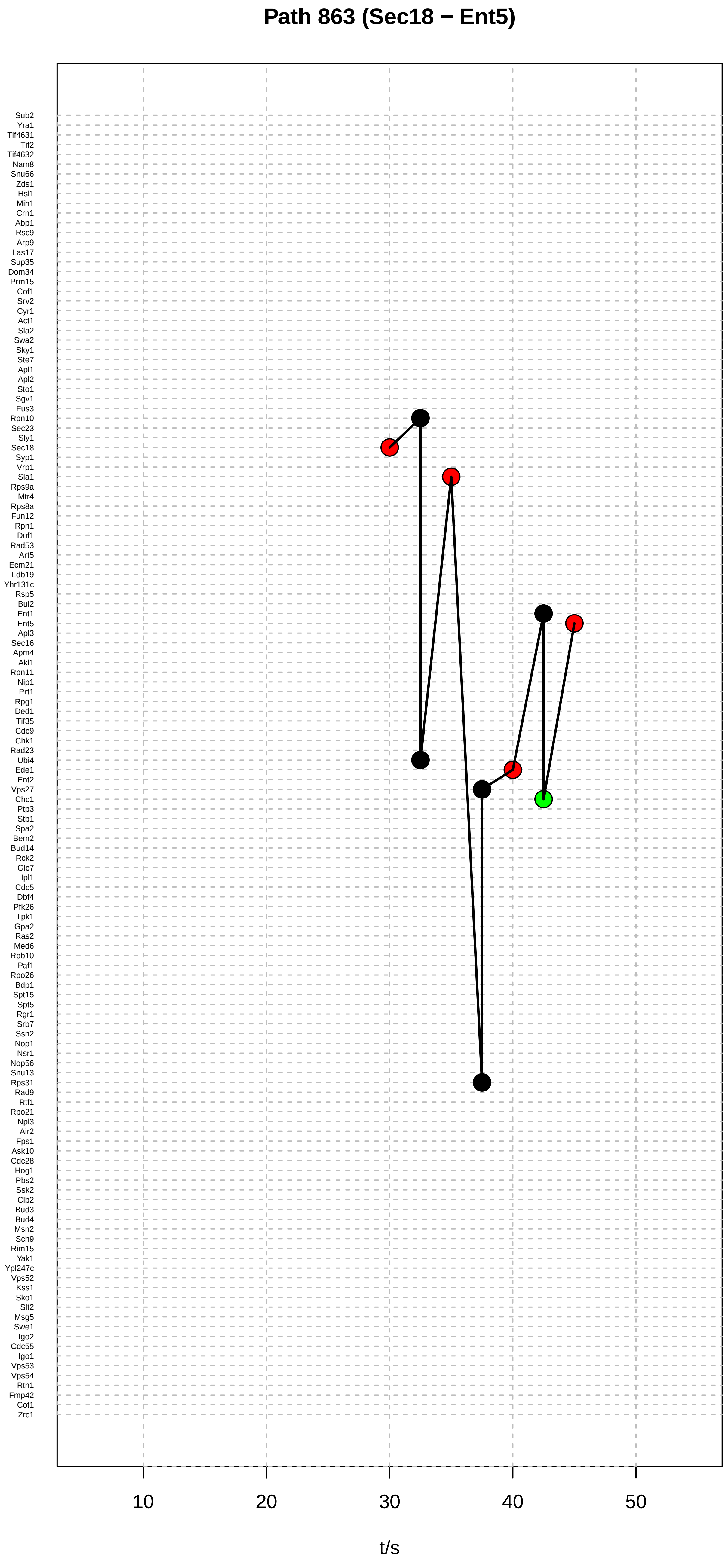

Path 863 (Sec18 − Ent5)
Sub2
Yra1
Tif4631
Tif2
Tif4632
Nam8
Snu66
Zds1
Hsl1
Mih1
Crn1
Abp1
Rsc9
Arp9
Las17
Sup35
Dom34
Prm15
Cof1
Srv2
Cyr1
Act1
Sla2
Swa2
Sky1
Ste7
Apl1
Apl2
Sto1
Sgv1
Fus3
Rpn10
Sec23
Sly1
Sec18
Syp1
Vrp1
Sla1
Rps9a
Mtr4
Rps8a
Fun12
Rpn1
Duf1
Rad53
Art5
Ecm21
Ldb19
Yhr131c
Rsp5
Bul2
Ent1
Ent5
Apl3
Sec16
Apm4
Akl1
Rpn11
Nip1
Prt1
Rpg1
Ded1
Tif35
Cdc9
Chk1
Rad23
Ubi4
Ede1
Ent2
Vps27
Chc1
Ptp3
Stb1
Spa2
Bem2
Bud14
Rck2
Glc7
Ipl1
Cdc5
Dbf4
Pfk26
Tpk1
Gpa2
Ras2
Med6
Rpb10
Paf1
Rpo26
Bdp1
Spt15
Spt5
Rgr1
Srb7
Ssn2
Nop1
Nsr1
Nop56
Snu13
Rps31
Rad9
Rtf1
Rpo21
Npl3
Air2
Fps1
Ask10
Cdc28
Hog1
Pbs2
Ssk2
Clb2
Bud3
Bud4
Msn2
Sch9
Rim15
Yak1
Ypl247c
Vps52
Kss1
Sko1
Slt2
Msg5
Swe1
Igo2
Cdc55
Igo1
Vps53
Vps54
Rtn1
Fmp42
Cot1
Zrc1
10
20
30
40
50
t/s

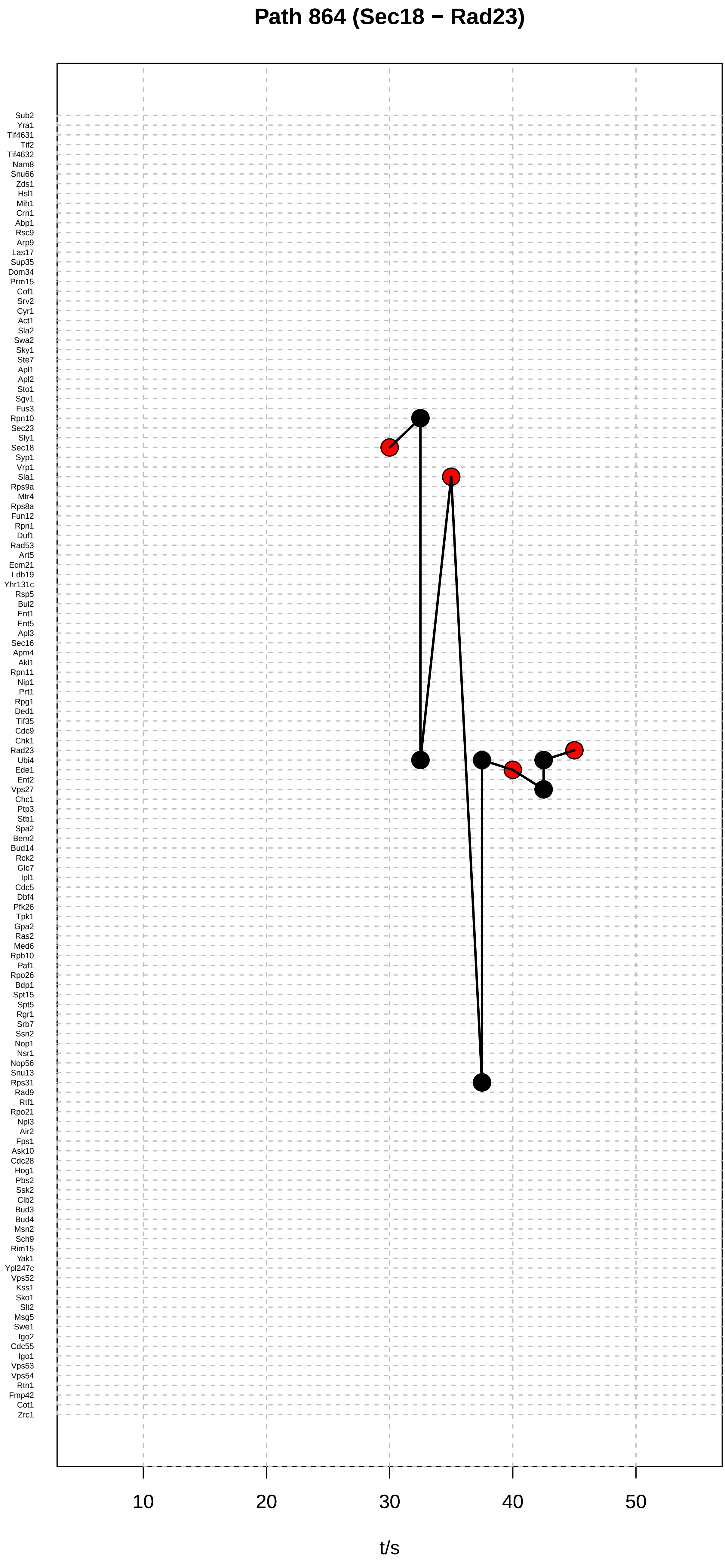

Path 864 (Sec18 − Rad23)
Sub2
Yra1
Tif4631
Tif2
Tif4632
Nam8
Snu66
Zds1
Hsl1
Mih1
Crn1
Abp1
Rsc9
Arp9
Las17
Sup35
Dom34
Prm15
Cof1
Srv2
Cyr1
Act1
Sla2
Swa2
Sky1
Ste7
Apl1
Apl2
Sto1
Sgv1
Fus3
Rpn10
Sec23
Sly1
Sec18
Syp1
Vrp1
Sla1
Rps9a
Mtr4
Rps8a
Fun12
Rpn1
Duf1
Rad53
Art5
Ecm21
Ldb19
Yhr131c
Rsp5
Bul2
Ent1
Ent5
Apl3
Sec16
Apm4
Akl1
Rpn11
Nip1
Prt1
Rpg1
Ded1
Tif35
Cdc9
Chk1
Rad23
Ubi4
Ede1
Ent2
Vps27
Chc1
Ptp3
Stb1
Spa2
Bem2
Bud14
Rck2
Glc7
Ipl1
Cdc5
Dbf4
Pfk26
Tpk1
Gpa2
Ras2
Med6
Rpb10
Paf1
Rpo26
Bdp1
Spt15
Spt5
Rgr1
Srb7
Ssn2
Nop1
Nsr1
Nop56
Snu13
Rps31
Rad9
Rtf1
Rpo21
Npl3
Air2
Fps1
Ask10
Cdc28
Hog1
Pbs2
Ssk2
Clb2
Bud3
Bud4
Msn2
Sch9
Rim15
Yak1
Ypl247c
Vps52
Kss1
Sko1
Slt2
Msg5
Swe1
Igo2
Cdc55
Igo1
Vps53
Vps54
Rtn1
Fmp42
Cot1
Zrc1
10
20
30
40
50
t/s

### Path 865 (Sec18 − Rad23)

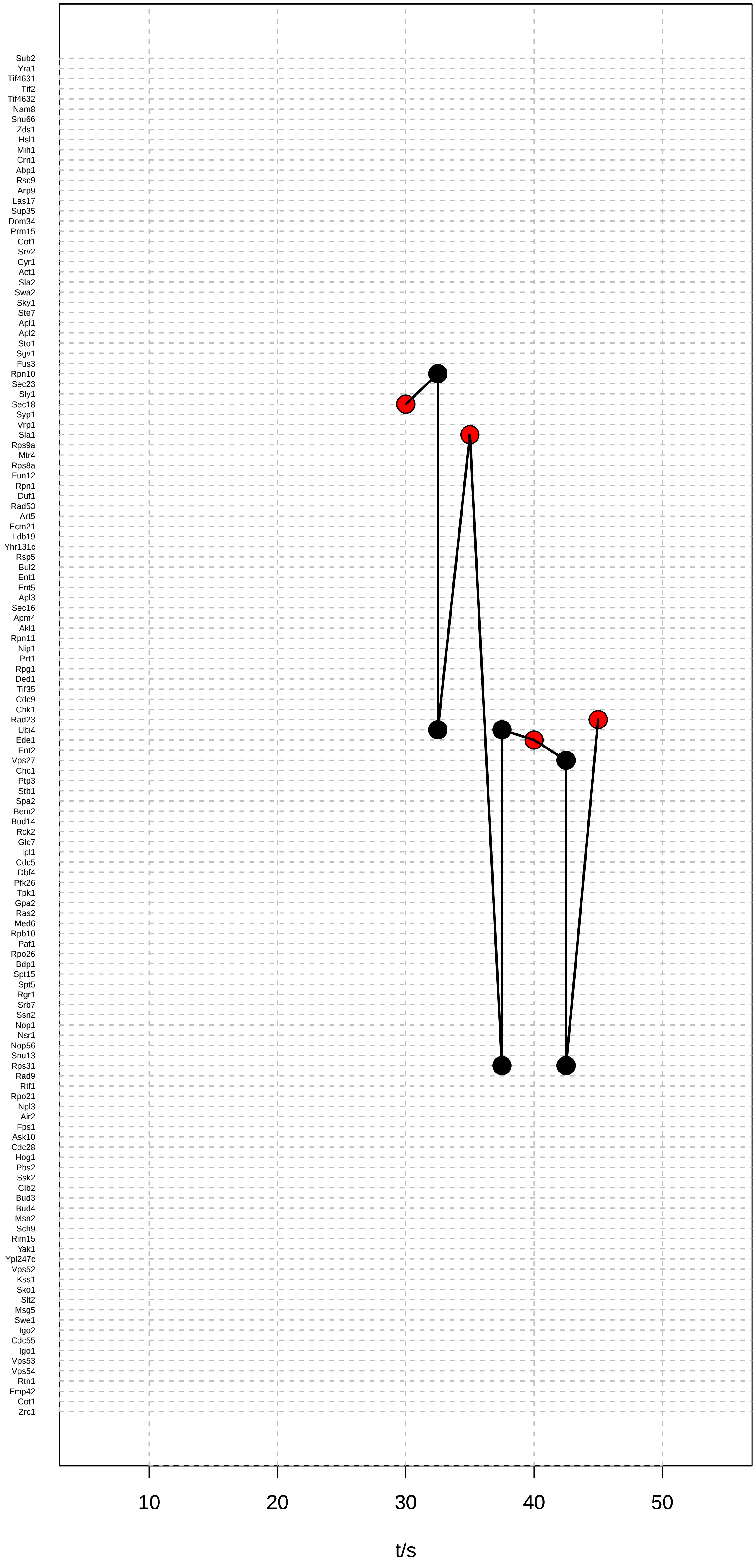

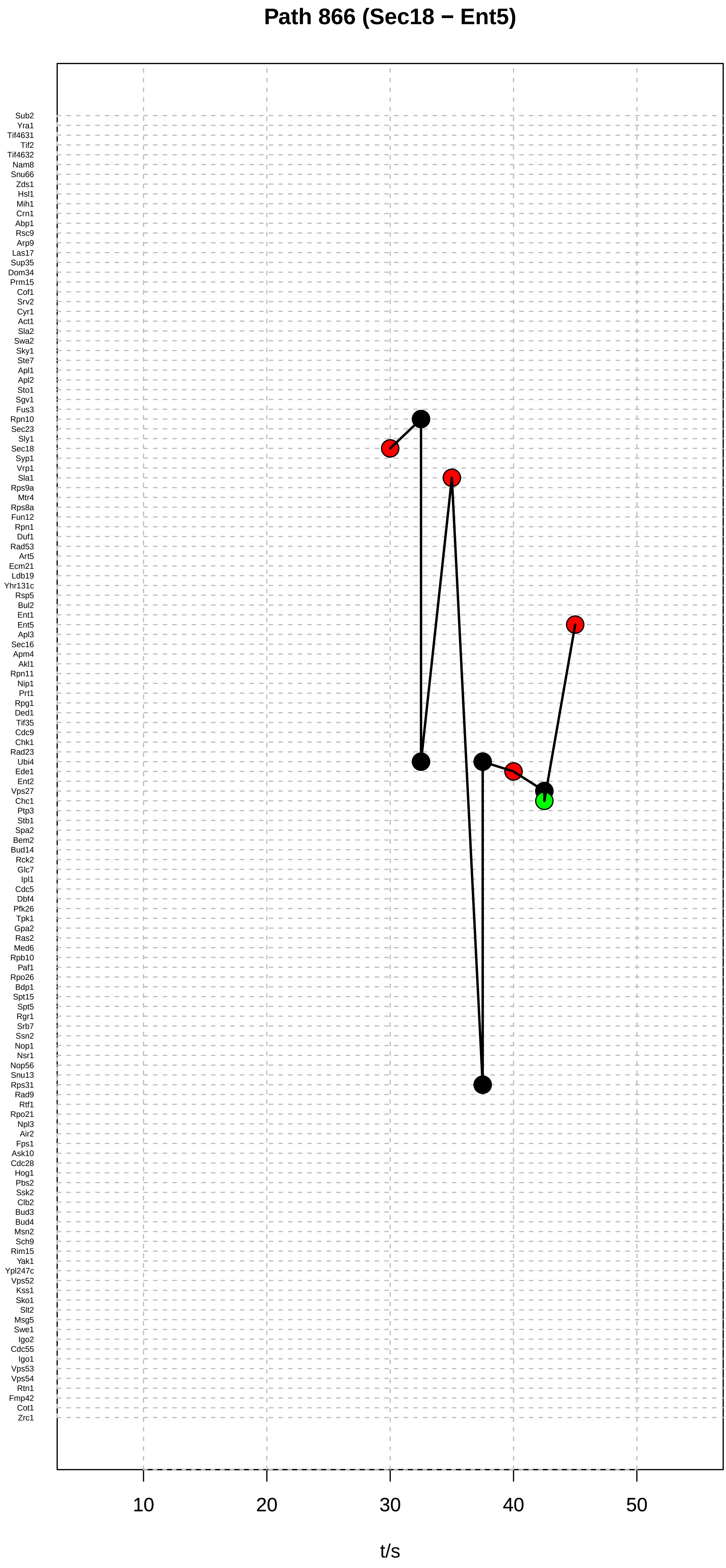
Path 866 (Sec18 − Ent5)
Sub2
Yra1
Tif4631
Tif2
Tif4632
Nam8
Snu66
Zds1
Hsl1
Mih1
Crn1
Abp1
Rsc9
Arp9
Las17
Sup35
Dom34
Prm15
Cof1
Srv2
Cyr1
Act1
Sla2
Swa2
Sky1
Ste7
Apl1
Apl2
Sto1
Sgv1
Fus3
Rpn10
Sec23
Sly1
Sec18
Syp1
Vrp1
Sla1
Rps9a
Mtr4
Rps8a
Fun12
Rpn1
Duf1
Rad53
Art5
Ecm21
Ldb19
Yhr131c
Rsp5
Bul2
Ent1
Ent5
Apl3
Sec16
Apm4
Akl1
Rpn11
Nip1
Prt1
Rpg1
Ded1
Tif35
Cdc9
Chk1
Rad23
Ubi4
Ede1
Ent2
Vps27
Chc1
Ptp3
Stb1
Spa2
Bem2
Bud14
Rck2
Glc7
Ipl1
Cdc5
Dbf4
Pfk26
Tpk1
Gpa2
Ras2
Med6
Rpb10
Paf1
Rpo26
Bdp1
Spt15
Spt5
Rgr1
Srb7
Ssn2
Nop1
Nsr1
Nop56
Snu13
Rps31
Rad9
Rtf1
Rpo21
Npl3
Air2
Fps1
Ask10
Cdc28
Hog1
Pbs2
Ssk2
Clb2
Bud3
Bud4
Msn2
Sch9
Rim15
Yak1
Ypl247c
Vps52
Kss1
Sko1
Slt2
Msg5
Swe1
Igo2
Cdc55
Igo1
Vps53
Vps54
Rtn1
Fmp42
Cot1
Zrc1
10
20
30
40
50
t/s

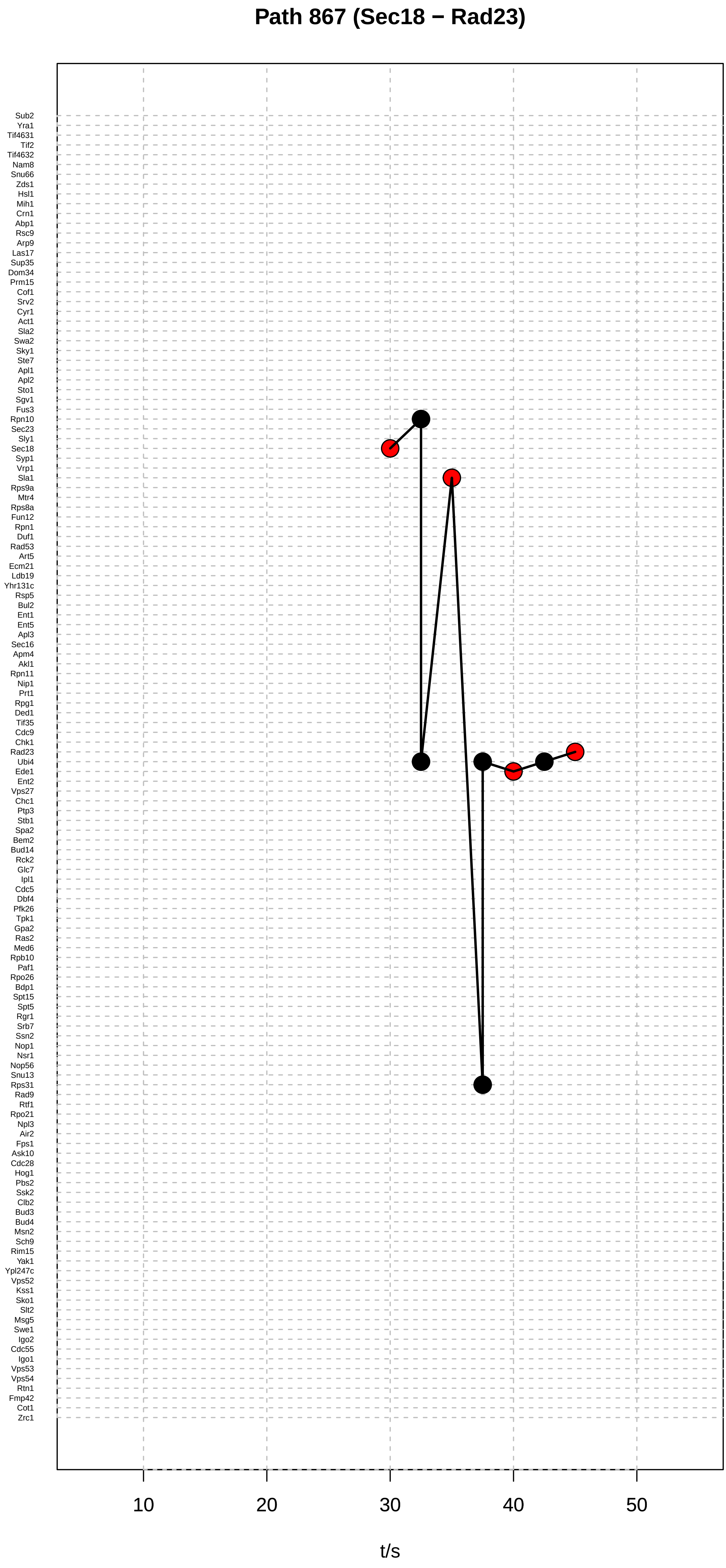
Path 867 (Sec18 − Rad23)
Sub2
Yra1
Tif4631
Tif2
Tif4632
Nam8
Snu66
Zds1
Hsl1
Mih1
Crn1
Abp1
Rsc9
Arp9
Las17
Sup35
Dom34
Prm15
Cof1
Srv2
Cyr1
Act1
Sla2
Swa2
Sky1
Ste7
Apl1
Apl2
Sto1
Sgv1
Fus3
Rpn10
Sec23
Sly1
Sec18
Syp1
Vrp1
Sla1
Rps9a
Mtr4
Rps8a
Fun12
Rpn1
Duf1
Rad53
Art5
Ecm21
Ldb19
Yhr131c
Rsp5
Bul2
Ent1
Ent5
Apl3
Sec16
Apm4
Akl1
Rpn11
Nip1
Prt1
Rpg1
Ded1
Tif35
Cdc9
Chk1
Rad23
Ubi4
Ede1
Ent2
Vps27
Chc1
Ptp3
Stb1
Spa2
Bem2
Bud14
Rck2
Glc7
Ipl1
Cdc5
Dbf4
Pfk26
Tpk1
Gpa2
Ras2
Med6
Rpb10
Paf1
Rpo26
Bdp1
Spt15
Spt5
Rgr1
Srb7
Ssn2
Nop1
Nsr1
Nop56
Snu13
Rps31
Rad9
Rtf1
Rpo21
Npl3
Air2
Fps1
Ask10
Cdc28
Hog1
Pbs2
Ssk2
Clb2
Bud3
Bud4
Msn2
Sch9
Rim15
Yak1
Ypl247c
Vps52
Kss1
Sko1
Slt2
Msg5
Swe1
Igo2
Cdc55
Igo1
Vps53
Vps54
Rtn1
Fmp42
Cot1
Zrc1
10
20
30
40
50
t/s

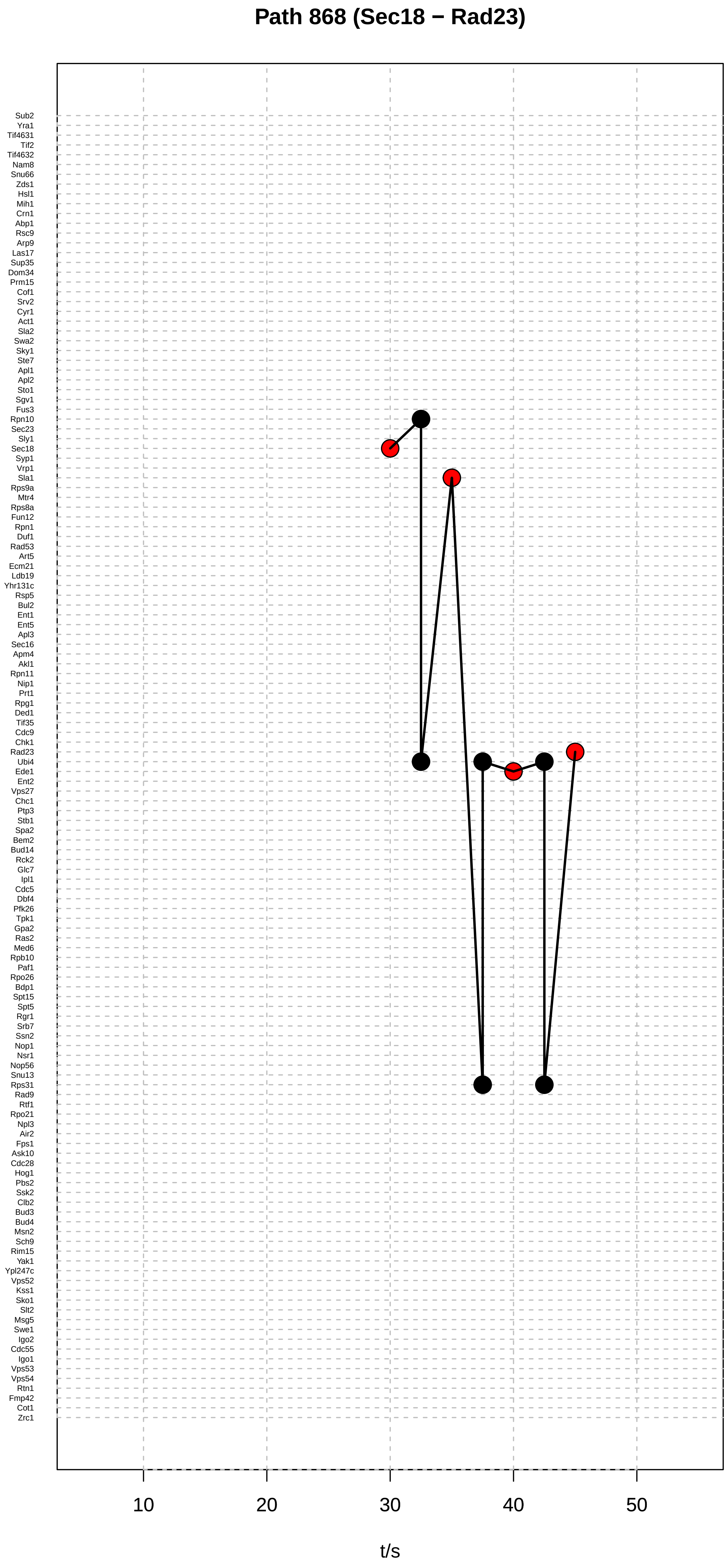
Path 868 (Sec18 − Rad23)
Sub2
Yra1
Tif4631
Tif2
Tif4632
Nam8
Snu66
Zds1
Hsl1
Mih1
Crn1
Abp1
Rsc9
Arp9
Las17
Sup35
Dom34
Prm15
Cof1
Srv2
Cyr1
Act1
Sla2
Swa2
Sky1
Ste7
Apl1
Apl2
Sto1
Sgv1
Fus3
Rpn10
Sec23
Sly1
Sec18
Syp1
Vrp1
Sla1
Rps9a
Mtr4
Rps8a
Fun12
Rpn1
Duf1
Rad53
Art5
Ecm21
Ldb19
Yhr131c
Rsp5
Bul2
Ent1
Ent5
Apl3
Sec16
Apm4
Akl1
Rpn11
Nip1
Prt1
Rpg1
Ded1
Tif35
Cdc9
Chk1
Rad23
Ubi4
Ede1
Ent2
Vps27
Chc1
Ptp3
Stb1
Spa2
Bem2
Bud14
Rck2
Glc7
Ipl1
Cdc5
Dbf4
Pfk26
Tpk1
Gpa2
Ras2
Med6
Rpb10
Paf1
Rpo26
Bdp1
Spt15
Spt5
Rgr1
Srb7
Ssn2
Nop1
Nsr1
Nop56
Snu13
Rps31
Rad9
Rtf1
Rpo21
Npl3
Air2
Fps1
Ask10
Cdc28
Hog1
Pbs2
Ssk2
Clb2
Bud3
Bud4
Msn2
Sch9
Rim15
Yak1
Ypl247c
Vps52
Kss1
Sko1
Slt2
Msg5
Swe1
Igo2
Cdc55
Igo1
Vps53
Vps54
Rtn1
Fmp42
Cot1
Zrc1
10
20
30
40
50
t/s

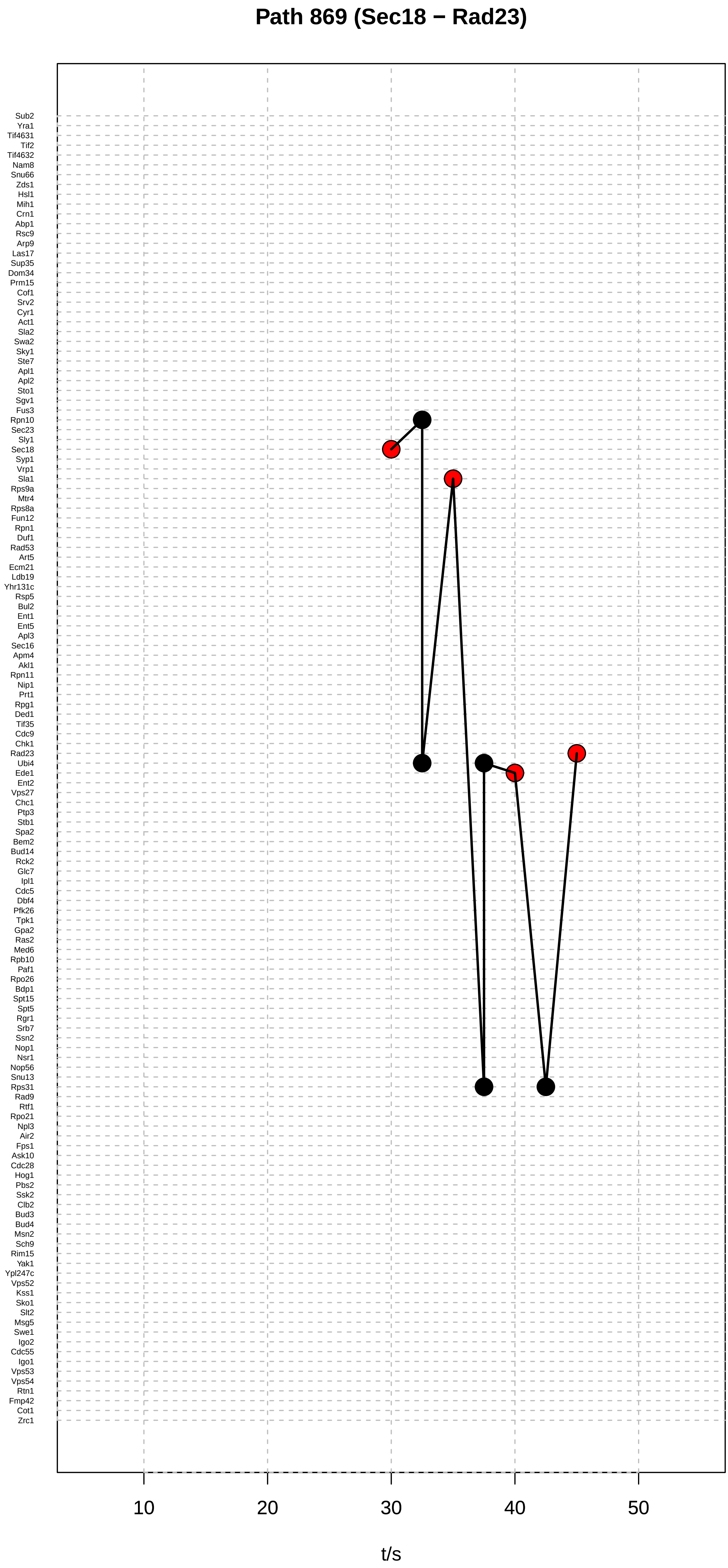

Path 869 (Sec18 − Rad23)
Sub2
Yra1
Tif4631
Tif2
Tif4632
Nam8
Snu66
Zds1
Hsl1
Mih1
Crn1
Abp1
Rsc9
Arp9
Las17
Sup35
Dom34
Prm15
Cof1
Srv2
Cyr1
Act1
Sla2
Swa2
Sky1
Ste7
Apl1
Apl2
Sto1
Sgv1
Fus3
Rpn10
Sec23
Sly1
Sec18
Syp1
Vrp1
Sla1
Rps9a
Mtr4
Rps8a
Fun12
Rpn1
Duf1
Rad53
Art5
Ecm21
Ldb19
Yhr131c
Rsp5
Bul2
Ent1
Ent5
Apl3
Sec16
Apm4
Akl1
Rpn11
Nip1
Prt1
Rpg1
Ded1
Tif35
Cdc9
Chk1
Rad23
Ubi4
Ede1
Ent2
Vps27
Chc1
Ptp3
Stb1
Spa2
Bem2
Bud14
Rck2
Glc7
Ipl1
Cdc5
Dbf4
Pfk26
Tpk1
Gpa2
Ras2
Med6
Rpb10
Paf1
Rpo26
Bdp1
Spt15
Spt5
Rgr1
Srb7
Ssn2
Nop1
Nsr1
Nop56
Snu13
Rps31
Rad9
Rtf1
Rpo21
Npl3
Air2
Fps1
Ask10
Cdc28
Hog1
Pbs2
Ssk2
Clb2
Bud3
Bud4
Msn2
Sch9
Rim15
Yak1
Ypl247c
Vps52
Kss1
Sko1
Slt2
Msg5
Swe1
Igo2
Cdc55
Igo1
Vps53
Vps54
Rtn1
Fmp42
Cot1
Zrc1
10
20
30
40
50
t/s

#### Path 870 (Sec18 − Rad23)

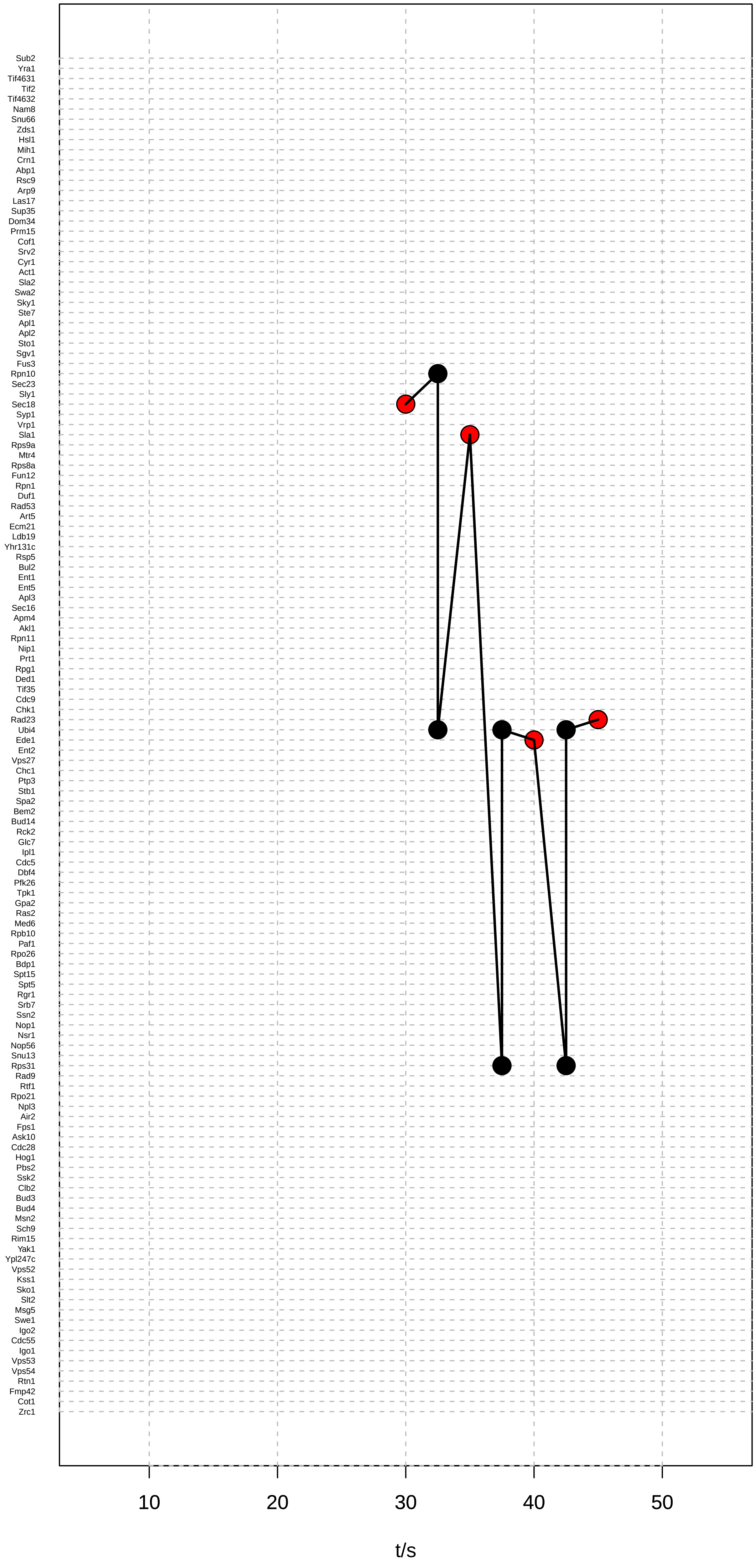

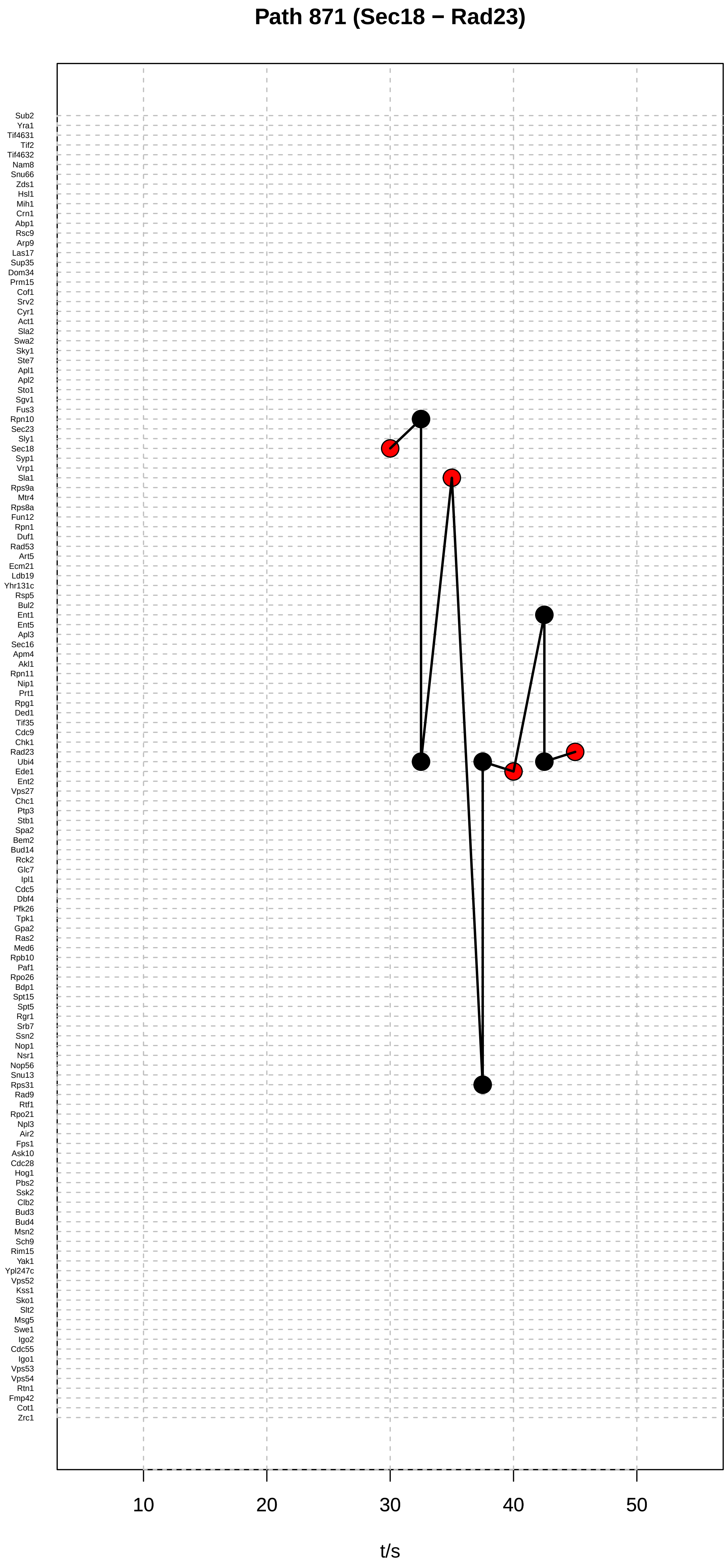

Path 871 (Sec18 − Rad23)
Sub2
Yra1
Tif4631
Tif2
Tif4632
Nam8
Snu66
Zds1
Hsl1
Mih1
Crn1
Abp1
Rsc9
Arp9
Las17
Sup35
Dom34
Prm15
Cof1
Srv2
Cyr1
Act1
Sla2
Swa2
Sky1
Ste7
Apl1
Apl2
Sto1
Sgv1
Fus3
Rpn10
Sec23
Sly1
Sec18
Syp1
Vrp1
Sla1
Rps9a
Mtr4
Rps8a
Fun12
Rpn1
Duf1
Rad53
Art5
Ecm21
Ldb19
Yhr131c
Rsp5
Bul2
Ent1
Ent5
Apl3
Sec16
Apm4
Akl1
Rpn11
Nip1
Prt1
Rpg1
Ded1
Tif35
Cdc9
Chk1
Rad23
Ubi4
Ede1
Ent2
Vps27
Chc1
Ptp3
Stb1
Spa2
Bem2
Bud14
Rck2
Glc7
Ipl1
Cdc5
Dbf4
Pfk26
Tpk1
Gpa2
Ras2
Med6
Rpb10
Paf1
Rpo26
Bdp1
Spt15
Spt5
Rgr1
Srb7
Ssn2
Nop1
Nsr1
Nop56
Snu13
Rps31
Rad9
Rtf1
Rpo21
Npl3
Air2
Fps1
Ask10
Cdc28
Hog1
Pbs2
Ssk2
Clb2
Bud3
Bud4
Msn2
Sch9
Rim15
Yak1
Ypl247c
Vps52
Kss1
Sko1
Slt2
Msg5
Swe1
Igo2
Cdc55
Igo1
Vps53
Vps54
Rtn1
Fmp42
Cot1
Zrc1
10
20
30
40
50
t/s

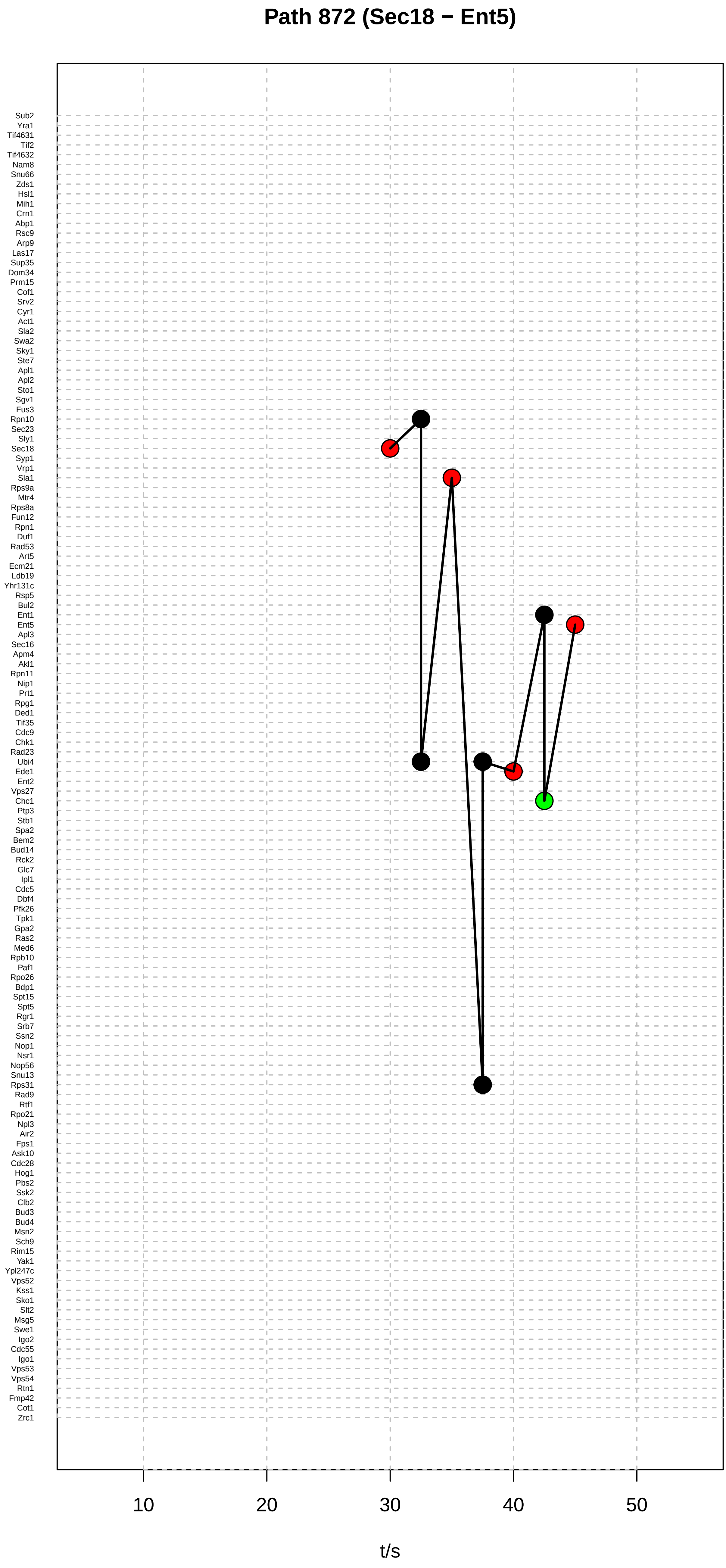

Path 872 (Sec18 − Ent5)
Sub2
Yra1
Tif4631
Tif2
Tif4632
Nam8
Snu66
Zds1
Hsl1
Mih1
Crn1
Abp1
Rsc9
Arp9
Las17
Sup35
Dom34
Prm15
Cof1
Srv2
Cyr1
Act1
Sla2
Swa2
Sky1
Ste7
Apl1
Apl2
Sto1
Sgv1
Fus3
Rpn10
Sec23
Sly1
Sec18
Syp1
Vrp1
Sla1
Rps9a
Mtr4
Rps8a
Fun12
Rpn1
Duf1
Rad53
Art5
Ecm21
Ldb19
Yhr131c
Rsp5
Bul2
Ent1
Ent5
Apl3
Sec16
Apm4
Akl1
Rpn11
Nip1
Prt1
Rpg1
Ded1
Tif35
Cdc9
Chk1
Rad23
Ubi4
Ede1
Ent2
Vps27
Chc1
Ptp3
Stb1
Spa2
Bem2
Bud14
Rck2
Glc7
Ipl1
Cdc5
Dbf4
Pfk26
Tpk1
Gpa2
Ras2
Med6
Rpb10
Paf1
Rpo26
Bdp1
Spt15
Spt5
Rgr1
Srb7
Ssn2
Nop1
Nsr1
Nop56
Snu13
Rps31
Rad9
Rtf1
Rpo21
Npl3
Air2
Fps1
Ask10
Cdc28
Hog1
Pbs2
Ssk2
Clb2
Bud3
Bud4
Msn2
Sch9
Rim15
Yak1
Ypl247c
Vps52
Kss1
Sko1
Slt2
Msg5
Swe1
Igo2
Cdc55
Igo1
Vps53
Vps54
Rtn1
Fmp42
Cot1
Zrc1
10
20
30
40
50
t/s

### Path 873 (Sec18 − Rad23)

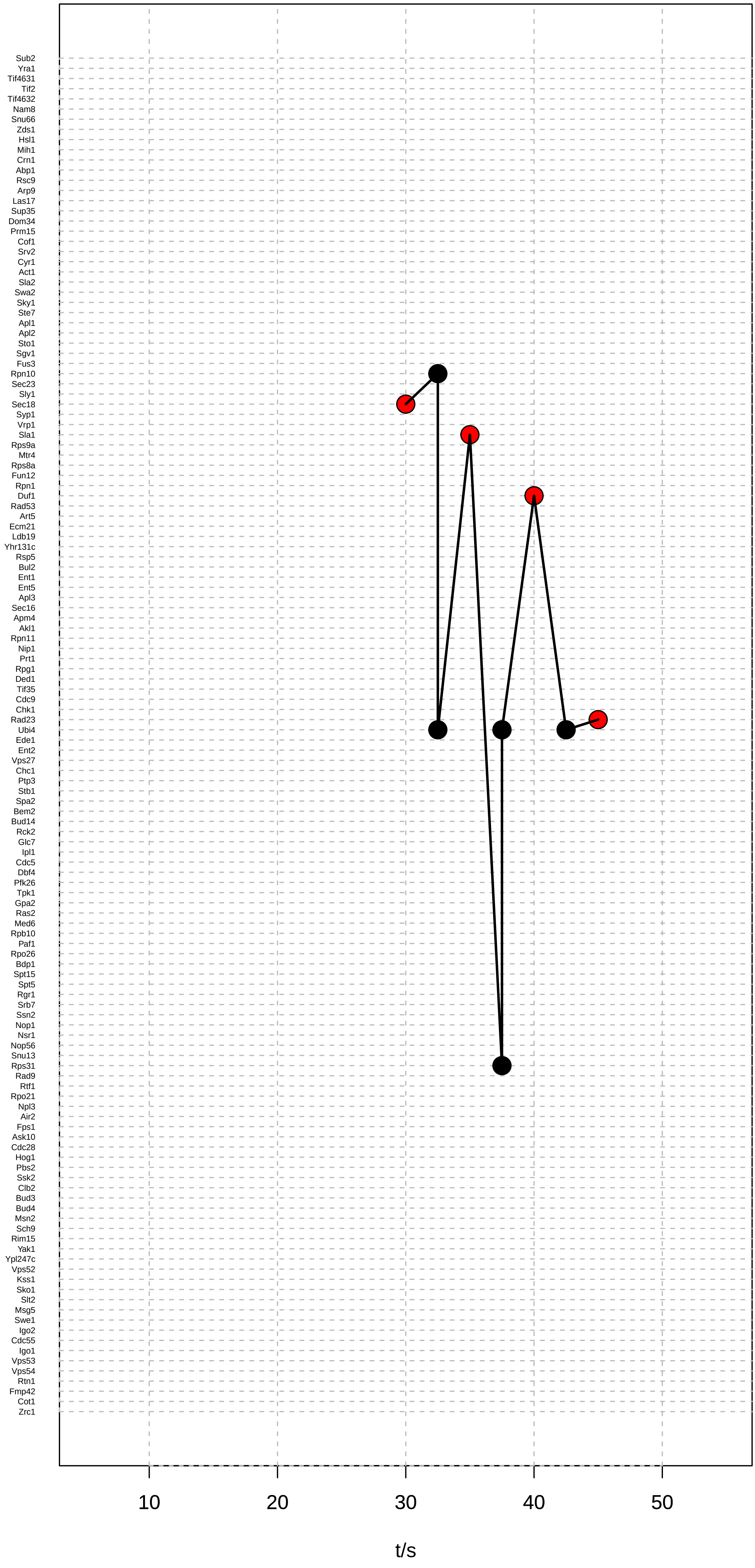

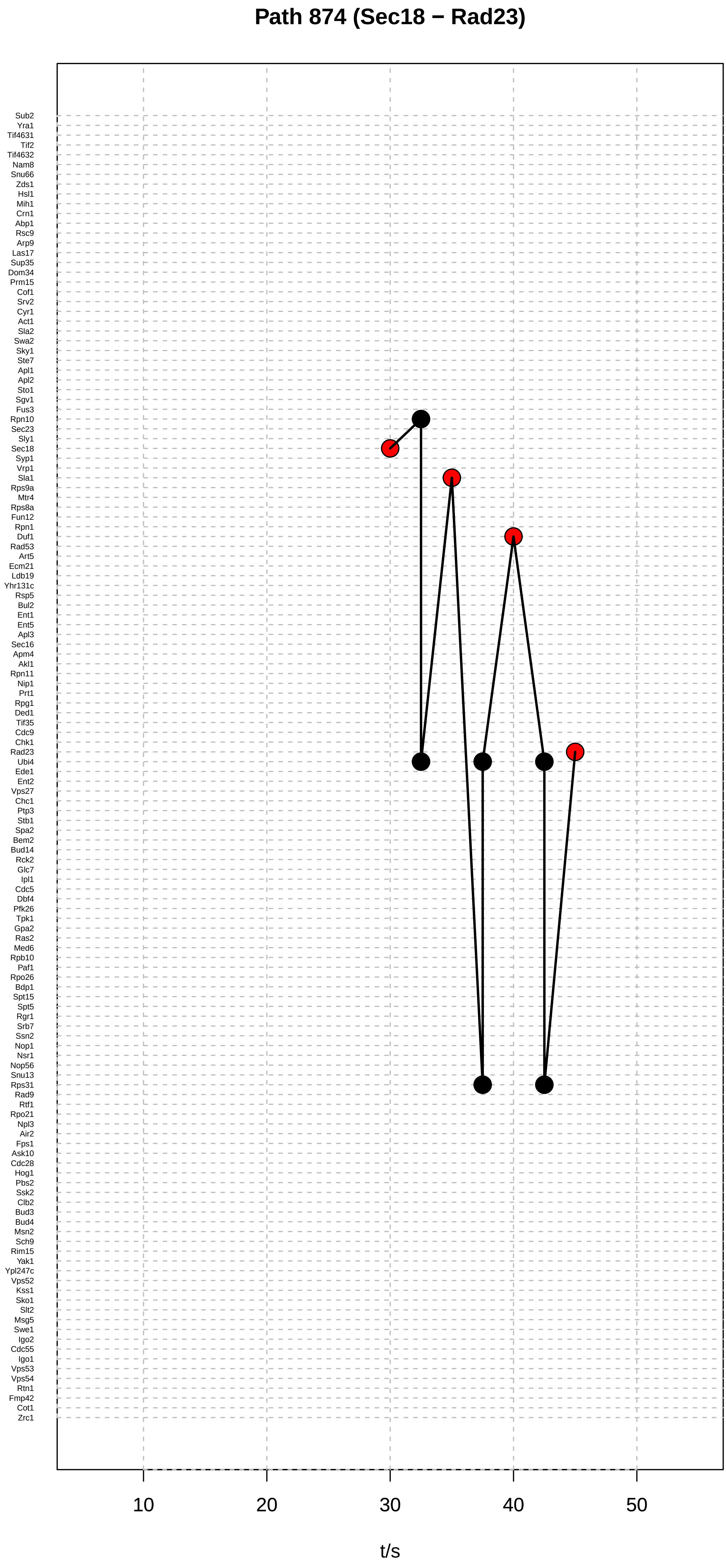

Path 874 (Sec18 − Rad23)
Sub2
Yra1
Tif4631
Tif2
Tif4632
Nam8
Snu66
Zds1
Hsl1
Mih1
Crn1
Abp1
Rsc9
Arp9
Las17
Sup35
Dom34
Prm15
Cof1
Srv2
Cyr1
Act1
Sla2
Swa2
Sky1
Ste7
Apl1
Apl2
Sto1
Sgv1
Fus3
Rpn10
Sec23
Sly1
Sec18
Syp1
Vrp1
Sla1
Rps9a
Mtr4
Rps8a
Fun12
Rpn1
Duf1
Rad53
Art5
Ecm21
Ldb19
Yhr131c
Rsp5
Bul2
Ent1
Ent5
Apl3
Sec16
Apm4
Akl1
Rpn11
Nip1
Prt1
Rpg1
Ded1
Tif35
Cdc9
Chk1
Rad23
Ubi4
Ede1
Ent2
Vps27
Chc1
Ptp3
Stb1
Spa2
Bem2
Bud14
Rck2
Glc7
Ipl1
Cdc5
Dbf4
Pfk26
Tpk1
Gpa2
Ras2
Med6
Rpb10
Paf1
Rpo26
Bdp1
Spt15
Spt5
Rgr1
Srb7
Ssn2
Nop1
Nsr1
Nop56
Snu13
Rps31
Rad9
Rtf1
Rpo21
Npl3
Air2
Fps1
Ask10
Cdc28
Hog1
Pbs2
Ssk2
Clb2
Bud3
Bud4
Msn2
Sch9
Rim15
Yak1
Ypl247c
Vps52
Kss1
Sko1
Slt2
Msg5
Swe1
Igo2
Cdc55
Igo1
Vps53
Vps54
Rtn1
Fmp42
Cot1
Zrc1
10
20
30
40
50
t/s

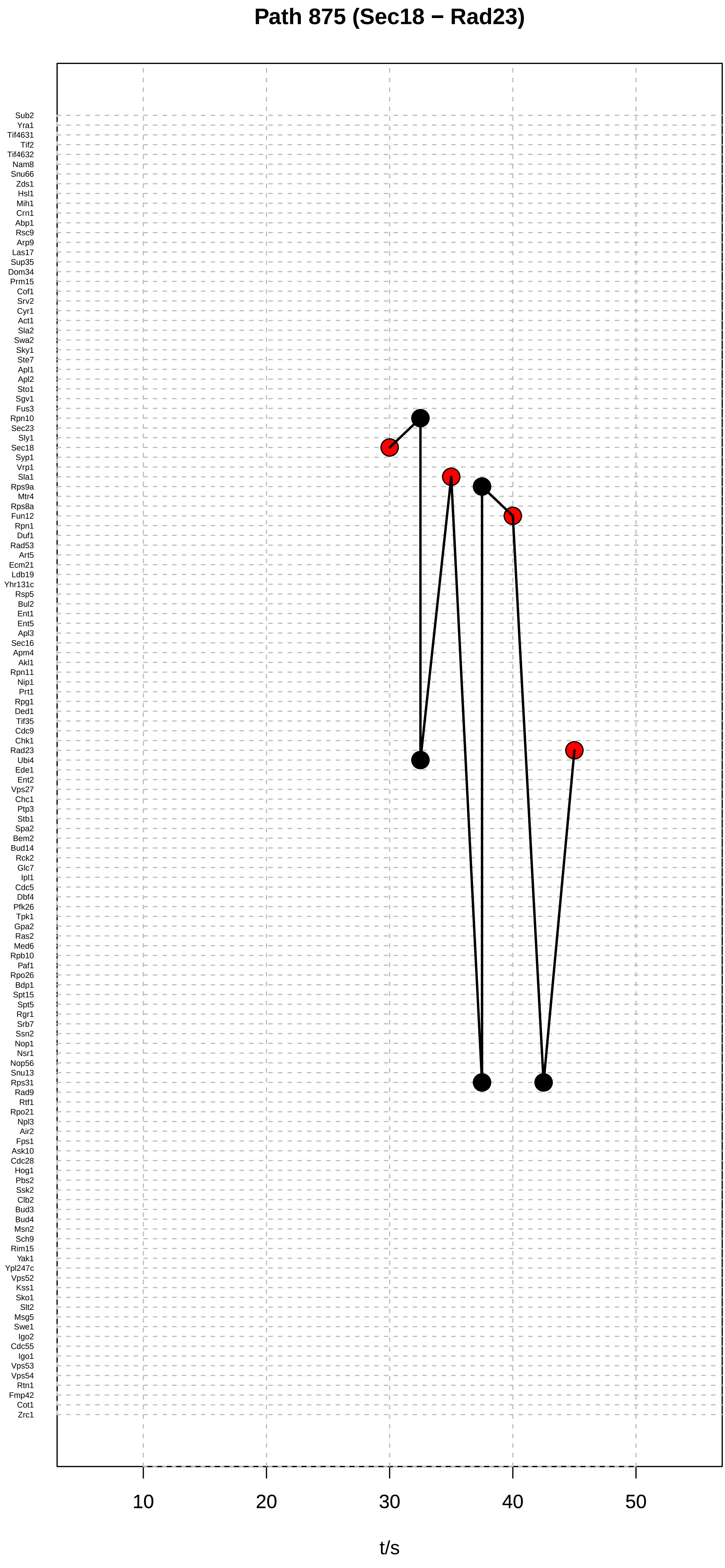
Path 875 (Sec18 − Rad23)
Sub2
Yra1
Tif4631
Tif2
Tif4632
Nam8
Snu66
Zds1
Hsl1
Mih1
Crn1
Abp1
Rsc9
Arp9
Las17
Sup35
Dom34
Prm15
Cof1
Srv2
Cyr1
Act1
Sla2
Swa2
Sky1
Ste7
Apl1
Apl2
Sto1
Sgv1
Fus3
Rpn10
Sec23
Sly1
Sec18
Syp1
Vrp1
Sla1
Rps9a
Mtr4
Rps8a
Fun12
Rpn1
Duf1
Rad53
Art5
Ecm21
Ldb19
Yhr131c
Rsp5
Bul2
Ent1
Ent5
Apl3
Sec16
Apm4
Akl1
Rpn11
Nip1
Prt1
Rpg1
Ded1
Tif35
Cdc9
Chk1
Rad23
Ubi4
Ede1
Ent2
Vps27
Chc1
Ptp3
Stb1
Spa2
Bem2
Bud14
Rck2
Glc7
Ipl1
Cdc5
Dbf4
Pfk26
Tpk1
Gpa2
Ras2
Med6
Rpb10
Paf1
Rpo26
Bdp1
Spt15
Spt5
Rgr1
Srb7
Ssn2
Nop1
Nsr1
Nop56
Snu13
Rps31
Rad9
Rtf1
Rpo21
Npl3
Air2
Fps1
Ask10
Cdc28
Hog1
Pbs2
Ssk2
Clb2
Bud3
Bud4
Msn2
Sch9
Rim15
Yak1
Ypl247c
Vps52
Kss1
Sko1
Slt2
Msg5
Swe1
Igo2
Cdc55
Igo1
Vps53
Vps54
Rtn1
Fmp42
Cot1
Zrc1
10
20
30
40
50
t/s

#### Path 876 (Sec18 − Rad23)

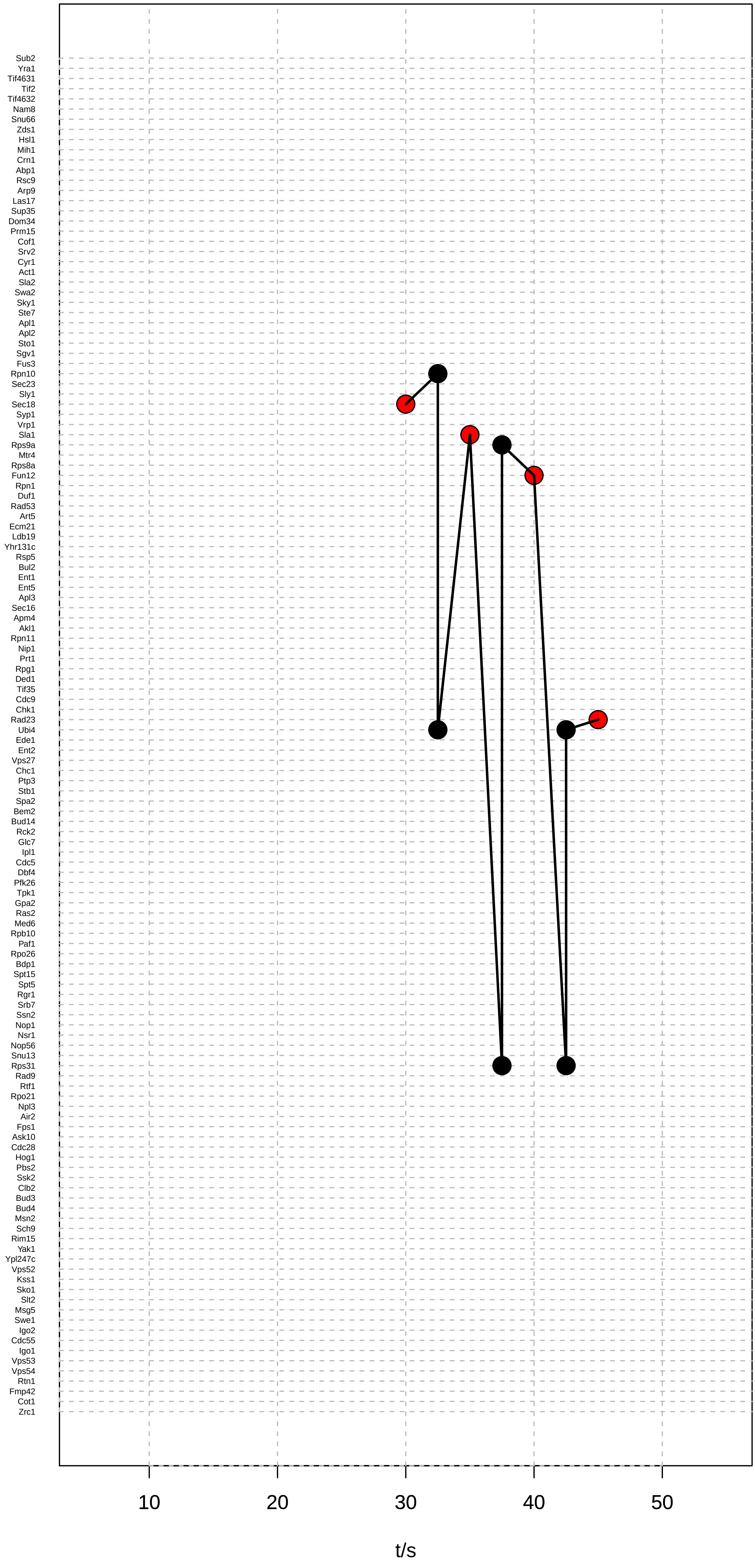

#### Path 877 (Sec18 − Rad23)

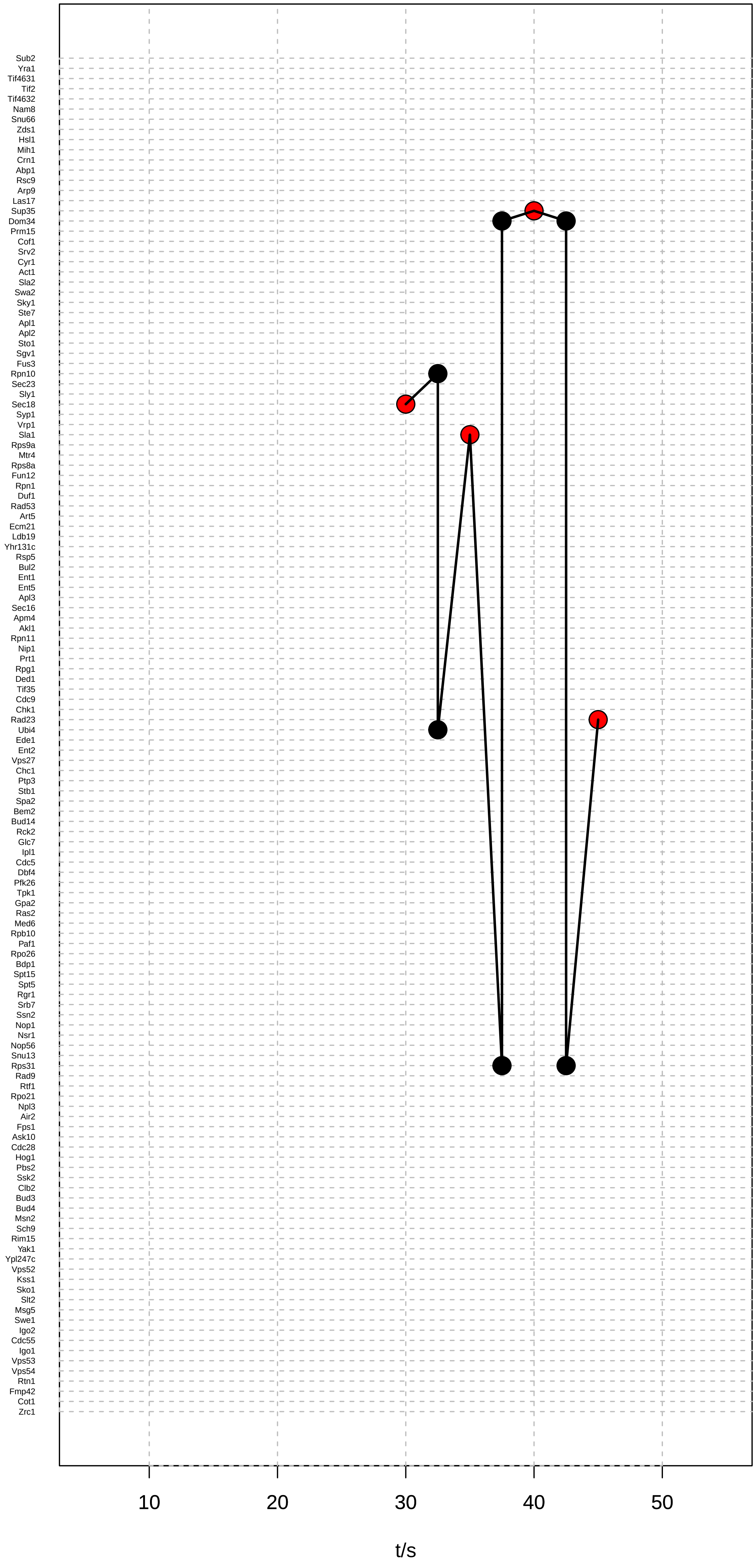

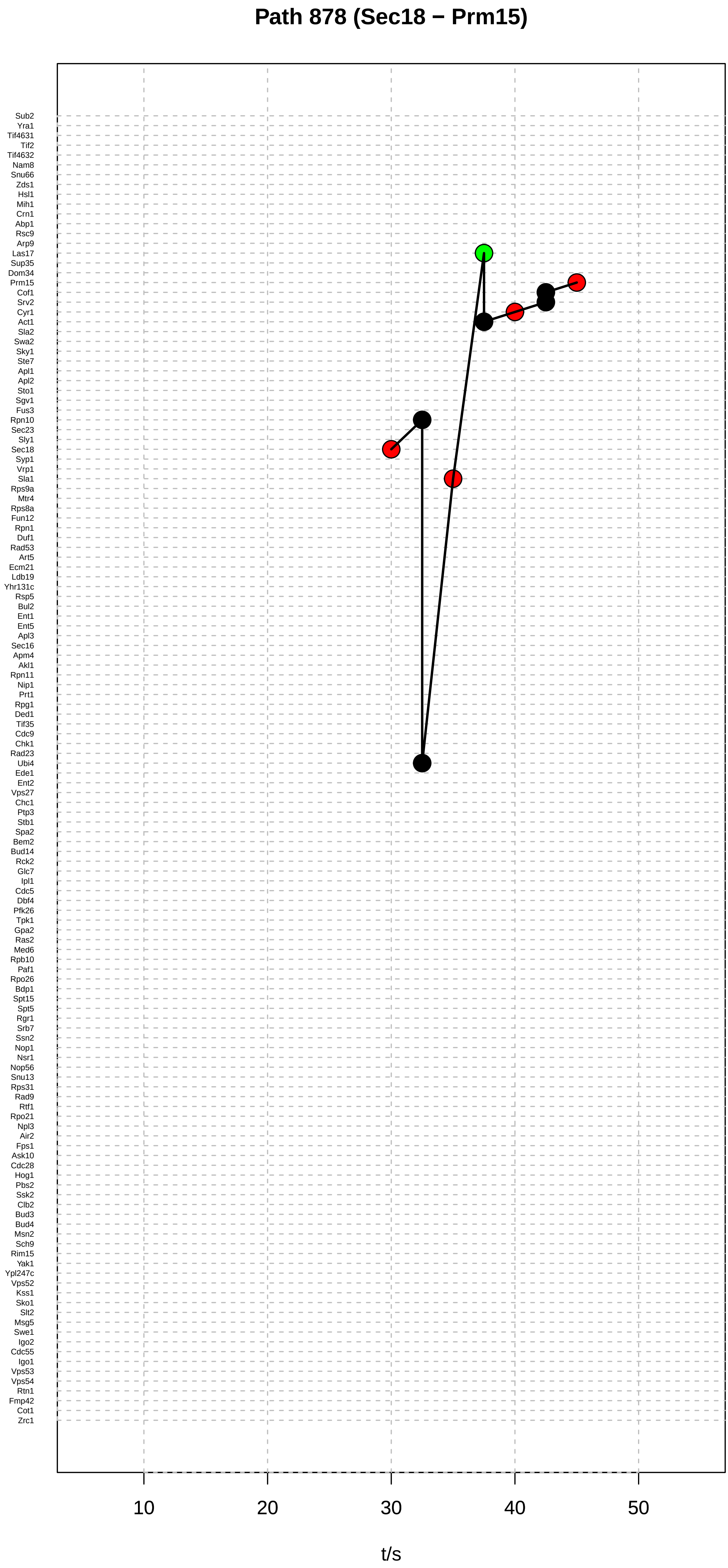

Path 878 (Sec18 − Prm15)
Sub2
Yra1
Tif4631
Tif2
Tif4632
Nam8
Snu66
Zds1
Hsl1
Mih1
Crn1
Abp1
Rsc9
Arp9
Las17
Sup35
Dom34
Prm15
Cof1
Srv2
Cyr1
Act1
Sla2
Swa2
Sky1
Ste7
Apl1
Apl2
Sto1
Sgv1
Fus3
Rpn10
Sec23
Sly1
Sec18
Syp1
Vrp1
Sla1
Rps9a
Mtr4
Rps8a
Fun12
Rpn1
Duf1
Rad53
Art5
Ecm21
Ldb19
Yhr131c
Rsp5
Bul2
Ent1
Ent5
Apl3
Sec16
Apm4
Akl1
Rpn11
Nip1
Prt1
Rpg1
Ded1
Tif35
Cdc9
Chk1
Rad23
Ubi4
Ede1
Ent2
Vps27
Chc1
Ptp3
Stb1
Spa2
Bem2
Bud14
Rck2
Glc7
Ipl1
Cdc5
Dbf4
Pfk26
Tpk1
Gpa2
Ras2
Med6
Rpb10
Paf1
Rpo26
Bdp1
Spt15
Spt5
Rgr1
Srb7
Ssn2
Nop1
Nsr1
Nop56
Snu13
Rps31
Rad9
Rtf1
Rpo21
Npl3
Air2
Fps1
Ask10
Cdc28
Hog1
Pbs2
Ssk2
Clb2
Bud3
Bud4
Msn2
Sch9
Rim15
Yak1
Ypl247c
Vps52
Kss1
Sko1
Slt2
Msg5
Swe1
Igo2
Cdc55
Igo1
Vps53
Vps54
Rtn1
Fmp42
Cot1
Zrc1
10
20
30
40
50
t/s

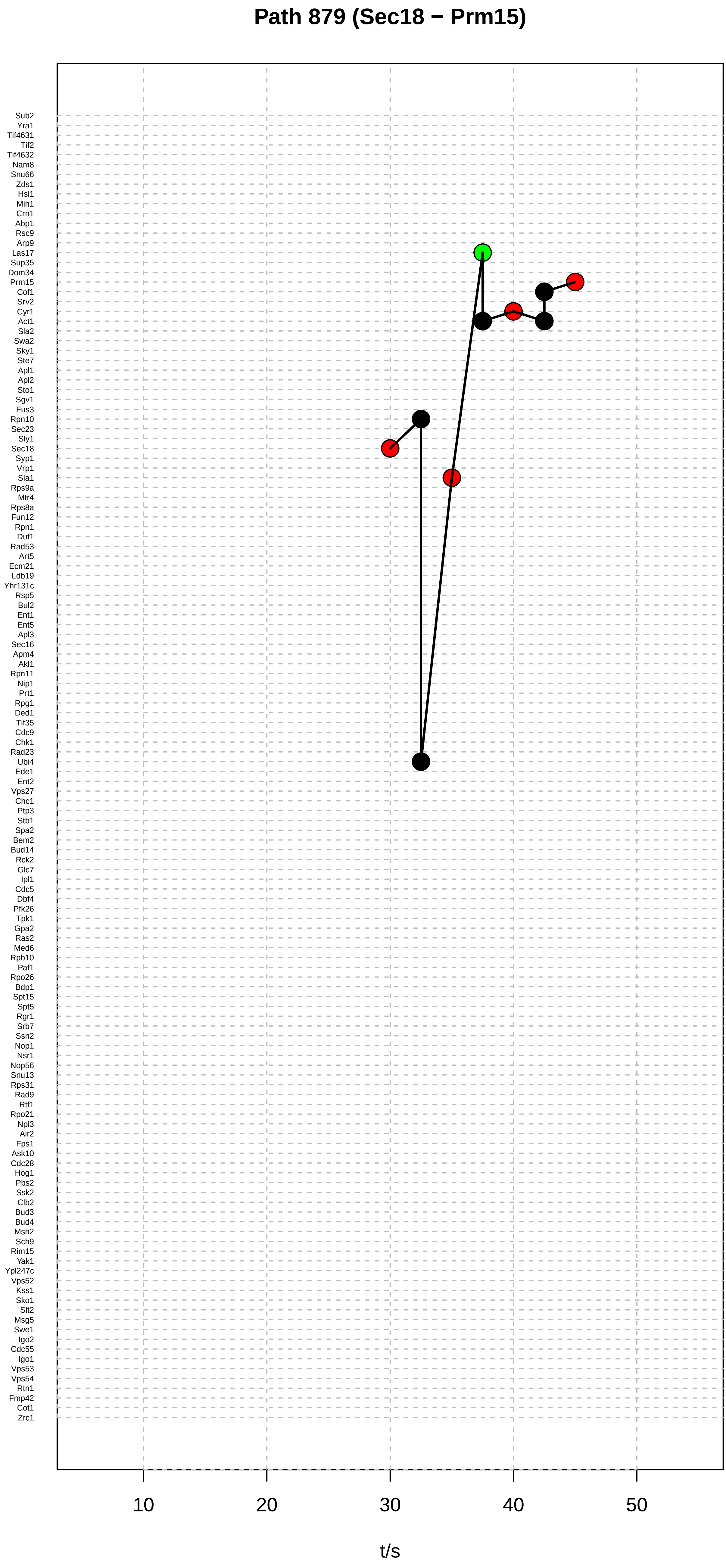

Path 879 (Sec18 − Prm15)
Sub2
Yra1
Tif4631
Tif2
Tif4632
Nam8
Snu66
Zds1
Hsl1
Mih1
Crn1
Abp1
Rsc9
Arp9
Las17
Sup35
Dom34
Prm15
Cof1
Srv2
Cyr1
Act1
Sla2
Swa2
Sky1
Ste7
Apl1
Apl2
Sto1
Sgv1
Fus3
Rpn10
Sec23
Sly1
Sec18
Syp1
Vrp1
Sla1
Rps9a
Mtr4
Rps8a
Fun12
Rpn1
Duf1
Rad53
Art5
Ecm21
Ldb19
Yhr131c
Rsp5
Bul2
Ent1
Ent5
Apl3
Sec16
Apm4
Akl1
Rpn11
Nip1
Prt1
Rpg1
Ded1
Tif35
Cdc9
Chk1
Rad23
Ubi4
Ede1
Ent2
Vps27
Chc1
Ptp3
Stb1
Spa2
Bem2
Bud14
Rck2
Glc7
Ipl1
Cdc5
Dbf4
Pfk26
Tpk1
Gpa2
Ras2
Med6
Rpb10
Paf1
Rpo26
Bdp1
Spt15
Spt5
Rgr1
Srb7
Ssn2
Nop1
Nsr1
Nop56
Snu13
Rps31
Rad9
Rtf1
Rpo21
Npl3
Air2
Fps1
Ask10
Cdc28
Hog1
Pbs2
Ssk2
Clb2
Bud3
Bud4
Msn2
Sch9
Rim15
Yak1
Ypl247c
Vps52
Kss1
Sko1
Slt2
Msg5
Swe1
Igo2
Cdc55
Igo1
Vps53
Vps54
Rtn1
Fmp42
Cot1
Zrc1
10
20
30
40
50
t/s

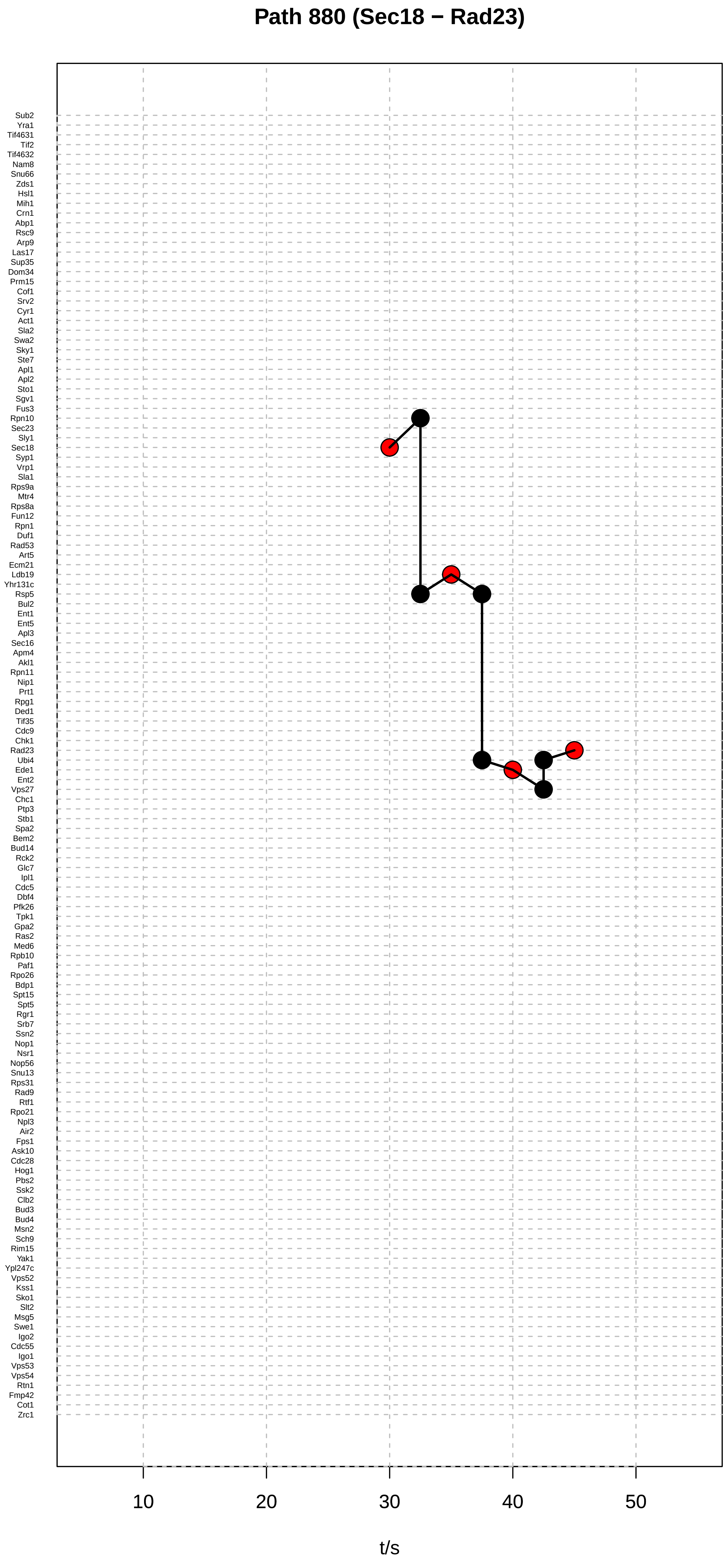
Path 880 (Sec18 − Rad23)
Sub2
Yra1
Tif4631
Tif2
Tif4632
Nam8
Snu66
Zds1
Hsl1
Mih1
Crn1
Abp1
Rsc9
Arp9
Las17
Sup35
Dom34
Prm15
Cof1
Srv2
Cyr1
Act1
Sla2
Swa2
Sky1
Ste7
Apl1
Apl2
Sto1
Sgv1
Fus3
Rpn10
Sec23
Sly1
Sec18
Syp1
Vrp1
Sla1
Rps9a
Mtr4
Rps8a
Fun12
Rpn1
Duf1
Rad53
Art5
Ecm21
Ldb19
Yhr131c
Rsp5
Bul2
Ent1
Ent5
Apl3
Sec16
Apm4
Akl1
Rpn11
Nip1
Prt1
Rpg1
Ded1
Tif35
Cdc9
Chk1
Rad23
Ubi4
Ede1
Ent2
Vps27
Chc1
Ptp3
Stb1
Spa2
Bem2
Bud14
Rck2
Glc7
Ipl1
Cdc5
Dbf4
Pfk26
Tpk1
Gpa2
Ras2
Med6
Rpb10
Paf1
Rpo26
Bdp1
Spt15
Spt5
Rgr1
Srb7
Ssn2
Nop1
Nsr1
Nop56
Snu13
Rps31
Rad9
Rtf1
Rpo21
Npl3
Air2
Fps1
Ask10
Cdc28
Hog1
Pbs2
Ssk2
Clb2
Bud3
Bud4
Msn2
Sch9
Rim15
Yak1
Ypl247c
Vps52
Kss1
Sko1
Slt2
Msg5
Swe1
Igo2
Cdc55
Igo1
Vps53
Vps54
Rtn1
Fmp42
Cot1
Zrc1
10
20
30
40
50
t/s

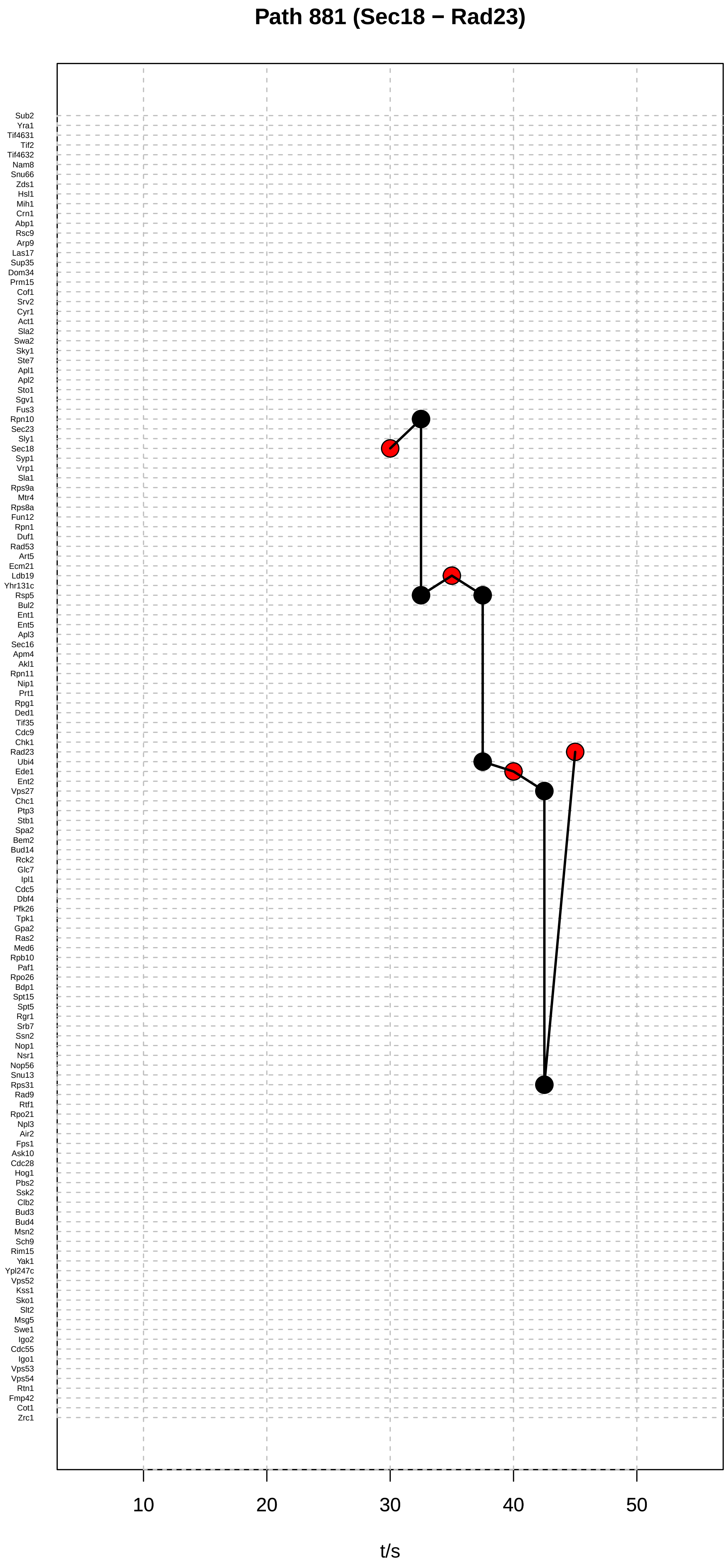
Path 881 (Sec18 − Rad23)
Sub2
Yra1
Tif4631
Tif2
Tif4632
Nam8
Snu66
Zds1
Hsl1
Mih1
Crn1
Abp1
Rsc9
Arp9
Las17
Sup35
Dom34
Prm15
Cof1
Srv2
Cyr1
Act1
Sla2
Swa2
Sky1
Ste7
Apl1
Apl2
Sto1
Sgv1
Fus3
Rpn10
Sec23
Sly1
Sec18
Syp1
Vrp1
Sla1
Rps9a
Mtr4
Rps8a
Fun12
Rpn1
Duf1
Rad53
Art5
Ecm21
Ldb19
Yhr131c
Rsp5
Bul2
Ent1
Ent5
Apl3
Sec16
Apm4
Akl1
Rpn11
Nip1
Prt1
Rpg1
Ded1
Tif35
Cdc9
Chk1
Rad23
Ubi4
Ede1
Ent2
Vps27
Chc1
Ptp3
Stb1
Spa2
Bem2
Bud14
Rck2
Glc7
Ipl1
Cdc5
Dbf4
Pfk26
Tpk1
Gpa2
Ras2
Med6
Rpb10
Paf1
Rpo26
Bdp1
Spt15
Spt5
Rgr1
Srb7
Ssn2
Nop1
Nsr1
Nop56
Snu13
Rps31
Rad9
Rtf1
Rpo21
Npl3
Air2
Fps1
Ask10
Cdc28
Hog1
Pbs2
Ssk2
Clb2
Bud3
Bud4
Msn2
Sch9
Rim15
Yak1
Ypl247c
Vps52
Kss1
Sko1
Slt2
Msg5
Swe1
Igo2
Cdc55
Igo1
Vps53
Vps54
Rtn1
Fmp42
Cot1
Zrc1
10
20
30
40
50
t/s

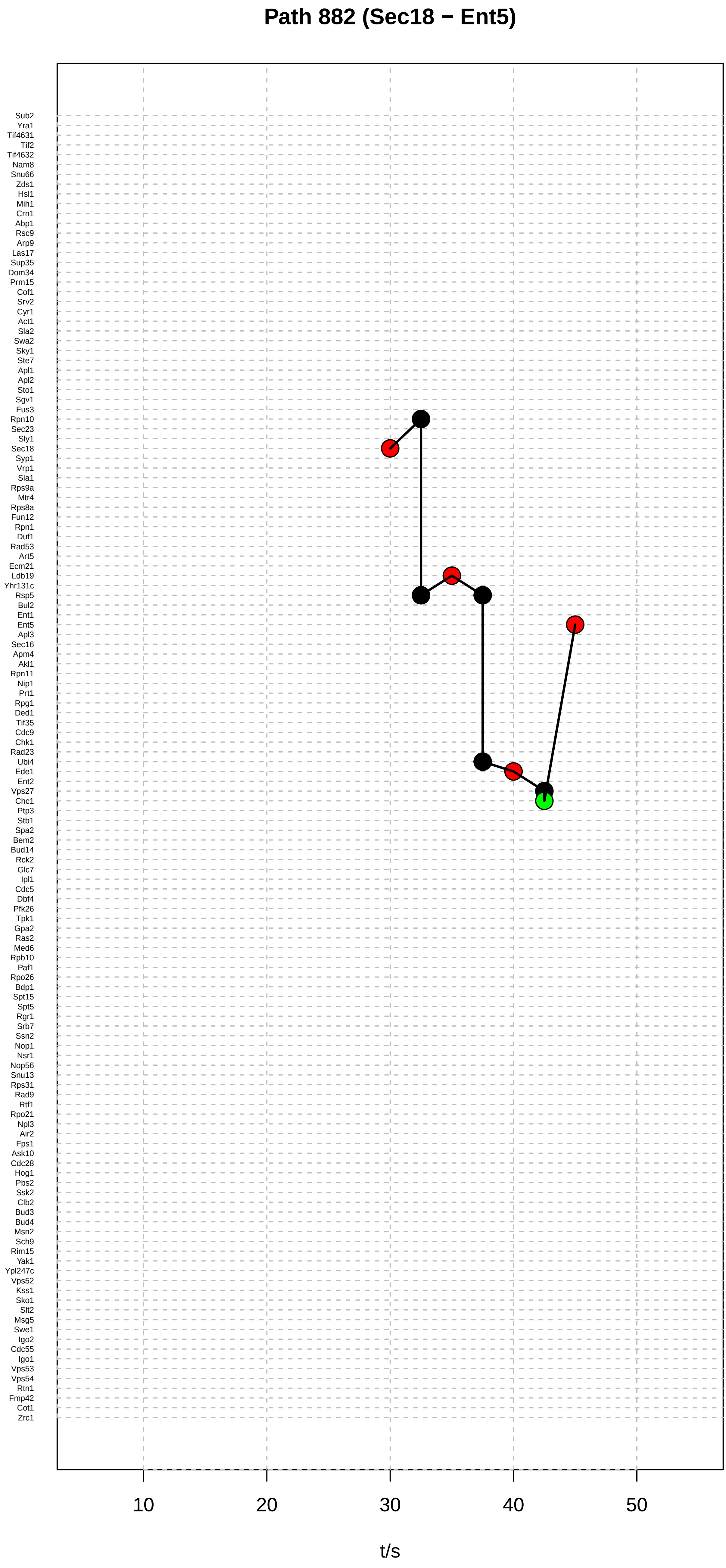

Path 882 (Sec18 − Ent5)
Sub2
Yra1
Tif4631
Tif2
Tif4632
Nam8
Snu66
Zds1
Hsl1
Mih1
Crn1
Abp1
Rsc9
Arp9
Las17
Sup35
Dom34
Prm15
Cof1
Srv2
Cyr1
Act1
Sla2
Swa2
Sky1
Ste7
Apl1
Apl2
Sto1
Sgv1
Fus3
Rpn10
Sec23
Sly1
Sec18
Syp1
Vrp1
Sla1
Rps9a
Mtr4
Rps8a
Fun12
Rpn1
Duf1
Rad53
Art5
Ecm21
Ldb19
Yhr131c
Rsp5
Bul2
Ent1
Ent5
Apl3
Sec16
Apm4
Akl1
Rpn11
Nip1
Prt1
Rpg1
Ded1
Tif35
Cdc9
Chk1
Rad23
Ubi4
Ede1
Ent2
Vps27
Chc1
Ptp3
Stb1
Spa2
Bem2
Bud14
Rck2
Glc7
Ipl1
Cdc5
Dbf4
Pfk26
Tpk1
Gpa2
Ras2
Med6
Rpb10
Paf1
Rpo26
Bdp1
Spt15
Spt5
Rgr1
Srb7
Ssn2
Nop1
Nsr1
Nop56
Snu13
Rps31
Rad9
Rtf1
Rpo21
Npl3
Air2
Fps1
Ask10
Cdc28
Hog1
Pbs2
Ssk2
Clb2
Bud3
Bud4
Msn2
Sch9
Rim15
Yak1
Ypl247c
Vps52
Kss1
Sko1
Slt2
Msg5
Swe1
Igo2
Cdc55
Igo1
Vps53
Vps54
Rtn1
Fmp42
Cot1
Zrc1
10
20
30
40
50
t/s

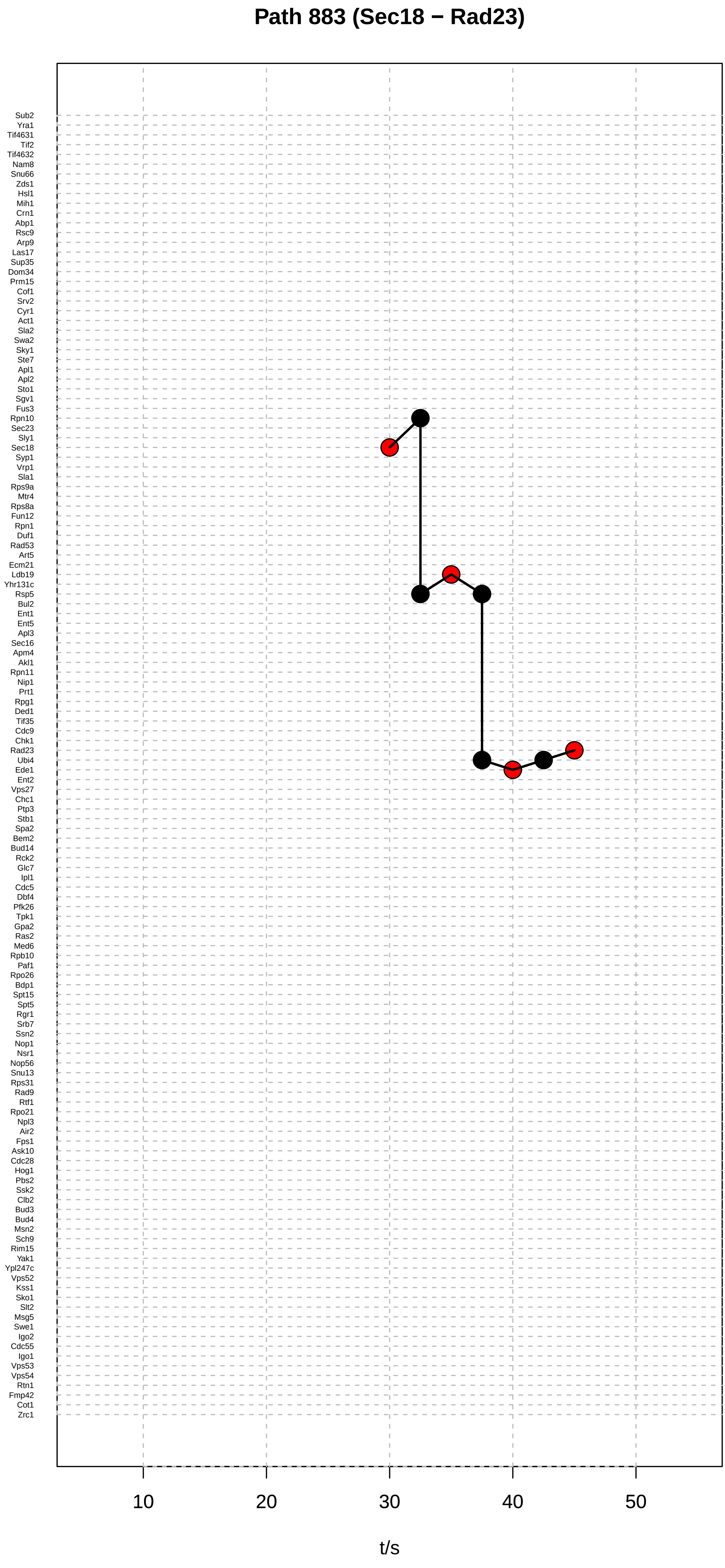

Path 883 (Sec18 − Rad23)
Sub2
Yra1
Tif4631
Tif2
Tif4632
Nam8
Snu66
Zds1
Hsl1
Mih1
Crn1
Abp1
Rsc9
Arp9
Las17
Sup35
Dom34
Prm15
Cof1
Srv2
Cyr1
Act1
Sla2
Swa2
Sky1
Ste7
Apl1
Apl2
Sto1
Sgv1
Fus3
Rpn10
Sec23
Sly1
Sec18
Syp1
Vrp1
Sla1
Rps9a
Mtr4
Rps8a
Fun12
Rpn1
Duf1
Rad53
Art5
Ecm21
Ldb19
Yhr131c
Rsp5
Bul2
Ent1
Ent5
Apl3
Sec16
Apm4
Akl1
Rpn11
Nip1
Prt1
Rpg1
Ded1
Tif35
Cdc9
Chk1
Rad23
Ubi4
Ede1
Ent2
Vps27
Chc1
Ptp3
Stb1
Spa2
Bem2
Bud14
Rck2
Glc7
Ipl1
Cdc5
Dbf4
Pfk26
Tpk1
Gpa2
Ras2
Med6
Rpb10
Paf1
Rpo26
Bdp1
Spt15
Spt5
Rgr1
Srb7
Ssn2
Nop1
Nsr1
Nop56
Snu13
Rps31
Rad9
Rtf1
Rpo21
Npl3
Air2
Fps1
Ask10
Cdc28
Hog1
Pbs2
Ssk2
Clb2
Bud3
Bud4
Msn2
Sch9
Rim15
Yak1
Ypl247c
Vps52
Kss1
Sko1
Slt2
Msg5
Swe1
Igo2
Cdc55
Igo1
Vps53
Vps54
Rtn1
Fmp42
Cot1
Zrc1
10
20
30
40
50
t/s

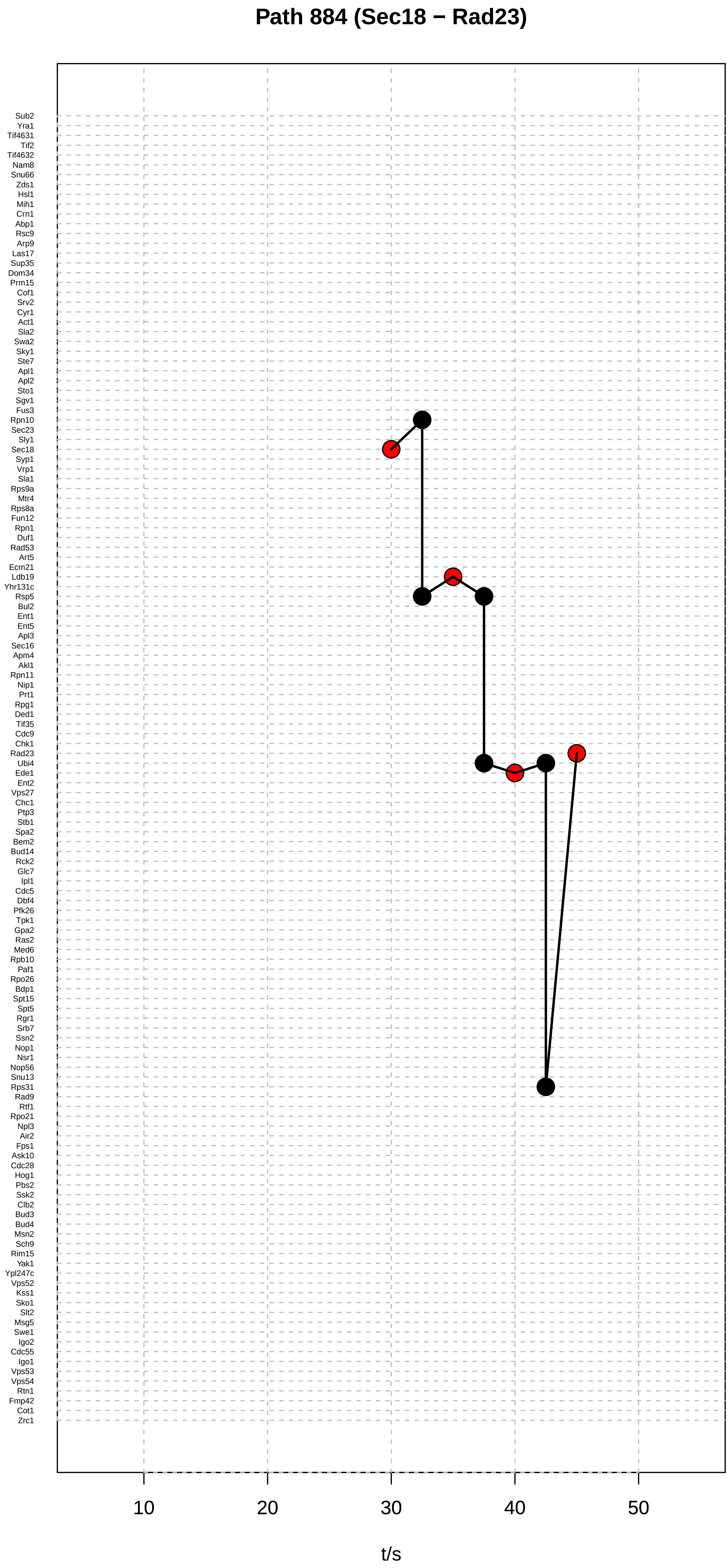

Path 884 (Sec18 − Rad23)
Sub2
Yra1
Tif4631
Tif2
Tif4632
Nam8
Snu66
Zds1
Hsl1
Mih1
Crn1
Abp1
Rsc9
Arp9
Las17
Sup35
Dom34
Prm15
Cof1
Srv2
Cyr1
Act1
Sla2
Swa2
Sky1
Ste7
Apl1
Apl2
Sto1
Sgv1
Fus3
Rpn10
Sec23
Sly1
Sec18
Syp1
Vrp1
Sla1
Rps9a
Mtr4
Rps8a
Fun12
Rpn1
Duf1
Rad53
Art5
Ecm21
Ldb19
Yhr131c
Rsp5
Bul2
Ent1
Ent5
Apl3
Sec16
Apm4
Akl1
Rpn11
Nip1
Prt1
Rpg1
Ded1
Tif35
Cdc9
Chk1
Rad23
Ubi4
Ede1
Ent2
Vps27
Chc1
Ptp3
Stb1
Spa2
Bem2
Bud14
Rck2
Glc7
Ipl1
Cdc5
Dbf4
Pfk26
Tpk1
Gpa2
Ras2
Med6
Rpb10
Paf1
Rpo26
Bdp1
Spt15
Spt5
Rgr1
Srb7
Ssn2
Nop1
Nsr1
Nop56
Snu13
Rps31
Rad9
Rtf1
Rpo21
Npl3
Air2
Fps1
Ask10
Cdc28
Hog1
Pbs2
Ssk2
Clb2
Bud3
Bud4
Msn2
Sch9
Rim15
Yak1
Ypl247c
Vps52
Kss1
Sko1
Slt2
Msg5
Swe1
Igo2
Cdc55
Igo1
Vps53
Vps54
Rtn1
Fmp42
Cot1
Zrc1
10
20
30
40
50
t/s

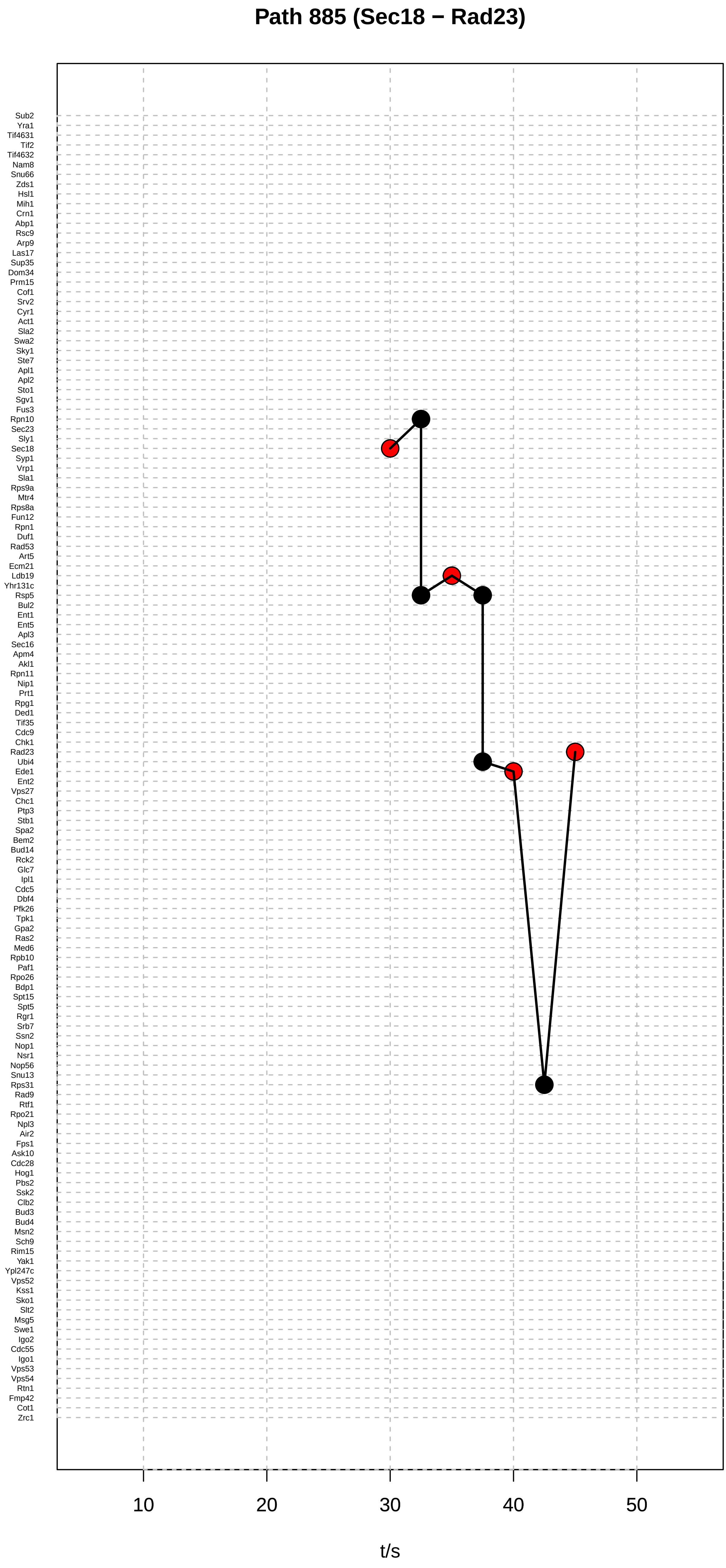
Path 885 (Sec18 − Rad23)
Sub2
Yra1
Tif4631
Tif2
Tif4632
Nam8
Snu66
Zds1
Hsl1
Mih1
Crn1
Abp1
Rsc9
Arp9
Las17
Sup35
Dom34
Prm15
Cof1
Srv2
Cyr1
Act1
Sla2
Swa2
Sky1
Ste7
Apl1
Apl2
Sto1
Sgv1
Fus3
Rpn10
Sec23
Sly1
Sec18
Syp1
Vrp1
Sla1
Rps9a
Mtr4
Rps8a
Fun12
Rpn1
Duf1
Rad53
Art5
Ecm21
Ldb19
Yhr131c
Rsp5
Bul2
Ent1
Ent5
Apl3
Sec16
Apm4
Akl1
Rpn11
Nip1
Prt1
Rpg1
Ded1
Tif35
Cdc9
Chk1
Rad23
Ubi4
Ede1
Ent2
Vps27
Chc1
Ptp3
Stb1
Spa2
Bem2
Bud14
Rck2
Glc7
Ipl1
Cdc5
Dbf4
Pfk26
Tpk1
Gpa2
Ras2
Med6
Rpb10
Paf1
Rpo26
Bdp1
Spt15
Spt5
Rgr1
Srb7
Ssn2
Nop1
Nsr1
Nop56
Snu13
Rps31
Rad9
Rtf1
Rpo21
Npl3
Air2
Fps1
Ask10
Cdc28
Hog1
Pbs2
Ssk2
Clb2
Bud3
Bud4
Msn2
Sch9
Rim15
Yak1
Ypl247c
Vps52
Kss1
Sko1
Slt2
Msg5
Swe1
Igo2
Cdc55
Igo1
Vps53
Vps54
Rtn1
Fmp42
Cot1
Zrc1
10
20
30
40
50
t/s

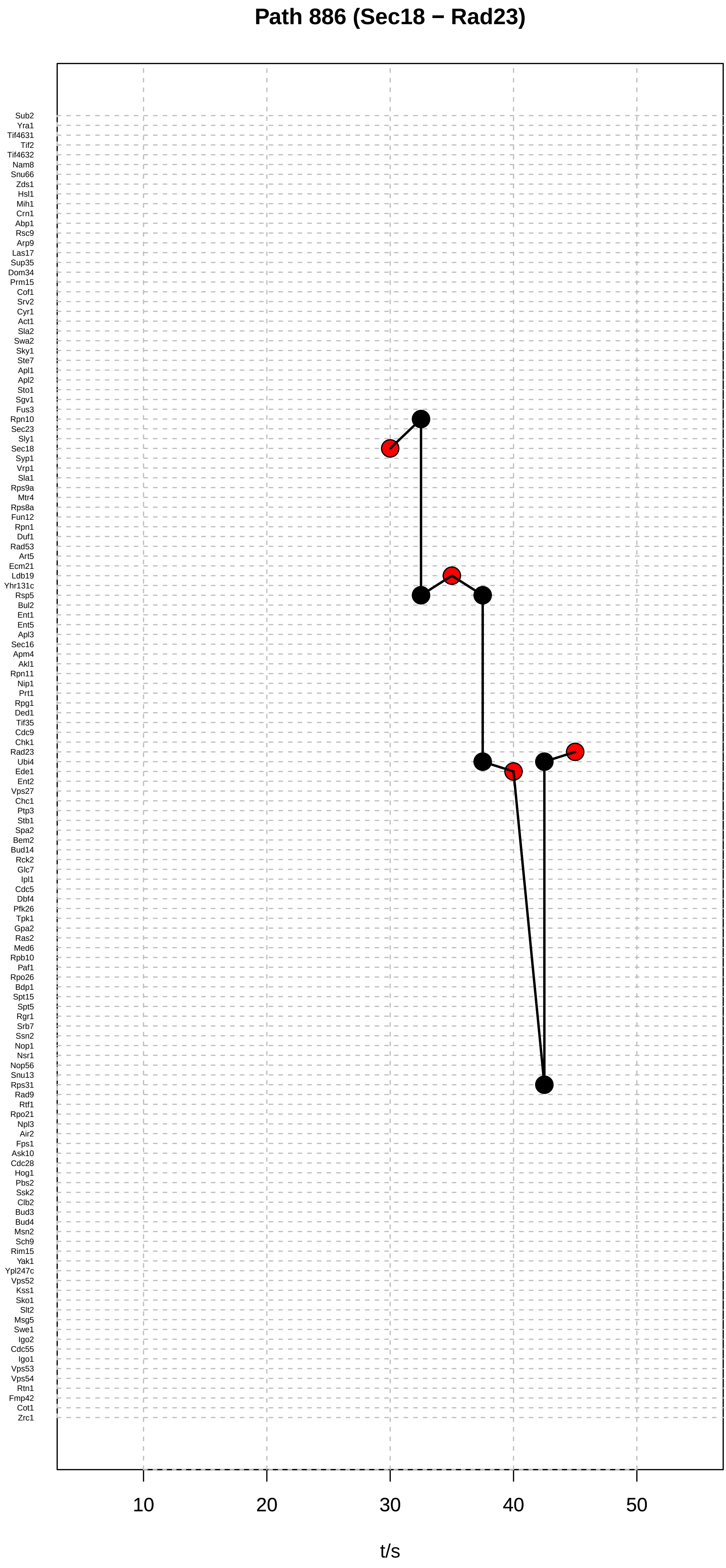
Path 886 (Sec18 − Rad23)
Sub2
Yra1
Tif4631
Tif2
Tif4632
Nam8
Snu66
Zds1
Hsl1
Mih1
Crn1
Abp1
Rsc9
Arp9
Las17
Sup35
Dom34
Prm15
Cof1
Srv2
Cyr1
Act1
Sla2
Swa2
Sky1
Ste7
Apl1
Apl2
Sto1
Sgv1
Fus3
Rpn10
Sec23
Sly1
Sec18
Syp1
Vrp1
Sla1
Rps9a
Mtr4
Rps8a
Fun12
Rpn1
Duf1
Rad53
Art5
Ecm21
Ldb19
Yhr131c
Rsp5
Bul2
Ent1
Ent5
Apl3
Sec16
Apm4
Akl1
Rpn11
Nip1
Prt1
Rpg1
Ded1
Tif35
Cdc9
Chk1
Rad23
Ubi4
Ede1
Ent2
Vps27
Chc1
Ptp3
Stb1
Spa2
Bem2
Bud14
Rck2
Glc7
Ipl1
Cdc5
Dbf4
Pfk26
Tpk1
Gpa2
Ras2
Med6
Rpb10
Paf1
Rpo26
Bdp1
Spt15
Spt5
Rgr1
Srb7
Ssn2
Nop1
Nsr1
Nop56
Snu13
Rps31
Rad9
Rtf1
Rpo21
Npl3
Air2
Fps1
Ask10
Cdc28
Hog1
Pbs2
Ssk2
Clb2
Bud3
Bud4
Msn2
Sch9
Rim15
Yak1
Ypl247c
Vps52
Kss1
Sko1
Slt2
Msg5
Swe1
Igo2
Cdc55
Igo1
Vps53
Vps54
Rtn1
Fmp42
Cot1
Zrc1
10
20
30
40
50
t/s

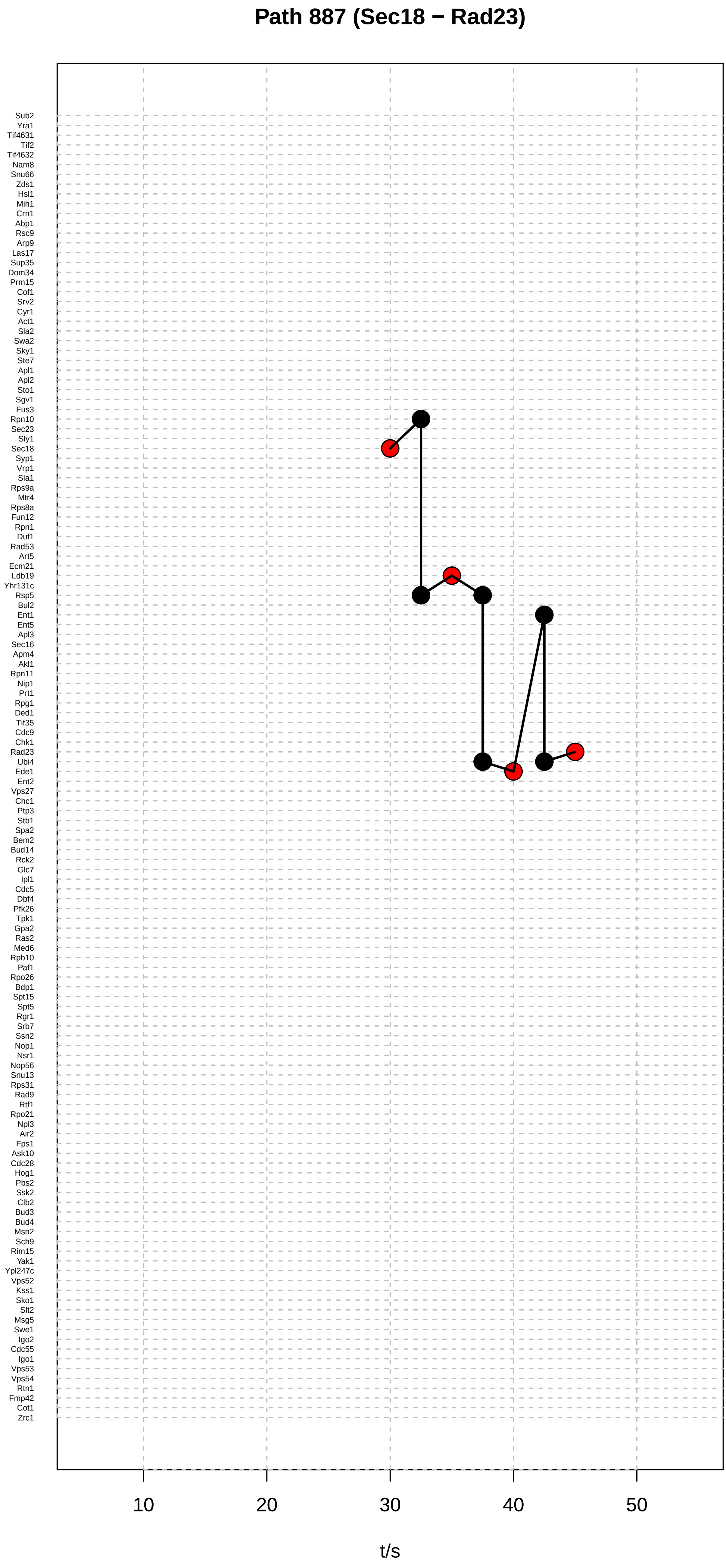
Path 887 (Sec18 − Rad23)
Sub2
Yra1
Tif4631
Tif2
Tif4632
Nam8
Snu66
Zds1
Hsl1
Mih1
Crn1
Abp1
Rsc9
Arp9
Las17
Sup35
Dom34
Prm15
Cof1
Srv2
Cyr1
Act1
Sla2
Swa2
Sky1
Ste7
Apl1
Apl2
Sto1
Sgv1
Fus3
Rpn10
Sec23
Sly1
Sec18
Syp1
Vrp1
Sla1
Rps9a
Mtr4
Rps8a
Fun12
Rpn1
Duf1
Rad53
Art5
Ecm21
Ldb19
Yhr131c
Rsp5
Bul2
Ent1
Ent5
Apl3
Sec16
Apm4
Akl1
Rpn11
Nip1
Prt1
Rpg1
Ded1
Tif35
Cdc9
Chk1
Rad23
Ubi4
Ede1
Ent2
Vps27
Chc1
Ptp3
Stb1
Spa2
Bem2
Bud14
Rck2
Glc7
Ipl1
Cdc5
Dbf4
Pfk26
Tpk1
Gpa2
Ras2
Med6
Rpb10
Paf1
Rpo26
Bdp1
Spt15
Spt5
Rgr1
Srb7
Ssn2
Nop1
Nsr1
Nop56
Snu13
Rps31
Rad9
Rtf1
Rpo21
Npl3
Air2
Fps1
Ask10
Cdc28
Hog1
Pbs2
Ssk2
Clb2
Bud3
Bud4
Msn2
Sch9
Rim15
Yak1
Ypl247c
Vps52
Kss1
Sko1
Slt2
Msg5
Swe1
Igo2
Cdc55
Igo1
Vps53
Vps54
Rtn1
Fmp42
Cot1
Zrc1
10
20
30
40
50
t/s

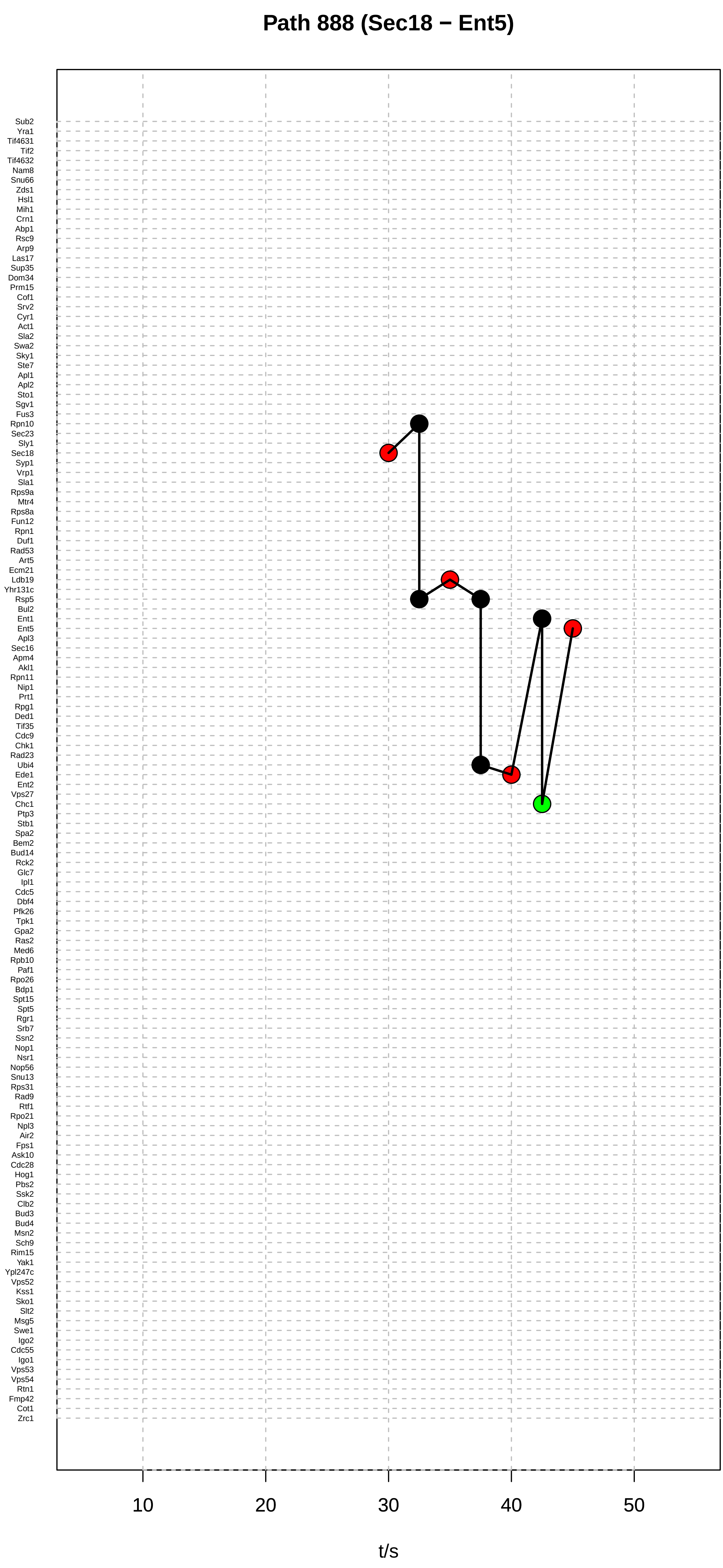

Path 888 (Sec18 − Ent5)
Sub2
Yra1
Tif4631
Tif2
Tif4632
Nam8
Snu66
Zds1
Hsl1
Mih1
Crn1
Abp1
Rsc9
Arp9
Las17
Sup35
Dom34
Prm15
Cof1
Srv2
Cyr1
Act1
Sla2
Swa2
Sky1
Ste7
Apl1
Apl2
Sto1
Sgv1
Fus3
Rpn10
Sec23
Sly1
Sec18
Syp1
Vrp1
Sla1
Rps9a
Mtr4
Rps8a
Fun12
Rpn1
Duf1
Rad53
Art5
Ecm21
Ldb19
Yhr131c
Rsp5
Bul2
Ent1
Ent5
Apl3
Sec16
Apm4
Akl1
Rpn11
Nip1
Prt1
Rpg1
Ded1
Tif35
Cdc9
Chk1
Rad23
Ubi4
Ede1
Ent2
Vps27
Chc1
Ptp3
Stb1
Spa2
Bem2
Bud14
Rck2
Glc7
Ipl1
Cdc5
Dbf4
Pfk26
Tpk1
Gpa2
Ras2
Med6
Rpb10
Paf1
Rpo26
Bdp1
Spt15
Spt5
Rgr1
Srb7
Ssn2
Nop1
Nsr1
Nop56
Snu13
Rps31
Rad9
Rtf1
Rpo21
Npl3
Air2
Fps1
Ask10
Cdc28
Hog1
Pbs2
Ssk2
Clb2
Bud3
Bud4
Msn2
Sch9
Rim15
Yak1
Ypl247c
Vps52
Kss1
Sko1
Slt2
Msg5
Swe1
Igo2
Cdc55
Igo1
Vps53
Vps54
Rtn1
Fmp42
Cot1
Zrc1
10
20
30
40
50
t/s

#### Path 889 (Sec18 − Rad23)

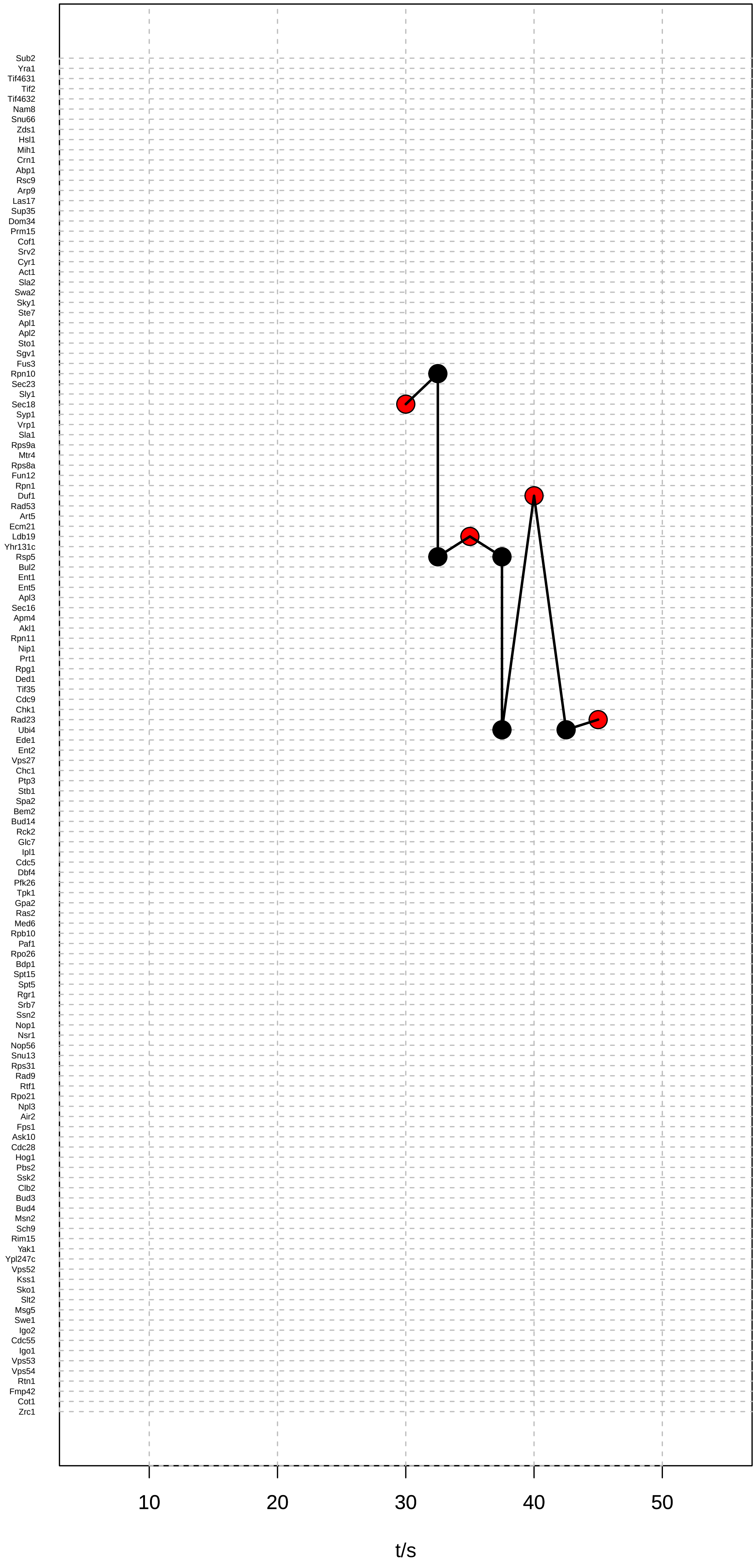

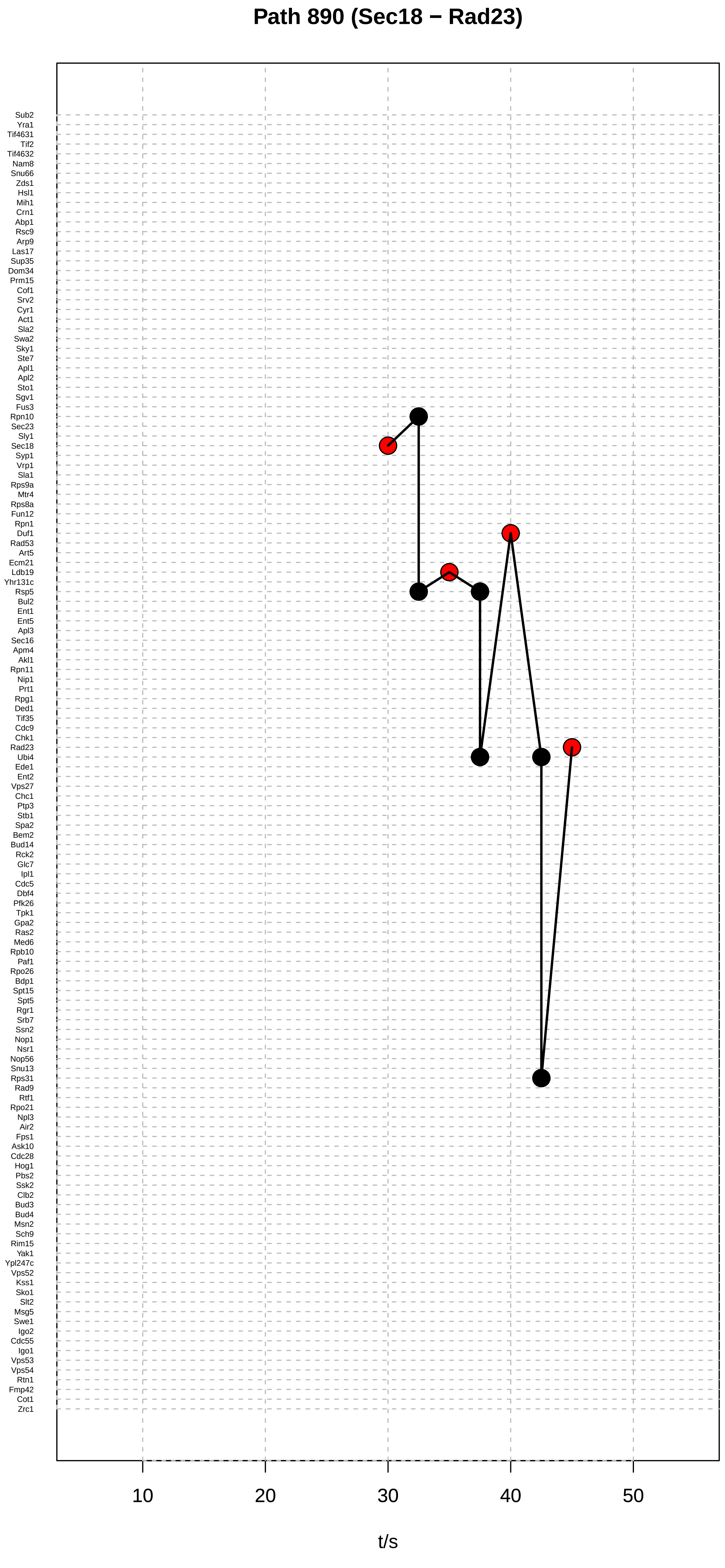
Path 890 (Sec18 − Rad23)
Sub2
Yra1
Tif4631
Tif2
Tif4632
Nam8
Snu66
Zds1
Hsl1
Mih1
Crn1
Abp1
Rsc9
Arp9
Las17
Sup35
Dom34
Prm15
Cof1
Srv2
Cyr1
Act1
Sla2
Swa2
Sky1
Ste7
Apl1
Apl2
Sto1
Sgv1
Fus3
Rpn10
Sec23
Sly1
Sec18
Syp1
Vrp1
Sla1
Rps9a
Mtr4
Rps8a
Fun12
Rpn1
Duf1
Rad53
Art5
Ecm21
Ldb19
Yhr131c
Rsp5
Bul2
Ent1
Ent5
Apl3
Sec16
Apm4
Akl1
Rpn11
Nip1
Prt1
Rpg1
Ded1
Tif35
Cdc9
Chk1
Rad23
Ubi4
Ede1
Ent2
Vps27
Chc1
Ptp3
Stb1
Spa2
Bem2
Bud14
Rck2
Glc7
Ipl1
Cdc5
Dbf4
Pfk26
Tpk1
Gpa2
Ras2
Med6
Rpb10
Paf1
Rpo26
Bdp1
Spt15
Spt5
Rgr1
Srb7
Ssn2
Nop1
Nsr1
Nop56
Snu13
Rps31
Rad9
Rtf1
Rpo21
Npl3
Air2
Fps1
Ask10
Cdc28
Hog1
Pbs2
Ssk2
Clb2
Bud3
Bud4
Msn2
Sch9
Rim15
Yak1
Ypl247c
Vps52
Kss1
Sko1
Slt2
Msg5
Swe1
Igo2
Cdc55
Igo1
Vps53
Vps54
Rtn1
Fmp42
Cot1
Zrc1
10
20
30
40
50
t/s

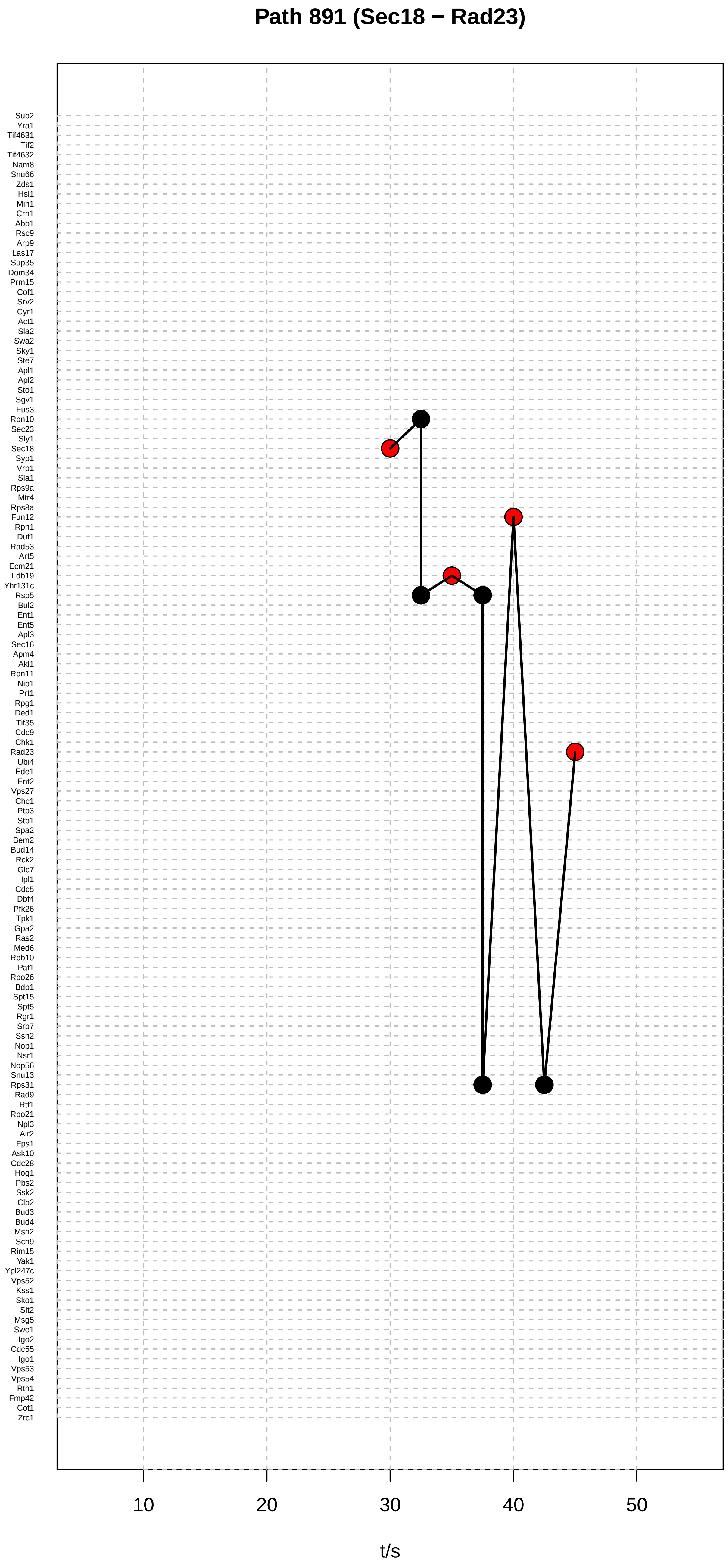
Path 891 (Sec18 − Rad23)
Sub2
Yra1
Tif4631
Tif2
Tif4632
Nam8
Snu66
Zds1
Hsl1
Mih1
Crn1
Abp1
Rsc9
Arp9
Las17
Sup35
Dom34
Prm15
Cof1
Srv2
Cyr1
Act1
Sla2
Swa2
Sky1
Ste7
Apl1
Apl2
Sto1
Sgv1
Fus3
Rpn10
Sec23
Sly1
Sec18
Syp1
Vrp1
Sla1
Rps9a
Mtr4
Rps8a
Fun12
Rpn1
Duf1
Rad53
Art5
Ecm21
Ldb19
Yhr131c
Rsp5
Bul2
Ent1
Ent5
Apl3
Sec16
Apm4
Akl1
Rpn11
Nip1
Prt1
Rpg1
Ded1
Tif35
Cdc9
Chk1
Rad23
Ubi4
Ede1
Ent2
Vps27
Chc1
Ptp3
Stb1
Spa2
Bem2
Bud14
Rck2
Glc7
Ipl1
Cdc5
Dbf4
Pfk26
Tpk1
Gpa2
Ras2
Med6
Rpb10
Paf1
Rpo26
Bdp1
Spt15
Spt5
Rgr1
Srb7
Ssn2
Nop1
Nsr1
Nop56
Snu13
Rps31
Rad9
Rtf1
Rpo21
Npl3
Air2
Fps1
Ask10
Cdc28
Hog1
Pbs2
Ssk2
Clb2
Bud3
Bud4
Msn2
Sch9
Rim15
Yak1
Ypl247c
Vps52
Kss1
Sko1
Slt2
Msg5
Swe1
Igo2
Cdc55
Igo1
Vps53
Vps54
Rtn1
Fmp42
Cot1
Zrc1
10
20
30
40
50
t/s

### Path 892 (Sec18 − Rad23)

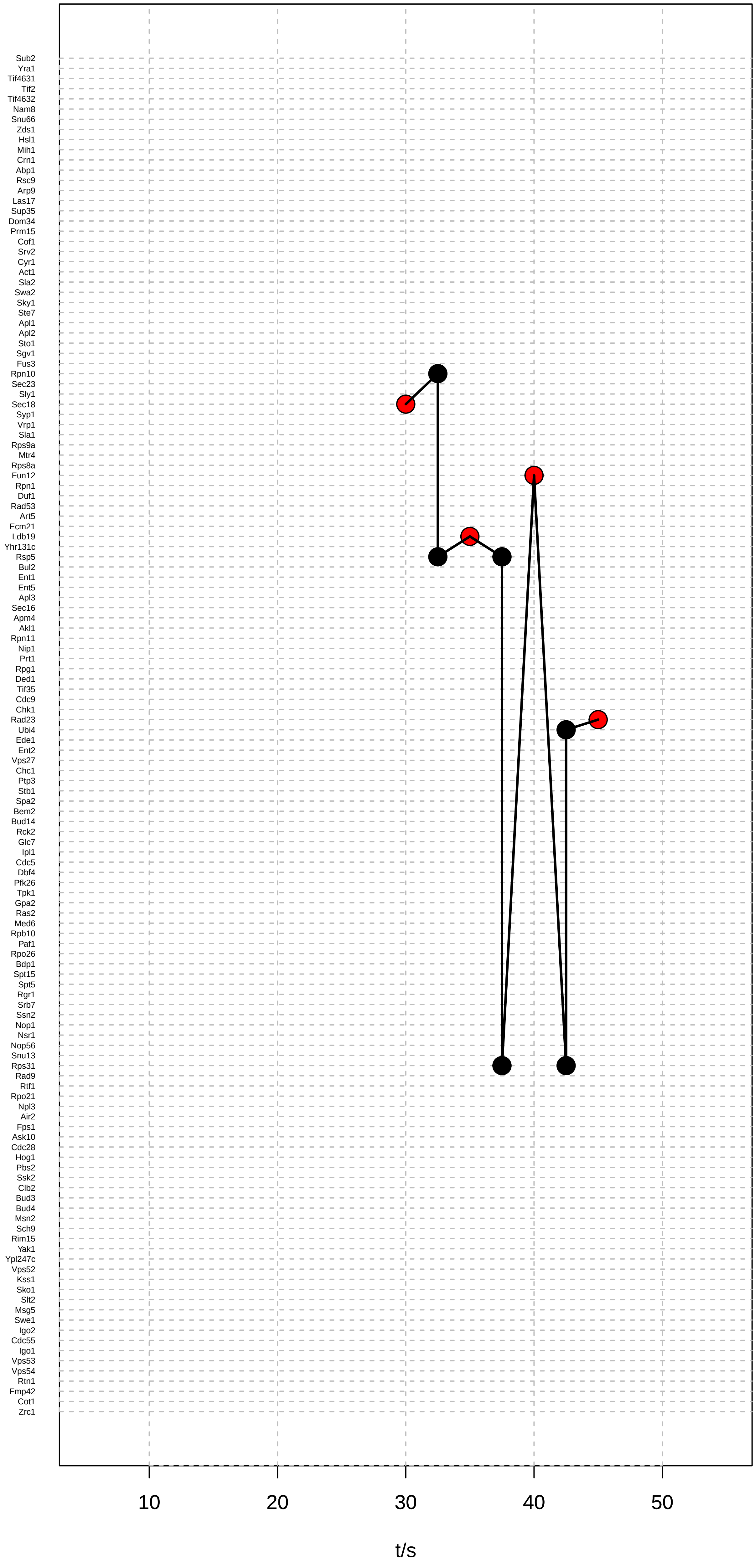

### Path 893 (Sec18 − Rad23)

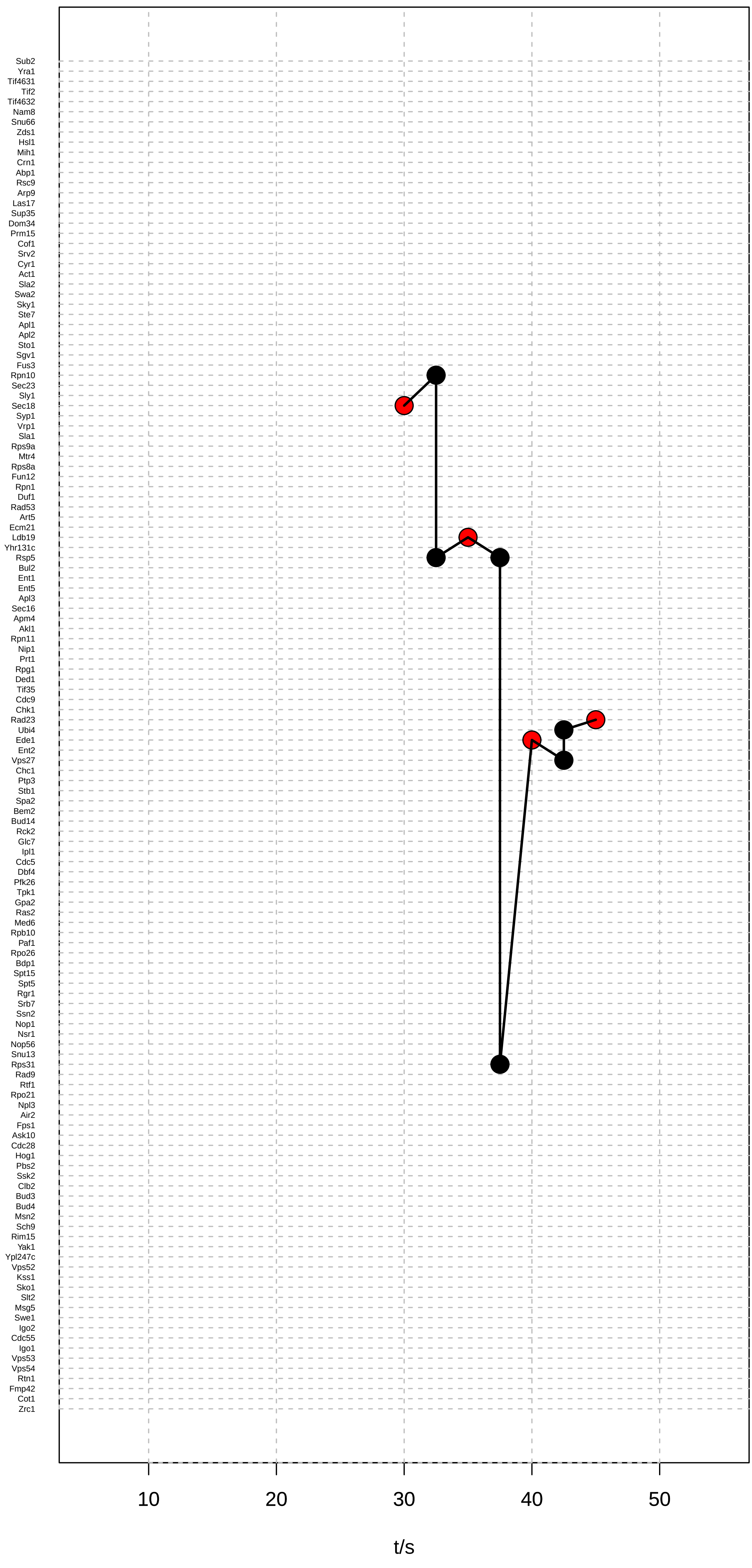

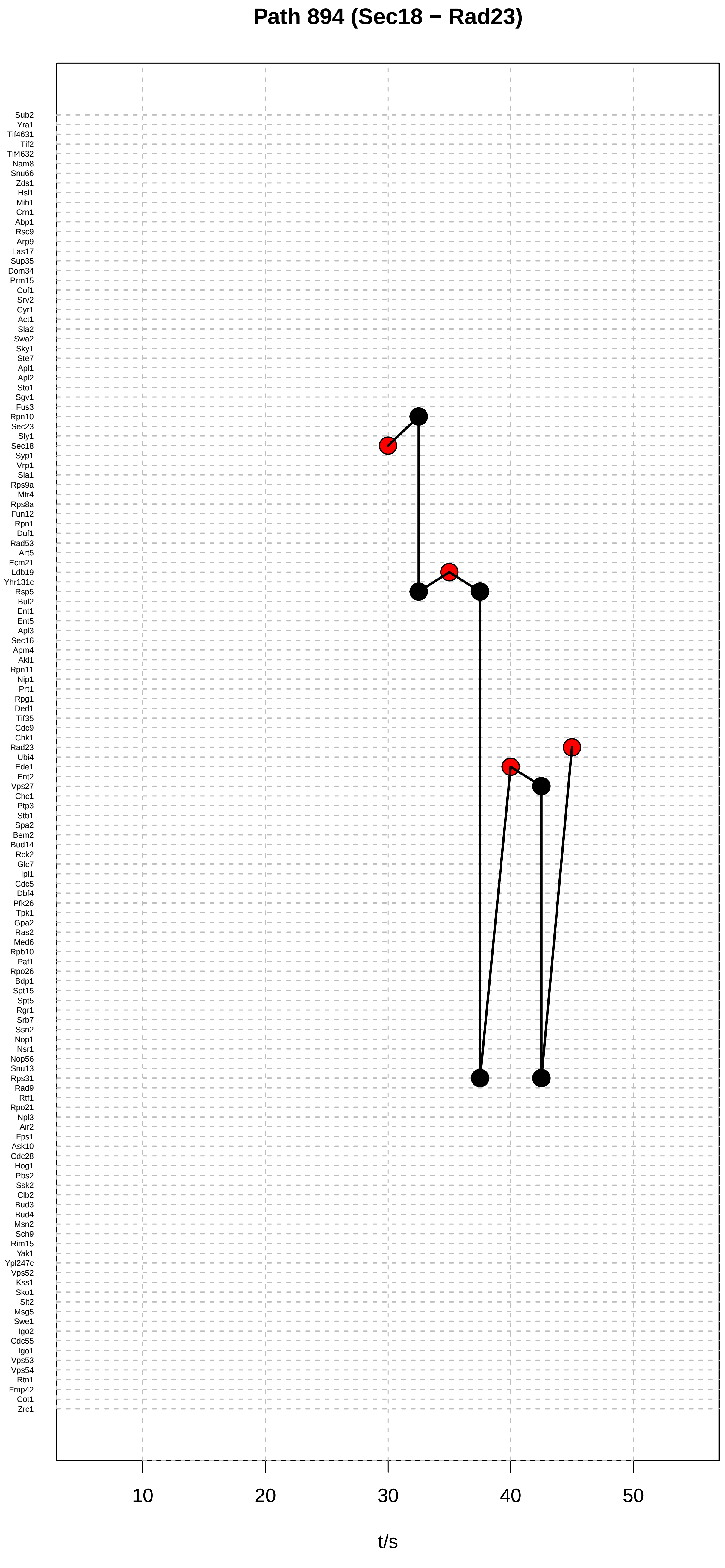

Path 894 (Sec18 − Rad23)
Sub2
Yra1
Tif4631
Tif2
Tif4632
Nam8
Snu66
Zds1
Hsl1
Mih1
Crn1
Abp1
Rsc9
Arp9
Las17
Sup35
Dom34
Prm15
Cof1
Srv2
Cyr1
Act1
Sla2
Swa2
Sky1
Ste7
Apl1
Apl2
Sto1
Sgv1
Fus3
Rpn10
Sec23
Sly1
Sec18
Syp1
Vrp1
Sla1
Rps9a
Mtr4
Rps8a
Fun12
Rpn1
Duf1
Rad53
Art5
Ecm21
Ldb19
Yhr131c
Rsp5
Bul2
Ent1
Ent5
Apl3
Sec16
Apm4
Akl1
Rpn11
Nip1
Prt1
Rpg1
Ded1
Tif35
Cdc9
Chk1
Rad23
Ubi4
Ede1
Ent2
Vps27
Chc1
Ptp3
Stb1
Spa2
Bem2
Bud14
Rck2
Glc7
Ipl1
Cdc5
Dbf4
Pfk26
Tpk1
Gpa2
Ras2
Med6
Rpb10
Paf1
Rpo26
Bdp1
Spt15
Spt5
Rgr1
Srb7
Ssn2
Nop1
Nsr1
Nop56
Snu13
Rps31
Rad9
Rtf1
Rpo21
Npl3
Air2
Fps1
Ask10
Cdc28
Hog1
Pbs2
Ssk2
Clb2
Bud3
Bud4
Msn2
Sch9
Rim15
Yak1
Ypl247c
Vps52
Kss1
Sko1
Slt2
Msg5
Swe1
Igo2
Cdc55
Igo1
Vps53
Vps54
Rtn1
Fmp42
Cot1
Zrc1
10
20
30
40
50
t/s

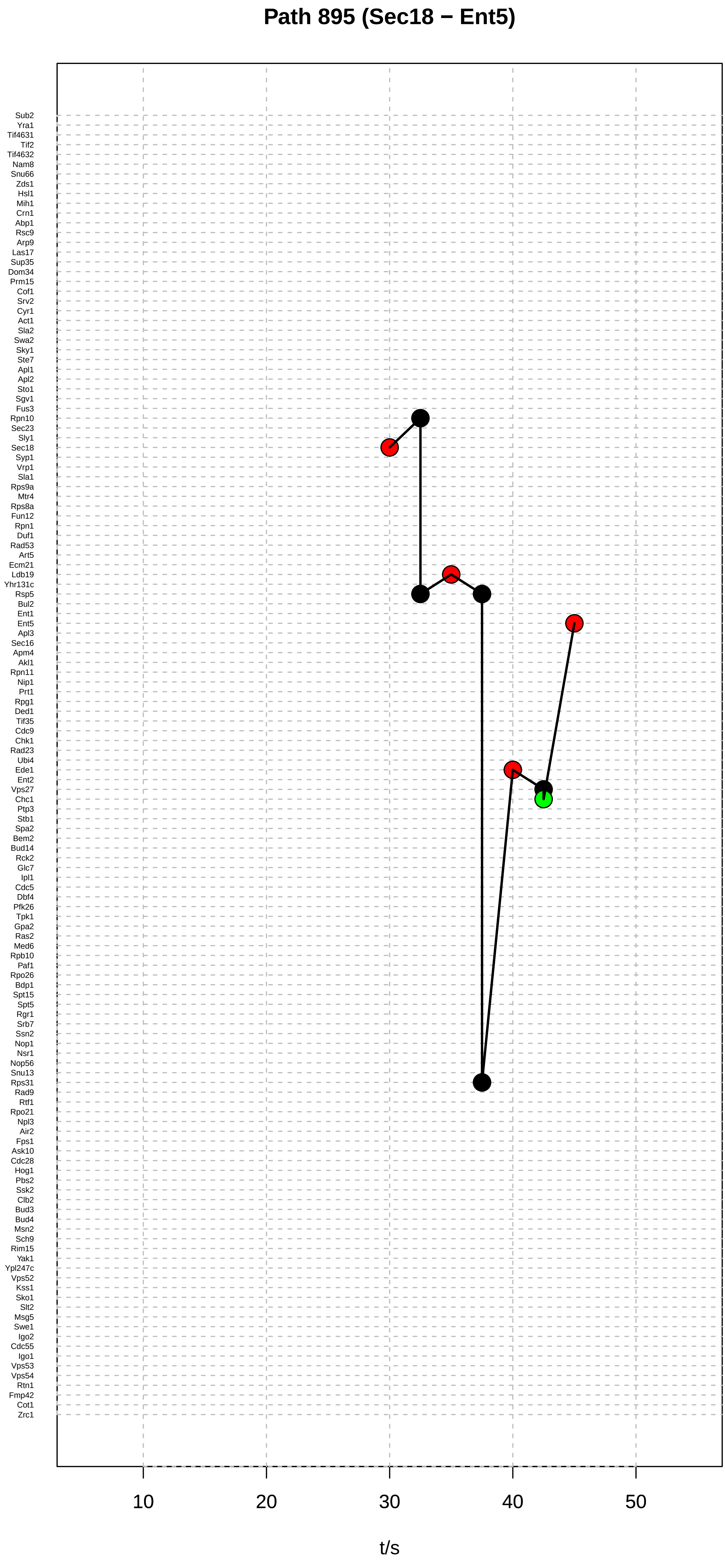

Path 895 (Sec18 − Ent5)
Sub2
Yra1
Tif4631
Tif2
Tif4632
Nam8
Snu66
Zds1
Hsl1
Mih1
Crn1
Abp1
Rsc9
Arp9
Las17
Sup35
Dom34
Prm15
Cof1
Srv2
Cyr1
Act1
Sla2
Swa2
Sky1
Ste7
Apl1
Apl2
Sto1
Sgv1
Fus3
Rpn10
Sec23
Sly1
Sec18
Syp1
Vrp1
Sla1
Rps9a
Mtr4
Rps8a
Fun12
Rpn1
Duf1
Rad53
Art5
Ecm21
Ldb19
Yhr131c
Rsp5
Bul2
Ent1
Ent5
Apl3
Sec16
Apm4
Akl1
Rpn11
Nip1
Prt1
Rpg1
Ded1
Tif35
Cdc9
Chk1
Rad23
Ubi4
Ede1
Ent2
Vps27
Chc1
Ptp3
Stb1
Spa2
Bem2
Bud14
Rck2
Glc7
Ipl1
Cdc5
Dbf4
Pfk26
Tpk1
Gpa2
Ras2
Med6
Rpb10
Paf1
Rpo26
Bdp1
Spt15
Spt5
Rgr1
Srb7
Ssn2
Nop1
Nsr1
Nop56
Snu13
Rps31
Rad9
Rtf1
Rpo21
Npl3
Air2
Fps1
Ask10
Cdc28
Hog1
Pbs2
Ssk2
Clb2
Bud3
Bud4
Msn2
Sch9
Rim15
Yak1
Ypl247c
Vps52
Kss1
Sko1
Slt2
Msg5
Swe1
Igo2
Cdc55
Igo1
Vps53
Vps54
Rtn1
Fmp42
Cot1
Zrc1
10
20
30
40
50
t/s

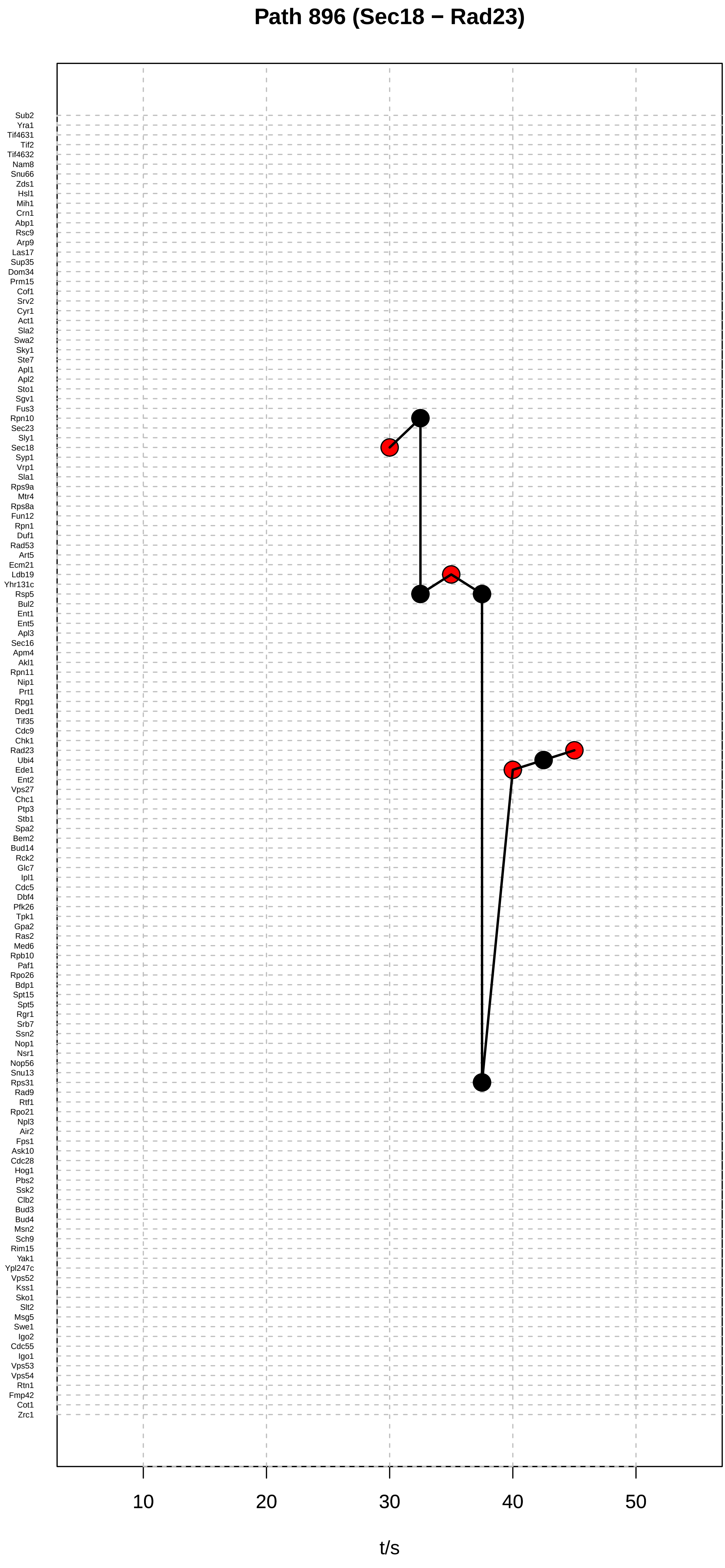
Path 896 (Sec18 − Rad23)
Sub2
Yra1
Tif4631
Tif2
Tif4632
Nam8
Snu66
Zds1
Hsl1
Mih1
Crn1
Abp1
Rsc9
Arp9
Las17
Sup35
Dom34
Prm15
Cof1
Srv2
Cyr1
Act1
Sla2
Swa2
Sky1
Ste7
Apl1
Apl2
Sto1
Sgv1
Fus3
Rpn10
Sec23
Sly1
Sec18
Syp1
Vrp1
Sla1
Rps9a
Mtr4
Rps8a
Fun12
Rpn1
Duf1
Rad53
Art5
Ecm21
Ldb19
Yhr131c
Rsp5
Bul2
Ent1
Ent5
Apl3
Sec16
Apm4
Akl1
Rpn11
Nip1
Prt1
Rpg1
Ded1
Tif35
Cdc9
Chk1
Rad23
Ubi4
Ede1
Ent2
Vps27
Chc1
Ptp3
Stb1
Spa2
Bem2
Bud14
Rck2
Glc7
Ipl1
Cdc5
Dbf4
Pfk26
Tpk1
Gpa2
Ras2
Med6
Rpb10
Paf1
Rpo26
Bdp1
Spt15
Spt5
Rgr1
Srb7
Ssn2
Nop1
Nsr1
Nop56
Snu13
Rps31
Rad9
Rtf1
Rpo21
Npl3
Air2
Fps1
Ask10
Cdc28
Hog1
Pbs2
Ssk2
Clb2
Bud3
Bud4
Msn2
Sch9
Rim15
Yak1
Ypl247c
Vps52
Kss1
Sko1
Slt2
Msg5
Swe1
Igo2
Cdc55
Igo1
Vps53
Vps54
Rtn1
Fmp42
Cot1
Zrc1
10
20
30
40
50
t/s

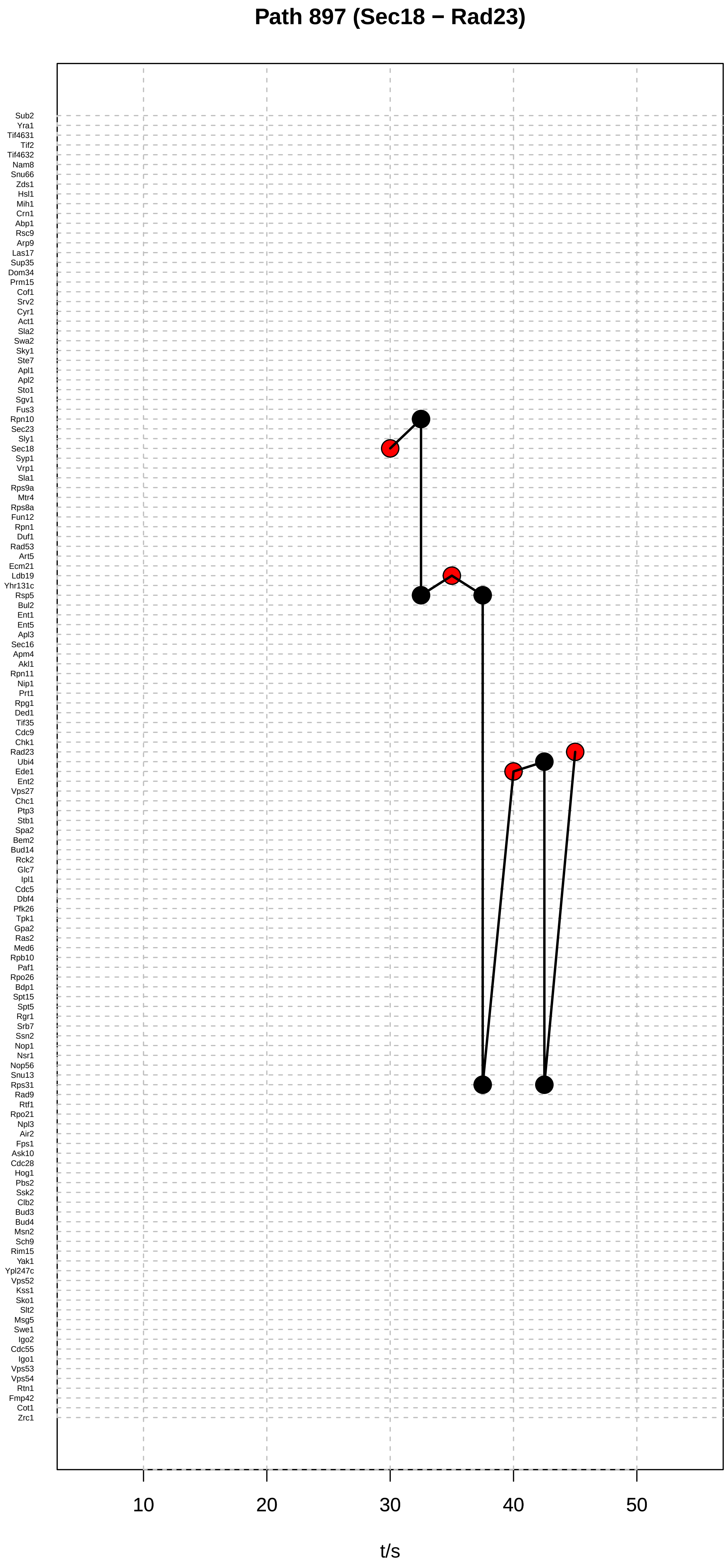
Path 897 (Sec18 − Rad23)
Sub2
Yra1
Tif4631
Tif2
Tif4632
Nam8
Snu66
Zds1
Hsl1
Mih1
Crn1
Abp1
Rsc9
Arp9
Las17
Sup35
Dom34
Prm15
Cof1
Srv2
Cyr1
Act1
Sla2
Swa2
Sky1
Ste7
Apl1
Apl2
Sto1
Sgv1
Fus3
Rpn10
Sec23
Sly1
Sec18
Syp1
Vrp1
Sla1
Rps9a
Mtr4
Rps8a
Fun12
Rpn1
Duf1
Rad53
Art5
Ecm21
Ldb19
Yhr131c
Rsp5
Bul2
Ent1
Ent5
Apl3
Sec16
Apm4
Akl1
Rpn11
Nip1
Prt1
Rpg1
Ded1
Tif35
Cdc9
Chk1
Rad23
Ubi4
Ede1
Ent2
Vps27
Chc1
Ptp3
Stb1
Spa2
Bem2
Bud14
Rck2
Glc7
Ipl1
Cdc5
Dbf4
Pfk26
Tpk1
Gpa2
Ras2
Med6
Rpb10
Paf1
Rpo26
Bdp1
Spt15
Spt5
Rgr1
Srb7
Ssn2
Nop1
Nsr1
Nop56
Snu13
Rps31
Rad9
Rtf1
Rpo21
Npl3
Air2
Fps1
Ask10
Cdc28
Hog1
Pbs2
Ssk2
Clb2
Bud3
Bud4
Msn2
Sch9
Rim15
Yak1
Ypl247c
Vps52
Kss1
Sko1
Slt2
Msg5
Swe1
Igo2
Cdc55
Igo1
Vps53
Vps54
Rtn1
Fmp42
Cot1
Zrc1
10
20
30
40
50
t/s

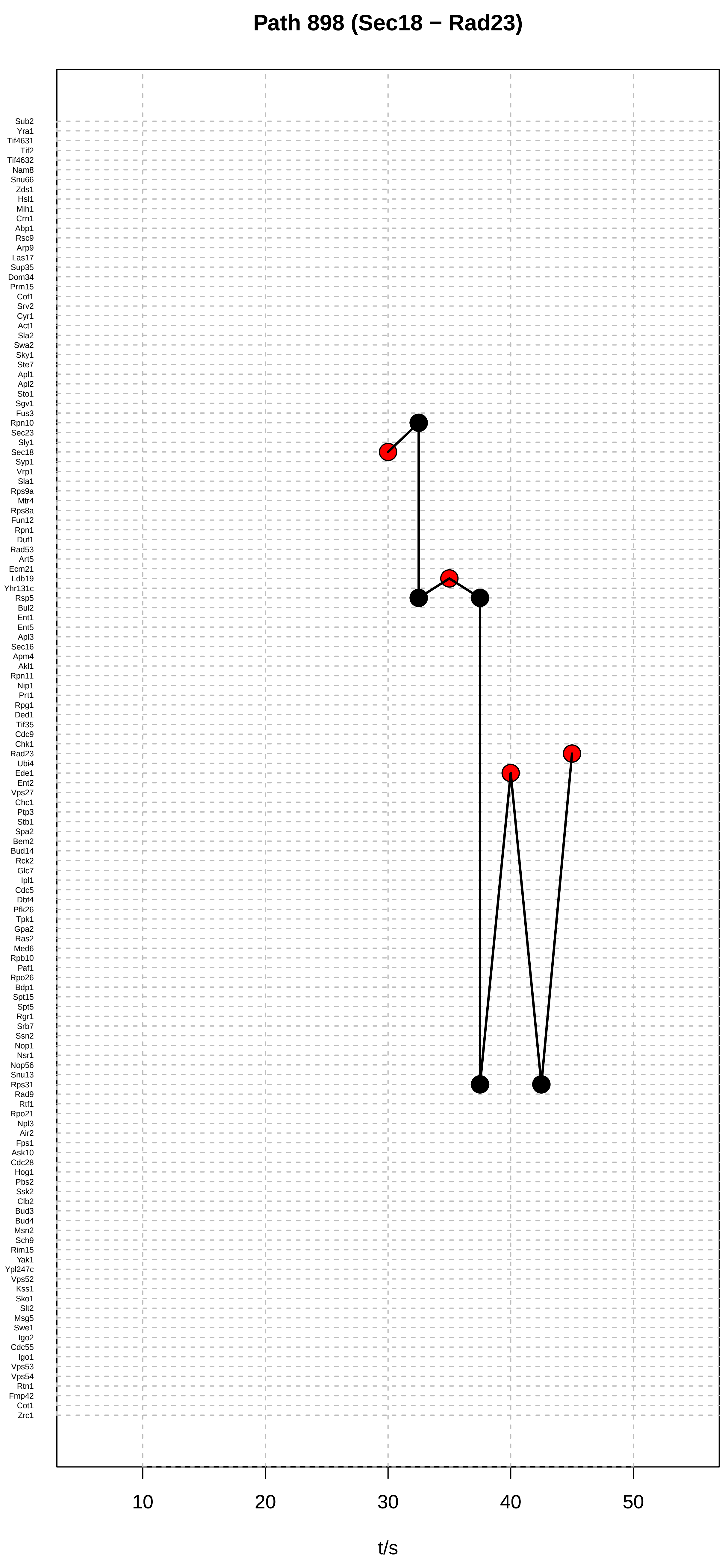

Path 898 (Sec18 − Rad23)
Sub2
Yra1
Tif4631
Tif2
Tif4632
Nam8
Snu66
Zds1
Hsl1
Mih1
Crn1
Abp1
Rsc9
Arp9
Las17
Sup35
Dom34
Prm15
Cof1
Srv2
Cyr1
Act1
Sla2
Swa2
Sky1
Ste7
Apl1
Apl2
Sto1
Sgv1
Fus3
Rpn10
Sec23
Sly1
Sec18
Syp1
Vrp1
Sla1
Rps9a
Mtr4
Rps8a
Fun12
Rpn1
Duf1
Rad53
Art5
Ecm21
Ldb19
Yhr131c
Rsp5
Bul2
Ent1
Ent5
Apl3
Sec16
Apm4
Akl1
Rpn11
Nip1
Prt1
Rpg1
Ded1
Tif35
Cdc9
Chk1
Rad23
Ubi4
Ede1
Ent2
Vps27
Chc1
Ptp3
Stb1
Spa2
Bem2
Bud14
Rck2
Glc7
Ipl1
Cdc5
Dbf4
Pfk26
Tpk1
Gpa2
Ras2
Med6
Rpb10
Paf1
Rpo26
Bdp1
Spt15
Spt5
Rgr1
Srb7
Ssn2
Nop1
Nsr1
Nop56
Snu13
Rps31
Rad9
Rtf1
Rpo21
Npl3
Air2
Fps1
Ask10
Cdc28
Hog1
Pbs2
Ssk2
Clb2
Bud3
Bud4
Msn2
Sch9
Rim15
Yak1
Ypl247c
Vps52
Kss1
Sko1
Slt2
Msg5
Swe1
Igo2
Cdc55
Igo1
Vps53
Vps54
Rtn1
Fmp42
Cot1
Zrc1
10
20
30
40
50
t/s

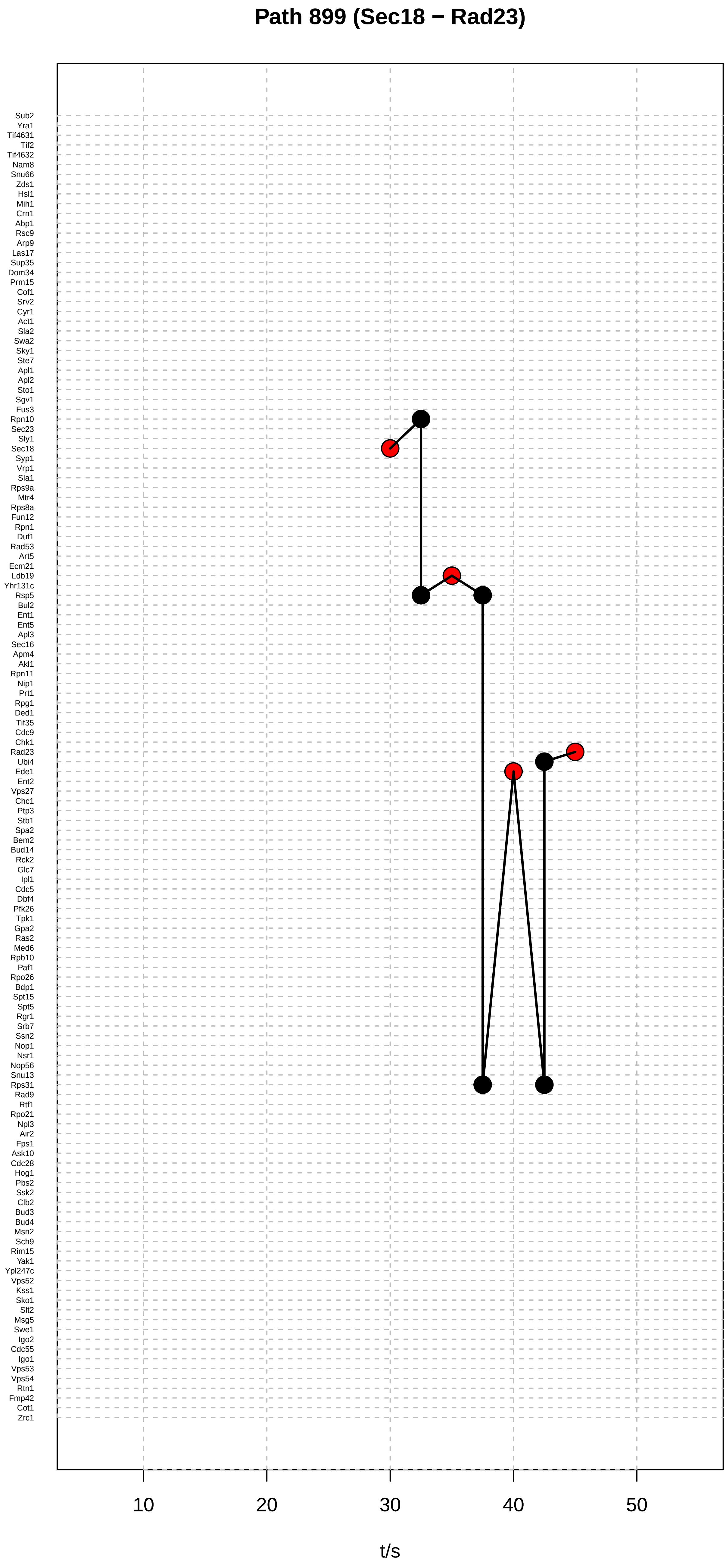

Path 899 (Sec18 − Rad23)
Sub2
Yra1
Tif4631
Tif2
Tif4632
Nam8
Snu66
Zds1
Hsl1
Mih1
Crn1
Abp1
Rsc9
Arp9
Las17
Sup35
Dom34
Prm15
Cof1
Srv2
Cyr1
Act1
Sla2
Swa2
Sky1
Ste7
Apl1
Apl2
Sto1
Sgv1
Fus3
Rpn10
Sec23
Sly1
Sec18
Syp1
Vrp1
Sla1
Rps9a
Mtr4
Rps8a
Fun12
Rpn1
Duf1
Rad53
Art5
Ecm21
Ldb19
Yhr131c
Rsp5
Bul2
Ent1
Ent5
Apl3
Sec16
Apm4
Akl1
Rpn11
Nip1
Prt1
Rpg1
Ded1
Tif35
Cdc9
Chk1
Rad23
Ubi4
Ede1
Ent2
Vps27
Chc1
Ptp3
Stb1
Spa2
Bem2
Bud14
Rck2
Glc7
Ipl1
Cdc5
Dbf4
Pfk26
Tpk1
Gpa2
Ras2
Med6
Rpb10
Paf1
Rpo26
Bdp1
Spt15
Spt5
Rgr1
Srb7
Ssn2
Nop1
Nsr1
Nop56
Snu13
Rps31
Rad9
Rtf1
Rpo21
Npl3
Air2
Fps1
Ask10
Cdc28
Hog1
Pbs2
Ssk2
Clb2
Bud3
Bud4
Msn2
Sch9
Rim15
Yak1
Ypl247c
Vps52
Kss1
Sko1
Slt2
Msg5
Swe1
Igo2
Cdc55
Igo1
Vps53
Vps54
Rtn1
Fmp42
Cot1
Zrc1
10
20
30
40
50
t/s

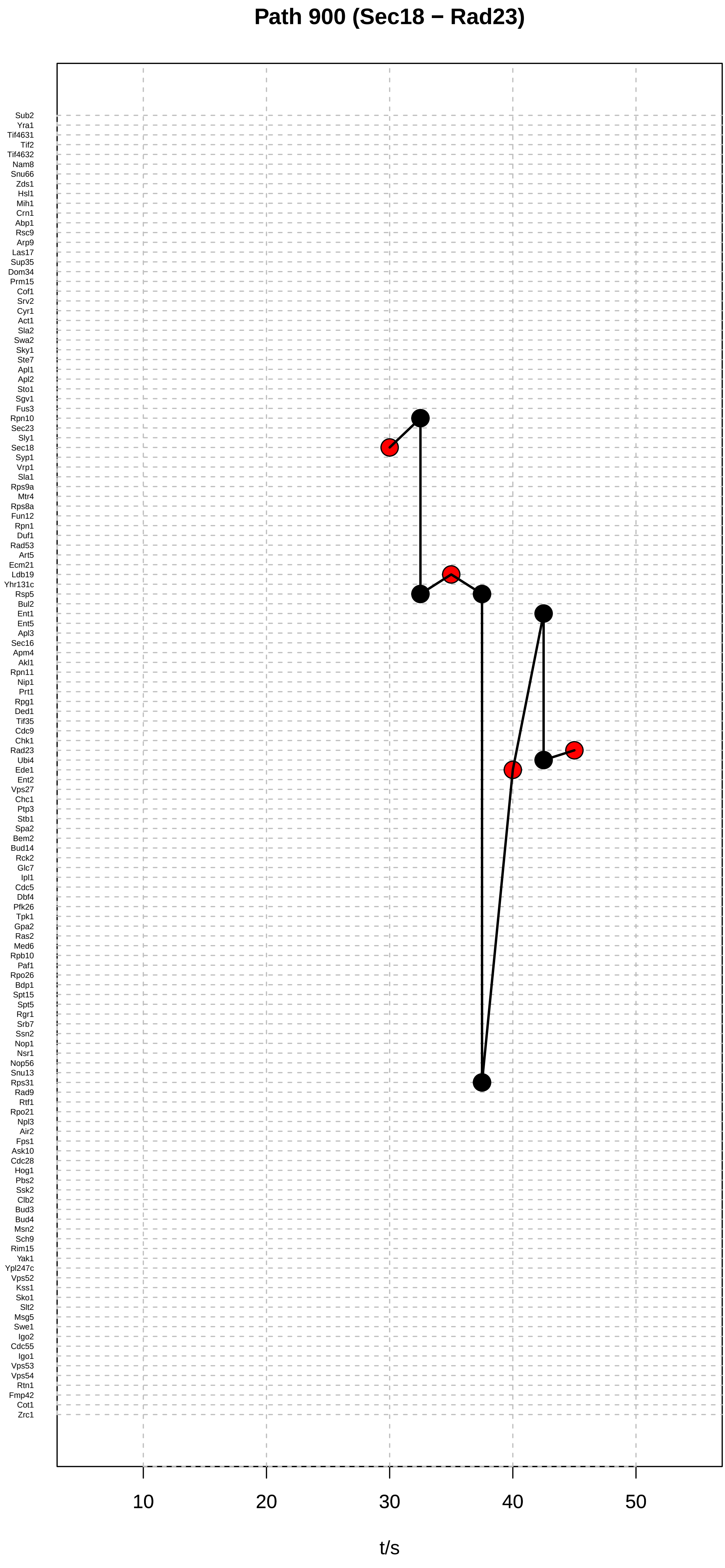
Path 900 (Sec18 − Rad23)
Sub2
Yra1
Tif4631
Tif2
Tif4632
Nam8
Snu66
Zds1
Hsl1
Mih1
Crn1
Abp1
Rsc9
Arp9
Las17
Sup35
Dom34
Prm15
Cof1
Srv2
Cyr1
Act1
Sla2
Swa2
Sky1
Ste7
Apl1
Apl2
Sto1
Sgv1
Fus3
Rpn10
Sec23
Sly1
Sec18
Syp1
Vrp1
Sla1
Rps9a
Mtr4
Rps8a
Fun12
Rpn1
Duf1
Rad53
Art5
Ecm21
Ldb19
Yhr131c
Rsp5
Bul2
Ent1
Ent5
Apl3
Sec16
Apm4
Akl1
Rpn11
Nip1
Prt1
Rpg1
Ded1
Tif35
Cdc9
Chk1
Rad23
Ubi4
Ede1
Ent2
Vps27
Chc1
Ptp3
Stb1
Spa2
Bem2
Bud14
Rck2
Glc7
Ipl1
Cdc5
Dbf4
Pfk26
Tpk1
Gpa2
Ras2
Med6
Rpb10
Paf1
Rpo26
Bdp1
Spt15
Spt5
Rgr1
Srb7
Ssn2
Nop1
Nsr1
Nop56
Snu13
Rps31
Rad9
Rtf1
Rpo21
Npl3
Air2
Fps1
Ask10
Cdc28
Hog1
Pbs2
Ssk2
Clb2
Bud3
Bud4
Msn2
Sch9
Rim15
Yak1
Ypl247c
Vps52
Kss1
Sko1
Slt2
Msg5
Swe1
Igo2
Cdc55
Igo1
Vps53
Vps54
Rtn1
Fmp42
Cot1
Zrc1
10
20
30
40
50
t/s

### Path 901 (Sec18 − Ent5)

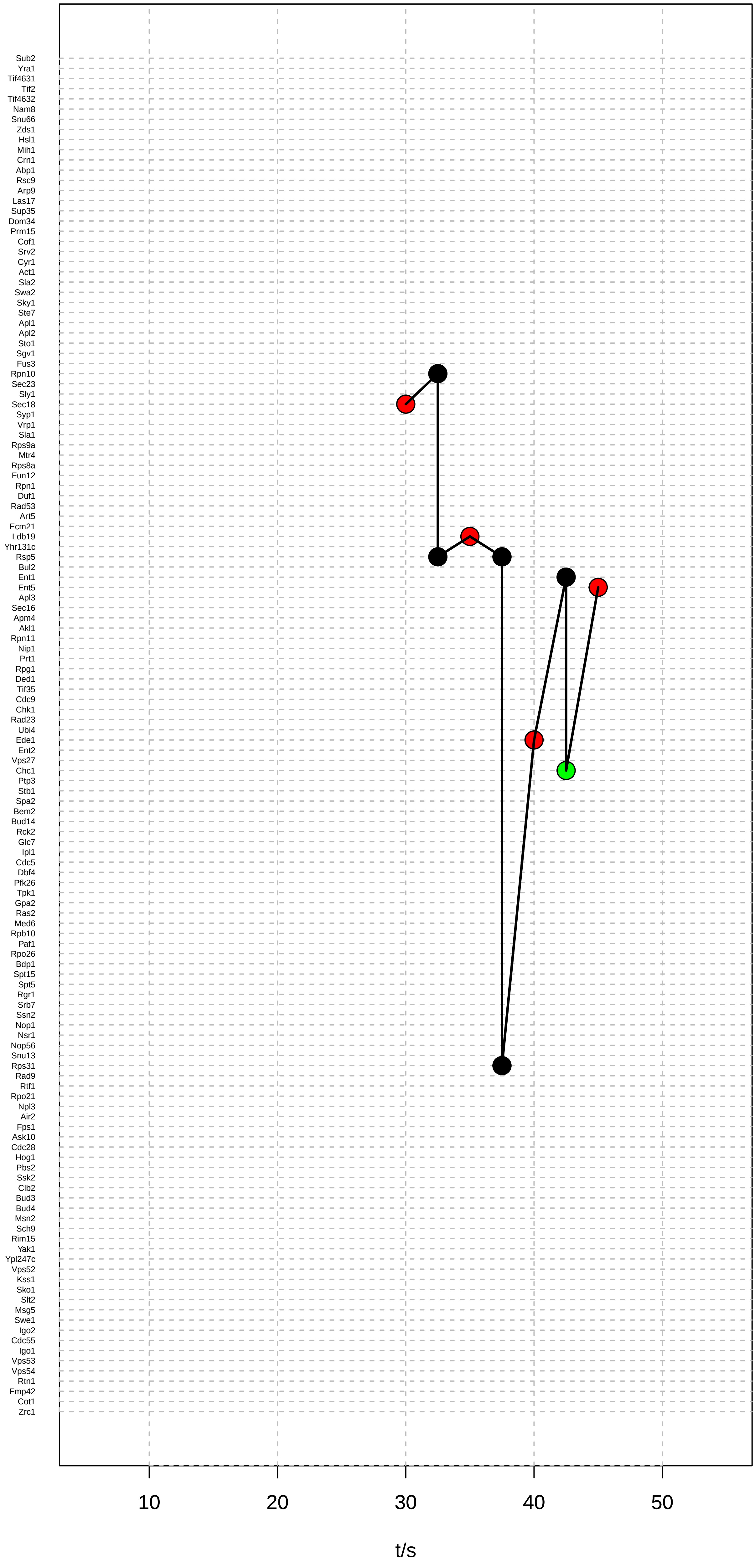

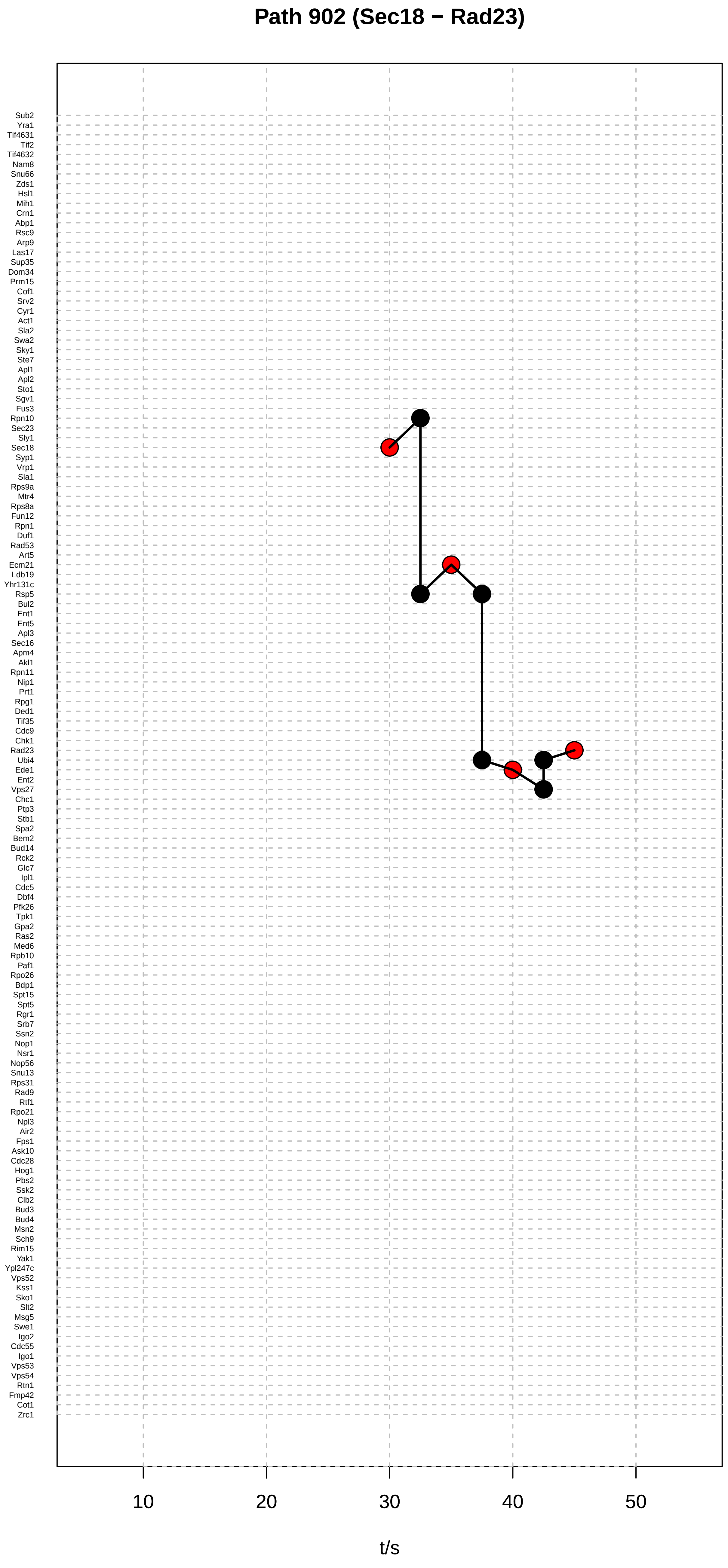
Path 902 (Sec18 − Rad23)
Sub2
Yra1
Tif4631
Tif2
Tif4632
Nam8
Snu66
Zds1
Hsl1
Mih1
Crn1
Abp1
Rsc9
Arp9
Las17
Sup35
Dom34
Prm15
Cof1
Srv2
Cyr1
Act1
Sla2
Swa2
Sky1
Ste7
Apl1
Apl2
Sto1
Sgv1
Fus3
Rpn10
Sec23
Sly1
Sec18
Syp1
Vrp1
Sla1
Rps9a
Mtr4
Rps8a
Fun12
Rpn1
Duf1
Rad53
Art5
Ecm21
Ldb19
Yhr131c
Rsp5
Bul2
Ent1
Ent5
Apl3
Sec16
Apm4
Akl1
Rpn11
Nip1
Prt1
Rpg1
Ded1
Tif35
Cdc9
Chk1
Rad23
Ubi4
Ede1
Ent2
Vps27
Chc1
Ptp3
Stb1
Spa2
Bem2
Bud14
Rck2
Glc7
Ipl1
Cdc5
Dbf4
Pfk26
Tpk1
Gpa2
Ras2
Med6
Rpb10
Paf1
Rpo26
Bdp1
Spt15
Spt5
Rgr1
Srb7
Ssn2
Nop1
Nsr1
Nop56
Snu13
Rps31
Rad9
Rtf1
Rpo21
Npl3
Air2
Fps1
Ask10
Cdc28
Hog1
Pbs2
Ssk2
Clb2
Bud3
Bud4
Msn2
Sch9
Rim15
Yak1
Ypl247c
Vps52
Kss1
Sko1
Slt2
Msg5
Swe1
Igo2
Cdc55
Igo1
Vps53
Vps54
Rtn1
Fmp42
Cot1
Zrc1
10
20
30
40
50
t/s

Path 903 (Sec18 − Rad23)

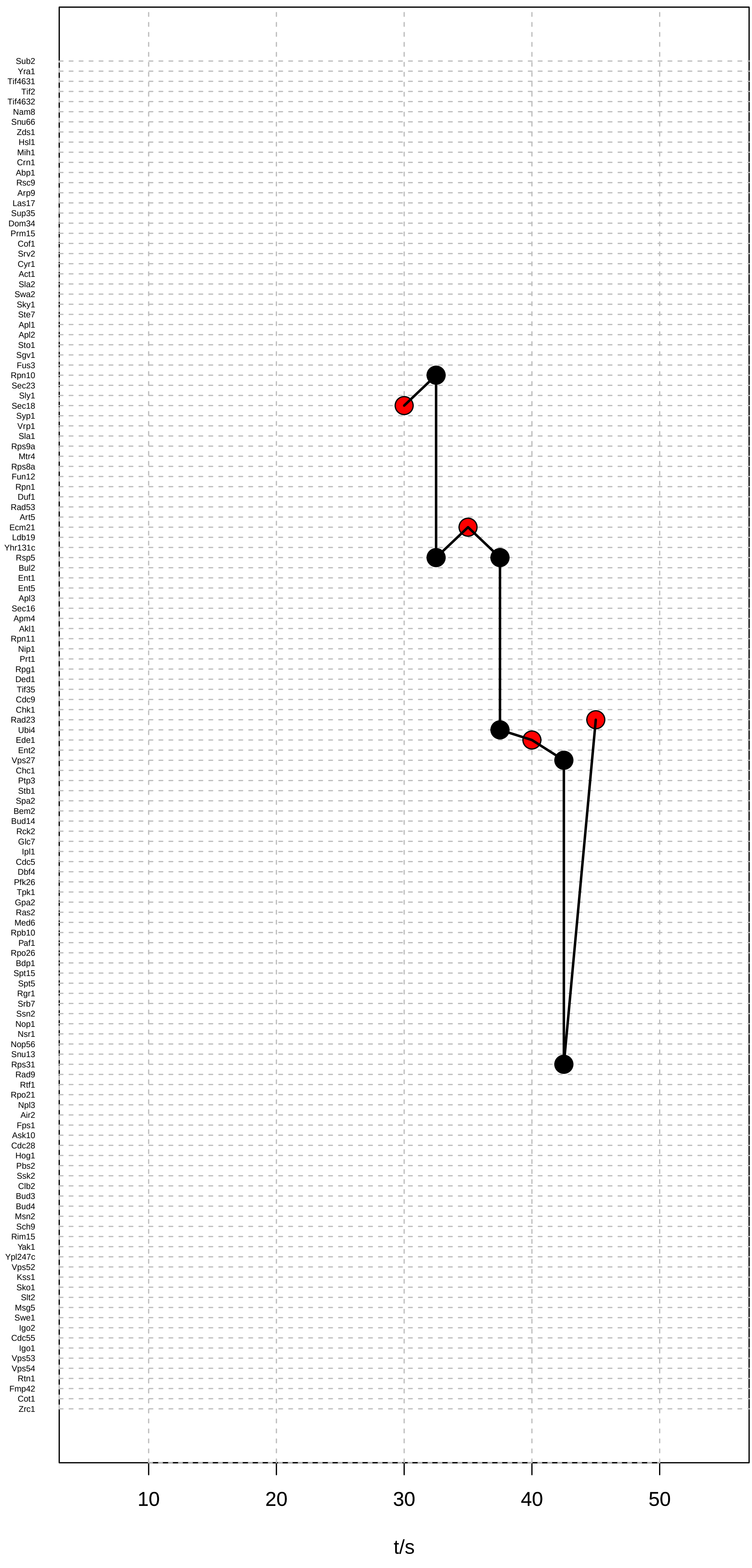

#### Path 904 (Sec18 − Ent5)

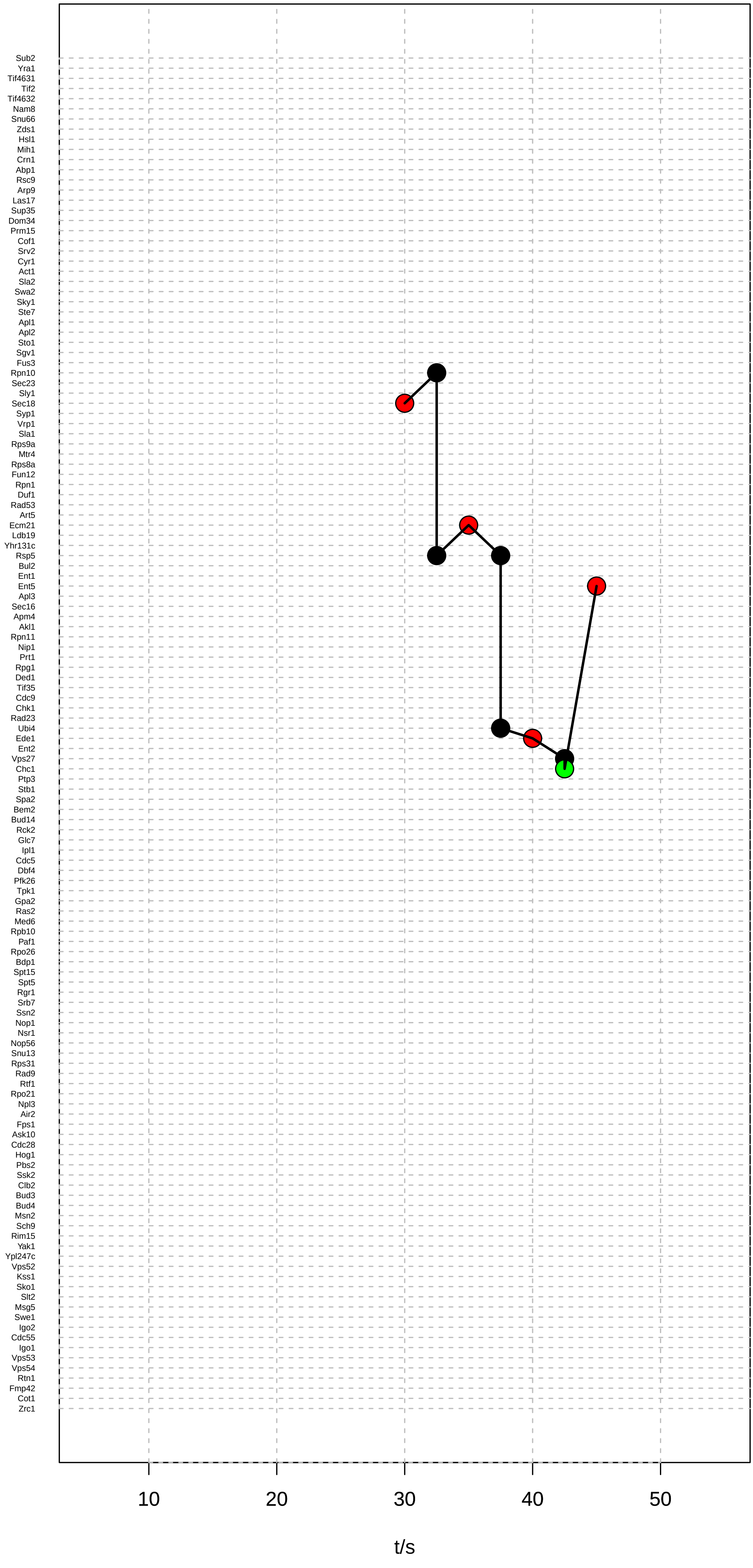

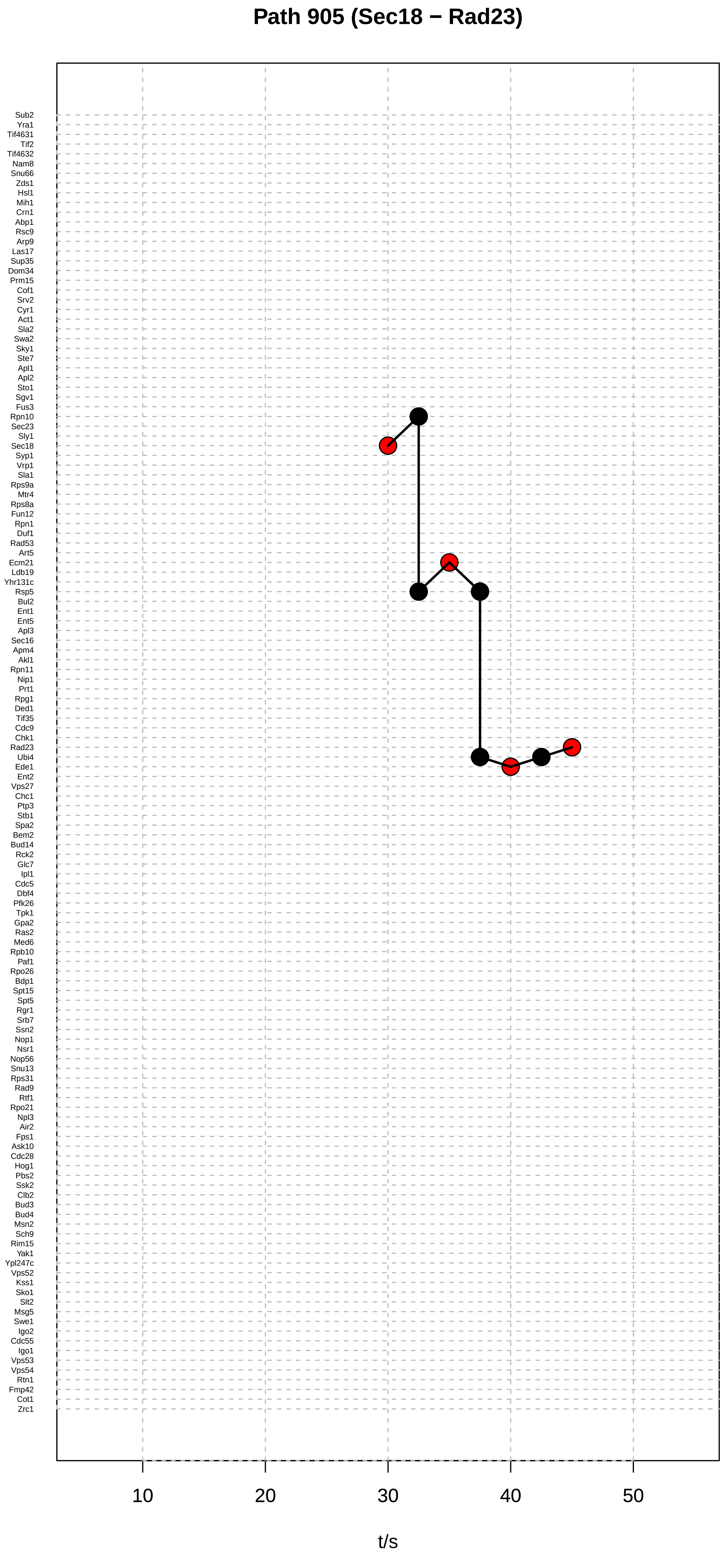

Path 905 (Sec18 − Rad23)
Sub2
Yra1
Tif4631
Tif2
Tif4632
Nam8
Snu66
Zds1
Hsl1
Mih1
Crn1
Abp1
Rsc9
Arp9
Las17
Sup35
Dom34
Prm15
Cof1
Srv2
Cyr1
Act1
Sla2
Swa2
Sky1
Ste7
Apl1
Apl2
Sto1
Sgv1
Fus3
Rpn10
Sec23
Sly1
Sec18
Syp1
Vrp1
Sla1
Rps9a
Mtr4
Rps8a
Fun12
Rpn1
Duf1
Rad53
Art5
Ecm21
Ldb19
Yhr131c
Rsp5
Bul2
Ent1
Ent5
Apl3
Sec16
Apm4
Akl1
Rpn11
Nip1
Prt1
Rpg1
Ded1
Tif35
Cdc9
Chk1
Rad23
Ubi4
Ede1
Ent2
Vps27
Chc1
Ptp3
Stb1
Spa2
Bem2
Bud14
Rck2
Glc7
Ipl1
Cdc5
Dbf4
Pfk26
Tpk1
Gpa2
Ras2
Med6
Rpb10
Paf1
Rpo26
Bdp1
Spt15
Spt5
Rgr1
Srb7
Ssn2
Nop1
Nsr1
Nop56
Snu13
Rps31
Rad9
Rtf1
Rpo21
Npl3
Air2
Fps1
Ask10
Cdc28
Hog1
Pbs2
Ssk2
Clb2
Bud3
Bud4
Msn2
Sch9
Rim15
Yak1
Ypl247c
Vps52
Kss1
Sko1
Slt2
Msg5
Swe1
Igo2
Cdc55
Igo1
Vps53
Vps54
Rtn1
Fmp42
Cot1
Zrc1
10
20
30
40
50
t/s

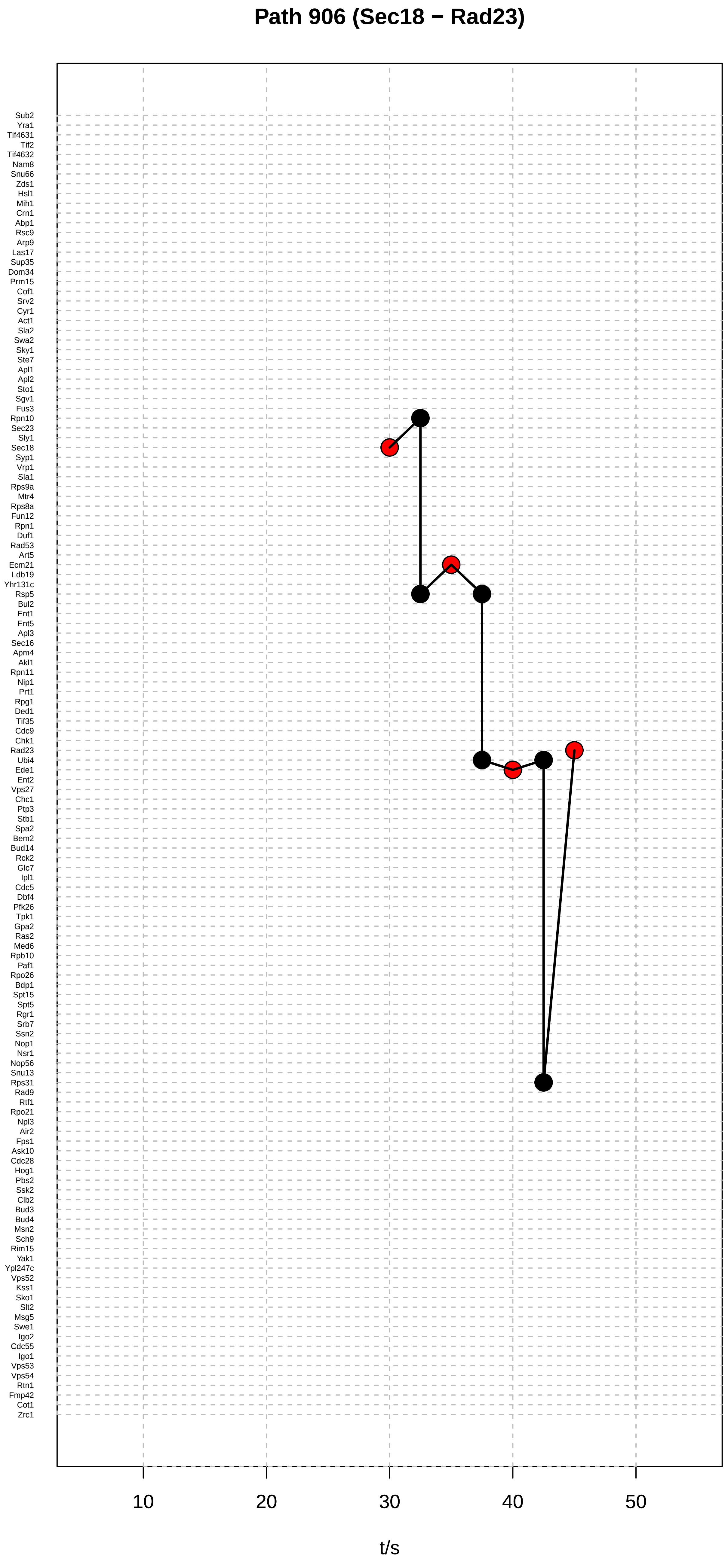
Path 906 (Sec18 − Rad23)
Sub2
Yra1
Tif4631
Tif2
Tif4632
Nam8
Snu66
Zds1
Hsl1
Mih1
Crn1
Abp1
Rsc9
Arp9
Las17
Sup35
Dom34
Prm15
Cof1
Srv2
Cyr1
Act1
Sla2
Swa2
Sky1
Ste7
Apl1
Apl2
Sto1
Sgv1
Fus3
Rpn10
Sec23
Sly1
Sec18
Syp1
Vrp1
Sla1
Rps9a
Mtr4
Rps8a
Fun12
Rpn1
Duf1
Rad53
Art5
Ecm21
Ldb19
Yhr131c
Rsp5
Bul2
Ent1
Ent5
Apl3
Sec16
Apm4
Akl1
Rpn11
Nip1
Prt1
Rpg1
Ded1
Tif35
Cdc9
Chk1
Rad23
Ubi4
Ede1
Ent2
Vps27
Chc1
Ptp3
Stb1
Spa2
Bem2
Bud14
Rck2
Glc7
Ipl1
Cdc5
Dbf4
Pfk26
Tpk1
Gpa2
Ras2
Med6
Rpb10
Paf1
Rpo26
Bdp1
Spt15
Spt5
Rgr1
Srb7
Ssn2
Nop1
Nsr1
Nop56
Snu13
Rps31
Rad9
Rtf1
Rpo21
Npl3
Air2
Fps1
Ask10
Cdc28
Hog1
Pbs2
Ssk2
Clb2
Bud3
Bud4
Msn2
Sch9
Rim15
Yak1
Ypl247c
Vps52
Kss1
Sko1
Slt2
Msg5
Swe1
Igo2
Cdc55
Igo1
Vps53
Vps54
Rtn1
Fmp42
Cot1
Zrc1
10
20
30
40
50
t/s

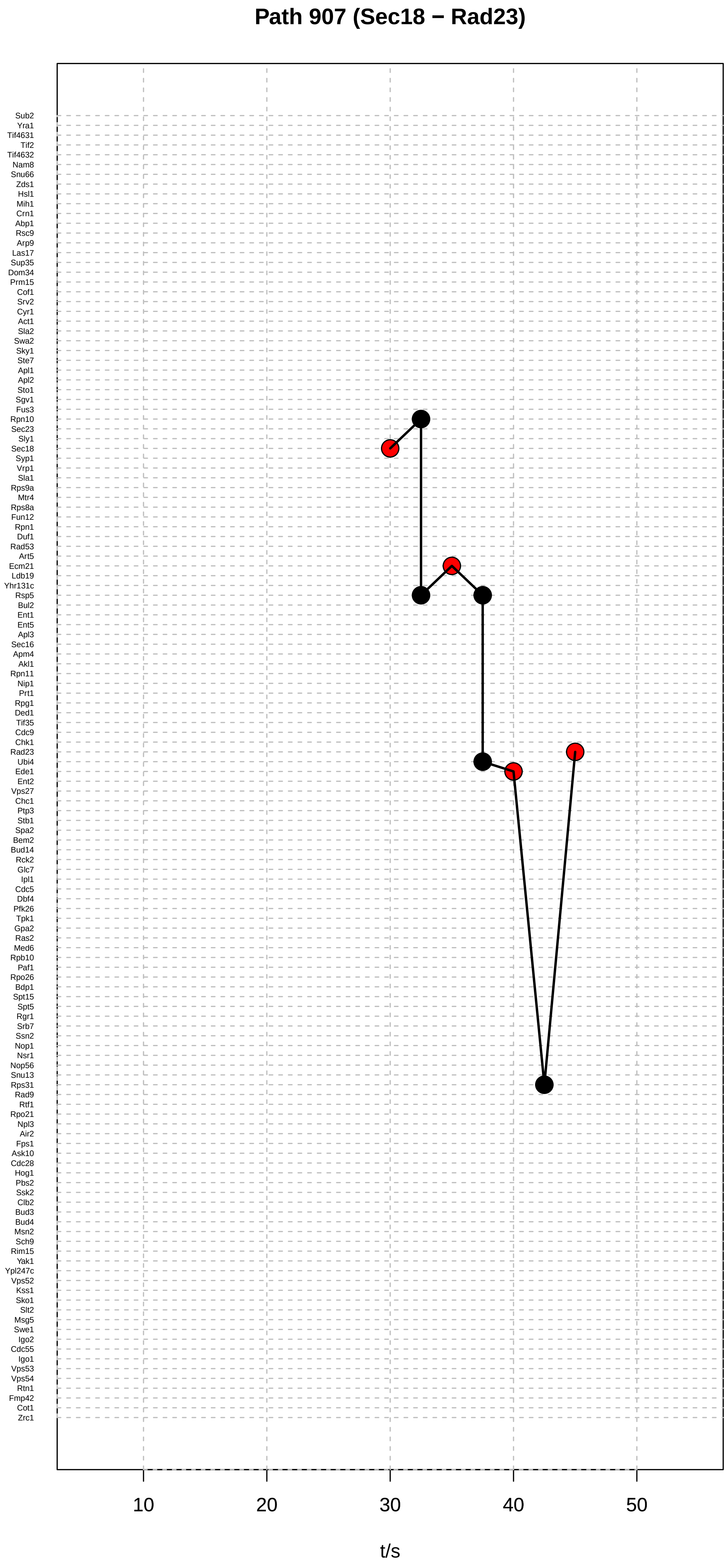

Path 907 (Sec18 − Rad23)
Sub2
Yra1
Tif4631
Tif2
Tif4632
Nam8
Snu66
Zds1
Hsl1
Mih1
Crn1
Abp1
Rsc9
Arp9
Las17
Sup35
Dom34
Prm15
Cof1
Srv2
Cyr1
Act1
Sla2
Swa2
Sky1
Ste7
Apl1
Apl2
Sto1
Sgv1
Fus3
Rpn10
Sec23
Sly1
Sec18
Syp1
Vrp1
Sla1
Rps9a
Mtr4
Rps8a
Fun12
Rpn1
Duf1
Rad53
Art5
Ecm21
Ldb19
Yhr131c
Rsp5
Bul2
Ent1
Ent5
Apl3
Sec16
Apm4
Akl1
Rpn11
Nip1
Prt1
Rpg1
Ded1
Tif35
Cdc9
Chk1
Rad23
Ubi4
Ede1
Ent2
Vps27
Chc1
Ptp3
Stb1
Spa2
Bem2
Bud14
Rck2
Glc7
Ipl1
Cdc5
Dbf4
Pfk26
Tpk1
Gpa2
Ras2
Med6
Rpb10
Paf1
Rpo26
Bdp1
Spt15
Spt5
Rgr1
Srb7
Ssn2
Nop1
Nsr1
Nop56
Snu13
Rps31
Rad9
Rtf1
Rpo21
Npl3
Air2
Fps1
Ask10
Cdc28
Hog1
Pbs2
Ssk2
Clb2
Bud3
Bud4
Msn2
Sch9
Rim15
Yak1
Ypl247c
Vps52
Kss1
Sko1
Slt2
Msg5
Swe1
Igo2
Cdc55
Igo1
Vps53
Vps54
Rtn1
Fmp42
Cot1
Zrc1
10
20
30
40
50
t/s

Path 908 (Sec18 − Rad23)

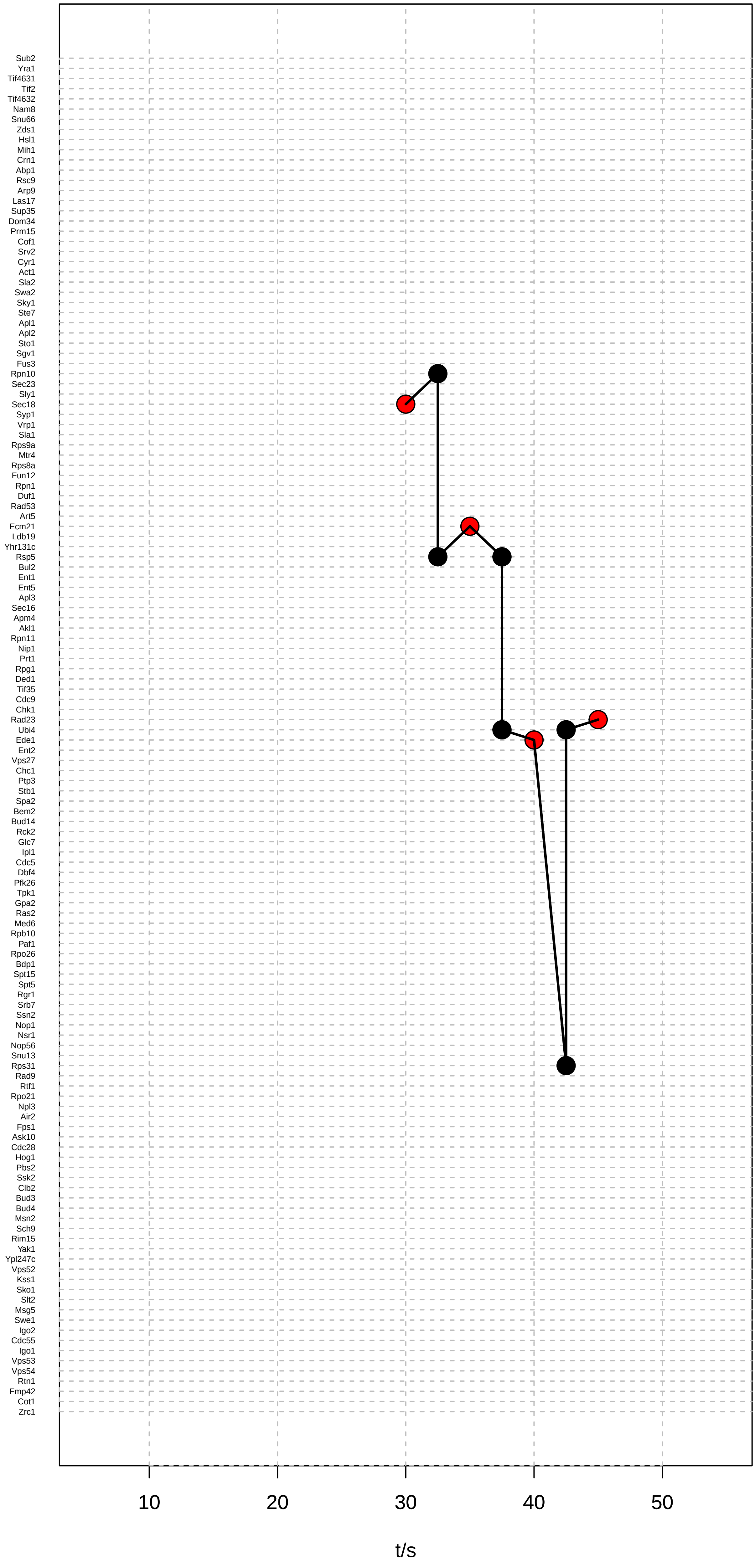

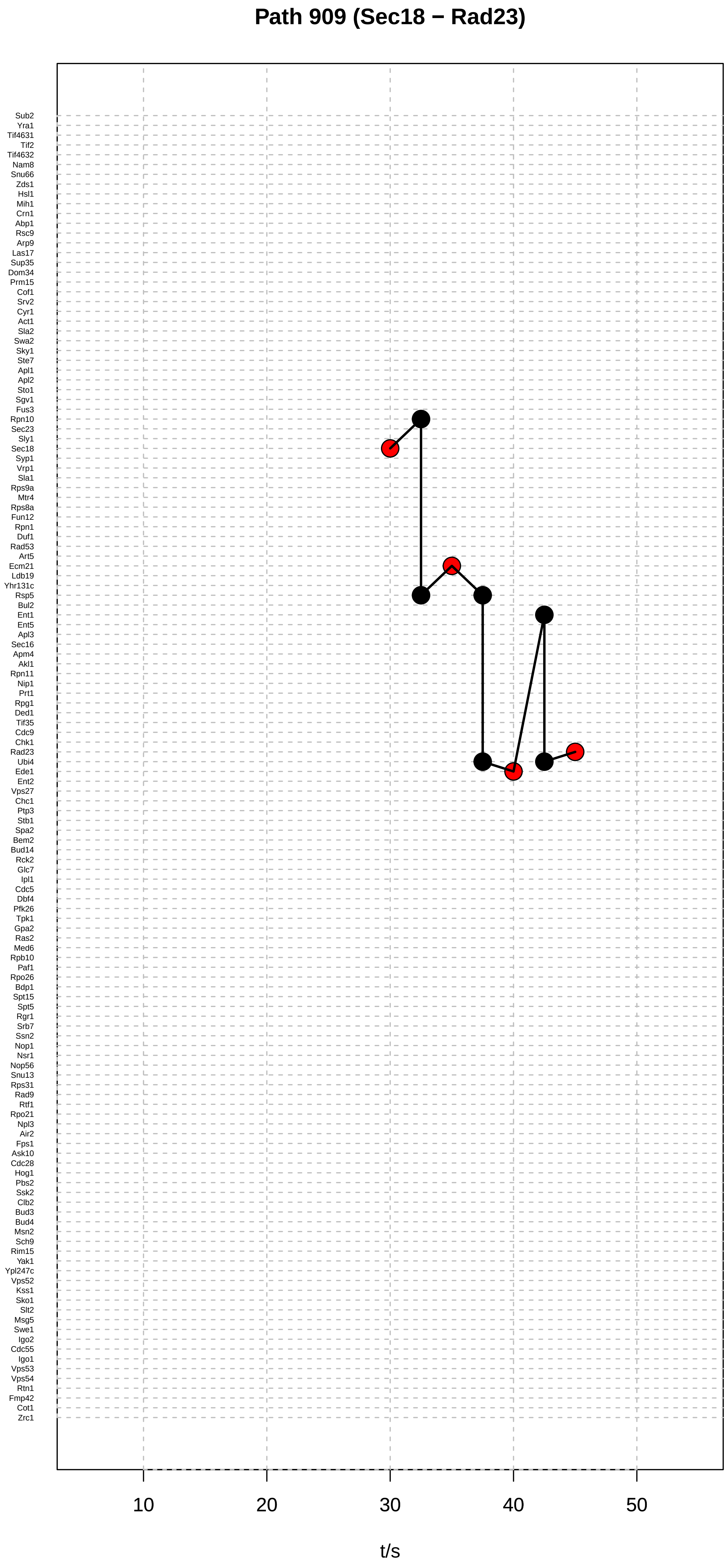

Path 909 (Sec18 − Rad23)
Sub2
Yra1
Tif4631
Tif2
Tif4632
Nam8
Snu66
Zds1
Hsl1
Mih1
Crn1
Abp1
Rsc9
Arp9
Las17
Sup35
Dom34
Prm15
Cof1
Srv2
Cyr1
Act1
Sla2
Swa2
Sky1
Ste7
Apl1
Apl2
Sto1
Sgv1
Fus3
Rpn10
Sec23
Sly1
Sec18
Syp1
Vrp1
Sla1
Rps9a
Mtr4
Rps8a
Fun12
Rpn1
Duf1
Rad53
Art5
Ecm21
Ldb19
Yhr131c
Rsp5
Bul2
Ent1
Ent5
Apl3
Sec16
Apm4
Akl1
Rpn11
Nip1
Prt1
Rpg1
Ded1
Tif35
Cdc9
Chk1
Rad23
Ubi4
Ede1
Ent2
Vps27
Chc1
Ptp3
Stb1
Spa2
Bem2
Bud14
Rck2
Glc7
Ipl1
Cdc5
Dbf4
Pfk26
Tpk1
Gpa2
Ras2
Med6
Rpb10
Paf1
Rpo26
Bdp1
Spt15
Spt5
Rgr1
Srb7
Ssn2
Nop1
Nsr1
Nop56
Snu13
Rps31
Rad9
Rtf1
Rpo21
Npl3
Air2
Fps1
Ask10
Cdc28
Hog1
Pbs2
Ssk2
Clb2
Bud3
Bud4
Msn2
Sch9
Rim15
Yak1
Ypl247c
Vps52
Kss1
Sko1
Slt2
Msg5
Swe1
Igo2
Cdc55
Igo1
Vps53
Vps54
Rtn1
Fmp42
Cot1
Zrc1
10
20
30
40
50
t/s

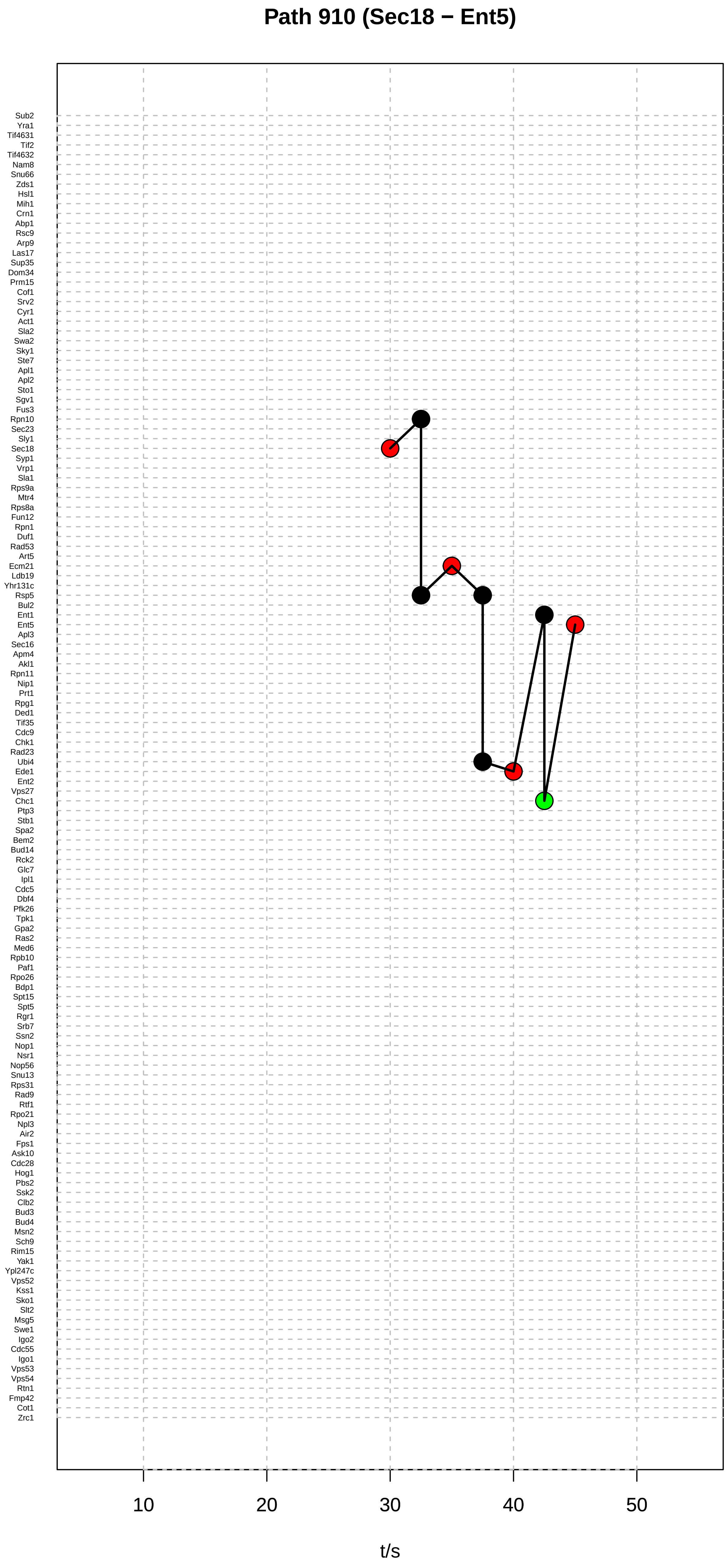
Path 910 (Sec18 − Ent5)
Sub2
Yra1
Tif4631
Tif2
Tif4632
Nam8
Snu66
Zds1
Hsl1
Mih1
Crn1
Abp1
Rsc9
Arp9
Las17
Sup35
Dom34
Prm15
Cof1
Srv2
Cyr1
Act1
Sla2
Swa2
Sky1
Ste7
Apl1
Apl2
Sto1
Sgv1
Fus3
Rpn10
Sec23
Sly1
Sec18
Syp1
Vrp1
Sla1
Rps9a
Mtr4
Rps8a
Fun12
Rpn1
Duf1
Rad53
Art5
Ecm21
Ldb19
Yhr131c
Rsp5
Bul2
Ent1
Ent5
Apl3
Sec16
Apm4
Akl1
Rpn11
Nip1
Prt1
Rpg1
Ded1
Tif35
Cdc9
Chk1
Rad23
Ubi4
Ede1
Ent2
Vps27
Chc1
Ptp3
Stb1
Spa2
Bem2
Bud14
Rck2
Glc7
Ipl1
Cdc5
Dbf4
Pfk26
Tpk1
Gpa2
Ras2
Med6
Rpb10
Paf1
Rpo26
Bdp1
Spt15
Spt5
Rgr1
Srb7
Ssn2
Nop1
Nsr1
Nop56
Snu13
Rps31
Rad9
Rtf1
Rpo21
Npl3
Air2
Fps1
Ask10
Cdc28
Hog1
Pbs2
Ssk2
Clb2
Bud3
Bud4
Msn2
Sch9
Rim15
Yak1
Ypl247c
Vps52
Kss1
Sko1
Slt2
Msg5
Swe1
Igo2
Cdc55
Igo1
Vps53
Vps54
Rtn1
Fmp42
Cot1
Zrc1
10
20
30
40
50
t/s

### Path 911 (Sec18 − Rad23)

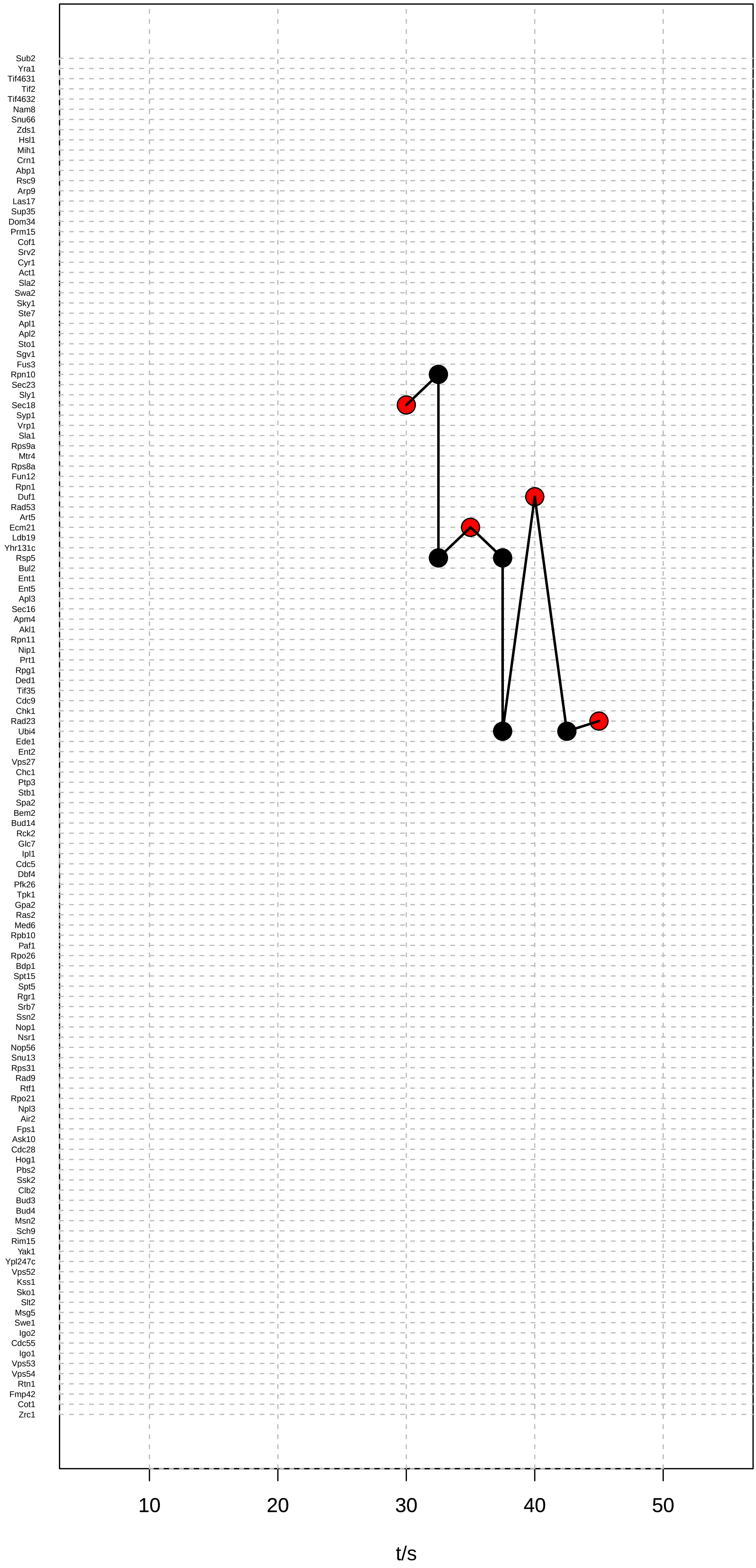

### Path 912 (Sec18 − Rad23)

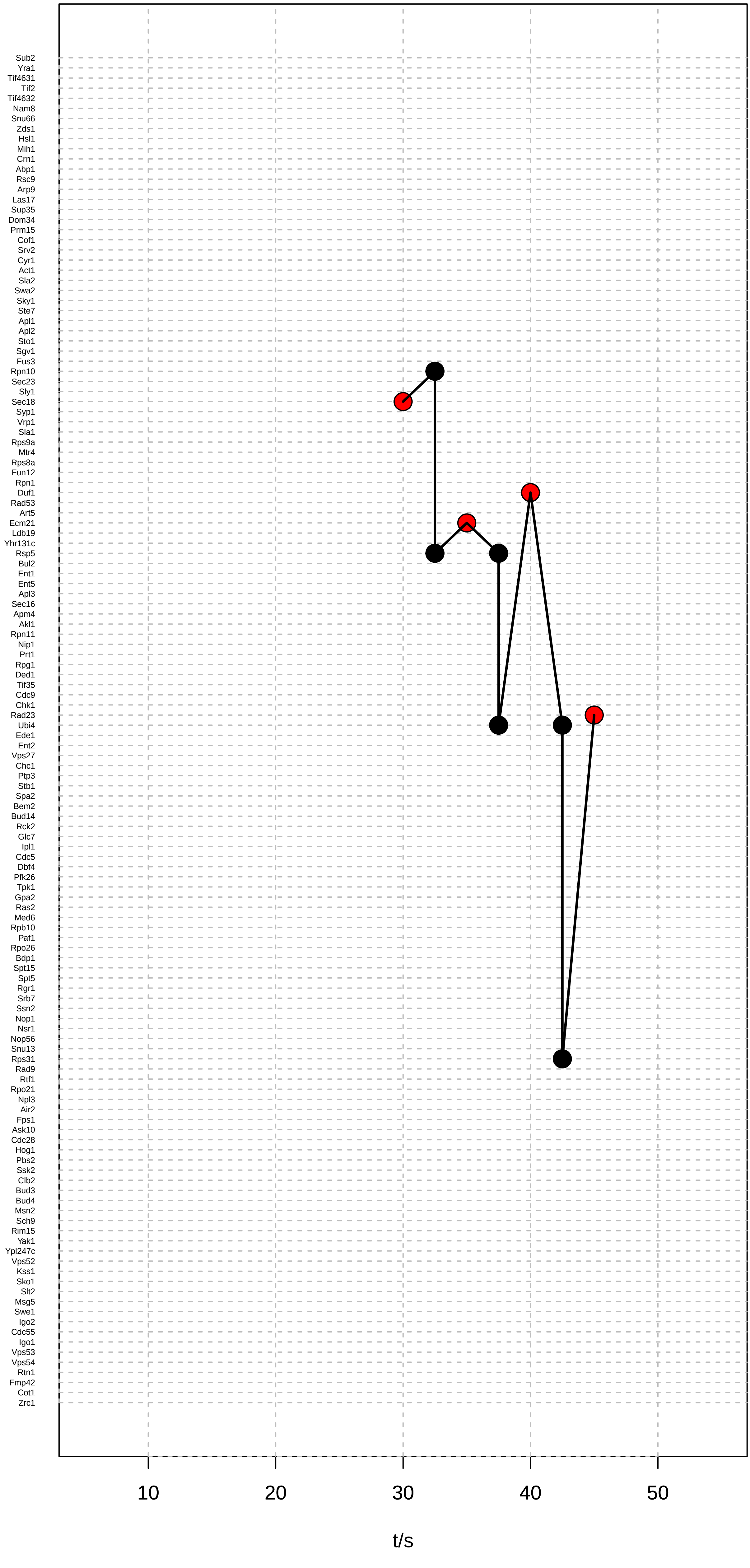

#### Path 913 (Sec18 − Rad23)

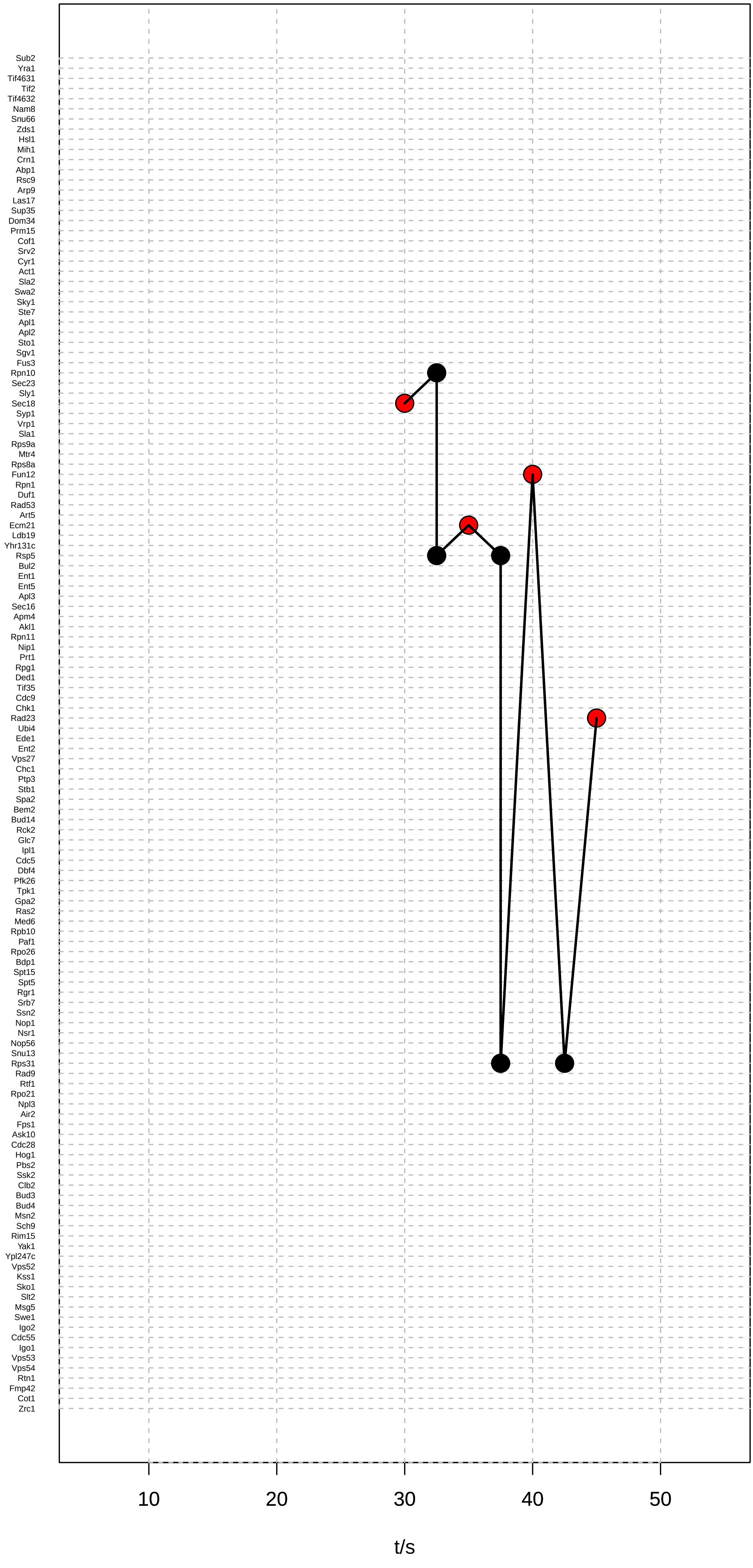

#### Path 914 (Sec18 − Rad23)

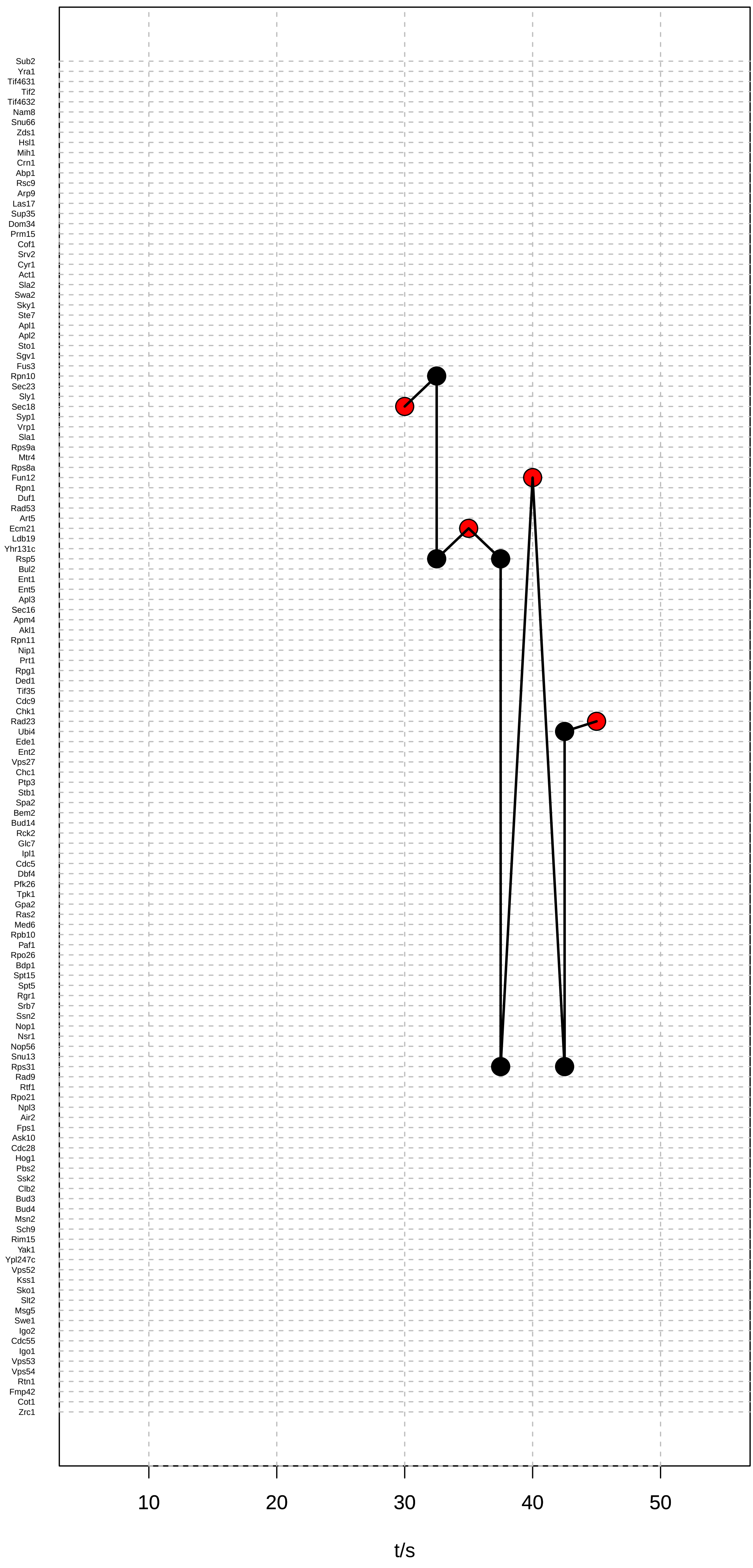

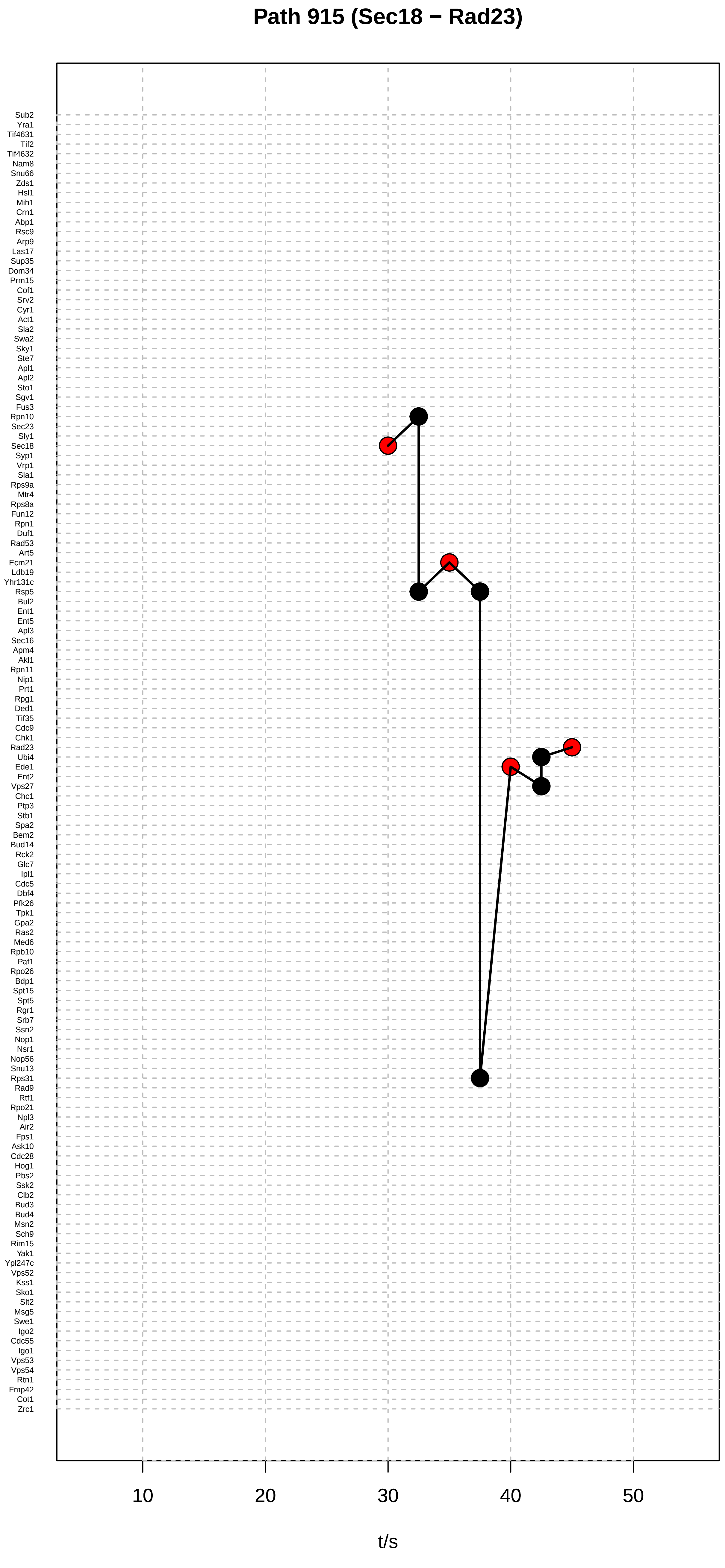
Path 915 (Sec18 − Rad23)
Sub2
Yra1
Tif4631
Tif2
Tif4632
Nam8
Snu66
Zds1
Hsl1
Mih1
Crn1
Abp1
Rsc9
Arp9
Las17
Sup35
Dom34
Prm15
Cof1
Srv2
Cyr1
Act1
Sla2
Swa2
Sky1
Ste7
Apl1
Apl2
Sto1
Sgv1
Fus3
Rpn10
Sec23
Sly1
Sec18
Syp1
Vrp1
Sla1
Rps9a
Mtr4
Rps8a
Fun12
Rpn1
Duf1
Rad53
Art5
Ecm21
Ldb19
Yhr131c
Rsp5
Bul2
Ent1
Ent5
Apl3
Sec16
Apm4
Akl1
Rpn11
Nip1
Prt1
Rpg1
Ded1
Tif35
Cdc9
Chk1
Rad23
Ubi4
Ede1
Ent2
Vps27
Chc1
Ptp3
Stb1
Spa2
Bem2
Bud14
Rck2
Glc7
Ipl1
Cdc5
Dbf4
Pfk26
Tpk1
Gpa2
Ras2
Med6
Rpb10
Paf1
Rpo26
Bdp1
Spt15
Spt5
Rgr1
Srb7
Ssn2
Nop1
Nsr1
Nop56
Snu13
Rps31
Rad9
Rtf1
Rpo21
Npl3
Air2
Fps1
Ask10
Cdc28
Hog1
Pbs2
Ssk2
Clb2
Bud3
Bud4
Msn2
Sch9
Rim15
Yak1
Ypl247c
Vps52
Kss1
Sko1
Slt2
Msg5
Swe1
Igo2
Cdc55
Igo1
Vps53
Vps54
Rtn1
Fmp42
Cot1
Zrc1
10
20
30
40
50
t/s

#### Path 916 (Sec18 − Rad23)

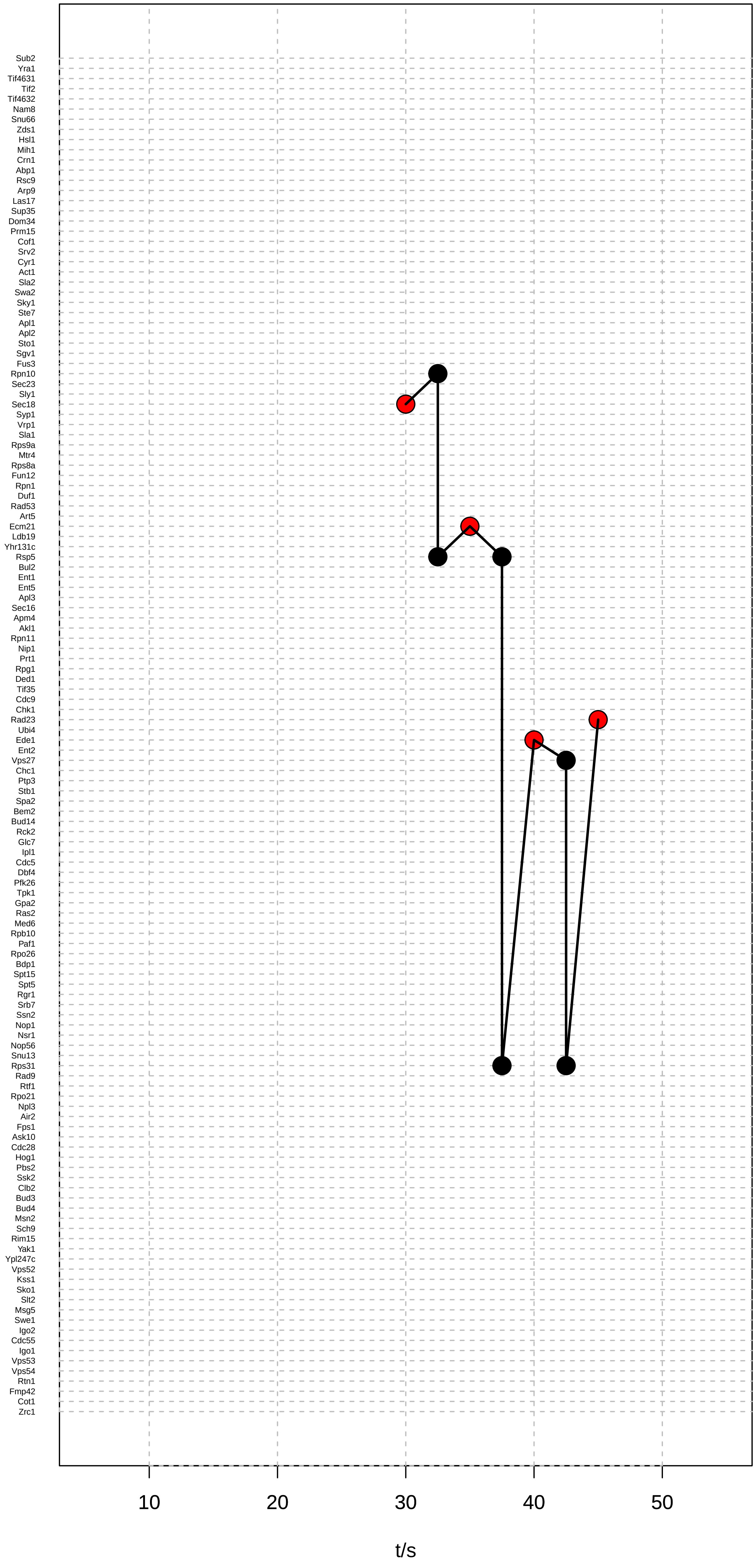

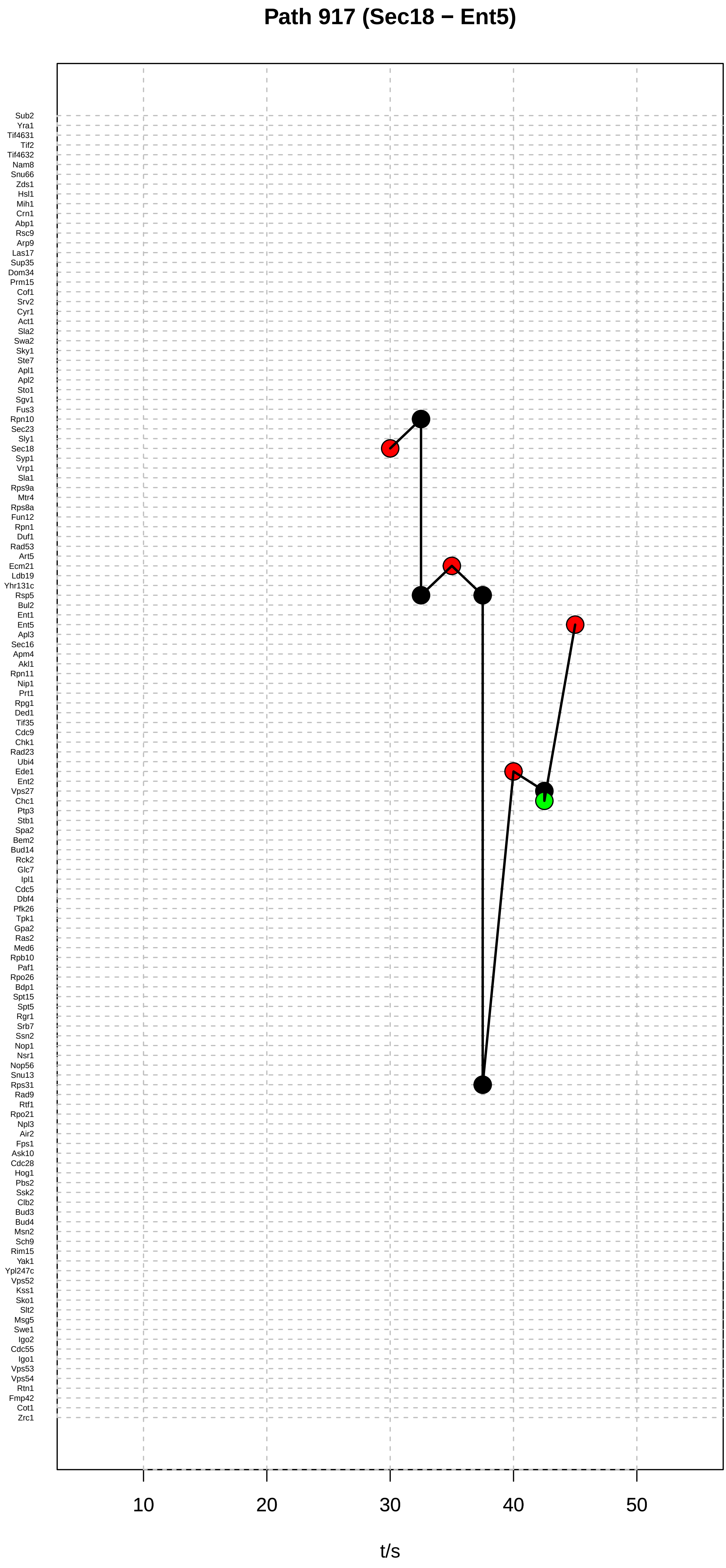

Path 917 (Sec18 − Ent5)
Sub2
Yra1
Tif4631
Tif2
Tif4632
Nam8
Snu66
Zds1
Hsl1
Mih1
Crn1
Abp1
Rsc9
Arp9
Las17
Sup35
Dom34
Prm15
Cof1
Srv2
Cyr1
Act1
Sla2
Swa2
Sky1
Ste7
Apl1
Apl2
Sto1
Sgv1
Fus3
Rpn10
Sec23
Sly1
Sec18
Syp1
Vrp1
Sla1
Rps9a
Mtr4
Rps8a
Fun12
Rpn1
Duf1
Rad53
Art5
Ecm21
Ldb19
Yhr131c
Rsp5
Bul2
Ent1
Ent5
Apl3
Sec16
Apm4
Akl1
Rpn11
Nip1
Prt1
Rpg1
Ded1
Tif35
Cdc9
Chk1
Rad23
Ubi4
Ede1
Ent2
Vps27
Chc1
Ptp3
Stb1
Spa2
Bem2
Bud14
Rck2
Glc7
Ipl1
Cdc5
Dbf4
Pfk26
Tpk1
Gpa2
Ras2
Med6
Rpb10
Paf1
Rpo26
Bdp1
Spt15
Spt5
Rgr1
Srb7
Ssn2
Nop1
Nsr1
Nop56
Snu13
Rps31
Rad9
Rtf1
Rpo21
Npl3
Air2
Fps1
Ask10
Cdc28
Hog1
Pbs2
Ssk2
Clb2
Bud3
Bud4
Msn2
Sch9
Rim15
Yak1
Ypl247c
Vps52
Kss1
Sko1
Slt2
Msg5
Swe1
Igo2
Cdc55
Igo1
Vps53
Vps54
Rtn1
Fmp42
Cot1
Zrc1
10
20
30
40
50
t/s

### Path 918 (Sec18 − Rad23)

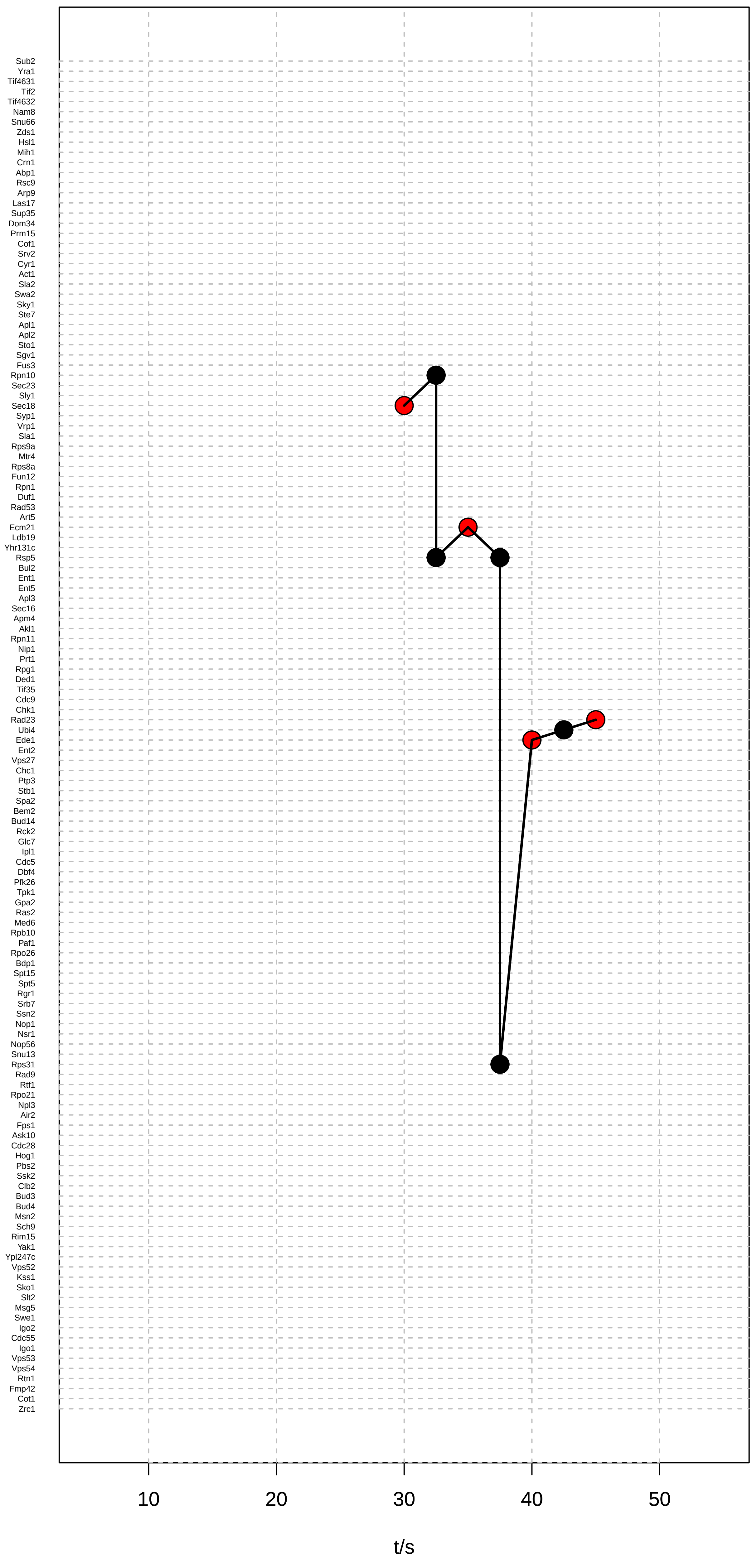

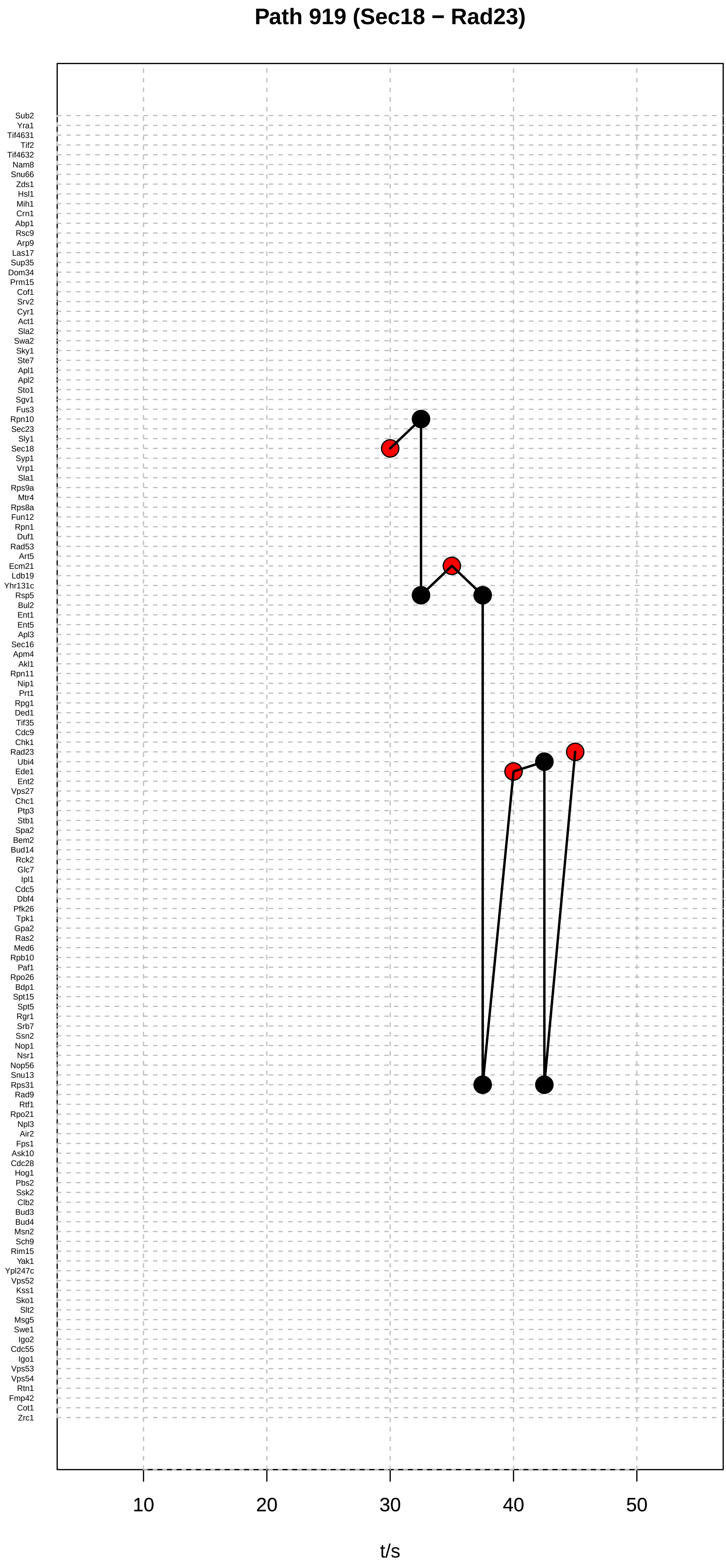

Path 919 (Sec18 − Rad23)
Sub2
Yra1
Tif4631
Tif2
Tif4632
Nam8
Snu66
Zds1
Hsl1
Mih1
Crn1
Abp1
Rsc9
Arp9
Las17
Sup35
Dom34
Prm15
Cof1
Srv2
Cyr1
Act1
Sla2
Swa2
Sky1
Ste7
Apl1
Apl2
Sto1
Sgv1
Fus3
Rpn10
Sec23
Sly1
Sec18
Syp1
Vrp1
Sla1
Rps9a
Mtr4
Rps8a
Fun12
Rpn1
Duf1
Rad53
Art5
Ecm21
Ldb19
Yhr131c
Rsp5
Bul2
Ent1
Ent5
Apl3
Sec16
Apm4
Akl1
Rpn11
Nip1
Prt1
Rpg1
Ded1
Tif35
Cdc9
Chk1
Rad23
Ubi4
Ede1
Ent2
Vps27
Chc1
Ptp3
Stb1
Spa2
Bem2
Bud14
Rck2
Glc7
Ipl1
Cdc5
Dbf4
Pfk26
Tpk1
Gpa2
Ras2
Med6
Rpb10
Paf1
Rpo26
Bdp1
Spt15
Spt5
Rgr1
Srb7
Ssn2
Nop1
Nsr1
Nop56
Snu13
Rps31
Rad9
Rtf1
Rpo21
Npl3
Air2
Fps1
Ask10
Cdc28
Hog1
Pbs2
Ssk2
Clb2
Bud3
Bud4
Msn2
Sch9
Rim15
Yak1
Ypl247c
Vps52
Kss1
Sko1
Slt2
Msg5
Swe1
Igo2
Cdc55
Igo1
Vps53
Vps54
Rtn1
Fmp42
Cot1
Zrc1
10
20
30
40
50
t/s

#### Path 920 (Sec18 − Rad23)

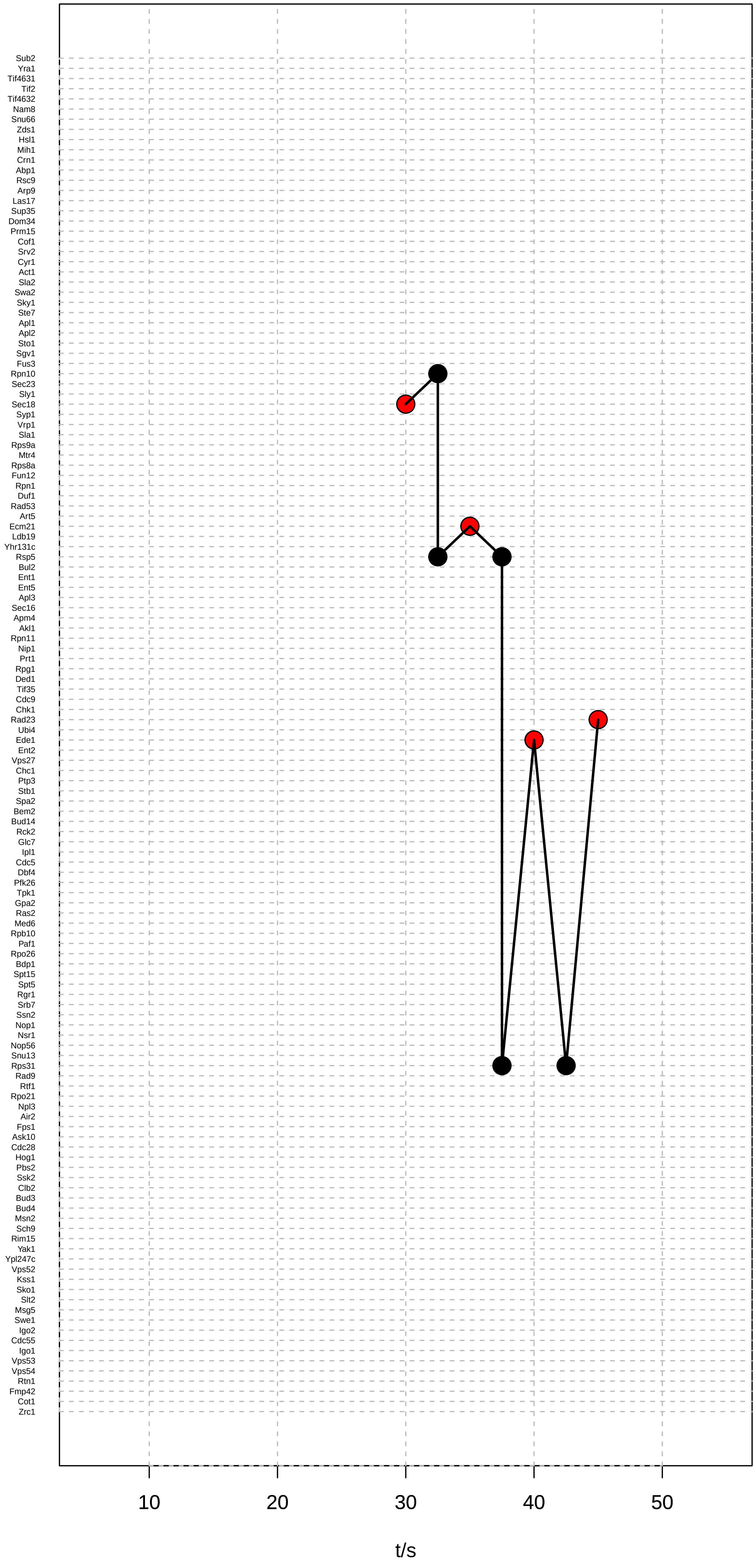

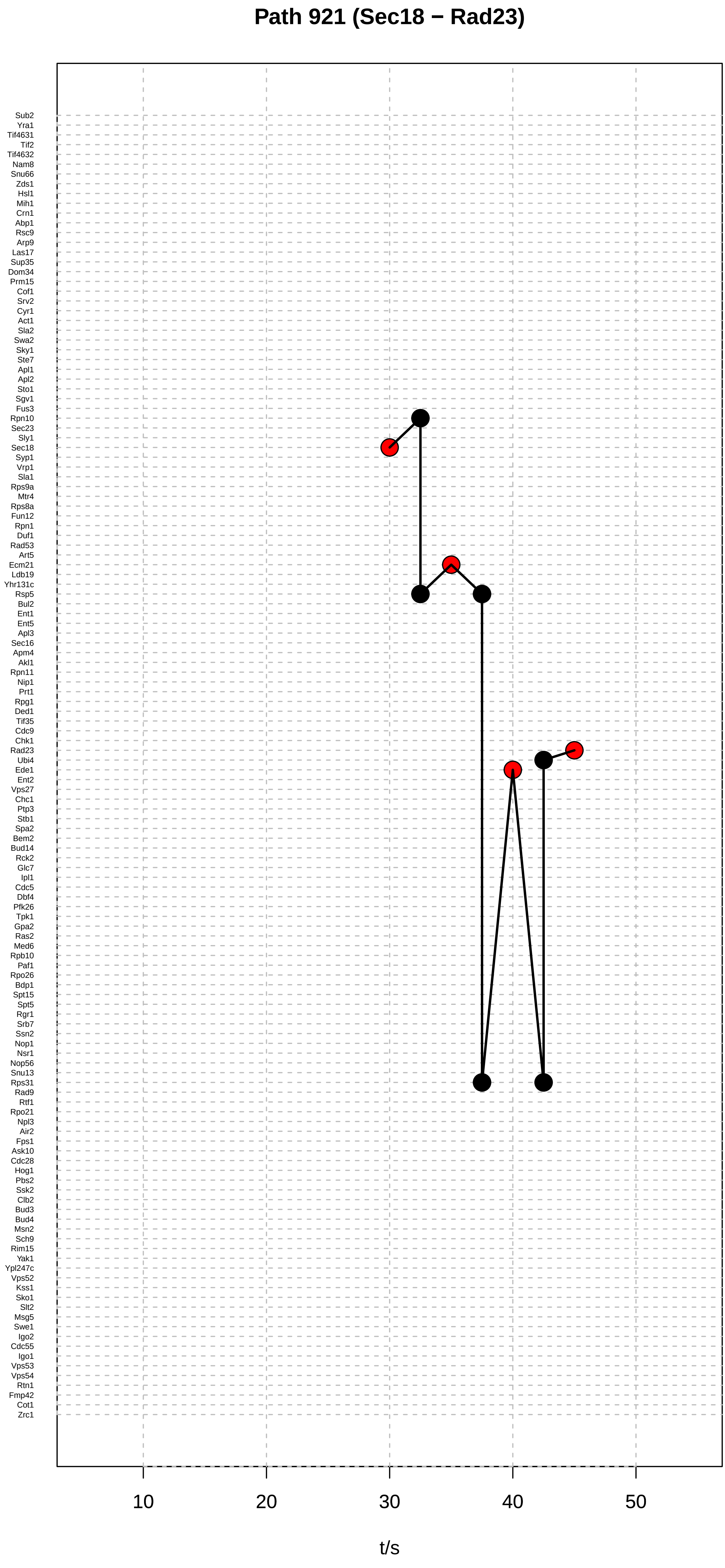
Path 921 (Sec18 − Rad23)
Sub2
Yra1
Tif4631
Tif2
Tif4632
Nam8
Snu66
Zds1
Hsl1
Mih1
Crn1
Abp1
Rsc9
Arp9
Las17
Sup35
Dom34
Prm15
Cof1
Srv2
Cyr1
Act1
Sla2
Swa2
Sky1
Ste7
Apl1
Apl2
Sto1
Sgv1
Fus3
Rpn10
Sec23
Sly1
Sec18
Syp1
Vrp1
Sla1
Rps9a
Mtr4
Rps8a
Fun12
Rpn1
Duf1
Rad53
Art5
Ecm21
Ldb19
Yhr131c
Rsp5
Bul2
Ent1
Ent5
Apl3
Sec16
Apm4
Akl1
Rpn11
Nip1
Prt1
Rpg1
Ded1
Tif35
Cdc9
Chk1
Rad23
Ubi4
Ede1
Ent2
Vps27
Chc1
Ptp3
Stb1
Spa2
Bem2
Bud14
Rck2
Glc7
Ipl1
Cdc5
Dbf4
Pfk26
Tpk1
Gpa2
Ras2
Med6
Rpb10
Paf1
Rpo26
Bdp1
Spt15
Spt5
Rgr1
Srb7
Ssn2
Nop1
Nsr1
Nop56
Snu13
Rps31
Rad9
Rtf1
Rpo21
Npl3
Air2
Fps1
Ask10
Cdc28
Hog1
Pbs2
Ssk2
Clb2
Bud3
Bud4
Msn2
Sch9
Rim15
Yak1
Ypl247c
Vps52
Kss1
Sko1
Slt2
Msg5
Swe1
Igo2
Cdc55
Igo1
Vps53
Vps54
Rtn1
Fmp42
Cot1
Zrc1
10
20
30
40
50
t/s

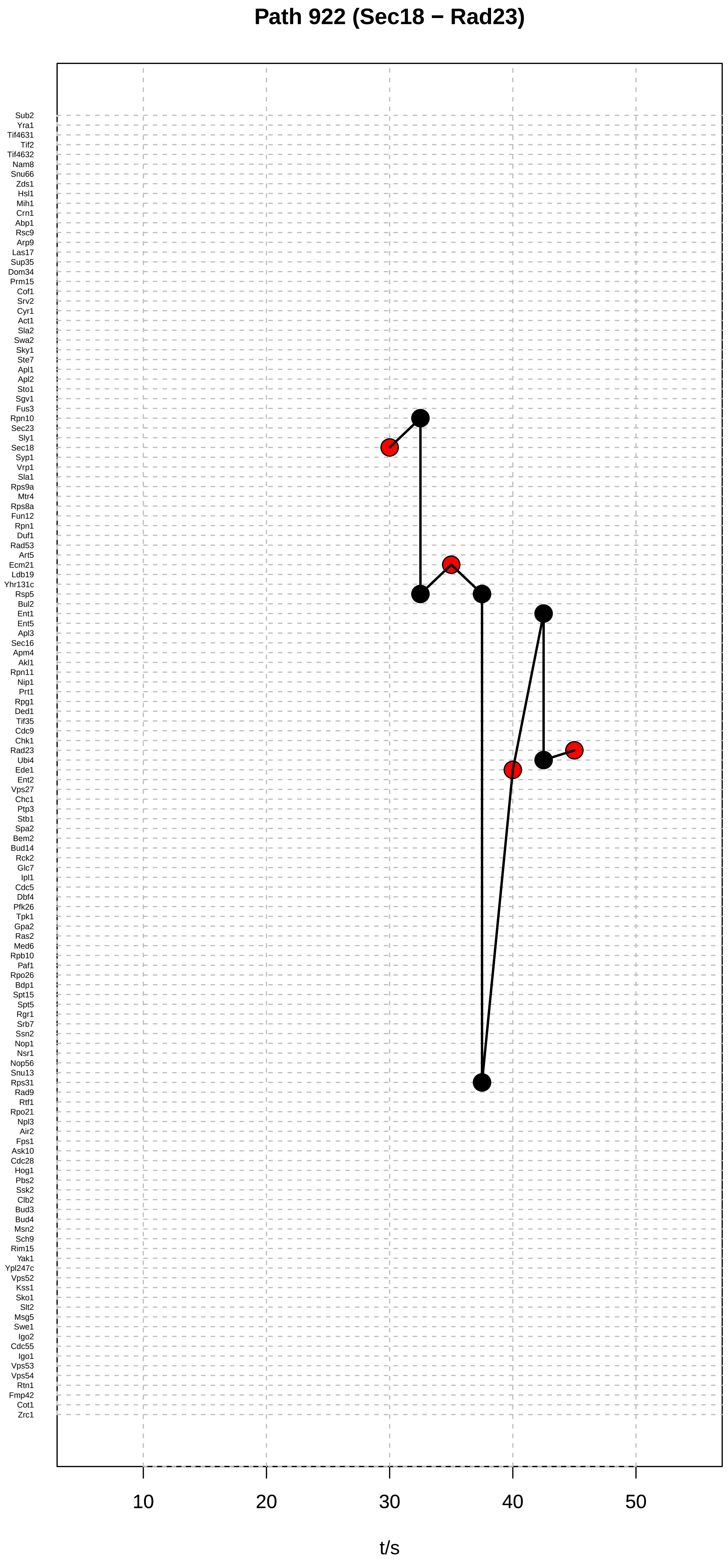
Path 922 (Sec18 − Rad23)
Sub2
Yra1
Tif4631
Tif2
Tif4632
Nam8
Snu66
Zds1
Hsl1
Mih1
Crn1
Abp1
Rsc9
Arp9
Las17
Sup35
Dom34
Prm15
Cof1
Srv2
Cyr1
Act1
Sla2
Swa2
Sky1
Ste7
Apl1
Apl2
Sto1
Sgv1
Fus3
Rpn10
Sec23
Sly1
Sec18
Syp1
Vrp1
Sla1
Rps9a
Mtr4
Rps8a
Fun12
Rpn1
Duf1
Rad53
Art5
Ecm21
Ldb19
Yhr131c
Rsp5
Bul2
Ent1
Ent5
Apl3
Sec16
Apm4
Akl1
Rpn11
Nip1
Prt1
Rpg1
Ded1
Tif35
Cdc9
Chk1
Rad23
Ubi4
Ede1
Ent2
Vps27
Chc1
Ptp3
Stb1
Spa2
Bem2
Bud14
Rck2
Glc7
Ipl1
Cdc5
Dbf4
Pfk26
Tpk1
Gpa2
Ras2
Med6
Rpb10
Paf1
Rpo26
Bdp1
Spt15
Spt5
Rgr1
Srb7
Ssn2
Nop1
Nsr1
Nop56
Snu13
Rps31
Rad9
Rtf1
Rpo21
Npl3
Air2
Fps1
Ask10
Cdc28
Hog1
Pbs2
Ssk2
Clb2
Bud3
Bud4
Msn2
Sch9
Rim15
Yak1
Ypl247c
Vps52
Kss1
Sko1
Slt2
Msg5
Swe1
Igo2
Cdc55
Igo1
Vps53
Vps54
Rtn1
Fmp42
Cot1
Zrc1
10
20
30
40
50
t/s

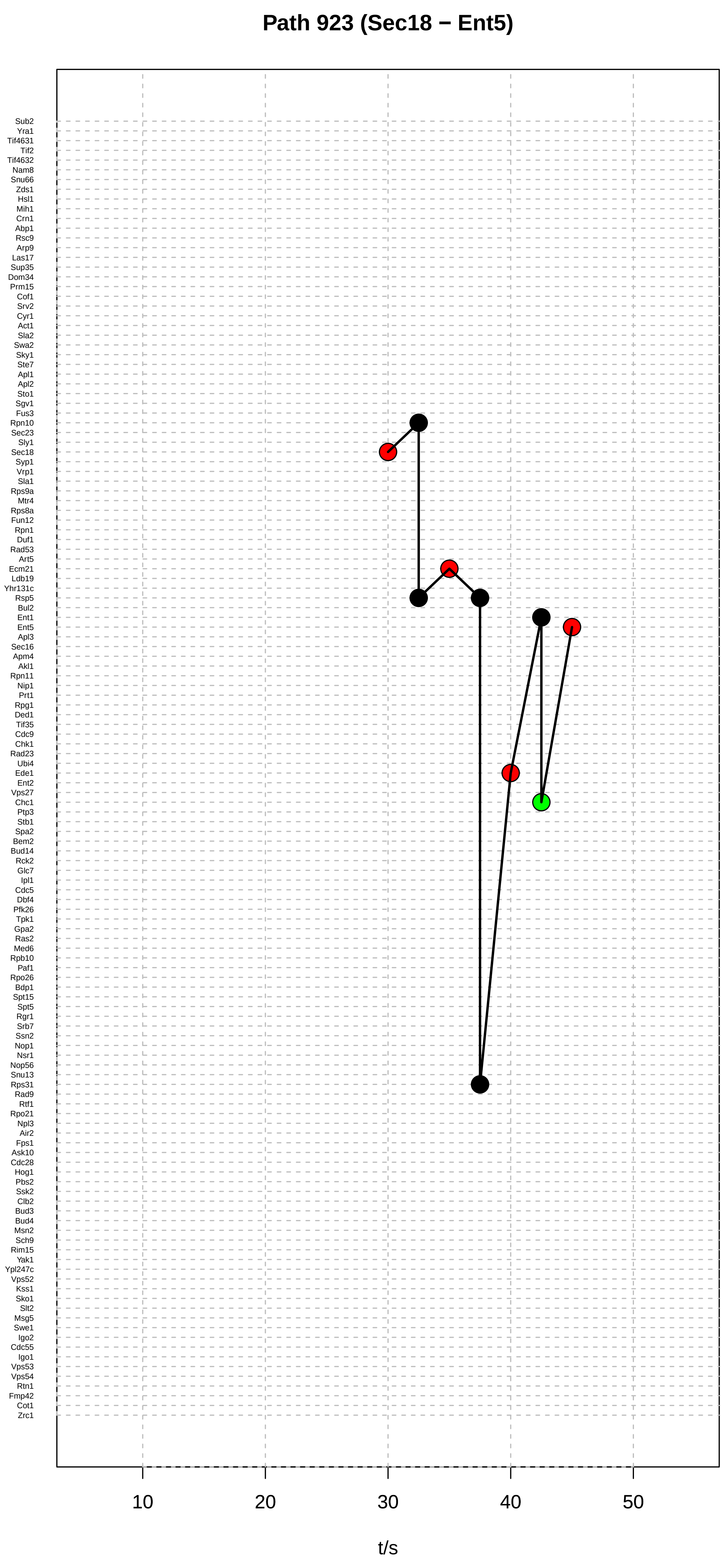
Path 923 (Sec18 − Ent5)
Sub2
Yra1
Tif4631
Tif2
Tif4632
Nam8
Snu66
Zds1
Hsl1
Mih1
Crn1
Abp1
Rsc9
Arp9
Las17
Sup35
Dom34
Prm15
Cof1
Srv2
Cyr1
Act1
Sla2
Swa2
Sky1
Ste7
Apl1
Apl2
Sto1
Sgv1
Fus3
Rpn10
Sec23
Sly1
Sec18
Syp1
Vrp1
Sla1
Rps9a
Mtr4
Rps8a
Fun12
Rpn1
Duf1
Rad53
Art5
Ecm21
Ldb19
Yhr131c
Rsp5
Bul2
Ent1
Ent5
Apl3
Sec16
Apm4
Akl1
Rpn11
Nip1
Prt1
Rpg1
Ded1
Tif35
Cdc9
Chk1
Rad23
Ubi4
Ede1
Ent2
Vps27
Chc1
Ptp3
Stb1
Spa2
Bem2
Bud14
Rck2
Glc7
Ipl1
Cdc5
Dbf4
Pfk26
Tpk1
Gpa2
Ras2
Med6
Rpb10
Paf1
Rpo26
Bdp1
Spt15
Spt5
Rgr1
Srb7
Ssn2
Nop1
Nsr1
Nop56
Snu13
Rps31
Rad9
Rtf1
Rpo21
Npl3
Air2
Fps1
Ask10
Cdc28
Hog1
Pbs2
Ssk2
Clb2
Bud3
Bud4
Msn2
Sch9
Rim15
Yak1
Ypl247c
Vps52
Kss1
Sko1
Slt2
Msg5
Swe1
Igo2
Cdc55
Igo1
Vps53
Vps54
Rtn1
Fmp42
Cot1
Zrc1
10
20
30
40
50
t/s

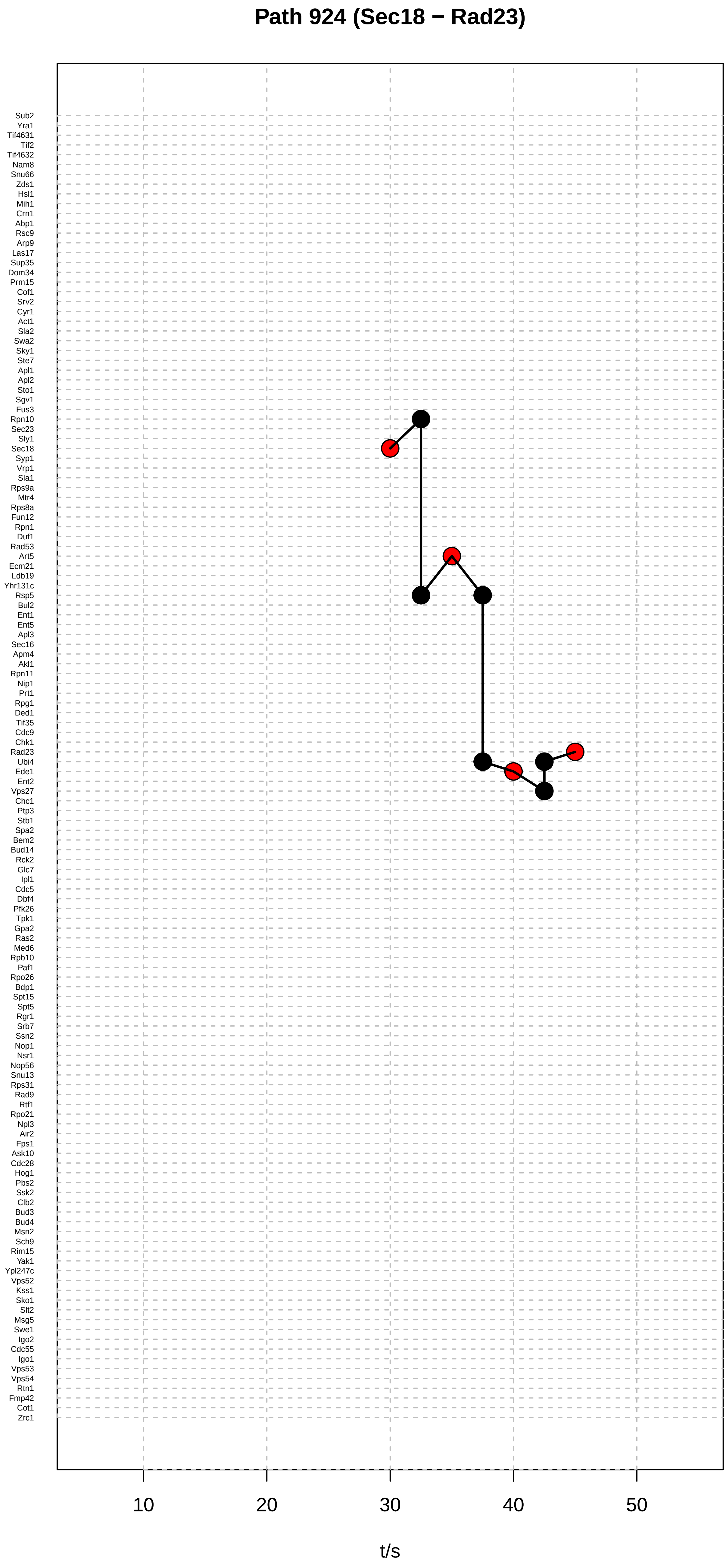
Path 924 (Sec18 − Rad23)
Sub2
Yra1
Tif4631
Tif2
Tif4632
Nam8
Snu66
Zds1
Hsl1
Mih1
Crn1
Abp1
Rsc9
Arp9
Las17
Sup35
Dom34
Prm15
Cof1
Srv2
Cyr1
Act1
Sla2
Swa2
Sky1
Ste7
Apl1
Apl2
Sto1
Sgv1
Fus3
Rpn10
Sec23
Sly1
Sec18
Syp1
Vrp1
Sla1
Rps9a
Mtr4
Rps8a
Fun12
Rpn1
Duf1
Rad53
Art5
Ecm21
Ldb19
Yhr131c
Rsp5
Bul2
Ent1
Ent5
Apl3
Sec16
Apm4
Akl1
Rpn11
Nip1
Prt1
Rpg1
Ded1
Tif35
Cdc9
Chk1
Rad23
Ubi4
Ede1
Ent2
Vps27
Chc1
Ptp3
Stb1
Spa2
Bem2
Bud14
Rck2
Glc7
Ipl1
Cdc5
Dbf4
Pfk26
Tpk1
Gpa2
Ras2
Med6
Rpb10
Paf1
Rpo26
Bdp1
Spt15
Spt5
Rgr1
Srb7
Ssn2
Nop1
Nsr1
Nop56
Snu13
Rps31
Rad9
Rtf1
Rpo21
Npl3
Air2
Fps1
Ask10
Cdc28
Hog1
Pbs2
Ssk2
Clb2
Bud3
Bud4
Msn2
Sch9
Rim15
Yak1
Ypl247c
Vps52
Kss1
Sko1
Slt2
Msg5
Swe1
Igo2
Cdc55
Igo1
Vps53
Vps54
Rtn1
Fmp42
Cot1
Zrc1
10
20
30
40
50
t/s

### Path 925 (Sec18 − Rad23)

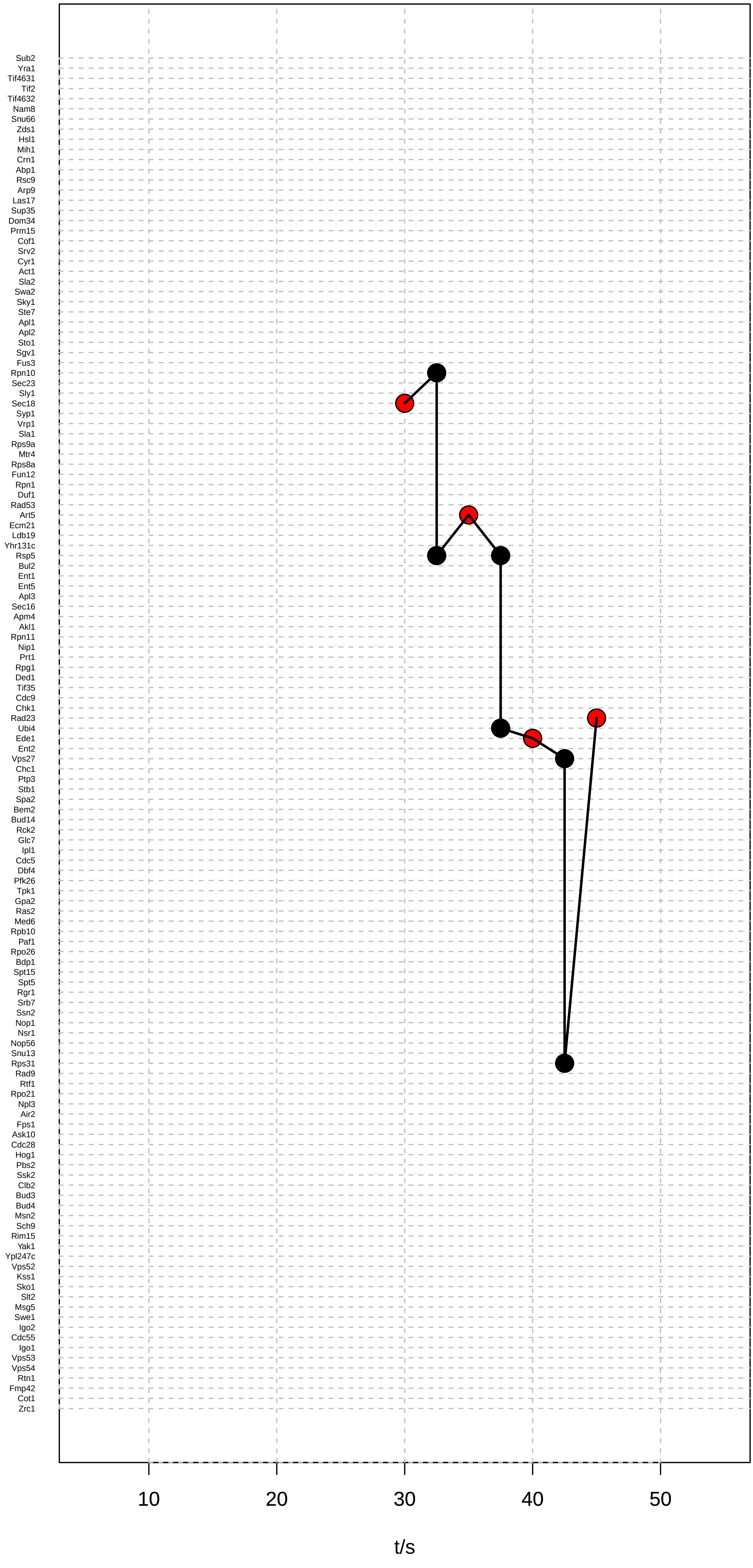

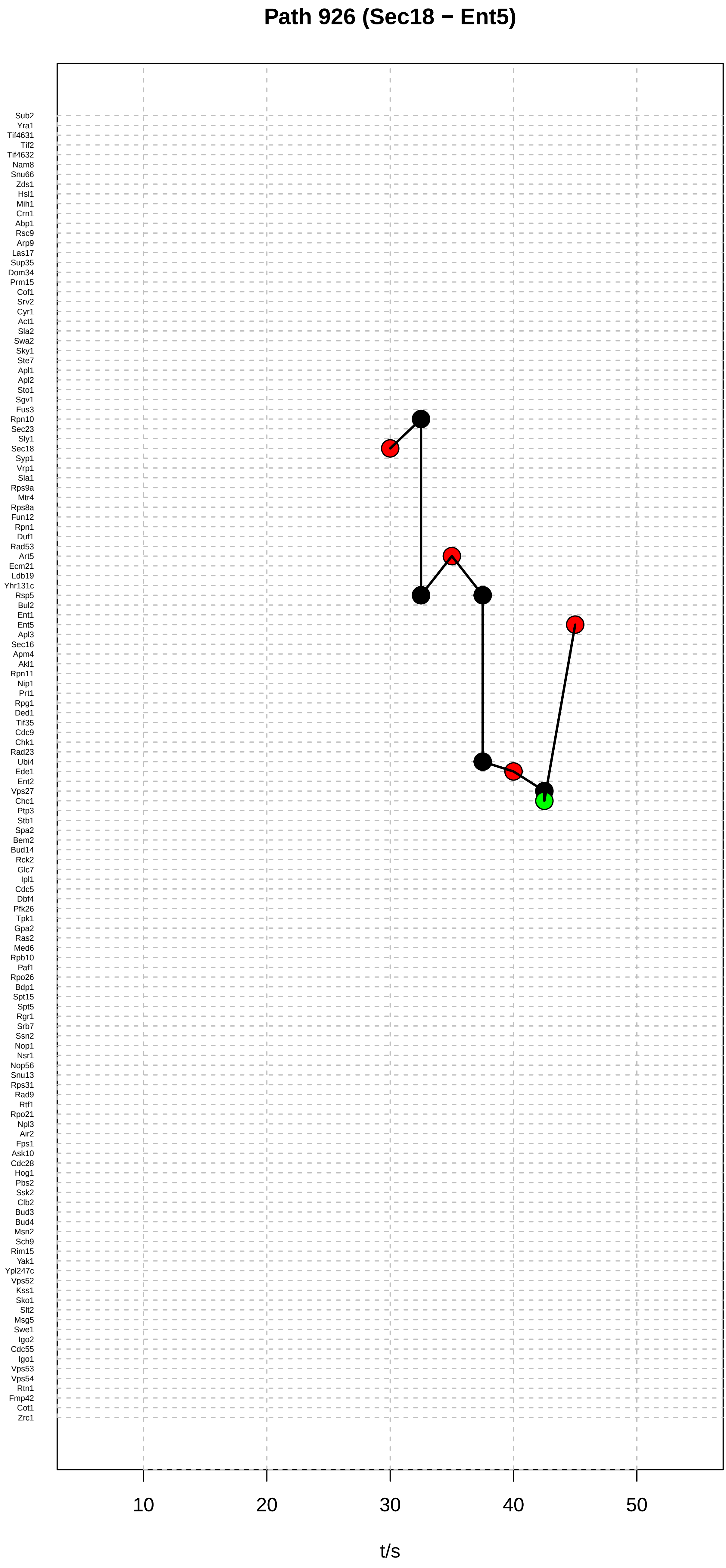

Path 926 (Sec18 − Ent5)
Sub2
Yra1
Tif4631
Tif2
Tif4632
Nam8
Snu66
Zds1
Hsl1
Mih1
Crn1
Abp1
Rsc9
Arp9
Las17
Sup35
Dom34
Prm15
Cof1
Srv2
Cyr1
Act1
Sla2
Swa2
Sky1
Ste7
Apl1
Apl2
Sto1
Sgv1
Fus3
Rpn10
Sec23
Sly1
Sec18
Syp1
Vrp1
Sla1
Rps9a
Mtr4
Rps8a
Fun12
Rpn1
Duf1
Rad53
Art5
Ecm21
Ldb19
Yhr131c
Rsp5
Bul2
Ent1
Ent5
Apl3
Sec16
Apm4
Akl1
Rpn11
Nip1
Prt1
Rpg1
Ded1
Tif35
Cdc9
Chk1
Rad23
Ubi4
Ede1
Ent2
Vps27
Chc1
Ptp3
Stb1
Spa2
Bem2
Bud14
Rck2
Glc7
Ipl1
Cdc5
Dbf4
Pfk26
Tpk1
Gpa2
Ras2
Med6
Rpb10
Paf1
Rpo26
Bdp1
Spt15
Spt5
Rgr1
Srb7
Ssn2
Nop1
Nsr1
Nop56
Snu13
Rps31
Rad9
Rtf1
Rpo21
Npl3
Air2
Fps1
Ask10
Cdc28
Hog1
Pbs2
Ssk2
Clb2
Bud3
Bud4
Msn2
Sch9
Rim15
Yak1
Ypl247c
Vps52
Kss1
Sko1
Slt2
Msg5
Swe1
Igo2
Cdc55
Igo1
Vps53
Vps54
Rtn1
Fmp42
Cot1
Zrc1
10
20
30
40
50
t/s

### Path 927 (Sec18 − Rad23)

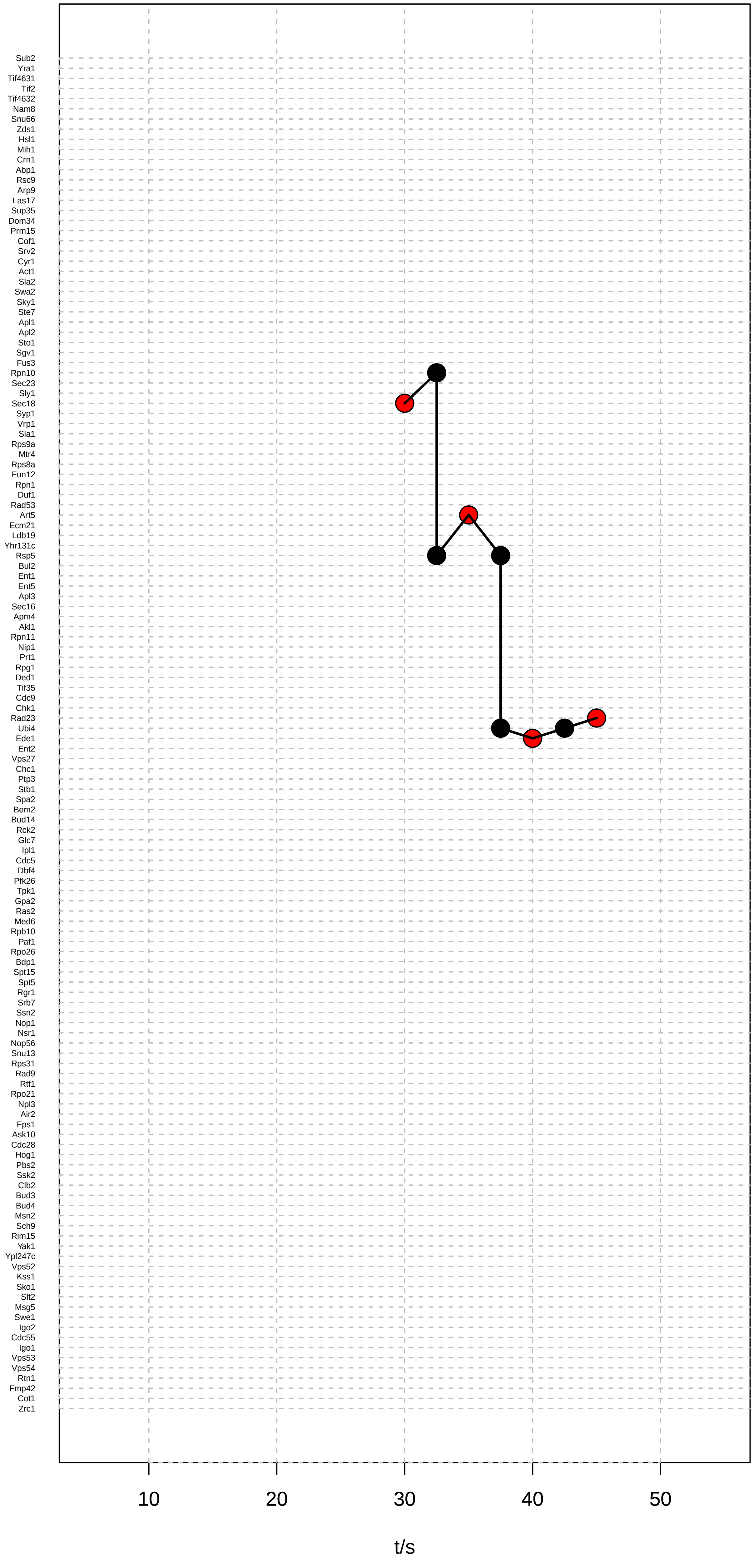

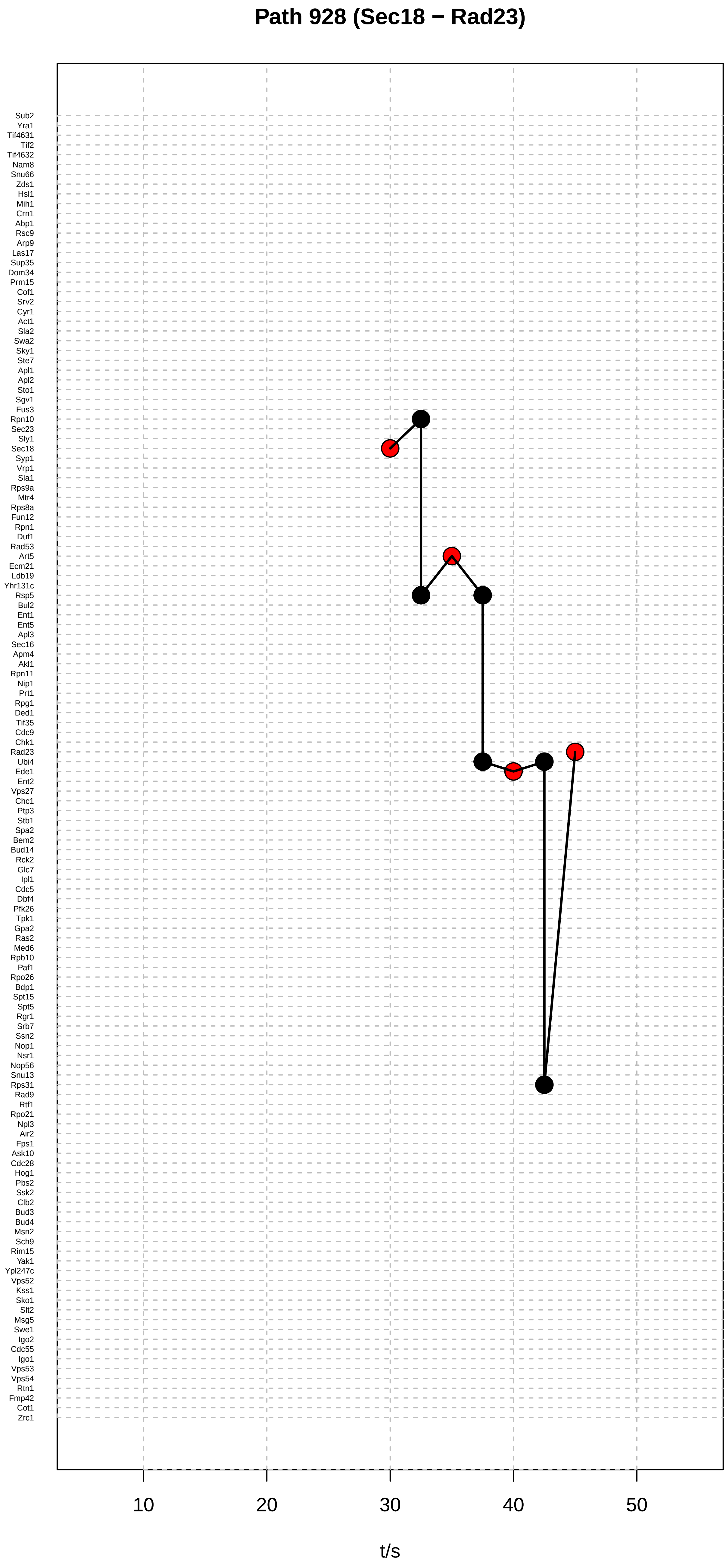
Path 928 (Sec18 − Rad23)
Sub2
Yra1
Tif4631
Tif2
Tif4632
Nam8
Snu66
Zds1
Hsl1
Mih1
Crn1
Abp1
Rsc9
Arp9
Las17
Sup35
Dom34
Prm15
Cof1
Srv2
Cyr1
Act1
Sla2
Swa2
Sky1
Ste7
Apl1
Apl2
Sto1
Sgv1
Fus3
Rpn10
Sec23
Sly1
Sec18
Syp1
Vrp1
Sla1
Rps9a
Mtr4
Rps8a
Fun12
Rpn1
Duf1
Rad53
Art5
Ecm21
Ldb19
Yhr131c
Rsp5
Bul2
Ent1
Ent5
Apl3
Sec16
Apm4
Akl1
Rpn11
Nip1
Prt1
Rpg1
Ded1
Tif35
Cdc9
Chk1
Rad23
Ubi4
Ede1
Ent2
Vps27
Chc1
Ptp3
Stb1
Spa2
Bem2
Bud14
Rck2
Glc7
Ipl1
Cdc5
Dbf4
Pfk26
Tpk1
Gpa2
Ras2
Med6
Rpb10
Paf1
Rpo26
Bdp1
Spt15
Spt5
Rgr1
Srb7
Ssn2
Nop1
Nsr1
Nop56
Snu13
Rps31
Rad9
Rtf1
Rpo21
Npl3
Air2
Fps1
Ask10
Cdc28
Hog1
Pbs2
Ssk2
Clb2
Bud3
Bud4
Msn2
Sch9
Rim15
Yak1
Ypl247c
Vps52
Kss1
Sko1
Slt2
Msg5
Swe1
Igo2
Cdc55
Igo1
Vps53
Vps54
Rtn1
Fmp42
Cot1
Zrc1
10
20
30
40
50
t/s

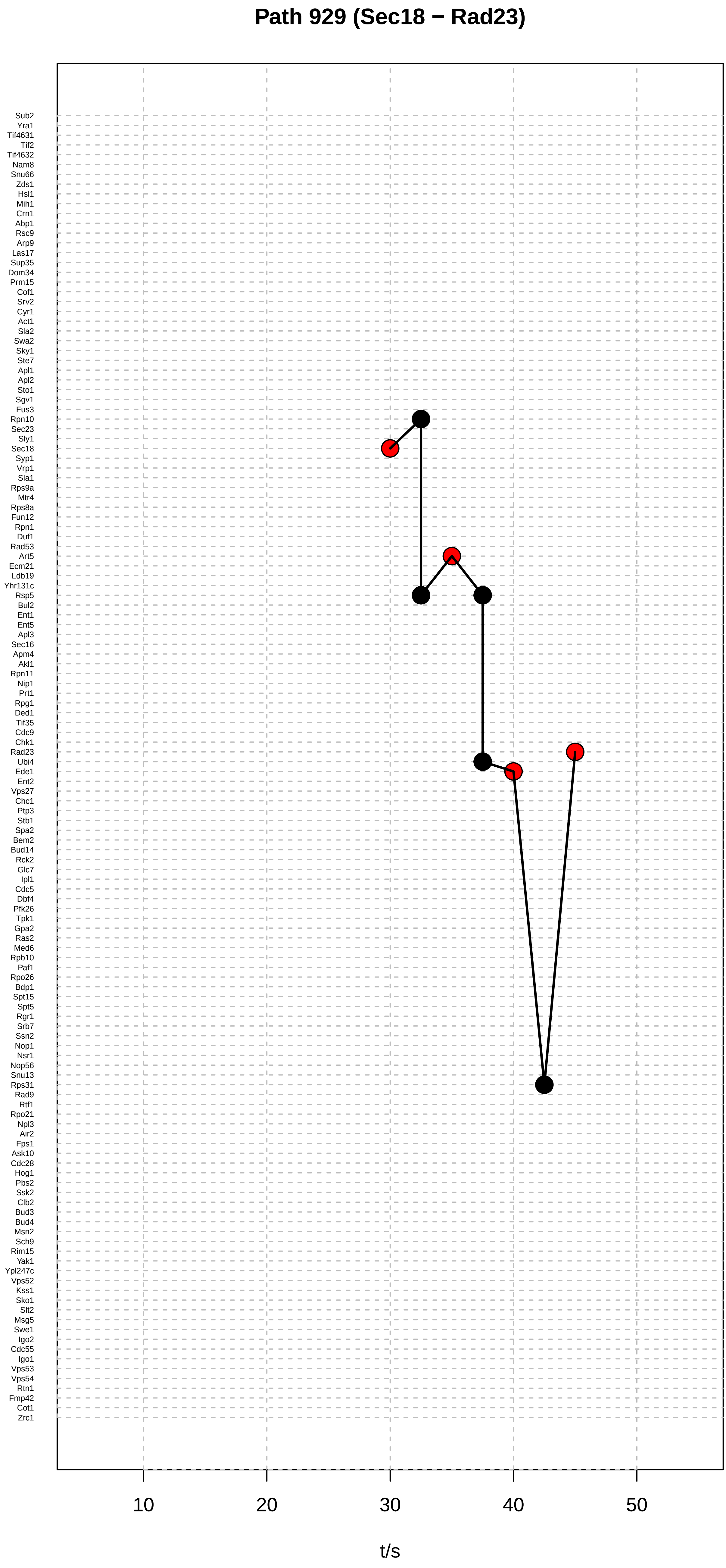

Path 929 (Sec18 − Rad23)
Sub2
Yra1
Tif4631
Tif2
Tif4632
Nam8
Snu66
Zds1
Hsl1
Mih1
Crn1
Abp1
Rsc9
Arp9
Las17
Sup35
Dom34
Prm15
Cof1
Srv2
Cyr1
Act1
Sla2
Swa2
Sky1
Ste7
Apl1
Apl2
Sto1
Sgv1
Fus3
Rpn10
Sec23
Sly1
Sec18
Syp1
Vrp1
Sla1
Rps9a
Mtr4
Rps8a
Fun12
Rpn1
Duf1
Rad53
Art5
Ecm21
Ldb19
Yhr131c
Rsp5
Bul2
Ent1
Ent5
Apl3
Sec16
Apm4
Akl1
Rpn11
Nip1
Prt1
Rpg1
Ded1
Tif35
Cdc9
Chk1
Rad23
Ubi4
Ede1
Ent2
Vps27
Chc1
Ptp3
Stb1
Spa2
Bem2
Bud14
Rck2
Glc7
Ipl1
Cdc5
Dbf4
Pfk26
Tpk1
Gpa2
Ras2
Med6
Rpb10
Paf1
Rpo26
Bdp1
Spt15
Spt5
Rgr1
Srb7
Ssn2
Nop1
Nsr1
Nop56
Snu13
Rps31
Rad9
Rtf1
Rpo21
Npl3
Air2
Fps1
Ask10
Cdc28
Hog1
Pbs2
Ssk2
Clb2
Bud3
Bud4
Msn2
Sch9
Rim15
Yak1
Ypl247c
Vps52
Kss1
Sko1
Slt2
Msg5
Swe1
Igo2
Cdc55
Igo1
Vps53
Vps54
Rtn1
Fmp42
Cot1
Zrc1
10
20
30
40
50
t/s

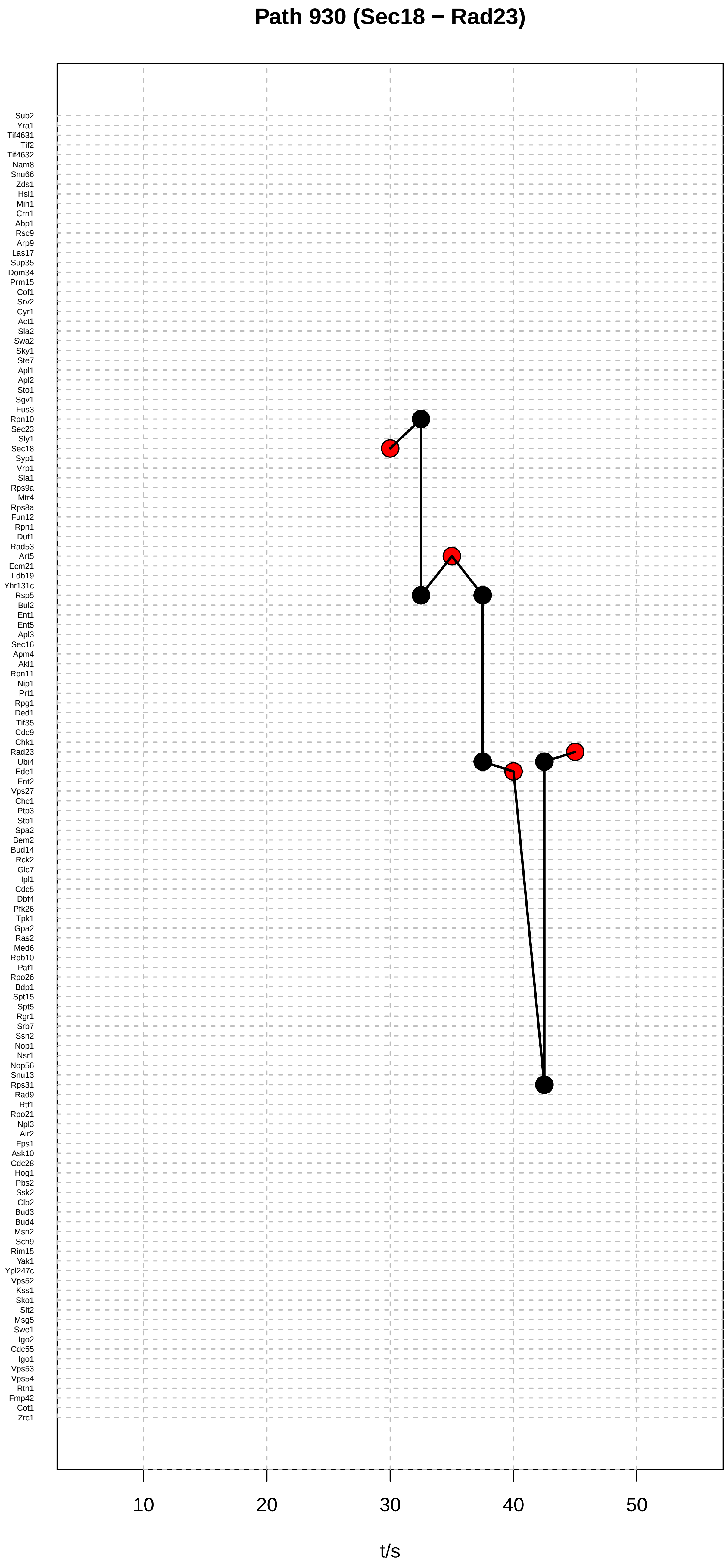
Path 930 (Sec18 − Rad23)
Sub2
Yra1
Tif4631
Tif2
Tif4632
Nam8
Snu66
Zds1
Hsl1
Mih1
Crn1
Abp1
Rsc9
Arp9
Las17
Sup35
Dom34
Prm15
Cof1
Srv2
Cyr1
Act1
Sla2
Swa2
Sky1
Ste7
Apl1
Apl2
Sto1
Sgv1
Fus3
Rpn10
Sec23
Sly1
Sec18
Syp1
Vrp1
Sla1
Rps9a
Mtr4
Rps8a
Fun12
Rpn1
Duf1
Rad53
Art5
Ecm21
Ldb19
Yhr131c
Rsp5
Bul2
Ent1
Ent5
Apl3
Sec16
Apm4
Akl1
Rpn11
Nip1
Prt1
Rpg1
Ded1
Tif35
Cdc9
Chk1
Rad23
Ubi4
Ede1
Ent2
Vps27
Chc1
Ptp3
Stb1
Spa2
Bem2
Bud14
Rck2
Glc7
Ipl1
Cdc5
Dbf4
Pfk26
Tpk1
Gpa2
Ras2
Med6
Rpb10
Paf1
Rpo26
Bdp1
Spt15
Spt5
Rgr1
Srb7
Ssn2
Nop1
Nsr1
Nop56
Snu13
Rps31
Rad9
Rtf1
Rpo21
Npl3
Air2
Fps1
Ask10
Cdc28
Hog1
Pbs2
Ssk2
Clb2
Bud3
Bud4
Msn2
Sch9
Rim15
Yak1
Ypl247c
Vps52
Kss1
Sko1
Slt2
Msg5
Swe1
Igo2
Cdc55
Igo1
Vps53
Vps54
Rtn1
Fmp42
Cot1
Zrc1
10
20
30
40
50
t/s

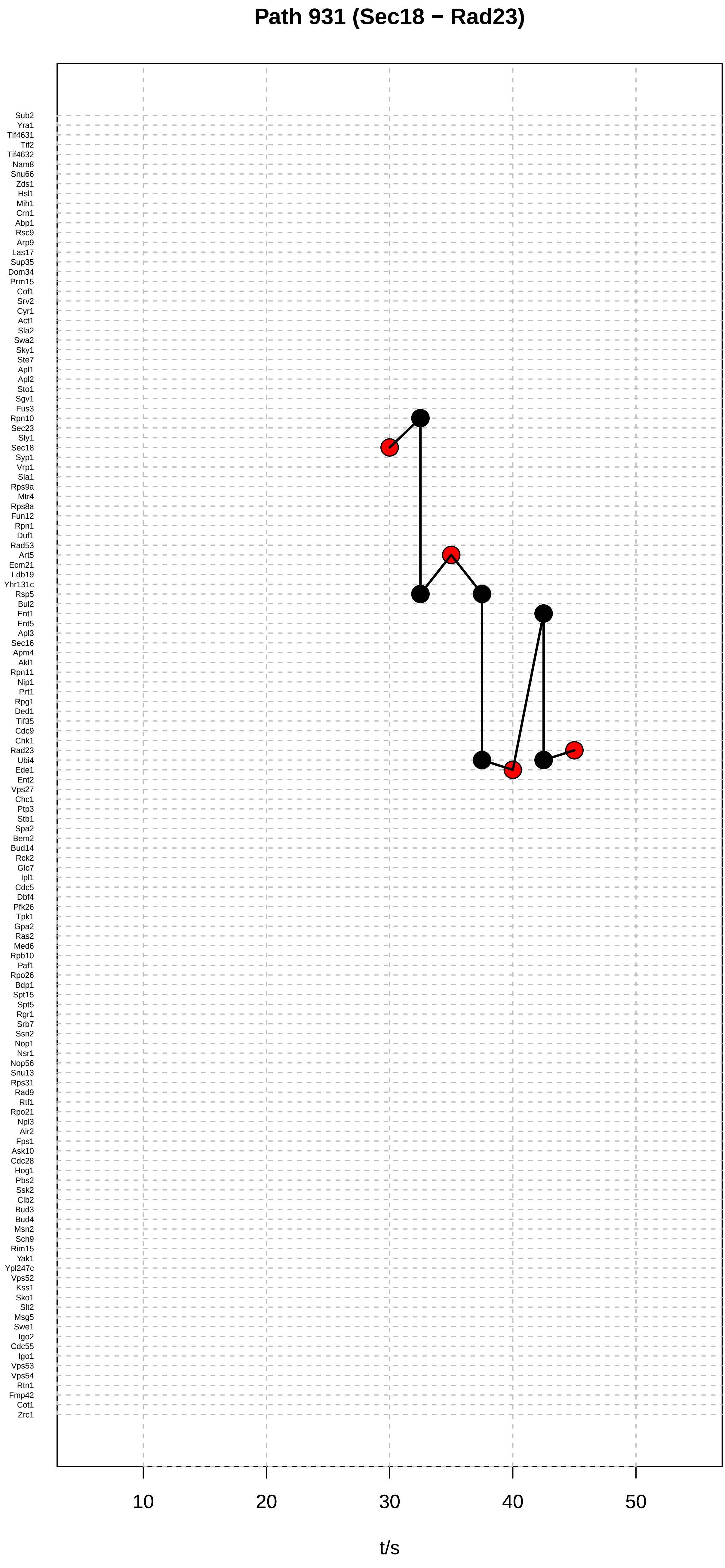

Path 931 (Sec18 − Rad23)
Sub2
Yra1
Tif4631
Tif2
Tif4632
Nam8
Snu66
Zds1
Hsl1
Mih1
Crn1
Abp1
Rsc9
Arp9
Las17
Sup35
Dom34
Prm15
Cof1
Srv2
Cyr1
Act1
Sla2
Swa2
Sky1
Ste7
Apl1
Apl2
Sto1
Sgv1
Fus3
Rpn10
Sec23
Sly1
Sec18
Syp1
Vrp1
Sla1
Rps9a
Mtr4
Rps8a
Fun12
Rpn1
Duf1
Rad53
Art5
Ecm21
Ldb19
Yhr131c
Rsp5
Bul2
Ent1
Ent5
Apl3
Sec16
Apm4
Akl1
Rpn11
Nip1
Prt1
Rpg1
Ded1
Tif35
Cdc9
Chk1
Rad23
Ubi4
Ede1
Ent2
Vps27
Chc1
Ptp3
Stb1
Spa2
Bem2
Bud14
Rck2
Glc7
Ipl1
Cdc5
Dbf4
Pfk26
Tpk1
Gpa2
Ras2
Med6
Rpb10
Paf1
Rpo26
Bdp1
Spt15
Spt5
Rgr1
Srb7
Ssn2
Nop1
Nsr1
Nop56
Snu13
Rps31
Rad9
Rtf1
Rpo21
Npl3
Air2
Fps1
Ask10
Cdc28
Hog1
Pbs2
Ssk2
Clb2
Bud3
Bud4
Msn2
Sch9
Rim15
Yak1
Ypl247c
Vps52
Kss1
Sko1
Slt2
Msg5
Swe1
Igo2
Cdc55
Igo1
Vps53
Vps54
Rtn1
Fmp42
Cot1
Zrc1
10
20
30
40
50
t/s

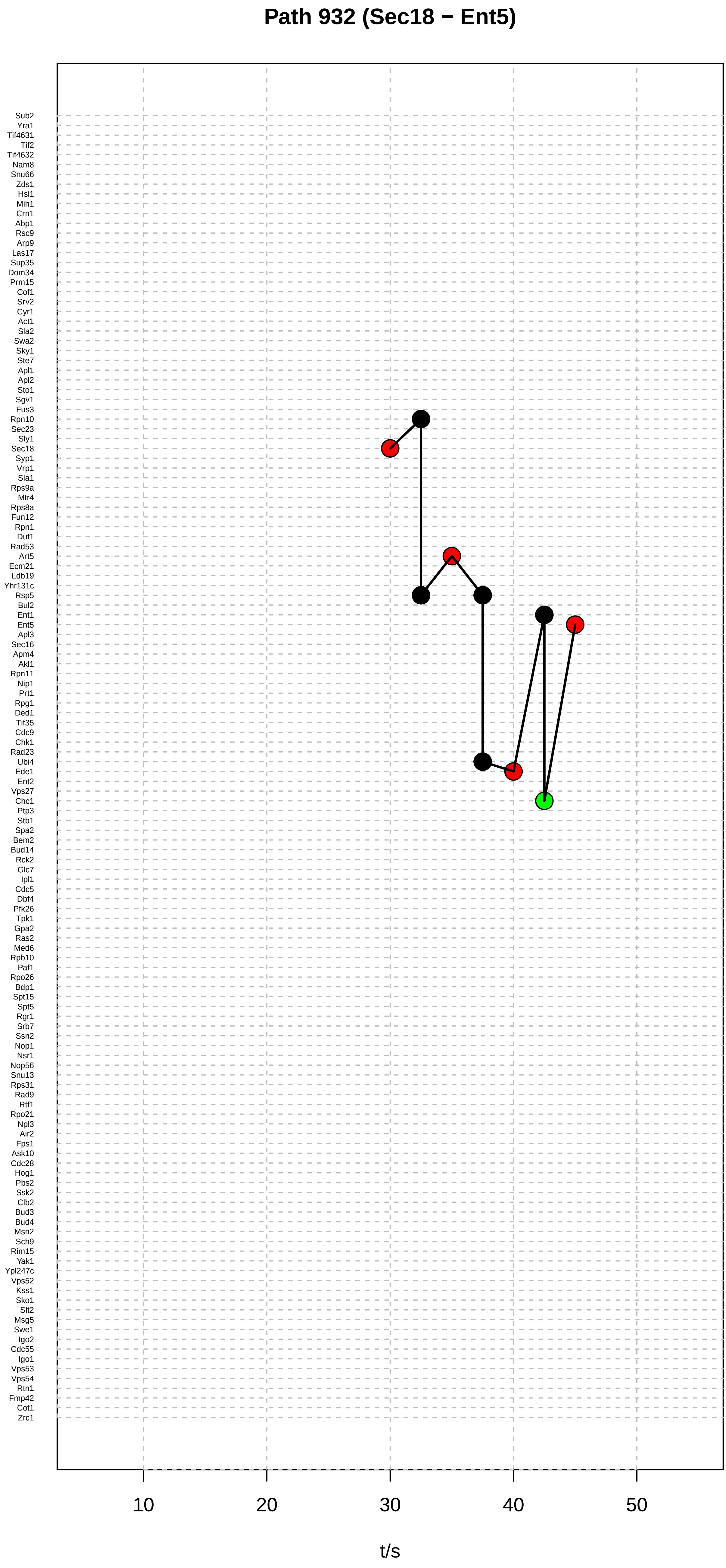

Path 932 (Sec18 − Ent5)
Sub2
Yra1
Tif4631
Tif2
Tif4632
Nam8
Snu66
Zds1
Hsl1
Mih1
Crn1
Abp1
Rsc9
Arp9
Las17
Sup35
Dom34
Prm15
Cof1
Srv2
Cyr1
Act1
Sla2
Swa2
Sky1
Ste7
Apl1
Apl2
Sto1
Sgv1
Fus3
Rpn10
Sec23
Sly1
Sec18
Syp1
Vrp1
Sla1
Rps9a
Mtr4
Rps8a
Fun12
Rpn1
Duf1
Rad53
Art5
Ecm21
Ldb19
Yhr131c
Rsp5
Bul2
Ent1
Ent5
Apl3
Sec16
Apm4
Akl1
Rpn11
Nip1
Prt1
Rpg1
Ded1
Tif35
Cdc9
Chk1
Rad23
Ubi4
Ede1
Ent2
Vps27
Chc1
Ptp3
Stb1
Spa2
Bem2
Bud14
Rck2
Glc7
Ipl1
Cdc5
Dbf4
Pfk26
Tpk1
Gpa2
Ras2
Med6
Rpb10
Paf1
Rpo26
Bdp1
Spt15
Spt5
Rgr1
Srb7
Ssn2
Nop1
Nsr1
Nop56
Snu13
Rps31
Rad9
Rtf1
Rpo21
Npl3
Air2
Fps1
Ask10
Cdc28
Hog1
Pbs2
Ssk2
Clb2
Bud3
Bud4
Msn2
Sch9
Rim15
Yak1
Ypl247c
Vps52
Kss1
Sko1
Slt2
Msg5
Swe1
Igo2
Cdc55
Igo1
Vps53
Vps54
Rtn1
Fmp42
Cot1
Zrc1
10
20
30
40
50
t/s

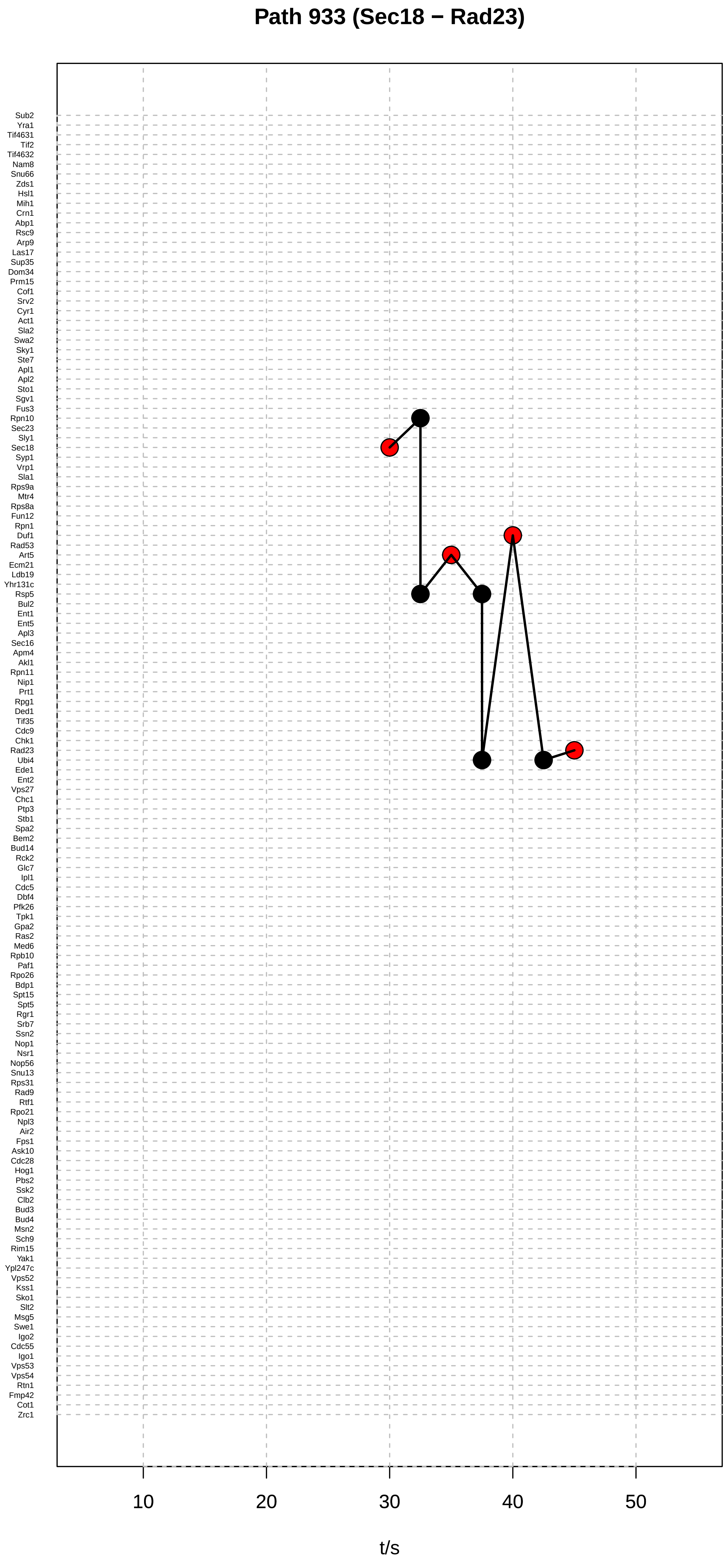

Path 933 (Sec18 − Rad23)
Sub2
Yra1
Tif4631
Tif2
Tif4632
Nam8
Snu66
Zds1
Hsl1
Mih1
Crn1
Abp1
Rsc9
Arp9
Las17
Sup35
Dom34
Prm15
Cof1
Srv2
Cyr1
Act1
Sla2
Swa2
Sky1
Ste7
Apl1
Apl2
Sto1
Sgv1
Fus3
Rpn10
Sec23
Sly1
Sec18
Syp1
Vrp1
Sla1
Rps9a
Mtr4
Rps8a
Fun12
Rpn1
Duf1
Rad53
Art5
Ecm21
Ldb19
Yhr131c
Rsp5
Bul2
Ent1
Ent5
Apl3
Sec16
Apm4
Akl1
Rpn11
Nip1
Prt1
Rpg1
Ded1
Tif35
Cdc9
Chk1
Rad23
Ubi4
Ede1
Ent2
Vps27
Chc1
Ptp3
Stb1
Spa2
Bem2
Bud14
Rck2
Glc7
Ipl1
Cdc5
Dbf4
Pfk26
Tpk1
Gpa2
Ras2
Med6
Rpb10
Paf1
Rpo26
Bdp1
Spt15
Spt5
Rgr1
Srb7
Ssn2
Nop1
Nsr1
Nop56
Snu13
Rps31
Rad9
Rtf1
Rpo21
Npl3
Air2
Fps1
Ask10
Cdc28
Hog1
Pbs2
Ssk2
Clb2
Bud3
Bud4
Msn2
Sch9
Rim15
Yak1
Ypl247c
Vps52
Kss1
Sko1
Slt2
Msg5
Swe1
Igo2
Cdc55
Igo1
Vps53
Vps54
Rtn1
Fmp42
Cot1
Zrc1
10
20
30
40
50
t/s

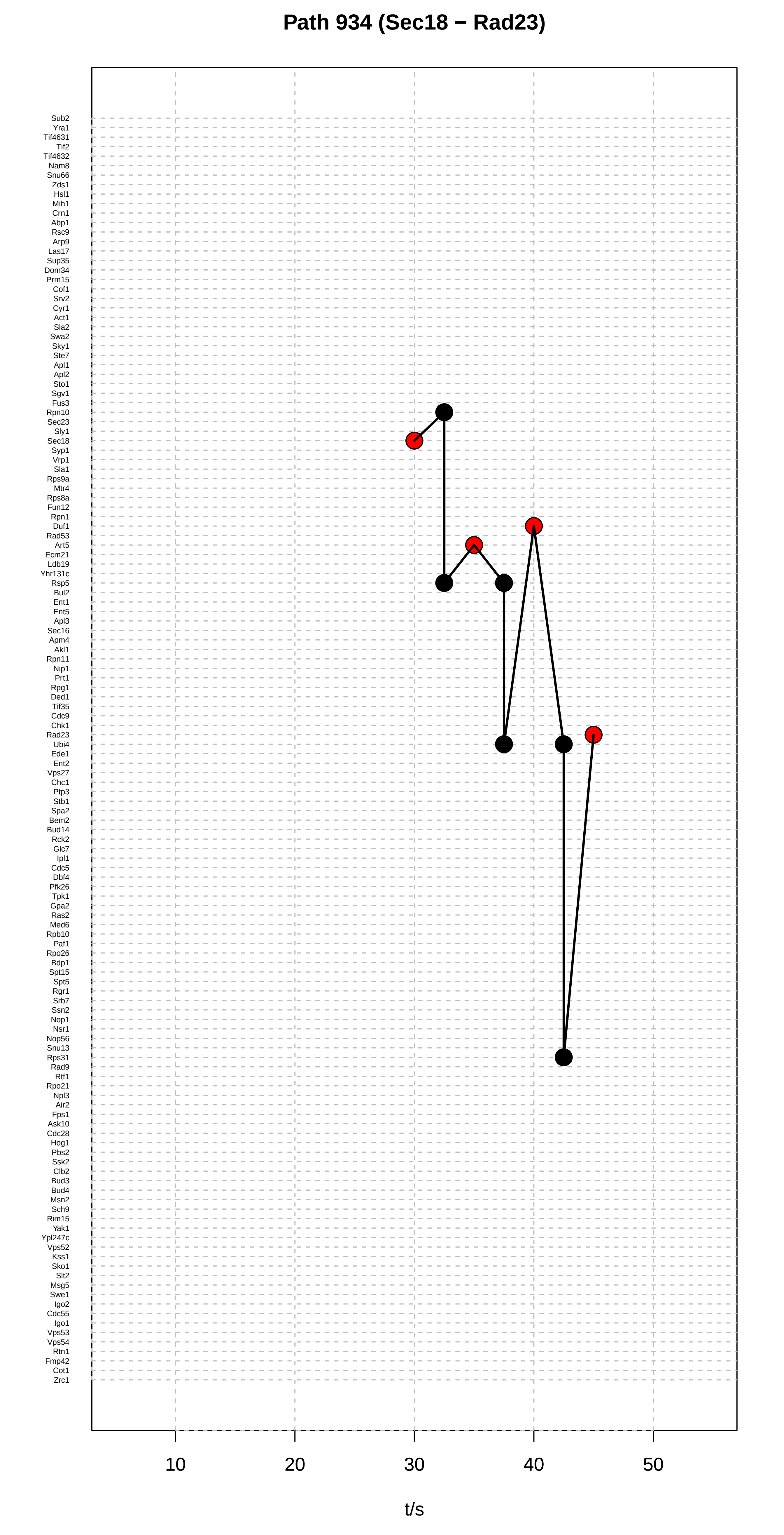

Path 934 (Sec18 − Rad23)
Sub2
Yra1
Tif4631
Tif2
Tif4632
Nam8
Snu66
Zds1
Hsl1
Mih1
Crn1
Abp1
Rsc9
Arp9
Las17
Sup35
Dom34
Prm15
Cof1
Srv2
Cyr1
Act1
Sla2
Swa2
Sky1
Ste7
Apl1
Apl2
Sto1
Sgv1
Fus3
Rpn10
Sec23
Sly1
Sec18
Syp1
Vrp1
Sla1
Rps9a
Mtr4
Rps8a
Fun12
Rpn1
Duf1
Rad53
Art5
Ecm21
Ldb19
Yhr131c
Rsp5
Bul2
Ent1
Ent5
Apl3
Sec16
Apm4
Akl1
Rpn11
Nip1
Prt1
Rpg1
Ded1
Tif35
Cdc9
Chk1
Rad23
Ubi4
Ede1
Ent2
Vps27
Chc1
Ptp3
Stb1
Spa2
Bem2
Bud14
Rck2
Glc7
Ipl1
Cdc5
Dbf4
Pfk26
Tpk1
Gpa2
Ras2
Med6
Rpb10
Paf1
Rpo26
Bdp1
Spt15
Spt5
Rgr1
Srb7
Ssn2
Nop1
Nsr1
Nop56
Snu13
Rps31
Rad9
Rtf1
Rpo21
Npl3
Air2
Fps1
Ask10
Cdc28
Hog1
Pbs2
Ssk2
Clb2
Bud3
Bud4
Msn2
Sch9
Rim15
Yak1
Ypl247c
Vps52
Kss1
Sko1
Slt2
Msg5
Swe1
Igo2
Cdc55
Igo1
Vps53
Vps54
Rtn1
Fmp42
Cot1
Zrc1
10
20
30
40
50
t/s

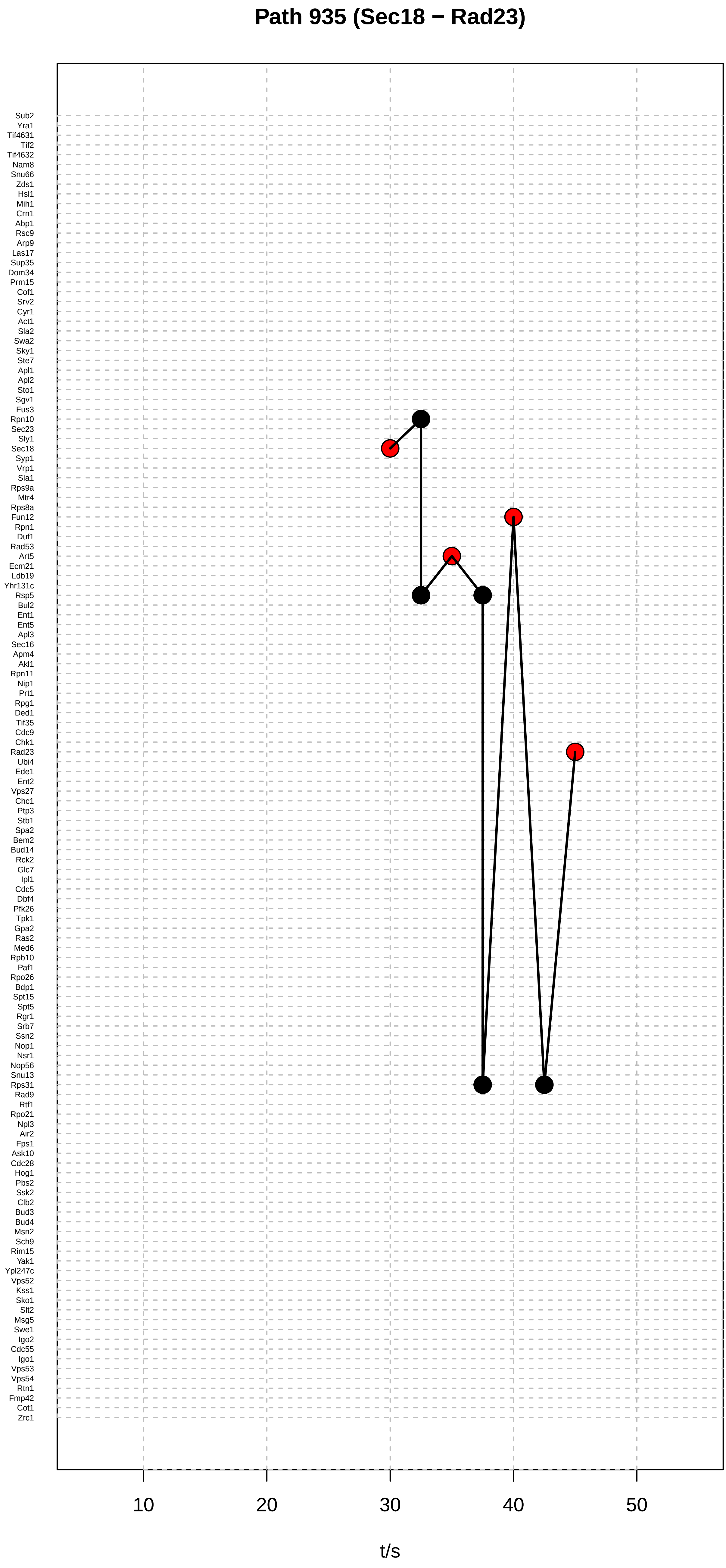
Path 935 (Sec18 − Rad23)
Sub2
Yra1
Tif4631
Tif2
Tif4632
Nam8
Snu66
Zds1
Hsl1
Mih1
Crn1
Abp1
Rsc9
Arp9
Las17
Sup35
Dom34
Prm15
Cof1
Srv2
Cyr1
Act1
Sla2
Swa2
Sky1
Ste7
Apl1
Apl2
Sto1
Sgv1
Fus3
Rpn10
Sec23
Sly1
Sec18
Syp1
Vrp1
Sla1
Rps9a
Mtr4
Rps8a
Fun12
Rpn1
Duf1
Rad53
Art5
Ecm21
Ldb19
Yhr131c
Rsp5
Bul2
Ent1
Ent5
Apl3
Sec16
Apm4
Akl1
Rpn11
Nip1
Prt1
Rpg1
Ded1
Tif35
Cdc9
Chk1
Rad23
Ubi4
Ede1
Ent2
Vps27
Chc1
Ptp3
Stb1
Spa2
Bem2
Bud14
Rck2
Glc7
Ipl1
Cdc5
Dbf4
Pfk26
Tpk1
Gpa2
Ras2
Med6
Rpb10
Paf1
Rpo26
Bdp1
Spt15
Spt5
Rgr1
Srb7
Ssn2
Nop1
Nsr1
Nop56
Snu13
Rps31
Rad9
Rtf1
Rpo21
Npl3
Air2
Fps1
Ask10
Cdc28
Hog1
Pbs2
Ssk2
Clb2
Bud3
Bud4
Msn2
Sch9
Rim15
Yak1
Ypl247c
Vps52
Kss1
Sko1
Slt2
Msg5
Swe1
Igo2
Cdc55
Igo1
Vps53
Vps54
Rtn1
Fmp42
Cot1
Zrc1
10
20
30
40
50
t/s

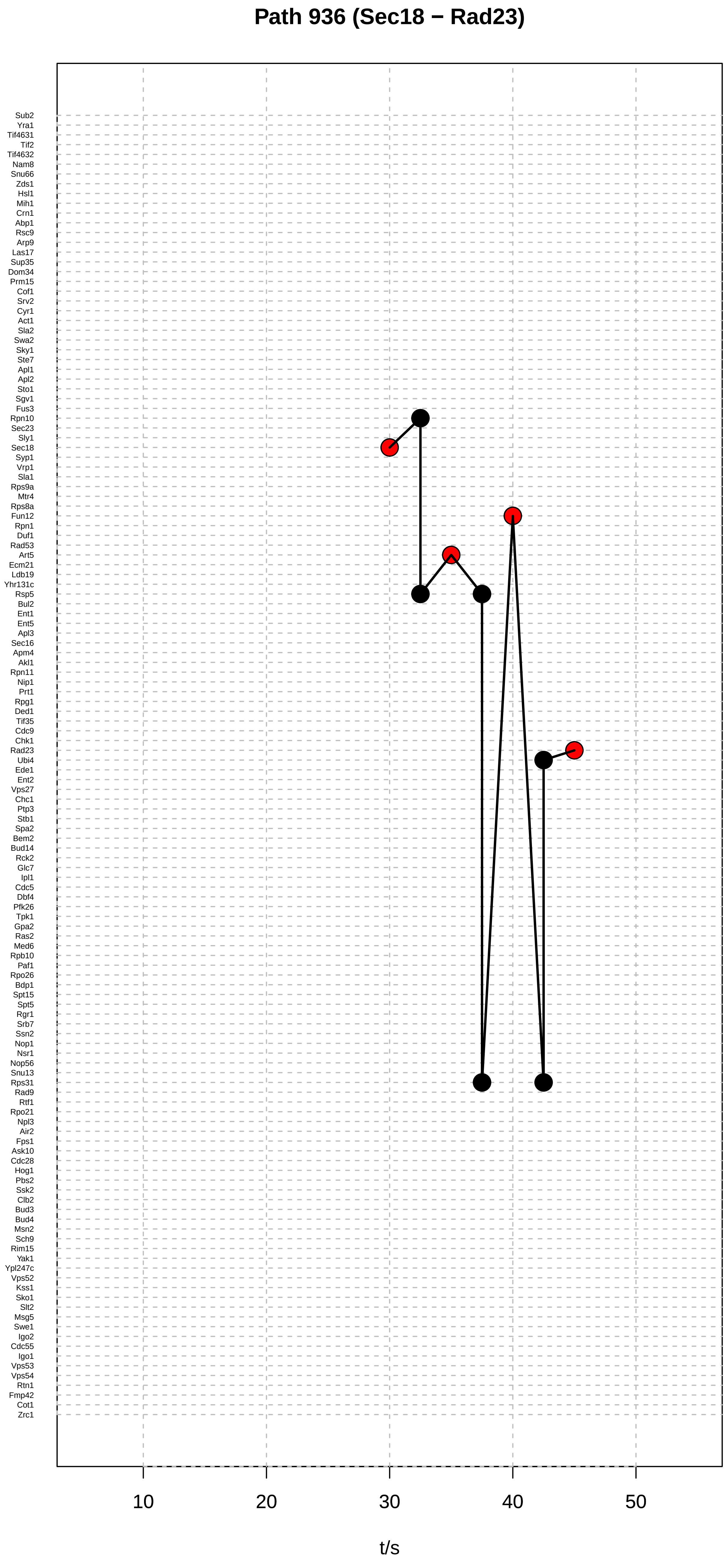

Path 936 (Sec18 − Rad23)
Sub2
Yra1
Tif4631
Tif2
Tif4632
Nam8
Snu66
Zds1
Hsl1
Mih1
Crn1
Abp1
Rsc9
Arp9
Las17
Sup35
Dom34
Prm15
Cof1
Srv2
Cyr1
Act1
Sla2
Swa2
Sky1
Ste7
Apl1
Apl2
Sto1
Sgv1
Fus3
Rpn10
Sec23
Sly1
Sec18
Syp1
Vrp1
Sla1
Rps9a
Mtr4
Rps8a
Fun12
Rpn1
Duf1
Rad53
Art5
Ecm21
Ldb19
Yhr131c
Rsp5
Bul2
Ent1
Ent5
Apl3
Sec16
Apm4
Akl1
Rpn11
Nip1
Prt1
Rpg1
Ded1
Tif35
Cdc9
Chk1
Rad23
Ubi4
Ede1
Ent2
Vps27
Chc1
Ptp3
Stb1
Spa2
Bem2
Bud14
Rck2
Glc7
Ipl1
Cdc5
Dbf4
Pfk26
Tpk1
Gpa2
Ras2
Med6
Rpb10
Paf1
Rpo26
Bdp1
Spt15
Spt5
Rgr1
Srb7
Ssn2
Nop1
Nsr1
Nop56
Snu13
Rps31
Rad9
Rtf1
Rpo21
Npl3
Air2
Fps1
Ask10
Cdc28
Hog1
Pbs2
Ssk2
Clb2
Bud3
Bud4
Msn2
Sch9
Rim15
Yak1
Ypl247c
Vps52
Kss1
Sko1
Slt2
Msg5
Swe1
Igo2
Cdc55
Igo1
Vps53
Vps54
Rtn1
Fmp42
Cot1
Zrc1
10
20
30
40
50
t/s

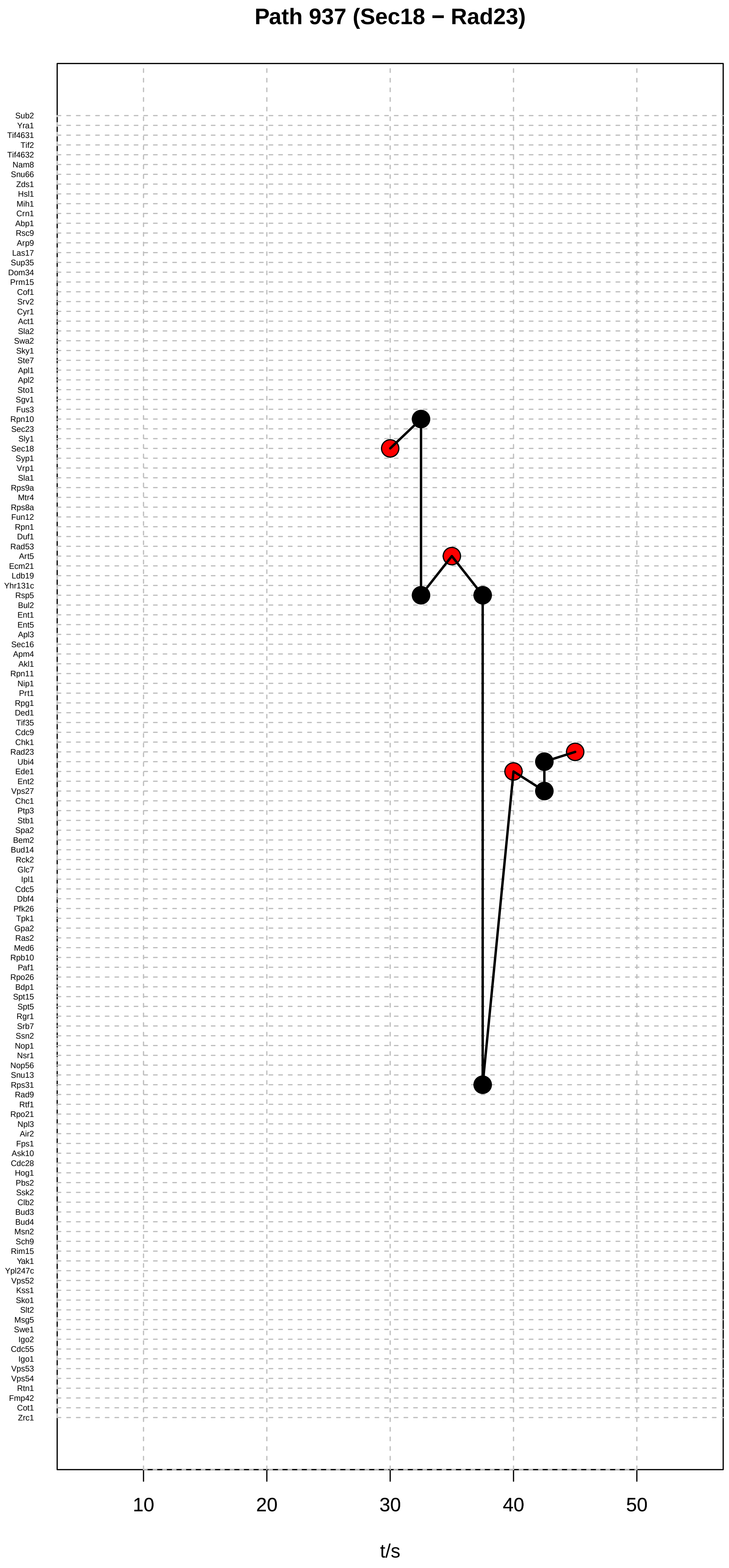
Path 937 (Sec18 − Rad23)
Sub2
Yra1
Tif4631
Tif2
Tif4632
Nam8
Snu66
Zds1
Hsl1
Mih1
Crn1
Abp1
Rsc9
Arp9
Las17
Sup35
Dom34
Prm15
Cof1
Srv2
Cyr1
Act1
Sla2
Swa2
Sky1
Ste7
Apl1
Apl2
Sto1
Sgv1
Fus3
Rpn10
Sec23
Sly1
Sec18
Syp1
Vrp1
Sla1
Rps9a
Mtr4
Rps8a
Fun12
Rpn1
Duf1
Rad53
Art5
Ecm21
Ldb19
Yhr131c
Rsp5
Bul2
Ent1
Ent5
Apl3
Sec16
Apm4
Akl1
Rpn11
Nip1
Prt1
Rpg1
Ded1
Tif35
Cdc9
Chk1
Rad23
Ubi4
Ede1
Ent2
Vps27
Chc1
Ptp3
Stb1
Spa2
Bem2
Bud14
Rck2
Glc7
Ipl1
Cdc5
Dbf4
Pfk26
Tpk1
Gpa2
Ras2
Med6
Rpb10
Paf1
Rpo26
Bdp1
Spt15
Spt5
Rgr1
Srb7
Ssn2
Nop1
Nsr1
Nop56
Snu13
Rps31
Rad9
Rtf1
Rpo21
Npl3
Air2
Fps1
Ask10
Cdc28
Hog1
Pbs2
Ssk2
Clb2
Bud3
Bud4
Msn2
Sch9
Rim15
Yak1
Ypl247c
Vps52
Kss1
Sko1
Slt2
Msg5
Swe1
Igo2
Cdc55
Igo1
Vps53
Vps54
Rtn1
Fmp42
Cot1
Zrc1
10
20
30
40
50
t/s

#### Path 938 (Sec18 − Rad23)

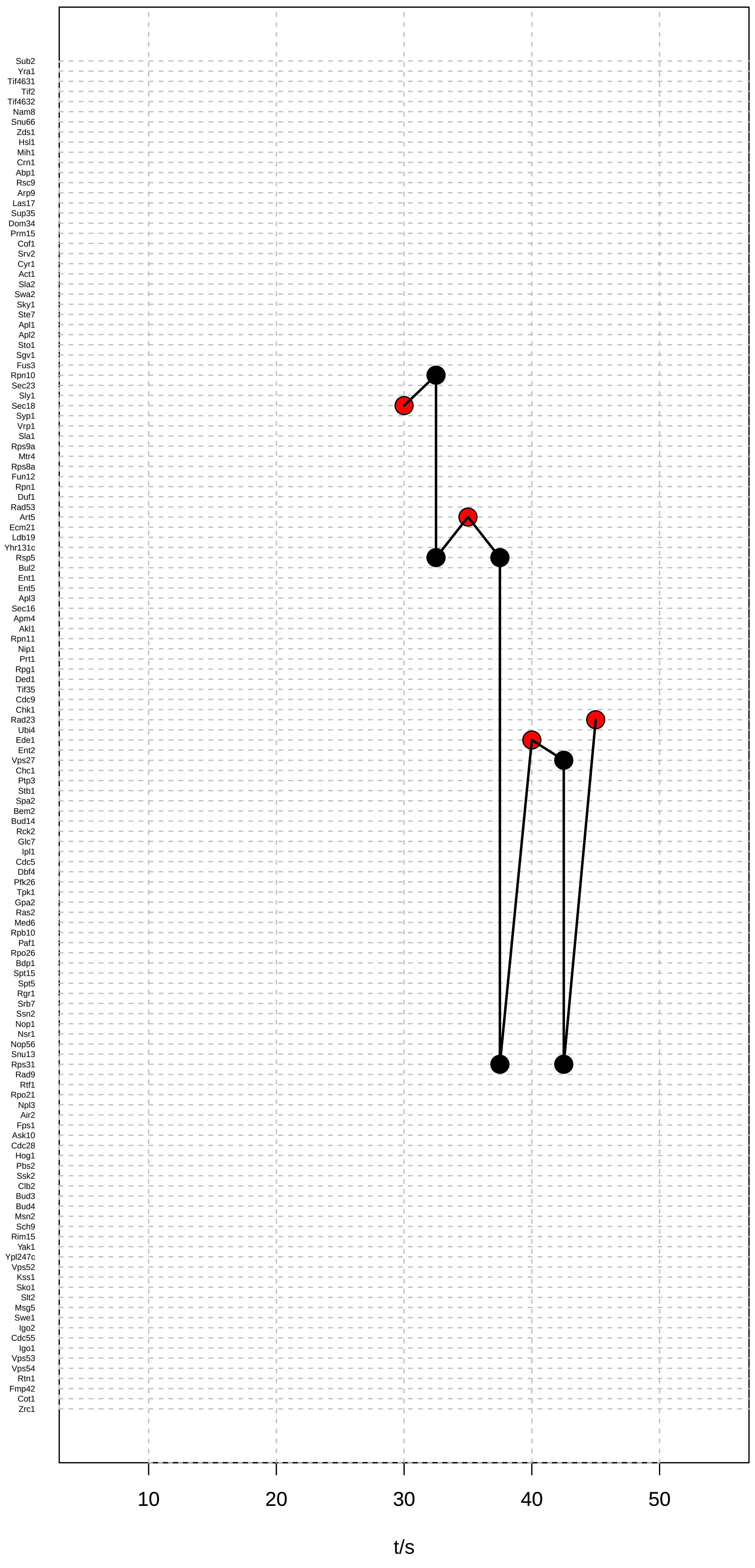

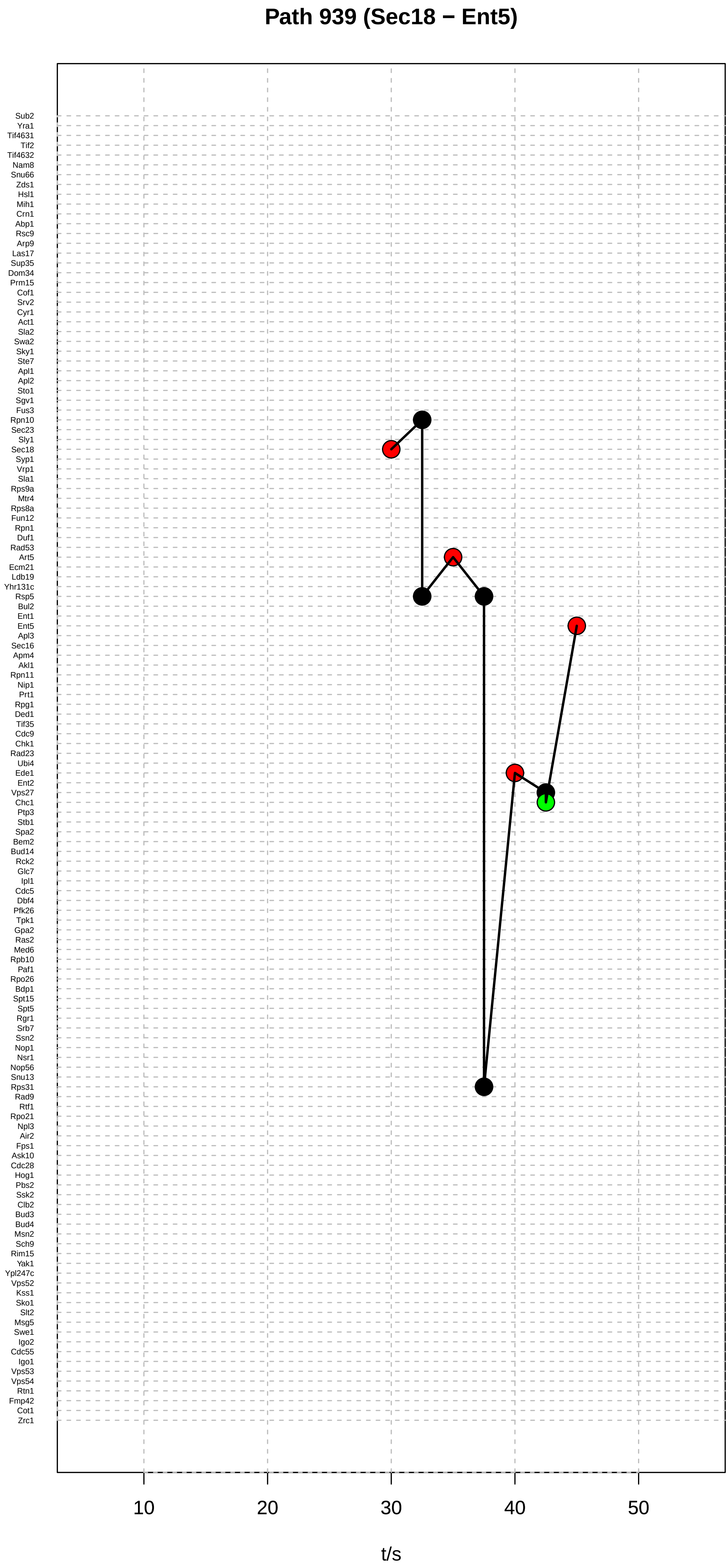

Path 939 (Sec18 − Ent5)
Sub2
Yra1
Tif4631
Tif2
Tif4632
Nam8
Snu66
Zds1
Hsl1
Mih1
Crn1
Abp1
Rsc9
Arp9
Las17
Sup35
Dom34
Prm15
Cof1
Srv2
Cyr1
Act1
Sla2
Swa2
Sky1
Ste7
Apl1
Apl2
Sto1
Sgv1
Fus3
Rpn10
Sec23
Sly1
Sec18
Syp1
Vrp1
Sla1
Rps9a
Mtr4
Rps8a
Fun12
Rpn1
Duf1
Rad53
Art5
Ecm21
Ldb19
Yhr131c
Rsp5
Bul2
Ent1
Ent5
Apl3
Sec16
Apm4
Akl1
Rpn11
Nip1
Prt1
Rpg1
Ded1
Tif35
Cdc9
Chk1
Rad23
Ubi4
Ede1
Ent2
Vps27
Chc1
Ptp3
Stb1
Spa2
Bem2
Bud14
Rck2
Glc7
Ipl1
Cdc5
Dbf4
Pfk26
Tpk1
Gpa2
Ras2
Med6
Rpb10
Paf1
Rpo26
Bdp1
Spt15
Spt5
Rgr1
Srb7
Ssn2
Nop1
Nsr1
Nop56
Snu13
Rps31
Rad9
Rtf1
Rpo21
Npl3
Air2
Fps1
Ask10
Cdc28
Hog1
Pbs2
Ssk2
Clb2
Bud3
Bud4
Msn2
Sch9
Rim15
Yak1
Ypl247c
Vps52
Kss1
Sko1
Slt2
Msg5
Swe1
Igo2
Cdc55
Igo1
Vps53
Vps54
Rtn1
Fmp42
Cot1
Zrc1
10
20
30
40
50
t/s

Path 940 (Sec18 − Rad23)

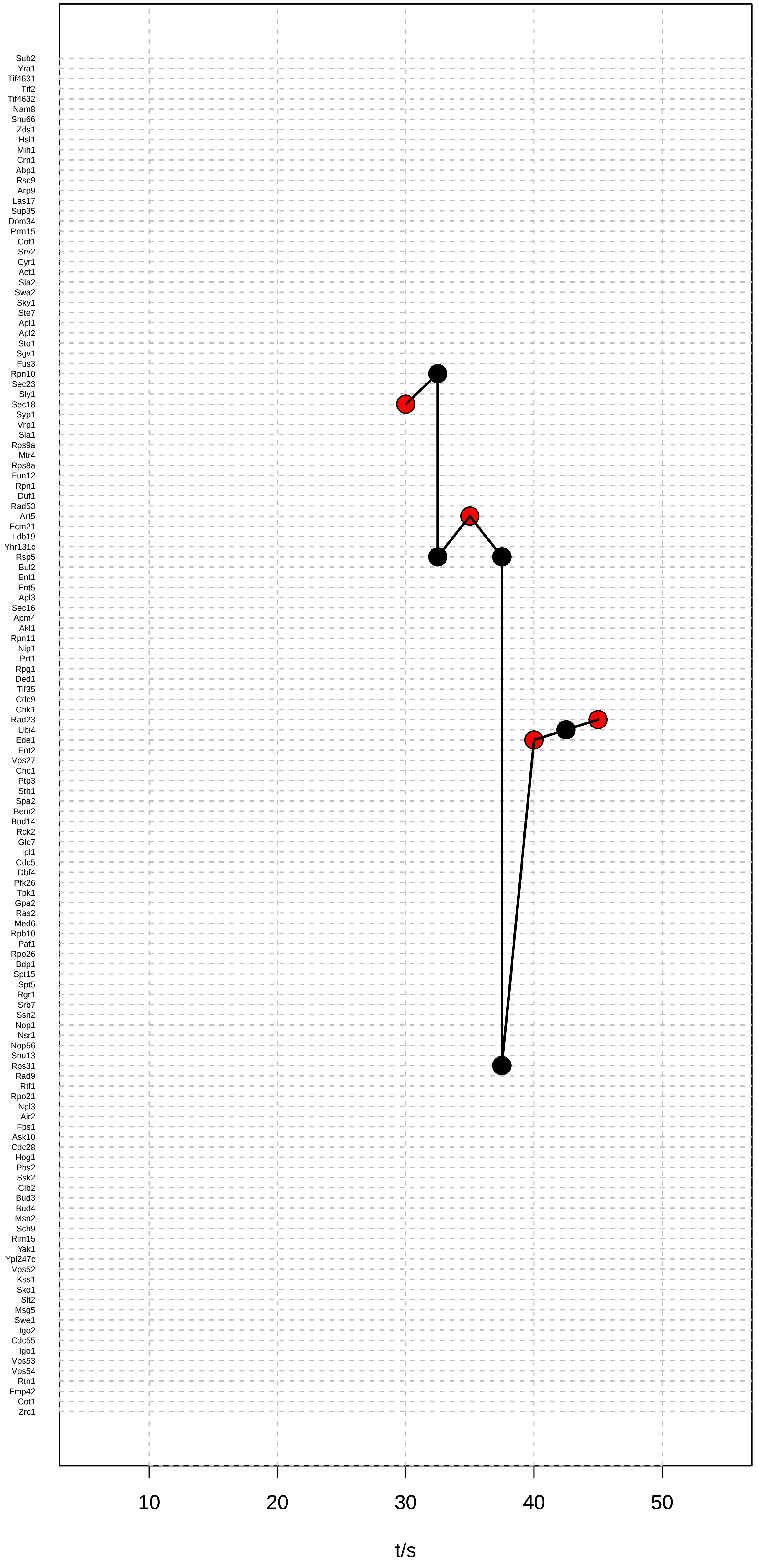

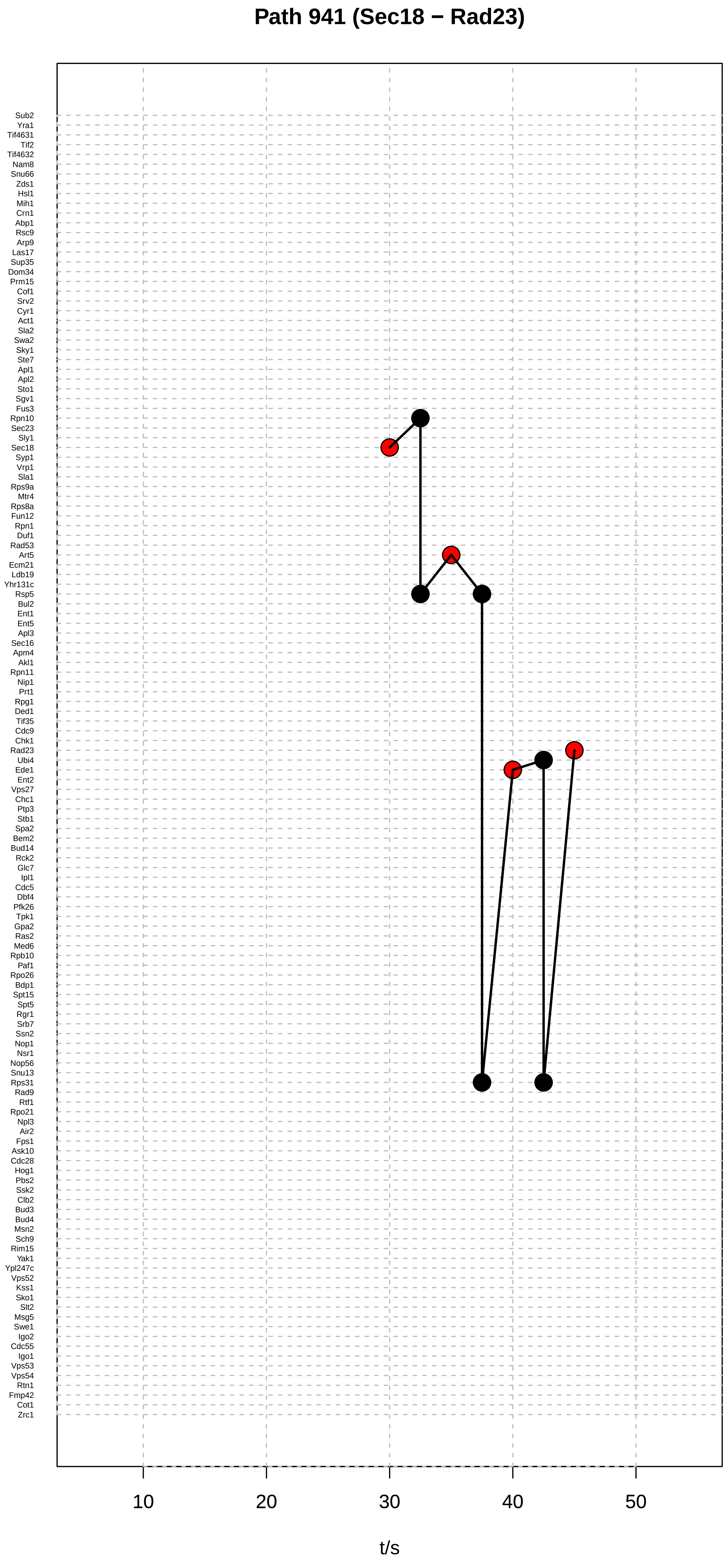

Path 941 (Sec18 − Rad23)
Sub2
Yra1
Tif4631
Tif2
Tif4632
Nam8
Snu66
Zds1
Hsl1
Mih1
Crn1
Abp1
Rsc9
Arp9
Las17
Sup35
Dom34
Prm15
Cof1
Srv2
Cyr1
Act1
Sla2
Swa2
Sky1
Ste7
Apl1
Apl2
Sto1
Sgv1
Fus3
Rpn10
Sec23
Sly1
Sec18
Syp1
Vrp1
Sla1
Rps9a
Mtr4
Rps8a
Fun12
Rpn1
Duf1
Rad53
Art5
Ecm21
Ldb19
Yhr131c
Rsp5
Bul2
Ent1
Ent5
Apl3
Sec16
Apm4
Akl1
Rpn11
Nip1
Prt1
Rpg1
Ded1
Tif35
Cdc9
Chk1
Rad23
Ubi4
Ede1
Ent2
Vps27
Chc1
Ptp3
Stb1
Spa2
Bem2
Bud14
Rck2
Glc7
Ipl1
Cdc5
Dbf4
Pfk26
Tpk1
Gpa2
Ras2
Med6
Rpb10
Paf1
Rpo26
Bdp1
Spt15
Spt5
Rgr1
Srb7
Ssn2
Nop1
Nsr1
Nop56
Snu13
Rps31
Rad9
Rtf1
Rpo21
Npl3
Air2
Fps1
Ask10
Cdc28
Hog1
Pbs2
Ssk2
Clb2
Bud3
Bud4
Msn2
Sch9
Rim15
Yak1
Ypl247c
Vps52
Kss1
Sko1
Slt2
Msg5
Swe1
Igo2
Cdc55
Igo1
Vps53
Vps54
Rtn1
Fmp42
Cot1
Zrc1
10
20
30
40
50
t/s

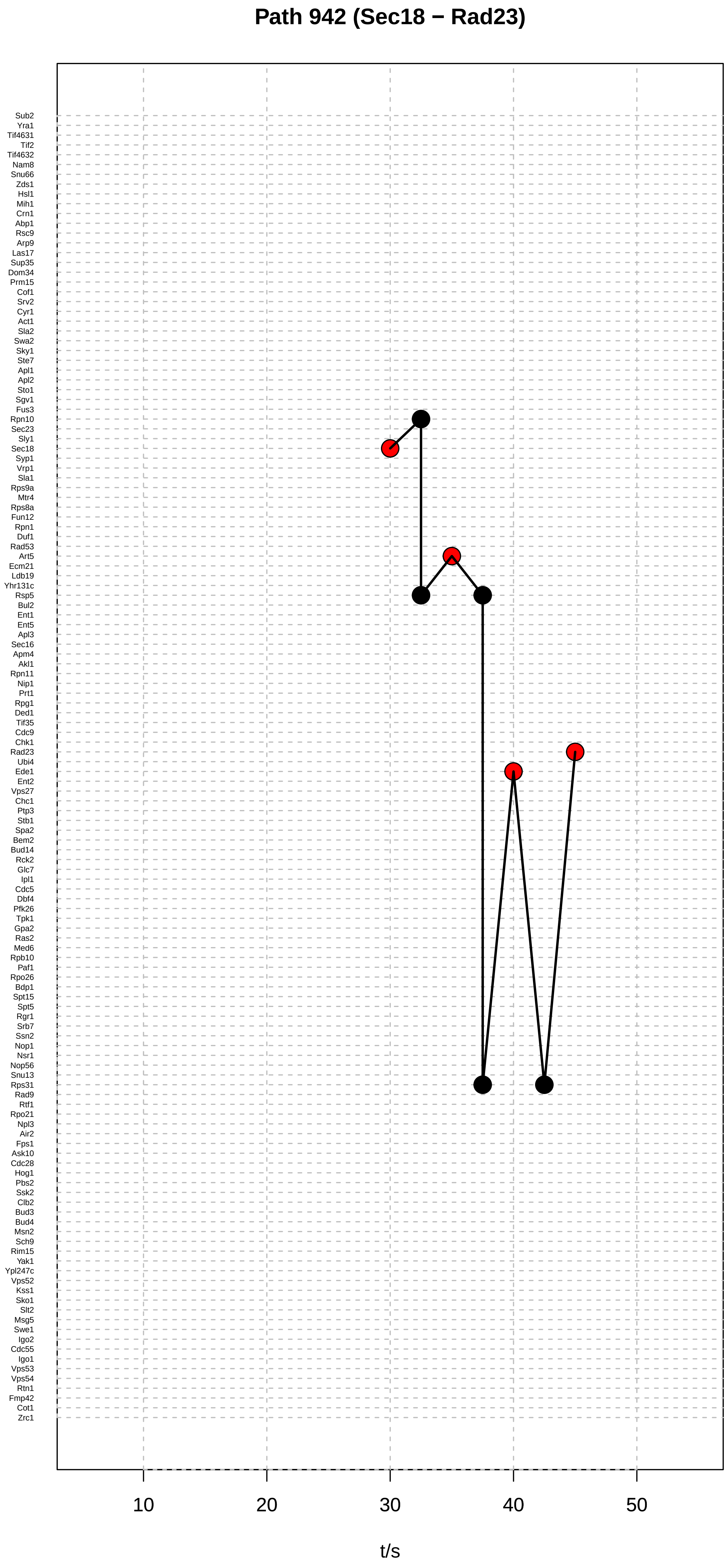
Path 942 (Sec18 − Rad23)
Sub2
Yra1
Tif4631
Tif2
Tif4632
Nam8
Snu66
Zds1
Hsl1
Mih1
Crn1
Abp1
Rsc9
Arp9
Las17
Sup35
Dom34
Prm15
Cof1
Srv2
Cyr1
Act1
Sla2
Swa2
Sky1
Ste7
Apl1
Apl2
Sto1
Sgv1
Fus3
Rpn10
Sec23
Sly1
Sec18
Syp1
Vrp1
Sla1
Rps9a
Mtr4
Rps8a
Fun12
Rpn1
Duf1
Rad53
Art5
Ecm21
Ldb19
Yhr131c
Rsp5
Bul2
Ent1
Ent5
Apl3
Sec16
Apm4
Akl1
Rpn11
Nip1
Prt1
Rpg1
Ded1
Tif35
Cdc9
Chk1
Rad23
Ubi4
Ede1
Ent2
Vps27
Chc1
Ptp3
Stb1
Spa2
Bem2
Bud14
Rck2
Glc7
Ipl1
Cdc5
Dbf4
Pfk26
Tpk1
Gpa2
Ras2
Med6
Rpb10
Paf1
Rpo26
Bdp1
Spt15
Spt5
Rgr1
Srb7
Ssn2
Nop1
Nsr1
Nop56
Snu13
Rps31
Rad9
Rtf1
Rpo21
Npl3
Air2
Fps1
Ask10
Cdc28
Hog1
Pbs2
Ssk2
Clb2
Bud3
Bud4
Msn2
Sch9
Rim15
Yak1
Ypl247c
Vps52
Kss1
Sko1
Slt2
Msg5
Swe1
Igo2
Cdc55
Igo1
Vps53
Vps54
Rtn1
Fmp42
Cot1
Zrc1
10
20
30
40
50
t/s

#### Path 943 (Sec18 − Rad23)

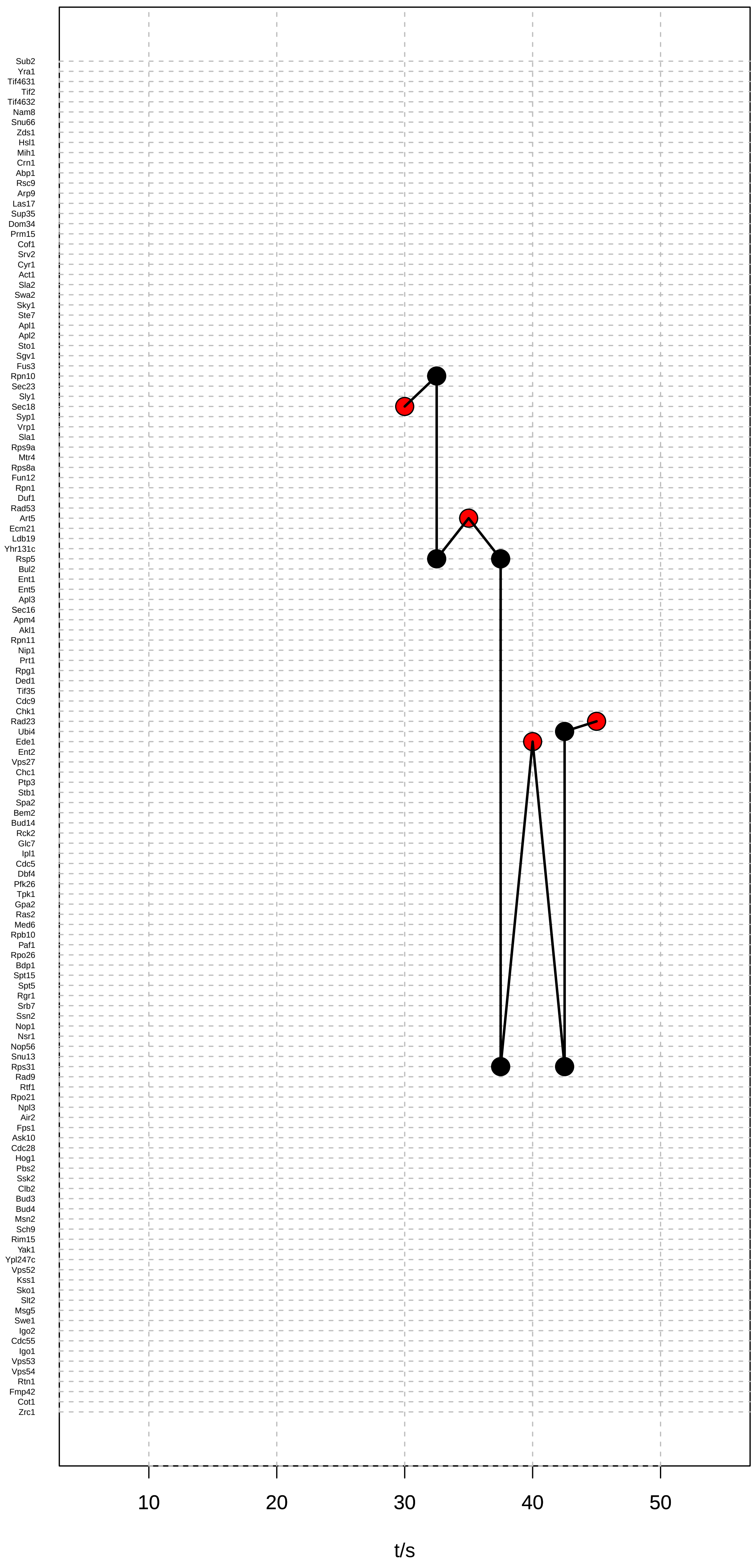

#### Path 944 (Sec18 − Rad23)

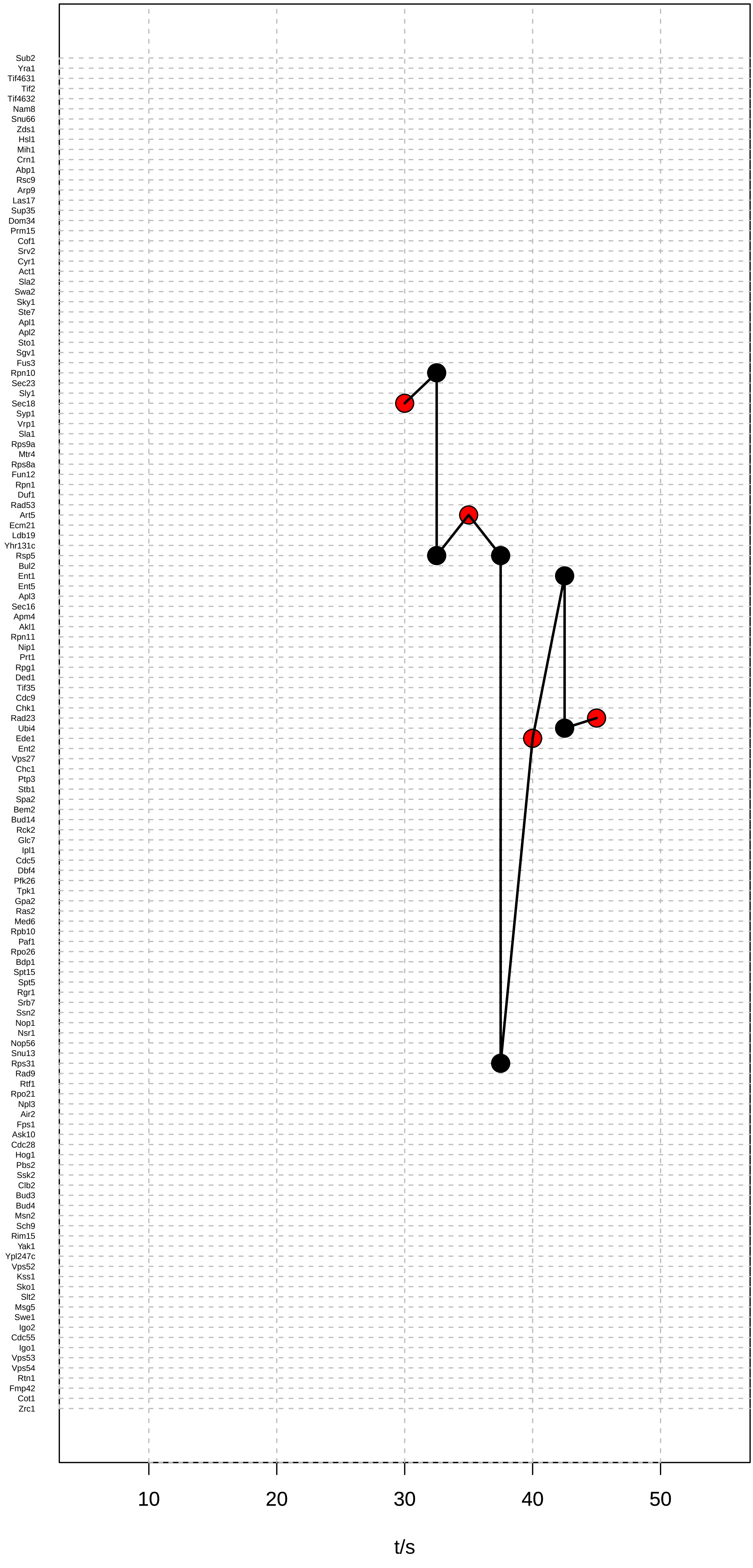

Path 945 (Sec18 − Ent5)

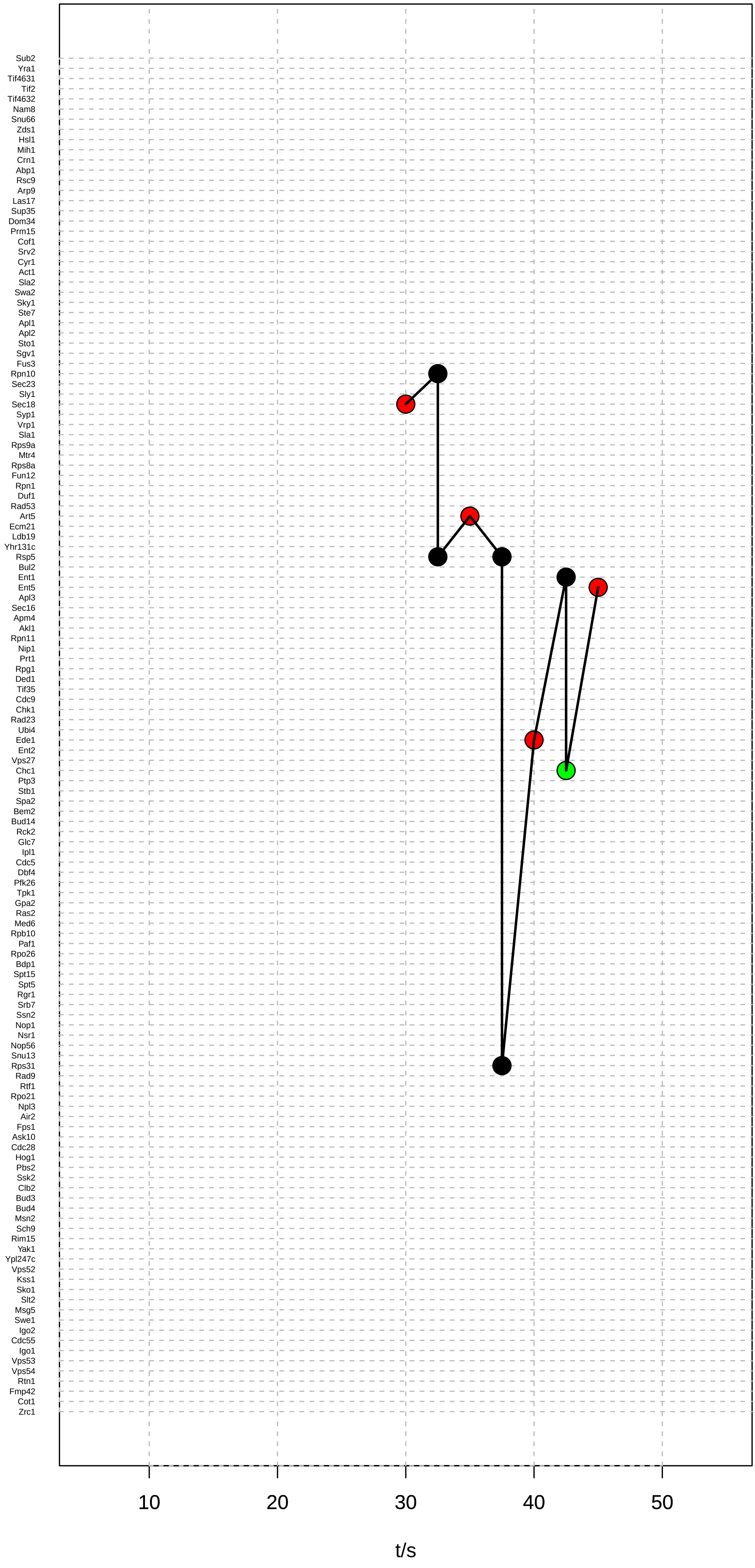

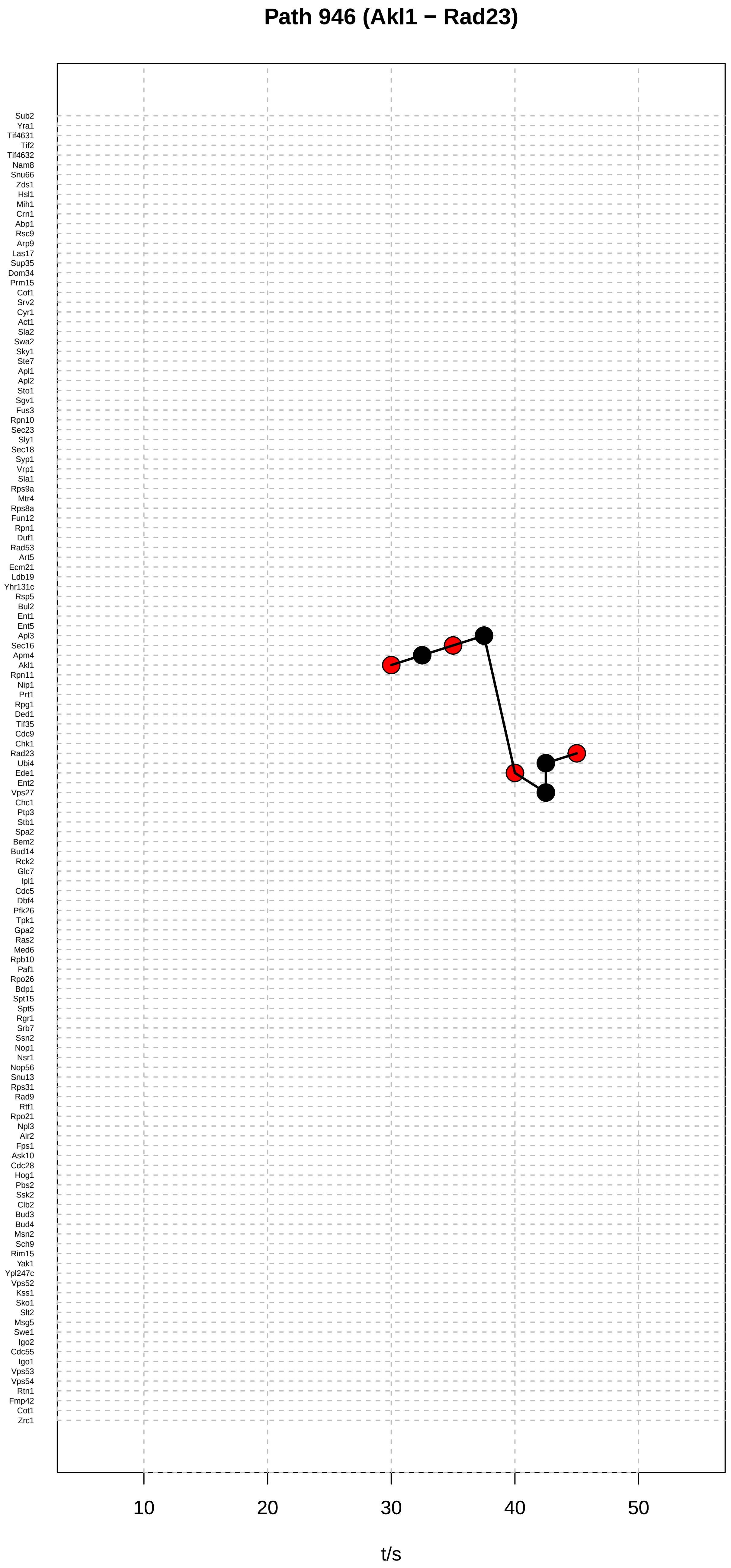

Path 946 (Akl1 − Rad23)
Sub2
Yra1
Tif4631
Tif2
Tif4632
Nam8
Snu66
Zds1
Hsl1
Mih1
Crn1
Abp1
Rsc9
Arp9
Las17
Sup35
Dom34
Prm15
Cof1
Srv2
Cyr1
Act1
Sla2
Swa2
Sky1
Ste7
Apl1
Apl2
Sto1
Sgv1
Fus3
Rpn10
Sec23
Sly1
Sec18
Syp1
Vrp1
Sla1
Rps9a
Mtr4
Rps8a
Fun12
Rpn1
Duf1
Rad53
Art5
Ecm21
Ldb19
Yhr131c
Rsp5
Bul2
Ent1
Ent5
Apl3
Sec16
Apm4
Akl1
Rpn11
Nip1
Prt1
Rpg1
Ded1
Tif35
Cdc9
Chk1
Rad23
Ubi4
Ede1
Ent2
Vps27
Chc1
Ptp3
Stb1
Spa2
Bem2
Bud14
Rck2
Glc7
Ipl1
Cdc5
Dbf4
Pfk26
Tpk1
Gpa2
Ras2
Med6
Rpb10
Paf1
Rpo26
Bdp1
Spt15
Spt5
Rgr1
Srb7
Ssn2
Nop1
Nsr1
Nop56
Snu13
Rps31
Rad9
Rtf1
Rpo21
Npl3
Air2
Fps1
Ask10
Cdc28
Hog1
Pbs2
Ssk2
Clb2
Bud3
Bud4
Msn2
Sch9
Rim15
Yak1
Ypl247c
Vps52
Kss1
Sko1
Slt2
Msg5
Swe1
Igo2
Cdc55
Igo1
Vps53
Vps54
Rtn1
Fmp42
Cot1
Zrc1
10
20
30
40
50
t/s

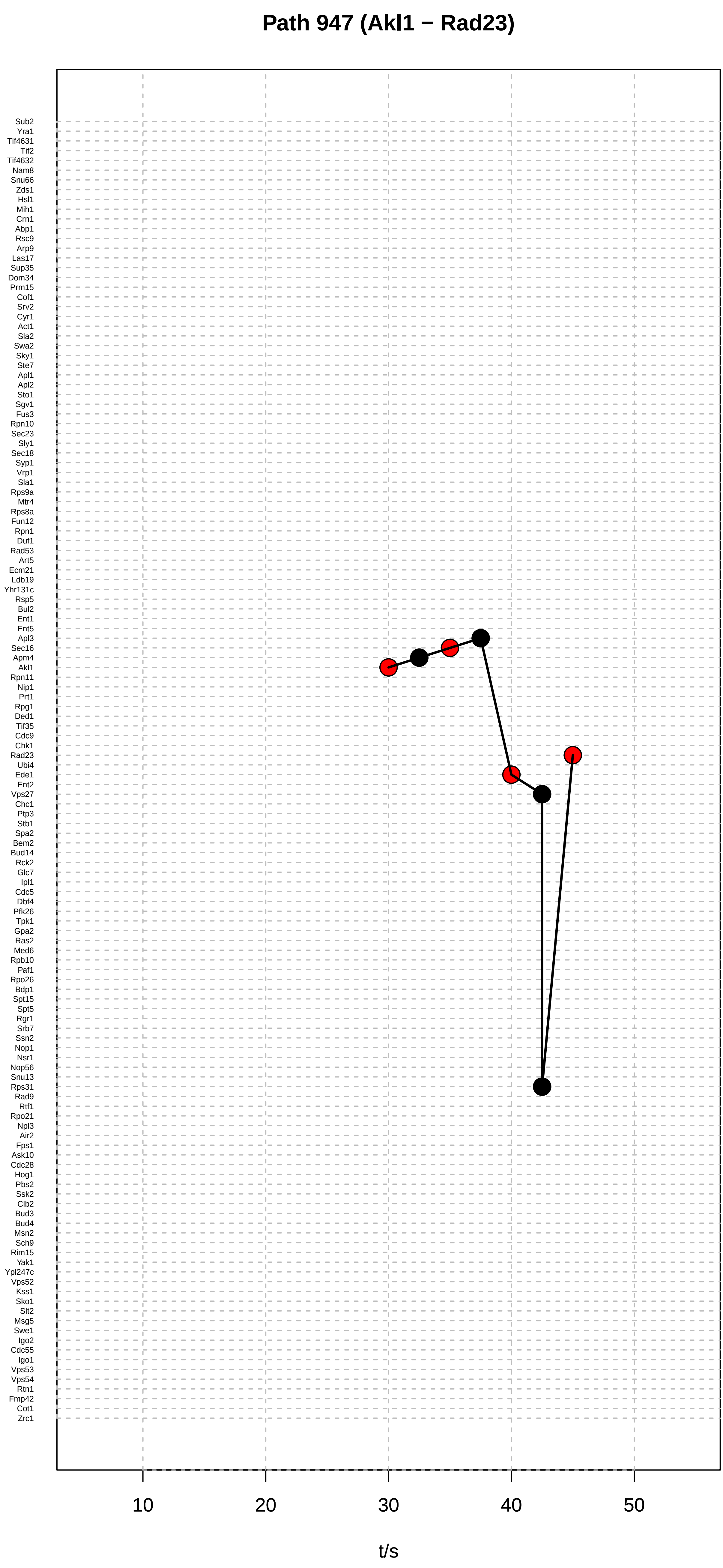
Path 947 (Akl1 − Rad23)
Sub2
Yra1
Tif4631
Tif2
Tif4632
Nam8
Snu66
Zds1
Hsl1
Mih1
Crn1
Abp1
Rsc9
Arp9
Las17
Sup35
Dom34
Prm15
Cof1
Srv2
Cyr1
Act1
Sla2
Swa2
Sky1
Ste7
Apl1
Apl2
Sto1
Sgv1
Fus3
Rpn10
Sec23
Sly1
Sec18
Syp1
Vrp1
Sla1
Rps9a
Mtr4
Rps8a
Fun12
Rpn1
Duf1
Rad53
Art5
Ecm21
Ldb19
Yhr131c
Rsp5
Bul2
Ent1
Ent5
Apl3
Sec16
Apm4
Akl1
Rpn11
Nip1
Prt1
Rpg1
Ded1
Tif35
Cdc9
Chk1
Rad23
Ubi4
Ede1
Ent2
Vps27
Chc1
Ptp3
Stb1
Spa2
Bem2
Bud14
Rck2
Glc7
Ipl1
Cdc5
Dbf4
Pfk26
Tpk1
Gpa2
Ras2
Med6
Rpb10
Paf1
Rpo26
Bdp1
Spt15
Spt5
Rgr1
Srb7
Ssn2
Nop1
Nsr1
Nop56
Snu13
Rps31
Rad9
Rtf1
Rpo21
Npl3
Air2
Fps1
Ask10
Cdc28
Hog1
Pbs2
Ssk2
Clb2
Bud3
Bud4
Msn2
Sch9
Rim15
Yak1
Ypl247c
Vps52
Kss1
Sko1
Slt2
Msg5
Swe1
Igo2
Cdc55
Igo1
Vps53
Vps54
Rtn1
Fmp42
Cot1
Zrc1
10
20
30
40
50
t/s

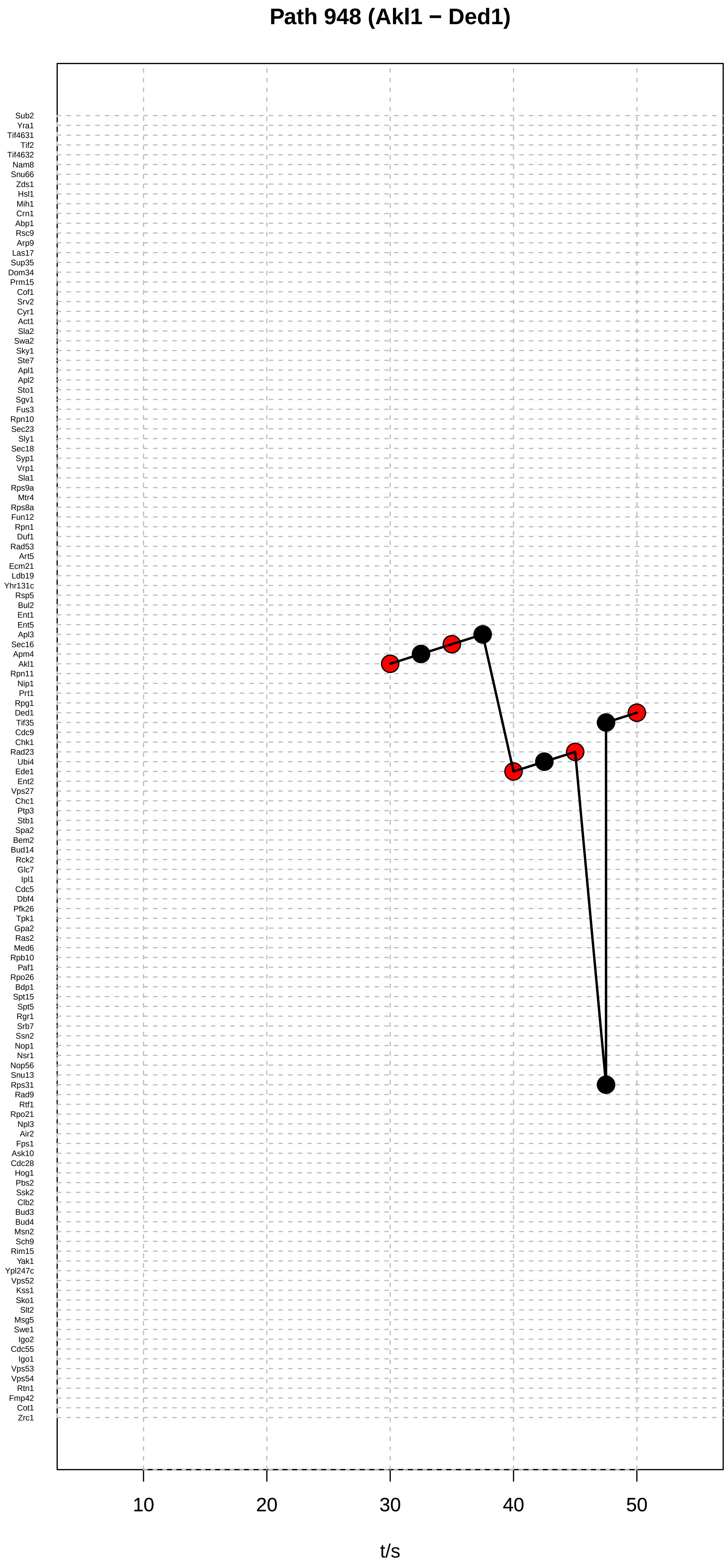
Path 948 (Akl1 − Ded1)
Sub2
Yra1
Tif4631
Tif2
Tif4632
Nam8
Snu66
Zds1
Hsl1
Mih1
Crn1
Abp1
Rsc9
Arp9
Las17
Sup35
Dom34
Prm15
Cof1
Srv2
Cyr1
Act1
Sla2
Swa2
Sky1
Ste7
Apl1
Apl2
Sto1
Sgv1
Fus3
Rpn10
Sec23
Sly1
Sec18
Syp1
Vrp1
Sla1
Rps9a
Mtr4
Rps8a
Fun12
Rpn1
Duf1
Rad53
Art5
Ecm21
Ldb19
Yhr131c
Rsp5
Bul2
Ent1
Ent5
Apl3
Sec16
Apm4
Akl1
Rpn11
Nip1
Prt1
Rpg1
Ded1
Tif35
Cdc9
Chk1
Rad23
Ubi4
Ede1
Ent2
Vps27
Chc1
Ptp3
Stb1
Spa2
Bem2
Bud14
Rck2
Glc7
Ipl1
Cdc5
Dbf4
Pfk26
Tpk1
Gpa2
Ras2
Med6
Rpb10
Paf1
Rpo26
Bdp1
Spt15
Spt5
Rgr1
Srb7
Ssn2
Nop1
Nsr1
Nop56
Snu13
Rps31
Rad9
Rtf1
Rpo21
Npl3
Air2
Fps1
Ask10
Cdc28
Hog1
Pbs2
Ssk2
Clb2
Bud3
Bud4
Msn2
Sch9
Rim15
Yak1
Ypl247c
Vps52
Kss1
Sko1
Slt2
Msg5
Swe1
Igo2
Cdc55
Igo1
Vps53
Vps54
Rtn1
Fmp42
Cot1
Zrc1
10
20
30
40
50
t/s

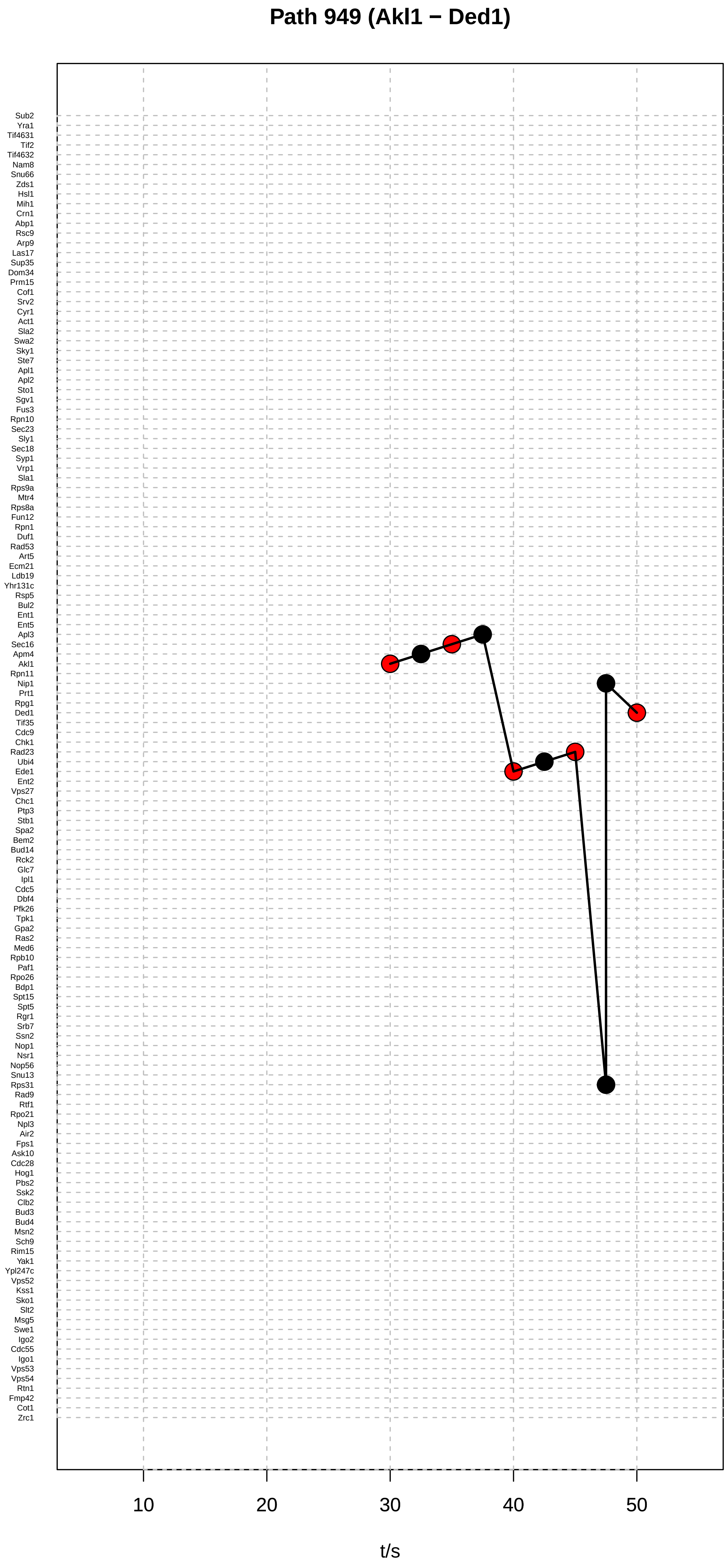

Path 949 (Akl1 − Ded1)
Sub2
Yra1
Tif4631
Tif2
Tif4632
Nam8
Snu66
Zds1
Hsl1
Mih1
Crn1
Abp1
Rsc9
Arp9
Las17
Sup35
Dom34
Prm15
Cof1
Srv2
Cyr1
Act1
Sla2
Swa2
Sky1
Ste7
Apl1
Apl2
Sto1
Sgv1
Fus3
Rpn10
Sec23
Sly1
Sec18
Syp1
Vrp1
Sla1
Rps9a
Mtr4
Rps8a
Fun12
Rpn1
Duf1
Rad53
Art5
Ecm21
Ldb19
Yhr131c
Rsp5
Bul2
Ent1
Ent5
Apl3
Sec16
Apm4
Akl1
Rpn11
Nip1
Prt1
Rpg1
Ded1
Tif35
Cdc9
Chk1
Rad23
Ubi4
Ede1
Ent2
Vps27
Chc1
Ptp3
Stb1
Spa2
Bem2
Bud14
Rck2
Glc7
Ipl1
Cdc5
Dbf4
Pfk26
Tpk1
Gpa2
Ras2
Med6
Rpb10
Paf1
Rpo26
Bdp1
Spt15
Spt5
Rgr1
Srb7
Ssn2
Nop1
Nsr1
Nop56
Snu13
Rps31
Rad9
Rtf1
Rpo21
Npl3
Air2
Fps1
Ask10
Cdc28
Hog1
Pbs2
Ssk2
Clb2
Bud3
Bud4
Msn2
Sch9
Rim15
Yak1
Ypl247c
Vps52
Kss1
Sko1
Slt2
Msg5
Swe1
Igo2
Cdc55
Igo1
Vps53
Vps54
Rtn1
Fmp42
Cot1
Zrc1
10
20
30
40
50
t/s

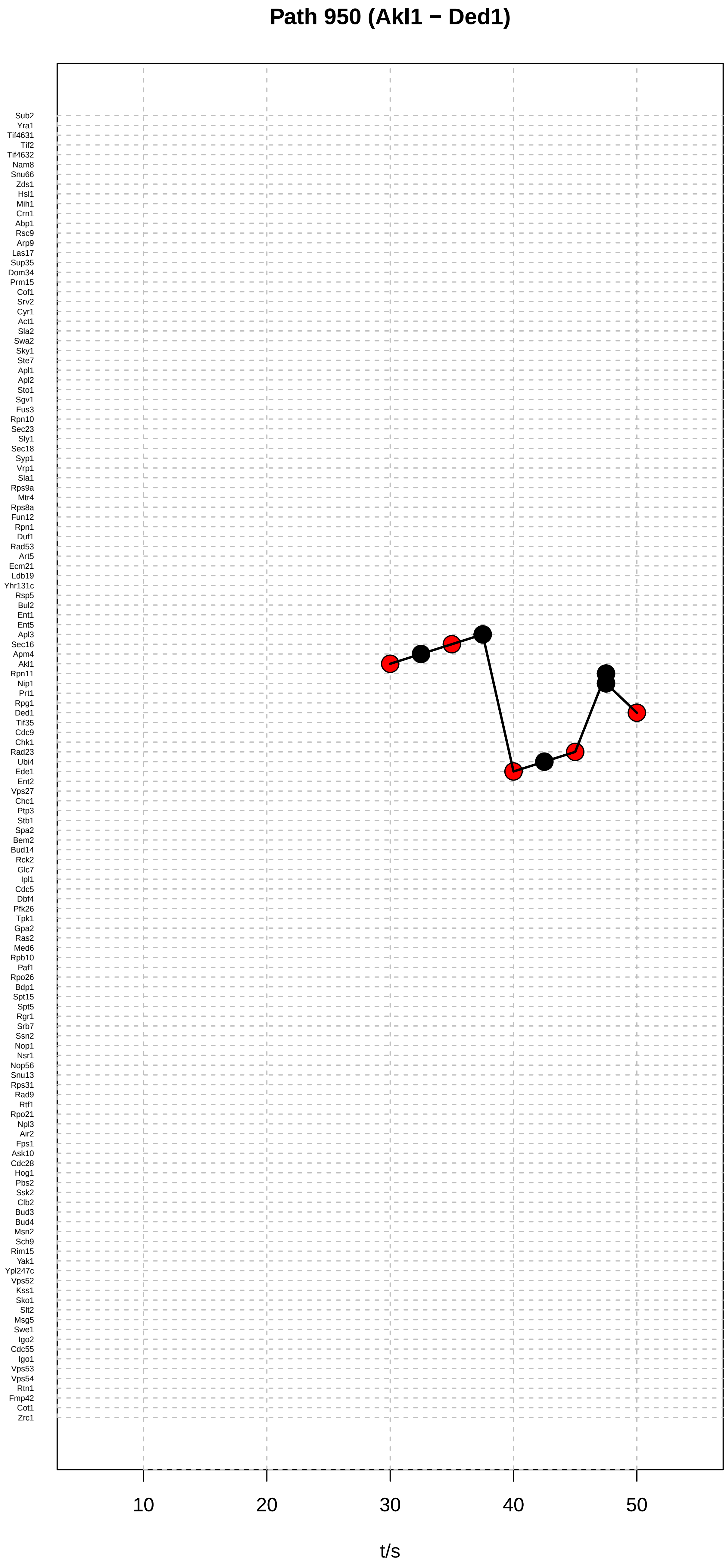

Path 950 (Akl1 − Ded1)
Sub2
Yra1
Tif4631
Tif2
Tif4632
Nam8
Snu66
Zds1
Hsl1
Mih1
Crn1
Abp1
Rsc9
Arp9
Las17
Sup35
Dom34
Prm15
Cof1
Srv2
Cyr1
Act1
Sla2
Swa2
Sky1
Ste7
Apl1
Apl2
Sto1
Sgv1
Fus3
Rpn10
Sec23
Sly1
Sec18
Syp1
Vrp1
Sla1
Rps9a
Mtr4
Rps8a
Fun12
Rpn1
Duf1
Rad53
Art5
Ecm21
Ldb19
Yhr131c
Rsp5
Bul2
Ent1
Ent5
Apl3
Sec16
Apm4
Akl1
Rpn11
Nip1
Prt1
Rpg1
Ded1
Tif35
Cdc9
Chk1
Rad23
Ubi4
Ede1
Ent2
Vps27
Chc1
Ptp3
Stb1
Spa2
Bem2
Bud14
Rck2
Glc7
Ipl1
Cdc5
Dbf4
Pfk26
Tpk1
Gpa2
Ras2
Med6
Rpb10
Paf1
Rpo26
Bdp1
Spt15
Spt5
Rgr1
Srb7
Ssn2
Nop1
Nsr1
Nop56
Snu13
Rps31
Rad9
Rtf1
Rpo21
Npl3
Air2
Fps1
Ask10
Cdc28
Hog1
Pbs2
Ssk2
Clb2
Bud3
Bud4
Msn2
Sch9
Rim15
Yak1
Ypl247c
Vps52
Kss1
Sko1
Slt2
Msg5
Swe1
Igo2
Cdc55
Igo1
Vps53
Vps54
Rtn1
Fmp42
Cot1
Zrc1
10
20
30
40
50
t/s

### Path 951 (Akl1 − Rad23)

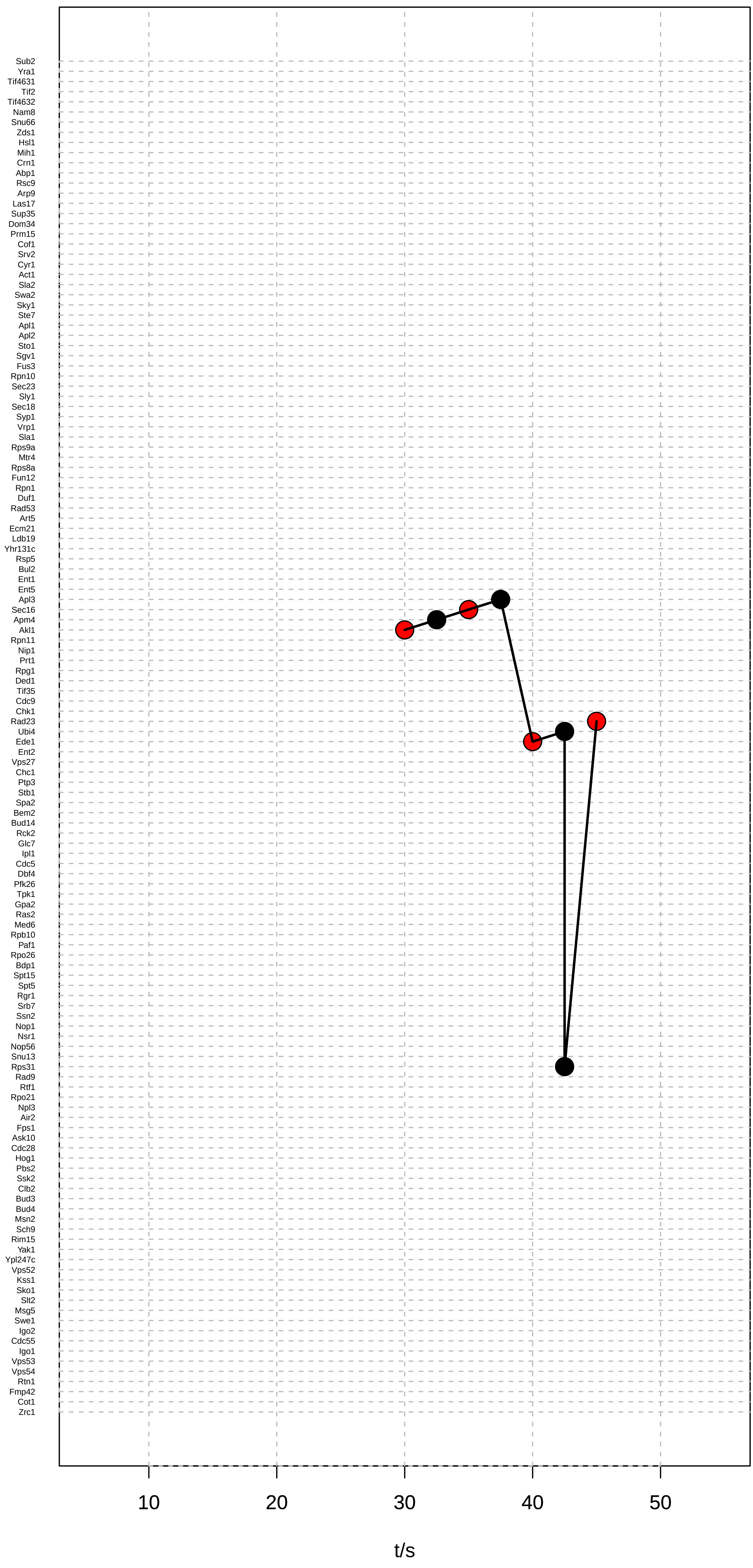

### Path 952 (Akl1 − Ded1)

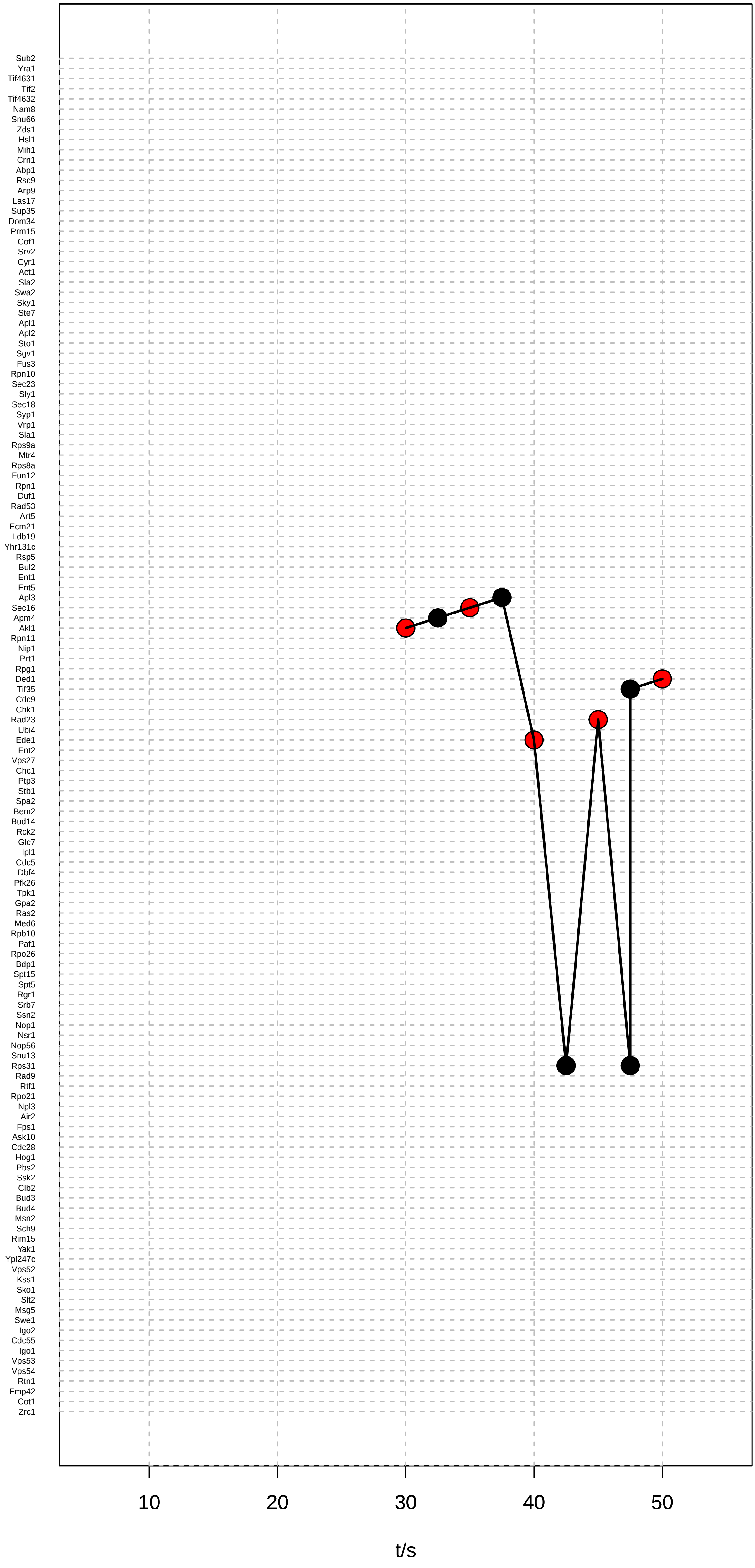

### Path 953 (Akl1 − Ded1)

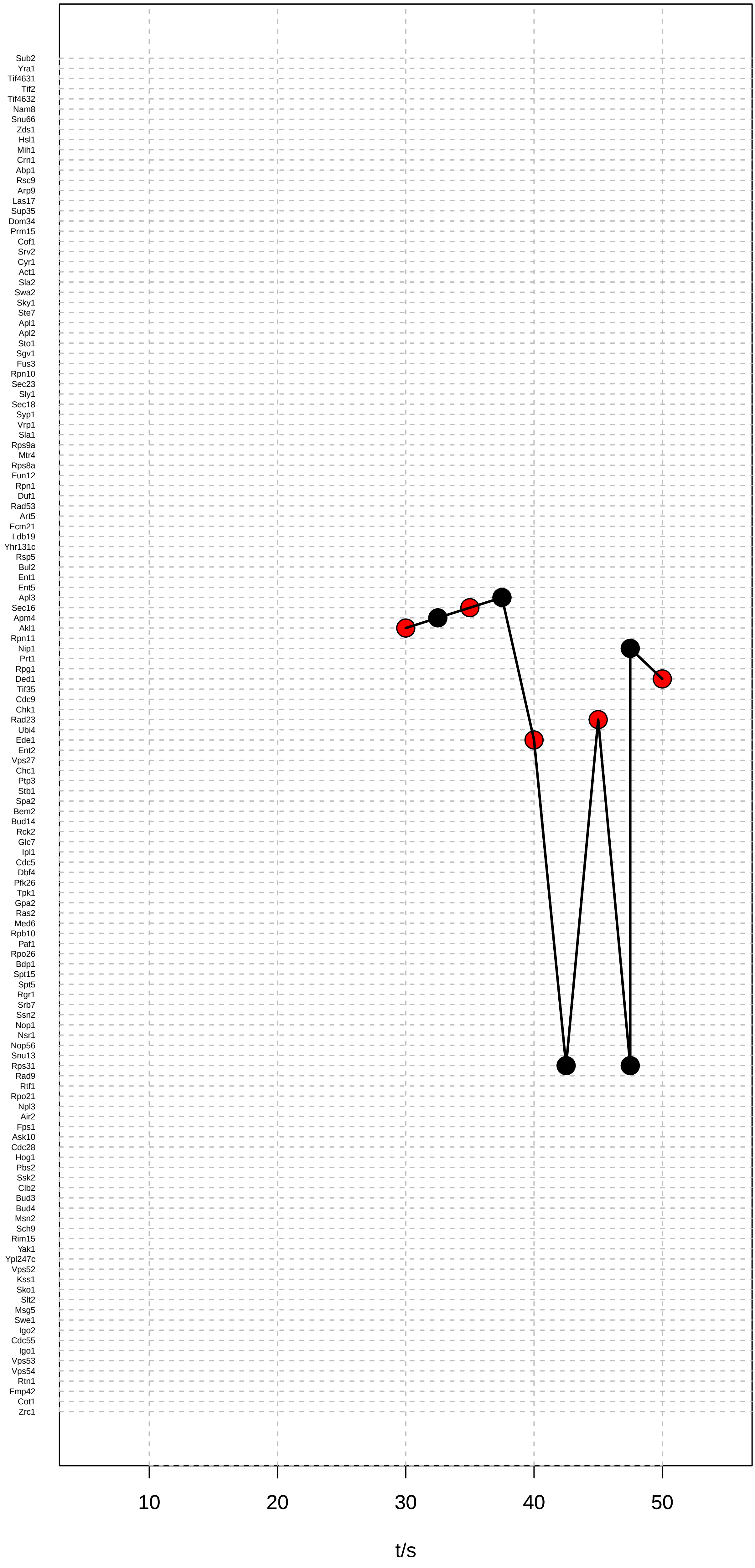

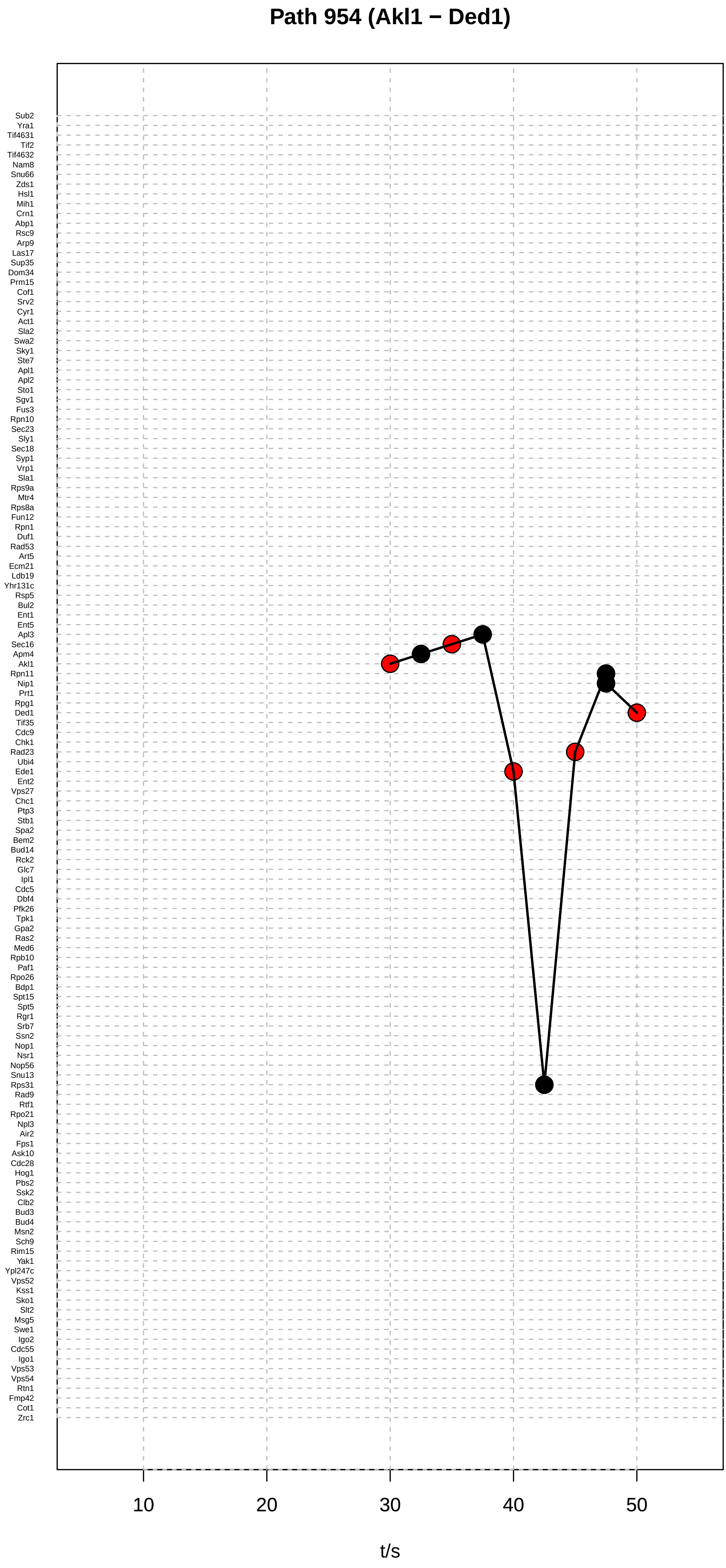
Path 954 (Akl1 − Ded1)
Sub2
Yra1
Tif4631
Tif2
Tif4632
Nam8
Snu66
Zds1
Hsl1
Mih1
Crn1
Abp1
Rsc9
Arp9
Las17
Sup35
Dom34
Prm15
Cof1
Srv2
Cyr1
Act1
Sla2
Swa2
Sky1
Ste7
Apl1
Apl2
Sto1
Sgv1
Fus3
Rpn10
Sec23
Sly1
Sec18
Syp1
Vrp1
Sla1
Rps9a
Mtr4
Rps8a
Fun12
Rpn1
Duf1
Rad53
Art5
Ecm21
Ldb19
Yhr131c
Rsp5
Bul2
Ent1
Ent5
Apl3
Sec16
Apm4
Akl1
Rpn11
Nip1
Prt1
Rpg1
Ded1
Tif35
Cdc9
Chk1
Rad23
Ubi4
Ede1
Ent2
Vps27
Chc1
Ptp3
Stb1
Spa2
Bem2
Bud14
Rck2
Glc7
Ipl1
Cdc5
Dbf4
Pfk26
Tpk1
Gpa2
Ras2
Med6
Rpb10
Paf1
Rpo26
Bdp1
Spt15
Spt5
Rgr1
Srb7
Ssn2
Nop1
Nsr1
Nop56
Snu13
Rps31
Rad9
Rtf1
Rpo21
Npl3
Air2
Fps1
Ask10
Cdc28
Hog1
Pbs2
Ssk2
Clb2
Bud3
Bud4
Msn2
Sch9
Rim15
Yak1
Ypl247c
Vps52
Kss1
Sko1
Slt2
Msg5
Swe1
Igo2
Cdc55
Igo1
Vps53
Vps54
Rtn1
Fmp42
Cot1
Zrc1
10
20
30
40
50
t/s

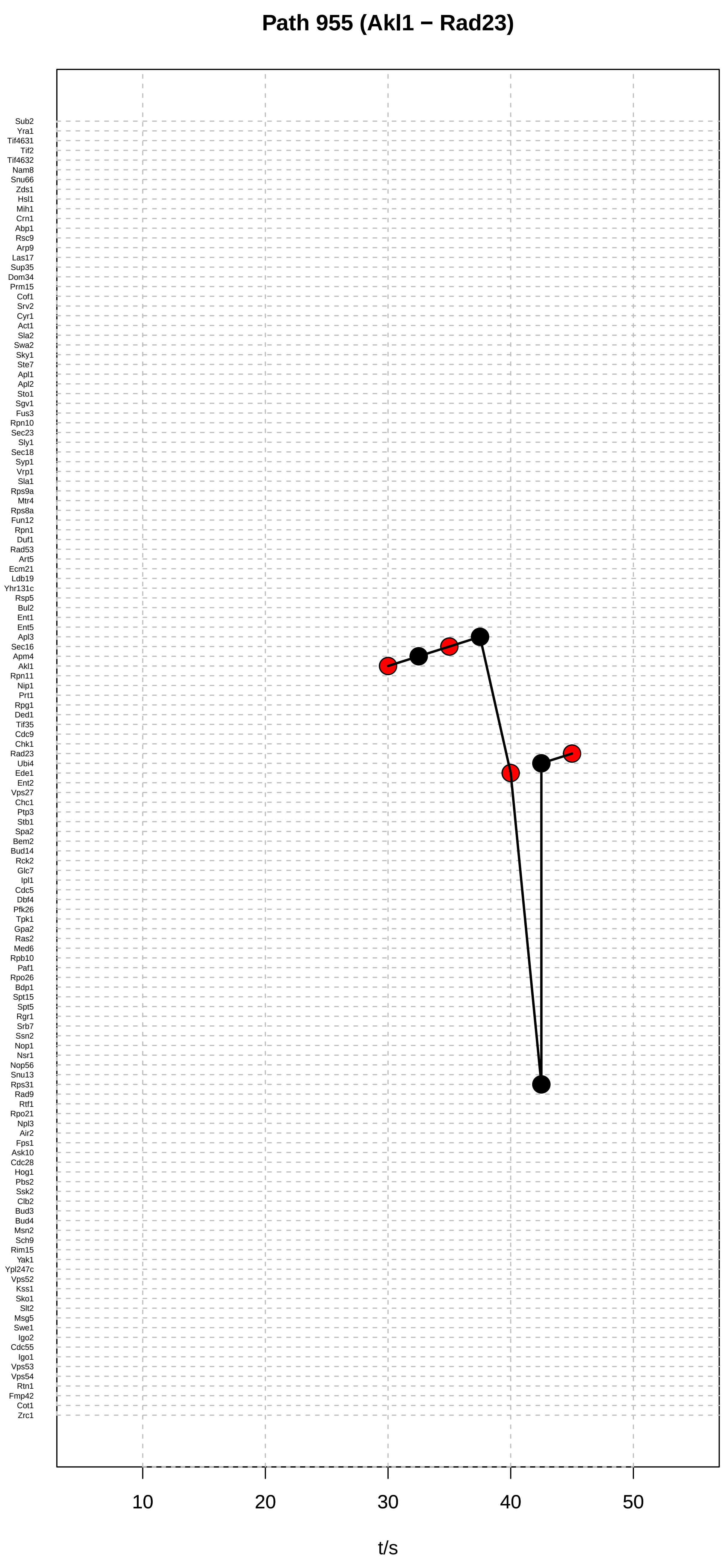

Path 955 (Akl1 − Rad23)
Sub2
Yra1
Tif4631
Tif2
Tif4632
Nam8
Snu66
Zds1
Hsl1
Mih1
Crn1
Abp1
Rsc9
Arp9
Las17
Sup35
Dom34
Prm15
Cof1
Srv2
Cyr1
Act1
Sla2
Swa2
Sky1
Ste7
Apl1
Apl2
Sto1
Sgv1
Fus3
Rpn10
Sec23
Sly1
Sec18
Syp1
Vrp1
Sla1
Rps9a
Mtr4
Rps8a
Fun12
Rpn1
Duf1
Rad53
Art5
Ecm21
Ldb19
Yhr131c
Rsp5
Bul2
Ent1
Ent5
Apl3
Sec16
Apm4
Akl1
Rpn11
Nip1
Prt1
Rpg1
Ded1
Tif35
Cdc9
Chk1
Rad23
Ubi4
Ede1
Ent2
Vps27
Chc1
Ptp3
Stb1
Spa2
Bem2
Bud14
Rck2
Glc7
Ipl1
Cdc5
Dbf4
Pfk26
Tpk1
Gpa2
Ras2
Med6
Rpb10
Paf1
Rpo26
Bdp1
Spt15
Spt5
Rgr1
Srb7
Ssn2
Nop1
Nsr1
Nop56
Snu13
Rps31
Rad9
Rtf1
Rpo21
Npl3
Air2
Fps1
Ask10
Cdc28
Hog1
Pbs2
Ssk2
Clb2
Bud3
Bud4
Msn2
Sch9
Rim15
Yak1
Ypl247c
Vps52
Kss1
Sko1
Slt2
Msg5
Swe1
Igo2
Cdc55
Igo1
Vps53
Vps54
Rtn1
Fmp42
Cot1
Zrc1
10
20
30
40
50
t/s

### Path 956 (Akl1 − Rad23)

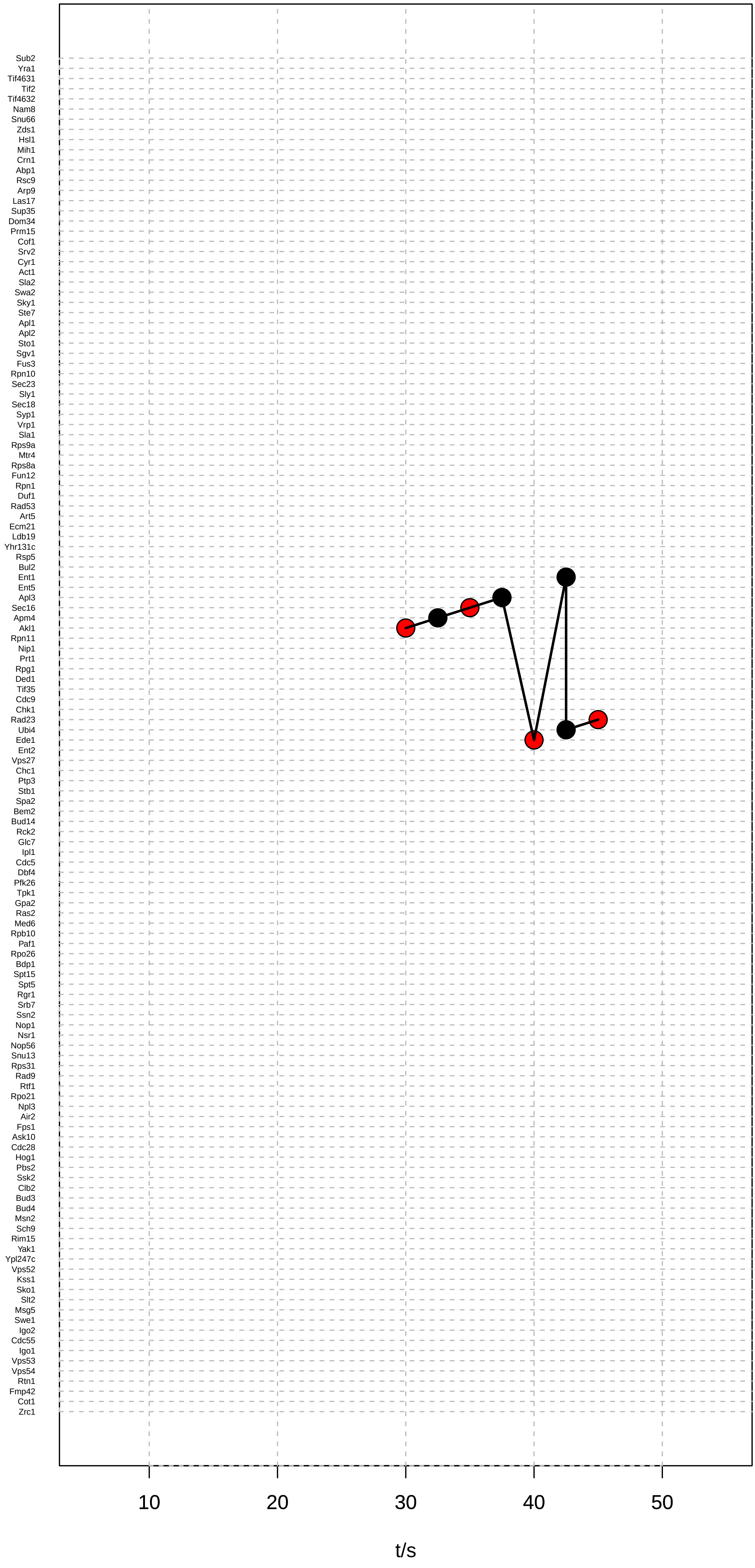

#### Path 957 (Akl1 − Rad23)

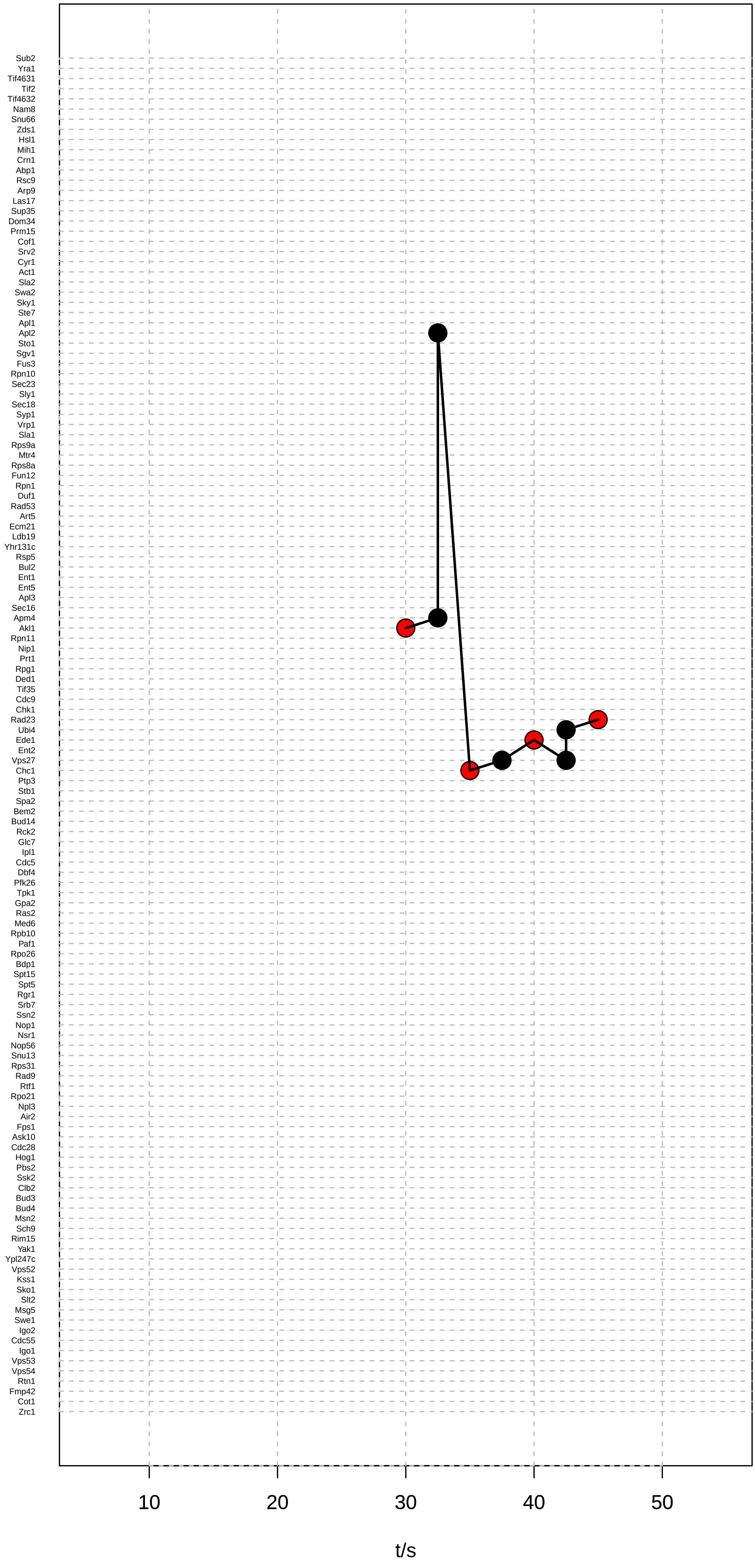

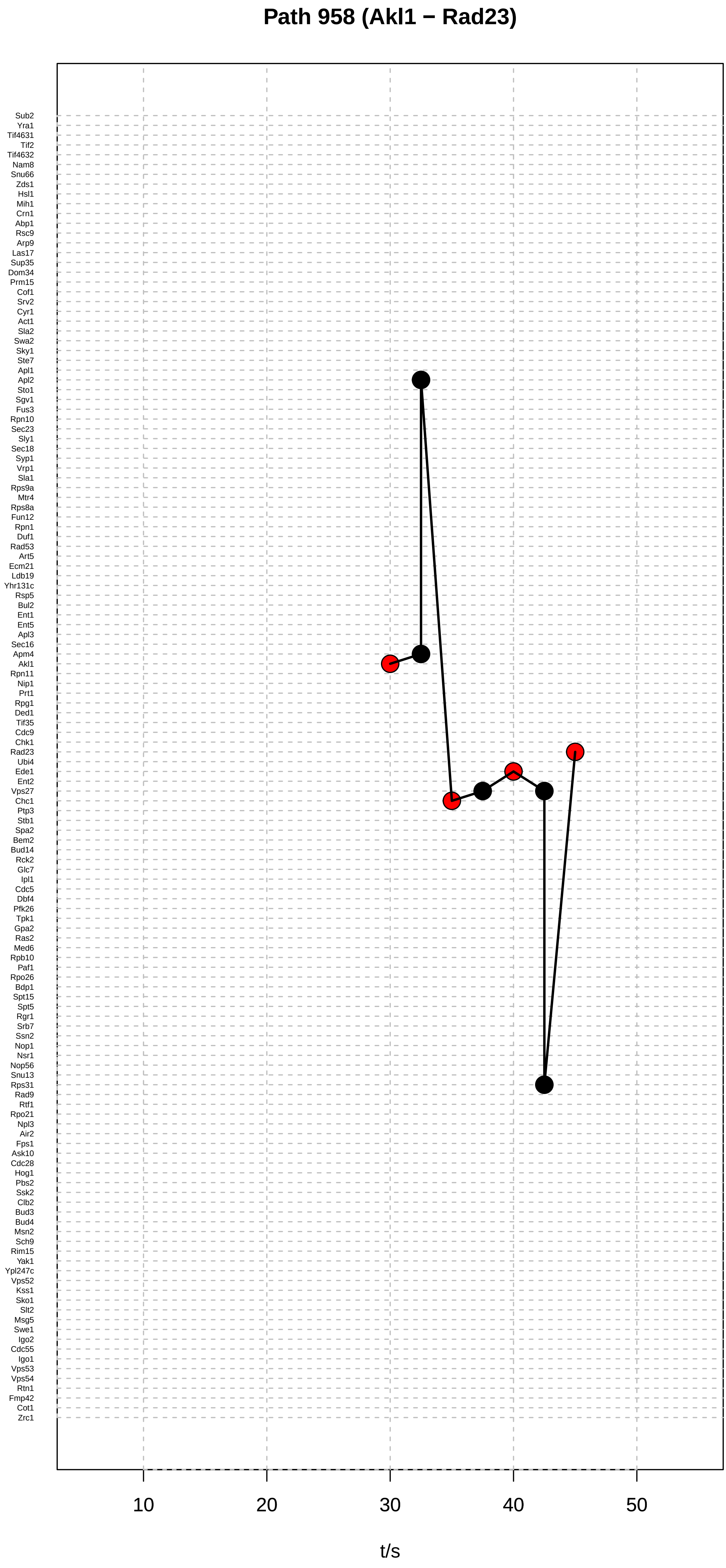
Path 958 (Akl1 − Rad23)
Sub2
Yra1
Tif4631
Tif2
Tif4632
Nam8
Snu66
Zds1
Hsl1
Mih1
Crn1
Abp1
Rsc9
Arp9
Las17
Sup35
Dom34
Prm15
Cof1
Srv2
Cyr1
Act1
Sla2
Swa2
Sky1
Ste7
Apl1
Apl2
Sto1
Sgv1
Fus3
Rpn10
Sec23
Sly1
Sec18
Syp1
Vrp1
Sla1
Rps9a
Mtr4
Rps8a
Fun12
Rpn1
Duf1
Rad53
Art5
Ecm21
Ldb19
Yhr131c
Rsp5
Bul2
Ent1
Ent5
Apl3
Sec16
Apm4
Akl1
Rpn11
Nip1
Prt1
Rpg1
Ded1
Tif35
Cdc9
Chk1
Rad23
Ubi4
Ede1
Ent2
Vps27
Chc1
Ptp3
Stb1
Spa2
Bem2
Bud14
Rck2
Glc7
Ipl1
Cdc5
Dbf4
Pfk26
Tpk1
Gpa2
Ras2
Med6
Rpb10
Paf1
Rpo26
Bdp1
Spt15
Spt5
Rgr1
Srb7
Ssn2
Nop1
Nsr1
Nop56
Snu13
Rps31
Rad9
Rtf1
Rpo21
Npl3
Air2
Fps1
Ask10
Cdc28
Hog1
Pbs2
Ssk2
Clb2
Bud3
Bud4
Msn2
Sch9
Rim15
Yak1
Ypl247c
Vps52
Kss1
Sko1
Slt2
Msg5
Swe1
Igo2
Cdc55
Igo1
Vps53
Vps54
Rtn1
Fmp42
Cot1
Zrc1
10
20
30
40
50
t/s

Path 959 (Akl1 − Rad23)

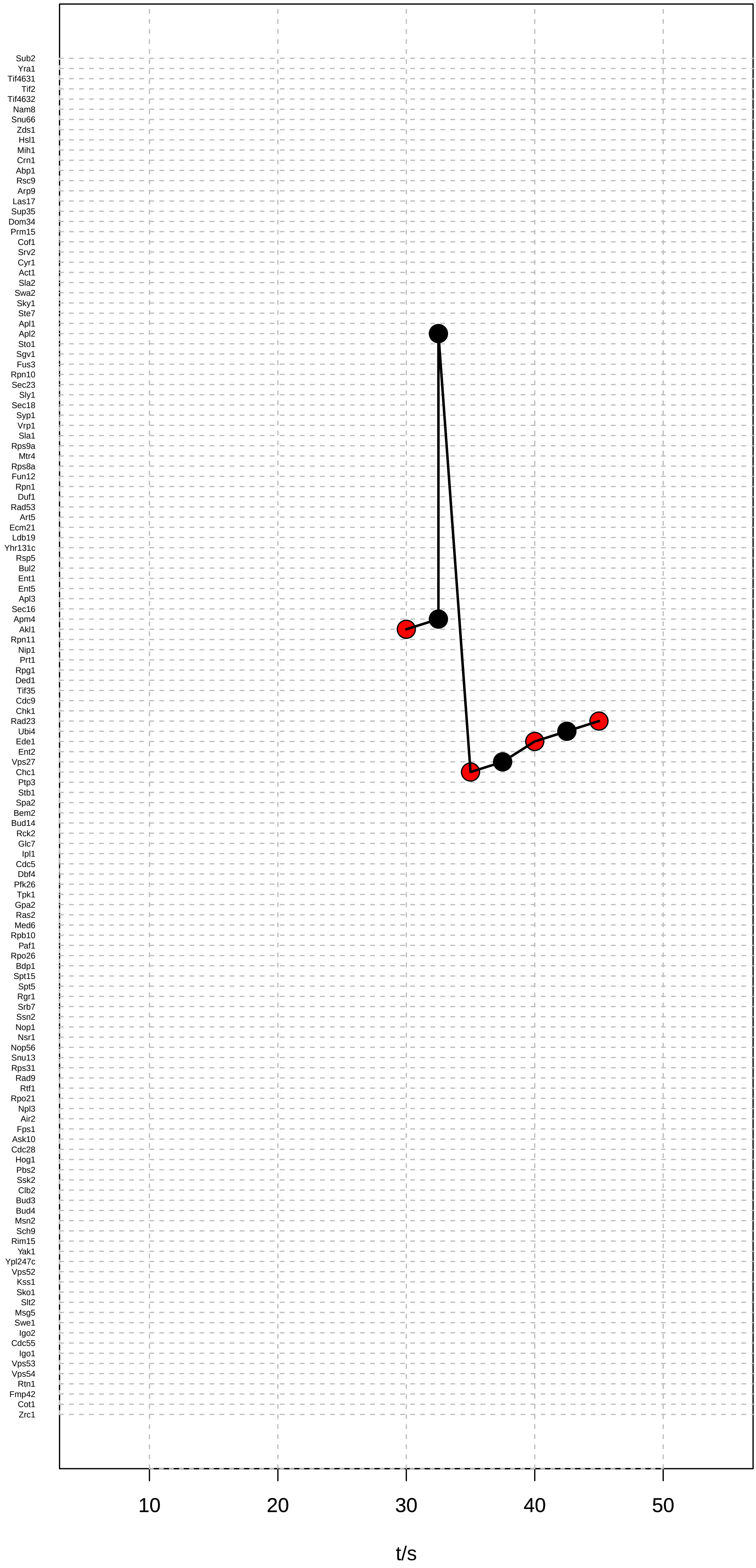

#### Path 960 (Akl1 − Rad23)

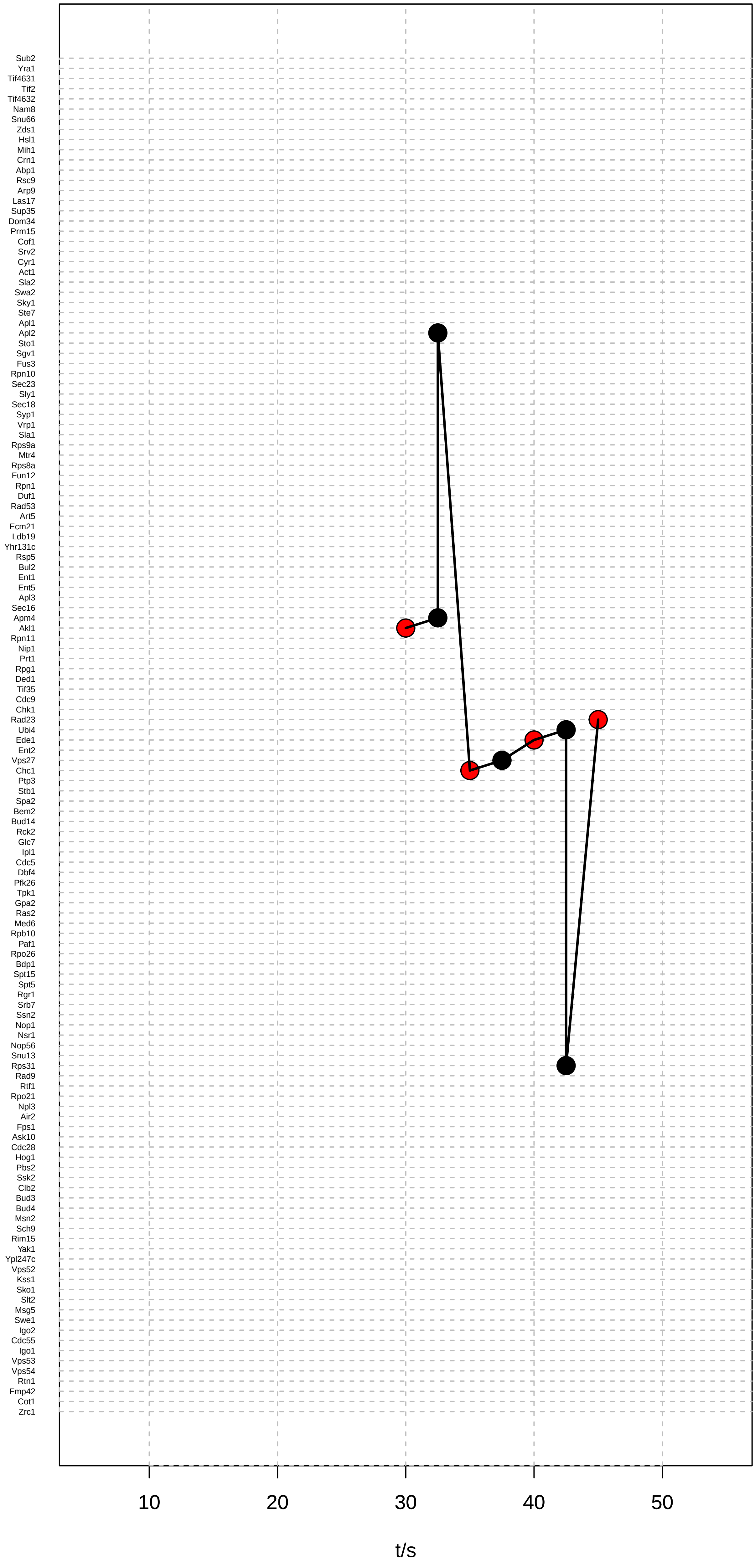

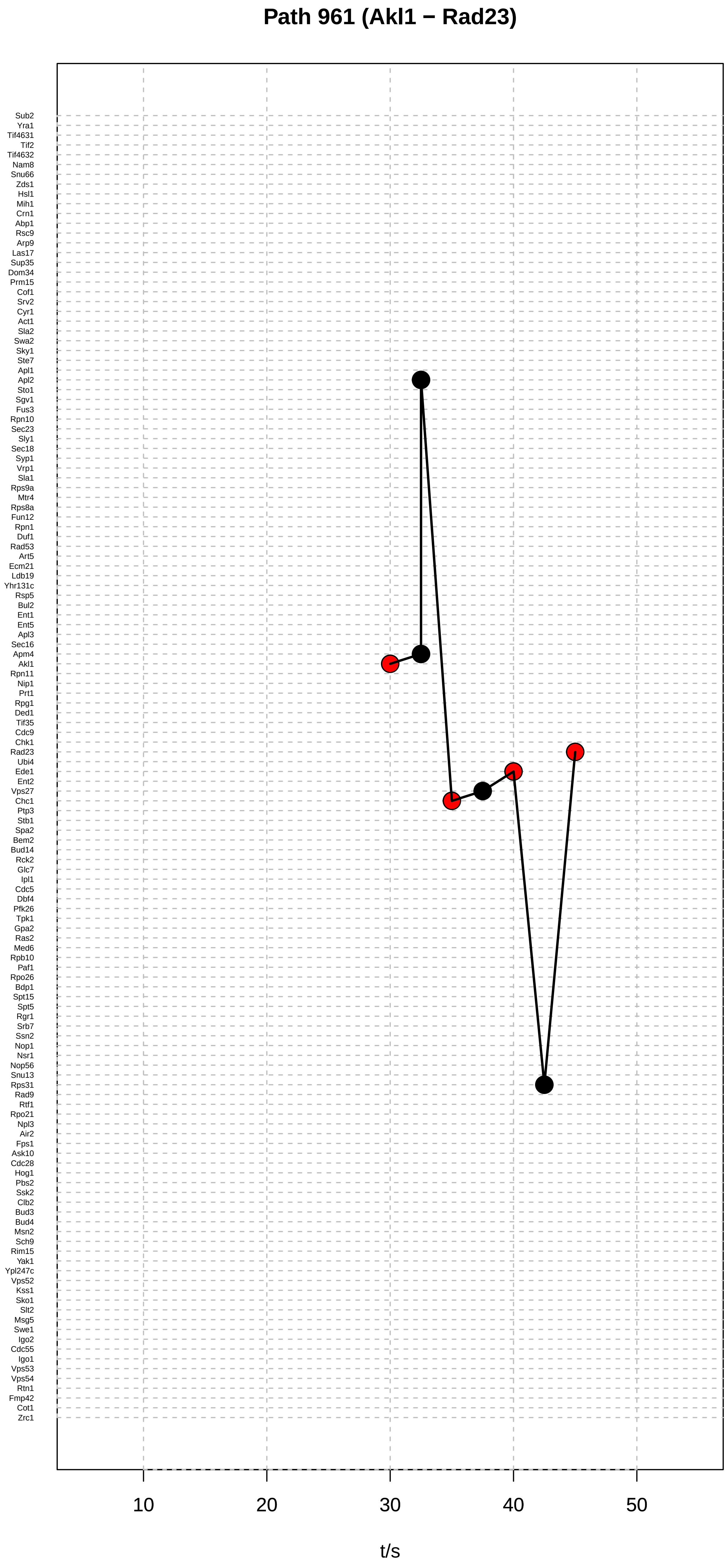

Path 961 (Akl1 − Rad23)
Sub2
Yra1
Tif4631
Tif2
Tif4632
Nam8
Snu66
Zds1
Hsl1
Mih1
Crn1
Abp1
Rsc9
Arp9
Las17
Sup35
Dom34
Prm15
Cof1
Srv2
Cyr1
Act1
Sla2
Swa2
Sky1
Ste7
Apl1
Apl2
Sto1
Sgv1
Fus3
Rpn10
Sec23
Sly1
Sec18
Syp1
Vrp1
Sla1
Rps9a
Mtr4
Rps8a
Fun12
Rpn1
Duf1
Rad53
Art5
Ecm21
Ldb19
Yhr131c
Rsp5
Bul2
Ent1
Ent5
Apl3
Sec16
Apm4
Akl1
Rpn11
Nip1
Prt1
Rpg1
Ded1
Tif35
Cdc9
Chk1
Rad23
Ubi4
Ede1
Ent2
Vps27
Chc1
Ptp3
Stb1
Spa2
Bem2
Bud14
Rck2
Glc7
Ipl1
Cdc5
Dbf4
Pfk26
Tpk1
Gpa2
Ras2
Med6
Rpb10
Paf1
Rpo26
Bdp1
Spt15
Spt5
Rgr1
Srb7
Ssn2
Nop1
Nsr1
Nop56
Snu13
Rps31
Rad9
Rtf1
Rpo21
Npl3
Air2
Fps1
Ask10
Cdc28
Hog1
Pbs2
Ssk2
Clb2
Bud3
Bud4
Msn2
Sch9
Rim15
Yak1
Ypl247c
Vps52
Kss1
Sko1
Slt2
Msg5
Swe1
Igo2
Cdc55
Igo1
Vps53
Vps54
Rtn1
Fmp42
Cot1
Zrc1
10
20
30
40
50
t/s

Path 962 (Akl1 − Rad23)

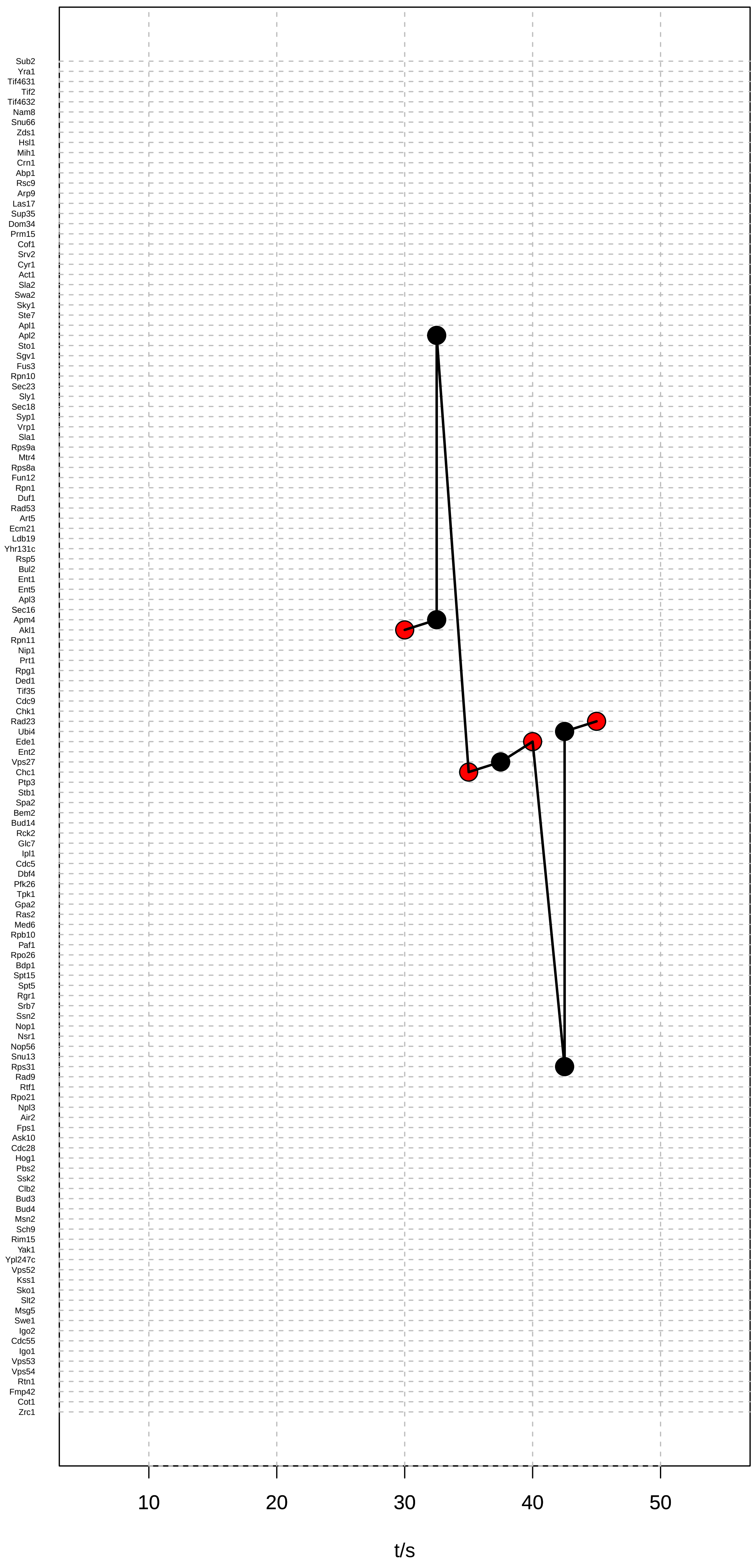

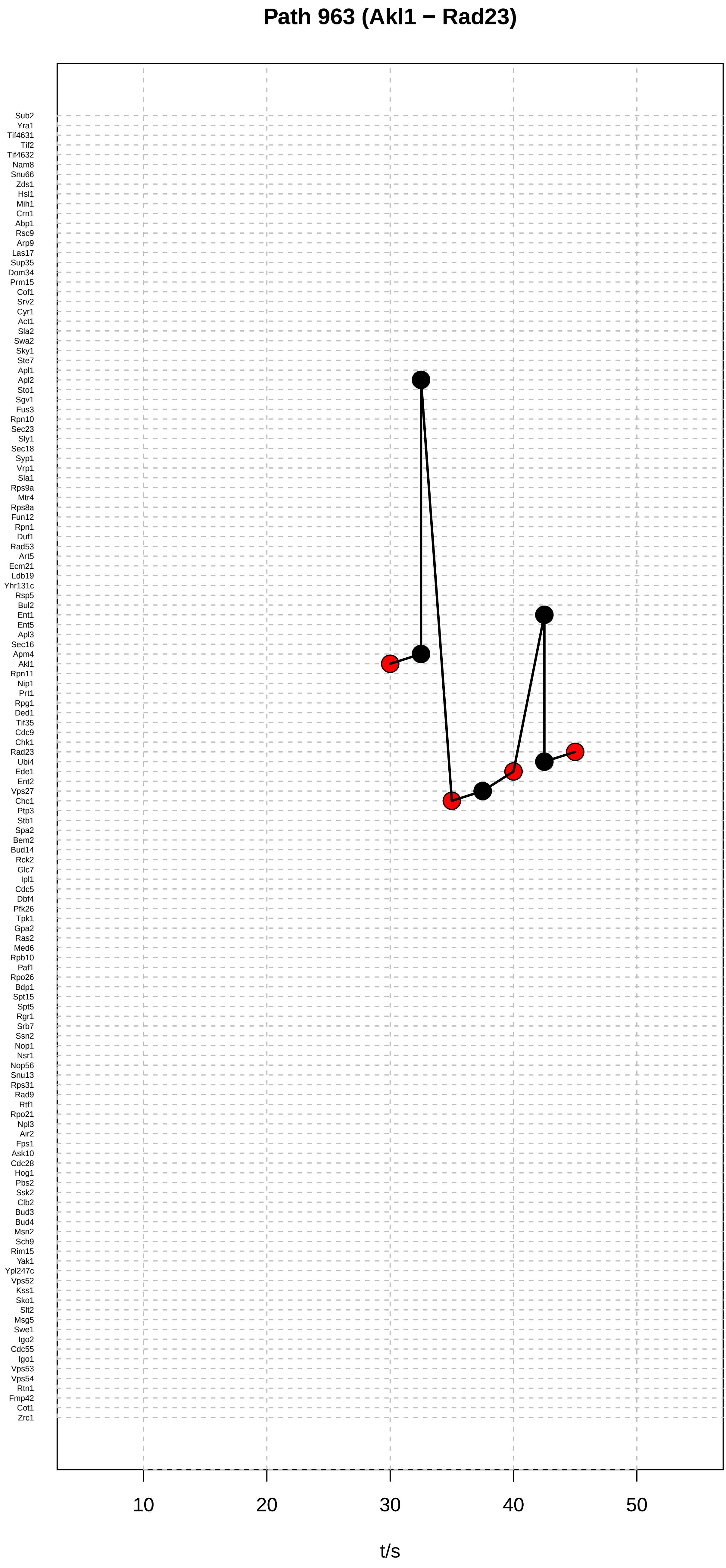
Path 963 (Akl1 − Rad23)
Sub2
Yra1
Tif4631
Tif2
Tif4632
Nam8
Snu66
Zds1
Hsl1
Mih1
Crn1
Abp1
Rsc9
Arp9
Las17
Sup35
Dom34
Prm15
Cof1
Srv2
Cyr1
Act1
Sla2
Swa2
Sky1
Ste7
Apl1
Apl2
Sto1
Sgv1
Fus3
Rpn10
Sec23
Sly1
Sec18
Syp1
Vrp1
Sla1
Rps9a
Mtr4
Rps8a
Fun12
Rpn1
Duf1
Rad53
Art5
Ecm21
Ldb19
Yhr131c
Rsp5
Bul2
Ent1
Ent5
Apl3
Sec16
Apm4
Akl1
Rpn11
Nip1
Prt1
Rpg1
Ded1
Tif35
Cdc9
Chk1
Rad23
Ubi4
Ede1
Ent2
Vps27
Chc1
Ptp3
Stb1
Spa2
Bem2
Bud14
Rck2
Glc7
Ipl1
Cdc5
Dbf4
Pfk26
Tpk1
Gpa2
Ras2
Med6
Rpb10
Paf1
Rpo26
Bdp1
Spt15
Spt5
Rgr1
Srb7
Ssn2
Nop1
Nsr1
Nop56
Snu13
Rps31
Rad9
Rtf1
Rpo21
Npl3
Air2
Fps1
Ask10
Cdc28
Hog1
Pbs2
Ssk2
Clb2
Bud3
Bud4
Msn2
Sch9
Rim15
Yak1
Ypl247c
Vps52
Kss1
Sko1
Slt2
Msg5
Swe1
Igo2
Cdc55
Igo1
Vps53
Vps54
Rtn1
Fmp42
Cot1
Zrc1
10
20
30
40
50
t/s

#### Path 964 (Akl1 − Rad23)

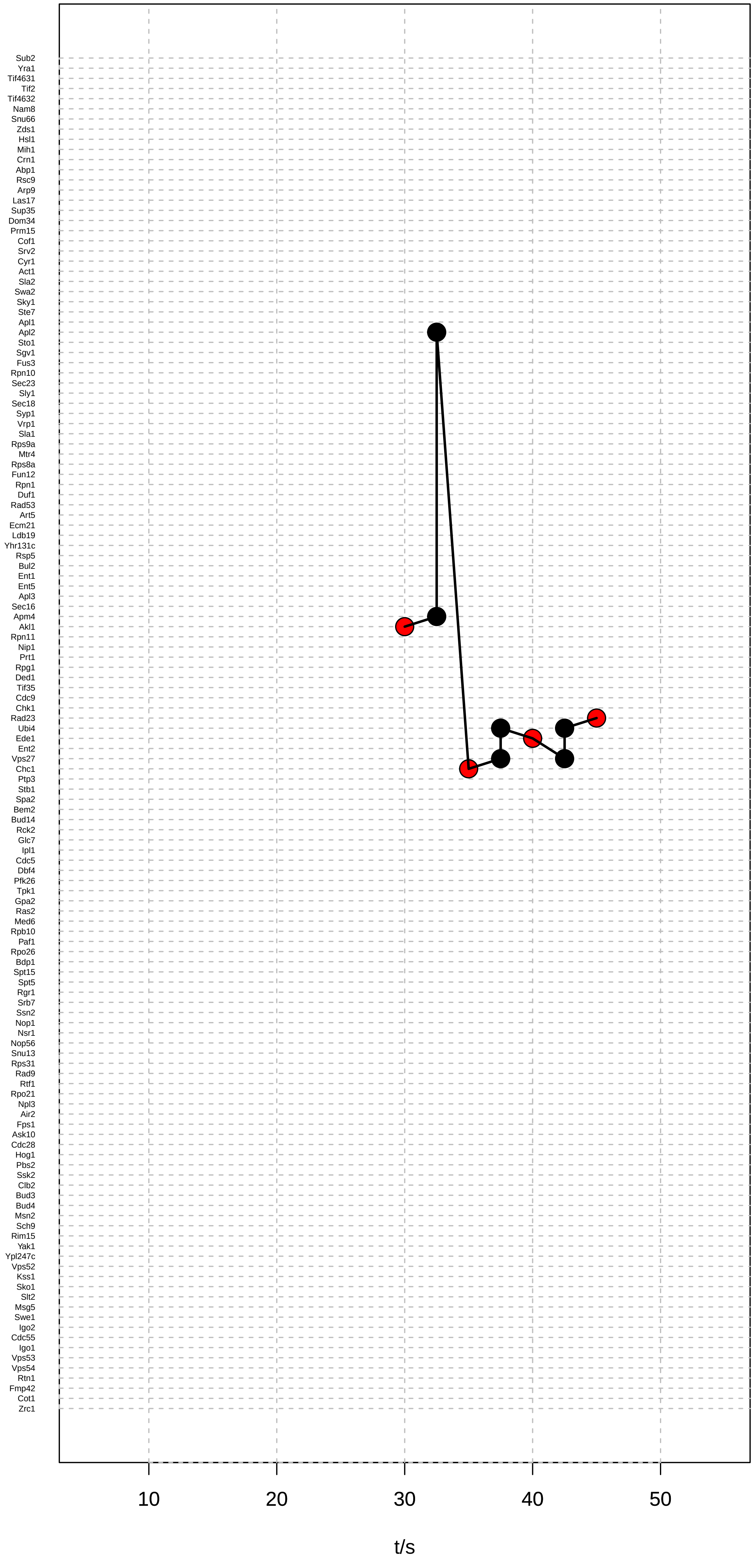

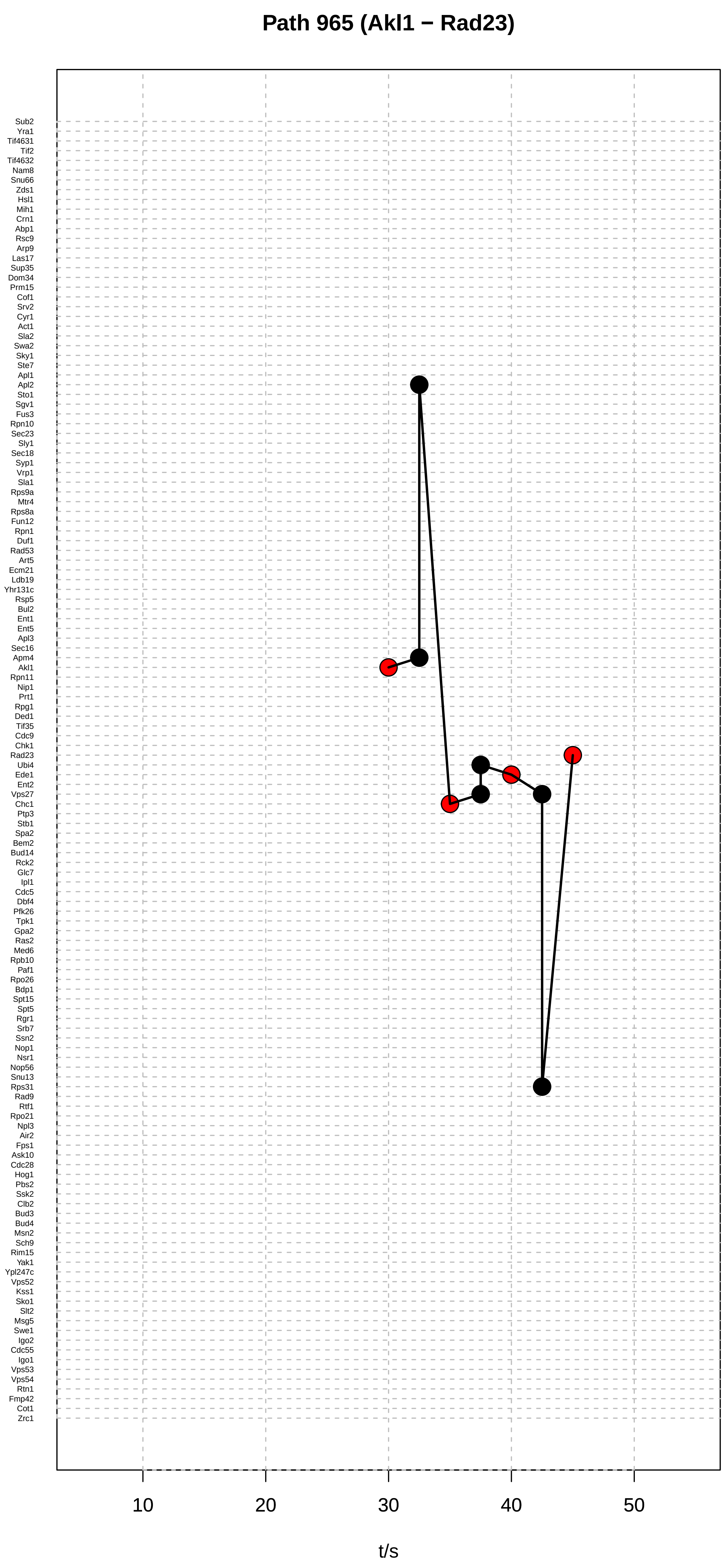
Path 965 (Akl1 − Rad23)
Sub2
Yra1
Tif4631
Tif2
Tif4632
Nam8
Snu66
Zds1
Hsl1
Mih1
Crn1
Abp1
Rsc9
Arp9
Las17
Sup35
Dom34
Prm15
Cof1
Srv2
Cyr1
Act1
Sla2
Swa2
Sky1
Ste7
Apl1
Apl2
Sto1
Sgv1
Fus3
Rpn10
Sec23
Sly1
Sec18
Syp1
Vrp1
Sla1
Rps9a
Mtr4
Rps8a
Fun12
Rpn1
Duf1
Rad53
Art5
Ecm21
Ldb19
Yhr131c
Rsp5
Bul2
Ent1
Ent5
Apl3
Sec16
Apm4
Akl1
Rpn11
Nip1
Prt1
Rpg1
Ded1
Tif35
Cdc9
Chk1
Rad23
Ubi4
Ede1
Ent2
Vps27
Chc1
Ptp3
Stb1
Spa2
Bem2
Bud14
Rck2
Glc7
Ipl1
Cdc5
Dbf4
Pfk26
Tpk1
Gpa2
Ras2
Med6
Rpb10
Paf1
Rpo26
Bdp1
Spt15
Spt5
Rgr1
Srb7
Ssn2
Nop1
Nsr1
Nop56
Snu13
Rps31
Rad9
Rtf1
Rpo21
Npl3
Air2
Fps1
Ask10
Cdc28
Hog1
Pbs2
Ssk2
Clb2
Bud3
Bud4
Msn2
Sch9
Rim15
Yak1
Ypl247c
Vps52
Kss1
Sko1
Slt2
Msg5
Swe1
Igo2
Cdc55
Igo1
Vps53
Vps54
Rtn1
Fmp42
Cot1
Zrc1
10
20
30
40
50
t/s

#### Path 966 (Akl1 − Ent5)

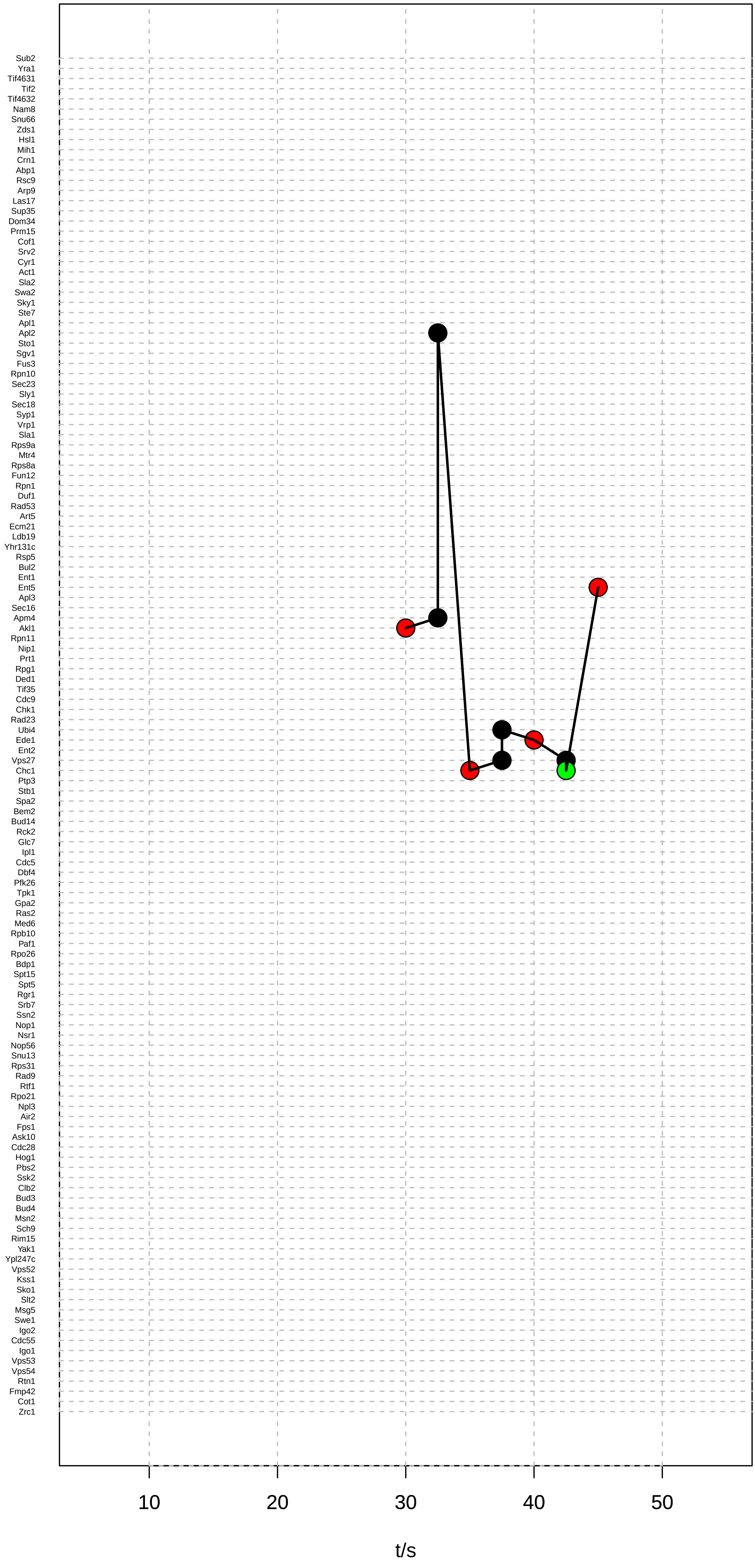

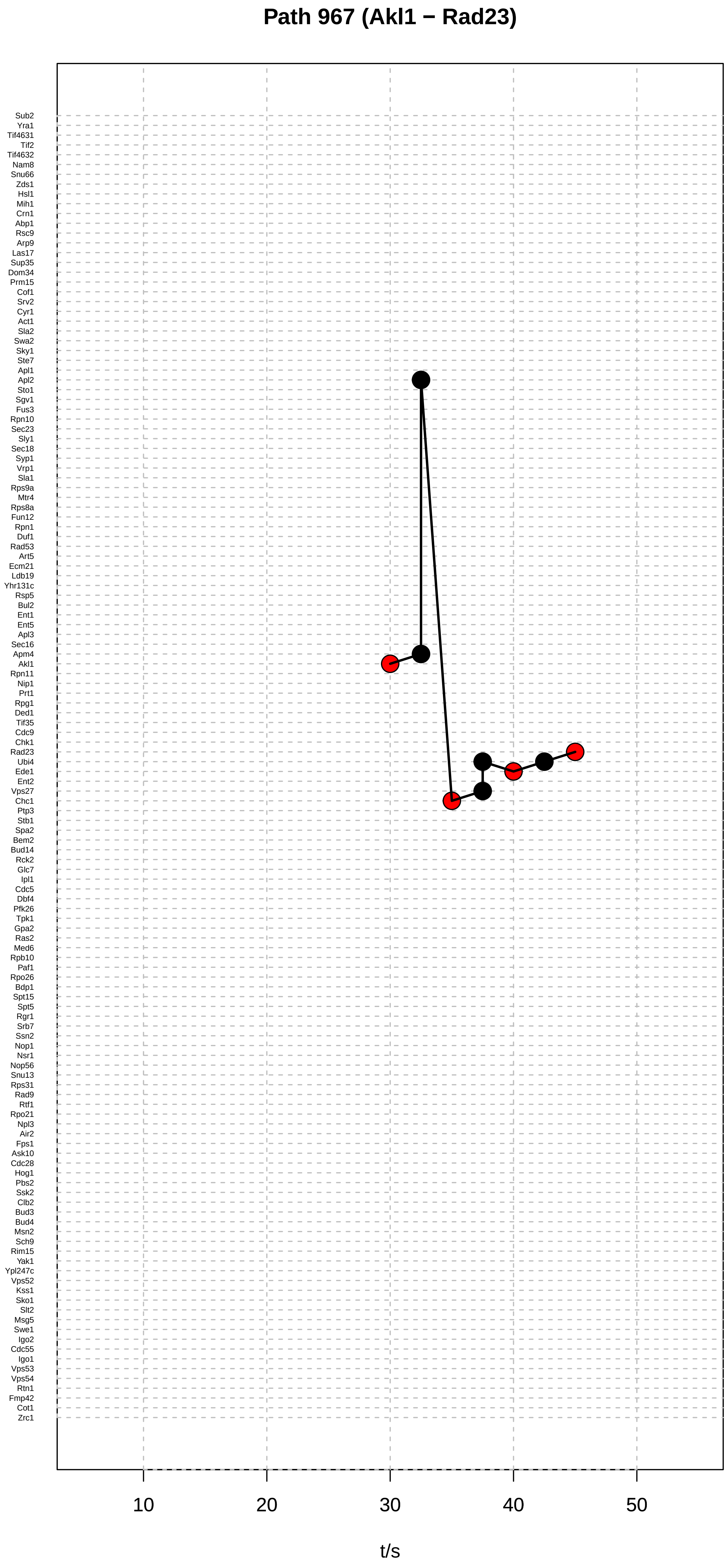

Path 967 (Akl1 − Rad23)
Sub2
Yra1
Tif4631
Tif2
Tif4632
Nam8
Snu66
Zds1
Hsl1
Mih1
Crn1
Abp1
Rsc9
Arp9
Las17
Sup35
Dom34
Prm15
Cof1
Srv2
Cyr1
Act1
Sla2
Swa2
Sky1
Ste7
Apl1
Apl2
Sto1
Sgv1
Fus3
Rpn10
Sec23
Sly1
Sec18
Syp1
Vrp1
Sla1
Rps9a
Mtr4
Rps8a
Fun12
Rpn1
Duf1
Rad53
Art5
Ecm21
Ldb19
Yhr131c
Rsp5
Bul2
Ent1
Ent5
Apl3
Sec16
Apm4
Akl1
Rpn11
Nip1
Prt1
Rpg1
Ded1
Tif35
Cdc9
Chk1
Rad23
Ubi4
Ede1
Ent2
Vps27
Chc1
Ptp3
Stb1
Spa2
Bem2
Bud14
Rck2
Glc7
Ipl1
Cdc5
Dbf4
Pfk26
Tpk1
Gpa2
Ras2
Med6
Rpb10
Paf1
Rpo26
Bdp1
Spt15
Spt5
Rgr1
Srb7
Ssn2
Nop1
Nsr1
Nop56
Snu13
Rps31
Rad9
Rtf1
Rpo21
Npl3
Air2
Fps1
Ask10
Cdc28
Hog1
Pbs2
Ssk2
Clb2
Bud3
Bud4
Msn2
Sch9
Rim15
Yak1
Ypl247c
Vps52
Kss1
Sko1
Slt2
Msg5
Swe1
Igo2
Cdc55
Igo1
Vps53
Vps54
Rtn1
Fmp42
Cot1
Zrc1
10
20
30
40
50
t/s

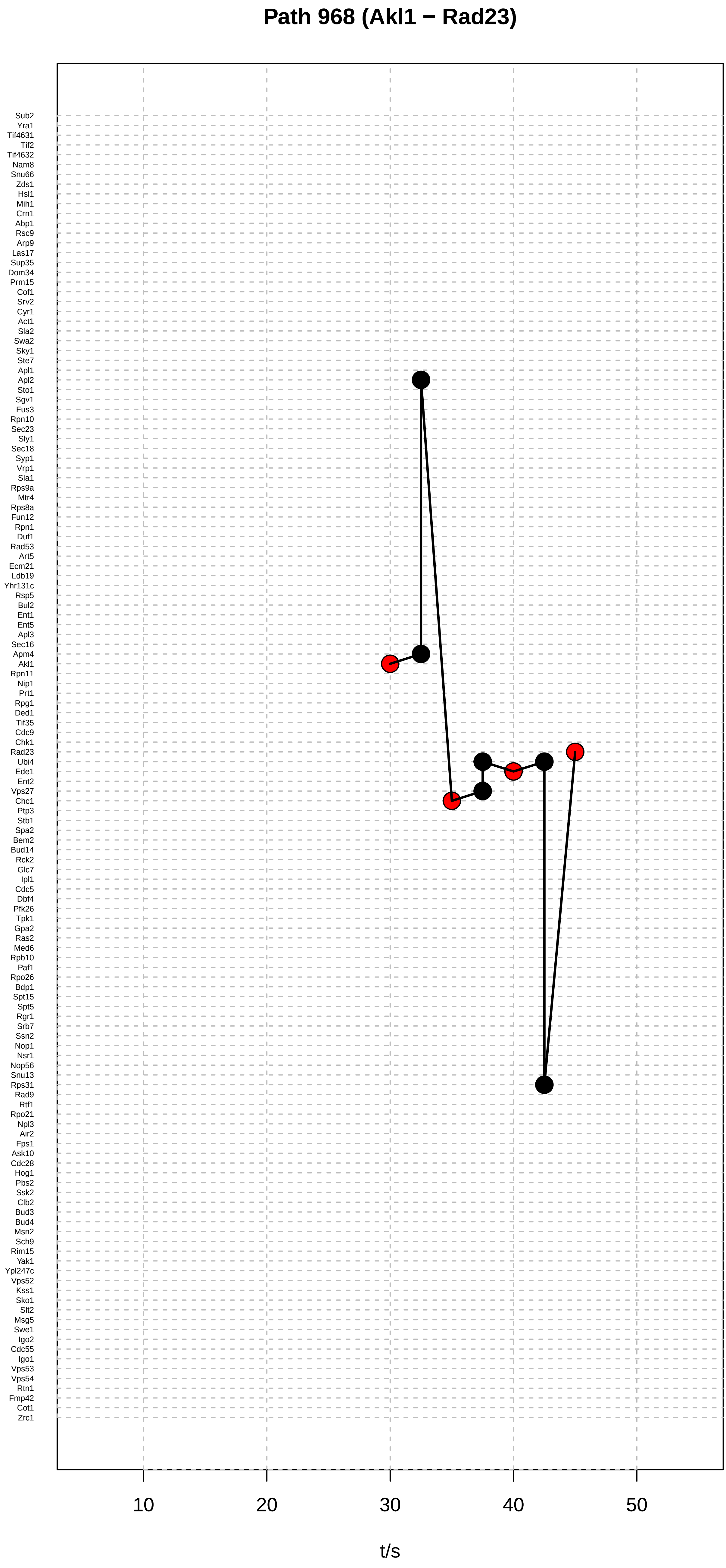

Path 968 (Akl1 − Rad23)
Sub2
Yra1
Tif4631
Tif2
Tif4632
Nam8
Snu66
Zds1
Hsl1
Mih1
Crn1
Abp1
Rsc9
Arp9
Las17
Sup35
Dom34
Prm15
Cof1
Srv2
Cyr1
Act1
Sla2
Swa2
Sky1
Ste7
Apl1
Apl2
Sto1
Sgv1
Fus3
Rpn10
Sec23
Sly1
Sec18
Syp1
Vrp1
Sla1
Rps9a
Mtr4
Rps8a
Fun12
Rpn1
Duf1
Rad53
Art5
Ecm21
Ldb19
Yhr131c
Rsp5
Bul2
Ent1
Ent5
Apl3
Sec16
Apm4
Akl1
Rpn11
Nip1
Prt1
Rpg1
Ded1
Tif35
Cdc9
Chk1
Rad23
Ubi4
Ede1
Ent2
Vps27
Chc1
Ptp3
Stb1
Spa2
Bem2
Bud14
Rck2
Glc7
Ipl1
Cdc5
Dbf4
Pfk26
Tpk1
Gpa2
Ras2
Med6
Rpb10
Paf1
Rpo26
Bdp1
Spt15
Spt5
Rgr1
Srb7
Ssn2
Nop1
Nsr1
Nop56
Snu13
Rps31
Rad9
Rtf1
Rpo21
Npl3
Air2
Fps1
Ask10
Cdc28
Hog1
Pbs2
Ssk2
Clb2
Bud3
Bud4
Msn2
Sch9
Rim15
Yak1
Ypl247c
Vps52
Kss1
Sko1
Slt2
Msg5
Swe1
Igo2
Cdc55
Igo1
Vps53
Vps54
Rtn1
Fmp42
Cot1
Zrc1
10
20
30
40
50
t/s

#### Path 969 (Akl1 − Rad23)

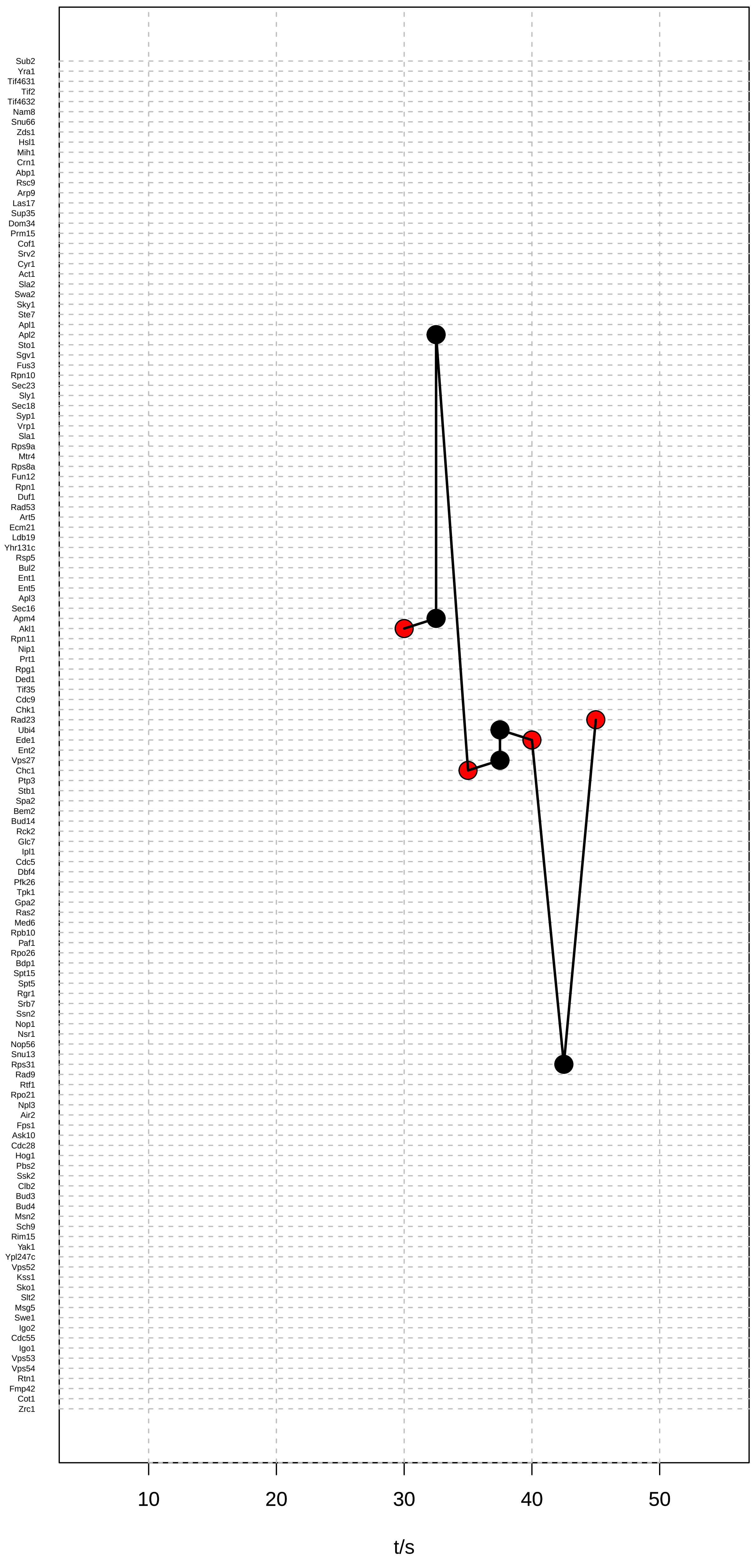

Path 970 (Akl1 − Rad23)

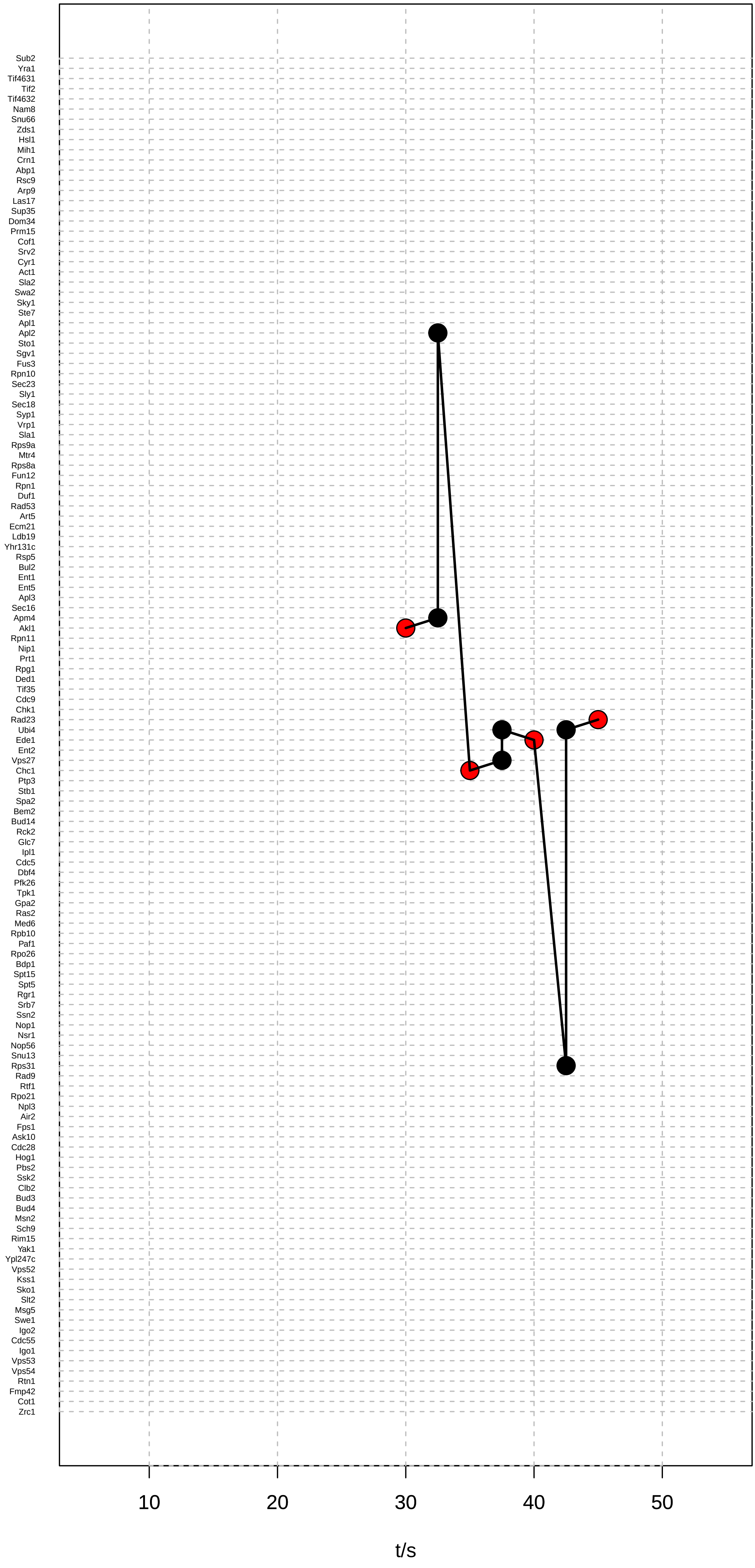

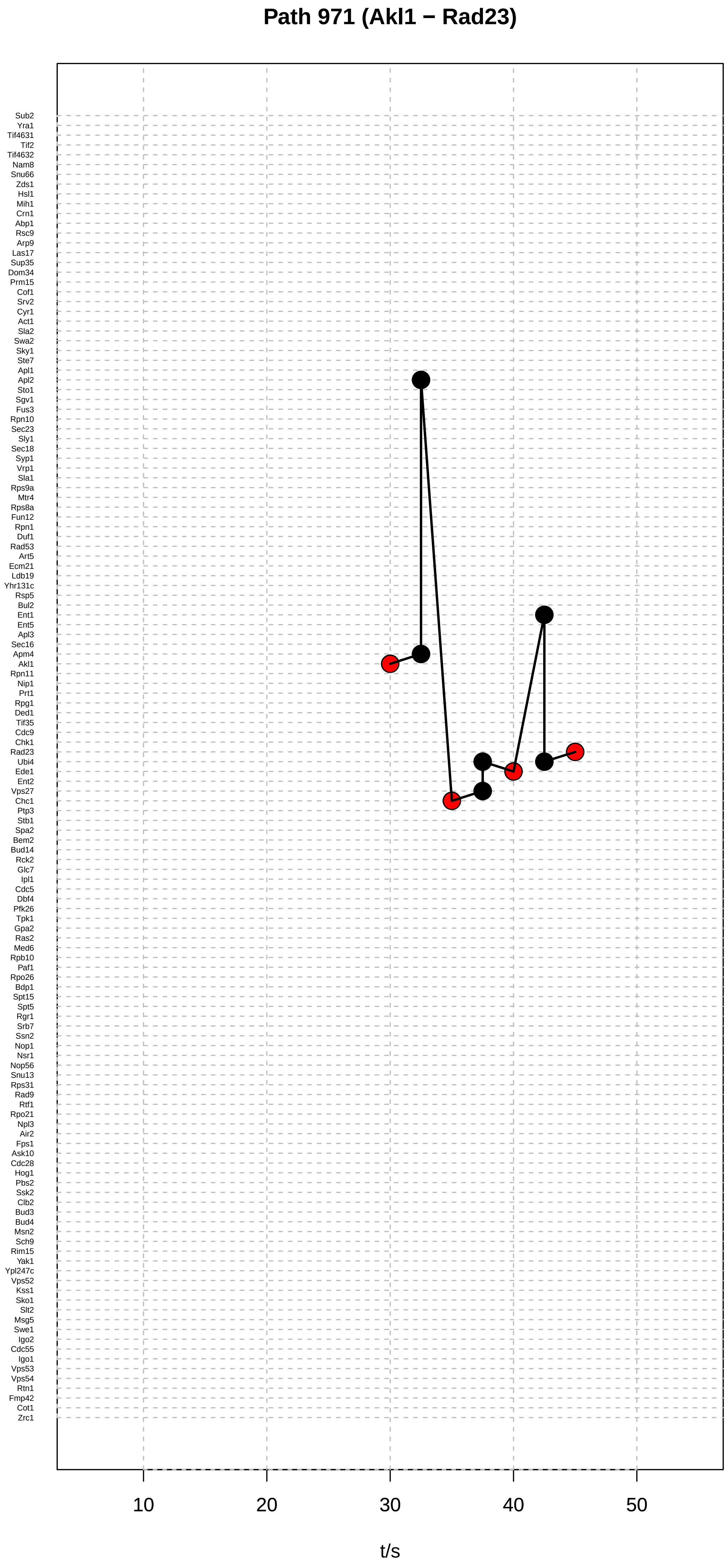
Path 971 (Akl1 − Rad23)
Sub2
Yra1
Tif4631
Tif2
Tif4632
Nam8
Snu66
Zds1
Hsl1
Mih1
Crn1
Abp1
Rsc9
Arp9
Las17
Sup35
Dom34
Prm15
Cof1
Srv2
Cyr1
Act1
Sla2
Swa2
Sky1
Ste7
Apl1
Apl2
Sto1
Sgv1
Fus3
Rpn10
Sec23
Sly1
Sec18
Syp1
Vrp1
Sla1
Rps9a
Mtr4
Rps8a
Fun12
Rpn1
Duf1
Rad53
Art5
Ecm21
Ldb19
Yhr131c
Rsp5
Bul2
Ent1
Ent5
Apl3
Sec16
Apm4
Akl1
Rpn11
Nip1
Prt1
Rpg1
Ded1
Tif35
Cdc9
Chk1
Rad23
Ubi4
Ede1
Ent2
Vps27
Chc1
Ptp3
Stb1
Spa2
Bem2
Bud14
Rck2
Glc7
Ipl1
Cdc5
Dbf4
Pfk26
Tpk1
Gpa2
Ras2
Med6
Rpb10
Paf1
Rpo26
Bdp1
Spt15
Spt5
Rgr1
Srb7
Ssn2
Nop1
Nsr1
Nop56
Snu13
Rps31
Rad9
Rtf1
Rpo21
Npl3
Air2
Fps1
Ask10
Cdc28
Hog1
Pbs2
Ssk2
Clb2
Bud3
Bud4
Msn2
Sch9
Rim15
Yak1
Ypl247c
Vps52
Kss1
Sko1
Slt2
Msg5
Swe1
Igo2
Cdc55
Igo1
Vps53
Vps54
Rtn1
Fmp42
Cot1
Zrc1
10
20
30
40
50
t/s

#### Path 972 (Akl1 − Ent5)

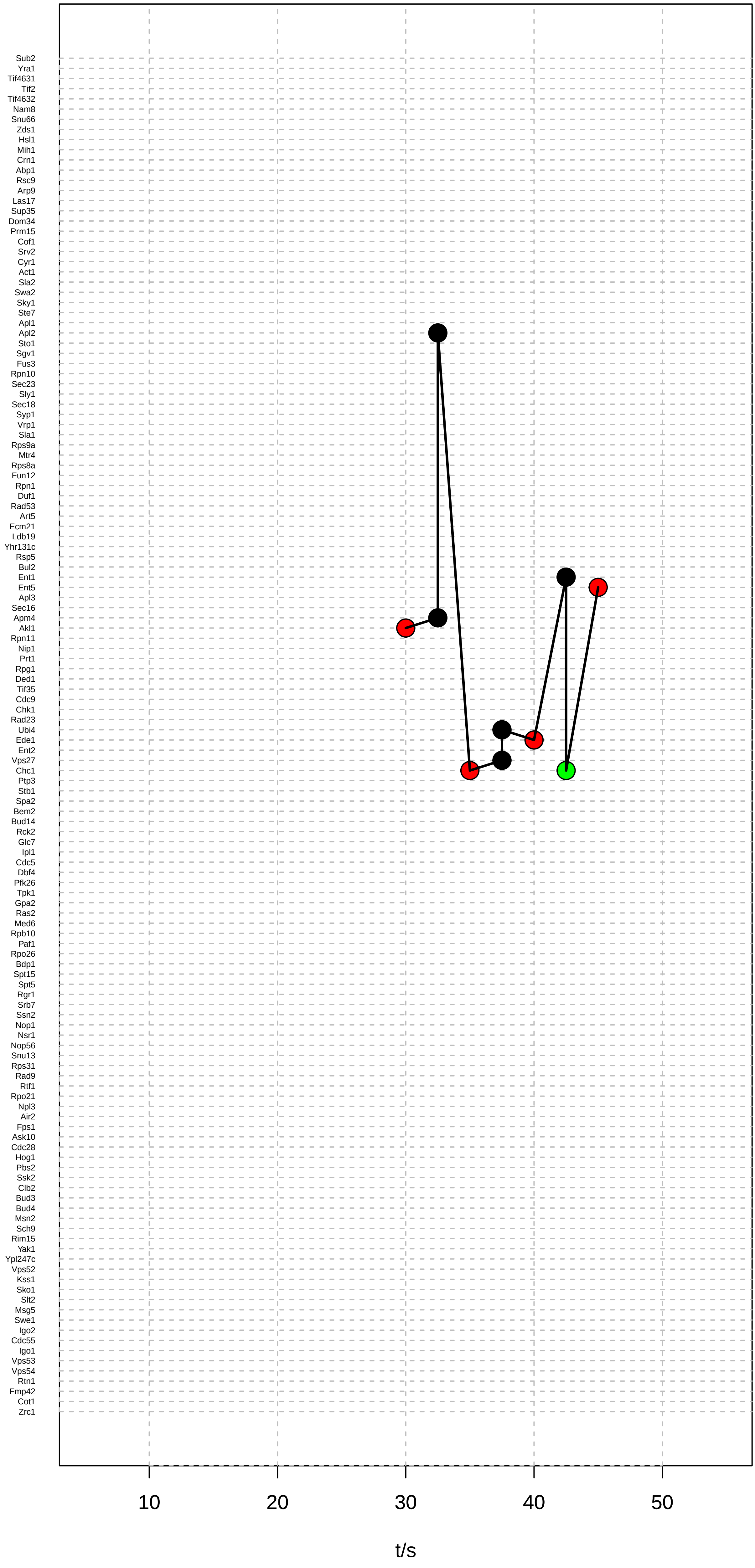

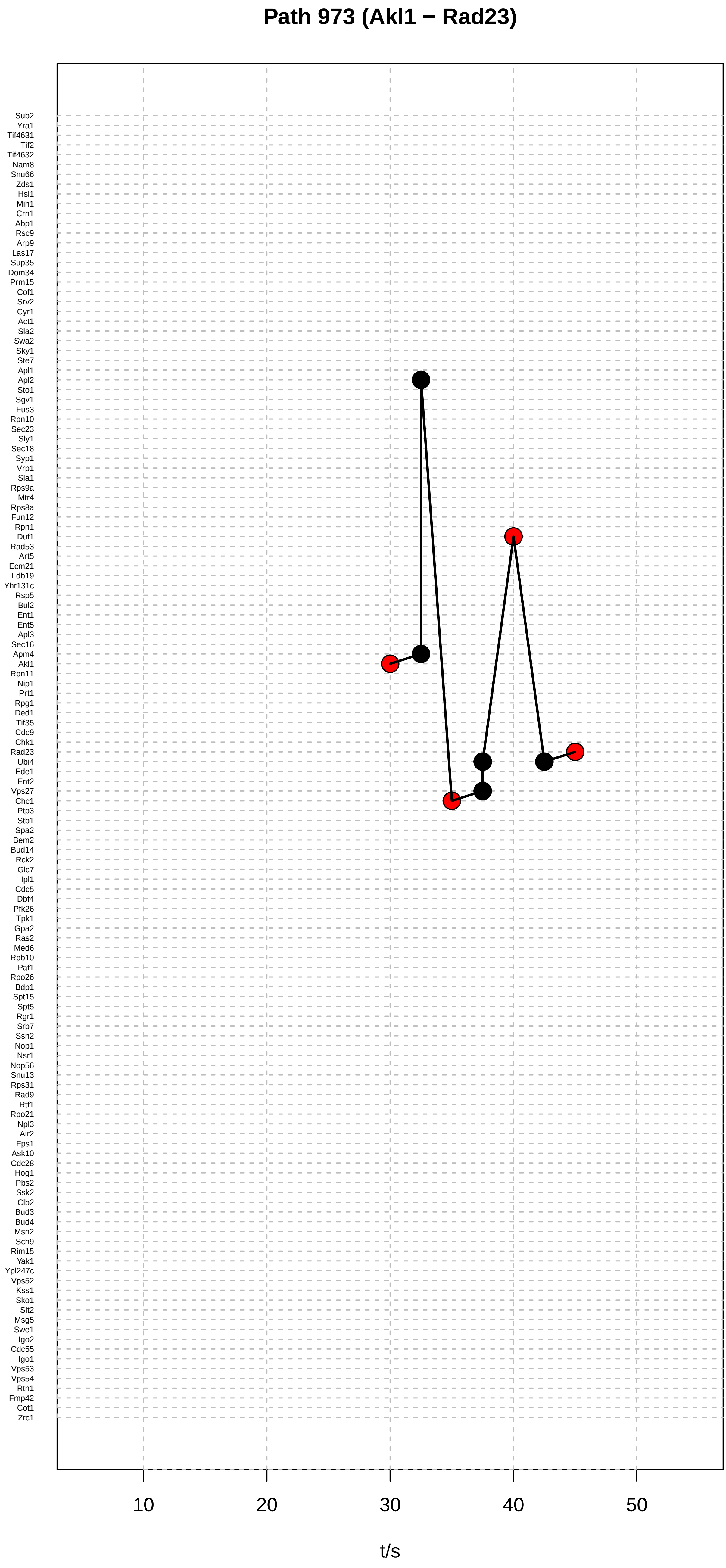
Path 973 (Akl1 − Rad23)
Sub2
Yra1
Tif4631
Tif2
Tif4632
Nam8
Snu66
Zds1
Hsl1
Mih1
Crn1
Abp1
Rsc9
Arp9
Las17
Sup35
Dom34
Prm15
Cof1
Srv2
Cyr1
Act1
Sla2
Swa2
Sky1
Ste7
Apl1
Apl2
Sto1
Sgv1
Fus3
Rpn10
Sec23
Sly1
Sec18
Syp1
Vrp1
Sla1
Rps9a
Mtr4
Rps8a
Fun12
Rpn1
Duf1
Rad53
Art5
Ecm21
Ldb19
Yhr131c
Rsp5
Bul2
Ent1
Ent5
Apl3
Sec16
Apm4
Akl1
Rpn11
Nip1
Prt1
Rpg1
Ded1
Tif35
Cdc9
Chk1
Rad23
Ubi4
Ede1
Ent2
Vps27
Chc1
Ptp3
Stb1
Spa2
Bem2
Bud14
Rck2
Glc7
Ipl1
Cdc5
Dbf4
Pfk26
Tpk1
Gpa2
Ras2
Med6
Rpb10
Paf1
Rpo26
Bdp1
Spt15
Spt5
Rgr1
Srb7
Ssn2
Nop1
Nsr1
Nop56
Snu13
Rps31
Rad9
Rtf1
Rpo21
Npl3
Air2
Fps1
Ask10
Cdc28
Hog1
Pbs2
Ssk2
Clb2
Bud3
Bud4
Msn2
Sch9
Rim15
Yak1
Ypl247c
Vps52
Kss1
Sko1
Slt2
Msg5
Swe1
Igo2
Cdc55
Igo1
Vps53
Vps54
Rtn1
Fmp42
Cot1
Zrc1
10
20
30
40
50
t/s

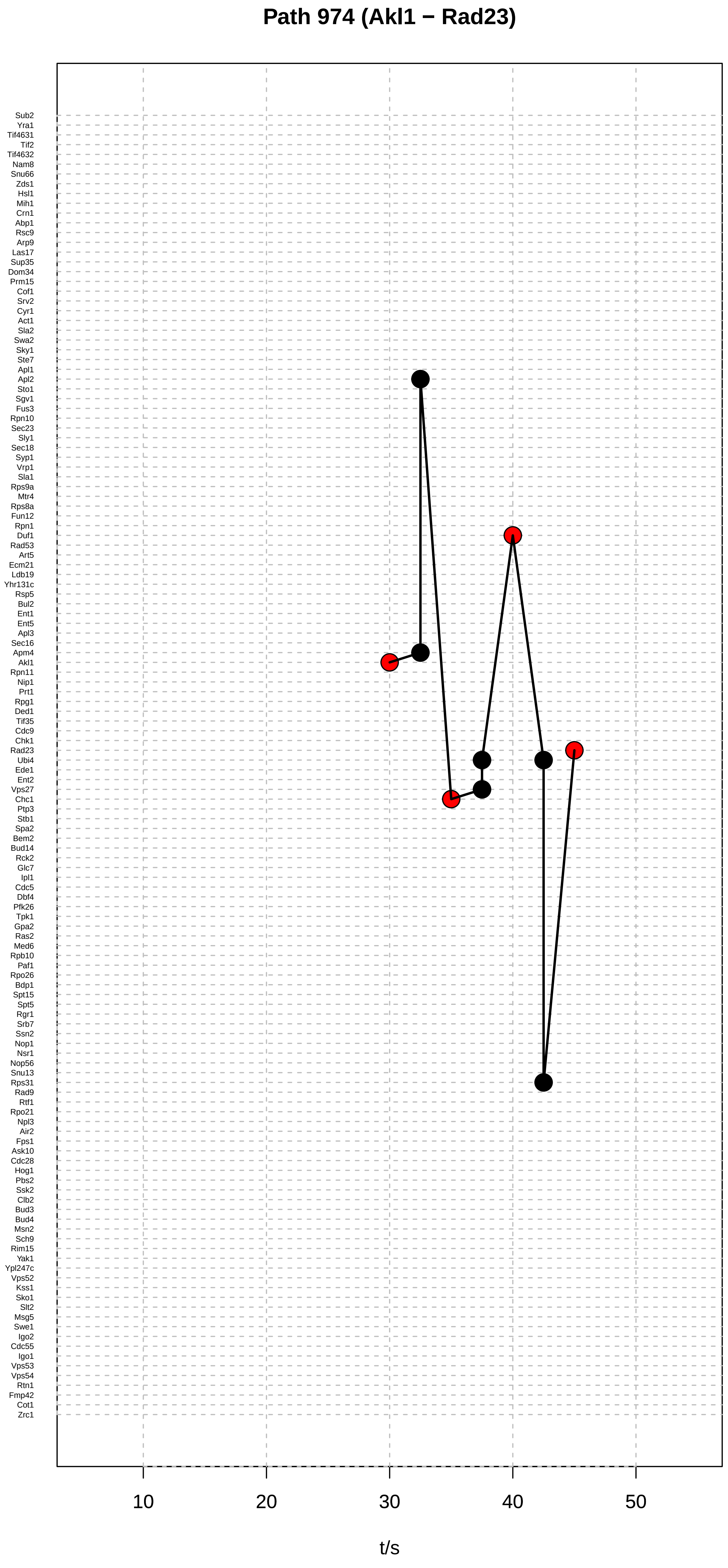
Path 974 (Akl1 − Rad23)
Sub2
Yra1
Tif4631
Tif2
Tif4632
Nam8
Snu66
Zds1
Hsl1
Mih1
Crn1
Abp1
Rsc9
Arp9
Las17
Sup35
Dom34
Prm15
Cof1
Srv2
Cyr1
Act1
Sla2
Swa2
Sky1
Ste7
Apl1
Apl2
Sto1
Sgv1
Fus3
Rpn10
Sec23
Sly1
Sec18
Syp1
Vrp1
Sla1
Rps9a
Mtr4
Rps8a
Fun12
Rpn1
Duf1
Rad53
Art5
Ecm21
Ldb19
Yhr131c
Rsp5
Bul2
Ent1
Ent5
Apl3
Sec16
Apm4
Akl1
Rpn11
Nip1
Prt1
Rpg1
Ded1
Tif35
Cdc9
Chk1
Rad23
Ubi4
Ede1
Ent2
Vps27
Chc1
Ptp3
Stb1
Spa2
Bem2
Bud14
Rck2
Glc7
Ipl1
Cdc5
Dbf4
Pfk26
Tpk1
Gpa2
Ras2
Med6
Rpb10
Paf1
Rpo26
Bdp1
Spt15
Spt5
Rgr1
Srb7
Ssn2
Nop1
Nsr1
Nop56
Snu13
Rps31
Rad9
Rtf1
Rpo21
Npl3
Air2
Fps1
Ask10
Cdc28
Hog1
Pbs2
Ssk2
Clb2
Bud3
Bud4
Msn2
Sch9
Rim15
Yak1
Ypl247c
Vps52
Kss1
Sko1
Slt2
Msg5
Swe1
Igo2
Cdc55
Igo1
Vps53
Vps54
Rtn1
Fmp42
Cot1
Zrc1
10
20
30
40
50
t/s

**Path 975 (Akl1 − Rad23)**

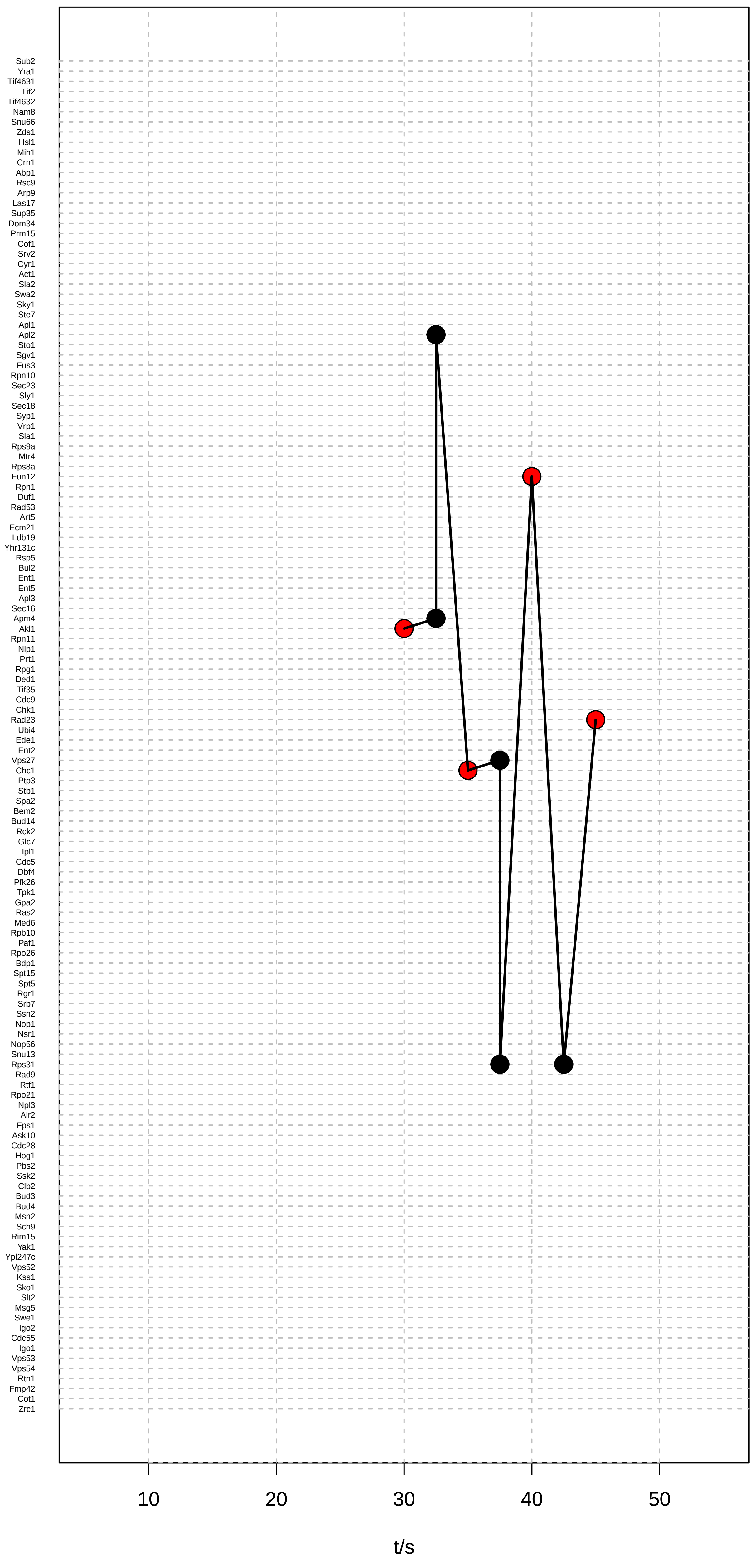

### Path 976 (Akl1 − Rad23)

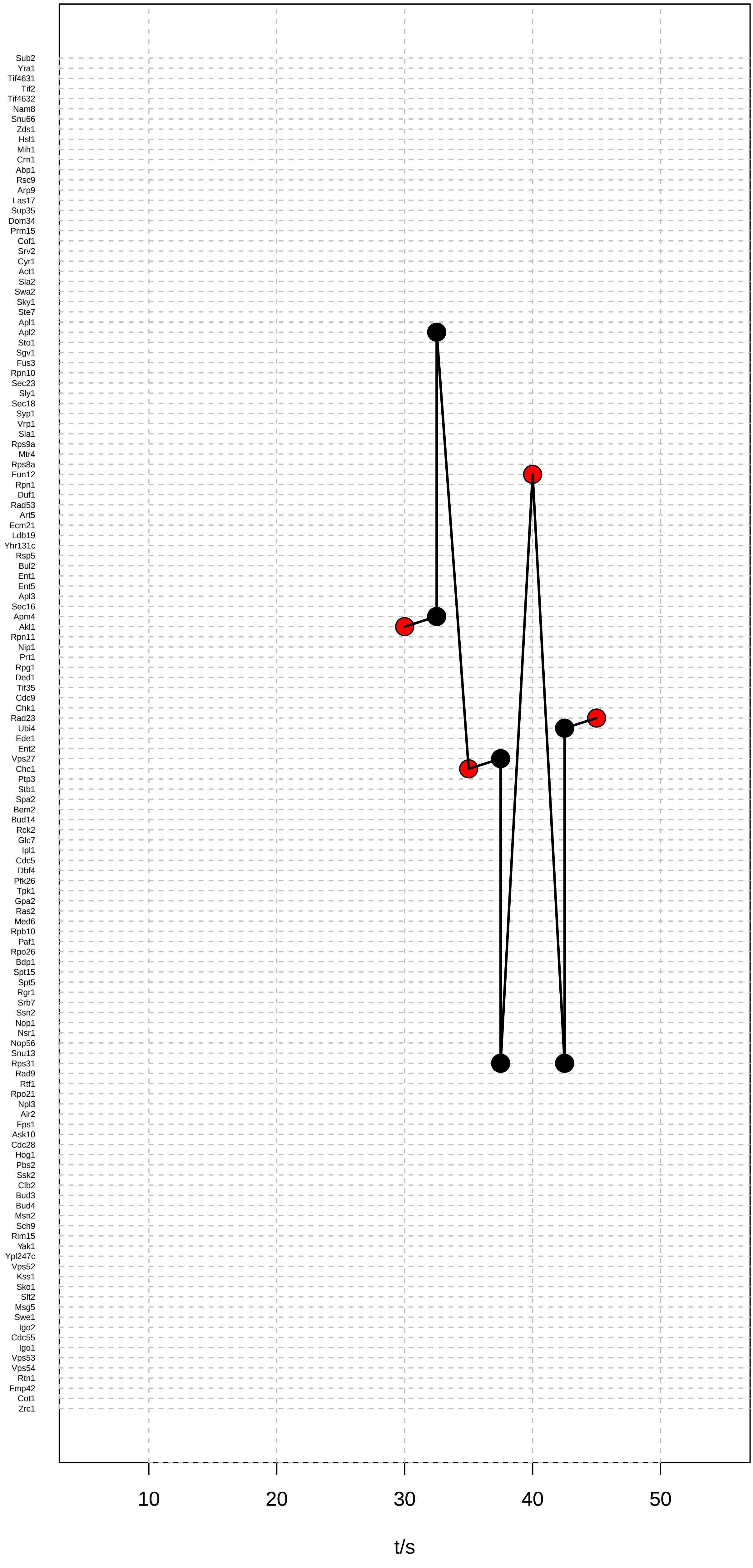

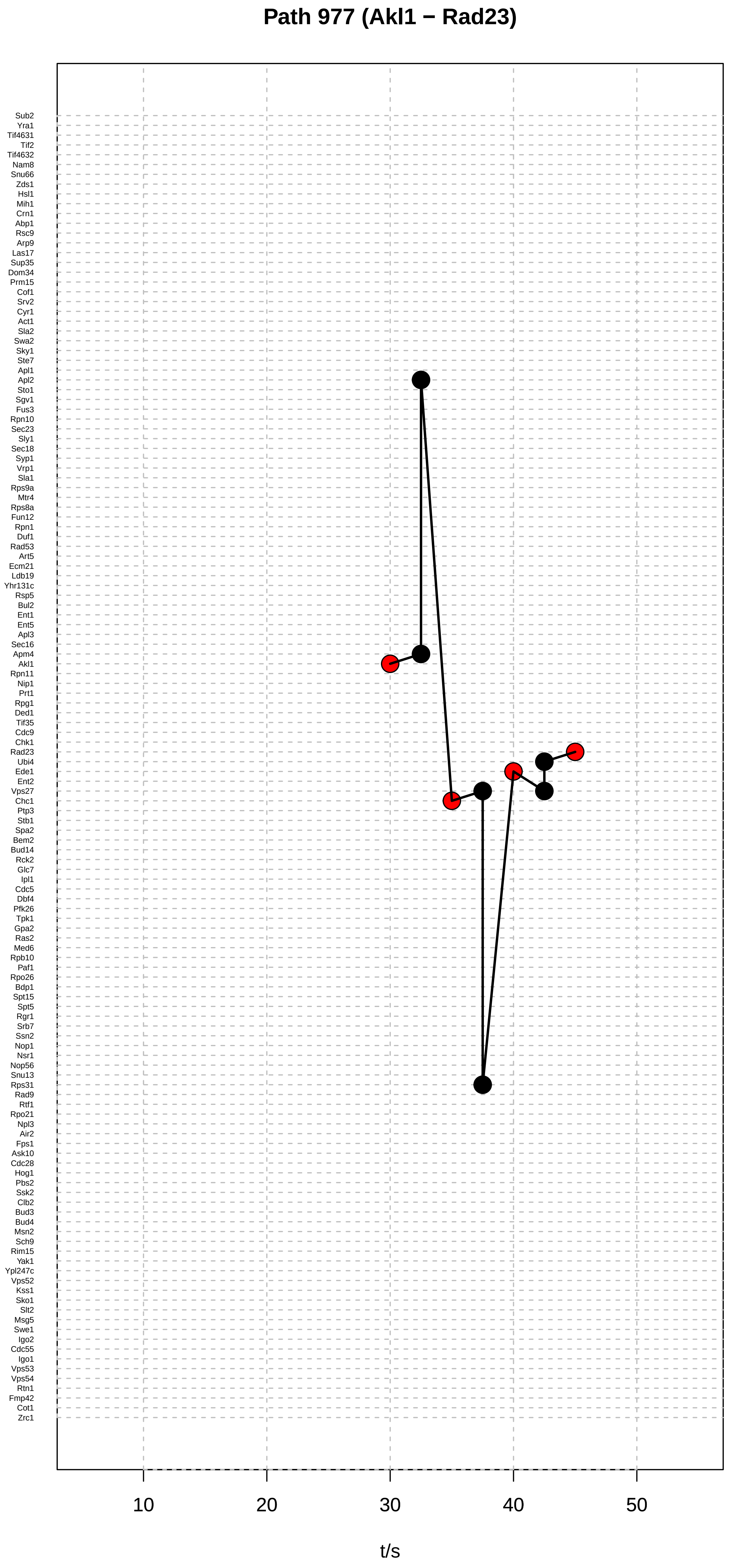

Path 977 (Akl1 − Rad23)
Sub2
Yra1
Tif4631
Tif2
Tif4632
Nam8
Snu66
Zds1
Hsl1
Mih1
Crn1
Abp1
Rsc9
Arp9
Las17
Sup35
Dom34
Prm15
Cof1
Srv2
Cyr1
Act1
Sla2
Swa2
Sky1
Ste7
Apl1
Apl2
Sto1
Sgv1
Fus3
Rpn10
Sec23
Sly1
Sec18
Syp1
Vrp1
Sla1
Rps9a
Mtr4
Rps8a
Fun12
Rpn1
Duf1
Rad53
Art5
Ecm21
Ldb19
Yhr131c
Rsp5
Bul2
Ent1
Ent5
Apl3
Sec16
Apm4
Akl1
Rpn11
Nip1
Prt1
Rpg1
Ded1
Tif35
Cdc9
Chk1
Rad23
Ubi4
Ede1
Ent2
Vps27
Chc1
Ptp3
Stb1
Spa2
Bem2
Bud14
Rck2
Glc7
Ipl1
Cdc5
Dbf4
Pfk26
Tpk1
Gpa2
Ras2
Med6
Rpb10
Paf1
Rpo26
Bdp1
Spt15
Spt5
Rgr1
Srb7
Ssn2
Nop1
Nsr1
Nop56
Snu13
Rps31
Rad9
Rtf1
Rpo21
Npl3
Air2
Fps1
Ask10
Cdc28
Hog1
Pbs2
Ssk2
Clb2
Bud3
Bud4
Msn2
Sch9
Rim15
Yak1
Ypl247c
Vps52
Kss1
Sko1
Slt2
Msg5
Swe1
Igo2
Cdc55
Igo1
Vps53
Vps54
Rtn1
Fmp42
Cot1
Zrc1
10
20
30
40
50
t/s

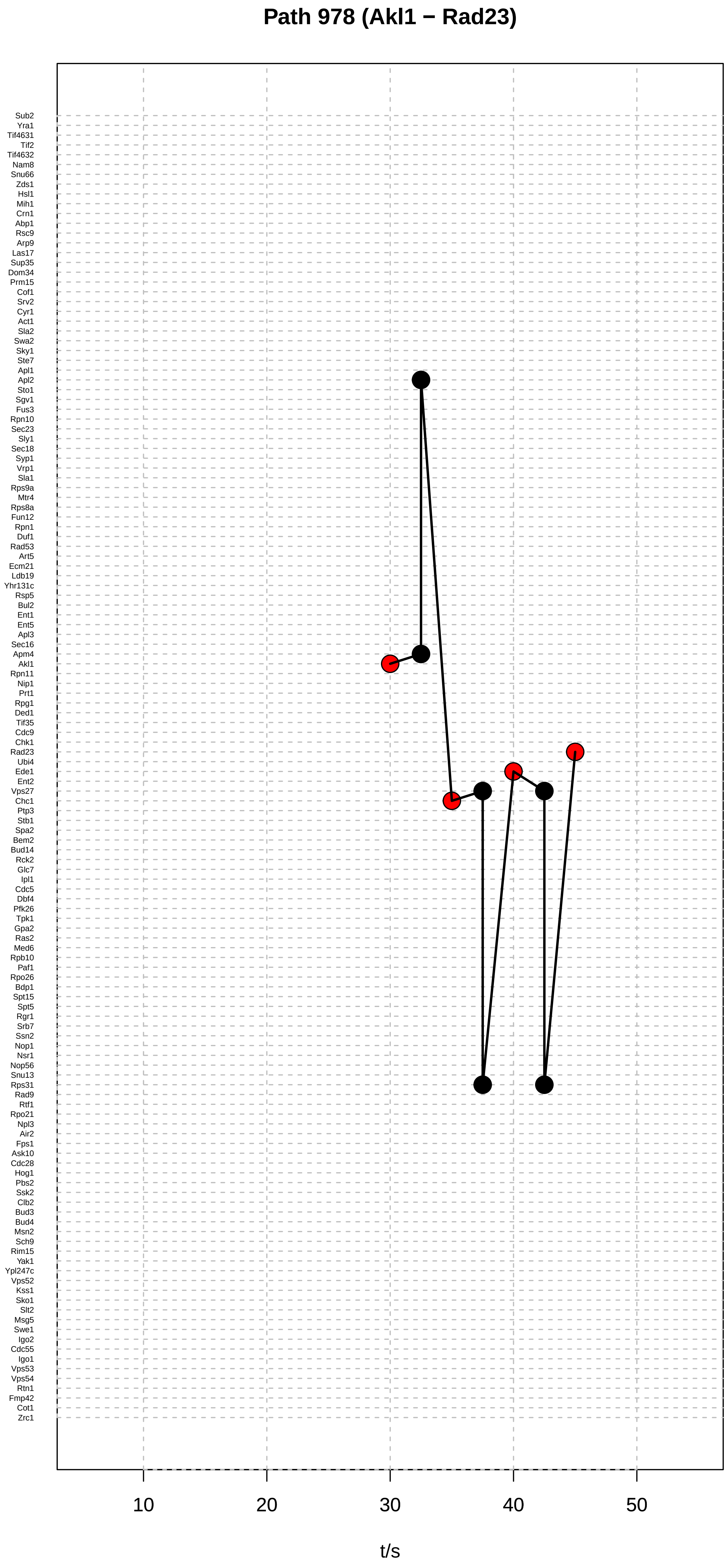

Path 978 (Akl1 − Rad23)
Sub2
Yra1
Tif4631
Tif2
Tif4632
Nam8
Snu66
Zds1
Hsl1
Mih1
Crn1
Abp1
Rsc9
Arp9
Las17
Sup35
Dom34
Prm15
Cof1
Srv2
Cyr1
Act1
Sla2
Swa2
Sky1
Ste7
Apl1
Apl2
Sto1
Sgv1
Fus3
Rpn10
Sec23
Sly1
Sec18
Syp1
Vrp1
Sla1
Rps9a
Mtr4
Rps8a
Fun12
Rpn1
Duf1
Rad53
Art5
Ecm21
Ldb19
Yhr131c
Rsp5
Bul2
Ent1
Ent5
Apl3
Sec16
Apm4
Akl1
Rpn11
Nip1
Prt1
Rpg1
Ded1
Tif35
Cdc9
Chk1
Rad23
Ubi4
Ede1
Ent2
Vps27
Chc1
Ptp3
Stb1
Spa2
Bem2
Bud14
Rck2
Glc7
Ipl1
Cdc5
Dbf4
Pfk26
Tpk1
Gpa2
Ras2
Med6
Rpb10
Paf1
Rpo26
Bdp1
Spt15
Spt5
Rgr1
Srb7
Ssn2
Nop1
Nsr1
Nop56
Snu13
Rps31
Rad9
Rtf1
Rpo21
Npl3
Air2
Fps1
Ask10
Cdc28
Hog1
Pbs2
Ssk2
Clb2
Bud3
Bud4
Msn2
Sch9
Rim15
Yak1
Ypl247c
Vps52
Kss1
Sko1
Slt2
Msg5
Swe1
Igo2
Cdc55
Igo1
Vps53
Vps54
Rtn1
Fmp42
Cot1
Zrc1
10
20
30
40
50
t/s

#### Path 979 (Akl1 − Ent5)

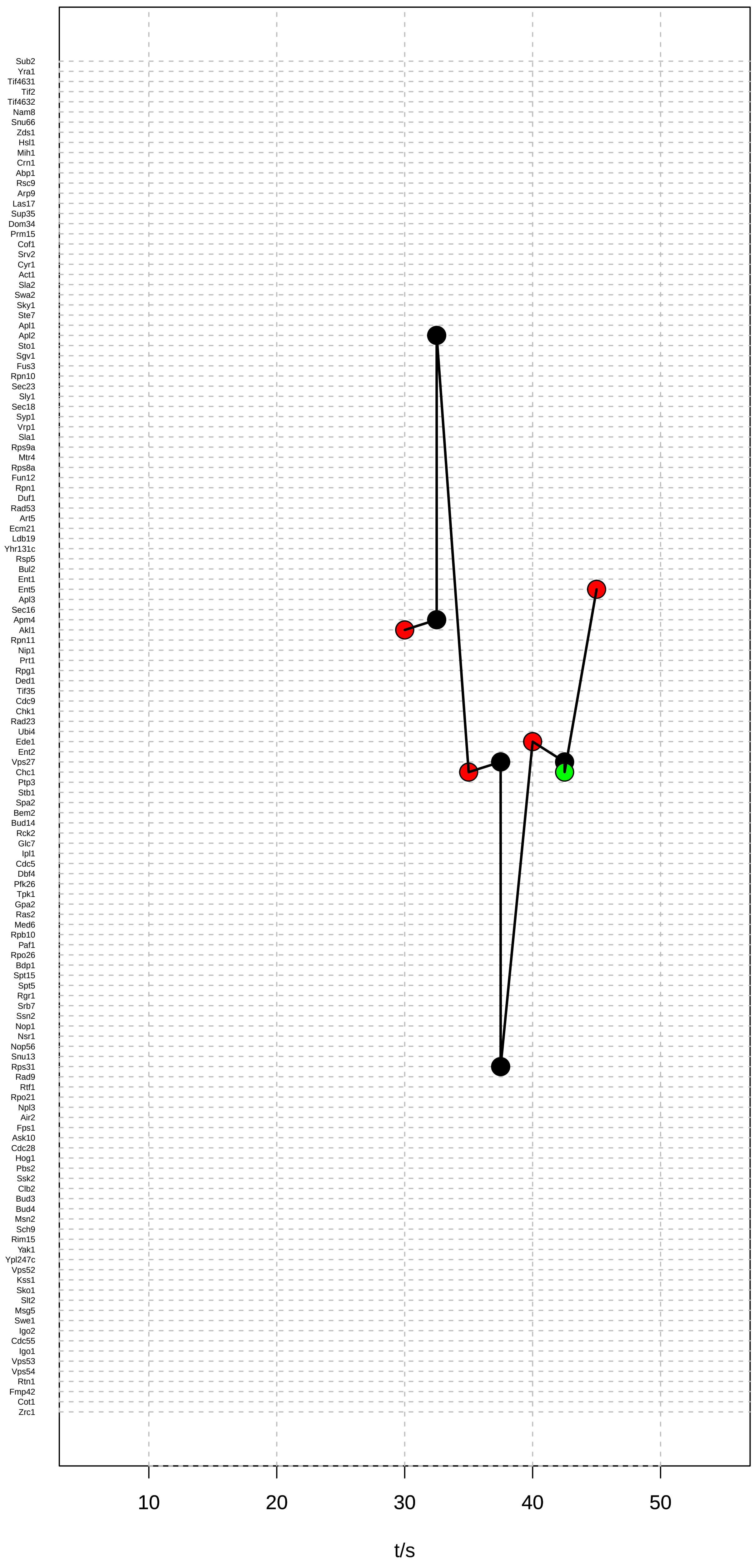

### Path 980 (Akl1 − Rad23)

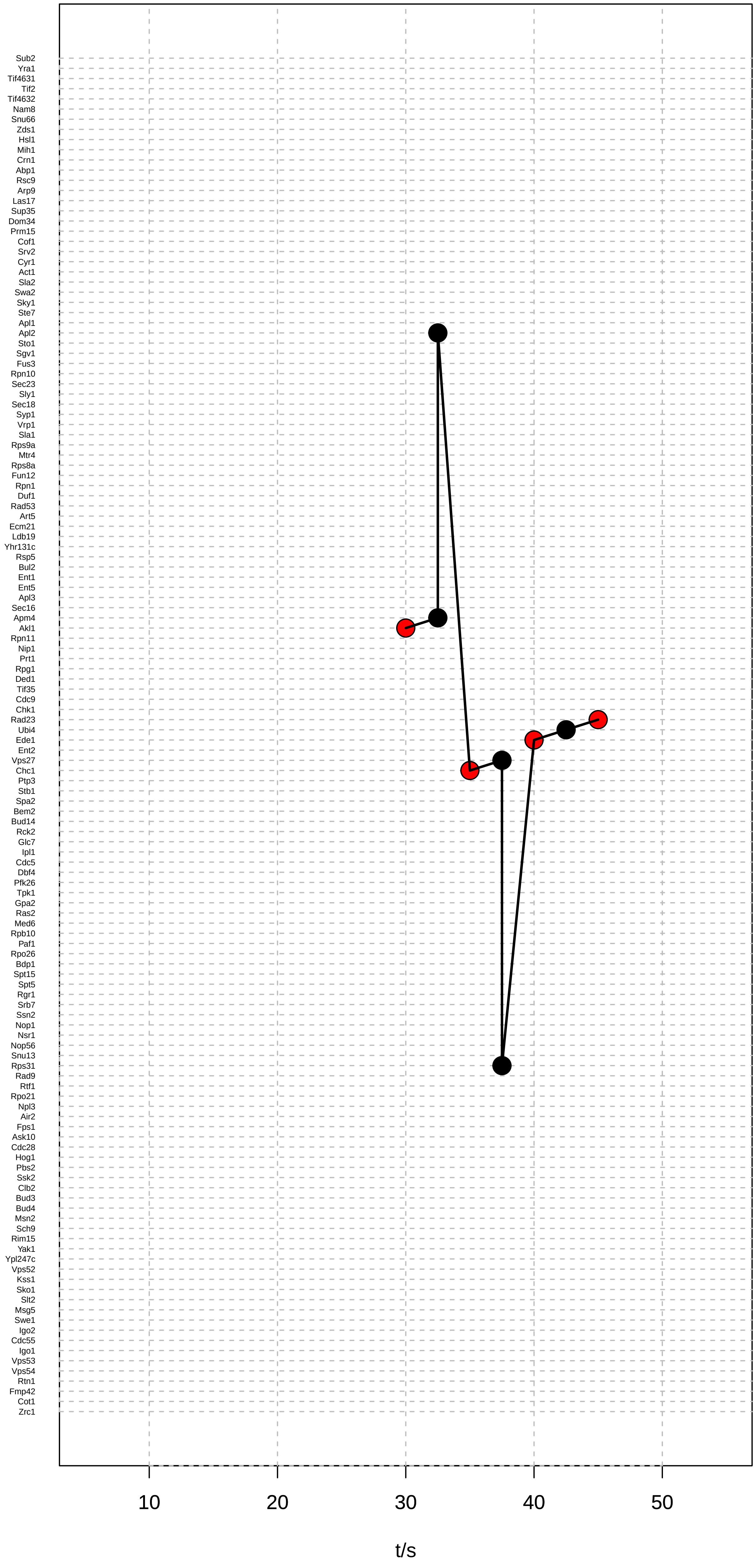

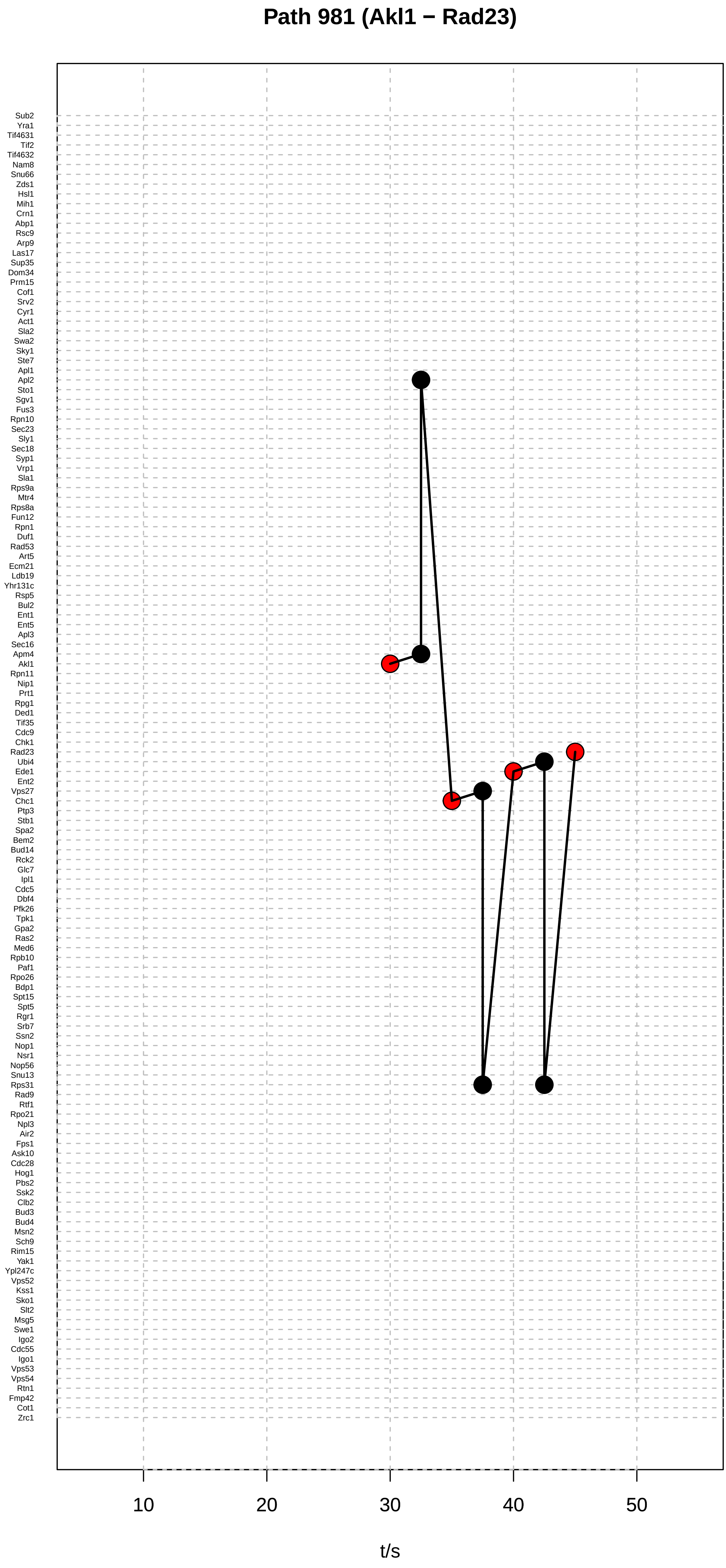
Path 981 (Akl1 − Rad23)
Sub2
Yra1
Tif4631
Tif2
Tif4632
Nam8
Snu66
Zds1
Hsl1
Mih1
Crn1
Abp1
Rsc9
Arp9
Las17
Sup35
Dom34
Prm15
Cof1
Srv2
Cyr1
Act1
Sla2
Swa2
Sky1
Ste7
Apl1
Apl2
Sto1
Sgv1
Fus3
Rpn10
Sec23
Sly1
Sec18
Syp1
Vrp1
Sla1
Rps9a
Mtr4
Rps8a
Fun12
Rpn1
Duf1
Rad53
Art5
Ecm21
Ldb19
Yhr131c
Rsp5
Bul2
Ent1
Ent5
Apl3
Sec16
Apm4
Akl1
Rpn11
Nip1
Prt1
Rpg1
Ded1
Tif35
Cdc9
Chk1
Rad23
Ubi4
Ede1
Ent2
Vps27
Chc1
Ptp3
Stb1
Spa2
Bem2
Bud14
Rck2
Glc7
Ipl1
Cdc5
Dbf4
Pfk26
Tpk1
Gpa2
Ras2
Med6
Rpb10
Paf1
Rpo26
Bdp1
Spt15
Spt5
Rgr1
Srb7
Ssn2
Nop1
Nsr1
Nop56
Snu13
Rps31
Rad9
Rtf1
Rpo21
Npl3
Air2
Fps1
Ask10
Cdc28
Hog1
Pbs2
Ssk2
Clb2
Bud3
Bud4
Msn2
Sch9
Rim15
Yak1
Ypl247c
Vps52
Kss1
Sko1
Slt2
Msg5
Swe1
Igo2
Cdc55
Igo1
Vps53
Vps54
Rtn1
Fmp42
Cot1
Zrc1
10
20
30
40
50
t/s

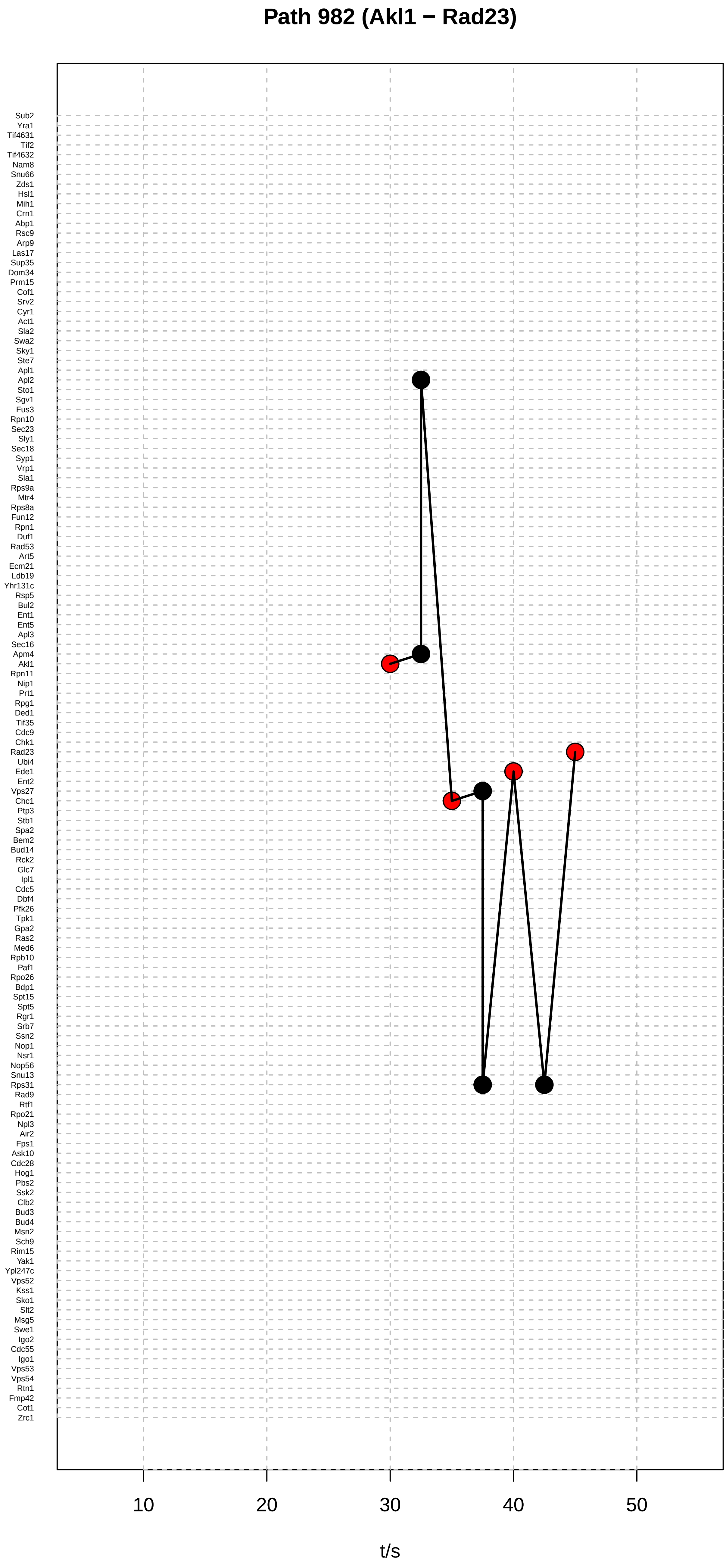
Path 982 (Akl1 − Rad23)
Sub2
Yra1
Tif4631
Tif2
Tif4632
Nam8
Snu66
Zds1
Hsl1
Mih1
Crn1
Abp1
Rsc9
Arp9
Las17
Sup35
Dom34
Prm15
Cof1
Srv2
Cyr1
Act1
Sla2
Swa2
Sky1
Ste7
Apl1
Apl2
Sto1
Sgv1
Fus3
Rpn10
Sec23
Sly1
Sec18
Syp1
Vrp1
Sla1
Rps9a
Mtr4
Rps8a
Fun12
Rpn1
Duf1
Rad53
Art5
Ecm21
Ldb19
Yhr131c
Rsp5
Bul2
Ent1
Ent5
Apl3
Sec16
Apm4
Akl1
Rpn11
Nip1
Prt1
Rpg1
Ded1
Tif35
Cdc9
Chk1
Rad23
Ubi4
Ede1
Ent2
Vps27
Chc1
Ptp3
Stb1
Spa2
Bem2
Bud14
Rck2
Glc7
Ipl1
Cdc5
Dbf4
Pfk26
Tpk1
Gpa2
Ras2
Med6
Rpb10
Paf1
Rpo26
Bdp1
Spt15
Spt5
Rgr1
Srb7
Ssn2
Nop1
Nsr1
Nop56
Snu13
Rps31
Rad9
Rtf1
Rpo21
Npl3
Air2
Fps1
Ask10
Cdc28
Hog1
Pbs2
Ssk2
Clb2
Bud3
Bud4
Msn2
Sch9
Rim15
Yak1
Ypl247c
Vps52
Kss1
Sko1
Slt2
Msg5
Swe1
Igo2
Cdc55
Igo1
Vps53
Vps54
Rtn1
Fmp42
Cot1
Zrc1
10
20
30
40
50
t/s

#### Path 983 (Akl1 − Rad23)

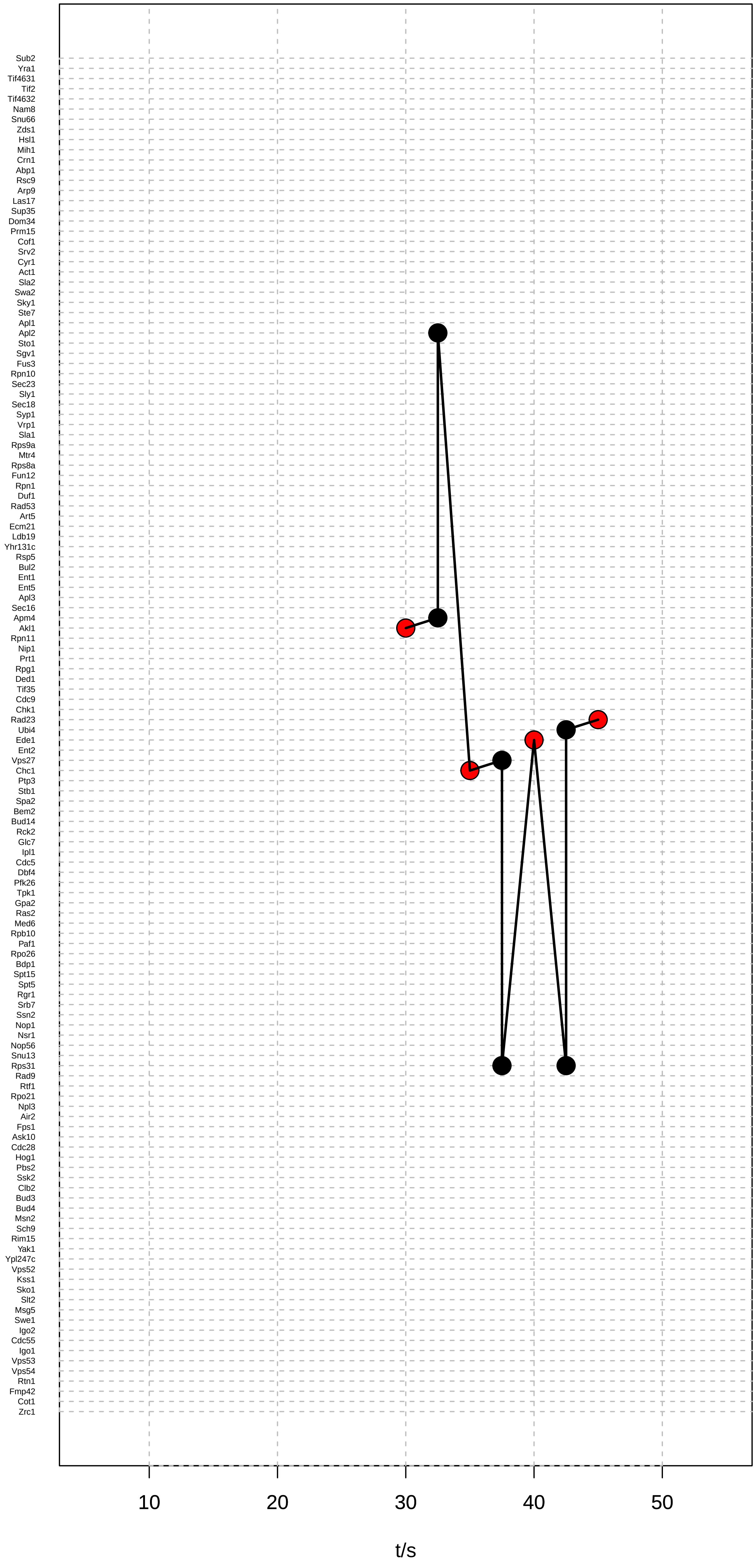

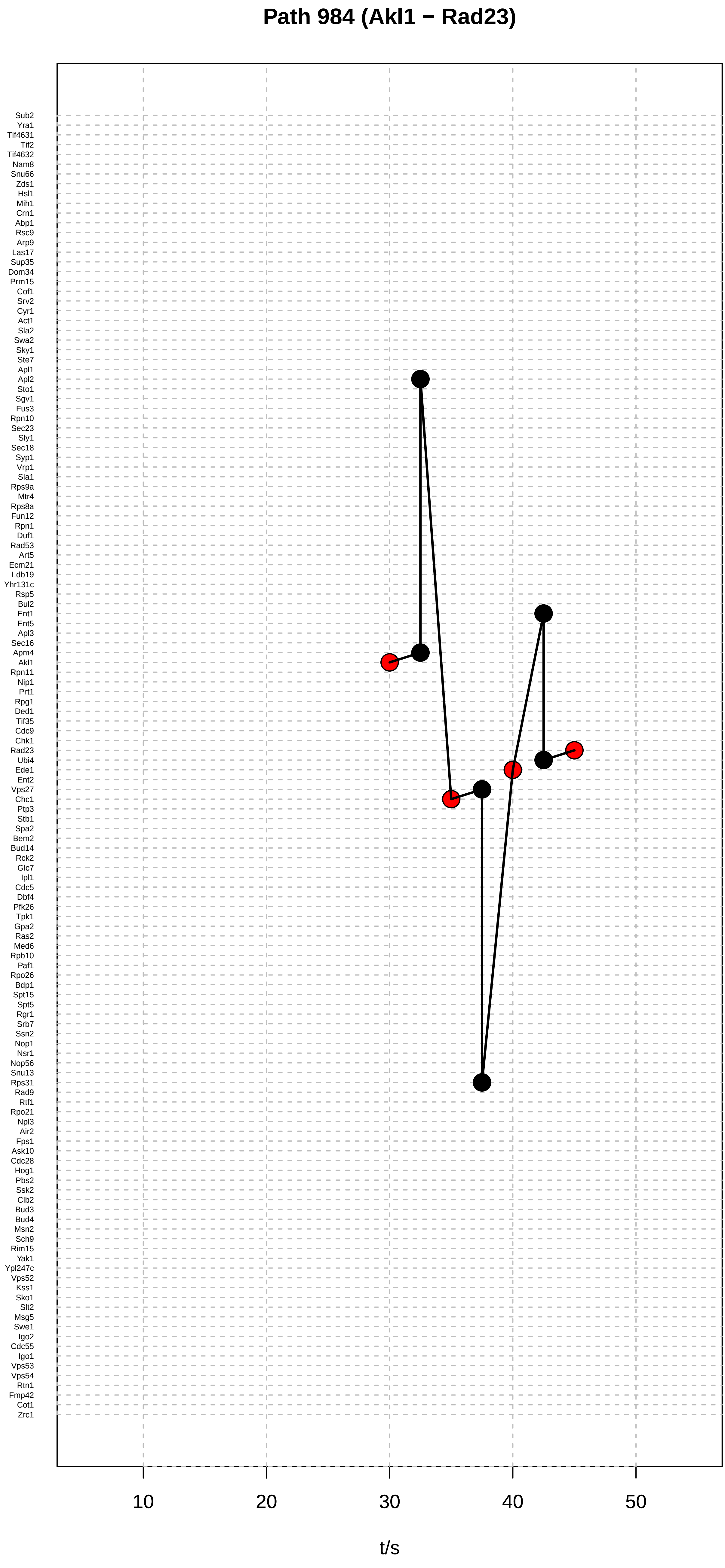

Path 984 (Akl1 − Rad23)
Sub2
Yra1
Tif4631
Tif2
Tif4632
Nam8
Snu66
Zds1
Hsl1
Mih1
Crn1
Abp1
Rsc9
Arp9
Las17
Sup35
Dom34
Prm15
Cof1
Srv2
Cyr1
Act1
Sla2
Swa2
Sky1
Ste7
Apl1
Apl2
Sto1
Sgv1
Fus3
Rpn10
Sec23
Sly1
Sec18
Syp1
Vrp1
Sla1
Rps9a
Mtr4
Rps8a
Fun12
Rpn1
Duf1
Rad53
Art5
Ecm21
Ldb19
Yhr131c
Rsp5
Bul2
Ent1
Ent5
Apl3
Sec16
Apm4
Akl1
Rpn11
Nip1
Prt1
Rpg1
Ded1
Tif35
Cdc9
Chk1
Rad23
Ubi4
Ede1
Ent2
Vps27
Chc1
Ptp3
Stb1
Spa2
Bem2
Bud14
Rck2
Glc7
Ipl1
Cdc5
Dbf4
Pfk26
Tpk1
Gpa2
Ras2
Med6
Rpb10
Paf1
Rpo26
Bdp1
Spt15
Spt5
Rgr1
Srb7
Ssn2
Nop1
Nsr1
Nop56
Snu13
Rps31
Rad9
Rtf1
Rpo21
Npl3
Air2
Fps1
Ask10
Cdc28
Hog1
Pbs2
Ssk2
Clb2
Bud3
Bud4
Msn2
Sch9
Rim15
Yak1
Ypl247c
Vps52
Kss1
Sko1
Slt2
Msg5
Swe1
Igo2
Cdc55
Igo1
Vps53
Vps54
Rtn1
Fmp42
Cot1
Zrc1
10
20
30
40
50
t/s

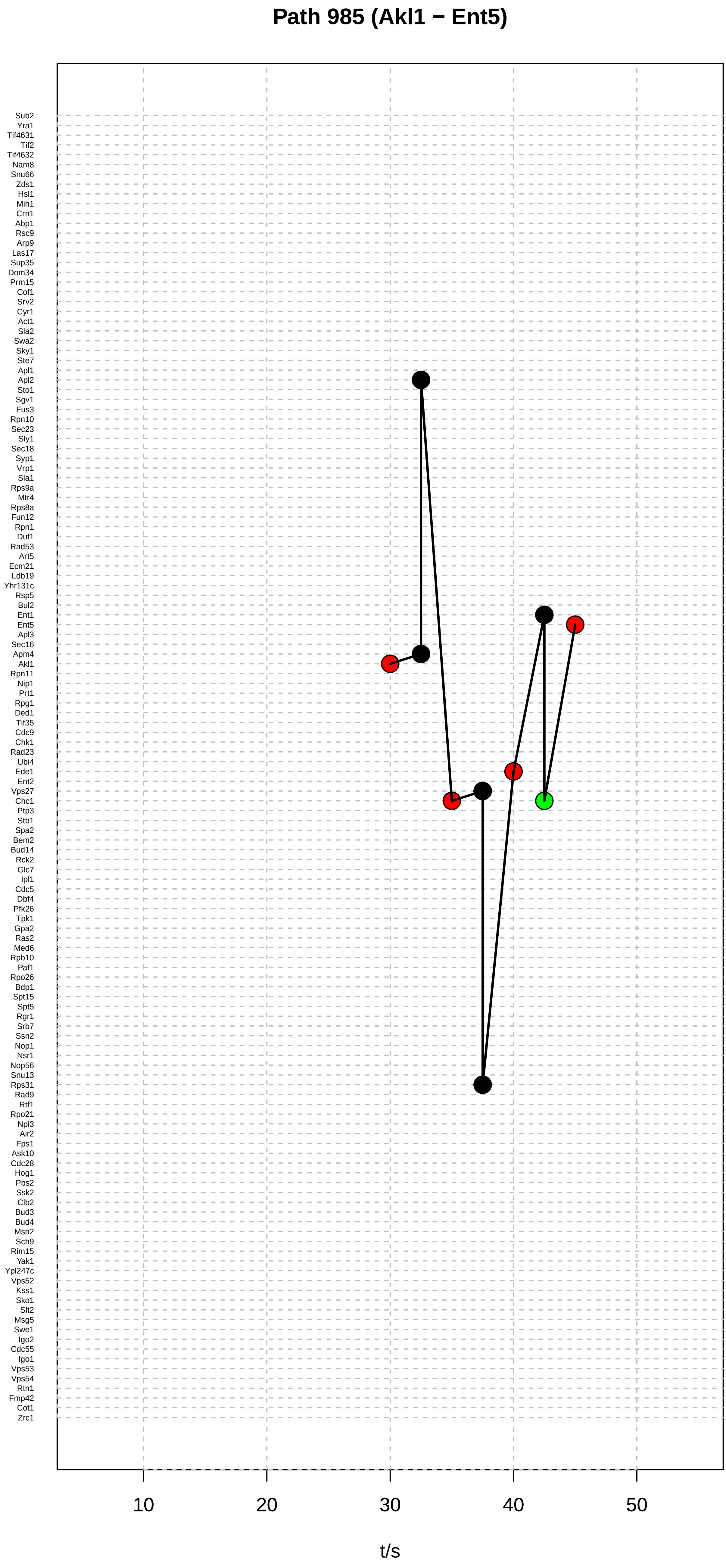

Path 985 (Akl1 − Ent5)
Sub2
Yra1
Tif4631
Tif2
Tif4632
Nam8
Snu66
Zds1
Hsl1
Mih1
Crn1
Abp1
Rsc9
Arp9
Las17
Sup35
Dom34
Prm15
Cof1
Srv2
Cyr1
Act1
Sla2
Swa2
Sky1
Ste7
Apl1
Apl2
Sto1
Sgv1
Fus3
Rpn10
Sec23
Sly1
Sec18
Syp1
Vrp1
Sla1
Rps9a
Mtr4
Rps8a
Fun12
Rpn1
Duf1
Rad53
Art5
Ecm21
Ldb19
Yhr131c
Rsp5
Bul2
Ent1
Ent5
Apl3
Sec16
Apm4
Akl1
Rpn11
Nip1
Prt1
Rpg1
Ded1
Tif35
Cdc9
Chk1
Rad23
Ubi4
Ede1
Ent2
Vps27
Chc1
Ptp3
Stb1
Spa2
Bem2
Bud14
Rck2
Glc7
Ipl1
Cdc5
Dbf4
Pfk26
Tpk1
Gpa2
Ras2
Med6
Rpb10
Paf1
Rpo26
Bdp1
Spt15
Spt5
Rgr1
Srb7
Ssn2
Nop1
Nsr1
Nop56
Snu13
Rps31
Rad9
Rtf1
Rpo21
Npl3
Air2
Fps1
Ask10
Cdc28
Hog1
Pbs2
Ssk2
Clb2
Bud3
Bud4
Msn2
Sch9
Rim15
Yak1
Ypl247c
Vps52
Kss1
Sko1
Slt2
Msg5
Swe1
Igo2
Cdc55
Igo1
Vps53
Vps54
Rtn1
Fmp42
Cot1
Zrc1
10
20
30
40
50
t/s

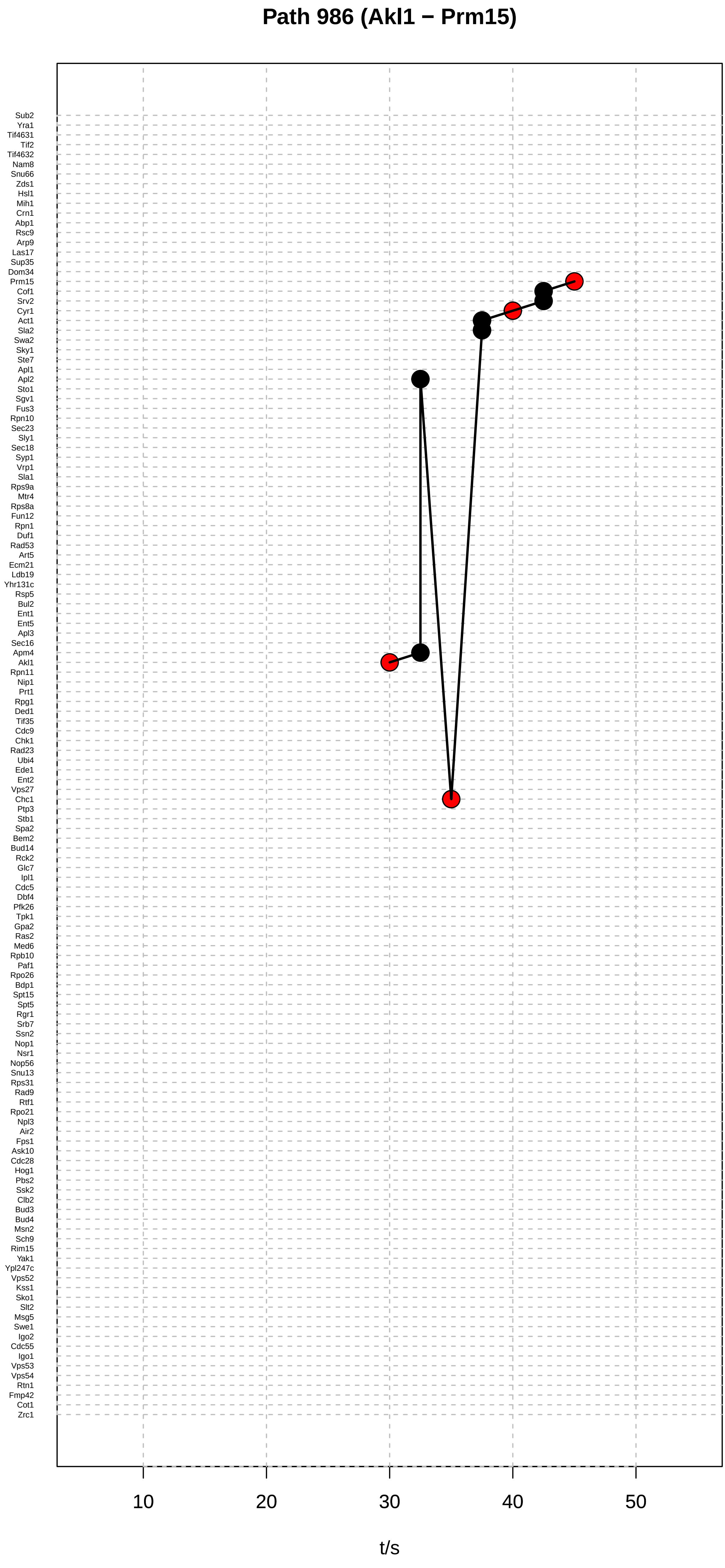
Path 986 (Akl1 − Prm15)
Sub2
Yra1
Tif4631
Tif2
Tif4632
Nam8
Snu66
Zds1
Hsl1
Mih1
Crn1
Abp1
Rsc9
Arp9
Las17
Sup35
Dom34
Prm15
Cof1
Srv2
Cyr1
Act1
Sla2
Swa2
Sky1
Ste7
Apl1
Apl2
Sto1
Sgv1
Fus3
Rpn10
Sec23
Sly1
Sec18
Syp1
Vrp1
Sla1
Rps9a
Mtr4
Rps8a
Fun12
Rpn1
Duf1
Rad53
Art5
Ecm21
Ldb19
Yhr131c
Rsp5
Bul2
Ent1
Ent5
Apl3
Sec16
Apm4
Akl1
Rpn11
Nip1
Prt1
Rpg1
Ded1
Tif35
Cdc9
Chk1
Rad23
Ubi4
Ede1
Ent2
Vps27
Chc1
Ptp3
Stb1
Spa2
Bem2
Bud14
Rck2
Glc7
Ipl1
Cdc5
Dbf4
Pfk26
Tpk1
Gpa2
Ras2
Med6
Rpb10
Paf1
Rpo26
Bdp1
Spt15
Spt5
Rgr1
Srb7
Ssn2
Nop1
Nsr1
Nop56
Snu13
Rps31
Rad9
Rtf1
Rpo21
Npl3
Air2
Fps1
Ask10
Cdc28
Hog1
Pbs2
Ssk2
Clb2
Bud3
Bud4
Msn2
Sch9
Rim15
Yak1
Ypl247c
Vps52
Kss1
Sko1
Slt2
Msg5
Swe1
Igo2
Cdc55
Igo1
Vps53
Vps54
Rtn1
Fmp42
Cot1
Zrc1
10
20
30
40
50
t/s

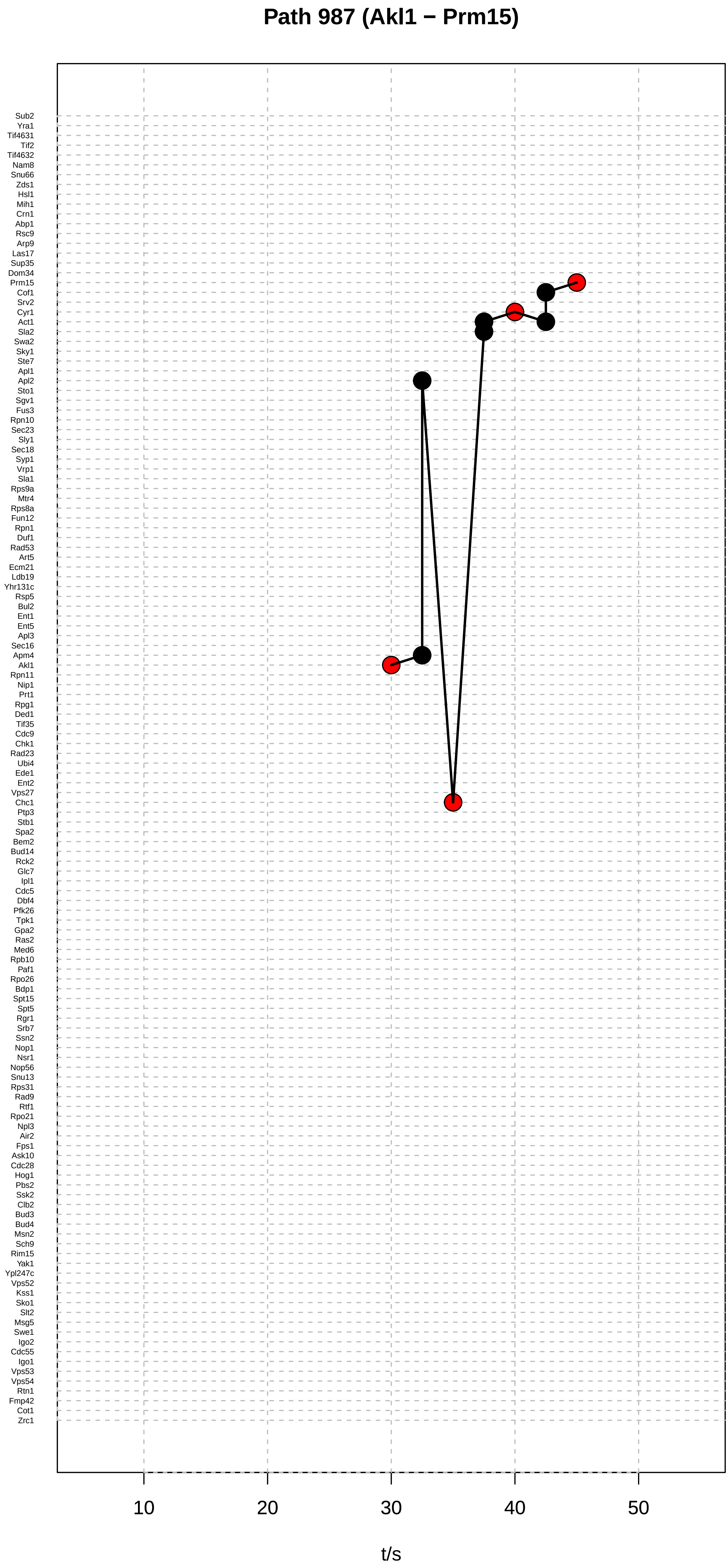
Path 987 (Akl1 − Prm15)
Sub2
Yra1
Tif4631
Tif2
Tif4632
Nam8
Snu66
Zds1
Hsl1
Mih1
Crn1
Abp1
Rsc9
Arp9
Las17
Sup35
Dom34
Prm15
Cof1
Srv2
Cyr1
Act1
Sla2
Swa2
Sky1
Ste7
Apl1
Apl2
Sto1
Sgv1
Fus3
Rpn10
Sec23
Sly1
Sec18
Syp1
Vrp1
Sla1
Rps9a
Mtr4
Rps8a
Fun12
Rpn1
Duf1
Rad53
Art5
Ecm21
Ldb19
Yhr131c
Rsp5
Bul2
Ent1
Ent5
Apl3
Sec16
Apm4
Akl1
Rpn11
Nip1
Prt1
Rpg1
Ded1
Tif35
Cdc9
Chk1
Rad23
Ubi4
Ede1
Ent2
Vps27
Chc1
Ptp3
Stb1
Spa2
Bem2
Bud14
Rck2
Glc7
Ipl1
Cdc5
Dbf4
Pfk26
Tpk1
Gpa2
Ras2
Med6
Rpb10
Paf1
Rpo26
Bdp1
Spt15
Spt5
Rgr1
Srb7
Ssn2
Nop1
Nsr1
Nop56
Snu13
Rps31
Rad9
Rtf1
Rpo21
Npl3
Air2
Fps1
Ask10
Cdc28
Hog1
Pbs2
Ssk2
Clb2
Bud3
Bud4
Msn2
Sch9
Rim15
Yak1
Ypl247c
Vps52
Kss1
Sko1
Slt2
Msg5
Swe1
Igo2
Cdc55
Igo1
Vps53
Vps54
Rtn1
Fmp42
Cot1
Zrc1
10
20
30
40
50
t/s

#### Path 988 (Akl1 − Prm15)

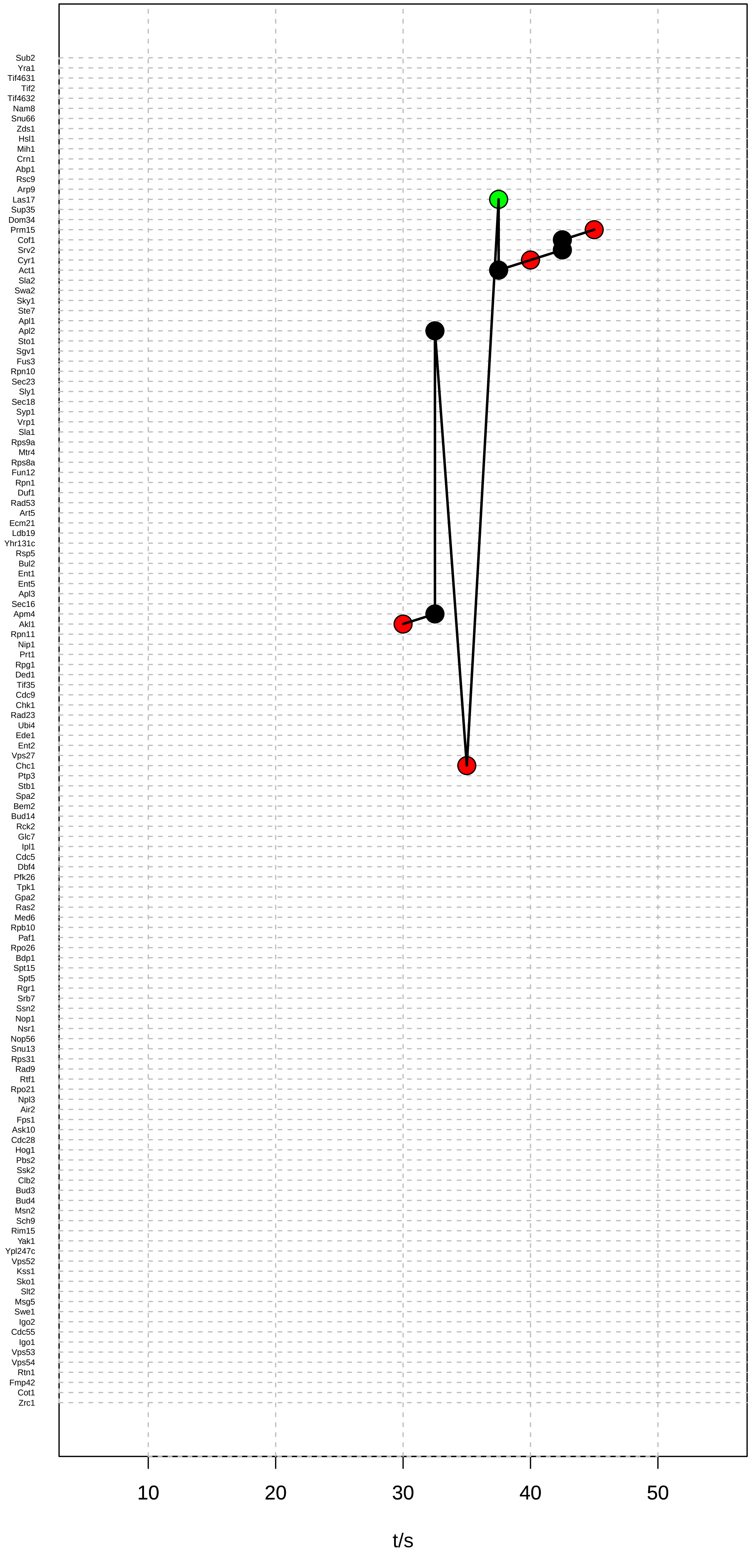

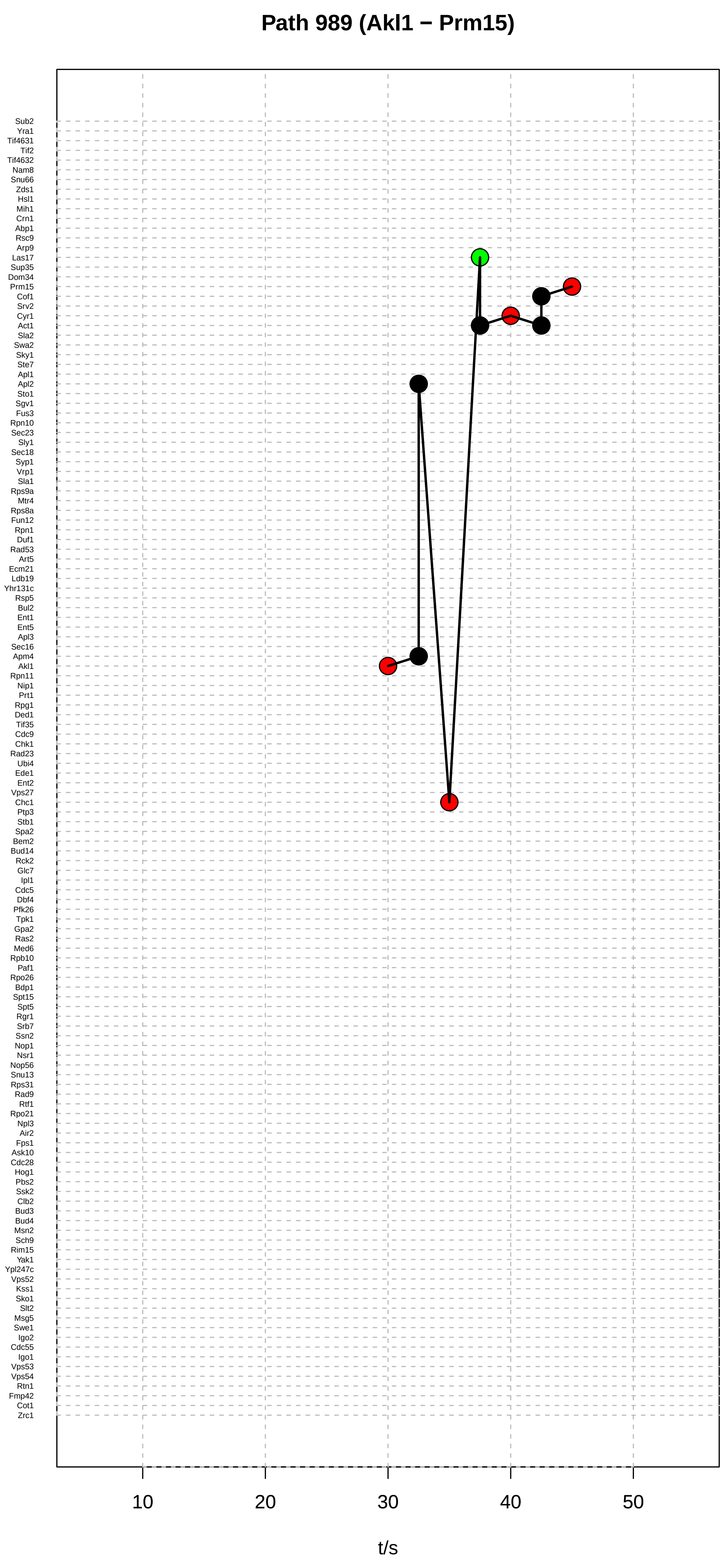

Path 989 (Akl1 − Prm15)
Sub2
Yra1
Tif4631
Tif2
Tif4632
Nam8
Snu66
Zds1
Hsl1
Mih1
Crn1
Abp1
Rsc9
Arp9
Las17
Sup35
Dom34
Prm15
Cof1
Srv2
Cyr1
Act1
Sla2
Swa2
Sky1
Ste7
Apl1
Apl2
Sto1
Sgv1
Fus3
Rpn10
Sec23
Sly1
Sec18
Syp1
Vrp1
Sla1
Rps9a
Mtr4
Rps8a
Fun12
Rpn1
Duf1
Rad53
Art5
Ecm21
Ldb19
Yhr131c
Rsp5
Bul2
Ent1
Ent5
Apl3
Sec16
Apm4
Akl1
Rpn11
Nip1
Prt1
Rpg1
Ded1
Tif35
Cdc9
Chk1
Rad23
Ubi4
Ede1
Ent2
Vps27
Chc1
Ptp3
Stb1
Spa2
Bem2
Bud14
Rck2
Glc7
Ipl1
Cdc5
Dbf4
Pfk26
Tpk1
Gpa2
Ras2
Med6
Rpb10
Paf1
Rpo26
Bdp1
Spt15
Spt5
Rgr1
Srb7
Ssn2
Nop1
Nsr1
Nop56
Snu13
Rps31
Rad9
Rtf1
Rpo21
Npl3
Air2
Fps1
Ask10
Cdc28
Hog1
Pbs2
Ssk2
Clb2
Bud3
Bud4
Msn2
Sch9
Rim15
Yak1
Ypl247c
Vps52
Kss1
Sko1
Slt2
Msg5
Swe1
Igo2
Cdc55
Igo1
Vps53
Vps54
Rtn1
Fmp42
Cot1
Zrc1
10
20
30
40
50
t/s

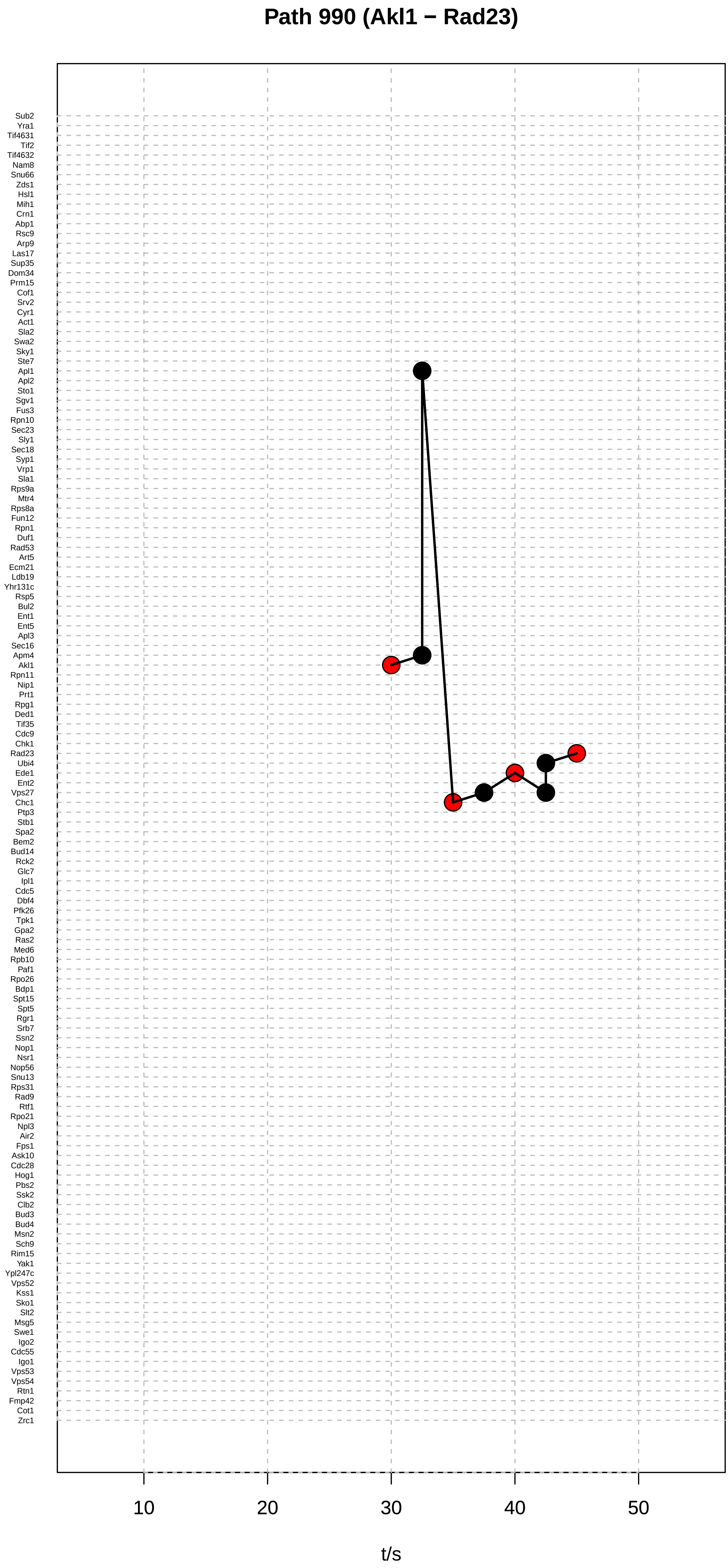

Path 990 (Akl1 − Rad23)
Sub2
Yra1
Tif4631
Tif2
Tif4632
Nam8
Snu66
Zds1
Hsl1
Mih1
Crn1
Abp1
Rsc9
Arp9
Las17
Sup35
Dom34
Prm15
Cof1
Srv2
Cyr1
Act1
Sla2
Swa2
Sky1
Ste7
Apl1
Apl2
Sto1
Sgv1
Fus3
Rpn10
Sec23
Sly1
Sec18
Syp1
Vrp1
Sla1
Rps9a
Mtr4
Rps8a
Fun12
Rpn1
Duf1
Rad53
Art5
Ecm21
Ldb19
Yhr131c
Rsp5
Bul2
Ent1
Ent5
Apl3
Sec16
Apm4
Akl1
Rpn11
Nip1
Prt1
Rpg1
Ded1
Tif35
Cdc9
Chk1
Rad23
Ubi4
Ede1
Ent2
Vps27
Chc1
Ptp3
Stb1
Spa2
Bem2
Bud14
Rck2
Glc7
Ipl1
Cdc5
Dbf4
Pfk26
Tpk1
Gpa2
Ras2
Med6
Rpb10
Paf1
Rpo26
Bdp1
Spt15
Spt5
Rgr1
Srb7
Ssn2
Nop1
Nsr1
Nop56
Snu13
Rps31
Rad9
Rtf1
Rpo21
Npl3
Air2
Fps1
Ask10
Cdc28
Hog1
Pbs2
Ssk2
Clb2
Bud3
Bud4
Msn2
Sch9
Rim15
Yak1
Ypl247c
Vps52
Kss1
Sko1
Slt2
Msg5
Swe1
Igo2
Cdc55
Igo1
Vps53
Vps54
Rtn1
Fmp42
Cot1
Zrc1
10
20
30
40
50
t/s

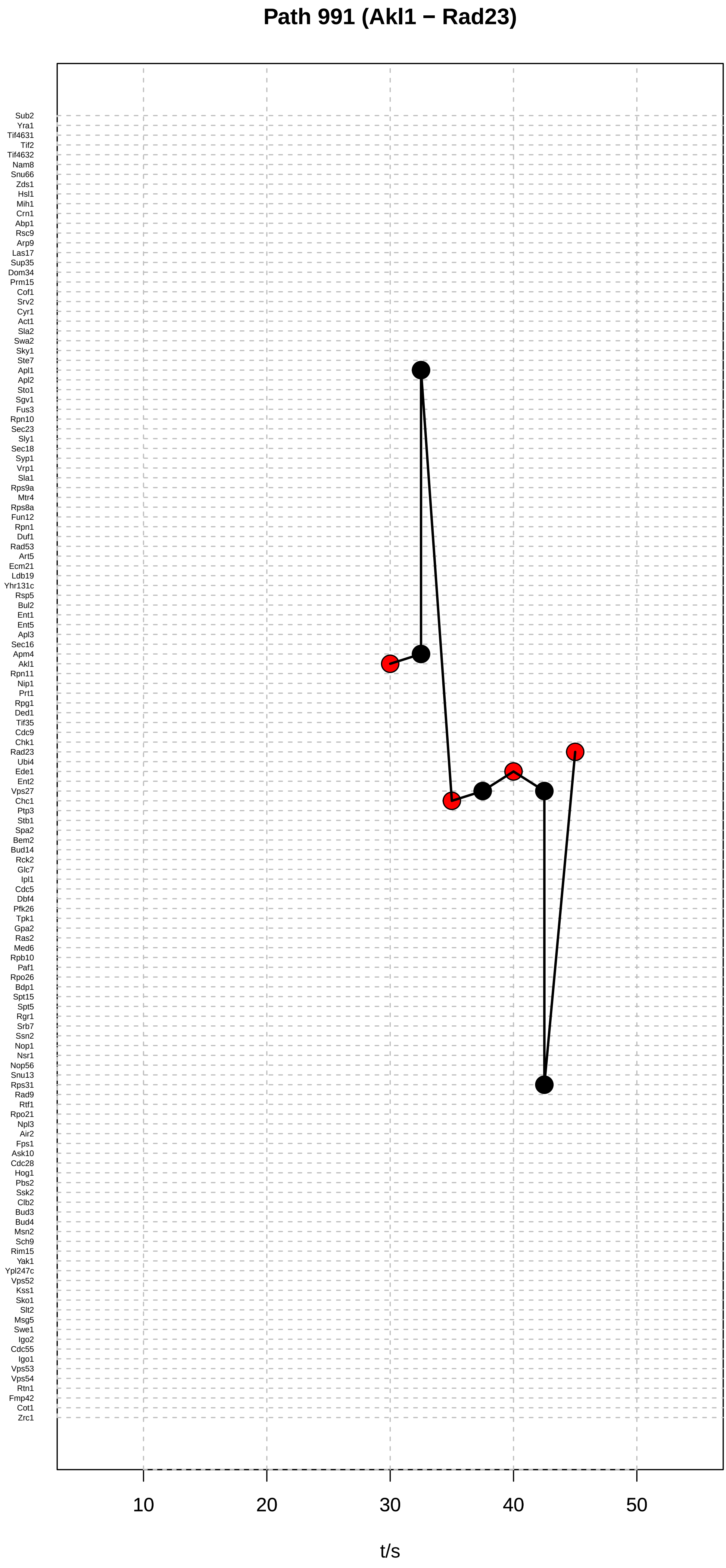
Path 991 (Akl1 − Rad23)
Sub2
Yra1
Tif4631
Tif2
Tif4632
Nam8
Snu66
Zds1
Hsl1
Mih1
Crn1
Abp1
Rsc9
Arp9
Las17
Sup35
Dom34
Prm15
Cof1
Srv2
Cyr1
Act1
Sla2
Swa2
Sky1
Ste7
Apl1
Apl2
Sto1
Sgv1
Fus3
Rpn10
Sec23
Sly1
Sec18
Syp1
Vrp1
Sla1
Rps9a
Mtr4
Rps8a
Fun12
Rpn1
Duf1
Rad53
Art5
Ecm21
Ldb19
Yhr131c
Rsp5
Bul2
Ent1
Ent5
Apl3
Sec16
Apm4
Akl1
Rpn11
Nip1
Prt1
Rpg1
Ded1
Tif35
Cdc9
Chk1
Rad23
Ubi4
Ede1
Ent2
Vps27
Chc1
Ptp3
Stb1
Spa2
Bem2
Bud14
Rck2
Glc7
Ipl1
Cdc5
Dbf4
Pfk26
Tpk1
Gpa2
Ras2
Med6
Rpb10
Paf1
Rpo26
Bdp1
Spt15
Spt5
Rgr1
Srb7
Ssn2
Nop1
Nsr1
Nop56
Snu13
Rps31
Rad9
Rtf1
Rpo21
Npl3
Air2
Fps1
Ask10
Cdc28
Hog1
Pbs2
Ssk2
Clb2
Bud3
Bud4
Msn2
Sch9
Rim15
Yak1
Ypl247c
Vps52
Kss1
Sko1
Slt2
Msg5
Swe1
Igo2
Cdc55
Igo1
Vps53
Vps54
Rtn1
Fmp42
Cot1
Zrc1
10
20
30
40
50
t/s

### Path 992 (Akl1 − Rad23)

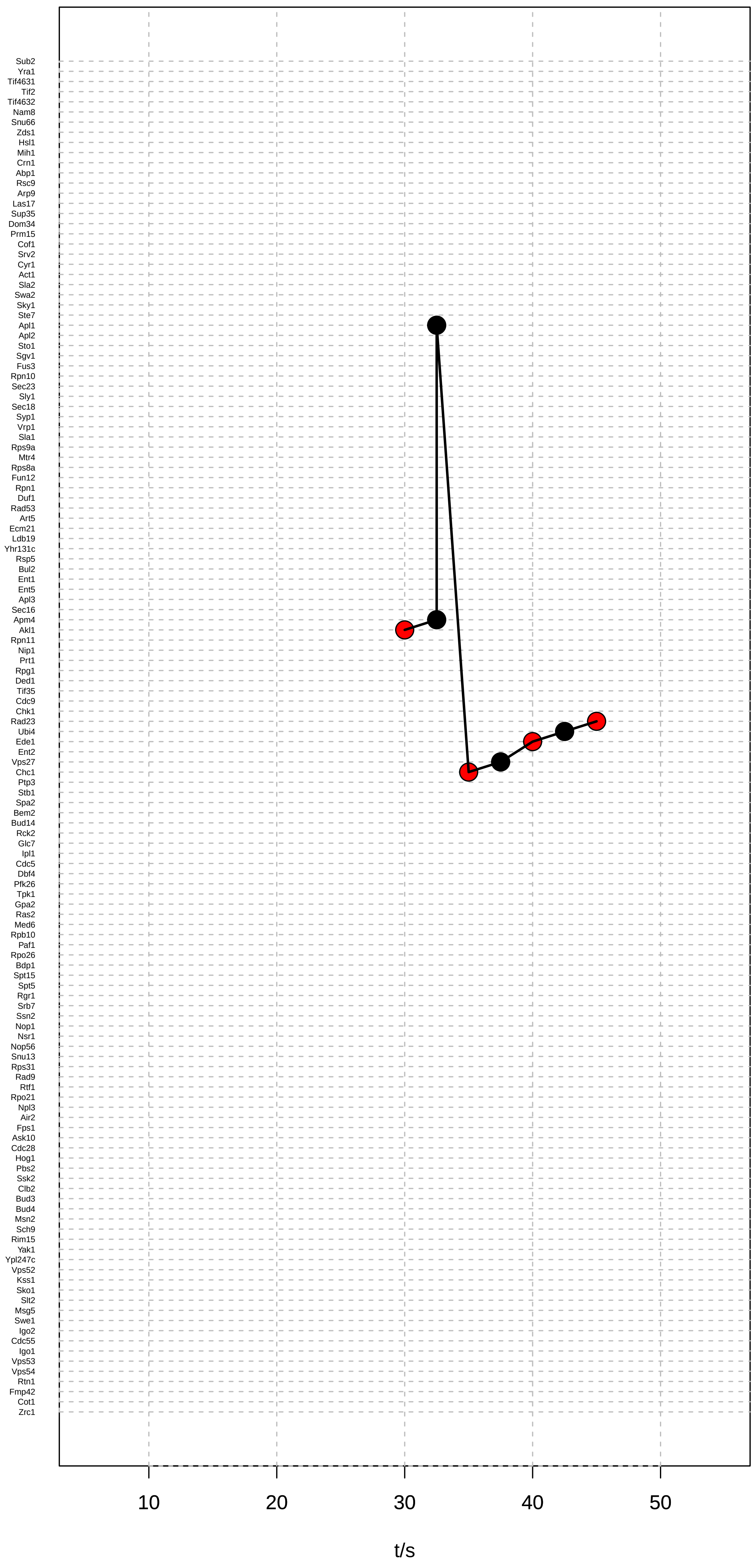

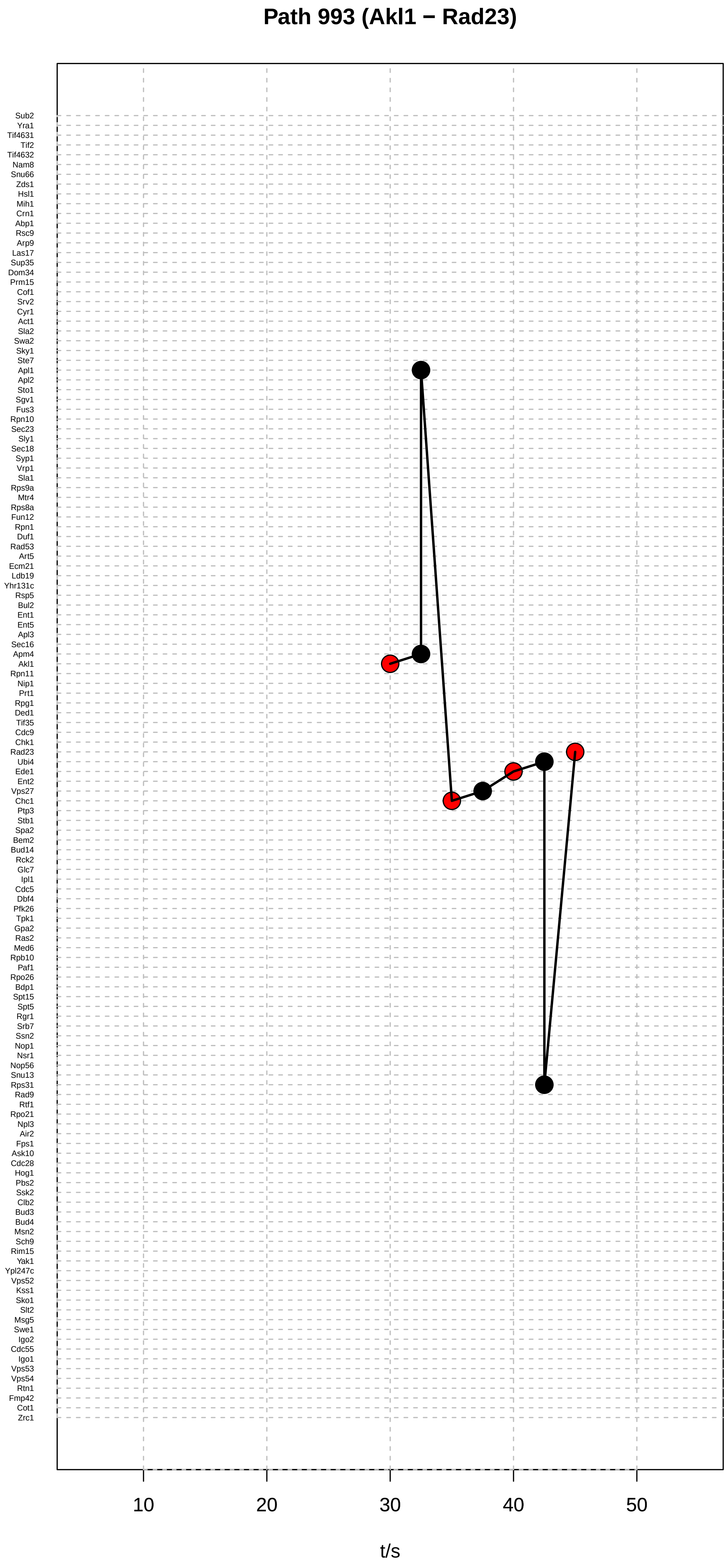
Path 993 (Akl1 − Rad23)
Sub2
Yra1
Tif4631
Tif2
Tif4632
Nam8
Snu66
Zds1
Hsl1
Mih1
Crn1
Abp1
Rsc9
Arp9
Las17
Sup35
Dom34
Prm15
Cof1
Srv2
Cyr1
Act1
Sla2
Swa2
Sky1
Ste7
Apl1
Apl2
Sto1
Sgv1
Fus3
Rpn10
Sec23
Sly1
Sec18
Syp1
Vrp1
Sla1
Rps9a
Mtr4
Rps8a
Fun12
Rpn1
Duf1
Rad53
Art5
Ecm21
Ldb19
Yhr131c
Rsp5
Bul2
Ent1
Ent5
Apl3
Sec16
Apm4
Akl1
Rpn11
Nip1
Prt1
Rpg1
Ded1
Tif35
Cdc9
Chk1
Rad23
Ubi4
Ede1
Ent2
Vps27
Chc1
Ptp3
Stb1
Spa2
Bem2
Bud14
Rck2
Glc7
Ipl1
Cdc5
Dbf4
Pfk26
Tpk1
Gpa2
Ras2
Med6
Rpb10
Paf1
Rpo26
Bdp1
Spt15
Spt5
Rgr1
Srb7
Ssn2
Nop1
Nsr1
Nop56
Snu13
Rps31
Rad9
Rtf1
Rpo21
Npl3
Air2
Fps1
Ask10
Cdc28
Hog1
Pbs2
Ssk2
Clb2
Bud3
Bud4
Msn2
Sch9
Rim15
Yak1
Ypl247c
Vps52
Kss1
Sko1
Slt2
Msg5
Swe1
Igo2
Cdc55
Igo1
Vps53
Vps54
Rtn1
Fmp42
Cot1
Zrc1
10
20
30
40
50
t/s

#### Path 994 (Akl1 − Rad23)

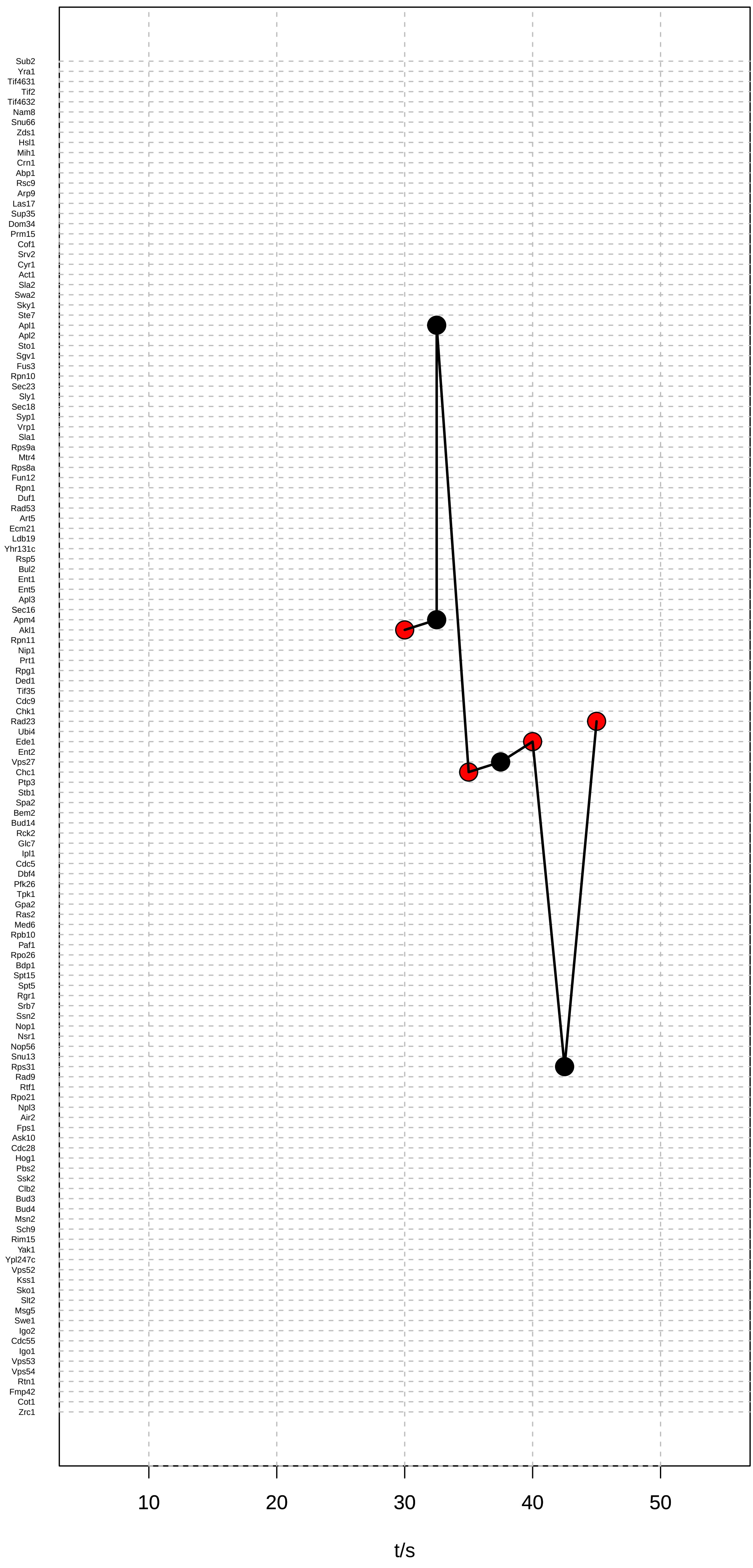

### Path 995 (Akl1 − Rad23)

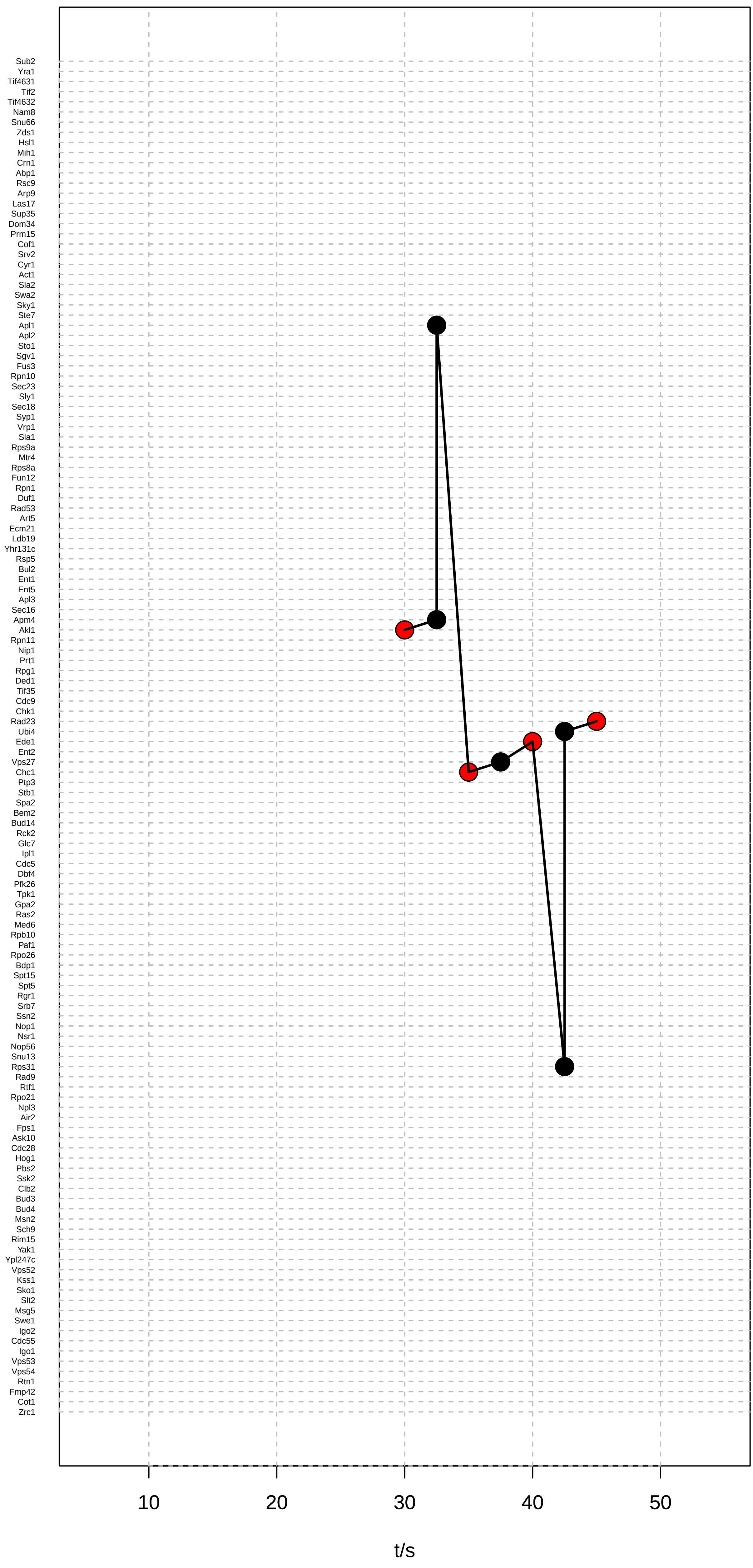

#### Path 996 (Akl1 − Rad23)

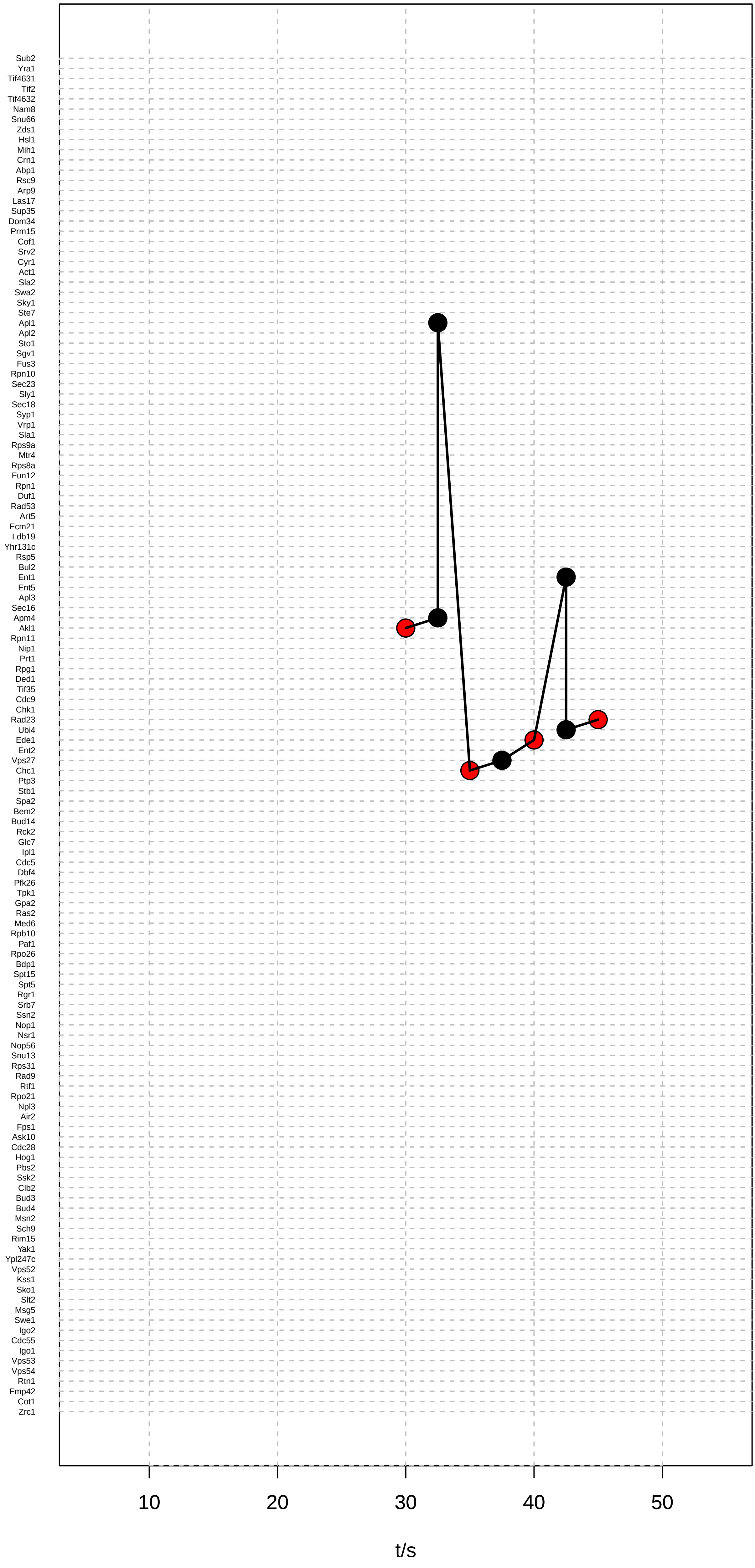

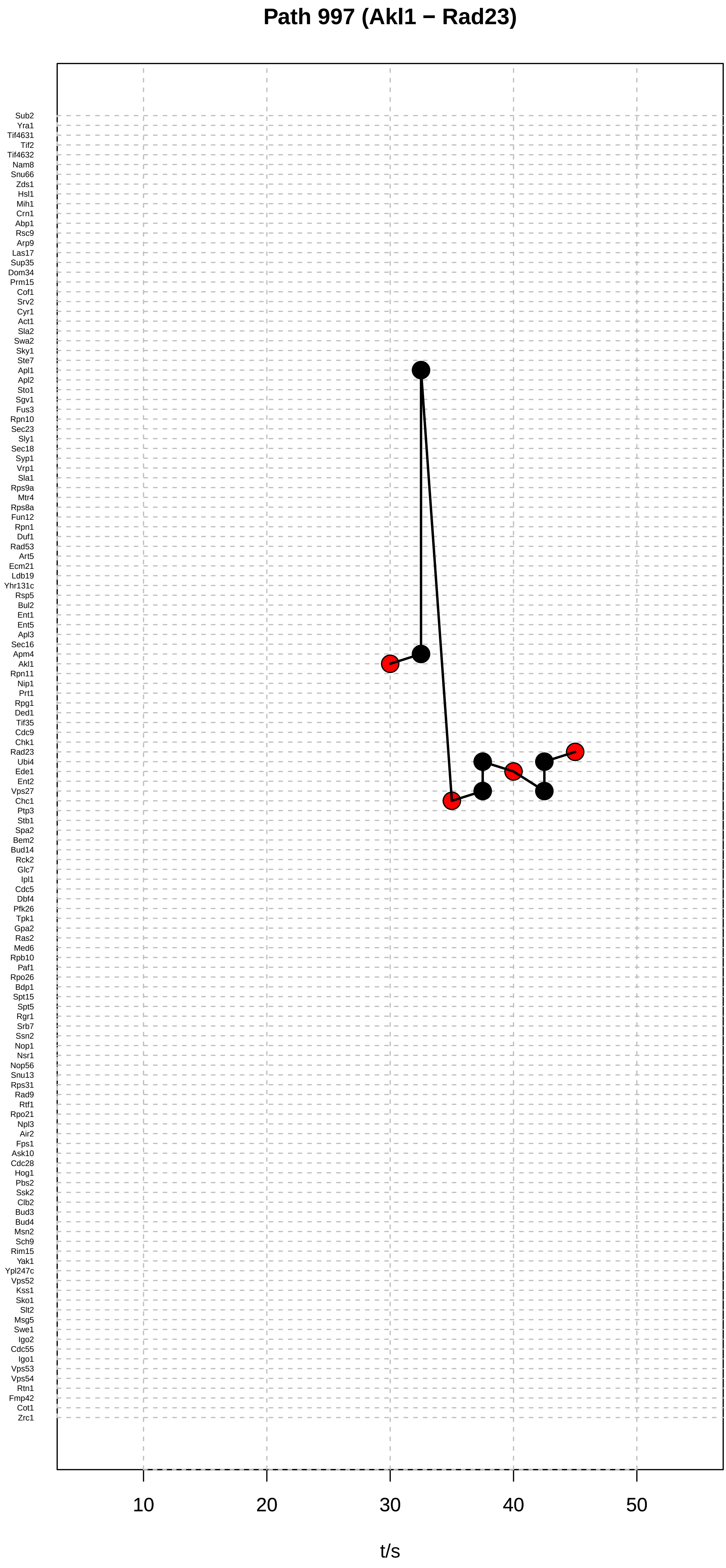

Path 997 (Akl1 − Rad23)
Sub2
Yra1
Tif4631
Tif2
Tif4632
Nam8
Snu66
Zds1
Hsl1
Mih1
Crn1
Abp1
Rsc9
Arp9
Las17
Sup35
Dom34
Prm15
Cof1
Srv2
Cyr1
Act1
Sla2
Swa2
Sky1
Ste7
Apl1
Apl2
Sto1
Sgv1
Fus3
Rpn10
Sec23
Sly1
Sec18
Syp1
Vrp1
Sla1
Rps9a
Mtr4
Rps8a
Fun12
Rpn1
Duf1
Rad53
Art5
Ecm21
Ldb19
Yhr131c
Rsp5
Bul2
Ent1
Ent5
Apl3
Sec16
Apm4
Akl1
Rpn11
Nip1
Prt1
Rpg1
Ded1
Tif35
Cdc9
Chk1
Rad23
Ubi4
Ede1
Ent2
Vps27
Chc1
Ptp3
Stb1
Spa2
Bem2
Bud14
Rck2
Glc7
Ipl1
Cdc5
Dbf4
Pfk26
Tpk1
Gpa2
Ras2
Med6
Rpb10
Paf1
Rpo26
Bdp1
Spt15
Spt5
Rgr1
Srb7
Ssn2
Nop1
Nsr1
Nop56
Snu13
Rps31
Rad9
Rtf1
Rpo21
Npl3
Air2
Fps1
Ask10
Cdc28
Hog1
Pbs2
Ssk2
Clb2
Bud3
Bud4
Msn2
Sch9
Rim15
Yak1
Ypl247c
Vps52
Kss1
Sko1
Slt2
Msg5
Swe1
Igo2
Cdc55
Igo1
Vps53
Vps54
Rtn1
Fmp42
Cot1
Zrc1
10
20
30
40
50
t/s

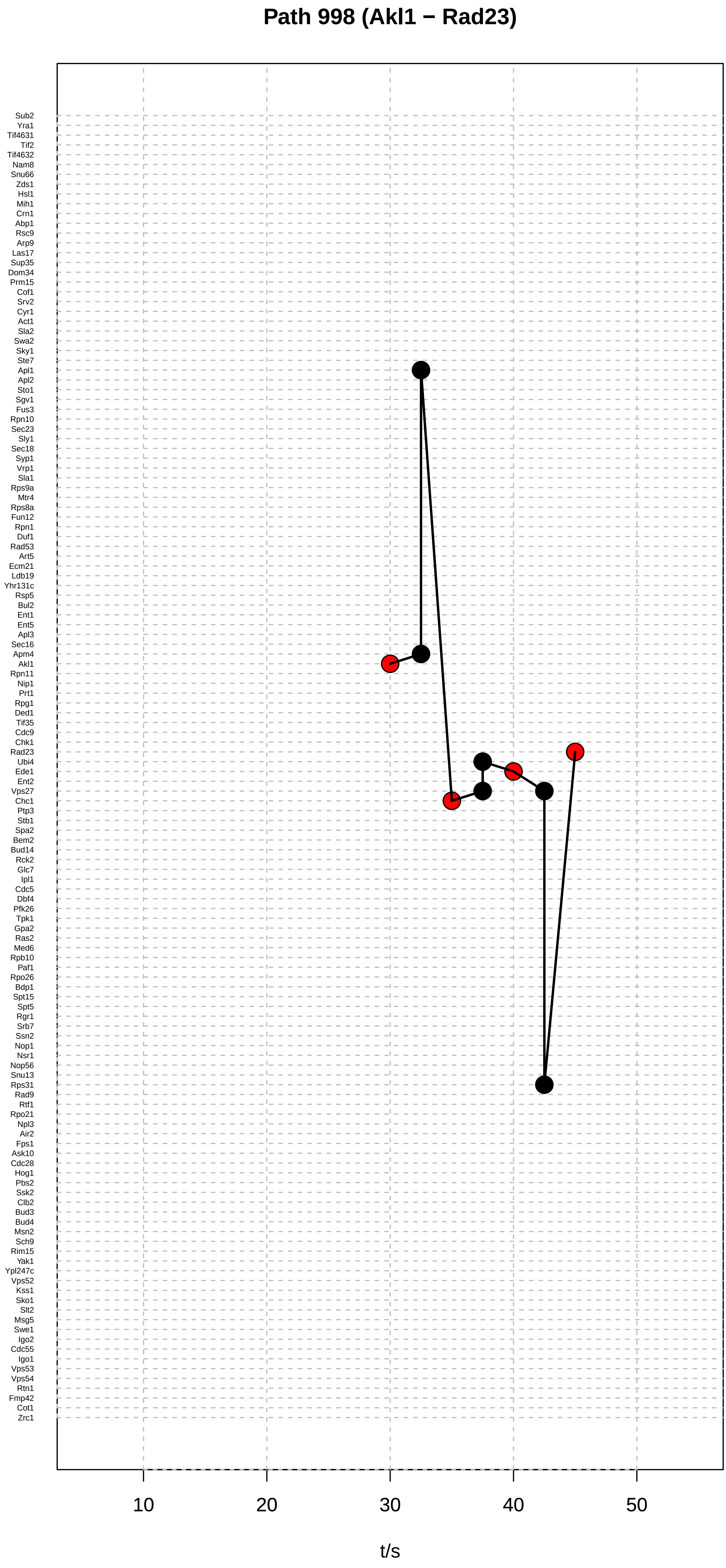
Path 998 (Akl1 − Rad23)
Sub2
Yra1
Tif4631
Tif2
Tif4632
Nam8
Snu66
Zds1
Hsl1
Mih1
Crn1
Abp1
Rsc9
Arp9
Las17
Sup35
Dom34
Prm15
Cof1
Srv2
Cyr1
Act1
Sla2
Swa2
Sky1
Ste7
Apl1
Apl2
Sto1
Sgv1
Fus3
Rpn10
Sec23
Sly1
Sec18
Syp1
Vrp1
Sla1
Rps9a
Mtr4
Rps8a
Fun12
Rpn1
Duf1
Rad53
Art5
Ecm21
Ldb19
Yhr131c
Rsp5
Bul2
Ent1
Ent5
Apl3
Sec16
Apm4
Akl1
Rpn11
Nip1
Prt1
Rpg1
Ded1
Tif35
Cdc9
Chk1
Rad23
Ubi4
Ede1
Ent2
Vps27
Chc1
Ptp3
Stb1
Spa2
Bem2
Bud14
Rck2
Glc7
Ipl1
Cdc5
Dbf4
Pfk26
Tpk1
Gpa2
Ras2
Med6
Rpb10
Paf1
Rpo26
Bdp1
Spt15
Spt5
Rgr1
Srb7
Ssn2
Nop1
Nsr1
Nop56
Snu13
Rps31
Rad9
Rtf1
Rpo21
Npl3
Air2
Fps1
Ask10
Cdc28
Hog1
Pbs2
Ssk2
Clb2
Bud3
Bud4
Msn2
Sch9
Rim15
Yak1
Ypl247c
Vps52
Kss1
Sko1
Slt2
Msg5
Swe1
Igo2
Cdc55
Igo1
Vps53
Vps54
Rtn1
Fmp42
Cot1
Zrc1
10
20
30
40
50
t/s

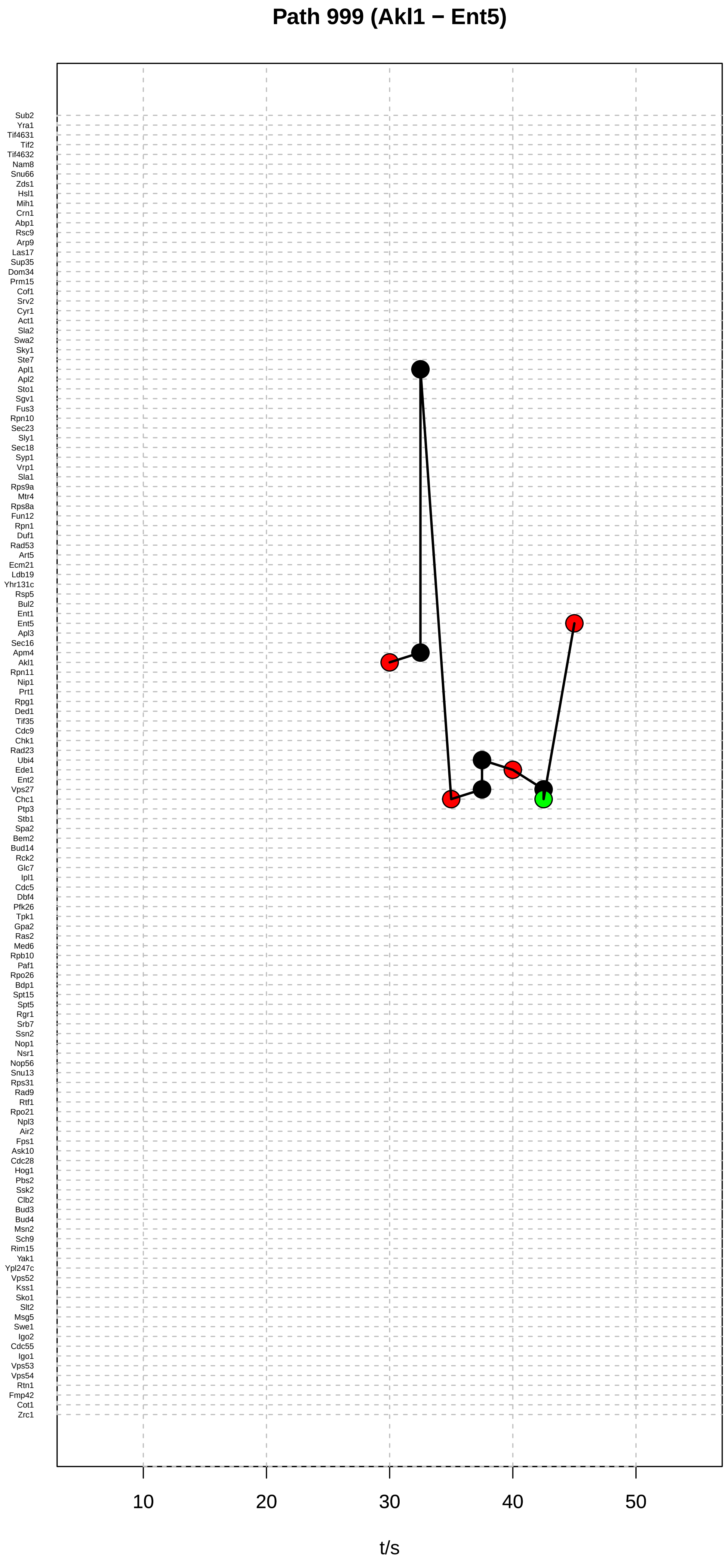
Path 999 (Akl1 − Ent5)
Sub2
Yra1
Tif4631
Tif2
Tif4632
Nam8
Snu66
Zds1
Hsl1
Mih1
Crn1
Abp1
Rsc9
Arp9
Las17
Sup35
Dom34
Prm15
Cof1
Srv2
Cyr1
Act1
Sla2
Swa2
Sky1
Ste7
Apl1
Apl2
Sto1
Sgv1
Fus3
Rpn10
Sec23
Sly1
Sec18
Syp1
Vrp1
Sla1
Rps9a
Mtr4
Rps8a
Fun12
Rpn1
Duf1
Rad53
Art5
Ecm21
Ldb19
Yhr131c
Rsp5
Bul2
Ent1
Ent5
Apl3
Sec16
Apm4
Akl1
Rpn11
Nip1
Prt1
Rpg1
Ded1
Tif35
Cdc9
Chk1
Rad23
Ubi4
Ede1
Ent2
Vps27
Chc1
Ptp3
Stb1
Spa2
Bem2
Bud14
Rck2
Glc7
Ipl1
Cdc5
Dbf4
Pfk26
Tpk1
Gpa2
Ras2
Med6
Rpb10
Paf1
Rpo26
Bdp1
Spt15
Spt5
Rgr1
Srb7
Ssn2
Nop1
Nsr1
Nop56
Snu13
Rps31
Rad9
Rtf1
Rpo21
Npl3
Air2
Fps1
Ask10
Cdc28
Hog1
Pbs2
Ssk2
Clb2
Bud3
Bud4
Msn2
Sch9
Rim15
Yak1
Ypl247c
Vps52
Kss1
Sko1
Slt2
Msg5
Swe1
Igo2
Cdc55
Igo1
Vps53
Vps54
Rtn1
Fmp42
Cot1
Zrc1
10
20
30
40
50
t/s

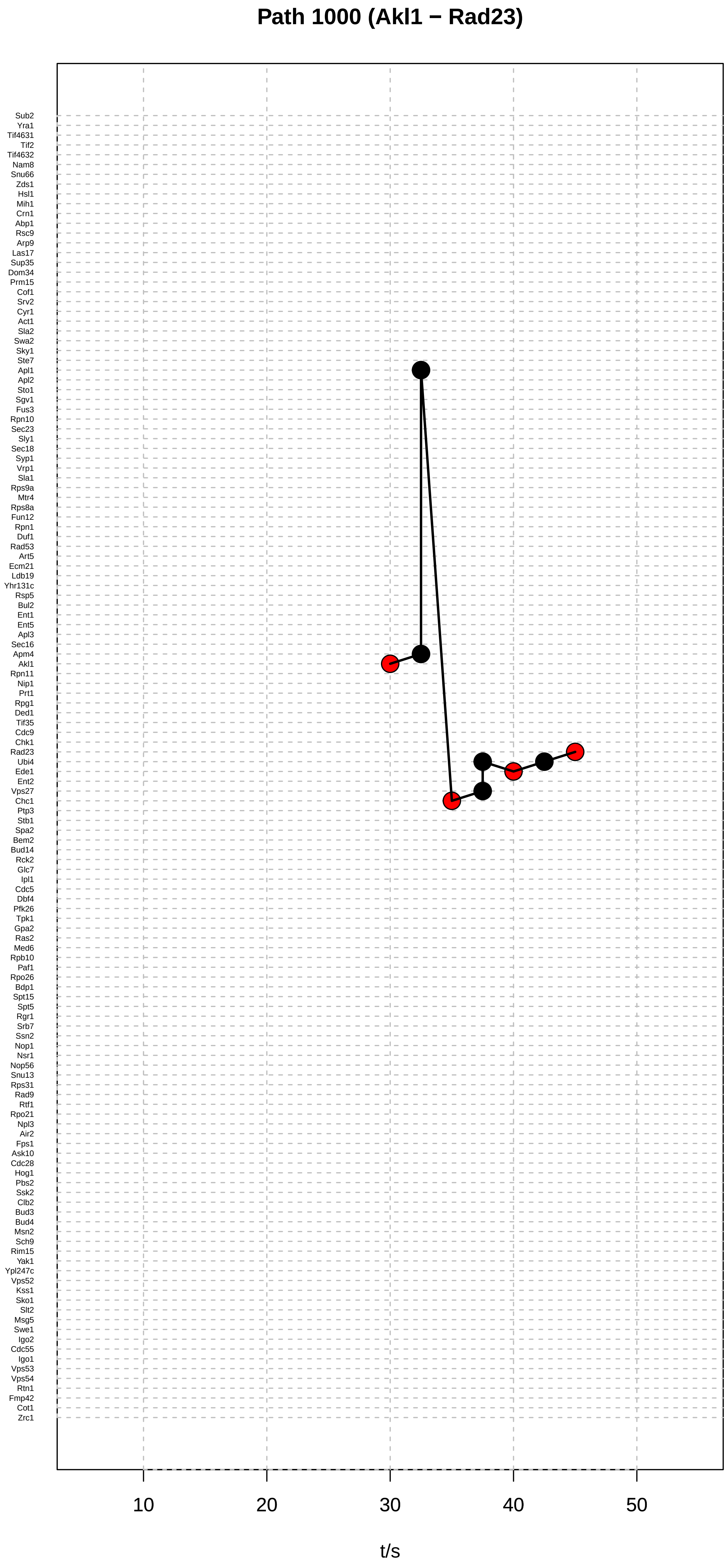

Path 1000 (Akl1 − Rad23)
Sub2
Yra1
Tif4631
Tif2
Tif4632
Nam8
Snu66
Zds1
Hsl1
Mih1
Crn1
Abp1
Rsc9
Arp9
Las17
Sup35
Dom34
Prm15
Cof1
Srv2
Cyr1
Act1
Sla2
Swa2
Sky1
Ste7
Apl1
Apl2
Sto1
Sgv1
Fus3
Rpn10
Sec23
Sly1
Sec18
Syp1
Vrp1
Sla1
Rps9a
Mtr4
Rps8a
Fun12
Rpn1
Duf1
Rad53
Art5
Ecm21
Ldb19
Yhr131c
Rsp5
Bul2
Ent1
Ent5
Apl3
Sec16
Apm4
Akl1
Rpn11
Nip1
Prt1
Rpg1
Ded1
Tif35
Cdc9
Chk1
Rad23
Ubi4
Ede1
Ent2
Vps27
Chc1
Ptp3
Stb1
Spa2
Bem2
Bud14
Rck2
Glc7
Ipl1
Cdc5
Dbf4
Pfk26
Tpk1
Gpa2
Ras2
Med6
Rpb10
Paf1
Rpo26
Bdp1
Spt15
Spt5
Rgr1
Srb7
Ssn2
Nop1
Nsr1
Nop56
Snu13
Rps31
Rad9
Rtf1
Rpo21
Npl3
Air2
Fps1
Ask10
Cdc28
Hog1
Pbs2
Ssk2
Clb2
Bud3
Bud4
Msn2
Sch9
Rim15
Yak1
Ypl247c
Vps52
Kss1
Sko1
Slt2
Msg5
Swe1
Igo2
Cdc55
Igo1
Vps53
Vps54
Rtn1
Fmp42
Cot1
Zrc1
10
20
30
40
50
t/s

#### Path 1001 (Akl1 − Rad23)

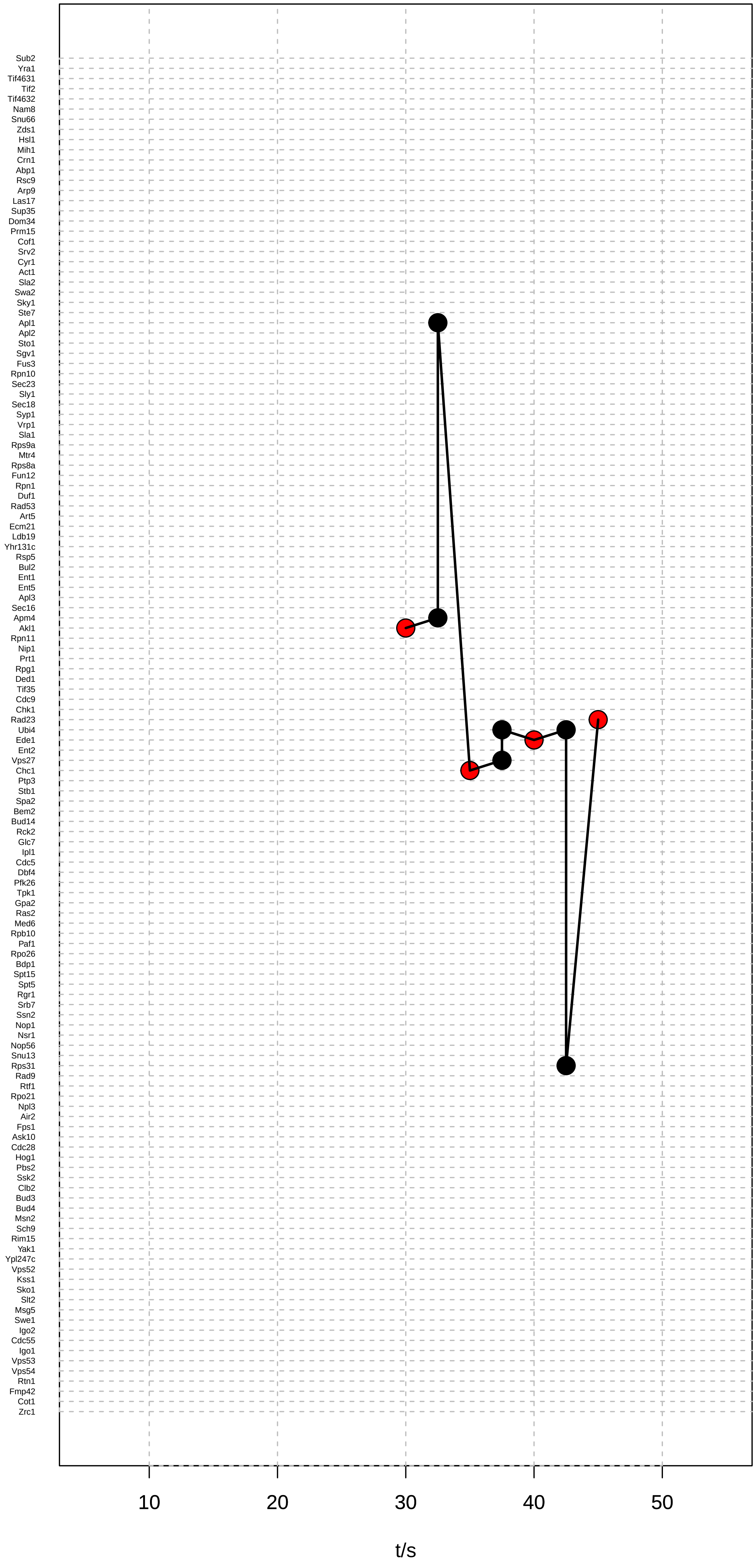

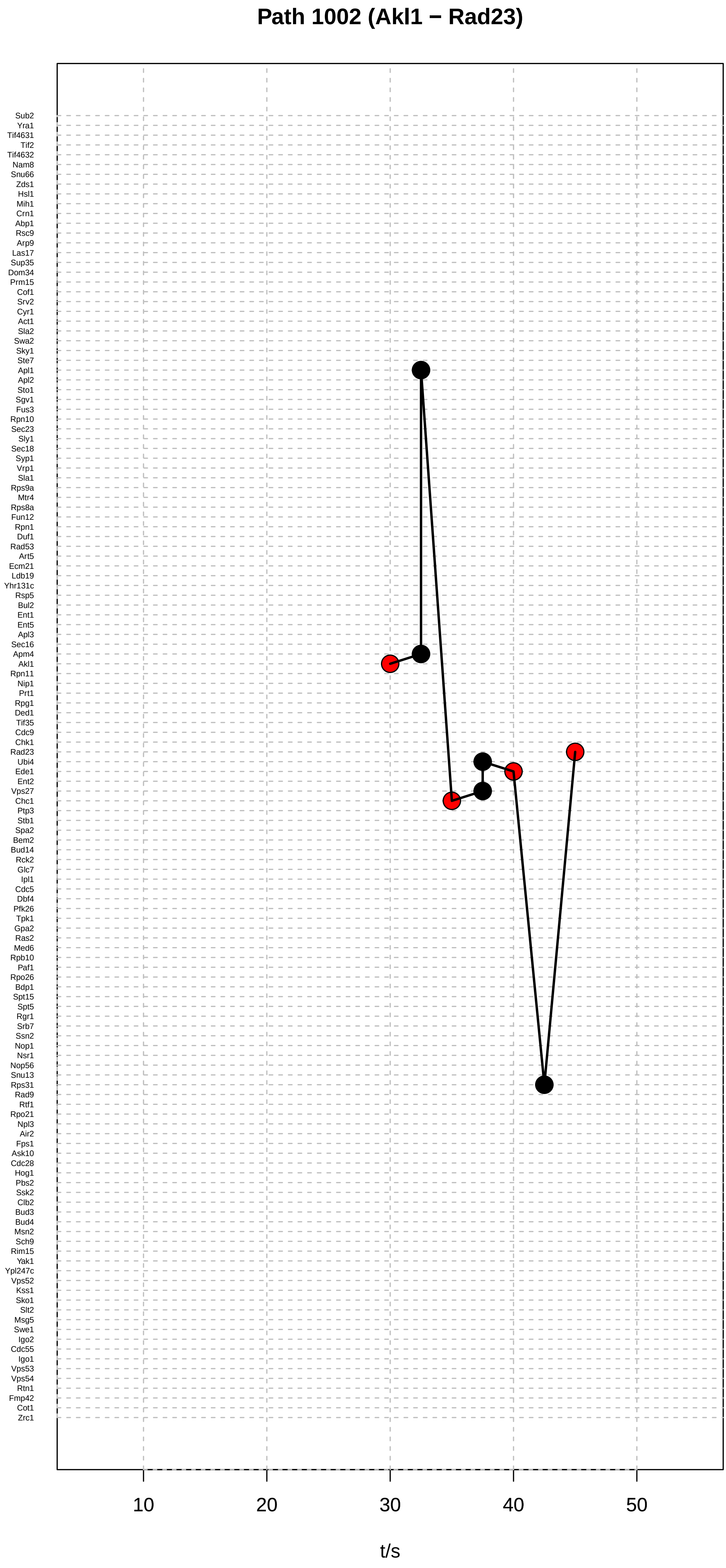
Path 1002 (Akl1 − Rad23)
Sub2
Yra1
Tif4631
Tif2
Tif4632
Nam8
Snu66
Zds1
Hsl1
Mih1
Crn1
Abp1
Rsc9
Arp9
Las17
Sup35
Dom34
Prm15
Cof1
Srv2
Cyr1
Act1
Sla2
Swa2
Sky1
Ste7
Apl1
Apl2
Sto1
Sgv1
Fus3
Rpn10
Sec23
Sly1
Sec18
Syp1
Vrp1
Sla1
Rps9a
Mtr4
Rps8a
Fun12
Rpn1
Duf1
Rad53
Art5
Ecm21
Ldb19
Yhr131c
Rsp5
Bul2
Ent1
Ent5
Apl3
Sec16
Apm4
Akl1
Rpn11
Nip1
Prt1
Rpg1
Ded1
Tif35
Cdc9
Chk1
Rad23
Ubi4
Ede1
Ent2
Vps27
Chc1
Ptp3
Stb1
Spa2
Bem2
Bud14
Rck2
Glc7
Ipl1
Cdc5
Dbf4
Pfk26
Tpk1
Gpa2
Ras2
Med6
Rpb10
Paf1
Rpo26
Bdp1
Spt15
Spt5
Rgr1
Srb7
Ssn2
Nop1
Nsr1
Nop56
Snu13
Rps31
Rad9
Rtf1
Rpo21
Npl3
Air2
Fps1
Ask10
Cdc28
Hog1
Pbs2
Ssk2
Clb2
Bud3
Bud4
Msn2
Sch9
Rim15
Yak1
Ypl247c
Vps52
Kss1
Sko1
Slt2
Msg5
Swe1
Igo2
Cdc55
Igo1
Vps53
Vps54
Rtn1
Fmp42
Cot1
Zrc1
10
20
30
40
50
t/s

### Path 1003 (Akl1 − Rad23)

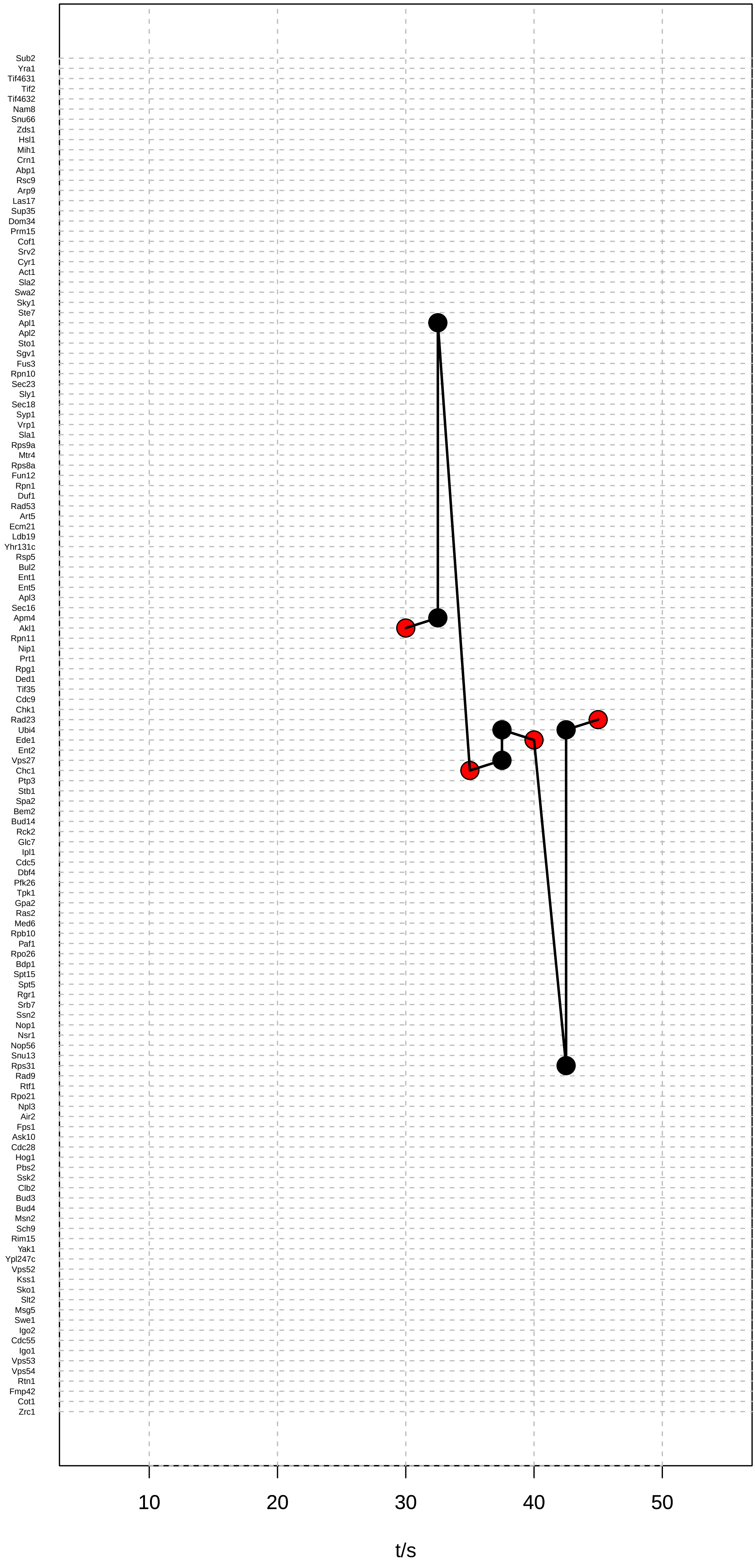

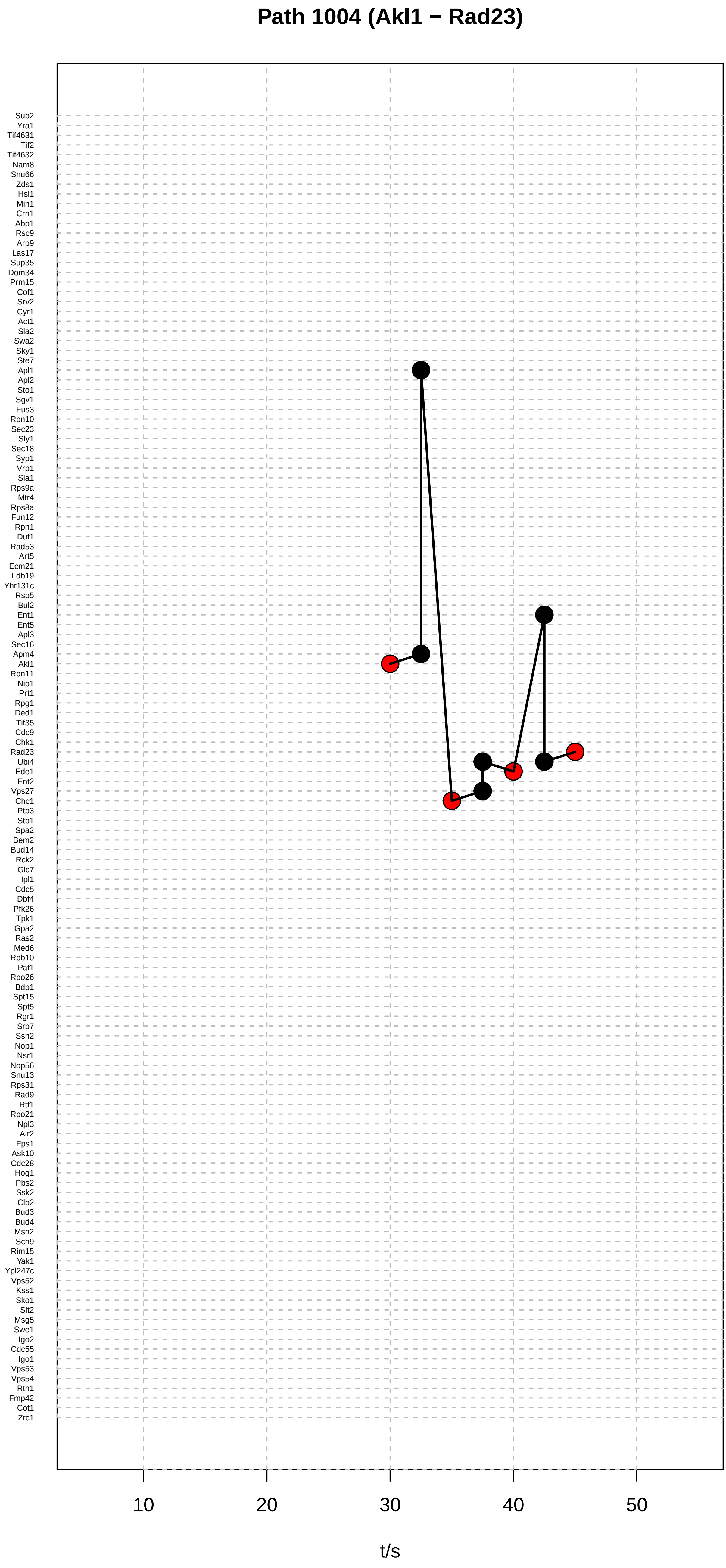
Path 1004 (Akl1 − Rad23)
Sub2
Yra1
Tif4631
Tif2
Tif4632
Nam8
Snu66
Zds1
Hsl1
Mih1
Crn1
Abp1
Rsc9
Arp9
Las17
Sup35
Dom34
Prm15
Cof1
Srv2
Cyr1
Act1
Sla2
Swa2
Sky1
Ste7
Apl1
Apl2
Sto1
Sgv1
Fus3
Rpn10
Sec23
Sly1
Sec18
Syp1
Vrp1
Sla1
Rps9a
Mtr4
Rps8a
Fun12
Rpn1
Duf1
Rad53
Art5
Ecm21
Ldb19
Yhr131c
Rsp5
Bul2
Ent1
Ent5
Apl3
Sec16
Apm4
Akl1
Rpn11
Nip1
Prt1
Rpg1
Ded1
Tif35
Cdc9
Chk1
Rad23
Ubi4
Ede1
Ent2
Vps27
Chc1
Ptp3
Stb1
Spa2
Bem2
Bud14
Rck2
Glc7
Ipl1
Cdc5
Dbf4
Pfk26
Tpk1
Gpa2
Ras2
Med6
Rpb10
Paf1
Rpo26
Bdp1
Spt15
Spt5
Rgr1
Srb7
Ssn2
Nop1
Nsr1
Nop56
Snu13
Rps31
Rad9
Rtf1
Rpo21
Npl3
Air2
Fps1
Ask10
Cdc28
Hog1
Pbs2
Ssk2
Clb2
Bud3
Bud4
Msn2
Sch9
Rim15
Yak1
Ypl247c
Vps52
Kss1
Sko1
Slt2
Msg5
Swe1
Igo2
Cdc55
Igo1
Vps53
Vps54
Rtn1
Fmp42
Cot1
Zrc1
10
20
30
40
50
t/s

### Path 1005 (Akl1 − Ent5)

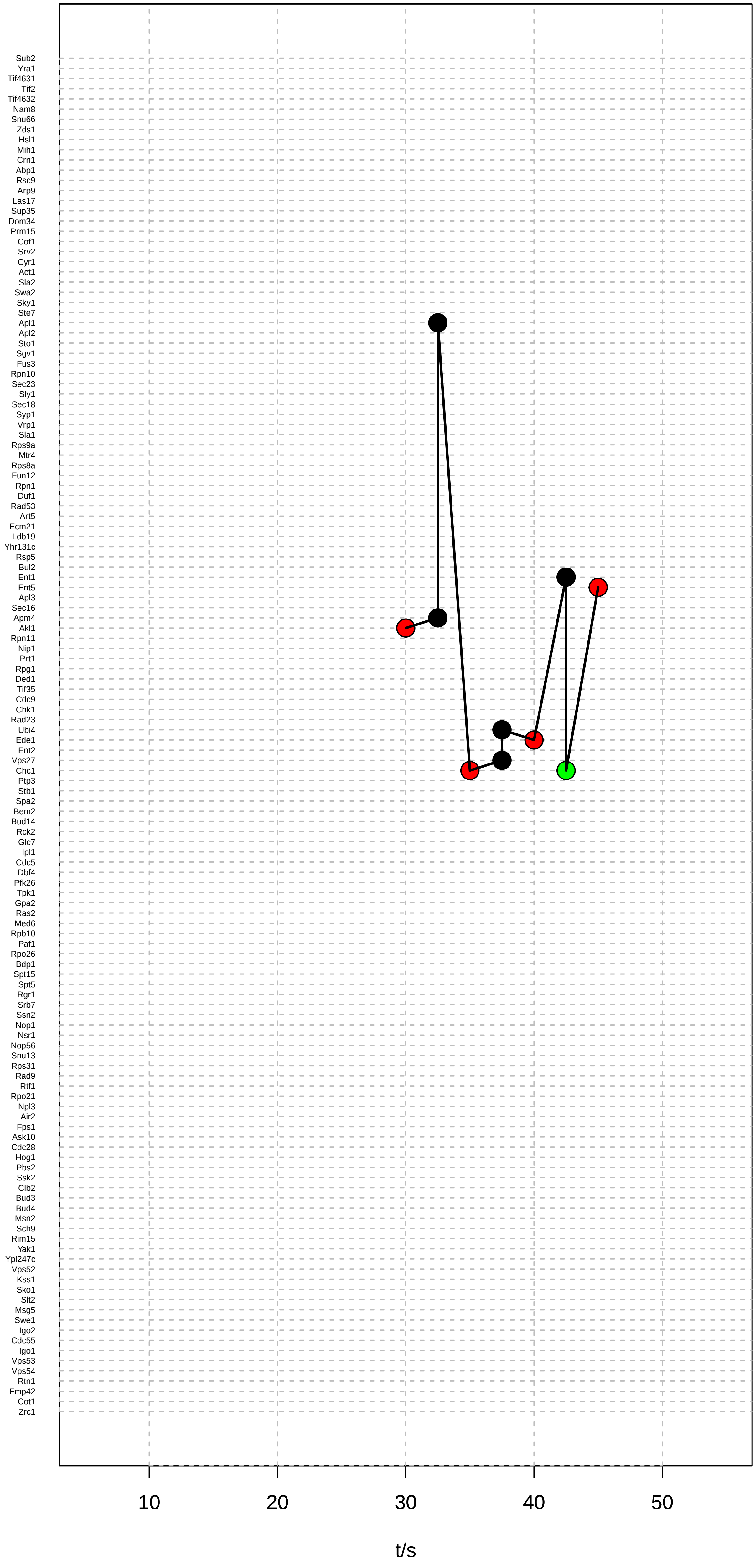

### Path 1006 (Akl1 − Rad23)

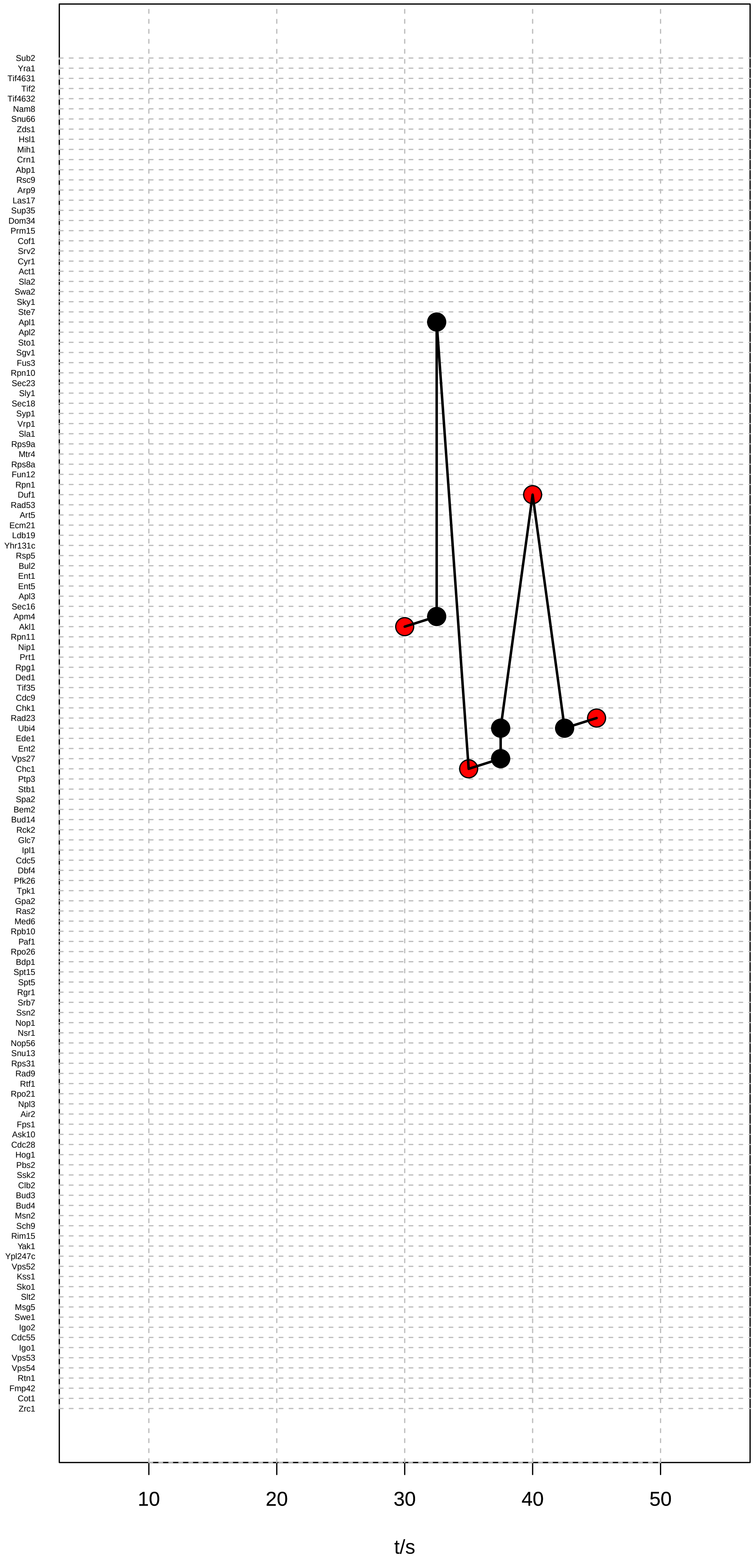

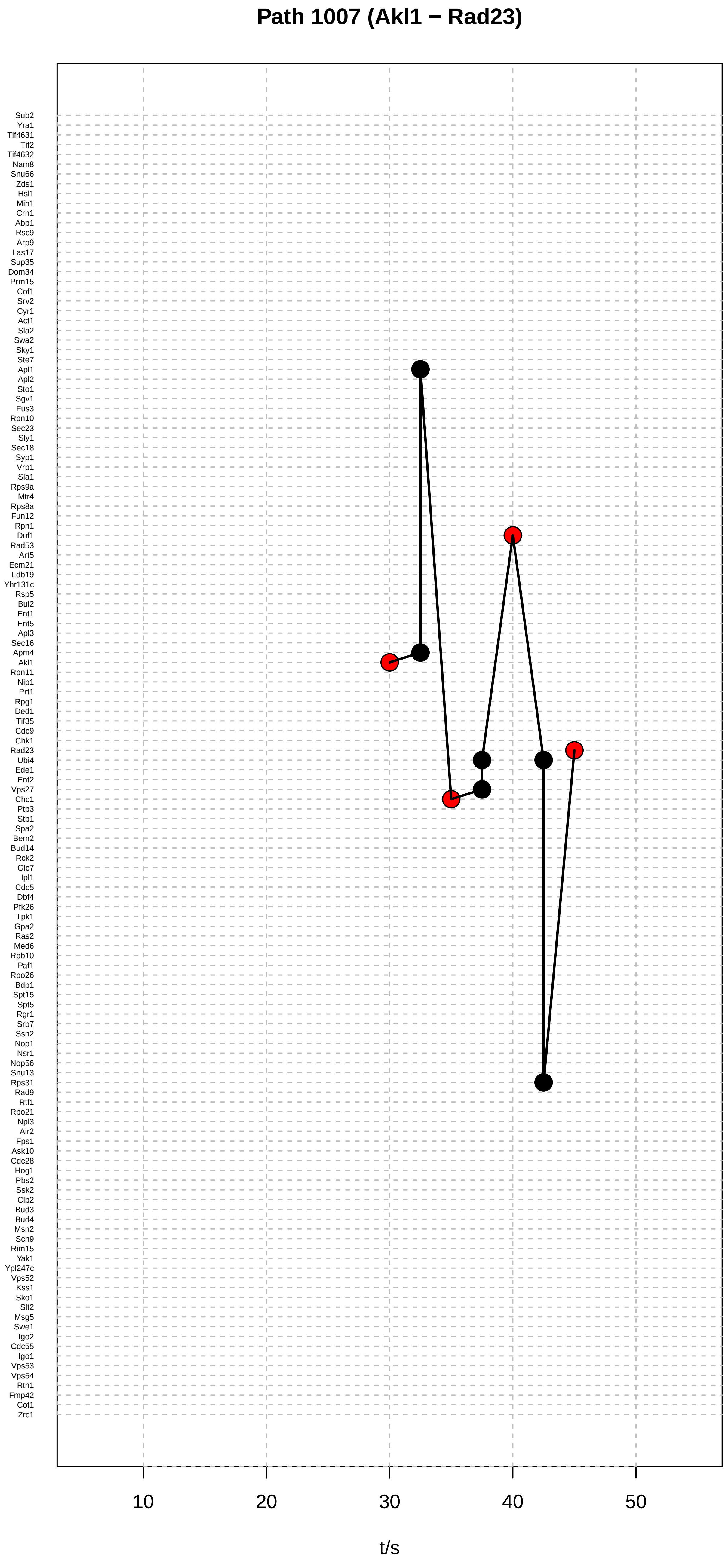
Path 1007 (Akl1 − Rad23)
Sub2
Yra1
Tif4631
Tif2
Tif4632
Nam8
Snu66
Zds1
Hsl1
Mih1
Crn1
Abp1
Rsc9
Arp9
Las17
Sup35
Dom34
Prm15
Cof1
Srv2
Cyr1
Act1
Sla2
Swa2
Sky1
Ste7
Apl1
Apl2
Sto1
Sgv1
Fus3
Rpn10
Sec23
Sly1
Sec18
Syp1
Vrp1
Sla1
Rps9a
Mtr4
Rps8a
Fun12
Rpn1
Duf1
Rad53
Art5
Ecm21
Ldb19
Yhr131c
Rsp5
Bul2
Ent1
Ent5
Apl3
Sec16
Apm4
Akl1
Rpn11
Nip1
Prt1
Rpg1
Ded1
Tif35
Cdc9
Chk1
Rad23
Ubi4
Ede1
Ent2
Vps27
Chc1
Ptp3
Stb1
Spa2
Bem2
Bud14
Rck2
Glc7
Ipl1
Cdc5
Dbf4
Pfk26
Tpk1
Gpa2
Ras2
Med6
Rpb10
Paf1
Rpo26
Bdp1
Spt15
Spt5
Rgr1
Srb7
Ssn2
Nop1
Nsr1
Nop56
Snu13
Rps31
Rad9
Rtf1
Rpo21
Npl3
Air2
Fps1
Ask10
Cdc28
Hog1
Pbs2
Ssk2
Clb2
Bud3
Bud4
Msn2
Sch9
Rim15
Yak1
Ypl247c
Vps52
Kss1
Sko1
Slt2
Msg5
Swe1
Igo2
Cdc55
Igo1
Vps53
Vps54
Rtn1
Fmp42
Cot1
Zrc1
10
20
30
40
50
t/s

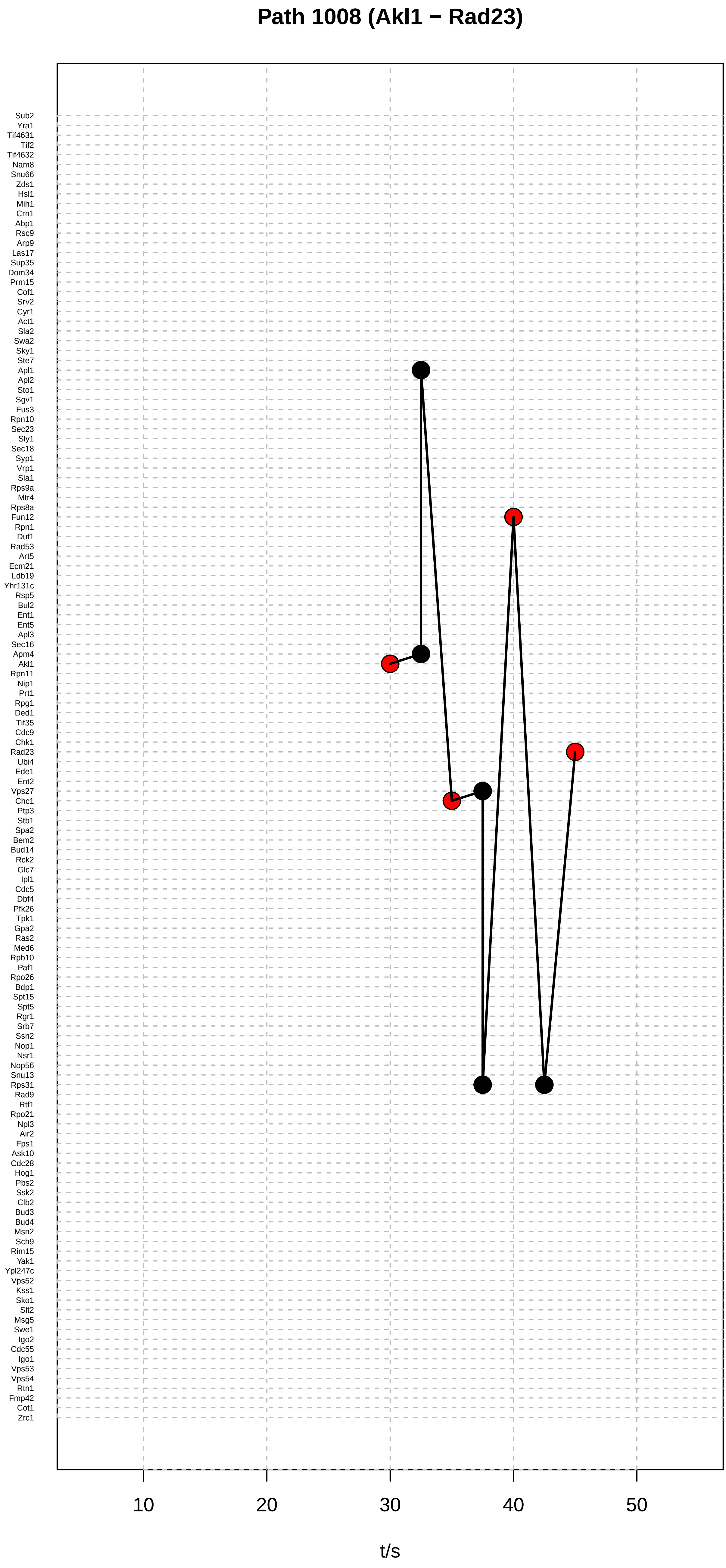
Path 1008 (Akl1 − Rad23)
Sub2
Yra1
Tif4631
Tif2
Tif4632
Nam8
Snu66
Zds1
Hsl1
Mih1
Crn1
Abp1
Rsc9
Arp9
Las17
Sup35
Dom34
Prm15
Cof1
Srv2
Cyr1
Act1
Sla2
Swa2
Sky1
Ste7
Apl1
Apl2
Sto1
Sgv1
Fus3
Rpn10
Sec23
Sly1
Sec18
Syp1
Vrp1
Sla1
Rps9a
Mtr4
Rps8a
Fun12
Rpn1
Duf1
Rad53
Art5
Ecm21
Ldb19
Yhr131c
Rsp5
Bul2
Ent1
Ent5
Apl3
Sec16
Apm4
Akl1
Rpn11
Nip1
Prt1
Rpg1
Ded1
Tif35
Cdc9
Chk1
Rad23
Ubi4
Ede1
Ent2
Vps27
Chc1
Ptp3
Stb1
Spa2
Bem2
Bud14
Rck2
Glc7
Ipl1
Cdc5
Dbf4
Pfk26
Tpk1
Gpa2
Ras2
Med6
Rpb10
Paf1
Rpo26
Bdp1
Spt15
Spt5
Rgr1
Srb7
Ssn2
Nop1
Nsr1
Nop56
Snu13
Rps31
Rad9
Rtf1
Rpo21
Npl3
Air2
Fps1
Ask10
Cdc28
Hog1
Pbs2
Ssk2
Clb2
Bud3
Bud4
Msn2
Sch9
Rim15
Yak1
Ypl247c
Vps52
Kss1
Sko1
Slt2
Msg5
Swe1
Igo2
Cdc55
Igo1
Vps53
Vps54
Rtn1
Fmp42
Cot1
Zrc1
10
20
30
40
50
t/s

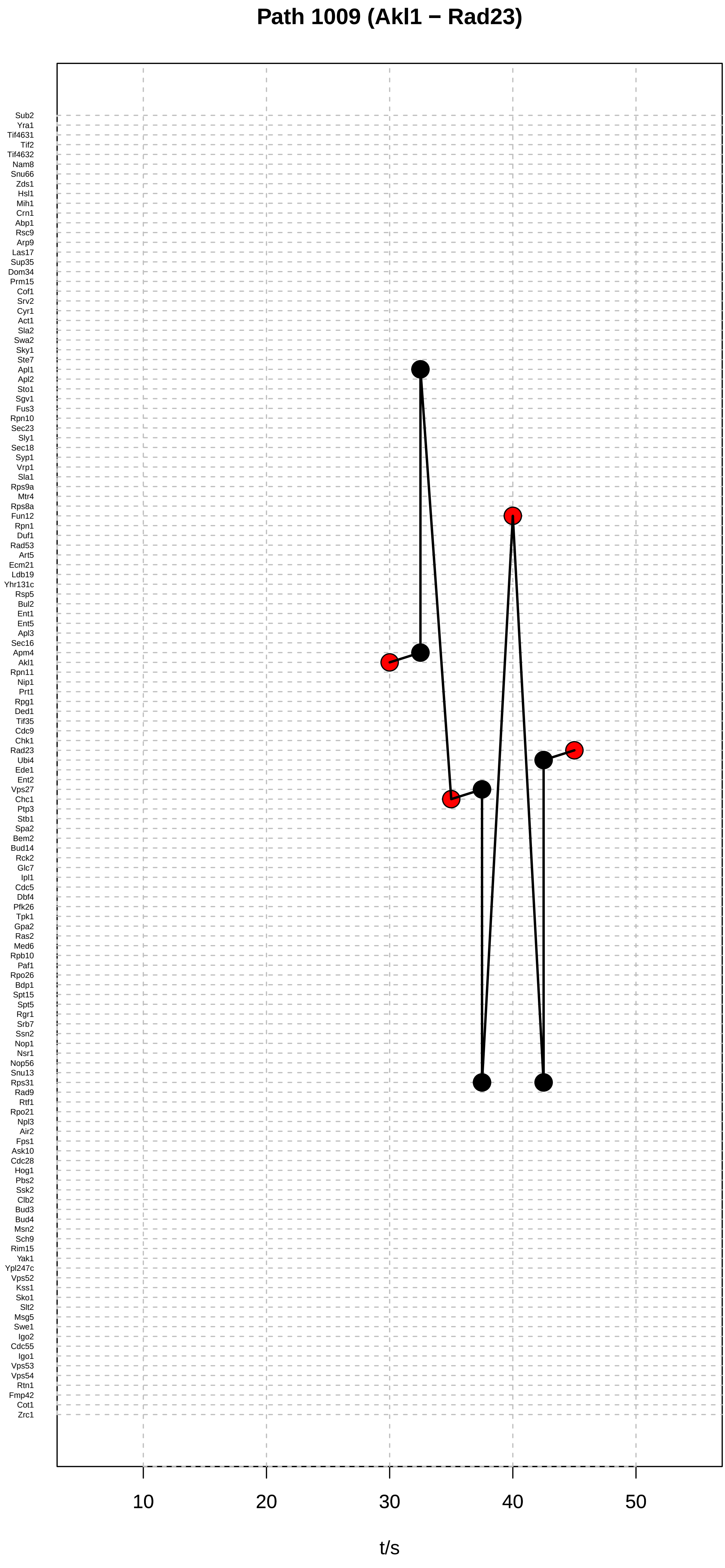

Path 1009 (Akl1 − Rad23)
Sub2
Yra1
Tif4631
Tif2
Tif4632
Nam8
Snu66
Zds1
Hsl1
Mih1
Crn1
Abp1
Rsc9
Arp9
Las17
Sup35
Dom34
Prm15
Cof1
Srv2
Cyr1
Act1
Sla2
Swa2
Sky1
Ste7
Apl1
Apl2
Sto1
Sgv1
Fus3
Rpn10
Sec23
Sly1
Sec18
Syp1
Vrp1
Sla1
Rps9a
Mtr4
Rps8a
Fun12
Rpn1
Duf1
Rad53
Art5
Ecm21
Ldb19
Yhr131c
Rsp5
Bul2
Ent1
Ent5
Apl3
Sec16
Apm4
Akl1
Rpn11
Nip1
Prt1
Rpg1
Ded1
Tif35
Cdc9
Chk1
Rad23
Ubi4
Ede1
Ent2
Vps27
Chc1
Ptp3
Stb1
Spa2
Bem2
Bud14
Rck2
Glc7
Ipl1
Cdc5
Dbf4
Pfk26
Tpk1
Gpa2
Ras2
Med6
Rpb10
Paf1
Rpo26
Bdp1
Spt15
Spt5
Rgr1
Srb7
Ssn2
Nop1
Nsr1
Nop56
Snu13
Rps31
Rad9
Rtf1
Rpo21
Npl3
Air2
Fps1
Ask10
Cdc28
Hog1
Pbs2
Ssk2
Clb2
Bud3
Bud4
Msn2
Sch9
Rim15
Yak1
Ypl247c
Vps52
Kss1
Sko1
Slt2
Msg5
Swe1
Igo2
Cdc55
Igo1
Vps53
Vps54
Rtn1
Fmp42
Cot1
Zrc1
10
20
30
40
50
t/s

#### Path 1010 (Akl1 − Rad23)

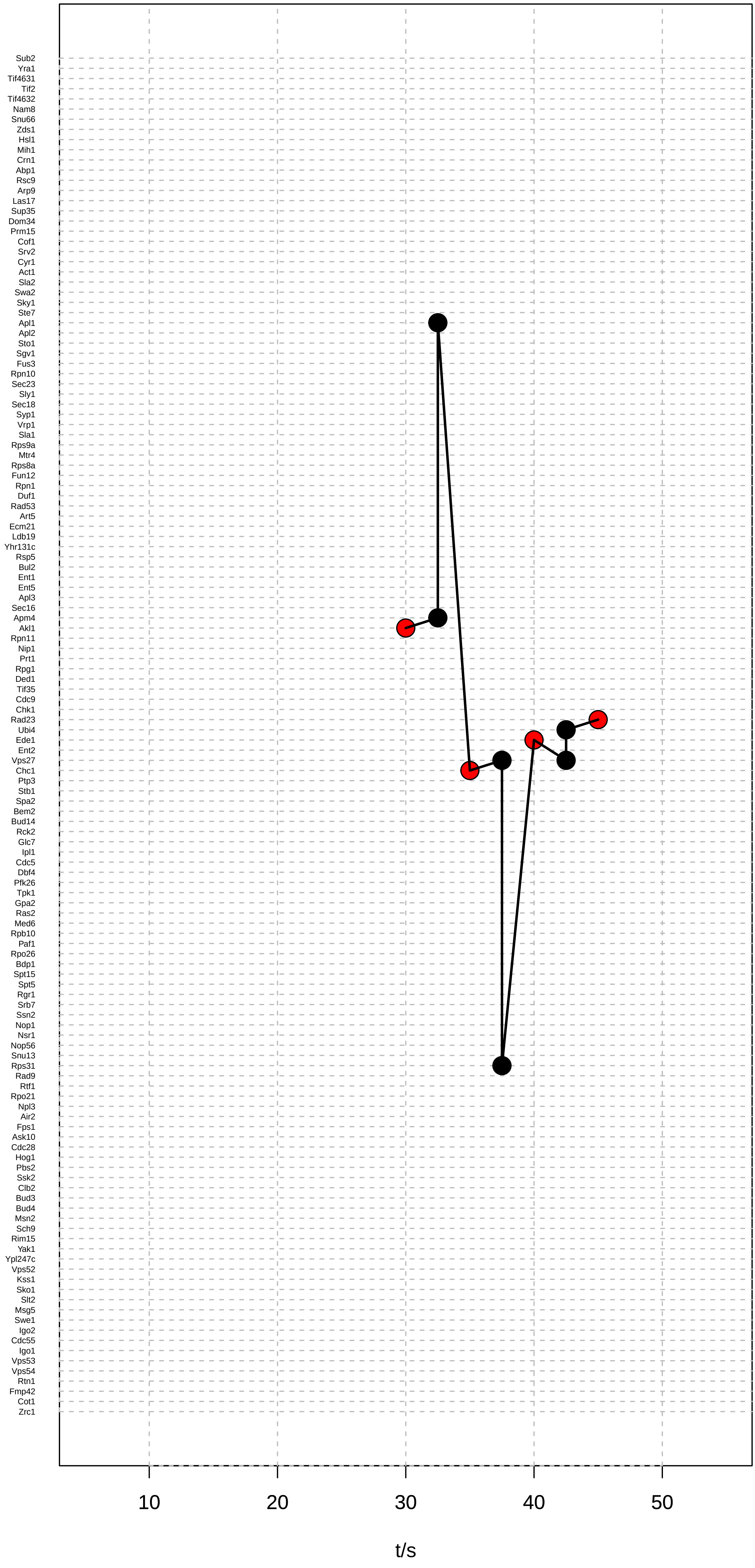

### Path 1011 (Akl1 − Rad23)

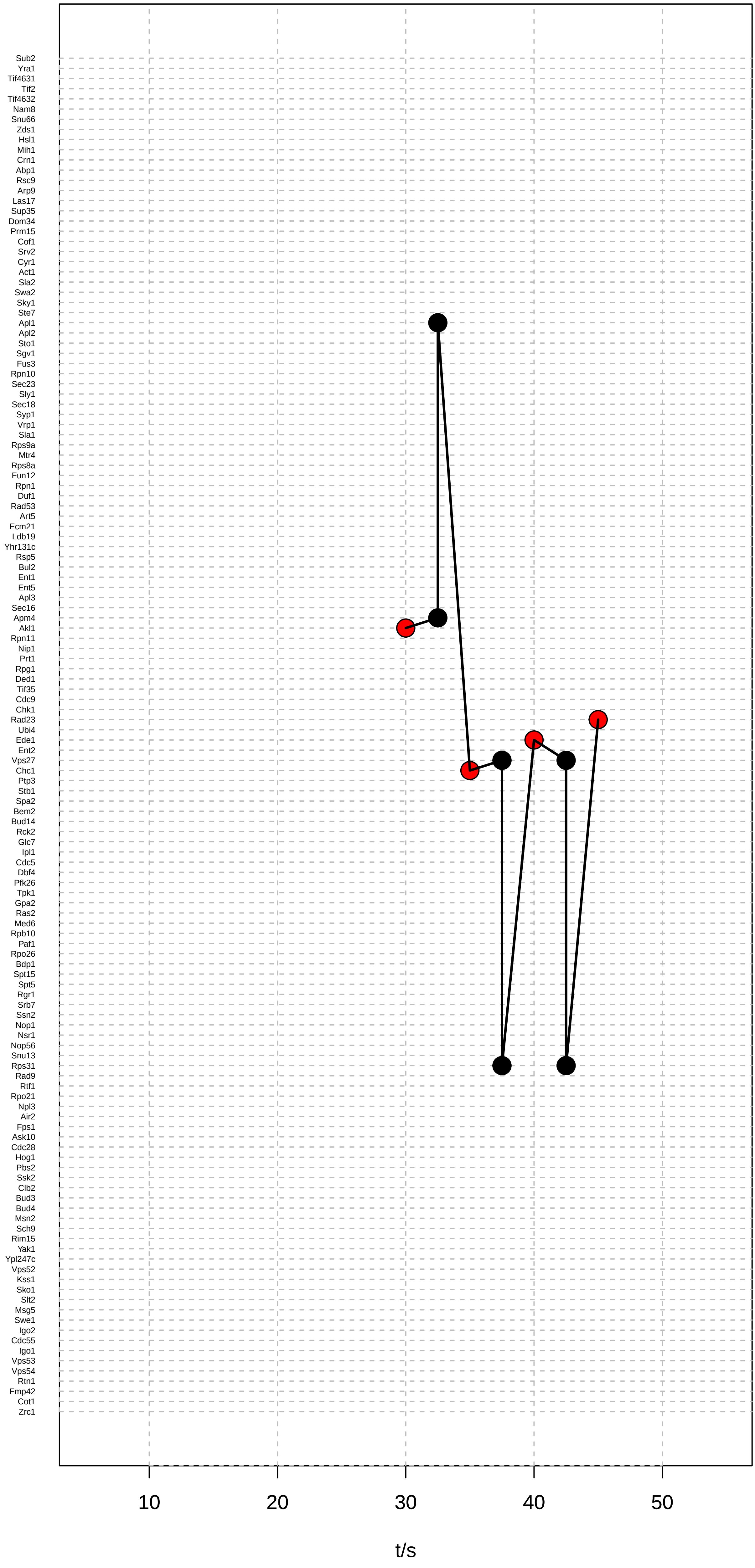

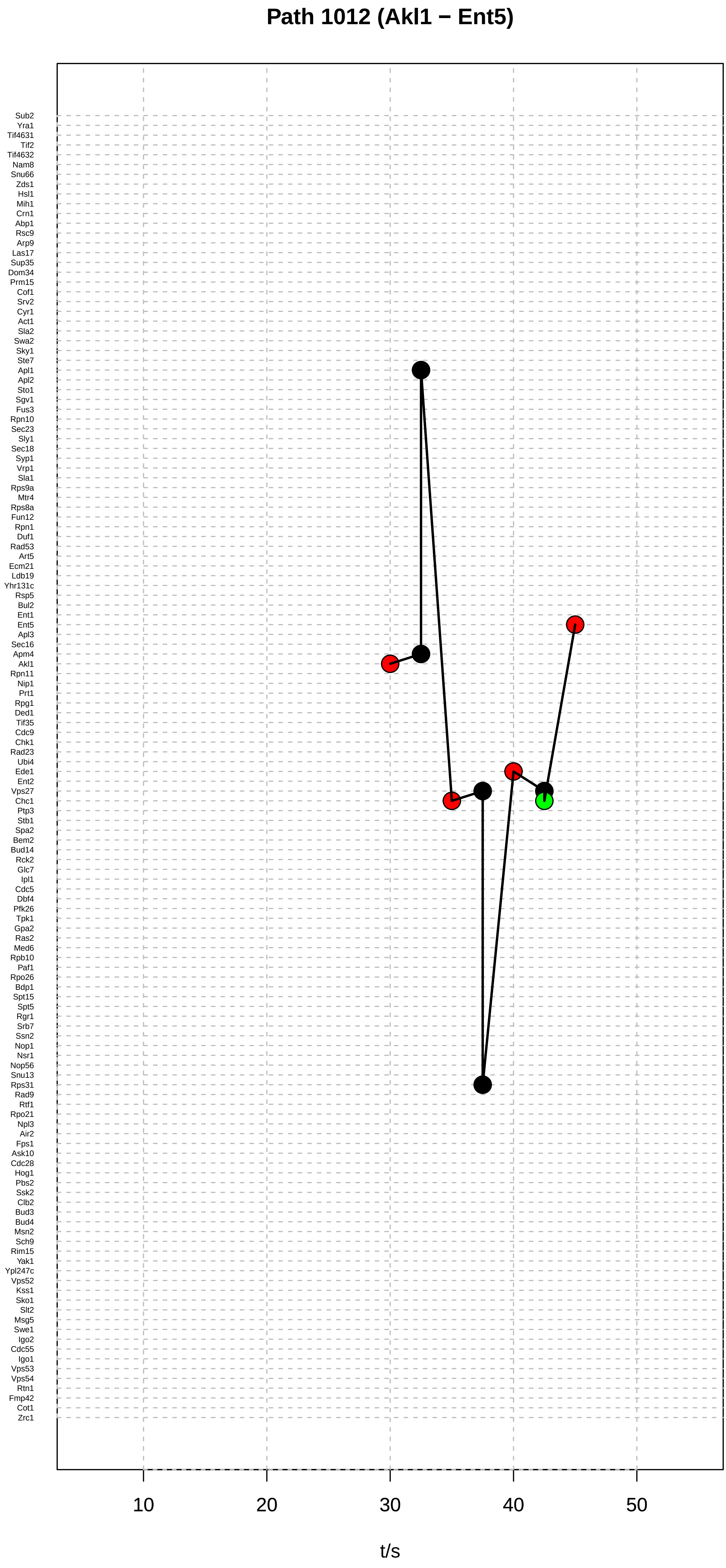
Path 1012 (Akl1 − Ent5)
Sub2
Yra1
Tif4631
Tif2
Tif4632
Nam8
Snu66
Zds1
Hsl1
Mih1
Crn1
Abp1
Rsc9
Arp9
Las17
Sup35
Dom34
Prm15
Cof1
Srv2
Cyr1
Act1
Sla2
Swa2
Sky1
Ste7
Apl1
Apl2
Sto1
Sgv1
Fus3
Rpn10
Sec23
Sly1
Sec18
Syp1
Vrp1
Sla1
Rps9a
Mtr4
Rps8a
Fun12
Rpn1
Duf1
Rad53
Art5
Ecm21
Ldb19
Yhr131c
Rsp5
Bul2
Ent1
Ent5
Apl3
Sec16
Apm4
Akl1
Rpn11
Nip1
Prt1
Rpg1
Ded1
Tif35
Cdc9
Chk1
Rad23
Ubi4
Ede1
Ent2
Vps27
Chc1
Ptp3
Stb1
Spa2
Bem2
Bud14
Rck2
Glc7
Ipl1
Cdc5
Dbf4
Pfk26
Tpk1
Gpa2
Ras2
Med6
Rpb10
Paf1
Rpo26
Bdp1
Spt15
Spt5
Rgr1
Srb7
Ssn2
Nop1
Nsr1
Nop56
Snu13
Rps31
Rad9
Rtf1
Rpo21
Npl3
Air2
Fps1
Ask10
Cdc28
Hog1
Pbs2
Ssk2
Clb2
Bud3
Bud4
Msn2
Sch9
Rim15
Yak1
Ypl247c
Vps52
Kss1
Sko1
Slt2
Msg5
Swe1
Igo2
Cdc55
Igo1
Vps53
Vps54
Rtn1
Fmp42
Cot1
Zrc1
10
20
30
40
50
t/s

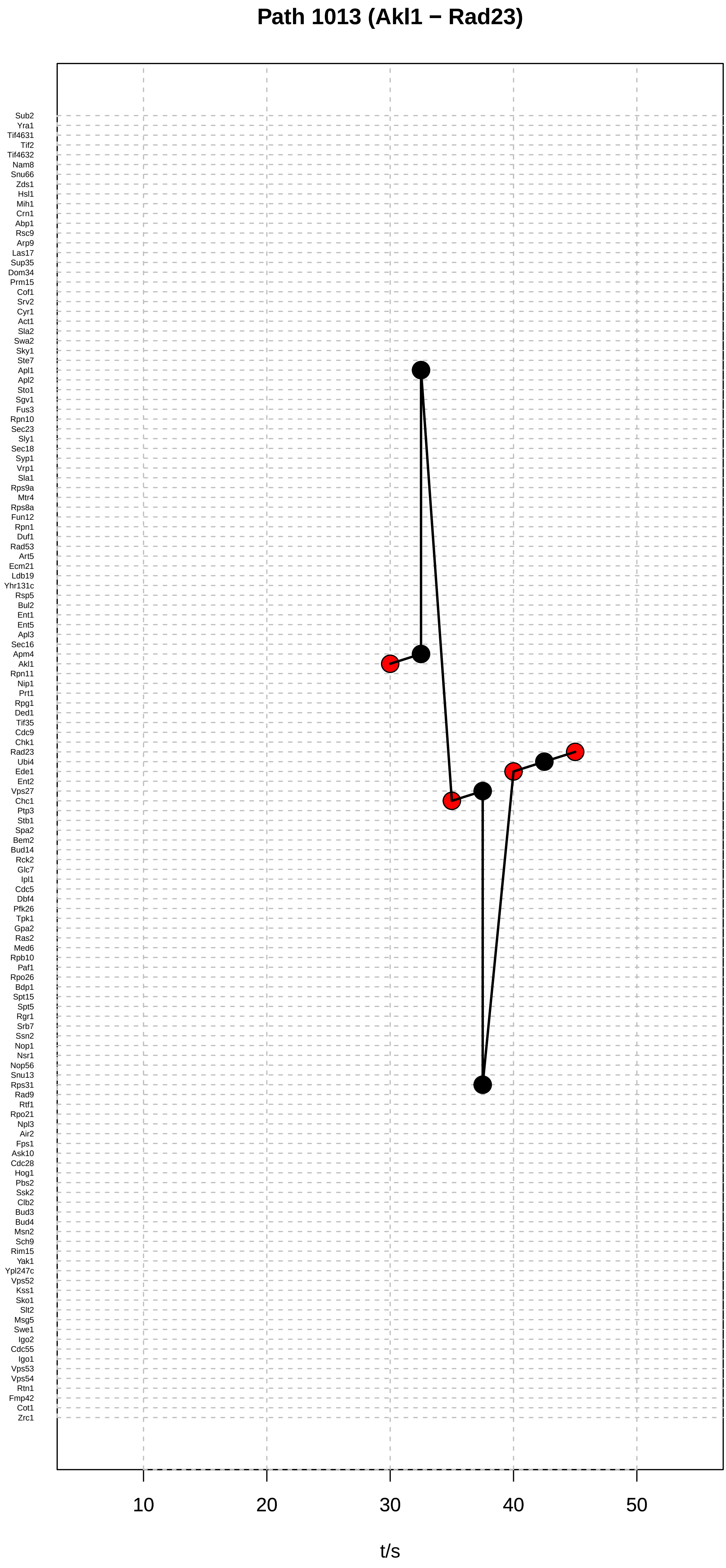

Path 1013 (Akl1 − Rad23)
Sub2
Yra1
Tif4631
Tif2
Tif4632
Nam8
Snu66
Zds1
Hsl1
Mih1
Crn1
Abp1
Rsc9
Arp9
Las17
Sup35
Dom34
Prm15
Cof1
Srv2
Cyr1
Act1
Sla2
Swa2
Sky1
Ste7
Apl1
Apl2
Sto1
Sgv1
Fus3
Rpn10
Sec23
Sly1
Sec18
Syp1
Vrp1
Sla1
Rps9a
Mtr4
Rps8a
Fun12
Rpn1
Duf1
Rad53
Art5
Ecm21
Ldb19
Yhr131c
Rsp5
Bul2
Ent1
Ent5
Apl3
Sec16
Apm4
Akl1
Rpn11
Nip1
Prt1
Rpg1
Ded1
Tif35
Cdc9
Chk1
Rad23
Ubi4
Ede1
Ent2
Vps27
Chc1
Ptp3
Stb1
Spa2
Bem2
Bud14
Rck2
Glc7
Ipl1
Cdc5
Dbf4
Pfk26
Tpk1
Gpa2
Ras2
Med6
Rpb10
Paf1
Rpo26
Bdp1
Spt15
Spt5
Rgr1
Srb7
Ssn2
Nop1
Nsr1
Nop56
Snu13
Rps31
Rad9
Rtf1
Rpo21
Npl3
Air2
Fps1
Ask10
Cdc28
Hog1
Pbs2
Ssk2
Clb2
Bud3
Bud4
Msn2
Sch9
Rim15
Yak1
Ypl247c
Vps52
Kss1
Sko1
Slt2
Msg5
Swe1
Igo2
Cdc55
Igo1
Vps53
Vps54
Rtn1
Fmp42
Cot1
Zrc1
10
20
30
40
50
t/s

#### Path 1014 (Akl1 − Rad23)

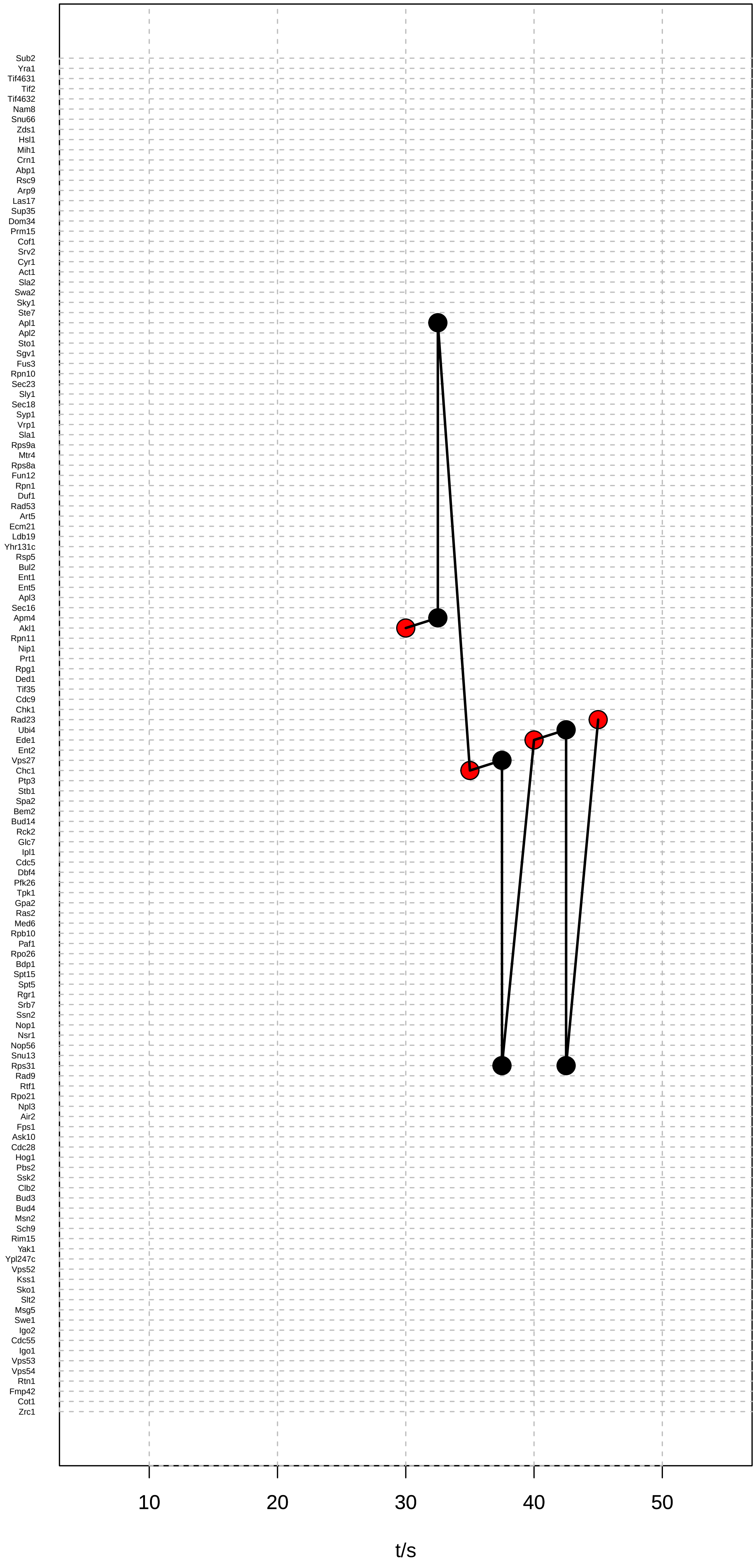

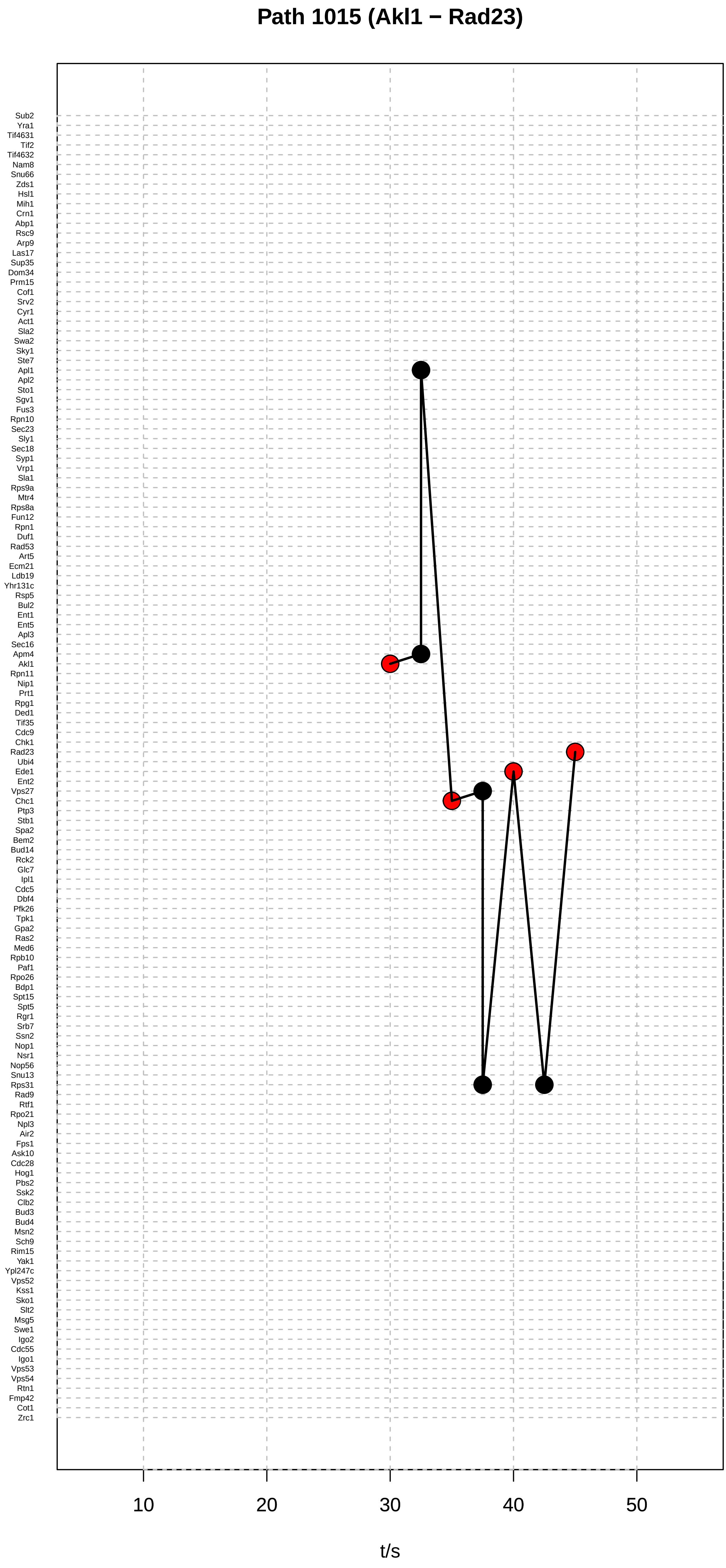
Path 1015 (Akl1 − Rad23)
Sub2
Yra1
Tif4631
Tif2
Tif4632
Nam8
Snu66
Zds1
Hsl1
Mih1
Crn1
Abp1
Rsc9
Arp9
Las17
Sup35
Dom34
Prm15
Cof1
Srv2
Cyr1
Act1
Sla2
Swa2
Sky1
Ste7
Apl1
Apl2
Sto1
Sgv1
Fus3
Rpn10
Sec23
Sly1
Sec18
Syp1
Vrp1
Sla1
Rps9a
Mtr4
Rps8a
Fun12
Rpn1
Duf1
Rad53
Art5
Ecm21
Ldb19
Yhr131c
Rsp5
Bul2
Ent1
Ent5
Apl3
Sec16
Apm4
Akl1
Rpn11
Nip1
Prt1
Rpg1
Ded1
Tif35
Cdc9
Chk1
Rad23
Ubi4
Ede1
Ent2
Vps27
Chc1
Ptp3
Stb1
Spa2
Bem2
Bud14
Rck2
Glc7
Ipl1
Cdc5
Dbf4
Pfk26
Tpk1
Gpa2
Ras2
Med6
Rpb10
Paf1
Rpo26
Bdp1
Spt15
Spt5
Rgr1
Srb7
Ssn2
Nop1
Nsr1
Nop56
Snu13
Rps31
Rad9
Rtf1
Rpo21
Npl3
Air2
Fps1
Ask10
Cdc28
Hog1
Pbs2
Ssk2
Clb2
Bud3
Bud4
Msn2
Sch9
Rim15
Yak1
Ypl247c
Vps52
Kss1
Sko1
Slt2
Msg5
Swe1
Igo2
Cdc55
Igo1
Vps53
Vps54
Rtn1
Fmp42
Cot1
Zrc1
10
20
30
40
50
t/s

### Path 1016 (Akl1 − Rad23)

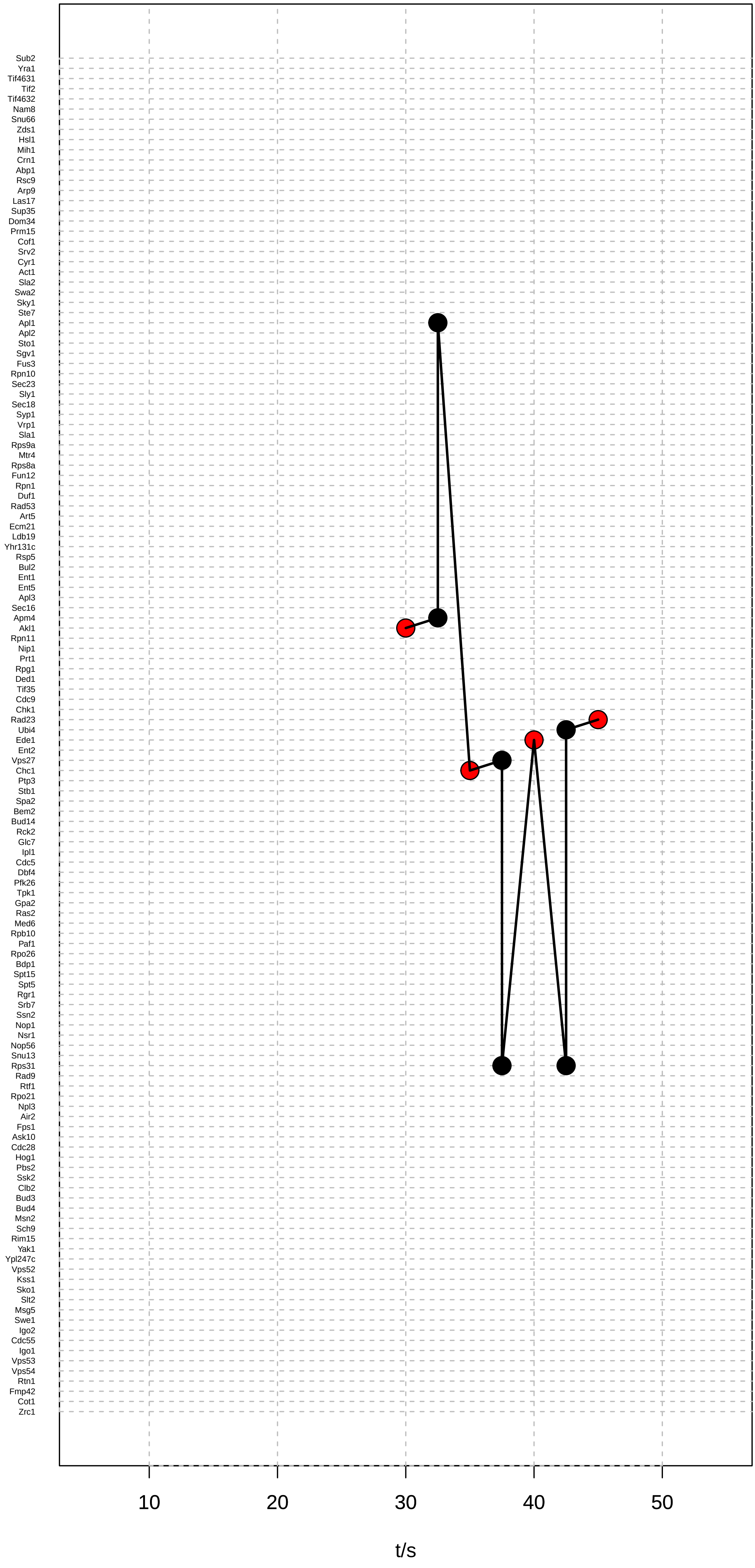

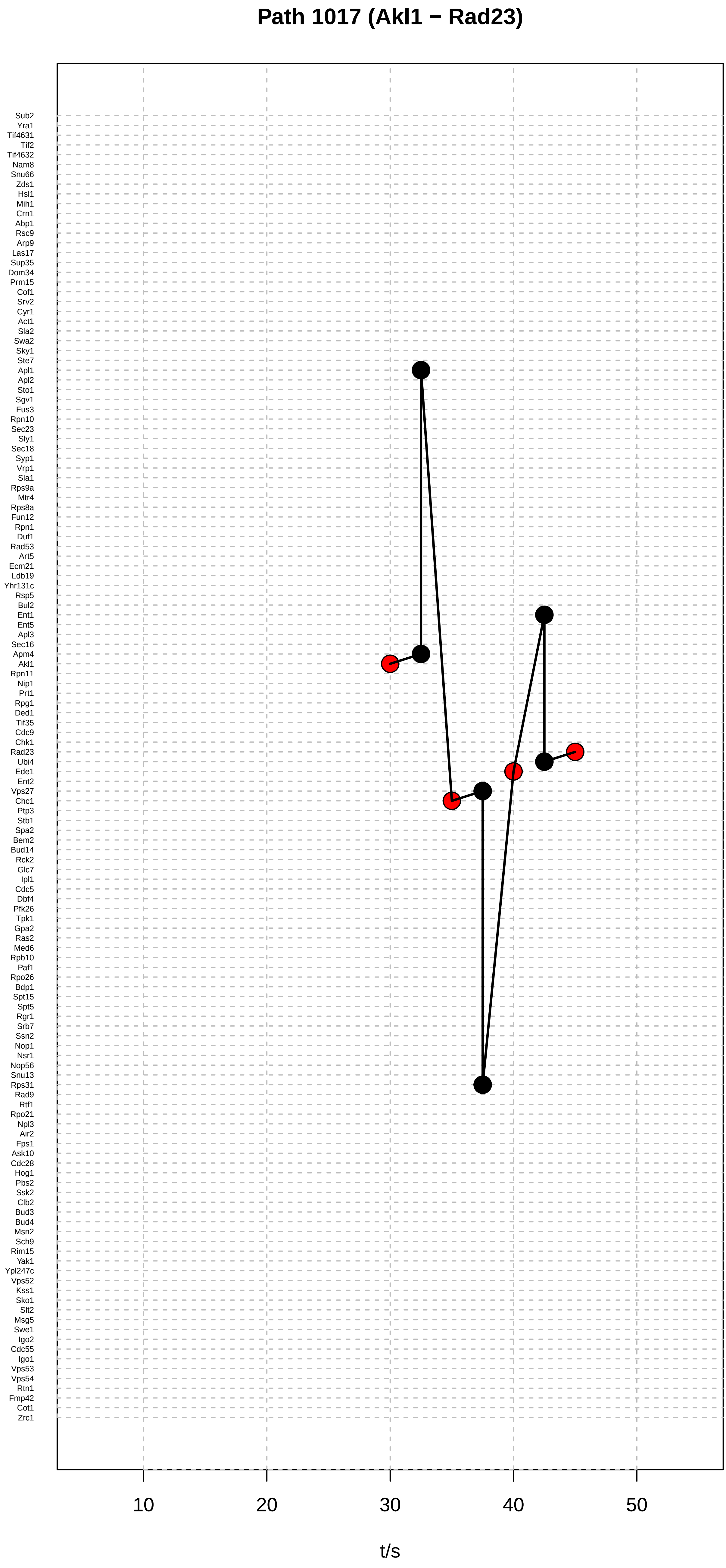
Path 1017 (Akl1 − Rad23)
Sub2
Yra1
Tif4631
Tif2
Tif4632
Nam8
Snu66
Zds1
Hsl1
Mih1
Crn1
Abp1
Rsc9
Arp9
Las17
Sup35
Dom34
Prm15
Cof1
Srv2
Cyr1
Act1
Sla2
Swa2
Sky1
Ste7
Apl1
Apl2
Sto1
Sgv1
Fus3
Rpn10
Sec23
Sly1
Sec18
Syp1
Vrp1
Sla1
Rps9a
Mtr4
Rps8a
Fun12
Rpn1
Duf1
Rad53
Art5
Ecm21
Ldb19
Yhr131c
Rsp5
Bul2
Ent1
Ent5
Apl3
Sec16
Apm4
Akl1
Rpn11
Nip1
Prt1
Rpg1
Ded1
Tif35
Cdc9
Chk1
Rad23
Ubi4
Ede1
Ent2
Vps27
Chc1
Ptp3
Stb1
Spa2
Bem2
Bud14
Rck2
Glc7
Ipl1
Cdc5
Dbf4
Pfk26
Tpk1
Gpa2
Ras2
Med6
Rpb10
Paf1
Rpo26
Bdp1
Spt15
Spt5
Rgr1
Srb7
Ssn2
Nop1
Nsr1
Nop56
Snu13
Rps31
Rad9
Rtf1
Rpo21
Npl3
Air2
Fps1
Ask10
Cdc28
Hog1
Pbs2
Ssk2
Clb2
Bud3
Bud4
Msn2
Sch9
Rim15
Yak1
Ypl247c
Vps52
Kss1
Sko1
Slt2
Msg5
Swe1
Igo2
Cdc55
Igo1
Vps53
Vps54
Rtn1
Fmp42
Cot1
Zrc1
10
20
30
40
50
t/s

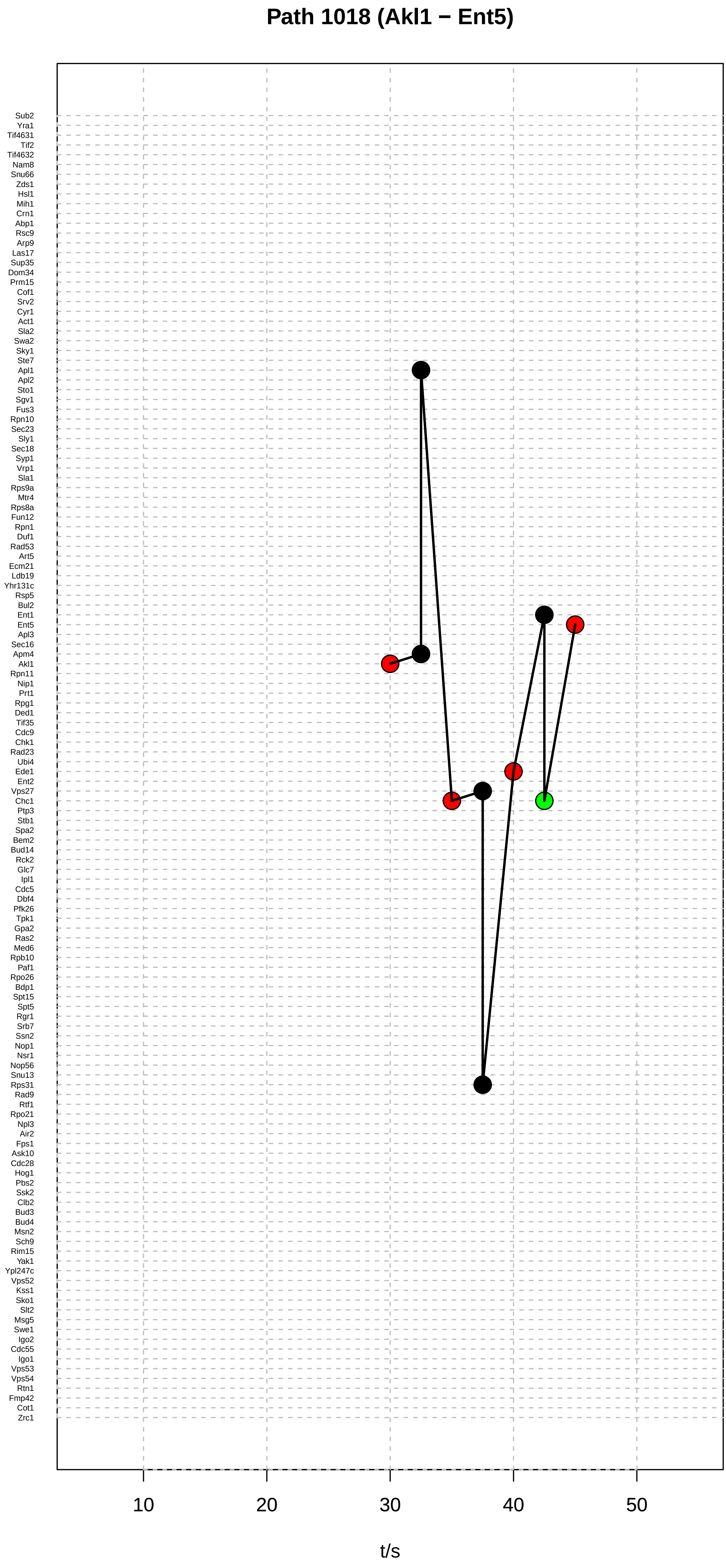
Path 1018 (Akl1 − Ent5)
Sub2
Yra1
Tif4631
Tif2
Tif4632
Nam8
Snu66
Zds1
Hsl1
Mih1
Crn1
Abp1
Rsc9
Arp9
Las17
Sup35
Dom34
Prm15
Cof1
Srv2
Cyr1
Act1
Sla2
Swa2
Sky1
Ste7
Apl1
Apl2
Sto1
Sgv1
Fus3
Rpn10
Sec23
Sly1
Sec18
Syp1
Vrp1
Sla1
Rps9a
Mtr4
Rps8a
Fun12
Rpn1
Duf1
Rad53
Art5
Ecm21
Ldb19
Yhr131c
Rsp5
Bul2
Ent1
Ent5
Apl3
Sec16
Apm4
Akl1
Rpn11
Nip1
Prt1
Rpg1
Ded1
Tif35
Cdc9
Chk1
Rad23
Ubi4
Ede1
Ent2
Vps27
Chc1
Ptp3
Stb1
Spa2
Bem2
Bud14
Rck2
Glc7
Ipl1
Cdc5
Dbf4
Pfk26
Tpk1
Gpa2
Ras2
Med6
Rpb10
Paf1
Rpo26
Bdp1
Spt15
Spt5
Rgr1
Srb7
Ssn2
Nop1
Nsr1
Nop56
Snu13
Rps31
Rad9
Rtf1
Rpo21
Npl3
Air2
Fps1
Ask10
Cdc28
Hog1
Pbs2
Ssk2
Clb2
Bud3
Bud4
Msn2
Sch9
Rim15
Yak1
Ypl247c
Vps52
Kss1
Sko1
Slt2
Msg5
Swe1
Igo2
Cdc55
Igo1
Vps53
Vps54
Rtn1
Fmp42
Cot1
Zrc1
10
20
30
40
50
t/s

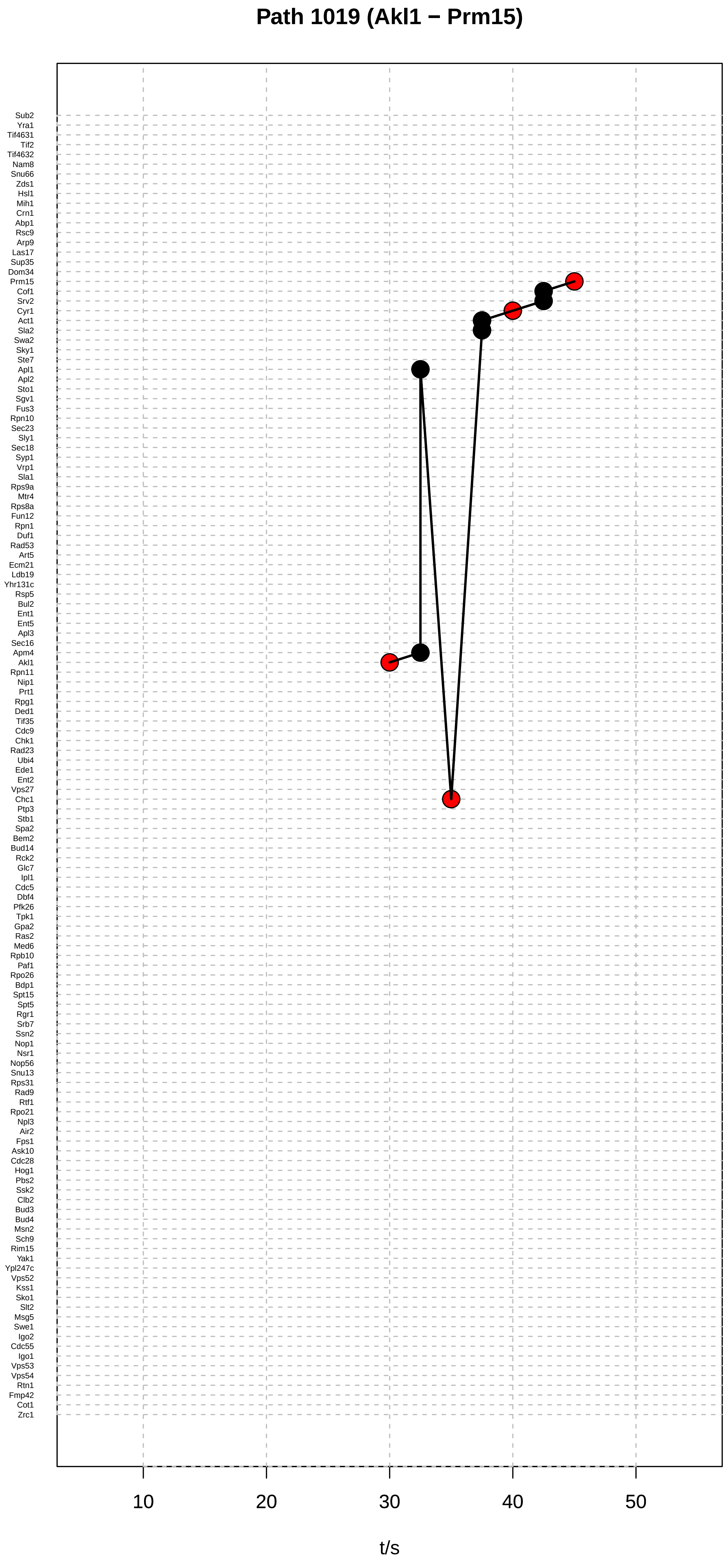
Path 1019 (Akl1 − Prm15)
Sub2
Yra1
Tif4631
Tif2
Tif4632
Nam8
Snu66
Zds1
Hsl1
Mih1
Crn1
Abp1
Rsc9
Arp9
Las17
Sup35
Dom34
Prm15
Cof1
Srv2
Cyr1
Act1
Sla2
Swa2
Sky1
Ste7
Apl1
Apl2
Sto1
Sgv1
Fus3
Rpn10
Sec23
Sly1
Sec18
Syp1
Vrp1
Sla1
Rps9a
Mtr4
Rps8a
Fun12
Rpn1
Duf1
Rad53
Art5
Ecm21
Ldb19
Yhr131c
Rsp5
Bul2
Ent1
Ent5
Apl3
Sec16
Apm4
Akl1
Rpn11
Nip1
Prt1
Rpg1
Ded1
Tif35
Cdc9
Chk1
Rad23
Ubi4
Ede1
Ent2
Vps27
Chc1
Ptp3
Stb1
Spa2
Bem2
Bud14
Rck2
Glc7
Ipl1
Cdc5
Dbf4
Pfk26
Tpk1
Gpa2
Ras2
Med6
Rpb10
Paf1
Rpo26
Bdp1
Spt15
Spt5
Rgr1
Srb7
Ssn2
Nop1
Nsr1
Nop56
Snu13
Rps31
Rad9
Rtf1
Rpo21
Npl3
Air2
Fps1
Ask10
Cdc28
Hog1
Pbs2
Ssk2
Clb2
Bud3
Bud4
Msn2
Sch9
Rim15
Yak1
Ypl247c
Vps52
Kss1
Sko1
Slt2
Msg5
Swe1
Igo2
Cdc55
Igo1
Vps53
Vps54
Rtn1
Fmp42
Cot1
Zrc1
10
20
30
40
50
t/s

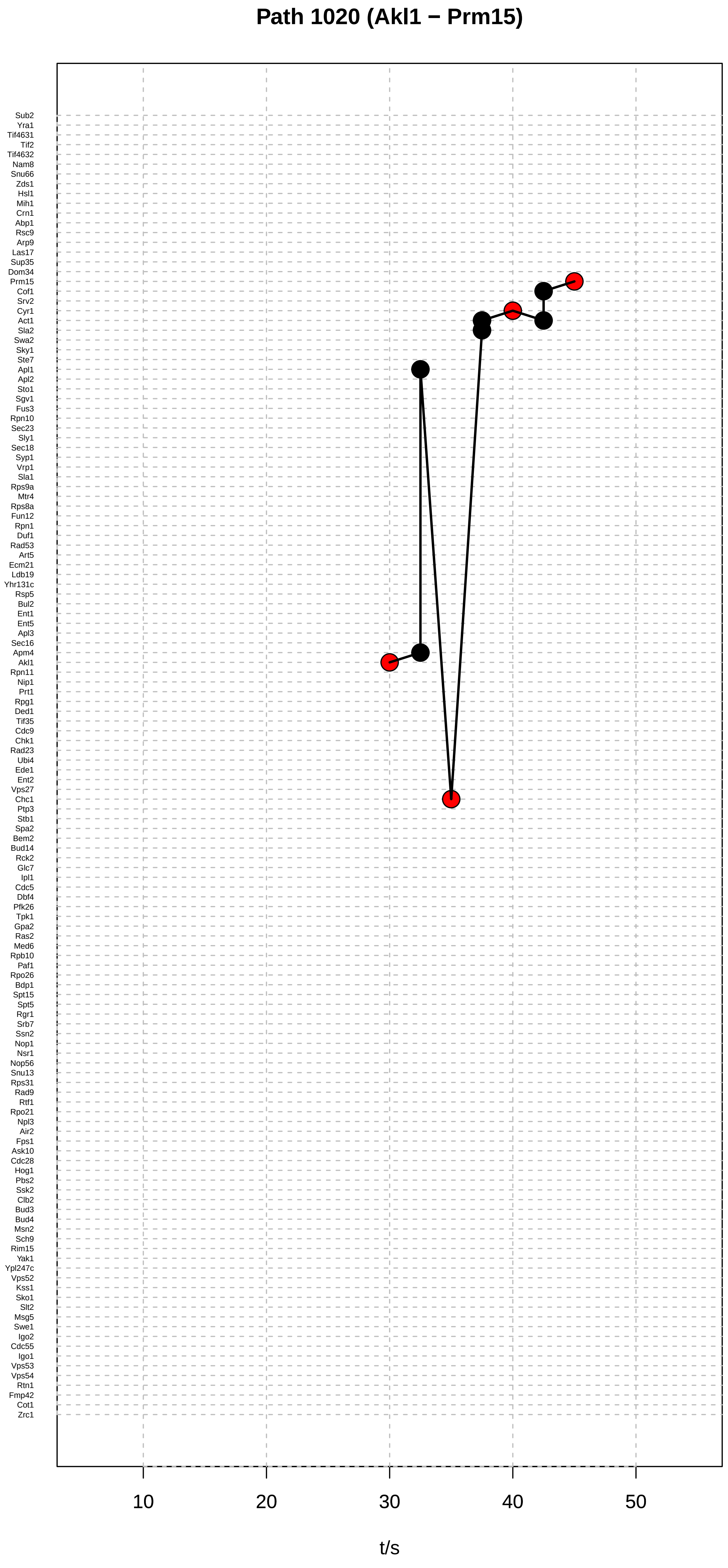

Path 1020 (Akl1 − Prm15)
Sub2
Yra1
Tif4631
Tif2
Tif4632
Nam8
Snu66
Zds1
Hsl1
Mih1
Crn1
Abp1
Rsc9
Arp9
Las17
Sup35
Dom34
Prm15
Cof1
Srv2
Cyr1
Act1
Sla2
Swa2
Sky1
Ste7
Apl1
Apl2
Sto1
Sgv1
Fus3
Rpn10
Sec23
Sly1
Sec18
Syp1
Vrp1
Sla1
Rps9a
Mtr4
Rps8a
Fun12
Rpn1
Duf1
Rad53
Art5
Ecm21
Ldb19
Yhr131c
Rsp5
Bul2
Ent1
Ent5
Apl3
Sec16
Apm4
Akl1
Rpn11
Nip1
Prt1
Rpg1
Ded1
Tif35
Cdc9
Chk1
Rad23
Ubi4
Ede1
Ent2
Vps27
Chc1
Ptp3
Stb1
Spa2
Bem2
Bud14
Rck2
Glc7
Ipl1
Cdc5
Dbf4
Pfk26
Tpk1
Gpa2
Ras2
Med6
Rpb10
Paf1
Rpo26
Bdp1
Spt15
Spt5
Rgr1
Srb7
Ssn2
Nop1
Nsr1
Nop56
Snu13
Rps31
Rad9
Rtf1
Rpo21
Npl3
Air2
Fps1
Ask10
Cdc28
Hog1
Pbs2
Ssk2
Clb2
Bud3
Bud4
Msn2
Sch9
Rim15
Yak1
Ypl247c
Vps52
Kss1
Sko1
Slt2
Msg5
Swe1
Igo2
Cdc55
Igo1
Vps53
Vps54
Rtn1
Fmp42
Cot1
Zrc1
10
20
30
40
50
t/s

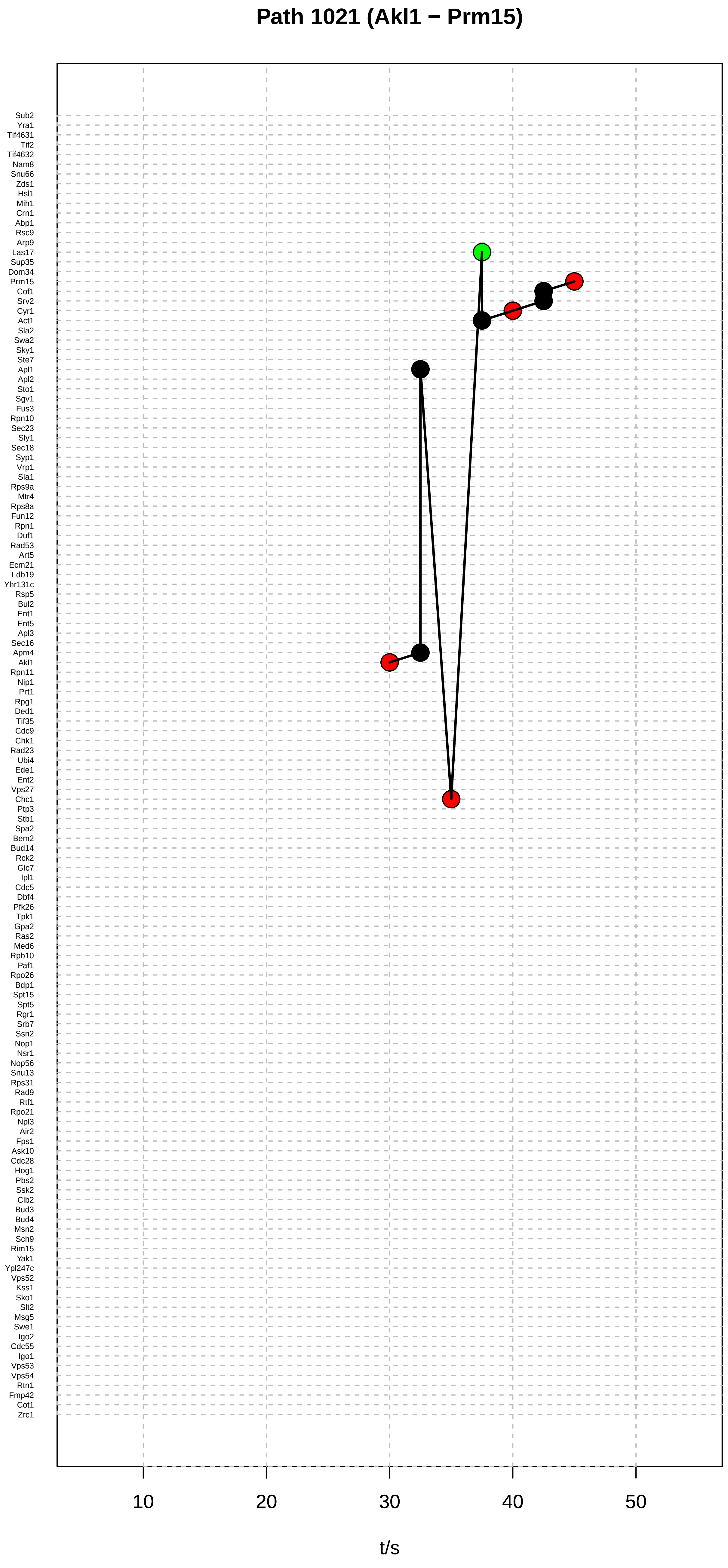

Path 1021 (Akl1 − Prm15)
Sub2
Yra1
Tif4631
Tif2
Tif4632
Nam8
Snu66
Zds1
Hsl1
Mih1
Crn1
Abp1
Rsc9
Arp9
Las17
Sup35
Dom34
Prm15
Cof1
Srv2
Cyr1
Act1
Sla2
Swa2
Sky1
Ste7
Apl1
Apl2
Sto1
Sgv1
Fus3
Rpn10
Sec23
Sly1
Sec18
Syp1
Vrp1
Sla1
Rps9a
Mtr4
Rps8a
Fun12
Rpn1
Duf1
Rad53
Art5
Ecm21
Ldb19
Yhr131c
Rsp5
Bul2
Ent1
Ent5
Apl3
Sec16
Apm4
Akl1
Rpn11
Nip1
Prt1
Rpg1
Ded1
Tif35
Cdc9
Chk1
Rad23
Ubi4
Ede1
Ent2
Vps27
Chc1
Ptp3
Stb1
Spa2
Bem2
Bud14
Rck2
Glc7
Ipl1
Cdc5
Dbf4
Pfk26
Tpk1
Gpa2
Ras2
Med6
Rpb10
Paf1
Rpo26
Bdp1
Spt15
Spt5
Rgr1
Srb7
Ssn2
Nop1
Nsr1
Nop56
Snu13
Rps31
Rad9
Rtf1
Rpo21
Npl3
Air2
Fps1
Ask10
Cdc28
Hog1
Pbs2
Ssk2
Clb2
Bud3
Bud4
Msn2
Sch9
Rim15
Yak1
Ypl247c
Vps52
Kss1
Sko1
Slt2
Msg5
Swe1
Igo2
Cdc55
Igo1
Vps53
Vps54
Rtn1
Fmp42
Cot1
Zrc1
10
20
30
40
50
t/s

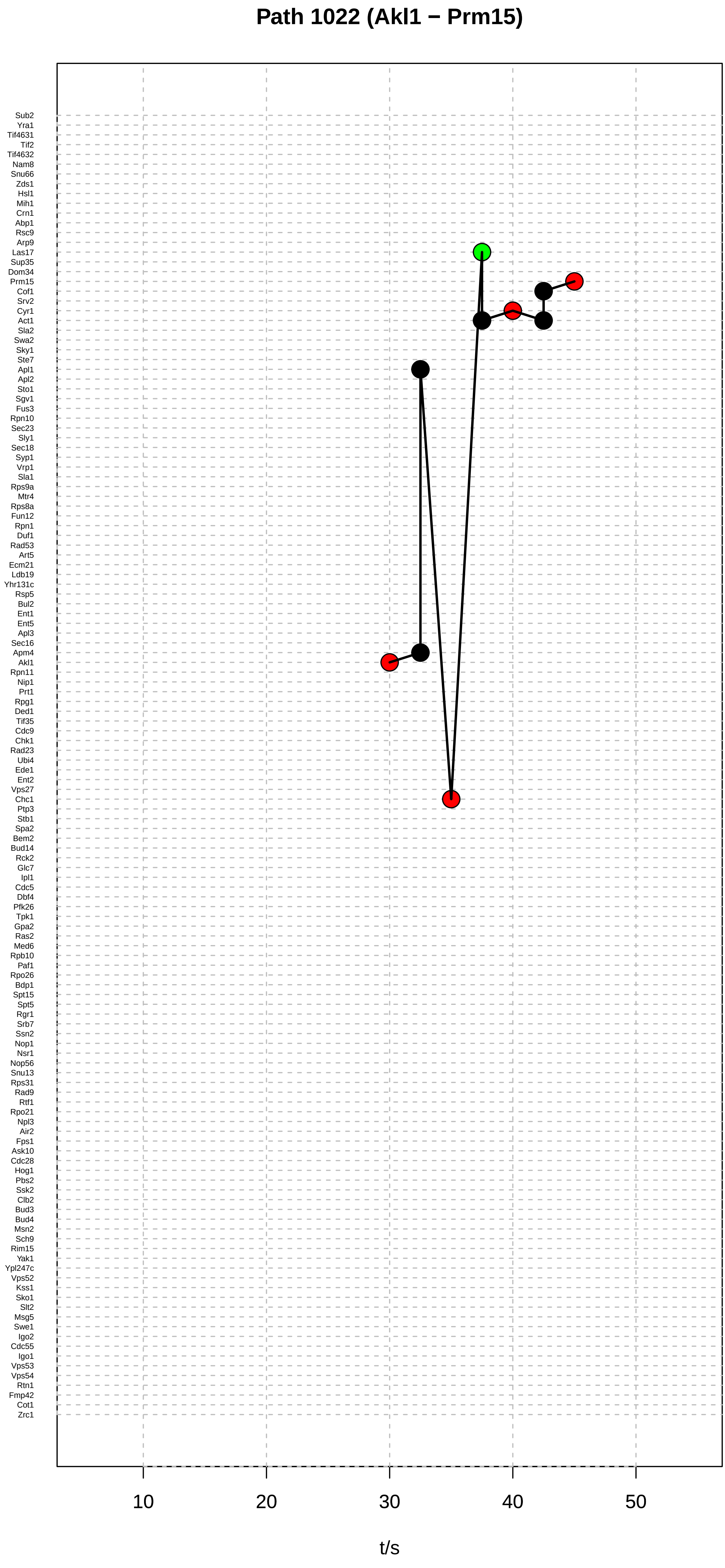

Path 1022 (Akl1 − Prm15)
Sub2
Yra1
Tif4631
Tif2
Tif4632
Nam8
Snu66
Zds1
Hsl1
Mih1
Crn1
Abp1
Rsc9
Arp9
Las17
Sup35
Dom34
Prm15
Cof1
Srv2
Cyr1
Act1
Sla2
Swa2
Sky1
Ste7
Apl1
Apl2
Sto1
Sgv1
Fus3
Rpn10
Sec23
Sly1
Sec18
Syp1
Vrp1
Sla1
Rps9a
Mtr4
Rps8a
Fun12
Rpn1
Duf1
Rad53
Art5
Ecm21
Ldb19
Yhr131c
Rsp5
Bul2
Ent1
Ent5
Apl3
Sec16
Apm4
Akl1
Rpn11
Nip1
Prt1
Rpg1
Ded1
Tif35
Cdc9
Chk1
Rad23
Ubi4
Ede1
Ent2
Vps27
Chc1
Ptp3
Stb1
Spa2
Bem2
Bud14
Rck2
Glc7
Ipl1
Cdc5
Dbf4
Pfk26
Tpk1
Gpa2
Ras2
Med6
Rpb10
Paf1
Rpo26
Bdp1
Spt15
Spt5
Rgr1
Srb7
Ssn2
Nop1
Nsr1
Nop56
Snu13
Rps31
Rad9
Rtf1
Rpo21
Npl3
Air2
Fps1
Ask10
Cdc28
Hog1
Pbs2
Ssk2
Clb2
Bud3
Bud4
Msn2
Sch9
Rim15
Yak1
Ypl247c
Vps52
Kss1
Sko1
Slt2
Msg5
Swe1
Igo2
Cdc55
Igo1
Vps53
Vps54
Rtn1
Fmp42
Cot1
Zrc1
10
20
30
40
50
t/s

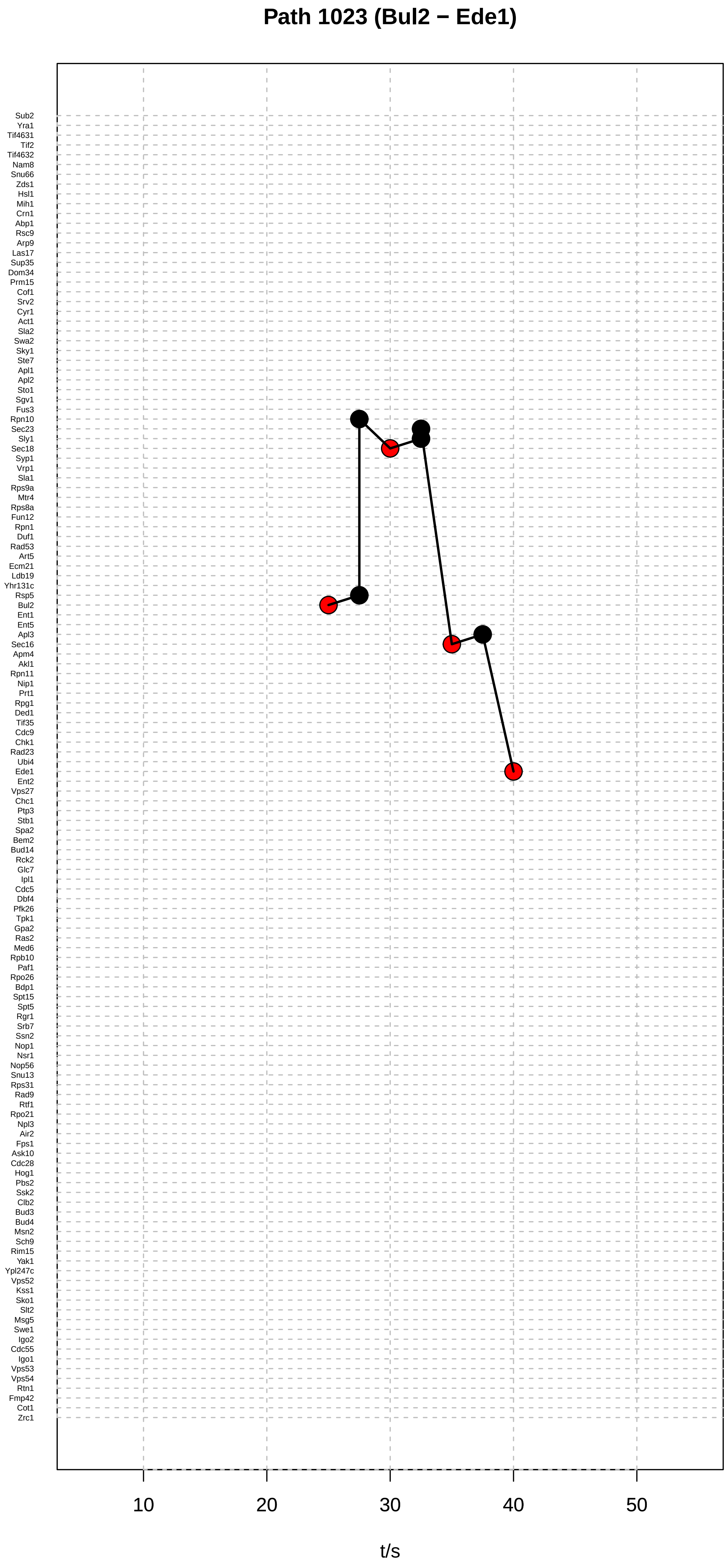

Path 1023 (Bul2 − Ede1)
Sub2
Yra1
Tif4631
Tif2
Tif4632
Nam8
Snu66
Zds1
Hsl1
Mih1
Crn1
Abp1
Rsc9
Arp9
Las17
Sup35
Dom34
Prm15
Cof1
Srv2
Cyr1
Act1
Sla2
Swa2
Sky1
Ste7
Apl1
Apl2
Sto1
Sgv1
Fus3
Rpn10
Sec23
Sly1
Sec18
Syp1
Vrp1
Sla1
Rps9a
Mtr4
Rps8a
Fun12
Rpn1
Duf1
Rad53
Art5
Ecm21
Ldb19
Yhr131c
Rsp5
Bul2
Ent1
Ent5
Apl3
Sec16
Apm4
Akl1
Rpn11
Nip1
Prt1
Rpg1
Ded1
Tif35
Cdc9
Chk1
Rad23
Ubi4
Ede1
Ent2
Vps27
Chc1
Ptp3
Stb1
Spa2
Bem2
Bud14
Rck2
Glc7
Ipl1
Cdc5
Dbf4
Pfk26
Tpk1
Gpa2
Ras2
Med6
Rpb10
Paf1
Rpo26
Bdp1
Spt15
Spt5
Rgr1
Srb7
Ssn2
Nop1
Nsr1
Nop56
Snu13
Rps31
Rad9
Rtf1
Rpo21
Npl3
Air2
Fps1
Ask10
Cdc28
Hog1
Pbs2
Ssk2
Clb2
Bud3
Bud4
Msn2
Sch9
Rim15
Yak1
Ypl247c
Vps52
Kss1
Sko1
Slt2
Msg5
Swe1
Igo2
Cdc55
Igo1
Vps53
Vps54
Rtn1
Fmp42
Cot1
Zrc1
10
20
30
40
50
t/s

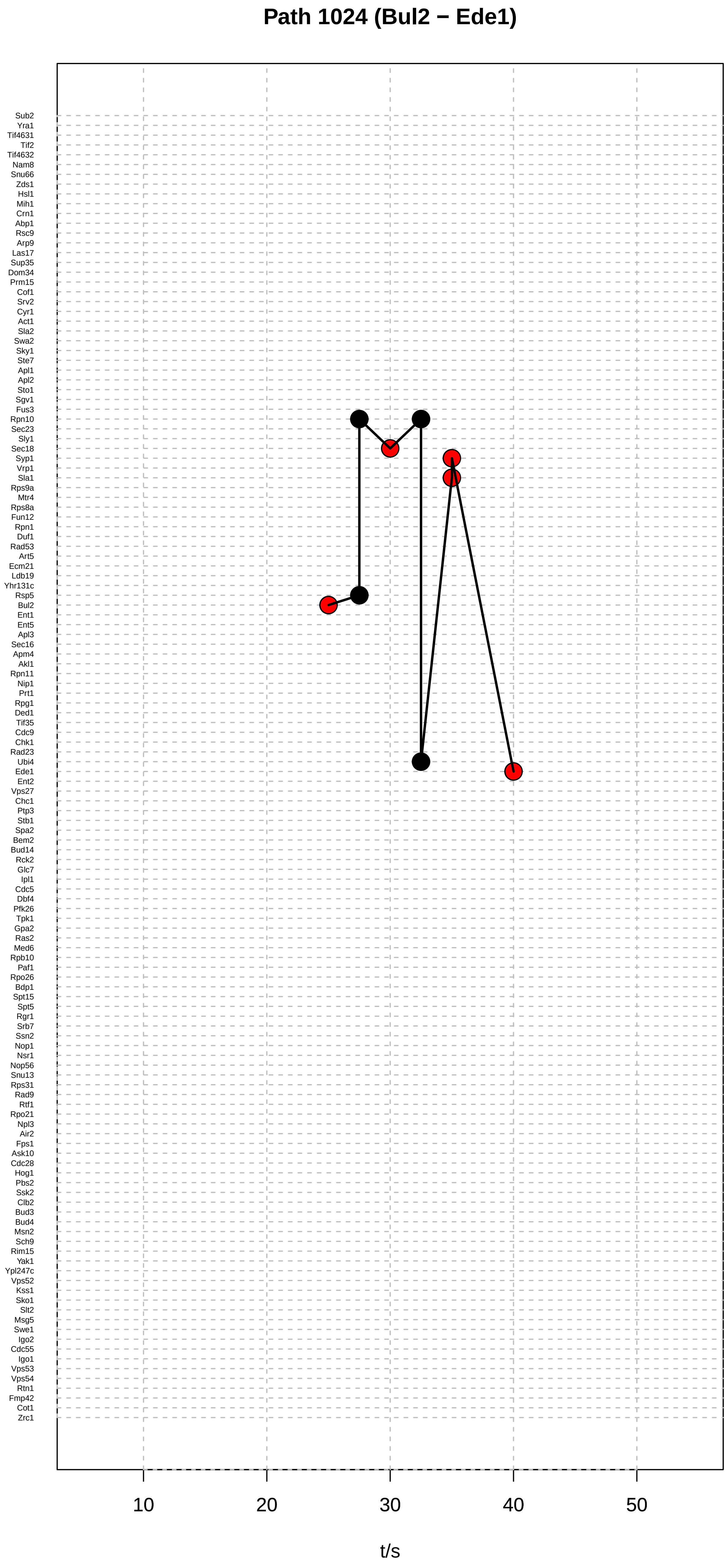

Path 1024 (Bul2 − Ede1)
Sub2
Yra1
Tif4631
Tif2
Tif4632
Nam8
Snu66
Zds1
Hsl1
Mih1
Crn1
Abp1
Rsc9
Arp9
Las17
Sup35
Dom34
Prm15
Cof1
Srv2
Cyr1
Act1
Sla2
Swa2
Sky1
Ste7
Apl1
Apl2
Sto1
Sgv1
Fus3
Rpn10
Sec23
Sly1
Sec18
Syp1
Vrp1
Sla1
Rps9a
Mtr4
Rps8a
Fun12
Rpn1
Duf1
Rad53
Art5
Ecm21
Ldb19
Yhr131c
Rsp5
Bul2
Ent1
Ent5
Apl3
Sec16
Apm4
Akl1
Rpn11
Nip1
Prt1
Rpg1
Ded1
Tif35
Cdc9
Chk1
Rad23
Ubi4
Ede1
Ent2
Vps27
Chc1
Ptp3
Stb1
Spa2
Bem2
Bud14
Rck2
Glc7
Ipl1
Cdc5
Dbf4
Pfk26
Tpk1
Gpa2
Ras2
Med6
Rpb10
Paf1
Rpo26
Bdp1
Spt15
Spt5
Rgr1
Srb7
Ssn2
Nop1
Nsr1
Nop56
Snu13
Rps31
Rad9
Rtf1
Rpo21
Npl3
Air2
Fps1
Ask10
Cdc28
Hog1
Pbs2
Ssk2
Clb2
Bud3
Bud4
Msn2
Sch9
Rim15
Yak1
Ypl247c
Vps52
Kss1
Sko1
Slt2
Msg5
Swe1
Igo2
Cdc55
Igo1
Vps53
Vps54
Rtn1
Fmp42
Cot1
Zrc1
10
20
30
40
50
t/s

### Path 1025 (Bul2 − Ede1)

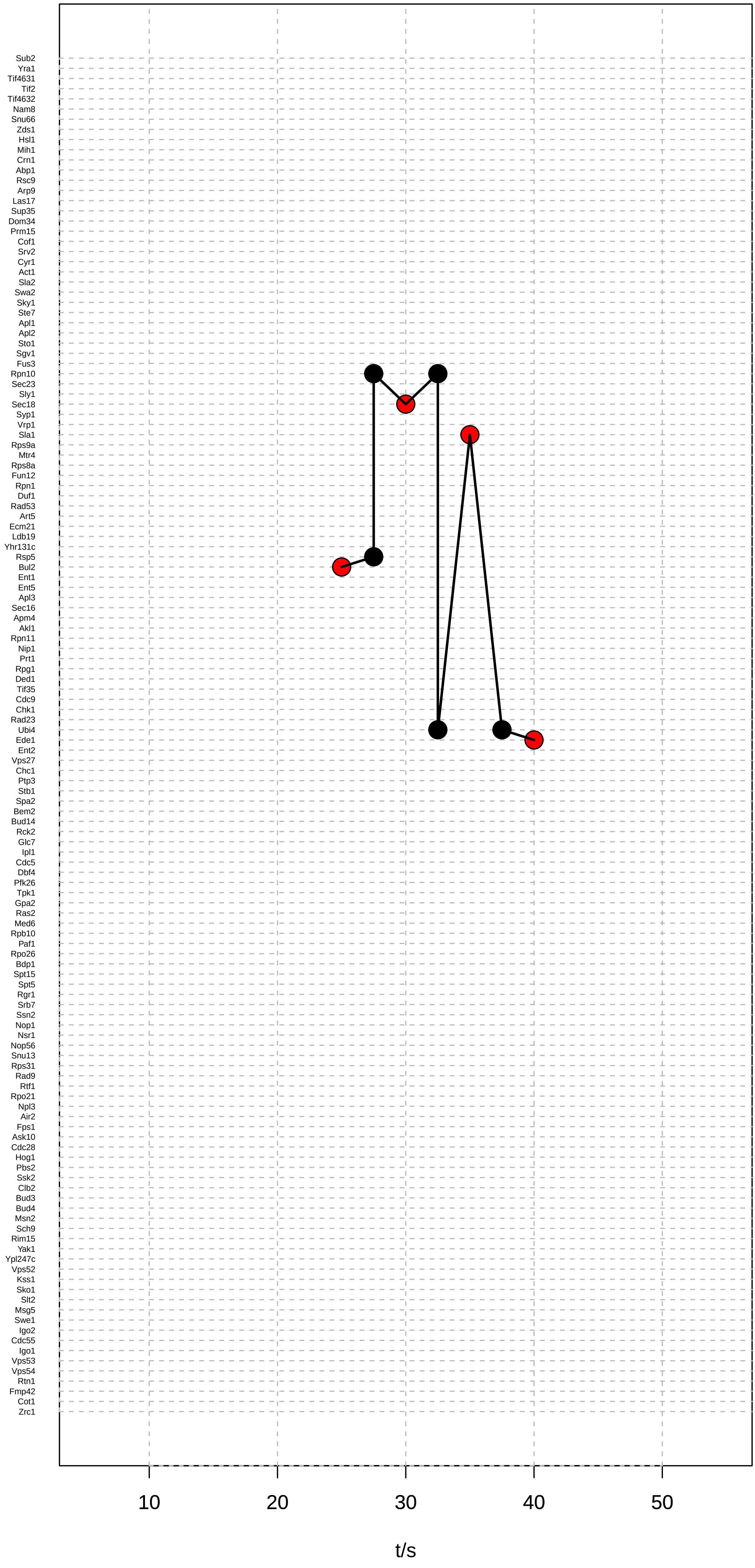

#### Path 1026 (Bul2 − Duf1)

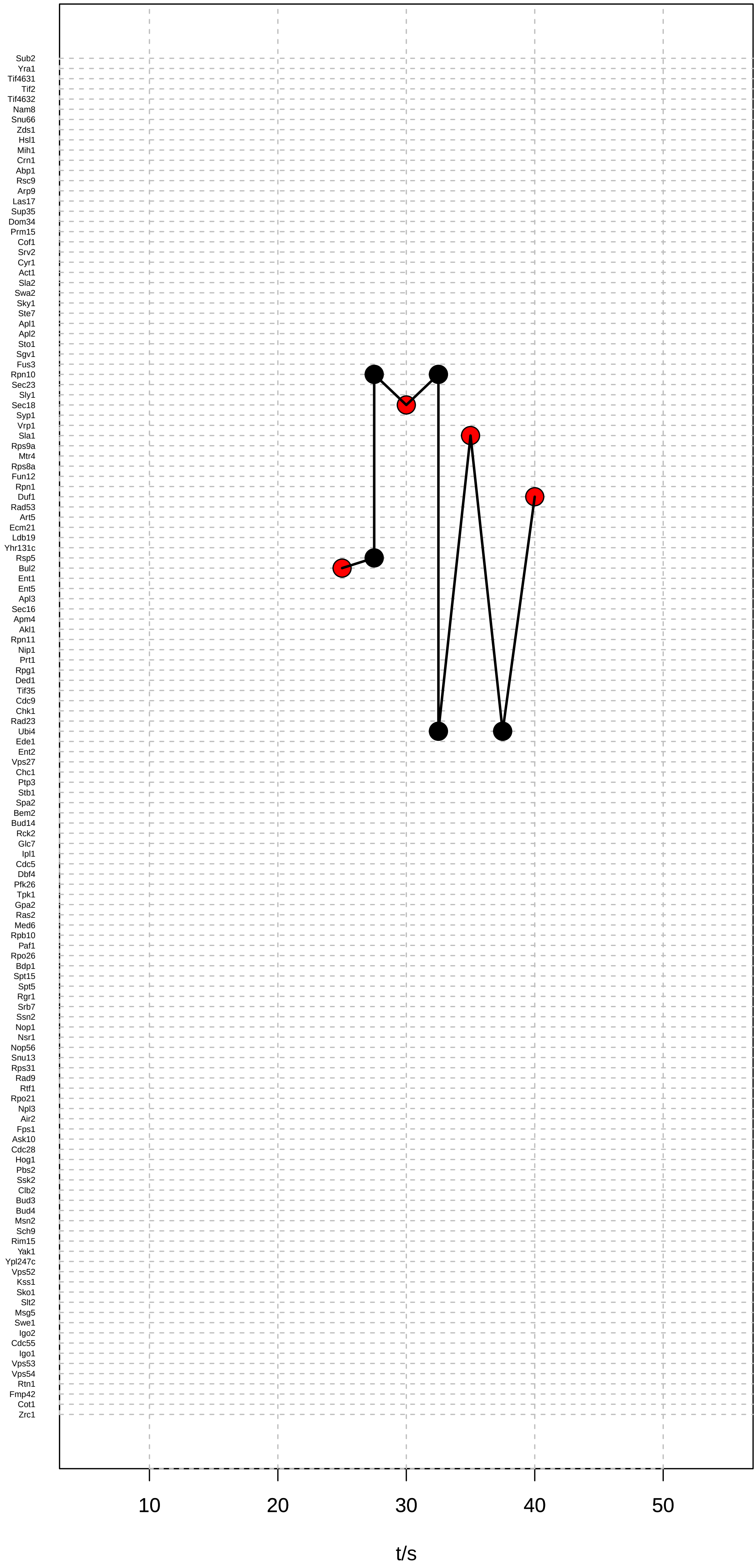

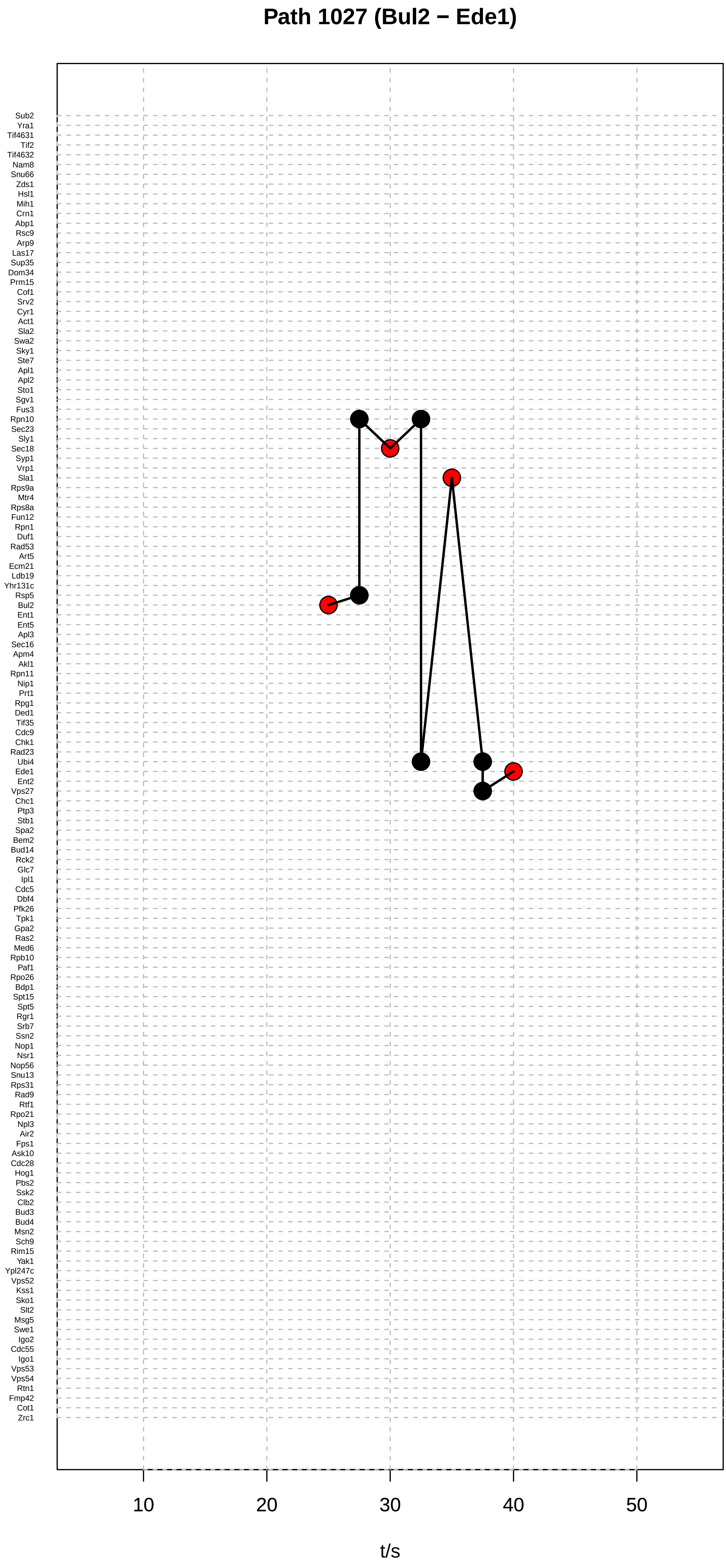

Path 1027 (Bul2 − Ede1)
Sub2
Yra1
Tif4631
Tif2
Tif4632
Nam8
Snu66
Zds1
Hsl1
Mih1
Crn1
Abp1
Rsc9
Arp9
Las17
Sup35
Dom34
Prm15
Cof1
Srv2
Cyr1
Act1
Sla2
Swa2
Sky1
Ste7
Apl1
Apl2
Sto1
Sgv1
Fus3
Rpn10
Sec23
Sly1
Sec18
Syp1
Vrp1
Sla1
Rps9a
Mtr4
Rps8a
Fun12
Rpn1
Duf1
Rad53
Art5
Ecm21
Ldb19
Yhr131c
Rsp5
Bul2
Ent1
Ent5
Apl3
Sec16
Apm4
Akl1
Rpn11
Nip1
Prt1
Rpg1
Ded1
Tif35
Cdc9
Chk1
Rad23
Ubi4
Ede1
Ent2
Vps27
Chc1
Ptp3
Stb1
Spa2
Bem2
Bud14
Rck2
Glc7
Ipl1
Cdc5
Dbf4
Pfk26
Tpk1
Gpa2
Ras2
Med6
Rpb10
Paf1
Rpo26
Bdp1
Spt15
Spt5
Rgr1
Srb7
Ssn2
Nop1
Nsr1
Nop56
Snu13
Rps31
Rad9
Rtf1
Rpo21
Npl3
Air2
Fps1
Ask10
Cdc28
Hog1
Pbs2
Ssk2
Clb2
Bud3
Bud4
Msn2
Sch9
Rim15
Yak1
Ypl247c
Vps52
Kss1
Sko1
Slt2
Msg5
Swe1
Igo2
Cdc55
Igo1
Vps53
Vps54
Rtn1
Fmp42
Cot1
Zrc1
10
20
30
40
50
t/s

### Path 1028 (Bul2 − Fun12)

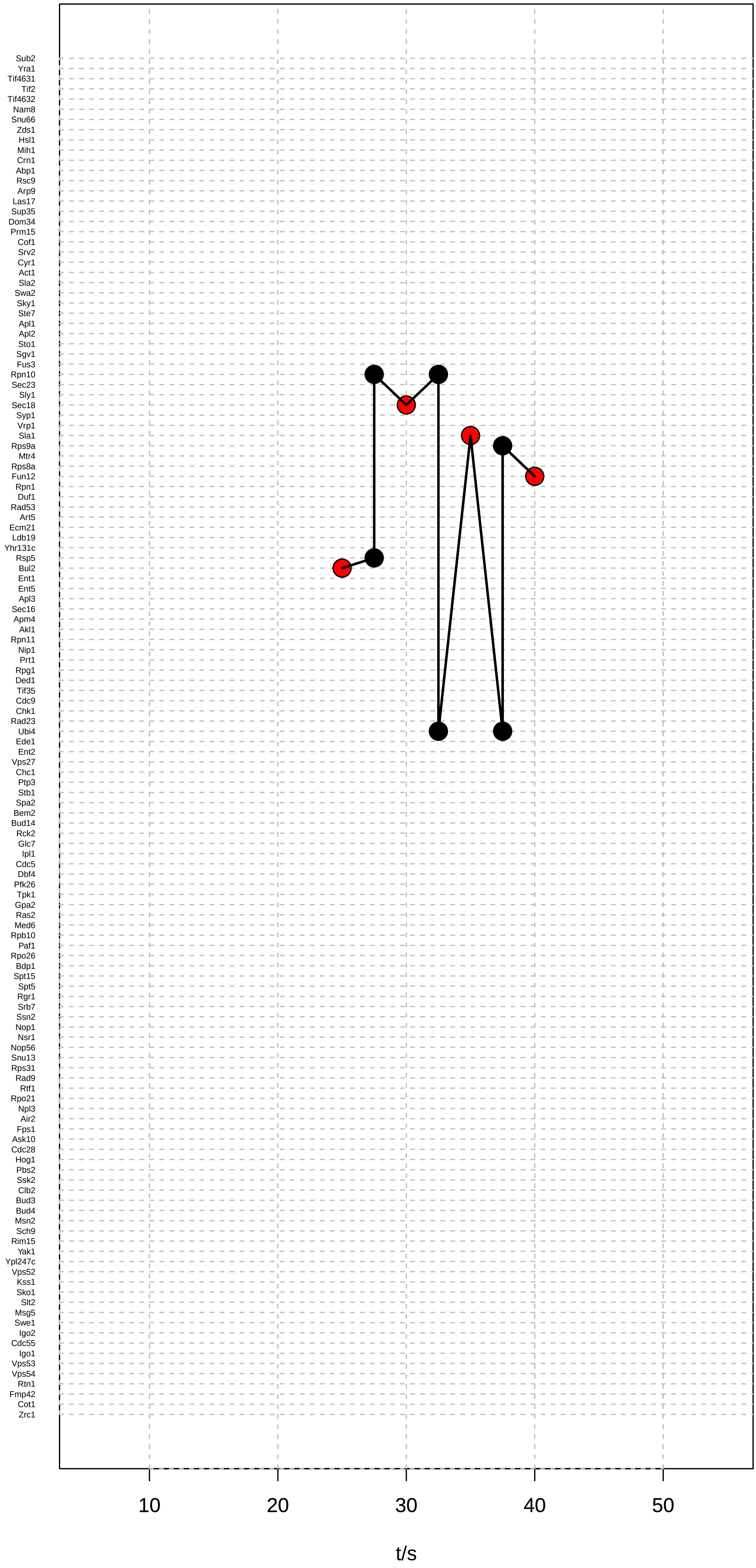

### Path 1029 (Bul2 − Fun12)

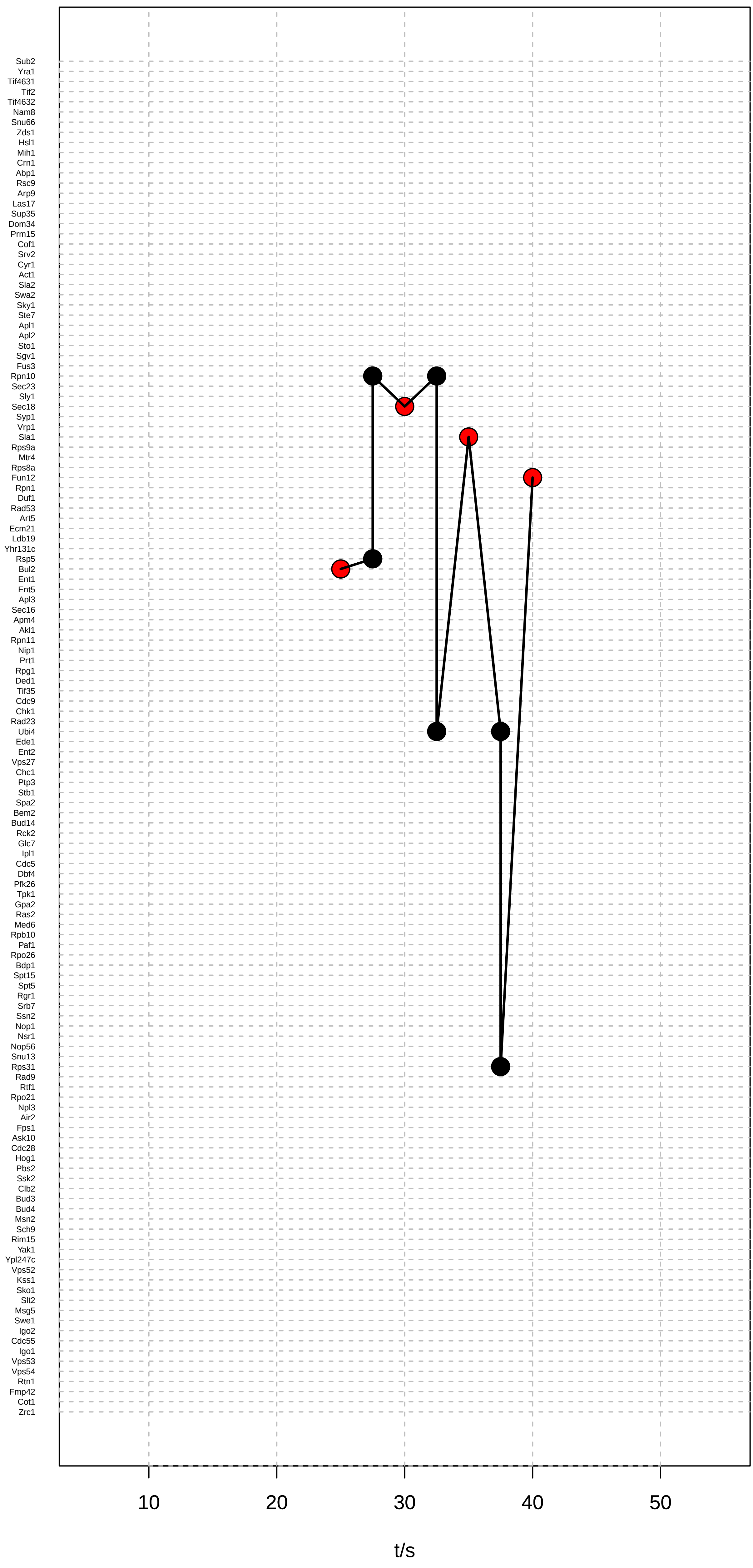

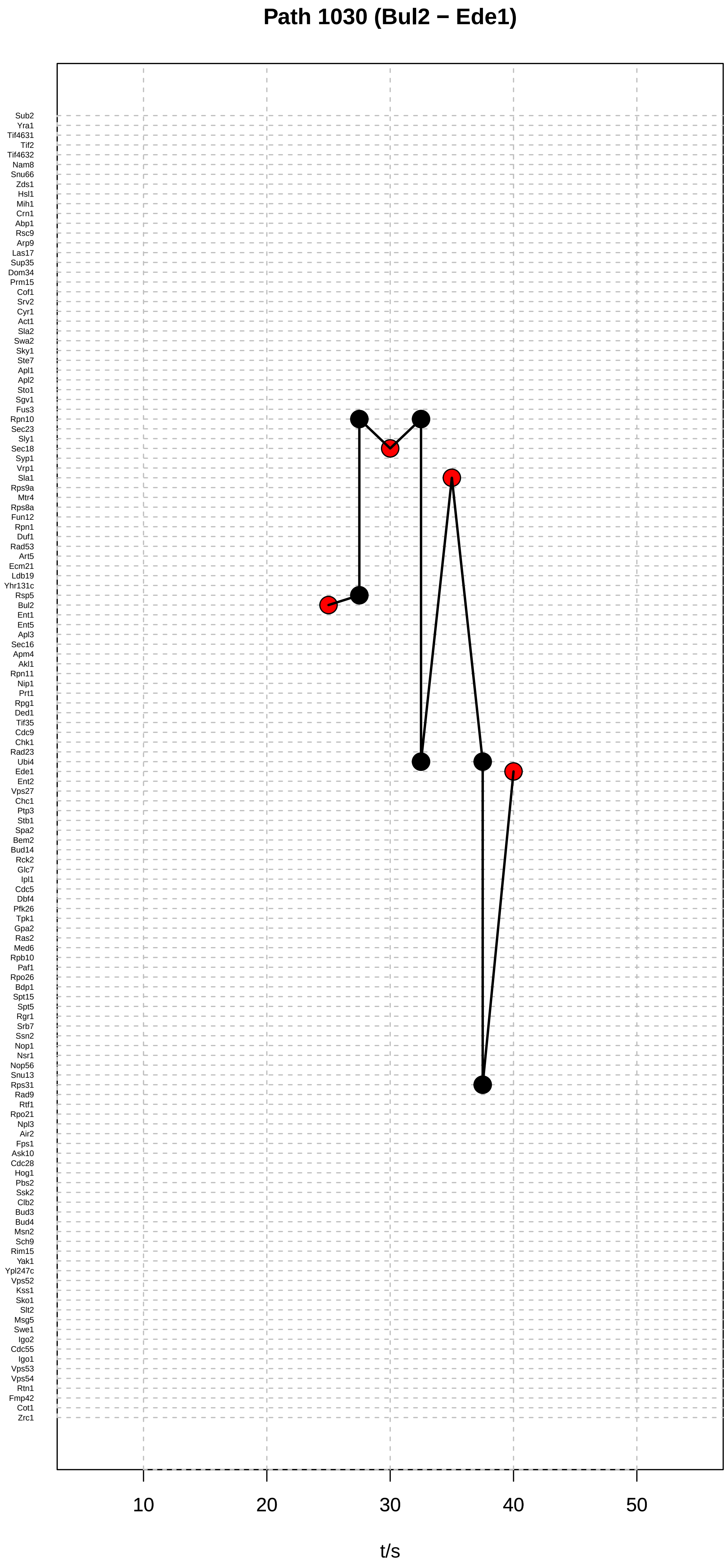
Path 1030 (Bul2 − Ede1)
Sub2
Yra1
Tif4631
Tif2
Tif4632
Nam8
Snu66
Zds1
Hsl1
Mih1
Crn1
Abp1
Rsc9
Arp9
Las17
Sup35
Dom34
Prm15
Cof1
Srv2
Cyr1
Act1
Sla2
Swa2
Sky1
Ste7
Apl1
Apl2
Sto1
Sgv1
Fus3
Rpn10
Sec23
Sly1
Sec18
Syp1
Vrp1
Sla1
Rps9a
Mtr4
Rps8a
Fun12
Rpn1
Duf1
Rad53
Art5
Ecm21
Ldb19
Yhr131c
Rsp5
Bul2
Ent1
Ent5
Apl3
Sec16
Apm4
Akl1
Rpn11
Nip1
Prt1
Rpg1
Ded1
Tif35
Cdc9
Chk1
Rad23
Ubi4
Ede1
Ent2
Vps27
Chc1
Ptp3
Stb1
Spa2
Bem2
Bud14
Rck2
Glc7
Ipl1
Cdc5
Dbf4
Pfk26
Tpk1
Gpa2
Ras2
Med6
Rpb10
Paf1
Rpo26
Bdp1
Spt15
Spt5
Rgr1
Srb7
Ssn2
Nop1
Nsr1
Nop56
Snu13
Rps31
Rad9
Rtf1
Rpo21
Npl3
Air2
Fps1
Ask10
Cdc28
Hog1
Pbs2
Ssk2
Clb2
Bud3
Bud4
Msn2
Sch9
Rim15
Yak1
Ypl247c
Vps52
Kss1
Sko1
Slt2
Msg5
Swe1
Igo2
Cdc55
Igo1
Vps53
Vps54
Rtn1
Fmp42
Cot1
Zrc1
10
20
30
40
50
t/s

### Path 1031 (Bul2 − Cyr1)

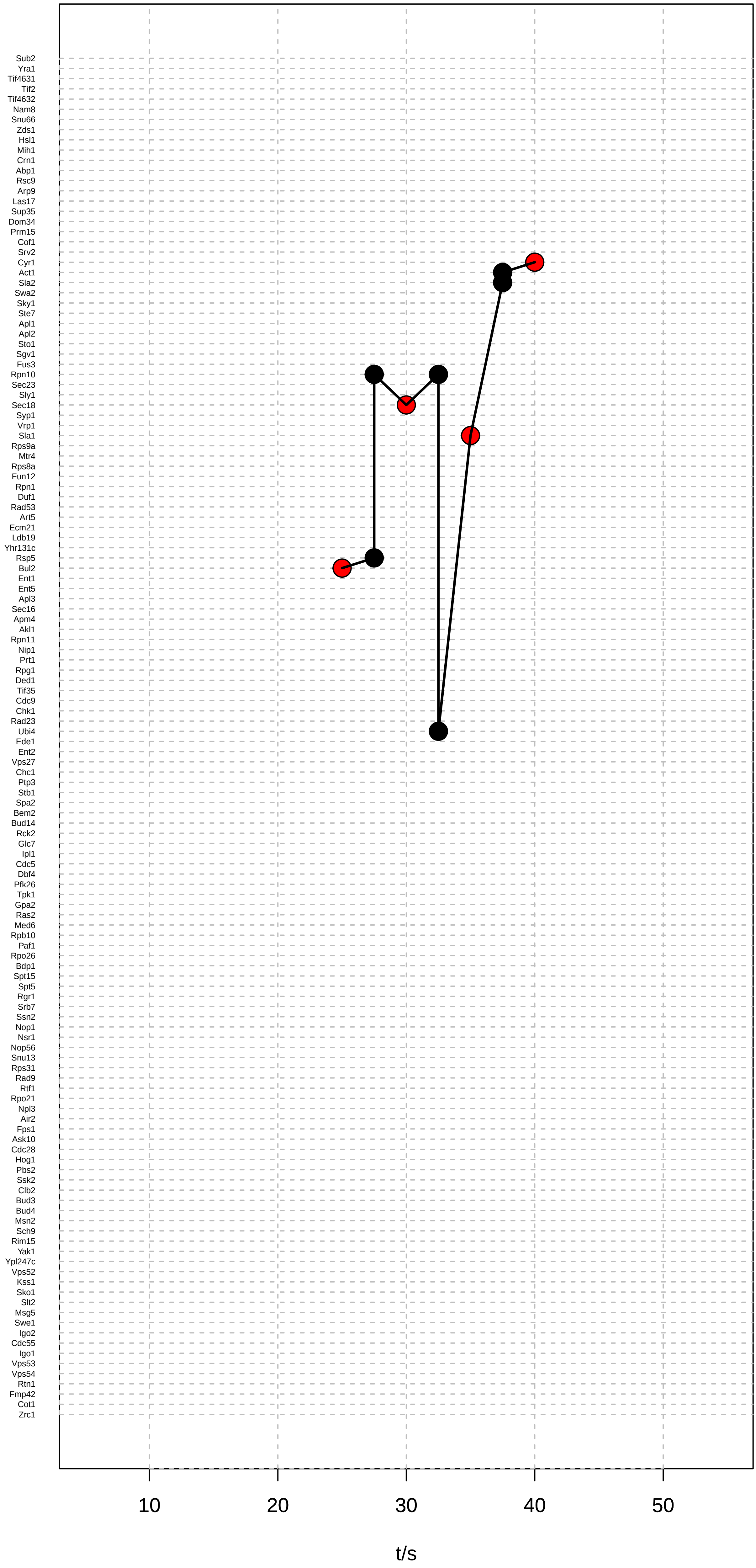

#### Path 1032 (Bul2 − Fun12)

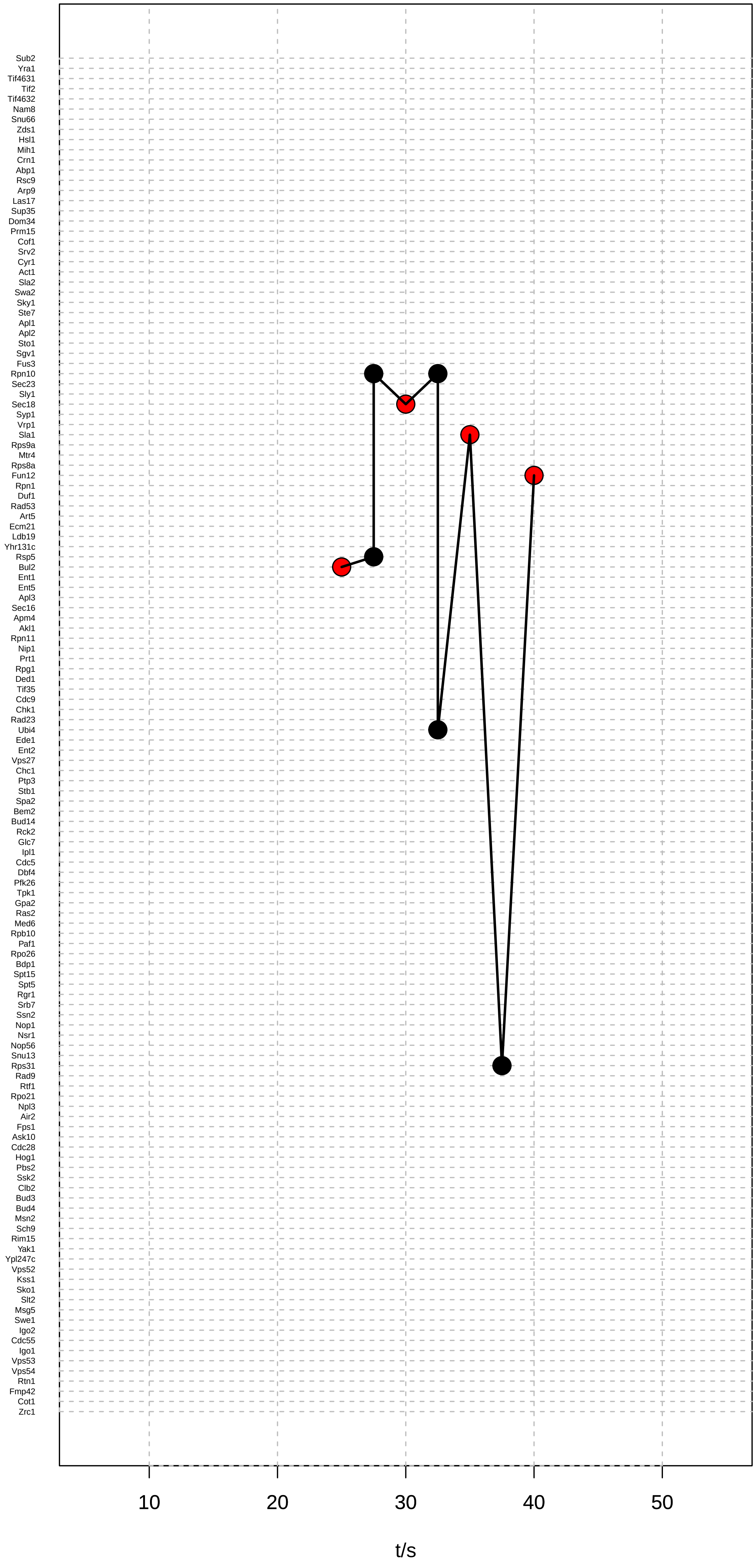

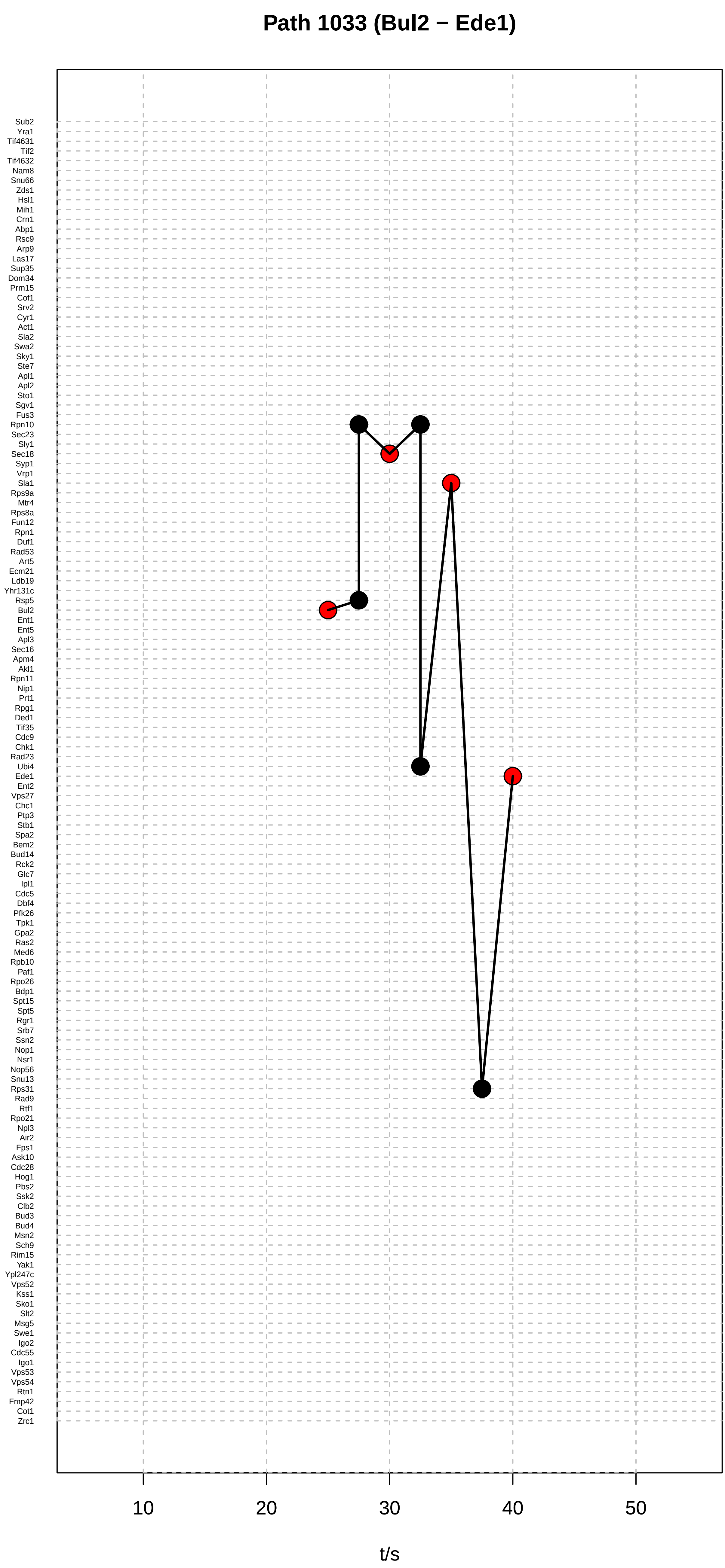

Path 1033 (Bul2 − Ede1)
Sub2
Yra1
Tif4631
Tif2
Tif4632
Nam8
Snu66
Zds1
Hsl1
Mih1
Crn1
Abp1
Rsc9
Arp9
Las17
Sup35
Dom34
Prm15
Cof1
Srv2
Cyr1
Act1
Sla2
Swa2
Sky1
Ste7
Apl1
Apl2
Sto1
Sgv1
Fus3
Rpn10
Sec23
Sly1
Sec18
Syp1
Vrp1
Sla1
Rps9a
Mtr4
Rps8a
Fun12
Rpn1
Duf1
Rad53
Art5
Ecm21
Ldb19
Yhr131c
Rsp5
Bul2
Ent1
Ent5
Apl3
Sec16
Apm4
Akl1
Rpn11
Nip1
Prt1
Rpg1
Ded1
Tif35
Cdc9
Chk1
Rad23
Ubi4
Ede1
Ent2
Vps27
Chc1
Ptp3
Stb1
Spa2
Bem2
Bud14
Rck2
Glc7
Ipl1
Cdc5
Dbf4
Pfk26
Tpk1
Gpa2
Ras2
Med6
Rpb10
Paf1
Rpo26
Bdp1
Spt15
Spt5
Rgr1
Srb7
Ssn2
Nop1
Nsr1
Nop56
Snu13
Rps31
Rad9
Rtf1
Rpo21
Npl3
Air2
Fps1
Ask10
Cdc28
Hog1
Pbs2
Ssk2
Clb2
Bud3
Bud4
Msn2
Sch9
Rim15
Yak1
Ypl247c
Vps52
Kss1
Sko1
Slt2
Msg5
Swe1
Igo2
Cdc55
Igo1
Vps53
Vps54
Rtn1
Fmp42
Cot1
Zrc1
10
20
30
40
50
t/s

### Path 1034 (Bul2 − Ede1)

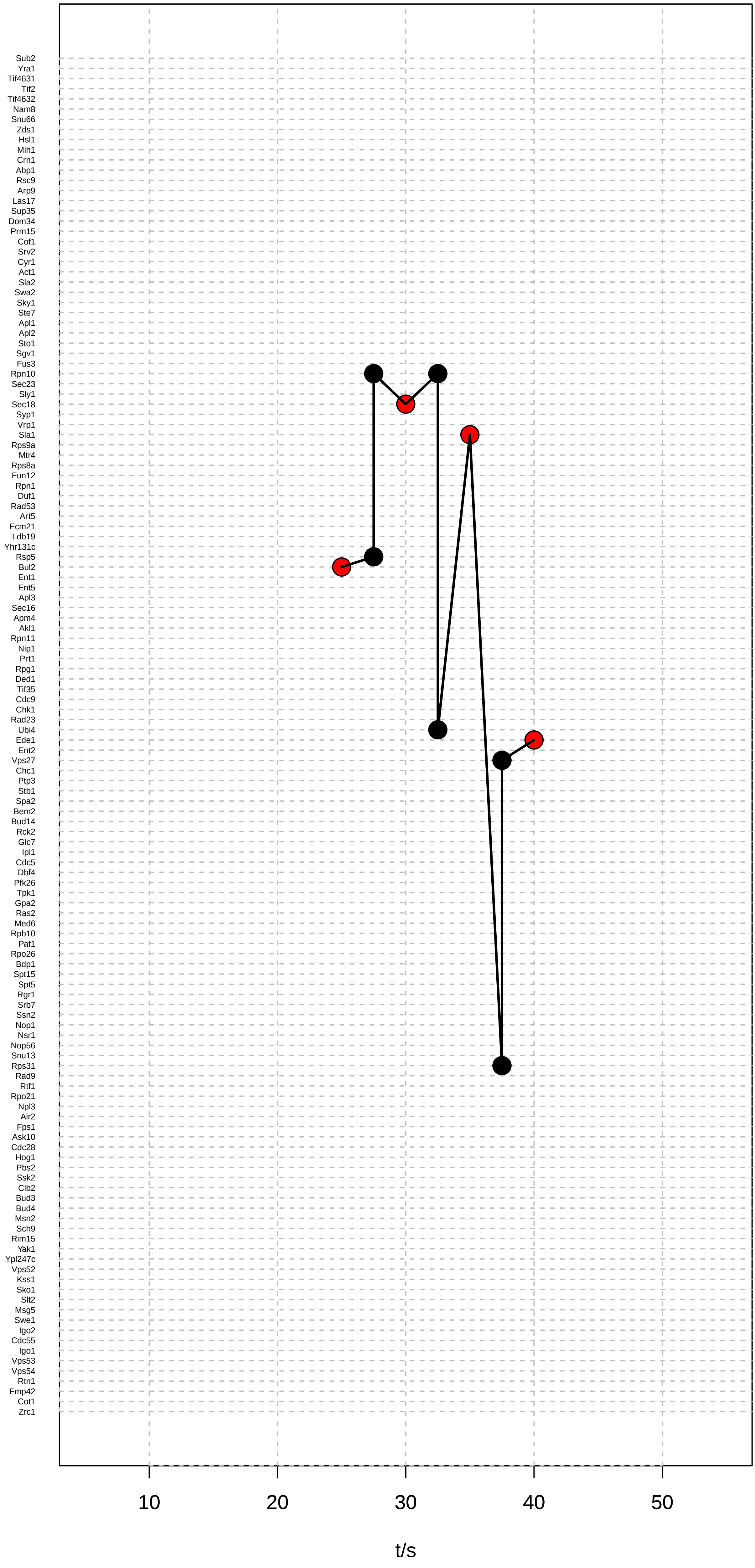

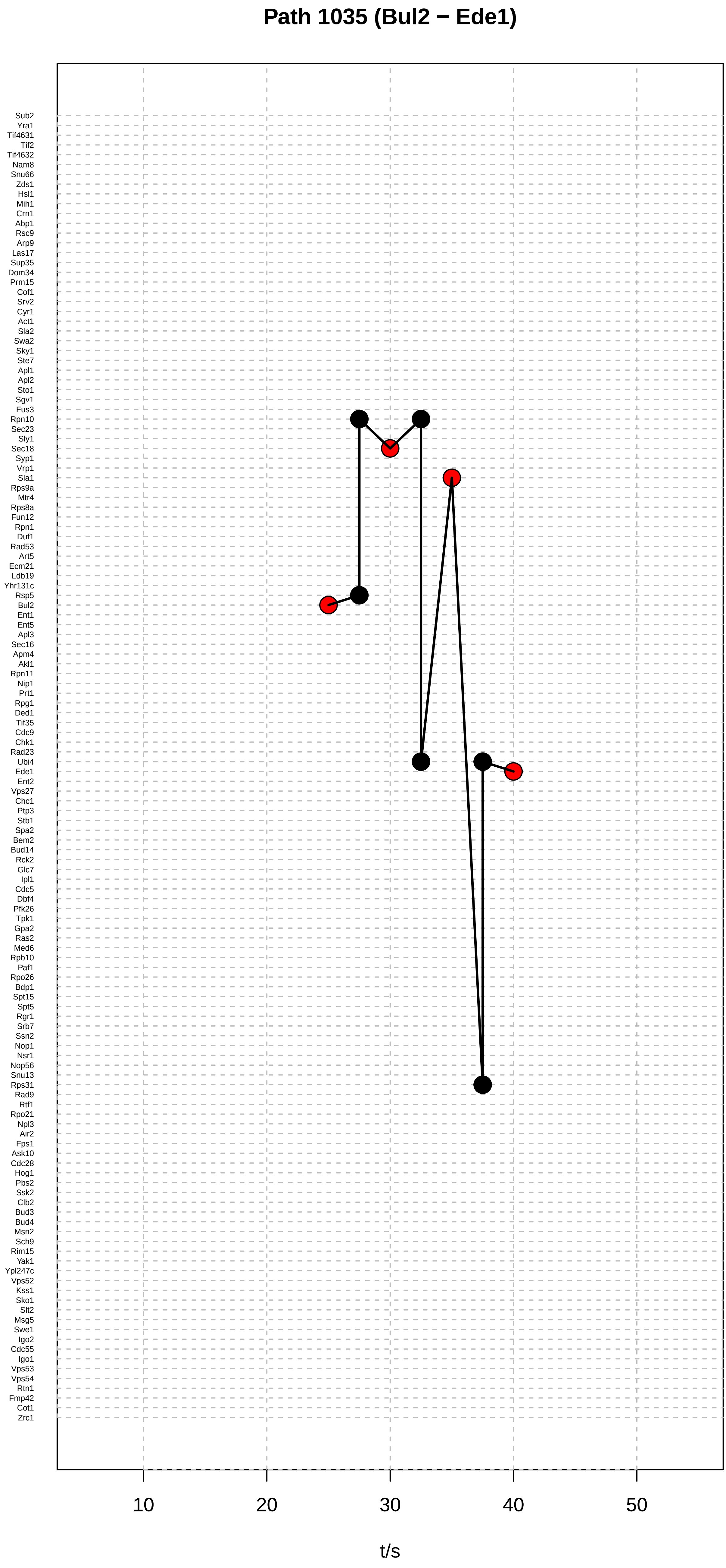

Path 1035 (Bul2 − Ede1)
Sub2
Yra1
Tif4631
Tif2
Tif4632
Nam8
Snu66
Zds1
Hsl1
Mih1
Crn1
Abp1
Rsc9
Arp9
Las17
Sup35
Dom34
Prm15
Cof1
Srv2
Cyr1
Act1
Sla2
Swa2
Sky1
Ste7
Apl1
Apl2
Sto1
Sgv1
Fus3
Rpn10
Sec23
Sly1
Sec18
Syp1
Vrp1
Sla1
Rps9a
Mtr4
Rps8a
Fun12
Rpn1
Duf1
Rad53
Art5
Ecm21
Ldb19
Yhr131c
Rsp5
Bul2
Ent1
Ent5
Apl3
Sec16
Apm4
Akl1
Rpn11
Nip1
Prt1
Rpg1
Ded1
Tif35
Cdc9
Chk1
Rad23
Ubi4
Ede1
Ent2
Vps27
Chc1
Ptp3
Stb1
Spa2
Bem2
Bud14
Rck2
Glc7
Ipl1
Cdc5
Dbf4
Pfk26
Tpk1
Gpa2
Ras2
Med6
Rpb10
Paf1
Rpo26
Bdp1
Spt15
Spt5
Rgr1
Srb7
Ssn2
Nop1
Nsr1
Nop56
Snu13
Rps31
Rad9
Rtf1
Rpo21
Npl3
Air2
Fps1
Ask10
Cdc28
Hog1
Pbs2
Ssk2
Clb2
Bud3
Bud4
Msn2
Sch9
Rim15
Yak1
Ypl247c
Vps52
Kss1
Sko1
Slt2
Msg5
Swe1
Igo2
Cdc55
Igo1
Vps53
Vps54
Rtn1
Fmp42
Cot1
Zrc1
10
20
30
40
50
t/s

#### Path 1036 (Bul2 − Duf1)

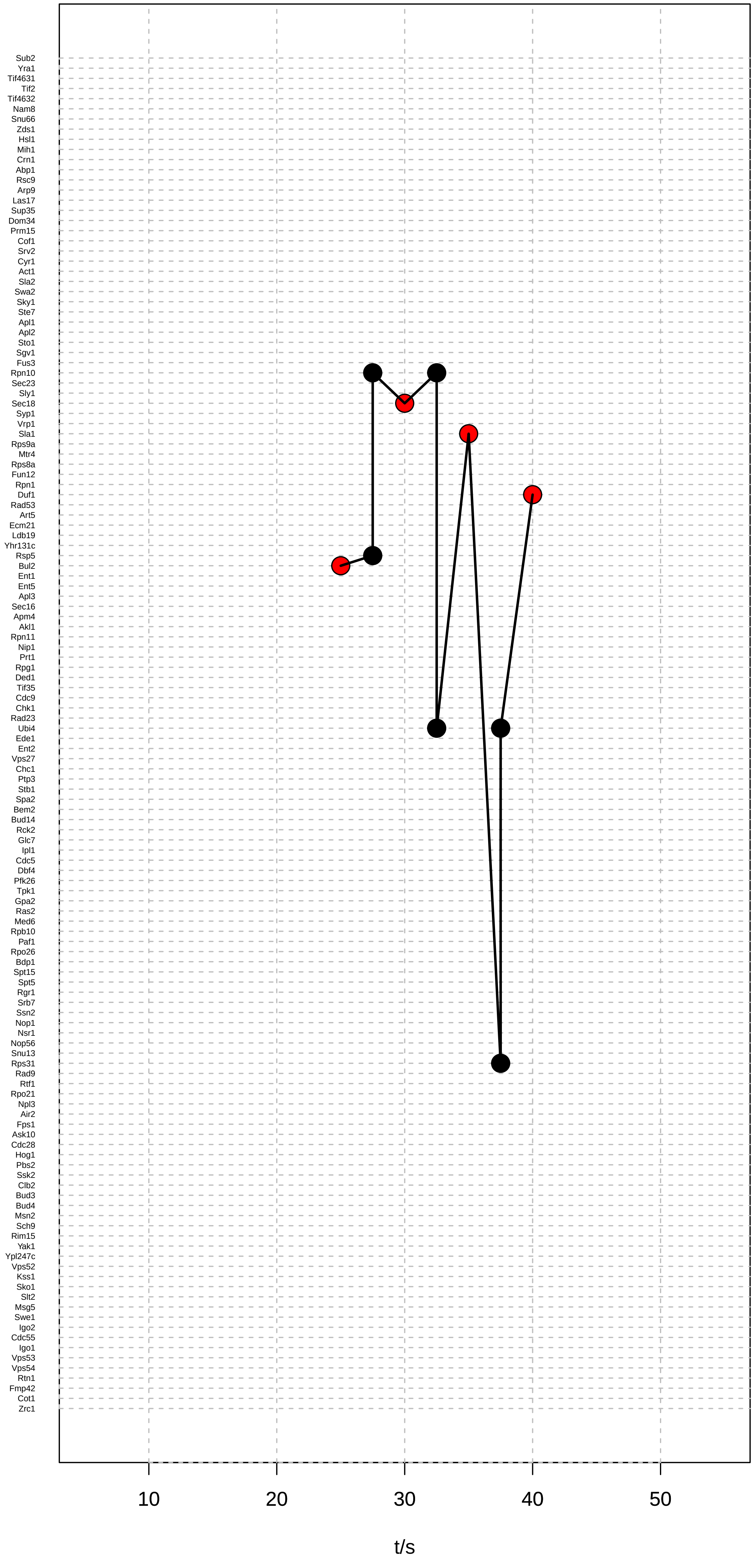

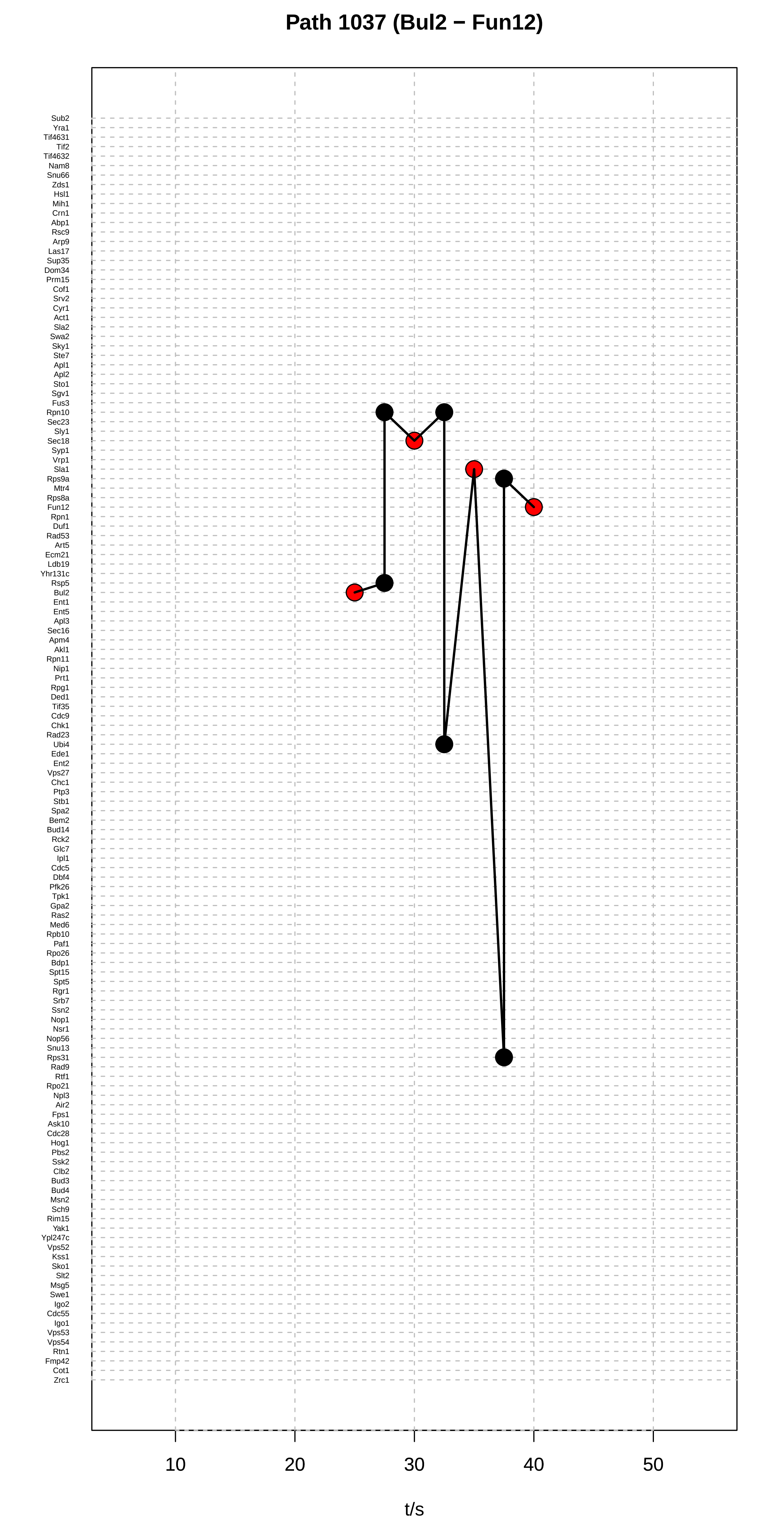

Path 1037 (Bul2 − Fun12)
Sub2
Yra1
Tif4631
Tif2
Tif4632
Nam8
Snu66
Zds1
Hsl1
Mih1
Crn1
Abp1
Rsc9
Arp9
Las17
Sup35
Dom34
Prm15
Cof1
Srv2
Cyr1
Act1
Sla2
Swa2
Sky1
Ste7
Apl1
Apl2
Sto1
Sgv1
Fus3
Rpn10
Sec23
Sly1
Sec18
Syp1
Vrp1
Sla1
Rps9a
Mtr4
Rps8a
Fun12
Rpn1
Duf1
Rad53
Art5
Ecm21
Ldb19
Yhr131c
Rsp5
Bul2
Ent1
Ent5
Apl3
Sec16
Apm4
Akl1
Rpn11
Nip1
Prt1
Rpg1
Ded1
Tif35
Cdc9
Chk1
Rad23
Ubi4
Ede1
Ent2
Vps27
Chc1
Ptp3
Stb1
Spa2
Bem2
Bud14
Rck2
Glc7
Ipl1
Cdc5
Dbf4
Pfk26
Tpk1
Gpa2
Ras2
Med6
Rpb10
Paf1
Rpo26
Bdp1
Spt15
Spt5
Rgr1
Srb7
Ssn2
Nop1
Nsr1
Nop56
Snu13
Rps31
Rad9
Rtf1
Rpo21
Npl3
Air2
Fps1
Ask10
Cdc28
Hog1
Pbs2
Ssk2
Clb2
Bud3
Bud4
Msn2
Sch9
Rim15
Yak1
Ypl247c
Vps52
Kss1
Sko1
Slt2
Msg5
Swe1
Igo2
Cdc55
Igo1
Vps53
Vps54
Rtn1
Fmp42
Cot1
Zrc1
10
20
30
40
50
t/s

#### Path 1038 (Bul2 − Sup35)

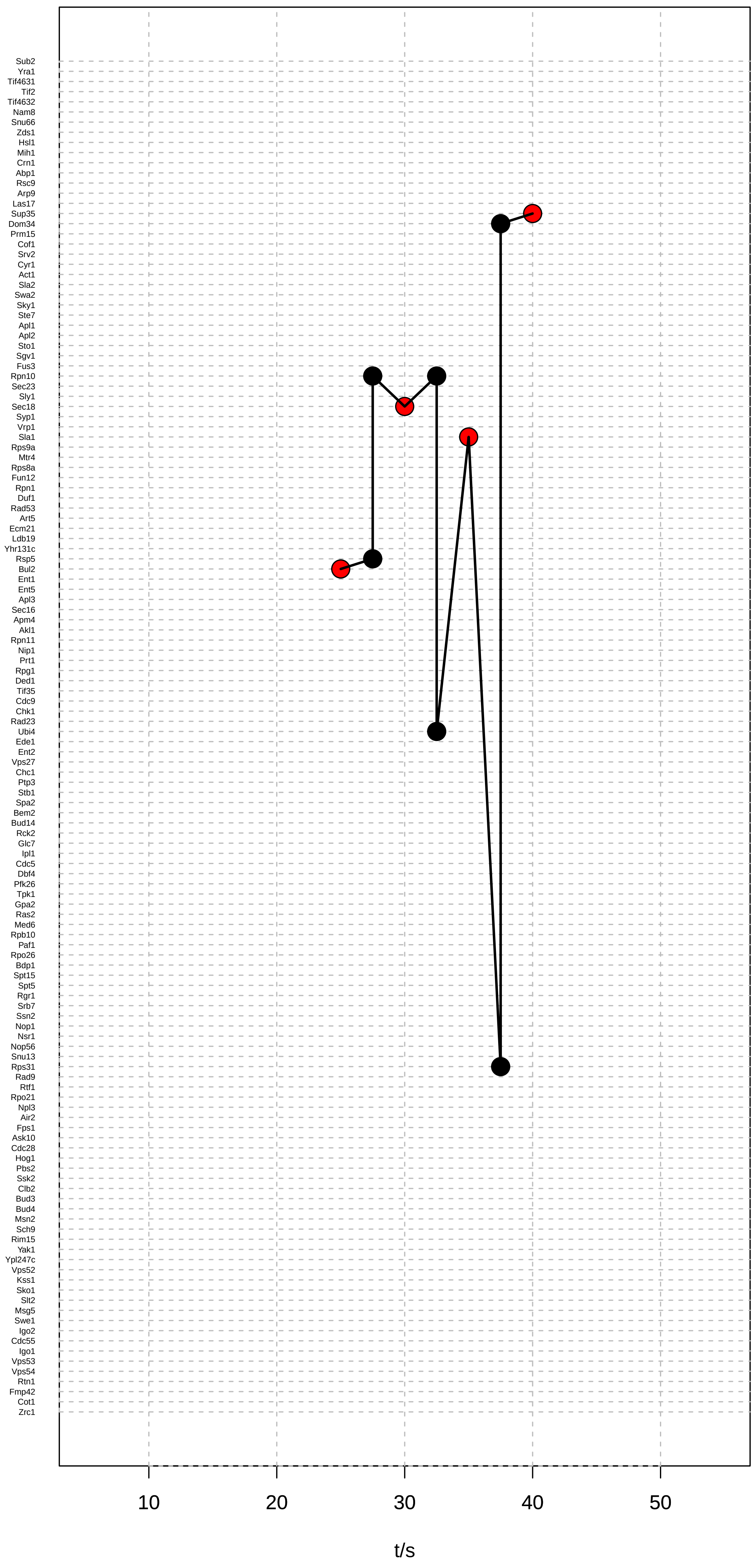

#### Path 1039 (Bul2 − Cyr1)

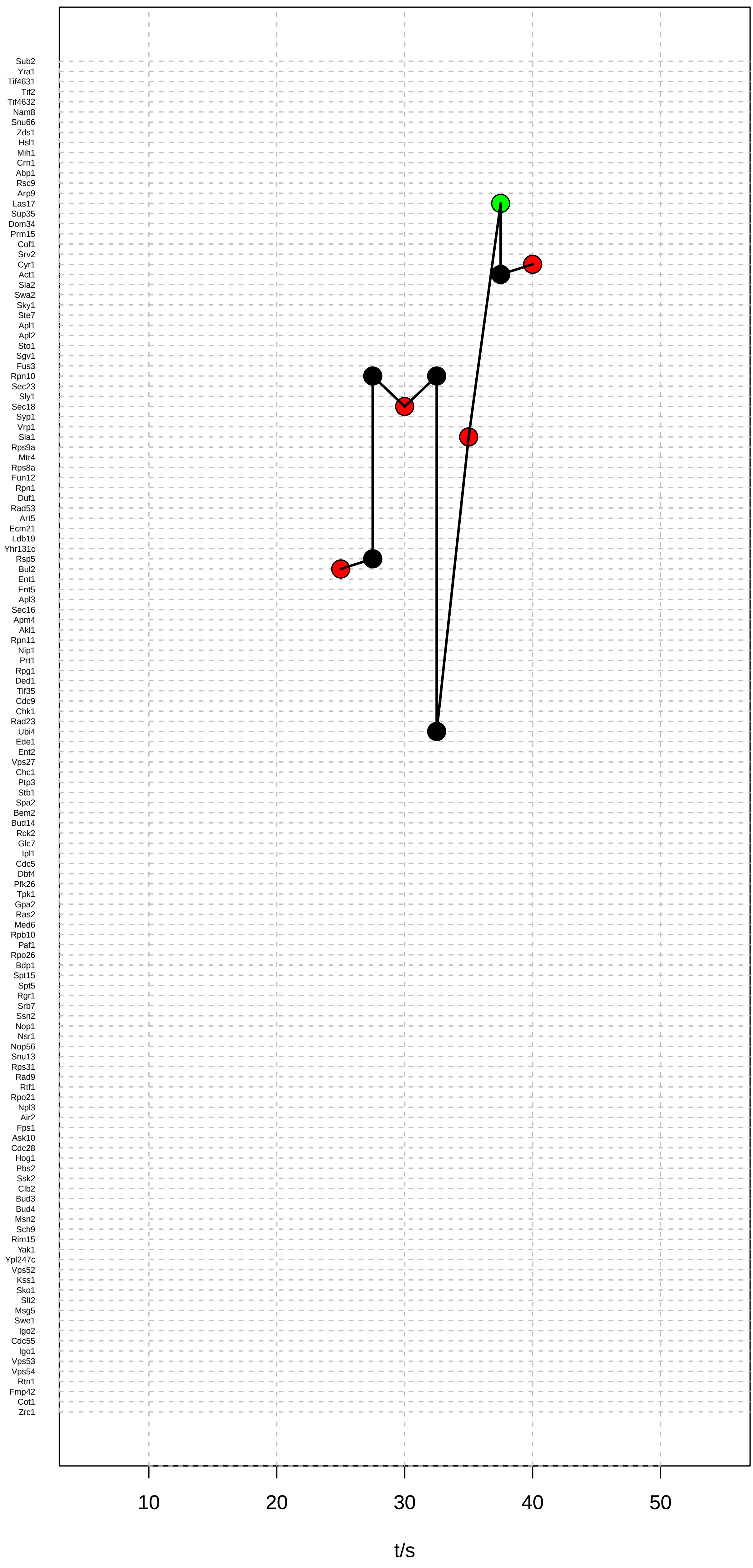

### Path 1040 (Bul2 − Ede1)

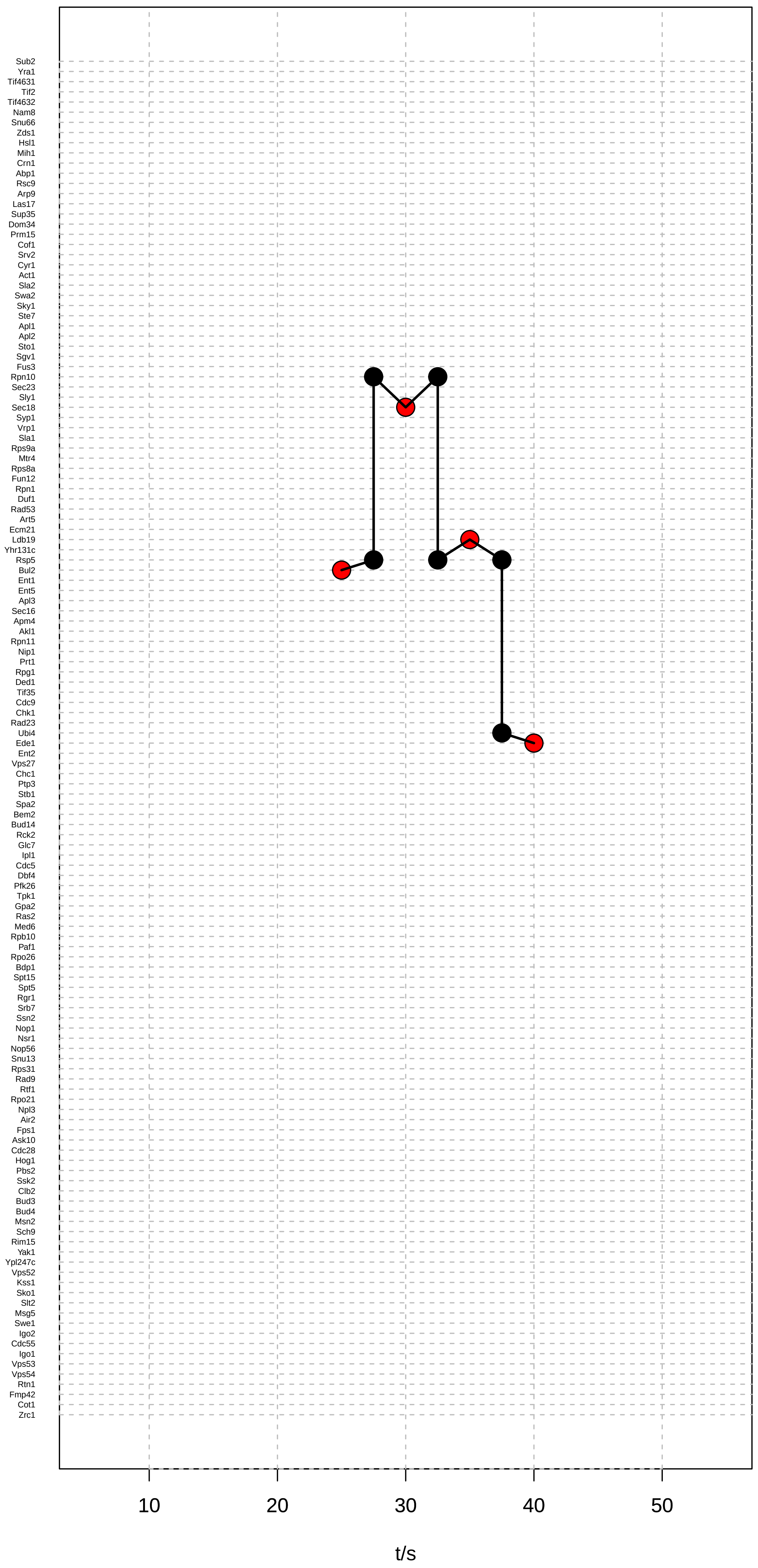

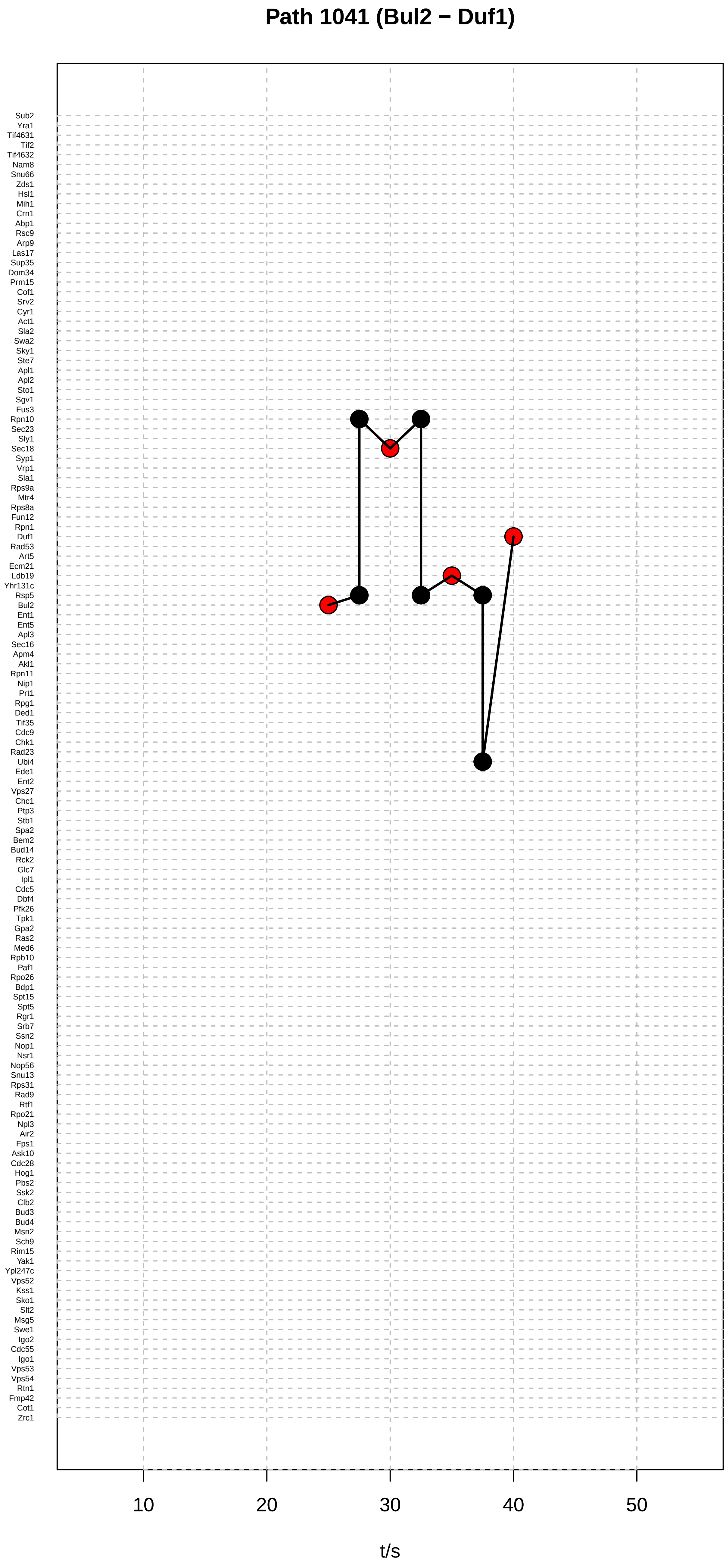

Path 1041 (Bul2 − Duf1)
Sub2
Yra1
Tif4631
Tif2
Tif4632
Nam8
Snu66
Zds1
Hsl1
Mih1
Crn1
Abp1
Rsc9
Arp9
Las17
Sup35
Dom34
Prm15
Cof1
Srv2
Cyr1
Act1
Sla2
Swa2
Sky1
Ste7
Apl1
Apl2
Sto1
Sgv1
Fus3
Rpn10
Sec23
Sly1
Sec18
Syp1
Vrp1
Sla1
Rps9a
Mtr4
Rps8a
Fun12
Rpn1
Duf1
Rad53
Art5
Ecm21
Ldb19
Yhr131c
Rsp5
Bul2
Ent1
Ent5
Apl3
Sec16
Apm4
Akl1
Rpn11
Nip1
Prt1
Rpg1
Ded1
Tif35
Cdc9
Chk1
Rad23
Ubi4
Ede1
Ent2
Vps27
Chc1
Ptp3
Stb1
Spa2
Bem2
Bud14
Rck2
Glc7
Ipl1
Cdc5
Dbf4
Pfk26
Tpk1
Gpa2
Ras2
Med6
Rpb10
Paf1
Rpo26
Bdp1
Spt15
Spt5
Rgr1
Srb7
Ssn2
Nop1
Nsr1
Nop56
Snu13
Rps31
Rad9
Rtf1
Rpo21
Npl3
Air2
Fps1
Ask10
Cdc28
Hog1
Pbs2
Ssk2
Clb2
Bud3
Bud4
Msn2
Sch9
Rim15
Yak1
Ypl247c
Vps52
Kss1
Sko1
Slt2
Msg5
Swe1
Igo2
Cdc55
Igo1
Vps53
Vps54
Rtn1
Fmp42
Cot1
Zrc1
10
20
30
40
50
t/s

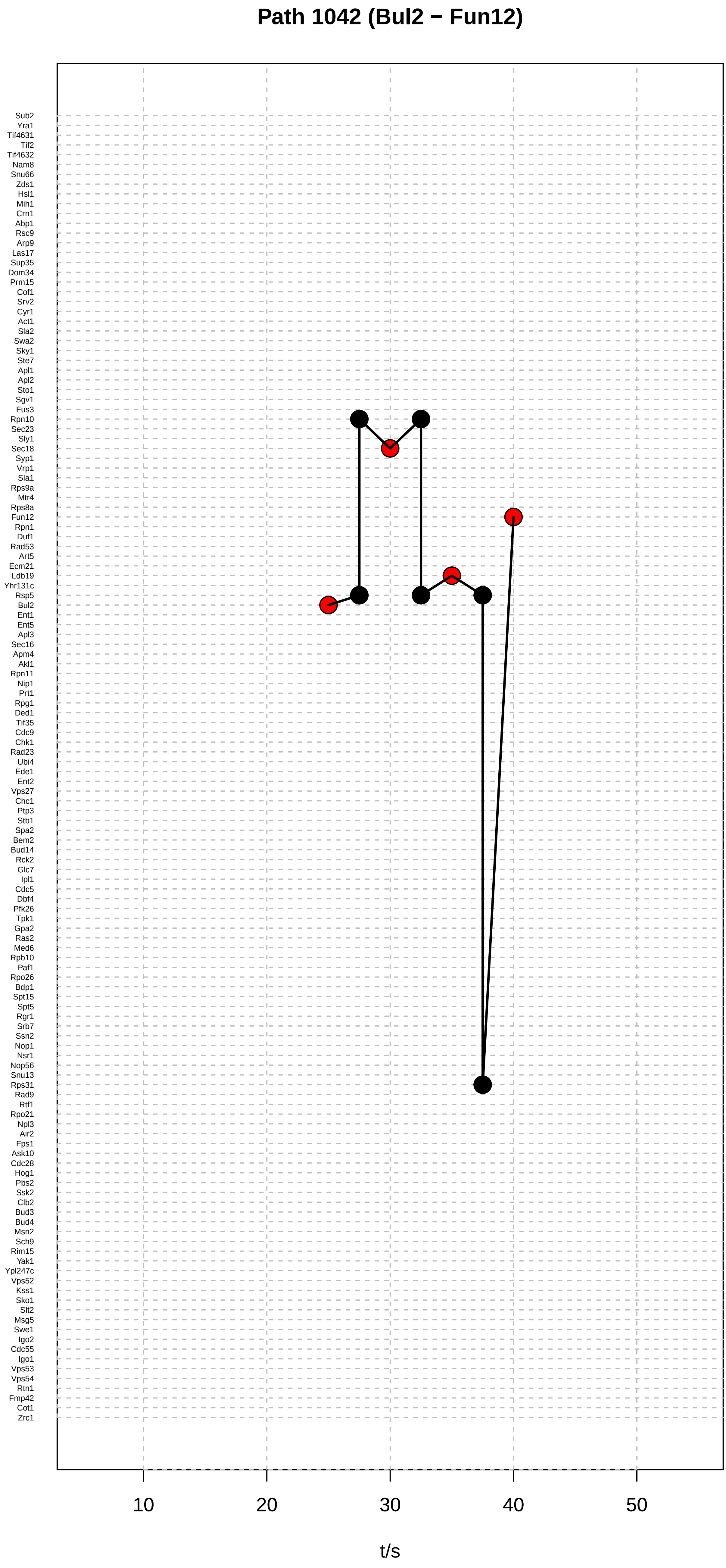

Path 1042 (Bul2 − Fun12)
Sub2
Yra1
Tif4631
Tif2
Tif4632
Nam8
Snu66
Zds1
Hsl1
Mih1
Crn1
Abp1
Rsc9
Arp9
Las17
Sup35
Dom34
Prm15
Cof1
Srv2
Cyr1
Act1
Sla2
Swa2
Sky1
Ste7
Apl1
Apl2
Sto1
Sgv1
Fus3
Rpn10
Sec23
Sly1
Sec18
Syp1
Vrp1
Sla1
Rps9a
Mtr4
Rps8a
Fun12
Rpn1
Duf1
Rad53
Art5
Ecm21
Ldb19
Yhr131c
Rsp5
Bul2
Ent1
Ent5
Apl3
Sec16
Apm4
Akl1
Rpn11
Nip1
Prt1
Rpg1
Ded1
Tif35
Cdc9
Chk1
Rad23
Ubi4
Ede1
Ent2
Vps27
Chc1
Ptp3
Stb1
Spa2
Bem2
Bud14
Rck2
Glc7
Ipl1
Cdc5
Dbf4
Pfk26
Tpk1
Gpa2
Ras2
Med6
Rpb10
Paf1
Rpo26
Bdp1
Spt15
Spt5
Rgr1
Srb7
Ssn2
Nop1
Nsr1
Nop56
Snu13
Rps31
Rad9
Rtf1
Rpo21
Npl3
Air2
Fps1
Ask10
Cdc28
Hog1
Pbs2
Ssk2
Clb2
Bud3
Bud4
Msn2
Sch9
Rim15
Yak1
Ypl247c
Vps52
Kss1
Sko1
Slt2
Msg5
Swe1
Igo2
Cdc55
Igo1
Vps53
Vps54
Rtn1
Fmp42
Cot1
Zrc1
10
20
30
40
50
t/s

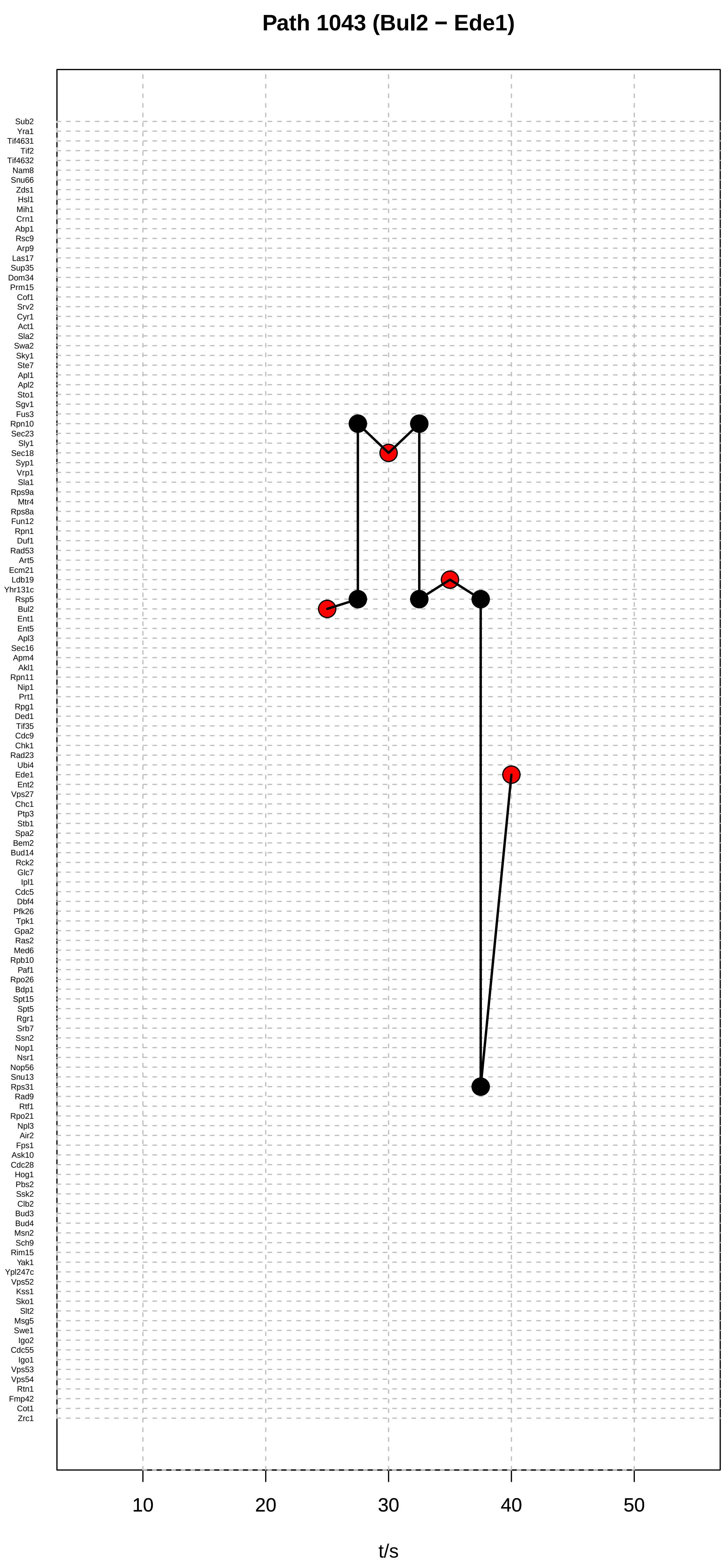
Path 1043 (Bul2 − Ede1)
Sub2
Yra1
Tif4631
Tif2
Tif4632
Nam8
Snu66
Zds1
Hsl1
Mih1
Crn1
Abp1
Rsc9
Arp9
Las17
Sup35
Dom34
Prm15
Cof1
Srv2
Cyr1
Act1
Sla2
Swa2
Sky1
Ste7
Apl1
Apl2
Sto1
Sgv1
Fus3
Rpn10
Sec23
Sly1
Sec18
Syp1
Vrp1
Sla1
Rps9a
Mtr4
Rps8a
Fun12
Rpn1
Duf1
Rad53
Art5
Ecm21
Ldb19
Yhr131c
Rsp5
Bul2
Ent1
Ent5
Apl3
Sec16
Apm4
Akl1
Rpn11
Nip1
Prt1
Rpg1
Ded1
Tif35
Cdc9
Chk1
Rad23
Ubi4
Ede1
Ent2
Vps27
Chc1
Ptp3
Stb1
Spa2
Bem2
Bud14
Rck2
Glc7
Ipl1
Cdc5
Dbf4
Pfk26
Tpk1
Gpa2
Ras2
Med6
Rpb10
Paf1
Rpo26
Bdp1
Spt15
Spt5
Rgr1
Srb7
Ssn2
Nop1
Nsr1
Nop56
Snu13
Rps31
Rad9
Rtf1
Rpo21
Npl3
Air2
Fps1
Ask10
Cdc28
Hog1
Pbs2
Ssk2
Clb2
Bud3
Bud4
Msn2
Sch9
Rim15
Yak1
Ypl247c
Vps52
Kss1
Sko1
Slt2
Msg5
Swe1
Igo2
Cdc55
Igo1
Vps53
Vps54
Rtn1
Fmp42
Cot1
Zrc1
10
20
30
40
50
t/s

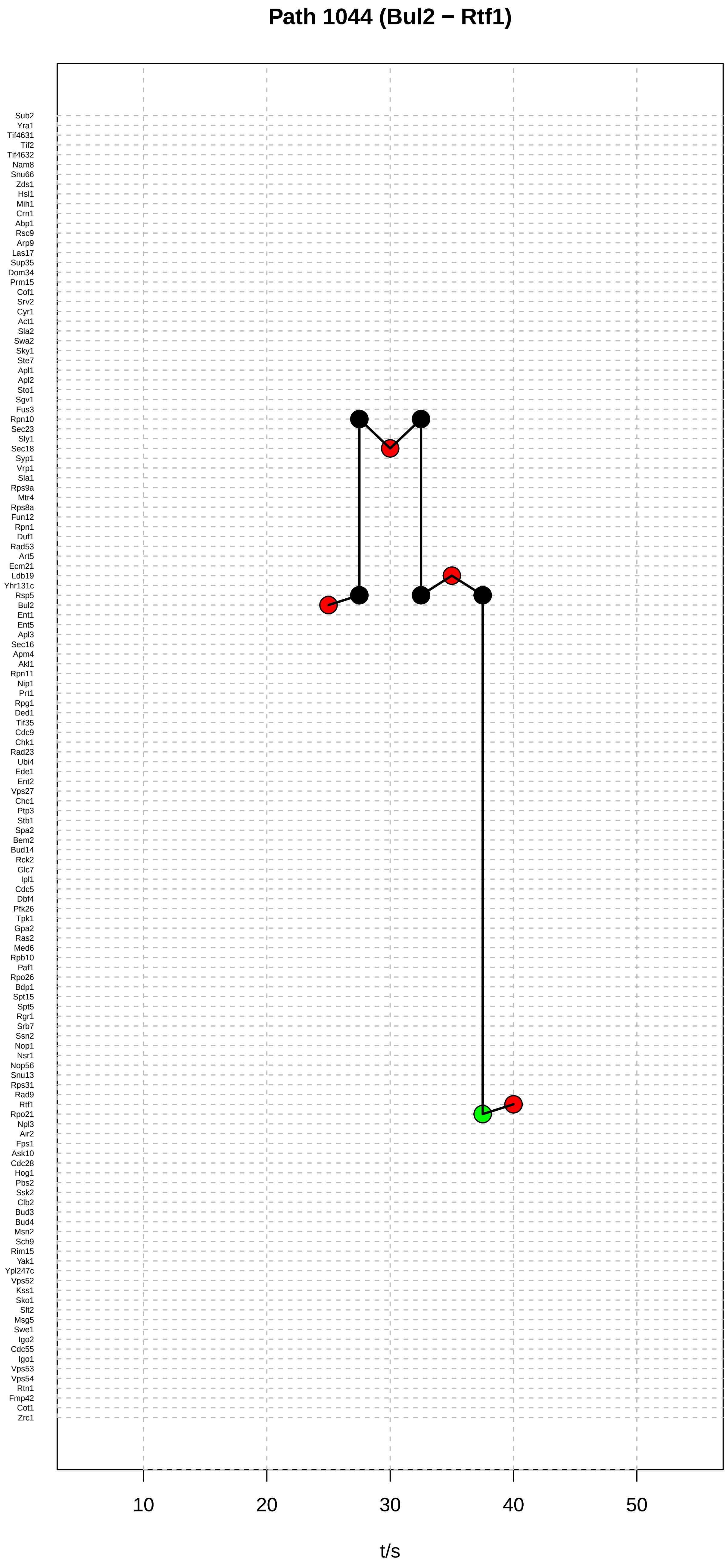

Path 1044 (Bul2 − Rtf1)
Sub2
Yra1
Tif4631
Tif2
Tif4632
Nam8
Snu66
Zds1
Hsl1
Mih1
Crn1
Abp1
Rsc9
Arp9
Las17
Sup35
Dom34
Prm15
Cof1
Srv2
Cyr1
Act1
Sla2
Swa2
Sky1
Ste7
Apl1
Apl2
Sto1
Sgv1
Fus3
Rpn10
Sec23
Sly1
Sec18
Syp1
Vrp1
Sla1
Rps9a
Mtr4
Rps8a
Fun12
Rpn1
Duf1
Rad53
Art5
Ecm21
Ldb19
Yhr131c
Rsp5
Bul2
Ent1
Ent5
Apl3
Sec16
Apm4
Akl1
Rpn11
Nip1
Prt1
Rpg1
Ded1
Tif35
Cdc9
Chk1
Rad23
Ubi4
Ede1
Ent2
Vps27
Chc1
Ptp3
Stb1
Spa2
Bem2
Bud14
Rck2
Glc7
Ipl1
Cdc5
Dbf4
Pfk26
Tpk1
Gpa2
Ras2
Med6
Rpb10
Paf1
Rpo26
Bdp1
Spt15
Spt5
Rgr1
Srb7
Ssn2
Nop1
Nsr1
Nop56
Snu13
Rps31
Rad9
Rtf1
Rpo21
Npl3
Air2
Fps1
Ask10
Cdc28
Hog1
Pbs2
Ssk2
Clb2
Bud3
Bud4
Msn2
Sch9
Rim15
Yak1
Ypl247c
Vps52
Kss1
Sko1
Slt2
Msg5
Swe1
Igo2
Cdc55
Igo1
Vps53
Vps54
Rtn1
Fmp42
Cot1
Zrc1
10
20
30
40
50
t/s

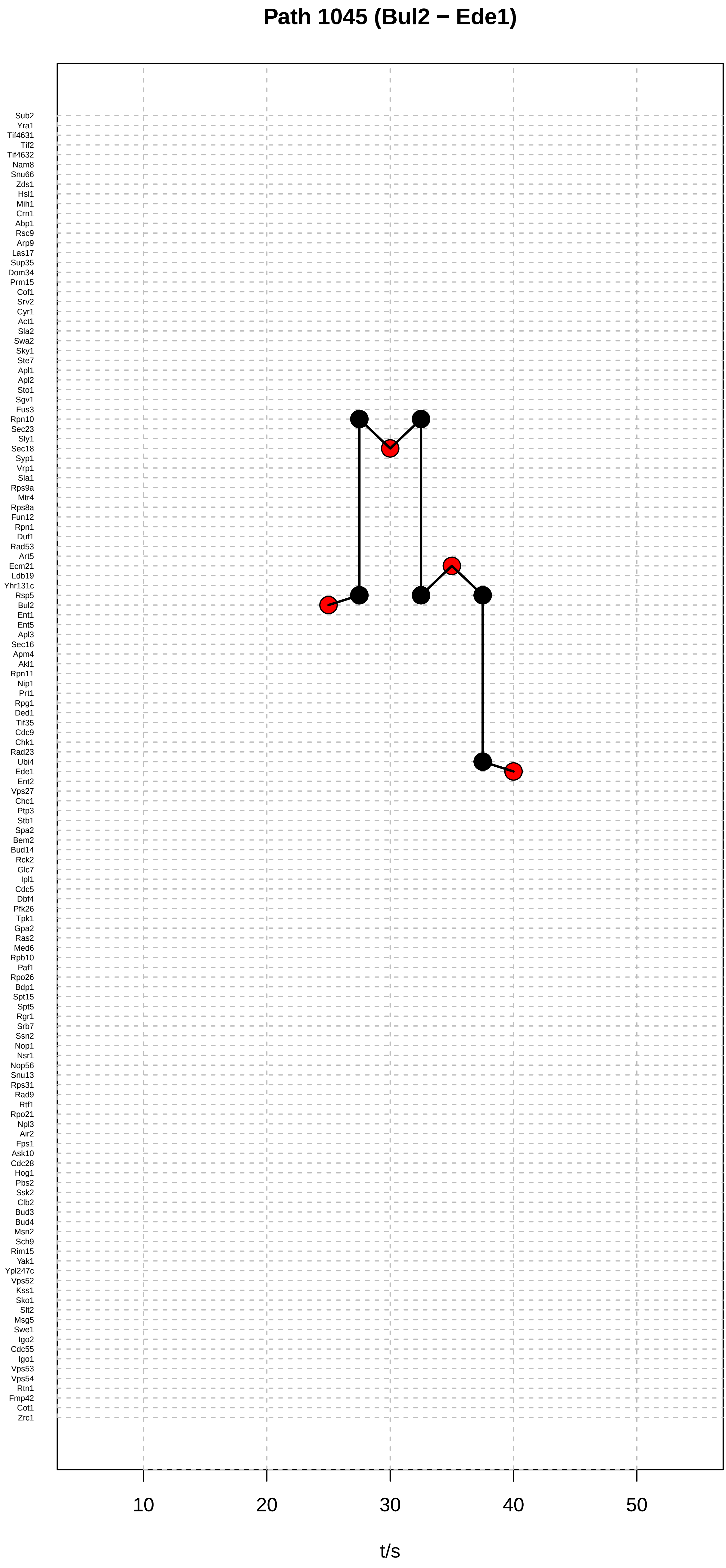

Path 1045 (Bul2 − Ede1)
Sub2
Yra1
Tif4631
Tif2
Tif4632
Nam8
Snu66
Zds1
Hsl1
Mih1
Crn1
Abp1
Rsc9
Arp9
Las17
Sup35
Dom34
Prm15
Cof1
Srv2
Cyr1
Act1
Sla2
Swa2
Sky1
Ste7
Apl1
Apl2
Sto1
Sgv1
Fus3
Rpn10
Sec23
Sly1
Sec18
Syp1
Vrp1
Sla1
Rps9a
Mtr4
Rps8a
Fun12
Rpn1
Duf1
Rad53
Art5
Ecm21
Ldb19
Yhr131c
Rsp5
Bul2
Ent1
Ent5
Apl3
Sec16
Apm4
Akl1
Rpn11
Nip1
Prt1
Rpg1
Ded1
Tif35
Cdc9
Chk1
Rad23
Ubi4
Ede1
Ent2
Vps27
Chc1
Ptp3
Stb1
Spa2
Bem2
Bud14
Rck2
Glc7
Ipl1
Cdc5
Dbf4
Pfk26
Tpk1
Gpa2
Ras2
Med6
Rpb10
Paf1
Rpo26
Bdp1
Spt15
Spt5
Rgr1
Srb7
Ssn2
Nop1
Nsr1
Nop56
Snu13
Rps31
Rad9
Rtf1
Rpo21
Npl3
Air2
Fps1
Ask10
Cdc28
Hog1
Pbs2
Ssk2
Clb2
Bud3
Bud4
Msn2
Sch9
Rim15
Yak1
Ypl247c
Vps52
Kss1
Sko1
Slt2
Msg5
Swe1
Igo2
Cdc55
Igo1
Vps53
Vps54
Rtn1
Fmp42
Cot1
Zrc1
10
20
30
40
50
t/s

### Path 1046 (Bul2 − Duf1)

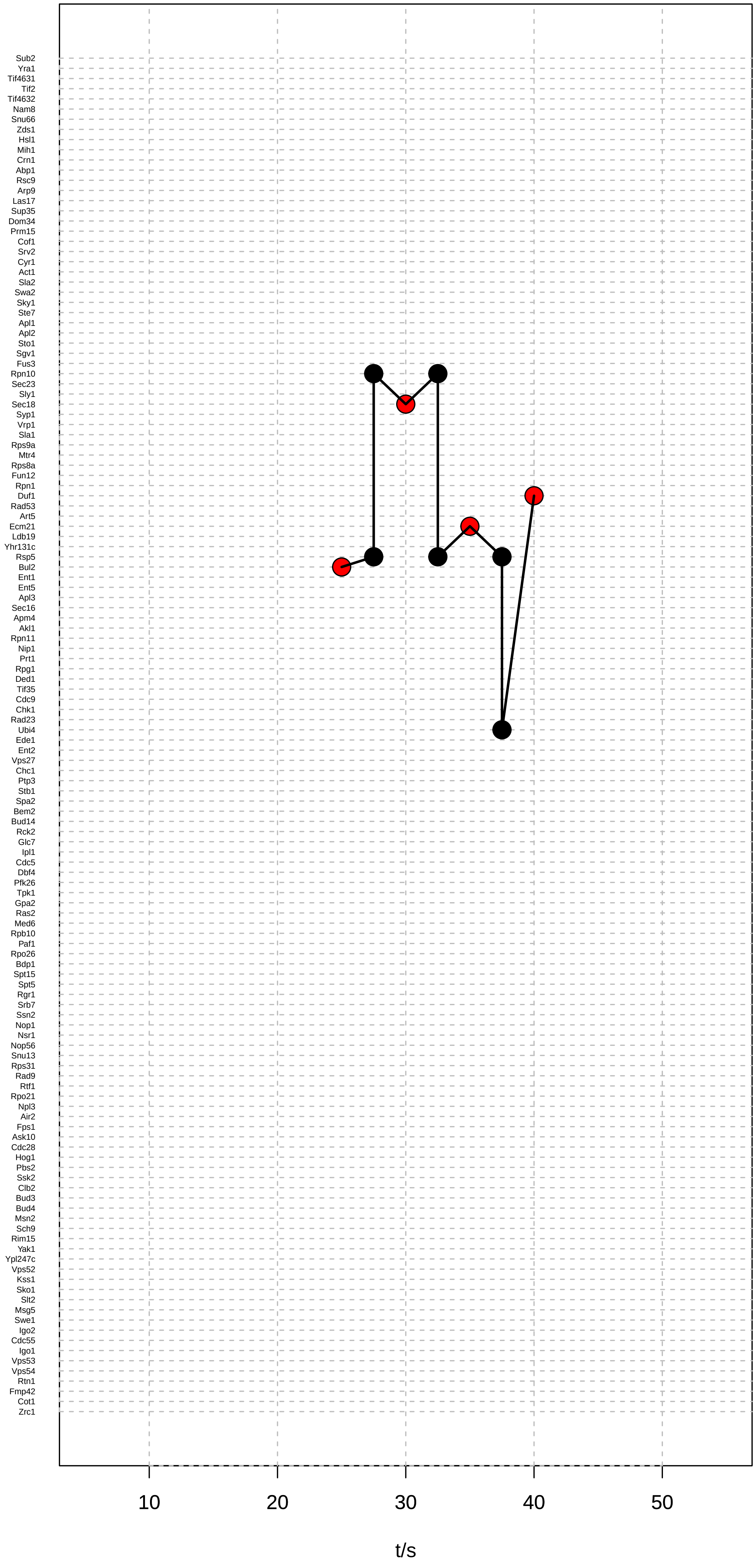

Path 1047 (Bul2 − Fun12)

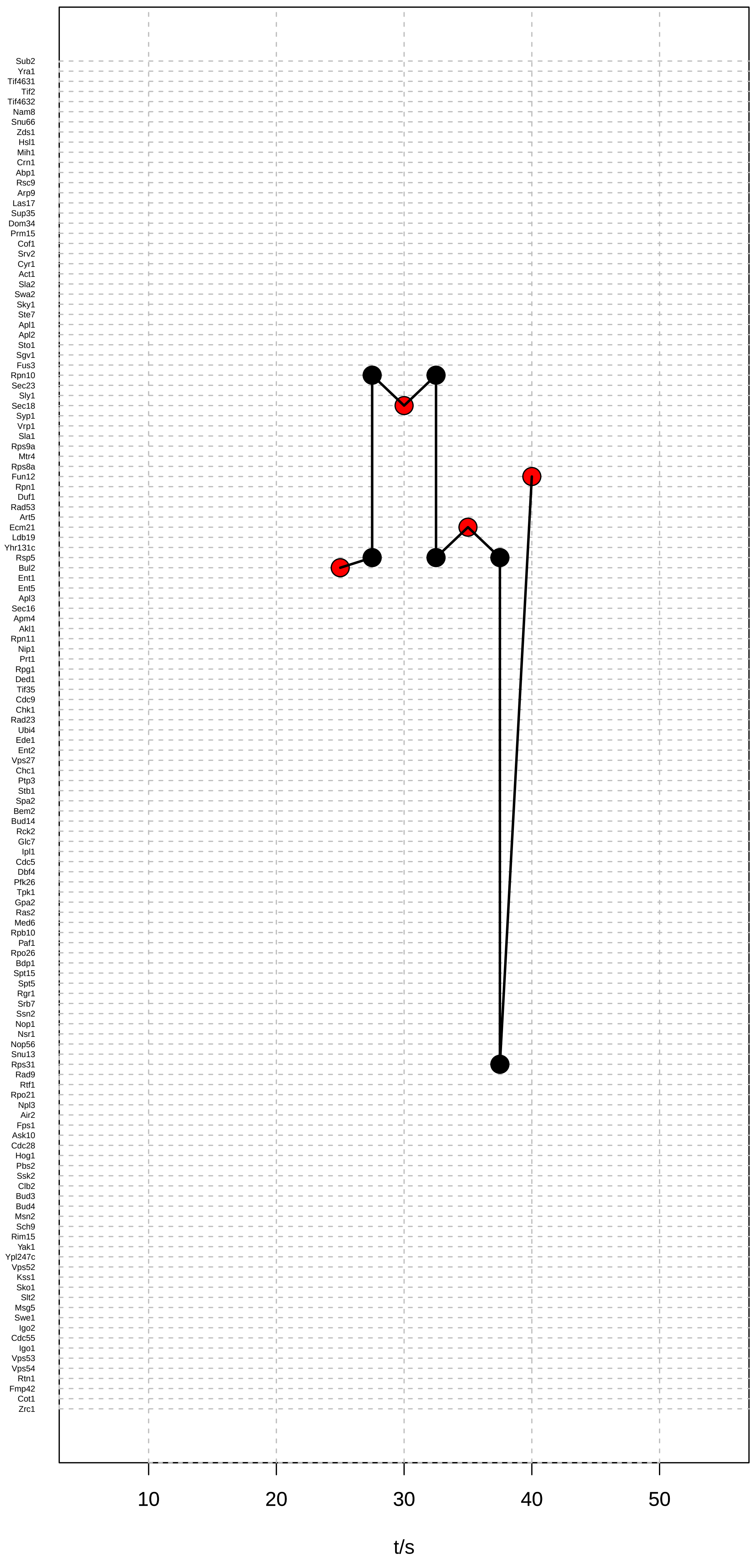

### Path 1048 (Bul2 − Ede1)

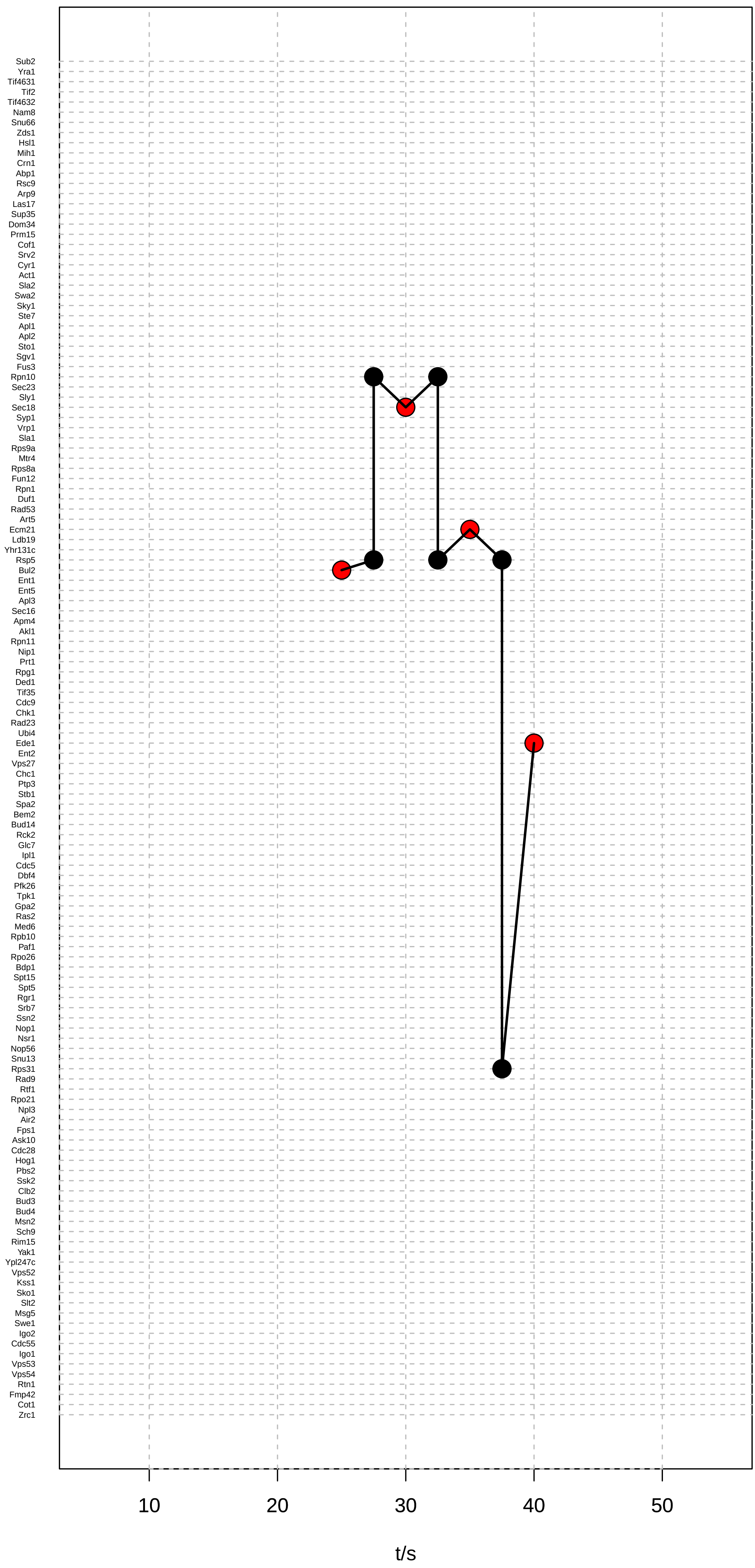

### Path 1049 (Bul2 − Rtf1)

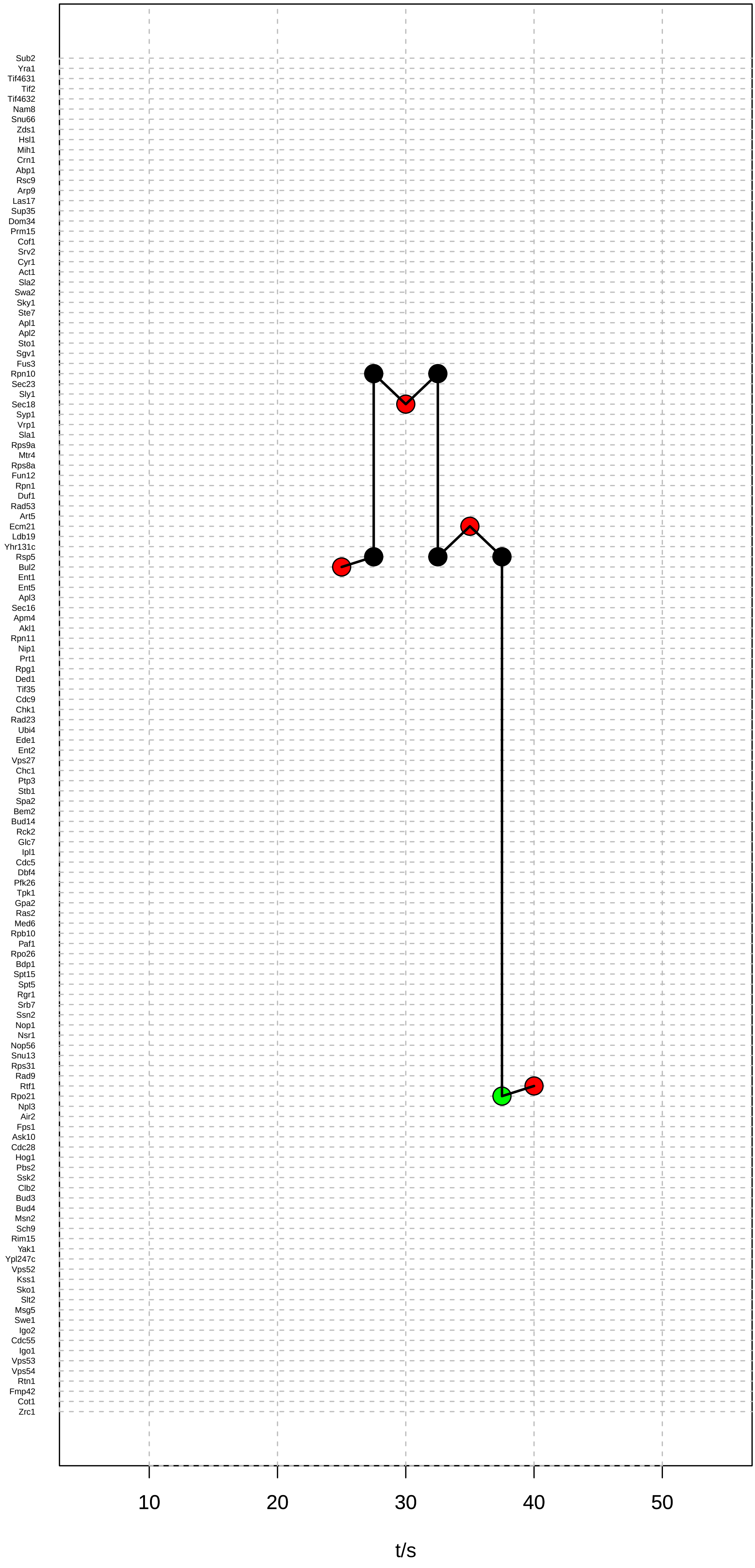

#### Path 1050 (Bul2 − Ede1)

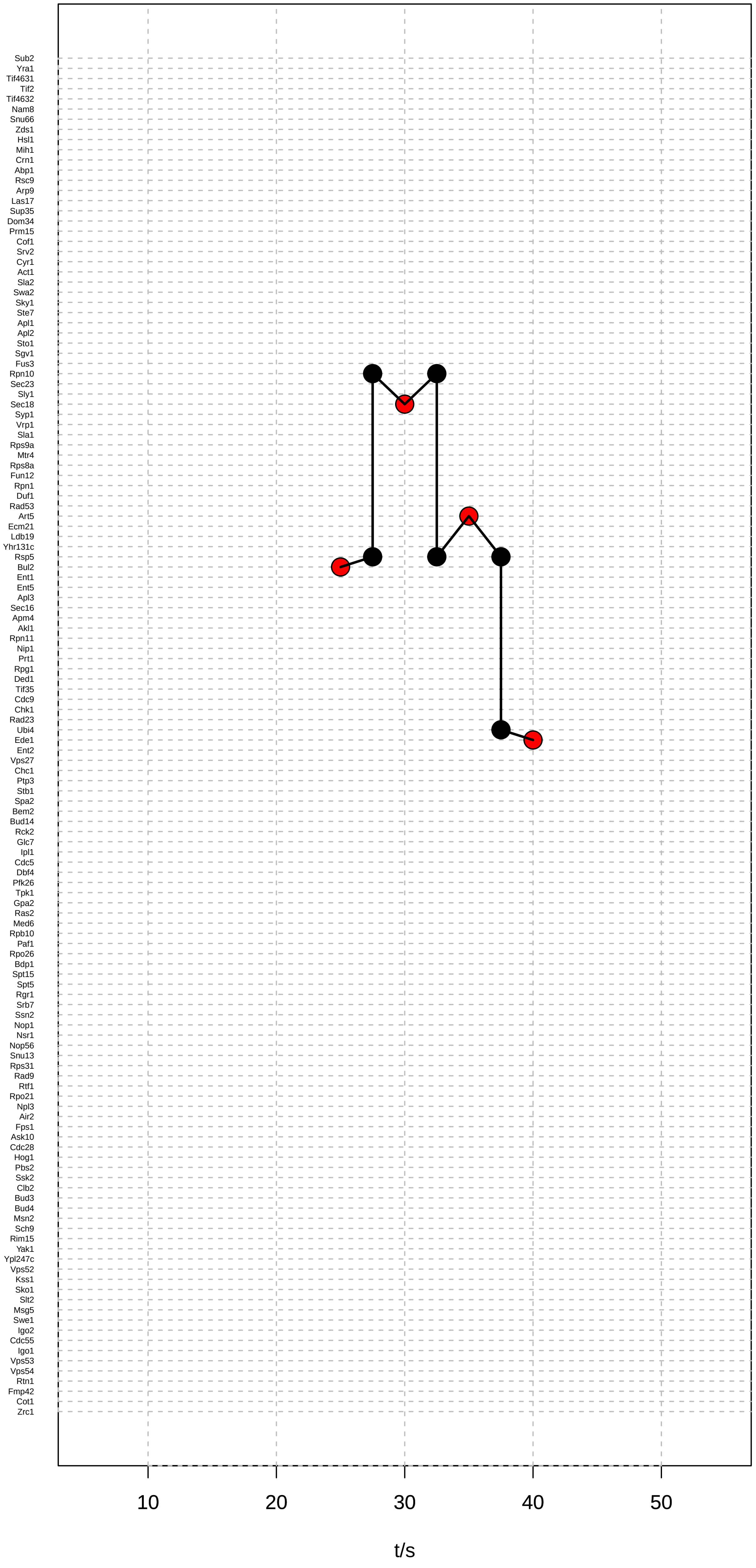

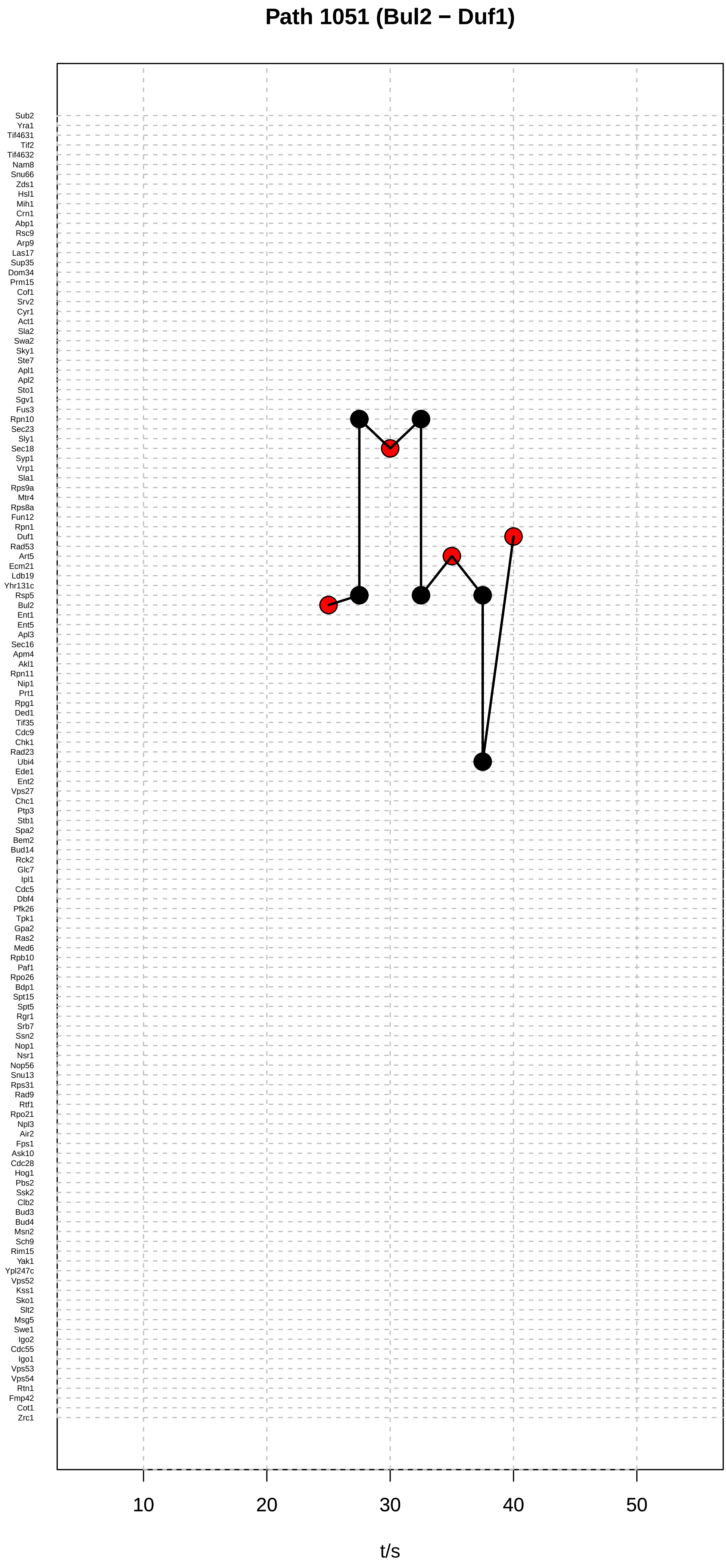

Path 1051 (Bul2 − Duf1)
Sub2
Yra1
Tif4631
Tif2
Tif4632
Nam8
Snu66
Zds1
Hsl1
Mih1
Crn1
Abp1
Rsc9
Arp9
Las17
Sup35
Dom34
Prm15
Cof1
Srv2
Cyr1
Act1
Sla2
Swa2
Sky1
Ste7
Apl1
Apl2
Sto1
Sgv1
Fus3
Rpn10
Sec23
Sly1
Sec18
Syp1
Vrp1
Sla1
Rps9a
Mtr4
Rps8a
Fun12
Rpn1
Duf1
Rad53
Art5
Ecm21
Ldb19
Yhr131c
Rsp5
Bul2
Ent1
Ent5
Apl3
Sec16
Apm4
Akl1
Rpn11
Nip1
Prt1
Rpg1
Ded1
Tif35
Cdc9
Chk1
Rad23
Ubi4
Ede1
Ent2
Vps27
Chc1
Ptp3
Stb1
Spa2
Bem2
Bud14
Rck2
Glc7
Ipl1
Cdc5
Dbf4
Pfk26
Tpk1
Gpa2
Ras2
Med6
Rpb10
Paf1
Rpo26
Bdp1
Spt15
Spt5
Rgr1
Srb7
Ssn2
Nop1
Nsr1
Nop56
Snu13
Rps31
Rad9
Rtf1
Rpo21
Npl3
Air2
Fps1
Ask10
Cdc28
Hog1
Pbs2
Ssk2
Clb2
Bud3
Bud4
Msn2
Sch9
Rim15
Yak1
Ypl247c
Vps52
Kss1
Sko1
Slt2
Msg5
Swe1
Igo2
Cdc55
Igo1
Vps53
Vps54
Rtn1
Fmp42
Cot1
Zrc1
10
20
30
40
50
t/s

### Path 1052 (Bul2 − Fun12)

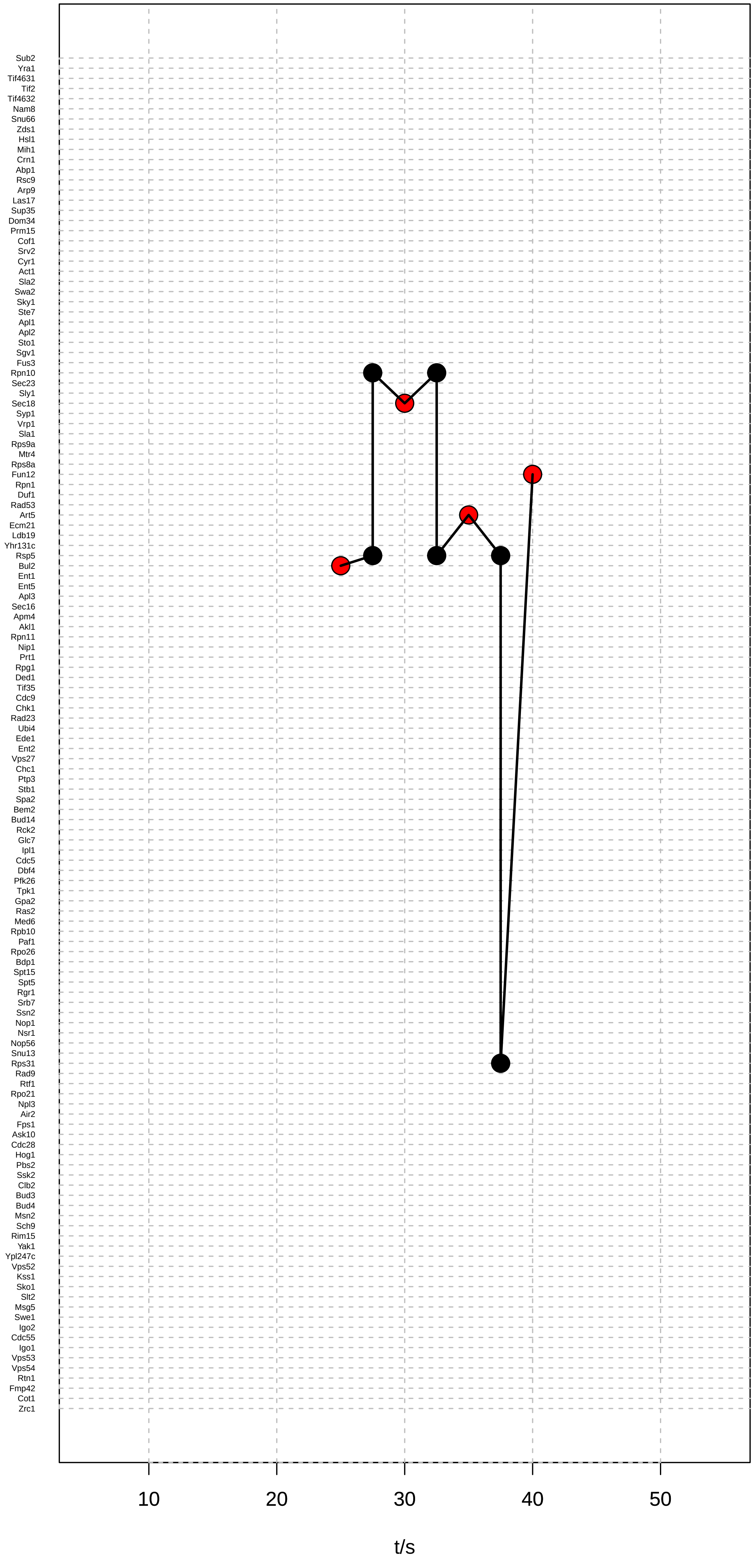

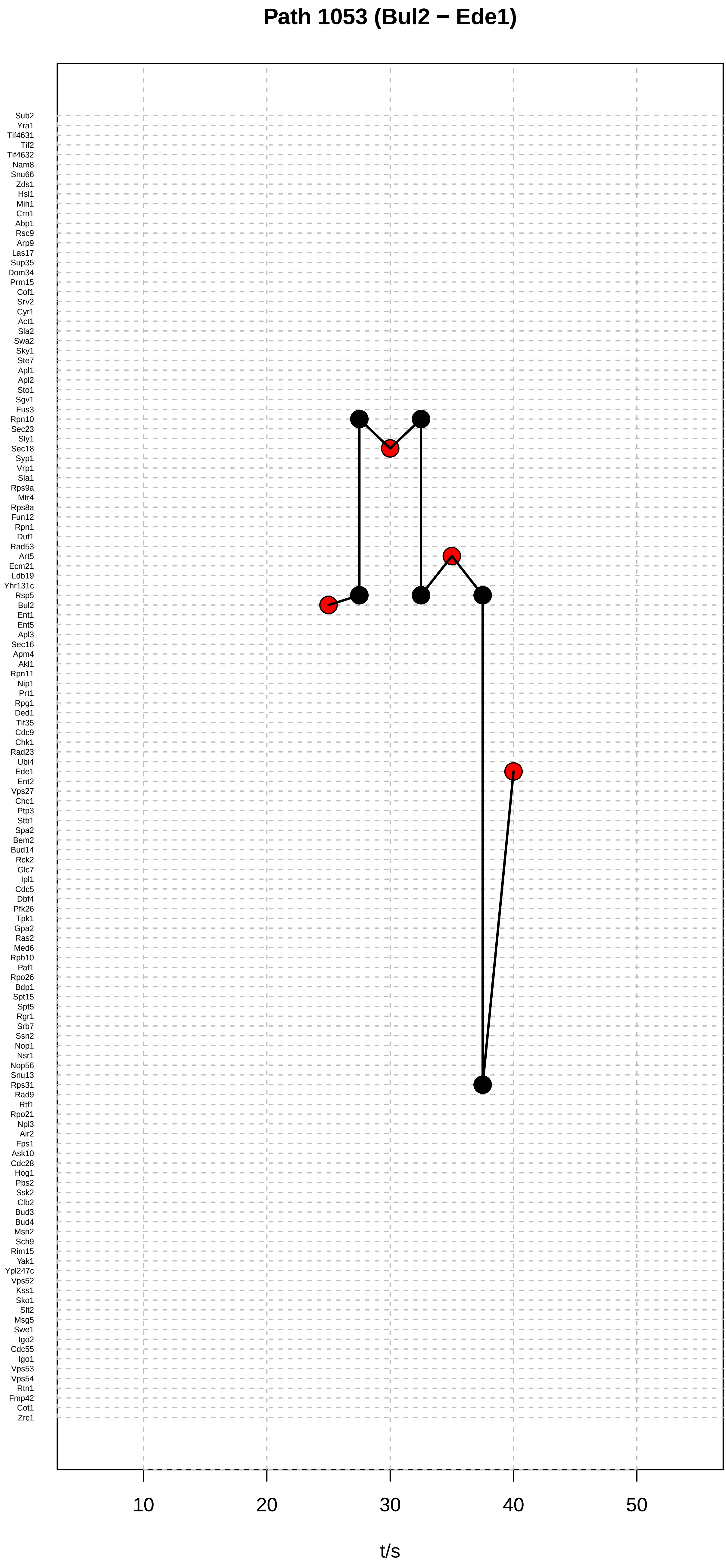

Path 1053 (Bul2 − Ede1)
Sub2
Yra1
Tif4631
Tif2
Tif4632
Nam8
Snu66
Zds1
Hsl1
Mih1
Crn1
Abp1
Rsc9
Arp9
Las17
Sup35
Dom34
Prm15
Cof1
Srv2
Cyr1
Act1
Sla2
Swa2
Sky1
Ste7
Apl1
Apl2
Sto1
Sgv1
Fus3
Rpn10
Sec23
Sly1
Sec18
Syp1
Vrp1
Sla1
Rps9a
Mtr4
Rps8a
Fun12
Rpn1
Duf1
Rad53
Art5
Ecm21
Ldb19
Yhr131c
Rsp5
Bul2
Ent1
Ent5
Apl3
Sec16
Apm4
Akl1
Rpn11
Nip1
Prt1
Rpg1
Ded1
Tif35
Cdc9
Chk1
Rad23
Ubi4
Ede1
Ent2
Vps27
Chc1
Ptp3
Stb1
Spa2
Bem2
Bud14
Rck2
Glc7
Ipl1
Cdc5
Dbf4
Pfk26
Tpk1
Gpa2
Ras2
Med6
Rpb10
Paf1
Rpo26
Bdp1
Spt15
Spt5
Rgr1
Srb7
Ssn2
Nop1
Nsr1
Nop56
Snu13
Rps31
Rad9
Rtf1
Rpo21
Npl3
Air2
Fps1
Ask10
Cdc28
Hog1
Pbs2
Ssk2
Clb2
Bud3
Bud4
Msn2
Sch9
Rim15
Yak1
Ypl247c
Vps52
Kss1
Sko1
Slt2
Msg5
Swe1
Igo2
Cdc55
Igo1
Vps53
Vps54
Rtn1
Fmp42
Cot1
Zrc1
10
20
30
40
50
t/s

### Path 1054 (Bul2 − Rtf1)

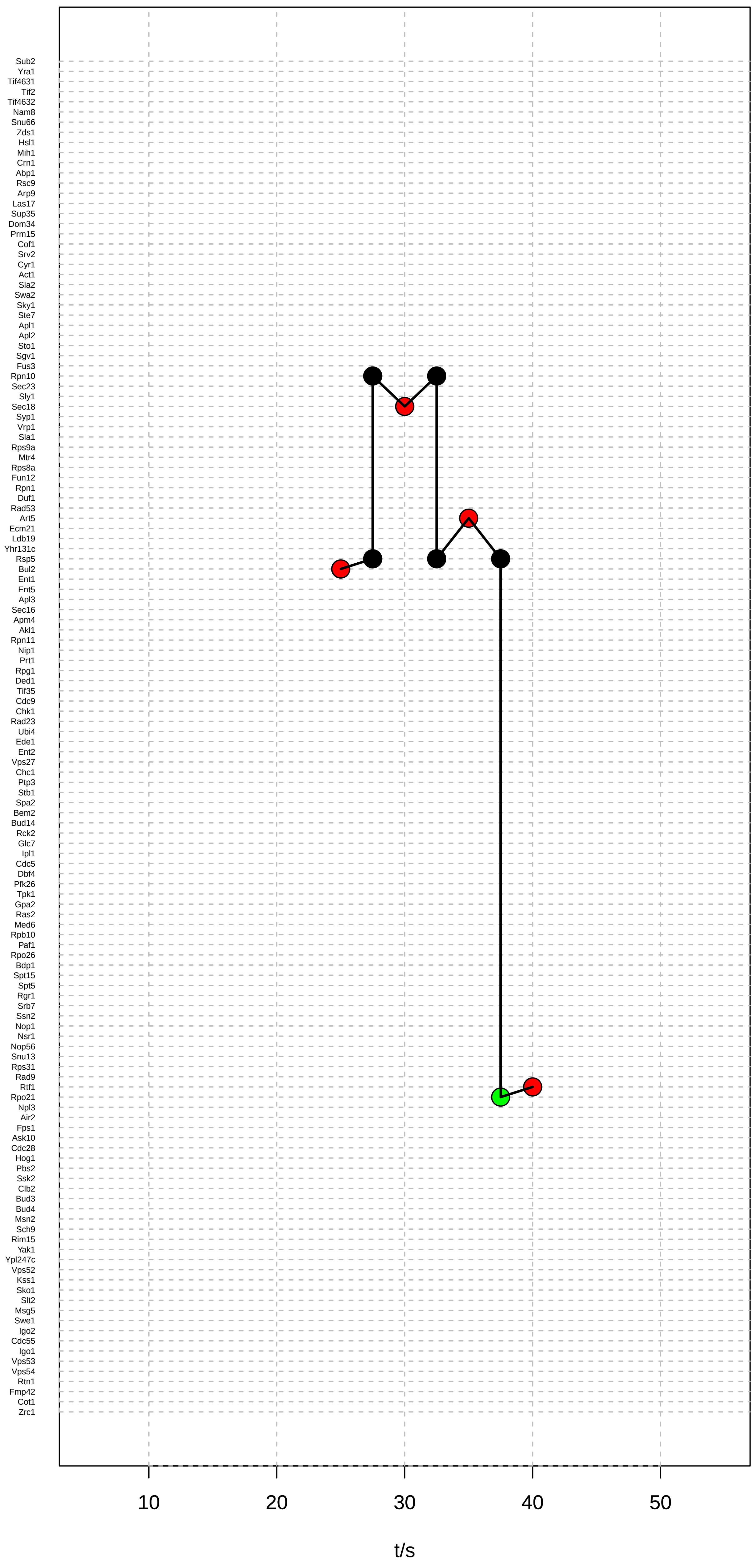

### Path 1055 (Bul2 − Rad23)

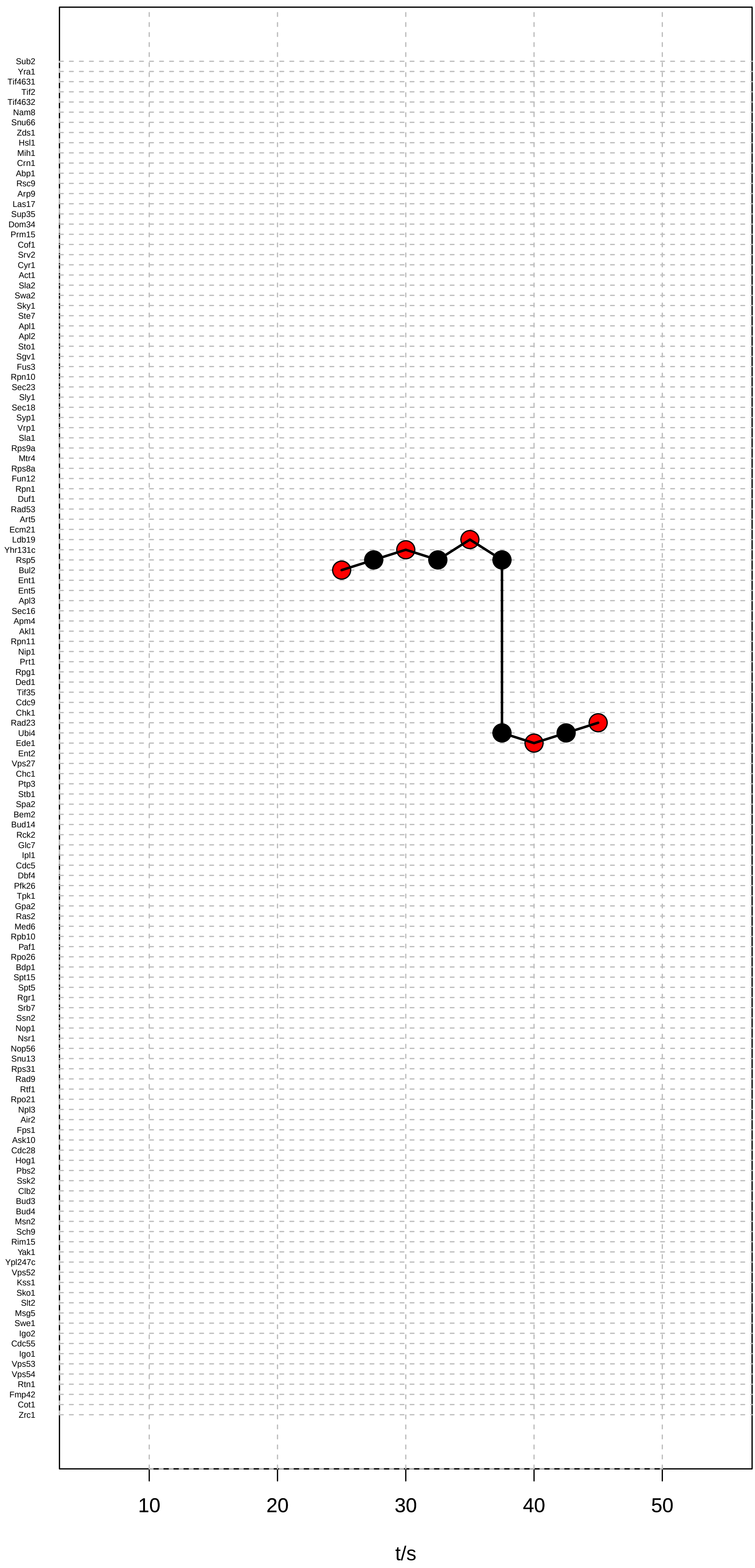

#### Path 1056 (Bul2 − Rad23)

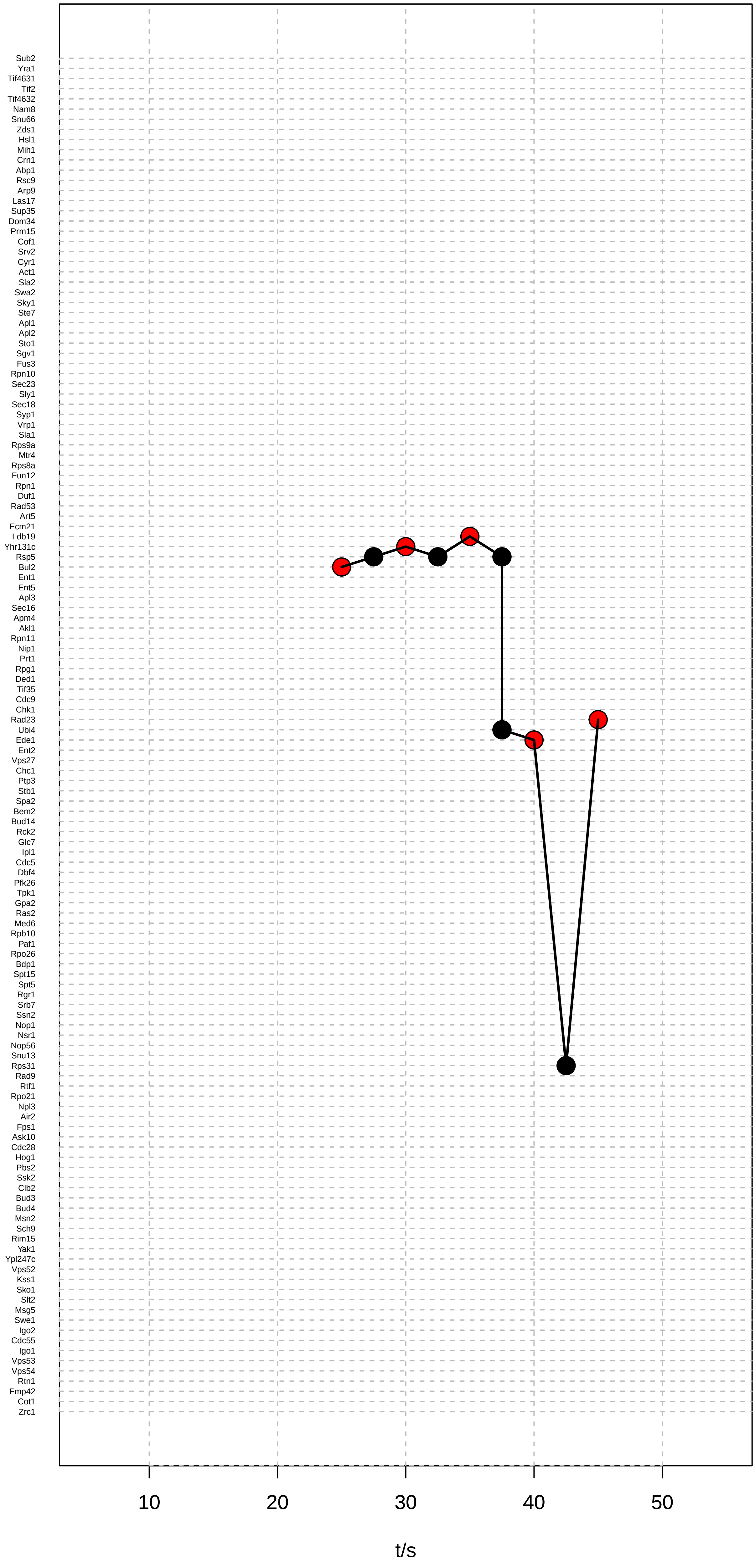

### Path 1057 (Bul2 − Rad23)

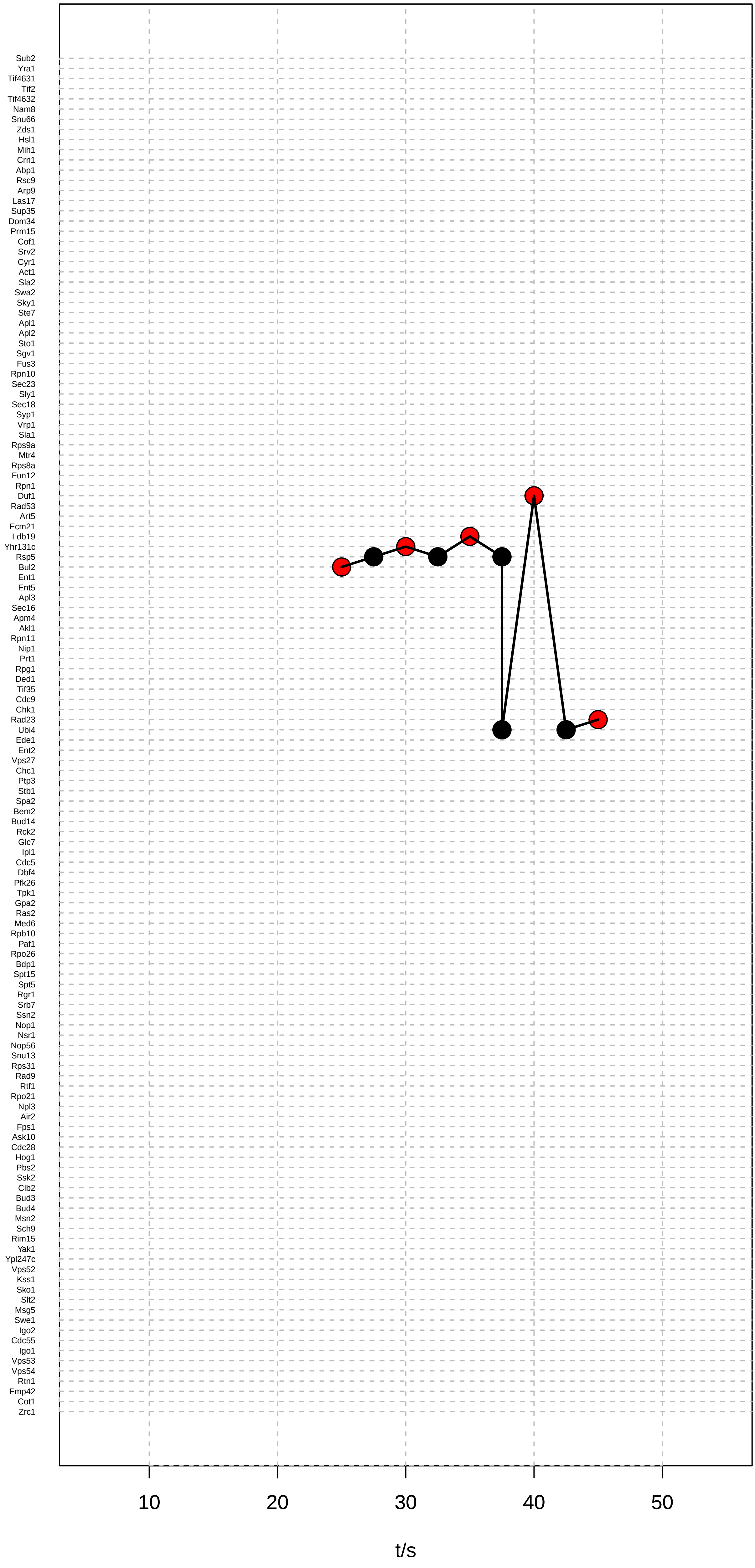

### Path 1058 (Bul2 − Rad23)

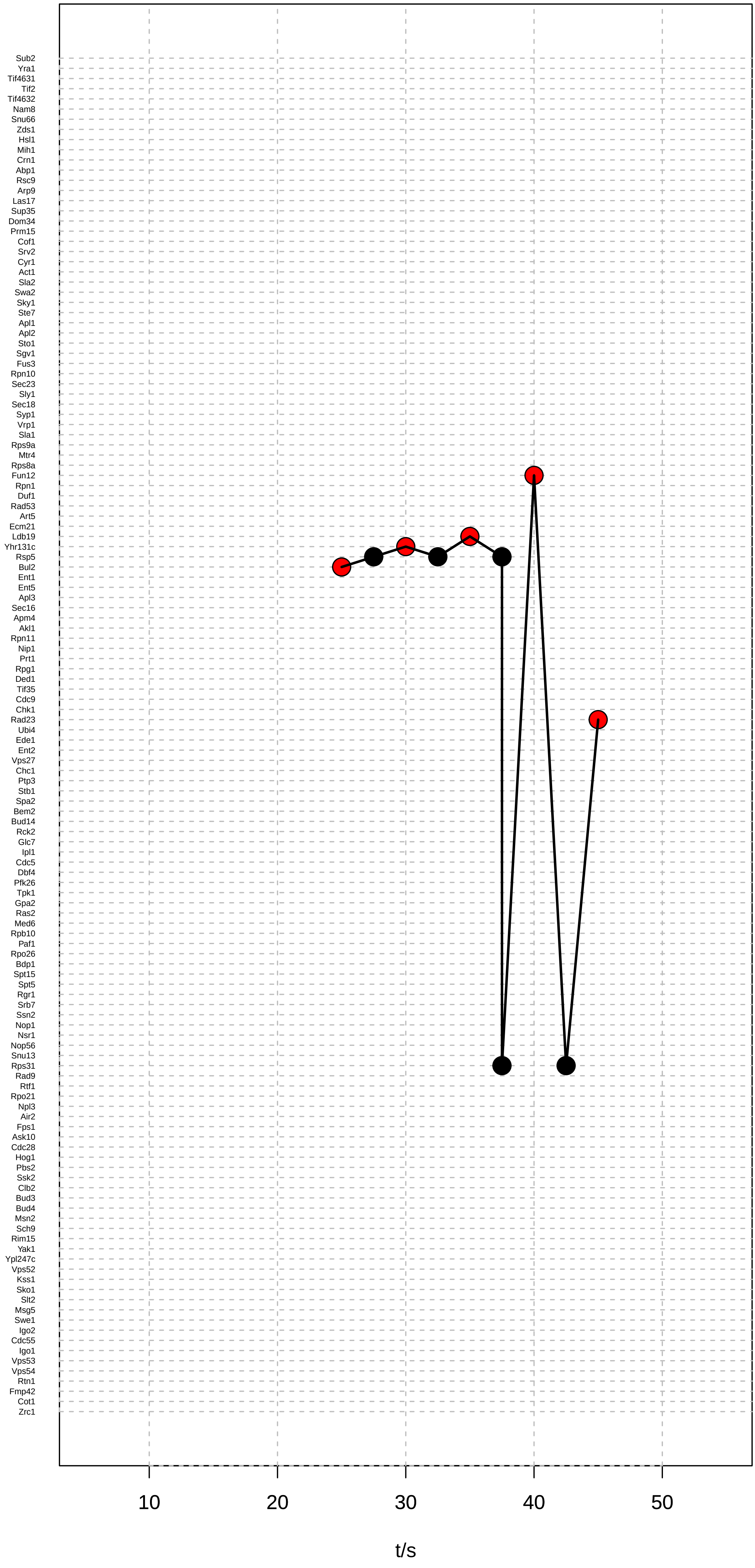

#### Path 1059 (Bul2 − Rad23)

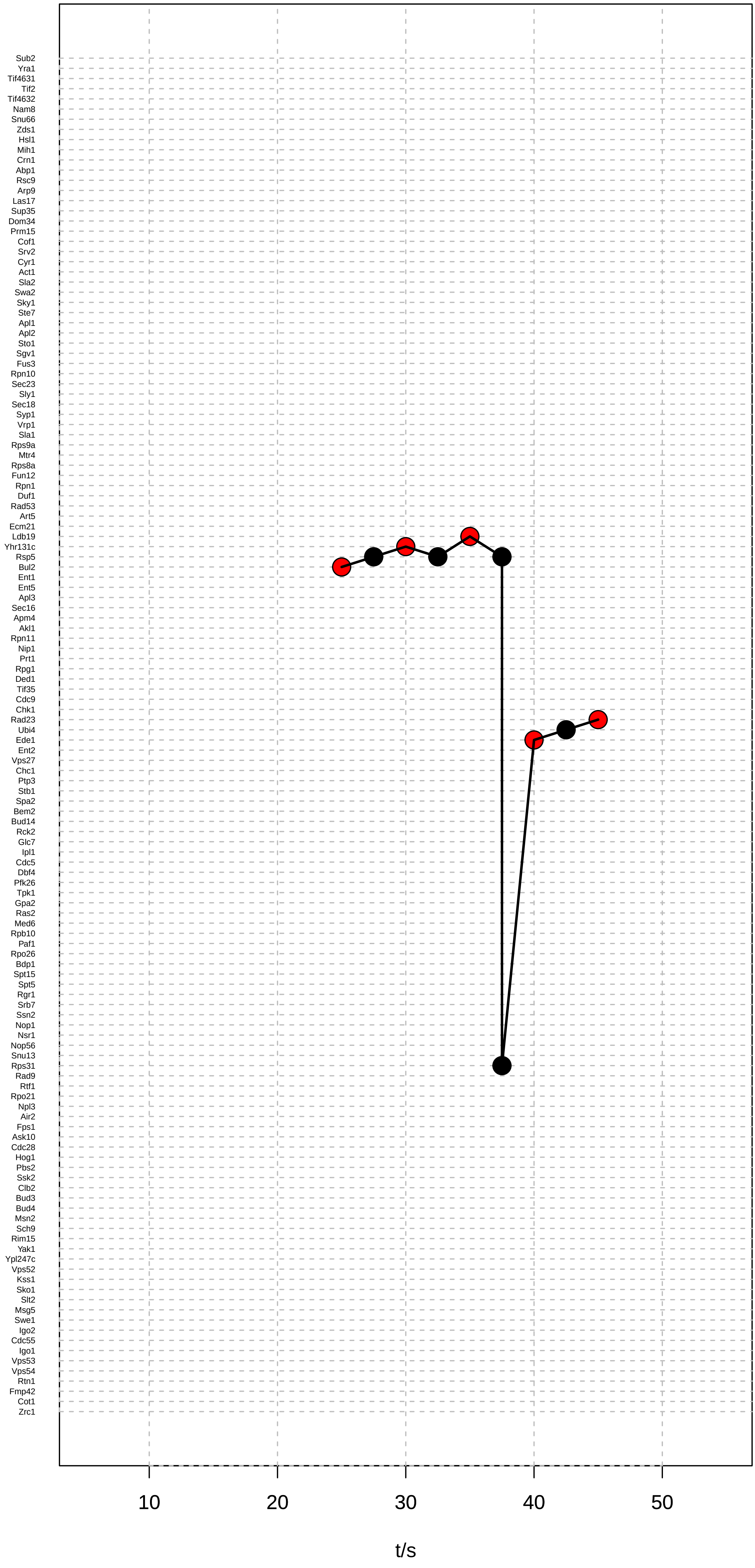

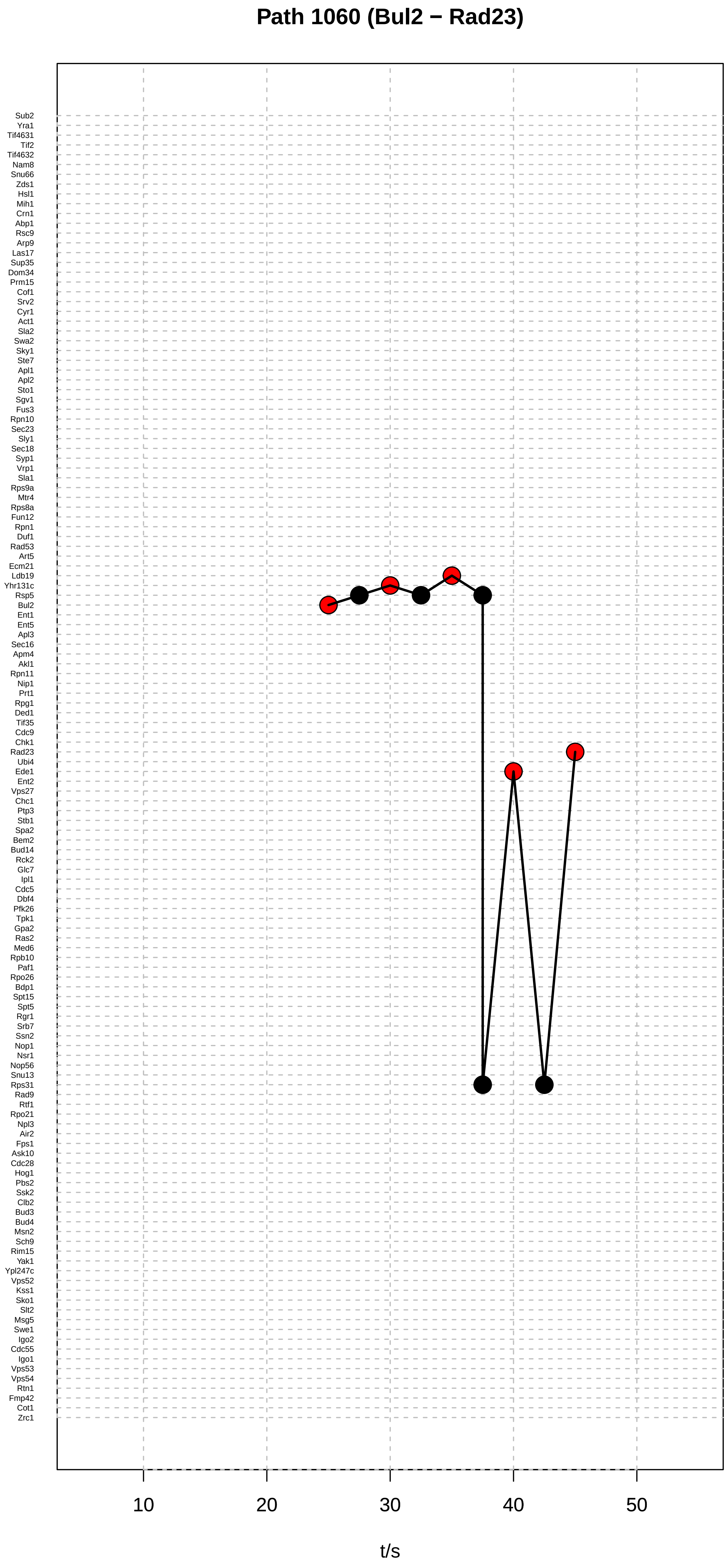

Path 1060 (Bul2 − Rad23)
Sub2
Yra1
Tif4631
Tif2
Tif4632
Nam8
Snu66
Zds1
Hsl1
Mih1
Crn1
Abp1
Rsc9
Arp9
Las17
Sup35
Dom34
Prm15
Cof1
Srv2
Cyr1
Act1
Sla2
Swa2
Sky1
Ste7
Apl1
Apl2
Sto1
Sgv1
Fus3
Rpn10
Sec23
Sly1
Sec18
Syp1
Vrp1
Sla1
Rps9a
Mtr4
Rps8a
Fun12
Rpn1
Duf1
Rad53
Art5
Ecm21
Ldb19
Yhr131c
Rsp5
Bul2
Ent1
Ent5
Apl3
Sec16
Apm4
Akl1
Rpn11
Nip1
Prt1
Rpg1
Ded1
Tif35
Cdc9
Chk1
Rad23
Ubi4
Ede1
Ent2
Vps27
Chc1
Ptp3
Stb1
Spa2
Bem2
Bud14
Rck2
Glc7
Ipl1
Cdc5
Dbf4
Pfk26
Tpk1
Gpa2
Ras2
Med6
Rpb10
Paf1
Rpo26
Bdp1
Spt15
Spt5
Rgr1
Srb7
Ssn2
Nop1
Nsr1
Nop56
Snu13
Rps31
Rad9
Rtf1
Rpo21
Npl3
Air2
Fps1
Ask10
Cdc28
Hog1
Pbs2
Ssk2
Clb2
Bud3
Bud4
Msn2
Sch9
Rim15
Yak1
Ypl247c
Vps52
Kss1
Sko1
Slt2
Msg5
Swe1
Igo2
Cdc55
Igo1
Vps53
Vps54
Rtn1
Fmp42
Cot1
Zrc1
10
20
30
40
50
t/s

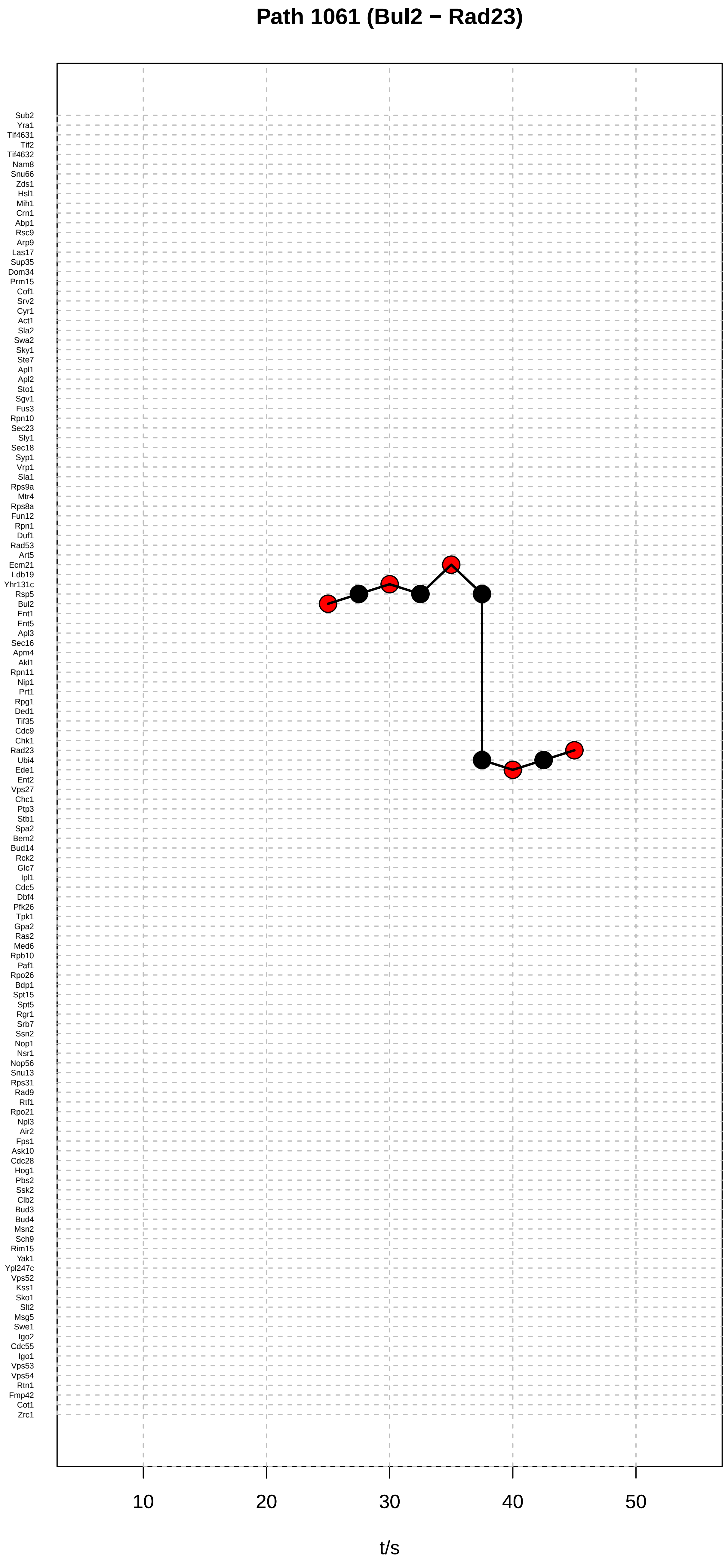

Path 1061 (Bul2 − Rad23)
Sub2
Yra1
Tif4631
Tif2
Tif4632
Nam8
Snu66
Zds1
Hsl1
Mih1
Crn1
Abp1
Rsc9
Arp9
Las17
Sup35
Dom34
Prm15
Cof1
Srv2
Cyr1
Act1
Sla2
Swa2
Sky1
Ste7
Apl1
Apl2
Sto1
Sgv1
Fus3
Rpn10
Sec23
Sly1
Sec18
Syp1
Vrp1
Sla1
Rps9a
Mtr4
Rps8a
Fun12
Rpn1
Duf1
Rad53
Art5
Ecm21
Ldb19
Yhr131c
Rsp5
Bul2
Ent1
Ent5
Apl3
Sec16
Apm4
Akl1
Rpn11
Nip1
Prt1
Rpg1
Ded1
Tif35
Cdc9
Chk1
Rad23
Ubi4
Ede1
Ent2
Vps27
Chc1
Ptp3
Stb1
Spa2
Bem2
Bud14
Rck2
Glc7
Ipl1
Cdc5
Dbf4
Pfk26
Tpk1
Gpa2
Ras2
Med6
Rpb10
Paf1
Rpo26
Bdp1
Spt15
Spt5
Rgr1
Srb7
Ssn2
Nop1
Nsr1
Nop56
Snu13
Rps31
Rad9
Rtf1
Rpo21
Npl3
Air2
Fps1
Ask10
Cdc28
Hog1
Pbs2
Ssk2
Clb2
Bud3
Bud4
Msn2
Sch9
Rim15
Yak1
Ypl247c
Vps52
Kss1
Sko1
Slt2
Msg5
Swe1
Igo2
Cdc55
Igo1
Vps53
Vps54
Rtn1
Fmp42
Cot1
Zrc1
10
20
30
40
50
t/s

### Path 1062 (Bul2 − Rad23)

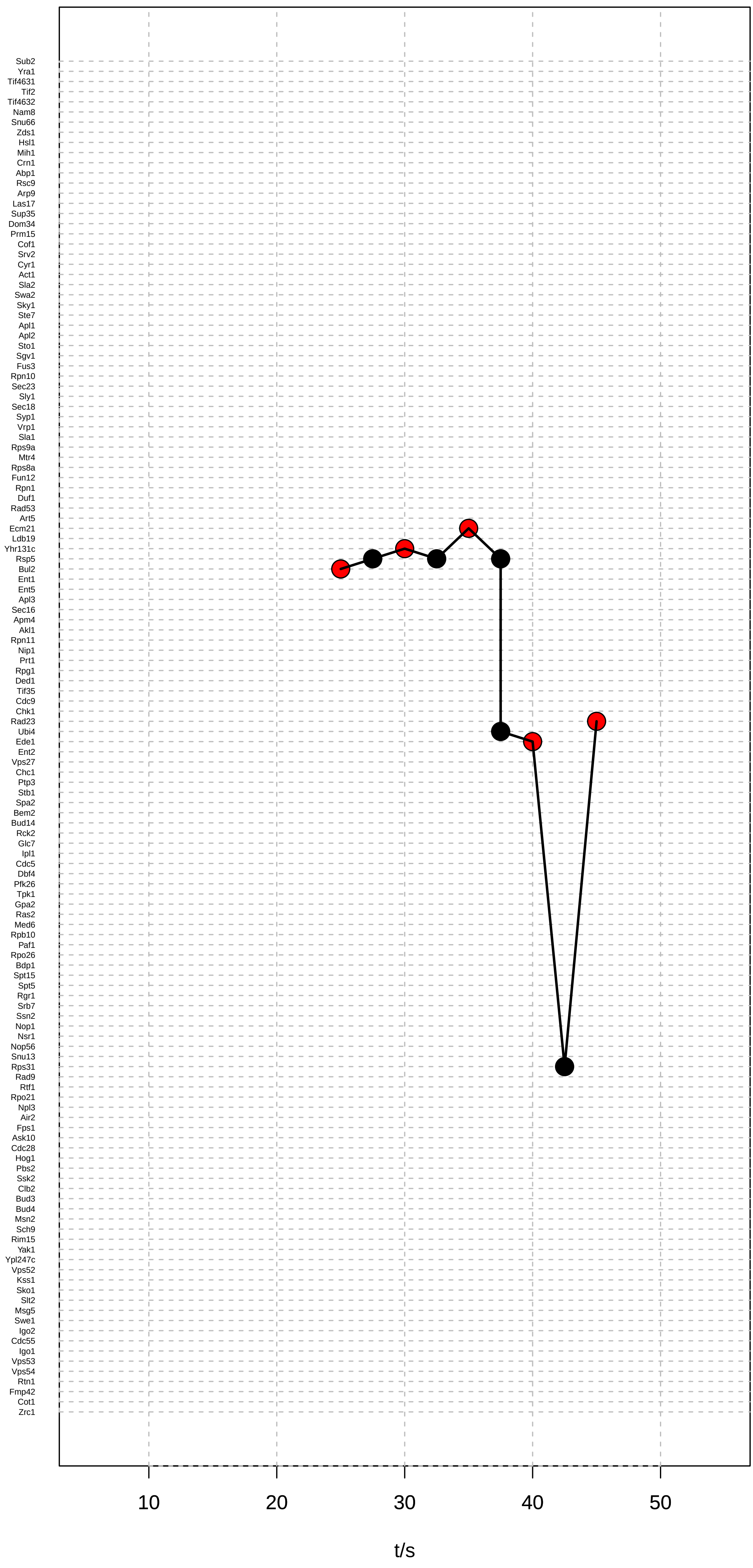

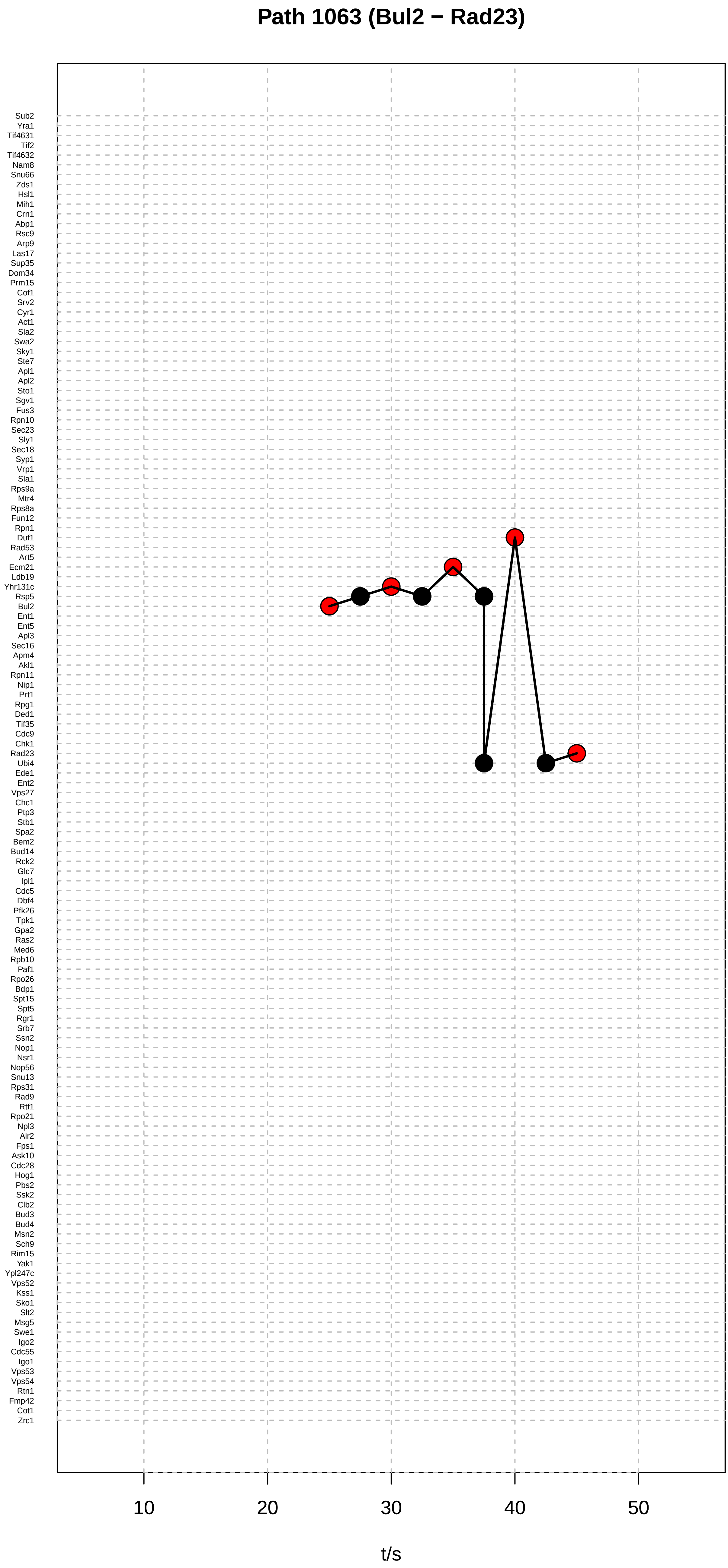

Path 1063 (Bul2 − Rad23)
Sub2
Yra1
Tif4631
Tif2
Tif4632
Nam8
Snu66
Zds1
Hsl1
Mih1
Crn1
Abp1
Rsc9
Arp9
Las17
Sup35
Dom34
Prm15
Cof1
Srv2
Cyr1
Act1
Sla2
Swa2
Sky1
Ste7
Apl1
Apl2
Sto1
Sgv1
Fus3
Rpn10
Sec23
Sly1
Sec18
Syp1
Vrp1
Sla1
Rps9a
Mtr4
Rps8a
Fun12
Rpn1
Duf1
Rad53
Art5
Ecm21
Ldb19
Yhr131c
Rsp5
Bul2
Ent1
Ent5
Apl3
Sec16
Apm4
Akl1
Rpn11
Nip1
Prt1
Rpg1
Ded1
Tif35
Cdc9
Chk1
Rad23
Ubi4
Ede1
Ent2
Vps27
Chc1
Ptp3
Stb1
Spa2
Bem2
Bud14
Rck2
Glc7
Ipl1
Cdc5
Dbf4
Pfk26
Tpk1
Gpa2
Ras2
Med6
Rpb10
Paf1
Rpo26
Bdp1
Spt15
Spt5
Rgr1
Srb7
Ssn2
Nop1
Nsr1
Nop56
Snu13
Rps31
Rad9
Rtf1
Rpo21
Npl3
Air2
Fps1
Ask10
Cdc28
Hog1
Pbs2
Ssk2
Clb2
Bud3
Bud4
Msn2
Sch9
Rim15
Yak1
Ypl247c
Vps52
Kss1
Sko1
Slt2
Msg5
Swe1
Igo2
Cdc55
Igo1
Vps53
Vps54
Rtn1
Fmp42
Cot1
Zrc1
10
20
30
40
50
t/s

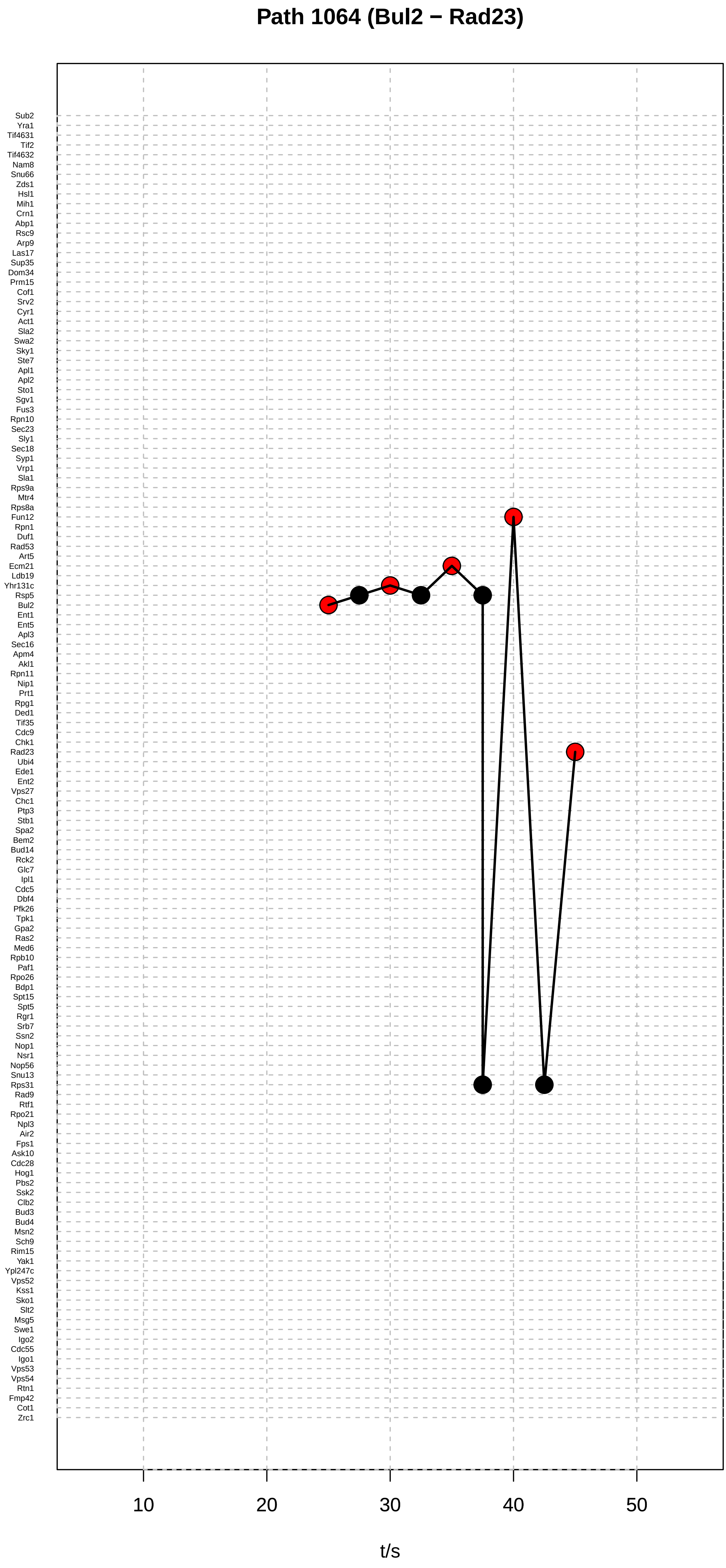

Path 1064 (Bul2 − Rad23)
Sub2
Yra1
Tif4631
Tif2
Tif4632
Nam8
Snu66
Zds1
Hsl1
Mih1
Crn1
Abp1
Rsc9
Arp9
Las17
Sup35
Dom34
Prm15
Cof1
Srv2
Cyr1
Act1
Sla2
Swa2
Sky1
Ste7
Apl1
Apl2
Sto1
Sgv1
Fus3
Rpn10
Sec23
Sly1
Sec18
Syp1
Vrp1
Sla1
Rps9a
Mtr4
Rps8a
Fun12
Rpn1
Duf1
Rad53
Art5
Ecm21
Ldb19
Yhr131c
Rsp5
Bul2
Ent1
Ent5
Apl3
Sec16
Apm4
Akl1
Rpn11
Nip1
Prt1
Rpg1
Ded1
Tif35
Cdc9
Chk1
Rad23
Ubi4
Ede1
Ent2
Vps27
Chc1
Ptp3
Stb1
Spa2
Bem2
Bud14
Rck2
Glc7
Ipl1
Cdc5
Dbf4
Pfk26
Tpk1
Gpa2
Ras2
Med6
Rpb10
Paf1
Rpo26
Bdp1
Spt15
Spt5
Rgr1
Srb7
Ssn2
Nop1
Nsr1
Nop56
Snu13
Rps31
Rad9
Rtf1
Rpo21
Npl3
Air2
Fps1
Ask10
Cdc28
Hog1
Pbs2
Ssk2
Clb2
Bud3
Bud4
Msn2
Sch9
Rim15
Yak1
Ypl247c
Vps52
Kss1
Sko1
Slt2
Msg5
Swe1
Igo2
Cdc55
Igo1
Vps53
Vps54
Rtn1
Fmp42
Cot1
Zrc1
10
20
30
40
50
t/s

### Path 1065 (Bul2 − Rad23)

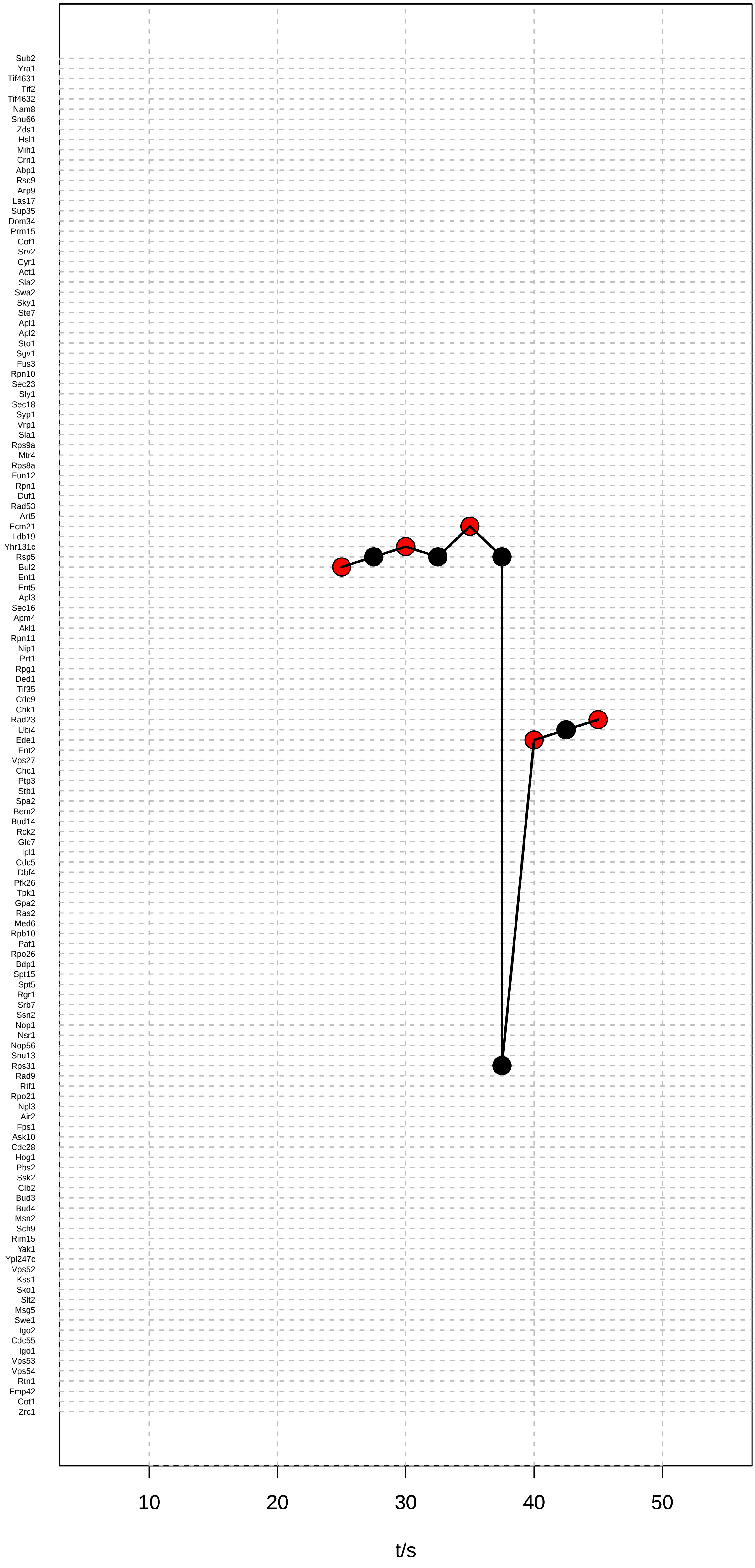

#### Path 1066 (Bul2 − Rad23)

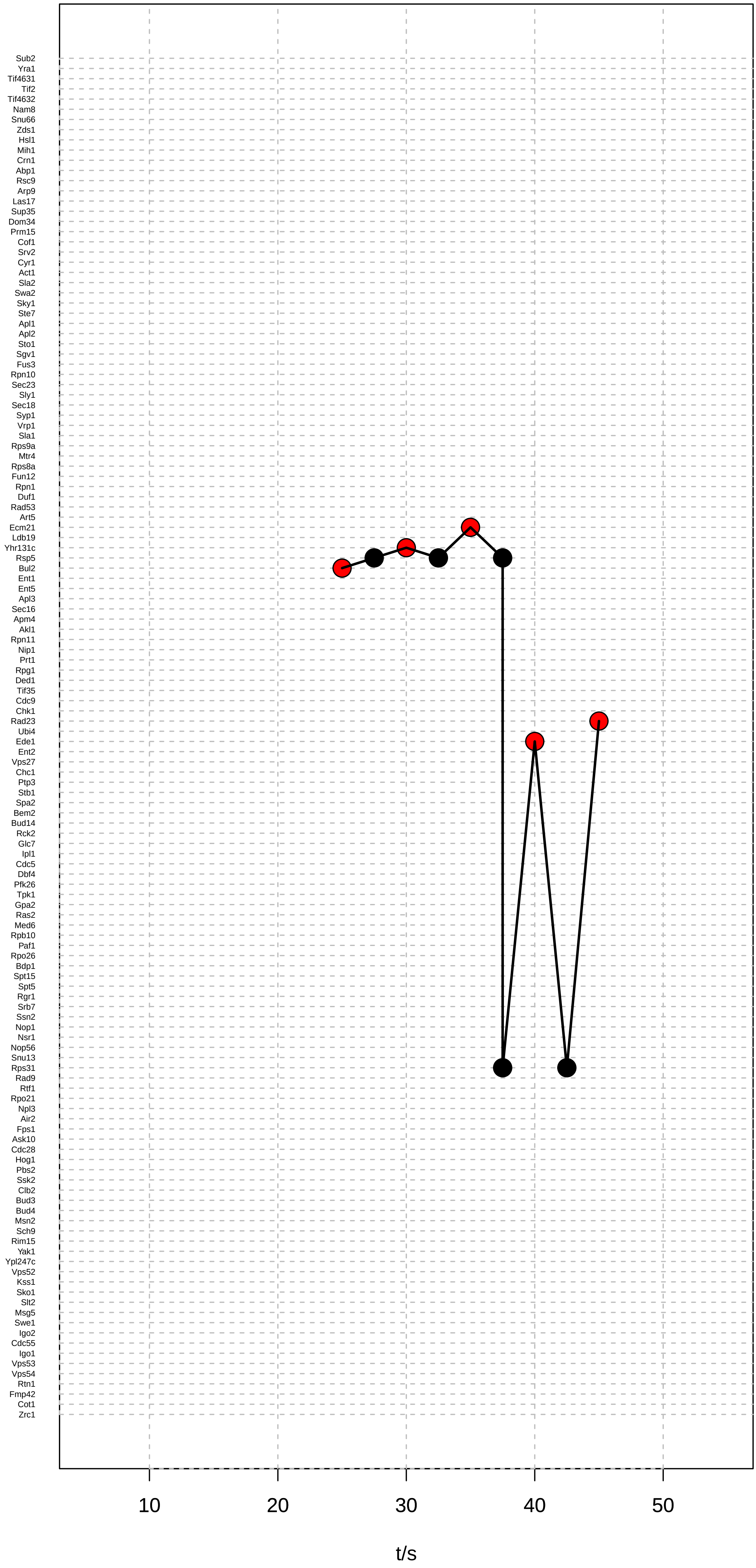

Path 1067 (Bul2 − Rad23)

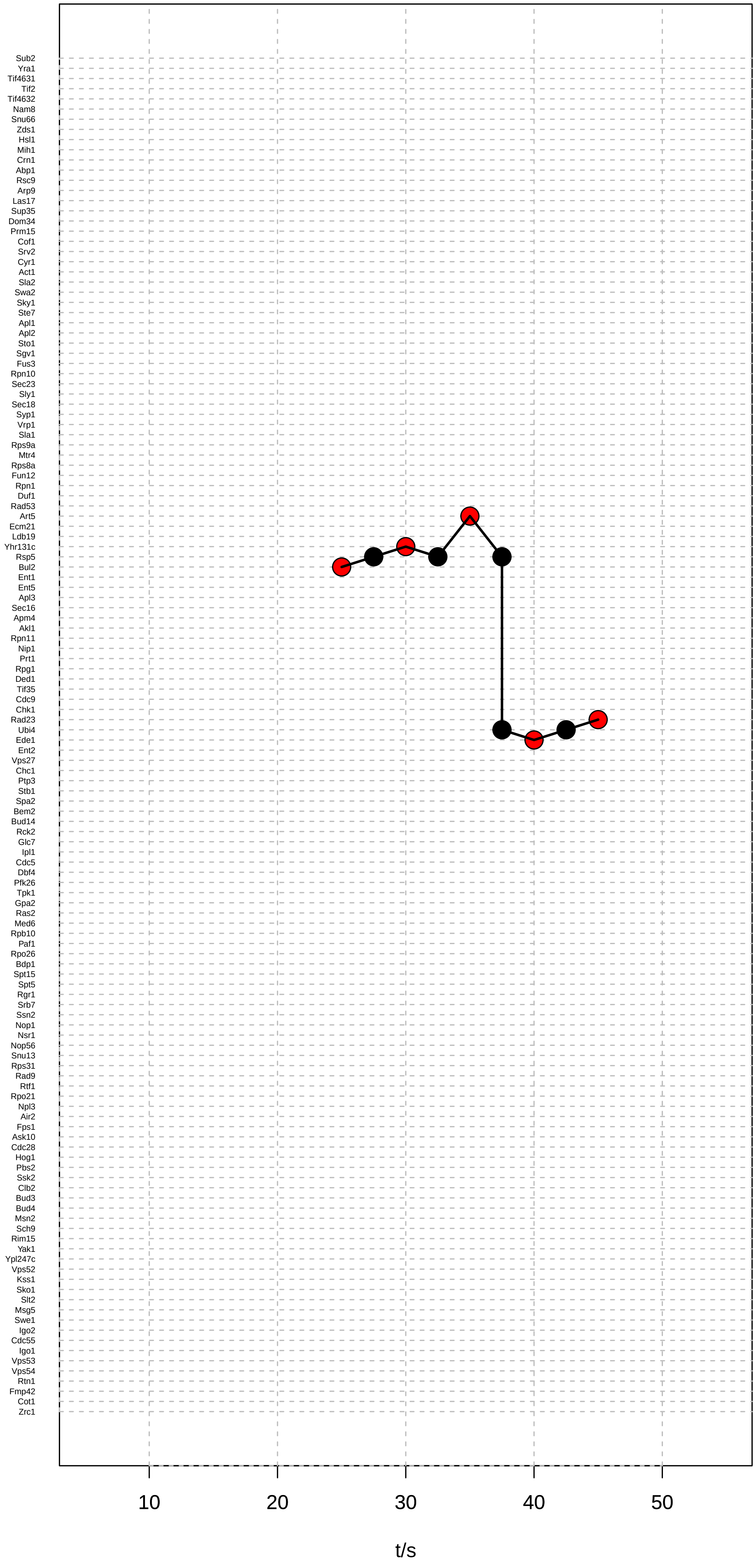

#### Path 1068 (Bul2 − Rad23)

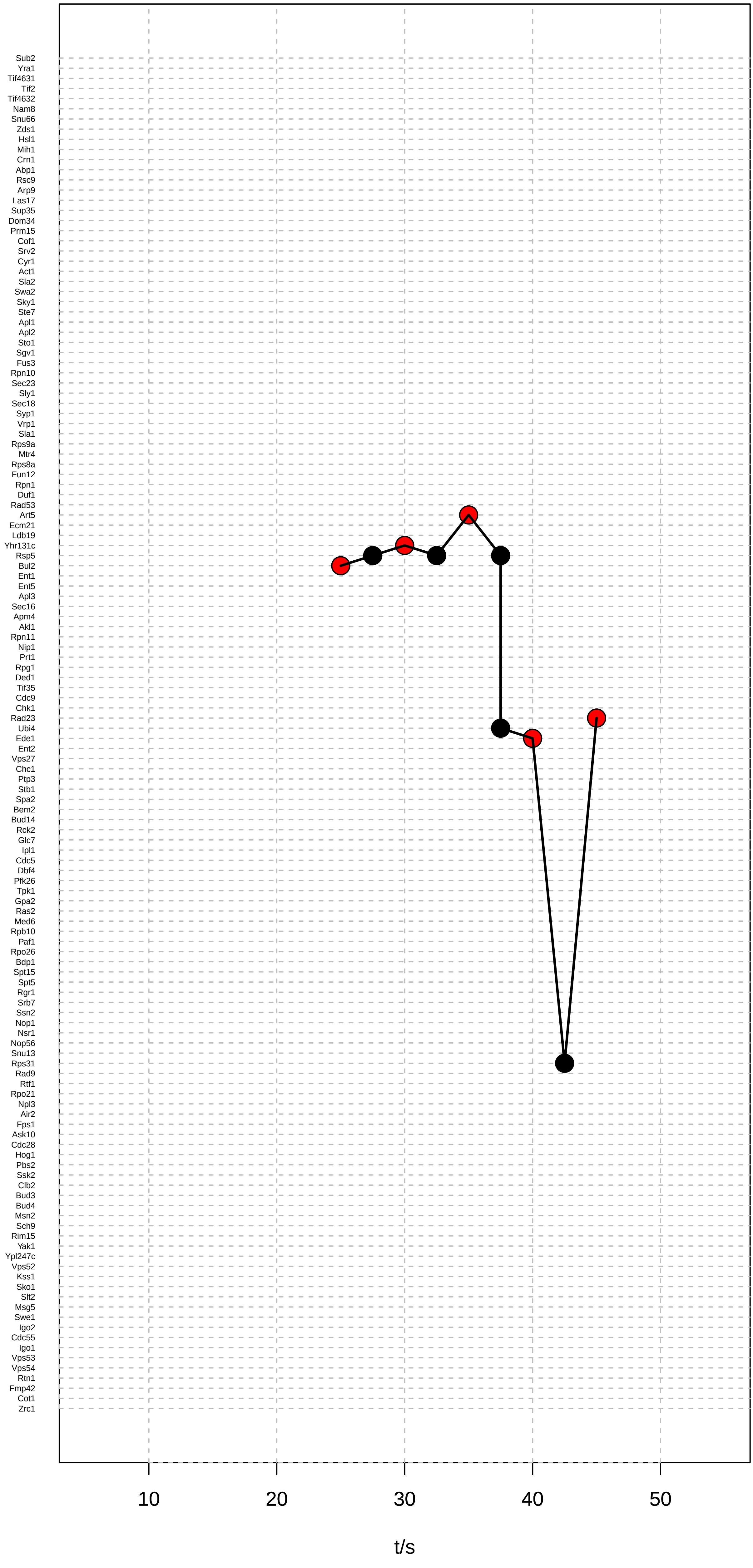

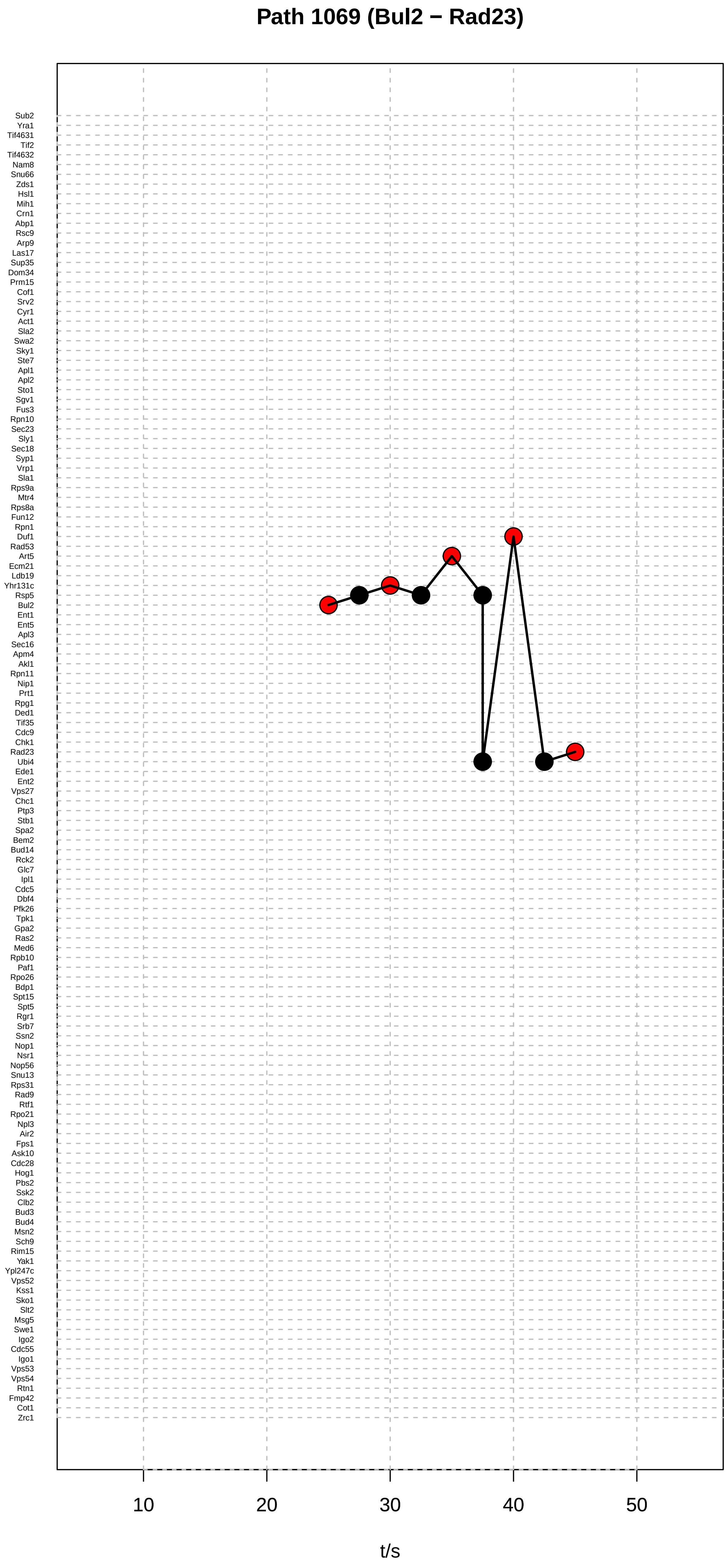
Path 1069 (Bul2 − Rad23)
Sub2
Yra1
Tif4631
Tif2
Tif4632
Nam8
Snu66
Zds1
Hsl1
Mih1
Crn1
Abp1
Rsc9
Arp9
Las17
Sup35
Dom34
Prm15
Cof1
Srv2
Cyr1
Act1
Sla2
Swa2
Sky1
Ste7
Apl1
Apl2
Sto1
Sgv1
Fus3
Rpn10
Sec23
Sly1
Sec18
Syp1
Vrp1
Sla1
Rps9a
Mtr4
Rps8a
Fun12
Rpn1
Duf1
Rad53
Art5
Ecm21
Ldb19
Yhr131c
Rsp5
Bul2
Ent1
Ent5
Apl3
Sec16
Apm4
Akl1
Rpn11
Nip1
Prt1
Rpg1
Ded1
Tif35
Cdc9
Chk1
Rad23
Ubi4
Ede1
Ent2
Vps27
Chc1
Ptp3
Stb1
Spa2
Bem2
Bud14
Rck2
Glc7
Ipl1
Cdc5
Dbf4
Pfk26
Tpk1
Gpa2
Ras2
Med6
Rpb10
Paf1
Rpo26
Bdp1
Spt15
Spt5
Rgr1
Srb7
Ssn2
Nop1
Nsr1
Nop56
Snu13
Rps31
Rad9
Rtf1
Rpo21
Npl3
Air2
Fps1
Ask10
Cdc28
Hog1
Pbs2
Ssk2
Clb2
Bud3
Bud4
Msn2
Sch9
Rim15
Yak1
Ypl247c
Vps52
Kss1
Sko1
Slt2
Msg5
Swe1
Igo2
Cdc55
Igo1
Vps53
Vps54
Rtn1
Fmp42
Cot1
Zrc1
10
20
30
40
50
t/s

### Path 1070 (Bul2 − Rad23)

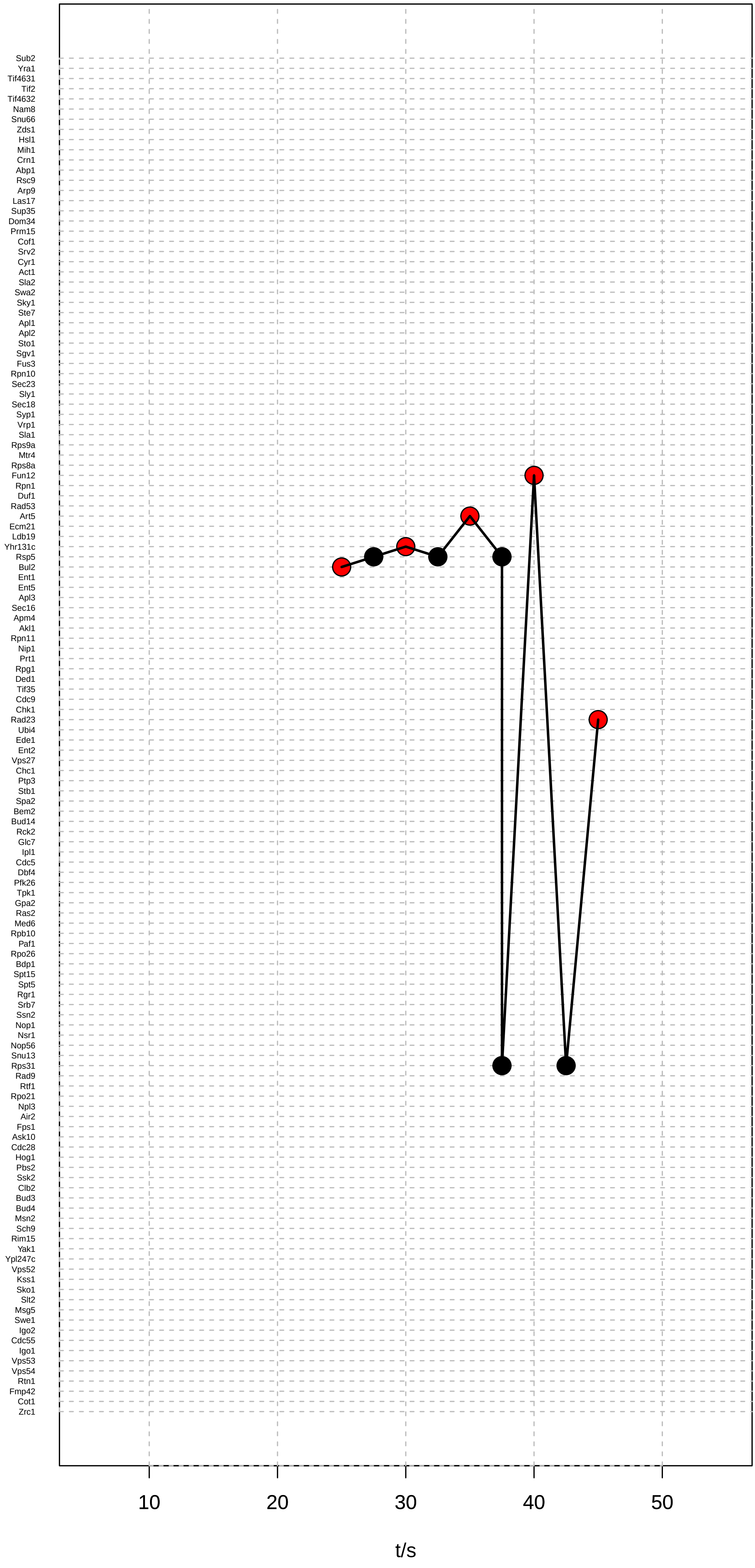

### Path 1071 (Bul2 − Rad23)

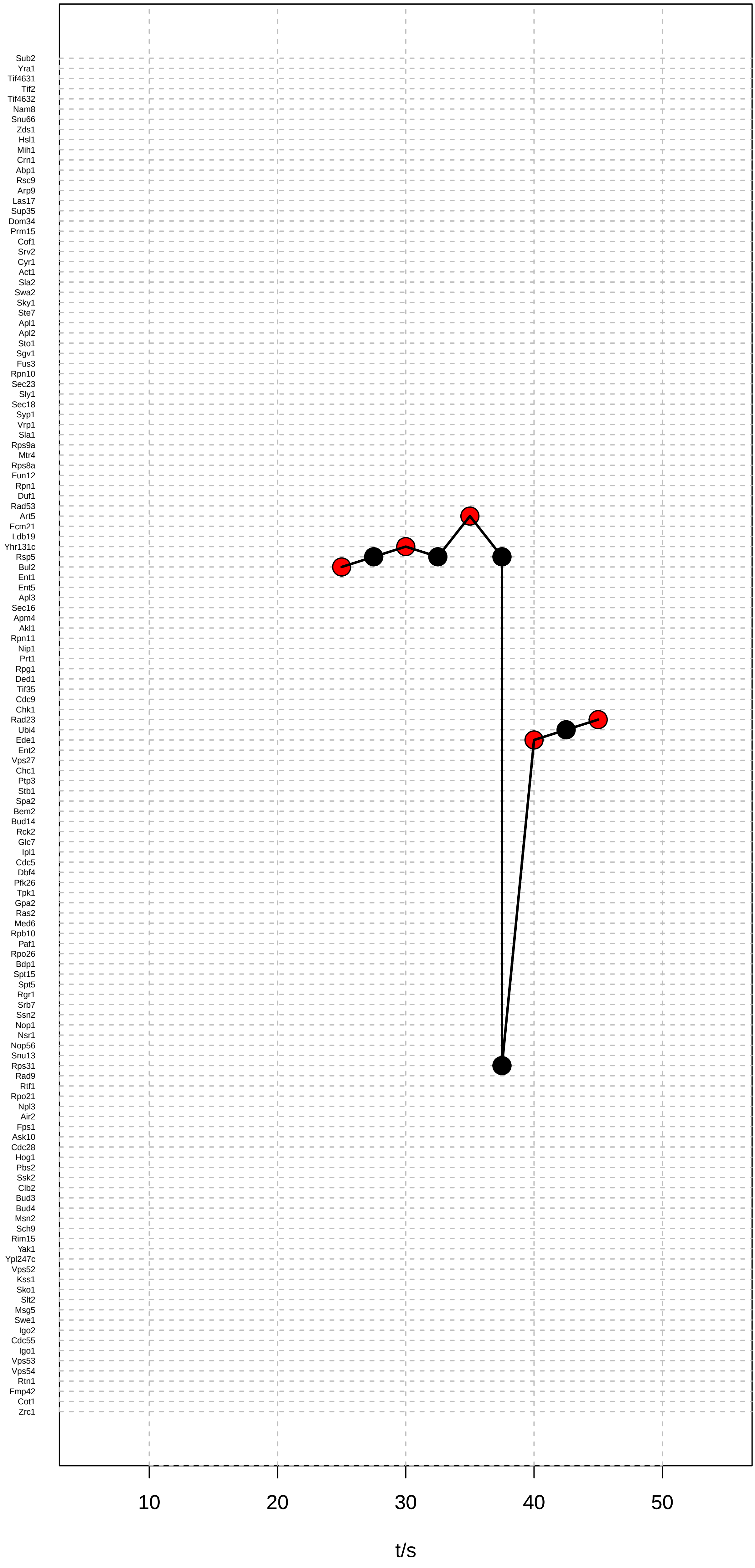

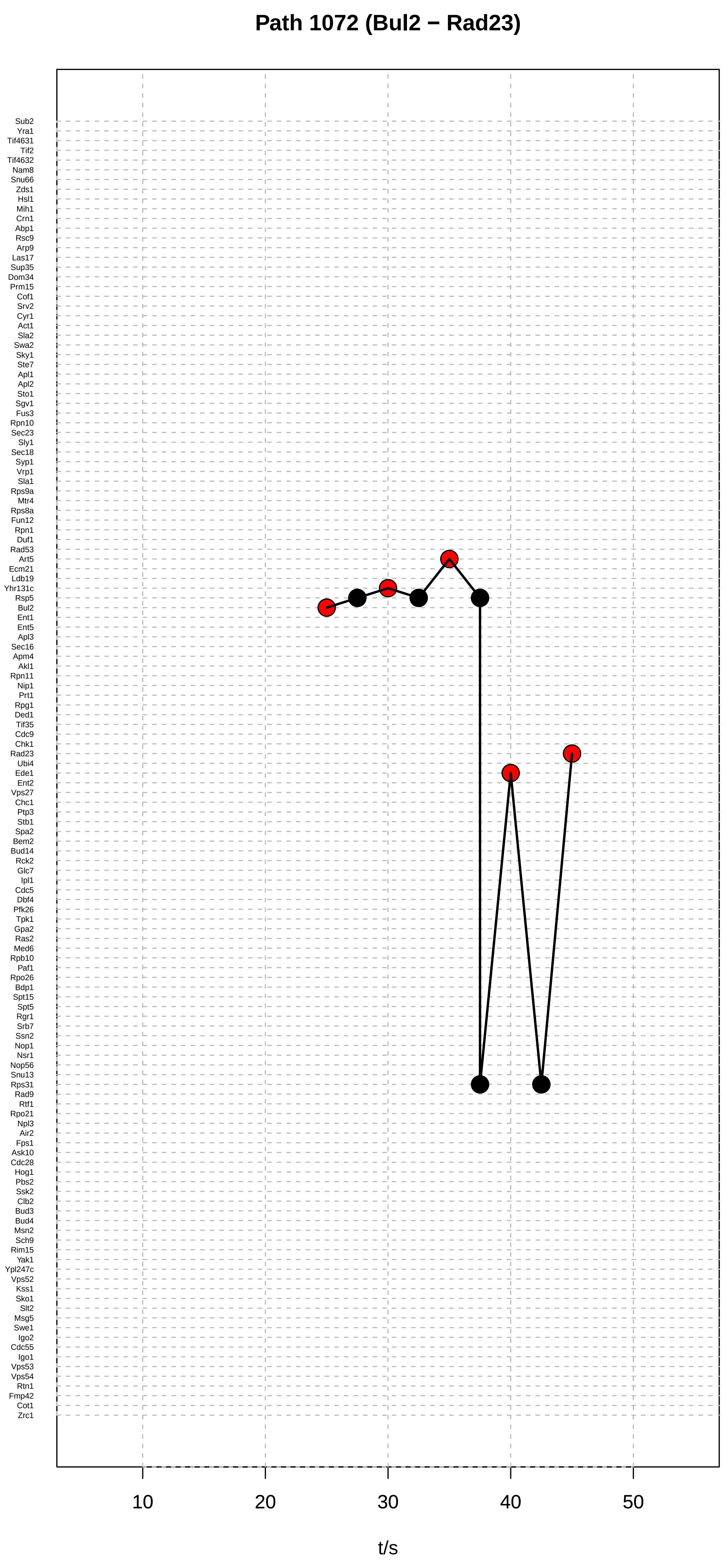

Path 1072 (Bul2 − Rad23)
Sub2
Yra1
Tif4631
Tif2
Tif4632
Nam8
Snu66
Zds1
Hsl1
Mih1
Crn1
Abp1
Rsc9
Arp9
Las17
Sup35
Dom34
Prm15
Cof1
Srv2
Cyr1
Act1
Sla2
Swa2
Sky1
Ste7
Apl1
Apl2
Sto1
Sgv1
Fus3
Rpn10
Sec23
Sly1
Sec18
Syp1
Vrp1
Sla1
Rps9a
Mtr4
Rps8a
Fun12
Rpn1
Duf1
Rad53
Art5
Ecm21
Ldb19
Yhr131c
Rsp5
Bul2
Ent1
Ent5
Apl3
Sec16
Apm4
Akl1
Rpn11
Nip1
Prt1
Rpg1
Ded1
Tif35
Cdc9
Chk1
Rad23
Ubi4
Ede1
Ent2
Vps27
Chc1
Ptp3
Stb1
Spa2
Bem2
Bud14
Rck2
Glc7
Ipl1
Cdc5
Dbf4
Pfk26
Tpk1
Gpa2
Ras2
Med6
Rpb10
Paf1
Rpo26
Bdp1
Spt15
Spt5
Rgr1
Srb7
Ssn2
Nop1
Nsr1
Nop56
Snu13
Rps31
Rad9
Rtf1
Rpo21
Npl3
Air2
Fps1
Ask10
Cdc28
Hog1
Pbs2
Ssk2
Clb2
Bud3
Bud4
Msn2
Sch9
Rim15
Yak1
Ypl247c
Vps52
Kss1
Sko1
Slt2
Msg5
Swe1
Igo2
Cdc55
Igo1
Vps53
Vps54
Rtn1
Fmp42
Cot1
Zrc1
10
20
30
40
50
t/s

### Path 1073 (Bul2 − Ede1)

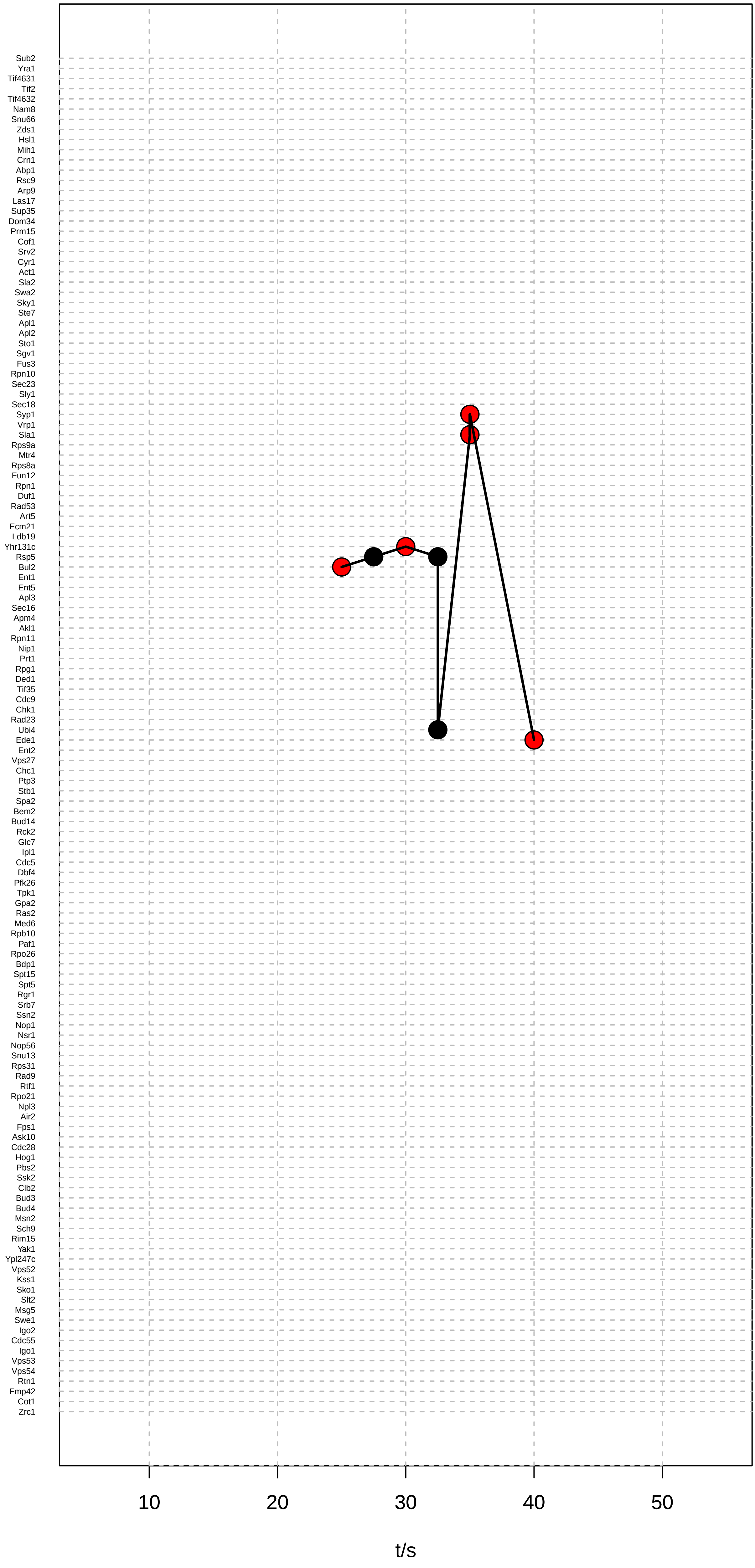

#### Path 1074 (Bul2 − Ede1)

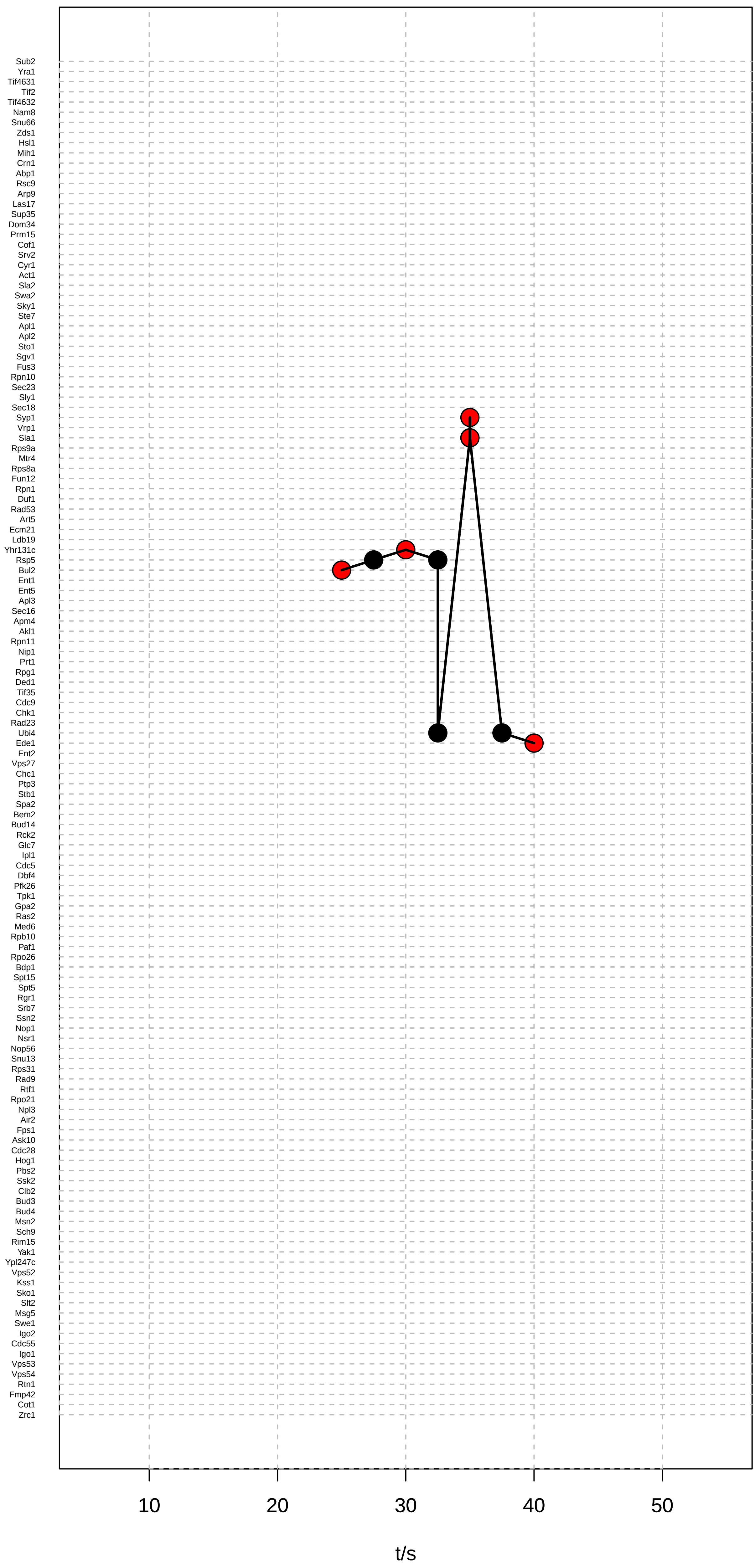

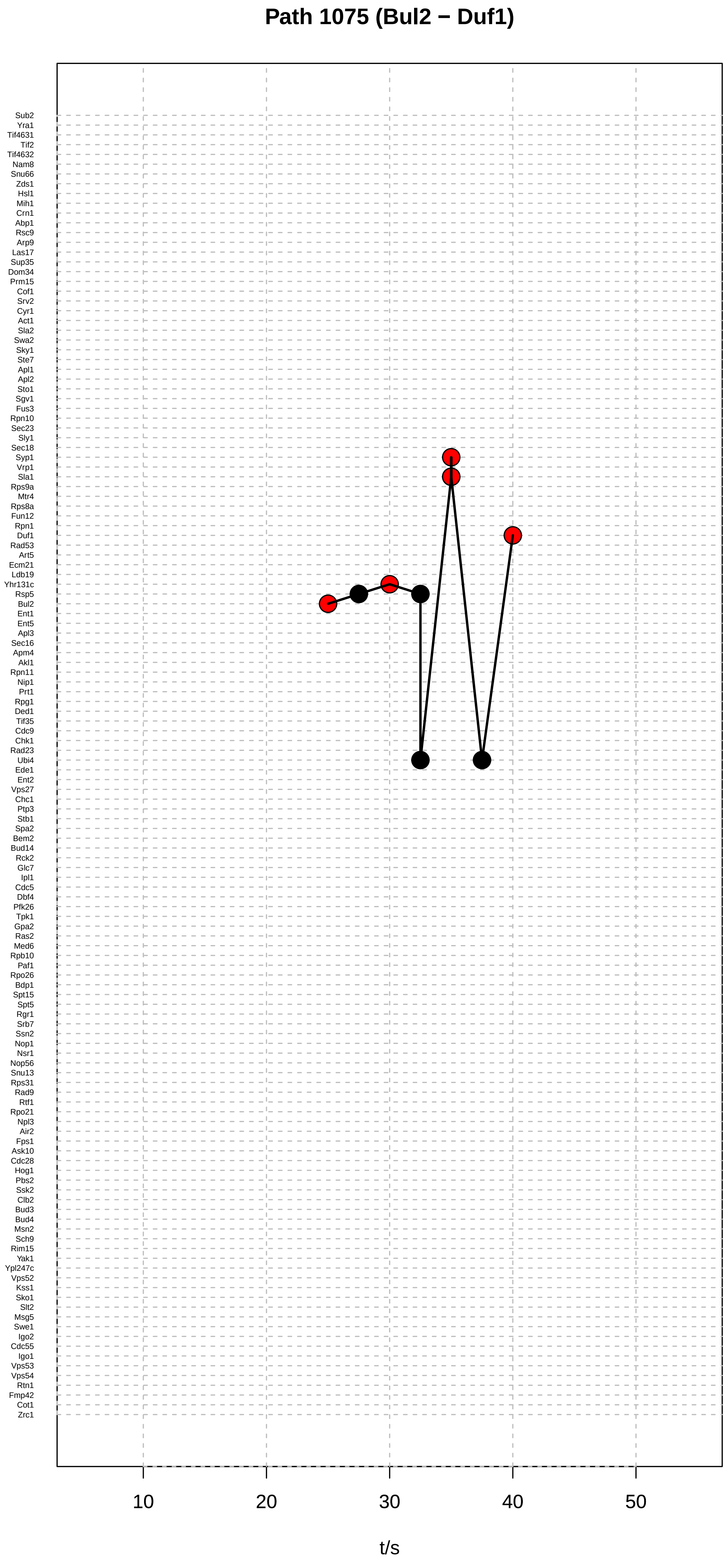
Path 1075 (Bul2 − Duf1)
Sub2
Yra1
Tif4631
Tif2
Tif4632
Nam8
Snu66
Zds1
Hsl1
Mih1
Crn1
Abp1
Rsc9
Arp9
Las17
Sup35
Dom34
Prm15
Cof1
Srv2
Cyr1
Act1
Sla2
Swa2
Sky1
Ste7
Apl1
Apl2
Sto1
Sgv1
Fus3
Rpn10
Sec23
Sly1
Sec18
Syp1
Vrp1
Sla1
Rps9a
Mtr4
Rps8a
Fun12
Rpn1
Duf1
Rad53
Art5
Ecm21
Ldb19
Yhr131c
Rsp5
Bul2
Ent1
Ent5
Apl3
Sec16
Apm4
Akl1
Rpn11
Nip1
Prt1
Rpg1
Ded1
Tif35
Cdc9
Chk1
Rad23
Ubi4
Ede1
Ent2
Vps27
Chc1
Ptp3
Stb1
Spa2
Bem2
Bud14
Rck2
Glc7
Ipl1
Cdc5
Dbf4
Pfk26
Tpk1
Gpa2
Ras2
Med6
Rpb10
Paf1
Rpo26
Bdp1
Spt15
Spt5
Rgr1
Srb7
Ssn2
Nop1
Nsr1
Nop56
Snu13
Rps31
Rad9
Rtf1
Rpo21
Npl3
Air2
Fps1
Ask10
Cdc28
Hog1
Pbs2
Ssk2
Clb2
Bud3
Bud4
Msn2
Sch9
Rim15
Yak1
Ypl247c
Vps52
Kss1
Sko1
Slt2
Msg5
Swe1
Igo2
Cdc55
Igo1
Vps53
Vps54
Rtn1
Fmp42
Cot1
Zrc1
10
20
30
40
50
t/s

### Path 1076 (Bul2 − Fun12)

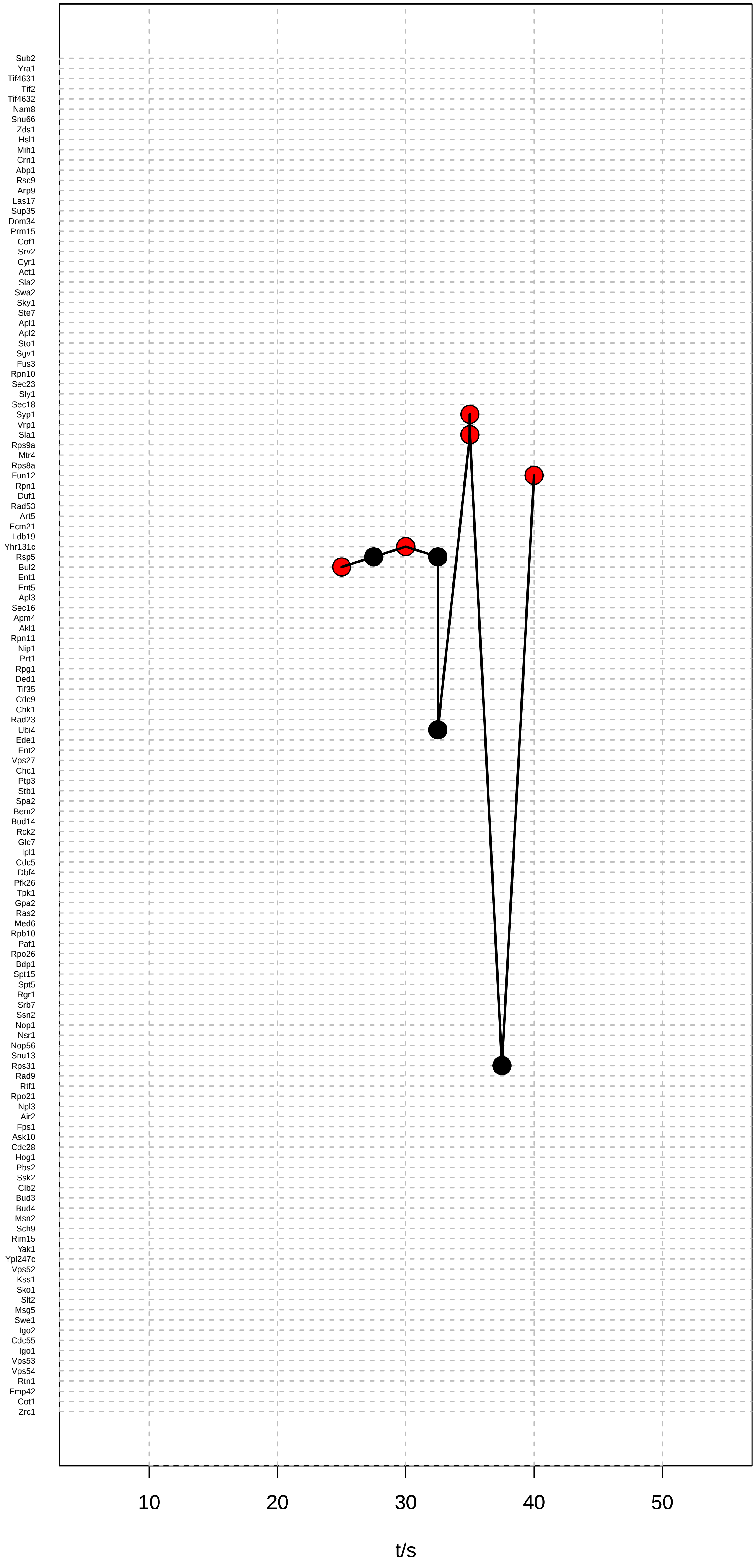

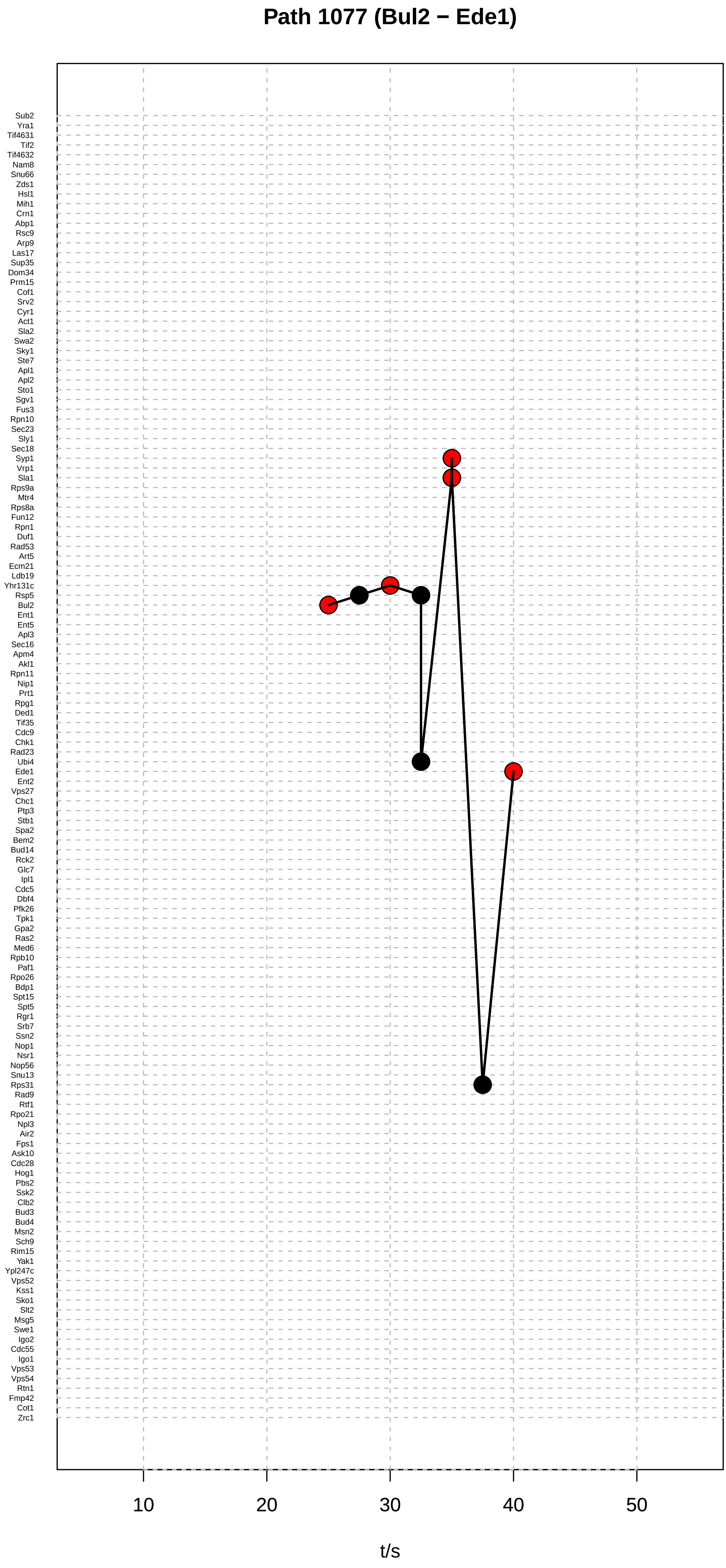
Path 1077 (Bul2 − Ede1)
Sub2
Yra1
Tif4631
Tif2
Tif4632
Nam8
Snu66
Zds1
Hsl1
Mih1
Crn1
Abp1
Rsc9
Arp9
Las17
Sup35
Dom34
Prm15
Cof1
Srv2
Cyr1
Act1
Sla2
Swa2
Sky1
Ste7
Apl1
Apl2
Sto1
Sgv1
Fus3
Rpn10
Sec23
Sly1
Sec18
Syp1
Vrp1
Sla1
Rps9a
Mtr4
Rps8a
Fun12
Rpn1
Duf1
Rad53
Art5
Ecm21
Ldb19
Yhr131c
Rsp5
Bul2
Ent1
Ent5
Apl3
Sec16
Apm4
Akl1
Rpn11
Nip1
Prt1
Rpg1
Ded1
Tif35
Cdc9
Chk1
Rad23
Ubi4
Ede1
Ent2
Vps27
Chc1
Ptp3
Stb1
Spa2
Bem2
Bud14
Rck2
Glc7
Ipl1
Cdc5
Dbf4
Pfk26
Tpk1
Gpa2
Ras2
Med6
Rpb10
Paf1
Rpo26
Bdp1
Spt15
Spt5
Rgr1
Srb7
Ssn2
Nop1
Nsr1
Nop56
Snu13
Rps31
Rad9
Rtf1
Rpo21
Npl3
Air2
Fps1
Ask10
Cdc28
Hog1
Pbs2
Ssk2
Clb2
Bud3
Bud4
Msn2
Sch9
Rim15
Yak1
Ypl247c
Vps52
Kss1
Sko1
Slt2
Msg5
Swe1
Igo2
Cdc55
Igo1
Vps53
Vps54
Rtn1
Fmp42
Cot1
Zrc1
10
20
30
40
50
t/s

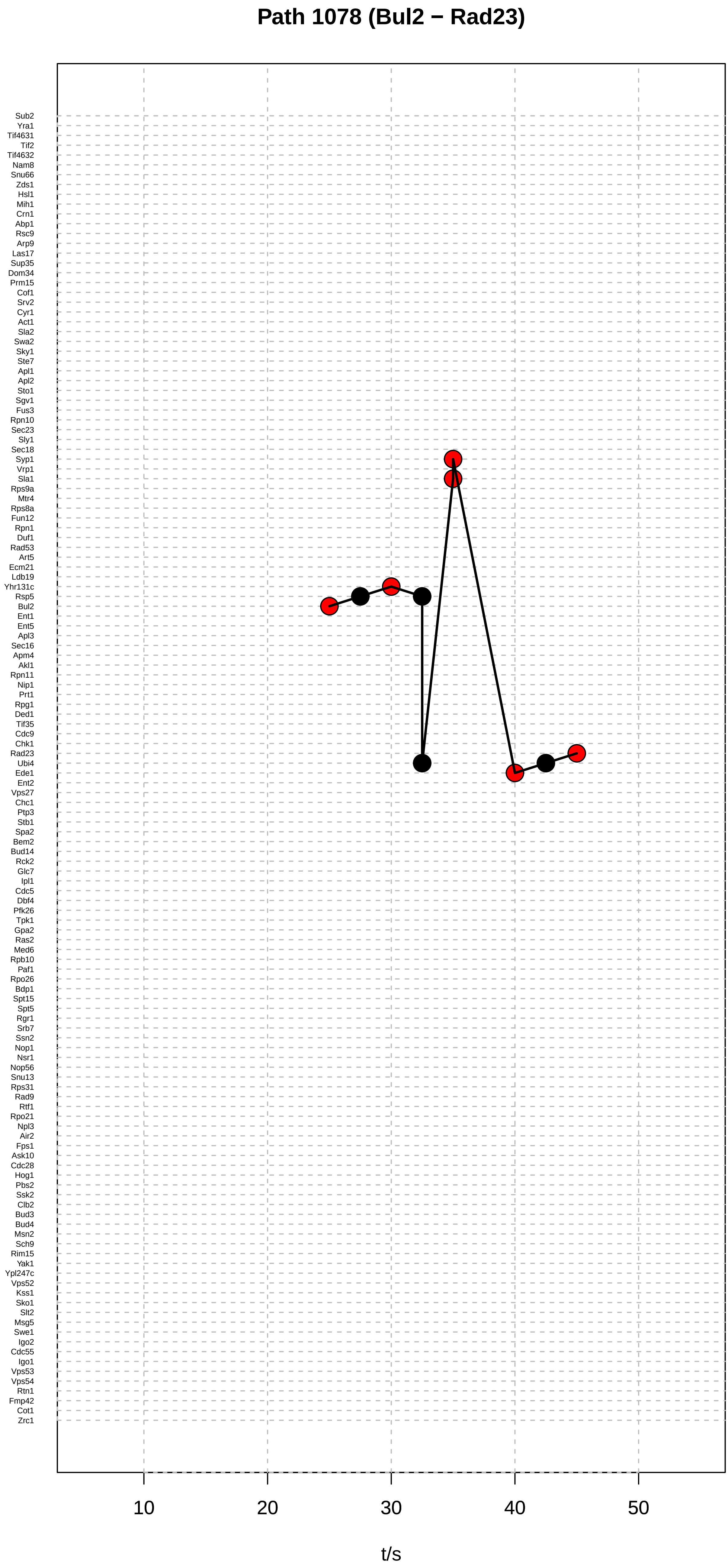

Path 1078 (Bul2 − Rad23)
Sub2
Yra1
Tif4631
Tif2
Tif4632
Nam8
Snu66
Zds1
Hsl1
Mih1
Crn1
Abp1
Rsc9
Arp9
Las17
Sup35
Dom34
Prm15
Cof1
Srv2
Cyr1
Act1
Sla2
Swa2
Sky1
Ste7
Apl1
Apl2
Sto1
Sgv1
Fus3
Rpn10
Sec23
Sly1
Sec18
Syp1
Vrp1
Sla1
Rps9a
Mtr4
Rps8a
Fun12
Rpn1
Duf1
Rad53
Art5
Ecm21
Ldb19
Yhr131c
Rsp5
Bul2
Ent1
Ent5
Apl3
Sec16
Apm4
Akl1
Rpn11
Nip1
Prt1
Rpg1
Ded1
Tif35
Cdc9
Chk1
Rad23
Ubi4
Ede1
Ent2
Vps27
Chc1
Ptp3
Stb1
Spa2
Bem2
Bud14
Rck2
Glc7
Ipl1
Cdc5
Dbf4
Pfk26
Tpk1
Gpa2
Ras2
Med6
Rpb10
Paf1
Rpo26
Bdp1
Spt15
Spt5
Rgr1
Srb7
Ssn2
Nop1
Nsr1
Nop56
Snu13
Rps31
Rad9
Rtf1
Rpo21
Npl3
Air2
Fps1
Ask10
Cdc28
Hog1
Pbs2
Ssk2
Clb2
Bud3
Bud4
Msn2
Sch9
Rim15
Yak1
Ypl247c
Vps52
Kss1
Sko1
Slt2
Msg5
Swe1
Igo2
Cdc55
Igo1
Vps53
Vps54
Rtn1
Fmp42
Cot1
Zrc1
10
20
30
40
50
t/s

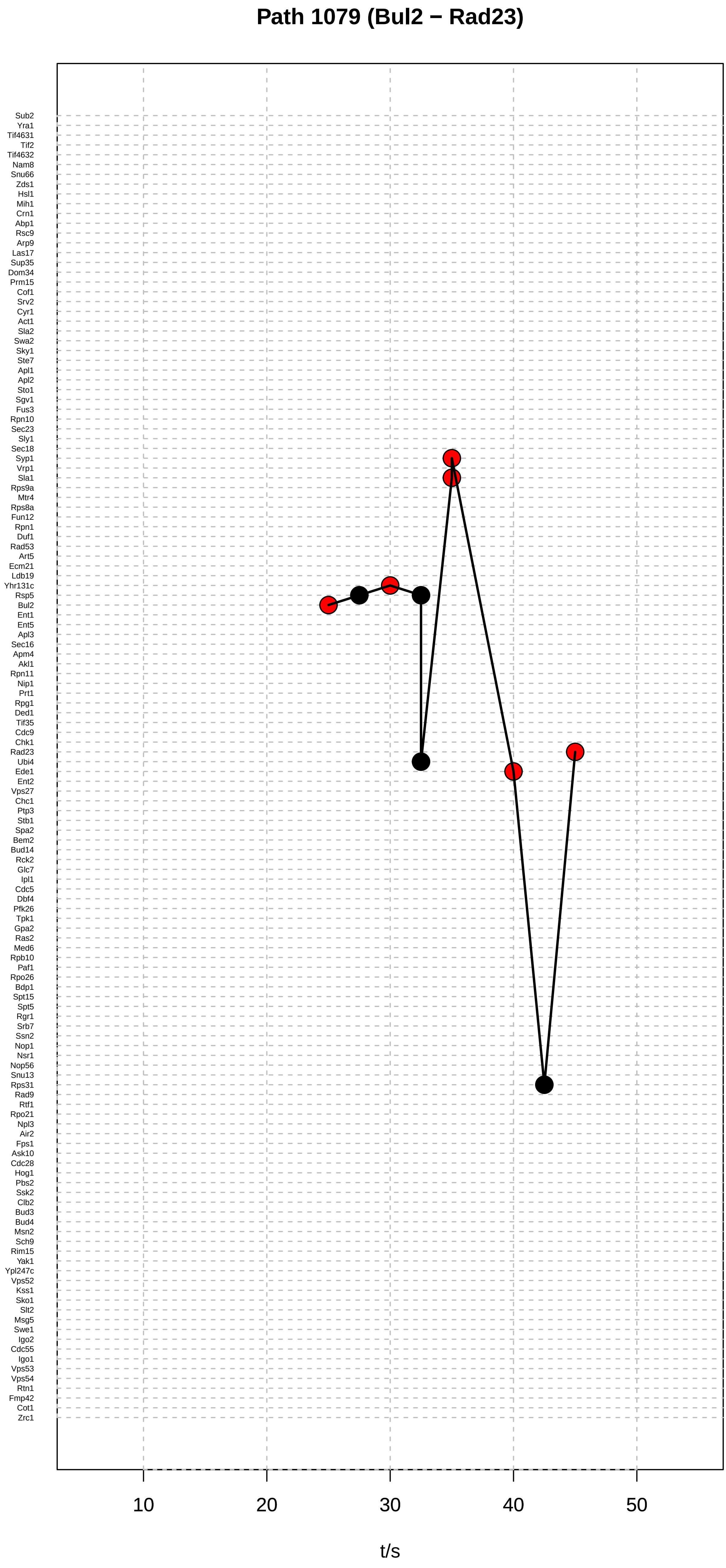

Path 1079 (Bul2 − Rad23)
Sub2
Yra1
Tif4631
Tif2
Tif4632
Nam8
Snu66
Zds1
Hsl1
Mih1
Crn1
Abp1
Rsc9
Arp9
Las17
Sup35
Dom34
Prm15
Cof1
Srv2
Cyr1
Act1
Sla2
Swa2
Sky1
Ste7
Apl1
Apl2
Sto1
Sgv1
Fus3
Rpn10
Sec23
Sly1
Sec18
Syp1
Vrp1
Sla1
Rps9a
Mtr4
Rps8a
Fun12
Rpn1
Duf1
Rad53
Art5
Ecm21
Ldb19
Yhr131c
Rsp5
Bul2
Ent1
Ent5
Apl3
Sec16
Apm4
Akl1
Rpn11
Nip1
Prt1
Rpg1
Ded1
Tif35
Cdc9
Chk1
Rad23
Ubi4
Ede1
Ent2
Vps27
Chc1
Ptp3
Stb1
Spa2
Bem2
Bud14
Rck2
Glc7
Ipl1
Cdc5
Dbf4
Pfk26
Tpk1
Gpa2
Ras2
Med6
Rpb10
Paf1
Rpo26
Bdp1
Spt15
Spt5
Rgr1
Srb7
Ssn2
Nop1
Nsr1
Nop56
Snu13
Rps31
Rad9
Rtf1
Rpo21
Npl3
Air2
Fps1
Ask10
Cdc28
Hog1
Pbs2
Ssk2
Clb2
Bud3
Bud4
Msn2
Sch9
Rim15
Yak1
Ypl247c
Vps52
Kss1
Sko1
Slt2
Msg5
Swe1
Igo2
Cdc55
Igo1
Vps53
Vps54
Rtn1
Fmp42
Cot1
Zrc1
10
20
30
40
50
t/s

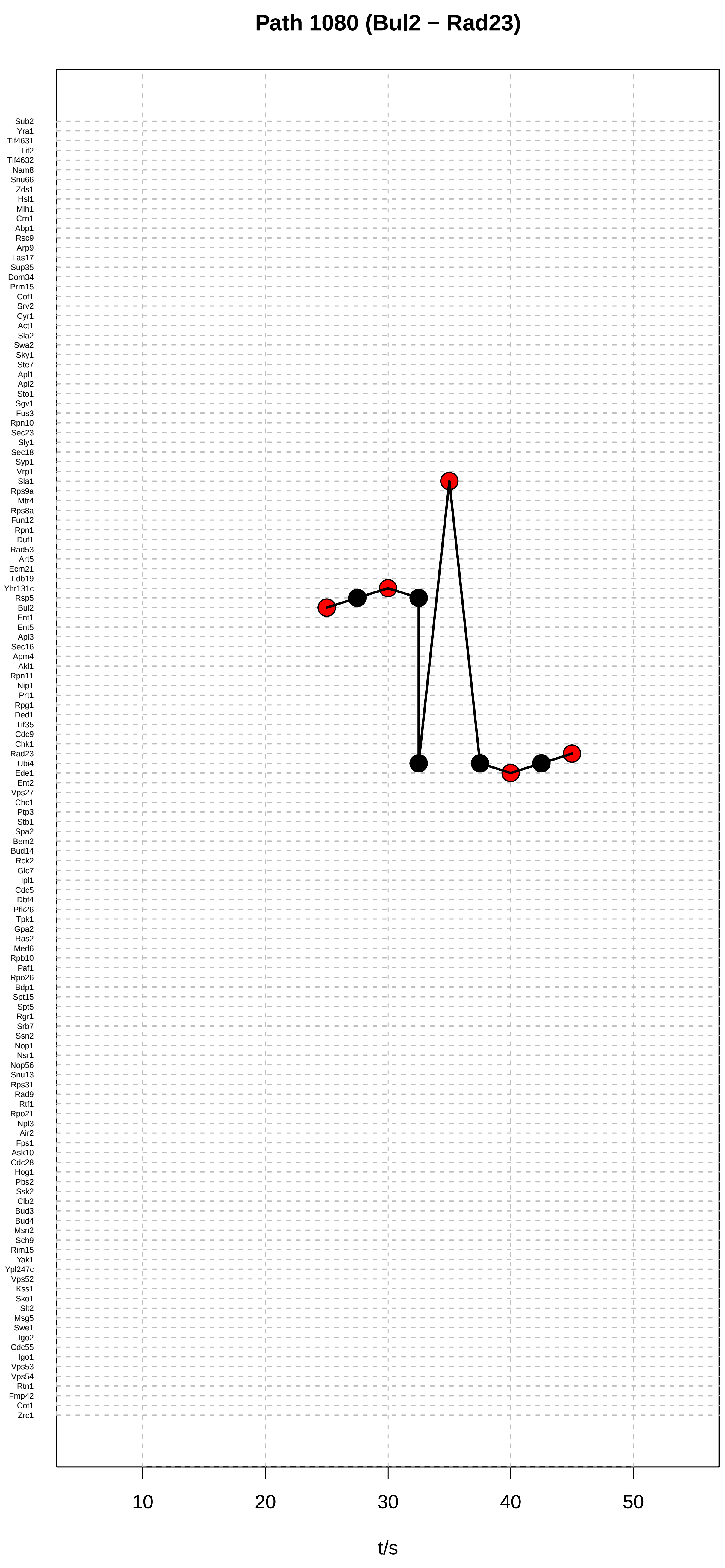

Path 1080 (Bul2 − Rad23)
Sub2
Yra1
Tif4631
Tif2
Tif4632
Nam8
Snu66
Zds1
Hsl1
Mih1
Crn1
Abp1
Rsc9
Arp9
Las17
Sup35
Dom34
Prm15
Cof1
Srv2
Cyr1
Act1
Sla2
Swa2
Sky1
Ste7
Apl1
Apl2
Sto1
Sgv1
Fus3
Rpn10
Sec23
Sly1
Sec18
Syp1
Vrp1
Sla1
Rps9a
Mtr4
Rps8a
Fun12
Rpn1
Duf1
Rad53
Art5
Ecm21
Ldb19
Yhr131c
Rsp5
Bul2
Ent1
Ent5
Apl3
Sec16
Apm4
Akl1
Rpn11
Nip1
Prt1
Rpg1
Ded1
Tif35
Cdc9
Chk1
Rad23
Ubi4
Ede1
Ent2
Vps27
Chc1
Ptp3
Stb1
Spa2
Bem2
Bud14
Rck2
Glc7
Ipl1
Cdc5
Dbf4
Pfk26
Tpk1
Gpa2
Ras2
Med6
Rpb10
Paf1
Rpo26
Bdp1
Spt15
Spt5
Rgr1
Srb7
Ssn2
Nop1
Nsr1
Nop56
Snu13
Rps31
Rad9
Rtf1
Rpo21
Npl3
Air2
Fps1
Ask10
Cdc28
Hog1
Pbs2
Ssk2
Clb2
Bud3
Bud4
Msn2
Sch9
Rim15
Yak1
Ypl247c
Vps52
Kss1
Sko1
Slt2
Msg5
Swe1
Igo2
Cdc55
Igo1
Vps53
Vps54
Rtn1
Fmp42
Cot1
Zrc1
10
20
30
40
50
t/s

### Path 1081 (Bul2 − Rad23)

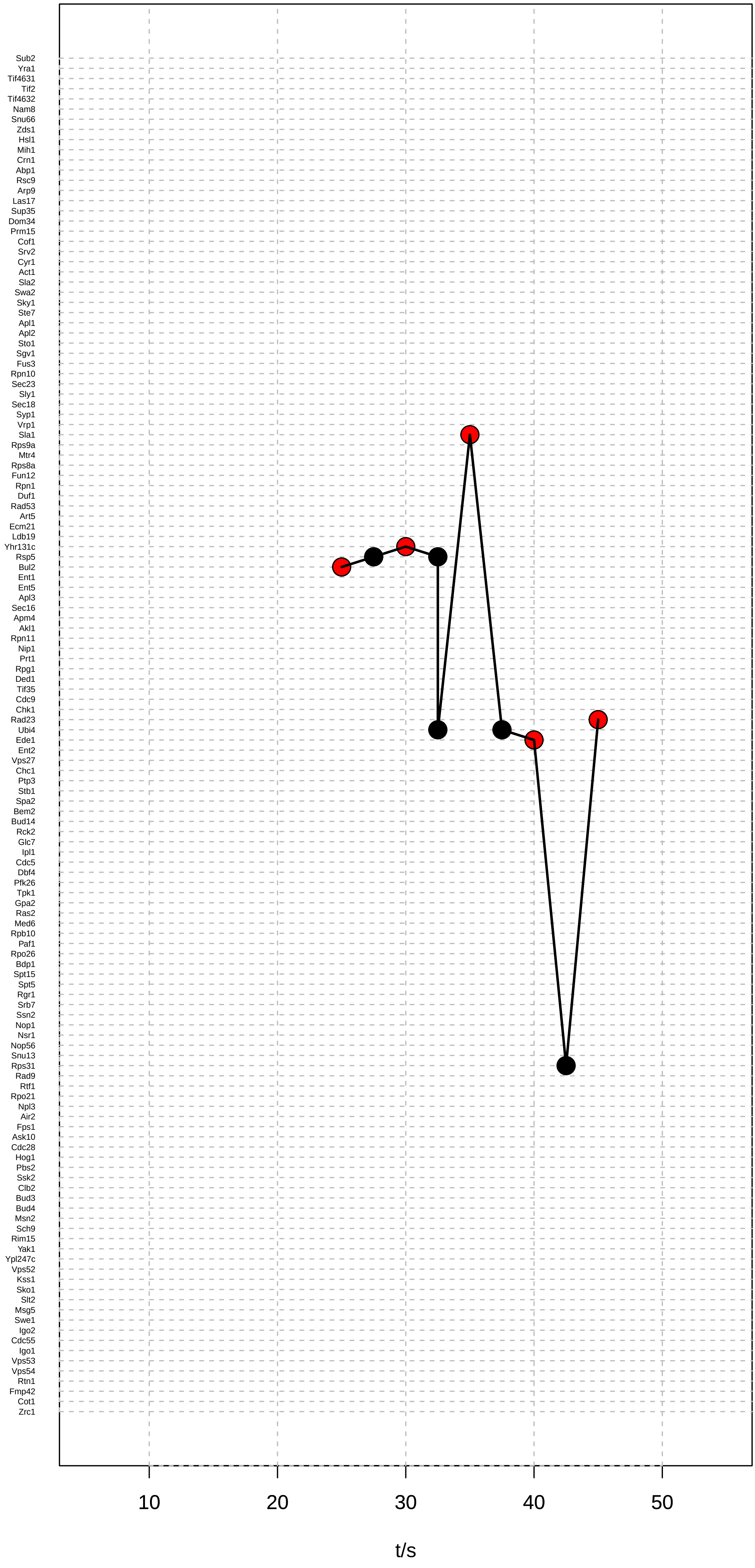

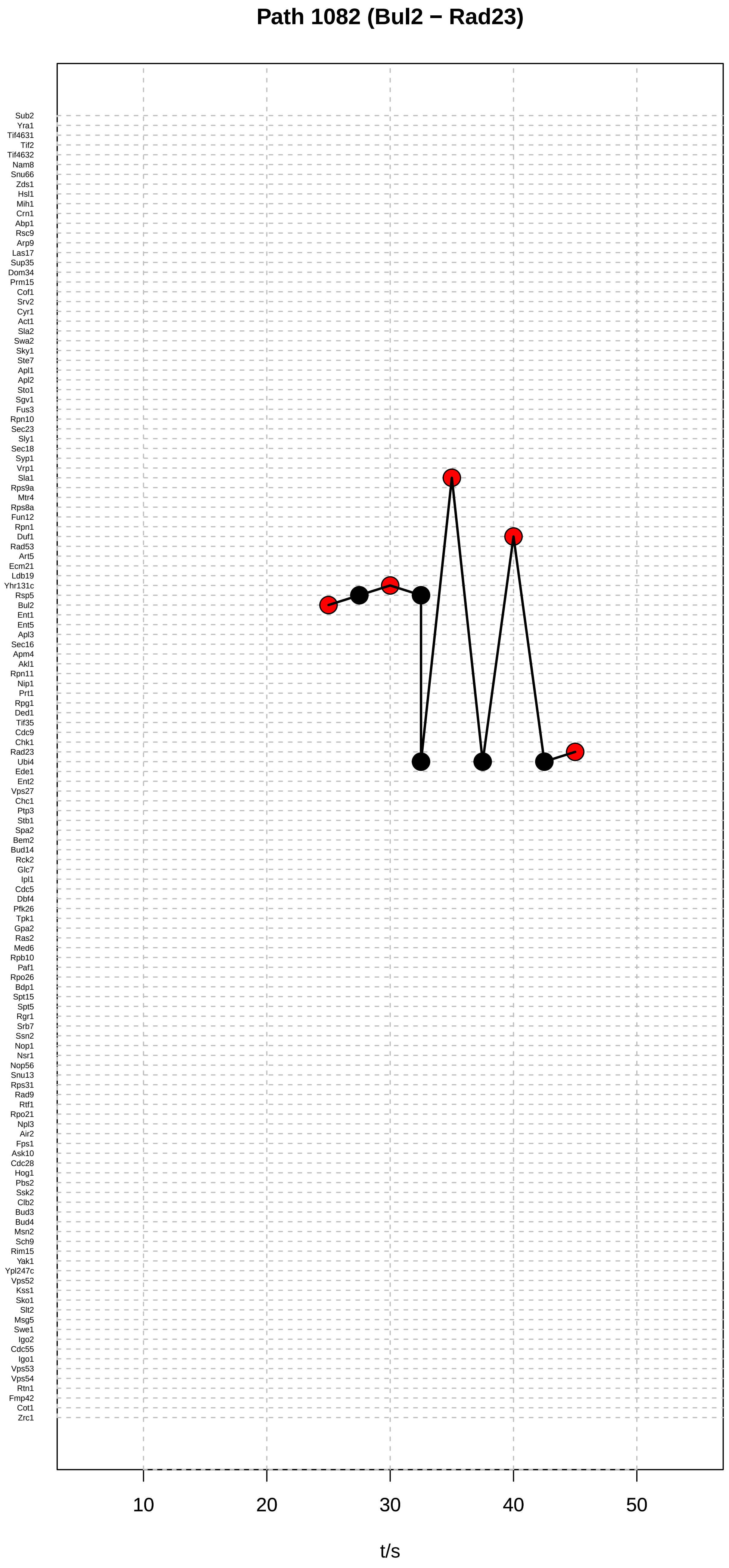
Path 1082 (Bul2 − Rad23)
Sub2
Yra1
Tif4631
Tif2
Tif4632
Nam8
Snu66
Zds1
Hsl1
Mih1
Crn1
Abp1
Rsc9
Arp9
Las17
Sup35
Dom34
Prm15
Cof1
Srv2
Cyr1
Act1
Sla2
Swa2
Sky1
Ste7
Apl1
Apl2
Sto1
Sgv1
Fus3
Rpn10
Sec23
Sly1
Sec18
Syp1
Vrp1
Sla1
Rps9a
Mtr4
Rps8a
Fun12
Rpn1
Duf1
Rad53
Art5
Ecm21
Ldb19
Yhr131c
Rsp5
Bul2
Ent1
Ent5
Apl3
Sec16
Apm4
Akl1
Rpn11
Nip1
Prt1
Rpg1
Ded1
Tif35
Cdc9
Chk1
Rad23
Ubi4
Ede1
Ent2
Vps27
Chc1
Ptp3
Stb1
Spa2
Bem2
Bud14
Rck2
Glc7
Ipl1
Cdc5
Dbf4
Pfk26
Tpk1
Gpa2
Ras2
Med6
Rpb10
Paf1
Rpo26
Bdp1
Spt15
Spt5
Rgr1
Srb7
Ssn2
Nop1
Nsr1
Nop56
Snu13
Rps31
Rad9
Rtf1
Rpo21
Npl3
Air2
Fps1
Ask10
Cdc28
Hog1
Pbs2
Ssk2
Clb2
Bud3
Bud4
Msn2
Sch9
Rim15
Yak1
Ypl247c
Vps52
Kss1
Sko1
Slt2
Msg5
Swe1
Igo2
Cdc55
Igo1
Vps53
Vps54
Rtn1
Fmp42
Cot1
Zrc1
10
20
30
40
50
t/s

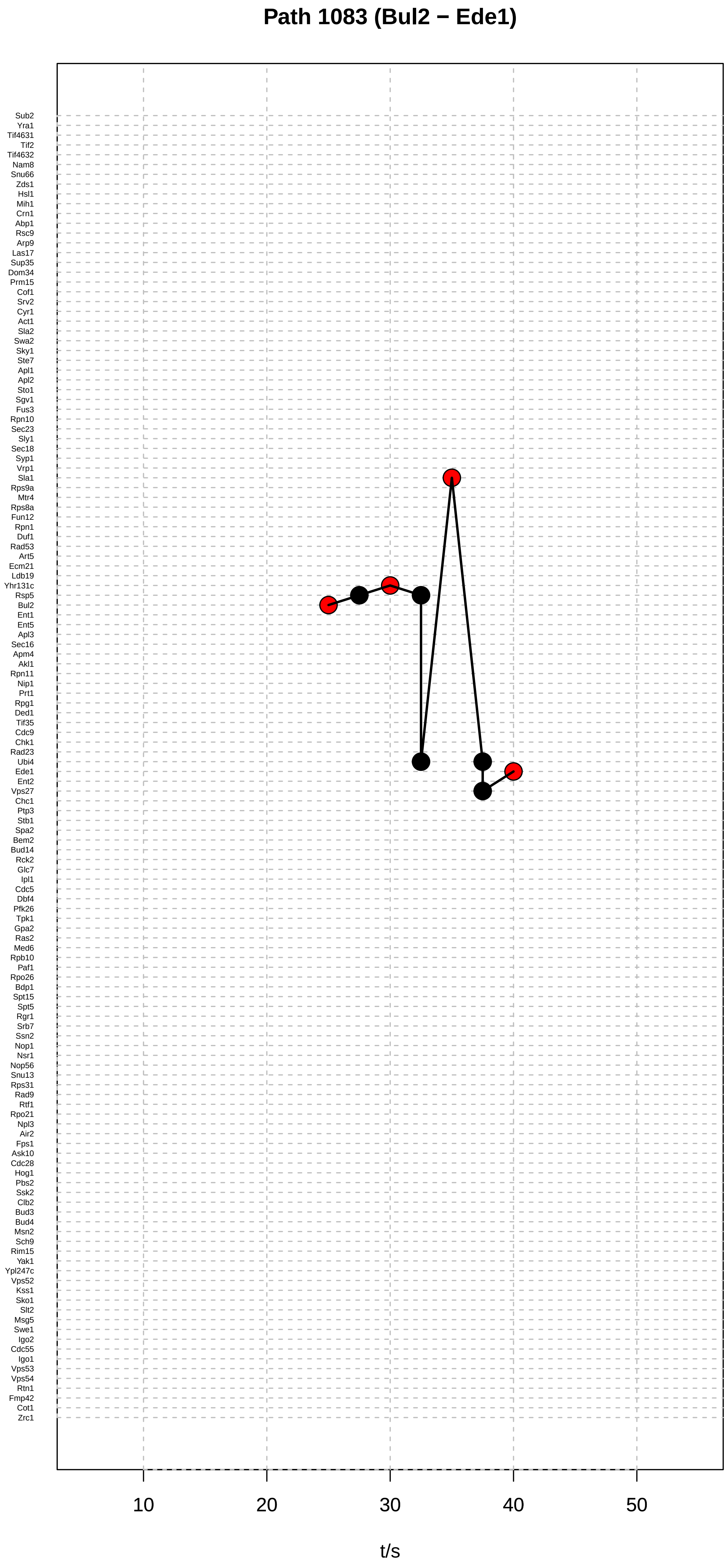

Path 1083 (Bul2 − Ede1)
Sub2
Yra1
Tif4631
Tif2
Tif4632
Nam8
Snu66
Zds1
Hsl1
Mih1
Crn1
Abp1
Rsc9
Arp9
Las17
Sup35
Dom34
Prm15
Cof1
Srv2
Cyr1
Act1
Sla2
Swa2
Sky1
Ste7
Apl1
Apl2
Sto1
Sgv1
Fus3
Rpn10
Sec23
Sly1
Sec18
Syp1
Vrp1
Sla1
Rps9a
Mtr4
Rps8a
Fun12
Rpn1
Duf1
Rad53
Art5
Ecm21
Ldb19
Yhr131c
Rsp5
Bul2
Ent1
Ent5
Apl3
Sec16
Apm4
Akl1
Rpn11
Nip1
Prt1
Rpg1
Ded1
Tif35
Cdc9
Chk1
Rad23
Ubi4
Ede1
Ent2
Vps27
Chc1
Ptp3
Stb1
Spa2
Bem2
Bud14
Rck2
Glc7
Ipl1
Cdc5
Dbf4
Pfk26
Tpk1
Gpa2
Ras2
Med6
Rpb10
Paf1
Rpo26
Bdp1
Spt15
Spt5
Rgr1
Srb7
Ssn2
Nop1
Nsr1
Nop56
Snu13
Rps31
Rad9
Rtf1
Rpo21
Npl3
Air2
Fps1
Ask10
Cdc28
Hog1
Pbs2
Ssk2
Clb2
Bud3
Bud4
Msn2
Sch9
Rim15
Yak1
Ypl247c
Vps52
Kss1
Sko1
Slt2
Msg5
Swe1
Igo2
Cdc55
Igo1
Vps53
Vps54
Rtn1
Fmp42
Cot1
Zrc1
10
20
30
40
50
t/s

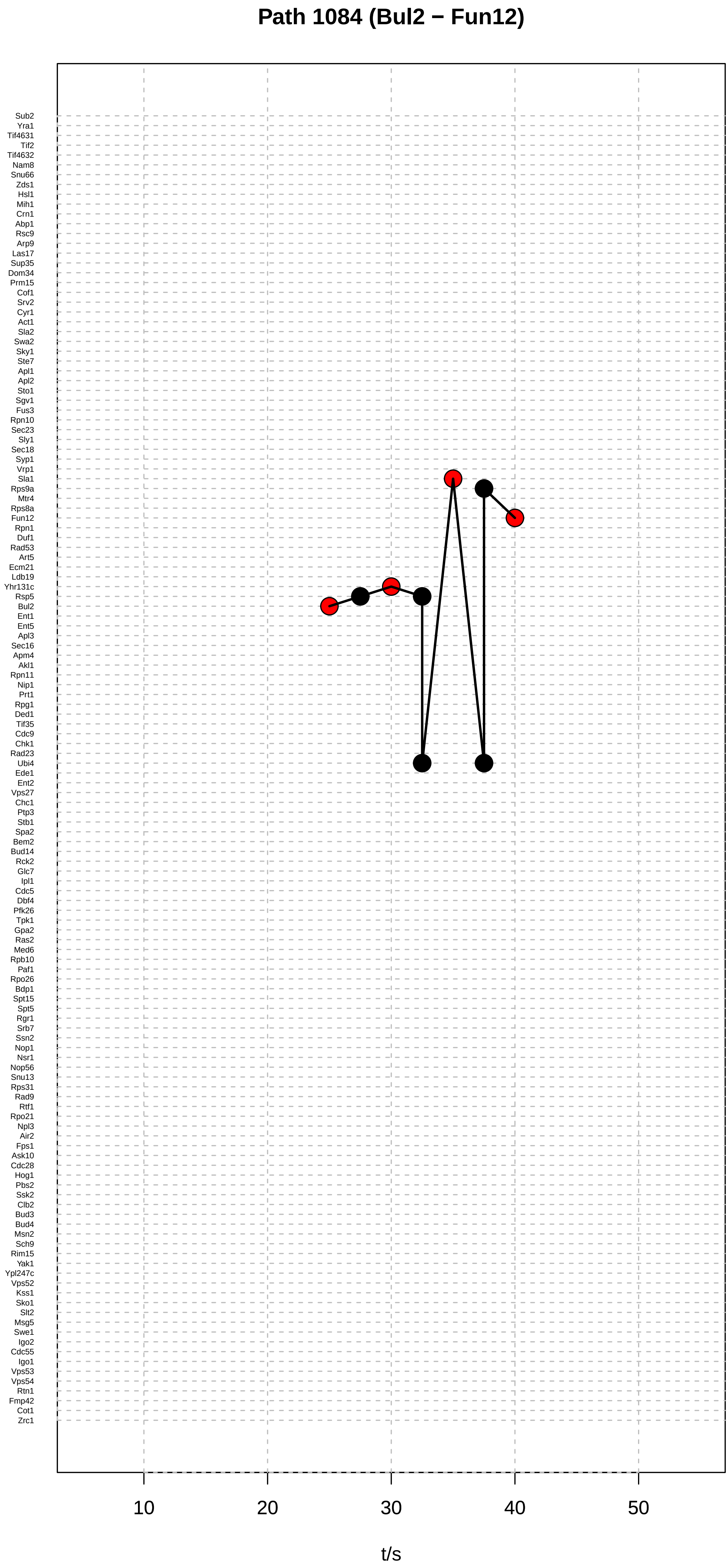

Path 1084 (Bul2 − Fun12)
Sub2
Yra1
Tif4631
Tif2
Tif4632
Nam8
Snu66
Zds1
Hsl1
Mih1
Crn1
Abp1
Rsc9
Arp9
Las17
Sup35
Dom34
Prm15
Cof1
Srv2
Cyr1
Act1
Sla2
Swa2
Sky1
Ste7
Apl1
Apl2
Sto1
Sgv1
Fus3
Rpn10
Sec23
Sly1
Sec18
Syp1
Vrp1
Sla1
Rps9a
Mtr4
Rps8a
Fun12
Rpn1
Duf1
Rad53
Art5
Ecm21
Ldb19
Yhr131c
Rsp5
Bul2
Ent1
Ent5
Apl3
Sec16
Apm4
Akl1
Rpn11
Nip1
Prt1
Rpg1
Ded1
Tif35
Cdc9
Chk1
Rad23
Ubi4
Ede1
Ent2
Vps27
Chc1
Ptp3
Stb1
Spa2
Bem2
Bud14
Rck2
Glc7
Ipl1
Cdc5
Dbf4
Pfk26
Tpk1
Gpa2
Ras2
Med6
Rpb10
Paf1
Rpo26
Bdp1
Spt15
Spt5
Rgr1
Srb7
Ssn2
Nop1
Nsr1
Nop56
Snu13
Rps31
Rad9
Rtf1
Rpo21
Npl3
Air2
Fps1
Ask10
Cdc28
Hog1
Pbs2
Ssk2
Clb2
Bud3
Bud4
Msn2
Sch9
Rim15
Yak1
Ypl247c
Vps52
Kss1
Sko1
Slt2
Msg5
Swe1
Igo2
Cdc55
Igo1
Vps53
Vps54
Rtn1
Fmp42
Cot1
Zrc1
10
20
30
40
50
t/s

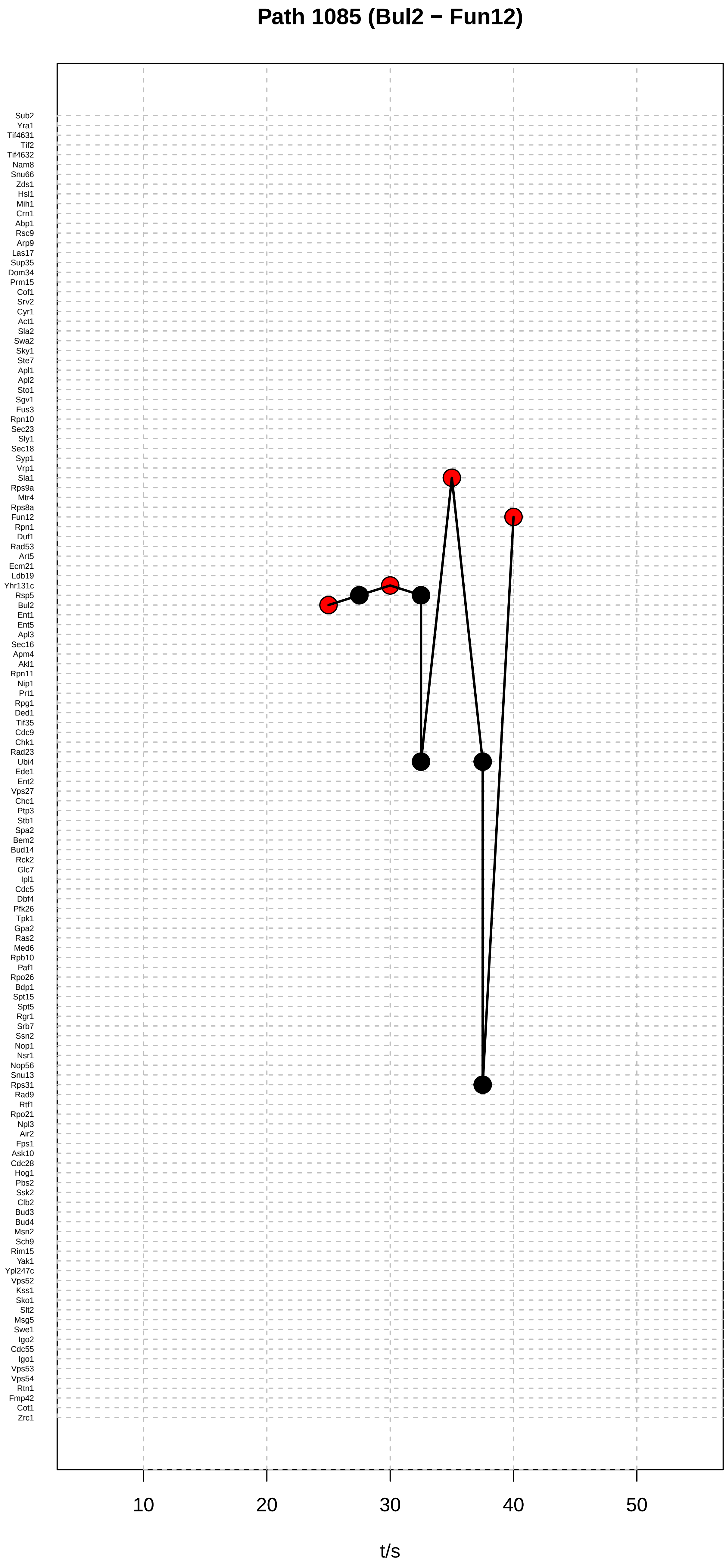

Path 1085 (Bul2 − Fun12)
Sub2
Yra1
Tif4631
Tif2
Tif4632
Nam8
Snu66
Zds1
Hsl1
Mih1
Crn1
Abp1
Rsc9
Arp9
Las17
Sup35
Dom34
Prm15
Cof1
Srv2
Cyr1
Act1
Sla2
Swa2
Sky1
Ste7
Apl1
Apl2
Sto1
Sgv1
Fus3
Rpn10
Sec23
Sly1
Sec18
Syp1
Vrp1
Sla1
Rps9a
Mtr4
Rps8a
Fun12
Rpn1
Duf1
Rad53
Art5
Ecm21
Ldb19
Yhr131c
Rsp5
Bul2
Ent1
Ent5
Apl3
Sec16
Apm4
Akl1
Rpn11
Nip1
Prt1
Rpg1
Ded1
Tif35
Cdc9
Chk1
Rad23
Ubi4
Ede1
Ent2
Vps27
Chc1
Ptp3
Stb1
Spa2
Bem2
Bud14
Rck2
Glc7
Ipl1
Cdc5
Dbf4
Pfk26
Tpk1
Gpa2
Ras2
Med6
Rpb10
Paf1
Rpo26
Bdp1
Spt15
Spt5
Rgr1
Srb7
Ssn2
Nop1
Nsr1
Nop56
Snu13
Rps31
Rad9
Rtf1
Rpo21
Npl3
Air2
Fps1
Ask10
Cdc28
Hog1
Pbs2
Ssk2
Clb2
Bud3
Bud4
Msn2
Sch9
Rim15
Yak1
Ypl247c
Vps52
Kss1
Sko1
Slt2
Msg5
Swe1
Igo2
Cdc55
Igo1
Vps53
Vps54
Rtn1
Fmp42
Cot1
Zrc1
10
20
30
40
50
t/s

### Path 1086 (Bul2 − Ede1)

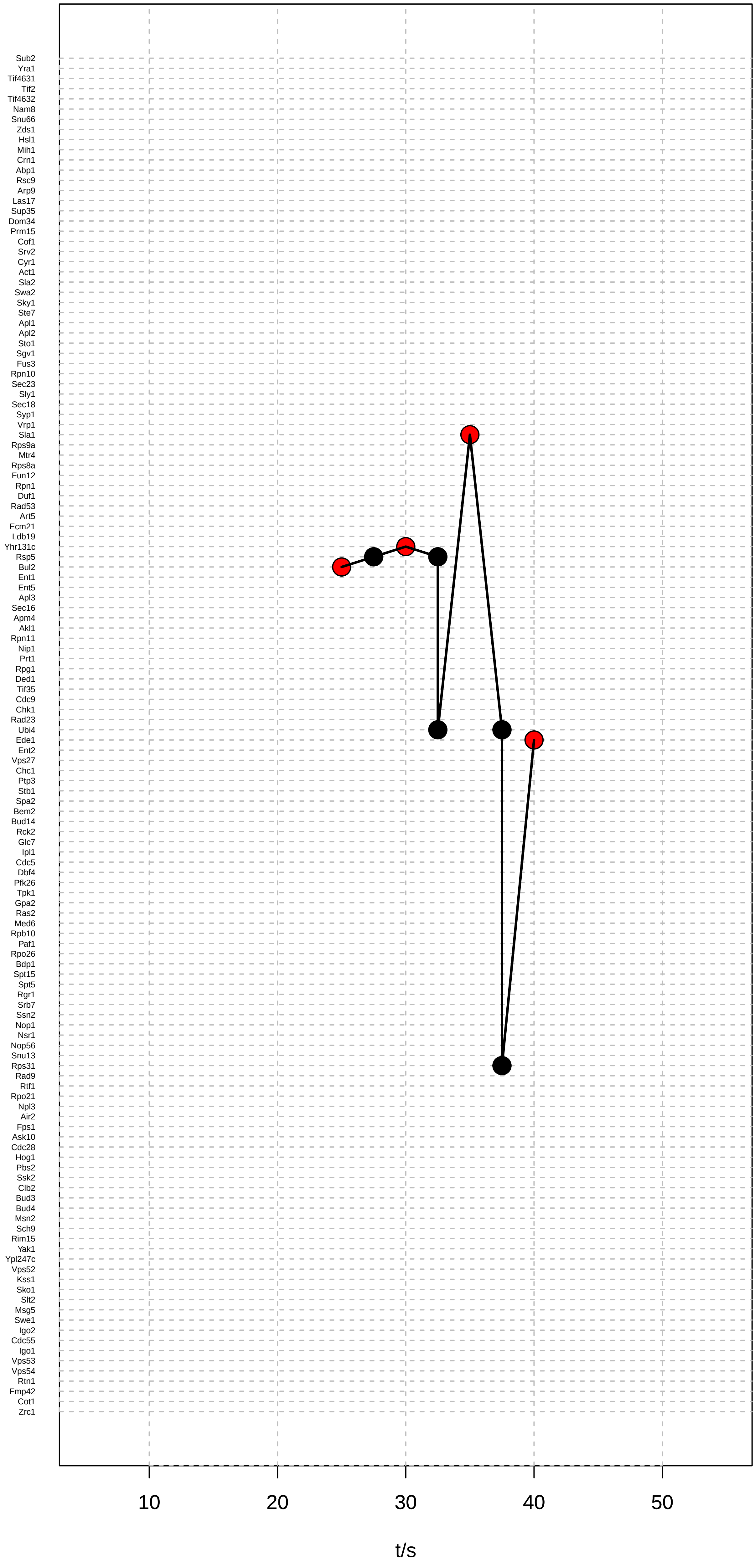

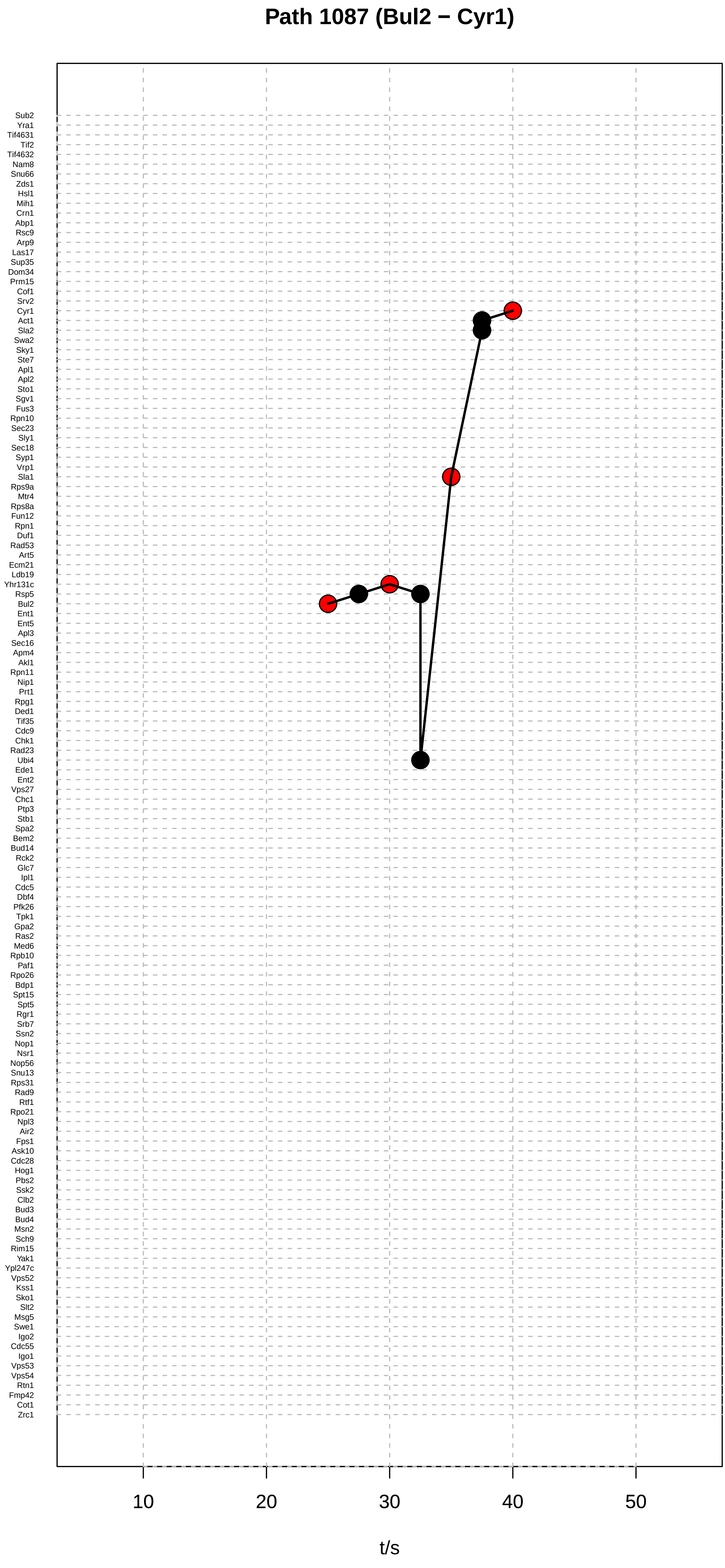

Path 1087 (Bul2 − Cyr1)
Sub2
Yra1
Tif4631
Tif2
Tif4632
Nam8
Snu66
Zds1
Hsl1
Mih1
Crn1
Abp1
Rsc9
Arp9
Las17
Sup35
Dom34
Prm15
Cof1
Srv2
Cyr1
Act1
Sla2
Swa2
Sky1
Ste7
Apl1
Apl2
Sto1
Sgv1
Fus3
Rpn10
Sec23
Sly1
Sec18
Syp1
Vrp1
Sla1
Rps9a
Mtr4
Rps8a
Fun12
Rpn1
Duf1
Rad53
Art5
Ecm21
Ldb19
Yhr131c
Rsp5
Bul2
Ent1
Ent5
Apl3
Sec16
Apm4
Akl1
Rpn11
Nip1
Prt1
Rpg1
Ded1
Tif35
Cdc9
Chk1
Rad23
Ubi4
Ede1
Ent2
Vps27
Chc1
Ptp3
Stb1
Spa2
Bem2
Bud14
Rck2
Glc7
Ipl1
Cdc5
Dbf4
Pfk26
Tpk1
Gpa2
Ras2
Med6
Rpb10
Paf1
Rpo26
Bdp1
Spt15
Spt5
Rgr1
Srb7
Ssn2
Nop1
Nsr1
Nop56
Snu13
Rps31
Rad9
Rtf1
Rpo21
Npl3
Air2
Fps1
Ask10
Cdc28
Hog1
Pbs2
Ssk2
Clb2
Bud3
Bud4
Msn2
Sch9
Rim15
Yak1
Ypl247c
Vps52
Kss1
Sko1
Slt2
Msg5
Swe1
Igo2
Cdc55
Igo1
Vps53
Vps54
Rtn1
Fmp42
Cot1
Zrc1
10
20
30
40
50
t/s

### Path 1088 (Bul2 − Rad23)

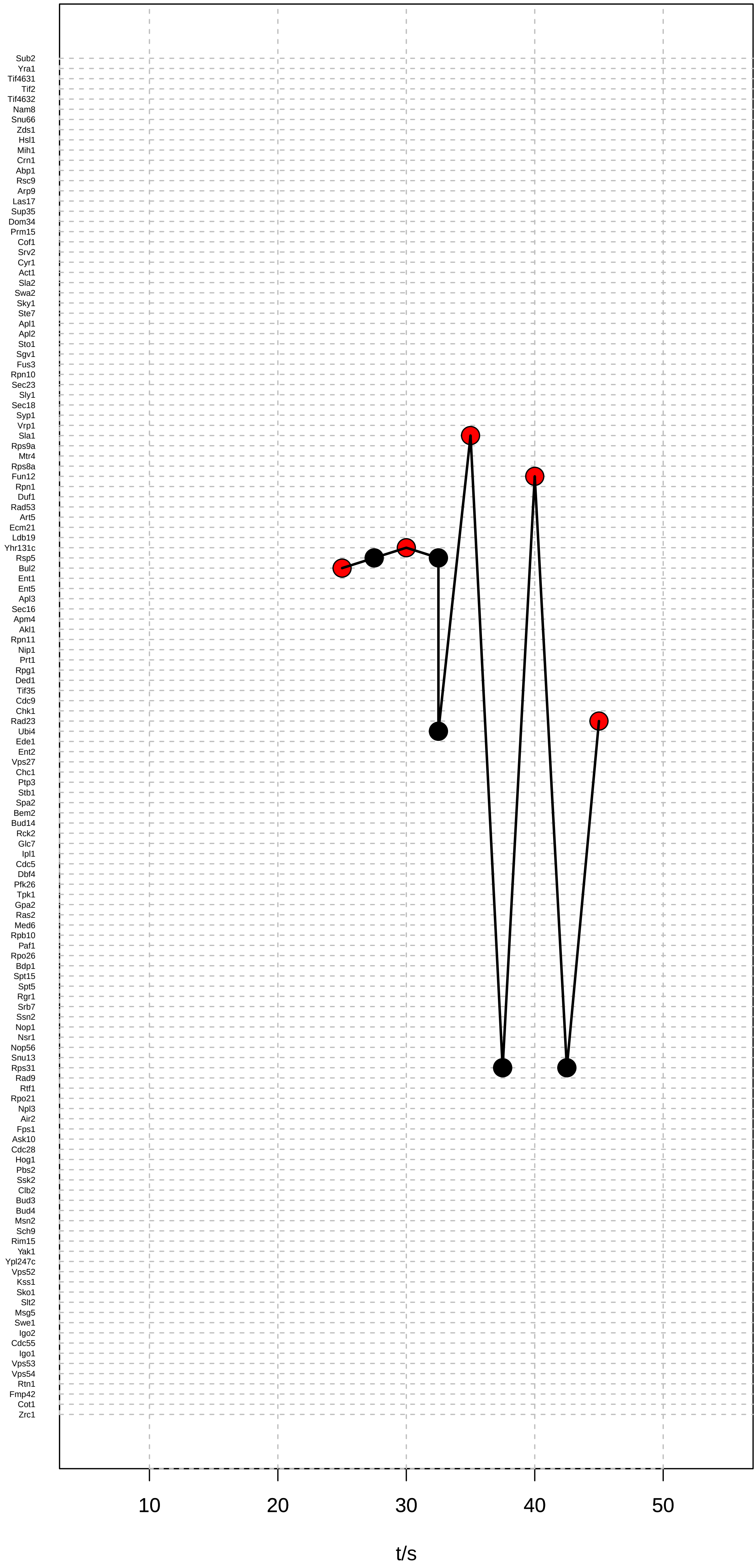

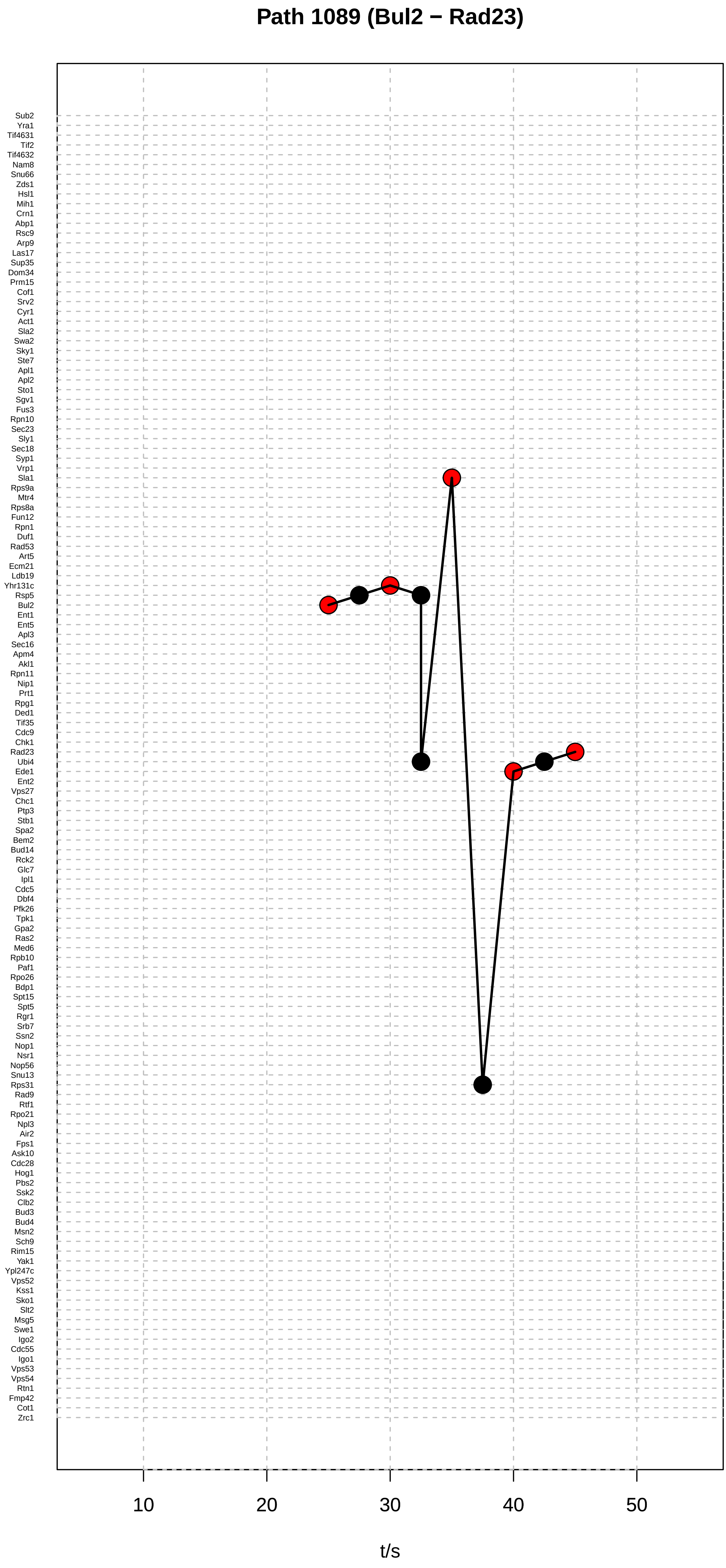

Path 1089 (Bul2 − Rad23)
Sub2
Yra1
Tif4631
Tif2
Tif4632
Nam8
Snu66
Zds1
Hsl1
Mih1
Crn1
Abp1
Rsc9
Arp9
Las17
Sup35
Dom34
Prm15
Cof1
Srv2
Cyr1
Act1
Sla2
Swa2
Sky1
Ste7
Apl1
Apl2
Sto1
Sgv1
Fus3
Rpn10
Sec23
Sly1
Sec18
Syp1
Vrp1
Sla1
Rps9a
Mtr4
Rps8a
Fun12
Rpn1
Duf1
Rad53
Art5
Ecm21
Ldb19
Yhr131c
Rsp5
Bul2
Ent1
Ent5
Apl3
Sec16
Apm4
Akl1
Rpn11
Nip1
Prt1
Rpg1
Ded1
Tif35
Cdc9
Chk1
Rad23
Ubi4
Ede1
Ent2
Vps27
Chc1
Ptp3
Stb1
Spa2
Bem2
Bud14
Rck2
Glc7
Ipl1
Cdc5
Dbf4
Pfk26
Tpk1
Gpa2
Ras2
Med6
Rpb10
Paf1
Rpo26
Bdp1
Spt15
Spt5
Rgr1
Srb7
Ssn2
Nop1
Nsr1
Nop56
Snu13
Rps31
Rad9
Rtf1
Rpo21
Npl3
Air2
Fps1
Ask10
Cdc28
Hog1
Pbs2
Ssk2
Clb2
Bud3
Bud4
Msn2
Sch9
Rim15
Yak1
Ypl247c
Vps52
Kss1
Sko1
Slt2
Msg5
Swe1
Igo2
Cdc55
Igo1
Vps53
Vps54
Rtn1
Fmp42
Cot1
Zrc1
10
20
30
40
50
t/s

### Path 1090 (Bul2 − Rad23)

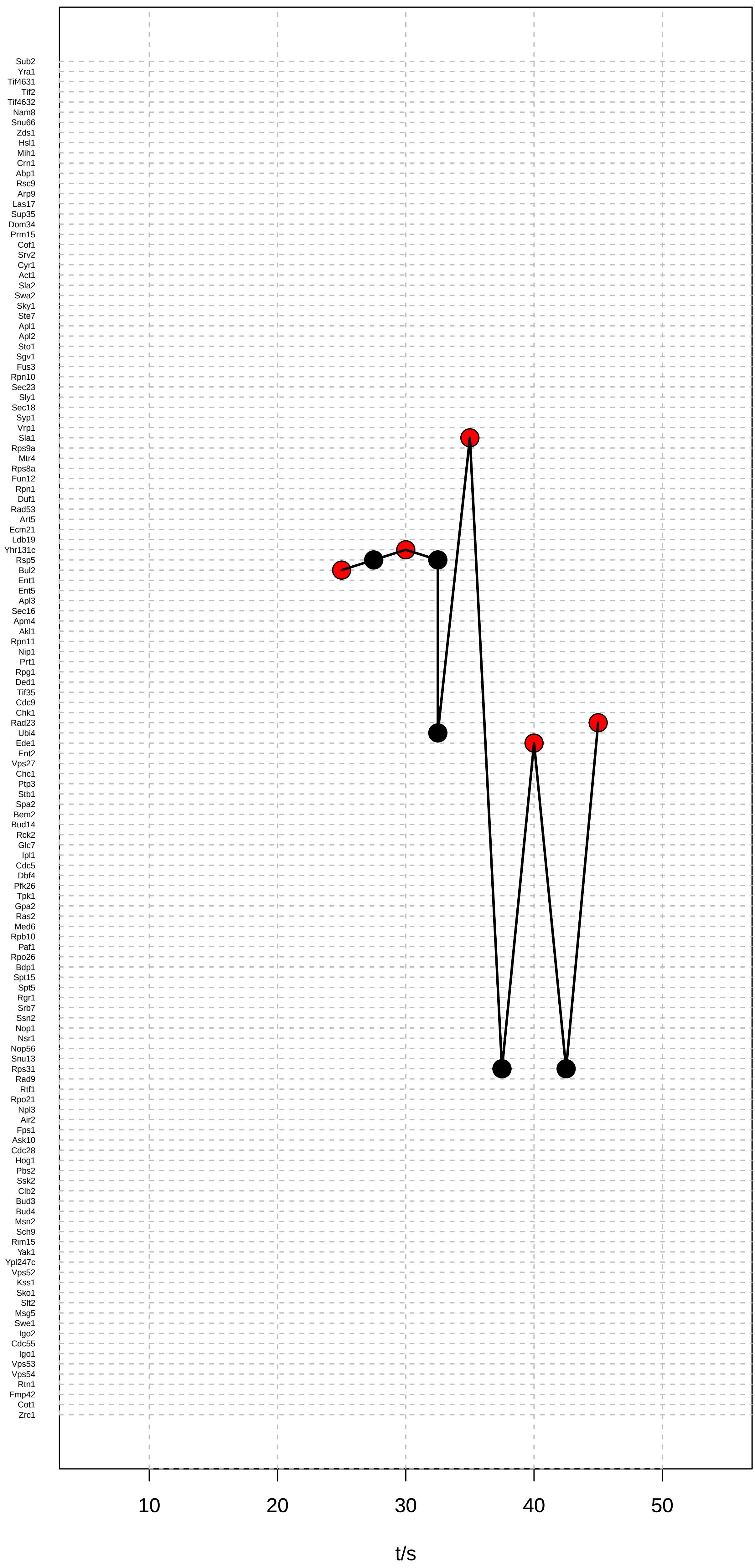

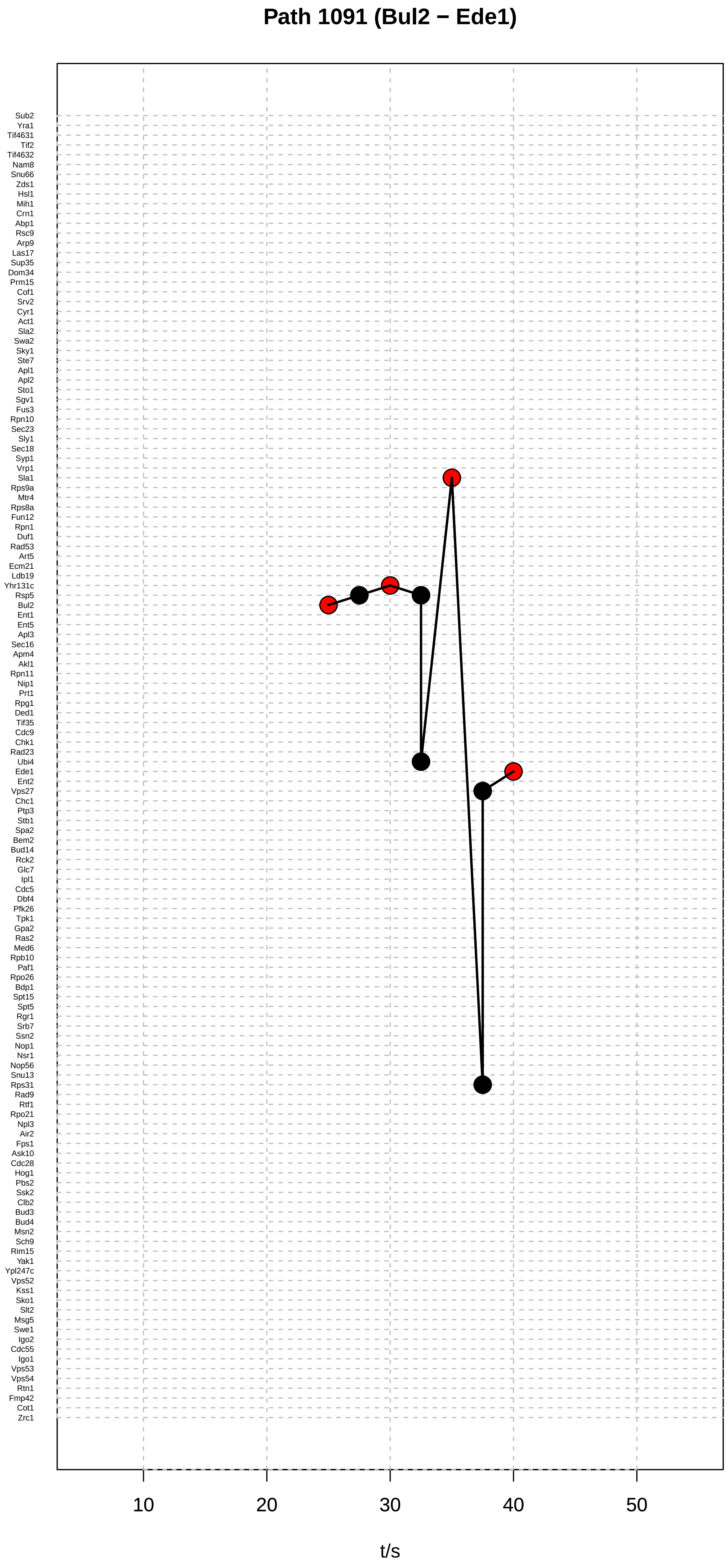

Path 1091 (Bul2 − Ede1)
Sub2
Yra1
Tif4631
Tif2
Tif4632
Nam8
Snu66
Zds1
Hsl1
Mih1
Crn1
Abp1
Rsc9
Arp9
Las17
Sup35
Dom34
Prm15
Cof1
Srv2
Cyr1
Act1
Sla2
Swa2
Sky1
Ste7
Apl1
Apl2
Sto1
Sgv1
Fus3
Rpn10
Sec23
Sly1
Sec18
Syp1
Vrp1
Sla1
Rps9a
Mtr4
Rps8a
Fun12
Rpn1
Duf1
Rad53
Art5
Ecm21
Ldb19
Yhr131c
Rsp5
Bul2
Ent1
Ent5
Apl3
Sec16
Apm4
Akl1
Rpn11
Nip1
Prt1
Rpg1
Ded1
Tif35
Cdc9
Chk1
Rad23
Ubi4
Ede1
Ent2
Vps27
Chc1
Ptp3
Stb1
Spa2
Bem2
Bud14
Rck2
Glc7
Ipl1
Cdc5
Dbf4
Pfk26
Tpk1
Gpa2
Ras2
Med6
Rpb10
Paf1
Rpo26
Bdp1
Spt15
Spt5
Rgr1
Srb7
Ssn2
Nop1
Nsr1
Nop56
Snu13
Rps31
Rad9
Rtf1
Rpo21
Npl3
Air2
Fps1
Ask10
Cdc28
Hog1
Pbs2
Ssk2
Clb2
Bud3
Bud4
Msn2
Sch9
Rim15
Yak1
Ypl247c
Vps52
Kss1
Sko1
Slt2
Msg5
Swe1
Igo2
Cdc55
Igo1
Vps53
Vps54
Rtn1
Fmp42
Cot1
Zrc1
10
20
30
40
50
t/s

### Path 1092 (Bul2 − Ede1)

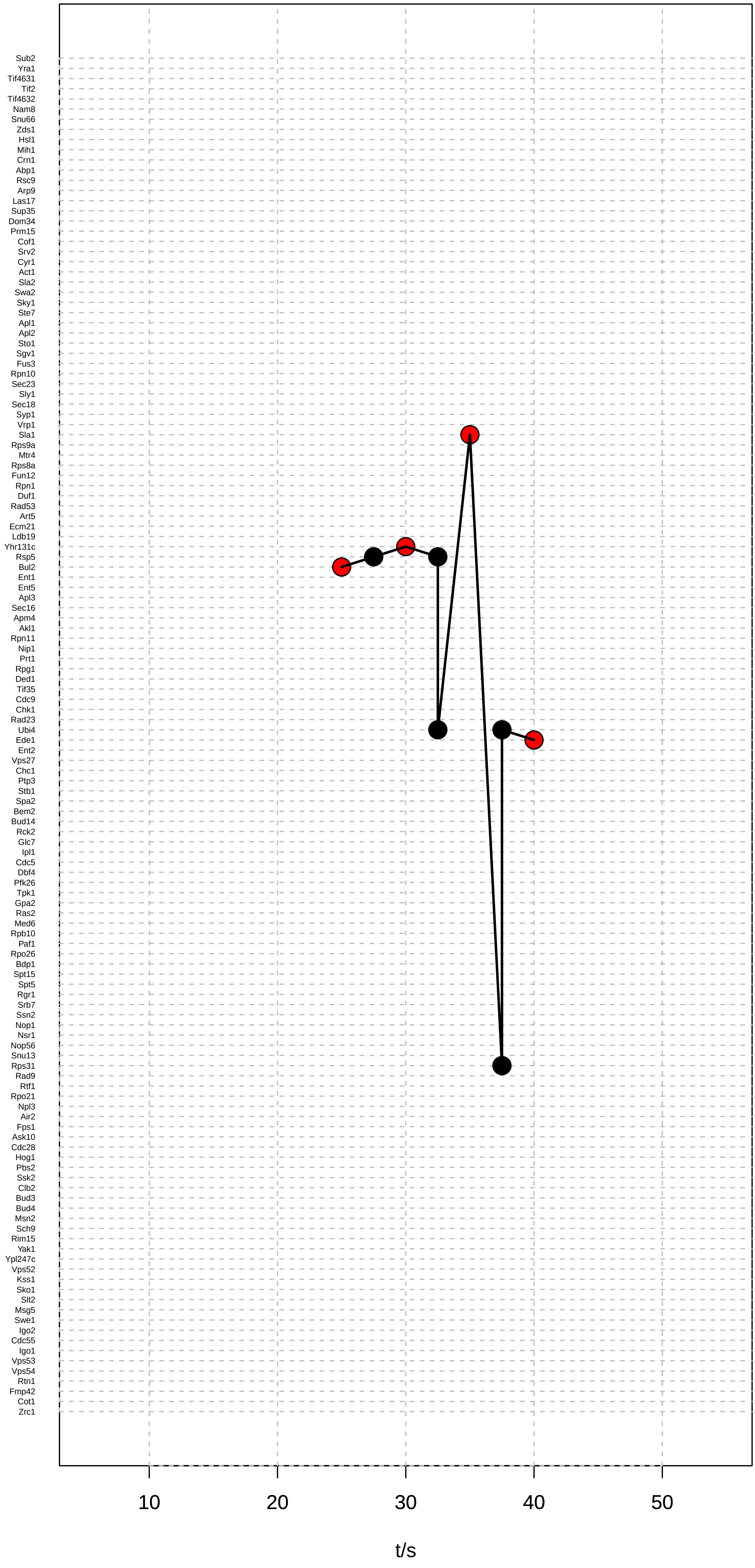

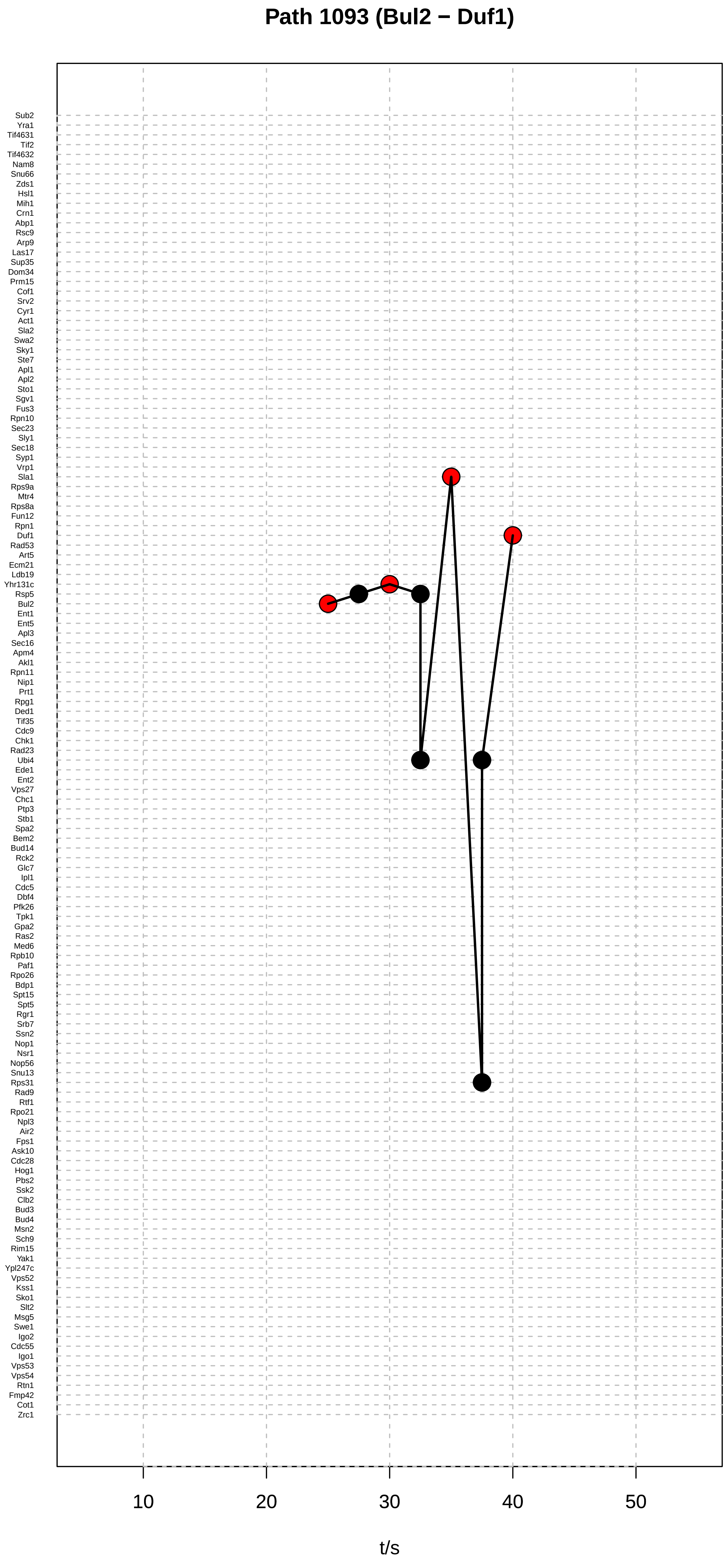
Path 1093 (Bul2 − Duf1)
Sub2
Yra1
Tif4631
Tif2
Tif4632
Nam8
Snu66
Zds1
Hsl1
Mih1
Crn1
Abp1
Rsc9
Arp9
Las17
Sup35
Dom34
Prm15
Cof1
Srv2
Cyr1
Act1
Sla2
Swa2
Sky1
Ste7
Apl1
Apl2
Sto1
Sgv1
Fus3
Rpn10
Sec23
Sly1
Sec18
Syp1
Vrp1
Sla1
Rps9a
Mtr4
Rps8a
Fun12
Rpn1
Duf1
Rad53
Art5
Ecm21
Ldb19
Yhr131c
Rsp5
Bul2
Ent1
Ent5
Apl3
Sec16
Apm4
Akl1
Rpn11
Nip1
Prt1
Rpg1
Ded1
Tif35
Cdc9
Chk1
Rad23
Ubi4
Ede1
Ent2
Vps27
Chc1
Ptp3
Stb1
Spa2
Bem2
Bud14
Rck2
Glc7
Ipl1
Cdc5
Dbf4
Pfk26
Tpk1
Gpa2
Ras2
Med6
Rpb10
Paf1
Rpo26
Bdp1
Spt15
Spt5
Rgr1
Srb7
Ssn2
Nop1
Nsr1
Nop56
Snu13
Rps31
Rad9
Rtf1
Rpo21
Npl3
Air2
Fps1
Ask10
Cdc28
Hog1
Pbs2
Ssk2
Clb2
Bud3
Bud4
Msn2
Sch9
Rim15
Yak1
Ypl247c
Vps52
Kss1
Sko1
Slt2
Msg5
Swe1
Igo2
Cdc55
Igo1
Vps53
Vps54
Rtn1
Fmp42
Cot1
Zrc1
10
20
30
40
50
t/s

Path 1094 (Bul2 − Fun12)

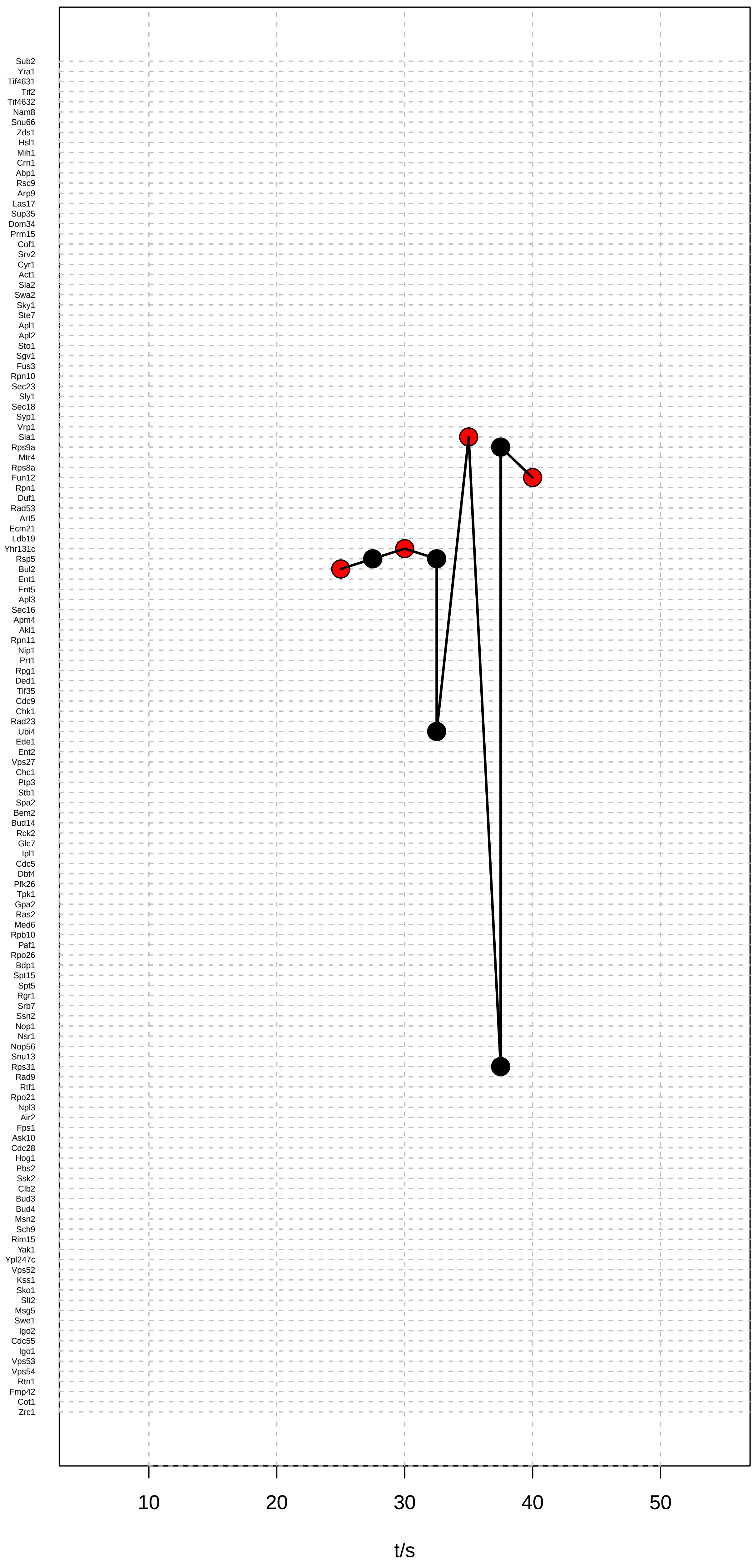

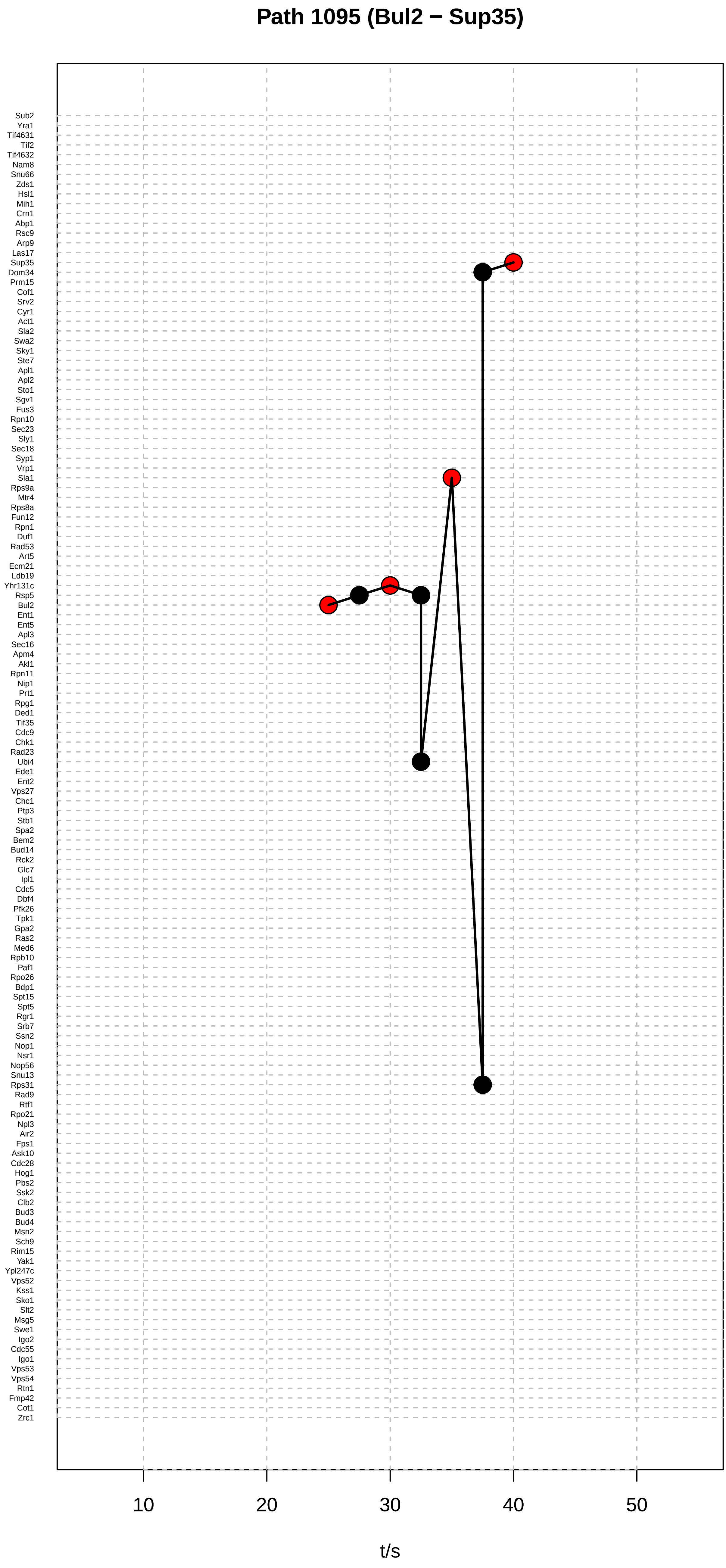

Path 1095 (Bul2 − Sup35)
Sub2
Yra1
Tif4631
Tif2
Tif4632
Nam8
Snu66
Zds1
Hsl1
Mih1
Crn1
Abp1
Rsc9
Arp9
Las17
Sup35
Dom34
Prm15
Cof1
Srv2
Cyr1
Act1
Sla2
Swa2
Sky1
Ste7
Apl1
Apl2
Sto1
Sgv1
Fus3
Rpn10
Sec23
Sly1
Sec18
Syp1
Vrp1
Sla1
Rps9a
Mtr4
Rps8a
Fun12
Rpn1
Duf1
Rad53
Art5
Ecm21
Ldb19
Yhr131c
Rsp5
Bul2
Ent1
Ent5
Apl3
Sec16
Apm4
Akl1
Rpn11
Nip1
Prt1
Rpg1
Ded1
Tif35
Cdc9
Chk1
Rad23
Ubi4
Ede1
Ent2
Vps27
Chc1
Ptp3
Stb1
Spa2
Bem2
Bud14
Rck2
Glc7
Ipl1
Cdc5
Dbf4
Pfk26
Tpk1
Gpa2
Ras2
Med6
Rpb10
Paf1
Rpo26
Bdp1
Spt15
Spt5
Rgr1
Srb7
Ssn2
Nop1
Nsr1
Nop56
Snu13
Rps31
Rad9
Rtf1
Rpo21
Npl3
Air2
Fps1
Ask10
Cdc28
Hog1
Pbs2
Ssk2
Clb2
Bud3
Bud4
Msn2
Sch9
Rim15
Yak1
Ypl247c
Vps52
Kss1
Sko1
Slt2
Msg5
Swe1
Igo2
Cdc55
Igo1
Vps53
Vps54
Rtn1
Fmp42
Cot1
Zrc1
10
20
30
40
50
t/s

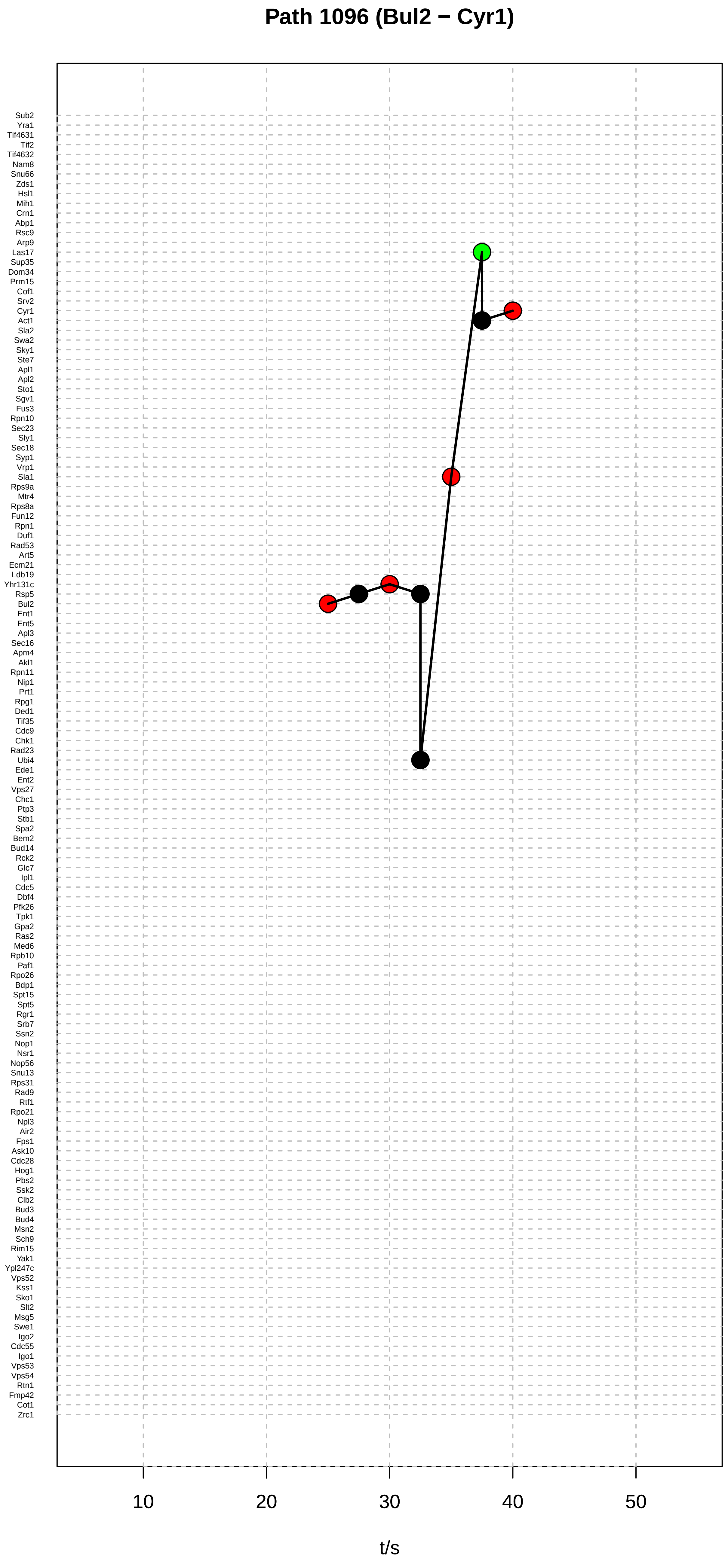

Path 1096 (Bul2 − Cyr1)
Sub2
Yra1
Tif4631
Tif2
Tif4632
Nam8
Snu66
Zds1
Hsl1
Mih1
Crn1
Abp1
Rsc9
Arp9
Las17
Sup35
Dom34
Prm15
Cof1
Srv2
Cyr1
Act1
Sla2
Swa2
Sky1
Ste7
Apl1
Apl2
Sto1
Sgv1
Fus3
Rpn10
Sec23
Sly1
Sec18
Syp1
Vrp1
Sla1
Rps9a
Mtr4
Rps8a
Fun12
Rpn1
Duf1
Rad53
Art5
Ecm21
Ldb19
Yhr131c
Rsp5
Bul2
Ent1
Ent5
Apl3
Sec16
Apm4
Akl1
Rpn11
Nip1
Prt1
Rpg1
Ded1
Tif35
Cdc9
Chk1
Rad23
Ubi4
Ede1
Ent2
Vps27
Chc1
Ptp3
Stb1
Spa2
Bem2
Bud14
Rck2
Glc7
Ipl1
Cdc5
Dbf4
Pfk26
Tpk1
Gpa2
Ras2
Med6
Rpb10
Paf1
Rpo26
Bdp1
Spt15
Spt5
Rgr1
Srb7
Ssn2
Nop1
Nsr1
Nop56
Snu13
Rps31
Rad9
Rtf1
Rpo21
Npl3
Air2
Fps1
Ask10
Cdc28
Hog1
Pbs2
Ssk2
Clb2
Bud3
Bud4
Msn2
Sch9
Rim15
Yak1
Ypl247c
Vps52
Kss1
Sko1
Slt2
Msg5
Swe1
Igo2
Cdc55
Igo1
Vps53
Vps54
Rtn1
Fmp42
Cot1
Zrc1
10
20
30
40
50
t/s

### Path 1097 (Ipl1 − Pfk26)

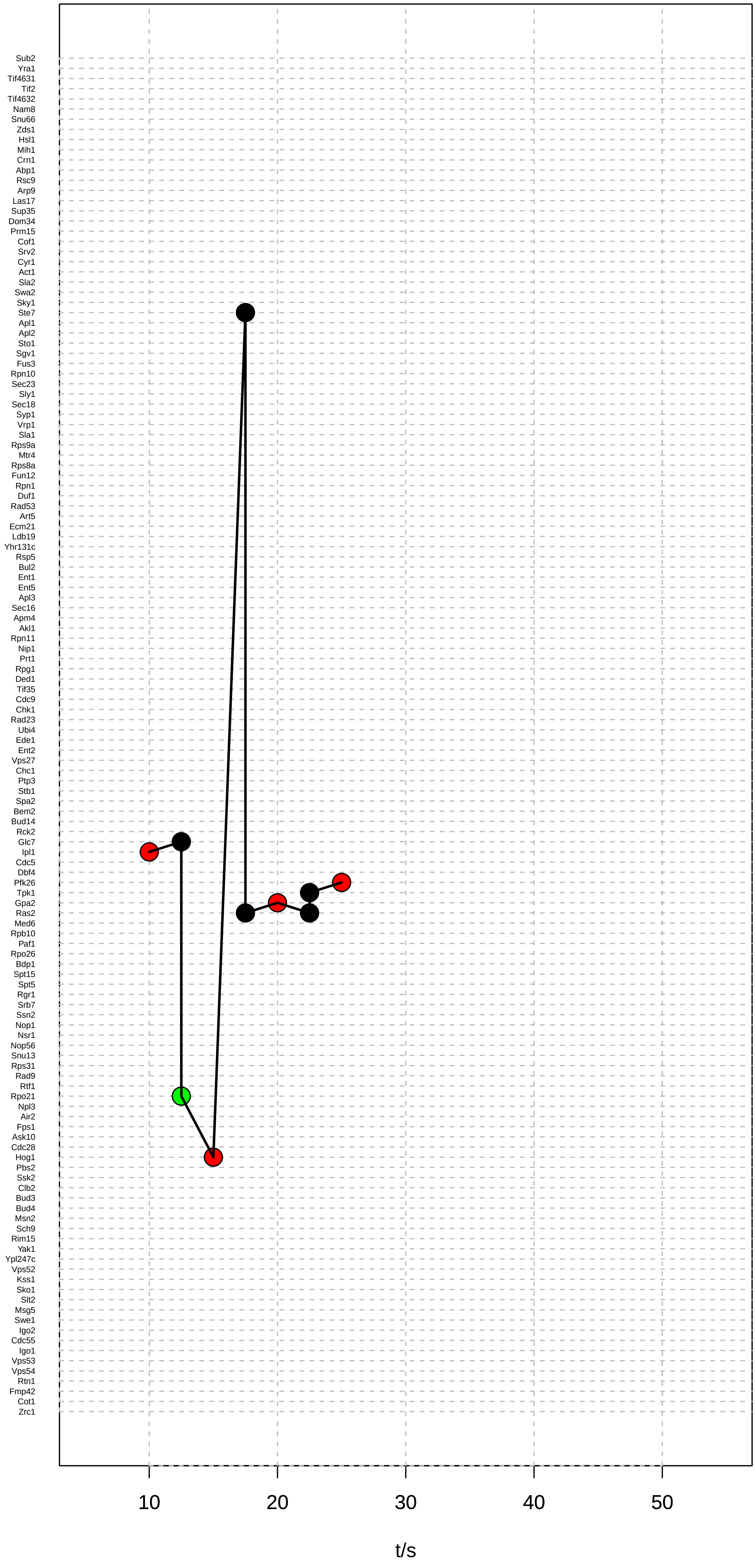

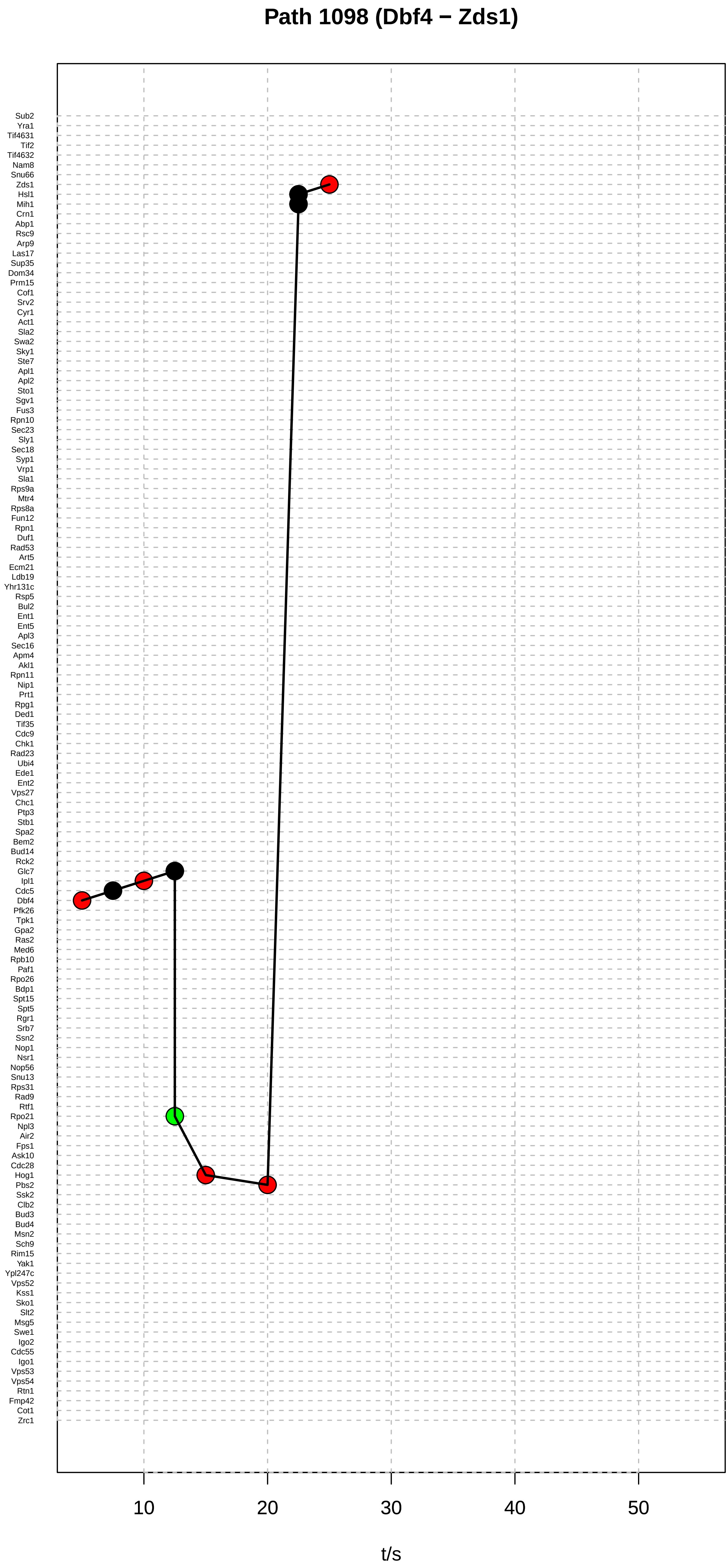

Path 1098 (Dbf4 − Zds1)
Sub2
Yra1
Tif4631
Tif2
Tif4632
Nam8
Snu66
Zds1
Hsl1
Mih1
Crn1
Abp1
Rsc9
Arp9
Las17
Sup35
Dom34
Prm15
Cof1
Srv2
Cyr1
Act1
Sla2
Swa2
Sky1
Ste7
Apl1
Apl2
Sto1
Sgv1
Fus3
Rpn10
Sec23
Sly1
Sec18
Syp1
Vrp1
Sla1
Rps9a
Mtr4
Rps8a
Fun12
Rpn1
Duf1
Rad53
Art5
Ecm21
Ldb19
Yhr131c
Rsp5
Bul2
Ent1
Ent5
Apl3
Sec16
Apm4
Akl1
Rpn11
Nip1
Prt1
Rpg1
Ded1
Tif35
Cdc9
Chk1
Rad23
Ubi4
Ede1
Ent2
Vps27
Chc1
Ptp3
Stb1
Spa2
Bem2
Bud14
Rck2
Glc7
Ipl1
Cdc5
Dbf4
Pfk26
Tpk1
Gpa2
Ras2
Med6
Rpb10
Paf1
Rpo26
Bdp1
Spt15
Spt5
Rgr1
Srb7
Ssn2
Nop1
Nsr1
Nop56
Snu13
Rps31
Rad9
Rtf1
Rpo21
Npl3
Air2
Fps1
Ask10
Cdc28
Hog1
Pbs2
Ssk2
Clb2
Bud3
Bud4
Msn2
Sch9
Rim15
Yak1
Ypl247c
Vps52
Kss1
Sko1
Slt2
Msg5
Swe1
Igo2
Cdc55
Igo1
Vps53
Vps54
Rtn1
Fmp42
Cot1
Zrc1
10
20
30
40
50
t/s

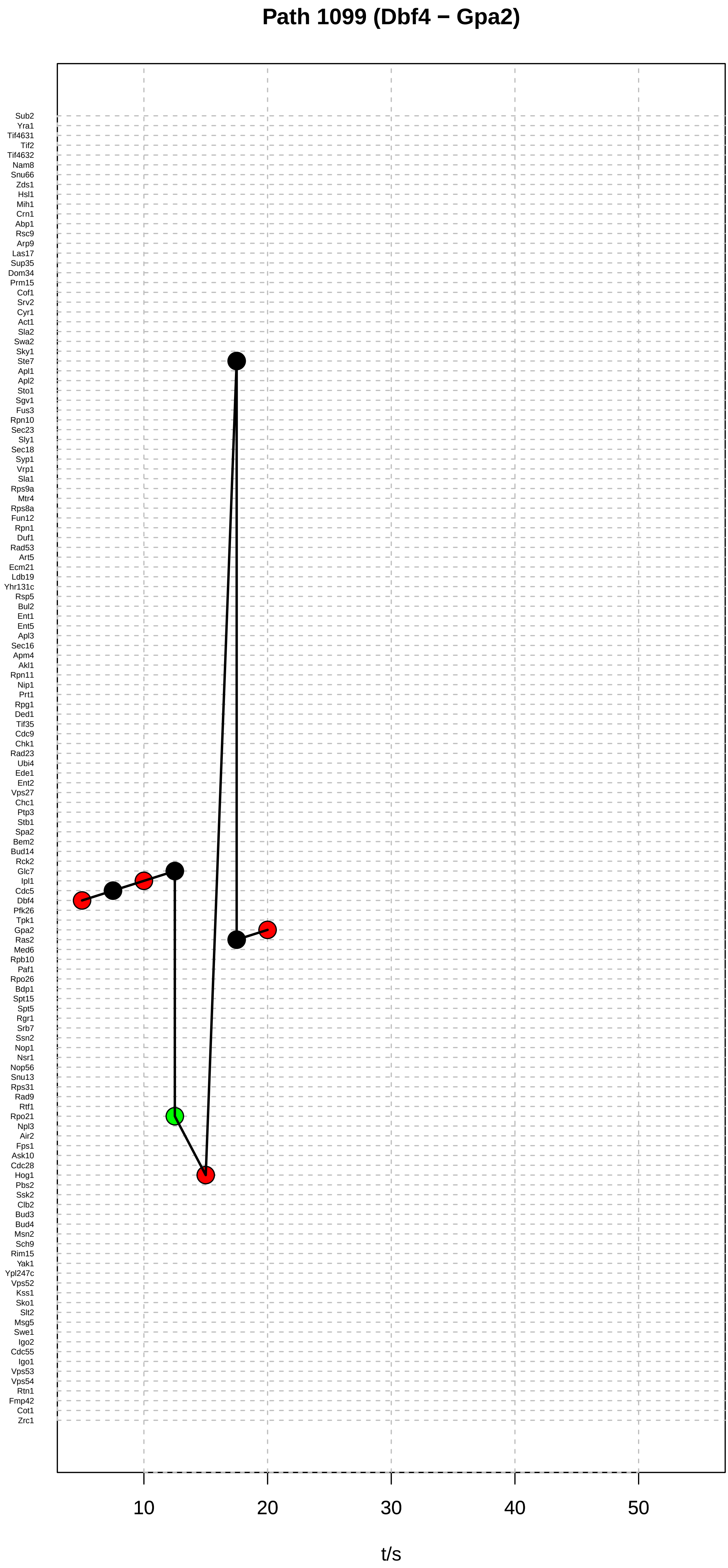

Path 1099 (Dbf4 − Gpa2)
Sub2
Yra1
Tif4631
Tif2
Tif4632
Nam8
Snu66
Zds1
Hsl1
Mih1
Crn1
Abp1
Rsc9
Arp9
Las17
Sup35
Dom34
Prm15
Cof1
Srv2
Cyr1
Act1
Sla2
Swa2
Sky1
Ste7
Apl1
Apl2
Sto1
Sgv1
Fus3
Rpn10
Sec23
Sly1
Sec18
Syp1
Vrp1
Sla1
Rps9a
Mtr4
Rps8a
Fun12
Rpn1
Duf1
Rad53
Art5
Ecm21
Ldb19
Yhr131c
Rsp5
Bul2
Ent1
Ent5
Apl3
Sec16
Apm4
Akl1
Rpn11
Nip1
Prt1
Rpg1
Ded1
Tif35
Cdc9
Chk1
Rad23
Ubi4
Ede1
Ent2
Vps27
Chc1
Ptp3
Stb1
Spa2
Bem2
Bud14
Rck2
Glc7
Ipl1
Cdc5
Dbf4
Pfk26
Tpk1
Gpa2
Ras2
Med6
Rpb10
Paf1
Rpo26
Bdp1
Spt15
Spt5
Rgr1
Srb7
Ssn2
Nop1
Nsr1
Nop56
Snu13
Rps31
Rad9
Rtf1
Rpo21
Npl3
Air2
Fps1
Ask10
Cdc28
Hog1
Pbs2
Ssk2
Clb2
Bud3
Bud4
Msn2
Sch9
Rim15
Yak1
Ypl247c
Vps52
Kss1
Sko1
Slt2
Msg5
Swe1
Igo2
Cdc55
Igo1
Vps53
Vps54
Rtn1
Fmp42
Cot1
Zrc1
10
20
30
40
50
t/s

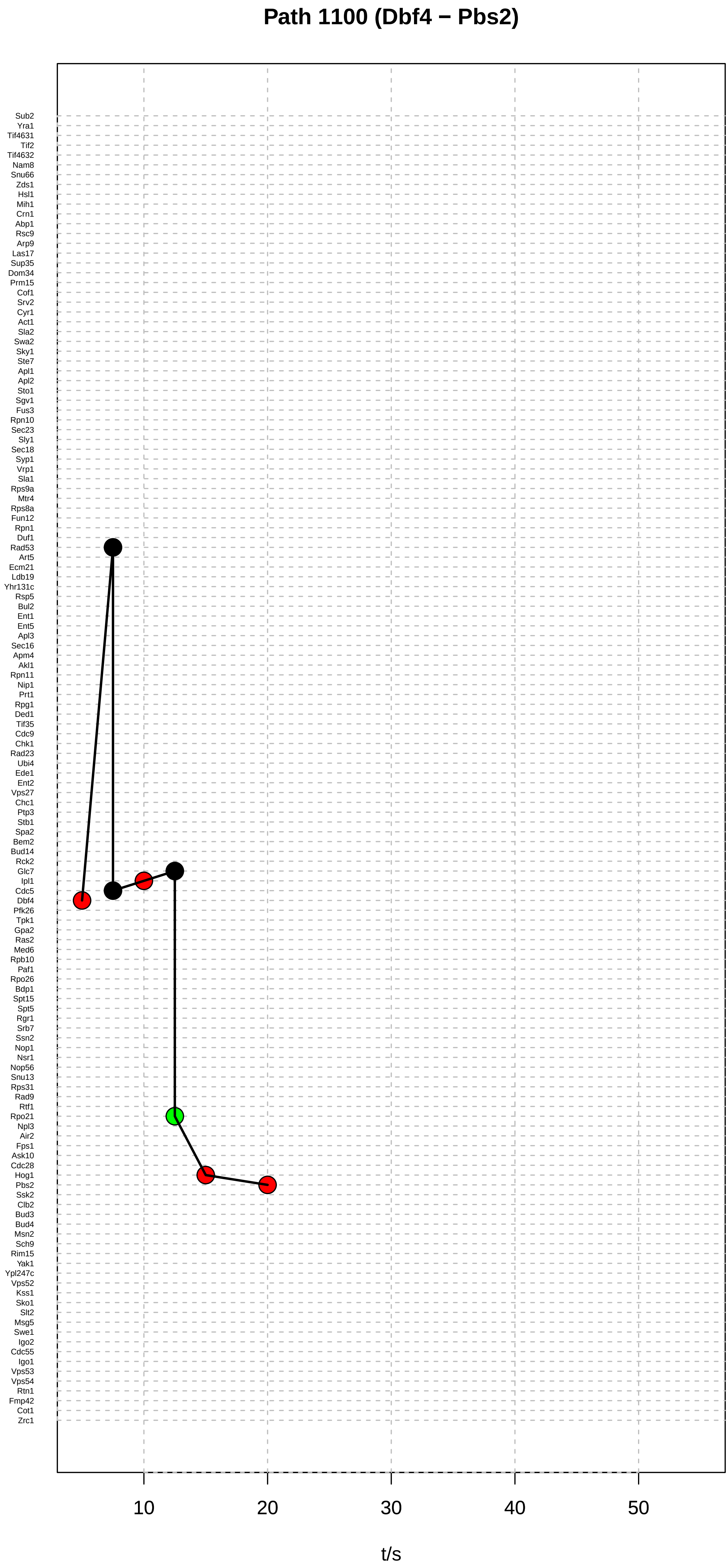
Path 1100 (Dbf4 − Pbs2)
Sub2
Yra1
Tif4631
Tif2
Tif4632
Nam8
Snu66
Zds1
Hsl1
Mih1
Crn1
Abp1
Rsc9
Arp9
Las17
Sup35
Dom34
Prm15
Cof1
Srv2
Cyr1
Act1
Sla2
Swa2
Sky1
Ste7
Apl1
Apl2
Sto1
Sgv1
Fus3
Rpn10
Sec23
Sly1
Sec18
Syp1
Vrp1
Sla1
Rps9a
Mtr4
Rps8a
Fun12
Rpn1
Duf1
Rad53
Art5
Ecm21
Ldb19
Yhr131c
Rsp5
Bul2
Ent1
Ent5
Apl3
Sec16
Apm4
Akl1
Rpn11
Nip1
Prt1
Rpg1
Ded1
Tif35
Cdc9
Chk1
Rad23
Ubi4
Ede1
Ent2
Vps27
Chc1
Ptp3
Stb1
Spa2
Bem2
Bud14
Rck2
Glc7
Ipl1
Cdc5
Dbf4
Pfk26
Tpk1
Gpa2
Ras2
Med6
Rpb10
Paf1
Rpo26
Bdp1
Spt15
Spt5
Rgr1
Srb7
Ssn2
Nop1
Nsr1
Nop56
Snu13
Rps31
Rad9
Rtf1
Rpo21
Npl3
Air2
Fps1
Ask10
Cdc28
Hog1
Pbs2
Ssk2
Clb2
Bud3
Bud4
Msn2
Sch9
Rim15
Yak1
Ypl247c
Vps52
Kss1
Sko1
Slt2
Msg5
Swe1
Igo2
Cdc55
Igo1
Vps53
Vps54
Rtn1
Fmp42
Cot1
Zrc1
10
20
30
40
50
t/s

#### Path 1101 (Dbf4 − Sky1)

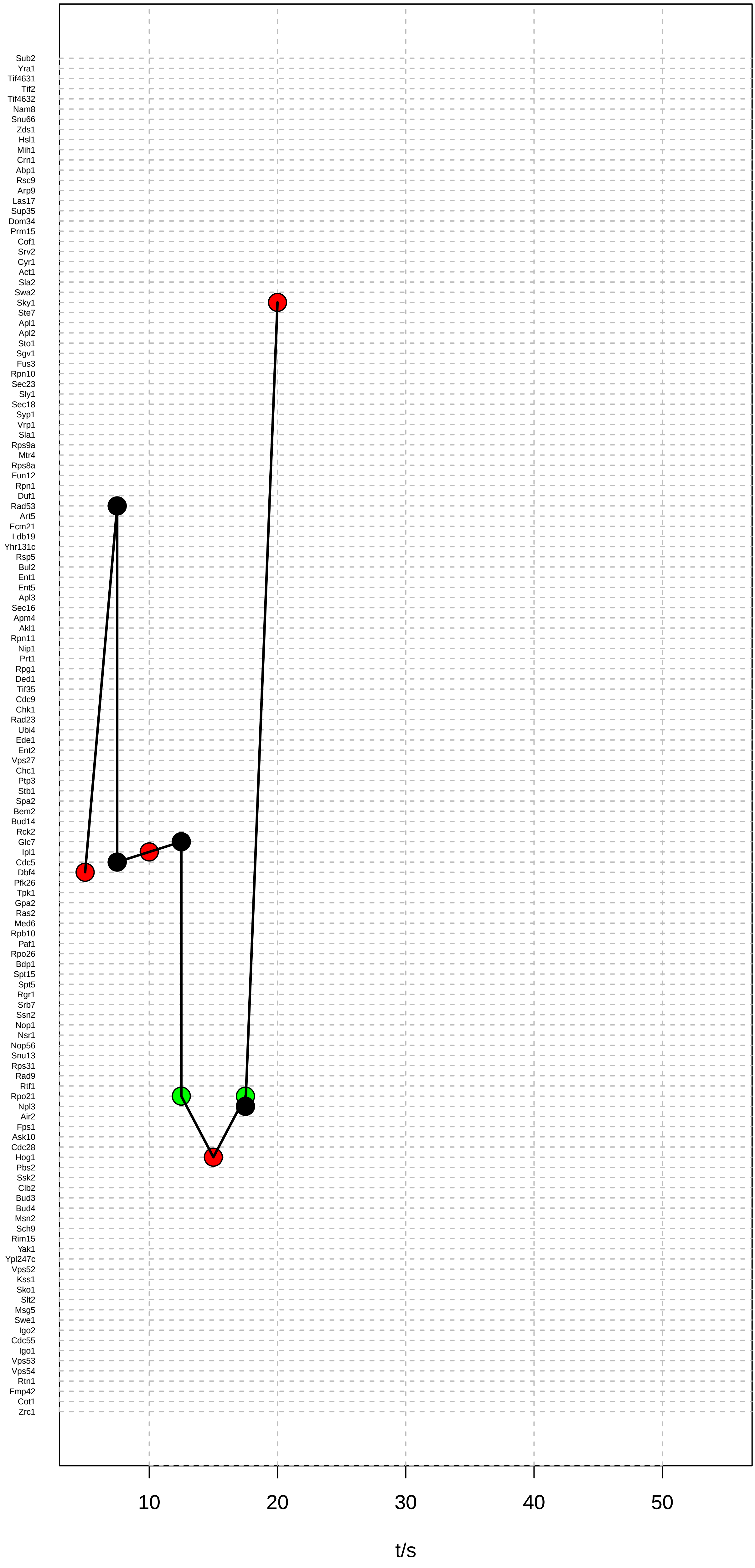

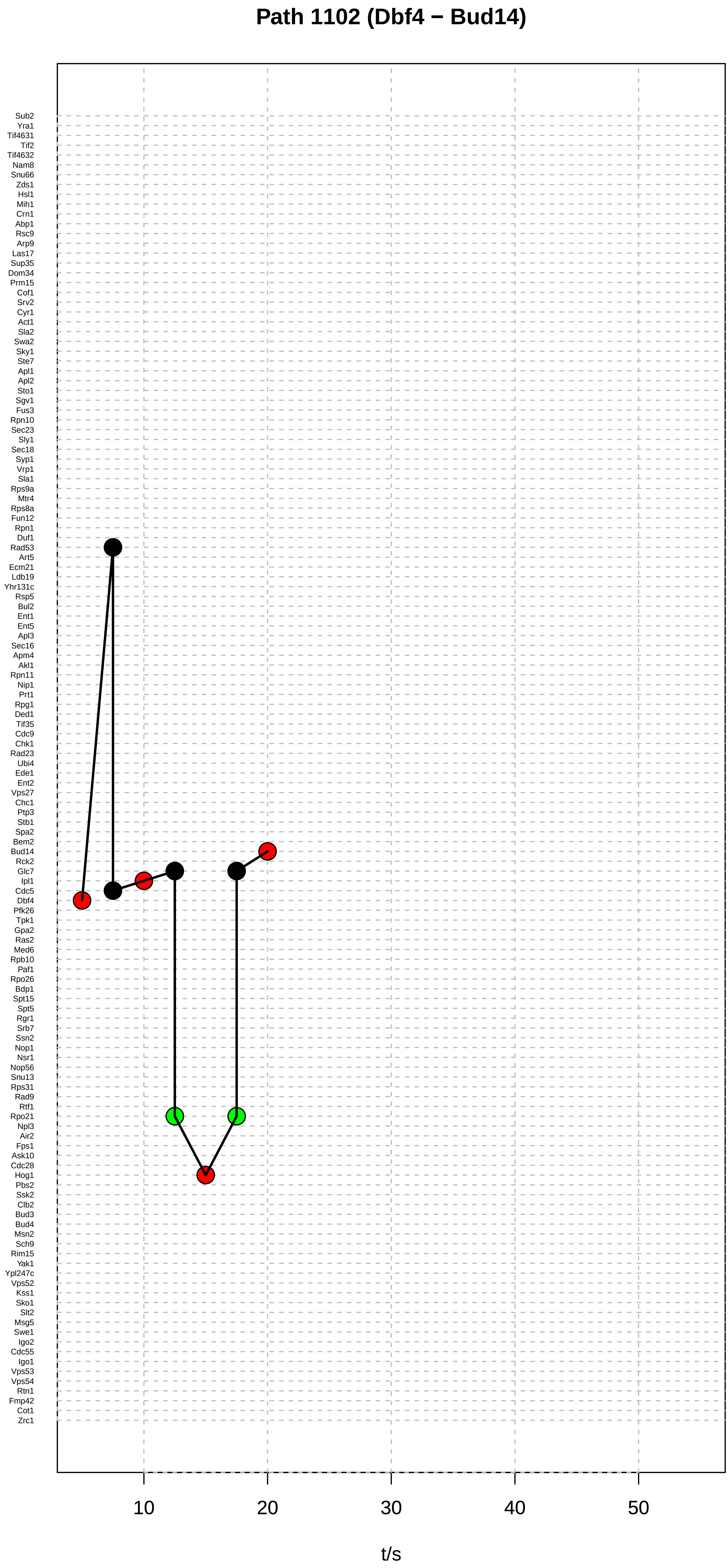

Path 1102 (Dbf4 − Bud14)
Sub2
Yra1
Tif4631
Tif2
Tif4632
Nam8
Snu66
Zds1
Hsl1
Mih1
Crn1
Abp1
Rsc9
Arp9
Las17
Sup35
Dom34
Prm15
Cof1
Srv2
Cyr1
Act1
Sla2
Swa2
Sky1
Ste7
Apl1
Apl2
Sto1
Sgv1
Fus3
Rpn10
Sec23
Sly1
Sec18
Syp1
Vrp1
Sla1
Rps9a
Mtr4
Rps8a
Fun12
Rpn1
Duf1
Rad53
Art5
Ecm21
Ldb19
Yhr131c
Rsp5
Bul2
Ent1
Ent5
Apl3
Sec16
Apm4
Akl1
Rpn11
Nip1
Prt1
Rpg1
Ded1
Tif35
Cdc9
Chk1
Rad23
Ubi4
Ede1
Ent2
Vps27
Chc1
Ptp3
Stb1
Spa2
Bem2
Bud14
Rck2
Glc7
Ipl1
Cdc5
Dbf4
Pfk26
Tpk1
Gpa2
Ras2
Med6
Rpb10
Paf1
Rpo26
Bdp1
Spt15
Spt5
Rgr1
Srb7
Ssn2
Nop1
Nsr1
Nop56
Snu13
Rps31
Rad9
Rtf1
Rpo21
Npl3
Air2
Fps1
Ask10
Cdc28
Hog1
Pbs2
Ssk2
Clb2
Bud3
Bud4
Msn2
Sch9
Rim15
Yak1
Ypl247c
Vps52
Kss1
Sko1
Slt2
Msg5
Swe1
Igo2
Cdc55
Igo1
Vps53
Vps54
Rtn1
Fmp42
Cot1
Zrc1
10
20
30
40
50
t/s

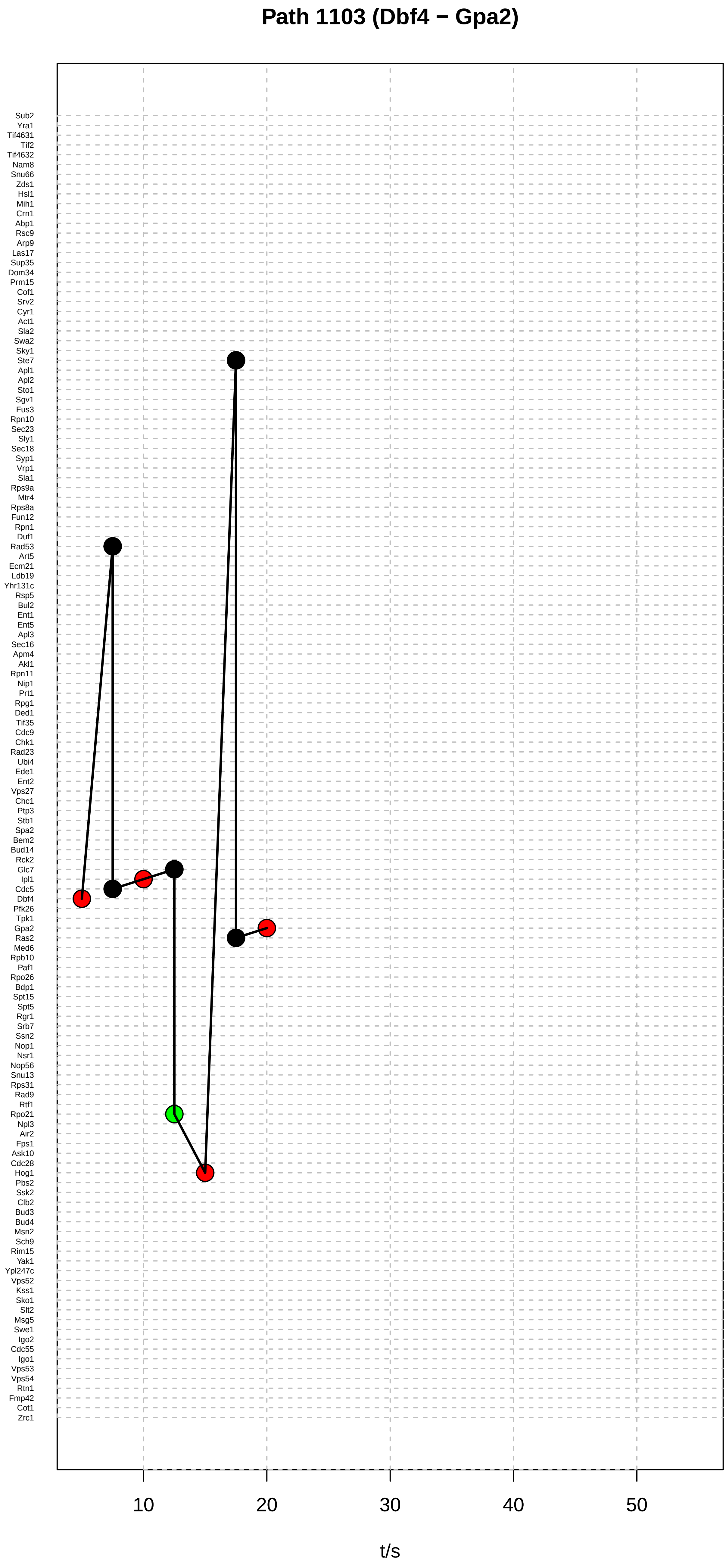
Path 1103 (Dbf4 − Gpa2)
Sub2
Yra1
Tif4631
Tif2
Tif4632
Nam8
Snu66
Zds1
Hsl1
Mih1
Crn1
Abp1
Rsc9
Arp9
Las17
Sup35
Dom34
Prm15
Cof1
Srv2
Cyr1
Act1
Sla2
Swa2
Sky1
Ste7
Apl1
Apl2
Sto1
Sgv1
Fus3
Rpn10
Sec23
Sly1
Sec18
Syp1
Vrp1
Sla1
Rps9a
Mtr4
Rps8a
Fun12
Rpn1
Duf1
Rad53
Art5
Ecm21
Ldb19
Yhr131c
Rsp5
Bul2
Ent1
Ent5
Apl3
Sec16
Apm4
Akl1
Rpn11
Nip1
Prt1
Rpg1
Ded1
Tif35
Cdc9
Chk1
Rad23
Ubi4
Ede1
Ent2
Vps27
Chc1
Ptp3
Stb1
Spa2
Bem2
Bud14
Rck2
Glc7
Ipl1
Cdc5
Dbf4
Pfk26
Tpk1
Gpa2
Ras2
Med6
Rpb10
Paf1
Rpo26
Bdp1
Spt15
Spt5
Rgr1
Srb7
Ssn2
Nop1
Nsr1
Nop56
Snu13
Rps31
Rad9
Rtf1
Rpo21
Npl3
Air2
Fps1
Ask10
Cdc28
Hog1
Pbs2
Ssk2
Clb2
Bud3
Bud4
Msn2
Sch9
Rim15
Yak1
Ypl247c
Vps52
Kss1
Sko1
Slt2
Msg5
Swe1
Igo2
Cdc55
Igo1
Vps53
Vps54
Rtn1
Fmp42
Cot1
Zrc1
10
20
30
40
50
t/s

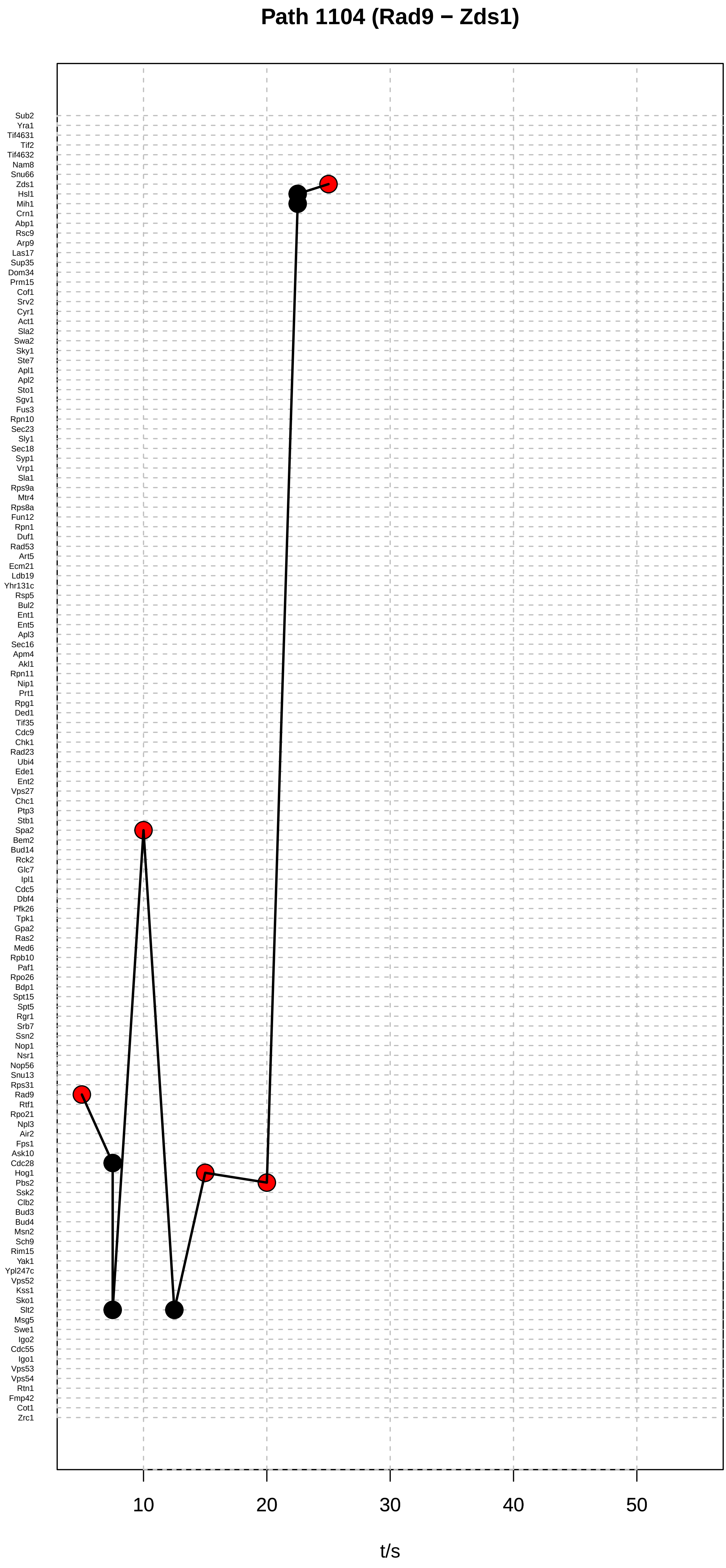
Path 1104 (Rad9 − Zds1)
Sub2
Yra1
Tif4631
Tif2
Tif4632
Nam8
Snu66
Zds1
Hsl1
Mih1
Crn1
Abp1
Rsc9
Arp9
Las17
Sup35
Dom34
Prm15
Cof1
Srv2
Cyr1
Act1
Sla2
Swa2
Sky1
Ste7
Apl1
Apl2
Sto1
Sgv1
Fus3
Rpn10
Sec23
Sly1
Sec18
Syp1
Vrp1
Sla1
Rps9a
Mtr4
Rps8a
Fun12
Rpn1
Duf1
Rad53
Art5
Ecm21
Ldb19
Yhr131c
Rsp5
Bul2
Ent1
Ent5
Apl3
Sec16
Apm4
Akl1
Rpn11
Nip1
Prt1
Rpg1
Ded1
Tif35
Cdc9
Chk1
Rad23
Ubi4
Ede1
Ent2
Vps27
Chc1
Ptp3
Stb1
Spa2
Bem2
Bud14
Rck2
Glc7
Ipl1
Cdc5
Dbf4
Pfk26
Tpk1
Gpa2
Ras2
Med6
Rpb10
Paf1
Rpo26
Bdp1
Spt15
Spt5
Rgr1
Srb7
Ssn2
Nop1
Nsr1
Nop56
Snu13
Rps31
Rad9
Rtf1
Rpo21
Npl3
Air2
Fps1
Ask10
Cdc28
Hog1
Pbs2
Ssk2
Clb2
Bud3
Bud4
Msn2
Sch9
Rim15
Yak1
Ypl247c
Vps52
Kss1
Sko1
Slt2
Msg5
Swe1
Igo2
Cdc55
Igo1
Vps53
Vps54
Rtn1
Fmp42
Cot1
Zrc1
10
20
30
40
50
t/s

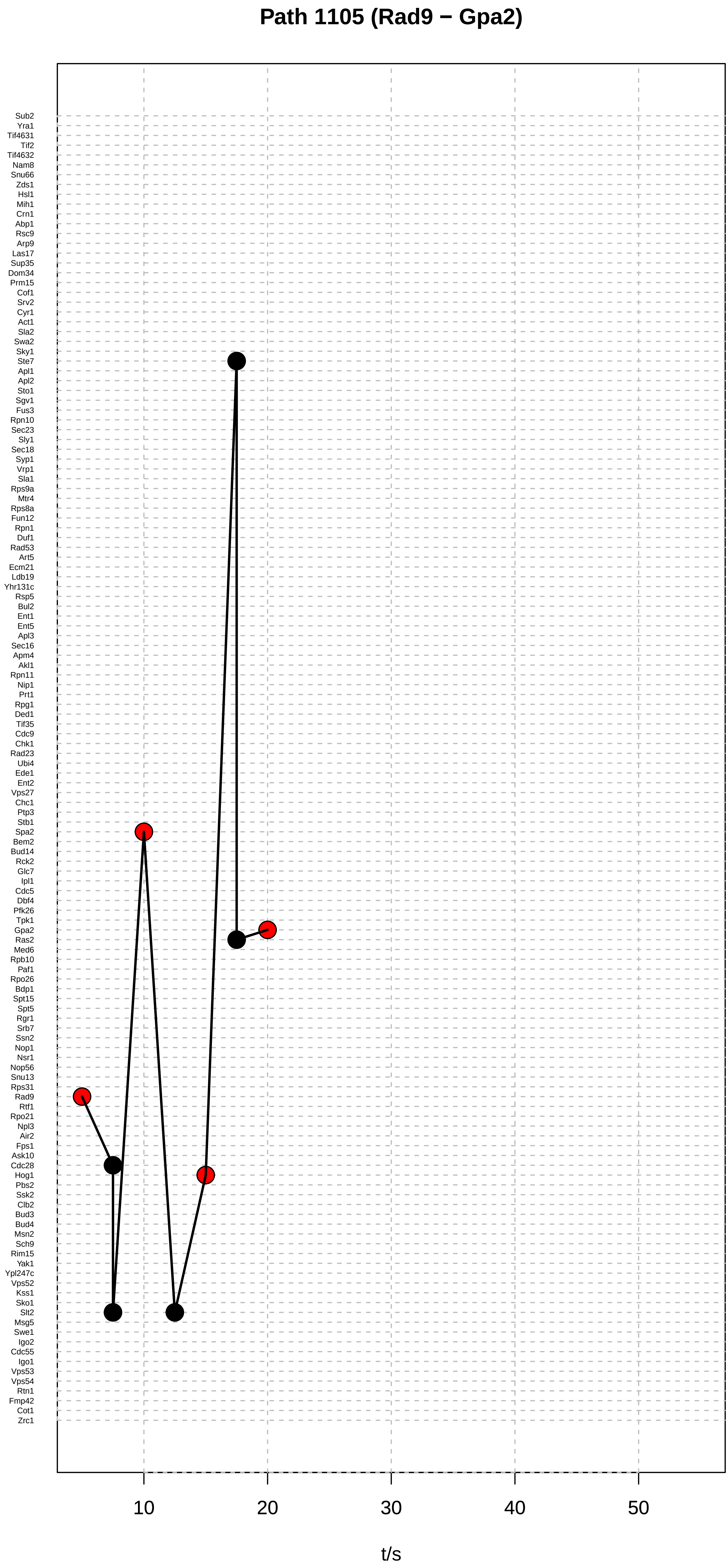
Path 1105 (Rad9 − Gpa2)
Sub2
Yra1
Tif4631
Tif2
Tif4632
Nam8
Snu66
Zds1
Hsl1
Mih1
Crn1
Abp1
Rsc9
Arp9
Las17
Sup35
Dom34
Prm15
Cof1
Srv2
Cyr1
Act1
Sla2
Swa2
Sky1
Ste7
Apl1
Apl2
Sto1
Sgv1
Fus3
Rpn10
Sec23
Sly1
Sec18
Syp1
Vrp1
Sla1
Rps9a
Mtr4
Rps8a
Fun12
Rpn1
Duf1
Rad53
Art5
Ecm21
Ldb19
Yhr131c
Rsp5
Bul2
Ent1
Ent5
Apl3
Sec16
Apm4
Akl1
Rpn11
Nip1
Prt1
Rpg1
Ded1
Tif35
Cdc9
Chk1
Rad23
Ubi4
Ede1
Ent2
Vps27
Chc1
Ptp3
Stb1
Spa2
Bem2
Bud14
Rck2
Glc7
Ipl1
Cdc5
Dbf4
Pfk26
Tpk1
Gpa2
Ras2
Med6
Rpb10
Paf1
Rpo26
Bdp1
Spt15
Spt5
Rgr1
Srb7
Ssn2
Nop1
Nsr1
Nop56
Snu13
Rps31
Rad9
Rtf1
Rpo21
Npl3
Air2
Fps1
Ask10
Cdc28
Hog1
Pbs2
Ssk2
Clb2
Bud3
Bud4
Msn2
Sch9
Rim15
Yak1
Ypl247c
Vps52
Kss1
Sko1
Slt2
Msg5
Swe1
Igo2
Cdc55
Igo1
Vps53
Vps54
Rtn1
Fmp42
Cot1
Zrc1
10
20
30
40
50
t/s

### Path 1106 (Rad9 − Pbs2)

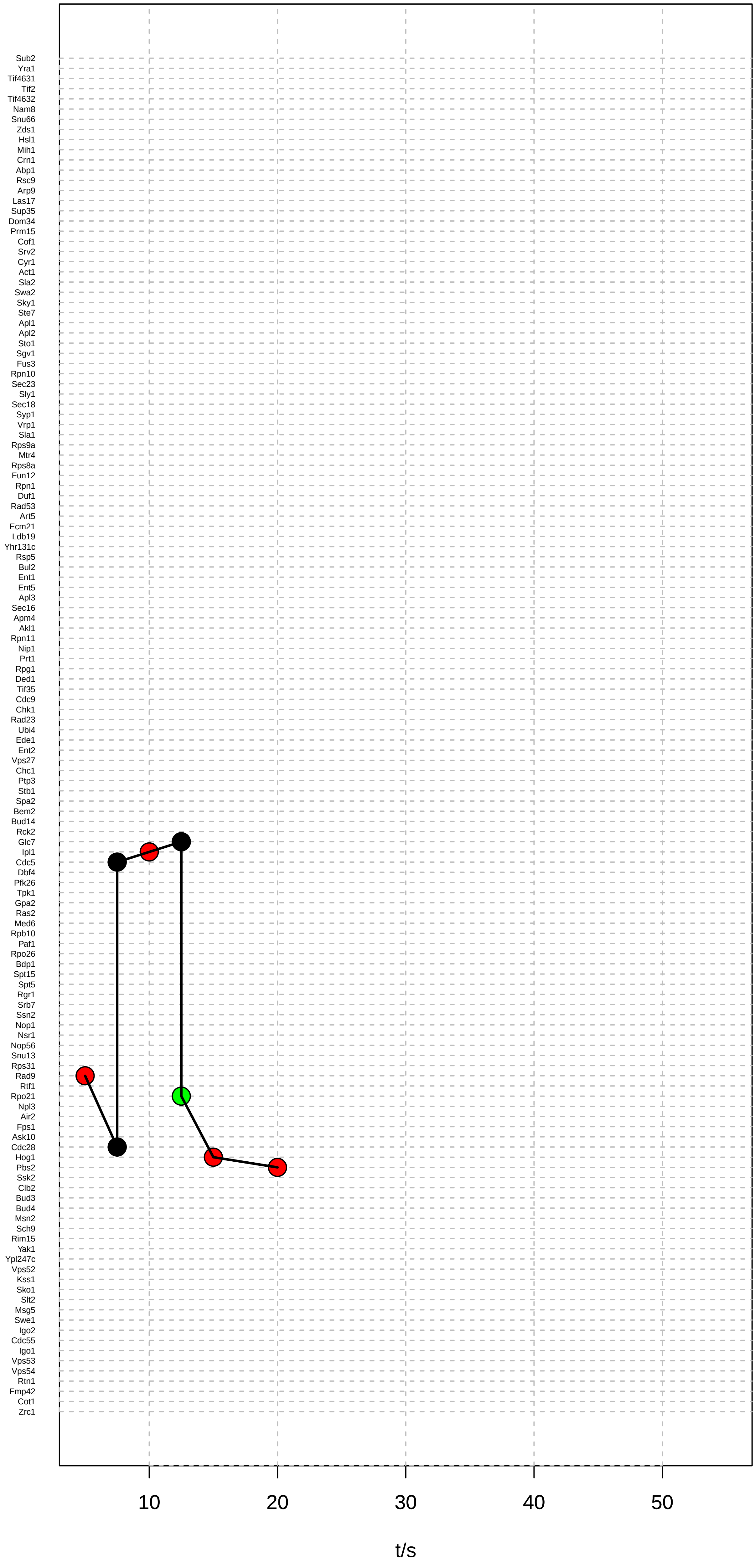

#### Path 1107 (Rad9 − Sky1)

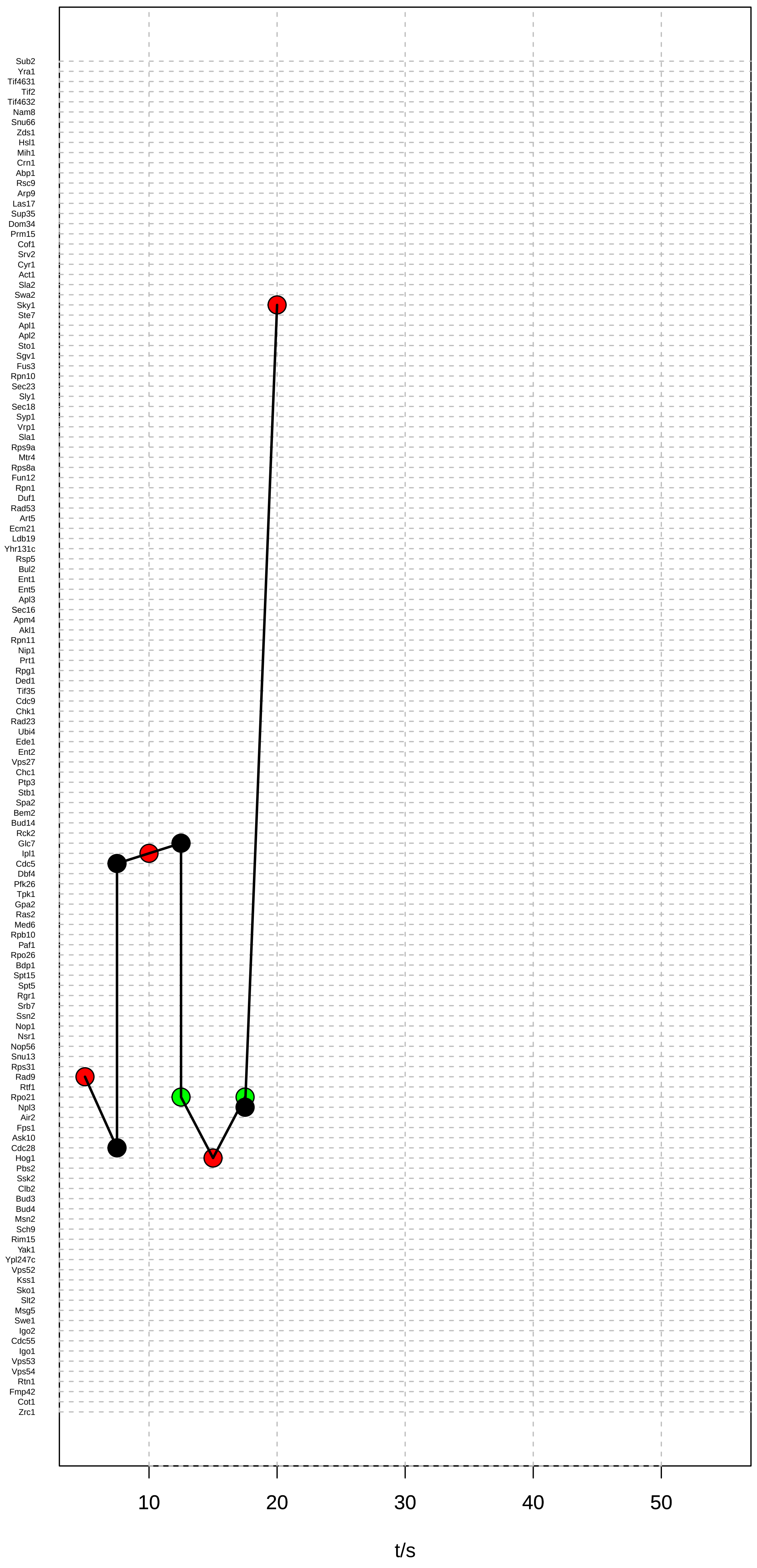

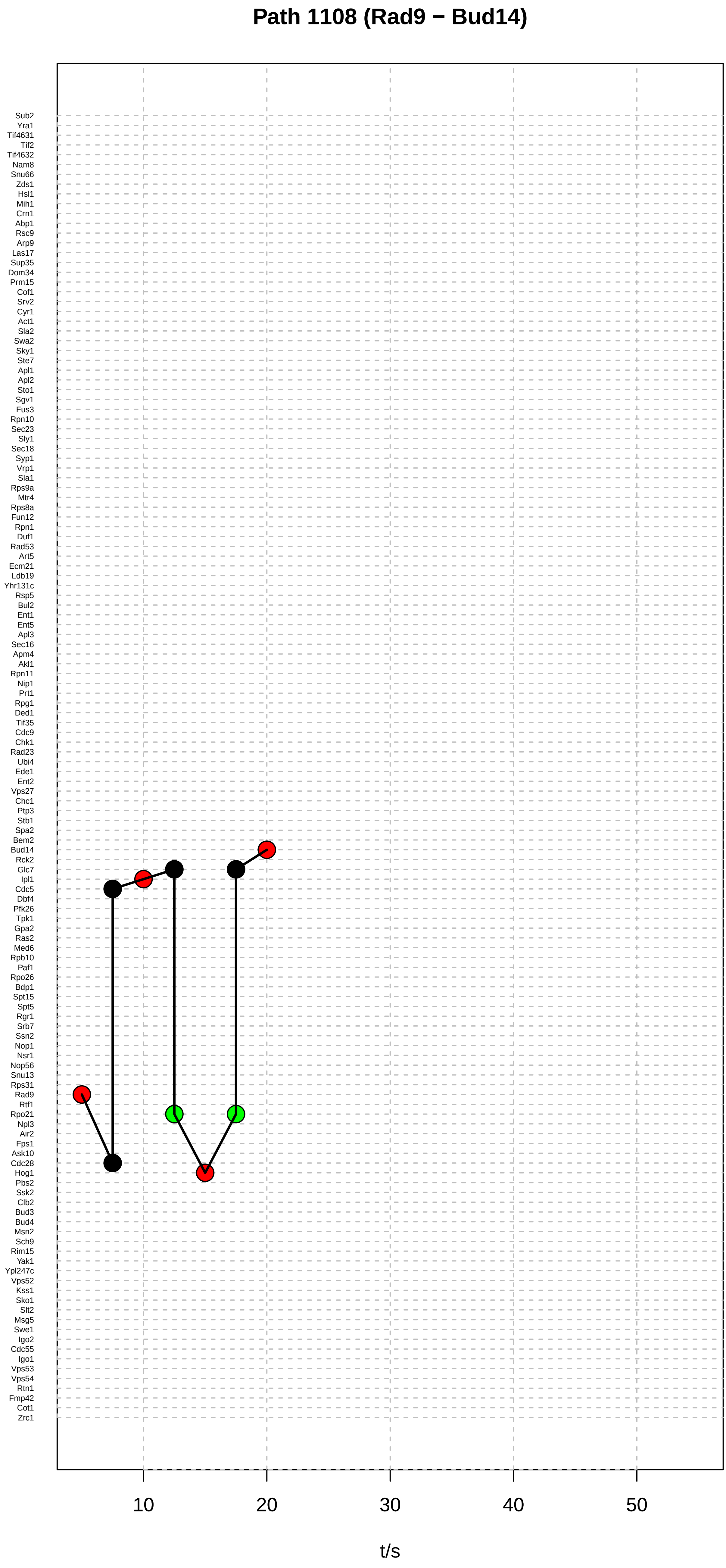

Path 1108 (Rad9 − Bud14)
Sub2
Yra1
Tif4631
Tif2
Tif4632
Nam8
Snu66
Zds1
Hsl1
Mih1
Crn1
Abp1
Rsc9
Arp9
Las17
Sup35
Dom34
Prm15
Cof1
Srv2
Cyr1
Act1
Sla2
Swa2
Sky1
Ste7
Apl1
Apl2
Sto1
Sgv1
Fus3
Rpn10
Sec23
Sly1
Sec18
Syp1
Vrp1
Sla1
Rps9a
Mtr4
Rps8a
Fun12
Rpn1
Duf1
Rad53
Art5
Ecm21
Ldb19
Yhr131c
Rsp5
Bul2
Ent1
Ent5
Apl3
Sec16
Apm4
Akl1
Rpn11
Nip1
Prt1
Rpg1
Ded1
Tif35
Cdc9
Chk1
Rad23
Ubi4
Ede1
Ent2
Vps27
Chc1
Ptp3
Stb1
Spa2
Bem2
Bud14
Rck2
Glc7
Ipl1
Cdc5
Dbf4
Pfk26
Tpk1
Gpa2
Ras2
Med6
Rpb10
Paf1
Rpo26
Bdp1
Spt15
Spt5
Rgr1
Srb7
Ssn2
Nop1
Nsr1
Nop56
Snu13
Rps31
Rad9
Rtf1
Rpo21
Npl3
Air2
Fps1
Ask10
Cdc28
Hog1
Pbs2
Ssk2
Clb2
Bud3
Bud4
Msn2
Sch9
Rim15
Yak1
Ypl247c
Vps52
Kss1
Sko1
Slt2
Msg5
Swe1
Igo2
Cdc55
Igo1
Vps53
Vps54
Rtn1
Fmp42
Cot1
Zrc1
10
20
30
40
50
t/s

### Path 1109 (Rad9 − Gpa2)

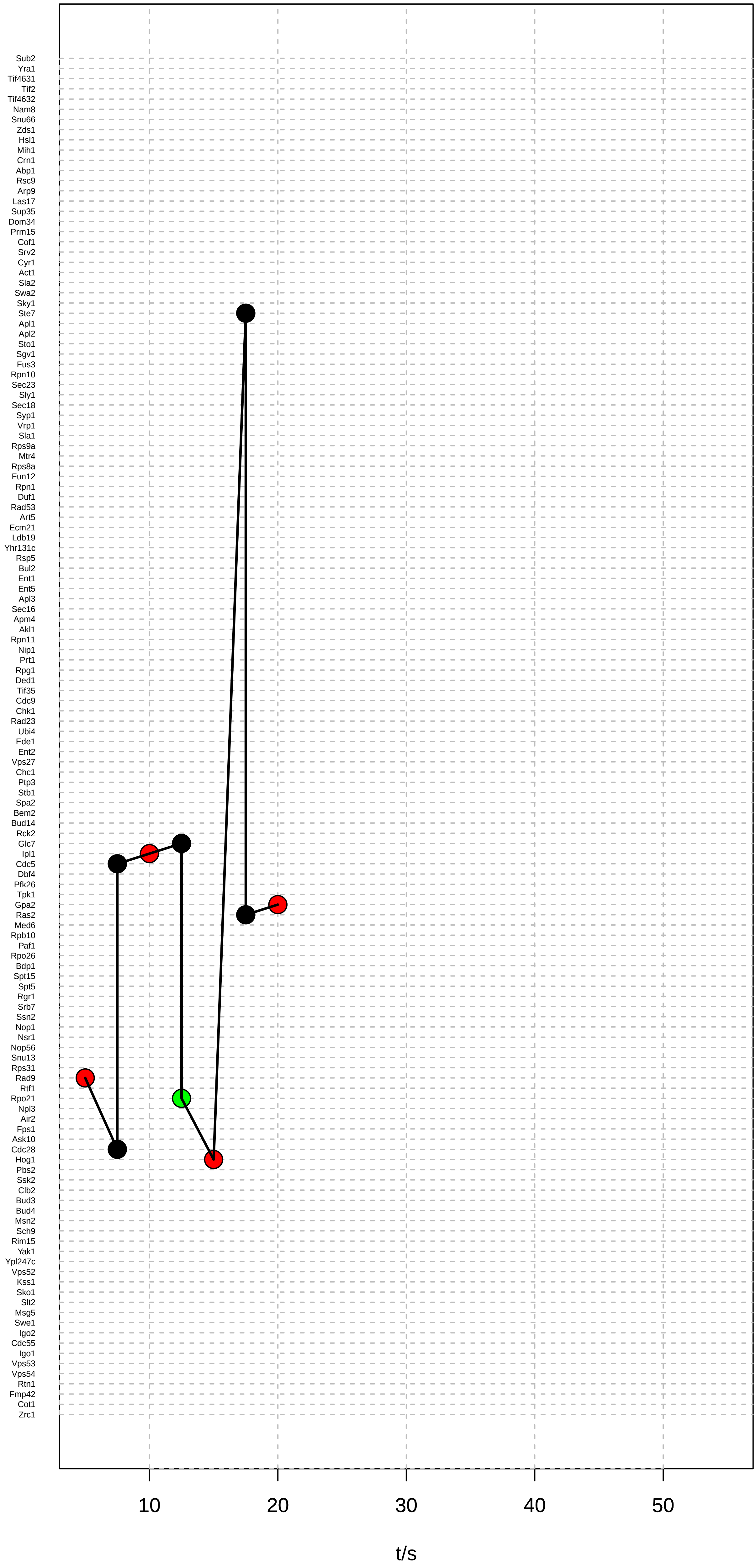

### Path 1110 (Rad9 − Pbs2)

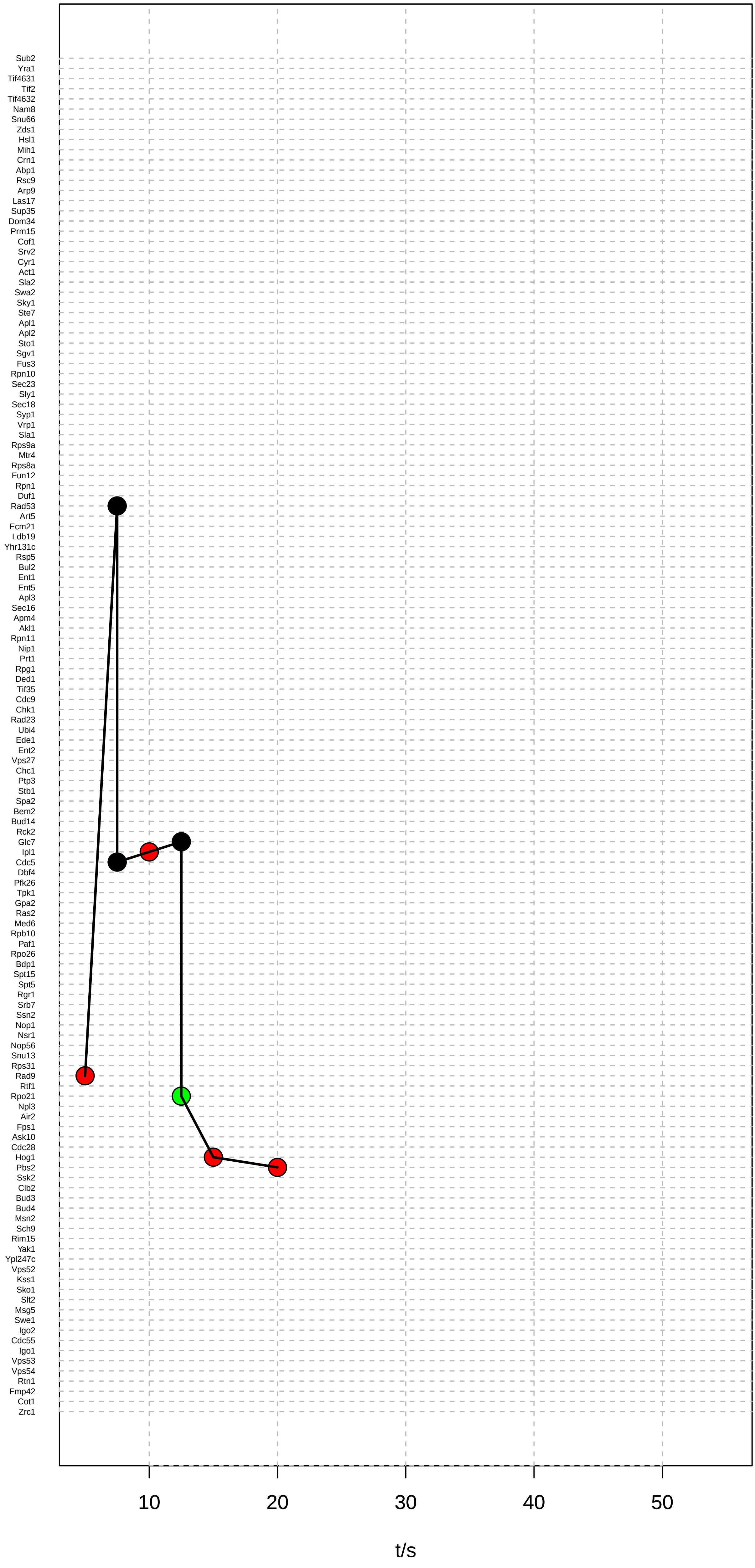

### Path 1111 (Rad9 − Sky1)

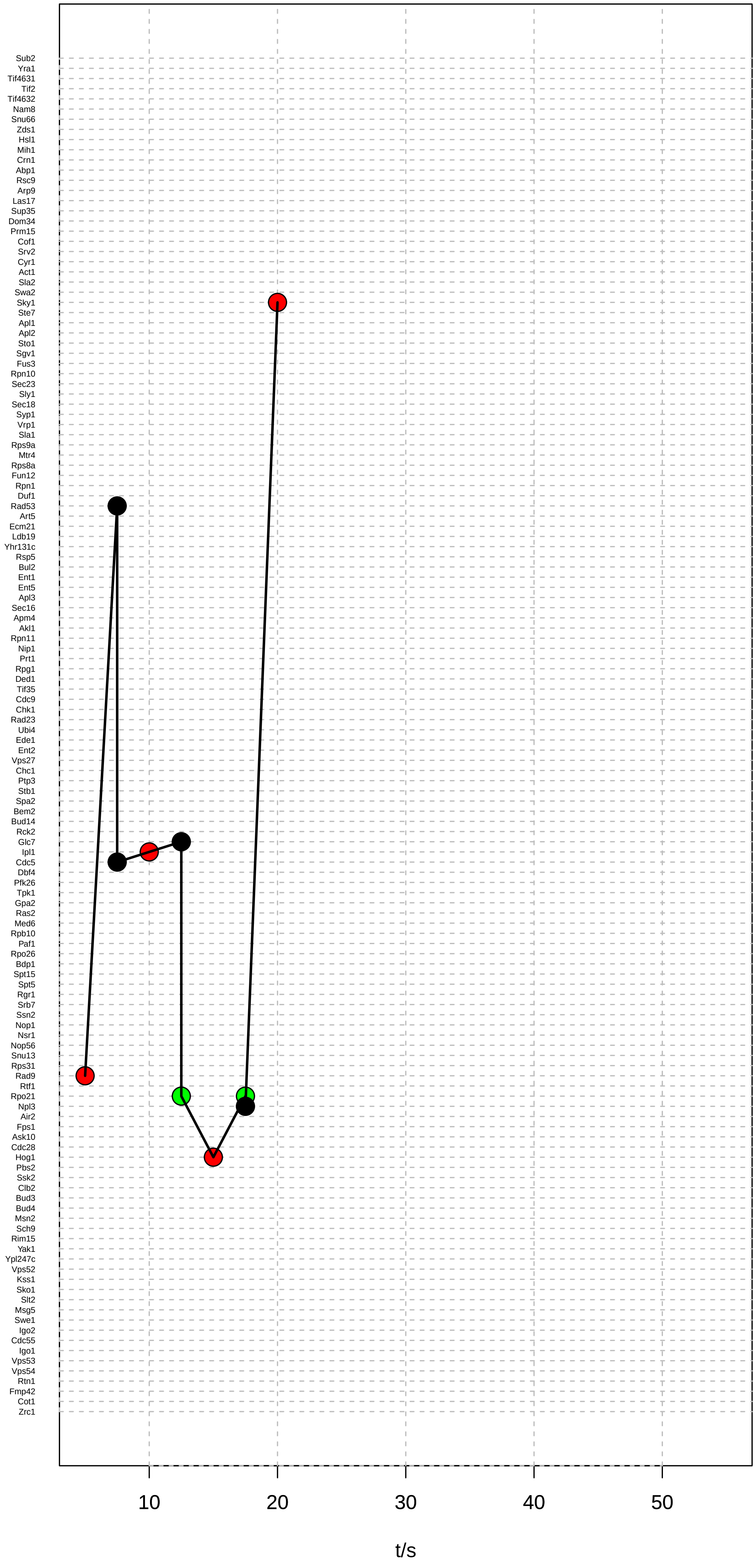

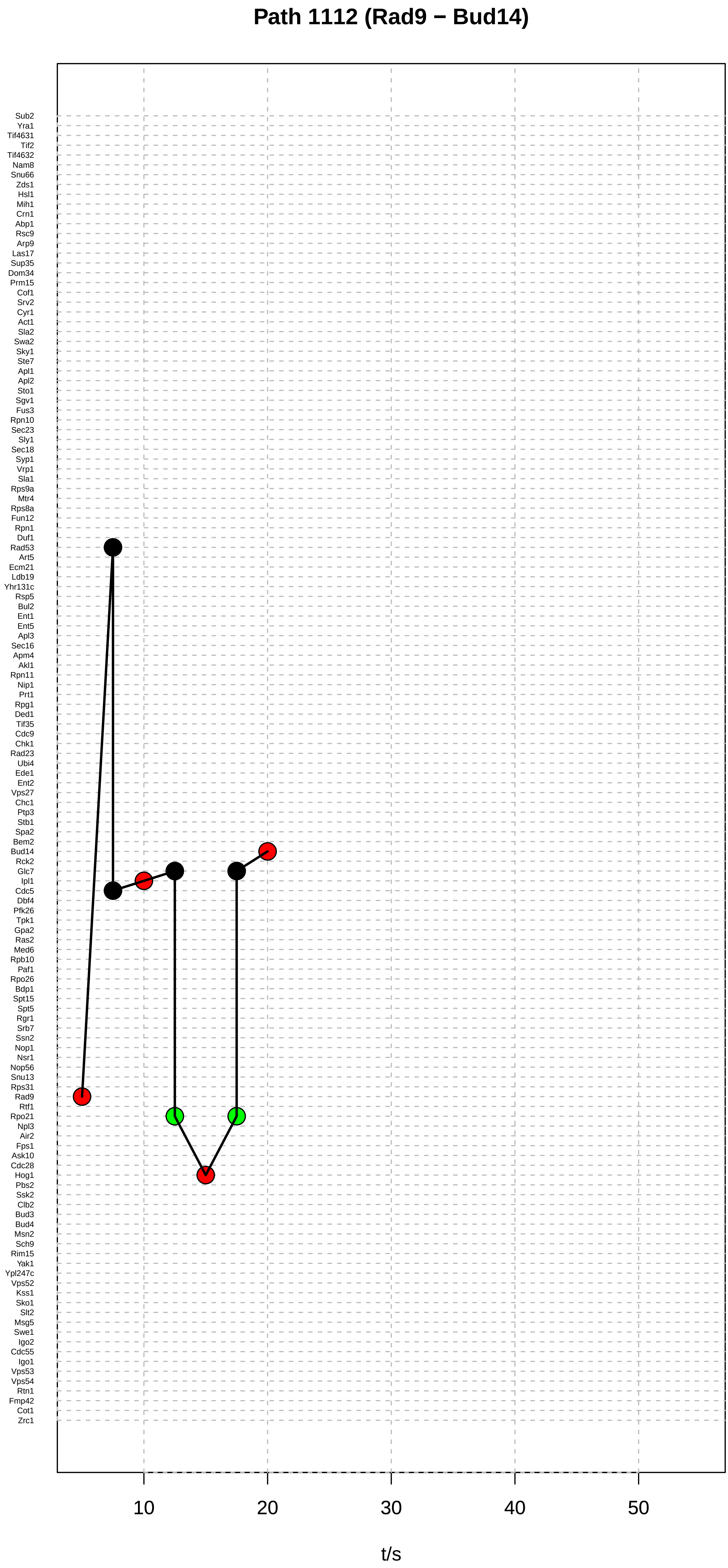
Path 1112 (Rad9 − Bud14)
Sub2
Yra1
Tif4631
Tif2
Tif4632
Nam8
Snu66
Zds1
Hsl1
Mih1
Crn1
Abp1
Rsc9
Arp9
Las17
Sup35
Dom34
Prm15
Cof1
Srv2
Cyr1
Act1
Sla2
Swa2
Sky1
Ste7
Apl1
Apl2
Sto1
Sgv1
Fus3
Rpn10
Sec23
Sly1
Sec18
Syp1
Vrp1
Sla1
Rps9a
Mtr4
Rps8a
Fun12
Rpn1
Duf1
Rad53
Art5
Ecm21
Ldb19
Yhr131c
Rsp5
Bul2
Ent1
Ent5
Apl3
Sec16
Apm4
Akl1
Rpn11
Nip1
Prt1
Rpg1
Ded1
Tif35
Cdc9
Chk1
Rad23
Ubi4
Ede1
Ent2
Vps27
Chc1
Ptp3
Stb1
Spa2
Bem2
Bud14
Rck2
Glc7
Ipl1
Cdc5
Dbf4
Pfk26
Tpk1
Gpa2
Ras2
Med6
Rpb10
Paf1
Rpo26
Bdp1
Spt15
Spt5
Rgr1
Srb7
Ssn2
Nop1
Nsr1
Nop56
Snu13
Rps31
Rad9
Rtf1
Rpo21
Npl3
Air2
Fps1
Ask10
Cdc28
Hog1
Pbs2
Ssk2
Clb2
Bud3
Bud4
Msn2
Sch9
Rim15
Yak1
Ypl247c
Vps52
Kss1
Sko1
Slt2
Msg5
Swe1
Igo2
Cdc55
Igo1
Vps53
Vps54
Rtn1
Fmp42
Cot1
Zrc1
10
20
30
40
50
t/s

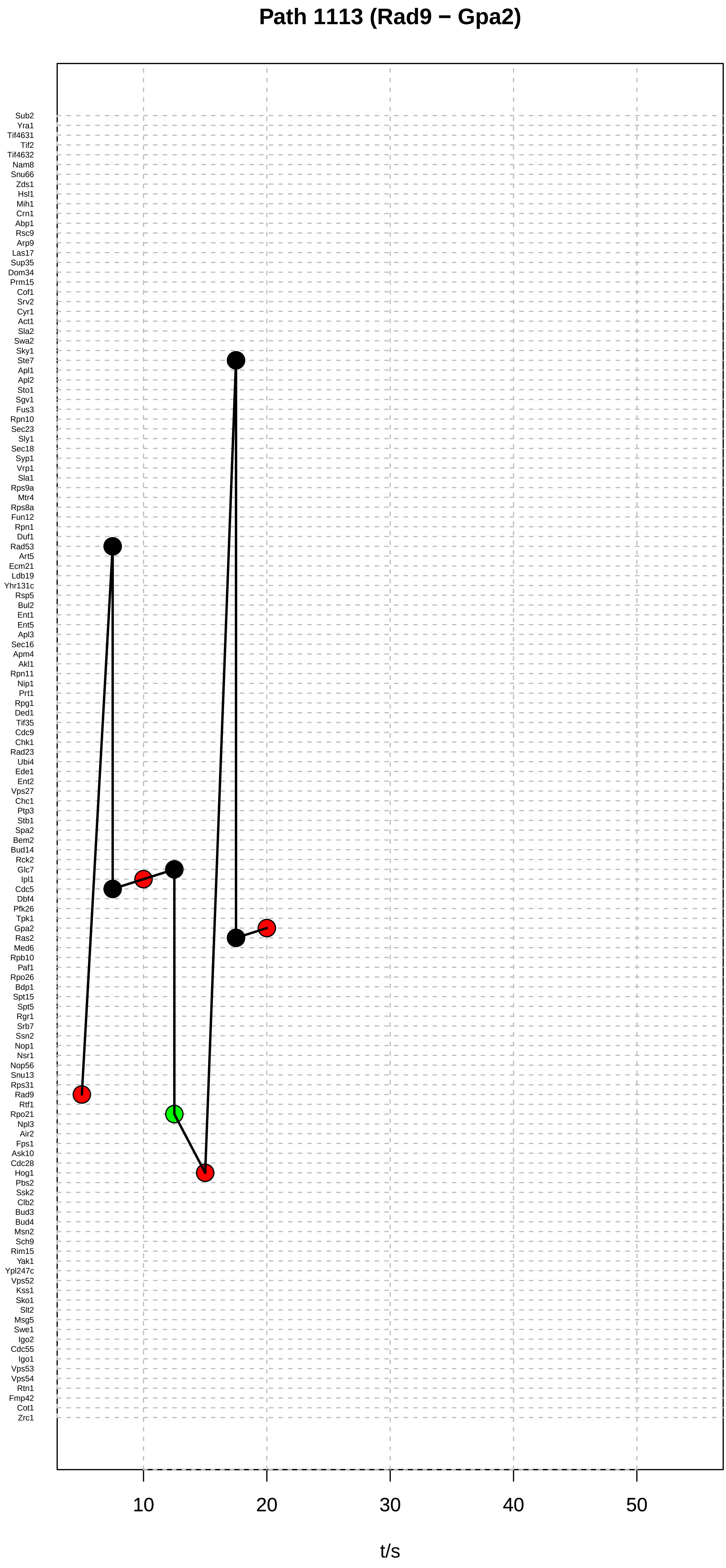

Path 1113 (Rad9 − Gpa2)
Sub2
Yra1
Tif4631
Tif2
Tif4632
Nam8
Snu66
Zds1
Hsl1
Mih1
Crn1
Abp1
Rsc9
Arp9
Las17
Sup35
Dom34
Prm15
Cof1
Srv2
Cyr1
Act1
Sla2
Swa2
Sky1
Ste7
Apl1
Apl2
Sto1
Sgv1
Fus3
Rpn10
Sec23
Sly1
Sec18
Syp1
Vrp1
Sla1
Rps9a
Mtr4
Rps8a
Fun12
Rpn1
Duf1
Rad53
Art5
Ecm21
Ldb19
Yhr131c
Rsp5
Bul2
Ent1
Ent5
Apl3
Sec16
Apm4
Akl1
Rpn11
Nip1
Prt1
Rpg1
Ded1
Tif35
Cdc9
Chk1
Rad23
Ubi4
Ede1
Ent2
Vps27
Chc1
Ptp3
Stb1
Spa2
Bem2
Bud14
Rck2
Glc7
Ipl1
Cdc5
Dbf4
Pfk26
Tpk1
Gpa2
Ras2
Med6
Rpb10
Paf1
Rpo26
Bdp1
Spt15
Spt5
Rgr1
Srb7
Ssn2
Nop1
Nsr1
Nop56
Snu13
Rps31
Rad9
Rtf1
Rpo21
Npl3
Air2
Fps1
Ask10
Cdc28
Hog1
Pbs2
Ssk2
Clb2
Bud3
Bud4
Msn2
Sch9
Rim15
Yak1
Ypl247c
Vps52
Kss1
Sko1
Slt2
Msg5
Swe1
Igo2
Cdc55
Igo1
Vps53
Vps54
Rtn1
Fmp42
Cot1
Zrc1
10
20
30
40
50
t/s

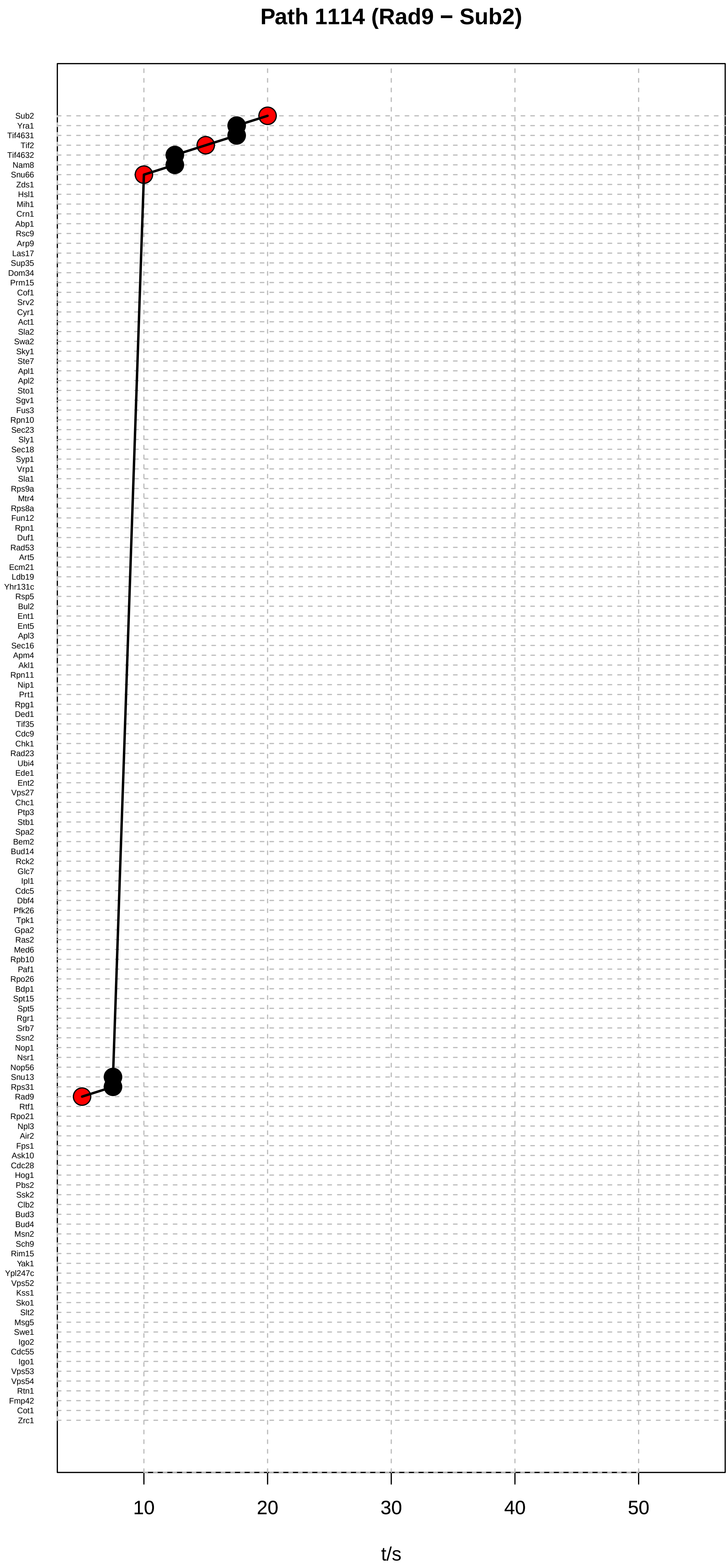

Path 1114 (Rad9 − Sub2)
Sub2
Yra1
Tif4631
Tif2
Tif4632
Nam8
Snu66
Zds1
Hsl1
Mih1
Crn1
Abp1
Rsc9
Arp9
Las17
Sup35
Dom34
Prm15
Cof1
Srv2
Cyr1
Act1
Sla2
Swa2
Sky1
Ste7
Apl1
Apl2
Sto1
Sgv1
Fus3
Rpn10
Sec23
Sly1
Sec18
Syp1
Vrp1
Sla1
Rps9a
Mtr4
Rps8a
Fun12
Rpn1
Duf1
Rad53
Art5
Ecm21
Ldb19
Yhr131c
Rsp5
Bul2
Ent1
Ent5
Apl3
Sec16
Apm4
Akl1
Rpn11
Nip1
Prt1
Rpg1
Ded1
Tif35
Cdc9
Chk1
Rad23
Ubi4
Ede1
Ent2
Vps27
Chc1
Ptp3
Stb1
Spa2
Bem2
Bud14
Rck2
Glc7
Ipl1
Cdc5
Dbf4
Pfk26
Tpk1
Gpa2
Ras2
Med6
Rpb10
Paf1
Rpo26
Bdp1
Spt15
Spt5
Rgr1
Srb7
Ssn2
Nop1
Nsr1
Nop56
Snu13
Rps31
Rad9
Rtf1
Rpo21
Npl3
Air2
Fps1
Ask10
Cdc28
Hog1
Pbs2
Ssk2
Clb2
Bud3
Bud4
Msn2
Sch9
Rim15
Yak1
Ypl247c
Vps52
Kss1
Sko1
Slt2
Msg5
Swe1
Igo2
Cdc55
Igo1
Vps53
Vps54
Rtn1
Fmp42
Cot1
Zrc1
10
20
30
40
50
t/s

Path 1115 (Rad9 − Sky1)

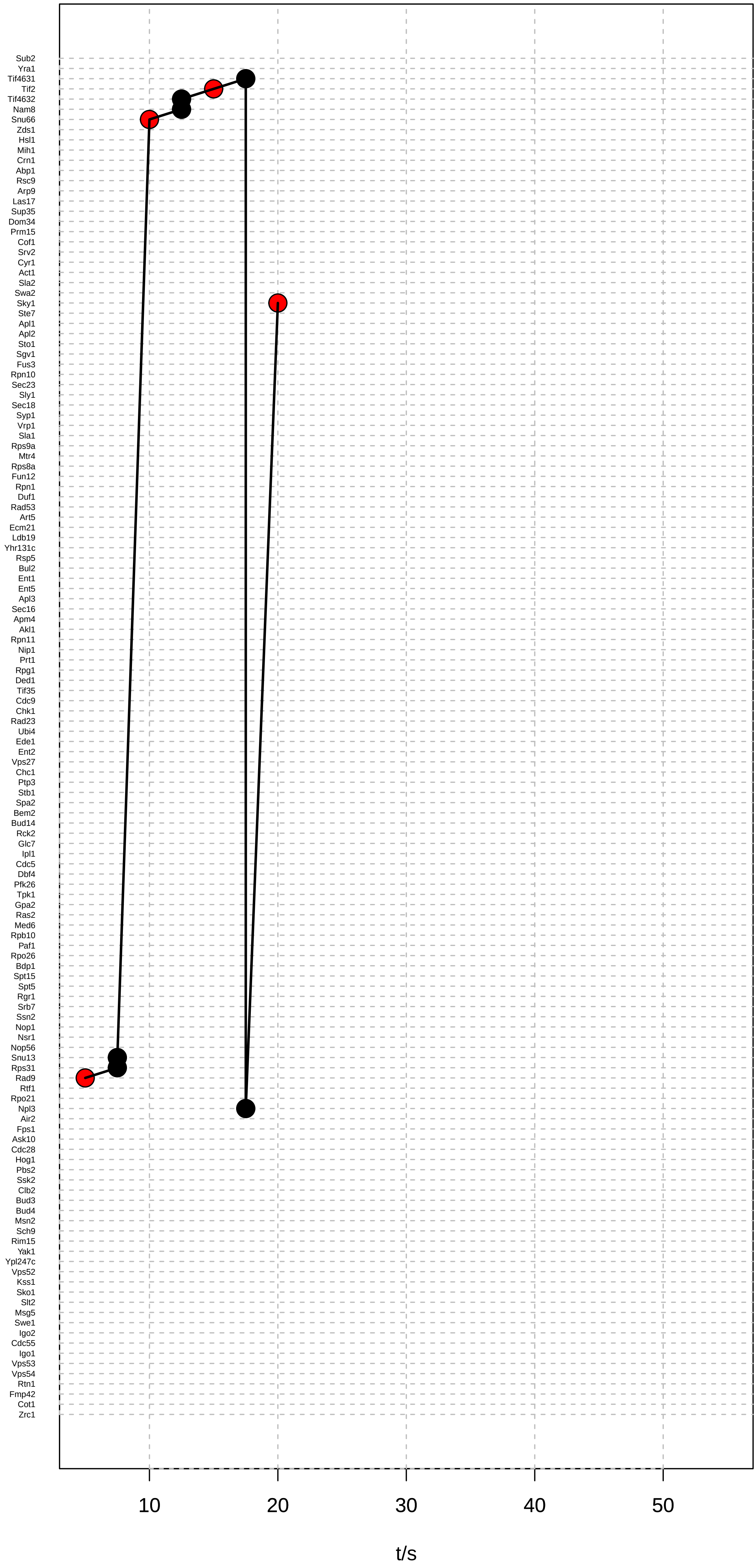

### Path 1116 (Rpo21 − Zds1)

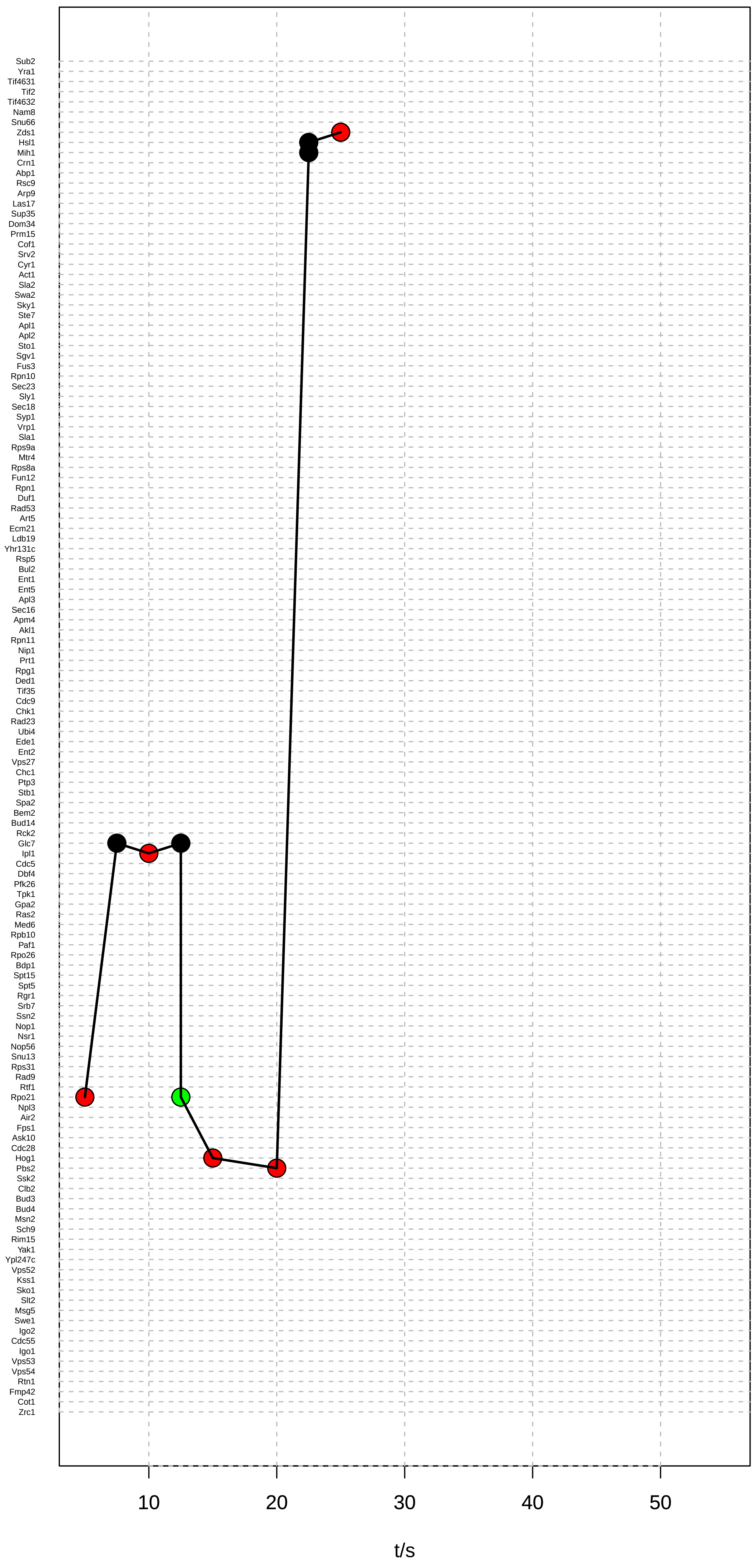

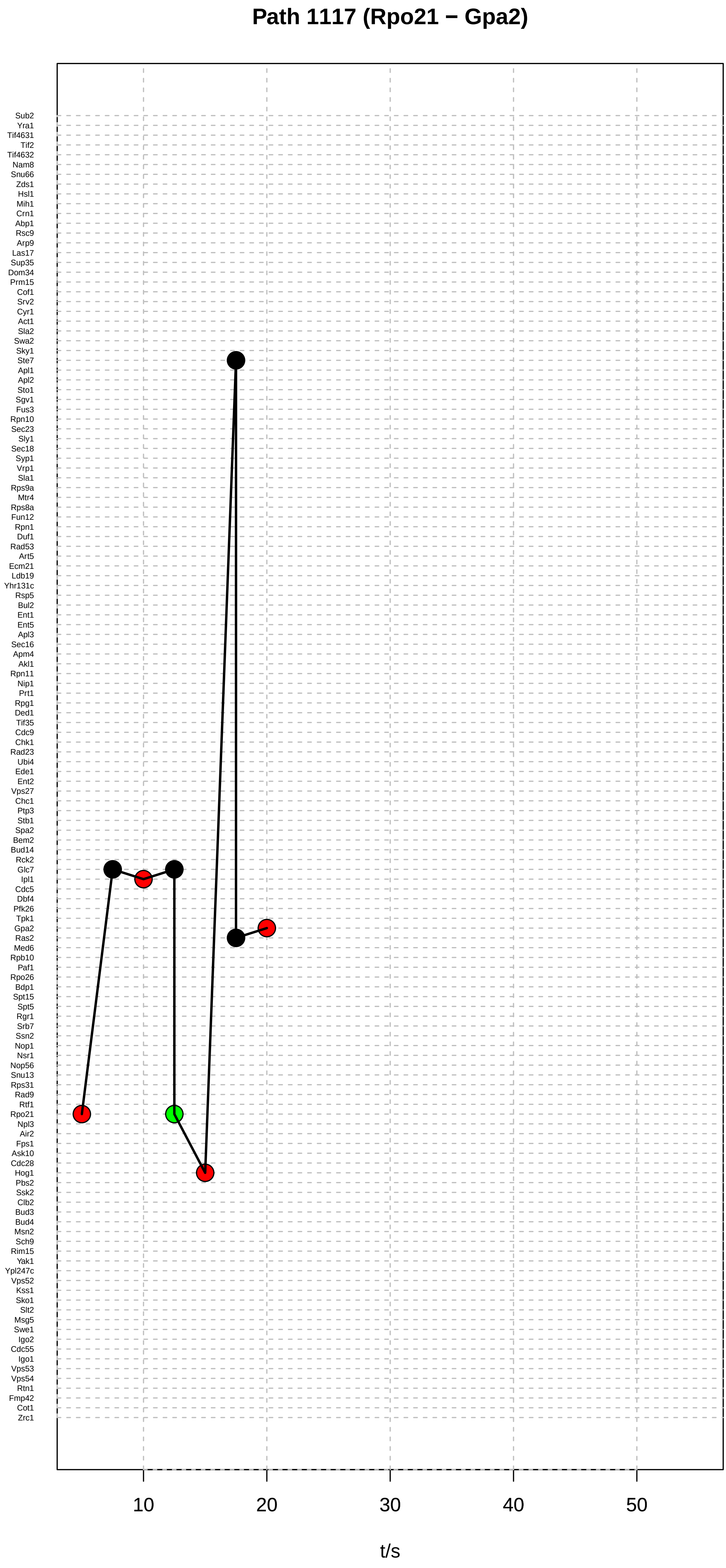
Path 1117 (Rpo21 − Gpa2)
Sub2
Yra1
Tif4631
Tif2
Tif4632
Nam8
Snu66
Zds1
Hsl1
Mih1
Crn1
Abp1
Rsc9
Arp9
Las17
Sup35
Dom34
Prm15
Cof1
Srv2
Cyr1
Act1
Sla2
Swa2
Sky1
Ste7
Apl1
Apl2
Sto1
Sgv1
Fus3
Rpn10
Sec23
Sly1
Sec18
Syp1
Vrp1
Sla1
Rps9a
Mtr4
Rps8a
Fun12
Rpn1
Duf1
Rad53
Art5
Ecm21
Ldb19
Yhr131c
Rsp5
Bul2
Ent1
Ent5
Apl3
Sec16
Apm4
Akl1
Rpn11
Nip1
Prt1
Rpg1
Ded1
Tif35
Cdc9
Chk1
Rad23
Ubi4
Ede1
Ent2
Vps27
Chc1
Ptp3
Stb1
Spa2
Bem2
Bud14
Rck2
Glc7
Ipl1
Cdc5
Dbf4
Pfk26
Tpk1
Gpa2
Ras2
Med6
Rpb10
Paf1
Rpo26
Bdp1
Spt15
Spt5
Rgr1
Srb7
Ssn2
Nop1
Nsr1
Nop56
Snu13
Rps31
Rad9
Rtf1
Rpo21
Npl3
Air2
Fps1
Ask10
Cdc28
Hog1
Pbs2
Ssk2
Clb2
Bud3
Bud4
Msn2
Sch9
Rim15
Yak1
Ypl247c
Vps52
Kss1
Sko1
Slt2
Msg5
Swe1
Igo2
Cdc55
Igo1
Vps53
Vps54
Rtn1
Fmp42
Cot1
Zrc1
10
20
30
40
50
t/s

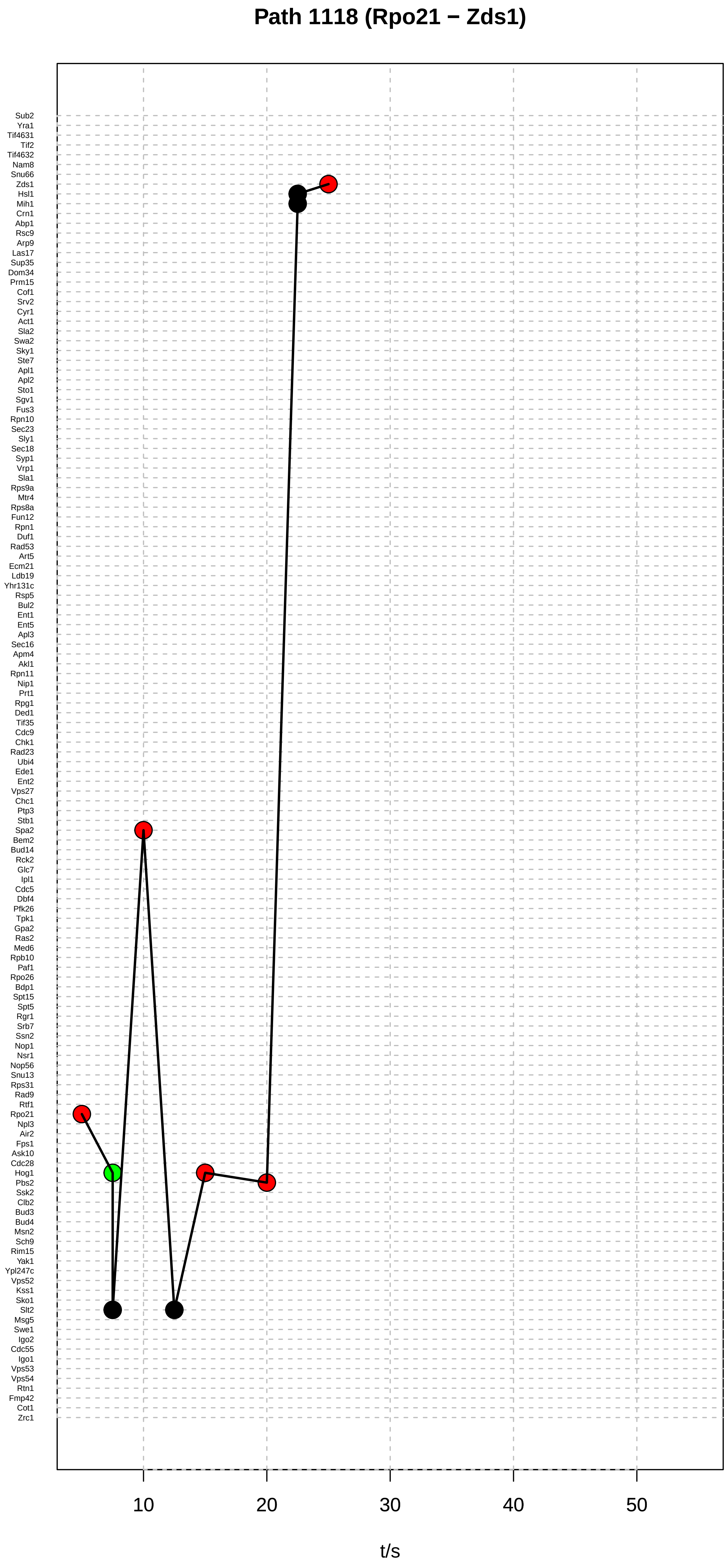

Path 1118 (Rpo21 − Zds1)
Sub2
Yra1
Tif4631
Tif2
Tif4632
Nam8
Snu66
Zds1
Hsl1
Mih1
Crn1
Abp1
Rsc9
Arp9
Las17
Sup35
Dom34
Prm15
Cof1
Srv2
Cyr1
Act1
Sla2
Swa2
Sky1
Ste7
Apl1
Apl2
Sto1
Sgv1
Fus3
Rpn10
Sec23
Sly1
Sec18
Syp1
Vrp1
Sla1
Rps9a
Mtr4
Rps8a
Fun12
Rpn1
Duf1
Rad53
Art5
Ecm21
Ldb19
Yhr131c
Rsp5
Bul2
Ent1
Ent5
Apl3
Sec16
Apm4
Akl1
Rpn11
Nip1
Prt1
Rpg1
Ded1
Tif35
Cdc9
Chk1
Rad23
Ubi4
Ede1
Ent2
Vps27
Chc1
Ptp3
Stb1
Spa2
Bem2
Bud14
Rck2
Glc7
Ipl1
Cdc5
Dbf4
Pfk26
Tpk1
Gpa2
Ras2
Med6
Rpb10
Paf1
Rpo26
Bdp1
Spt15
Spt5
Rgr1
Srb7
Ssn2
Nop1
Nsr1
Nop56
Snu13
Rps31
Rad9
Rtf1
Rpo21
Npl3
Air2
Fps1
Ask10
Cdc28
Hog1
Pbs2
Ssk2
Clb2
Bud3
Bud4
Msn2
Sch9
Rim15
Yak1
Ypl247c
Vps52
Kss1
Sko1
Slt2
Msg5
Swe1
Igo2
Cdc55
Igo1
Vps53
Vps54
Rtn1
Fmp42
Cot1
Zrc1
10
20
30
40
50
t/s

#### Path 1119 (Rpo21 − Gpa2)

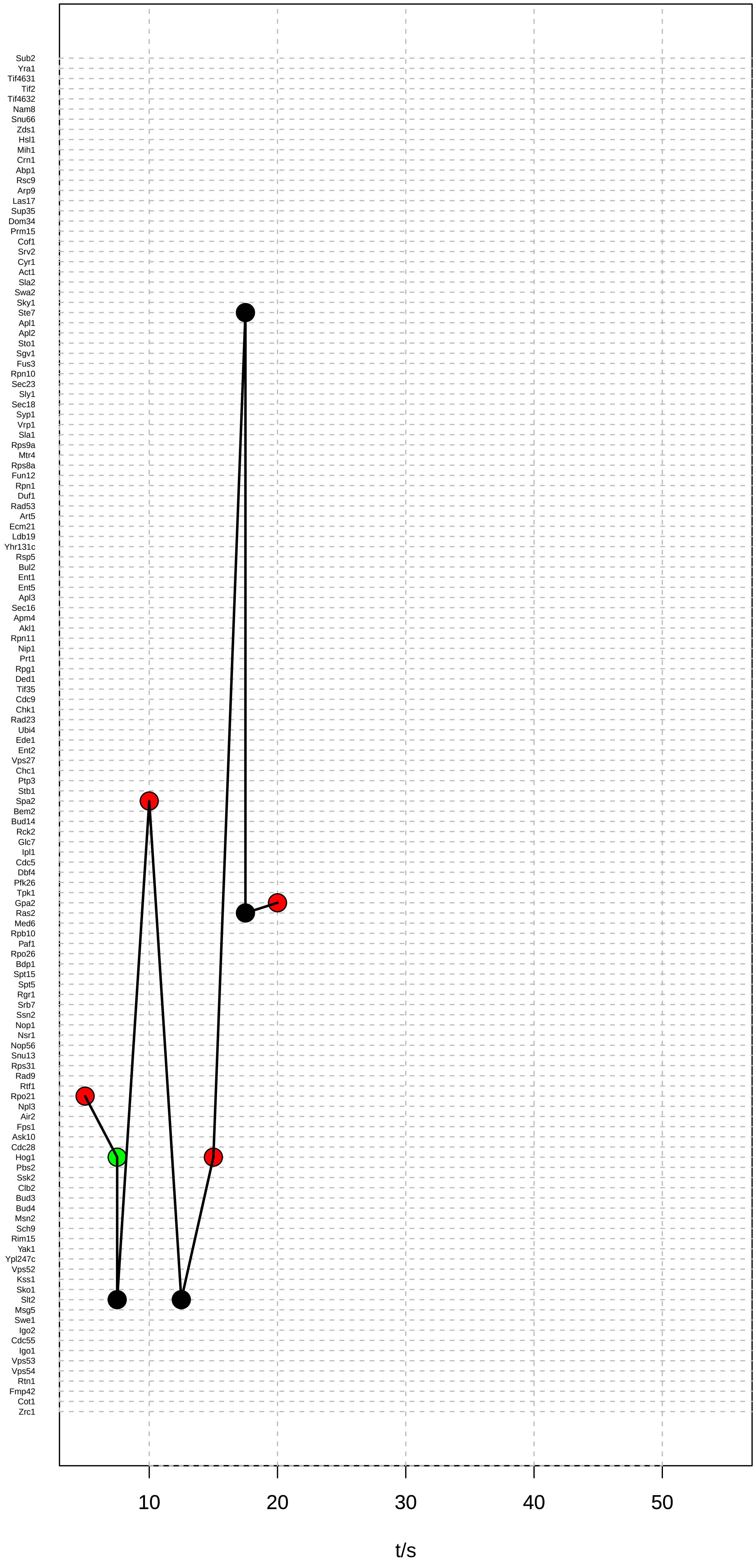

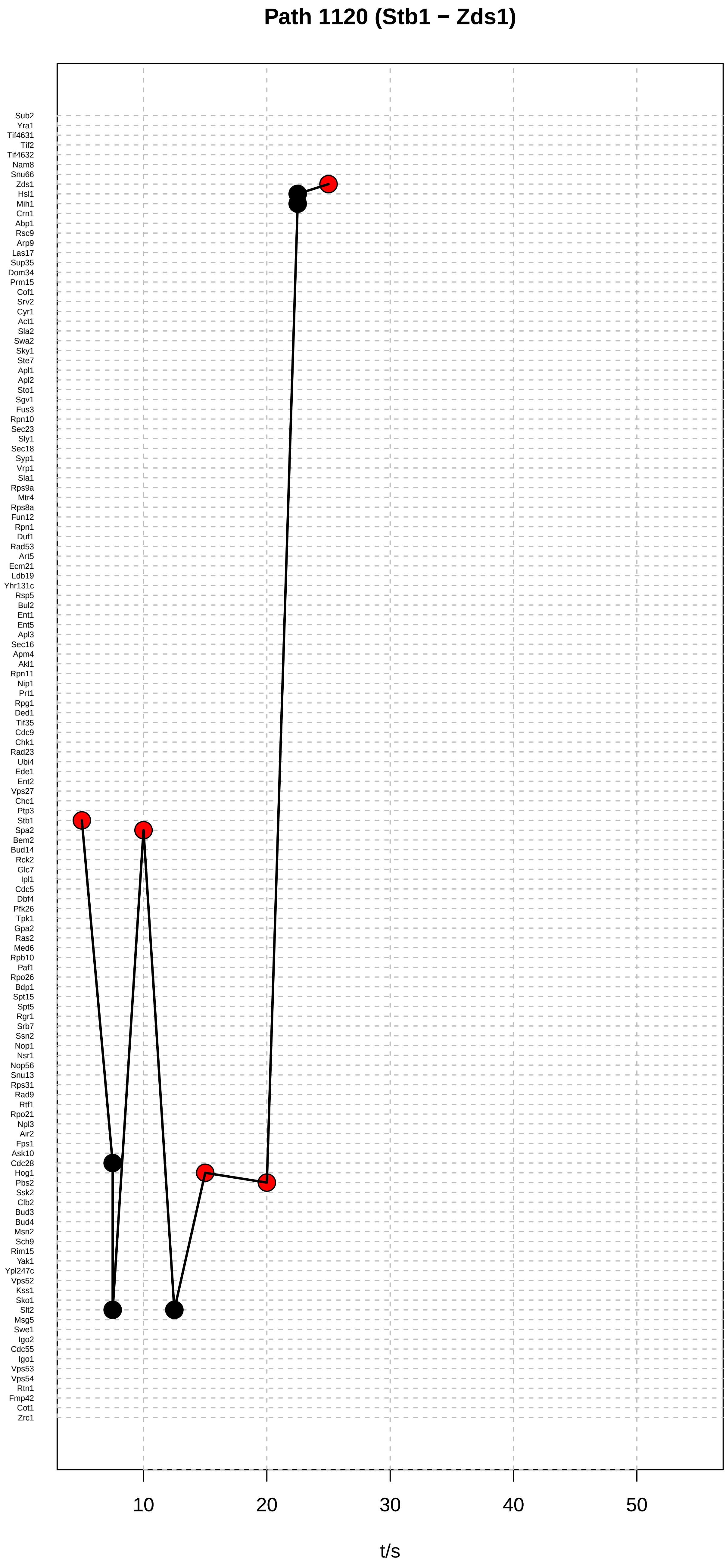
Path 1120 (Stb1 − Zds1)
Sub2
Yra1
Tif4631
Tif2
Tif4632
Nam8
Snu66
Zds1
Hsl1
Mih1
Crn1
Abp1
Rsc9
Arp9
Las17
Sup35
Dom34
Prm15
Cof1
Srv2
Cyr1
Act1
Sla2
Swa2
Sky1
Ste7
Apl1
Apl2
Sto1
Sgv1
Fus3
Rpn10
Sec23
Sly1
Sec18
Syp1
Vrp1
Sla1
Rps9a
Mtr4
Rps8a
Fun12
Rpn1
Duf1
Rad53
Art5
Ecm21
Ldb19
Yhr131c
Rsp5
Bul2
Ent1
Ent5
Apl3
Sec16
Apm4
Akl1
Rpn11
Nip1
Prt1
Rpg1
Ded1
Tif35
Cdc9
Chk1
Rad23
Ubi4
Ede1
Ent2
Vps27
Chc1
Ptp3
Stb1
Spa2
Bem2
Bud14
Rck2
Glc7
Ipl1
Cdc5
Dbf4
Pfk26
Tpk1
Gpa2
Ras2
Med6
Rpb10
Paf1
Rpo26
Bdp1
Spt15
Spt5
Rgr1
Srb7
Ssn2
Nop1
Nsr1
Nop56
Snu13
Rps31
Rad9
Rtf1
Rpo21
Npl3
Air2
Fps1
Ask10
Cdc28
Hog1
Pbs2
Ssk2
Clb2
Bud3
Bud4
Msn2
Sch9
Rim15
Yak1
Ypl247c
Vps52
Kss1
Sko1
Slt2
Msg5
Swe1
Igo2
Cdc55
Igo1
Vps53
Vps54
Rtn1
Fmp42
Cot1
Zrc1
10
20
30
40
50
t/s

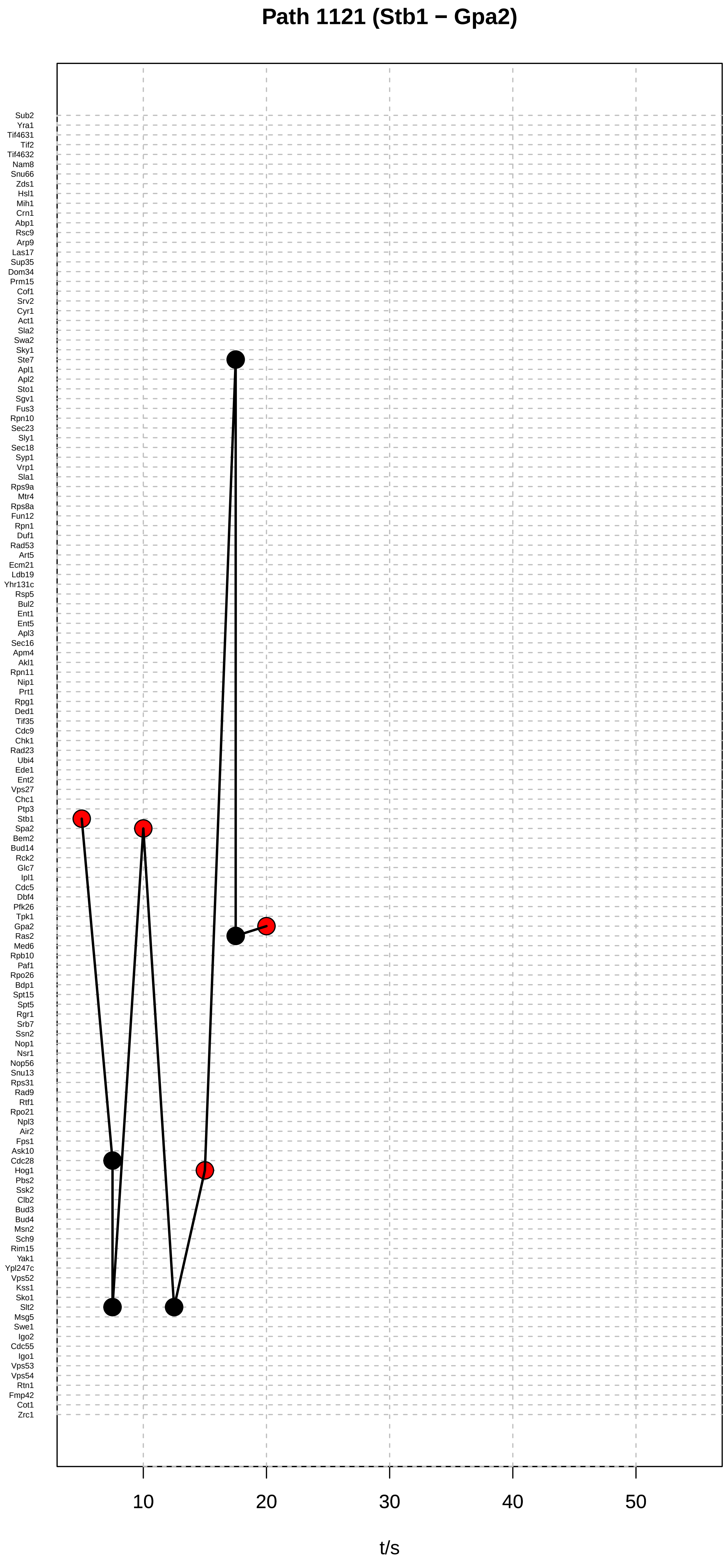

Path 1121 (Stb1 − Gpa2)
Sub2
Yra1
Tif4631
Tif2
Tif4632
Nam8
Snu66
Zds1
Hsl1
Mih1
Crn1
Abp1
Rsc9
Arp9
Las17
Sup35
Dom34
Prm15
Cof1
Srv2
Cyr1
Act1
Sla2
Swa2
Sky1
Ste7
Apl1
Apl2
Sto1
Sgv1
Fus3
Rpn10
Sec23
Sly1
Sec18
Syp1
Vrp1
Sla1
Rps9a
Mtr4
Rps8a
Fun12
Rpn1
Duf1
Rad53
Art5
Ecm21
Ldb19
Yhr131c
Rsp5
Bul2
Ent1
Ent5
Apl3
Sec16
Apm4
Akl1
Rpn11
Nip1
Prt1
Rpg1
Ded1
Tif35
Cdc9
Chk1
Rad23
Ubi4
Ede1
Ent2
Vps27
Chc1
Ptp3
Stb1
Spa2
Bem2
Bud14
Rck2
Glc7
Ipl1
Cdc5
Dbf4
Pfk26
Tpk1
Gpa2
Ras2
Med6
Rpb10
Paf1
Rpo26
Bdp1
Spt15
Spt5
Rgr1
Srb7
Ssn2
Nop1
Nsr1
Nop56
Snu13
Rps31
Rad9
Rtf1
Rpo21
Npl3
Air2
Fps1
Ask10
Cdc28
Hog1
Pbs2
Ssk2
Clb2
Bud3
Bud4
Msn2
Sch9
Rim15
Yak1
Ypl247c
Vps52
Kss1
Sko1
Slt2
Msg5
Swe1
Igo2
Cdc55
Igo1
Vps53
Vps54
Rtn1
Fmp42
Cot1
Zrc1
10
20
30
40
50
t/s

### Path 1122 (Stb1 − Pbs2)

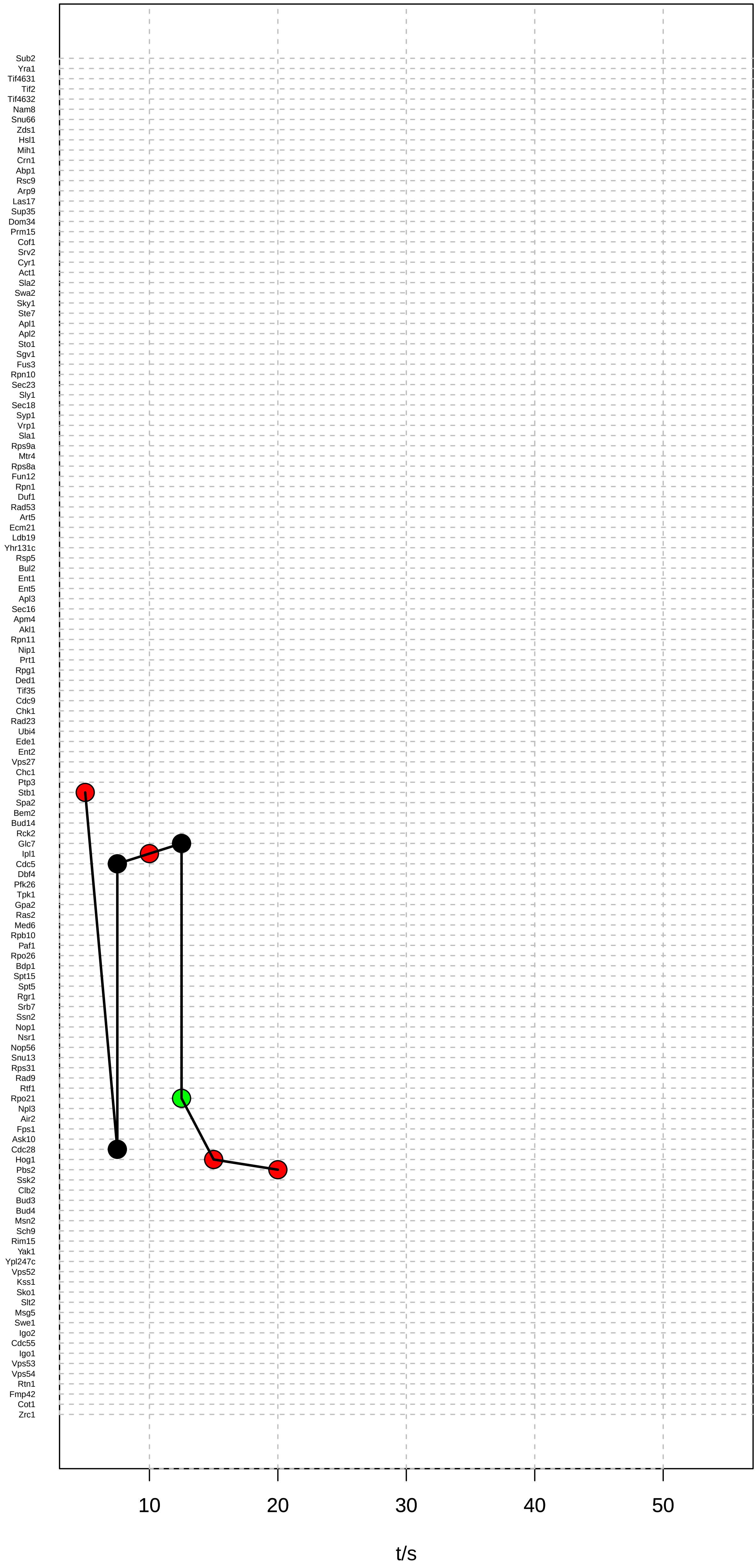

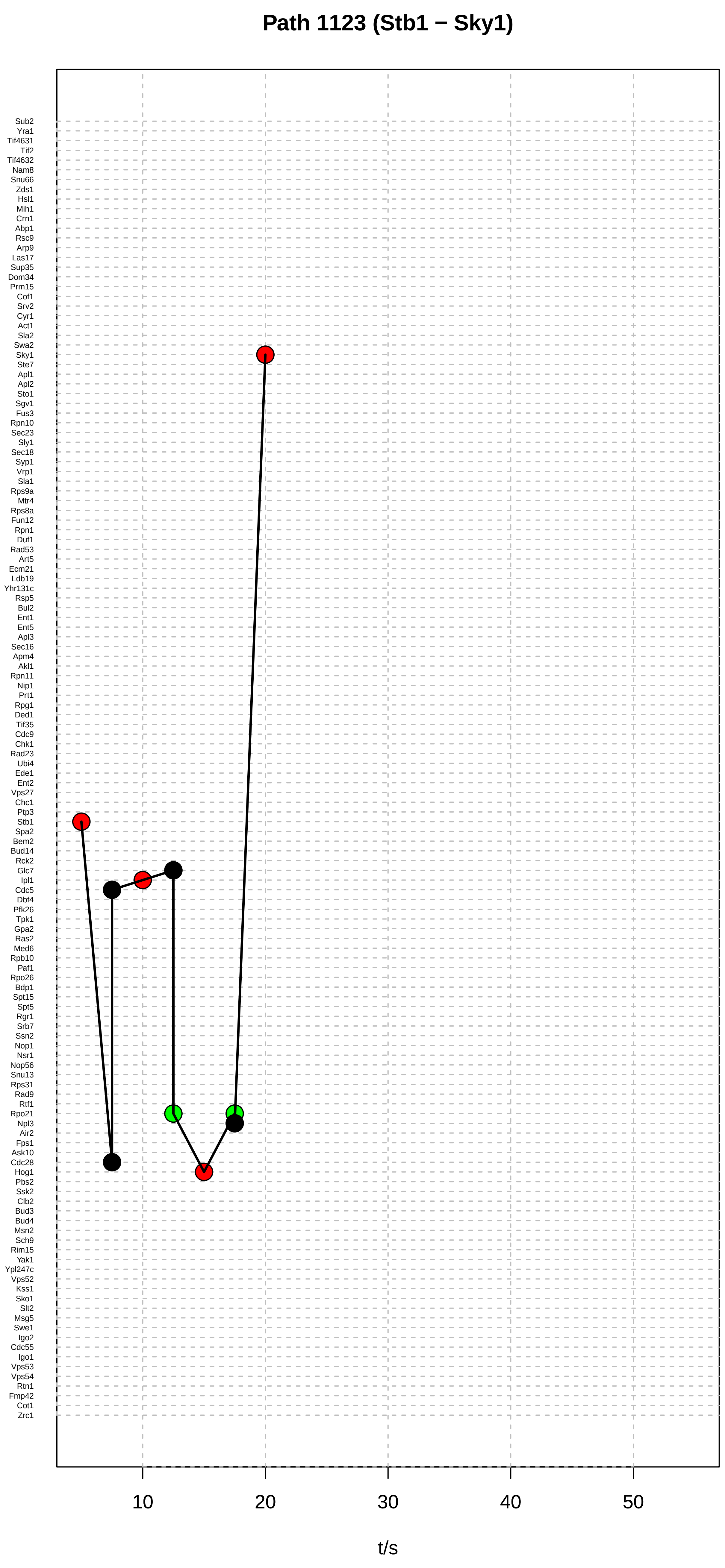

Path 1123 (Stb1 − Sky1)
Sub2
Yra1
Tif4631
Tif2
Tif4632
Nam8
Snu66
Zds1
Hsl1
Mih1
Crn1
Abp1
Rsc9
Arp9
Las17
Sup35
Dom34
Prm15
Cof1
Srv2
Cyr1
Act1
Sla2
Swa2
Sky1
Ste7
Apl1
Apl2
Sto1
Sgv1
Fus3
Rpn10
Sec23
Sly1
Sec18
Syp1
Vrp1
Sla1
Rps9a
Mtr4
Rps8a
Fun12
Rpn1
Duf1
Rad53
Art5
Ecm21
Ldb19
Yhr131c
Rsp5
Bul2
Ent1
Ent5
Apl3
Sec16
Apm4
Akl1
Rpn11
Nip1
Prt1
Rpg1
Ded1
Tif35
Cdc9
Chk1
Rad23
Ubi4
Ede1
Ent2
Vps27
Chc1
Ptp3
Stb1
Spa2
Bem2
Bud14
Rck2
Glc7
Ipl1
Cdc5
Dbf4
Pfk26
Tpk1
Gpa2
Ras2
Med6
Rpb10
Paf1
Rpo26
Bdp1
Spt15
Spt5
Rgr1
Srb7
Ssn2
Nop1
Nsr1
Nop56
Snu13
Rps31
Rad9
Rtf1
Rpo21
Npl3
Air2
Fps1
Ask10
Cdc28
Hog1
Pbs2
Ssk2
Clb2
Bud3
Bud4
Msn2
Sch9
Rim15
Yak1
Ypl247c
Vps52
Kss1
Sko1
Slt2
Msg5
Swe1
Igo2
Cdc55
Igo1
Vps53
Vps54
Rtn1
Fmp42
Cot1
Zrc1
10
20
30
40
50
t/s

#### Path 1124 (Stb1 − Bud14)

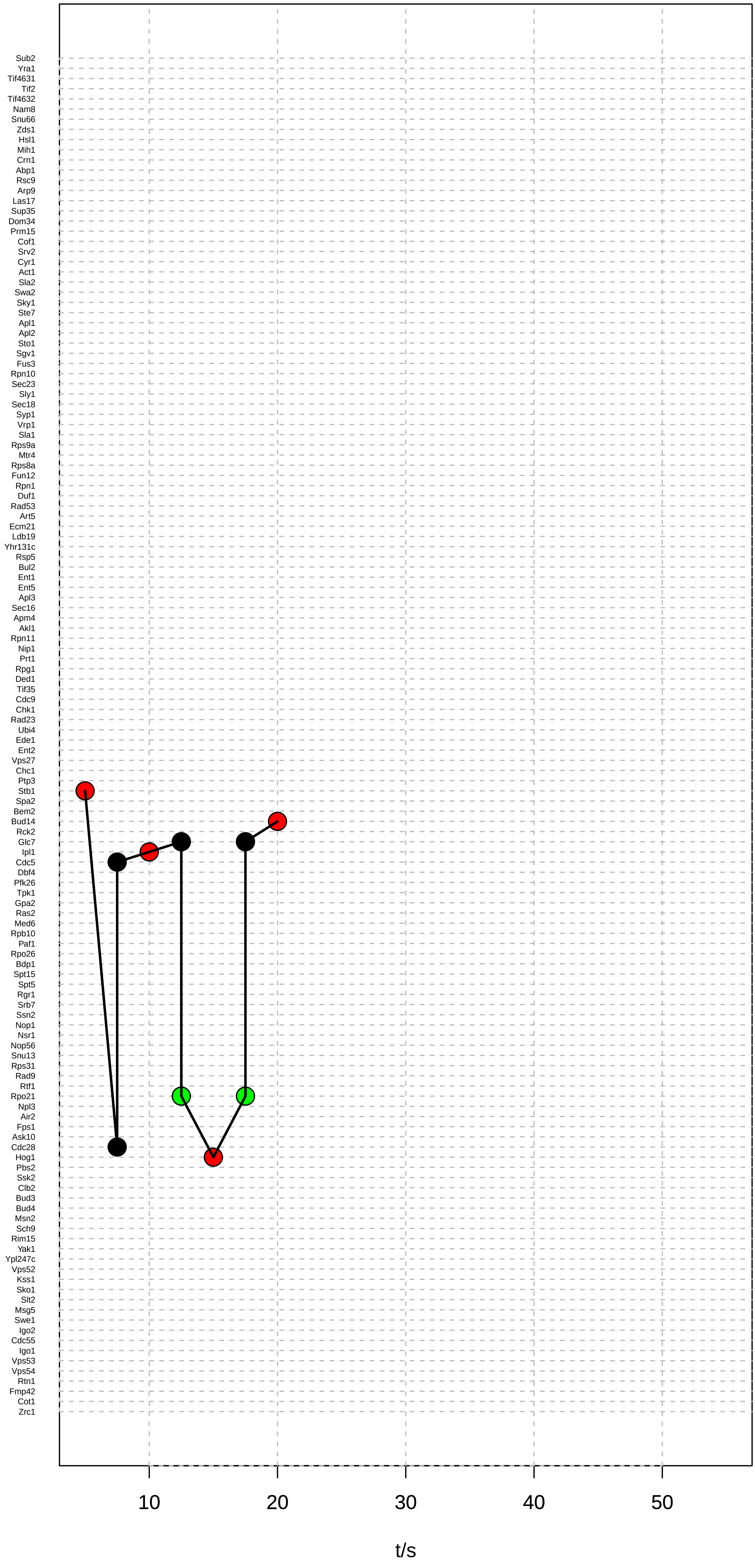

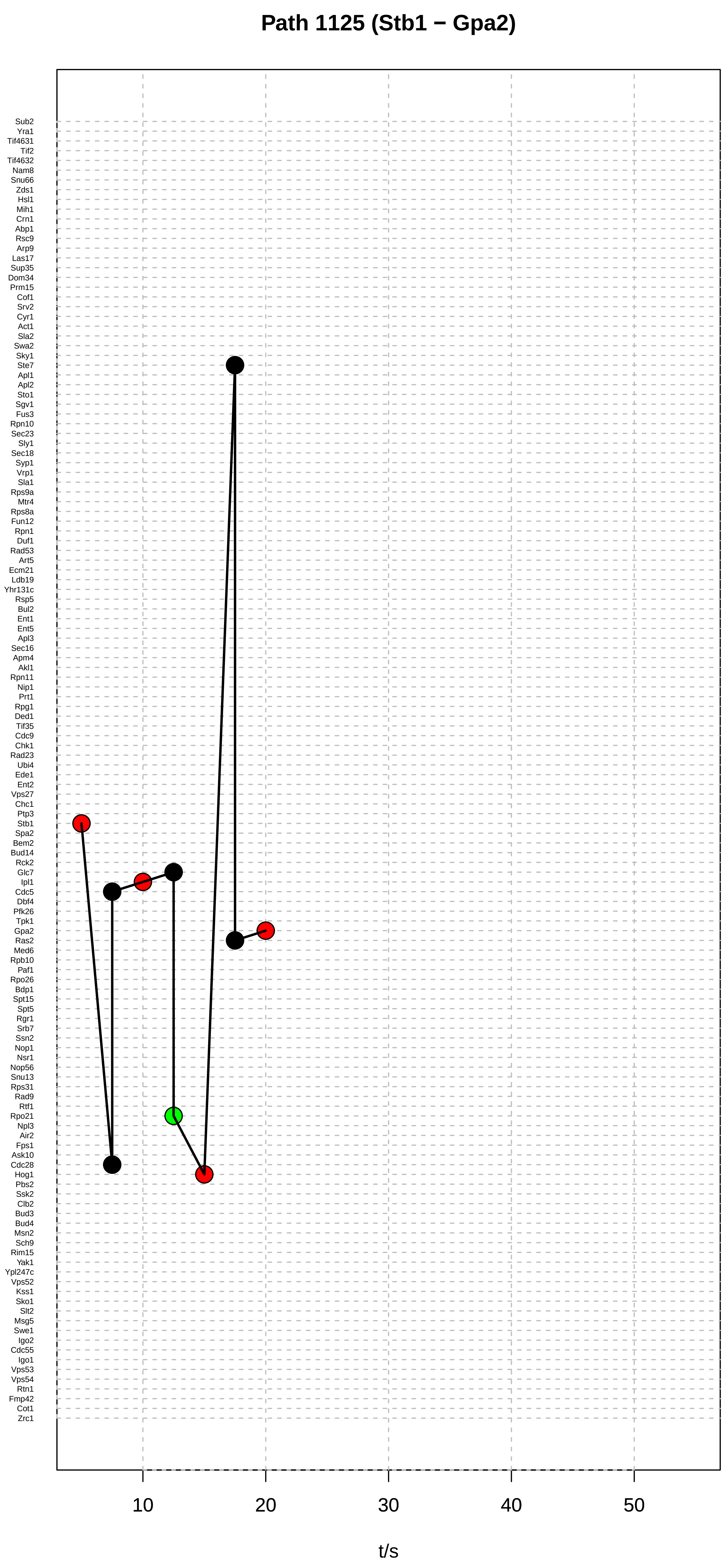

Path 1125 (Stb1 − Gpa2)
Sub2
Yra1
Tif4631
Tif2
Tif4632
Nam8
Snu66
Zds1
Hsl1
Mih1
Crn1
Abp1
Rsc9
Arp9
Las17
Sup35
Dom34
Prm15
Cof1
Srv2
Cyr1
Act1
Sla2
Swa2
Sky1
Ste7
Apl1
Apl2
Sto1
Sgv1
Fus3
Rpn10
Sec23
Sly1
Sec18
Syp1
Vrp1
Sla1
Rps9a
Mtr4
Rps8a
Fun12
Rpn1
Duf1
Rad53
Art5
Ecm21
Ldb19
Yhr131c
Rsp5
Bul2
Ent1
Ent5
Apl3
Sec16
Apm4
Akl1
Rpn11
Nip1
Prt1
Rpg1
Ded1
Tif35
Cdc9
Chk1
Rad23
Ubi4
Ede1
Ent2
Vps27
Chc1
Ptp3
Stb1
Spa2
Bem2
Bud14
Rck2
Glc7
Ipl1
Cdc5
Dbf4
Pfk26
Tpk1
Gpa2
Ras2
Med6
Rpb10
Paf1
Rpo26
Bdp1
Spt15
Spt5
Rgr1
Srb7
Ssn2
Nop1
Nsr1
Nop56
Snu13
Rps31
Rad9
Rtf1
Rpo21
Npl3
Air2
Fps1
Ask10
Cdc28
Hog1
Pbs2
Ssk2
Clb2
Bud3
Bud4
Msn2
Sch9
Rim15
Yak1
Ypl247c
Vps52
Kss1
Sko1
Slt2
Msg5
Swe1
Igo2
Cdc55
Igo1
Vps53
Vps54
Rtn1
Fmp42
Cot1
Zrc1
10
20
30
40
50
t/s

### Path 1126 (Ptp3 − Zds1)

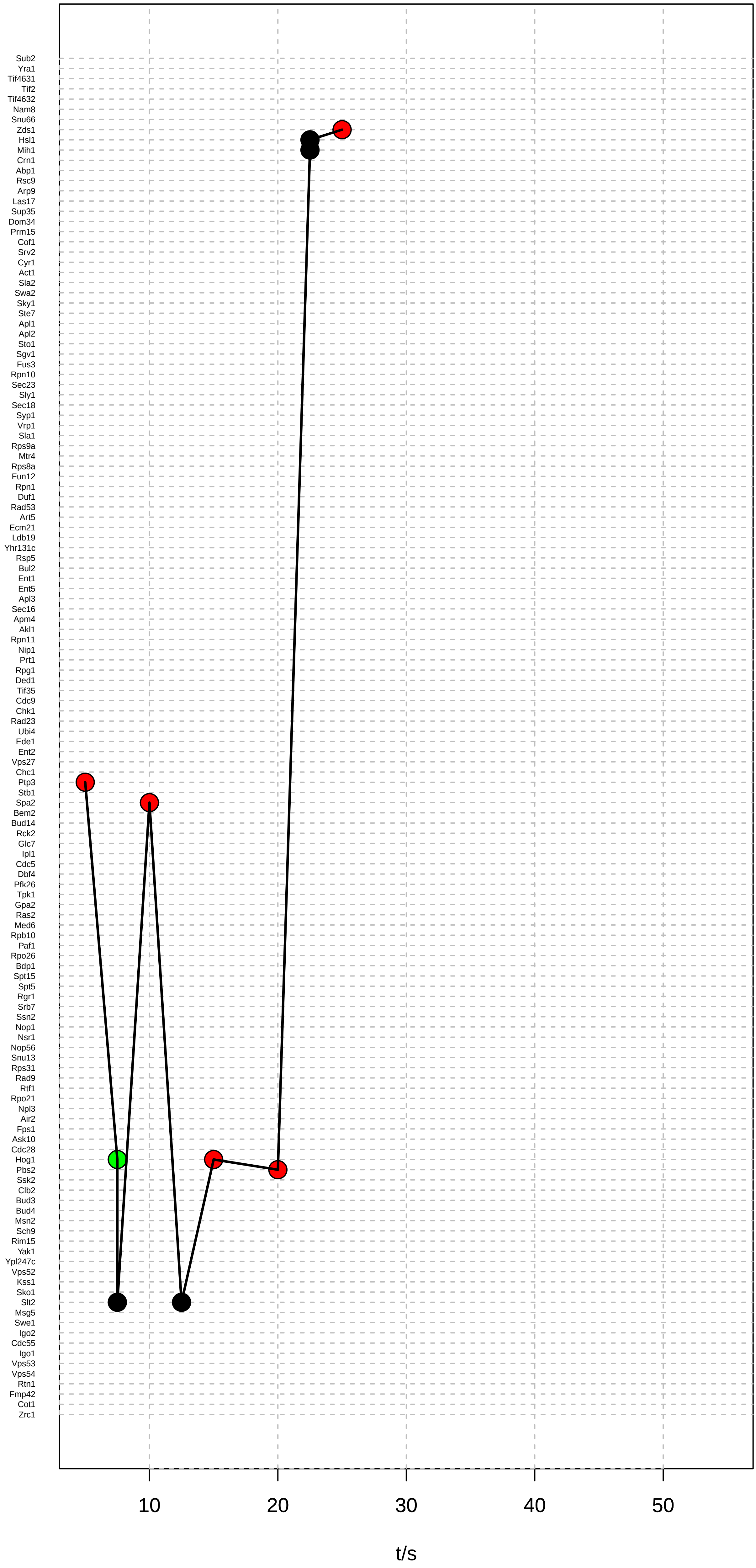

### Path 1127 (Ptp3 − Gpa2)

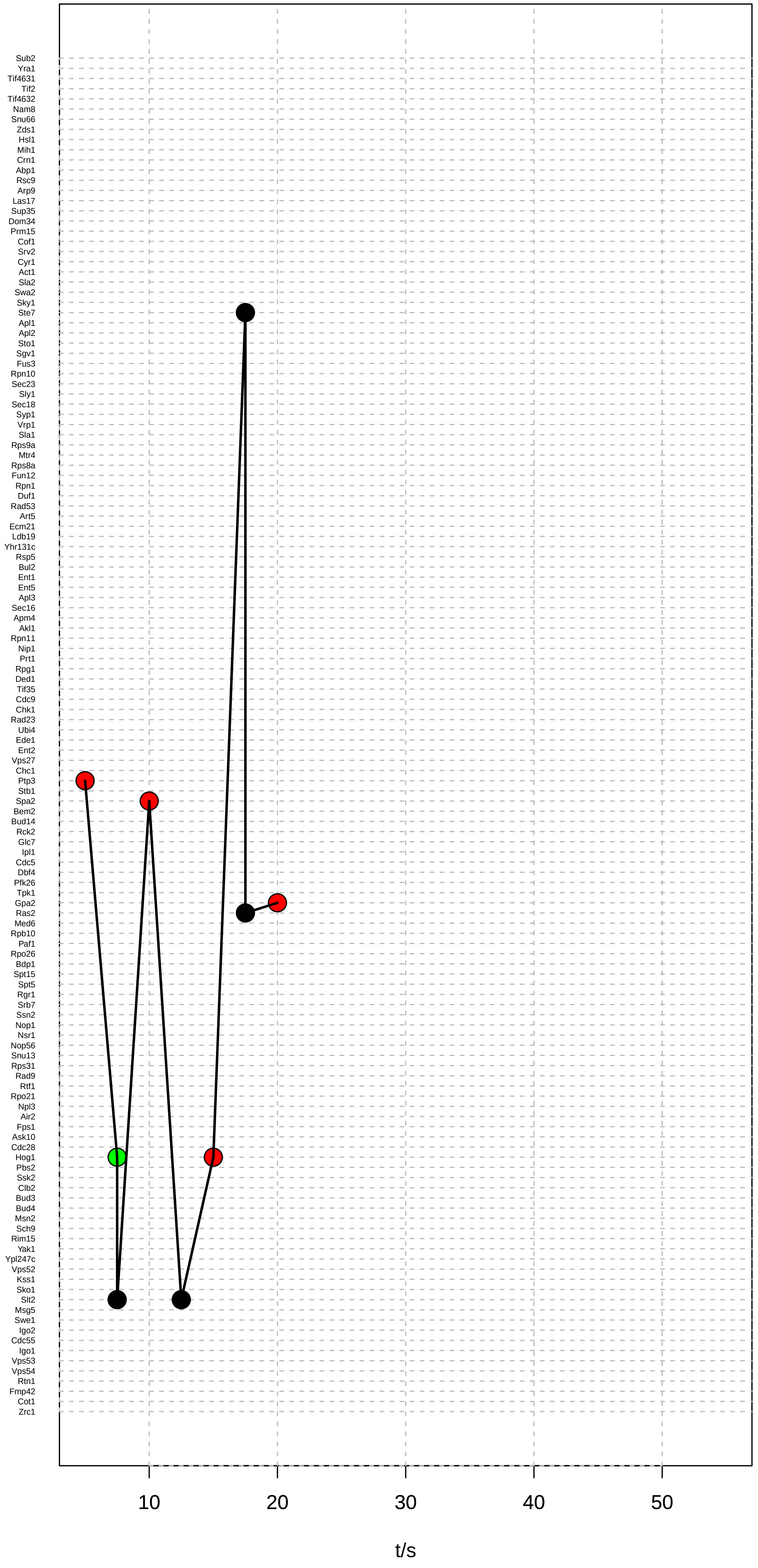
