## Supplemental File 6 for "Network inference from temporal phosphoproteomics informed by protein-protein interactions"

#### Path 1 (Zrc1 − Fmp42)

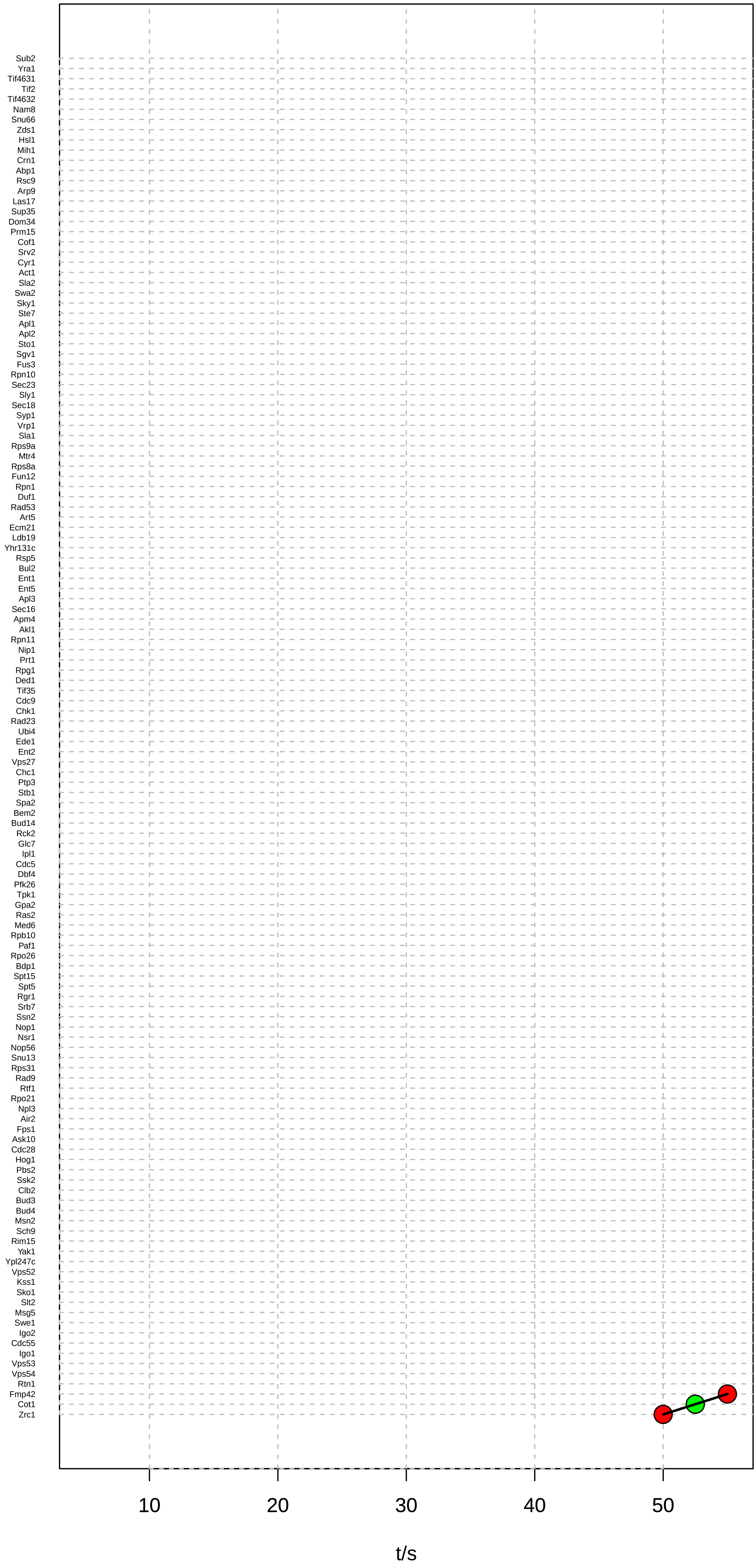

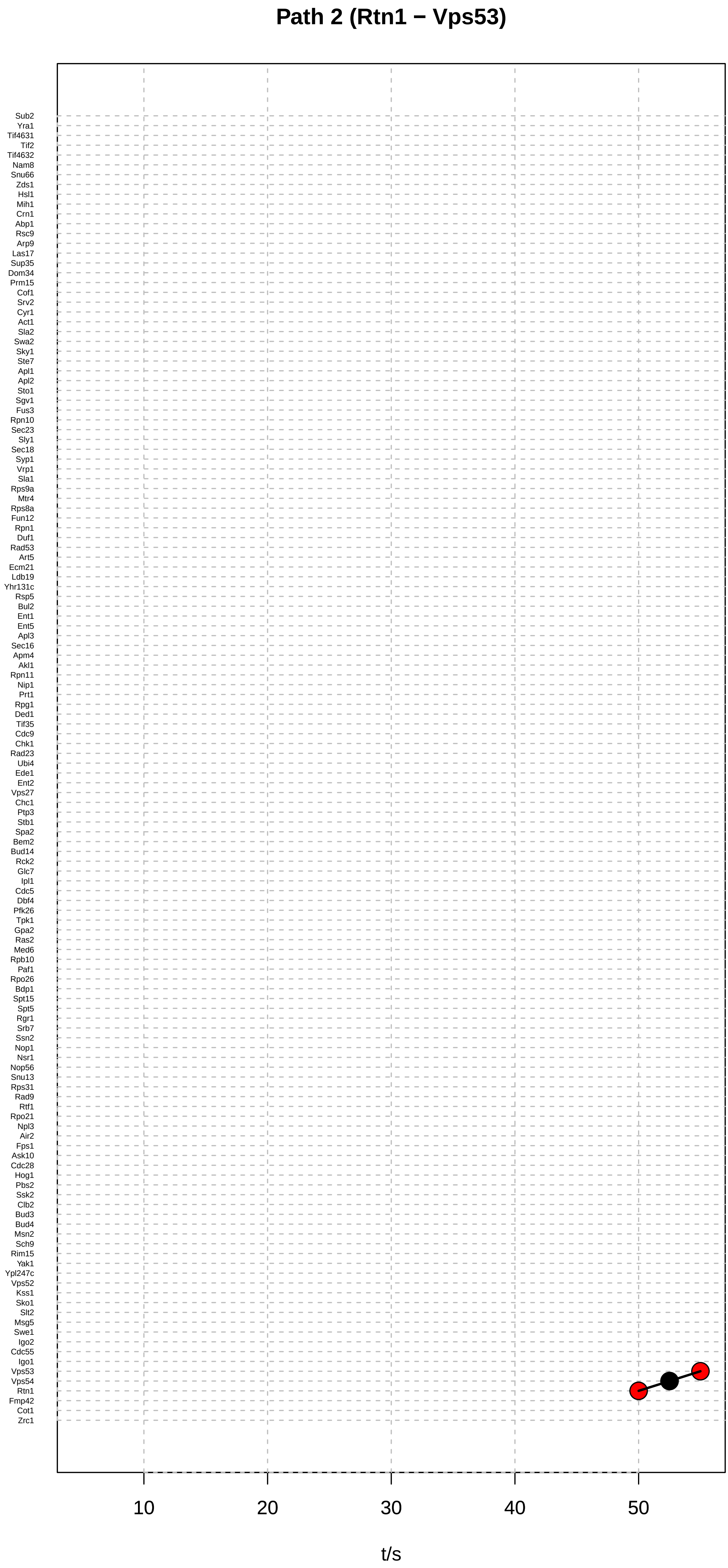
Path 2 (Rtn1 − Vps53)
Sub2
Yra1
Tif4631
Tif2
Tif4632
Nam8
Snu66
Zds1
Hsl1
Mih1
Crn1
Abp1
Rsc9
Arp9
Las17
Sup35
Dom34
Prm15
Cof1
Srv2
Cyr1
Act1
Sla2
Swa2
Sky1
Ste7
Apl1
Apl2
Sto1
Sgv1
Fus3
Rpn10
Sec23
Sly1
Sec18
Syp1
Vrp1
Sla1
Rps9a
Mtr4
Rps8a
Fun12
Rpn1
Duf1
Rad53
Art5
Ecm21
Ldb19
Yhr131c
Rsp5
Bul2
Ent1
Ent5
Apl3
Sec16
Apm4
Akl1
Rpn11
Nip1
Prt1
Rpg1
Ded1
Tif35
Cdc9
Chk1
Rad23
Ubi4
Ede1
Ent2
Vps27
Chc1
Ptp3
Stb1
Spa2
Bem2
Bud14
Rck2
Glc7
Ipl1
Cdc5
Dbf4
Pfk26
Tpk1
Gpa2
Ras2
Med6
Rpb10
Paf1
Rpo26
Bdp1
Spt15
Spt5
Rgr1
Srb7
Ssn2
Nop1
Nsr1
Nop56
Snu13
Rps31
Rad9
Rtf1
Rpo21
Npl3
Air2
Fps1
Ask10
Cdc28
Hog1
Pbs2
Ssk2
Clb2
Bud3
Bud4
Msn2
Sch9
Rim15
Yak1
Ypl247c
Vps52
Kss1
Sko1
Slt2
Msg5
Swe1
Igo2
Cdc55
Igo1
Vps53
Vps54
Rtn1
Fmp42
Cot1
Zrc1
10
20
30
40
50
t/s

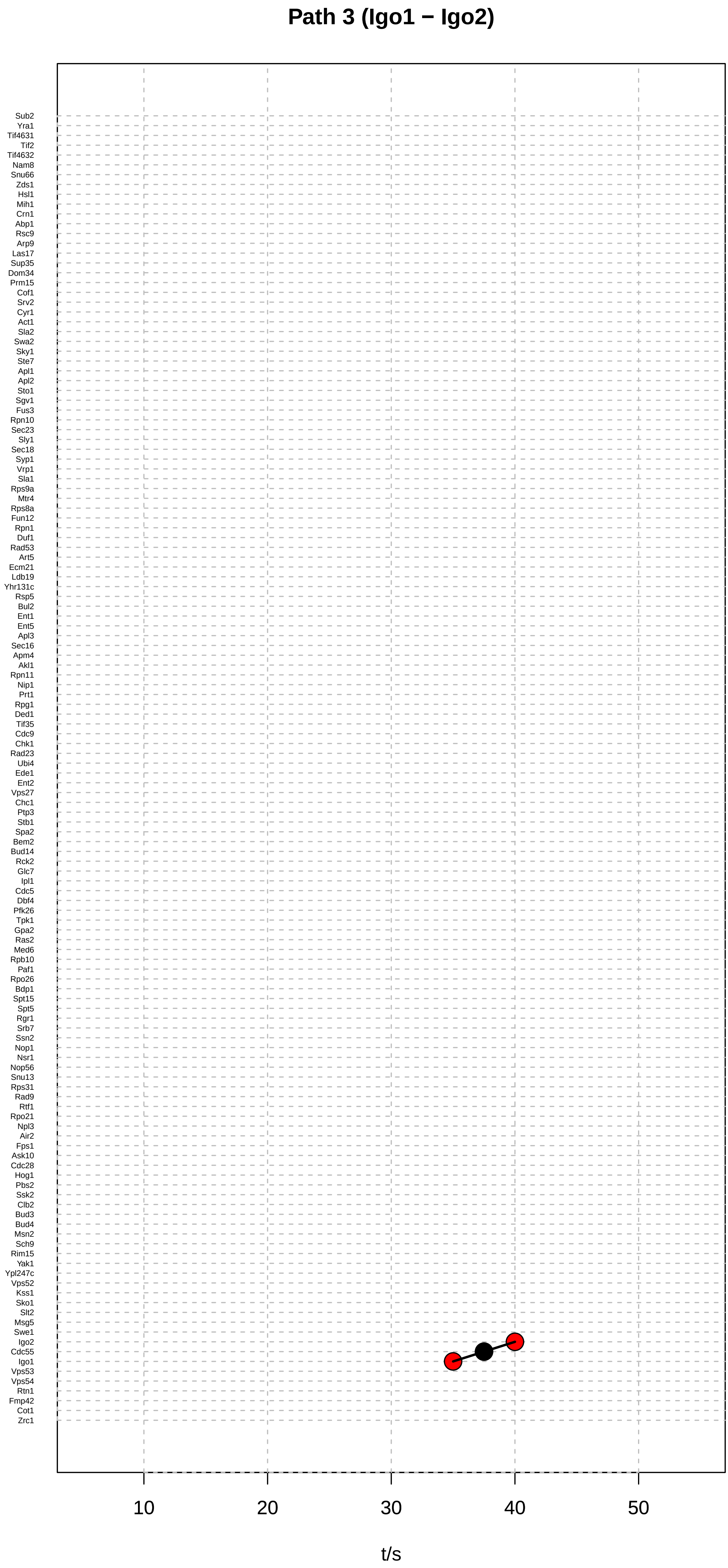

Path 3 (Igo1 − Igo2)
Sub2
Yra1
Tif4631
Tif2
Tif4632
Nam8
Snu66
Zds1
Hsl1
Mih1
Crn1
Abp1
Rsc9
Arp9
Las17
Sup35
Dom34
Prm15
Cof1
Srv2
Cyr1
Act1
Sla2
Swa2
Sky1
Ste7
Apl1
Apl2
Sto1
Sgv1
Fus3
Rpn10
Sec23
Sly1
Sec18
Syp1
Vrp1
Sla1
Rps9a
Mtr4
Rps8a
Fun12
Rpn1
Duf1
Rad53
Art5
Ecm21
Ldb19
Yhr131c
Rsp5
Bul2
Ent1
Ent5
Apl3
Sec16
Apm4
Akl1
Rpn11
Nip1
Prt1
Rpg1
Ded1
Tif35
Cdc9
Chk1
Rad23
Ubi4
Ede1
Ent2
Vps27
Chc1
Ptp3
Stb1
Spa2
Bem2
Bud14
Rck2
Glc7
Ipl1
Cdc5
Dbf4
Pfk26
Tpk1
Gpa2
Ras2
Med6
Rpb10
Paf1
Rpo26
Bdp1
Spt15
Spt5
Rgr1
Srb7
Ssn2
Nop1
Nsr1
Nop56
Snu13
Rps31
Rad9
Rtf1
Rpo21
Npl3
Air2
Fps1
Ask10
Cdc28
Hog1
Pbs2
Ssk2
Clb2
Bud3
Bud4
Msn2
Sch9
Rim15
Yak1
Ypl247c
Vps52
Kss1
Sko1
Slt2
Msg5
Swe1
Igo2
Cdc55
Igo1
Vps53
Vps54
Rtn1
Fmp42
Cot1
Zrc1
10
20
30
40
50
t/s

### Path 4 (Ypl247c − Sch9)

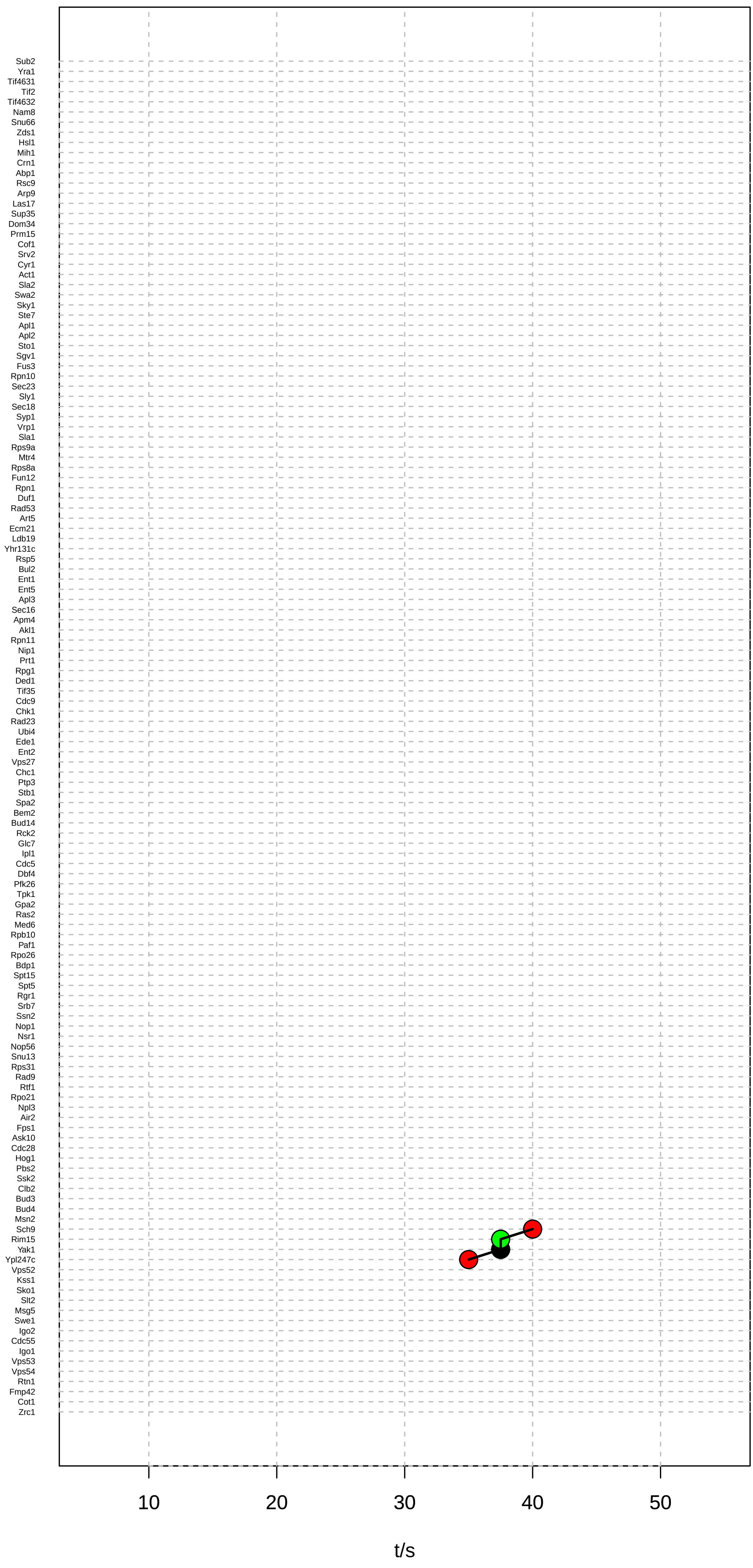

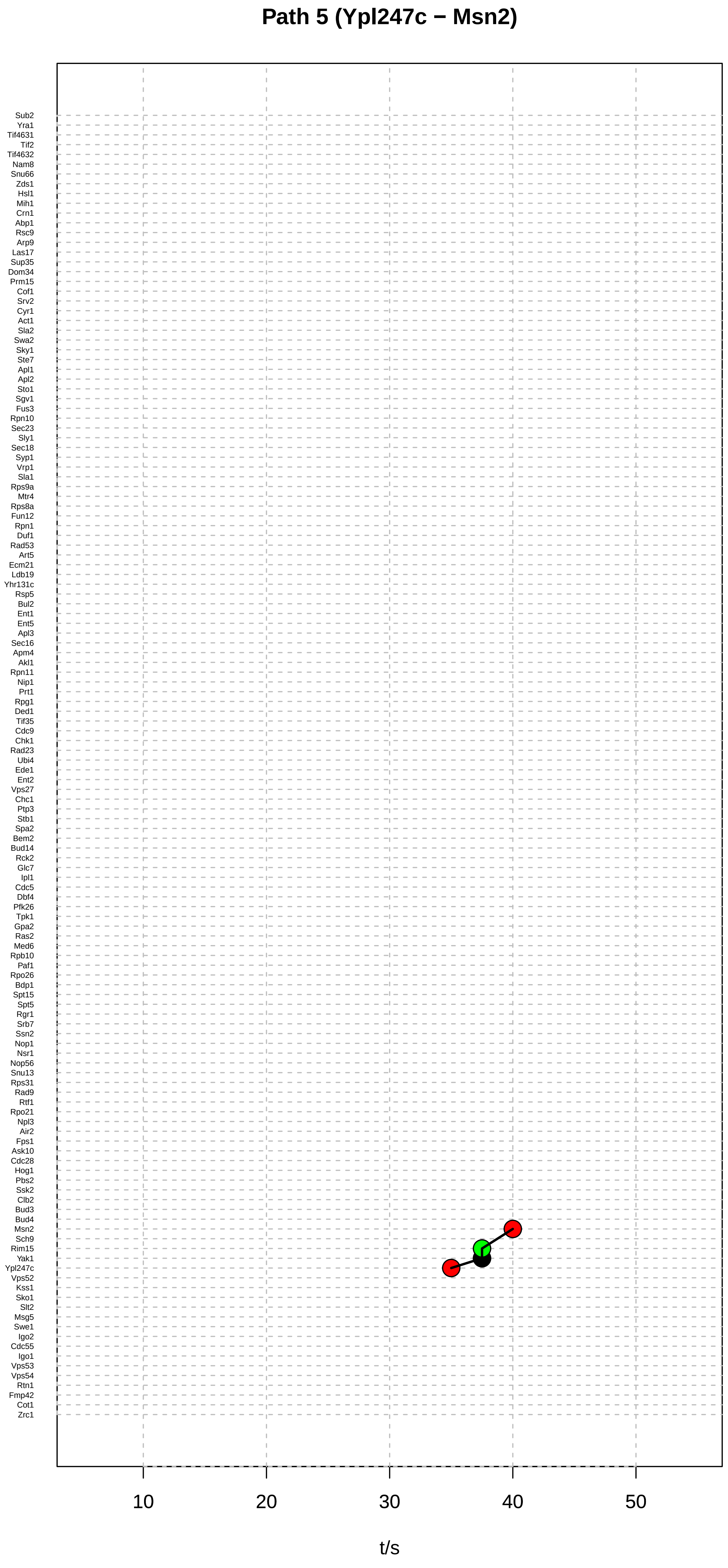
Path 5 (Ypl247c − Msn2)
Sub2
Yra1
Tif4631
Tif2
Tif4632
Nam8
Snu66
Zds1
Hsl1
Mih1
Crn1
Abp1
Rsc9
Arp9
Las17
Sup35
Dom34
Prm15
Cof1
Srv2
Cyr1
Act1
Sla2
Swa2
Sky1
Ste7
Apl1
Apl2
Sto1
Sgv1
Fus3
Rpn10
Sec23
Sly1
Sec18
Syp1
Vrp1
Sla1
Rps9a
Mtr4
Rps8a
Fun12
Rpn1
Duf1
Rad53
Art5
Ecm21
Ldb19
Yhr131c
Rsp5
Bul2
Ent1
Ent5
Apl3
Sec16
Apm4
Akl1
Rpn11
Nip1
Prt1
Rpg1
Ded1
Tif35
Cdc9
Chk1
Rad23
Ubi4
Ede1
Ent2
Vps27
Chc1
Ptp3
Stb1
Spa2
Bem2
Bud14
Rck2
Glc7
Ipl1
Cdc5
Dbf4
Pfk26
Tpk1
Gpa2
Ras2
Med6
Rpb10
Paf1
Rpo26
Bdp1
Spt15
Spt5
Rgr1
Srb7
Ssn2
Nop1
Nsr1
Nop56
Snu13
Rps31
Rad9
Rtf1
Rpo21
Npl3
Air2
Fps1
Ask10
Cdc28
Hog1
Pbs2
Ssk2
Clb2
Bud3
Bud4
Msn2
Sch9
Rim15
Yak1
Ypl247c
Vps52
Kss1
Sko1
Slt2
Msg5
Swe1
Igo2
Cdc55
Igo1
Vps53
Vps54
Rtn1
Fmp42
Cot1
Zrc1
10
20
30
40
50
t/s

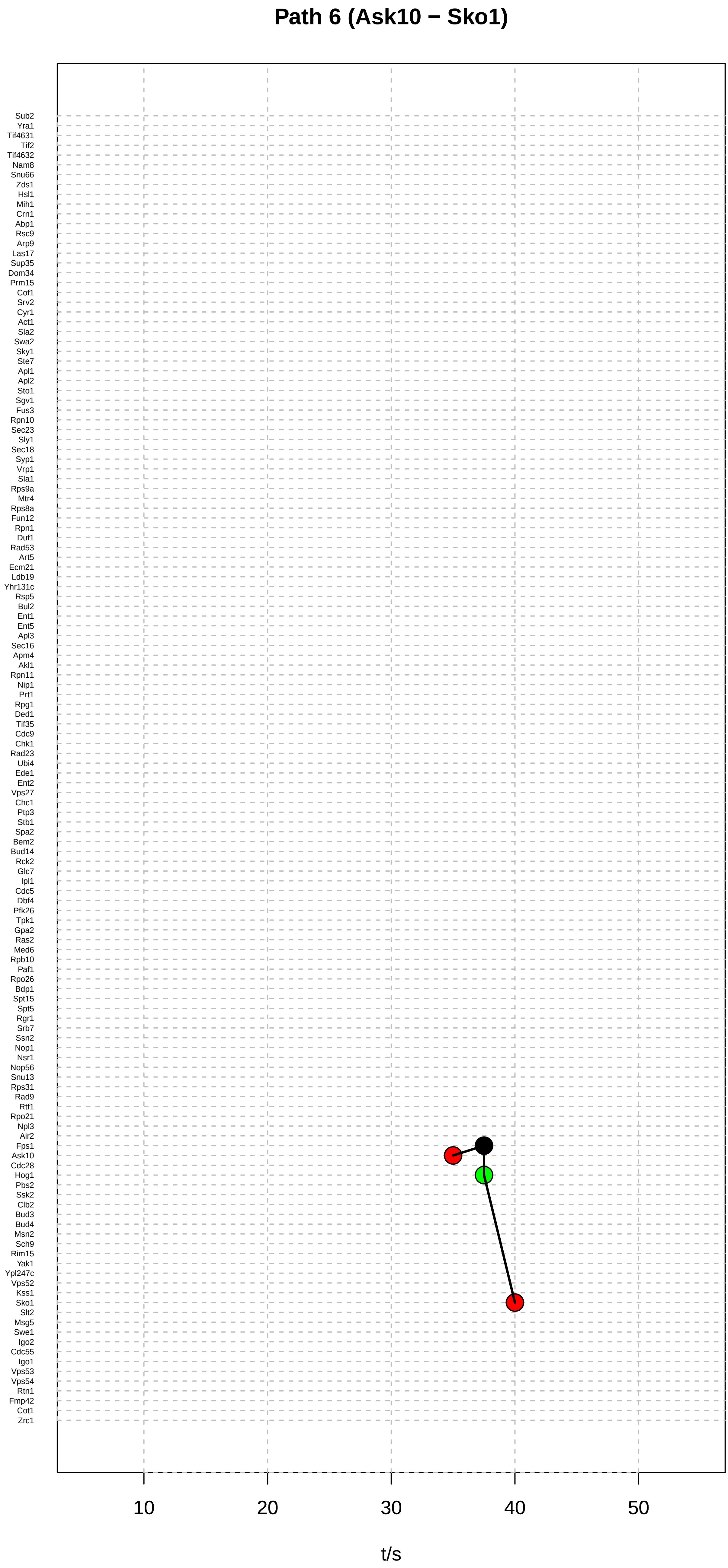

Path 6 (Ask10 − Sko1)
Sub2
Yra1
Tif4631
Tif2
Tif4632
Nam8
Snu66
Zds1
Hsl1
Mih1
Crn1
Abp1
Rsc9
Arp9
Las17
Sup35
Dom34
Prm15
Cof1
Srv2
Cyr1
Act1
Sla2
Swa2
Sky1
Ste7
Apl1
Apl2
Sto1
Sgv1
Fus3
Rpn10
Sec23
Sly1
Sec18
Syp1
Vrp1
Sla1
Rps9a
Mtr4
Rps8a
Fun12
Rpn1
Duf1
Rad53
Art5
Ecm21
Ldb19
Yhr131c
Rsp5
Bul2
Ent1
Ent5
Apl3
Sec16
Apm4
Akl1
Rpn11
Nip1
Prt1
Rpg1
Ded1
Tif35
Cdc9
Chk1
Rad23
Ubi4
Ede1
Ent2
Vps27
Chc1
Ptp3
Stb1
Spa2
Bem2
Bud14
Rck2
Glc7
Ipl1
Cdc5
Dbf4
Pfk26
Tpk1
Gpa2
Ras2
Med6
Rpb10
Paf1
Rpo26
Bdp1
Spt15
Spt5
Rgr1
Srb7
Ssn2
Nop1
Nsr1
Nop56
Snu13
Rps31
Rad9
Rtf1
Rpo21
Npl3
Air2
Fps1
Ask10
Cdc28
Hog1
Pbs2
Ssk2
Clb2
Bud3
Bud4
Msn2
Sch9
Rim15
Yak1
Ypl247c
Vps52
Kss1
Sko1
Slt2
Msg5
Swe1
Igo2
Cdc55
Igo1
Vps53
Vps54
Rtn1
Fmp42
Cot1
Zrc1
10
20
30
40
50
t/s

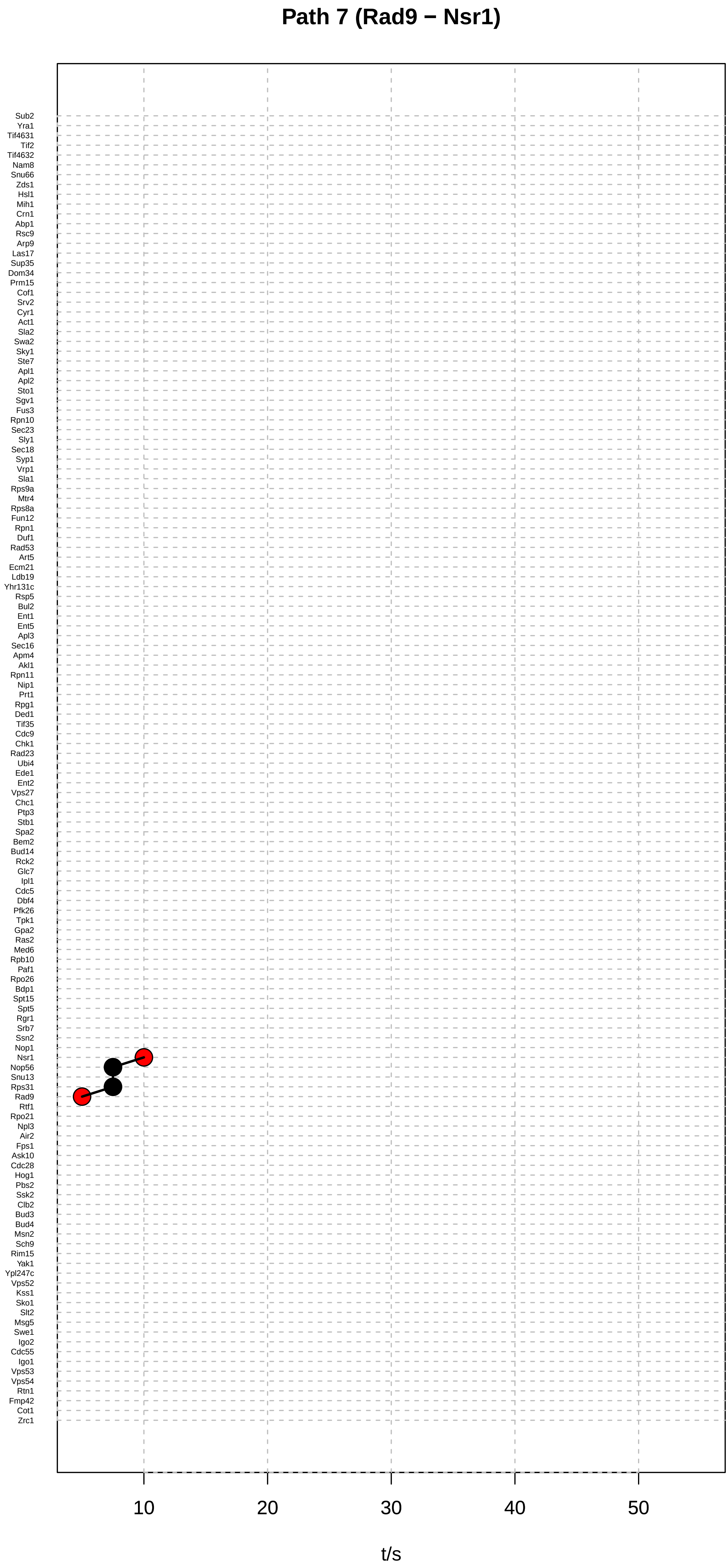
Path 7 (Rad9 − Nsr1)
Sub2
Yra1
Tif4631
Tif2
Tif4632
Nam8
Snu66
Zds1
Hsl1
Mih1
Crn1
Abp1
Rsc9
Arp9
Las17
Sup35
Dom34
Prm15
Cof1
Srv2
Cyr1
Act1
Sla2
Swa2
Sky1
Ste7
Apl1
Apl2
Sto1
Sgv1
Fus3
Rpn10
Sec23
Sly1
Sec18
Syp1
Vrp1
Sla1
Rps9a
Mtr4
Rps8a
Fun12
Rpn1
Duf1
Rad53
Art5
Ecm21
Ldb19
Yhr131c
Rsp5
Bul2
Ent1
Ent5
Apl3
Sec16
Apm4
Akl1
Rpn11
Nip1
Prt1
Rpg1
Ded1
Tif35
Cdc9
Chk1
Rad23
Ubi4
Ede1
Ent2
Vps27
Chc1
Ptp3
Stb1
Spa2
Bem2
Bud14
Rck2
Glc7
Ipl1
Cdc5
Dbf4
Pfk26
Tpk1
Gpa2
Ras2
Med6
Rpb10
Paf1
Rpo26
Bdp1
Spt15
Spt5
Rgr1
Srb7
Ssn2
Nop1
Nsr1
Nop56
Snu13
Rps31
Rad9
Rtf1
Rpo21
Npl3
Air2
Fps1
Ask10
Cdc28
Hog1
Pbs2
Ssk2
Clb2
Bud3
Bud4
Msn2
Sch9
Rim15
Yak1
Ypl247c
Vps52
Kss1
Sko1
Slt2
Msg5
Swe1
Igo2
Cdc55
Igo1
Vps53
Vps54
Rtn1
Fmp42
Cot1
Zrc1
10
20
30
40
50
t/s

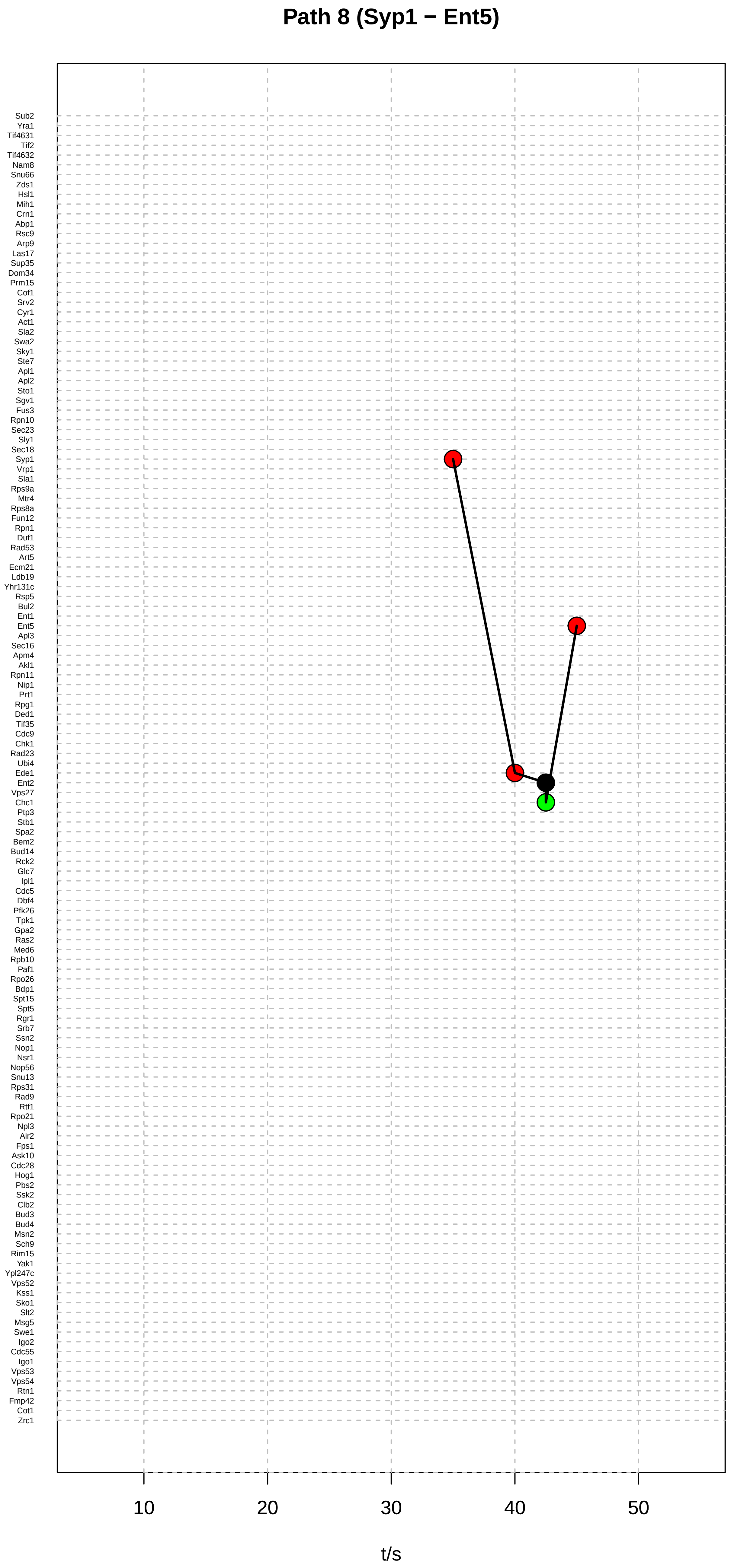
Path 8 (Syp1 − Ent5)
Sub2
Yra1
Tif4631
Tif2
Tif4632
Nam8
Snu66
Zds1
Hsl1
Mih1
Crn1
Abp1
Rsc9
Arp9
Las17
Sup35
Dom34
Prm15
Cof1
Srv2
Cyr1
Act1
Sla2
Swa2
Sky1
Ste7
Apl1
Apl2
Sto1
Sgv1
Fus3
Rpn10
Sec23
Sly1
Sec18
Syp1
Vrp1
Sla1
Rps9a
Mtr4
Rps8a
Fun12
Rpn1
Duf1
Rad53
Art5
Ecm21
Ldb19
Yhr131c
Rsp5
Bul2
Ent1
Ent5
Apl3
Sec16
Apm4
Akl1
Rpn11
Nip1
Prt1
Rpg1
Ded1
Tif35
Cdc9
Chk1
Rad23
Ubi4
Ede1
Ent2
Vps27
Chc1
Ptp3
Stb1
Spa2
Bem2
Bud14
Rck2
Glc7
Ipl1
Cdc5
Dbf4
Pfk26
Tpk1
Gpa2
Ras2
Med6
Rpb10
Paf1
Rpo26
Bdp1
Spt15
Spt5
Rgr1
Srb7
Ssn2
Nop1
Nsr1
Nop56
Snu13
Rps31
Rad9
Rtf1
Rpo21
Npl3
Air2
Fps1
Ask10
Cdc28
Hog1
Pbs2
Ssk2
Clb2
Bud3
Bud4
Msn2
Sch9
Rim15
Yak1
Ypl247c
Vps52
Kss1
Sko1
Slt2
Msg5
Swe1
Igo2
Cdc55
Igo1
Vps53
Vps54
Rtn1
Fmp42
Cot1
Zrc1
10
20
30
40
50
t/s

### Path 9 (Syp1 − Ent5)

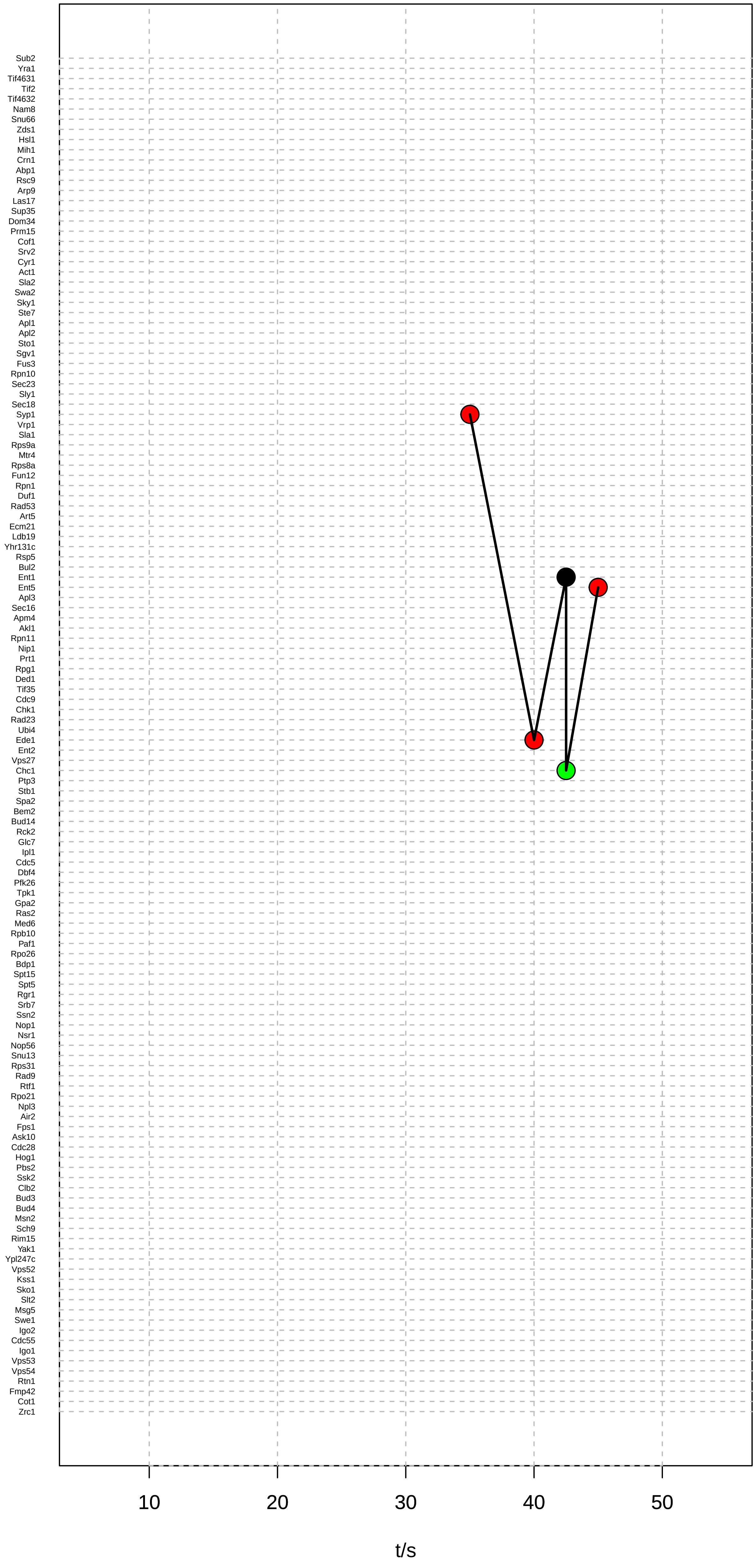

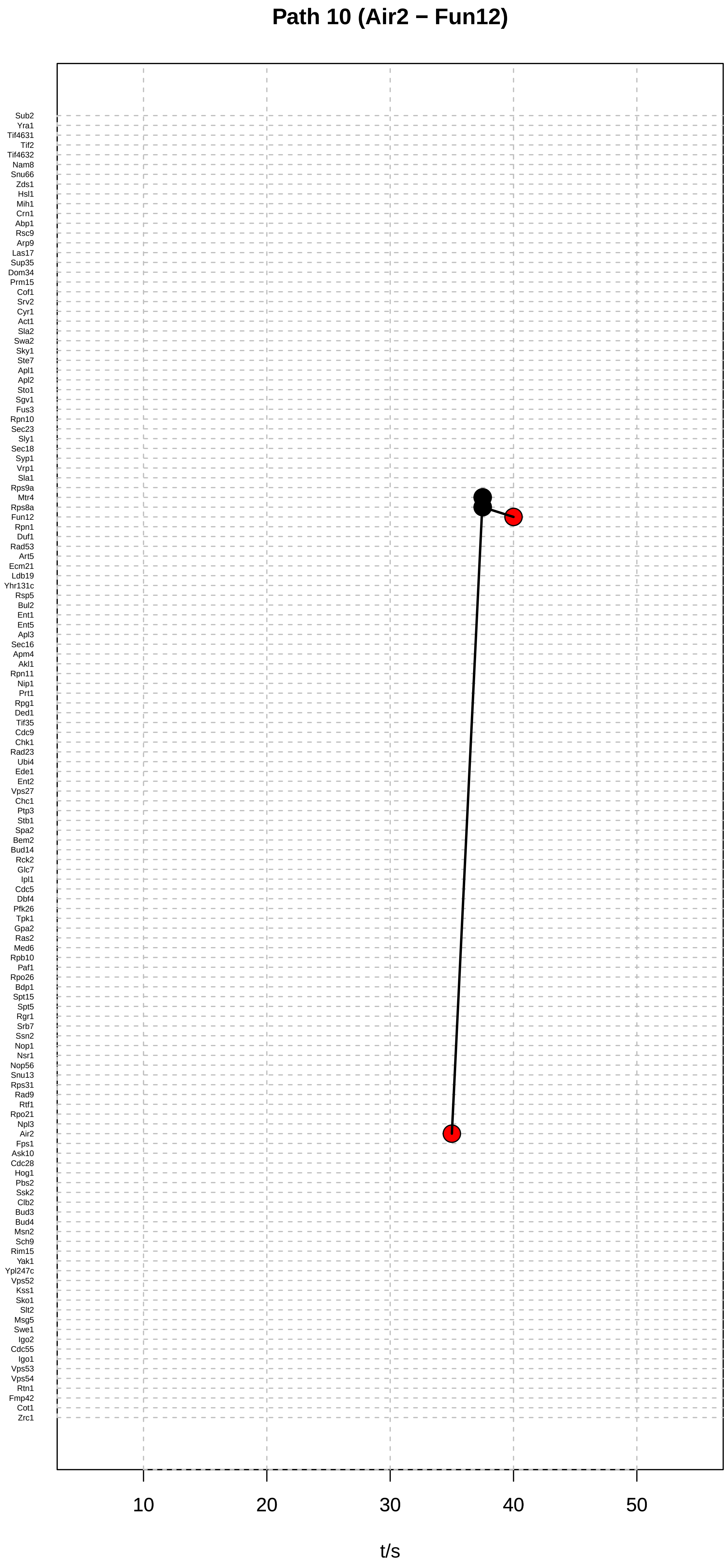

Path 10 (Air2 − Fun12)
Sub2
Yra1
Tif4631
Tif2
Tif4632
Nam8
Snu66
Zds1
Hsl1
Mih1
Crn1
Abp1
Rsc9
Arp9
Las17
Sup35
Dom34
Prm15
Cof1
Srv2
Cyr1
Act1
Sla2
Swa2
Sky1
Ste7
Apl1
Apl2
Sto1
Sgv1
Fus3
Rpn10
Sec23
Sly1
Sec18
Syp1
Vrp1
Sla1
Rps9a
Mtr4
Rps8a
Fun12
Rpn1
Duf1
Rad53
Art5
Ecm21
Ldb19
Yhr131c
Rsp5
Bul2
Ent1
Ent5
Apl3
Sec16
Apm4
Akl1
Rpn11
Nip1
Prt1
Rpg1
Ded1
Tif35
Cdc9
Chk1
Rad23
Ubi4
Ede1
Ent2
Vps27
Chc1
Ptp3
Stb1
Spa2
Bem2
Bud14
Rck2
Glc7
Ipl1
Cdc5
Dbf4
Pfk26
Tpk1
Gpa2
Ras2
Med6
Rpb10
Paf1
Rpo26
Bdp1
Spt15
Spt5
Rgr1
Srb7
Ssn2
Nop1
Nsr1
Nop56
Snu13
Rps31
Rad9
Rtf1
Rpo21
Npl3
Air2
Fps1
Ask10
Cdc28
Hog1
Pbs2
Ssk2
Clb2
Bud3
Bud4
Msn2
Sch9
Rim15
Yak1
Ypl247c
Vps52
Kss1
Sko1
Slt2
Msg5
Swe1
Igo2
Cdc55
Igo1
Vps53
Vps54
Rtn1
Fmp42
Cot1
Zrc1
10
20
30
40
50
t/s

#### Path 11 (Ssn2 − Rtf1)

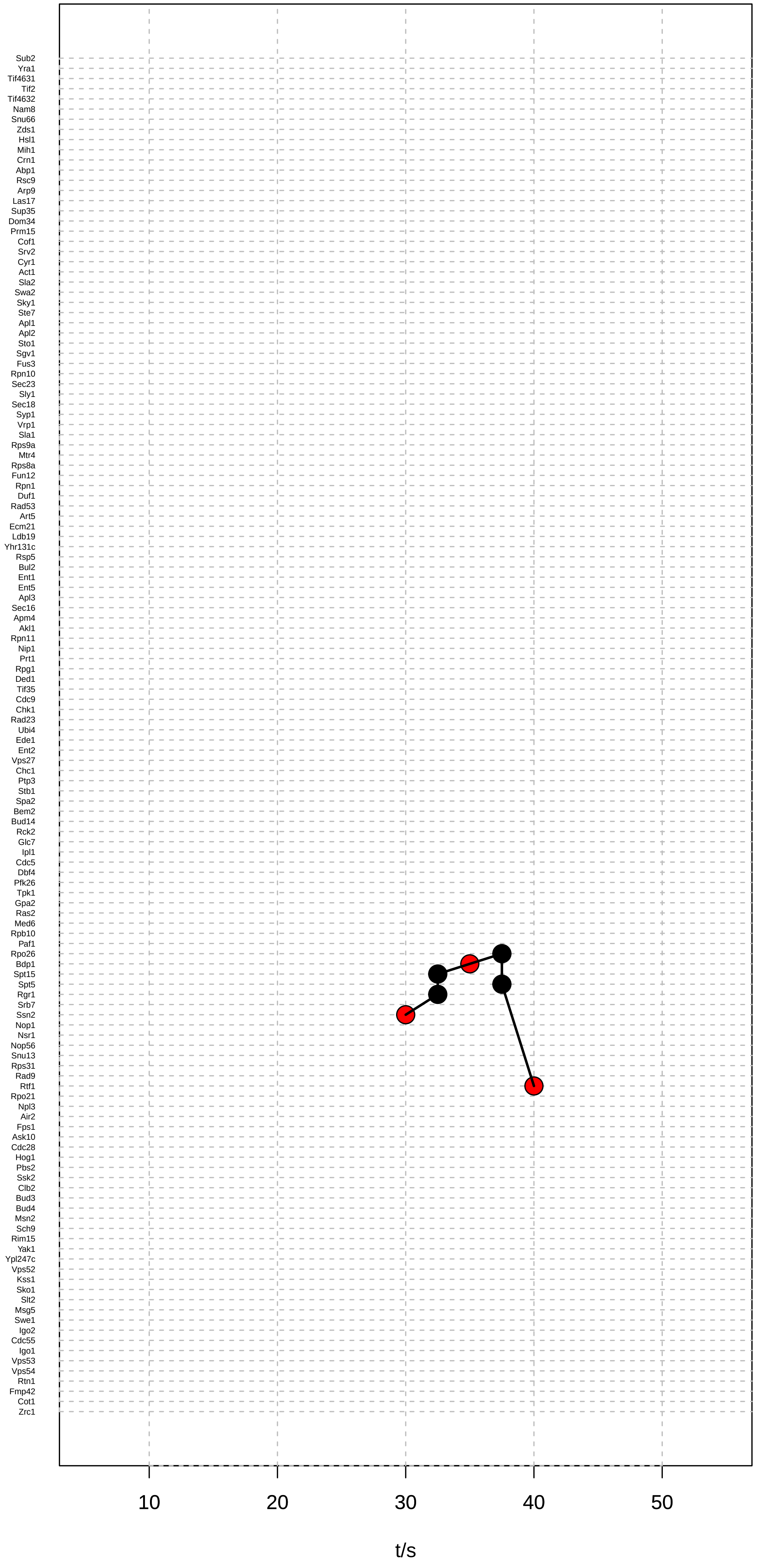

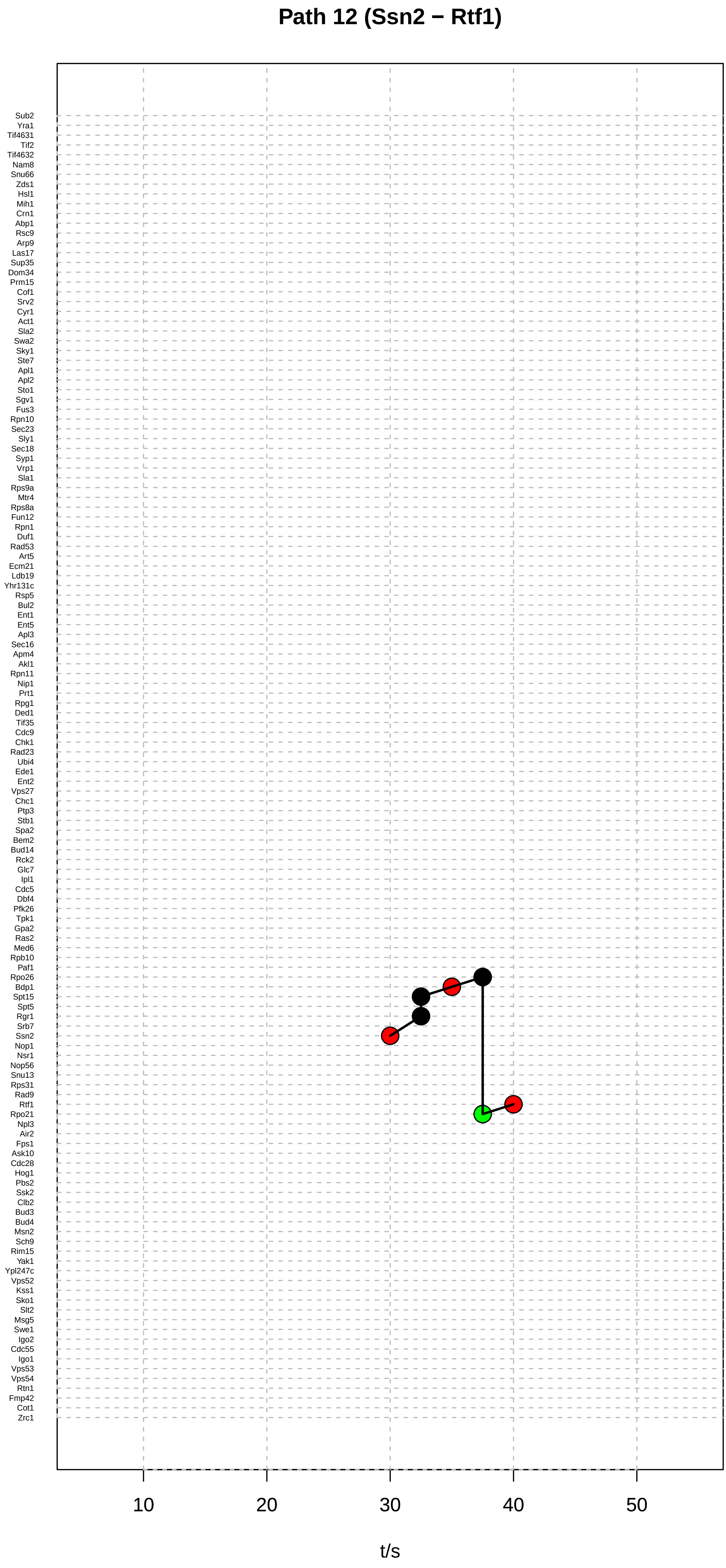

Path 12 (Ssn2 − Rtf1)
Sub2
Yra1
Tif4631
Tif2
Tif4632
Nam8
Snu66
Zds1
Hsl1
Mih1
Crn1
Abp1
Rsc9
Arp9
Las17
Sup35
Dom34
Prm15
Cof1
Srv2
Cyr1
Act1
Sla2
Swa2
Sky1
Ste7
Apl1
Apl2
Sto1
Sgv1
Fus3
Rpn10
Sec23
Sly1
Sec18
Syp1
Vrp1
Sla1
Rps9a
Mtr4
Rps8a
Fun12
Rpn1
Duf1
Rad53
Art5
Ecm21
Ldb19
Yhr131c
Rsp5
Bul2
Ent1
Ent5
Apl3
Sec16
Apm4
Akl1
Rpn11
Nip1
Prt1
Rpg1
Ded1
Tif35
Cdc9
Chk1
Rad23
Ubi4
Ede1
Ent2
Vps27
Chc1
Ptp3
Stb1
Spa2
Bem2
Bud14
Rck2
Glc7
Ipl1
Cdc5
Dbf4
Pfk26
Tpk1
Gpa2
Ras2
Med6
Rpb10
Paf1
Rpo26
Bdp1
Spt15
Spt5
Rgr1
Srb7
Ssn2
Nop1
Nsr1
Nop56
Snu13
Rps31
Rad9
Rtf1
Rpo21
Npl3
Air2
Fps1
Ask10
Cdc28
Hog1
Pbs2
Ssk2
Clb2
Bud3
Bud4
Msn2
Sch9
Rim15
Yak1
Ypl247c
Vps52
Kss1
Sko1
Slt2
Msg5
Swe1
Igo2
Cdc55
Igo1
Vps53
Vps54
Rtn1
Fmp42
Cot1
Zrc1
10
20
30
40
50
t/s

Path 13 (Bul2 − Ldb19)

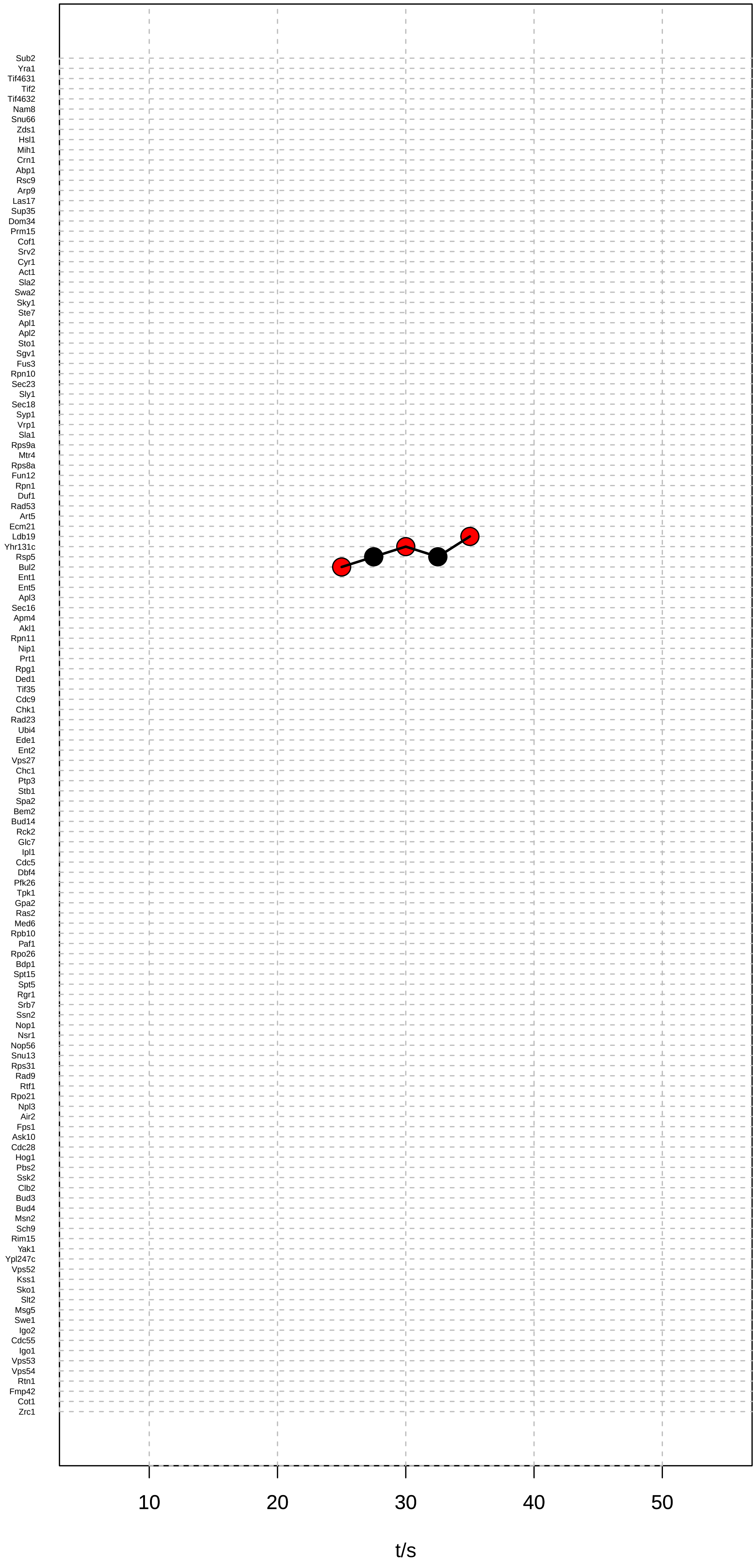

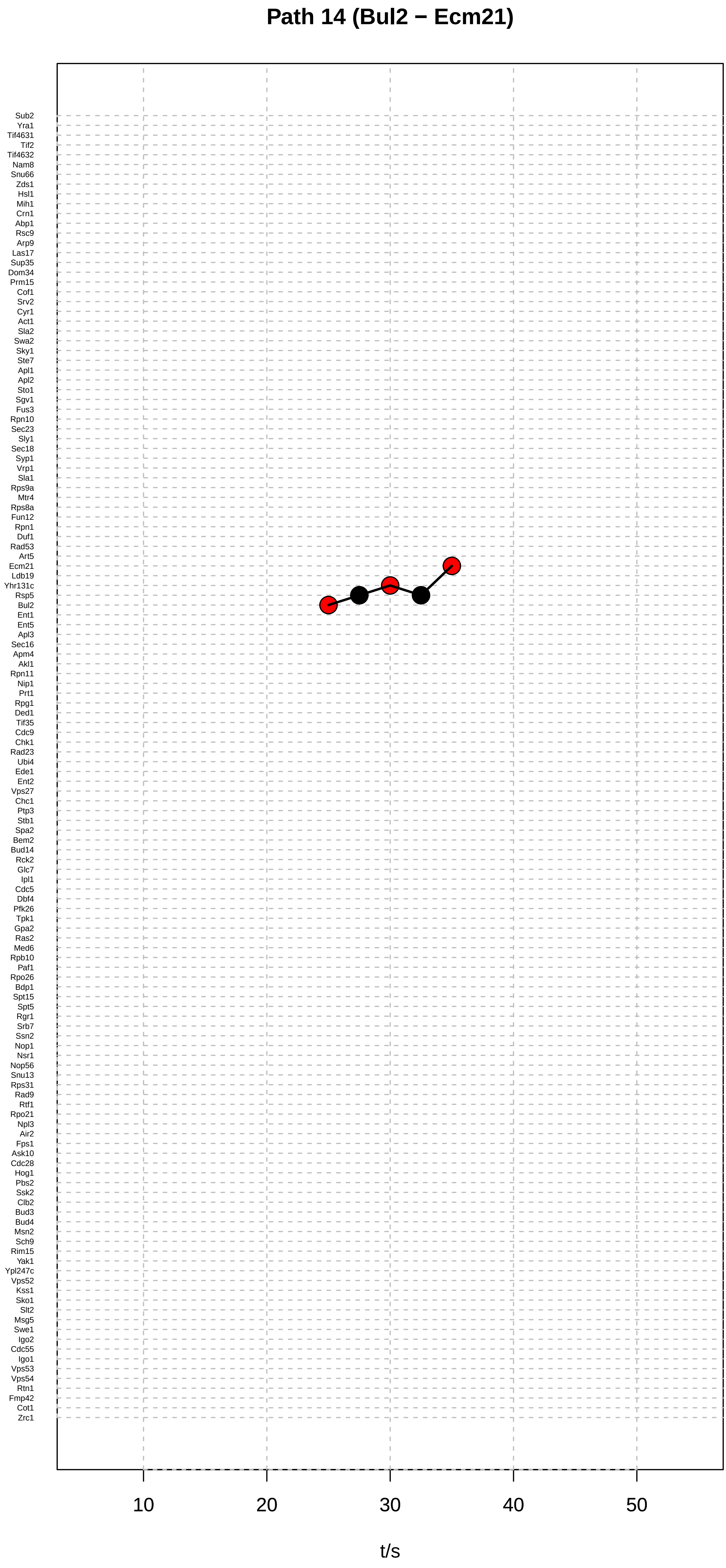

Path 14 (Bul2 − Ecm21)
Sub2
Yra1
Tif4631
Tif2
Tif4632
Nam8
Snu66
Zds1
Hsl1
Mih1
Crn1
Abp1
Rsc9
Arp9
Las17
Sup35
Dom34
Prm15
Cof1
Srv2
Cyr1
Act1
Sla2
Swa2
Sky1
Ste7
Apl1
Apl2
Sto1
Sgv1
Fus3
Rpn10
Sec23
Sly1
Sec18
Syp1
Vrp1
Sla1
Rps9a
Mtr4
Rps8a
Fun12
Rpn1
Duf1
Rad53
Art5
Ecm21
Ldb19
Yhr131c
Rsp5
Bul2
Ent1
Ent5
Apl3
Sec16
Apm4
Akl1
Rpn11
Nip1
Prt1
Rpg1
Ded1
Tif35
Cdc9
Chk1
Rad23
Ubi4
Ede1
Ent2
Vps27
Chc1
Ptp3
Stb1
Spa2
Bem2
Bud14
Rck2
Glc7
Ipl1
Cdc5
Dbf4
Pfk26
Tpk1
Gpa2
Ras2
Med6
Rpb10
Paf1
Rpo26
Bdp1
Spt15
Spt5
Rgr1
Srb7
Ssn2
Nop1
Nsr1
Nop56
Snu13
Rps31
Rad9
Rtf1
Rpo21
Npl3
Air2
Fps1
Ask10
Cdc28
Hog1
Pbs2
Ssk2
Clb2
Bud3
Bud4
Msn2
Sch9
Rim15
Yak1
Ypl247c
Vps52
Kss1
Sko1
Slt2
Msg5
Swe1
Igo2
Cdc55
Igo1
Vps53
Vps54
Rtn1
Fmp42
Cot1
Zrc1
10
20
30
40
50
t/s

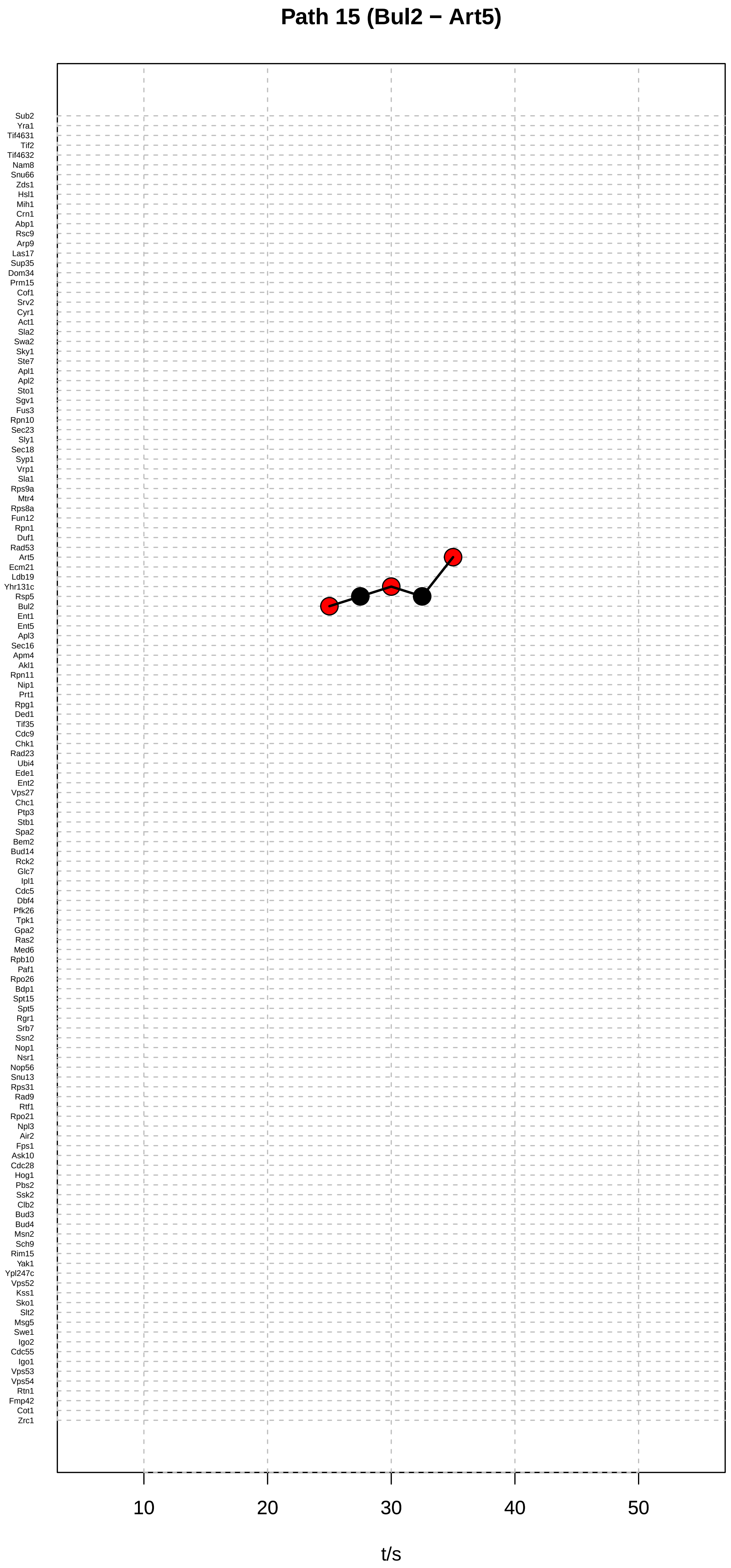
Path 15 (Bul2 − Art5)
Sub2
Yra1
Tif4631
Tif2
Tif4632
Nam8
Snu66
Zds1
Hsl1
Mih1
Crn1
Abp1
Rsc9
Arp9
Las17
Sup35
Dom34
Prm15
Cof1
Srv2
Cyr1
Act1
Sla2
Swa2
Sky1
Ste7
Apl1
Apl2
Sto1
Sgv1
Fus3
Rpn10
Sec23
Sly1
Sec18
Syp1
Vrp1
Sla1
Rps9a
Mtr4
Rps8a
Fun12
Rpn1
Duf1
Rad53
Art5
Ecm21
Ldb19
Yhr131c
Rsp5
Bul2
Ent1
Ent5
Apl3
Sec16
Apm4
Akl1
Rpn11
Nip1
Prt1
Rpg1
Ded1
Tif35
Cdc9
Chk1
Rad23
Ubi4
Ede1
Ent2
Vps27
Chc1
Ptp3
Stb1
Spa2
Bem2
Bud14
Rck2
Glc7
Ipl1
Cdc5
Dbf4
Pfk26
Tpk1
Gpa2
Ras2
Med6
Rpb10
Paf1
Rpo26
Bdp1
Spt15
Spt5
Rgr1
Srb7
Ssn2
Nop1
Nsr1
Nop56
Snu13
Rps31
Rad9
Rtf1
Rpo21
Npl3
Air2
Fps1
Ask10
Cdc28
Hog1
Pbs2
Ssk2
Clb2
Bud3
Bud4
Msn2
Sch9
Rim15
Yak1
Ypl247c
Vps52
Kss1
Sko1
Slt2
Msg5
Swe1
Igo2
Cdc55
Igo1
Vps53
Vps54
Rtn1
Fmp42
Cot1
Zrc1
10
20
30
40
50
t/s

Path 16 (Rim15 − Pfk26)

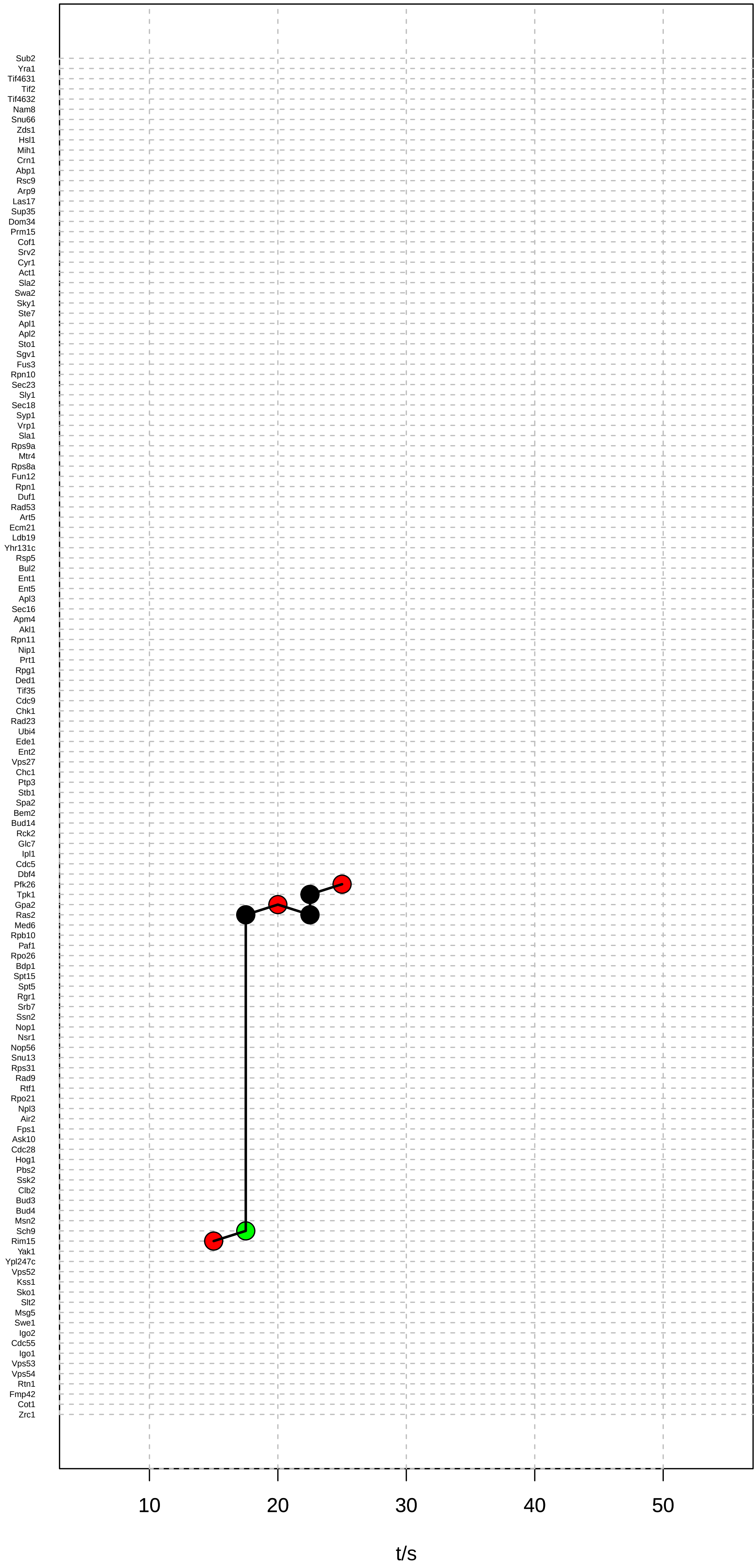

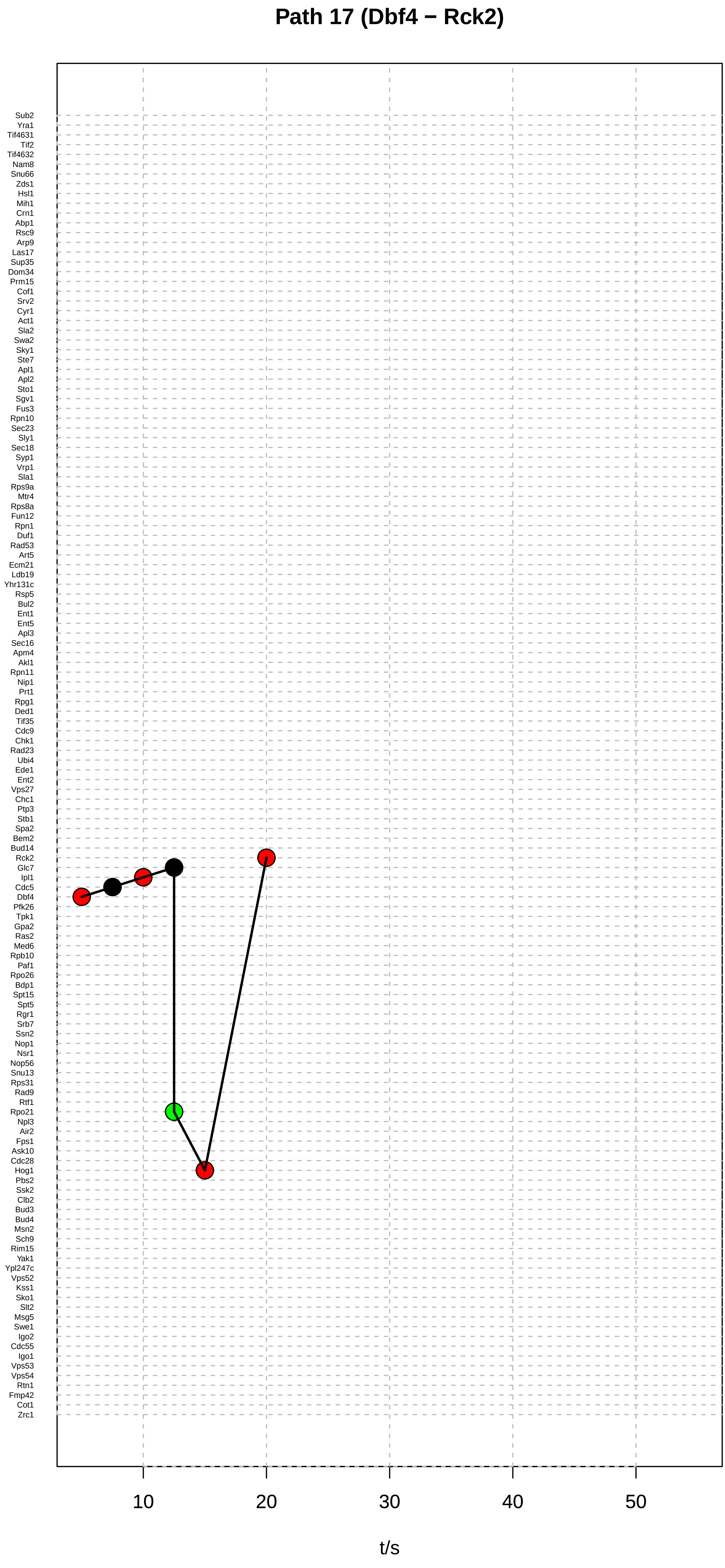

Path 17 (Dbf4 − Rck2)
Sub2
Yra1
Tif4631
Tif2
Tif4632
Nam8
Snu66
Zds1
Hsl1
Mih1
Crn1
Abp1
Rsc9
Arp9
Las17
Sup35
Dom34
Prm15
Cof1
Srv2
Cyr1
Act1
Sla2
Swa2
Sky1
Ste7
Apl1
Apl2
Sto1
Sgv1
Fus3
Rpn10
Sec23
Sly1
Sec18
Syp1
Vrp1
Sla1
Rps9a
Mtr4
Rps8a
Fun12
Rpn1
Duf1
Rad53
Art5
Ecm21
Ldb19
Yhr131c
Rsp5
Bul2
Ent1
Ent5
Apl3
Sec16
Apm4
Akl1
Rpn11
Nip1
Prt1
Rpg1
Ded1
Tif35
Cdc9
Chk1
Rad23
Ubi4
Ede1
Ent2
Vps27
Chc1
Ptp3
Stb1
Spa2
Bem2
Bud14
Rck2
Glc7
Ipl1
Cdc5
Dbf4
Pfk26
Tpk1
Gpa2
Ras2
Med6
Rpb10
Paf1
Rpo26
Bdp1
Spt15
Spt5
Rgr1
Srb7
Ssn2
Nop1
Nsr1
Nop56
Snu13
Rps31
Rad9
Rtf1
Rpo21
Npl3
Air2
Fps1
Ask10
Cdc28
Hog1
Pbs2
Ssk2
Clb2
Bud3
Bud4
Msn2
Sch9
Rim15
Yak1
Ypl247c
Vps52
Kss1
Sko1
Slt2
Msg5
Swe1
Igo2
Cdc55
Igo1
Vps53
Vps54
Rtn1
Fmp42
Cot1
Zrc1
10
20
30
40
50
t/s

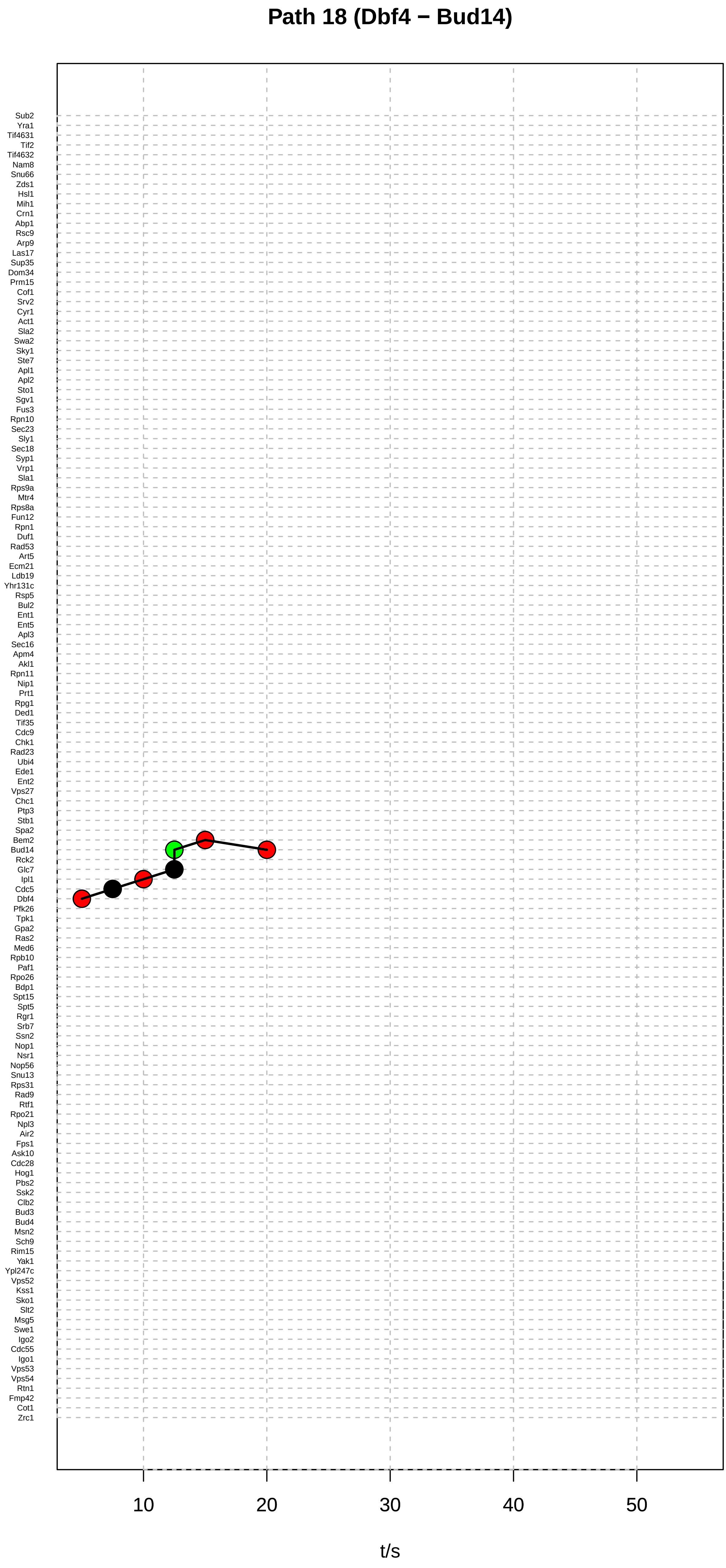
Path 18 (Dbf4 − Bud14)
Sub2
Yra1
Tif4631
Tif2
Tif4632
Nam8
Snu66
Zds1
Hsl1
Mih1
Crn1
Abp1
Rsc9
Arp9
Las17
Sup35
Dom34
Prm15
Cof1
Srv2
Cyr1
Act1
Sla2
Swa2
Sky1
Ste7
Apl1
Apl2
Sto1
Sgv1
Fus3
Rpn10
Sec23
Sly1
Sec18
Syp1
Vrp1
Sla1
Rps9a
Mtr4
Rps8a
Fun12
Rpn1
Duf1
Rad53
Art5
Ecm21
Ldb19
Yhr131c
Rsp5
Bul2
Ent1
Ent5
Apl3
Sec16
Apm4
Akl1
Rpn11
Nip1
Prt1
Rpg1
Ded1
Tif35
Cdc9
Chk1
Rad23
Ubi4
Ede1
Ent2
Vps27
Chc1
Ptp3
Stb1
Spa2
Bem2
Bud14
Rck2
Glc7
Ipl1
Cdc5
Dbf4
Pfk26
Tpk1
Gpa2
Ras2
Med6
Rpb10
Paf1
Rpo26
Bdp1
Spt15
Spt5
Rgr1
Srb7
Ssn2
Nop1
Nsr1
Nop56
Snu13
Rps31
Rad9
Rtf1
Rpo21
Npl3
Air2
Fps1
Ask10
Cdc28
Hog1
Pbs2
Ssk2
Clb2
Bud3
Bud4
Msn2
Sch9
Rim15
Yak1
Ypl247c
Vps52
Kss1
Sko1
Slt2
Msg5
Swe1
Igo2
Cdc55
Igo1
Vps53
Vps54
Rtn1
Fmp42
Cot1
Zrc1
10
20
30
40
50
t/s

### Path 19 (Dbf4 − Rck2)

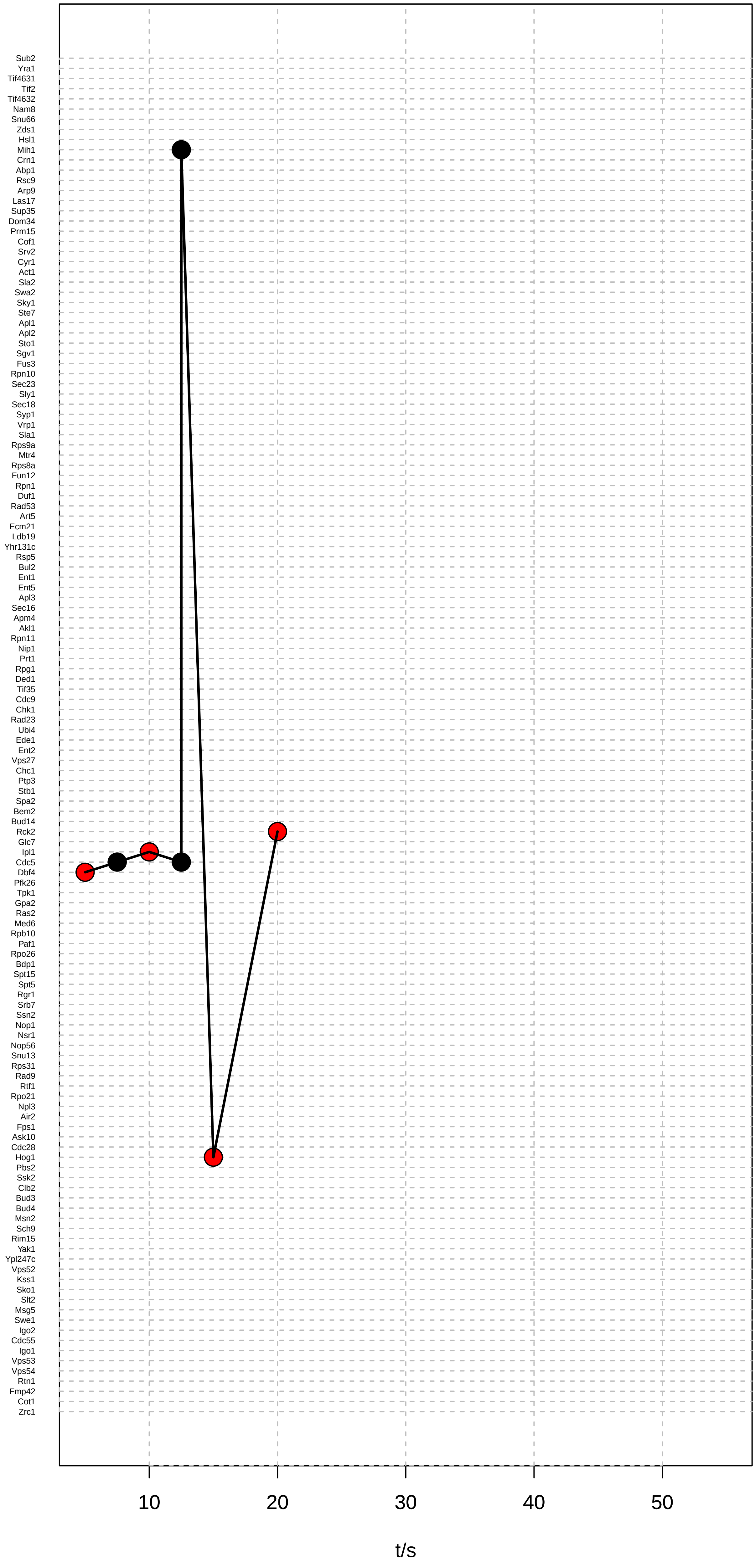

### Path 20 (Ptp3 − Rck2)

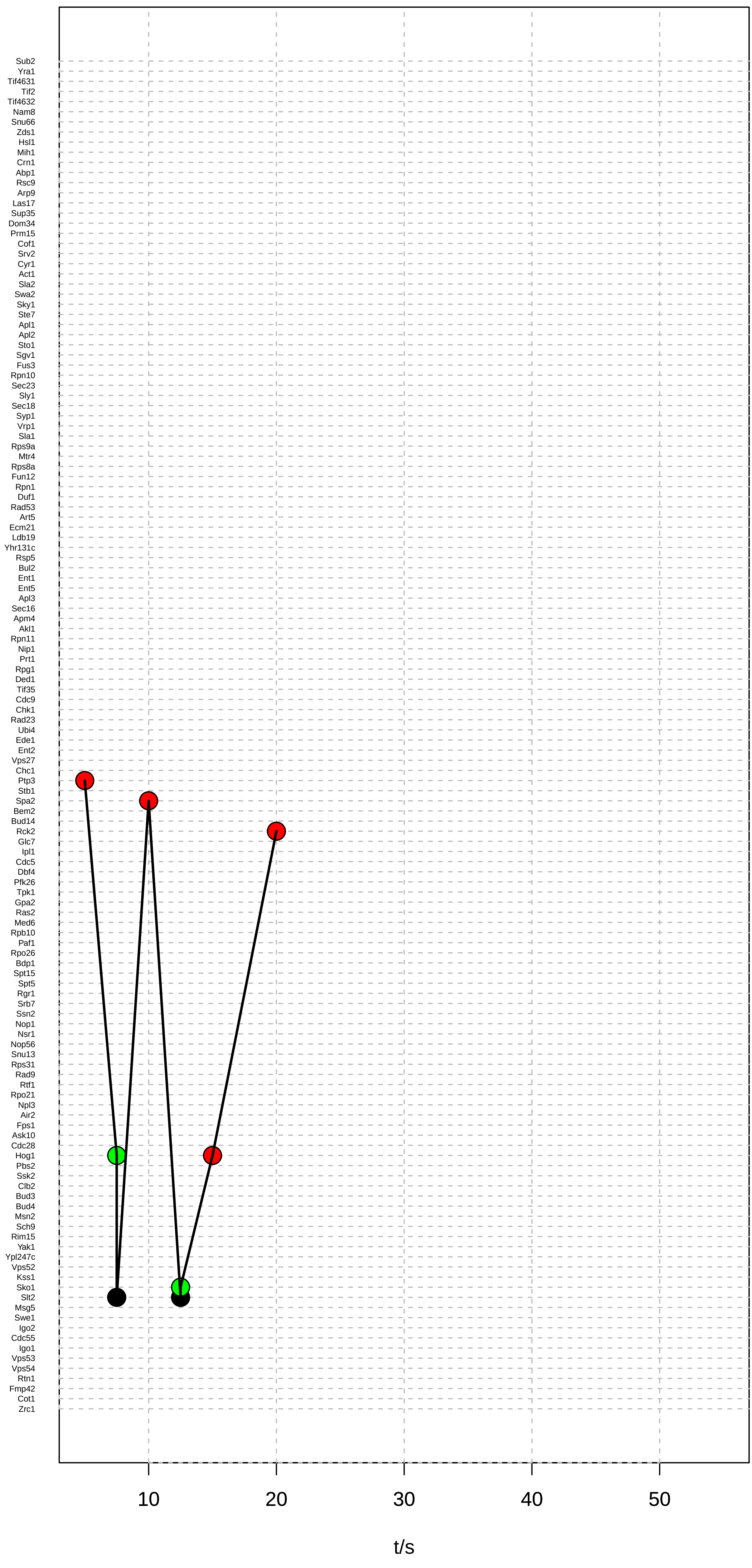

### Path 21 (Ptp3 − Rck2)

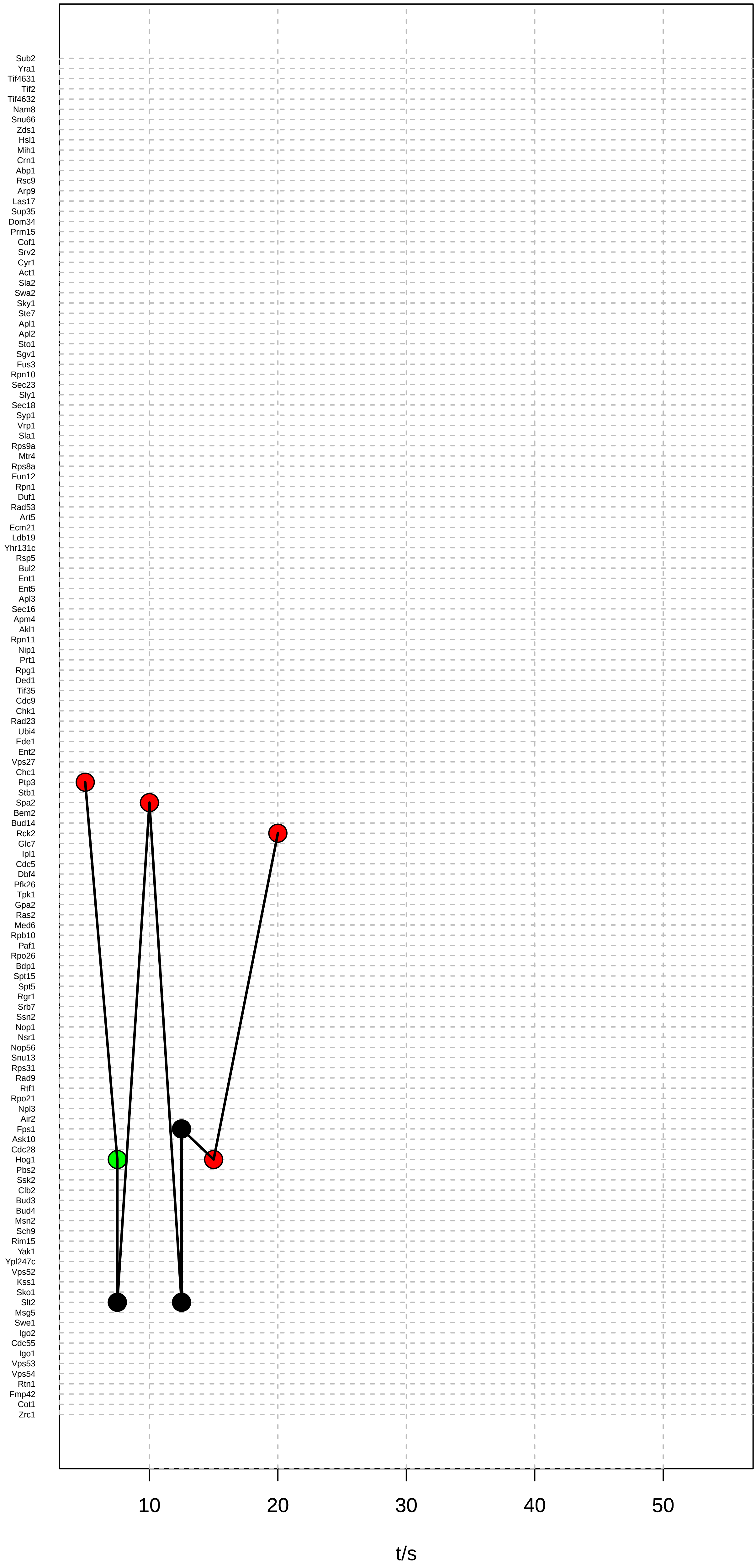

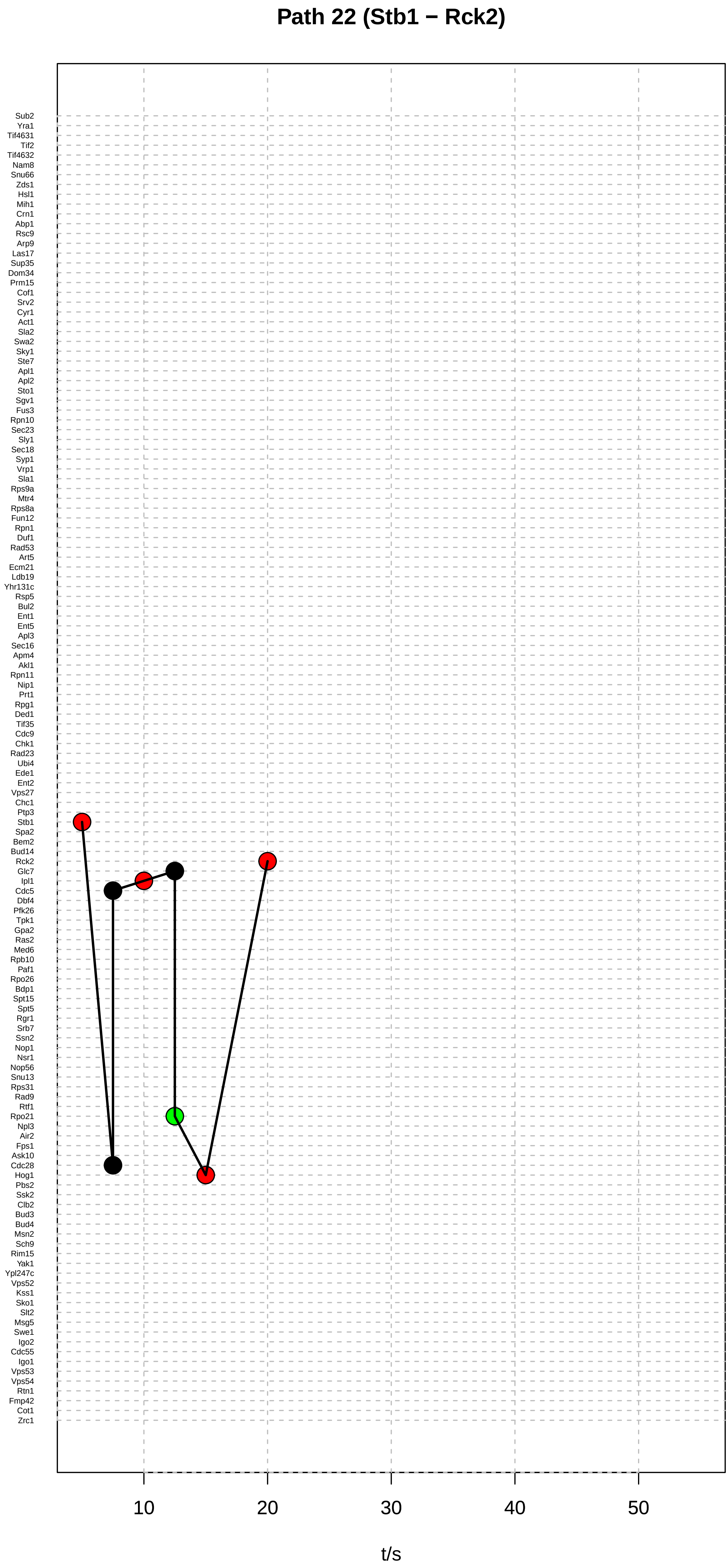
Path 22 (Stb1 − Rck2)
Sub2
Yra1
Tif4631
Tif2
Tif4632
Nam8
Snu66
Zds1
Hsl1
Mih1
Crn1
Abp1
Rsc9
Arp9
Las17
Sup35
Dom34
Prm15
Cof1
Srv2
Cyr1
Act1
Sla2
Swa2
Sky1
Ste7
Apl1
Apl2
Sto1
Sgv1
Fus3
Rpn10
Sec23
Sly1
Sec18
Syp1
Vrp1
Sla1
Rps9a
Mtr4
Rps8a
Fun12
Rpn1
Duf1
Rad53
Art5
Ecm21
Ldb19
Yhr131c
Rsp5
Bul2
Ent1
Ent5
Apl3
Sec16
Apm4
Akl1
Rpn11
Nip1
Prt1
Rpg1
Ded1
Tif35
Cdc9
Chk1
Rad23
Ubi4
Ede1
Ent2
Vps27
Chc1
Ptp3
Stb1
Spa2
Bem2
Bud14
Rck2
Glc7
Ipl1
Cdc5
Dbf4
Pfk26
Tpk1
Gpa2
Ras2
Med6
Rpb10
Paf1
Rpo26
Bdp1
Spt15
Spt5
Rgr1
Srb7
Ssn2
Nop1
Nsr1
Nop56
Snu13
Rps31
Rad9
Rtf1
Rpo21
Npl3
Air2
Fps1
Ask10
Cdc28
Hog1
Pbs2
Ssk2
Clb2
Bud3
Bud4
Msn2
Sch9
Rim15
Yak1
Ypl247c
Vps52
Kss1
Sko1
Slt2
Msg5
Swe1
Igo2
Cdc55
Igo1
Vps53
Vps54
Rtn1
Fmp42
Cot1
Zrc1
10
20
30
40
50
t/s

#### Path 23 (Stb1 − Bud14)

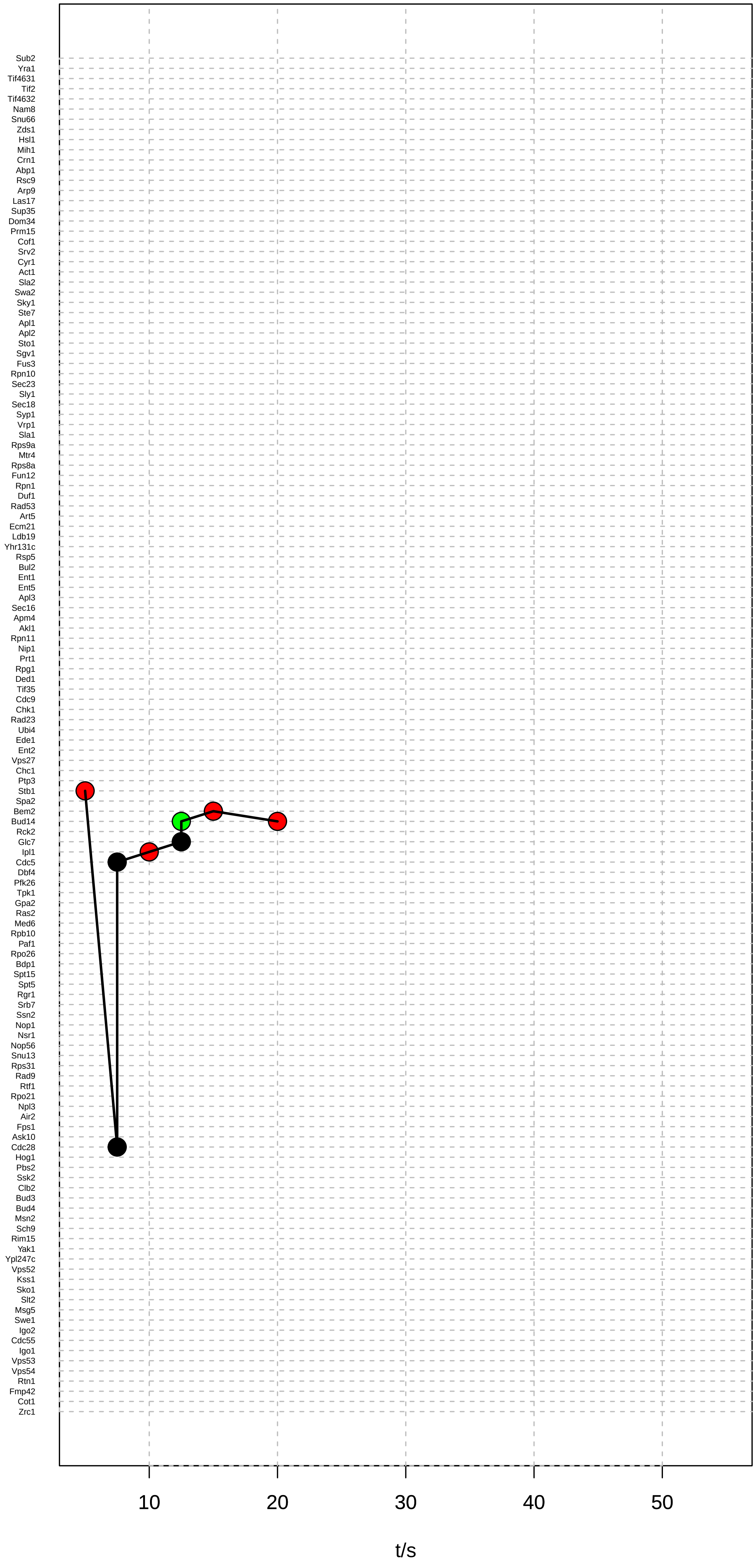

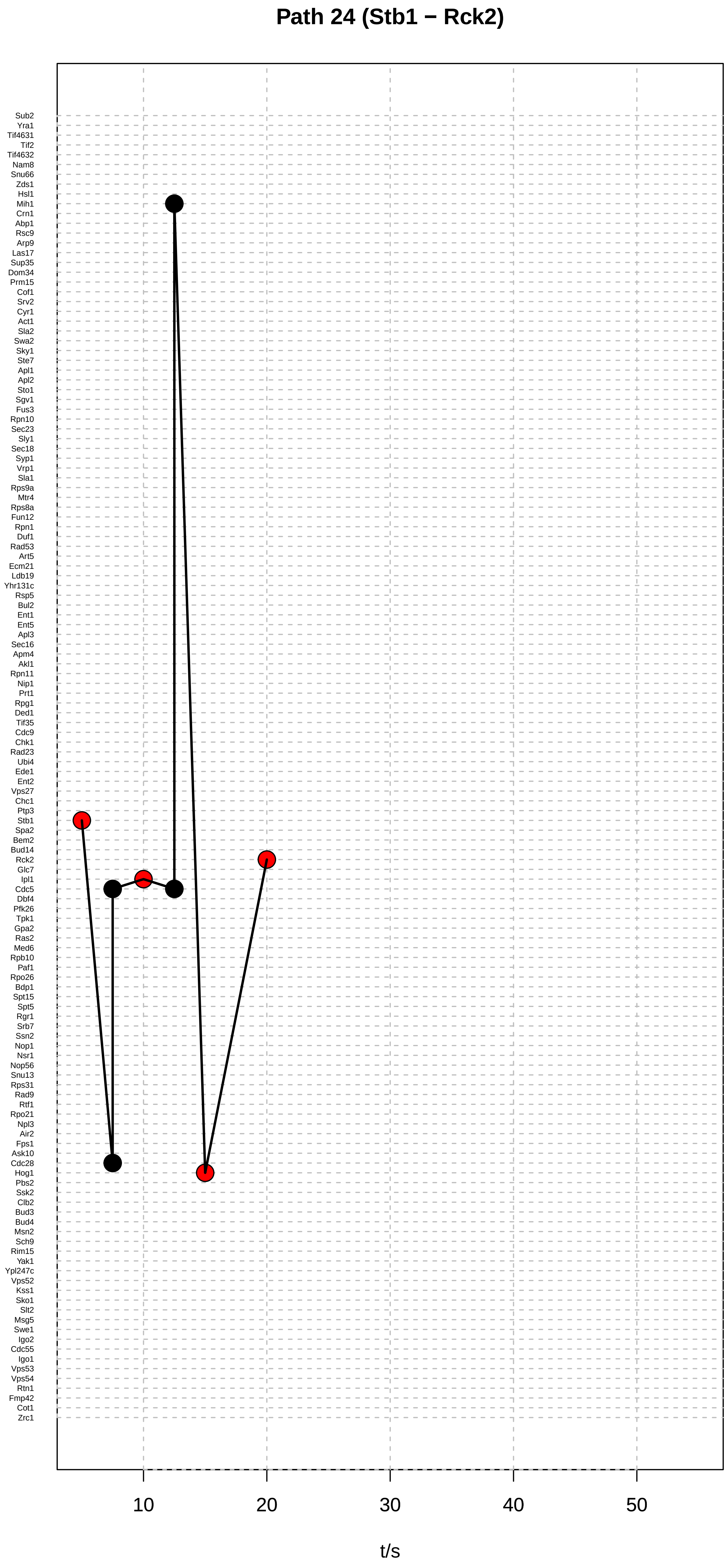
Path 24 (Stb1 − Rck2)
Sub2
Yra1
Tif4631
Tif2
Tif4632
Nam8
Snu66
Zds1
Hsl1
Mih1
Crn1
Abp1
Rsc9
Arp9
Las17
Sup35
Dom34
Prm15
Cof1
Srv2
Cyr1
Act1
Sla2
Swa2
Sky1
Ste7
Apl1
Apl2
Sto1
Sgv1
Fus3
Rpn10
Sec23
Sly1
Sec18
Syp1
Vrp1
Sla1
Rps9a
Mtr4
Rps8a
Fun12
Rpn1
Duf1
Rad53
Art5
Ecm21
Ldb19
Yhr131c
Rsp5
Bul2
Ent1
Ent5
Apl3
Sec16
Apm4
Akl1
Rpn11
Nip1
Prt1
Rpg1
Ded1
Tif35
Cdc9
Chk1
Rad23
Ubi4
Ede1
Ent2
Vps27
Chc1
Ptp3
Stb1
Spa2
Bem2
Bud14
Rck2
Glc7
Ipl1
Cdc5
Dbf4
Pfk26
Tpk1
Gpa2
Ras2
Med6
Rpb10
Paf1
Rpo26
Bdp1
Spt15
Spt5
Rgr1
Srb7
Ssn2
Nop1
Nsr1
Nop56
Snu13
Rps31
Rad9
Rtf1
Rpo21
Npl3
Air2
Fps1
Ask10
Cdc28
Hog1
Pbs2
Ssk2
Clb2
Bud3
Bud4
Msn2
Sch9
Rim15
Yak1
Ypl247c
Vps52
Kss1
Sko1
Slt2
Msg5
Swe1
Igo2
Cdc55
Igo1
Vps53
Vps54
Rtn1
Fmp42
Cot1
Zrc1
10
20
30
40
50
t/s

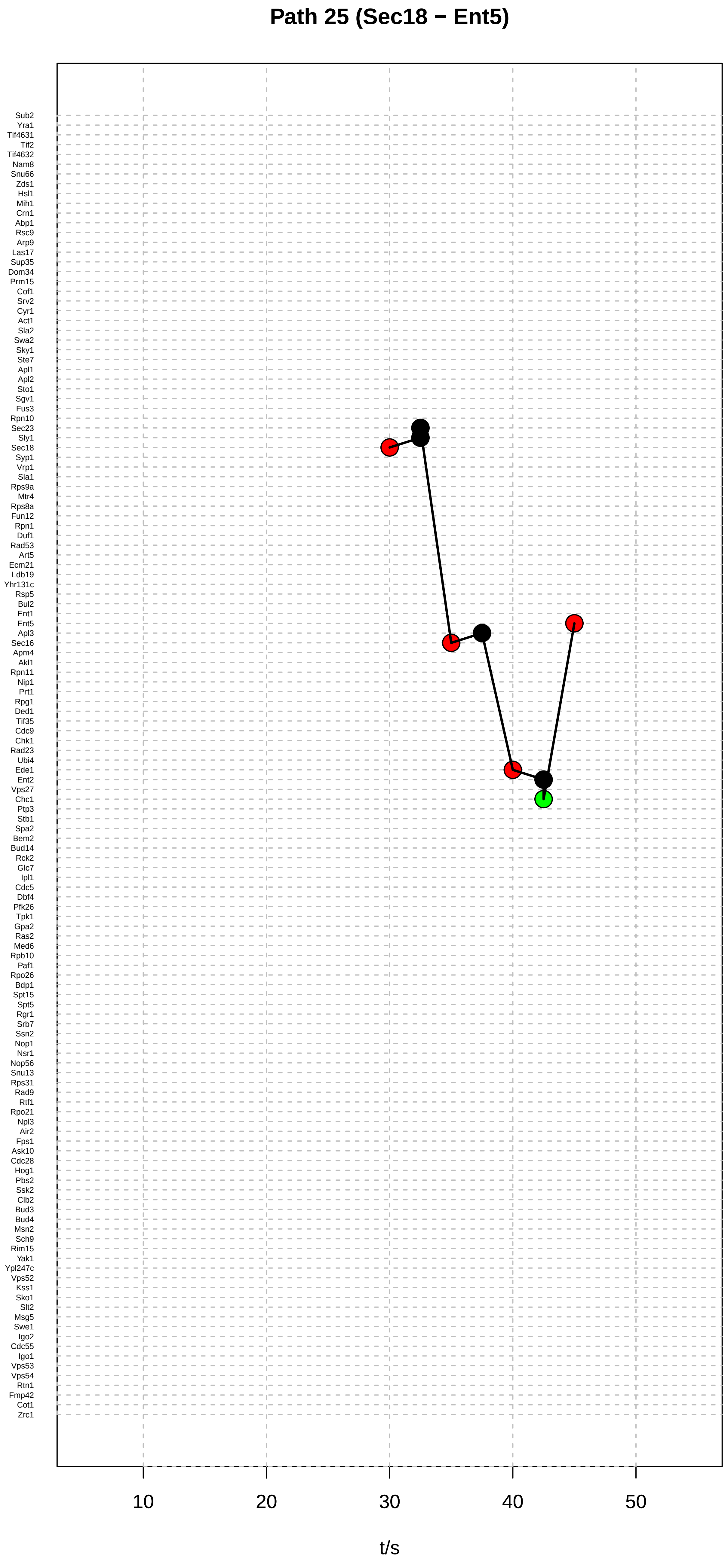
Path 25 (Sec18 − Ent5)
Sub2
Yra1
Tif4631
Tif2
Tif4632
Nam8
Snu66
Zds1
Hsl1
Mih1
Crn1
Abp1
Rsc9
Arp9
Las17
Sup35
Dom34
Prm15
Cof1
Srv2
Cyr1
Act1
Sla2
Swa2
Sky1
Ste7
Apl1
Apl2
Sto1
Sgv1
Fus3
Rpn10
Sec23
Sly1
Sec18
Syp1
Vrp1
Sla1
Rps9a
Mtr4
Rps8a
Fun12
Rpn1
Duf1
Rad53
Art5
Ecm21
Ldb19
Yhr131c
Rsp5
Bul2
Ent1
Ent5
Apl3
Sec16
Apm4
Akl1
Rpn11
Nip1
Prt1
Rpg1
Ded1
Tif35
Cdc9
Chk1
Rad23
Ubi4
Ede1
Ent2
Vps27
Chc1
Ptp3
Stb1
Spa2
Bem2
Bud14
Rck2
Glc7
Ipl1
Cdc5
Dbf4
Pfk26
Tpk1
Gpa2
Ras2
Med6
Rpb10
Paf1
Rpo26
Bdp1
Spt15
Spt5
Rgr1
Srb7
Ssn2
Nop1
Nsr1
Nop56
Snu13
Rps31
Rad9
Rtf1
Rpo21
Npl3
Air2
Fps1
Ask10
Cdc28
Hog1
Pbs2
Ssk2
Clb2
Bud3
Bud4
Msn2
Sch9
Rim15
Yak1
Ypl247c
Vps52
Kss1
Sko1
Slt2
Msg5
Swe1
Igo2
Cdc55
Igo1
Vps53
Vps54
Rtn1
Fmp42
Cot1
Zrc1
10
20
30
40
50
t/s

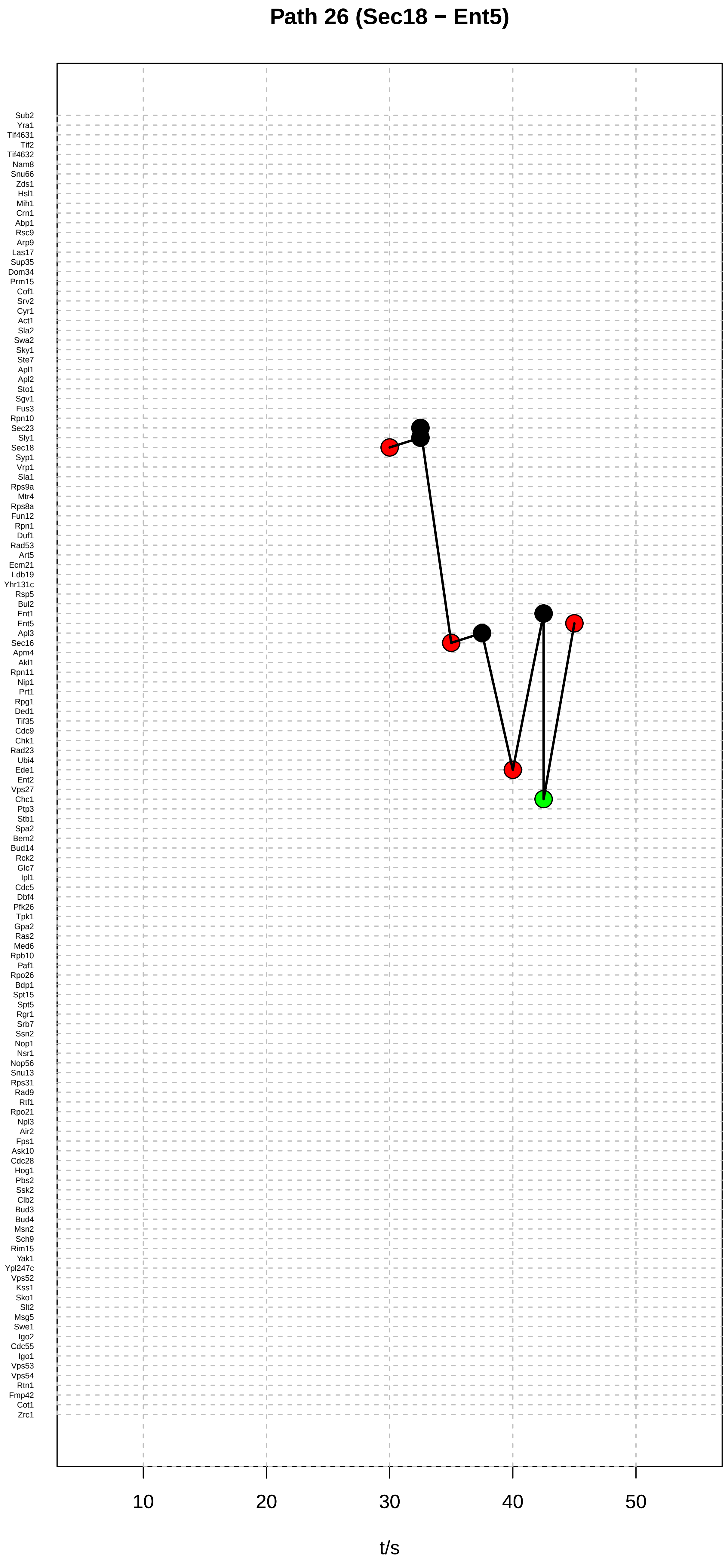
Path 26 (Sec18 − Ent5)
Sub2
Yra1
Tif4631
Tif2
Tif4632
Nam8
Snu66
Zds1
Hsl1
Mih1
Crn1
Abp1
Rsc9
Arp9
Las17
Sup35
Dom34
Prm15
Cof1
Srv2
Cyr1
Act1
Sla2
Swa2
Sky1
Ste7
Apl1
Apl2
Sto1
Sgv1
Fus3
Rpn10
Sec23
Sly1
Sec18
Syp1
Vrp1
Sla1
Rps9a
Mtr4
Rps8a
Fun12
Rpn1
Duf1
Rad53
Art5
Ecm21
Ldb19
Yhr131c
Rsp5
Bul2
Ent1
Ent5
Apl3
Sec16
Apm4
Akl1
Rpn11
Nip1
Prt1
Rpg1
Ded1
Tif35
Cdc9
Chk1
Rad23
Ubi4
Ede1
Ent2
Vps27
Chc1
Ptp3
Stb1
Spa2
Bem2
Bud14
Rck2
Glc7
Ipl1
Cdc5
Dbf4
Pfk26
Tpk1
Gpa2
Ras2
Med6
Rpb10
Paf1
Rpo26
Bdp1
Spt15
Spt5
Rgr1
Srb7
Ssn2
Nop1
Nsr1
Nop56
Snu13
Rps31
Rad9
Rtf1
Rpo21
Npl3
Air2
Fps1
Ask10
Cdc28
Hog1
Pbs2
Ssk2
Clb2
Bud3
Bud4
Msn2
Sch9
Rim15
Yak1
Ypl247c
Vps52
Kss1
Sko1
Slt2
Msg5
Swe1
Igo2
Cdc55
Igo1
Vps53
Vps54
Rtn1
Fmp42
Cot1
Zrc1
10
20
30
40
50
t/s

#### Path 27 (Akl1 − Ent5)

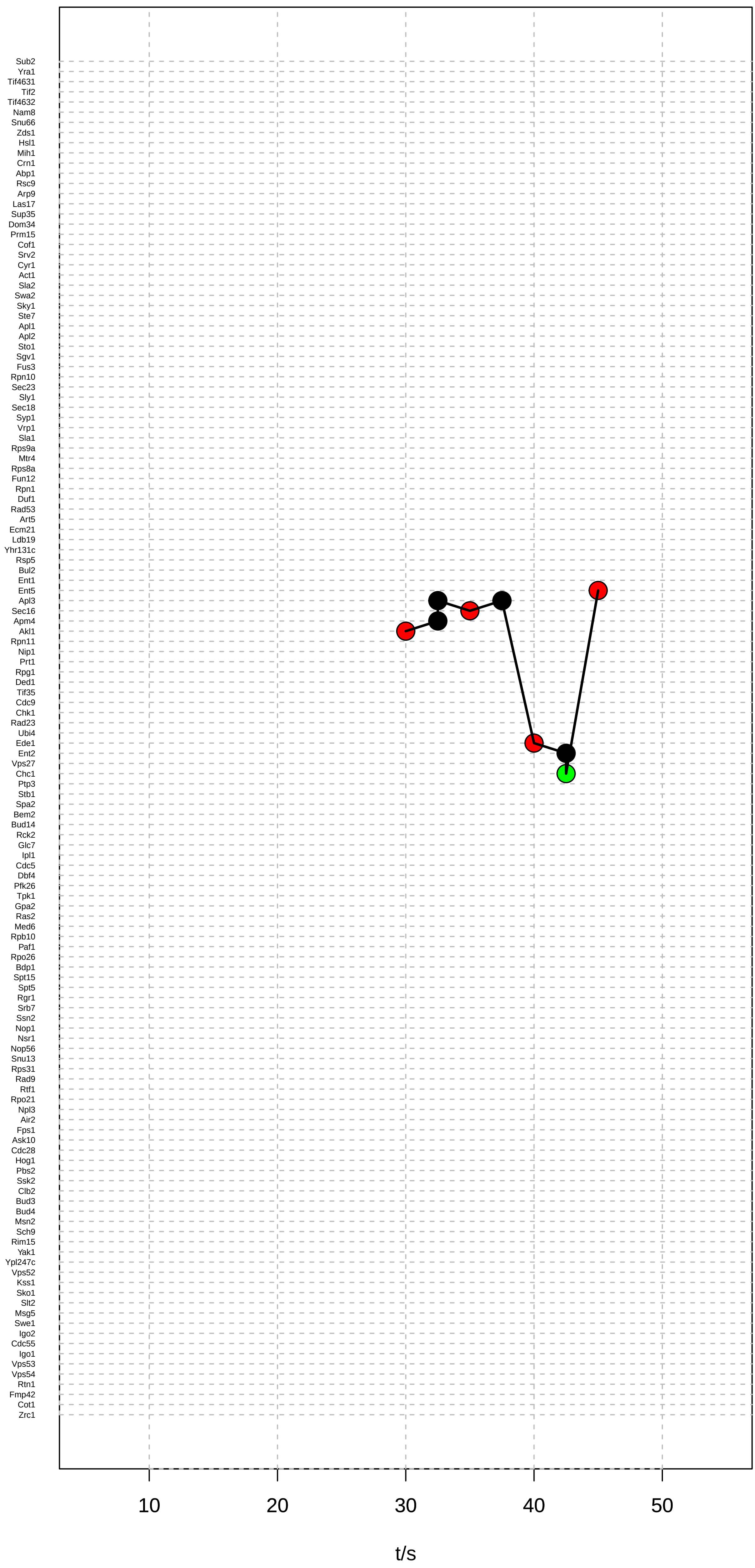

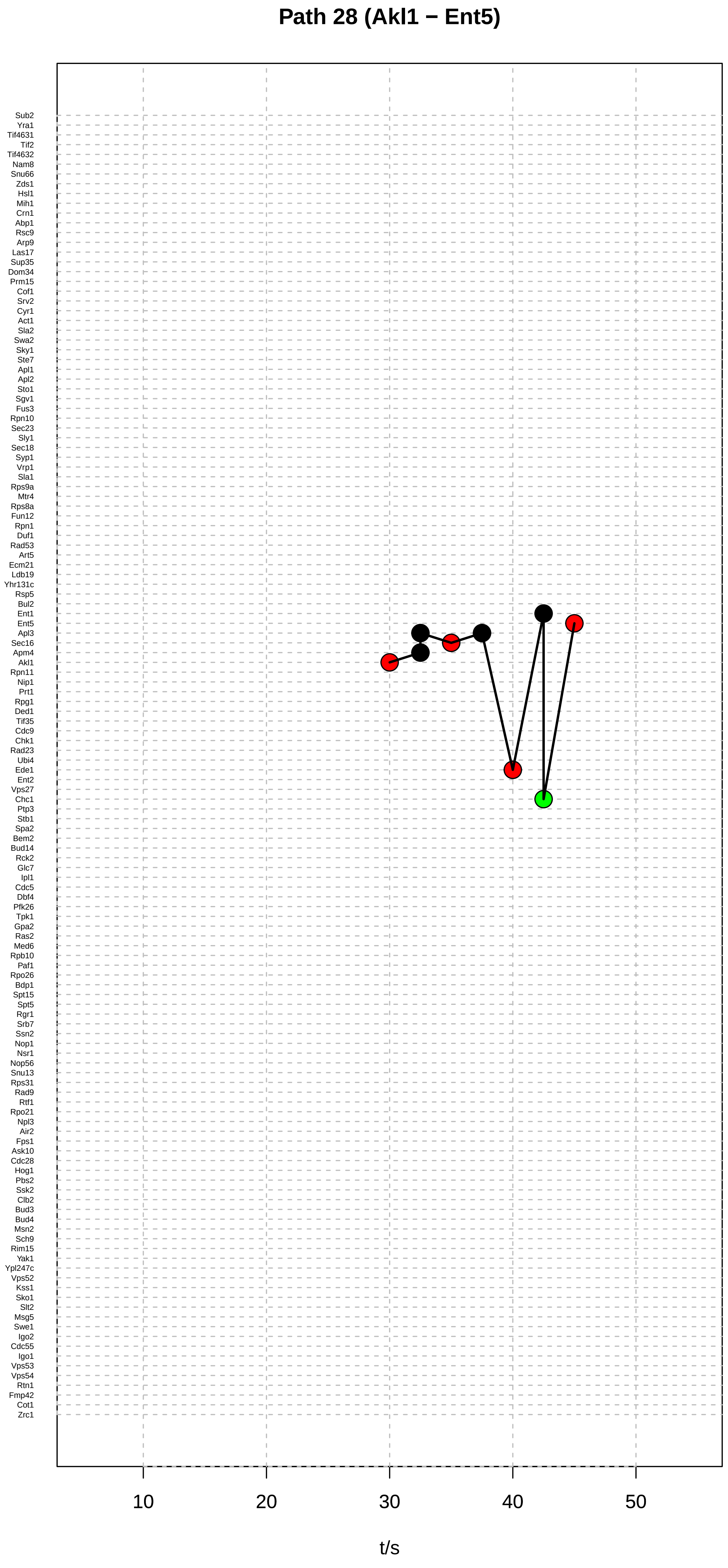

Path 28 (Akl1 − Ent5)
Sub2
Yra1
Tif4631
Tif2
Tif4632
Nam8
Snu66
Zds1
Hsl1
Mih1
Crn1
Abp1
Rsc9
Arp9
Las17
Sup35
Dom34
Prm15
Cof1
Srv2
Cyr1
Act1
Sla2
Swa2
Sky1
Ste7
Apl1
Apl2
Sto1
Sgv1
Fus3
Rpn10
Sec23
Sly1
Sec18
Syp1
Vrp1
Sla1
Rps9a
Mtr4
Rps8a
Fun12
Rpn1
Duf1
Rad53
Art5
Ecm21
Ldb19
Yhr131c
Rsp5
Bul2
Ent1
Ent5
Apl3
Sec16
Apm4
Akl1
Rpn11
Nip1
Prt1
Rpg1
Ded1
Tif35
Cdc9
Chk1
Rad23
Ubi4
Ede1
Ent2
Vps27
Chc1
Ptp3
Stb1
Spa2
Bem2
Bud14
Rck2
Glc7
Ipl1
Cdc5
Dbf4
Pfk26
Tpk1
Gpa2
Ras2
Med6
Rpb10
Paf1
Rpo26
Bdp1
Spt15
Spt5
Rgr1
Srb7
Ssn2
Nop1
Nsr1
Nop56
Snu13
Rps31
Rad9
Rtf1
Rpo21
Npl3
Air2
Fps1
Ask10
Cdc28
Hog1
Pbs2
Ssk2
Clb2
Bud3
Bud4
Msn2
Sch9
Rim15
Yak1
Ypl247c
Vps52
Kss1
Sko1
Slt2
Msg5
Swe1
Igo2
Cdc55
Igo1
Vps53
Vps54
Rtn1
Fmp42
Cot1
Zrc1
10
20
30
40
50
t/s

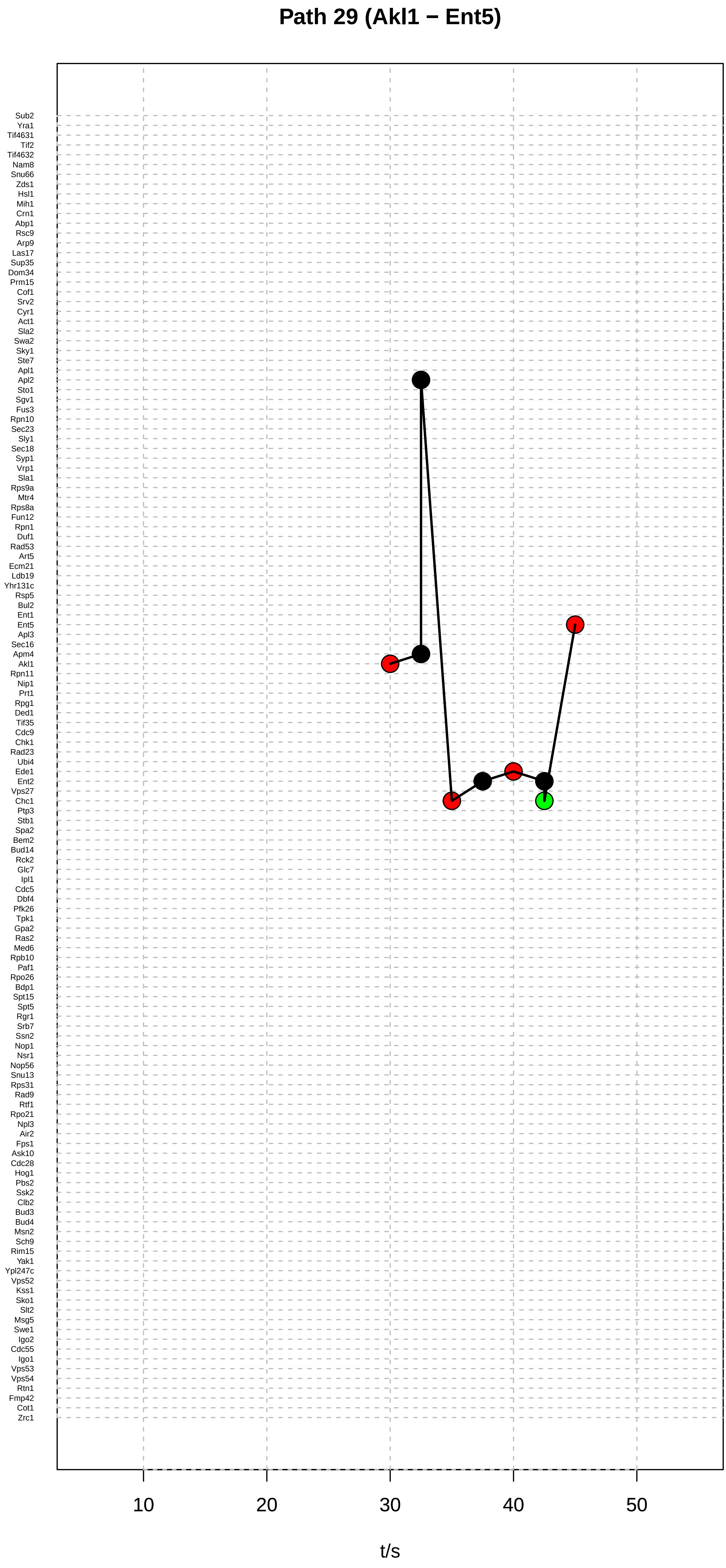

Path 29 (Akl1 − Ent5)
Sub2
Yra1
Tif4631
Tif2
Tif4632
Nam8
Snu66
Zds1
Hsl1
Mih1
Crn1
Abp1
Rsc9
Arp9
Las17
Sup35
Dom34
Prm15
Cof1
Srv2
Cyr1
Act1
Sla2
Swa2
Sky1
Ste7
Apl1
Apl2
Sto1
Sgv1
Fus3
Rpn10
Sec23
Sly1
Sec18
Syp1
Vrp1
Sla1
Rps9a
Mtr4
Rps8a
Fun12
Rpn1
Duf1
Rad53
Art5
Ecm21
Ldb19
Yhr131c
Rsp5
Bul2
Ent1
Ent5
Apl3
Sec16
Apm4
Akl1
Rpn11
Nip1
Prt1
Rpg1
Ded1
Tif35
Cdc9
Chk1
Rad23
Ubi4
Ede1
Ent2
Vps27
Chc1
Ptp3
Stb1
Spa2
Bem2
Bud14
Rck2
Glc7
Ipl1
Cdc5
Dbf4
Pfk26
Tpk1
Gpa2
Ras2
Med6
Rpb10
Paf1
Rpo26
Bdp1
Spt15
Spt5
Rgr1
Srb7
Ssn2
Nop1
Nsr1
Nop56
Snu13
Rps31
Rad9
Rtf1
Rpo21
Npl3
Air2
Fps1
Ask10
Cdc28
Hog1
Pbs2
Ssk2
Clb2
Bud3
Bud4
Msn2
Sch9
Rim15
Yak1
Ypl247c
Vps52
Kss1
Sko1
Slt2
Msg5
Swe1
Igo2
Cdc55
Igo1
Vps53
Vps54
Rtn1
Fmp42
Cot1
Zrc1
10
20
30
40
50
t/s

#### Path 30 (Akl1 − Ent5)

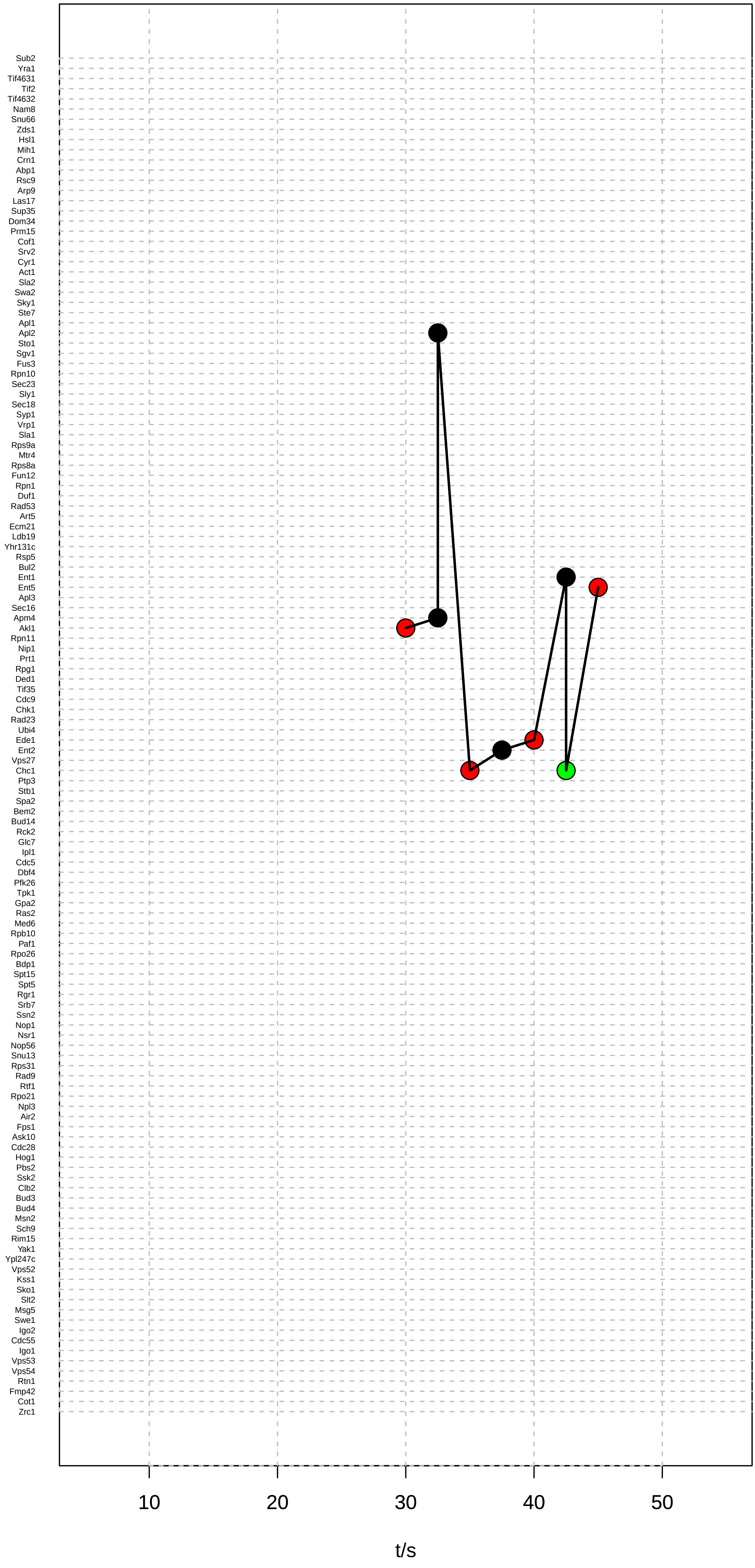

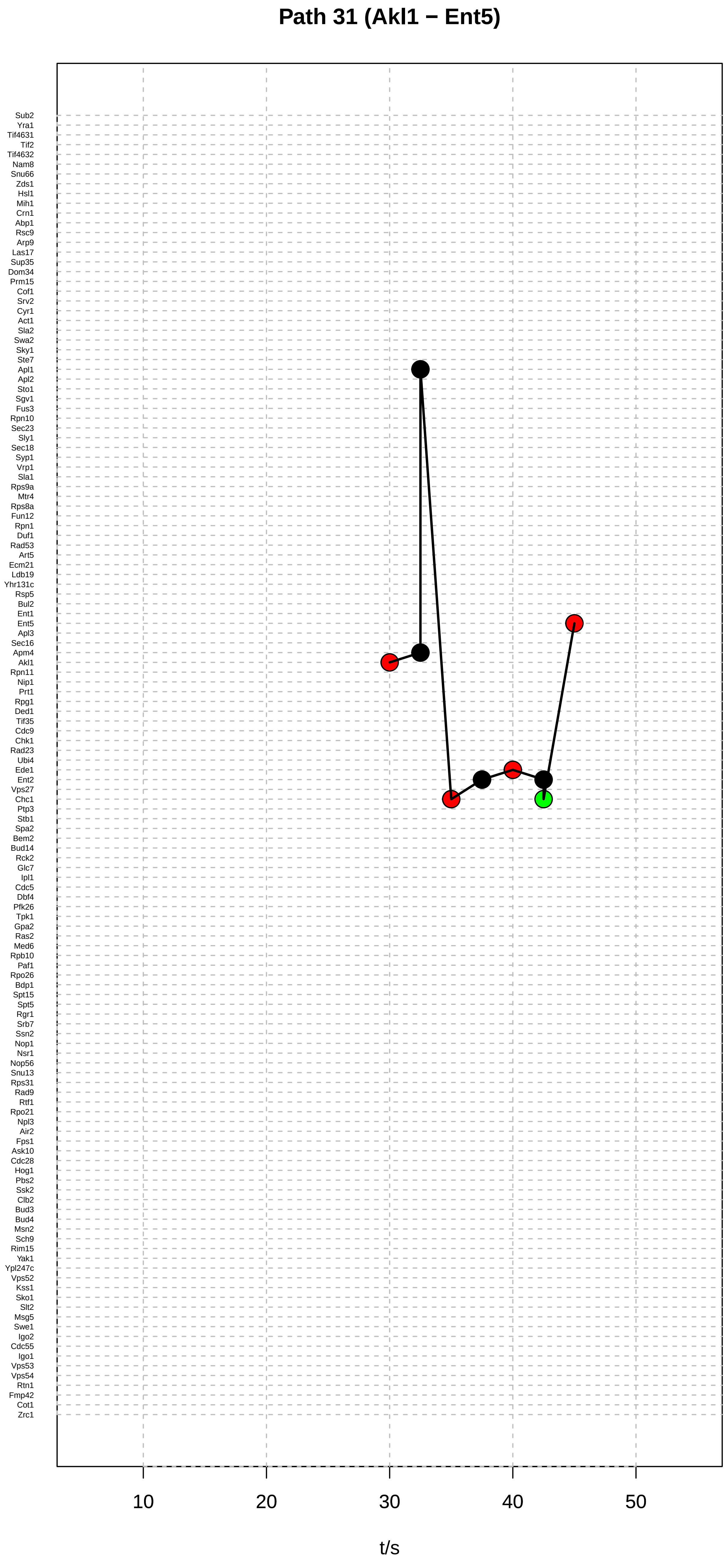
Path 31 (Akl1 − Ent5)
Sub2
Yra1
Tif4631
Tif2
Tif4632
Nam8
Snu66
Zds1
Hsl1
Mih1
Crn1
Abp1
Rsc9
Arp9
Las17
Sup35
Dom34
Prm15
Cof1
Srv2
Cyr1
Act1
Sla2
Swa2
Sky1
Ste7
Apl1
Apl2
Sto1
Sgv1
Fus3
Rpn10
Sec23
Sly1
Sec18
Syp1
Vrp1
Sla1
Rps9a
Mtr4
Rps8a
Fun12
Rpn1
Duf1
Rad53
Art5
Ecm21
Ldb19
Yhr131c
Rsp5
Bul2
Ent1
Ent5
Apl3
Sec16
Apm4
Akl1
Rpn11
Nip1
Prt1
Rpg1
Ded1
Tif35
Cdc9
Chk1
Rad23
Ubi4
Ede1
Ent2
Vps27
Chc1
Ptp3
Stb1
Spa2
Bem2
Bud14
Rck2
Glc7
Ipl1
Cdc5
Dbf4
Pfk26
Tpk1
Gpa2
Ras2
Med6
Rpb10
Paf1
Rpo26
Bdp1
Spt15
Spt5
Rgr1
Srb7
Ssn2
Nop1
Nsr1
Nop56
Snu13
Rps31
Rad9
Rtf1
Rpo21
Npl3
Air2
Fps1
Ask10
Cdc28
Hog1
Pbs2
Ssk2
Clb2
Bud3
Bud4
Msn2
Sch9
Rim15
Yak1
Ypl247c
Vps52
Kss1
Sko1
Slt2
Msg5
Swe1
Igo2
Cdc55
Igo1
Vps53
Vps54
Rtn1
Fmp42
Cot1
Zrc1
10
20
30
40
50
t/s

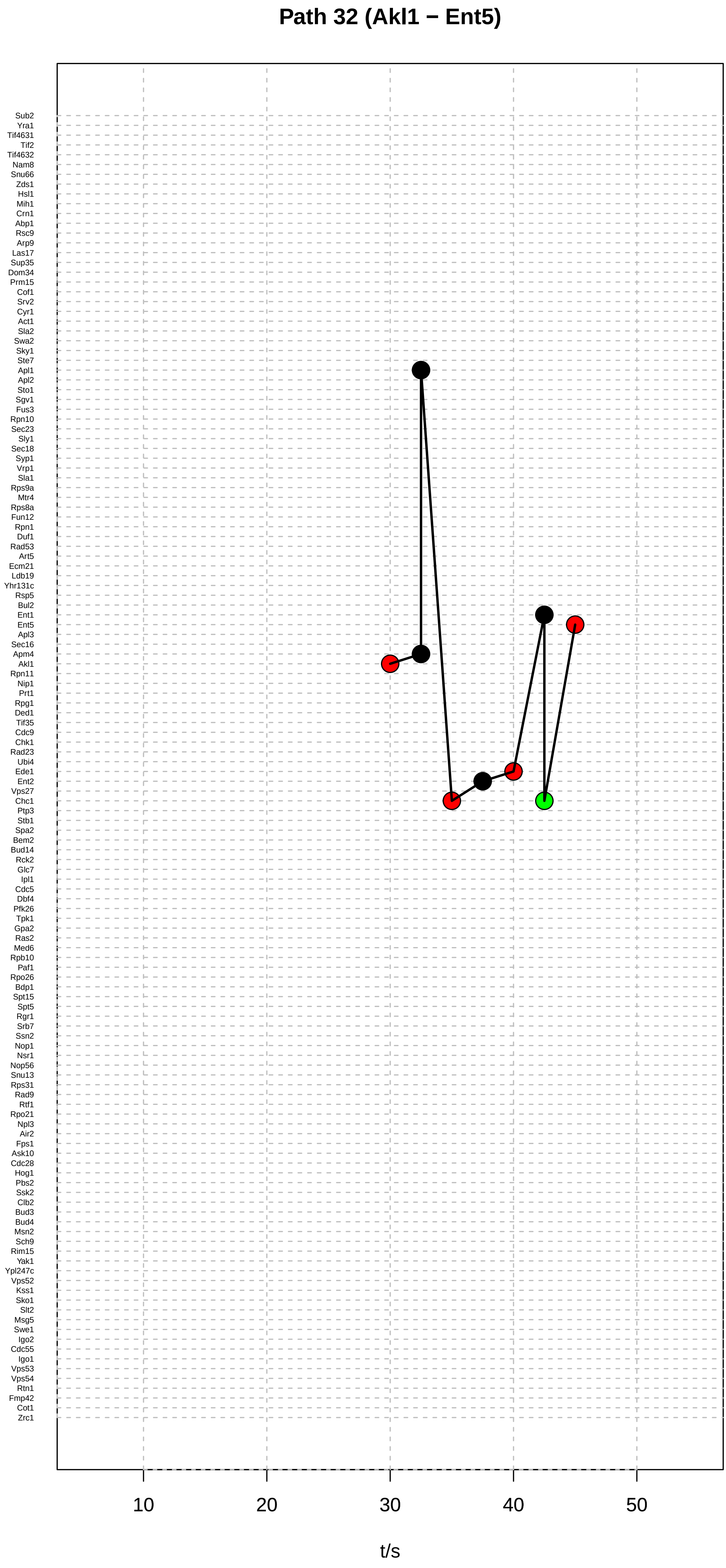

Path 32 (Akl1 − Ent5)
Sub2
Yra1
Tif4631
Tif2
Tif4632
Nam8
Snu66
Zds1
Hsl1
Mih1
Crn1
Abp1
Rsc9
Arp9
Las17
Sup35
Dom34
Prm15
Cof1
Srv2
Cyr1
Act1
Sla2
Swa2
Sky1
Ste7
Apl1
Apl2
Sto1
Sgv1
Fus3
Rpn10
Sec23
Sly1
Sec18
Syp1
Vrp1
Sla1
Rps9a
Mtr4
Rps8a
Fun12
Rpn1
Duf1
Rad53
Art5
Ecm21
Ldb19
Yhr131c
Rsp5
Bul2
Ent1
Ent5
Apl3
Sec16
Apm4
Akl1
Rpn11
Nip1
Prt1
Rpg1
Ded1
Tif35
Cdc9
Chk1
Rad23
Ubi4
Ede1
Ent2
Vps27
Chc1
Ptp3
Stb1
Spa2
Bem2
Bud14
Rck2
Glc7
Ipl1
Cdc5
Dbf4
Pfk26
Tpk1
Gpa2
Ras2
Med6
Rpb10
Paf1
Rpo26
Bdp1
Spt15
Spt5
Rgr1
Srb7
Ssn2
Nop1
Nsr1
Nop56
Snu13
Rps31
Rad9
Rtf1
Rpo21
Npl3
Air2
Fps1
Ask10
Cdc28
Hog1
Pbs2
Ssk2
Clb2
Bud3
Bud4
Msn2
Sch9
Rim15
Yak1
Ypl247c
Vps52
Kss1
Sko1
Slt2
Msg5
Swe1
Igo2
Cdc55
Igo1
Vps53
Vps54
Rtn1
Fmp42
Cot1
Zrc1
10
20
30
40
50
t/s

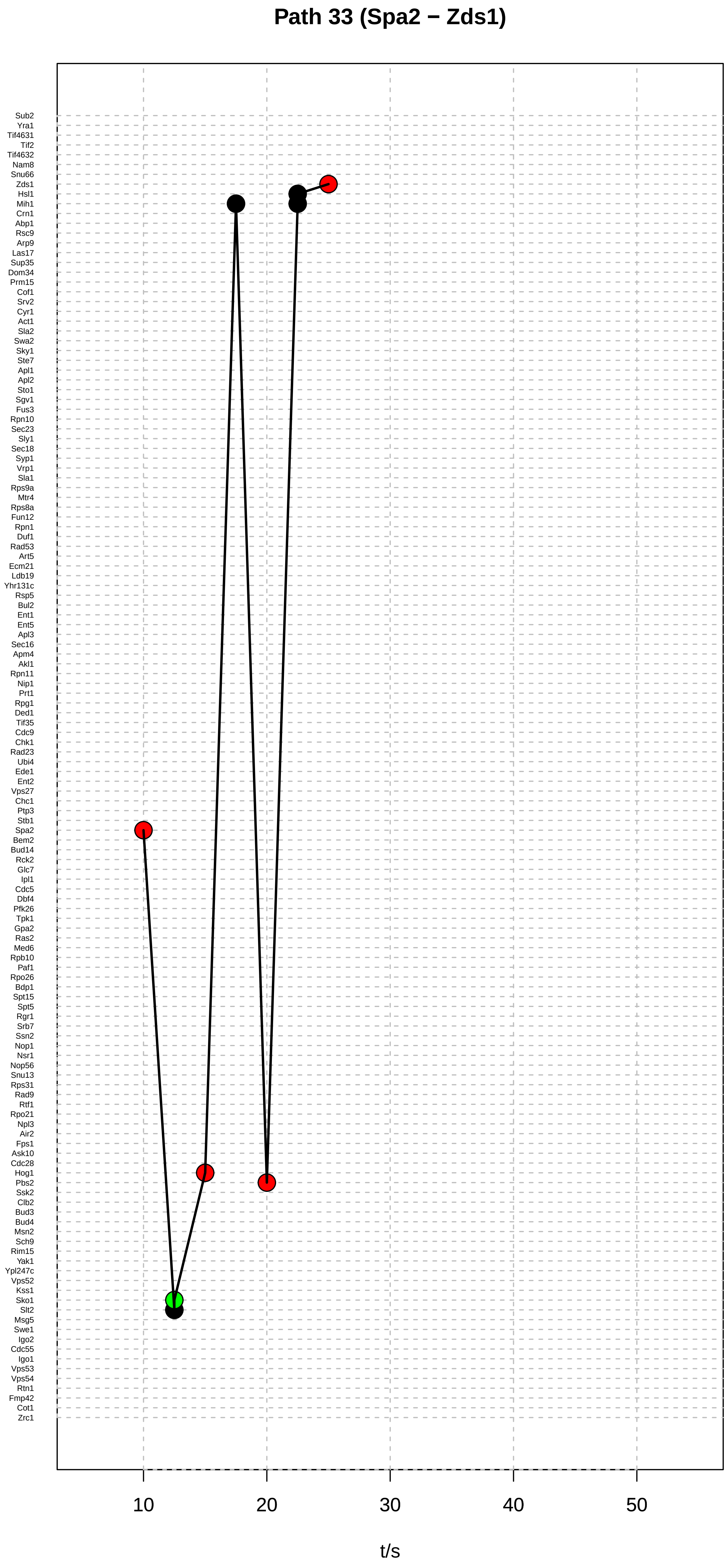

Path 33 (Spa2 − Zds1)
Sub2
Yra1
Tif4631
Tif2
Tif4632
Nam8
Snu66
Zds1
Hsl1
Mih1
Crn1
Abp1
Rsc9
Arp9
Las17
Sup35
Dom34
Prm15
Cof1
Srv2
Cyr1
Act1
Sla2
Swa2
Sky1
Ste7
Apl1
Apl2
Sto1
Sgv1
Fus3
Rpn10
Sec23
Sly1
Sec18
Syp1
Vrp1
Sla1
Rps9a
Mtr4
Rps8a
Fun12
Rpn1
Duf1
Rad53
Art5
Ecm21
Ldb19
Yhr131c
Rsp5
Bul2
Ent1
Ent5
Apl3
Sec16
Apm4
Akl1
Rpn11
Nip1
Prt1
Rpg1
Ded1
Tif35
Cdc9
Chk1
Rad23
Ubi4
Ede1
Ent2
Vps27
Chc1
Ptp3
Stb1
Spa2
Bem2
Bud14
Rck2
Glc7
Ipl1
Cdc5
Dbf4
Pfk26
Tpk1
Gpa2
Ras2
Med6
Rpb10
Paf1
Rpo26
Bdp1
Spt15
Spt5
Rgr1
Srb7
Ssn2
Nop1
Nsr1
Nop56
Snu13
Rps31
Rad9
Rtf1
Rpo21
Npl3
Air2
Fps1
Ask10
Cdc28
Hog1
Pbs2
Ssk2
Clb2
Bud3
Bud4
Msn2
Sch9
Rim15
Yak1
Ypl247c
Vps52
Kss1
Sko1
Slt2
Msg5
Swe1
Igo2
Cdc55
Igo1
Vps53
Vps54
Rtn1
Fmp42
Cot1
Zrc1
10
20
30
40
50
t/s

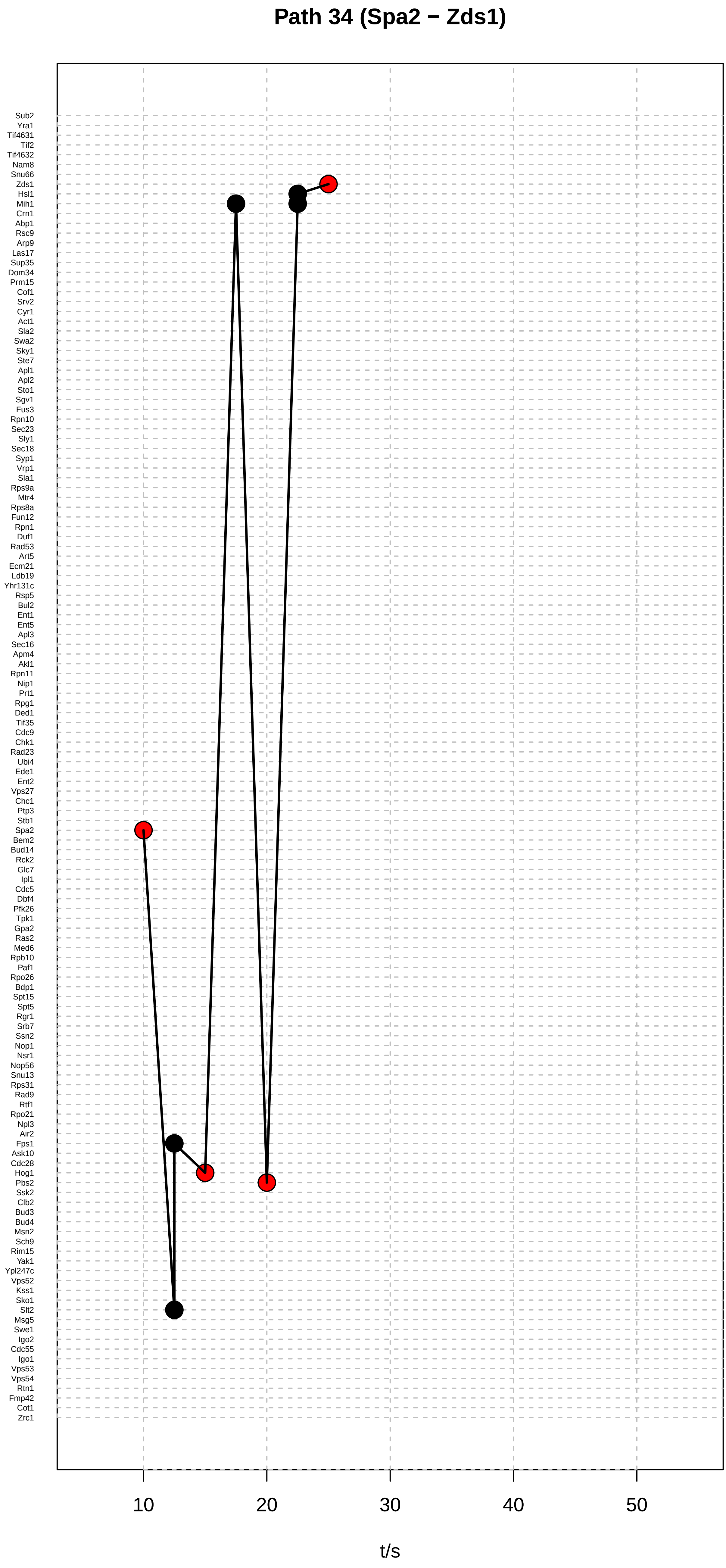
Path 34 (Spa2 − Zds1)
Sub2
Yra1
Tif4631
Tif2
Tif4632
Nam8
Snu66
Zds1
Hsl1
Mih1
Crn1
Abp1
Rsc9
Arp9
Las17
Sup35
Dom34
Prm15
Cof1
Srv2
Cyr1
Act1
Sla2
Swa2
Sky1
Ste7
Apl1
Apl2
Sto1
Sgv1
Fus3
Rpn10
Sec23
Sly1
Sec18
Syp1
Vrp1
Sla1
Rps9a
Mtr4
Rps8a
Fun12
Rpn1
Duf1
Rad53
Art5
Ecm21
Ldb19
Yhr131c
Rsp5
Bul2
Ent1
Ent5
Apl3
Sec16
Apm4
Akl1
Rpn11
Nip1
Prt1
Rpg1
Ded1
Tif35
Cdc9
Chk1
Rad23
Ubi4
Ede1
Ent2
Vps27
Chc1
Ptp3
Stb1
Spa2
Bem2
Bud14
Rck2
Glc7
Ipl1
Cdc5
Dbf4
Pfk26
Tpk1
Gpa2
Ras2
Med6
Rpb10
Paf1
Rpo26
Bdp1
Spt15
Spt5
Rgr1
Srb7
Ssn2
Nop1
Nsr1
Nop56
Snu13
Rps31
Rad9
Rtf1
Rpo21
Npl3
Air2
Fps1
Ask10
Cdc28
Hog1
Pbs2
Ssk2
Clb2
Bud3
Bud4
Msn2
Sch9
Rim15
Yak1
Ypl247c
Vps52
Kss1
Sko1
Slt2
Msg5
Swe1
Igo2
Cdc55
Igo1
Vps53
Vps54
Rtn1
Fmp42
Cot1
Zrc1
10
20
30
40
50
t/s

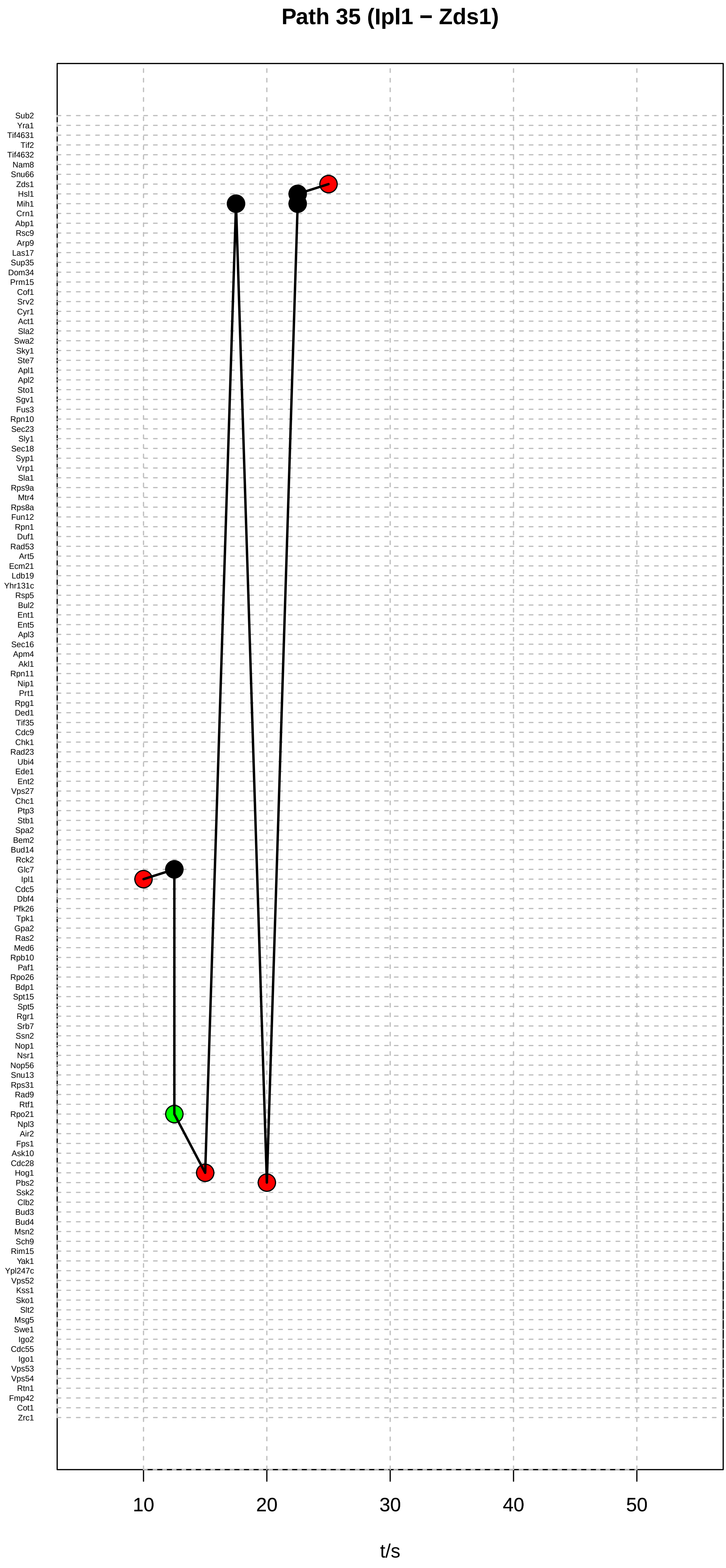

Path 35 (Ipl1 − Zds1)
Sub2
Yra1
Tif4631
Tif2
Tif4632
Nam8
Snu66
Zds1
Hsl1
Mih1
Crn1
Abp1
Rsc9
Arp9
Las17
Sup35
Dom34
Prm15
Cof1
Srv2
Cyr1
Act1
Sla2
Swa2
Sky1
Ste7
Apl1
Apl2
Sto1
Sgv1
Fus3
Rpn10
Sec23
Sly1
Sec18
Syp1
Vrp1
Sla1
Rps9a
Mtr4
Rps8a
Fun12
Rpn1
Duf1
Rad53
Art5
Ecm21
Ldb19
Yhr131c
Rsp5
Bul2
Ent1
Ent5
Apl3
Sec16
Apm4
Akl1
Rpn11
Nip1
Prt1
Rpg1
Ded1
Tif35
Cdc9
Chk1
Rad23
Ubi4
Ede1
Ent2
Vps27
Chc1
Ptp3
Stb1
Spa2
Bem2
Bud14
Rck2
Glc7
Ipl1
Cdc5
Dbf4
Pfk26
Tpk1
Gpa2
Ras2
Med6
Rpb10
Paf1
Rpo26
Bdp1
Spt15
Spt5
Rgr1
Srb7
Ssn2
Nop1
Nsr1
Nop56
Snu13
Rps31
Rad9
Rtf1
Rpo21
Npl3
Air2
Fps1
Ask10
Cdc28
Hog1
Pbs2
Ssk2
Clb2
Bud3
Bud4
Msn2
Sch9
Rim15
Yak1
Ypl247c
Vps52
Kss1
Sko1
Slt2
Msg5
Swe1
Igo2
Cdc55
Igo1
Vps53
Vps54
Rtn1
Fmp42
Cot1
Zrc1
10
20
30
40
50
t/s

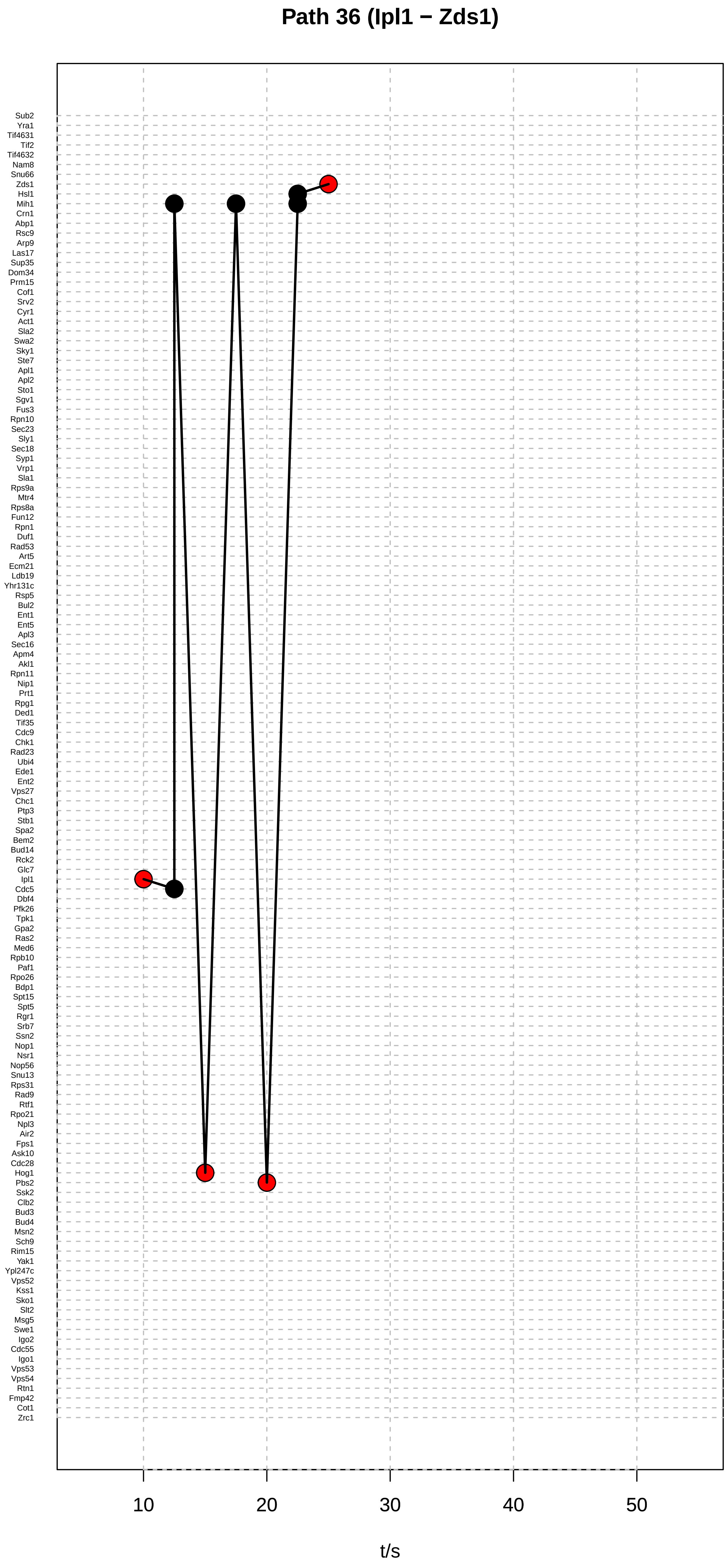
Path 36 (Ipl1 − Zds1)
Sub2
Yra1
Tif4631
Tif2
Tif4632
Nam8
Snu66
Zds1
Hsl1
Mih1
Crn1
Abp1
Rsc9
Arp9
Las17
Sup35
Dom34
Prm15
Cof1
Srv2
Cyr1
Act1
Sla2
Swa2
Sky1
Ste7
Apl1
Apl2
Sto1
Sgv1
Fus3
Rpn10
Sec23
Sly1
Sec18
Syp1
Vrp1
Sla1
Rps9a
Mtr4
Rps8a
Fun12
Rpn1
Duf1
Rad53
Art5
Ecm21
Ldb19
Yhr131c
Rsp5
Bul2
Ent1
Ent5
Apl3
Sec16
Apm4
Akl1
Rpn11
Nip1
Prt1
Rpg1
Ded1
Tif35
Cdc9
Chk1
Rad23
Ubi4
Ede1
Ent2
Vps27
Chc1
Ptp3
Stb1
Spa2
Bem2
Bud14
Rck2
Glc7
Ipl1
Cdc5
Dbf4
Pfk26
Tpk1
Gpa2
Ras2
Med6
Rpb10
Paf1
Rpo26
Bdp1
Spt15
Spt5
Rgr1
Srb7
Ssn2
Nop1
Nsr1
Nop56
Snu13
Rps31
Rad9
Rtf1
Rpo21
Npl3
Air2
Fps1
Ask10
Cdc28
Hog1
Pbs2
Ssk2
Clb2
Bud3
Bud4
Msn2
Sch9
Rim15
Yak1
Ypl247c
Vps52
Kss1
Sko1
Slt2
Msg5
Swe1
Igo2
Cdc55
Igo1
Vps53
Vps54
Rtn1
Fmp42
Cot1
Zrc1
10
20
30
40
50
t/s

### Path 37 (Dbf4 − Bud14)

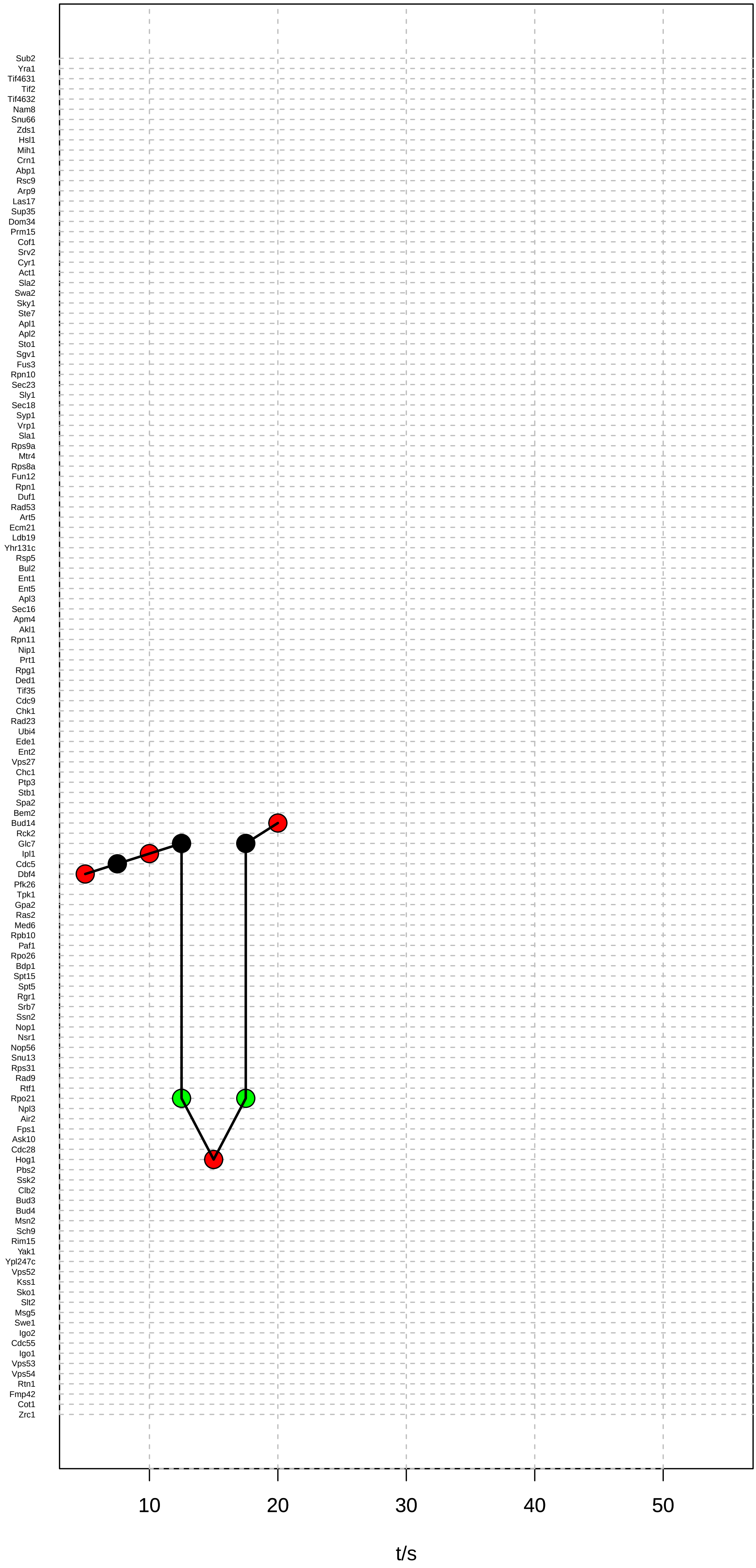

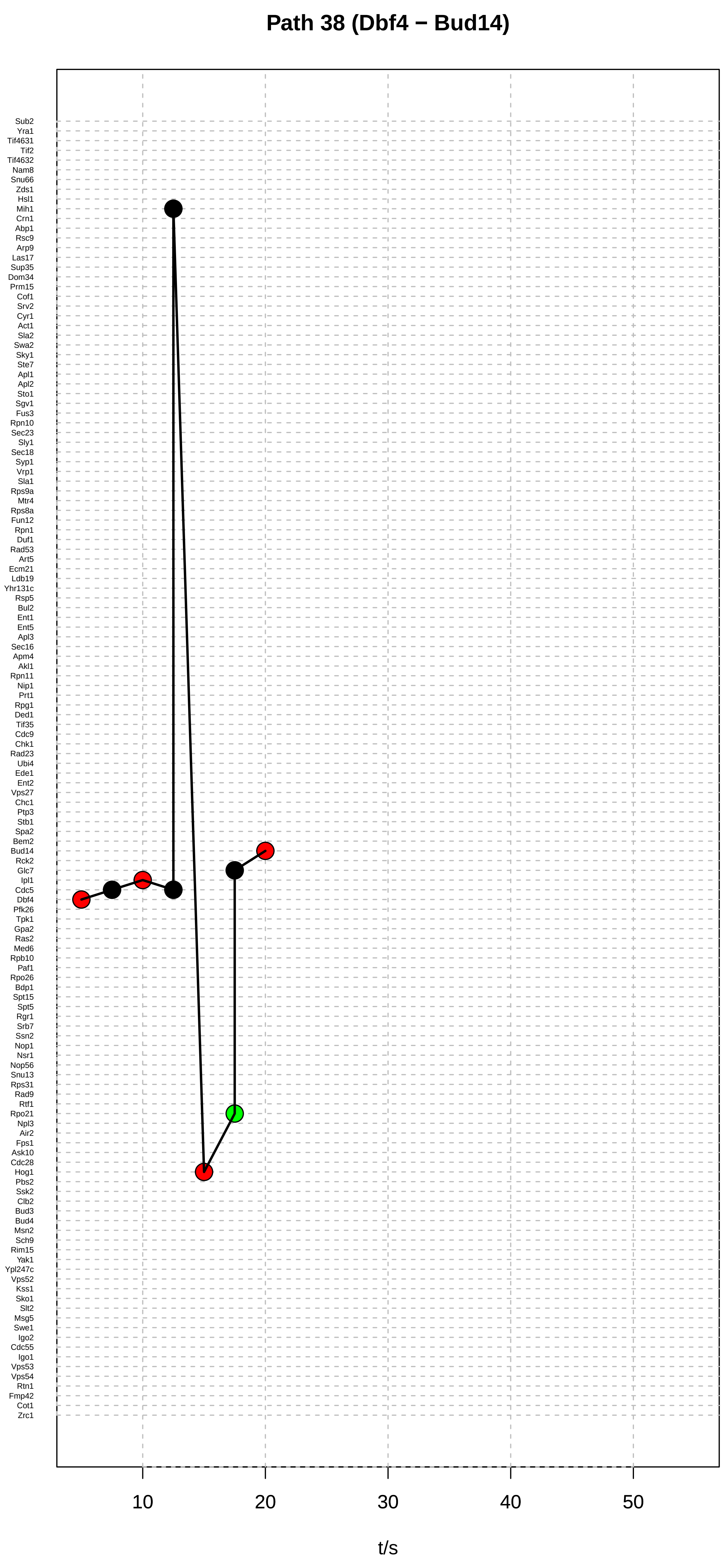

Path 38 (Dbf4 − Bud14)
Sub2
Yra1
Tif4631
Tif2
Tif4632
Nam8
Snu66
Zds1
Hsl1
Mih1
Crn1
Abp1
Rsc9
Arp9
Las17
Sup35
Dom34
Prm15
Cof1
Srv2
Cyr1
Act1
Sla2
Swa2
Sky1
Ste7
Apl1
Apl2
Sto1
Sgv1
Fus3
Rpn10
Sec23
Sly1
Sec18
Syp1
Vrp1
Sla1
Rps9a
Mtr4
Rps8a
Fun12
Rpn1
Duf1
Rad53
Art5
Ecm21
Ldb19
Yhr131c
Rsp5
Bul2
Ent1
Ent5
Apl3
Sec16
Apm4
Akl1
Rpn11
Nip1
Prt1
Rpg1
Ded1
Tif35
Cdc9
Chk1
Rad23
Ubi4
Ede1
Ent2
Vps27
Chc1
Ptp3
Stb1
Spa2
Bem2
Bud14
Rck2
Glc7
Ipl1
Cdc5
Dbf4
Pfk26
Tpk1
Gpa2
Ras2
Med6
Rpb10
Paf1
Rpo26
Bdp1
Spt15
Spt5
Rgr1
Srb7
Ssn2
Nop1
Nsr1
Nop56
Snu13
Rps31
Rad9
Rtf1
Rpo21
Npl3
Air2
Fps1
Ask10
Cdc28
Hog1
Pbs2
Ssk2
Clb2
Bud3
Bud4
Msn2
Sch9
Rim15
Yak1
Ypl247c
Vps52
Kss1
Sko1
Slt2
Msg5
Swe1
Igo2
Cdc55
Igo1
Vps53
Vps54
Rtn1
Fmp42
Cot1
Zrc1
10
20
30
40
50
t/s

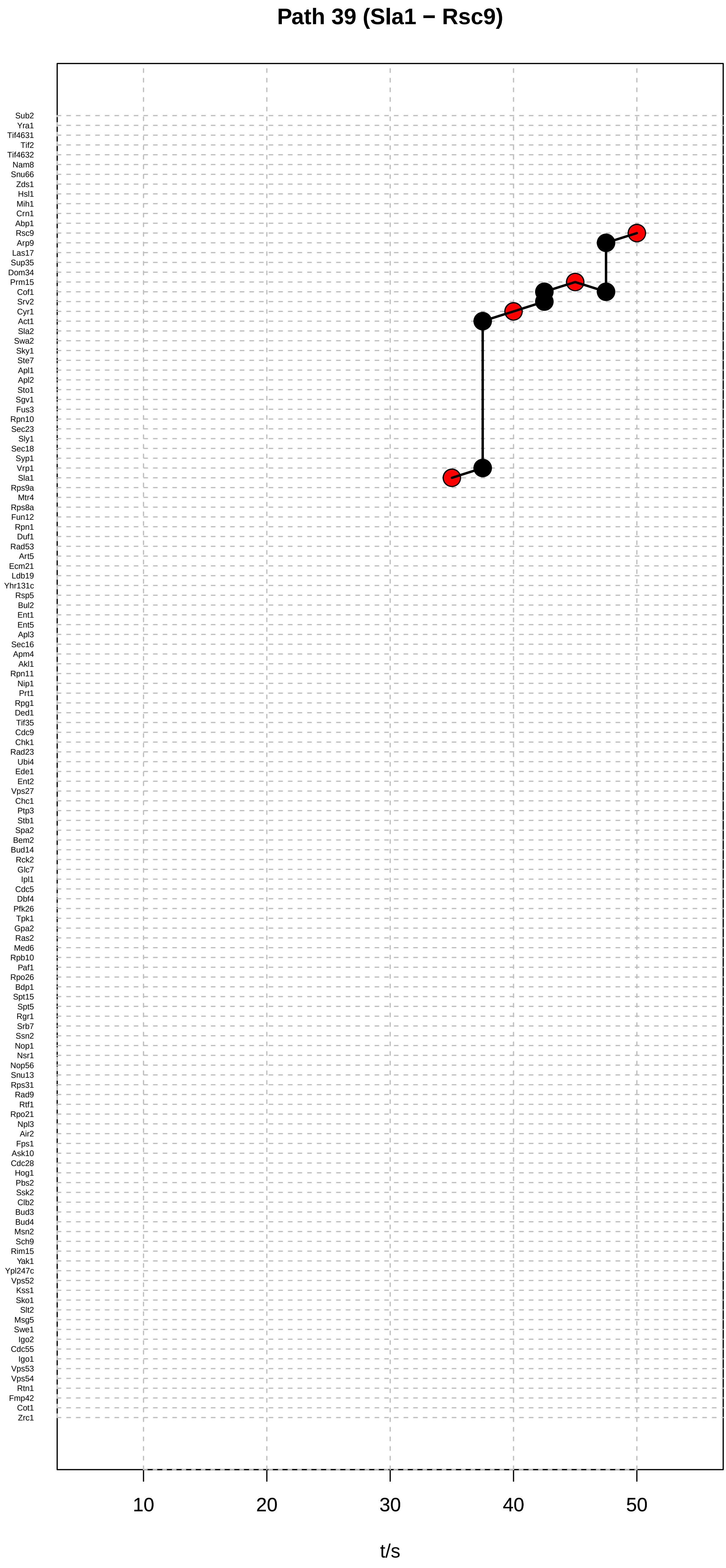

Path 39 (Sla1 − Rsc9)
Sub2
Yra1
Tif4631
Tif2
Tif4632
Nam8
Snu66
Zds1
Hsl1
Mih1
Crn1
Abp1
Rsc9
Arp9
Las17
Sup35
Dom34
Prm15
Cof1
Srv2
Cyr1
Act1
Sla2
Swa2
Sky1
Ste7
Apl1
Apl2
Sto1
Sgv1
Fus3
Rpn10
Sec23
Sly1
Sec18
Syp1
Vrp1
Sla1
Rps9a
Mtr4
Rps8a
Fun12
Rpn1
Duf1
Rad53
Art5
Ecm21
Ldb19
Yhr131c
Rsp5
Bul2
Ent1
Ent5
Apl3
Sec16
Apm4
Akl1
Rpn11
Nip1
Prt1
Rpg1
Ded1
Tif35
Cdc9
Chk1
Rad23
Ubi4
Ede1
Ent2
Vps27
Chc1
Ptp3
Stb1
Spa2
Bem2
Bud14
Rck2
Glc7
Ipl1
Cdc5
Dbf4
Pfk26
Tpk1
Gpa2
Ras2
Med6
Rpb10
Paf1
Rpo26
Bdp1
Spt15
Spt5
Rgr1
Srb7
Ssn2
Nop1
Nsr1
Nop56
Snu13
Rps31
Rad9
Rtf1
Rpo21
Npl3
Air2
Fps1
Ask10
Cdc28
Hog1
Pbs2
Ssk2
Clb2
Bud3
Bud4
Msn2
Sch9
Rim15
Yak1
Ypl247c
Vps52
Kss1
Sko1
Slt2
Msg5
Swe1
Igo2
Cdc55
Igo1
Vps53
Vps54
Rtn1
Fmp42
Cot1
Zrc1
10
20
30
40
50
t/s

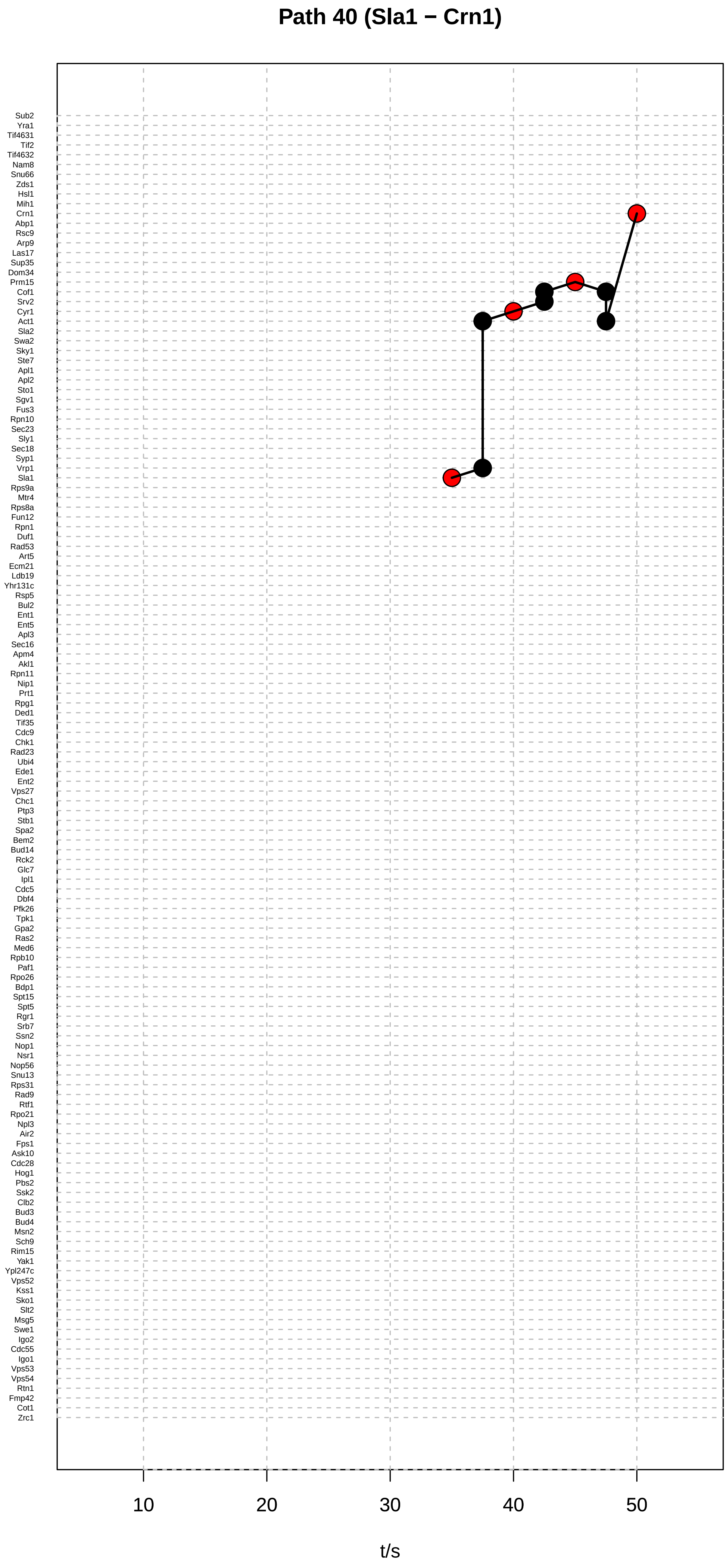

Path 40 (Sla1 − Crn1)
Sub2
Yra1
Tif4631
Tif2
Tif4632
Nam8
Snu66
Zds1
Hsl1
Mih1
Crn1
Abp1
Rsc9
Arp9
Las17
Sup35
Dom34
Prm15
Cof1
Srv2
Cyr1
Act1
Sla2
Swa2
Sky1
Ste7
Apl1
Apl2
Sto1
Sgv1
Fus3
Rpn10
Sec23
Sly1
Sec18
Syp1
Vrp1
Sla1
Rps9a
Mtr4
Rps8a
Fun12
Rpn1
Duf1
Rad53
Art5
Ecm21
Ldb19
Yhr131c
Rsp5
Bul2
Ent1
Ent5
Apl3
Sec16
Apm4
Akl1
Rpn11
Nip1
Prt1
Rpg1
Ded1
Tif35
Cdc9
Chk1
Rad23
Ubi4
Ede1
Ent2
Vps27
Chc1
Ptp3
Stb1
Spa2
Bem2
Bud14
Rck2
Glc7
Ipl1
Cdc5
Dbf4
Pfk26
Tpk1
Gpa2
Ras2
Med6
Rpb10
Paf1
Rpo26
Bdp1
Spt15
Spt5
Rgr1
Srb7
Ssn2
Nop1
Nsr1
Nop56
Snu13
Rps31
Rad9
Rtf1
Rpo21
Npl3
Air2
Fps1
Ask10
Cdc28
Hog1
Pbs2
Ssk2
Clb2
Bud3
Bud4
Msn2
Sch9
Rim15
Yak1
Ypl247c
Vps52
Kss1
Sko1
Slt2
Msg5
Swe1
Igo2
Cdc55
Igo1
Vps53
Vps54
Rtn1
Fmp42
Cot1
Zrc1
10
20
30
40
50
t/s

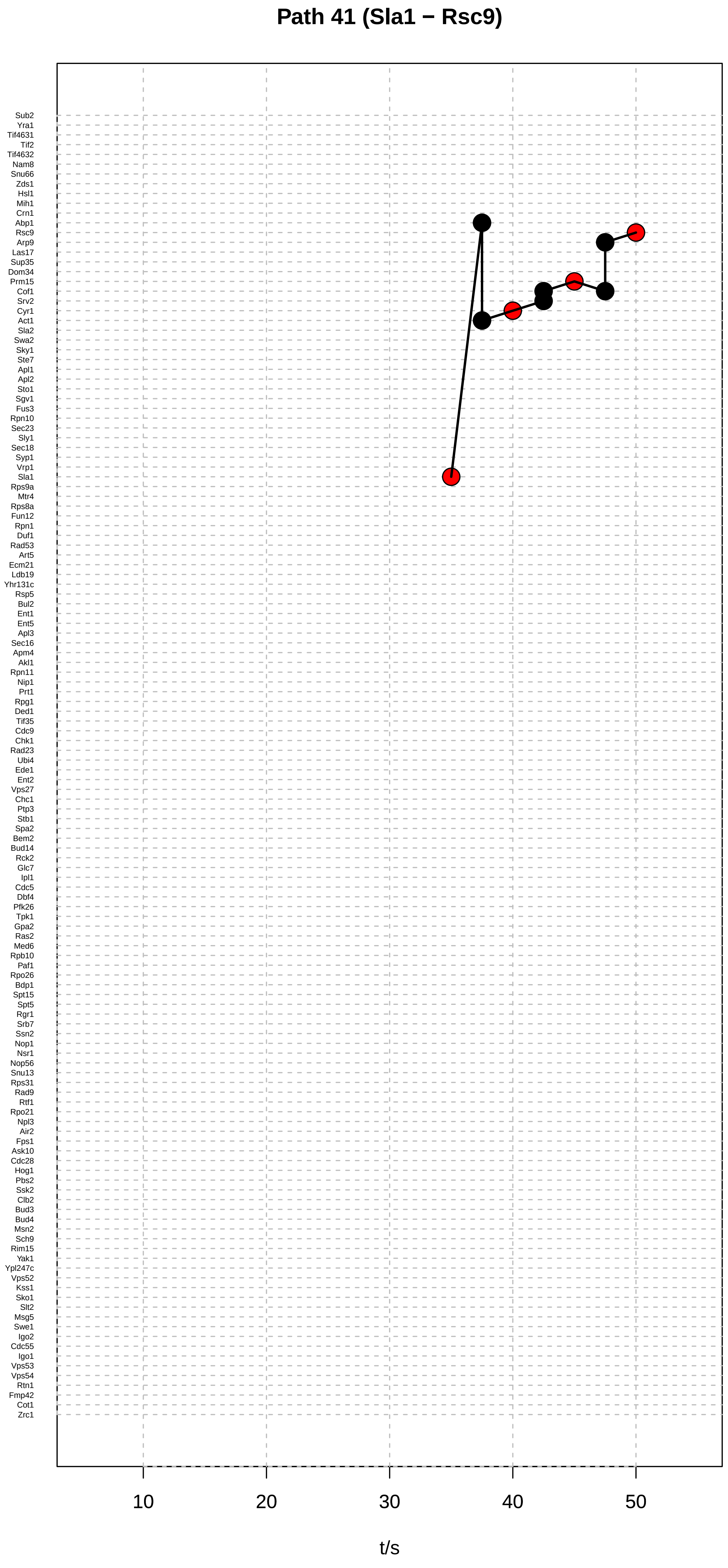

Path 41 (Sla1 − Rsc9)
Sub2
Yra1
Tif4631
Tif2
Tif4632
Nam8
Snu66
Zds1
Hsl1
Mih1
Crn1
Abp1
Rsc9
Arp9
Las17
Sup35
Dom34
Prm15
Cof1
Srv2
Cyr1
Act1
Sla2
Swa2
Sky1
Ste7
Apl1
Apl2
Sto1
Sgv1
Fus3
Rpn10
Sec23
Sly1
Sec18
Syp1
Vrp1
Sla1
Rps9a
Mtr4
Rps8a
Fun12
Rpn1
Duf1
Rad53
Art5
Ecm21
Ldb19
Yhr131c
Rsp5
Bul2
Ent1
Ent5
Apl3
Sec16
Apm4
Akl1
Rpn11
Nip1
Prt1
Rpg1
Ded1
Tif35
Cdc9
Chk1
Rad23
Ubi4
Ede1
Ent2
Vps27
Chc1
Ptp3
Stb1
Spa2
Bem2
Bud14
Rck2
Glc7
Ipl1
Cdc5
Dbf4
Pfk26
Tpk1
Gpa2
Ras2
Med6
Rpb10
Paf1
Rpo26
Bdp1
Spt15
Spt5
Rgr1
Srb7
Ssn2
Nop1
Nsr1
Nop56
Snu13
Rps31
Rad9
Rtf1
Rpo21
Npl3
Air2
Fps1
Ask10
Cdc28
Hog1
Pbs2
Ssk2
Clb2
Bud3
Bud4
Msn2
Sch9
Rim15
Yak1
Ypl247c
Vps52
Kss1
Sko1
Slt2
Msg5
Swe1
Igo2
Cdc55
Igo1
Vps53
Vps54
Rtn1
Fmp42
Cot1
Zrc1
10
20
30
40
50
t/s

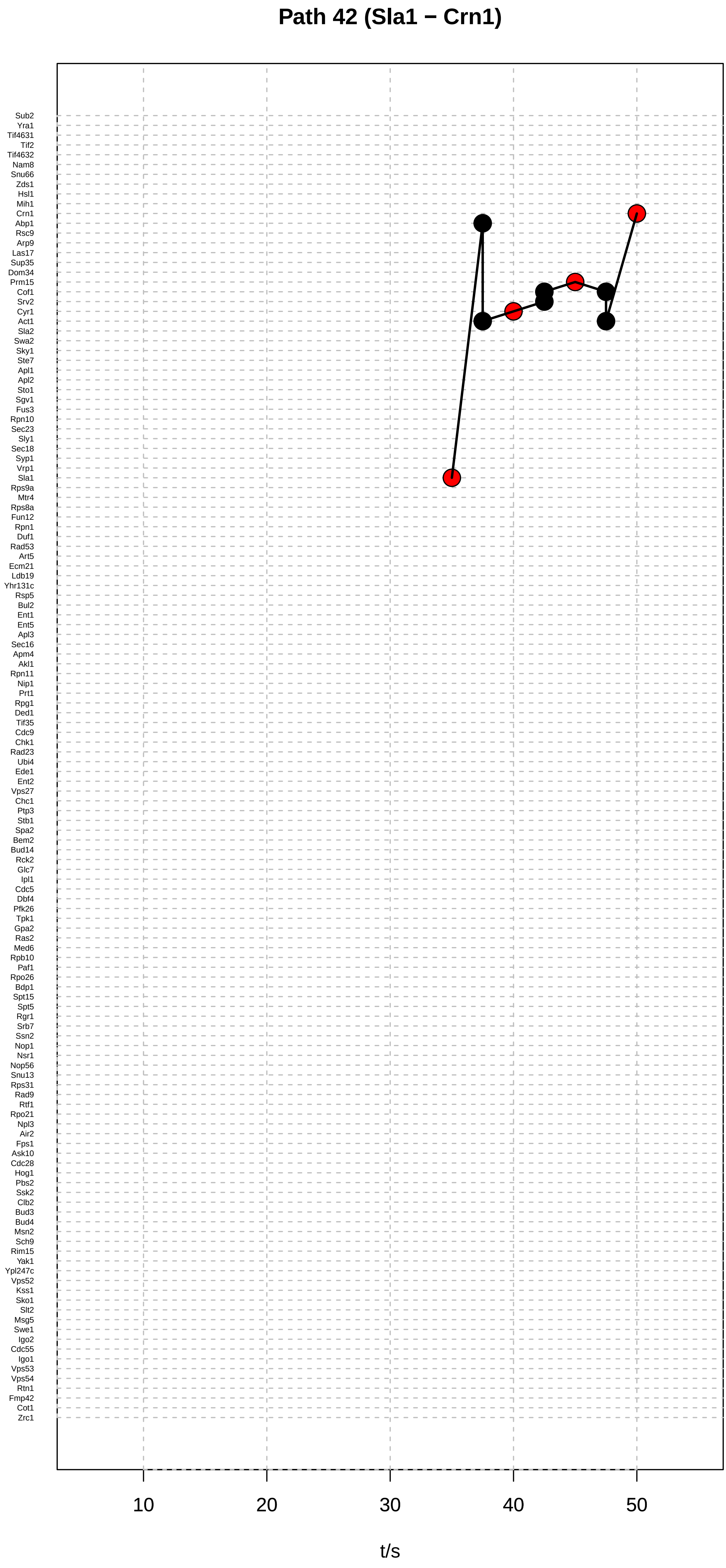

Path 42 (Sla1 − Crn1)
Sub2
Yra1
Tif4631
Tif2
Tif4632
Nam8
Snu66
Zds1
Hsl1
Mih1
Crn1
Abp1
Rsc9
Arp9
Las17
Sup35
Dom34
Prm15
Cof1
Srv2
Cyr1
Act1
Sla2
Swa2
Sky1
Ste7
Apl1
Apl2
Sto1
Sgv1
Fus3
Rpn10
Sec23
Sly1
Sec18
Syp1
Vrp1
Sla1
Rps9a
Mtr4
Rps8a
Fun12
Rpn1
Duf1
Rad53
Art5
Ecm21
Ldb19
Yhr131c
Rsp5
Bul2
Ent1
Ent5
Apl3
Sec16
Apm4
Akl1
Rpn11
Nip1
Prt1
Rpg1
Ded1
Tif35
Cdc9
Chk1
Rad23
Ubi4
Ede1
Ent2
Vps27
Chc1
Ptp3
Stb1
Spa2
Bem2
Bud14
Rck2
Glc7
Ipl1
Cdc5
Dbf4
Pfk26
Tpk1
Gpa2
Ras2
Med6
Rpb10
Paf1
Rpo26
Bdp1
Spt15
Spt5
Rgr1
Srb7
Ssn2
Nop1
Nsr1
Nop56
Snu13
Rps31
Rad9
Rtf1
Rpo21
Npl3
Air2
Fps1
Ask10
Cdc28
Hog1
Pbs2
Ssk2
Clb2
Bud3
Bud4
Msn2
Sch9
Rim15
Yak1
Ypl247c
Vps52
Kss1
Sko1
Slt2
Msg5
Swe1
Igo2
Cdc55
Igo1
Vps53
Vps54
Rtn1
Fmp42
Cot1
Zrc1
10
20
30
40
50
t/s

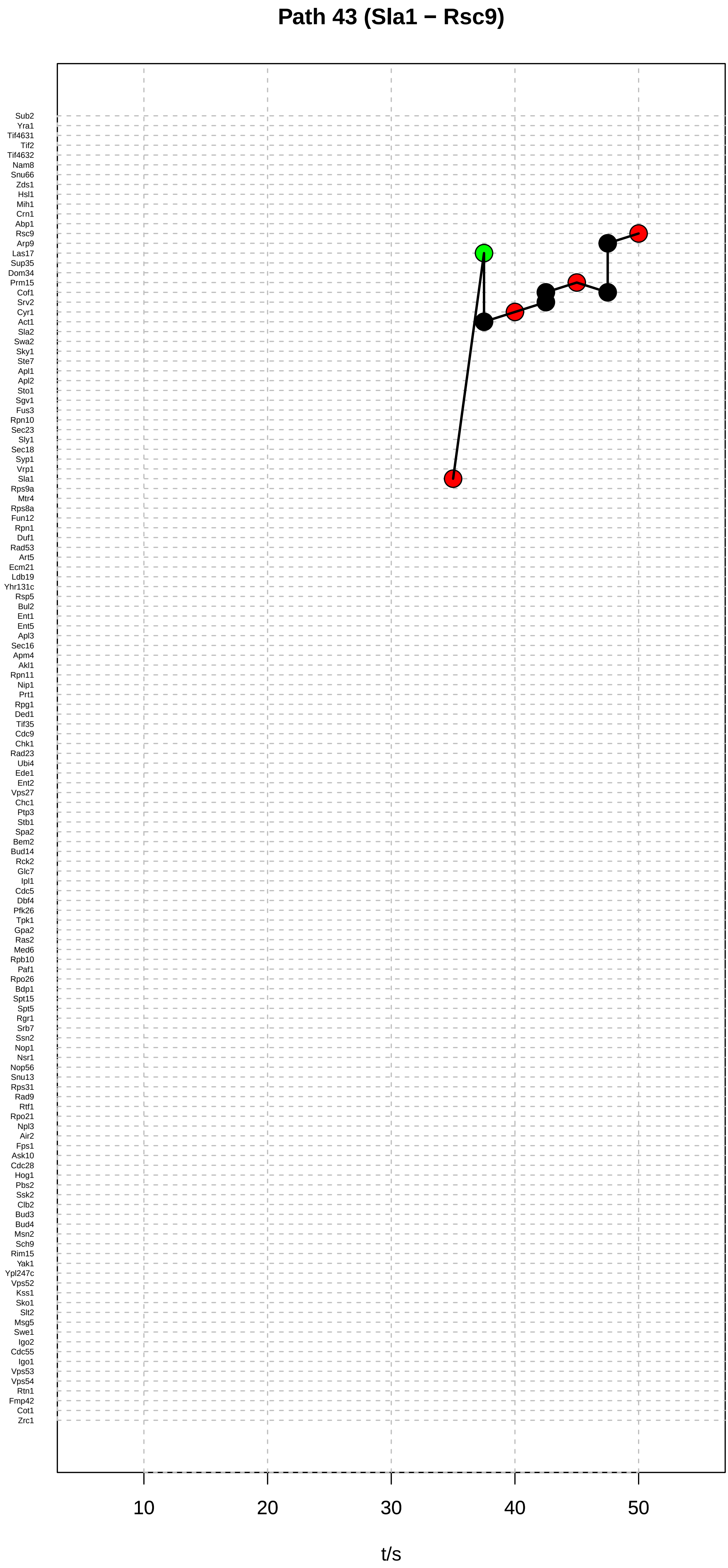

Path 43 (Sla1 − Rsc9)
Sub2
Yra1
Tif4631
Tif2
Tif4632
Nam8
Snu66
Zds1
Hsl1
Mih1
Crn1
Abp1
Rsc9
Arp9
Las17
Sup35
Dom34
Prm15
Cof1
Srv2
Cyr1
Act1
Sla2
Swa2
Sky1
Ste7
Apl1
Apl2
Sto1
Sgv1
Fus3
Rpn10
Sec23
Sly1
Sec18
Syp1
Vrp1
Sla1
Rps9a
Mtr4
Rps8a
Fun12
Rpn1
Duf1
Rad53
Art5
Ecm21
Ldb19
Yhr131c
Rsp5
Bul2
Ent1
Ent5
Apl3
Sec16
Apm4
Akl1
Rpn11
Nip1
Prt1
Rpg1
Ded1
Tif35
Cdc9
Chk1
Rad23
Ubi4
Ede1
Ent2
Vps27
Chc1
Ptp3
Stb1
Spa2
Bem2
Bud14
Rck2
Glc7
Ipl1
Cdc5
Dbf4
Pfk26
Tpk1
Gpa2
Ras2
Med6
Rpb10
Paf1
Rpo26
Bdp1
Spt15
Spt5
Rgr1
Srb7
Ssn2
Nop1
Nsr1
Nop56
Snu13
Rps31
Rad9
Rtf1
Rpo21
Npl3
Air2
Fps1
Ask10
Cdc28
Hog1
Pbs2
Ssk2
Clb2
Bud3
Bud4
Msn2
Sch9
Rim15
Yak1
Ypl247c
Vps52
Kss1
Sko1
Slt2
Msg5
Swe1
Igo2
Cdc55
Igo1
Vps53
Vps54
Rtn1
Fmp42
Cot1
Zrc1
10
20
30
40
50
t/s

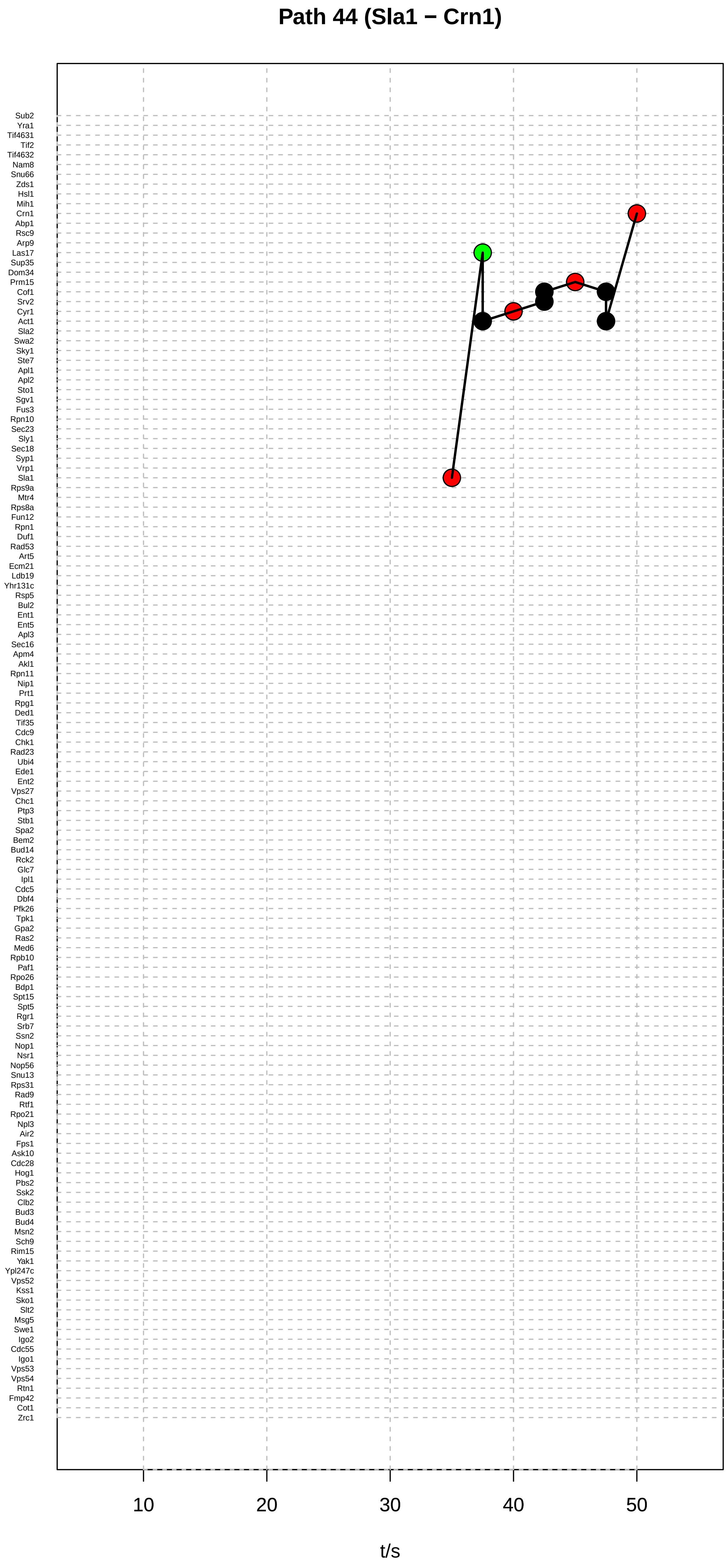

Path 44 (Sla1 − Crn1)
Sub2
Yra1
Tif4631
Tif2
Tif4632
Nam8
Snu66
Zds1
Hsl1
Mih1
Crn1
Abp1
Rsc9
Arp9
Las17
Sup35
Dom34
Prm15
Cof1
Srv2
Cyr1
Act1
Sla2
Swa2
Sky1
Ste7
Apl1
Apl2
Sto1
Sgv1
Fus3
Rpn10
Sec23
Sly1
Sec18
Syp1
Vrp1
Sla1
Rps9a
Mtr4
Rps8a
Fun12
Rpn1
Duf1
Rad53
Art5
Ecm21
Ldb19
Yhr131c
Rsp5
Bul2
Ent1
Ent5
Apl3
Sec16
Apm4
Akl1
Rpn11
Nip1
Prt1
Rpg1
Ded1
Tif35
Cdc9
Chk1
Rad23
Ubi4
Ede1
Ent2
Vps27
Chc1
Ptp3
Stb1
Spa2
Bem2
Bud14
Rck2
Glc7
Ipl1
Cdc5
Dbf4
Pfk26
Tpk1
Gpa2
Ras2
Med6
Rpb10
Paf1
Rpo26
Bdp1
Spt15
Spt5
Rgr1
Srb7
Ssn2
Nop1
Nsr1
Nop56
Snu13
Rps31
Rad9
Rtf1
Rpo21
Npl3
Air2
Fps1
Ask10
Cdc28
Hog1
Pbs2
Ssk2
Clb2
Bud3
Bud4
Msn2
Sch9
Rim15
Yak1
Ypl247c
Vps52
Kss1
Sko1
Slt2
Msg5
Swe1
Igo2
Cdc55
Igo1
Vps53
Vps54
Rtn1
Fmp42
Cot1
Zrc1
10
20
30
40
50
t/s

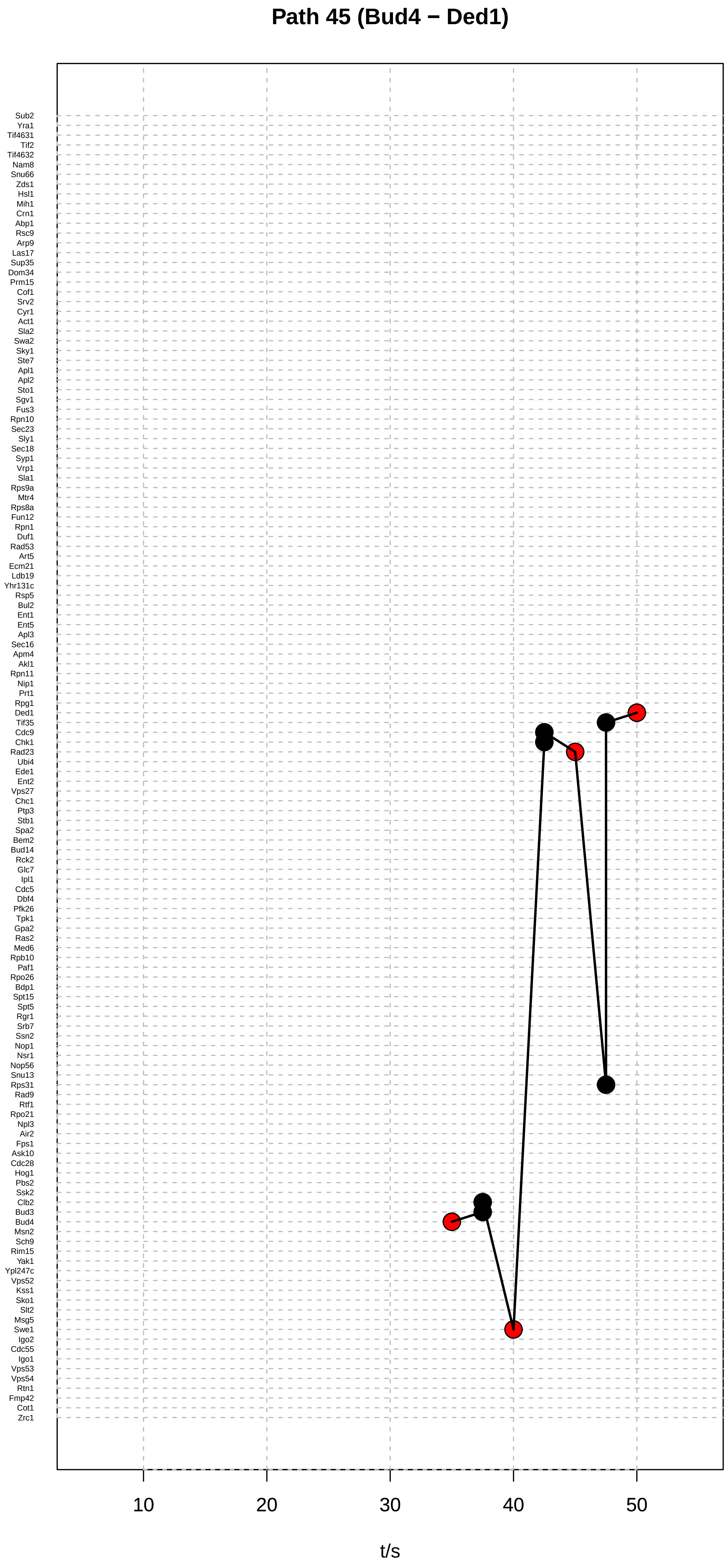
Path 45 (Bud4 − Ded1)
Sub2
Yra1
Tif4631
Tif2
Tif4632
Nam8
Snu66
Zds1
Hsl1
Mih1
Crn1
Abp1
Rsc9
Arp9
Las17
Sup35
Dom34
Prm15
Cof1
Srv2
Cyr1
Act1
Sla2
Swa2
Sky1
Ste7
Apl1
Apl2
Sto1
Sgv1
Fus3
Rpn10
Sec23
Sly1
Sec18
Syp1
Vrp1
Sla1
Rps9a
Mtr4
Rps8a
Fun12
Rpn1
Duf1
Rad53
Art5
Ecm21
Ldb19
Yhr131c
Rsp5
Bul2
Ent1
Ent5
Apl3
Sec16
Apm4
Akl1
Rpn11
Nip1
Prt1
Rpg1
Ded1
Tif35
Cdc9
Chk1
Rad23
Ubi4
Ede1
Ent2
Vps27
Chc1
Ptp3
Stb1
Spa2
Bem2
Bud14
Rck2
Glc7
Ipl1
Cdc5
Dbf4
Pfk26
Tpk1
Gpa2
Ras2
Med6
Rpb10
Paf1
Rpo26
Bdp1
Spt15
Spt5
Rgr1
Srb7
Ssn2
Nop1
Nsr1
Nop56
Snu13
Rps31
Rad9
Rtf1
Rpo21
Npl3
Air2
Fps1
Ask10
Cdc28
Hog1
Pbs2
Ssk2
Clb2
Bud3
Bud4
Msn2
Sch9
Rim15
Yak1
Ypl247c
Vps52
Kss1
Sko1
Slt2
Msg5
Swe1
Igo2
Cdc55
Igo1
Vps53
Vps54
Rtn1
Fmp42
Cot1
Zrc1
10
20
30
40
50
t/s

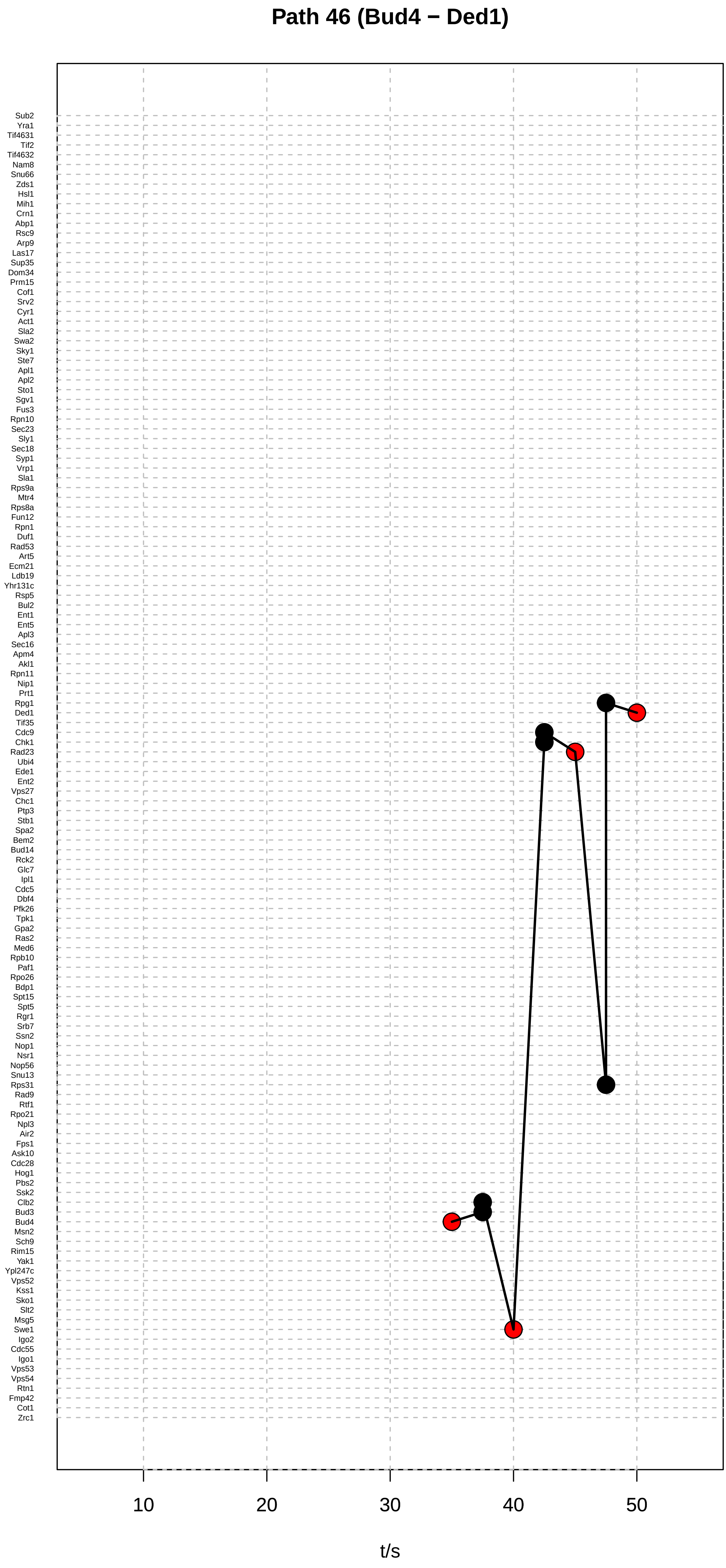

Path 46 (Bud4 − Ded1)
Sub2
Yra1
Tif4631
Tif2
Tif4632
Nam8
Snu66
Zds1
Hsl1
Mih1
Crn1
Abp1
Rsc9
Arp9
Las17
Sup35
Dom34
Prm15
Cof1
Srv2
Cyr1
Act1
Sla2
Swa2
Sky1
Ste7
Apl1
Apl2
Sto1
Sgv1
Fus3
Rpn10
Sec23
Sly1
Sec18
Syp1
Vrp1
Sla1
Rps9a
Mtr4
Rps8a
Fun12
Rpn1
Duf1
Rad53
Art5
Ecm21
Ldb19
Yhr131c
Rsp5
Bul2
Ent1
Ent5
Apl3
Sec16
Apm4
Akl1
Rpn11
Nip1
Prt1
Rpg1
Ded1
Tif35
Cdc9
Chk1
Rad23
Ubi4
Ede1
Ent2
Vps27
Chc1
Ptp3
Stb1
Spa2
Bem2
Bud14
Rck2
Glc7
Ipl1
Cdc5
Dbf4
Pfk26
Tpk1
Gpa2
Ras2
Med6
Rpb10
Paf1
Rpo26
Bdp1
Spt15
Spt5
Rgr1
Srb7
Ssn2
Nop1
Nsr1
Nop56
Snu13
Rps31
Rad9
Rtf1
Rpo21
Npl3
Air2
Fps1
Ask10
Cdc28
Hog1
Pbs2
Ssk2
Clb2
Bud3
Bud4
Msn2
Sch9
Rim15
Yak1
Ypl247c
Vps52
Kss1
Sko1
Slt2
Msg5
Swe1
Igo2
Cdc55
Igo1
Vps53
Vps54
Rtn1
Fmp42
Cot1
Zrc1
10
20
30
40
50
t/s

### Path 47 (Bud4 − Ded1)

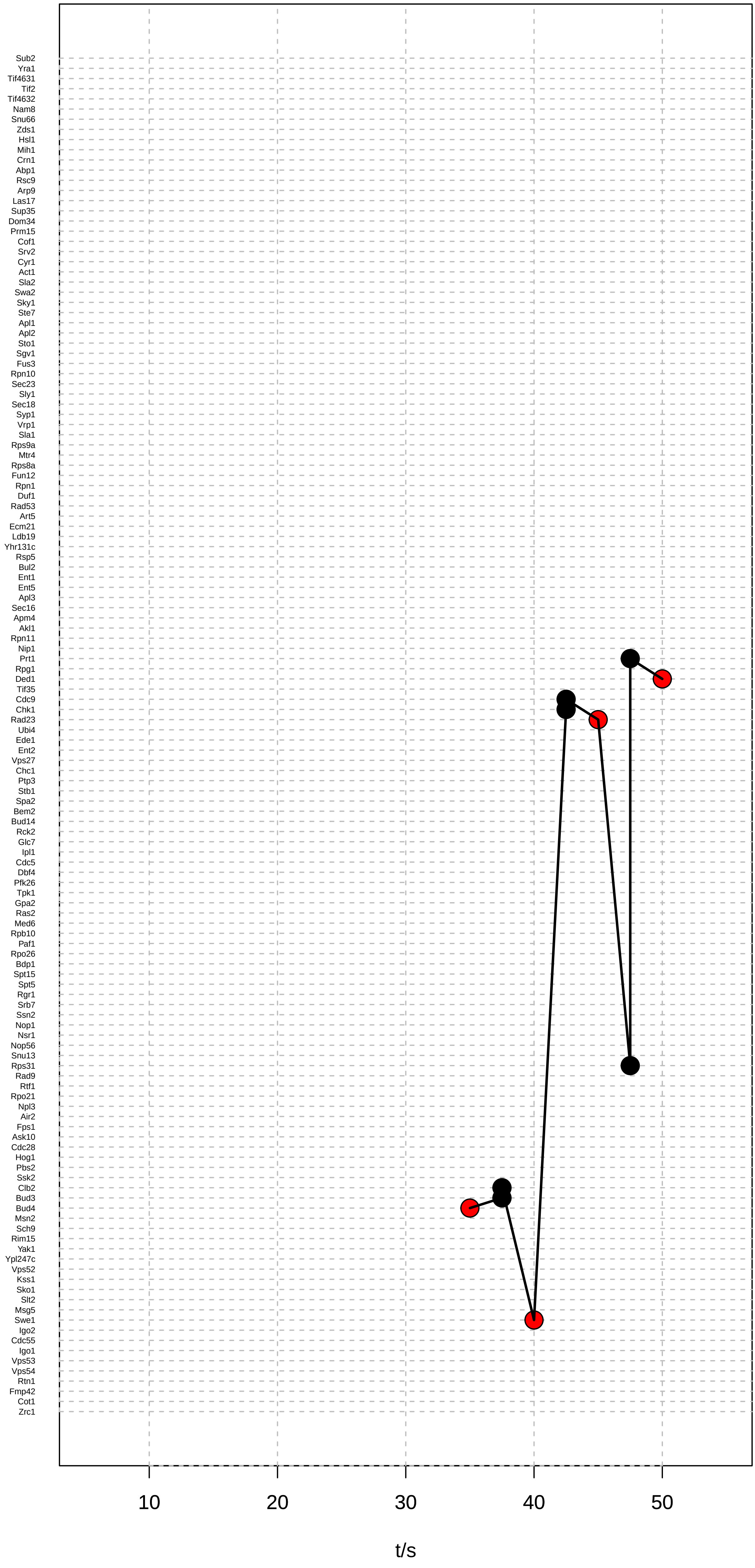

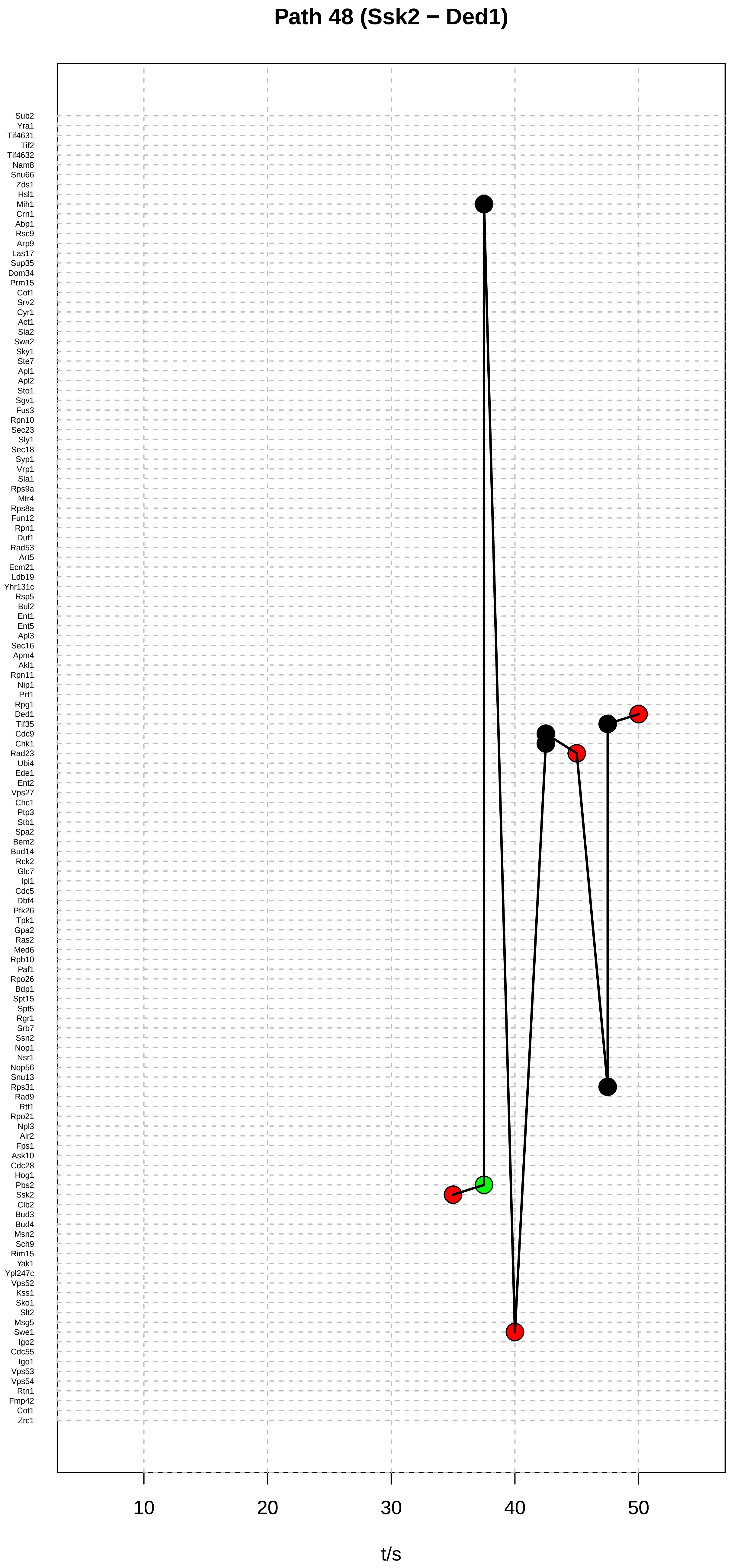
Path 48 (Ssk2 − Ded1)
Sub2
Yra1
Tif4631
Tif2
Tif4632
Nam8
Snu66
Zds1
Hsl1
Mih1
Crn1
Abp1
Rsc9
Arp9
Las17
Sup35
Dom34
Prm15
Cof1
Srv2
Cyr1
Act1
Sla2
Swa2
Sky1
Ste7
Apl1
Apl2
Sto1
Sgv1
Fus3
Rpn10
Sec23
Sly1
Sec18
Syp1
Vrp1
Sla1
Rps9a
Mtr4
Rps8a
Fun12
Rpn1
Duf1
Rad53
Art5
Ecm21
Ldb19
Yhr131c
Rsp5
Bul2
Ent1
Ent5
Apl3
Sec16
Apm4
Akl1
Rpn11
Nip1
Prt1
Rpg1
Ded1
Tif35
Cdc9
Chk1
Rad23
Ubi4
Ede1
Ent2
Vps27
Chc1
Ptp3
Stb1
Spa2
Bem2
Bud14
Rck2
Glc7
Ipl1
Cdc5
Dbf4
Pfk26
Tpk1
Gpa2
Ras2
Med6
Rpb10
Paf1
Rpo26
Bdp1
Spt15
Spt5
Rgr1
Srb7
Ssn2
Nop1
Nsr1
Nop56
Snu13
Rps31
Rad9
Rtf1
Rpo21
Npl3
Air2
Fps1
Ask10
Cdc28
Hog1
Pbs2
Ssk2
Clb2
Bud3
Bud4
Msn2
Sch9
Rim15
Yak1
Ypl247c
Vps52
Kss1
Sko1
Slt2
Msg5
Swe1
Igo2
Cdc55
Igo1
Vps53
Vps54
Rtn1
Fmp42
Cot1
Zrc1
10
20
30
40
50
t/s

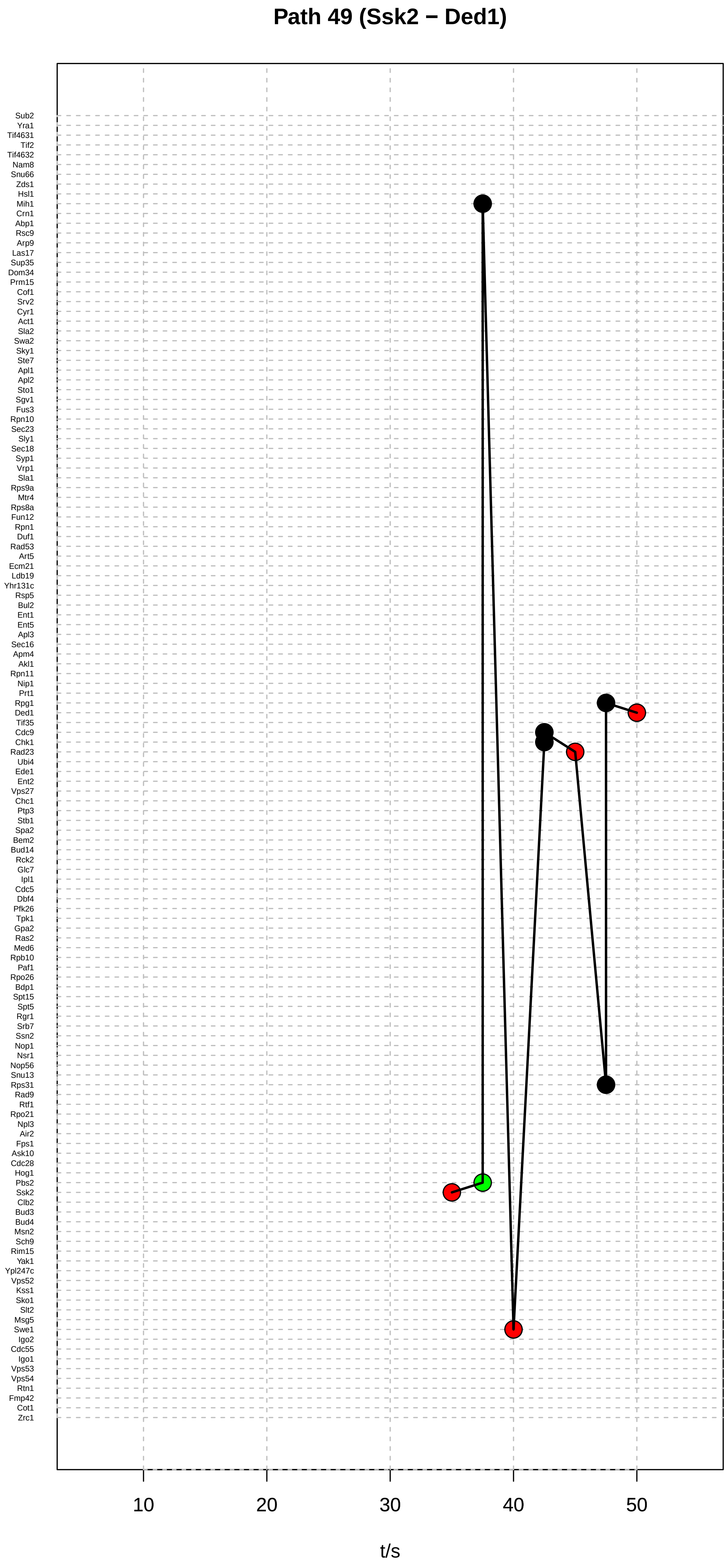
Path 49 (Ssk2 − Ded1)
Sub2
Yra1
Tif4631
Tif2
Tif4632
Nam8
Snu66
Zds1
Hsl1
Mih1
Crn1
Abp1
Rsc9
Arp9
Las17
Sup35
Dom34
Prm15
Cof1
Srv2
Cyr1
Act1
Sla2
Swa2
Sky1
Ste7
Apl1
Apl2
Sto1
Sgv1
Fus3
Rpn10
Sec23
Sly1
Sec18
Syp1
Vrp1
Sla1
Rps9a
Mtr4
Rps8a
Fun12
Rpn1
Duf1
Rad53
Art5
Ecm21
Ldb19
Yhr131c
Rsp5
Bul2
Ent1
Ent5
Apl3
Sec16
Apm4
Akl1
Rpn11
Nip1
Prt1
Rpg1
Ded1
Tif35
Cdc9
Chk1
Rad23
Ubi4
Ede1
Ent2
Vps27
Chc1
Ptp3
Stb1
Spa2
Bem2
Bud14
Rck2
Glc7
Ipl1
Cdc5
Dbf4
Pfk26
Tpk1
Gpa2
Ras2
Med6
Rpb10
Paf1
Rpo26
Bdp1
Spt15
Spt5
Rgr1
Srb7
Ssn2
Nop1
Nsr1
Nop56
Snu13
Rps31
Rad9
Rtf1
Rpo21
Npl3
Air2
Fps1
Ask10
Cdc28
Hog1
Pbs2
Ssk2
Clb2
Bud3
Bud4
Msn2
Sch9
Rim15
Yak1
Ypl247c
Vps52
Kss1
Sko1
Slt2
Msg5
Swe1
Igo2
Cdc55
Igo1
Vps53
Vps54
Rtn1
Fmp42
Cot1
Zrc1
10
20
30
40
50
t/s

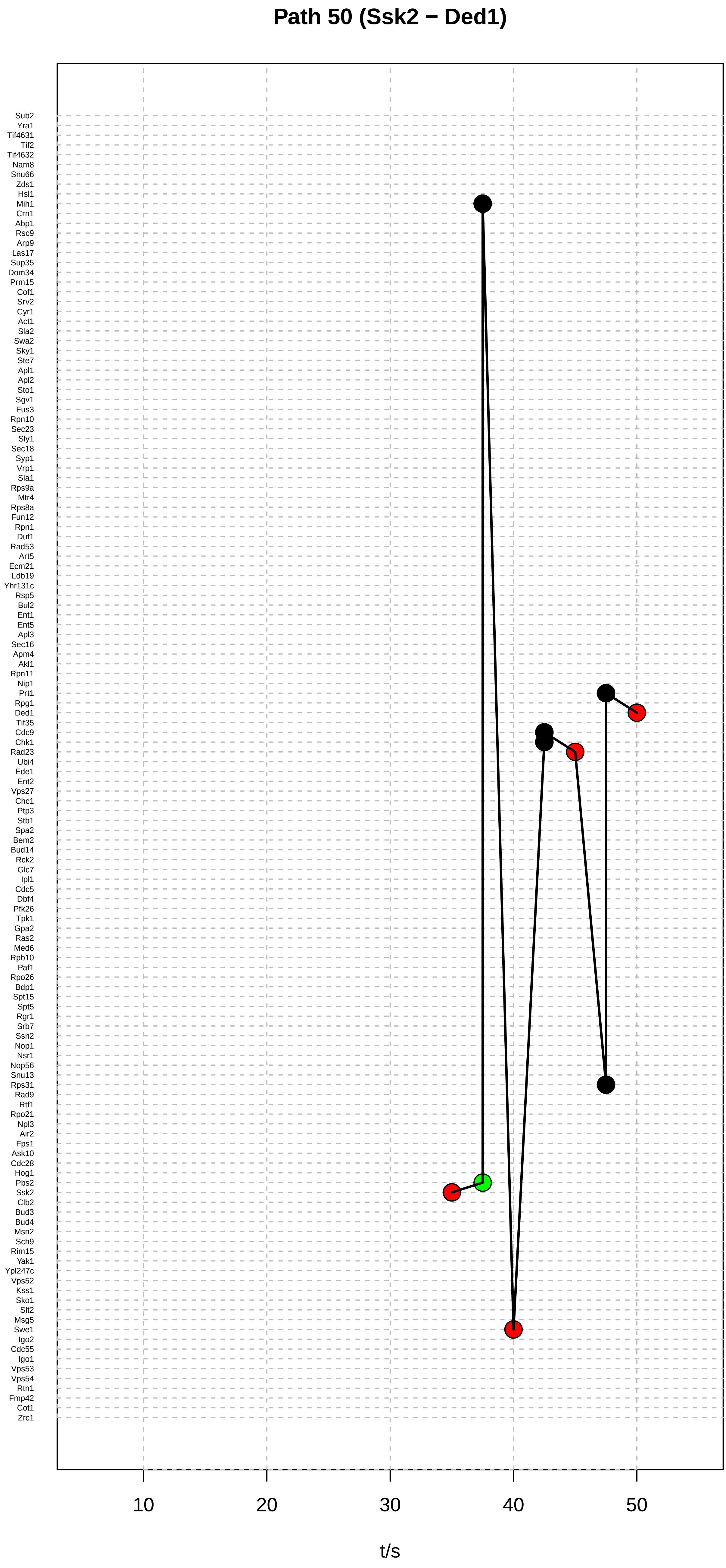
Path 50 (Ssk2 − Ded1)
Sub2
Yra1
Tif4631
Tif2
Tif4632
Nam8
Snu66
Zds1
Hsl1
Mih1
Crn1
Abp1
Rsc9
Arp9
Las17
Sup35
Dom34
Prm15
Cof1
Srv2
Cyr1
Act1
Sla2
Swa2
Sky1
Ste7
Apl1
Apl2
Sto1
Sgv1
Fus3
Rpn10
Sec23
Sly1
Sec18
Syp1
Vrp1
Sla1
Rps9a
Mtr4
Rps8a
Fun12
Rpn1
Duf1
Rad53
Art5
Ecm21
Ldb19
Yhr131c
Rsp5
Bul2
Ent1
Ent5
Apl3
Sec16
Apm4
Akl1
Rpn11
Nip1
Prt1
Rpg1
Ded1
Tif35
Cdc9
Chk1
Rad23
Ubi4
Ede1
Ent2
Vps27
Chc1
Ptp3
Stb1
Spa2
Bem2
Bud14
Rck2
Glc7
Ipl1
Cdc5
Dbf4
Pfk26
Tpk1
Gpa2
Ras2
Med6
Rpb10
Paf1
Rpo26
Bdp1
Spt15
Spt5
Rgr1
Srb7
Ssn2
Nop1
Nsr1
Nop56
Snu13
Rps31
Rad9
Rtf1
Rpo21
Npl3
Air2
Fps1
Ask10
Cdc28
Hog1
Pbs2
Ssk2
Clb2
Bud3
Bud4
Msn2
Sch9
Rim15
Yak1
Ypl247c
Vps52
Kss1
Sko1
Slt2
Msg5
Swe1
Igo2
Cdc55
Igo1
Vps53
Vps54
Rtn1
Fmp42
Cot1
Zrc1
10
20
30
40
50
t/s

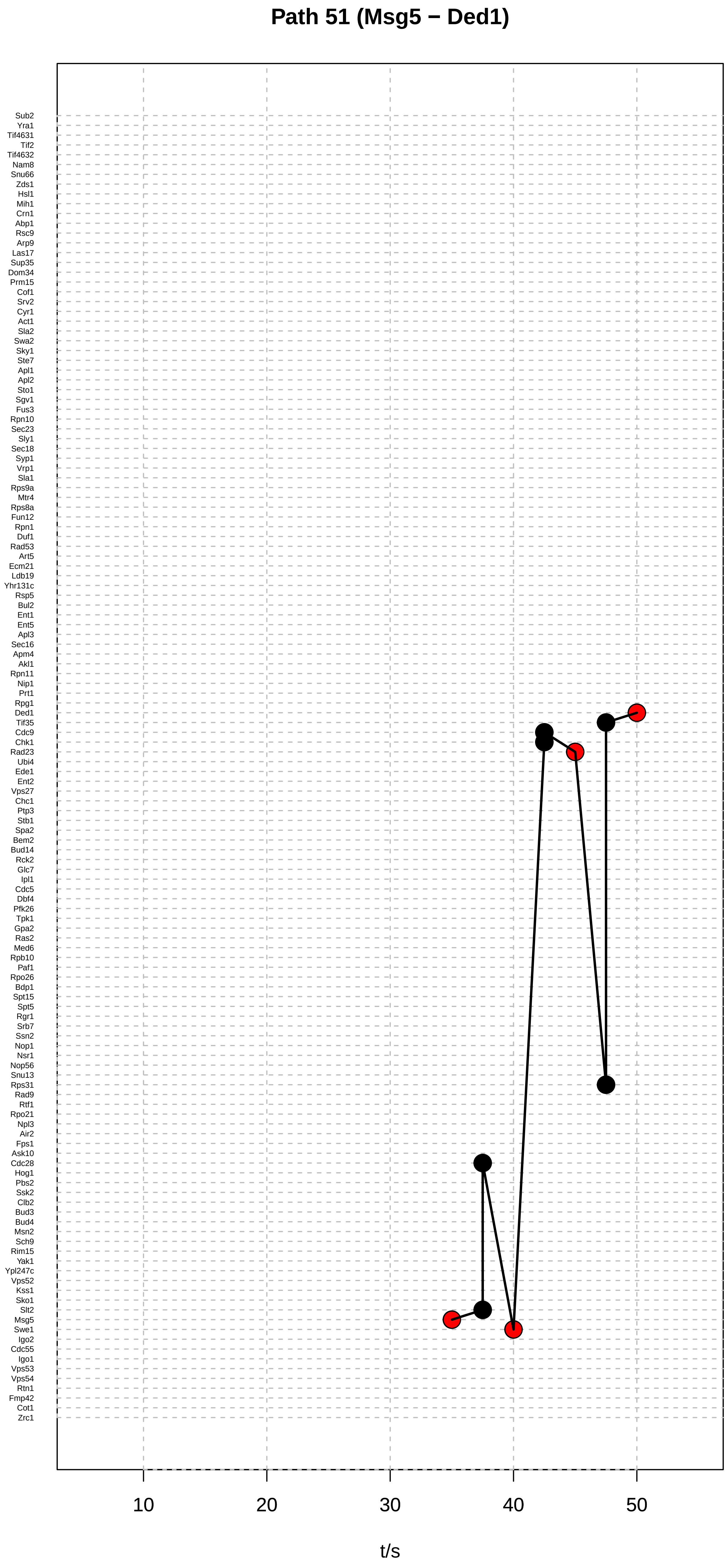

Path 51 (Msg5 − Ded1)
Sub2
Yra1
Tif4631
Tif2
Tif4632
Nam8
Snu66
Zds1
Hsl1
Mih1
Crn1
Abp1
Rsc9
Arp9
Las17
Sup35
Dom34
Prm15
Cof1
Srv2
Cyr1
Act1
Sla2
Swa2
Sky1
Ste7
Apl1
Apl2
Sto1
Sgv1
Fus3
Rpn10
Sec23
Sly1
Sec18
Syp1
Vrp1
Sla1
Rps9a
Mtr4
Rps8a
Fun12
Rpn1
Duf1
Rad53
Art5
Ecm21
Ldb19
Yhr131c
Rsp5
Bul2
Ent1
Ent5
Apl3
Sec16
Apm4
Akl1
Rpn11
Nip1
Prt1
Rpg1
Ded1
Tif35
Cdc9
Chk1
Rad23
Ubi4
Ede1
Ent2
Vps27
Chc1
Ptp3
Stb1
Spa2
Bem2
Bud14
Rck2
Glc7
Ipl1
Cdc5
Dbf4
Pfk26
Tpk1
Gpa2
Ras2
Med6
Rpb10
Paf1
Rpo26
Bdp1
Spt15
Spt5
Rgr1
Srb7
Ssn2
Nop1
Nsr1
Nop56
Snu13
Rps31
Rad9
Rtf1
Rpo21
Npl3
Air2
Fps1
Ask10
Cdc28
Hog1
Pbs2
Ssk2
Clb2
Bud3
Bud4
Msn2
Sch9
Rim15
Yak1
Ypl247c
Vps52
Kss1
Sko1
Slt2
Msg5
Swe1
Igo2
Cdc55
Igo1
Vps53
Vps54
Rtn1
Fmp42
Cot1
Zrc1
10
20
30
40
50
t/s

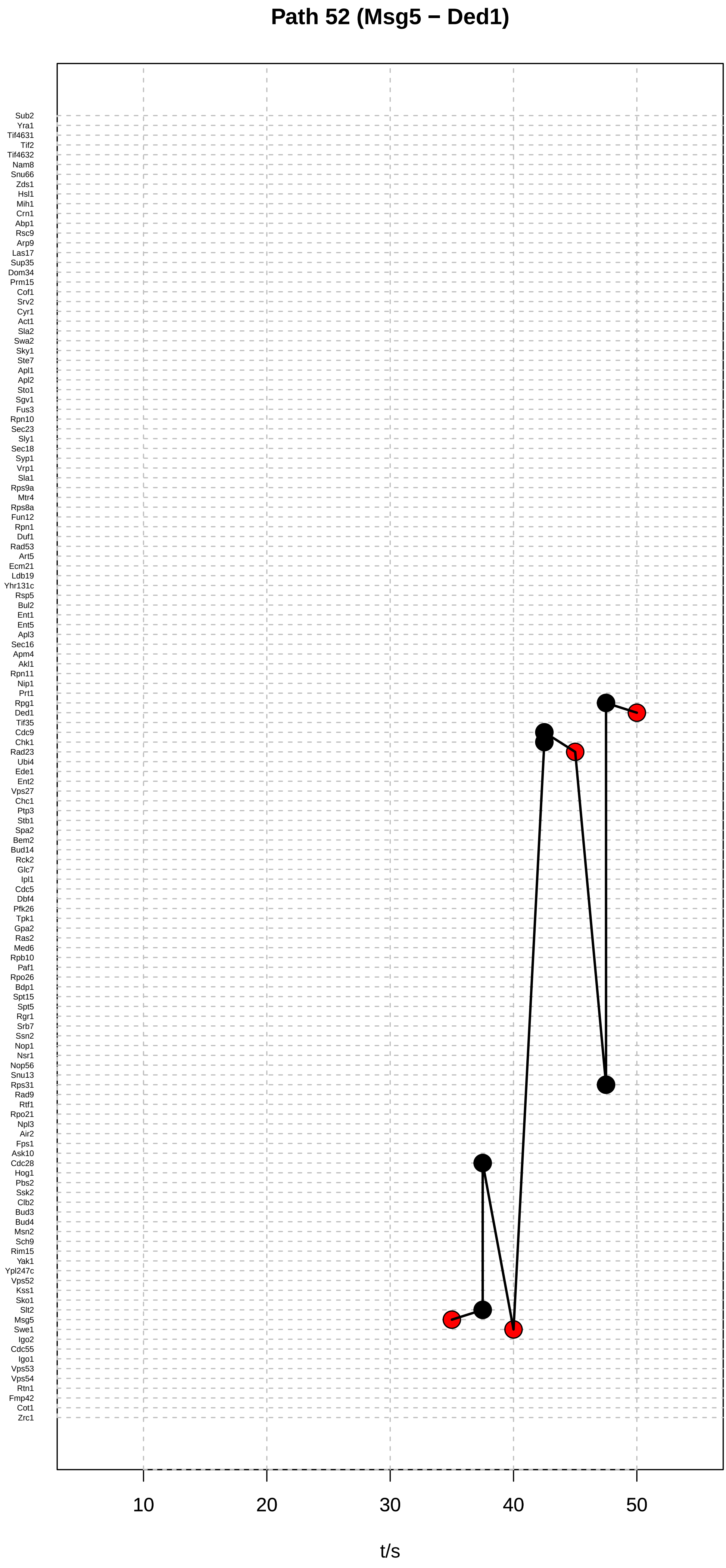
Path 52 (Msg5 − Ded1)
Sub2
Yra1
Tif4631
Tif2
Tif4632
Nam8
Snu66
Zds1
Hsl1
Mih1
Crn1
Abp1
Rsc9
Arp9
Las17
Sup35
Dom34
Prm15
Cof1
Srv2
Cyr1
Act1
Sla2
Swa2
Sky1
Ste7
Apl1
Apl2
Sto1
Sgv1
Fus3
Rpn10
Sec23
Sly1
Sec18
Syp1
Vrp1
Sla1
Rps9a
Mtr4
Rps8a
Fun12
Rpn1
Duf1
Rad53
Art5
Ecm21
Ldb19
Yhr131c
Rsp5
Bul2
Ent1
Ent5
Apl3
Sec16
Apm4
Akl1
Rpn11
Nip1
Prt1
Rpg1
Ded1
Tif35
Cdc9
Chk1
Rad23
Ubi4
Ede1
Ent2
Vps27
Chc1
Ptp3
Stb1
Spa2
Bem2
Bud14
Rck2
Glc7
Ipl1
Cdc5
Dbf4
Pfk26
Tpk1
Gpa2
Ras2
Med6
Rpb10
Paf1
Rpo26
Bdp1
Spt15
Spt5
Rgr1
Srb7
Ssn2
Nop1
Nsr1
Nop56
Snu13
Rps31
Rad9
Rtf1
Rpo21
Npl3
Air2
Fps1
Ask10
Cdc28
Hog1
Pbs2
Ssk2
Clb2
Bud3
Bud4
Msn2
Sch9
Rim15
Yak1
Ypl247c
Vps52
Kss1
Sko1
Slt2
Msg5
Swe1
Igo2
Cdc55
Igo1
Vps53
Vps54
Rtn1
Fmp42
Cot1
Zrc1
10
20
30
40
50
t/s

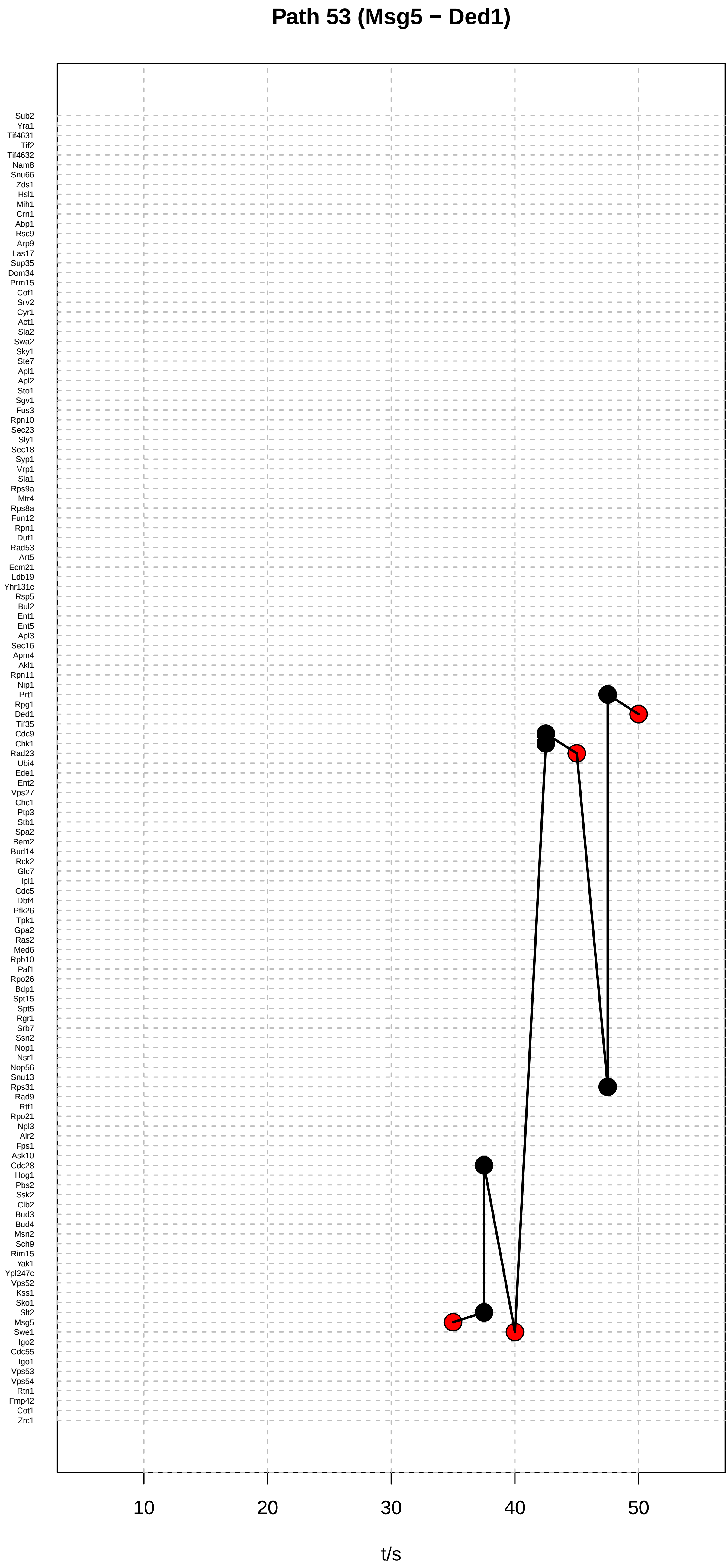
Path 53 (Msg5 − Ded1)
Sub2
Yra1
Tif4631
Tif2
Tif4632
Nam8
Snu66
Zds1
Hsl1
Mih1
Crn1
Abp1
Rsc9
Arp9
Las17
Sup35
Dom34
Prm15
Cof1
Srv2
Cyr1
Act1
Sla2
Swa2
Sky1
Ste7
Apl1
Apl2
Sto1
Sgv1
Fus3
Rpn10
Sec23
Sly1
Sec18
Syp1
Vrp1
Sla1
Rps9a
Mtr4
Rps8a
Fun12
Rpn1
Duf1
Rad53
Art5
Ecm21
Ldb19
Yhr131c
Rsp5
Bul2
Ent1
Ent5
Apl3
Sec16
Apm4
Akl1
Rpn11
Nip1
Prt1
Rpg1
Ded1
Tif35
Cdc9
Chk1
Rad23
Ubi4
Ede1
Ent2
Vps27
Chc1
Ptp3
Stb1
Spa2
Bem2
Bud14
Rck2
Glc7
Ipl1
Cdc5
Dbf4
Pfk26
Tpk1
Gpa2
Ras2
Med6
Rpb10
Paf1
Rpo26
Bdp1
Spt15
Spt5
Rgr1
Srb7
Ssn2
Nop1
Nsr1
Nop56
Snu13
Rps31
Rad9
Rtf1
Rpo21
Npl3
Air2
Fps1
Ask10
Cdc28
Hog1
Pbs2
Ssk2
Clb2
Bud3
Bud4
Msn2
Sch9
Rim15
Yak1
Ypl247c
Vps52
Kss1
Sko1
Slt2
Msg5
Swe1
Igo2
Cdc55
Igo1
Vps53
Vps54
Rtn1
Fmp42
Cot1
Zrc1
10
20
30
40
50
t/s

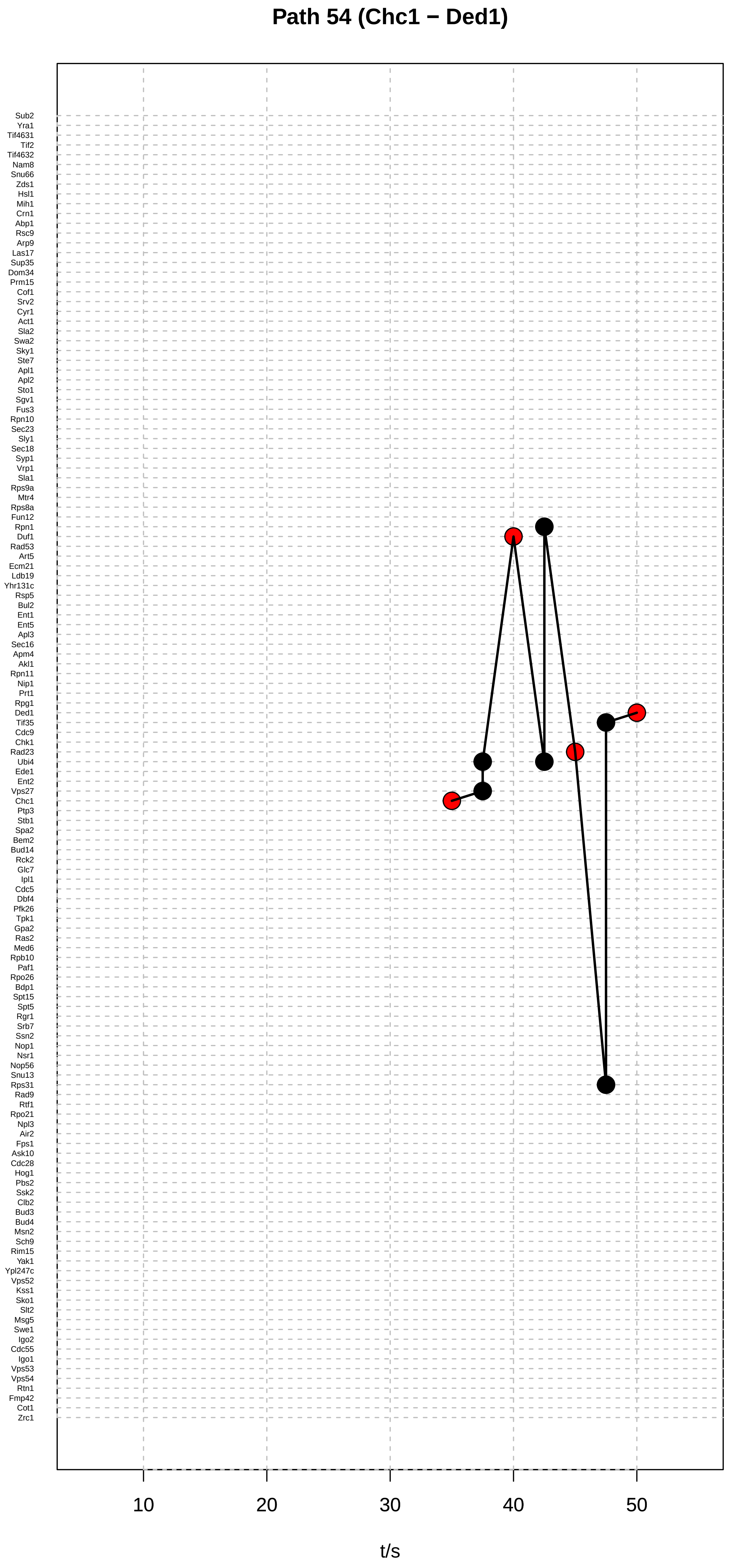

Path 54 (Chc1 − Ded1)
Sub2
Yra1
Tif4631
Tif2
Tif4632
Nam8
Snu66
Zds1
Hsl1
Mih1
Crn1
Abp1
Rsc9
Arp9
Las17
Sup35
Dom34
Prm15
Cof1
Srv2
Cyr1
Act1
Sla2
Swa2
Sky1
Ste7
Apl1
Apl2
Sto1
Sgv1
Fus3
Rpn10
Sec23
Sly1
Sec18
Syp1
Vrp1
Sla1
Rps9a
Mtr4
Rps8a
Fun12
Rpn1
Duf1
Rad53
Art5
Ecm21
Ldb19
Yhr131c
Rsp5
Bul2
Ent1
Ent5
Apl3
Sec16
Apm4
Akl1
Rpn11
Nip1
Prt1
Rpg1
Ded1
Tif35
Cdc9
Chk1
Rad23
Ubi4
Ede1
Ent2
Vps27
Chc1
Ptp3
Stb1
Spa2
Bem2
Bud14
Rck2
Glc7
Ipl1
Cdc5
Dbf4
Pfk26
Tpk1
Gpa2
Ras2
Med6
Rpb10
Paf1
Rpo26
Bdp1
Spt15
Spt5
Rgr1
Srb7
Ssn2
Nop1
Nsr1
Nop56
Snu13
Rps31
Rad9
Rtf1
Rpo21
Npl3
Air2
Fps1
Ask10
Cdc28
Hog1
Pbs2
Ssk2
Clb2
Bud3
Bud4
Msn2
Sch9
Rim15
Yak1
Ypl247c
Vps52
Kss1
Sko1
Slt2
Msg5
Swe1
Igo2
Cdc55
Igo1
Vps53
Vps54
Rtn1
Fmp42
Cot1
Zrc1
10
20
30
40
50
t/s

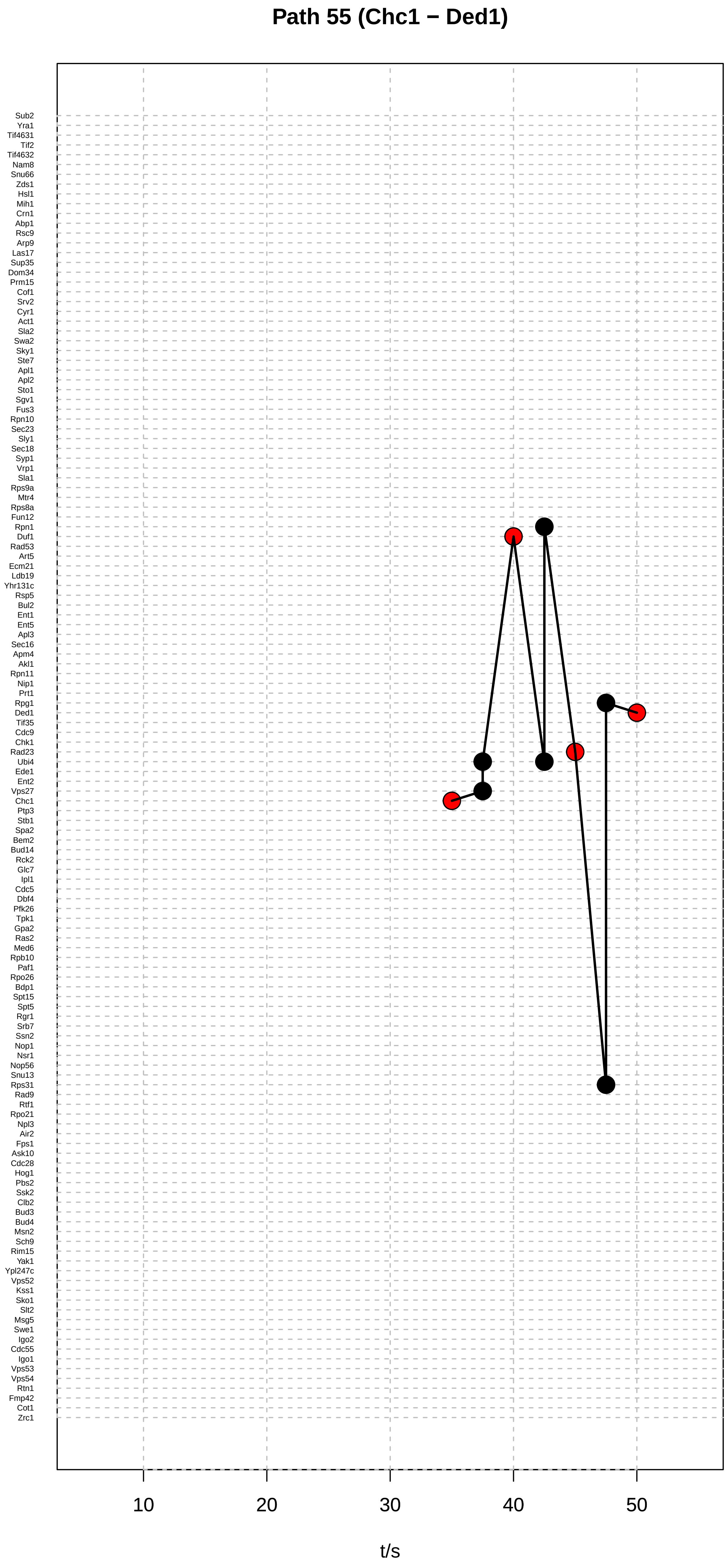

Path 55 (Chc1 − Ded1)
Sub2
Yra1
Tif4631
Tif2
Tif4632
Nam8
Snu66
Zds1
Hsl1
Mih1
Crn1
Abp1
Rsc9
Arp9
Las17
Sup35
Dom34
Prm15
Cof1
Srv2
Cyr1
Act1
Sla2
Swa2
Sky1
Ste7
Apl1
Apl2
Sto1
Sgv1
Fus3
Rpn10
Sec23
Sly1
Sec18
Syp1
Vrp1
Sla1
Rps9a
Mtr4
Rps8a
Fun12
Rpn1
Duf1
Rad53
Art5
Ecm21
Ldb19
Yhr131c
Rsp5
Bul2
Ent1
Ent5
Apl3
Sec16
Apm4
Akl1
Rpn11
Nip1
Prt1
Rpg1
Ded1
Tif35
Cdc9
Chk1
Rad23
Ubi4
Ede1
Ent2
Vps27
Chc1
Ptp3
Stb1
Spa2
Bem2
Bud14
Rck2
Glc7
Ipl1
Cdc5
Dbf4
Pfk26
Tpk1
Gpa2
Ras2
Med6
Rpb10
Paf1
Rpo26
Bdp1
Spt15
Spt5
Rgr1
Srb7
Ssn2
Nop1
Nsr1
Nop56
Snu13
Rps31
Rad9
Rtf1
Rpo21
Npl3
Air2
Fps1
Ask10
Cdc28
Hog1
Pbs2
Ssk2
Clb2
Bud3
Bud4
Msn2
Sch9
Rim15
Yak1
Ypl247c
Vps52
Kss1
Sko1
Slt2
Msg5
Swe1
Igo2
Cdc55
Igo1
Vps53
Vps54
Rtn1
Fmp42
Cot1
Zrc1
10
20
30
40
50
t/s

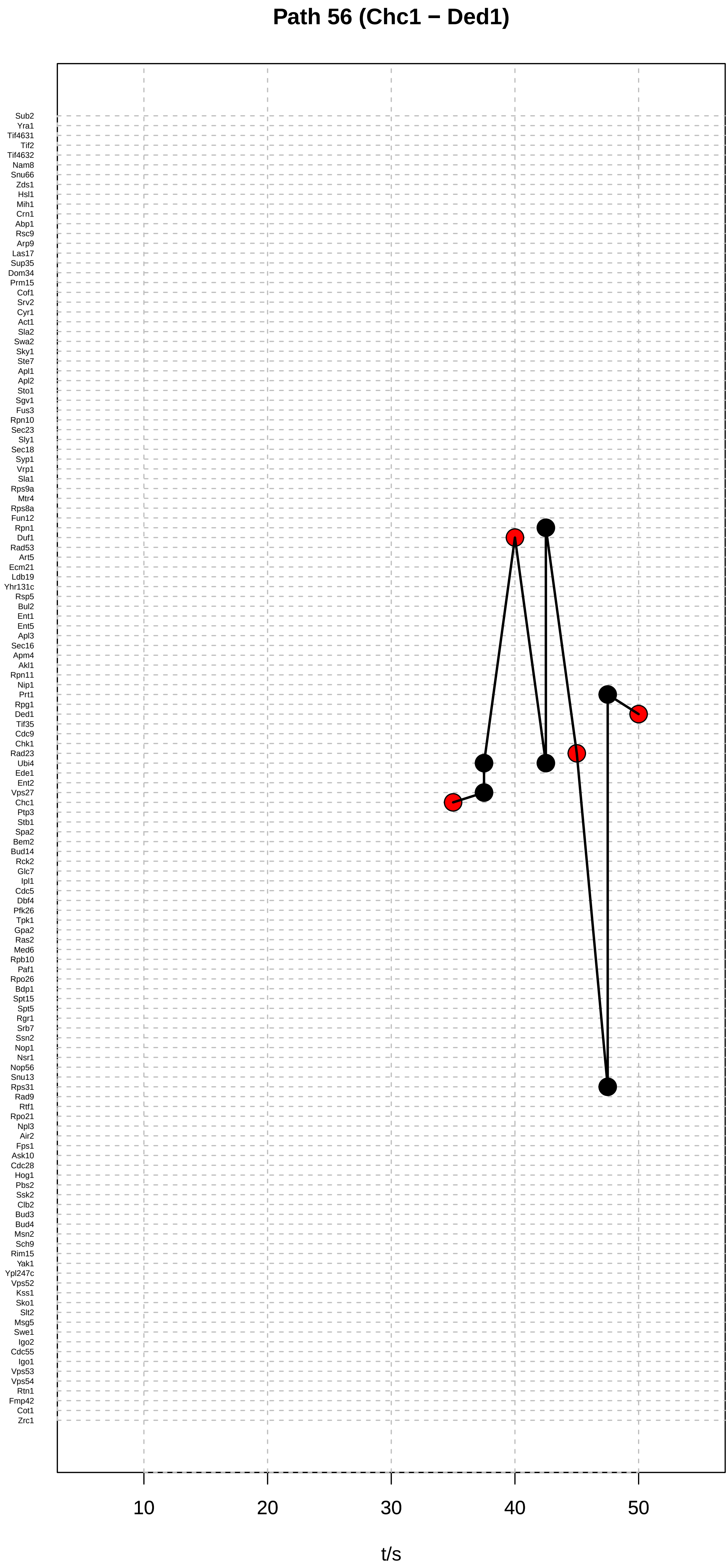
Path 56 (Chc1 − Ded1)
Sub2
Yra1
Tif4631
Tif2
Tif4632
Nam8
Snu66
Zds1
Hsl1
Mih1
Crn1
Abp1
Rsc9
Arp9
Las17
Sup35
Dom34
Prm15
Cof1
Srv2
Cyr1
Act1
Sla2
Swa2
Sky1
Ste7
Apl1
Apl2
Sto1
Sgv1
Fus3
Rpn10
Sec23
Sly1
Sec18
Syp1
Vrp1
Sla1
Rps9a
Mtr4
Rps8a
Fun12
Rpn1
Duf1
Rad53
Art5
Ecm21
Ldb19
Yhr131c
Rsp5
Bul2
Ent1
Ent5
Apl3
Sec16
Apm4
Akl1
Rpn11
Nip1
Prt1
Rpg1
Ded1
Tif35
Cdc9
Chk1
Rad23
Ubi4
Ede1
Ent2
Vps27
Chc1
Ptp3
Stb1
Spa2
Bem2
Bud14
Rck2
Glc7
Ipl1
Cdc5
Dbf4
Pfk26
Tpk1
Gpa2
Ras2
Med6
Rpb10
Paf1
Rpo26
Bdp1
Spt15
Spt5
Rgr1
Srb7
Ssn2
Nop1
Nsr1
Nop56
Snu13
Rps31
Rad9
Rtf1
Rpo21
Npl3
Air2
Fps1
Ask10
Cdc28
Hog1
Pbs2
Ssk2
Clb2
Bud3
Bud4
Msn2
Sch9
Rim15
Yak1
Ypl247c
Vps52
Kss1
Sko1
Slt2
Msg5
Swe1
Igo2
Cdc55
Igo1
Vps53
Vps54
Rtn1
Fmp42
Cot1
Zrc1
10
20
30
40
50
t/s

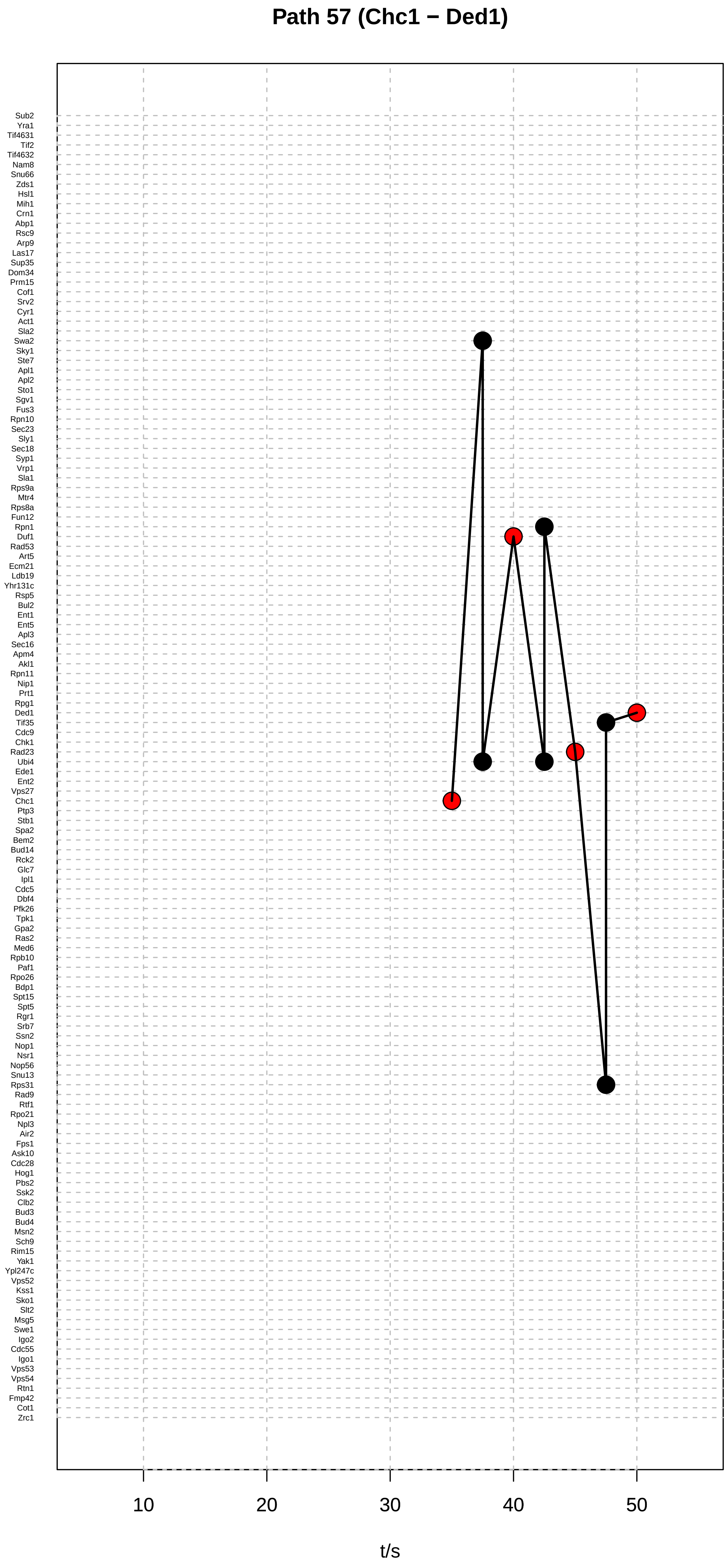

Path 57 (Chc1 − Ded1)
Sub2
Yra1
Tif4631
Tif2
Tif4632
Nam8
Snu66
Zds1
Hsl1
Mih1
Crn1
Abp1
Rsc9
Arp9
Las17
Sup35
Dom34
Prm15
Cof1
Srv2
Cyr1
Act1
Sla2
Swa2
Sky1
Ste7
Apl1
Apl2
Sto1
Sgv1
Fus3
Rpn10
Sec23
Sly1
Sec18
Syp1
Vrp1
Sla1
Rps9a
Mtr4
Rps8a
Fun12
Rpn1
Duf1
Rad53
Art5
Ecm21
Ldb19
Yhr131c
Rsp5
Bul2
Ent1
Ent5
Apl3
Sec16
Apm4
Akl1
Rpn11
Nip1
Prt1
Rpg1
Ded1
Tif35
Cdc9
Chk1
Rad23
Ubi4
Ede1
Ent2
Vps27
Chc1
Ptp3
Stb1
Spa2
Bem2
Bud14
Rck2
Glc7
Ipl1
Cdc5
Dbf4
Pfk26
Tpk1
Gpa2
Ras2
Med6
Rpb10
Paf1
Rpo26
Bdp1
Spt15
Spt5
Rgr1
Srb7
Ssn2
Nop1
Nsr1
Nop56
Snu13
Rps31
Rad9
Rtf1
Rpo21
Npl3
Air2
Fps1
Ask10
Cdc28
Hog1
Pbs2
Ssk2
Clb2
Bud3
Bud4
Msn2
Sch9
Rim15
Yak1
Ypl247c
Vps52
Kss1
Sko1
Slt2
Msg5
Swe1
Igo2
Cdc55
Igo1
Vps53
Vps54
Rtn1
Fmp42
Cot1
Zrc1
10
20
30
40
50
t/s

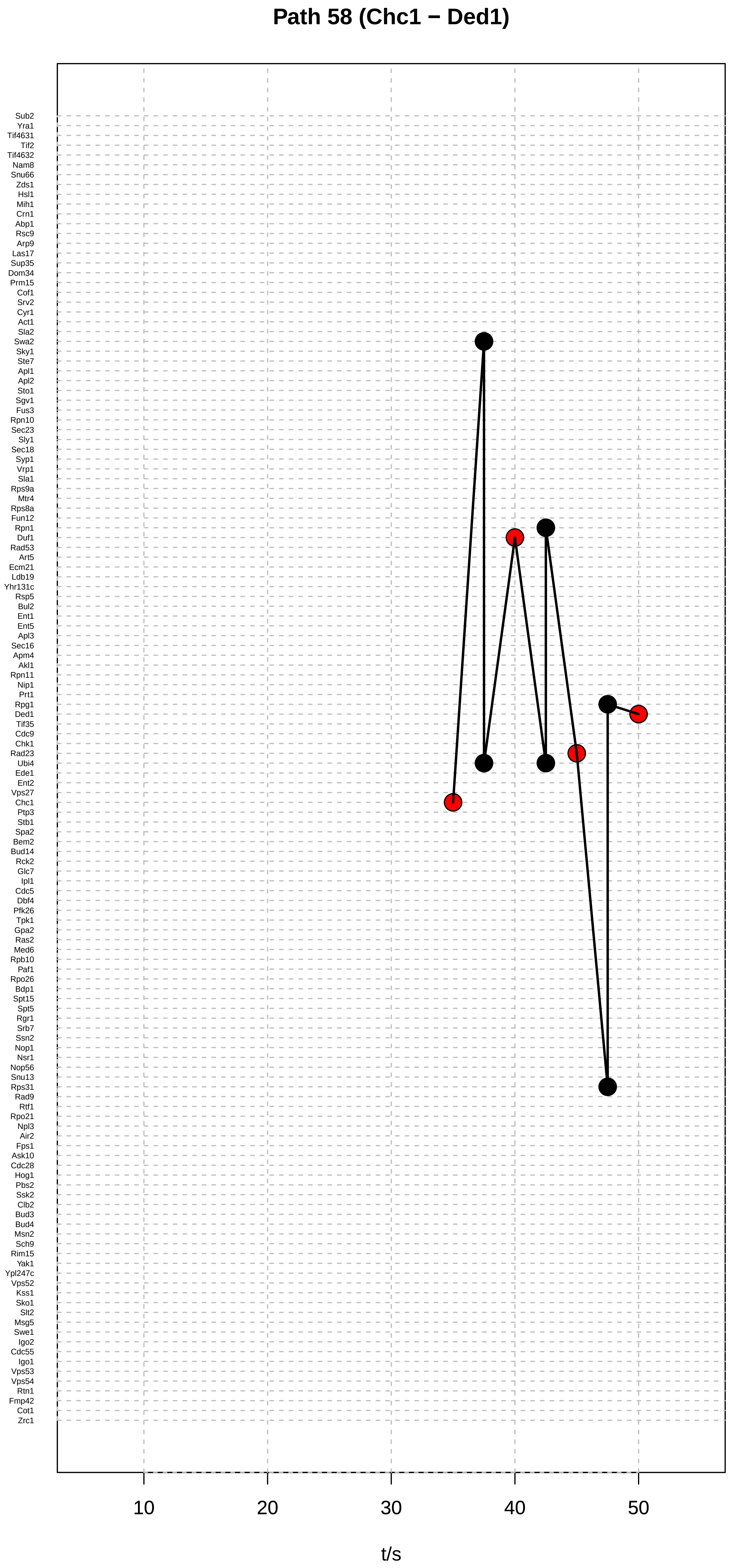

Path 58 (Chc1 − Ded1)
Sub2
Yra1
Tif4631
Tif2
Tif4632
Nam8
Snu66
Zds1
Hsl1
Mih1
Crn1
Abp1
Rsc9
Arp9
Las17
Sup35
Dom34
Prm15
Cof1
Srv2
Cyr1
Act1
Sla2
Swa2
Sky1
Ste7
Apl1
Apl2
Sto1
Sgv1
Fus3
Rpn10
Sec23
Sly1
Sec18
Syp1
Vrp1
Sla1
Rps9a
Mtr4
Rps8a
Fun12
Rpn1
Duf1
Rad53
Art5
Ecm21
Ldb19
Yhr131c
Rsp5
Bul2
Ent1
Ent5
Apl3
Sec16
Apm4
Akl1
Rpn11
Nip1
Prt1
Rpg1
Ded1
Tif35
Cdc9
Chk1
Rad23
Ubi4
Ede1
Ent2
Vps27
Chc1
Ptp3
Stb1
Spa2
Bem2
Bud14
Rck2
Glc7
Ipl1
Cdc5
Dbf4
Pfk26
Tpk1
Gpa2
Ras2
Med6
Rpb10
Paf1
Rpo26
Bdp1
Spt15
Spt5
Rgr1
Srb7
Ssn2
Nop1
Nsr1
Nop56
Snu13
Rps31
Rad9
Rtf1
Rpo21
Npl3
Air2
Fps1
Ask10
Cdc28
Hog1
Pbs2
Ssk2
Clb2
Bud3
Bud4
Msn2
Sch9
Rim15
Yak1
Ypl247c
Vps52
Kss1
Sko1
Slt2
Msg5
Swe1
Igo2
Cdc55
Igo1
Vps53
Vps54
Rtn1
Fmp42
Cot1
Zrc1
10
20
30
40
50
t/s

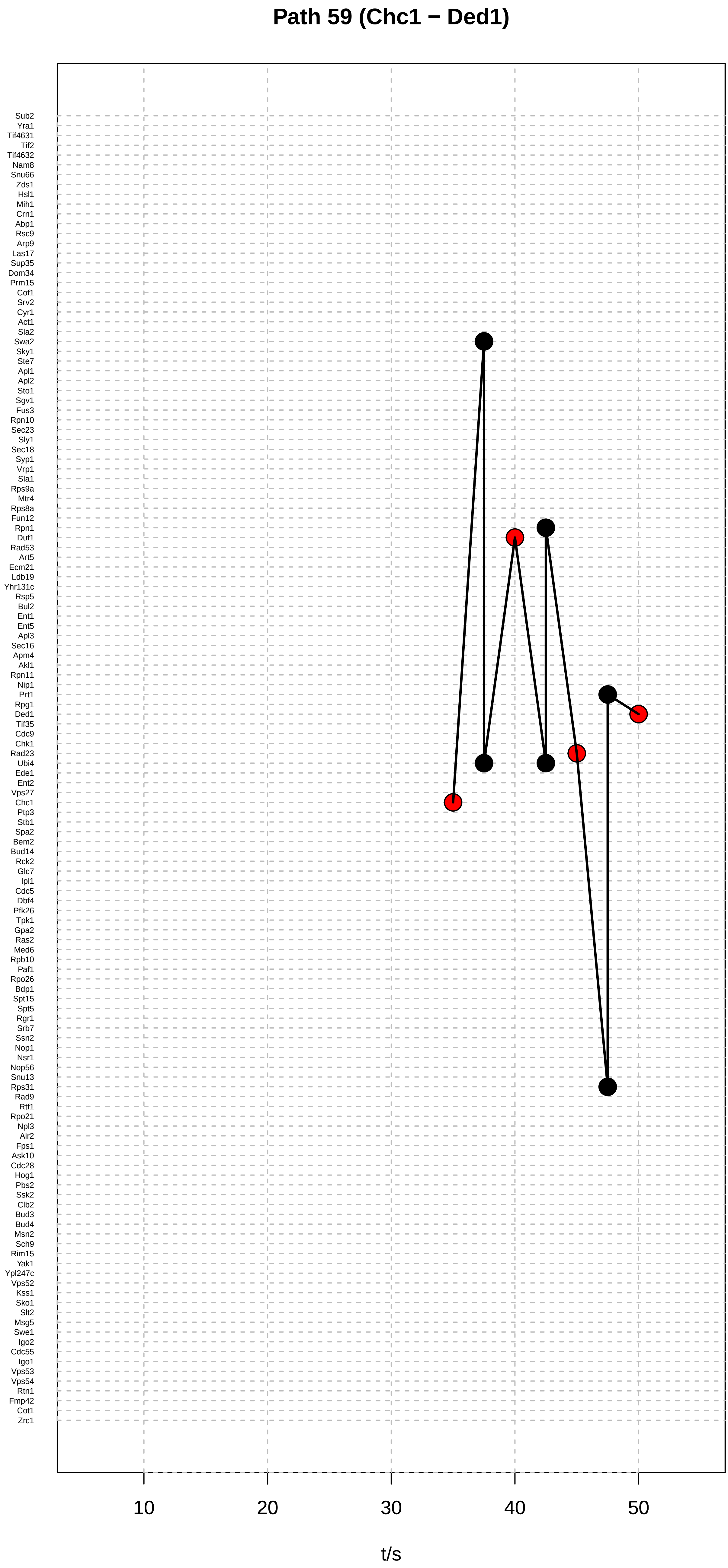

Path 59 (Chc1 − Ded1)
Sub2
Yra1
Tif4631
Tif2
Tif4632
Nam8
Snu66
Zds1
Hsl1
Mih1
Crn1
Abp1
Rsc9
Arp9
Las17
Sup35
Dom34
Prm15
Cof1
Srv2
Cyr1
Act1
Sla2
Swa2
Sky1
Ste7
Apl1
Apl2
Sto1
Sgv1
Fus3
Rpn10
Sec23
Sly1
Sec18
Syp1
Vrp1
Sla1
Rps9a
Mtr4
Rps8a
Fun12
Rpn1
Duf1
Rad53
Art5
Ecm21
Ldb19
Yhr131c
Rsp5
Bul2
Ent1
Ent5
Apl3
Sec16
Apm4
Akl1
Rpn11
Nip1
Prt1
Rpg1
Ded1
Tif35
Cdc9
Chk1
Rad23
Ubi4
Ede1
Ent2
Vps27
Chc1
Ptp3
Stb1
Spa2
Bem2
Bud14
Rck2
Glc7
Ipl1
Cdc5
Dbf4
Pfk26
Tpk1
Gpa2
Ras2
Med6
Rpb10
Paf1
Rpo26
Bdp1
Spt15
Spt5
Rgr1
Srb7
Ssn2
Nop1
Nsr1
Nop56
Snu13
Rps31
Rad9
Rtf1
Rpo21
Npl3
Air2
Fps1
Ask10
Cdc28
Hog1
Pbs2
Ssk2
Clb2
Bud3
Bud4
Msn2
Sch9
Rim15
Yak1
Ypl247c
Vps52
Kss1
Sko1
Slt2
Msg5
Swe1
Igo2
Cdc55
Igo1
Vps53
Vps54
Rtn1
Fmp42
Cot1
Zrc1
10
20
30
40
50
t/s

### Path 60 (Yhr131c − Prm15)

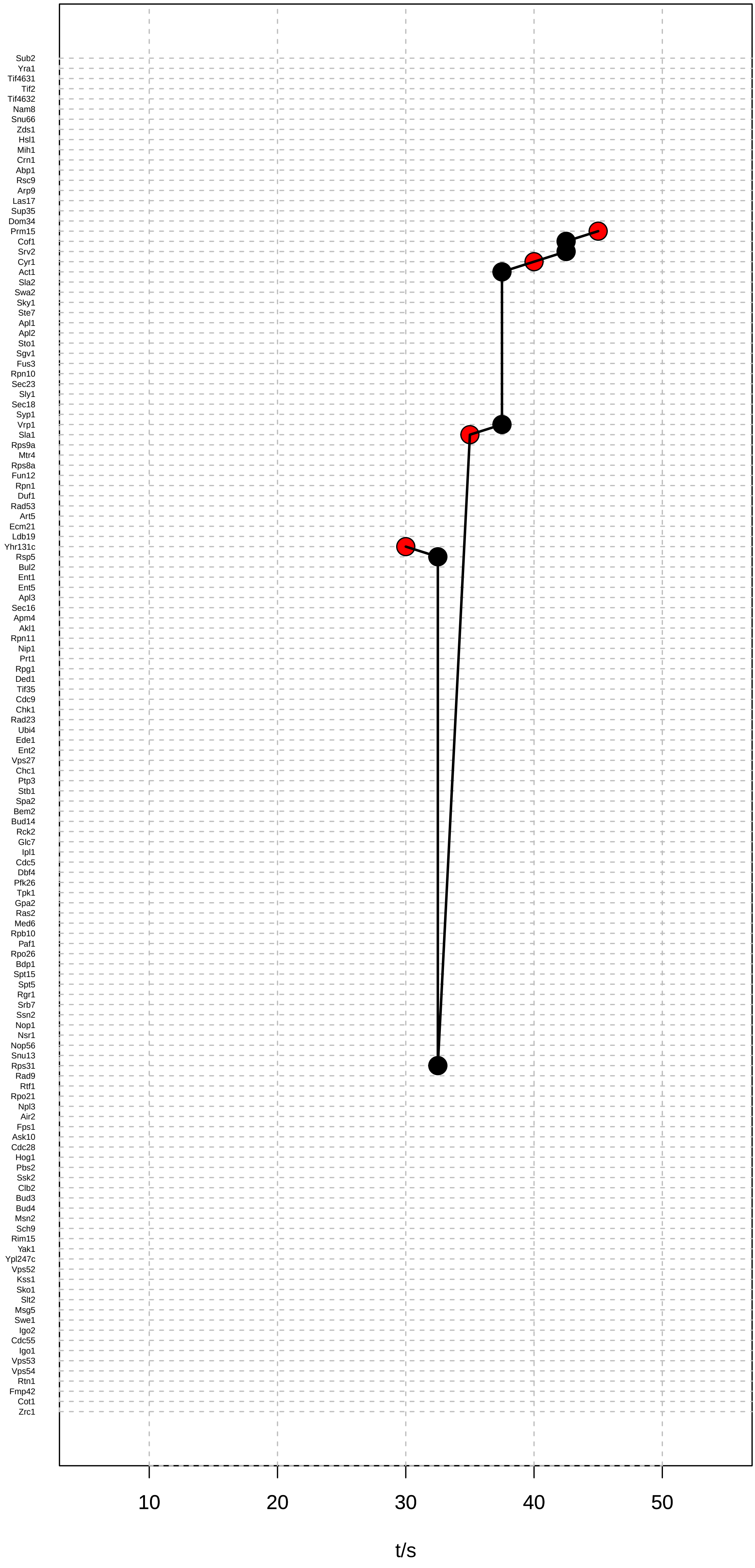

#### Path 61 (Yhr131c − Prm15)

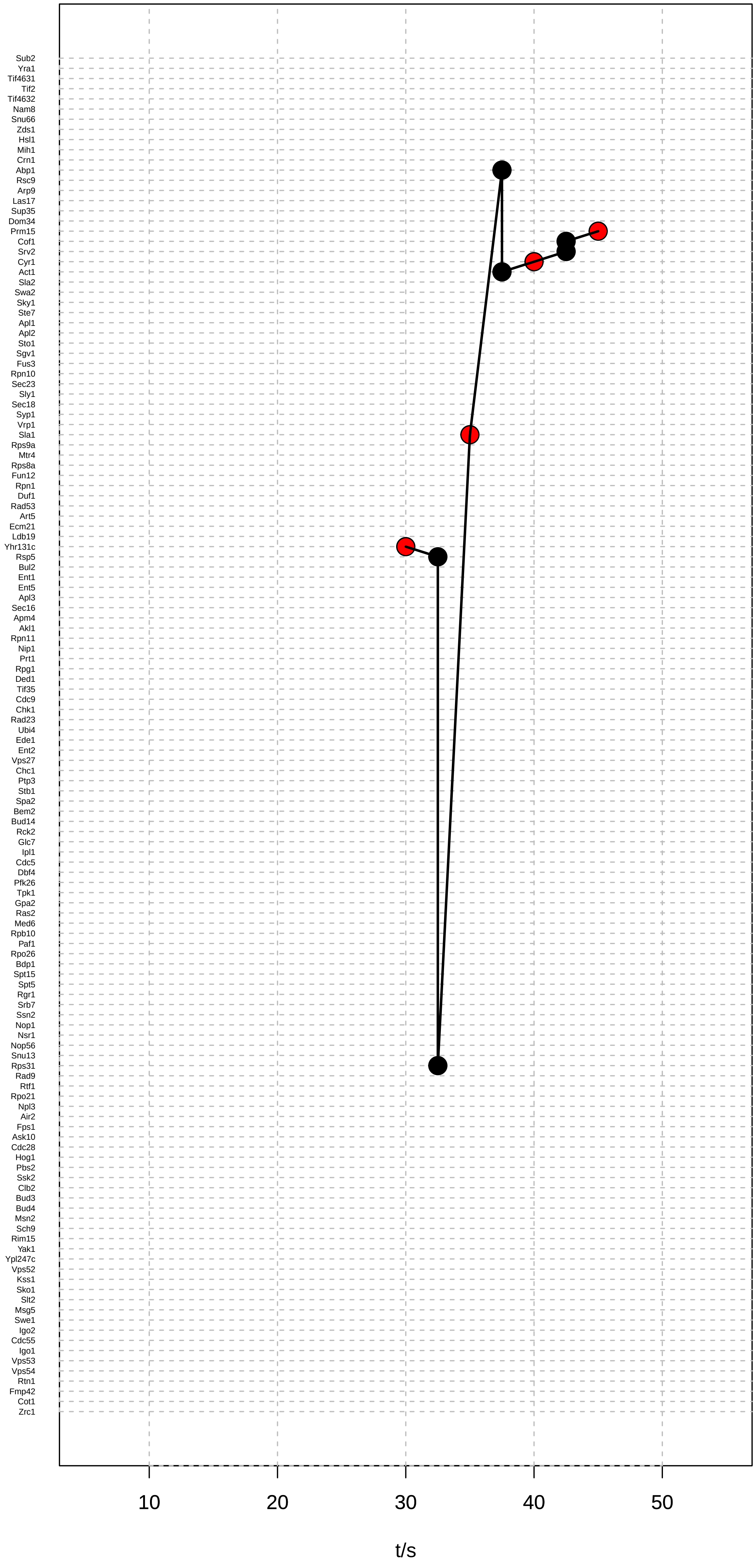

### Path 62 (Yhr131c − Prm15)

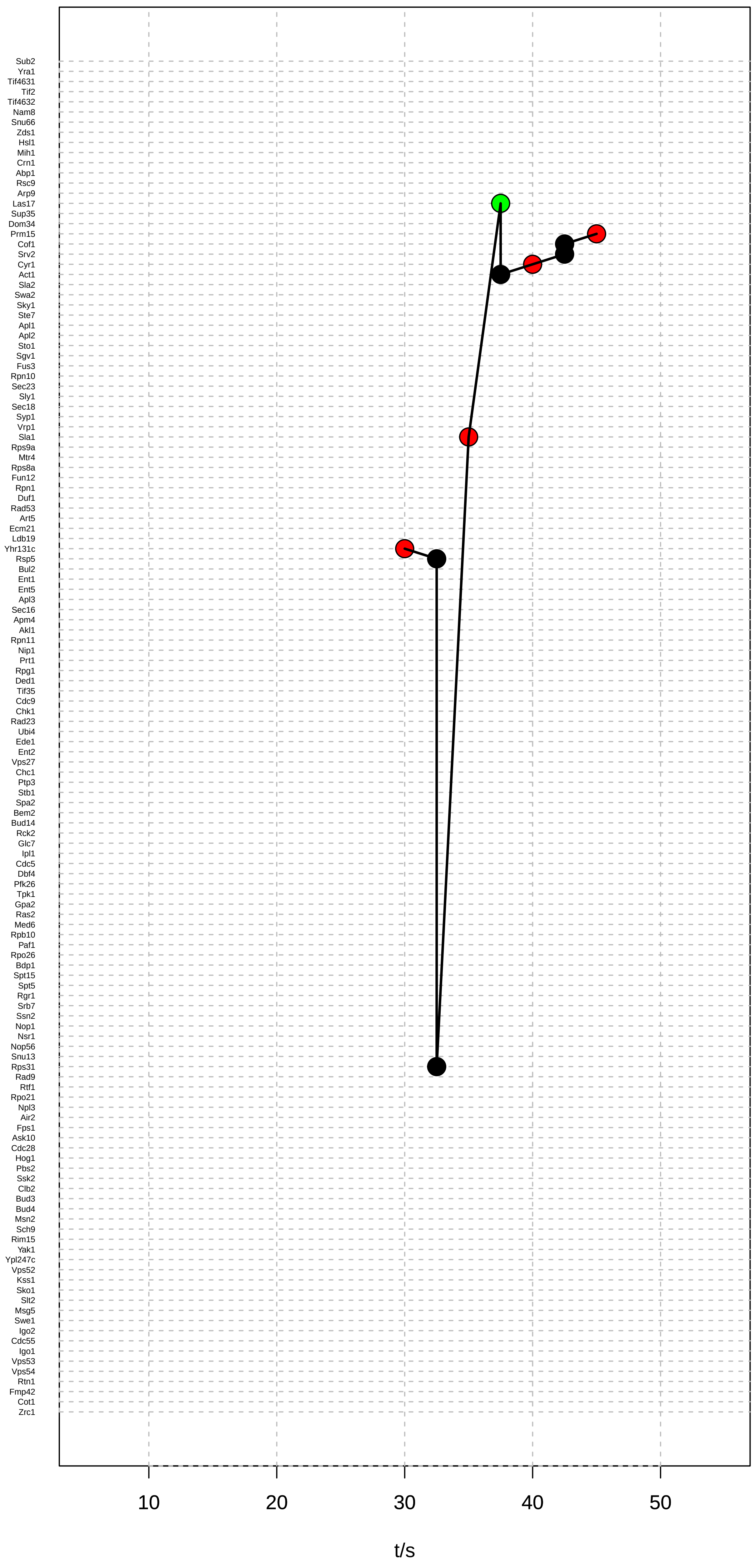

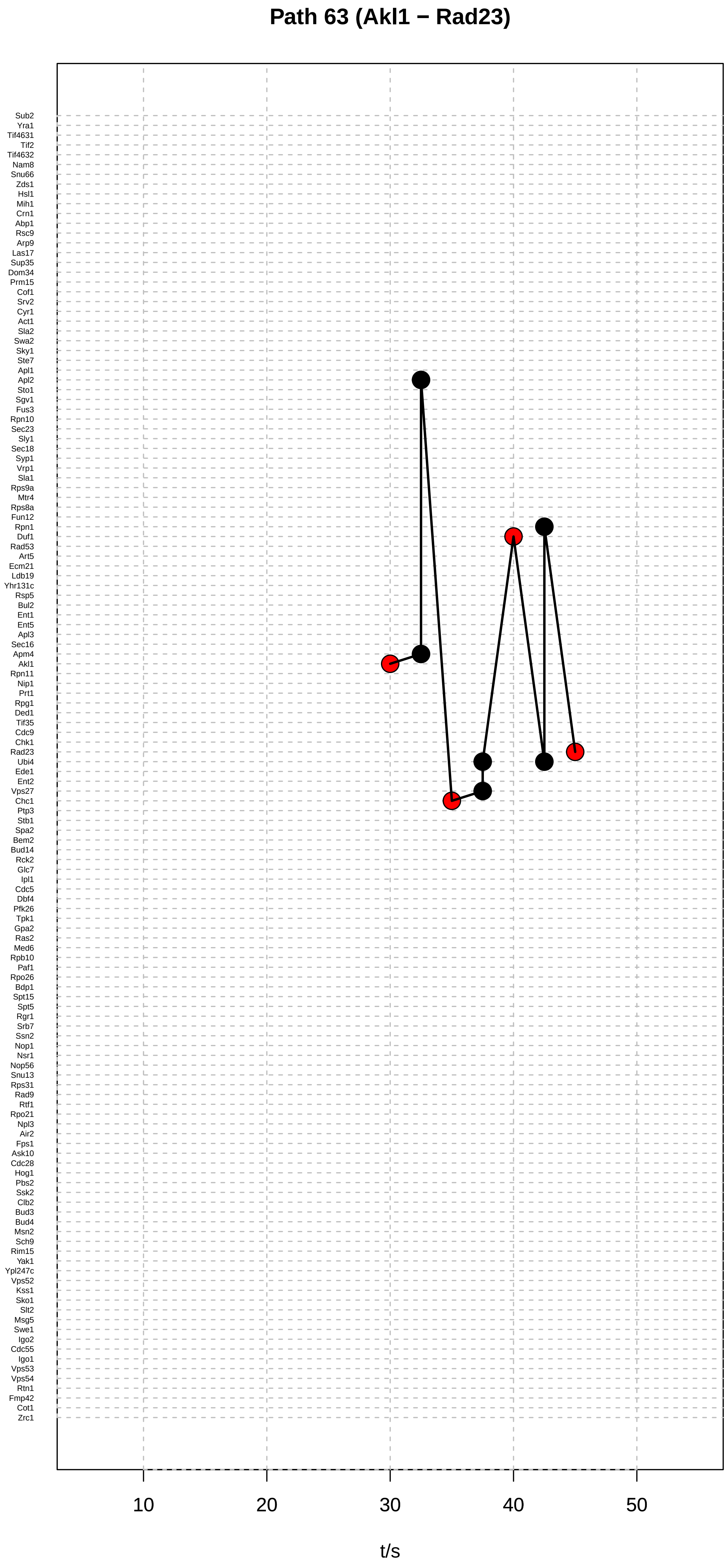
Path 63 (Akl1 − Rad23)
Sub2
Yra1
Tif4631
Tif2
Tif4632
Nam8
Snu66
Zds1
Hsl1
Mih1
Crn1
Abp1
Rsc9
Arp9
Las17
Sup35
Dom34
Prm15
Cof1
Srv2
Cyr1
Act1
Sla2
Swa2
Sky1
Ste7
Apl1
Apl2
Sto1
Sgv1
Fus3
Rpn10
Sec23
Sly1
Sec18
Syp1
Vrp1
Sla1
Rps9a
Mtr4
Rps8a
Fun12
Rpn1
Duf1
Rad53
Art5
Ecm21
Ldb19
Yhr131c
Rsp5
Bul2
Ent1
Ent5
Apl3
Sec16
Apm4
Akl1
Rpn11
Nip1
Prt1
Rpg1
Ded1
Tif35
Cdc9
Chk1
Rad23
Ubi4
Ede1
Ent2
Vps27
Chc1
Ptp3
Stb1
Spa2
Bem2
Bud14
Rck2
Glc7
Ipl1
Cdc5
Dbf4
Pfk26
Tpk1
Gpa2
Ras2
Med6
Rpb10
Paf1
Rpo26
Bdp1
Spt15
Spt5
Rgr1
Srb7
Ssn2
Nop1
Nsr1
Nop56
Snu13
Rps31
Rad9
Rtf1
Rpo21
Npl3
Air2
Fps1
Ask10
Cdc28
Hog1
Pbs2
Ssk2
Clb2
Bud3
Bud4
Msn2
Sch9
Rim15
Yak1
Ypl247c
Vps52
Kss1
Sko1
Slt2
Msg5
Swe1
Igo2
Cdc55
Igo1
Vps53
Vps54
Rtn1
Fmp42
Cot1
Zrc1
10
20
30
40
50
t/s

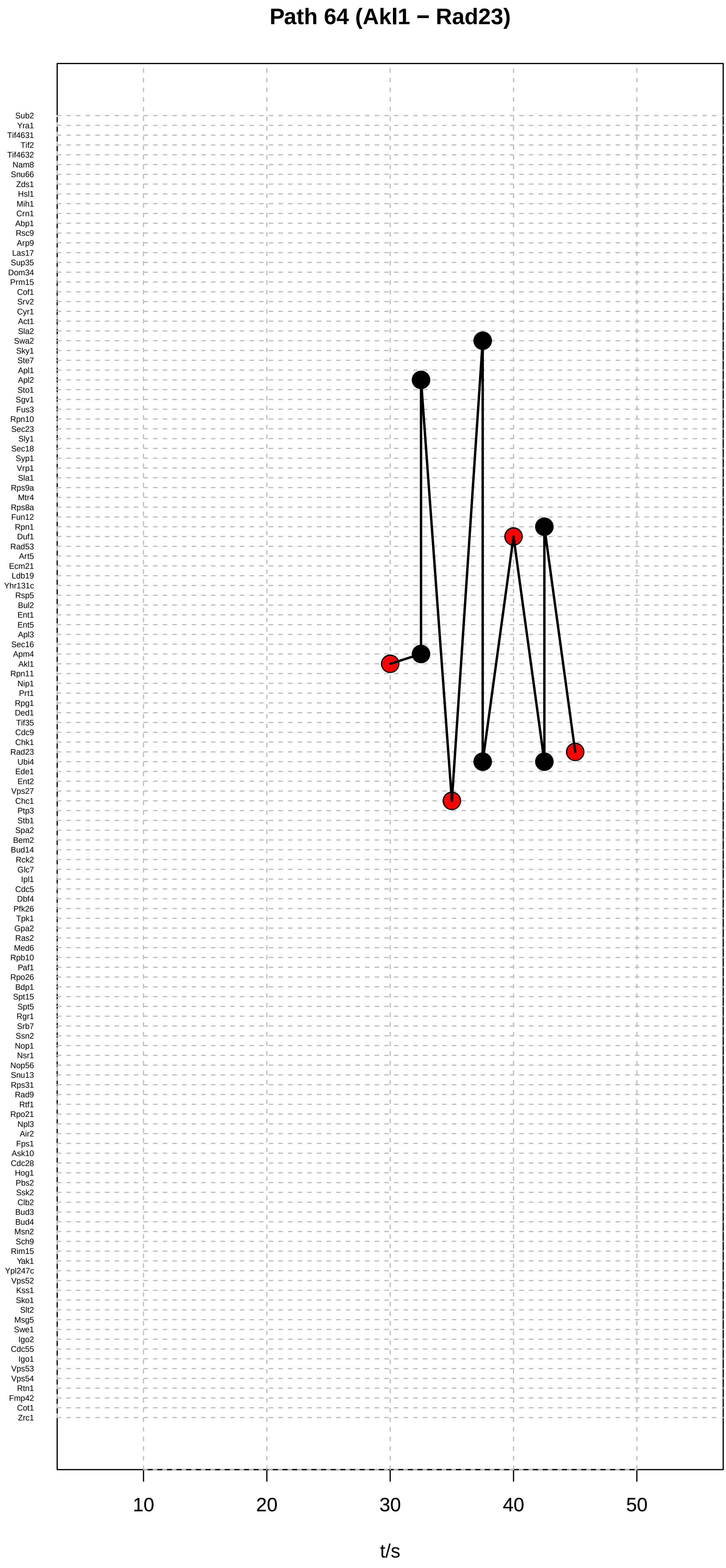
Path 64 (Akl1 − Rad23)
Sub2
Yra1
Tif4631
Tif2
Tif4632
Nam8
Snu66
Zds1
Hsl1
Mih1
Crn1
Abp1
Rsc9
Arp9
Las17
Sup35
Dom34
Prm15
Cof1
Srv2
Cyr1
Act1
Sla2
Swa2
Sky1
Ste7
Apl1
Apl2
Sto1
Sgv1
Fus3
Rpn10
Sec23
Sly1
Sec18
Syp1
Vrp1
Sla1
Rps9a
Mtr4
Rps8a
Fun12
Rpn1
Duf1
Rad53
Art5
Ecm21
Ldb19
Yhr131c
Rsp5
Bul2
Ent1
Ent5
Apl3
Sec16
Apm4
Akl1
Rpn11
Nip1
Prt1
Rpg1
Ded1
Tif35
Cdc9
Chk1
Rad23
Ubi4
Ede1
Ent2
Vps27
Chc1
Ptp3
Stb1
Spa2
Bem2
Bud14
Rck2
Glc7
Ipl1
Cdc5
Dbf4
Pfk26
Tpk1
Gpa2
Ras2
Med6
Rpb10
Paf1
Rpo26
Bdp1
Spt15
Spt5
Rgr1
Srb7
Ssn2
Nop1
Nsr1
Nop56
Snu13
Rps31
Rad9
Rtf1
Rpo21
Npl3
Air2
Fps1
Ask10
Cdc28
Hog1
Pbs2
Ssk2
Clb2
Bud3
Bud4
Msn2
Sch9
Rim15
Yak1
Ypl247c
Vps52
Kss1
Sko1
Slt2
Msg5
Swe1
Igo2
Cdc55
Igo1
Vps53
Vps54
Rtn1
Fmp42
Cot1
Zrc1
10
20
30
40
50
t/s

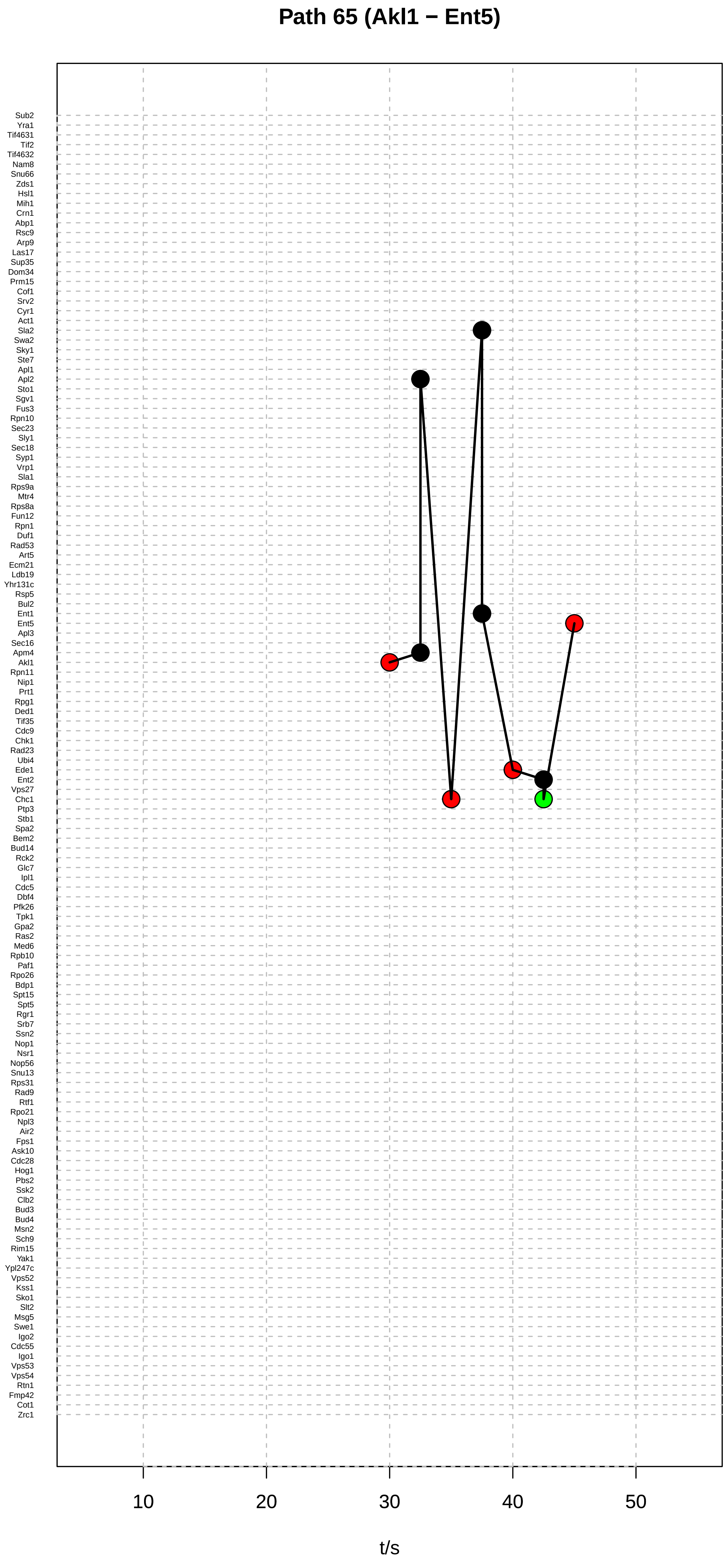

Path 65 (Akl1 − Ent5)
Sub2
Yra1
Tif4631
Tif2
Tif4632
Nam8
Snu66
Zds1
Hsl1
Mih1
Crn1
Abp1
Rsc9
Arp9
Las17
Sup35
Dom34
Prm15
Cof1
Srv2
Cyr1
Act1
Sla2
Swa2
Sky1
Ste7
Apl1
Apl2
Sto1
Sgv1
Fus3
Rpn10
Sec23
Sly1
Sec18
Syp1
Vrp1
Sla1
Rps9a
Mtr4
Rps8a
Fun12
Rpn1
Duf1
Rad53
Art5
Ecm21
Ldb19
Yhr131c
Rsp5
Bul2
Ent1
Ent5
Apl3
Sec16
Apm4
Akl1
Rpn11
Nip1
Prt1
Rpg1
Ded1
Tif35
Cdc9
Chk1
Rad23
Ubi4
Ede1
Ent2
Vps27
Chc1
Ptp3
Stb1
Spa2
Bem2
Bud14
Rck2
Glc7
Ipl1
Cdc5
Dbf4
Pfk26
Tpk1
Gpa2
Ras2
Med6
Rpb10
Paf1
Rpo26
Bdp1
Spt15
Spt5
Rgr1
Srb7
Ssn2
Nop1
Nsr1
Nop56
Snu13
Rps31
Rad9
Rtf1
Rpo21
Npl3
Air2
Fps1
Ask10
Cdc28
Hog1
Pbs2
Ssk2
Clb2
Bud3
Bud4
Msn2
Sch9
Rim15
Yak1
Ypl247c
Vps52
Kss1
Sko1
Slt2
Msg5
Swe1
Igo2
Cdc55
Igo1
Vps53
Vps54
Rtn1
Fmp42
Cot1
Zrc1
10
20
30
40
50
t/s

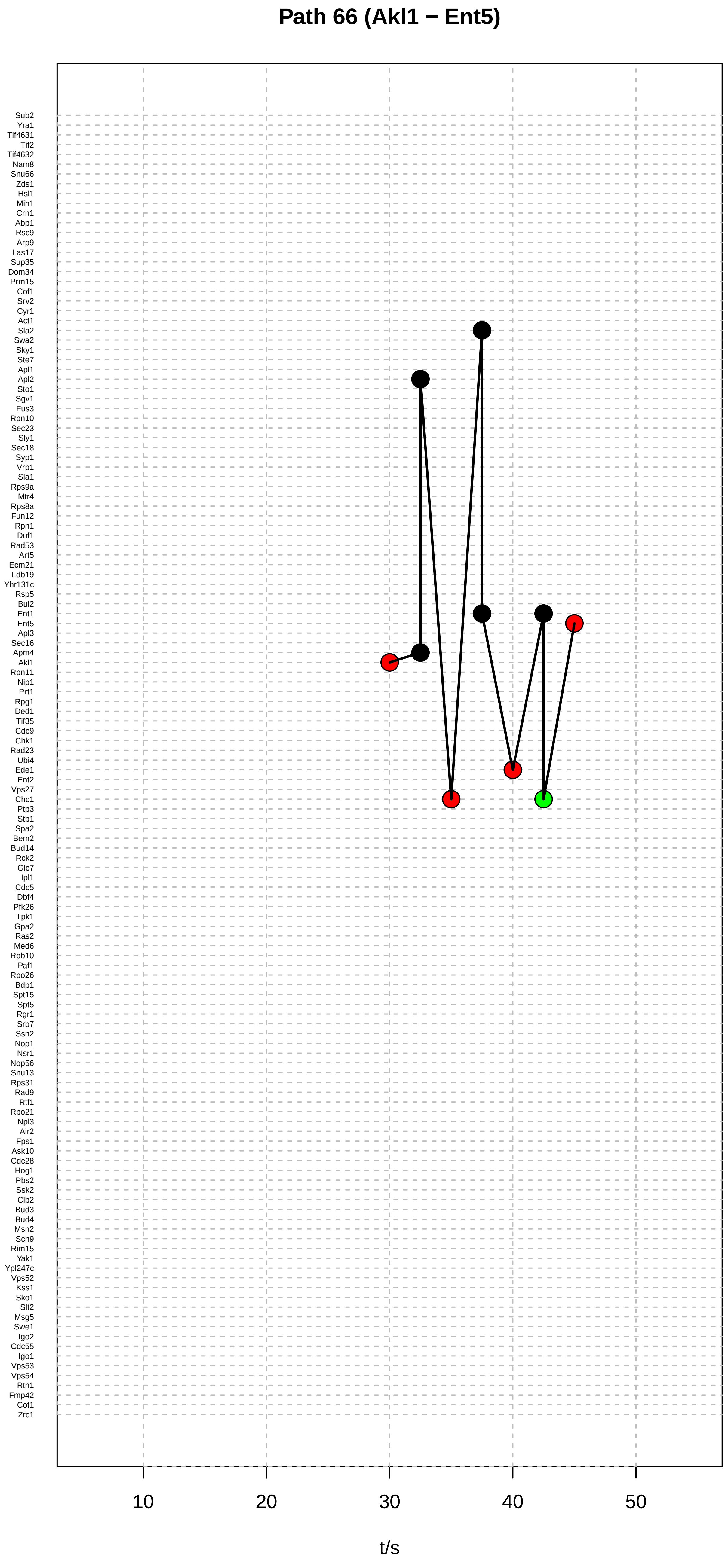

Path 66 (Akl1 − Ent5)
Sub2
Yra1
Tif4631
Tif2
Tif4632
Nam8
Snu66
Zds1
Hsl1
Mih1
Crn1
Abp1
Rsc9
Arp9
Las17
Sup35
Dom34
Prm15
Cof1
Srv2
Cyr1
Act1
Sla2
Swa2
Sky1
Ste7
Apl1
Apl2
Sto1
Sgv1
Fus3
Rpn10
Sec23
Sly1
Sec18
Syp1
Vrp1
Sla1
Rps9a
Mtr4
Rps8a
Fun12
Rpn1
Duf1
Rad53
Art5
Ecm21
Ldb19
Yhr131c
Rsp5
Bul2
Ent1
Ent5
Apl3
Sec16
Apm4
Akl1
Rpn11
Nip1
Prt1
Rpg1
Ded1
Tif35
Cdc9
Chk1
Rad23
Ubi4
Ede1
Ent2
Vps27
Chc1
Ptp3
Stb1
Spa2
Bem2
Bud14
Rck2
Glc7
Ipl1
Cdc5
Dbf4
Pfk26
Tpk1
Gpa2
Ras2
Med6
Rpb10
Paf1
Rpo26
Bdp1
Spt15
Spt5
Rgr1
Srb7
Ssn2
Nop1
Nsr1
Nop56
Snu13
Rps31
Rad9
Rtf1
Rpo21
Npl3
Air2
Fps1
Ask10
Cdc28
Hog1
Pbs2
Ssk2
Clb2
Bud3
Bud4
Msn2
Sch9
Rim15
Yak1
Ypl247c
Vps52
Kss1
Sko1
Slt2
Msg5
Swe1
Igo2
Cdc55
Igo1
Vps53
Vps54
Rtn1
Fmp42
Cot1
Zrc1
10
20
30
40
50
t/s

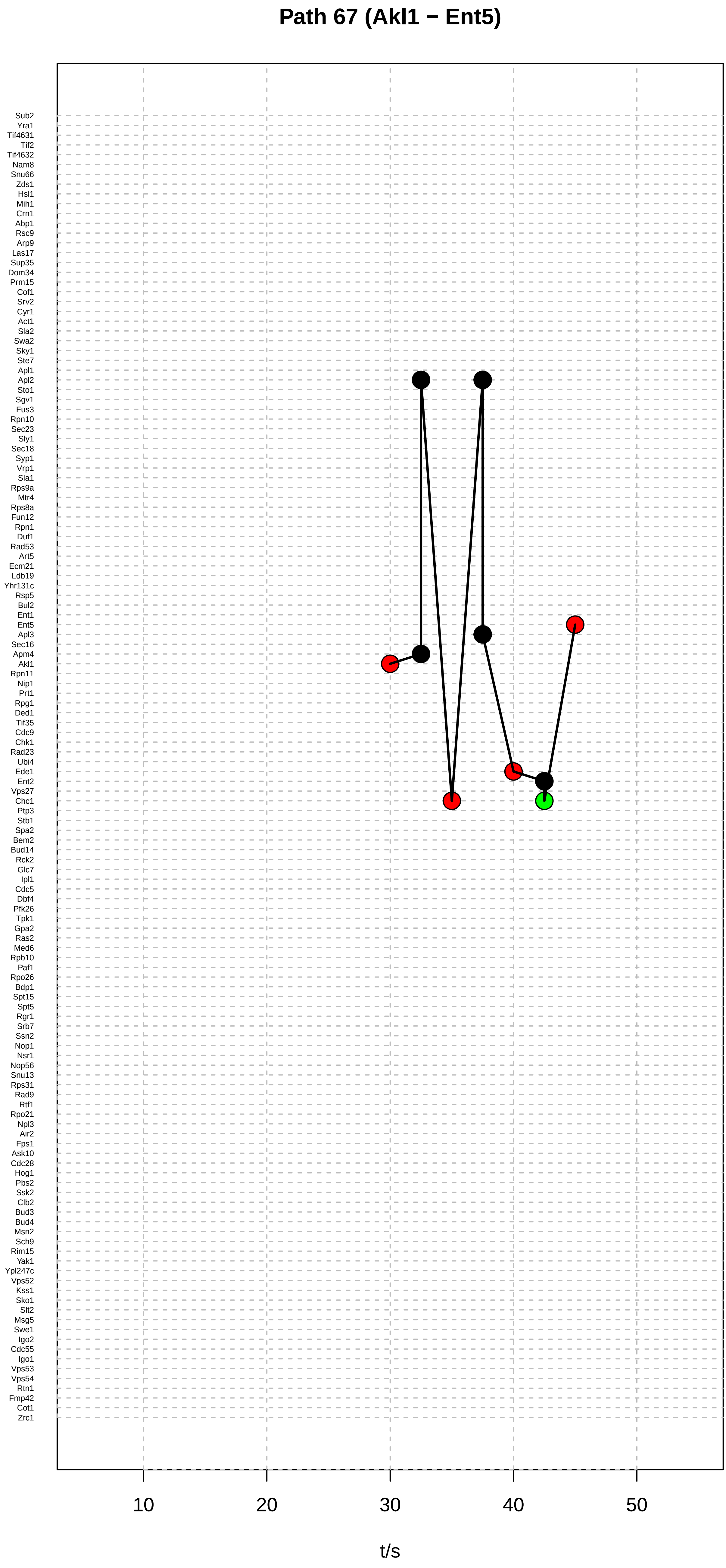

Path 67 (Akl1 − Ent5)
Sub2
Yra1
Tif4631
Tif2
Tif4632
Nam8
Snu66
Zds1
Hsl1
Mih1
Crn1
Abp1
Rsc9
Arp9
Las17
Sup35
Dom34
Prm15
Cof1
Srv2
Cyr1
Act1
Sla2
Swa2
Sky1
Ste7
Apl1
Apl2
Sto1
Sgv1
Fus3
Rpn10
Sec23
Sly1
Sec18
Syp1
Vrp1
Sla1
Rps9a
Mtr4
Rps8a
Fun12
Rpn1
Duf1
Rad53
Art5
Ecm21
Ldb19
Yhr131c
Rsp5
Bul2
Ent1
Ent5
Apl3
Sec16
Apm4
Akl1
Rpn11
Nip1
Prt1
Rpg1
Ded1
Tif35
Cdc9
Chk1
Rad23
Ubi4
Ede1
Ent2
Vps27
Chc1
Ptp3
Stb1
Spa2
Bem2
Bud14
Rck2
Glc7
Ipl1
Cdc5
Dbf4
Pfk26
Tpk1
Gpa2
Ras2
Med6
Rpb10
Paf1
Rpo26
Bdp1
Spt15
Spt5
Rgr1
Srb7
Ssn2
Nop1
Nsr1
Nop56
Snu13
Rps31
Rad9
Rtf1
Rpo21
Npl3
Air2
Fps1
Ask10
Cdc28
Hog1
Pbs2
Ssk2
Clb2
Bud3
Bud4
Msn2
Sch9
Rim15
Yak1
Ypl247c
Vps52
Kss1
Sko1
Slt2
Msg5
Swe1
Igo2
Cdc55
Igo1
Vps53
Vps54
Rtn1
Fmp42
Cot1
Zrc1
10
20
30
40
50
t/s

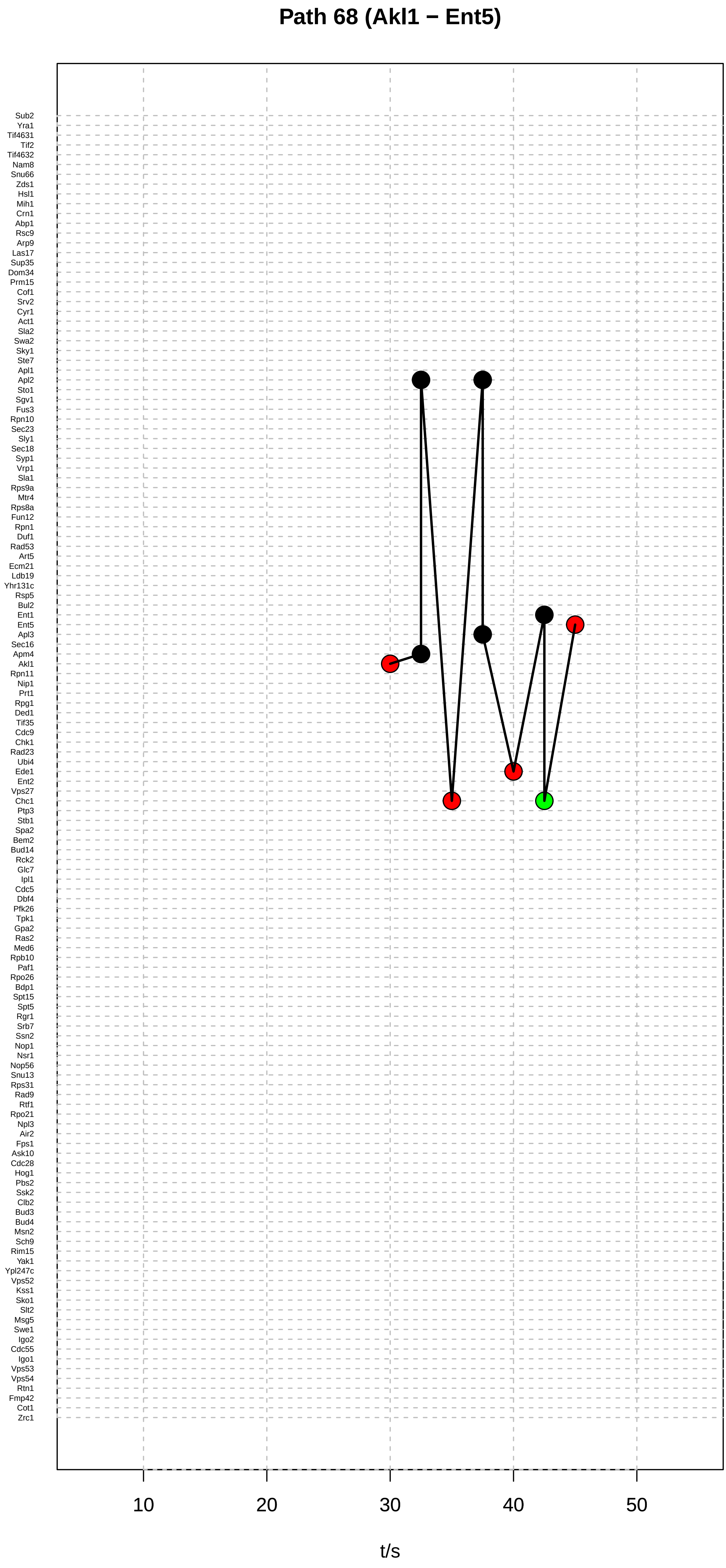

Path 68 (Akl1 − Ent5)
Sub2
Yra1
Tif4631
Tif2
Tif4632
Nam8
Snu66
Zds1
Hsl1
Mih1
Crn1
Abp1
Rsc9
Arp9
Las17
Sup35
Dom34
Prm15
Cof1
Srv2
Cyr1
Act1
Sla2
Swa2
Sky1
Ste7
Apl1
Apl2
Sto1
Sgv1
Fus3
Rpn10
Sec23
Sly1
Sec18
Syp1
Vrp1
Sla1
Rps9a
Mtr4
Rps8a
Fun12
Rpn1
Duf1
Rad53
Art5
Ecm21
Ldb19
Yhr131c
Rsp5
Bul2
Ent1
Ent5
Apl3
Sec16
Apm4
Akl1
Rpn11
Nip1
Prt1
Rpg1
Ded1
Tif35
Cdc9
Chk1
Rad23
Ubi4
Ede1
Ent2
Vps27
Chc1
Ptp3
Stb1
Spa2
Bem2
Bud14
Rck2
Glc7
Ipl1
Cdc5
Dbf4
Pfk26
Tpk1
Gpa2
Ras2
Med6
Rpb10
Paf1
Rpo26
Bdp1
Spt15
Spt5
Rgr1
Srb7
Ssn2
Nop1
Nsr1
Nop56
Snu13
Rps31
Rad9
Rtf1
Rpo21
Npl3
Air2
Fps1
Ask10
Cdc28
Hog1
Pbs2
Ssk2
Clb2
Bud3
Bud4
Msn2
Sch9
Rim15
Yak1
Ypl247c
Vps52
Kss1
Sko1
Slt2
Msg5
Swe1
Igo2
Cdc55
Igo1
Vps53
Vps54
Rtn1
Fmp42
Cot1
Zrc1
10
20
30
40
50
t/s

### Path 69 (Akl1 − Ent5)

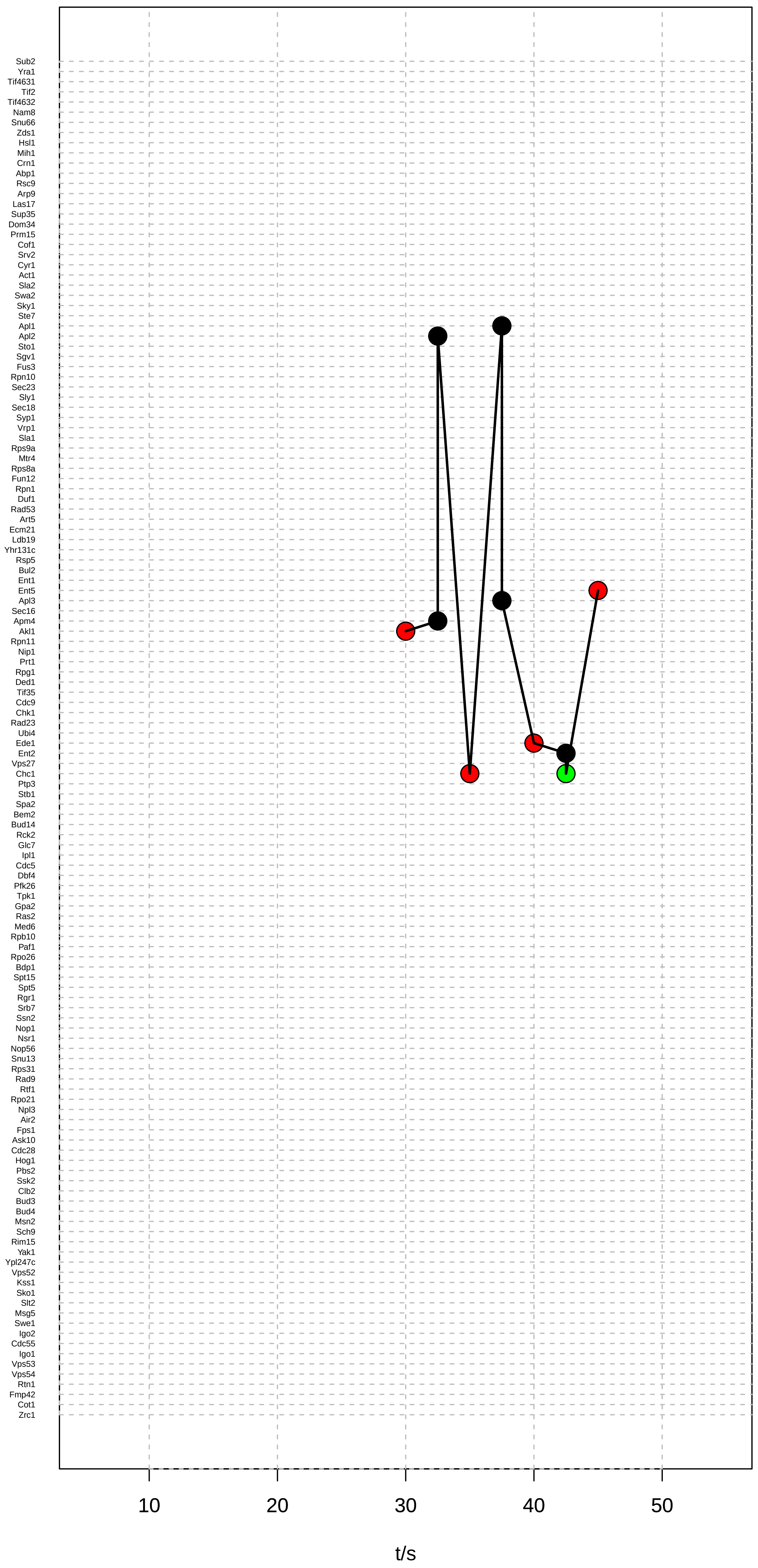

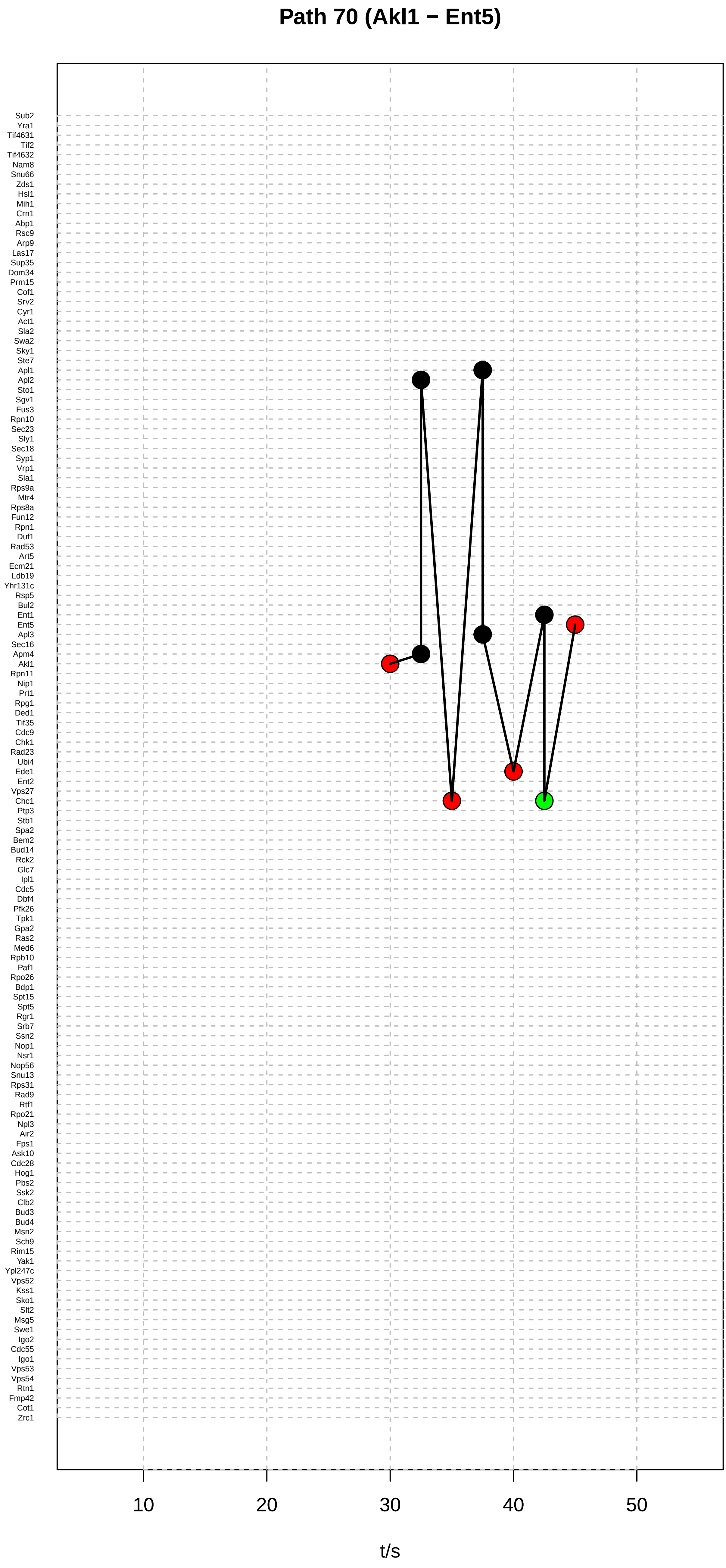

Path 70 (Akl1 − Ent5)
Sub2
Yra1
Tif4631
Tif2
Tif4632
Nam8
Snu66
Zds1
Hsl1
Mih1
Crn1
Abp1
Rsc9
Arp9
Las17
Sup35
Dom34
Prm15
Cof1
Srv2
Cyr1
Act1
Sla2
Swa2
Sky1
Ste7
Apl1
Apl2
Sto1
Sgv1
Fus3
Rpn10
Sec23
Sly1
Sec18
Syp1
Vrp1
Sla1
Rps9a
Mtr4
Rps8a
Fun12
Rpn1
Duf1
Rad53
Art5
Ecm21
Ldb19
Yhr131c
Rsp5
Bul2
Ent1
Ent5
Apl3
Sec16
Apm4
Akl1
Rpn11
Nip1
Prt1
Rpg1
Ded1
Tif35
Cdc9
Chk1
Rad23
Ubi4
Ede1
Ent2
Vps27
Chc1
Ptp3
Stb1
Spa2
Bem2
Bud14
Rck2
Glc7
Ipl1
Cdc5
Dbf4
Pfk26
Tpk1
Gpa2
Ras2
Med6
Rpb10
Paf1
Rpo26
Bdp1
Spt15
Spt5
Rgr1
Srb7
Ssn2
Nop1
Nsr1
Nop56
Snu13
Rps31
Rad9
Rtf1
Rpo21
Npl3
Air2
Fps1
Ask10
Cdc28
Hog1
Pbs2
Ssk2
Clb2
Bud3
Bud4
Msn2
Sch9
Rim15
Yak1
Ypl247c
Vps52
Kss1
Sko1
Slt2
Msg5
Swe1
Igo2
Cdc55
Igo1
Vps53
Vps54
Rtn1
Fmp42
Cot1
Zrc1
10
20
30
40
50
t/s

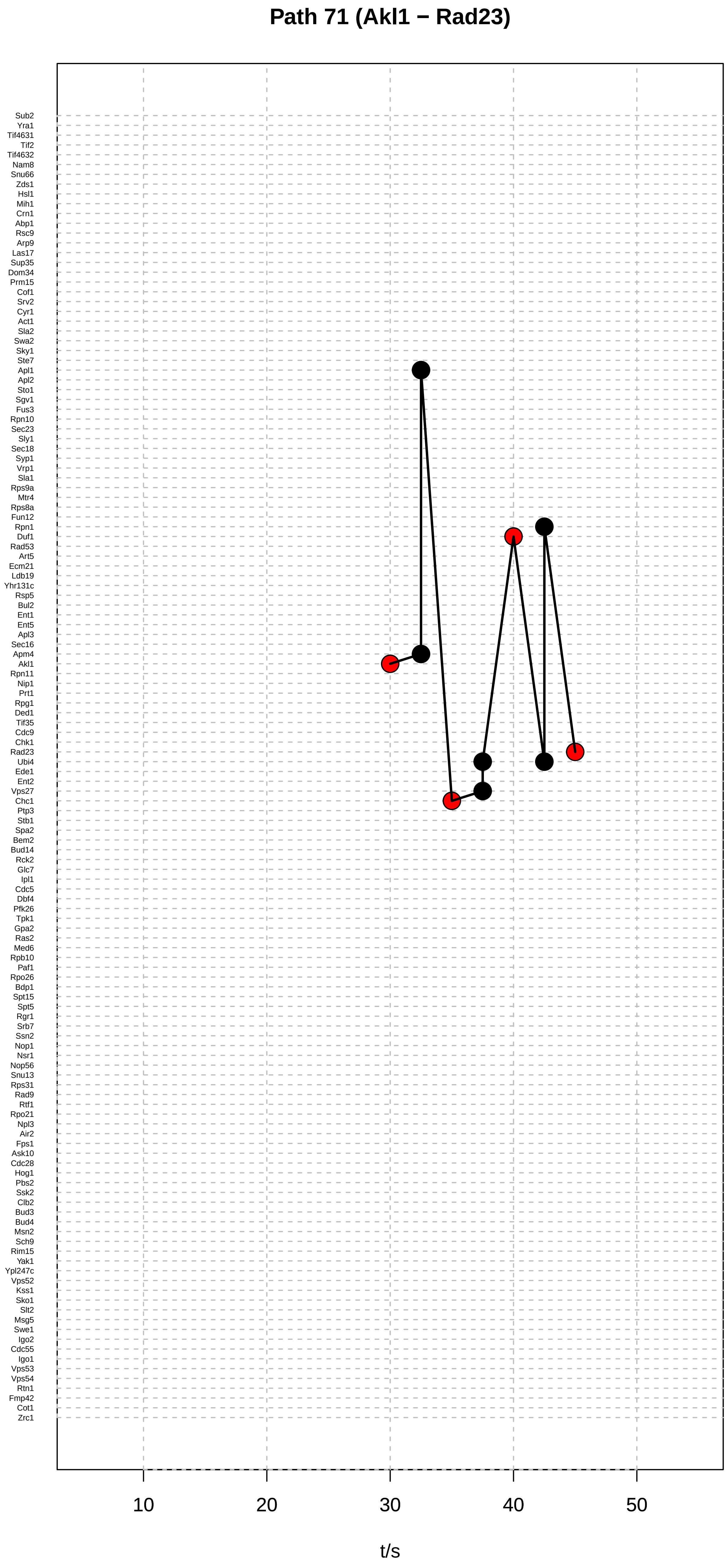

Path 71 (Akl1 − Rad23)
Sub2
Yra1
Tif4631
Tif2
Tif4632
Nam8
Snu66
Zds1
Hsl1
Mih1
Crn1
Abp1
Rsc9
Arp9
Las17
Sup35
Dom34
Prm15
Cof1
Srv2
Cyr1
Act1
Sla2
Swa2
Sky1
Ste7
Apl1
Apl2
Sto1
Sgv1
Fus3
Rpn10
Sec23
Sly1
Sec18
Syp1
Vrp1
Sla1
Rps9a
Mtr4
Rps8a
Fun12
Rpn1
Duf1
Rad53
Art5
Ecm21
Ldb19
Yhr131c
Rsp5
Bul2
Ent1
Ent5
Apl3
Sec16
Apm4
Akl1
Rpn11
Nip1
Prt1
Rpg1
Ded1
Tif35
Cdc9
Chk1
Rad23
Ubi4
Ede1
Ent2
Vps27
Chc1
Ptp3
Stb1
Spa2
Bem2
Bud14
Rck2
Glc7
Ipl1
Cdc5
Dbf4
Pfk26
Tpk1
Gpa2
Ras2
Med6
Rpb10
Paf1
Rpo26
Bdp1
Spt15
Spt5
Rgr1
Srb7
Ssn2
Nop1
Nsr1
Nop56
Snu13
Rps31
Rad9
Rtf1
Rpo21
Npl3
Air2
Fps1
Ask10
Cdc28
Hog1
Pbs2
Ssk2
Clb2
Bud3
Bud4
Msn2
Sch9
Rim15
Yak1
Ypl247c
Vps52
Kss1
Sko1
Slt2
Msg5
Swe1
Igo2
Cdc55
Igo1
Vps53
Vps54
Rtn1
Fmp42
Cot1
Zrc1
10
20
30
40
50
t/s

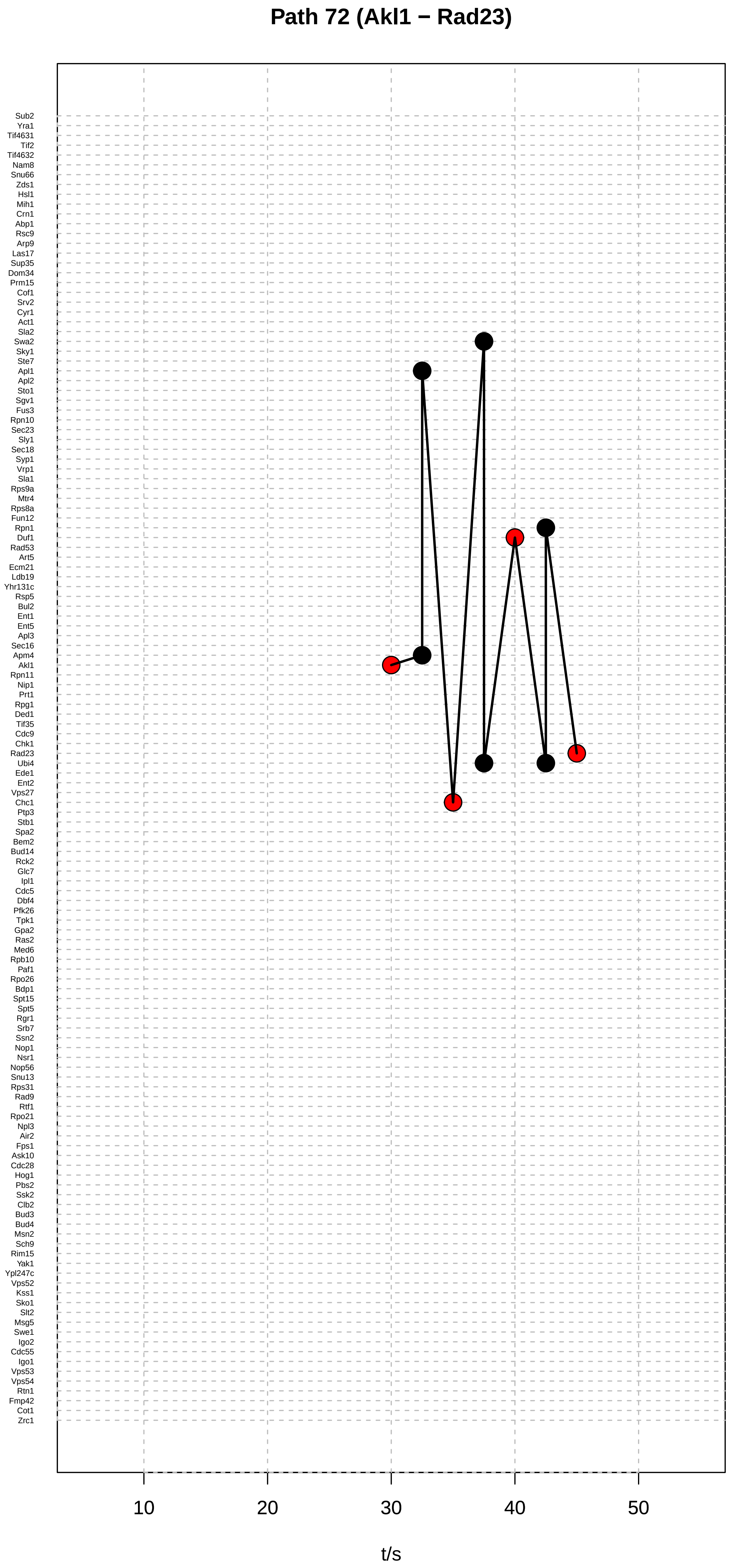

Path 72 (Akl1 − Rad23)
Sub2
Yra1
Tif4631
Tif2
Tif4632
Nam8
Snu66
Zds1
Hsl1
Mih1
Crn1
Abp1
Rsc9
Arp9
Las17
Sup35
Dom34
Prm15
Cof1
Srv2
Cyr1
Act1
Sla2
Swa2
Sky1
Ste7
Apl1
Apl2
Sto1
Sgv1
Fus3
Rpn10
Sec23
Sly1
Sec18
Syp1
Vrp1
Sla1
Rps9a
Mtr4
Rps8a
Fun12
Rpn1
Duf1
Rad53
Art5
Ecm21
Ldb19
Yhr131c
Rsp5
Bul2
Ent1
Ent5
Apl3
Sec16
Apm4
Akl1
Rpn11
Nip1
Prt1
Rpg1
Ded1
Tif35
Cdc9
Chk1
Rad23
Ubi4
Ede1
Ent2
Vps27
Chc1
Ptp3
Stb1
Spa2
Bem2
Bud14
Rck2
Glc7
Ipl1
Cdc5
Dbf4
Pfk26
Tpk1
Gpa2
Ras2
Med6
Rpb10
Paf1
Rpo26
Bdp1
Spt15
Spt5
Rgr1
Srb7
Ssn2
Nop1
Nsr1
Nop56
Snu13
Rps31
Rad9
Rtf1
Rpo21
Npl3
Air2
Fps1
Ask10
Cdc28
Hog1
Pbs2
Ssk2
Clb2
Bud3
Bud4
Msn2
Sch9
Rim15
Yak1
Ypl247c
Vps52
Kss1
Sko1
Slt2
Msg5
Swe1
Igo2
Cdc55
Igo1
Vps53
Vps54
Rtn1
Fmp42
Cot1
Zrc1
10
20
30
40
50
t/s

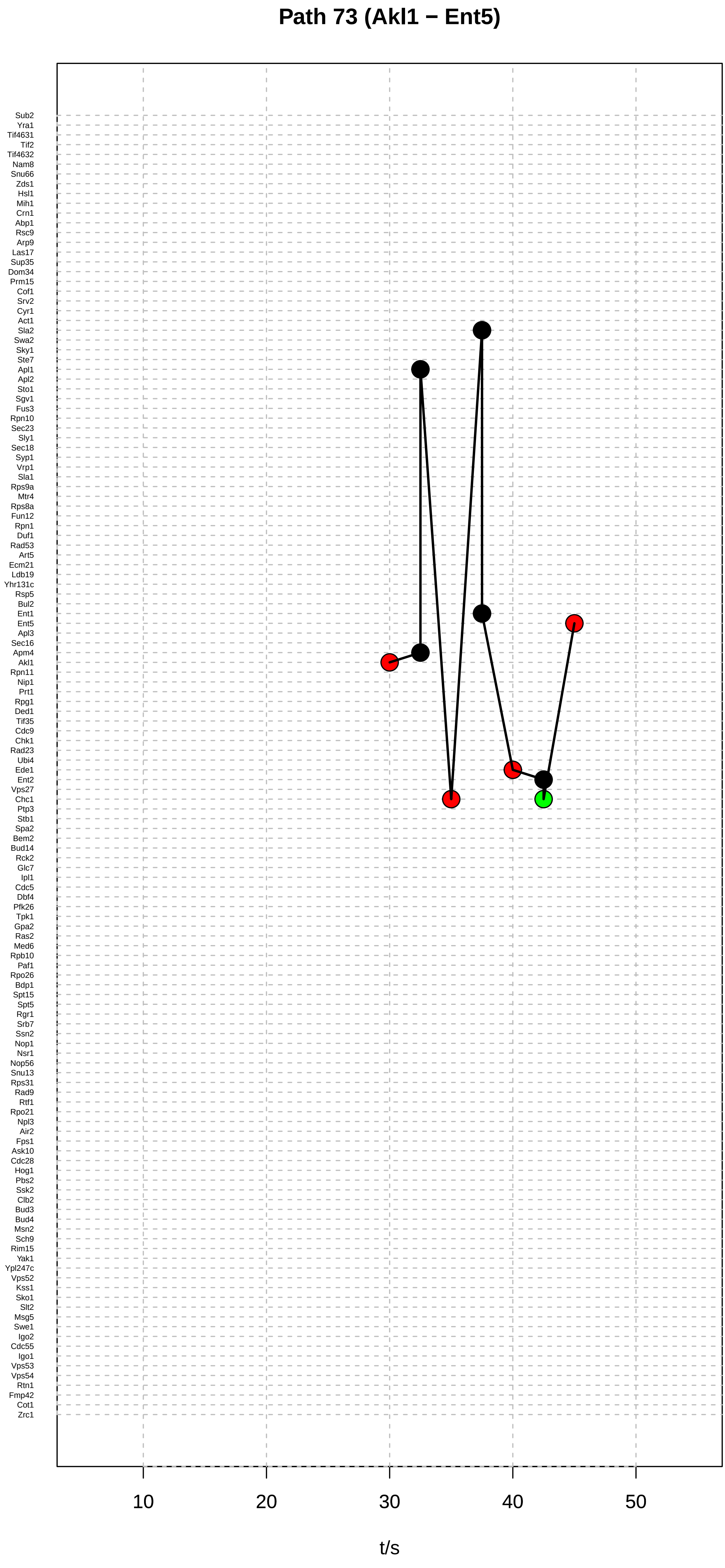

Path 73 (Akl1 − Ent5)
Sub2
Yra1
Tif4631
Tif2
Tif4632
Nam8
Snu66
Zds1
Hsl1
Mih1
Crn1
Abp1
Rsc9
Arp9
Las17
Sup35
Dom34
Prm15
Cof1
Srv2
Cyr1
Act1
Sla2
Swa2
Sky1
Ste7
Apl1
Apl2
Sto1
Sgv1
Fus3
Rpn10
Sec23
Sly1
Sec18
Syp1
Vrp1
Sla1
Rps9a
Mtr4
Rps8a
Fun12
Rpn1
Duf1
Rad53
Art5
Ecm21
Ldb19
Yhr131c
Rsp5
Bul2
Ent1
Ent5
Apl3
Sec16
Apm4
Akl1
Rpn11
Nip1
Prt1
Rpg1
Ded1
Tif35
Cdc9
Chk1
Rad23
Ubi4
Ede1
Ent2
Vps27
Chc1
Ptp3
Stb1
Spa2
Bem2
Bud14
Rck2
Glc7
Ipl1
Cdc5
Dbf4
Pfk26
Tpk1
Gpa2
Ras2
Med6
Rpb10
Paf1
Rpo26
Bdp1
Spt15
Spt5
Rgr1
Srb7
Ssn2
Nop1
Nsr1
Nop56
Snu13
Rps31
Rad9
Rtf1
Rpo21
Npl3
Air2
Fps1
Ask10
Cdc28
Hog1
Pbs2
Ssk2
Clb2
Bud3
Bud4
Msn2
Sch9
Rim15
Yak1
Ypl247c
Vps52
Kss1
Sko1
Slt2
Msg5
Swe1
Igo2
Cdc55
Igo1
Vps53
Vps54
Rtn1
Fmp42
Cot1
Zrc1
10
20
30
40
50
t/s

#### Path 74 (Akl1 − Ent5)

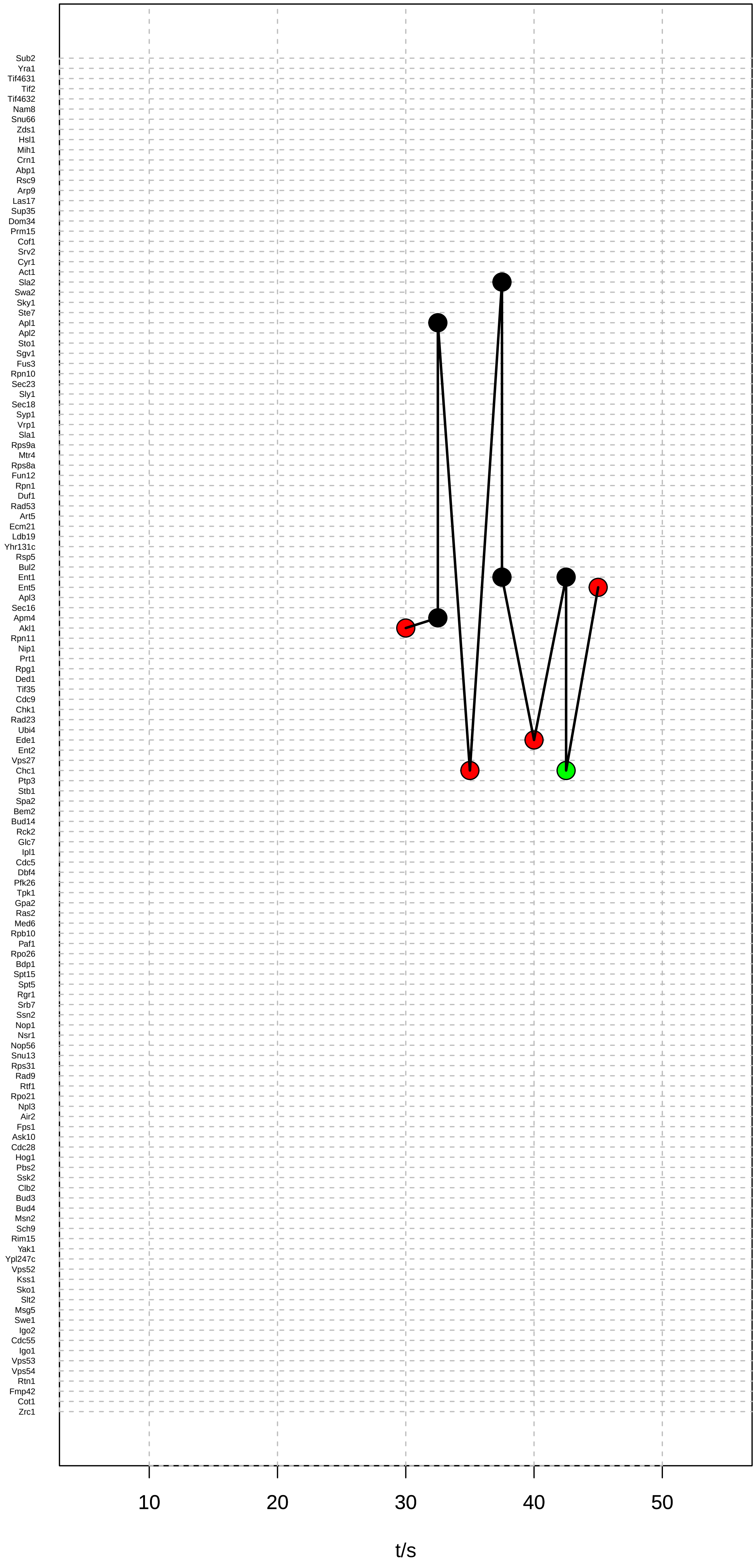

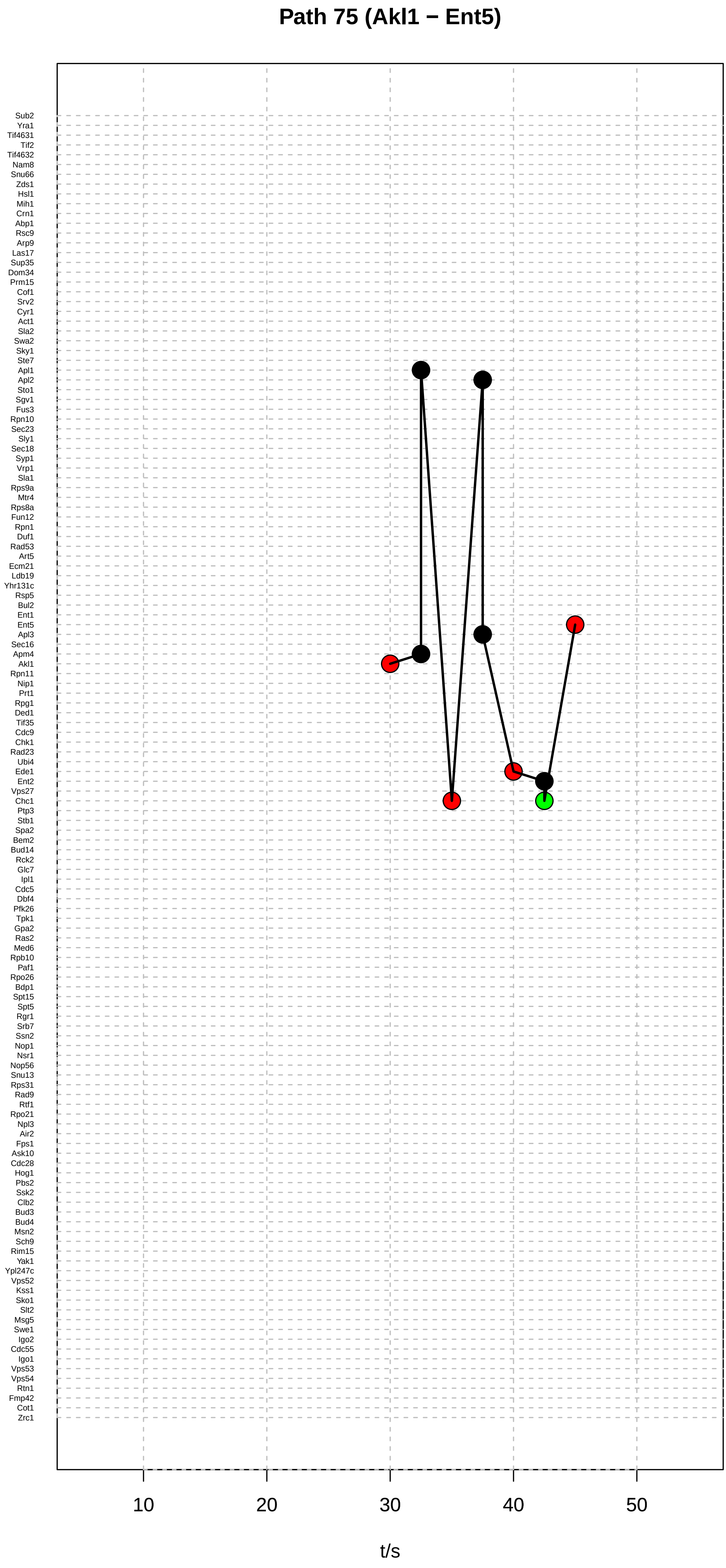

Path 75 (Akl1 − Ent5)
Sub2
Yra1
Tif4631
Tif2
Tif4632
Nam8
Snu66
Zds1
Hsl1
Mih1
Crn1
Abp1
Rsc9
Arp9
Las17
Sup35
Dom34
Prm15
Cof1
Srv2
Cyr1
Act1
Sla2
Swa2
Sky1
Ste7
Apl1
Apl2
Sto1
Sgv1
Fus3
Rpn10
Sec23
Sly1
Sec18
Syp1
Vrp1
Sla1
Rps9a
Mtr4
Rps8a
Fun12
Rpn1
Duf1
Rad53
Art5
Ecm21
Ldb19
Yhr131c
Rsp5
Bul2
Ent1
Ent5
Apl3
Sec16
Apm4
Akl1
Rpn11
Nip1
Prt1
Rpg1
Ded1
Tif35
Cdc9
Chk1
Rad23
Ubi4
Ede1
Ent2
Vps27
Chc1
Ptp3
Stb1
Spa2
Bem2
Bud14
Rck2
Glc7
Ipl1
Cdc5
Dbf4
Pfk26
Tpk1
Gpa2
Ras2
Med6
Rpb10
Paf1
Rpo26
Bdp1
Spt15
Spt5
Rgr1
Srb7
Ssn2
Nop1
Nsr1
Nop56
Snu13
Rps31
Rad9
Rtf1
Rpo21
Npl3
Air2
Fps1
Ask10
Cdc28
Hog1
Pbs2
Ssk2
Clb2
Bud3
Bud4
Msn2
Sch9
Rim15
Yak1
Ypl247c
Vps52
Kss1
Sko1
Slt2
Msg5
Swe1
Igo2
Cdc55
Igo1
Vps53
Vps54
Rtn1
Fmp42
Cot1
Zrc1
10
20
30
40
50
t/s

### Path 76 (Akl1 − Ent5)

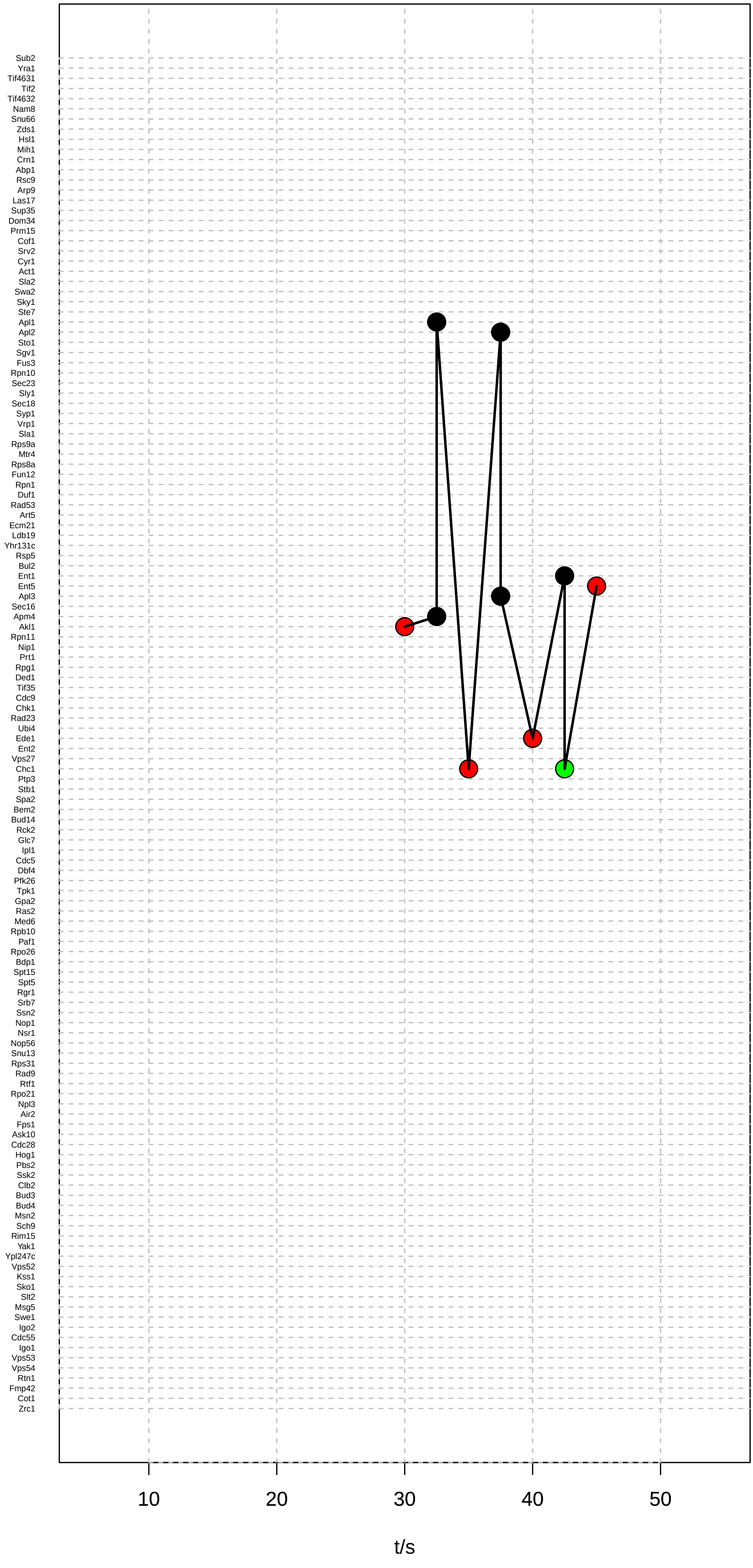

### Path 77 (Akl1 − Ent5)

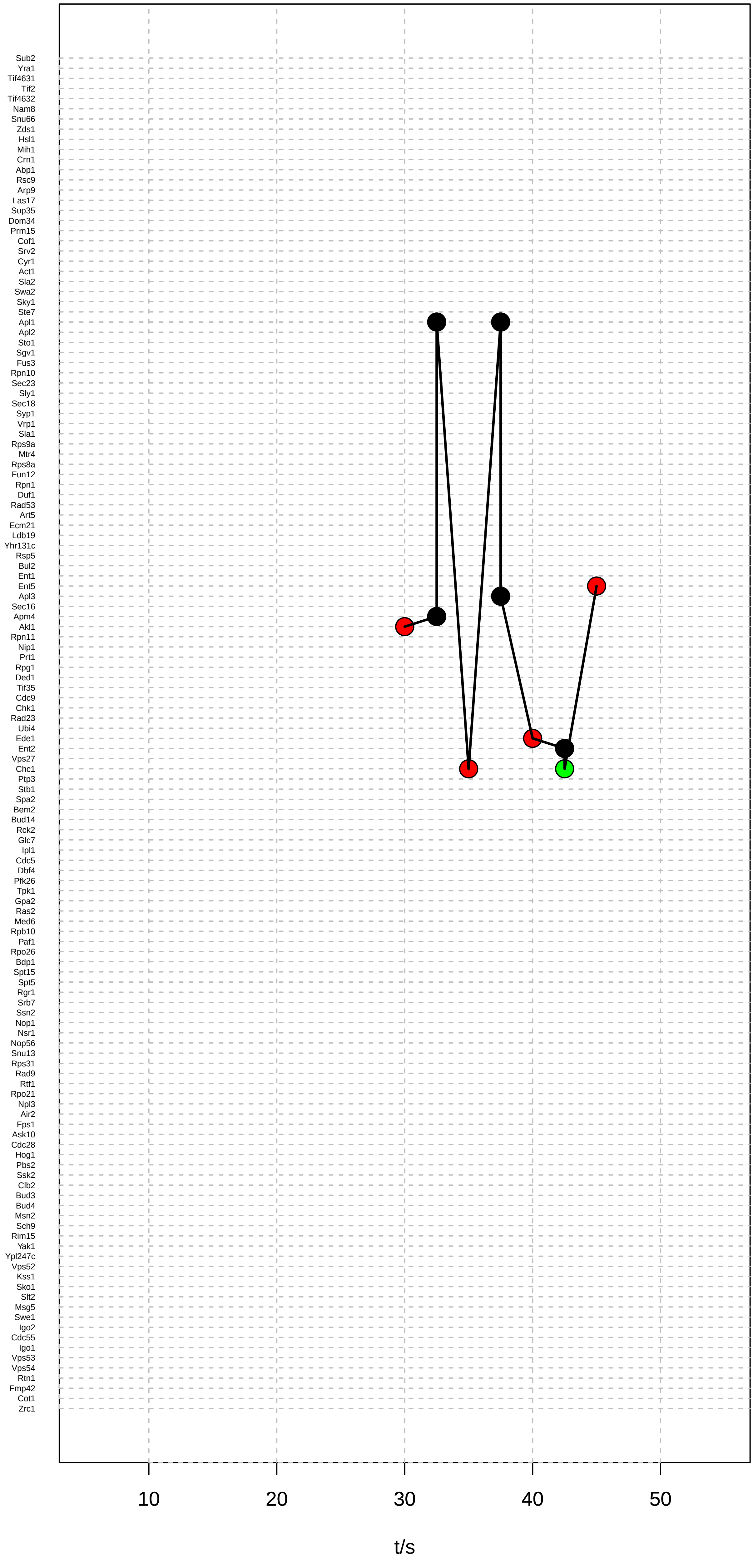

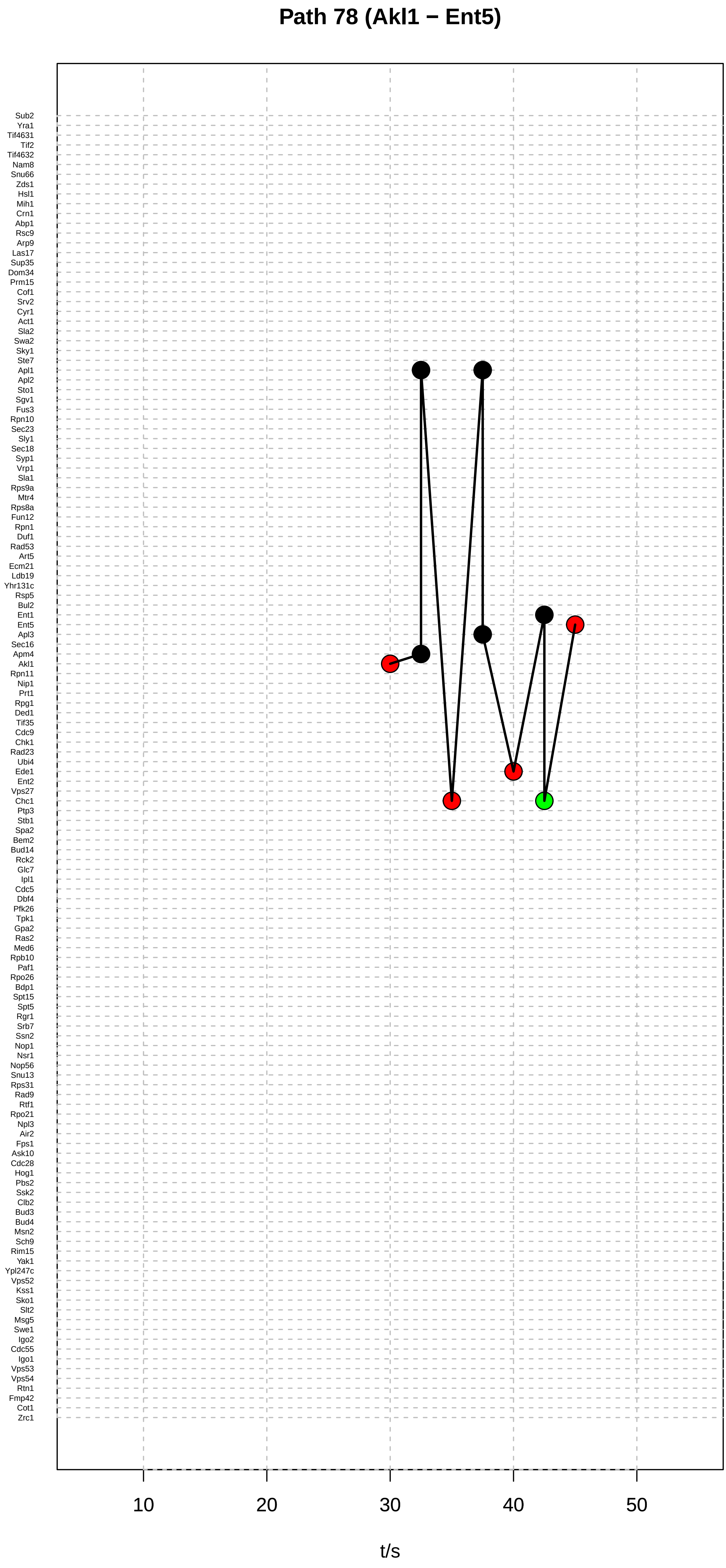

Path 78 (Akl1 − Ent5)
Sub2
Yra1
Tif4631
Tif2
Tif4632
Nam8
Snu66
Zds1
Hsl1
Mih1
Crn1
Abp1
Rsc9
Arp9
Las17
Sup35
Dom34
Prm15
Cof1
Srv2
Cyr1
Act1
Sla2
Swa2
Sky1
Ste7
Apl1
Apl2
Sto1
Sgv1
Fus3
Rpn10
Sec23
Sly1
Sec18
Syp1
Vrp1
Sla1
Rps9a
Mtr4
Rps8a
Fun12
Rpn1
Duf1
Rad53
Art5
Ecm21
Ldb19
Yhr131c
Rsp5
Bul2
Ent1
Ent5
Apl3
Sec16
Apm4
Akl1
Rpn11
Nip1
Prt1
Rpg1
Ded1
Tif35
Cdc9
Chk1
Rad23
Ubi4
Ede1
Ent2
Vps27
Chc1
Ptp3
Stb1
Spa2
Bem2
Bud14
Rck2
Glc7
Ipl1
Cdc5
Dbf4
Pfk26
Tpk1
Gpa2
Ras2
Med6
Rpb10
Paf1
Rpo26
Bdp1
Spt15
Spt5
Rgr1
Srb7
Ssn2
Nop1
Nsr1
Nop56
Snu13
Rps31
Rad9
Rtf1
Rpo21
Npl3
Air2
Fps1
Ask10
Cdc28
Hog1
Pbs2
Ssk2
Clb2
Bud3
Bud4
Msn2
Sch9
Rim15
Yak1
Ypl247c
Vps52
Kss1
Sko1
Slt2
Msg5
Swe1
Igo2
Cdc55
Igo1
Vps53
Vps54
Rtn1
Fmp42
Cot1
Zrc1
10
20
30
40
50
t/s

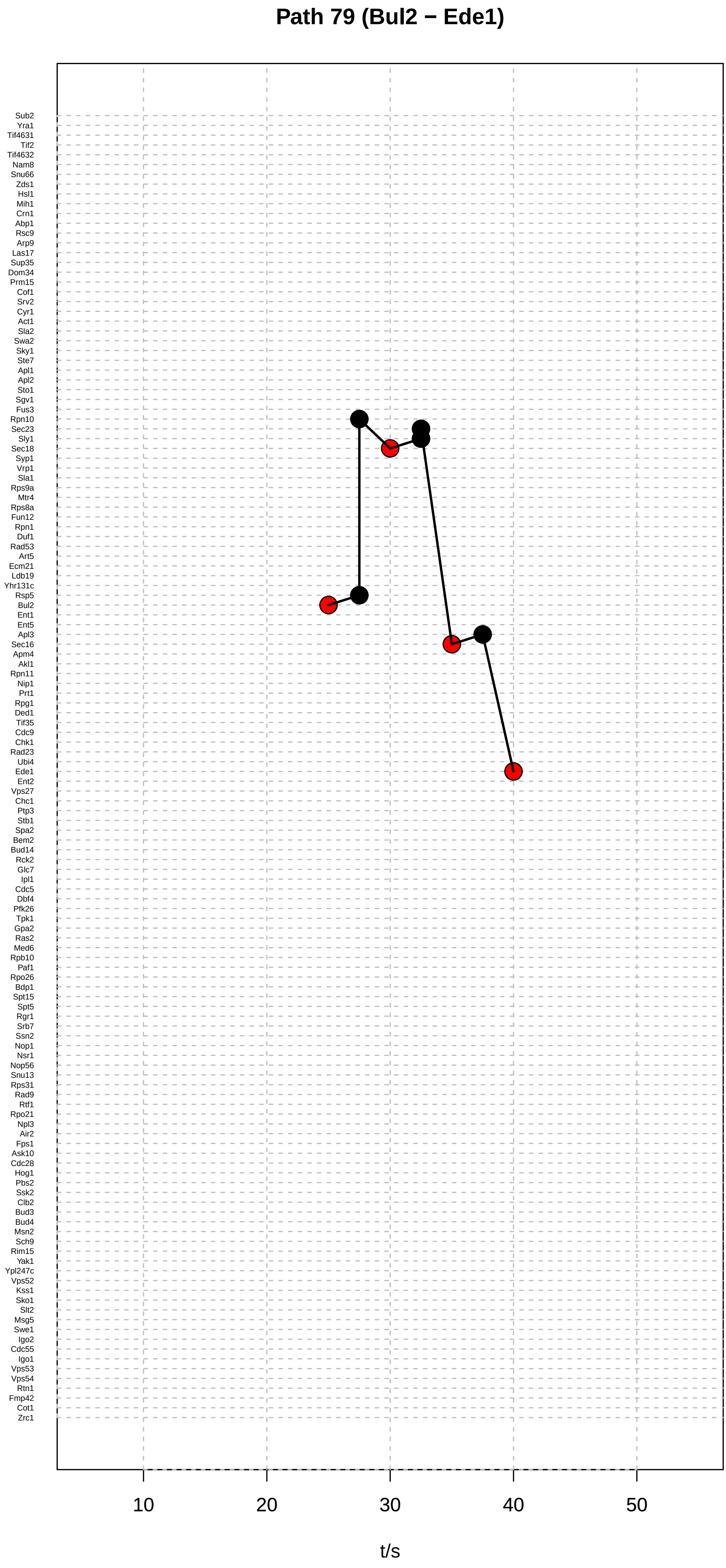

Path 79 (Bul2 − Ede1)
Sub2
Yra1
Tif4631
Tif2
Tif4632
Nam8
Snu66
Zds1
Hsl1
Mih1
Crn1
Abp1
Rsc9
Arp9
Las17
Sup35
Dom34
Prm15
Cof1
Srv2
Cyr1
Act1
Sla2
Swa2
Sky1
Ste7
Apl1
Apl2
Sto1
Sgv1
Fus3
Rpn10
Sec23
Sly1
Sec18
Syp1
Vrp1
Sla1
Rps9a
Mtr4
Rps8a
Fun12
Rpn1
Duf1
Rad53
Art5
Ecm21
Ldb19
Yhr131c
Rsp5
Bul2
Ent1
Ent5
Apl3
Sec16
Apm4
Akl1
Rpn11
Nip1
Prt1
Rpg1
Ded1
Tif35
Cdc9
Chk1
Rad23
Ubi4
Ede1
Ent2
Vps27
Chc1
Ptp3
Stb1
Spa2
Bem2
Bud14
Rck2
Glc7
Ipl1
Cdc5
Dbf4
Pfk26
Tpk1
Gpa2
Ras2
Med6
Rpb10
Paf1
Rpo26
Bdp1
Spt15
Spt5
Rgr1
Srb7
Ssn2
Nop1
Nsr1
Nop56
Snu13
Rps31
Rad9
Rtf1
Rpo21
Npl3
Air2
Fps1
Ask10
Cdc28
Hog1
Pbs2
Ssk2
Clb2
Bud3
Bud4
Msn2
Sch9
Rim15
Yak1
Ypl247c
Vps52
Kss1
Sko1
Slt2
Msg5
Swe1
Igo2
Cdc55
Igo1
Vps53
Vps54
Rtn1
Fmp42
Cot1
Zrc1
10
20
30
40
50
t/s

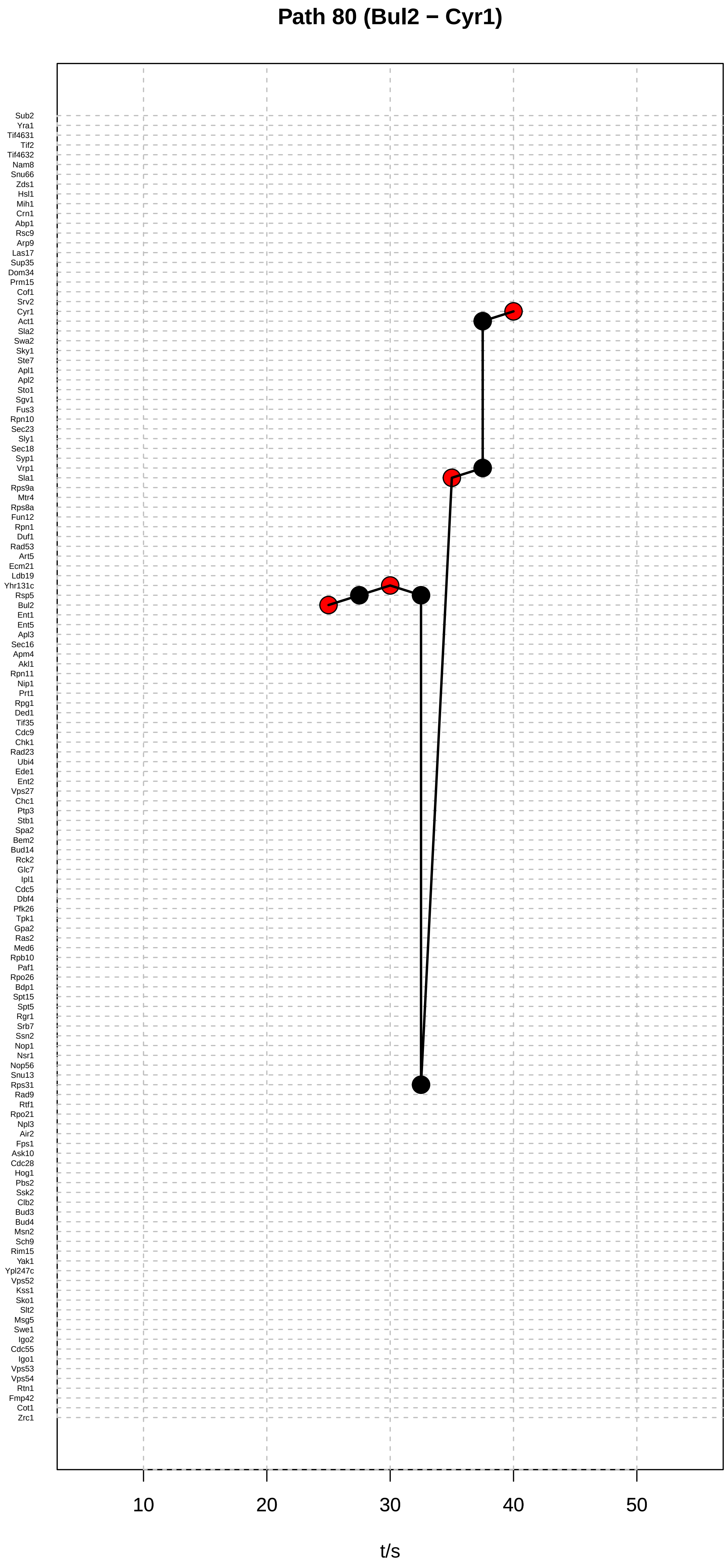

Path 80 (Bul2 − Cyr1)
Sub2
Yra1
Tif4631
Tif2
Tif4632
Nam8
Snu66
Zds1
Hsl1
Mih1
Crn1
Abp1
Rsc9
Arp9
Las17
Sup35
Dom34
Prm15
Cof1
Srv2
Cyr1
Act1
Sla2
Swa2
Sky1
Ste7
Apl1
Apl2
Sto1
Sgv1
Fus3
Rpn10
Sec23
Sly1
Sec18
Syp1
Vrp1
Sla1
Rps9a
Mtr4
Rps8a
Fun12
Rpn1
Duf1
Rad53
Art5
Ecm21
Ldb19
Yhr131c
Rsp5
Bul2
Ent1
Ent5
Apl3
Sec16
Apm4
Akl1
Rpn11
Nip1
Prt1
Rpg1
Ded1
Tif35
Cdc9
Chk1
Rad23
Ubi4
Ede1
Ent2
Vps27
Chc1
Ptp3
Stb1
Spa2
Bem2
Bud14
Rck2
Glc7
Ipl1
Cdc5
Dbf4
Pfk26
Tpk1
Gpa2
Ras2
Med6
Rpb10
Paf1
Rpo26
Bdp1
Spt15
Spt5
Rgr1
Srb7
Ssn2
Nop1
Nsr1
Nop56
Snu13
Rps31
Rad9
Rtf1
Rpo21
Npl3
Air2
Fps1
Ask10
Cdc28
Hog1
Pbs2
Ssk2
Clb2
Bud3
Bud4
Msn2
Sch9
Rim15
Yak1
Ypl247c
Vps52
Kss1
Sko1
Slt2
Msg5
Swe1
Igo2
Cdc55
Igo1
Vps53
Vps54
Rtn1
Fmp42
Cot1
Zrc1
10
20
30
40
50
t/s

### Path 81 (Bul2 − Cyr1)

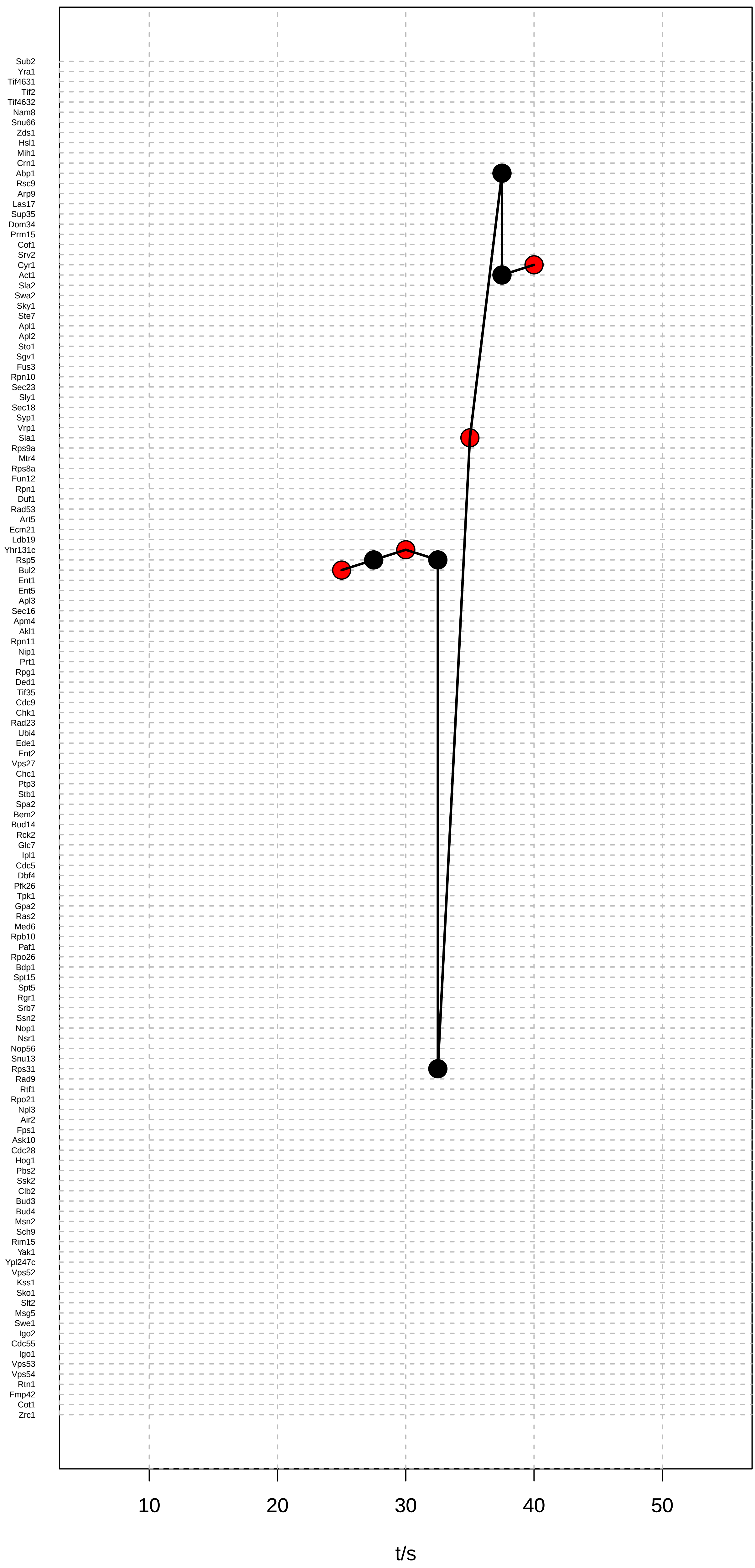

### Path 82 (Bul2 − Cyr1)

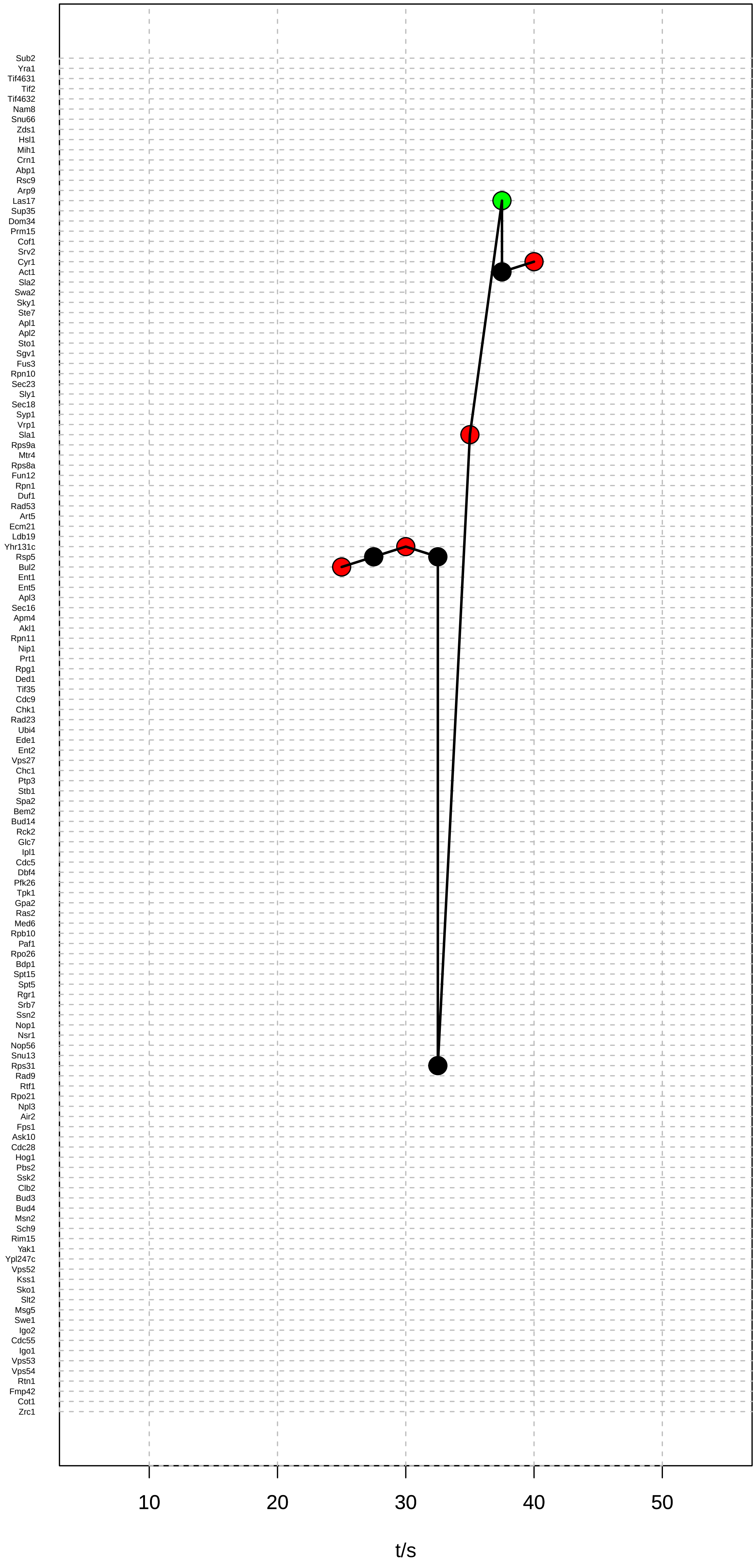

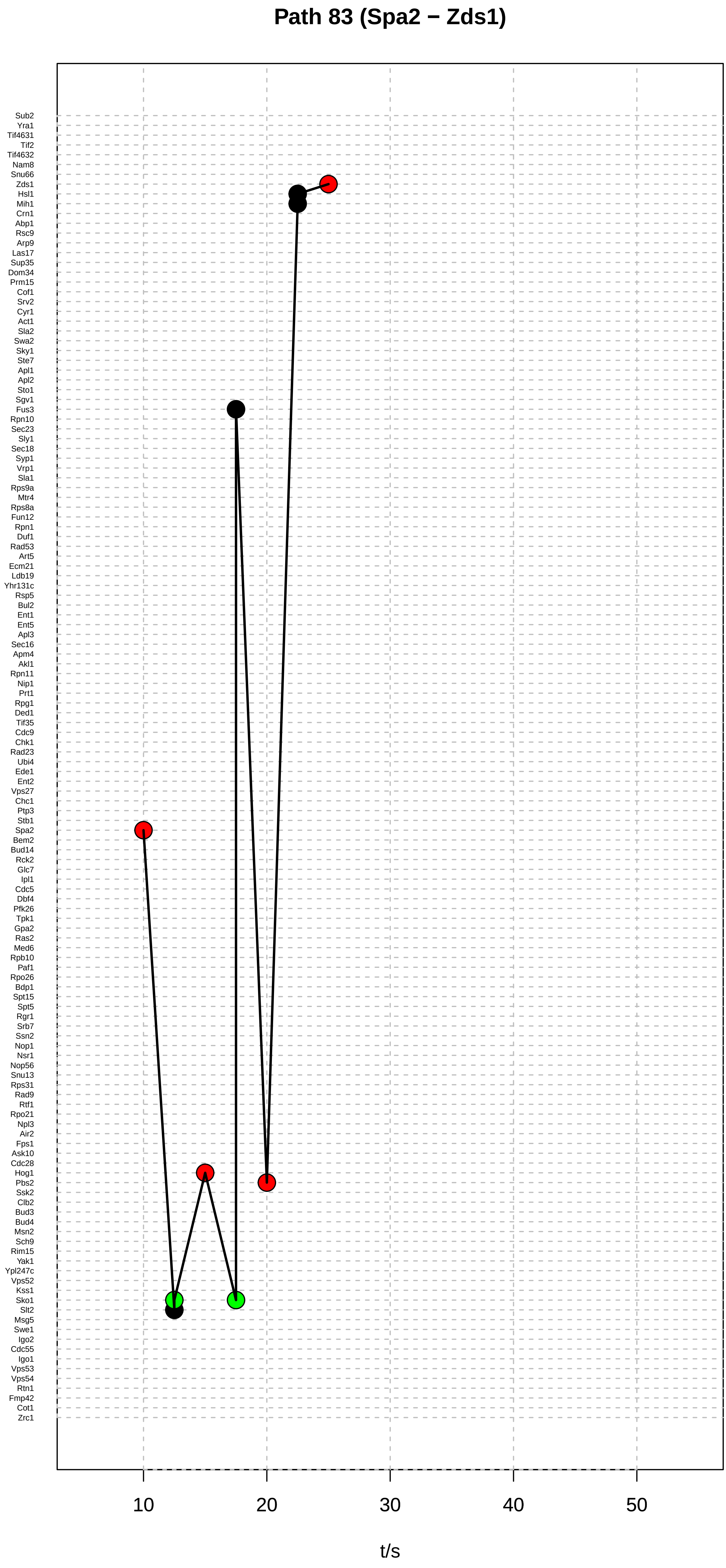
Path 83 (Spa2 − Zds1)
Sub2
Yra1
Tif4631
Tif2
Tif4632
Nam8
Snu66
Zds1
Hsl1
Mih1
Crn1
Abp1
Rsc9
Arp9
Las17
Sup35
Dom34
Prm15
Cof1
Srv2
Cyr1
Act1
Sla2
Swa2
Sky1
Ste7
Apl1
Apl2
Sto1
Sgv1
Fus3
Rpn10
Sec23
Sly1
Sec18
Syp1
Vrp1
Sla1
Rps9a
Mtr4
Rps8a
Fun12
Rpn1
Duf1
Rad53
Art5
Ecm21
Ldb19
Yhr131c
Rsp5
Bul2
Ent1
Ent5
Apl3
Sec16
Apm4
Akl1
Rpn11
Nip1
Prt1
Rpg1
Ded1
Tif35
Cdc9
Chk1
Rad23
Ubi4
Ede1
Ent2
Vps27
Chc1
Ptp3
Stb1
Spa2
Bem2
Bud14
Rck2
Glc7
Ipl1
Cdc5
Dbf4
Pfk26
Tpk1
Gpa2
Ras2
Med6
Rpb10
Paf1
Rpo26
Bdp1
Spt15
Spt5
Rgr1
Srb7
Ssn2
Nop1
Nsr1
Nop56
Snu13
Rps31
Rad9
Rtf1
Rpo21
Npl3
Air2
Fps1
Ask10
Cdc28
Hog1
Pbs2
Ssk2
Clb2
Bud3
Bud4
Msn2
Sch9
Rim15
Yak1
Ypl247c
Vps52
Kss1
Sko1
Slt2
Msg5
Swe1
Igo2
Cdc55
Igo1
Vps53
Vps54
Rtn1
Fmp42
Cot1
Zrc1
10
20
30
40
50
t/s

#### Path 84 (Spa2 − Zds1)

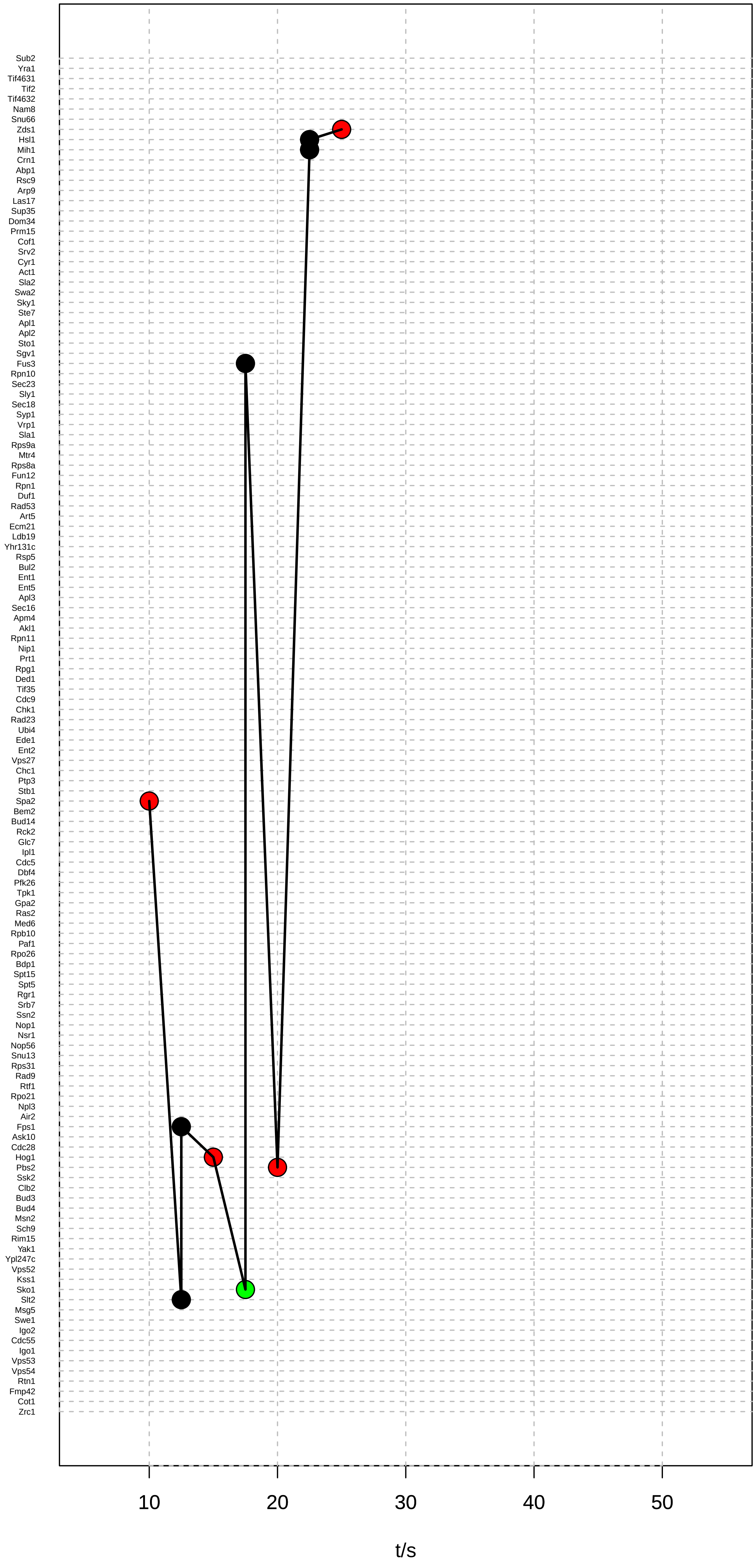

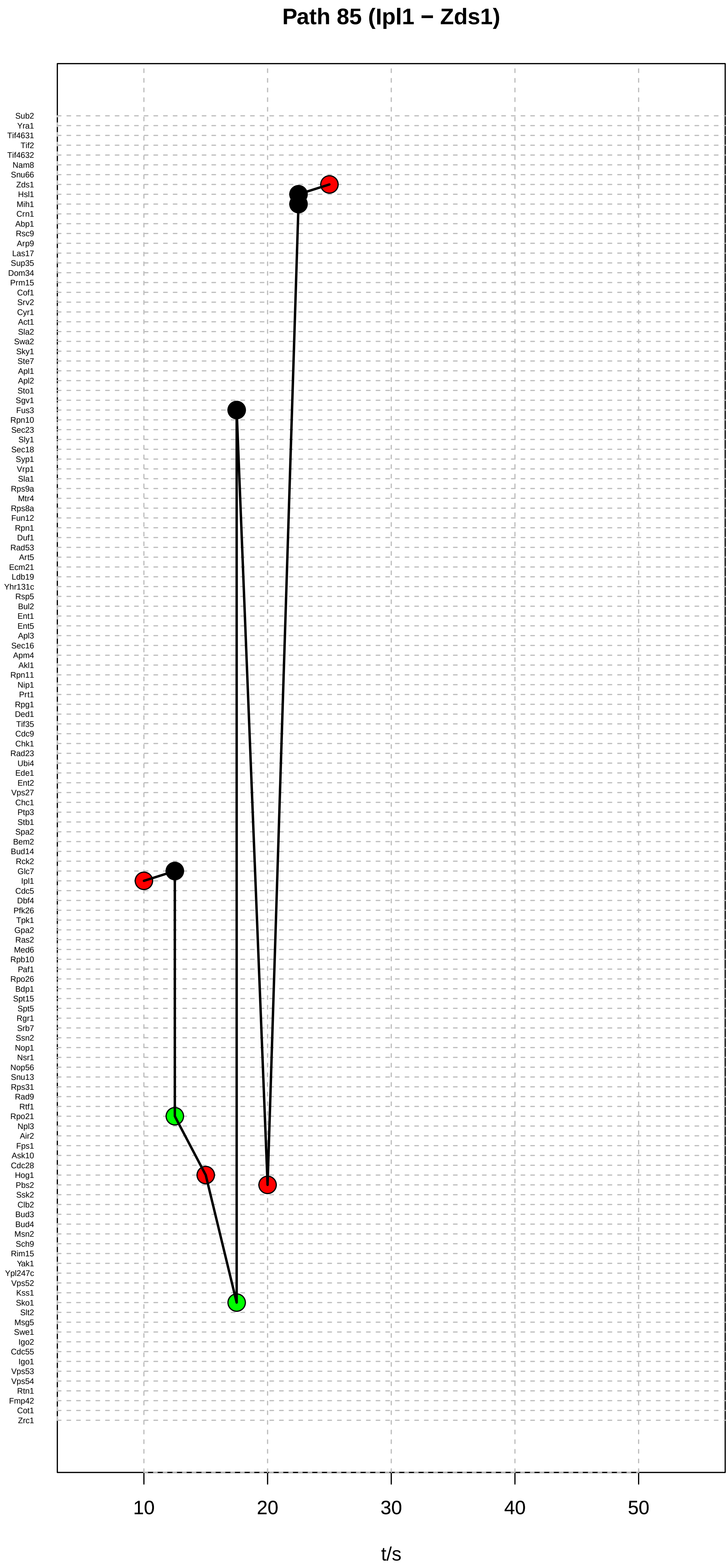
Path 85 (Ipl1 − Zds1)
Sub2
Yra1
Tif4631
Tif2
Tif4632
Nam8
Snu66
Zds1
Hsl1
Mih1
Crn1
Abp1
Rsc9
Arp9
Las17
Sup35
Dom34
Prm15
Cof1
Srv2
Cyr1
Act1
Sla2
Swa2
Sky1
Ste7
Apl1
Apl2
Sto1
Sgv1
Fus3
Rpn10
Sec23
Sly1
Sec18
Syp1
Vrp1
Sla1
Rps9a
Mtr4
Rps8a
Fun12
Rpn1
Duf1
Rad53
Art5
Ecm21
Ldb19
Yhr131c
Rsp5
Bul2
Ent1
Ent5
Apl3
Sec16
Apm4
Akl1
Rpn11
Nip1
Prt1
Rpg1
Ded1
Tif35
Cdc9
Chk1
Rad23
Ubi4
Ede1
Ent2
Vps27
Chc1
Ptp3
Stb1
Spa2
Bem2
Bud14
Rck2
Glc7
Ipl1
Cdc5
Dbf4
Pfk26
Tpk1
Gpa2
Ras2
Med6
Rpb10
Paf1
Rpo26
Bdp1
Spt15
Spt5
Rgr1
Srb7
Ssn2
Nop1
Nsr1
Nop56
Snu13
Rps31
Rad9
Rtf1
Rpo21
Npl3
Air2
Fps1
Ask10
Cdc28
Hog1
Pbs2
Ssk2
Clb2
Bud3
Bud4
Msn2
Sch9
Rim15
Yak1
Ypl247c
Vps52
Kss1
Sko1
Slt2
Msg5
Swe1
Igo2
Cdc55
Igo1
Vps53
Vps54
Rtn1
Fmp42
Cot1
Zrc1
10
20
30
40
50
t/s

### Path 86 (Ipl1 − Zds1)

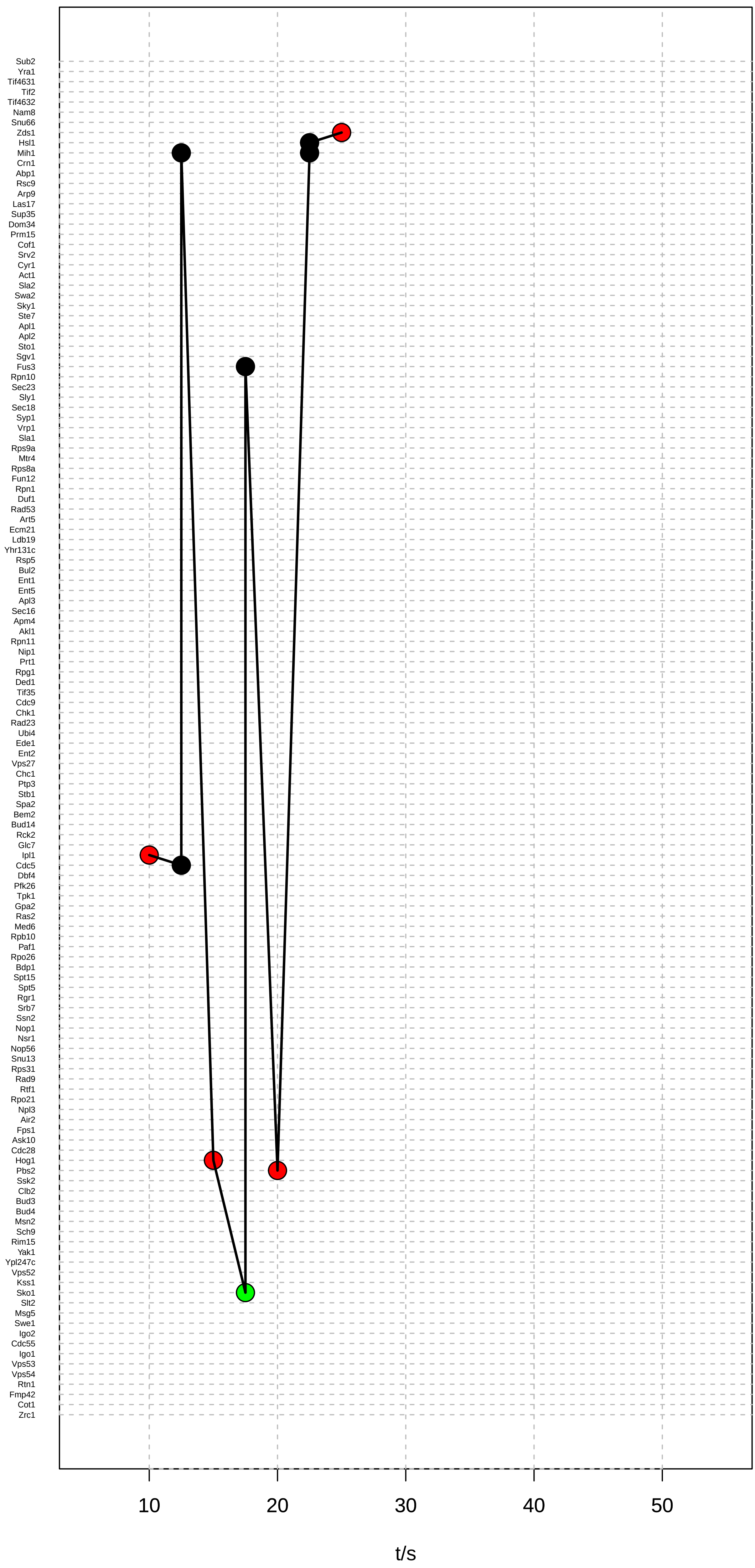

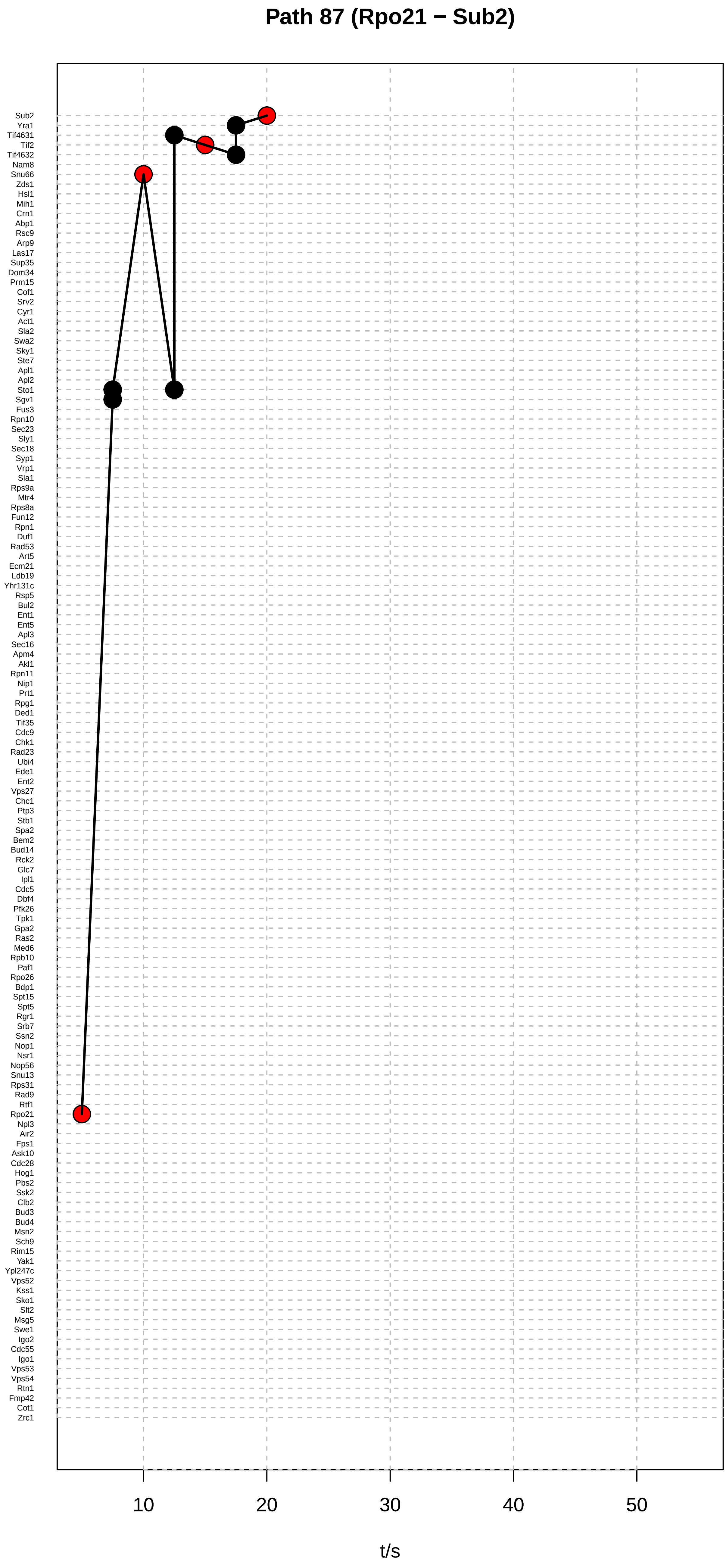

Path 87 (Rpo21 − Sub2)
Sub2
Yra1
Tif4631
Tif2
Tif4632
Nam8
Snu66
Zds1
Hsl1
Mih1
Crn1
Abp1
Rsc9
Arp9
Las17
Sup35
Dom34
Prm15
Cof1
Srv2
Cyr1
Act1
Sla2
Swa2
Sky1
Ste7
Apl1
Apl2
Sto1
Sgv1
Fus3
Rpn10
Sec23
Sly1
Sec18
Syp1
Vrp1
Sla1
Rps9a
Mtr4
Rps8a
Fun12
Rpn1
Duf1
Rad53
Art5
Ecm21
Ldb19
Yhr131c
Rsp5
Bul2
Ent1
Ent5
Apl3
Sec16
Apm4
Akl1
Rpn11
Nip1
Prt1
Rpg1
Ded1
Tif35
Cdc9
Chk1
Rad23
Ubi4
Ede1
Ent2
Vps27
Chc1
Ptp3
Stb1
Spa2
Bem2
Bud14
Rck2
Glc7
Ipl1
Cdc5
Dbf4
Pfk26
Tpk1
Gpa2
Ras2
Med6
Rpb10
Paf1
Rpo26
Bdp1
Spt15
Spt5
Rgr1
Srb7
Ssn2
Nop1
Nsr1
Nop56
Snu13
Rps31
Rad9
Rtf1
Rpo21
Npl3
Air2
Fps1
Ask10
Cdc28
Hog1
Pbs2
Ssk2
Clb2
Bud3
Bud4
Msn2
Sch9
Rim15
Yak1
Ypl247c
Vps52
Kss1
Sko1
Slt2
Msg5
Swe1
Igo2
Cdc55
Igo1
Vps53
Vps54
Rtn1
Fmp42
Cot1
Zrc1
10
20
30
40
50
t/s

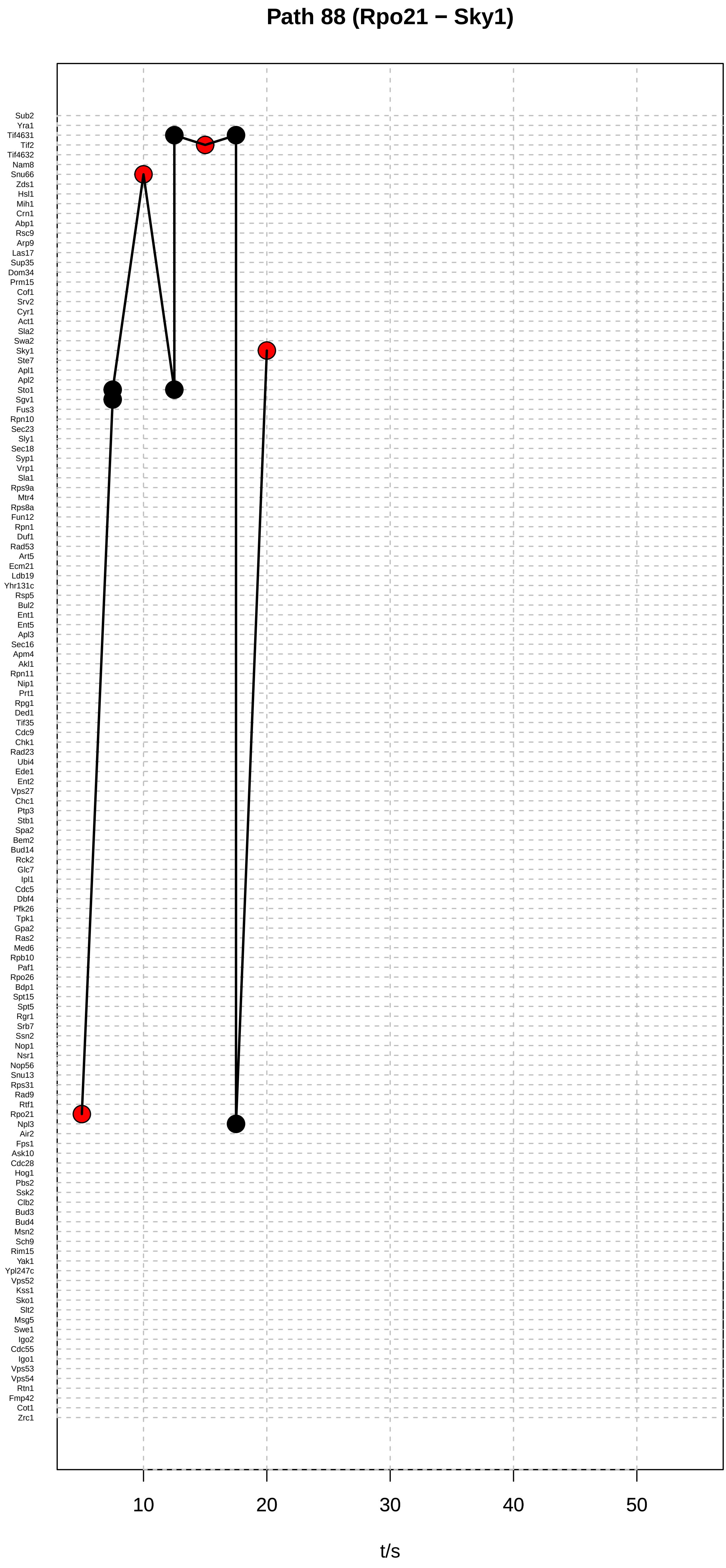
Path 88 (Rpo21 − Sky1)
Sub2
Yra1
Tif4631
Tif2
Tif4632
Nam8
Snu66
Zds1
Hsl1
Mih1
Crn1
Abp1
Rsc9
Arp9
Las17
Sup35
Dom34
Prm15
Cof1
Srv2
Cyr1
Act1
Sla2
Swa2
Sky1
Ste7
Apl1
Apl2
Sto1
Sgv1
Fus3
Rpn10
Sec23
Sly1
Sec18
Syp1
Vrp1
Sla1
Rps9a
Mtr4
Rps8a
Fun12
Rpn1
Duf1
Rad53
Art5
Ecm21
Ldb19
Yhr131c
Rsp5
Bul2
Ent1
Ent5
Apl3
Sec16
Apm4
Akl1
Rpn11
Nip1
Prt1
Rpg1
Ded1
Tif35
Cdc9
Chk1
Rad23
Ubi4
Ede1
Ent2
Vps27
Chc1
Ptp3
Stb1
Spa2
Bem2
Bud14
Rck2
Glc7
Ipl1
Cdc5
Dbf4
Pfk26
Tpk1
Gpa2
Ras2
Med6
Rpb10
Paf1
Rpo26
Bdp1
Spt15
Spt5
Rgr1
Srb7
Ssn2
Nop1
Nsr1
Nop56
Snu13
Rps31
Rad9
Rtf1
Rpo21
Npl3
Air2
Fps1
Ask10
Cdc28
Hog1
Pbs2
Ssk2
Clb2
Bud3
Bud4
Msn2
Sch9
Rim15
Yak1
Ypl247c
Vps52
Kss1
Sko1
Slt2
Msg5
Swe1
Igo2
Cdc55
Igo1
Vps53
Vps54
Rtn1
Fmp42
Cot1
Zrc1
10
20
30
40
50
t/s

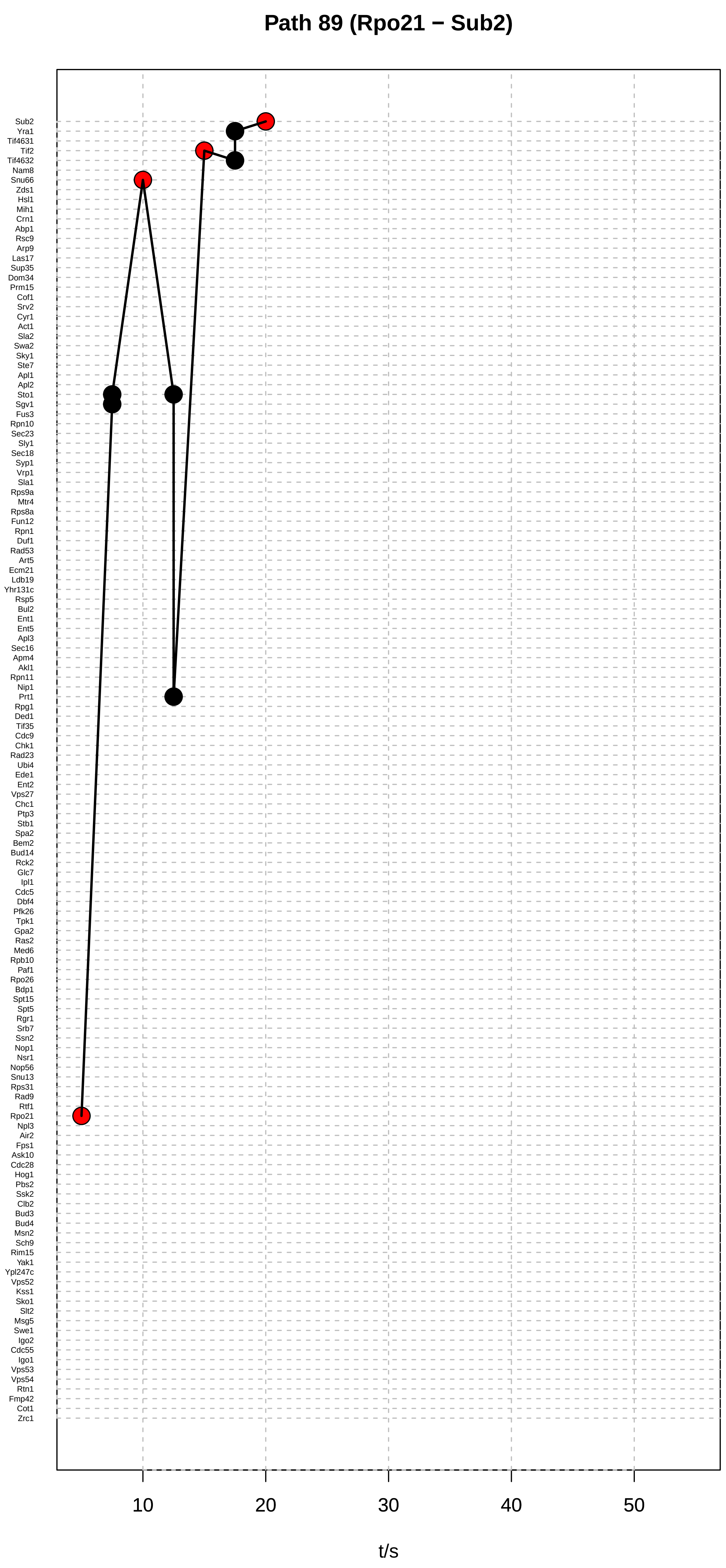

Path 89 (Rpo21 − Sub2)
Sub2
Yra1
Tif4631
Tif2
Tif4632
Nam8
Snu66
Zds1
Hsl1
Mih1
Crn1
Abp1
Rsc9
Arp9
Las17
Sup35
Dom34
Prm15
Cof1
Srv2
Cyr1
Act1
Sla2
Swa2
Sky1
Ste7
Apl1
Apl2
Sto1
Sgv1
Fus3
Rpn10
Sec23
Sly1
Sec18
Syp1
Vrp1
Sla1
Rps9a
Mtr4
Rps8a
Fun12
Rpn1
Duf1
Rad53
Art5
Ecm21
Ldb19
Yhr131c
Rsp5
Bul2
Ent1
Ent5
Apl3
Sec16
Apm4
Akl1
Rpn11
Nip1
Prt1
Rpg1
Ded1
Tif35
Cdc9
Chk1
Rad23
Ubi4
Ede1
Ent2
Vps27
Chc1
Ptp3
Stb1
Spa2
Bem2
Bud14
Rck2
Glc7
Ipl1
Cdc5
Dbf4
Pfk26
Tpk1
Gpa2
Ras2
Med6
Rpb10
Paf1
Rpo26
Bdp1
Spt15
Spt5
Rgr1
Srb7
Ssn2
Nop1
Nsr1
Nop56
Snu13
Rps31
Rad9
Rtf1
Rpo21
Npl3
Air2
Fps1
Ask10
Cdc28
Hog1
Pbs2
Ssk2
Clb2
Bud3
Bud4
Msn2
Sch9
Rim15
Yak1
Ypl247c
Vps52
Kss1
Sko1
Slt2
Msg5
Swe1
Igo2
Cdc55
Igo1
Vps53
Vps54
Rtn1
Fmp42
Cot1
Zrc1
10
20
30
40
50
t/s

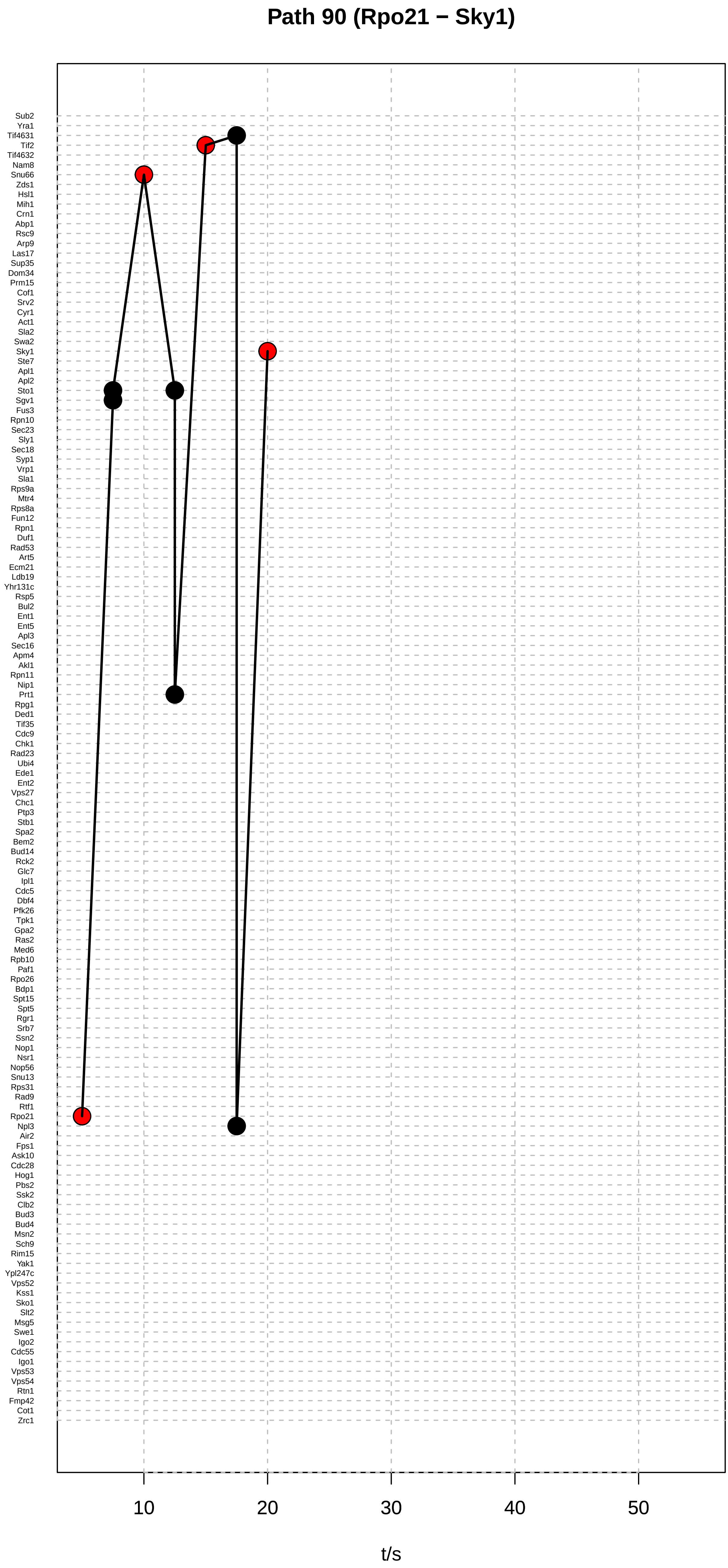

Path 90 (Rpo21 − Sky1)
Sub2
Yra1
Tif4631
Tif2
Tif4632
Nam8
Snu66
Zds1
Hsl1
Mih1
Crn1
Abp1
Rsc9
Arp9
Las17
Sup35
Dom34
Prm15
Cof1
Srv2
Cyr1
Act1
Sla2
Swa2
Sky1
Ste7
Apl1
Apl2
Sto1
Sgv1
Fus3
Rpn10
Sec23
Sly1
Sec18
Syp1
Vrp1
Sla1
Rps9a
Mtr4
Rps8a
Fun12
Rpn1
Duf1
Rad53
Art5
Ecm21
Ldb19
Yhr131c
Rsp5
Bul2
Ent1
Ent5
Apl3
Sec16
Apm4
Akl1
Rpn11
Nip1
Prt1
Rpg1
Ded1
Tif35
Cdc9
Chk1
Rad23
Ubi4
Ede1
Ent2
Vps27
Chc1
Ptp3
Stb1
Spa2
Bem2
Bud14
Rck2
Glc7
Ipl1
Cdc5
Dbf4
Pfk26
Tpk1
Gpa2
Ras2
Med6
Rpb10
Paf1
Rpo26
Bdp1
Spt15
Spt5
Rgr1
Srb7
Ssn2
Nop1
Nsr1
Nop56
Snu13
Rps31
Rad9
Rtf1
Rpo21
Npl3
Air2
Fps1
Ask10
Cdc28
Hog1
Pbs2
Ssk2
Clb2
Bud3
Bud4
Msn2
Sch9
Rim15
Yak1
Ypl247c
Vps52
Kss1
Sko1
Slt2
Msg5
Swe1
Igo2
Cdc55
Igo1
Vps53
Vps54
Rtn1
Fmp42
Cot1
Zrc1
10
20
30
40
50
t/s

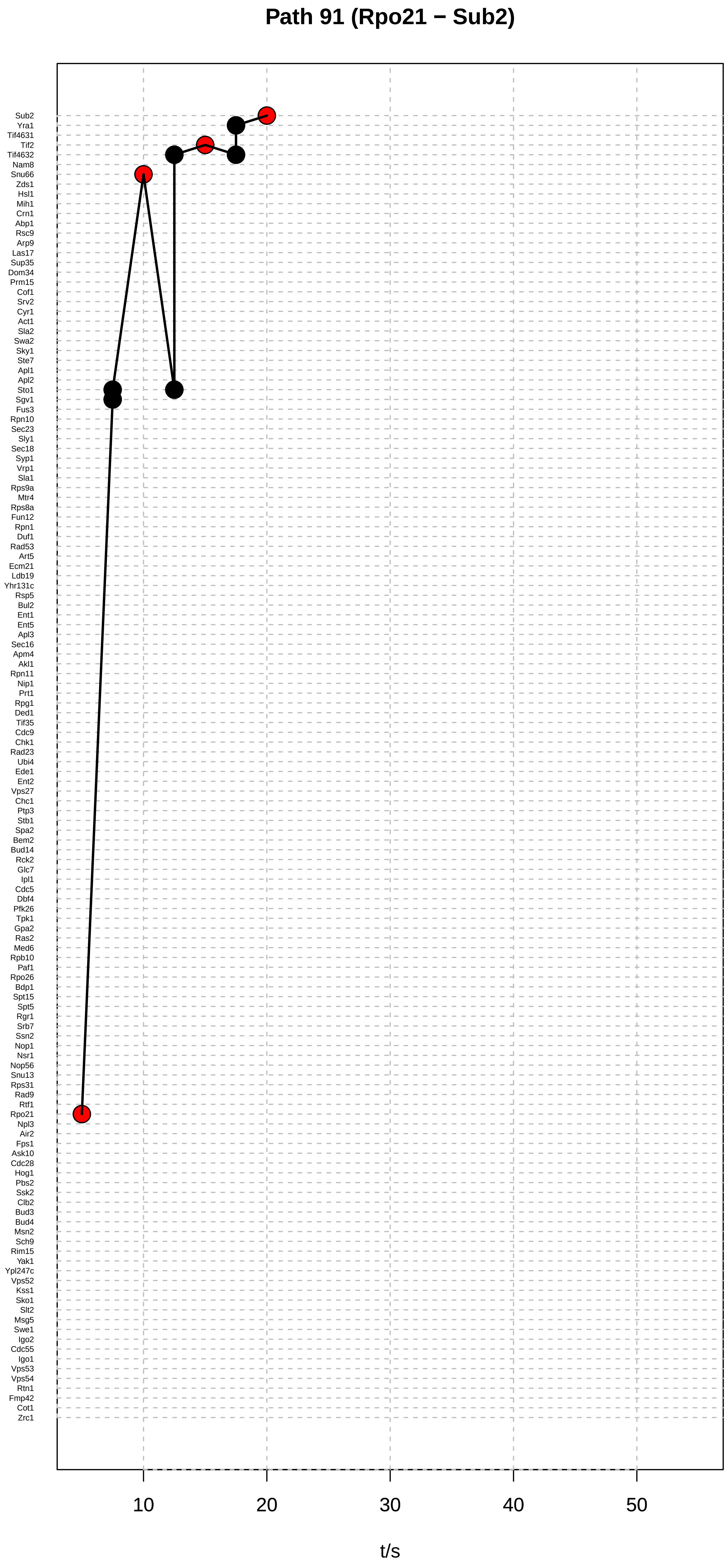

Path 91 (Rpo21 − Sub2)
Sub2
Yra1
Tif4631
Tif2
Tif4632
Nam8
Snu66
Zds1
Hsl1
Mih1
Crn1
Abp1
Rsc9
Arp9
Las17
Sup35
Dom34
Prm15
Cof1
Srv2
Cyr1
Act1
Sla2
Swa2
Sky1
Ste7
Apl1
Apl2
Sto1
Sgv1
Fus3
Rpn10
Sec23
Sly1
Sec18
Syp1
Vrp1
Sla1
Rps9a
Mtr4
Rps8a
Fun12
Rpn1
Duf1
Rad53
Art5
Ecm21
Ldb19
Yhr131c
Rsp5
Bul2
Ent1
Ent5
Apl3
Sec16
Apm4
Akl1
Rpn11
Nip1
Prt1
Rpg1
Ded1
Tif35
Cdc9
Chk1
Rad23
Ubi4
Ede1
Ent2
Vps27
Chc1
Ptp3
Stb1
Spa2
Bem2
Bud14
Rck2
Glc7
Ipl1
Cdc5
Dbf4
Pfk26
Tpk1
Gpa2
Ras2
Med6
Rpb10
Paf1
Rpo26
Bdp1
Spt15
Spt5
Rgr1
Srb7
Ssn2
Nop1
Nsr1
Nop56
Snu13
Rps31
Rad9
Rtf1
Rpo21
Npl3
Air2
Fps1
Ask10
Cdc28
Hog1
Pbs2
Ssk2
Clb2
Bud3
Bud4
Msn2
Sch9
Rim15
Yak1
Ypl247c
Vps52
Kss1
Sko1
Slt2
Msg5
Swe1
Igo2
Cdc55
Igo1
Vps53
Vps54
Rtn1
Fmp42
Cot1
Zrc1
10
20
30
40
50
t/s

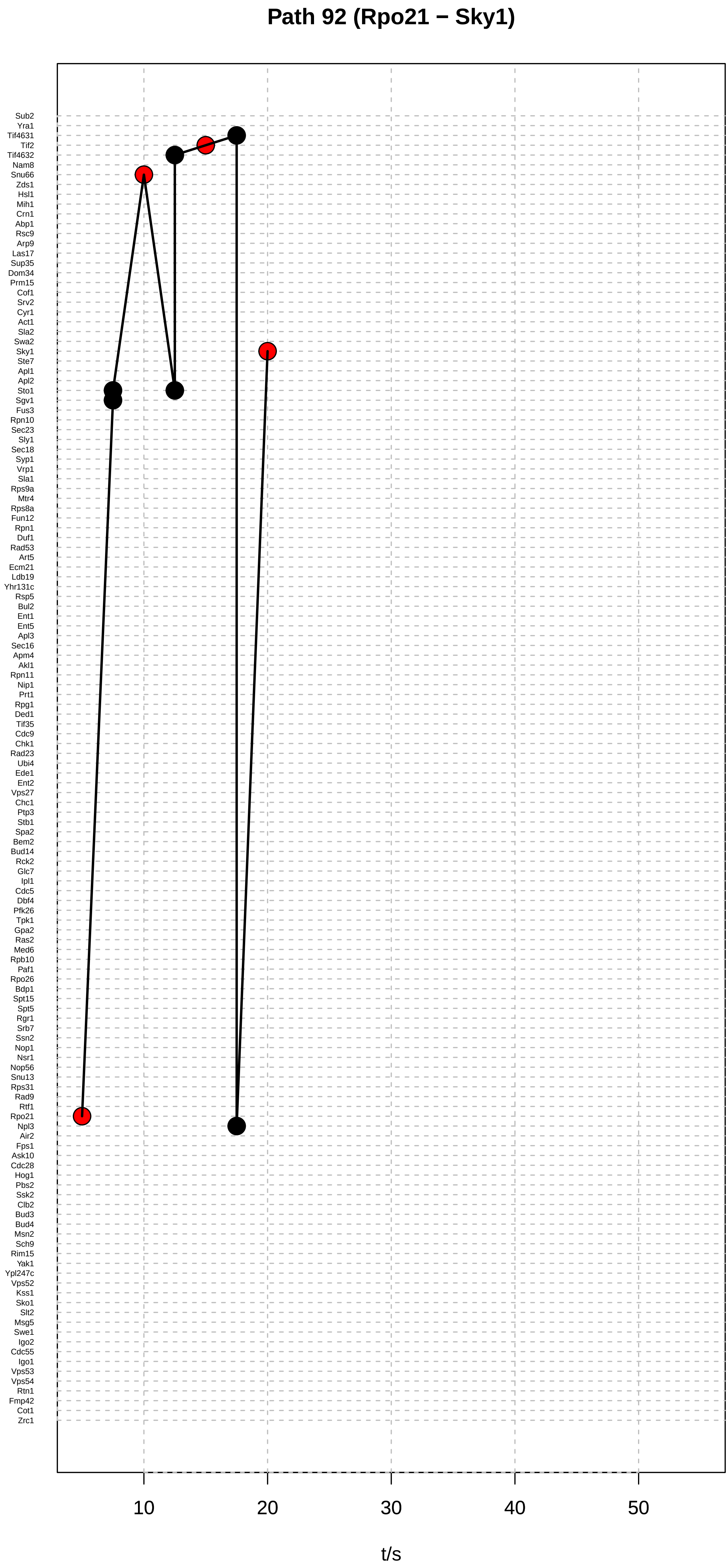

Path 92 (Rpo21 − Sky1)
Sub2
Yra1
Tif4631
Tif2
Tif4632
Nam8
Snu66
Zds1
Hsl1
Mih1
Crn1
Abp1
Rsc9
Arp9
Las17
Sup35
Dom34
Prm15
Cof1
Srv2
Cyr1
Act1
Sla2
Swa2
Sky1
Ste7
Apl1
Apl2
Sto1
Sgv1
Fus3
Rpn10
Sec23
Sly1
Sec18
Syp1
Vrp1
Sla1
Rps9a
Mtr4
Rps8a
Fun12
Rpn1
Duf1
Rad53
Art5
Ecm21
Ldb19
Yhr131c
Rsp5
Bul2
Ent1
Ent5
Apl3
Sec16
Apm4
Akl1
Rpn11
Nip1
Prt1
Rpg1
Ded1
Tif35
Cdc9
Chk1
Rad23
Ubi4
Ede1
Ent2
Vps27
Chc1
Ptp3
Stb1
Spa2
Bem2
Bud14
Rck2
Glc7
Ipl1
Cdc5
Dbf4
Pfk26
Tpk1
Gpa2
Ras2
Med6
Rpb10
Paf1
Rpo26
Bdp1
Spt15
Spt5
Rgr1
Srb7
Ssn2
Nop1
Nsr1
Nop56
Snu13
Rps31
Rad9
Rtf1
Rpo21
Npl3
Air2
Fps1
Ask10
Cdc28
Hog1
Pbs2
Ssk2
Clb2
Bud3
Bud4
Msn2
Sch9
Rim15
Yak1
Ypl247c
Vps52
Kss1
Sko1
Slt2
Msg5
Swe1
Igo2
Cdc55
Igo1
Vps53
Vps54
Rtn1
Fmp42
Cot1
Zrc1
10
20
30
40
50
t/s

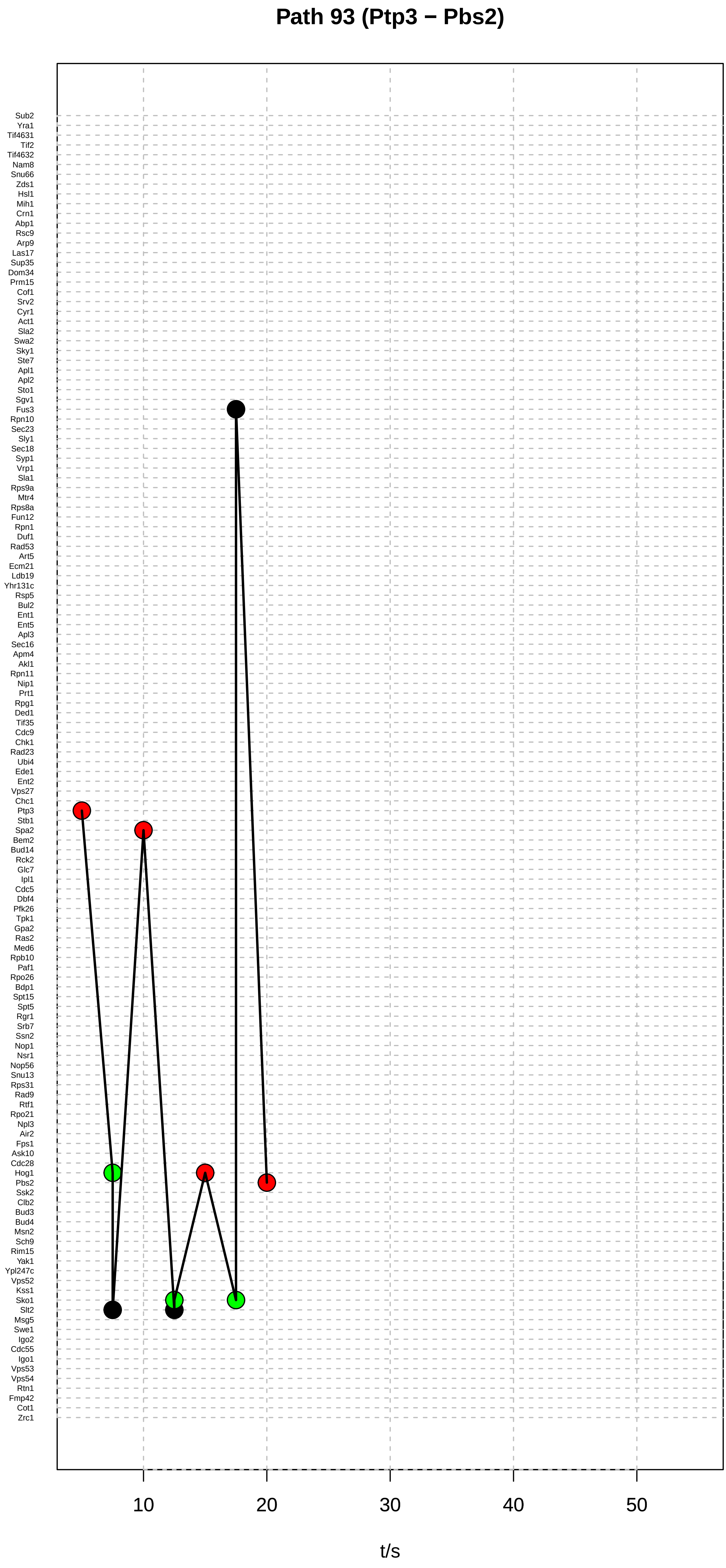

Path 93 (Ptp3 − Pbs2)
Sub2
Yra1
Tif4631
Tif2
Tif4632
Nam8
Snu66
Zds1
Hsl1
Mih1
Crn1
Abp1
Rsc9
Arp9
Las17
Sup35
Dom34
Prm15
Cof1
Srv2
Cyr1
Act1
Sla2
Swa2
Sky1
Ste7
Apl1
Apl2
Sto1
Sgv1
Fus3
Rpn10
Sec23
Sly1
Sec18
Syp1
Vrp1
Sla1
Rps9a
Mtr4
Rps8a
Fun12
Rpn1
Duf1
Rad53
Art5
Ecm21
Ldb19
Yhr131c
Rsp5
Bul2
Ent1
Ent5
Apl3
Sec16
Apm4
Akl1
Rpn11
Nip1
Prt1
Rpg1
Ded1
Tif35
Cdc9
Chk1
Rad23
Ubi4
Ede1
Ent2
Vps27
Chc1
Ptp3
Stb1
Spa2
Bem2
Bud14
Rck2
Glc7
Ipl1
Cdc5
Dbf4
Pfk26
Tpk1
Gpa2
Ras2
Med6
Rpb10
Paf1
Rpo26
Bdp1
Spt15
Spt5
Rgr1
Srb7
Ssn2
Nop1
Nsr1
Nop56
Snu13
Rps31
Rad9
Rtf1
Rpo21
Npl3
Air2
Fps1
Ask10
Cdc28
Hog1
Pbs2
Ssk2
Clb2
Bud3
Bud4
Msn2
Sch9
Rim15
Yak1
Ypl247c
Vps52
Kss1
Sko1
Slt2
Msg5
Swe1
Igo2
Cdc55
Igo1
Vps53
Vps54
Rtn1
Fmp42
Cot1
Zrc1
10
20
30
40
50
t/s

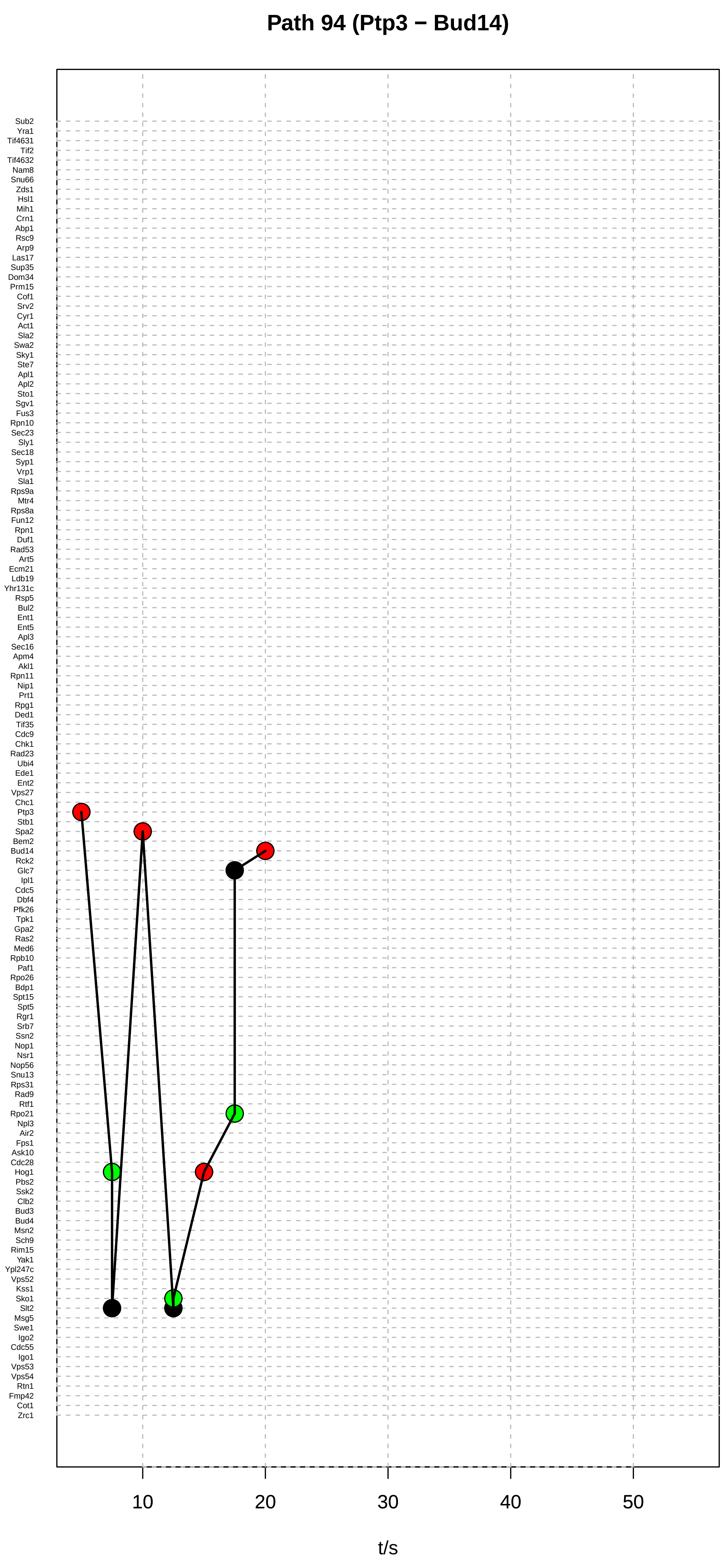

Path 94 (Ptp3 − Bud14)
Sub2
Yra1
Tif4631
Tif2
Tif4632
Nam8
Snu66
Zds1
Hsl1
Mih1
Crn1
Abp1
Rsc9
Arp9
Las17
Sup35
Dom34
Prm15
Cof1
Srv2
Cyr1
Act1
Sla2
Swa2
Sky1
Ste7
Apl1
Apl2
Sto1
Sgv1
Fus3
Rpn10
Sec23
Sly1
Sec18
Syp1
Vrp1
Sla1
Rps9a
Mtr4
Rps8a
Fun12
Rpn1
Duf1
Rad53
Art5
Ecm21
Ldb19
Yhr131c
Rsp5
Bul2
Ent1
Ent5
Apl3
Sec16
Apm4
Akl1
Rpn11
Nip1
Prt1
Rpg1
Ded1
Tif35
Cdc9
Chk1
Rad23
Ubi4
Ede1
Ent2
Vps27
Chc1
Ptp3
Stb1
Spa2
Bem2
Bud14
Rck2
Glc7
Ipl1
Cdc5
Dbf4
Pfk26
Tpk1
Gpa2
Ras2
Med6
Rpb10
Paf1
Rpo26
Bdp1
Spt15
Spt5
Rgr1
Srb7
Ssn2
Nop1
Nsr1
Nop56
Snu13
Rps31
Rad9
Rtf1
Rpo21
Npl3
Air2
Fps1
Ask10
Cdc28
Hog1
Pbs2
Ssk2
Clb2
Bud3
Bud4
Msn2
Sch9
Rim15
Yak1
Ypl247c
Vps52
Kss1
Sko1
Slt2
Msg5
Swe1
Igo2
Cdc55
Igo1
Vps53
Vps54
Rtn1
Fmp42
Cot1
Zrc1
10
20
30
40
50
t/s

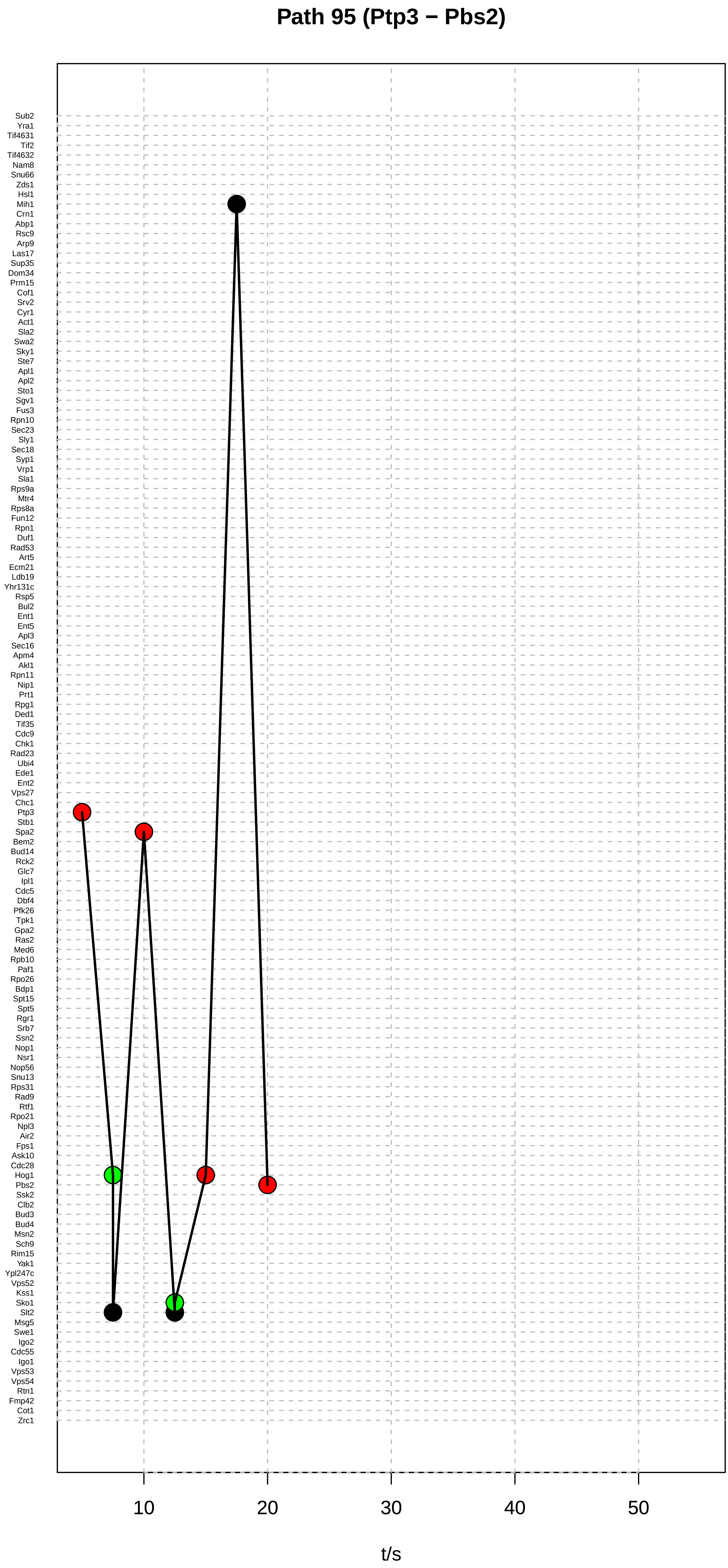
Path 95 (Ptp3 − Pbs2)
Sub2
Yra1
Tif4631
Tif2
Tif4632
Nam8
Snu66
Zds1
Hsl1
Mih1
Crn1
Abp1
Rsc9
Arp9
Las17
Sup35
Dom34
Prm15
Cof1
Srv2
Cyr1
Act1
Sla2
Swa2
Sky1
Ste7
Apl1
Apl2
Sto1
Sgv1
Fus3
Rpn10
Sec23
Sly1
Sec18
Syp1
Vrp1
Sla1
Rps9a
Mtr4
Rps8a
Fun12
Rpn1
Duf1
Rad53
Art5
Ecm21
Ldb19
Yhr131c
Rsp5
Bul2
Ent1
Ent5
Apl3
Sec16
Apm4
Akl1
Rpn11
Nip1
Prt1
Rpg1
Ded1
Tif35
Cdc9
Chk1
Rad23
Ubi4
Ede1
Ent2
Vps27
Chc1
Ptp3
Stb1
Spa2
Bem2
Bud14
Rck2
Glc7
Ipl1
Cdc5
Dbf4
Pfk26
Tpk1
Gpa2
Ras2
Med6
Rpb10
Paf1
Rpo26
Bdp1
Spt15
Spt5
Rgr1
Srb7
Ssn2
Nop1
Nsr1
Nop56
Snu13
Rps31
Rad9
Rtf1
Rpo21
Npl3
Air2
Fps1
Ask10
Cdc28
Hog1
Pbs2
Ssk2
Clb2
Bud3
Bud4
Msn2
Sch9
Rim15
Yak1
Ypl247c
Vps52
Kss1
Sko1
Slt2
Msg5
Swe1
Igo2
Cdc55
Igo1
Vps53
Vps54
Rtn1
Fmp42
Cot1
Zrc1
10
20
30
40
50
t/s

#### Path 96 (Ptp3 − Pbs2)

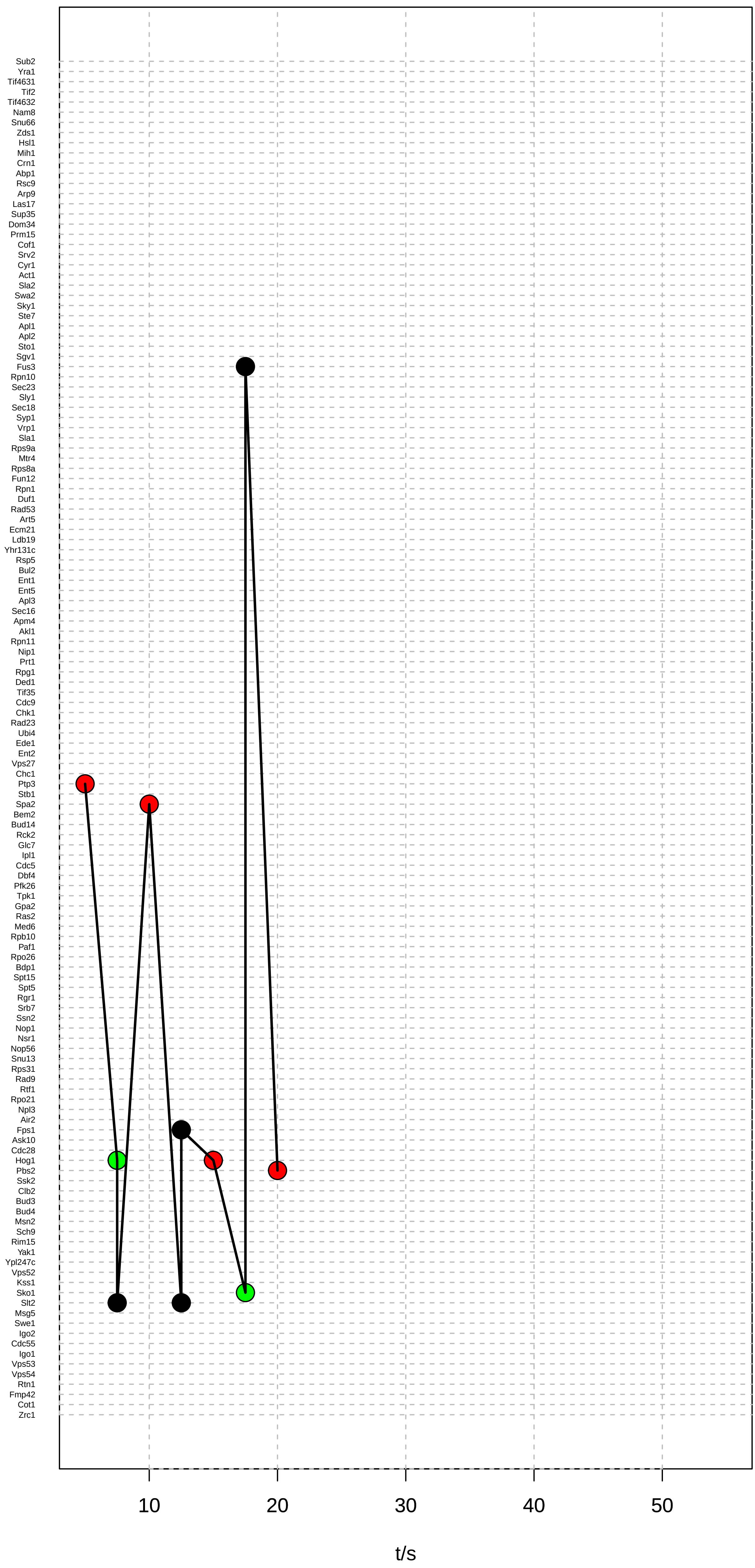

#### Path 97 (Ptp3 − Bud14)

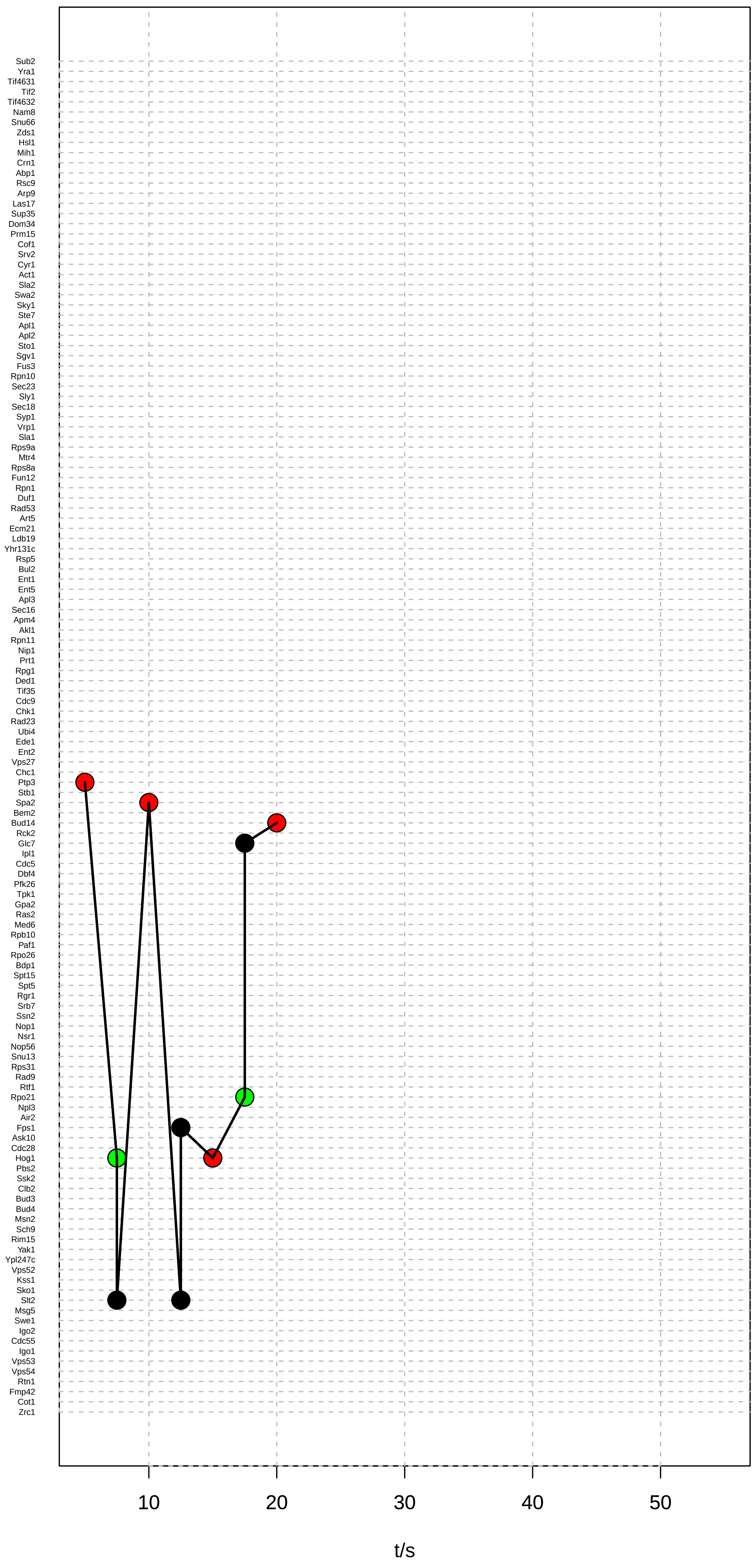

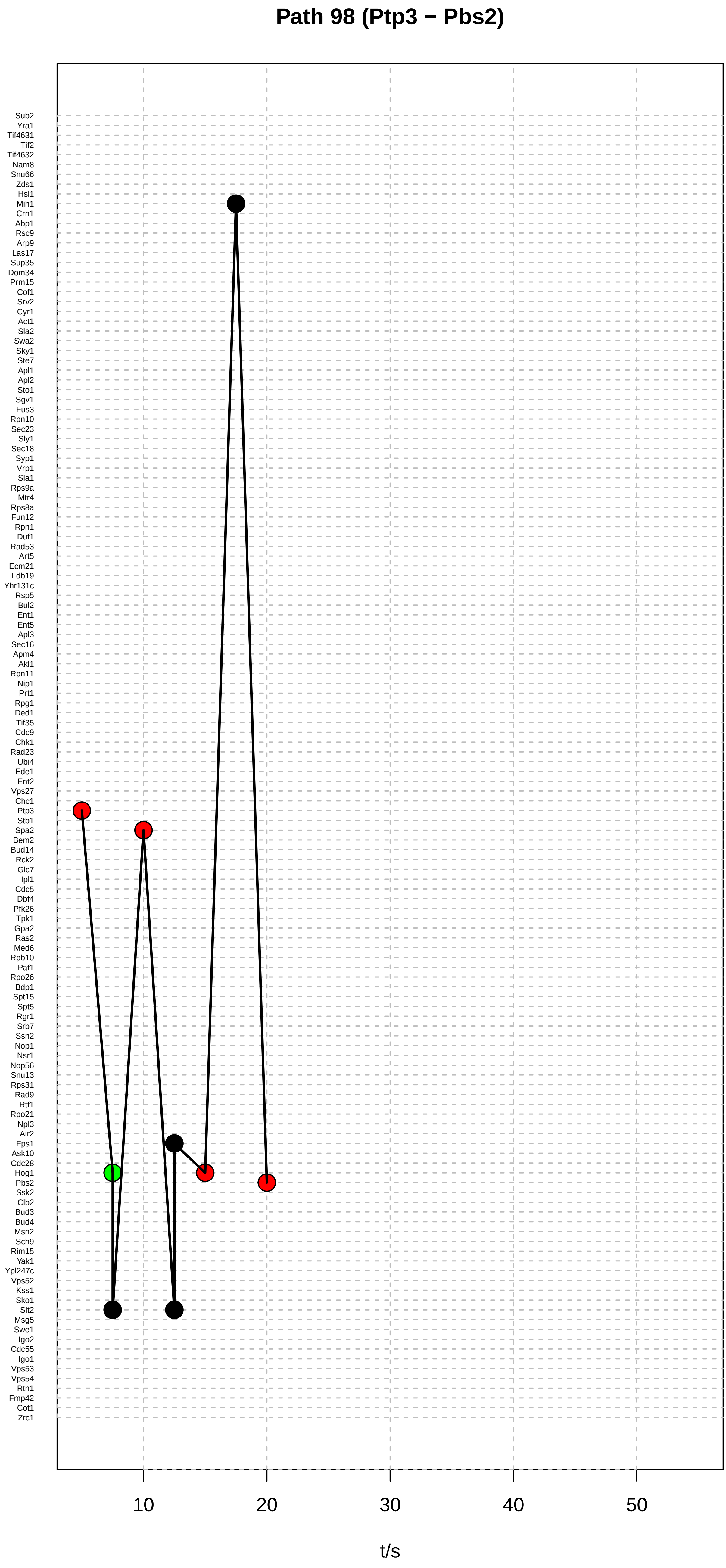

Path 98 (Ptp3 − Pbs2)
Sub2
Yra1
Tif4631
Tif2
Tif4632
Nam8
Snu66
Zds1
Hsl1
Mih1
Crn1
Abp1
Rsc9
Arp9
Las17
Sup35
Dom34
Prm15
Cof1
Srv2
Cyr1
Act1
Sla2
Swa2
Sky1
Ste7
Apl1
Apl2
Sto1
Sgv1
Fus3
Rpn10
Sec23
Sly1
Sec18
Syp1
Vrp1
Sla1
Rps9a
Mtr4
Rps8a
Fun12
Rpn1
Duf1
Rad53
Art5
Ecm21
Ldb19
Yhr131c
Rsp5
Bul2
Ent1
Ent5
Apl3
Sec16
Apm4
Akl1
Rpn11
Nip1
Prt1
Rpg1
Ded1
Tif35
Cdc9
Chk1
Rad23
Ubi4
Ede1
Ent2
Vps27
Chc1
Ptp3
Stb1
Spa2
Bem2
Bud14
Rck2
Glc7
Ipl1
Cdc5
Dbf4
Pfk26
Tpk1
Gpa2
Ras2
Med6
Rpb10
Paf1
Rpo26
Bdp1
Spt15
Spt5
Rgr1
Srb7
Ssn2
Nop1
Nsr1
Nop56
Snu13
Rps31
Rad9
Rtf1
Rpo21
Npl3
Air2
Fps1
Ask10
Cdc28
Hog1
Pbs2
Ssk2
Clb2
Bud3
Bud4
Msn2
Sch9
Rim15
Yak1
Ypl247c
Vps52
Kss1
Sko1
Slt2
Msg5
Swe1
Igo2
Cdc55
Igo1
Vps53
Vps54
Rtn1
Fmp42
Cot1
Zrc1
10
20
30
40
50
t/s

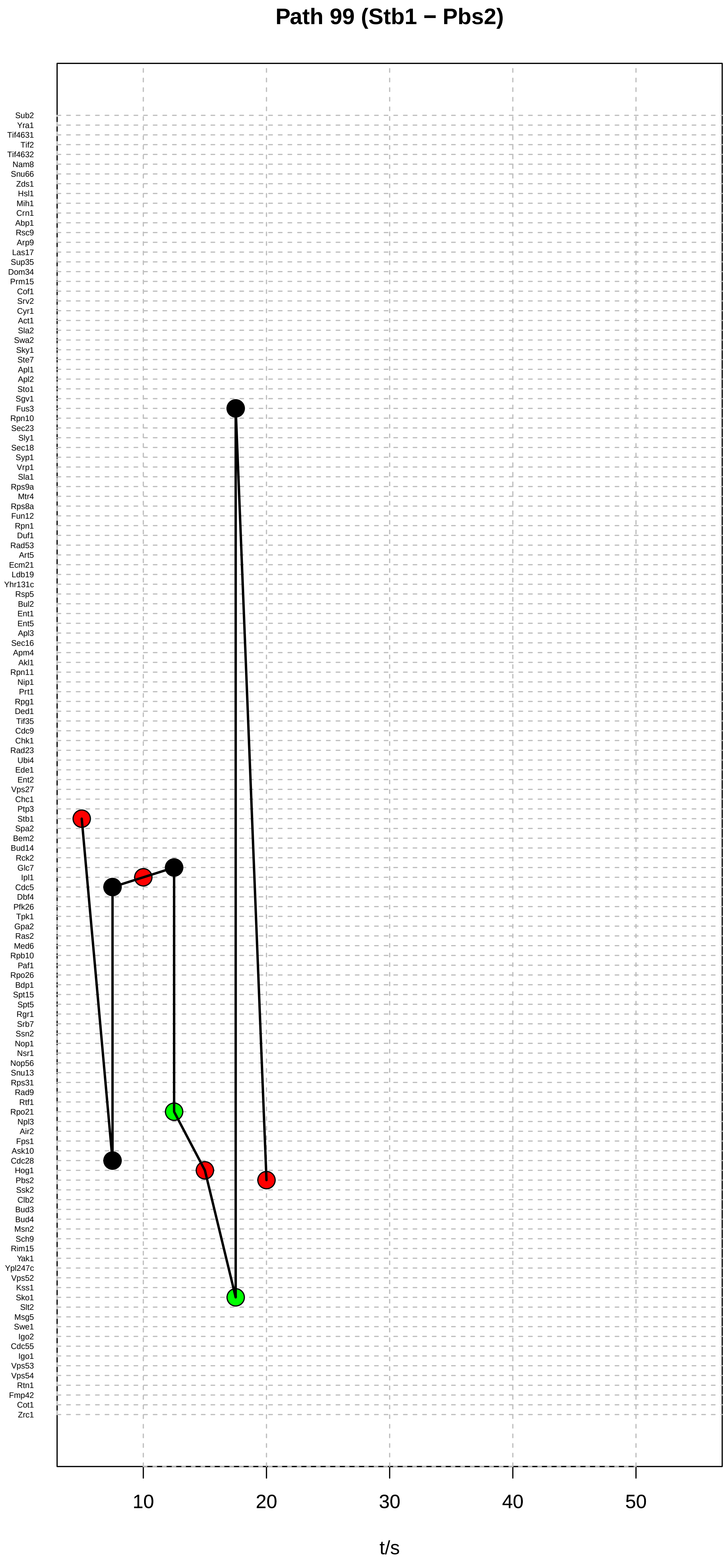
Path 99 (Stb1 − Pbs2)
Sub2
Yra1
Tif4631
Tif2
Tif4632
Nam8
Snu66
Zds1
Hsl1
Mih1
Crn1
Abp1
Rsc9
Arp9
Las17
Sup35
Dom34
Prm15
Cof1
Srv2
Cyr1
Act1
Sla2
Swa2
Sky1
Ste7
Apl1
Apl2
Sto1
Sgv1
Fus3
Rpn10
Sec23
Sly1
Sec18
Syp1
Vrp1
Sla1
Rps9a
Mtr4
Rps8a
Fun12
Rpn1
Duf1
Rad53
Art5
Ecm21
Ldb19
Yhr131c
Rsp5
Bul2
Ent1
Ent5
Apl3
Sec16
Apm4
Akl1
Rpn11
Nip1
Prt1
Rpg1
Ded1
Tif35
Cdc9
Chk1
Rad23
Ubi4
Ede1
Ent2
Vps27
Chc1
Ptp3
Stb1
Spa2
Bem2
Bud14
Rck2
Glc7
Ipl1
Cdc5
Dbf4
Pfk26
Tpk1
Gpa2
Ras2
Med6
Rpb10
Paf1
Rpo26
Bdp1
Spt15
Spt5
Rgr1
Srb7
Ssn2
Nop1
Nsr1
Nop56
Snu13
Rps31
Rad9
Rtf1
Rpo21
Npl3
Air2
Fps1
Ask10
Cdc28
Hog1
Pbs2
Ssk2
Clb2
Bud3
Bud4
Msn2
Sch9
Rim15
Yak1
Ypl247c
Vps52
Kss1
Sko1
Slt2
Msg5
Swe1
Igo2
Cdc55
Igo1
Vps53
Vps54
Rtn1
Fmp42
Cot1
Zrc1
10
20
30
40
50
t/s

#### Path 100 (Stb1 − Bud14)

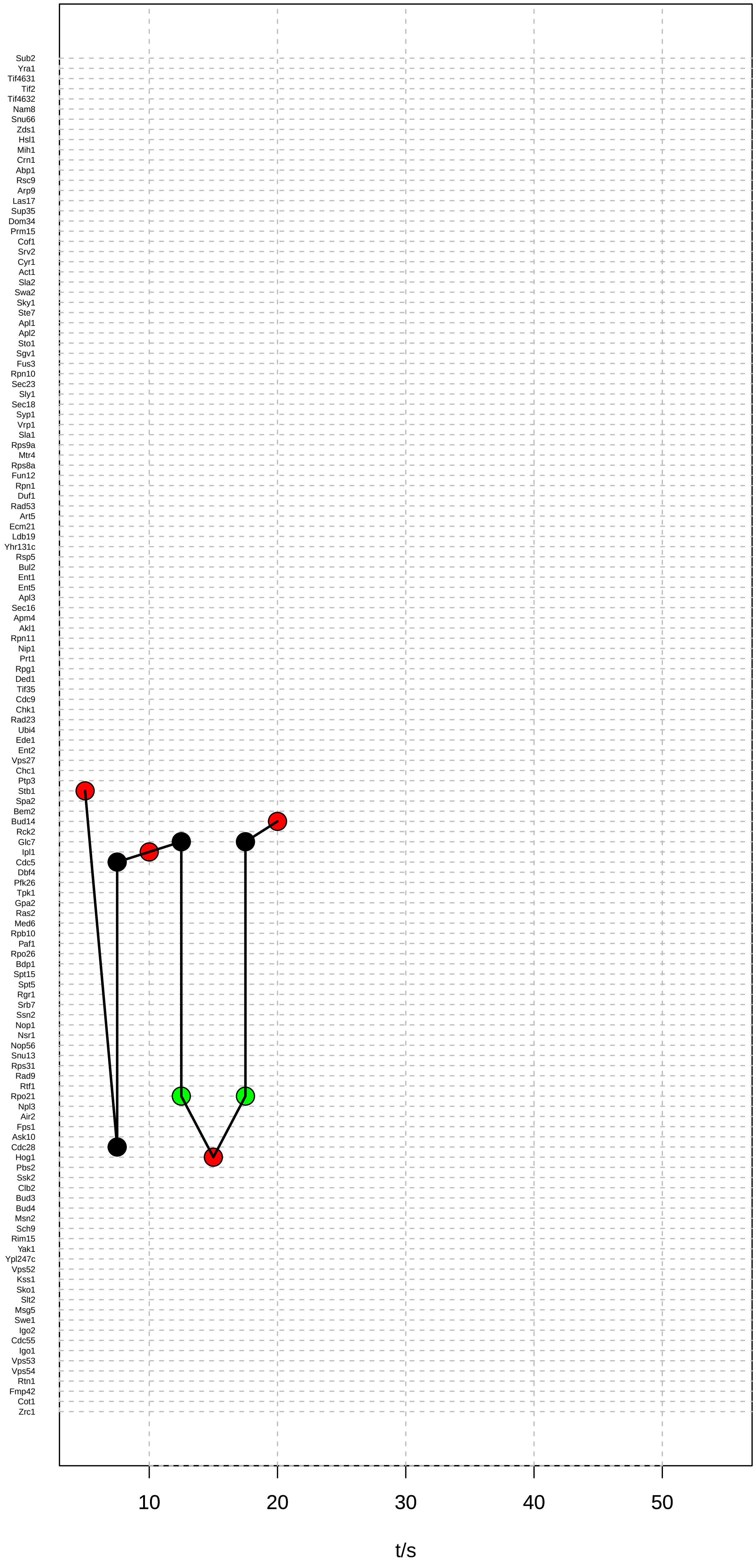

### Path 101 (Stb1 − Pbs2)

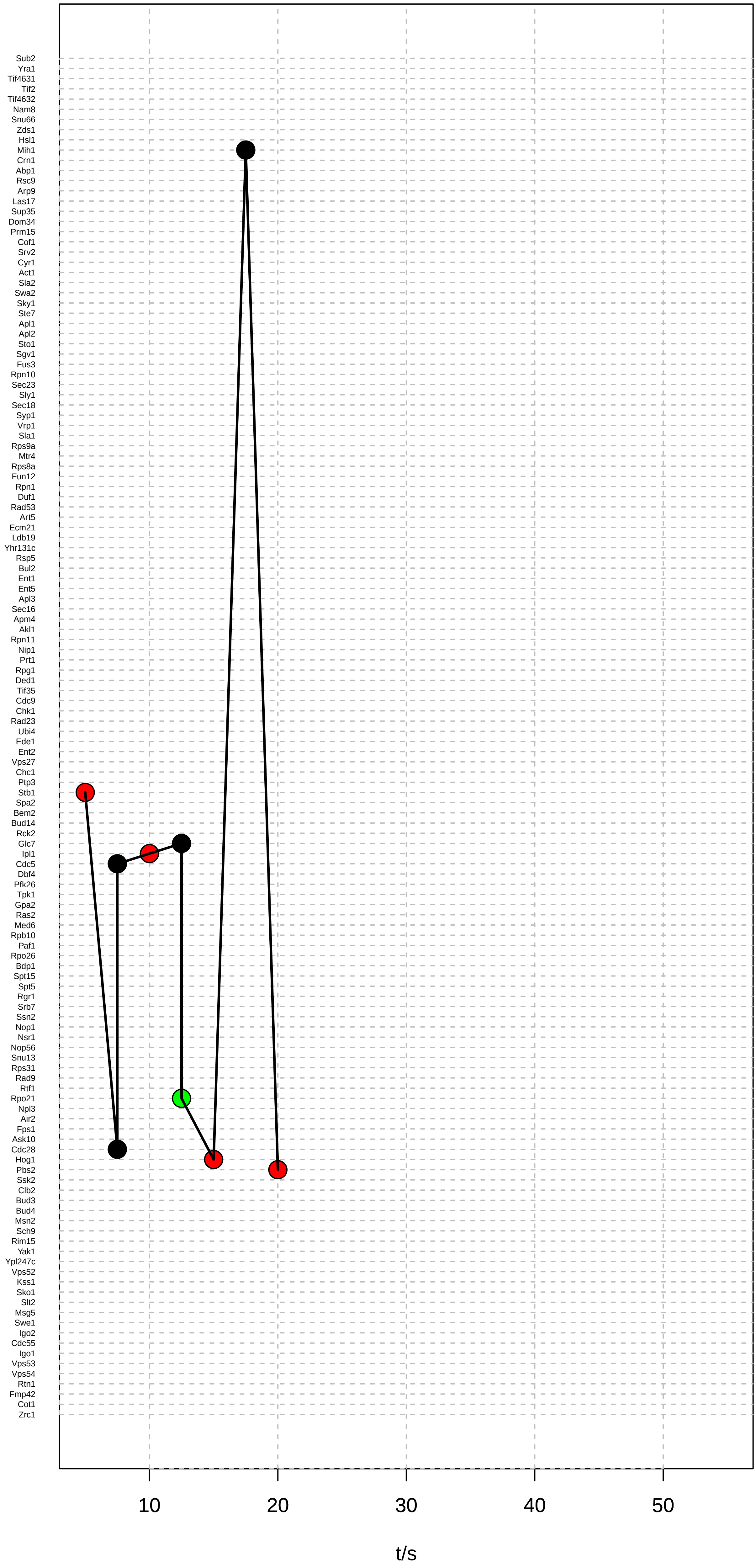

### Path 102 (Stb1 − Pbs2)

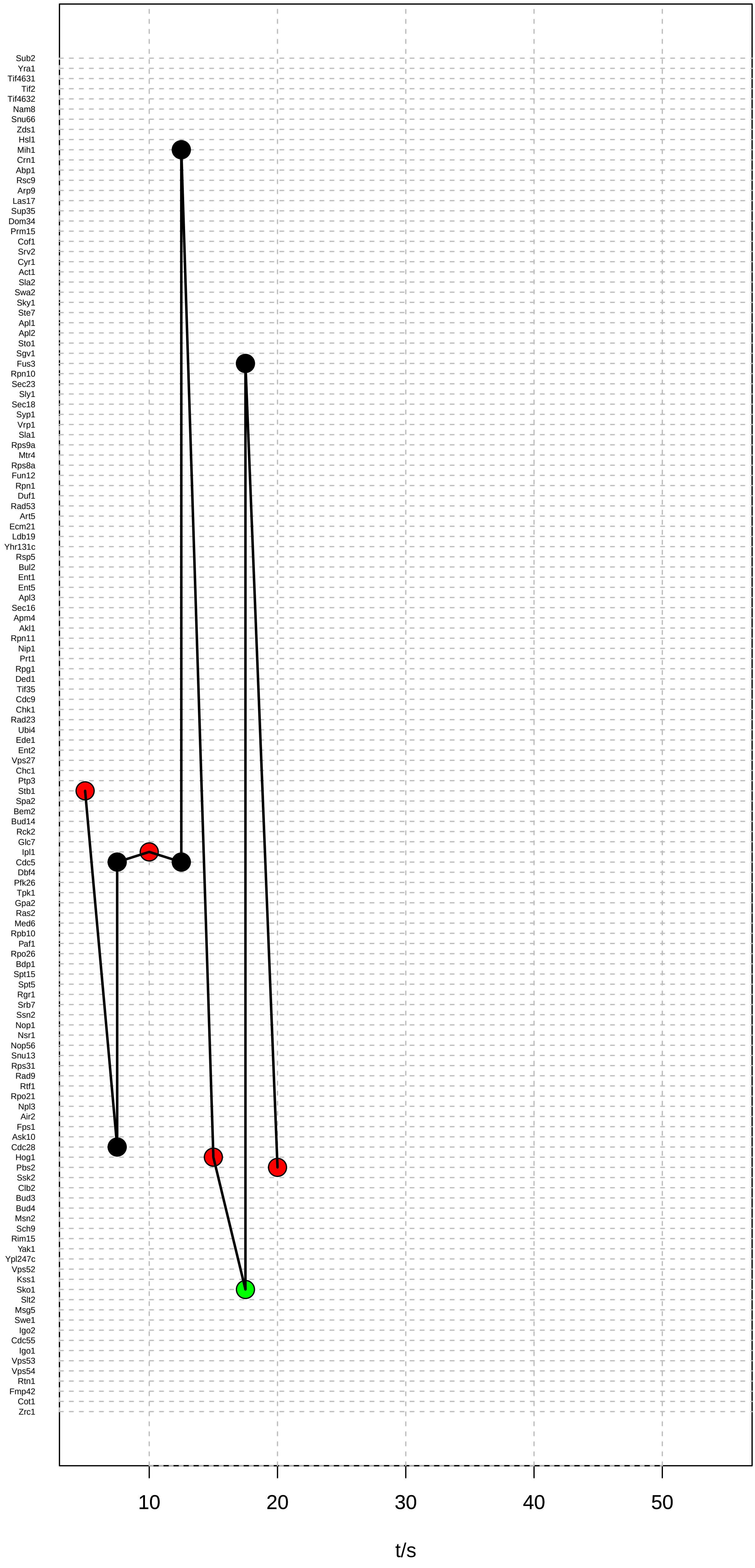

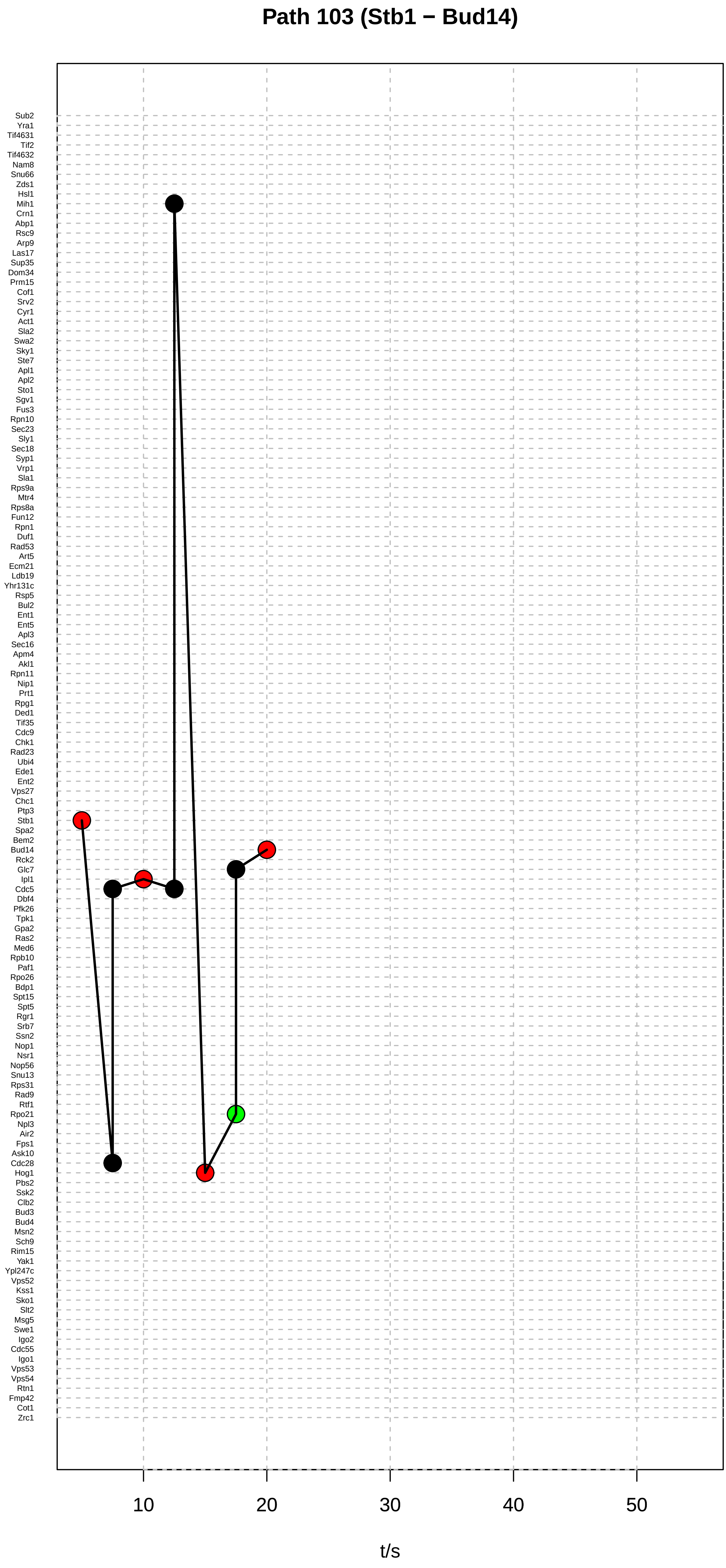

Path 103 (Stb1 − Bud14)
Sub2
Yra1
Tif4631
Tif2
Tif4632
Nam8
Snu66
Zds1
Hsl1
Mih1
Crn1
Abp1
Rsc9
Arp9
Las17
Sup35
Dom34
Prm15
Cof1
Srv2
Cyr1
Act1
Sla2
Swa2
Sky1
Ste7
Apl1
Apl2
Sto1
Sgv1
Fus3
Rpn10
Sec23
Sly1
Sec18
Syp1
Vrp1
Sla1
Rps9a
Mtr4
Rps8a
Fun12
Rpn1
Duf1
Rad53
Art5
Ecm21
Ldb19
Yhr131c
Rsp5
Bul2
Ent1
Ent5
Apl3
Sec16
Apm4
Akl1
Rpn11
Nip1
Prt1
Rpg1
Ded1
Tif35
Cdc9
Chk1
Rad23
Ubi4
Ede1
Ent2
Vps27
Chc1
Ptp3
Stb1
Spa2
Bem2
Bud14
Rck2
Glc7
Ipl1
Cdc5
Dbf4
Pfk26
Tpk1
Gpa2
Ras2
Med6
Rpb10
Paf1
Rpo26
Bdp1
Spt15
Spt5
Rgr1
Srb7
Ssn2
Nop1
Nsr1
Nop56
Snu13
Rps31
Rad9
Rtf1
Rpo21
Npl3
Air2
Fps1
Ask10
Cdc28
Hog1
Pbs2
Ssk2
Clb2
Bud3
Bud4
Msn2
Sch9
Rim15
Yak1
Ypl247c
Vps52
Kss1
Sko1
Slt2
Msg5
Swe1
Igo2
Cdc55
Igo1
Vps53
Vps54
Rtn1
Fmp42
Cot1
Zrc1
10
20
30
40
50
t/s

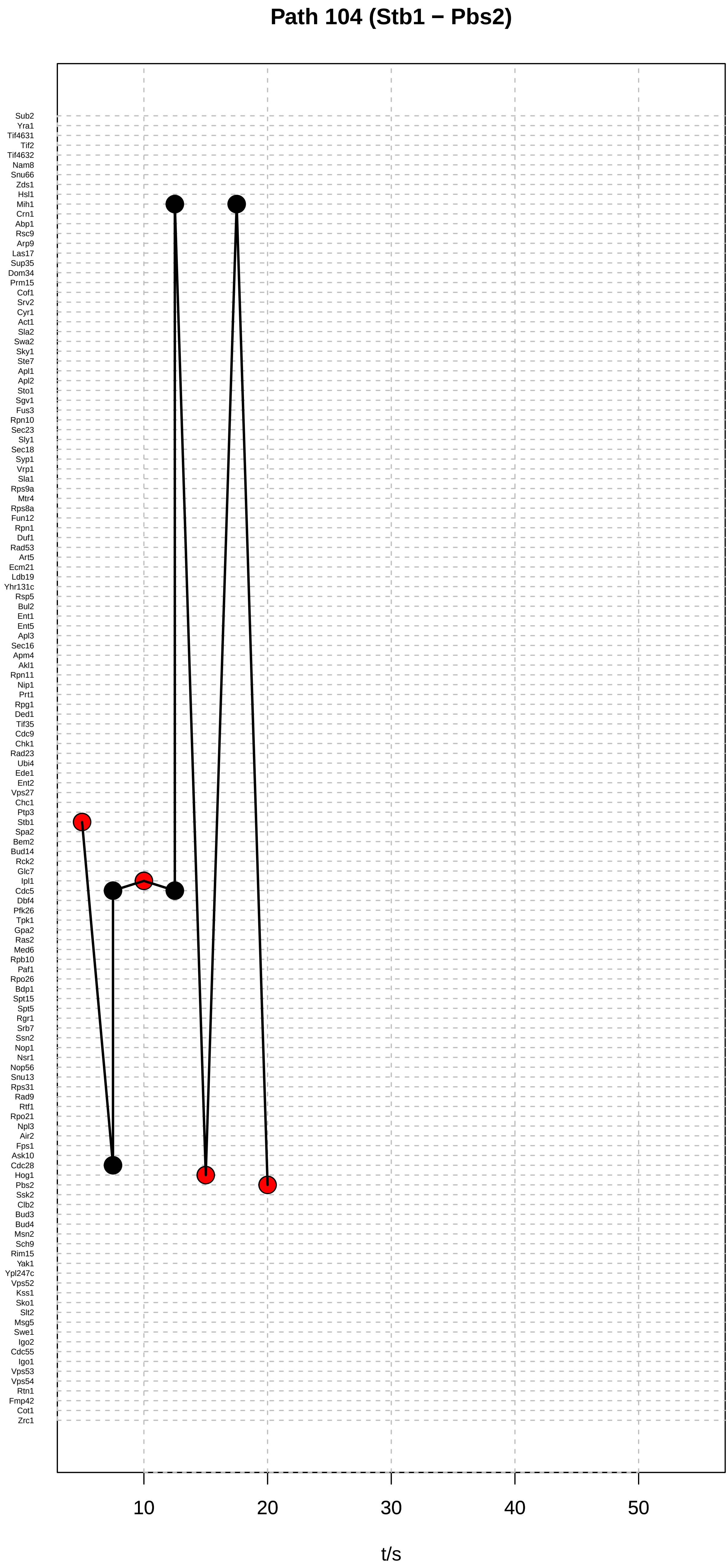

Path 104 (Stb1 − Pbs2)
Sub2
Yra1
Tif4631
Tif2
Tif4632
Nam8
Snu66
Zds1
Hsl1
Mih1
Crn1
Abp1
Rsc9
Arp9
Las17
Sup35
Dom34
Prm15
Cof1
Srv2
Cyr1
Act1
Sla2
Swa2
Sky1
Ste7
Apl1
Apl2
Sto1
Sgv1
Fus3
Rpn10
Sec23
Sly1
Sec18
Syp1
Vrp1
Sla1
Rps9a
Mtr4
Rps8a
Fun12
Rpn1
Duf1
Rad53
Art5
Ecm21
Ldb19
Yhr131c
Rsp5
Bul2
Ent1
Ent5
Apl3
Sec16
Apm4
Akl1
Rpn11
Nip1
Prt1
Rpg1
Ded1
Tif35
Cdc9
Chk1
Rad23
Ubi4
Ede1
Ent2
Vps27
Chc1
Ptp3
Stb1
Spa2
Bem2
Bud14
Rck2
Glc7
Ipl1
Cdc5
Dbf4
Pfk26
Tpk1
Gpa2
Ras2
Med6
Rpb10
Paf1
Rpo26
Bdp1
Spt15
Spt5
Rgr1
Srb7
Ssn2
Nop1
Nsr1
Nop56
Snu13
Rps31
Rad9
Rtf1
Rpo21
Npl3
Air2
Fps1
Ask10
Cdc28
Hog1
Pbs2
Ssk2
Clb2
Bud3
Bud4
Msn2
Sch9
Rim15
Yak1
Ypl247c
Vps52
Kss1
Sko1
Slt2
Msg5
Swe1
Igo2
Cdc55
Igo1
Vps53
Vps54
Rtn1
Fmp42
Cot1
Zrc1
10
20
30
40
50
t/s

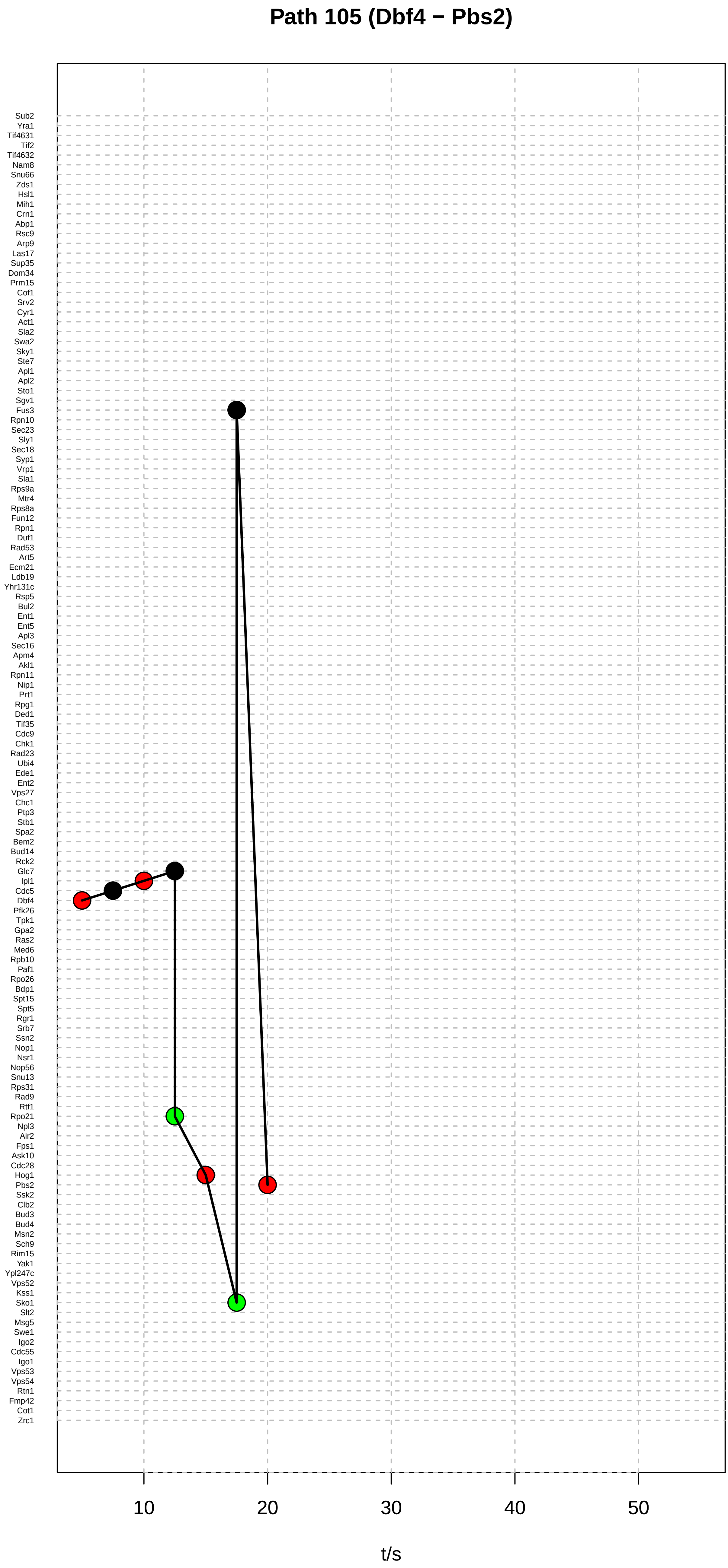

Path 105 (Dbf4 − Pbs2)
Sub2
Yra1
Tif4631
Tif2
Tif4632
Nam8
Snu66
Zds1
Hsl1
Mih1
Crn1
Abp1
Rsc9
Arp9
Las17
Sup35
Dom34
Prm15
Cof1
Srv2
Cyr1
Act1
Sla2
Swa2
Sky1
Ste7
Apl1
Apl2
Sto1
Sgv1
Fus3
Rpn10
Sec23
Sly1
Sec18
Syp1
Vrp1
Sla1
Rps9a
Mtr4
Rps8a
Fun12
Rpn1
Duf1
Rad53
Art5
Ecm21
Ldb19
Yhr131c
Rsp5
Bul2
Ent1
Ent5
Apl3
Sec16
Apm4
Akl1
Rpn11
Nip1
Prt1
Rpg1
Ded1
Tif35
Cdc9
Chk1
Rad23
Ubi4
Ede1
Ent2
Vps27
Chc1
Ptp3
Stb1
Spa2
Bem2
Bud14
Rck2
Glc7
Ipl1
Cdc5
Dbf4
Pfk26
Tpk1
Gpa2
Ras2
Med6
Rpb10
Paf1
Rpo26
Bdp1
Spt15
Spt5
Rgr1
Srb7
Ssn2
Nop1
Nsr1
Nop56
Snu13
Rps31
Rad9
Rtf1
Rpo21
Npl3
Air2
Fps1
Ask10
Cdc28
Hog1
Pbs2
Ssk2
Clb2
Bud3
Bud4
Msn2
Sch9
Rim15
Yak1
Ypl247c
Vps52
Kss1
Sko1
Slt2
Msg5
Swe1
Igo2
Cdc55
Igo1
Vps53
Vps54
Rtn1
Fmp42
Cot1
Zrc1
10
20
30
40
50
t/s

### Path 106 (Dbf4 − Pbs2)

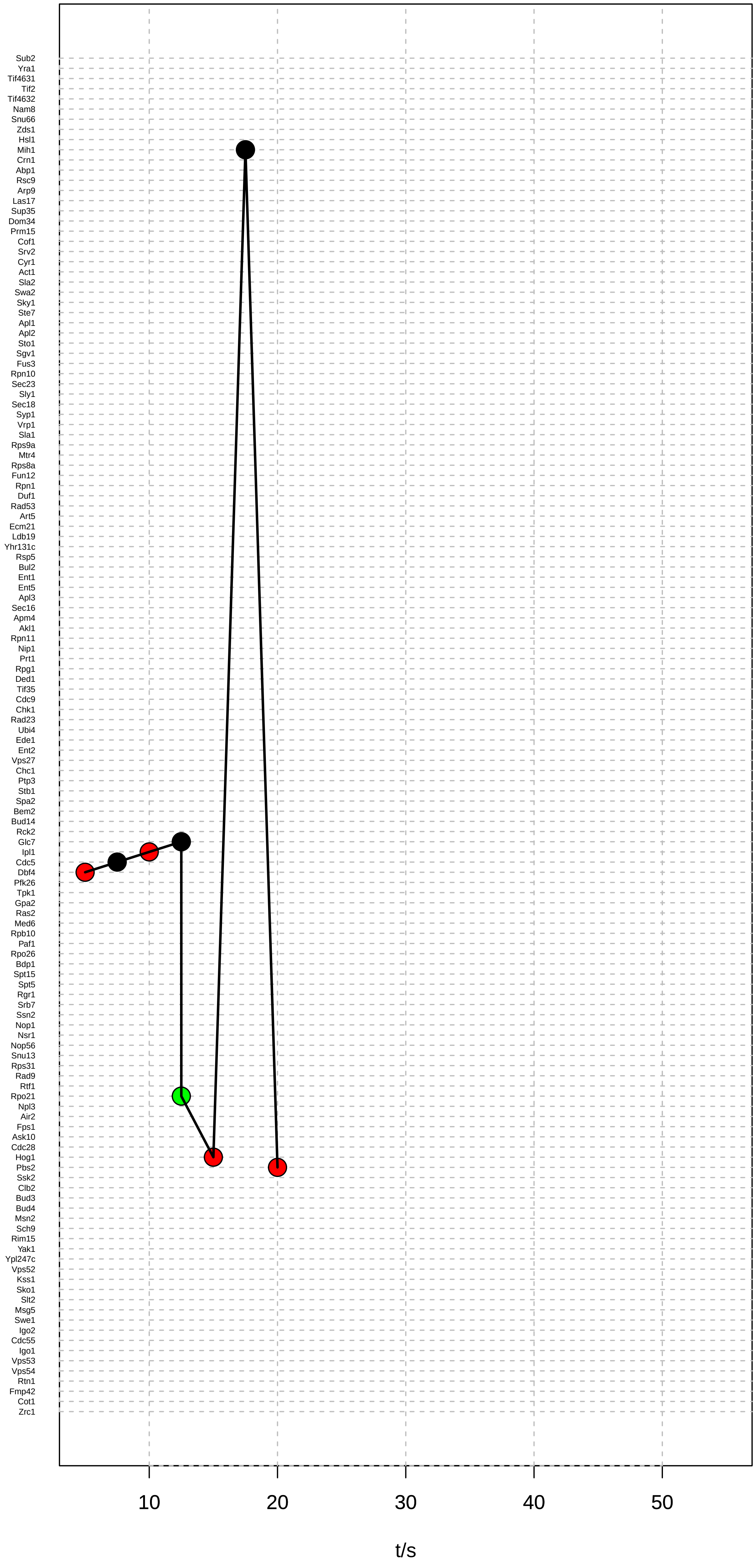

### Path 107 (Dbf4 − Pbs2)

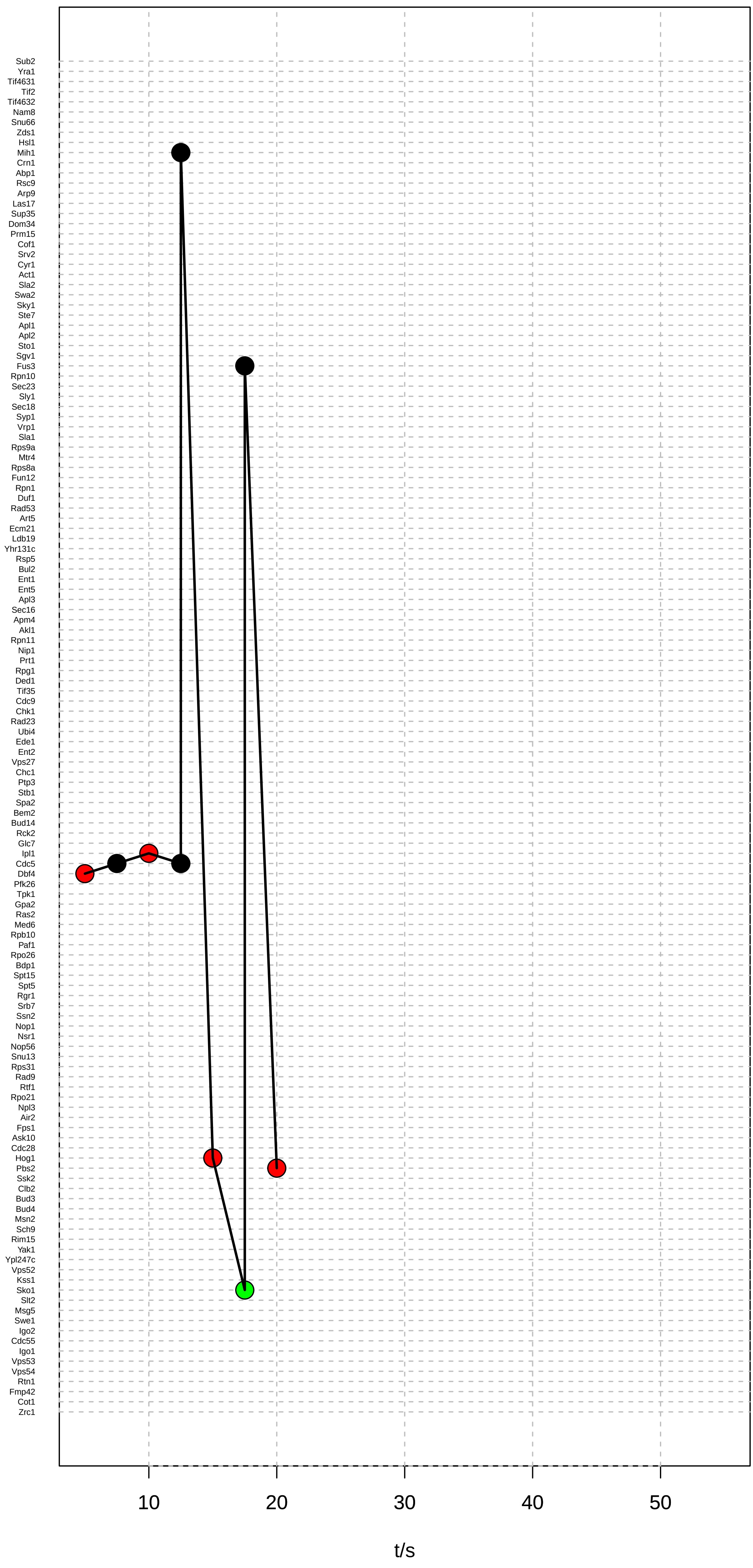

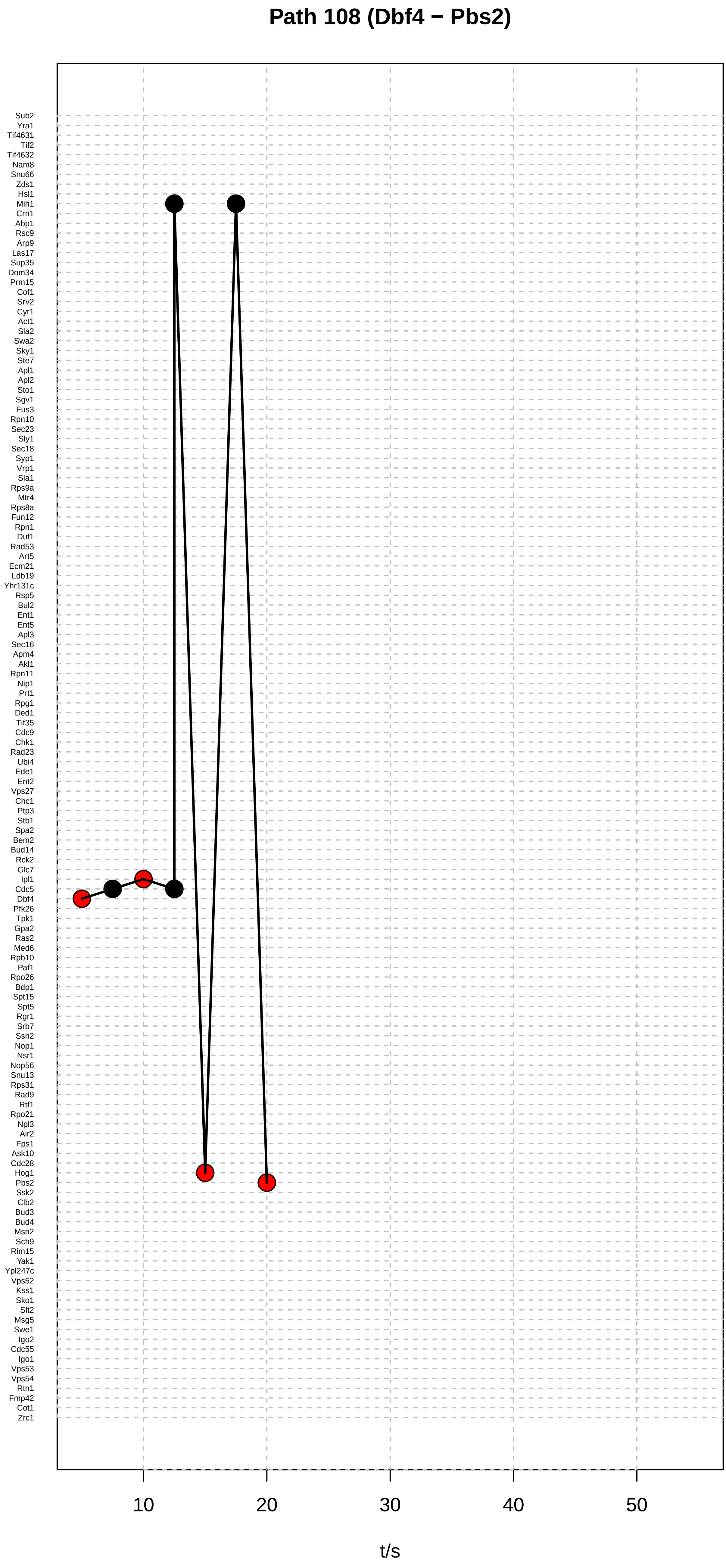

Path 108 (Dbf4 − Pbs2)
Sub2
Yra1
Tif4631
Tif2
Tif4632
Nam8
Snu66
Zds1
Hsl1
Mih1
Crn1
Abp1
Rsc9
Arp9
Las17
Sup35
Dom34
Prm15
Cof1
Srv2
Cyr1
Act1
Sla2
Swa2
Sky1
Ste7
Apl1
Apl2
Sto1
Sgv1
Fus3
Rpn10
Sec23
Sly1
Sec18
Syp1
Vrp1
Sla1
Rps9a
Mtr4
Rps8a
Fun12
Rpn1
Duf1
Rad53
Art5
Ecm21
Ldb19
Yhr131c
Rsp5
Bul2
Ent1
Ent5
Apl3
Sec16
Apm4
Akl1
Rpn11
Nip1
Prt1
Rpg1
Ded1
Tif35
Cdc9
Chk1
Rad23
Ubi4
Ede1
Ent2
Vps27
Chc1
Ptp3
Stb1
Spa2
Bem2
Bud14
Rck2
Glc7
Ipl1
Cdc5
Dbf4
Pfk26
Tpk1
Gpa2
Ras2
Med6
Rpb10
Paf1
Rpo26
Bdp1
Spt15
Spt5
Rgr1
Srb7
Ssn2
Nop1
Nsr1
Nop56
Snu13
Rps31
Rad9
Rtf1
Rpo21
Npl3
Air2
Fps1
Ask10
Cdc28
Hog1
Pbs2
Ssk2
Clb2
Bud3
Bud4
Msn2
Sch9
Rim15
Yak1
Ypl247c
Vps52
Kss1
Sko1
Slt2
Msg5
Swe1
Igo2
Cdc55
Igo1
Vps53
Vps54
Rtn1
Fmp42
Cot1
Zrc1
10
20
30
40
50
t/s
